# Novel Adomaviruses Associated with Blotchy Bass Syndrome in Black Basses (*Micropterus spp.)*

**DOI:** 10.1101/2025.06.01.657292

**Authors:** Luke R. Iwanowicz, Clayton D. Raines, Kelsey T. Young, Vicki S. Blazer, Heather L. Walsh, Geoff Smith, Cynthia Holt, John Odenkirk, Tom Jones, Jan-Michael Hessenauer, Morgan A. Biggs, Christopher B. Buck, Justin Greer, R. Scott Cornman

## Abstract

Black bass (*Micropterus spp.*) are the most important warmwater game fishes in the United States. They have high socioeconomic and recreational value and support an important aquaculture industry. Since 2008, fisheries managers have been reporting the observation of hyperpigmented melanistic lesions (HPMLs) on smallmouth bass (*M. dolomieu*) in different ecoregions of the United States. Similar HPMLs have been observed in largemouth bass (*M. nigricans*) since the 1980’s. Here, we report a close association between novel adomaviruses and the hallmark blotchy clinical presentation of hyperpigmented lesions on the skin smallmouth and largemouth black bass and provide evidence that satisfies Rivers’ postulates. The two adomaviruses are structurally and phylogenetically similar but share only 68.0% identity at aligned nucleotide sites and each has been found in only one host species to date. The manifestation of this skin disease appears to be seasonal in both species, primarily affects adults and is of unknown health consequence. Although the significance of infection to fish health remains unclear, understanding the disease ecology of these can inform biosecurity and the interjurisdictional movement of individuals. Moreover, as hyperpigmentation in other fish species is often idiopathic, our findings reframe perspectives for future investigations into this clinical presentation in other species.

## Introduction

Black bass (*Micropterus spp.*) are iconic North American sportfishes that support the most economically valuable freshwater sport fishery in the United States [1]. They are associated with a multibillion-dollar industry annually represented by recreational angling and aquaculture [2, 3]. Black bass are keystone predators within freshwater ecosystems and represent the most prized warmwater recreational sportfish in North America [4]. While indigenous to North America, largemouth bass (*M. nigricans,* LMB) and smallmouth bass (*M. dolomieu,* SMB) now have a cosmopolitan distribution as they have been introduced throughout the world due to their desirability as a sportfish [5]. Broodstock for black basses are maintained for private and public stocking. In addition, LMB production as a food fish is an emerging market in the United States and part of a thriving aquaculture industry in Asia [3, 6].

Between 2003 and 2010 there was an increased frequency of mortality events, low relative abundance, and poor year classes of SMB in specific areas of the Susquehanna and Potomac River Basins of the eastern United States [7, 8]. Coincident with these reports were increased observations of macroscopic lesions of the integument associated with opportunistic microbes [9]. These prolonged mortality events drew public attention to fish and ecosystem health. Notably, as these widespread mortality events abated in the Susquehanna River system during the latter end of this timeline (ca. 2010), hyperpigmented melanistic skin lesions (HPMLs) in SMB previously unreported in this watershed were increasingly observed with a springtime prevalence as high as 55% [10]. Interestingly, fisheries managers started observing or receiving reports of HPMLs on SMB in disparate watersheds in the United States around this same time (Lake St. Clair, MI in 2008, Lake Champlain, VT in 2009; rivers of the Great Lakes Region in 2010; Potomac River, WV in 2012)[10, 11]. The appearance of HPMLs among black bass has become so commonplace that the condition is colloquially referred to as “blotchy bass syndrome” (BBS) by anglers and resource managers. The first official report of BBS in LMB dates to a case in 1984 from the Hudson River, NY (S1 Fig) [12]. Although anecdotal evidence suggests sporadic observations of BBS in LMB for the last 40 years or more, documented accounts primarily exist in online message boards, angler-supported social media resources, or non-interpretive reports [13].

HPMLs associated with the integument have been reported in marine and freshwater fishes worldwide [10, 14–18]. These lesions often manifest as macroscopic, discrete, focal or multifocal areas of hypermelanization on the external surface of the fish. Histological descriptions of these lesions typically include an atypical increased density of melanocytes within the epidermal and dermal strata of the skin, disruption of the basement membrane, or pleiomorphic cell morphology [15, 19]. Other than black spot disease that is caused by parasitic infections in fishes and is clinically distinguishable from BBS, modest efforts have been made to identify the etiology of HPMLs [20–22]. Diagnostic analysis including histopathology, culture-based virology, molecular and classical bacteriology and electron microscopy have been attempted to identify putative microbial agents in many other instances; however, such lesions are typically idiopathic in nature. Hypermelanization has been ascribed to possible exposure to environmental contaminants, exposure to ultraviolet radiation or other unspecified environmental stressors, or genetics, yet there is a paucity of experimental evidence to support these candidate causes [14, 23].

Despite nearly a half century of BBS observations in black basses, little progress has been made to resolve the etiology of this idiopathic syndrome. We recently found evidence of a putative adomavirus associated with HPMLs in SMB [10]. To shed further light on a possible causal relationship, we searched for similar viruses in LMB with HPMLs. Here, we present complete genomes of two phylogenetically related, but clearly different species of adomaviruses associated with the HPMLs hallmark of BBS and provide phylogenetic evidence that the emergence of this disease in SMB may not be the result of recent introductions.

## Materials and Methods

### Specimen Sampling

All live animal handling and use procedures were reviewed and approved by the U.S. Geological Survey, Eastern Ecological Science Center Animal Care and Use Committee (IACUC 07001, 2020-06 and 2021-17L). Samples were acquired from both LMB and SMB from a variety of natural aquatic settings including the Susquehanna River Basin (PA), the Potomac River Basin (VA), Lake St. Clair (MI), Lake Champlain (VT) and western Texas. Alternatively, fish maintained in aquatic exhibits at Bass Pro Shops and Cabela’s with HPMLs were sampled (S1 Table).

Most of the samples utilized for this research were collected with non-lethal, minimally invasive approaches. Archived samples collected from tandem research efforts in 2017 and 2019 were also used [10, 24, 25]. Those fish were euthanized with an overdose of tricaine methanesulfonate (MS-222, Argent Labs, Redmond, WA). Otoliths were removed for ageing and phenotypic sex was determined in lethally sampled fishes. Age and sex could not be determined for non-lethally sampled individuals. Depending on the sample, clinically affected or normal scales were collected for analysis, and nucleic acids were preserved in RNA*later* (Thermo Fisher Scientific, Waltham, MA) or DNA/RNA Shield (Zymo Research, Irvine, CA). Most non-lethal sampling included the use of swabs and collection tubes containing DNA/ RNA Shield. The HPMLs or normal skin were sampled by rubbing the swab back and forth with gentle, yet firm pressure for 10-15 seconds. The swab head was then rotated such that the entire surface contacted the sample area. Transfer of melanin to the swab confirmed tissue transfer in the case of HPMLs (S1 Video). Swabs were then inserted in the collection tube for subsequent nucleic acid extraction. Details regarding specific methods are provided as supplemental material (S1 Text)

### Nucleic acid extraction, sequencing and *de novo* assembly of viral genomes

Nucleic acid extraction, sequencing and *de novo* assembly methods differed across the sample sets utilized for downstream analysis and are included as supplemental materials (S1 Text).

### Identification of virus positive samples and resolving repetitive regions

PCR primers were designed to amplify a loci of the RepE1 gene in each virus. The forward primers were anchored in the parvovirus non-structural protein NS1 conserved domain for MdA-1 and MnA-1 to minimize the likelihood of target sequence mismatch across samples given uncertainly in sequence variability. Primers were designed using the modified version of Primer3 2.3.7 bundled within Geneious Prime v2020.0.5 [26]. Primer sequences are listed in Table 1.

**Table 1.**
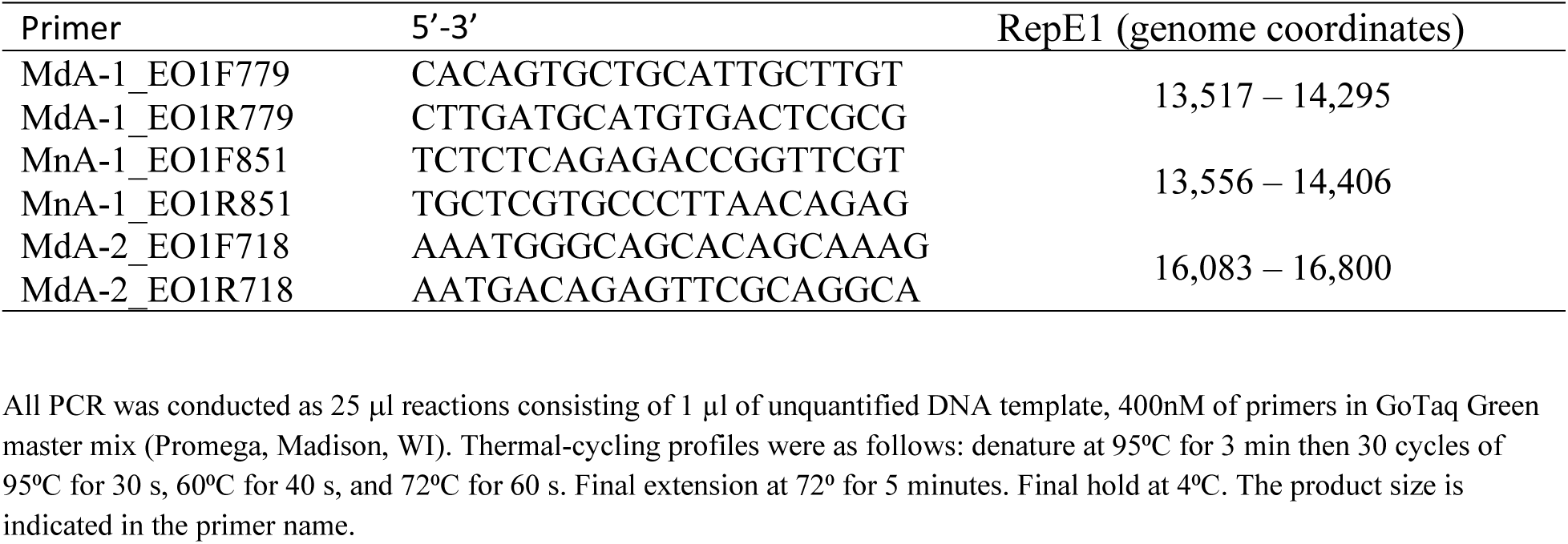
Primers and thermal cycling conditions for the amplification of bass adomavirus helicase genes.

Conventional PCR was conducted as 25 μl reactions consisting of 1 µl of unquantified DNA template, 400nM of primers in GoTaq Green master mix (Promega, Madison, WI). Thermal-cycling profiles were as follows: denature at 95⁰C for 3 min then 30 cycles of 95⁰C for 30 s, 60⁰C for 40 s, and 72⁰C for 60 s. Final extension at 72⁰ for 5 minutes. Final hold at 4⁰C. The product size is indicated in the primer name. A subset of RepE1 products PCR products were purified using a QIAquick PCR Purification Kit (Qiagen, Hilden, Germany) and confirmed via Sanger sequencing.

### Comparison of adomaviruses from large and smallmouth bass

Complete genomes of the black bass adomaviruses were ordinated such that the linearized sequence terminated at the beginning of the SET ORF. Open reading frames were predicted using Geneious Prime. Candidate protein sequences without clear hits via BLASTP were analyzed with HHpred using the NCBI Conserved Domains, PDB_mmCIF70, Pfam-A, and UniProtSwissProt-viral70 databases [27]. Sequences without clear hits in HHpred searches were queried via Dali using predicted protein structure from AlphaFold2 or RoseTTAfold predictions [28–30]. In some instances, we assigned protein names based on synteny even when not supported by prediction queries as not to further infer function in the absence of experimental data [31, 32].

Genomes were aligned using MUSCLE v5.1 bundled with Geneious Prime software v2024.0.2 to determine pairwise identity [33]. All homologous proteins were aligned to determine protein identity in a similar fashion.

### Visualizing viral nucleic acid (RNAscope)

In order to visualize viral nucleic acids in HPMLs we developed an *in situ* hybridization RNAscope® assay that utilized a 15ZZ probe targeting nucleotides 4 - 778 of the MdA-1 adenain gene ([UFQ21633](https://www.ncbi.nlm.nih.gov/nuccore/MZ673484.1?from=9800&to=10561)). A 12ZZ probe targeting nucleotides 2 – 684) of the MnA-1 adenain gene ([XQZ12369](https://www.ncbi.nlm.nih.gov/nuccore/PV430023.2?from=9524&to=10366)) was also developed. Because of the small, low complexity dsDNA genome, this probe effectively targets both viral mRNA and genomic DNA. We analyzed SMB samples from a health assessment study in the Susquehanna River, PA collected during fall 2013 and spring 2024 [24, 34]. Samples from LMB were collected from Raystown Lake, PA during the spring of 2024. Normal skin tissue as well as skin with HPMLs stabilized in PAXgene® Tissue FIX (PreAnalytiX, Switzerland) were decalcified with 0.5M EDTA for 48 hr, embedded into paraffin, and sectioned at 5 μm. RNAscope (Advanced Cell Diagnostics, Newark, California) was used to observe the localization of the target gene nucleic acids. Selected sections with HPMLs were deparaffinized with three changes of Pro-Par clearant (Anatech Ltd., Michigan) for 5 min each and rehydrated with a graded ethanol series of 100%, 95%, 80%, 70%, and 50% for 3 min each and air dried. RNAscope was conducted according to the manufacturer’s protocols for the RNAscope Multiplex Fluorescent Reagent Kit v2 Assay. The RNAscope Protease Plus incubation step was omitted given that assay optimization identified that PAXgene preserved samples were overly digested. The target probes were heated to 40°C for 10 min, cooled to room temperature and hybridization was carried out at 40°C for 2 h in an InSlide Out hybridization oven (Boekel, Pennsylvania). A negative control consisted of the probe diluent only. Sample controls included normal no HPML) skin. Slides were counterstained with Sudan Black B for 30 s, rinsed clear with deionised water and mounted with ProLong Gold Antifade Mountant (Thermo Fisher Scientific, Carlsbad, California). Slides were imaged with a Keyence BZ-X810 (Itasca, IL). Fluorescent signal from viral nucleic acids was visualized (ex: 470 ± 40nm; em: 525 ± 50nm). Digital images were imported into GIMP 2.10.32 to construct composite figures and correct for autofluorescence [24].

### Viral gene expression (RNA-seq)

We evaluated the expression of viral transcripts in HPMLs of smallmouth bass collected in 2017 (BioProject PRJNA530557) and 2022 (BioProject PRJNA1254126) as well as largemouth bass collected in 2022 (BioProject PRJNA1253685). Adapter removal and quality control of the raw sequencing reads was performed in trimmomatic with the parameters set as LEADING:5 TRAILING:5 SLIDINGWINDOW:4:15 MINLEN:25. Cleaned reads passing quality control were first aligned to the host transcriptome obtained from the NCBI (LMB: accession [GCF_014851395.1_ASM1485139v1](https://www.ncbi.nlm.nih.gov/datasets/genome/GCF_014851395.1/), SMB: accession [GCF_021292245.1_ASM2129224v1](https://www.ncbi.nlm.nih.gov/datasets/genome/GCF_021292245.1/)) to remove reads originating from the host. Reads were aligned to the appropriate transcriptome using salmon (v1.4.0) in mapping mode with GC bias and selective alignment [35]. Parameter settings were maintained for both paired-end (2022) and single-end (2017) sequencing datasets. Reads that did not align to the host transcriptome were obtained using –writeUnmappedNames in salmon followed by sequence extraction using subseq from the seqtk package [36]. These reads were then aligned all non-overlapping ORFs of MdA-1 or MnA-1 using salmon to quantify the abundance of viral transcripts.

Viral transcript quantification was imported into R using tximport and counts were normalized in DESeq2 using the counts function with normalized = TRUE [37, 38]. Viral transcripts were not detected in clinically normal fish or regions not presenting hyperpigmented lesions and these samples were therefore removed prior to downstream visualization in order to appropriately calculate dispersion and count variance.

### Phylogenetic analysis

We recovered adomavirus protein sequences of the replicase gene (RepE1, RepLT or CressRep), Adenain, and Hexon from NCBI (S2 Table). Sequences were aligned using MUSCLE v5.1 bundled within Geneious Prime software v2024.0.2 [33]. Alignment files were used to infer phylogenetic trees in IQ-Tree 1.6.12 for each of the viral proteins [39].

Phylogenetic trees were visualized using iTOL v6 [40]. Pairwise identity of translated amino acid sequences for these proteins were determined and visualized using the Sequence Demarcation Tool [41].

## Results

### Gross presentation of blotchy bass syndrome and site-specific prevalence

The HPMLs observed on both bass species investigated here shared an indistinguishable gross presentation (Fig 1). These areas of hyperpigmentation presented on the external surface of fish including the fins and margins of the oral cavity. Lesions consisted of sharply circumscribed, melanistic macules of variable size with irregular, asymmetric borders surrounded by normal-appearing skin. The gross morphology of discrete, melanistic blotches was visually suggestive of melanin irregularly radiating from a central focal point. In some instances, these blotches coalesced into more intricate, amorphous patterns. Of note, during the late spring these HPMLs were often fragile and the integument in these focal areas was disrupted with gentle pressure (S1 Video). HMPL fragility was not observed in the fall. Gross and histological descriptions in SMB have been reported previously [10].

**Fig 1.**
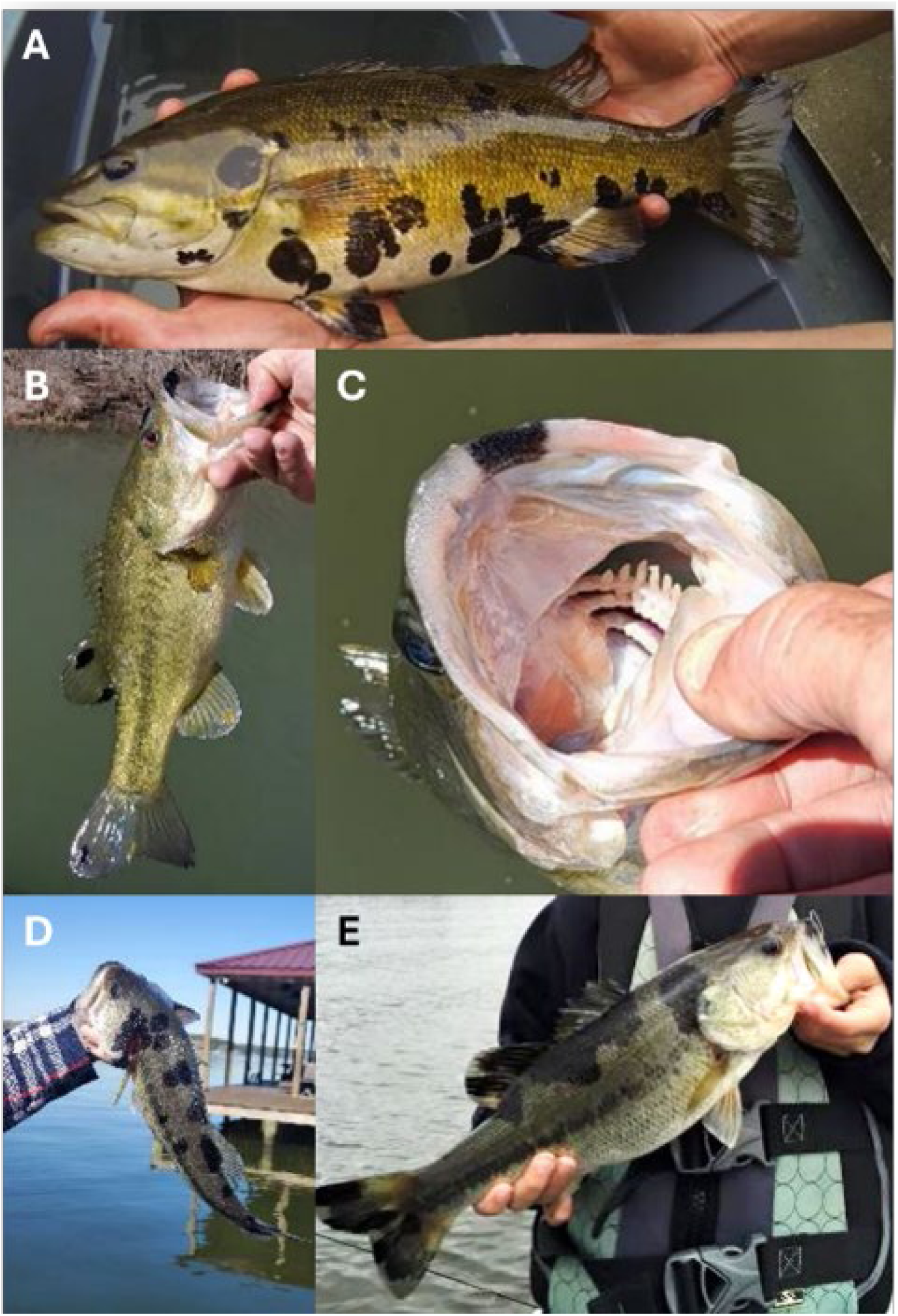
Gross presentation of blotchy bass syndrome in smallmouth and largemouth bass. Hyperpigmented lesions (HPMLs) on the lateral surface of a smallmouth bass (A), HPMLs on the dorsal and caudal fins of a largemouth bass (B), HPML along the premaxilla (C), HMPLs along the dorsolateral surface of a largemouth bass (D), and HPMLs coalescing into a significant area of hyperpigmentation (E).

Comprehensive size-distribution surveys of SMB were initiated in Lake St. Clair, MI during 2002. These surveys were always conducted during May prior to spawning. In 2008 the hallmark presentation of BBS was first observed at a frequency such that fisheries managers began documenting this external anomaly (Fig 2A). Since 2008, the prevalence of blotchy SMB ranged from 1.7 - 9.2% (x̅=5.1%; n=5,893). These surveys primarily targeted adults (180 - 560mm) and the smallest blotchy bass was 320 mm. Although blotchy LMB are observed in Lake St. Clair, a similar prevalence dataset is not available.

**Fig 2.**
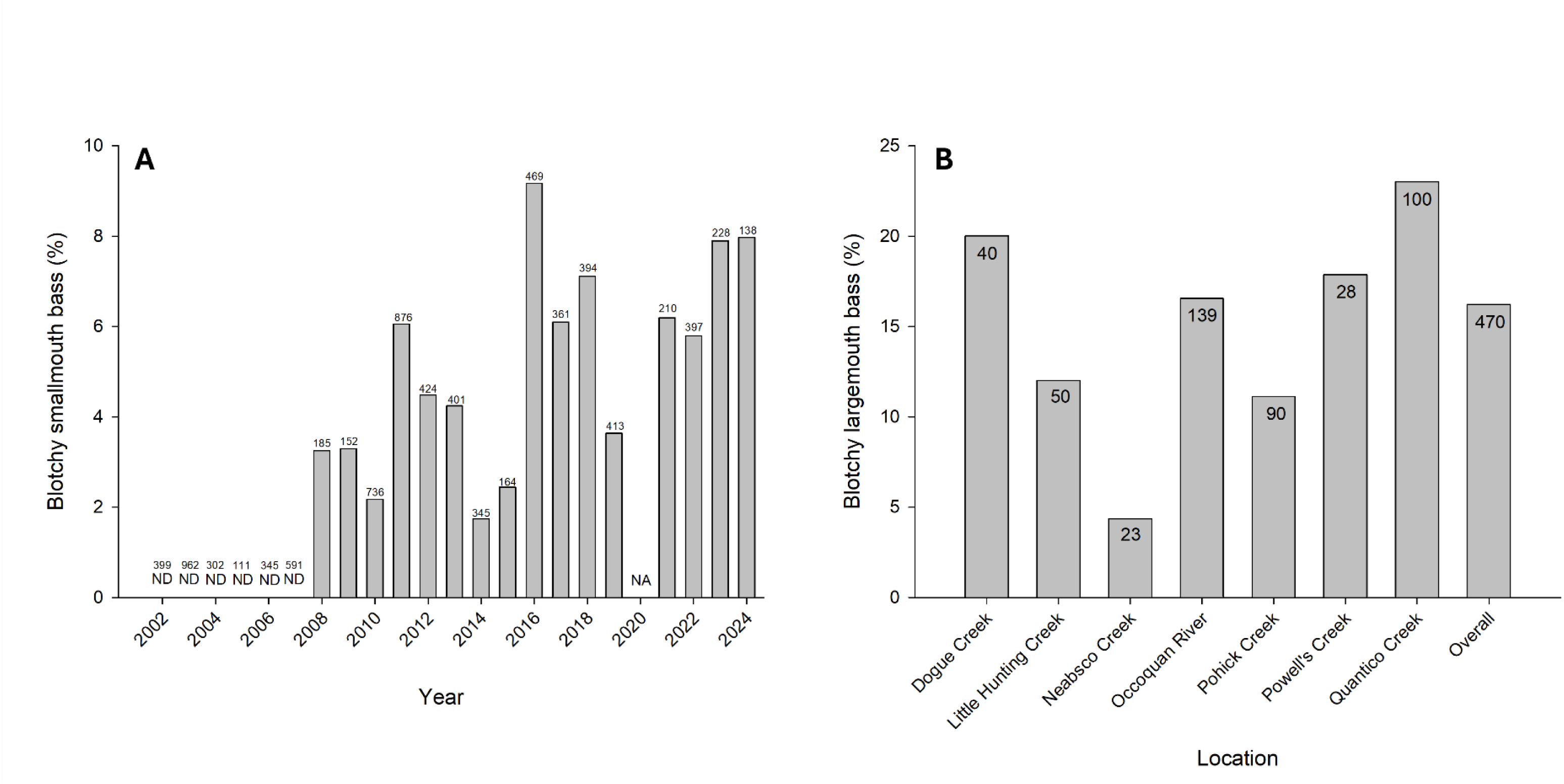
Prevalence of blotchy bass in different geographic locations. (A) Smallmouth bass were samples from Lake St Clair, Minnesota and (B) largemouth bass were sampled from tributaries of the Potomac River in Virginia. Sample size is indicated for each collection. ND indicates not detected, NA indicates data not collected.

The Virginia Department of Wildlife Resources performed mark-recapture surveys between March and April 2021 to estimate LMB population size structure across 7 tributaries in the Potomac River Watershed. The prevalence of BBS was also documented during this pre-spawn sampling. Blotchy LMB were observed at all sample locations (Fig 2B). The smallest blotchy fish was 278 mm. Prevalence of blotchy LMB ranged from 4.3 - 23.0% (x̅=16.2%; n=470).

### Association of adomaviruses with HPMLs

Initial transcriptome assemblies generated during a previous study identified a papillomavirus helicase-like sequence (MT010627) in HPMLs of SMB, motivating additional DNA sequencing to complete a reference genome (10). The genome model presented here proposes a circular dsDNA of 16,002 bp (MZ673484) with gene content and phylogenetic position consistent with other fish-associated adomaviruses (S2 Fig). We refer to this virus as *Micropterus dolomieu adomavirus 1* (MdA-1). In addition, a variant of MdA-1 was recovered from DNA sequencing of an HPML-affected SMB sampled from the mainstem of the Susquehanna River (SUSHM8; PV454363). We recovered a complete genome of an additional novel adomavirus (*Micropterus nigricans adomavirus 1*; PV430023) with a circularized length of 15,989 bp from a HPML lesions affecting an LMB specimen from Lake St. Clair, MI (S2 Fig).

Although the intent of this research was not to evaluate the geospatial distribution of BBS, the external presentation of pathognomonic HPMLs, hallmark of this disease, facilitated observation and collection of samples from geographically disparate regions of the United States. To confirm an association of adomaviruses with HPMLs we screened samples from clinically normal skin and HPMLs for the presence of viral DNA. We detected the viral helicase gene RepE1 in 100% and 95.2% of HPMLs from samples screened for MdA-1 (n=49) and MnA-1 (n=63), respectively (S1 Table). PCR product for RepE1 was never detected in normal skin samples (n=14) from clinically affected fish nor clinically normal fish (n=20). We screened two raised mucoid skin lesions (RMSLs) on SMB skin (S3 Fig), a lesion-type which we have previously ascribed to a different adomavirus, MdA-2 [42]. They were both PCR positive for MdA-2, but negative for MdA-1. Of note, both MdA-1 and MdA-2 were present in metagenomic and metatranscriptomic libraries of RMSLs (SRR10512798 and SRR10540651, respectively) deposited in the Sequence Read Archive (SRA) based on PebbleScout queries and local *in silico* mapping [43]. Albeit, MdA-1 was present orders of magnitude lower than MdA-2 (29x and 57260x coverage respectively). Further, we never detected MdA-1 in LMB nor MnA-1 in SMB.

### Relative expression of viral genes in HPMLs

We evaluated the relative expression of viral genes in skin samples from LMB and SMB. The intention was to verify active infection and survey expression patterns of viral genes.

Analyses included SMB collected from the Susquehanna River Watershed (PA) in late April 2017 and May 2022, and LMB from Little Hunting and Dogue Creek (VA) in April 2022. Low variability in expression profiles across temporally matched samples (Fig 3A and B) suggested a seasonal synchronicity of the viral lifecycle in naturally infected fish. Notably, expression patterns were similar between MdA-1 and MnA-1 in the different bass species collected during 2022 suggesting similar lifecycle strategies between these viruses. Across samples, the viral helicase gene (RepE1) presumed to be an early expressed gene was among the least abundant viral transcripts suggesting that fish were in late-stage infection.

**Fig 3.**
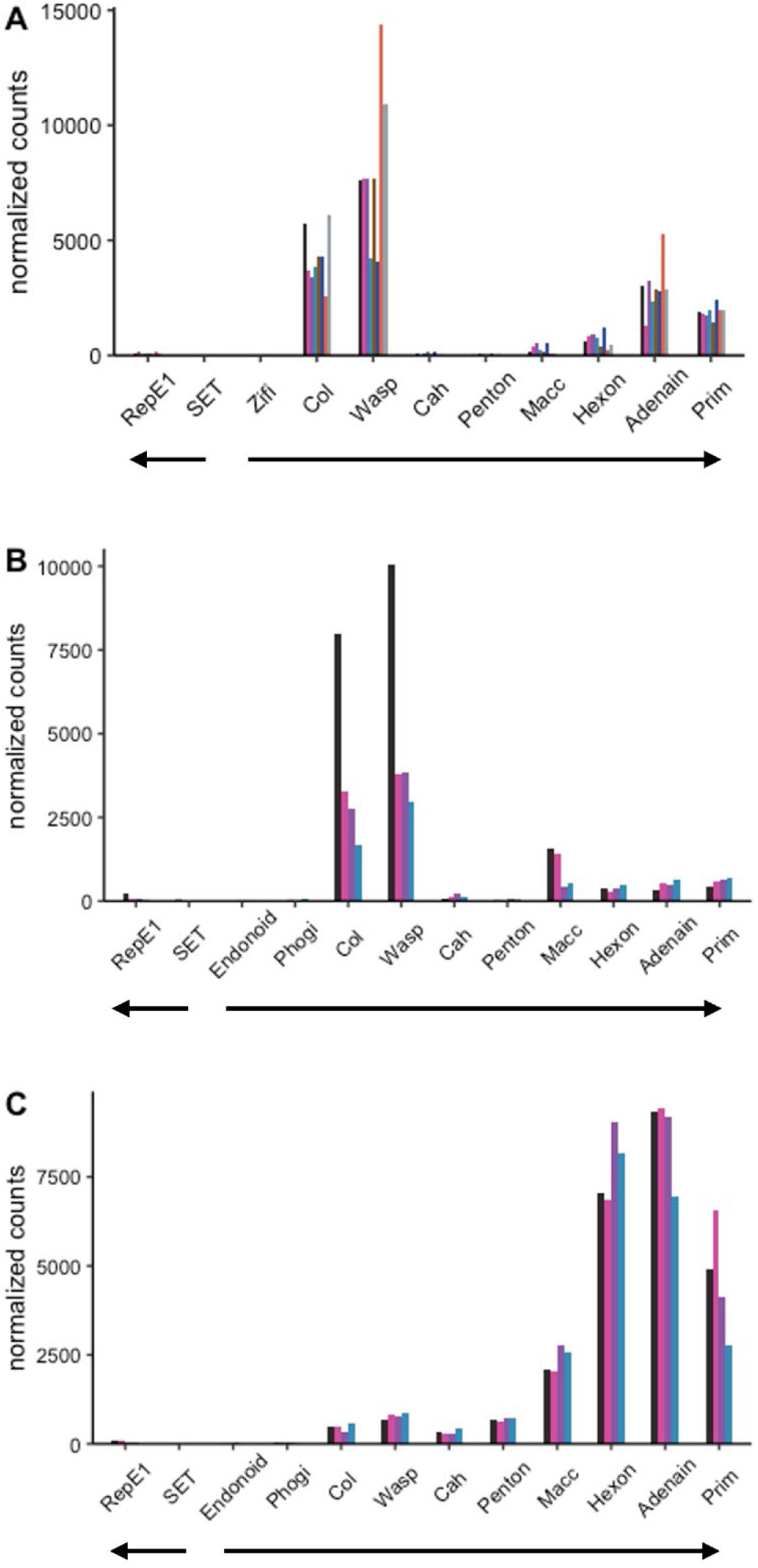
Expression of viral genes in hyperpigmented lesions in large and smallmouth bass. Viral transcript expression was detected in all swabs collected from areas presenting hyperpigmented lesions. Viral transcripts were not detected in swabs from skin regions without hyperpigmentation or in clinical normal fish. (A) MnA-1 in largemouth bass from VA sampled in 2022 (B) MnA-1 from smallmouth bass sampled in PA during 2022 (C) MdA-1 from smallmouth bass sampled in PA during 2017. Genes are arranged in genomic order of open reading frames. Bar colors within each panel indicate transcripts for specific individual fish. Block arrows indicate coding genomic strand.

In contrast, MdA-1 expression profiles from SMB samples collected in 2022 were quantitatively dissimilar from fish collected in 2017 (Fig 3B and C). Although this may highlight annual variation (April 2017 vs May 2022), expression patterns between fish from the same site during the same sampling season were remarkably similar. These differences across years, yet time and site-specific similarities further suggests a coordinated seasonal progression in the differential expression of viral genes.

### Localization of viral RNA within HPMLs

In an effort to identify the cell-type(s) infected by these adomaviruses, we developed RNAScope probes specific to the adenain transcripts of MdA-1 and MnA-1. Adenain was targeted given that RNA-seq results from HPMLs sampled in 2017 identified it as the most abundant viral transcript. Given the small dsDNA genome of these viruses, RNAScope probes also effectively targeted viral DNA which was evident in the nucleus of virus positive epithelial cells. In spring-collected samples from LMB and SMB, perinuclear staining was observed in melanocytes that had migrated to the epidermis indicating viral replication in this cell type (Fig 4). A cytoplasmic signal was not observed in melanocytes. Virus positive cells were also observed within the basement membrane in LMB infected with MnA-1 (S4 Fig). Notably, we identified viral transcripts in the cytoplasm of epithelial cells within the epidermis of a fall-collected SMB (S5 Fig). Nuclear staining of viral nucleic acids was evident in these cells as well. In fall-collected SMB, virus negative melanocytes were often observed proximate to virus positive cells in HPMLs. This suggests that the proliferation and infiltration of melanocytes from the dermis to the epidermis during the fall was part of a host response to viral infection. Positive signal was not observed in clinically normal skin.

**Fig 4.**
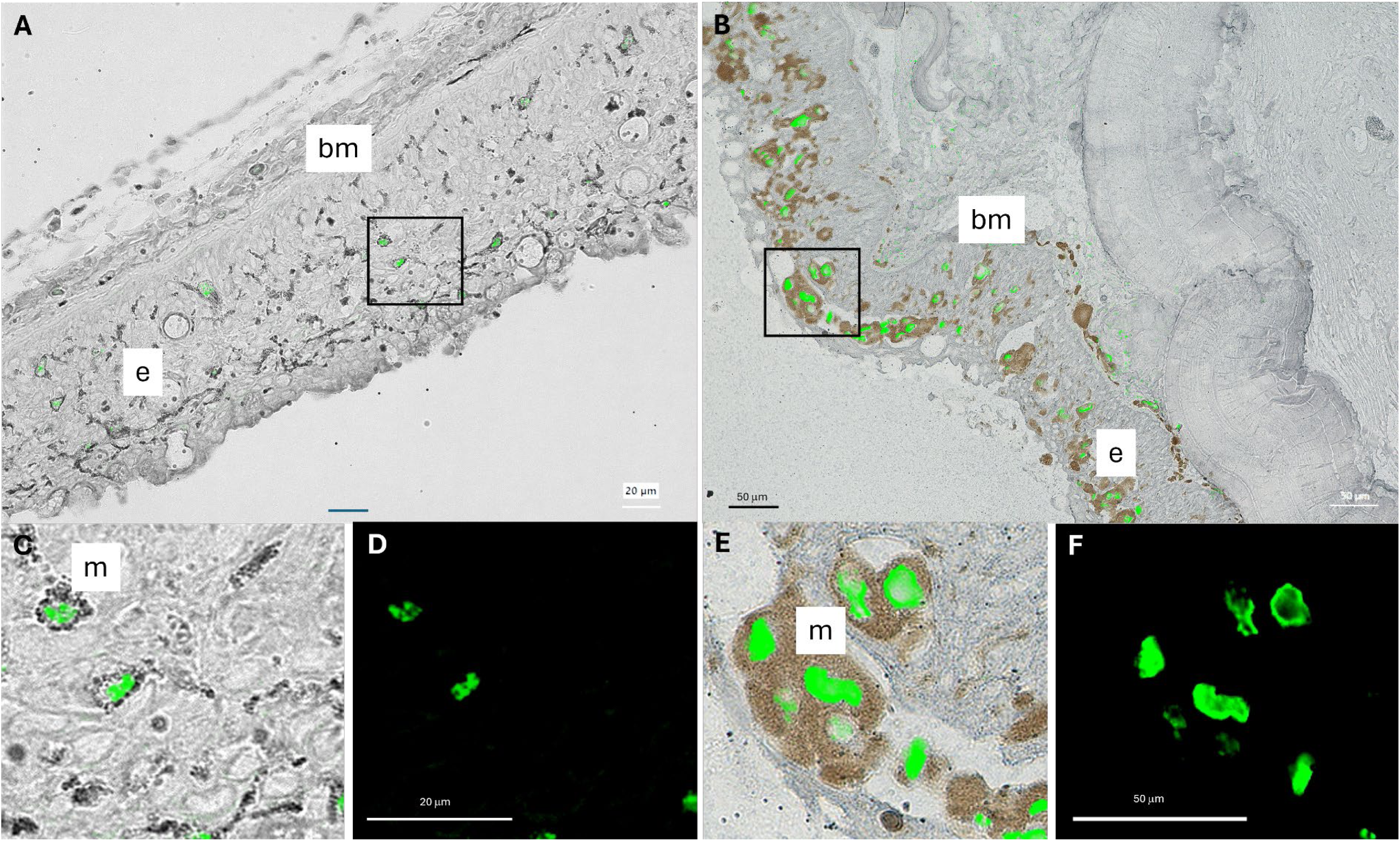
Composite images of histological sections of hyperpigmented lesions on large (A) and smallmouth bass (B). Images represent an overlay of RNAScope probes targeting the adenain sequence of these bass adomaviruses (green) and a brightfield image depicting melanocytes within the integument. Insets (C; overlay) and (D; darkfield); (E; overlay) and (F; darkfield) provide a high magnification view of nuclear staining within melanocytes located in the indicated regions. The epidermis (e), basement membrane (bm) and melanocytes (m) of the integument are indicated.

### Phylogenetically related, but different adomaviruses are associated with HPMLs in largemouth and smallmouth bass

Consistent with other adomaviruses, MdA-1 and MnA-1 encode two cassettes of bidirectionally transcribed genes (Fig 5) [31, 44, 45]. The annotated genomes included core genes associated with adomaviruses including Cah (hypothetical <u>c</u>apsid-surface protein with predicted <u>a</u>lpha <u>h</u>elical [particularly coiled-coil] character), Penton (<u>pent</u>americ single β-jellyroll capsid protein), Macc (hypothetical <u>m</u>embrane-<u>a</u>ctive <u>c</u>apsid <u>c</u>ore protein), Hexon (double β-jellyroll major capsid protein that trimerizes to form virion facets), Adenain (papain-like cysteine protease involved in virion maturation), Prim (homolog of archaeal-eukaryotic <u>prim</u>ase small catalytic subunits), Rep (superfamily 3 ATP-dependent <u>rep</u>licative DNA helicase with N-terminal nicking endonuclease-like domain) and SET (<u>S</u>u(var)3-9 <u>E</u>nhancer-of-Zeste and <u>T</u>rithorax homologue). In addition to these core genes, there were predicted open reading frames (ORFs) encoding genes with no clear homologues in GenBank or HHPred searches. These included: Endonoid (<u>Endon</u>uclease like protein), Phogi (remote similarity to <u>pho</u>spho<u>g</u>lucose isomerase), <u>Herpeto</u> (remote similarity to Herpeto_peptide), Zifi (Nab2-type <u>zi</u>nc <u>fi</u>nger), Col (predicted <u>col</u>lagen-like fold), and Wasp (WASP actin regulatory protein-like). Neologisms were assigned based on HHpred and AlphaFold/DALI best hits or by synteny adopted from previous adomavirus annotation efforts [31]. Predicted protein structures differed across these syntenic ORFs as well as HHpred and DALI predictions, but we retained syntenic naming assignments for simplicity.

**Fig 5.**
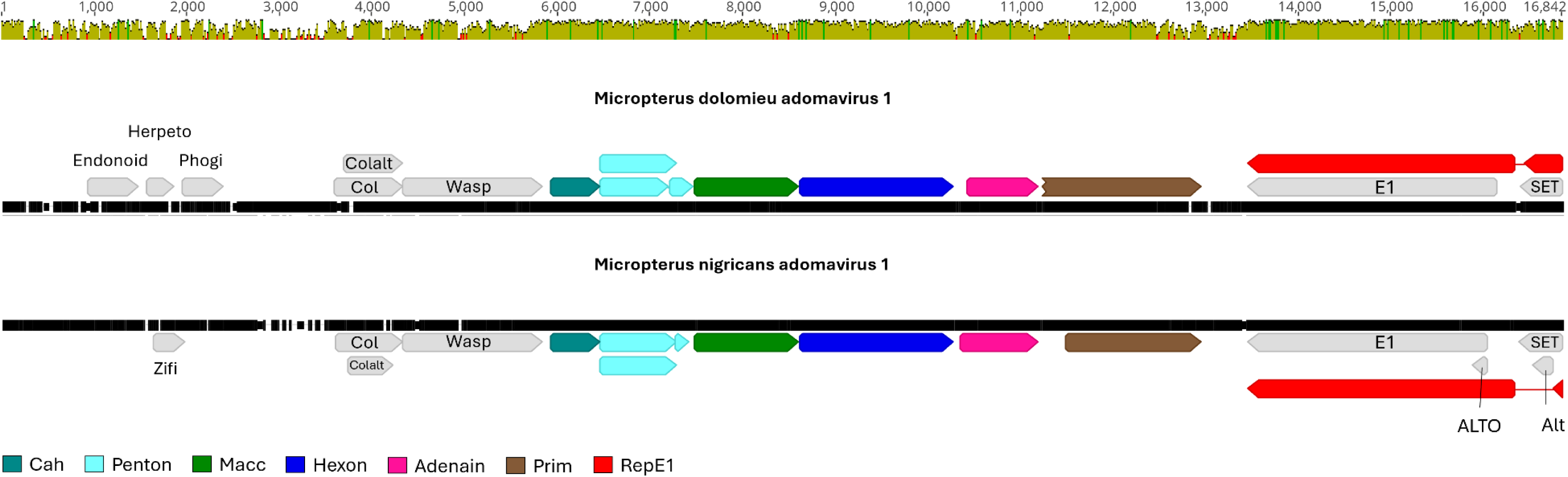
Genome alignment of the smallmouth bass adomavirus 1 (MdA-1) and largemouth bass adomavirus 1 (MnA-1). Core adomavirus open reading frames are colored in non-gray. Identity graph depicts similarity/dissimilarity between the genomes (sliding window = 1).

Genome alignment of MdA-1 and MnA-1 indicated 68.0% nucleotide identity. Predicted coding sequences (CDS) were similar across the genomes, yet there were a few distinctive differences. Open reading frames encoding homologs of MdA-1 Endonoid and Phogi were not predicted in MnA-1. In this genomic region upstream of Col, another ORF (Zifi) was identified in MnA-1 encoding a predicted Nab2-type <u>zi</u>nc <u>fi</u>nger. The MnA-1 genome also included an overprinted ALTO ORF within RepE1. A possible ALTO-like ORF in MdA-1 is interrupted by a nonsense mutation. Amino acid identity across shared predicted proteins ranged from 41.0 (Colalt) to 92.2% (Hexon; Table 2). In general, there was a notable decrease in protein identity in ORFs upstream of Cah. Predicted proteins in this region shared 63.3% identity or less between the two viruses. This included Col, Colalt and Wasp which we named based on synteny alone.

**Table 2.** Pairwise protein identity between predicted proteins of MdA-1 and MnA-1. Proteins are sorted by % identity. ALTO, Endonoid, Herpeto, Phogi and Zifi were not included in pairwise comparisons given that they were unique.

| Protein | % Identity |
| --- | --- |
| Hexon | 92.2 |
| Penton | 86.8 |
| Penton+ | 83.4 |
| Cah | 82.5 |
| RepE1 | 78.9 |
| Prim | 76.9 |
| Macc | 74.0 |
| Adenain | 72.4 |
| Col | 63.3 |
| SET | 62.0 |
| Wasp | 51.7 |
| Alt | 49.3 |
| Colalt | 41.0 |

This region also included divergent protein coding regions. Predicted three-dimensional protein structures of these proteins, and results of HHpred and DALI further emphasize the dissimilarity (Fig 6, S6 Fig, S7 Fig, S3 Table, S4 Table). Notably, HHpred and Dali predictions for Wasp from both MdA-1 and MnA-1 suggested similarity to porphyrins and myoglobin (S7 Fig). DALI yielded no results for MnA-1 Col. A predicted retinoblastoma pAB groove interacting motif was identified within RepE1 of both viruses, suggesting potential interactions with host tumor suppressing genes.

**Fig 6.**
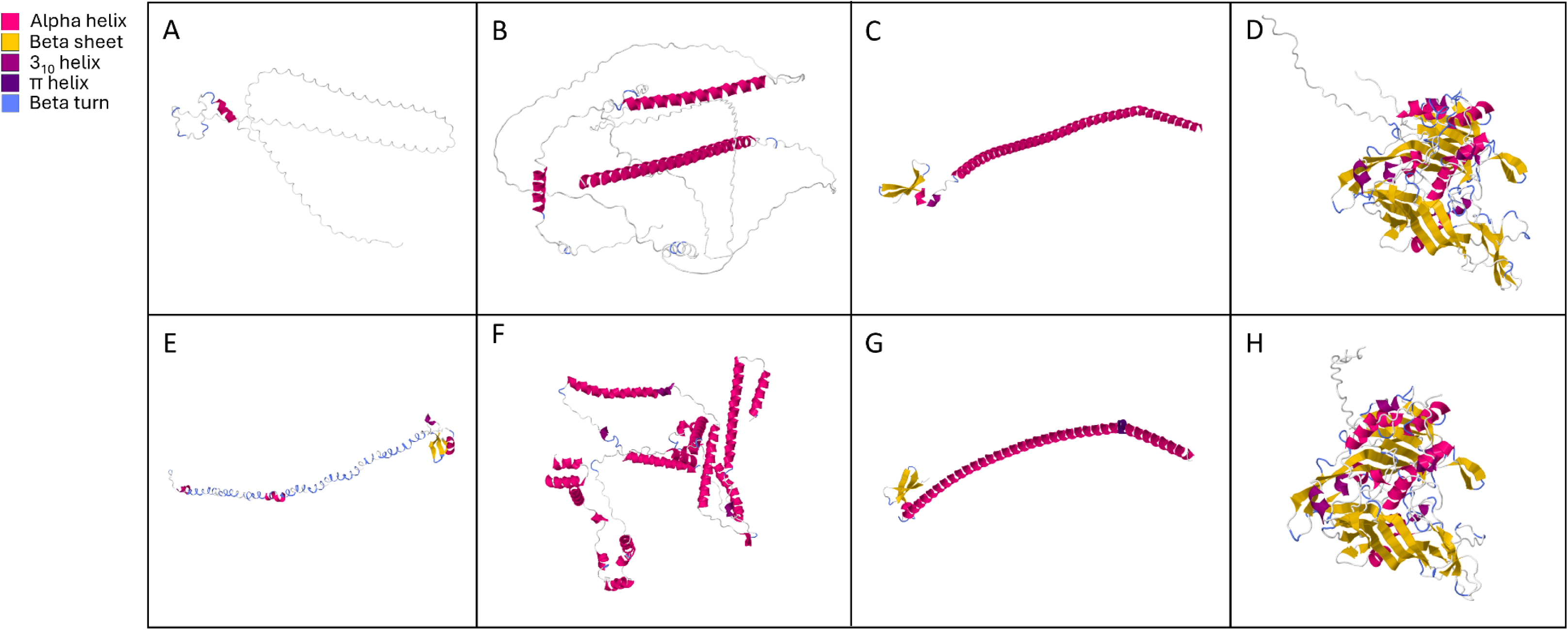
Predicted protein structures of Col (A, E), Wasp (B, F), Cah C, G) and Hexon (D, H) of MdA-1 and MnA-1, respectively. The predicted structures differ markedly between the Col and Wasp proteins from the different viruses whereas Cah and Hexon share similar structures.

### Genetic diversity across bass adomaviruses

Given the recent, concurrent emergence of BBS in disparate watersheds, we compared partial sequences of RepE1 across MdA-1 samples from SMB collected from Michigan, Pennsylvania and Vermont to gain insights into genetic diversity across samples as a preliminary epidemiological survey (S1 Table). Sequences obtained from the same location were always identical across the 699 bp PCR amplicon (Fig 7A, Fig 8A), whereas four base substitutions were noted between sites. All single nucleotide polymorphisms (SNPs) were synonymous transversions; A↔C (1) or C**↔**G (4). Complete genome alignments of the two MdA-1 isolates from PA during different years identified only 14 nucleotide differences across the genomes (99.9% identical) and no difference in the RepE1 amplicon region (S7 Fig), suggesting *prima facie* an overall low rate of accepted mutations within populations. Notably, of these SNPs only those within the Col and Wasp ORFs led to protein coding changes (Col; 99.5% and Wasp; 99.4% protein identity).

**Fig 7.**
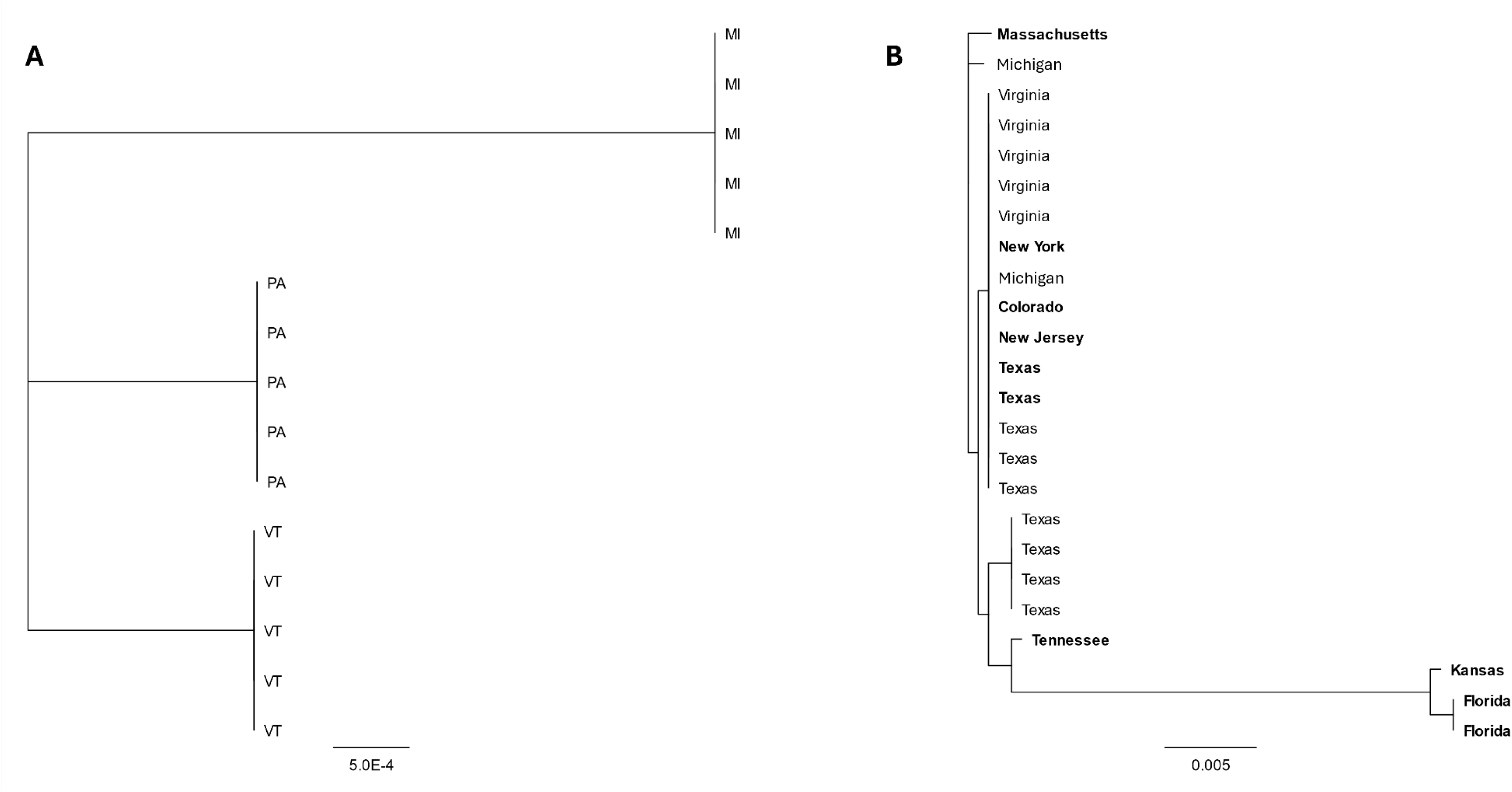
Neighbor joining trees depicting the relationships between partial sequences RepE1 of MdA-1 (A) and MnA-1 (B) from bass sampled from different locations in the United States. Samples identified in bold were collected from largemouth bass maintained in Bass Pro Shop live exhibits.

**Fig 8.**
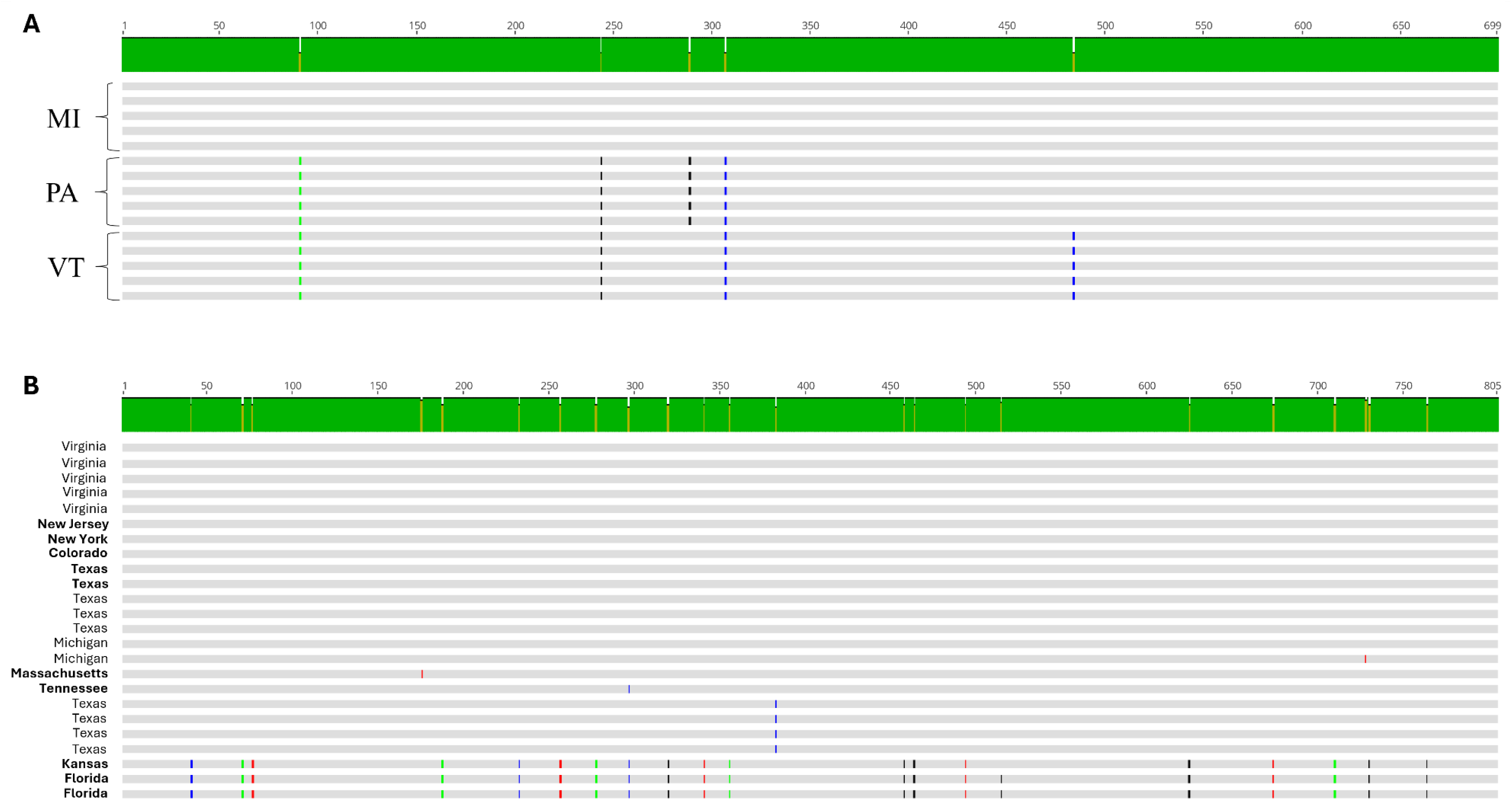
Alignment of RepE1 nucleotide sequences from samples collected from smallmouth bass (A) inhabiting waterbodies in Lake St Clair, MI; Susquehanna River, PA, and Lake Champlain (VT) or largemouth bass (B) represented in Figure 7A or 7B, respectively. Single nucleotide polymorphisms are identified in color to identify the base. Black = G, blue = C, green = A, red = T. Samples identified in bold were collected from largemouth bass maintained in Bass Pro Shop live exhibits

We compared a similar region of MnA-1 RepE1 from clinically affected LMB housed in live exhibits at Bass Pro Shops (BPS) located in different states across the USA as well as wild-caught fish from MI, VA, and TX. We observed up to 21 nucleotide differences (97.4% identity) across some samples. These were restricted to three samples that likely represent a different subtype of MnA-1 (Fig 7B). Whole-genome reconstructions of MnA-1 from these RepE1 distinctive samples are not available. Across this 805 bp locus, samples from six different locations were identical (Fig 8B). Identified SNPs were represented by both transitions (C↔T [9] and A↔G [5]) and transversions (G↔T [3], A↔C [2] and A↔T [2]). Notably, the nucleotide differences in the most divergent clade only led to two amino acid changes. When sequences from this distinct clade were not considered, we only observed at most two nucleotide differences between samples. Unfortunately, the origin of fish from BPS was not always known, and phylogeographic inferences for MnA-1 in wild fish are limited. Nonetheless, it can be concluded that the MnA-1 does not show the strong geographic structure apparent in MdA-1 based on this sampling.

### Phylogenetic placement of novel adomaviruses

Phylogenetic analysis of the replicase genes of the bass adomaviruses and other publicly available adomaviruses placed these viruses in a clade of RepE1 adomaviruses associated with a subgroup of African cichlids (subfamily Pseudocrenilabrinae; Fig 9). In general, the RepE1, RepLT and CressRep proteins resolved to distinct groups or to a clade represented by family-level taxa. CressRep was restricted to invertebrate hosts (specifically Bivalvia), but adomavirus replicases are not monophyletic. Subclades of RepE1 or RepLT were best predicted by host taxa. Similar phylogenetic placement was observed using other core adomavirus genes. Greatest sequence homology was observed across hexon proteins (Fig 10). Based on sequence similarity across these three proteins, representative adomaviruses from this clade are more similar to those from reptile hosts (leatherback sea turtle; *Dermochelys coriacea*) than fish hosts. Sequence similarity of adenain or hexon was not a predictor of replicase type which supports the proposal of horizontal gene transfer within this consortium of adomaviruses [46]. Although the three adomaviruses with cichlid hosts that resolved to the clade including MdA-1 and MnA-1 share the same genome architecture, similar to these viruses there is considerable variability in the number and putative identity of ORFs upstream of Col (S8 Fig).

**Fig 9.**
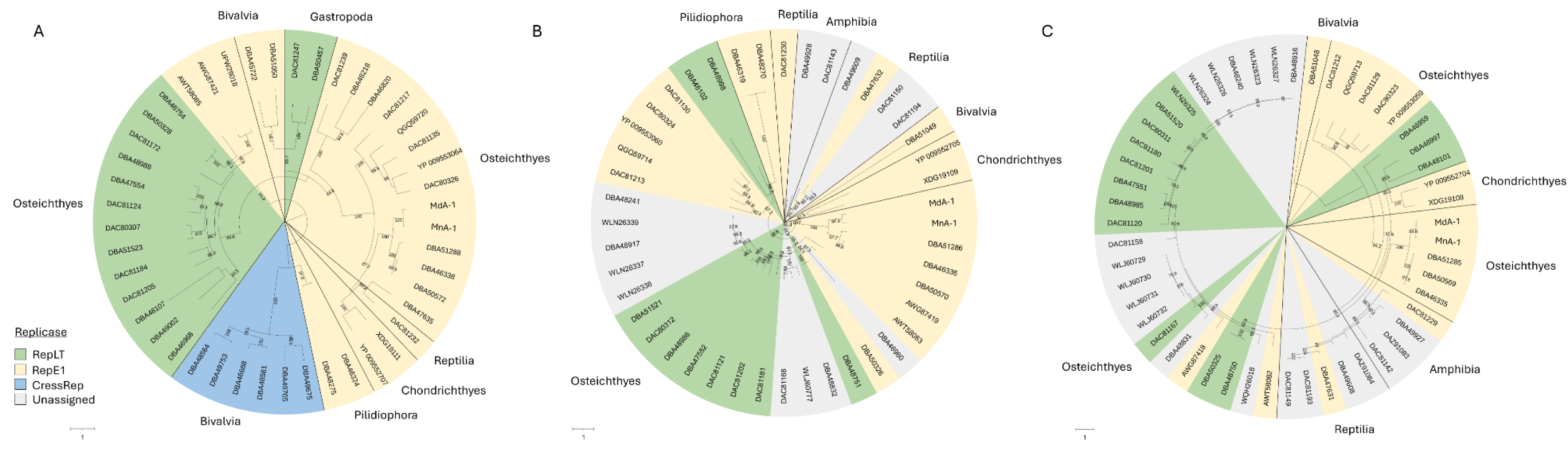
Circular maximum likelihood trees of replicase (A), adenain (B) and hexon (C) proteins of adomaviruses. Colors represent the replicase type. Class of the host organism is indicated along the margins of the tree. Bootstrap values are indicated for all branches where support is >75%.

**Fig 10.**
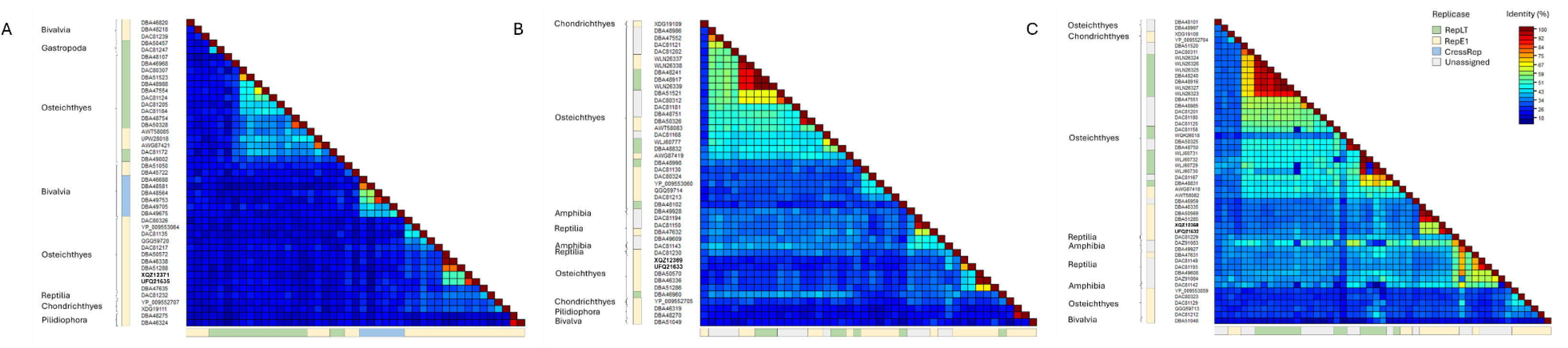
Pairwise protein identity matrix of replicase (A), adenain (B) and (C) hexon proteins. Replicase type is indicated along the margins as well as the Linnean class of the host organism. If known the replicase-type of each virus is indicated.

## Discussion

Adomaviruses are an enigmatic group of viruses with a circular dsDNA genome related to the small DNA tumor viruses [45, 47]. Their name is a chimeric portmanteau derived from the <u>ad</u>eno, papill<u>oma</u> and poly<u>oma</u>viruses for which they share protein homology and mixed elements of syntenic genome architecture. At present this group is represented by nearly 60 complete or partial genomes from a host range that includes invertebrates, fishes, and herptiles, but with greatest representation in fishes [31, 48]. They are broadly defined by a circular genome that is 9-20 kilobase pairs (kb) in length that encodes a superfamily 3 helicase, a candidate Penton, and a candidate Cah-class gene [31]. This group of viruses consists of three classes defined by the polyomavirus-like replicase (RepLT), papillomavirus-like replicase (RepE1), or CRESS-like replicase (CressRep). Notably, most of these viral genomes have been mined from metagenomic projects, and only a few have been identified in clinically diseased fishes and ascribed as the causative pathogen [31, 45, 48–55]. Although the host-microbe relationships of many adomaviruses are unclear, some representatives of this group lead to substantial disease in fishes and are associated with considerable economic losses in commercial aquaculture settings particularly in the case of anguillids [52–54, 56]. At present the adomaviruses are not formally recognized by the International Committee on Taxonomy of Viruses (ICTV), and are indexed as unclassified dsDNA viruses in the [NCBI taxonomy database](https://www.ncbi.nlm.nih.gov/datasets/taxonomy/tree/?taxon=2219222) [57].

Our investigation here was a deliberate attempt to clarify the relationship between a putative, novel adomavirus sequence identified in a SMB deep sequencing project and HPMLs in black basses [10]. Initial screening attempts to confirm virus in HPMLs from SMB routinely verified the presence of adomavirus DNA; however, primer sets for this virus always failed to amplify specific product from the same lesion type on LMB. Deep sequencing and *de novo* assembly approaches were required to identify the adomavirus associated with HPMLs in LMB. The adomaviruses herein described from bass exhibiting BBS are members of the RepE1 class of adomaviruses. While the gross clinical signs of BBS are indistinguishable between LMB and SMB, they appear to be the result of different but related viruses that likely co-evolved with their host upon the divergence of these species around six million years ago [1]. These genomes are only 68.0% identical with notable coding differences affecting the number of predicted viral proteins and identity across cognate proteins. Given these differences and despite the same clinical presentation across these two host species, BBS is caused by two different viruses in these distinct hosts and outcome(s) of viral infection may differ.

While we have not attempted to fulfill Koch’s postulates, we feel that Rivers’ postulates have been satisfied such that (a) the specific adomaviruses have been found to be associated with the clinical signs of disease with a degree of regularity, and (b) the adomaviruses were demonstrated to occur in clinically affected individuals not as an incidental or accidental finding but as the cause of the disease under investigation [58]. Moreover, the fact that two related but genetically distinct (68.0% identical) adomaviruses are associated with the hallmark clinical signs of BBS provides indubitable evidence that these viruses are indeed causative agents of this seasonal disease.

The emergence of BBS in SMB from geographically separated watersheds in Michigan, Pennsylvania and Vermont prompted us to hypothesize that perhaps there were recent, sequential introductions of this virus. While our data sets are not comprehensive, the viruses are clearly not identical across these watersheds and more informatively they share identical SNPs in the RepE1 loci specific to geographical collection sites suggestive of phylogeographic differences. Given that viruses with dsDNA genomes accumulate mutations more slowly than other virus-types, we anticipate that future analyses with larger datasets may provide further evidence that these viruses have been evolving in geographic isolation for hundreds to thousands of years and are not the result of recent introductions [59–62]. As an example, evolution rates for polyomaviruses are 0.5% per million years (5×10^-9^) and 1.3 × 10^−7^ spontaneous mutations per base per infection cycle for adenoviruses under minimal selection [62, 63]. That stated, it should be mentioned that SMB were introduced to the Susquehanna River watershed (PA) during the 1870’s but are indigenous to the other sampled locations [64]. Due to the frequent movement of LMB across the country, we anticipate more homologous viral genomes, and this is supported by our current dataset. Our sequencing analyses were limited to a small locus within RepE1 which is a conserved viral helicase gene [65]. Insights into phylodynamics may be better achieved by targeting loci between the SET and Cah ORFs where we observe the greatest predicted protein coding differences between the MdA-1 and MnA-1 genomes, as well as the two MdA-1 genomes [66]. Perhaps this region of the genome is similar to the hypervariable regions of adenoviruses [60]. If indeed new introductions of MdA-1 do not explain the emergence of BBS in these geographically distant watersheds, perhaps changes in thermal profiles at the ecosystem-scale has affected host-microbe balance in a fashion predicted by evolutionary mismatch theory [67].

BBS exhibits strong seasonality and the gross, seasonal manifestation has recently been modelled for SMB in the Susquehana River watershed [24]. Perhaps most unexpected in these data sets is the similarity in expression patterns between individuals at a given collection site considering that these fish were all naturally infected. This coordination suggests that these fish were either exposed to the virus at a similar time, or that environmental triggers such as temperature drive this response. In some regards BBS, resembles the retroviral diseases of walleye (*Stizostedion vitreus*) integument in that it affects adults, is seasonal (fall through spring), and resolves when water temperatures increase [68, 69]. Fish are poikilothermic vertebrates and seasonality of disease is well established and often ascribed to the downregulation of adaptive immune responses in cooler temperature [70–73]. As a result, fishes tend to rely on innate immune responses during these times and perhaps the recruitment of melanocytes is part of this innate immune response [72]. Of particular note, the clinical manifestation of BBS is observed in adults and the observation of clinical signs coincides with the seasonal timing of gonadal recrudescence [24]. This condition is observed in males and females, but it is unknown if there is a sex bias in respect to prevalence or severity. We anticipate that there is a physiological sex steroid background profile that influences this disease presentation. Elevated androgens and estrogens are part of the normal adult physiology during gonadal recrudescence which occurs during the fall through late spring in these temperate basses depending on latitude [74, 75]. This may explain the anecdotal observation that large healthy-looking fish of high condition factor seem to be disproportionally affected given that these fish typically have higher steroid hormone concentrations. Although HPMLs are not melanomas, they are composed of the same, but non-transformed cell-type for which androgens have been demonstrated to drive melanoma invasiveness [76]. Perhaps a similar steroid-mediated mechanism drives melanocyte migration in BBS.

Based on RNAScope evidence, it appears that the development of the hallmark melanistic blotches shared across the bass species here is likely an orchestrated host response to viral infection, rather than the direct result of melanocyte infection. Epithelial cells in the integument appear to be the primary cell-type infected with virus during the early phase (fall) of grossly observable disease. Proliferation and infiltration of melanocytes into the epidermis is observed at this stage. Melanocytes are clearly permissive to infection evinced by perinuclear staining of this cell type during a later stage of natural infection (spring). Similar to the reports of adomavirus infection in giant guitarfish (*Rhynchobatus djiddensis*) and sand tiger sharks (*Carcharias taurus*), epithelial cells of the epidermis appear to be the initial target cell of infection [45, 55]. Melanization is also associated with lesions in guitarfish and clinical disease appears to be transient [77]. Interestingly, there is some evidence that cichlids, including [*Perissodus microlepis*,](https://www.tanganyika.si/Tanganjika/Perissodus/Perissodus%20microlepis/slides/Perissodus%20microlepis%20%27Isanga%20Bay%27%20%28Kushangaza%29.html) exhibit HPMLs reminiscent of lesions observed in BBS. If melanization is the result of a host response to viral infection, there must be a stage of this disease when tissues are virus positive prior to the clinical observation of HPMLs.

Although melanocytes are commonly recognized for their photoprotective role to ultraviolet electromagnetic radiation by the melanin they produce, this cell type is also an antigen presenting cell (APC) similar to macrophages and dendritic cells - although they originate from the neural crest rather than from hematopoietic tissues [78]. While melanocytes are not classified as professional APCs, they do share functional similarly in that they are phagocytic and express the class II major histocompatibility complex upon IFNγ stimulation. They are involved in host defenses via phagocytosis and antigen presentation to T cells, secrete pro- or anti-inflammatory cytokines to regulate immune responses and are an important cell type in innate immunity to multiple pathogen types including viruses [79–83]. Inflammatory mediators have been demonstrated to promote melanogenesis [80]. Evidence supports that the M2 macrophage phenotype associated with post-infection tissue remodeling are responsible for providing an anti-inflammatory cytokine environment that promotes skin pigmentation [84]. Intermediates formed during melanogenesis are have potent antimicrobial properties [85]. Given the clinical interest in melanin and its association with diseases, perhaps BBS may serve as a model for therapeutic research in humans as specific viral proteins are likely associated with the melanistic response [86].

Although BBS is associated with viral infection, it is unclear if it has detrimental health effects. The syndrome has been observed in LMB for nearly 40 years and there have been no clear, negative health effects. During this time frame, if these highly visible markings of hyperpigmented lesions on the surface of bass were associated with substantial mortality events anglers and resource managers would likely have noticed. In the case of SMB, it is noteworthy that the emergence of this clinical presentation was observed after years of chronic mortality events that impacted the population structure [87]. It is also notable that ulcerative skin lesions are often infected with opportunistic bacterial pathogens and evidence suggests that LMB virus leads to similar lesions. However, molecular diagnostics were not available or employed to screen for largemouth bass virus (LMBV) nor these novel adomaviruses in this work during fish kills [9, 88, 89]. Fish affected in these mortality events were also typically of a size class (young-of-the-year) in which clinical BBS is not observed - this may not be indicative of lack of infection but rather lack of a host response). The seemingly simultaneous emergence of the clinical signs of BBS in SMB during the last 15 years in disparate watersheds prompts the question, “What has changed?”. The 22-year data set from Lake St. Clair serendipitously captured the emergence of this condition in a world class fishery. Mortality events similar to those in the Susquehanna River have not been reported in Lake St. Clair nor in Lake Champlain, VT. In the case of Lake St. Clair, fisheries managers had been monitoring fish for six years prior to observing blotchy bass at a prevalence that warranted inclusion on their datasheets. Perhaps the most intriguing observation in this data set is that no blotchy fish were reported prior to 2008. Although this does not suggest that blotchy bass were not present, it likely indicates that something may have changed to an extent that this observation became noteworthy of documentation. Since that time, springtime annual variation in blotchy bass has been reported with an average prevalence of 5.1% and as high as 9.2%. This is similar to the prevalence and annual variation observed in the Susquehanna River basin. One unifying commonality between these three regions is that they tout world-class smallmouth bass fisheries and respectively receive extensive angling pressure [24].

Transmission of bass adomaviruses is a black box. Given the fragility of HPMLs during the spring and viral gene expression data suggesting that this is a late phase of viral infection, it is possible that transmission occurs during spawning which involves physical contact such as contact nips during courtship [90]. During the winter months, these fish also congregate in deeper waters at densities that increase the likelihood of physical contact. Angler handling of infected fish may also pose a mechanism of transmission. Unlike that of terrestrial vertebrates, the skin of fishes is not keratinized and the outermost layer is a mucosal epithelium resembling that of the gut [91]. If indeed epithelial cells are the initial target cell of adomaviruses, minimal disruption to this mucosal layer may be sufficient to facilitate infection. In this instance, handling fish with BBS is likely a risk factor for transmission.

While simply conjecture, one possible explanation could be that hyperpigmentation during spawning season is advantageous and these adomaviruses are beneficial to the host particularly when turbidity increases due to spring run-off. Although less commonly considered in the context of viruses, examples of viral mutualistic symbioses have been documented [92–95]. Melanism and variation in color patterns influence mate selection in fishes [96]. Melanic side-spotting in poecilid fishes has been well investigated particularly in swordtails (*Xiphophorus spp*.). These side-spotting patterns are of low frequency (<30%), and while genetic factors are associated with this phenotype the selection mechanisms are largely unknown [97].

Adomaviruses have recently been identified in swordtails, but no research has been published that evaluates the outcome of infection or whether there is a non-host genetic association with side-spotting. There may be an unexplored relationship between seasonal side-spotting in some fishes associated with adomaviruses.

Hyperpigmented lesions grossly similar to those in the black basses described here have been reported in other fish species. This observation is sometimes reported in the published literature, but more often is communicated by fisheries managers. We have evidence that adomaviruses are associated with those HPMLs in some instances (unpublished data), suggesting that an adomavirus etiology could be investigated in efforts to identify causative agent for instances when metazoan parasites are ruled out. Given the level of divergence in nucleotide sequence across these viruses, such efforts will inevitably require *de novo* sequencing methods coupled with virus discovery bioinformatic workflows to identify them if indeed present [98–101].

Although the significance of adomavirus infection in black basses is not clear, these viruses are a substantial problem in eel aquaculture that could be abated with the development of an efficacious vaccine development. Due to the external and macroscopically observable clinical signs of adomavirus infection in black basses, development of an infection model for BBS could be developed to test antigen candidates for adomavirus vaccines. Disease outcomes of infectious microbes are often exacerbated in intensive aquaculture conditions [102]. LMB are increasingly cultured for stocking for recreational purposes as well as a commercial food fish. It is prudent that managers take notice of viral diseases such as this with unknown outcomes under different environmental settings. In the instance of blotchy bass syndrome, fish managers can now inform the angling public of the cause of HPMLs based on scientific findings. Additionally, given the microbial nature of this disease more research can be conducted in the context of a causative agent to support the development of management decisions.

## Supporting information

S1 PDB Files

## Acknowledgements

This research was funded by the Biological Threats and Invasive Species Research Program of the US Geological Survey, Ecosystems Mission Area. We wish to thank Blayk Michaels of Bass Pro Shops and Cabela’s for coordinating sample collection efforts from blotchy bass housed in live exhibits. This research would not have been possible without the in-kind support and sampling assistance by the Michigan Department of Natural Resources, Pennsylvania Fish & Boat Commission, Vermont Fish & Wildlife Department, and the Virginia Department of Inland Game and Fisheries. We wish to thank Meghan Kepler-Shall for providing the image of a clinically affected smallmouth bass and anglers for submitting images to the Texas Parks and Wildlife Department. Any use of trade, firm, or product names is for descriptive purposes only and does not imply endorsement by the U.S. Government. The USDA is an equal opportunity employer. Supporting data are archived in [FigShare.](https://doi.org/10.6084/m9.figshare.c.7760105)

