## Supplementary material for "Novel Adomaviruses Associated with Blotchy Bass Syndrome in Black Basses (*Micropterus spp.)*": S1 Fig

**Supplemental Figure 1**: Diagnostic case report of the first recorded observation of blotchy bass syndrome in largemouth bass.

| Date Submitted | ANIMAL PATHOLOGY RECORD | Accession No. |
|---|---|---|
| 11-20-84 | DEPARTMENT OF ANIMAL PATHOLOGY<br>COLLEGE OF RESOURCE DEVELOPMENT<br>UNIVERSITY OF RHODE ISLAND<br>KINGSTON, R. I. 02881 | 0 293<br>Date Received |

| Veterinarian | Address | Phone |
|---|---|---|
| Dr. Richard Wolke | 25 Main Street, Carolina, RI | 792-2334 (work) |

| Owner | Address |
|---|---|
| Marty Wencek | 811 Middlebridge Rd., Narragansett, RI 02882 |

| Ident. | Animal | Breed | Color | Age | Sex | Weight | Prev. Acces. |
|---|---|---|---|---|---|---|---|
| Micropterus | salmoides | | | | female | | |

Clinical Diagnosis

History and Clinical Summary:
Fish collected by means of angling. Caught in Roundout Creek, tributary to the Hudson River Kingston, NY. A total of 81 bass were taken over an 8 hour period (7:30 a.m. - 3:30 p.m.). Dark external patches occurred only on fish over approx. 30 cm in length. The larger the specimen, the more numerous the black patches. Not all fish over 30 cm exhibited these markings. They occurred on approx. 33% of fish between 30 cm and 35 cm. Specimens greater than 35 cm showed these markings approx. 50%+ of the time. The number of patches varied from fish to fish, some more covered than others. Patches were ~~xxx~~ most numerous on either side of the body, the fins, particularly the dorsal, anal

| Specimen Submitted | Preservation | Condition of Specimen When Received at Lab: |
|---|---|---|
| Live Animal ☐ Dead Animal ☒ Tissues ☐ | Fresh ☒ Frozen ☐ Fixed ☐ | |

| Biopsy Data Exact Location | Size | Duration | Encapsulated | Lymph node involvement |
|---|---|---|---|---|
| | 42 cm | | YES ☐ NO ☐ | YES ☐ NO ☐ |

| Autopsy Data Natural Death ☐ | Mode of Euthanasia | Time and Date of Death | Time and Date of Autopsy |
|---|---|---|---|

Tissues Submitted:

Findings:
and caudal, and on the lips. Fish were taken from 1 meter to 5 meters deep on 3" plastic tail jig lures and 4" plastic worms -- from weedy areas (submergents) to rocky shorelines - also adjacent to sunken barges. Water temp. ~ 9°C. Clarity <1 meter. Tidal amplitude approx. 1 meter. Weather conditions overcast, 38°, northerly breeze.

There are round to rectangular pigmented (black) areas above lateral line and on dorsal fin. Areas are 4 x 2 to 0.5 x 1 cm and do not extend into dermis. They number 8. The anal fin also has such an area. Mesentary was adhering to the peritoneum.

SEE OVER

| Diagnoses | Classification | |
|---|---|---|
| Peritonitis, Granulomatous, Verminous, Melanosis | P8 2a<br>U3C | Tissues saved ☐<br>Photographs ☐<br>Radiographs ☐ |

| Pathologist | Date Prepared | Fee |
|---|---|---|
| R. E. Wolke | 12-14084 | |
