## Supplementary material for "Novel Adomaviruses Associated with Blotchy Bass Syndrome in Black Basses (*Micropterus spp.)*": S2 Fig

**Supplemental Figure 2**: Circular genome maps of Micropterus dolomieu adomavirus 1 (MdA-1) and Micropterus nigricans adomavirus 1 (MnA-1). Core adomavirus ORFs are colored in non-gray.

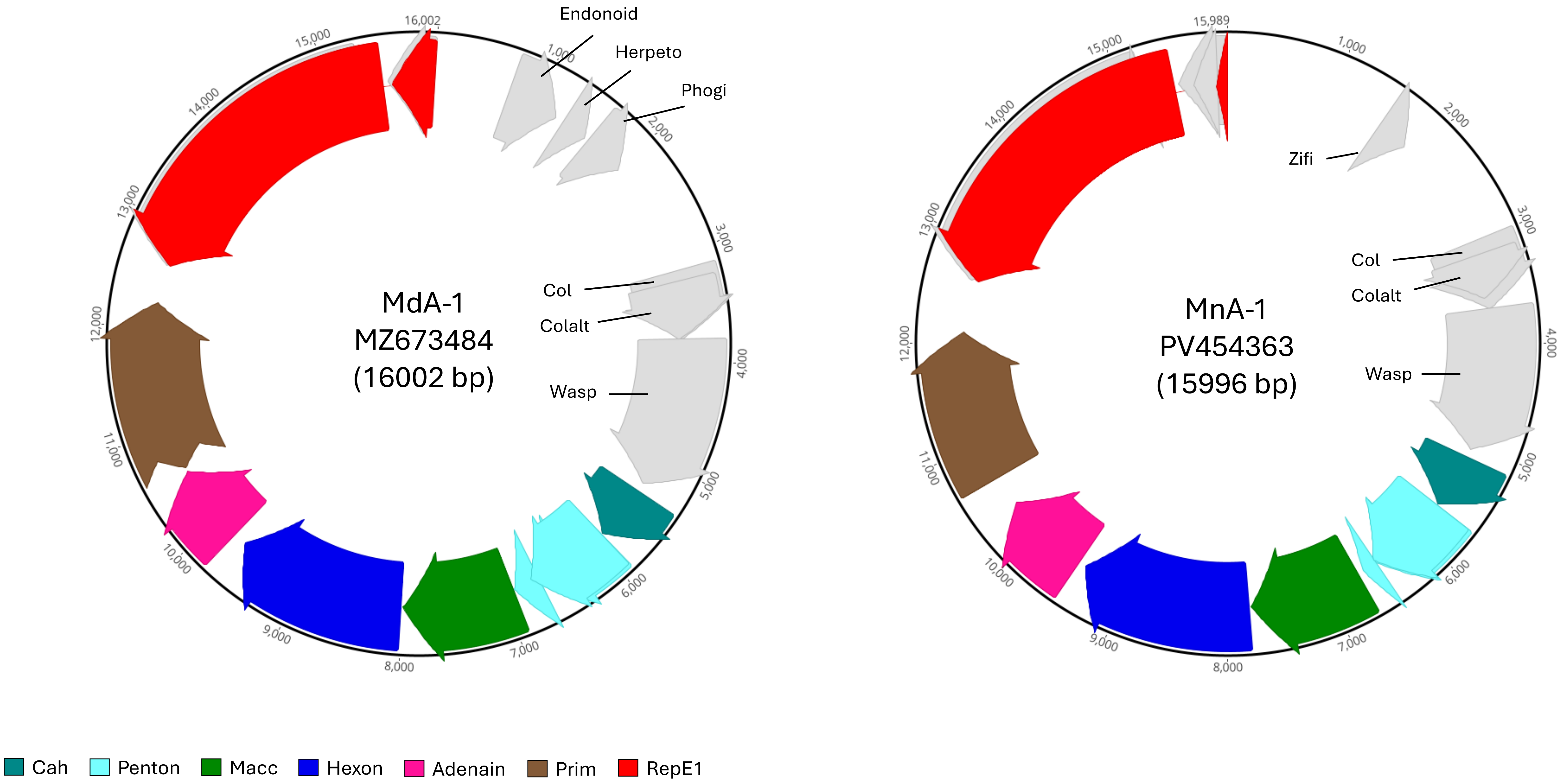
