## Supplementary material for "Novel Adomaviruses Associated with Blotchy Bass Syndrome in Black Basses (*Micropterus spp.)*": S3 Fig

**Supplemental Figure 3**: Gross and histological presentation of the hyperpigmented melanistic lesions (HPML) associated with MdA-1 infection and the mucoid lesions associated with MdA-2 in smallmouth bass.

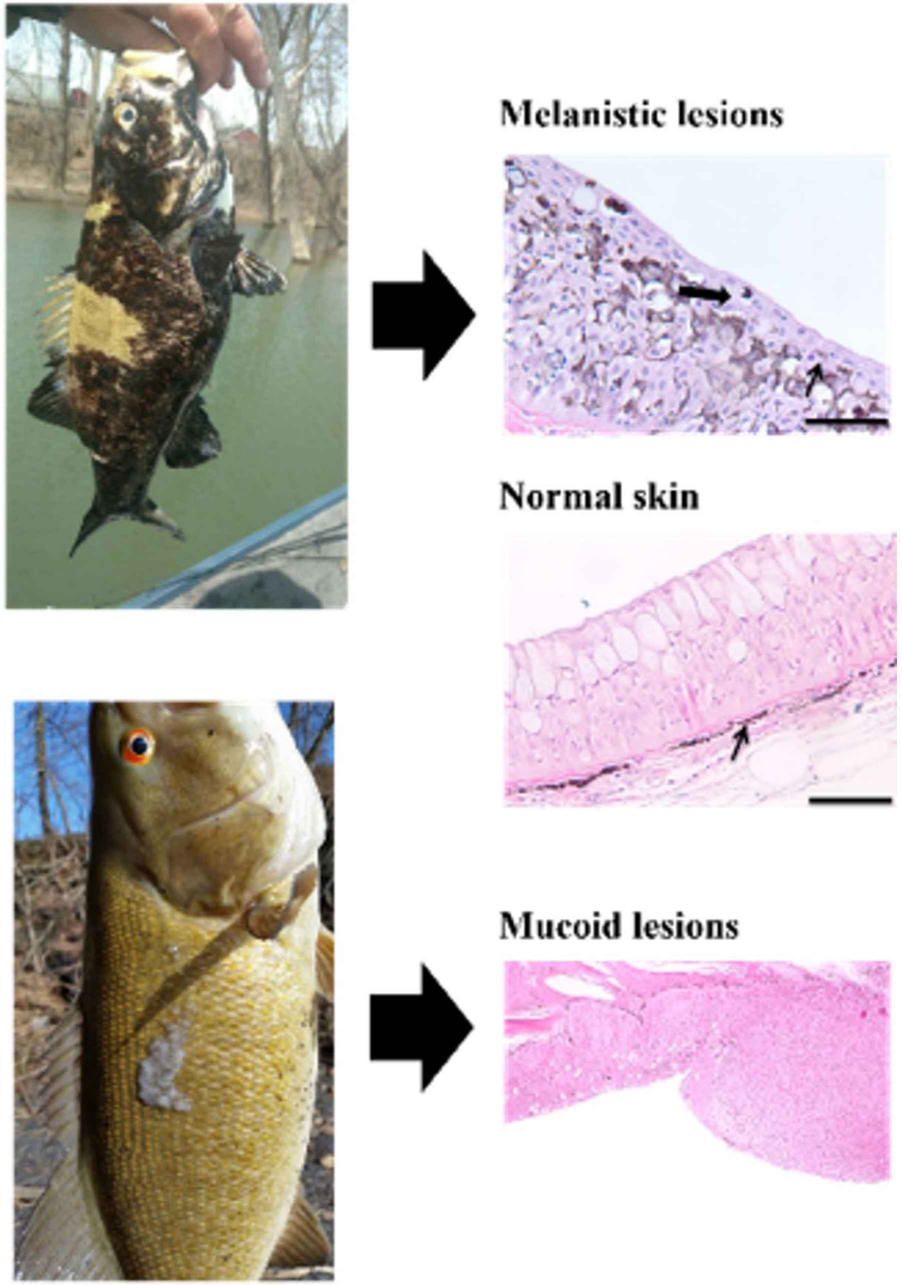
