## Supplementary material for "Novel Adomaviruses Associated with Blotchy Bass Syndrome in Black Basses (*Micropterus spp.)*": S4 Fig

**Supplemental Figure 4**. RNAScope analysis of HPMLs in infected smallmouth bass skin sampled during the fall. The adenain transcript of MdA-1 was targeted. Cells positive for adomavirus nucleic acids are restricted to the epidermis. While positive signal was sometimes observed in melanocytes, it was more commonly observed in non-pigmented, epithelial cells of the epidermis.

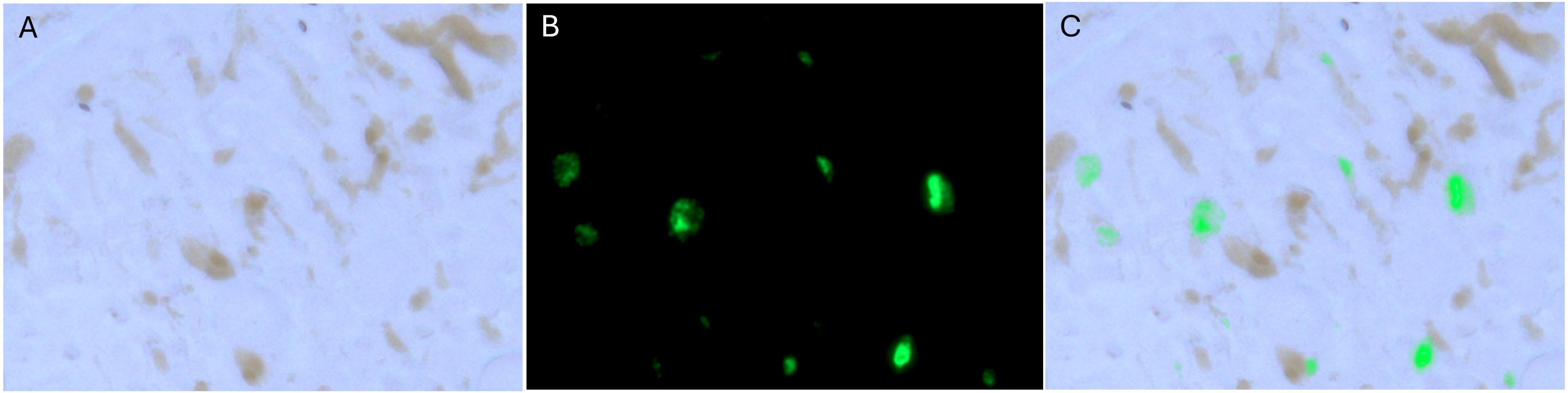
