## Supplementary material for "Novel Adomaviruses Associated with Blotchy Bass Syndrome in Black Basses (*Micropterus spp.)*": S5 Fig

**Supplemental Figure 5**. RNAScope analysis of HPMLS in infected largemouth bass sampled in the spring. The adenain transcript of MnA-1 was targeted. Cells positive for adomavirus nucleic acids are observed in the epidermis, but most are observed in the basement membrane.

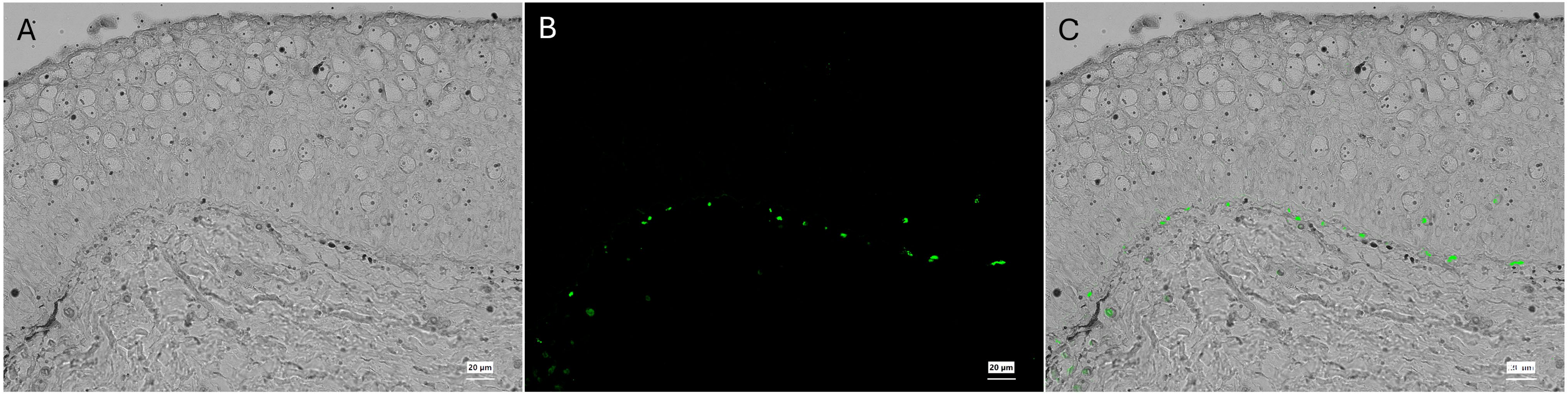
