## Supplementary material for "Novel Adomaviruses Associated with Blotchy Bass Syndrome in Black Basses (*Micropterus spp.)*": S6 Fig

### Col

Pairwise Identity = 63.4%

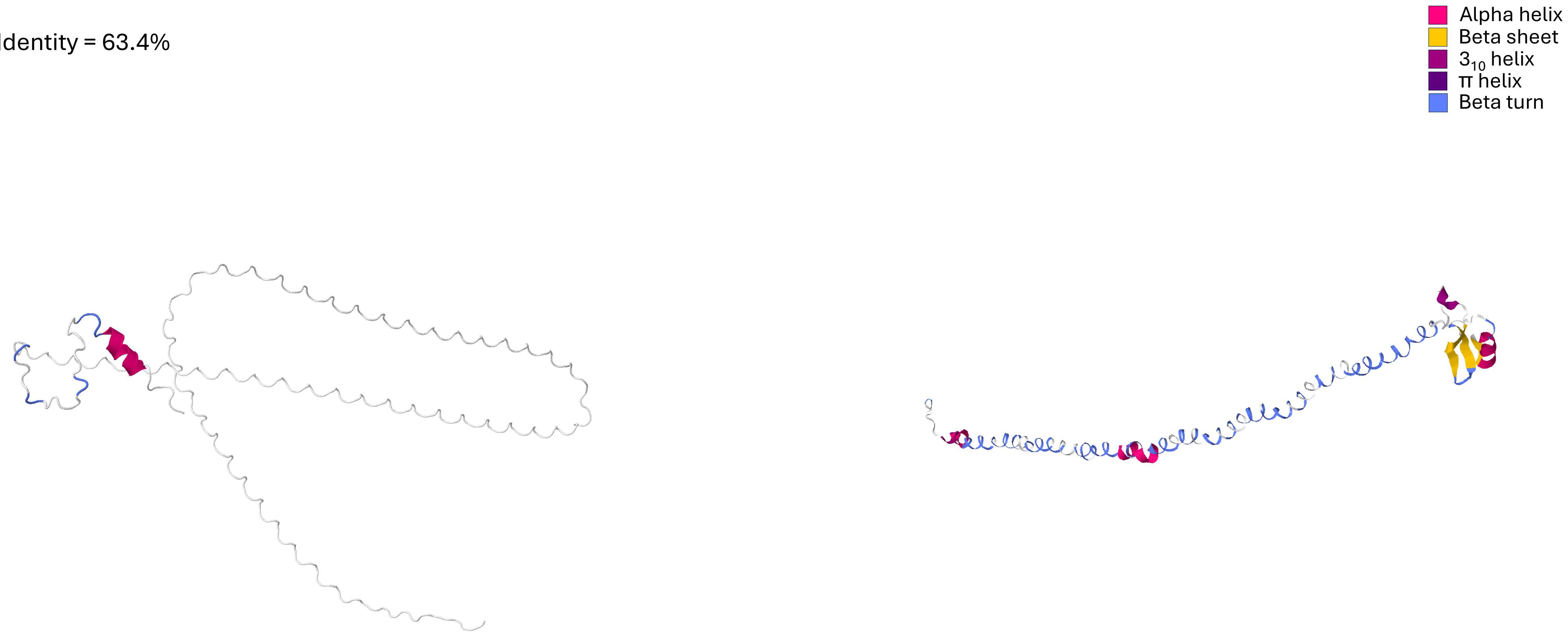


MdA-1
(UFQ21626)

MnA-1
(XQZ12361)

### Colalt

Pairwise Identity = 41.0%

- Alpha helix
- Beta sheet
- $3_{10}$ helix
- π helix
- Beta turn

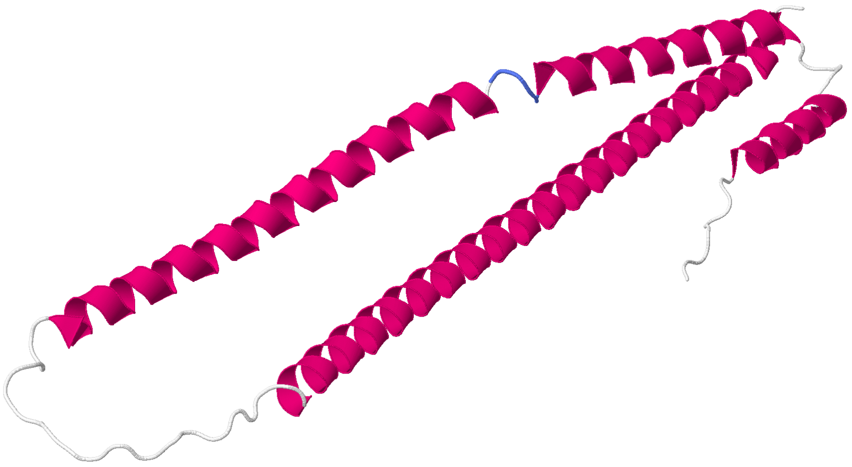

MdA-1
(UFQ21627)

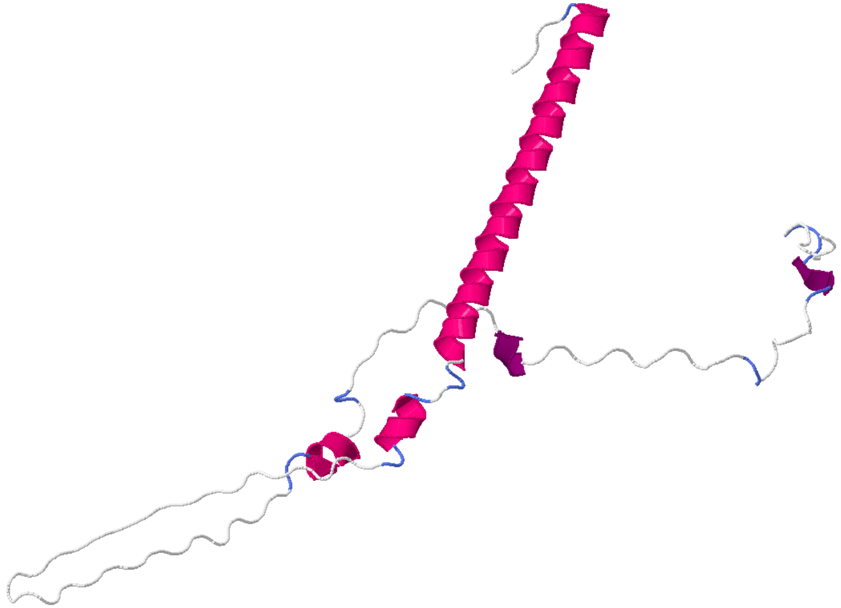

MnA-1
(XQZ12362)

### Wasp

Pairwise Identity = 51.7%

Alpha helix
Beta sheet
$3_{10}$ helix
π helix
Beta turn

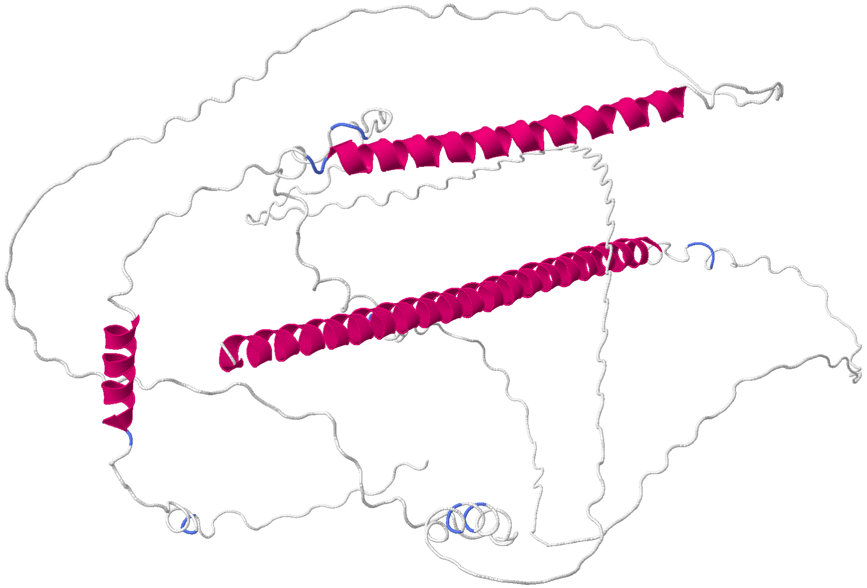

MdA-1
(UFQ21628)

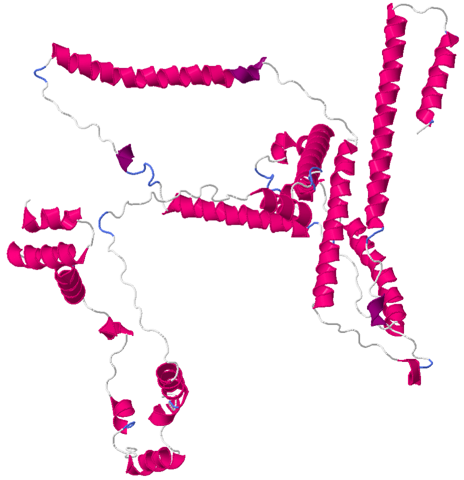

MnA-1
(XQZ12363)

#### Cah

Pairwise Identity = 82.5%

Alpha helix
Beta sheet
$3_{10}$ helix
π helix
Beta turn

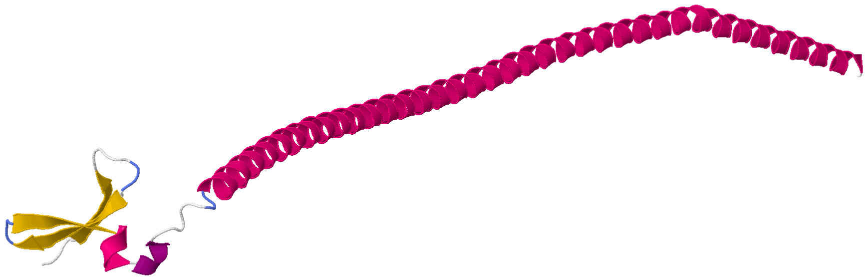

MdA-1
(UFQ21629)

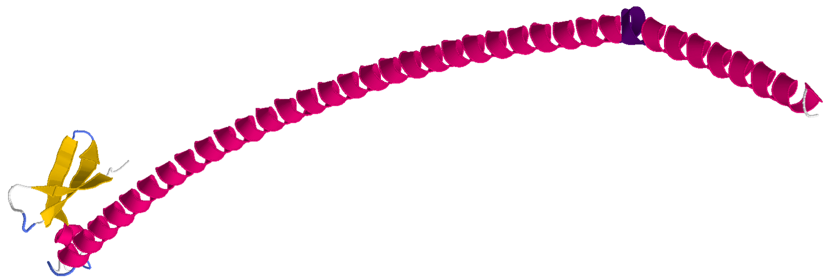

MnA-1
(XQZ12364)

### Macc

Pairwise Identity = 74.0%

- Alpha helix
- Beta sheet
- $3_{10}$ helix
- π helix
- Beta turn

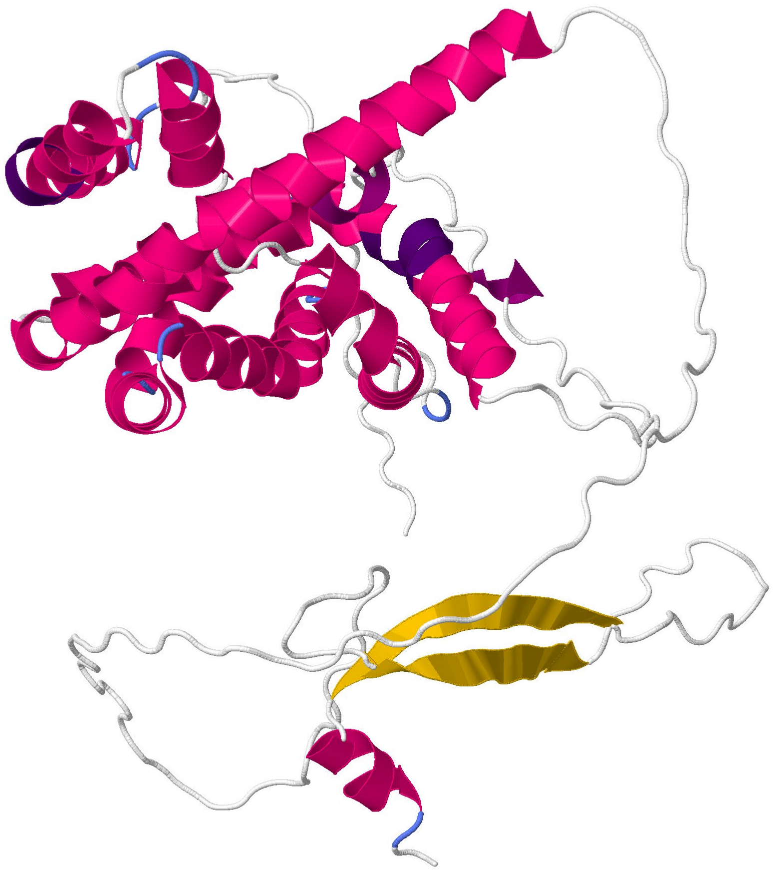

MdA-1
(UFQ21631)

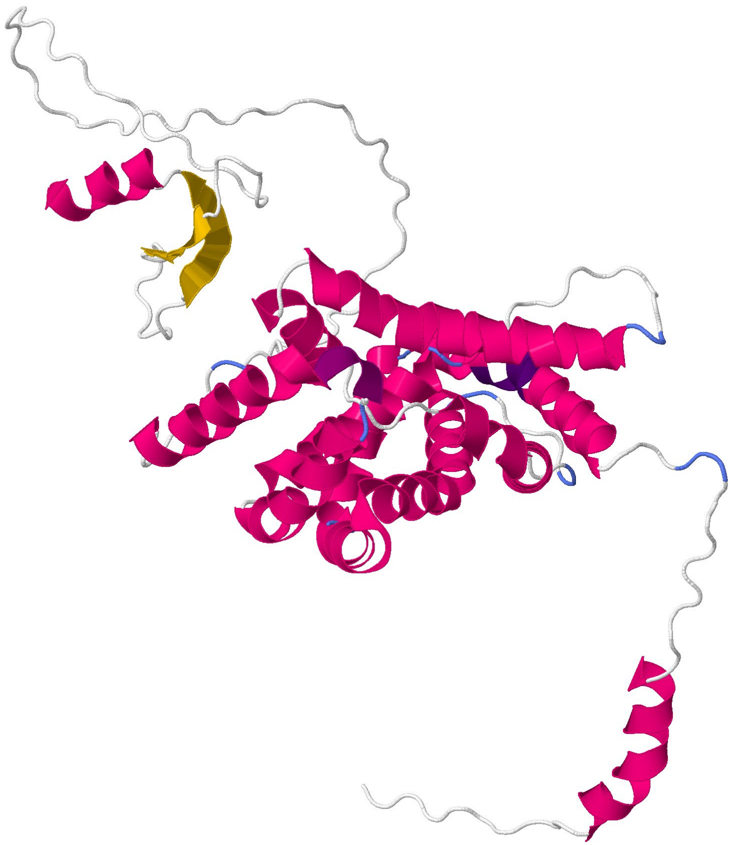

MnA-1
(XQZ12367)

### Penton

Pairwise Identity = 86.8%

Alpha helix
Beta sheet
$3_{10}$ helix
π helix
Beta turn

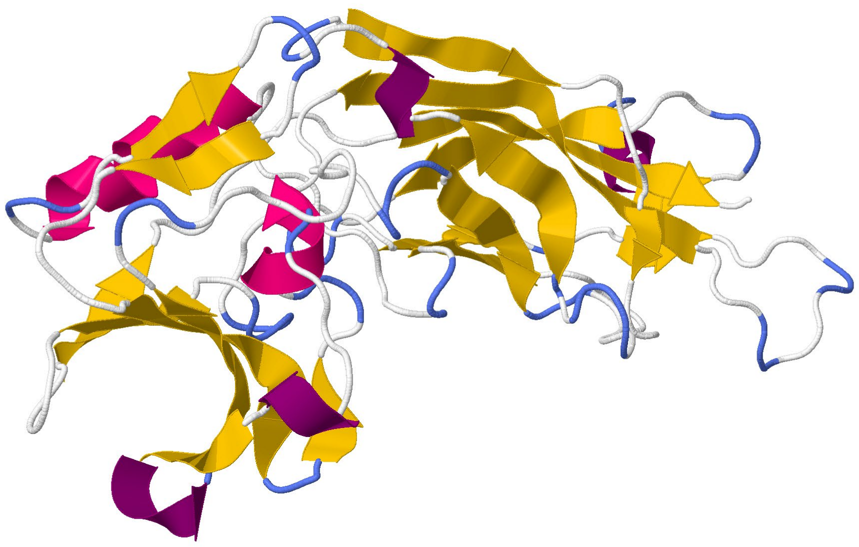

MdA-1
(UFQ21630)

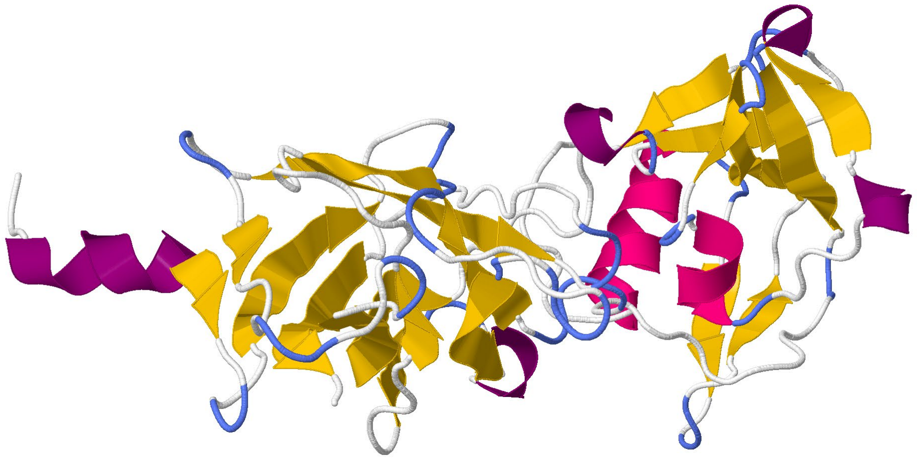

MnA-1
(XQZ12365)

### Penton+

Pairwise Identity = 83.4%

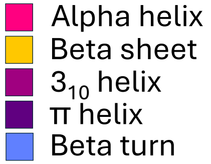


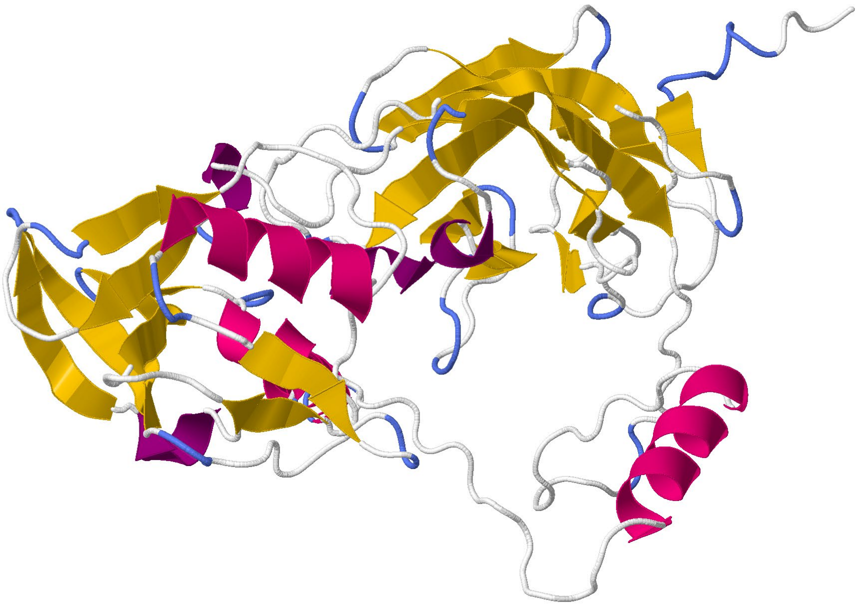

MdA-1
(XQU54337)

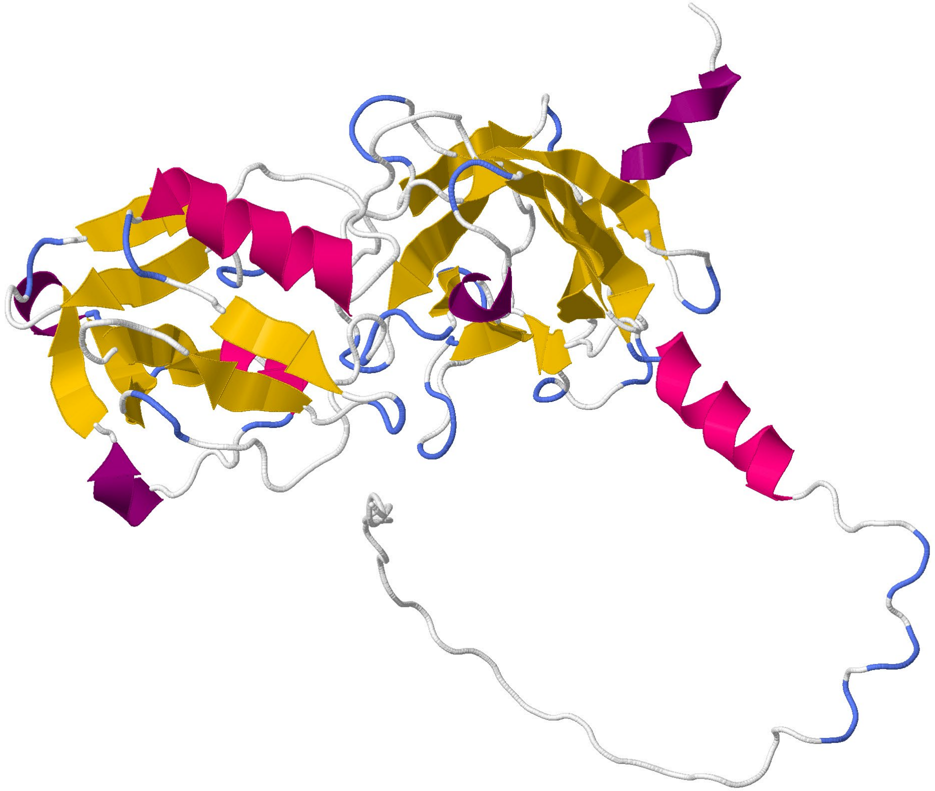

MnA-1
(XQZ12366)

### Hexon

Pairwise Identity = 92.2%

Alpha helix
Beta sheet
$3_{10}$ helix
π helix
Beta turn

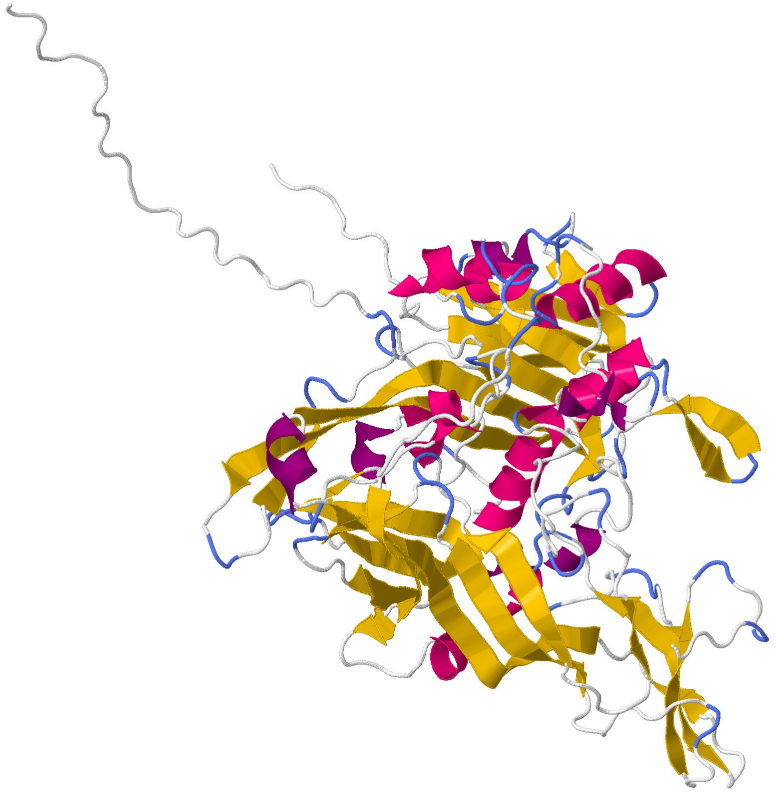

MdA-1
(UFQ21632)

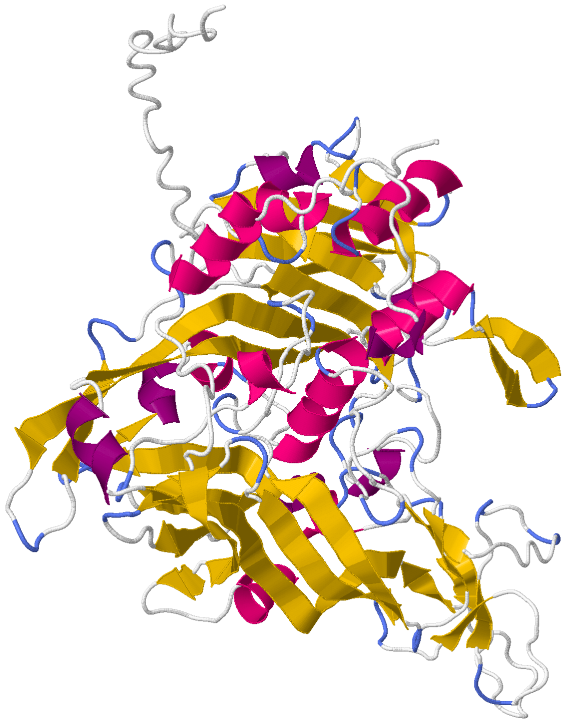

MnA-1
(XQZ1236)

### Adenain

Pairwise Identity = 72.4%

Alpha helix
Beta sheet
$3_{10}$ helix
π helix
Beta turn

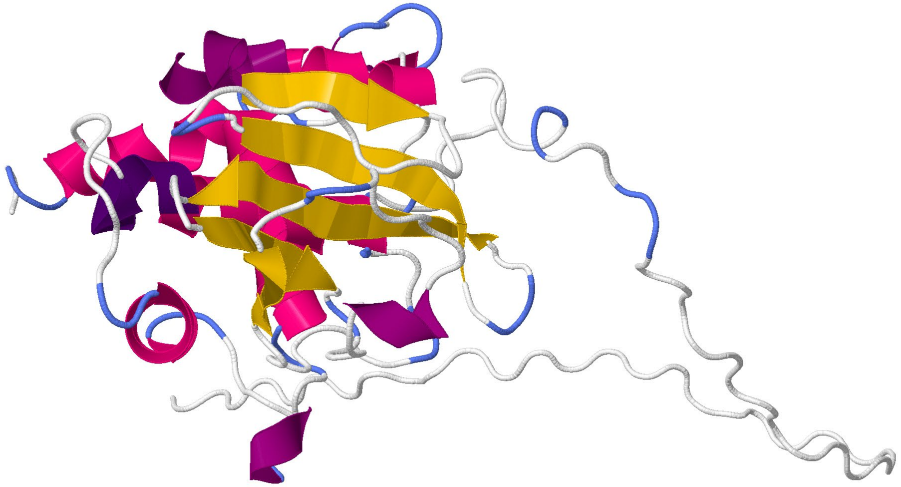

MdA-1
(UFQ21633)

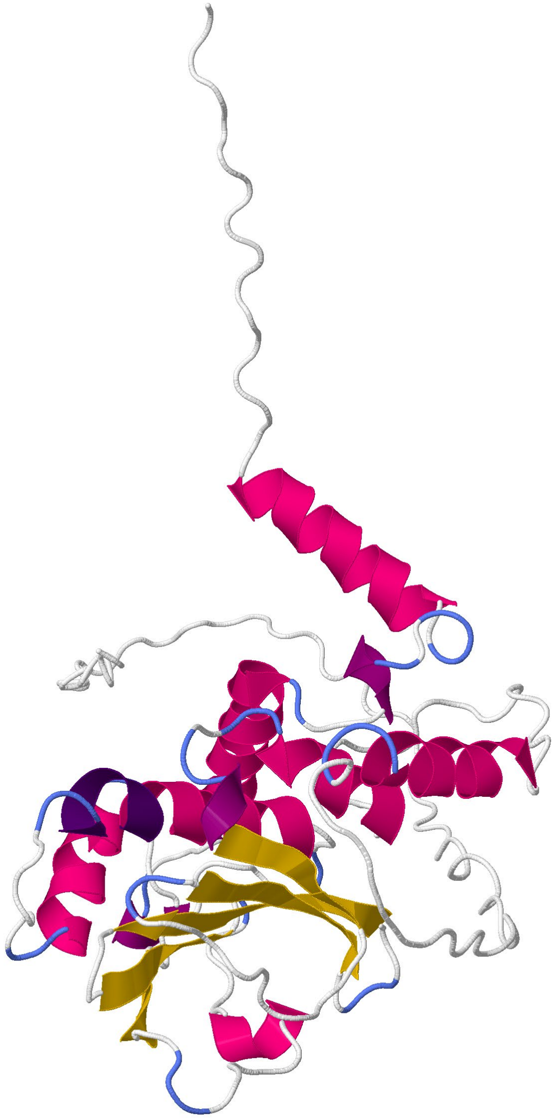

MnA-1
(XQZ12369)

### Prim

Pairwise Identity = 76.9%

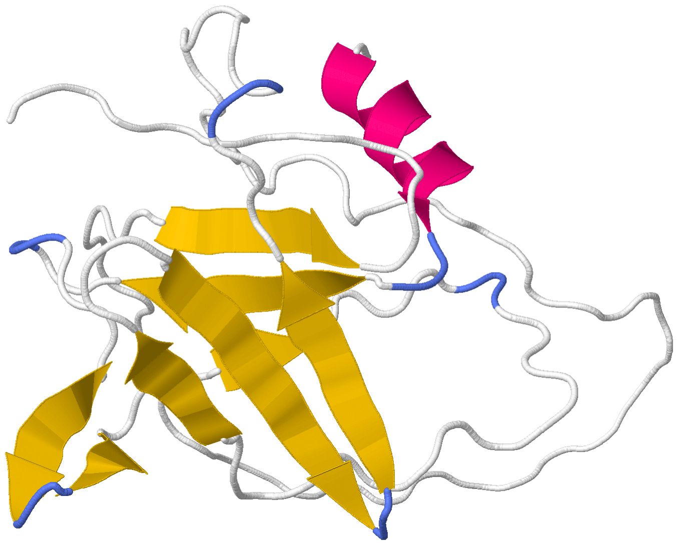

MdA-1
(UFQ21634)

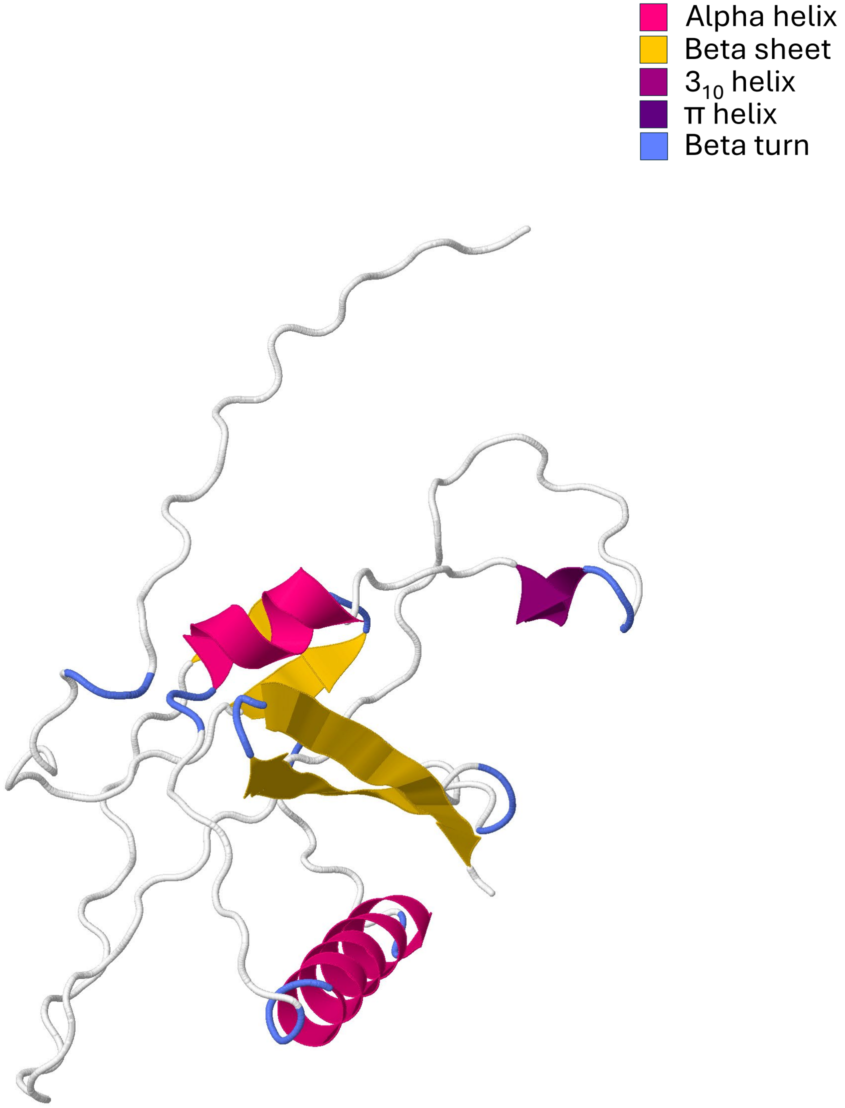


MnA-1
(XQZ12370)

### RepE1

Pairwise Identity = 78.9%

- Alpha helix
- Beta sheet
- $3_{10}$ helix
- π helix
- Beta turn

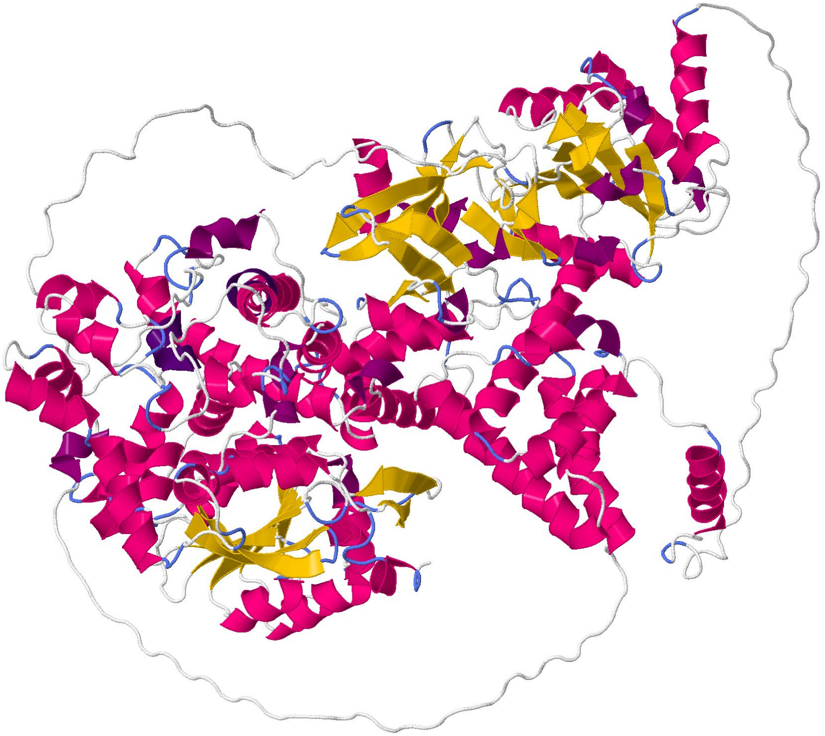

MdA-1
(UFQ21635)

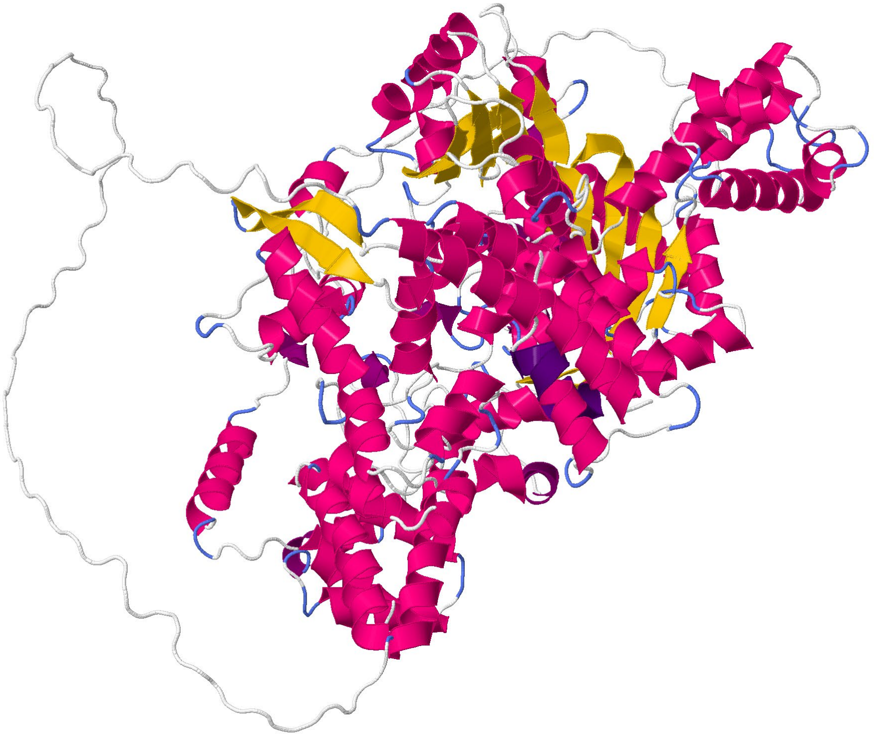

MnA-1
(XQZ12371)

#### SET

Pairwise Identity = 62.0%

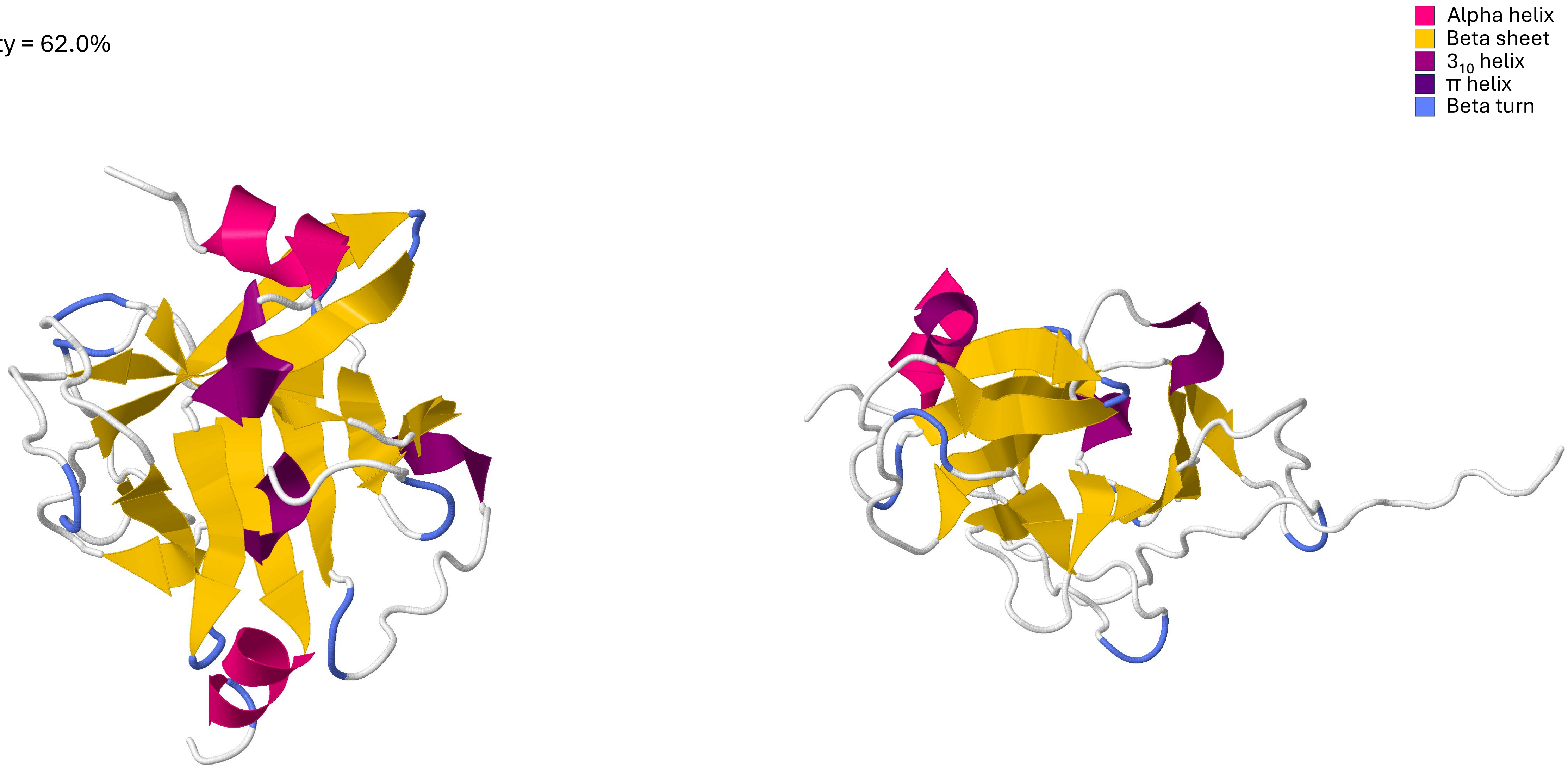


MdA-1
(UFQ21636)

MnA-1
(XQZ12374)

Alpha helix
Beta sheet
$3_{10}$ helix
π helix
Beta turn

**Zifi**

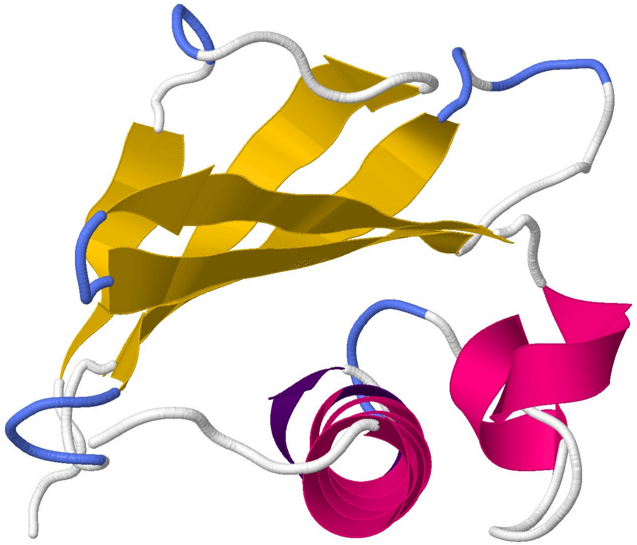

MnA-1
(XQZ12360)

Alpha helix
Beta sheet
$3_{10}$ helix
π helix
Beta turn

**Endonoid**

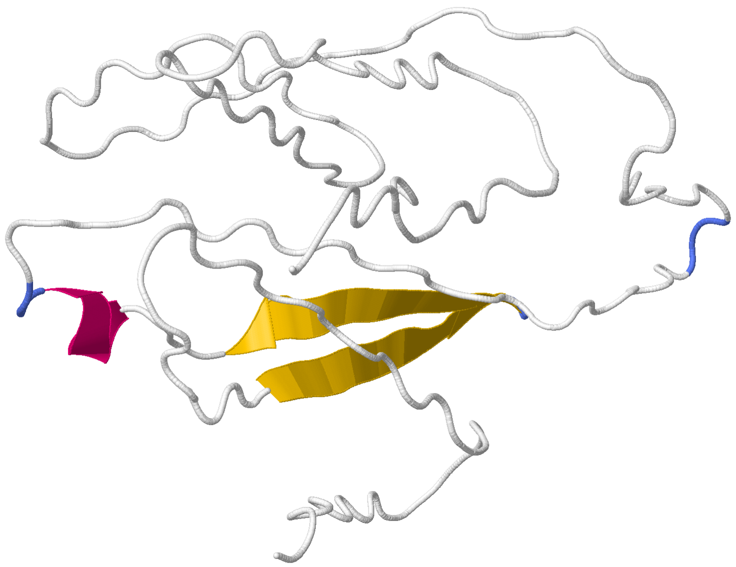

MdA-1
(UFQ21624)

**Herpeto**

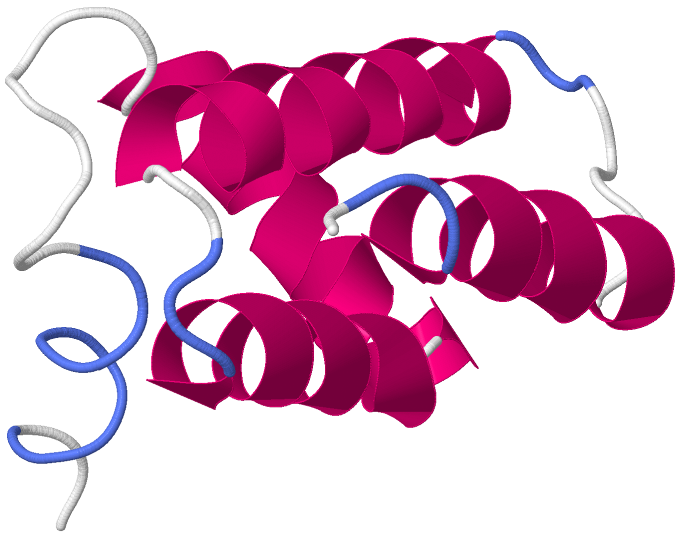

MdA-1
(XQU54336)

**Phogi**

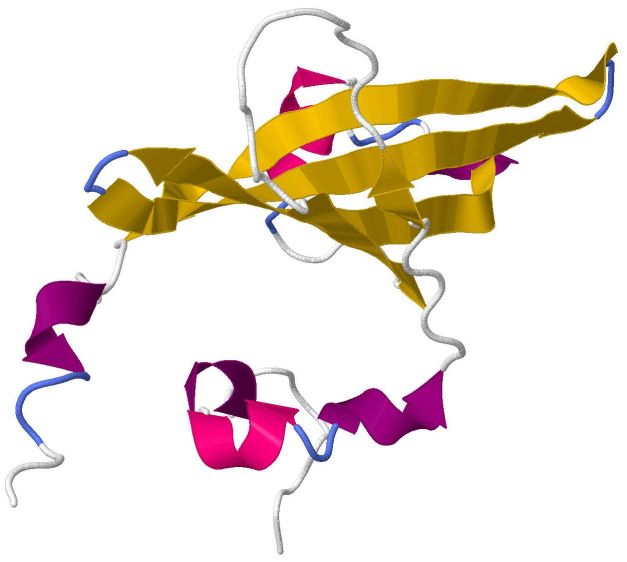

MdA-1
(UFQ21625)
