## Supplementary material for "Novel Adomaviruses Associated with Blotchy Bass Syndrome in Black Basses (*Micropterus spp.)*": S7 Fig

**Supplemental Figure 7**: MAFFT alignment of two MdA-1 genomes originating from different Pennsylvania rivers sampled during different years. SNPs are indicated with vertical lines. Open reading frames are identified.

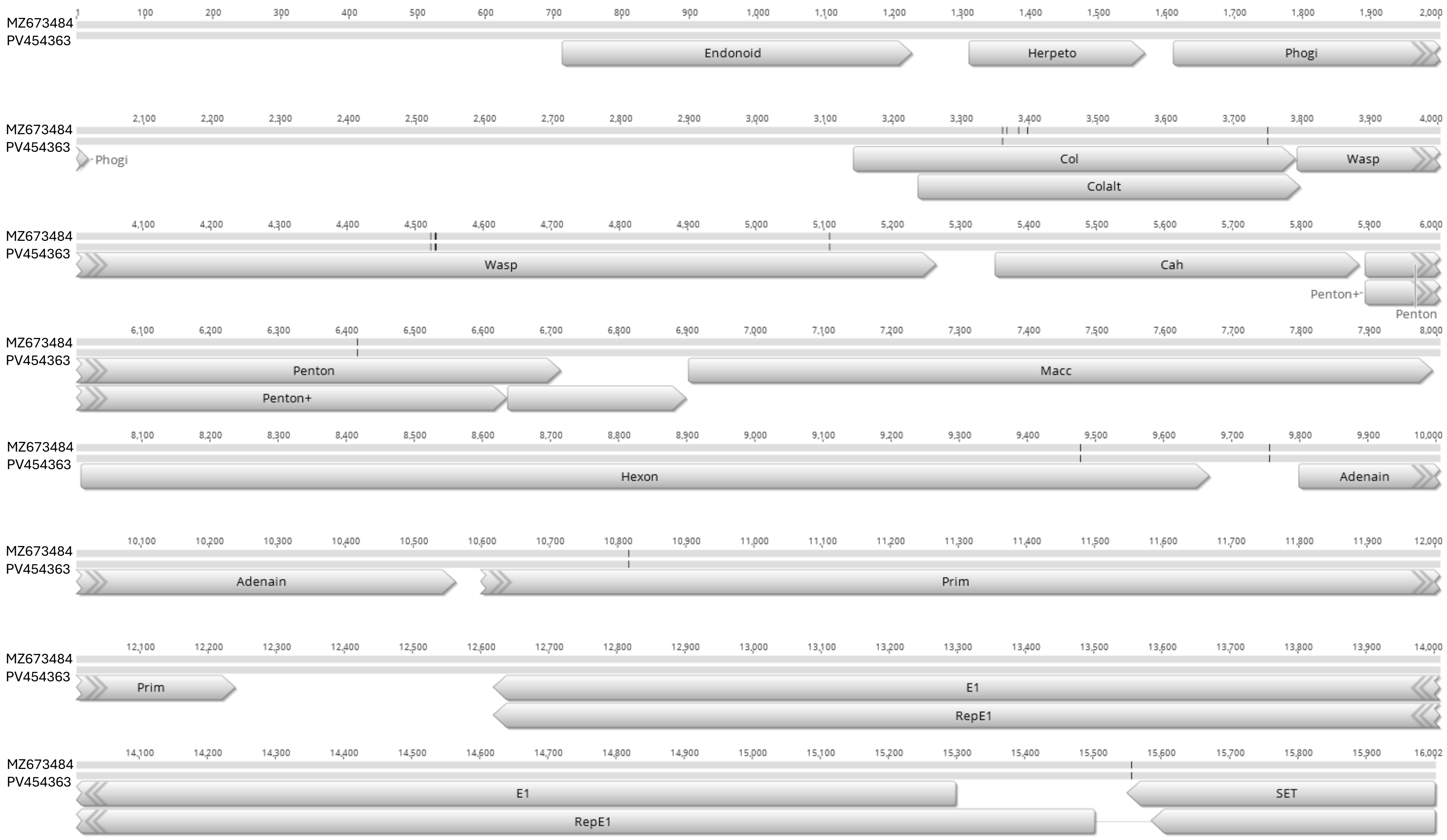
