## Supplementary material for "Novel Adomaviruses Associated with Blotchy Bass Syndrome in Black Basses (*Micropterus spp.)*": S8 Fig

**Supplemental Figure 8**: MAFFT genome alignment of adomaviruses in the same phylogenetic clade as MdA-1 and MnA-1. Core adomavirus ORFs are colored in non-gray. Identity graph depicts similarity/dissimilarity across the genomes (sliding window = 1).

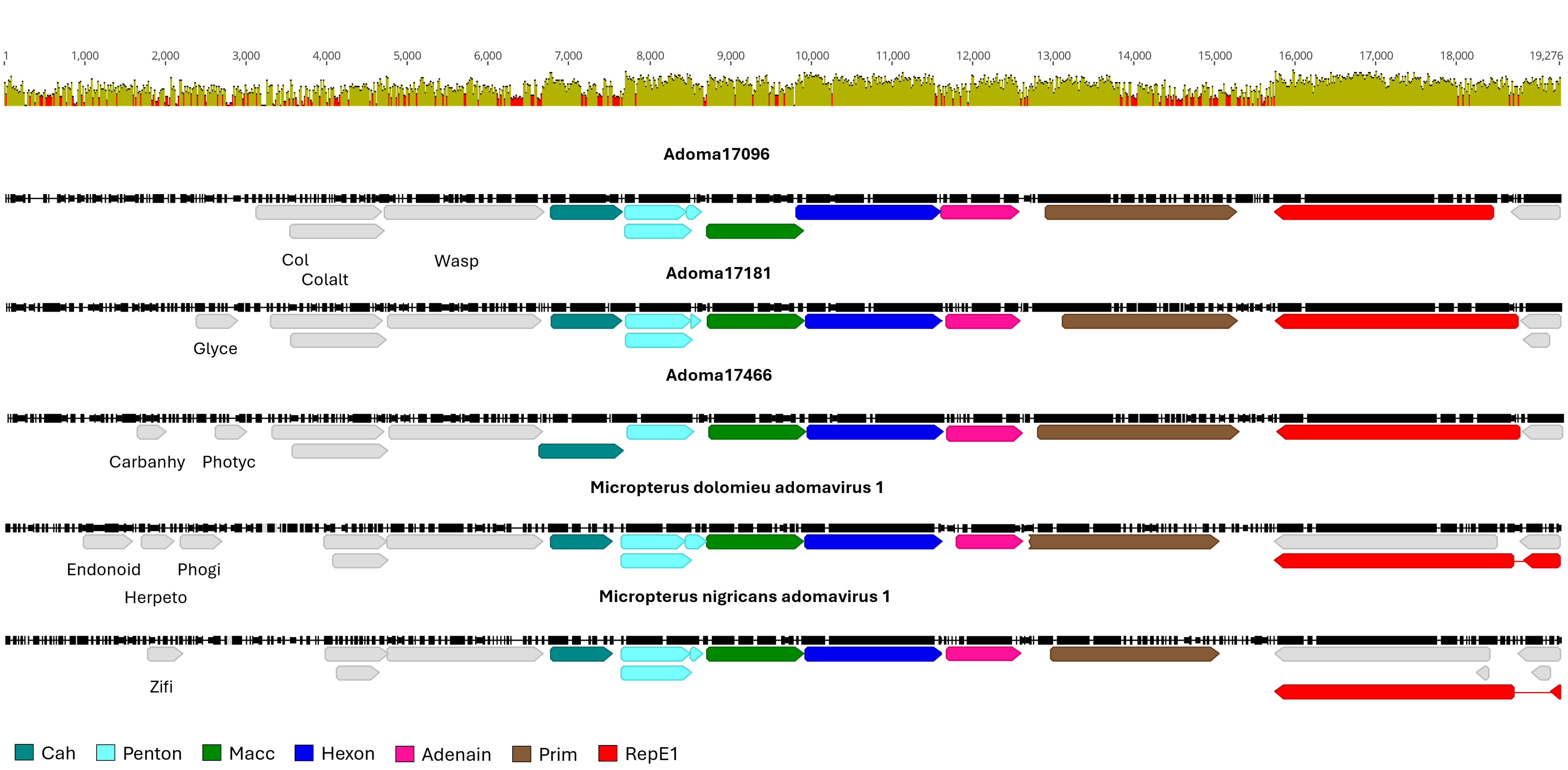
