## Supplementary material for "Novel Adomaviruses Associated with Blotchy Bass Syndrome in Black Basses (*Micropterus spp.)*": S3 Table

| Virus | Protein | Best hit | Name | Probablility | E-Value | Score | Target Length |
|---|---|---|---|---|---|---|---|
| MdA-1 | Prim | 6RB4_A | DNA primase small subunit; Primase, DNA-dependent RNA polymerase, ATP, priming, replication; HET: EDO; 1.5A {Homo sapien | 100 | 1.4-66 | 541.6 | 410 |
| MdA-1 | RepE1 | P26543 | VE1_HPV58 Replication protein E1 OS=Human papillomavirus 58 OX=10598 GN=E1 PE=3 SV=1 | 100 | 1.7-58 | 547.5 | 644 |
| MnA-1 | Prim | 6RB4_A | DNA primase small subunit; Primase, DNA-dependent RNA polymerase, ATP, priming, replication; HET: EDO; 1.5A | 100 | 1.40E-62 | 515.8 | 410 |
| MnA-1 | RepE1 | P26543 | VE1_HPV58 Replication protein E1 OS=Human papillomavirus 58 OX=10598 GN=E1 PE=3 SV=1 | 100 | 1.30E-65 | 602.7 | 644 |
| MdA-1 | Adenain | 4EKF_A | Adenain; alpha and beta protein (a+b), Hydrolase; HET: CSD; 0.98A {Human adenovirus 2} SCOP: d.3.1.7 | 99.96 | 1.50E-28 | 157 | 204 |
| MnA-1 | Adenain | P19119 | PRO_ADEB3 Protease OS=Bovine adenovirus B serotype 3 OX=10510 GN=L3 PE=3 SV=1 | 99.95 | 1.60E-27 | 214 | 204 |
| MnA-1 | SET | 9EH2_I | Histone-lysine N-methyltransferase SETD2; SETD2, Transcription, H3K36me3, TRANSFERASE-RNA-DNA complex; HET: TPO, SEP; 3. | 99.44 | 1.50E-12 | 90.57 | 1133 |
| MdA-1 | SET | cd10545 | SET_AtSUVH-like; SET domain found in Arabidopsis thaliana histone H3-K9 methyltransferases (SUVHs) and similar proteins. | 99.41 | 5.00E-11 | 68.52 | 236 |
| MnA-1 | Penton | Q5UQU9 | YL356_MIMIV Uncharacterized protein L356 OS=Acanthamoeba polyphaga mimivirus OX=212035 GN=MIMI_L356 PE=4 SV=1 | 99.36 | 1.40E-10 | 117.4 | 621 |
| MdA-1 | Penton | Q5UQU9 | P1v1; giant virus, nucleocytoplasmic large DNA viruses (NCLDVs), viral assembly, Paramecium bursaria chlorella virus 1 | 98.72 | 1.10E-06 | 91.48 | 621 |
| MdA-1 | Hexon | PF21738.1 | DJR_capsid ; Double jelly roll capsid-like protein | 97.35 | 0.045 | 57.29 | 316 |
| MnA-1 | Col | PF13717.11 | zinc_ribbon_4; zinc-ribbon domain | 97.2 | 0.00016 | 0.3 | 37 |
| MnA-1 | Hexon | PF21738.1 | DJR_capsid ; Double jelly roll capsid-like protein | 97.11 | 0.1 | 54.79 | 316 |
| MnA-1 | Macc | PF05829.16 | Adeno_PX ; Adenovirus late L2 mu core protein (Protein X) | 97.1 | 0.0014 | 49.75 | 42 |
| MdA-1 | Wasp | PF02514.20 | CobN-Mg_chel ; CobN/Magnesium Chelatase | 96.46 | 0.0031 | 71.18 | 1189 |
| MdA-1 | Macc | PF05829.16 | Adeno_PX ; Adenovirus late L2 mu core protein (Protein X) | 96.14 | 0.019 | 44.2 | 42 |
| MnA-1 | Cah | P04507 | SIGM1_REOVJ Outer capsid protein sigma-1 OS=Reovirus type 2 (strain D5/Jones) OX=10885 GN=S1 PE=3 SV=3 | 91.58 | 13 | 35.5 | 462 |
| MdA-1 | Cah | P04507 | SIGM1_REOVJ Outer capsid protein sigma-1 OS=Reovirus type 2 (strain D5/Jones) OX=10885 GN=S1 PE=3 SV=3 | 90.5 | 26 | 33.57 | 462 |
| MdA-1 | Phogi | cd05016 | SIS_PGI_2; Phosphoglucose isomerase (PGI) contains two SIS (Sugar ISomerase) domains. | 60.54 | 20 | 24.22 | 192 |
| MnA-1 | Zifi | PF14608.10 | zf-CCCH_2 ; RNA-binding, Nab2-type zinc finger | 54.44 | 9.2 | 18.61 | 18 |
| MnA-1 | Wasp | PF14357.10 | DUF4404 ; Domain of unknown function (DUF4404) | 51.82 | 26 | 28.47 | 85 |
| MnA-1 | Colalt | 7TOP_PR | PR20; Ribosomal binding peptide, ALS/FTD-associated dipeptide repeat protein, RIBOSOME; 2.4A {Saccharomyces cerevisiae} | 48.29 | 100 | 21.11 | 40 |
| MdA-1 | Herpeto | PF14409.11 | Herpeto_peptide ; Ribosomally synthesized peptide in Herpetosiphon | 36.76 | 32 | 23.38 | 62 |
| MdA-1 | Col | 2KLZ_A | Ataxin-3; UIM, Ataxin-3, Ubiquitin-binding, Hydrolase, Neurodegeneration, Nucleus, Phosphoprotein, Spinocerebellar ataxi | 27.89 | 110 | 17.3 | 52 |
| MdA-1 | Endonoid | PF17785.5 | PUA_3 ; PUA-like domain | 27.53 | 41 | 19.62 | 64 |
| MdA-1 | Colalt | 8Q7N_T | Transcription elongation regulator 1; spliceosome, pre-catalytic spliceosome, spliceosomal B complex, SPLICING; 3.1A | 7.74 | 810 | 23.03 | 1095 |
