## Supplementary material for "Novel Adomaviruses Associated with Blotchy Bass Syndrome in Black Basses (*Micropterus spp.)*": S4 Table

**DALI results: MdA-1 Endonoid**

| No: | Chain | Z | rmsd | lali | nres | %id | PDB_Description |
|---|---|---|---|---|---|---|---|
| 1:00 | s001-A | 22.3 | 20.5 | 170 | 0 | 0 | NA |

**DALI results: MdA-1 Herpeto**

| *No:* | *Chain* | *Z* | *rmsd* | *lali* | *nres* | *%id* | *PDB* | *Description* |
|---|---|---|---|---|---|---|---|---|
| 1:00 | 7ok0-C | 6 | 2 | 55 | 393 | 5 | MOLECULE: | DNA-DIRECTED RNA POLYMERASE SUBUNIT A' |
| 2:00 | 7oqy-C | 5.8 | 2 | 54 | 383 | 6 | MOLECULE: | DNA-DIRECTED RNA POLYMERASE SUBUNIT A' |
| 3:00 | 8p2i-C | 5.7 | 2 | 52 | 390 | 8 | MOLECULE: | DNA-DIRECTED RNA POLYMERASE SUBUNIT RPO1N |
| 4:00 | 8cro-C | 5.6 | 1.9 | 52 | 390 | 8 | MOLECULE: | DNA-DIRECTED RNA POLYMERASE SUBUNIT RPO1N |
| 5:00 | 7oq4-C | 5.5 | 1.9 | 54 | 393 | 6 | MOLECULE: | DNA-DIRECTED RNA POLYMERASE SUBUNIT A' |
| 6:00 | 4ayb-C | 5.4 | 1.7 | 52 | 376 | 12 | MOLECULE: | DNA-DIRECTED RNA POLYMERASE |
| 7:00 | 8rbo-C | 5.4 | 2 | 53 | 390 | 8 | MOLECULE: | DNA-DIRECTED RNA POLYMERASE SUBUNIT RPO1N |
| 8:00 | 8orq-C | 5.3 | 2 | 52 | 388 | 8 | MOLECULE: | DNA-DIRECTED RNA POLYMERASE SUBUNIT RPO1N |
| 9:00 | 6kf9-C | 5 | 2 | 51 | 388 | 12 | MOLECULE: | DNA-DIRECTED RNA POLYMERASE SUBUNIT |
| 10:00 | 6kf3-C | 5 | 1.9 | 51 | 388 | 12 | MOLECULE: | DNA-DIRECTED RNA POLYMERASE SUBUNIT |
| 11:00 | 8oki-C | 5 | 1.9 | 52 | 390 | 8 | MOLECULE: | DNA-DIRECTED RNA POLYMERASE SUBUNIT RPO1N |
| 12:00 | 2pmz-G | 5 | 1.9 | 51 | 279 | 4 | MOLECULE: | DNA-DIRECTED RNA POLYMERASE SUBUNIT A |
| 13:00 | 1k30-A | 5 | 1.9 | 54 | 363 | 9 | MOLECULE: | GLYCEROL-3-PHOSPHATE ACYLTRANSFERASE |
| 14:00 | 6kf4-C | 4.9 | 2.1 | 51 | 388 | 12 | MOLECULE: | DNA-DIRECTED RNA POLYMERASE SUBUNIT |
| 15:00 | 6swr-E | 4.9 | 2.1 | 55 | 222 | 11 | MOLECULE: | NANOBODY, MALTOSE/MALTODEXTRIN-BINDING PERIPLASMI |
| 16:00 | 4qiw-C | 4.8 | 1.7 | 47 | 369 | 13 | MOLECULE: | DNA-DIRECTED RNA POLYMERASE |
| 17:00 | 1iuq-A | 4.8 | 1.9 | 54 | 357 | 9 | MOLECULE: | GLYCEROL-3-PHOSPHATE ACYLTRANSFERASE |
| 18:00 | 7jgs-C | 4.7 | 3.3 | 60 | 614 | 13 | MOLECULE: | ORIGIN RECOGNITION COMPLEX SUBUNIT 2 |
| 19:00 | 3hkz-M | 4.7 | 1.8 | 51 | 370 | 4 | MOLECULE: | DNA-DIRECTED RNA POLYMERASE SUBUNIT A' |
| 20:00 | 2px0-B | 4.7 | 3.3 | 54 | 258 | 11 | MOLECULE: | FLAGELLAR BIOSYNTHESIS PROTEIN FLHF |
| 21:00 | 2waq-C | 4.6 | 1.8 | 49 | 367 | 6 | MOLECULE: | DNA-DIRECTED RNA POLYMERASE RPO1N SUBUNIT |
| 22:00 | 2y0s-Y | 4.6 | 1.9 | 49 | 368 | 8 | MOLECULE: | DNA-DIRECTED RNA POLYMERASE |
| 23:00 | 2y0s-C | 4.6 | 1.9 | 49 | 368 | 8 | MOLECULE: | DNA-DIRECTED RNA POLYMERASE |
| 24:00:00 | 8vej-A | 4.6 | 2.2 | 54 | 240 | 7 | MOLECULE: | CHD_BUTTRESS |
| 25:00:00 | 6hd8-B | 4.5 | 2.1 | 55 | 232 | 11 | MOLECULE: | NANOBODY,MALTOSE/MALTODEXTRIN-BINDING PERIPLASMIC |
| 26:00:00 | 6hd9-B | 4.5 | 2.1 | 55 | 232 | 11 | MOLECULE: | NANOBODY,MALTOSE/MALTODEXTRIN-BINDING PERIPLASMIC |
| 27:00:00 | 2wb1-C | 4.5 | 1.8 | 49 | 367 | 6 | MOLECULE: | DNA-DIRECTED RNA POLYMERASE RPO1N SUBUNIT |
| 28:00:00 | 6xj1-A | 4.5 | 3.2 | 58 | 266 | 3 | MOLECULE: | CELL DIVISION CONTROL PROTEIN 15 |
| 29:00:00 | 2pmz-C | 4.5 | 1.9 | 51 | 279 | 4 | MOLECULE: | DNA-DIRECTED RNA POLYMERASE SUBUNIT A |
| 30:00:00 | 4qiw-M | 4.5 | 1.7 | 47 | 369 | 13 | MOLECULE: | DNA-DIRECTED RNA POLYMERASE |
| 31:00:00 | 8v2d-c | 4.5 | 3.1 | 75 | 314 | 5 | MOLECULE: | O43_129 COMPONENT B |
| 32:00:00 | 8v3b-V | 4.5 | 3.1 | 75 | 314 | 5 | MOLECULE: | O43_129_+4 COMPONENT A |
| 33:00:00 | 8v3b-I | 4.5 | 3.2 | 77 | 314 | 5 | MOLECULE: | O43_129_+4 COMPONENT A |
| 34:00:00 | 6hdc-B | 4.5 | 2.1 | 55 | 232 | 11 | MOLECULE: | NANOBODY,MALTOSE/MALTODEXTRIN-BINDING PERIPLASMIC |
| 35:00:00 | 6swr-B | 4.5 | 2.1 | 55 | 230 | 11 | MOLECULE: | NANOBODY, MALTOSE/MALTODEXTRIN-BINDING PERIPLASMI |
| 36:00:00 | 2px0-C | 4.5 | 3.3 | 54 | 258 | 11 | MOLECULE: | FLAGELLAR BIOSYNTHESIS PROTEIN FLHF |
| 37:00:00 | 7yh8-A | 4.5 | 2.6 | 53 | 62 | 11 | MOLECULE: | L-19437 |
| 38:00:00 | 3adb-B | 4.5 | 3 | 57 | 247 | 12 | MOLECULE: | L-SERYL-TRNA(SEC) KINASE |
| 39:00:00 | 7yh8-C | 4.5 | 2.6 | 53 | 62 | 11 | MOLECULE: | L-19437 |
| 40:00:00 | 1nxh-A | 4.5 | 2.5 | 57 | 124 | 7 | MOLECULE: | MTH396 PROTEIN |
| 41:00:00 | 1nxh-B | 4.5 | 2.5 | 57 | 118 | 7 | MOLECULE: | MTH396 PROTEIN |
| 42:00:00 | 8v2d-k | 4.4 | 3.1 | 75 | 314 | 5 | MOLECULE: | O43_129 COMPONENT B |
| 43:00:00 | 8v2d-Z | 4.4 | 3 | 75 | 314 | 5 | MOLECULE: | O43_129 COMPONENT B |
| 44:00:00 | 3syn-C | 4.4 | 3.3 | 54 | 258 | 11 | MOLECULE: | FLAGELLAR BIOSYNTHESIS PROTEIN FLHF |
| 45:00:00 | 3syn-D | 4.4 | 3.3 | 54 | 258 | 11 | MOLECULE: | FLAGELLAR BIOSYNTHESIS PROTEIN FLHF |
| 46:00:00 | 3syn-B | 4.4 | 3.3 | 54 | 258 | 11 | MOLECULE: | FLAGELLAR BIOSYNTHESIS PROTEIN FLHF |
| 47:00:00 | 3syn-A | 4.4 | 3.3 | 54 | 258 | 11 | MOLECULE: | FLAGELLAR BIOSYNTHESIS PROTEIN FLHF |
| 48:00:00 | 7jk3-C | 4.4 | 3.1 | 60 | 614 | 10 | MOLECULE: | ORIGIN RECOGNITION COMPLEX SUBUNIT 2 |
| 49:00:00 | 7bve-A | 4.4 | 2.4 | 58 | 1034 | 5 | MOLECULE: | INTEGRAL MEMBRANE INDOLYLACETYLINOSITOL |
| 50:00:00 | 8v3b-U | 4.4 | 3.5 | 62 | 314 | 10 | MOLECULE: | O43_129_+4 COMPONENT A |
| 51:00:00 | 8v2d-f | 4.4 | 3.5 | 69 | 314 | 1 | MOLECULE: | O43_129 COMPONENT B |
| 52:00:00 | 8v3b-L | 4.4 | 2.9 | 59 | 314 | 5 | MOLECULE: | O43_129_+4 COMPONENT A |
| 53:00:00 | 7jzm-A | 4.3 | 3.8 | 55 | 64 | 13 | MOLECULE: | LCB3 |
| 54:00:00 | 2px0-A | 4.3 | 3.3 | 54 | 258 | 11 | MOLECULE: | FLAGELLAR BIOSYNTHESIS PROTEIN FLHF |

| | | | | | | | | |
|---|---|---|---|---|---|---|---|---|
| 55:00:00 | 3add-A | 4.3 | 3.5 | 57 | 251 | 5 | MOLECULE: | L-SERYL-TRNA(SEC) KINASE |
| 56:00:00 | 2wb1-Y | 4.3 | 1.8 | 49 | 367 | 6 | MOLECULE: | DNA-DIRECTED RNA POLYMERASE RPO1N SUBUNIT |
| 57:00:00 | 8v3b-R | 4.3 | 3.1 | 75 | 314 | 5 | MOLECULE: | O43_129_+4 COMPONENT A |
| 58:00:00 | 1r4g-A | 4.3 | 2.5 | 49 | 53 | 6 | MOLECULE: | RNA POLYMERASE ALPHA SUBUNIT |
| 59:00:00 | 8v3b-C | 4.3 | 3.1 | 75 | 314 | 5 | MOLECULE: | O43_129_+4 COMPONENT A |
| 60:00:00 | 8v2d-X | 4.3 | 3.1 | 76 | 314 | 5 | MOLECULE: | O43_129 COMPONENT B |
| 61:00:00 | 2px0-F | 4.3 | 3.4 | 54 | 258 | 11 | MOLECULE: | FLAGELLAR BIOSYNTHESIS PROTEIN FLHF |
| 62:00:00 | 3add-B | 4.3 | 2.8 | 52 | 239 | 12 | MOLECULE: | L-SERYL-TRNA(SEC) KINASE |
| 63:00:00 | 7ctg-C | 4.3 | 3.5 | 61 | 592 | 7 | MOLECULE: | ORIGIN RECOGNITION COMPLEX SUBUNIT 1 |
| 64:00:00 | 3a4m-A | 4.3 | 3.6 | 68 | 236 | 9 | MOLECULE: | L-SERYL-TRNA(SEC) KINASE |
| 65:00:00 | 8v3b-H | 4.3 | 3.7 | 65 | 314 | 9 | MOLECULE: | O43_129_+4 COMPONENT A |
| 66:00:00 | 7upq-G | 4.3 | 3.9 | 74 | 198 | 3 | MOLECULE: | DHT03 PROTEIN A |
| 67:00:00 | 8v3b-J | 4.3 | 3.3 | 60 | 314 | 5 | MOLECULE: | O43_129_+4 COMPONENT A |
| 68:00:00 | 8fbi-A | 4.3 | 3.1 | 64 | 274 | 8 | MOLECULE: | KWOCA_39 |
| 69:00:00 | 3a4m-B | 4.3 | 3.8 | 68 | 236 | 12 | MOLECULE: | L-SERYL-TRNA(SEC) KINASE |
| 70:00:00 | 8vej-B | 4.3 | 2.3 | 54 | 242 | 7 | MOLECULE: | CHD_BUTTRESS |
| 71:00:00 | 7jk2-C | 4.3 | 2.7 | 56 | 609 | 11 | MOLECULE: | ORIGIN RECOGNITION COMPLEX SUBUNIT 2 |
| 72:00:00 | 8v3b-W | 4.2 | 3.1 | 76 | 314 | 5 | MOLECULE: | O43_129_+4 COMPONENT A |
| 73:00:00 | 8v2d-g | 4.2 | 3.1 | 75 | 314 | 5 | MOLECULE: | O43_129 COMPONENT B |
| 74:00:00 | 4xgc-C | 4.2 | 3.2 | 61 | 567 | 11 | MOLECULE: | ORIGIN RECOGNITION COMPLEX SUBUNIT 2 |
| 75:00:00 | 3adc-A | 4.2 | 2.8 | 53 | 246 | 11 | MOLECULE: | L-SERYL-TRNA(SEC) KINASE |
| 76:00:00 | 2px0-E | 4.2 | 3.2 | 53 | 258 | 11 | MOLECULE: | FLAGELLAR BIOSYNTHESIS PROTEIN FLHF |
| 77:00:00 | 8v3b-X | 4.2 | 3.7 | 63 | 314 | 10 | MOLECULE: | O43_129_+4 COMPONENT A |
| 78:00:00 | 8v3b-P | 4.2 | 3.8 | 64 | 314 | 9 | MOLECULE: | O43_129_+4 COMPONENT A |
| 79:00:00 | 8v2d-m | 4.2 | 2.9 | 59 | 176 | 12 | MOLECULE: | O43_129 COMPONENT B |
| 80:00:00 | 8v3b-T | 4.2 | 2.7 | 56 | 314 | 5 | MOLECULE: | O43_129_+4 COMPONENT A |
| 81:00:00 | 3a4n-A | 4.2 | 3.4 | 66 | 237 | 11 | MOLECULE: | L-SERYL-TRNA(SEC) KINASE |
| 82:00:00 | 8v3b-N | 4.2 | 3.5 | 61 | 314 | 11 | MOLECULE: | O43_129_+4 COMPONENT A |
| 83:00:00 | 8v3b-K | 4.1 | 3.2 | 76 | 314 | 5 | MOLECULE: | O43_129_+4 COMPONENT A |
| 84:00:00 | 8v2d-a | 4.1 | 3.2 | 75 | 314 | 5 | MOLECULE: | O43_129 COMPONENT B |
| 85:00:00 | 2px3-A | 4.1 | 3.2 | 54 | 256 | 11 | MOLECULE: | FLAGELLAR BIOSYNTHESIS PROTEIN FLHF |
| 86:00:00 | 5uj8-C | 4.1 | 3.1 | 62 | 553 | 6 | MOLECULE: | ORIGIN RECOGNITION COMPLEX SUBUNIT 3 |
| 87:00:00 | 7bvf-B | 4.1 | 2.3 | 55 | 1054 | 4 | MOLECULE: | PROBABLE ARABINOSYLTRANSFERASE B |
| 88:00:00 | 8v2d-b | 4.1 | 3.1 | 77 | 314 | 6 | MOLECULE: | O43_129 COMPONENT B |
| 89:00:00 | 8v3b-S | 4.1 | 3.2 | 78 | 314 | 6 | MOLECULE: | O43_129_+4 COMPONENT A |
| 90:00:00 | 3am1-A | 4.1 | 2.8 | 52 | 236 | 12 | MOLECULE: | L-SERYL-TRNA(SEC) KINASE |
| 91:00:00 | 3adc-B | 4.1 | 2.8 | 51 | 239 | 12 | MOLECULE: | L-SERYL-TRNA(SEC) KINASE |
| 92:00:00 | 8v3b-G | 4.1 | 3.8 | 65 | 314 | 9 | MOLECULE: | O43_129_+4 COMPONENT A |
| 93:00:00 | 8v2d-e | 4.1 | 2.8 | 55 | 314 | 5 | MOLECULE: | O43_129 COMPONENT B |
| 94:00:00 | 3vr5-A | 4.1 | 3.3 | 67 | 588 | 9 | MOLECULE: | V-TYPE SODIUM ATPASE CATALYTIC SUBUNIT A |
| 95:00:00 | 7tmp-C | 4.1 | 4.5 | 66 | 592 | 5 | MOLECULE: | H(+)-TRANSPORTING TWO-SECTOR ATPASE |
| 96:00:00 | 3vr4-A | 4.1 | 3.1 | 69 | 588 | 10 | MOLECULE: | V-TYPE SODIUM ATPASE CATALYTIC SUBUNIT A |
| 97:00:00 | 8v2d-J | 4.1 | 3.6 | 64 | 314 | 9 | MOLECULE: | O43_129 COMPONENT B |
| 98:00:00 | 8v3b-M | 4.1 | 2.9 | 64 | 314 | 2 | MOLECULE: | O43_129_+4 COMPONENT A |
| 99:00:00 | 8v3b-O | 4.1 | 3.5 | 63 | 314 | 10 | MOLECULE: | O43_129_+4 COMPONENT A |
| 0:00 | 4p1w-C | 4 | 3.9 | 58 | 399 | 10 | MOLECULE: | ATG29 |
| 1:00 | 7jzn-E | 4 | 3.7 | 54 | 64 | 13 | MOLECULE: | SPIKE GLYCOPROTEIN |
| 2:00 | 6vmt-A | 4 | 3.1 | 64 | 99 | 9 | MOLECULE: | PROGRAMMED CELL DEATH PROTEIN 5 HOMOLOG |
| 3:00 | 7jzn-F | 4 | 3.8 | 54 | 64 | 13 | MOLECULE: | SPIKE GLYCOPROTEIN |
| 4:00 | 8v3b-F | 4 | 3.1 | 75 | 314 | 5 | MOLECULE: | O43_129_+4 COMPONENT A |
| 5:00 | 8v2d-h | 4 | 3.1 | 75 | 314 | 5 | MOLECULE: | O43_129 COMPONENT B |
| 6:00 | 1vdz-A | 4 | 3 | 70 | 513 | 10 | MOLECULE: | A-TYPE ATPASE SUBUNIT A |
| 7:00 | 2v3c-C | 4 | 2.7 | 56 | 403 | 9 | MOLECULE: | SIGNAL RECOGNITION PARTICLE 19 KDA PROTEIN |
| 8:00 | 8ia1-A | 4 | 2.2 | 54 | 361 | 6 | MOLECULE: | GLYCEROL-3-PHOSPHATE ACYLTRANSFERASE, CHLOROPLAST |
| 9:00 | 8ia1-B | 4 | 2.2 | 54 | 361 | 6 | MOLECULE: | GLYCEROL-3-PHOSPHATE ACYLTRANSFERASE, CHLOROPLAST |
| 10:00 | 1j8m-F | 4 | 2.5 | 55 | 295 | 4 | MOLECULE: | SIGNAL RECOGNITION 54 KDA PROTEIN |

| | | | | | | | | |
|---|---|---|---|---|---|---|---|---|
| 11:00 | 2px0-G | 4 | 3.2 | 54 | 258 | 11 | MOLECULE: | FLAGELLAR BIOSYNTHESIS PROTEIN FLHF |
| 12:00 | 6x0o-A | 4 | 2.8 | 58 | 1037 | 3 | MOLECULE: | INTEGRAL MEMBRANE INDOLYLACETYLINOSITOL |
| 13:00 | 6ty9-A | 4 | 3.8 | 76 | 1208 | 7 | MOLECULE: | RNA-DEPENDENT RNA POLYMERASE |
| 14:00 | 3a4l-A | 4 | 3.6 | 67 | 236 | 9 | MOLECULE: | L-SERYL-TRNA(SEC) KINASE |
| 15:00 | 1oks-A | 4 | 2.1 | 48 | 53 | 8 | MOLECULE: | RNA POLYMERASE ALPHA SUBUNIT |
| 16:00 | 3a4l-B | 4 | 3.7 | 68 | 237 | 9 | MOLECULE: | L-SERYL-TRNA(SEC) KINASE |
| 17:00 | 2px0-D | 4 | 3.4 | 54 | 258 | 11 | MOLECULE: | FLAGELLAR BIOSYNTHESIS PROTEIN FLHF |
| 18:00 | 3a4n-B | 4 | 3.4 | 66 | 237 | 12 | MOLECULE: | L-SERYL-TRNA(SEC) KINASE |
| 19:00 | 7bvf-A | 4 | 2.7 | 57 | 1087 | 9 | MOLECULE: | PROBABLE ARABINOSYLTRANSFERASE B |
| 20:00 | 7tmr-A | 4 | 2.9 | 64 | 593 | 8 | MOLECULE: | H(+)-TRANSPORTING TWO-SECTOR ATPASE |
| 21:00 | 7c2n-A | 4 | 2.8 | 54 | 900 | 7 | MOLECULE: | DRUG EXPORTERS OF THE RND SUPERFAMILY-LIKE PROTEI |
| 22:00 | 8v2d-Y | 4 | 3.4 | 62 | 314 | 10 | MOLECULE: | O43_129 COMPONENT B |
| 23:00 | 8v3b-B | 4 | 2.9 | 64 | 314 | 2 | MOLECULE: | O43_129_+4 COMPONENT A |
| 24:00:00 | 7cos-I | 4 | 3 | 65 | 583 | 6 | MOLECULE: | V-TYPE SODIUM ATPASE CATALYTIC SUBUNIT A |
| 25:00:00 | 3hkz-C | 3.9 | 2.1 | 51 | 370 | 6 | MOLECULE: | DNA-DIRECTED RNA POLYMERASE SUBUNIT A' |
| 26:00:00 | 8v3b-D | 3.9 | 3.2 | 75 | 314 | 5 | MOLECULE: | O43_129_+4 COMPONENT A |
| 27:00:00 | 8v2d-l | 3.9 | 3.2 | 76 | 314 | 5 | MOLECULE: | O43_129 COMPONENT B |
| 28:00:00 | 5mj4-C | 3.9 | 4 | 56 | 81 | 2 | MOLECULE: | INTERLEUKIN-12 SUBUNIT BETA |
| 29:00:00 | 5lxj-A | 3.9 | 3.4 | 51 | 53 | 6 | MOLECULE: | PHOSPHOPROTEIN |
| 30:00:00 | 6em5-R | 3.9 | 3.3 | 55 | 120 | 7 | MOLECULE: | 5.8S RIBOSOMAL RNA |
| 31:00:00 | 4og9-C | 3.9 | 4 | 56 | 81 | 2 | MOLECULE: | INTERLEUKIN-12 SUBUNIT BETA |
| 32:00:00 | 7tcd-A | 3.9 | 3.4 | 66 | 436 | 11 | MOLECULE: | D13 LOV2-DARPIN FUSION |
| 33:00:00 | 3adb-A | 3.9 | 2.7 | 51 | 250 | 12 | MOLECULE: | L-SERYL-TRNA(SEC) KINASE |
| 34:00:00 | 7coq-A | 3.9 | 3 | 66 | 586 | 9 | MOLECULE: | V-TYPE SODIUM ATPASE CATALYTIC SUBUNIT A |
| 35:00:00 | 8v2d-j | 3.9 | 3.4 | 64 | 314 | 0 | MOLECULE: | O43_129 COMPONENT B |
| 36:00:00 | 7upq-J | 3.9 | 3.6 | 75 | 197 | 4 | MOLECULE: | DHT03 PROTEIN A |
| 37:00:00 | 7cop-B | 3.9 | 2.8 | 59 | 586 | 5 | MOLECULE: | V-TYPE SODIUM ATPASE CATALYTIC SUBUNIT A |
| 38:00:00 | 7qps-A | 3.9 | 2.3 | 54 | 322 | 6 | MOLECULE: | ACT DOMAIN PROTEIN |
| 39:00:00 | 7bv1-C | 3.8 | 3.7 | 53 | 63 | 4 | MOLECULE: | NSP12 |
| 40:00:00 | 8ce8-b | 3.8 | 3.4 | 70 | 219 | 11 | MOLECULE: | CYTOCHROME C BIOGENESIS ATP-BINDING EXPORT PROTEI |
| 41:00:00 | 4oe8-C | 3.8 | 3.8 | 57 | 87 | 2 | MOLECULE: | INTERLEUKIN-12 SUBUNIT BETA |
| 42:00:00 | 6zr2-n | 3.8 | 4.5 | 58 | 177 | 3 | MOLECULE: | NADH-UBIQUINONE OXIDOREDUCTASE CHAIN 3 |
| 43:00:00 | 8kdb-B | 3.8 | 2.5 | 51 | 65 | 12 | MOLECULE: | RNA-DIRECTED RNA POLYMERASE L |
| 44:00:00 | 8v9u-A | 3.8 | 2.5 | 52 | 81 | 8 | MOLECULE: | DNA (CYTOSINE-5)-METHYLTRANSFERASE 1 |
| 45:00:00 | 8kdc-B | 3.8 | 2.6 | 51 | 65 | 12 | MOLECULE: | RNA-DIRECTED RNA POLYMERASE L |
| 46:00:00 | 8etc-R | 3.8 | 3 | 51 | 124 | 6 | MOLECULE: | RNA (2151-MER) |
| 47:00:00 | 5f3p-B | 3.8 | 4.1 | 60 | 188 | 10 | MOLECULE: | PUTATIVE UNCHARACTERIZED PROTEIN |
| 48:00:00 | 7jgr-C | 3.8 | 3.1 | 59 | 612 | 8 | MOLECULE: | ORIGIN RECOGNITION COMPLEX SUBUNIT 2 |
| 49:00:00 | 6s2v-C | 3.8 | 2.8 | 58 | 336 | 3 | MOLECULE: | (P)PPGPP SYNTHETASE I, SPOT/RELA |
| 50:00:00 | 5vre-D | 3.8 | 3 | 54 | 188 | 6 | MOLECULE: | PUTATIVE INTEGRAL MEMBRANE PROTEIN |
| 51:00:00 | 7dqd-K | 3.8 | 4.2 | 61 | 584 | 7 | MOLECULE: | V-TYPE SODIUM ATPASE CATALYTIC SUBUNIT A |
| 52:00:00 | 5uj8-D | 3.8 | 3.1 | 62 | 553 | 6 | MOLECULE: | ORIGIN RECOGNITION COMPLEX SUBUNIT 3 |
| 53:00:00 | 6vq9-A | 3.8 | 4.5 | 65 | 600 | 5 | MOLECULE: | ATPASE H+-TRANSPORTING V1 SUBUNIT A |
| 54:00:00 | 8rgr-n | 3.8 | 4.4 | 56 | 178 | 4 | MOLECULE: | NADH DEHYDROGENASE [UBIQUINONE] IRON-SULFUR PROTE |
| 55:00:00 | 7jzn-G | 3.8 | 3.8 | 54 | 64 | 13 | MOLECULE: | SPIKE GLYCOPROTEIN |
| 56:00:00 | 7w1t-p | 3.8 | 4.5 | 58 | 178 | 3 | MOLECULE: | NADH DEHYDROGENASE [UBIQUINONE] FLAVOPROTEIN 1, |
| 57:00:00 | 5mj3-C | 3.8 | 3.6 | 57 | 87 | 0 | MOLECULE: | INTERLEUKIN-12 SUBUNIT BETA |
| 58:00:00 | 7jpq-C | 3.8 | 3.2 | 60 | 587 | 5 | MOLECULE: | ORIGIN RECOGNITION COMPLEX SUBUNIT 2 |
| 59:00:00 | 8rgq-n | 3.8 | 4.4 | 56 | 178 | 4 | MOLECULE: | NADH DEHYDROGENASE [UBIQUINONE] IRON-SULFUR PROTE |
| 60:00:00 | 8rgp-n | 3.8 | 4.5 | 57 | 178 | 4 | MOLECULE: | NADH DEHYDROGENASE [UBIQUINONE] IRON-SULFUR PROTE |
| 61:00:00 | 8v3b-E | 3.8 | 2 | 43 | 314 | 12 | MOLECULE: | O43_129_+4 COMPONENT A |
| 62:00:00 | 8v3b-A | 3.8 | 2.1 | 43 | 314 | 12 | MOLECULE: | O43_129_+4 COMPONENT A |
| 63:00:00 | 7fde-C | 3.8 | 2.8 | 66 | 594 | 8 | MOLECULE: | V-TYPE PROTON ATPASE SUBUNIT C |
| 64:00:00 | 5zea-H | 3.8 | 3.1 | 67 | 586 | 9 | MOLECULE: | V-TYPE SODIUM ATPASE CATALYTIC SUBUNIT A |
| 65:00:00 | 7dqd-J | 3.8 | 3.9 | 74 | 584 | 5 | MOLECULE: | V-TYPE SODIUM ATPASE CATALYTIC SUBUNIT A |
| 66:00:00 | 6qum-B | 3.8 | 3 | 67 | 578 | 10 | MOLECULE: | V-TYPE ATP SYNTHASE ALPHA CHAIN |

| | | | | | | | | |
|---|---|---|---|---|---|---|---|---|
| 67:00:00 | 6vq6-A | 3.7 | 4.5 | 65 | 600 | 5 | MOLECULE: | ATPASE H+-TRANSPORTING V1 SUBUNIT A |
| 68:00:00 | 4d7x-A | 3.7 | 2.2 | 50 | 86 | 6 | MOLECULE: | MEDIATOR OF RNA POLYMERASE II TRANSCRIPTION SUBUN |
| 69:00:00 | 6j73-A | 3.7 | 3 | 53 | 539 | 6 | MOLECULE: | ISONIAZID INDUCIBLE GENE PROTEIN INIA |
| 70:00:00 | 6ajf-A | 3.7 | 5.2 | 61 | 901 | 8 | MOLECULE: | DRUG EXPORTERS OF THE RND SUPERFAMILY-LIKE PROTEI |
| 71:00:00 | 7udl-A | 3.7 | 3.2 | 56 | 278 | 9 | MOLECULE: | DESIGNED HELICAL REPEAT PROTEIN (DHR) RPB_PLP1_R6 |
| 72:00:00 | 6dfk-G | 3.7 | 2.7 | 55 | 213 | 0 | MOLECULE: | SUBUNIT OF PROTEASEOME ACTIVATOR COMPLEX,PUTATIVE |
| 73:00:00 | 6dfk-C | 3.7 | 3.5 | 59 | 222 | 0 | MOLECULE: | SUBUNIT OF PROTEASEOME ACTIVATOR COMPLEX,PUTATIVE |
| 74:00:00 | 1t6o-A | 3.7 | 2.1 | 46 | 49 | 9 | MOLECULE: | PHOSPHOPROTEIN |
| 75:00:00 | 7qvb-A | 3.7 | 4.3 | 67 | 213 | 3 | MOLECULE: | DNA DAMAGE RESPONSE PROTEIN C |
| 76:00:00 | 8ovt-C | 3.7 | 4.3 | 72 | 286 | 8 | MOLECULE: | YEGT GLYCOSYLTRANSFERASE |
| 77:00:00 | 3ctd-A | 3.7 | 5.6 | 60 | 150 | 10 | MOLECULE: | PUTATIVE ATPASE, AAA FAMILY |
| 78:00:00 | 2qex-P | 3.7 | 2.9 | 50 | 143 | 6 | MOLECULE: | 23S RIBOSOMAL RNA |
| 79:00:00 | 7zd6-t | 3.7 | 5.1 | 54 | 177 | 6 | MOLECULE: | NADH-UBIQUINONE OXIDOREDUCTASE CHAIN 3 |
| 80:00:00 | 7w2y-p | 3.7 | 4.5 | 57 | 178 | 4 | MOLECULE: | NADH DEHYDROGENASE [UBIQUINONE] FLAVOPROTEIN 1, |
| 81:00:00 | 7r44-n | 3.7 | 3 | 52 | 172 | 6 | MOLECULE: | NADH-UBIQUINONE OXIDOREDUCTASE CHAIN 3 |
| 82:00:00 | 7r4f-n | 3.7 | 3 | 52 | 171 | 6 | MOLECULE: | NADH-UBIQUINONE OXIDOREDUCTASE CHAIN 3 |
| 83:00:00 | 7w1u-p | 3.7 | 4.4 | 58 | 178 | 3 | MOLECULE: | NADH DEHYDROGENASE [UBIQUINONE] FLAVOPROTEIN 1, |
| 84:00:00 | 7w2u-p | 3.7 | 4.5 | 57 | 178 | 4 | MOLECULE: | NADH DEHYDROGENASE [UBIQUINONE] FLAVOPROTEIN 1, |
| 85:00:00 | 7w0y-p | 3.7 | 4.5 | 58 | 178 | 3 | MOLECULE: | NADH DEHYDROGENASE [UBIQUINONE] FLAVOPROTEIN 1, |
| 86:00:00 | 7w1v-p | 3.7 | 4.3 | 56 | 178 | 4 | MOLECULE: | NADH DEHYDROGENASE [UBIQUINONE] FLAVOPROTEIN 1, |
| 87:00:00 | 7r4g-n | 3.7 | 3 | 52 | 171 | 6 | MOLECULE: | NADH-UBIQUINONE OXIDOREDUCTASE CHAIN 3 |
| 88:00:00 | 7r46-n | 3.7 | 3 | 52 | 171 | 6 | MOLECULE: | NADH-UBIQUINONE OXIDOREDUCTASE CHAIN 3 |
| 89:00:00 | 7w1z-p | 3.7 | 4.6 | 56 | 178 | 5 | MOLECULE: | NADH DEHYDROGENASE [UBIQUINONE] FLAVOPROTEIN 1, |
| 90:00:00 | 7v31-p | 3.7 | 4.4 | 56 | 178 | 4 | MOLECULE: | NADH DEHYDROGENASE [UBIQUINONE] FLAVOPROTEIN 1, |
| 91:00:00 | 8ow0-N | 3.7 | 2.6 | 50 | 391 | 14 | MOLECULE: | C0N3 DNA |
| 92:00:00 | 8q0o-n | 3.7 | 3.1 | 53 | 171 | 6 | MOLECULE: | NADH-UBIQUINONE OXIDOREDUCTASE CHAIN 3 |
| 93:00:00 | 7w4e-p | 3.7 | 4.4 | 57 | 178 | 4 | MOLECULE: | NADH DEHYDROGENASE [UBIQUINONE] FLAVOPROTEIN 1, |
| 94:00:00 | 8ca3-n | 3.7 | 4.5 | 57 | 177 | 4 | MOLECULE: | NADH-UBIQUINONE OXIDOREDUCTASE CHAIN 3 |
| 95:00:00 | 7b93-n | 3.7 | 4.5 | 57 | 178 | 4 | MOLECULE: | NADH-UBIQUINONE OXIDOREDUCTASE CHAIN 3 |
| 96:00:00 | 7vyg-p | 3.7 | 4.5 | 57 | 178 | 4 | MOLECULE: | NADH DEHYDROGENASE [UBIQUINONE] IRON-SULFUR PROTE |
| 97:00:00 | 5cbn-A | 3.7 | 3.2 | 56 | 176 | 5 | MOLECULE: | MALTOSE-BINDING PERIPLASMIC PROTEIN |
| 98:00:00 | 7w2r-p | 3.7 | 4.5 | 57 | 178 | 4 | MOLECULE: | NADH DEHYDROGENASE [UBIQUINONE] FLAVOPROTEIN 1, |
| 99:00:00 | 3x38-D | 3.7 | 3.3 | 56 | 77 | 5 | MOLECULE: | MITOCHONDRIAL MORPHOGENESIS PROTEIN SLD7 |
| 0:00 | 8v2d-d | 3.7 | 3.5 | 61 | 314 | 8 | MOLECULE: | O43_129 COMPONENT B |
| 1:00 | 7bvg-A | 3.7 | 2.6 | 56 | 1078 | 4 | MOLECULE: | INTEGRAL MEMBRANE INDOLYLACETYLINOSITOL |
| 2:00 | 7bvc-A | 3.7 | 2.7 | 57 | 1078 | 5 | MOLECULE: | INTEGRAL MEMBRANE INDOLYLACETYLINOSITOL |
| 3:00 | 5bw9-B | 3.7 | 3.7 | 65 | 594 | 6 | MOLECULE: | V-TYPE PROTON ATPASE CATALYTIC SUBUNIT A |
| 4:00 | 7bvc-B | 3.7 | 3 | 60 | 1063 | 2 | MOLECULE: | INTEGRAL MEMBRANE INDOLYLACETYLINOSITOL |
| 5:00 | 7upp-A | 3.7 | 3.8 | 80 | 312 | 5 | MOLECULE: | DHT03 PROTEIN A |
| 6:00 | 5bw9-b | 3.7 | 3.7 | 65 | 589 | 6 | MOLECULE: | V-TYPE PROTON ATPASE CATALYTIC SUBUNIT A |
| 7:00 | 7fdb-C | 3.7 | 3 | 63 | 594 | 8 | MOLECULE: | YEAST VACUOLAR ATPASE A SUBUNIT |
| 8:00 | 7qps-B | 3.7 | 2.3 | 53 | 303 | 6 | MOLECULE: | ACT DOMAIN PROTEIN |
| 9:00 | 6r10-C | 3.7 | 3.2 | 68 | 577 | 1 | MOLECULE: | V-TYPE ATP SYNTHASE ALPHA CHAIN |
| 10:00 | 8iwh-g | 3.6 | 3.7 | 55 | 120 | 5 | MOLECULE: | PHOTOSYSTEM II PROTEIN D1 |
| 11:00 | 2m6u-A | 3.6 | 2.8 | 56 | 82 | 13 | MOLECULE: | CHOLINE BINDING PROTEIN A |
| 12:00 | 4k12-B | 3.6 | 2.8 | 56 | 82 | 13 | MOLECULE: | COMPLEMENT FACTOR H |
| 13:00 | 7d4f-C | 3.6 | 3.6 | 53 | 63 | 4 | MOLECULE: | NON-STRUCTURAL PROTEIN 8 |
| 14:00 | 7nfu-B | 3.6 | 3.4 | 63 | 213 | 10 | MOLECULE: | NUCLEOID OCCLUSION PROTEIN |
| 15:00 | 3x38-A | 3.6 | 3.3 | 56 | 78 | 5 | MOLECULE: | MITOCHONDRIAL MORPHOGENESIS PROTEIN SLD7 |
| 16:00 | 6t6v-A | 3.6 | 2.5 | 52 | 742 | 6 | MOLECULE: | NITRIC OXIDE REDUCTASE SUBUNIT B |
| 17:00 | 8v2d-K | 3.6 | 3.2 | 76 | 314 | 5 | MOLECULE: | O43_129 COMPONENT B |
| 18:00 | 8ap8-d | 3.6 | 3 | 54 | 270 | 4 | MOLECULE: | OSCP |
| 19:00 | 7v2c-p | 3.6 | 4.5 | 57 | 178 | 4 | MOLECULE: | NADH DEHYDROGENASE [UBIQUINONE] FLAVOPROTEIN 1, |
| 20:00 | 7ak6-n | 3.6 | 4.5 | 57 | 176 | 4 | MOLECULE: | NADH-UBIQUINONE OXIDOREDUCTASE CHAIN 3 |
| 21:00 | 1st6-A | 3.6 | 3.9 | 59 | 1049 | 10 | MOLECULE: | VINCULIN |
| 22:00 | 5wlw-F | 3.6 | 2.6 | 52 | 83 | 6 | MOLECULE: | CYSTEINE DESULFURASE, MITOCHONDRIAL |

| | | | | | | | | |
|---|---|---|---|---|---|---|---|---|
| 23:00 | 6zkg-t | 3.6 | 5 | 53 | 177 | 6 | MOLECULE: | NADH DEHYDROGENASE [UBIQUINONE] FLAVOPROTEIN 1, |
| 24:00:00 | 6o7w-A | 3.6 | 4 | 62 | 593 | 3 | MOLECULE: | V-TYPE PROTON ATPASE SUBUNIT D |
| 25:00:00 | 3g71-P | 3.6 | 3 | 50 | 143 | 4 | MOLECULE: | 23S RIBOSOMAL RNA |
| 26:00:00 | 5uj8-A | 3.6 | 3 | 60 | 553 | 7 | MOLECULE: | ORIGIN RECOGNITION COMPLEX SUBUNIT 3 |
| 27:00:00 | 7qpr-C | 3.6 | 3 | 57 | 678 | 7 | MOLECULE: | ACT DOMAIN PROTEIN |
| 28:00:00 | 2otj-P | 3.6 | 2.9 | 50 | 143 | 6 | MOLECULE: | 23S RIBOSOMAL RNA |
| 29:00:00 | 7coq-B | 3.6 | 2.8 | 60 | 586 | 5 | MOLECULE: | V-TYPE SODIUM ATPASE CATALYTIC SUBUNIT A |
| 30:00:00 | 3ctd-B | 3.6 | 3.6 | 56 | 158 | 11 | MOLECULE: | PUTATIVE ATPASE, AAA FAMILY |
| 31:00:00 | 7vc0-p | 3.6 | 4.6 | 56 | 178 | 5 | MOLECULE: | NADH DEHYDROGENASE [UBIQUINONE] IRON-SULFUR PROTE |
| 32:00:00 | 7vzv-p | 3.6 | 4.5 | 57 | 178 | 4 | MOLECULE: | NADH DEHYDROGENASE [UBIQUINONE] FLAVOPROTEIN 1, |
| 33:00:00 | 7r42-n | 3.6 | 3 | 52 | 172 | 6 | MOLECULE: | NADH-UBIQUINONE OXIDOREDUCTASE CHAIN 3 |
| 34:00:00 | 3ccm-P | 3.6 | 3 | 50 | 143 | 4 | MOLECULE: | 50S RIBOSOMAL PROTEIN L2P |
| 35:00:00 | 7vys-p | 3.6 | 4.5 | 57 | 178 | 4 | MOLECULE: | NADH DEHYDROGENASE [UBIQUINONE] IRON-SULFUR PROTE |
| 36:00:00 | 7v2h-p | 3.6 | 4.4 | 56 | 178 | 4 | MOLECULE: | NADH DEHYDROGENASE [UBIQUINONE] FLAVOPROTEIN 1, |
| 37:00:00 | 7r48-n | 3.6 | 3 | 52 | 171 | 6 | MOLECULE: | NADH-UBIQUINONE OXIDOREDUCTASE CHAIN 3 |
| 38:00:00 | 1yhq-P | 3.6 | 2.9 | 50 | 143 | 6 | MOLECULE: | 23S RIBOSOMAL RNA |
| 39:00:00 | 7r45-n | 3.6 | 3 | 52 | 171 | 6 | MOLECULE: | NADH-UBIQUINONE OXIDOREDUCTASE CHAIN 3 |
| 40:00:00 | 7w0r-p | 3.6 | 4.4 | 56 | 178 | 4 | MOLECULE: | NADH DEHYDROGENASE [UBIQUINONE] FLAVOPROTEIN 1, |
| 41:00:00 | 3ccv-P | 3.6 | 3 | 50 | 143 | 4 | MOLECULE: | 50S RIBOSOMAL PROTEIN L2P |
| 42:00:00 | 7r4c-n | 3.6 | 3 | 52 | 171 | 6 | MOLECULE: | NADH-UBIQUINONE OXIDOREDUCTASE CHAIN 3 |
| 43:00:00 | 7vbp-p | 3.6 | 4.5 | 58 | 178 | 3 | MOLECULE: | NADH DEHYDROGENASE [UBIQUINONE] IRON-SULFUR PROTE |
| 44:00:00 | 8q49-n | 3.6 | 3 | 52 | 171 | 6 | MOLECULE: | NADH-UBIQUINONE OXIDOREDUCTASE CHAIN 3 |
| 45:00:00 | 7zdm-t | 3.6 | 5 | 54 | 177 | 6 | MOLECULE: | NADH DEHYDROGENASE [UBIQUINONE] FLAVOPROTEIN 1, |
| 46:00:00 | 8q0a-n | 3.6 | 2.9 | 52 | 171 | 6 | MOLECULE: | NADH-UBIQUINONE OXIDOREDUCTASE CHAIN 3 |
| 47:00:00 | 8rgt-n | 3.6 | 4.5 | 56 | 178 | 4 | MOLECULE: | NADH DEHYDROGENASE [UBIQUINONE] IRON-SULFUR PROTE |
| 48:00:00 | 7r47-n | 3.6 | 3 | 52 | 171 | 6 | MOLECULE: | NADH-UBIQUINONE OXIDOREDUCTASE CHAIN 3 |
| 49:00:00 | 7w1p-p | 3.6 | 4.5 | 57 | 178 | 4 | MOLECULE: | NADH DEHYDROGENASE [UBIQUINONE] FLAVOPROTEIN 1, |
| 50:00:00 | 8q46-n | 3.6 | 3 | 52 | 171 | 6 | MOLECULE: | NADH-UBIQUINONE OXIDOREDUCTASE CHAIN 3 |
| 51:00:00 | 1vql-P | 3.6 | 3 | 50 | 143 | 4 | MOLECULE: | 23S RIBOSOMAL RNA |
| 52:00:00 | 7w4g-p | 3.6 | 4.6 | 57 | 178 | 5 | MOLECULE: | NADH DEHYDROGENASE [UBIQUINONE] FLAVOPROTEIN 1, |
| 53:00:00 | 7v33-p | 3.6 | 4.5 | 57 | 178 | 4 | MOLECULE: | NADH DEHYDROGENASE [UBIQUINONE] FLAVOPROTEIN 1, |
| 54:00:00 | 7vxs-p | 3.6 | 4.5 | 57 | 178 | 4 | MOLECULE: | NADH DEHYDROGENASE [UBIQUINONE] IRON-SULFUR PROTE |
| 55:00:00 | 8c2s-n | 3.6 | 4.5 | 56 | 177 | 4 | MOLECULE: | NADH-UBIQUINONE OXIDOREDUCTASE CHAIN 3 |
| 56:00:00 | 7r41-n | 3.6 | 3 | 52 | 172 | 6 | MOLECULE: | NADH-UBIQUINONE OXIDOREDUCTASE CHAIN 3 |
| 57:00:00 | 8om1-n | 3.6 | 4.5 | 56 | 178 | 4 | MOLECULE: | NADH-UBIQUINONE OXIDOREDUCTASE CHAIN 3 |
| 58:00:00 | 5f3o-A | 3.6 | 3.9 | 56 | 194 | 7 | MOLECULE: | PUTATIVE UNCHARACTERIZED PROTEIN |
| 59:00:00 | 8q4a-n | 3.6 | 3.2 | 53 | 171 | 6 | MOLECULE: | NADH-UBIQUINONE OXIDOREDUCTASE CHAIN 3 |
| 60:00:00 | 8g28-B | 3.6 | 3.6 | 56 | 104 | 9 | MOLECULE: | ATPASE, AAA FAMILY |
| 61:00:00 | 3x38-C | 3.6 | 3.2 | 55 | 79 | 5 | MOLECULE: | MITOCHONDRIAL MORPHOGENESIS PROTEIN SLD7 |
| 62:00:00 | 7qsm-n | 3.6 | 3 | 52 | 171 | 6 | MOLECULE: | NADH-UBIQUINONE OXIDOREDUCTASE CHAIN 3 |
| 63:00:00 | 8g28-A | 3.6 | 3.7 | 56 | 108 | 9 | MOLECULE: | ATPASE, AAA FAMILY |
| 64:00:00 | 7bve-B | 3.6 | 2.7 | 57 | 1034 | 9 | MOLECULE: | INTEGRAL MEMBRANE INDOLYLACETYLINOSITOL |
| 65:00:00 | 7zdh-t | 3.6 | 5.1 | 54 | 177 | 6 | MOLECULE: | NADH-UBIQUINONE OXIDOREDUCTASE CHAIN 3 |
| 66:00:00 | 7r4d-n | 3.6 | 3 | 52 | 171 | 6 | MOLECULE: | NADH-UBIQUINONE OXIDOREDUCTASE CHAIN 3 |
| 67:00:00 | 8ca5-n | 3.6 | 4.4 | 58 | 177 | 3 | MOLECULE: | NADH DEHYDROGENASE [UBIQUINONE] 1 ALPHA SUBCOMPLE |
| 68:00:00 | 3i56-P | 3.6 | 2.9 | 50 | 143 | 6 | MOLECULE: | 50S RIBOSOMAL PROTEIN L2P |
| 69:00:00 | 7v2e-p | 3.6 | 4.5 | 58 | 178 | 3 | MOLECULE: | NADH DEHYDROGENASE [UBIQUINONE] FLAVOPROTEIN 1, |
| 70:00:00 | 7tmr-E | 3.6 | 3.5 | 69 | 593 | 7 | MOLECULE: | H(+)-TRANSPORTING TWO-SECTOR ATPASE |
| 71:00:00 | 3j9v-A | 3.6 | 3.1 | 68 | 593 | 7 | MOLECULE: | V-TYPE PROTON ATPASE SUBUNIT A, VACUOLAR ISOFORM |
| 72:00:00 | 7qpr-A | 3.6 | 2.6 | 56 | 687 | 5 | MOLECULE: | ACT DOMAIN PROTEIN |
| 73:00:00 | 5h78-A | 3.6 | 9.1 | 56 | 192 | 7 | MOLECULE: | CAMP-DEPENDENT PROTEIN KINASE TYPE II-ALPHA REGUL |
| 74:00:00 | 8ovt-A | 3.5 | 4.1 | 69 | 288 | 10 | MOLECULE: | YEGT GLYCOSYLTRANSFERASE |
| 75:00:00 | 8ovw-N | 3.5 | 2.7 | 50 | 391 | 14 | MOLECULE: | CENTROMERE-BINDING PROTEIN 1 |
| 76:00:00 | 7qpr-D | 3.5 | 3 | 56 | 689 | 9 | MOLECULE: | ACT DOMAIN PROTEIN |
| 77:00:00 | 3x38-B | 3.5 | 3.2 | 55 | 79 | 5 | MOLECULE: | MITOCHONDRIAL MORPHOGENESIS PROTEIN SLD7 |
| 78:00:00 | 6hdb-B | 3.5 | 2.3 | 56 | 232 | 11 | MOLECULE: | NANOBODY,MALTOSE/MALTODEXTRIN-BINDING PERIPLASMIC |

| | | | | | | | | |
|---|---|---|---|---|---|---|---|---|
| 79:00:00 | 6r3q-A | 3.5 | 3.5 | 56 | 841 | 11 | MOLECULE: | ADENYLATE CYCLASE 9 |
| 80:00:00 | 7ose-A | 3.5 | 3.3 | 55 | 503 | 9 | MOLECULE: | CYTOCHROME BD-II UBIQUINOL OXIDASE SUBUNIT 1 |
| 81:00:00 | 6hda-B | 3.5 | 3.1 | 58 | 232 | 10 | MOLECULE: | NANOBODY,MALTOSE/MALTODEXTRIN-BINDING PERIPLASMIC |
| 82:00:00 | 8ce8-B | 3.5 | 3.7 | 70 | 219 | 10 | MOLECULE: | CYTOCHROME C BIOGENESIS ATP-BINDING EXPORT PROTEI |
| 83:00:00 | 2qw6-C | 3.5 | 3.2 | 52 | 83 | 12 | MOLECULE: | AAA ATPASE, CENTRAL REGION |
| 84:00:00 | 7saz-A | 3.5 | 3.7 | 54 | 217 | 6 | MOLECULE: | GLDM |
| 85:00:00 | 6dfk-L | 3.5 | 2.6 | 55 | 233 | 0 | MOLECULE: | SUBUNIT OF PROTEASEOME ACTIVATOR COMPLEX,PUTATIVE |
| 86:00:00 | 1k73-Q | 3.5 | 2.9 | 50 | 143 | 6 | MOLECULE: | 23S RRNA |
| 87:00:00 | 1kd1-Q | 3.5 | 3.1 | 50 | 143 | 4 | MOLECULE: | 23S RRNA |
| 88:00:00 | 7bw4-C | 3.5 | 4.1 | 57 | 65 | 2 | MOLECULE: | REPLICASE POLYPROTEIN 1AB |
| 89:00:00 | 7udj-H | 3.5 | 3.4 | 64 | 192 | 6 | MOLECULE: | 4XPAW PEPTIDE |
| 90:00:00 | 1q86-Q | 3.5 | 2.9 | 50 | 143 | 6 | MOLECULE: | 23S RIBOSOMAL RNA |
| 91:00:00 | 1vq7-P | 3.5 | 2.9 | 50 | 143 | 6 | MOLECULE: | 23S RIBOSOMAL RNA |
| 92:00:00 | 1yjn-P | 3.5 | 3 | 50 | 143 | 4 | MOLECULE: | 23S RIBOSOMAL RNA |
| 93:00:00 | 8ovt-D | 3.5 | 4 | 68 | 286 | 9 | MOLECULE: | YEGT GLYCOSYLTRANSFERASE |
| 94:00:00 | 7ak5-n | 3.5 | 4.4 | 55 | 176 | 5 | MOLECULE: | NADH-UBIQUINONE OXIDOREDUCTASE CHAIN 3 |
| 95:00:00 | 7w1o-p | 3.5 | 4.5 | 57 | 178 | 4 | MOLECULE: | NADH DEHYDROGENASE [UBIQUINONE] FLAVOPROTEIN 1, |
| 96:00:00 | 6zkv-t | 3.5 | 5 | 53 | 177 | 6 | MOLECULE: | NADH DEHYDROGENASE [UBIQUINONE] FLAVOPROTEIN 1, |
| 97:00:00 | 7qsn-n | 3.5 | 3.1 | 52 | 171 | 8 | MOLECULE: | NADH-UBIQUINONE OXIDOREDUCTASE CHAIN 3 |
| 98:00:00 | 7vwl-p | 3.5 | 4.4 | 56 | 178 | 4 | MOLECULE: | NADH DEHYDROGENASE [UBIQUINONE] IRON-SULFUR PROTE |
| 99:00:00 | 7vy1-p | 3.5 | 4.6 | 57 | 178 | 5 | MOLECULE: | NADH DEHYDROGENASE [UBIQUINONE] IRON-SULFUR PROTE |
| 0:00 | 5xtd-p | 3.5 | 3.1 | 53 | 172 | 4 | MOLECULE: | NADH DEHYDROGENASE [UBIQUINONE] FLAVOPROTEIN 1, |
| 1:00 | 7qso-n | 3.5 | 3.3 | 53 | 171 | 6 | MOLECULE: | NADH-UBIQUINONE OXIDOREDUCTASE CHAIN 3 |
| 2:00 | 6zkd-t | 3.5 | 5 | 54 | 177 | 6 | MOLECULE: | NADH DEHYDROGENASE [UBIQUINONE] FLAVOPROTEIN 1, |
| 3:00 | 3cma-P | 3.5 | 3 | 50 | 142 | 6 | MOLECULE: | 50S RIBOSOMAL PROTEIN L2P |
| 4:00 | 7qsk-n | 3.5 | 3 | 52 | 172 | 6 | MOLECULE: | NADH-UBIQUINONE OXIDOREDUCTASE CHAIN 3 |
| 5:00 | 1kqs-O | 3.5 | 2.9 | 50 | 143 | 6 | MOLECULE: | 23S RRNA |
| 6:00 | 8q25-n | 3.5 | 3.2 | 53 | 171 | 6 | MOLECULE: | NADH-UBIQUINONE OXIDOREDUCTASE CHAIN 3 |
| 7:00 | 8q47-n | 3.5 | 3 | 51 | 171 | 6 | MOLECULE: | NADH-UBIQUINONE OXIDOREDUCTASE CHAIN 3 |
| 8:00 | 3i55-P | 3.5 | 2.9 | 50 | 143 | 6 | MOLECULE: | 23S RIBOSOMAL RNA |
| 9:00 | 6zki-t | 3.5 | 5 | 53 | 177 | 6 | MOLECULE: | NADH DEHYDROGENASE [UBIQUINONE] FLAVOPROTEIN 1, |
| 10:00 | 7bvg-B | 3.5 | 2.5 | 55 | 1063 | 4 | MOLECULE: | INTEGRAL MEMBRANE INDOLYLACETYLINOSITOL |
| 11:00 | 1q82-Q | 3.5 | 2.9 | 50 | 143 | 6 | MOLECULE: | 23S RIBOSOMAL RNA |
| 12:00 | 7w2k-p | 3.5 | 4.5 | 57 | 178 | 4 | MOLECULE: | NADH DEHYDROGENASE [UBIQUINONE] FLAVOPROTEIN 1, |
| 13:00 | 3cce-P | 3.5 | 2.9 | 50 | 143 | 6 | MOLECULE: | 50S RIBOSOMAL PROTEIN L2P |
| 14:00 | 7v2r-p | 3.5 | 4.4 | 56 | 178 | 4 | MOLECULE: | NADH DEHYDROGENASE [UBIQUINONE] FLAVOPROTEIN 1, |
| 15:00 | 1vqo-P | 3.5 | 3 | 50 | 143 | 4 | MOLECULE: | 23S RIBOSOMAL RNA |
| 16:00 | 7zdp-t | 3.5 | 2.9 | 52 | 177 | 6 | MOLECULE: | COMPLEX I-49KD |
| 17:00 | 7w4n-p | 3.5 | 4.4 | 57 | 178 | 4 | MOLECULE: | NADH DEHYDROGENASE [UBIQUINONE] FLAVOPROTEIN 1, |
| 18:00 | 1qvg-O | 3.5 | 2.9 | 50 | 143 | 6 | MOLECULE: | 23S RIBOSOMAL RNA |
| 19:00 | 1yij-P | 3.5 | 2.9 | 50 | 143 | 6 | MOLECULE: | 23S RIBOSOMAL RNA |
| 20:00 | 1vqk-P | 3.5 | 2.9 | 50 | 143 | 6 | MOLECULE: | 23S RIBOSOMAL RNA |
| 21:00 | 5d80-b | 3.5 | 3.4 | 72 | 598 | 4 | MOLECULE: | V-TYPE PROTON ATPASE CATALYTIC SUBUNIT A |
| 22:00 | 1nji-Q | 3.5 | 2.9 | 50 | 143 | 6 | MOLECULE: | 23S RIBOSOMAL RNA |
| 23:00 | 1yi2-P | 3.5 | 2.9 | 50 | 143 | 6 | MOLECULE: | 23S RIBOSOMAL RNA |
| 24:00:00 | 1s72-P | 3.5 | 2.9 | 50 | 143 | 6 | MOLECULE: | 23S RIBOSOMAL RNA |
| 25:00:00 | 1vqn-P | 3.5 | 3 | 50 | 143 | 4 | MOLECULE: | 23S RIBOSOMAL RNA |
| 26:00:00 | 8q45-n | 3.5 | 3 | 52 | 171 | 6 | MOLECULE: | NADH-UBIQUINONE OXIDOREDUCTASE CHAIN 3 |
| 27:00:00 | 8beh-n | 3.5 | 2.7 | 52 | 109 | 4 | MOLECULE: | NADH-UBIQUINONE OXIDOREDUCTASE CHAIN 5 |
| 28:00:00 | 3cd6-P | 3.5 | 2.9 | 50 | 143 | 6 | MOLECULE: | 50S RIBOSOMAL PROTEIN L2P |
| 29:00:00 | 6zkb-t | 3.5 | 5.1 | 53 | 177 | 6 | MOLECULE: | MITOCHONDRIAL COMPLEX I, 49 KDA SUBUNIT |
| 30:00:00 | 1qvf-O | 3.5 | 2.9 | 50 | 143 | 6 | MOLECULE: | 23S RIBOSOMAL RNA |
| 31:00:00 | 1m1k-Q | 3.5 | 2.9 | 50 | 143 | 6 | MOLECULE: | 23S RRNA |
| 32:00:00 | 1vq8-P | 3.5 | 2.9 | 50 | 143 | 6 | MOLECULE: | 23S RIBOSOMAL RNA |
| 33:00:00 | 7vyi-p | 3.5 | 4.6 | 57 | 178 | 5 | MOLECULE: | NADH DEHYDROGENASE [UBIQUINONE] IRON-SULFUR PROTE |
| 34:00:00 | 1jj2-O | 3.5 | 3.1 | 50 | 143 | 4 | MOLECULE: | 23S RRNA |

| | | | | | | | |
|---|---|---|---|---|---|---|---|
| 35:00:00 | 7zeb-t | 3.5 | 5 | 54 | 177 | 6 | MOLECULE: NADH DEHYDROGENASE [UBIQUINONE] IRON-SULFUR PROTE |
| 36:00:00 | 8q1p-n | 3.5 | 3.1 | 53 | 171 | 6 | MOLECULE: NADH-UBIQUINONE OXIDOREDUCTASE CHAIN 3 |
| 37:00:00 | 8q48-n | 3.5 | 3.1 | 52 | 171 | 8 | MOLECULE: NADH-UBIQUINONE OXIDOREDUCTASE CHAIN 3 |
| 38:00:00 | 1vqp-P | 3.5 | 2.9 | 50 | 143 | 6 | MOLECULE: 23S RIBOSOMAL RNA |
| 39:00:00 | 6wm4-B | 3.5 | 3 | 66 | 600 | 11 | MOLECULE: V-TYPE PROTON ATPASE 116 KDA SUBUNIT A ISOFORM 1 |
| 40:00:00 | 7upq-A | 3.5 | 3.7 | 67 | 198 | 9 | MOLECULE: DHT03 PROTEIN A |
| 41:00:00 | 6wm3-C | 3.5 | 3.1 | 61 | 600 | 5 | MOLECULE: V-TYPE PROTON ATPASE 116 KDA SUBUNIT A ISOFORM 1 |
| 42:00:00 | 7qpr-B | 3.5 | 2.6 | 56 | 687 | 5 | MOLECULE: ACT DOMAIN PROTEIN |
| 43:00:00 | 8v3b-Q | 3.5 | 3 | 62 | 314 | 2 | MOLECULE: O43_129_+4 COMPONENT A |
| 44:00:00 | 3j9t-C | 3.5 | 3.1 | 69 | 593 | 7 | MOLECULE: V-TYPE PROTON ATPASE SUBUNIT D |
| 45:00:00 | 8v2d-i | 3.5 | 3 | 60 | 314 | 0 | MOLECULE: O43_129 COMPONENT B |
| 46:00:00 | 3j9v-C | 3.5 | 3.3 | 69 | 593 | 7 | MOLECULE: V-TYPE PROTON ATPASE SUBUNIT A, VACUOLAR ISOFORM |
| 47:00:00 | 3jb9-R | 3.4 | 2.8 | 59 | 575 | 0 | MOLECULE: PRE-MRNA-SPLICING FACTOR SPP42 |
| 48:00:00 | 7xyq-B | 3.4 | 3.5 | 52 | 124 | 13 | MOLECULE: CD274 MOLECULE |
| 49:00:00 | 3bbz-A | 3.4 | 2.4 | 45 | 48 | 4 | MOLECULE: P PROTEIN |
| 50:00:00 | 2g8l-B | 3.4 | 2.7 | 52 | 286 | 6 | MOLECULE: 287AA LONG HYPOTHETICAL PROTEIN |
| 51:00:00 | 5gmk-d | 3.4 | 2.8 | 59 | 549 | 0 | MOLECULE: PRE-MRNA-SPLICING FACTOR 8 |
| 52:00:00 | 6ybw-z | 3.4 | 6 | 60 | 160 | 5 | MOLECULE: 40S RIBOSOMAL PROTEIN S4, X ISOFORM |
| 53:00:00 | 6qfo-A | 3.4 | 4 | 55 | 1348 | 4 | MOLECULE: PEGA DOMAIN-CONTAINING PROTEIN,PEGA DOMAIN-CONTAI |
| 54:00:00 | 7v2d-p | 3.4 | 4.6 | 57 | 178 | 5 | MOLECULE: NADH DEHYDROGENASE [UBIQUINONE] FLAVOPROTEIN 1, |
| 55:00:00 | 7cxm-C | 3.4 | 3.6 | 53 | 72 | 4 | MOLECULE: NSP12 |
| 56:00:00 | 3gn4-A | 3.4 | 3.1 | 51 | 129 | 6 | MOLECULE: MYOSIN-VI |
| 57:00:00 | 6m71-C | 3.4 | 3.8 | 55 | 70 | 4 | MOLECULE: SARS-COV-2 NSP 12 |
| 58:00:00 | 5h7a-F | 3.4 | 2.2 | 46 | 188 | 7 | MOLECULE: IMMUNOGLOBULIN G-BINDING PROTEIN A |
| 59:00:00 | 6k68-L | 3.4 | 3 | 49 | 50 | 4 | MOLECULE: 3MNZ VARIABLE HEAVY CHAIN |
| 60:00:00 | 7cxn-C | 3.4 | 3.5 | 53 | 72 | 4 | MOLECULE: NSP12 |
| 61:00:00 | 6zkn-t | 3.4 | 5 | 53 | 177 | 6 | MOLECULE: NADH DEHYDROGENASE [UBIQUINONE] FLAVOPROTEIN 1, |
| 62:00:00 | 6zkm-t | 3.4 | 5 | 53 | 177 | 6 | MOLECULE: NADH DEHYDROGENASE [UBIQUINONE] FLAVOPROTEIN 1, |
| 63:00:00 | 8tl7-R | 3.4 | 2.3 | 56 | 694 | 5 | MOLECULE: COMPUTATIONALLY DESIGNED PROTEIN |
| 64:00:00 | 6g72-n | 3.4 | 4.5 | 57 | 176 | 5 | MOLECULE: NADH-UBIQUINONE OXIDOREDUCTASE CHAIN 3 |
| 65:00:00 | 6g2j-n | 3.4 | 4.5 | 57 | 176 | 5 | MOLECULE: NADH-UBIQUINONE OXIDOREDUCTASE CHAIN 3 |
| 66:00:00 | 5nbz-B | 3.4 | 3.1 | 58 | 237 | 9 | MOLECULE: WZZB |
| 67:00:00 | 3ze5-B | 3.4 | 3.2 | 66 | 116 | 14 | MOLECULE: DIACYLGLYCEROL KINASE |
| 68:00:00 | 1k8a-Q | 3.4 | 2.9 | 50 | 143 | 6 | MOLECULE: 23S RRNA |
| 69:00:00 | 5ujm-C | 3.4 | 3 | 60 | 553 | 7 | MOLECULE: ORIGIN RECOGNITION COMPLEX SUBUNIT 1 |
| 70:00:00 | 7ohg-A | 3.4 | 2.5 | 53 | 351 | 4 | MOLECULE: GUANOSINE-3',5'-BIS(DIPHOSPHATE) 3'-PYROPHOSPHOHY |
| 71:00:00 | 8acu-A | 3.4 | 2.3 | 50 | 349 | 8 | MOLECULE: GTP PYROPHOSPHOKINASE |
| 72:00:00 | 1n8r-Q | 3.4 | 2.9 | 50 | 143 | 6 | MOLECULE: 23S RIBOSOMAL RNA |
| 73:00:00 | 7w32-p | 3.4 | 4.5 | 56 | 178 | 5 | MOLECULE: NADH DEHYDROGENASE [UBIQUINONE] FLAVOPROTEIN 1, |
| 74:00:00 | 8q1u-n | 3.4 | 2.5 | 50 | 171 | 6 | MOLECULE: NADH-UBIQUINONE OXIDOREDUCTASE CHAIN 3 |
| 75:00:00 | 7v3m-p | 3.4 | 4.6 | 56 | 178 | 5 | MOLECULE: NADH DEHYDROGENASE [UBIQUINONE] FLAVOPROTEIN 1, |
| 76:00:00 | 3cc7-P | 3.4 | 3 | 50 | 143 | 4 | MOLECULE: 50S RIBOSOMAL PROTEIN L2P |
| 77:00:00 | 8q0f-n | 3.4 | 3 | 52 | 171 | 6 | MOLECULE: NADH-UBIQUINONE OXIDOREDUCTASE CHAIN 3 |
| 78:00:00 | 7zdj-t | 3.4 | 4.9 | 54 | 177 | 6 | MOLECULE: COMPLEX I-49KD |
| 79:00:00 | 7w4k-p | 3.4 | 4.4 | 56 | 178 | 5 | MOLECULE: NADH DEHYDROGENASE [UBIQUINONE] FLAVOPROTEIN 1, |
| 80:00:00 | 7qsl-n | 3.4 | 3 | 52 | 172 | 6 | MOLECULE: NADH-UBIQUINONE OXIDOREDUCTASE CHAIN 3 |
| 81:00:00 | 6zke-t | 3.4 | 5.1 | 53 | 177 | 6 | MOLECULE: NADH DEHYDROGENASE [UBIQUINONE] FLAVOPROTEIN 1, |
| 82:00:00 | 7cos-H | 3.4 | 3.5 | 72 | 586 | 4 | MOLECULE: V-TYPE SODIUM ATPASE CATALYTIC SUBUNIT A |
| 83:00:00 | 7w2l-p | 3.4 | 4.5 | 57 | 178 | 4 | MOLECULE: NADH DEHYDROGENASE [UBIQUINONE] FLAVOPROTEIN 1, |
| 84:00:00 | 6zkc-t | 3.4 | 5.1 | 54 | 177 | 6 | MOLECULE: NADH DEHYDROGENASE [UBIQUINONE] FLAVOPROTEIN 1, |
| 85:00:00 | 6qev-B | 3.4 | 3 | 51 | 1346 | 2 | MOLECULE: PEGA DOMAIN-CONTAINING PROTEIN,PEGA DOMAIN-CONTAI |
| 86:00:00 | 7k8b-A | 3.4 | 3.2 | 58 | 726 | 7 | MOLECULE: DRUG EXPORTERS OF THE RND SUPERFAMILY-LIKE PROTEI |
| 87:00:00 | 7v32-p | 3.4 | 4.4 | 56 | 178 | 4 | MOLECULE: NADH DEHYDROGENASE [UBIQUINONE] FLAVOPROTEIN 1, |
| 88:00:00 | 7w35-p | 3.4 | 4.4 | 56 | 178 | 4 | MOLECULE: NADH DEHYDROGENASE [UBIQUINONE] FLAVOPROTEIN 1, |
| 89:00:00 | 7arb-n | 3.4 | 2.9 | 52 | 109 | 6 | MOLECULE: NADH-UBIQUINONE OXIDOREDUCTASE CHAIN 3 |
| 90:00:00 | 5xtc-p | 3.4 | 3.1 | 52 | 172 | 6 | MOLECULE: NADH DEHYDROGENASE [UBIQUINONE] IRON-SULFUR PROTE |

| | | | | | | | | |
|---|---|---|---|---|---|---|---|---|
| 91:00:00 | 7w4j-p | 3.4 | 4.4 | 56 | 178 | 5 | MOLECULE: | NADH DEHYDROGENASE [UBIQUINONE] FLAVOPROTEIN 1, |
| 92:00:00 | 1q81-Q | 3.4 | 2.9 | 50 | 143 | 6 | MOLECULE: | 23S RIBOSOMAL RNA |
| 93:00:00 | 3ccu-P | 3.4 | 2.9 | 50 | 143 | 6 | MOLECULE: | 50S RIBOSOMAL PROTEIN L2P |
| 94:00:00 | 3g6e-P | 3.4 | 3 | 50 | 143 | 4 | MOLECULE: | 23S RIBOSOMAL RNA |
| 95:00:00 | 7aqw-n | 3.4 | 2.9 | 52 | 109 | 6 | MOLECULE: | NADH-UBIQUINONE OXIDOREDUCTASE CHAIN 5 |
| 96:00:00 | 7w0h-p | 3.4 | 4.6 | 57 | 178 | 5 | MOLECULE: | NADH DEHYDROGENASE [UBIQUINONE] FLAVOPROTEIN 1, |
| 97:00:00 | 6vqb-B | 3.4 | 3.5 | 68 | 600 | 10 | MOLECULE: | ATPASE H+-TRANSPORTING V1 SUBUNIT A |
| 98:00:00 | 6r0z-B | 3.4 | 4 | 71 | 578 | 1 | MOLECULE: | V-TYPE ATP SYNTHASE ALPHA CHAIN |
| 99:00:00 | 5cwo-B | 3.4 | 3.4 | 67 | 219 | 9 | MOLECULE: | DESIGNED HELICAL REPEAT PROTEIN |
| 0:00 | 3j9v-E | 3.4 | 3.1 | 70 | 593 | 9 | MOLECULE: | V-TYPE PROTON ATPASE SUBUNIT A, VACUOLAR ISOFORM |
| 1:00 | 7tmq-C | 3.4 | 3.8 | 66 | 590 | 8 | MOLECULE: | H(+)-TRANSPORTING TWO-SECTOR ATPASE |
| 2:00 | 5knc-B | 3.4 | 3.2 | 68 | 588 | 10 | MOLECULE: | V-TYPE SODIUM ATPASE CATALYTIC SUBUNIT A |
| 3:00 | 7khr-B | 3.4 | 3.3 | 63 | 591 | 3 | MOLECULE: | V-TYPE PROTON ATPASE CATALYTIC SUBUNIT A |
| 4:00 | 5y5z-C | 3.4 | 3 | 67 | 576 | 3 | MOLECULE: | V-TYPE ATP SYNTHASE ALPHA CHAIN |
| 5:00 | 6ly8-C | 3.4 | 3.2 | 69 | 577 | 3 | MOLECULE: | V-TYPE ATP SYNTHASE ALPHA CHAIN |
| 6:00 | 5cwo-A | 3.4 | 3.2 | 61 | 219 | 10 | MOLECULE: | DESIGNED HELICAL REPEAT PROTEIN |
| 7:00 | 7fda-C | 3.4 | 3.3 | 62 | 594 | 6 | MOLECULE: | YEAST VACUOLAR ATPASE A SUBUNIT |
| 8:00 | 2zhg-A | 3.3 | 3 | 58 | 121 | 9 | MOLECULE: | REDOX-SENSITIVE TRANSCRIPTIONAL ACTIVATOR SOXR |
| 9:00 | 6l7r-A | 3.3 | 2.7 | 49 | 105 | 12 | MOLECULE: | PUTATIVE SPINDLE POLE BODY COMPONENT ALP6 PROTEIN |
| 10:00 | 8jpa-A | 3.3 | 3.2 | 61 | 139 | 7 | MOLECULE: | DE NOVO DESIGN CAVITATED PROTEIN |
| 11:00 | 4uxv-A | 3.3 | 3.5 | 53 | 522 | 9 | MOLECULE: | SEPTATION RING FORMATION REGULATOR EZRA |
| 12:00 | 6rie-A | 3.3 | 3.1 | 61 | 1285 | 7 | MOLECULE: | DNA-DEPENDENT RNA POLYMERASE SUBUNIT RPO147 |
| 13:00 | 6l1q-A | 3.3 | 5.3 | 56 | 267 | 13 | MOLECULE: | CBBQ PROTEIN |
| 14:00 | 7d2w-A | 3.3 | 2.6 | 58 | 125 | 10 | MOLECULE: | PRESAN DOMAIN-CONTAINING PROTEIN |
| 15:00 | 8bob-C | 3.3 | 3.6 | 57 | 412 | 0 | MOLECULE: | PROTEIN MALY |
| 16:00 | 2n5j-A | 3.3 | 3.2 | 44 | 49 | 18 | MOLECULE: | RIBONUCLEASE ZC3H12A |
| 17:00 | 7obq-y | 3.3 | 4.2 | 60 | 454 | 8 | MOLECULE: | SRP RNA |
| 18:00 | 2hjm-A | 3.3 | 2.1 | 50 | 91 | 4 | MOLECULE: | HYPOTHETICAL PROTEIN PF1176 |
| 19:00 | 7ng0-A | 3.3 | 3.4 | 61 | 199 | 10 | MOLECULE: | NUCLEOID OCCLUSION PROTEIN |
| 20:00 | 5u5q-A | 3.3 | 3.7 | 67 | 1434 | 6 | MOLECULE: | DNA-DIRECTED RNA POLYMERASE II SUBUNIT RPB1 |
| 21:00 | 7bzf-C | 3.3 | 3.8 | 55 | 68 | 4 | MOLECULE: | RNA-DIRECTED RNA POLYMERASE |
| 22:00 | 7bv2-C | 3.3 | 2.8 | 49 | 63 | 4 | MOLECULE: | NSP12 |
| 23:00 | 6cgh-A | 3.3 | 3 | 59 | 89 | 2 | MOLECULE: | DNAJ HOMOLOG SUBFAMILY C MEMBER 2 |
| 24:00:00 | 7rmx-A | 3.3 | 2.7 | 53 | 229 | 4 | MOLECULE: | TUNABLE SYMMETRIC PROTEIN, D_3_212 |
| 25:00:00 | 6uxe-B | 3.3 | 2.3 | 49 | 85 | 6 | MOLECULE: | CYSTEINE DESULFURASE, MITOCHONDRIAL |
| 26:00:00 | 5i6r-A | 3.3 | 3.2 | 57 | 419 | 2 | MOLECULE: | SLIT-ROBO RHO GTPASE-ACTIVATING PROTEIN 2 |
| 27:00:00 | 6zkh-t | 3.3 | 5 | 53 | 177 | 6 | MOLECULE: | NADH DEHYDROGENASE [UBIQUINONE] FLAVOPROTEIN 1, |
| 28:00:00 | 7d2w-B | 3.3 | 2.6 | 58 | 125 | 3 | MOLECULE: | PRESAN DOMAIN-CONTAINING PROTEIN |
| 29:00:00 | 2ffj-B | 3.3 | 2.9 | 51 | 271 | 6 | MOLECULE: | CONSERVED HYPOTHETICAL PROTEIN |
| 30:00:00 | 6dfk-N | 3.3 | 2.6 | 55 | 233 | 0 | MOLECULE: | SUBUNIT OF PROTEASEOME ACTIVATOR COMPLEX,PUTATIVE |
| 31:00:00 | 1w2b-O | 3.3 | 3.3 | 50 | 143 | 4 | MOLECULE: | 23S RRNA |
| 32:00:00 | 2g8l-A | 3.3 | 2.7 | 52 | 284 | 6 | MOLECULE: | 287AA LONG HYPOTHETICAL PROTEIN |
| 33:00:00 | 3vr6-A | 3.3 | 4.4 | 62 | 587 | 6 | MOLECULE: | V-TYPE SODIUM ATPASE CATALYTIC SUBUNIT A |
| 34:00:00 | 7bx8-B | 3.3 | 2.9 | 58 | 1038 | 3 | MOLECULE: | INTEGRAL MEMBRANE INDOLYLACETYLINOSITOL |
| 35:00:00 | 3ccq-P | 3.3 | 3.1 | 50 | 143 | 4 | MOLECULE: | 50S RIBOSOMAL PROTEIN L2P |
| 36:00:00 | 6vq8-B | 3.3 | 3.5 | 68 | 600 | 10 | MOLECULE: | ATPASE H+-TRANSPORTING V1 SUBUNIT A |
| 37:00:00 | 6zkf-t | 3.3 | 5.1 | 52 | 177 | 6 | MOLECULE: | NADH DEHYDROGENASE [UBIQUINONE] FLAVOPROTEIN 1, |
| 38:00:00 | 7w4q-p | 3.3 | 4.6 | 57 | 178 | 5 | MOLECULE: | NADH DEHYDROGENASE [UBIQUINONE] FLAVOPROTEIN 1, |
| 39:00:00 | 7w4d-p | 3.3 | 4.5 | 58 | 178 | 3 | MOLECULE: | NADH DEHYDROGENASE [UBIQUINONE] FLAVOPROTEIN 1, |
| 40:00:00 | 3ow2-O | 3.3 | 3 | 50 | 143 | 4 | MOLECULE: | 23S RIBOSOMAL RNA |
| 41:00:00 | 3jb7-A | 3.3 | 3.4 | 74 | 1198 | 4 | MOLECULE: | CPV RNA-DEPENDENT RNA POLYMERASE |
| 42:00:00 | 3ccj-P | 3.3 | 2.9 | 50 | 143 | 6 | MOLECULE: | 50S RIBOSOMAL PROTEIN L2P |
| 43:00:00 | 8q0m-n | 3.3 | 3.1 | 52 | 171 | 6 | MOLECULE: | NADH-UBIQUINONE OXIDOREDUCTASE CHAIN 3 |
| 44:00:00 | 3g4s-P | 3.3 | 3 | 50 | 143 | 4 | MOLECULE: | 23S RIBOSOMAL RNA |
| 45:00:00 | 7v30-p | 3.3 | 4.5 | 56 | 178 | 5 | MOLECULE: | NADH DEHYDROGENASE [UBIQUINONE] FLAVOPROTEIN 1, |
| 46:00:00 | 1yit-P | 3.3 | 3 | 50 | 143 | 4 | MOLECULE: | 23S RIBOSOMAL RNA |

| | | | | | | | | |
|---|---|---|---|---|---|---|---|---|
| 47:00:00 | 8olt-n | 3.3 | 4.5 | 57 | 177 | 4 | MOLECULE: | NADH-UBIQUINONE OXIDOREDUCTASE CHAIN 3 |
| 48:00:00 | 7jpo-C | 3.3 | 3.1 | 60 | 594 | 5 | MOLECULE: | ORIGIN RECOGNITION COMPLEX SUBUNIT 1 |
| 49:00:00 | 6ajj-A | 3.3 | 4.4 | 58 | 901 | 9 | MOLECULE: | DRUG EXPORTERS OF THE RND SUPERFAMILY-LIKE PROTEI |
| 50:00:00 | 6o7x-E | 3.3 | 3.1 | 67 | 593 | 7 | MOLECULE: | V-TYPE PROTON ATPASE SUBUNIT C |
| 51:00:00 | 7dqc-A | 3.3 | 3.8 | 63 | 586 | 8 | MOLECULE: | V-TYPE SODIUM ATPASE CATALYTIC SUBUNIT A |
| 52:00:00 | 7cor-A | 3.3 | 3.2 | 68 | 587 | 9 | MOLECULE: | V-TYPE SODIUM ATPASE CATALYTIC SUBUNIT A |
| 53:00:00 | 6ly8-A | 3.3 | 3.2 | 69 | 577 | 1 | MOLECULE: | V-TYPE ATP SYNTHASE ALPHA CHAIN |
| 54:00:00 | 6o7v-C | 3.3 | 3.3 | 67 | 593 | 7 | MOLECULE: | V-TYPE PROTON ATPASE SUBUNIT D |
| 55:00:00 | 3kdw-A | 3.2 | 3.2 | 61 | 206 | 11 | MOLECULE: | PUTATIVE SUGAR BINDING PROTEIN |
| 56:00:00 | 5mmj-c | 3.2 | 8.2 | 52 | 216 | 10 | MOLECULE: | 50S RIBOSOMAL PROTEIN L31 |
| 57:00:00 | 1xfi-A | 3.2 | 3.4 | 54 | 343 | 6 | MOLECULE: | UNKNOWN PROTEIN |
| 58:00:00 | 6zko-t | 3.2 | 5 | 54 | 177 | 6 | MOLECULE: | NADH DEHYDROGENASE [UBIQUINONE] FLAVOPROTEIN 1, |
| 59:00:00 | 6v85-F | 3.2 | 2.4 | 47 | 47 | 4 | MOLECULE: | RNA-DIRECTED RNA POLYMERASE L |
| 60:00:00 | 1sb0-A | 3.2 | 2.3 | 49 | 87 | 4 | MOLECULE: | PROTEIN CBP |
| 61:00:00 | 7jlt-C | 3.2 | 3.8 | 53 | 81 | 4 | MOLECULE: | NON-STRUCTURAL PROTEIN 7 |
| 62:00:00 | 8df7-A | 3.2 | 3.3 | 54 | 850 | 4 | MOLECULE: | TOPOISOMERASE V |
| 63:00:00 | 5oxf-C | 3.2 | 3.4 | 64 | 601 | 9 | MOLECULE: | GTP-BINDING PROTEIN |
| 64:00:00 | 8hra-E | 3.2 | 6.4 | 59 | 834 | 10 | MOLECULE: | ARCHAEAL ATPASE |
| 65:00:00 | 8hr9-J | 3.2 | 6.2 | 58 | 835 | 10 | MOLECULE: | ARCHAEAL ATPASE |
| 66:00:00 | 7v2k-p | 3.2 | 4.5 | 56 | 178 | 5 | MOLECULE: | NADH DEHYDROGENASE [UBIQUINONE] FLAVOPROTEIN 1, |
| 67:00:00 | 4uq8-W | 3.2 | 2.7 | 54 | 72 | 0 | MOLECULE: | NADH UBIQUINONE OXIDOREDUCTASE CHAIN 3 |
| 68:00:00 | 6poo-A | 3.2 | 3.2 | 54 | 273 | 4 | MOLECULE: | BIBA |
| 69:00:00 | 6rjx-A | 3.2 | 3.1 | 54 | 157 | 2 | MOLECULE: | LYSM DOMAIN PROTEIN |
| 70:00:00 | 4heo-B | 3.2 | 2.4 | 48 | 60 | 4 | MOLECULE: | PHOSPHOPROTEIN |
| 71:00:00 | 3cc4-P | 3.2 | 2.9 | 50 | 143 | 6 | MOLECULE: | 50S RIBOSOMAL PROTEIN L2P |
| 72:00:00 | 2qa4-P | 3.2 | 2.9 | 50 | 143 | 6 | MOLECULE: | 23S RIBOSOMAL RNA |
| 73:00:00 | 2px0-H | 3.2 | 3.3 | 54 | 258 | 11 | MOLECULE: | FLAGELLAR BIOSYNTHESIS PROTEIN FLHF |
| 74:00:00 | 6r0w-B | 3.2 | 4.6 | 60 | 577 | 7 | MOLECULE: | V-TYPE ATP SYNTHASE ALPHA CHAIN |
| 75:00:00 | 3vr2-A | 3.2 | 3 | 59 | 586 | 5 | MOLECULE: | V-TYPE SODIUM ATPASE CATALYTIC SUBUNIT A |
| 76:00:00 | 7upq-D | 3.2 | 3.6 | 78 | 198 | 10 | MOLECULE: | DHT03 PROTEIN A |
| 77:00:00 | 7vye-p | 3.2 | 4.5 | 57 | 178 | 4 | MOLECULE: | NADH DEHYDROGENASE [UBIQUINONE] IRON-SULFUR PROTE |
| 78:00:00 | 7w20-p | 3.2 | 4.4 | 56 | 178 | 4 | MOLECULE: | NADH DEHYDROGENASE [UBIQUINONE] FLAVOPROTEIN 1, |
| 79:00:00 | 7w31-p | 3.2 | 4.6 | 57 | 178 | 5 | MOLECULE: | NADH DEHYDROGENASE [UBIQUINONE] FLAVOPROTEIN 1, |
| 80:00:00 | 3ccr-P | 3.2 | 2.9 | 50 | 143 | 6 | MOLECULE: | 50S RIBOSOMAL PROTEIN L2P |
| 81:00:00 | 3a5c-B | 3.2 | 3.6 | 72 | 561 | 6 | MOLECULE: | V-TYPE ATP SYNTHASE ALPHA CHAIN |
| 82:00:00 | 1yj9-P | 3.2 | 2.9 | 50 | 143 | 6 | MOLECULE: | 23S RIBOSOMAL RNA |
| 83:00:00 | 7w4m-p | 3.2 | 4.5 | 56 | 178 | 5 | MOLECULE: | NADH DEHYDROGENASE [UBIQUINONE] FLAVOPROTEIN 1, |
| 84:00:00 | 7v2f-p | 3.2 | 4.5 | 57 | 178 | 4 | MOLECULE: | NADH DEHYDROGENASE [UBIQUINONE] FLAVOPROTEIN 1, |
| 85:00:00 | 3bge-A | 3.2 | 5.8 | 59 | 163 | 12 | MOLECULE: | PREDICTED ATPASE |
| 86:00:00 | 6r10-B | 3.2 | 3.3 | 69 | 577 | 1 | MOLECULE: | V-TYPE ATP SYNTHASE ALPHA CHAIN |
| 87:00:00 | 7fdb-A | 3.2 | 3.3 | 58 | 594 | 7 | MOLECULE: | YEAST VACUOLAR ATPASE A SUBUNIT |
| 88:00:00 | 6r0w-C | 3.2 | 3.2 | 67 | 577 | 1 | MOLECULE: | V-TYPE ATP SYNTHASE ALPHA CHAIN |
| 89:00:00 | 7nad-R | 3.2 | 3.3 | 52 | 134 | 8 | MOLECULE: | 25S RRNA |
| 90:00:00 | 6r0z-A | 3.2 | 3 | 67 | 577 | 3 | MOLECULE: | V-TYPE ATP SYNTHASE ALPHA CHAIN |
| 91:00:00 | 6o7w-C | 3.2 | 3.2 | 68 | 593 | 7 | MOLECULE: | V-TYPE PROTON ATPASE SUBUNIT D |
| 92:00:00 | 5y60-C | 3.2 | 4 | 65 | 576 | 3 | MOLECULE: | V-TYPE ATP SYNTHASE ALPHA CHAIN |
| 93:00:00 | 6r0y-C | 3.2 | 3.2 | 66 | 577 | 2 | MOLECULE: | V-TYPE ATP SYNTHASE ALPHA CHAIN |
| 94:00:00 | 7jk5-C | 3.2 | 3.2 | 59 | 590 | 12 | MOLECULE: | ORIGIN RECOGNITION COMPLEX SUBUNIT 2 |
| 95:00:00 | 3gqb-C | 3.2 | 2.9 | 66 | 578 | 3 | MOLECULE: | V-TYPE ATP SYNTHASE ALPHA CHAIN |
| 96:00:00 | 5knc-A | 3.2 | 3.4 | 70 | 588 | 9 | MOLECULE: | V-TYPE SODIUM ATPASE CATALYTIC SUBUNIT A |
| 97:00:00 | 6s2t-A | 3.2 | 2.5 | 52 | 350 | 4 | MOLECULE: | (P)PPGPP SYNTHETASE I, SPOT/RELA |
| 98:00:00 | 6xby-B | 3.2 | 3.6 | 72 | 577 | 3 | MOLECULE: | V-TYPE PROTON ATPASE CATALYTIC SUBUNIT A |
| 99:00:00 | 4zil-A | 3.1 | 3.5 | 63 | 210 | 2 | MOLECULE: | DSBA OXIDOREDUCTASE |
| 0:00 | 3u5z-B | 3.1 | 5.1 | 60 | 320 | 7 | MOLECULE: | DNA POLYMERASE ACCESSORY PROTEIN 44 |
| 1:00 | 5cwc-A | 3.1 | 2.5 | 54 | 196 | 2 | MOLECULE: | DESIGNED HELICAL REPEAT PROTEIN |
| 2:00 | 7sgz-A | 3.1 | 3.3 | 56 | 418 | 2 | MOLECULE: | CHECKPOINT PROTEIN RAD24 |

| | | | | | | | | |
|---|---|---|---|---|---|---|---|---|
| 3:00 | 5lc5-n | 3.1 | 2.7 | 52 | 166 | 6 | MOLECULE: | NADH-UBIQUINONE OXIDOREDUCTASE CHAIN 3 |
| 4:00 | 6wc9-A | 3.1 | 5.2 | 55 | 367 | 7 | MOLECULE: | ENDOSOMAL/LYSOSOMAL POTASSIUM CHANNEL TMEM175 |
| 5:00 | 5z8q-A | 3.1 | 3.6 | 56 | 101 | 9 | MOLECULE: | HEAT SHOCK PROTEIN SSA1 |
| 6:00 | 5ys9-A | 3.1 | 3.7 | 66 | 692 | 8 | MOLECULE: | ACYL-COENZYME A OXIDASE 3 |
| 7:00 | 6qyi-B | 3.1 | 5.6 | 57 | 519 | 4 | MOLECULE: | 4-HYDROXYPHENYLACETATE 3-MONOOXYGENASE OXYGENASE |
| 8:00 | 8hr9-L | 3.1 | 6.2 | 57 | 833 | 11 | MOLECULE: | ARCHAEAL ATPASE |
| 9:00 | 2hz8-A | 3.1 | 4.4 | 58 | 115 | 5 | MOLECULE: | DE NOVO DESIGNED DIIRON PROTEIN |
| 10:00 | 6nur-C | 3.1 | 3.7 | 53 | 70 | 4 | MOLECULE: | NSP12 |
| 11:00 | 8q0q-n | 3.1 | 3.1 | 53 | 171 | 6 | MOLECULE: | NADH-UBIQUINONE OXIDOREDUCTASE CHAIN 3 |
| 12:00 | 6m5i-A | 3.1 | 3.7 | 53 | 81 | 4 | MOLECULE: | SARS-COV-2 NSP8 |
| 13:00 | 5lfj-B | 3.1 | 4.4 | 52 | 113 | 0 | MOLECULE: | BACTERIAL PROTEASOME ACTIVATOR |
| 14:00 | 7rkc-A | 3.1 | 3.6 | 67 | 220 | 7 | MOLECULE: | D_3_633 |
| 15:00 | 7tdr-A | 3.1 | 2.4 | 58 | 289 | 10 | MOLECULE: | 34K2 SALIVARY PROTEIN |
| 16:00 | 2ewf-A | 3.1 | 3.9 | 54 | 587 | 6 | MOLECULE: | NICKING ENDONUCLEASE N.BSPD6I |
| 17:00 | 2vda-A | 3.1 | 3.8 | 75 | 828 | 5 | MOLECULE: | TRANSLOCASE SUBUNIT SECA |
| 18:00 | 8t03-A | 3.1 | 3 | 63 | 199 | 3 | MOLECULE: | PROTEIN MYOMAKER |
| 19:00 | 8buy-A | 3.1 | 2.7 | 59 | 134 | 5 | MOLECULE: | GRANULE ASSOCIATED RAC AND RHOG EFFECTOR PROTEIN |
| 20:00 | 7vfp-F | 3.1 | 3.7 | 68 | 220 | 10 | MOLECULE: | CYTOCHROME C BIOGENESIS ATP-BINDING EXPORT PROTEI |
| 21:00 | 2otl-P | 3.1 | 2.9 | 50 | 143 | 6 | MOLECULE: | 23S RIBOSOMAL RNA |
| 22:00 | 5zea-I | 3.1 | 4.2 | 60 | 584 | 7 | MOLECULE: | V-TYPE SODIUM ATPASE CATALYTIC SUBUNIT A |
| 23:00 | 5vre-B | 3.1 | 3.1 | 54 | 188 | 6 | MOLECULE: | PUTATIVE INTEGRAL MEMBRANE PROTEIN |
| 24:00:00 | 3cpw-O | 3.1 | 2.9 | 50 | 143 | 6 | MOLECULE: | 23S RIBOSOMAL RNA |
| 25:00:00 | 6o7v-E | 3.1 | 3.6 | 71 | 593 | 3 | MOLECULE: | V-TYPE PROTON ATPASE SUBUNIT D |
| 26:00:00 | 6zkj-t | 3.1 | 4.9 | 53 | 177 | 6 | MOLECULE: | NADH DEHYDROGENASE [UBIQUINONE] FLAVOPROTEIN 1, |
| 27:00:00 | 3cc2-P | 3.1 | 2.9 | 50 | 143 | 6 | MOLECULE: | 50S RIBOSOMAL PROTEIN L2P |
| 28:00:00 | 6zkl-t | 3.1 | 2.9 | 52 | 177 | 6 | MOLECULE: | NADH DEHYDROGENASE [UBIQUINONE] FLAVOPROTEIN 1, |
| 29:00:00 | 5cbo-G | 3.1 | 2.9 | 49 | 176 | 6 | MOLECULE: | MBP3-16,IMMUNOGLOBULIN G-BINDING PROTEIN A |
| 30:00:00 | 5ze9-A | 3.1 | 3.6 | 72 | 588 | 4 | MOLECULE: | V-TYPE SODIUM ATPASE CATALYTIC SUBUNIT A |
| 31:00:00 | 6vq7-B | 3.1 | 3.6 | 73 | 600 | 4 | MOLECULE: | ATPASE H+-TRANSPORTING V1 SUBUNIT A |
| 32:00:00 | 3a5d-B | 3.1 | 3.7 | 74 | 561 | 5 | MOLECULE: | V-TYPE ATP SYNTHASE ALPHA CHAIN |
| 33:00:00 | 3ccs-P | 3.1 | 3 | 50 | 143 | 4 | MOLECULE: | 50S RIBOSOMAL PROTEIN L2P |
| 34:00:00 | 1vqm-P | 3.1 | 3 | 50 | 143 | 4 | MOLECULE: | 23S RIBOSOMAL RNA |
| 35:00:00 | 6qep-A | 3.1 | 3.2 | 51 | 1341 | 2 | MOLECULE: | PEGA DOMAIN-CONTAINING PROTEIN,ENGBF-DARPIN FUSIO |
| 36:00:00 | 3ccl-P | 3.1 | 2.9 | 50 | 143 | 6 | MOLECULE: | 50S RIBOSOMAL PROTEIN L2P |
| 37:00:00 | 1vq4-P | 3.1 | 3.1 | 50 | 143 | 4 | MOLECULE: | 23S RIBOSOMAL RNA |
| 38:00:00 | 5voz-A | 3.1 | 3.6 | 69 | 592 | 3 | MOLECULE: | V-TYPE PROTON ATPASE CATALYTIC SUBUNIT A,V-TYPE P |
| 39:00:00 | 6sh9-B | 3.1 | 4 | 56 | 1339 | 4 | MOLECULE: | ENDO-ALPHA-N-ACETYLGALACTOSAMINIDASE,DARPIN 4B D1 |
| 40:00:00 | 7fda-E | 3.1 | 3.6 | 64 | 594 | 6 | MOLECULE: | YEAST VACUOLAR ATPASE A SUBUNIT |
| 41:00:00 | 5y5z-A | 3.1 | 3.1 | 66 | 576 | 2 | MOLECULE: | V-TYPE ATP SYNTHASE ALPHA CHAIN |
| 42:00:00 | 6qle-N | 3.1 | 2.2 | 47 | 371 | 15 | MOLECULE: | CENTRAL KINETOCHORE SUBUNIT MCM16,CENTRAL KINETOC |
| 43:00:00 | 7fda-A | 3.1 | 4.6 | 67 | 594 | 4 | MOLECULE: | YEAST VACUOLAR ATPASE A SUBUNIT |
| 44:00:00 | 5y5y-A | 3.1 | 3.1 | 69 | 577 | 7 | MOLECULE: | V-TYPE ATP SYNTHASE ALPHA CHAIN |
| 45:00:00 | 6tz2-A | 3.1 | 4.7 | 78 | 1208 | 6 | MOLECULE: | RNA-DEPENDENT RNA POLYMERASE |
| 46:00:00 | 6r0y-A | 3.1 | 3.3 | 69 | 577 | 1 | MOLECULE: | V-TYPE ATP SYNTHASE ALPHA CHAIN |
| 47:00:00 | 6wm3-A | 3.1 | 3.6 | 69 | 600 | 3 | MOLECULE: | V-TYPE PROTON ATPASE 116 KDA SUBUNIT A ISOFORM 1 |
| 48:00:00 | 3j9u-E | 3.1 | 2.8 | 63 | 593 | 8 | MOLECULE: | V-TYPE PROTON ATPASE SUBUNIT D |
| 49:00:00 | 6ly8-B | 3.1 | 3.1 | 69 | 577 | 4 | MOLECULE: | V-TYPE ATP SYNTHASE ALPHA CHAIN |
| 50:00:00 | 6r10-A | 3.1 | 3.4 | 68 | 577 | 3 | MOLECULE: | V-TYPE ATP SYNTHASE ALPHA CHAIN |
| 51:00:00 | 8utk-A | 3 | 3.3 | 48 | 55 | 6 | MOLECULE: | 23R-B04DSLF02IB |
| 52:00:00 | 2a3v-B | 3 | 2.9 | 50 | 320 | 4 | MOLECULE: | DNA (31-MER) |
| 53:00:00 | 8oyy-A | 3 | 4.2 | 62 | 233 | 5 | MOLECULE: | DE NOVO DESIGNED SOLUBLE GPCR-LIKE PROTEIN |
| 54:00:00 | 7m9a-G | 3 | 3.6 | 58 | 257 | 5 | MOLECULE: | TNSC |
| 55:00:00 | 7m9c-G | 3 | 3.6 | 58 | 257 | 5 | MOLECULE: | TNSC |
| 56:00:00 | 7m9b-G | 3 | 3.4 | 57 | 257 | 5 | MOLECULE: | TNSC |
| 57:00:00 | 7m9b-D | 3 | 3.9 | 61 | 257 | 7 | MOLECULE: | TNSC |
| 58:00:00 | 7m9b-C | 3 | 3.1 | 55 | 257 | 5 | MOLECULE: | TNSC |

| | | | | | | | | |
|---|---|---|---|---|---|---|---|---|
| 59:00:00 | 7m9a-D | 3 | 3.9 | 60 | 257 | 7 | MOLECULE: | TNSC |
| 60:00:00 | 7m9c-D | 3 | 3.9 | 60 | 257 | 7 | MOLECULE: | TNSC |
| 61:00:00 | 7ole-A | 3 | 3.4 | 60 | 433 | 3 | MOLECULE: | RUVB-LIKE 1 |
| 62:00:00 | 3a98-A | 3 | 3.7 | 58 | 157 | 7 | MOLECULE: | DEDICATOR OF CYTOKINESIS PROTEIN 2 |
| 63:00:00 | 7a24-X | 3 | 2.8 | 52 | 103 | 2 | MOLECULE: | 51KDA |
| 64:00:00 | 5y2g-A | 3 | 3.6 | 66 | 581 | 12 | MOLECULE: | MALTOSE-BINDING PERIPLASMIC PROTEIN,PROTEIN B |
| 65:00:00 | 6v86-F | 3 | 2.4 | 44 | 47 | 5 | MOLECULE: | RNA-DIRECTED RNA POLYMERASE L |
| 66:00:00 | 5h7b-A | 3 | 2.2 | 46 | 230 | 7 | MOLECULE: | IMMUNOGLOBULIN G-BINDING PROTEIN A |
| 67:00:00 | 5h7a-H | 3 | 2.4 | 46 | 187 | 7 | MOLECULE: | IMMUNOGLOBULIN G-BINDING PROTEIN A |
| 68:00:00 | 6dlu-P | 3 | 2.7 | 54 | 747 | 6 | MOLECULE: | DYNAMIN-1 |
| 69:00:00 | 6dlc-A | 3 | 4.3 | 55 | 105 | 5 | MOLECULE: | DESIGNED PROTEIN DHD1:234_A |
| 70:00:00 | 7a23-X | 3 | 2.8 | 52 | 103 | 2 | MOLECULE: | 51KDA |
| 71:00:00 | 8q0j-n | 3 | 2.5 | 50 | 171 | 6 | MOLECULE: | NADH-UBIQUINONE OXIDOREDUCTASE CHAIN 3 |
| 72:00:00 | 6hum-F | 3 | 3.5 | 57 | 611 | 5 | MOLECULE: | NAD(P)H-QUINONE OXIDOREDUCTASE SUBUNIT 1 |
| 73:00:00 | 6tgz-F | 3 | 2.6 | 56 | 352 | 2 | MOLECULE: | 55 KDA IMMEDIATE-EARLY PROTEIN 1 |
| 74:00:00 | 3ok8-A | 3 | 2.6 | 53 | 219 | 8 | MOLECULE: | BRAIN-SPECIFIC ANGIOGENESIS INHIBITOR 1-ASSOCIATE |
| 75:00:00 | 4dvz-A | 3 | 4.7 | 58 | 459 | 9 | MOLECULE: | CYTOTOXICITY-ASSOCIATED IMMUNODOMINANT ANTIGEN |
| 76:00:00 | 8q7n-L | 3 | 3.9 | 50 | 373 | 6 | MOLECULE: | U5 SNRNA |
| 77:00:00 | 8tl7-S | 3 | 3.3 | 66 | 694 | 8 | MOLECULE: | COMPUTATIONALLY DESIGNED PROTEIN |
| 78:00:00 | 2o3l-A | 3 | 2.5 | 48 | 83 | 6 | MOLECULE: | HYPOTHETICAL PROTEIN |
| 79:00:00 | 2qxl-B | 3 | 2.8 | 52 | 625 | 2 | MOLECULE: | HEAT SHOCK PROTEIN HOMOLOG SSE1 |
| 80:00:00 | 6bar-A | 3 | 4.6 | 64 | 299 | 11 | MOLECULE: | ROD SHAPE DETERMINING PROTEIN RODA |
| 81:00:00 | 2wpv-A | 3 | 9.3 | 50 | 281 | 6 | MOLECULE: | UPF0363 PROTEIN YOR164C |
| 82:00:00 | 6th1-R | 3 | 2.8 | 61 | 360 | 5 | MOLECULE: | IMMEDIATE EARLY PROTEIN 1 |
| 83:00:00 | 1vq6-P | 3 | 2.9 | 50 | 143 | 6 | MOLECULE: | 23S RIBOSOMAL RNA |
| 84:00:00 | 5knc-C | 3 | 4.4 | 62 | 586 | 6 | MOLECULE: | V-TYPE SODIUM ATPASE CATALYTIC SUBUNIT A |
| 85:00:00 | 3vr2-B | 3 | 3 | 59 | 586 | 5 | MOLECULE: | V-TYPE SODIUM ATPASE CATALYTIC SUBUNIT A |
| 86:00:00 | 6zks-t | 3 | 5 | 53 | 177 | 6 | MOLECULE: | NADH DEHYDROGENASE [UBIQUINONE] FLAVOPROTEIN 1, |
| 87:00:00 | 1m90-Q | 3 | 2.9 | 50 | 143 | 6 | MOLECULE: | 23S RRNA |
| 88:00:00 | 6vq8-A | 3 | 3.6 | 73 | 600 | 4 | MOLECULE: | ATPASE H+-TRANSPORTING V1 SUBUNIT A |
| 89:00:00 | 6zkt-t | 3 | 5.1 | 53 | 177 | 6 | MOLECULE: | NADH DEHYDROGENASE [UBIQUINONE] FLAVOPROTEIN 1, |
| 90:00:00 | 2zxq-A | 3 | 2.8 | 51 | 1178 | 0 | MOLECULE: | ENDO-ALPHA-N-ACETYLGALACTOSAMINIDASE |
| 91:00:00 | 3j9t-E | 3 | 3.6 | 71 | 593 | 3 | MOLECULE: | V-TYPE PROTON ATPASE SUBUNIT D |
| 92:00:00 | 7w00-p | 3 | 4.5 | 57 | 178 | 5 | MOLECULE: | NADH DEHYDROGENASE [UBIQUINONE] FLAVOPROTEIN 1, |
| 93:00:00 | 3cxc-O | 3 | 2.9 | 50 | 143 | 6 | MOLECULE: | 23S RIBOSOMAL RNA |
| 94:00:00 | 1vq5-P | 3 | 2.9 | 50 | 143 | 6 | MOLECULE: | 23S RIBOSOMAL RNA |
| 95:00:00 | 6vqa-B | 3 | 3.6 | 70 | 600 | 3 | MOLECULE: | ATPASE H+-TRANSPORTING V1 SUBUNIT A |
| 96:00:00 | 1yjw-P | 3 | 2.9 | 50 | 143 | 6 | MOLECULE: | 23S RIBOSOMAL RNA |
| 97:00:00 | 6vq6-C | 3 | 3.5 | 68 | 600 | 10 | MOLECULE: | ATPASE H+-TRANSPORTING V1 SUBUNIT A |
| 98:00:00 | 1k9m-Q | 3 | 3.1 | 50 | 143 | 4 | MOLECULE: | 23S RRNA |
| 99:00:00 | 1kc8-Q | 3 | 2.9 | 50 | 143 | 6 | MOLECULE: | 23S RRNA |
| 0:00 | 6vqb-A | 3 | 3.6 | 73 | 600 | 4 | MOLECULE: | ATPASE H+-TRANSPORTING V1 SUBUNIT A |
| 1:00 | 7oy2-C | 3 | 2.9 | 53 | 454 | 9 | MOLECULE: | APPB PROTEIN |
| 2:00 | 2qw6-D | 3 | 4.4 | 53 | 86 | 6 | MOLECULE: | AAA ATPASE, CENTRAL REGION |
| 3:00 | 6zkp-t | 3 | 5.1 | 54 | 177 | 6 | MOLECULE: | NADH DEHYDROGENASE [UBIQUINONE] FLAVOPROTEIN 1, |
| 4:00 | 5cbo-K | 3 | 4.3 | 49 | 176 | 4 | MOLECULE: | MBP3-16,IMMUNOGLOBULIN G-BINDING PROTEIN A |
| 5:00 | 8a1a-A | 3 | 3 | 51 | 1345 | 2 | MOLECULE: | L1F11V1 |
| 6:00 | 1vq9-P | 3 | 2.9 | 50 | 143 | 6 | MOLECULE: | 23S RIBOSOMAL RNA |
| 7:00 | 3jb6-A | 3 | 3.6 | 77 | 1196 | 6 | MOLECULE: | RNA-DEPENDENT RNA POLYMERASE |
| 8:00 | 6vq9-C | 3 | 3.5 | 68 | 600 | 10 | MOLECULE: | ATPASE H+-TRANSPORTING V1 SUBUNIT A |
| 9:00 | 7fde-E | 3 | 3.3 | 67 | 594 | 1 | MOLECULE: | V-TYPE PROTON ATPASE SUBUNIT C |
| 10:00 | 7bx8-A | 3 | 3.1 | 59 | 1038 | 2 | MOLECULE: | INTEGRAL MEMBRANE INDOLYLACETYLINOSITOL |
| 11:00 | 6o7v-A | 3 | 3.2 | 67 | 593 | 7 | MOLECULE: | V-TYPE PROTON ATPASE SUBUNIT D |
| 12:00 | 5voz-C | 3 | 2.9 | 68 | 592 | 7 | MOLECULE: | V-TYPE PROTON ATPASE CATALYTIC SUBUNIT A,V-TYPE P |
| 13:00 | 3bge-B | 3 | 5.5 | 57 | 163 | 9 | MOLECULE: | PREDICTED ATPASE |
| 14:00 | 6wm3-B | 3 | 3.3 | 68 | 600 | 3 | MOLECULE: | V-TYPE PROTON ATPASE 116 KDA SUBUNIT A ISOFORM 1 |

| | | | | | | | | |
|---|---|---|---|---|---|---|---|---|
| 15:00 | 8wqb-F | 3 | 9.3 | 67 | 627 | 9 | MOLECULE: | CULLIN-2 |
| 16:00 | 8c81-B | 2.9 | 2.7 | 49 | 539 | 4 | MOLECULE: | PROTEIN ORM1 |
| 17:00 | 7xqk-B | 2.9 | 3.6 | 59 | 276 | 0 | MOLECULE: | GLUTATHIONE S-TRANSFERASE CLASS-MU 26 KDA ISOZYME |
| 18:00 | 5iy6-U | 2.9 | 2.7 | 49 | 170 | 6 | MOLECULE: | DNA-DIRECTED RNA POLYMERASE II SUBUNIT RPB1 |
| 19:00 | 2ejq-B | 2.9 | 3.4 | 65 | 115 | 2 | MOLECULE: | HYPOTHETICAL PROTEIN TTHA0227 |
| 20:00 | 7mi8-A | 2.9 | 3.1 | 66 | 868 | 9 | MOLECULE: | FUSION PROTEIN OF DYNEIN AND ENDOLYSIN |
| 21:00 | 7m99-C | 2.9 | 3.8 | 59 | 257 | 7 | MOLECULE: | TNSC |
| 22:00 | 7n6i-E | 2.9 | 4.2 | 61 | 257 | 7 | MOLECULE: | TNIQ (HOMOLOGY MODEL) |
| 23:00 | 7jlv-D | 2.9 | 3.5 | 61 | 437 | 13 | MOLECULE: | DISEASE RESISTANCE PROTEIN ROQ1 |
| 24:00:00 | 7m9c-P | 2.9 | 3.6 | 57 | 257 | 5 | MOLECULE: | TNSC |
| 25:00:00 | 7m9a-C | 2.9 | 3.7 | 59 | 257 | 7 | MOLECULE: | TNSC |
| 26:00:00 | 7jlv-A | 2.9 | 3.5 | 61 | 437 | 13 | MOLECULE: | DISEASE RESISTANCE PROTEIN ROQ1 |
| 27:00:00 | 1xme-A | 2.9 | 4.2 | 67 | 557 | 4 | MOLECULE: | CYTOCHROME C OXIDASE POLYPEPTIDE I |
| 28:00:00 | 8ba0-n | 2.9 | 4.7 | 55 | 134 | 7 | MOLECULE: | NADH-UBIQUINONE OXIDOREDUCTASE CHAIN 3 |
| 29:00:00 | 8hra-B | 2.9 | 6.4 | 59 | 835 | 10 | MOLECULE: | ARCHAEAL ATPASE |
| 30:00:00 | 3cqz-A | 2.9 | 3.8 | 66 | 1349 | 9 | MOLECULE: | DNA-DIRECTED RNA POLYMERASE II SUBUNIT RPB1 |
| 31:00:00 | 6jbq-F | 2.9 | 3.7 | 57 | 186 | 7 | MOLECULE: | DNA-DIRECTED RNA POLYMERASE SUBUNIT ALPHA |
| 32:00:00 | 2d1l-A | 2.9 | 3.9 | 57 | 249 | 2 | MOLECULE: | METASTASIS SUPPRESSOR PROTEIN 1 |
| 33:00:00 | 6soy-A | 2.9 | 4.1 | 55 | 323 | 7 | MOLECULE: | ESAG6, SUBUNIT OF HETERODIMERIC TRANSFERRIN RECEP |
| 34:00:00 | 8afz-A | 2.9 | 3 | 58 | 381 | 5 | MOLECULE: | SORTING NEXIN-1 |
| 35:00:00 | 5tgw-A | 2.9 | 3.8 | 58 | 109 | 3 | MOLECULE: | PS1 |
| 36:00:00 | 5wau-A | 2.9 | 3.7 | 67 | 514 | 7 | MOLECULE: | CYTOCHROME C OXIDASE SUBUNIT 1 |
| 37:00:00 | 6peq-G | 2.9 | 4.3 | 55 | 132 | 2 | MOLECULE: | GLUTAMATE RECEPTOR 2 |
| 38:00:00 | 8djk-A | 2.9 | 3.8 | 56 | 245 | 7 | MOLECULE: | 3-HYDROXY-3-METHYLGLUTARYL-COENZYME A REDUCTASE |
| 39:00:00 | 6peq-E | 2.9 | 4.3 | 55 | 132 | 2 | MOLECULE: | GLUTAMATE RECEPTOR 2 |
| 40:00:00 | 2qff-A | 2.9 | 3.4 | 52 | 74 | 0 | MOLECULE: | HYPOTHETICAL PROTEIN |
| 41:00:00 | 1oed-A | 2.9 1 | 0.2 | 57 | 127 | 12 | MOLECULE: | ACETYLCHOLINE RECEPTOR PROTEIN, ALPHA CHAIN |
| 42:00:00 | 1j78-A | 2.9 | 3.4 | 61 | 433 | 5 | MOLECULE: | VITAMIN D BINDING PROTEIN |
| 43:00:00 | 1y2o-A | 2.9 | 2.6 | 53 | 248 | 8 | MOLECULE: | BAI1-ASSOCIATED PROTEIN 2 ISOFORM 1 |
| 44:00:00 | 8g6r-C | 2.9 | 3.9 | 55 | 62 | 4 | MOLECULE: | NSP12 |
| 45:00:00 | 2awy-A | 2.9 | 3.2 | 58 | 133 | 10 | MOLECULE: | HEMERYTHRIN-LIKE DOMAIN PROTEIN DCRH |
| 46:00:00 | 8evm-A | 2.9 | 3.1 | 65 | 235 | 3 | MOLECULE: | CHLOROPHYLL DIMER PROTEIN DESIGNS, SPECIAL PAIR 3 |
| 47:00:00 | 1oed-E | 2.9 1 | 0.1 | 56 | 128 | 7 | MOLECULE: | ACETYLCHOLINE RECEPTOR PROTEIN, ALPHA CHAIN |
| 48:00:00 | 7du0-A | 2.9 | 3.8 | 51 | 125 | 4 | MOLECULE: | ACRIF14 |
| 49:00:00 | 7o6v-A | 2.9 | 2.7 | 48 | 71 | 10 | MOLECULE: | VIN3-LIKE PROTEIN 2 |
| 50:00:00 | 7z6h-D | 2.9 | 3.7 | 62 | 324 | 6 | MOLECULE: | CELL CYCLE CHECKPOINT CONTROL PROTEIN RAD9A |
| 51:00:00 | 3waq-A | 2.9 | 3.7 | 59 | 134 | 12 | MOLECULE: | HEMERYTHRIN-LIKE DOMAIN PROTEIN DCRH |
| 52:00:00 | 1i4y-A | 2.9 | 3.5 | 57 | 113 | 0 | MOLECULE: | METHEMERYTHRIN |
| 53:00:00 | 7cop-A | 2.9 | 4.4 | 62 | 586 | 6 | MOLECULE: | V-TYPE SODIUM ATPASE CATALYTIC SUBUNIT A |
| 54:00:00 | 3j9u-C | 2.9 | 3.7 | 70 | 593 | 3 | MOLECULE: | V-TYPE PROTON ATPASE SUBUNIT D |
| 55:00:00 | 6nuw-E | 2.9 | 2 | 44 | 369 | 7 | MOLECULE: | INNER KINETOCHORE SUBUNIT IML3 |
| 56:00:00 | 6s2u-A | 2.9 | 2.5 | 51 | 341 | 4 | MOLECULE: | (P)PPGPP SYNTHETASE I, SPOT/RELA |
| 57:00:00 | 3vr3-C | 2.9 | 4.3 | 62 | 584 | 6 | MOLECULE: | V-TYPE SODIUM ATPASE CATALYTIC SUBUNIT A |
| 58:00:00 | 3vr5-B | 2.9 | 4.4 | 62 | 593 | 6 | MOLECULE: | V-TYPE SODIUM ATPASE CATALYTIC SUBUNIT A |
| 59:00:00 | 2qw6-A | 2.9 | 2.9 | 47 | 85 | 4 | MOLECULE: | AAA ATPASE, CENTRAL REGION |
| 60:00:00 | 6qfk-A | 2.9 | 3 | 51 | 1347 | 2 | MOLECULE: | PEGA DOMAIN-CONTAINING PROTEIN,PEGA DOMAIN-CONTAI |
| 61:00:00 | 3cme-P | 2.9 | 3.2 | 50 | 142 | 4 | MOLECULE: | 50S RIBOSOMAL PROTEIN L2P |
| 62:00:00 | 6qum-C | 2.9 | 3.8 | 75 | 578 | 5 | MOLECULE: | V-TYPE ATP SYNTHASE ALPHA CHAIN |
| 63:00:00 | 7bvh-B | 2.9 | 3.2 | 59 | 1035 | 5 | MOLECULE: | INTEGRAL MEMBRANE INDOLYLACETYLINOSITOL |
| 64:00:00 | 2k0n-A | 2.9 | 2.3 | 50 | 85 | 8 | MOLECULE: | MEDIATOR OF RNA POLYMERASE II TRANSCRIPTION |
| 65:00:00 | 2k9d-A | 2.9 | 2.3 | 43 | 44 | 7 | MOLECULE: | PHOSPHOPROTEIN |
| 66:00:00 | 6r0w-A | 2.9 | 3.6 | 74 | 577 | 5 | MOLECULE: | V-TYPE ATP SYNTHASE ALPHA CHAIN |
| 67:00:00 | 7zme-R | 2.9 | 2.4 | 49 | 98 | 8 | MOLECULE: | NADH-UBIQUINONE OXIDOREDUCTASE CHAIN 1 |
| 68:00:00 | 7jpp-C | 2.9 | 3.1 | 60 | 597 | 5 | MOLECULE: | ORIGIN RECOGNITION COMPLEX SUBUNIT 1 |
| 69:00:00 | 6zkr-t | 2.9 | 5.1 | 53 | 177 | 6 | MOLECULE: | NADH DEHYDROGENASE [UBIQUINONE] FLAVOPROTEIN 1, |
| 70:00:00 | 1q7y-Q | 2.9 | 2.9 | 50 | 143 | 6 | MOLECULE: | 23S RIBOSOMAL RNA |

| | | | | | | | | |
|---|---|---|---|---|---|---|---|---|
| 71:00:00 | 7tmm-A | 2.9 | 3.5 | 70 | 593 | 3 | MOLECULE: | H(+)-TRANSPORTING TWO-SECTOR ATPASE |
| 72:00:00 | 7ar8-n | 2.9 | 2.7 | 52 | 109 | 6 | MOLECULE: | NADH-UBIQUINONE OXIDOREDUCTASE CHAIN 3 |
| 73:00:00 | 6qld-N | 2.9 | 2.1 | 45 | 371 | 9 | MOLECULE: | INNER KINETOCHORE SUBUNIT MIF2 |
| 74:00:00 | 7tmq-A | 2.9 | 3.9 | 72 | 591 | 4 | MOLECULE: | H(+)-TRANSPORTING TWO-SECTOR ATPASE |
| 75:00:00 | 5bw9-c | 2.9 | 3 | 64 | 586 | 6 | MOLECULE: | V-TYPE PROTON ATPASE CATALYTIC SUBUNIT A |
| 76:00:00 | 7bvh-A | 2.9 | 2.8 | 58 | 1035 | 5 | MOLECULE: | INTEGRAL MEMBRANE INDOLYLACETYLINOSITOL |
| 77:00:00 | 7tmo-E | 2.9 | 4.5 | 66 | 590 | 5 | MOLECULE: | H(+)-TRANSPORTING TWO-SECTOR ATPASE |
| 78:00:00 | 5h7c-C | 2.9 | 2.9 | 45 | 397 | 11 | MOLECULE: | IMMUNOGLOBULIN G-BINDING PROTEIN A, DHR14 |
| 79:00:00 | 5bn5-A | 2.9 | 3.7 | 64 | 562 | 3 | MOLECULE: | V-TYPE ATP SYNTHASE ALPHA CHAIN |
| 80:00:00 | 6o7w-E | 2.9 | 3.1 | 69 | 593 | 7 | MOLECULE: | V-TYPE PROTON ATPASE SUBUNIT D |
| 81:00:00 | 5voz-E | 2.9 | 3.1 | 68 | 592 | 7 | MOLECULE: | V-TYPE PROTON ATPASE CATALYTIC SUBUNIT A,V-TYPE P |
| 82:00:00 | 7tmq-E | 2.9 | 4.5 | 66 | 583 | 5 | MOLECULE: | H(+)-TRANSPORTING TWO-SECTOR ATPASE |
| 83:00:00 | 8je2-D | 2.9 1 | 0.1 | 68 | 494 | 3 | MOLECULE: | CULLIN-2 |
| 84:00:00 | 7v1n-A | 2.9 | 4.6 | 52 | 2344 | 6 | MOLECULE: | TOXIN B |
| 85:00:00 | 6vq9-B | 2.9 | 3.5 | 68 | 600 | 10 | MOLECULE: | ATPASE H+-TRANSPORTING V1 SUBUNIT A |
| 86:00:00 | 8wqe-B | 2.9 | 8.4 | 61 | 627 | 10 | MOLECULE: | CULLIN-2 |
| 87:00:00 | 7nj0-C | 2.8 | 3.9 | 65 | 270 | 3 | MOLECULE: | SECURIN,SEPARIN |
| 88:00:00 | 5iqc-A | 2.8 | 3.5 | 59 | 301 | 2 | MOLECULE: | BIFUNCTIONAL AAC/APH |
| 89:00:00 | 8eov-A | 2.8 | 2.8 | 47 | 139 | 9 | MOLECULE: | C2HR1_4R |
| 90:00:00 | 4cc9-B | 2.8 | 2.8 | 50 | 98 | 6 | MOLECULE: | PROTEIN VPRBP |
| 91:00:00 | 4hlq-C | 2.8 | 3.4 | 57 | 281 | 16 | MOLECULE: | TBC1 DOMAIN FAMILY MEMBER 20 |
| 92:00:00 | 7m9a-A | 2.8 | 3.6 | 56 | 257 | 7 | MOLECULE: | TNSC |
| 93:00:00 | 7m9c-O | 2.8 | 3.8 | 57 | 257 | 7 | MOLECULE: | TNSC |
| 94:00:00 | 7m9b-A | 2.8 | 3.8 | 57 | 257 | 7 | MOLECULE: | TNSC |
| 95:00:00 | 7m9c-C | 2.8 | 3.9 | 60 | 257 | 7 | MOLECULE: | TNSC |
| 96:00:00 | 7m9c-A | 2.8 | 3.6 | 56 | 257 | 7 | MOLECULE: | TNSC |
| 97:00:00 | 7m9c-B | 2.8 | 4 | 61 | 257 | 7 | MOLECULE: | TNSC |
| 98:00:00 | 7m9b-B | 2.8 | 4 | 61 | 257 | 7 | MOLECULE: | TNSC |
| 99:00:00 | 7m9a-B | 2.8 | 4 | 61 | 257 | 7 | MOLECULE: | TNSC |
| 0:00 | 2qyw-A | 2.8 | 8.9 | 56 | 96 | 5 | MOLECULE: | VESICLE TRANSPORT THROUGH INTERACTION WITH T-SNAR |
| 1:00 | 6zka-t | 2.8 | 4.9 | 54 | 177 | 6 | MOLECULE: | NADH-UBIQUINONE OXIDOREDUCTASE CHAIN 3 |
| 2:00 | 8jzl-F | 2.8 | 3.7 | 55 | 498 | 9 | MOLECULE: | INOSITOL PHOSPHATE PHOSPHATASE SOPB |
| 3:00 | 8jzl-D | 2.8 | 3.7 | 55 | 498 | 9 | MOLECULE: | INOSITOL PHOSPHATE PHOSPHATASE SOPB |
| 4:00 | 3rip-A | 2.8 | 5.9 | 56 | 568 | 7 | MOLECULE: | GAMMA-TUBULIN COMPLEX COMPONENT 4 |
| 5:00 | 6rfq-F | 2.8 | 3.9 | 53 | 121 | 9 | MOLECULE: | SUBUNIT NUAM OF NADH:UBIQUINONE OXIDOREDUCTASE (C |
| 6:00 | 8jzl-C | 2.8 | 3.7 | 55 | 498 | 9 | MOLECULE: | INOSITOL PHOSPHATE PHOSPHATASE SOPB |
| 7:00 | 5jk7-H | 2.8 | 3.2 | 51 | 74 | 6 | MOLECULE: | DNA DAMAGE-BINDING PROTEIN 1 |
| 8:00 | 1ywm-A | 2.8 | 3 | 55 | 180 | 5 | MOLECULE: | C PROTEIN ALPHA-ANTIGEN |
| 9:00 | 6zhi-B | 2.8 | 3.9 | 62 | 222 | 11 | MOLECULE: | HEAT SHOCK PROTEIN 70 |
| 10:00 | 8hr8-E | 2.8 | 6.1 | 57 | 833 | 9 | MOLECULE: | ARCHAEAL ATPASE |
| 11:00 | 8jzl-A | 2.8 | 3.7 | 55 | 498 | 9 | MOLECULE: | INOSITOL PHOSPHATE PHOSPHATASE SOPB |
| 12:00 | 8bms-C | 2.8 | 5.4 | 62 | 263 | 6 | MOLECULE: | ENERGY-COUPLING FACTOR TRANSPORTER ATP-BINDING PR |
| 13:00 | 8jzl-B | 2.8 | 3.7 | 55 | 498 | 9 | MOLECULE: | INOSITOL PHOSPHATE PHOSPHATASE SOPB |
| 14:00 | 5oqk-A | 2.8 | 2.5 | 52 | 147 | 4 | MOLECULE: | VOLTAGE-GATED HYDROGEN CHANNEL 1 |
| 15:00 | 1k04-A | 2.8 | 3.5 | 58 | 142 | 9 | MOLECULE: | FOCAL ADHESION KINASE 1 |
| 16:00 | 3g6i-A | 2.8 | 3.4 | 62 | 200 | 8 | MOLECULE: | PUTATIVE OUTER MEMBRANE PROTEIN, PART OF CARBOHYD |
| 17:00 | 8hdv-A | 2.8 | 3.1 | 52 | 137 | 6 | MOLECULE: | DE NOVO DESIGN CAVITATED PROTEIN |
| 18:00 | 2hmz-A | 2.8 | 3.7 | 59 | 113 | 3 | MOLECULE: | HEMERYTHRIN |
| 19:00 | 5x5y-F | 2.8 | 3.7 | 64 | 351 | 3 | MOLECULE: | PROBABLE ATP-BINDING COMPONENT OF ABC TRANSPORTER |
| 20:00 | 1dow-A | 2.8 | 3.7 | 58 | 205 | 5 | MOLECULE: | ALPHA-CATENIN |
| 21:00 | 5lqx-Y | 2.8 | 2.9 | 55 | 122 | 9 | MOLECULE: | ATP SYNTHASE SUBUNIT F |
| 22:00 | 6v3f-A | 2.8 | 4.1 | 69 | 1234 | 3 | MOLECULE: | NPC1-LIKE INTRACELLULAR CHOLESTEROL TRANSPORTER 1 |
| 23:00 | 7qsd-n | 2.8 | 3 | 52 | 172 | 6 | MOLECULE: | NADH-UBIQUINONE OXIDOREDUCTASE CHAIN 3 |
| 24:00:00 | 8jzl-E | 2.8 | 3.7 | 55 | 498 | 9 | MOLECULE: | INOSITOL PHOSPHATE PHOSPHATASE SOPB |
| 25:00:00 | 8buy-B | 2.8 | 3.4 | 56 | 126 | 5 | MOLECULE: | GRANULE ASSOCIATED RAC AND RHOG EFFECTOR PROTEIN |
| 26:00:00 | 6xby-A | 2.8 | 4.4 | 64 | 597 | 5 | MOLECULE: | V-TYPE PROTON ATPASE CATALYTIC SUBUNIT A |

| | | | | | | | | |
|---|---|---|---|---|---|---|---|---|
| 27:00:00 | 6xbw-C | 2.8 | 4.3 | 63 | 597 | 5 | MOLECULE: | V-TYPE PROTON ATPASE CATALYTIC SUBUNIT A |
| 28:00:00 | 7oiw-A | 2.8 | 2.6 | 51 | 327 | 8 | MOLECULE: | GTP PYROPHOSPHOKINASE |
| 29:00:00 | 7fdc-E | 2.8 | 3.2 | 63 | 594 | 8 | MOLECULE: | YEAST VACUOLAR ATPASE A SUBUNIT |
| 30:00:00 | 7dqe-C | 2.8 | 4.4 | 62 | 584 | 6 | MOLECULE: | V-TYPE SODIUM ATPASE CATALYTIC SUBUNIT A |
| 31:00:00 | 5lnk-t | 2.8 | 3 | 52 | 166 | 6 | MOLECULE: | MITOCHONDRIAL COMPLEX I, 51 KDA SUBUNIT |
| 32:00:00 | 6wm2-A | 2.8 | 3.7 | 73 | 600 | 4 | MOLECULE: | V-TYPE PROTON ATPASE SUBUNIT E 1 |
| 33:00:00 | 5ze9-B | 2.8 | 3.6 | 72 | 592 | 4 | MOLECULE: | V-TYPE SODIUM ATPASE CATALYTIC SUBUNIT A |
| 34:00:00 | 6vqa-C | 2.8 | 3.1 | 65 | 600 | 11 | MOLECULE: | ATPASE H+-TRANSPORTING V1 SUBUNIT A |
| 35:00:00 | 8a19-A | 2.8 | 3 | 51 | 1345 | 2 | MOLECULE: | L1E4V1 |
| 36:00:00 | 8pql-H | 2.8 | 9.6 | 64 | 597 | 13 | MOLECULE: | UBIQUITIN-CONJUGATING ENZYME E2 R2 |
| 37:00:00 | 6wlz-C | 2.8 | 3.6 | 73 | 600 | 4 | MOLECULE: | V-TYPE PROTON ATPASE CATALYTIC SUBUNIT A |
| 38:00:00 | 5h76-A | 2.8 | 4.2 | 55 | 197 | 7 | MOLECULE: | DARPIN,IMMUNOGLOBULIN G-BINDING PROTEIN A |
| 39:00:00 | 6aji-A | 2.8 | 4.7 | 59 | 873 | 10 | MOLECULE: | DRUG EXPORTERS OF THE RND SUPERFAMILY-LIKE PROTEI |
| 40:00:00 | 6rx4-A | 2.8 | 4.3 | 55 | 474 | 5 | MOLECULE: | CYTOCHROME BD-I UBIQUINOL OXIDASE SUBUNIT 1 |
| 41:00:00 | 6zl1-C | 2.8 | 3.3 | 61 | 122 | 2 | MOLECULE: | ALBUMIN |
| 42:00:00 | 7k8a-A | 2.8 | 2.7 | 56 | 722 | 7 | MOLECULE: | DRUG EXPORTERS OF THE RND SUPERFAMILY-LIKE PROTEI |
| 43:00:00 | 3j0j-C | 2.8 | 3.6 | 71 | 561 | 6 | MOLECULE: | V-TYPE ATP SYNTHASE ALPHA CHAIN |
| 44:00:00 | 8b9z-n | 2.8 | 4.5 | 55 | 134 | 7 | MOLECULE: | NADH-UBIQUINONE OXIDOREDUCTASE CHAIN 3 |
| 45:00:00 | 6r0z-C | 2.8 | 3.7 | 73 | 577 | 4 | MOLECULE: | V-TYPE ATP SYNTHASE ALPHA CHAIN |
| 46:00:00 | 6wlz-A | 2.8 | 3.6 | 72 | 600 | 3 | MOLECULE: | V-TYPE PROTON ATPASE CATALYTIC SUBUNIT A |
| 47:00:00 | 6wlz-B | 2.8 | 3.3 | 68 | 600 | 4 | MOLECULE: | V-TYPE PROTON ATPASE CATALYTIC SUBUNIT A |
| 48:00:00 | 5d80-A | 2.8 | 3.1 | 67 | 591 | 6 | MOLECULE: | V-TYPE PROTON ATPASE CATALYTIC SUBUNIT A |
| 49:00:00 | 7tmo-A | 2.8 | 3.8 | 62 | 592 | 10 | MOLECULE: | H(+)-TRANSPORTING TWO-SECTOR ATPASE |
| 50:00:00 | 6s2v-B | 2.8 | 3.5 | 59 | 326 | 8 | MOLECULE: | (P)PPGPP SYNTHETASE I, SPOT/RELA |
| 51:00:00 | 7khr-A | 2.8 | 2.9 | 62 | 589 | 2 | MOLECULE: | V-TYPE PROTON ATPASE CATALYTIC SUBUNIT A |
| 52:00:00 | 7w4c-p | 2.8 | 4.7 | 57 | 178 | 0 | MOLECULE: | NADH DEHYDROGENASE [UBIQUINONE] FLAVOPROTEIN 1, |
| 53:00:00 | 5uql-A | 2.8 | 3.7 | 64 | 542 | 9 | MOLECULE: | TOXIN A |
| 54:00:00 | 7fdb-E | 2.8 | 3.8 | 73 | 594 | 3 | MOLECULE: | YEAST VACUOLAR ATPASE A SUBUNIT |
| 55:00:00 | 7d5i-A | 2.8 | 3.1 | 50 | 443 | 4 | MOLECULE: | CYTOCHROME D UBIQUINOL OXIDASE SUBUNIT 1 |
| 56:00:00 | 3j9t-A | 2.8 | 2.8 | 63 | 593 | 8 | MOLECULE: | V-TYPE PROTON ATPASE SUBUNIT D |
| 57:00:00 | 6xby-C | 2.8 | 3.3 | 67 | 583 | 1 | MOLECULE: | V-TYPE PROTON ATPASE CATALYTIC SUBUNIT A |
| 58:00:00 | 5h78-B | 2.8 | 9.1 | 70 | 192 | 7 | MOLECULE: | CAMP-DEPENDENT PROTEIN KINASE TYPE II-ALPHA REGUL |
| 59:00:00 | 7cor-C | 2.8 | 3.2 | 66 | 584 | 8 | MOLECULE: | V-TYPE SODIUM ATPASE CATALYTIC SUBUNIT A |
| 60:00:00 | 6xbw-A | 2.8 | 4.4 | 59 | 589 | 5 | MOLECULE: | V-TYPE PROTON ATPASE CATALYTIC SUBUNIT A |
| 61:00:00 | 5d80-a | 2.8 | 3.5 | 72 | 591 | 6 | MOLECULE: | V-TYPE PROTON ATPASE CATALYTIC SUBUNIT A |
| 62:00:00 | 6bzh-B | 2.7 | 1.9 | 38 | 192 | 8 | MOLECULE: | PROBABLE ATP-DEPENDENT RNA HELICASE DDX58 |
| 63:00:00 | 6dan-B | 2.7 | 3.3 | 55 | 328 | 7 | MOLECULE: | PHDJ |
| 64:00:00 | 7tjk-F | 2.7 | 2.7 | 52 | 262 | 8 | MOLECULE: | ORIGIN RECOGNITION COMPLEX SUBUNIT 1 |
| 65:00:00 | 8bmw-N | 2.7 | 3.6 | 55 | 251 | 7 | MOLECULE: | CRISPR-ASSOCIATED SMALL SUBUNIT PROTEIN (TYPE III |
| 66:00:00 | 7m9b-F | 2.7 | 3.8 | 58 | 257 | 7 | MOLECULE: | TNSC |
| 67:00:00 | 7m9c-F | 2.7 | 3.7 | 58 | 257 | 7 | MOLECULE: | TNSC |
| 68:00:00 | 7m99-G | 2.7 | 3.3 | 55 | 257 | 7 | MOLECULE: | TNSC |
| 69:00:00 | 7jlv-B | 2.7 | 3.6 | 61 | 437 | 11 | MOLECULE: | DISEASE RESISTANCE PROTEIN ROQ1 |
| 70:00:00 | 7m9a-F | 2.7 | 3.8 | 58 | 257 | 7 | MOLECULE: | TNSC |
| 71:00:00 | 7m99-H | 2.7 | 3.3 | 55 | 257 | 7 | MOLECULE: | TNSC |
| 72:00:00 | 7n6i-J | 2.7 | 3.3 | 55 | 257 | 7 | MOLECULE: | TNIQ (HOMOLOGY MODEL) |
| 73:00:00 | 7n6i-I | 2.7 | 3.3 | 55 | 257 | 7 | MOLECULE: | TNIQ (HOMOLOGY MODEL) |
| 74:00:00 | 7n6i-G | 2.7 | 3.3 | 55 | 257 | 7 | MOLECULE: | TNIQ (HOMOLOGY MODEL) |
| 75:00:00 | 7m99-B | 2.7 | 3.3 | 55 | 257 | 7 | MOLECULE: | TNSC |
| 76:00:00 | 7n6i-F | 2.7 | 3.8 | 58 | 257 | 7 | MOLECULE: | TNIQ (HOMOLOGY MODEL) |
| 77:00:00 | 8gyw-A | 2.7 | 3 | 61 | 380 | 7 | MOLECULE: | CHOLINE/ETHANOLAMINEPHOSPHOTRANSFERASE 1 |
| 78:00:00 | 5l1a-A | 2.7 | 2.8 | 54 | 108 | 7 | MOLECULE: | UNCHARACTERIZED PROTEIN |
| 79:00:00 | 7uni-C | 2.7 | 3.5 | 53 | 242 | 2 | MOLECULE: | SP2-ZNPPAM DESIGNED CHLOROPHYLL DIMER PROTEIN |
| 80:00:00 | 8j1x-A | 2.7 | 3.8 | 54 | 174 | 6 | MOLECULE: | NEAR-INFRARED FLUORESCENT PROTEIN |
| 81:00:00 | 3dpn-A | 2.7 | 1.9 | 48 | 537 | 13 | MOLECULE: | PROTEIN CT_858 |
| 82:00:00 | 6y93-A | 2.7 | 3.1 | 53 | 105 | 6 | MOLECULE: | NUCLEOID OCCLUSION PROTEIN |

| | | | | | | | | |
|---|---|---|---|---|---|---|---|---|
| 83:00:00 | 5h77-A | 2.7 | 5 | 51 | 84 | 4 | MOLECULE: | CAMP-DEPENDENT PROTEIN KINASE TYPE II-ALPHA REGUL |
| 84:00:00 | 7ar9-N | 2.7 | 4.3 | 60 | 375 | 13 | MOLECULE: | ND3 |
| 85:00:00 | 3h0g-A | 2.7 | 3.7 | 64 | 1496 | 5 | MOLECULE: | DNA-DIRECTED RNA POLYMERASE II SUBUNIT RPB1 |
| 86:00:00 | 4by1-A | 2.7 | 3.8 | 67 | 1427 | 9 | MOLECULE: | DNA-DIRECTED RNA POLYMERASE II SUBUNIT RPB1 |
| 87:00:00 | 7opb-D | 2.7 | 2.6 | 43 | 55 | 5 | MOLECULE: | INTERLEUKIN-7 RECEPTOR SUBUNIT ALPHA |
| 88:00:00 | 5ul2-A | 2.7 | 4 | 71 | 731 | 1 | MOLECULE: | OXSB PROTEIN |
| 89:00:00 | 1sfo-A | 2.7 | 3.5 | 66 | 1395 | 5 | MOLECULE: | RNA STRAND |
| 90:00:00 | 6xss-A | 2.7 | 3.6 | 70 | 260 | 4 | MOLECULE: | C4_NAT_HFUSE-7900 |
| 91:00:00 | 6rfr-C | 2.7 | 4.3 | 69 | 438 | 7 | MOLECULE: | SUBUNIT NUAM OF NADH:UBIQUINONE OXIDOREDUCTASE (C |
| 92:00:00 | 8ga6-A | 2.7 | 6.8 | 58 | 476 | 5 | MOLECULE: | THR6 |
| 93:00:00 | 6z5l-A | 2.7 | 5.8 | 63 | 251 | 5 | MOLECULE: | MATRIX PROTEIN 1 |
| 94:00:00 | 6zkq-t | 2.7 | 5.1 | 53 | 177 | 6 | MOLECULE: | NADH DEHYDROGENASE [UBIQUINONE] FLAVOPROTEIN 1, |
| 95:00:00 | 7ctt-C | 2.7 | 3.9 | 55 | 63 | 4 | MOLECULE: | NSP12 |
| 96:00:00 | 4h6h-B | 2.7 | 3.1 | 51 | 80 | 8 | MOLECULE: | FIBRINOGEN-BINDING PROTEIN |
| 97:00:00 | 5luf-W | 2.7 | 3 | 53 | 72 | 0 | MOLECULE: | CYTOCHROME B-C1 COMPLEX SUBUNIT 1, MITOCHONDRIAL |
| 98:00:00 | 8a1g-A | 2.7 | 4.2 | 53 | 218 | 8 | MOLECULE: | SORTING NEXIN-1 |
| 99:00:00 | 4up6-A | 2.7 | 3.4 | 57 | 114 | 7 | MOLECULE: | DIACYLGLYCEROL KINASE |
| 0:00 | 1wgw-A | 2.7 | 2.9 | 54 | 99 | 7 | MOLECULE: | SIGNAL RECOGINITION PARTICLE 54 |
| 1:00 | 2p61-A | 2.7 | 3.7 | 56 | 114 | 5 | MOLECULE: | HYPOTHETICAL PROTEIN TM_1646 |
| 2:00 | 8e55-A | 2.7 | 3.2 | 59 | 195 | 0 | MOLECULE: | SG135 |
| 3:00 | 3h74-A | 2.7 | 3.6 | 64 | 278 | 6 | MOLECULE: | PYRIDOXAL KINASE |
| 4:00 | 7dfh-C | 2.7 | 4.5 | 55 | 63 | 5 | MOLECULE: | RNA-DIRECTED RNA POLYMERAS |
| 5:00 | 4nsr-E | 2.7 | 3.1 | 56 | 89 | 4 | MOLECULE: | IMMUNITY PROTEIN |
| 6:00 | 4nsr-A | 2.7 | 3.1 | 56 | 89 | 4 | MOLECULE: | IMMUNITY PROTEIN |
| 7:00 | 8ub3-k | 2.7 | 4 | 58 | 224 | 3 | MOLECULE: | DPHF7 FILAMENT |
| 8:00 | 2ozn-B | 2.7 | 3.9 | 56 | 131 | 4 | MOLECULE: | O-GLCNACASE NAGJ |
| 9:00 | 4uis-C | 2.7 | 3.9 | 69 | 196 | 0 | MOLECULE: | GAMMA-SECRETASE |
| 10:00 | 7sb2-A | 2.7 | 3.8 | 57 | 299 | 5 | MOLECULE: | GLDM |
| 11:00 | 5nvs-A | 2.7 | 3.3 | 62 | 932 | 0 | MOLECULE: | DYNEIN HEAVY CHAIN |
| 12:00 | 1i5n-D | 2.7 | 3.6 | 60 | 129 | 3 | MOLECULE: | CHEMOTAXIS PROTEIN CHEA |
| 13:00 | 3o42-A | 2.7 | 3.8 | 55 | 197 | 9 | MOLECULE: | GP41-5 |
| 14:00 | 5i9k-A | 2.7 | 3.2 | 56 | 123 | 9 | MOLECULE: | MICROSOMAL GLUTATHIONE S-TRANSFERASE 1 |
| 15:00 | 2qup-A | 2.7 | 2.9 | 57 | 119 | 11 | MOLECULE: | BH1478 PROTEIN |
| 16:00 | 2e2h-A | 2.7 | 4 | 68 | 1398 | 7 | MOLECULE: | 5'-R(*AP*UP*CP*GP*AP*GP*AP*GP*GP*A)-3' |
| 17:00 | 5knb-B | 2.7 | 4.3 | 62 | 592 | 6 | MOLECULE: | V-TYPE SODIUM ATPASE CATALYTIC SUBUNIT A |
| 18:00 | 6o7x-A | 2.7 | 3.9 | 61 | 593 | 3 | MOLECULE: | V-TYPE PROTON ATPASE SUBUNIT C |
| 19:00 | 6vq7-C | 2.7 | 4.5 | 65 | 600 | 5 | MOLECULE: | ATPASE H+-TRANSPORTING V1 SUBUNIT A |
| 20:00 | 7cos-A | 2.7 | 3 | 59 | 586 | 5 | MOLECULE: | V-TYPE SODIUM ATPASE CATALYTIC SUBUNIT A |
| 21:00 | 7zm7-R | 2.7 | 2.6 | 50 | 98 | 8 | MOLECULE: | NADH-UBIQUINONE OXIDOREDUCTASE CHAIN 1 |
| 22:00 | 7zmh-R | 2.7 | 2.4 | 49 | 98 | 6 | MOLECULE: | NADH-UBIQUINONE OXIDOREDUCTASE CHAIN 1 |
| 23:00 | 6zie-A | 2.7 | 4 | 56 | 123 | 5 | MOLECULE: | CMPX-383B |
| 24:00:00 | 7khr-C | 2.7 | 3.6 | 72 | 600 | 4 | MOLECULE: | V-TYPE PROTON ATPASE CATALYTIC SUBUNIT A |
| 25:00:00 | 7fde-A | 2.7 | 3.6 | 72 | 594 | 1 | MOLECULE: | V-TYPE PROTON ATPASE SUBUNIT C |
| 26:00:00 | 5knd-C | 2.7 | 3.6 | 72 | 586 | 4 | MOLECULE: | V-TYPE SODIUM ATPASE CATALYTIC SUBUNIT A |
| 27:00:00 | 7dqc-B | 2.7 | 3.7 | 71 | 586 | 4 | MOLECULE: | V-TYPE SODIUM ATPASE CATALYTIC SUBUNIT A |
| 28:00:00 | 7zmg-R | 2.7 | 2.4 | 50 | 98 | 6 | MOLECULE: | NADH-UBIQUINONE OXIDOREDUCTASE CHAIN 1 |
| 29:00:00 | 6aun-A | 2.7 | 5.1 | 61 | 618 | 8 | MOLECULE: | PLA2G6, IPLA2BETA |
| 30:00:00 | 3ikj-A | 2.7 | 3.9 | 74 | 513 | 8 | MOLECULE: | V-TYPE ATP SYNTHASE ALPHA CHAIN |
| 31:00:00 | 7dh0-f | 2.7 | 3.5 | 52 | 167 | 8 | MOLECULE: | NADH-UBIQUINONE OXIDOREDUCTASE CHAIN 2 |
| 32:00:00 | 6xbw-B | 2.7 | 3.5 | 72 | 590 | 3 | MOLECULE: | V-TYPE PROTON ATPASE CATALYTIC SUBUNIT A |
| 33:00:00 | 3pvs-A | 2.7 | 3.4 | 57 | 414 | 11 | MOLECULE: | REPLICATION-ASSOCIATED RECOMBINATION PROTEIN A |
| 34:00:00 | 6zl1-D | 2.7 | 2.8 | 51 | 120 | 4 | MOLECULE: | ALBUMIN |
| 35:00:00 | 7dqd-B | 2.7 | 3.1 | 65 | 584 | 8 | MOLECULE: | V-TYPE SODIUM ATPASE CATALYTIC SUBUNIT A |
| 36:00:00 | 7vz8-p | 2.7 | 4.7 | 57 | 178 | 0 | MOLECULE: | NADH DEHYDROGENASE [UBIQUINONE] IRON-SULFUR PROTE |
| 37:00:00 | 7upp-B | 2.7 | 3.8 | 81 | 309 | 9 | MOLECULE: | DHT03 PROTEIN A |
| 38:00:00 | 6oq7-A | 2.7 | 4.3 | 59 | 541 | 7 | MOLECULE: | TOXIN B |

| | | | | | | | | |
|---|---|---|---|---|---|---|---|---|
| 39:00:00 | 7fdc-A | 2.7 | 3.1 | 66 | 594 | 5 | MOLECULE: | YEAST VACUOLAR ATPASE A SUBUNIT |
| 40:00:00 | 5d80-c | 2.7 | 3.6 | 62 | 589 | 10 | MOLECULE: | V-TYPE PROTON ATPASE CATALYTIC SUBUNIT A |
| 41:00:00 | 3i72-A | 2.7 | 3.1 | 65 | 514 | 2 | MOLECULE: | A-TYPE ATP SYNTHASE CATALYTIC SUBUNIT A |
| 42:00:00 | 5voy-A | 2.7 | 3 | 62 | 592 | 3 | MOLECULE: | V-TYPE PROTON ATPASE CATALYTIC SUBUNIT A,V-TYPE P |
| 43:00:00 | 6qum-A | 2.7 | 3.7 | 71 | 578 | 3 | MOLECULE: | V-TYPE ATP SYNTHASE ALPHA CHAIN |
| 44:00:00 | 5bw9-C | 2.7 | 3.1 | 61 | 589 | 5 | MOLECULE: | V-TYPE PROTON ATPASE CATALYTIC SUBUNIT A |
| 45:00:00 | 3vr4-B | 2.7 | 4 | 65 | 593 | 8 | MOLECULE: | V-TYPE SODIUM ATPASE CATALYTIC SUBUNIT A |
| 46:00:00 | 7arc-W | 2.6 | 3 | 54 | 127 | 7 | MOLECULE: | PSST |
| 47:00:00 | 8wo8-A | 2.6 | 3 | 53 | 463 | 2 | MOLECULE: | PROBABLE RIBONUCLEASE FAU-1 |
| 48:00:00 | 5cwf-A | 2.6 | 3.2 | 54 | 177 | 2 | MOLECULE: | DESIGNED HELICAL REPEAT PROTEIN |
| 49:00:00 | 5h75-D | 2.6 | 2.9 | 43 | 225 | 9 | MOLECULE: | MERSACIDIN DECARBOXYLASE,IMMUNOGLOBULIN G-BINDING |
| 50:00:00 | 1ixs-A | 2.6 | 3.3 | 45 | 50 | 11 | MOLECULE: | HOLLIDAY JUNCTION DNA HELICASE RUVA |
| 51:00:00 | 8jh7-D | 2.6 | 3.3 | 55 | 431 | 9 | MOLECULE: | ANTI-BRIL FAB HEAVY CHAIN |
| 52:00:00 | 2fm8-C | 2.6 | 3.4 | 62 | 220 | 3 | MOLECULE: | SURFACE PRESENTATION OF ANTIGENS PROTEIN SPAK |
| 53:00:00 | 7m9b-K | 2.6 | 3.7 | 59 | 257 | 7 | MOLECULE: | TNSC |
| 54:00:00 | 7n6i-C | 2.6 | 3.3 | 55 | 257 | 7 | MOLECULE: | TNIQ (HOMOLOGY MODEL) |
| 55:00:00 | 7n6i-D | 2.6 | 3.3 | 55 | 257 | 7 | MOLECULE: | TNIQ (HOMOLOGY MODEL) |
| 56:00:00 | 7m99-E | 2.6 | 3.3 | 55 | 257 | 7 | MOLECULE: | TNSC |
| 57:00:00 | 7m9c-K | 2.6 | 3.3 | 56 | 257 | 5 | MOLECULE: | TNSC |
| 58:00:00 | 7m9a-K | 2.6 | 3.7 | 59 | 257 | 7 | MOLECULE: | TNSC |
| 59:00:00 | 7jlv-G | 2.6 | 3.5 | 59 | 437 | 12 | MOLECULE: | DISEASE RESISTANCE PROTEIN ROQ1 |
| 60:00:00 | 6mbz-A | 2.6 | 9.7 | 56 | 536 | 5 | MOLECULE: | SIGNAL TRANSDUCER AND ACTIVATOR OF TRANSCRIPTION |
| 61:00:00 | 4v3i-A | 2.6 | 9.9 | 57 | 153 | 0 | MOLECULE: | VCA0115 |
| 62:00:00 | 7a23-d | 2.6 | 2.8 | 52 | 66 | 6 | MOLECULE: | 51KDA |
| 63:00:00 | 6orb-A | 2.6 | 3.2 | 64 | 3352 | 6 | MOLECULE: | MIDASIN |
| 64:00:00 | 2j68-A | 2.6 | 4.1 | 55 | 680 | 5 | MOLECULE: | BACTERIAL DYNAMIN-LIKE PROTEIN |
| 65:00:00 | 3c4a-A | 2.6 | 3.3 | 68 | 365 | 10 | MOLECULE: | PROBABLE TRYPTOPHAN HYDROXYLASE VIOD |
| 66:00:00 | 8tl7-Z | 2.6 | 2.8 | 63 | 694 | 8 | MOLECULE: | COMPUTATIONALLY DESIGNED PROTEIN |
| 67:00:00 | 5j0l-A | 2.6 | 2.7 | 53 | 130 | 2 | MOLECULE: | DESIGNED PROTEIN 3L6HC2_2 |
| 68:00:00 | 4roe-A | 2.6 | 3.1 | 57 | 307 | 7 | MOLECULE: | TRANSCRIPTION FACTOR IIIB 50 KDA SUBUNIT |
| 69:00:00 | 7os0-A | 2.6 | 3.9 | 60 | 1130 | 3 | MOLECULE: | CAS13A |
| 70:00:00 | 6fh1-A | 2.6 | 3.5 | 65 | 353 | 9 | MOLECULE: | PROTEIN-ARGININE KINASE |
| 71:00:00 | 8v2d-A | 2.6 | 9.8 | 62 | 307 | 8 | MOLECULE: | O43_129 COMPONENT B |
| 72:00:00 | 5vhx-E | 2.6 | 4.1 | 58 | 179 | 5 | MOLECULE: | GLUTAMATE RECEPTOR 2,GERM CELL-SPECIFIC GENE 1-LI |
| 73:00:00 | 8ub3-H | 2.6 | 4.1 | 58 | 224 | 3 | MOLECULE: | DPHF7 FILAMENT |
| 74:00:00 | 1l2f-A | 2.6 | 2.8 | 50 | 344 | 10 | MOLECULE: | N UTILIZATION SUBSTANCE PROTEIN A |
| 75:00:00 | 1lot-A | 2.6 | 3.3 | 60 | 436 | 5 | MOLECULE: | VITAMIN D-BINDING PROTEIN |
| 76:00:00 | 8x6g-H | 2.6 | 4.2 | 63 | 244 | 5 | MOLECULE: | DNA-DIRECTED RNA POLYMERASE SUBUNIT ALPHA |
| 77:00:00 | 3if8-B | 2.6 | 3.6 | 65 | 237 | 9 | MOLECULE: | PROTEIN ZWILCH HOMOLOG |
| 78:00:00 | 4nsr-D | 2.6 | 3.1 | 56 | 92 | 4 | MOLECULE: | IMMUNITY PROTEIN |
| 79:00:00 | 2y1d-A | 2.6 | 4.2 | 70 | 359 | 11 | MOLECULE: | 1-DEOXY-D-XYLULOSE 5-PHOSPHATE REDUCTOISOMERASE |
| 80:00:00 | 8ub3-C | 2.6 | 4.1 | 58 | 224 | 3 | MOLECULE: | DPHF7 FILAMENT |
| 81:00:00 | 2occ-A | 2.6 | 3.2 | 69 | 514 | 3 | MOLECULE: | CYTOCHROME C OXIDASE |
| 82:00:00 | 1nze-A | 2.6 | 4.7 | 60 | 112 | 5 | MOLECULE: | OXYGEN-EVOLVING ENHANCER PROTEIN 3 |
| 83:00:00 | 2fu2-A | 2.6 | 3.4 | 47 | 78 | 11 | MOLECULE: | HYPOTHETICAL PROTEIN SPY2152 |
| 84:00:00 | 6wiq-A | 2.6 | 3.6 | 54 | 80 | 4 | MOLECULE: | SARS-COV-2 NSP7 |
| 85:00:00 | 7tmp-E | 2.6 | 4.4 | 62 | 590 | 5 | MOLECULE: | H(+)-TRANSPORTING TWO-SECTOR ATPASE |
| 86:00:00 | 3vr6-B | 2.6 | 4.3 | 62 | 592 | 6 | MOLECULE: | V-TYPE SODIUM ATPASE CATALYTIC SUBUNIT A |
| 87:00:00 | 5o31-n | 2.6 | 3 | 53 | 166 | 8 | MOLECULE: | NADH-UBIQUINONE OXIDOREDUCTASE CHAIN 3 |
| 88:00:00 | 5tsj-B | 2.6 | 3.5 | 70 | 575 | 4 | MOLECULE: | V-TYPE ATP SYNTHASE ALPHA CHAIN |
| 89:00:00 | 3se0-A | 2.6 | 3.7 | 74 | 519 | 8 | MOLECULE: | V-TYPE ATP SYNTHASE ALPHA CHAIN |
| 90:00:00 | 5h76-C | 2.6 | 4.3 | 53 | 197 | 8 | MOLECULE: | DARPIN,IMMUNOGLOBULIN G-BINDING PROTEIN A |
| 91:00:00 | 7cor-B | 2.6 | 3.8 | 72 | 586 | 7 | MOLECULE: | V-TYPE SODIUM ATPASE CATALYTIC SUBUNIT A |
| 92:00:00 | 7ar7-n | 2.6 | 2.9 | 52 | 109 | 6 | MOLECULE: | NADH-UBIQUINONE OXIDOREDUCTASE CHAIN 3 |
| 93:00:00 | 6wm2-B | 2.6 | 3.7 | 73 | 600 | 4 | MOLECULE: | V-TYPE PROTON ATPASE SUBUNIT E 1 |
| 94:00:00 | 5gar-B | 2.6 | 3.7 | 73 | 575 | 4 | MOLECULE: | V-TYPE ATP SYNTHASE ALPHA CHAIN |

| | | | | | | | | |
|---|---|---|---|---|---|---|---|---|
| 95:00:00 | 6vq7-A | 2.6 | 3.6 | 72 | 600 | 3 | MOLECULE: | ATPASE H+-TRANSPORTING V1 SUBUNIT A |
| 96:00:00 | 7fdc-C | 2.6 | 3.8 | 73 | 594 | 3 | MOLECULE: | YEAST VACUOLAR ATPASE A SUBUNIT |
| 97:00:00 | 2r9g-N | 2.6 | 6 | 58 | 180 | 9 | MOLECULE: | AAA ATPASE, CENTRAL REGION |
| 98:00:00 | 5f3p-A | 2.6 | 4.3 | 56 | 193 | 7 | MOLECULE: | PUTATIVE UNCHARACTERIZED PROTEIN |
| 99:00:00 | 5cbo-F | 2.6 | 2.9 | 50 | 176 | 6 | MOLECULE: | MBP3-16,IMMUNOGLOBULIN G-BINDING PROTEIN A |
| 0:00 | 5cbo-H | 2.6 | 4.2 | 49 | 176 | 4 | MOLECULE: | MBP3-16,IMMUNOGLOBULIN G-BINDING PROTEIN A |
| 1:00 | 8q1y-n | 2.6 | 3 | 52 | 171 | 6 | MOLECULE: | NADH-UBIQUINONE OXIDOREDUCTASE CHAIN 3 |
| 2:00 | 6wm4-A | 2.6 | 3.6 | 73 | 600 | 4 | MOLECULE: | V-TYPE PROTON ATPASE 116 KDA SUBUNIT A ISOFORM 1 |
| 3:00 | 5bn4-A | 2.6 | 3.7 | 63 | 567 | 3 | MOLECULE: | V-TYPE ATP SYNTHASE ALPHA CHAIN |
| 4:00 | 7tmr-C | 2.6 | 3.9 | 73 | 593 | 4 | MOLECULE: | H(+)-TRANSPORTING TWO-SECTOR ATPASE |
| 5:00 | 3pvs-B | 2.6 | 5.1 | 58 | 417 | 10 | MOLECULE: | REPLICATION-ASSOCIATED RECOMBINATION PROTEIN A |
| 6:00 | 3m4y-A | 2.6 | 3.9 | 70 | 558 | 1 | MOLECULE: | V-TYPE ATP SYNTHASE ALPHA CHAIN |
| 7:00 | 6yxa-A | 2.6 | 2.6 | 48 | 536 | 8 | MOLECULE: | GTP PYROPHOSPHOKINASE |
| 8:00 | 5gas-C | 2.6 | 3.3 | 67 | 576 | 4 | MOLECULE: | V-TYPE ATP SYNTHASE ALPHA CHAIN |
| 9:00 | 5h7c-A | 2.6 | 2.7 | 41 | 400 | 12 | MOLECULE: | IMMUNOGLOBULIN G-BINDING PROTEIN A, DHR14 |
| 10:00 | 2r9g-B | 2.6 | 6 | 58 | 180 | 9 | MOLECULE: | AAA ATPASE, CENTRAL REGION |
| 11:00 | 3w3a-J | 2.6 | 3.2 | 68 | 577 | 4 | MOLECULE: | V-TYPE ATP SYNTHASE ALPHA CHAIN |
| 12:00 | 5voy-C | 2.6 | 3.9 | 72 | 592 | 3 | MOLECULE: | V-TYPE PROTON ATPASE CATALYTIC SUBUNIT A,V-TYPE P |
| 13:00 | 8q7r-H | 2.6 1 | 0.2 | 68 | 593 | 10 | MOLECULE: | CULLIN-2 |
| 14:00 | 4dmw-A | 2.6 | 3.5 | 67 | 535 | 6 | MOLECULE: | TOXIN A |
| 15:00 | 7dqe-A | 2.6 | 3.3 | 65 | 584 | 5 | MOLECULE: | V-TYPE SODIUM ATPASE CATALYTIC SUBUNIT A |
| 16:00 | 6wm4-C | 2.6 | 3.4 | 67 | 600 | 1 | MOLECULE: | V-TYPE PROTON ATPASE 116 KDA SUBUNIT A ISOFORM 1 |
| 17:00 | 6fes-A | 2.6 | 4.7 | 65 | 365 | 12 | MOLECULE: | D12_BRIC2, A SYNTHETIC PROTEIN,D12_BRIC2, A SYNTH |
| 18:00 | 8vei-A | 2.6 | 3.7 | 47 | 117 | 9 | MOLECULE: | CHD_R1 |
| 19:00 | 7deg-A | 2.5 | 3.8 | 72 | 581 | 3 | MOLECULE: | CYTOCHROME C OXIDASE SUBUNIT I |
| 20:00 | 7qe7-A | 2.5 | 4 | 69 | 1648 | 9 | MOLECULE: | ANAPHASE-PROMOTING COMPLEX SUBUNIT 10 |
| 21:00 | 2ve7-B | 2.5 | 2.2 | 36 | 303 | 6 | MOLECULE: | KINETOCHORE PROTEIN HEC1, KINETOCHORE PROTEIN SPC |
| 22:00 | 7but-A | 2.5 | 6 | 57 | 142 | 2 | MOLECULE: | ACINIFORM SPIDROIN |
| 23:00 | 7m9c-E | 2.5 | 3.4 | 58 | 257 | 5 | MOLECULE: | TNSC |
| 24:00:00 | 4wvm-B | 2.5 | 4.2 | 67 | 616 | 4 | MOLECULE: | STONUSTOXIN SUBUNIT ALPHA |
| 25:00:00 | 7m99-A | 2.5 | 3.3 | 54 | 257 | 7 | MOLECULE: | TNSC |
| 26:00:00 | 7l7v-A | 2.5 | 4.4 | 52 | 121 | 0 | MOLECULE: | PROBABLE DISEASE RESISTANCE PROTEIN AT5G66900 |
| 27:00:00 | 5wd8-A | 2.5 | 9.8 | 57 | 90 | 0 | MOLECULE: | UNCHARACTERIZED PROTEIN |
| 28:00:00 | 8bcs-A | 2.5 | 4.5 | 41 | 49 | 7 | MOLECULE: | CC-HP1.0 |
| 29:00:00 | 8vjn-B | 2.5 | 3.6 | 39 | 51 | 0 | MOLECULE: | ENCAPSULIN NANOCOMPARTMENT CARGO PROTEIN ENCD |
| 30:00:00 | 6w2r-A | 2.5 | 4.1 | 70 | 218 | 7 | MOLECULE: | JUNCTION 19 DHR54-DHR79 |
| 31:00:00 | 7lsv-B | 2.5 | 3.5 | 60 | 158 | 7 | MOLECULE: | CALMODULIN-DEPENDENT PROTEIN KINASE |
| 32:00:00 | 3gtg-A | 2.5 | 4 | 68 | 1442 | 9 | MOLECULE: | DNA-DIRECTED RNA POLYMERASE II SUBUNIT RPB1 |
| 33:00:00 | 6wbf-A | 2.5 | 3.3 | 66 | 344 | 6 | MOLECULE: | PANNEXIN-1 |
| 34:00:00 | 1nu7-D | 2.5 | 3.1 | 54 | 282 | 13 | MOLECULE: | THROMBIN LIGHT CHAIN |
| 35:00:00 | 1pc6-A | 2.5 | 4.3 | 56 | 141 | 4 | MOLECULE: | PROTEIN NINB |
| 36:00:00 | 6sdk-D | 2.5 | 3.4 | 58 | 199 | 7 | MOLECULE: | STAGE 0 SPORULATION PROTEIN J |
| 37:00:00 | 6hra-B | 2.5 | 4 | 68 | 679 | 4 | MOLECULE: | POTASSIUM-TRANSPORTING ATPASE POTASSIUM-BINDING S |
| 38:00:00 | 1sxj-D | 2.5 | 4 | 57 | 328 | 9 | MOLECULE: | ACTIVATOR 1 95 KDA SUBUNIT |
| 39:00:00 | 4bzj-C | 2.5 | 8.5 | 55 | 693 | 2 | MOLECULE: | PROTEIN TRANSPORT PROTEIN SEC31 |
| 40:00:00 | 2p5t-E | 2.5 | 3.8 | 54 | 95 | 6 | MOLECULE: | FRAGMENT OF PEZA HELIX-TURN-HELIX MOTIF |
| 41:00:00 | 7wtu-K | 2.5 1 | 0.6 | 56 | 1022 | 14 | MOLECULE: | 18S RRNA |
| 42:00:00 | 7ytj-B | 2.5 | 3.2 | 55 | 102 | 7 | MOLECULE: | VACUOLAR TRANSPORTER CHAPERONE 4 |
| 43:00:00 | 8i13-H | 2.5 | 9.7 | 53 | 280 | 6 | MOLECULE: | SMC6 ISOFORM 1 |
| 44:00:00 | 1lq7-A | 2.5 | 3 | 48 | 67 | 6 | MOLECULE: | ALPHA3W |
| 45:00:00 | 7jzu-A | 2.5 | 2.6 | 45 | 55 | 4 | MOLECULE: | LCB1 |
| 46:00:00 | 4y9j-B | 2.5 | 5.5 | 61 | 593 | 10 | MOLECULE: | PROTEIN ACDH-11, ISOFORM B |
| 47:00:00 | 6rfl-C | 2.5 1 | 0.8 | 53 | 304 | 4 | MOLECULE: | DNA-DEPENDENT RNA POLYMERASE SUBUNIT RPO132 |
| 48:00:00 | 2c5i-T | 2.5 | 9.7 | 55 | 94 | 9 | MOLECULE: | T-SNARE AFFECTING A LATE GOLGI COMPARTMENT |
| 49:00:00 | 7jzl-E | 2.5 | 2.6 | 45 | 55 | 4 | MOLECULE: | SPIKE GLYCOPROTEIN |
| 50:00:00 | 7wtt-K | 2.5 1 | 0.4 | 70 | 1022 | 11 | MOLECULE: | 18S RRNA |

| | | | | | | | | |
|---|---|---|---|---|---|---|---|---|
| 51:00:00 | 7jzl-G | 2.5 | 2.6 | 45 | 55 | 4 | MOLECULE: | SPIKE GLYCOPROTEIN |
| 52:00:00 | 5e85-A | 2.5 | 5.5 | 53 | 235 | 6 | MOLECULE: | 78 KDA GLUCOSE-REGULATED PROTEIN |
| 53:00:00 | 8ub3-A | 2.5 | 4.1 | 58 | 224 | 3 | MOLECULE: | DPHF7 FILAMENT |
| 54:00:00 | 8ub3-J | 2.5 | 4.1 | 58 | 224 | 3 | MOLECULE: | DPHF7 FILAMENT |
| 55:00:00 | 5j4o-A | 2.5 | 9.9 | 57 | 222 | 11 | MOLECULE: | SPECTRIN ALPHA CHAIN, ERYTHROCYTIC 1 |
| 56:00:00 | 2pno-C | 2.5 | 3.5 | 54 | 146 | 2 | MOLECULE: | LEUKOTRIENE C4 SYNTHASE |
| 57:00:00 | 7p14-A | 2.5 | 3.2 | 53 | 329 | 9 | MOLECULE: | XK-RELATED PROTEIN |
| 58:00:00 | 5wuq-A | 2.5 | 3.3 | 56 | 158 | 5 | MOLECULE: | ECF RNA POLYMERASE SIGMA FACTOR SIGW |
| 59:00:00 | 1kw2-A | 2.5 | 4 | 59 | 455 | 8 | MOLECULE: | VITAMIN D-BINDING PROTEIN |
| 60:00:00 | 8j8y-A | 2.5 | 3.3 | 56 | 129 | 7 | MOLECULE: | IMMUNODOMINANT MEMBRANE PROTEIN |
| 61:00:00 | 7nd2-A | 2.5 | 3.9 | 57 | 330 | 5 | MOLECULE: | PROTEIN PHOSPHATASE 1 REGULATORY SUBUNIT 21 |
| 62:00:00 | 6l82-A | 2.5 | 2.9 | 49 | 98 | 2 | MOLECULE: | SPINDLE POLE BODY COMPONENT |
| 63:00:00 | 8vjn-D | 2.5 | 3.8 | 40 | 52 | 0 | MOLECULE: | ENCAPSULIN NANOCOMPARTMENT CARGO PROTEIN ENCD |
| 64:00:00 | 7ytj-C | 2.5 | 3.4 | 55 | 102 | 7 | MOLECULE: | VACUOLAR TRANSPORTER CHAPERONE 4 |
| 65:00:00 | 1au1-A | 2.5 | 4.3 | 41 | 166 | 7 | MOLECULE: | INTERFERON-BETA |
| 66:00:00 | 8euy-b | 2.5 | 2.9 | 52 | 120 | 12 | MOLECULE: | RNA (1095-MER) |
| 67:00:00 | 6rth-A | 2.5 | 4 | 56 | 504 | 4 | MOLECULE: | RTX TOXIN AND CA2+-BINDING PROTEIN |
| 68:00:00 | 5y5x-C | 2.5 | 4.5 | 60 | 576 | 7 | MOLECULE: | V-TYPE ATP SYNTHASE ALPHA CHAIN |
| 69:00:00 | 2r9g-G | 2.5 | 6 | 58 | 179 | 9 | MOLECULE: | AAA ATPASE, CENTRAL REGION |
| 70:00:00 | 2r9g-K | 2.5 | 6 | 61 | 180 | 8 | MOLECULE: | AAA ATPASE, CENTRAL REGION |
| 71:00:00 | 2r9g-F | 2.5 | 6 | 58 | 180 | 9 | MOLECULE: | AAA ATPASE, CENTRAL REGION |
| 72:00:00 | 7r43-n | 2.5 | 3 | 52 | 172 | 6 | MOLECULE: | NADH-UBIQUINONE OXIDOREDUCTASE CHAIN 3 |
| 73:00:00 | 2r9g-J | 2.5 | 6.6 | 60 | 181 | 8 | MOLECULE: | AAA ATPASE, CENTRAL REGION |
| 74:00:00 | 8ovt-B | 2.5 | 3.6 | 68 | 284 | 7 | MOLECULE: | YEGT GLYCOSYLTRANSFERASE |
| 75:00:00 | 2r9g-I | 2.5 | 6 | 58 | 178 | 9 | MOLECULE: | AAA ATPASE, CENTRAL REGION |
| 76:00:00 | 6zkk-t | 2.5 | 4.9 | 53 | 177 | 6 | MOLECULE: | NADH DEHYDROGENASE [UBIQUINONE] FLAVOPROTEIN 1, |
| 77:00:00 | 6r0y-B | 2.5 | 4 | 75 | 577 | 7 | MOLECULE: | V-TYPE ATP SYNTHASE ALPHA CHAIN |
| 78:00:00 | 2r9g-A | 2.5 | 6 | 58 | 180 | 9 | MOLECULE: | AAA ATPASE, CENTRAL REGION |
| 79:00:00 | 3qia-A | 2.5 | 3.8 | 71 | 564 | 7 | MOLECULE: | V-TYPE ATP SYNTHASE ALPHA CHAIN |
| 80:00:00 | 2r9g-M | 2.5 | 6 | 58 | 179 | 9 | MOLECULE: | AAA ATPASE, CENTRAL REGION |
| 81:00:00 | 5y5y-B | 2.5 | 3.5 | 61 | 577 | 7 | MOLECULE: | V-TYPE ATP SYNTHASE ALPHA CHAIN |
| 82:00:00 | 4k7b-A | 2.5 | 3.6 | 52 | 111 | 8 | MOLECULE: | EXTRINSIC PROTEIN IN PHOTOSYSTEM II |
| 83:00:00 | 8iwh-G | 2.5 | 3.6 | 54 | 120 | 6 | MOLECULE: | PHOTOSYSTEM II PROTEIN D1 |
| 84:00:00 | 2r9g-D | 2.5 | 6 | 58 | 180 | 9 | MOLECULE: | AAA ATPASE, CENTRAL REGION |
| 85:00:00 | 3sdz-A | 2.5 | 3.1 | 65 | 548 | 2 | MOLECULE: | V-TYPE ATP SYNTHASE ALPHA CHAIN |
| 86:00:00 | 3pvs-C | 2.5 | 3.7 | 52 | 420 | 12 | MOLECULE: | REPLICATION-ASSOCIATED RECOMBINATION PROTEIN A |
| 87:00:00 | 6fes-D | 2.5 | 4.9 | 63 | 363 | 8 | MOLECULE: | D12_BRIC2, A SYNTHETIC PROTEIN,D12_BRIC2, A SYNTH |
| 88:00:00 | 2vl8-B | 2.5 | 3.6 | 66 | 533 | 6 | MOLECULE: | CYTOTOXIN L |
| 89:00:00 | 5voy-E | 2.5 | 3.5 | 71 | 592 | 4 | MOLECULE: | V-TYPE PROTON ATPASE CATALYTIC SUBUNIT A,V-TYPE P |
| 90:00:00 | 6vq6-B | 2.5 | 3.5 | 69 | 600 | 1 | MOLECULE: | ATPASE H+-TRANSPORTING V1 SUBUNIT A |
| 91:00:00 | 5d80-C | 2.5 | 3.5 | 58 | 589 | 7 | MOLECULE: | V-TYPE PROTON ATPASE CATALYTIC SUBUNIT A |
| 92:00:00 | 3pvs-D | 2.5 | 6.1 | 55 | 424 | 11 | MOLECULE: | REPLICATION-ASSOCIATED RECOMBINATION PROTEIN A |
| 93:00:00 | 5cwh-A | 2.5 | 9.1 | 50 | 159 | 6 | MOLECULE: | DESIGNED HELICAL REPEAT PROTEIN |
| 94:00:00 | 7or1-D | 2.5 | 3.8 | 54 | 596 | 6 | MOLECULE: | TRANSIENT RECEPTOR POTENTIAL CATION CHANNEL SUBFA |
| 95:00:00 | 2vkh-C | 2.5 | 3.4 | 64 | 537 | 6 | MOLECULE: | CYTOTOXIN L |
| 96:00:00 | 5y81-H | 2.4 | 3 | 66 | 154 | 14 | MOLECULE: | TRANSCRIPTION-ASSOCIATED PROTEIN 1 |
| 97:00:00 | 7m99-F | 2.4 | 4.2 | 60 | 257 | 7 | MOLECULE: | TNSC |
| 98:00:00 | 7m9b-E | 2.4 | 3.2 | 56 | 257 | 5 | MOLECULE: | TNSC |
| 99:00:00 | 7l7b-F | 2.4 | 4.1 | 62 | 272 | 6 | MOLECULE: | DNA-DIRECTED RNA POLYMERASE SUBUNIT ALPHA |
| 0:00 | 8f6q-B | 2.4 | 8.3 | 56 | 199 | 7 | MOLECULE: | C8-71 |
| 1:00 | 7m9a-E | 2.4 | 3.2 | 56 | 257 | 5 | MOLECULE: | TNSC |
| 2:00 | 8vjn-A | 2.4 | 3.8 | 40 | 52 | 0 | MOLECULE: | ENCAPSULIN NANOCOMPARTMENT CARGO PROTEIN ENCD |
| 3:00 | 7uhb-K | 2.4 | 4.3 | 56 | 75 | 2 | MOLECULE: | SPIKE GLYCOPROTEIN |
| 4:00 | 7udc-B | 2.4 | 9.7 | 58 | 215 | 5 | MOLECULE: | SYNTAXIN-BINDING PROTEIN 1 |
| 5:00 | 7uni-B | 2.4 | 3.8 | 54 | 245 | 4 | MOLECULE: | SP2-ZNPPAM DESIGNED CHLOROPHYLL DIMER PROTEIN |
| 6:00 | 4u5a-B | 2.4 | 2.8 | 55 | 184 | 9 | MOLECULE: | SPOROZOITE MICRONEME PROTEIN ESSENTIAL FOR CELL T |

| | | | | | | | | |
|---|---|---|---|---|---|---|---|---|
| 7:00 | 6sp2-A | 2.4 | 3.8 | 58 | 366 | 5 | MOLECULE: | MEMBRANE PROTEIN TMS1D |
| 8:00 | 3ucq-A | 2.4 | 3.5 | 53 | 651 | 6 | MOLECULE: | AMYLOSUCRASE |
| 9:00 | 8hr8-D | 2.4 | 6 | 58 | 836 | 7 | MOLECULE: | ARCHAEAL ATPASE |
| 10:00 | 1k83-A | 2.4 | 3.8 | 65 | 1366 | 9 | MOLECULE: | DNA-DIRECTED RNA POLYMERASE II LARGEST SUBUNIT |
| 11:00 | 3zjc-E | 2.4 | 2.9 | 39 | 286 | 5 | MOLECULE: | GTPASE IMAP FAMILY MEMBER 7 |
| 12:00 | 1y1v-A | 2.4 | 4 | 66 | 1426 | 9 | MOLECULE: | DNA-DIRECTED RNA POLYMERASE II LARGEST SUBUNIT |
| 13:00 | 6nbq-B | 2.4 | 2.7 | 52 | 483 | 19 | MOLECULE: | NAD(P)H-QUINONE OXIDOREDUCTASE SUBUNIT H |
| 14:00 | 6n1l-A | 2.4 | 4 | 58 | 141 | 2 | MOLECULE: | FIBRONECTIN-BINDING PROTEIN BBK32 |
| 15:00 | 6tgb-A | 2.4 | 3.6 | 60 | 1450 | 13 | MOLECULE: | DEDICATOR OF CYTOKINESIS PROTEIN 2 |
| 16:00 | 6wca-A | 2.4 | 5.9 | 60 | 367 | 5 | MOLECULE: | ENDOSOMAL/LYSOSOMAL POTASSIUM CHANNEL TMEM175 |
| 17:00 | 6nbx-F | 2.4 | 3.5 | 57 | 655 | 2 | MOLECULE: | NAD(P)H-QUINONE OXIDOREDUCTASE SUBUNIT 1 |
| 18:00 | 3k7d-A | 2.4 | 2.6 | 57 | 497 | 5 | MOLECULE: | GLUTAMATE-AMMONIA-LIGASE ADENYLYLTRANSFERASE |
| 19:00 | 7lh7-B | 2.4 | 4 | 47 | 135 | 9 | MOLECULE: | BCL-2-LIKE PROTEIN 1 |
| 20:00 | 8ugc-A | 2.4 | 3.5 | 73 | 384 | 4 | MOLECULE: | FD15 |
| 21:00 | 5l09-B | 2.4 | 2.1 | 35 | 164 | 6 | MOLECULE: | QUORUM-SENSING TRANSCRIPTIONAL ACTIVATOR |
| 22:00 | 6e6t-A | 2.4 | 4.5 | 55 | 266 | 2 | MOLECULE: | NCMC |
| 23:00 | 3c7n-A | 2.4 | 4.4 | 54 | 648 | 4 | MOLECULE: | HEAT SHOCK PROTEIN HOMOLOG SSE1 |
| 24:00:00 | 3fki-A | 2.4 | 3.8 | 65 | 1427 | 9 | MOLECULE: | DNA-DIRECTED RNA POLYMERASE II SUBUNIT RPB1 |
| 25:00:00 | 2y31-A | 2.4 | 3.5 | 55 | 242 | 5 | MOLECULE: | PUTATIVE REPRESSOR SIMREG2 |
| 26:00:00 | 6fh2-B | 2.4 | 2.9 | 62 | 304 | 10 | MOLECULE: | PROTEIN-ARGININE KINASE |
| 27:00:00 | 3l8i-A | 2.4 | 3.6 | 59 | 198 | 2 | MOLECULE: | PROGRAMMED CELL DEATH PROTEIN 10 |
| 28:00:00 | 3o1j-A | 2.4 | 3.6 | 58 | 273 | 3 | MOLECULE: | SENSOR PROTEIN TORS |
| 29:00:00 | 6in7-B | 2.4 | 4 | 58 | 174 | 7 | MOLECULE: | SIGMA FACTOR ALGU NEGATIVE REGULATORY PROTEIN |
| 30:00:00 | 5uz5-G | 2.4 | 2.8 | 51 | 229 | 8 | MOLECULE: | U1 SMALL NUCLEAR RIBONUCLEOPROTEIN 70 KDA HOMOLOG |
| 31:00:00 | 2apl-A | 2.4 | 3.5 | 59 | 149 | 10 | MOLECULE: | HYPOTHETICAL PROTEIN PG0816 |
| 32:00:00 | 4nsr-C | 2.4 | 3.2 | 56 | 93 | 2 | MOLECULE: | IMMUNITY PROTEIN |
| 33:00:00 | 2ns7-D | 2.4 | 3.4 | 53 | 178 | 13 | MOLECULE: | TETRACYCLINE REPRESSOR PROTEIN |
| 34:00:00 | 6zyv-A | 2.4 1 | 0.1 | 59 | 238 | 7 | MOLECULE: | CIR PROTEIN |
| 35:00:00 | 5vre-A | 2.4 | 3 | 56 | 188 | 5 | MOLECULE: | PUTATIVE INTEGRAL MEMBRANE PROTEIN |
| 36:00:00 | 6gdj-A | 2.4 | 3.8 | 51 | 71 | 8 | MOLECULE: | MTO2 |
| 37:00:00 | 8ve7-A | 2.4 | 3.9 | 57 | 294 | 9 | MOLECULE: | DARPIN PROTEIN SCAFFOLD |
| 38:00:00 | 6tz0-A | 2.4 | 3.8 | 75 | 1208 | 7 | MOLECULE: | RNA-DEPENDENT RNA POLYMERASE |
| 39:00:00 | 3vr2-C | 2.4 | 3.6 | 72 | 583 | 6 | MOLECULE: | V-TYPE SODIUM ATPASE CATALYTIC SUBUNIT A |
| 40:00:00 | 2r9g-L | 2.4 | 6.6 | 62 | 188 | 8 | MOLECULE: | AAA ATPASE, CENTRAL REGION |
| 41:00:00 | 6qlf-N | 2.4 | 2.2 | 45 | 371 | 9 | MOLECULE: | INNER KINETOCHORE SUBUNIT IML3 |
| 42:00:00 | 2r9g-H | 2.4 | 6 | 58 | 179 | 9 | MOLECULE: | AAA ATPASE, CENTRAL REGION |
| 43:00:00 | 3i73-A | 2.4 | 3.8 | 72 | 512 | 8 | MOLECULE: | A-TYPE ATP SYNTHASE CATALYTIC SUBUNIT A |
| 44:00:00 | 6ty8-A | 2.4 | 3.7 | 76 | 1196 | 7 | MOLECULE: | RNA-DEPENDENT RNA POLYMERASE |
| 45:00:00 | 5knb-A | 2.4 | 3.9 | 73 | 587 | 4 | MOLECULE: | V-TYPE SODIUM ATPASE CATALYTIC SUBUNIT A |
| 46:00:00 | 7dqc-C | 2.4 | 3.7 | 73 | 584 | 5 | MOLECULE: | V-TYPE SODIUM ATPASE CATALYTIC SUBUNIT A |
| 47:00:00 | 6rko-A | 2.4 | 3.2 | 55 | 444 | 11 | MOLECULE: | CYTOCHROME BD-I UBIQUINOL OXIDASE SUBUNIT 2 |
| 48:00:00 | 6tz1-A | 2.4 | 3.8 | 75 | 1208 | 7 | MOLECULE: | RNA-DEPENDENT RNA POLYMERASE |
| 49:00:00 | 7ctf-C | 2.4 | 3.3 | 61 | 592 | 5 | MOLECULE: | ORIGIN RECOGNITION COMPLEX SUBUNIT 1 |
| 50:00:00 | 7vy9-p | 2.4 | 4.5 | 58 | 178 | 3 | MOLECULE: | NADH DEHYDROGENASE [UBIQUINONE] IRON-SULFUR PROTE |
| 51:00:00 | 5uj8-B | 2.4 | 3 | 60 | 553 | 7 | MOLECULE: | ORIGIN RECOGNITION COMPLEX SUBUNIT 3 |
| 52:00:00 | 8v9o-E | 2.4 | 3.5 | 60 | 299 | 5 | MOLECULE: | TETRAHEDRAL NANOCAGE CAGE, NON-FUSION COMPONENT |
| 53:00:00 | 2r9g-P | 2.4 | 6.6 | 60 | 187 | 8 | MOLECULE: | AAA ATPASE, CENTRAL REGION |
| 54:00:00 | 5bn3-A | 2.4 | 3.4 | 59 | 563 | 2 | MOLECULE: | V-TYPE ATP SYNTHASE ALPHA CHAIN |
| 55:00:00 | 8ffq-B | 2.4 | 3.1 | 51 | 602 | 4 | MOLECULE: | TRANSIENT RECEPTOR POTENTIAL CATION CHANNEL SUBFA |
| 56:00:00 | 7cos-C | 2.4 | 4.7 | 62 | 585 | 10 | MOLECULE: | V-TYPE SODIUM ATPASE CATALYTIC SUBUNIT A |
| 57:00:00 | 8ffq-A | 2.4 | 3.1 | 51 | 602 | 4 | MOLECULE: | TRANSIENT RECEPTOR POTENTIAL CATION CHANNEL SUBFA |
| 58:00:00 | 3nd8-A | 2.4 | 3.6 | 68 | 510 | 1 | MOLECULE: | V-TYPE ATP SYNTHASE ALPHA CHAIN |
| 59:00:00 | 6lu0-A | 2.3 | 2.5 | 48 | 937 | 8 | MOLECULE: | CAS12I2 |
| 60:00:00 | 7m9b-L | 2.3 | 3.5 | 55 | 257 | 5 | MOLECULE: | TNSC |
| 61:00:00 | 7mbw-A | 2.3 | 5.2 | 63 | 475 | 8 | MOLECULE: | TRANSPOSON TN7 TRANSPOSITION PROTEIN TNSC |
| 62:00:00 | 7m9c-J | 2.3 | 3.5 | 58 | 257 | 5 | MOLECULE: | TNSC |

| | | | | | | | | |
|---|---|---|---|---|---|---|---|---|
| 63:00:00 | 7m9a-L | 2.3 | 3.6 | 56 | 257 | 5 | MOLECULE: | TNSC |
| 64:00:00 | 7m9a-J | 2.3 | 3.5 | 58 | 257 | 5 | MOLECULE: | TNSC |
| 65:00:00 | 6wcc-B | 2.3 | 2.8 | 55 | 367 | 7 | MOLECULE: | ENDOSOMAL/LYSOSOMAL POTASSIUM CHANNEL TMEM175 |
| 66:00:00 | 7m9c-L | 2.3 | 3.5 | 55 | 257 | 5 | MOLECULE: | TNSC |
| 67:00:00 | 7m9b-J | 2.3 | 3.5 | 58 | 257 | 5 | MOLECULE: | TNSC |
| 68:00:00 | 2uui-A | 2.3 | 2.9 | 57 | 155 | 5 | MOLECULE: | LEUKOTRIENE C4 SYNTHASE |
| 69:00:00 | 7unh-A | 2.3 | 3.3 | 65 | 246 | 11 | MOLECULE: | SP2 DESIGNED CHLOROPHYLL DIMER PROTEIN |
| 70:00:00 | 7ag9-A | 2.3 | 9.8 | 56 | 375 | 4 | MOLECULE: | KAR9 |
| 71:00:00 | 8g9j-A | 2.3 | 8.9 | 54 | 201 | 4 | MOLECULE: | THR1 |
| 72:00:00 | 2lse-A | 2.3 | 3.7 | 57 | 101 | 2 | MOLECULE: | FOUR HELIX BUNDLE PROTEIN |
| 73:00:00 | 6ksa-B | 2.3 | 3.9 | 68 | 611 | 9 | MOLECULE: | ACYL-COA DEHYDROGENASE |
| 74:00:00 | 2lp4-A | 2.3 | 4.8 | 69 | 225 | 10 | MOLECULE: | CHEMOTAXIS PROTEIN CHEA |
| 75:00:00 | 2ncg-A | 2.3 | 4.4 | 59 | 118 | 7 | MOLECULE: | RGA1E |
| 76:00:00 | 5wah-A | 2.3 | 2.9 | 54 | 99 | 6 | MOLECULE: | IGA FC RECEPTOR |
| 77:00:00 | 6n4n-F | 2.3 | 3.5 | 63 | 191 | 6 | MOLECULE: | NS3 PROTEASE |
| 78:00:00 | 8a64-A | 2.3 | 4.7 | 61 | 940 | 10 | MOLECULE: | ENDO-BETA-N-ACETYLGLUCOSAMINIDASE F2 |
| 79:00:00 | 8tl7-X | 2.3 | 2.6 | 62 | 694 | 10 | MOLECULE: | COMPUTATIONALLY DESIGNED PROTEIN |
| 80:00:00 | 5z2c-A | 2.3 | 3.7 | 58 | 445 | 12 | MOLECULE: | ALPHA-PROTEIN KINASE 1 |
| 81:00:00 | 7wtz-K | 2.3 1 | 0.5 | 70 | 909 | 11 | MOLECULE: | 18S RRNA |
| 82:00:00 | 6tjv-F | 2.3 | 3 | 53 | 600 | 6 | MOLECULE: | NAD(P)H-QUINONE OXIDOREDUCTASE SUBUNIT 1 |
| 83:00:00 | 6xke-A | 2.3 | 2.8 | 54 | 116 | 4 | MOLECULE: | ALBICIN |
| 84:00:00 | 3h0d-A | 2.3 | 2.9 | 52 | 155 | 6 | MOLECULE: | CTSR |
| 85:00:00 | 1vbg-A | 2.3 | 6.1 | 59 | 874 | 10 | MOLECULE: | PYRUVATE,ORTHOPHOSPHATE DIKINASE |
| 86:00:00 | 6n4n-C | 2.3 | 3.6 | 65 | 184 | 3 | MOLECULE: | NS3 PROTEASE |
| 87:00:00 | 1t6u-A | 2.3 | 4.7 | 60 | 117 | 3 | MOLECULE: | SUPEROXIDE DISMUTASE [NI] |
| 88:00:00 | 4fyf-A | 2.3 | 2.4 | 58 | 739 | 9 | MOLECULE: | SIDF, INHIBITOR OF GROWTH FAMILY, MEMBER 3 |
| 89:00:00 | 7b5n-H | 2.3 | 4.9 | 66 | 423 | 5 | MOLECULE: | CULLIN-1 |
| 90:00:00 | 5uha-F | 2.3 | 4.2 | 64 | 322 | 8 | MOLECULE: | DNA-DIRECTED RNA POLYMERASE SUBUNIT ALPHA |
| 91:00:00 | 6a2j-A | 2.3 | 4.2 | 51 | 309 | 6 | MOLECULE: | HEME A SYNTHASE |
| 92:00:00 | 7qho-V | 2.3 | 3.8 | 38 | 129 | 18 | MOLECULE: | CYTOCHROME BC1 COMPLEX RIESKE IRON-SULFUR SUBUNIT |
| 93:00:00 | 2v0o-A | 2.3 | 3.4 | 55 | 273 | 7 | MOLECULE: | FCH DOMAIN ONLY PROTEIN 2 |
| 94:00:00 | 7k8c-A | 2.3 | 2.9 | 58 | 722 | 7 | MOLECULE: | TREHALOSE MONOMYCOLATE EXPORTER MMPL3 |
| 95:00:00 | 5cbo-E | 2.3 | 2.9 | 46 | 176 | 7 | MOLECULE: | MBP3-16,IMMUNOGLOBULIN G-BINDING PROTEIN A |
| 96:00:00 | 6c9k-D | 2.3 | 4 | 59 | 294 | 7 | MOLECULE: | DARP14 - SUBUNIT A WITH DARPIN |
| 97:00:00 | 7vbl-p | 2.3 | 4.4 | 56 | 178 | 4 | MOLECULE: | NADH DEHYDROGENASE [UBIQUINONE] IRON-SULFUR PROTE |
| 98:00:00 | 7k8d-A | 2.3 | 4.7 | 62 | 724 | 10 | MOLECULE: | DRUG EXPORTERS OF THE RND SUPERFAMILY-LIKE PROTEI |
| 99:00:00 | 5cbo-L | 2.3 | 3 | 50 | 176 | 6 | MOLECULE: | MBP3-16,IMMUNOGLOBULIN G-BINDING PROTEIN A |
| 0:00 | 8v9o-F | 2.3 | 3.6 | 60 | 300 | 5 | MOLECULE: | TETRAHEDRAL NANOCAGE CAGE, NON-FUSION COMPONENT |
| 1:00 | 8v9o-D | 2.3 | 3.6 | 60 | 301 | 5 | MOLECULE: | TETRAHEDRAL NANOCAGE CAGE, NON-FUSION COMPONENT |
| 2:00 | 7jps-C | 2.3 | 3.2 | 61 | 591 | 5 | MOLECULE: | ORIGIN RECOGNITION COMPLEX SUBUNIT 1 |
| 3:00 | 5h76-B | 2.3 | 4.6 | 53 | 197 | 8 | MOLECULE: | DARPIN,IMMUNOGLOBULIN G-BINDING PROTEIN A |
| 4:00 | 6c9k-J | 2.3 | 4.1 | 59 | 292 | 7 | MOLECULE: | DARP14 - SUBUNIT A WITH DARPIN |
| 5:00 | 5cbo-C | 2.3 | 2.8 | 43 | 176 | 9 | MOLECULE: | MBP3-16,IMMUNOGLOBULIN G-BINDING PROTEIN A |
| 6:00 | 6v9v-A | 2.3 | 3.7 | 53 | 597 | 6 | MOLECULE: | TRANSIENT RECEPTOR POTENTIAL CATION CHANNEL SUBFA |
| 7:00 | 7tmm-C | 2.3 | 4 | 67 | 592 | 3 | MOLECULE: | H(+)-TRANSPORTING TWO-SECTOR ATPASE |
| 8:00 | 6v9v-D | 2.3 | 3.7 | 53 | 597 | 6 | MOLECULE: | TRANSIENT RECEPTOR POTENTIAL CATION CHANNEL SUBFA |
| 9:00 | 3oe7-X | 2.3 | 3.1 | 57 | 469 | 9 | MOLECULE: | ATP SYNTHASE SUBUNIT ALPHA |
| 10:00 | 5cbo-B | 2.3 | 2.8 | 43 | 176 | 9 | MOLECULE: | MBP3-16,IMMUNOGLOBULIN G-BINDING PROTEIN A |
| 11:00 | 8fin-A | 2.3 | 9.5 | 53 | 174 | 9 | MOLECULE: | CS207A |
| 12:00 | 6fes-B | 2.3 | 4.9 | 66 | 363 | 12 | MOLECULE: | D12_BRIC2, A SYNTHETIC PROTEIN,D12_BRIC2, A SYNTH |
| 13:00 | 6ztq-n | 2.3 | 4.6 | 56 | 177 | 0 | MOLECULE: | NADH-UBIQUINONE OXIDOREDUCTASE CHAIN 3 |
| 14:00 | 7bwr-A | 2.3 | 5.3 | 54 | 1058 | 4 | MOLECULE: | INTEGRAL MEMBRANE INDOLYLACETYLINOSITOL |
| 15:00 | 7w4f-p | 2.3 | 4.7 | 57 | 178 | 0 | MOLECULE: | NADH DEHYDROGENASE [UBIQUINONE] FLAVOPROTEIN 1, |
| 16:00 | 6fes-C | 2.3 | 3.8 | 58 | 363 | 14 | MOLECULE: | D12_BRIC2, A SYNTHETIC PROTEIN,D12_BRIC2, A SYNTH |
| 17:00 | 4dmv-A | 2.3 | 4 | 50 | 537 | 6 | MOLECULE: | TOXIN A |
| 18:00 | 7n6b-A | 2.3 | 3.2 | 49 | 757 | 6 | MOLECULE: | MMPL3 TRANSPORTER |

| | | | | | | | | |
|---|---|---|---|---|---|---|---|---|
| 19:00 | 5bw9-a | 2.3 | 4.3 | 64 | 591 | 5 | MOLECULE: | V-TYPE PROTON ATPASE CATALYTIC SUBUNIT A |
| 20:00 | 3qjy-A | 2.3 | 3.1 | 65 | 556 | 2 | MOLECULE: | V-TYPE ATP SYNTHASE ALPHA CHAIN |
| 21:00 | 6ej7-A | 2.2 | 3.2 | 47 | 706 | 11 | MOLECULE: | XYLOSYLTRANSFERASE 1 |
| 22:00 | 2kob-A | 2.2 | 2.9 | 43 | 108 | 5 | MOLECULE: | UNCHARACTERIZED PROTEIN |
| 23:00 | 7o8c-A | 2.2 | 2.3 | 43 | 582 | 2 | MOLECULE: | SURFACE GLYCAN-BINDING PROTEIN BO2743 |
| 24:00:00 | 7n6i-H | 2.2 | 3.3 | 54 | 257 | 7 | MOLECULE: | TNIQ (HOMOLOGY MODEL) |
| 25:00:00 | 6ahx-A | 2.2 | 3.9 | 40 | 60 | 8 | MOLECULE: | PUTATIVE CYTOSOLIC PROTEIN |
| 26:00:00 | 6gcs-F | 2.2 | 3.9 | 53 | 119 | 9 | MOLECULE: | 75-KDA PROTEIN (NUAM) |
| 27:00:00 | 6irr-A | 2.2 | 8.9 | 48 | 133 | 8 | MOLECULE: | DISRUPTED IN SCHIZOPHRENIA 1 HOMOLOG,CYCLIC AMP-D |
| 28:00:00 | 3qwe-A | 2.2 | 3.7 | 56 | 260 | 2 | MOLECULE: | GEM-INTERACTING PROTEIN |
| 29:00:00 | 3p8c-F | 2.2 | 4.8 | 43 | 156 | 5 | MOLECULE: | CYTOPLASMIC FMR1-INTERACTING PROTEIN 1 |
| 30:00:00 | 7vqo-A | 2.2 | 4.2 | 65 | 1180 | 6 | MOLECULE: | AMS1, NBR1 AND MALE FUSION PROTEIN |
| 31:00:00 | 3s4w-A | 2.2 1 | 0.2 | 64 | 1206 | 2 | MOLECULE: | FANCONI ANEMIA GROUP I PROTEIN HOMOLOG |
| 32:00:00 | 7aqr-V | 2.2 | 3.9 | 53 | 140 | 4 | MOLECULE: | NADH DEHYDROGENASE [UBIQUINONE] IRON-SULFUR PROTE |
| 33:00:00 | 2uxw-A | 2.2 | 3.7 | 66 | 567 | 8 | MOLECULE: | VERY-LONG-CHAIN SPECIFIC ACYL-COA DEHYDROGENASE |
| 34:00:00 | 5ijj-A | 2.2 | 9 | 54 | 175 | 4 | MOLECULE: | SPX DOMAIN |
| 35:00:00 | 7s0r-A | 2.2 | 3.1 | 49 | 85 | 12 | MOLECULE: | C PROTEIN BETA ANTIGEN |
| 36:00:00 | 6r9r-A | 2.2 | 4.8 | 55 | 454 | 7 | MOLECULE: | DNA (5'-R(P*AP*AP*AP*A)-3') |
| 37:00:00 | 4m0m-A | 2.2 | 3.4 | 67 | 721 | 7 | MOLECULE: | PUTATIVE UNCHARACTERIZED PROTEIN |
| 38:00:00 | 2dgj-A | 2.2 | 3.3 | 48 | 246 | 4 | MOLECULE: | HYPOTHETICAL PROTEIN EBHA |
| 39:00:00 | 8ebt-C | 2.2 1 | 1.3 | 60 | 191 | 5 | MOLECULE: | GENERAL TRANSCRIPTION AND DNA REPAIR FACTOR IIH H |
| 40:00:00 | 7wu0-K | 2.2 1 | 0.3 | 60 | 909 | 13 | MOLECULE: | 18S RRNA |
| 41:00:00 | 7mkk-B | 2.2 | 4.6 | 54 | 73 | 7 | MOLECULE: | SMALL OVARY, ISOFORM A |
| 42:00:00 | 4dyl-A | 2.2 | 3.5 | 57 | 376 | 4 | MOLECULE: | TYROSINE-PROTEIN KINASE FES/FPS |
| 43:00:00 | 6xj1-B | 2.2 | 3.1 | 58 | 277 | 3 | MOLECULE: | CELL DIVISION CONTROL PROTEIN 15 |
| 44:00:00 | 4he8-A | 2.2 | 3.9 | 41 | 92 | 5 | MOLECULE: | NADH-QUINONE OXIDOREDUCTASE SUBUNIT 7 |
| 45:00:00 | 1k05-B | 2.2 | 3.4 | 56 | 142 | 9 | MOLECULE: | FOCAL ADHESION KINASE 1 |
| 46:00:00 | 6oeh-A | 2.2 | 3 | 57 | 470 | 0 | MOLECULE: | POLYALA MODEL OF OMCC I-LAYER |
| 47:00:00 | 6z35-A | 2.2 | 3.9 | 56 | 109 | 2 | MOLECULE: | MAQUETTE 2-1IP |
| 48:00:00 | 3t38-A | 2.2 | 3.7 | 52 | 199 | 6 | MOLECULE: | ARSENATE REDUCTASE |
| 49:00:00 | 6f1v-m | 2.2 | 3.6 | 54 | 217 | 7 | MOLECULE: | CYTOPLASMIC DYNEIN 1 HEAVY CHAIN 1 |
| 50:00:00 | 4nsr-B | 2.2 | 2.7 | 51 | 90 | 2 | MOLECULE: | IMMUNITY PROTEIN |
| 51:00:00 | 4nsr-F | 2.2 | 2.8 | 51 | 89 | 2 | MOLECULE: | IMMUNITY PROTEIN |
| 52:00:00 | 2hxo-A | 2.2 | 3 | 51 | 207 | 10 | MOLECULE: | PUTATIVE TETR-FAMILY TRANSCRIPTIONAL REGULATOR |
| 53:00:00 | 1q0k-A | 2.2 | 3.9 | 56 | 117 | 0 | MOLECULE: | SUPEROXIDE DISMUTASE [NI] |
| 54:00:00 | 1i4d-A | 2.2 | 3.4 | 53 | 188 | 6 | MOLECULE: | ARFAPTIN 2 |
| 55:00:00 | 8c5v-Q | 2.2 | 3.6 | 42 | 516 | 5 | MOLECULE: | CHEMOTAXIS PROTEIN CHEA |
| 56:00:00 | 2pmr-A | 2.2 | 4.1 | 49 | 76 | 10 | MOLECULE: | UNCHARACTERIZED PROTEIN |
| 57:00:00 | 1rke-A | 2.2 | 6.4 | 61 | 262 | 3 | MOLECULE: | VINCULIN |
| 58:00:00 | 2ip6-A | 2.2 | 2.9 | 50 | 87 | 8 | MOLECULE: | PAPB |
| 59:00:00 | 6dfk-I | 2.2 | 3.8 | 54 | 236 | 7 | MOLECULE: | SUBUNIT OF PROTEASEOME ACTIVATOR COMPLEX,PUTATIVE |
| 60:00:00 | 2hye-C | 2.2 1 | 0.6 | 56 | 719 | 11 | MOLECULE: | DNA DAMAGE-BINDING PROTEIN 1 |
| 61:00:00 | 3les-A | 2.2 | 4.2 | 62 | 179 | 5 | MOLECULE: | RNA POLYMERASE SIGMA FACTOR |
| 62:00:00 | 6xh5-F | 2.2 | 8.6 | 59 | 348 | 5 | MOLECULE: | HELICAL FUSION DESIGN |
| 63:00:00 | 6c9k-T | 2.2 | 3.9 | 57 | 294 | 9 | MOLECULE: | DARP14 - SUBUNIT A WITH DARPIN |
| 64:00:00 | 7cop-C | 2.2 | 3.7 | 72 | 584 | 4 | MOLECULE: | V-TYPE SODIUM ATPASE CATALYTIC SUBUNIT A |
| 65:00:00 | 5vre-C | 2.2 | 3.1 | 56 | 188 | 5 | MOLECULE: | PUTATIVE INTEGRAL MEMBRANE PROTEIN |
| 66:00:00 | 6c9k-I | 2.2 | 4 | 59 | 293 | 7 | MOLECULE: | DARP14 - SUBUNIT A WITH DARPIN |
| 67:00:00 | 3i4l-A | 2.2 | 3.8 | 74 | 524 | 8 | MOLECULE: | A-TYPE ATP SYNTHASE CATALYTIC SUBUNIT A |
| 68:00:00 | 6c9k-B | 2.2 | 4 | 57 | 292 | 9 | MOLECULE: | DARP14 - SUBUNIT A WITH DARPIN |
| 69:00:00 | 5knd-B | 2.2 | 3.6 | 72 | 592 | 4 | MOLECULE: | V-TYPE SODIUM ATPASE CATALYTIC SUBUNIT A |
| 70:00:00 | 6c9k-Q | 2.2 | 4.5 | 60 | 293 | 8 | MOLECULE: | DARP14 - SUBUNIT A WITH DARPIN |
| 71:00:00 | 7w4l-p | 2.2 | 4.5 | 56 | 178 | 5 | MOLECULE: | NADH DEHYDROGENASE [UBIQUINONE] FLAVOPROTEIN 1, |
| 72:00:00 | 6c9k-K | 2.2 | 4 | 57 | 291 | 9 | MOLECULE: | DARP14 - SUBUNIT A WITH DARPIN |
| 73:00:00 | 6c9k-S | 2.2 | 4 | 57 | 291 | 9 | MOLECULE: | DARP14 - SUBUNIT A WITH DARPIN |
| 74:00:00 | 7pty-B | 2.2 1 | 1 | 67 | 865 | 6 | MOLECULE: | DELTA-LATROINSECTOTOXIN-LT1A |

| | | | | | | | | |
|---|---|---|---|---|---|---|---|---|
| 75:00:00 | 6c9k-A | 2.2 | 4.5 | 60 | 293 | 8 | MOLECULE: | DARP14 - SUBUNIT A WITH DARPIN |
| 76:00:00 | 5cwl-A | 2.2 1 | 0.1 | 53 | 177 | 8 | MOLECULE: | DESIGNED HELICAL REPEAT PROTEIN |
| 77:00:00 | 5x09-A | 2.2 | 3.6 | 68 | 530 | 1 | MOLECULE: | V-TYPE ATP SYNTHASE ALPHA CHAIN,V-TYPE ATP SYNTHA |
| 78:00:00 | 7yks-D | 2.2 | 3.1 | 52 | 591 | 10 | MOLECULE: | TRANSIENT RECEPTOR POTENTIAL CATION CHANNEL SUBFA |
| 79:00:00 | 2bvm-A | 2.2 | 3.7 | 50 | 541 | 10 | MOLECULE: | TOXIN B |
| 80:00:00 | 7wnx-A | 2.2 | 3 | 56 | 734 | 2 | MOLECULE: | TREHALOSE MONOMYCOLATE EXPORTER MMPL3 |
| 81:00:00 | 7dqd-C | 2.2 | 4.2 | 74 | 584 | 7 | MOLECULE: | V-TYPE SODIUM ATPASE CATALYTIC SUBUNIT A |
| 82:00:00 | 7ose-D | 2.2 | 4.4 | 52 | 503 | 13 | MOLECULE: | CYTOCHROME BD-II UBIQUINOL OXIDASE SUBUNIT 1 |
| 83:00:00 | 7jk4-C | 2.2 | 6.9 | 59 | 612 | 3 | MOLECULE: | ORIGIN RECOGNITION COMPLEX SUBUNIT 2 |
| 84:00:00 | 6v9v-C | 2.2 | 3.7 | 53 | 597 | 6 | MOLECULE: | TRANSIENT RECEPTOR POTENTIAL CATION CHANNEL SUBFA |
| 85:00:00 | 5cbo-I | 2.2 | 2.7 | 43 | 176 | 9 | MOLECULE: | MBP3-16,IMMUNOGLOBULIN G-BINDING PROTEIN A |
| 86:00:00 | 7lkm-A | 2.1 | 2.5 | 39 | 149 | 10 | MOLECULE: | PILUS BIOGENESIS PROTEIN |
| 87:00:00 | 7m9c-H | 2.1 | 3.9 | 59 | 257 | 7 | MOLECULE: | TNSC |
| 88:00:00 | 7m9a-H | 2.1 | 3.3 | 54 | 257 | 4 | MOLECULE: | TNSC |
| 89:00:00 | 7m9b-H | 2.1 | 3.3 | 54 | 257 | 4 | MOLECULE: | TNSC |
| 90:00:00 | 7p2p-C | 2.1 | 4.3 | 40 | 174 | 8 | MOLECULE: | SIGNAL PEPTIDASE COMPLEX CATALYTIC SUBUNIT SEC11A |
| 91:00:00 | 8e1e-B | 2.1 | 4.3 | 50 | 232 | 4 | MOLECULE: | SG122_C3 |
| 92:00:00 | 6gvw-B | 2.1 | 6.6 | 44 | 262 | 5 | MOLECULE: | BRCA1-A COMPLEX SUBUNIT ABRAXAS 1 |
| 93:00:00 | 6l7o-G | 2.1 | 3 | 39 | 192 | 13 | MOLECULE: | NAD(P)H-QUINONE OXIDOREDUCTASE SUBUNIT 1 |
| 94:00:00 | 7phk-E | 2.1 | 7.7 | 47 | 393 | 6 | MOLECULE: | POTASSIUM VOLTAGE-GATED CHANNEL, SHAW-RELATED SUB |
| 95:00:00 | 8tl7-Y | 2.1 | 2.6 | 62 | 694 | 10 | MOLECULE: | COMPUTATIONALLY DESIGNED PROTEIN |
| 96:00:00 | 7e1w-B | 2.1 | 4.6 | 52 | 524 | 8 | MOLECULE: | CYTOCHROME C OXIDASE SUBUNIT 2 |
| 97:00:00 | 7zmb-F | 2.1 | 4 | 53 | 238 | 6 | MOLECULE: | NADH-UBIQUINONE OXIDOREDUCTASE CHAIN 1 |
| 98:00:00 | 5flz-B | 2.1 | 4.9 | 61 | 565 | 11 | MOLECULE: | SPINDLE POLE BODY COMPONENT SPC97 |
| 99:00:00 | 1r9t-A | 2.1 | 4 | 65 | 1395 | 9 | MOLECULE: | RNA STRAND |
| 0:00 | 8j0h-A | 2.1 | 8.9 | 56 | 134 | 9 | MOLECULE: | UNCHARACTERIZED PROTEIN C4H3.06 |
| 1:00 | 3rgu-A | 2.1 | 4.7 | 53 | 88 | 8 | MOLECULE: | FIMBRIAE-ASSOCIATED PROTEIN FAP1 |
| 2:00 | 5jbr-A | 2.1 | 8.5 | 45 | 149 | 4 | MOLECULE: | UNCHARACTERIZED PROTEIN BCAV_2135 |
| 3:00 | 4uy3-A | 2.1 | 2.9 | 53 | 194 | 8 | MOLECULE: | SEPTATION RING FORMATION REGULATOR EZRA |
| 4:00 | 4noo-B | 2.1 | 2.7 | 50 | 95 | 2 | MOLECULE: | VGRG PROTEIN |
| 5:00 | 4nqi-D | 2.1 | 3.3 | 53 | 232 | 0 | MOLECULE: | SH3 DOMAIN-CONTAINING PROTEIN |
| 6:00 | 8bpr-G | 2.1 | 3.5 | 65 | 193 | 5 | MOLECULE: | DNA REPLICATION AND REPAIR PROTEIN RECF |
| 7:00 | 7e1x-B | 2.1 | 4.6 | 52 | 524 | 8 | MOLECULE: | CYTOCHROME C OXIDASE SUBUNIT 2 |
| 8:00 | 3ats-A | 2.1 | 6.3 | 58 | 352 | 5 | MOLECULE: | PUTATIVE UNCHARACTERIZED PROTEIN |
| 9:00 | 8w41-A | 2.1 | 4 | 55 | 1162 | 0 | MOLECULE: | CALMODULIN-1 |
| 10:00 | 3hai-A | 2.1 | 3.2 | 51 | 294 | 4 | MOLECULE: | HUMAN PACSIN1 F-BAR |
| 11:00 | 6irw-A | 2.1 | 4.5 | 56 | 495 | 11 | MOLECULE: | PHOSPHORYLATED CTD-INTERACTING FACTOR 1 |
| 12:00 | 8e7s-O | 2.1 | 9.5 | 49 | 269 | 2 | MOLECULE: | CYTOCHROME B-C1 COMPLEX SUBUNIT 1, MITOCHONDRIAL |
| 13:00 | 8pmq-2 | 2.1 | 7.8 | 55 | 355 | 11 | MOLECULE: | E3 UBIQUITIN-PROTEIN LIGASE RMD5 |
| 14:00 | 6tjv-C | 2.1 | 3.4 | 39 | 114 | 13 | MOLECULE: | NAD(P)H-QUINONE OXIDOREDUCTASE SUBUNIT 1 |
| 15:00 | 3t46-A | 2.1 | 4.2 | 53 | 75 | 9 | MOLECULE: | SCIN-D |
| 16:00 | 6q8x-A | 2.1 | 4 | 41 | 117 | 5 | MOLECULE: | NADH-QUINONE OXIDOREDUCTASE SUBUNIT 1 |
| 17:00 | 4bpd-F | 2.1 | 2.7 | 46 | 92 | 15 | MOLECULE: | DIACYLGLYCEROL KINASE |
| 18:00 | 8ia1-C | 2.1 | 2.3 | 54 | 361 | 6 | MOLECULE: | GLYCEROL-3-PHOSPHATE ACYLTRANSFERASE, CHLOROPLAST |
| 19:00 | 3ckd-B | 2.1 | 4.5 | 55 | 244 | 4 | MOLECULE: | INVASION PLASMID ANTIGEN, SECRETED BY THE MXI-SPA |
| 20:00 | 7wej-A | 2.1 | 3.3 | 49 | 270 | 8 | MOLECULE: | WD REPEAT-CONTAINING PROTEIN 47 |
| 21:00 | 1n5u-A | 2.1 | 3.7 | 63 | 583 | 2 | MOLECULE: | SERUM ALBUMIN |
| 22:00 | 1k40-A | 2.1 | 3.3 | 57 | 126 | 9 | MOLECULE: | ADHESION KINASE |
| 23:00 | 4brb-F | 2.1 | 3 | 47 | 92 | 15 | MOLECULE: | DIACYLGLYCEROL KINASE |
| 24:00:00 | 7khc-F | 2.1 | 3.7 | 55 | 447 | 9 | MOLECULE: | DNA-DIRECTED RNA POLYMERASE SUBUNIT ALPHA |
| 25:00:00 | 2z0v-A | 2.1 | 3.8 | 52 | 223 | 4 | MOLECULE: | SH3-CONTAINING GRB2-LIKE PROTEIN 3 |
| 26:00:00 | 6fbv-F | 2.1 | 3.6 | 60 | 293 | 3 | MOLECULE: | DNA-DIRECTED RNA POLYMERASE SUBUNIT ALPHA |
| 27:00:00 | 1z23-A | 2.1 | 3.9 | 57 | 163 | 7 | MOLECULE: | CRK-ASSOCIATED SUBSTRATE |
| 28:00:00 | 4by9-C | 2.1 | 3.2 | 51 | 366 | 2 | MOLECULE: | SSR26 |
| 29:00:00 | 1lvf-A | 2.1 | 9.5 | 53 | 106 | 8 | MOLECULE: | SYNTAXIN 6 |
| 30:00:00 | 1m6t-A | 2.1 | 3.7 | 55 | 106 | 4 | MOLECULE: | SOLUBLE CYTOCHROME B562 |

| | | | | | | | | |
|---|---|---|---|---|---|---|---|---|
| 31:00:00 | 7rmy-A | 2.1 | 3.8 | 75 | 266 | 4 | MOLECULE: | DE NOVO DESIGNED TUNABLE HOMODIMER, D_3-337 |
| 32:00:00 | 3kyj-A | 2.1 | 4.1 | 58 | 129 | 3 | MOLECULE: | PUTATIVE HISTIDINE PROTEIN KINASE |
| 33:00:00 | 2c2l-A | 2.1 | 2.7 | 42 | 281 | 0 | MOLECULE: | CARBOXY TERMINUS OF HSP70-INTERACTING PROTEIN |
| 34:00:00 | 8swd-D | 2.1 | 3 | 39 | 53 | 10 | MOLECULE: | 2-OXOGLUTARATE:ACCEPTOR OXIDOREDUCTASE |
| 35:00:00 | 1ysm-A | 2.1 | 2.2 | 38 | 55 | 8 | MOLECULE: | CALCYCLIN-BINDING PROTEIN |
| 36:00:00 | 4jza-A | 2.1 | 3.9 | 61 | 817 | 10 | MOLECULE: | UNCHARACTERIZED PROTEIN |
| 37:00:00 | 7m4m-A | 2.1 | 6.9 | 42 | 173 | 5 | MOLECULE: | E3 UBIQUITIN-PROTEIN LIGASE RNF216 |
| 38:00:00 | 3l32-B | 2.1 | 1.6 | 35 | 45 | 3 | MOLECULE: | PHOSPHOPROTEIN |
| 39:00:00 | 6vq8-C | 2.1 | 3.7 | 73 | 600 | 4 | MOLECULE: | ATPASE H+-TRANSPORTING V1 SUBUNIT A |
| 40:00:00 | 2qw6-B | 2.1 | 3.7 | 48 | 85 | 6 | MOLECULE: | AAA ATPASE, CENTRAL REGION |
| 41:00:00 | 7r72-R | 2.1 | 3.2 | 52 | 134 | 8 | MOLECULE: | 25S RRNA |
| 42:00:00 | 5f3q-B | 2.1 | 2.9 | 53 | 176 | 6 | MOLECULE: | PUTATIVE UNCHARACTERIZED PROTEIN |
| 43:00:00 | 5f3o-B | 2.1 | 3.1 | 55 | 189 | 5 | MOLECULE: | PUTATIVE UNCHARACTERIZED PROTEIN |
| 44:00:00 | 7tts-F | 2.1 | 4.9 | 59 | 433 | 15 | MOLECULE: | CASEINOLYTIC PEPTIDASE B PROTEIN HOMOLOG |
| 45:00:00 | 7cte-C | 2.1 | 3.2 | 60 | 592 | 5 | MOLECULE: | ORIGIN RECOGNITION COMPLEX SUBUNIT 2 |
| 46:00:00 | 6xh5-E | 2.1 | 8.5 | 61 | 348 | 5 | MOLECULE: | HELICAL FUSION DESIGN |
| 47:00:00 | 3vr4-C | 2.1 | 3.3 | 64 | 586 | 5 | MOLECULE: | V-TYPE SODIUM ATPASE CATALYTIC SUBUNIT A |
| 48:00:00 | 5zea-A | 2.1 | 3.3 | 65 | 587 | 5 | MOLECULE: | V-TYPE SODIUM ATPASE CATALYTIC SUBUNIT A |
| 49:00:00 | 7uby-B | 2.1 | 3.8 | 51 | 529 | 6 | MOLECULE: | GLUCOSYLTRANSFERASE TCDA |
| 50:00:00 | 5zea-G | 2.1 | 3.6 | 72 | 587 | 4 | MOLECULE: | V-TYPE SODIUM ATPASE CATALYTIC SUBUNIT A |
| 51:00:00 | 7or0-B | 2.1 | 3.8 | 54 | 596 | 6 | MOLECULE: | TRANSIENT RECEPTOR POTENTIAL CATION CHANNEL SUBFA |
| 52:00:00 | 7dqe-B | 2.1 | 3.3 | 66 | 584 | 6 | MOLECULE: | V-TYPE SODIUM ATPASE CATALYTIC SUBUNIT A |
| 53:00:00 | 7zmb-R | 2.1 | 2.9 | 49 | 98 | 8 | MOLECULE: | NADH-UBIQUINONE OXIDOREDUCTASE CHAIN 1 |
| 54:00:00 | 1pjq-B | 2 | 4 | 60 | 456 | 8 | MOLECULE: | SIROHEME SYNTHASE |
| 55:00:00 | 7p61-A | 2 | 4.9 | 43 | 106 | 14 | MOLECULE: | NADH-QUINONE OXIDOREDUCTASE SUBUNIT F |
| 56:00:00 | 4a4z-A | 2 | 3.8 | 37 | 874 | 8 | MOLECULE: | ANTIVIRAL HELICASE SKI2 |
| 57:00:00 | 7upp-C | 2 | 3.9 | 42 | 75 | 5 | MOLECULE: | DHT03 PROTEIN A |
| 58:00:00 | 7d0i-F | 2 | 4.6 | 45 | 424 | 7 | MOLECULE: | AUTOPHAGY-RELATED PROTEIN 9 |
| 59:00:00 | 5lnk-A | 2 | 4.8 | 44 | 115 | 9 | MOLECULE: | MITOCHONDRIAL COMPLEX I, 51 KDA SUBUNIT |
| 60:00:00 | 6ziy-A | 2 | 4 | 41 | 117 | 5 | MOLECULE: | NADH-QUINONE OXIDOREDUCTASE SUBUNIT 1 |
| 61:00:00 | 8ebs-C | 2.0 1 | 0.7 | 67 | 358 | 4 | MOLECULE: | TFIIH BASAL TRANSCRIPTION FACTOR COMPLEX HELICASE |
| 62:00:00 | 6jhm-A | 2 | 4.5 | 66 | 494 | 8 | MOLECULE: | CHLOROPHENOL MONOOXYGENASE |
| 63:00:00 | 8ab0-G | 2 | 3.7 | 64 | 193 | 6 | MOLECULE: | RECOMBINATION PROTEIN RECR |
| 64:00:00 | 6khi-C | 2 | 3.8 | 41 | 120 | 12 | MOLECULE: | NAD(P)H-QUINONE OXIDOREDUCTASE SUBUNIT 1 |
| 65:00:00 | 6l80-C | 2 | 2.9 | 47 | 107 | 4 | MOLECULE: | GAMMA-TUBULIN COMPLEX SUBUNIT MOD21 |
| 66:00:00 | 7nyh-A | 2 | 4.7 | 43 | 86 | 14 | MOLECULE: | NADH-QUINONE OXIDOREDUCTASE SUBUNIT A |
| 67:00:00 | 6l7p-C | 2 | 3.7 | 41 | 121 | 12 | MOLECULE: | NAD(P)H-QUINONE OXIDOREDUCTASE SUBUNIT 1 |
| 68:00:00 | 6vp0-E | 2 | 7.8 | 52 | 406 | 10 | MOLECULE: | DIACYLGLYCEROL O-ACYLTRANSFERASE 1 |
| 69:00:00 | 6ptq-A | 2 | 8.2 | 48 | 482 | 6 | MOLECULE: | PHOTORECEPTOR-HISTIDINE KINASE BPHP |
| 70:00:00 | 8tl7-V | 2 | 3 | 64 | 694 | 8 | MOLECULE: | COMPUTATIONALLY DESIGNED PROTEIN |
| 71:00:00 | 8f6q-A | 2 | 9 | 66 | 199 | 9 | MOLECULE: | C8-71 |
| 72:00:00 | 1eo0-A | 2 | 2.4 | 47 | 77 | 13 | MOLECULE: | TRANSCRIPTION ELONGATION FACTOR S-II |
| 73:00:00 | 8om1-A | 2 | 4.1 | 42 | 115 | 14 | MOLECULE: | NADH-UBIQUINONE OXIDOREDUCTASE CHAIN 3 |
| 74:00:00 | 8swd-C | 2 | 3.1 | 39 | 53 | 10 | MOLECULE: | 2-OXOGLUTARATE:ACCEPTOR OXIDOREDUCTASE |
| 75:00:00 | 7tad-C | 2.0 1 | 0 | 53 | 210 | 9 | MOLECULE: | REGULATORY PROTEIN NPR1 |
| 76:00:00 | 7d0i-L | 2 | 4.6 | 45 | 424 | 7 | MOLECULE: | AUTOPHAGY-RELATED PROTEIN 9 |
| 77:00:00 | 6umr-A | 2 | 2.9 | 56 | 418 | 5 | MOLECULE: | DAMAGE-CONTROL PHOSPHATASE DUF89 |
| 78:00:00 | 6zku-t | 2 | 5.1 | 53 | 177 | 6 | MOLECULE: | NADH DEHYDROGENASE [UBIQUINONE] FLAVOPROTEIN 1, |
| 79:00:00 | 7jpr-C | 2 | 3 | 58 | 593 | 5 | MOLECULE: | ORIGIN RECOGNITION COMPLEX SUBUNIT 1 |
| 80:00:00 | 4mix-A | 2 | 3.6 | 68 | 276 | 3 | MOLECULE: | PUTATIVE INSECTICIDAL TOXIN |
| 81:00:00 | 3mfy-A | 2 | 3.6 | 70 | 512 | 3 | MOLECULE: | V-TYPE ATP SYNTHASE ALPHA CHAIN |
| 82:00:00 | 8pwl-A | 2.0 1 | 0.8 | 66 | 830 | 6 | MOLECULE: | E3 UBIQUITIN-PROTEIN LIGASE HACE1 |
| 83:00:00 | 7tmp-A | 2 | 3.2 | 64 | 593 | 5 | MOLECULE: | H(+)-TRANSPORTING TWO-SECTOR ATPASE |
| 84:00:00 | 3srz-A | 2 | 3.7 | 64 | 537 | 9 | MOLECULE: | TOXIN A |
| 85:00:00 | 7yks-B | 2 | 2.9 | 52 | 591 | 10 | MOLECULE: | TRANSIENT RECEPTOR POTENTIAL CATION CHANNEL SUBFA |

**DALI results: MdA-1 Phogi**

| No: | Chain | Z | rmsd | lali | nres | %id | PDB | Description |
|---|---|---|---|---|---|---|---|---|
| 1:00 | 7r5s-P | 5.8 | 5.7 | 72 | 224 | 1 | MOLECULE: | CENTROMERE PROTEIN H |
| 2:00 | 7qoo-P | 5.6 | 7.1 | 83 | 233 | 1 | MOLECULE: | CENTROMERE PROTEIN C |
| 3:00 | 7pkn-P | 5.6 | 6.8 | 74 | 224 | 3 | MOLECULE: | CENTROMERE PROTEIN H |
| 4:00 | 7uik-p | 5.3 | 5.2 | 73 | 895 | 4 | MOLECULE: | DNA (38-MER) |
| 5:00 | 1wzn-A | 5.2 | 4.2 | 70 | 245 | 7 | MOLECULE: | SAM-DEPENDENT METHYLTRANSFERASE |
| 6:00 | 7ywx-P | 5.2 | 5.6 | 72 | 224 | 1 | MOLECULE: | CENTROMERE PROTEIN H |
| 7:00 | 6i7s-G | 5 | 5.4 | 88 | 863 | 6 | MOLECULE: | PROTEIN DISULFIDE-ISOMERASE |
| 8:00 | 3u1w-C | 5 | 4.7 | 78 | 250 | 6 | MOLECULE: | HYPOTHETICAL PERIPLASMIC PROTEIN |
| 9:00 | 5d16-A | 4.9 | 3 | 64 | 161 | 6 | MOLECULE: | TRANSPOSON TN7 TRANSPOSITION PROTEI |
| 10:00 | 7xhn-P | 4.9 | 7.8 | 79 | 224 | 4 | MOLECULE: | CENTROMERE PROTEIN O |
| 11:00 | 6lk8-J | 4.8 | 5.2 | 72 | 1026 | 11 | MOLECULE: | MGC83295 PROTEIN |
| 12:00 | 7uic-p | 4.8 | 5.1 | 72 | 903 | 4 | MOLECULE: | MEDIATOR OF RNA POLYMERASE II TRANS |
| 13:00 | 8eoj-B | 4.8 | 6.9 | 92 | 781 | 4 | MOLECULE: | PROTEIN DISULFIDE-ISOMERASE |
| 14:00 | 5o68-D | 4.8 | 7.1 | 90 | 271 | 7 | MOLECULE: | FAPF |
| 15:00 | 7yyh-P | 4.8 | 7.3 | 68 | 219 | 7 | MOLECULE: | HISTONE H3-LIKE CENTROMERIC PROTEIN |
| 16:00 | 7r5v-P | 4.8 | 7.4 | 69 | 219 | 7 | MOLECULE: | CENTROMERE PROTEIN H |
| 17:00 | 7uil-6 | 4.8 | 5.2 | 73 | 903 | 4 | MOLECULE: | MEDIATOR OF RNA POLYMERASE II TRANS |
| 18:00 | 7px8-A | 4.7 | 4.9 | 74 | 700 | 5 | MOLECULE: | ACYLAMINO-ACID-RELEASING ENZYME |
| 19:00 | 5o68-H | 4.7 | 7.1 | 88 | 272 | 6 | MOLECULE: | FAPF |
| 20:00 | 5o68-K | 4.7 | 7.1 | 89 | 272 | 6 | MOLECULE: | FAPF |
| 21:00 | 7qun-A | 4.7 | 4.9 | 73 | 695 | 5 | MOLECULE: | ACYLAMINO-ACID-RELEASING ENZYME |
| 22:00 | 7qun-C | 4.7 | 4.6 | 73 | 694 | 5 | MOLECULE: | ACYLAMINO-ACID-RELEASING ENZYME |
| 23:00 | 7px8-B | 4.7 | 4.9 | 72 | 700 | 6 | MOLECULE: | ACYLAMINO-ACID-RELEASING ENZYME |
| 24:00:00 | 7qun-B | 4.7 | 5 | 73 | 694 | 5 | MOLECULE: | ACYLAMINO-ACID-RELEASING ENZYME |
| 25:00:00 | 7qun-D | 4.7 | 4.6 | 73 | 694 | 5 | MOLECULE: | ACYLAMINO-ACID-RELEASING ENZYME |
| 26:00:00 | 3cnx-C | 4.6 | 5.3 | 65 | 148 | 9 | MOLECULE: | UNCHARACTERIZED PROTEIN |
| 27:00:00 | 6zce-p | 4.6 | 6.4 | 76 | 646 | 8 | MOLECULE: | EUKARYOTIC TRANSLATION INITIATION F |
| 28:00:00 | 5o65-B | 4.6 | 6 | 89 | 296 | 4 | MOLECULE: | FAPF |
| 29:00:00 | 7s7u-A | 4.6 | 6.7 | 83 | 511 | 4 | MOLECULE: | INICSNFR 3.0 FLUORESCENT NICOTINE S |
| 30:00:00 | 8cas-p | 4.6 | 4.9 | 70 | 646 | 9 | MOLECULE: | 40S RIBOSOMAL PROTEIN S0-A |
| 31:00:00 | 5o68-I | 4.6 | 5.4 | 76 | 256 | 12 | MOLECULE: | FAPF |
| 32:00:00 | 4ik8-A | 4.6 | 6.7 | 83 | 395 | 2 | MOLECULE: | RCAMP, GREEN FLUORESCENT PROTEIN |
| 33:00:00 | 3wlc-A | 4.6 | 6.7 | 82 | 392 | 2 | MOLECULE: | MYOSIN LIGHT CHAIN KINASE, GREEN FL |
| 34:00:00 | 4it7-A | 4.5 | 3.7 | 65 | 107 | 5 | MOLECULE: | CPI |
| 35:00:00 | 6qvi-A | 4.5 | 7.4 | 76 | 518 | 8 | MOLECULE: | COMZ |
| 36:00:00 | 5tgn-A | 4.5 | 3.8 | 57 | 109 | 12 | MOLECULE: | UNCHARACTERIZED PROTEIN |
| 37:00:00 | 3d9r-B | 4.5 | 6.8 | 74 | 134 | 7 | MOLECULE: | KETOSTEROID ISOMERASE-LIKE PROTEIN |
| 38:00:00 | 6i7s-H | 4.5 | 5.2 | 86 | 862 | 6 | MOLECULE: | PROTEIN DISULFIDE-ISOMERASE |
| 39:00:00 | 7s7v-C | 4.5 | 8 | 87 | 506 | 3 | MOLECULE: | INICSNFR 3.0 FLUORESCENT NICOTINE S |
| 40:00:00 | 6e8a-A | 4.4 | 2.8 | 56 | 148 | 4 | MOLECULE: | DUF1795 DOMAIN-CONTAINING PROTEIN |
| 41:00:00 | 7wb4-E | 4.4 | 4.8 | 76 | 1363 | 4 | MOLECULE: | OUTER NUP133 |
| 42:00:00 | 2o5p-A | 4.4 | 12.2 | 100 | 772 | 6 | MOLECULE: | FERRIPYOVERDINE RECEPTOR |
| 43:00:00 | 5dl5-A | 4.4 | 7.7 | 79 | 416 | 9 | MOLECULE: | MEMBRANE PROTEIN |
| 44:00:00 | 5tgn-C | 4.4 | 4.1 | 56 | 107 | 11 | MOLECULE: | UNCHARACTERIZED PROTEIN |

| | | | | | | | | |
|---|---|---|---|---|---|---|---|---|
| 45:00:00 | 3blz-A | 4.4 | 3.6 | 57 | 124 | 9 | MOLECULE: | NTF2-LIKE PROTEIN OF UNKNOWN FUNCTI |
| 46:00:00 | 7jta-B | 4.4 | 3.3 | 52 | 56 | 4 | MOLECULE: | NTF2-LIKE NUCLEASE/ANTI-CRISPR |
| 47:00:00 | 7pb8-P | 4.4 | 7.7 | 75 | 224 | 8 | MOLECULE: | CENTROMERE PROTEIN O |
| 48:00:00 | 6zu9-p | 4.4 | 5.4 | 70 | 646 | 9 | MOLECULE: | 18S RIBOSOMAL RNA |
| 49:00:00 | 5o67-C | 4.4 | 6.4 | 84 | 284 | 6 | MOLECULE: | FAPF |
| 50:00:00 | 3u1w-B | 4.4 | 3.2 | 74 | 248 | 1 | MOLECULE: | HYPOTHETICAL PERIPLASMIC PROTEIN |
| 51:00:00 | 5o65-A | 4.4 | 7.1 | 89 | 276 | 6 | MOLECULE: | FAPF |
| 52:00:00 | 7s7t-A | 4.4 | 6.7 | 83 | 509 | 4 | MOLECULE: | INICSNFR 3.0 FLUORESCENT NICOTINE S |
| 53:00:00 | 3ek4-A | 4.4 | 6.7 | 82 | 348 | 2 | MOLECULE: | MYOSIN LIGHT CHAIN KINASE, GREEN FL |
| 54:00:00 | 7jta-A | 4.3 | 3.6 | 52 | 56 | 4 | MOLECULE: | NTF2-LIKE NUCLEASE/ANTI-CRISPR |
| 55:00:00 | 3gzr-A | 4.3 | 4.5 | 68 | 143 | 6 | MOLECULE: | UNCHARACTERIZED PROTEIN WITH A NTF2 |
| 56:00:00 | 2w4y-A | 4.3 | 4 | 69 | 122 | 4 | MOLECULE: | CAULOBACTER 5 VIRUS-LIKE PARTICLE |
| 57:00:00 | 2d73-A | 4.3 | 3 | 62 | 717 | 11 | MOLECULE: | ALPHA-GLUCOSIDASE SUSB |
| 58:00:00 | 7zkh-A | 4.3 | 5.6 | 74 | 267 | 9 | MOLECULE: | METHYLTRANSFERASE |
| 59:00:00 | 6nd4-H | 4.3 | 5 | 74 | 834 | 7 | MOLECULE: | ETS RRNA |
| 60:00:00 | 6bjt-B | 4.3 | 2.9 | 55 | 130 | 4 | MOLECULE: | DUF4440 DOMAIN-CONTAINING PROTEIN |
| 61:00:00 | 7syf-A | 4.3 | 9.7 | 80 | 1391 | 9 | MOLECULE: | PHOSPHATIDYLINOSITOL 3,4,5-TRISPHOS |
| 62:00:00 | 8cah-p | 4.3 | 5.5 | 70 | 646 | 6 | MOLECULE: | EUKARYOTIC TRANSLATION INITIATION F |
| 63:00:00 | 3cnx-A | 4.3 | 5.3 | 65 | 138 | 9 | MOLECULE: | UNCHARACTERIZED PROTEIN |
| 64:00:00 | 3wld-A | 4.3 | 6.8 | 83 | 393 | 2 | MOLECULE: | MYOSIN LIGHT CHAIN KINASE, GREEN FL |
| 65:00:00 | 7aug-A | 4.3 | 6.8 | 82 | 398 | 2 | MOLECULE: | RSGCAMP1.3 |
| 66:00:00 | 3u1w-A | 4.3 | 3.1 | 74 | 249 | 1 | MOLECULE: | HYPOTHETICAL PERIPLASMIC PROTEIN |
| 67:00:00 | 4pr7-A | 4.2 | 6.8 | 73 | 171 | 7 | MOLECULE: | OLIGOGALACTURONATE-SPECIFIC PORIN K |
| 68:00:00 | 8feh-D | 4.2 | 6.5 | 70 | 268 | 7 | MOLECULE: | MINOR CAPSID PROTEIN A1 FUSION |
| 69:00:00 | 5yvf-A | 4.2 | 3.7 | 73 | 322 | 4 | MOLECULE: | BFA1 |
| 70:00:00 | 6i1m-A | 4.2 | 4 | 61 | 93 | 3 | MOLECULE: | CYSTATIN |
| 71:00:00 | 5fq6-M | 4.2 | 9.5 | 95 | 948 | 9 | MOLECULE: | PUTATIVE LIPOPROTEIN |
| 72:00:00 | 7sjy-A | 4.2 | 8.3 | 78 | 313 | 4 | MOLECULE: | ANTI-SIGMA-I FACTOR RSGI9 |
| 73:00:00 | 6iqt-A | 4.2 | 5.3 | 68 | 135 | 10 | MOLECULE: | CAG PATHOGENICITY ISLAND PROTEIN (C |
| 74:00:00 | 6r2q-B | 4.2 | 7.4 | 91 | 649 | 4 | MOLECULE: | CYSTATHIONINE BETA-SYNTHASE |
| 75:00:00 | 7oqe-P | 4.2 | 5.5 | 74 | 1186 | 8 | MOLECULE: | PROTEIN NAM8 |
| 76:00:00 | 4lzi-A | 4.1 | 7.1 | 69 | 277 | 10 | MOLECULE: | MULTICYSTATIN |
| 77:00:00 | 8cli-B | 4.1 | 4 | 66 | 689 | 8 | MOLECULE: | GENERAL TRANSCRIPTION FACTOR 3C POL |
| 78:00:00 | 6dbj-B | 4.1 | 6 | 77 | 352 | 12 | MOLECULE: | RECOMBINATION ACTIVATING GENE 1 - M |
| 79:00:00 | 4p42-A | 4.1 | 7.4 | 85 | 455 | 8 | MOLECULE: | EXTENDED SYNAPTOTAGMIN-2 |
| 80:00:00 | 3f8x-B | 4.1 | 2.8 | 49 | 132 | 4 | MOLECULE: | PUTATIVE DELTA-5-3-KETOSTEROID ISOM |
| 81:00:00 | 6zrd-A | 4.1 | 6.3 | 77 | 411 | 5 | MOLECULE: | HISTONE-BINDING PROTEIN RBBP4 |
| 82:00:00 | 6jp6-A | 4.1 | 4.7 | 72 | 998 | 4 | MOLECULE: | TRNA (GUANOSINE(34)-2'-O)-METHYLTRA |
| 83:00:00 | 6m63-A | 4.1 | 6.8 | 81 | 359 | 2 | MOLECULE: | CHIMERA OF CYCLIC NUCLEOTIDE-GATED |
| 84:00:00 | 3cu3-A | 4.1 | 6.4 | 74 | 162 | 7 | MOLECULE: | DOMAIN OF UNKNOWN FUNCTION WITH A C |
| 85:00:00 | 3fsd-A | 4.1 | 6 | 67 | 121 | 4 | MOLECULE: | NTF2-LIKE PROTEIN OF UNKNOWN FUNCTI |
| 86:00:00 | 5d17-I | 4.1 | 3.1 | 65 | 158 | 5 | MOLECULE: | TRANSPOSON TN7 TRANSPOSITION PROTEI |
| 87:00:00 | 8bys-A | 4 | 9.8 | 85 | 323 | 5 | MOLECULE: | S-LAYER HOMOLOGY DOMAIN-CONTAINING |
| 88:00:00 | 7qe7-I | 4 | 6.4 | 69 | 742 | 6 | MOLECULE: | ANAPHASE-PROMOTING COMPLEX SUBUNIT |
| 89:00:00 | 2ags-A | 4 | 7 | 66 | 634 | 6 | MOLECULE: | SIALIDASE |
| 90:00:00 | 7oc9-A | 4 | 5.1 | 70 | 133 | 7 | MOLECULE: | BD0675 |

| | | | | | | | | |
|---|---|---|---|---|---|---|---|---|
| 91:00:00 | 7cd7-A | 4 | 3.6 | 64 | 99 | 8 | MOLECULE: | GREEN FLUORESCENT PROTEIN |
| 92:00:00 | 2obd-A | 4 | 5 | 79 | 472 | 8 | MOLECULE: | CHOLESTERYL ESTER TRANSFER PROTEIN |
| 93:00:00 | 1uyn-X | 4 | 6.9 | 94 | 279 | 7 | MOLECULE: | NALP |
| 94:00:00 | 8sy3-A | 4 | 5.7 | 69 | 209 | 7 | MOLECULE: | BURP DOMAIN-CONTAINING PROTEIN |
| 95:00:00 | 4xzv-B | 4 | 4.1 | 65 | 142 | 6 | MOLECULE: | MALTOSE-BINDING PERIPLASMIC PROTEIN |
| 96:00:00 | 3vu4-A | 4 | 4.1 | 66 | 319 | 5 | MOLECULE: | KMHSV2 |
| 97:00:00 | 3cnx-B | 4 | 4.9 | 61 | 146 | 10 | MOLECULE: | UNCHARACTERIZED PROTEIN |
| 98:00:00 | 5o68-B | 4 | 6.4 | 84 | 274 | 6 | MOLECULE: | FAPF |
| 99:00:00 | 7px8-D | 4 | 4.8 | 71 | 699 | 4 | MOLECULE: | ACYLAMINO-ACID-RELEASING ENZYME |
| 0:00 | 7px8-C | 4 | 4.5 | 70 | 699 | 4 | MOLECULE: | ACYLAMINO-ACID-RELEASING ENZYME |
| 1:00 | 7whm-A | 3.9 | 4.7 | 96 | 1214 | 5 | MOLECULE: | PPPDE DOMAIN-CONTAINING PROTEIN |
| 2:00 | 6qp8-A | 3.9 | 4.5 | 82 | 622 | 7 | MOLECULE: | IP13724P |
| 3:00 | 6zmq-A | 3.9 | 6 | 61 | 589 | 5 | MOLECULE: | CYTOCHROME C-TYPE BIOGENESIS PROTEI |
| 4:00 | 7agv-F | 3.9 | 9.8 | 51 | 163 | 8 | MOLECULE: | K(+)/H(+) ANTIPORTER SUBUNIT KHTT |
| 5:00 | 3csl-A | 3.9 | 7 | 97 | 753 | 4 | MOLECULE: | HASR PROTEIN |
| 6:00 | 1ww1-A | 3.9 | 7.6 | 73 | 250 | 8 | MOLECULE: | TRNASE Z |
| 7:00 | 8rf0-A | 3.9 | 4.4 | 71 | 2780 | 4 | MOLECULE: | CYCLIC BETA-(1,2)-GLUCAN SYNTHASE N |
| 8:00 | 4aip-B | 3.9 | 7.1 | 97 | 665 | 6 | MOLECULE: | FE-REGULATED PROTEIN B |
| 9:00 | 5gyd-C | 3.9 | 7.6 | 84 | 228 | 10 | MOLECULE: | MITOCHONDRIAL DISTRIBUTION AND MORP |
| 10:00 | 2gjv-A | 3.9 | 2.9 | 71 | 136 | 7 | MOLECULE: | PUTATIVE CYTOPLASMIC PROTEIN |
| 11:00 | 4epa-A | 3.9 | 11.8 | 97 | 632 | 3 | MOLECULE: | PESTICIN RECEPTOR |
| 12:00 | 8asw-A | 3.9 | 5.6 | 58 | 1255 | 5 | MOLECULE: | ELONGATOR COMPLEX PROTEIN 1 |
| 13:00 | 3nm7-A | 3.9 | 2.3 | 52 | 84 | 6 | MOLECULE: | UNCHARACTERIZED PROTEIN |
| 14:00 | 8e2g-A | 3.9 | 6.2 | 66 | 523 | 8 | MOLECULE: | BACULOVIRAL IAP REPEAT-CONTAINING P |
| 15:00 | 7uil-p | 3.9 | 9.1 | 73 | 903 | 8 | MOLECULE: | MEDIATOR OF RNA POLYMERASE II TRANS |
| 16:00 | 7xho-P | 3.9 | 7.1 | 80 | 224 | 1 | MOLECULE: | CENTROMERE PROTEIN C |
| 17:00 | 6ht9-B | 3.8 | 4.5 | 70 | 316 | 7 | MOLECULE: | ASTACIN |
| 18:00 | 3lh4-A | 3.8 | 4.6 | 64 | 115 | 5 | MOLECULE: | SECRETED CYSTATIN |
| 19:00 | 2pff-B | 3.8 | 4.3 | 76 | 2006 | 0 | MOLECULE: | FATTY ACID SYNTHASE SUBUNIT ALPHA |
| 20:00 | 3v89-A | 3.8 | 9.3 | 95 | 853 | 5 | MOLECULE: | TRANSFERRIN-BINDING PROTEIN A |
| 21:00 | 1jmx-B | 3.8 | 5.3 | 73 | 339 | 8 | MOLECULE: | AMINE DEHYDROGENASE |
| 22:00 | 6w3g-A | 3.8 | 3.8 | 58 | 109 | 10 | MOLECULE: | RD1NTF2_04 |
| 23:00 | 8ovw-O | 3.8 | 7.3 | 78 | 241 | 5 | MOLECULE: | CENTROMERE-BINDING PROTEIN 1 |
| 24:00:00 | 5jrk-A | 3.8 | 4.2 | 63 | 695 | 6 | MOLECULE: | DIPEPTIDYL AMINOPEPTIDASES/ACYLAMIN |
| 25:00:00 | 3e99-A | 3.8 | 5.7 | 74 | 149 | 7 | MOLECULE: | BENZOATE 1,2-DIOXYGENASE BETA SUBUN |
| 26:00:00 | 7qog-A | 3.8 | 4.4 | 97 | 657 | 4 | MOLECULE: | PORTAL PROTEIN GP20 |
| 27:00:00 | 3os7-A | 3.8 | 4.3 | 82 | 338 | 5 | MOLECULE: | GALACTOSE MUTAROTASE-LIKE PROTEIN |
| 28:00:00 | 7apk-m | 3.8 | 10.7 | 84 | 549 | 11 | MOLECULE: | THO COMPLEX SUBUNIT 1 |
| 29:00:00 | 5mq0-t | 3.8 | 2.8 | 58 | 438 | 7 | MOLECULE: | YEAST UBC4 GENE FOR UBIQUITIN-CONJU |
| 30:00:00 | 7t5q-I | 3.8 | 6.1 | 90 | 283 | 2 | MOLECULE: | ACTIN-RELATED PROTEIN 3 |
| 31:00:00 | 4hbr-A | 3.8 | 4.1 | 78 | 140 | 5 | MOLECULE: | PUTATIVE PERIPLASMIC PROTEIN |
| 32:00:00 | 6baq-F | 3.8 | 4.8 | 89 | 211 | 8 | MOLECULE: | BPI FOLD-CONTAINING FAMILY A MEMBER |
| 33:00:00 | 7el9-A | 3.8 | 6.1 | 75 | 1731 | 0 | MOLECULE: | RNA-DIRECTED RNA POLYMERASE L |
| 34:00:00 | 8ovw-P | 3.8 | 7.5 | 66 | 257 | 11 | MOLECULE: | CENTROMERE-BINDING PROTEIN 1 |
| 35:00:00 | 8g8v-A | 3.8 | 4.5 | 77 | 254 | 8 | MOLECULE: | GTP CYCLOHYDROLASE FOLE2 |
| 36:00:00 | 7zb1-B | 3.8 | 4.3 | 70 | 725 | 4 | MOLECULE: | PROLYL ENDOPEPTIDASE |

| | | | | | | | | |
|---|---|---|---|---|---|---|---|---|
| 37:00:00 | 5tgn-B | 3.8 | 3.7 | 56 | 108 | 9 | MOLECULE: | UNCHARACTERIZED PROTEIN |
| 38:00:00 | 7oqb-P | 3.8 | 5.6 | 72 | 1186 | 8 | MOLECULE: | U2 SNRNP COMPONENT HSH155 |
| 39:00:00 | 3ke7-B | 3.7 | 4.4 | 62 | 132 | 16 | MOLECULE: | PUTATIVE KETOSTEROID ISOMERASE |
| 40:00:00 | 3dcz-A | 3.7 | 3.3 | 70 | 170 | 7 | MOLECULE: | PUTATIVE RNFG SUBUNIT OF ELECTRON T |
| 41:00:00 | 6cb1-s | 3.7 | 3.4 | 69 | 512 | 7 | MOLECULE: | 35S PRE-RIBOSOMAL RNA MISCRNA |
| 42:00:00 | 6h04-I | 3.7 | 8 | 79 | 501 | 1 | MOLECULE: | COMPLEMENT COMPONENT C9 |
| 43:00:00 | 3db7-A | 3.7 | 2.8 | 52 | 127 | 2 | MOLECULE: | PUTATIVE CALCIUM-REGULATED PERIPLAS |
| 44:00:00 | 1mg2-A | 3.7 | 4 | 55 | 382 | 7 | MOLECULE: | METHYLAMINE DEHYDROGENASE, HEAVY CH |
| 45:00:00 | 2o3o-I | 3.7 | 8 | 70 | 244 | 9 | MOLECULE: | YYCI PROTEIN |
| 46:00:00 | 7ep9-A | 3.7 | 4 | 65 | 660 | 3 | MOLECULE: | S9 FAMILY PEPTIDASE |
| 47:00:00 | 1yem-B | 3.7 | 5.9 | 56 | 166 | 0 | MOLECULE: | CONSERVED HYPOTHETICAL PROTEIN PFU- |
| 48:00:00 | 7n3v-A | 3.7 | 4.5 | 67 | 266 | 10 | MOLECULE: | LMCA |
| 49:00:00 | 8c7g-A | 3.7 | 5.3 | 67 | 607 | 3 | MOLECULE: | MIC1 DOMAIN-CONTAINING PROTEIN |
| 50:00:00 | 3wpv-A | 3.7 | 4.9 | 80 | 538 | 8 | MOLECULE: | BETA-FRUCTOFURANOSIDASE |
| 51:00:00 | 4zkq-A | 3.7 | 9.6 | 68 | 388 | 1 | MOLECULE: | PUTATIVE UNCHARACTERIZED PROTEIN |
| 52:00:00 | 7vud-A | 3.7 | 4.8 | 74 | 535 | 7 | MOLECULE: | CAROTENOID CLEAVAGE DIOXYGENASE 1 |
| 53:00:00 | 2qkd-A | 3.7 | 8.4 | 66 | 384 | 5 | MOLECULE: | ZINC FINGER PROTEIN ZPR1 |
| 54:00:00 | 7esn-A | 3.7 | 3.6 | 64 | 435 | 2 | MOLECULE: | L-RHAMNOSE-ALPHA-1,4-D-GLUCURONATE |
| 55:00:00 | 6aig-A | 3.7 | 7 | 64 | 258 | 8 | MOLECULE: | DNA POLYMERASE SLIDING CLAMP 1 |
| 56:00:00 | 7wb4-f | 3.7 | 5.5 | 61 | 321 | 7 | MOLECULE: | OUTER NUP133 |
| 57:00:00 | 4jf7-A | 3.7 | 4.7 | 67 | 501 | 6 | MOLECULE: | HEMAGGLUTININ-NEURAMINIDASE |
| 58:00:00 | 2oaj-A | 3.7 | 6.5 | 77 | 875 | 5 | MOLECULE: | PROTEIN SNI1 |
| 59:00:00 | 4h5j-B | 3.7 | 5.5 | 73 | 347 | 3 | MOLECULE: | GUANINE NUCLEOTIDE-EXCHANGE FACTOR |
| 60:00:00 | 6h3i-A | 3.7 | 6 | 70 | 2124 | 10 | MOLECULE: | PROTEIN INVOLVED IN GLIDING MOTILIT |
| 61:00:00 | 4q35-A | 3.7 | 8.8 | 99 | 758 | 9 | MOLECULE: | LPS-ASSEMBLY PROTEIN LPTD |
| 62:00:00 | 5tf2-A | 3.7 | 4.7 | 55 | 338 | 5 | MOLECULE: | PROLACTIN REGULATORY ELEMENT-BINDIN |
| 63:00:00 | 3en8-A | 3.7 | 7.9 | 64 | 128 | 11 | MOLECULE: | UNCHARACTERIZED NTF-2 LIKE PROTEIN |
| 64:00:00 | 2wjq-A | 3.7 | 6 | 61 | 205 | 0 | MOLECULE: | PROBABLE N-ACETYLNEURAMINIC ACID OU |
| 65:00:00 | 7btx-L | 3.7 | 10.2 | 75 | 314 | 7 | MOLECULE: | MITOCHONDRIAL OUTER MEMBRANE BETA-B |
| 66:00:00 | 8xcj-F | 3.7 | 8 | 62 | 420 | 6 | MOLECULE: | MALTOPORIN |
| 67:00:00 | 6ehd-A | 3.7 | 6.3 | 89 | 322 | 4 | MOLECULE: | OMPT PROTEIN |
| 68:00:00 | 6lk8-E | 3.7 | 13.6 | 65 | 1030 | 2 | MOLECULE: | MGC83295 PROTEIN |
| 69:00:00 | 8dte-A | 3.7 | 4.7 | 62 | 172 | 16 | MOLECULE: | PERIPLASMIC TRANSPORT PROTEIN |
| 70:00:00 | 7e7w-C | 3.7 | 3.2 | 66 | 264 | 9 | MOLECULE: | FUCOSE-BINDING LECTIN PROTEIN,FUCOS |
| 71:00:00 | 5kuy-G | 3.7 | 3 | 57 | 84 | 7 | MOLECULE: | HEMAGGLUTININ HA1 |
| 72:00:00 | 5bka-B | 3.7 | 5.3 | 71 | 131 | 11 | MOLECULE: | HYDROXYLACYL-COA DEHYDROGENASE |
| 73:00:00 | 5ml9-B | 3.7 | 4 | 62 | 103 | 3 | MOLECULE: | LOW AFFINITY IMMUNOGLOBULIN GAMMA F |
| 74:00:00 | 7wb4-J | 3.7 | 5.2 | 73 | 1021 | 5 | MOLECULE: | OUTER NUP133 |
| 75:00:00 | 5d17-C | 3.7 | 3.1 | 67 | 158 | 4 | MOLECULE: | TRANSPOSON TN7 TRANSPOSITION PROTEI |
| 76:00:00 | 8p0w-A | 3.6 | 3.2 | 48 | 190 | 4 | MOLECULE: | COMM DOMAIN-CONTAINING PROTEIN 1 |
| 77:00:00 | 7mhu-A | 3.6 | 7.1 | 68 | 374 | 13 | MOLECULE: | EXO-ALPHA-SIALIDASE |
| 78:00:00 | 6sg8-B | 3.6 | 5.6 | 70 | 381 | 6 | MOLECULE: | HEMAGGLUTININ-NEURAMINIDASE |
| 79:00:00 | 5hax-A | 3.6 | 5.9 | 74 | 698 | 11 | MOLECULE: | NUCLEOPORIN NUP170 |
| 80:00:00 | 6sgo-A | 3.6 | 5.5 | 67 | 136 | 3 | MOLECULE: | SECRETED PROTEIN |
| 81:00:00 | 6l77-A | 3.6 | 3.5 | 67 | 214 | 7 | MOLECULE: | MS5A |
| 82:00:00 | 5bw0-F | 3.6 | 2.7 | 53 | 91 | 6 | MOLECULE: | TYPE II SECRETION SYSTEM PROTEIN J |

| | | | | | | | | |
|---|---|---|---|---|---|---|---|---|
| 83:00:00 | 1rn7-A | 3.6 | 4.4 | 68 | 112 | 7 | MOLECULE: | CYSTATIN D |
| 84:00:00 | 6c0f-A | 3.6 | 5.6 | 70 | 394 | 9 | MOLECULE: | SACCHAROMYCES CEREVISIAE S288C 35S |
| 85:00:00 | 9c4o-A | 3.6 | 7.8 | 93 | 661 | 4 | MOLECULE: | PYRROLOQUINOLINE QUINONE TRANSPORTE |
| 86:00:00 | 6em5-m | 3.6 | 3.4 | 72 | 645 | 11 | MOLECULE: | 5.8S RIBOSOMAL RNA |
| 87:00:00 | 5own-A | 3.6 | 7.9 | 82 | 295 | 4 | MOLECULE: | PERFORIN-LIKE PROTEIN 1 |
| 88:00:00 | 4rhz-A | 3.6 | 3.7 | 79 | 259 | 8 | MOLECULE: | CRY23AA1 |
| 89:00:00 | 5vob-D | 3.6 | 8.7 | 62 | 170 | 6 | MOLECULE: | ENVELOPE GLYCOPROTEIN H |
| 90:00:00 | 6z37-A | 3.6 | 6.7 | 82 | 418 | 5 | MOLECULE: | TODX |
| 91:00:00 | 6w4q-C | 3.6 | 4.3 | 62 | 758 | 10 | MOLECULE: | TAIL FIBER |
| 92:00:00 | 7e4h-A | 3.6 | 6.8 | 76 | 439 | 4 | MOLECULE: | SORTING ASSEMBLY MACHINERY 50 KDA S |
| 93:00:00 | 2h36-X | 3.6 | 2.5 | 49 | 108 | 6 | MOLECULE: | HYPOTHETICAL PROTEIN SIFV0014 |
| 94:00:00 | 7eeb-F | 3.6 | 5 | 67 | 1042 | 9 | MOLECULE: | ENHANCED GREEN FLUORESCENT PROTEIN, |
| 95:00:00 | 8cpz-B | 3.6 | 10.4 | 69 | 2122 | 9 | MOLECULE: | TCDA1 |
| 96:00:00 | 7eeb-H | 3.6 | 6.1 | 75 | 916 | 5 | MOLECULE: | ENHANCED GREEN FLUORESCENT PROTEIN, |
| 97:00:00 | 8ppr-K | 3.6 | 6.5 | 62 | 246 | 5 | MOLECULE: | KINETOCHORE-ASSOCIATED PROTEIN DSN1 |
| 98:00:00 | 6cp9-E | 3.6 | 8.6 | 62 | 119 | 10 | MOLECULE: | CDIA |
| 99:00:00 | 5nqz-A | 3.6 | 5.2 | 70 | 267 | 9 | MOLECULE: | FACTOR H BINDING PROTEIN,MAJOR OUTE |
| 0:00 | 4hh4-C | 3.6 | 5.9 | 65 | 257 | 5 | MOLECULE: | CCBJ |
| 1:00 | 8tie-g | 3.6 | 6.7 | 76 | 1157 | 5 | MOLECULE: | NUCLEOPORIN NUP120 |
| 2:00 | 2rjz-A | 3.6 | 4.2 | 71 | 135 | 4 | MOLECULE: | PILO PROTEIN |
| 3:00 | 6zlt-B | 3.6 | 8.3 | 90 | 933 | 4 | MOLECULE: | SUSD HOMOLOG |
| 4:00 | 3kvn-X | 3.6 | 4.5 | 65 | 628 | 3 | MOLECULE: | ESTERASE ESTA |
| 5:00 | 7bc4-B | 3.6 | 7.4 | 70 | 2054 | 4 | MOLECULE: | FATTY ACID SYNTHASE SUBUNIT ALPHA |
| 6:00 | 7st9-H | 3.6 | 8.6 | 59 | 316 | 10 | MOLECULE: | CHECKPOINT PROTEIN RAD24 |
| 7:00 | 2x2i-D | 3.6 | 5.5 | 57 | 1025 | 9 | MOLECULE: | ALPHA-1,4-GLUCAN LYASE ISOZYME 1 |
| 8:00 | 7kd9-F | 3.6 | 6.3 | 83 | 225 | 7 | MOLECULE: | GALLATE DECARBOXYLASE |
| 9:00 | 7e7w-B | 3.6 | 3.2 | 64 | 264 | 13 | MOLECULE: | FUCOSE-BINDING LECTIN PROTEIN,FUCOS |
| 10:00 | 6w40-B | 3.6 | 2.4 | 48 | 115 | 2 | MOLECULE: | DENOVO NTF2 |
| 11:00 | 5d17-A | 3.6 | 3.2 | 67 | 159 | 4 | MOLECULE: | TRANSPOSON TN7 TRANSPOSITION PROTEI |
| 12:00 | 5d17-L | 3.6 | 3.2 | 67 | 158 | 4 | MOLECULE: | TRANSPOSON TN7 TRANSPOSITION PROTEI |
| 13:00 | 6gel-B | 3.6 | 7 | 68 | 528 | 3 | MOLECULE: | GREEN FLUORESCENT PROTEIN,OPTIMIZED |
| 14:00 | 5d17-F | 3.6 | 3.3 | 67 | 158 | 4 | MOLECULE: | TRANSPOSON TN7 TRANSPOSITION PROTEI |
| 15:00 | 8j1g-B | 3.5 | 5.3 | 76 | 313 | 7 | MOLECULE: | ORNITHINE CYCLODEAMINASE FAMILY PRO |
| 16:00 | 6hsy-A | 3.5 | 4.5 | 59 | 190 | 5 | MOLECULE: | TOLUENE TOLERANCE PROTEIN TTG2D |
| 17:00 | 1y8c-A | 3.5 | 7.6 | 61 | 246 | 7 | MOLECULE: | S-ADENOSYLMETHIONINE-DEPENDENT METH |
| 18:00 | 2gsk-A | 3.5 | 12.7 | 96 | 590 | 7 | MOLECULE: | VITAMIN B12 TRANSPORTER BTUB |
| 19:00 | 8dml-B | 3.5 | 3.8 | 61 | 133 | 3 | MOLECULE: | VTRA |
| 20:00 | 7krw-A | 3.5 | 6.1 | 74 | 608 | 12 | MOLECULE: | CHAPERONE PROTEIN DNAK FUSED WITH S |
| 21:00 | 8y7e-3 | 3.5 | 3.5 | 57 | 1193 | 9 | MOLECULE: | PRE-MRNA |
| 22:00 | 3boy-A | 3.5 | 4.9 | 83 | 150 | 4 | MOLECULE: | 5'-R(*UP*UP*UP*AP*GP*UP*UP*UP*UP*UP |
| 23:00 | 4zn4-B | 3.5 | 4.9 | 68 | 411 | 6 | MOLECULE: | SQT1 |
| 24:00:00 | 2gtl-N | 3.5 | 3.8 | 69 | 220 | 1 | MOLECULE: | EXTRACELLULAR GLOBIN 4 |
| 25:00:00 | 5efv-B | 3.5 | 3.9 | 70 | 635 | 7 | MOLECULE: | PHI ETA ORF 56-LIKE PROTEIN |
| 26:00:00 | 2gvh-B | 3.5 | 3.6 | 74 | 250 | 1 | MOLECULE: | AGR_L_2016P |
| 27:00:00 | 2h1t-A | 3.5 | 12 | 61 | 186 | 3 | MOLECULE: | HYPOTHETICAL PROTEIN |
| 28:00:00 | 3h51-A | 3.5 | 5.1 | 64 | 142 | 9 | MOLECULE: | PUTATIVE CALCIUM/CALMODULIN DEPENDE |

| | | | | | | | | |
|---|---|---|---|---|---|---|---|---|
| 29:00:00 | 6ud7-A | 3.5 | 5.4 | 75 | 426 | 3 | MOLECULE: | DDB1- AND CUL4-ASSOCIATED FACTOR 15 |
| 30:00:00 | 7s00-c | 3.5 | 6.3 | 66 | 484 | 6 | MOLECULE: | DNA-DIRECTED RNA POLYMERASE BETA SU |
| 31:00:00 | 5ldt-A | 3.5 | 8.2 | 84 | 403 | 2 | MOLECULE: | MOMP PORIN |
| 32:00:00 | 5h3z-A | 3.5 | 6.5 | 76 | 1113 | 1 | MOLECULE: | UNCHARACTERIZED PROTEIN |
| 33:00:00 | 5n1a-B | 3.5 | 4 | 65 | 706 | 3 | MOLECULE: | UTP4 |
| 34:00:00 | 5z8o-B | 3.5 | 4.8 | 71 | 146 | 6 | MOLECULE: | CYCLASE/DEHYDRASE |
| 35:00:00 | 2n2l-A | 3.5 | 3.1 | 59 | 156 | 3 | MOLECULE: | OUTER MEMBRANE PROTEIN X |
| 36:00:00 | 4uvq-A | 3.5 | 5.6 | 69 | 263 | 4 | MOLECULE: | THIAZOLINE OXIDASE/SUBTILISIN-LIKE |
| 37:00:00 | 7e85-A | 3.5 | 4.2 | 52 | 114 | 4 | MOLECULE: | SNOAL-LIKE DOMAIN-CONTAINING PROTEI |
| 38:00:00 | 4hvt-A | 3.5 | 4.8 | 78 | 691 | 8 | MOLECULE: | POST-PROLINE CLEAVING ENZYME |
| 39:00:00 | 3k7c-A | 3.5 | 4 | 58 | 108 | 7 | MOLECULE: | PUTATIVE NTF2-LIKE TRANSPEPTIDASE |
| 40:00:00 | 6abo-A | 3.5 | 5.8 | 51 | 213 | 8 | MOLECULE: | DNA REPAIR PROTEIN XRCC4 |
| 41:00:00 | 6h0f-J | 3.5 | 6.2 | 74 | 842 | 4 | MOLECULE: | DNA DAMAGE-BINDING PROTEIN 1,DNA DA |
| 42:00:00 | 7t5q-J | 3.5 | 6.4 | 92 | 291 | 5 | MOLECULE: | ACTIN-RELATED PROTEIN 3 |
| 43:00:00 | 8ptx-B | 3.5 | 5.1 | 76 | 800 | 7 | MOLECULE: | ELONGATOR COMPLEX PROTEIN 1 |
| 44:00:00 | 7q4p-C | 3.5 | 3.6 | 54 | 878 | 4 | MOLECULE: | SPLICING FACTOR 3A SUBUNIT 2 |
| 45:00:00 | 6tzk-A | 3.5 | 6.7 | 77 | 451 | 4 | MOLECULE: | CELLULOSE SYNTHASE OPERON PROTEIN C |
| 46:00:00 | 5mt2-A | 3.5 | 4.8 | 84 | 352 | 7 | MOLECULE: | BETA-GALACTOSIDASE |
| 47:00:00 | 7zsk-A | 3.5 | 5.2 | 72 | 1482 | 3 | MOLECULE: | PUTATIVE POLYKETIDE SYNTHASE |
| 48:00:00 | 3qv0-A | 3.5 | 3.3 | 61 | 179 | 3 | MOLECULE: | MITOCHONDRIAL ACIDIC PROTEIN MAM33 |
| 49:00:00 | 6z8y-A | 3.5 | 10.7 | 86 | 658 | 6 | MOLECULE: | PUTATIVE COPPER TRANSPORT OUTER MEM |
| 50:00:00 | 3grd-A | 3.5 | 5.5 | 71 | 133 | 4 | MOLECULE: | UNCHARACTERIZED NTF2-SUPERFAMILY PR |
| 51:00:00 | 6rw6-B | 3.5 | 6.3 | 64 | 2485 | 2 | MOLECULE: | TCDA1 |
| 52:00:00 | 7qe7-A | 3.5 | 5.4 | 71 | 1648 | 10 | MOLECULE: | ANAPHASE-PROMOTING COMPLEX SUBUNIT |
| 53:00:00 | 3k4x-A | 3.5 | 5.2 | 60 | 758 | 5 | MOLECULE: | PROLIFERATING CELL NUCLEAR ANTIGEN |
| 54:00:00 | 7ne0-D | 3.5 | 4.9 | 60 | 161 | 8 | MOLECULE: | NETRIN-1 |
| 55:00:00 | 5bun-C | 3.5 | 8.9 | 72 | 433 | 6 | MOLECULE: | OUTER MEMBRANE PROTEIN |
| 56:00:00 | 3hx8-A | 3.5 | 5.2 | 66 | 128 | 6 | MOLECULE: | PUTATIVE KETOSTEROID ISOMERASE |
| 57:00:00 | 7e7n-A | 3.5 | 3.7 | 61 | 259 | 10 | MOLECULE: | FUCOSE-BINDING LECTIN PROTEIN,FUCOS |
| 58:00:00 | 6q0r-C | 3.5 | 2.9 | 58 | 183 | 5 | MOLECULE: | DNA DAMAGE-BINDING PROTEIN 1 |
| 59:00:00 | 7c5w-B | 3.5 | 4.8 | 63 | 163 | 3 | MOLECULE: | IOTA-CARBONIC ANHYDRASE |
| 60:00:00 | 3rwa-F | 3.5 | 5.3 | 62 | 232 | 2 | MOLECULE: | FLUORESCENT PROTEIN FP480 |
| 61:00:00 | 3rwa-B | 3.5 | 5.3 | 62 | 232 | 2 | MOLECULE: | FLUORESCENT PROTEIN FP480 |
| 62:00:00 | 6ogb-B | 3.5 | 9 | 57 | 225 | 5 | MOLECULE: | CRYSTAL STRUCTURE OF GREEN FLUORESC |
| 63:00:00 | 6gel-A | 3.5 | 7.1 | 68 | 530 | 3 | MOLECULE: | GREEN FLUORESCENT PROTEIN,OPTIMIZED |
| 64:00:00 | 4w6g-B | 3.5 | 8.5 | 55 | 217 | 5 | MOLECULE: | FLUORESCENT PROTEIN D190C |
| 65:00:00 | 4wvm-B | 3.4 | 6.3 | 80 | 616 | 6 | MOLECULE: | STONUSTOXIN SUBUNIT ALPHA |
| 66:00:00 | 7nyw-B | 3.4 | 12.2 | 72 | 858 | 7 | MOLECULE: | CHROMOSOME PARTITION PROTEIN MUKB |
| 67:00:00 | 7kzn-A | 3.4 | 4.1 | 66 | 315 | 5 | MOLECULE: | HEAVY CHAIN ALPHA |
| 68:00:00 | 5nj5-A | 3.4 | 10.6 | 89 | 480 | 4 | MOLECULE: | METALLOPROTEASE TLDD |
| 69:00:00 | 5mtz-A | 3.4 | 6.1 | 75 | 768 | 5 | MOLECULE: | RIBONUCLEASE Z |
| 70:00:00 | 5kpe-A | 3.4 | 4.3 | 58 | 120 | 5 | MOLECULE: | DE NOVO BETA SHEET DESIGN PROTEIN O |
| 71:00:00 | 7pd7-B | 3.4 | 5.1 | 75 | 251 | 3 | MOLECULE: | METHYLTRANSFERASE |
| 72:00:00 | 6gie-A | 3.4 | 11.1 | 77 | 218 | 6 | MOLECULE: | 33-36 KDA OUTER MEMBRANE PROTEIN |
| 73:00:00 | 2mc8-A | 3.4 | 3.7 | 63 | 114 | 5 | MOLECULE: | UNCHARACTERIZED PROTEIN |
| 74:00:00 | 4irt-A | 3.4 | 4.4 | 65 | 398 | 5 | MOLECULE: | UNCHARACTERIZED PROTEIN |

| | | | | | | | | |
|---|---|---|---|---|---|---|---|---|
| 75:00:00 | 6lvb-A | 3.4 | 4.2 | 68 | 762 | 3 | MOLECULE: | N,N-DIMETHYLFORMAMIDASE LARGE SUBUN |
| 76:00:00 | 6tyd-V | 3.4 | 7.5 | 89 | 219 | 10 | MOLECULE: | LIM DOMAIN-BINDING PROTEIN 1 |
| 77:00:00 | 6h0a-A | 3.4 | 5.6 | 77 | 339 | 5 | MOLECULE: | SERUM PARAOXONASE-1 BY DIRECTED EVO |
| 78:00:00 | 5tc1-M | 3.4 | 5 | 77 | 330 | 4 | MOLECULE: | CAPSID PROTEIN |
| 79:00:00 | 7zao-A | 3.4 | 7.6 | 75 | 429 | 8 | MOLECULE: | SIALIDASE (NEURAMINIDASE) FAMILY PR |
| 80:00:00 | 3kg7-C | 3.4 | 5.7 | 80 | 287 | 5 | MOLECULE: | CURH |
| 81:00:00 | 6jzf-A | 3.4 | 5.7 | 65 | 153 | 5 | MOLECULE: | PLASTID DIVISION PROTEIN CDP1, CHLO |
| 82:00:00 | 7nyw-A | 3.4 | 12 | 72 | 685 | 7 | MOLECULE: | CHROMOSOME PARTITION PROTEIN MUKB |
| 83:00:00 | 7czf-B | 3.4 | 4.1 | 62 | 644 | 2 | MOLECULE: | EPHRIN TYPE-A RECEPTOR 2 |
| 84:00:00 | 6cd2-C | 3.4 | 4.6 | 77 | 727 | 8 | MOLECULE: | CHAPERONE PROTEIN PAPD |
| 85:00:00 | 5a1u-E | 3.4 | 5.3 | 69 | 822 | 3 | MOLECULE: | ADP-RIBOSYLATION FACTOR 1 |
| 86:00:00 | 7azn-A | 3.4 | 4.6 | 71 | 177 | 6 | MOLECULE: | PROTEIN ASTER-C |
| 87:00:00 | 4n58-A | 3.4 | 4.4 | 64 | 266 | 6 | MOLECULE: | PECTOCIN M2 |
| 88:00:00 | 4y25-A | 3.4 | 7.5 | 85 | 290 | 7 | MOLECULE: | POLY-BETA-1,6-N-ACETYL-D-GLUCOSAMIN |
| 89:00:00 | 7y22-Y | 3.4 | 4.3 | 65 | 794 | 8 | MOLECULE: | PHAGE CONNECTOR PROTEIN |
| 90:00:00 | 3pgb-A | 3.4 | 4.8 | 93 | 740 | 4 | MOLECULE: | PUTATIVE UNCHARACTERIZED PROTEIN |
| 91:00:00 | 4o3v-B | 3.4 | 5.5 | 71 | 140 | 7 | MOLECULE: | VIRB8-LIKE PROTEIN OF TYPE IV SECRE |
| 92:00:00 | 3h3h-B | 3.4 | 4.3 | 53 | 120 | 2 | MOLECULE: | UNCHARACTERIZED SNOAL-LIKE PROTEIN |
| 93:00:00 | 3hsa-A | 3.4 | 3.9 | 53 | 124 | 8 | MOLECULE: | PLECKSTRIN HOMOLOGY DOMAIN |
| 94:00:00 | 4zsv-A | 3.4 | 11.3 | 65 | 294 | 2 | MOLECULE: | UNCHARACTERIZED PROTEIN |
| 95:00:00 | 3kvp-D | 3.4 | 4.2 | 46 | 49 | 2 | MOLECULE: | UNCHARACTERIZED PROTEIN YMZC |
| 96:00:00 | 4rdr-A | 3.4 | 6.6 | 77 | 706 | 5 | MOLECULE: | ZNUD |
| 97:00:00 | 1u1i-A | 3.4 | 4.3 | 65 | 392 | 9 | MOLECULE: | MYO-INOSITOL-1-PHOSPHATE SYNTHASE |
| 98:00:00 | 6hxv-A | 3.4 | 5.6 | 54 | 163 | 2 | MOLECULE: | COILED-COIL DOMAIN-CONTAINING PROTE |
| 99:00:00 | 8chv-B | 3.4 | 2.1 | 47 | 147 | 6 | MOLECULE: | TRANSCRIPTIONAL ACTIVATOR PROTEIN P |
| 0:00 | 4qtq-A | 3.4 | 3.7 | 62 | 209 | 6 | MOLECULE: | XAC2610 PROTEIN |
| 1:00 | 7e7u-B | 3.4 | 4 | 62 | 270 | 10 | MOLECULE: | FUCOSE-BINDING LECTIN PROTEIN,FUCOS |
| 2:00 | 7kd9-I | 3.4 | 6.5 | 83 | 226 | 7 | MOLECULE: | GALLATE DECARBOXYLASE |
| 3:00 | 6w3d-A | 3.4 | 3.7 | 57 | 114 | 12 | MOLECULE: | RD1NTF2_05 |
| 4:00 | 5bka-D | 3.4 | 5.3 | 65 | 130 | 12 | MOLECULE: | HYDROXYLACYL-COA DEHYDROGENASE |
| 5:00 | 7kd9-C | 3.4 | 6 | 83 | 227 | 7 | MOLECULE: | GALLATE DECARBOXYLASE |
| 6:00 | 3sg2-A | 3.4 | 7.1 | 78 | 394 | 5 | MOLECULE: | MYOSIN LIGHT CHAIN KINASE, GREEN FL |
| 7:00 | 7rrk-C | 3.4 | 4.9 | 62 | 213 | 10 | MOLECULE: | FLUORESCENT PROTEIN DRONPA |
| 8:00 | 3rwa-C | 3.4 | 6.6 | 65 | 232 | 3 | MOLECULE: | FLUORESCENT PROTEIN FP480 |
| 9:00 | 5o68-A | 3.4 | 6.4 | 83 | 272 | 6 | MOLECULE: | FAPF |
| 10:00 | 5d16-B | 3.4 | 3.2 | 65 | 157 | 5 | MOLECULE: | TRANSPOSON TN7 TRANSPOSITION PROTEI |
| 11:00 | 5d17-K | 3.4 | 3.1 | 63 | 157 | 5 | MOLECULE: | TRANSPOSON TN7 TRANSPOSITION PROTEI |
| 12:00 | 8f2r-B | 3.3 | 3.6 | 48 | 199 | 2 | MOLECULE: | COMM DOMAIN-CONTAINING PROTEIN 1 |
| 13:00 | 5k36-F | 3.3 | 9.5 | 61 | 215 | 7 | MOLECULE: | EXOSOME COMPLEX COMPONENT RRP45 |
| 14:00 | 7kd9-A | 3.3 | 6.9 | 88 | 231 | 9 | MOLECULE: | GALLATE DECARBOXYLASE |
| 15:00 | 6kaw-A | 3.3 | 7.3 | 75 | 381 | 4 | MOLECULE: | CGHA |
| 16:00 | 6uio-C | 3.3 | 4.2 | 67 | 112 | 7 | MOLECULE: | CYSTATIN-8 |
| 17:00 | 3rob-A | 3.3 | 5.4 | 71 | 131 | 4 | MOLECULE: | UNCHARACTERIZED CONSERVED PROTEIN |
| 18:00 | 5e1v-B | 3.3 | 3.6 | 69 | 274 | 1 | MOLECULE: | POLYKETIDE SYNTHASE PKSL |
| 19:00 | 1eut-A | 3.3 | 4.3 | 64 | 601 | 6 | MOLECULE: | SIALIDASE |
| 20:00 | 2yfr-A | 3.3 | 5.7 | 80 | 533 | 5 | MOLECULE: | LEVANSUCRASE |

| | | | | | | | | |
|---|---|---|---|---|---|---|---|---|
| 21:00 | 8ih8-D | 3.3 | 2.6 | 56 | 130 | 7 | MOLECULE: | ANTI-SIGMA-F FACTOR ANTAGONIST RSFB |
| 22:00 | 3ci0-K | 3.3 | 3.5 | 70 | 280 | 9 | MOLECULE: | PSEUDOPILIN GSPI |
| 23:00 | 8cyk-A | 3.3 | 4.9 | 67 | 131 | 6 | MOLECULE: | HALC1_878 |
| 24:00:00 | 8qca-C | 3.3 | 4.3 | 71 | 384 | 6 | MOLECULE: | ANTIVIRAL HELICASE SKI2 |
| 25:00:00 | 8jbq-A | 3.3 | 5.6 | 79 | 692 | 3 | MOLECULE: | HEMOLYSIN |
| 26:00:00 | 6myv-C | 3.3 | 9.1 | 67 | 526 | 1 | MOLECULE: | SIALIDASE26 |
| 27:00:00 | 6urg-A | 3.3 | 4.1 | 70 | 1202 | 6 | MOLECULE: | CLEAVAGE AND POLYADENYLATION SPECIF |
| 28:00:00 | 5hal-A | 3.3 | 3.4 | 60 | 103 | 7 | MOLECULE: | UNCHARACTERIZED PROTEIN |
| 29:00:00 | 4i0o-A | 3.3 | 5 | 77 | 463 | 5 | MOLECULE: | PROTEIN ELYS |
| 30:00:00 | 3emo-A | 3.3 | 3.4 | 53 | 126 | 4 | MOLECULE: | HIA (ADHESIN) |
| 31:00:00 | 4ebr-A | 3.3 | 4 | 83 | 157 | 7 | MOLECULE: | UBIQUITIN-LIKE-CONJUGATING ENZYME A |
| 32:00:00 | 2gia-G | 3.3 | 4.9 | 63 | 155 | 6 | MOLECULE: | MITOCHONDRIAL RNA-BINDING PROTEIN 2 |
| 33:00:00 | 4ccv-A | 3.3 | 4.2 | 62 | 115 | 3 | MOLECULE: | HISTIDINE-RICH GLYCOPROTEIN |
| 34:00:00 | 1nkg-A | 3.3 | 9.7 | 58 | 508 | 10 | MOLECULE: | RHAMNOGALACTURONASE B |
| 35:00:00 | 3a76-B | 3.3 | 6.3 | 79 | 153 | 8 | MOLECULE: | GAMMA-HEXACHLOROCYCLOHEXANE DEHYDRO |
| 36:00:00 | 8f2r-I | 3.3 | 3.4 | 48 | 198 | 10 | MOLECULE: | COMM DOMAIN-CONTAINING PROTEIN 1 |
| 37:00:00 | 2rgq-B | 3.3 | 5.5 | 67 | 134 | 7 | MOLECULE: | DOMAIN OF UNKNOWN FUNCTION WITH A C |
| 38:00:00 | 6r8f-E | 3.3 | 4.1 | 64 | 210 | 0 | MOLECULE: | LYS-63-SPECIFIC DEUBIQUITINASE BRCC |
| 39:00:00 | 4i4k-A | 3.3 | 5.3 | 63 | 138 | 2 | MOLECULE: | UNCHARACTERIZED PROTEIN SGCJ |
| 40:00:00 | 6j6a-A | 3.3 | 9.7 | 87 | 442 | 3 | MOLECULE: | ZINC-DEPENDENT PROTEASE, TLDD/PMBA |
| 41:00:00 | 5h7z-B | 3.3 | 6.5 | 66 | 302 | 3 | MOLECULE: | UNCHARACTERIZED PROTEIN |
| 42:00:00 | 5ogs-A | 3.3 | 5.5 | 63 | 403 | 10 | MOLECULE: | WD REPEAT AND HMG-BOX DNA-BINDING P |
| 43:00:00 | 2kxg-A | 3.3 | 3.8 | 68 | 95 | 7 | MOLECULE: | ASPARTIC PROTEASE INHIBITOR |
| 44:00:00 | 6w3w-A | 3.3 | 5.4 | 54 | 107 | 11 | MOLECULE: | DENOVO NTF2 |
| 45:00:00 | 6hdv-A | 3.3 | 9 | 81 | 136 | 6 | MOLECULE: | AFIFAVIDIN |
| 46:00:00 | 8bmx-C | 3.3 | 11.1 | 92 | 633 | 4 | MOLECULE: | PUTATIVE TONB-LINKED OUTER MEMBRANE |
| 47:00:00 | 8adl-J | 3.3 | 4 | 52 | 801 | 2 | MOLECULE: | MTC5 ISOFORM 1 |
| 48:00:00 | 7svm-A | 3.3 | 5.3 | 89 | 852 | 7 | MOLECULE: | DIPEPTIDYL PEPTIDASE 8 |
| 49:00:00 | 5dfz-B | 3.3 | 6 | 70 | 1224 | 1 | MOLECULE: | VACUOLAR PROTEIN SORTING-ASSOCIATED |
| 50:00:00 | 6igb-A | 3.3 | 4.2 | 68 | 363 | 4 | MOLECULE: | PERIPLASMIC GLUCONOLACTONASE, PPGL |
| 51:00:00 | 4ui9-A | 3.3 | 5.8 | 75 | 1441 | 7 | MOLECULE: | ANAPHASE-PROMOTING COMPLEX SUBUNIT |
| 52:00:00 | 8xcj-A | 3.3 | 10.3 | 77 | 422 | 4 | MOLECULE: | MALTOPORIN |
| 53:00:00 | 7pi3-A | 3.3 | 5.9 | 76 | 328 | 1 | MOLECULE: | CYSTEINE-RICH PROTECTIVE ANTIGEN |
| 54:00:00 | 6q50-A | 3.3 | 5.2 | 68 | 318 | 7 | MOLECULE: | MPT-4 |
| 55:00:00 | 7s5n-B | 3.3 | 5.7 | 65 | 128 | 2 | MOLECULE: | MYMAA.17060.A |
| 56:00:00 | 6ewz-A | 3.3 | 9.3 | 62 | 202 | 6 | MOLECULE: | GTP PYROPHOSPHOKINASE |
| 57:00:00 | 4acv-B | 3.3 | 3.1 | 61 | 122 | 2 | MOLECULE: | PROPHAGE LAMBDALM01, ANTIGEN B |
| 58:00:00 | 3f40-A | 3.3 | 5.6 | 63 | 112 | 5 | MOLECULE: | UNCHARACTERIZED NTF2-LIKE PROTEIN |
| 59:00:00 | 6gn5-A | 3.3 | 5.2 | 81 | 177 | 5 | MOLECULE: | GRAM DOMAIN-CONTAINING PROTEIN 1C |
| 60:00:00 | 4i8e-X | 3.3 | 2.3 | 51 | 359 | 10 | MOLECULE: | PLATELET BINDING PROTEIN GSPB |
| 61:00:00 | 8fs6-G | 3.3 | 3.3 | 59 | 298 | 3 | MOLECULE: | CHECKPOINT PROTEIN RAD24 |
| 62:00:00 | 8jxe-A | 3.3 | 7 | 68 | 1469 | 12 | MOLECULE: | LDL RECEPTOR RELATED PROTEIN 2 |
| 63:00:00 | 1wnh-A | 3.3 | 4.4 | 68 | 220 | 6 | MOLECULE: | LATEXIN |
| 64:00:00 | 6f37-A | 3.3 | 3.2 | 54 | 113 | 9 | MOLECULE: | NANO3,FUCOSE-BINDING LECTIN PROTEIN |
| 65:00:00 | 8j0n-A | 3.3 | 4.6 | 62 | 949 | 5 | MOLECULE: | ER MEMBRANE PROTEIN COMPLEX SUBUNIT |
| 66:00:00 | 1r5m-A | 3.3 | 8.6 | 53 | 351 | 6 | MOLECULE: | SIR4-INTERACTING PROTEIN SIF2 |

| | | | | | | | | |
|---|---|---|---|---|---|---|---|---|
| 67:00:00 | 8ck0-A | 3.3 | 4.2 | 90 | 478 | 7 | MOLECULE: | PORTAL PROTEIN |
| 68:00:00 | 8igd-A | 3.3 | 2.9 | 54 | 82 | 4 | MOLECULE: | DOUBLE-STRANDED RNA-BINDING DOMAIN |
| 69:00:00 | 5xta-A | 3.3 | 3.4 | 71 | 120 | 1 | MOLECULE: | VIRK PROTEIN |
| 70:00:00 | 7kpr-A | 3.3 | 7.9 | 89 | 403 | 3 | MOLECULE: | PROTEIN PHOSPHATASE 1H |
| 71:00:00 | 4hxg-F | 3.3 | 5.9 | 64 | 614 | 3 | MOLECULE: | PUTATIVE UNCHARACTERIZED PROTEIN PH |
| 72:00:00 | 7qok-A | 3.3 | 6 | 51 | 1361 | 6 | MOLECULE: | MUZZLE PROTEIN GP44 |
| 73:00:00 | 4rlc-A | 3.3 | 6.9 | 75 | 135 | 4 | MOLECULE: | OUTER MEMBRANE PORIN F |
| 74:00:00 | 5uac-C | 3.3 | 5.6 | 62 | 1342 | 6 | MOLECULE: | DNA-DIRECTED RNA POLYMERASE SUBUNIT |
| 75:00:00 | 5l8s-A | 3.3 | 4.1 | 68 | 604 | 6 | MOLECULE: | AMINO ACYL PEPTIDASE |
| 76:00:00 | 7d9b-A | 3.3 | 4.9 | 64 | 588 | 5 | MOLECULE: | ALPHA-GLYCOSIDASE |
| 77:00:00 | 4zgv-A | 3.3 | 12.5 | 89 | 809 | 1 | MOLECULE: | FERREDOXIN RECEPTOR |
| 78:00:00 | 8xci-Z | 3.3 | 7 | 57 | 383 | 7 | MOLECULE: | TIP ATTACHMENT PROTEIN J |
| 79:00:00 | 6jql-A | 3.3 | 3.8 | 69 | 678 | 6 | MOLECULE: | BIFUNCTIONAL PROTEIN PAAZ |
| 80:00:00 | 8ghl-C | 3.3 | 9.9 | 51 | 377 | 6 | MOLECULE: | PROTEIN HIR1 |
| 81:00:00 | 6hpv-A | 3.3 | 5.4 | 70 | 299 | 6 | MOLECULE: | FETUIN-B |
| 82:00:00 | 8ghn-C | 3.3 | 9.9 | 51 | 377 | 6 | MOLECULE: | PROTEIN HIR1 |
| 83:00:00 | 7e85-C | 3.3 | 4 | 53 | 114 | 9 | MOLECULE: | SNOAL-LIKE DOMAIN-CONTAINING PROTEI |
| 84:00:00 | 7e85-B | 3.3 | 4.7 | 55 | 114 | 11 | MOLECULE: | SNOAL-LIKE DOMAIN-CONTAINING PROTEI |
| 85:00:00 | 4w6r-P | 3.3 | 8.7 | 56 | 204 | 7 | MOLECULE: | FLUORESCENT PROTEIN D102C |
| 86:00:00 | 5o67-A | 3.3 | 7.7 | 80 | 282 | 9 | MOLECULE: | FAPF |
| 87:00:00 | 3rwa-E | 3.3 | 5.2 | 62 | 232 | 2 | MOLECULE: | FLUORESCENT PROTEIN FP480 |
| 88:00:00 | 4u1f-A | 3.3 | 6 | 73 | 474 | 4 | MOLECULE: | EUKARYOTIC TRANSLATION INITIATION F |
| 89:00:00 | 6mxw-H | 3.3 | 4.7 | 61 | 215 | 11 | MOLECULE: | PHOTOSWITCHABLE PROTEIN TETDRON |
| 90:00:00 | 5o68-J | 3.3 | 7 | 93 | 268 | 4 | MOLECULE: | FAPF |
| 91:00:00 | 5o68-L | 3.3 | 5.4 | 76 | 258 | 12 | MOLECULE: | FAPF |
| 92:00:00 | 3sg7-A | 3.3 | 6.7 | 83 | 398 | 2 | MOLECULE: | MYOSIN LIGHT CHAIN KINASE, GREEN FL |
| 93:00:00 | 5exb-L | 3.3 | 4.4 | 61 | 225 | 5 | MOLECULE: | GREEN FLUORESCENT PROTEIN |
| 94:00:00 | 7t4e-A | 3.2 | 5.6 | 72 | 307 | 6 | MOLECULE: | EPX1 |
| 95:00:00 | 6cb1-k | 3.2 | 3.2 | 52 | 77 | 8 | MOLECULE: | 35S PRE-RIBOSOMAL RNA MISCRNA |
| 96:00:00 | 6imj-A | 3.2 | 3.7 | 70 | 412 | 3 | MOLECULE: | DNA LIGASE |
| 97:00:00 | 5ien-A | 3.2 | 5.8 | 71 | 129 | 4 | MOLECULE: | D30H_F14 |
| 98:00:00 | 6s6z-A | 3.2 | 6.2 | 67 | 1083 | 4 | MOLECULE: | BETA-GALACTOSIDASE |
| 99:00:00 | 5u6y-A | 3.2 | 5.3 | 66 | 459 | 9 | MOLECULE: | CALCIUM/CALMODULIN-DEPENDENT PROTEI |
| 0:00 | 8b61-B | 3.2 | 4.4 | 60 | 200 | 7 | MOLECULE: | CONSERVED HYPOTHETICAL LIPOPROTEIN |
| 1:00 | 7zei-A | 3.2 | 4.9 | 71 | 310 | 7 | MOLECULE: | CH_GAF159A |
| 2:00 | 1aq3-A | 3.2 | 4.6 | 78 | 129 | 6 | MOLECULE: | RNA (5'- |
| 3:00 | 7jmn-P | 3.2 | 7.3 | 91 | 810 | 5 | MOLECULE: | MEDIATOR OF RNA POLYMERASE II TRANS |
| 4:00 | 8gdl-A | 3.2 | 2.4 | 60 | 355 | 7 | MOLECULE: | SECRETED SALIVARY ACID PHOSPHATASE |
| 5:00 | 3a1j-A | 3.2 | 7.2 | 65 | 265 | 8 | MOLECULE: | CELL CYCLE CHECKPOINT CONTROL PROTE |
| 6:00 | 7a0h-A | 3.2 | 5.9 | 79 | 285 | 3 | MOLECULE: | F-ACTIN-CAPPING PROTEIN SUBUNIT ALP |
| 7:00 | 7st9-G | 3.2 | 7.5 | 67 | 340 | 3 | MOLECULE: | CHECKPOINT PROTEIN RAD24 |
| 8:00 | 5zbt-A | 3.2 | 3.5 | 63 | 212 | 8 | MOLECULE: | LECTIN-LIKE PROTEIN |
| 9:00 | 5tse-A | 3.2 | 3.6 | 74 | 136 | 3 | MOLECULE: | LPS-ASSEMBLY LIPOPROTEIN LPTE |
| 10:00 | 5w3s-A | 3.2 | 5.3 | 81 | 485 | 6 | MOLECULE: | MUCOLIPIN-3 ISOFORM 1 |
| 11:00 | 4aee-A | 3.2 | 6.2 | 70 | 684 | 7 | MOLECULE: | ALPHA AMYLASE, CATALYTIC REGION |
| 12:00 | 7zgr-A | 3.2 | 6 | 80 | 1253 | 5 | MOLECULE: | PROTEIN CFT1 |

| | | | | | | | | |
|---|---|---|---|---|---|---|---|---|
| 13:00 | 6sl1-A | 3.2 | 6.7 | 65 | 2651 | 3 | MOLECULE: | SERINE/THREONINE-PROTEIN KINASE TEL |
| 14:00 | 6r6u-A | 3.2 | 7.5 | 68 | 458 | 6 | MOLECULE: | CIS-ACONITATE DECARBOXYLASE |
| 15:00 | 6lyq-A | 3.2 | 8.9 | 91 | 787 | 5 | MOLECULE: | OUTER MEMBRANE PROTEIN ASSEMBLY FAC |
| 16:00 | 6m1u-A | 3.2 | 3.6 | 58 | 122 | 9 | MOLECULE: | RNA N6-ADENOSINE-METHYLTRANSFERASE |
| 17:00 | 8gqq-A | 3.2 | 4.1 | 61 | 99 | 3 | MOLECULE: | CYSTATIN-A2 |
| 18:00 | 7o5y-D | 3.2 | 9 | 71 | 131 | 11 | MOLECULE: | TYPE IV PILUS BIOGENESIS PROTEIN PI |
| 19:00 | 6hmj-A | 3.2 | 9 | 77 | 359 | 6 | MOLECULE: | PUTATIVE PAS/PAC SENSOR PROTEIN |
| 20:00 | 7vcf-I | 3.2 | 7.5 | 76 | 363 | 8 | MOLECULE: | TIC214 |
| 21:00 | 3qqz-A | 3.2 | 4.5 | 63 | 243 | 2 | MOLECULE: | PUTATIVE UNCHARACTERIZED PROTEIN YJ |
| 22:00 | 4q51-B | 3.2 | 3.2 | 73 | 265 | 8 | MOLECULE: | UNCHARACTERIZED PROTEIN |
| 23:00 | 5ft0-A | 3.2 | 8.7 | 77 | 256 | 5 | MOLECULE: | GP37 |
| 24:00:00 | 7rxu-A | 3.2 | 3.3 | 71 | 134 | 7 | MOLECULE: | LIPOPROTEIN |
| 25:00:00 | 2yeq-A | 3.2 | 4.8 | 72 | 522 | 7 | MOLECULE: | ALKALINE PHOSPHATASE D |
| 26:00:00 | 5wlz-C | 3.2 | 9.3 | 51 | 206 | 6 | MOLECULE: | DNA REPAIR PROTEIN XRCC4,MYOSIN-7 |
| 27:00:00 | 7ype-A | 3.2 | 8.3 | 77 | 286 | 8 | MOLECULE: | E301R |
| 28:00:00 | 2xe4-A | 3.2 | 5.3 | 70 | 721 | 3 | MOLECULE: | OLIGOPEPTIDASE B |
| 29:00:00 | 3vsf-C | 3.2 | 3.3 | 71 | 482 | 3 | MOLECULE: | RICIN B LECTIN |
| 30:00:00 | 6dhx-A | 3.2 | 5.8 | 69 | 182 | 7 | MOLECULE: | TIPC2 |
| 31:00:00 | 5nxh-A | 3.2 | 9.8 | 69 | 546 | 3 | MOLECULE: | LONG-TAIL FIBER PROXIMAL SUBUNIT |
| 32:00:00 | 6h3i-F | 3.2 | 5.8 | 81 | 345 | 4 | MOLECULE: | PROTEIN INVOLVED IN GLIDING MOTILIT |
| 33:00:00 | 8pjn-2 | 3.2 | 3.8 | 56 | 156 | 5 | MOLECULE: | E3 UBIQUITIN-PROTEIN TRANSFERASE RM |
| 34:00:00 | 6ekv-A | 3.2 | 3.6 | 72 | 718 | 6 | MOLECULE: | TOXIN COMPLEX COMPONENT ORF-X2 |
| 35:00:00 | 7obm-A | 3.2 | 4.5 | 67 | 624 | 9 | MOLECULE: | PROLYL ENDOPEPTIDASE-LIKE |
| 36:00:00 | 6sb5-J | 3.2 | 6.5 | 76 | 543 | 7 | MOLECULE: | MACROPHAGE-EXPRESSED GENE 1 PROTEIN |
| 37:00:00 | 7nri-E | 3.2 | 4.4 | 56 | 84 | 7 | MOLECULE: | OUTER MEMBRANE PROTEIN ASSEMBLY FAC |
| 38:00:00 | 6iys-O | 3.2 | 8.3 | 59 | 245 | 7 | MOLECULE: | OUTER SURFACE PROTEIN A |
| 39:00:00 | 6w3f-A | 3.2 | 5.3 | 59 | 115 | 8 | MOLECULE: | RD1NTF2_05_I64F_A80G_T94P_D101K_L10 |
| 40:00:00 | 5d17-H | 3.2 | 3.4 | 61 | 156 | 5 | MOLECULE: | TRANSPOSON TN7 TRANSPOSITION PROTEI |
| 41:00:00 | 4u1f-B | 3.2 | 6 | 73 | 475 | 4 | MOLECULE: | EUKARYOTIC TRANSLATION INITIATION F |
| 42:00:00 | 8es4-E | 3.1 | 4.3 | 69 | 772 | 6 | MOLECULE: | GP35 |
| 43:00:00 | 6bym-A | 3.1 | 4.4 | 80 | 200 | 11 | MOLECULE: | STEROL-BINDING PROTEIN |
| 44:00:00 | 3ggr-C | 3.1 | 6.2 | 72 | 264 | 3 | MOLECULE: | CELL CYCLE CHECKPOINT CONTROL PROTE |
| 45:00:00 | 4w78-F | 3.1 | 2.9 | 56 | 127 | 4 | MOLECULE: | HYDRATASE CHSH1 |
| 46:00:00 | 2vf9-A | 3.1 | 5 | 76 | 131 | 4 | MOLECULE: | COAT PROTEIN |
| 47:00:00 | 8bd7-S | 3.1 | 5.2 | 62 | 1104 | 3 | MOLECULE: | IFT88 |
| 48:00:00 | 7qpg-R | 3.1 | 5.2 | 65 | 2208 | 8 | MOLECULE: | PROTEIN ZWILCH HOMOLOG |
| 49:00:00 | 7aed-B | 3.1 | 5.7 | 60 | 138 | 5 | MOLECULE: | PRGL |
| 50:00:00 | 3hk4-A | 3.1 | 5.2 | 60 | 118 | 5 | MOLECULE: | MLR7391 PROTEIN |
| 51:00:00 | 8vwr-A | 3.1 | 5.4 | 71 | 154 | 3 | MOLECULE: | ACYL DEHYDRATASE |
| 52:00:00 | 5mdq-A | 3.1 | 9.7 | 89 | 350 | 7 | MOLECULE: | CHITOPORIN |
| 53:00:00 | 1jki-A | 3.1 | 10.8 | 59 | 525 | 3 | MOLECULE: | MYO-INOSITOL-1-PHOSPHATE SYNTHASE |
| 54:00:00 | 1l0o-A | 3.1 | 3.2 | 62 | 141 | 5 | MOLECULE: | ANTI-SIGMA F FACTOR |
| 55:00:00 | 8alo-B | 3.1 | 5.1 | 79 | 148 | 5 | MOLECULE: | CHOLERA TOXIN TRANSCRIPTIONAL ACTIV |
| 56:00:00 | 2zwa-B | 3.1 | 3.8 | 57 | 684 | 2 | MOLECULE: | LEUCINE CARBOXYL METHYLTRANSFERASE |
| 57:00:00 | 7q3e-A | 3.1 | 6.9 | 71 | 551 | 7 | MOLECULE: | WD REPEAT-CONTAINING AND PLANAR CEL |
| 58:00:00 | 8f2r-F | 3.1 | 3.5 | 49 | 77 | 6 | MOLECULE: | COMM DOMAIN-CONTAINING PROTEIN 1 |

| | | | | | | | | |
|---|---|---|---|---|---|---|---|---|
| 59:00:00 | 6ekt-A | 3.1 | 5.9 | 81 | 419 | 4 | MOLECULE: | P-47 PROTEIN |
| 60:00:00 | 6a70-B | 3.1 | 5.3 | 78 | 704 | 6 | MOLECULE: | POLYCYSTIN-2 |
| 61:00:00 | 8tie-a | 3.1 | 4.3 | 64 | 1012 | 3 | MOLECULE: | NUCLEOPORIN NUP120 |
| 62:00:00 | 1fi1-A | 3.1 | 11.8 | 94 | 707 | 6 | MOLECULE: | FERRICHROME-IRON RECEPTOR |
| 63:00:00 | 2rau-A | 3.1 | 6.7 | 62 | 350 | 2 | MOLECULE: | PUTATIVE ESTERASE |
| 64:00:00 | 7uyx-B | 3.1 | 5.2 | 68 | 169 | 6 | MOLECULE: | BACTERIOPHAGE PA1C GP2 |
| 65:00:00 | 5k47-A | 3.1 | 8.2 | 82 | 484 | 7 | MOLECULE: | POLYCYSTIN-2 |
| 66:00:00 | 3nqz-A | 3.1 | 6.5 | 70 | 175 | 10 | MOLECULE: | SECRETED METALLOPROTEASE MCP02 |
| 67:00:00 | 4ddg-A | 3.1 | 6.9 | 60 | 399 | 7 | MOLECULE: | UBIQUITIN-CONJUGATING ENZYME E2 D2, |
| 68:00:00 | 7yls-A | 3.1 | 5.9 | 74 | 158 | 8 | MOLECULE: | AROMATIC-RING-HYDROXYLATING DIOXYGE |
| 69:00:00 | 4l9h-A | 3.1 | 3.3 | 75 | 155 | 8 | MOLECULE: | F-BOX ONLY PROTEIN 7 |
| 70:00:00 | 7kra-A | 3.1 | 9.8 | 70 | 694 | 4 | MOLECULE: | ER MEMBRANE PROTEIN COMPLEX SUBUNIT |
| 71:00:00 | 2wyh-A | 3.1 | 9.7 | 75 | 905 | 7 | MOLECULE: | ALPHA-MANNOSIDASE |
| 72:00:00 | 4o2h-A | 3.1 | 8.4 | 69 | 137 | 6 | MOLECULE: | PROTEIN BCAM1869 |
| 73:00:00 | 4a7z-A | 3.1 | 5.2 | 75 | 881 | 9 | MOLECULE: | ALDOS-2-ULOSE DEHYDRATASE |
| 74:00:00 | 7fh6-A | 3.1 | 9.4 | 77 | 653 | 3 | MOLECULE: | CYLK |
| 75:00:00 | 7rag-A | 3.1 | 7.6 | 63 | 170 | 3 | MOLECULE: | LIPOPROTEIN |
| 76:00:00 | 5kph-A | 3.1 | 3.7 | 55 | 85 | 7 | MOLECULE: | DE NOVO BETA SHEET DESIGN PROTEIN O |
| 77:00:00 | 6wlw-U | 3.1 | 5.8 | 79 | 205 | 6 | MOLECULE: | V-TYPE PROTON ATPASE 21 KDA PROTEOL |
| 78:00:00 | 2w5n-A | 3.1 | 4.8 | 74 | 365 | 1 | MOLECULE: | ALPHA-L-ARABINOFURANOSIDASE |
| 79:00:00 | 8tum-a | 3.1 | 4.7 | 55 | 143 | 0 | MOLECULE: | TYPE IV MAJOR PILIN PROTEIN PILA |
| 80:00:00 | 5edf-A | 3.1 | 7.1 | 77 | 224 | 1 | MOLECULE: | FRPC OPERON PROTEIN |
| 81:00:00 | 7z8b-F | 3.1 | 5.6 | 63 | 457 | 6 | MOLECULE: | CULLIN-7 |
| 82:00:00 | 8qc6-A | 3.1 | 7.1 | 74 | 366 | 7 | MOLECULE: | OXIDOREDUCTASE |
| 83:00:00 | 3o61-A | 3.1 | 8.3 | 51 | 187 | 6 | MOLECULE: | GDP-MANNOSE PYROPHOSPHATASE NUDK |
| 84:00:00 | 5u0p-Q | 3.1 | 7 | 66 | 508 | 6 | MOLECULE: | MEDIATOR COMPLEX SUBUNIT 14 |
| 85:00:00 | 8thf-A | 3.1 | 5.4 | 59 | 900 | 5 | MOLECULE: | SPIKE PROTEIN S1,SPIKE GLYCOPROTEIN |
| 86:00:00 | 6eus-A | 3.1 | 7.8 | 71 | 346 | 4 | MOLECULE: | DCAP-LIKE PROTEIN |
| 87:00:00 | 2x3j-A | 3.1 | 11 | 67 | 586 | 7 | MOLECULE: | ACSD |
| 88:00:00 | 6yhk-A | 3.1 | 3.3 | 81 | 994 | 6 | MOLECULE: | CYTOTOXIC NECROTIZING FACTOR |
| 89:00:00 | 3duk-A | 3.1 | 3.6 | 56 | 125 | 5 | MOLECULE: | NTF2-LIKE PROTEIN OF UNKNOWN FUNCTI |
| 90:00:00 | 3ebw-A | 3.1 | 4.8 | 61 | 157 | 7 | MOLECULE: | PER A 4 ALLERGEN |
| 91:00:00 | 8ae1-A | 3.1 | 7.4 | 91 | 953 | 5 | MOLECULE: | S-LAYER PROTEIN SLPA |
| 92:00:00 | 5wd6-A | 3.1 | 4 | 68 | 195 | 10 | MOLECULE: | SHORT PALATE, LUNG AND NASAL EPITHE |
| 93:00:00 | 5chx-A | 3.1 | 5.5 | 55 | 210 | 4 | MOLECULE: | XRCC4-MYH7-1590-1657 |
| 94:00:00 | 3muu-A | 3.1 | 6.5 | 65 | 623 | 5 | MOLECULE: | STRUCTURAL POLYPROTEIN |
| 95:00:00 | 4pj2-B | 3.1 | 3.6 | 59 | 123 | 0 | MOLECULE: | PUTATIVE EXPORTED PROTEIN |
| 96:00:00 | 6ohh-B | 3.1 | 5.3 | 64 | 128 | 6 | MOLECULE: | EF1P2_MFAP2B |
| 97:00:00 | 8a9y-E | 3.1 | 4.3 | 72 | 497 | 8 | MOLECULE: | GLYCOSIDE HYDROLASE FAMILY 32 |
| 98:00:00 | 7okx-O | 3.1 | 4.5 | 53 | 106 | 4 | MOLECULE: | DNA-DIRECTED RNA POLYMERASE II SUBU |
| 99:00:00 | 6te9-F | 3.1 | 2.9 | 61 | 134 | 8 | MOLECULE: | ADAPTOR PROTEIN RCC01688 |
| 0:00 | 1oh1-A | 3.1 | 3.7 | 63 | 109 | 13 | MOLECULE: | STAPHOSTATIN A |
| 1:00 | 6w90-A | 3.1 | 3.7 | 53 | 130 | 6 | MOLECULE: | NTF2 FOLD PROTEIN LOOP-HELIX-LOOP D |
| 2:00 | 5xqi-B | 3.1 | 4.5 | 55 | 276 | 2 | MOLECULE: | PROTEIN ROGDI HOMOLOG |
| 3:00 | 6jsh-B | 3.1 | 4.5 | 70 | 1904 | 0 | MOLECULE: | FATTY ACID SYNTHASE SUBUNIT BETA |
| 4:00 | 8ghn-A | 3.1 | 4.8 | 55 | 667 | 7 | MOLECULE: | PROTEIN HIR1 |

| | | | | | | | | |
|---|---|---|---|---|---|---|---|---|
| 5:00 | 5bka-C | 3.1 | 5.1 | 63 | 126 | 11 | MOLECULE: | HYDROXYLACYL-COA DEHYDROGENASE |
| 6:00 | 7f0o-A | 3.1 | 10 | 75 | 144 | 8 | MOLECULE: | NSRQ |
| 7:00 | 6isl-A | 3.1 | 4.2 | 65 | 120 | 12 | MOLECULE: | XIME, SNOAL-LIKE DOMAIN PROTEIN |
| 8:00 | 3dxo-A | 3.1 | 4.6 | 58 | 117 | 2 | MOLECULE: | UNCHARACTERIZED SNOAL-LIKE PROTEIN |
| 9:00 | 7e7w-A | 3.1 | 3.3 | 68 | 264 | 9 | MOLECULE: | FUCOSE-BINDING LECTIN PROTEIN,FUCOS |
| 10:00 | 4nox-A | 3.1 | 7.4 | 79 | 460 | 9 | MOLECULE: | EUKARYOTIC TRANSLATION INITIATION F |
| 11:00 | 5d17-B | 3.1 | 3.2 | 60 | 158 | 5 | MOLECULE: | TRANSPOSON TN7 TRANSPOSITION PROTEI |
| 12:00 | 7dmx-B | 3.1 | 5.1 | 57 | 209 | 7 | MOLECULE: | PHOCL GREEN |
| 13:00 | 3tmt-B | 3.1 | 4.1 | 59 | 220 | 5 | MOLECULE: | GREEN TO RED PHOTOCONVERTIBLE GPF-L |
| 14:00 | 2ib6-A | 3.1 | 4.2 | 62 | 225 | 6 | MOLECULE: | YELLOW MUTANT CHROMO PROTEIN |
| 15:00 | 6k96-B | 3 | 8.5 | 75 | 343 | 3 | MOLECULE: | FIVE-MEMBERED-CYCLITOL-PHOSPHATE SY |
| 16:00 | 7pkn-O | 3 | 4.7 | 56 | 210 | 4 | MOLECULE: | CENTROMERE PROTEIN H |
| 17:00 | 7rtn-A | 3 | 11.6 | 62 | 270 | 5 | MOLECULE: | OUTER CAPSID PROTEIN VP5 |
| 18:00 | 8g7w-B | 3 | 4.6 | 81 | 1059 | 1 | MOLECULE: | TYPE I PKS MODULE 4, MODULE 5 |
| 19:00 | 8s9t-B | 3 | 11.5 | 71 | 185 | 7 | MOLECULE: | CAS7-CAS5-CAS11 |
| 20:00 | 6e14-D | 3 | 9.1 | 83 | 805 | 7 | MOLECULE: | TYPE 1 FIMBRIN D-MANNOSE SPECIFIC A |
| 21:00 | 7w63-B | 3 | 7 | 77 | 395 | 6 | MOLECULE: | TOXIN-COREGULATED PILUS BIOSYNTHESI |
| 22:00 | 4ec6-A | 3 | 5.5 | 61 | 109 | 7 | MOLECULE: | PUTATIVE UNCHARACTERIZED PROTEIN |
| 23:00 | 7aoh-B | 3 | 5.8 | 77 | 1134 | 4 | MOLECULE: | DNA-DIRECTED RNA POLYMERASE 147 KDA |
| 24:00:00 | 8f2r-D | 3 | 2.9 | 49 | 199 | 4 | MOLECULE: | COMM DOMAIN-CONTAINING PROTEIN 1 |
| 25:00:00 | 2hhi-A | 3 | 3.1 | 65 | 204 | 6 | MOLECULE: | IMMUNOGENIC PROTEIN MPT64 |
| 26:00:00 | 6xd4-A | 3 | 6.1 | 87 | 458 | 9 | MOLECULE: | HEMOLYSIN |
| 27:00:00 | 7pkq-N | 3 | 4.2 | 69 | 195 | 3 | MOLECULE: | MS35 |
| 28:00:00 | 3nja-D | 3 | 4.1 | 51 | 105 | 8 | MOLECULE: | PROBABLE GGDEF FAMILY PROTEIN |
| 29:00:00 | 2hye-A | 3 | 5.8 | 72 | 1140 | 10 | MOLECULE: | DNA DAMAGE-BINDING PROTEIN 1 |
| 30:00:00 | 8p0w-J | 3 | 3.2 | 48 | 202 | 2 | MOLECULE: | COMM DOMAIN-CONTAINING PROTEIN 1 |
| 31:00:00 | 5x04-A | 3 | 4.6 | 67 | 276 | 7 | MOLECULE: | DODECANOYL-[ACYL-CARRIER-PROTEIN] H |
| 32:00:00 | 6mrk-B | 3 | 5.8 | 68 | 188 | 6 | MOLECULE: | NUCLEAR RNA EXPORT FACTOR 2 |
| 33:00:00 | 6m36-I | 3 | 2.6 | 48 | 104 | 4 | MOLECULE: | SERINE-PROTEIN KINASE RSBW |
| 34:00:00 | 5v1d-D | 3 | 5.4 | 71 | 247 | 6 | MOLECULE: | EIF2AK3 PROTEIN |
| 35:00:00 | 6eu4-A | 3 | 6.1 | 68 | 587 | 6 | MOLECULE: | TAIL SPIKE PROTEIN |
| 36:00:00 | 7kbj-I | 3 | 5.4 | 59 | 153 | 8 | MOLECULE: | NEUTRAL ALPHA-GLUCOSIDASE AB TRYPSI |
| 37:00:00 | 4mkq-A | 3 | 4.9 | 76 | 169 | 8 | MOLECULE: | MONALYSIN |
| 38:00:00 | 2ch9-A | 3 | 5.9 | 73 | 126 | 3 | MOLECULE: | CYSTATIN F |
| 39:00:00 | 3esi-A | 3 | 3.1 | 68 | 124 | 3 | MOLECULE: | UNCHARACTERIZED PROTEIN |
| 40:00:00 | 4h5b-A | 3 | 2.4 | 67 | 152 | 3 | MOLECULE: | DR_1245 PROTEIN |
| 41:00:00 | 1k1x-A | 3 | 5.1 | 72 | 636 | 7 | MOLECULE: | 4-ALPHA-GLUCANOTRANSFERASE |
| 42:00:00 | 7bvj-A | 3 | 7.1 | 68 | 313 | 4 | MOLECULE: | OXIDOREDUCTASE, NAD-BINDING |
| 43:00:00 | 7c5x-A | 3 | 6.2 | 70 | 506 | 6 | MOLECULE: | IOTA-CARBONIC ANHYDRASE |
| 44:00:00 | 5wi2-B | 3 | 3.6 | 54 | 93 | 9 | MOLECULE: | CDNA FLJ56409, HIGHLY SIMILAR TO SE |
| 45:00:00 | 8t0b-A | 3 | 5.7 | 61 | 121 | 7 | MOLECULE: | DUF1842 DOMAIN-CONTAINING PROTEIN |
| 46:00:00 | 3put-B | 3 | 3.2 | 71 | 155 | 4 | MOLECULE: | HYPOTHETICAL CONSERVED PROTEIN |
| 47:00:00 | 2lyx-A | 3 | 4 | 60 | 87 | 15 | MOLECULE: | UNCHARACTERIZED PROTEIN YQZG |
| 48:00:00 | 8q72-A | 3 | 13 | 71 | 750 | 6 | MOLECULE: | JETC |
| 49:00:00 | 4d0v-D | 3 | 6 | 67 | 116 | 1 | MOLECULE: | FIBER PROTEIN |
| 50:00:00 | 8p0w-H | 3 | 3.3 | 47 | 183 | 2 | MOLECULE: | COMM DOMAIN-CONTAINING PROTEIN 1 |

| | | | | | | | | |
|---|---|---|---|---|---|---|---|---|
| 51:00:00 | 1dwn-A | 3 | 4.5 | 64 | 127 | 6 | MOLECULE: | PHAGE COAT PROTEIN |
| 52:00:00 | 1mm4-A | 3 | 5.5 | 82 | 170 | 4 | MOLECULE: | CRCA PROTEIN |
| 53:00:00 | 3b8l-A | 3 | 5.7 | 77 | 147 | 9 | MOLECULE: | UNCHARACTERIZED PROTEIN |
| 54:00:00 | 6dn4-A | 3 | 7.4 | 62 | 275 | 6 | MOLECULE: | BETA-LACTAMASE |
| 55:00:00 | 3df6-B | 3 | 3.1 | 51 | 99 | 2 | MOLECULE: | ORF99 |
| 56:00:00 | 7eep-A | 3 | 4.5 | 87 | 529 | 7 | MOLECULE: | PAM1 PORTAL PROTEINS |
| 57:00:00 | 3but-A | 3 | 3.5 | 69 | 125 | 6 | MOLECULE: | UNCHARACTERIZED PROTEIN AF_0446 |
| 58:00:00 | 6oh1-A | 3 | 4.9 | 54 | 84 | 9 | MOLECULE: | IMMUNOGLOBULIN A1 PROTEASE |
| 59:00:00 | 6exp-F | 3 | 2.9 | 53 | 103 | 9 | MOLECULE: | SIRV3 ACRID1 (GP02) ANTI-CRISPR PRO |
| 60:00:00 | 3mmy-C | 3 | 4.2 | 55 | 355 | 7 | MOLECULE: | MRNA EXPORT FACTOR |
| 61:00:00 | 8p26-J | 3 | 5.4 | 56 | 147 | 0 | MOLECULE: | U2 SMALL NUCLEAR RIBONUCLEOPROTEIN |
| 62:00:00 | 6u3r-A | 3 | 3.4 | 51 | 190 | 2 | MOLECULE: | DNAJ HOMOLOG SUBFAMILY B MEMBER 6,D |
| 63:00:00 | 8gex-A | 3 | 3.8 | 62 | 411 | 8 | MOLECULE: | DUF4374 DOMAIN-CONTAINING PROTEIN |
| 64:00:00 | 4agi-A | 3 | 4 | 68 | 314 | 4 | MOLECULE: | FUCOSE-SPECIFIC LECTIN FLEA |
| 65:00:00 | 6yle-B | 3 | 7.5 | 71 | 415 | 8 | MOLECULE: | PRE-RRNA-PROCESSING PROTEIN IPI3 |
| 66:00:00 | 4j5t-A | 3 | 12.6 | 58 | 788 | 9 | MOLECULE: | MANNOSYL-OLIGOSACCHARIDE GLUCOSIDAS |
| 67:00:00 | 5jtl-A | 3 | 6 | 58 | 155 | 12 | MOLECULE: | PROTEIN-EXPORT PROTEIN SECB |
| 68:00:00 | 5cd6-A | 3 | 4.6 | 80 | 575 | 6 | MOLECULE: | TPR-DOMAIN CONTAINING PROTEIN |
| 69:00:00 | 8jto-A | 3 | 8.6 | 76 | 343 | 4 | MOLECULE: | OUTER MEMBRANE PORIN (FRAGMENT) |
| 70:00:00 | 3fz2-E | 3 | 3.1 | 63 | 133 | 8 | MOLECULE: | MINOR TAIL PROTEIN U |
| 71:00:00 | 5y58-A | 3 | 6.2 | 73 | 548 | 5 | MOLECULE: | ATP-DEPENDENT DNA HELICASE II SUBUN |
| 72:00:00 | 7uea-U | 3 | 12.4 | 78 | 364 | 6 | MOLECULE: | PHOTOSYSTEM P840 REACTION CENTER, L |
| 73:00:00 | 8fn4-2 | 3 | 5.6 | 58 | 357 | 9 | MOLECULE: | RNA-EDITING SUBSTRATE-BINDING COMPL |
| 74:00:00 | 8bbg-B | 3 | 5.9 | 68 | 1426 | 1 | MOLECULE: | WD REPEAT-CONTAINING PROTEIN 19 |
| 75:00:00 | 6e8e-A | 3 | 7.4 | 64 | 468 | 3 | MOLECULE: | BETA SLIDING CLAMP,DNA MISMATCH REP |
| 76:00:00 | 1sli-A | 3 | 5.5 | 75 | 679 | 7 | MOLECULE: | INTRAMOLECULAR TRANS-SIALIDASE |
| 77:00:00 | 1xsi-A | 3 | 4.2 | 53 | 773 | 8 | MOLECULE: | PUTATIVE FAMILY 31 GLUCOSIDASE YICI |
| 78:00:00 | 1mdc-A | 3 | 3.1 | 53 | 131 | 6 | MOLECULE: | INSECT FATTY ACID BINDING PROTEIN |
| 79:00:00 | 8r6y-A | 3 | 12.3 | 63 | 1991 | 14 | MOLECULE: | RNA-DIRECTED RNA POLYMERASE L |
| 80:00:00 | 4u8u-M | 3 | 3.7 | 68 | 222 | 4 | MOLECULE: | GLOBIN A CHAIN |
| 81:00:00 | 8evk-A | 3 | 4.9 | 63 | 115 | 8 | MOLECULE: | DIHYDRONEOPTERIN ALDOLASE |
| 82:00:00 | 7vwk-A | 3 | 3.6 | 70 | 260 | 7 | MOLECULE: | POLYKETIDE SYNTHASE |
| 83:00:00 | 4qvh-A | 3 | 7.8 | 57 | 596 | 2 | MOLECULE: | MALTOSE-BINDING PERIPLASMIC PROTEIN |
| 84:00:00 | 7elb-A | 3 | 6.7 | 73 | 1937 | 0 | MOLECULE: | RNA-DIRECTED RNA POLYMERASE L |
| 85:00:00 | 7vd7-A | 3 | 3.5 | 48 | 94 | 8 | MOLECULE: | TOXIN |
| 86:00:00 | 8ye0-C | 3 | 5.1 | 73 | 290 | 8 | MOLECULE: | LYNF/TRUF/PATF FAMILY PEPTIDE O-PRE |
| 87:00:00 | 4xrw-A | 3 | 11.4 | 64 | 305 | 6 | MOLECULE: | BEXL |
| 88:00:00 | 6nd4-T | 3 | 4.8 | 69 | 812 | 7 | MOLECULE: | ETS RRNA |
| 89:00:00 | 4g75-A | 3 | 4.5 | 66 | 292 | 3 | MOLECULE: | PHOSPHODIESTERASE |
| 90:00:00 | 7qwt-A | 3 | 3.2 | 80 | 352 | 6 | MOLECULE: | RIESKE (2FE-2S) DOMAIN PROTEIN |
| 91:00:00 | 2g8s-A | 3 | 4.3 | 70 | 347 | 9 | MOLECULE: | GLUCOSE/SORBOSONE DEHYDROGENASES |
| 92:00:00 | 4neh-A | 3 | 7.4 | 78 | 1083 | 6 | MOLECULE: | INTEGRIN ALPHA-X |
| 93:00:00 | 6yai-E | 3 | 4.4 | 64 | 726 | 6 | MOLECULE: | CLATHRIN HEAVY CHAIN |
| 94:00:00 | 6yfi-A | 3 | 4.6 | 66 | 151 | 3 | MOLECULE: | COAT PROTEIN |
| 95:00:00 | 4gq2-P | 3 | 3.9 | 64 | 371 | 6 | MOLECULE: | NUCLEOPORIN NUP120 |
| 96:00:00 | 8pz4-A | 3 | 9.6 | 81 | 466 | 4 | MOLECULE: | ALGINATE PRODUCTION PROTEIN ALGE |

| | | | | | | | | |
|---|---|---|---|---|---|---|---|---|
| 97:00:00 | 3av0-B | 3 | 12 | 64 | 365 | 5 | MOLECULE: | DNA DOUBLE-STRAND BREAK REPAIR PROT |
| 98:00:00 | 1dp4-C | 3 | 5.6 | 63 | 429 | 6 | MOLECULE: | ATRIAL NATRIURETIC PEPTIDE RECEPTOR |
| 99:00:00 | 8uzl-A | 3 | 4.3 | 68 | 143 | 6 | MOLECULE: | DESIGNED TRANSMEMBRANE BETA-BARREL |
| 0:00 | 6gxs-B | 3 | 3.6 | 63 | 363 | 6 | MOLECULE: | CV39L LECTIN |
| 1:00 | 6h7f-A | 3 | 5.5 | 76 | 674 | 9 | MOLECULE: | BAUA |
| 2:00 | 4n4r-B | 3 | 3.2 | 70 | 151 | 9 | MOLECULE: | LPS-ASSEMBLY PROTEIN LPTD |
| 3:00 | 6isk-A | 3 | 4.4 | 67 | 120 | 12 | MOLECULE: | XIME, SNOAL-LIKE DOMAIN PROTEIN |
| 4:00 | 7q3d-A | 3 | 6.9 | 68 | 545 | 6 | MOLECULE: | WD REPEAT-CONTAINING AND PLANAR CEL |
| 5:00 | 7cd8-B | 3 | 11.3 | 60 | 228 | 5 | MOLECULE: | GREEN FLUORESCENT PROTEIN |
| 6:00 | 4zf4-B | 3 | 4.7 | 71 | 223 | 10 | MOLECULE: | GREEN FLUORESCENT PROTEIN |
| 7:00 | 2rh7-A | 3 | 8.3 | 52 | 211 | 4 | MOLECULE: | GREEN FLUORESCENT PROTEIN |
| 8:00 | 5o67-B | 3 | 3.9 | 66 | 290 | 2 | MOLECULE: | FAPF |
| 9:00 | 6ofm-A | 3 | 8.7 | 56 | 229 | 7 | MOLECULE: | GREEN FLUORESCENT PROTEIN (GFP) S6 |
| 10:00 | 2g16-A | 3 | 6.5 | 49 | 62 | 10 | MOLECULE: | GREEN FLUORESCENT PROTEIN |
| 11:00 | 3evr-A | 3 | 8.4 | 81 | 397 | 5 | MOLECULE: | MYOSIN LIGHT CHAIN KINASE, GREEN FL |
| 12:00 | 4pa0-A | 3 | 8.3 | 54 | 974 | 6 | MOLECULE: | MYOSIN-7,GREEN FLUORESCENT PROTEIN |
| 13:00 | 5d74-B | 2.9 | 4.2 | 57 | 234 | 7 | MOLECULE: | PUTATIVE PHAGE LYSIN |
| 14:00 | 5jk2-A | 2.9 | 3.4 | 61 | 128 | 3 | MOLECULE: | TP0751 |
| 15:00 | 8f2r-G | 2.9 | 3.6 | 50 | 200 | 12 | MOLECULE: | COMM DOMAIN-CONTAINING PROTEIN 1 |
| 16:00 | 4mtu-A | 2.9 | 4.7 | 74 | 143 | 5 | MOLECULE: | BETA-ALANYL-COA:AMMONIA LYASE 2 |
| 17:00 | 6c9j-A | 2.9 | 5.3 | 63 | 386 | 2 | MOLECULE: | 5'-AMP-ACTIVATED PROTEIN KINASE CAT |
| 18:00 | 8ptx-C | 2.9 | 4.9 | 67 | 539 | 4 | MOLECULE: | ELONGATOR COMPLEX PROTEIN 1 |
| 19:00 | 4y4v-A | 2.9 | 6.4 | 72 | 318 | 8 | MOLECULE: | CONSERVED HYPOTHETICAL SECRETED PRO |
| 20:00 | 2kcd-A | 2.9 | 3.7 | 62 | 120 | 0 | MOLECULE: | UNCHARACTERIZED PROTEIN SSP0047 |
| 21:00 | 3zpm-A | 2.9 | 6 | 92 | 211 | 13 | MOLECULE: | LATHERIN |
| 22:00 | 3h6j-A | 2.9 | 7.2 | 68 | 438 | 7 | MOLECULE: | NEURAMINIDASE |
| 23:00 | 4nlm-A | 2.9 | 5.7 | 58 | 326 | 2 | MOLECULE: | LMO1340 PROTEIN |
| 24:00:00 | 7k10-A | 2.9 | 7.8 | 69 | 1259 | 3 | MOLECULE: | DNA-DEPENDENT PROTEIN KINASE CATALY |
| 25:00:00 | 8pbz-A | 2.9 | 4.7 | 80 | 1003 | 6 | MOLECULE: | MGP-OPERON PROTEIN 3 |
| 26:00:00 | 4j0x-B | 2.9 | 4.7 | 53 | 366 | 11 | MOLECULE: | RIBOSOMAL RNA-PROCESSING PROTEIN 9 |
| 27:00:00 | 3j9e-D | 2.9 | 8.3 | 74 | 520 | 7 | MOLECULE: | VP5 |
| 28:00:00 | 8cvo-E | 2.9 | 2.2 | 46 | 64 | 7 | MOLECULE: | 16S RIBOSOMAL RNA |
| 29:00:00 | 6rwb-A | 2.9 | 6.2 | 63 | 1873 | 5 | MOLECULE: | TOXIN,TOXIN COMPLEX SUBUNIT TCAB,PU |
| 30:00:00 | 7qh3-B | 2.9 | 7.6 | 58 | 148 | 5 | MOLECULE: | RSFG |
| 31:00:00 | 4v12-A | 2.9 | 4.9 | 70 | 337 | 3 | MOLECULE: | MAOC LIKE DOMAIN PROTEIN |
| 32:00:00 | 8deo-A | 2.9 | 5.1 | 57 | 454 | 9 | MOLECULE: | ACCUMULATION ASSOCIATED PROTEIN |
| 33:00:00 | 5c9p-A | 2.9 | 7.4 | 71 | 354 | 3 | MOLECULE: | PLL LECTIN |
| 34:00:00 | 5mmj-y | 2.9 | 3.8 | 60 | 116 | 12 | MOLECULE: | 50S RIBOSOMAL PROTEIN L31 |
| 35:00:00 | 8hpo-K | 2.9 | 8.4 | 62 | 413 | 2 | MOLECULE: | TRANSCRIPTIONAL REGULATORY PROTEIN |
| 36:00:00 | 8oml-A | 2.9 | 7.9 | 68 | 325 | 6 | MOLECULE: | CELL WALL SURFACE ANCHOR FAMILY PRO |
| 37:00:00 | 6qzh-A | 2.9 | 4.4 | 69 | 744 | 7 | MOLECULE: | HUMAN CHEMOKINE RECEPTOR 7 |
| 38:00:00 | 4gcz-B | 2.9 | 7.1 | 59 | 378 | 8 | MOLECULE: | BLUE-LIGHT PHOTORECEPTOR, SENSOR PR |
| 39:00:00 | 1tp6-A | 2.9 | 5.3 | 69 | 126 | 3 | MOLECULE: | HYPOTHETICAL PROTEIN PA1314 |
| 40:00:00 | 8ovo-A | 2.9 | 4.1 | 63 | 504 | 10 | MOLECULE: | PUTATIVE PERIPLASMIC BINDING TRANSP |
| 41:00:00 | 4lgq-A | 2.9 | 5.2 | 64 | 134 | 11 | MOLECULE: | PUTATIVE POLYKETIDE CYCLASE |
| 42:00:00 | 3ua3-A | 2.9 | 5.7 | 86 | 663 | 2 | MOLECULE: | PROTEIN ARGININE N-METHYLTRANSFERAS |

| | | | | | | | | |
|---|---|---|---|---|---|---|---|---|
| 43:00:00 | 8f2r-C | 2.9 | 3.6 | 48 | 195 | 4 | MOLECULE: | COMM DOMAIN-CONTAINING PROTEIN 1 |
| 44:00:00 | 5iw9-A | 2.9 | 3.4 | 50 | 126 | 8 | MOLECULE: | BASEPLATE WEDGE PROTEIN GP25 |
| 45:00:00 | 1s28-A | 2.9 | 2.2 | 43 | 130 | 9 | MOLECULE: | ORF1 |
| 46:00:00 | 6p59-B | 2.9 | 7.8 | 62 | 196 | 5 | MOLECULE: | CORE-BINDING FACTOR SUBUNIT BETA |
| 47:00:00 | 8cb6-F | 2.9 | 4.8 | 57 | 110 | 4 | MOLECULE: | LYSOSOMAL ALPHA-GLUCOSIDASE (76 KDA |
| 48:00:00 | 1l0q-A | 2.9 | 7.1 | 65 | 391 | 9 | MOLECULE: | SURFACE LAYER PROTEIN |
| 49:00:00 | 6pnq-B | 2.9 | 5.2 | 64 | 133 | 2 | MOLECULE: | NEUREXIN-1 |
| 50:00:00 | 5yk6-A | 2.9 | 5.2 | 87 | 239 | 10 | MOLECULE: | MAINTENANCE OF MITOCHONDRIAL MORPHO |
| 51:00:00 | 6x1i-B | 2.9 | 6.4 | 65 | 146 | 0 | MOLECULE: | COB_ADENO_TRANS DOMAIN-CONTAINING P |
| 52:00:00 | 7r2x-A | 2.9 | 5.7 | 72 | 762 | 8 | MOLECULE: | MANNURONIC ACID SPECIFIC LYASE |
| 53:00:00 | 2k54-A | 2.9 | 3.6 | 67 | 123 | 6 | MOLECULE: | PROTEIN ATU0742 |
| 54:00:00 | 7uhy-C | 2.9 | 5.3 | 69 | 578 | 1 | MOLECULE: | GATOR COMPLEX PROTEIN MIOS |
| 55:00:00 | 8f5o-E | 2.9 | 6.9 | 66 | 1076 | 15 | MOLECULE: | INTRAFLAGELLAR TRANSPORT PROTEIN 12 |
| 56:00:00 | 6u5j-A | 2.9 | 5.1 | 72 | 164 | 8 | MOLECULE: | COLLAR PA0615 |
| 57:00:00 | 5bp3-A | 2.9 | 5.7 | 74 | 286 | 8 | MOLECULE: | MYCOCEROSIC ACID SYNTHASE-LIKE POLY |
| 58:00:00 | 6icl-A | 2.9 | 3.7 | 62 | 338 | 3 | MOLECULE: | METHYLXANTHINE N3-DEMETHYLASE NDMB |
| 59:00:00 | 2ns9-A | 2.9 | 5.3 | 72 | 152 | 7 | MOLECULE: | HYPOTHETICAL PROTEIN APE2225 |
| 60:00:00 | 6sw9-F | 2.9 | 2 | 47 | 229 | 0 | MOLECULE: | 16S RIBOSOMAL RNA |
| 61:00:00 | 7vcf-B | 2.9 | 8.5 | 94 | 651 | 10 | MOLECULE: | TIC214 |
| 62:00:00 | 8d8k-E | 2.9 | 2.9 | 50 | 291 | 8 | MOLECULE: | PROBABLE S-ADENOSYL-L-METHIONINE-DE |
| 63:00:00 | 6kyb-B | 2.9 | 3.9 | 72 | 330 | 6 | MOLECULE: | AUTOPHAGY-RELATED PROTEIN 18 |
| 64:00:00 | 6b7l-A | 2.9 | 3.8 | 72 | 290 | 1 | MOLECULE: | IMMUNE MODULATOR A |
| 65:00:00 | 3c8w-A | 2.9 | 4.8 | 69 | 250 | 6 | MOLECULE: | ACETOACETATE DECARBOXYLASE ADC |
| 66:00:00 | 7pge-B | 2.9 | 8.6 | 77 | 198 | 3 | MOLECULE: | COPPER RESISTANCE PROTEIN B |
| 67:00:00 | 5nzr-K | 2.9 | 7 | 70 | 559 | 4 | MOLECULE: | COATOMER SUBUNIT ALPHA |
| 68:00:00 | 7ktr-C | 2.9 | 3.9 | 44 | 407 | 5 | MOLECULE: | TRANSFORMATION/TRANSCRIPTION DOMAIN |
| 69:00:00 | 3jro-A | 2.9 | 4.6 | 71 | 701 | 0 | MOLECULE: | FUSION PROTEIN OF PROTEIN TRANSPORT |
| 70:00:00 | 6k7f-A | 2.9 | 4.6 | 71 | 499 | 1 | MOLECULE: | MALTOSE/MALTODEXTRIN-BINDING PERIPL |
| 71:00:00 | 3eu7-A | 2.9 | 7.2 | 63 | 313 | 5 | MOLECULE: | PARTNER AND LOCALIZER OF BRCA2 |
| 72:00:00 | 6nau-A | 2.9 | 3.7 | 59 | 334 | 2 | MOLECULE: | 6-PHOSPHOGLUCONOLACTONASE |
| 73:00:00 | 7mo7-B | 2.9 | 4.6 | 65 | 699 | 3 | MOLECULE: | HEPATOCYTE GROWTH FACTOR |
| 74:00:00 | 2zpa-B | 2.9 | 7.8 | 62 | 662 | 3 | MOLECULE: | UNCHARACTERIZED PROTEIN YPFI |
| 75:00:00 | 8an0-A | 2.9 | 5.7 | 53 | 282 | 6 | MOLECULE: | PUTATIVE SECRETED PROTEIN |
| 76:00:00 | 3g8z-A | 2.9 | 4.5 | 62 | 129 | 6 | MOLECULE: | PROTEIN OF UNKNOWN FUNCTION WITH CY |
| 77:00:00 | 6rw8-B | 2.9 | 10 | 69 | 2337 | 9 | MOLECULE: | A COMPONENT OF INSECTICIDAL TOXIN C |
| 78:00:00 | 2du7-A | 2.9 | 7.5 | 70 | 539 | 11 | MOLECULE: | O-PHOSPHOSERYL-TRNA SYNTHETASE |
| 79:00:00 | 4zel-A | 2.9 | 4.1 | 51 | 550 | 4 | MOLECULE: | DOPAMINE BETA-HYDROXYLASE |
| 80:00:00 | 4bg7-A | 2.9 | 4 | 52 | 97 | 4 | MOLECULE: | PUTATIVE TRANSCRIPTIONAL COACTIVATO |
| 81:00:00 | 7jrd-A | 2.9 | 4.6 | 66 | 566 | 2 | MOLECULE: | LACTOFERRIN-BINDING PROTEIN B |
| 82:00:00 | 2ia7-A | 2.9 | 3.9 | 59 | 111 | 7 | MOLECULE: | TAIL LYSOZYME, PUTATIVE |
| 83:00:00 | 8rtd-S | 2.9 | 5.7 | 63 | 223 | 10 | MOLECULE: | TRWJ PROTEIN |
| 84:00:00 | 4n6l-A | 2.9 | 3.8 | 61 | 114 | 5 | MOLECULE: | CYSTATIN-M |
| 85:00:00 | 2xgl-A | 2.9 | 3.6 | 55 | 90 | 5 | MOLECULE: | COLICIN-M IMMUNITY PROTEIN |
| 86:00:00 | 3lhn-A | 2.9 | 4.2 | 53 | 108 | 8 | MOLECULE: | LIPOPROTEIN |
| 87:00:00 | 4qa8-A | 2.9 | 4.3 | 59 | 210 | 3 | MOLECULE: | PUTATIVE LIPOPROTEIN LPRF |
| 88:00:00 | 3b33-A | 2.9 | 4.8 | 49 | 109 | 6 | MOLECULE: | SENSOR PROTEIN |

| | | | | | | | | |
|---|---|---|---|---|---|---|---|---|
| 89:00:00 | 1kmo-A | 2.9 | 7.1 | 94 | 661 | 6 | MOLECULE: | IRON(III) DICITRATE TRANSPORT PROTE |
| 90:00:00 | 6l4i-B | 2.9 | 3.3 | 58 | 90 | 0 | MOLECULE: | SINGLE CHAIN MONELLIN |
| 91:00:00 | 6l44-A | 2.9 | 3.5 | 59 | 89 | 0 | MOLECULE: | SINGLE CHAIN MONELLIN |
| 92:00:00 | 6l44-B | 2.9 | 3.5 | 59 | 90 | 2 | MOLECULE: | SINGLE CHAIN MONELLIN |
| 93:00:00 | 4mhc-A | 2.9 | 6.9 | 68 | 671 | 10 | MOLECULE: | NUCLEOPORIN NUP157 |
| 94:00:00 | 7xsi-A | 2.9 | 5.3 | 63 | 130 | 6 | MOLECULE: | SORDARIN/HYPOXYSORDARIN BIOSYNTHESI |
| 95:00:00 | 3dxo-B | 2.9 | 4.6 | 61 | 118 | 2 | MOLECULE: | UNCHARACTERIZED SNOAL-LIKE PROTEIN |
| 96:00:00 | 3ksp-A | 2.9 | 7.9 | 70 | 129 | 7 | MOLECULE: | CALCIUM/CALMODULIN-DEPENDENT KINASE |
| 97:00:00 | 5zc1-E | 2.9 | 3.6 | 57 | 98 | 14 | MOLECULE: | CYSTATIN-1 |
| 98:00:00 | 1sq9-A | 2.9 | 4.5 | 71 | 378 | 3 | MOLECULE: | ANTIVIRAL PROTEIN SKI8 |
| 99:00:00 | 4ouq-A | 2.9 | 3.4 | 53 | 105 | 8 | MOLECULE: | UNCHARACTERIZED PROTEIN |
| 0:00 | 6mwq-G | 2.9 | 6.3 | 57 | 226 | 4 | MOLECULE: | DARPIN, MUSCLE-TYPE ALDOLASE CHIMER |
| 1:00 | 3o78-B | 2.9 | 6 | 71 | 400 | 11 | MOLECULE: | MYOSIN LIGHT CHAIN KINASE, SMOOTH M |
| 2:00 | 8ill-B | 2.9 | 8.6 | 54 | 212 | 2 | MOLECULE: | GREEN FLUORESCENT PROTEIN |
| 3:00 | 7a7v-A | 2.9 | 8.3 | 55 | 225 | 5 | MOLECULE: | GREEN FLUORESCENT PROTEIN |
| 4:00 | 4w6i-A | 2.9 | 7.8 | 52 | 219 | 13 | MOLECULE: | FLUORESCENT PROTEIN D190C |
| 5:00 | 4w7d-B | 2.9 | 8.5 | 55 | 223 | 5 | MOLECULE: | FLUORESCENT PROTEIN D21H/K26H |
| 6:00 | 5o68-F | 2.9 | 5.4 | 93 | 254 | 9 | MOLECULE: | FAPF |
| 7:00 | 6uhl-A | 2.9 | 8.3 | 55 | 224 | 4 | MOLECULE: | C148 MGFP-SCDNA-1 |
| 8:00 | 8b6t-B | 2.9 | 8.3 | 55 | 516 | 4 | MOLECULE: | GREEN FLUORESCENT PROTEIN,HALOALKAN |
| 9:00 | 7a86-A | 2.9 | 8.9 | 58 | 224 | 5 | MOLECULE: | GREEN FLUORESCENT PROTEIN |
| 10:00 | 2g5z-A | 2.9 | 6.5 | 49 | 62 | 10 | MOLECULE: | GREEN FLUORESCENT PROTEIN |
| 11:00 | 6hzn-A | 2.8 | 4 | 57 | 742 | 7 | MOLECULE: | DERMATAN-SULFATE EPIMERASE |
| 12:00 | 2erv-A | 2.8 | 6.1 | 80 | 150 | 9 | MOLECULE: | HYPOTHETICAL PROTEIN PAER03002360 |
| 13:00 | 5a5y-A | 2.8 | 6.9 | 64 | 211 | 2 | MOLECULE: | TRIPHOSPHATE TUNEL METALLOENZYME 3 |
| 14:00 | 2e1q-A | 2.8 | 7.5 | 82 | 1307 | 4 | MOLECULE: | XANTHINE DEHYDROGENASE/OXIDASE |
| 15:00 | 3vb8-A | 2.8 | 3.9 | 68 | 153 | 6 | MOLECULE: | ENGINEERED PROTEIN |
| 16:00 | 7wkk-D | 2.8 | 3.7 | 60 | 1115 | 7 | MOLECULE: | MGC83295 PROTEIN |
| 17:00 | 7yqh-B | 2.8 | 11.6 | 72 | 1066 | 6 | MOLECULE: | STRUCTURAL MAINTENANCE OF CHROMOSOM |
| 18:00 | 7x80-A | 2.8 | 6.3 | 68 | 148 | 1 | MOLECULE: | PLOI4 |
| 19:00 | 6n8s-A | 2.8 | 6.2 | 77 | 834 | 9 | MOLECULE: | LETHAL(2) GIANT LARVAE PROTEIN HOMO |
| 20:00 | 8abv-A | 2.8 | 5.1 | 78 | 244 | 3 | MOLECULE: | SNOAL-LIKE DOMAIN-CONTAINING PROTEI |
| 21:00 | 6al2-A | 2.8 | 4.9 | 85 | 484 | 2 | MOLECULE: | MEMBRANE PROTEIN INSERTASE YIDC |
| 22:00 | 3g0k-A | 2.8 | 4.7 | 65 | 129 | 3 | MOLECULE: | PUTATIVE MEMBRANE PROTEIN |
| 23:00 | 5njb-B | 2.8 | 9.9 | 85 | 445 | 2 | MOLECULE: | METALLOPROTEASE TLDD |
| 24:00:00 | 6nu9-A | 2.8 | 3.9 | 50 | 168 | 10 | MOLECULE: | ZINC-BINDING NON-STRUCTURAL PROTEIN |
| 25:00:00 | 5nfi-B | 2.8 | 4.3 | 68 | 277 | 7 | MOLECULE: | MINOR FIMBRIUM ANCHORING SUBUNIT MF |
| 26:00:00 | 1bp1-A | 2.8 | 5.6 | 81 | 456 | 4 | MOLECULE: | BACTERICIDAL/PERMEABILITY-INCREASIN |
| 27:00:00 | 3sbu-A | 2.8 | 4.9 | 63 | 230 | 6 | MOLECULE: | HYPOTHETICAL NTF2-LIKE PROTEIN |
| 28:00:00 | 8sv7-A | 2.8 | 5 | 66 | 272 | 2 | MOLECULE: | TRANSPORT AND GOLGI ORGANIZATION PR |
| 29:00:00 | 7nyx-A | 2.8 | 4.8 | 63 | 1467 | 13 | MOLECULE: | CHROMOSOME PARTITION PROTEIN MUKB |
| 30:00:00 | 6p74-A | 2.8 | 11.2 | 59 | 512 | 8 | MOLECULE: | PUTATIVE ATP-DEPENDENT ENDONUCLEASE |
| 31:00:00 | 6tlb-A | 2.8 | 3.3 | 72 | 185 | 8 | MOLECULE: | SERINE/THREONINE PROTEIN KINASE |
| 32:00:00 | 5n6u-A | 2.8 | 4.5 | 51 | 811 | 6 | MOLECULE: | BETA-MANNOSIDASE |
| 33:00:00 | 4pux-A | 2.8 | 5.8 | 79 | 157 | 11 | MOLECULE: | UNCHARACTERIZED PROTEIN |
| 34:00:00 | 5tod-D | 2.8 | 8.1 | 70 | 157 | 4 | MOLECULE: | TRANSMEMBRANE PROTEIN 24 |

| | | | | | | | | |
|---|---|---|---|---|---|---|---|---|
| 35:00:00 | 8cli-C | 2.8 | 8.1 | 82 | 579 | 2 | MOLECULE: | GENERAL TRANSCRIPTION FACTOR 3C POL |
| 36:00:00 | 8oxh-A | 2.8 | 2.7 | 51 | 93 | 6 | MOLECULE: | AVRA6 |
| 37:00:00 | 8u7i-A | 2.8 | 11.2 | 73 | 626 | 4 | MOLECULE: | ENDONUCLEASE GAJA |
| 38:00:00 | 7c8z-B | 2.8 | 6 | 71 | 161 | 4 | MOLECULE: | SALICYLATE 5-HYDROXYLASE, LARGE OXY |
| 39:00:00 | 8shj-A | 2.8 | 5 | 65 | 334 | 3 | MOLECULE: | WD REPEAT-CONTAINING PROTEIN 91 |
| 40:00:00 | 5wzr-A | 2.8 | 8.4 | 69 | 630 | 3 | MOLECULE: | ALPHA-N-ACETYLGALACTOSAMINIDASE |
| 41:00:00 | 3e1e-C | 2.8 | 3.4 | 67 | 141 | 6 | MOLECULE: | THIOESTERASE FAMILY PROTEIN |
| 42:00:00 | 8ptx-A | 2.8 | 4.1 | 70 | 1215 | 9 | MOLECULE: | ELONGATOR COMPLEX PROTEIN 1 |
| 43:00:00 | 8iha-A | 2.8 | 4.6 | 56 | 129 | 4 | MOLECULE: | POLYKETIDE CYCLASE / DEHYDRASE AND |
| 44:00:00 | 2cg8-C | 2.8 | 8 | 70 | 248 | 3 | MOLECULE: | DIHYDRONEOPTERIN ALDOLASE 6-HYDROXY |
| 45:00:00 | 4ue0-A | 2.8 | 4.1 | 54 | 115 | 6 | MOLECULE: | FIBER |
| 46:00:00 | 3wy4-B | 2.8 | 7.4 | 53 | 421 | 4 | MOLECULE: | ALPHA-GLUCOSIDASE |
| 47:00:00 | 1jkg-A | 2.8 | 5.5 | 67 | 139 | 16 | MOLECULE: | P15 |
| 48:00:00 | 1he8-A | 2.8 | 6.3 | 66 | 749 | 5 | MOLECULE: | PHOSPHATIDYLINOSITOL 3-KINASE CATAL |
| 49:00:00 | 5hd6-A | 2.8 | 6.4 | 81 | 175 | 5 | MOLECULE: | 3-HYDROXYDECANOYL-[ACYL-CARRIER-PRO |
| 50:00:00 | 3msw-A | 2.8 | 6.9 | 71 | 139 | 3 | MOLECULE: | UNCHARACTERIZED PROTEIN |
| 51:00:00 | 2qiy-A | 2.8 | 4.3 | 59 | 134 | 8 | MOLECULE: | UBP3-ASSOCIATED PROTEIN BRE5 |
| 52:00:00 | 6w6m-A | 2.8 | 5.5 | 64 | 412 | 5 | MOLECULE: | TYPE IV PILUS SECRETIN PILQ FAMILY |
| 53:00:00 | 7l5m-A | 2.8 | 4 | 62 | 87 | 6 | MOLECULE: | LIPOCALIN FAMILY PROTEIN |
| 54:00:00 | 8vwp-A | 2.8 | 7.8 | 67 | 502 | 9 | MOLECULE: | LANGYA VIRUS ATTACHMENT (G) PROTEIN |
| 55:00:00 | 8wxf-A | 2.8 | 10.7 | 63 | 730 | 3 | MOLECULE: | BIFUNCTIONAL GUANOSINE PENTAPHOSPHA |
| 56:00:00 | 4rki-A | 2.8 | 6.9 | 53 | 374 | 4 | MOLECULE: | DNA POLYMERASE III SUBUNIT BETA |
| 57:00:00 | 6nd4-L | 2.8 | 5.3 | 58 | 473 | 3 | MOLECULE: | ETS RRNA |
| 58:00:00 | 7xxa-E | 2.8 | 2.5 | 48 | 144 | 4 | MOLECULE: | VP1 |
| 59:00:00 | 3q6k-B | 2.8 | 5.8 | 68 | 381 | 10 | MOLECULE: | 43.2 KDA SALIVARY PROTEIN |
| 60:00:00 | 7zty-A | 2.8 | 7.8 | 58 | 345 | 5 | MOLECULE: | CNH DOMAIN-CONTAINING PROTEIN |
| 61:00:00 | 6s9u-A | 2.8 | 3.6 | 64 | 520 | 5 | MOLECULE: | PUTATIVE SUCROSE PHOSPHORYLASE |
| 62:00:00 | 2ooj-B | 2.8 | 6.1 | 69 | 134 | 4 | MOLECULE: | HYPOTHETICAL PROTEIN |
| 63:00:00 | 6nd4-W | 2.8 | 4.6 | 61 | 385 | 2 | MOLECULE: | ETS RRNA |
| 64:00:00 | 6qhg-A | 2.8 | 5.6 | 52 | 119 | 10 | MOLECULE: | POLYMERASE |
| 65:00:00 | 5mnt-B | 2.8 | 6.1 | 74 | 421 | 4 | MOLECULE: | A2 MATURATION PROTEIN |
| 66:00:00 | 6xt9-I | 2.8 | 8.3 | 62 | 767 | 5 | MOLECULE: | BARDET-BIEDL SYNDROME 1 PROTEIN |
| 67:00:00 | 4r5o-A | 2.8 | 3.6 | 59 | 426 | 5 | MOLECULE: | QUINONPROTEIN ALCOHOL DEHYDROGENASE |
| 68:00:00 | 8r8q-B | 2.8 | 9.3 | 54 | 341 | 6 | MOLECULE: | MAJOR FACILITATOR SUPERFAMILY DOMAI |
| 69:00:00 | 8dit-B | 2.8 | 8 | 62 | 834 | 3 | MOLECULE: | VACUOLAR PROTEIN SORTING-ASSOCIATED |
| 70:00:00 | 5iwd-A | 2.8 | 2.9 | 53 | 261 | 9 | MOLECULE: | DNA POLYMERASE PROCESSIVITY FACTOR |
| 71:00:00 | 6ozd-B | 2.8 | 3.4 | 61 | 523 | 10 | MOLECULE: | PUTATIVE EXPORTED PROTEIN |
| 72:00:00 | 6eys-B | 2.8 | 4.4 | 71 | 478 | 6 | MOLECULE: | PVDP |
| 73:00:00 | 6mly-C | 2.8 | 3.6 | 73 | 774 | 7 | MOLECULE: | BIFUNCTIONAL GH43-CE PROTEIN |
| 74:00:00 | 3hrp-A | 2.8 | 4.2 | 61 | 400 | 7 | MOLECULE: | UNCHARACTERIZED PROTEIN |
| 75:00:00 | 4gwp-B | 2.8 | 3.6 | 73 | 492 | 5 | MOLECULE: | MEDIATOR OF RNA POLYMERASE II TRANS |
| 76:00:00 | 4exr-A | 2.8 | 3 | 52 | 152 | 0 | MOLECULE: | PUTATIVE LIPOPROTEIN |
| 77:00:00 | 7sfn-B | 2.8 | 3 | 68 | 153 | 4 | MOLECULE: | OLMO - OLIGOMYCIN SPIROCYCLASE |
| 78:00:00 | 8es5-A | 2.8 | 4.2 | 60 | 146 | 3 | MOLECULE: | ACTIVATOR OF HSP90 ATPASE |
| 79:00:00 | 7wmp-a | 2.8 | 4.7 | 99 | 559 | 5 | MOLECULE: | PORTAL PROTEIN |
| 80:00:00 | 6jq9-B | 2.8 | 3.9 | 79 | 483 | 5 | MOLECULE: | SHORT ULVAN LYASE |

| | | | | | | | | |
|---|---|---|---|---|---|---|---|---|
| 81:00:00 | 5n9j-A | 2.8 | 3.1 | 58 | 566 | 5 | MOLECULE: | MEDIATOR OF RNA POLYMERASE II TRANS |
| 82:00:00 | 8chw-A | 2.8 | 2.9 | 50 | 67 | 6 | MOLECULE: | TRANSCRIPTIONAL ACTIVATOR PROTEIN P |
| 83:00:00 | 5f75-C | 2.8 | 3.9 | 67 | 475 | 4 | MOLECULE: | THIOCYANATE DEHYDROGENASE |
| 84:00:00 | 3was-A | 2.8 | 4.5 | 69 | 389 | 9 | MOLECULE: | 4-O-BETA-D-MANNOSYL-D-GLUCOSE PHOSP |
| 85:00:00 | 3s5m-A | 2.8 | 8.8 | 58 | 1094 | 5 | MOLECULE: | FALCILYSIN |
| 86:00:00 | 5gqt-A | 2.8 | 8.8 | 63 | 470 | 3 | MOLECULE: | NITRILE-SPECIFIER PROTEIN 1 |
| 87:00:00 | 6rfy-D | 2.8 | 12 | 65 | 407 | 6 | MOLECULE: | EIS2 |
| 88:00:00 | 3w7s-A | 2.8 | 8.9 | 71 | 760 | 6 | MOLECULE: | UNCHARACTERIZED PROTEIN YGJK |
| 89:00:00 | 6vbv-7 | 2.8 | 4.4 | 74 | 706 | 5 | MOLECULE: | BARDET-BIEDL SYNDROME 18 PROTEIN |
| 90:00:00 | 7tom-A | 2.8 | 5.7 | 53 | 499 | 0 | MOLECULE: | GLYCEROL DIBIPHYTANYL GLYCEROL TETR |
| 91:00:00 | 7bgh-A | 2.8 | 9.7 | 80 | 177 | 3 | MOLECULE: | OUTER ENVELOPE PORE PROTEIN 21, CHL |
| 92:00:00 | 3s9j-A | 2.8 | 8.6 | 57 | 362 | 7 | MOLECULE: | MEMBER OF DUF4221 FAMILY |
| 93:00:00 | 4ew7-A | 2.8 | 4.8 | 52 | 113 | 2 | MOLECULE: | CONJUGATIVE TRANSFER: REGULATION |
| 94:00:00 | 8h8b-A | 2.8 | 12.5 | 68 | 1027 | 6 | MOLECULE: | PUTATIVE RHS-FAMILY PROTEIN |
| 95:00:00 | 2p4f-A | 2.8 | 3.8 | 62 | 191 | 8 | MOLECULE: | SIMILAR TO SP\|P32453 SACCHAROMYCES |
| 96:00:00 | 8y6p-Q | 2.8 | 7.7 | 65 | 1228 | 2 | MOLECULE: | APAF-1 RELATED KILLER DARK |
| 97:00:00 | 4xi5-A | 2.8 | 9.8 | 67 | 728 | 3 | MOLECULE: | ENVELOPE GLYCOPROTEIN H |
| 98:00:00 | 3nqh-A | 2.8 | 3.5 | 54 | 439 | 6 | MOLECULE: | GLYCOSYL HYDROLASE |
| 99:00:00 | 4bp8-A | 2.8 | 5.6 | 62 | 712 | 10 | MOLECULE: | OLIGOPEPTIDASE B |
| 0:00 | 1xks-A | 2.8 | 4.6 | 73 | 374 | 7 | MOLECULE: | NUCLEAR PORE COMPLEX PROTEIN NUP133 |
| 1:00 | 5bka-A | 2.8 | 5.3 | 67 | 115 | 12 | MOLECULE: | HYDROXYLACYL-COA DEHYDROGENASE |
| 2:00 | 3eby-A | 2.8 | 5.8 | 68 | 153 | 7 | MOLECULE: | BETA SUBUNIT OF A PUTATIVE AROMATIC |
| 3:00 | 5tph-A | 2.8 | 5.7 | 61 | 120 | 10 | MOLECULE: | DE NOVO NTF2 HOMODIMER |
| 4:00 | 7pkq-L | 2.8 | 4.5 | 72 | 175 | 7 | MOLECULE: | MS35 |
| 5:00 | 5i97-A | 2.8 | 5.9 | 64 | 143 | 8 | MOLECULE: | CONJUGAL TRANSFER PROTEIN |
| 6:00 | 8h2t-H | 2.8 | 5.5 | 71 | 157 | 7 | MOLECULE: | RIESKE (2FE-2S) DOMAIN PROTEIN |
| 7:00 | 6zgn-A | 2.8 | 5 | 58 | 101 | 3 | MOLECULE: | PUTATIVE TRANSFER PROTEIN |
| 8:00 | 8f5p-E | 2.8 | 5.8 | 60 | 1057 | 3 | MOLECULE: | NET DOMAIN-CONTAINING PROTEIN |
| 9:00 | 5o65-C | 2.8 | 7.3 | 94 | 278 | 4 | MOLECULE: | FAPF |
| 10:00 | 5o68-G | 2.8 | 5.8 | 86 | 271 | 6 | MOLECULE: | FAPF |
| 11:00 | 5d17-G | 2.8 | 3.3 | 61 | 158 | 5 | MOLECULE: | TRANSPOSON TN7 TRANSPOSITION PROTEI |
| 12:00 | 5wj3-B | 2.8 | 8.4 | 55 | 225 | 5 | MOLECULE: | GREEN FLUORESCENT PROTEIN |
| 13:00 | 1rrx-A | 2.8 | 8.9 | 58 | 225 | 5 | MOLECULE: | SIGF1-GFP FUSION PROTEIN |
| 14:00 | 6xu4-A | 2.8 | 6.2 | 82 | 389 | 2 | MOLECULE: | FGCAMP |
| 15:00 | 3osr-A | 2.8 | 6.3 | 82 | 603 | 2 | MOLECULE: | MALTOSE-BINDING PERIPLASMIC PROTEIN |
| 16:00 | 4w6r-I | 2.8 | 9 | 52 | 209 | 10 | MOLECULE: | FLUORESCENT PROTEIN D102C |
| 17:00 | 4w75-B | 2.8 | 8 | 50 | 187 | 8 | MOLECULE: | FLUORESCENT PROTEIN D21H/K26C |
| 18:00 | 2gu3-A | 2.7 | 3.3 | 49 | 128 | 10 | MOLECULE: | YPMB PROTEIN |
| 19:00 | 2gey-A | 2.7 | 5.9 | 71 | 157 | 7 | MOLECULE: | ACLR PROTEIN |
| 20:00 | 3cyg-A | 2.7 | 5.9 | 66 | 215 | 3 | MOLECULE: | UNCHARACTERIZED PROTEIN |
| 21:00 | 1lqm-H | 2.7 | 3.5 | 51 | 84 | 12 | MOLECULE: | URACIL-DNA GLYCOSYLASE |
| 22:00 | 2ffs-A | 2.7 | 2.9 | 63 | 146 | 3 | MOLECULE: | HYPOTHETICAL PROTEIN PA1206 |
| 23:00 | 1u5o-A | 2.7 | 5.9 | 62 | 125 | 10 | MOLECULE: | NUCLEAR TRANSPORT FACTOR 2 |
| 24:00:00 | 7eu3-7 | 2.7 | 3.9 | 69 | 317 | 6 | MOLECULE: | NAD(P)H-QUINONE OXIDOREDUCTASE SUBU |
| 25:00:00 | 4ffg-A | 2.7 | 4.4 | 68 | 480 | 7 | MOLECULE: | LEVAN FRUCTOTRANSFERASE |
| 26:00:00 | 8hqo-a | 2.7 | 3.4 | 60 | 160 | 8 | MOLECULE: | PORTAL PROTEIN |

| | | | | | | | | |
|---|---|---|---|---|---|---|---|---|
| 27:00:00 | 5mtw-A | 2.7 | 4.2 | 67 | 138 | 6 | MOLECULE: | SECB-LIKE CHAPERONE RV1957 |
| 28:00:00 | 1vl4-A | 2.7 | 10.7 | 87 | 428 | 11 | MOLECULE: | PMBA-RELATED PROTEIN |
| 29:00:00 | 8dk3-A | 2.7 | 9.3 | 64 | 455 | 3 | MOLECULE: | JETC |
| 30:00:00 | 7cpx-A | 2.7 | 9.3 | 78 | 2262 | 5 | MOLECULE: | LOVASTATIN NONAKETIDE SYNTHASE, POL |
| 31:00:00 | 6r5w-A | 2.7 | 6 | 66 | 171 | 3 | MOLECULE: | GP15 PROTEIN |
| 32:00:00 | 6stx-C | 2.7 | 4.9 | 61 | 663 | 11 | MOLECULE: | KELCH DOMAIN-CONTAINING PROTEIN |
| 33:00:00 | 2ogf-B | 2.7 | 4.4 | 60 | 122 | 2 | MOLECULE: | HYPOTHETICAL PROTEIN MJ0408 |
| 34:00:00 | 6y6k-A | 2.7 | 4 | 58 | 1299 | 10 | MOLECULE: | RNA-DEPENDENT RNA POLYMERASE |
| 35:00:00 | 3f7s-A | 2.7 | 4.4 | 69 | 142 | 6 | MOLECULE: | UNCHARACTERIZED NTF2-LIKE PROTEIN |
| 36:00:00 | 1nbu-A | 2.7 | 4.3 | 62 | 118 | 5 | MOLECULE: | PROBABLE DIHYDRONEOPTERIN ALDOLASE |
| 37:00:00 | 8tnn-A | 2.7 | 11.2 | 64 | 658 | 5 | MOLECULE: | ENVELOPE GLYCOPROTEIN H |
| 38:00:00 | 3c1a-A | 2.7 | 4.9 | 62 | 307 | 5 | MOLECULE: | PUTATIVE OXIDOREDUCTASE |
| 39:00:00 | 4ghb-B | 2.7 | 5 | 63 | 257 | 3 | MOLECULE: | HYPOTHETICAL PROTEIN |
| 40:00:00 | 2nyk-A | 2.7 | 11.5 | 67 | 237 | 9 | MOLECULE: | M157 |
| 41:00:00 | 6mlt-A | 2.7 | 7.1 | 60 | 603 | 3 | MOLECULE: | HEMOLYSIN-RELATED PROTEIN |
| 42:00:00 | 5hi8-A | 2.7 | 3.8 | 53 | 136 | 4 | MOLECULE: | ANTENNA PROTEIN |
| 43:00:00 | 1oac-B | 2.7 | 4.4 | 88 | 723 | 3 | MOLECULE: | COPPER AMINE OXIDASE |
| 44:00:00 | 4r03-A | 2.7 | 3.9 | 65 | 109 | 5 | MOLECULE: | UNCHARACTERIZED PROTEIN |
| 45:00:00 | 8i0a-A | 2.7 | 6.9 | 67 | 524 | 6 | MOLECULE: | ALPHA-L-ARABINOFURANOSIDASE |
| 46:00:00 | 4p5i-A | 2.7 | 6.5 | 69 | 240 | 12 | MOLECULE: | CHEMOKINE BINDING PROTEIN |
| 47:00:00 | 8bsh-C | 2.7 | 5.6 | 68 | 794 | 9 | MOLECULE: | PROTEIN TRANSPORT PROTEIN SEC23 |
| 48:00:00 | 3dm8-A | 2.7 | 6.6 | 66 | 135 | 6 | MOLECULE: | UNCHARACTERIZED PROTEIN RPA4348 |
| 49:00:00 | 2vnv-B | 2.7 | 3.1 | 60 | 128 | 3 | MOLECULE: | BCLA |
| 50:00:00 | 1b9l-A | 2.7 | 5.1 | 63 | 119 | 2 | MOLECULE: | PROTEIN (EPIMERASE) |
| 51:00:00 | 7vma-A | 2.7 | 3.5 | 67 | 620 | 4 | MOLECULE: | AMYLO-ALPHA-1,6-GLUCOSIDASE, PUTATI |
| 52:00:00 | 8hmc-B | 2.7 | 3.5 | 61 | 1195 | 7 | MOLECULE: | INTRAFLAGELLAR TRANSPORT PROTEIN 12 |
| 53:00:00 | 6s8f-F | 2.7 | 6.1 | 62 | 2571 | 3 | MOLECULE: | SERINE/THREONINE-PROTEIN KINASE TEL |
| 54:00:00 | 5k8g-A | 2.7 | 5.6 | 64 | 511 | 5 | MOLECULE: | ANTIFREEZE PROTEIN |
| 55:00:00 | 3er7-B | 2.7 | 4.8 | 67 | 123 | 6 | MOLECULE: | UNCHARACTERIZED NTF2-LIKE PROTEIN |
| 56:00:00 | 6nnw-A | 2.7 | 8.2 | 74 | 208 | 7 | MOLECULE: | TETRONASIN |
| 57:00:00 | 6v04-A | 2.7 | 5 | 71 | 268 | 4 | MOLECULE: | UNCHARACTERIZED SRPBCC DOMAIN-CONTA |
| 58:00:00 | 3slj-A | 2.7 | 10.2 | 98 | 305 | 7 | MOLECULE: | SERINE PROTEASE ESPP |
| 59:00:00 | 7fse-A | 2.7 | 7.8 | 55 | 1040 | 15 | MOLECULE: | REVERSE GYRASE |
| 60:00:00 | 3alx-B | 2.7 | 4.3 | 58 | 526 | 2 | MOLECULE: | HEMAGGLUTININ,LINKER,CDW150 |
| 61:00:00 | 4l3a-A | 2.7 | 3.4 | 56 | 469 | 4 | MOLECULE: | INTERNALIN K |
| 62:00:00 | 6sum-A | 2.7 | 11.2 | 49 | 335 | 10 | MOLECULE: | AMICOUMACIN KINASE |
| 63:00:00 | 6kko-A | 2.7 | 4.2 | 63 | 170 | 8 | MOLECULE: | PUTATIVE SERINE PHOSPHATASE |
| 64:00:00 | 5jzr-A | 2.7 | 6.5 | 74 | 131 | 9 | MOLECULE: | COAT PROTEIN |
| 65:00:00 | 5hzv-A | 2.7 | 5.1 | 66 | 598 | 8 | MOLECULE: | MALTOSE-BINDING PERIPLASMIC PROTEIN |
| 66:00:00 | 8aim-B | 2.7 | 4.1 | 52 | 86 | 4 | MOLECULE: | URACIL-DNA GLYCOSYLASE INHIBITOR |
| 67:00:00 | 5d9g-A | 2.7 | 4 | 80 | 246 | 1 | MOLECULE: | TIP41-LIKE PROTEIN |
| 68:00:00 | 2in5-A | 2.7 | 4.8 | 69 | 195 | 10 | MOLECULE: | HYPOTHETICAL LIPOPROTEIN YMCC |
| 69:00:00 | 4v1a-i | 2.7 | 3.6 | 70 | 242 | 6 | MOLECULE: | MITORIBOSOMAL PROTEIN ML37, MRPL37 |
| 70:00:00 | 7p8s-A | 2.7 | 3.6 | 71 | 293 | 11 | MOLECULE: | LEUCOTOXIN LUKEV |
| 71:00:00 | 7ohi-A | 2.7 | 6.7 | 67 | 250 | 4 | MOLECULE: | AP-2 COMPLEX SUBUNIT ALPHA-2 |
| 72:00:00 | 7okx-M | 2.7 | 10.3 | 56 | 139 | 5 | MOLECULE: | DNA-DIRECTED RNA POLYMERASE II SUBU |

| | | | | | | | | |
|---|---|---|---|---|---|---|---|---|
| 73:00:00 | 3n7n-B | 2.7 | 5.8 | 58 | 164 | 3 | MOLECULE: | MONOPOLIN COMPLEX SUBUNIT CSM1 |
| 74:00:00 | 7eni-C | 2.7 | 3.3 | 59 | 126 | 7 | MOLECULE: | CRISPR-ASSOCIATED ENDONUCLEASE CAS9 |
| 75:00:00 | 3klq-A | 2.7 | 4.9 | 70 | 137 | 4 | MOLECULE: | PUTATIVE PILUS ANCHORING PROTEIN |
| 76:00:00 | 4u3v-A | 2.7 | 6.1 | 67 | 242 | 3 | MOLECULE: | POLYKETIDE SYNTHASE PKSR |
| 77:00:00 | 7erl-A | 2.7 | 5.2 | 69 | 545 | 6 | MOLECULE: | BETA-XYLANASE |
| 78:00:00 | 2hqs-A | 2.7 | 3.6 | 55 | 412 | 13 | MOLECULE: | PROTEIN TOLB |
| 79:00:00 | 6h6g-A | 2.7 | 5.1 | 61 | 2139 | 2 | MOLECULE: | TCDB2,TCCC3 |
| 80:00:00 | 1qjs-A | 2.7 | 5.8 | 60 | 408 | 5 | MOLECULE: | HEMOPEXIN |
| 81:00:00 | 8h36-A | 2.7 | 7.2 | 69 | 567 | 6 | MOLECULE: | E3 UBIQUITIN-PROTEIN LIGASE RBX1 |
| 82:00:00 | 5ts4-B | 2.7 | 4.6 | 57 | 106 | 7 | MOLECULE: | DENOVO NTF2 |
| 83:00:00 | 8a3t-C | 2.7 | 5.9 | 65 | 1410 | 8 | MOLECULE: | ANAPHASE-PROMOTING COMPLEX SUBUNIT |
| 84:00:00 | 4mob-A | 2.7 | 3 | 64 | 319 | 11 | MOLECULE: | ACYL-COENZYME A THIOESTERASE 12 |
| 85:00:00 | 6wil-A | 2.7 | 8.1 | 93 | 536 | 6 | MOLECULE: | HEMOLYSIN ACTIVATOR PROTEIN CDIB |
| 86:00:00 | 6goc-A | 2.7 | 5.5 | 70 | 444 | 9 | MOLECULE: | DUF3826 DOMAIN-CONTAINING PROTEIN |
| 87:00:00 | 7to3-A | 2.7 | 11.1 | 73 | 568 | 4 | MOLECULE: | CAP2 |
| 88:00:00 | 7dna-A | 2.7 | 8.4 | 55 | 213 | 5 | MOLECULE: | GREEN-TO-RED PHOTOCONVERTIBLE GFP-L |
| 89:00:00 | 7x2n-A | 2.7 | 4.4 | 66 | 340 | 3 | MOLECULE: | PYCR1 |
| 90:00:00 | 6vxk-B | 2.7 | 5.2 | 69 | 747 | 7 | MOLECULE: | SEMAPHORIN-LIKE PROTEIN 139 |
| 91:00:00 | 2z5b-A | 2.7 | 4.4 | 41 | 127 | 10 | MOLECULE: | PROTEIN YPL144W |
| 92:00:00 | 1sa8-A | 2.7 | 3.3 | 59 | 106 | 5 | MOLECULE: | FATTY ACID-BINDING PROTEIN, INTESTI |
| 93:00:00 | 6wxu-B | 2.7 | 3.9 | 58 | 273 | 3 | MOLECULE: | DUAL OXIDASE 1 |
| 94:00:00 | 6jcy-C | 2.7 | 5.3 | 62 | 1139 | 6 | MOLECULE: | DNA-DIRECTED RNA POLYMERASE SUBUNIT |
| 95:00:00 | 3ats-A | 2.7 | 3.8 | 57 | 352 | 18 | MOLECULE: | PUTATIVE UNCHARACTERIZED PROTEIN |
| 96:00:00 | 6vs7-A | 2.7 | 3.2 | 54 | 409 | 9 | MOLECULE: | ADHESIN |
| 97:00:00 | 7eeb-E | 2.7 | 4.6 | 64 | 1053 | 5 | MOLECULE: | ENHANCED GREEN FLUORESCENT PROTEIN, |
| 98:00:00 | 8axs-A | 2.7 | 3.4 | 73 | 573 | 3 | MOLECULE: | EXO-ALPHA-SIALIDASE |
| 99:00:00 | 7lxf-A | 2.7 | 3.1 | 56 | 134 | 9 | MOLECULE: | PROTEIN ENABLED HOMOLOG,PHOTORECEPT |
| 0:00 | 7mjp-A | 2.7 | 4.7 | 59 | 366 | 5 | MOLECULE: | ATP-SENSITIVE INWARD RECTIFIER POTA |
| 1:00 | 8c7i-A | 2.7 | 8.4 | 55 | 404 | 11 | MOLECULE: | GREEN FLUORESCENT PROTEIN,TLL0404 P |
| 2:00 | 3oan-A | 2.7 | 3.3 | 51 | 116 | 0 | MOLECULE: | ABR034WP |
| 3:00 | 8h7g-C | 2.7 | 5.3 | 68 | 3261 | 4 | MOLECULE: | TRANSFORMATION/TRANSCRIPTION DOMAIN |
| 4:00 | 2j04-A | 2.7 | 6 | 70 | 587 | 10 | MOLECULE: | HYPOTHETICAL PROTEIN YPL007C |
| 5:00 | 3kkg-A | 2.7 | 5 | 68 | 144 | 12 | MOLECULE: | PUTATIVE SNOAL-LIKE POLYKETIDE CYCL |
| 6:00 | 3s25-A | 2.7 | 4.6 | 54 | 295 | 9 | MOLECULE: | HYPOTHETICAL 7-BLADED BETA-PROPELLE |
| 7:00 | 5a4o-A | 2.7 | 3.3 | 47 | 74 | 2 | MOLECULE: | BPSL1147 |
| 8:00 | 8env-a | 2.7 | 3.1 | 49 | 102 | 2 | MOLECULE: | SHEATH PROTEIN GP31 |
| 9:00 | 6dj8-A | 2.7 | 11.8 | 62 | 384 | 8 | MOLECULE: | BETA SLIDING CLAMP |
| 10:00 | 8us3-A | 2.7 | 5.9 | 88 | 552 | 10 | MOLECULE: | TRANSLOCATION AND ASSEMBLY MODULE S |
| 11:00 | 6p29-A | 2.7 | 4 | 49 | 134 | 8 | MOLECULE: | N-DEMETHYLINDOLMYCIN SYNTHASE (PLUN |
| 12:00 | 3ehc-B | 2.7 | 4.1 | 62 | 128 | 2 | MOLECULE: | SNOAL-LIKE POLYKETIDE CYCLASE |
| 13:00 | 4w4l-C | 2.7 | 5 | 60 | 297 | 7 | MOLECULE: | PE FAMILY PROTEIN PE25 |
| 14:00 | 3azo-A | 2.7 | 6.1 | 83 | 661 | 6 | MOLECULE: | AMINOPEPTIDASE |
| 15:00 | 2yzy-A | 2.7 | 5.2 | 55 | 163 | 2 | MOLECULE: | PUTATIVE UNCHARACTERIZED PROTEIN TT |
| 16:00 | 4ktp-A | 2.7 | 8.2 | 68 | 767 | 7 | MOLECULE: | GLYCOSIDE HYDROLASE FAMILY 65 CENTR |
| 17:00 | 1k32-A | 2.7 | 4.5 | 60 | 1023 | 3 | MOLECULE: | TRICORN PROTEASE |
| 18:00 | 3i2n-A | 2.7 | 5.3 | 68 | 343 | 10 | MOLECULE: | WD REPEAT-CONTAINING PROTEIN 92 |

| | | | | | | | | |
|---|---|---|---|---|---|---|---|---|
| 19:00 | 5hx0-A | 2.7 | 7.8 | 64 | 365 | 6 | MOLECULE: | UNCHARACTERIZED PROTEIN DFER_1899 |
| 20:00 | 7emf-P | 2.7 | 6 | 74 | 766 | 14 | MOLECULE: | MEDIATOR OF RNA POLYMERASE II TRANS |
| 21:00 | 7qrd-A | 2.7 | 3.6 | 64 | 364 | 8 | MOLECULE: | HISTONE-ARGININE METHYLTRANSFERASE |
| 22:00 | 2ojh-A | 2.7 | 5 | 54 | 277 | 7 | MOLECULE: | UNCHARACTERIZED PROTEIN ATU1656/AGR |
| 23:00 | 7v2w-H | 2.7 | 6.9 | 71 | 379 | 4 | MOLECULE: | THO COMPLEX SUBUNIT HPR1 |
| 24:00:00 | 7z4w-A | 2.7 | 4.3 | 74 | 447 | 4 | MOLECULE: | PORTAL PROTEIN |
| 25:00:00 | 1yli-B | 2.7 | 4.2 | 65 | 147 | 9 | MOLECULE: | PUTATIVE ACYL-COA THIOESTER HYDROLA |
| 26:00:00 | 4gxb-A | 2.7 | 6 | 69 | 263 | 9 | MOLECULE: | SORTING NEXIN-17 |
| 27:00:00 | 6icz-A | 2.7 | 9.3 | 71 | 2253 | 8 | MOLECULE: | PROTEIN MAGO NASHI HOMOLOG 2 |
| 28:00:00 | 4o3v-A | 2.7 | 4.4 | 61 | 138 | 5 | MOLECULE: | VIRB8-LIKE PROTEIN OF TYPE IV SECRE |
| 29:00:00 | 6fej-B | 2.7 | 4 | 60 | 118 | 5 | MOLECULE: | ALL4940 PROTEIN |
| 30:00:00 | 3d9r-A | 2.7 | 4.8 | 66 | 132 | 14 | MOLECULE: | KETOSTEROID ISOMERASE-LIKE PROTEIN |
| 31:00:00 | 6isk-B | 2.7 | 4 | 62 | 120 | 11 | MOLECULE: | XIME, SNOAL-LIKE DOMAIN PROTEIN |
| 32:00:00 | 5o7s-B | 2.7 | 2.9 | 56 | 90 | 0 | MOLECULE: | MONELLIN CHAIN B,MONELLIN CHAIN A |
| 33:00:00 | 4q9t-A | 2.7 | 4.9 | 62 | 384 | 6 | MOLECULE: | NUCLEOPORIN NUP133 |
| 34:00:00 | 8abt-A | 2.7 | 4.2 | 73 | 235 | 1 | MOLECULE: | SNOAL-LIKE DOMAIN-CONTAINING PROTEI |
| 35:00:00 | 5o68-E | 2.7 | 6.3 | 88 | 273 | 7 | MOLECULE: | FAPF |
| 36:00:00 | 6og8-B | 2.7 | 6.8 | 60 | 227 | 3 | MOLECULE: | CRYSTAL STRUCTURE OF GREEN FLUORESC |
| 37:00:00 | 6zsm-A | 2.7 | 7 | 78 | 396 | 4 | MOLECULE: | GREEN FLUORESCENT PROTEIN,GREEN FLU |
| 38:00:00 | 3sg4-A | 2.7 | 7.1 | 78 | 394 | 4 | MOLECULE: | MYOSIN LIGHT CHAIN KINASE, GREEN FL |
| 39:00:00 | 7lug-A | 2.7 | 6 | 75 | 223 | 5 | MOLECULE: | RED FLUORESCENT PNRFP B30Y MUTANT |
| 40:00:00 | 5d17-D | 2.7 | 3.3 | 61 | 157 | 5 | MOLECULE: | TRANSPOSON TN7 TRANSPOSITION PROTEI |
| 41:00:00 | 4w6k-B | 2.7 | 3.4 | 51 | 182 | 4 | MOLECULE: | FLUORESCENT PROTEIN D117C |
| 42:00:00 | 6uhr-A | 2.7 | 10.4 | 58 | 222 | 12 | MOLECULE: | C148 MGFP-SCDNA-2 |
| 43:00:00 | 6hr1-B | 2.7 | 8.3 | 55 | 401 | 4 | MOLECULE: | MYOSIN LIGHT CHAIN KINASE 2, SKELET |
| 44:00:00 | 7cd7-D | 2.7 | 8.4 | 55 | 235 | 11 | MOLECULE: | GREEN FLUORESCENT PROTEIN |
| 45:00:00 | 4w6p-F | 2.7 | 10.3 | 57 | 214 | 9 | MOLECULE: | FLUORESCENT PROTEIN D102C |
| 46:00:00 | 8yq4-A | 2.7 | 8.6 | 54 | 213 | 2 | MOLECULE: | MBAOJIN2 - BRIGHT AND STABLE GREEN |
| 47:00:00 | 4jfg-F | 2.7 | 10.2 | 55 | 217 | 13 | MOLECULE: | GREEN FLUORESCENT PROTEIN |
| 48:00:00 | 3ztf-A | 2.7 | 8.4 | 55 | 236 | 11 | MOLECULE: | GREEN FLUORESCENT PROTEIN |
| 49:00:00 | 8uzh-B | 2.6 | 3.3 | 52 | 626 | 4 | MOLECULE: | SUMO FUSED TREHALOSE SYNTHASE (TRES |
| 50:00:00 | 3hzp-A | 2.6 | 5.2 | 67 | 128 | 7 | MOLECULE: | NTF2-LIKE PROTEIN OF UNKNOWN FUNCTI |
| 51:00:00 | 4eae-B | 2.6 | 4.1 | 65 | 188 | 8 | MOLECULE: | LMO1068 PROTEIN |
| 52:00:00 | 7ty0-C | 2.6 | 6.7 | 67 | 488 | 3 | MOLECULE: | GLYCOPROTEIN G |
| 53:00:00 | 6cvz-A | 2.6 | 5.6 | 66 | 337 | 6 | MOLECULE: | E3 UBIQUITIN-PROTEIN LIGASE RFWD3 |
| 54:00:00 | 4mp6-A | 2.6 | 4.9 | 81 | 334 | 2 | MOLECULE: | PUTATIVE ORNITHINE CYCLODEAMINASE |
| 55:00:00 | 2chc-A | 2.6 | 6.1 | 74 | 169 | 5 | MOLECULE: | PROTEIN RV3472 |
| 56:00:00 | 8fcj-I | 2.6 | 9.5 | 74 | 494 | 1 | MOLECULE: | TYPE I-B CRISPR-ASSOCIATED PROTEIN |
| 57:00:00 | 1z94-B | 2.6 | 4.2 | 71 | 144 | 3 | MOLECULE: | CONSERVED HYPOTHETICAL PROTEIN |
| 58:00:00 | 2rfr-A | 2.6 | 5.3 | 71 | 154 | 7 | MOLECULE: | UNCHARACTERIZED PROTEIN |
| 59:00:00 | 6dfl-A | 2.6 | 10.2 | 58 | 241 | 3 | MOLECULE: | LIPOPOLYSACCHARIDE CORE HEPTOSE(I) |
| 60:00:00 | 5fw5-A | 2.6 | 4.7 | 62 | 139 | 8 | MOLECULE: | RAS GTPASE-ACTIVATING PROTEIN-BINDI |
| 61:00:00 | 7ojn-L | 2.6 | 7.3 | 93 | 2010 | 5 | MOLECULE: | RNA-DIRECTED RNA POLYMERASE L |
| 62:00:00 | 3lyd-B | 2.6 | 3.2 | 59 | 160 | 3 | MOLECULE: | UNCHARACTERIZED PROTEIN |
| 63:00:00 | 6nob-A | 2.6 | 6.5 | 76 | 634 | 3 | MOLECULE: | BETA-FRUCTOFURANOSIDASE |
| 64:00:00 | 3ef8-A | 2.6 | 5.2 | 70 | 149 | 6 | MOLECULE: | PUTATIVE SCYALONE DEHYDRATASE |

| | | | | | | | | |
|---|---|---|---|---|---|---|---|---|
| 65:00:00 | 1fw9-A | 2.6 | 2.9 | 62 | 164 | 8 | MOLECULE: | CHORISMATE LYASE |
| 66:00:00 | 6mit-G | 2.6 | 6.9 | 64 | 355 | 3 | MOLECULE: | LIPOPOLYSACCHARIDE EXPORT SYSTEM AT |
| 67:00:00 | 4zv3-B | 2.6 | 3.5 | 63 | 300 | 3 | MOLECULE: | CYTOSOLIC ACYL COENZYME A THIOESTER |
| 68:00:00 | 3b18-A | 2.6 | 4.4 | 72 | 167 | 7 | MOLECULE: | UNCHARACTERIZED PROTEIN RV0098/MT01 |
| 69:00:00 | 6itd-A | 2.6 | 5 | 65 | 328 | 5 | MOLECULE: | SLR0355 PROTEIN |
| 70:00:00 | 8jyx-A | 2.6 | 6.2 | 90 | 565 | 6 | MOLECULE: | MALTODEXTRIN-BINDING PROTEIN,GASDER |
| 71:00:00 | 7nie-C | 2.6 | 6.1 | 68 | 294 | 7 | MOLECULE: | GLYCEROL KINASE |
| 72:00:00 | 8u01-A | 2.6 | 5.2 | 56 | 909 | 2 | MOLECULE: | GLYCOSYL HYDROLASE FAMILY 2, SUGAR |
| 73:00:00 | 8suc-A | 2.6 | 5.1 | 62 | 282 | 10 | MOLECULE: | NHL (RING FINGER B-BOX COILED COIL) |
| 74:00:00 | 7alc-C | 2.6 | 4.4 | 62 | 162 | 3 | MOLECULE: | GTP CYCLOHYDROLASE 1 |
| 75:00:00 | 3db2-A | 2.6 | 6.2 | 71 | 347 | 3 | MOLECULE: | PUTATIVE NADPH-DEPENDENT OXIDOREDUC |
| 76:00:00 | 4r7k-A | 2.6 | 5.8 | 74 | 168 | 4 | MOLECULE: | HYPOTHETICAL PROTEIN JHP0584 |
| 77:00:00 | 6xux-A | 2.6 | 9 | 72 | 879 | 6 | MOLECULE: | NANOBODY,GLUCOSIDASE YGJK,GLUCOSIDA |
| 78:00:00 | 7s9w-B | 2.6 | 5.6 | 68 | 529 | 6 | MOLECULE: | DRMA |
| 79:00:00 | 7wug-4 | 2.6 | 6 | 69 | 230 | 10 | MOLECULE: | VACUOLAR IMPORT AND DEGRADATION PRO |
| 80:00:00 | 6e3k-E | 2.6 | 3.1 | 58 | 214 | 3 | MOLECULE: | INTERFERON GAMMA |
| 81:00:00 | 8h0h-A | 2.6 | 8.6 | 75 | 145 | 8 | MOLECULE: | UNCHARACTERIZED PROTEIN RV1546 |
| 82:00:00 | 7e2c-K | 2.6 | 4.7 | 71 | 251 | 1 | MOLECULE: | TRAPP-ASSOCIATED PROTEIN TCA17 |
| 83:00:00 | 8adl-C | 2.6 | 4.5 | 68 | 782 | 4 | MOLECULE: | MTC5 ISOFORM 1 |
| 84:00:00 | 3g16-A | 2.6 | 4.4 | 60 | 156 | 2 | MOLECULE: | UNCHARACTERIZED PROTEIN WITH CYSTAT |
| 85:00:00 | 3t8n-B | 2.6 | 4.5 | 72 | 130 | 8 | MOLECULE: | STEROID DELTA-ISOMERASE |
| 86:00:00 | 1zc4-B | 2.6 | 3.7 | 57 | 113 | 5 | MOLECULE: | RAS-RELATED PROTEIN RAL-A |
| 87:00:00 | 4boe-A | 2.6 | 4.3 | 58 | 154 | 9 | MOLECULE: | JAPANIN |
| 88:00:00 | 2fs2-B | 2.6 | 4.2 | 73 | 138 | 7 | MOLECULE: | PHENYLACETIC ACID DEGRADATION PROTE |
| 89:00:00 | 5img-A | 2.6 | 7.6 | 60 | 467 | 8 | MOLECULE: | DIPEPTIDASE |
| 90:00:00 | 8iex-A | 2.6 | 4.8 | 54 | 88 | 4 | MOLECULE: | PROBABLE WRKY TRANSCRIPTION FACTOR |
| 91:00:00 | 6xlf-C | 2.6 | 4.5 | 70 | 316 | 11 | MOLECULE: | ATP-DEPENDENT MOLECULAR CHAPERONE H |
| 92:00:00 | 8dek-A | 2.6 | 4.8 | 69 | 440 | 3 | MOLECULE: | NPCBM/NEW2 DOMAIN-CONTAINING PROTEI |
| 93:00:00 | 4gb5-A | 2.6 | 5.5 | 72 | 148 | 6 | MOLECULE: | UNCHARACTERIZED PROTEIN |
| 94:00:00 | 4rzm-B | 2.6 | 5.1 | 64 | 283 | 0 | MOLECULE: | EPOXIDE HYDROLASE LASB |
| 95:00:00 | 8ath-A | 2.6 | 3.9 | 68 | 168 | 7 | MOLECULE: | LYSOSOME-ASSOCIATED MEMBRANE GLYCOP |
| 96:00:00 | 5iri-A | 2.6 | 3 | 52 | 105 | 8 | MOLECULE: | SERINE/THREONINE-PROTEIN KINASE BRS |
| 97:00:00 | 7tpr-A | 2.6 | 10 | 75 | 1145 | 7 | MOLECULE: | SPIKE GLYCOPROTEIN |
| 98:00:00 | 7aqc-R | 2.6 | 3.6 | 73 | 558 | 5 | MOLECULE: | 23S RIBOSOMAL RNA |
| 99:00:00 | 7pkw-A | 2.6 | 8.4 | 70 | 240 | 6 | MOLECULE: | PUTATIVE TRANSFER PROTEIN |
| 0:00 | 7jsr-A | 2.6 | 4.6 | 66 | 1496 | 9 | MOLECULE: | NAD-SPECIFIC GLUTAMATE DEHYDROGENAS |
| 1:00 | 6gc1-C | 2.6 | 4.6 | 68 | 551 | 6 | MOLECULE: | NHL REPEAT-CONTAINING PROTEIN 2 |
| 2:00 | 6lwz-A | 2.6 | 3.9 | 44 | 53 | 9 | MOLECULE: | BACTERIOCIN |
| 3:00 | 8e5f-A | 2.6 | 9.6 | 73 | 293 | 10 | MOLECULE: | C-TYPE CYTOCHROME |
| 4:00 | 2ret-A | 2.6 | 3.4 | 53 | 84 | 6 | MOLECULE: | PSEUDOPILIN EPSI |
| 5:00 | 6ppb-k | 2.6 | 3.9 | 50 | 344 | 10 | MOLECULE: | CAPSID VERTEX COMPONENT 1 |
| 6:00 | 8veh-C | 2.6 | 4.9 | 55 | 194 | 5 | MOLECULE: | OUTER MEMBRANE LIPOPROTEIN-SORTING |
| 7:00 | 6gez-B | 2.6 | 7 | 68 | 532 | 3 | MOLECULE: | GREEN FLUORESCENT PROTEIN,OPTIMIZED |
| 8:00 | 3adj-A | 2.6 | 3.1 | 48 | 72 | 2 | MOLECULE: | F21M12.9 PROTEIN |
| 9:00 | 3sn0-A | 2.6 | 2.4 | 46 | 401 | 4 | MOLECULE: | PUTATIVE L-ALANINE-DL-GLUTAMATE EPI |
| 10:00 | 6dzk-Y | 2.6 | 3.8 | 59 | 103 | 12 | MOLECULE: | 16S RRNA |

| | | | | | | | | |
|---|---|---|---|---|---|---|---|---|
| 11:00 | 8dh8-A | 2.6 | 5.2 | 69 | 700 | 6 | MOLECULE: | LEPTIN RECEPTOR |
| 12:00 | 6jx7-A | 2.6 | 6.3 | 61 | 1245 | 7 | MOLECULE: | FELINE INFECTIOUS PERITONITIS VIRUS |
| 13:00 | 6bpm-A | 2.6 | 8 | 87 | 711 | 6 | MOLECULE: | CATECHOLATE SIDEROPHORE RECEPTOR FI |
| 14:00 | 3o8e-D | 2.6 | 5.7 | 58 | 248 | 0 | MOLECULE: | FIBER 36.1 KDA PROTEIN |
| 15:00 | 8t1l-I | 2.6 | 4.8 | 65 | 1111 | 8 | MOLECULE: | MEDIATOR OF RNA POLYMERASE II TRANS |
| 16:00 | 6d72-C | 2.6 | 3.9 | 59 | 177 | 3 | MOLECULE: | SPERMIDINE N1-ACETYLTRANSFERASE |
| 17:00 | 7z32-A | 2.6 | 4.9 | 74 | 412 | 7 | MOLECULE: | ACIDPHOSPHATASE |
| 18:00 | 8umd-C | 2.6 | 4.6 | 61 | 290 | 8 | MOLECULE: | FLAGELLAR M-RING PROTEIN |
| 19:00 | 4or8-B | 2.6 | 3.8 | 61 | 226 | 8 | MOLECULE: | MEMBRANE-ASSOCIATED PROTEIN VP24 |
| 20:00 | 7riz-A | 2.6 | 6.6 | 62 | 306 | 2 | MOLECULE: | BETA-PROPELLER LACTONASE |
| 21:00 | 3cmb-B | 2.6 | 6.8 | 78 | 266 | 8 | MOLECULE: | ACETOACETATE DECARBOXYLASE |
| 22:00 | 4fhn-B | 2.6 | 4.1 | 77 | 1022 | 1 | MOLECULE: | NUCLEOPORIN NUP37 |
| 23:00 | 3uc2-B | 2.6 | 2.8 | 43 | 126 | 2 | MOLECULE: | HYPOTHETICAL PROTEIN WITH IMMUNOGLO |
| 24:00:00 | 4yhc-A | 2.6 | 5.2 | 68 | 439 | 6 | MOLECULE: | STEROL REGULATORY ELEMENT-BINDING P |
| 25:00:00 | 8w4j-J | 2.6 | 4 | 64 | 574 | 6 | MOLECULE: | GLUTAMATE DEHYDROGENASE 1, MITOCHON |
| 26:00:00 | 2bvb-A | 2.6 | 4 | 66 | 137 | 9 | MOLECULE: | MICRONEMAL PROTEIN 1 |
| 27:00:00 | 7uaw-A | 2.6 | 4.1 | 55 | 125 | 7 | MOLECULE: | ABC TRANSPORTER ATPASE |
| 28:00:00 | 5x7o-A | 2.6 | 7.6 | 59 | 1247 | 8 | MOLECULE: | GLYCOSIDE HYDROLASE FAMILY 31 ALPHA |
| 29:00:00 | 5isv-A | 2.6 | 3.2 | 56 | 156 | 4 | MOLECULE: | RIBOSOMAL-PROTEIN-ALANINE ACETYLTRA |
| 30:00:00 | 8tar-R | 2.6 | 4.6 | 63 | 440 | 2 | MOLECULE: | ANAPHASE-PROMOTING COMPLEX SUBUNIT |
| 31:00:00 | 8v8b-A | 2.6 | 4.7 | 83 | 476 | 7 | MOLECULE: | PORTAL PROTEIN |
| 32:00:00 | 4qxz-A | 2.6 | 3 | 53 | 146 | 8 | MOLECULE: | UNCHARACTERIZED PROTEIN |
| 33:00:00 | 1ei5-A | 2.6 | 4 | 47 | 518 | 2 | MOLECULE: | D-AMINOPEPTIDASE |
| 34:00:00 | 5e6m-A | 2.6 | 4.9 | 78 | 520 | 4 | MOLECULE: | GLYCINE--TRNA LIGASE |
| 35:00:00 | 1xsz-A | 2.6 | 12 | 62 | 356 | 10 | MOLECULE: | GUANINE NUCLEOTIDE EXCHANGE PROTEIN |
| 36:00:00 | 2r4i-A | 2.6 | 5.4 | 69 | 123 | 7 | MOLECULE: | UNCHARACTERIZED PROTEIN |
| 37:00:00 | 6sy0-A | 2.6 | 3.2 | 50 | 137 | 2 | MOLECULE: | TRANSCRIPTION FACTOR WITH AP2 DOMAI |
| 38:00:00 | 2n8x-A | 2.6 | 3.5 | 68 | 166 | 7 | MOLECULE: | LPS-ASSEMBLY LIPOPROTEIN LPTE |
| 39:00:00 | 7eeb-G | 2.6 | 5.4 | 67 | 714 | 9 | MOLECULE: | ENHANCED GREEN FLUORESCENT PROTEIN, |
| 40:00:00 | 8sh2-D | 2.6 | 5.9 | 70 | 384 | 10 | MOLECULE: | KELCH DOMAIN-CONTAINING PROTEIN 2 |
| 41:00:00 | 8ssm-C | 2.6 | 3.3 | 47 | 89 | 2 | MOLECULE: | CONTACT-DEPENDENT INHIBITOR A |
| 42:00:00 | 3tzg-A | 2.6 | 5.6 | 76 | 239 | 0 | MOLECULE: | HYPOTHETICAL PROTEIN BVU_2266 |
| 43:00:00 | 2q7a-A | 2.6 | 2.9 | 54 | 152 | 7 | MOLECULE: | CELL SURFACE HEME-BINDING PROTEIN |
| 44:00:00 | 5ftx-A | 2.6 | 3 | 61 | 651 | 5 | MOLECULE: | SURFACE LAYER PROTEIN |
| 45:00:00 | 5lql-A | 2.6 | 3.8 | 62 | 155 | 8 | MOLECULE: | 4-HYDROXYBENZOYL-COA THIOESTERASE |
| 46:00:00 | 6p2k-B | 2.6 | 5.1 | 62 | 778 | 6 | MOLECULE: | FIBRONECTIN TYPE III DOMAIN-CONTAIN |
| 47:00:00 | 3kya-A | 2.6 | 4.4 | 65 | 471 | 6 | MOLECULE: | PUTATIVE PHOSPHATASE |
| 48:00:00 | 5kku-B | 2.6 | 4.3 | 65 | 289 | 3 | MOLECULE: | POLYKETIDE SYNTHASE TYPE I |
| 49:00:00 | 3wz4-G | 2.6 | 5.1 | 64 | 140 | 11 | MOLECULE: | DOTI |
| 50:00:00 | 7jmn-N | 2.6 | 6.4 | 63 | 198 | 10 | MOLECULE: | MEDIATOR OF RNA POLYMERASE II TRANS |
| 51:00:00 | 4akx-B | 2.6 | 9.4 | 56 | 534 | 7 | MOLECULE: | SPCU |
| 52:00:00 | 6cbc-B | 2.6 | 8 | 93 | 270 | 8 | MOLECULE: | VACUOLAR PROTEIN SORTING-ASSOCIATED |
| 53:00:00 | 7fhr-A | 2.6 | 3.5 | 63 | 439 | 3 | MOLECULE: | PUTATIVE PHTHALATE 4,5-DIOXYGENASE, |
| 54:00:00 | 5boi-A | 2.6 | 4.4 | 55 | 226 | 9 | MOLECULE: | GERMINATION PROTEIN YPEB |
| 55:00:00 | 3dmc-A | 2.6 | 4.1 | 62 | 134 | 10 | MOLECULE: | NTF2-LIKE PROTEIN |
| 56:00:00 | 2p4o-A | 2.6 | 7.6 | 54 | 302 | 9 | MOLECULE: | HYPOTHETICAL PROTEIN |

| | | | | | | | | |
|---|---|---|---|---|---|---|---|---|
| 57:00:00 | 1vla-A | 2.6 | 3.9 | 61 | 144 | 3 | MOLECULE: | HYDROPEROXIDE RESISTANCE PROTEIN OS |
| 58:00:00 | 3w5g-A | 2.6 | 4.6 | 56 | 705 | 5 | MOLECULE: | BETA-GALACTOSIDASE |
| 59:00:00 | 8r1o-F | 2.6 | 12.9 | 70 | 281 | 4 | MOLECULE: | RRP45 |
| 60:00:00 | 3wz3-A | 2.6 | 5.1 | 71 | 137 | 8 | MOLECULE: | TRAM PROTEIN |
| 61:00:00 | 8avg-A | 2.6 | 4.3 | 62 | 603 | 3 | MOLECULE: | ELONGATOR COMPLEX PROTEIN 1 |
| 62:00:00 | 8cuk-B | 2.6 | 7.1 | 67 | 334 | 9 | MOLECULE: | E3 UBIQUITIN-PROTEIN LIGASE PEP5 |
| 63:00:00 | 5u35-A | 2.6 | 3.9 | 57 | 124 | 9 | MOLECULE: | DE NOVO NTF2 WITH LARGE CAVITY |
| 64:00:00 | 6isl-B | 2.6 | 4.2 | 63 | 120 | 8 | MOLECULE: | XIME, SNOAL-LIKE DOMAIN PROTEIN |
| 65:00:00 | 3gwr-A | 2.6 | 5.2 | 65 | 126 | 2 | MOLECULE: | PUTATIVE CALCIUM/CALMODULIN-DEPENDE |
| 66:00:00 | 8cuk-C | 2.6 | 7 | 64 | 330 | 14 | MOLECULE: | E3 UBIQUITIN-PROTEIN LIGASE PEP5 |
| 67:00:00 | 5a5u-B | 2.6 | 8.1 | 65 | 613 | 8 | MOLECULE: | EUKARYOTIC INITIATION FACTOR 3 |
| 68:00:00 | 6l4j-A | 2.6 | 2.8 | 52 | 90 | 0 | MOLECULE: | SINGLE CHAIN MONELLIN |
| 69:00:00 | 5tpj-A | 2.6 | 4.4 | 55 | 122 | 11 | MOLECULE: | DENOVO NTF2 |
| 70:00:00 | 5d17-E | 2.6 | 3.4 | 63 | 158 | 5 | MOLECULE: | TRANSPOSON TN7 TRANSPOSITION PROTEI |
| 71:00:00 | 6quh-C | 2.6 | 9.8 | 59 | 228 | 8 | MOLECULE: | GREEN FLUORESCENT PROTEIN |
| 72:00:00 | 5b61-F | 2.6 | 9 | 58 | 223 | 5 | MOLECULE: | GREEN FLUORESCENT PROTEIN |
| 73:00:00 | 2ib6-C | 2.6 | 10.8 | 56 | 225 | 11 | MOLECULE: | YELLOW MUTANT CHROMO PROTEIN |
| 74:00:00 | 5wj2-B | 2.6 | 8.7 | 57 | 226 | 7 | MOLECULE: | CYAN FLUORESCENT PROTEIN |
| 75:00:00 | 4w76-B | 2.6 | 9.8 | 59 | 226 | 7 | MOLECULE: | FLUORESCENT PROTEIN D21H/K26C |
| 76:00:00 | 7sws-C | 2.6 | 7.6 | 54 | 217 | 6 | MOLECULE: | CHROMOPROTEIN AMILCP |
| 77:00:00 | 7amb-D | 2.6 | 8.7 | 57 | 229 | 7 | MOLECULE: | GREEN FLUORESCENT PROTEIN |
| 78:00:00 | 4tzg-E | 2.6 | 4.4 | 63 | 217 | 8 | MOLECULE: | FLUORESCENT PROTEIN |
| 79:00:00 | 7rrh-D | 2.6 | 9.7 | 63 | 215 | 5 | MOLECULE: | FLUORESCENT PROTEIN DRONPA |
| 80:00:00 | 6og9-B | 2.6 | 5.9 | 53 | 226 | 6 | MOLECULE: | CRYSTAL STRUCTURE OF GREEN FLUORESC |
| 81:00:00 | 6fll-A | 2.6 | 10.3 | 60 | 223 | 5 | MOLECULE: | GREEN FLUORESCENT PROTEIN |
| 82:00:00 | 3ls3-B | 2.6 | 4.4 | 62 | 214 | 8 | MOLECULE: | PADRON0.9 |
| 83:00:00 | 5hzu-D | 2.6 | 8.4 | 52 | 212 | 6 | MOLECULE: | FLUORESCENT PROTEIN DRONPA |
| 84:00:00 | 5dpi-F | 2.6 | 10 | 58 | 224 | 5 | MOLECULE: | GREEN FLUORESCENT PROTEIN |
| 85:00:00 | 2ib6-E | 2.6 | 10.8 | 55 | 225 | 11 | MOLECULE: | YELLOW MUTANT CHROMO PROTEIN |
| 86:00:00 | 5dtl-H | 2.6 | 9.7 | 63 | 219 | 2 | MOLECULE: | GREEN TO RED PHOTOCONVERTIBLE GFP-L |
| 87:00:00 | 5cm8-A | 2.5 | 3.5 | 44 | 419 | 5 | MOLECULE: | RAL GUANINE NUCLEOTIDE DISSOCIATION |
| 88:00:00 | 3zwl-D | 2.5 | 4.7 | 63 | 352 | 10 | MOLECULE: | EUKARYOTIC TRANSLATION INITIATION F |
| 89:00:00 | 3ge2-A | 2.5 | 3 | 46 | 89 | 2 | MOLECULE: | LIPOPROTEIN, PUTATIVE |
| 90:00:00 | 3ub1-D | 2.5 | 6.6 | 65 | 253 | 11 | MOLECULE: | ORF13-LIKE PROTEIN |
| 91:00:00 | 8h93-B | 2.5 | 6.5 | 65 | 367 | 6 | MOLECULE: | NACHT, LRR AND PYD DOMAINS-CONTAINI |
| 92:00:00 | 8bo9-A | 2.5 | 3.3 | 66 | 174 | 3 | MOLECULE: | NON STRUCTURAL POLYPROTEIN |
| 93:00:00 | 4ps6-A | 2.5 | 3.8 | 60 | 130 | 3 | MOLECULE: | INHIBITOR OF VERTEBRATE LYSOZYME |
| 94:00:00 | 2l4v-A | 2.5 | 3.2 | 65 | 135 | 8 | MOLECULE: | CYSTATIN |
| 95:00:00 | 2r5x-B | 2.5 | 2.9 | 48 | 118 | 4 | MOLECULE: | UNCHARACTERIZED CONSERVED PROTEIN |
| 96:00:00 | 3edp-A | 2.5 | 8.2 | 83 | 223 | 4 | MOLECULE: | LIN2111 PROTEIN |
| 97:00:00 | 8ju3-A | 2.5 | 10.5 | 72 | 290 | 6 | MOLECULE: | TAIL FIBER PROTEIN S,TAIL FIBER PRO |
| 98:00:00 | 3hdp-A | 2.5 | 11.7 | 53 | 132 | 8 | MOLECULE: | GLYOXALASE-I |
| 99:00:00 | 7myj-C | 2.5 | 5 | 63 | 464 | 3 | MOLECULE: | 5'-AMP-ACTIVATED PROTEIN KINASE CAT |
| 0:00 | 4ava-A | 2.5 | 4.5 | 77 | 327 | 6 | MOLECULE: | LYSINE ACETYLTRANSFERASE |
| 1:00 | 5c9i-A | 2.5 | 4.1 | 59 | 768 | 5 | MOLECULE: | PROTEIN RELATED TO PENICILLIN ACYLA |
| 2:00 | 8a43-B | 2.5 | 3.5 | 65 | 1131 | 2 | MOLECULE: | DNA-DIRECTED RNA POLYMERASE I SUBUN |

| | | | | | | | | |
|---|---|---|---|---|---|---|---|---|
| 3:00 | 4rl8-C | 2.5 | 7.2 | 89 | 268 | 7 | MOLECULE: | PROTEIN INVOLVED IN META-PATHWAY OF |
| 4:00 | 8d82-A | 2.5 | 3.5 | 53 | 589 | 0 | MOLECULE: | SOLUBLE INTERLEUKIN-6 RECEPTOR SUBU |
| 5:00 | 7acv-B | 2.5 | 3.4 | 50 | 127 | 8 | MOLECULE: | LMW SLP |
| 6:00 | 1xs0-C | 2.5 | 3.2 | 63 | 129 | 5 | MOLECULE: | INHIBITOR OF VERTEBRATE LYSOZYME |
| 7:00 | 3exz-B | 2.5 | 4.3 | 64 | 147 | 6 | MOLECULE: | MAOC-LIKE DEHYDRATASE |
| 8:00 | 7cbl-A | 2.5 | 7.6 | 69 | 211 | 4 | MOLECULE: | FLAGELLAR L-RING PROTEIN |
| 9:00 | 8gun-A | 2.5 | 5.4 | 70 | 166 | 1 | MOLECULE: | TYPE VII SECRETION SYSTEM PROTEIN E |
| 10:00 | 6zwm-C | 2.5 | 5.4 | 71 | 319 | 13 | MOLECULE: | SERINE/THREONINE-PROTEIN KINASE MTO |
| 11:00 | 2rqf-A | 2.5 | 7.8 | 86 | 227 | 5 | MOLECULE: | HEMOLYMPH JUVENILE HORMONE BINDING |
| 12:00 | 8k44-A | 2.5 | 6.7 | 72 | 280 | 6 | MOLECULE: | VP9 |
| 13:00 | 3fgy-A | 2.5 | 4.3 | 57 | 135 | 12 | MOLECULE: | UNCHARACTERIZED NTF2-LIKE PROTEIN |
| 14:00 | 8t2v-B | 2.5 | 9.7 | 66 | 718 | 11 | MOLECULE: | INTEGRIN ALPHA-IIB |
| 15:00 | 3q63-F | 2.5 | 4.3 | 69 | 139 | 6 | MOLECULE: | MLL2253 PROTEIN |
| 16:00 | 6xp5-Q | 2.5 | 7.2 | 72 | 476 | 6 | MOLECULE: | MEDIATOR OF RNA POLYMERASE II TRANS |
| 17:00 | 4xb6-C | 2.5 | 7.5 | 78 | 353 | 8 | MOLECULE: | ALPHA-D-RIBOSE 1-METHYLPHOSPHONATE |
| 18:00 | 7ml9-A | 2.5 | 5 | 81 | 295 | 5 | MOLECULE: | INSECTICIDAL PROTEIN |
| 19:00 | 6gut-B | 2.5 | 5.3 | 58 | 239 | 3 | MOLECULE: | 23S RRNA PSEUDOURIDINE SYNTHASE D,P |
| 20:00 | 5b2o-A | 2.5 | 5.2 | 58 | 1455 | 3 | MOLECULE: | CRISPR-ASSOCIATED ENDONUCLEASE CAS9 |
| 21:00 | 5awh-A | 2.5 | 6.4 | 68 | 759 | 4 | MOLECULE: | UNCHARACTERIZED PROTEIN |
| 22:00 | 5vgl-A | 2.5 | 5.4 | 57 | 154 | 2 | MOLECULE: | LACHRYMATORY-FACTOR SYNTHASE |
| 23:00 | 8cgm-A | 2.5 | 11.8 | 57 | 190 | 7 | MOLECULE: | OUTER MEMBRANE LIPOPROTEIN CARRIER |
| 24:00:00 | 2wgo-A | 2.5 | 3.9 | 55 | 98 | 5 | MOLECULE: | RANASPUMIN-2 |
| 25:00:00 | 4i60-A | 2.5 | 4.7 | 74 | 128 | 3 | MOLECULE: | AVIDIN |
| 26:00:00 | 6s50-A | 2.5 | 4 | 70 | 248 | 4 | MOLECULE: | STREPTAVIDIN |
| 27:00:00 | 4fvs-A | 2.5 | 4.8 | 76 | 214 | 5 | MOLECULE: | PUTATIVE LIPOPROTEIN |
| 28:00:00 | 4jhy-A | 2.5 | 3.9 | 61 | 156 | 7 | MOLECULE: | HYPOTHETICAL PROTEIN |
| 29:00:00 | 1tlw-A | 2.5 | 11.7 | 93 | 251 | 6 | MOLECULE: | NUCLEOSIDE-SPECIFIC CHANNEL-FORMING |
| 30:00:00 | 1lo7-A | 2.5 | 5.6 | 65 | 140 | 6 | MOLECULE: | 4-HYDROXYBENZOYL-COA THIOESTERASE |
| 31:00:00 | 4ak1-A | 2.5 | 5.4 | 58 | 600 | 5 | MOLECULE: | BT_4661 |
| 32:00:00 | 2joe-A | 2.5 | 4.3 | 59 | 139 | 7 | MOLECULE: | HYPOTHETICAL LIPOPROTEIN YEHR |
| 33:00:00 | 3qoo-A | 2.5 | 6.8 | 72 | 135 | 7 | MOLECULE: | UNCHARACTERIZED PROTEIN |
| 34:00:00 | 5dyv-A | 2.5 | 6.4 | 64 | 135 | 6 | MOLECULE: | YD REPEAT-CONTAINING PROTEIN |
| 35:00:00 | 6kme-A | 2.5 | 5.1 | 68 | 284 | 4 | MOLECULE: | PHYTOCHROMOBILIN SYNTHASE |
| 36:00:00 | 6qzk-A | 2.5 | 5.2 | 75 | 746 | 15 | MOLECULE: | CLOSTRIDIUM BUTYRICUM ARGONAUTE |
| 37:00:00 | 8i6v-D | 2.5 | 5.9 | 65 | 678 | 2 | MOLECULE: | VACUOLAR TRANSPORTER CHAPERONE COMP |
| 38:00:00 | 4rul-A | 2.5 | 9.8 | 60 | 821 | 5 | MOLECULE: | DNA TOPOISOMERASE 1 |
| 39:00:00 | 6gvw-C | 2.5 | 4.1 | 77 | 383 | 1 | MOLECULE: | BRCA1-A COMPLEX SUBUNIT ABRAXAS 1 |
| 40:00:00 | 7omm-B | 2.5 | 2.7 | 55 | 128 | 7 | MOLECULE: | LPS-ASSEMBLY PROTEIN LPTD |
| 41:00:00 | 1nbh-A | 2.5 | 8.6 | 70 | 293 | 7 | MOLECULE: | GLYCINE N-METHYLTRANSFERASE |
| 42:00:00 | 4uw8-A | 2.5 | 6.2 | 72 | 401 | 6 | MOLECULE: | L-SHAPED TAIL FIBER PROTEIN |
| 43:00:00 | 5h3k-A | 2.5 | 9.6 | 79 | 535 | 5 | MOLECULE: | SLR0280 PROTEIN |
| 44:00:00 | 3fbq-A | 2.5 | 4.7 | 68 | 283 | 4 | MOLECULE: | CONSERVED DOMAIN PROTEIN |
| 45:00:00 | 8th8-H | 2.5 | 7.5 | 67 | 820 | 3 | MOLECULE: | DYNEIN REGULATORY COMPLEX PROTEIN 1 |
| 46:00:00 | 5w5y-O | 2.5 | 3.7 | 76 | 640 | 7 | MOLECULE: | DNA-DIRECTED RNA POLYMERASE I SUBUN |
| 47:00:00 | 7qb6-A | 2.5 | 4.3 | 74 | 164 | 4 | MOLECULE: | NODULIN-13 |
| 48:00:00 | 1yfq-A | 2.5 | 5.8 | 55 | 342 | 2 | MOLECULE: | CELL CYCLE ARREST PROTEIN BUB3 |

| | | | | | | | | |
|---|---|---|---|---|---|---|---|---|
| 49:00:00 | 4e72-A | 2.5 | 7 | 62 | 218 | 5 | MOLECULE: | UNCHARACTERIZED PROTEIN |
| 50:00:00 | 3ewk-A | 2.5 | 5.2 | 54 | 227 | 0 | MOLECULE: | SENSOR PROTEIN |
| 51:00:00 | 3wyb-A | 2.5 | 4.7 | 66 | 327 | 8 | MOLECULE: | MESO-DIAMINOPIMELATE D-DEHYDROGENAS |
| 52:00:00 | 6jp4-A | 2.5 | 9 | 75 | 770 | 1 | MOLECULE: | ALGINATE LYASE |
| 53:00:00 | 6m15-A | 2.5 | 5.6 | 71 | 965 | 7 | MOLECULE: | SPIKE GLYCOPROTEIN |
| 54:00:00 | 3f14-A | 2.5 | 2.9 | 53 | 112 | 13 | MOLECULE: | UNCHARACTERIZED NTF2-LIKE PROTEIN |
| 55:00:00 | 5h2d-A | 2.5 | 10.1 | 84 | 418 | 7 | MOLECULE: | KLLA0C04147P |
| 56:00:00 | 7bi2-A | 2.5 | 5.4 | 59 | 1021 | 7 | MOLECULE: | PHOSPHATIDYLINOSITOL 4-PHOSPHATE 3- |
| 57:00:00 | 3icy-A | 2.5 | 9.8 | 56 | 118 | 4 | MOLECULE: | SENSOR PROTEIN |
| 58:00:00 | 6xzq-C | 2.5 | 8.8 | 65 | 772 | 8 | MOLECULE: | RNA (5'- |
| 59:00:00 | 4tmd-A | 2.5 | 3.8 | 69 | 190 | 4 | MOLECULE: | UNCHARACTERIZED PROTEIN |
| 60:00:00 | 7dsk-A | 2.5 | 3 | 53 | 471 | 2 | MOLECULE: | 4F2 CELL-SURFACE ANTIGEN HEAVY CHAI |
| 61:00:00 | 2x12-A | 2.5 | 2.8 | 55 | 200 | 0 | MOLECULE: | FIMBRIAE-ASSOCIATED PROTEIN FAP1 |
| 62:00:00 | 1lsh-B | 2.5 | 6.8 | 70 | 174 | 4 | MOLECULE: | LIPOVITELLIN (LV-1N, LV-1C) |
| 63:00:00 | 7sg7-S | 2.5 | 3.7 | 65 | 471 | 6 | MOLECULE: | GENE 14 PROTEIN |
| 64:00:00 | 3o77-A | 2.5 | 8.5 | 81 | 388 | 4 | MOLECULE: | MYOSIN LIGHT CHAIN KINASE, SMOOTH M |
| 65:00:00 | 1bh3-A | 2.5 | 6.2 | 80 | 289 | 5 | MOLECULE: | PORIN |
| 66:00:00 | 6r9t-A | 2.5 | 5.5 | 53 | 2185 | 8 | MOLECULE: | TALIN-1 |
| 67:00:00 | 6b12-B | 2.5 | 3.7 | 52 | 153 | 2 | MOLECULE: | TNE2 |
| 68:00:00 | 5h1j-A | 2.5 | 3.4 | 55 | 693 | 11 | MOLECULE: | GEM-ASSOCIATED PROTEIN 5 |
| 69:00:00 | 5wox-A | 2.5 | 7.2 | 53 | 145 | 8 | MOLECULE: | UNCHARACTERIZED PROTEIN |
| 70:00:00 | 7djl-A | 2.5 | 4.4 | 61 | 473 | 8 | MOLECULE: | PROTEIN SUPPRESSOR OF QUENCHING 1, |
| 71:00:00 | 2ia9-D | 2.5 | 3.7 | 44 | 99 | 5 | MOLECULE: | PUTATIVE SEPTATION PROTEIN SPOVG |
| 72:00:00 | 5mqr-A | 2.5 | 3.6 | 61 | 1082 | 2 | MOLECULE: | BETA-L-ARABINOBIOSIDASE |
| 73:00:00 | 4id2-A | 2.5 | 3.6 | 64 | 136 | 8 | MOLECULE: | UNCHARACTERIZED PROTEIN |
| 74:00:00 | 5o8o-A | 2.5 | 7 | 75 | 332 | 3 | MOLECULE: | MITOCHONDRIAL IMPORT RECEPTOR SUBUN |
| 75:00:00 | 3k1l-A | 2.5 | 7 | 57 | 376 | 0 | MOLECULE: | FANCL |
| 76:00:00 | 7su6-B | 2.5 | 4.6 | 56 | 111 | 5 | MOLECULE: | 7,8-DIHYDRONEOPTERIN ALDOLASE |
| 77:00:00 | 6vbu-2 | 2.5 | 3 | 47 | 659 | 6 | MOLECULE: | BARDET-BIEDL SYNDROME 18 PROTEIN |
| 78:00:00 | 2iv9-A | 2.5 | 3.6 | 67 | 236 | 4 | MOLECULE: | AP-2 COMPLEX SUBUNIT BETA-2 |
| 79:00:00 | 6u9f-A | 2.5 | 4.7 | 51 | 717 | 4 | MOLECULE: | PDPA |
| 80:00:00 | 5xfm-A | 2.5 | 5.6 | 93 | 632 | 8 | MOLECULE: | ALPHA-GLUCOSIDASE |
| 81:00:00 | 7lc1-B | 2.5 | 4.4 | 56 | 218 | 4 | MOLECULE: | ISOFORM 2B OF GTPASE KRAS |
| 82:00:00 | 6f2p-A | 2.5 | 6.5 | 74 | 1040 | 7 | MOLECULE: | PAENIBACILLUS XANTHAN LYASE |
| 83:00:00 | 1cvm-A | 2.5 | 3.9 | 67 | 353 | 4 | MOLECULE: | PHYTASE |
| 84:00:00 | 3dzm-A | 2.5 | 7 | 81 | 208 | 2 | MOLECULE: | HYPOTHETICAL CONSERVED PROTEIN |
| 85:00:00 | 3u97-A | 2.5 | 3.1 | 45 | 77 | 11 | MOLECULE: | RIBONUCLEASE TOXIN BRNT |
| 86:00:00 | 6a9e-A | 2.5 | 7.2 | 55 | 339 | 7 | MOLECULE: | ENDOLYSIN,AUTOPHAGY-RELATED PROTEIN |
| 87:00:00 | 4lxr-A | 2.5 | 3.8 | 42 | 755 | 7 | MOLECULE: | PROTEIN TOLL |
| 88:00:00 | 5uwa-A | 2.5 | 5 | 67 | 185 | 12 | MOLECULE: | PROBABLE PHOSPHOLIPID-BINDING PROTE |
| 89:00:00 | 2bib-A | 2.5 | 5.1 | 59 | 540 | 2 | MOLECULE: | TEICHOIC ACID PHOSPHORYLCHOLINE EST |
| 90:00:00 | 5ho2-A | 2.5 | 4.4 | 80 | 803 | 4 | MOLECULE: | EXTRACELLULAR ARABINANASE |
| 91:00:00 | 8d2j-B | 2.5 | 5.3 | 48 | 483 | 6 | MOLECULE: | IPD113_COW |
| 92:00:00 | 7qzq-A | 2.5 | 3.6 | 50 | 344 | 14 | MOLECULE: | KELCH REPEAT AND BTB DOMAIN-CONTAIN |
| 93:00:00 | 8g6e-b | 2.5 | 3.8 | 56 | 256 | 7 | MOLECULE: | PROTEASOME SUBUNIT ALPHA TYPE-6 |
| 94:00:00 | 1tu1-A | 2.5 | 3.2 | 55 | 144 | 4 | MOLECULE: | HYPOTHETICAL PROTEIN PA0094 |

| | | | | | | | | |
|---|---|---|---|---|---|---|---|---|
| 95:00:00 | 5fr8-A | 2.5 | 13.8 | 93 | 707 | 4 | MOLECULE: | TONB-DEPENDENT SIDEROPHORE RECEPTOR |
| 96:00:00 | 3q9d-A | 2.5 | 1.9 | 38 | 175 | 11 | MOLECULE: | PROTEIN CPN_0803/CP_1068/CPJ0803/CP |
| 97:00:00 | 3k5j-A | 2.5 | 5.1 | 51 | 182 | 2 | MOLECULE: | SUPPRESSOR OF FUSED FAMILY PROTEIN |
| 98:00:00 | 5if6-A | 2.5 | 5.7 | 72 | 134 | 7 | MOLECULE: | OHP9_1C |
| 99:00:00 | 8qrl-D | 2.5 | 4 | 62 | 343 | 2 | MOLECULE: | 12S MITOCHONDRIAL RRNA |
| 0:00 | 7qrj-B | 2.5 | 6.5 | 66 | 182 | 6 | MOLECULE: | ZAV_19 PROTEIN |
| 1:00 | 6ulg-N | 2.5 | 7.9 | 56 | 483 | 2 | MOLECULE: | FOLLICULIN |
| 2:00 | 1nr0-A | 2.5 | 9.7 | 63 | 610 | 5 | MOLECULE: | ACTIN INTERACTING PROTEIN 1 |
| 3:00 | 2fkj-A | 2.5 | 2.6 | 40 | 361 | 5 | MOLECULE: | OUTER SURFACE PROTEIN A |
| 4:00 | 6yug-A | 2.5 | 4 | 59 | 143 | 5 | MOLECULE: | DIAMINE ACETYLTRANSFERASE |
| 5:00 | 5m1i-A | 2.5 | 7 | 68 | 539 | 9 | MOLECULE: | ALPHA-GALACTOSIDASE |
| 6:00 | 6n7p-M | 2.5 | 3.3 | 54 | 107 | 2 | MOLECULE: | U1 SMALL NUCLEAR RIBONUCLEOPROTEIN |
| 7:00 | 3o4j-C | 2.5 | 4.6 | 64 | 579 | 6 | MOLECULE: | ACYLAMINO-ACID-RELEASING ENZYME |
| 8:00 | 4fyg-A | 2.5 | 4.4 | 83 | 743 | 5 | MOLECULE: | SIDF, INHIBITOR OF GROWTH FAMILY, M |
| 9:00 | 6cl5-A | 2.5 | 9.1 | 69 | 374 | 4 | MOLECULE: | TAIL FIBER PROTEIN |
| 10:00 | 5w0p-C | 2.5 | 5 | 76 | 850 | 7 | MOLECULE: | ENDOLYSIN,RHODOPSIN,S-ARRESTIN |
| 11:00 | 7wug-Y | 2.5 | 4.8 | 67 | 508 | 9 | MOLECULE: | VACUOLAR IMPORT AND DEGRADATION PRO |
| 12:00 | 8smu-A | 2.5 | 9.1 | 59 | 393 | 10 | MOLECULE: | HTAACR2 INTEGRAL FUSION WITHIN ENHA |
| 13:00 | 6itw-A | 2.5 | 3.2 | 58 | 221 | 7 | MOLECULE: | TYPE VI IMMUNITY PROTEIN ATU4351 |
| 14:00 | 8jnx-A | 2.5 | 5.3 | 64 | 662 | 2 | MOLECULE: | ALPHA-AMYLASE |
| 15:00 | 8g9t-K | 2.5 | 4 | 50 | 582 | 8 | MOLECULE: | ACRIC9 |
| 16:00 | 5ztm-A | 2.5 | 10.4 | 54 | 168 | 7 | MOLECULE: | DOSAGE COMPENSATION REGULATOR |
| 17:00 | 6qam-A | 2.5 | 7.3 | 65 | 211 | 6 | MOLECULE: | OUTER MEMBRANE PROTEIN ALKL |
| 18:00 | 6rlb-C | 2.5 | 6 | 61 | 494 | 5 | MOLECULE: | O6-ALKYLGUANINE-DNA ALKYLTRANSFERAS |
| 19:00 | 3afl-A | 2.5 | 4.9 | 73 | 766 | 7 | MOLECULE: | OLIGO ALGINATE LYASE |
| 20:00 | 8ewi-A | 2.5 | 4.7 | 71 | 1775 | 6 | MOLECULE: | E3 UBIQUITIN-PROTEIN LIGASE UBR5 |
| 21:00 | 5mwv-A | 2.5 | 5.1 | 72 | 281 | 4 | MOLECULE: | OUTER MEMBRANE PROTEIN G |
| 22:00 | 6nd4-Q | 2.5 | 4.5 | 64 | 862 | 6 | MOLECULE: | ETS RRNA |
| 23:00 | 6le0-A | 2.5 | 5.9 | 67 | 209 | 3 | MOLECULE: | ABMU |
| 24:00:00 | 8p98-A | 2.5 | 10.4 | 90 | 621 | 3 | MOLECULE: | VITAMIN B12 TRANSPORTER BTUB |
| 25:00:00 | 7agv-A | 2.5 | 2.9 | 56 | 160 | 5 | MOLECULE: | K(+)/H(+) ANTIPORTER SUBUNIT KHTT |
| 26:00:00 | 8h2t-E | 2.5 | 5.6 | 70 | 157 | 7 | MOLECULE: | RIESKE (2FE-2S) DOMAIN PROTEIN |
| 27:00:00 | 8f5p-C | 2.5 | 6.1 | 62 | 1176 | 8 | MOLECULE: | NET DOMAIN-CONTAINING PROTEIN |
| 28:00:00 | 3gwr-B | 2.5 | 5.4 | 66 | 128 | 2 | MOLECULE: | PUTATIVE CALCIUM/CALMODULIN-DEPENDE |
| 29:00:00 | 5trv-A | 2.5 | 3.6 | 56 | 118 | 9 | MOLECULE: | DENOVO NTF2 |
| 30:00:00 | 7qh3-C | 2.5 | 9.9 | 63 | 137 | 6 | MOLECULE: | RSFG |
| 31:00:00 | 8h2t-A | 2.5 | 5.5 | 71 | 157 | 7 | MOLECULE: | RIESKE (2FE-2S) DOMAIN PROTEIN |
| 32:00:00 | 5u9o-D | 2.5 | 5.1 | 67 | 146 | 4 | MOLECULE: | PLASTID DIVISION PROTEIN CDP1, CHLO |
| 33:00:00 | 8aha-A | 2.5 | 11.1 | 65 | 237 | 2 | MOLECULE: | GREEN FLUORESCENT PROTEIN |
| 34:00:00 | 5o68-C | 2.5 | 6.4 | 84 | 262 | 6 | MOLECULE: | FAPF |
| 35:00:00 | 4w7x-B | 2.5 | 6.1 | 57 | 220 | 4 | MOLECULE: | FLUORESCENT PROTEIN E115C/T118H |
| 36:00:00 | 4uer-b | 2.5 | 7.6 | 81 | 480 | 2 | MOLECULE: | EIF1A |
| 37:00:00 | 5k1h-B | 2.5 | 5.9 | 61 | 554 | 3 | MOLECULE: | EUKARYOTIC TRANSLATION INITIATION F |
| 38:00:00 | 8am4-A | 2.5 | 8.2 | 54 | 229 | 9 | MOLECULE: | GREEN FLUORESCENT PROTEIN |
| 39:00:00 | 4w7f-A | 2.5 | 8.2 | 53 | 213 | 9 | MOLECULE: | FLUORESCENT PROTEIN E124H/K126H |
| 40:00:00 | 5exb-J | 2.5 | 4 | 73 | 221 | 8 | MOLECULE: | GREEN FLUORESCENT PROTEIN |

| | | | | | | | | |
|---|---|---|---|---|---|---|---|---|
| 41:00:00 | 4jfg-A | 2.5 | 8.9 | 57 | 225 | 5 | MOLECULE: | GREEN FLUORESCENT PROTEIN |
| 42:00:00 | 3ssy-A | 2.5 | 10.7 | 69 | 227 | 6 | MOLECULE: | GREEN FLUORESCENT PROTEIN |
| 43:00:00 | 1emc-A | 2.5 | 8.2 | 54 | 224 | 9 | MOLECULE: | GREEN FLUORESCENT PROTEIN |
| 44:00:00 | 3bpq-D | 2.4 | 2.9 | 44 | 86 | 0 | MOLECULE: | ANTITOXIN RELB3 |
| 45:00:00 | 5yx4-A | 2.4 | 4.8 | 65 | 232 | 6 | MOLECULE: | CHALCONE-FLAVONONE ISOMERASE FAMILY |
| 46:00:00 | 7aak-A | 2.4 | 4.9 | 71 | 345 | 6 | MOLECULE: | PORPHOBILINOGEN DEAMINASE |
| 47:00:00 | 2imj-A | 2.4 | 4.3 | 68 | 158 | 9 | MOLECULE: | HYPOTHETICAL PROTEIN DUF1348 |
| 48:00:00 | 1lsh-A | 2.4 | 8.1 | 93 | 955 | 11 | MOLECULE: | LIPOVITELLIN (LV-1N, LV-1C) |
| 49:00:00 | 4lee-A | 2.4 | 5.1 | 64 | 313 | 3 | MOLECULE: | AGGLUTININ-LIKE PROTEIN 3 |
| 50:00:00 | 8wkl-G | 2.4 | 4.2 | 66 | 307 | 8 | MOLECULE: | STRICTOSIDINE SYNTHASE |
| 51:00:00 | 2ztb-B | 2.4 | 3.7 | 71 | 248 | 1 | MOLECULE: | CRYSTAL PROTEIN |
| 52:00:00 | 3dgc-S | 2.4 | 3.2 | 54 | 207 | 6 | MOLECULE: | INTERLEUKIN-22 |
| 53:00:00 | 3bb9-B | 2.4 | 4.7 | 60 | 125 | 5 | MOLECULE: | PUTATIVE ORPHAN PROTEIN |
| 54:00:00 | 4l8p-A | 2.4 | 6.1 | 73 | 168 | 1 | MOLECULE: | BILE ACID 7A-DEHYDRATASE, BAIE |
| 55:00:00 | 6td9-B | 2.4 | 3.4 | 71 | 247 | 6 | MOLECULE: | PA1624 |
| 56:00:00 | 3ty1-A | 2.4 | 5.3 | 87 | 384 | 1 | MOLECULE: | HYPOTHETICAL ALDOSE 1-EPIMERASE |
| 57:00:00 | 3rby-A | 2.4 | 4.1 | 69 | 245 | 4 | MOLECULE: | UNCHARACTERIZED PROTEIN YLR301W |
| 58:00:00 | 5jrb-K | 2.4 | 8.1 | 66 | 188 | 6 | MOLECULE: | DNA REPAIR PROTEIN RAD52 HOMOLOG |
| 59:00:00 | 3g1j-A | 2.4 | 3.2 | 49 | 90 | 6 | MOLECULE: | INTEGRON CASSETTE PROTEIN |
| 60:00:00 | 3pu2-B | 2.4 | 4.6 | 69 | 153 | 4 | MOLECULE: | UNCHARACTERIZED PROTEIN |
| 61:00:00 | 2wb7-A | 2.4 | 5.3 | 74 | 525 | 4 | MOLECULE: | PT26-6P |
| 62:00:00 | 5gv0-A | 2.4 | 4.8 | 68 | 162 | 7 | MOLECULE: | LYSOSOME-ASSOCIATED MEMBRANE GLYCOP |
| 63:00:00 | 2qzi-A | 2.4 | 3 | 49 | 101 | 10 | MOLECULE: | UNCHARACTERIZED PROTEIN |
| 64:00:00 | 9be2-A | 2.4 | 10.1 | 70 | 335 | 6 | MOLECULE: | NUCLEOID-ASSOCIATED PROTEIN YEJK |
| 65:00:00 | 8r54-A | 2.4 | 4.4 | 79 | 1982 | 8 | MOLECULE: | TENEURIN-3 |
| 66:00:00 | 1l3a-C | 2.4 | 6.6 | 60 | 182 | 2 | MOLECULE: | P24: PLANT TRANSCRIPTIONAL REGULATO |
| 67:00:00 | 7vqo-A | 2.4 | 9 | 77 | 1180 | 4 | MOLECULE: | AMS1, NBR1 AND MALE FUSION PROTEIN |
| 68:00:00 | 3k0z-A | 2.4 | 5.6 | 72 | 149 | 7 | MOLECULE: | PUTATIVE POLYKETIDE CYCLASE |
| 69:00:00 | 7f13-A | 2.4 | 10.4 | 78 | 154 | 9 | MOLECULE: | DCR3 |
| 70:00:00 | 7zr1-D | 2.4 | 10.2 | 72 | 780 | 7 | MOLECULE: | DOUBLE-STRAND BREAK REPAIR PROTEIN |
| 71:00:00 | 4esq-A | 2.4 | 4.8 | 72 | 194 | 4 | MOLECULE: | SERINE/THREONINE PROTEIN KINASE |
| 72:00:00 | 6xrb-A | 2.4 | 2.9 | 60 | 150 | 10 | MOLECULE: | SCIW |
| 73:00:00 | 3mi6-D | 2.4 | 5.8 | 84 | 743 | 11 | MOLECULE: | ALPHA-GALACTOSIDASE |
| 74:00:00 | 8rkh-A | 2.4 | 8.1 | 66 | 219 | 6 | MOLECULE: | ZONA PELLUCIDA SPERM-BINDING PROTEI |
| 75:00:00 | 6ipv-A | 2.4 | 3 | 68 | 221 | 6 | MOLECULE: | CQSB2 |
| 76:00:00 | 2il5-A | 2.4 | 6.7 | 72 | 162 | 7 | MOLECULE: | HYPOTHETICAL PROTEIN |
| 77:00:00 | 3ggr-B | 2.4 | 7.3 | 61 | 278 | 3 | MOLECULE: | CELL CYCLE CHECKPOINT CONTROL PROTE |
| 78:00:00 | 4ql0-A | 2.4 | 7.4 | 89 | 520 | 4 | MOLECULE: | FILAMENTOUS HEMAGGLUTININ TRANSPORT |
| 79:00:00 | 6e5u-A | 2.4 | 4.7 | 70 | 339 | 10 | MOLECULE: | NUCLEAR RNA EXPORT FACTOR 1 |
| 80:00:00 | 5da9-A | 2.4 | 10.9 | 71 | 433 | 7 | MOLECULE: | PUTATIVE UNCHARACTERIZED PROTEIN,PU |
| 81:00:00 | 4kc5-A | 2.4 | 5.5 | 83 | 914 | 2 | MOLECULE: | RHIE PROTEIN |
| 82:00:00 | 3c5p-A | 2.4 | 9.5 | 78 | 194 | 14 | MOLECULE: | PROTEIN BAS0735 OF UNKNOWN FUNCTION |
| 83:00:00 | 4kwy-A | 2.4 | 4.6 | 67 | 137 | 9 | MOLECULE: | PUTATIVE UNCHARACTERIZED PROTEIN |
| 84:00:00 | 5lal-B | 2.4 | 6.1 | 69 | 158 | 7 | MOLECULE: | DIRIGENT PROTEIN 6 |
| 85:00:00 | 3bcy-A | 2.4 | 2.9 | 57 | 146 | 5 | MOLECULE: | PROTEIN YER067W |
| 86:00:00 | 4dkk-A | 2.4 | 2.1 | 45 | 104 | 2 | MOLECULE: | DOUBLE-STRANDED RNA-BINDING PROTEIN |

| | | | | | | | | |
|---|---|---|---|---|---|---|---|---|
| 87:00:00 | 6zu9-Q | 2.4 | 9.9 | 70 | 226 | 4 | MOLECULE: | 18S RIBOSOMAL RNA |
| 88:00:00 | 2fpn-A | 2.4 | 5.9 | 64 | 205 | 5 | MOLECULE: | YWMB |
| 89:00:00 | 3v9o-A | 2.4 | 6.6 | 63 | 121 | 5 | MOLECULE: | DIHYDRONEOPTERIN ALDOLASE |
| 90:00:00 | 2w9j-B | 2.4 | 3.4 | 51 | 82 | 4 | MOLECULE: | SIGNAL RECOGNITION PARTICLE SUBUNIT |
| 91:00:00 | 4qan-A | 2.4 | 4.4 | 65 | 385 | 2 | MOLECULE: | HYPOTHETICAL PROTEIN |
| 92:00:00 | 5k21-B | 2.4 | 4.6 | 63 | 123 | 11 | MOLECULE: | PYOCYANIN DEMETHYLASE |
| 93:00:00 | 1vjh-A | 2.4 | 4.3 | 59 | 120 | 5 | MOLECULE: | BET V I ALLERGEN FAMILY |
| 94:00:00 | 4ol4-A | 2.4 | 3.5 | 63 | 177 | 8 | MOLECULE: | PROLINE-RICH 28 KDA ANTIGEN |
| 95:00:00 | 1t6a-A | 2.4 | 4.2 | 52 | 124 | 10 | MOLECULE: | RBSTP2229 GENE PRODUCT |
| 96:00:00 | 7pw9-A | 2.4 | 8.2 | 63 | 1952 | 6 | MOLECULE: | SERINE/THREONINE-PROTEIN KINASE SMG |
| 97:00:00 | 8kg3-A | 2.4 | 5.4 | 67 | 311 | 7 | MOLECULE: | OS06G0623700 PROTEIN |
| 98:00:00 | 3gek-A | 2.4 | 4.6 | 60 | 132 | 8 | MOLECULE: | PUTATIVE THIOESTERASE YHDA |
| 99:00:00 | 7w7i-B | 2.4 | 7.4 | 80 | 342 | 5 | MOLECULE: | FCTA DOMAIN-CONTAINING PROTEIN |
| 0:00 | 5e4m-A | 2.4 | 4.5 | 61 | 177 | 2 | MOLECULE: | HYDROXYNITRILE LYASE |
| 1:00 | 4fnv-A | 2.4 | 6 | 76 | 659 | 9 | MOLECULE: | HEPARINASE III PROTEIN, HEPARITIN S |
| 2:00 | 7bcz-A | 2.4 | 4.1 | 59 | 242 | 3 | MOLECULE: | 4'-PHOSPHOPANTETHEINYL TRANSFERASE |
| 3:00 | 3qq2-B | 2.4 | 7.6 | 89 | 250 | 6 | MOLECULE: | BRKA AUTOTRANSPORTER |
| 4:00 | 8e4g-P | 2.4 | 7.5 | 76 | 793 | 5 | MOLECULE: | PORTAL PROTEIN |
| 5:00 | 7brm-A | 2.4 | 6.6 | 84 | 257 | 5 | MOLECULE: | CURLI PRODUCTION ASSEMBLY/TRANSPORT |
| 6:00 | 3gzb-A | 2.4 | 5.3 | 78 | 154 | 6 | MOLECULE: | PUTATIVE SNOAL-LIKE POLYKETIDE CYCL |
| 7:00 | 6z9c-A | 2.4 | 3.6 | 70 | 258 | 4 | MOLECULE: | POLYMERASE DELTA-INTERACTING PROTEI |
| 8:00 | 3n91-A | 2.4 | 5.5 | 74 | 315 | 9 | MOLECULE: | UNCHARACTERIZED PROTEIN |
| 9:00 | 4wp5-A | 2.4 | 8.5 | 81 | 199 | 4 | MOLECULE: | MRNA EXPORT PROTEIN |
| 10:00 | 4dk6-C | 2.4 | 4.6 | 38 | 87 | 8 | MOLECULE: | SINGLE DOMAIN ANTIBODY VHH |
| 11:00 | 1bqs-A | 2.4 | 3.7 | 58 | 209 | 9 | MOLECULE: | PROTEIN (MUCOSAL ADDRESSIN CELL ADH |
| 12:00 | 4hg6-A | 2.4 | 5.8 | 81 | 747 | 9 | MOLECULE: | CELLULOSE SYNTHASE SUBUNIT A |
| 13:00 | 4u3q-B | 2.4 | 3.4 | 48 | 99 | 13 | MOLECULE: | 17 KDA LIPOPROTEIN |
| 14:00 | 6em5-d | 2.4 | 4.7 | 53 | 107 | 2 | MOLECULE: | 5.8S RIBOSOMAL RNA |
| 15:00 | 3aki-A | 2.4 | 6.5 | 72 | 448 | 3 | MOLECULE: | PUTATIVE SECRETED ALPHA L-ARABINOFU |
| 16:00 | 1dml-E | 2.4 | 6.6 | 53 | 275 | 13 | MOLECULE: | DNA POLYMERASE PROCESSIVITY FACTOR |
| 17:00 | 5yje-B | 2.4 | 11.5 | 59 | 318 | 7 | MOLECULE: | PROTEIN HIRA |
| 18:00 | 4ll4-C | 2.4 | 5 | 65 | 281 | 8 | MOLECULE: | THIOREDOXIN-INTERACTING PROTEIN |
| 19:00 | 5a1v-K | 2.4 | 8 | 50 | 1125 | 10 | MOLECULE: | ADP-RIBOSYLATION FACTOR 1 |
| 20:00 | 8a9b-B | 2.4 | 6.3 | 84 | 809 | 4 | MOLECULE: | LIPID BINDING PROTEIN P116 (MPN213) |
| 21:00 | 7z6e-C | 2.4 | 6.8 | 63 | 534 | 8 | MOLECULE: | SERINE/THREONINE-PROTEIN KINASE MRC |
| 22:00 | 6orj-A | 2.4 | 12.1 | 61 | 243 | 3 | MOLECULE: | PHIKZ164 |
| 23:00 | 3af5-A | 2.4 | 4.3 | 58 | 638 | 5 | MOLECULE: | PUTATIVE UNCHARACTERIZED PROTEIN PH |
| 24:00:00 | 5hl8-A | 2.4 | 3.2 | 48 | 81 | 2 | MOLECULE: | TYPE II SECRETION SYSTEM PROTEIN L |
| 25:00:00 | 6gj1-D | 2.4 | 4.1 | 59 | 121 | 3 | MOLECULE: | PUTATIVE TYPE VI SECRETION PROTEIN |
| 26:00:00 | 6z3b-A | 2.4 | 8.1 | 59 | 371 | 7 | MOLECULE: | GFO/IDH/MOCA FAMILY OXIDOREDUCTASE |
| 27:00:00 | 8cm1-A | 2.4 | 7 | 63 | 179 | 3 | MOLECULE: | OUTER-MEMBRANE LIPOPROTEIN LOLB |
| 28:00:00 | 8y19-A | 2.4 | 6.3 | 61 | 1208 | 7 | MOLECULE: | SPIKE GLYCOPROTEIN |
| 29:00:00 | 7rpy-A | 2.4 | 3.2 | 66 | 241 | 6 | MOLECULE: | COHESIN-CONTAINING PROTEIN |
| 30:00:00 | 7oix-A | 2.4 | 7.2 | 81 | 265 | 5 | MOLECULE: | SYNCYTIN-2 |
| 31:00:00 | 3u4y-A | 2.4 | 4.4 | 63 | 319 | 11 | MOLECULE: | UNCHARACTERIZED PROTEIN |
| 32:00:00 | 5nzu-D | 2.4 | 8.3 | 61 | 425 | 11 | MOLECULE: | COATOMER SUBUNIT ALPHA |

| | | | | | | | | |
|---|---|---|---|---|---|---|---|---|
| 33:00:00 | 3hw2-A | 2.4 | 5.1 | 62 | 308 | 3 | MOLECULE: | PROTEIN SIFA |
| 34:00:00 | 8ahx-G | 2.4 | 5.1 | 74 | 197 | 4 | MOLECULE: | ION-TRANSLOCATING OXIDOREDUCTASE CO |
| 35:00:00 | 8uci-A | 2.4 | 7.4 | 52 | 381 | 4 | MOLECULE: | ATP DEPENDENT DNA LIGASE |
| 36:00:00 | 1n7u-A | 2.4 | 6.1 | 75 | 554 | 7 | MOLECULE: | ADSORPTION PROTEIN P2 |
| 37:00:00 | 7z4f-J | 2.4 | 5.3 | 92 | 603 | 9 | MOLECULE: | PUTATIVE STRUCTURAL PROTEIN |
| 38:00:00 | 7bjt-A | 2.4 | 7.8 | 69 | 727 | 4 | MOLECULE: | ALGINATE LYASE, FAMILY PL17 |
| 39:00:00 | 7ylz-A | 2.4 | 14.8 | 78 | 576 | 4 | MOLECULE: | HYDROXYAMIDOTRANSFERASE |
| 40:00:00 | 8dys-A | 2.4 | 5.2 | 62 | 409 | 5 | MOLECULE: | EUKARYOTIC TRANSLATION INITIATION F |
| 41:00:00 | 2pmq-B | 2.4 | 6.1 | 52 | 374 | 6 | MOLECULE: | MANDELATE RACEMASE/MUCONATE LACTONI |
| 42:00:00 | 3fvz-A | 2.4 | 5.1 | 56 | 329 | 9 | MOLECULE: | PEPTIDYL-GLYCINE ALPHA-AMIDATING MO |
| 43:00:00 | 6rkw-B | 2.4 | 3.1 | 47 | 781 | 6 | MOLECULE: | DNA GYRASE SUBUNIT A |
| 44:00:00 | 4ozq-B | 2.4 | 10 | 78 | 687 | 6 | MOLECULE: | CHIMERA OF MALTOSE-BINDING PERIPLAS |
| 45:00:00 | 4k15-A | 2.4 | 9.9 | 60 | 138 | 2 | MOLECULE: | LMO2686 PROTEIN |
| 46:00:00 | 7eu9-A | 2.4 | 6.3 | 70 | 1078 | 4 | MOLECULE: | CAS12I1 D647A MUTANT |
| 47:00:00 | 6trq-C | 2.4 | 9.6 | 52 | 336 | 6 | MOLECULE: | M7GPPPX DIPHOSPHATASE |
| 48:00:00 | 4asc-A | 2.4 | 4.3 | 68 | 308 | 0 | MOLECULE: | KELCH REPEAT AND BTB DOMAIN-CONTAIN |
| 49:00:00 | 5jyx-B | 2.4 | 6.8 | 63 | 109 | 2 | MOLECULE: | ARCHEAOSINE SYNTHASE QUEF-LIKE |
| 50:00:00 | 4fvk-A | 2.4 | 5.3 | 71 | 367 | 3 | MOLECULE: | NEURAMINIDASE |
| 51:00:00 | 6chg-A | 2.4 | 4.1 | 61 | 305 | 2 | MOLECULE: | KLLA0E24487P |
| 52:00:00 | 5oa3-0 | 2.4 | 9.1 | 63 | 479 | 5 | MOLECULE: | EUKARYOTIC TRANSLATION INITIATION F |
| 53:00:00 | 3b7c-A | 2.4 | 4.8 | 66 | 121 | 5 | MOLECULE: | UNCHARACTERIZED PROTEIN |
| 54:00:00 | 8pel-E | 2.4 | 10 | 67 | 304 | 4 | MOLECULE: | RRP45 |
| 55:00:00 | 5vzt-B | 2.4 | 5.9 | 69 | 418 | 4 | MOLECULE: | S-PHASE KINASE-ASSOCIATED PROTEIN 1 |
| 56:00:00 | 8t8s-B | 2.4 | 6.7 | 66 | 664 | 8 | MOLECULE: | SORTILIN |
| 57:00:00 | 5c0p-A | 2.4 | 3.8 | 65 | 284 | 5 | MOLECULE: | ENDO-ARABINASE |
| 58:00:00 | 5y6q-C | 2.4 | 6.3 | 82 | 748 | 9 | MOLECULE: | ALDEHYDE OXIDASE SMALL SUBUNIT |
| 59:00:00 | 8etf-C | 2.4 | 7.2 | 62 | 330 | 8 | MOLECULE: | CHOLOYLGLYCINE HYDROLASE |
| 60:00:00 | 1lqs-R | 2.4 | 3.4 | 61 | 207 | 7 | MOLECULE: | INTERLEUKIN-10 RECEPTOR ALPHA CHAIN |
| 61:00:00 | 6m76-A | 2.4 | 8.5 | 64 | 923 | 3 | MOLECULE: | LPXTG-MOTIF CELL WALL ANCHOR DOMAIN |
| 62:00:00 | 6zyg-A | 2.4 | 3.1 | 43 | 113 | 2 | MOLECULE: | PROTEALYSIN-ASSOCIATED PROTEIN |
| 63:00:00 | 4fmb-A | 2.4 | 4.4 | 67 | 351 | 1 | MOLECULE: | CYSTEINE PROTEASE-LIKE VIRA |
| 64:00:00 | 6mfi-A | 2.4 | 5.2 | 51 | 264 | 6 | MOLECULE: | METALLO-BETA-LACTAMASE |
| 65:00:00 | 5lox-1 | 2.4 | 6.6 | 75 | 241 | 7 | MOLECULE: | PEPTIDASE |
| 66:00:00 | 7vnx-A | 2.4 | 7.5 | 48 | 216 | 8 | MOLECULE: | TKARKI |
| 67:00:00 | 7uwf-A | 2.4 | 3.3 | 54 | 358 | 13 | MOLECULE: | WD REPEAT-CONTAINING PROTEIN 18 |
| 68:00:00 | 7bkb-k | 2.4 | 3.8 | 45 | 386 | 4 | MOLECULE: | COB--COM HETERODISULFIDE REDUCTASE |
| 69:00:00 | 7mge-A | 2.4 | 5.5 | 61 | 349 | 3 | MOLECULE: | WD REPEAT-CONTAINING PROTEIN 41 |
| 70:00:00 | 5dky-A | 2.4 | 3.2 | 64 | 928 | 3 | MOLECULE: | ALPHA GLUCOSIDASE-LIKE PROTEIN |
| 71:00:00 | 7c5w-A | 2.4 | 4.9 | 61 | 164 | 5 | MOLECULE: | IOTA-CARBONIC ANHYDRASE |
| 72:00:00 | 7plc-D | 2.4 | 4.6 | 64 | 286 | 2 | MOLECULE: | SMP-30/CGR1 FAMILY PROTEIN |
| 73:00:00 | 2oit-A | 2.4 | 5.5 | 74 | 434 | 4 | MOLECULE: | NUCLEOPORIN 214KDA |
| 74:00:00 | 6uio-B | 2.4 | 4.8 | 55 | 110 | 7 | MOLECULE: | CYSTATIN-8 |
| 75:00:00 | 7epn-A | 2.4 | 4.7 | 61 | 133 | 7 | MOLECULE: | SNOAL-LIKE DOMAIN-CONTAINING PROTEI |
| 76:00:00 | 8ghl-G | 2.4 | 6.2 | 57 | 276 | 7 | MOLECULE: | PROTEIN HIR1 |
| 77:00:00 | 6h01-C | 2.4 | 4.9 | 57 | 230 | 7 | MOLECULE: | GREEN FLUORESCENT PROTEIN |
| 78:00:00 | 4w6l-A | 2.4 | 5.8 | 51 | 204 | 4 | MOLECULE: | FLUORESCENT PROTEIN D117C |

| | | | | | | | | |
|---|---|---|---|---|---|---|---|---|
| 79:00:00 | 3adf-B | 2.4 | 5 | 60 | 209 | 5 | MOLECULE: | MONOMERIC AZAMI GREEN |
| 80:00:00 | 2whu-A | 2.4 | 10.8 | 58 | 215 | 9 | MOLECULE: | LARGE STOKES SHIFT FLUORESCENT PROT |
| 81:00:00 | 2wht-C | 2.4 | 10.9 | 59 | 215 | 8 | MOLECULE: | LARGE STOKES SHIFT FLUORESCENT PROT |
| 82:00:00 | 5hzo-A | 2.4 | 8.4 | 54 | 228 | 9 | MOLECULE: | GREEN FLUORESCENT PROTEIN |
| 83:00:00 | 3evv-A | 2.4 | 6.9 | 76 | 385 | 5 | MOLECULE: | MYOSIN LIGHT CHAIN KINASE, GREEN FL |
| 84:00:00 | 6fp8-A | 2.4 | 8.1 | 54 | 216 | 11 | MOLECULE: | GFP-LIKE FLUORESCENT CHROMOPROTEIN |
| 85:00:00 | 8ill-A | 2.4 | 8.5 | 54 | 213 | 2 | MOLECULE: | GREEN FLUORESCENT PROTEIN |
| 86:00:00 | 3svs-C | 2.4 | 11 | 57 | 222 | 11 | MOLECULE: | MKATE S158A/S143C |
| 87:00:00 | 6wv5-A | 2.4 | 10.4 | 56 | 370 | 9 | MOLECULE: | VITAMIN K EPOXIDE REDUCTASE CYS43SE |
| 88:00:00 | 8i4o-C | 2.4 | 5.2 | 75 | 206 | 7 | MOLECULE: | SPLIT GREEN FLOURESCENT PROTEIN |
| 89:00:00 | 6y1g-C | 2.4 | 7.6 | 45 | 221 | 4 | MOLECULE: | GFP-LIKE NON-FLUORESCENT CHROMOPROT |
| 90:00:00 | 3mgf-B | 2.4 | 4.7 | 61 | 209 | 5 | MOLECULE: | FLUORESCENT PROTEIN |
| 91:00:00 | 4w77-B | 2.4 | 8.2 | 54 | 217 | 9 | MOLECULE: | FLUORESCENT PROTEIN D21H/K26C |
| 92:00:00 | 4w6r-N | 2.4 | 5.6 | 50 | 174 | 4 | MOLECULE: | FLUORESCENT PROTEIN D102C |
| 93:00:00 | 5f9g-A | 2.4 | 6.9 | 77 | 228 | 5 | MOLECULE: | PNGFP1.5-Y.CRO,GREEN FLUORESCENT PR |
| 94:00:00 | 5exb-C | 2.4 | 8.2 | 54 | 223 | 6 | MOLECULE: | GREEN FLUORESCENT PROTEIN |
| 95:00:00 | 6nql-C | 2.4 | 8.4 | 52 | 216 | 6 | MOLECULE: | FLUORESCENT PROTEIN DRONPA |
| 96:00:00 | 8i4o-I | 2.4 | 5.1 | 75 | 204 | 7 | MOLECULE: | SPLIT GREEN FLOURESCENT PROTEIN |
| 97:00:00 | 6uru-A | 2.4 | 10 | 82 | 500 | 5 | MOLECULE: | IACHSNFR PRECURSOR |
| 98:00:00 | 4jfg-H | 2.4 | 8.4 | 55 | 225 | 5 | MOLECULE: | GREEN FLUORESCENT PROTEIN |
| 99:00:00 | 2ib5-F | 2.4 | 10.8 | 56 | 225 | 11 | MOLECULE: | CHROMO PROTEIN |
| 0:00 | 5dpi-E | 2.4 | 11.5 | 47 | 225 | 6 | MOLECULE: | GREEN FLUORESCENT PROTEIN |
| 1:00 | 5ox9-A | 2.4 | 9.6 | 60 | 226 | 5 | MOLECULE: | GREEN FLUORESCENT PROTEIN |
| 2:00 | 8i4o-E | 2.4 | 5.2 | 74 | 203 | 5 | MOLECULE: | SPLIT GREEN FLOURESCENT PROTEIN |
| 3:00 | 6un4-A | 2.4 | 9.5 | 62 | 245 | 6 | MOLECULE: | GREEN FLUORESCENT PROTEIN |
| 4:00 | 8i4o-A | 2.4 | 5 | 74 | 208 | 5 | MOLECULE: | SPLIT GREEN FLOURESCENT PROTEIN |
| 5:00 | 8jxg-A | 2.3 | 4.7 | 63 | 1114 | 3 | MOLECULE: | RAT RAP |
| 6:00 | 2kie-A | 2.3 | 5.1 | 56 | 124 | 2 | MOLECULE: | INOSITOL POLYPHOSPHATE 5-PHOSPHATAS |
| 7:00 | 2hjs-A | 2.3 | 5.7 | 67 | 334 | 6 | MOLECULE: | USG-1 PROTEIN HOMOLOG |
| 8:00 | 1yax-A | 2.3 | 6 | 66 | 153 | 8 | MOLECULE: | VIRULENCE SENSOR PROTEIN PHOQ, SENS |
| 9:00 | 2jgs-B | 2.3 | 4.2 | 70 | 116 | 11 | MOLECULE: | CIRCULAR PERMUTANT OF AVIDIN |
| 10:00 | 8dl1-A | 2.3 | 6 | 53 | 713 | 8 | MOLECULE: | ALPHA AMYLASE, CATALYTIC DOMAIN PRO |
| 11:00 | 4f8b-B | 2.3 | 4.4 | 63 | 145 | 0 | MOLECULE: | NADPH-DEPENDENT 7-CYANO-7-DEAZAGUAN |
| 12:00 | 7kra-G | 2.3 | 5.4 | 74 | 151 | 7 | MOLECULE: | ER MEMBRANE PROTEIN COMPLEX SUBUNIT |
| 13:00 | 5coz-A | 2.3 | 4.9 | 85 | 347 | 7 | MOLECULE: | UNCHARACTERIZED PROTEIN |
| 14:00 | 5k36-C | 2.3 | 10.1 | 67 | 340 | 6 | MOLECULE: | EXOSOME COMPLEX COMPONENT RRP45 |
| 15:00 | 5bsz-A | 2.3 | 4.2 | 68 | 248 | 6 | MOLECULE: | N-METHYLTRANSFERASE |
| 16:00 | 2vzs-A | 2.3 | 6.2 | 63 | 857 | 5 | MOLECULE: | EXO-BETA-D-GLUCOSAMINIDASE |
| 17:00 | 3en1-B | 2.3 | 6.5 | 74 | 180 | 8 | MOLECULE: | BENZENE 1,2-DIOXYGENASE SUBUNIT ALP |
| 18:00 | 7dpy-A | 2.3 | 3 | 53 | 90 | 4 | MOLECULE: | BRUCELLA ABORTUS PHIA |
| 19:00 | 4imh-A | 2.3 | 9.1 | 61 | 347 | 10 | MOLECULE: | HEMIN DEGRADING FACTOR |
| 20:00 | 1tbu-B | 2.3 | 3 | 52 | 98 | 6 | MOLECULE: | PEROXISOMAL ACYL-COENZYME A THIOEST |
| 21:00 | 4x2m-A | 2.3 | 5.8 | 73 | 177 | 11 | MOLECULE: | MTR2 |
| 22:00 | 3m83-A | 2.3 | 6.5 | 70 | 325 | 0 | MOLECULE: | ACETYL XYLAN ESTERASE |
| 23:00 | 5imm-A | 2.3 | 3.8 | 53 | 120 | 8 | MOLECULE: | NANOBODY |
| 24:00:00 | 3l5z-A | 2.3 | 3.3 | 60 | 143 | 3 | MOLECULE: | TRANSCRIPTIONAL REGULATOR, GNTR FAM |

| | | | | | | | | |
|---|---|---|---|---|---|---|---|---|
| 25:00:00 | 8b7d-A | 2.3 | 4.4 | 62 | 147 | 3 | MOLECULE: | TRANSMEMBRANE PROTEIN 106B |
| 26:00:00 | 7rpo-A | 2.3 | 5.6 | 58 | 252 | 7 | MOLECULE: | DNA POLYMERASE SLIDING CLAMP 1 |
| 27:00:00 | 7tmu-C | 2.3 | 5 | 70 | 146 | 6 | MOLECULE: | SRPBCC FAMILY PROTEIN |
| 28:00:00 | 8gy2-A | 2.3 | 4.6 | 79 | 723 | 4 | MOLECULE: | ALCOHOL DEHYDROGENASE (QUINONE), DE |
| 29:00:00 | 7ns9-A | 2.3 | 6.5 | 58 | 178 | 5 | MOLECULE: | TRIPHOSPHATE TUNNEL METALLOENZYME S |
| 30:00:00 | 6f7k-B | 2.3 | 3.9 | 61 | 547 | 7 | MOLECULE: | TAILSPIKE |
| 31:00:00 | 3bws-A | 2.3 | 4 | 70 | 407 | 3 | MOLECULE: | PROTEIN LP49 |
| 32:00:00 | 4mis-A | 2.3 | 3.7 | 49 | 98 | 2 | MOLECULE: | PUTATIVE UNCHARACTERIZED PROTEIN |
| 33:00:00 | 6k4e-B | 2.3 | 10.9 | 68 | 246 | 7 | MOLECULE: | HAMP DOMAIN-CONTAINING PROTEIN |
| 34:00:00 | 7b2a-A | 2.3 | 4.9 | 65 | 162 | 5 | MOLECULE: | CIRPA5 |
| 35:00:00 | 5eq7-A | 2.3 | 4.8 | 55 | 258 | 4 | MOLECULE: | INOSITOL MONOPHOSPHATASE |
| 36:00:00 | 8ts6-A | 2.3 | 5.8 | 85 | 500 | 2 | MOLECULE: | PORTAL PROTEIN |
| 37:00:00 | 7cu9-B | 2.3 | 4.5 | 62 | 193 | 11 | MOLECULE: | TUBE-FORMING PROTEIN IN MYCOBACTERI |
| 38:00:00 | 4e6f-B | 2.3 | 6.1 | 64 | 173 | 11 | MOLECULE: | UNCHARACTERIZED PROTEIN |
| 39:00:00 | 8e1m-C | 2.3 | 5.1 | 74 | 264 | 7 | MOLECULE: | MITOCHONDRIAL IMPORT INNER MEMBRANE |
| 40:00:00 | 7mo0-B | 2.3 | 2.6 | 45 | 126 | 4 | MOLECULE: | GTP-BINDING NUCLEAR PROTEIN RAN |
| 41:00:00 | 5g25-A | 2.3 | 2.9 | 45 | 82 | 2 | MOLECULE: | TYPE-IV LIKE COMPETENCE PILIN TTHA1 |
| 42:00:00 | 7d1b-A | 2.3 | 9.3 | 62 | 302 | 6 | MOLECULE: | LEUCINE AMINOPEPTIDASE |
| 43:00:00 | 2wzp-A | 2.3 | 5.4 | 59 | 266 | 3 | MOLECULE: | PUTATIVE RECEPTOR BINDING PROTEIN |
| 44:00:00 | 4jp0-A | 2.3 | 4.5 | 83 | 378 | 5 | MOLECULE: | 43.8 KDA INSECTICIDAL CRYSTAL PROTE |
| 45:00:00 | 2k4v-A | 2.3 | 4.9 | 58 | 125 | 2 | MOLECULE: | UNCHARACTERIZED PROTEIN PA1076 |
| 46:00:00 | 3nwz-A | 2.3 | 5.3 | 62 | 155 | 5 | MOLECULE: | BH2602 PROTEIN |
| 47:00:00 | 7zlv-h | 2.3 | 3.6 | 69 | 527 | 3 | MOLECULE: | PROBABLE CENTRAL STRAIGHT FIBER |
| 48:00:00 | 5nwm-A | 2.3 | 10.6 | 65 | 132 | 3 | MOLECULE: | NUCLEAR RECEPTOR COACTIVATOR 1 |
| 49:00:00 | 2xgr-A | 2.3 | 4.9 | 78 | 197 | 5 | MOLECULE: | SPD1 NUCLEASE |
| 50:00:00 | 3gvj-A | 2.3 | 3.7 | 80 | 670 | 8 | MOLECULE: | ENDO-N-ACETYLNEURAMINIDASE |
| 51:00:00 | 3rqb-B | 2.3 | 6.9 | 68 | 265 | 9 | MOLECULE: | UNCHARACTERIZED PROTEIN |
| 52:00:00 | 1hpw-A | 2.3 | 9.5 | 65 | 129 | 5 | MOLECULE: | FIMBRIAL PROTEIN |
| 53:00:00 | 4eg9-A | 2.3 | 5.4 | 81 | 231 | 4 | MOLECULE: | UNCHARACTERIZED PROTEIN SAOUHSC_027 |
| 54:00:00 | 3hfi-A | 2.3 | 2.8 | 55 | 137 | 5 | MOLECULE: | PUTATIVE REGULATOR |
| 55:00:00 | 2d42-A | 2.3 | 2.6 | 48 | 249 | 2 | MOLECULE: | NON-TOXIC CRYSTAL PROTEIN |
| 56:00:00 | 4ffu-K | 2.3 | 3.2 | 59 | 135 | 10 | MOLECULE: | OXIDASE |
| 57:00:00 | 2m7o-A | 2.3 | 3.3 | 43 | 70 | 14 | MOLECULE: | UNCHARACTERIZED PROTEIN |
| 58:00:00 | 2a8i-A | 2.3 | 7.6 | 91 | 341 | 5 | MOLECULE: | THREONINE ASPARTASE 1 |
| 59:00:00 | 4fzq-A | 2.3 | 4.3 | 52 | 79 | 2 | MOLECULE: | UNCHARACTERIZED PROTEIN CONSERVED I |
| 60:00:00 | 2wbv-B | 2.3 | 5 | 60 | 186 | 8 | MOLECULE: | FIBER PROTEIN |
| 61:00:00 | 8wzb-G | 2.3 | 8.6 | 59 | 387 | 3 | MOLECULE: | DPY30 DOMAIN CONTAINING 2 |
| 62:00:00 | 2gxf-A | 2.3 | 3.2 | 53 | 119 | 2 | MOLECULE: | HYPOTHETICAL PROTEIN YYBH |
| 63:00:00 | 4q28-A | 2.3 | 3.6 | 51 | 110 | 6 | MOLECULE: | PERIPLAKIN |
| 64:00:00 | 5e27-A | 2.3 | 8.9 | 58 | 226 | 7 | MOLECULE: | RESUSCITATION-PROMOTING FACTOR RPFB |
| 65:00:00 | 1su3-B | 2.3 | 4 | 55 | 416 | 5 | MOLECULE: | INTERSTITIAL COLLAGENASE |
| 66:00:00 | 7kqq-B | 2.3 | 6.5 | 60 | 306 | 5 | MOLECULE: | WD REPEAT-CONTAINING PROTEIN 55 |
| 67:00:00 | 4ggt-B | 2.3 | 4.1 | 63 | 106 | 5 | MOLECULE: | BRADAVIDIN 2 |
| 68:00:00 | 5f0j-B | 2.3 | 6.2 | 75 | 306 | 4 | MOLECULE: | VACUOLAR PROTEIN SORTING-ASSOCIATED |
| 69:00:00 | 7vmb-A | 2.3 | 6.1 | 67 | 359 | 7 | MOLECULE: | IQ MOTIF AND SEC7 DOMAIN-CONTAINING |
| 70:00:00 | 5ukg-A | 2.3 | 9.1 | 79 | 393 | 3 | MOLECULE: | K-GECO |

| | | | | | | | | |
|---|---|---|---|---|---|---|---|---|
| 71:00:00 | 5gzl-A | 2.3 | 3.9 | 77 | 358 | 3 | MOLECULE: | LYSINE CYCLODEAMINASE |
| 72:00:00 | 3lju-X | 2.3 | 3.8 | 56 | 373 | 9 | MOLECULE: | ARF-GAP WITH DUAL PH DOMAIN-CONTAIN |
| 73:00:00 | 8ffz-E | 2.3 | 4.8 | 69 | 518 | 7 | MOLECULE: | TRANSCRIPTION FACTOR IIIA |
| 74:00:00 | 3ijc-A | 2.3 | 4.9 | 60 | 357 | 0 | MOLECULE: | POLYCOMB PROTEIN EED |
| 75:00:00 | 5gyy-B | 2.3 | 7.2 | 67 | 405 | 3 | MOLECULE: | S-RECEPTOR KINASE SRK9 |
| 76:00:00 | 7pfp-A | 2.3 | 6.9 | 63 | 563 | 5 | MOLECULE: | UROMODULIN |
| 77:00:00 | 5wru-E | 2.3 | 5.2 | 56 | 311 | 4 | MOLECULE: | PROBABLE INORGANIC PYROPHOSPHATASE |
| 78:00:00 | 8ey4-A | 2.3 | 2 | 39 | 88 | 10 | MOLECULE: | CYS_RICH_CPCC DOMAIN-CONTAINING PRO |
| 79:00:00 | 6upk-G | 2.3 | 3 | 45 | 423 | 4 | MOLECULE: | HISTONE H3.1 |
| 80:00:00 | 3kk7-A | 2.3 | 7.5 | 77 | 502 | 1 | MOLECULE: | PUTATIVE CELL INVASION PROTEIN WITH |
| 81:00:00 | 2dsb-A | 2.3 | 10.6 | 47 | 206 | 11 | MOLECULE: | ADP-SUGAR PYROPHOSPHATASE |
| 82:00:00 | 8dit-C | 2.3 | 5.3 | 69 | 736 | 4 | MOLECULE: | VACUOLAR PROTEIN SORTING-ASSOCIATED |
| 83:00:00 | 6um1-A | 2.3 | 5 | 53 | 2208 | 8 | MOLECULE: | CATION-INDEPENDENT MANNOSE-6-PHOSPH |
| 84:00:00 | 8eec-A | 2.3 | 5.1 | 67 | 317 | 4 | MOLECULE: | ISOFORM 2 OF MITOGEN-ACTIVATED PROT |
| 85:00:00 | 5hr7-A | 2.3 | 8.2 | 63 | 361 | 11 | MOLECULE: | TRNA GLU |
| 86:00:00 | 5ch1-A | 2.3 | 5.2 | 62 | 481 | 5 | MOLECULE: | PUTATIVE POLYCOMB PROTEIN EED |
| 87:00:00 | 3rri-A | 2.3 | 3.9 | 51 | 132 | 4 | MOLECULE: | GLYOXALASE/BLEOMYCIN RESISTANCE PRO |
| 88:00:00 | 5fql-A | 2.3 | 5.3 | 83 | 507 | 7 | MOLECULE: | IDURONATE-2-SULFATASE |
| 89:00:00 | 8c8g-B | 2.3 | 6.5 | 57 | 1195 | 7 | MOLECULE: | PUTATIVE BOTULINUM-LIKE TOXIN WO |
| 90:00:00 | 7w1m-B | 2.3 | 12 | 61 | 528 | 7 | MOLECULE: | STRUCTURAL MAINTENANCE OF CHROMOSOM |
| 91:00:00 | 1r0u-A | 2.3 | 4.6 | 60 | 142 | 5 | MOLECULE: | PROTEIN YWIB |
| 92:00:00 | 1kit-A | 2.3 | 5.3 | 69 | 757 | 3 | MOLECULE: | SIALIDASE |
| 93:00:00 | 8jxc-A | 2.3 | 4.6 | 65 | 1337 | 11 | MOLECULE: | LDL RECEPTOR RELATED PROTEIN 2 |
| 94:00:00 | 7usl-C | 2.3 | 17.3 | 64 | 715 | 6 | MOLECULE: | INTEGRIN ALPHA-M |
| 95:00:00 | 4fzm-A | 2.3 | 4.7 | 65 | 276 | 2 | MOLECULE: | BACTERIOCIN |
| 96:00:00 | 3jbr-F | 2.3 | 5.9 | 58 | 872 | 0 | MOLECULE: | VOLTAGE-DEPENDENT L-TYPE CALCIUM CH |
| 97:00:00 | 5fzp-A | 2.3 | 3.7 | 57 | 344 | 2 | MOLECULE: | DISPASE AUTOLYSIS-INDUCING PROTEIN |
| 98:00:00 | 3dm0-A | 2.3 | 5 | 64 | 675 | 5 | MOLECULE: | MALTOSE-BINDING PERIPLASMIC PROTEIN |
| 99:00:00 | 7klj-A | 2.3 | 4.3 | 48 | 324 | 8 | MOLECULE: | ISOFORM 2 OF KINESIN-LIKE PROTEIN K |
| 0:00 | 6bxr-A | 2.3 | 5.7 | 57 | 273 | 0 | MOLECULE: | MITOCHONDRIAL ASSOCIATION FACTOR 1 |
| 1:00 | 7woo-D | 2.3 | 5.9 | 69 | 1398 | 12 | MOLECULE: | NUCLEOPORIN NIC96 |
| 2:00 | 3mlq-A | 2.3 | 5 | 74 | 186 | 8 | MOLECULE: | DNA-DIRECTED RNA POLYMERASE SUBUNIT |
| 3:00 | 4orl-A | 2.3 | 2.5 | 52 | 110 | 8 | MOLECULE: | UNCHARACTERIZED PROTEIN |
| 4:00 | 8t1l-L | 2.3 | 3.2 | 58 | 559 | 2 | MOLECULE: | MEDIATOR OF RNA POLYMERASE II TRANS |
| 5:00 | 6nd4-O | 2.3 | 4.4 | 60 | 832 | 10 | MOLECULE: | ETS RRNA |
| 6:00 | 5k1c-C | 2.3 | 8.3 | 65 | 407 | 8 | MOLECULE: | UBIQUITIN CARBOXYL-TERMINAL HYDROLA |
| 7:00 | 5o63-B | 2.3 | 3 | 54 | 161 | 7 | MOLECULE: | RESTRICTION ENDONUCLEASE UBALAI |
| 8:00 | 3f5o-A | 2.3 | 2.9 | 65 | 138 | 5 | MOLECULE: | THIOESTERASE SUPERFAMILY MEMBER 2 |
| 9:00 | 1gju-A | 2.3 | 8 | 51 | 636 | 4 | MOLECULE: | MALTODEXTRIN GLYCOSYLTRANSFERASE |
| 10:00 | 8of7-A | 2.3 | 4.5 | 70 | 143 | 14 | MOLECULE: | RHS FAMILY PROTEIN |
| 11:00 | 4i27-A | 2.3 | 10.5 | 68 | 361 | 10 | MOLECULE: | DNA NUCLEOTIDYLEXOTRANSFERASE |
| 12:00 | 8f5o-C | 2.3 | 4.8 | 55 | 1179 | 0 | MOLECULE: | INTRAFLAGELLAR TRANSPORT PROTEIN 12 |
| 13:00 | 5ixh-B | 2.3 | 7.2 | 59 | 161 | 8 | MOLECULE: | YCEI-LIKE DOMAIN PROTEIN |
| 14:00 | 3no2-A | 2.3 | 3.7 | 64 | 274 | 3 | MOLECULE: | UNCHARACTERIZED PROTEIN |
| 15:00 | 5i8u-C | 2.3 | 7.7 | 46 | 203 | 0 | MOLECULE: | ADP-RIBOSE PYROPHOSPHATASE |
| 16:00 | 6nd4-S | 2.3 | 7.6 | 68 | 481 | 1 | MOLECULE: | ETS RRNA |

| | | | | | | | | |
|---|---|---|---|---|---|---|---|---|
| 17:00 | 8pbz-B | 2.3 | 5.7 | 56 | 1380 | 5 | MOLECULE: | MGP-OPERON PROTEIN 3 |
| 18:00 | 2y3u-A | 2.3 | 4.5 | 67 | 679 | 4 | MOLECULE: | COLLAGENASE |
| 19:00 | 6fpt-A | 2.3 | 5.7 | 58 | 391 | 5 | MOLECULE: | E3 UBIQUITIN-PROTEIN LIGASE TRIM71 |
| 20:00 | 3htx-D | 2.3 | 4.4 | 55 | 796 | 5 | MOLECULE: | HEN1 |
| 21:00 | 2z2o-C | 2.3 | 5.7 | 59 | 299 | 3 | MOLECULE: | VIRGINIAMYCIN B LYASE |
| 22:00 | 8glo-A | 2.3 | 3.8 | 60 | 255 | 7 | MOLECULE: | HEMOPHILIN |
| 23:00 | 6w17-F | 2.3 | 5.3 | 55 | 168 | 4 | MOLECULE: | ACTIN-RELATED PROTEIN 3 |
| 24:00:00 | 3kny-A | 2.3 | 8.8 | 76 | 194 | 7 | MOLECULE: | HYPOTHETICAL PROTEIN BT_3535 |
| 25:00:00 | 6ah0-W | 2.3 | 6 | 58 | 463 | 3 | MOLECULE: | U5SNRNA |
| 26:00:00 | 8wqr-B | 2.3 | 7.4 | 59 | 356 | 5 | MOLECULE: | ACTIVATING MOLECULE IN BECN1-REGULA |
| 27:00:00 | 6rlb-D | 2.3 | 5.1 | 66 | 446 | 9 | MOLECULE: | O6-ALKYLGUANINE-DNA ALKYLTRANSFERAS |
| 28:00:00 | 3w0s-A | 2.3 | 2.8 | 45 | 298 | 2 | MOLECULE: | HYGROMYCIN-B 4-O-KINASE |
| 29:00:00 | 8ttb-B | 2.3 | 3.4 | 58 | 426 | 9 | MOLECULE: | SERINE/THREONINE-PROTEIN PHOSPHATAS |
| 30:00:00 | 6wg3-A | 2.3 | 10.6 | 61 | 561 | 5 | MOLECULE: | STRUCTURAL MAINTENANCE OF CHROMOSOM |
| 31:00:00 | 8q7n-F | 2.3 | 6 | 60 | 431 | 8 | MOLECULE: | U5 SNRNA |
| 32:00:00 | 5ncl-A | 2.3 | 13.5 | 58 | 418 | 5 | MOLECULE: | SERINE/THREONINE-PROTEIN KINASE CBK |
| 33:00:00 | 4ci8-A | 2.3 | 9.2 | 56 | 640 | 4 | MOLECULE: | ECHINODERM MICROTUBULE-ASSOCIATED P |
| 34:00:00 | 7xp3-A | 2.3 | 4.4 | 58 | 153 | 2 | MOLECULE: | NAC DOMAIN-CONTAINING PROTEIN 92 |
| 35:00:00 | 2por-A | 2.3 | 7.9 | 78 | 301 | 5 | MOLECULE: | PORIN |
| 36:00:00 | 7wh0-A | 2.3 | 5.7 | 66 | 529 | 6 | MOLECULE: | BETA-CAROTENE 15,15'-MONOOXYGENASE |
| 37:00:00 | 8h6h-A | 2.3 | 5.2 | 70 | 986 | 4 | MOLECULE: | CELLODEXTRIN PHOSPHORYLASE |
| 38:00:00 | 8boz-J | 2.3 | 5.5 | 54 | 216 | 9 | MOLECULE: | TRANSMEMBRANE PROTEIN |
| 39:00:00 | 2cwz-A | 2.3 | 4.5 | 67 | 138 | 6 | MOLECULE: | THIOESTERASE FAMILY PROTEIN |
| 40:00:00 | 4gk9-A | 2.3 | 4.2 | 54 | 279 | 4 | MOLECULE: | AGGLUTININ (BOA) |
| 41:00:00 | 3i0y-A | 2.3 | 4.6 | 67 | 138 | 4 | MOLECULE: | PUTATIVE POLYKETIDE CYCLASE |
| 42:00:00 | 2p9w-A | 2.3 | 5.4 | 66 | 333 | 8 | MOLECULE: | MAL S 1 ALLERGENIC PROTEIN |
| 43:00:00 | 8ajm-B | 2.3 | 5.2 | 60 | 395 | 3 | MOLECULE: | DNA DAMAGE-BINDING PROTEIN 1 |
| 44:00:00 | 2jox-A | 2.3 | 3.5 | 54 | 106 | 4 | MOLECULE: | CHURCHILL PROTEIN |
| 45:00:00 | 4g1e-A | 2.3 | 7.8 | 75 | 964 | 15 | MOLECULE: | INTEGRIN ALPHA-V |
| 46:00:00 | 7kpz-B | 2.3 | 5.7 | 58 | 126 | 7 | MOLECULE: | HUPZ |
| 47:00:00 | 1sjw-A | 2.3 | 7.5 | 64 | 142 | 5 | MOLECULE: | NOGALONIC ACID METHYL ESTER CYCLASE |
| 48:00:00 | 8ghn-B | 2.3 | 5.7 | 65 | 800 | 5 | MOLECULE: | PROTEIN HIR1 |
| 49:00:00 | 4po7-A | 2.3 | 7 | 59 | 658 | 2 | MOLECULE: | SORTILIN |
| 50:00:00 | 4r1k-A | 2.3 | 5.8 | 62 | 135 | 3 | MOLECULE: | UNCHARACTERIZED PROTEIN |
| 51:00:00 | 5tgn-D | 2.3 | 4 | 56 | 109 | 11 | MOLECULE: | UNCHARACTERIZED PROTEIN |
| 52:00:00 | 7c5v-A | 2.3 | 5.2 | 59 | 164 | 0 | MOLECULE: | IOTA-CARBONIC ANHYDRASE |
| 53:00:00 | 1mwt-A | 2.3 | 6.3 | 72 | 635 | 11 | MOLECULE: | PBP2A |
| 54:00:00 | 7c5v-B | 2.3 | 4.9 | 62 | 163 | 6 | MOLECULE: | IOTA-CARBONIC ANHYDRASE |
| 55:00:00 | 7aqo-E | 2.3 | 5.5 | 59 | 348 | 7 | MOLECULE: | THO COMPLEX SUBUNIT 2 |
| 56:00:00 | 7kd9-E | 2.3 | 6.6 | 82 | 219 | 7 | MOLECULE: | GALLATE DECARBOXYLASE |
| 57:00:00 | 5ts4-A | 2.3 | 4.9 | 48 | 101 | 6 | MOLECULE: | DENOVO NTF2 |
| 58:00:00 | 3ec9-A | 2.3 | 5.4 | 63 | 130 | 3 | MOLECULE: | UNCHARACTERIZED NTF2-LIKE PROTEIN |
| 59:00:00 | 7uhy-B | 2.3 | 3.8 | 58 | 761 | 5 | MOLECULE: | GATOR COMPLEX PROTEIN MIOS |
| 60:00:00 | 7plc-C | 2.3 | 6 | 64 | 286 | 2 | MOLECULE: | SMP-30/CGR1 FAMILY PROTEIN |
| 61:00:00 | 4xzz-B | 2.3 | 5.4 | 70 | 316 | 9 | MOLECULE: | CONSERVED HYPOTHETICAL SECRETED PRO |
| 62:00:00 | 5d17-M | 2.3 | 3.3 | 62 | 159 | 5 | MOLECULE: | TRANSPOSON TN7 TRANSPOSITION PROTEI |

| | | | | | | | | |
|---|---|---|---|---|---|---|---|---|
| 63:00:00 | 4w6p-B | 2.3 | 5.7 | 50 | 214 | 4 | MOLECULE: | FLUORESCENT PROTEIN D102C |
| 64:00:00 | 4w6g-A | 2.3 | 6.8 | 58 | 218 | 3 | MOLECULE: | FLUORESCENT PROTEIN D190C |
| 65:00:00 | 7lqo-A | 2.3 | 5.9 | 75 | 229 | 5 | MOLECULE: | RED FLUORESCENT PEROXYNITRITE BIOSE |
| 66:00:00 | 5ukg-B | 2.3 | 8.9 | 81 | 389 | 2 | MOLECULE: | K-GECO |
| 67:00:00 | 4w6p-A | 2.3 | 10.6 | 57 | 217 | 4 | MOLECULE: | FLUORESCENT PROTEIN D102C |
| 68:00:00 | 7a7n-A | 2.3 | 6.6 | 61 | 227 | 3 | MOLECULE: | GREEN FLUORESCENT PROTEIN |
| 69:00:00 | 4w7a-C | 2.3 | 10.7 | 58 | 220 | 9 | MOLECULE: | FLUORESCENT PROTEIN D21H/K26C |
| 70:00:00 | 1yhg-B | 2.3 | 8.1 | 53 | 224 | 13 | MOLECULE: | GREEN FLUORESCENT PROTEIN |
| 71:00:00 | 6un6-B | 2.3 | 4.7 | 71 | 230 | 10 | MOLECULE: | GREEN FLUORESCENT PROTEIN |
| 72:00:00 | 6jc6-D | 2.3 | 4.4 | 63 | 220 | 2 | MOLECULE: | SHCP |
| 73:00:00 | 4ik3-A | 2.3 | 11.9 | 80 | 393 | 5 | MOLECULE: | RCAMP, GREEN FLUORESCENT PROTEIN |
| 74:00:00 | 4emq-F | 2.3 | 4.4 | 62 | 213 | 6 | MOLECULE: | FLUORESCENT PROTEIN DRONPA |
| 75:00:00 | 5exb-D | 2.3 | 9.3 | 58 | 221 | 7 | MOLECULE: | GREEN FLUORESCENT PROTEIN |
| 76:00:00 | 3lvc-B | 2.3 | 8.3 | 55 | 226 | 9 | MOLECULE: | GREEN FLUORESCENT PROTEIN |
| 77:00:00 | 7y96-A | 2.3 | 10.3 | 57 | 309 | 9 | MOLECULE: | GREEN FLUORESCENT PROTEIN,MEMBRANE |
| 78:00:00 | 8bxt-A | 2.3 | 8.5 | 54 | 214 | 2 | MOLECULE: | STAYGOLD |
| 79:00:00 | 7qlj-A | 2.3 | 4.7 | 62 | 219 | 5 | MOLECULE: | RSKIIRO |
| 80:00:00 | 4w6n-C | 2.3 | 8.7 | 57 | 218 | 7 | MOLECULE: | FLUORESCENT PROTEIN D117C |
| 81:00:00 | 8i4o-G | 2.3 | 5.8 | 72 | 205 | 7 | MOLECULE: | SPLIT GREEN FLOURESCENT PROTEIN |
| 82:00:00 | 6lnp-D | 2.3 | 10.4 | 56 | 362 | 7 | MOLECULE: | FUSION PROTEIN OF GREEN FLUORESCENT |
| 83:00:00 | 6ofk-A | 2.3 | 8.9 | 58 | 225 | 5 | MOLECULE: | GREEN FLUORESCENT PROTEIN (GFP) S6 |
| 84:00:00 | 6ofn-B | 2.3 | 5.3 | 70 | 236 | 10 | MOLECULE: | GREEN FLUORESCENT PROTEIN (GFP) S6 |
| 85:00:00 | 6ya9-A | 2.3 | 6.4 | 75 | 400 | 1 | MOLECULE: | RSCGAMP |
| 86:00:00 | 4w6r-F | 2.3 | 8.4 | 53 | 201 | 9 | MOLECULE: | FLUORESCENT PROTEIN D102C |
| 87:00:00 | 2ib5-H | 2.3 | 11.1 | 57 | 225 | 11 | MOLECULE: | CHROMO PROTEIN |
| 88:00:00 | 3mgf-D | 2.3 | 5.1 | 60 | 212 | 5 | MOLECULE: | FLUORESCENT PROTEIN |
| 89:00:00 | 3wck-A | 2.3 | 4.7 | 67 | 217 | 3 | MOLECULE: | MONOMERIC PHOTOSENITIZING FLUORESCE |
| 90:00:00 | 2odr-A | 2.2 | 7.7 | 65 | 491 | 6 | MOLECULE: | PHOSPHOSERYL-TRNA SYNTHETASE |
| 91:00:00 | 1s5a-B | 2.2 | 5.2 | 68 | 143 | 9 | MOLECULE: | HYPOTHETICAL PROTEIN YESE |
| 92:00:00 | 3d4e-A | 2.2 | 7.3 | 45 | 162 | 2 | MOLECULE: | PUTATIVE BETA-LACTAMASE INHIBITOR P |
| 93:00:00 | 2amu-A | 2.2 | 2.7 | 52 | 132 | 6 | MOLECULE: | PUTATIVE SUPEROXIDE REDUCTASE |
| 94:00:00 | 2jmu-A | 2.2 | 3.3 | 62 | 224 | 2 | MOLECULE: | THIAMINE-TRIPHOSPHATASE |
| 95:00:00 | 7lhg-C | 2.2 | 6.9 | 76 | 805 | 1 | MOLECULE: | P FIMBRIAL USHER PROTEIN PAPC |
| 96:00:00 | 1qab-E | 2.2 | 4.4 | 69 | 180 | 12 | MOLECULE: | PROTEIN (TRANSTHYRETIN) |
| 97:00:00 | 3j2m-A | 2.2 | 4 | 65 | 211 | 3 | MOLECULE: | TAIL CONNECTOR PROTEIN GP15 |
| 98:00:00 | 2q78-F | 2.2 | 8.5 | 71 | 136 | 6 | MOLECULE: | UNCHARACTERIZED PROTEIN |
| 99:00:00 | 4gah-B | 2.2 | 4 | 66 | 195 | 6 | MOLECULE: | THIOESTERASE SUPERFAMILY MEMBER 4 |
| 0:00 | 8ih7-B | 2.2 | 4 | 63 | 317 | 3 | MOLECULE: | 4-HYDROXY-2-OXOVALERATE ALDOLASE |
| 1:00 | 3bc9-A | 2.2 | 4.3 | 54 | 585 | 7 | MOLECULE: | ALPHA AMYLASE, CATALYTIC REGION |
| 2:00 | 7orl-A | 2.2 | 7.3 | 82 | 2183 | 7 | MOLECULE: | RNA (5'-D(*(GTG))-R(P*AP*AP*UP*AP*C |
| 3:00 | 4rqo-B | 2.2 | 4.2 | 68 | 448 | 6 | MOLECULE: | L-SERINE DEHYDRATASE |
| 4:00 | 4q6u-B | 2.2 | 5 | 64 | 324 | 3 | MOLECULE: | UNCHARACTERIZED PROTEIN |
| 5:00 | 8bad-B | 2.2 | 4.9 | 88 | 351 | 6 | MOLECULE: | BINARY TOXIN A-LIKE PROTEIN |
| 6:00 | 8k49-T | 2.2 | 5.7 | 74 | 627 | 5 | MOLECULE: | VP2 |
| 7:00 | 7wb4-c | 2.2 | 3.8 | 59 | 346 | 2 | MOLECULE: | OUTER NUP133 |
| 8:00 | 1f13-A | 2.2 | 12.2 | 68 | 721 | 4 | MOLECULE: | CELLULAR COAGULATION FACTOR XIII ZY |

| | | | | | | | | |
|---|---|---|---|---|---|---|---|---|
| 9:00 | 8r1o-D | 2.2 | 10.4 | 63 | 233 | 5 | MOLECULE: | RRP45 |
| 10:00 | 5jxl-A | 2.2 | 3.4 | 74 | 851 | 7 | MOLECULE: | FLAGELLAR HOOK PROTEIN FLGE |
| 11:00 | 5zx9-A | 2.2 | 2.7 | 57 | 165 | 2 | MOLECULE: | ALANINE AND PROLINE-RICH SECRETED P |
| 12:00 | 6ihc-F | 2.2 | 4.1 | 67 | 153 | 4 | MOLECULE: | 3-HYDROXYACYL-[ACYL-CARRIER-PROTEIN |
| 13:00 | 8x6f-C | 2.2 | 6.6 | 65 | 1153 | 3 | MOLECULE: | DNA-DIRECTED RNA POLYMERASE SUBUNIT |
| 14:00 | 7p2p-C | 2.2 | 4.3 | 72 | 174 | 6 | MOLECULE: | SIGNAL PEPTIDASE COMPLEX CATALYTIC |
| 15:00 | 3ceb-A | 2.2 | 6.8 | 47 | 194 | 6 | MOLECULE: | D-AMINOACID AMINOTRANSFERASE-LIKE P |
| 16:00 | 4doh-B | 2.2 | 3.5 | 51 | 193 | 2 | MOLECULE: | INTERLEUKIN-20 |
| 17:00 | 4uoj-A | 2.2 | 5.3 | 74 | 917 | 7 | MOLECULE: | BETA-MANNOSIDASE GH2 |
| 18:00 | 7k4o-A | 2.2 | 6.2 | 78 | 554 | 10 | MOLECULE: | CARBOXYLIC ESTER HYDROLASE |
| 19:00 | 1jmo-A | 2.2 | 3.4 | 67 | 427 | 7 | MOLECULE: | THROMBIN, LIGHT CHAIN |
| 20:00 | 4pw1-B | 2.2 | 5.9 | 56 | 215 | 5 | MOLECULE: | UNCHARACTERIZED PROTEIN |
| 21:00 | 5fmv-A | 2.2 | 3.3 | 57 | 351 | 7 | MOLECULE: | RECEPTOR-TYPE TYROSINE-PROTEIN PHOS |
| 22:00 | 2xem-D | 2.2 | 3.4 | 60 | 146 | 13 | MOLECULE: | DYNE7 |
| 23:00 | 2bmo-B | 2.2 | 5.9 | 82 | 194 | 7 | MOLECULE: | OXYGENASE-ALPHA NBDO |
| 24:00:00 | 3bnv-D | 2.2 | 4.3 | 64 | 143 | 9 | MOLECULE: | CJ0977 |
| 25:00:00 | 6wum-a | 2.2 | 11.3 | 90 | 445 | 7 | MOLECULE: | SAM35 |
| 26:00:00 | 4l8n-A | 2.2 | 9.2 | 72 | 448 | 6 | MOLECULE: | PDZ DOMAIN PROTEIN |
| 27:00:00 | 3r87-A | 2.2 | 3.7 | 58 | 132 | 7 | MOLECULE: | PUTATIVE UNCHARACTERIZED PROTEIN |
| 28:00:00 | 3fig-B | 2.2 | 4.1 | 64 | 578 | 5 | MOLECULE: | 2-ISOPROPYLMALATE SYNTHASE |
| 29:00:00 | 6ged-A | 2.2 | 3.8 | 76 | 295 | 4 | MOLECULE: | PRGB |
| 30:00:00 | 2v5y-A | 2.2 | 3.4 | 59 | 564 | 5 | MOLECULE: | RECEPTOR-TYPE TYROSINE-PROTEIN PHOS |
| 31:00:00 | 6zyx-Y | 2.2 | 8.9 | 90 | 1067 | 4 | MOLECULE: | DYNEIN HEAVY CHAIN, OUTER ARM PROTE |
| 32:00:00 | 2ve7-B | 2.2 | 6.5 | 61 | 303 | 5 | MOLECULE: | KINETOCHORE PROTEIN HEC1, KINETOCHO |
| 33:00:00 | 6hiu-A | 2.2 | 7 | 65 | 143 | 3 | MOLECULE: | CYTOCHROME P460 |
| 34:00:00 | 8oe4-D | 2.2 | 3.7 | 56 | 510 | 2 | MOLECULE: | INTERLEUKIN-12 SUBUNIT BETA |
| 35:00:00 | 2kd2-A | 2.2 | 2.8 | 47 | 94 | 4 | MOLECULE: | FAS APOPTOTIC INHIBITORY MOLECULE 1 |
| 36:00:00 | 3rd4-B | 2.2 | 4.3 | 50 | 82 | 6 | MOLECULE: | UNCHARACTERIZED PROTEIN |
| 37:00:00 | 2ymw-B | 2.2 | 5.5 | 77 | 685 | 9 | MOLECULE: | L-LYSINE 6-OXIDASE |
| 38:00:00 | 2dyt-A | 2.2 | 5.6 | 82 | 213 | 1 | MOLECULE: | AUTOPHAGY-RELATED PROTEIN 3 |
| 39:00:00 | 5cn2-A | 2.2 | 3.1 | 53 | 115 | 11 | MOLECULE: | ADP-RIBOSYLATION FACTOR-BINDING PRO |
| 40:00:00 | 4e0s-B | 2.2 | 7.5 | 85 | 898 | 6 | MOLECULE: | COMPLEMENT C5 |
| 41:00:00 | 3to3-B | 2.2 | 9.2 | 53 | 595 | 8 | MOLECULE: | PETROBACTIN BIOSYNTHESIS PROTEIN AS |
| 42:00:00 | 8uq9-a | 2.2 | 3.6 | 66 | 433 | 6 | MOLECULE: | E3 UBIQUITIN-PROTEIN LIGASE RNF168, |
| 43:00:00 | 8pel-C | 2.2 | 11.3 | 69 | 311 | 10 | MOLECULE: | RRP45 |
| 44:00:00 | 8hbn-A | 2.2 | 4.1 | 66 | 371 | 5 | MOLECULE: | MRNA EXPORT FACTOR MEX67 |
| 45:00:00 | 3w15-A | 2.2 | 6.4 | 55 | 336 | 5 | MOLECULE: | PEROXISOMAL TARGETING SIGNAL 2 RECE |
| 46:00:00 | 1z24-A | 2.2 | 4.2 | 59 | 189 | 8 | MOLECULE: | INSECTICYANIN A FORM |
| 47:00:00 | 4ycz-A | 2.2 | 4.3 | 58 | 713 | 5 | MOLECULE: | FUSION PROTEIN OF SEC13 AND NUP145C |
| 48:00:00 | 6nqz-B | 2.2 | 4.5 | 60 | 215 | 2 | MOLECULE: | FLAGELLAR COILING PROTEIN B |
| 49:00:00 | 5j6q-A | 2.2 | 10.1 | 63 | 595 | 6 | MOLECULE: | CELL WALL BINDING PROTEIN CWP8 |
| 50:00:00 | 6j6q-F | 2.2 | 4.8 | 65 | 161 | 2 | MOLECULE: | PRE-MRNA-SPLICING FACTOR 8 |
| 51:00:00 | 6fwv-A | 2.2 | 4.3 | 68 | 522 | 3 | MOLECULE: | COLLAGEN ADHESION PROTEIN |
| 52:00:00 | 3fka-B | 2.2 | 4.8 | 58 | 120 | 3 | MOLECULE: | UNCHARACTERIZED NTF-2 LIKE PROTEIN |
| 53:00:00 | 7uic-n | 2.2 | 10 | 82 | 220 | 5 | MOLECULE: | MEDIATOR OF RNA POLYMERASE II TRANS |
| 54:00:00 | 3qs3-H | 2.2 | 3.4 | 65 | 177 | 12 | MOLECULE: | FIMBRILLIN MATB HOMOLOG, ECPD |

| | | | | | | | | |
|---|---|---|---|---|---|---|---|---|
| 55:00:00 | 5u8o-B | 2.2 | 3.8 | 55 | 360 | 0 | MOLECULE: | ZN-DEPENDENT HYDROLASE |
| 56:00:00 | 2ooi-B | 2.2 | 9.1 | 67 | 158 | 6 | MOLECULE: | SA0254 PROTEIN |
| 57:00:00 | 6wo0-A | 2.2 | 5.7 | 67 | 359 | 7 | MOLECULE: | PROTEIN ARTEMIS |
| 58:00:00 | 2f1z-B | 2.2 | 2.9 | 52 | 481 | 10 | MOLECULE: | UBIQUITIN CARBOXYL-TERMINAL HYDROLA |
| 59:00:00 | 3i0o-A | 2.2 | 4.2 | 51 | 329 | 2 | MOLECULE: | SPECTINOMYCIN PHOSPHOTRANSFERASE |
| 60:00:00 | 6s62-A | 2.2 | 4.9 | 59 | 485 | 3 | MOLECULE: | PROPIONATE CATABOLIC PROTEIN PRPD |
| 61:00:00 | 8dfv-K | 2.2 | 5.6 | 57 | 253 | 7 | MOLECULE: | ENDORIBONUCLEASE DCR-1 |
| 62:00:00 | 4ckn-C | 2.2 | 3.2 | 57 | 181 | 7 | MOLECULE: | SAS-6 |
| 63:00:00 | 4q1z-A | 2.2 | 3.7 | 48 | 389 | 8 | MOLECULE: | PUTATIVE LIPOPROTEIN |
| 64:00:00 | 3c6k-D | 2.2 | 8.3 | 52 | 348 | 4 | MOLECULE: | SPERMINE SYNTHASE |
| 65:00:00 | 6wes-A | 2.2 | 3.8 | 63 | 158 | 8 | MOLECULE: | TOX3 |
| 66:00:00 | 4jgl-A | 2.2 | 4.6 | 67 | 152 | 7 | MOLECULE: | HYPOTHETICAL PROTEIN |
| 67:00:00 | 7vcf-F | 2.2 | 6.5 | 79 | 402 | 9 | MOLECULE: | TIC214 |
| 68:00:00 | 1ss4-A | 2.2 | 3 | 44 | 149 | 9 | MOLECULE: | GLYOXALASE FAMILY PROTEIN |
| 69:00:00 | 1im0-A | 2.2 | 7.2 | 90 | 262 | 3 | MOLECULE: | OUTER MEMBRANE PHSOPHOLIPASE A |
| 70:00:00 | 6rap-E | 2.2 | 5.1 | 73 | 275 | 4 | MOLECULE: | AFP1 |
| 71:00:00 | 4cg4-A | 2.2 | 4 | 62 | 376 | 8 | MOLECULE: | PYRIN |
| 72:00:00 | 8khg-A | 2.2 | 4.5 | 63 | 271 | 3 | MOLECULE: | ITACONYL-COA HYDRATASE |
| 73:00:00 | 5c0w-B | 2.2 | 11.9 | 70 | 244 | 4 | MOLECULE: | EXOSOME COMPLEX COMPONENT RRP45 |
| 74:00:00 | 7e5c-A | 2.2 | 6.7 | 79 | 439 | 3 | MOLECULE: | XAA-PRO DIPEPTIDASE |
| 75:00:00 | 3soy-A | 2.2 | 5.1 | 65 | 142 | 8 | MOLECULE: | NTF2-LIKE SUPERFAMILY PROTEIN |
| 76:00:00 | 3osm-A | 2.2 | 3.4 | 59 | 119 | 3 | MOLECULE: | SERINE/THREONINE-PROTEIN KINASE KCC |
| 77:00:00 | 6w40-A | 2.2 | 4.6 | 57 | 120 | 11 | MOLECULE: | DENOVO NTF2 |
| 78:00:00 | 7zbo-A | 2.2 | 5.2 | 75 | 348 | 8 | MOLECULE: | AMINE DEHYDROGENASE |
| 79:00:00 | 6vbu-5 | 2.2 | 4.2 | 59 | 300 | 7 | MOLECULE: | BARDET-BIEDL SYNDROME 18 PROTEIN |
| 80:00:00 | 3b7f-A | 2.2 | 4.5 | 63 | 368 | 8 | MOLECULE: | GLYCOSYL HYDROLASE, BNR REPEAT |
| 81:00:00 | 1rwi-A | 2.2 | 4.8 | 63 | 256 | 11 | MOLECULE: | SERINE/THREONINE-PROTEIN KINASE PKN |
| 82:00:00 | 7e38-B | 2.2 | 7.3 | 65 | 305 | 5 | MOLECULE: | DEOXYPODOPHYLLOTOXIN SYNTHASE |
| 83:00:00 | 5g24-A | 2.2 | 4.8 | 63 | 200 | 5 | MOLECULE: | TYPE-IV LIKE PILIN TTHA1219 |
| 84:00:00 | 7pgr-F | 2.2 | 9.1 | 64 | 2423 | 9 | MOLECULE: | NEUROFIBROMIN |
| 85:00:00 | 8hsb-B | 2.2 | 3.2 | 59 | 159 | 10 | MOLECULE: | CDNG |
| 86:00:00 | 7lyt-A | 2.2 | 3.2 | 59 | 712 | 3 | MOLECULE: | CASPHI |
| 87:00:00 | 3azv-A | 2.2 | 4.4 | 66 | 418 | 3 | MOLECULE: | D/C MOSAIC NEUROTOXIN |
| 88:00:00 | 7zm6-B | 2.2 | 6 | 80 | 433 | 5 | MOLECULE: | ATTACHMENT PROTEIN |
| 89:00:00 | 6qm7-H | 2.2 | 3.8 | 58 | 229 | 5 | MOLECULE: | PROTEASOME ALPHA1 CHAIN |
| 90:00:00 | 3e8p-A | 2.2 | 3.6 | 70 | 153 | 7 | MOLECULE: | UNCHARACTERIZED PROTEIN |
| 91:00:00 | 6qmu-A | 2.2 | 5.6 | 68 | 184 | 4 | MOLECULE: | NEUTROPHIL GELATINASE-ASSOCIATED LI |
| 92:00:00 | 4p07-A | 2.2 | 3.8 | 71 | 569 | 8 | MOLECULE: | ARYLSULFATE SULFOTRANSFERASE ASST |
| 93:00:00 | 5aiw-A | 2.2 | 4.5 | 69 | 127 | 7 | MOLECULE: | TRAH |
| 94:00:00 | 8hbn-B | 2.2 | 5.1 | 69 | 165 | 10 | MOLECULE: | MRNA EXPORT FACTOR MEX67 |
| 95:00:00 | 8fne-A | 2.2 | 7.6 | 68 | 525 | 4 | MOLECULE: | MALTOSE/MALTODEXTRIN-BINDING PERIPL |
| 96:00:00 | 8ee7-A | 2.2 | 10 | 64 | 456 | 11 | MOLECULE: | PTUA |
| 97:00:00 | 8cp6-B | 2.2 | 16.1 | 73 | 1084 | 10 | MOLECULE: | TOXIN PROTEIN TSE5 |
| 98:00:00 | 7q05-E | 2.2 | 5.2 | 64 | 409 | 5 | MOLECULE: | TEREPHTHALATE 1,2-DIOXYGENASE, TERM |
| 99:00:00 | 7mni-C | 2.2 | 4.1 | 61 | 412 | 5 | MOLECULE: | NUCLEAR PORE COMPLEX PROTEIN NUP88 |
| 0:00 | 6qz4-A | 2.2 | 7.9 | 80 | 568 | 6 | MOLECULE: | MONO(2-HYDROXYETHYL) TEREPHTHALATE |

| | | | | | | | | |
|---|---|---|---|---|---|---|---|---|
| 1:00 | 5mzv-C | 2.2 | 6.5 | 61 | 293 | 3 | MOLECULE: | INTERLEUKIN-12 SUBUNIT BETA |
| 2:00 | 5je1-A | 2.2 | 9.1 | 54 | 245 | 6 | MOLECULE: | METHYL TRANSFERASE |
| 3:00 | 4wl2-A | 2.2 | 6.1 | 61 | 346 | 2 | MOLECULE: | PUTATIVE EXPORTED CHOLOYLGLYCINE HY |
| 4:00 | 6t9i-D | 2.2 | 4.9 | 59 | 566 | 7 | MOLECULE: | TRANSCRIPTION FACTOR SPT20 |
| 5:00 | 4xnn-A | 2.2 | 5.4 | 66 | 445 | 8 | MOLECULE: | CELLOBIOHYDROLASE CHBI |
| 6:00 | 4fi1-A | 2.2 | 12.8 | 54 | 371 | 6 | MOLECULE: | CASEIN KINASE II SUBUNIT ALPHA |
| 7:00 | 4v0n-D | 2.2 | 5.5 | 59 | 304 | 5 | MOLECULE: | BARDET-BIEDL SYNDROME 1 PROTEIN |
| 8:00 | 4nkq-A | 2.2 | 5.3 | 47 | 412 | 11 | MOLECULE: | CYTOKINE RECEPTOR COMMON SUBUNIT BE |
| 9:00 | 7qpg-B | 2.2 | 9.5 | 59 | 591 | 5 | MOLECULE: | PROTEIN ZWILCH HOMOLOG |
| 10:00 | 2p3p-B | 2.2 | 3.1 | 63 | 198 | 6 | MOLECULE: | HYPOTHETICAL PROTEIN |
| 11:00 | 4epc-A | 2.2 | 3.5 | 50 | 152 | 2 | MOLECULE: | N-ACETYLMURAMOYL-L-ALANINE AMIDASE |
| 12:00 | 5o9z-L | 2.2 | 4.5 | 56 | 459 | 4 | MOLECULE: | PRE-MRNA-PROCESSING-SPLICING FACTOR |
| 13:00 | 2vdu-D | 2.2 | 4.9 | 62 | 376 | 0 | MOLECULE: | TRNA (GUANINE-N(7)-)-METHYLTRANSFER |
| 14:00 | 8onf-A | 2.2 | 7.5 | 66 | 258 | 5 | MOLECULE: | CELL WALL SURFACE ANCHOR FAMILY PRO |
| 15:00 | 6zxf-z | 2.2 | 3.8 | 52 | 373 | 10 | MOLECULE: | PRE-18S RIBOSOMAL RNA |
| 16:00 | 6a5g-B | 2.2 | 6.1 | 65 | 157 | 5 | MOLECULE: | [4+2] AND [4+6] CYCLASE STMD |
| 17:00 | 6i9f-A | 2.2 | 4.6 | 51 | 155 | 4 | MOLECULE: | FATTY ACID-BINDING PROTEIN HOMOLOG |
| 18:00 | 8day-A | 2.2 | 4.2 | 58 | 399 | 5 | MOLECULE: | DIMETHYLALLYLTRYPTOPHAN SYNTHASE 1 |
| 19:00 | 4dvy-P | 2.2 | 5.7 | 61 | 656 | 5 | MOLECULE: | CYTOTOXICITY-ASSOCIATED IMMUNODOMIN |
| 20:00 | 6yvd-D | 2.2 | 9.9 | 65 | 414 | 3 | MOLECULE: | CONDENSIN COMPLEX SUBUNIT 2 |
| 21:00 | 6dgv-A | 2.2 | 10.4 | 89 | 553 | 3 | MOLECULE: | FLUORESCENT GABA SENSOR PRECURSOR |
| 22:00 | 3cje-A | 2.2 | 7 | 67 | 150 | 9 | MOLECULE: | OSMC-LIKE PROTEIN |
| 23:00 | 8qx8-B | 2.2 | 8.4 | 57 | 728 | 2 | MOLECULE: | VACUOLAR PROTEIN SORTING-ASSOCIATED |
| 24:00:00 | 3s5t-A | 2.2 | 7.3 | 61 | 253 | 2 | MOLECULE: | DUF3298 FAMILY PROTEIN |
| 25:00:00 | 6pon-A | 2.2 | 3.2 | 66 | 255 | 8 | MOLECULE: | ADHERENCE AND VIRULENCE PROTEIN A |
| 26:00:00 | 8c47-B | 2.2 | 4.9 | 47 | 151 | 9 | MOLECULE: | PROFILIN |
| 27:00:00 | 5oc7-D | 2.2 | 4.1 | 47 | 112 | 11 | MOLECULE: | BREAKPOINT CLUSTER REGION PROTEIN,P |
| 28:00:00 | 2pso-B | 2.2 | 9.3 | 57 | 198 | 11 | MOLECULE: | STAR-RELATED LIPID TRANSFER PROTEIN |
| 29:00:00 | 1k4n-A | 2.2 | 9 | 54 | 183 | 2 | MOLECULE: | PROTEIN EC4020 |
| 30:00:00 | 8opr-A | 2.2 | 7.5 | 62 | 651 | 6 | MOLECULE: | S-LAYER PROTEIN EA1 |
| 31:00:00 | 5cl2-A | 2.2 | 3.7 | 60 | 246 | 2 | MOLECULE: | SPORULATION-CONTROL PROTEIN SPO0M |
| 32:00:00 | 5k3w-A | 2.2 | 5.1 | 64 | 298 | 5 | MOLECULE: | CPUTA1 |
| 33:00:00 | 7c3c-B | 2.2 | 4.2 | 74 | 346 | 1 | MOLECULE: | AOFLEA |
| 34:00:00 | 5yww-A | 2.2 | 8.9 | 59 | 485 | 3 | MOLECULE: | NUCLEOTIDE BINDING PROTEIN PINC |
| 35:00:00 | 1eqr-A | 2.2 | 4.3 | 64 | 590 | 13 | MOLECULE: | ASPARTYL-TRNA SYNTHETASE |
| 36:00:00 | 6eke-C | 2.2 | 4 | 40 | 43 | 5 | MOLECULE: | LECTIN |
| 37:00:00 | 8qyn-N | 2.2 | 3.6 | 57 | 216 | 7 | MOLECULE: | PROTEASOME SUBUNIT ALPHA TYPE-2 |
| 38:00:00 | 5cww-B | 2.2 | 6 | 71 | 538 | 6 | MOLECULE: | NUCLEOPORIN NUP145N |
| 39:00:00 | 8beq-A | 2.2 | 4.1 | 66 | 534 | 3 | MOLECULE: | FRUCTOFURANOSIDASE FROM RHODOTORULA |
| 40:00:00 | 2o62-A | 2.2 | 6.3 | 60 | 269 | 8 | MOLECULE: | HYPOTHETICAL PROTEIN |
| 41:00:00 | 6wg3-B | 2.2 | 11.4 | 62 | 693 | 8 | MOLECULE: | STRUCTURAL MAINTENANCE OF CHROMOSOM |
| 42:00:00 | 3e29-B | 2.2 | 3.5 | 69 | 135 | 3 | MOLECULE: | UNCHARACTERIZED PROTEIN Q7WE92_BORB |
| 43:00:00 | 1ns0-B | 2.2 | 3.6 | 48 | 346 | 4 | MOLECULE: | GALACTOSE MUTAROTASE |
| 44:00:00 | 5ejq-A | 2.2 | 5.8 | 69 | 501 | 6 | MOLECULE: | MYOSIN-I HEAVY CHAIN |
| 45:00:00 | 6b3y-A | 2.2 | 4.6 | 49 | 225 | 2 | MOLECULE: | DENN DOMAIN-CONTAINING PROTEIN 3 |
| 46:00:00 | 4z48-A | 2.2 | 8 | 69 | 240 | 1 | MOLECULE: | UNCHARACTERIZED PROTEIN |

| | | | | | | | | |
|---|---|---|---|---|---|---|---|---|
| 47:00:00 | 5ine-A | 2.2 | 6.1 | 51 | 348 | 6 | MOLECULE: | PRE-GLYCOPROTEIN POLYPROTEIN GP COM |
| 48:00:00 | 2fwv-A | 2.2 | 3.8 | 65 | 190 | 5 | MOLECULE: | HYPOTHETICAL PROTEIN MTUBF_01000852 |
| 49:00:00 | 8osv-A | 2.2 | 5.2 | 53 | 248 | 8 | MOLECULE: | GDMF |
| 50:00:00 | 4yy8-B | 2.2 | 6.5 | 71 | 377 | 11 | MOLECULE: | KELCH PROTEIN |
| 51:00:00 | 7zgr-D | 2.2 | 4.7 | 58 | 395 | 3 | MOLECULE: | PROTEIN CFT1 |
| 52:00:00 | 6qx6-A | 2.2 | 5.5 | 63 | 264 | 11 | MOLECULE: | FERREDOXIN BILIN REDUCTASE PLASTID |
| 53:00:00 | 7pab-A | 2.2 | 3.2 | 54 | 415 | 7 | MOLECULE: | NUCLEAR EGRESS PROTEIN 2,NUCLEAR EG |
| 54:00:00 | 5mus-B | 2.2 | 8 | 64 | 328 | 6 | MOLECULE: | L PROTEIN |
| 55:00:00 | 5xn7-B | 2.2 | 10.1 | 55 | 551 | 2 | MOLECULE: | PUTATIVE RTX-TOXIN |
| 56:00:00 | 4je3-A | 2.2 | 5.3 | 67 | 244 | 10 | MOLECULE: | CENTRAL KINETOCHORE SUBUNIT IML3 |
| 57:00:00 | 6n90-A | 2.2 | 8.2 | 52 | 88 | 6 | MOLECULE: | UNCHARACTERIZED PROTEIN |
| 58:00:00 | 1ve3-B | 2.2 | 8.7 | 66 | 227 | 5 | MOLECULE: | HYPOTHETICAL PROTEIN PH0226 |
| 59:00:00 | 3u0s-A | 2.2 | 7.6 | 75 | 327 | 0 | MOLECULE: | DIISOPROPYL-FLUOROPHOSPHATASE |
| 60:00:00 | 1vhz-A | 2.2 | 8.5 | 44 | 186 | 9 | MOLECULE: | ADP COMPOUNDS HYDROLASE NUDE |
| 61:00:00 | 7ztq-A | 2.2 | 6.3 | 55 | 236 | 5 | MOLECULE: | CAROTENOID-BINDING PROTEIN |
| 62:00:00 | 6y86-A | 2.2 | 9.1 | 53 | 425 | 6 | MOLECULE: | MEMBRANE PROTEIN INSERTASE YIDC |
| 63:00:00 | 6f3a-h | 2.2 | 6.5 | 53 | 401 | 11 | MOLECULE: | ARP1 ACTIN RELATED PROTEIN 1 HOMOLO |
| 64:00:00 | 4wp5-B | 2.2 | 6.1 | 73 | 171 | 10 | MOLECULE: | MRNA EXPORT PROTEIN |
| 65:00:00 | 7jmv-A | 2.2 | 6.2 | 85 | 223 | 6 | MOLECULE: | PEA PATHOGENICITY PROTEIN 2 |
| 66:00:00 | 7f0o-B | 2.2 | 8.9 | 70 | 149 | 9 | MOLECULE: | NSRQ |
| 67:00:00 | 4dki-A | 2.2 | 5.3 | 68 | 642 | 9 | MOLECULE: | PENICILLIN-BINDING PROTEIN 2' |
| 68:00:00 | 7jmr-A | 2.2 | 6.4 | 87 | 224 | 5 | MOLECULE: | PEA PATHOGENICITY PROTEIN 2 |
| 69:00:00 | 7z6e-A | 2.2 | 7.6 | 60 | 519 | 7 | MOLECULE: | SERINE/THREONINE-PROTEIN KINASE MRC |
| 70:00:00 | 5bka-F | 2.2 | 5 | 68 | 134 | 10 | MOLECULE: | HYDROXYLACYL-COA DEHYDROGENASE |
| 71:00:00 | 8ghl-I | 2.2 | 9.4 | 51 | 377 | 6 | MOLECULE: | PROTEIN HIR1 |
| 72:00:00 | 6chg-D | 2.2 | 5.1 | 61 | 408 | 3 | MOLECULE: | KLLA0E24487P |
| 73:00:00 | 8boz-N | 2.2 | 5.6 | 52 | 206 | 10 | MOLECULE: | TRANSMEMBRANE PROTEIN |
| 74:00:00 | 8ghn-I | 2.2 | 9.4 | 51 | 377 | 6 | MOLECULE: | PROTEIN HIR1 |
| 75:00:00 | 4q53-B | 2.2 | 2.4 | 51 | 109 | 8 | MOLECULE: | UNCHARACTERIZED PROTEIN |
| 76:00:00 | 8boz-L | 2.2 | 5.5 | 51 | 206 | 8 | MOLECULE: | TRANSMEMBRANE PROTEIN |
| 77:00:00 | 2gex-A | 2.2 | 4.8 | 65 | 146 | 5 | MOLECULE: | SNOL |
| 78:00:00 | 7f0z-A | 2.2 | 8.8 | 69 | 139 | 7 | MOLECULE: | NSRQ |
| 79:00:00 | 8t1m-B | 2.2 | 6 | 66 | 116 | 8 | MOLECULE: | DUF1842 DOMAIN-CONTAINING PROTEIN |
| 80:00:00 | 7yao-B | 2.2 | 3.6 | 52 | 213 | 0 | MOLECULE: | OXSTAYGOLD |
| 81:00:00 | 3st0-A | 2.2 | 10.3 | 57 | 226 | 2 | MOLECULE: | GREEN FLUORESCENT PROTEIN |
| 82:00:00 | 6xu4-B | 2.2 | 8.6 | 78 | 382 | 5 | MOLECULE: | FGCAMP |
| 83:00:00 | 3osr-B | 2.2 | 6.5 | 82 | 600 | 4 | MOLECULE: | MALTOSE-BINDING PERIPLASMIC PROTEIN |
| 84:00:00 | 5b61-A | 2.2 | 8.4 | 55 | 222 | 5 | MOLECULE: | GREEN FLUORESCENT PROTEIN |
| 85:00:00 | 2ah8-A | 2.2 | 6.6 | 62 | 232 | 3 | MOLECULE: | GREEN FLUORESCENT PROTEIN |
| 86:00:00 | 5yr2-B | 2.2 | 6.1 | 74 | 228 | 4 | MOLECULE: | GREEN FLUORESCENT PROTEIN |
| 87:00:00 | 4w73-B | 2.2 | 11.7 | 58 | 214 | 3 | MOLECULE: | FLUORESCENT PROTEIN E115C/T118H |
| 88:00:00 | 3rwa-G | 2.2 | 5.2 | 62 | 232 | 2 | MOLECULE: | FLUORESCENT PROTEIN FP480 |
| 89:00:00 | 4jfg-B | 2.2 | 5.5 | 52 | 222 | 4 | MOLECULE: | GREEN FLUORESCENT PROTEIN |
| 90:00:00 | 5yr2-A | 2.2 | 6 | 74 | 226 | 5 | MOLECULE: | GREEN FLUORESCENT PROTEIN |
| 91:00:00 | 6gqh-A | 2.2 | 7.2 | 59 | 226 | 7 | MOLECULE: | GREEN FLUORESCENT PROTEIN |
| 92:00:00 | 5du0-A | 2.2 | 9.8 | 59 | 227 | 7 | MOLECULE: | GREEN FLUORESCENT PROTEIN |

| | | | | | | | | |
|---|---|---|---|---|---|---|---|---|
| 93:00:00 | 6nqq-F | 2.2 | 4.3 | 59 | 219 | 8 | MOLECULE: | FLUORESCENT PROTEIN DRONPA |
| 94:00:00 | 7y40-B | 2.2 | 7.2 | 47 | 213 | 6 | MOLECULE: | STAYGOLD |
| 95:00:00 | 3ek8-A | 2.2 | 6.7 | 76 | 393 | 5 | MOLECULE: | MYOSIN LIGHT CHAIN KINASE, GREEN FL |
| 96:00:00 | 2ib5-C | 2.2 | 11 | 57 | 225 | 11 | MOLECULE: | CHROMO PROTEIN |
| 97:00:00 | 7dna-F | 2.2 | 5 | 56 | 205 | 7 | MOLECULE: | GREEN-TO-RED PHOTOCONVERTIBLE GFP-L |
| 98:00:00 | 8gyf-A | 2.2 | 7.2 | 47 | 213 | 6 | MOLECULE: | STAYGOLD(K192Y) |
| 99:00:00 | 4w7x-C | 2.2 | 8.7 | 57 | 223 | 7 | MOLECULE: | FLUORESCENT PROTEIN E115C/T118H |
| 0:00 | 5wj2-A | 2.2 | 8.2 | 55 | 234 | 5 | MOLECULE: | CYAN FLUORESCENT PROTEIN |
| 1:00 | 8i4o-K | 2.2 | 5.8 | 71 | 204 | 7 | MOLECULE: | SPLIT GREEN FLOURESCENT PROTEIN |
| 2:00 | 3bxc-G | 2.2 | 9.9 | 59 | 218 | 0 | MOLECULE: | FAR-RED FLUORESCENT PROTEIN MKATE |
| 3:00 | 5jzl-A | 2.2 | 10.6 | 68 | 231 | 9 | MOLECULE: | GREEN FLUORESCENT PROTEIN |
| 4:00 | 8j2k-B | 2.2 | 8.7 | 47 | 213 | 6 | MOLECULE: | STAYGOLD(N137A, Q140S) |
| 5:00 | 6un7-A | 2.2 | 6 | 68 | 226 | 10 | MOLECULE: | GREEN FLUORESCENT PROTEIN |
| 6:00 | 8ovp-A | 2.2 | 4.6 | 71 | 500 | 4 | MOLECULE: | PUTATIVE PERIPLASMIC BINDING TRANSP |
| 7:00 | 8j2h-B | 2.2 | 7.5 | 50 | 214 | 6 | MOLECULE: | STAYGOLD(N137A) |
| 8:00 | 5ktg-A | 2.2 | 8.4 | 54 | 308 | 4 | MOLECULE: | GREEN FLUORESCENT PROTEIN, BCL-2 HO |
| 9:00 | 8xld-A | 2.2 | 7.3 | 48 | 213 | 6 | MOLECULE: | STAYGOLD |
| 10:00 | 2p4m-G | 2.2 | 8.5 | 53 | 218 | 6 | MOLECULE: | GFP-LIKE NON-FLUORESCENT CHROMOPROT |
| 11:00 | 4gio-A | 2.1 | 2.7 | 51 | 96 | 6 | MOLECULE: | PUTATIVE LIPOPROTEIN |
| 12:00 | 3k67-A | 2.1 | 4.1 | 63 | 156 | 3 | MOLECULE: | PUTATIVE DEHYDRATASE AF1124 |
| 13:00 | 5dl7-A | 2.1 | 7.9 | 91 | 402 | 4 | MOLECULE: | PORIN |
| 14:00 | 5xev-A | 2.1 | 9.1 | 79 | 409 | 8 | MOLECULE: | XAA-PRO DIPEPTIDASE,PEPTIDASE-RELAT |
| 15:00 | 7n34-A | 2.1 | 6.8 | 60 | 108 | 5 | MOLECULE: | CAP15 |
| 16:00 | 6pwv-D | 2.1 | 5.8 | 70 | 357 | 4 | MOLECULE: | RETINOBLASTOMA-BINDING PROTEIN 5 |
| 17:00 | 5mr3-B | 2.1 | 4.1 | 63 | 117 | 5 | MOLECULE: | EGG-LYSIN |
| 18:00 | 4ruw-A | 2.1 | 7 | 63 | 386 | 3 | MOLECULE: | ENDONUCLEASE/EXONUCLEASE/PHOSPHATAS |
| 19:00 | 1ln1-A | 2.1 | 4.1 | 54 | 203 | 6 | MOLECULE: | PHOSPHATIDYLCHOLINE TRANSFER PROTEI |
| 20:00 | 7jjn-A | 2.1 | 4.8 | 54 | 514 | 2 | MOLECULE: | GLYCOSIDASES |
| 21:00 | 6e57-A | 2.1 | 3.6 | 53 | 399 | 2 | MOLECULE: | SURFACE GLYCAN BINDING PROTEIN B |
| 22:00 | 3mes-B | 2.1 | 4.5 | 50 | 358 | 6 | MOLECULE: | CHOLINE KINASE |
| 23:00 | 8oq3-D | 2.1 | 6.8 | 63 | 1556 | 8 | MOLECULE: | COMPLEMENT C3 |
| 24:00:00 | 5flv-M | 2.1 | 4.9 | 65 | 229 | 12 | MOLECULE: | HOMEOBOX PROTEIN NKX-2.5, T-BOX TRA |
| 25:00:00 | 4e1t-A | 2.1 | 8.6 | 82 | 245 | 6 | MOLECULE: | INVASIN |
| 26:00:00 | 2pyw-A | 2.1 | 4 | 55 | 417 | 5 | MOLECULE: | UNCHARACTERIZED PROTEIN |
| 27:00:00 | 4tq2-A | 2.1 | 4.9 | 72 | 177 | 7 | MOLECULE: | PUTATIVE PHYCOERYTHRIN LYASE |
| 28:00:00 | 1i78-A | 2.1 | 7.4 | 92 | 297 | 5 | MOLECULE: | PROTEASE VII |
| 29:00:00 | 2onf-A | 2.1 | 3.4 | 59 | 140 | 7 | MOLECULE: | HYPOTHETICAL PROTEIN TA0195 |
| 30:00:00 | 3og6-B | 2.1 | 3.5 | 60 | 200 | 5 | MOLECULE: | INTERLEUKIN-29 |
| 31:00:00 | 6k7l-A | 2.1 | 6.2 | 67 | 992 | 7 | MOLECULE: | PHOSPHOLIPID-TRANSPORTING ATPASE |
| 32:00:00 | 1zxf-A | 2.1 | 5.7 | 67 | 155 | 7 | MOLECULE: | CALC |
| 33:00:00 | 8ew9-A | 2.1 | 6.3 | 73 | 241 | 5 | MOLECULE: | ALTERED INHERITANCE OF MITOCHONDRIA |
| 34:00:00 | 5npy-A | 2.1 | 4.3 | 75 | 472 | 5 | MOLECULE: | FLAGELLAR BASAL BODY PROTEIN |
| 35:00:00 | 3n72-B | 2.1 | 5.2 | 61 | 152 | 5 | MOLECULE: | PUTATIVE ACTIVATOR OF HSP90 |
| 36:00:00 | 6ph4-B | 2.1 | 5.2 | 56 | 460 | 4 | MOLECULE: | BLUE-LIGHT-ACTIVATED HISTIDINE KINA |
| 37:00:00 | 4mjt-A | 2.1 | 6.6 | 82 | 236 | 7 | MOLECULE: | MONALYSIN |
| 38:00:00 | 8p1l-A | 2.1 | 10.6 | 71 | 2036 | 7 | MOLECULE: | RNA-DIRECTED RNA POLYMERASE L |

| | | | | | | | | |
|---|---|---|---|---|---|---|---|---|
| 39:00:00 | 3uv1-A | 2.1 | 5.9 | 82 | 190 | 6 | MOLECULE: | DER F 7 ALLERGEN |
| 40:00:00 | 5zki-B | 2.1 | 4.6 | 77 | 304 | 8 | MOLECULE: | NUCLEASE EXOG, MITOCHONDRIAL |
| 41:00:00 | 5yz0-A | 2.1 | 3 | 57 | 2362 | 4 | MOLECULE: | SERINE/THREONINE-PROTEIN KINASE ATR |
| 42:00:00 | 6egt-A | 2.1 | 4.3 | 57 | 445 | 5 | MOLECULE: | GLYCOPROTEIN |
| 43:00:00 | 4nzd-A | 2.1 | 6.4 | 66 | 210 | 2 | MOLECULE: | INTERLEUKIN-21 RECEPTOR |
| 44:00:00 | 3rwx-A | 2.1 | 7.5 | 75 | 254 | 3 | MOLECULE: | HYPOTHETICAL BACTERIAL OUTER MEMBRA |
| 45:00:00 | 3p51-A | 2.1 | 4.5 | 68 | 145 | 1 | MOLECULE: | UNCHARACTERIZED PROTEIN |
| 46:00:00 | 3tw5-A | 2.1 | 5.5 | 80 | 344 | 1 | MOLECULE: | TRANSGLUTAMINASE ELICITOR |
| 47:00:00 | 2o5c-A | 2.1 | 9.5 | 61 | 634 | 3 | MOLECULE: | 5'-D(*CP*GP*CP*AP*AP*CP*TP*T)-3' |
| 48:00:00 | 2j8f-A | 2.1 | 4.5 | 51 | 341 | 10 | MOLECULE: | LYSOZYME |
| 49:00:00 | 3oml-A | 2.1 | 10 | 72 | 532 | 7 | MOLECULE: | PEROXISOMAL MULTIFUNCTIONAL ENZYME |
| 50:00:00 | 8hgn-A | 2.1 | 4.6 | 70 | 353 | 6 | MOLECULE: | BETA-METHYLMALYL-COA DEHYDRATASE |
| 51:00:00 | 3tek-A | 2.1 | 3.9 | 58 | 139 | 0 | MOLECULE: | THERMODBP-SINGLE STRANDED DNA BINDI |
| 52:00:00 | 6agf-B | 2.1 | 8.5 | 64 | 173 | 6 | MOLECULE: | SODIUM CHANNEL PROTEIN TYPE 4 SUBUN |
| 53:00:00 | 4qni-A | 2.1 | 5.7 | 73 | 321 | 5 | MOLECULE: | UNCHARACTERIZED PROTEIN |
| 54:00:00 | 6jjt-D | 2.1 | 7.3 | 73 | 133 | 4 | MOLECULE: | PHNH |
| 55:00:00 | 3zt9-A | 2.1 | 5.7 | 65 | 192 | 6 | MOLECULE: | SERINE PHOSPHATASE |
| 56:00:00 | 1j6r-A | 2.1 | 3.8 | 68 | 197 | 16 | MOLECULE: | METHIONINE SYNTHASE |
| 57:00:00 | 4p2i-A | 2.1 | 4 | 50 | 117 | 0 | MOLECULE: | MKIAA0254 PROTEIN |
| 58:00:00 | 8a16-B | 2.1 | 4.1 | 53 | 244 | 8 | MOLECULE: | RECEPTOR-TYPE TYROSINE-PROTEIN PHOS |
| 59:00:00 | 6zgd-A | 2.1 | 5.6 | 58 | 288 | 9 | MOLECULE: | PROTON-GATED ION CHANNEL |
| 60:00:00 | 2wpw-A | 2.1 | 4 | 59 | 328 | 3 | MOLECULE: | ORF14 |
| 61:00:00 | 2ntk-A | 2.1 | 5.9 | 51 | 206 | 2 | MOLECULE: | IMP CYCLOHYDROLASE |
| 62:00:00 | 5jpq-i | 2.1 | 5.4 | 57 | 659 | 7 | MOLECULE: | WD40 DOMAIN PROTEINS |
| 63:00:00 | 4i2x-E | 2.1 | 3.4 | 51 | 314 | 8 | MOLECULE: | FABOX117 LIGHT CHAIN |
| 64:00:00 | 3ie5-B | 2.1 | 3.6 | 72 | 164 | 7 | MOLECULE: | PHENOLIC OXIDATIVE COUPLING PROTEIN |
| 65:00:00 | 2cuh-A | 2.1 | 5.3 | 59 | 115 | 7 | MOLECULE: | TENASCIN-X |
| 66:00:00 | 5yq0-C | 2.1 | 7.2 | 99 | 304 | 8 | MOLECULE: | COFJ |
| 67:00:00 | 1zwy-B | 2.1 | 4.3 | 61 | 180 | 5 | MOLECULE: | HYPOTHETICAL UPF0244 PROTEIN VC0702 |
| 68:00:00 | 2j2z-A | 2.1 | 7.9 | 64 | 216 | 11 | MOLECULE: | CHAPERONE PROTEIN PAPD |
| 69:00:00 | 6hsw-A | 2.1 | 8.5 | 70 | 422 | 9 | MOLECULE: | CARBOHYDRATE ESTERASE FAMILY 15 DOM |
| 70:00:00 | 5v77-A | 2.1 | 3.3 | 46 | 106 | 0 | MOLECULE: | UNCHARACTERIZED PROTEIN |
| 71:00:00 | 4hrv-B | 2.1 | 4.4 | 63 | 144 | 8 | MOLECULE: | PUTATIVE LIPOPROTEIN GNA1162 |
| 72:00:00 | 7pvh-A | 2.1 | 4.4 | 77 | 195 | 1 | MOLECULE: | POR SECRETION SYSTEM PROTEIN PORN/G |
| 73:00:00 | 5ymx-A | 2.1 | 4.2 | 51 | 202 | 4 | MOLECULE: | MUTUAL GLIDING-MOTILITY PROTEIN MGL |
| 74:00:00 | 2ehb-D | 2.1 | 3.7 | 53 | 126 | 6 | MOLECULE: | CALCINEURIN B-LIKE PROTEIN 4 |
| 75:00:00 | 5dz8-A | 2.1 | 5.7 | 69 | 167 | 1 | MOLECULE: | BSPA (BSPA_V) |
| 76:00:00 | 1vi8-B | 2.1 | 4.5 | 60 | 146 | 7 | MOLECULE: | HYPOTHETICAL PROTEIN YDII |
| 77:00:00 | 2mvb-A | 2.1 | 7.6 | 67 | 165 | 3 | MOLECULE: | UNCHARACTERIZED PROTEIN |
| 78:00:00 | 4gbs-B | 2.1 | 6.5 | 58 | 186 | 5 | MOLECULE: | PUTATIVE LIPOPROTEIN |
| 79:00:00 | 4mjg-A | 2.1 | 3.2 | 61 | 177 | 2 | MOLECULE: | HYPOTHETICAL PROTEIN |
| 80:00:00 | 3pcs-B | 2.1 | 4.5 | 86 | 355 | 2 | MOLECULE: | ESPG |
| 81:00:00 | 4lsd-A | 2.1 | 3.6 | 51 | 98 | 4 | MOLECULE: | FIBRONECTIN TYPE III DOMAIN-CONTAIN |
| 82:00:00 | 6ycq-A | 2.1 | 8.2 | 65 | 347 | 2 | MOLECULE: | AUXIN RESPONSE FACTOR 1 |
| 83:00:00 | 4qdc-A | 2.1 | 4.1 | 78 | 369 | 6 | MOLECULE: | 3-KETOSTEROID 9ALPHA-HYDROXYLASE OX |
| 84:00:00 | 3osq-A | 2.1 | 8.8 | 78 | 607 | 4 | MOLECULE: | MALTOSE-BINDING PERIPLASMIC PROTEIN |

| | | | | | | | | |
|---|---|---|---|---|---|---|---|---|
| 85:00:00 | 8f3h-A | 2.1 | 5.5 | 70 | 643 | 3 | MOLECULE: | PENICILLIN BINDING PROTEIN 5 |
| 86:00:00 | 7bny-B | 2.1 | 7.8 | 68 | 140 | 6 | MOLECULE: | GENOME POLYPROTEIN |
| 87:00:00 | 4aq1-A | 2.1 | 4.8 | 57 | 721 | 7 | MOLECULE: | SBSB PROTEIN |
| 88:00:00 | 3fid-A | 2.1 | 6.1 | 82 | 296 | 2 | MOLECULE: | PUTATIVE OUTER MEMBRANE PROTEIN (LP |
| 89:00:00 | 3nz3-A | 2.1 | 4.5 | 56 | 100 | 5 | MOLECULE: | PUTATIVE UNCHARACTERIZED PROTEIN |
| 90:00:00 | 3kog-A | 2.1 | 4.7 | 75 | 226 | 5 | MOLECULE: | PUTATIVE PORE-FORMING TOXIN |
| 91:00:00 | 6hq9-A | 2.1 | 2.9 | 43 | 60 | 7 | MOLECULE: | DNA EXCISION REPAIR PROTEIN ERCC-6- |
| 92:00:00 | 8fn9-F | 2.1 | 3.9 | 55 | 103 | 5 | MOLECULE: | TENASCIN-R |
| 93:00:00 | 2ikk-A | 2.1 | 4.2 | 58 | 146 | 12 | MOLECULE: | HYPOTHETICAL TRANSCRIPTIONAL REGULA |
| 94:00:00 | 5m1u-A | 2.1 | 6.4 | 49 | 127 | 10 | MOLECULE: | CURLI PRODUCTION ASSEMBLY/TRANSPORT |
| 95:00:00 | 5mob-A | 2.1 | 5.9 | 77 | 193 | 6 | MOLECULE: | SLPYL1_ABA |
| 96:00:00 | 6cfw-K | 2.1 | 5.9 | 68 | 166 | 4 | MOLECULE: | MONOVALENT CATION/H+ ANTIPORTER SUB |
| 97:00:00 | 6e0a-B | 2.1 | 4.7 | 68 | 262 | 3 | MOLECULE: | METHYL-ACCEPTING CHEMOTAXIS PROTEIN |
| 98:00:00 | 4les-A | 2.1 | 3.1 | 61 | 142 | 7 | MOLECULE: | PROTEIN - CONSERVED HYPOTHETICAL |
| 99:00:00 | 7vgr-A | 2.1 | 3.8 | 57 | 198 | 4 | MOLECULE: | YN7756_1 FAB LIGHT CHAIN |
| 0:00 | 5a1u-G | 2.1 | 4.1 | 61 | 811 | 3 | MOLECULE: | ADP-RIBOSYLATION FACTOR 1 |
| 1:00 | 5lth-A | 2.1 | 3.3 | 64 | 334 | 5 | MOLECULE: | HEME DEPENDENT OXIDATIVE N-DEMETHYL |
| 2:00 | 8bl4-e | 2.1 | 4.4 | 59 | 149 | 7 | MOLECULE: | PHAGE TAIL SHEATH FAMILY PROTEIN |
| 3:00 | 5yvq-A | 2.1 | 4 | 56 | 358 | 7 | MOLECULE: | TAIL FIBER PROTEIN S |
| 4:00 | 7ct1-A | 2.1 | 5.9 | 74 | 263 | 3 | MOLECULE: | FUSION PROTEIN SORTING NEXIN-27 AND |
| 5:00 | 7xyz-B | 2.1 | 3.2 | 61 | 456 | 5 | MOLECULE: | TRIPARTITE MOTIF-CONTAINING PROTEIN |
| 6:00 | 5mje-A | 2.1 | 5.9 | 45 | 111 | 4 | MOLECULE: | CYTOTOXIC TRANSLATIONAL REPRESSOR O |
| 7:00 | 5af0-C | 2.1 | 6 | 57 | 239 | 5 | MOLECULE: | MAELSTROM |
| 8:00 | 7kyx-A | 2.1 | 4.5 | 63 | 397 | 6 | MOLECULE: | CORONIN-6 |
| 9:00 | 2x2h-A | 2.1 | 4.4 | 65 | 1025 | 2 | MOLECULE: | ALPHA-1,4-GLUCAN LYASE ISOZYME 1 |
| 10:00 | 7np8-B | 2.1 | 4.1 | 67 | 620 | 10 | MOLECULE: | COENZYME F420-DEPENDENT SULFITE RED |
| 11:00 | 6zls-A | 2.1 | 3.6 | 56 | 292 | 5 | MOLECULE: | HISTIDINE KINASE |
| 12:00 | 5u30-A | 2.1 | 4.3 | 67 | 1085 | 4 | MOLECULE: | CRISPR-ASSOCIATED ENDONUCLEASE C2C1 |
| 13:00 | 8wb2-A | 2.1 | 2.8 | 57 | 179 | 4 | MOLECULE: | TEMPERATURE-INDUCED LIPOCALIN-1 |
| 14:00 | 3j1w-A | 2.1 | 3.9 | 50 | 106 | 4 | MOLECULE: | PROTEIN PRGH |
| 15:00 | 6ef3-A | 2.1 | 5.5 | 70 | 247 | 7 | MOLECULE: | PROTEASOME SUBUNIT BETA TYPE-1 |
| 16:00 | 2dso-C | 2.1 | 4.6 | 69 | 323 | 6 | MOLECULE: | DRP35 |
| 17:00 | 4qvs-A | 2.1 | 9.3 | 60 | 217 | 7 | MOLECULE: | S-LAYER DOMAIN-CONTAINING PROTEIN |
| 18:00 | 5yyl-A | 2.1 | 6.4 | 64 | 405 | 5 | MOLECULE: | MAJOR ROYAL JELLY PROTEIN 1 |
| 19:00 | 5opq-A | 2.1 | 4.3 | 64 | 659 | 9 | MOLECULE: | 3,6-ANHYDRO-D-GALACTOSIDASE |
| 20:00 | 4fca-A | 2.1 | 5.5 | 64 | 482 | 6 | MOLECULE: | CONSERVED DOMAIN PROTEIN |
| 21:00 | 3acp-A | 2.1 | 8.6 | 52 | 417 | 8 | MOLECULE: | WD REPEAT-CONTAINING PROTEIN YGL004 |
| 22:00 | 1ecg-B | 2.1 | 14.6 | 75 | 501 | 11 | MOLECULE: | GLUTAMINE PHOSPHORIBOSYLPYROPHOSPHA |
| 23:00 | 4bq6-A | 2.1 | 3 | 61 | 205 | 3 | MOLECULE: | NEOGENIN |
| 24:00:00 | 7dqx-E | 2.1 | 8.7 | 58 | 293 | 7 | MOLECULE: | 6-HYDROXYPSEUDOOXYNICOTINE DEHYDROG |
| 25:00:00 | 6peu-B | 2.1 | 9.3 | 54 | 385 | 2 | MOLECULE: | UNCHARACTERIZED PROTEIN MJ1094 |
| 26:00:00 | 4c2m-I | 2.1 | 7.3 | 43 | 124 | 2 | MOLECULE: | DNA-DIRECTED RNA POLYMERASE I SUBUN |
| 27:00:00 | 5z4g-A | 2.1 | 3.5 | 67 | 145 | 6 | MOLECULE: | SAHS4 |
| 28:00:00 | 7w2t-A | 2.1 | 7.5 | 73 | 773 | 4 | MOLECULE: | GLUCOSYLCERAMIDASE |
| 29:00:00 | 3v7d-B | 2.1 | 3.9 | 47 | 450 | 4 | MOLECULE: | SUPPRESSOR OF KINETOCHORE PROTEIN 1 |
| 30:00:00 | 8kei-D | 2.1 | 3.6 | 62 | 164 | 10 | MOLECULE: | CYTOCHROME B-245 LIGHT CHAIN |

| | | | | | | | | |
|---|---|---|---|---|---|---|---|---|
| 31:00:00 | 8f2r-E | 2.1 | 3.1 | 46 | 205 | 13 | MOLECULE: | COMM DOMAIN-CONTAINING PROTEIN 1 |
| 32:00:00 | 5hx2-A | 2.1 | 6.5 | 57 | 1030 | 5 | MOLECULE: | BASEPLATE WEDGE PROTEIN GP7 |
| 33:00:00 | 8e98-D | 2.1 | 12.2 | 67 | 638 | 7 | MOLECULE: | GLUTAMATE RECEPTOR IONOTROPIC, NMDA |
| 34:00:00 | 5mc9-A | 2.1 | 4.4 | 51 | 597 | 8 | MOLECULE: | LAMININ SUBUNIT ALPHA-1 |
| 35:00:00 | 6nd4-I | 2.1 | 4.2 | 49 | 487 | 0 | MOLECULE: | ETS RRNA |
| 36:00:00 | 2h9u-A | 2.1 | 3 | 51 | 101 | 4 | MOLECULE: | DNA/RNA-BINDING PROTEIN ALBA 2 |
| 37:00:00 | 1fwx-A | 2.1 | 4.8 | 59 | 591 | 5 | MOLECULE: | NITROUS OXIDE REDUCTASE |
| 38:00:00 | 3b77-A | 2.1 | 7.4 | 60 | 188 | 12 | MOLECULE: | UNCHARACTERIZED PROTEIN |
| 39:00:00 | 4jmh-A | 2.1 | 2.6 | 47 | 200 | 6 | MOLECULE: | CLAMP SHC1_PY239/240 |
| 40:00:00 | 2x5h-D | 2.1 | 3.6 | 45 | 94 | 7 | MOLECULE: | ORF 131 |
| 41:00:00 | 8bn0-A | 2.1 | 6.6 | 62 | 342 | 0 | MOLECULE: | PUTATIVE SURFACE LAYER PROTEIN |
| 42:00:00 | 6nd4-J | 2.1 | 3.7 | 69 | 493 | 9 | MOLECULE: | ETS RRNA |
| 43:00:00 | 6il9-A | 2.1 | 3 | 43 | 490 | 7 | MOLECULE: | FRUCTURONATE-TAGATURONATE EPIMERASE |
| 44:00:00 | 5oj3-A | 2.1 | 5.3 | 55 | 322 | 4 | MOLECULE: | PHOTOSYSTEM II STABILITY/ASSEMBLY F |
| 45:00:00 | 8p4k-A | 2.1 | 3.9 | 47 | 599 | 2 | MOLECULE: | CORE PROTEIN OPG136 |
| 46:00:00 | 6rtv-A | 2.1 | 8.2 | 49 | 386 | 8 | MOLECULE: | 4-O-METHYL-GLUCURONOYL METHYLESTERA |
| 47:00:00 | 4wju-A | 2.1 | 5.8 | 62 | 457 | 3 | MOLECULE: | RIBOSOME ASSEMBLY PROTEIN 4 |
| 48:00:00 | 5i08-A | 2.1 | 6.2 | 66 | 958 | 9 | MOLECULE: | SPIKE GLYCOPROTEIN, ENVELOPE GLYCOP |
| 49:00:00 | 5c0x-K | 2.1 | 6.4 | 55 | 350 | 7 | MOLECULE: | EXOSOME COMPLEX COMPONENT RRP45 |
| 50:00:00 | 1g6o-A | 2.1 | 4.9 | 49 | 323 | 8 | MOLECULE: | CAG-ALPHA |
| 51:00:00 | 4kha-A | 2.1 | 7.4 | 53 | 360 | 4 | MOLECULE: | SPT16M-HISTONE H2B 1.1 CHIMERA |
| 52:00:00 | 5cvo-D | 2.1 | 4.7 | 54 | 612 | 2 | MOLECULE: | WD REPEAT-CONTAINING PROTEIN 48 |
| 53:00:00 | 6rte-A | 2.1 | 3.5 | 56 | 464 | 7 | MOLECULE: | CYTOCHROME C |
| 54:00:00 | 7wj7-A | 2.1 | 10 | 57 | 242 | 11 | MOLECULE: | SERINE/THREONINE PROTEIN KINASE |
| 55:00:00 | 8y6b-D | 2.1 | 4.7 | 68 | 517 | 4 | MOLECULE: | DISINTEGRIN AND METALLOPROTEINASE D |
| 56:00:00 | 6muq-A | 2.1 | 4.3 | 64 | 267 | 8 | MOLECULE: | MUREIN-DD-ENDOPEPTIDASE |
| 57:00:00 | 6gkw-A | 2.1 | 4.5 | 50 | 351 | 4 | MOLECULE: | PUTATIVE PHAGE XKDK-LIKE PROTEIN |
| 58:00:00 | 5o60-H | 2.1 | 2.8 | 48 | 151 | 2 | MOLECULE: | 50S RIBOSOMAL PROTEIN BL37 |
| 59:00:00 | 8ide-A | 2.1 | 6.4 | 46 | 929 | 4 | MOLECULE: | N(6)-L-THREONYLCARBAMOYLADENINE SYN |
| 60:00:00 | 8v1k-A | 2.1 | 5.5 | 72 | 185 | 3 | MOLECULE: | OUTER-MEMBRANE LIPOPROTEIN CARRIER |
| 61:00:00 | 7s69-A | 2.1 | 4.4 | 52 | 174 | 6 | MOLECULE: | N-ACETYLGLUCOSAMINE-1-PHOSPHOTRANSF |
| 62:00:00 | 5uc2-C | 2.1 | 4 | 56 | 242 | 0 | MOLECULE: | DOMAIN OF UNKNOWN FUNCTION DUF1849 |
| 63:00:00 | 2hqv-A | 2.1 | 8 | 56 | 172 | 2 | MOLECULE: | AGR_C_4470P |
| 64:00:00 | 8h8p-B | 2.1 | 4.9 | 80 | 355 | 5 | MOLECULE: | THIOMORPHOLINE-CARBOXYLATE DEHYDROG |
| 65:00:00 | 6h41-A | 2.1 | 4.7 | 69 | 309 | 4 | MOLECULE: | INTERLEUKIN-5 RECEPTOR SUBUNIT ALPH |
| 66:00:00 | 6sno-A | 2.1 | 5.7 | 66 | 573 | 5 | MOLECULE: | PHOSPHOGLUCOMUTASE-1 |
| 67:00:00 | 8k2v-A | 2.1 | 9.3 | 56 | 214 | 5 | MOLECULE: | METHYLCROTONOYL-COA CARBOXYLASE SUB |
| 68:00:00 | 6cp9-D | 2.1 | 4.7 | 57 | 116 | 12 | MOLECULE: | CDIA |
| 69:00:00 | 7w42-B | 2.1 | 10.7 | 61 | 402 | 8 | MOLECULE: | UNCHARACTERIZED ATPASE YJOB |
| 70:00:00 | 7krj-C | 2.1 | 8.8 | 50 | 133 | 10 | MOLECULE: | HEAT SHOCK PROTEIN HSP 90-ALPHA |
| 71:00:00 | 5w5p-A | 2.1 | 7.3 | 92 | 610 | 4 | MOLECULE: | TAIL FIBER PROTEIN |
| 72:00:00 | 6ahr-F | 2.1 | 5.2 | 59 | 129 | 7 | MOLECULE: | H1 RNA |
| 73:00:00 | 5owg-A | 2.1 | 4.4 | 56 | 181 | 5 | MOLECULE: | PCYX_EBK42635 |
| 74:00:00 | 7kzn-D | 2.1 | 4.3 | 65 | 456 | 6 | MOLECULE: | HEAVY CHAIN ALPHA |
| 75:00:00 | 8acs-A | 2.1 | 6.4 | 65 | 598 | 14 | MOLECULE: | FAD-DEPENDENT OXIDOREDUCTASE |
| 76:00:00 | 5ltw-D | 2.1 | 2.4 | 44 | 98 | 7 | MOLECULE: | 14-3-3 PROTEIN SIGMA |

| | | | | | | | | |
|---|---|---|---|---|---|---|---|---|
| 77:00:00 | 1w94-A | 2.1 | 3.2 | 48 | 155 | 6 | MOLECULE: | PROBABLE BRIX-DOMAIN RIBOSOMAL BIOG |
| 78:00:00 | 5ejr-A | 2.1 | 4.7 | 58 | 500 | 3 | MOLECULE: | MYOSIN-I HEAVY CHAIN |
| 79:00:00 | 5ek5-B | 2.1 | 2.5 | 38 | 120 | 5 | MOLECULE: | IRMA |
| 80:00:00 | 7apk-F | 2.1 | 5.3 | 67 | 337 | 0 | MOLECULE: | THO COMPLEX SUBUNIT 1 |
| 81:00:00 | 6fhm-A | 2.1 | 6.9 | 58 | 193 | 9 | MOLECULE: | OUTER-MEMBRANE LIPOPROTEIN CARRIER |
| 82:00:00 | 1qqg-B | 2.1 | 6.2 | 56 | 210 | 5 | MOLECULE: | INSULIN RECEPTOR SUBSTRATE 1 |
| 83:00:00 | 7et0-C | 2.1 | 5.4 | 66 | 684 | 3 | MOLECULE: | BACTERIA FACTOR B |
| 84:00:00 | 8orn-B | 2.1 | 6.6 | 57 | 180 | 7 | MOLECULE: | OUTER-MEMBRANE LIPOPROTEIN CARRIER |
| 85:00:00 | 6jt5-A | 2.1 | 7.5 | 57 | 409 | 4 | MOLECULE: | EXTRACELLULAR PQQ-DEPENDENT SUGAR D |
| 86:00:00 | 6y9c-A | 2.1 | 3.4 | 60 | 365 | 8 | MOLECULE: | CARNITINE MONOOXYGENASE OXYGENASE S |
| 87:00:00 | 5ezu-A | 2.1 | 3 | 47 | 76 | 4 | MOLECULE: | PROTEIN A46 |
| 88:00:00 | 1x1g-A | 2.1 | 3.8 | 53 | 129 | 8 | MOLECULE: | PLECKSTRIN 2 |
| 89:00:00 | 3ejv-A | 2.1 | 6 | 65 | 159 | 2 | MOLECULE: | UNCHARACTERIZED PROTEIN WITH CYSTAT |
| 90:00:00 | 8bl3-A | 2.1 | 4.7 | 65 | 141 | 3 | MOLECULE: | SCIG12 |
| 91:00:00 | 3zfo-A | 2.1 | 3.6 | 53 | 150 | 2 | MOLECULE: | N-TERMINAL PROTEASE NPRO |
| 92:00:00 | 6wqc-A | 2.1 | 5 | 54 | 293 | 7 | MOLECULE: | N-TERMINAL ACETYLTRANSFERASE, GNAT |
| 93:00:00 | 6yzg-A | 2.1 | 4.6 | 58 | 279 | 2 | MOLECULE: | SURFACE-ASSOCIATED PROTEIN CSHB |
| 94:00:00 | 5gt1-A | 2.1 | 3.3 | 46 | 153 | 9 | MOLECULE: | CHOLINE BINDING PROTEIN A |
| 95:00:00 | 1m1g-C | 2.1 | 3.8 | 50 | 244 | 2 | MOLECULE: | TRANSCRIPTION ANTITERMINATION PROTE |
| 96:00:00 | 7lt1-A | 2.1 | 8 | 62 | 403 | 3 | MOLECULE: | PROTEIN MB21D2 |
| 97:00:00 | 6ffy-A | 2.1 | 6.8 | 56 | 918 | 2 | MOLECULE: | VPS10 DOMAIN-CONTAINING RECEPTOR SO |
| 98:00:00 | 4tqj-A | 2.1 | 6.8 | 61 | 403 | 8 | MOLECULE: | LECTIN 2 |
| 99:00:00 | 7nnl-B | 2.1 | 5 | 65 | 682 | 11 | MOLECULE: | POTASSIUM-TRANSPORTING ATPASE POTAS |
| 0:00 | 8oml-B | 2.1 | 4.1 | 66 | 317 | 0 | MOLECULE: | CELL WALL SURFACE ANCHOR FAMILY PRO |
| 1:00 | 7f10-B | 2.1 | 9.5 | 68 | 140 | 4 | MOLECULE: | NSRQ |
| 2:00 | 2w2c-A | 2.1 | 5.2 | 61 | 128 | 7 | MOLECULE: | CALCIUM/CALMODULIN-DEPENDENT PROTEI |
| 3:00 | 7f0y-A | 2.1 | 9.6 | 67 | 141 | 3 | MOLECULE: | NSRQ |
| 4:00 | 3ff0-A | 2.1 | 6.7 | 76 | 162 | 4 | MOLECULE: | PHENAZINE BIOSYNTHESIS PROTEIN PHZB |
| 5:00 | 4y4v-B | 2.1 | 5.1 | 70 | 317 | 10 | MOLECULE: | CONSERVED HYPOTHETICAL SECRETED PRO |
| 6:00 | 8jxd-A | 2.1 | 6.2 | 62 | 493 | 2 | MOLECULE: | LDL RECEPTOR RELATED PROTEIN 2 |
| 7:00 | 4x2h-B | 2.1 | 5.6 | 69 | 175 | 12 | MOLECULE: | PUTATIVE MRNA EXPORT PROTEIN |
| 8:00 | 5yzn-A | 2.1 | 4.9 | 57 | 649 | 7 | MOLECULE: | ACYL-PEPTIDE HYDROLASE, PUTATIVE |
| 9:00 | 7f14-A | 2.1 | 9.3 | 70 | 148 | 4 | MOLECULE: | DCR3 |
| 10:00 | 5evh-A | 2.1 | 6.8 | 64 | 121 | 3 | MOLECULE: | UNCHARACTERIZED PROTEIN |
| 11:00 | 6swy-4 | 2.1 | 4.6 | 68 | 217 | 4 | MOLECULE: | VACUOLAR IMPORT AND DEGRADATION PRO |
| 12:00 | 8ovn-A | 2.1 | 9.6 | 78 | 503 | 5 | MOLECULE: | PUTATIVE PERIPLASMIC BINDING TRANSP |
| 13:00 | 6vio-D | 2.1 | 6.5 | 59 | 225 | 3 | MOLECULE: | GREEN FLUORESCENT PROTEIN GFP |
| 14:00 | 3evu-A | 2.1 | 12.5 | 79 | 393 | 5 | MOLECULE: | MYOSIN LIGHT CHAIN KINASE, GREEN FL |
| 15:00 | 4w7e-A | 2.1 | 6.8 | 59 | 221 | 3 | MOLECULE: | FLUORESCENT PROTEIN D21H/K26H |
| 16:00 | 4bdu-B | 2.1 | 7.8 | 63 | 295 | 3 | MOLECULE: | GREEN FLUORESCENT PROTEIN, APOPTOSI |
| 17:00 | 3ek7-A | 2.1 | 12.3 | 79 | 389 | 6 | MOLECULE: | MYOSIN LIGHT CHAIN KINASE, GREEN FL |
| 18:00 | 5dpi-D | 2.1 | 5.9 | 56 | 225 | 4 | MOLECULE: | GREEN FLUORESCENT PROTEIN |
| 19:00 | 3sg5-A | 2.1 | 12.3 | 78 | 387 | 5 | MOLECULE: | MYOSIN LIGHT CHAIN KINASE, GREEN FL |
| 20:00 | 6uru-B | 2.1 | 8.8 | 74 | 500 | 4 | MOLECULE: | IACHSNFR PRECURSOR |
| 21:00 | 3sg6-A | 2.1 | 12.1 | 78 | 383 | 4 | MOLECULE: | MYOSIN LIGHT CHAIN KINASE, GREEN FL |
| 22:00 | 7bwn-M | 2.1 | 8.4 | 55 | 280 | 5 | MOLECULE: | CHIMERA OF GREEN FLUORESCENT PROTEI |

| | | | | | | | | |
|---|---|---|---|---|---|---|---|---|
| 23:00 | 7bwn-J | 2.1 | 10 | 57 | 280 | 5 | MOLECULE: | CHIMERA OF GREEN FLUORESCENT PROTEI |
| 24:00:00 | 6wva-A | 2.1 | 6.8 | 68 | 411 | 3 | MOLECULE: | VITAMIN K EPOXIDE REDUCTASE-LIKE PR |
| 25:00:00 | 3k1k-B | 2.1 | 6.4 | 58 | 224 | 3 | MOLECULE: | GREEN FLUORESCENT PROTEIN |
| 26:00:00 | 8j3j-A | 2.1 | 8.6 | 57 | 214 | 9 | MOLECULE: | STAYGOLD(Q140S, Y187F) |
| 27:00:00 | 6efr-A | 2.1 | 9.3 | 78 | 503 | 4 | MOLECULE: | INICSNFR 1.0, A GENETICALLY ENCODED |
| 28:00:00 | 4ik9-A | 2.1 | 12.3 | 78 | 393 | 5 | MOLECULE: | RCAMP, GREEN FLUORESCENT PROTEIN |
| 29:00:00 | 8ovp-B | 2.1 | 6.1 | 74 | 497 | 4 | MOLECULE: | PUTATIVE PERIPLASMIC BINDING TRANSP |
| 30:00:00 | 3sg3-A | 2.1 | 7.1 | 78 | 394 | 4 | MOLECULE: | MYOSIN LIGHT CHAIN KINASE, GREEN FL |
| 31:00:00 | 4ik4-A | 2.1 | 13.3 | 74 | 395 | 7 | MOLECULE: | RCAMP, GREEN FLUORESCENT PROTEIN |
| 32:00:00 | 6zsn-A | 2.1 | 7 | 78 | 397 | 4 | MOLECULE: | GREEN FLUORESCENT PROTEIN,GREEN FLU |
| 33:00:00 | 7st4-A | 2.1 | 7.8 | 79 | 385 | 5 | MOLECULE: | JGCAMP8.410.80 |
| 34:00:00 | 4zf4-A | 2.1 | 4.7 | 71 | 226 | 10 | MOLECULE: | GREEN FLUORESCENT PROTEIN |
| 35:00:00 | 2wiq-A | 2.1 | 10 | 57 | 225 | 9 | MOLECULE: | KILLERRED |
| 36:00:00 | 7amf-C | 2.1 | 11.2 | 72 | 234 | 7 | MOLECULE: | GREEN FLUORESCENT PROTEIN |
| 37:00:00 | 3vic-D | 2.1 | 9.4 | 55 | 218 | 2 | MOLECULE: | GFP-LIKE NON-FLUORESCENT CHROMOPROT |
| 38:00:00 | 4ndj-A | 2.1 | 11.6 | 47 | 279 | 9 | MOLECULE: | GREEN FLUORESCENT PROTEIN, CHIMERIC |
| 39:00:00 | 8bxt-B | 2.1 | 7.1 | 48 | 213 | 6 | MOLECULE: | STAYGOLD |
| 40:00:00 | 7rrh-F | 2.1 | 10.2 | 63 | 213 | 2 | MOLECULE: | FLUORESCENT PROTEIN DRONPA |
| 41:00:00 | 7bwn-C | 2.1 | 8.2 | 55 | 280 | 5 | MOLECULE: | CHIMERA OF GREEN FLUORESCENT PROTEI |
| 42:00:00 | 4w7r-D | 2.1 | 8.8 | 57 | 225 | 9 | MOLECULE: | FLUORESCENT PROTEIN E124H/K126C |
| 43:00:00 | 2ote-B | 2.1 | 11.8 | 49 | 213 | 6 | MOLECULE: | GFP-LIKE FLUORESCENT CHROMOPROTEIN |
| 44:00:00 | 3cgl-E | 2.1 | 10.4 | 65 | 212 | 0 | MOLECULE: | GFP-LIKE FLUORESCENT CHROMOPROTEIN |
| 45:00:00 | 7rri-D | 2.1 | 10.3 | 65 | 219 | 2 | MOLECULE: | FLUORESCENT PROTEIN DRONPA |
| 46:00:00 | 5mfc-D | 2.1 | 8.9 | 58 | 237 | 5 | MOLECULE: | YIIIM5AII |
| 47:00:00 | 6ofo-A | 2.1 | 8.9 | 61 | 220 | 3 | MOLECULE: | GREEN FLUORESCENT PROTEIN (GFP) S1 |
| 48:00:00 | 5hzt-I | 2.1 | 11.6 | 51 | 212 | 6 | MOLECULE: | FLUORESCENT PROTEIN DRONPA |
| 49:00:00 | 7o7c-A | 2.1 | 10.7 | 71 | 232 | 3 | MOLECULE: | GREEN FLUORESCENT PROTEIN |
| 50:00:00 | 4w6k-A | 2.1 | 10.9 | 47 | 194 | 11 | MOLECULE: | FLUORESCENT PROTEIN D117C |
| 51:00:00 | 2oiw-A | 2 | 3 | 58 | 133 | 7 | MOLECULE: | PUTATIVE 4-HYDROXYBENZOYL-COA THIOE |
| 52:00:00 | 6uhd-A | 2 | 13.4 | 86 | 322 | 2 | MOLECULE: | HISTONE ACETYLTRANSFERASE TYPE B CA |
| 53:00:00 | 7v9a-B | 2 | 4.5 | 68 | 377 | 4 | MOLECULE: | TELOMERASE CAJAL BODY PROTEIN 1 |
| 54:00:00 | 2hq7-B | 2 | 8.6 | 64 | 142 | 9 | MOLECULE: | PROTEIN, RELATED TO GENERAL STRESS |
| 55:00:00 | 1a87-A | 2 | 4.1 | 54 | 297 | 6 | MOLECULE: | COLICIN N |
| 56:00:00 | 6lix-B | 2 | 11 | 69 | 196 | 7 | MOLECULE: | CHROMOPHORE LYASE CRL, CHLOROPLASTI |
| 57:00:00 | 6shc-A | 2 | 5.7 | 62 | 273 | 8 | MOLECULE: | SERINE/THREONINE-PROTEIN KINASE/END |
| 58:00:00 | 2yrl-A | 2 | 4.3 | 55 | 102 | 5 | MOLECULE: | KIAA1837 PROTEIN |
| 59:00:00 | 4lmi-B | 2 | 5.5 | 62 | 136 | 8 | MOLECULE: | UNCHARACTERIZED PROTEIN |
| 60:00:00 | 4ge1-A | 2 | 3.8 | 68 | 195 | 10 | MOLECULE: | BIOGENIC AMINE-BINDING PROTEIN |
| 61:00:00 | 8ok9-C | 2 | 5.6 | 65 | 236 | 9 | MOLECULE: | PIWI PROTEIN AF_1318 |
| 62:00:00 | 3jvn-A | 2 | 3.6 | 54 | 122 | 4 | MOLECULE: | ACETYLTRANSFERASE |
| 63:00:00 | 3w81-A | 2 | 10.3 | 76 | 616 | 5 | MOLECULE: | ALPHA-L-IDURONIDASE |
| 64:00:00 | 5oj2-B | 2 | 2.6 | 47 | 692 | 9 | MOLECULE: | MAM DOMAIN-CONTAINING GLYCOSYLPHOSP |
| 65:00:00 | 5hdf-B | 2 | 8.7 | 46 | 320 | 9 | MOLECULE: | HYDROLASE |
| 66:00:00 | 4v3g-A | 2 | 11 | 90 | 330 | 3 | MOLECULE: | CYMA PROTEIN |
| 67:00:00 | 2dc1-A | 2 | 4.5 | 62 | 236 | 3 | MOLECULE: | L-ASPARTATE DEHYDROGENASE |
| 68:00:00 | 2qml-A | 2 | 4.4 | 59 | 193 | 3 | MOLECULE: | BH2621 PROTEIN |

| | | | | | | | | |
|---|---|---|---|---|---|---|---|---|
| 69:00:00 | 6dv3-A | 2 | 7.3 | 77 | 500 | 4 | MOLECULE: | PROTEIN INVG |
| 70:00:00 | 2qp2-A | 2 | 8.1 | 73 | 498 | 10 | MOLECULE: | UNKNOWN PROTEIN |
| 71:00:00 | 2r41-C | 2 | 2.8 | 55 | 106 | 9 | MOLECULE: | UNCHARACTERIZED PROTEIN |
| 72:00:00 | 3ubg-B | 2 | 4.1 | 52 | 310 | 6 | MOLECULE: | NEURAL-CADHERIN |
| 73:00:00 | 6gef-A | 2 | 6.9 | 60 | 388 | 12 | MOLECULE: | TYPE IV SECRETION SYSTEM PROTEIN DO |
| 74:00:00 | 2mpo-A | 2 | 4.6 | 75 | 182 | 9 | MOLECULE: | MIC2-ASSOCIATED PROTEIN |
| 75:00:00 | 7cun-G | 2 | 4 | 64 | 895 | 6 | MOLECULE: | INTEGRATOR COMPLEX SUBUNIT 1 |
| 76:00:00 | 3c5m-B | 2 | 9 | 65 | 378 | 3 | MOLECULE: | OLIGOGALACTURONATE LYASE |
| 77:00:00 | 7qof-d | 2 | 4.1 | 56 | 333 | 2 | MOLECULE: | MAJOR CAPSID PROTEIN GP32 |
| 78:00:00 | 7nky-O | 2 | 1.9 | 42 | 427 | 10 | MOLECULE: | RNA (5'- |
| 79:00:00 | 7s8v-B | 2 | 8.6 | 75 | 518 | 3 | MOLECULE: | INSULIN-LIKE GROWTH FACTOR 1 RECEPT |
| 80:00:00 | 8as8-A | 2 | 10 | 70 | 597 | 4 | MOLECULE: | JETC |
| 81:00:00 | 6e5d-A | 2 | 5.6 | 59 | 173 | 5 | MOLECULE: | LIPID BINDING PROTEIN LPQN |
| 82:00:00 | 2fkc-A | 2 | 4.2 | 65 | 247 | 6 | MOLECULE: | 5'-D(*CP*CP*AP*GP*CP*GP*CP*TP*GP*G) |
| 83:00:00 | 5n83-C | 2 | 6.9 | 69 | 196 | 0 | MOLECULE: | FIBER |
| 84:00:00 | 7e2c-I | 2 | 9.6 | 64 | 799 | 6 | MOLECULE: | TRAPP-ASSOCIATED PROTEIN TCA17 |
| 85:00:00 | 4kh6-B | 2 | 5 | 58 | 596 | 7 | MOLECULE: | NUCLEOSIDE-TRIPHOSPHATASE 2 |
| 86:00:00 | 5nbs-A | 2 | 8.1 | 62 | 842 | 5 | MOLECULE: | BETA-GLUCOSIDASE |
| 87:00:00 | 3e8w-A | 2 | 7.5 | 82 | 219 | 5 | MOLECULE: | TAKEOUT-LIKE PROTEIN 1 |
| 88:00:00 | 8fbe-A | 2 | 4.5 | 73 | 144 | 3 | MOLECULE: | NEUROTOXIN COMPLEX COMPONENT ORF-X1 |
| 89:00:00 | 4m8r-A | 2 | 3.9 | 64 | 406 | 8 | MOLECULE: | HYPOTHETICAL PROTEIN |
| 90:00:00 | 2d7v-A | 2 | 3.5 | 60 | 155 | 13 | MOLECULE: | HYPOTHETICAL PROTEIN VCA0330 |
| 91:00:00 | 8i5f-A | 2 | 8.3 | 62 | 433 | 2 | MOLECULE: | DEDICATOR OF CYTOKINESIS PROTEIN 10 |
| 92:00:00 | 4r1k-B | 2 | 4.8 | 58 | 136 | 2 | MOLECULE: | UNCHARACTERIZED PROTEIN |
| 93:00:00 | 3fh1-A | 2 | 3.4 | 56 | 122 | 2 | MOLECULE: | UNCHARACTERIZED NTF2-LIKE PROTEIN |
| 94:00:00 | 6uf2-A | 2 | 3.1 | 50 | 125 | 6 | MOLECULE: | BIOFILM-RELATED PROTEIN |
| 95:00:00 | 2eab-A | 2 | 7.4 | 66 | 888 | 6 | MOLECULE: | ALPHA-FUCOSIDASE |
| 96:00:00 | 6v4n-M | 2 | 4.6 | 69 | 391 | 1 | MOLECULE: | NEURAMINIDASE |
| 97:00:00 | 6f3h-B | 2 | 10.2 | 69 | 790 | 4 | MOLECULE: | EXORIBONUCLEASE II, MITOCHONDRIAL |
| 98:00:00 | 6k9k-A | 2 | 9.5 | 70 | 2501 | 4 | MOLECULE: | SERINE-PROTEIN KINASE ATM |
| 99:00:00 | 5aor-A | 2 | 19.6 | 66 | 1009 | 3 | MOLECULE: | DOSAGE COMPENSATION REGULATOR |
| 0:00 | 3nqn-A | 2 | 6.3 | 58 | 151 | 10 | MOLECULE: | UNCHARACTERIZED PROTEIN |
| 1:00 | 7vgh-B | 2 | 4.3 | 73 | 1182 | 7 | MOLECULE: | CELL CYCLE CONTROL PROTEIN 50B |
| 2:00 | 6j7n-A | 2 | 7.4 | 64 | 290 | 5 | MOLECULE: | GUANYLYLTRANSFERASE-LIKE TOXIN |
| 3:00 | 1jeq-A | 2 | 5.7 | 68 | 548 | 9 | MOLECULE: | KU70 |
| 4:00 | 6u4k-A | 2 | 3.8 | 52 | 391 | 10 | MOLECULE: | TALIN-2 |
| 5:00 | 2crf-A | 2 | 4.9 | 67 | 150 | 6 | MOLECULE: | RAN BINDING PROTEIN 3 |
| 6:00 | 4ruq-B | 2 | 5 | 60 | 237 | 0 | MOLECULE: | FISH-EGG LECTIN |
| 7:00 | 4l1n-A | 2 | 5.1 | 55 | 161 | 4 | MOLECULE: | CONSERVED LIPOPROTEIN, PUTATIVE |
| 8:00 | 5dcj-A | 2 | 6.9 | 50 | 118 | 4 | MOLECULE: | OSMOLARITY SENSOR PROTEIN ENVZ |
| 9:00 | 8ck1-A | 2 | 9.7 | 51 | 826 | 4 | MOLECULE: | TAIL NOZZLE |
| 10:00 | 2fg9-A | 2 | 5.9 | 63 | 166 | 6 | MOLECULE: | 5-NITROIMIDAZOLE ANTIBIOTIC RESISTA |
| 11:00 | 2m89-A | 2 | 3.7 | 63 | 134 | 5 | MOLECULE: | AHA1 DOMAIN PROTEIN |
| 12:00 | 3c7g-A | 2 | 5.2 | 74 | 488 | 7 | MOLECULE: | ENDO-1,4-BETA-XYLANASE |
| 13:00 | 4k90-B | 2 | 5.2 | 62 | 207 | 11 | MOLECULE: | EXTRACELLULAR METALLOPROTEINASE MEP |
| 14:00 | 1d09-B | 2 | 3.8 | 56 | 153 | 4 | MOLECULE: | ASPARTATE CARBAMOYLTRANSFERASE CATA |

| | | | | | | | | |
|---|---|---|---|---|---|---|---|---|
| 15:00 | 5dfz-C | 2 | 6.8 | 65 | 783 | 8 | MOLECULE: | VACUOLAR PROTEIN SORTING-ASSOCIATED |
| 16:00 | 4std-A | 2 | 5.9 | 68 | 164 | 6 | MOLECULE: | SCYTALONE DEHYDRATASE |
| 17:00 | 5hww-A | 2 | 4.5 | 54 | 128 | 9 | MOLECULE: | SENSOR HISTIDINE KINASE TODS |
| 18:00 | 4qxa-B | 2 | 6.5 | 56 | 147 | 7 | MOLECULE: | RAS-RELATED PROTEIN RAB-9A |
| 19:00 | 3tvv-B | 2 | 4.3 | 51 | 235 | 2 | MOLECULE: | HISTONE CHAPERONE RTT106 |
| 20:00 | 5csk-A | 2 | 5.6 | 49 | 1996 | 4 | MOLECULE: | ACETYL-COA CARBOXYLASE |
| 21:00 | 8owu-E | 2 | 3 | 46 | 115 | 4 | MOLECULE: | TRANSPOSASE |
| 22:00 | 8t1l-A | 2 | 6.7 | 66 | 472 | 6 | MOLECULE: | MEDIATOR OF RNA POLYMERASE II TRANS |
| 23:00 | 5txc-A | 2 | 5.1 | 74 | 653 | 9 | MOLECULE: | ATXE2 |
| 24:00:00 | 5d9b-A | 2 | 6.8 | 61 | 307 | 8 | MOLECULE: | LUCIFERIN REGENERATING ENZYME |
| 25:00:00 | 5b2g-C | 2 | 5.5 | 50 | 350 | 8 | MOLECULE: | ENDOLYSIN,CLAUDIN-4 |
| 26:00:00 | 8adl-A | 2 | 4.4 | 64 | 633 | 5 | MOLECULE: | MTC5 ISOFORM 1 |
| 27:00:00 | 8a43-A | 2 | 2.9 | 49 | 1549 | 8 | MOLECULE: | DNA-DIRECTED RNA POLYMERASE I SUBUN |
| 28:00:00 | 7s3l-A | 2 | 3.6 | 42 | 254 | 5 | MOLECULE: | AMINOGLYCOSIDE PHOSPHOTRANSFERASE F |
| 29:00:00 | 8usw-B | 2 | 13 | 72 | 745 | 3 | MOLECULE: | GLUTAMATE RECEPTOR IONOTROPIC, NMDA |
| 30:00:00 | 5xmz-A | 2 | 4.9 | 64 | 122 | 8 | MOLECULE: | EFFECTOR PROTEIN PEVD1 |
| 31:00:00 | 2pcs-A | 2 | 5.1 | 70 | 152 | 4 | MOLECULE: | CONSERVED PROTEIN |
| 32:00:00 | 6od1-B | 2 | 3.4 | 54 | 106 | 4 | MOLECULE: | REGULATOR OF RPOS |
| 33:00:00 | 2ow6-A | 2 | 8.6 | 75 | 1016 | 3 | MOLECULE: | ALPHA-MANNOSIDASE 2 |
| 34:00:00 | 3tkn-A | 2 | 5 | 48 | 451 | 10 | MOLECULE: | NUCLEOPORIN NUP82 |
| 35:00:00 | 7jrl-A | 2 | 3.9 | 57 | 457 | 5 | MOLECULE: | F5/8 TYPE C DOMAIN PROTEIN |
| 36:00:00 | 8a3t-B | 2 | 4.7 | 58 | 440 | 10 | MOLECULE: | ANAPHASE-PROMOTING COMPLEX SUBUNIT |
| 37:00:00 | 9b21-A | 2 | 7.9 | 49 | 205 | 0 | MOLECULE: | ADP-RIBOSE PYROPHOSPHATASE |
| 38:00:00 | 6qvf-B | 2 | 3.4 | 62 | 173 | 6 | MOLECULE: | PREPILIN-LIKE PROTEIN |
| 39:00:00 | 8ae6-W | 2 | 3.7 | 49 | 941 | 8 | MOLECULE: | NITROGEN PERMEASE REGULATOR 3 |
| 40:00:00 | 4yu6-A | 2 | 12.1 | 81 | 754 | 9 | MOLECULE: | IMMUNE INHIBITOR A, METALLOPROTEASE |
| 41:00:00 | 1ofd-A | 2 | 7.2 | 64 | 1491 | 8 | MOLECULE: | FERREDOXIN-DEPENDENT GLUTAMATE SYNT |
| 42:00:00 | 6do4-A | 2 | 4.9 | 43 | 319 | 7 | MOLECULE: | KELCH DOMAIN-CONTAINING PROTEIN 2 |
| 43:00:00 | 5wqh-F | 2 | 6.9 | 74 | 140 | 7 | MOLECULE: | ISOMERASE TRT14 |
| 44:00:00 | 7z4f-G | 2 | 5.5 | 69 | 922 | 3 | MOLECULE: | PUTATIVE STRUCTURAL PROTEIN |
| 45:00:00 | 3wj9-B | 2 | 7.3 | 64 | 402 | 9 | MOLECULE: | EUKARYOTIC TRANSLATION INITIATION F |
| 46:00:00 | 7aed-A | 2 | 4.3 | 55 | 128 | 5 | MOLECULE: | PRGL |
| 47:00:00 | 7f13-B | 2 | 9.2 | 70 | 148 | 4 | MOLECULE: | DCR3 |
| 48:00:00 | 6xzf-A | 2 | 6 | 57 | 225 | 4 | MOLECULE: | EGFP |
| 49:00:00 | 2whs-D | 2 | 8.9 | 55 | 226 | 4 | MOLECULE: | LARGE STOKES SHIFT FLUORESCENT PROT |
| 50:00:00 | 6nqs-D | 2 | 4.3 | 59 | 214 | 8 | MOLECULE: | FLUORESCENT PROTEIN DRONPA |
| 51:00:00 | 4w75-A | 2 | 12.2 | 54 | 209 | 2 | MOLECULE: | FLUORESCENT PROTEIN D21H/K26C |
| 52:00:00 | 3ekj-A | 2 | 11.4 | 74 | 301 | 4 | MOLECULE: | MYOSIN LIGHT CHAIN KINASE, GREEN FL |
| 53:00:00 | 4pfe-B | 2 | 9.4 | 60 | 340 | 3 | MOLECULE: | GREEN FLUORESCENT PROTEIN |
| 54:00:00 | 6tv7-A | 2 | 7 | 78 | 397 | 4 | MOLECULE: | RSGCAMP |
| 55:00:00 | 4w6u-C | 2 | 10.1 | 60 | 221 | 7 | MOLECULE: | FLUORESCENT PROTEIN E115H/T118H |
| 56:00:00 | 4dxp-A | 2 | 10.4 | 65 | 215 | 2 | MOLECULE: | LEA X121 GFP-LIKE PROTEINS |
| 57:00:00 | 5eb7-A | 2 | 9.7 | 63 | 213 | 2 | MOLECULE: | REVERSIBLY PHOTOSWITCHING CHROMOPRO |
| 58:00:00 | 7a8i-D | 2 | 11 | 58 | 224 | 5 | MOLECULE: | GREEN FLUORESCENT PROTEIN |

**DALI results: MdA-1 Col**

| No: | Chain | Z | rmsd | lali | nres | %id | PDB Description |
|---|---|---|---|---|---|---|---|
| 1:00 | s001A | 7.6 | 62.4 | 215 | 0 | 0 | |

**DALI results: MdA-1 Colalt**

| No: | Chain | Z | rmsd | lali | nres | %id | PDB | Description |
|---|---|---|---|---|---|---|---|---|
| 1:00 | 8th8-s | 6.3 | 11 | 116 | 187 | 9 | MOLECULE: | DYNEIN REGULATORY COMPLEX PROTEIN 1/2 N-TERMINAL |
| 2:00 | 6j9r-B | 6.1 | 4.2 | 98 | 126 | 8 | MOLECULE: | BRAIN TUMOR PROTEIN |
| 3:00 | 7wxz-D | 6 | 6.5 | 90 | 187 | 14 | MOLECULE: | SPIKE PROTEIN S2' |
| 4:00 | 8ga8-G | 6 | 3.5 | 78 | 97 | 5 | MOLECULE: | TRANSCRIPTIONAL REGULATORY PROTEIN SDS3 |
| 5:00 | 8fxf-A | 6 | 2.1 | 78 | 78 | 10 | MOLECULE: | E3 UBIQUITIN-PROTEIN LIGASE TRIM56 |
| 6:00 | 1pl5-S | 6 | 3 | 76 | 76 | 11 | MOLECULE: | REGULATORY PROTEIN SIR4 |
| 7:00 | 5fiy-C | 6 | 3.6 | 78 | 78 | 8 | MOLECULE: | PRKC APOPTOSIS WT1 REGULATOR PROTEIN |
| 8:00 | 6hk9-B | 5.9 | 2.3 | 78 | 78 | 5 | MOLECULE: | TESTIS-EXPRESSED PROTEIN 12 |
| 9:00 | 3vp8-D | 5.9 | 2.5 | 78 | 78 | 5 | MOLECULE: | GENERAL TRANSCRIPTIONAL COREPRESSOR TUP1 |
| 10:00 | 3vp8-C | 5.9 | 2.1 | 77 | 77 | 6 | MOLECULE: | GENERAL TRANSCRIPTIONAL COREPRESSOR TUP1 |
| 11:00 | 8sjj-A | 5.9 | 2.9 | 74 | 74 | 5 | MOLECULE: | SEPTIN-14 |
| 12:00 | 8sjj-B | 5.9 | 3.3 | 74 | 74 | 5 | MOLECULE: | SEPTIN-14 |
| 13:00 | 8t9d-D | 5.9 | 6.5 | 87 | 161 | 8 | MOLECULE: | MEDIATOR OF RNA POLYMERASE II TRANSCRIPTION SUBUN |
| 14:00 | 8czi-A | 5.9 | 2.2 | 71 | 71 | 11 | MOLECULE: | SCAFFOLDED SPIKE PROTEIN S2' HR1 |
| 15:00 | 8fcf-D | 5.9 | 3.6 | 79 | 79 | 4 | MOLECULE: | PLASMALEMMA VESICLE-ASSOCIATED PROTEIN |
| 16:00 | 8fxf-D | 5.9 | 2.1 | 78 | 78 | 10 | MOLECULE: | E3 UBIQUITIN-PROTEIN LIGASE TRIM56 |
| 17:00 | 8tgt-A | 5.9 | 1.6 | 76 | 76 | 8 | MOLECULE: | M PROTEIN |
| 18:00 | 8czi-C | 5.9 | 2.2 | 71 | 71 | 11 | MOLECULE: | SCAFFOLDED SPIKE PROTEIN S2' HR1 |
| 19:00 | 8afh-P | 5.9 | 4.1 | 79 | 79 | 14 | MOLECULE: | CRESCENTIN |
| 20:00 | 8fxf-B | 5.9 | 2 | 78 | 78 | 10 | MOLECULE: | E3 UBIQUITIN-PROTEIN LIGASE TRIM56 |
| 21:00 | 1pl5-A | 5.9 | 2.9 | 75 | 75 | 11 | MOLECULE: | REGULATORY PROTEIN SIR4 |
| 22:00 | 8afe-F | 5.9 | 4.1 | 79 | 79 | 14 | MOLECULE: | CRESCENTIN |
| 23:00 | 8afh-F | 5.9 | 4.1 | 79 | 79 | 14 | MOLECULE: | CRESCENTIN |
| 24:00:00 | 8csp-U | 5.9 | 9.8 | 98 | 174 | 6 | MOLECULE: | 28S RIBOSOMAL PROTEIN S34, MITOCHONDRIAL |
| 25:00:00 | 1gk4-D | 5.9 | 3.6 | 78 | 78 | 8 | MOLECULE: | VIMENTIN |
| 26:00:00 | 1gk4-B | 5.9 | 4.2 | 79 | 79 | 5 | MOLECULE: | VIMENTIN |
| 27:00:00 | 2bez-C | 5.9 | 2.5 | 77 | 77 | 9 | MOLECULE: | E2 GLYCOPROTEIN |
| 28:00:00 | 5jxc-C | 5.9 | 3.5 | 78 | 78 | 10 | MOLECULE: | RAS/RAP GTPASE-ACTIVATING PROTEIN SYNGAP |
| 29:00:00 | 1n7s-C | 5.9 | 5.4 | 79 | 79 | 8 | MOLECULE: | VESICLE-ASSOCIATED MEMBRANE PROTEIN 2 |
| 30:00:00 | 5djn-A | 5.9 | 2.2 | 77 | 77 | 10 | MOLECULE: | KINESIN-LIKE PROTEIN |
| 31:00:00 | 1gk4-A | 5.9 | 3 | 78 | 79 | 8 | MOLECULE: | VIMENTIN |
| 32:00:00 | 6aoz-A | 5.9 | 3.8 | 74 | 74 | 7 | MOLECULE: | CASP8-ASSOCIATED PROTEIN 2 |
| 33:00:00 | 8x7z-A | 5.8 | 3.6 | 81 | 83 | 12 | MOLECULE: | HR1 |
| 34:00:00 | 8v0y-G | 5.8 | 3.3 | 81 | 82 | 5 | MOLECULE: | ACTIN, ALPHA CARDIAC MUSCLE 1 |
| 35:00:00 | 8t1i-B | 5.8 | 7.6 | 82 | 158 | 9 | MOLECULE: | MEDIATOR OF RNA POLYMERASE II TRANSCRIPTION SUBUN |
| 36:00:00 | 1nyh-A | 5.8 | 2.9 | 76 | 76 | 11 | MOLECULE: | REGULATORY PROTEIN SIR4 |
| 37:00:00 | 7tik-B | 5.8 | 1.7 | 71 | 71 | 10 | MOLECULE: | FERRITIN, DPS FAMILY PROTEIN AND SPIKE PROTEIN S2 |
| 38:00:00 | 8fa2-A | 5.8 | 2.2 | 71 | 71 | 10 | MOLECULE: | SCAFFOLDED SPIKE PROTEIN S2' HR1 |
| 39:00:00 | 8fxf-C | 5.8 | 2 | 78 | 78 | 10 | MOLECULE: | E3 UBIQUITIN-PROTEIN LIGASE TRIM56 |
| 40:00:00 | 8fa1-C | 5.8 | 1.8 | 71 | 71 | 10 | MOLECULE: | FERRITIN, DPS FAMILY PROTEIN AND SPIKE PROTEIN S2 |
| 41:00:00 | 8fa2-C | 5.8 | 2.2 | 71 | 71 | 10 | MOLECULE: | SCAFFOLDED SPIKE PROTEIN S2' HR1 |
| 42:00:00 | 8fa1-A | 5.8 | 1.8 | 71 | 71 | 10 | MOLECULE: | FERRITIN, DPS FAMILY PROTEIN AND SPIKE PROTEIN S2 |
| 43:00:00 | 8czi-B | 5.8 | 2.2 | 71 | 71 | 11 | MOLECULE: | SCAFFOLDED SPIKE PROTEIN S2' HR1 |
| 44:00:00 | 7tik-C | 5.8 | 1.7 | 71 | 71 | 10 | MOLECULE: | FERRITIN, DPS FAMILY PROTEIN AND SPIKE PROTEIN S2 |
| 45:00:00 | 8fa1-B | 5.8 | 1.8 | 71 | 71 | 10 | MOLECULE: | FERRITIN, DPS FAMILY PROTEIN AND SPIKE PROTEIN S2 |
| 46:00:00 | 8sdi-A | 5.8 | 1.9 | 80 | 80 | 6 | MOLECULE: | TRIPARTITE MOTIF-CONTAINING PROTEIN 45 |
| 47:00:00 | 8fa2-B | 5.8 | 2.2 | 71 | 71 | 10 | MOLECULE: | SCAFFOLDED SPIKE PROTEIN S2' HR1 |
| 48:00:00 | 7z44-A | 5.8 | 6.9 | 100 | 598 | 5 | MOLECULE: | PORTAL PROTEIN |
| 49:00:00 | 7tik-A | 5.8 | 1.7 | 71 | 71 | 10 | MOLECULE: | FERRITIN, DPS FAMILY PROTEIN AND SPIKE PROTEIN S2 |
| 50:00:00 | 6h9l-B | 5.8 | 7.5 | 93 | 129 | 4 | MOLECULE: | UNCHARACTERIZED PROTEIN |
| 51:00:00 | 8sdi-D | 5.8 | 1.9 | 80 | 80 | 6 | MOLECULE: | TRIPARTITE MOTIF-CONTAINING PROTEIN 45 |
| 52:00:00 | 2wps-B | 5.8 | 5.2 | 86 | 98 | 10 | MOLECULE: | TRIMERIC AUTOTRANSPORTER ADHESIN FRAGMENT |
| 53:00:00 | 2wps-C | 5.8 | 5.2 | 86 | 98 | 9 | MOLECULE: | TRIMERIC AUTOTRANSPORTER ADHESIN FRAGMENT |
| 54:00:00 | 3vp8-A | 5.8 | 2.3 | 75 | 75 | 8 | MOLECULE: | GENERAL TRANSCRIPTIONAL COREPRESSOR TUP1 |

| | | | | | | | | |
|---|---|---|---|---|---|---|---|---|
| 55:00:00 | 5djn-C | 5.8 | 4 | 78 | 81 | 10 | MOLECULE: | KINESIN-LIKE PROTEIN |
| 56:00:00 | 3fwc-N | 5.8 | 3.1 | 82 | 84 | 5 | MOLECULE: | CELL DIVISION CONTROL PROTEIN 31 |
| 57:00:00 | 3fwc-J | 5.8 | 1.3 | 77 | 78 | 5 | MOLECULE: | CELL DIVISION CONTROL PROTEIN 31 |
| 58:00:00 | 1x8y-A | 5.8 | 3.7 | 74 | 74 | 5 | MOLECULE: | LAMIN A/C |
| 59:00:00 | 7apk-m | 5.7 | 3.5 | 71 | 549 | 3 | MOLECULE: | THO COMPLEX SUBUNIT 1 |
| 60:00:00 | 3iv1-A | 5.7 | 2.9 | 76 | 78 | 5 | MOLECULE: | TUMOR SUSCEPTIBILITY GENE 101 PROTEIN |
| 61:00:00 | 6eun-A | 5.7 | 6.8 | 89 | 147 | 8 | MOLECULE: | ADHESIN |
| 62:00:00 | 3vp8-B | 5.7 | 2.8 | 79 | 80 | 6 | MOLECULE: | GENERAL TRANSCRIPTIONAL COREPRESSOR TUP1 |
| 63:00:00 | 2jee-A | 5.7 | 2.7 | 77 | 78 | 8 | MOLECULE: | CELL DIVISION PROTEIN ZAPB |
| 64:00:00 | 3iv1-E | 5.7 | 2.9 | 76 | 78 | 5 | MOLECULE: | TUMOR SUSCEPTIBILITY GENE 101 PROTEIN |
| 65:00:00 | 3iv1-F | 5.7 | 3.3 | 77 | 78 | 5 | MOLECULE: | TUMOR SUSCEPTIBILITY GENE 101 PROTEIN |
| 66:00:00 | 3iv1-B | 5.7 | 2.9 | 76 | 78 | 5 | MOLECULE: | TUMOR SUSCEPTIBILITY GENE 101 PROTEIN |
| 67:00:00 | 3iv1-H | 5.7 | 3.1 | 77 | 78 | 5 | MOLECULE: | TUMOR SUSCEPTIBILITY GENE 101 PROTEIN |
| 68:00:00 | 2jee-D | 5.7 | 2.7 | 77 | 78 | 8 | MOLECULE: | CELL DIVISION PROTEIN ZAPB |
| 69:00:00 | 7x2e-A | 5.7 | 6.1 | 94 | 163 | 6 | MOLECULE: | HARMONIN |
| 70:00:00 | 2jee-B | 5.7 | 2.7 | 77 | 78 | 8 | MOLECULE: | CELL DIVISION PROTEIN ZAPB |
| 71:00:00 | 7say-B | 5.7 | 2.6 | 70 | 70 | 6 | MOLECULE: | GENERAL CONTROL TRANSCRIPTION FACTOR GCN4/M PROTE |
| 72:00:00 | 8v0i-F | 5.7 | 4.6 | 81 | 82 | 9 | MOLECULE: | ACTIN, ALPHA CARDIAC MUSCLE 1 |
| 73:00:00 | 8uzx-G | 5.7 | 3.3 | 81 | 82 | 5 | MOLECULE: | ACTIN, ALPHA CARDIAC MUSCLE 1 |
| 74:00:00 | 8fcf-C | 5.7 | 2.4 | 81 | 83 | 9 | MOLECULE: | PLASMALEMMA VESICLE-ASSOCIATED PROTEIN |
| 75:00:00 | 8uz5-G | 5.7 | 3.9 | 81 | 82 | 5 | MOLECULE: | ACTIN, ALPHA CARDIAC MUSCLE 1 |
| 76:00:00 | 8sdi-B | 5.7 | 1.3 | 78 | 80 | 10 | MOLECULE: | TRIPARTITE MOTIF-CONTAINING PROTEIN 45 |
| 77:00:00 | 8uz6-G | 5.7 | 4.6 | 80 | 82 | 6 | MOLECULE: | ACTIN, ALPHA CARDIAC MUSCLE 1 |
| 78:00:00 | 8uwx-G | 5.7 | 4.6 | 80 | 82 | 6 | MOLECULE: | ACTIN, ALPHA CARDIAC MUSCLE 1 |
| 79:00:00 | 8v0i-G | 5.7 | 3.8 | 80 | 82 | 6 | MOLECULE: | ACTIN, ALPHA CARDIAC MUSCLE 1 |
| 80:00:00 | 8sdi-C | 5.7 | 1.5 | 79 | 80 | 10 | MOLECULE: | TRIPARTITE MOTIF-CONTAINING PROTEIN 45 |
| 81:00:00 | 8x7x-C | 5.7 | 2.1 | 72 | 72 | 13 | MOLECULE: | HR1 |
| 82:00:00 | 8sjj-D | 5.7 | 2.2 | 83 | 86 | 5 | MOLECULE: | SEPTIN-14 |
| 83:00:00 | 8sor-C | 5.7 | 6.9 | 96 | 264 | 6 | MOLECULE: | PHOSPHATIDYLINOSITOL 3-KINASE CATALYTIC SUBUNIT T |
| 84:00:00 | 7zd6-q | 5.7 | 11.8 | 101 | 139 | 9 | MOLECULE: | NADH-UBIQUINONE OXIDOREDUCTASE CHAIN 3 |
| 85:00:00 | 7y5d-b | 5.7 | 6.2 | 95 | 145 | 2 | MOLECULE: | ATP SYNTHASE SUBUNIT A |
| 86:00:00 | 1sfc-J | 5.7 | 2.6 | 74 | 74 | 5 | MOLECULE: | PROTEIN (SYNAPTOBREVIN 2) |
| 87:00:00 | 3vem-D | 5.7 | 1.9 | 78 | 80 | 5 | MOLECULE: | HELICASE PROTEIN MOM1 |
| 88:00:00 | 8uzx-F | 5.7 | 4.4 | 81 | 82 | 9 | MOLECULE: | ACTIN, ALPHA CARDIAC MUSCLE 1 |
| 89:00:00 | 5wq4-D | 5.7 | 3.8 | 72 | 72 | 11 | MOLECULE: | UBIQUITIN |
| 90:00:00 | 8v0y-F | 5.7 | 4.4 | 80 | 82 | 11 | MOLECULE: | ACTIN, ALPHA CARDIAC MUSCLE 1 |
| 91:00:00 | 1uix-B | 5.7 | 2.9 | 68 | 68 | 12 | MOLECULE: | RHO-ASSOCIATED KINASE |
| 92:00:00 | 4e61-A | 5.7 | 3.5 | 84 | 91 | 7 | MOLECULE: | PROTEIN BIM1 |
| 93:00:00 | 4mvd-B | 5.6 | 14 | 81 | 253 | 2 | MOLECULE: | CHOLINE-PHOSPHATE CYTIDYLYLTRANSFERASE A |
| 94:00:00 | 3trt-A | 5.6 | 1.6 | 75 | 75 | 8 | MOLECULE: | VIMENTIN |
| 95:00:00 | 3iv1-C | 5.6 | 3.3 | 78 | 78 | 5 | MOLECULE: | TUMOR SUSCEPTIBILITY GENE 101 PROTEIN |
| 96:00:00 | 3iv1-G | 5.6 | 3 | 76 | 78 | 5 | MOLECULE: | TUMOR SUSCEPTIBILITY GENE 101 PROTEIN |
| 97:00:00 | 3iv1-D | 5.6 | 3.2 | 77 | 78 | 5 | MOLECULE: | TUMOR SUSCEPTIBILITY GENE 101 PROTEIN |
| 98:00:00 | 5h07-D | 5.6 | 3.8 | 78 | 78 | 6 | MOLECULE: | POLYUBIQUITIN-C |
| 99:00:00 | 8sjj-C | 5.6 | 3.2 | 81 | 86 | 5 | MOLECULE: | SEPTIN-14 |
| 0:00 | 8j9s-A | 5.6 | 3.3 | 71 | 71 | 10 | MOLECULE: | AMINOACYL TRNA SYNTHASE COMPLEX-INTERACTING MULTI |
| 1:00 | 8fby-D | 5.6 | 3.4 | 82 | 86 | 4 | MOLECULE: | PLASMALEMMA VESICLE-ASSOCIATED PROTEIN |
| 2:00 | 8t1l-B | 5.6 | 8.7 | 89 | 158 | 3 | MOLECULE: | MEDIATOR OF RNA POLYMERASE II TRANSCRIPTION SUBUN |
| 3:00 | 8v0k-G | 5.6 | 3.4 | 81 | 82 | 9 | MOLECULE: | ACTIN, ALPHA CARDIAC MUSCLE 1 |
| 4:00 | 4wy4-C | 5.6 | 3 | 75 | 78 | 7 | MOLECULE: | VESICLE-ASSOCIATED MEMBRANE PROTEIN 8 |
| 5:00 | 5xau-E | 5.6 | 3.1 | 72 | 72 | 8 | MOLECULE: | LAMININ SUBUNIT ALPHA-5 |
| 6:00 | 8afh-E | 5.6 | 3.7 | 79 | 79 | 14 | MOLECULE: | CRESCENTIN |
| 7:00 | 1fav-A | 5.6 | 3.9 | 78 | 78 | 9 | MOLECULE: | HIV-1 ENVELOPE PROTEIN CHIMERA |
| 8:00 | 8afe-E | 5.6 | 3.7 | 79 | 79 | 14 | MOLECULE: | CRESCENTIN |
| 9:00 | 8dd0-O | 5.6 | 2 | 70 | 70 | 6 | MOLECULE: | ACTIN, ALPHA CARDIAC MUSCLE 1 |
| 10:00 | 8sde-A | 5.6 | 2.3 | 83 | 89 | 8 | MOLECULE: | TRIPARTITE MOTIF-CONTAINING PROTEIN 29 |

| | | | | | | | | |
|---|---|---|---|---|---|---|---|---|
| 11:00 | 8gt9-B | 5.6 | 4.9 | 83 | 88 | 7 | MOLECULE: | BECLIN-2 |
| 12:00 | 8uyd-G | 5.6 | 4.5 | 80 | 82 | 11 | MOLECULE: | ACTIN, ALPHA CARDIAC MUSCLE 1 |
| 13:00 | 8dd0-N | 5.6 | 4 | 70 | 70 | 6 | MOLECULE: | ACTIN, ALPHA CARDIAC MUSCLE 1 |
| 14:00 | 8i3e-B | 5.6 | 6.7 | 93 | 135 | 6 | MOLECULE: | ELKS/RAB6-INTERACTING/CAST FAMILY MEMBER 1 |
| 15:00 | 8fby-B | 5.6 | 3.2 | 84 | 88 | 4 | MOLECULE: | PLASMALEMMA VESICLE-ASSOCIATED PROTEIN |
| 16:00 | 3cvf-C | 5.6 | 3.4 | 74 | 74 | 8 | MOLECULE: | HOMER PROTEIN HOMOLOG 3 |
| 17:00 | 8afh-O | 5.6 | 3.7 | 79 | 79 | 14 | MOLECULE: | CRESCENTIN |
| 18:00 | 7ucc-J | 5.6 | 1.7 | 68 | 68 | 9 | MOLECULE: | PROTEIN FOSB |
| 19:00 | 8dd0-J | 5.6 | 1.8 | 70 | 70 | 13 | MOLECULE: | ACTIN, ALPHA CARDIAC MUSCLE 1 |
| 20:00 | 1gk4-F | 5.6 | 2.6 | 74 | 74 | 7 | MOLECULE: | VIMENTIN |
| 21:00 | 1sfc-F | 5.6 | 3.6 | 73 | 73 | 4 | MOLECULE: | PROTEIN (SYNAPTOBREVIN 2) |
| 22:00 | 3tyy-B | 5.6 | 2.2 | 71 | 71 | 13 | MOLECULE: | LAMIN-B1 |
| 23:00 | 3thf-A | 5.6 | 7.1 | 129 | 175 | 9 | MOLECULE: | PROTEIN SHROOM |
| 24:00:00 | 3klt-D | 5.6 | 1.4 | 70 | 70 | 7 | MOLECULE: | VIMENTIN |
| 25:00:00 | 3bas-B | 5.6 | 4.6 | 79 | 87 | 9 | MOLECULE: | MYOSIN HEAVY CHAIN, STRIATED MUSCLE/GENERAL CONTR |
| 26:00:00 | 3vem-B | 5.6 | 1.7 | 79 | 82 | 6 | MOLECULE: | HELICASE PROTEIN MOM1 |
| 27:00:00 | 3fwc-F | 5.6 | 2.3 | 82 | 84 | 5 | MOLECULE: | CELL DIVISION CONTROL PROTEIN 31 |
| 28:00:00 | 4dhx-A | 5.6 | 2.3 | 72 | 72 | 6 | MOLECULE: | 80 KDA MCM3-ASSOCIATED PROTEIN |
| 29:00:00 | 4bl6-A | 5.6 | 2.4 | 79 | 82 | 14 | MOLECULE: | PROTEIN BICAUDAL D |
| 30:00:00 | 4dhx-D | 5.6 | 2.2 | 69 | 69 | 6 | MOLECULE: | 80 KDA MCM3-ASSOCIATED PROTEIN |
| 31:00:00 | 5whf-D | 5.6 | 3.7 | 82 | 85 | 10 | MOLECULE: | VIMENTIN |
| 32:00:00 | 3ni0-B | 5.6 | 2.9 | 80 | 87 | 10 | MOLECULE: | BONE MARROW STROMAL ANTIGEN 2 |
| 33:00:00 | 1uix-A | 5.6 | 3.6 | 69 | 69 | 12 | MOLECULE: | RHO-ASSOCIATED KINASE |
| 34:00:00 | 5oi7-A | 5.6 | 4.9 | 83 | 88 | 17 | MOLECULE: | CENTROSOMAL PROTEIN OF 85 KDA |
| 35:00:00 | 3v4q-A | 5.6 | 3.5 | 74 | 74 | 5 | MOLECULE: | PRELAMIN-A/C |
| 36:00:00 | 3vp9-B | 5.6 | 5.1 | 77 | 77 | 6 | MOLECULE: | GENERAL TRANSCRIPTIONAL COREPRESSOR TUP1 |
| 37:00:00 | 3htk-B | 5.6 | 4 | 73 | 73 | 11 | MOLECULE: | STRUCTURAL MAINTENANCE OF CHROMOSOMES PROTEIN 5 |
| 38:00:00 | 7znl-E | 5.6 | 2.3 | 65 | 538 | 8 | MOLECULE: | THO COMPLEX SUBUNIT 1 |
| 39:00:00 | 4mvd-H | 5.6 | 14 | 81 | 253 | 2 | MOLECULE: | CHOLINE-PHOSPHATE CYTIDYLYLTRANSFERASE A |
| 40:00:00 | 8fwj-A | 5.5 | 4.8 | 101 | 552 | 6 | MOLECULE: | CIRCADIAN CLOCK PROTEIN KAIC |
| 41:00:00 | 7znk-o | 5.5 | 8.2 | 91 | 164 | 10 | MOLECULE: | RNA |
| 42:00:00 | 6ap0-B | 5.5 | 3.3 | 81 | 88 | 4 | MOLECULE: | CASP8-ASSOCIATED PROTEIN 2 |
| 43:00:00 | 6oqa-G | 5.5 | 3.3 | 81 | 87 | 11 | MOLECULE: | PEPTIDYL-PROLYL CIS-TRANS ISOMERASE FKBP1A |
| 44:00:00 | 7fde-G | 5.5 | 3.5 | 83 | 231 | 7 | MOLECULE: | V-TYPE PROTON ATPASE SUBUNIT C |
| 45:00:00 | 7yfj-B | 5.5 | 4.4 | 84 | 92 | 6 | MOLECULE: | PRE-MRNA-SPLICING REGULATOR WTAP |
| 46:00:00 | 8uz6-F | 5.5 | 4 | 80 | 82 | 5 | MOLECULE: | ACTIN, ALPHA CARDIAC MUSCLE 1 |
| 47:00:00 | 7z8i-x | 5.5 | 3 | 83 | 93 | 7 | MOLECULE: | ARP1 ACTIN RELATED PROTEIN 1 HOMOLOG A |
| 48:00:00 | 8uwy-F | 5.5 | 4 | 80 | 82 | 5 | MOLECULE: | ACTIN, ALPHA CARDIAC MUSCLE 1 |
| 49:00:00 | 8dd0-I | 5.5 | 3.6 | 70 | 70 | 13 | MOLECULE: | ACTIN, ALPHA CARDIAC MUSCLE 1 |
| 50:00:00 | 4ll7-G | 5.5 | 4.2 | 83 | 87 | 16 | MOLECULE: | SWI5-DEPENDENT HO EXPRESSION PROTEIN 3 |
| 51:00:00 | 8uzy-F | 5.5 | 4.2 | 79 | 82 | 6 | MOLECULE: | ACTIN, ALPHA CARDIAC MUSCLE 1 |
| 52:00:00 | 6cfz-D | 5.5 | 2.7 | 72 | 73 | 3 | MOLECULE: | ASK1 |
| 53:00:00 | 8uww-F | 5.5 | 4 | 80 | 82 | 5 | MOLECULE: | ACTIN, ALPHA CARDIAC MUSCLE 1 |
| 54:00:00 | 1gk4-C | 5.5 | 2.8 | 70 | 70 | 3 | MOLECULE: | VIMENTIN |
| 55:00:00 | 8uz5-F | 5.5 | 4 | 80 | 82 | 5 | MOLECULE: | ACTIN, ALPHA CARDIAC MUSCLE 1 |
| 56:00:00 | 7z8i-X | 5.5 | 4.7 | 83 | 93 | 7 | MOLECULE: | ARP1 ACTIN RELATED PROTEIN 1 HOMOLOG A |
| 57:00:00 | 8uwx-F | 5.5 | 4 | 80 | 82 | 5 | MOLECULE: | ACTIN, ALPHA CARDIAC MUSCLE 1 |
| 58:00:00 | 8gt9-C | 5.5 | 5.1 | 82 | 90 | 11 | MOLECULE: | BECLIN-2 |
| 59:00:00 | 8v01-F | 5.5 | 4.4 | 79 | 82 | 6 | MOLECULE: | ACTIN, ALPHA CARDIAC MUSCLE 1 |
| 60:00:00 | 1x79-C | 5.5 | 3.4 | 80 | 88 | 9 | MOLECULE: | ADP-RIBOSYLATION FACTOR BINDING PROTEIN GGA1 |
| 61:00:00 | 4n3y-B | 5.5 | 3.2 | 81 | 83 | 6 | MOLECULE: | RAB5 GDP/GTP EXCHANGE FACTOR |
| 62:00:00 | 5jxc-A | 5.5 | 4.5 | 81 | 86 | 11 | MOLECULE: | RAS/RAP GTPASE-ACTIVATING PROTEIN SYNGAP |
| 63:00:00 | 3vem-C | 5.5 | 2 | 78 | 83 | 5 | MOLECULE: | HELICASE PROTEIN MOM1 |
| 64:00:00 | 3klt-B | 5.5 | 2.5 | 71 | 71 | 7 | MOLECULE: | VIMENTIN |
| 65:00:00 | 3vem-A | 5.5 | 2.6 | 80 | 82 | 9 | MOLECULE: | HELICASE PROTEIN MOM1 |
| 66:00:00 | 5mq4-B | 5.5 | 4.3 | 79 | 111 | 4 | MOLECULE: | PROTEIN KINASE C-BINDING PROTEIN 1 |

| | | | | | | | | |
|---|---|---|---|---|---|---|---|---|
| 67:00:00 | 4xa6-D | 5.5 | 4 | 93 | 105 | 4 | MOLECULE: | GP7-MYH7(1777-1855)-EB1 CHIMERA PROTEIN |
| 68:00:00 | 4bl6-C | 5.5 | 7 | 83 | 86 | 13 | MOLECULE: | PROTEIN BICAUDAL D |
| 69:00:00 | 5whf-A | 5.5 | 5.8 | 83 | 86 | 7 | MOLECULE: | VIMENTIN |
| 70:00:00 | 5hda-C | 5.5 | 4.9 | 83 | 117 | 4 | MOLECULE: | ZINC FINGER MYND DOMAIN-CONTAINING PROTEIN 11 |
| 71:00:00 | 6ap0-A | 5.5 | 3.2 | 78 | 82 | 8 | MOLECULE: | CASP8-ASSOCIATED PROTEIN 2 |
| 72:00:00 | 4lhz-E | 5.5 | 5.2 | 76 | 76 | 4 | MOLECULE: | RAS-RELATED PROTEIN RAB-8A |
| 73:00:00 | 5k7b-D | 5.5 | 3.5 | 80 | 88 | 4 | MOLECULE: | BECLIN-2 |
| 74:00:00 | 4lhy-E | 5.5 | 5.2 | 76 | 76 | 4 | MOLECULE: | RAS-RELATED PROTEIN RAB-8A |
| 75:00:00 | 4n5b-A | 5.4 | 4.6 | 89 | 104 | 1 | MOLECULE: | PHOSPHOPROTEIN |
| 76:00:00 | 8gt9-A | 5.4 | 6.3 | 82 | 91 | 12 | MOLECULE: | BECLIN-2 |
| 77:00:00 | 7vf2-C | 5.4 | 6.8 | 86 | 184 | 7 | MOLECULE: | PROTEIN VIRILIZER HOMOLOG |
| 78:00:00 | 3n7n-D | 5.4 | 3.1 | 79 | 158 | 8 | MOLECULE: | MONOPOLIN COMPLEX SUBUNIT CSM1 |
| 79:00:00 | 3n7n-C | 5.4 | 4 | 81 | 158 | 9 | MOLECULE: | MONOPOLIN COMPLEX SUBUNIT CSM1 |
| 80:00:00 | 4wij-B | 5.4 | 7.5 | 106 | 301 | 4 | MOLECULE: | SPLICING FACTOR, PROLINE- AND GLUTAMINE-RICH |
| 81:00:00 | 3q8t-A | 5.4 | 4.4 | 86 | 94 | 7 | MOLECULE: | BECLIN-1 |
| 82:00:00 | 8ims-A | 5.4 | 3.7 | 85 | 93 | 11 | MOLECULE: | E3 UBIQUITIN-PROTEIN LIGASE TRAF7 |
| 83:00:00 | 6y09-C | 5.4 | 5.3 | 82 | 86 | 5 | MOLECULE: | RAS-RELATED PROTEIN RAB-33B |
| 84:00:00 | 6otn-B | 5.4 | 2.3 | 67 | 67 | 6 | MOLECULE: | TROPOMYOSIN ALPHA-3 CHAIN |
| 85:00:00 | 2jee-C | 5.4 | 2.5 | 74 | 78 | 8 | MOLECULE: | CELL DIVISION PROTEIN ZAPB |
| 86:00:00 | 3cve-B | 5.4 | 2.1 | 66 | 66 | 11 | MOLECULE: | HOMER PROTEIN HOMOLOG 1 |
| 87:00:00 | 8fby-C | 5.4 | 3.2 | 85 | 93 | 6 | MOLECULE: | PLASMALEMMA VESICLE-ASSOCIATED PROTEIN |
| 88:00:00 | 8uyd-F | 5.4 | 3.5 | 79 | 82 | 6 | MOLECULE: | ACTIN, ALPHA CARDIAC MUSCLE 1 |
| 89:00:00 | 7ucd-J | 5.4 | 1.7 | 68 | 68 | 9 | MOLECULE: | PROTEIN FOSB |
| 90:00:00 | 7saf-B | 5.4 | 2.7 | 67 | 70 | 10 | MOLECULE: | GENERAL CONTROL TRANSCRIPTION FACTOR GCN4/M PROTE |
| 91:00:00 | 4ll7-B | 5.4 | 4.1 | 82 | 88 | 16 | MOLECULE: | SWI5-DEPENDENT HO EXPRESSION PROTEIN 3 |
| 92:00:00 | 8uzy-G | 5.4 | 3.9 | 79 | 82 | 5 | MOLECULE: | ACTIN, ALPHA CARDIAC MUSCLE 1 |
| 93:00:00 | 7say-D | 5.4 | 2.9 | 67 | 67 | 6 | MOLECULE: | GENERAL CONTROL TRANSCRIPTION FACTOR GCN4/M PROTE |
| 94:00:00 | 7apk-o | 5.4 | 6.2 | 82 | 155 | 6 | MOLECULE: | THO COMPLEX SUBUNIT 1 |
| 95:00:00 | 7eqb-A | 5.4 | 2.9 | 68 | 68 | 3 | MOLECULE: | KINESIN-LIKE PROTEIN |
| 96:00:00 | 1sfc-B | 5.4 | 3.2 | 72 | 72 | 4 | MOLECULE: | PROTEIN (SYNAPTOBREVIN 2) |
| 97:00:00 | 4ll7-C | 5.4 | 4 | 83 | 90 | 19 | MOLECULE: | SWI5-DEPENDENT HO EXPRESSION PROTEIN 3 |
| 98:00:00 | 8bs7-B | 5.4 | 2.3 | 76 | 76 | 7 | MOLECULE: | PHOSPHOPROTEIN |
| 99:00:00 | 1n7s-B | 5.4 | 3.6 | 68 | 68 | 7 | MOLECULE: | VESICLE-ASSOCIATED MEMBRANE PROTEIN 2 |
| 0:00 | 8gt9-D | 5.4 | 5.2 | 83 | 91 | 11 | MOLECULE: | BECLIN-2 |
| 1:00 | 3cvf-D | 5.4 | 3 | 72 | 72 | 8 | MOLECULE: | HOMER PROTEIN HOMOLOG 3 |
| 2:00 | 7yg4-E | 5.4 | 2.3 | 67 | 67 | 7 | MOLECULE: | PROTEIN VIRILIZER HOMOLOG |
| 3:00 | 1gk4-E | 5.4 | 2.7 | 69 | 70 | 3 | MOLECULE: | VIMENTIN |
| 4:00 | 8srq-Y | 5.4 | 5.2 | 86 | 252 | 7 | MOLECULE: | RB1-INDUCIBLE COILED-COIL PROTEIN 1 |
| 5:00 | 1x79-B | 5.4 | 5.1 | 85 | 90 | 8 | MOLECULE: | ADP-RIBOSYLATION FACTOR BINDING PROTEIN GGA1 |
| 6:00 | 4ll7-F | 5.4 | 4.5 | 83 | 91 | 16 | MOLECULE: | SWI5-DEPENDENT HO EXPRESSION PROTEIN 3 |
| 7:00 | 7z8j-X | 5.4 | 5.2 | 87 | 93 | 7 | MOLECULE: | BICD FAMILY-LIKE CARGO ADAPTER 1 |
| 8:00 | 5k9l-D | 5.4 | 5.4 | 83 | 89 | 10 | MOLECULE: | BECLIN-2 |
| 9:00 | 4zku-A | 5.4 | 5.9 | 85 | 179 | 9 | MOLECULE: | TAIL NEEDLE PROTEIN GP26 |
| 10:00 | 2xdj-B | 5.4 | 4 | 67 | 67 | 12 | MOLECULE: | UNCHARACTERIZED PROTEIN YBGF |
| 11:00 | 6tt7-K | 5.4 | 7.8 | 82 | 208 | 5 | MOLECULE: | ATP SYNTHASE SUBUNIT ALPHA |
| 12:00 | 5vpb-D | 5.4 | 3.2 | 66 | 66 | 9 | MOLECULE: | PROTEIN FOSB |
| 13:00 | 3vir-A | 5.4 | 2 | 69 | 69 | 13 | MOLECULE: | MATING-TYPE SWITCHING PROTEIN SWI5 |
| 14:00 | 5tby-A | 5.4 | 11.3 | 99 | 954 | 6 | MOLECULE: | MYOSIN-7 |
| 15:00 | 5m88-A | 5.4 | 5.4 | 81 | 121 | 5 | MOLECULE: | CORE |
| 16:00 | 5lxn-A | 5.4 | 3.9 | 78 | 81 | 10 | MOLECULE: | TRANSFORMING ACIDIC COILED-COIL-CONTAINING PROTEI |
| 17:00 | 1nkn-C | 5.4 | 3.9 | 73 | 75 | 5 | MOLECULE: | S2N51-GCN4 |
| 18:00 | 4i6m-C | 5.4 | 2.5 | 69 | 69 | 6 | MOLECULE: | ACTIN-RELATED PROTEIN 7 |
| 19:00 | 1nkn-A | 5.4 | 3.9 | 73 | 74 | 5 | MOLECULE: | S2N51-GCN4 |
| 20:00 | 5cch-B | 5.4 | 3 | 67 | 67 | 6 | MOLECULE: | VESICLE-ASSOCIATED MEMBRANE PROTEIN 2 |
| 21:00 | 5ccg-B | 5.4 | 3 | 66 | 66 | 5 | MOLECULE: | VESICLE-ASSOCIATED MEMBRANE PROTEIN 2 |
| 22:00 | 5cci-B | 5.4 | 3 | 67 | 67 | 3 | MOLECULE: | VESICLE-ASSOCIATED MEMBRANE PROTEIN 2 |

| | | | | | | | | |
|---|---|---|---|---|---|---|---|---|
| 23:00 | 5lob-D | 5.4 | 5.6 | 84 | 85 | 4 | MOLECULE: | RABPHILIN-3A |
| 24:00:00 | 3tnu-A | 5.4 | 3 | 83 | 90 | 10 | MOLECULE: | KERATIN, TYPE I CYTOSKELETAL 14 |
| 25:00:00 | 1l4a-C | 5.4 | 5.1 | 73 | 73 | 10 | MOLECULE: | SYNAPTOBREVIN |
| 26:00:00 | 6mti-B | 5.4 | 3 | 67 | 67 | 3 | MOLECULE: | SYNAPTOTAGMIN-1 |
| 27:00:00 | 4lhx-E | 5.4 | 5.2 | 76 | 76 | 4 | MOLECULE: | RAS-RELATED PROTEIN RAB-8A |
| 28:00:00 | 6mti-F | 5.4 | 3.3 | 67 | 67 | 4 | MOLECULE: | SYNAPTOTAGMIN-1 |
| 29:00:00 | 6mti-J | 5.4 | 3 | 67 | 67 | 3 | MOLECULE: | SYNAPTOTAGMIN-1 |
| 30:00:00 | 5eof-A | 5.4 | 5.5 | 72 | 74 | 7 | MOLECULE: | OPTINEURIN |
| 31:00:00 | 8e4g-a | 5.4 | 2.2 | 90 | 127 | 10 | MOLECULE: | PORTAL PROTEIN |
| 32:00:00 | 3v5b-A | 5.4 | 3 | 74 | 74 | 8 | MOLECULE: | PRELAMIN-A/C |
| 33:00:00 | 6mti-N | 5.4 | 3 | 67 | 67 | 3 | MOLECULE: | SYNAPTOTAGMIN-1 |
| 34:00:00 | 6mti-R | 5.4 | 3.3 | 67 | 67 | 4 | MOLECULE: | SYNAPTOTAGMIN-1 |
| 35:00:00 | 6mti-V | 5.4 | 3 | 67 | 67 | 3 | MOLECULE: | SYNAPTOTAGMIN-1 |
| 36:00:00 | 4mvd-F | 5.4 | 5.6 | 84 | 253 | 7 | MOLECULE: | CHOLINE-PHOSPHATE CYTIDYLYLTRANSFERASE A |
| 37:00:00 | 4om3-D | 5.3 | 23.6 | 98 | 122 | 6 | MOLECULE: | TRANSDUCIN-LIKE ENHANCER PROTEIN 1 |
| 38:00:00 | 7jts-s | 5.3 | 9.4 | 97 | 290 | 2 | MOLECULE: | RADIAL SPOKE PROTEIN 3 |
| 39:00:00 | 3q8t-B | 5.3 | 4.1 | 87 | 95 | 7 | MOLECULE: | BECLIN-1 |
| 40:00:00 | 5wsu-D | 5.3 | 3.7 | 87 | 91 | 3 | MOLECULE: | CALMODULIN |
| 41:00:00 | 7xuz-A | 5.3 | 2.9 | 84 | 120 | 10 | MOLECULE: | HISTONE DEACETYLASE 4 |
| 42:00:00 | 2doq-D | 5.3 | 3.9 | 81 | 83 | 6 | MOLECULE: | CELL DIVISION CONTROL PROTEIN 31 |
| 43:00:00 | 5wwl-N | 5.3 | 7.7 | 107 | 155 | 6 | MOLECULE: | CENTROMERE PROTEIN MIS12 |
| 44:00:00 | 6wmz-C | 5.3 | 6 | 95 | 292 | 3 | MOLECULE: | SPLICING FACTOR, PROLINE- AND GLUTAMINE-RICH |
| 45:00:00 | 5bw9-g | 5.3 | 5.5 | 83 | 141 | 5 | MOLECULE: | V-TYPE PROTON ATPASE CATALYTIC SUBUNIT A |
| 46:00:00 | 5kiu-A | 5.3 | 6.4 | 75 | 76 | 5 | MOLECULE: | SELENOPROTEIN S |
| 47:00:00 | 8arf-B | 5.3 | 4.1 | 83 | 97 | 10 | MOLECULE: | PROTEIN SPINDLY |
| 48:00:00 | 6iak-F | 5.3 | 4.9 | 71 | 71 | 11 | MOLECULE: | UNCHARACTERIZED PROTEIN |
| 49:00:00 | 8bs7-F | 5.3 | 2.4 | 74 | 74 | 5 | MOLECULE: | PHOSPHOPROTEIN |
| 50:00:00 | 7znl-g | 5.3 | 8.1 | 84 | 161 | 5 | MOLECULE: | THO COMPLEX SUBUNIT 1 |
| 51:00:00 | 7wxz-B | 5.3 | 4.4 | 88 | 185 | 9 | MOLECULE: | SPIKE PROTEIN S2' |
| 52:00:00 | 7znk-g | 5.3 | 8.1 | 84 | 161 | 5 | MOLECULE: | RNA |
| 53:00:00 | 8v01-G | 5.3 | 3.7 | 77 | 82 | 8 | MOLECULE: | ACTIN, ALPHA CARDIAC MUSCLE 1 |
| 54:00:00 | 5toh-B | 5.3 | 4 | 70 | 70 | 7 | MOLECULE: | POLYMERASE COFACTOR VP35 |
| 55:00:00 | 7znl-G | 5.3 | 8.1 | 84 | 161 | 5 | MOLECULE: | THO COMPLEX SUBUNIT 1 |
| 56:00:00 | 7z8j-x | 5.3 | 2.5 | 83 | 93 | 7 | MOLECULE: | BICD FAMILY-LIKE CARGO ADAPTER 1 |
| 57:00:00 | 7znk-G | 5.3 | 8.1 | 84 | 161 | 5 | MOLECULE: | RNA |
| 58:00:00 | 5c9n-A | 5.3 | 2.6 | 64 | 64 | 20 | MOLECULE: | GEMININ COILED-COIL DOMAIN-CONTAINING PROTEIN 1 |
| 59:00:00 | 4ll7-D | 5.3 | 4 | 83 | 91 | 19 | MOLECULE: | SWI5-DEPENDENT HO EXPRESSION PROTEIN 3 |
| 60:00:00 | 7yes-E | 5.3 | 4.9 | 67 | 67 | 9 | MOLECULE: | RNA-DIRECTED RNA POLYMERASE L |
| 61:00:00 | 7yfj-A | 5.3 | 4.1 | 83 | 97 | 11 | MOLECULE: | PRE-MRNA-SPLICING REGULATOR WTAP |
| 62:00:00 | 3cve-C | 5.3 | 3.2 | 66 | 66 | 17 | MOLECULE: | HOMER PROTEIN HOMOLOG 1 |
| 63:00:00 | 8ims-B | 5.3 | 3.5 | 84 | 92 | 6 | MOLECULE: | E3 UBIQUITIN-PROTEIN LIGASE TRAF7 |
| 64:00:00 | 8p0v-K | 5.3 | 6.9 | 101 | 304 | 4 | MOLECULE: | COILED-COIL DOMAIN-CONTAINING PROTEIN 93 |
| 65:00:00 | 6obi-A | 5.3 | 1.8 | 68 | 68 | 3 | MOLECULE: | MYOSIN-VI |
| 66:00:00 | 8arf-A | 5.3 | 4.1 | 83 | 96 | 5 | MOLECULE: | PROTEIN SPINDLY |
| 67:00:00 | 1jth-B | 5.3 | 4.7 | 67 | 67 | 6 | MOLECULE: | SNAP25 |
| 68:00:00 | 7puz-A | 5.3 | 7 | 109 | 148 | 8 | MOLECULE: | MICOS COMPLEX SUBUNIT MIC60 |
| 69:00:00 | 8jsm-E | 5.3 | 5 | 67 | 67 | 10 | MOLECULE: | RNA-DIRECTED RNA POLYMERASE L |
| 70:00:00 | 5l1x-L | 5.3 | 4.8 | 89 | 370 | 4 | MOLECULE: | HMPV F2 SUBUNIT |
| 71:00:00 | 5l1x-B | 5.3 | 3.3 | 86 | 369 | 5 | MOLECULE: | HMPV F2 SUBUNIT |
| 72:00:00 | 8jsn-E | 5.3 | 4.4 | 67 | 67 | 10 | MOLECULE: | RNA-DIRECTED RNA POLYMERASE L |
| 73:00:00 | 5d3a-A | 5.3 | 4.2 | 79 | 82 | 4 | MOLECULE: | KINESIN-LIKE PROTEIN KIF21A |
| 74:00:00 | 3s4r-B | 5.3 | 3.1 | 85 | 91 | 11 | MOLECULE: | VIMENTIN |
| 75:00:00 | 2wpr-C | 5.3 | 5.4 | 85 | 97 | 6 | MOLECULE: | TRIMERIC AUTOTRANSPORTER ADHESIN FRAGMENT |
| 76:00:00 | 3klt-A | 5.3 | 2.1 | 70 | 71 | 7 | MOLECULE: | VIMENTIN |
| 77:00:00 | 2xdj-E | 5.3 | 2.8 | 67 | 67 | 16 | MOLECULE: | UNCHARACTERIZED PROTEIN YBGF |
| 78:00:00 | 2xdj-D | 5.3 | 3.8 | 66 | 66 | 12 | MOLECULE: | UNCHARACTERIZED PROTEIN YBGF |

| | | | | | | | | |
|---|---|---|---|---|---|---|---|---|
| 79:00:00 | 7x5g-E | 5.3 | 4.7 | 80 | 95 | 10 | MOLECULE: | TRANSCRIPTION FACTOR MAFG |
| 80:00:00 | 3vir-B | 5.3 | 1.1 | 66 | 66 | 14 | MOLECULE: | MATING-TYPE SWITCHING PROTEIN SWI5 |
| 81:00:00 | 5oag-D | 5.3 | 4.7 | 87 | 98 | 8 | MOLECULE: | RAB-3A-INTERACTING PROTEIN |
| 82:00:00 | 3h6p-B | 5.3 | 3.2 | 67 | 67 | 7 | MOLECULE: | ESAT-6 LIKE PROTEIN ESXS |
| 83:00:00 | 6ano-B | 5.3 | 3 | 70 | 70 | 13 | MOLECULE: | CASP8-ASSOCIATED PROTEIN 2 |
| 84:00:00 | 6f6p-D | 5.3 | 4.9 | 87 | 98 | 6 | MOLECULE: | RAB-3A-INTERACTING PROTEIN |
| 85:00:00 | 5cj1-C | 5.3 | 3.1 | 80 | 98 | 13 | MOLECULE: | GP7-MYH7-(1526-1571) CHIMERA PROTEIN |
| 86:00:00 | 3b5n-B | 5.3 | 3.8 | 69 | 69 | 3 | MOLECULE: | SYNAPTOBREVIN HOMOLOG 1 |
| 87:00:00 | 2q6q-B | 5.3 | 3 | 65 | 65 | 8 | MOLECULE: | SPINDLE POLE BODY COMPONENT SPC42 |
| 88:00:00 | 3klt-C | 5.3 | 2.4 | 71 | 71 | 7 | MOLECULE: | VIMENTIN |
| 89:00:00 | 4ytd-B | 5.3 | 4.5 | 87 | 99 | 7 | MOLECULE: | PROTEIN BICAUDAL D HOMOLOG 1 |
| 90:00:00 | 4c46-C | 5.3 | 3.1 | 74 | 74 | 8 | MOLECULE: | GENERAL CONTROL PROTEIN GCN4, GENERAL CONTROL PRO |
| 91:00:00 | 5ccg-H | 5.3 | 3.3 | 65 | 65 | 5 | MOLECULE: | VESICLE-ASSOCIATED MEMBRANE PROTEIN 2 |
| 92:00:00 | 3u1a-B | 5.3 | 3.5 | 77 | 84 | 3 | MOLECULE: | SMOOTH MUSCLE TROPOMYOSIN ALPHA |
| 93:00:00 | 7upz-B | 5.3 | 4.1 | 68 | 68 | 9 | MOLECULE: | CCAAT/ENHANCER-BINDING PROTEIN BETA |
| 94:00:00 | 8ims-C | 5.3 | 4.2 | 87 | 92 | 9 | MOLECULE: | E3 UBIQUITIN-PROTEIN LIGASE TRAF7 |
| 95:00:00 | 3u1a-D | 5.3 | 2.8 | 75 | 80 | 3 | MOLECULE: | SMOOTH MUSCLE TROPOMYOSIN ALPHA |
| 96:00:00 | 3vp9-A | 5.3 | 2.2 | 70 | 70 | 11 | MOLECULE: | GENERAL TRANSCRIPTIONAL COREPRESSOR TUP1 |
| 97:00:00 | 7tv4-D | 5.3 | 3.4 | 79 | 79 | 5 | MOLECULE: | NF-KAPPA-B ESSENTIAL MODULATOR |
| 98:00:00 | 4wbn-E | 5.3 | 4.4 | 88 | 120 | 3 | MOLECULE: | TUBULIN ALPHA-1B CHAIN |
| 99:00:00 | 3v4w-A | 5.3 | 3.4 | 74 | 74 | 5 | MOLECULE: | PRELAMIN-A/C |
| 0:00 | 5cch-D | 5.3 | 3.9 | 64 | 64 | 3 | MOLECULE: | VESICLE-ASSOCIATED MEMBRANE PROTEIN 2 |
| 1:00 | 2e43-B | 5.3 | 3.2 | 64 | 64 | 5 | MOLECULE: | CCAAT/ENHANCER-BINDING PROTEIN BETA |
| 2:00 | 4tuy-E | 5.3 | 4.7 | 88 | 120 | 3 | MOLECULE: | TUBULIN ALPHA-1B CHAIN |
| 3:00 | 2yo3-A | 5.3 | 5.7 | 80 | 257 | 6 | MOLECULE: | GENERAL CONTROL PROTEIN GCN4, PUTATIVE INNER MEMB |
| 4:00 | 7vhp-G | 5.2 | 8.1 | 109 | 299 | 4 | MOLECULE: | ATP-DEPENDENT ZINC METALLOPROTEASE FTSH |
| 5:00 | 5hda-A | 5.2 | 1.9 | 71 | 121 | 11 | MOLECULE: | ZINC FINGER MYND DOMAIN-CONTAINING PROTEIN 11 |
| 6:00 | 7yhn-E | 5.2 | 4.5 | 91 | 123 | 8 | MOLECULE: | TUBULIN ALPHA-1B CHAIN |
| 7:00 | 7uw5-A | 5.2 | 7.6 | 84 | 723 | 10 | MOLECULE: | MECHANOSENSITIVE CHANNEL MSCK |
| 8:00 | 3trt-B | 5.2 | 5.4 | 73 | 73 | 10 | MOLECULE: | VIMENTIN |
| 9:00 | 5oag-A | 5.2 | 3.2 | 87 | 98 | 8 | MOLECULE: | RAB-3A-INTERACTING PROTEIN |
| 10:00 | 8r67-E | 5.2 | 3.1 | 84 | 122 | 6 | MOLECULE: | DETYROSINATED TUBULIN ALPHA-1B CHAIN |
| 11:00 | 8b7a-E | 5.2 | 4.5 | 87 | 121 | 7 | MOLECULE: | TUBULIN ALPHA-1B CHAIN |
| 12:00 | 5cci-D | 5.2 | 3.9 | 64 | 64 | 3 | MOLECULE: | VESICLE-ASSOCIATED MEMBRANE PROTEIN 2 |
| 13:00 | 4n78-E | 5.2 | 2.2 | 66 | 67 | 5 | MOLECULE: | CYTOPLASMIC FMR1-INTERACTING PROTEIN 1 |
| 14:00 | 7yg4-B | 5.2 | 3.6 | 67 | 67 | 9 | MOLECULE: | PROTEIN VIRILIZER HOMOLOG |
| 15:00 | 8xwx-D | 5.2 | 2.7 | 65 | 65 | 8 | MOLECULE: | B-CELL RECEPTOR-ASSOCIATED PROTEIN 31 |
| 16:00 | 7znk-O | 5.2 | 7.9 | 90 | 164 | 3 | MOLECULE: | RNA |
| 17:00 | 8j9s-B | 5.2 | 4.1 | 67 | 67 | 9 | MOLECULE: | AMINOACYL TRNA SYNTHASE COMPLEX-INTERACTING MULTI |
| 18:00 | 8j0s-d | 5.2 | 6.4 | 89 | 445 | 9 | MOLECULE: | ATP SYNTHASE SUBUNIT A |
| 19:00 | 7znl-o | 5.2 | 7.9 | 90 | 164 | 3 | MOLECULE: | THO COMPLEX SUBUNIT 1 |
| 20:00 | 8bs7-H | 5.2 | 2.1 | 71 | 71 | 6 | MOLECULE: | PHOSPHOPROTEIN |
| 21:00 | 8rc1-E | 5.2 | 2.3 | 84 | 89 | 0 | MOLECULE: | TUBULIN ALPHA-1B CHAIN |
| 22:00 | 8bs7-E | 5.2 | 1.8 | 71 | 71 | 7 | MOLECULE: | PHOSPHOPROTEIN |
| 23:00 | 9b8q-N | 5.2 | 2.7 | 88 | 106 | 6 | MOLECULE: | V-TYPE PROTON ATPASE SUBUNIT C 1 |
| 24:00:00 | 7xfr-D | 5.2 | 2.2 | 62 | 62 | 11 | MOLECULE: | ISOFORM 2 OF WD REPEAT DOMAIN PHOSPHOINOSITIDE-IN |
| 25:00:00 | 4ll7-A | 5.2 | 3.8 | 80 | 94 | 13 | MOLECULE: | SWI5-DEPENDENT HO EXPRESSION PROTEIN 3 |
| 26:00:00 | 3b5n-F | 5.2 | 4.1 | 68 | 68 | 6 | MOLECULE: | SYNAPTOBREVIN HOMOLOG 1 |
| 27:00:00 | 8ppr-P | 5.2 | 19.7 | 95 | 176 | 7 | MOLECULE: | KINETOCHORE-ASSOCIATED PROTEIN DSN1 HOMOLOG |
| 28:00:00 | 7x5e-A | 5.2 | 2.4 | 77 | 99 | 4 | MOLECULE: | TRANSCRIPTION FACTOR MAFG |
| 29:00:00 | 7x5f-E | 5.2 | 4.2 | 79 | 95 | 9 | MOLECULE: | TRANSCRIPTION FACTOR MAFG |
| 30:00:00 | 7x5f-A | 5.2 | 2.8 | 81 | 100 | 10 | MOLECULE: | TRANSCRIPTION FACTOR MAFG |
| 31:00:00 | 6mti-D | 5.2 | 4 | 64 | 64 | 3 | MOLECULE: | SYNAPTOTAGMIN-1 |
| 32:00:00 | 6mti-H | 5.2 | 4 | 64 | 64 | 3 | MOLECULE: | SYNAPTOTAGMIN-1 |
| 33:00:00 | 6mti-L | 5.2 | 4 | 64 | 64 | 3 | MOLECULE: | SYNAPTOTAGMIN-1 |
| 34:00:00 | 1gtw-B | 5.2 | 2.8 | 65 | 67 | 6 | MOLECULE: | CAAT/ENHANCER BINDING PROTEIN BETA |

| | | | | | | | | |
|---|---|---|---|---|---|---|---|---|
| 35:00:00 | 2wpq-A | 5.2 | 5.2 | 87 | 99 | 6 | MOLECULE: | TRIMERIC AUTOTRANSPORTER ADHESIN FRAGMENT |
| 36:00:00 | 3mq7-C | 5.2 | 3.8 | 87 | 99 | 6 | MOLECULE: | BONE MARROW STROMAL ANTIGEN 2 |
| 37:00:00 | 3mq7-E | 5.2 | 4 | 87 | 99 | 9 | MOLECULE: | BONE MARROW STROMAL ANTIGEN 2 |
| 38:00:00 | 6bl7-B | 5.2 | 4.1 | 87 | 98 | 9 | MOLECULE: | PROTEIN STU2 |
| 39:00:00 | 2wpq-B | 5.2 | 5 | 87 | 99 | 6 | MOLECULE: | TRIMERIC AUTOTRANSPORTER ADHESIN FRAGMENT |
| 40:00:00 | 3mqc-D | 5.2 | 4 | 86 | 100 | 5 | MOLECULE: | BONE MARROW STROMAL ANTIGEN 2 |
| 41:00:00 | 5bu8-A | 5.2 | 5.8 | 83 | 199 | 6 | MOLECULE: | DNA STABILIZATION PROTEIN |
| 42:00:00 | 3mq7-I | 5.2 | 3.2 | 87 | 99 | 7 | MOLECULE: | BONE MARROW STROMAL ANTIGEN 2 |
| 43:00:00 | 5d80-M | 5.2 | 5.8 | 94 | 203 | 6 | MOLECULE: | V-TYPE PROTON ATPASE CATALYTIC SUBUNIT A |
| 44:00:00 | 3mq7-D | 5.2 | 4.9 | 87 | 99 | 9 | MOLECULE: | BONE MARROW STROMAL ANTIGEN 2 |
| 45:00:00 | 3mq7-A | 5.2 | 4.2 | 87 | 99 | 8 | MOLECULE: | BONE MARROW STROMAL ANTIGEN 2 |
| 46:00:00 | 3mq7-G | 5.2 | 3.1 | 87 | 99 | 8 | MOLECULE: | BONE MARROW STROMAL ANTIGEN 2 |
| 47:00:00 | 6mti-P | 5.2 | 3.9 | 64 | 64 | 3 | MOLECULE: | SYNAPTOTAGMIN-1 |
| 48:00:00 | 7x5g-A | 5.2 | 3.5 | 80 | 99 | 10 | MOLECULE: | TRANSCRIPTION FACTOR MAFG |
| 49:00:00 | 2wt7-A | 5.2 | 2.4 | 63 | 63 | 6 | MOLECULE: | PROTO-ONCOGENE PROTEIN C-FOS |
| 50:00:00 | 2ba2-A | 5.2 | 5.6 | 76 | 81 | 8 | MOLECULE: | HYPOTHETICAL UPF0134 PROTEIN MPN010 |
| 51:00:00 | 3ni0-A | 5.2 | 5.1 | 86 | 95 | 10 | MOLECULE: | BONE MARROW STROMAL ANTIGEN 2 |
| 52:00:00 | 2e42-B | 5.2 | 3.1 | 65 | 67 | 6 | MOLECULE: | CCAAT/ENHANCER-BINDING PROTEIN BETA |
| 53:00:00 | 4owf-B | 5.2 | 4.8 | 82 | 86 | 5 | MOLECULE: | NF-KAPPA-B ESSENTIAL MODULATOR |
| 54:00:00 | 7xuz-B | 5.2 | 3.6 | 86 | 120 | 9 | MOLECULE: | HISTONE DEACETYLASE 4 |
| 55:00:00 | 6mti-T | 5.2 | 4 | 64 | 64 | 3 | MOLECULE: | SYNAPTOTAGMIN-1 |
| 56:00:00 | 6mti-X | 5.2 | 4 | 64 | 64 | 3 | MOLECULE: | SYNAPTOTAGMIN-1 |
| 57:00:00 | 4bry-B | 5.2 | 3.8 | 69 | 69 | 6 | MOLECULE: | GEMININ |
| 58:00:00 | 3mq7-K | 5.2 | 3.8 | 86 | 99 | 8 | MOLECULE: | BONE MARROW STROMAL ANTIGEN 2 |
| 59:00:00 | 5cj1-A | 5.2 | 3.6 | 81 | 98 | 14 | MOLECULE: | GP7-MYH7-(1526-1571) CHIMERA PROTEIN |
| 60:00:00 | 4c46-A | 5.2 | 2.9 | 72 | 72 | 7 | MOLECULE: | GENERAL CONTROL PROTEIN GCN4, GENERAL CONTROL PRO |
| 61:00:00 | 6fia-E | 5.2 | 5.8 | 82 | 88 | 4 | MOLECULE: | LINE-1 RETROTRANSPOSABLE ELEMENT ORF1 PROTEIN |
| 62:00:00 | 1sfc-A | 5.2 | 4.7 | 69 | 69 | 10 | MOLECULE: | PROTEIN (SYNAPTOBREVIN 2) |
| 63:00:00 | 4o4h-E | 5.2 | 4.5 | 88 | 123 | 3 | MOLECULE: | TUBULIN ALPHA-1B CHAIN |
| 64:00:00 | 7bjg-A | 5.2 | 5.5 | 87 | 98 | 6 | MOLECULE: | SD21996P |
| 65:00:00 | 6abo-A | 5.2 | 4.9 | 104 | 213 | 12 | MOLECULE: | DNA REPAIR PROTEIN XRCC4 |
| 66:00:00 | 3hd7-E | 5.2 | 5.6 | 83 | 90 | 2 | MOLECULE: | VESICLE-ASSOCIATED MEMBRANE PROTEIN 2 |
| 67:00:00 | 4i4t-E | 5.2 | 4.4 | 88 | 123 | 3 | MOLECULE: | TUBULIN ALPHA-1B CHAIN |
| 68:00:00 | 4o2b-E | 5.2 | 4.5 | 88 | 121 | 3 | MOLECULE: | TUBULIN ALPHA-1B CHAIN |
| 69:00:00 | 1gu5-B | 5.2 | 3.1 | 65 | 67 | 8 | MOLECULE: | CAAT/ENHANCER BINDING PROTEIN BETA |
| 70:00:00 | 4tv9-E | 5.2 | 4.5 | 88 | 122 | 3 | MOLECULE: | TUBULIN ALPHA-1B CHAIN |
| 71:00:00 | 4i50-E | 5.2 | 4.4 | 88 | 123 | 3 | MOLECULE: | TUBULIN ALPHA-1B CHAIN |
| 72:00:00 | 4ihj-E | 5.2 | 4.4 | 88 | 121 | 3 | MOLECULE: | TUBULIN ALPHA-1B CHAIN |
| 73:00:00 | 1gu4-B | 5.2 | 3.3 | 65 | 67 | 8 | MOLECULE: | CAAT/ENHANCER BINDING PROTEIN BETA |
| 74:00:00 | 5c8y-E | 5.2 | 4.6 | 88 | 121 | 3 | MOLECULE: | TUBULIN ALPHA |
| 75:00:00 | 4o4j-E | 5.2 | 4.5 | 88 | 123 | 3 | MOLECULE: | TUBULIN ALPHA-1B CHAIN |
| 76:00:00 | 4tv8-E | 5.2 | 4.5 | 88 | 121 | 3 | MOLECULE: | TUBULIN ALPHA-1B CHAIN |
| 77:00:00 | 5ccg-D | 5.2 | 4.1 | 64 | 64 | 3 | MOLECULE: | VESICLE-ASSOCIATED MEMBRANE PROTEIN 2 |
| 78:00:00 | 4nqj-A | 5.1 | 37.2 | 126 | 177 | 8 | MOLECULE: | E3 UBIQUITIN-PROTEIN LIGASE TRIM69 |
| 79:00:00 | 6fia-D | 5.1 | 23 | 93 | 104 | 2 | MOLECULE: | LINE-1 RETROTRANSPOSABLE ELEMENT ORF1 PROTEIN |
| 80:00:00 | 4bt9-B | 5.1 | 4.3 | 81 | 238 | 7 | MOLECULE: | PROLYL 4-HYDROXYLASE SUBUNIT ALPHA-1 |
| 81:00:00 | 8ppr-F | 5.1 | 8.4 | 78 | 112 | 8 | MOLECULE: | KINETOCHORE-ASSOCIATED PROTEIN DSN1 HOMOLOG |
| 82:00:00 | 5mq4-C | 5.1 | 5.5 | 78 | 117 | 5 | MOLECULE: | PROTEIN KINASE C-BINDING PROTEIN 1 |
| 83:00:00 | 8hpo-D | 5.1 | 6.2 | 92 | 165 | 7 | MOLECULE: | TRANSCRIPTIONAL REGULATORY PROTEIN UME1 |
| 84:00:00 | 8rgq-Z | 5.1 | 13.8 | 109 | 141 | 6 | MOLECULE: | NADH DEHYDROGENASE [UBIQUINONE] IRON-SULFUR PROTE |
| 85:00:00 | 3cve-A | 5.1 | 2.8 | 64 | 68 | 9 | MOLECULE: | HOMER PROTEIN HOMOLOG 1 |
| 86:00:00 | 4aj5-Y | 5.1 | 3.2 | 78 | 99 | 8 | MOLECULE: | SPINDLE AND KINETOCHORE-ASSOCIATED PROTEIN 1 |
| 87:00:00 | 4aj5-1 | 5.1 | 3.2 | 78 | 99 | 8 | MOLECULE: | SPINDLE AND KINETOCHORE-ASSOCIATED PROTEIN 1 |
| 88:00:00 | 7x5d-A | 5.1 | 4 | 87 | 99 | 7 | MOLECULE: | LAMIN-A/C |
| 89:00:00 | 3cve-D | 5.1 | 1.8 | 61 | 61 | 16 | MOLECULE: | HOMER PROTEIN HOMOLOG 1 |
| 90:00:00 | 3rk2-D | 5.1 | 4.1 | 65 | 65 | 3 | MOLECULE: | VESICLE-ASSOCIATED MEMBRANE PROTEIN 2 |

| | | | | | | | | |
|---|---|---|---|---|---|---|---|---|
| 91:00:00 | 7tv4-B | 5.1 | 4 | 79 | 81 | 8 | MOLECULE: | NF-KAPPA-B ESSENTIAL MODULATOR |
| 92:00:00 | 7saf-A | 5.1 | 2 | 62 | 62 | 6 | MOLECULE: | GENERAL CONTROL TRANSCRIPTION FACTOR GCN4/M PROTE |
| 93:00:00 | 9b8q-O | 5.1 | 4 | 84 | 103 | 5 | MOLECULE: | V-TYPE PROTON ATPASE SUBUNIT C 1 |
| 94:00:00 | 7yet-E | 5.1 | 5.3 | 67 | 67 | 9 | MOLECULE: | RNA-DIRECTED RNA POLYMERASE L |
| 95:00:00 | 8q85-W | 5.1 | 5.8 | 81 | 106 | 5 | MOLECULE: | KINETOCHORE PROTEIN NDC80 |
| 96:00:00 | 8k8d-A | 5.1 | 2.1 | 66 | 68 | 12 | MOLECULE: | CCAAT/ENHANCER-BINDING PROTEIN BETA |
| 97:00:00 | 7znl-O | 5.1 | 6.9 | 86 | 164 | 9 | MOLECULE: | THO COMPLEX SUBUNIT 1 |
| 98:00:00 | 8bs7-G | 5.1 | 1.6 | 68 | 68 | 6 | MOLECULE: | PHOSPHOPROTEIN |
| 99:00:00 | 8ard-A | 5.1 | 1.7 | 62 | 62 | 13 | MOLECULE: | UNCONVENTIONAL MYOSIN-VI |
| 0:00 | 8h9g-K | 5.1 | 5.7 | 96 | 198 | 2 | MOLECULE: | ATP SYNTHASE F(0) COMPLEX SUBUNIT B1, MITOCHONDRI |
| 1:00 | 8bs7-D | 5.1 | 2.3 | 78 | 80 | 5 | MOLECULE: | PHOSPHOPROTEIN |
| 2:00 | 7x5d-B | 5.1 | 4.2 | 83 | 94 | 8 | MOLECULE: | LAMIN-A/C |
| 3:00 | 4ll7-E | 5.1 | 4.3 | 82 | 90 | 18 | MOLECULE: | SWI5-DEPENDENT HO EXPRESSION PROTEIN 3 |
| 4:00 | 1n7s-A | 5.1 | 4.4 | 63 | 63 | 13 | MOLECULE: | VESICLE-ASSOCIATED MEMBRANE PROTEIN 2 |
| 5:00 | 1urq-B | 5.1 | 2.7 | 63 | 63 | 5 | MOLECULE: | M-TOMOSYN ISOFORM |
| 6:00 | 4e61-B | 5.1 | 5.6 | 78 | 84 | 8 | MOLECULE: | PROTEIN BIM1 |
| 7:00 | 4ll7-H | 5.1 | 3.8 | 81 | 91 | 10 | MOLECULE: | SWI5-DEPENDENT HO EXPRESSION PROTEIN 3 |
| 8:00 | 8t1l-D | 5.1 | 8 | 98 | 161 | 8 | MOLECULE: | MEDIATOR OF RNA POLYMERASE II TRANSCRIPTION SUBUN |
| 9:00 | 8b7c-E | 5.1 | 5.5 | 90 | 123 | 3 | MOLECULE: | TUBULIN ALPHA-1B CHAIN |
| 10:00 | 1hjb-D | 5.1 | 3.2 | 68 | 68 | 12 | MOLECULE: | CCAAT/ENHANCER BINDING PROTEIN BETA |
| 11:00 | 3rk2-H | 5.1 | 4.1 | 65 | 65 | 3 | MOLECULE: | VESICLE-ASSOCIATED MEMBRANE PROTEIN 2 |
| 12:00 | 7c4j-H | 5.1 | 6.2 | 93 | 242 | 3 | MOLECULE: | TRANSCRIPTION REGULATORY PROTEIN SNF12 |
| 13:00 | 1h8a-B | 5.1 | 2.8 | 67 | 67 | 7 | MOLECULE: | CAAT/ENHANCER BINDING PROTEIN BETA |
| 14:00 | 3mq7-L | 5.1 | 4.8 | 87 | 99 | 9 | MOLECULE: | BONE MARROW STROMAL ANTIGEN 2 |
| 15:00 | 3pp5-A | 5.1 | 3.4 | 63 | 63 | 13 | MOLECULE: | BRK1 |
| 16:00 | 6vol-I | 5.1 | 3 | 82 | 150 | 6 | MOLECULE: | ATP SYNTHASE SUBUNIT ALPHA, CHLOROPLASTIC |
| 17:00 | 3hd7-A | 5.1 | 5.6 | 83 | 91 | 2 | MOLECULE: | VESICLE-ASSOCIATED MEMBRANE PROTEIN 2 |
| 18:00 | 3mqc-A | 5.1 | 3.3 | 87 | 100 | 9 | MOLECULE: | BONE MARROW STROMAL ANTIGEN 2 |
| 19:00 | 3mqc-C | 5.1 | 3.5 | 84 | 100 | 10 | MOLECULE: | BONE MARROW STROMAL ANTIGEN 2 |
| 20:00 | 3mq7-H | 5.1 | 4.8 | 87 | 99 | 9 | MOLECULE: | BONE MARROW STROMAL ANTIGEN 2 |
| 21:00 | 3s9g-A | 5.1 | 4.8 | 78 | 82 | 18 | MOLECULE: | PROTEIN HEXIM1 |
| 22:00 | 2xdj-C | 5.1 | 3.1 | 64 | 67 | 14 | MOLECULE: | UNCHARACTERIZED PROTEIN YBGF |
| 23:00 | 3tnu-B | 5.1 | 4.9 | 86 | 95 | 9 | MOLECULE: | KERATIN, TYPE I CYTOSKELETAL 14 |
| 24:00:00 | 3u1c-A | 5.1 | 6.6 | 88 | 101 | 10 | MOLECULE: | TROPOMYOSIN ALPHA-1 CHAIN |
| 25:00:00 | 3b5n-A | 5.1 | 3.9 | 61 | 61 | 5 | MOLECULE: | SYNAPTOBREVIN HOMOLOG 1 |
| 26:00:00 | 5wme-D | 5.1 | 2.3 | 75 | 89 | 5 | MOLECULE: | CAPSID ASSEMBLY SCAFFOLDING PROTEIN,MYOSIN-7 |
| 27:00:00 | 3mqc-B | 5.1 | 4.4 | 86 | 100 | 5 | MOLECULE: | BONE MARROW STROMAL ANTIGEN 2 |
| 28:00:00 | 1l2p-A | 5.1 | 1.2 | 61 | 61 | 7 | MOLECULE: | ATP SYNTHASE B CHAIN |
| 29:00:00 | 2nps-B | 5.1 | 2.8 | 68 | 68 | 4 | MOLECULE: | VESICLE-ASSOCIATED MEMBRANE PROTEIN 4 |
| 30:00:00 | 7x5e-E | 5.1 | 3.4 | 80 | 101 | 10 | MOLECULE: | TRANSCRIPTION FACTOR MAFG |
| 31:00:00 | 5toi-B | 5.1 | 4.7 | 70 | 70 | 6 | MOLECULE: | POLYMERASE COFACTOR VP35 |
| 32:00:00 | 4ytd-A | 5.1 | 6.4 | 88 | 96 | 9 | MOLECULE: | PROTEIN BICAUDAL D HOMOLOG 1 |
| 33:00:00 | 2ba2-B | 5.1 | 4.5 | 74 | 80 | 7 | MOLECULE: | HYPOTHETICAL UPF0134 PROTEIN MPN010 |
| 34:00:00 | 5dol-A | 5.1 | 2 | 61 | 61 | 7 | MOLECULE: | INITIATION-CONTROL PROTEIN YABA |
| 35:00:00 | 2ba2-C | 5.1 | 4.8 | 74 | 81 | 7 | MOLECULE: | HYPOTHETICAL UPF0134 PROTEIN MPN010 |
| 36:00:00 | 5ca0-E | 5.1 | 4.7 | 88 | 121 | 3 | MOLECULE: | TUBULIN ALPHA-1B CHAIN |
| 37:00:00 | 4lhx-F | 5.1 | 6.1 | 75 | 76 | 4 | MOLECULE: | RAS-RELATED PROTEIN RAB-8A |
| 38:00:00 | 4i55-E | 5.1 | 4.4 | 88 | 124 | 3 | MOLECULE: | TUBULIN ALPHA-1B CHAIN |
| 39:00:00 | 4o4l-E | 5.1 | 4.4 | 88 | 123 | 3 | MOLECULE: | TUBULIN ALPHA-1B CHAIN |
| 40:00:00 | 4o4i-E | 5.1 | 4.4 | 88 | 121 | 3 | MOLECULE: | TUBULIN ALPHA-1B CHAIN |
| 41:00:00 | 5cb4-E | 5.1 | 4.7 | 88 | 121 | 3 | MOLECULE: | TUBULIN ALPHA |
| 42:00:00 | 4x20-E | 5.1 | 4.6 | 88 | 123 | 3 | MOLECULE: | TUBULIN ALPHA CHAIN |
| 43:00:00 | 4o2a-E | 5.1 | 5 | 91 | 124 | 3 | MOLECULE: | TUBULIN ALPHA-1B CHAIN |
| 44:00:00 | 1kil-D | 5.1 | 4.5 | 65 | 66 | 3 | MOLECULE: | SYNAPTOBREVIN SNARE MOTIF |
| 45:00:00 | 1hjb-B | 5.1 | 2.1 | 65 | 67 | 12 | MOLECULE: | CCAAT/ENHANCER BINDING PROTEIN BETA |
| 46:00:00 | 6uci-B | 5.1 | 1.5 | 61 | 61 | 7 | MOLECULE: | PROTEIN FOSB |

| | | | | | | | | |
|---|---|---|---|---|---|---|---|---|
| 47:00:00 | 5ca1-E | 5.1 | 4.6 | 88 | 121 | 3 | MOLECULE: | TUBULIN ALPHA |
| 48:00:00 | 3i00-B | 5.1 | 2.4 | 74 | 80 | 7 | MOLECULE: | HUNTINGTIN-INTERACTING PROTEIN 1 |
| 49:00:00 | 1n7s-D | 5.1 | 4.8 | 66 | 66 | 3 | MOLECULE: | VESICLE-ASSOCIATED MEMBRANE PROTEIN 2 |
| 50:00:00 | 2qa7-B | 5.1 | 2.7 | 69 | 70 | 9 | MOLECULE: | HUNTINGTIN-INTERACTING PROTEIN 1 |
| 51:00:00 | 4x1i-E | 5.1 | 4.6 | 88 | 123 | 3 | MOLECULE: | TUBULIN ALPHA CHAIN |
| 52:00:00 | 3b5n-H | 5.1 | 3.7 | 63 | 63 | 3 | MOLECULE: | SYNAPTOBREVIN HOMOLOG 1 |
| 53:00:00 | 2qa7-D | 5.1 | 2.8 | 69 | 70 | 9 | MOLECULE: | HUNTINGTIN-INTERACTING PROTEIN 1 |
| 54:00:00 | 4x1y-E | 5.1 | 4.5 | 87 | 123 | 3 | MOLECULE: | TUBULIN ALPHA CHAIN |
| 55:00:00 | 1h88-A | 5.1 | 2.2 | 67 | 70 | 9 | MOLECULE: | CCAAT/ENHANCER BINDING PROTEIN BETA |
| 56:00:00 | 1ebo-B | 5.1 | 4.3 | 92 | 112 | 5 | MOLECULE: | EBOLA VIRUS ENVELOPE PROTEIN CHIMERA CONSISTING |
| 57:00:00 | 7znk-e | 5.1 | 2.4 | 65 | 527 | 8 | MOLECULE: | RNA |
| 58:00:00 | 7znl-e | 5.1 | 2.4 | 65 | 527 | 8 | MOLECULE: | THO COMPLEX SUBUNIT 1 |
| 59:00:00 | 7znk-E | 5.1 | 2.4 | 65 | 538 | 8 | MOLECULE: | RNA |
| 60:00:00 | 3opc-B | 5 | 2.8 | 87 | 131 | 9 | MOLECULE: | UNCHARACTERIZED PROTEIN |
| 61:00:00 | 7woo-G | 5 | 13.3 | 89 | 200 | 12 | MOLECULE: | NUCLEOPORIN NIC96 |
| 62:00:00 | 4he8-A | 5 | 25.4 | 84 | 92 | 10 | MOLECULE: | NADH-QUINONE OXIDOREDUCTASE SUBUNIT 7 |
| 63:00:00 | 7sqk-C | 5 | 31.1 | 122 | 600 | 11 | MOLECULE: | HAUS AUGMIN-LIKE COMPLEX SUBUNIT 1 |
| 64:00:00 | 5oxf-A | 5 | 7.3 | 115 | 703 | 5 | MOLECULE: | GTP-BINDING PROTEIN |
| 65:00:00 | 6ian-C | 5 | 26.7 | 93 | 432 | 9 | MOLECULE: | INTRAFLAGELLAR TRANSPORT PROTEIN 74 |
| 66:00:00 | 4aj5-2 | 5 | 4.4 | 78 | 100 | 5 | MOLECULE: | SPINDLE AND KINETOCHORE-ASSOCIATED PROTEIN 1 |
| 67:00:00 | 3n7n-B | 5 | 3.7 | 85 | 164 | 7 | MOLECULE: | MONOPOLIN COMPLEX SUBUNIT CSM1 |
| 68:00:00 | 3b5n-D | 5 | 4.1 | 64 | 64 | 3 | MOLECULE: | SYNAPTOBREVIN HOMOLOG 1 |
| 69:00:00 | 4aj5-4 | 5 | 3.3 | 77 | 98 | 8 | MOLECULE: | SPINDLE AND KINETOCHORE-ASSOCIATED PROTEIN 1 |
| 70:00:00 | 4aj5-V | 5 | 3.4 | 77 | 98 | 8 | MOLECULE: | SPINDLE AND KINETOCHORE-ASSOCIATED PROTEIN 1 |
| 71:00:00 | 4aj5-X | 5 | 3.8 | 78 | 98 | 8 | MOLECULE: | SPINDLE AND KINETOCHORE-ASSOCIATED PROTEIN 1 |
| 72:00:00 | 3m0d-A | 5 | 3.2 | 63 | 63 | 5 | MOLECULE: | TNF RECEPTOR-ASSOCIATED FACTOR 2 |
| 73:00:00 | 6z26-B | 5 | 4.4 | 87 | 103 | 8 | MOLECULE: | SPINDLE ASSEMBLY ABNORMAL PROTEIN 6 HOMOLOG |
| 74:00:00 | 6zpm-A | 5 | 3.2 | 85 | 103 | 5 | MOLECULE: | TRYPANOSOMA CRUZI KKT4 117-218 |
| 75:00:00 | 7y5b-b | 5 | 10.8 | 104 | 145 | 3 | MOLECULE: | ATP SYNTHASE SUBUNIT ALPHA |
| 76:00:00 | 4m3l-A | 5 | 1.5 | 60 | 60 | 8 | MOLECULE: | E3 UBIQUITIN-PROTEIN LIGASE TRIM63 |
| 77:00:00 | 8a5a-G | 5 | 7.1 | 91 | 110 | 7 | MOLECULE: | CHROMATIN-REMODELING ATPASE INO80 |
| 78:00:00 | 3zdo-F | 5 | 3.1 | 65 | 65 | 9 | MOLECULE: | PHOSPHOPROTEIN |
| 79:00:00 | 7wjt-B | 5 | 1.5 | 60 | 60 | 8 | MOLECULE: | ISOFORM 2 OF TRANSMEMBRANE PROTEIN 266 |
| 80:00:00 | 3rl0-X | 5 | 3.8 | 63 | 63 | 2 | MOLECULE: | VESICLE-ASSOCIATED MEMBRANE PROTEIN 2 |
| 81:00:00 | 7w2u-W | 5 | 13.3 | 105 | 142 | 10 | MOLECULE: | NADH DEHYDROGENASE [UBIQUINONE] FLAVOPROTEIN 1, |
| 82:00:00 | 3rl0-L | 5 | 3.9 | 64 | 64 | 2 | MOLECULE: | VESICLE-ASSOCIATED MEMBRANE PROTEIN 2 |
| 83:00:00 | 7yer-E | 5 | 5.4 | 67 | 67 | 9 | MOLECULE: | RNA-DIRECTED RNA POLYMERASE L |
| 84:00:00 | 1qbz-C | 5 | 3.1 | 70 | 120 | 7 | MOLECULE: | PROTEIN (SIV GP41 ECTODOMAIN) |
| 85:00:00 | 8g0c-b | 5 | 6.2 | 94 | 141 | 3 | MOLECULE: | ATP SYNTHASE SUBUNIT ALPHA |
| 86:00:00 | 8ap7-L | 5 | 2 | 65 | 65 | 5 | MOLECULE: | ATP SYNTHASE SUBUNIT A |
| 87:00:00 | 1h89-A | 5 | 1.9 | 64 | 64 | 5 | MOLECULE: | CAAT/ENHANCER BINDING PROTEIN BETA |
| 88:00:00 | 1gtw-A | 5 | 2.9 | 65 | 65 | 8 | MOLECULE: | CAAT/ENHANCER BINDING PROTEIN BETA |
| 89:00:00 | 2yo3-C | 5 | 4.1 | 75 | 257 | 5 | MOLECULE: | GENERAL CONTROL PROTEIN GCN4, PUTATIVE INNER MEMB |
| 90:00:00 | 8k8d-B | 5 | 2.7 | 66 | 66 | 11 | MOLECULE: | CCAAT/ENHANCER-BINDING PROTEIN BETA |
| 91:00:00 | 8bs7-C | 5 | 2.7 | 77 | 81 | 8 | MOLECULE: | PHOSPHOPROTEIN |
| 92:00:00 | 8at4-E | 5 | 23.9 | 106 | 217 | 7 | MOLECULE: | HAUS AUGMIN-LIKE COMPLEX SUBUNIT 1 |
| 93:00:00 | 8bd7-M | 5 | 8.1 | 95 | 164 | 6 | MOLECULE: | IFT88 |
| 94:00:00 | 6zpj-A | 5 | 7.1 | 90 | 110 | 11 | MOLECULE: | LEISHMANIA MEXICANA KKT4 |
| 95:00:00 | 6r0z-I | 5 | 2.2 | 77 | 103 | 5 | MOLECULE: | V-TYPE ATP SYNTHASE ALPHA CHAIN |
| 96:00:00 | 8bs7-A | 5 | 1.7 | 68 | 68 | 6 | MOLECULE: | PHOSPHOPROTEIN |
| 97:00:00 | 8afh-B | 5 | 4 | 84 | 97 | 6 | MOLECULE: | CRESCENTIN |
| 98:00:00 | 1hvv-B | 5 | 2.2 | 62 | 62 | 3 | MOLECULE: | SYNTAXIN 1A |
| 99:00:00 | 8afe-B | 5 | 4 | 84 | 97 | 6 | MOLECULE: | CRESCENTIN |
| 0:00 | 1gd2-F | 5 | 4.3 | 64 | 64 | 9 | MOLECULE: | DNA (5'- |
| 1:00 | 8afh-L | 5 | 4 | 84 | 97 | 6 | MOLECULE: | CRESCENTIN |
| 2:00 | 4q9u-D | 5 | 3.5 | 74 | 75 | 11 | MOLECULE: | RAB5 GDP/GTP EXCHANGE FACTOR |

| | | | | | | | | |
|---|---|---|---|---|---|---|---|---|
| 3:00 | 6yjd-A | 5 | 3.7 | 83 | 104 | 8 | MOLECULE: | CAPSID ASSEMBLY SCAFFOLDING PROTEIN,PRELAMIN-A/C |
| 4:00 | 4wpx-B | 5 | 9.7 | 78 | 81 | 5 | MOLECULE: | CELL DIVISION CONTROL PROTEIN 31-LIKE PROTEIN |
| 5:00 | 2e43-A | 5 | 2.2 | 62 | 62 | 8 | MOLECULE: | CCAAT/ENHANCER-BINDING PROTEIN BETA |
| 6:00 | 7f3t-A | 5 | 4.6 | 85 | 330 | 11 | MOLECULE: | TRANSMEMBRANE PROTEIN 120A |
| 7:00 | 5l1x-F | 5 | 5.1 | 88 | 369 | 3 | MOLECULE: | HMPV F2 SUBUNIT |
| 8:00 | 2no2-A | 5 | 4.5 | 87 | 102 | 10 | MOLECULE: | HUNTINGTIN-INTERACTING PROTEIN 1 |
| 9:00 | 4wpx-E | 5 | 9.8 | 78 | 81 | 5 | MOLECULE: | CELL DIVISION CONTROL PROTEIN 31-LIKE PROTEIN |
| 10:00 | 5ed9-B | 5 | 4.4 | 72 | 72 | 10 | MOLECULE: | SUN DOMAIN-CONTAINING PROTEIN 2 |
| 11:00 | 1hjb-E | 5 | 4.7 | 66 | 68 | 9 | MOLECULE: | CCAAT/ENHANCER BINDING PROTEIN BETA |
| 12:00 | 5j9t-K | 5 | 23.9 | 123 | 269 | 11 | MOLECULE: | HISTONE ACETYLTRANSFERASE ESA1 |
| 13:00 | 4n21-C | 5 | 3.8 | 91 | 124 | 10 | MOLECULE: | GP2 ECTODOMAIN |
| 14:00 | 5j9w-G | 5 | 11.7 | 101 | 275 | 10 | MOLECULE: | HISTONE ACETYLTRANSFERASE ESA1 |
| 15:00 | 2xdj-A | 5 | 4.3 | 66 | 66 | 14 | MOLECULE: | UNCHARACTERIZED PROTEIN YBGF |
| 16:00 | 6ys4-F | 5 | 3.5 | 87 | 101 | 6 | MOLECULE: | SPINDLE ASSEMBLY ABNORMAL PROTEIN 6 HOMOLOG |
| 17:00 | 5vpf-C | 5 | 2.3 | 64 | 64 | 6 | MOLECULE: | PROTEIN FOSB |
| 18:00 | 6a9p-E | 5 | 6.1 | 87 | 102 | 11 | MOLECULE: | GLIAL FIBRILLARY ACIDIC PROTEIN |
| 19:00 | 2xu6-A | 5 | 2.7 | 63 | 63 | 8 | MOLECULE: | MDV1 COILED COIL |
| 20:00 | 5wsu-C | 5 | 2.4 | 82 | 90 | 4 | MOLECULE: | CALMODULIN |
| 21:00 | 3ryc-E | 5 | 5.2 | 91 | 136 | 3 | MOLECULE: | TUBULIN ALPHA CHAIN |
| 22:00 | 3h6p-A | 5 | 2.7 | 60 | 60 | 5 | MOLECULE: | ESAT-6 LIKE PROTEIN ESXS |
| 23:00 | 4jzl-A | 5 | 1.8 | 62 | 62 | 5 | MOLECULE: | B-CELL RECEPTOR-ASSOCIATED PROTEIN 31 |
| 24:00:00 | 5wme-B | 5 | 2.1 | 74 | 92 | 8 | MOLECULE: | CAPSID ASSEMBLY SCAFFOLDING PROTEIN,MYOSIN-7 |
| 25:00:00 | 1t3j-A | 5 | 1.9 | 62 | 62 | 13 | MOLECULE: | MITOFUSIN 1 |
| 26:00:00 | 3lt6-D | 5 | 2.9 | 62 | 62 | 13 | MOLECULE: | ADHESIN YADA |
| 27:00:00 | 1gu4-A | 5 | 2.8 | 65 | 65 | 14 | MOLECULE: | CAAT/ENHANCER BINDING PROTEIN BETA |
| 28:00:00 | 5d60-D | 5 | 3 | 61 | 61 | 13 | MOLECULE: | PUTATIVE TRANSCRIPTION FACTOR |
| 29:00:00 | 3mq7-B | 5 | 4.7 | 87 | 99 | 9 | MOLECULE: | BONE MARROW STROMAL ANTIGEN 2 |
| 30:00:00 | 2wt7-B | 5 | 4 | 75 | 90 | 7 | MOLECULE: | PROTO-ONCOGENE PROTEIN C-FOS |
| 31:00:00 | 2xu6-B | 5 | 2.7 | 64 | 64 | 8 | MOLECULE: | MDV1 COILED COIL |
| 32:00:00 | 2q6q-A | 5 | 2.7 | 62 | 62 | 11 | MOLECULE: | SPINDLE POLE BODY COMPONENT SPC42 |
| 33:00:00 | 2wps-A | 5 | 5.2 | 87 | 98 | 3 | MOLECULE: | TRIMERIC AUTOTRANSPORTER ADHESIN FRAGMENT |
| 34:00:00 | 6znl-N | 5 | 5.9 | 84 | 280 | 11 | MOLECULE: | ARP1 ACTIN RELATED PROTEIN 1 HOMOLOG A |
| 35:00:00 | 2qa7-C | 5 | 5 | 87 | 102 | 7 | MOLECULE: | HUNTINGTIN-INTERACTING PROTEIN 1 |
| 36:00:00 | 3u59-D | 5 | 5.1 | 89 | 98 | 4 | MOLECULE: | TROPOMYOSIN BETA CHAIN |
| 37:00:00 | 4n21-F | 5 | 3.9 | 92 | 124 | 11 | MOLECULE: | GP2 ECTODOMAIN |
| 38:00:00 | 3u1a-A | 5 | 3.3 | 76 | 82 | 4 | MOLECULE: | SMOOTH MUSCLE TROPOMYOSIN ALPHA |
| 39:00:00 | 4lhx-D | 5 | 4.6 | 71 | 71 | 4 | MOLECULE: | RAS-RELATED PROTEIN RAB-8A |
| 40:00:00 | 1sfc-I | 5 | 4.9 | 69 | 69 | 13 | MOLECULE: | PROTEIN (SYNAPTOBREVIN 2) |
| 41:00:00 | 4li0-D | 5 | 3.9 | 69 | 69 | 10 | MOLECULE: | RAS-RELATED PROTEIN RAB-8A |
| 42:00:00 | 4jzl-D | 5 | 2 | 61 | 61 | 5 | MOLECULE: | B-CELL RECEPTOR-ASSOCIATED PROTEIN 31 |
| 43:00:00 | 4iij-E | 5 | 4.6 | 88 | 122 | 3 | MOLECULE: | TUBULIN ALPHA-1B CHAIN |
| 44:00:00 | 4n21-E | 5 | 3.9 | 91 | 121 | 9 | MOLECULE: | GP2 ECTODOMAIN |
| 45:00:00 | 4jzl-B | 5 | 2 | 61 | 61 | 5 | MOLECULE: | B-CELL RECEPTOR-ASSOCIATED PROTEIN 31 |
| 46:00:00 | 2qa7-A | 5 | 4.6 | 87 | 99 | 7 | MOLECULE: | HUNTINGTIN-INTERACTING PROTEIN 1 |
| 47:00:00 | 5ccg-J | 5 | 3.7 | 63 | 63 | 3 | MOLECULE: | VESICLE-ASSOCIATED MEMBRANE PROTEIN 2 |
| 48:00:00 | 6yf5-A | 5 | 5.9 | 84 | 89 | 6 | MOLECULE: | PRELAMIN-A/C,MICROTUBULE-ASSOCIATED PROTEIN RP/EB |
| 49:00:00 | 3rl0-P | 5 | 3.8 | 64 | 64 | 2 | MOLECULE: | VESICLE-ASSOCIATED MEMBRANE PROTEIN 2 |
| 50:00:00 | 2e42-A | 5 | 2.6 | 65 | 65 | 8 | MOLECULE: | CCAAT/ENHANCER-BINDING PROTEIN BETA |
| 51:00:00 | 1ebo-C | 5 | 5.5 | 92 | 111 | 9 | MOLECULE: | EBOLA VIRUS ENVELOPE PROTEIN CHIMERA CONSISTING |
| 52:00:00 | 4g2k-B | 5 | 5.2 | 86 | 108 | 12 | MOLECULE: | GENERAL CONTROL PROTEIN GCN4, ENVELOPE GLYCOPROTE |
| 53:00:00 | 4aj5-T | 4.9 | 27.1 | 101 | 110 | 6 | MOLECULE: | SPINDLE AND KINETOCHORE-ASSOCIATED PROTEIN 1 |
| 54:00:00 | 8wjo-B | 4.9 | 42.1 | 84 | 283 | 8 | MOLECULE: | STRUCTURAL MAINTENANCE OF CHROMOSOMES PROTEIN 5 |
| 55:00:00 | 8tek-P | 4.9 | 3.1 | 78 | 138 | 5 | MOLECULE: | DYNEIN REGULATORY COMPLEX PROTEIN 1/2 N-TERMINAL |
| 56:00:00 | 8idc-E | 4.9 | 4.3 | 95 | 300 | 9 | MOLECULE: | CELL DIVISION ATP-BINDING PROTEIN FTSE |
| 57:00:00 | 1m1j-D | 4.9 | 10.8 | 112 | 194 | 5 | MOLECULE: | FIBRINOGEN ALPHA SUBUNIT |
| 58:00:00 | 4r3z-A | 4.9 | 3.6 | 73 | 76 | 10 | MOLECULE: | AMINOACYL TRNA SYNTHASE COMPLEX-INTERACTING MULTI |

| | | | | | | | | |
|---|---|---|---|---|---|---|---|---|
| 59:00:00 | 2p2u-B | 4.9 | 5.7 | 75 | 153 | 8 | MOLECULE: | HOST-NUCLEASE INHIBITOR PROTEIN GAM, PUTATIVE |
| 60:00:00 | 4hl8-A | 4.9 | 6.6 | 101 | 782 | 3 | MOLECULE: | MAJOR VAULT PROTEIN |
| 61:00:00 | 5wlz-C | 4.9 | 7.8 | 100 | 206 | 7 | MOLECULE: | DNA REPAIR PROTEIN XRCC4,MYOSIN-7 |
| 62:00:00 | 1s1c-Y | 4.9 | 3.3 | 70 | 70 | 4 | MOLECULE: | TRANSFORMING PROTEIN RHOA |
| 63:00:00 | 4aj5-3 | 4.9 | 3.5 | 76 | 98 | 8 | MOLECULE: | SPINDLE AND KINETOCHORE-ASSOCIATED PROTEIN 1 |
| 64:00:00 | 4emc-B | 4.9 | 4 | 75 | 158 | 7 | MOLECULE: | MONOPOLIN COMPLEX SUBUNIT CSM1 |
| 65:00:00 | 5d60-B | 4.9 | 2.7 | 61 | 61 | 16 | MOLECULE: | PUTATIVE TRANSCRIPTION FACTOR |
| 66:00:00 | 4h22-B | 4.9 | 3.2 | 75 | 83 | 4 | MOLECULE: | LEUCINE-RICH REPEAT FLIGHTLESS-INTERACTING PROTEI |
| 67:00:00 | 5ajs-B | 4.9 | 2.5 | 64 | 64 | 6 | MOLECULE: | THAP DOMAIN-CONTAINING PROTEIN 11 |
| 68:00:00 | 8q85-b | 4.9 | 5.3 | 69 | 70 | 12 | MOLECULE: | KINETOCHORE PROTEIN NDC80 |
| 69:00:00 | 4aj5-B | 4.9 | 11.3 | 80 | 87 | 14 | MOLECULE: | SPINDLE AND KINETOCHORE-ASSOCIATED PROTEIN 1 |
| 70:00:00 | 1wt6-D | 4.9 | 3.3 | 63 | 63 | 5 | MOLECULE: | MYOTONIN-PROTEIN KINASE |
| 71:00:00 | 3b5n-L | 4.9 | 3.5 | 62 | 62 | 3 | MOLECULE: | SYNAPTOBREVIN HOMOLOG 1 |
| 72:00:00 | 7yes-D | 4.9 | 2.7 | 68 | 68 | 13 | MOLECULE: | RNA-DIRECTED RNA POLYMERASE L |
| 73:00:00 | 7upz-A | 4.9 | 2.3 | 64 | 64 | 9 | MOLECULE: | CCAAT/ENHANCER-BINDING PROTEIN BETA |
| 74:00:00 | 8pr4-x | 4.9 | 2 | 62 | 92 | 8 | MOLECULE: | ARP11 |
| 75:00:00 | 8xwx-F | 4.9 | 2.5 | 62 | 66 | 6 | MOLECULE: | B-CELL RECEPTOR-ASSOCIATED PROTEIN 31 |
| 76:00:00 | 7usd-C | 4.9 | 5.1 | 74 | 191 | 8 | MOLECULE: | CYTOPLASMIC FMR1-INTERACTING PROTEIN 1 |
| 77:00:00 | 7xho-K | 4.9 | 15.7 | 105 | 230 | 7 | MOLECULE: | CENTROMERE PROTEIN C |
| 78:00:00 | 8uwy-G | 4.9 | 4.6 | 73 | 82 | 5 | MOLECULE: | ACTIN, ALPHA CARDIAC MUSCLE 1 |
| 79:00:00 | 8p1u-C | 4.9 | 5.6 | 80 | 93 | 10 | MOLECULE: | CELL DIVISION PROTEIN FTSL |
| 80:00:00 | 8ap7-l | 4.9 | 1.8 | 65 | 65 | 6 | MOLECULE: | ATP SYNTHASE SUBUNIT A |
| 81:00:00 | 7ucd-F | 4.9 | 1.9 | 64 | 64 | 8 | MOLECULE: | PROTEIN FOSB |
| 82:00:00 | 8jsm-C | 4.9 | 5.3 | 70 | 99 | 7 | MOLECULE: | RNA-DIRECTED RNA POLYMERASE L |
| 83:00:00 | 8pr2-B | 4.9 | 2.8 | 87 | 106 | 8 | MOLECULE: | C-JUN-AMINO-TERMINAL KINASE-INTERACTING PROTEIN 3 |
| 84:00:00 | 8q85-F | 4.9 | 7.5 | 95 | 230 | 8 | MOLECULE: | KINETOCHORE PROTEIN NDC80 |
| 85:00:00 | 3hrn-A | 4.9 | 2.3 | 63 | 63 | 8 | MOLECULE: | TRANSIENT RECEPTOR POTENTIAL (TRP) CHANNEL |
| 86:00:00 | 6gbo-A | 4.9 | 6.2 | 72 | 73 | 7 | MOLECULE: | POLYMERASE COFACTOR VP35 |
| 87:00:00 | 3jb9-U | 4.9 | 3.8 | 89 | 430 | 9 | MOLECULE: | PRE-MRNA-SPLICING FACTOR SPP42 |
| 88:00:00 | 1c1g-D | 4.9 | 11.8 | 105 | 284 | 6 | MOLECULE: | TROPOMYOSIN |
| 89:00:00 | 7xhn-U | 4.9 | 5.7 | 83 | 140 | 4 | MOLECULE: | CENTROMERE PROTEIN O |
| 90:00:00 | 1hvv-D | 4.9 | 3.6 | 61 | 61 | 3 | MOLECULE: | SYNTAXIN 1A |
| 91:00:00 | 3b5n-J | 4.9 | 3.3 | 64 | 64 | 6 | MOLECULE: | SYNAPTOBREVIN HOMOLOG 1 |
| 92:00:00 | 8bd7-X | 4.9 | 7.4 | 87 | 106 | 2 | MOLECULE: | IFT88 |
| 93:00:00 | 7usc-D | 4.9 | 1.8 | 60 | 60 | 7 | MOLECULE: | CYTOPLASMIC FMR1-INTERACTING PROTEIN 1 |
| 94:00:00 | 7usd-D | 4.9 | 2.3 | 62 | 62 | 5 | MOLECULE: | CYTOPLASMIC FMR1-INTERACTING PROTEIN 1 |
| 95:00:00 | 4q9u-H | 4.9 | 3.9 | 77 | 82 | 10 | MOLECULE: | RAB5 GDP/GTP EXCHANGE FACTOR |
| 96:00:00 | 4gkw-A | 4.9 | 3.4 | 84 | 158 | 6 | MOLECULE: | SPINDLE ASSEMBLY ABNORMAL PROTEIN 6 |
| 97:00:00 | 5ylz-t | 4.9 | 7 | 82 | 128 | 4 | MOLECULE: | PRE-MRNA-SPLICING FACTOR 8 |
| 98:00:00 | 5omd-A | 4.9 | 1.8 | 61 | 61 | 7 | MOLECULE: | DNA DAMAGE CHECKPOINT PROTEIN LCD1 |
| 99:00:00 | 3rl0-f | 4.9 | 3.6 | 62 | 62 | 0 | MOLECULE: | VESICLE-ASSOCIATED MEMBRANE PROTEIN 2 |
| 0:00 | 3m0d-B | 4.9 | 3.4 | 63 | 63 | 5 | MOLECULE: | TNF RECEPTOR-ASSOCIATED FACTOR 2 |
| 1:00 | 2b9c-B | 4.9 | 8.6 | 94 | 142 | 6 | MOLECULE: | STRIATED-MUSCLE ALPHA TROPOMYOSIN |
| 2:00 | 6dd9-D | 4.9 | 3.3 | 87 | 121 | 9 | MOLECULE: | SYNAPTONEMAL COMPLEX PROTEIN 3 |
| 3:00 | 5c3l-A | 4.9 | 8.6 | 93 | 134 | 8 | MOLECULE: | NUP54 |
| 4:00 | 3zdo-C | 4.9 | 3.1 | 63 | 63 | 10 | MOLECULE: | PHOSPHOPROTEIN |
| 5:00 | 6v8o-J | 4.9 | 7.2 | 97 | 115 | 5 | MOLECULE: | HIGH TEMPERATURE LETHAL PROTEIN 1 |
| 6:00 | 3u59-B | 4.9 | 5.1 | 91 | 101 | 3 | MOLECULE: | TROPOMYOSIN BETA CHAIN |
| 7:00 | 5w5d-C | 4.9 | 3.8 | 61 | 61 | 7 | MOLECULE: | VESICLE-ASSOCIATED MEMBRANE PROTEIN 2 |
| 8:00 | 1sa0-E | 4.9 | 3.8 | 85 | 124 | 4 | MOLECULE: | TUBULIN ALPHA CHAIN |
| 9:00 | 2wz7-A | 4.9 | 4.4 | 66 | 66 | 14 | MOLECULE: | UNCHARACTERIZED PROTEIN YBGF |
| 10:00 | 2wpr-A | 4.9 | 5.6 | 87 | 97 | 7 | MOLECULE: | TRIMERIC AUTOTRANSPORTER ADHESIN FRAGMENT |
| 11:00 | 4b2q-t | 4.9 | 4.6 | 83 | 129 | 11 | MOLECULE: | ATP SYNTHASE SUBUNIT ALPHA, MITOCHONDRIAL |
| 12:00 | 4zdw-B | 4.9 | 5.9 | 84 | 91 | 7 | MOLECULE: | RAS-RELATED PROTEIN SEC4 |
| 13:00 | 3vir-D | 4.9 | 1.2 | 63 | 63 | 8 | MOLECULE: | MATING-TYPE SWITCHING PROTEIN SWI5 |
| 14:00 | 2d3e-D | 4.9 | 4 | 86 | 130 | 3 | MOLECULE: | GENERAL CONTROL PROTEIN GCN4 AND TROPOMYOSIN 1 AL |

| | | | | | | | | |
|---|---|---|---|---|---|---|---|---|
| 15:00 | 3p8c-E | 4.9 | 3.5 | 64 | 64 | 6 | MOLECULE: | CYTOPLASMIC FMR1-INTERACTING PROTEIN 1 |
| 16:00 | 3thf-B | 4.9 | 8 | 139 | 174 | 11 | MOLECULE: | PROTEIN SHROOM |
| 17:00 | 1sfc-D | 4.9 | 4.7 | 68 | 74 | 4 | MOLECULE: | PROTEIN (SYNAPTOBREVIN 2) |
| 18:00 | 2kz9-A | 4.9 | 5.7 | 69 | 69 | 6 | MOLECULE: | V-TYPE PROTON ATPASE SUBUNIT E |
| 19:00 | 3a7o-E | 4.9 | 1.9 | 60 | 60 | 8 | MOLECULE: | AUTOPHAGY PROTEIN 16 |
| 20:00 | 3lt7-B | 4.9 | 2.9 | 60 | 60 | 12 | MOLECULE: | ADHESIN YADA |
| 21:00 | 2xdj-F | 4.9 | 2.7 | 61 | 61 | 8 | MOLECULE: | UNCHARACTERIZED PROTEIN YBGF |
| 22:00 | 8aia-B | 4.9 | 3.7 | 82 | 85 | 5 | MOLECULE: | CRESCENTIN |
| 23:00 | 3j97-L | 4.9 | 3.8 | 66 | 66 | 5 | MOLECULE: | VESICLE-FUSING ATPASE |
| 24:00:00 | 5wlq-A | 4.9 | 4.4 | 82 | 104 | 10 | MOLECULE: | CAPSID ASSEMBLY SCAFFOLDING PROTEIN,MYOSIN-7,MICR |
| 25:00:00 | 6r0w-K | 4.9 | 5.1 | 85 | 98 | 4 | MOLECULE: | V-TYPE ATP SYNTHASE ALPHA CHAIN |
| 26:00:00 | 7yes-C | 4.9 | 4.8 | 70 | 99 | 11 | MOLECULE: | RNA-DIRECTED RNA POLYMERASE L |
| 27:00:00 | 3j96-L | 4.9 | 3.7 | 66 | 66 | 5 | MOLECULE: | VESICLE-FUSING ATPASE |
| 28:00:00 | 3mqb-E | 4.9 | 5.5 | 91 | 106 | 3 | MOLECULE: | BONE MARROW STROMAL ANTIGEN 2 |
| 29:00:00 | 3hd7-B | 4.9 | 4.5 | 87 | 98 | 1 | MOLECULE: | VESICLE-ASSOCIATED MEMBRANE PROTEIN 2 |
| 30:00:00 | 5ccg-G | 4.9 | 4.1 | 62 | 63 | 15 | MOLECULE: | VESICLE-ASSOCIATED MEMBRANE PROTEIN 2 |
| 31:00:00 | 5apv-B | 4.9 | 2.9 | 67 | 67 | 10 | MOLECULE: | GENERAL CONTROL PROTEIN GCN4 |
| 32:00:00 | 3mqb-B | 4.9 | 6.2 | 91 | 106 | 4 | MOLECULE: | BONE MARROW STROMAL ANTIGEN 2 |
| 33:00:00 | 3fx0-B | 4.9 | 3.7 | 71 | 71 | 8 | MOLECULE: | NF-KAPPA-B ESSENTIAL MODULATOR |
| 34:00:00 | 4qkv-A | 4.9 | 6.2 | 80 | 101 | 6 | MOLECULE: | POLYMERASE I AND TRANSCRIPT RELEASE FACTOR |
| 35:00:00 | 4n21-D | 4.9 | 3.8 | 91 | 123 | 9 | MOLECULE: | GP2 ECTODOMAIN |
| 36:00:00 | 3ipd-A | 4.9 | 5.6 | 85 | 91 | 4 | MOLECULE: | VESICLE-ASSOCIATED MEMBRANE PROTEIN 2 |
| 37:00:00 | 5cch-A | 4.9 | 3.9 | 63 | 63 | 14 | MOLECULE: | VESICLE-ASSOCIATED MEMBRANE PROTEIN 2 |
| 38:00:00 | 4li0-F | 4.9 | 3.6 | 69 | 69 | 4 | MOLECULE: | RAS-RELATED PROTEIN RAB-8A |
| 39:00:00 | 5ccg-A | 4.9 | 4 | 63 | 63 | 14 | MOLECULE: | VESICLE-ASSOCIATED MEMBRANE PROTEIN 2 |
| 40:00:00 | 5cci-A | 4.9 | 3.9 | 62 | 62 | 15 | MOLECULE: | VESICLE-ASSOCIATED MEMBRANE PROTEIN 2 |
| 41:00:00 | 4w7z-A | 4.9 | 3 | 64 | 64 | 9 | MOLECULE: | B-CELL RECEPTOR-ASSOCIATED PROTEIN 29 |
| 42:00:00 | 7xho-U | 4.9 | 7 | 83 | 140 | 6 | MOLECULE: | CENTROMERE PROTEIN C |
| 43:00:00 | 4w7z-C | 4.9 | 1.6 | 59 | 59 | 10 | MOLECULE: | B-CELL RECEPTOR-ASSOCIATED PROTEIN 29 |
| 44:00:00 | 4w7z-D | 4.9 | 1.7 | 60 | 60 | 10 | MOLECULE: | B-CELL RECEPTOR-ASSOCIATED PROTEIN 29 |
| 45:00:00 | 4w7z-B | 4.9 | 3 | 64 | 64 | 9 | MOLECULE: | B-CELL RECEPTOR-ASSOCIATED PROTEIN 29 |
| 46:00:00 | 3rl0-T | 4.9 | 3.7 | 62 | 62 | 0 | MOLECULE: | VESICLE-ASSOCIATED MEMBRANE PROTEIN 2 |
| 47:00:00 | 4w80-A | 4.9 | 2.9 | 64 | 64 | 9 | MOLECULE: | B-CELL RECEPTOR-ASSOCIATED PROTEIN 29 |
| 48:00:00 | 1ebo-A | 4.9 | 5.4 | 92 | 112 | 5 | MOLECULE: | EBOLA VIRUS ENVELOPE PROTEIN CHIMERA CONSISTING |
| 49:00:00 | 8fed-E | 4.8 | 17.6 | 97 | 358 | 5 | MOLECULE: | VIRULENCE FACTOR MCE FAMILY PROTEIN |
| 50:00:00 | 6ian-A | 4.8 | 9.8 | 79 | 306 | 9 | MOLECULE: | INTRAFLAGELLAR TRANSPORT PROTEIN 74 |
| 51:00:00 | 4orh-H | 4.8 | 4 | 89 | 142 | 8 | MOLECULE: | UBIQUITIN-CONJUGATING ENZYME E2 VARIANT 2 |
| 52:00:00 | 8g0q-B | 4.8 | 12.1 | 103 | 230 | 8 | MOLECULE: | KINETOCHORE PROTEIN NDC80 |
| 53:00:00 | 5opt-P | 4.8 | 6.8 | 89 | 249 | 8 | MOLECULE: | ACTIVATED PROTEIN KINASE C RECEPTOR, PUTATIVE |
| 54:00:00 | 2ocy-A | 4.8 | 8.8 | 83 | 149 | 12 | MOLECULE: | RAB GUANINE NUCLEOTIDE EXCHANGE FACTOR SEC2 |
| 55:00:00 | 6ff7-G | 4.8 | 6.9 | 92 | 132 | 5 | MOLECULE: | RNA-BINDING MOTIF PROTEIN, X-LINKED 2 |
| 56:00:00 | 4aj5-Z | 4.8 | 3.3 | 76 | 97 | 7 | MOLECULE: | SPINDLE AND KINETOCHORE-ASSOCIATED PROTEIN 1 |
| 57:00:00 | 8amr-A | 4.8 | 5 | 97 | 155 | 4 | MOLECULE: | TRIPARTITE MOTIF-CONTAINING PROTEIN 3 |
| 58:00:00 | 7syg-H | 4.8 | 3.8 | 71 | 237 | 4 | MOLECULE: | 18S RRNA |
| 59:00:00 | 5jvp-B | 4.8 | 5.1 | 84 | 90 | 5 | MOLECULE: | CHIMERA PROTEIN OF CENTROMERE-ASSOCIATED PROTEIN |
| 60:00:00 | 6ys4-A | 4.8 | 4.4 | 87 | 103 | 11 | MOLECULE: | SPINDLE ASSEMBLY ABNORMAL PROTEIN 6 HOMOLOG |
| 61:00:00 | 7bjs-A | 4.8 | 2.8 | 64 | 69 | 5 | MOLECULE: | KINESIN HEAVY CHAIN |
| 62:00:00 | 2v4h-A | 4.8 | 5.9 | 84 | 98 | 5 | MOLECULE: | NF-KAPPA-B ESSENTIAL MODULATOR |
| 63:00:00 | 6cfz-B | 4.8 | 5.3 | 65 | 65 | 9 | MOLECULE: | ASK1 |
| 64:00:00 | 1qbz-B | 4.8 | 2.7 | 63 | 114 | 6 | MOLECULE: | PROTEIN (SIV GP41 ECTODOMAIN) |
| 65:00:00 | 8h9v-K | 4.8 | 5.9 | 90 | 198 | 3 | MOLECULE: | ATP SYNTHASE F(0) COMPLEX SUBUNIT C1, MITOCHONDRI |
| 66:00:00 | 8akr-Q | 4.8 | 4.7 | 79 | 196 | 9 | MOLECULE: | CHLOROPLAST MEMBRANE-ASSOCIATED 30 KD PROTEIN |
| 67:00:00 | 3brt-D | 4.8 | 4 | 62 | 62 | 6 | MOLECULE: | INHIBITOR OF NUCLEAR FACTOR KAPPA-B KINASE SUBUNI |
| 68:00:00 | 3gn4-A | 4.8 | 4.1 | 78 | 129 | 8 | MOLECULE: | MYOSIN-VI |
| 69:00:00 | 8bh1-D | 4.8 | 7.2 | 71 | 85 | 8 | MOLECULE: | PROBABLE PEPTIDOGLYCAN GLYCOSYLTRANSFERASE FTSW |
| 70:00:00 | 1qbz-A | 4.8 | 3.1 | 62 | 102 | 5 | MOLECULE: | PROTEIN (SIV GP41 ECTODOMAIN) |

| | | | | | | | | |
|---|---|---|---|---|---|---|---|---|
| 71:00:00 | 7wjt-C | 4.8 | 1.7 | 59 | 59 | 7 | MOLECULE: | ISOFORM 2 OF TRANSMEMBRANE PROTEIN 266 |
| 72:00:00 | 5mq0-t | 4.8 | 8.8 | 89 | 438 | 9 | MOLECULE: | YEAST UBC4 GENE FOR UBIQUITIN-CONJUGATING ENZYME |
| 73:00:00 | 7yer-D | 4.8 | 2.8 | 68 | 68 | 13 | MOLECULE: | RNA-DIRECTED RNA POLYMERASE L |
| 74:00:00 | 7y8r-Q | 4.8 | 10 | 76 | 91 | 11 | MOLECULE: | HISTONE H3 |
| 75:00:00 | 8fby-A | 4.8 | 3.5 | 75 | 89 | 4 | MOLECULE: | PLASMALEMMA VESICLE-ASSOCIATED PROTEIN |
| 76:00:00 | 8q85-V | 4.8 | 3.9 | 69 | 117 | 4 | MOLECULE: | KINETOCHORE PROTEIN NDC80 |
| 77:00:00 | 6cfz-A | 4.8 | 5.7 | 66 | 66 | 8 | MOLECULE: | ASK1 |
| 78:00:00 | 1gmj-C | 4.8 | 1.3 | 59 | 59 | 2 | MOLECULE: | ATPASE INHIBITOR |
| 79:00:00 | 7syo-H | 4.8 | 6.1 | 73 | 237 | 4 | MOLECULE: | 18S RRNA |
| 80:00:00 | 7naf-5 | 4.8 | 15.6 | 84 | 105 | 5 | MOLECULE: | 25S RRNA |
| 81:00:00 | 6lth-Q | 4.8 | 21.7 | 92 | 105 | 5 | MOLECULE: | TRANSCRIPTION ACTIVATOR BRG1 |
| 82:00:00 | 8sdj-A | 4.8 | 4.3 | 87 | 112 | 7 | MOLECULE: | PYRIN |
| 83:00:00 | 8sdj-D | 4.8 | 4.3 | 88 | 112 | 7 | MOLECULE: | PYRIN |
| 84:00:00 | 6h9m-A | 4.8 | 4.4 | 81 | 94 | 11 | MOLECULE: | COILED-COIL DOMAIN-CONTAINING PROTEIN 90B, MITOCH |
| 85:00:00 | 5xau-C | 4.8 | 4.7 | 70 | 70 | 4 | MOLECULE: | LAMININ SUBUNIT ALPHA-5 |
| 86:00:00 | 7yet-C | 4.8 | 4.4 | 68 | 99 | 13 | MOLECULE: | RNA-DIRECTED RNA POLYMERASE L |
| 87:00:00 | 8sdj-C | 4.8 | 4.3 | 88 | 112 | 7 | MOLECULE: | PYRIN |
| 88:00:00 | 5c9n-B | 4.8 | 3.6 | 61 | 61 | 20 | MOLECULE: | GEMININ COILED-COIL DOMAIN-CONTAINING PROTEIN 1 |
| 89:00:00 | 1io4-A | 4.8 | 2.4 | 63 | 63 | 11 | MOLECULE: | CSF-1R PROMOTER |
| 90:00:00 | 7use-D | 4.8 | 1.9 | 61 | 62 | 5 | MOLECULE: | CYTOPLASMIC FMR1-INTERACTING PROTEIN 1 |
| 91:00:00 | 8sdj-B | 4.8 | 4.3 | 87 | 112 | 7 | MOLECULE: | PYRIN |
| 92:00:00 | 8uww-G | 4.8 | 3.9 | 76 | 82 | 7 | MOLECULE: | ACTIN, ALPHA CARDIAC MUSCLE 1 |
| 93:00:00 | 8pr2-C | 4.8 | 5.7 | 87 | 106 | 8 | MOLECULE: | C-JUN-AMINO-TERMINAL KINASE-INTERACTING PROTEIN 3 |
| 94:00:00 | 7xr0-E | 4.8 | 5.8 | 90 | 123 | 4 | MOLECULE: | TUBULIN ALPHA-1B CHAIN |
| 95:00:00 | 7xqy-E | 4.8 | 5.8 | 90 | 123 | 3 | MOLECULE: | TUBULIN ALPHA-1B CHAIN |
| 96:00:00 | 3b5n-I | 4.8 | 3.2 | 58 | 58 | 2 | MOLECULE: | SYNAPTOBREVIN HOMOLOG 1 |
| 97:00:00 | 3rl0-H | 4.8 | 3.6 | 61 | 61 | 0 | MOLECULE: | VESICLE-ASSOCIATED MEMBRANE PROTEIN 2 |
| 98:00:00 | 2xzr-A | 4.8 | 4.7 | 87 | 108 | 6 | MOLECULE: | IMMUNOGLOBULIN-BINDING PROTEIN EIBD |
| 99:00:00 | 3uf1-B | 4.8 | 4.6 | 89 | 106 | 8 | MOLECULE: | VIMENTIN |
| 0:00 | 3mq7-J | 4.8 | 6.5 | 87 | 99 | 5 | MOLECULE: | BONE MARROW STROMAL ANTIGEN 2 |
| 1:00 | 6fia-C | 4.8 | 6.2 | 80 | 89 | 4 | MOLECULE: | LINE-1 RETROTRANSPOSABLE ELEMENT ORF1 PROTEIN |
| 2:00 | 3q0x-A | 4.8 | 10.2 | 83 | 206 | 6 | MOLECULE: | CENTRIOLE PROTEIN |
| 3:00 | 5y60-K | 4.8 | 3.6 | 87 | 99 | 7 | MOLECULE: | V-TYPE ATP SYNTHASE ALPHA CHAIN |
| 4:00 | 2d3e-C | 4.8 | 5.1 | 81 | 130 | 10 | MOLECULE: | GENERAL CONTROL PROTEIN GCN4 AND TROPOMYOSIN 1 AL |
| 5:00 | 3hd7-F | 4.8 | 4.2 | 83 | 99 | 1 | MOLECULE: | VESICLE-ASSOCIATED MEMBRANE PROTEIN 2 |
| 6:00 | 4eot-A | 4.8 | 2.5 | 73 | 92 | 12 | MOLECULE: | TRANSCRIPTION FACTOR MAFA |
| 7:00 | 5toi-A | 4.8 | 5.2 | 68 | 68 | 7 | MOLECULE: | POLYMERASE COFACTOR VP35 |
| 8:00 | 6f63-A | 4.8 | 5.7 | 85 | 96 | 7 | MOLECULE: | SYNAPTONEMAL COMPLEX PROTEIN 1 |
| 9:00 | 3m0a-A | 4.8 | 3 | 62 | 62 | 5 | MOLECULE: | TNF RECEPTOR-ASSOCIATED FACTOR 2 |
| 10:00 | 4bry-A | 4.8 | 3.1 | 68 | 69 | 6 | MOLECULE: | GEMININ |
| 11:00 | 3zdo-D | 4.8 | 3 | 64 | 64 | 11 | MOLECULE: | PHOSPHOPROTEIN |
| 12:00 | 3zdo-B | 4.8 | 2.7 | 62 | 62 | 13 | MOLECULE: | PHOSPHOPROTEIN |
| 13:00 | 3lt7-E | 4.8 | 2.9 | 61 | 61 | 8 | MOLECULE: | ADHESIN YADA |
| 14:00 | 2q2f-A | 4.8 | 5.5 | 71 | 72 | 7 | MOLECULE: | SELENOPROTEIN S |
| 15:00 | 5gmk-q | 4.8 | 7.4 | 88 | 387 | 8 | MOLECULE: | PRE-MRNA-SPLICING FACTOR 8 |
| 16:00 | 6zqm-b | 4.8 | 6.6 | 103 | 209 | 7 | MOLECULE: | ATP SYNTHASE SUBUNIT ALPHA, MITOCHONDRIAL |
| 17:00 | 3a7o-A | 4.8 | 2.1 | 59 | 59 | 12 | MOLECULE: | AUTOPHAGY PROTEIN 16 |
| 18:00 | 8b8b-D | 4.8 | 3.7 | 85 | 97 | 11 | MOLECULE: | MUNIA BORNAVIRUS 1 PHOSPHOPROTEIN |
| 19:00 | 3rk2-B | 4.8 | 2.6 | 59 | 59 | 3 | MOLECULE: | VESICLE-ASSOCIATED MEMBRANE PROTEIN 2 |
| 20:00 | 8aix-F | 4.8 | 4.7 | 80 | 85 | 5 | MOLECULE: | CRESCENTIN |
| 21:00 | 3zdo-A | 4.8 | 3.2 | 63 | 63 | 13 | MOLECULE: | PHOSPHOPROTEIN |
| 22:00 | 2n1t-B | 4.8 | 5 | 71 | 72 | 4 | MOLECULE: | VESICLE-ASSOCIATED MEMBRANE PROTEIN 2 |
| 23:00 | 4aj5-A | 4.8 | 11 | 73 | 79 | 10 | MOLECULE: | SPINDLE AND KINETOCHORE-ASSOCIATED PROTEIN 1 |
| 24:00:00 | 3rl0-F | 4.8 | 3 | 61 | 61 | 5 | MOLECULE: | VESICLE-ASSOCIATED MEMBRANE PROTEIN 2 |
| 25:00:00 | 3rl0-R | 4.8 | 3 | 61 | 61 | 5 | MOLECULE: | VESICLE-ASSOCIATED MEMBRANE PROTEIN 2 |
| 26:00:00 | 8aix-B | 4.8 | 4.7 | 80 | 85 | 5 | MOLECULE: | CRESCENTIN |

| | | | | | | | | |
|---|---|---|---|---|---|---|---|---|
| 27:00:00 | 2nps-A | 4.8 | 3.3 | 63 | 63 | 5 | MOLECULE: | VESICLE-ASSOCIATED MEMBRANE PROTEIN 4 |
| 28:00:00 | 3rk2-F | 4.8 | 2.6 | 59 | 59 | 3 | MOLECULE: | VESICLE-ASSOCIATED MEMBRANE PROTEIN 2 |
| 29:00:00 | 1kil-A | 4.8 | 5 | 65 | 65 | 14 | MOLECULE: | SYNAPTOBREVIN SNARE MOTIF |
| 30:00:00 | 2n1t-A | 4.8 | 4.8 | 69 | 69 | 13 | MOLECULE: | VESICLE-ASSOCIATED MEMBRANE PROTEIN 2 |
| 31:00:00 | 3v6i-A | 4.8 | 5 | 104 | 186 | 6 | MOLECULE: | V-TYPE ATP SYNTHASE SUBUNIT E |
| 32:00:00 | 1h8a-A | 4.8 | 2.7 | 65 | 68 | 9 | MOLECULE: | CAAT/ENHANCER BINDING PROTEIN BETA |
| 33:00:00 | 4li0-E | 4.8 | 3.5 | 64 | 64 | 5 | MOLECULE: | RAS-RELATED PROTEIN RAB-8A |
| 34:00:00 | 3u1a-C | 4.8 | 3.9 | 76 | 84 | 4 | MOLECULE: | SMOOTH MUSCLE TROPOMYOSIN ALPHA |
| 35:00:00 | 3i00-A | 4.8 | 3.4 | 73 | 76 | 5 | MOLECULE: | HUNTINGTIN-INTERACTING PROTEIN 1 |
| 36:00:00 | 2zxx-A | 4.8 | 7.9 | 69 | 70 | 6 | MOLECULE: | GEMININ |
| 37:00:00 | 2v66-D | 4.8 | 6.3 | 91 | 111 | 10 | MOLECULE: | NUCLEAR DISTRIBUTION PROTEIN NUDE-LIKE 1 |
| 38:00:00 | 4w80-B | 4.8 | 3 | 64 | 64 | 9 | MOLECULE: | B-CELL RECEPTOR-ASSOCIATED PROTEIN 29 |
| 39:00:00 | 4w80-D | 4.8 | 1.6 | 58 | 58 | 10 | MOLECULE: | B-CELL RECEPTOR-ASSOCIATED PROTEIN 29 |
| 40:00:00 | 2v66-C | 4.8 | 6 | 93 | 111 | 10 | MOLECULE: | NUCLEAR DISTRIBUTION PROTEIN NUDE-LIKE 1 |
| 41:00:00 | 1sfc-L | 4.8 | 4.4 | 67 | 73 | 3 | MOLECULE: | PROTEIN (SYNAPTOBREVIN 2) |
| 42:00:00 | 3rl0-D | 4.8 | 3.7 | 62 | 62 | 0 | MOLECULE: | VESICLE-ASSOCIATED MEMBRANE PROTEIN 2 |
| 43:00:00 | 3rl0-b | 4.8 | 3.6 | 61 | 61 | 0 | MOLECULE: | VESICLE-ASSOCIATED MEMBRANE PROTEIN 2 |
| 44:00:00 | 3zdo-G | 4.8 | 2.8 | 64 | 64 | 5 | MOLECULE: | PHOSPHOPROTEIN |
| 45:00:00 | 1urq-D | 4.8 | 4.1 | 62 | 62 | 0 | MOLECULE: | M-TOMOSYN ISOFORM |
| 46:00:00 | 3rk3-D | 4.8 | 4.3 | 65 | 65 | 3 | MOLECULE: | VAMP2 |
| 47:00:00 | 1h89-B | 4.8 | 4.9 | 64 | 64 | 6 | MOLECULE: | CAAT/ENHANCER BINDING PROTEIN BETA |
| 48:00:00 | 5jst-A | 4.8 | 6.2 | 86 | 428 | 7 | MOLECULE: | MALTOSE-BINDING PERIPLASMIC PROTEIN,MITOCHONDRIAL |
| 49:00:00 | 8xwx-G | 4.8 | 2.2 | 57 | 57 | 9 | MOLECULE: | B-CELL RECEPTOR-ASSOCIATED PROTEIN 31 |
| 50:00:00 | 4w80-C | 4.8 | 1.7 | 59 | 59 | 10 | MOLECULE: | B-CELL RECEPTOR-ASSOCIATED PROTEIN 29 |
| 51:00:00 | 7apk-E | 4.8 | 4.7 | 61 | 523 | 11 | MOLECULE: | THO COMPLEX SUBUNIT 1 |
| 52:00:00 | 1joc-A | 4.7 | 3.3 | 75 | 123 | 9 | MOLECULE: | EARLY ENDOSOMAL AUTOANTIGEN 1 |
| 53:00:00 | 1aa0-A | 4.7 | 4 | 74 | 113 | 8 | MOLECULE: | FIBRITIN |
| 54:00:00 | 7eeb-C | 4.7 | 7.6 | 104 | 278 | 6 | MOLECULE: | ENHANCED GREEN FLUORESCENT PROTEIN,CATION CHANNEL |
| 55:00:00 | 6kn7-T | 4.7 | 15.3 | 67 | 138 | 9 | MOLECULE: | ACTIN, ALPHA SKELETAL MUSCLE |
| 56:00:00 | 8at3-A | 4.7 | 8.2 | 133 | 286 | 8 | MOLECULE: | HAUS AUGMIN-LIKE COMPLEX SUBUNIT 1 |
| 57:00:00 | 4aj5-W | 4.7 | 4.8 | 85 | 98 | 8 | MOLECULE: | SPINDLE AND KINETOCHORE-ASSOCIATED PROTEIN 1 |
| 58:00:00 | 6v92-R | 4.7 | 4.5 | 79 | 325 | 11 | MOLECULE: | ACTIN-RELATED PROTEIN 7 |
| 59:00:00 | 4emc-A | 4.7 | 3.6 | 70 | 151 | 6 | MOLECULE: | MONOPOLIN COMPLEX SUBUNIT CSM1 |
| 60:00:00 | 8bd7-W | 4.7 | 6.3 | 86 | 114 | 8 | MOLECULE: | IFT88 |
| 61:00:00 | 7eaa-C | 4.7 | 6 | 91 | 114 | 12 | MOLECULE: | RB1-INDUCIBLE COILED-COIL PROTEIN 1 |
| 62:00:00 | 3uf1-D | 4.7 | 4.6 | 90 | 112 | 7 | MOLECULE: | VIMENTIN |
| 63:00:00 | 3b5n-C | 4.7 | 3.2 | 65 | 70 | 9 | MOLECULE: | SYNAPTOBREVIN HOMOLOG 1 |
| 64:00:00 | 7s5u-B | 4.7 | 7.8 | 96 | 180 | 6 | MOLECULE: | KINESIN-LIKE PROTEIN KLP61F |
| 65:00:00 | 5hmo-A | 4.7 | 8 | 88 | 131 | 9 | MOLECULE: | UNCONVENTIONAL MYOSIN-X |
| 66:00:00 | 6xor-B | 4.7 | 5.9 | 71 | 71 | 10 | MOLECULE: | PROTEIN SWALLOW |
| 67:00:00 | 4m3l-B | 4.7 | 1.9 | 58 | 58 | 14 | MOLECULE: | E3 UBIQUITIN-PROTEIN LIGASE TRIM63 |
| 68:00:00 | 2w6a-B | 4.7 | 1.8 | 56 | 56 | 5 | MOLECULE: | ARF GTPASE-ACTIVATING PROTEIN GIT1 |
| 69:00:00 | 8b4i-D | 4.7 | 3.7 | 60 | 60 | 5 | MOLECULE: | MITOCHONDRIAL IMPORT RECEPTOR SUBUNIT TOM40 |
| 70:00:00 | 7z47-F | 4.7 | 5.9 | 67 | 68 | 9 | MOLECULE: | ADAPTOR PROTEIN |
| 71:00:00 | 6xor-A | 4.7 | 6.2 | 71 | 71 | 8 | MOLECULE: | PROTEIN SWALLOW |
| 72:00:00 | 4i1l-A | 4.7 | 2.5 | 62 | 62 | 5 | MOLECULE: | FORKHEAD BOX PROTEIN P3 |
| 73:00:00 | 8jsm-D | 4.7 | 4.2 | 68 | 68 | 12 | MOLECULE: | RNA-DIRECTED RNA POLYMERASE L |
| 74:00:00 | 5kc1-J | 4.7 | 1.6 | 58 | 58 | 3 | MOLECULE: | AUTOPHAGY-RELATED PROTEIN 38 |
| 75:00:00 | 7b93-Z | 4.7 | 13 | 101 | 143 | 7 | MOLECULE: | NADH-UBIQUINONE OXIDOREDUCTASE CHAIN 3 |
| 76:00:00 | 3p8c-D | 4.7 | 4.3 | 77 | 204 | 8 | MOLECULE: | CYTOPLASMIC FMR1-INTERACTING PROTEIN 1 |
| 77:00:00 | 8afl-E | 4.7 | 3.3 | 68 | 68 | 13 | MOLECULE: | CRESCENTIN |
| 78:00:00 | 6cp3-Z | 4.7 | 29.2 | 118 | 155 | 6 | MOLECULE: | ATP SYNTHASE SUBUNIT 9, MITOCHONDRIAL |
| 79:00:00 | 6cfz-E | 4.7 | 6.8 | 66 | 68 | 8 | MOLECULE: | ASK1 |
| 80:00:00 | 8q85-c | 4.7 | 4.3 | 62 | 64 | 5 | MOLECULE: | KINETOCHORE PROTEIN NDC80 |
| 81:00:00 | 7yet-D | 4.7 | 2.7 | 68 | 68 | 13 | MOLECULE: | RNA-DIRECTED RNA POLYMERASE L |
| 82:00:00 | 8e2j-C | 4.7 | 2.4 | 57 | 57 | 0 | MOLECULE: | DIABLO IAP-BINDING MITOCHONDRIAL PROTEIN |

| | | | | | | | | |
|---|---|---|---|---|---|---|---|---|
| 83:00:00 | 8afm-K | 4.7 | 3.1 | 68 | 68 | 9 | MOLECULE: | CRESCENTIN |
| 84:00:00 | 8afm-E | 4.7 | 3.3 | 68 | 68 | 13 | MOLECULE: | CRESCENTIN |
| 85:00:00 | 8b4i-C | 4.7 | 3.6 | 60 | 60 | 7 | MOLECULE: | MITOCHONDRIAL IMPORT RECEPTOR SUBUNIT TOM40 |
| 86:00:00 | 7bv6-B | 4.7 | 2.4 | 61 | 61 | 5 | MOLECULE: | VESICLE-ASSOCIATED MEMBRANE PROTEIN 8 |
| 87:00:00 | 1io4-B | 4.7 | 3.4 | 65 | 70 | 12 | MOLECULE: | CSF-1R PROMOTER |
| 88:00:00 | 4bwd-A | 4.7 | 4.4 | 61 | 61 | 8 | MOLECULE: | SHORT COILED-COIL PROTEIN |
| 89:00:00 | 7y5c-b | 4.7 | 4.8 | 93 | 145 | 4 | MOLECULE: | ATP SYNTHASE SUBUNIT ALPHA |
| 90:00:00 | 1hvv-C | 4.7 | 3.5 | 66 | 66 | 3 | MOLECULE: | SYNTAXIN 1A |
| 91:00:00 | 8b8b-B | 4.7 | 3 | 81 | 95 | 7 | MOLECULE: | MUNIA BORNAVIRUS 1 PHOSPHOPROTEIN |
| 92:00:00 | 8afm-B | 4.7 | 3.4 | 77 | 77 | 8 | MOLECULE: | CRESCENTIN |
| 93:00:00 | 3iox-A | 4.7 | 2.7 | 80 | 489 | 4 | MOLECULE: | AGI/II |
| 94:00:00 | 8b8b-C | 4.7 | 3.2 | 85 | 95 | 7 | MOLECULE: | MUNIA BORNAVIRUS 1 PHOSPHOPROTEIN |
| 95:00:00 | 8at3-E | 4.7 | 13 | 111 | 217 | 7 | MOLECULE: | HAUS AUGMIN-LIKE COMPLEX SUBUNIT 1 |
| 96:00:00 | 8dbs-X | 4.7 | 4.9 | 85 | 156 | 2 | MOLECULE: | ATP SYNTHASE SUBUNIT ALPHA |
| 97:00:00 | 3b5n-E | 4.7 | 4.2 | 58 | 58 | 10 | MOLECULE: | SYNAPTOBREVIN HOMOLOG 1 |
| 98:00:00 | 6dd8-D | 4.7 | 4.2 | 87 | 117 | 3 | MOLECULE: | SYNAPTONEMAL COMPLEX PROTEIN 3 |
| 99:00:00 | 1ik9-A | 4.7 | 4.8 | 85 | 207 | 8 | MOLECULE: | DNA REPAIR PROTEIN XRCC4 |
| 0:00 | 3mq7-F | 4.7 | 5.6 | 86 | 99 | 2 | MOLECULE: | BONE MARROW STROMAL ANTIGEN 2 |
| 1:00 | 3v6i-B | 4.7 | 7.7 | 89 | 101 | 3 | MOLECULE: | V-TYPE ATP SYNTHASE SUBUNIT E |
| 2:00 | 3m06-D | 4.7 | 2.8 | 62 | 62 | 6 | MOLECULE: | TNF RECEPTOR-ASSOCIATED FACTOR 2 |
| 3:00 | 3zdo-E | 4.7 | 2.7 | 62 | 62 | 13 | MOLECULE: | PHOSPHOPROTEIN |
| 4:00 | 2v66-B | 4.7 | 5.8 | 91 | 111 | 12 | MOLECULE: | NUCLEAR DISTRIBUTION PROTEIN NUDE-LIKE 1 |
| 5:00 | 3m0a-B | 4.7 | 2.9 | 62 | 62 | 6 | MOLECULE: | TNF RECEPTOR-ASSOCIATED FACTOR 2 |
| 6:00 | 3m06-F | 4.7 | 3.5 | 62 | 62 | 5 | MOLECULE: | TNF RECEPTOR-ASSOCIATED FACTOR 2 |
| 7:00 | 3h7z-A | 4.7 | 3.3 | 61 | 61 | 10 | MOLECULE: | ADHESIN YADA |
| 8:00 | 3rl0-J | 4.7 | 2.7 | 59 | 59 | 3 | MOLECULE: | VESICLE-ASSOCIATED MEMBRANE PROTEIN 2 |
| 9:00 | 5apv-F | 4.7 | 6.7 | 67 | 67 | 3 | MOLECULE: | GENERAL CONTROL PROTEIN GCN4 |
| 10:00 | 3rl0-d | 4.7 | 2.7 | 59 | 59 | 3 | MOLECULE: | VESICLE-ASSOCIATED MEMBRANE PROTEIN 2 |
| 11:00 | 4c46-B | 4.7 | 3.5 | 71 | 73 | 1 | MOLECULE: | GENERAL CONTROL PROTEIN GCN4, GENERAL CONTROL PRO |
| 12:00 | 3m06-A | 4.7 | 3.6 | 62 | 62 | 3 | MOLECULE: | TNF RECEPTOR-ASSOCIATED FACTOR 2 |
| 13:00 | 2wvr-B | 4.7 | 6.7 | 70 | 74 | 6 | MOLECULE: | GEMININ |
| 14:00 | 2x7a-A | 4.7 | 1.8 | 59 | 59 | 7 | MOLECULE: | BONE MARROW STROMAL ANTIGEN 2 |
| 15:00 | 3m06-E | 4.7 | 3.1 | 62 | 62 | 6 | MOLECULE: | TNF RECEPTOR-ASSOCIATED FACTOR 2 |
| 16:00 | 7z47-D | 4.7 | 5.1 | 76 | 87 | 9 | MOLECULE: | ADAPTOR PROTEIN |
| 17:00 | 5f5p-E | 4.7 | 1.8 | 57 | 57 | 7 | MOLECULE: | PROTEIN SHROOM2 |
| 18:00 | 4efa-E | 4.7 | 4 | 74 | 225 | 4 | MOLECULE: | V-TYPE PROTON ATPASE SUBUNIT C |
| 19:00 | 1t2k-D | 4.7 | 1.6 | 61 | 61 | 5 | MOLECULE: | 31-MER |
| 20:00 | 4eb6-E | 4.7 | 4.4 | 91 | 132 | 8 | MOLECULE: | TUBULIN ALPHA CHAIN |
| 21:00 | 4x1k-E | 4.7 | 4.8 | 87 | 123 | 5 | MOLECULE: | TUBULIN ALPHA CHAIN |
| 22:00 | 2yo3-B | 4.7 | 6.4 | 80 | 258 | 6 | MOLECULE: | GENERAL CONTROL PROTEIN GCN4, PUTATIVE INNER MEMB |
| 23:00 | 5d80-n | 4.7 | 3.4 | 71 | 105 | 7 | MOLECULE: | V-TYPE PROTON ATPASE CATALYTIC SUBUNIT A |
| 24:00:00 | 3lt6-C | 4.7 | 2.4 | 58 | 58 | 10 | MOLECULE: | ADHESIN YADA |
| 25:00:00 | 8pr3-B | 4.7 | 4.1 | 82 | 105 | 11 | MOLECULE: | C-JUN-AMINO-TERMINAL KINASE-INTERACTING PROTEIN 3 |
| 26:00:00 | 4m3l-C | 4.7 | 1.9 | 57 | 57 | 14 | MOLECULE: | E3 UBIQUITIN-PROTEIN LIGASE TRIM63 |
| 27:00:00 | 2v66-E | 4.7 | 6.5 | 92 | 111 | 10 | MOLECULE: | NUCLEAR DISTRIBUTION PROTEIN NUDE-LIKE 1 |
| 28:00:00 | 5apw-B | 4.7 | 4.4 | 64 | 64 | 11 | MOLECULE: | GENERAL CONTROL PROTEIN GCN4 |
| 29:00:00 | 8xwx-E | 4.7 | 2.3 | 58 | 58 | 9 | MOLECULE: | B-CELL RECEPTOR-ASSOCIATED PROTEIN 31 |
| 30:00:00 | 3fwc-B | 4.7 | 3.9 | 75 | 83 | 4 | MOLECULE: | CELL DIVISION CONTROL PROTEIN 31 |
| 31:00:00 | 4xa1-C | 4.7 | 9.5 | 90 | 130 | 4 | MOLECULE: | GP7-MYH7(1173-1238)-EB1 CHIMERA PROTEIN |
| 32:00:00 | 6fln-B | 4.6 | 23.4 | 98 | 367 | 5 | MOLECULE: | E3 UBIQUITIN/ISG15 LIGASE TRIM25 |
| 33:00:00 | 6fkf-p | 4.6 | 21.7 | 104 | 143 | 7 | MOLECULE: | ATP SYNTHASE SUBUNIT ALPHA, CHLOROPLASTIC |
| 34:00:00 | 6oei-A | 4.6 | 5.9 | 96 | 243 | 9 | MOLECULE: | SPINDLE POLE BODY COMPONENT SPC42,SIGMA-54-DEPEND |
| 35:00:00 | 7uic-c | 4.6 | 3.1 | 65 | 110 | 9 | MOLECULE: | MEDIATOR OF RNA POLYMERASE II TRANSCRIPTION SUBUN |
| 36:00:00 | 7w20-W | 4.6 | 10 | 86 | 142 | 7 | MOLECULE: | NADH DEHYDROGENASE [UBIQUINONE] FLAVOPROTEIN 1, |
| 37:00:00 | 3jb9-i | 4.6 | 10.3 | 97 | 161 | 5 | MOLECULE: | PRE-MRNA-SPLICING FACTOR SPP42 |
| 38:00:00 | 6ppb-l | 4.6 | 7.2 | 80 | 83 | 9 | MOLECULE: | CAPSID VERTEX COMPONENT 1 |

| | | | | | | | | |
|---|---|---|---|---|---|---|---|---|
| 39:00:00 | 7bgf-A | 4.6 | 4.2 | 87 | 112 | 5 | MOLECULE: | DNA ENDONUCLEASE RBBP8,CTIP/RBBP8 |
| 40:00:00 | 8tgt-B | 4.6 | 4.7 | 82 | 90 | 5 | MOLECULE: | M PROTEIN |
| 41:00:00 | 8g0p-B | 4.6 | 3.2 | 86 | 98 | 7 | MOLECULE: | KINETOCHORE PROTEIN NDC80 HOMOLOG |
| 42:00:00 | 3zdo-H | 4.6 | 3.1 | 62 | 62 | 6 | MOLECULE: | PHOSPHOPROTEIN |
| 43:00:00 | 3hd7-H | 4.6 | 4.1 | 62 | 62 | 0 | MOLECULE: | VESICLE-ASSOCIATED MEMBRANE PROTEIN 2 |
| 44:00:00 | 5omb-C | 4.6 | 9.5 | 85 | 98 | 11 | MOLECULE: | REPLICATION FACTOR A PROTEIN 1 |
| 45:00:00 | 6id0-s | 4.6 | 3.2 | 67 | 67 | 7 | MOLECULE: | PRE-MRNA-PROCESSING-SPLICING FACTOR 8 |
| 46:00:00 | 5mqf-L | 4.6 | 17.1 | 87 | 336 | 7 | MOLECULE: | PRE-MRNA-PROCESSING-SPLICING FACTOR 8 |
| 47:00:00 | 8e2j-D | 4.6 | 2.2 | 57 | 57 | 0 | MOLECULE: | DIABLO IAP-BINDING MITOCHONDRIAL PROTEIN |
| 48:00:00 | 4dl0-K | 4.6 | 2.8 | 68 | 97 | 6 | MOLECULE: | V-TYPE PROTON ATPASE SUBUNIT C |
| 49:00:00 | 7wjt-D | 4.6 | 1.7 | 56 | 56 | 7 | MOLECULE: | ISOFORM 2 OF TRANSMEMBRANE PROTEIN 266 |
| 50:00:00 | 8yxr-B | 4.6 | 5.6 | 68 | 69 | 1 | MOLECULE: | PHOSPHOPROTEIN |
| 51:00:00 | 6un9-A | 4.6 | 2.6 | 66 | 69 | 11 | MOLECULE: | UNCHARACTERIZED PROTEIN |
| 52:00:00 | 5mw9-H | 4.6 | 2.4 | 57 | 57 | 9 | MOLECULE: | CENTROSOMIN |
| 53:00:00 | 3pn7-D | 4.6 | 5.8 | 68 | 69 | 6 | MOLECULE: | MYOSIN HEAVY CHAIN |
| 54:00:00 | 8swd-A | 4.6 | 1.3 | 54 | 54 | 13 | MOLECULE: | 2-OXOGLUTARATE:ACCEPTOR OXIDOREDUCTASE |
| 55:00:00 | 7sqk-A | 4.6 | 10.9 | 88 | 278 | 9 | MOLECULE: | HAUS AUGMIN-LIKE COMPLEX SUBUNIT 1 |
| 56:00:00 | 8owi-A | 4.6 | 1.2 | 55 | 55 | 4 | MOLECULE: | CENTROMERE-ASSOCIATED PROTEIN E |
| 57:00:00 | 8aix-A | 4.6 | 4 | 81 | 84 | 5 | MOLECULE: | CRESCENTIN |
| 58:00:00 | 7xfr-B | 4.6 | 4.9 | 66 | 69 | 6 | MOLECULE: | ISOFORM 2 OF WD REPEAT DOMAIN PHOSPHOINOSITIDE-IN |
| 59:00:00 | 7wjt-A | 4.6 | 1.4 | 55 | 55 | 9 | MOLECULE: | ISOFORM 2 OF TRANSMEMBRANE PROTEIN 266 |
| 60:00:00 | 8aix-E | 4.6 | 4 | 81 | 84 | 5 | MOLECULE: | CRESCENTIN |
| 61:00:00 | 3mqb-A | 4.6 | 6.6 | 95 | 109 | 9 | MOLECULE: | BONE MARROW STROMAL ANTIGEN 2 |
| 62:00:00 | 7nad-5 | 4.6 | 4.9 | 69 | 123 | 6 | MOLECULE: | 25S RRNA |
| 63:00:00 | 8aia-A | 4.6 | 4 | 81 | 84 | 5 | MOLECULE: | CRESCENTIN |
| 64:00:00 | 7uo4-B | 4.6 | 2.7 | 78 | 187 | 6 | MOLECULE: | RNA-DIRECTED RNA POLYMERASE |
| 65:00:00 | 8afm-H | 4.6 | 2.7 | 77 | 77 | 9 | MOLECULE: | CRESCENTIN |
| 66:00:00 | 8k86-B | 4.6 | 2 | 57 | 58 | 11 | MOLECULE: | NUCLEAR FACTOR INTERLEUKIN-3-REGULATED PROTEIN |
| 67:00:00 | 8swd-B | 4.6 | 1.1 | 54 | 54 | 13 | MOLECULE: | 2-OXOGLUTARATE:ACCEPTOR OXIDOREDUCTASE |
| 68:00:00 | 8ahl-J | 4.6 | 5.2 | 87 | 120 | 3 | MOLECULE: | CRESCENTIN |
| 69:00:00 | 8q3v-W | 4.6 | 6 | 74 | 75 | 8 | MOLECULE: | TETRAHYDROMETHANOPTERIN S-METHYLTRANSFERASE SUBUN |
| 70:00:00 | 8afl-B | 4.6 | 3.5 | 77 | 77 | 8 | MOLECULE: | CRESCENTIN |
| 71:00:00 | 1kil-B | 4.6 | 2.5 | 59 | 59 | 3 | MOLECULE: | SYNAPTOBREVIN SNARE MOTIF |
| 72:00:00 | 8ajb-V | 4.6 | 5.4 | 90 | 120 | 3 | MOLECULE: | CRESCENTIN |
| 73:00:00 | 5f5p-F | 4.6 | 1.9 | 58 | 58 | 9 | MOLECULE: | PROTEIN SHROOM2 |
| 74:00:00 | 5t58-B | 4.6 | 4.2 | 73 | 190 | 4 | MOLECULE: | KLLA0F02343P |
| 75:00:00 | 4yv4-C | 4.6 | 1.7 | 57 | 57 | 4 | MOLECULE: | SPINDLE ASSEMBLY ABNORMAL PROTEIN 5 |
| 76:00:00 | 8pr3-C | 4.6 | 3.4 | 81 | 106 | 2 | MOLECULE: | C-JUN-AMINO-TERMINAL KINASE-INTERACTING PROTEIN 3 |
| 77:00:00 | 3ipd-D | 4.6 | 4.5 | 63 | 63 | 5 | MOLECULE: | VESICLE-ASSOCIATED MEMBRANE PROTEIN 2 |
| 78:00:00 | 3n7n-A | 4.6 | 6.4 | 85 | 163 | 0 | MOLECULE: | MONOPOLIN COMPLEX SUBUNIT CSM1 |
| 79:00:00 | 3k5b-B | 4.6 | 5.3 | 86 | 100 | 5 | MOLECULE: | V-TYPE ATP SYNTHASE, SUBUNIT (VAPC-THERM) |
| 80:00:00 | 7jg7-b | 4.6 | 6.2 | 95 | 138 | 5 | MOLECULE: | ATP SYNTHASE SUBUNIT ALPHA |
| 81:00:00 | 3uf1-C | 4.6 | 5.5 | 87 | 109 | 8 | MOLECULE: | VIMENTIN |
| 82:00:00 | 6dd9-C | 4.6 | 6.6 | 94 | 125 | 11 | MOLECULE: | SYNAPTONEMAL COMPLEX PROTEIN 3 |
| 83:00:00 | 5y5z-K | 4.6 | 8.1 | 88 | 99 | 5 | MOLECULE: | V-TYPE ATP SYNTHASE ALPHA CHAIN |
| 84:00:00 | 5wjb-C | 4.6 | 4.7 | 82 | 112 | 13 | MOLECULE: | CAPSID ASSEMBLY SCAFFOLDING PROTEIN,MYOSIN-7 |
| 85:00:00 | 2xv5-B | 4.6 | 1.7 | 57 | 57 | 9 | MOLECULE: | LAMIN-A/C |
| 86:00:00 | 3n4x-B | 4.6 | 3.6 | 69 | 151 | 7 | MOLECULE: | MONOPOLIN COMPLEX SUBUNIT CSM1 |
| 87:00:00 | 6dd9-A | 4.6 | 3.3 | 86 | 128 | 3 | MOLECULE: | SYNAPTONEMAL COMPLEX PROTEIN 3 |
| 88:00:00 | 1t2k-C | 4.6 | 3 | 62 | 62 | 3 | MOLECULE: | 31-MER |
| 89:00:00 | 5y60-I | 4.6 | 4.6 | 87 | 99 | 6 | MOLECULE: | V-TYPE ATP SYNTHASE ALPHA CHAIN |
| 90:00:00 | 3rl0-Z | 4.6 | 2.5 | 58 | 58 | 3 | MOLECULE: | VESICLE-ASSOCIATED MEMBRANE PROTEIN 2 |
| 91:00:00 | 3rl0-B | 4.6 | 2.7 | 59 | 59 | 3 | MOLECULE: | VESICLE-ASSOCIATED MEMBRANE PROTEIN 2 |
| 92:00:00 | 3rk3-B | 4.6 | 2.9 | 62 | 62 | 3 | MOLECULE: | VAMP2 |
| 93:00:00 | 2fyz-E | 4.6 | 2.3 | 62 | 62 | 8 | MOLECULE: | FUSION GLYCOPROTEIN F0 |
| 94:00:00 | 1l4a-B | 4.6 | 4 | 79 | 82 | 4 | MOLECULE: | SYNAPTOBREVIN |

| | | | | | | | | |
|---|---|---|---|---|---|---|---|---|
| 95:00:00 | 3rl0-V | 4.6 | 2.7 | 58 | 58 | 3 | MOLECULE: | VESICLE-ASSOCIATED MEMBRANE PROTEIN 2 |
| 96:00:00 | 3j99-L | 4.6 | 4 | 66 | 66 | 6 | MOLECULE: | VESICLE-FUSING ATPASE |
| 97:00:00 | 1fos-E | 4.6 | 1.9 | 60 | 60 | 3 | MOLECULE: | DNA (5'- |
| 98:00:00 | 3u59-A | 4.6 | 4.7 | 86 | 100 | 7 | MOLECULE: | TROPOMYOSIN BETA CHAIN |
| 99:00:00 | 3m0a-C | 4.6 | 3.2 | 62 | 62 | 6 | MOLECULE: | TNF RECEPTOR-ASSOCIATED FACTOR 2 |
| 0:00 | 3mqb-F | 4.6 | 4.8 | 86 | 102 | 8 | MOLECULE: | BONE MARROW STROMAL ANTIGEN 2 |
| 1:00 | 1z56-A | 4.6 | 3.4 | 76 | 77 | 8 | MOLECULE: | LIGASE INTERACTING FACTOR 1 |
| 2:00 | 3lt6-E | 4.6 | 2.2 | 57 | 57 | 9 | MOLECULE: | ADHESIN YADA |
| 3:00 | 1nwq-A | 4.6 | 2.2 | 60 | 60 | 10 | MOLECULE: | 5'- |
| 4:00 | 2zxx-D | 4.6 | 7.2 | 65 | 65 | 5 | MOLECULE: | GEMININ |
| 5:00 | 3u59-C | 4.6 | 4.5 | 87 | 101 | 8 | MOLECULE: | TROPOMYOSIN BETA CHAIN |
| 6:00 | 3m06-B | 4.6 | 2.9 | 62 | 62 | 6 | MOLECULE: | TNF RECEPTOR-ASSOCIATED FACTOR 2 |
| 7:00 | 3vh7-E | 4.6 | 1.9 | 56 | 56 | 9 | MOLECULE: | ENVELOPE GLYCOPROTEIN GP160 |
| 8:00 | 3lt7-F | 4.6 | 2.5 | 59 | 59 | 12 | MOLECULE: | ADHESIN YADA |
| 9:00 | 2xv5-A | 4.6 | 2.7 | 57 | 57 | 9 | MOLECULE: | LAMIN-A/C |
| 10:00 | 3a7o-F | 4.6 | 2.3 | 58 | 58 | 5 | MOLECULE: | AUTOPHAGY PROTEIN 16 |
| 11:00 | 2guv-A | 4.6 | 2.7 | 56 | 56 | 4 | MOLECULE: | MAJOR OUTER MEMBRANE LIPOPROTEIN |
| 12:00 | 3ipd-H | 4.6 | 4.5 | 63 | 63 | 5 | MOLECULE: | VESICLE-ASSOCIATED MEMBRANE PROTEIN 2 |
| 13:00 | 3lt7-C | 4.6 | 2.5 | 59 | 59 | 12 | MOLECULE: | ADHESIN YADA |
| 14:00 | 7bv6-G | 4.6 | 3.8 | 68 | 74 | 9 | MOLECULE: | VESICLE-ASSOCIATED MEMBRANE PROTEIN 8 |
| 15:00 | 3bat-B | 4.6 | 5 | 71 | 81 | 4 | MOLECULE: | MYOSIN HEAVY CHAIN, STRIATED MUSCLE/GENERAL CONTR |
| 16:00 | 8th8-S | 4.6 | 43.4 | 94 | 187 | 3 | MOLECULE: | DYNEIN REGULATORY COMPLEX PROTEIN 1/2 N-TERMINAL |
| 17:00 | 1kmi-Z | 4.5 | 23.4 | 109 | 177 | 6 | MOLECULE: | CHEMOTAXIS PROTEIN CHEY |
| 18:00 | 6znl-n | 4.5 | 16.2 | 144 | 343 | 10 | MOLECULE: | ARP1 ACTIN RELATED PROTEIN 1 HOMOLOG A |
| 19:00 | 8gb3-F | 4.5 | 3.3 | 79 | 90 | 9 | MOLECULE: | CHAPERONE PROTEIN DNAK |
| 20:00 | 7emf-D | 4.5 | 6.7 | 86 | 158 | 9 | MOLECULE: | MEDIATOR OF RNA POLYMERASE II TRANSCRIPTION SUBUN |
| 21:00 | 8u95-A | 4.5 | 98.5 | 103 | 1009 | 9 | MOLECULE: | MYOSIN HEAVY CHAIN, ISOFORM U |
| 22:00 | 4jle-B | 4.5 | 4.5 | 64 | 159 | 2 | MOLECULE: | PHIST |
| 23:00 | 1m1j-B | 4.5 | 8 | 101 | 402 | 8 | MOLECULE: | FIBRINOGEN ALPHA SUBUNIT |
| 24:00:00 | 8h36-G | 4.5 | 6 | 103 | 379 | 9 | MOLECULE: | E3 UBIQUITIN-PROTEIN LIGASE RBX1 |
| 25:00:00 | 8q3v-g | 4.5 | 9.1 | 76 | 76 | 5 | MOLECULE: | TETRAHYDROMETHANOPTERIN S-METHYLTRANSFERASE SUBUN |
| 26:00:00 | 3zx6-B | 4.5 | 13.8 | 116 | 303 | 4 | MOLECULE: | HAMP, METHYL-ACCEPTING CHEMOTAXIS PROTEIN I |
| 27:00:00 | 8q85-X | 4.5 | 9.5 | 68 | 68 | 10 | MOLECULE: | KINETOCHORE PROTEIN NDC80 |
| 28:00:00 | 2b9b-B | 4.5 | 10 | 82 | 482 | 7 | MOLECULE: | FUSION GLYCOPROTEIN F0 |
| 29:00:00 | 7eqc-G | 4.5 | 6.3 | 89 | 107 | 2 | MOLECULE: | CYTOKINESIS DEFECT |
| 30:00:00 | 8p0v-L | 4.5 | 20.6 | 96 | 416 | 9 | MOLECULE: | COILED-COIL DOMAIN-CONTAINING PROTEIN 93 |
| 31:00:00 | 5wq4-E | 4.5 | 3.6 | 66 | 69 | 3 | MOLECULE: | UBIQUITIN |
| 32:00:00 | 8cra-D | 4.5 | 24.6 | 75 | 93 | 16 | MOLECULE: | FLORAL HOMEOTIC PROTEIN AGAMOUS |
| 33:00:00 | 5a5t-A | 4.5 | 21.4 | 93 | 599 | 8 | MOLECULE: | EUKARYOTIC TRANSLATION INITIATION FACTOR 3 SUBUNI |
| 34:00:00 | 8izl-E | 4.5 | 2.7 | 58 | 58 | 9 | MOLECULE: | PHOSPHOPROTEIN |
| 35:00:00 | 6jn2-B | 4.5 | 9.8 | 67 | 67 | 12 | MOLECULE: | PROTEIN AF-10 |
| 36:00:00 | 8q3v-G | 4.5 | 5.9 | 73 | 74 | 8 | MOLECULE: | TETRAHYDROMETHANOPTERIN S-METHYLTRANSFERASE SUBUN |
| 37:00:00 | 8yxo-E | 4.5 | 2.7 | 58 | 58 | 9 | MOLECULE: | PHOSPHOPROTEIN |
| 38:00:00 | 8k8a-B | 4.5 | 2 | 57 | 59 | 11 | MOLECULE: | NUCLEAR FACTOR INTERLEUKIN-3-REGULATED PROTEIN |
| 39:00:00 | 8k8a-A | 4.5 | 4.2 | 61 | 62 | 11 | MOLECULE: | NUCLEAR FACTOR INTERLEUKIN-3-REGULATED PROTEIN |
| 40:00:00 | 2ix7-C | 4.5 | 1.7 | 58 | 59 | 2 | MOLECULE: | MYOSIN-5A |
| 41:00:00 | 5kc1-F | 4.5 | 5.7 | 61 | 61 | 5 | MOLECULE: | AUTOPHAGY-RELATED PROTEIN 38 |
| 42:00:00 | 8hpo-H | 4.5 | 8.4 | 95 | 233 | 8 | MOLECULE: | TRANSCRIPTIONAL REGULATORY PROTEIN UME1 |
| 43:00:00 | 8sq9-B | 4.5 | 2.6 | 80 | 186 | 6 | MOLECULE: | RNA-DIRECTED RNA POLYMERASE |
| 44:00:00 | 3f6n-B | 4.5 | 5.9 | 71 | 74 | 11 | MOLECULE: | VIRION-ASSOCIATED PROTEIN |
| 45:00:00 | 5ed9-C | 4.5 | 2.2 | 63 | 63 | 11 | MOLECULE: | SUN DOMAIN-CONTAINING PROTEIN 2 |
| 46:00:00 | 6fkh-b | 4.5 | 6.6 | 92 | 161 | 2 | MOLECULE: | ATP SYNTHASE SUBUNIT A, CHLOROPLASTIC |
| 47:00:00 | 8i3e-C | 4.5 | 2.3 | 56 | 56 | 11 | MOLECULE: | ELKS/RAB6-INTERACTING/CAST FAMILY MEMBER 1 |
| 48:00:00 | 8aix-V | 4.5 | 4.3 | 79 | 84 | 6 | MOLECULE: | CRESCENTIN |
| 49:00:00 | 5zuv-A | 4.5 | 4.6 | 80 | 130 | 6 | MOLECULE: | SPIKE GLYCOPROTEIN,INHIBITOR EK1 |
| 50:00:00 | 3uf1-A | 4.5 | 6.3 | 88 | 104 | 5 | MOLECULE: | VIMENTIN |

| | | | | | | | | |
|---|---|---|---|---|---|---|---|---|
| 51:00:00 | 6dd8-C | 4.5 | 3.8 | 67 | 123 | 7 | MOLECULE: | SYNAPTONEMAL COMPLEX PROTEIN 3 |
| 52:00:00 | 1sa1-E | 4.5 | 6.8 | 95 | 134 | 4 | MOLECULE: | TUBULIN ALPHA CHAIN |
| 53:00:00 | 2zxx-E | 4.5 | 5.9 | 69 | 78 | 6 | MOLECULE: | GEMININ |
| 54:00:00 | 1wyy-B | 4.5 | 2.6 | 79 | 124 | 10 | MOLECULE: | E2 GLYCOPROTEIN |
| 55:00:00 | 3rl0-N | 4.5 | 2.6 | 57 | 57 | 4 | MOLECULE: | VESICLE-ASSOCIATED MEMBRANE PROTEIN 2 |
| 56:00:00 | 8afl-A | 4.5 | 4.5 | 76 | 77 | 9 | MOLECULE: | CRESCENTIN |
| 57:00:00 | 3uux-B | 4.5 | 9 | 91 | 132 | 12 | MOLECULE: | MITOCHONDRIA FISSION 1 PROTEIN |
| 58:00:00 | 4cjd-A | 4.5 | 18.5 | 78 | 123 | 6 | MOLECULE: | NADA |
| 59:00:00 | 3m06-C | 4.5 | 3.2 | 62 | 62 | 10 | MOLECULE: | TNF RECEPTOR-ASSOCIATED FACTOR 2 |
| 60:00:00 | 4kht-A | 4.5 | 3.6 | 66 | 66 | 6 | MOLECULE: | GP41 HELIX |
| 61:00:00 | 8b8b-A | 4.5 | 5.2 | 84 | 99 | 7 | MOLECULE: | MUNIA BORNAVIRUS 1 PHOSPHOPROTEIN |
| 62:00:00 | 7oz3-A | 4.5 | 7.1 | 82 | 202 | 2 | MOLECULE: | GNTR FAMILY TRANSCRIPTIONAL REGULATOR |
| 63:00:00 | 4pxj-C | 4.5 | 2.6 | 60 | 61 | 8 | MOLECULE: | C-JUN-AMINO-TERMINAL KINASE-INTERACTING PROTEIN 3 |
| 64:00:00 | 2ahm-G | 4.5 | 4 | 86 | 191 | 5 | MOLECULE: | REPLICASE POLYPROTEIN 1AB, LIGHT CHAIN |
| 65:00:00 | 3a7o-D | 4.5 | 2.6 | 58 | 58 | 12 | MOLECULE: | AUTOPHAGY PROTEIN 16 |
| 66:00:00 | 8aia-L | 4.5 | 4.3 | 79 | 84 | 6 | MOLECULE: | CRESCENTIN |
| 67:00:00 | 3htk-A | 4.5 | 1.8 | 60 | 60 | 3 | MOLECULE: | STRUCTURAL MAINTENANCE OF CHROMOSOMES PROTEIN 5 |
| 68:00:00 | 6j9r-A | 4.5 | 6.8 | 89 | 125 | 9 | MOLECULE: | BRAIN TUMOR PROTEIN |
| 69:00:00 | 7aej-B | 4.5 | 2.8 | 84 | 145 | 12 | MOLECULE: | ENVELOPE GLYCOPROTEIN GP160,ENVELOPE GLYCOPROTEIN |
| 70:00:00 | 7xyz-B | 4.4 | 29.6 | 117 | 456 | 7 | MOLECULE: | TRIPARTITE MOTIF-CONTAINING PROTEIN 72 |
| 71:00:00 | 5cws-D | 4.4 | 26.5 | 99 | 180 | 4 | MOLECULE: | SAB-158 FAB LIGHT CHAIN |
| 72:00:00 | 6xp5-G | 4.4 | 6.2 | 86 | 122 | 6 | MOLECULE: | MEDIATOR OF RNA POLYMERASE II TRANSCRIPTION SUBUN |
| 73:00:00 | 8tek-B | 4.4 | 22.8 | 101 | 208 | 10 | MOLECULE: | DYNEIN REGULATORY COMPLEX PROTEIN 1/2 N-TERMINAL |
| 74:00:00 | 5d80-G | 4.4 | 6.1 | 87 | 223 | 8 | MOLECULE: | V-TYPE PROTON ATPASE CATALYTIC SUBUNIT A |
| 75:00:00 | 4tql-B | 4.4 | 3.2 | 100 | 235 | 6 | MOLECULE: | THREE HELIX BUNDLE |
| 76:00:00 | 8fef-B | 4.4 | 17 | 93 | 335 | 4 | MOLECULE: | VIRULENCE FACTOR MCE FAMILY PROTEIN |
| 77:00:00 | 6zls-A | 4.4 | 3.2 | 80 | 292 | 18 | MOLECULE: | HISTIDINE KINASE |
| 78:00:00 | 8ixk-E | 4.4 | 3.2 | 81 | 109 | 5 | MOLECULE: | ATTACHMENT PROTEIN G3P |
| 79:00:00 | 8fed-F | 4.4 | 18.7 | 101 | 399 | 9 | MOLECULE: | VIRULENCE FACTOR MCE FAMILY PROTEIN |
| 80:00:00 | 6h3a-A | 4.4 | 46.1 | 84 | 266 | 12 | MOLECULE: | SWI/SNF-RELATED MATRIX-ASSOCIATED ACTIN-DEPENDENT |
| 81:00:00 | 3hnw-A | 4.4 | 4.5 | 91 | 128 | 7 | MOLECULE: | UNCHARACTERIZED PROTEIN |
| 82:00:00 | 4f61-I | 4.4 | 6.6 | 91 | 234 | 5 | MOLECULE: | TUBULIN ALPHA CHAIN |
| 83:00:00 | 4jpp-D | 4.4 | 29.9 | 120 | 129 | 13 | MOLECULE: | MINOR SPIKE PROTEIN H |
| 84:00:00 | 8w6b-H | 4.4 | 2.7 | 56 | 57 | 14 | MOLECULE: | TAX1-BINDING PROTEIN 1 |
| 85:00:00 | 8pqw-F | 4.4 | 6.4 | 89 | 150 | 3 | MOLECULE: | CYTOPLASMIC DYNEIN 1 HEAVY CHAIN 1 |
| 86:00:00 | 7xho-Q | 4.4 | 5.2 | 74 | 164 | 3 | MOLECULE: | CENTROMERE PROTEIN C |
| 87:00:00 | 5odw-D | 4.4 | 2.7 | 64 | 198 | 9 | MOLECULE: | FERRIPYOVERDINE RECEPTOR |
| 88:00:00 | 7vc4-H | 4.4 | 10.4 | 76 | 107 | 5 | MOLECULE: | MITOCHONDRIAL IMPORT RECEPTOR SUBUNIT TOM6 HOMOLO |
| 89:00:00 | 8rtd-B | 4.4 | 3.7 | 66 | 197 | 11 | MOLECULE: | TRWJ PROTEIN |
| 90:00:00 | 8k86-A | 4.4 | 4.5 | 61 | 63 | 11 | MOLECULE: | NUCLEAR FACTOR INTERLEUKIN-3-REGULATED PROTEIN |
| 91:00:00 | 8owi-B | 4.4 | 2.1 | 54 | 54 | 6 | MOLECULE: | CENTROMERE-ASSOCIATED PROTEIN E |
| 92:00:00 | 8rtd-A | 4.4 | 3.6 | 67 | 197 | 10 | MOLECULE: | TRWJ PROTEIN |
| 93:00:00 | 7say-A | 4.4 | 2.4 | 56 | 56 | 11 | MOLECULE: | GENERAL CONTROL TRANSCRIPTION FACTOR GCN4/M PROTE |
| 94:00:00 | 8b8a-A | 4.4 | 7.5 | 85 | 95 | 7 | MOLECULE: | PHOSPHOPROTEIN |
| 95:00:00 | 8w6b-C | 4.4 | 3 | 57 | 57 | 14 | MOLECULE: | TAX1-BINDING PROTEIN 1 |
| 96:00:00 | 6wbp-A | 4.4 | 2.2 | 56 | 56 | 7 | MOLECULE: | SEPTIN-6 |
| 97:00:00 | 6qle-Q | 4.4 | 10.9 | 94 | 224 | 11 | MOLECULE: | CENTRAL KINETOCHORE SUBUNIT MCM16,CENTRAL KINETOC |
| 98:00:00 | 8aix-K | 4.4 | 4.1 | 72 | 72 | 3 | MOLECULE: | CRESCENTIN |
| 99:00:00 | 7use-E | 4.4 | 4.2 | 79 | 155 | 9 | MOLECULE: | CYTOPLASMIC FMR1-INTERACTING PROTEIN 1 |
| 0:00 | 8ftk-A | 4.4 | 30.1 | 103 | 1584 | 2 | MOLECULE: | 5'-3' RNA HELICASE-LIKE PROTEIN |
| 1:00 | 7w35-W | 4.4 | 13.4 | 108 | 142 | 8 | MOLECULE: | NADH DEHYDROGENASE [UBIQUINONE] FLAVOPROTEIN 1, |
| 2:00 | 7uo4-D | 4.4 | 8.8 | 82 | 187 | 5 | MOLECULE: | RNA-DIRECTED RNA POLYMERASE |
| 3:00 | 6zr2-Z | 4.4 | 13 | 90 | 141 | 2 | MOLECULE: | NADH-UBIQUINONE OXIDOREDUCTASE CHAIN 3 |
| 4:00 | 7tkd-U | 4.4 | 7.1 | 92 | 155 | 2 | MOLECULE: | ATP SYNTHASE SUBUNIT 9, MITOCHONDRIAL |
| 5:00 | 2fxo-B | 4.4 | 9.5 | 100 | 125 | 10 | MOLECULE: | MYOSIN HEAVY CHAIN, CARDIAC MUSCLE BETA ISOFORM |
| 6:00 | 3hkb-E | 4.4 | 4.6 | 86 | 124 | 6 | MOLECULE: | TUBULIN ALPHA CHAIN |

| | | | | | | | | |
|---|---|---|---|---|---|---|---|---|
| 7:00 | 3j98-L | 4.4 | 4.1 | 66 | 66 | 6 | MOLECULE: | VESICLE-FUSING ATPASE |
| 8:00 | 3j98-K | 4.4 | 3.8 | 61 | 61 | 15 | MOLECULE: | VESICLE-FUSING ATPASE |
| 9:00 | 3j96-K | 4.4 | 3.9 | 61 | 61 | 15 | MOLECULE: | VESICLE-FUSING ATPASE |
| 10:00 | 3j99-K | 4.4 | 3.8 | 61 | 61 | 15 | MOLECULE: | VESICLE-FUSING ATPASE |
| 11:00 | 7lsy-F | 4.4 | 4.6 | 92 | 201 | 7 | MOLECULE: | X-RAY REPAIR CROSS-COMPLEMENTING PROTEIN 6 |
| 12:00 | 1fos-H | 4.4 | 4 | 58 | 58 | 3 | MOLECULE: | DNA (5'- |
| 13:00 | 3hhm-B | 4.4 | 5.1 | 100 | 247 | 6 | MOLECULE: | PHOSPHATIDYLINOSITOL-4,5-BISPHOSPHATE 3-KINASE |
| 14:00 | 7xhn-K | 4.4 | 13.9 | 101 | 230 | 10 | MOLECULE: | CENTROMERE PROTEIN O |
| 15:00 | 4m3l-D | 4.4 | 1.7 | 53 | 53 | 9 | MOLECULE: | E3 UBIQUITIN-PROTEIN LIGASE TRIM63 |
| 16:00 | 8aix-L | 4.4 | 4.3 | 79 | 84 | 6 | MOLECULE: | CRESCENTIN |
| 17:00 | 4g2k-A | 4.4 | 21.6 | 87 | 110 | 9 | MOLECULE: | GENERAL CONTROL PROTEIN GCN4, ENVELOPE GLYCOPROTE |
| 18:00 | 2p22-A | 4.3 | 6.8 | 79 | 168 | 6 | MOLECULE: | SUPPRESSOR PROTEIN STP22 OF TEMPERATURE-SENSITIVE |
| 19:00 | 5vpe-A | 4.3 | 4.1 | 67 | 67 | 4 | MOLECULE: | PROTEIN FOSB |
| 20:00 | 8i03-G | 4.3 | 7.4 | 111 | 166 | 5 | MOLECULE: | PAIRED AMPHIPATHIC HELIX PROTEIN PST1 |
| 21:00 | 5oqm-h | 4.3 | 7.9 | 79 | 131 | 5 | MOLECULE: | DNA-DIRECTED RNA POLYMERASE II SUBUNIT RPB1 |
| 22:00 | 3e98-A | 4.3 | 2.6 | 71 | 178 | 10 | MOLECULE: | GAF DOMAIN OF UNKNOWN FUNCTION |
| 23:00 | 7x5e-B | 4.3 | 7.5 | 75 | 106 | 7 | MOLECULE: | TRANSCRIPTION FACTOR MAFG |
| 24:00:00 | 5j9u-G | 4.3 | 25.8 | 91 | 297 | 7 | MOLECULE: | HISTONE ACETYLTRANSFERASE ESA1 |
| 25:00:00 | 8q7e-J | 4.3 | 6 | 71 | 895 | 6 | MOLECULE: | CULLIN-9 |
| 26:00:00 | 4xa6-A | 4.3 | 6.6 | 78 | 168 | 8 | MOLECULE: | GP7-MYH7(1777-1855)-EB1 CHIMERA PROTEIN |
| 27:00:00 | 3viq-B | 4.3 | 10 | 74 | 85 | 11 | MOLECULE: | SWI5-DEPENDENT RECOMBINATION DNA REPAIR PROTEIN 1 |
| 28:00:00 | 8aix-Q | 4.3 | 6.4 | 95 | 183 | 7 | MOLECULE: | CRESCENTIN |
| 29:00:00 | 8b8d-D | 4.3 | 4 | 85 | 103 | 8 | MOLECULE: | PHOSPHOPROTEIN |
| 30:00:00 | 8q0j-Z | 4.3 | 11.3 | 99 | 142 | 11 | MOLECULE: | NADH-UBIQUINONE OXIDOREDUCTASE CHAIN 3 |
| 31:00:00 | 6v85-C | 4.3 | 4.2 | 68 | 74 | 10 | MOLECULE: | RNA-DIRECTED RNA POLYMERASE L |
| 32:00:00 | 8ovw-B | 4.3 | 7.3 | 86 | 107 | 8 | MOLECULE: | CENTROMERE-BINDING PROTEIN 1 |
| 33:00:00 | 7tmp-H | 4.3 | 9.2 | 79 | 86 | 8 | MOLECULE: | H(+)-TRANSPORTING TWO-SECTOR ATPASE |
| 34:00:00 | 6j6g-c | 4.3 | 7.9 | 89 | 436 | 8 | MOLECULE: | PRE-MRNA-SPLICING FACTOR 8 |
| 35:00:00 | 2wss-T | 4.3 | 9.1 | 79 | 86 | 1 | MOLECULE: | ATP SYNTHASE SUBUNIT ALPHA, MITOCHONDRIAL |
| 36:00:00 | 3hd7-D | 4.3 | 3.4 | 58 | 63 | 2 | MOLECULE: | VESICLE-ASSOCIATED MEMBRANE PROTEIN 2 |
| 37:00:00 | 8urb-B | 4.3 | 3.6 | 78 | 184 | 9 | MOLECULE: | NSP12 |
| 38:00:00 | 8gwb-D | 4.3 | 8.3 | 80 | 186 | 6 | MOLECULE: | RNA-DIRECTED RNA POLYMERASE |
| 39:00:00 | 7uo7-D | 4.3 | 9.3 | 84 | 186 | 6 | MOLECULE: | RNA-DIRECTED RNA POLYMERASE |
| 40:00:00 | 7r7c-J | 4.3 | 8.5 | 69 | 96 | 6 | MOLECULE: | 25S RRNA |
| 41:00:00 | 8t0p-B | 4.3 | 2.1 | 70 | 107 | 9 | MOLECULE: | INNER KINETOCHORE SUBUNIT AME1 |
| 42:00:00 | 8x01-E | 4.3 | 2.5 | 54 | 54 | 9 | MOLECULE: | RNA-DIRECTED RNA POLYMERASE L |
| 43:00:00 | 7qog-A | 4.3 | 9.9 | 97 | 657 | 6 | MOLECULE: | PORTAL PROTEIN GP20 |
| 44:00:00 | 7uv8-U | 4.3 | 6.5 | 93 | 161 | 8 | MOLECULE: | UBIQUITIN-CONJUGATING ENZYME E2 2 |
| 45:00:00 | 8cra-F | 4.3 | 13.6 | 68 | 93 | 16 | MOLECULE: | FLORAL HOMEOTIC PROTEIN AGAMOUS |
| 46:00:00 | 5gna-B | 4.3 | 2.9 | 56 | 56 | 5 | MOLECULE: | FLAGELLAR PROTEIN FLIT |
| 47:00:00 | 3ghg-D | 4.3 | 8.2 | 84 | 174 | 8 | MOLECULE: | FIBRINOGEN ALPHA CHAIN |
| 48:00:00 | 2x7r-C | 4.3 | 3.7 | 58 | 58 | 7 | MOLECULE: | TRANSMEMBRANE PROTEIN GP41 |
| 49:00:00 | 7w2l-W | 4.3 | 13.2 | 107 | 142 | 8 | MOLECULE: | NADH DEHYDROGENASE [UBIQUINONE] FLAVOPROTEIN 1, |
| 50:00:00 | 6dd9-B | 4.3 | 8.5 | 98 | 127 | 4 | MOLECULE: | SYNAPTONEMAL COMPLEX PROTEIN 3 |
| 51:00:00 | 2ocy-B | 4.3 | 6.5 | 91 | 149 | 8 | MOLECULE: | RAB GUANINE NUCLEOTIDE EXCHANGE FACTOR SEC2 |
| 52:00:00 | 2fxo-C | 4.3 | 5.6 | 91 | 127 | 10 | MOLECULE: | MYOSIN HEAVY CHAIN, CARDIAC MUSCLE BETA ISOFORM |
| 53:00:00 | 3hke-E | 4.3 | 4.6 | 89 | 124 | 8 | MOLECULE: | TUBULIN ALPHA CHAIN |
| 54:00:00 | 5noj-H | 4.3 | 6.9 | 91 | 136 | 0 | MOLECULE: | ACTIN, ALPHA SKELETAL MUSCLE |
| 55:00:00 | 6gak-C | 4.3 | 6.4 | 95 | 131 | 16 | MOLECULE: | OUTER CAPSID PROTEIN SIGMA-1 |
| 56:00:00 | 6gak-A | 4.3 | 7.2 | 95 | 129 | 12 | MOLECULE: | OUTER CAPSID PROTEIN SIGMA-1 |
| 57:00:00 | 2e7s-C | 4.3 | 4 | 84 | 118 | 8 | MOLECULE: | RAB GUANINE NUCLEOTIDE EXCHANGE FACTOR SEC2 |
| 58:00:00 | 8hhg-L | 4.3 | 10.3 | 78 | 81 | 13 | MOLECULE: | CELL DIVISION PROTEIN FTSQ |
| 59:00:00 | 2wty-A | 4.3 | 4.4 | 77 | 94 | 4 | MOLECULE: | TRANSCRIPTION FACTOR MAFB |
| 60:00:00 | 7vby-H | 4.3 | 5.9 | 56 | 56 | 9 | MOLECULE: | TRANSLOCASE OF THE OUTER MEMBRANE |
| 61:00:00 | 4bhv-C | 4.3 | 1.7 | 57 | 57 | 5 | MOLECULE: | PHOSPHOPROTEIN |
| 62:00:00 | 7w4d-W | 4.3 | 14.7 | 112 | 142 | 9 | MOLECULE: | NADH DEHYDROGENASE [UBIQUINONE] FLAVOPROTEIN 1, |

| | | | | | | | | |
|---|---|---|---|---|---|---|---|---|
| 63:00:00 | 3h01-A | 4.3 | 1.3 | 54 | 54 | 6 | MOLECULE: | ENVELOPE GLYCOPROTEIN GP160 |
| 64:00:00 | 8h9r-K | 4.3 | 5.8 | 97 | 198 | 2 | MOLECULE: | ATP SYNTHASE F(0) COMPLEX SUBUNIT B1, MITOCHONDRI |
| 65:00:00 | 8afm-A | 4.3 | 3 | 72 | 77 | 6 | MOLECULE: | CRESCENTIN |
| 66:00:00 | 6e95-B | 4.3 | 19.3 | 94 | 229 | 5 | MOLECULE: | STAPHYLOCOCCUS AUREUS AGRC HISTIDINE KINASE MODUL |
| 67:00:00 | 6e52-B | 4.3 | 20.1 | 94 | 229 | 6 | MOLECULE: | STAPHYLOCOCCUS AUREUS AGRC HISTIDINE KINASE MODUL |
| 68:00:00 | 6h9l-A | 4.3 | 6.7 | 92 | 127 | 3 | MOLECULE: | UNCHARACTERIZED PROTEIN |
| 69:00:00 | 6ewy-A | 4.2 | 26 | 104 | 201 | 9 | MOLECULE: | PEPTIDOGLYCAN ENDOPEPTIDASE RIPA |
| 70:00:00 | 6lk8-a | 4.2 | 23.6 | 103 | 1272 | 8 | MOLECULE: | MGC83295 PROTEIN |
| 71:00:00 | 6b85-A | 4.2 | 3.3 | 70 | 215 | 16 | MOLECULE: | TMHC4_R |
| 72:00:00 | 5yfp-C | 4.2 | 5.7 | 84 | 790 | 10 | MOLECULE: | EXOCYST COMPLEX COMPONENT SEC3 |
| 73:00:00 | 6wuc-K | 4.2 | 18.5 | 105 | 235 | 14 | MOLECULE: | INNER KINETOCHORE SUBUNIT MCM16 |
| 74:00:00 | 5wkq-A | 4.2 | 4.8 | 92 | 166 | 10 | MOLECULE: | INVASIN IPAB |
| 75:00:00 | 8srm-B | 4.2 | 10 | 90 | 235 | 8 | MOLECULE: | RB1-INDUCIBLE COILED-COIL PROTEIN 1 |
| 76:00:00 | 8r1a-A | 4.2 | 48.2 | 142 | 587 | 7 | MOLECULE: | GUANYLATE BINDING PROTEIN 1 |
| 77:00:00 | 3hgf-A | 4.2 | 3.7 | 82 | 98 | 9 | MOLECULE: | RHOPTRY PROTEIN FRAGMENT |
| 78:00:00 | 6exn-D | 4.2 | 9.4 | 86 | 97 | 10 | MOLECULE: | U2 SNRNA |
| 79:00:00 | 4aj5-U | 4.2 | 3.3 | 79 | 100 | 8 | MOLECULE: | SPINDLE AND KINETOCHORE-ASSOCIATED PROTEIN 1 |
| 80:00:00 | 6id0-L | 4.2 | 5.4 | 91 | 475 | 13 | MOLECULE: | PRE-MRNA-PROCESSING-SPLICING FACTOR 8 |
| 81:00:00 | 2wz7-B | 4.2 | 3.3 | 61 | 67 | 15 | MOLECULE: | UNCHARACTERIZED PROTEIN YBGF |
| 82:00:00 | 6wmz-A | 4.2 | 3.6 | 97 | 299 | 4 | MOLECULE: | SPLICING FACTOR, PROLINE- AND GLUTAMINE-RICH |
| 83:00:00 | 5gmk-c | 4.2 | 8.1 | 90 | 436 | 7 | MOLECULE: | PRE-MRNA-SPLICING FACTOR 8 |
| 84:00:00 | 8ahl-C | 4.2 | 71.9 | 152 | 234 | 7 | MOLECULE: | CRESCENTIN |
| 85:00:00 | 8x01-B | 4.2 | 3.5 | 63 | 69 | 8 | MOLECULE: | RNA-DIRECTED RNA POLYMERASE L |
| 86:00:00 | 8afl-F | 4.2 | 3.2 | 59 | 59 | 17 | MOLECULE: | CRESCENTIN |
| 87:00:00 | 8izm-B | 4.2 | 5.8 | 68 | 69 | 1 | MOLECULE: | PHOSPHOPROTEIN |
| 88:00:00 | 8afm-F | 4.2 | 3.2 | 59 | 59 | 17 | MOLECULE: | CRESCENTIN |
| 89:00:00 | 8ow0-U | 4.2 | 5.5 | 75 | 184 | 8 | MOLECULE: | C0N3 DNA |
| 90:00:00 | 1eq7-A | 4.2 | 2.5 | 56 | 56 | 13 | MOLECULE: | OUTER MEMBRANE LIPOPROTEIN |
| 91:00:00 | 8gwi-B | 4.2 | 4.5 | 82 | 187 | 1 | MOLECULE: | RNA-DIRECTED RNA POLYMERASE |
| 92:00:00 | 6vq6-M | 4.2 | 4.8 | 86 | 114 | 5 | MOLECULE: | ATPASE H+-TRANSPORTING V1 SUBUNIT A |
| 93:00:00 | 8tes-E | 4.2 | 8 | 79 | 84 | 6 | MOLECULE: | LARGE TEGUMENT PROTEIN DENEDDYLASE |
| 94:00:00 | 6exn-O | 4.2 | 17.4 | 82 | 320 | 7 | MOLECULE: | U2 SNRNA |
| 95:00:00 | 5dfz-D | 4.2 | 6.3 | 96 | 341 | 11 | MOLECULE: | VACUOLAR PROTEIN SORTING-ASSOCIATED PROTEIN 38 |
| 96:00:00 | 8yxr-E | 4.2 | 2.5 | 54 | 54 | 9 | MOLECULE: | PHOSPHOPROTEIN |
| 97:00:00 | 8gwg-B | 4.2 | 4.5 | 82 | 187 | 1 | MOLECULE: | RNA-DIRECTED RNA POLYMERASE |
| 98:00:00 | 7jtk-F | 4.2 | 8.9 | 97 | 287 | 3 | MOLECULE: | FLAGELLAR RADIAL SPOKE PROTEIN 1 |
| 99:00:00 | 8gwf-B | 4.2 | 4.5 | 82 | 187 | 1 | MOLECULE: | RNA-DIRECTED RNA POLYMERASE |
| 0:00 | 8cra-E | 4.2 | 3.7 | 63 | 96 | 6 | MOLECULE: | FLORAL HOMEOTIC PROTEIN AGAMOUS |
| 1:00 | 3a5t-B | 4.2 | 2.3 | 68 | 93 | 4 | MOLECULE: | TRANSCRIPTION FACTOR MAFG |
| 2:00 | 8cra-B | 4.2 | 2.9 | 61 | 91 | 7 | MOLECULE: | FLORAL HOMEOTIC PROTEIN AGAMOUS |
| 3:00 | 8afm-L | 4.2 | 3.1 | 59 | 59 | 5 | MOLECULE: | CRESCENTIN |
| 4:00 | 2xnx-M | 4.2 | 6.6 | 90 | 107 | 8 | MOLECULE: | FIBRINOGEN ALPHA CHAIN |
| 5:00 | 6j5i-b | 4.2 | 9.4 | 99 | 209 | 6 | MOLECULE: | ATP SYNTHASE SUBUNIT ALPHA, MITOCHONDRIAL |
| 6:00 | 7nju-b | 4.2 | 4.8 | 63 | 63 | 6 | MOLECULE: | ATP SYNTHASE SUBUNIT C |
| 7:00 | 7vpx-J | 4.2 | 5.3 | 54 | 54 | 4 | MOLECULE: | SPLICING FACTOR 3A SUBUNIT 2 |
| 8:00 | 4a7h-H | 4.2 | 8.4 | 98 | 136 | 4 | MOLECULE: | ACTIN, ALPHA SKELETAL MUSCLE |
| 9:00 | 5noj-F | 4.2 | 9.5 | 99 | 136 | 0 | MOLECULE: | ACTIN, ALPHA SKELETAL MUSCLE |
| 10:00 | 3j4k-G | 4.2 | 8.4 | 98 | 136 | 0 | MOLECULE: | ACTIN, ALPHA SKELETAL MUSCLE |
| 11:00 | 6xby-N | 4.2 | 3.4 | 82 | 108 | 2 | MOLECULE: | V-TYPE PROTON ATPASE CATALYTIC SUBUNIT A |
| 12:00 | 4a7f-H | 4.2 | 8.4 | 98 | 136 | 4 | MOLECULE: | ACTIN, ALPHA SKELETAL MUSCLE |
| 13:00 | 4a7l-H | 4.2 | 8.4 | 98 | 136 | 4 | MOLECULE: | ACTIN, ALPHA SKELETON MUSCLE |
| 14:00 | 4jol-A | 4.2 | 2.5 | 57 | 60 | 7 | MOLECULE: | PROTEIN CBFA2T1 |
| 15:00 | 8akr-v | 4.2 | 4.9 | 79 | 196 | 11 | MOLECULE: | CHLOROPLAST MEMBRANE-ASSOCIATED 30 KD PROTEIN |
| 16:00 | 4bhv-A | 4.2 | 1.7 | 57 | 57 | 5 | MOLECULE: | PHOSPHOPROTEIN |
| 17:00 | 8dd0-P | 4.2 | 5.8 | 70 | 75 | 4 | MOLECULE: | ACTIN, ALPHA CARDIAC MUSCLE 1 |
| 18:00 | 7znk-M | 4.2 | 49.5 | 80 | 549 | 8 | MOLECULE: | RNA |

| | | | | | | | | |
|---|---|---|---|---|---|---|---|---|
| 19:00 | 7znl-M | 4.2 | 53.3 | 82 | 549 | 6 | MOLECULE: | THO COMPLEX SUBUNIT 1 |
| 20:00 | 4xa1-B | 4.2 | 30.7 | 115 | 155 | 8 | MOLECULE: | GP7-MYH7(1173-1238)-EB1 CHIMERA PROTEIN |
| 21:00 | 2yny-C | 4.2 | 5.3 | 76 | 105 | 5 | MOLECULE: | GENERAL CONTROL PROTEIN GCN4, PUTATIVE INNER MEMB |
| 22:00 | 6cfz-C | 4.1 | 6.4 | 76 | 79 | 7 | MOLECULE: | ASK1 |
| 23:00 | 5y06-A | 4.1 | 25.3 | 99 | 235 | 9 | MOLECULE: | MSMEG_4306 |
| 24:00:00 | 3na7-A | 4.1 | 24.8 | 106 | 237 | 7 | MOLECULE: | HP0958 |
| 25:00:00 | 4jio-B | 4.1 | 4.7 | 71 | 320 | 8 | MOLECULE: | BRO1 |
| 26:00:00 | 5oqj-1 | 4.1 | 20.7 | 137 | 491 | 4 | MOLECULE: | DNA-DIRECTED RNA POLYMERASE II SUBUNIT RPB1 |
| 27:00:00 | 7q83-D | 4.1 | 3.9 | 64 | 156 | 11 | MOLECULE: | EXOCYST COMPLEX COMPONENT SEC3 |
| 28:00:00 | 6zsi-D | 4.1 | 4.3 | 79 | 133 | 8 | MOLECULE: | RAS-RELATED PROTEIN RAB-8A |
| 29:00:00 | 6b3o-A | 4.1 | 8 | 89 | 344 | 9 | MOLECULE: | SPIKE GLYCOPROTEIN |
| 30:00:00 | 8rgr-Z | 4.1 | 14.4 | 106 | 141 | 6 | MOLECULE: | NADH DEHYDROGENASE [UBIQUINONE] IRON-SULFUR PROTE |
| 31:00:00 | 8rgp-Z | 4.1 | 14.4 | 108 | 141 | 6 | MOLECULE: | NADH DEHYDROGENASE [UBIQUINONE] IRON-SULFUR PROTE |
| 32:00:00 | 8jsn-D | 4.1 | 3.8 | 63 | 68 | 10 | MOLECULE: | RNA-DIRECTED RNA POLYMERASE L |
| 33:00:00 | 6ucv-J | 4.1 | 4.5 | 55 | 55 | 7 | MOLECULE: | MITOCHONDRIAL IMPORT RECEPTOR SUBUNIT TOM40 |
| 34:00:00 | 8gwo-D | 4.1 | 8.2 | 82 | 186 | 4 | MOLECULE: | RNA-DIRECTED RNA POLYMERASE |
| 35:00:00 | 7sxk-j | 4.1 | 5.8 | 81 | 578 | 7 | MOLECULE: | PORTAL PROTEIN |
| 36:00:00 | 8ajb-I | 4.1 | 5.5 | 90 | 120 | 8 | MOLECULE: | CRESCENTIN |
| 37:00:00 | 7w91-A | 4.1 | 2.3 | 53 | 53 | 6 | MOLECULE: | CENTROSOMAL PROTEIN OF 63 KDA |
| 38:00:00 | 8i03-E | 4.1 | 12.9 | 97 | 169 | 4 | MOLECULE: | PAIRED AMPHIPATHIC HELIX PROTEIN PST1 |
| 39:00:00 | 8ahl-I | 4.1 | 5.5 | 90 | 120 | 9 | MOLECULE: | CRESCENTIN |
| 40:00:00 | 7usc-C | 4.1 | 3.7 | 78 | 190 | 8 | MOLECULE: | CYTOPLASMIC FMR1-INTERACTING PROTEIN 1 |
| 41:00:00 | 8ajb-U | 4.1 | 5.5 | 90 | 120 | 9 | MOLECULE: | CRESCENTIN |
| 42:00:00 | 6od2-A | 4.1 | 1.8 | 53 | 53 | 9 | MOLECULE: | SPINDLE POLE BODY COMPONENT SPC42 |
| 43:00:00 | 1kfm-A | 4.1 | 2.2 | 50 | 50 | 8 | MOLECULE: | MAJOR OUTER MEMBRANE LIPOPROTEIN |
| 44:00:00 | 7w91-H | 4.1 | 1.8 | 52 | 52 | 6 | MOLECULE: | CENTROSOMAL PROTEIN OF 63 KDA |
| 45:00:00 | 6f1t-M | 4.1 | 14.2 | 146 | 587 | 0 | MOLECULE: | ARP1 ACTIN RELATED PROTEIN 1 HOMOLOG A |
| 46:00:00 | 5to7-A | 4.1 | 9.7 | 99 | 141 | 10 | MOLECULE: | NUCLEOPROTEIN TPR |
| 47:00:00 | 4zxq-D | 4.1 | 5.7 | 87 | 140 | 6 | MOLECULE: | TAIL NEEDLE PROTEIN GP26 |
| 48:00:00 | 6gak-B | 4.1 | 7.6 | 93 | 131 | 5 | MOLECULE: | OUTER CAPSID PROTEIN SIGMA-1 |
| 49:00:00 | 6snz-C | 4.1 | 6.5 | 92 | 146 | 4 | MOLECULE: | PRELAMIN-A/C |
| 50:00:00 | 5to6-A | 4.1 | 6.7 | 96 | 141 | 8 | MOLECULE: | NUCLEOPROTEIN TPR |
| 51:00:00 | 3lt6-B | 4.1 | 3.1 | 58 | 61 | 7 | MOLECULE: | ADHESIN YADA |
| 52:00:00 | 5tvb-B | 4.1 | 7.9 | 98 | 140 | 9 | MOLECULE: | NUCLEOPROTEIN TPR |
| 53:00:00 | 8hhh-L | 4.1 | 10.4 | 76 | 81 | 12 | MOLECULE: | CELL DIVISION PROTEIN FTSQ |
| 54:00:00 | 3rrr-H | 4.1 | 5.6 | 89 | 359 | 7 | MOLECULE: | FUSION GLYCOPROTEIN F0 |
| 55:00:00 | 5nol-G | 4.1 | 7.3 | 90 | 135 | 0 | MOLECULE: | CARDIAC MUSCLE ALPHA ACTIN 1 |
| 56:00:00 | 5nol-F | 4.1 | 7.4 | 95 | 135 | 0 | MOLECULE: | CARDIAC MUSCLE ALPHA ACTIN 1 |
| 57:00:00 | 5to7-D | 4.1 | 5 | 86 | 141 | 10 | MOLECULE: | NUCLEOPROTEIN TPR |
| 58:00:00 | 6tda-L | 4.1 | 7.4 | 86 | 596 | 9 | MOLECULE: | HISTONE H3.2 |
| 59:00:00 | 3swf-B | 4.1 | 3.3 | 56 | 58 | 5 | MOLECULE: | CGMP-GATED CATION CHANNEL ALPHA-1 |
| 60:00:00 | 3j97-K | 4.1 | 3.8 | 61 | 61 | 15 | MOLECULE: | VESICLE-FUSING ATPASE |
| 61:00:00 | 6o7x-J | 4.1 | 2.8 | 79 | 105 | 4 | MOLECULE: | V-TYPE PROTON ATPASE SUBUNIT C |
| 62:00:00 | 1fe6-C | 4.1 | 1.1 | 49 | 49 | 8 | MOLECULE: | TETRABRACHION |
| 63:00:00 | 8bef-u | 4.1 | 5.5 | 55 | 55 | 4 | MOLECULE: | NADH-UBIQUINONE OXIDOREDUCTASE CHAIN 3 |
| 64:00:00 | 2w6a-A | 4.1 | 3.4 | 57 | 61 | 16 | MOLECULE: | ARF GTPASE-ACTIVATING PROTEIN GIT1 |
| 65:00:00 | 7w91-G | 4.1 | 2.3 | 53 | 53 | 6 | MOLECULE: | CENTROSOMAL PROTEIN OF 63 KDA |
| 66:00:00 | 1ik7-A | 4.1 | 1.5 | 52 | 52 | 10 | MOLECULE: | PROBABLE SERINE/THREONINE-PROTEIN KINASE PELLE |
| 67:00:00 | 1ebo-E | 4.1 | 19.2 | 89 | 111 | 6 | MOLECULE: | EBOLA VIRUS ENVELOPE PROTEIN CHIMERA CONSISTING |
| 68:00:00 | 1ebo-D | 4.1 | 21 | 90 | 112 | 4 | MOLECULE: | EBOLA VIRUS ENVELOPE PROTEIN CHIMERA CONSISTING |
| 69:00:00 | 6pfp-D | 4.1 | 41 | 116 | 156 | 7 | MOLECULE: | MYOSIN-7 FUSED TO GP7 AND EB1 |
| 70:00:00 | 2yny-B | 4.1 | 4.7 | 75 | 104 | 5 | MOLECULE: | GENERAL CONTROL PROTEIN GCN4, PUTATIVE INNER MEMB |
| 71:00:00 | 6a68-A | 4 | 19.4 | 90 | 162 | 7 | MOLECULE: | CALCIUM-DEPENDENT SECRETION ACTIVATOR 1 |
| 72:00:00 | 6y1y-B | 4 | 4.5 | 79 | 128 | 4 | MOLECULE: | CHEA |
| 73:00:00 | 8ovw-U | 4 | 6.9 | 78 | 184 | 8 | MOLECULE: | CENTROMERE-BINDING PROTEIN 1 |
| 74:00:00 | 5twv-B | 4 | 27.3 | 134 | 1375 | 9 | MOLECULE: | ATP-SENSITIVE INWARD RECTIFIER POTASSIUM CHANNEL |

| | | | | | | | | |
|---|---|---|---|---|---|---|---|---|
| 75:00:00 | 1jad-A | 4 | 3.2 | 109 | 242 | 6 | MOLECULE: | PHOSPHOLIPASE C BETA |
| 76:00:00 | 8pmq-9 | 4 | 4.2 | 79 | 412 | 14 | MOLECULE: | E3 UBIQUITIN-PROTEIN LIGASE RMD5 |
| 77:00:00 | 7qoo-H | 4 | 21.4 | 109 | 210 | 8 | MOLECULE: | CENTROMERE PROTEIN C |
| 78:00:00 | 4p1m-B | 4 | 3.7 | 74 | 107 | 12 | MOLECULE: | CELL DIVISION PROTEIN ZAPA |
| 79:00:00 | 7mge-B | 4 | 6.4 | 82 | 442 | 9 | MOLECULE: | WD REPEAT-CONTAINING PROTEIN 41 |
| 80:00:00 | 8wpe-D | 4 | 4.1 | 63 | 69 | 10 | MOLECULE: | DNA POLYMERASE |
| 81:00:00 | 8afz-B | 4 | 9 | 104 | 376 | 9 | MOLECULE: | SORTING NEXIN-1 |
| 82:00:00 | 6z6o-C | 4 | 14.7 | 104 | 548 | 6 | MOLECULE: | HISTONE DEACETYLASE HDA1 |
| 83:00:00 | 2xqh-A | 4 | 8.7 | 101 | 258 | 4 | MOLECULE: | IMMUNOGLOBULIN-BINDING PROTEIN EIBD |
| 84:00:00 | 5gai-B | 4 | 6.5 | 89 | 721 | 9 | MOLECULE: | PORTAL PROTEIN |
| 85:00:00 | 8aia-K | 4 | 3.9 | 68 | 72 | 3 | MOLECULE: | CRESCENTIN |
| 86:00:00 | 8yxo-C | 4 | 3.3 | 58 | 91 | 10 | MOLECULE: | PHOSPHOPROTEIN |
| 87:00:00 | 8aix-U | 4 | 3.9 | 68 | 72 | 3 | MOLECULE: | CRESCENTIN |
| 88:00:00 | 3jb9-W | 4 | 6.4 | 93 | 426 | 9 | MOLECULE: | PRE-MRNA-SPLICING FACTOR SPP42 |
| 89:00:00 | 7z4f-F | 4 | 7 | 68 | 68 | 10 | MOLECULE: | PUTATIVE STRUCTURAL PROTEIN |
| 90:00:00 | 8ajb-J | 4 | 8.2 | 85 | 120 | 8 | MOLECULE: | CRESCENTIN |
| 91:00:00 | 7e9t-A | 4 | 22.6 | 98 | 371 | 8 | MOLECULE: | SPIKE PROTEIN S2 |
| 92:00:00 | 7w91-K | 4 | 1.6 | 51 | 51 | 6 | MOLECULE: | CENTROSOMAL PROTEIN OF 63 KDA |
| 93:00:00 | 7w91-B | 4 | 1.8 | 52 | 52 | 6 | MOLECULE: | CENTROSOMAL PROTEIN OF 63 KDA |
| 94:00:00 | 1jth-D | 4 | 2.6 | 51 | 51 | 4 | MOLECULE: | SNAP25 |
| 95:00:00 | 5jlf-F | 4 | 7.5 | 96 | 135 | 0 | MOLECULE: | ACTIN, ALPHA SKELETAL MUSCLE |
| 96:00:00 | 6eb8-H | 4 | 4.2 | 83 | 101 | 7 | MOLECULE: | PHOSPHOPROTEIN |
| 97:00:00 | 2fxm-A | 4 | 6.7 | 95 | 126 | 4 | MOLECULE: | MYOSIN HEAVY CHAIN, CARDIAC MUSCLE BETA ISOFORM |
| 98:00:00 | 6id1-K | 4 | 11.1 | 97 | 152 | 7 | MOLECULE: | PRE-MRNA-PROCESSING-SPLICING FACTOR 8 |
| 99:00:00 | 4eot-B | 4 | 4.4 | 67 | 92 | 4 | MOLECULE: | TRANSCRIPTION FACTOR MAFA |
| 0:00 | 5j3d-F | 4 | 4 | 84 | 358 | 6 | MOLECULE: | 14N4 HEAVY CHAIN |
| 1:00 | 6eup-A | 4 | 4 | 82 | 147 | 4 | MOLECULE: | ADHESIN A |
| 2:00 | 3u1c-B | 4 | 5.1 | 82 | 101 | 10 | MOLECULE: | TROPOMYOSIN ALPHA-1 CHAIN |
| 3:00 | 6ign-A | 4 | 6.7 | 90 | 123 | 9 | MOLECULE: | KINESIN-1 HEAVY CHAIN |
| 4:00 | 3ii6-C | 4 | 4.6 | 89 | 201 | 4 | MOLECULE: | DNA REPAIR PROTEIN XRCC4 |
| 5:00 | 6gbp-J | 4 | 4.4 | 63 | 66 | 14 | MOLECULE: | POLYMERASE COFACTOR VP35 |
| 6:00 | 4c5q-B | 4 | 2.9 | 55 | 55 | 7 | MOLECULE: | PHOSPHOPROTEIN |
| 7:00 | 6h9m-B | 4 | 5.1 | 74 | 94 | 4 | MOLECULE: | COILED-COIL DOMAIN-CONTAINING PROTEIN 90B, MITOCH |
| 8:00 | 6h9m-C | 4 | 4.9 | 73 | 94 | 12 | MOLECULE: | COILED-COIL DOMAIN-CONTAINING PROTEIN 90B, MITOCH |
| 9:00 | 7woo-I | 3.9 | 10.2 | 100 | 187 | 9 | MOLECULE: | NUCLEOPORIN NIC96 |
| 10:00 | 4c47-A | 3.9 | 7.8 | 87 | 191 | 8 | MOLECULE: | INNER MEMBRANE LIPOPROTEIN |
| 11:00 | 3edv-A | 3.9 | 6.9 | 108 | 322 | 6 | MOLECULE: | SPECTRIN BETA CHAIN, BRAIN 1 |
| 12:00 | 1ezj-A | 3.9 | 7.2 | 72 | 114 | 10 | MOLECULE: | NUCLEOCAPSID PHOSPHOPROTEIN |
| 13:00 | 8etc-F | 3.9 | 4.7 | 53 | 218 | 2 | MOLECULE: | RNA (2151-MER) |
| 14:00 | 8pda-A | 3.9 | 10.2 | 108 | 973 | 8 | MOLECULE: | ERAD-ASSOCIATED E3 UBIQUITIN-PROTEIN LIGASE DOA10 |
| 15:00 | 6xby-a | 3.9 | 21.8 | 89 | 494 | 2 | MOLECULE: | V-TYPE PROTON ATPASE CATALYTIC SUBUNIT A |
| 16:00 | 8hhf-B | 3.9 | 4 | 72 | 89 | 7 | MOLECULE: | CELL DIVISION PROTEIN FTSQ |
| 17:00 | 7xzj-G | 3.9 | 2.5 | 63 | 73 | 6 | MOLECULE: | CTAP3 |
| 18:00 | 8auc-B | 3.9 | 14.3 | 107 | 493 | 3 | MOLECULE: | CELL WALL-ASSOCIATED HYDROLASES (INVASION-ASSOCIA |
| 19:00 | 6gmh-Q | 3.9 | 9.8 | 103 | 884 | 5 | MOLECULE: | RPB1 |
| 20:00 | 5l4k-S | 3.9 | 16.2 | 135 | 491 | 5 | MOLECULE: | 26S PROTEASOME NON-ATPASE REGULATORY SUBUNIT 4 |
| 21:00 | 8bd7-G | 3.9 | 20.4 | 93 | 205 | 6 | MOLECULE: | IFT88 |
| 22:00 | 4abx-A | 3.9 | 8.8 | 95 | 167 | 7 | MOLECULE: | DNA REPAIR PROTEIN RECN |
| 23:00 | 8cr1-C | 3.9 | 8.2 | 93 | 266 | 4 | MOLECULE: | ATPASE ASNA1 |
| 24:00:00 | 4ilo-A | 3.9 | 3.5 | 104 | 236 | 6 | MOLECULE: | CT398 |
| 25:00:00 | 6u1s-A | 3.9 | 7.7 | 127 | 310 | 7 | MOLECULE: | DE NOVO DESIGNED 16-HELIX TRANSMEMBRANE NANOPORE, |
| 26:00:00 | 8ajb-G | 3.9 | 6.9 | 93 | 155 | 6 | MOLECULE: | CRESCENTIN |
| 27:00:00 | 2efr-A | 3.9 | 8.7 | 97 | 155 | 6 | MOLECULE: | GENERAL CONTROL PROTEIN GCN4 AND TROPOMYOSIN 1 AL |
| 28:00:00 | 4n5b-C | 3.9 | 6 | 69 | 100 | 9 | MOLECULE: | PHOSPHOPROTEIN |
| 29:00:00 | 2n64-A | 3.9 | 3.6 | 72 | 75 | 7 | MOLECULE: | SH3 DOMAIN-CONTAINING KINASE-BINDING PROTEIN 1 |
| 30:00:00 | 1urq-A | 3.9 | 3.3 | 52 | 57 | 8 | MOLECULE: | M-TOMOSYN ISOFORM |

| | | | | | | | | |
|---|---|---|---|---|---|---|---|---|
| 31:00:00 | 4cpc-G | 3.9 | 7.3 | 90 | 152 | 10 | MOLECULE: | SYNAPTONEMAL COMPLEX PROTEIN 3 |
| 32:00:00 | 4bhv-B | 3.9 | 1.8 | 56 | 56 | 5 | MOLECULE: | PHOSPHOPROTEIN |
| 33:00:00 | 8ahl-G | 3.9 | 16.6 | 105 | 155 | 9 | MOLECULE: | CRESCENTIN |
| 34:00:00 | 8ajb-S | 3.9 | 16.2 | 103 | 155 | 8 | MOLECULE: | CRESCENTIN |
| 35:00:00 | 8qu5-q | 3.9 | 7.7 | 86 | 128 | 6 | MOLECULE: | 39S RIBOSOMAL PROTEIN L32, MITOCHONDRIAL |
| 36:00:00 | 6qp4-A | 3.9 | 7.4 | 89 | 157 | 7 | MOLECULE: | USPA1 |
| 37:00:00 | 6w1s-W | 3.9 | 19.8 | 89 | 118 | 7 | MOLECULE: | MEDIATOR OF RNA POLYMERASE II TRANSCRIPTION SUBUN |
| 38:00:00 | 3h7x-E | 3.9 | 2.3 | 53 | 53 | 4 | MOLECULE: | ADHESIN YADA |
| 39:00:00 | 2f8x-M | 3.9 | 4.7 | 55 | 55 | 9 | MOLECULE: | 5'- |
| 40:00:00 | 5wjb-D | 3.9 | 6.1 | 88 | 121 | 7 | MOLECULE: | CAPSID ASSEMBLY SCAFFOLDING PROTEIN,MYOSIN-7 |
| 41:00:00 | 3n4x-D | 3.9 | 4.7 | 77 | 159 | 9 | MOLECULE: | MONOPOLIN COMPLEX SUBUNIT CSM1 |
| 42:00:00 | 5vox-J | 3.9 | 3.7 | 86 | 104 | 5 | MOLECULE: | V-TYPE PROTON ATPASE CATALYTIC SUBUNIT A,V-TYPE P |
| 43:00:00 | 7oad-C | 3.9 | 6.9 | 78 | 80 | 5 | MOLECULE: | GENERAL CONTROL TRANSCRIPTION FACTOR GCN4,CONSERV |
| 44:00:00 | 5to6-B | 3.9 | 6.5 | 97 | 139 | 6 | MOLECULE: | NUCLEOPROTEIN TPR |
| 45:00:00 | 1ysa-D | 3.9 | 1.5 | 55 | 57 | 9 | MOLECULE: | DNA (5'- |
| 46:00:00 | 2efs-D | 3.9 | 7.8 | 90 | 154 | 7 | MOLECULE: | GENERAL CONTROL PROTEIN GCN4 AND TROPOMYOSIN 1 AL |
| 47:00:00 | 2efr-D | 3.9 | 7.6 | 89 | 155 | 4 | MOLECULE: | GENERAL CONTROL PROTEIN GCN4 AND TROPOMYOSIN 1 AL |
| 48:00:00 | 7oac-B | 3.9 | 6.7 | 78 | 81 | 5 | MOLECULE: | GENERAL CONTROL TRANSCRIPTION FACTOR GCN4,CONSERV |
| 49:00:00 | 5t4o-I | 3.9 | 5.5 | 92 | 155 | 4 | MOLECULE: | ATP SYNTHASE SUBUNIT ALPHA |
| 50:00:00 | 5lqy-G | 3.9 | 3.7 | 73 | 267 | 10 | MOLECULE: | ATP SYNTHASE SUBUNIT F |
| 51:00:00 | 5n9j-A | 3.9 | 11.2 | 76 | 566 | 8 | MOLECULE: | MEDIATOR OF RNA POLYMERASE II TRANSCRIPTION SUBUN |
| 52:00:00 | 6dd8-A | 3.9 | 4.2 | 87 | 131 | 10 | MOLECULE: | SYNAPTONEMAL COMPLEX PROTEIN 3 |
| 53:00:00 | 7apk-e | 3.9 | 5.8 | 59 | 512 | 8 | MOLECULE: | THO COMPLEX SUBUNIT 1 |
| 54:00:00 | 7aej-A | 3.9 | 4.9 | 77 | 129 | 10 | MOLECULE: | ENVELOPE GLYCOPROTEIN GP160,ENVELOPE GLYCOPROTEIN |
| 55:00:00 | 2b9c-A | 3.9 | 6.5 | 85 | 136 | 9 | MOLECULE: | STRIATED-MUSCLE ALPHA TROPOMYOSIN |
| 56:00:00 | 7aej-C | 3.9 | 4.3 | 77 | 156 | 12 | MOLECULE: | ENVELOPE GLYCOPROTEIN GP160,ENVELOPE GLYCOPROTEIN |
| 57:00:00 | 3hh0-A | 3.8 | 10.9 | 73 | 134 | 4 | MOLECULE: | TRANSCRIPTIONAL REGULATOR, MERR FAMILY |
| 58:00:00 | 8q85-G | 3.8 | 53.1 | 87 | 178 | 9 | MOLECULE: | KINETOCHORE PROTEIN NDC80 |
| 59:00:00 | 7zu0-C | 3.8 | 3.6 | 53 | 842 | 0 | MOLECULE: | E3 UBIQUITIN-PROTEIN LIGASE PEP5 |
| 60:00:00 | 8q7n-X | 3.8 | 10.5 | 69 | 81 | 3 | MOLECULE: | U5 SNRNA |
| 61:00:00 | 5szj-B | 3.8 | 3.3 | 63 | 145 | 5 | MOLECULE: | RAS-RELATED PROTEIN RAB-10 |
| 62:00:00 | 3viq-A | 3.8 | 11.3 | 84 | 122 | 2 | MOLECULE: | SWI5-DEPENDENT RECOMBINATION DNA REPAIR PROTEIN 1 |
| 63:00:00 | 1owa-A | 3.8 | 20.7 | 96 | 156 | 8 | MOLECULE: | SPECTRIN ALPHA CHAIN, ERYTHROCYTE |
| 64:00:00 | 7egm-H | 3.8 | 11.5 | 88 | 388 | 9 | MOLECULE: | TRANSCRIPTION REGULATORY PROTEIN SNF2 |
| 65:00:00 | 7tdq-A | 3.8 | 11.7 | 76 | 85 | 7 | MOLECULE: | TEGUMENT PROTEIN ORF52 |
| 66:00:00 | 7uh4-A | 3.8 | 2.8 | 74 | 110 | 5 | MOLECULE: | LXG-ASSOCIATED ALPHA-HELICAL PROTEIN D2 |
| 67:00:00 | 4n5b-B | 3.8 | 5.5 | 67 | 100 | 6 | MOLECULE: | PHOSPHOPROTEIN |
| 68:00:00 | 4n5b-E | 3.8 | 5.3 | 68 | 103 | 9 | MOLECULE: | PHOSPHOPROTEIN |
| 69:00:00 | 6wm2-K | 3.8 | 4.8 | 86 | 114 | 5 | MOLECULE: | V-TYPE PROTON ATPASE SUBUNIT E 1 |
| 70:00:00 | 7yv9-A | 3.8 | 8.9 | 96 | 1067 | 7 | MOLECULE: | UNCONVENTIONAL MYOSIN-VA |
| 71:00:00 | 4wij-A | 3.8 | 5.2 | 97 | 298 | 4 | MOLECULE: | SPLICING FACTOR, PROLINE- AND GLUTAMINE-RICH |
| 72:00:00 | 6xra-A | 3.8 | 26.1 | 97 | 348 | 8 | MOLECULE: | SPIKE GLYCOPROTEIN |
| 73:00:00 | 1sfc-H | 3.8 | 3.6 | 61 | 72 | 3 | MOLECULE: | PROTEIN (SYNAPTOBREVIN 2) |
| 74:00:00 | 1g2c-O | 3.8 | 1.7 | 50 | 50 | 8 | MOLECULE: | FUSION PROTEIN (F) |
| 75:00:00 | 1ifk-A | 3.8 | 2.9 | 51 | 51 | 14 | MOLECULE: | INOVIRUS |
| 76:00:00 | 1g2c-I | 3.8 | 1.7 | 50 | 50 | 10 | MOLECULE: | FUSION PROTEIN (F) |
| 77:00:00 | 2d3e-B | 3.8 | 4 | 82 | 130 | 4 | MOLECULE: | GENERAL CONTROL PROTEIN GCN4 AND TROPOMYOSIN 1 AL |
| 78:00:00 | 3j9v-J | 3.8 | 3 | 71 | 105 | 10 | MOLECULE: | V-TYPE PROTON ATPASE SUBUNIT A, VACUOLAR ISOFORM |
| 79:00:00 | 6g2t-N | 3.8 | 8.2 | 94 | 135 | 0 | MOLECULE: | ACTIN, CYTOPLASMIC 2 |
| 80:00:00 | 7jh7-J | 3.8 | 8 | 94 | 135 | 0 | MOLECULE: | ACTIN, ALPHA CARDIAC MUSCLE 1 |
| 81:00:00 | 7tj7-T | 3.8 | 8 | 93 | 135 | 0 | MOLECULE: | CARDIAC ACTIN |
| 82:00:00 | 7tj7-V | 3.8 | 6.9 | 92 | 135 | 0 | MOLECULE: | CARDIAC ACTIN |
| 83:00:00 | 2efs-A | 3.8 | 6.1 | 92 | 154 | 8 | MOLECULE: | GENERAL CONTROL PROTEIN GCN4 AND TROPOMYOSIN 1 AL |
| 84:00:00 | 5vr2-A | 3.8 | 2.3 | 49 | 50 | 10 | MOLECULE: | MYOCILIN |
| 85:00:00 | 4w7y-B | 3.8 | 3.2 | 57 | 64 | 4 | MOLECULE: | B-CELL RECEPTOR-ASSOCIATED PROTEIN 29 |
| 86:00:00 | 6snz-A | 3.8 | 6.8 | 90 | 160 | 9 | MOLECULE: | PRELAMIN-A/C |

| | | | | | | | | |
|---|---|---|---|---|---|---|---|---|
| 87:00:00 | 2oto-C | 3.8 | 8 | 96 | 138 | 7 | MOLECULE: | M PROTEIN |
| 88:00:00 | 2oto-A | 3.8 | 6.7 | 91 | 137 | 8 | MOLECULE: | M PROTEIN |
| 89:00:00 | 3gjo-C | 3.8 | 3.4 | 59 | 65 | 7 | MOLECULE: | MICROTUBULE-ASSOCIATED PROTEIN RP/EB FAMILY MEMBE |
| 90:00:00 | 6eun-B | 3.8 | 2.9 | 75 | 145 | 4 | MOLECULE: | ADHESIN |
| 91:00:00 | 6eun-C | 3.8 | 4 | 76 | 145 | 1 | MOLECULE: | ADHESIN |
| 92:00:00 | 7d7n-A | 3.7 | 6.8 | 70 | 704 | 9 | MOLECULE: | ATP-BINDING CASSETTE SUB-FAMILY B MEMBER 6, MITOC |
| 93:00:00 | 3o1j-A | 3.7 | 6.1 | 106 | 273 | 7 | MOLECULE: | SENSOR PROTEIN TORS |
| 94:00:00 | 8gel-A | 3.7 | 23.6 | 93 | 498 | 2 | MOLECULE: | SC4 |
| 95:00:00 | 8tvh-A | 3.7 | 8.5 | 82 | 86 | 11 | MOLECULE: | 4G5 LIGHT CHAIN |
| 96:00:00 | 6tmi-B | 3.7 | 8.2 | 104 | 246 | 10 | MOLECULE: | ATP SYNTHASE SUBUNIT ALPHA |
| 97:00:00 | 6gvw-F | 3.7 | 9.6 | 97 | 320 | 4 | MOLECULE: | BRCA1-A COMPLEX SUBUNIT ABRAXAS 1 |
| 98:00:00 | 4dyl-A | 3.7 | 2.7 | 82 | 376 | 12 | MOLECULE: | TYROSINE-PROTEIN KINASE FES/FPS |
| 99:00:00 | 3p8c-F | 3.7 | 3.4 | 77 | 156 | 5 | MOLECULE: | CYTOPLASMIC FMR1-INTERACTING PROTEIN 1 |
| 0:00 | 5vo5-A | 3.7 | 5.2 | 90 | 179 | 6 | MOLECULE: | COILED-COIL AND C2 DOMAIN-CONTAINING PROTEIN 1-LI |
| 1:00 | 2fo1-D | 3.7 | 6.1 | 60 | 63 | 10 | MOLECULE: | 5'- |
| 2:00 | 5lc5-Z | 3.7 | 13.4 | 104 | 138 | 3 | MOLECULE: | NADH-UBIQUINONE OXIDOREDUCTASE CHAIN 3 |
| 3:00 | 4r8g-E | 3.7 | 5 | 73 | 324 | 10 | MOLECULE: | UNCONVENTIONAL MYOSIN-IC |
| 4:00 | 5to5-A | 3.7 | 39 | 112 | 141 | 6 | MOLECULE: | NUCLEOPROTEIN TPR |
| 5:00 | 4n5b-G | 3.7 | 5.6 | 67 | 101 | 3 | MOLECULE: | PHOSPHOPROTEIN |
| 6:00 | 2v71-B | 3.7 | 38.2 | 108 | 160 | 8 | MOLECULE: | NUCLEAR DISTRIBUTION PROTEIN NUDE-LIKE 1 |
| 7:00 | 9b8q-I | 3.7 | 6.9 | 94 | 148 | 2 | MOLECULE: | V-TYPE PROTON ATPASE SUBUNIT C 1 |
| 8:00 | 8tek-M | 3.7 | 3.9 | 69 | 164 | 3 | MOLECULE: | DYNEIN REGULATORY COMPLEX PROTEIN 1/2 N-TERMINAL |
| 9:00 | 1h88-B | 3.7 | 3.8 | 60 | 71 | 5 | MOLECULE: | CCAAT/ENHANCER BINDING PROTEIN BETA |
| 10:00 | 7use-C | 3.7 | 3.7 | 72 | 112 | 8 | MOLECULE: | CYTOPLASMIC FMR1-INTERACTING PROTEIN 1 |
| 11:00 | 8t1i-I | 3.7 | 9.3 | 80 | 1086 | 6 | MOLECULE: | MEDIATOR OF RNA POLYMERASE II TRANSCRIPTION SUBUN |
| 12:00 | 2z5h-A | 3.7 | 2.8 | 49 | 49 | 2 | MOLECULE: | GENERAL CONTROL PROTEIN GCN4 AND TROPOMYOSIN ALPH |
| 13:00 | 8ovw-Y | 3.7 | 7.9 | 94 | 223 | 5 | MOLECULE: | CENTROMERE-BINDING PROTEIN 1 |
| 14:00 | 2v71-A | 3.7 | 8.7 | 97 | 160 | 6 | MOLECULE: | NUCLEAR DISTRIBUTION PROTEIN NUDE-LIKE 1 |
| 15:00 | 4c7n-B | 3.7 | 2.3 | 51 | 51 | 6 | MOLECULE: | MICROPHTHALMIA ASSOCIATED TRANSCRIPTION FACTOR |
| 16:00 | 8q3v-A | 3.7 | 4.6 | 56 | 61 | 16 | MOLECULE: | TETRAHYDROMETHANOPTERIN S-METHYLTRANSFERASE SUBUN |
| 17:00 | 5mqf-J | 3.7 | 6.9 | 76 | 135 | 9 | MOLECULE: | PRE-MRNA-PROCESSING-SPLICING FACTOR 8 |
| 18:00 | 6cxj-T | 3.7 | 7.8 | 93 | 127 | 0 | MOLECULE: | ACTIN, CYTOPLASMIC 2 |
| 19:00 | 6cxi-T | 3.7 | 7.8 | 93 | 127 | 0 | MOLECULE: | ACTIN, CYTOPLASMIC 2 |
| 20:00 | 6xbw-O | 3.7 | 4.6 | 82 | 108 | 2 | MOLECULE: | V-TYPE PROTON ATPASE CATALYTIC SUBUNIT A |
| 21:00 | 4dl0-G | 3.7 | 3.4 | 84 | 105 | 5 | MOLECULE: | V-TYPE PROTON ATPASE SUBUNIT C |
| 22:00 | 8q3v-Q | 3.7 | 5 | 57 | 61 | 16 | MOLECULE: | TETRAHYDROMETHANOPTERIN S-METHYLTRANSFERASE SUBUN |
| 23:00 | 5jx1-A | 3.7 | 2.3 | 55 | 64 | 4 | MOLECULE: | CHIMERA PROTEIN OF KINESIN-LIKE PROTEIN KIF3A AND |
| 24:00:00 | 2n1t-D | 3.7 | 4 | 62 | 74 | 2 | MOLECULE: | VESICLE-ASSOCIATED MEMBRANE PROTEIN 2 |
| 25:00:00 | 6mi3-A | 3.7 | 6.7 | 80 | 122 | 8 | MOLECULE: | NF-KB ESSENTIAL MODULATOR,NF-KAPPA-B ESSENTIAL MO |
| 26:00:00 | 7oah-C | 3.7 | 8 | 74 | 82 | 4 | MOLECULE: | GENERAL CONTROL TRANSCRIPTION FACTOR GCN4,CONSERV |
| 27:00:00 | 3wpr-C | 3.7 | 2.8 | 54 | 218 | 6 | MOLECULE: | TRIMERIC AUTOTRANSPORTER ADHESIN |
| 28:00:00 | 4mvd-C | 3.7 | 14.7 | 93 | 253 | 6 | MOLECULE: | CHOLINE-PHOSPHATE CYTIDYLYLTRANSFERASE A |
| 29:00:00 | 8es7-B | 3.6 | 2.2 | 41 | 287 | 7 | MOLECULE: | T-CELL SURFACE GLYCOPROTEIN CD3 ZETA CHAIN |
| 30:00:00 | 3a6m-A | 3.6 | 24.2 | 101 | 168 | 8 | MOLECULE: | PROTEIN GRPE |
| 31:00:00 | 8th8-B | 3.6 | 50 | 86 | 276 | 3 | MOLECULE: | DYNEIN REGULATORY COMPLEX PROTEIN 1/2 N-TERMINAL |
| 32:00:00 | 7aw7-D | 3.6 | 4.3 | 67 | 93 | 3 | MOLECULE: | HAPB |
| 33:00:00 | 6zz6-B | 3.6 | 36.8 | 83 | 423 | 7 | MOLECULE: | STRUCTURAL MAINTENANCE OF CHROMOSOMES PROTEIN 1,S |
| 34:00:00 | 5t96-B | 3.6 | 2.1 | 41 | 330 | 0 | MOLECULE: | HE PROTEIN |
| 35:00:00 | 5gox-A | 3.6 | 6.8 | 95 | 181 | 11 | MOLECULE: | DNA REPAIR PROTEIN RAD50 |
| 36:00:00 | 3ub0-D | 3.6 | 12.5 | 83 | 194 | 6 | MOLECULE: | NON-STRUCTURAL PROTEIN 6, NSP6, |
| 37:00:00 | 6sp2-A | 3.6 | 7.7 | 71 | 366 | 4 | MOLECULE: | MEMBRANE PROTEIN TMS1D |
| 38:00:00 | 8b9z-Z | 3.6 | 14.4 | 97 | 146 | 4 | MOLECULE: | NADH-UBIQUINONE OXIDOREDUCTASE CHAIN 3 |
| 39:00:00 | 5gw1-C | 3.6 | 10.5 | 86 | 176 | 10 | MOLECULE: | SORTING NEXIN-16 |
| 40:00:00 | 5nmo-A | 3.6 | 6.4 | 84 | 162 | 6 | MOLECULE: | CHROMOSOME PARTITION PROTEIN SMC,CHROMOSOME PARTI |
| 41:00:00 | 6wcj-B | 3.6 | 1.3 | 51 | 109 | 4 | MOLECULE: | CLATHRIN HEAVY CHAIN 1 |
| 42:00:00 | 4m8m-B | 3.6 | 5.5 | 84 | 574 | 5 | MOLECULE: | GCN4 COILED-COIL FUSED ZEBRAFISH PLEXINC1 |

| | | | | | | | | |
|---|---|---|---|---|---|---|---|---|
| 43:00:00 | 7zmg-n | 3.6 | 12.1 | 75 | 136 | 9 | MOLECULE: | NADH-UBIQUINONE OXIDOREDUCTASE CHAIN 1 |
| 44:00:00 | 8x5f-A | 3.6 | 7.9 | 80 | 616 | 5 | MOLECULE: | SOLUTE CARRIER FAMILY 53 MEMBER 1 |
| 45:00:00 | 5o31-Z | 3.6 | 12.9 | 104 | 137 | 3 | MOLECULE: | NADH-UBIQUINONE OXIDOREDUCTASE CHAIN 3 |
| 46:00:00 | 6ch2-E | 3.6 | 4.2 | 62 | 147 | 8 | MOLECULE: | FLAGELLAR BIOSYNTHESIS PROTEIN FLHA |
| 47:00:00 | 7tit-N | 3.6 | 55.7 | 119 | 135 | 0 | MOLECULE: | CARDIAC ACTIN |
| 48:00:00 | 7qoo-K | 3.6 | 17.2 | 90 | 253 | 9 | MOLECULE: | CENTROMERE PROTEIN C |
| 49:00:00 | 8hhh-B | 3.6 | 4.6 | 74 | 89 | 9 | MOLECULE: | CELL DIVISION PROTEIN FTSQ |
| 50:00:00 | 7x6g-D | 3.6 | 4.8 | 80 | 152 | 14 | MOLECULE: | QUORUM-SENSING REGULATOR PROTEIN G |
| 51:00:00 | 7jg8-d | 3.6 | 9.2 | 107 | 428 | 10 | MOLECULE: | ATP SYNTHASE SUBUNIT ALPHA |
| 52:00:00 | 2e7s-A | 3.6 | 3.3 | 76 | 114 | 9 | MOLECULE: | RAB GUANINE NUCLEOTIDE EXCHANGE FACTOR SEC2 |
| 53:00:00 | 3lj5-A | 3.6 | 4 | 91 | 692 | 5 | MOLECULE: | PORTAL PROTEIN |
| 54:00:00 | 5jlh-K | 3.6 | 7.2 | 91 | 135 | 0 | MOLECULE: | ACTIN, CYTOPLASMIC 2 |
| 55:00:00 | 7pkq-w | 3.6 | 7.7 | 88 | 155 | 7 | MOLECULE: | MS35 |
| 56:00:00 | 3o0z-A | 3.6 | 7.8 | 97 | 152 | 10 | MOLECULE: | RHO-ASSOCIATED PROTEIN KINASE 1 |
| 57:00:00 | 4jgs-C | 3.6 | 2.9 | 62 | 88 | 10 | MOLECULE: | MLV-RELATED PROVIRAL ENV POLYPROTEIN |
| 58:00:00 | 7oaf-A | 3.6 | 6.8 | 72 | 81 | 6 | MOLECULE: | GENERAL CONTROL TRANSCRIPTION FACTOR GCN4,CONSERV |
| 59:00:00 | 3wpr-B | 3.6 | 5.4 | 67 | 207 | 3 | MOLECULE: | TRIMERIC AUTOTRANSPORTER ADHESIN |
| 60:00:00 | 3gjo-A | 3.6 | 3.3 | 60 | 66 | 7 | MOLECULE: | MICROTUBULE-ASSOCIATED PROTEIN RP/EB FAMILY MEMBE |
| 61:00:00 | 4egx-A | 3.6 | 2 | 54 | 180 | 7 | MOLECULE: | KINESIN-LIKE PROTEIN KIF1A |
| 62:00:00 | 4mh6-A | 3.5 | 20.8 | 92 | 159 | 4 | MOLECULE: | PUTATIVE TYPE III SECRETION PROTEIN YSCO |
| 63:00:00 | 8qfs-A | 3.5 | 30.4 | 99 | 1183 | 4 | MOLECULE: | ELONGATION FACTOR TU |
| 64:00:00 | 5ncl-A | 3.5 | 9.9 | 73 | 418 | 7 | MOLECULE: | SERINE/THREONINE-PROTEIN KINASE CBK1 |
| 65:00:00 | 6em5-t | 3.5 | 16 | 84 | 290 | 8 | MOLECULE: | 5.8S RIBOSOMAL RNA |
| 66:00:00 | 6kn7-U | 3.5 | 8.3 | 114 | 170 | 2 | MOLECULE: | ACTIN, ALPHA SKELETAL MUSCLE |
| 67:00:00 | 7yoj-A | 3.5 | 3.1 | 66 | 867 | 3 | MOLECULE: | CASPI |
| 68:00:00 | 5n9j-E | 3.5 | 16.1 | 98 | 201 | 6 | MOLECULE: | MEDIATOR OF RNA POLYMERASE II TRANSCRIPTION SUBUN |
| 69:00:00 | 6ted-R | 3.5 | 11.2 | 81 | 244 | 9 | MOLECULE: | DNA-DIRECTED RNA POLYMERASE SUBUNIT |
| 70:00:00 | 5che-A | 3.5 | 14.5 | 112 | 427 | 6 | MOLECULE: | GLUTAMYL-TRNA REDUCTASE 1, CHLOROPLASTIC |
| 71:00:00 | 5cd4-I | 3.5 | 3.6 | 75 | 494 | 7 | MOLECULE: | CRISPR SYSTEM CASCADE SUBUNIT CASE |
| 72:00:00 | 6nyi-A | 3.5 | 3.2 | 65 | 97 | 5 | MOLECULE: | DESIGN CONSTRUCT XXA |
| 73:00:00 | 6wuc-H | 3.5 | 13.3 | 85 | 179 | 6 | MOLECULE: | INNER KINETOCHORE SUBUNIT MCM16 |
| 74:00:00 | 5n9j-Z | 3.5 | 3.6 | 58 | 119 | 7 | MOLECULE: | MEDIATOR OF RNA POLYMERASE II TRANSCRIPTION SUBUN |
| 75:00:00 | 7wkk-A | 3.5 | 10.4 | 95 | 1684 | 8 | MOLECULE: | MGC83295 PROTEIN |
| 76:00:00 | 4abn-B | 3.5 | 2.4 | 56 | 426 | 11 | MOLECULE: | TETRATRICOPEPTIDE REPEAT PROTEIN 5 |
| 77:00:00 | 6ynw-H | 3.5 | 1.8 | 60 | 75 | 10 | MOLECULE: | SUBUNIT C |
| 78:00:00 | 5yfp-H | 3.5 | 21.4 | 75 | 518 | 11 | MOLECULE: | EXOCYST COMPLEX COMPONENT SEC3 |
| 79:00:00 | 5vk5-B | 3.5 | 6.8 | 79 | 282 | 10 | MOLECULE: | POTASSIUM CHANNEL SUBFAMILY K MEMBER 2 |
| 80:00:00 | 7uus-Q | 3.5 | 9.2 | 64 | 170 | 6 | MOLECULE: | HYDROGENASE-2, LARGE SUBUNIT |
| 81:00:00 | 6iac-E | 3.5 | 11.9 | 98 | 146 | 6 | MOLECULE: | PORTAL PROTEIN |
| 82:00:00 | 7jw1-e | 3.5 | 10.6 | 123 | 235 | 13 | MOLECULE: | CAPSID PROTEINS |
| 83:00:00 | 8w8q-C | 3.5 | 8.2 | 74 | 287 | 5 | MOLECULE: | SOLUBLE CYTOCHROME B562,PROBABLE G-PROTEIN COUPLE |
| 84:00:00 | 8exh-a | 3.5 | 3.9 | 60 | 69 | 5 | MOLECULE: | PROTEIN VIRB2 |
| 85:00:00 | 3o0z-D | 3.5 | 8 | 95 | 160 | 6 | MOLECULE: | RHO-ASSOCIATED PROTEIN KINASE 1 |
| 86:00:00 | 8q4g-H | 3.5 | 8.9 | 97 | 179 | 4 | MOLECULE: | ACTIN, ALPHA CARDIAC MUSCLE 1 |
| 87:00:00 | 2oto-B | 3.5 | 57 | 121 | 138 | 7 | MOLECULE: | M PROTEIN |
| 88:00:00 | 4tt1-B | 3.5 | 7.6 | 95 | 129 | 3 | MOLECULE: | DENEDDYLASE |
| 89:00:00 | 8q4g-G | 3.5 | 9.1 | 96 | 179 | 5 | MOLECULE: | ACTIN, ALPHA CARDIAC MUSCLE 1 |
| 90:00:00 | 3eff-K | 3.5 | 4.7 | 76 | 139 | 7 | MOLECULE: | FAB |
| 91:00:00 | 1czq-A | 3.5 | 1.8 | 45 | 45 | 2 | MOLECULE: | FUSION PROTEIN BETWEEN THE HYDROPHOBIC POCKET OF |
| 92:00:00 | 3vmx-A | 3.5 | 1.1 | 45 | 48 | 4 | MOLECULE: | VOLTAGE-GATED HYDROGEN CHANNEL 1 |
| 93:00:00 | 5yr0-B | 3.5 | 2 | 44 | 44 | 7 | MOLECULE: | BECLIN-1 |
| 94:00:00 | 8gwe-B | 3.5 | 4.3 | 79 | 190 | 6 | MOLECULE: | RNA-DIRECTED RNA POLYMERASE |
| 95:00:00 | 8t1i-W | 3.5 | 20.8 | 85 | 118 | 12 | MOLECULE: | MEDIATOR OF RNA POLYMERASE II TRANSCRIPTION SUBUN |
| 96:00:00 | 5bvz-A | 3.5 | 5 | 83 | 180 | 6 | MOLECULE: | DNA STABILIZATION PROTEIN |
| 97:00:00 | 5jlh-I | 3.5 | 7.2 | 91 | 135 | 0 | MOLECULE: | ACTIN, CYTOPLASMIC 2 |
| 98:00:00 | 5nog-G | 3.5 | 7.2 | 91 | 135 | 0 | MOLECULE: | CARDIAC MUSCLE ALPHA ACTIN 1 |

| | | | | | | | | |
|---|---|---|---|---|---|---|---|---|
| 99:00:00 | 4a7l-B | 3.5 | 7.2 | 91 | 136 | 10 | MOLECULE: | ACTIN, ALPHA SKELETON MUSCLE |
| 0:00 | 3j4k-F | 3.5 | 7.2 | 91 | 136 | 0 | MOLECULE: | ACTIN, ALPHA SKELETAL MUSCLE |
| 1:00 | 4a7f-B | 3.5 | 7.2 | 91 | 136 | 10 | MOLECULE: | ACTIN, ALPHA SKELETAL MUSCLE |
| 2:00 | 8q3v-a | 3.5 | 4.8 | 58 | 61 | 2 | MOLECULE: | TETRAHYDROMETHANOPTERIN S-METHYLTRANSFERASE SUBUN |
| 3:00 | 4tko-B | 3.5 | 9.4 | 105 | 326 | 8 | MOLECULE: | EMRA |
| 4:00 | 5nen-A | 3.5 | 6.5 | 100 | 266 | 8 | MOLECULE: | LIPASE C |
| 5:00 | 7apk-M | 3.5 | 59.2 | 93 | 549 | 8 | MOLECULE: | THO COMPLEX SUBUNIT 1 |
| 6:00 | 7oaf-B | 3.5 | 6.5 | 72 | 81 | 6 | MOLECULE: | GENERAL CONTROL TRANSCRIPTION FACTOR GCN4,CONSERV |
| 7:00 | 7oaf-C | 3.5 | 6.3 | 74 | 81 | 3 | MOLECULE: | GENERAL CONTROL TRANSCRIPTION FACTOR GCN4,CONSERV |
| 8:00 | 6mi3-B | 3.5 | 6.4 | 78 | 124 | 4 | MOLECULE: | NF-KB ESSENTIAL MODULATOR,NF-KAPPA-B ESSENTIAL MO |
| 9:00 | 7oac-C | 3.5 | 7.9 | 72 | 81 | 4 | MOLECULE: | GENERAL CONTROL TRANSCRIPTION FACTOR GCN4,CONSERV |
| 10:00 | 2pms-C | 3.4 | 21.1 | 88 | 109 | 10 | MOLECULE: | LACTOTRANSFERRIN |
| 11:00 | 8cra-A | 3.4 | 15.8 | 76 | 96 | 7 | MOLECULE: | FLORAL HOMEOTIC PROTEIN AGAMOUS |
| 12:00 | 6w1s-Q | 3.4 | 38 | 76 | 131 | 5 | MOLECULE: | MEDIATOR OF RNA POLYMERASE II TRANSCRIPTION SUBUN |
| 13:00 | 8vxq-A | 3.4 | 17.7 | 79 | 155 | 5 | MOLECULE: | GP72 |
| 14:00 | 6gap-A | 3.4 | 61.3 | 121 | 218 | 5 | MOLECULE: | OUTER CAPSID PROTEIN SIGMA-1 |
| 15:00 | 7qoo-Q | 3.4 | 10.6 | 75 | 209 | 8 | MOLECULE: | CENTROMERE PROTEIN C |
| 16:00 | 7wkk-I | 3.4 | 29.3 | 101 | 171 | 12 | MOLECULE: | MGC83295 PROTEIN |
| 17:00 | 7emf-G | 3.4 | 7.5 | 88 | 161 | 7 | MOLECULE: | MEDIATOR OF RNA POLYMERASE II TRANSCRIPTION SUBUN |
| 18:00 | 4cq4-A | 3.4 | 3.1 | 73 | 106 | 11 | MOLECULE: | ENGINEERED VERSION OF TRANSMEMBRANE RECEPTOR AF15 |
| 19:00 | 4cgk-A | 3.4 | 9.4 | 101 | 351 | 6 | MOLECULE: | SECRETED 45 KDA PROTEIN |
| 20:00 | 7c4j-F | 3.4 | 4.5 | 75 | 383 | 9 | MOLECULE: | TRANSCRIPTION REGULATORY PROTEIN SNF12 |
| 21:00 | 4whj-A | 3.4 | 6.7 | 88 | 565 | 10 | MOLECULE: | INTERFERON-INDUCED GTP-BINDING PROTEIN MX2 |
| 22:00 | 3nr7-A | 3.4 | 4.9 | 64 | 81 | 9 | MOLECULE: | DNA-BINDING PROTEIN H-NS |
| 23:00 | 8hqo-x | 3.4 | 8 | 75 | 80 | 7 | MOLECULE: | PORTAL PROTEIN |
| 24:00:00 | 8d8k-V | 3.4 | 6 | 83 | 233 | 10 | MOLECULE: | PROBABLE S-ADENOSYL-L-METHIONINE-DEPENDENT RNA |
| 25:00:00 | 3k66-A | 3.4 | 3.7 | 91 | 217 | 5 | MOLECULE: | BETA-AMYLOID-LIKE PROTEIN |
| 26:00:00 | 8cqn-B | 3.4 | 7.3 | 97 | 187 | 13 | MOLECULE: | LIPOPROTEIN, PUTATIVE |
| 27:00:00 | 7vv6-R | 3.4 | 5.8 | 78 | 271 | 5 | MOLECULE: | MAS-RELATED G-PROTEIN COUPLED RECEPTOR MEMBER X2 |
| 28:00:00 | 2xeq-C | 3.4 | 2.7 | 57 | 241 | 7 | MOLECULE: | PAT1 HOMOLOG 1, |
| 29:00:00 | 7v2w-I | 3.4 | 41.9 | 124 | 239 | 7 | MOLECULE: | THO COMPLEX SUBUNIT HPR1 |
| 30:00:00 | 9b8o-U | 3.4 | 7.1 | 60 | 214 | 7 | MOLECULE: | ATPASE H+-TRANSPORTING V1 SUBUNIT D |
| 31:00:00 | 7sc0-A | 3.4 | 13.1 | 75 | 129 | 3 | MOLECULE: | CAVEOLIN-1 |
| 32:00:00 | 6qm5-A | 3.4 | 17.7 | 100 | 673 | 4 | MOLECULE: | PREDICTED PROTEIN |
| 33:00:00 | 7zmb-3 | 3.4 | 20.9 | 79 | 130 | 8 | MOLECULE: | NADH-UBIQUINONE OXIDOREDUCTASE CHAIN 1 |
| 34:00:00 | 3oja-A | 3.4 | 11.7 | 98 | 482 | 4 | MOLECULE: | LEUCINE-RICH IMMUNE MOLECULE 1 |
| 35:00:00 | 7w01-A | 3.4 | 15.4 | 105 | 1598 | 8 | MOLECULE: | PHOSPHOLIPID-TRANSPORTING ATPASE ABCA3 |
| 36:00:00 | 7m2w-F | 3.4 | 2.7 | 55 | 674 | 9 | MOLECULE: | TUBULIN GAMMA CHAIN |
| 37:00:00 | 6c48-D | 3.4 | 3.1 | 61 | 87 | 13 | MOLECULE: | PROTEIN LIN-9 HOMOLOG |
| 38:00:00 | 7cun-D | 3.4 | 15.3 | 93 | 827 | 5 | MOLECULE: | INTEGRATOR COMPLEX SUBUNIT 1 |
| 39:00:00 | 4n5b-F | 3.4 | 6 | 70 | 101 | 4 | MOLECULE: | PHOSPHOPROTEIN |
| 40:00:00 | 6w4f-D | 3.4 | 10.9 | 77 | 198 | 10 | MOLECULE: | APOLIPOPROTEIN A-I |
| 41:00:00 | 7fdb-G | 3.4 | 7.2 | 92 | 225 | 4 | MOLECULE: | YEAST VACUOLAR ATPASE A SUBUNIT |
| 42:00:00 | 5uky-A | 3.4 | 2.8 | 62 | 74 | 8 | MOLECULE: | ATP-BINDING PROTEIN |
| 43:00:00 | 7tit-P | 3.4 | 55.7 | 117 | 135 | 0 | MOLECULE: | CARDIAC ACTIN |
| 44:00:00 | 6cfz-I | 3.4 | 11.4 | 87 | 106 | 6 | MOLECULE: | ASK1 |
| 45:00:00 | 3vyi-F | 3.4 | 1.4 | 45 | 49 | 2 | MOLECULE: | VOLTAGE-GATED HYDROGEN CHANNEL 1 |
| 46:00:00 | 1w2e-A | 3.4 | 2.9 | 54 | 91 | 6 | MOLECULE: | ZAPA |
| 47:00:00 | 5nvu-A | 3.4 | 6.6 | 99 | 3169 | 0 | MOLECULE: | DYNEIN MOTOR DOMAIN |
| 48:00:00 | 8aud-A | 3.4 | 6 | 91 | 350 | 9 | MOLECULE: | CELL WALL-ASSOCIATED HYDROLASES (INVASION-ASSOCIA |
| 49:00:00 | 8fed-A | 3.4 | 14.5 | 83 | 392 | 4 | MOLECULE: | VIRULENCE FACTOR MCE FAMILY PROTEIN |
| 50:00:00 | 1ebo-F | 3.4 | 19.4 | 89 | 113 | 6 | MOLECULE: | EBOLA VIRUS ENVELOPE PROTEIN CHIMERA CONSISTING |
| 51:00:00 | 3wpo-A | 3.4 | 26.9 | 96 | 193 | 6 | MOLECULE: | TRIMERIC AUTOTRANSPORTER ADHESIN |
| 52:00:00 | 4xa1-A | 3.4 | 17.2 | 94 | 153 | 9 | MOLECULE: | GP7-MYH7(1173-1238)-EB1 CHIMERA PROTEIN |
| 53:00:00 | 4egx-C | 3.4 | 2.6 | 55 | 174 | 4 | MOLECULE: | KINESIN-LIKE PROTEIN KIF1A |
| 54:00:00 | 6a70-B | 3.3 | 12.9 | 73 | 704 | 3 | MOLECULE: | POLYCYSTIN-2 |

| | | | | | | | | |
|---|---|---|---|---|---|---|---|---|
| 55:00:00 | 5ijn-G | 3.3 | 26.9 | 99 | 171 | 8 | MOLECULE: | NUCLEAR PORE COMPLEX PROTEIN NUP155 |
| 56:00:00 | 8k9t-A | 3.3 | 31.7 | 106 | 961 | 5 | MOLECULE: | GPI INOSITOL-DEACYLASE,MCHERRY PROTEIN |
| 57:00:00 | 7qo4-J | 3.3 | 4.2 | 60 | 405 | 7 | MOLECULE: | 26S PROTEASOME REGULATORY SUBUNIT RPN1 |
| 58:00:00 | 8bjw-F | 3.3 | 5 | 69 | 97 | 12 | MOLECULE: | PROTEIN C |
| 59:00:00 | 2m67-A | 3.3 | 3.2 | 61 | 81 | 8 | MOLECULE: | MERF |
| 60:00:00 | 8tdl-A | 3.3 | 10.1 | 76 | 364 | 11 | MOLECULE: | MECHANOSENSITIVE ION CHANNEL PROTEIN 10 |
| 61:00:00 | 1t7s-A | 3.3 | 3.2 | 63 | 129 | 13 | MOLECULE: | BAG-1 COCHAPERONE |
| 62:00:00 | 7xzi-A | 3.3 | 46.6 | 118 | 1598 | 9 | MOLECULE: | CTAP3 |
| 63:00:00 | 6c14-B | 3.3 | 2.7 | 55 | 164 | 7 | MOLECULE: | PROTOCADHERIN-15 |
| 64:00:00 | 7t6d-A | 3.3 | 3.4 | 58 | 389 | 7 | MOLECULE: | LIPOPOLYSACCHARIDE ASSEMBLY PROTEIN B |
| 65:00:00 | 7v2c-d | 3.3 | 9.7 | 66 | 175 | 5 | MOLECULE: | NADH DEHYDROGENASE [UBIQUINONE] FLAVOPROTEIN 1, |
| 66:00:00 | 4gcz-B | 3.3 | 7.8 | 90 | 378 | 9 | MOLECULE: | BLUE-LIGHT PHOTORECEPTOR, SENSOR PROTEIN FIXL |
| 67:00:00 | 6xf1-A | 3.3 | 4.8 | 96 | 230 | 6 | MOLECULE: | NESPRIN-2 |
| 68:00:00 | 7zn7-D | 3.3 | 3.1 | 58 | 128 | 9 | MOLECULE: | DNA DAMAGE-BINDING PROTEIN 1 |
| 69:00:00 | 7kzn-B | 3.3 | 15.6 | 106 | 683 | 6 | MOLECULE: | HEAVY CHAIN ALPHA |
| 70:00:00 | 4mer-D | 3.3 | 5.2 | 68 | 97 | 10 | MOLECULE: | STREPTOCOCCAL HISTIDINE-RICH GLYCOPROTEIN INTERAC |
| 71:00:00 | 8qfc-C | 3.3 | 4.9 | 101 | 380 | 13 | MOLECULE: | 60S RIBOSOMAL PROTEIN L10A |
| 72:00:00 | 7pgg-A | 3.3 | 7.6 | 81 | 147 | 9 | MOLECULE: | ION TRANSPORT PROTEIN |
| 73:00:00 | 8f7c-A | 3.3 | 9.7 | 72 | 287 | 6 | MOLECULE: | PANNEXIN-2, SOLUBLE CYTOCHROME B562 FUSION |
| 74:00:00 | 7v2w-J | 3.3 | 6.4 | 80 | 236 | 5 | MOLECULE: | THO COMPLEX SUBUNIT HPR1 |
| 75:00:00 | 6hgc-A | 3.3 | 10.5 | 78 | 304 | 1 | MOLECULE: | UBIQUITIN CARBOXYL-TERMINAL HYDROLASE CALYPSO,UBI |
| 76:00:00 | 6wg3-B | 3.3 | 12.4 | 108 | 693 | 5 | MOLECULE: | STRUCTURAL MAINTENANCE OF CHROMOSOMES PROTEIN 1A |
| 77:00:00 | 6rlb-A | 3.3 | 18 | 129 | 969 | 6 | MOLECULE: | O6-ALKYLGUANINE-DNA ALKYLTRANSFERASE MUTANT,DYNC2 |
| 78:00:00 | 6ogd-E | 3.3 | 13.7 | 121 | 1078 | 7 | MOLECULE: | TOXIN SUBUNIT YENA1 |
| 79:00:00 | 8aia-G | 3.3 | 6.3 | 95 | 183 | 7 | MOLECULE: | CRESCENTIN |
| 80:00:00 | 8c5v-I | 3.3 | 53.2 | 107 | 516 | 7 | MOLECULE: | CHEMOTAXIS PROTEIN CHEA |
| 81:00:00 | 8ovw-Q | 3.3 | 26.5 | 112 | 258 | 8 | MOLECULE: | CENTROMERE-BINDING PROTEIN 1 |
| 82:00:00 | 7z4a-J | 3.3 | 40.7 | 89 | 588 | 10 | MOLECULE: | ADAPTOR PROTEIN |
| 83:00:00 | 4mer-A | 3.3 | 7.4 | 76 | 96 | 8 | MOLECULE: | STREPTOCOCCAL HISTIDINE-RICH GLYCOPROTEIN INTERAC |
| 84:00:00 | 8ovw-Z | 3.3 | 24.2 | 101 | 151 | 10 | MOLECULE: | CENTROMERE-BINDING PROTEIN 1 |
| 85:00:00 | 1gmj-D | 3.3 | 1.8 | 52 | 56 | 10 | MOLECULE: | ATPASE INHIBITOR |
| 86:00:00 | 3oja-B | 3.3 | 5.8 | 123 | 534 | 11 | MOLECULE: | LEUCINE-RICH IMMUNE MOLECULE 1 |
| 87:00:00 | 5cws-E | 3.3 | 26.6 | 111 | 241 | 7 | MOLECULE: | SAB-158 FAB LIGHT CHAIN |
| 88:00:00 | 2e7s-E | 3.3 | 6.9 | 82 | 124 | 5 | MOLECULE: | RAB GUANINE NUCLEOTIDE EXCHANGE FACTOR SEC2 |
| 89:00:00 | 5z7g-C | 3.3 | 1.5 | 43 | 43 | 7 | MOLECULE: | TAX1-BINDING PROTEIN 1 |
| 90:00:00 | 4nqi-D | 3.3 | 4 | 87 | 232 | 5 | MOLECULE: | SH3 DOMAIN-CONTAINING PROTEIN |
| 91:00:00 | 7oaa-A | 3.3 | 9.2 | 76 | 82 | 7 | MOLECULE: | GENERAL CONTROL TRANSCRIPTION FACTOR GCN4,CONSERV |
| 92:00:00 | 8th8-A | 3.2 | 23.9 | 103 | 290 | 6 | MOLECULE: | DYNEIN REGULATORY COMPLEX PROTEIN 1/2 N-TERMINAL |
| 93:00:00 | 6jy0-A | 3.2 | 25 | 109 | 410 | 6 | MOLECULE: | FLAGELLIN |
| 94:00:00 | 2d1l-A | 3.2 | 4.7 | 93 | 249 | 6 | MOLECULE: | METASTASIS SUPPRESSOR PROTEIN 1 |
| 95:00:00 | 6tpi-A | 3.2 | 23.4 | 94 | 380 | 14 | MOLECULE: | MUREIN HYDROLASE ACTIVATOR ENVC |
| 96:00:00 | 6oqw-Y | 3.2 | 52.1 | 90 | 155 | 8 | MOLECULE: | ATP SYNTHASE SUBUNIT DELTA |
| 97:00:00 | 6gh5-M | 3.2 | 5.5 | 50 | 338 | 8 | MOLECULE: | DNA-DIRECTED RNA POLYMERASE SUBUNIT ALPHA |
| 98:00:00 | 8eja-A | 3.2 | 3.1 | 65 | 113 | 8 | MOLECULE: | ABAK |
| 99:00:00 | 6xxv-C | 3.2 | 3.2 | 62 | 111 | 8 | MOLECULE: | ANTIBODY C57, HEAVY CHAIN |
| 0:00 | 7edr-D | 3.2 | 9.5 | 71 | 122 | 8 | MOLECULE: | MOESIN/EZRIN/RADIXIN HOMOLOG 2 |
| 1:00 | 4wfc-B | 3.2 | 3.9 | 57 | 118 | 4 | MOLECULE: | EXOSOME COMPLEX EXONUCLEASE RRP6 |
| 2:00 | 6xdc-A | 3.2 | 3.5 | 57 | 193 | 12 | MOLECULE: | PROTEIN 3A |
| 3:00 | 5td8-C | 3.2 | 10.4 | 77 | 111 | 9 | MOLECULE: | KINETOCHORE PROTEIN NDC80 |
| 4:00 | 5c21-A | 3.2 | 3 | 83 | 267 | 7 | MOLECULE: | CHROMOSOMAL HEMOLYSIN D |
| 5:00 | 4gx2-B | 3.2 | 2.1 | 56 | 548 | 9 | MOLECULE: | TRKA DOMAIN PROTEIN |
| 6:00 | 6gmh-M | 3.2 | 6.4 | 68 | 991 | 7 | MOLECULE: | RPB1 |
| 7:00 | 8oyy-A | 3.2 | 11.9 | 67 | 233 | 1 | MOLECULE: | DE NOVO DESIGNED SOLUBLE GPCR-LIKE PROTEIN |
| 8:00 | 1s35-A | 3.2 | 4.8 | 89 | 211 | 10 | MOLECULE: | SPECTRIN BETA CHAIN, ERYTHROCYTE |
| 9:00 | 6k7x-C | 3.2 | 15.1 | 99 | 276 | 6 | MOLECULE: | CALCIUM UNIPORTER PROTEIN, MITOCHONDRIAL |
| 10:00 | 8hmy-B | 3.2 | 31.1 | 88 | 264 | 5 | MOLECULE: | TRNA-SPLICING ENDONUCLEASE SUBUNIT SEN2 |

| | | | | | | | | |
|---|---|---|---|---|---|---|---|---|
| 11:00 | 4u6u-D | 3.2 | 8.7 | 86 | 283 | 12 | MOLECULE: | COG7 |
| 12:00 | 7cgp-C | 3.2 | 12.3 | 95 | 223 | 11 | MOLECULE: | MITOCHONDRIAL IMPORT INNER MEMBRANE TRANSLOCASE S |
| 13:00 | 7khw-a | 3.2 | 3.4 | 55 | 174 | 4 | MOLECULE: | TRANSLOCON ESPA |
| 14:00 | 5yfp-B | 3.2 | 10.5 | 101 | 927 | 8 | MOLECULE: | EXOCYST COMPLEX COMPONENT SEC3 |
| 15:00 | 7pv1-D | 3.2 | 3.9 | 71 | 141 | 4 | MOLECULE: | MICOS COMPLEX SUBUNIT MIC60 FUSED TO MIC19 |
| 16:00 | 4rp5-A | 3.2 | 12.2 | 71 | 94 | 6 | MOLECULE: | DISKS LARGE 1 TUMOR SUPPRESSOR PROTEIN |
| 17:00 | 6yvu-A | 3.2 | 15.3 | 118 | 1127 | 8 | MOLECULE: | STRUCTURAL MAINTENANCE OF CHROMOSOMES PROTEIN 2,S |
| 18:00 | 3cqx-D | 3.2 | 2.8 | 62 | 84 | 6 | MOLECULE: | HEAT SHOCK COGNATE 71 KDA PROTEIN |
| 19:00 | 8ij1-D | 3.2 | 6.7 | 68 | 627 | 3 | MOLECULE: | CULLIN-2 |
| 20:00 | 6k9k-A | 3.2 | 31 | 72 | 2501 | 8 | MOLECULE: | SERINE-PROTEIN KINASE ATM |
| 21:00 | 7pl9-A | 3.2 | 16.5 | 117 | 994 | 9 | MOLECULE: | RHODOPSIN |
| 22:00 | 7z8b-C | 3.2 | 8.7 | 106 | 1224 | 10 | MOLECULE: | CULLIN-7 |
| 23:00 | 8smr-G | 3.2 | 3.1 | 54 | 312 | 11 | MOLECULE: | UBIQUINOL-CYTOCHROME C REDUCTASE IRON-SULFUR SUBU |
| 24:00:00 | 8z9a-A | 3.2 | 5.4 | 66 | 406 | 6 | MOLECULE: | ODORANT RECEPTOR, APISORCO |
| 25:00:00 | 6lqo-L | 3.2 | 4.1 | 66 | 94 | 8 | MOLECULE: | CYTOPLASMIC ENVELOPMENT PROTEIN 1 |
| 26:00:00 | 7vcf-C | 3.2 | 4.7 | 80 | 217 | 8 | MOLECULE: | TIC214 |
| 27:00:00 | 8rz0-E | 3.2 | 3.3 | 66 | 113 | 5 | MOLECULE: | RH5-34EM |
| 28:00:00 | 6rd4-5 | 3.2 | 5.4 | 70 | 123 | 10 | MOLECULE: | ASA-10: POLYTOMELLA F-ATP SYNTHASE ASSOCIATED SUB |
| 29:00:00 | 8a3t-Q | 3.2 | 11.6 | 86 | 623 | 10 | MOLECULE: | ANAPHASE-PROMOTING COMPLEX SUBUNIT CDC27 |
| 30:00:00 | 6qaj-B | 3.2 | 7.1 | 82 | 441 | 6 | MOLECULE: | ENDOLYSIN,TRANSCRIPTION INTERMEDIARY FACTOR 1-BET |
| 31:00:00 | 6z6o-D | 3.2 | 9.1 | 95 | 542 | 7 | MOLECULE: | HISTONE DEACETYLASE HDA1 |
| 32:00:00 | 1yig-B | 3.2 | 4.4 | 51 | 61 | 6 | MOLECULE: | MICROTUBULE-ASSOCIATED PROTEIN RP/EB FAMILY |
| 33:00:00 | 1ifm-A | 3.2 | 1.9 | 46 | 46 | 7 | MOLECULE: | INOVIRUS |
| 34:00:00 | 7oac-A | 3.2 | 8.6 | 73 | 81 | 8 | MOLECULE: | GENERAL CONTROL TRANSCRIPTION FACTOR GCN4,CONSERV |
| 35:00:00 | 2efr-C | 3.2 | 62.1 | 93 | 154 | 4 | MOLECULE: | GENERAL CONTROL PROTEIN GCN4 AND TROPOMYOSIN 1 AL |
| 36:00:00 | 6pfp-A | 3.2 | 11 | 74 | 156 | 4 | MOLECULE: | MYOSIN-7 FUSED TO GP7 AND EB1 |
| 37:00:00 | 6n6r-D | 3.2 | 8.7 | 54 | 67 | 6 | MOLECULE: | UBIQUITIN |
| 38:00:00 | 5djo-A | 3.2 | 2.8 | 59 | 158 | 0 | MOLECULE: | KINESIN-LIKE PROTEIN |
| 39:00:00 | 5j8v-A | 3.1 | 25.4 | 102 | 3398 | 6 | MOLECULE: | RYANODINE RECEPTOR 1 |
| 40:00:00 | 8t1l-W | 3.1 | 16.5 | 76 | 119 | 4 | MOLECULE: | MEDIATOR OF RNA POLYMERASE II TRANSCRIPTION SUBUN |
| 41:00:00 | 7z4f-J | 3.1 | 29.1 | 78 | 603 | 5 | MOLECULE: | PUTATIVE STRUCTURAL PROTEIN |
| 42:00:00 | 7vdv-A | 3.1 | 21.2 | 77 | 809 | 6 | MOLECULE: | HISTONE H4 |
| 43:00:00 | 7bw0-R | 3.1 | 17.9 | 52 | 280 | 4 | MOLECULE: | SOLUBLE CYTOCHROME B562,G-PROTEIN COUPLED BILE AC |
| 44:00:00 | 6c0f-8 | 3.1 | 8.8 | 65 | 98 | 6 | MOLECULE: | SACCHAROMYCES CEREVISIAE S288C 35S PRE-RIBOSOMAL |
| 45:00:00 | 3zcj-A | 3.1 | 22.9 | 95 | 205 | 8 | MOLECULE: | CAGL |
| 46:00:00 | 5ew5-A | 3.1 | 22.2 | 116 | 490 | 3 | MOLECULE: | COLICIN-E9 |
| 47:00:00 | 6r1j-J | 3.1 | 9.1 | 102 | 264 | 8 | MOLECULE: | UNCHARACTERIZED PROTEIN |
| 48:00:00 | 7vc4-C | 3.1 | 8.2 | 66 | 107 | 6 | MOLECULE: | MITOCHONDRIAL IMPORT RECEPTOR SUBUNIT TOM6 HOMOLO |
| 49:00:00 | 6m3q-F | 3.1 | 3.5 | 88 | 301 | 8 | MOLECULE: | ANKYRIN-2 |
| 50:00:00 | 6qeq-A | 3.1 | 10.7 | 78 | 111 | 6 | MOLECULE: | PCFF |
| 51:00:00 | 6bog-A | 3.1 | 30.1 | 101 | 967 | 7 | MOLECULE: | RNA POLYMERASE-ASSOCIATED PROTEIN RAPA |
| 52:00:00 | 7oqz-A | 3.1 | 2.4 | 51 | 237 | 8 | MOLECULE: | TRANSMEMBRANE PROTEIN 45A |
| 53:00:00 | 8hvj-A | 3.1 | 10.2 | 114 | 281 | 10 | MOLECULE: | UPF0701 PROTEIN YICC |
| 54:00:00 | 2kn8-A | 3.1 | 10.8 | 66 | 68 | 8 | MOLECULE: | DNA CLEAVAGE AND PACKAGING PROTEIN LARGE SUBUNIT, |
| 55:00:00 | 6vls-D | 3.1 | 19.8 | 93 | 963 | 10 | MOLECULE: | MALTOSE/MALTODEXTRIN-BINDING PERIPLASMIC PROTEIN, |
| 56:00:00 | 8th8-J | 3.1 | 51.8 | 103 | 372 | 5 | MOLECULE: | DYNEIN REGULATORY COMPLEX PROTEIN 1/2 N-TERMINAL |
| 57:00:00 | 7e7i-A | 3.1 | 7.4 | 71 | 2011 | 8 | MOLECULE: | RETINAL-SPECIFIC PHOSPHOLIPID-TRANSPORTING ATPASE |
| 58:00:00 | 6q0x-A | 3.1 | 7.3 | 113 | 373 | 11 | MOLECULE: | SORTING NEXIN MVP1 |
| 59:00:00 | 6cp3-7 | 3.1 | 14.3 | 114 | 171 | 8 | MOLECULE: | ATP SYNTHASE SUBUNIT 9, MITOCHONDRIAL |
| 60:00:00 | 3qwe-A | 3.1 | 47.3 | 123 | 260 | 7 | MOLECULE: | GEM-INTERACTING PROTEIN |
| 61:00:00 | 7oqe-K | 3.1 | 6.4 | 89 | 406 | 7 | MOLECULE: | PROTEIN NAM8 |
| 62:00:00 | 7xdi-D | 3.1 | 4.8 | 71 | 124 | 18 | MOLECULE: | VP1 |
| 63:00:00 | 5vjx-J | 3.1 | 4.2 | 59 | 64 | 10 | MOLECULE: | CLOCK-INTERACTING PACEMAKER |
| 64:00:00 | 6pl5-A | 3.1 | 3.9 | 53 | 336 | 8 | MOLECULE: | PEPTIDOGLYCAN GLYCOSYLTRANSFERASE RODA |
| 65:00:00 | 7wkk-B | 3.1 | 5.5 | 66 | 1482 | 6 | MOLECULE: | MGC83295 PROTEIN |
| 66:00:00 | 8ceg-B | 3.1 | 3.2 | 76 | 214 | 7 | MOLECULE: | CBY1-INTERACTING BAR DOMAIN-CONTAINING PROTEIN 1 |

| | | | | | | | | |
|---|---|---|---|---|---|---|---|---|
| 67:00:00 | 6d5f-a | 3.1 | 15 | 77 | 131 | 5 | MOLECULE: | FIMBRIAL PROTEIN |
| 68:00:00 | 7xc2-A | 3.1 | 3.2 | 53 | 858 | 9 | MOLECULE: | CNL9 |
| 69:00:00 | 4dci-B | 3.1 | 3.8 | 72 | 148 | 10 | MOLECULE: | UNCHARACTERIZED PROTEIN |
| 70:00:00 | 8qfc-D | 3.1 | 2.6 | 75 | 185 | 4 | MOLECULE: | 60S RIBOSOMAL PROTEIN L10A |
| 71:00:00 | 3s4w-A | 3.1 | 4.2 | 71 | 1206 | 3 | MOLECULE: | FANCONI ANEMIA GROUP I PROTEIN HOMOLOG |
| 72:00:00 | 7ty0-C | 3.1 | 2.8 | 57 | 488 | 7 | MOLECULE: | GLYCOPROTEIN G |
| 73:00:00 | 6gy6-A | 3.1 | 7.2 | 104 | 365 | 13 | MOLECULE: | XAXA |
| 74:00:00 | 3rip-A | 3.1 | 3.1 | 54 | 568 | 6 | MOLECULE: | GAMMA-TUBULIN COMPLEX COMPONENT 4 |
| 75:00:00 | 5mpd-U | 3.1 | 24.5 | 113 | 298 | 6 | MOLECULE: | 26S PROTEASOME REGULATORY SUBUNIT RPN10 |
| 76:00:00 | 8tl7-A | 3.1 | 3.4 | 74 | 694 | 0 | MOLECULE: | COMPUTATIONALLY DESIGNED PROTEIN |
| 77:00:00 | 6fze-A | 3.1 | 11.2 | 71 | 209 | 7 | MOLECULE: | PUTATIVE SURFACE PROTEIN |
| 78:00:00 | 6nbx-C | 3.1 | 10.3 | 68 | 100 | 10 | MOLECULE: | NAD(P)H-QUINONE OXIDOREDUCTASE SUBUNIT 1 |
| 79:00:00 | 7v2c-j | 3.1 | 21.4 | 74 | 115 | 16 | MOLECULE: | NADH DEHYDROGENASE [UBIQUINONE] FLAVOPROTEIN 1, |
| 80:00:00 | 5oxf-C | 3.1 | 12.9 | 126 | 601 | 6 | MOLECULE: | GTP-BINDING PROTEIN |
| 81:00:00 | 5dlq-B | 3.1 | 7.3 | 74 | 1052 | 11 | MOLECULE: | EXPORTIN-4 |
| 82:00:00 | 6oqv-Y | 3.1 | 5.5 | 78 | 156 | 5 | MOLECULE: | ATP SYNTHASE SUBUNIT DELTA |
| 83:00:00 | 4aj5-L | 3.1 | 9.2 | 74 | 106 | 7 | MOLECULE: | SPINDLE AND KINETOCHORE-ASSOCIATED PROTEIN 1 |
| 84:00:00 | 3jax-A | 3.1 | 4.9 | 50 | 938 | 16 | MOLECULE: | MYOSIN 2 HEAVY CHAIN |
| 85:00:00 | 2efr-B | 3.1 | 6.6 | 79 | 154 | 3 | MOLECULE: | GENERAL CONTROL PROTEIN GCN4 AND TROPOMYOSIN 1 AL |
| 86:00:00 | 7eao-C | 3.1 | 3.9 | 46 | 62 | 9 | MOLECULE: | POLYUBIQUITIN-C |
| 87:00:00 | 3dtp-B | 3.1 | 26.3 | 112 | 940 | 4 | MOLECULE: | MYOSIN 2 HEAVY CHAIN CHIMERA OF SMOOTH AND CARDIA |
| 88:00:00 | 5yma-A | 3 | 6.5 | 85 | 179 | 2 | MOLECULE: | PUTATIVE RRNA PROCESSING PROTEIN |
| 89:00:00 | 3uux-D | 3 | 4.5 | 73 | 135 | 7 | MOLECULE: | MITOCHONDRIA FISSION 1 PROTEIN |
| 90:00:00 | 8dt0-A | 3 | 28.1 | 62 | 140 | 2 | MOLECULE: | SCAFFOLDING PROTEIN FUNCTIONAL SITES |
| 91:00:00 | 7wb4-E | 3 | 31.9 | 92 | 1363 | 5 | MOLECULE: | OUTER NUP133 |
| 92:00:00 | 5dfz-A | 3 | 52 | 85 | 343 | 5 | MOLECULE: | VACUOLAR PROTEIN SORTING-ASSOCIATED PROTEIN 38 |
| 93:00:00 | 7wkk-D | 3 | 22.8 | 54 | 1115 | 9 | MOLECULE: | MGC83295 PROTEIN |
| 94:00:00 | 4mu6-A | 3 | 5.9 | 100 | 274 | 11 | MOLECULE: | KINECTIN 1 (KINESIN RECEPTOR) |
| 95:00:00 | 7wji-A | 3 | 4.6 | 57 | 1763 | 9 | MOLECULE: | PROTEIN UNC-80 HOMOLOG |
| 96:00:00 | 6jfk-A | 3 | 11.8 | 111 | 428 | 5 | MOLECULE: | MITOFUSIN-2,CDNA FLJ57997, HIGHLY SIMILAR TO TRAN |
| 97:00:00 | 8ag8-A | 3 | 2.9 | 60 | 106 | 7 | MOLECULE: | FLUORESCENCE RECOVERY-LIKE PROTEIN |
| 98:00:00 | 7o3x-B | 3 | 28.4 | 80 | 215 | 10 | MOLECULE: | PROTEIN SLL0617 |
| 99:00:00 | 7nyw-C | 3 | 9.8 | 88 | 335 | 6 | MOLECULE: | CHROMOSOME PARTITION PROTEIN MUKB |
| 0:00 | 8bd7-L | 3 | 31.3 | 97 | 303 | 7 | MOLECULE: | IFT88 |
| 1:00 | 3plt-A | 3 | 5.7 | 88 | 214 | 10 | MOLECULE: | SPHINGOLIPID LONG CHAIN BASE-RESPONSIVE PROTEIN L |
| 2:00 | 8h2j-A | 3 | 6.4 | 67 | 92 | 7 | MOLECULE: | P26 |
| 3:00 | 6r7l-E | 3 | 17.1 | 78 | 98 | 6 | MOLECULE: | SECG |
| 4:00 | 4bk0-B | 3 | 7.2 | 74 | 97 | 7 | MOLECULE: | ATP-DEPENDENT DNA HELICASE Q5 |
| 5:00 | 1lj2-B | 3 | 4.7 | 66 | 110 | 6 | MOLECULE: | NONSTRUCTURAL RNA-BINDING PROTEIN 34 |
| 6:00 | 4lws-A | 3 | 8.7 | 53 | 100 | 15 | MOLECULE: | UNCHARACTERIZED PROTEIN |
| 7:00 | 7loi-A | 3 | 20.7 | 103 | 175 | 5 | MOLECULE: | TRANSMEMBRANE PROTEIN GP41 |
| 8:00 | 8fhc-B | 3 | 2.9 | 55 | 215 | 9 | MOLECULE: | TRNA-(MS[2]IO[6]A)-HYDROXYLASE |
| 9:00 | 7nh9-A | 3 | 5.1 | 114 | 327 | 4 | MOLECULE: | CMAX PROTEIN |
| 10:00 | 7pkq-C | 3 | 13.4 | 91 | 369 | 11 | MOLECULE: | MS35 |
| 11:00 | 6arz-A | 3 | 4.9 | 64 | 96 | 6 | MOLECULE: | UNCHARACTERIZED PROTEIN |
| 12:00 | 7zu0-F | 3 | 5.9 | 63 | 112 | 13 | MOLECULE: | E3 UBIQUITIN-PROTEIN LIGASE PEP5 |
| 13:00 | 8vc1-A | 3 | 5.7 | 65 | 391 | 5 | MOLECULE: | GUSTATORY RECEPTOR |
| 14:00 | 8e1m-C | 3 | 13.4 | 65 | 264 | 3 | MOLECULE: | MITOCHONDRIAL IMPORT INNER MEMBRANE TRANSLOCASE S |
| 15:00 | 8bhw-A | 3 | 4.4 | 106 | 332 | 5 | MOLECULE: | ECA POLYSACCHARIDE CHAIN LENGTH MODULATION PROTEI |
| 16:00 | 5da9-A | 3 | 5.1 | 90 | 433 | 3 | MOLECULE: | PUTATIVE UNCHARACTERIZED PROTEIN,PUTATIVE UNCHARA |
| 17:00 | 7p2y-a | 3 | 11.9 | 75 | 277 | 17 | MOLECULE: | ATP SYNTHASE SUBUNIT ALPHA |
| 18:00 | 6tmh-H | 3 | 2.2 | 55 | 71 | 11 | MOLECULE: | INHIBITOR OF F1 |
| 19:00 | 7tch-A | 3 | 13.8 | 73 | 645 | 3 | MOLECULE: | BACITRACIN EXPORT PERMEASE PROTEIN BCEB |
| 20:00 | 4zm1-B | 3 | 5.1 | 91 | 238 | 10 | MOLECULE: | CHAIN LENGTH DETERMINANT PROTEIN |
| 21:00 | 4x0j-B | 3 | 4.8 | 90 | 262 | 2 | MOLECULE: | HAPTOGLOBIN-HEMOGLOBIN RECEPTOR |
| 22:00 | 8ap7-c | 3 | 4.7 | 57 | 64 | 5 | MOLECULE: | ATP SYNTHASE SUBUNIT A |

| | | | | | | | |
|---|---|---|---|---|---|---|---|
| 23:00 | 6jho-A | 3 | 3.6 | 55 | 200 | 4 MOLECULE: | CAG PATHOGENICITY ISLAND PROTEIN (CAG6) |
| 24:00:00 | 6ar7-C | 3 | 2.8 | 57 | 207 | 5 MOLECULE: | UNCHARACTERIZED PROTEIN |
| 25:00:00 | 8b9z-A | 3 | 21.6 | 76 | 117 | 12 MOLECULE: | NADH-UBIQUINONE OXIDOREDUCTASE CHAIN 3 |
| 26:00:00 | 6qld-Y | 3 | 27.3 | 99 | 223 | 12 MOLECULE: | INNER KINETOCHORE SUBUNIT MIF2 |
| 27:00:00 | 7jsv-a | 3 | 3.6 | 57 | 66 | 5 MOLECULE: | PILIN |
| 28:00:00 | 5n9j-D | 3 | 14.6 | 84 | 135 | 7 MOLECULE: | MEDIATOR OF RNA POLYMERASE II TRANSCRIPTION SUBUN |
| 29:00:00 | 8d2s-A | 3 | 13.5 | 83 | 475 | 5 MOLECULE: | SODIUM-DEPENDENT LYSOPHOSPHATIDYLCHOLINE SYMPORTE |
| 30:00:00 | 7pw9-A | 3 | 30.9 | 69 | 1952 | 6 MOLECULE: | SERINE/THREONINE-PROTEIN KINASE SMG1,SERINE/THREO |
| 31:00:00 | 3fey-A | 3 | 7.6 | 91 | 761 | 3 MOLECULE: | NUCLEAR CAP-BINDING PROTEIN SUBUNIT 1 |
| 32:00:00 | 7woo-H | 3 | 6.4 | 80 | 246 | 13 MOLECULE: | NUCLEOPORIN NIC96 |
| 33:00:00 | 8r5i-F | 3 | 2.9 | 43 | 43 | 7 MOLECULE: | CORE PROTEIN A10 |
| 34:00:00 | 4aj5-M | 3 | 10.7 | 75 | 109 | 7 MOLECULE: | SPINDLE AND KINETOCHORE-ASSOCIATED PROTEIN 1 |
| 35:00:00 | 4aj5-R | 3 | 9.4 | 74 | 108 | 7 MOLECULE: | SPINDLE AND KINETOCHORE-ASSOCIATED PROTEIN 1 |
| 36:00:00 | 4aj5-O | 3 | 9 | 74 | 109 | 7 MOLECULE: | SPINDLE AND KINETOCHORE-ASSOCIATED PROTEIN 1 |
| 37:00:00 | 4aj5-Q | 3 | 9.4 | 73 | 102 | 5 MOLECULE: | SPINDLE AND KINETOCHORE-ASSOCIATED PROTEIN 1 |
| 38:00:00 | 7a48-B | 3 | 1.9 | 40 | 40 | 8 MOLECULE: | NANOBODY 49 |
| 39:00:00 | 3kyp-A | 3 | 3.4 | 55 | 171 | 5 MOLECULE: | NUCLEOSOME ASSEMBLY PROTEIN |
| 40:00:00 | 5oqm-k | 3 | 10.8 | 86 | 127 | 3 MOLECULE: | DNA-DIRECTED RNA POLYMERASE II SUBUNIT RPB1 |
| 41:00:00 | 4bk0-A | 3 | 5.3 | 65 | 92 | 11 MOLECULE: | ATP-DEPENDENT DNA HELICASE Q5 |
| 42:00:00 | 6l5j-A | 3 | 9.1 | 100 | 208 | 9 MOLECULE: | ROOTLETIN |
| 43:00:00 | 2efl-A | 3 | 3.6 | 95 | 281 | 4 MOLECULE: | FORMIN-BINDING PROTEIN 1 |
| 44:00:00 | 6eup-C | 3 | 2.7 | 67 | 147 | 9 MOLECULE: | ADHESIN A |
| 45:00:00 | 6eup-B | 3 | 2.7 | 67 | 147 | 9 MOLECULE: | ADHESIN A |
| 46:00:00 | 2yny-A | 3 | 4.8 | 65 | 106 | 12 MOLECULE: | GENERAL CONTROL PROTEIN GCN4, PUTATIVE INNER MEMB |
| 47:00:00 | 7y8r-P | 2.9 | 40.7 | 78 | 385 | 8 MOLECULE: | HISTONE H3 |
| 48:00:00 | 5xg2-A | 2.9 | 24 | 96 | 237 | 4 MOLECULE: | CHROMOSOME PARTITION PROTEIN SMC |
| 49:00:00 | 7nna-A | 2.9 | 30.1 | 92 | 194 | 1 MOLECULE: | KLEBICIN C ACTIVITY |
| 50:00:00 | 3kkd-C | 2.9 | 4.9 | 76 | 206 | 8 MOLECULE: | TRANSCRIPTIONAL REGULATOR |
| 51:00:00 | 8ia3-E | 2.9 | 7.7 | 78 | 111 | 6 MOLECULE: | UPSTREAM STIMULATORY FACTOR 2 |
| 52:00:00 | 8th8-L | 2.9 | 13.8 | 89 | 862 | 4 MOLECULE: | DYNEIN REGULATORY COMPLEX PROTEIN 1/2 N-TERMINAL |
| 53:00:00 | 7d8t-A | 2.9 | 15.7 | 87 | 199 | 2 MOLECULE: | DNA (5'- |
| 54:00:00 | 7yx4-A | 2.9 | 30.3 | 95 | 650 | 8 MOLECULE: | PUTATIVE GLUCOSE-METHANOL-CHOLINE OXIDOREDUCTASE |
| 55:00:00 | 8iw0-A | 2.9 | 3.6 | 61 | 84 | 5 MOLECULE: | LIPRIN-BETA-1,KN MOTIF AND ANKYRIN REPEAT DOMAIN- |
| 56:00:00 | 5v7p-A | 2.9 | 3.1 | 53 | 281 | 0 MOLECULE: | PROTEIN-S-ISOPRENYLCYSTEINE O-METHYLTRANSFERASE |
| 57:00:00 | 6yrf-A | 2.9 | 4 | 56 | 777 | 13 MOLECULE: | VEGETATIVE INSECTICIDAL PROTEIN |
| 58:00:00 | 6lo8-D | 2.9 | 4.5 | 57 | 119 | 9 MOLECULE: | MITOCHONDRIAL IMPORT INNER MEMBRANE TRANSLOCASE S |
| 59:00:00 | 6mdm-H | 2.9 | 18.7 | 84 | 147 | 4 MOLECULE: | VESICLE-FUSING ATPASE |
| 60:00:00 | 7pgs-N | 2.9 | 5.6 | 65 | 1787 | 6 MOLECULE: | NEUROFIBROMIN |
| 61:00:00 | 7emf-J | 2.9 | 5.2 | 70 | 122 | 13 MOLECULE: | MEDIATOR OF RNA POLYMERASE II TRANSCRIPTION SUBUN |
| 62:00:00 | 6ikn-A | 2.9 | 5.2 | 88 | 305 | 7 MOLECULE: | GROWTH ARREST-SPECIFIC PROTEIN 7 |
| 63:00:00 | 8j8p-R | 2.9 | 16.8 | 66 | 69 | 3 MOLECULE: | CTR9-LIKE PROTEIN |
| 64:00:00 | 4wzi-B | 2.9 | 9.3 | 71 | 424 | 8 MOLECULE: | CAMP-SPECIFIC 3',5'-CYCLIC PHOSPHODIESTERASE 4B |
| 65:00:00 | 4cz8-A | 2.9 | 8.6 | 74 | 422 | 8 MOLECULE: | NA+/H+ ANTIPORTER, PUTATIVE |
| 66:00:00 | 8g3a-D | 2.9 | 3.7 | 51 | 334 | 6 MOLECULE: | BACITRACIN EXPORT PERMEASE PROTEIN BCEB |
| 67:00:00 | 8i8c-K | 2.9 | 24.2 | 85 | 227 | 6 MOLECULE: | P40 |
| 68:00:00 | 2p4w-B | 2.9 | 9.4 | 85 | 198 | 1 MOLECULE: | TRANSCRIPTIONAL REGULATORY PROTEIN ARSR FAMILY |
| 69:00:00 | 3lvg-D | 2.9 | 6.9 | 84 | 180 | 5 MOLECULE: | CLATHRIN HEAVY CHAIN 1 |
| 70:00:00 | 5j1i-A | 2.9 | 7.3 | 128 | 346 | 6 MOLECULE: | PLECTIN |
| 71:00:00 | 5sva-V | 2.9 | 8.4 | 59 | 85 | 14 MOLECULE: | DNA-DIRECTED RNA POLYMERASE II SUBUNIT RPB1 |
| 72:00:00 | 8eki-C | 2.9 | 8.8 | 101 | 701 | 8 MOLECULE: | PROTEIN TRANSPORT PROTEIN SEC20 |
| 73:00:00 | 2aze-A | 2.9 | 5 | 66 | 149 | 6 MOLECULE: | TRANSCRIPTION FACTOR DP-1 |
| 74:00:00 | 8txr-C | 2.9 | 31.8 | 93 | 431 | 8 MOLECULE: | EXODEOXYRIBONUCLEASE 7 LARGE SUBUNIT |
| 75:00:00 | 5ztc-A | 2.9 | 2.3 | 49 | 197 | 4 MOLECULE: | LMO2088 PROTEIN |
| 76:00:00 | 1wgz-A | 2.9 | 2.4 | 59 | 510 | 12 MOLECULE: | CARBOXYPEPTIDASE 1 |
| 77:00:00 | 7dpa-A | 2.9 | 3.9 | 59 | 1642 | 10 MOLECULE: | DEDICATOR OF CYTOKINESIS PROTEIN 5 |
| 78:00:00 | 8soi-B | 2.9 | 7.6 | 118 | 515 | 7 MOLECULE: | RB1-INDUCIBLE COILED-COIL PROTEIN 1 |

| | | | | | | | | |
|---|---|---|---|---|---|---|---|---|
| 79:00:00 | 7lhe-A | 2.9 | 12.1 | 89 | 2222 | 11 | MOLECULE: | INOSITOL 1,4,5-TRISPHOSPHATE RECEPTOR TYPE 1 |
| 80:00:00 | 2wwb-B | 2.9 | 10.9 | 60 | 68 | 2 | MOLECULE: | PROTEIN TRANSPORT PROTEIN SEC61 SUBUNIT ALPHA ISO |
| 81:00:00 | 7y7q-A | 2.9 | 19.5 | 65 | 951 | 6 | MOLECULE: | RNA-DEPENDENT RNA POLYMERASE |
| 82:00:00 | 5xe3-E | 2.9 | 12 | 64 | 81 | 6 | MOLECULE: | ENDORIBONUCLEASE MAZF4 |
| 83:00:00 | 7z1n-O | 2.9 | 14.2 | 85 | 570 | 7 | MOLECULE: | DNA-DIRECTED RNA POLYMERASE III SUBUNIT RPC1 |
| 84:00:00 | 6ln2-A | 2.9 | 13.6 | 98 | 437 | 9 | MOLECULE: | GLUCAGON-LIKE PEPTIDE 1 RECEPTOR,RUBREDOXIN,GLUCA |
| 85:00:00 | 7ab4-F | 2.9 | 9.2 | 69 | 340 | 6 | MOLECULE: | PREDICTED TRANSCRIPTIONAL REGULATOR, XRE FAMILY |
| 86:00:00 | 4yxw-G | 2.9 | 6.9 | 90 | 213 | 8 | MOLECULE: | ATP SYNTHASE SUBUNIT ALPHA, MITOCHONDRIAL |
| 87:00:00 | 6j72-A | 2.9 | 6.2 | 112 | 561 | 6 | MOLECULE: | ISONIAZID INDUCIBLE GENE PROTEIN INIA |
| 88:00:00 | 5oqm-n | 2.9 | 14.2 | 85 | 136 | 9 | MOLECULE: | DNA-DIRECTED RNA POLYMERASE II SUBUNIT RPB1 |
| 89:00:00 | 6idp-A | 2.9 | 8.9 | 85 | 438 | 9 | MOLECULE: | MATE FAMILY EFFLUX TRANSPORTER |
| 90:00:00 | 8dft-A | 2.9 | 5.1 | 69 | 112 | 10 | MOLECULE: | PILIN PROTEIN |
| 91:00:00 | 8dkc-E | 2.9 | 3.4 | 59 | 103 | 5 | MOLECULE: | DNA-DIRECTED RNA POLYMERASE SUBUNIT ALPHA |
| 92:00:00 | 3l9f-A | 2.9 | 6 | 61 | 170 | 11 | MOLECULE: | PUTATIVE UNCHARACTERIZED PROTEIN SMU.1604C |
| 93:00:00 | 4pi0-A | 2.9 | 15 | 88 | 390 | 6 | MOLECULE: | UNKNOWN PEPTIDE |
| 94:00:00 | 3zni-A | 2.9 | 4.4 | 61 | 390 | 8 | MOLECULE: | E3 UBIQUITIN-PROTEIN LIGASE CBL-B |
| 95:00:00 | 8ih5-A | 2.9 | 22.1 | 105 | 689 | 4 | MOLECULE: | SYN-COPALYL DIPHOSPHATE SYNTHASE, CHLOROPLASTIC |
| 96:00:00 | 3aei-A | 2.9 | 3.7 | 63 | 94 | 11 | MOLECULE: | PREFOLDIN BETA SUBUNIT 2 |
| 97:00:00 | 6ird-B | 2.9 | 4.5 | 80 | 254 | 6 | MOLECULE: | 1-PHOSPHATIDYLINOSITOL 4,5-BISPHOSPHATE PHOSPHODI |
| 98:00:00 | 4lin-A | 2.9 | 7.1 | 87 | 289 | 9 | MOLECULE: | TAIL NEEDLE PROTEIN GP26 |
| 99:00:00 | 8i8b-C | 2.9 | 8.7 | 77 | 229 | 5 | MOLECULE: | MAJOR VIRAL CAPSID PROTEIN |
| 0:00 | 5apv-A | 2.9 | 7.3 | 64 | 70 | 3 | MOLECULE: | GENERAL CONTROL PROTEIN GCN4 |
| 1:00 | 8ajb-N | 2.9 | 39.9 | 99 | 238 | 7 | MOLECULE: | CRESCENTIN |
| 2:00 | 4aj5-S | 2.9 | 8.7 | 72 | 107 | 6 | MOLECULE: | SPINDLE AND KINETOCHORE-ASSOCIATED PROTEIN 1 |
| 3:00 | 4aj5-K | 2.9 | 8.4 | 72 | 108 | 8 | MOLECULE: | SPINDLE AND KINETOCHORE-ASSOCIATED PROTEIN 1 |
| 4:00 | 7z4f-K | 2.9 | 5.3 | 90 | 603 | 8 | MOLECULE: | PUTATIVE STRUCTURAL PROTEIN |
| 5:00 | 1qce-C | 2.9 | 3.2 | 68 | 123 | 15 | MOLECULE: | PROTEIN (GP41) |
| 6:00 | 1qce-B | 2.9 | 3.2 | 68 | 123 | 15 | MOLECULE: | PROTEIN (GP41) |
| 7:00 | 3uia-C | 2.9 | 1.5 | 39 | 52 | 13 | MOLECULE: | GLYCOPROTEIN 41 |
| 8:00 | 3wpo-C | 2.9 | 3.6 | 50 | 193 | 10 | MOLECULE: | TRIMERIC AUTOTRANSPORTER ADHESIN |
| 9:00 | 6h9l-C | 2.9 | 4.8 | 75 | 128 | 5 | MOLECULE: | UNCHARACTERIZED PROTEIN |
| 10:00 | 3wqa-C | 2.9 | 4.9 | 53 | 201 | 4 | MOLECULE: | TRIMERIC AUTOTRANSPORTER ADHESIN |
| 11:00 | 3zmf-B | 2.9 | 6.4 | 53 | 111 | 4 | MOLECULE: | GENERAL CONTROL PROTEIN GCN4, PUTATIVE INNER MEMB |
| 12:00 | 8pr4-X | 2.8 | 5.2 | 74 | 128 | 7 | MOLECULE: | ARP11 |
| 13:00 | 7p5j-A | 2.8 | 31.2 | 83 | 399 | 6 | MOLECULE: | PROTEIN TWEETY HOMOLOG 1 |
| 14:00 | 7rz4-A | 2.8 | 11.9 | 71 | 993 | 6 | MOLECULE: | GLUTAMATE RECEPTOR 2 |
| 15:00 | 5dfz-C | 2.8 | 8 | 80 | 783 | 6 | MOLECULE: | VACUOLAR PROTEIN SORTING-ASSOCIATED PROTEIN 38 |
| 16:00 | 7usd-A | 2.8 | 15.2 | 85 | 1185 | 7 | MOLECULE: | CYTOPLASMIC FMR1-INTERACTING PROTEIN 1 |
| 17:00 | 6zz6-C | 2.8 | 30.5 | 63 | 83 | 6 | MOLECULE: | STRUCTURAL MAINTENANCE OF CHROMOSOMES PROTEIN 1,S |
| 18:00 | 8ip4-A | 2.8 | 4.4 | 109 | 320 | 6 | MOLECULE: | MAGNESIUM TRANSPORTER MRS2 HOMOLOG, MITOCHONDRIAL |
| 19:00 | 2lck-A | 2.8 | 17.1 | 102 | 296 | 1 | MOLECULE: | MITOCHONDRIAL UNCOUPLING PROTEIN 2 |
| 20:00 | 4tsh-A | 2.8 | 8.5 | 69 | 121 | 4 | MOLECULE: | SURFACE PROTEIN ADHESIN |
| 21:00 | 6o8b-B | 2.8 | 4.2 | 105 | 657 | 9 | MOLECULE: | STIMULATOR OF INTERFERON GENES PROTEIN |
| 22:00 | 5tuu-B | 2.8 | 4 | 49 | 106 | 6 | MOLECULE: | TRANSCRIPTION FACTOR DP1 |
| 23:00 | 3b77-A | 2.8 | 10.3 | 87 | 188 | 10 | MOLECULE: | UNCHARACTERIZED PROTEIN |
| 24:00:00 | 5xbt-A | 2.8 | 10 | 84 | 271 | 10 | MOLECULE: | PROBABLE TRANSCRIPTIONAL REGULATOR |
| 25:00:00 | 5vj4-A | 2.8 | 2.8 | 57 | 275 | 7 | MOLECULE: | UNCHARACTERIZED PROTEIN |
| 26:00:00 | 8t1l-Y | 2.8 | 3.3 | 56 | 154 | 11 | MOLECULE: | MEDIATOR OF RNA POLYMERASE II TRANSCRIPTION SUBUN |
| 27:00:00 | 5nnv-D | 2.8 | 6.2 | 93 | 252 | 10 | MOLECULE: | CHROMOSOME PARTITION PROTEIN SMC,CHROMOSOME PARTI |
| 28:00:00 | 4bpm-A | 2.8 | 6 | 64 | 162 | 5 | MOLECULE: | PROSTAGLANDIN E SYNTHASE, FUSION PEPTIDE |
| 29:00:00 | 5t58-A | 2.8 | 13 | 104 | 233 | 6 | MOLECULE: | KLLA0F02343P |
| 30:00:00 | 8teu-E | 2.8 | 14.2 | 74 | 495 | 7 | MOLECULE: | LARGE TEGUMENT PROTEIN DENEDDYLASE |
| 31:00:00 | 8p62-D | 2.8 | 4.9 | 61 | 245 | 11 | MOLECULE: | DNA REPLICATION LICENSING FACTOR MCM2 |
| 32:00:00 | 8j22-C | 2.8 | 13 | 83 | 285 | 6 | MOLECULE: | GUANINE NUCLEOTIDE-BINDING PROTEIN G(I)/G(S)/G(T) |
| 33:00:00 | 6sny-A | 2.8 | 3.5 | 53 | 107 | 4 | MOLECULE: | SYNTHETIC EPCR BINDING PROTEIN |
| 34:00:00 | 8rt9-A | 2.8 | 3.9 | 69 | 207 | 6 | MOLECULE: | TRWJ PROTEIN |

| | | | | | | | | |
|---|---|---|---|---|---|---|---|---|
| 35:00:00 | 8ebt-K | 2.8 | 9 | 78 | 172 | 3 | MOLECULE: | GENERAL TRANSCRIPTION AND DNA REPAIR FACTOR IIH H |
| 36:00:00 | 8ptx-A | 2.8 | 31.1 | 103 | 1215 | 5 | MOLECULE: | ELONGATOR COMPLEX PROTEIN 1 |
| 37:00:00 | 8hir-A | 2.8 | 3.8 | 62 | 918 | 6 | MOLECULE: | POTASSIUM CHANNEL SUBFAMILY T MEMBER 1 |
| 38:00:00 | 6ye4-A | 2.8 | 5.4 | 82 | 236 | 4 | MOLECULE: | BIOPOLYMER TRANSPORT PROTEIN EXBB |
| 39:00:00 | 7kpx-C | 2.8 | 4 | 70 | 1352 | 7 | MOLECULE: | MEIOTIC MRNA STABILITY PROTEIN KINASE SSN3 |
| 40:00:00 | 6v35-A | 2.8 | 7.5 | 79 | 914 | 11 | MOLECULE: | CALCIUM-ACTIVATED POTASSIUM CHANNEL SUBUNIT ALPHA |
| 41:00:00 | 1y2o-A | 2.8 | 3.1 | 80 | 248 | 5 | MOLECULE: | BAI1-ASSOCIATED PROTEIN 2 ISOFORM 1 |
| 42:00:00 | 6dlu-P | 2.8 | 39.2 | 77 | 747 | 1 | MOLECULE: | DYNAMIN-1 |
| 43:00:00 | 4uxv-A | 2.8 | 3.1 | 61 | 522 | 5 | MOLECULE: | SEPTATION RING FORMATION REGULATOR EZRA |
| 44:00:00 | 7uxc-R | 2.8 | 10.9 | 51 | 107 | 2 | MOLECULE: | SERINE/THREONINE-PROTEIN KINASE MTOR |
| 45:00:00 | 2e50-P | 2.8 | 8.5 | 70 | 186 | 7 | MOLECULE: | PROTEIN SET |
| 46:00:00 | 7w76-D | 2.8 | 7.2 | 78 | 168 | 5 | MOLECULE: | UBIQUITIN-CONJUGATING ENZYME E2 2 |
| 47:00:00 | 6t5a-A | 2.8 | 6 | 71 | 96 | 10 | MOLECULE: | TEGUMENT PROTEIN UL51 |
| 48:00:00 | 1llm-C | 2.8 | 4.1 | 55 | 87 | 7 | MOLECULE: | 5'-D(*TP*CP*CP*CP*AP*CP*GP*CP*GP*TP*GP*GP*G)-3' |
| 49:00:00 | 6s3l-K | 2.8 | 7.9 | 86 | 193 | 9 | MOLECULE: | FLAGELLAR BIOSYNTHETIC PROTEIN FLIP |
| 50:00:00 | 7xad-C | 2.8 | 3 | 59 | 96 | 10 | MOLECULE: | PROGRAMMED CELL DEATH 1 LIGAND 1 |
| 51:00:00 | 8b6j-i | 2.8 | 12.5 | 85 | 117 | 7 | MOLECULE: | PEPTIDASE M16 INACTIVE DOMAIN PROTEIN |
| 52:00:00 | 8wcn-A | 2.8 | 6.6 | 71 | 379 | 11 | MOLECULE: | DIGUANYLATE CYCLASE |
| 53:00:00 | 4uy3-A | 2.8 | 4.6 | 80 | 194 | 9 | MOLECULE: | SEPTATION RING FORMATION REGULATOR EZRA |
| 54:00:00 | 8kde-G | 2.8 | 8.2 | 67 | 145 | 10 | MOLECULE: | PHOTOSYSTEM II CP47 REACTION CENTER PROTEIN |
| 55:00:00 | 8fef-D | 2.8 | 32.7 | 89 | 385 | 7 | MOLECULE: | VIRULENCE FACTOR MCE FAMILY PROTEIN |
| 56:00:00 | 6xxd-A | 2.8 | 4.5 | 71 | 125 | 7 | MOLECULE: | PILA |
| 57:00:00 | 6b2z-M | 2.8 | 10.5 | 85 | 249 | 5 | MOLECULE: | ATP SYNTHASE SUBUNIT C, MITOCHONDRIAL |
| 58:00:00 | 7wze-A | 2.8 | 4.5 | 63 | 161 | 11 | MOLECULE: | UNCHARACTERIZED HTH-TYPE TRANSCRIPTIONAL REGULATO |
| 59:00:00 | 6em5-b | 2.8 | 4.1 | 66 | 421 | 12 | MOLECULE: | 5.8S RIBOSOMAL RNA |
| 60:00:00 | 8sgi-A | 2.8 | 38.7 | 101 | 751 | 7 | MOLECULE: | SODIUM/CALCIUM EXCHANGER 1 |
| 61:00:00 | 2kjy-A | 2.8 | 4.2 | 56 | 60 | 4 | MOLECULE: | PROTEIN PHOSPHATASE 1 REGULATORY SUBUNIT 12A |
| 62:00:00 | 5j6f-A | 2.8 | 4 | 55 | 352 | 7 | MOLECULE: | 3-DEOXY-D-ARABINO-HEPTULOSONATE 7-PHOSPHATE SYNTH |
| 63:00:00 | 1u2m-C | 2.8 | 2.3 | 63 | 143 | 5 | MOLECULE: | HISTONE-LIKE PROTEIN HLP-1 |
| 64:00:00 | 3g9g-A | 2.8 | 9.1 | 93 | 250 | 5 | MOLECULE: | SUPPRESSOR OF YEAST PROFILIN DELETION |
| 65:00:00 | 8sl3-A | 2.8 | 12.6 | 87 | 578 | 9 | MOLECULE: | ADENYLATE CYCLASE TYPE 5 |
| 66:00:00 | 3jc5-A | 2.8 | 5.5 | 63 | 208 | 8 | MOLECULE: | DNA REPLICATION LICENSING FACTOR MCM2 |
| 67:00:00 | 5ip0-D | 2.8 | 7.1 | 69 | 108 | 7 | MOLECULE: | PHA GRANULE-ASSOCIATED PROTEIN |
| 68:00:00 | 8pm4-A | 2.8 | 10.8 | 73 | 604 | 3 | MOLECULE: | TRANSPOSASE |
| 69:00:00 | 7etm-A | 2.8 | 12.3 | 91 | 498 | 9 | MOLECULE: | PORTAL PROTEIN |
| 70:00:00 | 4tt0-B | 2.8 | 10.6 | 79 | 131 | 9 | MOLECULE: | DENEDDYLASE |
| 71:00:00 | 8hnc-A | 2.8 | 20.5 | 92 | 570 | 5 | MOLECULE: | SOLUTE CARRIER ORGANIC ANION TRANSPORTER FAMILY M |
| 72:00:00 | 7n15-A | 2.8 | 13.5 | 73 | 520 | 3 | MOLECULE: | CYCLIC NUCLEOTIDE-GATED CATION CHANNEL |
| 73:00:00 | 4n5b-D | 2.8 | 5.8 | 61 | 101 | 3 | MOLECULE: | PHOSPHOPROTEIN |
| 74:00:00 | 6jlb-C | 2.8 | 8 | 93 | 256 | 3 | MOLECULE: | LAMIN A/C |
| 75:00:00 | 7yfv-B | 2.8 | 4.1 | 47 | 50 | 9 | MOLECULE: | PROTEIN FANTOM |
| 76:00:00 | 1fze-A | 2.8 | 2.9 | 54 | 81 | 7 | MOLECULE: | FIBRINOGEN |
| 77:00:00 | 8pew-a | 2.8 | 3.2 | 64 | 187 | 3 | MOLECULE: | TRANSCRIPTION TERMINATION FACTOR RHO |
| 78:00:00 | 6tgz-F | 2.8 | 8.2 | 128 | 352 | 5 | MOLECULE: | 55 KDA IMMEDIATE-EARLY PROTEIN 1 |
| 79:00:00 | 1fzb-A | 2.8 | 3 | 54 | 81 | 7 | MOLECULE: | FIBRINOGEN |
| 80:00:00 | 8tek-N | 2.8 | 7.4 | 96 | 187 | 8 | MOLECULE: | DYNEIN REGULATORY COMPLEX PROTEIN 1/2 N-TERMINAL |
| 81:00:00 | 5voy-K | 2.8 | 8.8 | 96 | 216 | 6 | MOLECULE: | V-TYPE PROTON ATPASE CATALYTIC SUBUNIT A,V-TYPE P |
| 82:00:00 | 5tby-B | 2.8 | 10.6 | 117 | 950 | 6 | MOLECULE: | MYOSIN-7 |
| 83:00:00 | 8ek4-B | 2.8 | 2.5 | 58 | 135 | 5 | MOLECULE: | ICE-BINDING PROTEIN TIP-99A |
| 84:00:00 | 3bat-D | 2.8 | 5.2 | 50 | 80 | 4 | MOLECULE: | MYOSIN HEAVY CHAIN, STRIATED MUSCLE/GENERAL CONTR |
| 85:00:00 | 6pfp-B | 2.8 | 6.7 | 75 | 156 | 8 | MOLECULE: | MYOSIN-7 FUSED TO GP7 AND EB1 |
| 86:00:00 | 6mi4-A | 2.8 | 6.8 | 72 | 123 | 6 | MOLECULE: | NF-KB ESSENTIAL MODULATOR |
| 87:00:00 | 5jvu-B | 2.8 | 3.1 | 43 | 69 | 5 | MOLECULE: | CHIMERA PROTEIN OF KINESIN HEAVY CHAIN AND MICROT |
| 88:00:00 | 1am9-C | 2.7 | 7 | 60 | 82 | 5 | MOLECULE: | DNA (5'- |
| 89:00:00 | 6wkr-C | 2.7 | 32.8 | 77 | 607 | 9 | MOLECULE: | UBIQUITIN |
| 90:00:00 | 7rtn-A | 2.7 | 11.1 | 76 | 270 | 5 | MOLECULE: | OUTER CAPSID PROTEIN VP5 |

| | | | | | | | | |
|---|---|---|---|---|---|---|---|---|
| 91:00:00 | 8fck-E | 2.7 | 9.4 | 71 | 222 | 8 | MOLECULE: | HAUS AUGMIN-LIKE COMPLEX SUBUNIT 1 |
| 92:00:00 | 7c4j-E | 2.7 | 6.6 | 51 | 288 | 2 | MOLECULE: | TRANSCRIPTION REGULATORY PROTEIN SNF12 |
| 93:00:00 | 4uig-A | 2.7 | 4 | 59 | 91 | 7 | MOLECULE: | COPPER SENSITIVE OPERON REPRESSOR |
| 94:00:00 | 8jw0-b | 2.7 | 27.9 | 128 | 617 | 8 | MOLECULE: | PHOTOSYSTEM I PSAA |
| 95:00:00 | 6ibl-A | 2.7 | 16.3 | 95 | 397 | 8 | MOLECULE: | THIOREDOXIN 1,BETA-1 ADRENERGIC RECEPTOR |
| 96:00:00 | 6pwn-A | 2.7 | 3.7 | 71 | 280 | 10 | MOLECULE: | SMALL-CONDUCTANCE MECHANOSENSITIVE CHANNEL |
| 97:00:00 | 8gyz-B | 2.7 | 8.6 | 74 | 215 | 11 | MOLECULE: | TRANSCRIPTION FACTOR TGA7 |
| 98:00:00 | 6qti-A | 2.7 | 15.8 | 96 | 1038 | 6 | MOLECULE: | NICOTINAMIDE NUCLEOTIDE TRANSHYDROGENASE |
| 99:00:00 | 7tmw-R | 2.7 | 11.8 | 60 | 499 | 7 | MOLECULE: | RELAXIN RECEPTOR 1, GUANINE NUCLEOTIDE-BINDING PR |
| 0:00 | 1na6-B | 2.7 | 12.7 | 75 | 395 | 7 | MOLECULE: | RESTRICTION ENDONUCLEASE ECORII |
| 1:00 | 5yfp-G | 2.7 | 20.4 | 70 | 615 | 13 | MOLECULE: | EXOCYST COMPLEX COMPONENT SEC3 |
| 2:00 | 2odu-A | 2.7 | 5.7 | 81 | 217 | 10 | MOLECULE: | PLECTIN 1 |
| 3:00 | 8qq7-A | 2.7 | 5.5 | 67 | 399 | 9 | MOLECULE: | FAD-BINDING FR-TYPE DOMAIN-CONTAINING PROTEIN |
| 4:00 | 6v3f-A | 2.7 | 27.7 | 82 | 1234 | 10 | MOLECULE: | NPC1-LIKE INTRACELLULAR CHOLESTEROL TRANSPORTER 1 |
| 5:00 | 6z16-d | 2.7 | 4.1 | 57 | 489 | 7 | MOLECULE: | MULTISUBUNIT NA+/H+ ANTIPORTER, A SUBUNIT |
| 6:00 | 5td8-D | 2.7 | 9.1 | 77 | 128 | 8 | MOLECULE: | KINETOCHORE PROTEIN NDC80 |
| 7:00 | 8ik3-A | 2.7 | 5.4 | 65 | 328 | 9 | MOLECULE: | STIMULATOR OF INTERFERON GENES PROTEIN,IMMUNE PRO |
| 8:00 | 5ziy-A | 2.7 | 3.8 | 72 | 195 | 13 | MOLECULE: | FLAGELLAR HOOK-ASSOCIATED PROTEIN 3 |
| 9:00 | 6djl-E | 2.7 | 5.4 | 75 | 226 | 4 | MOLECULE: | RAS-RELATED PROTEIN RAB-11A |
| 10:00 | 8ap7-m | 2.7 | 11.9 | 70 | 129 | 4 | MOLECULE: | ATP SYNTHASE SUBUNIT A |
| 11:00 | 8hub-B | 2.7 | 17.2 | 88 | 545 | 5 | MOLECULE: | AMP DEAMINASE 2 |
| 12:00 | 5tgz-A | 2.7 | 8.9 | 97 | 439 | 1 | MOLECULE: | CANNABINOID RECEPTOR 1,FLAVODOXIN,CANNABINOID REC |
| 13:00 | 3mhh-C | 2.7 | 8.3 | 51 | 93 | 8 | MOLECULE: | UBIQUITIN CARBOXYL-TERMINAL HYDROLASE 8 |
| 14:00 | 2oeq-C | 2.7 | 3.4 | 51 | 117 | 12 | MOLECULE: | PROTEIN OF UNKNOWN FUNCTION, DUF964 |
| 15:00 | 3sjr-A | 2.7 | 3.7 | 63 | 126 | 5 | MOLECULE: | UNCHARACTERIZED PROTEIN |
| 16:00 | 8ebw-H | 2.7 | 4.3 | 89 | 532 | 8 | MOLECULE: | TFIIH BASAL TRANSCRIPTION FACTOR COMPLEX HELICASE |
| 17:00 | 2lhk-A | 2.7 | 14.4 | 89 | 107 | 6 | MOLECULE: | L0052 |
| 18:00 | 6zyx-C | 2.7 | 16.5 | 107 | 278 | 8 | MOLECULE: | DYNEIN HEAVY CHAIN, OUTER ARM PROTEIN |
| 19:00 | 6ah0-w | 2.7 | 33.4 | 99 | 443 | 5 | MOLECULE: | U5SNRNA |
| 20:00 | 5w82-B | 2.7 | 9 | 67 | 101 | 4 | MOLECULE: | PROTEIN DELTA |
| 21:00 | 7o4i-W | 2.7 | 5.4 | 73 | 312 | 8 | MOLECULE: | GENERAL TRANSCRIPTION AND DNA REPAIR FACTOR IIH H |
| 22:00 | 3pwx-A | 2.7 | 4 | 85 | 224 | 9 | MOLECULE: | PUTATIVE FLAGELLAR HOOK-ASSOCIATED PROTEIN |
| 23:00 | 6weg-A | 2.7 | 7.5 | 71 | 204 | 8 | MOLECULE: | STRINGENT STARVATION PROTEIN A, REGULATOR OF TRAN |
| 24:00:00 | 7b93-L | 2.7 | 17.5 | 129 | 607 | 5 | MOLECULE: | NADH-UBIQUINONE OXIDOREDUCTASE CHAIN 3 |
| 25:00:00 | 6zie-A | 2.7 | 2.6 | 50 | 123 | 4 | MOLECULE: | CMPX-383B |
| 26:00:00 | 7p64-L | 2.7 | 8.7 | 106 | 606 | 4 | MOLECULE: | NADH-QUINONE OXIDOREDUCTASE SUBUNIT F |
| 27:00:00 | 2ve7-D | 2.7 | 5 | 76 | 242 | 5 | MOLECULE: | KINETOCHORE PROTEIN HEC1, KINETOCHORE PROTEIN SPC |
| 28:00:00 | 3cwz-B | 2.7 | 3.4 | 54 | 310 | 9 | MOLECULE: | RAS-RELATED PROTEIN RAB-6A |
| 29:00:00 | 1few-A | 2.7 | 4.9 | 70 | 173 | 4 | MOLECULE: | SECOND MITOCHONDRIA-DERIVED ACTIVATOR OF CASPASES |
| 30:00:00 | 6x1g-A | 2.7 | 3.9 | 59 | 216 | 8 | MOLECULE: | ULP_PROTEASE DOMAIN-CONTAINING PROTEIN |
| 31:00:00 | 6ezn-B | 2.7 | 4.3 | 56 | 110 | 13 | MOLECULE: | DOLICHYL-DIPHOSPHOOLIGOSACCHARIDE--PROTEIN |
| 32:00:00 | 5aj3-d | 2.7 | 23.6 | 96 | 177 | 6 | MOLECULE: | MITORIBOSOMAL 12S RRNA |
| 33:00:00 | 7f3x-A | 2.7 | 3.8 | 53 | 449 | 8 | MOLECULE: | LPCAT3 |
| 34:00:00 | 8hfs-Z | 2.7 | 8.7 | 73 | 303 | 5 | MOLECULE: | MANNOSE-SPECIFIC PTS SYSTEM, IIC COMPONENT |
| 35:00:00 | 2pih-A | 2.7 | 11.4 | 62 | 123 | 3 | MOLECULE: | PROTEIN YMCA |
| 36:00:00 | 8eki-A | 2.7 | 5 | 75 | 170 | 4 | MOLECULE: | PROTEIN TRANSPORT PROTEIN SEC20 |
| 37:00:00 | 4j2c-A | 2.7 | 4.2 | 61 | 108 | 3 | MOLECULE: | SYNTAXIN-6 |
| 38:00:00 | 6ejq-B | 2.7 | 14.6 | 83 | 141 | 6 | MOLECULE: | TERMINASE SMALL SUBUNIT |
| 39:00:00 | 8j0h-A | 2.7 | 2.8 | 57 | 134 | 9 | MOLECULE: | UNCHARACTERIZED PROTEIN C4H3.06 |
| 40:00:00 | 1e7p-C | 2.7 | 3.4 | 62 | 254 | 6 | MOLECULE: | FUMARATE REDUCTASE FLAVOPROTEIN SUBUNIT |
| 41:00:00 | 5x5y-F | 2.7 | 5.4 | 65 | 351 | 15 | MOLECULE: | PROBABLE ATP-BINDING COMPONENT OF ABC TRANSPORTER |
| 42:00:00 | 2pjw-H | 2.7 | 5.6 | 63 | 88 | 8 | MOLECULE: | UNCHARACTERIZED PROTEIN YHL002W |
| 43:00:00 | 5dw7-A | 2.7 | 3.1 | 70 | 309 | 3 | MOLECULE: | GERMACRADIENOL/GEOSMIN SYNTHASE |
| 44:00:00 | 6ybt-u | 2.7 | 36 | 113 | 134 | 4 | MOLECULE: | EUKARYOTIC TRANSLATION INITIATION FACTOR 3 SUBUNI |
| 45:00:00 | 1fnt-c | 2.7 | 3.8 | 61 | 198 | 5 | MOLECULE: | PROTEASOME COMPONENT C7-ALPHA |
| 46:00:00 | 7yfv-C | 2.7 | 3.7 | 45 | 48 | 4 | MOLECULE: | PROTEIN FANTOM |

| | | | | | | | | |
|---|---|---|---|---|---|---|---|---|
| 47:00:00 | 6id1-q | 2.7 | 13.4 | 64 | 132 | 9 | MOLECULE: | PRE-MRNA-PROCESSING-SPLICING FACTOR 8 |
| 48:00:00 | 2ezo-B | 2.7 | 2.4 | 68 | 123 | 13 | MOLECULE: | GP41 |
| 49:00:00 | 2ezp-B | 2.7 | 2.4 | 68 | 123 | 12 | MOLECULE: | GP41 |
| 50:00:00 | 2ezr-C | 2.7 | 2.5 | 68 | 123 | 15 | MOLECULE: | GP41 |
| 51:00:00 | 2ezq-C | 2.7 | 2.5 | 68 | 123 | 13 | MOLECULE: | GP41 |
| 52:00:00 | 2ezp-C | 2.7 | 2.4 | 68 | 123 | 12 | MOLECULE: | GP41 |
| 53:00:00 | 8dk2-C | 2.7 | 7 | 85 | 723 | 7 | MOLECULE: | JETA |
| 54:00:00 | 2ezr-B | 2.7 | 2.6 | 68 | 123 | 13 | MOLECULE: | GP41 |
| 55:00:00 | 2ezp-A | 2.7 | 2.4 | 68 | 123 | 12 | MOLECULE: | GP41 |
| 56:00:00 | 2ezo-C | 2.7 | 2.4 | 68 | 123 | 10 | MOLECULE: | GP41 |
| 57:00:00 | 2q7c-C | 2.7 | 2.3 | 36 | 45 | 3 | MOLECULE: | FUSION PROTEIN BETWEEN YEAST VARIANT GCN4 AND HIV |
| 58:00:00 | 3jax-B | 2.7 | 26.3 | 112 | 940 | 4 | MOLECULE: | MYOSIN 2 HEAVY CHAIN |
| 59:00:00 | 8f3b-A | 2.7 | 15.4 | 46 | 50 | 9 | MOLECULE: | IQN22 |
| 60:00:00 | 7xe4-F | 2.6 | 35 | 112 | 1503 | 6 | MOLECULE: | 1,3-BETA-GLUCAN SYNTHASE COMPONENT FKS1 |
| 61:00:00 | 8ulg-A | 2.6 | 6.1 | 50 | 827 | 2 | MOLECULE: | ROD CGMP-SPECIFIC 3',5'-CYCLIC PHOSPHODIESTERASE |
| 62:00:00 | 8ozh-A | 2.6 | 13.3 | 47 | 58 | 2 | MOLECULE: | ENVELOPE GLYCOPROTEIN |
| 63:00:00 | 6jni-A | 2.6 | 10.3 | 68 | 145 | 7 | MOLECULE: | CADR |
| 64:00:00 | 7d60-B | 2.6 | 34.4 | 83 | 288 | 11 | MOLECULE: | CALCIUM HOMEOSTASIS MODULATOR PROTEIN 5 |
| 65:00:00 | 7a23-Z | 2.6 | 6.2 | 77 | 140 | 3 | MOLECULE: | 51KDA |
| 66:00:00 | 5n9j-U | 2.6 | 7.6 | 102 | 198 | 7 | MOLECULE: | MEDIATOR OF RNA POLYMERASE II TRANSCRIPTION SUBUN |
| 67:00:00 | 5jdo-A | 2.6 | 5.4 | 84 | 249 | 2 | MOLECULE: | HAPTOGLOBIN-HAEMOGLOBIN RECEPTOR |
| 68:00:00 | 8d8k-Q | 2.6 | 8.5 | 71 | 205 | 8 | MOLECULE: | PROBABLE S-ADENOSYL-L-METHIONINE-DEPENDENT RNA |
| 69:00:00 | 1s7o-C | 2.6 | 6 | 64 | 108 | 6 | MOLECULE: | HYPOTHETICAL UPF0122 PROTEIN |
| 70:00:00 | 6lo8-C | 2.6 | 3.3 | 55 | 108 | 13 | MOLECULE: | MITOCHONDRIAL IMPORT INNER MEMBRANE TRANSLOCASE S |
| 71:00:00 | 5y5a-A | 2.6 | 2.4 | 54 | 514 | 19 | MOLECULE: | KLLA0F20702P |
| 72:00:00 | 6en8-A | 2.6 | 14.1 | 52 | 191 | 13 | MOLECULE: | TRANSCRIPTIONAL REGULATOR TETR FAMILY |
| 73:00:00 | 6tmi-A | 2.6 | 5.9 | 67 | 109 | 1 | MOLECULE: | ATP SYNTHASE SUBUNIT ALPHA |
| 74:00:00 | 1gnc-A | 2.6 | 4.9 | 67 | 175 | 9 | MOLECULE: | GRANULOCYTE COLONY-STIMULATING FACTOR |
| 75:00:00 | 4wxa-E | 2.6 | 5 | 50 | 69 | 2 | MOLECULE: | EKC/KEOPS COMPLEX SUBUNIT PCC1 |
| 76:00:00 | 8ttf-A | 2.6 | 3.5 | 53 | 388 | 13 | MOLECULE: | QUINOLONE RESISTANCE PROTEIN NORA |
| 77:00:00 | 8q7h-A | 2.6 | 8.6 | 104 | 1403 | 5 | MOLECULE: | CULLIN-9 |
| 78:00:00 | 2v0o-A | 2.6 | 6 | 84 | 273 | 5 | MOLECULE: | FCH DOMAIN ONLY PROTEIN 2 |
| 79:00:00 | 7mbz-A | 2.6 | 7.2 | 75 | 222 | 5 | MOLECULE: | ABC TRANSPORTER, PERMEASE PROTEIN |
| 80:00:00 | 7x0a-B | 2.6 | 7.3 | 63 | 429 | 5 | MOLECULE: | T-COMPLEX PROTEIN 1 SUBUNIT ALPHA |
| 81:00:00 | 6hyd-A | 2.6 | 24.9 | 62 | 1574 | 13 | MOLECULE: | MIDASIN,MIDASIN,MIDASIN |
| 82:00:00 | 8wjn-A | 2.6 | 10.3 | 95 | 491 | 15 | MOLECULE: | STRUCTURAL MAINTENANCE OF CHROMOSOMES PROTEIN 6 |
| 83:00:00 | 7yzp-C | 2.6 | 3.7 | 79 | 448 | 9 | MOLECULE: | DNA HAIRPIN (59-MER) |
| 84:00:00 | 8bsb-B | 2.6 | 2.8 | 56 | 162 | 4 | MOLECULE: | METHYL-ACCEPTING CHEMOTAXIS PROTEIN |
| 85:00:00 | 8eoi-K | 2.6 | 12 | 85 | 1116 | 8 | MOLECULE: | ER MEMBRANE PROTEIN COMPLEX SUBUNIT 1 |
| 86:00:00 | 7ozs-F | 2.6 | 15.8 | 98 | 571 | 6 | MOLECULE: | PUTATIVE RIBOSOMAL PROTEIN |
| 87:00:00 | 5i6r-A | 2.6 | 3.8 | 92 | 419 | 7 | MOLECULE: | SLIT-ROBO RHO GTPASE-ACTIVATING PROTEIN 2 |
| 88:00:00 | 7oca-E | 2.6 | 4.2 | 67 | 158 | 10 | MOLECULE: | GLUTAMATE RECEPTOR 1 |
| 89:00:00 | 8qah-B | 2.6 | 4.5 | 69 | 426 | 1 | MOLECULE: | SC-CC-8-58 |
| 90:00:00 | 8imi-0 | 2.6 | 10.4 | 79 | 1045 | 9 | MOLECULE: | APCE |
| 91:00:00 | 5yfp-A | 2.6 | 22.5 | 86 | 672 | 12 | MOLECULE: | EXOCYST COMPLEX COMPONENT SEC3 |
| 92:00:00 | 7drt-B | 2.6 | 12.6 | 70 | 496 | 3 | MOLECULE: | PROTEIN WNT-3A |
| 93:00:00 | 7jtk-C | 2.6 | 5.7 | 84 | 427 | 5 | MOLECULE: | FLAGELLAR RADIAL SPOKE PROTEIN 1 |
| 94:00:00 | 5n9j-F | 2.6 | 9.3 | 58 | 79 | 12 | MOLECULE: | MEDIATOR OF RNA POLYMERASE II TRANSCRIPTION SUBUN |
| 95:00:00 | 7ura-A | 2.6 | 12.8 | 87 | 432 | 11 | MOLECULE: | ISOFORM 2 OF PROTEIN-SERINE O-PALMITOLEOYLTRANSFE |
| 96:00:00 | 6s8g-G | 2.6 | 5.4 | 66 | 244 | 6 | MOLECULE: | LIPOPOLYSACCHARIDE ABC TRANSPORTER, ATP-BINDING P |
| 97:00:00 | 3nbx-X | 2.6 | 6 | 59 | 481 | 8 | MOLECULE: | ATPASE RAVA |
| 98:00:00 | 7yxx-A | 2.6 | 6.2 | 67 | 1754 | 15 | MOLECULE: | PROBABLE UBIQUITIN CARBOXYL-TERMINAL HYDROLASE FA |
| 99:00:00 | 8tn1-A | 2.6 | 3.7 | 60 | 147 | 12 | MOLECULE: | DE NOVO DESIGNED 4 HELIX BUNDLES |
| 0:00 | 7yp0-B | 2.6 | 7.2 | 68 | 278 | 6 | MOLECULE: | GLUTATHIONE S-TRANSFERASE |
| 1:00 | 4n5c-D | 2.6 | 5.9 | 62 | 757 | 5 | MOLECULE: | CARGO-TRANSPORT PROTEIN YPP1 |
| 2:00 | 6kkk-A | 2.6 | 4.5 | 54 | 380 | 9 | MOLECULE: | SUGAR EFFLUX TRANSPORTER |

| | | | | | | | | |
|---|---|---|---|---|---|---|---|---|
| 3:00 | 5dfz-B | 2.6 | 7.3 | 73 | 1224 | 8 | MOLECULE: | VACUOLAR PROTEIN SORTING-ASSOCIATED PROTEIN 38 |
| 4:00 | 5img-A | 2.6 | 19.7 | 92 | 467 | 2 | MOLECULE: | DIPEPTIDASE |
| 5:00 | 7byl-A | 2.6 | 27 | 101 | 354 | 3 | MOLECULE: | GREEN FLUORESCENT PROTEIN,POTASSIUM VOLTAGE-GATED |
| 6:00 | 7emf-K | 2.6 | 13.3 | 69 | 112 | 1 | MOLECULE: | MEDIATOR OF RNA POLYMERASE II TRANSCRIPTION SUBUN |
| 7:00 | 8oyv-A | 2.6 | 10.6 | 68 | 182 | 9 | MOLECULE: | DE NOVO DESIGNED SOLUBLE CLAUDIN |
| 8:00 | 6v4a-C | 2.6 | 7.7 | 95 | 593 | 13 | MOLECULE: | NEUR_CHAN_LBD DOMAIN-CONTAINING PROTEIN |
| 9:00 | 4xal-A | 2.6 | 5.2 | 57 | 90 | 9 | MOLECULE: | TEGUMENT PROTEIN VP22 |
| 10:00 | 5ijo-J | 2.6 | 6.2 | 74 | 1256 | 8 | MOLECULE: | NUCLEAR PORE COMPLEX PROTEIN NUP155 |
| 11:00 | 3oa7-A | 2.6 | 8.9 | 85 | 193 | 15 | MOLECULE: | HEAD MORPHOGENESIS PROTEIN, CHAOTIC NUCLEAR MIGRA |
| 12:00 | 8a3t-C | 2.6 | 27.1 | 74 | 1410 | 5 | MOLECULE: | ANAPHASE-PROMOTING COMPLEX SUBUNIT CDC27 |
| 13:00 | 7elb-A | 2.6 | 32.3 | 79 | 1937 | 9 | MOLECULE: | RNA-DIRECTED RNA POLYMERASE L |
| 14:00 | 5j9q-D | 2.6 | 6.6 | 66 | 120 | 14 | MOLECULE: | HISTONE ACETYLTRANSFERASE ESA1 |
| 15:00 | 6dql-A | 2.6 | 4.8 | 48 | 227 | 6 | MOLECULE: | REGULATOR OF PROTEINASE B ROPB |
| 16:00 | 7jqe-A | 2.6 | 3 | 70 | 357 | 7 | MOLECULE: | ESAT-6/WXG100 SECRETION SYSTEM PROTEIN |
| 17:00 | 7kak-E | 2.6 | 26.5 | 67 | 172 | 6 | MOLECULE: | PROTEIN TRANSPORT CHANNEL SEC61 COMPLEX, ALPHA SU |
| 18:00 | 6v6d-A | 2.6 | 5.9 | 74 | 209 | 8 | MOLECULE: | PANNEXIN-1 |
| 19:00 | 5vgz-D | 2.6 | 3.8 | 54 | 107 | 6 | MOLECULE: | 26S PROTEASOME REGULATORY SUBUNIT 7 |
| 20:00 | 3m62-A | 2.6 | 5.6 | 57 | 955 | 14 | MOLECULE: | UBIQUITIN CONJUGATION FACTOR E4 |
| 21:00 | 6rax-N | 2.6 | 3.4 | 51 | 207 | 10 | MOLECULE: | DNA REPLICATION LICENSING FACTOR MCM2 |
| 22:00 | 6yle-D | 2.6 | 10.1 | 57 | 547 | 11 | MOLECULE: | PRE-RRNA-PROCESSING PROTEIN IPI3 |
| 23:00 | 8b7b-E | 2.6 | 51.5 | 93 | 123 | 6 | MOLECULE: | TUBULIN ALPHA-1B CHAIN |
| 24:00:00 | 7xqx-E | 2.6 | 50.3 | 93 | 123 | 0 | MOLECULE: | TUBULIN ALPHA-1B CHAIN |
| 25:00:00 | 7xr1-E | 2.6 | 50.2 | 93 | 123 | 0 | MOLECULE: | TUBULIN ALPHA-1B CHAIN |
| 26:00:00 | 6znm-x | 2.6 | 45.8 | 81 | 142 | 9 | MOLECULE: | ARP1 ACTIN RELATED PROTEIN 1 HOMOLOG A |
| 27:00:00 | 2e7s-K | 2.6 | 9 | 77 | 117 | 4 | MOLECULE: | RAB GUANINE NUCLEOTIDE EXCHANGE FACTOR SEC2 |
| 28:00:00 | 6b3r-A | 2.6 | 22.8 | 70 | 1502 | 4 | MOLECULE: | PIEZO-TYPE MECHANOSENSITIVE ION CHANNEL COMPONENT |
| 29:00:00 | 8pew-i | 2.6 | 5.4 | 67 | 187 | 3 | MOLECULE: | TRANSCRIPTION TERMINATION FACTOR RHO |
| 30:00:00 | 2ezs-B | 2.6 | 2.4 | 68 | 123 | 15 | MOLECULE: | GP41 |
| 31:00:00 | 2yo1-A | 2.6 | 8.7 | 83 | 273 | 6 | MOLECULE: | GENERAL CONTROL PROTEIN GCN4, PUTATIVE INNER MEMB |
| 32:00:00 | 2ezq-B | 2.6 | 2.6 | 68 | 123 | 15 | MOLECULE: | GP41 |
| 33:00:00 | 2ezq-A | 2.6 | 2.6 | 68 | 123 | 15 | MOLECULE: | GP41 |
| 34:00:00 | 2ezs-C | 2.6 | 2.4 | 68 | 123 | 16 | MOLECULE: | GP41 |
| 35:00:00 | 2d3e-A | 2.6 | 13.8 | 74 | 130 | 8 | MOLECULE: | GENERAL CONTROL PROTEIN GCN4 AND TROPOMYOSIN 1 AL |
| 36:00:00 | 2wpr-B | 2.6 | 9.7 | 62 | 97 | 13 | MOLECULE: | TRIMERIC AUTOTRANSPORTER ADHESIN FRAGMENT |
| 37:00:00 | 3mtu-A | 2.6 | 4.4 | 47 | 75 | 6 | MOLECULE: | TROPOMYOSIN ALPHA-1 CHAIN,MICROTUBULE-ASSOCIATED |
| 38:00:00 | 4mvd-D | 2.6 | 13.6 | 93 | 253 | 2 | MOLECULE: | CHOLINE-PHOSPHATE CYTIDYLYLTRANSFERASE A |
| 39:00:00 | 6bua-A | 2.5 | 39.9 | 62 | 942 | 5 | MOLECULE: | DICER-2, ISOFORM A |
| 40:00:00 | 6xky-A | 2.5 | 25.3 | 104 | 271 | 8 | MOLECULE: | FLAGELLIN |
| 41:00:00 | 6t9i-G | 2.5 | 8.1 | 75 | 140 | 8 | MOLECULE: | TRANSCRIPTION FACTOR SPT20 |
| 42:00:00 | 7xym-A | 2.5 | 4.6 | 54 | 488 | 11 | MOLECULE: | ENVELOPE GLYCOPROTEIN |
| 43:00:00 | 8ewi-A | 2.5 | 28.2 | 101 | 1775 | 5 | MOLECULE: | E3 UBIQUITIN-PROTEIN LIGASE UBR5 |
| 44:00:00 | 6zh3-A | 2.5 | 36.7 | 86 | 170 | 3 | MOLECULE: | VACUOLAR PROTEIN-SORTING-ASSOCIATED PROTEIN 24 |
| 45:00:00 | 3jac-A | 2.5 | 31.6 | 122 | 918 | 4 | MOLECULE: | PIEZO-TYPE MECHANOSENSITIVE ION CHANNEL COMPONENT |
| 46:00:00 | 8s7c-A | 2.5 | 21.1 | 96 | 1030 | 7 | MOLECULE: | VWFA AND CACHE DOMAIN-CONTAINING PROTEIN 1 |
| 47:00:00 | 6rjw-A | 2.5 | 19.7 | 94 | 161 | 11 | MOLECULE: | LYSM DOMAIN PROTEIN |
| 48:00:00 | 1sc7-A | 2.5 | 22.9 | 63 | 567 | 6 | MOLECULE: | 5'-D(*AP*AP*AP*AP*AP*GP*AP*CP*TP*T)-3' |
| 49:00:00 | 8v46-A | 2.5 | 10.5 | 74 | 412 | 5 | MOLECULE: | ARIA ANTITOXIN |
| 50:00:00 | 6xt9-A | 2.5 | 37.6 | 60 | 567 | 5 | MOLECULE: | BARDET-BIEDL SYNDROME 1 PROTEIN |
| 51:00:00 | 4wid-A | 2.5 | 3.3 | 96 | 353 | 7 | MOLECULE: | RHUL123 |
| 52:00:00 | 7p3x-B | 2.5 | 2.6 | 48 | 621 | 8 | MOLECULE: | AP-3 COMPLEX SUBUNIT DELTA |
| 53:00:00 | 7nyw-A | 2.5 | 9.3 | 113 | 685 | 10 | MOLECULE: | CHROMOSOME PARTITION PROTEIN MUKB |
| 54:00:00 | 2ncg-A | 2.5 | 4.3 | 58 | 118 | 7 | MOLECULE: | RGA1E |
| 55:00:00 | 8arl-A | 2.5 | 8.1 | 87 | 232 | 5 | MOLECULE: | TRYPTOPHAN-RICH ANTIGEN |
| 56:00:00 | 7rmy-A | 2.5 | 4.8 | 61 | 266 | 11 | MOLECULE: | DE NOVO DESIGNED TUNABLE HOMODIMER, D_3-337 |
| 57:00:00 | 8w41-A | 2.5 | 7.9 | 92 | 1162 | 10 | MOLECULE: | CALMODULIN-1 |
| 58:00:00 | 5c1f-A | 2.5 | 20.4 | 99 | 298 | 5 | MOLECULE: | SEPTATION PROTEIN IMP2 |

| | | | | | | | | |
|---|---|---|---|---|---|---|---|---|
| 59:00:00 | 8h6u-A | 2.5 | 3.2 | 69 | 338 | 4 | MOLECULE: | PRESILPHIPERFOLAN-8-BETA-OL SYNTHASE |
| 60:00:00 | 8ap7-a | 2.5 | 11 | 100 | 231 | 12 | MOLECULE: | ATP SYNTHASE SUBUNIT A |
| 61:00:00 | 7qlr-A | 2.5 | 27.1 | 94 | 603 | 10 | MOLECULE: | CDHS1_22 PUTATIVE TAIL FIBER PROTEIN |
| 62:00:00 | 6ny2-Y | 2.5 | 23.2 | 65 | 915 | 6 | MOLECULE: | DNA TARGET STRAND |
| 63:00:00 | 6its-A | 2.5 | 3.6 | 59 | 147 | 2 | MOLECULE: | METHYL-ACCEPTING CHEMOTAXIS SENSORY TRANSDUCER |
| 64:00:00 | 1wa8-A | 2.5 | 3.9 | 62 | 99 | 8 | MOLECULE: | ESAT-6 LIKE PROTEIN ESXB |
| 65:00:00 | 8jec-B | 2.5 | 8.6 | 74 | 663 | 7 | MOLECULE: | POTASSIUM CHANNEL SKOR |
| 66:00:00 | 3vou-A | 2.5 | 6.7 | 68 | 139 | 15 | MOLECULE: | ION TRANSPORT 2 DOMAIN PROTEIN, VOLTAGE-GATED SOD |
| 67:00:00 | 7nyw-B | 2.5 | 10 | 113 | 858 | 7 | MOLECULE: | CHROMOSOME PARTITION PROTEIN MUKB |
| 68:00:00 | 6uo8-B | 2.5 | 13.3 | 75 | 696 | 5 | MOLECULE: | GAMMA-AMINOBUTYRIC ACID TYPE B RECEPTOR SUBUNIT 1 |
| 69:00:00 | 7uhb-K | 2.5 | 2.8 | 49 | 75 | 2 | MOLECULE: | SPIKE GLYCOPROTEIN |
| 70:00:00 | 4myy-B | 2.5 | 7.7 | 60 | 84 | 8 | MOLECULE: | CURG, CURH FUSION PROTEIN |
| 71:00:00 | 6wu0-B | 2.5 | 8 | 91 | 854 | 8 | MOLECULE: | HOPANOID BIOSYNTHESIS ASSOCIATED RND TRANSPORTER |
| 72:00:00 | 3sae-A | 2.5 | 7 | 65 | 780 | 8 | MOLECULE: | ALPHA-BISABOLENE SYNTHASE |
| 73:00:00 | 6e1k-A | 2.5 | 9.3 | 88 | 536 | 9 | MOLECULE: | TWO PORE CALCIUM CHANNEL PROTEIN 1 |
| 74:00:00 | 4gwp-A | 2.5 | 7.1 | 54 | 112 | 9 | MOLECULE: | MEDIATOR OF RNA POLYMERASE II TRANSCRIPTION SUBUN |
| 75:00:00 | 8d8j-0 | 2.5 | 11.6 | 57 | 493 | 7 | MOLECULE: | PROBABLE S-ADENOSYL-L-METHIONINE-DEPENDENT RNA |
| 76:00:00 | 2p01-A | 2.5 | 9.7 | 58 | 323 | 14 | MOLECULE: | ALPHA-2-MACROGLOBULIN RECEPTOR-ASSOCIATED |
| 77:00:00 | 2c5i-T | 2.5 | 3 | 54 | 94 | 4 | MOLECULE: | T-SNARE AFFECTING A LATE GOLGI COMPARTMENT |
| 78:00:00 | 8hk0-B | 2.5 | 12 | 69 | 379 | 4 | MOLECULE: | DEHYDROGENASE |
| 79:00:00 | 5b86-A | 2.5 | 19.2 | 57 | 579 | 9 | MOLECULE: | TUMOR NECROSIS FACTOR ALPHA-INDUCED PROTEIN 2 |
| 80:00:00 | 7ewp-A | 2.5 | 32.6 | 68 | 577 | 7 | MOLECULE: | PROBABLE G-PROTEIN COUPLED RECEPTOR 158 |
| 81:00:00 | 6yj4-o | 2.5 | 4.3 | 53 | 83 | 4 | MOLECULE: | NADH-UBIQUINONE OXIDOREDUCTASE CHAIN 3 |
| 82:00:00 | 8jwd-A | 2.5 | 1.5 | 53 | 123 | 4 | MOLECULE: | HISTIDINE KINASE |
| 83:00:00 | 8ikg-R | 2.5 | 9.3 | 66 | 275 | 9 | MOLECULE: | GUANINE NUCLEOTIDE-BINDING PROTEIN G(I) SUBUNIT A |
| 84:00:00 | 3f0c-A | 2.5 | 3.4 | 50 | 193 | 18 | MOLECULE: | TRANSCRIPTIONAL REGULATOR |
| 85:00:00 | 3bm3-A | 2.5 | 3.8 | 60 | 259 | 5 | MOLECULE: | DNA (5'-D(*CP*AP*TP*CP*CP*AP*GP*GP*TP*AP*C)-3') |
| 86:00:00 | 3lf9-A | 2.5 | 3 | 66 | 120 | 8 | MOLECULE: | 4E10_D0_1IS1A_001_C (T161) |
| 87:00:00 | 8hti-R | 2.5 | 3.1 | 63 | 303 | 5 | MOLECULE: | GUANINE NUCLEOTIDE-BINDING PROTEIN G(S) SUBUNIT A |
| 88:00:00 | 4cem-B | 2.5 | 15.6 | 114 | 309 | 5 | MOLECULE: | REGULATOR OF NONSENSE TRANSCRIPTS 2 |
| 89:00:00 | 6uz3-A | 2.5 | 6.1 | 54 | 1126 | 7 | MOLECULE: | SODIUM CHANNEL PROTEIN TYPE 5 SUBUNIT ALPHA,GREEN |
| 90:00:00 | 3kkb-B | 2.5 | 2.2 | 57 | 128 | 12 | MOLECULE: | SENSOR PROTEIN |
| 91:00:00 | 8yxl-A | 2.5 | 9.7 | 71 | 699 | 4 | MOLECULE: | RNA-DIRECTED RNA POLYMERASE L |
| 92:00:00 | 7oi8-q | 2.5 | 9.8 | 91 | 164 | 7 | MOLECULE: | 39S RIBOSOMAL PROTEIN L2, MITOCHONDRIAL |
| 93:00:00 | 8hc0-A | 2.5 | 2.5 | 42 | 360 | 10 | MOLECULE: | ADHESION G-PROTEIN COUPLED RECEPTOR F1 |
| 94:00:00 | 5wd8-A | 2.5 | 4.5 | 53 | 90 | 9 | MOLECULE: | UNCHARACTERIZED PROTEIN |
| 95:00:00 | 8erc-A | 2.5 | 15.9 | 88 | 441 | 3 | MOLECULE: | LYSOPHOSPHOLIPID ACYLTRANSFERASE 7 |
| 96:00:00 | 2lor-A | 2.5 | 7.8 | 70 | 108 | 9 | MOLECULE: | TRANSMEMBRANE PROTEIN 141 |
| 97:00:00 | 3caz-B | 2.5 | 4.9 | 80 | 210 | 6 | MOLECULE: | BAR PROTEIN |
| 98:00:00 | 7emf-I | 2.5 | 5.4 | 58 | 73 | 5 | MOLECULE: | MEDIATOR OF RNA POLYMERASE II TRANSCRIPTION SUBUN |
| 99:00:00 | 8tek-E | 2.5 | 5.9 | 64 | 219 | 3 | MOLECULE: | DYNEIN REGULATORY COMPLEX PROTEIN 1/2 N-TERMINAL |
| 0:00 | 7r5s-S | 2.5 | 3 | 48 | 120 | 6 | MOLECULE: | CENTROMERE PROTEIN H |
| 1:00 | 3n1e-B | 2.5 | 5.8 | 66 | 141 | 3 | MOLECULE: | VACUOLAR PROTEIN SORTING-ASSOCIATED PROTEIN 54 |
| 2:00 | 2jmh-A | 2.5 | 4.4 | 61 | 117 | 2 | MOLECULE: | MITE ALLERGEN BLO T 5 |
| 3:00 | 7qoo-U | 2.5 | 18.7 | 85 | 186 | 9 | MOLECULE: | CENTROMERE PROTEIN C |
| 4:00 | 3rx6-A | 2.5 | 9.7 | 78 | 187 | 3 | MOLECULE: | POLARITY SUPPRESSION PROTEIN |
| 5:00 | 6ugm-X | 2.5 | 3.1 | 54 | 73 | 6 | MOLECULE: | HISTONE H3 |
| 6:00 | 6u9w-A | 2.5 | 9.5 | 61 | 562 | 5 | MOLECULE: | P2X PURINOCEPTOR 7 |
| 7:00 | 6xy6-C | 2.5 | 3.4 | 45 | 138 | 13 | MOLECULE: | ANTI-APOPTOTIC MEMBRANE PROTEIN |
| 8:00 | 4wpe-A | 2.5 | 20.1 | 100 | 275 | 3 | MOLECULE: | CYTOKINESIS PROTEIN 2 |
| 9:00 | 8dd7-B | 2.5 | 4.2 | 66 | 759 | 9 | MOLECULE: | METHYLATED-DNA--PROTEIN-CYSTEINE METHYLTRANSFERAS |
| 10:00 | 6cnn-A | 2.5 | 8.5 | 83 | 360 | 7 | MOLECULE: | INTERMEDIATE CONDUCTANCE CALCIUM-ACTIVATED POTASS |
| 11:00 | 7pc1-A | 2.5 | 4.2 | 50 | 72 | 10 | MOLECULE: | STBA |
| 12:00 | 8jjb-E | 2.5 | 51.9 | 93 | 123 | 8 | MOLECULE: | TUBULIN ALPHA-1B CHAIN |
| 13:00 | 8clg-E | 2.5 | 51.9 | 94 | 123 | 6 | MOLECULE: | TUBULIN ALPHA-1B CHAIN |
| 14:00 | 6ff7-K | 2.5 | 32.7 | 94 | 213 | 7 | MOLECULE: | RNA-BINDING MOTIF PROTEIN, X-LINKED 2 |

| | | | | | | | | |
|---|---|---|---|---|---|---|---|---|
| 15:00 | 4n5b-H | 2.5 | 5.6 | 68 | 103 | 4 | MOLECULE: | PHOSPHOPROTEIN |
| 16:00 | 8clc-E | 2.5 | 51.8 | 93 | 123 | 6 | MOLECULE: | TUBULIN ALPHA-1B CHAIN |
| 17:00 | 7jfr-E | 2.5 | 50.7 | 92 | 123 | 3 | MOLECULE: | TUBULIN ALPHA-1B CHAIN |
| 18:00 | 8jjc-E | 2.5 | 52 | 94 | 123 | 4 | MOLECULE: | TUBULIN ALPHA-1B CHAIN |
| 19:00 | 5j2u-E | 2.5 | 51 | 92 | 123 | 7 | MOLECULE: | TUBULIN ALPHA-1B CHAIN |
| 20:00 | 6sgz-H | 2.5 | 9.7 | 87 | 442 | 6 | MOLECULE: | ESX-3 SECRETION SYSTEM PROTEIN ECCD3 |
| 21:00 | 8c5v-K | 2.5 | 55.2 | 134 | 516 | 5 | MOLECULE: | CHEMOTAXIS PROTEIN CHEA |
| 22:00 | 8q0f-p | 2.5 | 10.1 | 75 | 174 | 7 | MOLECULE: | NADH-UBIQUINONE OXIDOREDUCTASE CHAIN 3 |
| 23:00 | 2xnx-A | 2.5 | 2.7 | 48 | 74 | 8 | MOLECULE: | FIBRINOGEN ALPHA CHAIN |
| 24:00:00 | 8cok-B | 2.5 | 6 | 58 | 101 | 2 | MOLECULE: | INHIBITOR OF GROWTH PROTEIN 3 |
| 25:00:00 | 8pew-k | 2.5 | 5 | 69 | 187 | 4 | MOLECULE: | TRANSCRIPTION TERMINATION FACTOR RHO |
| 26:00:00 | 3c9i-B | 2.5 | 8.1 | 84 | 242 | 7 | MOLECULE: | TAIL NEEDLE PROTEIN GP26 |
| 27:00:00 | 4ckg-A | 2.5 | 3.2 | 80 | 368 | 9 | MOLECULE: | ARF-GAP WITH COILED-COIL, ANK REPEAT AND PH DOMAI |
| 28:00:00 | 5m9e-C | 2.5 | 1.4 | 40 | 68 | 3 | MOLECULE: | MICROTUBULE INTEGRITY PROTEIN MAL3 |
| 29:00:00 | 6j5e-K | 2.5 | 14.5 | 38 | 42 | 8 | MOLECULE: | ENVELOPE GLYCOPROTEIN |
| 30:00:00 | 4mbe-E | 2.5 | 15.6 | 46 | 51 | 0 | MOLECULE: | CELL DIVISION CONTROL PROTEIN 31 |
| 31:00:00 | 3zmf-C | 2.5 | 8.7 | 64 | 111 | 8 | MOLECULE: | GENERAL CONTROL PROTEIN GCN4, PUTATIVE INNER MEMB |
| 32:00:00 | 1llm-D | 2.5 | 6 | 59 | 85 | 7 | MOLECULE: | 5'-D(*TP*CP*CP*CP*AP*CP*GP*CP*GP*TP*GP*GP*G)-3' |
| 33:00:00 | 7zet-A | 2.4 | 18.2 | 96 | 377 | 6 | MOLECULE: | CLUSTERIN |
| 34:00:00 | 4tn3-B | 2.4 | 11.8 | 128 | 362 | 5 | MOLECULE: | TRIM5/CYCLOPHILIN A FUSION PROTEIN/T4 LYSOZYME CH |
| 35:00:00 | 4gwp-B | 2.4 | 35.7 | 95 | 492 | 7 | MOLECULE: | MEDIATOR OF RNA POLYMERASE II TRANSCRIPTION SUBUN |
| 36:00:00 | 8ifg-A | 2.4 | 17.6 | 84 | 879 | 5 | MOLECULE: | RBAP48-RELATED WD40 REPEAT-CONTAINING PROTEIN PRW |
| 37:00:00 | 7qtt-Y | 2.4 | 24.8 | 75 | 306 | 1 | MOLECULE: | SPLICING FACTOR 3B SUBUNIT 3 |
| 38:00:00 | 8tie-c | 2.4 | 23.8 | 55 | 610 | 2 | MOLECULE: | NUCLEOPORIN NUP120 |
| 39:00:00 | 3tul-A | 2.4 | 4.2 | 61 | 136 | 11 | MOLECULE: | CELL INVASION PROTEIN SIPB |
| 40:00:00 | 3wxx-B | 2.4 | 22.3 | 73 | 183 | 4 | MOLECULE: | ACRH |
| 41:00:00 | 6yj4-g | 2.4 | 12.6 | 78 | 198 | 5 | MOLECULE: | NADH-UBIQUINONE OXIDOREDUCTASE CHAIN 3 |
| 42:00:00 | 8hpo-C | 2.4 | 39.6 | 93 | 138 | 5 | MOLECULE: | TRANSCRIPTIONAL REGULATORY PROTEIN UME1 |
| 43:00:00 | 7qhm-I | 2.4 | 4.5 | 61 | 135 | 8 | MOLECULE: | CYTOCHROME BC1 COMPLEX RIESKE IRON-SULFUR SUBUNIT |
| 44:00:00 | 6fes-A | 2.4 | 11.8 | 95 | 365 | 5 | MOLECULE: | D12_BRIC2, A SYNTHETIC PROTEIN,D12_BRIC2, A SYNTH |
| 45:00:00 | 6vz1-A | 2.4 | 20.7 | 103 | 411 | 7 | MOLECULE: | DIACYLGLYCEROL O-ACYLTRANSFERASE 1 |
| 46:00:00 | 7nb6-A | 2.4 | 10.5 | 92 | 340 | 9 | MOLECULE: | AI-2 TRANSPORT PROTEIN TQSA |
| 47:00:00 | 3nrg-A | 2.4 | 13.2 | 55 | 217 | 13 | MOLECULE: | TETR FAMILY TRANSCRIPTIONAL REGULATOR |
| 48:00:00 | 2qfc-A | 2.4 | 4.5 | 54 | 284 | 4 | MOLECULE: | PLCR PROTEIN |
| 49:00:00 | 4fz4-A | 2.4 | 3.6 | 53 | 154 | 11 | MOLECULE: | UNCHARACTERIZED PROTEIN CONSERVED IN BACTERIA |
| 50:00:00 | 2yfb-B | 2.4 | 4.9 | 84 | 238 | 8 | MOLECULE: | METHYL-ACCEPTING CHEMOTAXIS TRANSDUCER |
| 51:00:00 | 7woo-F | 2.4 | 8.4 | 79 | 1622 | 10 | MOLECULE: | NUCLEOPORIN NIC96 |
| 52:00:00 | 5wjt-A | 2.4 | 5.4 | 101 | 302 | 10 | MOLECULE: | FLAGELLIN |
| 53:00:00 | 7uwf-C | 2.4 | 3.4 | 73 | 517 | 8 | MOLECULE: | WD REPEAT-CONTAINING PROTEIN 18 |
| 54:00:00 | 6irr-A | 2.4 | 4.6 | 59 | 133 | 8 | MOLECULE: | DISRUPTED IN SCHIZOPHRENIA 1 HOMOLOG,CYCLIC AMP-D |
| 55:00:00 | 7wiu-B | 2.4 | 7.6 | 91 | 576 | 7 | MOLECULE: | MYCOBACTIN IMPORT ATP-BINDING/PERMEASE PROTEIN IR |
| 56:00:00 | 6lul-A | 2.4 | 5.7 | 67 | 223 | 13 | MOLECULE: | RESPIRATORY SUPERCOMPLEX FACTOR 2, MITOCHONDRIAL |
| 57:00:00 | 7mx2-B | 2.4 | 9.4 | 61 | 648 | 7 | MOLECULE: | N-ALPHA-ACETYLTRANSFERASE 35, NATC AUXILIARY SUBU |
| 58:00:00 | 8eup-3 | 2.4 | 8.2 | 68 | 194 | 7 | MOLECULE: | RNA (1422-MER) |
| 59:00:00 | 8dql-A | 2.4 | 11.6 | 67 | 257 | 6 | MOLECULE: | SECRETION SYSTEM PROTEIN |
| 60:00:00 | 2a79-B | 2.4 | 16.5 | 89 | 259 | 7 | MOLECULE: | VOLTAGE-GATED POTASSIUM CHANNEL BETA-2 SUBUNIT |
| 61:00:00 | 8jh7-D | 2.4 | 14.3 | 96 | 431 | 4 | MOLECULE: | ANTI-BRIL FAB HEAVY CHAIN |
| 62:00:00 | 7qpg-W | 2.4 | 6.4 | 69 | 779 | 9 | MOLECULE: | PROTEIN ZWILCH HOMOLOG |
| 63:00:00 | 3no6-A | 2.4 | 3.5 | 58 | 235 | 5 | MOLECULE: | TRANSCRIPTIONAL ACTIVATOR TENA |
| 64:00:00 | 8b9z-K | 2.4 | 2.9 | 53 | 96 | 6 | MOLECULE: | NADH-UBIQUINONE OXIDOREDUCTASE CHAIN 3 |
| 65:00:00 | 6ayi-A | 2.4 | 12 | 43 | 183 | 14 | MOLECULE: | HTH-TYPE TRANSCRIPTIONAL REGULATOR UIDR |
| 66:00:00 | 4wlp-B | 2.4 | 15.7 | 72 | 114 | 6 | MOLECULE: | UBIQUITIN CARBOXYL-TERMINAL HYDROLASE ISOZYME L5 |
| 67:00:00 | 5mnt-B | 2.4 | 14.9 | 115 | 421 | 9 | MOLECULE: | A2 MATURATION PROTEIN |
| 68:00:00 | 7swj-A | 2.4 | 8.2 | 76 | 300 | 14 | MOLECULE: | INWARD RECTIFIER POTASSIUM CHANNEL |
| 69:00:00 | 8buy-A | 2.4 | 2.9 | 61 | 134 | 10 | MOLECULE: | GRANULE ASSOCIATED RAC AND RHOG EFFECTOR PROTEIN |
| 70:00:00 | 6vxm-A | 2.4 | 3.1 | 57 | 277 | 5 | MOLECULE: | MECHANOSENSITIVE ION CHANNEL PROTEIN 1, MITOCHOND |

| | | | | | | | | |
|---|---|---|---|---|---|---|---|---|
| 71:00:00 | 4xng-D | 2.4 | 3.9 | 65 | 144 | 12 | MOLECULE: | UNCHARACTERIZED PROTEIN MG218.1 |
| 72:00:00 | 6ajf-A | 2.4 | 7.1 | 78 | 901 | 3 | MOLECULE: | DRUG EXPORTERS OF THE RND SUPERFAMILY-LIKE PROTEI |
| 73:00:00 | 8t53-A | 2.4 | 7.9 | 63 | 398 | 8 | MOLECULE: | UNDECAPRENYL-PHOSPHATE GALACTOSE PHOSPHOTRANSFERA |
| 74:00:00 | 3p01-A | 2.4 | 2.7 | 49 | 178 | 14 | MOLECULE: | TWO-COMPONENT RESPONSE REGULATOR |
| 75:00:00 | 7r5s-T | 2.4 | 4.8 | 50 | 112 | 10 | MOLECULE: | CENTROMERE PROTEIN H |
| 76:00:00 | 3afl-A | 2.4 | 7.5 | 66 | 766 | 9 | MOLECULE: | OLIGO ALGINATE LYASE |
| 77:00:00 | 3tix-A | 2.4 | 6.2 | 60 | 153 | 10 | MOLECULE: | UBIQUITIN-LIKE PROTEIN SMT3,RNA-INDUCED TRANSCRIP |
| 78:00:00 | 5x3q-A | 2.4 | 2.7 | 58 | 313 | 9 | MOLECULE: | ENVELOPE GLYCOPROTEIN |
| 79:00:00 | 5lbm-C | 2.4 | 4 | 56 | 90 | 4 | MOLECULE: | TRANSCRIPTIONAL REPRESSOR FRMR |
| 80:00:00 | 2qup-A | 2.4 | 2.3 | 53 | 119 | 9 | MOLECULE: | BH1478 PROTEIN |
| 81:00:00 | 7f9l-A | 2.4 | 3.2 | 69 | 162 | 3 | MOLECULE: | RIFIN |
| 82:00:00 | 8hij-A | 2.4 | 20.3 | 103 | 499 | 8 | MOLECULE: | BRIL-SLC19A1 CHIMERA |
| 83:00:00 | 5ijj-B | 2.4 | 3.3 | 66 | 157 | 11 | MOLECULE: | SPX DOMAIN |
| 84:00:00 | 3lic-A | 2.4 | 3 | 70 | 265 | 10 | MOLECULE: | SENSOR PROTEIN |
| 85:00:00 | 6zce-q | 2.4 | 19.6 | 78 | 636 | 4 | MOLECULE: | EUKARYOTIC TRANSLATION INITIATION FACTOR 3 SUBUNI |
| 86:00:00 | 6o84-A | 2.4 | 7.4 | 68 | 415 | 9 | MOLECULE: | LOC100127796 PROTEIN,LOC100127796 PROTEIN,OTOP3, |
| 87:00:00 | 4f52-E | 2.4 | 6.5 | 78 | 523 | 6 | MOLECULE: | CULLIN-1 |
| 88:00:00 | 8gxl-B | 2.4 | 3.8 | 58 | 122 | 7 | MOLECULE: | SURP AND G-PATCH DOMAIN-CONTAINING PROTEIN 1 |
| 89:00:00 | 6e67-B | 2.4 | 8.9 | 72 | 476 | 8 | MOLECULE: | BETA-2 ADRENERGIC RECEPTOR,ENDOLYSIN,GUANINE NUCL |
| 90:00:00 | 8jhk-A | 2.4 | 5.4 | 63 | 727 | 8 | MOLECULE: | ENGULFMENT AND CELL MOTILITY PROTEIN 1 |
| 91:00:00 | 8gju-F | 2.4 | 11.5 | 84 | 332 | 7 | MOLECULE: | METHYLMALONIC ACIDURIA TYPE A PROTEIN, MITOCHONDR |
| 92:00:00 | 4y66-C | 2.4 | 21.3 | 94 | 197 | 7 | MOLECULE: | MND1 |
| 93:00:00 | 5xtc-j | 2.4 | 14.3 | 53 | 115 | 19 | MOLECULE: | NADH DEHYDROGENASE [UBIQUINONE] IRON-SULFUR PROTE |
| 94:00:00 | 8ebt-C | 2.4 | 18.5 | 61 | 191 | 10 | MOLECULE: | GENERAL TRANSCRIPTION AND DNA REPAIR FACTOR IIH H |
| 95:00:00 | 2qqy-A | 2.4 | 2.7 | 55 | 139 | 9 | MOLECULE: | SIGMA B OPERON |
| 96:00:00 | 7p5v-B | 2.4 | 4.3 | 65 | 732 | 6 | MOLECULE: | VOLUME-REGULATED ANION CHANNEL SUBUNIT LRRC8A |
| 97:00:00 | 8dqk-A | 2.4 | 5.2 | 65 | 733 | 5 | MOLECULE: | CELLULOSE SYNTHASE-LIKE CSLF6 |
| 98:00:00 | 8ugc-A | 2.4 | 15.3 | 72 | 384 | 3 | MOLECULE: | FD15 |
| 99:00:00 | 3nmz-A | 2.4 | 6.3 | 50 | 406 | 2 | MOLECULE: | APC VARIANT PROTEIN |
| 0:00 | 2py8-C | 2.4 | 10.8 | 71 | 123 | 7 | MOLECULE: | HYPOTHETICAL PROTEIN RBCX |
| 1:00 | 6vtk-A | 2.4 | 4.4 | 62 | 446 | 13 | MOLECULE: | ACID-SENSING ION CHANNEL 1 |
| 2:00 | 6uak-A | 2.4 | 7.6 | 59 | 298 | 3 | MOLECULE: | SAM DEPENDENT METHYLTRANSFERASE LAHSB |
| 3:00 | 3hj1-B | 2.4 | 21 | 98 | 387 | 6 | MOLECULE: | MINOR EDITOSOME-ASSOCIATED TUTASE |
| 4:00 | 8z1e-A | 2.4 | 7.8 | 64 | 278 | 6 | MOLECULE: | UNCHARACTERIZED PROTEIN UL78 |
| 5:00 | 6wq0-A | 2.4 | 14.5 | 75 | 131 | 11 | MOLECULE: | DNA (301-MER) |
| 6:00 | 1zbd-B | 2.4 | 4.5 | 66 | 124 | 5 | MOLECULE: | RABPHILIN-3A |
| 7:00 | 6wqz-A | 2.4 | 7.1 | 62 | 536 | 5 | MOLECULE: | AUTOPHAGY-RELATED PROTEIN 9A |
| 8:00 | 6poo-A | 2.4 | 9.8 | 104 | 273 | 1 | MOLECULE: | BIBA |
| 9:00 | 4hkr-A | 2.4 | 2.2 | 54 | 165 | 7 | MOLECULE: | CALCIUM RELEASE-ACTIVATED CALCIUM CHANNEL PROTEIN |
| 10:00 | 7pp2-A | 2.4 | 4.3 | 56 | 442 | 9 | MOLECULE: | EXOCYST SUBUNIT EXO70 FAMILY PROTEIN |
| 11:00 | 7c79-L | 2.4 | 6.7 | 60 | 131 | 7 | MOLECULE: | RIBONUCLEASE MRP RNA SUBUNIT NME1 |
| 12:00 | 8tlq-A | 2.4 | 7.5 | 70 | 1283 | 6 | MOLECULE: | DNA POLYMERASE ZETA CATALYTIC SUBUNIT |
| 13:00 | 7tuk-A | 2.4 | 5.1 | 49 | 142 | 6 | MOLECULE: | SAG PROTEIN |
| 14:00 | 7zyw-E | 2.4 | 49.8 | 88 | 119 | 0 | MOLECULE: | TUBULIN ALPHA-1B CHAIN |
| 15:00 | 8clf-E | 2.4 | 50.8 | 91 | 121 | 8 | MOLECULE: | TUBULIN ALPHA-1B CHAIN |
| 16:00 | 8clb-E | 2.4 | 50.8 | 91 | 121 | 7 | MOLECULE: | TUBULIN ALPHA-1B CHAIN |
| 17:00 | 8cld-E | 2.4 | 50.6 | 91 | 121 | 8 | MOLECULE: | DETYROSINATED TUBULIN ALPHA-1B CHAIN |
| 18:00 | 8huh-E | 2.4 | 50.6 | 91 | 121 | 8 | MOLECULE: | TUBULIN ALPHA-1B CHAIN |
| 19:00 | 6zkn-Z | 2.4 | 9.7 | 75 | 171 | 4 | MOLECULE: | NADH DEHYDROGENASE [UBIQUINONE] FLAVOPROTEIN 1, |
| 20:00 | 6zkm-Z | 2.4 | 9.7 | 75 | 171 | 4 | MOLECULE: | NADH DEHYDROGENASE [UBIQUINONE] FLAVOPROTEIN 1, |
| 21:00 | 5cw3-A | 2.4 | 4.8 | 69 | 248 | 4 | MOLECULE: | BRCA1/BRCA2-CONTAINING COMPLEX SUBUNIT 3 |
| 22:00 | 8cpz-B | 2.4 | 7.1 | 99 | 2122 | 9 | MOLECULE: | TCDA1 |
| 23:00 | 1kql-A | 2.4 | 14.2 | 46 | 55 | 9 | MOLECULE: | FUSION PROTEIN OF AND STRIATED MUSCLE ALPHA-TROPO |
| 24:00:00 | 5h0n-K | 2.4 | 12.3 | 37 | 43 | 16 | MOLECULE: | HIV-1 GP41 NHR |
| 25:00:00 | 2z5h-F | 2.4 | 15.9 | 48 | 51 | 4 | MOLECULE: | GENERAL CONTROL PROTEIN GCN4 AND TROPOMYOSIN ALPH |
| 26:00:00 | 2r5b-C | 2.4 | 16 | 42 | 45 | 2 | MOLECULE: | GP41 N-PEPTIDE |

| | | | | | | | | |
|---|---|---|---|---|---|---|---|---|
| 27:00:00 | 6n5m-D | 2.4 | 14.5 | 51 | 62 | 8 | MOLECULE: | POLYUBIQUITIN-C |
| 28:00:00 | 3vgy-C | 2.4 | 15.3 | 44 | 49 | 9 | MOLECULE: | ENVELOPE GLYCOPROTEIN GP160 |
| 29:00:00 | 5cmz-C | 2.4 | 14.9 | 39 | 45 | 8 | MOLECULE: | ENVELOPE GLYCOPROTEIN |
| 30:00:00 | 5m9e-B | 2.4 | 1.6 | 40 | 71 | 3 | MOLECULE: | MICROTUBULE INTEGRITY PROTEIN MAL3 |
| 31:00:00 | 5jvp-F | 2.4 | 9.6 | 58 | 84 | 5 | MOLECULE: | CHIMERA PROTEIN OF CENTROMERE-ASSOCIATED PROTEIN |
| 32:00:00 | 2q7c-B | 2.4 | 15.4 | 42 | 45 | 10 | MOLECULE: | FUSION PROTEIN BETWEEN YEAST VARIANT GCN4 AND HIV |
| 33:00:00 | 7l8y-F | 2.4 | 4.5 | 56 | 143 | 7 | MOLECULE: | RH.33311 PABC-5 - HEAVY CHAIN |
| 34:00:00 | 7l8t-D | 2.4 | 4.5 | 56 | 144 | 7 | MOLECULE: | BG505 SOSIP.V5.2 N241/N289 - GP120 |
| 35:00:00 | 7rai-D | 2.4 | 4.1 | 45 | 128 | 13 | MOLECULE: | ENVELOPE GLYCOPROTEIN GP160 |
| 36:00:00 | 6yvd-D | 2.3 | 38.8 | 78 | 414 | 6 | MOLECULE: | CONDENSIN COMPLEX SUBUNIT 2 |
| 37:00:00 | 8pbz-B | 2.3 | 6.6 | 55 | 1380 | 4 | MOLECULE: | MGP-OPERON PROTEIN 3 |
| 38:00:00 | 5ofb-B | 2.3 | 17.9 | 85 | 541 | 2 | MOLECULE: | MORC FAMILY CW-TYPE ZINC FINGER PROTEIN 2 |
| 39:00:00 | 8qae-A | 2.3 | 29.6 | 68 | 211 | 6 | MOLECULE: | SC-APCC-6-SLLA |
| 40:00:00 | 7zr1-D | 2.3 | 18 | 110 | 780 | 2 | MOLECULE: | DOUBLE-STRAND BREAK REPAIR PROTEIN |
| 41:00:00 | 3ter-B | 2.3 | 5.3 | 68 | 121 | 7 | MOLECULE: | MAMMALIAN STROMAL INTERACTION MOLECULE-1 |
| 42:00:00 | 8ipr-C | 2.3 | 7.4 | 89 | 515 | 8 | MOLECULE: | COMPONENT LINKED WITH THE ASSEMBLY OF CYTOCHROME' |
| 43:00:00 | 6d5f-A | 2.3 | 12 | 79 | 193 | 8 | MOLECULE: | FIMBRIAL PROTEIN |
| 44:00:00 | 5e5w-B | 2.3 | 3.9 | 67 | 149 | 9 | MOLECULE: | HEMAGGLUTININ-ESTERASE |
| 45:00:00 | 7yfu-A | 2.3 | 12.1 | 37 | 44 | 14 | MOLECULE: | PROTEIN FANTOM |
| 46:00:00 | 7y22-e | 2.3 | 3.8 | 58 | 172 | 14 | MOLECULE: | PHAGE CONNECTOR PROTEIN |
| 47:00:00 | 7rvb-A | 2.3 | 3.1 | 56 | 209 | 2 | MOLECULE: | FERRITIN |
| 48:00:00 | 6qg0-G | 2.3 | 5.9 | 58 | 355 | 10 | MOLECULE: | TRANSLATION INITIATION FACTOR EIF-2B SUBUNIT ALPH |
| 49:00:00 | 1jfi-B | 2.3 | 11.2 | 83 | 135 | 8 | MOLECULE: | 5'- |
| 50:00:00 | 5tr1-A | 2.3 | 23.3 | 65 | 606 | 3 | MOLECULE: | CHLORIDE CHANNEL PROTEIN |
| 51:00:00 | 7ykr-A | 2.3 | 28.1 | 79 | 1035 | 6 | MOLECULE: | TRANSIENT RECEPTOR POTENTIAL CATION CHANNEL SUBFA |
| 52:00:00 | 8rhn-A | 2.3 | 4.2 | 56 | 198 | 7 | MOLECULE: | ATPASE FAMILY GENE 2 PROTEIN HOMOLOG A |
| 53:00:00 | 2lqt-A | 2.3 | 5.3 | 50 | 72 | 0 | MOLECULE: | COILED-COIL-HELIX-COILED-COIL-HELIX DOMAIN-CONTAI |
| 54:00:00 | 7b9f-D | 2.3 | 15.1 | 78 | 485 | 5 | MOLECULE: | ECCE5 |
| 55:00:00 | 8cqr-A | 2.3 | 4.2 | 59 | 103 | 10 | MOLECULE: | NINJURIN-1 |
| 56:00:00 | 2r44-A | 2.3 | 3.4 | 51 | 330 | 4 | MOLECULE: | UNCHARACTERIZED PROTEIN |
| 57:00:00 | 6xp5-D | 2.3 | 4.2 | 68 | 134 | 7 | MOLECULE: | MEDIATOR OF RNA POLYMERASE II TRANSCRIPTION SUBUN |
| 58:00:00 | 7d5p-B | 2.3 | 14.7 | 77 | 413 | 3 | MOLECULE: | DRUG TRANSPORTER, PUTATIVE |
| 59:00:00 | 6vz0-B | 2.3 | 3.1 | 56 | 76 | 11 | MOLECULE: | PULMONARY SURFACTANT-ASSOCIATED PROTEIN B |
| 60:00:00 | 7dgy-A | 2.3 | 3.6 | 48 | 99 | 4 | MOLECULE: | C2 |
| 61:00:00 | 2g8l-B | 2.3 | 8 | 57 | 286 | 4 | MOLECULE: | 287AA LONG HYPOTHETICAL PROTEIN |
| 62:00:00 | 7dwb-A | 2.3 | 10.5 | 99 | 421 | 4 | MOLECULE: | PANNEXIN-1 |
| 63:00:00 | 8b9z-h | 2.3 | 17.5 | 78 | 145 | 8 | MOLECULE: | NADH-UBIQUINONE OXIDOREDUCTASE CHAIN 3 |
| 64:00:00 | 6z5l-A | 2.3 | 16.8 | 91 | 251 | 1 | MOLECULE: | MATRIX PROTEIN 1 |
| 65:00:00 | 6icz-A | 2.3 | 14.2 | 83 | 2253 | 5 | MOLECULE: | PROTEIN MAGO NASHI HOMOLOG 2 |
| 66:00:00 | 7a6h-O | 2.3 | 13.2 | 69 | 512 | 7 | MOLECULE: | DNA-DIRECTED RNA POLYMERASE III SUBUNIT RPC1 |
| 67:00:00 | 2q83-A | 2.3 | 18.8 | 70 | 332 | 9 | MOLECULE: | YTAA PROTEIN |
| 68:00:00 | 8z9a-D | 2.3 | 7.1 | 88 | 357 | 5 | MOLECULE: | ODORANT RECEPTOR, APISORCO |
| 69:00:00 | 6q6e-A | 2.3 | 9.5 | 88 | 221 | 8 | MOLECULE: | CONDENSIN COMPLEX SUBUNIT 2,STRUCTURAL MAINTENANC |
| 70:00:00 | 4gzu-A | 2.3 | 2.6 | 56 | 436 | 7 | MOLECULE: | FERM, RHOGEF AND PLECKSTRIN DOMAIN-CONTAINING PRO |
| 71:00:00 | 6yj4-V | 2.3 | 3.4 | 49 | 126 | 0 | MOLECULE: | NADH-UBIQUINONE OXIDOREDUCTASE CHAIN 3 |
| 72:00:00 | 6bfi-B | 2.3 | 4.1 | 80 | 806 | 5 | MOLECULE: | VIN1 |
| 73:00:00 | 8bh8-A | 2.3 | 3.7 | 55 | 523 | 7 | MOLECULE: | PCIF1_WW DOMAIN-CONTAINING PROTEIN |
| 74:00:00 | 8fbi-A | 2.3 | 2.7 | 51 | 274 | 6 | MOLECULE: | KWOCA_39 |
| 75:00:00 | 7cun-B | 2.3 | 11.7 | 67 | 1058 | 7 | MOLECULE: | INTEGRATOR COMPLEX SUBUNIT 1 |
| 76:00:00 | 4mk6-A | 2.3 | 12.8 | 62 | 188 | 10 | MOLECULE: | PROBABLE DIHYDROXYACETONE KINASE REGULATOR DHSK_R |
| 77:00:00 | 5x11-E | 2.3 | 7.1 | 63 | 181 | 8 | MOLECULE: | DNA (28-MER) |
| 78:00:00 | 8wlj-A | 2.3 | 13.7 | 68 | 409 | 3 | MOLECULE: | SYNAPTIC VESICULAR AMINE TRANSPORTER |
| 79:00:00 | 8k0b-A | 2.3 | 7.2 | 86 | 689 | 7 | MOLECULE: | CALCIUM PERMEABLE STRESS-GATED CATION CHANNEL 1 |
| 80:00:00 | 6li2-A | 2.3 | 8.4 | 73 | 347 | 8 | MOLECULE: | CHIMERA OF G-PROTEIN COUPLED RECEPTOR 52 AND RUBR |
| 81:00:00 | 6whp-A | 2.3 | 12.1 | 80 | 428 | 6 | MOLECULE: | CHOLINE KINASE |
| 82:00:00 | 3wvo-A | 2.3 | 5.4 | 81 | 543 | 4 | MOLECULE: | CRISPR-ASSOCIATED PROTEIN, CSE1 FAMILY |

| | | | | | | | | |
|---|---|---|---|---|---|---|---|---|
| 83:00:00 | 8ftw-A | 2.3 | 3.1 | 61 | 166 | 15 | MOLECULE: | FLAGELLAR FLIT PROTEIN, FLAGELLAR FLIJ PROTEIN FU |
| 84:00:00 | 3kwo-A | 2.3 | 2.5 | 58 | 149 | 10 | MOLECULE: | PUTATIVE BACTERIOFERRITIN |
| 85:00:00 | 8ity-4 | 2.3 | 13.2 | 92 | 365 | 3 | MOLECULE: | SNRNA-ACTIVATING PROTEIN COMPLEX SUBUNIT 1 |
| 86:00:00 | 6xss-A | 2.3 | 18.2 | 56 | 260 | 9 | MOLECULE: | C4_NAT_HFUSE-7900 |
| 87:00:00 | 7ux0-A | 2.3 | 3.2 | 44 | 115 | 7 | MOLECULE: | SPERM-EGG FUSION PROTEIN TMEM95 |
| 88:00:00 | 2d6y-B | 2.3 | 14.9 | 57 | 188 | 7 | MOLECULE: | PUTATIVE TETR FAMILY REGULATORY PROTEIN |
| 89:00:00 | 7rcy-A | 2.3 | 5.4 | 69 | 829 | 13 | MOLECULE: | PENICILLIN-BINDING PROTEIN |
| 90:00:00 | 8e9g-L | 2.3 | 12.9 | 69 | 625 | 7 | MOLECULE: | TWO-COMPONENT SYSTEM RESPONSE REGULATOR |
| 91:00:00 | 1uus-A | 2.3 | 3.4 | 66 | 465 | 8 | MOLECULE: | STAT PROTEIN |
| 92:00:00 | 8sl4-A | 2.3 | 7.6 | 99 | 970 | 6 | MOLECULE: | ADENYLATE CYCLASE TYPE 5 |
| 93:00:00 | 2qyw-A | 2.3 | 3.4 | 54 | 96 | 7 | MOLECULE: | VESICLE TRANSPORT THROUGH INTERACTION WITH T-SNAR |
| 94:00:00 | 7woo-K | 2.3 | 11.7 | 84 | 254 | 13 | MOLECULE: | NUCLEOPORIN NIC96 |
| 95:00:00 | 1ku9-A | 2.3 | 6.9 | 72 | 151 | 6 | MOLECULE: | HYPOTHETICAL PROTEIN MJ223 |
| 96:00:00 | 4dlq-A | 2.3 | 5.6 | 68 | 353 | 10 | MOLECULE: | LATROPHILIN-1 |
| 97:00:00 | 7nad-8 | 2.3 | 8.7 | 58 | 58 | 10 | MOLECULE: | 25S RRNA |
| 98:00:00 | 8hf2-A | 2.3 | 2.7 | 52 | 120 | 10 | MOLECULE: | PRA1 FAMILY PROTEIN |
| 99:00:00 | 5n8k-C | 2.3 | 4.8 | 58 | 80 | 10 | MOLECULE: | GALACTOCEREBROSIDASE |
| 0:00 | 7w5a-2 | 2.3 | 2.2 | 57 | 123 | 2 | MOLECULE: | PRE-MRNA-PROCESSING-SPLICING FACTOR 8 |
| 1:00 | 8fck-H | 2.3 | 5.7 | 68 | 218 | 12 | MOLECULE: | HAUS AUGMIN-LIKE COMPLEX SUBUNIT 1 |
| 2:00 | 8q3v-b | 2.3 | 3.4 | 56 | 70 | 9 | MOLECULE: | TETRAHYDROMETHANOPTERIN S-METHYLTRANSFERASE SUBUN |
| 3:00 | 7n84-b | 2.3 | 3.2 | 60 | 675 | 10 | MOLECULE: | NUCLEOPORIN NUP188 |
| 4:00 | 8jlf-A | 2.3 | 3.8 | 71 | 592 | 8 | MOLECULE: | SYNAPTIC VESICLE GLYCOPROTEIN 2A |
| 5:00 | 8b9z-m | 2.3 | 12.7 | 81 | 108 | 6 | MOLECULE: | NADH-UBIQUINONE OXIDOREDUCTASE CHAIN 3 |
| 6:00 | 1xdo-A | 2.3 | 3.7 | 53 | 687 | 11 | MOLECULE: | POLYPHOSPHATE KINASE |
| 7:00 | 8ap7-f | 2.3 | 14.7 | 78 | 135 | 6 | MOLECULE: | ATP SYNTHASE SUBUNIT A |
| 8:00 | 6fj3-A | 2.3 | 8.4 | 71 | 562 | 6 | MOLECULE: | PARATHYROID HORMONE/PARATHYROID HORMONE-RELATED P |
| 9:00 | 6f1y-f | 2.3 | 3.6 | 46 | 280 | 7 | MOLECULE: | CYTOPLASMIC DYNEIN 1 HEAVY CHAIN 1,DYNEIN HEAVY C |
| 10:00 | 3c64-A | 2.3 | 3.9 | 52 | 152 | 4 | MOLECULE: | PFEMP1 VARIANT 2 OF STRAIN MC |
| 11:00 | 1tjl-A | 2.3 | 3.7 | 61 | 145 | 3 | MOLECULE: | DNAK SUPPRESSOR PROTEIN |
| 12:00 | 5ebz-A | 2.3 | 8.9 | 115 | 655 | 5 | MOLECULE: | INHIBITOR OF NUCLEAR FACTOR KAPPA-B KINASE SUBUNI |
| 13:00 | 7qj2-G | 2.3 | 31.7 | 70 | 640 | 4 | MOLECULE: | GAMMA-TUBULIN COMPLEX COMPONENT 5 |
| 14:00 | 4p1n-B | 2.3 | 6.3 | 56 | 207 | 11 | MOLECULE: | ATG1 TMIT |
| 15:00 | 6xz6-A | 2.3 | 6 | 75 | 213 | 5 | MOLECULE: | GARP DOMAIN-CONTAINING PROTEIN |
| 16:00 | 8hf3-A | 2.3 | 9.3 | 101 | 295 | 5 | MOLECULE: | PALMITOYLTRANSFERASE ZDHHC9 |
| 17:00 | 5a7d-R | 2.3 | 18.5 | 76 | 311 | 11 | MOLECULE: | PINS |
| 18:00 | 8ear-A | 2.3 | 8.4 | 102 | 2389 | 7 | MOLECULE: | INOSITOL 1,4,5-TRISPHOSPHATE RECEPTOR TYPE 1 |
| 19:00 | 8qa2-A | 2.3 | 6.1 | 70 | 215 | 4 | MOLECULE: | GAP JUNCTION BETA-2 PROTEIN |
| 20:00 | 6lcn-F | 2.3 | 8.6 | 68 | 309 | 10 | MOLECULE: | SERINE O-ACETYLTRANSFERASE |
| 21:00 | 8ikj-R | 2.3 | 2.8 | 51 | 532 | 8 | MOLECULE: | ADHESION G PROTEIN-COUPLED RECEPTOR E5,SOLUBLE CY |
| 22:00 | 7w1m-B | 2.3 | 11.8 | 117 | 528 | 5 | MOLECULE: | STRUCTURAL MAINTENANCE OF CHROMOSOMES PROTEIN 1A |
| 23:00 | 6br8-A | 2.3 | 4.3 | 47 | 249 | 4 | MOLECULE: | PROTEIN A6 HOMOLOG |
| 24:00:00 | 6nsj-A | 2.3 | 12.1 | 59 | 182 | 10 | MOLECULE: | ACID-ACTIVATED UREA CHANNEL |
| 25:00:00 | 7d2w-A | 2.3 | 2.8 | 57 | 125 | 4 | MOLECULE: | PRESAN DOMAIN-CONTAINING PROTEIN |
| 26:00:00 | 2uxx-B | 2.3 | 11.6 | 73 | 134 | 5 | MOLECULE: | LYSINE-SPECIFIC HISTONE DEMETHYLASE 1 |
| 27:00:00 | 2oh3-A | 2.3 | 3.5 | 55 | 149 | 5 | MOLECULE: | COG1633: UNCHARACTERIZED CONSERVED PROTEIN |
| 28:00:00 | 2fbq-A | 2.3 | 10.4 | 61 | 213 | 7 | MOLECULE: | PROBABLE TRANSCRIPTIONAL REGULATOR |
| 29:00:00 | 8kih-A | 2.3 | 8.3 | 67 | 291 | 1 | MOLECULE: | DITERPENE SYNTHASE, PHMA |
| 30:00:00 | 6e7e-A | 2.3 | 5.1 | 65 | 169 | 12 | MOLECULE: | INCLUSION MEMBRANE PROTEIN A |
| 31:00:00 | 7wji-C | 2.3 | 17.2 | 117 | 1394 | 6 | MOLECULE: | PROTEIN UNC-80 HOMOLOG |
| 32:00:00 | 3mxz-A | 2.3 | 4.3 | 60 | 107 | 8 | MOLECULE: | TUBULIN-SPECIFIC CHAPERONE A |
| 33:00:00 | 8uc3-C | 2.3 | 2.7 | 47 | 98 | 9 | MOLECULE: | ALBONOURSIN SYNTHASE |
| 34:00:00 | 6yxq-B | 2.3 | 4.9 | 54 | 396 | 6 | MOLECULE: | ACTIVATING SIGNAL COINTEGRATOR 1 COMPLEX SUBUNIT |
| 35:00:00 | 8cle-E | 2.3 | 50.1 | 90 | 120 | 8 | MOLECULE: | TUBULIN ALPHA-1B CHAIN |
| 36:00:00 | 7w2y-d | 2.3 | 8.7 | 69 | 175 | 7 | MOLECULE: | NADH DEHYDROGENASE [UBIQUINONE] FLAVOPROTEIN 1, |
| 37:00:00 | 5kht-D | 2.3 | 15.4 | 41 | 47 | 2 | MOLECULE: | TROPOMYOSIN ALPHA-1 CHAIN,GENERAL CONTROL PROTEIN |
| 38:00:00 | 1gzl-B | 2.3 | 15.5 | 42 | 45 | 10 | MOLECULE: | FUSION PROTEIN BETWEEN THE HYDROPHOBIC POCKET OF |

| | | | | | | | | |
|---|---|---|---|---|---|---|---|---|
| 39:00:00 | 5h0n-C | 2.3 | 13 | 36 | 42 | 8 | MOLECULE: | HIV-1 GP41 NHR |
| 40:00:00 | 3jbh-H | 2.3 | 26.8 | 104 | 964 | 2 | MOLECULE: | MYOSIN 2 HEAVY CHAIN STRIATED MUSCLE |
| 41:00:00 | 5jv3-C | 2.3 | 14 | 45 | 64 | 0 | MOLECULE: | CHIMERA PROTEIN OF KINESIN-LIKE PROTEIN KIF11 AND |
| 42:00:00 | 2efs-C | 2.3 | 11.4 | 72 | 154 | 8 | MOLECULE: | GENERAL CONTROL PROTEIN GCN4 AND TROPOMYOSIN 1 AL |
| 43:00:00 | 3cvf-B | 2.3 | 9.5 | 55 | 70 | 4 | MOLECULE: | HOMER PROTEIN HOMOLOG 3 |
| 44:00:00 | 3l35-C | 2.3 | 16.5 | 43 | 45 | 9 | MOLECULE: | GP41 N-PEPTIDE |
| 45:00:00 | 8fis-B | 2.3 | 10 | 47 | 125 | 6 | MOLECULE: | ENVELOPE GLYCOPROTEIN GP41 |
| 46:00:00 | 7ur6-B | 2.3 | 12.2 | 50 | 132 | 4 | MOLECULE: | GP120 |
| 47:00:00 | 6vo3-B | 2.3 | 4.1 | 43 | 124 | 7 | MOLECULE: | PGV04 HEAVY CHAIN |
| 48:00:00 | 2yo1-C | 2.3 | 6.9 | 84 | 273 | 4 | MOLECULE: | GENERAL CONTROL PROTEIN GCN4, PUTATIVE INNER MEMB |
| 49:00:00 | 7rai-B | 2.3 | 5.8 | 53 | 128 | 11 | MOLECULE: | ENVELOPE GLYCOPROTEIN GP160 |
| 50:00:00 | 6vo3-F | 2.3 | 4.1 | 43 | 124 | 7 | MOLECULE: | PGV04 HEAVY CHAIN |
| 51:00:00 | 6vo3-E | 2.3 | 4.1 | 43 | 124 | 7 | MOLECULE: | PGV04 HEAVY CHAIN |
| 52:00:00 | 4avm-A | 2.2 | 4.9 | 77 | 230 | 4 | MOLECULE: | BRIDGING INTEGRATOR 2 |
| 53:00:00 | 7sn7-a | 2.2 | 34.5 | 108 | 546 | 6 | MOLECULE: | FLAGELLIN |
| 54:00:00 | 6h2x-A | 2.2 | 22 | 95 | 352 | 5 | MOLECULE: | CHROMOSOME PARTITION PROTEIN MUKB,CHROMOSOME PART |
| 55:00:00 | 5zb2-A | 2.2 | 20.9 | 59 | 402 | 2 | MOLECULE: | DNA REPAIR PROTEIN RAD7 |
| 56:00:00 | 5gao-E | 2.2 | 5.1 | 48 | 338 | 0 | MOLECULE: | SMALL NUCLEAR RIBONUCLEOPROTEIN-ASSOCIATED PROTEI |
| 57:00:00 | 5y88-U | 2.2 | 15.4 | 56 | 488 | 9 | MOLECULE: | PRE-MRNA-SPLICING FACTOR 8 |
| 58:00:00 | 5lnc-B | 2.2 | 13.6 | 94 | 367 | 6 | MOLECULE: | VACUOLAR TRANSPORTER CHAPERONE 4,CORE HISTONE MAC |
| 59:00:00 | 6z2w-E | 2.2 | 31.5 | 90 | 2325 | 3 | MOLECULE: | DNA DAMAGE CHECKPOINT PROTEIN LCD1 |
| 60:00:00 | 7yqh-A | 2.2 | 12.4 | 97 | 1069 | 9 | MOLECULE: | STRUCTURAL MAINTENANCE OF CHROMOSOMES PROTEIN 5 |
| 61:00:00 | 8tek-A | 2.2 | 38.6 | 83 | 200 | 4 | MOLECULE: | DYNEIN REGULATORY COMPLEX PROTEIN 1/2 N-TERMINAL |
| 62:00:00 | 8j2f-A | 2.2 | 2.8 | 44 | 384 | 5 | MOLECULE: | SPHINGOMYELIN PHOSPHODIESTERASE 2 |
| 63:00:00 | 8g0l-B | 2.2 | 28.5 | 68 | 925 | 4 | MOLECULE: | N-ALPHA-ACETYLTRANSFERASE 20 |
| 64:00:00 | 5c8j-L | 2.2 | 3.8 | 50 | 137 | 10 | MOLECULE: | ANTIBODY FRAGMENT, HEAVY CHAIN |
| 65:00:00 | 8xgr-R | 2.2 | 14.4 | 67 | 305 | 7 | MOLECULE: | GUANINE NUCLEOTIDE-BINDING PROTEIN G(I)/G(S)/G(O) |
| 66:00:00 | 8k7t-B | 2.2 | 3.6 | 60 | 160 | 10 | MOLECULE: | HIGH AFFINITY IMMUNOGLOBULIN EPSILON RECEPTOR SUB |
| 67:00:00 | 8uza-A | 2.2 | 6.8 | 73 | 910 | 5 | MOLECULE: | CRISPR-ASSOCIATED ENDONUCLEASE CAS9 |
| 68:00:00 | 5oqk-A | 2.2 | 4.3 | 66 | 147 | 9 | MOLECULE: | VOLTAGE-GATED HYDROGEN CHANNEL 1 |
| 69:00:00 | 3c98-B | 2.2 | 12.8 | 85 | 230 | 11 | MOLECULE: | SYNTAXIN-BINDING PROTEIN 1 |
| 70:00:00 | 6yj4-L | 2.2 | 9.4 | 102 | 655 | 3 | MOLECULE: | NADH-UBIQUINONE OXIDOREDUCTASE CHAIN 3 |
| 71:00:00 | 8bf9-G | 2.2 | 9.4 | 62 | 158 | 6 | MOLECULE: | RNA (1766) |
| 72:00:00 | 8f72-B | 2.2 | 10.3 | 58 | 595 | 12 | MOLECULE: | TPR_REGION DOMAIN-CONTAINING PROTEIN |
| 73:00:00 | 7oci-C | 2.2 | 4.7 | 62 | 119 | 10 | MOLECULE: | DOLICHYL-DIPHOSPHOOLIGOSACCHARIDE--PROTEIN |
| 74:00:00 | 3k1s-A | 2.2 | 4.6 | 58 | 109 | 3 | MOLECULE: | PTS SYSTEM, CELLOBIOSE-SPECIFIC IIA COMPONENT |
| 75:00:00 | 6c66-A | 2.2 | 20.5 | 73 | 511 | 8 | MOLECULE: | CRISPR-ASSOCIATED HELICASE, CAS3 FAMILY |
| 76:00:00 | 2wzk-A | 2.2 | 23.5 | 62 | 369 | 10 | MOLECULE: | CULLIN-5 |
| 77:00:00 | 5tth-A | 2.2 | 12.8 | 70 | 671 | 9 | MOLECULE: | C-TERMINAL SPYCATCHER FUSION OF WILDTYPE ZEBRAFIS |
| 78:00:00 | 8on7-A | 2.2 | 4.1 | 57 | 525 | 9 | MOLECULE: | FMRFAMIDE-GATED SODIUM CHANNEL 1 (FANAC1) |
| 79:00:00 | 4h54-A | 2.2 | 2.2 | 50 | 274 | 8 | MOLECULE: | DIGUANYLATE CYCLASE YDEH |
| 80:00:00 | 8i6v-D | 2.2 | 29.5 | 76 | 678 | 8 | MOLECULE: | VACUOLAR TRANSPORTER CHAPERONE COMPLEX SUBUNIT 1 |
| 81:00:00 | 6aay-A | 2.2 | 21.1 | 107 | 1199 | 8 | MOLECULE: | BERGEYELLA ZOOHELCUM CAS13B (R1177A) MUTANT |
| 82:00:00 | 3o6q-A | 2.2 | 4.1 | 51 | 145 | 14 | MOLECULE: | STAGE II SPORULATION PROTEIN SA |
| 83:00:00 | 4d6k-A | 2.2 | 6.2 | 50 | 76 | 4 | MOLECULE: | DEOXYNUCLEOTIDYLTRANSFERASE TERMINAL-INTERACTING |
| 84:00:00 | 7e40-D | 2.2 | 2.7 | 67 | 345 | 9 | MOLECULE: | PROTEIN PHOSPHATE STARVATION RESPONSE 2 |
| 85:00:00 | 5f5p-A | 2.2 | 5.2 | 86 | 181 | 10 | MOLECULE: | PROTEIN SHROOM2 |
| 86:00:00 | 8r8q-A | 2.2 | 11.4 | 77 | 391 | 8 | MOLECULE: | MAJOR FACILITATOR SUPERFAMILY DOMAIN-CONTAINING P |
| 87:00:00 | 4o8w-D | 2.2 | 11.7 | 69 | 120 | 6 | MOLECULE: | SPORE GERMINATION PROTEIN |
| 88:00:00 | 6vrc-A | 2.2 | 14.3 | 68 | 1007 | 10 | MOLECULE: | CRISPR-ASSOCIATED ENDORIBONUCLEASE CAS13A |
| 89:00:00 | 5kua-A | 2.2 | 3.8 | 63 | 161 | 10 | MOLECULE: | PILIN |
| 90:00:00 | 2zet-C | 2.2 | 4.7 | 70 | 141 | 6 | MOLECULE: | RAS-RELATED PROTEIN RAB-27B |
| 91:00:00 | 3aai-A | 2.2 | 4.4 | 50 | 78 | 2 | MOLECULE: | COPPER HOMEOSTASIS OPERON REGULATORY PROTEIN |
| 92:00:00 | 7vrc-C | 2.2 | 2.5 | 57 | 109 | 14 | MOLECULE: | TRANSCRIPTION REGULATORY PROTEIN SNF11 |
| 93:00:00 | 6y92-A | 2.2 | 9.5 | 67 | 178 | 12 | MOLECULE: | B-LYMPHOCYTE ANTIGEN CD20 |
| 94:00:00 | 8d07-A | 2.2 | 3 | 45 | 65 | 13 | MOLECULE: | HALC3_109 |

| | | | | | | | | |
|---|---|---|---|---|---|---|---|---|
| 95:00:00 | 7k10-A | 2.2 | 3.4 | 44 | 1259 | 20 | MOLECULE: | DNA-DEPENDENT PROTEIN KINASE CATALYTIC SUBUNIT |
| 96:00:00 | 8fvt-A | 2.2 | 3 | 53 | 96 | 8 | MOLECULE: | 3HB12 |
| 97:00:00 | 8pop-A | 2.2 | 6.9 | 71 | 122 | 1 | MOLECULE: | TERMINASE SMALL SUBUNIT |
| 98:00:00 | 6rwb-A | 2.2 | 11 | 102 | 1873 | 7 | MOLECULE: | TOXIN,TOXIN COMPLEX SUBUNIT TCAB,PUTATIVE TOXIN S |
| 99:00:00 | 7yim-A | 2.2 | 13.8 | 77 | 592 | 8 | MOLECULE: | ALPHA-FETOPROTEIN |
| 0:00 | 3q9d-A | 2.2 | 22.7 | 64 | 175 | 11 | MOLECULE: | PROTEIN CPN_0803/CP_1068/CPJ0803/CPB0832 |
| 1:00 | 8wcs-A | 2.2 | 10.9 | 66 | 1044 | 2 | MOLECULE: | CAS13H1 |
| 2:00 | 4h8s-C | 2.2 | 6.8 | 84 | 382 | 6 | MOLECULE: | DCC-INTERACTING PROTEIN 13-BETA |
| 3:00 | 7f4u-A | 2.2 | 6.1 | 69 | 440 | 10 | MOLECULE: | TELOMERE LENGTH REGULATION PROTEIN TEL2 HOMOLOG |
| 4:00 | 2oap-1 | 2.2 | 14.3 | 64 | 498 | 5 | MOLECULE: | TYPE II SECRETION SYSTEM PROTEIN |
| 5:00 | 7eu3-D | 2.2 | 3.6 | 54 | 499 | 11 | MOLECULE: | NAD(P)H-QUINONE OXIDOREDUCTASE SUBUNIT 1, CHLOROP |
| 6:00 | 7vcf-H | 2.2 | 2.4 | 52 | 101 | 13 | MOLECULE: | TIC214 |
| 7:00 | 8t1i-F | 2.2 | 7.2 | 66 | 73 | 11 | MOLECULE: | MEDIATOR OF RNA POLYMERASE II TRANSCRIPTION SUBUN |
| 8:00 | 5wb2-A | 2.2 | 9.1 | 68 | 421 | 7 | MOLECULE: | ENVELOPE PROTEIN US28, NANOBODY 7 FUSION PROTEIN |
| 9:00 | 5lc5-h | 2.2 | 30.8 | 87 | 134 | 0 | MOLECULE: | NADH-UBIQUINONE OXIDOREDUCTASE CHAIN 3 |
| 10:00 | 1vcs-A | 2.2 | 7.2 | 64 | 102 | 8 | MOLECULE: | VESICLE TRANSPORT THROUGH INTERACTION WITH T- |
| 11:00 | 8jxo-A | 2.2 | 8.6 | 55 | 262 | 5 | MOLECULE: | BLCHR2 |
| 12:00 | 7bmh-A | 2.2 | 5.4 | 78 | 247 | 8 | MOLECULE: | OPSIN |
| 13:00 | 6o1q-A | 2.2 | 3.8 | 58 | 119 | 10 | MOLECULE: | NEPHROCYSTIN-1 |
| 14:00 | 7zpo-M | 2.2 | 7.9 | 74 | 438 | 8 | MOLECULE: | KTR SYSTEM POTASSIUM UPTAKE PROTEIN A |
| 15:00 | 2ewg-A | 2.2 | 6.3 | 49 | 367 | 6 | MOLECULE: | FARNESYL PYROPHOSPHATE SYNTHASE |
| 16:00 | 1u4q-A | 2.2 | 4.9 | 87 | 318 | 5 | MOLECULE: | SPECTRIN ALPHA CHAIN, BRAIN |
| 17:00 | 3h36-A | 2.2 | 3.1 | 48 | 78 | 10 | MOLECULE: | POLYRIBONUCLEOTIDE NUCLEOTIDYLTRANSFERASE |
| 18:00 | 3f46-A | 2.2 | 13.1 | 58 | 345 | 9 | MOLECULE: | 5,10-METHENYLTETRAHYDROMETHANOPTERIN HYDROGENASE |
| 19:00 | 5ce3-B | 2.2 | 18.6 | 81 | 598 | 7 | MOLECULE: | ACTIN |
| 20:00 | 4ad8-A | 2.2 | 13.6 | 111 | 452 | 4 | MOLECULE: | DNA REPAIR PROTEIN RECN |
| 21:00 | 6ulg-N | 2.2 | 11.9 | 59 | 483 | 3 | MOLECULE: | FOLLICULIN |
| 22:00 | 6eo1-A | 2.2 | 6.2 | 61 | 355 | 10 | MOLECULE: | CYCLIC NUCLEOTIDE-GATED POTASSIUM CHANNEL MLL3241 |
| 23:00 | 4y9j-B | 2.2 | 17.2 | 79 | 593 | 9 | MOLECULE: | PROTEIN ACDH-11, ISOFORM B |
| 24:00:00 | 3j83-A | 2.2 | 4.2 | 72 | 276 | 10 | MOLECULE: | ESX-1 SECRETION-ASSOCIATED PROTEIN ESPB |
| 25:00:00 | 4k2u-A | 2.2 | 4.4 | 71 | 230 | 8 | MOLECULE: | ERYTHROCYTE BINDING ANTIGEN 175 |
| 26:00:00 | 5loi-A | 2.2 | 17.7 | 75 | 391 | 8 | MOLECULE: | RAD26 |
| 27:00:00 | 3emo-A | 2.2 | 5.4 | 68 | 126 | 7 | MOLECULE: | HIA (ADHESIN) |
| 28:00:00 | 7rps-A | 2.2 | 3.2 | 52 | 154 | 8 | MOLECULE: | FIBRONECTIN-BINDING LIPOPROTEIN FBPB |
| 29:00:00 | 7lzh-A | 2.2 | 11 | 84 | 799 | 5 | MOLECULE: | GLUTAMATE RECEPTOR 3.4 |
| 30:00:00 | 7t71-A | 2.2 | 15 | 68 | 348 | 6 | MOLECULE: | MEVALONATE 3,5-BISPHOSPHATE DECARBOXYLASE |
| 31:00:00 | 4aur-A | 2.2 | 5.6 | 123 | 564 | 6 | MOLECULE: | LEOA |
| 32:00:00 | 5wg6-A | 2.2 | 18.2 | 57 | 517 | 7 | MOLECULE: | HISTONE-LYSINE N-METHYLTRANSFERASE EZH2,POLYCOMB |
| 33:00:00 | 7jrg-G | 2.2 | 7.6 | 61 | 70 | 8 | MOLECULE: | MITOCHONDRIAL-PROCESSING PEPTIDASE SUBUNIT BETA, |
| 34:00:00 | 6e9n-A | 2.2 | 12 | 75 | 409 | 7 | MOLECULE: | D-GALACTONATE TRANSPORT |
| 35:00:00 | 7d5i-A | 2.2 | 12.4 | 94 | 443 | 11 | MOLECULE: | CYTOCHROME D UBIQUINOL OXIDASE SUBUNIT 1 |
| 36:00:00 | 6ka4-A | 2.2 | 4 | 47 | 534 | 11 | MOLECULE: | F22L4.1 PROTEIN |
| 37:00:00 | 3zhe-A | 2.2 | 2.7 | 51 | 399 | 8 | MOLECULE: | NONSENSE-MEDIATED MRNA DECAY PROTEIN |
| 38:00:00 | 5t58-N | 2.2 | 12.1 | 77 | 196 | 8 | MOLECULE: | KLLA0F02343P |
| 39:00:00 | 7etw-A | 2.2 | 5.8 | 60 | 197 | 10 | MOLECULE: | INSULIN-INDUCED GENE 2 PROTEIN |
| 40:00:00 | 6f1t-f | 2.2 | 23.6 | 79 | 929 | 6 | MOLECULE: | ARP1 ACTIN RELATED PROTEIN 1 HOMOLOG A |
| 41:00:00 | 7xzj-A | 2.2 | 18.5 | 77 | 423 | 8 | MOLECULE: | CTAP3 |
| 42:00:00 | 7qe5-A | 2.2 | 15.9 | 71 | 616 | 13 | MOLECULE: | SIALIC ACID TRAP TRANSPORTER PERMEASE PROTEIN SIA |
| 43:00:00 | 2xwu-B | 2.2 | 5.6 | 63 | 910 | 13 | MOLECULE: | SUMO-CONJUGATING ENZYME UBC9 |
| 44:00:00 | 2o8p-A | 2.2 | 3.6 | 62 | 220 | 6 | MOLECULE: | 14-3-3 DOMAIN CONTAINING PROTEIN |
| 45:00:00 | 6f2d-G | 2.2 | 5.8 | 59 | 89 | 5 | MOLECULE: | FLAGELLAR BIOSYNTHETIC PROTEIN FLIP |
| 46:00:00 | 4n9n-A | 2.2 | 4.7 | 54 | 428 | 9 | MOLECULE: | STEROL UPTAKE CONTROL PROTEIN 2, LYSOZYME |
| 47:00:00 | 3oop-A | 2.2 | 5 | 64 | 139 | 6 | MOLECULE: | LIN2960 PROTEIN |
| 48:00:00 | 5mmj-n | 2.2 | 3.5 | 50 | 99 | 0 | MOLECULE: | 50S RIBOSOMAL PROTEIN L31 |
| 49:00:00 | 4g6d-B | 2.2 | 6.3 | 57 | 198 | 5 | MOLECULE: | RNA POLYMERASE SIGMA FACTOR RPOD |
| 50:00:00 | 3ut5-E | 2.2 | 51 | 94 | 132 | 7 | MOLECULE: | TUBULIN ALPHA CHAIN |

| | | | | | | | | |
|---|---|---|---|---|---|---|---|---|
| 51:00:00 | 8clh-E | 2.2 | 44.6 | 90 | 123 | 6 | MOLECULE: | TUBULIN ALPHA-1B CHAIN |
| 52:00:00 | 7b9s-Z | 2.2 | 12.5 | 86 | 462 | 7 | MOLECULE: | ECCE5 |
| 53:00:00 | 8aud-B | 2.2 | 6.4 | 78 | 346 | 8 | MOLECULE: | CELL WALL-ASSOCIATED HYDROLASES (INVASION-ASSOCIA |
| 54:00:00 | 6o7v-a | 2.2 | 34 | 68 | 625 | 10 | MOLECULE: | V-TYPE PROTON ATPASE SUBUNIT D |
| 55:00:00 | 6zkh-q | 2.2 | 9.4 | 68 | 139 | 6 | MOLECULE: | NADH DEHYDROGENASE [UBIQUINONE] FLAVOPROTEIN 1, |
| 56:00:00 | 7z4a-K | 2.2 | 5.9 | 92 | 588 | 9 | MOLECULE: | ADAPTOR PROTEIN |
| 57:00:00 | 8t1l-G | 2.2 | 8.9 | 75 | 122 | 7 | MOLECULE: | MEDIATOR OF RNA POLYMERASE II TRANSCRIPTION SUBUN |
| 58:00:00 | 3jac-B | 2.2 | 3.8 | 73 | 918 | 0 | MOLECULE: | PIEZO-TYPE MECHANOSENSITIVE ION CHANNEL COMPONENT |
| 59:00:00 | 2z5i-B | 2.2 | 16.9 | 46 | 49 | 4 | MOLECULE: | GENERAL CONTROL PROTEIN GCN4 AND TROPOMYOSIN ALPH |
| 60:00:00 | 3dtp-A | 2.2 | 5.2 | 50 | 938 | 14 | MOLECULE: | MYOSIN 2 HEAVY CHAIN CHIMERA OF SMOOTH AND CARDIA |
| 61:00:00 | 3l35-A | 2.2 | 16.6 | 43 | 45 | 2 | MOLECULE: | GP41 N-PEPTIDE |
| 62:00:00 | 2q5u-B | 2.2 | 15.6 | 40 | 45 | 8 | MOLECULE: | FUSION PROTEIN BETWEEN YEAST VARIANT GCN4 AND |
| 63:00:00 | 2z5i-E | 2.2 | 15.9 | 45 | 49 | 4 | MOLECULE: | GENERAL CONTROL PROTEIN GCN4 AND TROPOMYOSIN ALPH |
| 64:00:00 | 2z5i-A | 2.2 | 15.9 | 45 | 49 | 4 | MOLECULE: | GENERAL CONTROL PROTEIN GCN4 AND TROPOMYOSIN ALPH |
| 65:00:00 | 8jtd-B | 2.2 | 6.7 | 58 | 126 | 7 | MOLECULE: | GP120 PROTEIN OF HIV ENVELOPE TRIMER |
| 66:00:00 | 8tni-F | 2.2 | 6 | 55 | 126 | 11 | MOLECULE: | HIV-1 BG505 DS-SOSIP GP120 |
| 67:00:00 | 6x98-I | 2.2 | 3.1 | 41 | 122 | 12 | MOLECULE: | BG505 HIV-1 ENV GP120 |
| 68:00:00 | 6u59-B | 2.2 | 3.8 | 43 | 120 | 7 | MOLECULE: | SOSIP.664 GP120,SOSIP.664 GP120 |
| 69:00:00 | 6rw6-B | 2.1 | 25.1 | 97 | 2485 | 3 | MOLECULE: | TCDA1 |
| 70:00:00 | 8qrn-7 | 2.1 | 24.2 | 85 | 571 | 4 | MOLECULE: | 12S MITOCHONDRIAL RRNA |
| 71:00:00 | 8yb7-C | 2.1 | 24.1 | 101 | 455 | 9 | MOLECULE: | PAPAIN-LIKE PROTEASE NSP3 |
| 72:00:00 | 7syf-A | 2.1 | 24.8 | 92 | 1391 | 7 | MOLECULE: | PHOSPHATIDYLINOSITOL 3,4,5-TRISPHOSPHATE-DEPENDEN |
| 73:00:00 | 6qd6-G | 2.1 | 40.5 | 89 | 469 | 6 | MOLECULE: | MB-CHOPQ-NB207,OUTER MEMBRANE PROTEIN,MB-CHOPQ-NB |
| 74:00:00 | 5z7b-B | 2.1 | 32.3 | 55 | 197 | 2 | MOLECULE: | PADR FAMILY TRANSCRIPTIONAL REGULATOR |
| 75:00:00 | 6gyp-E | 2.1 | 27.7 | 90 | 512 | 3 | MOLECULE: | CENTROMERE DNA-BINDING PROTEIN COMPLEX CBF3 SUBUN |
| 76:00:00 | 7qo4-M | 2.1 | 15.7 | 49 | 421 | 8 | MOLECULE: | 26S PROTEASOME REGULATORY SUBUNIT RPN1 |
| 77:00:00 | 5ucg-A | 2.1 | 24.7 | 75 | 345 | 5 | MOLECULE: | STAGE II SPORULATION PROTEIN E |
| 78:00:00 | 2oa5-B | 2.1 | 12.4 | 78 | 97 | 13 | MOLECULE: | HYPOTHETICAL PROTEIN BQLF2 |
| 79:00:00 | 7qj0-L | 2.1 | 9.9 | 89 | 566 | 8 | MOLECULE: | MITOTIC-SPINDLE ORGANIZING PROTEIN 1 |
| 80:00:00 | 6wq2-A | 2.1 | 21.9 | 80 | 154 | 6 | MOLECULE: | A-DNA |
| 81:00:00 | 7v2y-B | 2.1 | 35.5 | 117 | 1226 | 9 | MOLECULE: | THO COMPLEX SUBUNIT HPR1 |
| 82:00:00 | 5b52-B | 2.1 | 15.4 | 47 | 60 | 6 | MOLECULE: | H-NS FAMILY PROTEIN MVAT |
| 83:00:00 | 2qfa-B | 2.1 | 18 | 55 | 62 | 11 | MOLECULE: | BACULOVIRAL IAP REPEAT-CONTAINING PROTEIN 5 |
| 84:00:00 | 8d8j-d | 2.1 | 10.8 | 69 | 660 | 14 | MOLECULE: | PROBABLE S-ADENOSYL-L-METHIONINE-DEPENDENT RNA |
| 85:00:00 | 8t1l-X | 2.1 | 3.6 | 42 | 127 | 7 | MOLECULE: | MEDIATOR OF RNA POLYMERASE II TRANSCRIPTION SUBUN |
| 86:00:00 | 6lcc-A | 2.1 | 18.1 | 74 | 375 | 7 | MOLECULE: | AATPS |
| 87:00:00 | 5h11-A | 2.1 | 4.5 | 49 | 494 | 6 | MOLECULE: | UNCHARACTERIZED PROTEIN |
| 88:00:00 | 8yzk-A | 2.1 | 12.6 | 75 | 264 | 5 | MOLECULE: | SOLUBLE CYTOCHROME B562,G-PROTEIN COUPLED RECEPTO |
| 89:00:00 | 5ouz-A | 2.1 | 5.1 | 65 | 178 | 6 | MOLECULE: | FERRITIN |
| 90:00:00 | 8rc4-o | 2.1 | 16.5 | 68 | 372 | 9 | MOLECULE: | SERINE/THREONINE-PROTEIN PHOSPHATASE 2A 65 KDA RE |
| 91:00:00 | 5b49-B | 2.1 | 14.6 | 69 | 247 | 9 | MOLECULE: | UDP-2,3-DIACYLGLUCOSAMINE HYDROLASE |
| 92:00:00 | 3b09-A | 2.1 | 3.8 | 44 | 65 | 5 | MOLECULE: | PEPTIDYL-PROLYL CIS-TRANS ISOMERASE |
| 93:00:00 | 5ctq-A | 2.1 | 26.8 | 83 | 518 | 5 | MOLECULE: | SQUAMOUS CELL CARCINOMA ANTIGEN RECOGNIZED BY T-C |
| 94:00:00 | 8h1l-B | 2.1 | 7.3 | 53 | 423 | 9 | MOLECULE: | N-ACYLGLUCOSAMINE 2-EPIMERASE |
| 95:00:00 | 7znk-b | 2.1 | 31.3 | 115 | 919 | 5 | MOLECULE: | RNA |
| 96:00:00 | 6h95-A | 2.1 | 6.1 | 55 | 225 | 5 | MOLECULE: | ALBICIDIN RESISTANCE PROTEIN |
| 97:00:00 | 1wrd-A | 2.1 | 3.5 | 54 | 98 | 15 | MOLECULE: | TARGET OF MYB PROTEIN 1 |
| 98:00:00 | 4um2-A | 2.1 | 4.9 | 61 | 510 | 11 | MOLECULE: | TELOMERASE-BINDING PROTEIN EST1A |
| 99:00:00 | 8t1l-S | 2.1 | 7.1 | 63 | 913 | 5 | MOLECULE: | MEDIATOR OF RNA POLYMERASE II TRANSCRIPTION SUBUN |
| 0:00 | 6tgb-A | 2.1 | 8.3 | 62 | 1450 | 8 | MOLECULE: | DEDICATOR OF CYTOKINESIS PROTEIN 2 |
| 1:00 | 4f23-A | 2.1 | 10.9 | 126 | 509 | 10 | MOLECULE: | HEMAGGLUTININ |
| 2:00 | 6v9z-A | 2.1 | 32.4 | 106 | 715 | 8 | MOLECULE: | ABC-TYPE BACTERIOCIN TRANSPORTER |
| 3:00 | 4pl0-B | 2.1 | 9.5 | 108 | 576 | 6 | MOLECULE: | MICROCIN-J25 EXPORT ATP-BINDING/PERMEASE PROTEIN |
| 4:00 | 6h8q-A | 2.1 | 5.5 | 73 | 879 | 4 | MOLECULE: | COHESIN SUBUNIT SCC3 |
| 5:00 | 7ooc-S | 2.1 | 3.3 | 56 | 77 | 7 | MOLECULE: | 30S RIBOSOMAL PROTEIN S3 |
| 6:00 | 3ont-A | 2.1 | 8.2 | 74 | 112 | 3 | MOLECULE: | SPOT 14 PROTEIN |

| | | | | | | | | |
|---|---|---|---|---|---|---|---|---|
| 7:00 | 1ox3-A | 2.1 | 3.5 | 59 | 108 | 10 | MOLECULE: | FIBRITIN |
| 8:00 | 2zdi-C | 2.1 | 4.9 | 65 | 148 | 3 | MOLECULE: | PREFOLDIN SUBUNIT BETA |
| 9:00 | 6rfl-C | 2.1 | 3.5 | 54 | 304 | 4 | MOLECULE: | DNA-DEPENDENT RNA POLYMERASE SUBUNIT RPO132 |
| 10:00 | 2oev-A | 2.1 | 8 | 111 | 697 | 5 | MOLECULE: | PROGRAMMED CELL DEATH 6-INTERACTING PROTEIN |
| 11:00 | 6ds9-A | 2.1 | 3.3 | 53 | 93 | 8 | MOLECULE: | DE NOVO DESIGNED THREE HELIX BUNDLE GRA3D |
| 12:00 | 7xzi-B | 2.1 | 3.2 | 59 | 166 | 5 | MOLECULE: | CTAP3 |
| 13:00 | 8ioi-A | 2.1 | 11.2 | 72 | 190 | 6 | MOLECULE: | PADR FAMILY TRANSCRIPTIONAL REGULATOR |
| 14:00 | 2hxo-A | 2.1 | 12.5 | 53 | 207 | 2 | MOLECULE: | PUTATIVE TETR-FAMILY TRANSCRIPTIONAL REGULATOR |
| 15:00 | 7etr-A | 2.1 | 7.8 | 53 | 88 | 6 | MOLECULE: | TRANSCRIPTIONAL REGULATOR COPG FAMILY |
| 16:00 | 7t5p-B | 2.1 | 4.6 | 67 | 419 | 7 | MOLECULE: | SUMO-INTERACTING MOTIF-CONTAINING PROTEIN 1 |
| 17:00 | 2raj-A | 2.1 | 5.6 | 94 | 382 | 10 | MOLECULE: | SORTING NEXIN-9 |
| 18:00 | 8j0n-D | 2.1 | 23.7 | 60 | 156 | 7 | MOLECULE: | ER MEMBRANE PROTEIN COMPLEX SUBUNIT 1 |
| 19:00 | 8ppr-N | 2.1 | 7.8 | 90 | 217 | 6 | MOLECULE: | KINETOCHORE-ASSOCIATED PROTEIN DSN1 HOMOLOG |
| 20:00 | 8i5f-A | 2.1 | 12.7 | 74 | 433 | 9 | MOLECULE: | DEDICATOR OF CYTOKINESIS PROTEIN 10 |
| 21:00 | 8fbj-A | 2.1 | 3.5 | 82 | 307 | 6 | MOLECULE: | KWOCA_60 |
| 22:00 | 7t1s-A | 2.1 | 3.3 | 49 | 234 | 6 | MOLECULE: | BSTC |
| 23:00 | 6z0f-A | 2.1 | 8.8 | 93 | 384 | 3 | MOLECULE: | ESX SECRETION SYSTEM PROTEIN YUKC |
| 24:00:00 | 6c5w-A | 2.1 | 10.7 | 87 | 285 | 2 | MOLECULE: | CALCIUM UNIPORTER |
| 25:00:00 | 7lby-A | 2.1 | 23.8 | 98 | 755 | 7 | MOLECULE: | CELLULOSE SYNTHASE CATALYTIC SUBUNIT [UDP-FORMING |
| 26:00:00 | 8djk-B | 2.1 | 3.7 | 49 | 281 | 10 | MOLECULE: | 3-HYDROXY-3-METHYLGLUTARYL-COENZYME A REDUCTASE |
| 27:00:00 | 6al9-B | 2.1 | 4.4 | 55 | 91 | 7 | MOLECULE: | CHORISMATE MUTASE |
| 28:00:00 | 6t8d-X | 2.1 | 4.8 | 81 | 355 | 16 | MOLECULE: | MAKB |
| 29:00:00 | 3owa-C | 2.1 | 18.5 | 96 | 587 | 9 | MOLECULE: | ACYL-COA DEHYDROGENASE |
| 30:00:00 | 3fn2-A | 2.1 | 4.1 | 50 | 97 | 6 | MOLECULE: | PUTATIVE SENSOR HISTIDINE KINASE DOMAIN |
| 31:00:00 | 6hty-B | 2.1 | 5.7 | 60 | 302 | 8 | MOLECULE: | NUCLEAR RECEPTOR SUBFAMILY 1 GROUP I MEMBER 2,NUC |
| 32:00:00 | 6xu2-A | 2.1 | 7.2 | 89 | 1091 | 9 | MOLECULE: | IMPORTIN-5 |
| 33:00:00 | 6t1z-A | 2.1 | 3 | 53 | 393 | 8 | MOLECULE: | LMRP INTEGRAL MEMBRANE PROTEIN |
| 34:00:00 | 6upn-A | 2.1 | 2.8 | 52 | 245 | 8 | MOLECULE: | ENDOPHILIN-B1 |
| 35:00:00 | 6zyy-C | 2.1 | 50.9 | 110 | 2901 | 2 | MOLECULE: | DYNEIN HEAVY CHAIN, OUTER ARM PROTEIN |
| 36:00:00 | 7use-B | 2.1 | 5.3 | 67 | 1099 | 6 | MOLECULE: | CYTOPLASMIC FMR1-INTERACTING PROTEIN 1 |
| 37:00:00 | 7p34-A | 2.1 | 4.1 | 49 | 391 | 4 | MOLECULE: | PEPTIDE ANTIBIOTIC TRANSPORTER SBMA |
| 38:00:00 | 8k32-C | 2.1 | 32.1 | 80 | 333 | 10 | MOLECULE: | KETOL-ACID REDUCTOISOMERASE (NADP(+)) |
| 39:00:00 | 4qn1-A | 2.1 | 2.1 | 58 | 372 | 3 | MOLECULE: | E3 UBIQUITIN-PROTEIN LIGASE SHPRH |
| 40:00:00 | 8a43-A | 2.1 | 8.8 | 68 | 1549 | 13 | MOLECULE: | DNA-DIRECTED RNA POLYMERASE I SUBUNIT RPA1 |
| 41:00:00 | 5tpm-A | 2.1 | 2.1 | 44 | 155 | 2 | MOLECULE: | PYRUVATE DEHYDROGENASE COMPLEX REPRESSOR |
| 42:00:00 | 5c73-A | 2.1 | 5.4 | 79 | 564 | 6 | MOLECULE: | PROTEIN GLYCOSYLATION K |
| 43:00:00 | 6xrx-A | 2.1 | 19.6 | 50 | 551 | 6 | MOLECULE: | MALTOSE/MALTODEXTRIN-BINDING PERIPLASMIC PROTEIN, |
| 44:00:00 | 2o6i-B | 2.1 | 23.4 | 81 | 430 | 4 | MOLECULE: | HD DOMAIN PROTEIN |
| 45:00:00 | 9auc-E | 2.1 | 2.5 | 42 | 117 | 5 | MOLECULE: | GUANINE NUCLEOTIDE-BINDING PROTEIN G(S) SUBUNIT A |
| 46:00:00 | 8u7i-A | 2.1 | 28.7 | 70 | 626 | 4 | MOLECULE: | ENDONUCLEASE GAJA |
| 47:00:00 | 6tdv-f | 2.1 | 12.4 | 115 | 274 | 8 | MOLECULE: | ATPTB1 |
| 48:00:00 | 3bvo-A | 2.1 | 19 | 62 | 197 | 6 | MOLECULE: | CO-CHAPERONE PROTEIN HSCB, MITOCHONDRIAL PRECURSO |
| 49:00:00 | 7ekp-A | 2.1 | 7.5 | 76 | 396 | 5 | MOLECULE: | NEURONAL ACETYLCHOLINE RECEPTOR SUBUNIT ALPHA-7 |
| 50:00:00 | 8xej-X | 2.1 | 21.6 | 78 | 358 | 5 | MOLECULE: | ISOFORM 2 OF BASIGIN |
| 51:00:00 | 1lqs-L | 2.1 | 3.8 | 50 | 142 | 6 | MOLECULE: | INTERLEUKIN-10 RECEPTOR ALPHA CHAIN |
| 52:00:00 | 8jhu-A | 2.1 | 8.9 | 85 | 811 | 8 | MOLECULE: | LEGIONELLA PNEUMOPHILA EFFECTOR PROTEIN SIDI |
| 53:00:00 | 8ccr-A | 2.1 | 3.3 | 50 | 105 | 6 | MOLECULE: | 4D2 (MUTANT T19D) |
| 54:00:00 | 8b6l-M | 2.1 | 3.5 | 54 | 110 | 6 | MOLECULE: | PROTEIN TRANSPORT PROTEIN SEC61 SUBUNIT ALPHA ISO |
| 55:00:00 | 3zjc-E | 2.1 | 7.2 | 81 | 286 | 9 | MOLECULE: | GTPASE IMAP FAMILY MEMBER 7 |
| 56:00:00 | 4acl-A | 2.1 | 3.4 | 51 | 155 | 8 | MOLECULE: | TSSL |
| 57:00:00 | 1z2c-B | 2.1 | 3 | 43 | 346 | 7 | MOLECULE: | RHO-RELATED GTP-BINDING PROTEIN RHOC |
| 58:00:00 | 5n77-A | 2.1 | 7.5 | 74 | 257 | 9 | MOLECULE: | MAGNESIUM TRANSPORT PROTEIN CORA |
| 59:00:00 | 1d7m-A | 2.1 | 39.4 | 83 | 101 | 7 | MOLECULE: | CORTEXILLIN I |
| 60:00:00 | 5jqz-B | 2.1 | 2.8 | 51 | 75 | 8 | MOLECULE: | DE NOVO DESIGNED HOMOTETRAMER |
| 61:00:00 | 8at4-F | 2.1 | 11.4 | 88 | 387 | 2 | MOLECULE: | HAUS AUGMIN-LIKE COMPLEX SUBUNIT 1 |
| 62:00:00 | 2xnx-J | 2.1 | 2.6 | 46 | 72 | 9 | MOLECULE: | FIBRINOGEN ALPHA CHAIN |

| | | | | | | | | |
|---|---|---|---|---|---|---|---|---|
| 63:00:00 | 1ar1-B | 2.1 | 5.5 | 55 | 252 | 11 | MOLECULE: | CYTOCHROME C OXIDASE |
| 64:00:00 | 2z5h-C | 2.1 | 17.8 | 49 | 52 | 4 | MOLECULE: | GENERAL CONTROL PROTEIN GCN4 AND TROPOMYOSIN ALPH |
| 65:00:00 | 7bv6-O | 2.1 | 15.6 | 53 | 73 | 11 | MOLECULE: | VESICLE-ASSOCIATED MEMBRANE PROTEIN 8 |
| 66:00:00 | 2z5i-D | 2.1 | 16.8 | 46 | 49 | 4 | MOLECULE: | GENERAL CONTROL PROTEIN GCN4 AND TROPOMYOSIN ALPH |
| 67:00:00 | 2z5i-G | 2.1 | 19 | 47 | 50 | 2 | MOLECULE: | GENERAL CONTROL PROTEIN GCN4 AND TROPOMYOSIN ALPH |
| 68:00:00 | 3p7k-A | 2.1 | 16.3 | 42 | 45 | 10 | MOLECULE: | GP41 PEPTIDE |
| 69:00:00 | 2z5i-C | 2.1 | 19.8 | 47 | 50 | 2 | MOLECULE: | GENERAL CONTROL PROTEIN GCN4 AND TROPOMYOSIN ALPH |
| 70:00:00 | 6vy2-D | 2.1 | 5.6 | 52 | 121 | 15 | MOLECULE: | GLYCOPROTEIN 120 |
| 71:00:00 | 7txd-F | 2.1 | 4 | 49 | 122 | 8 | MOLECULE: | ENVELOPE GLYCOPROTEIN GP120 |
| 72:00:00 | 6vy2-F | 2.1 | 5.9 | 54 | 121 | 11 | MOLECULE: | GLYCOPROTEIN 120 |
| 73:00:00 | 7rai-F | 2.1 | 5.7 | 48 | 128 | 13 | MOLECULE: | ENVELOPE GLYCOPROTEIN GP160 |
| 74:00:00 | 6vy2-B | 2.1 | 6.2 | 53 | 121 | 13 | MOLECULE: | GLYCOPROTEIN 120 |
| 75:00:00 | 3o40-A | 2.1 | 3.6 | 63 | 198 | 14 | MOLECULE: | GP41-5 |
| 76:00:00 | 8tnh-F | 2.1 | 6.7 | 62 | 126 | 11 | MOLECULE: | HIV-1 BG505 DS-SOSIP GP120 |
| 77:00:00 | 3jbh-B | 2.1 | 28.4 | 69 | 964 | 9 | MOLECULE: | MYOSIN 2 HEAVY CHAIN STRIATED MUSCLE |
| 78:00:00 | 3zmf-A | 2.1 | 6.2 | 52 | 111 | 8 | MOLECULE: | GENERAL CONTROL PROTEIN GCN4, PUTATIVE INNER MEMB |
| 79:00:00 | 8jtm-B | 2.1 | 6.1 | 56 | 126 | 11 | MOLECULE: | GP120 PROTEIN OF HIV ENVELOPE TRIMER |
| 80:00:00 | 6u0n-Z | 2.1 | 4.2 | 58 | 124 | 5 | MOLECULE: | ENVELOPE GLYCOPROTEIN GP120 |
| 81:00:00 | 3o43-A | 2.1 | 10.6 | 80 | 198 | 9 | MOLECULE: | GP41-5 |
| 82:00:00 | 8euv-B | 2.1 | 5.6 | 54 | 130 | 13 | MOLECULE: | ENVELOPE GLYCOPROTEIN GP120 |
| 83:00:00 | 7sq1-D | 2.1 | 5.1 | 41 | 121 | 10 | MOLECULE: | TRANSMEMBRANE PROTEIN GP41 |
| 84:00:00 | 8jtm-F | 2.1 | 5.5 | 54 | 123 | 11 | MOLECULE: | GP120 PROTEIN OF HIV ENVELOPE TRIMER |
| 85:00:00 | 7l6o-f | 2.1 | 4.2 | 44 | 126 | 7 | MOLECULE: | CH848.3.D0949.10.17CHIM.6R.DS.SOSIP.664 - GP120 |
| 86:00:00 | 7qru-E | 2 | 12.2 | 79 | 158 | 11 | MOLECULE: | NA+/H+ ANTIPORTER SUBUNIT D |
| 87:00:00 | 7roq-A | 2 | 31.5 | 135 | 1831 | 10 | MOLECULE: | PHOSPHOLIPID-TRANSPORTING ATPASE ABCA1 |
| 88:00:00 | 6vbu-2 | 2 | 4.3 | 43 | 659 | 2 | MOLECULE: | BARDET-BIEDL SYNDROME 18 PROTEIN |
| 89:00:00 | 6th1-R | 2 | 34 | 92 | 360 | 4 | MOLECULE: | IMMEDIATE EARLY PROTEIN 1 |
| 90:00:00 | 6uen-E | 2 | 4 | 50 | 114 | 8 | MOLECULE: | RNA-DIRECTED RNA POLYMERASE L |
| 91:00:00 | 8t5e-A | 2 | 3.2 | 48 | 130 | 8 | MOLECULE: | BIM_FULLDIFF |
| 92:00:00 | 7w7t-A | 2 | 18.5 | 84 | 1021 | 10 | MOLECULE: | SARCOPLASMIC/ENDOPLASMIC RETICULUM CALCIUM ATPASE |
| 93:00:00 | 8kde-3 | 2 | 26.6 | 63 | 97 | 5 | MOLECULE: | PHOTOSYSTEM II CP47 REACTION CENTER PROTEIN |
| 94:00:00 | 6sz9-D | 2 | 10.6 | 102 | 282 | 8 | MOLECULE: | ICMO (DOTL) |
| 95:00:00 | 2odm-B | 2 | 2.3 | 48 | 83 | 6 | MOLECULE: | UPF0358 PROTEIN MW0995 |
| 96:00:00 | 8csz-D | 2 | 5.4 | 78 | 494 | 6 | MOLECULE: | ISCB |
| 97:00:00 | 1q6u-A | 2 | 2.9 | 45 | 213 | 18 | MOLECULE: | FKBP-TYPE PEPTIDYL-PROLYL CIS-TRANS ISOMERASE FKP |
| 98:00:00 | 5ktf-A | 2 | 8.3 | 58 | 73 | 14 | MOLECULE: | SCAVENGER RECEPTOR CLASS B MEMBER 1 |
| 99:00:00 | 8ppr-D | 2 | 17.9 | 86 | 248 | 6 | MOLECULE: | KINETOCHORE-ASSOCIATED PROTEIN DSN1 HOMOLOG |
| 0:00 | 8eki-B | 2 | 4.7 | 49 | 85 | 0 | MOLECULE: | PROTEIN TRANSPORT PROTEIN SEC20 |
| 1:00 | 7aqw-m | 2 | 10.3 | 57 | 70 | 2 | MOLECULE: | NADH-UBIQUINONE OXIDOREDUCTASE CHAIN 5 |
| 2:00 | 4m1p-A | 2 | 4.2 | 54 | 96 | 4 | MOLECULE: | COPPER-SENSITIVE OPERON REPRESSOR (CSOR) |
| 3:00 | 8g7m-G | 2 | 9.5 | 62 | 343 | 5 | MOLECULE: | 60 KDA HEAT SHOCK PROTEIN, MITOCHONDRIAL |
| 4:00 | 7e2c-I | 2 | 5.6 | 52 | 799 | 10 | MOLECULE: | TRAPP-ASSOCIATED PROTEIN TCA17 |
| 5:00 | 1ypy-A | 2 | 5.3 | 52 | 182 | 8 | MOLECULE: | VIRION MEMBRANE PROTEIN |
| 6:00 | 3lss-B | 2 | 3.6 | 62 | 468 | 6 | MOLECULE: | SERYL-TRNA SYNTHETASE |
| 7:00 | 7kuw-A | 2 | 3.2 | 47 | 62 | 11 | MOLECULE: | SEQUENCE-BASED DESIGNED PROTEIN NMT_0994_GUIDED_0 |
| 8:00 | 3cit-A | 2 | 2.9 | 48 | 155 | 13 | MOLECULE: | SENSOR HISTIDINE KINASE |
| 9:00 | 8qt5-A | 2 | 3.2 | 60 | 248 | 5 | MOLECULE: | 14-3-3-LIKE PROTEIN G-BOX FACTOR 14 LAMBDA,PROTEI |
| 10:00 | 7f1t-A | 2 | 8.5 | 86 | 423 | 5 | MOLECULE: | C-C MOTIF CHEMOKINE 3,C-C CHEMOKINE RECEPTOR TYPE |
| 11:00 | 6s4m-A | 2 | 2.6 | 56 | 426 | 11 | MOLECULE: | MAJOR FACILITATOR SUPERFAMILY DOMAIN-CONTAINING P |
| 12:00 | 7aqr-V | 2 | 3.4 | 49 | 140 | 4 | MOLECULE: | NADH DEHYDROGENASE [UBIQUINONE] IRON-SULFUR PROTE |
| 13:00 | 5ndd-A | 2 | 7 | 77 | 567 | 8 | MOLECULE: | LYSOZYME,PROTEINASE-ACTIVATED RECEPTOR 2,SOLUBLE |
| 14:00 | 6xm1-D | 2 | 6.2 | 66 | 181 | 6 | MOLECULE: | VPS45 |
| 15:00 | 7w6k-A | 2 | 10.1 | 73 | 410 | 8 | MOLECULE: | GMALMT12/QUAC1 |
| 16:00 | 7mvv-A | 2 | 15.9 | 92 | 1542 | 7 | MOLECULE: | NUCLEOPORIN NUP192 |
| 17:00 | 8pog-A | 2 | 6.6 | 56 | 156 | 14 | MOLECULE: | BCSD OF ENTEROBACTER SP. 638 |
| 18:00 | 3esl-B | 2 | 3.7 | 49 | 202 | 6 | MOLECULE: | CHECKPOINT SERINE/THREONINE-PROTEIN KINASE BUB1 |

| | | | | | | | | |
|---|---|---|---|---|---|---|---|---|
| 19:00 | 7d3u-D | 2 | 12.6 | 72 | 534 | 7 | MOLECULE: | MONOVALENT NA+/H+ ANTIPORTER SUBUNIT D |
| 20:00 | 8gji-A | 2 | 3.7 | 65 | 167 | 8 | MOLECULE: | GCG BINDER |
| 21:00 | 6tpk-A | 2 | 13.8 | 70 | 461 | 1 | MOLECULE: | OXYTOCIN RECEPTOR |
| 22:00 | 4r1i-A | 2 | 9 | 59 | 507 | 8 | MOLECULE: | AMINOBENZOYL-GLUTAMATE TRANSPORTER |
| 23:00 | 6f2d-F | 2 | 8.6 | 83 | 258 | 11 | MOLECULE: | FLAGELLAR BIOSYNTHETIC PROTEIN FLIP |
| 24:00:00 | 1qoy-A | 2 | 11.1 | 92 | 303 | 5 | MOLECULE: | HEMOLYSIN E |
| 25:00:00 | 3kkz-A | 2 | 4 | 59 | 257 | 7 | MOLECULE: | UNCHARACTERIZED PROTEIN Q5LES9 |
| 26:00:00 | 5sy1-A | 2 | 10 | 81 | 582 | 10 | MOLECULE: | CALMODULIN |
| 27:00:00 | 6sl2-A | 2 | 5 | 103 | 619 | 5 | MOLECULE: | CALPONIN HOMOLOGY DOMAIN PROTEIN PUTATIVE |
| 28:00:00 | 5d2s-A | 2 | 13.7 | 52 | 91 | 8 | MOLECULE: | FIBROIN-MODULATOR-BINDING PROTEIN-1 |
| 29:00:00 | 7rsl-A | 2 | 5.5 | 52 | 235 | 12 | MOLECULE: | SEIPIN |
| 30:00:00 | 7ahd-A | 2 | 4.8 | 64 | 547 | 6 | MOLECULE: | ABC-TYPE PROLINE/GLYCINE BETAINE TRANSPORT SYSTEM |
| 31:00:00 | 1ej6-D | 2 | 22.4 | 67 | 417 | 7 | MOLECULE: | LAMBDA2 |
| 32:00:00 | 1y9b-A | 2 | 9.6 | 59 | 81 | 10 | MOLECULE: | CONSERVED HYPOTHETICAL PROTEIN |
| 33:00:00 | 5ys9-A | 2 | 10.7 | 87 | 692 | 5 | MOLECULE: | ACYL-COENZYME A OXIDASE 3 |
| 34:00:00 | 7mi6-A | 2 | 28.3 | 119 | 2419 | 4 | MOLECULE: | FUSION PROTEIN OF DYNEIN AND ENDOLYSIN |
| 35:00:00 | 6v4l-A | 2 | 13 | 72 | 479 | 10 | MOLECULE: | TRK SYSTEM POTASSIUM UPTAKE PROTEIN TRKH |
| 36:00:00 | 8cwo-T | 2 | 3.1 | 54 | 87 | 7 | MOLECULE: | 16S RIBOSOMAL RNA |
| 37:00:00 | 6w2q-A | 2 | 8 | 65 | 195 | 5 | MOLECULE: | JUNCTION 34 |
| 38:00:00 | 6s8h-F | 2 | 5.2 | 58 | 239 | 16 | MOLECULE: | LIPOPOLYSACCHARIDE ABC TRANSPORTER, ATP-BINDING P |
| 39:00:00 | 7xn9-A | 2 | 7.2 | 70 | 476 | 1 | MOLECULE: | SOMATOSTATIN RECEPTOR TYPE 2,ENDO-1,4-BETA-XYLANA |
| 40:00:00 | 6d03-E | 2 | 3.8 | 55 | 466 | 4 | MOLECULE: | TRANSFERRIN RECEPTOR PROTEIN 1 |
| 41:00:00 | 7a5p-u | 2 | 12.1 | 43 | 43 | 2 | MOLECULE: | U2 SNRNA |
| 42:00:00 | 7nyx-A | 2 | 22.4 | 115 | 1467 | 3 | MOLECULE: | CHROMOSOME PARTITION PROTEIN MUKB |
| 43:00:00 | 1e1d-A | 2 | 7.8 | 55 | 553 | 5 | MOLECULE: | HYDROXYLAMINE REDUCTASE |
| 44:00:00 | 7xzi-E | 2 | 14.1 | 88 | 770 | 7 | MOLECULE: | CTAP3 |
| 45:00:00 | 8q72-A | 2 | 5.2 | 109 | 750 | 6 | MOLECULE: | JETC |
| 46:00:00 | 2z5i-F | 2 | 17.2 | 46 | 49 | 4 | MOLECULE: | GENERAL CONTROL PROTEIN GCN4 AND TROPOMYOSIN ALPH |
| 47:00:00 | 6osy-A | 2 | 8.3 | 59 | 132 | 12 | MOLECULE: | BG505 GP120 |
| 48:00:00 | 5c0r-A | 2 | 3.8 | 77 | 270 | 8 | MOLECULE: | HEMAGGLUTININ, ENVELOPE GLYCOPROTEIN, FIBRITIN FU |
| 49:00:00 | 4r61-A | 2 | 2.7 | 56 | 140 | 14 | MOLECULE: | GP41-BASED CONSTRUCT COVNHR3-ABC |

**DALI results: MdA-1 Wasp**

| No: | Chain | Z | rmsd | lali | nres | %id | PDB | Description |
|---|---|---|---|---|---|---|---|---|
| 1:00 | 8puz-A | 7.8 | 6.8 | 139 | 161 | 11 | MOLECULE: | TROPOMYOSIN |
| 2:00 | 6yvu-B | 7.6 | 10 | 151 | 1191 | 5 | MOLECULE: | STRUCTURAL MAINTENANCE OF CHROMOSOMES PROTEI |
| 3:00 | 6fkf-p | 7.3 | 8.8 | 142 | 143 | 4 | MOLECULE: | ATP SYNTHASE SUBUNIT ALPHA, CHLOROPLASTIC |
| 4:00 | 5app-B | 7.3 | 8.9 | 128 | 128 | 5 | MOLECULE: | GENERAL CONTROL PROTEIN GCN4, OUTER MEMBRANE |
| 5:00 | 2ocy-A | 7.1 | 9.2 | 141 | 149 | 9 | MOLECULE: | RAB GUANINE NUCLEOTIDE EXCHANGE FACTOR SEC2 |
| 6:00 | 4cpc-G | 7 | 8.5 | 144 | 152 | 6 | MOLECULE: | SYNAPTONEMAL COMPLEX PROTEIN 3 |
| 7:00 | 4gkw-A | 6.9 | 7.8 | 149 | 158 | 5 | MOLECULE: | SPINDLE ASSEMBLY ABNORMAL PROTEIN 6 |
| 8:00 | 2efr-A | 6.9 | 7.5 | 145 | 155 | 6 | MOLECULE: | GENERAL CONTROL PROTEIN GCN4 AND TROPOMYOSIN |
| 9:00 | 8bd7-M | 6.8 | 19.8 | 134 | 164 | 7 | MOLECULE: | IFT88 |
| 10:00 | 7znk-o | 6.8 | 8.5 | 133 | 164 | 9 | MOLECULE: | RNA |
| 11:00 | 6z9l-A | 6.8 | 6.9 | 142 | 744 | 10 | MOLECULE: | PRGA |
| 12:00 | 7pkq-w | 6.7 | 7.4 | 127 | 155 | 5 | MOLECULE: | MS35 |
| 13:00 | 3uf1-D | 6.7 | 6.2 | 112 | 112 | 11 | MOLECULE: | VIMENTIN |
| 14:00 | 8pqw-F | 6.7 | 12.9 | 146 | 150 | 9 | MOLECULE: | CYTOPLASMIC DYNEIN 1 HEAVY CHAIN 1 |
| 15:00 | 2fxo-A | 6.7 | 3.7 | 127 | 129 | 7 | MOLECULE: | MYOSIN HEAVY CHAIN, CARDIAC MUSCLE BETA ISOF |
| 16:00 | 5hmo-A | 6.6 | 3.7 | 121 | 131 | 11 | MOLECULE: | UNCONVENTIONAL MYOSIN-X |
| 17:00 | 4ytd-B | 6.6 | 4.2 | 99 | 99 | 7 | MOLECULE: | PROTEIN BICAUDAL D HOMOLOG 1 |
| 18:00 | 1d7m-A | 6.6 | 3.3 | 101 | 101 | 9 | MOLECULE: | CORTEXILLIN I |
| 19:00 | 8q85-F | 6.6 | 8.6 | 149 | 230 | 4 | MOLECULE: | KINETOCHORE PROTEIN NDC80 |
| 20:00 | 3o0z-D | 6.6 | 7.5 | 145 | 160 | 10 | MOLECULE: | RHO-ASSOCIATED PROTEIN KINASE 1 |
| 21:00 | 6yvu-A | 6.6 | 10.7 | 154 | 1127 | 6 | MOLECULE: | STRUCTURAL MAINTENANCE OF CHROMOSOMES PROTEI |
| 22:00 | 8xi2-L | 6.5 | 38.5 | 130 | 468 | 5 | MOLECULE: | MPN DOMAIN-CONTAINING PROTEIN |
| 23:00 | 4xa6-A | 6.5 | 11.5 | 138 | 168 | 9 | MOLECULE: | GP7-MYH7(1777-1855)-EB1 CHIMERA PROTEIN |
| 24:00:00 | 3mqb-A | 6.5 | 3 | 109 | 109 | 6 | MOLECULE: | BONE MARROW STROMAL ANTIGEN 2 |
| 25:00:00 | 2qa7-C | 6.4 | 2.9 | 102 | 102 | 7 | MOLECULE: | HUNTINGTIN-INTERACTING PROTEIN 1 |
| 26:00:00 | 5nvu-A | 6.3 | 10.1 | 144 | 3169 | 0 | MOLECULE: | DYNEIN MOTOR DOMAIN |
| 27:00:00 | 6qp4-A | 6.3 | 5.8 | 137 | 157 | 8 | MOLECULE: | USPA1 |
| 28:00:00 | 7bjg-A | 6.2 | 2.6 | 98 | 98 | 6 | MOLECULE: | SD21996P |
| 29:00:00 | 6abo-A | 6.2 | 10.8 | 111 | 213 | 11 | MOLECULE: | DNA REPAIR PROTEIN XRCC4 |
| 30:00:00 | 8i03-E | 6.2 | 5.1 | 113 | 169 | 8 | MOLECULE: | PAIRED AMPHIPATHIC HELIX PROTEIN PST1 |
| 31:00:00 | 8q85-G | 6.1 | 8.1 | 145 | 178 | 6 | MOLECULE: | KINETOCHORE PROTEIN NDC80 |
| 32:00:00 | 4f61-I | 6.1 | 6.3 | 148 | 234 | 5 | MOLECULE: | TUBULIN ALPHA CHAIN |
| 33:00:00 | 9mhh-D | 6.1 | 14.6 | 143 | 313 | 6 | MOLECULE: | PHOSPHOINOSITIDE 3-KINASE REGULATORY SUBUNIT |
| 34:00:00 | 6eun-A | 6 | 4.7 | 112 | 147 | 8 | MOLECULE: | ADHESIN |
| 35:00:00 | 7apk-m | 6 | 5.8 | 126 | 549 | 7 | MOLECULE: | THO COMPLEX SUBUNIT 1 |
| 36:00:00 | 7vf2-C | 6 | 35.8 | 134 | 184 | 3 | MOLECULE: | PROTEIN VIRILIZER HOMOLOG |
| 37:00:00 | 8i03-G | 6 | 5.7 | 116 | 166 | 14 | MOLECULE: | PAIRED AMPHIPATHIC HELIX PROTEIN PST1 |
| 38:00:00 | 3oja-A | 5.9 | 11.5 | 123 | 482 | 6 | MOLECULE: | LEUCINE-RICH IMMUNE MOLECULE 1 |
| 39:00:00 | 1x79-B | 5.9 | 4.7 | 90 | 90 | 9 | MOLECULE: | ADP-RIBOSYLATION FACTOR BINDING PROTEIN GGA1 |
| 40:00:00 | 4nqj-A | 5.9 | 5.8 | 124 | 177 | 12 | MOLECULE: | E3 UBIQUITIN-PROTEIN LIGASE TRIM69 |
| 41:00:00 | 8th8-B | 5.9 | 9.7 | 153 | 276 | 7 | MOLECULE: | DYNEIN REGULATORY COMPLEX PROTEIN 1/2 N-TERM |
| 42:00:00 | 5cws-C | 5.9 | 20.9 | 96 | 169 | 8 | MOLECULE: | SAB-158 FAB LIGHT CHAIN |
| 43:00:00 | 8th8-J | 5.8 | 43 | 149 | 372 | 5 | MOLECULE: | DYNEIN REGULATORY COMPLEX PROTEIN 1/2 N-TERM |
| 44:00:00 | 5aj3-d | 5.8 | 21.3 | 149 | 177 | 4 | MOLECULE: | MITORIBOSOMAL 12S RRNA |
| 45:00:00 | 5cff-B | 5.8 | 4 | 88 | 88 | 3 | MOLECULE: | MIRANDA |
| 46:00:00 | 5d80-G | 5.8 | 2.6 | 98 | 223 | 14 | MOLECULE: | V-TYPE PROTON ATPASE CATALYTIC SUBUNIT A |
| 47:00:00 | 6ff7-K | 5.8 | 36.3 | 132 | 213 | 7 | MOLECULE: | RNA-BINDING MOTIF PROTEIN, X-LINKED 2 |
| 48:00:00 | 8r1a-A | 5.8 | 5.1 | 134 | 587 | 9 | MOLECULE: | GUANYLATE BINDING PROTEIN 1 |

| | | | | | | | | |
|---|---|---|---|---|---|---|---|---|
| 49:00:00 | 3s4r-B | 5.8 | 3.8 | 91 | 91 | 13 | MOLECULE: | VIMENTIN |
| 50:00:00 | 6iac-E | 5.7 | 16.7 | 126 | 146 | 11 | MOLECULE: | PORTAL PROTEIN |
| 51:00:00 | 6j5i-b | 5.7 | 14.4 | 143 | 209 | 1 | MOLECULE: | ATP SYNTHASE SUBUNIT ALPHA, MITOCHONDRIAL |
| 52:00:00 | 3a7p-A | 5.7 | 3.4 | 88 | 88 | 8 | MOLECULE: | AUTOPHAGY PROTEIN 16 |
| 53:00:00 | 8tek-P | 5.6 | 4.9 | 119 | 138 | 7 | MOLECULE: | DYNEIN REGULATORY COMPLEX PROTEIN 1/2 N-TERM |
| 54:00:00 | 7qoo-H | 5.6 | 22.1 | 125 | 210 | 10 | MOLECULE: | CENTROMERE PROTEIN C |
| 55:00:00 | 8txr-C | 5.6 | 10.6 | 146 | 431 | 5 | MOLECULE: | EXODEOXYRIBONUCLEASE 7 LARGE SUBUNIT |
| 56:00:00 | 8fef-B | 5.6 | 22.5 | 146 | 335 | 13 | MOLECULE: | VIRULENCE FACTOR MCE FAMILY PROTEIN |
| 57:00:00 | 6fia-D | 5.6 | 9.3 | 104 | 104 | 7 | MOLECULE: | LINE-1 RETROTRANSPOSABLE ELEMENT ORF1 PROTEI |
| 58:00:00 | 7woo-I | 5.6 | 36.3 | 131 | 187 | 8 | MOLECULE: | NUCLEOPORIN NIC96 |
| 59:00:00 | 6h9l-B | 5.6 | 7.6 | 116 | 129 | 6 | MOLECULE: | UNCHARACTERIZED PROTEIN |
| 60:00:00 | 8ovw-Z | 5.6 | 8.9 | 123 | 151 | 6 | MOLECULE: | CENTROMERE-BINDING PROTEIN 1 |
| 61:00:00 | 5dfz-D | 5.6 | 11.9 | 136 | 341 | 10 | MOLECULE: | VACUOLAR PROTEIN SORTING-ASSOCIATED PROTEIN |
| 62:00:00 | 5bw9-g | 5.5 | 2.9 | 87 | 141 | 2 | MOLECULE: | V-TYPE PROTON ATPASE CATALYTIC SUBUNIT A |
| 63:00:00 | 8th8-A | 5.5 | 5.3 | 133 | 290 | 5 | MOLECULE: | DYNEIN REGULATORY COMPLEX PROTEIN 1/2 N-TERM |
| 64:00:00 | 3fwc-F | 5.5 | 2.6 | 84 | 84 | 5 | MOLECULE: | CELL DIVISION CONTROL PROTEIN 31 |
| 65:00:00 | 5mqf-L | 5.5 | 36.3 | 118 | 336 | 3 | MOLECULE: | PRE-MRNA-PROCESSING-SPLICING FACTOR 8 |
| 66:00:00 | 7e9t-A | 5.5 | 23 | 133 | 371 | 9 | MOLECULE: | SPIKE PROTEIN S2 |
| 67:00:00 | 6f1t-X | 5.5 | 8 | 154 | 285 | 8 | MOLECULE: | ARP1 ACTIN RELATED PROTEIN 1 HOMOLOG A |
| 68:00:00 | 8a5a-G | 5.5 | 6.8 | 105 | 110 | 7 | MOLECULE: | CHROMATIN-REMODELING ATPASE INO80 |
| 69:00:00 | 8ovw-U | 5.4 | 13.3 | 129 | 184 | 7 | MOLECULE: | CENTROMERE-BINDING PROTEIN 1 |
| 70:00:00 | 2xnx-M | 5.4 | 4.7 | 107 | 107 | 7 | MOLECULE: | FIBRINOGEN ALPHA CHAIN |
| 71:00:00 | 6exn-D | 5.4 | 8.9 | 96 | 97 | 9 | MOLECULE: | U2 SNRNA |
| 72:00:00 | 4h22-B | 5.4 | 5.1 | 83 | 83 | 7 | MOLECULE: | LEUCINE-RICH REPEAT FLIGHTLESS-INTERACTING P |
| 73:00:00 | 8p0v-L | 5.4 | 32.7 | 152 | 416 | 5 | MOLECULE: | COILED-COIL DOMAIN-CONTAINING PROTEIN 93 |
| 74:00:00 | 7puz-A | 5.3 | 3.6 | 102 | 148 | 8 | MOLECULE: | MICOS COMPLEX SUBUNIT MIC60 |
| 75:00:00 | 8fwj-A | 5.3 | 8.7 | 114 | 552 | 5 | MOLECULE: | CIRCADIAN CLOCK PROTEIN KAIC |
| 76:00:00 | 5cws-D | 5.3 | 5.1 | 90 | 180 | 7 | MOLECULE: | SAB-158 FAB LIGHT CHAIN |
| 77:00:00 | 6j9r-B | 5.3 | 4.2 | 102 | 126 | 10 | MOLECULE: | BRAIN TUMOR PROTEIN |
| 78:00:00 | 8q85-d | 5.3 | 19.1 | 104 | 108 | 3 | MOLECULE: | KINETOCHORE PROTEIN NDC80 |
| 79:00:00 | 5ijn-H | 5.3 | 3.1 | 94 | 169 | 7 | MOLECULE: | NUCLEAR PORE COMPLEX PROTEIN NUP155 |
| 80:00:00 | 5hda-A | 5.2 | 7.9 | 97 | 121 | 7 | MOLECULE: | ZINC FINGER MYND DOMAIN-CONTAINING PROTEIN 1 |
| 81:00:00 | 8fed-F | 5.2 | 20.9 | 153 | 399 | 9 | MOLECULE: | VIRULENCE FACTOR MCE FAMILY PROTEIN |
| 82:00:00 | 8bd7-L | 5.2 | 31.1 | 155 | 303 | 8 | MOLECULE: | IFT88 |
| 83:00:00 | 4lin-A | 5.2 | 9.3 | 149 | 289 | 3 | MOLECULE: | TAIL NEEDLE PROTEIN GP26 |
| 84:00:00 | 4y66-C | 5.2 | 20.5 | 128 | 197 | 9 | MOLECULE: | MND1 |
| 85:00:00 | 5gmk-c | 5.2 | 38.6 | 134 | 436 | 7 | MOLECULE: | PRE-MRNA-SPLICING FACTOR 8 |
| 86:00:00 | 8qbv-A | 5.2 | 11.7 | 129 | 217 | 11 | MOLECULE: | MEMBRANE-ASSOCIATED PROTEIN VIPP1 |
| 87:00:00 | 9dtr-D | 5.2 | 9.9 | 104 | 108 | 11 | MOLECULE: | U2 SNRNA |
| 88:00:00 | 6j6g-c | 5.2 | 47.2 | 143 | 436 | 9 | MOLECULE: | PRE-MRNA-SPLICING FACTOR 8 |
| 89:00:00 | 5lxn-A | 5.1 | 4.1 | 81 | 81 | 10 | MOLECULE: | TRANSFORMING ACIDIC COILED-COIL-CONTAINING P |
| 90:00:00 | 3vp8-B | 5.1 | 2.5 | 80 | 80 | 6 | MOLECULE: | GENERAL TRANSCRIPTIONAL COREPRESSOR TUP1 |
| 91:00:00 | 8hpo-D | 5.1 | 10.2 | 118 | 165 | 6 | MOLECULE: | TRANSCRIPTIONAL REGULATORY PROTEIN UME1 |
| 92:00:00 | 8cqn-B | 5 | 8.4 | 123 | 187 | 7 | MOLECULE: | LIPOPROTEIN, PUTATIVE |
| 93:00:00 | 2xqh-A | 5 | 10.1 | 125 | 258 | 7 | MOLECULE: | IMMUNOGLOBULIN-BINDING PROTEIN EIBD |
| 94:00:00 | 2p22-A | 5 | 13.5 | 123 | 168 | 6 | MOLECULE: | SUPPRESSOR PROTEIN STP22 OF TEMPERATURE-SENS |
| 95:00:00 | 4mvd-B | 5 | 6.2 | 116 | 253 | 6 | MOLECULE: | CHOLINE-PHOSPHATE CYTIDYLYLTRANSFERASE A |
| 96:00:00 | 3jb9-i | 5 | 46.6 | 125 | 161 | 7 | MOLECULE: | PRE-MRNA-SPLICING FACTOR SPP42 |
| 97:00:00 | 6yjd-A | 5 | 3.2 | 87 | 104 | 7 | MOLECULE: | CAPSID ASSEMBLY SCAFFOLDING PROTEIN,PRELAMIN |
| 98:00:00 | 7fde-H | 5 | 4.8 | 96 | 112 | 6 | MOLECULE: | V-TYPE PROTON ATPASE SUBUNIT C |

| | | | | | | | | |
|---|---|---|---|---|---|---|---|---|
| 99:00:00 | 8ppr-P | 5 | 7.8 | 124 | 176 | 5 | MOLECULE: | KINETOCHORE-ASSOCIATED PROTEIN DSN1 HOMOLOG |
| 0:00 | 3jb9-W | 5 | 33.9 | 135 | 426 | 7 | MOLECULE: | PRE-MRNA-SPLICING FACTOR SPP42 |
| 1:00 | 8tek-A | 5 | 9.2 | 131 | 200 | 6 | MOLECULE: | DYNEIN REGULATORY COMPLEX PROTEIN 1/2 N-TERM |
| 2:00 | 3v6i-A | 4.9 | 6.4 | 115 | 186 | 9 | MOLECULE: | V-TYPE ATP SYNTHASE SUBUNIT E |
| 3:00 | 6znl-n | 4.9 | 11.1 | 113 | 343 | 7 | MOLECULE: | ARP1 ACTIN RELATED PROTEIN 1 HOMOLOG A |
| 4:00 | 6cfz-I | 4.9 | 12 | 105 | 106 | 5 | MOLECULE: | ASK1 |
| 5:00 | 4aj5-2 | 4.9 | 11 | 99 | 100 | 4 | MOLECULE: | SPINDLE AND KINETOCHORE-ASSOCIATED PROTEIN 1 |
| 6:00 | 1nyh-A | 4.9 | 3.8 | 76 | 76 | 9 | MOLECULE: | REGULATORY PROTEIN SIR4 |
| 7:00 | 2jee-A | 4.9 | 3 | 78 | 78 | 6 | MOLECULE: | CELL DIVISION PROTEIN ZAPB |
| 8:00 | 3iv1-A | 4.9 | 2.9 | 77 | 78 | 10 | MOLECULE: | TUMOR SUSCEPTIBILITY GENE 101 PROTEIN |
| 9:00 | 8tek-B | 4.8 | 15.5 | 121 | 208 | 7 | MOLECULE: | DYNEIN REGULATORY COMPLEX PROTEIN 1/2 N-TERM |
| 10:00 | 6oei-A | 4.8 | 2.1 | 97 | 243 | 10 | MOLECULE: | SPINDLE POLE BODY COMPONENT SPC42,SIGMA-54-D |
| 11:00 | 9na8-E | 4.8 | 4.7 | 84 | 244 | 10 | MOLECULE: | AUGMIN SUBUNIT 1 |
| 12:00 | 3vem-C | 4.8 | 2.8 | 79 | 83 | 9 | MOLECULE: | HELICASE PROTEIN MOM1 |
| 13:00 | 8ppr-D | 4.8 | 11.2 | 138 | 248 | 7 | MOLECULE: | KINETOCHORE-ASSOCIATED PROTEIN DSN1 HOMOLOG |
| 14:00 | 8amr-A | 4.7 | 7.7 | 116 | 155 | 9 | MOLECULE: | TRIPARTITE MOTIF-CONTAINING PROTEIN 3 |
| 15:00 | 5nnv-D | 4.7 | 7.6 | 125 | 252 | 6 | MOLECULE: | CHROMOSOME PARTITION PROTEIN SMC,CHROMOSOME |
| 16:00 | 3b5n-C | 4.7 | 3.7 | 70 | 70 | 6 | MOLECULE: | SYNAPTOBREVIN HOMOLOG 1 |
| 17:00 | 3trt-A | 4.7 | 2.9 | 75 | 75 | 3 | MOLECULE: | VIMENTIN |
| 18:00 | 3thf-A | 4.7 | 3.8 | 85 | 175 | 8 | MOLECULE: | PROTEIN SHROOM |
| 19:00 | 6id0-L | 4.7 | 22.8 | 125 | 475 | 6 | MOLECULE: | PRE-MRNA-PROCESSING-SPLICING FACTOR 8 |
| 20:00 | 5mq4-C | 4.6 | 6.2 | 90 | 117 | 3 | MOLECULE: | PROTEIN KINASE C-BINDING PROTEIN 1 |
| 21:00 | 7jtk-F | 4.6 | 7.2 | 130 | 287 | 8 | MOLECULE: | FLAGELLAR RADIAL SPOKE PROTEIN 1 |
| 22:00 | 4dhx-A | 4.6 | 2.2 | 72 | 72 | 4 | MOLECULE: | 80 KDA MCM3-ASSOCIATED PROTEIN |
| 23:00 | 7qoo-U | 4.6 | 4.5 | 110 | 186 | 6 | MOLECULE: | CENTROMERE PROTEIN C |
| 24:00:00 | 6wcj-B | 4.5 | 8.5 | 81 | 109 | 4 | MOLECULE: | CLATHRIN HEAVY CHAIN 1 |
| 25:00:00 | 8q85-V | 4.5 | 9.3 | 93 | 117 | 9 | MOLECULE: | KINETOCHORE PROTEIN NDC80 |
| 26:00:00 | 2ba2-A | 4.5 | 5.8 | 81 | 81 | 12 | MOLECULE: | HYPOTHETICAL UPF0134 PROTEIN MPN010 |
| 27:00:00 | 8wjo-B | 4.5 | 7.8 | 129 | 283 | 8 | MOLECULE: | STRUCTURAL MAINTENANCE OF CHROMOSOMES PROTEI |
| 28:00:00 | 8a5p-G | 4.5 | 5.8 | 89 | 99 | 9 | MOLECULE: | INO80 ATPASE |
| 29:00:00 | 6yvv-A | 4.5 | 56.8 | 107 | 465 | 6 | MOLECULE: | STRUCTURAL MAINTENANCE OF CHROMOSOMES PROTEI |
| 30:00:00 | 7emf-D | 4.4 | 9.7 | 107 | 158 | 14 | MOLECULE: | MEDIATOR OF RNA POLYMERASE II TRANSCRIPTION |
| 31:00:00 | 8ppr-M | 4.4 | 18.3 | 124 | 205 | 10 | MOLECULE: | KINETOCHORE-ASSOCIATED PROTEIN DSN1 HOMOLOG |
| 32:00:00 | 3n7n-B | 4.4 | 6.2 | 92 | 164 | 4 | MOLECULE: | MONOPOLIN COMPLEX SUBUNIT CSM1 |
| 33:00:00 | 7jts-s | 4.4 | 18.4 | 113 | 290 | 5 | MOLECULE: | RADIAL SPOKE PROTEIN 3 |
| 34:00:00 | 5dfz-A | 4.4 | 21.3 | 125 | 343 | 8 | MOLECULE: | VACUOLAR PROTEIN SORTING-ASSOCIATED PROTEIN |
| 35:00:00 | 7uw5-A | 4.4 | 36.1 | 105 | 723 | 8 | MOLECULE: | MECHANOSENSITIVE CHANNEL MSCK |
| 36:00:00 | 4wpx-B | 4.4 | 7.9 | 78 | 81 | 12 | MOLECULE: | CELL DIVISION CONTROL PROTEIN 31-LIKE PROTEI |
| 37:00:00 | 5kiu-A | 4.4 | 5.9 | 76 | 76 | 5 | MOLECULE: | SELENOPROTEIN S |
| 38:00:00 | 5omb-C | 4.4 | 9.7 | 88 | 98 | 9 | MOLECULE: | REPLICATION FACTOR A PROTEIN 1 |
| 39:00:00 | 3qh9-A | 4.4 | 3.1 | 66 | 66 | 9 | MOLECULE: | LIPRIN-BETA-2 |
| 40:00:00 | 7vhp-G | 4.4 | 13.4 | 132 | 299 | 10 | MOLECULE: | ATP-DEPENDENT ZINC METALLOPROTEASE FTSH |
| 41:00:00 | 1gmj-A | 4.4 | 3.1 | 65 | 65 | 5 | MOLECULE: | ATPASE INHIBITOR |
| 42:00:00 | 2wz7-B | 4.4 | 4.2 | 67 | 67 | 6 | MOLECULE: | UNCHARACTERIZED PROTEIN YBGF |
| 43:00:00 | 5wwl-N | 4.4 | 5.6 | 73 | 155 | 4 | MOLECULE: | CENTROMERE PROTEIN MIS12 |
| 44:00:00 | 8hpo-H | 4.4 | 5.1 | 126 | 233 | 11 | MOLECULE: | TRANSCRIPTIONAL REGULATORY PROTEIN UME1 |
| 45:00:00 | 7woo-G | 4.3 | 3.8 | 91 | 200 | 13 | MOLECULE: | NUCLEOPORIN NIC96 |
| 46:00:00 | 4n78-E | 4.3 | 2.9 | 66 | 67 | 5 | MOLECULE: | CYTOPLASMIC FMR1-INTERACTING PROTEIN 1 |
| 47:00:00 | 5c9n-A | 4.3 | 1.7 | 64 | 64 | 9 | MOLECULE: | GEMININ COILED-COIL DOMAIN-CONTAINING PROTEI |
| 48:00:00 | 8fed-A | 4.3 | 21.3 | 137 | 392 | 4 | MOLECULE: | VIRULENCE FACTOR MCE FAMILY PROTEIN |

| | | | | | | | | |
|---|---|---|---|---|---|---|---|---|
| 49:00:00 | 5ed9-B | 4.2 | 2.7 | 72 | 72 | 10 | MOLECULE: | SUN DOMAIN-CONTAINING PROTEIN 2 |
| 50:00:00 | 8ppr-F | 4.2 | 4.2 | 71 | 112 | 4 | MOLECULE: | KINETOCHORE-ASSOCIATED PROTEIN DSN1 HOMOLOG |
| 51:00:00 | 8p0v-K | 4.2 | 26.4 | 129 | 304 | 13 | MOLECULE: | COILED-COIL DOMAIN-CONTAINING PROTEIN 93 |
| 52:00:00 | 5t58-B | 4.2 | 11 | 112 | 190 | 9 | MOLECULE: | KLLA0F02343P |
| 53:00:00 | 3b5n-D | 4.2 | 3.6 | 64 | 64 | 9 | MOLECULE: | SYNAPTOBREVIN HOMOLOG 1 |
| 54:00:00 | 4bry-B | 4.2 | 4.7 | 69 | 69 | 6 | MOLECULE: | GEMININ |
| 55:00:00 | 6cfz-C | 4.2 | 3.5 | 70 | 79 | 4 | MOLECULE: | ASK1 |
| 56:00:00 | 4c47-A | 4.1 | 8.6 | 81 | 191 | 7 | MOLECULE: | INNER MEMBRANE LIPOPROTEIN |
| 57:00:00 | 4r8g-E | 4.1 | 15.8 | 109 | 324 | 6 | MOLECULE: | UNCONVENTIONAL MYOSIN-IC |
| 58:00:00 | 9cgi-C | 4.1 | 11.8 | 107 | 201 | 4 | MOLECULE: | RNA-DIRECTED RNA POLYMERASE L |
| 59:00:00 | 7sqk-C | 4.1 | 8.7 | 136 | 600 | 4 | MOLECULE: | HAUS AUGMIN-LIKE COMPLEX SUBUNIT 1 |
| 60:00:00 | 4r3z-A | 4.1 | 2 | 71 | 76 | 3 | MOLECULE: | AMINOACYL TRNA SYNTHASE COMPLEX-INTERACTING |
| 61:00:00 | 8tek-M | 4.1 | 23.8 | 96 | 164 | 8 | MOLECULE: | DYNEIN REGULATORY COMPLEX PROTEIN 1/2 N-TERM |
| 62:00:00 | 8fed-E | 4.1 | 21.2 | 136 | 358 | 6 | MOLECULE: | VIRULENCE FACTOR MCE FAMILY PROTEIN |
| 63:00:00 | 1s1c-Y | 4.1 | 3.7 | 70 | 70 | 7 | MOLECULE: | TRANSFORMING PROTEIN RHOA |
| 64:00:00 | 3m0d-A | 4.1 | 2.3 | 63 | 63 | 11 | MOLECULE: | TNF RECEPTOR-ASSOCIATED FACTOR 2 |
| 65:00:00 | 3m0d-C | 4.1 | 3.7 | 63 | 63 | 11 | MOLECULE: | TNF RECEPTOR-ASSOCIATED FACTOR 2 |
| 66:00:00 | 3lt6-F | 4.1 | 4.1 | 63 | 63 | 2 | MOLECULE: | ADHESIN YADA |
| 67:00:00 | 3zdo-F | 4.1 | 3.7 | 65 | 65 | 9 | MOLECULE: | PHOSPHOPROTEIN |
| 68:00:00 | 5wlz-C | 4.1 | 14.4 | 93 | 206 | 8 | MOLECULE: | DNA REPAIR PROTEIN XRCC4,MYOSIN-7 |
| 69:00:00 | 5mq0-t | 4.1 | 12.2 | 104 | 438 | 5 | MOLECULE: | YEAST UBC4 GENE FOR UBIQUITIN-CONJUGATING EN |
| 70:00:00 | 5dol-B | 4.1 | 3.1 | 62 | 62 | 5 | MOLECULE: | INITIATION-CONTROL PROTEIN YABA |
| 71:00:00 | 8p1u-C | 4.1 | 4.2 | 77 | 93 | 8 | MOLECULE: | CELL DIVISION PROTEIN FTSL |
| 72:00:00 | 9io5-G | 4 | 5.2 | 89 | 260 | 6 | MOLECULE: | G1-ATPASE SUBUNIT BETA |
| 73:00:00 | 8q85-b | 4 | 3.9 | 69 | 70 | 7 | MOLECULE: | KINETOCHORE PROTEIN NDC80 |
| 74:00:00 | 8aud-A | 4 | 3.8 | 94 | 350 | 10 | MOLECULE: | CELL WALL-ASSOCIATED HYDROLASES (INVASION-AS |
| 75:00:00 | 4orh-H | 4 | 3 | 77 | 142 | 8 | MOLECULE: | UBIQUITIN-CONJUGATING ENZYME E2 VARIANT 2 |
| 76:00:00 | 6wuc-K | 4 | 23.1 | 115 | 235 | 4 | MOLECULE: | INNER KINETOCHORE SUBUNIT MCM16 |
| 77:00:00 | 8srm-B | 4 | 11.4 | 89 | 235 | 9 | MOLECULE: | RB1-INDUCIBLE COILED-COIL PROTEIN 1 |
| 78:00:00 | 9d80-C | 4 | 11.4 | 121 | 275 | 7 | MOLECULE: | PORTAL PROTEIN |
| 79:00:00 | 1m1j-B | 4 | 13.6 | 119 | 402 | 4 | MOLECULE: | FIBRINOGEN ALPHA SUBUNIT |
| 80:00:00 | 2zxx-B | 4 | 6.1 | 71 | 78 | 7 | MOLECULE: | GEMININ |
| 81:00:00 | 1gd2-E | 4 | 3 | 65 | 65 | 6 | MOLECULE: | DNA (5'- |
| 82:00:00 | 1h88-B | 4 | 5.8 | 70 | 71 | 13 | MOLECULE: | CCAAT/ENHANCER BINDING PROTEIN BETA |
| 83:00:00 | 3hrn-A | 4 | 2.9 | 63 | 63 | 10 | MOLECULE: | TRANSIENT RECEPTOR POTENTIAL (TRP) CHANNEL |
| 84:00:00 | 5chx-A | 4 | 13 | 110 | 210 | 6 | MOLECULE: | XRCC4-MYH7-1590-1657 |
| 85:00:00 | 1aa0-A | 4 | 4.4 | 84 | 113 | 5 | MOLECULE: | FIBRITIN |
| 86:00:00 | 6ian-A | 3.9 | 34.7 | 100 | 306 | 4 | MOLECULE: | INTRAFLAGELLAR TRANSPORT PROTEIN 74 |
| 87:00:00 | 8g0q-B | 3.9 | 19.5 | 99 | 230 | 7 | MOLECULE: | KINETOCHORE PROTEIN NDC80 |
| 88:00:00 | 5oqm-h | 3.9 | 9.2 | 104 | 131 | 7 | MOLECULE: | DNA-DIRECTED RNA POLYMERASE II SUBUNIT RPB1 |
| 89:00:00 | 3p8c-D | 3.9 | 31.5 | 92 | 204 | 8 | MOLECULE: | CYTOPLASMIC FMR1-INTERACTING PROTEIN 1 |
| 90:00:00 | 7eeb-C | 3.9 | 20.5 | 88 | 278 | 6 | MOLECULE: | ENHANCED GREEN FLUORESCENT PROTEIN,CATION CH |
| 91:00:00 | 8q85-X | 3.9 | 7.6 | 68 | 68 | 4 | MOLECULE: | KINETOCHORE PROTEIN NDC80 |
| 92:00:00 | 4cjd-A | 3.9 | 17.4 | 80 | 123 | 8 | MOLECULE: | NADA |
| 93:00:00 | 6kn7-T | 3.9 | 14.7 | 67 | 138 | 6 | MOLECULE: | ACTIN, ALPHA SKELETAL MUSCLE |
| 94:00:00 | 5n9j-D | 3.9 | 16.9 | 97 | 135 | 5 | MOLECULE: | MEDIATOR OF RNA POLYMERASE II TRANSCRIPTION |
| 95:00:00 | 6gmh-Q | 3.9 | 18.8 | 116 | 884 | 9 | MOLECULE: | RPB1 |
| 96:00:00 | 7x2e-A | 3.9 | 9.4 | 97 | 163 | 8 | MOLECULE: | HARMONIN |
| 97:00:00 | 1z56-A | 3.9 | 2 | 75 | 77 | 8 | MOLECULE: | LIGASE INTERACTING FACTOR 1 |
| 98:00:00 | 3wmi-A | 3.9 | 2.3 | 57 | 57 | 2 | MOLECULE: | EIAV GP45 WILD TYPE |

| | | | | | | | | |
|---|---|---|---|---|---|---|---|---|
| 99:00:00 | 7sqk-F | 3.9 | 28.5 | 128 | 385 | 5 | MOLECULE: | HAUS AUGMIN-LIKE COMPLEX SUBUNIT 1 |
| 0:00 | 4ke2-A | 3.9 | 3.1 | 94 | 196 | 4 | MOLECULE: | TYPE I HYPERACTIVE ANTIFREEZE PROTEIN |
| 1:00 | 7nad-5 | 3.9 | 16.1 | 73 | 123 | 12 | MOLECULE: | 25S RRNA |
| 2:00 | 4njl-A | 3.9 | 3.9 | 81 | 129 | 5 | MOLECULE: | S PROTEIN |
| 3:00 | 3b5n-A | 3.9 | 2.4 | 60 | 61 | 5 | MOLECULE: | SYNAPTOBREVIN HOMOLOG 1 |
| 4:00 | 5opt-P | 3.9 | 9.4 | 98 | 249 | 8 | MOLECULE: | ACTIVATED PROTEIN KINASE C RECEPTOR, PUTATIV |
| 5:00 | 4om3-D | 3.9 | 3.4 | 80 | 122 | 11 | MOLECULE: | TRANSDUCIN-LIKE ENHANCER PROTEIN 1 |
| 6:00 | 2p2u-B | 3.8 | 22.6 | 80 | 153 | 5 | MOLECULE: | HOST-NUCLEASE INHIBITOR PROTEIN GAM, PUTATIV |
| 7:00 | 2p22-C | 3.8 | 8.1 | 108 | 186 | 7 | MOLECULE: | SUPPRESSOR PROTEIN STP22 OF TEMPERATURE-SENS |
| 8:00 | 8u95-A | 3.8 | 8.9 | 154 | 1009 | 7 | MOLECULE: | MYOSIN HEAVY CHAIN, ISOFORM U |
| 9:00 | 6kn7-U | 3.8 | 8.5 | 107 | 170 | 7 | MOLECULE: | ACTIN, ALPHA SKELETAL MUSCLE |
| 10:00 | 6gvw-F | 3.8 | 32.5 | 110 | 320 | 5 | MOLECULE: | BRCA1-A COMPLEX SUBUNIT ABRAXAS 1 |
| 11:00 | 5c1f-A | 3.8 | 5.8 | 123 | 298 | 7 | MOLECULE: | SEPTATION PROTEIN IMP2 |
| 12:00 | 6qld-Y | 3.8 | 14 | 130 | 223 | 10 | MOLECULE: | INNER KINETOCHORE SUBUNIT MIF2 |
| 13:00 | 8qyd-A | 3.8 | 7.5 | 113 | 280 | 12 | MOLECULE: | ANTI-PHAGE DEFENSE ZORAB SYSTEM ZORA |
| 14:00 | 8d8k-V | 3.8 | 17.7 | 119 | 233 | 10 | MOLECULE: | PROBABLE S-ADENOSYL-L-METHIONINE-DEPENDENT R |
| 15:00 | 8wge-A | 3.8 | 1.4 | 57 | 348 | 2 | MOLECULE: | ZAC |
| 16:00 | 7zr1-D | 3.8 | 14.3 | 133 | 780 | 7 | MOLECULE: | DOUBLE-STRAND BREAK REPAIR PROTEIN |
| 17:00 | 4i1l-A | 3.8 | 2.3 | 62 | 62 | 8 | MOLECULE: | FORKHEAD BOX PROTEIN P3 |
| 18:00 | 4aqr-D | 3.8 | 2.4 | 57 | 57 | 4 | MOLECULE: | CALMODULIN-7 |
| 19:00 | 4bwd-A | 3.8 | 3.7 | 61 | 61 | 8 | MOLECULE: | SHORT COILED-COIL PROTEIN |
| 20:00 | 1joc-A | 3.8 | 6.1 | 74 | 123 | 7 | MOLECULE: | EARLY ENDOSOMAL AUTOANTIGEN 1 |
| 21:00 | 9cpo-B | 3.7 | 10.8 | 95 | 195 | 8 | MOLECULE: | RNA-DIRECTED RNA POLYMERASE NSP12 |
| 22:00 | 5y06-A | 3.7 | 12.8 | 101 | 235 | 10 | MOLECULE: | MSMEG_4306 |
| 23:00 | 4wpe-A | 3.7 | 5.9 | 112 | 275 | 7 | MOLECULE: | CYTOKINESIS PROTEIN 2 |
| 24:00:00 | 1urq-A | 3.7 | 3.4 | 57 | 57 | 5 | MOLECULE: | M-TOMOSYN ISOFORM |
| 25:00:00 | 7aw7-D | 3.7 | 18.6 | 74 | 93 | 4 | MOLECULE: | HAPB |
| 26:00:00 | 7kdf-A | 3.7 | 10 | 114 | 270 | 4 | MOLECULE: | NDC80 ISOFORM 1,NDC80 ISOFORM 1 |
| 27:00:00 | 8q3v-g | 3.6 | 23.6 | 76 | 76 | 9 | MOLECULE: | TETRAHYDROMETHANOPTERIN S-METHYLTRANSFERASE |
| 28:00:00 | 5odw-D | 3.6 | 2.5 | 66 | 198 | 8 | MOLECULE: | FERRIPYOVERDINE RECEPTOR |
| 29:00:00 | 2efl-A | 3.6 | 4.7 | 121 | 281 | 3 | MOLECULE: | FORMIN-BINDING PROTEIN 1 |
| 30:00:00 | 3cl3-D | 3.6 | 2.2 | 59 | 59 | 0 | MOLECULE: | ORF K13 |
| 31:00:00 | 3e7k-H | 3.6 | 3.1 | 56 | 56 | 7 | MOLECULE: | TRPM7 CHANNEL |
| 32:00:00 | 4jol-A | 3.6 | 3.7 | 60 | 60 | 8 | MOLECULE: | PROTEIN CBFA2T1 |
| 33:00:00 | 7qog-A | 3.5 | 3.8 | 91 | 657 | 8 | MOLECULE: | PORTAL PROTEIN GP20 |
| 34:00:00 | 9f63-A | 3.5 | 35.3 | 87 | 422 | 7 | MOLECULE: | PROTEIN PNS1 |
| 35:00:00 | 5yfp-C | 3.5 | 11.8 | 101 | 790 | 9 | MOLECULE: | EXOCYST COMPLEX COMPONENT SEC3 |
| 36:00:00 | 2v0o-A | 3.5 | 6.7 | 115 | 273 | 7 | MOLECULE: | FCH DOMAIN ONLY PROTEIN 2 |
| 37:00:00 | 7w5a-X | 3.5 | 8.4 | 77 | 87 | 3 | MOLECULE: | PRE-MRNA-PROCESSING-SPLICING FACTOR 8 |
| 38:00:00 | 8v2q-A | 3.5 | 15 | 103 | 184 | 6 | MOLECULE: | CHARGED MULTIVESICULAR BODY PROTEIN 1B |
| 39:00:00 | 7x5e-B | 3.5 | 5.6 | 71 | 106 | 8 | MOLECULE: | TRANSCRIPTION FACTOR MAFG |
| 40:00:00 | 5mg8-A | 3.5 | 20.8 | 85 | 282 | 8 | MOLECULE: | STRUCTURAL MAINTENANCE OF CHROMOSOMES PROTEI |
| 41:00:00 | 2x7r-C | 3.5 | 2.8 | 58 | 58 | 3 | MOLECULE: | TRANSMEMBRANE PROTEIN GP41 |
| 42:00:00 | 7sgs-A | 3.5 | 1.2 | 53 | 53 | 8 | MOLECULE: | TUBULIN ALPHA-1B CHAIN |
| 43:00:00 | 5oqm-n | 3.4 | 14.3 | 94 | 136 | 4 | MOLECULE: | DNA-DIRECTED RNA POLYMERASE II SUBUNIT RPB1 |
| 44:00:00 | 5n9j-U | 3.4 | 25.3 | 104 | 198 | 5 | MOLECULE: | MEDIATOR OF RNA POLYMERASE II TRANSCRIPTION |
| 45:00:00 | 8fck-F | 3.4 | 14.8 | 122 | 393 | 8 | MOLECULE: | HAUS AUGMIN-LIKE COMPLEX SUBUNIT 1 |
| 46:00:00 | 3p8c-F | 3.4 | 10.4 | 106 | 156 | 7 | MOLECULE: | CYTOPLASMIC FMR1-INTERACTING PROTEIN 1 |
| 47:00:00 | 7emf-G | 3.4 | 3.8 | 79 | 161 | 3 | MOLECULE: | MEDIATOR OF RNA POLYMERASE II TRANSCRIPTION |
| 48:00:00 | 3opc-B | 3.4 | 2.2 | 70 | 131 | 4 | MOLECULE: | UNCHARACTERIZED PROTEIN |

| | | | | | | | | |
|---|---|---|---|---|---|---|---|---|
| 49:00:00 | 8afz-B | 3.4 | 2.7 | 74 | 376 | 9 | MOLECULE: | SORTING NEXIN-1 |
| 50:00:00 | 3eff-K | 3.4 | 3.9 | 82 | 139 | 12 | MOLECULE: | FAB |
| 51:00:00 | 7emf-K | 3.4 | 14.4 | 81 | 112 | 5 | MOLECULE: | MEDIATOR OF RNA POLYMERASE II TRANSCRIPTION |
| 52:00:00 | 6b85-A | 3.4 | 10.3 | 78 | 215 | 10 | MOLECULE: | TMHC4_R |
| 53:00:00 | 3a5c-G | 3.4 | 2.3 | 73 | 129 | 7 | MOLECULE: | V-TYPE ATP SYNTHASE ALPHA CHAIN |
| 54:00:00 | 6z6o-C | 3.3 | 8.6 | 101 | 548 | 10 | MOLECULE: | HISTONE DEACETYLASE HDA1 |
| 55:00:00 | 8qfc-D | 3.3 | 13.1 | 98 | 185 | 10 | MOLECULE: | 60S RIBOSOMAL PROTEIN L10A |
| 56:00:00 | 9c57-G | 3.3 | 43.1 | 125 | 878 | 6 | MOLECULE: | RUVB-LIKE 1 |
| 57:00:00 | 5jeq-A | 3.3 | 4.8 | 86 | 227 | 7 | MOLECULE: | NITRATE/NITRITE SENSOR PROTEIN NARQ |
| 58:00:00 | 9na9-A | 3.3 | 7.5 | 71 | 103 | 17 | MOLECULE: | AUGMIN SUBUNIT 1 |
| 59:00:00 | 7vc4-C | 3.3 | 16.5 | 85 | 107 | 9 | MOLECULE: | MITOCHONDRIAL IMPORT RECEPTOR SUBUNIT TOM6 H |
| 60:00:00 | 4tt0-B | 3.3 | 4.6 | 84 | 131 | 7 | MOLECULE: | DENEDDYLASE |
| 61:00:00 | 7x6g-D | 3.3 | 7.8 | 81 | 152 | 9 | MOLECULE: | QUORUM-SENSING REGULATOR PROTEIN G |
| 62:00:00 | 6ikn-A | 3.3 | 5.5 | 115 | 305 | 10 | MOLECULE: | GROWTH ARREST-SPECIFIC PROTEIN 7 |
| 63:00:00 | 7m2w-K | 3.3 | 16.2 | 87 | 95 | 8 | MOLECULE: | TUBULIN GAMMA CHAIN |
| 64:00:00 | 5oxf-A | 3.3 | 12.4 | 120 | 703 | 7 | MOLECULE: | GTP-BINDING PROTEIN |
| 65:00:00 | 4p1m-B | 3.3 | 7.7 | 65 | 107 | 12 | MOLECULE: | CELL DIVISION PROTEIN ZAPA |
| 66:00:00 | 7f3t-A | 3.3 | 4 | 70 | 330 | 13 | MOLECULE: | TRANSMEMBRANE PROTEIN 120A |
| 67:00:00 | 8q85-Z | 3.3 | 10.8 | 83 | 94 | 8 | MOLECULE: | KINETOCHORE PROTEIN NDC80 |
| 68:00:00 | 7sc0-A | 3.2 | 24.4 | 101 | 129 | 5 | MOLECULE: | CAVEOLIN-1 |
| 69:00:00 | 8hhf-B | 3.2 | 3.8 | 67 | 89 | 9 | MOLECULE: | CELL DIVISION PROTEIN FTSQ |
| 70:00:00 | 7vdv-A | 3.2 | 16 | 113 | 809 | 6 | MOLECULE: | HISTONE H4 |
| 71:00:00 | 5ew5-A | 3.2 | 23.7 | 109 | 490 | 5 | MOLECULE: | COLICIN-E9 |
| 72:00:00 | 7v2w-J | 3.2 | 15.2 | 115 | 236 | 5 | MOLECULE: | THO COMPLEX SUBUNIT HPR1 |
| 73:00:00 | 7nvr-n | 3.2 | 12.4 | 91 | 132 | 7 | MOLECULE: | TFIIH BASAL TRANSCRIPTION FACTOR COMPLEX HEL |
| 74:00:00 | 4h8s-C | 3.2 | 11.1 | 116 | 382 | 5 | MOLECULE: | DCC-INTERACTING PROTEIN 13-BETA |
| 75:00:00 | 4jio-B | 3.2 | 5.7 | 59 | 320 | 10 | MOLECULE: | BRO1 |
| 76:00:00 | 1kmi-Z | 3.2 | 11.3 | 99 | 177 | 5 | MOLECULE: | CHEMOTAXIS PROTEIN CHEY |
| 77:00:00 | 7wkk-I | 3.2 | 3.6 | 88 | 171 | 10 | MOLECULE: | MGC83295 PROTEIN |
| 78:00:00 | 7mge-B | 3.2 | 10.1 | 68 | 442 | 15 | MOLECULE: | WD REPEAT-CONTAINING PROTEIN 41 |
| 79:00:00 | 5yfp-F | 3.2 | 16.1 | 121 | 725 | 10 | MOLECULE: | EXOCYST COMPLEX COMPONENT SEC3 |
| 80:00:00 | 6em5-t | 3.2 | 15.5 | 78 | 290 | 3 | MOLECULE: | 5.8S RIBOSOMAL RNA |
| 81:00:00 | 2d1l-A | 3.2 | 40.5 | 105 | 249 | 3 | MOLECULE: | METASTASIS SUPPRESSOR PROTEIN 1 |
| 82:00:00 | 4nad-A | 3.2 | 6.1 | 92 | 133 | 4 | MOLECULE: | REGULATION OF NUCLEAR PRE-MRNA DOMAIN-CONTAI |
| 83:00:00 | 5wkq-A | 3.1 | 3.2 | 83 | 166 | 8 | MOLECULE: | INVASIN IPAB |
| 84:00:00 | 4nqi-D | 3.1 | 4.1 | 97 | 232 | 6 | MOLECULE: | SH3 DOMAIN-CONTAINING PROTEIN |
| 85:00:00 | 1ezj-A | 3.1 | 7.3 | 71 | 114 | 7 | MOLECULE: | NUCLEOCAPSID PHOSPHOPROTEIN |
| 86:00:00 | 8tek-E | 3.1 | 6 | 111 | 219 | 5 | MOLECULE: | DYNEIN REGULATORY COMPLEX PROTEIN 1/2 N-TERM |
| 87:00:00 | 6r7l-E | 3.1 | 22 | 78 | 98 | 8 | MOLECULE: | SECG |
| 88:00:00 | 3hgf-A | 3.1 | 4.6 | 69 | 98 | 3 | MOLECULE: | RHOPTRY PROTEIN FRAGMENT |
| 89:00:00 | 7v2w-I | 3.1 | 18.1 | 123 | 239 | 9 | MOLECULE: | THO COMPLEX SUBUNIT HPR1 |
| 90:00:00 | 7d7n-A | 3.1 | 18.8 | 89 | 704 | 4 | MOLECULE: | ATP-BINDING CASSETTE SUB-FAMILY B MEMBER 6, |
| 91:00:00 | 6c0f-8 | 3.1 | 8.9 | 71 | 98 | 8 | MOLECULE: | SACCHAROMYCES CEREVISIAE S288C 35S PRE-RIBOS |
| 92:00:00 | 4ilo-A | 3.1 | 9.5 | 95 | 236 | 4 | MOLECULE: | CT398 |
| 93:00:00 | 6gmh-M | 3.1 | 49.1 | 108 | 991 | 6 | MOLECULE: | RPB1 |
| 94:00:00 | 8vvc-A | 3 | 43.4 | 100 | 1390 | 8 | MOLECULE: | MULTIDRUG RESISTANCE-ASSOCIATED PROTEIN 1 |
| 95:00:00 | 4m8m-B | 3 | 16.5 | 85 | 574 | 6 | MOLECULE: | GCN4 COILED-COIL FUSED ZEBRAFISH PLEXINC1 |
| 96:00:00 | 6oee-A | 3 | 54.7 | 104 | 243 | 3 | MOLECULE: | TYPE IV SECRETION SYSTEM APPARATUS PROTEIN C |
| 97:00:00 | 4tko-B | 3 | 11.6 | 111 | 326 | 11 | MOLECULE: | EMRA |
| 98:00:00 | 4mu6-A | 3 | 5.5 | 99 | 274 | 9 | MOLECULE: | KINECTIN 1 (KINESIN RECEPTOR) |

| | | | | | | | | |
|---|---|---|---|---|---|---|---|---|
| 99:00:00 | 7zmg-n | 3 | 12.6 | 94 | 136 | 11 | MOLECULE: | NADH-UBIQUINONE OXIDOREDUCTASE CHAIN 1 |
| 0:00 | 8x5f-A | 3 | 36.1 | 94 | 616 | 6 | MOLECULE: | SOLUTE CARRIER FAMILY 53 MEMBER 1 |
| 1:00 | 4cq4-A | 3 | 3.7 | 77 | 106 | 6 | MOLECULE: | ENGINEERED VERSION OF TRANSMEMBRANE RECEPTOR |
| 2:00 | 6rd4-5 | 3 | 5.2 | 65 | 123 | 3 | MOLECULE: | ASA-10: POLYTOMELLA F-ATP SYNTHASE ASSOCIATE |
| 3:00 | 8xks-M | 3 | 14.9 | 93 | 135 | 1 | MOLECULE: | CTAP1 |
| 4:00 | 9eri-G | 3 | 4.5 | 66 | 207 | 15 | MOLECULE: | NA(+)-TRANSLOCATING FERREDOXIN:NAD(+) OXIDOR |
| 5:00 | 1y2o-A | 3 | 5.3 | 99 | 248 | 7 | MOLECULE: | BAI1-ASSOCIATED PROTEIN 2 ISOFORM 1 |
| 6:00 | 7oi8-q | 3 | 16.7 | 108 | 164 | 8 | MOLECULE: | 39S RIBOSOMAL PROTEIN L2, MITOCHONDRIAL |
| 7:00 | 8fef-D | 3 | 17.9 | 144 | 385 | 8 | MOLECULE: | VIRULENCE FACTOR MCE FAMILY PROTEIN |
| 8:00 | 2fo1-D | 3 | 4.1 | 59 | 63 | 3 | MOLECULE: | 5'- |
| 9:00 | 8cpz-B | 3 | 10.1 | 158 | 2122 | 6 | MOLECULE: | TCDA1 |
| 10:00 | 5vo5-A | 3 | 2.9 | 76 | 179 | 4 | MOLECULE: | COILED-COIL AND C2 DOMAIN-CONTAINING PROTEIN |
| 11:00 | 8xva-K | 3 | 38.6 | 81 | 145 | 10 | MOLECULE: | MITOCHONDRIAL IMPORT RECEPTOR SUBUNIT TOM6 H |
| 12:00 | 7egm-H | 3 | 18.2 | 122 | 388 | 9 | MOLECULE: | TRANSCRIPTION REGULATORY PROTEIN SNF2 |
| 13:00 | 4dyl-A | 3 | 4.6 | 117 | 376 | 7 | MOLECULE: | TYROSINE-PROTEIN KINASE FES/FPS |
| 14:00 | 5n9j-E | 2.9 | 11.9 | 97 | 201 | 7 | MOLECULE: | MEDIATOR OF RNA POLYMERASE II TRANSCRIPTION |
| 15:00 | 5mpd-U | 2.9 | 29.3 | 102 | 298 | 6 | MOLECULE: | 26S PROTEASOME REGULATORY SUBUNIT RPN10 |
| 16:00 | 9na8-D | 2.9 | 8.5 | 102 | 269 | 11 | MOLECULE: | AUGMIN SUBUNIT 1 |
| 17:00 | 5nen-A | 2.9 | 4.5 | 95 | 266 | 3 | MOLECULE: | LIPASE C |
| 18:00 | 6tdv-M | 2.9 | 19.2 | 87 | 166 | 8 | MOLECULE: | ATPTB1 |
| 19:00 | 3viq-A | 2.9 | 16.6 | 75 | 122 | 11 | MOLECULE: | SWI5-DEPENDENT RECOMBINATION DNA REPAIR PROT |
| 20:00 | 5c21-A | 2.9 | 3.9 | 94 | 267 | 5 | MOLECULE: | CHROMOSOMAL HEMOLYSIN D |
| 21:00 | 8hqo-x | 2.9 | 7 | 77 | 80 | 4 | MOLECULE: | PORTAL PROTEIN |
| 22:00 | 6sl2-A | 2.9 | 5.8 | 126 | 619 | 10 | MOLECULE: | CALPONIN HOMOLOGY DOMAIN PROTEIN PUTATIVE |
| 23:00 | 1jad-A | 2.9 | 3.7 | 89 | 242 | 8 | MOLECULE: | PHOSPHOLIPASE C BETA |
| 24:00:00 | 6wuc-H | 2.9 | 11.7 | 77 | 179 | 10 | MOLECULE: | INNER KINETOCHORE SUBUNIT MCM16 |
| 25:00:00 | 7jtk-C | 2.9 | 10.7 | 125 | 427 | 2 | MOLECULE: | FLAGELLAR RADIAL SPOKE PROTEIN 1 |
| 26:00:00 | 8f7n-A | 2.9 | 8.6 | 110 | 364 | 9 | MOLECULE: | METHYL-ACCEPTING CHEMOTAXIS PROTEIN |
| 27:00:00 | 3na7-A | 2.9 | 13.2 | 99 | 237 | 8 | MOLECULE: | HP0958 |
| 28:00:00 | 1lj2-B | 2.8 | 7.3 | 66 | 110 | 6 | MOLECULE: | NONSTRUCTURAL RNA-BINDING PROTEIN 34 |
| 29:00:00 | 7c4j-F | 2.8 | 20.3 | 108 | 383 | 7 | MOLECULE: | TRANSCRIPTION REGULATORY PROTEIN SNF12 |
| 30:00:00 | 6ted-R | 2.8 | 11.7 | 77 | 244 | 5 | MOLECULE: | DNA-DIRECTED RNA POLYMERASE SUBUNIT |
| 31:00:00 | 8qby-L | 2.8 | 32.3 | 108 | 660 | 6 | MOLECULE: | NADH-QUINONE OXIDOREDUCTASE SUBUNIT K |
| 32:00:00 | 7uus-Q | 2.8 | 15.3 | 90 | 170 | 7 | MOLECULE: | HYDROGENASE-2, LARGE SUBUNIT |
| 33:00:00 | 8i3j-A | 2.8 | 5 | 99 | 277 | 8 | MOLECULE: | DYNEIN AXONEMAL HEAVY CHAIN 1 |
| 34:00:00 | 3wqa-A | 2.8 | 7.3 | 106 | 201 | 4 | MOLECULE: | TRIMERIC AUTOTRANSPORTER ADHESIN |
| 35:00:00 | 8iw0-A | 2.8 | 2.9 | 61 | 84 | 11 | MOLECULE: | LIPRIN-BETA-1,KN MOTIF AND ANKYRIN REPEAT DO |
| 36:00:00 | 3qwe-A | 2.8 | 5.4 | 93 | 260 | 2 | MOLECULE: | GEM-INTERACTING PROTEIN |
| 37:00:00 | 8i4t-R | 2.8 | 11.9 | 97 | 501 | 4 | MOLECULE: | ENVELOPMENT POLYPROTEIN |
| 38:00:00 | 7ty0-C | 2.7 | 3.2 | 60 | 488 | 0 | MOLECULE: | GLYCOPROTEIN G |
| 39:00:00 | 7jw1-e | 2.7 | 9.1 | 101 | 235 | 6 | MOLECULE: | CAPSID PROTEINS |
| 40:00:00 | 3lvg-D | 2.7 | 4.1 | 88 | 180 | 3 | MOLECULE: | CLATHRIN HEAVY CHAIN 1 |
| 41:00:00 | 6hgc-A | 2.7 | 7.6 | 63 | 304 | 8 | MOLECULE: | UBIQUITIN CARBOXYL-TERMINAL HYDROLASE CALYPS |
| 42:00:00 | 6rw8-B | 2.7 | 51.7 | 122 | 2337 | 11 | MOLECULE: | A COMPONENT OF INSECTICIDAL TOXIN COMPLEX (T |
| 43:00:00 | 9f5y-p | 2.7 | 8.6 | 90 | 129 | 6 | MOLECULE: | NADH:UBIQUINONE OXIDOREDUCTASE 24 KD SUBUNIT |
| 44:00:00 | 7w9l-A | 2.7 | 48.8 | 80 | 1422 | 4 | MOLECULE: | SODIUM CHANNEL PROTEIN TYPE 9 SUBUNIT ALPHA |
| 45:00:00 | 6tda-L | 2.7 | 15.6 | 79 | 596 | 8 | MOLECULE: | HISTONE H3.2 |
| 46:00:00 | 6tpi-A | 2.7 | 6.7 | 103 | 380 | 4 | MOLECULE: | MUREIN HYDROLASE ACTIVATOR ENVC |
| 47:00:00 | 2ve7-D | 2.7 | 15.1 | 103 | 242 | 6 | MOLECULE: | KINETOCHORE PROTEIN HEC1, KINETOCHORE PROTEI |
| 48:00:00 | 8ap7-c | 2.7 | 4.2 | 59 | 64 | 7 | MOLECULE: | ATP SYNTHASE SUBUNIT A |

| | | | | | | | | |
|---|---|---|---|---|---|---|---|---|
| 49:00:00 | 5lm2-B | 2.7 | 14.1 | 112 | 340 | 4 | MOLECULE: | TYROSINE-PROTEIN PHOSPHATASE NON-RECEPTOR TY |
| 50:00:00 | 5lc5-Z | 2.7 | 16.1 | 96 | 138 | 4 | MOLECULE: | NADH-UBIQUINONE OXIDOREDUCTASE CHAIN 3 |
| 51:00:00 | 8ebw-H | 2.7 | 26.1 | 94 | 532 | 12 | MOLECULE: | TFIIH BASAL TRANSCRIPTION FACTOR COMPLEX HEL |
| 52:00:00 | 6djl-E | 2.7 | 27.7 | 99 | 226 | 8 | MOLECULE: | RAS-RELATED PROTEIN RAB-11A |
| 53:00:00 | 9na8-C | 2.7 | 6.1 | 106 | 234 | 6 | MOLECULE: | AUGMIN SUBUNIT 1 |
| 54:00:00 | 5knc-G | 2.7 | 4.1 | 83 | 195 | 6 | MOLECULE: | V-TYPE SODIUM ATPASE CATALYTIC SUBUNIT A |
| 55:00:00 | 3w3a-G | 2.7 | 2.9 | 93 | 210 | 6 | MOLECULE: | V-TYPE ATP SYNTHASE ALPHA CHAIN |
| 56:00:00 | 6r1j-J | 2.6 | 4.2 | 90 | 264 | 12 | MOLECULE: | UNCHARACTERIZED PROTEIN |
| 57:00:00 | 8at3-A | 2.6 | 42.2 | 68 | 286 | 12 | MOLECULE: | HAUS AUGMIN-LIKE COMPLEX SUBUNIT 1 |
| 58:00:00 | 6ian-C | 2.6 | 34.2 | 86 | 432 | 5 | MOLECULE: | INTRAFLAGELLAR TRANSPORT PROTEIN 74 |
| 59:00:00 | 4u6u-D | 2.6 | 9.7 | 84 | 283 | 13 | MOLECULE: | COG7 |
| 60:00:00 | 8av6-H | 2.6 | 8.2 | 76 | 98 | 11 | MOLECULE: | RUVB-LIKE HELICASE |
| 61:00:00 | 4abx-A | 2.6 | 6.3 | 82 | 167 | 10 | MOLECULE: | DNA REPAIR PROTEIN RECN |
| 62:00:00 | 6xky-A | 2.6 | 6.1 | 101 | 271 | 7 | MOLECULE: | FLAGELLIN |
| 63:00:00 | 7bqx-G | 2.6 | 22.5 | 78 | 84 | 8 | MOLECULE: | TRIPLEX CAPSID PROTEIN 1 |
| 64:00:00 | 5t58-A | 2.6 | 23.3 | 113 | 233 | 9 | MOLECULE: | KLLA0F02343P |
| 65:00:00 | 8y7m-I | 2.6 | 3.8 | 74 | 117 | 4 | MOLECULE: | POLYMERASE ACIDIC PROTEIN |
| 66:00:00 | 8hpo-C | 2.6 | 14.2 | 83 | 138 | 5 | MOLECULE: | TRANSCRIPTIONAL REGULATORY PROTEIN UME1 |
| 67:00:00 | 8hmy-B | 2.6 | 14.5 | 77 | 264 | 14 | MOLECULE: | TRNA-SPLICING ENDONUCLEASE SUBUNIT SEN2 |
| 68:00:00 | 8xku-A | 2.6 | 37.3 | 81 | 730 | 11 | MOLECULE: | PROBABLE INACTIVE ATP-DEPENDENT ZINC METALLO |
| 69:00:00 | 5yfp-A | 2.6 | 48.6 | 111 | 672 | 6 | MOLECULE: | EXOCYST COMPLEX COMPONENT SEC3 |
| 70:00:00 | 4cgk-A | 2.6 | 5 | 84 | 351 | 10 | MOLECULE: | SECRETED 45 KDA PROTEIN |
| 71:00:00 | 8fck-E | 2.6 | 24 | 89 | 222 | 11 | MOLECULE: | HAUS AUGMIN-LIKE COMPLEX SUBUNIT 1 |
| 72:00:00 | 7nyx-A | 2.6 | 10.9 | 118 | 1467 | 8 | MOLECULE: | CHROMOSOME PARTITION PROTEIN MUKB |
| 73:00:00 | 9gk2-G | 2.6 | 13.6 | 102 | 432 | 4 | MOLECULE: | PS2 |
| 74:00:00 | 7zqy-B | 2.5 | 5.3 | 80 | 177 | 15 | MOLECULE: | DH DOMAIN-CONTAINING PROTEIN |
| 75:00:00 | 3vou-A | 2.5 | 5.2 | 68 | 139 | 7 | MOLECULE: | ION TRANSPORT 2 DOMAIN PROTEIN, VOLTAGE-GATE |
| 76:00:00 | 5td8-C | 2.5 | 9.8 | 74 | 111 | 9 | MOLECULE: | KINETOCHORE PROTEIN NDC80 |
| 77:00:00 | 8cr1-C | 2.5 | 7.1 | 73 | 266 | 7 | MOLECULE: | ATPASE ASNA1 |
| 78:00:00 | 4tql-B | 2.5 | 3 | 82 | 235 | 11 | MOLECULE: | THREE HELIX BUNDLE |
| 79:00:00 | 1y9b-A | 2.5 | 5.9 | 58 | 81 | 7 | MOLECULE: | CONSERVED HYPOTHETICAL PROTEIN |
| 80:00:00 | 8tmj-D | 2.5 | 12.8 | 95 | 352 | 4 | MOLECULE: | SAB C18 HEAVY CHAIN |
| 81:00:00 | 6bpz-A | 2.5 | 38.7 | 113 | 900 | 8 | MOLECULE: | PIEZO-TYPE MECHANOSENSITIVE ION CHANNEL COMP |
| 82:00:00 | 6ye4-A | 2.5 | 3.7 | 83 | 236 | 4 | MOLECULE: | BIOPOLYMER TRANSPORT PROTEIN EXBB |
| 83:00:00 | 7qoo-Q | 2.5 | 21 | 96 | 209 | 10 | MOLECULE: | CENTROMERE PROTEIN C |
| 84:00:00 | 7woo-K | 2.5 | 11.2 | 95 | 254 | 6 | MOLECULE: | NUCLEOPORIN NIC96 |
| 85:00:00 | 8tkp-A | 2.5 | 35.2 | 84 | 614 | 6 | MOLECULE: | TRANSMEMBRANE CHANNEL-LIKE PROTEIN 2 |
| 86:00:00 | 1few-A | 2.5 | 4.3 | 75 | 173 | 9 | MOLECULE: | SECOND MITOCHONDRIA-DERIVED ACTIVATOR OF CAS |
| 87:00:00 | 8rhn-A | 2.5 | 4.1 | 64 | 198 | 9 | MOLECULE: | ATPASE FAMILY GENE 2 PROTEIN HOMOLOG A |
| 88:00:00 | 5wjt-A | 2.5 | 5.3 | 101 | 302 | 9 | MOLECULE: | FLAGELLIN |
| 89:00:00 | 6sp2-A | 2.5 | 11.3 | 57 | 366 | 4 | MOLECULE: | MEMBRANE PROTEIN TMS1D |
| 90:00:00 | 7oqe-K | 2.5 | 16.9 | 103 | 406 | 6 | MOLECULE: | PROTEIN NAM8 |
| 91:00:00 | 7sn7-a | 2.5 | 4.2 | 101 | 546 | 9 | MOLECULE: | FLAGELLIN |
| 92:00:00 | 8qfc-C | 2.5 | 12.4 | 118 | 380 | 9 | MOLECULE: | 60S RIBOSOMAL PROTEIN L10A |
| 93:00:00 | 8ceg-B | 2.5 | 2.9 | 79 | 214 | 4 | MOLECULE: | CBY1-INTERACTING BAR DOMAIN-CONTAINING PROTE |
| 94:00:00 | 7n5d-A | 2.5 | 48.4 | 110 | 386 | 5 | MOLECULE: | MECHANOSENSITIVE ION CHANNEL FLYCATCHER1 |
| 95:00:00 | 3r6n-A | 2.5 | 5.9 | 124 | 450 | 6 | MOLECULE: | DESMOPLAKIN |
| 96:00:00 | 5sva-V | 2.5 | 8.6 | 62 | 85 | 5 | MOLECULE: | DNA-DIRECTED RNA POLYMERASE II SUBUNIT RPB1 |
| 97:00:00 | 8wjn-A | 2.4 | 4.2 | 92 | 491 | 11 | MOLECULE: | STRUCTURAL MAINTENANCE OF CHROMOSOMES PROTEI |
| 98:00:00 | 5gox-A | 2.4 | 44.1 | 82 | 181 | 9 | MOLECULE: | DNA REPAIR PROTEIN RAD50 |

| | | | | | | | | |
|---|---|---|---|---|---|---|---|---|
| 99:00:00 | 6xp5-D | 2.4 | 9.6 | 98 | 134 | 8 | MOLECULE: | MEDIATOR OF RNA POLYMERASE II TRANSCRIPTION |
| 0:00 | 7tch-A | 2.4 | 11.3 | 78 | 645 | 4 | MOLECULE: | BACITRACIN EXPORT PERMEASE PROTEIN BCEB |
| 1:00 | 8b9z-m | 2.4 | 18.8 | 82 | 108 | 10 | MOLECULE: | NADH-UBIQUINONE OXIDOREDUCTASE CHAIN 3 |
| 2:00 | 5j1i-A | 2.4 | 6.7 | 102 | 346 | 8 | MOLECULE: | PLECTIN |
| 3:00 | 4gcz-B | 2.4 | 7.5 | 78 | 378 | 5 | MOLECULE: | BLUE-LIGHT PHOTORECEPTOR, SENSOR PROTEIN FIX |
| 4:00 | 4he8-D | 2.4 | 59.2 | 86 | 160 | 7 | MOLECULE: | NADH-QUINONE OXIDOREDUCTASE SUBUNIT 7 |
| 5:00 | 8ixk-E | 2.4 | 3.4 | 62 | 109 | 5 | MOLECULE: | ATTACHMENT PROTEIN G3P |
| 6:00 | 6gy6-A | 2.4 | 6.7 | 102 | 365 | 10 | MOLECULE: | XAXA |
| 7:00 | 5j6f-A | 2.4 | 13.9 | 58 | 352 | 9 | MOLECULE: | 3-DEOXY-D-ARABINO-HEPTULOSONATE 7-PHOSPHATE |
| 8:00 | 8e1m-C | 2.4 | 17.4 | 66 | 264 | 11 | MOLECULE: | MITOCHONDRIAL IMPORT INNER MEMBRANE TRANSLOC |
| 9:00 | 7ctp-A | 2.4 | 43.4 | 123 | 553 | 6 | MOLECULE: | PROTEIN NIBAN 2 |
| 10:00 | 7uh4-A | 2.4 | 3.8 | 63 | 110 | 6 | MOLECULE: | LXG-ASSOCIATED ALPHA-HELICAL PROTEIN D2 |
| 11:00 | 8ymk-A | 2.4 | 5.3 | 50 | 226 | 14 | MOLECULE: | ISOFORM S OF LARGE ENVELOPE PROTEIN |
| 12:00 | 6c5w-A | 2.4 | 5.2 | 79 | 285 | 9 | MOLECULE: | CALCIUM UNIPORTER |
| 13:00 | 8qx8-A | 2.4 | 21.5 | 106 | 969 | 4 | MOLECULE: | VACUOLAR PROTEIN SORTING-ASSOCIATED PROTEIN |
| 14:00 | 5lc5-h | 2.4 | 20.7 | 92 | 134 | 1 | MOLECULE: | NADH-UBIQUINONE OXIDOREDUCTASE CHAIN 3 |
| 15:00 | 4o8w-D | 2.4 | 11.6 | 90 | 120 | 6 | MOLECULE: | SPORE GERMINATION PROTEIN |
| 16:00 | 2pjw-H | 2.4 | 6.8 | 64 | 88 | 6 | MOLECULE: | UNCHARACTERIZED PROTEIN YHL002W |
| 17:00 | 6j6q-F | 2.4 | 13.4 | 77 | 161 | 8 | MOLECULE: | PRE-MRNA-SPLICING FACTOR 8 |
| 18:00 | 8rt9-A | 2.4 | 10 | 97 | 207 | 11 | MOLECULE: | TRWJ PROTEIN |
| 19:00 | 6zyy-C | 2.4 | 38.2 | 106 | 2901 | 9 | MOLECULE: | DYNEIN HEAVY CHAIN, OUTER ARM PROTEIN |
| 20:00 | 8k9c-A | 2.4 | 6.9 | 82 | 207 | 5 | MOLECULE: | CAPRIN-2 |
| 21:00 | 7c4j-C | 2.4 | 20.2 | 83 | 154 | 12 | MOLECULE: | TRANSCRIPTION REGULATORY PROTEIN SNF12 |
| 22:00 | 6tmi-B | 2.4 | 6.9 | 88 | 246 | 5 | MOLECULE: | ATP SYNTHASE SUBUNIT ALPHA |
| 23:00 | 7zr2-A | 2.4 | 6.2 | 84 | 227 | 8 | MOLECULE: | SPIKE PROTEIN S2',CHIMERIC PROTEIN MIMIC OF |
| 24:00:00 | 4wpc-B | 2.4 | 8.6 | 114 | 288 | 11 | MOLECULE: | RHO GTPASE-ACTIVATING PROTEIN RGD1 |
| 25:00:00 | 3on0-A | 2.3 | 12.6 | 67 | 121 | 7 | MOLECULE: | PROTEIN TRAM |
| 26:00:00 | 5xe3-E | 2.3 | 11.1 | 57 | 81 | 5 | MOLECULE: | ENDORIBONUCLEASE MAZF4 |
| 27:00:00 | 8h36-G | 2.3 | 63.9 | 96 | 379 | 5 | MOLECULE: | E3 UBIQUITIN-PROTEIN LIGASE RBX1 |
| 28:00:00 | 6zbj-A | 2.3 | 33.7 | 119 | 666 | 4 | MOLECULE: | PRECURSOR OF THE MAJOR MEROZOITE SURFACE ANT |
| 29:00:00 | 6cfw-I | 2.3 | 36.3 | 93 | 114 | 8 | MOLECULE: | MONOVALENT CATION/H+ ANTIPORTER SUBUNIT D |
| 30:00:00 | 7zu0-F | 2.3 | 8.3 | 58 | 112 | 5 | MOLECULE: | E3 UBIQUITIN-PROTEIN LIGASE PEP5 |
| 31:00:00 | 8at3-D | 2.3 | 70.4 | 116 | 666 | 8 | MOLECULE: | HAUS AUGMIN-LIKE COMPLEX SUBUNIT 1 |
| 32:00:00 | 3bk6-A | 2.3 | 10.5 | 72 | 170 | 3 | MOLECULE: | PH STOMATIN |
| 33:00:00 | 8b9z-h | 2.3 | 20.8 | 90 | 145 | 8 | MOLECULE: | NADH-UBIQUINONE OXIDOREDUCTASE CHAIN 3 |
| 34:00:00 | 2pih-A | 2.3 | 31.9 | 68 | 123 | 9 | MOLECULE: | PROTEIN YMCA |
| 35:00:00 | 6gsi-J | 2.3 | 28.9 | 75 | 89 | 7 | MOLECULE: | CAPSID PROTEIN |
| 36:00:00 | 8dft-A | 2.3 | 2.4 | 46 | 112 | 4 | MOLECULE: | PILIN PROTEIN |
| 37:00:00 | 6tgz-F | 2.3 | 3.4 | 93 | 352 | 3 | MOLECULE: | 55 KDA IMMEDIATE-EARLY PROTEIN 1 |
| 38:00:00 | 5o31-Z | 2.3 | 5.7 | 70 | 137 | 9 | MOLECULE: | NADH-UBIQUINONE OXIDOREDUCTASE CHAIN 3 |
| 39:00:00 | 8x2q-A | 2.3 | 4.7 | 62 | 109 | 5 | MOLECULE: | ADENOMATOUS POLYPOSIS COLI PROTEIN |
| 40:00:00 | 7vxs-o | 2.3 | 21.9 | 88 | 128 | 7 | MOLECULE: | NADH DEHYDROGENASE [UBIQUINONE] IRON-SULFUR |
| 41:00:00 | 8vb2-M | 2.3 | 23.6 | 85 | 125 | 4 | MOLECULE: | TETRAMERIC EJECTION PROTEIN (GP48) |
| 42:00:00 | 6mdm-H | 2.3 | 20.1 | 83 | 147 | 5 | MOLECULE: | VESICLE-FUSING ATPASE |
| 43:00:00 | 9bax-C | 2.3 | 16.3 | 62 | 64 | 2 | MOLECULE: | PROTEIN EFR3 HOMOLOG A |
| 44:00:00 | 8xks-S | 2.3 | 7.7 | 60 | 114 | 7 | MOLECULE: | CTAP1 |
| 45:00:00 | 6z6o-D | 2.3 | 33 | 111 | 542 | 7 | MOLECULE: | HISTONE DEACETYLASE HDA1 |
| 46:00:00 | 5t58-N | 2.2 | 12.7 | 79 | 196 | 8 | MOLECULE: | KLLA0F02343P |
| 47:00:00 | 4gwp-D | 2.2 | 7.3 | 78 | 121 | 3 | MOLECULE: | MEDIATOR OF RNA POLYMERASE II TRANSCRIPTION |
| 48:00:00 | 3f6n-B | 2.2 | 5.6 | 67 | 74 | 6 | MOLECULE: | VIRION-ASSOCIATED PROTEIN |

| | | | | | | | | |
|---|---|---|---|---|---|---|---|---|
| 49:00:00 | 9ivk-A | 2.2 | 3.8 | 71 | 278 | 11 | MOLECULE: | HBC599 MEMBRANE PROTEIN BINDER |
| 50:00:00 | 6h2x-A | 2.2 | 8 | 105 | 352 | 7 | MOLECULE: | CHROMOSOME PARTITION PROTEIN MUKB,CHROMOSOME |
| 51:00:00 | 8e4c-C | 2.2 | 7.7 | 79 | 142 | 4 | MOLECULE: | ISOFORM 2 OF IMMUNOGLOBULIN HEAVY CONSTANT M |
| 52:00:00 | 1u4q-A | 2.2 | 5.8 | 103 | 318 | 9 | MOLECULE: | SPECTRIN ALPHA CHAIN, BRAIN |
| 53:00:00 | 7sgr-A | 2.2 | 12.8 | 63 | 700 | 6 | MOLECULE: | ALPHA-HEMOLYSIN TRANSLOCATION ATP-BINDING PR |
| 54:00:00 | 8pmq-9 | 2.2 | 10.5 | 77 | 412 | 5 | MOLECULE: | E3 UBIQUITIN-PROTEIN LIGASE RMD5 |
| 55:00:00 | 2odu-A | 2.2 | 2.3 | 74 | 217 | 12 | MOLECULE: | PLECTIN 1 |
| 56:00:00 | 8ip4-A | 2.2 | 11.3 | 97 | 320 | 7 | MOLECULE: | MAGNESIUM TRANSPORTER MRS2 HOMOLOG, MITOCHON |
| 57:00:00 | 7b93-L | 2.2 | 26.9 | 112 | 607 | 5 | MOLECULE: | NADH-UBIQUINONE OXIDOREDUCTASE CHAIN 3 |
| 58:00:00 | 3edv-A | 2.2 | 9.2 | 105 | 322 | 8 | MOLECULE: | SPECTRIN BETA CHAIN, BRAIN 1 |
| 59:00:00 | 4lws-A | 2.2 | 2.5 | 59 | 100 | 7 | MOLECULE: | UNCHARACTERIZED PROTEIN |
| 60:00:00 | 8bd7-G | 2.2 | 35.6 | 89 | 205 | 11 | MOLECULE: | IFT88 |
| 61:00:00 | 6zh3-A | 2.2 | 16.9 | 91 | 170 | 7 | MOLECULE: | VACUOLAR PROTEIN-SORTING-ASSOCIATED PROTEIN |
| 62:00:00 | 5xbt-A | 2.2 | 10 | 69 | 271 | 12 | MOLECULE: | PROBABLE TRANSCRIPTIONAL REGULATOR |
| 63:00:00 | 5vk5-B | 2.2 | 3.9 | 61 | 282 | 10 | MOLECULE: | POTASSIUM CHANNEL SUBFAMILY K MEMBER 2 |
| 64:00:00 | 1jfi-B | 2.2 | 19.2 | 78 | 135 | 10 | MOLECULE: | 5'- |
| 65:00:00 | 7uxc-R | 2.2 | 18.3 | 60 | 107 | 5 | MOLECULE: | SERINE/THREONINE-PROTEIN KINASE MTOR |
| 66:00:00 | 8t9l-B | 2.2 | 14.7 | 66 | 110 | 8 | MOLECULE: | NUCLEOPORIN POM152 |
| 67:00:00 | 4y66-D | 2.2 | 18.1 | 100 | 208 | 10 | MOLECULE: | MND1 |
| 68:00:00 | 6e52-A | 2.2 | 29.4 | 90 | 243 | 6 | MOLECULE: | STAPHYLOCOCCUS AUREUS AGRC HISTIDINE KINASE |
| 69:00:00 | 7xzj-G | 2.2 | 2.9 | 52 | 73 | 2 | MOLECULE: | CTAP3 |
| 70:00:00 | 2mi2-A | 2.2 | 16.2 | 79 | 104 | 9 | MOLECULE: | SEC-INDEPENDENT PROTEIN TRANSLOCASE PROTEIN |
| 71:00:00 | 4aur-A | 2.2 | 9.1 | 82 | 564 | 7 | MOLECULE: | LEOA |
| 72:00:00 | 5ik2-G | 2.2 | 2.6 | 82 | 285 | 9 | MOLECULE: | ATP SYNTHASE SUBUNIT ALPHA |
| 73:00:00 | 6zsi-D | 2.2 | 5.4 | 66 | 133 | 9 | MOLECULE: | RAS-RELATED PROTEIN RAB-8A |
| 74:00:00 | 6lk8-a | 2.2 | 16.3 | 93 | 1272 | 12 | MOLECULE: | MGC83295 PROTEIN |
| 75:00:00 | 2p4w-B | 2.1 | 21.8 | 73 | 198 | 8 | MOLECULE: | TRANSCRIPTIONAL REGULATORY PROTEIN ARSR FAMI |
| 76:00:00 | 5j9q-D | 2.1 | 6.1 | 63 | 120 | 0 | MOLECULE: | HISTONE ACETYLTRANSFERASE ESA1 |
| 77:00:00 | 3caz-B | 2.1 | 6.4 | 82 | 210 | 5 | MOLECULE: | BAR PROTEIN |
| 78:00:00 | 7vcf-C | 2.1 | 17.7 | 88 | 217 | 7 | MOLECULE: | TIC214 |
| 79:00:00 | 4gnk-E | 2.1 | 50.5 | 84 | 235 | 4 | MOLECULE: | GUANINE NUCLEOTIDE-BINDING PROTEIN G(Q) SUBU |
| 80:00:00 | 8q3v-b | 2.1 | 3.2 | 55 | 70 | 11 | MOLECULE: | TETRAHYDROMETHANOPTERIN S-METHYLTRANSFERASE |
| 81:00:00 | 6lom-A | 2.1 | 19.1 | 97 | 295 | 7 | MOLECULE: | CALCIUM HOMEOSTASIS MODULATOR PROTEIN |
| 82:00:00 | 8tsh-F | 2.1 | 10.2 | 93 | 265 | 4 | MOLECULE: | ABC TRANSPORTER ATP-BINDING PROTEIN |
| 83:00:00 | 6j72-A | 2.1 | 14.9 | 85 | 561 | 6 | MOLECULE: | ISONIAZID INDUCIBLE GENE PROTEIN INIA |
| 84:00:00 | 8cbk-F | 2.1 | 27.6 | 79 | 344 | 5 | MOLECULE: | 3-HYDROXYACYL-COA DEHYDROGENASE TYPE-2 |
| 85:00:00 | 2m0q-A | 2.1 | 41 | 85 | 123 | 4 | MOLECULE: | POTASSIUM VOLTAGE-GATED CHANNEL SUBFAMILY E |
| 86:00:00 | 6x0a-a | 2.1 | 31.3 | 64 | 87 | 6 | MOLECULE: | PLASMID STABILIZATION SYSTEM |
| 87:00:00 | 2w0c-S | 2.1 | 22.5 | 70 | 84 | 6 | MOLECULE: | MAJOR CAPSID PROTEIN P2 |
| 88:00:00 | 2ehw-C | 2.1 | 2.1 | 47 | 117 | 9 | MOLECULE: | HYPOTHETICAL PROTEIN TTHB059 |
| 89:00:00 | 6cnn-A | 2.1 | 37.4 | 73 | 360 | 11 | MOLECULE: | INTERMEDIATE CONDUCTANCE CALCIUM-ACTIVATED P |
| 90:00:00 | 8b9z-Z | 2.1 | 8.4 | 70 | 146 | 9 | MOLECULE: | NADH-UBIQUINONE OXIDOREDUCTASE CHAIN 3 |
| 91:00:00 | 7z7s-A | 2.1 | 34.6 | 82 | 131 | 5 | MOLECULE: | NADH-QUINONE OXIDOREDUCTASE SUBUNIT F |
| 92:00:00 | 8kde-3 | 2.1 | 10.7 | 88 | 97 | 13 | MOLECULE: | PHOTOSYSTEM II CP47 REACTION CENTER PROTEIN |
| 93:00:00 | 8b6j-i | 2.1 | 19.5 | 88 | 117 | 6 | MOLECULE: | PEPTIDASE M16 INACTIVE DOMAIN PROTEIN |
| 94:00:00 | 4ad8-A | 2.1 | 10.3 | 110 | 452 | 6 | MOLECULE: | DNA REPAIR PROTEIN RECN |
| 95:00:00 | 7zu0-E | 2.1 | 35.5 | 107 | 343 | 8 | MOLECULE: | E3 UBIQUITIN-PROTEIN LIGASE PEP5 |
| 96:00:00 | 8eki-A | 2.1 | 3.7 | 71 | 170 | 13 | MOLECULE: | PROTEIN TRANSPORT PROTEIN SEC20 |
| 97:00:00 | 6zka-p | 2.1 | 22.4 | 89 | 128 | 7 | MOLECULE: | NADH-UBIQUINONE OXIDOREDUCTASE CHAIN 3 |
| 98:00:00 | 7nb6-A | 2.1 | 31.8 | 75 | 340 | 9 | MOLECULE: | AI-2 TRANSPORT PROTEIN TQSA |

| | | | | | | | | |
|---|---|---|---|---|---|---|---|---|
| 99:00:00 | 3mhh-C | 2.1 | 10.2 | 64 | 93 | 14 | MOLECULE: | UBIQUITIN CARBOXYL-TERMINAL HYDROLASE 8 |
| 0:00 | 1llm-C | 2.1 | 3.9 | 55 | 87 | 13 | MOLECULE: | 5'-D(*TP*CP*CP*CP*AP*CP*GP*CP*GP*TP*GP*GP*G) |
| 1:00 | 3fx7-B | 2.1 | 45.7 | 62 | 87 | 5 | MOLECULE: | PUTATIVE UNCHARACTERIZED PROTEIN |
| 2:00 | 7mvv-A | 2.1 | 20.1 | 102 | 1542 | 7 | MOLECULE: | NUCLEOPORIN NUP192 |
| 3:00 | 2wwb-B | 2 | 10.3 | 61 | 68 | 10 | MOLECULE: | PROTEIN TRANSPORT PROTEIN SEC61 SUBUNIT ALPH |
| 4:00 | 7yzp-C | 2 | 8.1 | 94 | 448 | 9 | MOLECULE: | DNA HAIRPIN (59-MER) |
| 5:00 | 8z9y-F | 2 | 15.2 | 77 | 95 | 6 | MOLECULE: | PROTEIN TIC 214 |
| 6:00 | 7emf-J | 2 | 6.3 | 68 | 122 | 6 | MOLECULE: | MEDIATOR OF RNA POLYMERASE II TRANSCRIPTION |
| 7:00 | 6thk-A | 2 | 9 | 107 | 466 | 13 | MOLECULE: | PYOCIN S5 |
| 8:00 | 6rwb-A | 2 | 39.5 | 141 | 1873 | 10 | MOLECULE: | TOXIN,TOXIN COMPLEX SUBUNIT TCAB,PUTATIVE TO |
| 9:00 | 1u2m-C | 2 | 5.4 | 63 | 143 | 11 | MOLECULE: | HISTONE-LIKE PROTEIN HLP-1 |
| 10:00 | 6xp5-G | 2 | 4.3 | 70 | 122 | 9 | MOLECULE: | MEDIATOR OF RNA POLYMERASE II TRANSCRIPTION |
| 11:00 | 8rrh-B | 2 | 29.2 | 94 | 299 | 6 | MOLECULE: | PROHIBITIN 1 |
| 12:00 | 8t1i-F | 2 | 6.8 | 56 | 73 | 7 | MOLECULE: | MEDIATOR OF RNA POLYMERASE II TRANSCRIPTION |
| 13:00 | 9mli-B | 2 | 50.3 | 118 | 2519 | 8 | MOLECULE: | A COMPONENT OF INSECTICIDAL TOXIN COMPLEX (T |
| 14:00 | 5n9j-A | 2 | 26.1 | 118 | 566 | 5 | MOLECULE: | MEDIATOR OF RNA POLYMERASE II TRANSCRIPTION |
| 15:00 | 1i49-A | 2 | 3.4 | 71 | 201 | 8 | MOLECULE: | ARFAPTIN 2 |

**DALI results: MdA-1 Cah**

| No: | Chain | Z | rmsd | lali | nres | %id | PDB | Description | | |
|---|---|---|---|---|---|---|---|---|---|---|
| 1:00 | 6gao-B | 2.8 | 71.6 | 162 | 225 | 7 | MOLECULE: | OUTER CAPSID PROTEIN SIGMA-1 | | |
| 2:00 | 6gao-A | 2.8 | 68.3 | 156 | 223 | 5 | MOLECULE: | OUTER CAPSID PROTEIN SIGMA-1 | | |
| 3:00 | 8th8-B | 2.7 | 64.4 | 154 | 276 | 3 | MOLECULE: | DYNEIN REGULATORY COMPLEX PROTEIN 1/2 | N-TERMINAL | |
| 4:00 | 7dl2-C | 2.7 | 85.9 | 168 | 224 | 7 | MOLECULE: | HAMARTIN | | |
| 5:00 | 2v71-B | 2.7 | 60.3 | 137 | 160 | 8 | MOLECULE: | NUCLEAR DISTRIBUTION PROTEIN NUDE-LIK | E 1 | |
| 6:00 | 8at3-H | 2.7 | 42.5 | 160 | 203 | 4 | MOLECULE: | HAUS AUGMIN-LIKE COMPLEX SUBUNIT 1 | | |
| 7:00 | 7dl2-D | 2.6 | 89.5 | 173 | 226 | 9 | MOLECULE: | HAMARTIN | | |
| 8:00 | 8tek-B | 2.6 | 44.6 | 136 | 208 | 9 | MOLECULE: | DYNEIN REGULATORY COMPLEX PROTEIN 1/2 | N-TERMINAL | |
| 9:00 | 8tek-P | 2.6 | 40.9 | 114 | 138 | 8 | MOLECULE: | DYNEIN REGULATORY COMPLEX PROTEIN 1/2 | N-TERMINAL | |
| 10:00 | 8th8-A | 2.6 | 68.4 | 159 | 290 | 8 | MOLECULE: | DYNEIN REGULATORY COMPLEX PROTEIN 1/2 | N-TERMINAL | |
| 11:00 | 5cws-J | 2.6 | 22.4 | 130 | 180 | 6 | MOLECULE: | SAB-158 FAB LIGHT CHAIN | | |
| 12:00 | 8sre-D | 2.6 | 38.1 | 101 | 1377 | 6 | MOLECULE: | TRPM2 CHANZYME | | |
| 13:00 | 8at4-H | 2.6 | 42.1 | 161 | 203 | 5 | MOLECULE: | HAUS AUGMIN-LIKE COMPLEX SUBUNIT 1 | | |
| 14:00 | 6gak-A | 2.6 | 50.2 | 119 | 129 | 1 | MOLECULE: | OUTER CAPSID PROTEIN SIGMA-1 | | |
| 15:00 | 8sri-C | 2.6 | 36.4 | 94 | 1368 | 2 | MOLECULE: | TRPM2 CHANZYME | | |
| 16:00 | 8sr7-A | 2.6 | 37 | 100 | 1375 | 5 | MOLECULE: | TRPM2 CHANZYME | | |
| 17:00 | 5mq4-D | 2.6 | 5.1 | 83 | 111 | 7 | MOLECULE: | PROTEIN KINASE C-BINDING PROTEIN 1 | | |
| 18:00 | 4cpc-G | 2.5 | 60.7 | 130 | 152 | 6 | MOLECULE: | SYNAPTONEMAL COMPLEX PROTEIN 3 | | |
| 19:00 | 2v71-A | 2.5 | 61.1 | 133 | 160 | 5 | MOLECULE: | NUCLEAR DISTRIBUTION PROTEIN NUDE-LIK | E 1 | |
| 20:00 | 8sre-A | 2.5 | 38 | 98 | 1377 | 6 | MOLECULE: | TRPM2 CHANZYME | | |
| 21:00 | 8fck-H | 2.5 | 42.2 | 167 | 218 | 4 | MOLECULE: | HAUS AUGMIN-LIKE COMPLEX SUBUNIT 1 | | |
| 22:00 | 8p0v-L | 2.5 | 51.3 | 174 | 416 | 7 | MOLECULE: | COILED-COIL DOMAIN-CONTAINING PROTEIN | | 93 |
| 23:00 | 5cws-D | 2.5 | 23 | 134 | 180 | 3 | MOLECULE: | SAB-158 FAB LIGHT CHAIN | | |
| 24:00:00 | 3o0z-D | 2.5 | 59.5 | 135 | 160 | 7 | MOLECULE: | RHO-ASSOCIATED PROTEIN KINASE 1 | | |
| 25:00:00 | 8th8-J | 2.5 | 68.2 | 156 | 372 | 8 | MOLECULE: | DYNEIN REGULATORY COMPLEX PROTEIN 1/2 | N-TERMINAL | |
| 26:00:00 | 8t9l-A | 2.5 | 7.5 | 93 | 108 | 9 | MOLECULE: | NUCLEOPORIN POM152 | | |
| 27:00:00 | 8fwj-A | 2.5 | 2.4 | 82 | 552 | 5 | MOLECULE: | CIRCADIAN CLOCK PROTEIN KAIC | | |
| 28:00:00 | 3v6i-A | 2.5 | 25.2 | 104 | 186 | 2 | MOLECULE: | V-TYPE ATP SYNTHASE SUBUNIT E | | |
| 29:00:00 | 7qoo-H | 2.5 | 26 | 107 | 210 | 8 | MOLECULE: | CENTROMERE PROTEIN C | | |
| 30:00:00 | 5hda-A | 2.5 | 5.6 | 85 | 121 | 5 | MOLECULE: | ZINC FINGER MYND DOMAIN-CONTAINING PR | OTEIN 11 | |
| 31:00:00 | 7apk-m | 2.5 | 44.1 | 100 | 549 | 7 | MOLECULE: | THO COMPLEX SUBUNIT 1 | | |
| 32:00:00 | 7x2e-A | 2.5 | 2.3 | 75 | 163 | 8 | MOLECULE: | HARMONIN | | |
| 33:00:00 | 4lin-L | 2.5 | 84.9 | 181 | 289 | 9 | MOLECULE: | TAIL NEEDLE PROTEIN GP26 | | |
| 34:00:00 | 2ocy-B | 2.5 | 59 | 136 | 149 | 6 | MOLECULE: | RAB GUANINE NUCLEOTIDE EXCHANGE FACTO | R SEC2 | |
| 35:00:00 | 8sre-C | 2.5 | 38.3 | 100 | 1377 | 8 | MOLECULE: | TRPM2 CHANZYME | | |
| 36:00:00 | 4cpc-A | 2.5 | 59.8 | 127 | 143 | 6 | MOLECULE: | SYNAPTONEMAL COMPLEX PROTEIN 3 | | |
| 37:00:00 | 8srf-B | 2.5 | 38.4 | 107 | 1377 | 8 | MOLECULE: | TRPM2 CHANZYME | | |
| 38:00:00 | 8sr7-C | 2.5 | 38.2 | 103 | 1375 | 8 | MOLECULE: | TRPM2 CHANZYME | | |
| 39:00:00 | 8sr7-D | 2.5 | 37.5 | 98 | 1375 | 8 | MOLECULE: | TRPM2 CHANZYME | | |
| 40:00:00 | 6xer-E | 2.5 | 3 | 87 | 120 | 2 | MOLECULE: | TUBULIN ALPHA-1B CHAIN | | |
| 41:00:00 | 6i5c-E | 2.5 | 3.4 | 88 | 120 | 10 | MOLECULE: | TUBULIN ALPHA-1B CHAIN | | |
| 42:00:00 | 7znk-O | 2.5 | 39.9 | 102 | 164 | 7 | MOLECULE: | RNA | | |
| 43:00:00 | 7l05-E | 2.5 | 3.6 | 89 | 121 | 9 | MOLECULE: | TUBULIN ALPHA-1B CHAIN | | |
| 44:00:00 | 8bdf-E | 2.5 | 3.4 | 88 | 121 | 10 | MOLECULE: | TUBULIN ALPHA-1B CHAIN | | |
| 45:00:00 | 4mvd-F | 2.5 | 4.7 | 86 | 253 | 9 | MOLECULE: | CHOLINE-PHOSPHATE CYTIDYLYLTRANSFERAS | E A | |
| 46:00:00 | 7ttd-E | 2.5 | 3.3 | 88 | 124 | 9 | MOLECULE: | TUBULIN ALPHA-1B CHAIN | | |
| 47:00:00 | 4wbn-E | 2.5 | 3.5 | 89 | 120 | 9 | MOLECULE: | TUBULIN ALPHA-1B CHAIN | | |
| 48:00:00 | 5jcb-E | 2.5 | 3.4 | 88 | 121 | 10 | MOLECULE: | TUBULIN ALPHA-1B CHAIN | | |
| 49:00:00 | 8fwj-D | 2.5 | 2.4 | 82 | 552 | 5 | MOLECULE: | CIRCADIAN CLOCK PROTEIN KAIC | | |

| | | | | | | | | | |
|---|---|---|---|---|---|---|---|---|---|
| 50:00:00 | 5yl2-E | 2.5 | 3.6 | 88 | 121 | 10 | MOLECULE: | TUBULIN ALPHA-1B CHAIN | |
| 51:00:00 | 6tde-E | 2.5 | 3.5 | 91 | 131 | 8 | MOLECULE: | TUBULIN ALPHA CHAIN | |
| 52:00:00 | 7ttf-E | 2.5 | 3.3 | 88 | 123 | 9 | MOLECULE: | TUBULIN ALPHA-1B CHAIN | |
| 53:00:00 | 6pc4-E | 2.5 | 3.5 | 88 | 121 | 10 | MOLECULE: | TUBULIN ALPHA-1B CHAIN | |
| 54:00:00 | 7apk-O | 2.5 | 43.9 | 108 | 155 | 6 | MOLECULE: | THO COMPLEX SUBUNIT 1 | |
| 55:00:00 | 5o7a-E | 2.5 | 3.5 | 88 | 118 | 10 | MOLECULE: | TUBULIN ALPHA-1B CHAIN | |
| 56:00:00 | 7znl-o | 2.5 | 44.5 | 105 | 164 | 7 | MOLECULE: | THO COMPLEX SUBUNIT 1 | |
| 57:00:00 | 5xkf-E | 2.5 | 3.4 | 88 | 121 | 10 | MOLECULE: | TUBULIN ALPHA-1B CHAIN | |
| 58:00:00 | 5xke-E | 2.5 | 3.5 | 88 | 121 | 10 | MOLECULE: | TUBULIN ALPHA-1B CHAIN | |
| 59:00:00 | 6r10-L | 2.5 | 3.3 | 98 | 186 | 3 | MOLECULE: | V-TYPE ATP SYNTHASE ALPHA CHAIN | |
| 60:00:00 | 6th4-E | 2.5 | 3.4 | 88 | 128 | 10 | MOLECULE: | TUBULIN ALPHA CHAIN | |
| 61:00:00 | 5lxs-E | 2.5 | 4.9 | 89 | 123 | 4 | MOLECULE: | TUBULIN ALPHA-1B CHAIN | |
| 62:00:00 | 5lyj-E | 2.5 | 3.7 | 90 | 123 | 10 | MOLECULE: | TUBULIN ALPHA-1B CHAIN | |
| 63:00:00 | 5hda-C | 2.5 | 6.5 | 84 | 117 | 15 | MOLECULE: | ZINC FINGER MYND DOMAIN-CONTAINING PR | OTEIN 11 |
| 64:00:00 | 3v6i-Y | 2.5 | 26.4 | 104 | 185 | 5 | MOLECULE: | V-TYPE ATP SYNTHASE SUBUNIT E | |
| 65:00:00 | 7f1m-A | 2.4 | 19.2 | 71 | 394 | 13 | MOLECULE: | NUCLEOPROTEIN | |
| 66:00:00 | 4f61-I | 2.4 | 77.4 | 157 | 234 | 5 | MOLECULE: | TUBULIN ALPHA CHAIN | |
| 67:00:00 | 4hl8-A | 2.4 | 50.9 | 136 | 782 | 9 | MOLECULE: | MAJOR VAULT PROTEIN | |
| 68:00:00 | 4gkw-A | 2.4 | 59.3 | 133 | 158 | 8 | MOLECULE: | SPINDLE ASSEMBLY ABNORMAL PROTEIN 6 | |
| 69:00:00 | 4ke2-A | 2.4 | 24.4 | 120 | 196 | 7 | MOLECULE: | TYPE I HYPERACTIVE ANTIFREEZE PROTEIN | |
| 70:00:00 | 5mq4-C | 2.4 | 6.2 | 85 | 117 | 7 | MOLECULE: | PROTEIN KINASE C-BINDING PROTEIN 1 | |
| 71:00:00 | 8i03-G | 2.4 | 14.8 | 112 | 166 | 7 | MOLECULE: | PAIRED AMPHIPATHIC HELIX PROTEIN PST1 | |
| 72:00:00 | 7znk-o | 2.4 | 43.7 | 106 | 164 | 7 | MOLECULE: | RNA | |
| 73:00:00 | 6oei-A | 2.4 | 16.3 | 107 | 243 | 6 | MOLECULE: | SPINDLE POLE BODY COMPONENT SPC42,SIG | MA-54-DEPEND |
| 74:00:00 | 7woo-I | 2.4 | 17.2 | 107 | 187 | 7 | MOLECULE: | NUCLEOPORIN NIC96 | |
| 75:00:00 | 6abo-A | 2.4 | 3.8 | 87 | 213 | 7 | MOLECULE: | DNA REPAIR PROTEIN XRCC4 | |
| 76:00:00 | 4cpc-C | 2.4 | 57.9 | 125 | 144 | 9 | MOLECULE: | SYNAPTONEMAL COMPLEX PROTEIN 3 | |
| 77:00:00 | 7ypw-A | 2.4 | 17 | 68 | 388 | 13 | MOLECULE: | NUCLEOPROTEIN | |
| 78:00:00 | 8srd-B | 2.4 | 38.2 | 108 | 1368 | 5 | MOLECULE: | TRPM2 CHANZYME | |
| 79:00:00 | 2v66-B | 2.4 | 39.7 | 104 | 111 | 12 | MOLECULE: | NUCLEAR DISTRIBUTION PROTEIN NUDE-LIK | E 1 |
| 80:00:00 | 4cpc-E | 2.4 | 57.6 | 129 | 143 | 7 | MOLECULE: | SYNAPTONEMAL COMPLEX PROTEIN 3 | |
| 81:00:00 | 8sr7-B | 2.4 | 38.1 | 103 | 1375 | 9 | MOLECULE: | TRPM2 CHANZYME | |
| 82:00:00 | 5xag-E | 2.4 | 3.6 | 88 | 120 | 9 | MOLECULE: | TUBULIN ALPHA-1B CHAIN | |
| 83:00:00 | 5s5s-E | 2.4 | 3.4 | 89 | 123 | 9 | MOLECULE: | TUBULIN ALPHA-1B CHAIN | |
| 84:00:00 | 5bmv-E | 2.4 | 3.6 | 89 | 123 | 9 | MOLECULE: | TUBULIN ALPHA-1B CHAIN | |
| 85:00:00 | 6r0z-L | 2.4 | 26.7 | 108 | 185 | 2 | MOLECULE: | V-TYPE ATP SYNTHASE ALPHA CHAIN | |
| 86:00:00 | 7z7d-E | 2.4 | 3.5 | 86 | 122 | 9 | MOLECULE: | TUBULIN ALPHA-1B CHAIN | |
| 87:00:00 | 5s5u-E | 2.4 | 3.5 | 89 | 123 | 9 | MOLECULE: | TUBULIN ALPHA-1B CHAIN | |
| 88:00:00 | 4i55-E | 2.4 | 3.4 | 89 | 124 | 9 | MOLECULE: | TUBULIN ALPHA-1B CHAIN | |
| 89:00:00 | 6y4n-E | 2.4 | 3.5 | 89 | 123 | 9 | MOLECULE: | TUBULIN ALPHA-1B CHAIN | |
| 90:00:00 | 7dp8-E | 2.4 | 4.7 | 91 | 123 | 9 | MOLECULE: | TUBULIN ALPHA-1B CHAIN | |
| 91:00:00 | 5gon-E | 2.4 | 22.5 | 89 | 121 | 9 | MOLECULE: | TUBULIN ALPHA-1B CHAIN | |
| 92:00:00 | 5s62-E | 2.4 | 3.5 | 89 | 123 | 9 | MOLECULE: | TUBULIN ALPHA-1B CHAIN | |
| 93:00:00 | 7apk-o | 2.4 | 41.6 | 107 | 155 | 7 | MOLECULE: | THO COMPLEX SUBUNIT 1 | |
| 94:00:00 | 7cpd-E | 2.4 | 3.6 | 88 | 121 | 10 | MOLECULE: | TUBULIN ALPHA-1B CHAIN | |
| 95:00:00 | 7db9-E | 2.4 | 3.5 | 88 | 124 | 10 | MOLECULE: | TUBULIN ALPHA-1B CHAIN | |
| 96:00:00 | 7znk-G | 2.4 | 38 | 100 | 161 | 6 | MOLECULE: | RNA | |
| 97:00:00 | 5sba-E | 2.4 | 3.4 | 88 | 121 | 10 | MOLECULE: | TUBULIN ALPHA-1B CHAIN | |
| 98:00:00 | 5ezy-E | 2.4 | 3.2 | 86 | 121 | 8 | MOLECULE: | TUBULIN ALPHA-1B CHAIN | |
| 99:00:00 | 5s5o-E | 2.4 | 3.4 | 89 | 123 | 9 | MOLECULE: | TUBULIN ALPHA-1B CHAIN | |
| 0:00 | 7xr1-E | 2.4 | 3.4 | 88 | 123 | 10 | MOLECULE: | TUBULIN ALPHA-1B CHAIN | |

**DALI results: MdA-1 Macc**

| No: | Chain | Z | rmsd | lali | nres | %id | PDB | Description |
|---|---|---|---|---|---|---|---|---|
| 1:00 | 5jdo-C | 5.9 | 3.4 | 86 | 141 | 3 | MOLECULE: | HAPTOGLOBIN |
| 2:00 | 7kyr-A | 5.7 | 3.8 | 90 | 151 | 11 | MOLECULE: | MYOGLOBIN |
| 3:00 | 6g5b-A | 5.7 | 3.8 | 90 | 154 | 12 | MOLECULE: | MYOGLOBIN |
| 4:00 | 3e5o-A | 5.6 | 3.9 | 88 | 153 | 13 | MOLECULE: | MYOGLOBIN |
| 5:00 | 7a44-A | 5.6 | 4 | 93 | 154 | 11 | MOLECULE: | MYOGLOBIN |
| 6:00 | 7sph-A | 5.6 | 3.9 | 92 | 154 | 11 | MOLECULE: | MYOGLOBIN |
| 7:00 | 5yzf-A | 5.5 | 4 | 93 | 154 | 11 | MOLECULE: | SPERM WHALE MY( |
| 8:00 | 6f1a-A | 5.4 | 3.8 | 90 | 154 | 11 | MOLECULE: | MYOGLOBIN |
| 9:00 | 5xkv-A | 5.4 | 3.8 | 89 | 153 | 12 | MOLECULE: | MYOGLOBIN |
| 10:00 | 5b84-A | 5.4 | 3.8 | 89 | 153 | 12 | MOLECULE: | MYOGLOBIN |
| 11:00 | 5hav-A | 5.4 | 3.8 | 89 | 153 | 12 | MOLECULE: | MYOGLOBIN |
| 12:00 | 6e03-A | 5.4 | 3.9 | 88 | 153 | 11 | MOLECULE: | MYOGLOBIN |
| 13:00 | 4tyx-A | 5.4 | 3.8 | 89 | 153 | 12 | MOLECULE: | MYOGLOBIN |
| 14:00 | 2blh-A | 5.3 | 4 | 90 | 153 | 13 | MOLECULE: | MYOGLOBIN |
| 15:00 | 8eko-A | 5.3 | 3.9 | 88 | 153 | 11 | MOLECULE: | MYOGLOBIN |
| 16:00 | 7l3u-A | 5.3 | 3.9 | 91 | 153 | 11 | MOLECULE: | MYOGLOBIN |
| 17:00 | 6krc-A | 5.3 | 3.9 | 89 | 153 | 12 | MOLECULE: | MYOGLOBIN |
| 18:00 | 1ch9-A | 5.3 | 4 | 92 | 154 | 12 | MOLECULE: | PROTEIN (MY |
| 19:00 | 7xcq-A | 5.2 | 3.9 | 88 | 153 | 11 | MOLECULE: | MYOGLOBIN |
| 20:00 | 6az6-B | 5.2 | 7.4 | 99 | 205 | 10 | MOLECULE: | GNTR FAMILY |
| 21:00 | 8j4k-A | 5.2 | 3.9 | 91 | 153 | 11 | MOLECULE: | MYOGLOBIN |
| 22:00 | 7xc9-A | 5.2 | 4 | 91 | 153 | 11 | MOLECULE: | MYOGLOBIN |
| 23:00 | 7l3y-A | 5.2 | 3.8 | 90 | 153 | 11 | MOLECULE: | MYOGLOBIN |
| 24:00:00 | 2r4w-A | 5.2 | 3.3 | 84 | 145 | 10 | MOLECULE: | GLOBIN-1 |
| 25:00:00 | 1myt-A | 5.2 | 3.8 | 85 | 145 | 8 | MOLECULE: | MYOGLOBIN |
| 26:00:00 | 3a59-B | 5.2 | 3.6 | 85 | 146 | 8 | MOLECULE: | HEMOGLOBIN |
| 27:00:00 | 7spe-A | 5.2 | 4 | 90 | 154 | 11 | MOLECULE: | MYOGLOBIN |
| 28:00:00 | 6az6-A | 5.1 | 8 | 102 | 209 | 9 | MOLECULE: | GNTR FAMILY |
| 29:00:00 | 5azr-A | 5.1 | 3.9 | 88 | 153 | 14 | MOLECULE: | MYOGLOBIN |
| 30:00:00 | 5zzf-A | 5.1 | 4.1 | 90 | 153 | 11 | MOLECULE: | MYOGLOBIN |
| 31:00:00 | 3ecx-A | 5.1 | 3.9 | 88 | 153 | 13 | MOLECULE: | MYOGLOBIN |
| 32:00:00 | 3w4u-E | 5.1 | 3.1 | 85 | 138 | 4 | MOLECULE: | HEMOGLOBIN |
| 33:00:00 | 1dtm-A | 5.1 | 3.8 | 89 | 152 | 11 | MOLECULE: | RECOMBINANT |
| 34:00:00 | 5yce-A | 5.1 | 3.9 | 89 | 151 | 13 | MOLECULE: | MYOGLOBIN |
| 35:00:00 | 2bnl-C | 5 | 4 | 84 | 134 | 8 | MOLECULE: | MODULATOR P |
| 36:00:00 | 5yci-B | 5 | 3.9 | 94 | 157 | 14 | MOLECULE: | ANCESTRAL M |
| 37:00:00 | 7xzi-B | 5 | 4.4 | 93 | 166 | 6 | MOLECULE: | CTAP3 |
| 38:00:00 | 3qm9-A | 5 | 4 | 85 | 144 | 8 | MOLECULE: | MYOGLOBIN |
| 39:00:00 | 3w4u-C | 5 | 3.5 | 86 | 141 | 2 | MOLECULE: | HEMOGLOBIN |
| 40:00:00 | 5ycj-B | 5 | 4 | 94 | 157 | 14 | MOLECULE: | ANCESTRAL M |
| 41:00:00 | 6ls8-I | 5 | 3.8 | 91 | 153 | 13 | MOLECULE: | MYOGLOBIN |
| 42:00:00 | 3rgk-A | 5 | 3.9 | 90 | 149 | 12 | MOLECULE: | MYOGLOBIN |

| | | | | | | | | |
|---|---|---|---|---|---|---|---|---|
| 43:00:00 | 3w4u-A | 5 | 3.6 | 86 | 141 | 2 | MOLECULE: | HEMOGLOBIN |
| 44:00:00 | 4nxa-A | 5 | 3.8 | 89 | 153 | 12 | MOLECULE: | MYOGLOBIN |
| 45:00:00 | 1v75-A | 5 | 3.5 | 87 | 141 | 6 | MOLECULE: | HEMOGLOBIN |
| 46:00:00 | 1nz3-A | 5 | 4 | 90 | 153 | 14 | MOLECULE: | MYOGLOBIN |
| 47:00:00 | 7spg-A | 5 | 4 | 92 | 152 | 12 | MOLECULE: | MYOGLOBIN |
| 48:00:00 | 3s1i-A | 5 | 3.7 | 85 | 131 | 9 | MOLECULE: | HEMOGLOBIN- |
| 49:00:00 | 8oup-D | 5 | 4.2 | 92 | 151 | 7 | MOLECULE: | NERVE HEMOG |
| 50:00:00 | 1u7s-A | 4.9 | 3.9 | 91 | 153 | 11 | MOLECULE: | MYOGLOBIN |
| 51:00:00 | 1ebc-A | 4.9 | 3.9 | 91 | 153 | 11 | MOLECULE: | PROTEIN (MY |
| 52:00:00 | 2frk-X | 4.9 | 4 | 91 | 152 | 14 | MOLECULE: | MYOGLOBIN |
| 53:00:00 | 1wvp-A | 4.9 | 3.9 | 89 | 151 | 12 | MOLECULE: | MYOGLOBIN |
| 54:00:00 | 3lr7-A | 4.9 | 4 | 90 | 151 | 14 | MOLECULE: | MYOGLOBIN |
| 55:00:00 | 2frj-X | 4.9 | 4 | 90 | 152 | 14 | MOLECULE: | MYOGLOBIN |
| 56:00:00 | 1lht-A | 4.9 | 3.7 | 89 | 153 | 12 | MOLECULE: | MYOGLOBIN |
| 57:00:00 | 6q6p-B | 4.9 | 4.2 | 91 | 157 | 3 | MOLECULE: | CYTOGLOBIN |
| 58:00:00 | 2nsr-A | 4.9 | 4 | 90 | 152 | 14 | MOLECULE: | MYOGLOBIN |
| 59:00:00 | 1jeb-C | 4.9 | 3.9 | 88 | 141 | 2 | MOLECULE: | HEMOGLOBIN |
| 60:00:00 | 5zzg-A | 4.9 | 4.1 | 90 | 153 | 11 | MOLECULE: | MYOGLOBIN |
| 61:00:00 | 6ls8-G | 4.9 | 3.9 | 90 | 153 | 14 | MOLECULE: | MYOGLOBIN |
| 62:00:00 | 4dc7-A | 4.9 | 4 | 90 | 152 | 14 | MOLECULE: | MYOGLOBIN |
| 63:00:00 | 1lhs-A | 4.9 | 3.7 | 89 | 153 | 12 | MOLECULE: | MYOGLOBIN |
| 64:00:00 | 6ls8-K | 4.9 | 3.9 | 90 | 153 | 14 | MOLECULE: | MYOGLOBIN |
| 65:00:00 | 1nz4-A | 4.9 | 4 | 90 | 153 | 14 | MOLECULE: | MYOGLOBIN |
| 66:00:00 | 4nxc-A | 4.9 | 3.8 | 89 | 153 | 12 | MOLECULE: | MYOGLOBIN |
| 67:00:00 | 3qm7-A | 4.9 | 3.9 | 85 | 144 | 8 | MOLECULE: | MYOGLOBIN |
| 68:00:00 | 1l2k-A | 4.9 | 3.9 | 89 | 151 | 12 | MOLECULE: | MYOGLOBIN |
| 69:00:00 | 6ls8-C | 4.9 | 3.9 | 91 | 153 | 14 | MOLECULE: | MYOGLOBIN |
| 70:00:00 | 1jeb-A | 4.9 | 3.4 | 86 | 141 | 3 | MOLECULE: | HEMOGLOBIN |
| 71:00:00 | 5o41-A | 4.9 | 4 | 93 | 154 | 11 | MOLECULE: | MYOGLOBIN |
| 72:00:00 | 8oup-B | 4.9 | 3.8 | 91 | 151 | 8 | MOLECULE: | NERVE HEMOG |
| 73:00:00 | 8oup-A | 4.9 | 3.9 | 89 | 150 | 6 | MOLECULE: | NERVE HEMOG |
| 74:00:00 | 3s1j-A | 4.9 | 3.5 | 84 | 131 | 10 | MOLECULE: | HEMOGLOBIN- |
| 75:00:00 | 8oup-C | 4.9 | 3.9 | 91 | 151 | 8 | MOLECULE: | NERVE HEMOG |
| 76:00:00 | 3ubc-D | 4.9 | 3.7 | 84 | 131 | 7 | MOLECULE: | HEMOGLOBIN- |
| 77:00:00 | 3s1i-C | 4.9 | 3.7 | 85 | 131 | 8 | MOLECULE: | HEMOGLOBIN- |
| 78:00:00 | 2ri4-C | 4.9 | 3.7 | 86 | 138 | 7 | MOLECULE: | HEMOGLOBIN |
| 79:00:00 | 3pt7-B | 4.8 | 4.2 | 89 | 152 | 4 | MOLECULE: | HEMOGLOBIN |
| 80:00:00 | 3ase-A | 4.8 | 4 | 90 | 154 | 11 | MOLECULE: | MYOGLOBIN |
| 81:00:00 | 1hbr-C | 4.8 | 3.5 | 87 | 140 | 6 | MOLECULE: | PROTEIN (HE |
| 82:00:00 | 1hbr-A | 4.8 | 3.6 | 88 | 141 | 6 | MOLECULE: | PROTEIN (HE |
| 83:00:00 | 1do7-A | 4.8 | 3.9 | 88 | 153 | 13 | MOLECULE: | MYOGLOBIN |
| 84:00:00 | 105m-A | 4.8 | 3.9 | 90 | 153 | 11 | MOLECULE: | MYOGLOBIN |
| 85:00:00 | 2eb8-A | 4.8 | 3.8 | 91 | 154 | 11 | MOLECULE: | MYOGLOBIN |
| 86:00:00 | 3qm5-A | 4.8 | 3.9 | 84 | 145 | 8 | MOLECULE: | MYOGLOBIN |

| | | | | | | | | |
|---|---|---|---|---|---|---|---|---|
| 87:00:00 | 4rol-A | 4.8 | 3 | 85 | 141 | 4 | MOLECULE: | HEMOGLOBIN |
| 88:00:00 | 4pqb-A | 4.8 | 4 | 90 | 153 | 12 | MOLECULE: | MYOGLOBIN |
| 89:00:00 | 1rse-A | 4.8 | 4 | 90 | 153 | 14 | MOLECULE: | HORSE HEART |
| 90:00:00 | 7ddt-A | 4.8 | 3.9 | 90 | 153 | 12 | MOLECULE: | ANCESTRAL M |
| 91:00:00 | 5cnc-A | 4.8 | 3.9 | 90 | 152 | 14 | MOLECULE: | MYOGLOBIN |
| 92:00:00 | 1j3f-A | 4.8 | 3.5 | 88 | 152 | 13 | MOLECULE: | MYOGLOBIN |
| 93:00:00 | 5mbn-A | 4.8 | 4 | 91 | 153 | 11 | MOLECULE: | MYOGLOBIN |
| 94:00:00 | 6ls8-A | 4.8 | 3.9 | 91 | 153 | 13 | MOLECULE: | MYOGLOBIN |
| 95:00:00 | 1iop-A | 4.8 | 3.9 | 91 | 153 | 11 | MOLECULE: | MYOGLOBIN |
| 96:00:00 | 5cne-A | 4.8 | 4 | 90 | 152 | 14 | MOLECULE: | MYOGLOBIN |
| 97:00:00 | 1nz5-A | 4.8 | 4 | 90 | 153 | 14 | MOLECULE: | MYOGLOBIN |
| 98:00:00 | 1yhr-A | 4.8 | 3.1 | 85 | 141 | 4 | MOLECULE: | HEMOGLOBIN |
| 99:00:00 | 4dc8-A | 4.8 | 4 | 90 | 152 | 14 | MOLECULE: | MYOGLOBIN |
| 0:00 | 1yoi-A | 4.8 | 3.9 | 89 | 153 | 12 | MOLECULE: | MYOGLOBIN |
| 1:00 | 5cnb-A | 4.8 | 3.9 | 90 | 152 | 14 | MOLECULE: | MYOGLOBIN |
| 2:00 | 4fwy-A | 4.8 | 3.8 | 90 | 153 | 11 | MOLECULE: | MYOGLOBIN |
| 3:00 | 2o5o-X | 4.8 | 4 | 90 | 152 | 14 | MOLECULE: | MYOGLOBIN |
| 4:00 | 1swm-A | 4.8 | 3.9 | 89 | 153 | 12 | MOLECULE: | MYOGLOBIN |
| 5:00 | 1dwr-A | 4.8 | 4 | 91 | 152 | 14 | MOLECULE: | MYOGLOBIN |
| 6:00 | 2v1i-A | 4.8 | 4 | 90 | 153 | 14 | MOLECULE: | MYOGLOBIN |
| 7:00 | 2bw9-M | 4.8 | 3.9 | 90 | 153 | 12 | MOLECULE: | MYOGLOBIN |
| 8:00 | 1mbc-A | 4.8 | 3.9 | 89 | 153 | 12 | MOLECULE: | MYOGLOBIN |
| 9:00 | 2nss-A | 4.8 | 4 | 90 | 152 | 14 | MOLECULE: | MYOGLOBIN |
| 10:00 | 2o5b-X | 4.8 | 4 | 90 | 152 | 14 | MOLECULE: | MYOGLOBIN |
| 11:00 | 1m6m-A | 4.8 | 4 | 91 | 153 | 13 | MOLECULE: | PROTEIN (MY |
| 12:00 | 3vau-A | 4.8 | 4 | 90 | 151 | 14 | MOLECULE: | MYOGLOBIN |
| 13:00 | 5utc-A | 4.8 | 4 | 89 | 152 | 13 | MOLECULE: | MYOGLOBIN |
| 14:00 | 1bvc-A | 4.8 | 3.9 | 90 | 153 | 11 | MOLECULE: | APOMYOGLOBI |
| 15:00 | 1wla-A | 4.8 | 4 | 90 | 153 | 14 | MOLECULE: | MYOGLOBIN |
| 16:00 | 5cn5-A | 4.8 | 4 | 90 | 152 | 14 | MOLECULE: | MYOGLOBIN |
| 17:00 | 4rol-C | 4.8 | 3.4 | 86 | 141 | 3 | MOLECULE: | HEMOGLOBIN |
| 18:00 | 1npg-A | 4.8 | 4 | 90 | 152 | 14 | MOLECULE: | MYOGLOBIN |
| 19:00 | 8a3t-T | 4.8 | 16.6 | 97 | 645 | 8 | MOLECULE: | ANAPHASE-PR |
| 20:00 | 1vxf-A | 4.8 | 3.9 | 89 | 153 | 12 | MOLECULE: | MYOGLOBIN |
| 21:00 | 1dwt-A | 4.8 | 4 | 90 | 152 | 14 | MOLECULE: | MYOGLOBIN |
| 22:00 | 104m-A | 4.8 | 3.9 | 89 | 153 | 12 | MOLECULE: | MYOGLOBIN |
| 23:00 | 1vxe-A | 4.8 | 3.9 | 89 | 153 | 12 | MOLECULE: | MYOGLOBIN |
| 24:00:00 | 2v1h-A | 4.8 | 4 | 90 | 153 | 14 | MOLECULE: | MYOGLOBIN |
| 25:00:00 | 1vxg-A | 4.8 | 3.9 | 89 | 153 | 12 | MOLECULE: | MYOGLOBIN |
| 26:00:00 | 2myd-A | 4.8 | 3.9 | 89 | 153 | 12 | MOLECULE: | MYOGLOBIN ( |
| 27:00:00 | 5utd-A | 4.8 | 3.9 | 90 | 151 | 12 | MOLECULE: | MYOGLOBIN |
| 28:00:00 | 4twv-A | 4.8 | 4 | 90 | 153 | 14 | MOLECULE: | MYOGLOBIN |
| 29:00:00 | 2o5s-X | 4.8 | 4 | 90 | 152 | 14 | MOLECULE: | MYOGLOBIN |
| 30:00:00 | 3lr9-A | 4.8 | 4 | 90 | 152 | 14 | MOLECULE: | MYOGLOBIN |

| | | | | | | | |
|---|---|---|---|---|---|---|---|
| 31:00:00 | 3rj6-B | 4.8 | 4 | 90 | 152 | 14 | MOLECULE: MYOGLOBIN |
| 32:00:00 | 2frf-A | 4.8 | 4.1 | 90 | 152 | 16 | MOLECULE: MYOGLOBIN |
| 33:00:00 | 2vlz-A | 4.8 | 4 | 91 | 153 | 14 | MOLECULE: MYOGLOBIN |
| 34:00:00 | 1yog-A | 4.8 | 3.9 | 89 | 153 | 12 | MOLECULE: MYOGLOBIN |
| 35:00:00 | 4mbn-A | 4.8 | 3.9 | 90 | 153 | 12 | MOLECULE: MYOGLOBIN |
| 36:00:00 | 5cng-A | 4.8 | 3.9 | 90 | 152 | 14 | MOLECULE: MYOGLOBIN |
| 37:00:00 | 2myb-A | 4.8 | 3.9 | 89 | 153 | 12 | MOLECULE: MYOGLOBIN ( |
| 38:00:00 | 2mb5-A | 4.8 | 3.9 | 89 | 153 | 12 | MOLECULE: MYOGLOBIN |
| 39:00:00 | 1emy-A | 4.8 | 3.9 | 90 | 153 | 11 | MOLECULE: MYOGLOBIN |
| 40:00:00 | 1bje-A | 4.8 | 4 | 90 | 153 | 14 | MOLECULE: MYOGLOBIN |
| 41:00:00 | 2o58-X | 4.8 | 4 | 90 | 152 | 14 | MOLECULE: MYOGLOBIN |
| 42:00:00 | 2mye-A | 4.8 | 3.9 | 89 | 153 | 12 | MOLECULE: MYOGLOBIN ( |
| 43:00:00 | 1mbi-A | 4.8 | 3.9 | 89 | 153 | 12 | MOLECULE: MYOGLOBIN |
| 44:00:00 | 2fri-X | 4.8 | 4 | 90 | 152 | 14 | MOLECULE: MYOGLOBIN |
| 45:00:00 | 2o5m-X | 4.8 | 4 | 90 | 152 | 14 | MOLECULE: MYOGLOBIN |
| 46:00:00 | 6q6p-A | 4.8 | 4.1 | 90 | 156 | 3 | MOLECULE: CYTOGLOBIN |
| 47:00:00 | 3m3b-A | 4.8 | 3.9 | 91 | 153 | 11 | MOLECULE: MYOGLOBIN |
| 48:00:00 | 1bzp-A | 4.8 | 3.9 | 89 | 153 | 12 | MOLECULE: PROTEIN (MY |
| 49:00:00 | 5cn8-A | 4.8 | 3.9 | 90 | 152 | 14 | MOLECULE: MYOGLOBIN |
| 50:00:00 | 2vlx-A | 4.8 | 4 | 90 | 153 | 14 | MOLECULE: MYOGLOBIN |
| 51:00:00 | 2v1j-A | 4.8 | 4 | 90 | 153 | 14 | MOLECULE: MYOGLOBIN |
| 52:00:00 | 1vxh-A | 4.8 | 3.9 | 89 | 153 | 12 | MOLECULE: MYOGLOBIN |
| 53:00:00 | 5zze-A | 4.8 | 4 | 91 | 152 | 14 | MOLECULE: MYOGLOBIN |
| 54:00:00 | 2ekt-A | 4.8 | 4 | 91 | 153 | 12 | MOLECULE: MYOGLOBIN |
| 55:00:00 | 2o5t-X | 4.8 | 4 | 90 | 152 | 14 | MOLECULE: MYOGLOBIN |
| 56:00:00 | 1vxd-A | 4.8 | 3.9 | 89 | 153 | 12 | MOLECULE: MYOGLOBIN |
| 57:00:00 | 5cnd-A | 4.8 | 4 | 90 | 152 | 14 | MOLECULE: MYOGLOBIN |
| 58:00:00 | 1bz6-A | 4.8 | 3.9 | 89 | 153 | 12 | MOLECULE: PROTEIN (MY |
| 59:00:00 | 3at5-A | 4.8 | 3.4 | 86 | 141 | 6 | MOLECULE: ALPHAA-GLOB |
| 60:00:00 | 1yoh-A | 4.8 | 3.9 | 89 | 153 | 12 | MOLECULE: MYOGLOBIN |
| 61:00:00 | 5cmv-A | 4.8 | 3.9 | 90 | 152 | 14 | MOLECULE: MYOGLOBIN |
| 62:00:00 | 1gjn-A | 4.8 | 4 | 90 | 153 | 14 | MOLECULE: MYOGLOBIN |
| 63:00:00 | 5azq-A | 4.8 | 3.9 | 90 | 153 | 14 | MOLECULE: MYOGLOBIN |
| 64:00:00 | 5cn4-A | 4.8 | 3.9 | 90 | 152 | 14 | MOLECULE: MYOGLOBIN |
| 65:00:00 | 1npf-A | 4.8 | 4 | 90 | 152 | 14 | MOLECULE: MYOGLOBIN |
| 66:00:00 | 5vzn-A | 4.8 | 3.9 | 91 | 151 | 11 | MOLECULE: MYOGLOBIN |
| 67:00:00 | 5vzq-A | 4.8 | 3.9 | 90 | 153 | 12 | MOLECULE: MYOGLOBIN |
| 68:00:00 | 2v1g-A | 4.8 | 4 | 90 | 153 | 14 | MOLECULE: MYOGLOBIN |
| 69:00:00 | 5cn9-A | 4.8 | 4 | 90 | 152 | 14 | MOLECULE: MYOGLOBIN |
| 70:00:00 | 2v1k-A | 4.8 | 4 | 90 | 153 | 14 | MOLECULE: MYOGLOBIN |
| 71:00:00 | 2v1f-A | 4.8 | 4 | 90 | 153 | 14 | MOLECULE: MYOGLOBIN |
| 72:00:00 | 5z7f-A | 4.8 | 4 | 90 | 152 | 13 | MOLECULE: MYOGLOBIN |
| 73:00:00 | 6ls8-E | 4.8 | 3.9 | 91 | 153 | 14 | MOLECULE: MYOGLOBIN |
| 74:00:00 | 1jp6-A | 4.8 | 3.9 | 89 | 152 | 12 | MOLECULE: MYOGLOBIN |

| | | | | | | | | |
|---|---|---|---|---|---|---|---|---|
| 75:00:00 | 5z7e-A | 4.8 | 4 | 90 | 153 | 14 | MOLECULE: | MYOGLOBIN |
| 76:00:00 | 2vly-A | 4.8 | 4 | 90 | 153 | 14 | MOLECULE: | MYOGLOBIN |
| 77:00:00 | 5cnf-A | 4.8 | 4 | 90 | 152 | 14 | MOLECULE: | MYOGLOBIN |
| 78:00:00 | 8wf5-A | 4.8 | 4 | 90 | 152 | 14 | MOLECULE: | MYOGLOBIN |
| 79:00:00 | 5yl3-A | 4.8 | 4 | 90 | 151 | 14 | MOLECULE: | MYOGLOBIN |
| 80:00:00 | 1xch-A | 4.8 | 4 | 90 | 153 | 14 | MOLECULE: | MYOGLOBIN |
| 81:00:00 | 1vxc-A | 4.8 | 3.9 | 89 | 153 | 12 | MOLECULE: | MYOGLOBIN |
| 82:00:00 | 1cq2-A | 4.8 | 3.9 | 89 | 153 | 12 | MOLECULE: | MYOGLOBIN |
| 83:00:00 | 2v1e-A | 4.8 | 4 | 90 | 153 | 14 | MOLECULE: | MYOGLOBIN |
| 84:00:00 | 2o5q-X | 4.8 | 4 | 90 | 152 | 14 | MOLECULE: | MYOGLOBIN |
| 85:00:00 | 1nz2-A | 4.8 | 4 | 90 | 153 | 14 | MOLECULE: | MYOGLOBIN |
| 86:00:00 | 1bzr-A | 4.8 | 3.9 | 90 | 153 | 12 | MOLECULE: | PROTEIN (MY |
| 87:00:00 | 2blj-M | 4.8 | 3.9 | 90 | 153 | 12 | MOLECULE: | MYOGLOBIN |
| 88:00:00 | 1spe-A | 4.8 | 4 | 89 | 153 | 13 | MOLECULE: | MYOGLOBIN |
| 89:00:00 | 1dws-A | 4.8 | 4 | 91 | 152 | 14 | MOLECULE: | MYOGLOBIN |
| 90:00:00 | 1jp8-A | 4.8 | 3.9 | 89 | 152 | 12 | MOLECULE: | MYOGLOBIN |
| 91:00:00 | 2o5l-X | 4.8 | 4 | 90 | 152 | 14 | MOLECULE: | MYOGLOBIN |
| 92:00:00 | 3rj6-A | 4.8 | 4 | 90 | 152 | 14 | MOLECULE: | MYOGLOBIN |
| 93:00:00 | 2vm0-A | 4.8 | 4 | 90 | 153 | 14 | MOLECULE: | MYOGLOBIN |
| 94:00:00 | 2cmm-A | 4.8 | 3.8 | 88 | 153 | 11 | MOLECULE: | MYOGLOBIN |
| 95:00:00 | 2mya-A | 4.8 | 4 | 89 | 153 | 13 | MOLECULE: | MYOGLOBIN ( |
| 96:00:00 | 6oty-B | 4.8 | 4.2 | 88 | 152 | 5 | MOLECULE: | HEMOGLOBIN |
| 97:00:00 | 6hit-A | 4.8 | 3.3 | 86 | 142 | 3 | MOLECULE: | HEMOGLOBIN |
| 98:00:00 | 1xz4-C | 4.8 | 3 | 84 | 141 | 6 | MOLECULE: | HEMOGLOBIN |
| 99:00:00 | 3bom-C | 4.8 | 3.4 | 86 | 142 | 8 | MOLECULE: | HEMOGLOBIN |
| 0:00 | 1mll-A | 4.8 | 4 | 90 | 154 | 11 | MOLECULE: | MYOGLOBIN |
| 1:00 | 3ubv-D | 4.8 | 3.7 | 85 | 131 | 9 | MOLECULE: | HEMOGLOBIN- |
| 2:00 | 1o1p-A | 4.8 | 4.8 | 95 | 283 | 3 | MOLECULE: | HEMOGLOBIN |
| 3:00 | 1o1l-A | 4.8 | 5 | 94 | 283 | 4 | MOLECULE: | HEMOGLOBIN |
| 4:00 | 3s1j-C | 4.8 | 3.8 | 84 | 131 | 7 | MOLECULE: | HEMOGLOBIN- |
| 5:00 | 1ltw-A | 4.8 | 4 | 91 | 154 | 12 | MOLECULE: | MYOGLOBIN |
| 6:00 | 6n03-A | 4.8 | 3.8 | 89 | 154 | 12 | MOLECULE: | MYOGLOBIN, |
| 7:00 | 107m-A | 4.8 | 3.9 | 92 | 154 | 11 | MOLECULE: | MYOGLOBIN |
| 8:00 | 1jw8-A | 4.8 | 4.5 | 92 | 154 | 9 | MOLECULE: | MYOGLOBIN |
| 9:00 | 2r1h-A | 4.8 | 3.4 | 86 | 142 | 8 | MOLECULE: | HEMOGLOBIN |
| 10:00 | 3ubc-G | 4.8 | 3.8 | 84 | 131 | 8 | MOLECULE: | HEMOGLOBIN- |
| 11:00 | 106m-A | 4.8 | 4 | 90 | 154 | 11 | MOLECULE: | MYOGLOBIN |
| 12:00 | 1jdo-A | 4.8 | 4 | 91 | 154 | 12 | MOLECULE: | MYOGLOBIN |
| 13:00 | 5utb-A | 4.8 | 3.9 | 90 | 152 | 11 | MOLECULE: | MYOGLOBIN |
| 14:00 | 2r1h-C | 4.8 | 3.4 | 86 | 142 | 8 | MOLECULE: | HEMOGLOBIN |
| 15:00 | 3ogb-A | 4.8 | 4 | 92 | 154 | 12 | MOLECULE: | MYOGLOBIN |
| 16:00 | 3h58-A | 4.8 | 3.9 | 90 | 154 | 11 | MOLECULE: | MYOGLOBIN |
| 17:00 | 8f9i-B | 4.8 | 4.1 | 92 | 154 | 12 | MOLECULE: | MYOGLOBIN |
| 18:00 | 1o1j-A | 4.8 | 5.2 | 95 | 283 | 3 | MOLECULE: | HEMOGLOBIN |

| | | | | | | | | |
|---|---|---|---|---|---|---|---|---|
| 19:00 | 111m-A | 4.8 | 3.9 | 91 | 154 | 12 | MOLECULE: | MYOGLOBIN |
| 20:00 | 5m3s-A | 4.8 | 4.5 | 93 | 154 | 8 | MOLECULE: | MYOGLOBIN |
| 21:00 | 1dti-A | 4.8 | 4 | 92 | 154 | 11 | MOLECULE: | PROTEIN (MY |
| 22:00 | 1ofj-A | 4.8 | 3.9 | 92 | 154 | 11 | MOLECULE: | MYOGLOBIN |
| 23:00 | 108m-A | 4.8 | 3.9 | 91 | 154 | 12 | MOLECULE: | MYOGLOBIN |
| 24:00:00 | 3bom-A | 4.8 | 2.9 | 84 | 142 | 8 | MOLECULE: | HEMOGLOBIN |
| 25:00:00 | 1c7c-A | 4.8 | 4.8 | 95 | 283 | 3 | MOLECULE: | PROTEIN (DE |
| 26:00:00 | 6mv0-A | 4.8 | 3.9 | 90 | 154 | 12 | MOLECULE: | MYOGLOBIN |
| 27:00:00 | 2g0r-A | 4.8 | 4 | 91 | 154 | 12 | MOLECULE: | MYOGLOBIN |
| 28:00:00 | 1tes-A | 4.8 | 3.9 | 91 | 154 | 12 | MOLECULE: | MYOGLOBIN |
| 29:00:00 | 101m-A | 4.8 | 4 | 92 | 154 | 11 | MOLECULE: | MYOGLOBIN |
| 30:00:00 | 8a5y-T | 4.7 | 17.7 | 101 | 650 | 8 | MOLECULE: | ANAPHASE-PR |
| 31:00:00 | 1abw-A | 4.7 | 13.4 | 119 | 287 | 8 | MOLECULE: | HEMOGLOBIN- |
| 32:00:00 | 6otw-B | 4.7 | 4.1 | 87 | 152 | 5 | MOLECULE: | HEMOGLOBIN |
| 33:00:00 | 3pt8-B | 4.7 | 4.2 | 89 | 152 | 4 | MOLECULE: | HEMOGLOBIN |
| 34:00:00 | 6kye-A | 4.7 | 3.6 | 88 | 140 | 2 | MOLECULE: | HEMOGLOBIN |
| 35:00:00 | 6kye-K | 4.7 | 3.4 | 86 | 140 | 3 | MOLECULE: | HEMOGLOBIN |
| 36:00:00 | 6otx-B | 4.7 | 4.2 | 90 | 152 | 6 | MOLECULE: | HEMOGLOBIN |
| 37:00:00 | 5ilm-A | 4.7 | 4 | 88 | 153 | 13 | MOLECULE: | MYOGLOBIN |
| 38:00:00 | 4h07-A | 4.7 | 4 | 90 | 154 | 11 | MOLECULE: | MYOGLOBIN |
| 39:00:00 | 1do4-A | 4.7 | 3.8 | 87 | 153 | 11 | MOLECULE: | MYOGLOBIN |
| 40:00:00 | 3m3a-A | 4.7 | 4 | 90 | 153 | 12 | MOLECULE: | MYOGLOBIN |
| 41:00:00 | 5kkk-A | 4.7 | 4 | 90 | 154 | 12 | MOLECULE: | MYOGLOBIN |
| 42:00:00 | 1yen-A | 4.7 | 3.6 | 87 | 138 | 2 | MOLECULE: | HEMOGLOBIN |
| 43:00:00 | 1do1-A | 4.7 | 3.9 | 88 | 153 | 13 | MOLECULE: | MYOGLOBIN |
| 44:00:00 | 1dxc-A | 4.7 | 3.9 | 88 | 154 | 11 | MOLECULE: | MYOGLOBIN |
| 45:00:00 | 1lfl-P | 4.7 | 2.9 | 85 | 141 | 5 | MOLECULE: | HEMOGLOBIN |
| 46:00:00 | 3hxn-C | 4.7 | 3.1 | 85 | 141 | 4 | MOLECULE: | HEMOGLOBIN |
| 47:00:00 | 1dxd-A | 4.7 | 4 | 88 | 153 | 13 | MOLECULE: | MYOGLOBIN |
| 48:00:00 | 1yca-A | 4.7 | 3.9 | 90 | 153 | 13 | MOLECULE: | MYOGLOBIN |
| 49:00:00 | 1sdk-A | 4.7 | 3.5 | 87 | 141 | 2 | MOLECULE: | HEMOGLOBIN |
| 50:00:00 | 1aj9-A | 4.7 | 3.5 | 87 | 141 | 2 | MOLECULE: | HEMOGLOBIN |
| 51:00:00 | 2w6w-A | 4.7 | 4 | 89 | 153 | 11 | MOLECULE: | MYOGLOBIN |
| 52:00:00 | 7pcf-A | 4.7 | 3.6 | 87 | 141 | 3 | MOLECULE: | HEMOGLOBIN |
| 53:00:00 | 3v2v-A | 4.7 | 4 | 89 | 152 | 16 | MOLECULE: | MYOGLOBIN |
| 54:00:00 | 3u3e-A | 4.7 | 3.9 | 88 | 153 | 13 | MOLECULE: | MYOGLOBIN |
| 55:00:00 | 4twu-A | 4.7 | 3.8 | 88 | 153 | 15 | MOLECULE: | MYOGLOBIN |
| 56:00:00 | 4rom-A | 4.7 | 3.1 | 85 | 141 | 4 | MOLECULE: | HEMOGLOBIN |
| 57:00:00 | 1do3-A | 4.7 | 3.9 | 88 | 153 | 13 | MOLECULE: | MYOGLOBIN |
| 58:00:00 | 2bnl-F | 4.7 | 3.9 | 79 | 129 | 8 | MOLECULE: | MODULATOR P |
| 59:00:00 | 6bmg-B | 4.7 | 4 | 91 | 154 | 13 | MOLECULE: | MYOGLOBIN |
| 60:00:00 | 1hjt-A | 4.7 | 4 | 90 | 153 | 12 | MOLECULE: | MYOGLOBIN |
| 61:00:00 | 8u5a-A | 4.7 | 3.5 | 84 | 149 | 7 | MOLECULE: | DESIGNED MY |
| 62:00:00 | 1azi-A | 4.7 | 4 | 90 | 153 | 14 | MOLECULE: | MYOGLOBIN |

| | | | | | | | | |
|---|---|---|---|---|---|---|---|---|
| 63:00:00 | 1mnk-A | 4.7 | 4.1 | 91 | 151 | 13 | MOLECULE: | MYOGLOBIN |
| 64:00:00 | 2zsn-A | 4.7 | 3.9 | 88 | 153 | 11 | MOLECULE: | MYOGLOBIN |
| 65:00:00 | 3edb-A | 4.7 | 3.9 | 88 | 153 | 11 | MOLECULE: | MYOGLOBIN |
| 66:00:00 | 2hbd-A | 4.7 | 3.4 | 86 | 141 | 2 | MOLECULE: | HEMOGLOBIN |
| 67:00:00 | 1vwt-C | 4.7 | 3.3 | 85 | 141 | 5 | MOLECULE: | HEMOGLOBIN |
| 68:00:00 | 6cf0-A | 4.7 | 4 | 88 | 153 | 13 | MOLECULE: | MYOGLOBIN |
| 69:00:00 | 6f17-A | 4.7 | 3.8 | 89 | 153 | 11 | MOLECULE: | MYOGLOBIN |
| 70:00:00 | 1myh-A | 4.7 | 3.9 | 91 | 153 | 13 | MOLECULE: | MYOGLOBIN |
| 71:00:00 | 1sdl-C | 4.7 | 3.4 | 86 | 141 | 3 | MOLECULE: | HEMOGLOBIN |
| 72:00:00 | 1y45-A | 4.7 | 3 | 84 | 141 | 5 | MOLECULE: | HEMOGLOBIN |
| 73:00:00 | 5vrt-A | 4.7 | 3.9 | 88 | 153 | 11 | MOLECULE: | MYOGLOBIN |
| 74:00:00 | 5ut9-A | 4.7 | 4 | 88 | 153 | 13 | MOLECULE: | MYOGLOBIN |
| 75:00:00 | 8f9j-A | 4.7 | 4 | 88 | 153 | 13 | MOLECULE: | MYOGLOBIN |
| 76:00:00 | 4m4b-A | 4.7 | 3.4 | 85 | 140 | 5 | MOLECULE: | HEMOGLOBIN |
| 77:00:00 | 2jho-A | 4.7 | 3.9 | 89 | 153 | 12 | MOLECULE: | MYOGLOBIN |
| 78:00:00 | 1rq3-C | 4.7 | 3.1 | 85 | 141 | 4 | MOLECULE: | HEMOGLOBIN |
| 79:00:00 | 1yev-C | 4.7 | 2.9 | 83 | 138 | 5 | MOLECULE: | HEMOGLOBIN |
| 80:00:00 | 1duo-A | 4.7 | 3.8 | 90 | 152 | 11 | MOLECULE: | SPERM WHALE |
| 81:00:00 | 2zt2-A | 4.7 | 3.9 | 89 | 153 | 12 | MOLECULE: | MYOGLOBIN |
| 82:00:00 | 1c7b-A | 4.7 | 3 | 84 | 141 | 5 | MOLECULE: | PROTEIN (DE |
| 83:00:00 | 1xzv-C | 4.7 | 3.1 | 85 | 141 | 4 | MOLECULE: | HEMOGLOBIN |
| 84:00:00 | 1y7g-C | 4.7 | 3.1 | 85 | 141 | 4 | MOLECULE: | HEMOGLOBIN |
| 85:00:00 | 3sdn-A | 4.7 | 3.9 | 89 | 152 | 12 | MOLECULE: | MYOGLOBIN |
| 86:00:00 | 2myc-A | 4.7 | 3.9 | 89 | 153 | 12 | MOLECULE: | MYOGLOBIN ( |
| 87:00:00 | 2bnl-A | 4.7 | 4 | 79 | 129 | 8 | MOLECULE: | MODULATOR P |
| 88:00:00 | 1yeo-C | 4.7 | 3.1 | 85 | 141 | 4 | MOLECULE: | HEMOGLOBIN |
| 89:00:00 | 5vnu-A | 4.7 | 3.9 | 90 | 153 | 12 | MOLECULE: | MYOGLOBIN |
| 90:00:00 | 1y5j-C | 4.7 | 3.1 | 85 | 141 | 4 | MOLECULE: | HEMOGLOBIN |
| 91:00:00 | 1dxv-A | 4.7 | 3 | 84 | 141 | 5 | MOLECULE: | HEMOGLOBIN |
| 92:00:00 | 4of9-A | 4.7 | 3.9 | 88 | 153 | 11 | MOLECULE: | MYOGLOBIN |
| 93:00:00 | 5ilr-A | 4.7 | 3.9 | 88 | 151 | 11 | MOLECULE: | MYOGLOBIN |
| 94:00:00 | 5iks-A | 4.7 | 3.9 | 89 | 151 | 12 | MOLECULE: | MYOGLOBIN |
| 95:00:00 | 1mwc-A | 4.7 | 3.9 | 90 | 153 | 13 | MOLECULE: | PROTEIN (MY |
| 96:00:00 | 2hbf-A | 4.7 | 3.4 | 86 | 141 | 2 | MOLECULE: | HEMOGLOBIN |
| 97:00:00 | 3hhb-A | 4.7 | 3.5 | 86 | 141 | 3 | MOLECULE: | HEMOGLOBIN |
| 98:00:00 | 1ymc-A | 4.7 | 4 | 90 | 153 | 14 | MOLECULE: | CYANOMET-SU |
| 99:00:00 | 3mn0-A | 4.7 | 3.8 | 90 | 153 | 11 | MOLECULE: | MYOGLOBIN |
| 0:00 | 4it8-A | 4.7 | 3.8 | 89 | 153 | 12 | MOLECULE: | MYOGLOBIN |
| 1:00 | 1lfl-C | 4.7 | 3.1 | 85 | 141 | 4 | MOLECULE: | HEMOGLOBIN |
| 2:00 | 1a0z-C | 4.7 | 3 | 85 | 141 | 6 | MOLECULE: | HEMOGLOBIN |
| 3:00 | 2dn2-A | 4.7 | 3 | 84 | 141 | 5 | MOLECULE: | HEMOGLOBIN |
| 4:00 | 1y4f-C | 4.7 | 3.1 | 85 | 141 | 4 | MOLECULE: | HEMOGLOBIN |
| 5:00 | 1y4v-C | 4.7 | 3.5 | 87 | 141 | 2 | MOLECULE: | HEMOGLOBIN |
| 6:00 | 3wfu-A | 4.7 | 4 | 90 | 152 | 14 | MOLECULE: | MYOGLOBIN |

| | | | | | | | | |
|---|---|---|---|---|---|---|---|---|
| 7:00 | 1a6k-A | 4.7 | 3.9 | 89 | 151 | 13 | MOLECULE: | MYOGLOBIN |
| 8:00 | 5ilp-A | 4.7 | 3.9 | 88 | 151 | 11 | MOLECULE: | MYOGLOBIN |
| 9:00 | 1c7d-A | 4.7 | 12.9 | 116 | 284 | 6 | MOLECULE: | PROTEIN (DE |
| 10:00 | 1hgc-C | 4.7 | 3.1 | 85 | 141 | 4 | MOLECULE: | HEMOGLOBIN |
| 11:00 | 3k9z-A | 4.7 | 3.9 | 91 | 153 | 11 | MOLECULE: | MYOGLOBIN |
| 12:00 | 1yh9-C | 4.7 | 3 | 84 | 141 | 5 | MOLECULE: | HEMOGLOBIN |
| 13:00 | 1fsx-C | 4.7 | 3.1 | 84 | 141 | 6 | MOLECULE: | HEMOGLOBIN |
| 14:00 | 1y4g-C | 4.7 | 3.1 | 85 | 141 | 4 | MOLECULE: | HEMOGLOBIN |
| 15:00 | 1ye2-C | 4.7 | 3 | 84 | 141 | 5 | MOLECULE: | HEMOGLOBIN |
| 16:00 | 1xzu-C | 4.7 | 3.5 | 87 | 141 | 2 | MOLECULE: | HEMOGLOBIN |
| 17:00 | 1c7b-C | 4.7 | 3 | 84 | 141 | 5 | MOLECULE: | PROTEIN (DE |
| 18:00 | 1xy0-C | 4.7 | 3.1 | 85 | 141 | 4 | MOLECULE: | HEMOGLOBIN |
| 19:00 | 1y8w-C | 4.7 | 3.4 | 86 | 141 | 3 | MOLECULE: | HEMOGLOBIN |
| 20:00 | 1gbv-A | 4.7 | 3.3 | 85 | 141 | 5 | MOLECULE: | HEMOGLOBIN |
| 21:00 | 3d17-C | 4.7 | 3.5 | 87 | 141 | 3 | MOLECULE: | HEMOGLOBIN |
| 22:00 | 1myj-A | 4.7 | 4.1 | 90 | 153 | 14 | MOLECULE: | MYOGLOBIN |
| 23:00 | 1mno-A | 4.7 | 3.8 | 90 | 153 | 13 | MOLECULE: | PROTEIN (MY |
| 24:00:00 | 2d6c-B | 4.7 | 4 | 89 | 151 | 12 | MOLECULE: | MYOGLOBIN |
| 25:00:00 | 1gbv-C | 4.7 | 3.4 | 86 | 141 | 3 | MOLECULE: | HEMOGLOBIN |
| 26:00:00 | 1myg-B | 4.7 | 3.9 | 90 | 153 | 13 | MOLECULE: | MYOGLOBIN |
| 27:00:00 | 8fdj-A | 4.7 | 3.9 | 89 | 151 | 12 | MOLECULE: | MYOGLOBIN |
| 28:00:00 | 5xl0-A | 4.7 | 3.9 | 88 | 151 | 11 | MOLECULE: | MYOGLOBIN |
| 29:00:00 | 1ymb-A | 4.7 | 4.2 | 90 | 153 | 14 | MOLECULE: | METMYOGLOBI |
| 30:00:00 | 5c6y-A | 4.7 | 3.9 | 89 | 153 | 12 | MOLECULE: | MYOGLOBIN |
| 31:00:00 | 3wi8-A | 4.7 | 4 | 90 | 151 | 14 | MOLECULE: | MYOGLOBIN |
| 32:00:00 | 1xz2-C | 4.7 | 3 | 84 | 141 | 5 | MOLECULE: | HEMOGLOBIN |
| 33:00:00 | 1v4w-C | 4.7 | 3.4 | 86 | 143 | 7 | MOLECULE: | HEMOGLOBIN |
| 34:00:00 | 1m6c-B | 4.7 | 3.9 | 90 | 153 | 14 | MOLECULE: | PROTEIN (MY |
| 35:00:00 | 7sli-A | 4.7 | 3.9 | 88 | 152 | 11 | MOLECULE: | MYOGLOBIN |
| 36:00:00 | 1myj-B | 4.7 | 3.9 | 90 | 153 | 13 | MOLECULE: | MYOGLOBIN |
| 37:00:00 | 1rvw-A | 4.7 | 3.8 | 88 | 141 | 2 | MOLECULE: | HEMOGLOBIN |
| 38:00:00 | 3e4n-A | 4.7 | 3.9 | 88 | 153 | 13 | MOLECULE: | MYOGLOBIN |
| 39:00:00 | 3hhb-C | 4.7 | 3.5 | 86 | 141 | 3 | MOLECULE: | HEMOGLOBIN |
| 40:00:00 | 1qsh-A | 4.7 | 3 | 84 | 141 | 5 | MOLECULE: | PROTEIN (HE |
| 41:00:00 | 2zt4-A | 4.7 | 4 | 88 | 153 | 13 | MOLECULE: | MYOGLOBIN |
| 42:00:00 | 1b86-C | 4.7 | 3.6 | 87 | 141 | 2 | MOLECULE: | PROTEIN (HE |
| 43:00:00 | 2z6s-A | 4.7 | 3.9 | 88 | 151 | 13 | MOLECULE: | MYOGLOBIN |
| 44:00:00 | 1mni-B | 4.7 | 3.9 | 90 | 153 | 13 | MOLECULE: | MYOGLOBIN |
| 45:00:00 | 1ajh-A | 4.7 | 3.9 | 88 | 153 | 11 | MOLECULE: | MYOGLOBIN |
| 46:00:00 | 4m4a-A | 4.7 | 3.8 | 88 | 140 | 2 | MOLECULE: | HEMOGLOBIN |
| 47:00:00 | 1dxt-A | 4.7 | 3 | 84 | 141 | 5 | MOLECULE: | HEMOGLOBIN |
| 48:00:00 | 2hbc-A | 4.7 | 3.4 | 85 | 141 | 4 | MOLECULE: | HEMOGLOBIN |
| 49:00:00 | 1qsh-C | 4.7 | 3.5 | 86 | 141 | 3 | MOLECULE: | PROTEIN (HE |
| 50:00:00 | 2hhe-C | 4.7 | 2.9 | 84 | 141 | 6 | MOLECULE: | HEMOGLOBIN |

| | | | | | | | | |
|---|---|---|---|---|---|---|---|---|
| 51:00:00 | 2zst-A | 4.7 | 3.9 | 88 | 153 | 11 | MOLECULE: | MYOGLOBIN |
| 52:00:00 | 1myh-B | 4.7 | 3.9 | 90 | 153 | 13 | MOLECULE: | MYOGLOBIN |
| 53:00:00 | 1j7w-A | 4.7 | 3.5 | 86 | 141 | 3 | MOLECULE: | HEMOGLOBIN |
| 54:00:00 | 4pqc-A | 4.7 | 3.8 | 89 | 153 | 12 | MOLECULE: | MYOGLOBIN |
| 55:00:00 | 1yev-A | 4.7 | 3 | 84 | 138 | 4 | MOLECULE: | HEMOGLOBIN |
| 56:00:00 | 1vxa-A | 4.7 | 3.9 | 89 | 153 | 12 | MOLECULE: | MYOGLOBIN |
| 57:00:00 | 1xz7-A | 4.7 | 3.1 | 85 | 141 | 4 | MOLECULE: | HEMOGLOBIN |
| 58:00:00 | 3hc9-A | 4.7 | 3.8 | 88 | 153 | 14 | MOLECULE: | MYOGLOBIN |
| 59:00:00 | 2bwh-A | 4.7 | 3.9 | 90 | 153 | 12 | MOLECULE: | MYOGLOBIN |
| 60:00:00 | 1gli-C | 4.7 | 3.5 | 86 | 141 | 3 | MOLECULE: | DEOXYHEMOGL |
| 61:00:00 | 1dxu-C | 4.7 | 3.5 | 87 | 141 | 2 | MOLECULE: | HEMOGLOBIN |
| 62:00:00 | 4pq6-A | 4.7 | 3.8 | 89 | 153 | 12 | MOLECULE: | MYOGLOBIN |
| 63:00:00 | 3e5i-A | 4.7 | 3.9 | 88 | 153 | 11 | MOLECULE: | MYOGLOBIN |
| 64:00:00 | 2zsz-A | 4.7 | 4 | 88 | 153 | 13 | MOLECULE: | MYOGLOBIN |
| 65:00:00 | 1duk-A | 4.7 | 3.8 | 90 | 152 | 11 | MOLECULE: | WILD-TYPE R |
| 66:00:00 | 7xcf-A | 4.7 | 3.9 | 90 | 153 | 11 | MOLECULE: | MYOGLOBIN |
| 67:00:00 | 2zss-A | 4.7 | 4 | 88 | 153 | 13 | MOLECULE: | MYOGLOBIN |
| 68:00:00 | 2hhb-C | 4.7 | 3.5 | 87 | 141 | 2 | MOLECULE: | HEMOGLOBIN |
| 69:00:00 | 2zt0-A | 4.7 | 3.9 | 88 | 153 | 11 | MOLECULE: | MYOGLOBIN |
| 70:00:00 | 6d45-A | 4.7 | 3.9 | 89 | 153 | 12 | MOLECULE: | MYOGLOBIN |
| 71:00:00 | 1dxv-C | 4.7 | 3.5 | 86 | 141 | 3 | MOLECULE: | HEMOGLOBIN |
| 72:00:00 | 3onz-A | 4.7 | 3.4 | 85 | 140 | 5 | MOLECULE: | HEMOGLOBIN |
| 73:00:00 | 2eku-A | 4.7 | 3.9 | 88 | 153 | 13 | MOLECULE: | MYOGLOBIN |
| 74:00:00 | 6e02-A | 4.7 | 4 | 88 | 153 | 13 | MOLECULE: | MYOGLOBIN |
| 75:00:00 | 2zt3-A | 4.7 | 3.9 | 88 | 153 | 13 | MOLECULE: | MYOGLOBIN |
| 76:00:00 | 2w6y-A | 4.7 | 3.9 | 88 | 153 | 11 | MOLECULE: | MYOGLOBIN |
| 77:00:00 | 1y0a-C | 4.7 | 3 | 84 | 141 | 5 | MOLECULE: | HEMOGLOBIN |
| 78:00:00 | 1mnj-B | 4.7 | 4 | 90 | 151 | 13 | MOLECULE: | MYOGLOBIN |
| 79:00:00 | 1a6g-A | 4.7 | 3.9 | 88 | 151 | 11 | MOLECULE: | MYOGLOBIN |
| 80:00:00 | 3rjn-B | 4.7 | 3.8 | 88 | 152 | 14 | MOLECULE: | MYOGLOBIN |
| 81:00:00 | 1mbn-A | 4.7 | 4 | 91 | 153 | 11 | MOLECULE: | MYOGLOBIN |
| 82:00:00 | 3ecz-A | 4.7 | 3.9 | 88 | 153 | 11 | MOLECULE: | MYOGLOBIN |
| 83:00:00 | 1myi-B | 4.7 | 3.8 | 90 | 153 | 13 | MOLECULE: | MYOGLOBIN |
| 84:00:00 | 1y46-C | 4.7 | 3.6 | 87 | 141 | 2 | MOLECULE: | HEMOGLOBIN |
| 85:00:00 | 2zso-A | 4.7 | 3.9 | 88 | 153 | 11 | MOLECULE: | MYOGLOBIN |
| 86:00:00 | 1myg-A | 4.7 | 4 | 91 | 153 | 13 | MOLECULE: | MYOGLOBIN |
| 87:00:00 | 3m39-A | 4.7 | 3.9 | 89 | 153 | 11 | MOLECULE: | MYOGLOBIN |
| 88:00:00 | 1hga-C | 4.7 | 3.1 | 85 | 141 | 4 | MOLECULE: | HEMOGLOBIN |
| 89:00:00 | 1gzx-C | 4.7 | 3.1 | 85 | 141 | 4 | MOLECULE: | HEMOGLOBIN |
| 90:00:00 | 6nq5-A | 4.7 | 3.2 | 86 | 140 | 3 | MOLECULE: | HEMOGLOBIN |
| 91:00:00 | 4qau-A | 4.7 | 3.8 | 89 | 153 | 12 | MOLECULE: | MYOGLOBIN |
| 92:00:00 | 1hgb-A | 4.7 | 3.1 | 85 | 141 | 4 | MOLECULE: | HEMOGLOBIN |
| 93:00:00 | 1nqp-A | 4.7 | 3.4 | 86 | 141 | 2 | MOLECULE: | HEMOGLOBIN |
| 94:00:00 | 1y5f-C | 4.7 | 3.5 | 87 | 141 | 2 | MOLECULE: | HEMOGLOBIN |

| | | | | | | | | |
|---|---|---|---|---|---|---|---|---|
| 95:00:00 | 1hbb-C | 4.7 | 3.1 | 85 | 141 | 4 | MOLECULE: | HEMOGLOBIN |
| 96:00:00 | 3e55-A | 4.7 | 3.9 | 88 | 153 | 11 | MOLECULE: | MYOGLOBIN |
| 97:00:00 | 1yhe-C | 4.7 | 3 | 84 | 141 | 5 | MOLECULE: | HEMOGLOBIN |
| 98:00:00 | 4n7n-K | 4.7 | 3.8 | 88 | 141 | 2 | MOLECULE: | HEMOGLOBIN |
| 99:00:00 | 2zsx-A | 4.7 | 3.9 | 88 | 153 | 11 | MOLECULE: | MYOGLOBIN |
| 0:00 | 1qsi-A | 4.7 | 3.3 | 85 | 141 | 5 | MOLECULE: | PROTEIN (HE |
| 1:00 | 1yen-C | 4.7 | 3 | 84 | 138 | 4 | MOLECULE: | HEMOGLOBIN |
| 2:00 | 1wmu-A | 4.7 | 4.1 | 88 | 141 | 8 | MOLECULE: | HEMOGLOBIN |
| 3:00 | 1dxt-C | 4.7 | 3.5 | 86 | 141 | 3 | MOLECULE: | HEMOGLOBIN |
| 4:00 | 1y22-C | 4.7 | 3.1 | 85 | 141 | 4 | MOLECULE: | HEMOGLOBIN |
| 5:00 | 4mxk-A | 4.7 | 3.8 | 88 | 153 | 11 | MOLECULE: | MYOGLOBIN |
| 6:00 | 1yca-B | 4.7 | 4 | 90 | 153 | 14 | MOLECULE: | MYOGLOBIN |
| 7:00 | 1gli-A | 4.7 | 3.5 | 86 | 141 | 3 | MOLECULE: | DEOXYHEMOGL |
| 8:00 | 2bli-A | 4.7 | 4 | 90 | 153 | 13 | MOLECULE: | MYOGLOBIN |
| 9:00 | 8f9h-A | 4.7 | 3.9 | 90 | 151 | 12 | MOLECULE: | MYOGLOBIN |
| 10:00 | 1y0w-C | 4.7 | 3.1 | 85 | 141 | 4 | MOLECULE: | HEMOGLOBIN |
| 11:00 | 1xz4-A | 4.7 | 3.5 | 87 | 141 | 3 | MOLECULE: | HEMOGLOBIN |
| 12:00 | 4ns2-A | 4.7 | 3.8 | 88 | 153 | 14 | MOLECULE: | MYOGLOBIN |
| 13:00 | 1hdb-C | 4.7 | 3.1 | 85 | 141 | 4 | MOLECULE: | HEMOGLOBIN |
| 14:00 | 5e6e-A | 4.7 | 3.4 | 85 | 141 | 5 | MOLECULE: | HEMOGLOBIN |
| 15:00 | 4lpi-A | 4.7 | 3.9 | 89 | 153 | 12 | MOLECULE: | MYOGLOBIN |
| 16:00 | 1v4w-A | 4.7 | 3.4 | 86 | 143 | 7 | MOLECULE: | HEMOGLOBIN |
| 17:00 | 1k1k-A | 4.7 | 3.4 | 86 | 141 | 3 | MOLECULE: | HEMOGLOBIN |
| 18:00 | 5wjk-A | 4.7 | 3.9 | 90 | 153 | 12 | MOLECULE: | MYOGLOBIN |
| 19:00 | 5xkw-A | 4.7 | 3.9 | 89 | 153 | 12 | MOLECULE: | MYOGLOBIN |
| 20:00 | 1a00-C | 4.7 | 3 | 85 | 141 | 6 | MOLECULE: | HEMOGLOBIN |
| 21:00 | 3ecl-A | 4.7 | 3.9 | 88 | 153 | 13 | MOLECULE: | MYOGLOBIN |
| 22:00 | 6ihx-A | 4.7 | 3.6 | 87 | 140 | 5 | MOLECULE: | HEMOGLOBIN |
| 23:00 | 3whm-E | 4.7 | 3.4 | 86 | 140 | 2 | MOLECULE: | HEMOGLOBIN |
| 24:00:00 | 1yma-A | 4.7 | 4.2 | 90 | 153 | 14 | MOLECULE: | MYOGLOBIN |
| 25:00:00 | 1yih-C | 4.7 | 3.4 | 86 | 141 | 3 | MOLECULE: | HEMOGLOBIN |
| 26:00:00 | 1y0t-C | 4.7 | 3.5 | 87 | 141 | 2 | MOLECULE: | HEMOGLOBIN |
| 27:00:00 | 3ciu-A | 4.7 | 3.4 | 86 | 140 | 5 | MOLECULE: | HEMOGLOBIN |
| 28:00:00 | 2hbs-C | 4.7 | 3.1 | 85 | 141 | 6 | MOLECULE: | HEMOGLOBIN |
| 29:00:00 | 2zsy-A | 4.7 | 3.9 | 88 | 153 | 11 | MOLECULE: | MYOGLOBIN |
| 30:00:00 | 1y7z-C | 4.7 | 3.1 | 85 | 141 | 4 | MOLECULE: | HEMOGLOBIN |
| 31:00:00 | 1y0d-C | 4.7 | 3.1 | 85 | 140 | 4 | MOLECULE: | HEMOGLOBIN |
| 32:00:00 | 1jpb-A | 4.7 | 3.9 | 88 | 151 | 11 | MOLECULE: | MYOGLOBIN |
| 33:00:00 | 1g08-C | 4.7 | 3.1 | 85 | 141 | 6 | MOLECULE: | HEMOGLOBIN |
| 34:00:00 | 1bzz-C | 4.7 | 3.5 | 86 | 141 | 3 | MOLECULE: | PROTEIN (HE |
| 35:00:00 | 1yhr-C | 4.7 | 3.1 | 85 | 141 | 4 | MOLECULE: | HEMOGLOBIN |
| 36:00:00 | 2hhd-C | 4.7 | 3.3 | 85 | 141 | 5 | MOLECULE: | HEMOGLOBIN |
| 37:00:00 | 1y09-C | 4.7 | 3.1 | 85 | 141 | 4 | MOLECULE: | HEMOGLOBIN |
| 38:00:00 | 1ydz-C | 4.7 | 3.4 | 86 | 141 | 3 | MOLECULE: | HEMOGLOBIN |

| | | | | | | | | |
|---|---|---|---|---|---|---|---|---|
| 39:00:00 | 5vzo-A | 4.7 | 3.9 | 88 | 153 | 11 | MOLECULE: | MYOGLOBIN |
| 40:00:00 | 1j3z-C | 4.7 | 3 | 84 | 141 | 5 | MOLECULE: | HEMOGLOBIN |
| 41:00:00 | 2z6t-A | 4.7 | 3.9 | 88 | 151 | 11 | MOLECULE: | MYOGLOBIN |
| 42:00:00 | 1y4r-C | 4.7 | 3.5 | 87 | 141 | 2 | MOLECULE: | HEMOGLOBIN |
| 43:00:00 | 1g9v-C | 4.7 | 3.4 | 86 | 141 | 3 | MOLECULE: | HEMOGLOBIN |
| 44:00:00 | 2hhb-A | 4.7 | 3.5 | 86 | 141 | 3 | MOLECULE: | HEMOGLOBIN |
| 45:00:00 | 1vwt-A | 4.7 | 3 | 84 | 141 | 5 | MOLECULE: | HEMOGLOBIN |
| 46:00:00 | 1a0u-C | 4.7 | 3.5 | 87 | 141 | 2 | MOLECULE: | HEMOGLOBIN |
| 47:00:00 | 1jp9-A | 4.7 | 3.9 | 89 | 151 | 12 | MOLECULE: | MYOGLOBIN |
| 48:00:00 | 1a6n-A | 4.7 | 3.9 | 88 | 151 | 11 | MOLECULE: | MYOGLOBIN |
| 49:00:00 | 5uta-A | 4.7 | 4 | 88 | 153 | 13 | MOLECULE: | MYOGLOBIN |
| 50:00:00 | 1j3y-C | 4.7 | 3 | 84 | 141 | 5 | MOLECULE: | HEMOGLOBIN |
| 51:00:00 | 1o1n-A | 4.7 | 13.8 | 118 | 282 | 4 | MOLECULE: | HEMOGLOBIN |
| 52:00:00 | 1y35-C | 4.7 | 3.5 | 87 | 141 | 2 | MOLECULE: | HEMOGLOBIN |
| 53:00:00 | 1pmb-B | 4.7 | 3.9 | 90 | 153 | 13 | MOLECULE: | MYOGLOBIN |
| 54:00:00 | 2zsq-A | 4.7 | 4 | 88 | 153 | 13 | MOLECULE: | MYOGLOBIN |
| 55:00:00 | 1y7d-C | 4.7 | 3.6 | 87 | 141 | 2 | MOLECULE: | HEMOGLOBIN |
| 56:00:00 | 7cen-A | 4.7 | 3.9 | 90 | 153 | 11 | MOLECULE: | MYOGLOBIN |
| 57:00:00 | 1dke-C | 4.7 | 2.9 | 84 | 141 | 4 | MOLECULE: | HEMOGLOBIN: |
| 58:00:00 | 1y0d-A | 4.7 | 3.1 | 85 | 140 | 4 | MOLECULE: | HEMOGLOBIN |
| 59:00:00 | 7slh-A | 4.7 | 4 | 88 | 152 | 13 | MOLECULE: | MYOGLOBIN |
| 60:00:00 | 5kd1-A | 4.7 | 4 | 88 | 153 | 13 | MOLECULE: | MYOGLOBIN |
| 61:00:00 | 1y7c-C | 4.7 | 3.1 | 85 | 141 | 4 | MOLECULE: | HEMOGLOBIN |
| 62:00:00 | 7cez-A | 4.7 | 3.9 | 90 | 153 | 12 | MOLECULE: | MYOGLOBIN |
| 63:00:00 | 1k0y-A | 4.7 | 3 | 84 | 141 | 5 | MOLECULE: | HEMOGLOBIN |
| 64:00:00 | 1o1o-A | 4.7 | 3.5 | 86 | 141 | 3 | MOLECULE: | HEMOGLOBIN |
| 65:00:00 | 1a6m-A | 4.7 | 3.9 | 88 | 151 | 11 | MOLECULE: | MYOGLOBIN |
| 66:00:00 | 7vdn-A | 4.7 | 3.9 | 88 | 151 | 11 | MOLECULE: | MYOGLOBIN |
| 67:00:00 | 1bbb-C | 4.7 | 3.4 | 86 | 141 | 2 | MOLECULE: | HEMOGLOBIN |
| 68:00:00 | 1bbb-A | 4.7 | 3.4 | 86 | 141 | 2 | MOLECULE: | HEMOGLOBIN |
| 69:00:00 | 3m38-A | 4.7 | 4 | 91 | 153 | 11 | MOLECULE: | MYOGLOBIN |
| 70:00:00 | 1buw-A | 4.7 | 3.4 | 86 | 141 | 2 | MOLECULE: | PROTEIN (HE |
| 71:00:00 | 1mbd-A | 4.7 | 4 | 91 | 153 | 11 | MOLECULE: | MYOGLOBIN |
| 72:00:00 | 1mnj-A | 4.7 | 4.1 | 91 | 153 | 13 | MOLECULE: | MYOGLOBIN |
| 73:00:00 | 6hit-E | 4.7 | 3.4 | 86 | 142 | 3 | MOLECULE: | HEMOGLOBIN |
| 74:00:00 | 1bz0-C | 4.7 | 3.5 | 87 | 141 | 2 | MOLECULE: | PROTEIN (HE |
| 75:00:00 | 1hsy-A | 4.7 | 4 | 90 | 153 | 14 | MOLECULE: | MYOGLOBIN |
| 76:00:00 | 1coh-C | 4.7 | 3.5 | 86 | 141 | 3 | MOLECULE: | HEMOGLOBIN |
| 77:00:00 | 4mxl-A | 4.7 | 3.8 | 90 | 153 | 11 | MOLECULE: | MYOGLOBIN |
| 78:00:00 | 1myi-A | 4.7 | 4 | 91 | 153 | 13 | MOLECULE: | MYOGLOBIN |
| 79:00:00 | 3ed9-A | 4.7 | 3.9 | 88 | 153 | 13 | MOLECULE: | MYOGLOBIN |
| 80:00:00 | 3szk-D | 4.7 | 3.1 | 85 | 140 | 2 | MOLECULE: | HEMOGLOBIN |
| 81:00:00 | 1y4q-C | 4.7 | 3.5 | 87 | 141 | 2 | MOLECULE: | HEMOGLOBIN |
| 82:00:00 | 2zsp-A | 4.7 | 3.9 | 88 | 153 | 11 | MOLECULE: | MYOGLOBIN |

| | | | | | | | | |
|---|---|---|---|---|---|---|---|---|
| 83:00:00 | 3dut-A | 4.7 | 3 | 84 | 141 | 5 | MOLECULE: | HEMOGLOBIN |
| 84:00:00 | 1y0c-C | 4.7 | 3.1 | 85 | 141 | 4 | MOLECULE: | HEMOGLOBIN |
| 85:00:00 | 1ajg-A | 4.7 | 3.9 | 88 | 153 | 11 | MOLECULE: | MYOGLOBIN |
| 86:00:00 | 6krf-A | 4.7 | 3.8 | 90 | 153 | 11 | MOLECULE: | MYOGLOBIN |
| 87:00:00 | 8dov-G | 4.7 | 3.4 | 86 | 140 | 3 | MOLECULE: | HEMOGLOBIN |
| 88:00:00 | 8fb0-A | 4.7 | 3.9 | 88 | 153 | 11 | MOLECULE: | MYOGLOBIN |
| 89:00:00 | 5ut8-A | 4.7 | 4 | 88 | 153 | 13 | MOLECULE: | MYOGLOBIN |
| 90:00:00 | 1bz0-A | 4.7 | 3 | 84 | 141 | 5 | MOLECULE: | PROTEIN (HE |
| 91:00:00 | 1mnk-B | 4.7 | 3.9 | 90 | 151 | 13 | MOLECULE: | MYOGLOBIN |
| 92:00:00 | 2hbs-E | 4.7 | 3.5 | 87 | 141 | 2 | MOLECULE: | HEMOGLOBIN |
| 93:00:00 | 6nbd-A | 4.7 | 3.4 | 86 | 140 | 3 | MOLECULE: | HEMOGLOBIN |
| 94:00:00 | 1y4p-C | 4.7 | 3.1 | 85 | 141 | 4 | MOLECULE: | HEMOGLOBIN |
| 95:00:00 | 3oo5-A | 4.7 | 3.4 | 85 | 140 | 5 | MOLECULE: | HEMOGLOBIN |
| 96:00:00 | 2zt1-A | 4.7 | 3.9 | 88 | 153 | 11 | MOLECULE: | MYOGLOBIN |
| 97:00:00 | 1a01-C | 4.7 | 3.5 | 86 | 141 | 3 | MOLECULE: | HEMOGLOBIN |
| 98:00:00 | 1hrm-A | 4.7 | 4 | 90 | 153 | 14 | MOLECULE: | MYOGLOBIN |
| 99:00:00 | 1mbo-A | 4.7 | 3.9 | 89 | 153 | 12 | MOLECULE: | MYOGLOBIN |
| 0:00 | 1k0y-C | 4.7 | 3.5 | 87 | 141 | 2 | MOLECULE: | HEMOGLOBIN |
| 1:00 | 1qsi-C | 4.7 | 3.4 | 85 | 141 | 5 | MOLECULE: | PROTEIN (HE |
| 2:00 | 1y83-C | 4.7 | 3.5 | 87 | 141 | 2 | MOLECULE: | HEMOGLOBIN |
| 3:00 | 1bzz-A | 4.7 | 3.5 | 86 | 141 | 3 | MOLECULE: | PROTEIN (HE |
| 4:00 | 1m6c-A | 4.7 | 4 | 91 | 153 | 13 | MOLECULE: | PROTEIN (MY |
| 5:00 | 3eda-A | 4.7 | 3.9 | 88 | 153 | 13 | MOLECULE: | MYOGLOBIN |
| 6:00 | 2zsr-A | 4.7 | 3.9 | 88 | 153 | 13 | MOLECULE: | MYOGLOBIN |
| 7:00 | 6zmx-A | 4.7 | 3.3 | 86 | 138 | 5 | MOLECULE: | HEMOGLOBIN |
| 8:00 | 3at6-A | 4.7 | 2.9 | 84 | 141 | 7 | MOLECULE: | ALPHAA-GLOB |
| 9:00 | 1mlq-A | 4.7 | 4 | 90 | 154 | 11 | MOLECULE: | MYOGLOBIN |
| 10:00 | 1obm-A | 4.7 | 4 | 92 | 154 | 11 | MOLECULE: | MYOGLOBIN |
| 11:00 | 2mgi-A | 4.7 | 4 | 92 | 154 | 11 | MOLECULE: | MYOGLOBIN |
| 12:00 | 2mga-A | 4.7 | 4 | 91 | 154 | 12 | MOLECULE: | MYOGLOBIN |
| 13:00 | 2d5z-C | 4.7 | 3 | 84 | 141 | 5 | MOLECULE: | HEMOGLOBIN |
| 14:00 | 3szk-A | 4.7 | 3.4 | 85 | 140 | 4 | MOLECULE: | HEMOGLOBIN |
| 15:00 | 1abs-A | 4.7 | 3.9 | 90 | 154 | 11 | MOLECULE: | MYOGLOBIN |
| 16:00 | 6n02-A | 4.7 | 3.9 | 90 | 154 | 12 | MOLECULE: | MYOGLOBIN, |
| 17:00 | 8qbc-A | 4.7 | 3.9 | 89 | 156 | 11 | MOLECULE: | MYOGLOBIN |
| 18:00 | 5jnz-A | 4.7 | 3.1 | 85 | 142 | 4 | MOLECULE: | ALPHA CHAIN |
| 19:00 | 1moc-A | 4.7 | 4 | 91 | 154 | 12 | MOLECULE: | MYOGLOBIN |
| 20:00 | 4n7o-K | 4.7 | 3.4 | 86 | 141 | 3 | MOLECULE: | HEMOGLOBIN |
| 21:00 | 2mgg-A | 4.7 | 4 | 90 | 154 | 11 | MOLECULE: | MYOGLOBIN |
| 22:00 | 8esu-A | 4.7 | 4.1 | 90 | 154 | 12 | MOLECULE: | MYOGLOBIN |
| 23:00 | 2hbs-G | 4.7 | 3 | 84 | 141 | 5 | MOLECULE: | HEMOGLOBIN |
| 24:00:00 | 1xy0-A | 4.7 | 3 | 84 | 141 | 5 | MOLECULE: | HEMOGLOBIN |
| 25:00:00 | 4esa-C | 4.7 | 3.3 | 85 | 142 | 6 | MOLECULE: | HEMOGLOBIN |
| 26:00:00 | 1g9v-A | 4.7 | 3 | 84 | 141 | 5 | MOLECULE: | HEMOGLOBIN |

| | | | | | | | | |
|---|---|---|---|---|---|---|---|---|
| 27:00:00 | 1mob-A | 4.7 | 4 | 92 | 154 | 11 | MOLECULE: | MYOGLOBIN |
| 28:00:00 | 3ubv-G | 4.7 | 3.7 | 85 | 131 | 11 | MOLECULE: | HEMOGLOBIN- |
| 29:00:00 | 4n7o-I | 4.7 | 3.1 | 86 | 141 | 3 | MOLECULE: | HEMOGLOBIN |
| 30:00:00 | 1ufp-A | 4.7 | 3.9 | 89 | 154 | 11 | MOLECULE: | MYOGLOBIN |
| 31:00:00 | 3qje-A | 4.7 | 3.1 | 85 | 141 | 4 | MOLECULE: | HEMOGLOBIN |
| 32:00:00 | 1gbu-C | 4.7 | 3.5 | 87 | 141 | 2 | MOLECULE: | HEMOGLOBIN |
| 33:00:00 | 1mgn-A | 4.7 | 4 | 92 | 154 | 11 | MOLECULE: | METMYOGLOBI |
| 34:00:00 | 6kah-G | 4.7 | 3.1 | 85 | 141 | 4 | MOLECULE: | HEMOGLOBIN |
| 35:00:00 | 1mtj-A | 4.7 | 3.9 | 90 | 154 | 11 | MOLECULE: | MYOGLOBIN |
| 36:00:00 | 5ile-A | 4.7 | 4.1 | 90 | 154 | 12 | MOLECULE: | MYOGLOBIN |
| 37:00:00 | 3ubc-A | 4.7 | 3.8 | 84 | 131 | 8 | MOLECULE: | HEMOGLOBIN- |
| 38:00:00 | 5kdq-A | 4.7 | 3 | 84 | 141 | 5 | MOLECULE: | HEMOGLOBIN |
| 39:00:00 | 7k4m-G | 4.7 | 3.1 | 85 | 140 | 4 | MOLECULE: | HEMOGLOBIN |
| 40:00:00 | 1mlr-A | 4.7 | 4 | 90 | 154 | 11 | MOLECULE: | MYOGLOBIN |
| 41:00:00 | 8qba-A | 4.7 | 4 | 91 | 156 | 12 | MOLECULE: | MYOGLOBIN |
| 42:00:00 | 3ubv-A | 4.7 | 3.8 | 84 | 131 | 8 | MOLECULE: | HEMOGLOBIN- |
| 43:00:00 | 3s1i-B | 4.7 | 3.8 | 84 | 131 | 8 | MOLECULE: | HEMOGLOBIN- |
| 44:00:00 | 1moa-A | 4.7 | 4 | 90 | 154 | 11 | MOLECULE: | MYOGLOBIN |
| 45:00:00 | 1o16-A | 4.7 | 4 | 90 | 154 | 11 | MOLECULE: | MYOGLOBIN |
| 46:00:00 | 1j40-G | 4.7 | 3.1 | 85 | 141 | 4 | MOLECULE: | HEMOGLOBIN |
| 47:00:00 | 1v4x-A | 4.7 | 3.3 | 85 | 143 | 8 | MOLECULE: | HEMOGLOBIN |
| 48:00:00 | 6iyh-A | 4.7 | 3.4 | 86 | 142 | 3 | MOLECULE: | ALPHA CHAIN |
| 49:00:00 | 1fcs-A | 4.7 | 3.9 | 91 | 154 | 12 | MOLECULE: | MYOGLOBIN |
| 50:00:00 | 1ye0-A | 4.7 | 2.9 | 83 | 141 | 5 | MOLECULE: | HEMOGLOBIN |
| 51:00:00 | 110m-A | 4.7 | 3.8 | 89 | 154 | 12 | MOLECULE: | MYOGLOBIN |
| 52:00:00 | 2mbw-A | 4.7 | 4 | 91 | 154 | 12 | MOLECULE: | MYOGLOBIN |
| 53:00:00 | 2mgm-A | 4.7 | 4 | 90 | 154 | 11 | MOLECULE: | MYOGLOBIN |
| 54:00:00 | 1rps-C | 4.7 | 3.1 | 85 | 141 | 4 | MOLECULE: | HEMOGLOBIN |
| 55:00:00 | 1bab-A | 4.7 | 3 | 84 | 142 | 5 | MOLECULE: | HEMOGLOBIN |
| 56:00:00 | 1xz7-C | 4.7 | 3 | 84 | 141 | 5 | MOLECULE: | HEMOGLOBIN |
| 57:00:00 | 4fwz-A | 4.7 | 4 | 92 | 153 | 11 | MOLECULE: | MYOGLOBIN |
| 58:00:00 | 3bj1-C | 4.7 | 3.4 | 86 | 142 | 6 | MOLECULE: | HEMOGLOBIN |
| 59:00:00 | 1mod-A | 4.7 | 4 | 92 | 154 | 11 | MOLECULE: | MYOGLOBIN |
| 60:00:00 | 3bj1-A | 4.7 | 3.4 | 86 | 142 | 6 | MOLECULE: | HEMOGLOBIN |
| 61:00:00 | 103m-A | 4.7 | 4 | 91 | 154 | 12 | MOLECULE: | MYOGLOBIN |
| 62:00:00 | 8ess-A | 4.7 | 3.9 | 90 | 154 | 11 | MOLECULE: | MYOGLOBIN |
| 63:00:00 | 6bwp-A | 4.7 | 3 | 84 | 141 | 5 | MOLECULE: | HEMOGLOBIN |
| 64:00:00 | 3s1j-B | 4.7 | 3.8 | 84 | 131 | 8 | MOLECULE: | HEMOGLOBIN- |
| 65:00:00 | 6m8f-A | 4.7 | 4 | 90 | 154 | 11 | MOLECULE: | MYOGLOBIN |
| 66:00:00 | 1y5k-C | 4.7 | 3.1 | 85 | 141 | 4 | MOLECULE: | HEMOGLOBIN |
| 67:00:00 | 3bj3-C | 4.7 | 3.3 | 86 | 142 | 6 | MOLECULE: | HEMOGLOBIN |
| 68:00:00 | 2bnl-E | 4.7 | 4 | 79 | 129 | 8 | MOLECULE: | MODULATOR P |
| 69:00:00 | 1mln-A | 4.7 | 4 | 92 | 154 | 11 | MOLECULE: | MYOGLOBIN |
| 70:00:00 | 1xz2-A | 4.7 | 3.1 | 85 | 141 | 4 | MOLECULE: | HEMOGLOBIN |

| | | | | | | | | |
|---|---|---|---|---|---|---|---|---|
| 71:00:00 | 6iyi-A | 4.7 | 3.1 | 85 | 142 | 4 | MOLECULE: | ALPHA CHAIN |
| 72:00:00 | 5wog-A | 4.7 | 3 | 84 | 137 | 4 | MOLECULE: | HEMOGLOBIN |
| 73:00:00 | 1dxu-A | 4.7 | 3 | 84 | 141 | 5 | MOLECULE: | HEMOGLOBIN |
| 74:00:00 | 2mgf-A | 4.7 | 4 | 91 | 154 | 12 | MOLECULE: | MYOGLOBIN |
| 75:00:00 | 1mlj-A | 4.7 | 4 | 90 | 154 | 11 | MOLECULE: | MYOGLOBIN |
| 76:00:00 | 8vyl-A | 4.7 | 3.4 | 86 | 137 | 3 | MOLECULE: | HEMOGLOBIN |
| 77:00:00 | 1hba-C | 4.7 | 3.1 | 85 | 141 | 4 | MOLECULE: | HEMOGLOBIN |
| 78:00:00 | 1bz1-C | 4.7 | 3 | 84 | 142 | 5 | MOLECULE: | PROTEIN (HE |
| 79:00:00 | 1a3n-C | 4.7 | 3 | 84 | 141 | 5 | MOLECULE: | HEMOGLOBIN |
| 80:00:00 | 1hbg-A | 4.6 | 3.6 | 88 | 147 | 6 | MOLECULE: | HEMOGLOBIN |
| 81:00:00 | 6zca-H | 4.6 | 16.6 | 122 | 771 | 6 | MOLECULE: | PROBABLE DN |
| 82:00:00 | 5x2u-G | 4.6 | 3.5 | 87 | 140 | 2 | MOLECULE: | HEMOGLOBIN |
| 83:00:00 | 1rq4-A | 4.6 | 3 | 84 | 141 | 5 | MOLECULE: | HEMOGLOBIN |
| 84:00:00 | 1y09-A | 4.6 | 3.6 | 88 | 141 | 3 | MOLECULE: | HEMOGLOBIN |
| 85:00:00 | 1ut0-B | 4.6 | 3.9 | 89 | 154 | 11 | MOLECULE: | CYTOGLOBIN |
| 86:00:00 | 1ut0-A | 4.6 | 4.1 | 90 | 154 | 10 | MOLECULE: | CYTOGLOBIN |
| 87:00:00 | 7ddu-C | 4.6 | 4 | 89 | 154 | 13 | MOLECULE: | MYOGLOBIN |
| 88:00:00 | 2bnl-B | 4.6 | 4 | 78 | 128 | 8 | MOLECULE: | MODULATOR P |
| 89:00:00 | 2bnl-D | 4.6 | 4 | 79 | 129 | 8 | MOLECULE: | MODULATOR P |
| 90:00:00 | 1kd2-A | 4.6 | 3.1 | 85 | 141 | 4 | MOLECULE: | HEMOGLOBIN |
| 91:00:00 | 4mqi-A | 4.6 | 3.4 | 85 | 140 | 5 | MOLECULE: | HEMOGLOBIN |
| 92:00:00 | 1xq5-C | 4.6 | 3.4 | 86 | 142 | 8 | MOLECULE: | HEMOGLOBIN |
| 93:00:00 | 1lfv-A | 4.6 | 3.2 | 85 | 141 | 4 | MOLECULE: | HEMOGLOBIN |
| 94:00:00 | 1urv-B | 4.6 | 3.9 | 89 | 154 | 11 | MOLECULE: | CYTOGLOBIN |
| 95:00:00 | 1umo-B | 4.6 | 4.2 | 90 | 154 | 10 | MOLECULE: | CYTOGLOBIN |
| 96:00:00 | 1urv-A | 4.6 | 4.1 | 90 | 154 | 10 | MOLECULE: | CYTOGLOBIN |
| 97:00:00 | 3bj2-C | 4.6 | 3.4 | 86 | 142 | 6 | MOLECULE: | HEMOGLOBIN |
| 98:00:00 | 8u5a-B | 4.6 | 3.7 | 85 | 149 | 7 | MOLECULE: | DESIGNED MY |
| 99:00:00 | 1lft-A | 4.6 | 3.5 | 86 | 141 | 3 | MOLECULE: | HEMOGLOBIN |
| 0:00 | 1fdh-A | 4.6 | 3 | 85 | 141 | 4 | MOLECULE: | HEMOGLOBIN |
| 1:00 | 7k4m-C | 4.6 | 3.4 | 86 | 140 | 2 | MOLECULE: | HEMOGLOBIN |
| 2:00 | 1spg-A | 4.6 | 3.4 | 86 | 142 | 8 | MOLECULE: | HEMOGLOBIN |
| 3:00 | 1yie-A | 4.6 | 3.4 | 85 | 141 | 5 | MOLECULE: | HEMOGLOBIN |
| 4:00 | 1y5k-A | 4.6 | 3.1 | 85 | 141 | 4 | MOLECULE: | HEMOGLOBIN |
| 5:00 | 1hac-C | 4.6 | 3.8 | 88 | 141 | 2 | MOLECULE: | HEMOGLOBIN |
| 6:00 | 3bcq-A | 4.6 | 3.3 | 85 | 142 | 9 | MOLECULE: | ALPHA-CHAIN |
| 7:00 | 6nbc-C | 4.6 | 3.4 | 85 | 140 | 4 | MOLECULE: | HEMOGLOBIN |
| 8:00 | 6lcx-G | 4.6 | 3 | 85 | 141 | 5 | MOLECULE: | HEMOGLOBIN |
| 9:00 | 1nih-A | 4.6 | 3.1 | 85 | 141 | 6 | MOLECULE: | HEMOGLOBIN |
| 10:00 | 1vxb-A | 4.6 | 3.8 | 88 | 153 | 11 | MOLECULE: | MYOGLOBIN |
| 11:00 | 7uvb-A | 4.6 | 3.4 | 86 | 141 | 2 | MOLECULE: | HEMOGLOBIN |
| 12:00 | 1gzx-A | 4.6 | 3.1 | 85 | 141 | 4 | MOLECULE: | HEMOGLOBIN |
| 13:00 | 5e83-C | 4.6 | 3.4 | 86 | 141 | 2 | MOLECULE: | HEMOGLOBIN |
| 14:00 | 2hbe-A | 4.6 | 3 | 84 | 141 | 4 | MOLECULE: | HEMOGLOBIN |

| | | | | | | | | |
|---|---|---|---|---|---|---|---|---|
| 15:00 | 2d5z-A | 4.6 | 3.1 | 86 | 141 | 5 | MOLECULE: | HEMOGLOBIN |
| 16:00 | 3d17-A | 4.6 | 3.5 | 86 | 141 | 3 | MOLECULE: | HEMOGLOBIN |
| 17:00 | 3cy5-A | 4.6 | 3.5 | 87 | 141 | 3 | MOLECULE: | HEMOGLOBIN |
| 18:00 | 1j3y-G | 4.6 | 2.9 | 84 | 141 | 6 | MOLECULE: | HEMOGLOBIN |
| 19:00 | 3qjb-A | 4.6 | 3.4 | 85 | 141 | 5 | MOLECULE: | HEMOGLOBIN |
| 20:00 | 5urc-C | 4.6 | 3.4 | 86 | 141 | 2 | MOLECULE: | HEMOGLOBIN |
| 21:00 | 6nbd-C | 4.6 | 3.4 | 86 | 140 | 3 | MOLECULE: | HEMOGLOBIN |
| 22:00 | 1cls-A | 4.6 | 3 | 84 | 141 | 5 | MOLECULE: | HEMOGLOBIN |
| 23:00 | 1y4f-A | 4.6 | 3 | 84 | 141 | 5 | MOLECULE: | HEMOGLOBIN |
| 24:00:00 | 1hab-A | 4.6 | 3.1 | 85 | 141 | 4 | MOLECULE: | HEMOGLOBIN |
| 25:00:00 | 2yrs-M | 4.6 | 3.4 | 86 | 141 | 3 | MOLECULE: | HEMOGLOBIN |
| 26:00:00 | 4n7p-G | 4.6 | 3.1 | 85 | 141 | 4 | MOLECULE: | HEMOGLOBIN |
| 27:00:00 | 1y4b-A | 4.6 | 3.1 | 86 | 141 | 5 | MOLECULE: | HEMOGLOBIN |
| 28:00:00 | 6bwp-C | 4.6 | 2.9 | 84 | 141 | 6 | MOLECULE: | HEMOGLOBIN |
| 29:00:00 | 6di4-C | 4.6 | 3.4 | 86 | 141 | 2 | MOLECULE: | HEMOGLOBIN |
| 30:00:00 | 1kd2-C | 4.6 | 3.1 | 85 | 141 | 6 | MOLECULE: | HEMOGLOBIN |
| 31:00:00 | 1rq4-C | 4.6 | 3.6 | 87 | 141 | 2 | MOLECULE: | HEMOGLOBIN |
| 32:00:00 | 6ii1-C | 4.6 | 3.5 | 86 | 138 | 6 | MOLECULE: | HEMOGLOBIN |
| 33:00:00 | 7uvb-C | 4.6 | 3.4 | 85 | 141 | 4 | MOLECULE: | HEMOGLOBIN |
| 34:00:00 | 3ic0-A | 4.6 | 3.4 | 86 | 141 | 2 | MOLECULE: | HEMOGLOBIN |
| 35:00:00 | 1ygd-A | 4.6 | 3.1 | 85 | 141 | 4 | MOLECULE: | HEMOGLOBIN |
| 36:00:00 | 1g09-A | 4.6 | 3.6 | 88 | 141 | 5 | MOLECULE: | HEMOGLOBIN |
| 37:00:00 | 1yeo-A | 4.6 | 3.1 | 85 | 141 | 4 | MOLECULE: | HEMOGLOBIN |
| 38:00:00 | 7uf6-C | 4.6 | 3.4 | 86 | 141 | 2 | MOLECULE: | HEMOGLOBIN |
| 39:00:00 | 6z4t-A | 4.6 | 4 | 90 | 154 | 12 | MOLECULE: | MYOGLOBIN |
| 40:00:00 | 2dn2-C | 4.6 | 2.9 | 84 | 141 | 6 | MOLECULE: | HEMOGLOBIN |
| 41:00:00 | 1a3o-C | 4.6 | 3 | 85 | 141 | 6 | MOLECULE: | HEMOGLOBIN |
| 42:00:00 | 1j7y-A | 4.6 | 2.9 | 84 | 141 | 6 | MOLECULE: | HEMOGLOBIN |
| 43:00:00 | 1yeq-C | 4.6 | 3 | 84 | 141 | 4 | MOLECULE: | HEMOGLOBIN |
| 44:00:00 | 1rps-A | 4.6 | 3.1 | 86 | 141 | 5 | MOLECULE: | HEMOGLOBIN |
| 45:00:00 | 7ehx-A | 4.6 | 3.9 | 90 | 153 | 11 | MOLECULE: | MYOGLOBIN |
| 46:00:00 | 1g0a-A | 4.6 | 3.4 | 86 | 141 | 5 | MOLECULE: | HEMOGLOBIN |
| 47:00:00 | 8dov-A | 4.6 | 3.5 | 86 | 140 | 3 | MOLECULE: | HEMOGLOBIN |
| 48:00:00 | 1u7r-A | 4.6 | 3.8 | 89 | 153 | 12 | MOLECULE: | MYOGLOBIN |
| 49:00:00 | 2hbs-A | 4.6 | 3.1 | 85 | 141 | 4 | MOLECULE: | HEMOGLOBIN |
| 50:00:00 | 8fdm-C | 4.6 | 3.4 | 85 | 139 | 5 | MOLECULE: | HEMOGLOBIN |
| 51:00:00 | 7jy3-C | 4.6 | 3.4 | 85 | 141 | 4 | MOLECULE: | HEMOGLOBIN |
| 52:00:00 | 6lcw-C | 4.6 | 3.1 | 85 | 141 | 4 | MOLECULE: | HEMOGLOBIN |
| 53:00:00 | 1y22-A | 4.6 | 3.6 | 87 | 141 | 2 | MOLECULE: | HEMOGLOBIN |
| 54:00:00 | 2hco-A | 4.6 | 3.3 | 85 | 141 | 5 | MOLECULE: | HEMOGLOBIN |
| 55:00:00 | 1xzv-A | 4.6 | 2.9 | 83 | 141 | 5 | MOLECULE: | HEMOGLOBIN |
| 56:00:00 | 7pch-C | 4.6 | 3.8 | 88 | 141 | 2 | MOLECULE: | HEMOGLOBIN |
| 57:00:00 | 7uf7-C | 4.6 | 3.4 | 86 | 141 | 2 | MOLECULE: | HEMOGLOBIN |
| 58:00:00 | 6hbw-C | 4.6 | 2.9 | 84 | 141 | 6 | MOLECULE: | PROTEIN (HE |

| | | | | | | | | |
|---|---|---|---|---|---|---|---|---|
| 59:00:00 | 1y85-A | 4.6 | 3 | 84 | 141 | 5 | MOLECULE: | HEMOGLOBIN |
| 60:00:00 | 1xxt-A | 4.6 | 3.1 | 85 | 141 | 4 | MOLECULE: | HEMOGLOBIN |
| 61:00:00 | 3vrf-A | 4.6 | 3.2 | 86 | 141 | 2 | MOLECULE: | HEMOGLOBIN |
| 62:00:00 | 5ufj-A | 4.6 | 3.5 | 86 | 141 | 2 | MOLECULE: | HEMOGLOBIN |
| 63:00:00 | 7jy0-C | 4.6 | 3.4 | 86 | 141 | 2 | MOLECULE: | HEMOGLOBIN |
| 64:00:00 | 3hen-A | 4.6 | 3.9 | 89 | 153 | 16 | MOLECULE: | MYOGLOBIN |
| 65:00:00 | 1aby-A | 4.6 | 9.5 | 110 | 283 | 5 | MOLECULE: | HEMOGLOBIN |
| 66:00:00 | 1mni-A | 4.6 | 4 | 90 | 153 | 13 | MOLECULE: | MYOGLOBIN |
| 67:00:00 | 6hk2-A | 4.6 | 3.4 | 85 | 141 | 5 | MOLECULE: | HEMOGLOBIN |
| 68:00:00 | 1yhe-A | 4.6 | 3.6 | 87 | 141 | 2 | MOLECULE: | HEMOGLOBIN |
| 69:00:00 | 1xz5-A | 4.6 | 3.1 | 85 | 141 | 4 | MOLECULE: | HEMOGLOBIN |
| 70:00:00 | 1yff-C | 4.6 | 3.5 | 86 | 141 | 2 | MOLECULE: | HEMOGLOBIN |
| 71:00:00 | 1yvq-A | 4.6 | 3 | 84 | 141 | 4 | MOLECULE: | HEMOGLOBIN |
| 72:00:00 | 2qss-C | 4.6 | 3.4 | 85 | 141 | 6 | MOLECULE: | HEMOGLOBIN |
| 73:00:00 | 6ihx-C | 4.6 | 3.4 | 86 | 140 | 6 | MOLECULE: | HEMOGLOBIN |
| 74:00:00 | 1yg5-A | 4.6 | 3.6 | 87 | 141 | 2 | MOLECULE: | HEMOGLOBIN |
| 75:00:00 | 1r1y-A | 4.6 | 3.4 | 87 | 141 | 5 | MOLECULE: | HEMOGLOBIN |
| 76:00:00 | 5zeo-A | 4.6 | 4 | 89 | 153 | 11 | MOLECULE: | MYOGLOBIN |
| 77:00:00 | 1jy7-C | 4.6 | 3.5 | 86 | 141 | 2 | MOLECULE: | HEMOGLOBIN |
| 78:00:00 | 2evk-A | 4.6 | 3.9 | 89 | 153 | 12 | MOLECULE: | MYOGLOBIN |
| 79:00:00 | 3pi8-C | 4.6 | 3.5 | 85 | 141 | 5 | MOLECULE: | HEMOGLOBIN |
| 80:00:00 | 1xxt-C | 4.6 | 3.1 | 85 | 141 | 4 | MOLECULE: | HEMOGLOBIN |
| 81:00:00 | 6kao-A | 4.6 | 3.6 | 87 | 141 | 2 | MOLECULE: | HEMOGLOBIN |
| 82:00:00 | 3v2z-A | 4.6 | 4 | 88 | 152 | 15 | MOLECULE: | MYOGLOBIN |
| 83:00:00 | 1yie-C | 4.6 | 3 | 84 | 141 | 4 | MOLECULE: | HEMOGLOBIN |
| 84:00:00 | 6xd9-A | 4.6 | 3.5 | 86 | 141 | 2 | MOLECULE: | HEMOGLOBIN |
| 85:00:00 | 1ygd-C | 4.6 | 3.1 | 85 | 141 | 4 | MOLECULE: | HEMOGLOBIN |
| 86:00:00 | 6kai-C | 4.6 | 2.9 | 83 | 141 | 5 | MOLECULE: | HEMOGLOBIN |
| 87:00:00 | 1mdn-A | 4.6 | 4 | 90 | 153 | 13 | MOLECULE: | PROTEIN (MY |
| 88:00:00 | 4mqh-A | 4.6 | 3.1 | 85 | 139 | 4 | MOLECULE: | HEMOGLOBIN |
| 89:00:00 | 1y45-C | 4.6 | 3.1 | 85 | 141 | 4 | MOLECULE: | HEMOGLOBIN |
| 90:00:00 | 1y5j-A | 4.6 | 3.6 | 87 | 141 | 2 | MOLECULE: | HEMOGLOBIN |
| 91:00:00 | 1hda-C | 4.6 | 3.1 | 85 | 141 | 6 | MOLECULE: | HEMOGLOBIN |
| 92:00:00 | 6kai-G | 4.6 | 3.1 | 86 | 141 | 5 | MOLECULE: | HEMOGLOBIN |
| 93:00:00 | 6bnr-C | 4.6 | 3.4 | 86 | 141 | 2 | MOLECULE: | HEMOGLOBIN |
| 94:00:00 | 1mwd-A | 4.6 | 3.9 | 90 | 153 | 13 | MOLECULE: | PROTEIN (MY |
| 95:00:00 | 1y31-C | 4.6 | 3.5 | 87 | 141 | 2 | MOLECULE: | HEMOGLOBIN |
| 96:00:00 | 5hlx-A | 4.6 | 4 | 90 | 153 | 11 | MOLECULE: | MYOGLOBIN |
| 97:00:00 | 5oj9-A | 4.6 | 3.9 | 89 | 154 | 11 | MOLECULE: | MYOGLOBIN |
| 98:00:00 | 1mwd-B | 4.6 | 4 | 90 | 153 | 14 | MOLECULE: | PROTEIN (MY |
| 99:00:00 | 1y7g-A | 4.6 | 3.1 | 85 | 141 | 4 | MOLECULE: | HEMOGLOBIN |
| 0:00 | 1a0z-A | 4.6 | 2.9 | 84 | 141 | 6 | MOLECULE: | HEMOGLOBIN |
| 1:00 | 1r1x-A | 4.6 | 3.8 | 87 | 141 | 3 | MOLECULE: | HEMOGLOBIN |
| 2:00 | 4mqk-C | 4.6 | 3.5 | 86 | 141 | 3 | MOLECULE: | HEMOGLOBIN |

| | | | | | | | | |
|---|---|---|---|---|---|---|---|---|
| 3:00 | 3nmm-A | 4.6 | 3 | 84 | 141 | 5 | MOLECULE: | HEMOGLOBIN |
| 4:00 | 2ef2-A | 4.6 | 3.5 | 88 | 153 | 11 | MOLECULE: | MYOGLOBIN |
| 5:00 | 4hhb-C | 4.6 | 3.5 | 86 | 141 | 3 | MOLECULE: | HEMOGLOBIN |
| 6:00 | 6kah-C | 4.6 | 3 | 84 | 141 | 4 | MOLECULE: | HEMOGLOBIN |
| 7:00 | 1y7d-A | 4.6 | 3.6 | 87 | 141 | 2 | MOLECULE: | HEMOGLOBIN |
| 8:00 | 3heo-A | 4.6 | 3.8 | 88 | 153 | 14 | MOLECULE: | MYOGLOBIN |
| 9:00 | 1j3z-G | 4.6 | 2.9 | 84 | 141 | 6 | MOLECULE: | HEMOGLOBIN |
| 10:00 | 8fdl-A | 4.6 | 2.9 | 84 | 141 | 6 | MOLECULE: | HEMOGLOBIN |
| 11:00 | 2yrs-C | 4.6 | 3.3 | 86 | 141 | 6 | MOLECULE: | HEMOGLOBIN |
| 12:00 | 1j7w-C | 4.6 | 3 | 84 | 141 | 5 | MOLECULE: | HEMOGLOBIN |
| 13:00 | 4mqg-A | 4.6 | 3.9 | 88 | 141 | 2 | MOLECULE: | HEMOGLOBIN |
| 14:00 | 7ud7-A | 4.6 | 3 | 85 | 141 | 5 | MOLECULE: | HEMOGLOBIN |
| 15:00 | 1j3z-E | 4.6 | 3 | 84 | 141 | 5 | MOLECULE: | HEMOGLOBIN |
| 16:00 | 3nl7-A | 4.6 | 3.6 | 87 | 141 | 2 | MOLECULE: | HEMOGLOBIN |
| 17:00 | 6lcx-C | 4.6 | 3.1 | 86 | 141 | 5 | MOLECULE: | HEMOGLOBIN |
| 18:00 | 6kae-G | 4.6 | 3 | 85 | 141 | 5 | MOLECULE: | HEMOGLOBIN |
| 19:00 | 6l5v-A | 4.6 | 3.5 | 86 | 141 | 3 | MOLECULE: | HEMOGLOBIN |
| 20:00 | 1xz5-C | 4.6 | 3.6 | 87 | 141 | 2 | MOLECULE: | HEMOGLOBIN |
| 21:00 | 5ucu-A | 4.6 | 3.4 | 85 | 140 | 5 | MOLECULE: | HEMOGLOBIN |
| 22:00 | 7dy3-E | 4.6 | 3 | 85 | 141 | 5 | MOLECULE: | HEMOGLOBIN |
| 23:00 | 3dut-C | 4.6 | 3 | 84 | 141 | 5 | MOLECULE: | HEMOGLOBIN |
| 24:00:00 | 3s48-D | 4.6 | 3.5 | 87 | 140 | 2 | MOLECULE: | IRON-REGULA |
| 25:00:00 | 3gdj-A | 4.6 | 3.5 | 87 | 141 | 6 | MOLECULE: | HEMOGLOBIN |
| 26:00:00 | 3qje-C | 4.6 | 3.1 | 85 | 141 | 6 | MOLECULE: | HEMOGLOBIN |
| 27:00:00 | 1yeu-A | 4.6 | 3.1 | 85 | 141 | 4 | MOLECULE: | HEMOGLOBIN |
| 28:00:00 | 6hbw-A | 4.6 | 3 | 84 | 141 | 5 | MOLECULE: | PROTEIN (HE |
| 29:00:00 | 6kat-A | 4.6 | 3.4 | 86 | 141 | 2 | MOLECULE: | HEMOGLOBIN |
| 30:00:00 | 3p5q-A | 4.6 | 3.4 | 86 | 139 | 6 | MOLECULE: | HEMOGLOBIN |
| 31:00:00 | 1r1y-C | 4.6 | 3.3 | 85 | 141 | 5 | MOLECULE: | HEMOGLOBIN |
| 32:00:00 | 1rq3-A | 4.6 | 3.5 | 86 | 141 | 3 | MOLECULE: | HEMOGLOBIN |
| 33:00:00 | 4mqc-A | 4.6 | 3.1 | 84 | 141 | 5 | MOLECULE: | HEMOGLOBIN |
| 34:00:00 | 8f9n-A | 4.6 | 4.1 | 88 | 153 | 13 | MOLECULE: | MYOGLOBIN |
| 35:00:00 | 5sw7-A | 4.6 | 3.4 | 85 | 142 | 5 | MOLECULE: | HEMOGLOBIN |
| 36:00:00 | 6f18-A | 4.6 | 3.8 | 88 | 154 | 11 | MOLECULE: | MYOGLOBIN |
| 37:00:00 | 1hgc-A | 4.6 | 3.4 | 86 | 141 | 3 | MOLECULE: | HEMOGLOBIN |
| 38:00:00 | 1y83-A | 4.6 | 3.1 | 85 | 141 | 4 | MOLECULE: | HEMOGLOBIN |
| 39:00:00 | 8fdk-A | 4.6 | 3.6 | 88 | 141 | 3 | MOLECULE: | HEMOGLOBIN |
| 40:00:00 | 7jjq-C | 4.6 | 3.6 | 86 | 139 | 3 | MOLECULE: | HEMOGLOBIN |
| 41:00:00 | 1ljw-A | 4.6 | 3.5 | 87 | 141 | 2 | MOLECULE: | HEMOGLOBIN |
| 42:00:00 | 2z6n-A | 4.6 | 4.1 | 88 | 141 | 8 | MOLECULE: | HEMOGLOBIN |
| 43:00:00 | 3vrg-A | 4.6 | 3.2 | 85 | 141 | 2 | MOLECULE: | HEMOGLOBIN |
| 44:00:00 | 8fdk-C | 4.6 | 3.6 | 86 | 141 | 3 | MOLECULE: | HEMOGLOBIN |
| 45:00:00 | 1y4r-A | 4.6 | 3.6 | 87 | 141 | 2 | MOLECULE: | HEMOGLOBIN |
| 46:00:00 | 7cue-C | 4.6 | 3.5 | 86 | 141 | 3 | MOLECULE: | HEMOGLOBIN |

| | | | | | | | | |
|---|---|---|---|---|---|---|---|---|
| 47:00:00 | 1y31-A | 4.6 | 3.6 | 87 | 141 | 2 | MOLECULE: | HEMOGLOBIN |
| 48:00:00 | 1bvd-A | 4.6 | 4 | 88 | 153 | 11 | MOLECULE: | APOMYOGLOBI |
| 49:00:00 | 4hhb-A | 4.6 | 3 | 84 | 141 | 5 | MOLECULE: | HEMOGLOBIN |
| 50:00:00 | 1j7s-A | 4.6 | 3.1 | 85 | 141 | 4 | MOLECULE: | HEMOGLOBIN |
| 51:00:00 | 1y5f-A | 4.6 | 3.6 | 87 | 141 | 2 | MOLECULE: | HEMOGLOBIN |
| 52:00:00 | 3gys-G | 4.6 | 3 | 84 | 138 | 6 | MOLECULE: | HEMOGLOBIN |
| 53:00:00 | 1yvt-A | 4.6 | 3.5 | 86 | 141 | 3 | MOLECULE: | HEMOGLOBIN |
| 54:00:00 | 6l5x-A | 4.6 | 3.4 | 86 | 141 | 2 | MOLECULE: | HEMOGLOBIN |
| 55:00:00 | 6kae-C | 4.6 | 3 | 84 | 141 | 5 | MOLECULE: | HEMOGLOBIN |
| 56:00:00 | 3qjd-A | 4.6 | 2.9 | 84 | 141 | 6 | MOLECULE: | HEMOGLOBIN |
| 57:00:00 | 6kaq-A | 4.6 | 3.5 | 87 | 141 | 2 | MOLECULE: | HEMOGLOBIN |
| 58:00:00 | 4fwx-A | 4.6 | 3.9 | 90 | 153 | 11 | MOLECULE: | MYOGLOBIN |
| 59:00:00 | 1ygf-C | 4.6 | 3.6 | 87 | 141 | 2 | MOLECULE: | HEMOGLOBIN |
| 60:00:00 | 1y2z-C | 4.6 | 3.6 | 87 | 141 | 2 | MOLECULE: | HEMOGLOBIN |
| 61:00:00 | 2w72-C | 4.6 | 2.9 | 84 | 141 | 4 | MOLECULE: | HUMAN HEMOG |
| 62:00:00 | 5vzp-A | 4.6 | 3.9 | 88 | 153 | 11 | MOLECULE: | MYOGLOBIN |
| 63:00:00 | 3vre-C | 4.6 | 3.1 | 85 | 141 | 2 | MOLECULE: | HEMOGLOBIN |
| 64:00:00 | 1xye-A | 4.6 | 3.1 | 85 | 141 | 4 | MOLECULE: | HEMOGLOBIN |
| 65:00:00 | 6l5y-A | 4.6 | 3.4 | 86 | 141 | 2 | MOLECULE: | HEMOGLOBIN |
| 66:00:00 | 8fdn-C | 4.6 | 3.5 | 87 | 141 | 5 | MOLECULE: | HEMOGLOBIN |
| 67:00:00 | 5ufj-C | 4.6 | 3.4 | 86 | 141 | 2 | MOLECULE: | HEMOGLOBIN |
| 68:00:00 | 4x0i-A | 4.6 | 3 | 83 | 141 | 5 | MOLECULE: | HEMOGLOBIN |
| 69:00:00 | 1o1i-A | 4.6 | 3.1 | 84 | 141 | 4 | MOLECULE: | HEMOGLOBIN |
| 70:00:00 | 5b85-A | 4.6 | 3.9 | 88 | 153 | 11 | MOLECULE: | MYOGLOBIN |
| 71:00:00 | 1y4v-A | 4.6 | 3.6 | 87 | 141 | 2 | MOLECULE: | HEMOGLOBIN |
| 72:00:00 | 3oo4-A | 4.6 | 3.8 | 88 | 141 | 5 | MOLECULE: | HEMOGLOBIN |
| 73:00:00 | 1ycb-A | 4.6 | 4.1 | 91 | 153 | 13 | MOLECULE: | MYOGLOBIN |
| 74:00:00 | 5x2t-G | 4.6 | 3.9 | 88 | 140 | 2 | MOLECULE: | HEMOGLOBIN |
| 75:00:00 | 1j3y-E | 4.6 | 3 | 84 | 141 | 5 | MOLECULE: | HEMOGLOBIN |
| 76:00:00 | 6f19-A | 4.6 | 3.8 | 88 | 154 | 11 | MOLECULE: | MYOGLOBIN |
| 77:00:00 | 1mno-B | 4.6 | 4 | 90 | 153 | 14 | MOLECULE: | PROTEIN (MY |
| 78:00:00 | 1cls-C | 4.6 | 2.9 | 84 | 141 | 4 | MOLECULE: | HEMOGLOBIN |
| 79:00:00 | 1uiw-A | 4.6 | 2.9 | 83 | 141 | 5 | MOLECULE: | HEMOGLOBIN |
| 80:00:00 | 1y7z-A | 4.6 | 3.1 | 85 | 141 | 4 | MOLECULE: | HEMOGLOBIN |
| 81:00:00 | 6g5a-A | 4.6 | 3.7 | 88 | 153 | 13 | MOLECULE: | MYOGLOBIN |
| 82:00:00 | 1a0u-A | 4.6 | 3.1 | 85 | 141 | 4 | MOLECULE: | HEMOGLOBIN |
| 83:00:00 | 1hbb-A | 4.6 | 3.6 | 87 | 141 | 2 | MOLECULE: | HEMOGLOBIN |
| 84:00:00 | 4ni0-A | 4.6 | 3.6 | 87 | 141 | 2 | MOLECULE: | HEMOGLOBIN |
| 85:00:00 | 5ker-E | 4.6 | 3.4 | 86 | 140 | 5 | MOLECULE: | ALPHA-GLOBI |
| 86:00:00 | 2hhe-A | 4.6 | 3.1 | 86 | 141 | 5 | MOLECULE: | HEMOGLOBIN |
| 87:00:00 | 1y2z-A | 4.6 | 3 | 84 | 141 | 4 | MOLECULE: | HEMOGLOBIN |
| 88:00:00 | 2qsp-C | 4.6 | 3.6 | 87 | 141 | 5 | MOLECULE: | HEMOGLOBIN |
| 89:00:00 | 1ye2-A | 4.6 | 3 | 84 | 141 | 5 | MOLECULE: | HEMOGLOBIN |
| 90:00:00 | 6kye-I | 4.6 | 3.8 | 87 | 141 | 3 | MOLECULE: | HEMOGLOBIN |

| | | | | | | | | |
|---|---|---|---|---|---|---|---|---|
| 91:00:00 | 6ka9-A | 4.6 | 3.4 | 85 | 141 | 5 | MOLECULE: | HEMOGLOBIN |
| 92:00:00 | 1yeu-C | 4.6 | 3 | 84 | 141 | 4 | MOLECULE: | HEMOGLOBIN |
| 93:00:00 | 1yff-G | 4.6 | 3.4 | 86 | 141 | 2 | MOLECULE: | HEMOGLOBIN |
| 94:00:00 | 3nmm-C | 4.6 | 2.9 | 84 | 141 | 6 | MOLECULE: | HEMOGLOBIN |
| 95:00:00 | 5jdo-E | 4.6 | 3.4 | 86 | 140 | 3 | MOLECULE: | HAPTOGLOBIN |
| 96:00:00 | 1y4g-A | 4.6 | 3 | 84 | 141 | 5 | MOLECULE: | HEMOGLOBIN |
| 97:00:00 | 8fdm-A | 4.6 | 3.4 | 85 | 139 | 5 | MOLECULE: | HEMOGLOBIN |
| 98:00:00 | 1yih-A | 4.6 | 3.4 | 85 | 141 | 5 | MOLECULE: | HEMOGLOBIN |
| 99:00:00 | 1lfl-A | 4.6 | 3 | 85 | 141 | 5 | MOLECULE: | HEMOGLOBIN |
| 0:00 | 1ye1-C | 4.6 | 3.5 | 87 | 141 | 2 | MOLECULE: | HEMOGLOBIN |
| 1:00 | 1yg5-C | 4.6 | 3.1 | 85 | 141 | 4 | MOLECULE: | HEMOGLOBIN |
| 2:00 | 1y35-A | 4.6 | 3.1 | 85 | 141 | 4 | MOLECULE: | HEMOGLOBIN |
| 3:00 | 8a61-T | 4.6 | 23.1 | 112 | 650 | 12 | MOLECULE: | ANAPHASE-PR |
| 4:00 | 1sdk-C | 4.6 | 3.4 | 85 | 141 | 5 | MOLECULE: | HEMOGLOBIN |
| 5:00 | 1m6m-B | 4.6 | 4 | 90 | 153 | 14 | MOLECULE: | PROTEIN (MY |
| 6:00 | 6kav-A | 4.6 | 3.4 | 86 | 141 | 2 | MOLECULE: | HEMOGLOBIN |
| 7:00 | 7pcq-C | 4.6 | 3.9 | 88 | 141 | 2 | MOLECULE: | HEMOGLOBIN |
| 8:00 | 4n8t-A | 4.6 | 3.4 | 85 | 140 | 5 | MOLECULE: | HEMOGLOBIN |
| 9:00 | 2dn3-A | 4.6 | 3.6 | 87 | 141 | 2 | MOLECULE: | HEMOGLOBIN |
| 10:00 | 1nih-C | 4.6 | 3.1 | 85 | 141 | 4 | MOLECULE: | HEMOGLOBIN |
| 11:00 | 4ni1-A | 4.6 | 3.4 | 86 | 141 | 6 | MOLECULE: | HEMOGLOBIN |
| 12:00 | 1hab-C | 4.6 | 3.4 | 86 | 141 | 3 | MOLECULE: | HEMOGLOBIN |
| 13:00 | 1xzu-A | 4.6 | 3.1 | 85 | 141 | 4 | MOLECULE: | HEMOGLOBIN |
| 14:00 | 2w72-A | 4.6 | 3.4 | 85 | 141 | 5 | MOLECULE: | HUMAN HEMOG |
| 15:00 | 1ycb-B | 4.6 | 4 | 90 | 153 | 14 | MOLECULE: | MYOGLOBIN |
| 16:00 | 1hda-A | 4.6 | 3.1 | 85 | 141 | 6 | MOLECULE: | HEMOGLOBIN |
| 17:00 | 1qi8-A | 4.6 | 3.1 | 85 | 141 | 4 | MOLECULE: | HEMOGLOBIN |
| 18:00 | 1y85-C | 4.6 | 3.6 | 87 | 141 | 2 | MOLECULE: | HEMOGLOBIN |
| 19:00 | 2d60-C | 4.6 | 2.9 | 84 | 141 | 6 | MOLECULE: | HEMOGLOBIN |
| 20:00 | 1y0w-A | 4.6 | 3 | 84 | 141 | 5 | MOLECULE: | HEMOGLOBIN |
| 21:00 | 6kye-E | 4.6 | 3.4 | 86 | 140 | 3 | MOLECULE: | HEMOGLOBIN |
| 22:00 | 1y0c-A | 4.6 | 3 | 85 | 141 | 5 | MOLECULE: | HEMOGLOBIN |
| 23:00 | 4l7y-C | 4.6 | 3 | 84 | 141 | 5 | MOLECULE: | HEMOGLOBIN |
| 24:00:00 | 1fsx-A | 4.6 | 3.1 | 85 | 141 | 6 | MOLECULE: | HEMOGLOBIN |
| 25:00:00 | 6e04-A | 4.6 | 4 | 88 | 153 | 13 | MOLECULE: | MYOGLOBIN |
| 26:00:00 | 7dy4-E | 4.6 | 3.1 | 84 | 141 | 5 | MOLECULE: | HEMOGLOBIN |
| 27:00:00 | 6kap-A | 4.6 | 3.5 | 87 | 141 | 2 | MOLECULE: | HEMOGLOBIN |
| 28:00:00 | 6lcw-G | 4.6 | 3 | 85 | 141 | 5 | MOLECULE: | HEMOGLOBIN |
| 29:00:00 | 7jy3-A | 4.6 | 3.4 | 86 | 141 | 2 | MOLECULE: | HEMOGLOBIN |
| 30:00:00 | 1j41-C | 4.6 | 2.9 | 83 | 141 | 5 | MOLECULE: | HEMOGLOBIN |
| 31:00:00 | 1fhj-C | 4.6 | 3.5 | 86 | 141 | 5 | MOLECULE: | HEMOGLOBIN |
| 32:00:00 | 1fdh-B | 4.6 | 3 | 85 | 141 | 4 | MOLECULE: | HEMOGLOBIN |
| 33:00:00 | 1yh9-A | 4.6 | 3.1 | 85 | 141 | 4 | MOLECULE: | HEMOGLOBIN |
| 34:00:00 | 2yrs-I | 4.6 | 3 | 85 | 141 | 5 | MOLECULE: | HEMOGLOBIN |

| | | | | | | | | |
|---|---|---|---|---|---|---|---|---|
| 35:00:00 | 2w6v-A | 4.6 | 2.9 | 84 | 141 | 6 | MOLECULE: | HEMOGLOBIN |
| 36:00:00 | 1yeq-A | 4.6 | 3 | 84 | 141 | 4 | MOLECULE: | HEMOGLOBIN |
| 37:00:00 | 1hdb-A | 4.6 | 3.5 | 86 | 141 | 3 | MOLECULE: | HEMOGLOBIN |
| 38:00:00 | 5kdq-C | 4.6 | 3.1 | 85 | 141 | 6 | MOLECULE: | HEMOGLOBIN |
| 39:00:00 | 7pch-A | 4.6 | 3.9 | 88 | 141 | 2 | MOLECULE: | HEMOGLOBIN |
| 40:00:00 | 1y0a-A | 4.6 | 2.9 | 83 | 141 | 5 | MOLECULE: | HEMOGLOBIN |
| 41:00:00 | 2d6c-A | 4.6 | 4 | 88 | 153 | 13 | MOLECULE: | MYOGLOBIN |
| 42:00:00 | 1mwc-B | 4.6 | 3.9 | 90 | 153 | 13 | MOLECULE: | PROTEIN (MY |
| 43:00:00 | 6ka9-C | 4.6 | 3.3 | 85 | 141 | 5 | MOLECULE: | HEMOGLOBIN |
| 44:00:00 | 6kye-C | 4.6 | 3.6 | 87 | 140 | 2 | MOLECULE: | HEMOGLOBIN |
| 45:00:00 | 1pmb-A | 4.6 | 3.9 | 90 | 153 | 13 | MOLECULE: | MYOGLOBIN |
| 46:00:00 | 7dy4-C | 4.6 | 3.5 | 86 | 141 | 3 | MOLECULE: | HEMOGLOBIN |
| 47:00:00 | 1rqa-C | 4.6 | 3.4 | 86 | 141 | 2 | MOLECULE: | HEMOGLOBIN |
| 48:00:00 | 1mdn-B | 4.6 | 4 | 90 | 153 | 14 | MOLECULE: | PROTEIN (MY |
| 49:00:00 | 1m9p-A | 4.6 | 3.4 | 85 | 141 | 4 | MOLECULE: | HEMOGLOBIN |
| 50:00:00 | 7vde-C | 4.6 | 3.5 | 87 | 141 | 2 | MOLECULE: | HEMOGLOBIN |
| 51:00:00 | 5hu6-A | 4.6 | 3 | 83 | 141 | 5 | MOLECULE: | HEMOGLOBIN |
| 52:00:00 | 3o89-A | 4.6 | 4 | 89 | 153 | 12 | MOLECULE: | MYOGLOBIN |
| 53:00:00 | 6rp5-A | 4.6 | 3.1 | 84 | 142 | 6 | MOLECULE: | HEMOGLOBIN |
| 54:00:00 | 1ye1-A | 4.6 | 3.6 | 87 | 141 | 2 | MOLECULE: | HEMOGLOBIN |
| 55:00:00 | 1j7y-C | 4.6 | 2.9 | 84 | 141 | 6 | MOLECULE: | HEMOGLOBIN |
| 56:00:00 | 1j41-G | 4.6 | 3 | 85 | 141 | 6 | MOLECULE: | HEMOGLOBIN |
| 57:00:00 | 1y7c-A | 4.6 | 3.1 | 85 | 141 | 4 | MOLECULE: | HEMOGLOBIN |
| 58:00:00 | 1y4p-A | 4.6 | 3.1 | 85 | 141 | 4 | MOLECULE: | HEMOGLOBIN |
| 59:00:00 | 6bwu-A | 4.6 | 3.6 | 87 | 140 | 5 | MOLECULE: | HEMOGLOBIN |
| 60:00:00 | 6kau-A | 4.6 | 3.4 | 86 | 141 | 2 | MOLECULE: | HEMOGLOBIN |
| 61:00:00 | 1y4q-A | 4.6 | 3.1 | 85 | 141 | 4 | MOLECULE: | HEMOGLOBIN |
| 62:00:00 | 6kas-A | 4.6 | 3.4 | 86 | 141 | 2 | MOLECULE: | HEMOGLOBIN |
| 63:00:00 | 1g08-A | 4.6 | 3.5 | 86 | 141 | 5 | MOLECULE: | HEMOGLOBIN |
| 64:00:00 | 3ba2-A | 4.6 | 4 | 88 | 152 | 15 | MOLECULE: | MYOGLOBIN |
| 65:00:00 | 5u3i-A | 4.6 | 3.4 | 86 | 141 | 2 | MOLECULE: | HEMOGLOBIN |
| 66:00:00 | 3hep-A | 4.6 | 3.8 | 88 | 153 | 14 | MOLECULE: | MYOGLOBIN |
| 67:00:00 | 8fdn-A | 4.6 | 3.1 | 84 | 141 | 5 | MOLECULE: | HEMOGLOBIN |
| 68:00:00 | 6kae-A | 4.6 | 3 | 84 | 141 | 5 | MOLECULE: | HEMOGLOBIN |
| 69:00:00 | 1y46-A | 4.6 | 3.6 | 87 | 141 | 2 | MOLECULE: | HEMOGLOBIN |
| 70:00:00 | 3d7o-A | 4.6 | 3.6 | 86 | 141 | 3 | MOLECULE: | HEMOGLOBIN |
| 71:00:00 | 1gbu-A | 4.6 | 3 | 84 | 141 | 5 | MOLECULE: | HEMOGLOBIN |
| 72:00:00 | 8j4l-A | 4.6 | 3.9 | 88 | 153 | 11 | MOLECULE: | MYOGLOBIN |
| 73:00:00 | 1y0t-A | 4.6 | 3.1 | 85 | 141 | 4 | MOLECULE: | HEMOGLOBIN |
| 74:00:00 | 1hba-A | 4.6 | 3.1 | 85 | 141 | 4 | MOLECULE: | HEMOGLOBIN |
| 75:00:00 | 2evp-A | 4.6 | 3.8 | 90 | 153 | 11 | MOLECULE: | MYOGLOBIN |
| 76:00:00 | 1j7s-C | 4.6 | 3 | 84 | 141 | 5 | MOLECULE: | HEMOGLOBIN |
| 77:00:00 | 2hhd-A | 4.6 | 2.9 | 84 | 141 | 6 | MOLECULE: | HEMOGLOBIN |
| 78:00:00 | 1irc-A | 4.6 | 3.8 | 90 | 152 | 11 | MOLECULE: | MYOGLOBIN ( |

| | | | | | | | | |
|---|---|---|---|---|---|---|---|---|
| 79:00:00 | 8dov-C | 4.6 | 3.5 | 87 | 140 | 5 | MOLECULE: | HEMOGLOBIN |
| 80:00:00 | 1hgb-C | 4.6 | 3.1 | 86 | 141 | 5 | MOLECULE: | HEMOGLOBIN |
| 81:00:00 | 1xq5-A | 4.6 | 3.4 | 86 | 142 | 8 | MOLECULE: | HEMOGLOBIN |
| 82:00:00 | 7jxz-A | 4.6 | 3.4 | 86 | 141 | 2 | MOLECULE: | HEMOGLOBIN |
| 83:00:00 | 6zmy-A | 4.6 | 3.1 | 85 | 139 | 5 | MOLECULE: | HEMOGLOBIN |
| 84:00:00 | 1umo-A | 4.6 | 4.2 | 89 | 154 | 10 | MOLECULE: | CYTOGLOBIN |
| 85:00:00 | 3a59-C | 4.6 | 3.4 | 86 | 141 | 6 | MOLECULE: | HEMOGLOBIN |
| 86:00:00 | 7k4m-A | 4.6 | 3.1 | 85 | 141 | 4 | MOLECULE: | HEMOGLOBIN |
| 87:00:00 | 3odq-C | 4.6 | 3.4 | 86 | 141 | 3 | MOLECULE: | HEMOGLOBIN |
| 88:00:00 | 4rom-C | 4.6 | 3 | 84 | 141 | 5 | MOLECULE: | HEMOGLOBIN |
| 89:00:00 | 1qxd-A | 4.6 | 3 | 84 | 141 | 4 | MOLECULE: | HEMOGLOBIN |
| 90:00:00 | 3hxn-A | 4.6 | 3.4 | 86 | 141 | 3 | MOLECULE: | HEMOGLOBIN |
| 91:00:00 | 7ddu-A | 4.6 | 3.8 | 89 | 154 | 12 | MOLECULE: | MYOGLOBIN |
| 92:00:00 | 6xdt-C | 4.6 | 3 | 84 | 141 | 4 | MOLECULE: | HEMOGLOBIN |
| 93:00:00 | 3wtg-A | 4.6 | 3 | 85 | 138 | 11 | MOLECULE: | HEMOGLOBIN |
| 94:00:00 | 3bj3-A | 4.6 | 3.3 | 86 | 142 | 6 | MOLECULE: | HEMOGLOBIN |
| 95:00:00 | 4ij2-A | 4.6 | 3.5 | 86 | 138 | 3 | MOLECULE: | HEMOGLOBIN |
| 96:00:00 | 5ojb-A | 4.6 | 3.9 | 91 | 154 | 13 | MOLECULE: | MYOGLOBIN |
| 97:00:00 | 7vw4-A | 4.6 | 4 | 89 | 152 | 12 | MOLECULE: | MYOGLOBIN |
| 98:00:00 | 2aa1-C | 4.6 | 3.3 | 85 | 142 | 7 | MOLECULE: | HEMOGLOBIN |
| 99:00:00 | 5oja-A | 4.6 | 4.1 | 92 | 152 | 12 | MOLECULE: | MYOGLOBIN |
| 0:00 | 8dov-E | 4.6 | 3.5 | 86 | 138 | 3 | MOLECULE: | HEMOGLOBIN |
| 1:00 | 2yrs-A | 4.6 | 3.4 | 86 | 141 | 3 | MOLECULE: | HEMOGLOBIN |
| 2:00 | 1hga-A | 4.6 | 3.4 | 86 | 141 | 3 | MOLECULE: | HEMOGLOBIN |
| 3:00 | 1yzi-A | 4.6 | 3.5 | 87 | 141 | 2 | MOLECULE: | HEMOGLOBIN |
| 4:00 | 6l5w-A | 4.6 | 3.5 | 86 | 141 | 3 | MOLECULE: | HEMOGLOBIN |
| 5:00 | 4esa-A | 4.6 | 3.3 | 85 | 142 | 6 | MOLECULE: | HEMOGLOBIN |
| 6:00 | 2w6v-C | 4.6 | 3.5 | 87 | 141 | 5 | MOLECULE: | HEMOGLOBIN |
| 7:00 | 5jgg-A | 4.6 | 3.4 | 86 | 142 | 3 | MOLECULE: | ALPHA CHAIN |
| 8:00 | 2aa1-A | 4.6 | 3.3 | 85 | 142 | 7 | MOLECULE: | HEMOGLOBIN |
| 9:00 | 3ic2-A | 4.6 | 3 | 84 | 141 | 4 | MOLECULE: | HEMOGLOBIN |
| 10:00 | 8fdl-C | 4.6 | 3.1 | 85 | 141 | 6 | MOLECULE: | HEMOGLOBIN |
| 11:00 | 3r5i-A | 4.6 | 3 | 84 | 141 | 4 | MOLECULE: | HEMOGLOBIN |
| 12:00 | 1hac-A | 4.6 | 3.4 | 86 | 141 | 3 | MOLECULE: | HEMOGLOBIN |
| 13:00 | 5ee4-E | 4.6 | 3 | 84 | 141 | 4 | MOLECULE: | HPUA |
| 14:00 | 7pcq-A | 4.6 | 3.5 | 87 | 141 | 2 | MOLECULE: | HEMOGLOBIN |
| 15:00 | 4ij2-C | 4.6 | 3.6 | 87 | 138 | 2 | MOLECULE: | HEMOGLOBIN |
| 16:00 | 6hit-G | 4.6 | 3.1 | 84 | 142 | 5 | MOLECULE: | HEMOGLOBIN |
| 17:00 | 1rqa-A | 4.6 | 2.9 | 83 | 141 | 5 | MOLECULE: | HEMOGLOBIN |
| 18:00 | 5yci-A | 4.6 | 3.9 | 89 | 154 | 16 | MOLECULE: | ANCESTRAL M |
| 19:00 | 6ii1-A | 4.6 | 3.2 | 85 | 137 | 6 | MOLECULE: | HEMOGLOBIN |
| 20:00 | 2ri4-I | 4.6 | 3 | 84 | 137 | 5 | MOLECULE: | HEMOGLOBIN |
| 21:00 | 4l7y-A | 4.6 | 3.1 | 85 | 141 | 4 | MOLECULE: | HEMOGLOBIN |
| 22:00 | 3s66-A | 4.6 | 3.5 | 86 | 139 | 3 | MOLECULE: | HEMOGLOBIN |

| | | | | | | | | |
|---|---|---|---|---|---|---|---|---|
| 23:00 | 6l5y-C | 4.6 | 3 | 84 | 141 | 4 | MOLECULE: | HEMOGLOBIN |
| 24:00:00 | 5x2s-G | 4.6 | 3.5 | 86 | 140 | 3 | MOLECULE: | HEMOGLOBIN |
| 25:00:00 | 7dy3-G | 4.6 | 3 | 84 | 141 | 4 | MOLECULE: | HEMOGLOBIN |
| 26:00:00 | 5woh-A | 4.6 | 3 | 84 | 137 | 4 | MOLECULE: | HEMOGLOBIN |
| 27:00:00 | 1qxe-A | 4.6 | 3 | 84 | 141 | 4 | MOLECULE: | HEMOGLOBIN |
| 28:00:00 | 6hit-C | 4.6 | 3.4 | 86 | 142 | 3 | MOLECULE: | HEMOGLOBIN |
| 29:00:00 | 4b3w-A | 4.6 | 4.2 | 90 | 154 | 10 | MOLECULE: | CYTOGLOBIN |
| 30:00:00 | 5e29-A | 4.6 | 3.1 | 84 | 141 | 5 | MOLECULE: | HEMOGLOBIN |
| 31:00:00 | 6xe7-A | 4.6 | 3.5 | 86 | 141 | 2 | MOLECULE: | HEMOGLOBIN |
| 32:00:00 | 1a01-A | 4.6 | 3 | 85 | 141 | 6 | MOLECULE: | HEMOGLOBIN |
| 33:00:00 | 3bcq-C | 4.6 | 3.3 | 85 | 142 | 9 | MOLECULE: | ALPHA-CHAIN |
| 34:00:00 | 1v4u-A | 4.6 | 3.3 | 85 | 143 | 8 | MOLECULE: | HEMOGLOBIN |
| 35:00:00 | 6kae-E | 4.6 | 3.1 | 85 | 141 | 4 | MOLECULE: | HEMOGLOBIN |
| 36:00:00 | 1uiw-C | 4.6 | 2.9 | 83 | 141 | 5 | MOLECULE: | HEMOGLOBIN |
| 37:00:00 | 6kas-C | 4.6 | 3.2 | 86 | 141 | 2 | MOLECULE: | HEMOGLOBIN |
| 38:00:00 | 1lfl-R | 4.6 | 3.4 | 86 | 141 | 3 | MOLECULE: | HEMOGLOBIN |
| 39:00:00 | 3d1k-A | 4.6 | 3.4 | 85 | 142 | 7 | MOLECULE: | HEMOGLOBIN |
| 40:00:00 | 6bnr-A | 4.6 | 3 | 84 | 141 | 4 | MOLECULE: | HEMOGLOBIN |
| 41:00:00 | 7uf6-A | 4.6 | 3 | 84 | 141 | 4 | MOLECULE: | HEMOGLOBIN |
| 42:00:00 | 6ka9-E | 4.6 | 3.4 | 85 | 141 | 5 | MOLECULE: | HEMOGLOBIN |
| 43:00:00 | 5x2s-K | 4.6 | 3.5 | 86 | 140 | 3 | MOLECULE: | HEMOGLOBIN |
| 44:00:00 | 7dy4-G | 4.6 | 3.1 | 85 | 141 | 4 | MOLECULE: | HEMOGLOBIN |
| 45:00:00 | 6kau-C | 4.6 | 3 | 84 | 141 | 4 | MOLECULE: | HEMOGLOBIN |
| 46:00:00 | 1j40-C | 4.6 | 3 | 84 | 141 | 5 | MOLECULE: | HEMOGLOBIN |
| 47:00:00 | 4n7n-G | 4.6 | 3.4 | 87 | 141 | 5 | MOLECULE: | HEMOGLOBIN |
| 48:00:00 | 1bz1-A | 4.6 | 3 | 84 | 142 | 5 | MOLECULE: | PROTEIN (HE |
| 49:00:00 | 1fhj-A | 4.6 | 3.4 | 86 | 141 | 6 | MOLECULE: | HEMOGLOBIN |
| 50:00:00 | 6jp1-A | 4.6 | 3.9 | 89 | 151 | 12 | MOLECULE: | MYOGLOBIN |
| 51:00:00 | 1o1k-C | 4.6 | 3.6 | 87 | 140 | 2 | MOLECULE: | HEMOGLOBIN |
| 52:00:00 | 3qjc-A | 4.6 | 3.5 | 86 | 141 | 3 | MOLECULE: | HEMOGLOBIN |
| 53:00:00 | 1bab-C | 4.6 | 3.3 | 85 | 142 | 5 | MOLECULE: | HEMOGLOBIN |
| 54:00:00 | 5x2r-G | 4.6 | 3.5 | 86 | 140 | 3 | MOLECULE: | HEMOGLOBIN |
| 55:00:00 | 2vwa-A | 4.5 | 2.8 | 71 | 99 | 13 | MOLECULE: | PUTATIVE UN |
| 56:00:00 | 1d8u-A | 4.5 | 5.5 | 95 | 165 | 9 | MOLECULE: | NON-SYMBIOT |
| 57:00:00 | 5ziq-A | 4.5 | 4 | 89 | 155 | 7 | MOLECULE: | GLOBIN PROT |
| 58:00:00 | 1yhu-U | 4.5 | 3.3 | 84 | 145 | 11 | MOLECULE: | HEMOGLOBIN |
| 59:00:00 | 4mqj-D | 4.5 | 3.4 | 85 | 145 | 7 | MOLECULE: | HEMOGLOBIN |
| 60:00:00 | 3gou-C | 4.5 | 3.8 | 88 | 141 | 5 | MOLECULE: | HEMOGLOBIN |
| 61:00:00 | 1iwh-A | 4.5 | 3.2 | 85 | 141 | 4 | MOLECULE: | HEMOGLOBIN |
| 62:00:00 | 1qpw-A | 4.5 | 3.1 | 85 | 141 | 5 | MOLECULE: | PORCINE HEM |
| 63:00:00 | 5x2s-I | 4.5 | 3.6 | 87 | 140 | 2 | MOLECULE: | HEMOGLOBIN |
| 64:00:00 | 1ux9-A | 4.5 | 4.1 | 89 | 154 | 10 | MOLECULE: | CYTOGLOBIN |
| 65:00:00 | 1i3d-A | 4.5 | 3.4 | 85 | 146 | 7 | MOLECULE: | HEMOGLOBIN |
| 66:00:00 | 3gqr-E | 4.5 | 3.6 | 87 | 141 | 5 | MOLECULE: | HEMOGLOBIN |

| | | | | | | | | |
|---|---|---|---|---|---|---|---|---|
| 67:00:00 | 1hbr-B | 4.5 | 3.5 | 85 | 143 | 9 | MOLECULE: | PROTEIN (HE |
| 68:00:00 | 3mkb-A | 4.5 | 3.4 | 86 | 140 | 6 | MOLECULE: | HEMOGLOBIN |
| 69:00:00 | 1jy7-W | 4.5 | 3.5 | 86 | 141 | 2 | MOLECULE: | HEMOGLOBIN |
| 70:00:00 | 4mqk-B | 4.5 | 3.5 | 86 | 145 | 6 | MOLECULE: | HEMOGLOBIN |
| 71:00:00 | 1ury-A | 4.5 | 4.1 | 89 | 154 | 10 | MOLECULE: | CYTOGLOBIN |
| 72:00:00 | 1jy7-U | 4.5 | 3.3 | 86 | 141 | 2 | MOLECULE: | HEMOGLOBIN |
| 73:00:00 | 1j40-E | 4.5 | 3.1 | 86 | 141 | 5 | MOLECULE: | HEMOGLOBIN |
| 74:00:00 | 1lfq-A | 4.5 | 3.1 | 84 | 141 | 5 | MOLECULE: | HEMOGLOBIN |
| 75:00:00 | 1nej-A | 4.5 | 3.6 | 86 | 141 | 3 | MOLECULE: | HEMOGLOBIN |
| 76:00:00 | 3pi9-A | 4.5 | 3.5 | 86 | 141 | 5 | MOLECULE: | HEMOGLOBIN |
| 77:00:00 | 1lfy-A | 4.5 | 3.6 | 88 | 141 | 3 | MOLECULE: | HEMOGLOBIN |
| 78:00:00 | 2zlu-A | 4.5 | 3.6 | 87 | 140 | 2 | MOLECULE: | HEMOGLOBIN |
| 79:00:00 | 2qls-A | 4.5 | 3.5 | 87 | 141 | 6 | MOLECULE: | HEMOGLOBIN |
| 80:00:00 | 2mhb-A | 4.5 | 3.4 | 86 | 141 | 3 | MOLECULE: | HEMOGLOBIN |
| 81:00:00 | 1hco-A | 4.5 | 3.1 | 86 | 141 | 5 | MOLECULE: | HEMOGLOBIN |
| 82:00:00 | 1cmy-A | 4.5 | 3.5 | 86 | 141 | 3 | MOLECULE: | HEMOGLOBIN |
| 83:00:00 | 1hho-A | 4.5 | 3.6 | 87 | 141 | 2 | MOLECULE: | HEMOGLOBIN |
| 84:00:00 | 6hk2-C | 4.5 | 4 | 89 | 141 | 2 | MOLECULE: | HEMOGLOBIN |
| 85:00:00 | 6lcw-A | 4.5 | 3.1 | 85 | 141 | 6 | MOLECULE: | HEMOGLOBIN |
| 86:00:00 | 1ns9-A | 4.5 | 3.6 | 87 | 141 | 2 | MOLECULE: | HEMOGLOBIN |
| 87:00:00 | 5e29-C | 4.5 | 3.1 | 85 | 141 | 6 | MOLECULE: | HEMOGLOBIN |
| 88:00:00 | 7jy1-A | 4.5 | 3 | 85 | 141 | 5 | MOLECULE: | HEMOGLOBIN |
| 89:00:00 | 1g0b-A | 4.5 | 3.2 | 85 | 141 | 4 | MOLECULE: | HEMOGLOBIN |
| 90:00:00 | 6hal-A | 4.5 | 3.6 | 87 | 139 | 2 | MOLECULE: | HEMOGLOBIN |
| 91:00:00 | 3pia-C | 4.5 | 3.5 | 85 | 141 | 5 | MOLECULE: | HEMOGLOBIN |
| 92:00:00 | 7jy0-A | 4.5 | 3.4 | 86 | 141 | 2 | MOLECULE: | HEMOGLOBIN |
| 93:00:00 | 1ns6-A | 4.5 | 3.2 | 86 | 141 | 3 | MOLECULE: | HEMOGLOBIN |
| 94:00:00 | 3a0g-A | 4.5 | 3.5 | 86 | 139 | 6 | MOLECULE: | HEMOGLOBIN |
| 95:00:00 | 1uiw-G | 4.5 | 3.1 | 85 | 141 | 6 | MOLECULE: | HEMOGLOBIN |
| 96:00:00 | 6lcx-A | 4.5 | 3.1 | 85 | 141 | 6 | MOLECULE: | HEMOGLOBIN |
| 97:00:00 | 1uiw-E | 4.5 | 2.9 | 83 | 141 | 5 | MOLECULE: | HEMOGLOBIN |
| 98:00:00 | 2qsp-A | 4.5 | 3.5 | 85 | 141 | 5 | MOLECULE: | HEMOGLOBIN |
| 99:00:00 | 1j41-A | 4.5 | 3.2 | 86 | 141 | 5 | MOLECULE: | HEMOGLOBIN |
| 0:00 | 3d1a-C | 4.5 | 3.9 | 88 | 141 | 3 | MOLECULE: | HEMOGLOBIN |
| 1:00 | 3pel-A | 4.5 | 3.5 | 87 | 141 | 5 | MOLECULE: | HEMOGLOBIN |
| 2:00 | 1mko-C | 4.5 | 3.1 | 85 | 141 | 4 | MOLECULE: | HEMOGLOBIN |
| 3:00 | 1a9w-C | 4.5 | 3.1 | 85 | 141 | 2 | MOLECULE: | HEMOGLOBIN |
| 4:00 | 6r2o-C | 4.5 | 3.4 | 86 | 141 | 2 | MOLECULE: | HEMOGLOBIN |
| 5:00 | 5c6e-A | 4.5 | 3.2 | 85 | 141 | 4 | MOLECULE: | HEMOGLOBIN |
| 6:00 | 5ee4-C | 4.5 | 3 | 85 | 141 | 5 | MOLECULE: | HPUA |
| 7:00 | 3vre-A | 4.5 | 3.5 | 88 | 141 | 2 | MOLECULE: | HEMOGLOBIN |
| 8:00 | 1out-A | 4.5 | 3 | 84 | 142 | 10 | MOLECULE: | HEMOGLOBIN |
| 9:00 | 3d1a-A | 4.5 | 4 | 89 | 141 | 3 | MOLECULE: | HEMOGLOBIN |
| 10:00 | 5ksi-C | 4.5 | 2.9 | 83 | 141 | 5 | MOLECULE: | HEMOGLOBIN |

| | | | | | | | | |
|---|---|---|---|---|---|---|---|---|
| 11:00 | 3gys-C | 4.5 | 3.5 | 87 | 139 | 6 | MOLECULE: | HEMOGLOBIN |
| 12:00 | 3whm-A | 4.5 | 3 | 84 | 140 | 4 | MOLECULE: | HEMOGLOBIN |
| 13:00 | 3pi8-A | 4.5 | 3.4 | 85 | 141 | 5 | MOLECULE: | HEMOGLOBIN |
| 14:00 | 2qls-C | 4.5 | 3.5 | 87 | 141 | 5 | MOLECULE: | HEMOGLOBIN |
| 15:00 | 6kai-A | 4.5 | 3.1 | 86 | 141 | 5 | MOLECULE: | HEMOGLOBIN |
| 16:00 | 2qss-A | 4.5 | 3.4 | 85 | 141 | 5 | MOLECULE: | HEMOGLOBIN |
| 17:00 | 1fn3-A | 4.5 | 3.6 | 88 | 141 | 2 | MOLECULE: | HEMOGLOBIN |
| 18:00 | 1m9p-C | 4.5 | 3.5 | 87 | 141 | 3 | MOLECULE: | HEMOGLOBIN |
| 19:00 | 3wcp-A | 4.5 | 3.2 | 86 | 141 | 5 | MOLECULE: | HEMOGLOBIN |
| 20:00 | 5ksj-C | 4.5 | 3 | 84 | 141 | 4 | MOLECULE: | HEMOGLOBIN |
| 21:00 | 3cy5-C | 4.5 | 3.5 | 87 | 141 | 5 | MOLECULE: | HEMOGLOBIN |
| 22:00 | 4mqj-C | 4.5 | 3.5 | 86 | 141 | 3 | MOLECULE: | HEMOGLOBIN |
| 23:00 | 4mqj-A | 4.5 | 3.5 | 86 | 141 | 3 | MOLECULE: | HEMOGLOBIN |
| 24:00:00 | 1b86-A | 4.5 | 3.4 | 86 | 141 | 6 | MOLECULE: | PROTEIN (HE |
| 25:00:00 | 1cmy-C | 4.5 | 3.1 | 84 | 141 | 5 | MOLECULE: | HEMOGLOBIN |
| 26:00:00 | 3eu1-C | 4.5 | 3.5 | 87 | 141 | 5 | MOLECULE: | HEMOGLOBIN |
| 27:00:00 | 3hyu-A | 4.5 | 3.5 | 86 | 141 | 6 | MOLECULE: | HEMOGLOBIN |
| 28:00:00 | 6kye-G | 4.5 | 3.8 | 88 | 140 | 3 | MOLECULE: | HEMOGLOBIN |
| 29:00:00 | 4f4o-G | 4.5 | 3.4 | 86 | 141 | 5 | MOLECULE: | HEMOGLOBIN |
| 30:00:00 | 7vde-A | 4.5 | 3.5 | 87 | 141 | 2 | MOLECULE: | HEMOGLOBIN |
| 31:00:00 | 3r5i-C | 4.5 | 3.5 | 85 | 141 | 4 | MOLECULE: | HEMOGLOBIN |
| 32:00:00 | 2in4-A | 4.5 | 3.9 | 88 | 153 | 14 | MOLECULE: | MYOGLOBIN |
| 33:00:00 | 3s65-C | 4.5 | 3.4 | 86 | 141 | 2 | MOLECULE: | HEMOGLOBIN |
| 34:00:00 | 4mqk-E | 4.5 | 3.6 | 87 | 138 | 3 | MOLECULE: | HEMOGLOBIN |
| 35:00:00 | 7jjq-A | 4.5 | 3.4 | 85 | 140 | 5 | MOLECULE: | HEMOGLOBIN |
| 36:00:00 | 1g09-C | 4.5 | 3.5 | 85 | 141 | 5 | MOLECULE: | HEMOGLOBIN |
| 37:00:00 | 1ouu-A | 4.5 | 3.4 | 85 | 142 | 9 | MOLECULE: | HEMOGLOBIN |
| 38:00:00 | 1ibe-A | 4.5 | 3.4 | 86 | 141 | 5 | MOLECULE: | HEMOGLOBIN |
| 39:00:00 | 1sdl-A | 4.5 | 3.6 | 88 | 141 | 3 | MOLECULE: | HEMOGLOBIN |
| 40:00:00 | 1dke-A | 4.5 | 3 | 84 | 141 | 6 | MOLECULE: | HEMOGLOBIN: |
| 41:00:00 | 5e83-A | 4.5 | 3.5 | 86 | 141 | 2 | MOLECULE: | HEMOGLOBIN |
| 42:00:00 | 5x2u-K | 4.5 | 2.9 | 84 | 140 | 5 | MOLECULE: | HEMOGLOBIN |
| 43:00:00 | 1a3n-A | 4.5 | 2.9 | 84 | 141 | 6 | MOLECULE: | HEMOGLOBIN |
| 44:00:00 | 3pi9-C | 4.5 | 3.5 | 85 | 141 | 5 | MOLECULE: | HEMOGLOBIN |
| 45:00:00 | 7dy4-A | 4.5 | 3.1 | 85 | 141 | 6 | MOLECULE: | HEMOGLOBIN |
| 46:00:00 | 6tb2-A | 4.5 | 3.4 | 85 | 141 | 5 | MOLECULE: | HEMOGLOBIN |
| 47:00:00 | 2rao-A | 4.5 | 3.5 | 85 | 141 | 5 | MOLECULE: | HEMOGLOBIN |
| 48:00:00 | 1ydz-A | 4.5 | 3.4 | 86 | 141 | 3 | MOLECULE: | HEMOGLOBIN |
| 49:00:00 | 4wjg-Z | 4.5 | 3.5 | 86 | 141 | 2 | MOLECULE: | HEMOGLOBIN |
| 50:00:00 | 6ka9-G | 4.5 | 2.9 | 84 | 141 | 6 | MOLECULE: | HEMOGLOBIN |
| 51:00:00 | 4f4o-A | 4.5 | 3.5 | 86 | 141 | 3 | MOLECULE: | HEMOGLOBIN |
| 52:00:00 | 5ksj-A | 4.5 | 3.1 | 85 | 141 | 4 | MOLECULE: | HEMOGLOBIN |
| 53:00:00 | 4wjg-U | 4.5 | 3.6 | 87 | 141 | 2 | MOLECULE: | HEMOGLOBIN |
| 54:00:00 | 1shr-C | 4.5 | 3.5 | 86 | 141 | 2 | MOLECULE: | HEMOGLOBIN |

| | | | | | | | | |
|---|---|---|---|---|---|---|---|---|
| 55:00:00 | 1j3y-A | 4.5 | 2.9 | 84 | 141 | 6 | MOLECULE: | HEMOGLOBIN |
| 56:00:00 | 1fn3-C | 4.5 | 3.6 | 88 | 141 | 2 | MOLECULE: | HEMOGLOBIN |
| 57:00:00 | 1g0a-C | 4.5 | 3.5 | 86 | 141 | 5 | MOLECULE: | HEMOGLOBIN |
| 58:00:00 | 3dhr-E | 4.5 | 3.5 | 86 | 141 | 6 | MOLECULE: | HEMOGLOBIN |
| 59:00:00 | 7ud8-C | 4.5 | 3.5 | 85 | 141 | 4 | MOLECULE: | HEMOGLOBIN |
| 60:00:00 | 7ud7-C | 4.5 | 3.1 | 85 | 141 | 6 | MOLECULE: | HEMOGLOBIN |
| 61:00:00 | 7dy3-C | 4.5 | 3 | 84 | 141 | 6 | MOLECULE: | HEMOGLOBIN |
| 62:00:00 | 4mqk-A | 4.5 | 3.5 | 86 | 141 | 3 | MOLECULE: | HEMOGLOBIN |
| 63:00:00 | 2qu0-C | 4.5 | 3.6 | 87 | 141 | 3 | MOLECULE: | HEMOGLOBIN |
| 64:00:00 | 1thb-A | 4.5 | 3.1 | 85 | 141 | 6 | MOLECULE: | HEMOGLOBIN |
| 65:00:00 | 1y4b-C | 4.5 | 3.4 | 87 | 141 | 3 | MOLECULE: | HEMOGLOBIN |
| 66:00:00 | 4wjg-K | 4.5 | 3.6 | 87 | 141 | 2 | MOLECULE: | HEMOGLOBIN |
| 67:00:00 | 5ni1-C | 4.5 | 3.5 | 87 | 141 | 2 | MOLECULE: | HEMOGLOBIN |
| 68:00:00 | 3gqp-C | 4.5 | 3.5 | 87 | 141 | 6 | MOLECULE: | HEMOGLOBIN |
| 69:00:00 | 1y8k-C | 4.5 | 3 | 84 | 141 | 4 | MOLECULE: | HEMOGLOBIN |
| 70:00:00 | 1v4x-C | 4.5 | 3.3 | 85 | 143 | 8 | MOLECULE: | HEMOGLOBIN |
| 71:00:00 | 7k4m-I | 4.5 | 3 | 84 | 140 | 4 | MOLECULE: | HEMOGLOBIN |
| 72:00:00 | 4wjg-F | 4.5 | 3.6 | 87 | 141 | 2 | MOLECULE: | HEMOGLOBIN |
| 73:00:00 | 1j3z-A | 4.5 | 2.9 | 84 | 141 | 6 | MOLECULE: | HEMOGLOBIN |
| 74:00:00 | 1yff-A | 4.5 | 3.5 | 87 | 141 | 2 | MOLECULE: | HEMOGLOBIN |
| 75:00:00 | 7jxz-C | 4.5 | 3.1 | 85 | 141 | 2 | MOLECULE: | HEMOGLOBIN |
| 76:00:00 | 1a9w-A | 4.5 | 3.5 | 86 | 141 | 2 | MOLECULE: | HEMOGLOBIN |
| 77:00:00 | 2d60-A | 4.5 | 2.9 | 84 | 141 | 6 | MOLECULE: | HEMOGLOBIN |
| 78:00:00 | 1si4-A | 4.5 | 3.4 | 86 | 141 | 2 | MOLECULE: | HEMOGLOBIN |
| 79:00:00 | 4f4o-J | 4.5 | 3.5 | 86 | 141 | 3 | MOLECULE: | HEMOGLOBIN |
| 80:00:00 | 4yu4-A | 4.5 | 3.5 | 86 | 141 | 5 | MOLECULE: | HEMOGLOBIN |
| 81:00:00 | 3wcp-C | 4.5 | 3.2 | 86 | 141 | 5 | MOLECULE: | HEMOGLOBIN |
| 82:00:00 | 3gys-E | 4.5 | 3 | 84 | 138 | 6 | MOLECULE: | HEMOGLOBIN |
| 83:00:00 | 6hal-C | 4.5 | 3.7 | 87 | 139 | 2 | MOLECULE: | HEMOGLOBIN |
| 84:00:00 | 4wjg-A | 4.5 | 3.6 | 87 | 141 | 2 | MOLECULE: | HEMOGLOBIN |
| 85:00:00 | 6lcx-E | 4.5 | 3.1 | 86 | 141 | 5 | MOLECULE: | HEMOGLOBIN |
| 86:00:00 | 1j40-A | 4.5 | 3.2 | 86 | 141 | 5 | MOLECULE: | HEMOGLOBIN |
| 87:00:00 | 8pur-A | 4.5 | 3.6 | 87 | 140 | 3 | MOLECULE: | HEMOGLOBIN, |
| 88:00:00 | 6kah-A | 4.5 | 3.1 | 86 | 141 | 5 | MOLECULE: | HEMOGLOBIN |
| 89:00:00 | 6zmx-C | 4.5 | 3.4 | 85 | 138 | 7 | MOLECULE: | HEMOGLOBIN |
| 90:00:00 | 4f4o-D | 4.5 | 3.4 | 86 | 141 | 5 | MOLECULE: | HEMOGLOBIN |
| 91:00:00 | 1jy7-R | 4.5 | 3.5 | 86 | 141 | 2 | MOLECULE: | HEMOGLOBIN |
| 92:00:00 | 1thb-C | 4.5 | 3.6 | 88 | 141 | 3 | MOLECULE: | HEMOGLOBIN |
| 93:00:00 | 1ygf-A | 4.5 | 3.1 | 86 | 141 | 5 | MOLECULE: | HEMOGLOBIN |
| 94:00:00 | 4fc3-A | 4.5 | 3.4 | 86 | 140 | 2 | MOLECULE: | HEMOGLOBIN |
| 95:00:00 | 1y8i-C | 4.5 | 3 | 84 | 141 | 4 | MOLECULE: | HEMOGLOBIN |
| 96:00:00 | 5ksi-A | 4.5 | 3 | 84 | 141 | 6 | MOLECULE: | HEMOGLOBIN |
| 97:00:00 | 7jy1-C | 4.5 | 3 | 84 | 141 | 6 | MOLECULE: | HEMOGLOBIN |
| 98:00:00 | 6z4r-A | 4.5 | 4 | 89 | 154 | 11 | MOLECULE: | MYOGLOBIN |

| | | | | | | | | |
|---|---|---|---|---|---|---|---|---|
| 99:00:00 | 3odq-A | 4.5 | 3.4 | 86 | 141 | 3 | MOLECULE: | HEMOGLOBIN |
| 0:00 | 1jy7-A | 4.5 | 3 | 83 | 141 | 5 | MOLECULE: | HEMOGLOBIN |
| 1:00 | 3b75-S | 4.5 | 3.5 | 86 | 141 | 2 | MOLECULE: | HEMOGLOBIN |
| 2:00 | 4n7o-E | 4.5 | 3.5 | 86 | 141 | 3 | MOLECULE: | HEMOGLOBIN |
| 3:00 | 6nbc-A | 4.5 | 3 | 83 | 140 | 5 | MOLECULE: | HEMOGLOBIN |
| 4:00 | 3qjd-C | 4.5 | 3.5 | 87 | 141 | 5 | MOLECULE: | HEMOGLOBIN |
| 5:00 | 2eb9-A | 4.5 | 3.4 | 84 | 150 | 13 | MOLECULE: | MYOGLOBIN |
| 6:00 | 2h8f-A | 4.5 | 2.8 | 83 | 142 | 7 | MOLECULE: | HEMOGLOBIN |
| 7:00 | 1v4u-C | 4.5 | 3.1 | 85 | 143 | 7 | MOLECULE: | HEMOGLOBIN |
| 8:00 | 1y01-B | 4.5 | 3.2 | 85 | 126 | 4 | MOLECULE: | ALPHA-HEMOG |
| 9:00 | 7qu4-B | 4.5 | 3 | 83 | 141 | 5 | MOLECULE: | HEMOGLOBIN |
| 10:00 | 4odc-A | 4.5 | 3.4 | 85 | 142 | 7 | MOLECULE: | HEMOGLOBIN |
| 11:00 | 1nej-C | 4.5 | 3 | 83 | 141 | 5 | MOLECULE: | HEMOGLOBIN |
| 12:00 | 3nfe-A | 4.5 | 3 | 84 | 142 | 7 | MOLECULE: | HEMOGLOBIN |
| 13:00 | 1hbh-C | 4.5 | 2.8 | 83 | 142 | 7 | MOLECULE: | HEMOGLOBIN |
| 14:00 | 1mko-A | 4.5 | 3.5 | 86 | 141 | 3 | MOLECULE: | HEMOGLOBIN |
| 15:00 | 3gkv-A | 4.5 | 3.4 | 85 | 142 | 7 | MOLECULE: | HEMOGLOBIN |
| 16:00 | 2peg-A | 4.5 | 3.4 | 85 | 142 | 7 | MOLECULE: | HEMOGLOBIN |
| 17:00 | 4n7p-E | 4.5 | 3.4 | 86 | 141 | 3 | MOLECULE: | HEMOGLOBIN |
| 18:00 | 2h8d-A | 4.5 | 3.3 | 85 | 142 | 7 | MOLECULE: | HEMOGLOBIN |
| 19:00 | 3gqg-A | 4.5 | 3.3 | 85 | 142 | 7 | MOLECULE: | HEMOGLOBIN |
| 20:00 | 5ni1-A | 4.5 | 3.5 | 87 | 141 | 2 | MOLECULE: | HEMOGLOBIN |
| 21:00 | 5lfg-A | 4.5 | 3.4 | 86 | 142 | 6 | MOLECULE: | HEMOGLOBIN |
| 22:00 | 7qu4-A | 4.5 | 3 | 84 | 142 | 4 | MOLECULE: | HEMOGLOBIN |
| 23:00 | 5woh-C | 4.5 | 3.4 | 86 | 137 | 3 | MOLECULE: | HEMOGLOBIN |
| 24:00:00 | 5x2t-K | 4.5 | 3.2 | 85 | 140 | 4 | MOLECULE: | HEMOGLOBIN |
| 25:00:00 | 2dn1-A | 4.5 | 3.5 | 85 | 140 | 5 | MOLECULE: | HEMOGLOBIN |
| 26:00:00 | 1la6-A | 4.5 | 3.4 | 85 | 142 | 7 | MOLECULE: | HEMOGLOBIN |
| 27:00:00 | 1t1n-A | 4.5 | 3.3 | 85 | 142 | 7 | MOLECULE: | PROTEIN (HE |
| 28:00:00 | 3s48-C | 4.5 | 3.4 | 86 | 138 | 3 | MOLECULE: | IRON-REGULA |
| 29:00:00 | 3gqg-C | 4.5 | 3.3 | 85 | 142 | 7 | MOLECULE: | HEMOGLOBIN |
| 30:00:00 | 2h8d-C | 4.5 | 3.3 | 85 | 142 | 7 | MOLECULE: | HEMOGLOBIN |
| 31:00:00 | 1pbx-A | 4.5 | 3.3 | 85 | 142 | 7 | MOLECULE: | HEMOGLOBIN |
| 32:00:00 | 1hbh-A | 4.5 | 2.8 | 83 | 142 | 7 | MOLECULE: | HEMOGLOBIN |
| 33:00:00 | 1o1o-C | 4.5 | 3.5 | 86 | 141 | 5 | MOLECULE: | HEMOGLOBIN |
| 34:00:00 | 1ouu-C | 4.5 | 3.4 | 86 | 142 | 9 | MOLECULE: | HEMOGLOBIN |
| 35:00:00 | 1o1k-A | 4.5 | 3 | 84 | 140 | 6 | MOLECULE: | HEMOGLOBIN |
| 36:00:00 | 4mqk-H | 4.5 | 3.4 | 85 | 145 | 7 | MOLECULE: | HEMOGLOBIN |
| 37:00:00 | 4mqk-D | 4.5 | 3.5 | 86 | 145 | 6 | MOLECULE: | HEMOGLOBIN |
| 38:00:00 | 3ng6-C | 4.5 | 2.8 | 83 | 142 | 7 | MOLECULE: | HEMOGLOBIN |
| 39:00:00 | 5x2t-I | 4.5 | 3.2 | 85 | 140 | 4 | MOLECULE: | HEMOGLOBIN |
| 40:00:00 | 5x2u-C | 4.5 | 3.5 | 86 | 140 | 3 | MOLECULE: | HEMOGLOBIN |
| 41:00:00 | 5wog-B | 4.5 | 3.4 | 85 | 137 | 5 | MOLECULE: | HEMOGLOBIN |
| 42:00:00 | 3ng6-A | 4.5 | 3.3 | 85 | 142 | 7 | MOLECULE: | HEMOGLOBIN |

| | | | | | | | | |
|---|---|---|---|---|---|---|---|---|
| 43:00:00 | 1tu9-A | 4.4 | 4.2 | 83 | 131 | 8 | MOLECULE: | HYPOTHETICA |
| 44:00:00 | 5hv1-A | 4.4 | 4.4 | 93 | 847 | 8 | MOLECULE: | PHOSPHOENOL |
| 45:00:00 | 4hrr-A | 4.4 | 3.5 | 86 | 149 | 3 | MOLECULE: | GLOBIN-2 A |
| 46:00:00 | 6p5b-A | 4.4 | 4.7 | 93 | 388 | 2 | MOLECULE: | MAVC |
| 47:00:00 | 6bb5-A | 4.4 | 3.4 | 86 | 139 | 5 | MOLECULE: | HEMOGLOBIN |
| 48:00:00 | 3pt8-A | 4.4 | 3.8 | 85 | 151 | 6 | MOLECULE: | HEMOGLOBIN |
| 49:00:00 | 3mkb-C | 4.4 | 3.4 | 85 | 140 | 6 | MOLECULE: | HEMOGLOBIN |
| 50:00:00 | 2olp-B | 4.4 | 4 | 85 | 151 | 5 | MOLECULE: | HEMOGLOBIN |
| 51:00:00 | 3b75-G | 4.4 | 4 | 89 | 141 | 2 | MOLECULE: | HEMOGLOBIN |
| 52:00:00 | 1y8h-C | 4.4 | 3.5 | 85 | 141 | 4 | MOLECULE: | HEMOGLOBIN |
| 53:00:00 | 1hbr-D | 4.4 | 3.5 | 85 | 146 | 9 | MOLECULE: | PROTEIN (HE |
| 54:00:00 | 4iro-C | 4.4 | 3.3 | 85 | 142 | 7 | MOLECULE: | HEMOGLOBIN |
| 55:00:00 | 1lfz-A | 4.4 | 3.4 | 86 | 141 | 3 | MOLECULE: | HEMOGLOBIN |
| 56:00:00 | 2dhb-A | 4.4 | 3 | 83 | 141 | 4 | MOLECULE: | HEMOGLOBIN |
| 57:00:00 | 3s65-A | 4.4 | 3.5 | 85 | 141 | 4 | MOLECULE: | HEMOGLOBIN |
| 58:00:00 | 7qu4-G | 4.4 | 3.5 | 84 | 143 | 7 | MOLECULE: | HEMOGLOBIN |
| 59:00:00 | 4mqj-F | 4.4 | 3.5 | 86 | 146 | 7 | MOLECULE: | HEMOGLOBIN |
| 60:00:00 | 5vmm-C | 4.4 | 3.6 | 88 | 141 | 2 | MOLECULE: | HEMOGLOBIN |
| 61:00:00 | 4mqk-G | 4.4 | 4 | 89 | 140 | 2 | MOLECULE: | HEMOGLOBIN |
| 62:00:00 | 3hf4-E | 4.4 | 3.4 | 86 | 141 | 6 | MOLECULE: | HEMOGLOBIN |
| 63:00:00 | 1o1p-D | 4.4 | 3.6 | 86 | 146 | 6 | MOLECULE: | HEMOGLOBIN |
| 64:00:00 | 1yev-B | 4.4 | 3.5 | 85 | 146 | 7 | MOLECULE: | HEMOGLOBIN |
| 65:00:00 | 1a4f-A | 4.4 | 3.7 | 86 | 141 | 8 | MOLECULE: | HEMOGLOBIN |
| 66:00:00 | 1yeo-B | 4.4 | 3.5 | 86 | 146 | 7 | MOLECULE: | HEMOGLOBIN |
| 67:00:00 | 5x2r-C | 4.4 | 3 | 84 | 140 | 4 | MOLECULE: | HEMOGLOBIN |
| 68:00:00 | 1i3e-A | 4.4 | 3.5 | 86 | 146 | 7 | MOLECULE: | HEMOGLOBIN |
| 69:00:00 | 2d5x-A | 4.4 | 3.5 | 86 | 141 | 3 | MOLECULE: | HEMOGLOBIN |
| 70:00:00 | 3wtg-B | 4.4 | 3.4 | 83 | 146 | 10 | MOLECULE: | HEMOGLOBIN |
| 71:00:00 | 3b75-E | 4.4 | 3.8 | 88 | 141 | 3 | MOLECULE: | HEMOGLOBIN |
| 72:00:00 | 3gqr-G | 4.4 | 3.1 | 85 | 141 | 7 | MOLECULE: | HEMOGLOBIN |
| 73:00:00 | 3gou-A | 4.4 | 3.7 | 88 | 141 | 5 | MOLECULE: | HEMOGLOBIN |
| 74:00:00 | 1y8i-A | 4.4 | 3 | 84 | 141 | 4 | MOLECULE: | HEMOGLOBIN |
| 75:00:00 | 4yu3-A | 4.4 | 3.5 | 86 | 141 | 5 | MOLECULE: | HEMOGLOBIN |
| 76:00:00 | 3hf4-A | 4.4 | 3.4 | 86 | 141 | 6 | MOLECULE: | HEMOGLOBIN |
| 77:00:00 | 4wjg-P | 4.4 | 3.5 | 86 | 141 | 2 | MOLECULE: | HEMOGLOBIN |
| 78:00:00 | 5ker-C | 4.4 | 3.5 | 86 | 140 | 5 | MOLECULE: | ALPHA-GLOBI |
| 79:00:00 | 3gqr-C | 4.4 | 3.3 | 87 | 141 | 6 | MOLECULE: | HEMOGLOBIN |
| 80:00:00 | 4x0l-A | 4.4 | 3.5 | 86 | 141 | 3 | MOLECULE: | HEMOGLOBIN |
| 81:00:00 | 1qpw-C | 4.4 | 3.6 | 87 | 141 | 3 | MOLECULE: | PORCINE HEM |
| 82:00:00 | 1v9q-A | 4.4 | 4.8 | 80 | 154 | 11 | MOLECULE: | MYOGLOBIN |
| 83:00:00 | 2b7h-A | 4.4 | 3.5 | 87 | 141 | 7 | MOLECULE: | HEMOGLOBIN |
| 84:00:00 | 3eok-A | 4.4 | 3 | 84 | 141 | 6 | MOLECULE: | HEMOGLOBIN |
| 85:00:00 | 8puq-A | 4.4 | 3.5 | 86 | 139 | 2 | MOLECULE: | HEMOGLOBIN |
| 86:00:00 | 1a3o-A | 4.4 | 2.9 | 83 | 141 | 6 | MOLECULE: | HEMOGLOBIN |

| | | | | | | | | |
|---|---|---|---|---|---|---|---|---|
| 87:00:00 | 6kah-E | 4.4 | 3.1 | 83 | 141 | 4 | MOLECULE: | HEMOGLOBIN |
| 88:00:00 | 6sva-A | 4.4 | 3.2 | 85 | 139 | 4 | MOLECULE: | HEMOGLOBIN |
| 89:00:00 | 1y8k-A | 4.4 | 3 | 84 | 141 | 4 | MOLECULE: | HEMOGLOBIN |
| 90:00:00 | 2zlv-A | 4.4 | 3.1 | 84 | 141 | 4 | MOLECULE: | HEMOGLOBIN |
| 91:00:00 | 3wr1-A | 4.4 | 3.5 | 87 | 141 | 9 | MOLECULE: | HEMOGLOBIN |
| 92:00:00 | 4mqj-E | 4.4 | 3.9 | 87 | 141 | 2 | MOLECULE: | HEMOGLOBIN |
| 93:00:00 | 1s0h-A | 4.4 | 3.4 | 86 | 141 | 5 | MOLECULE: | HEMOGLOBIN |
| 94:00:00 | 7cue-A | 4.4 | 3.6 | 86 | 141 | 2 | MOLECULE: | HEMOGLOBIN |
| 95:00:00 | 4mqj-G | 4.4 | 3.5 | 86 | 141 | 3 | MOLECULE: | HEMOGLOBIN |
| 96:00:00 | 2rao-C | 4.4 | 3.5 | 86 | 141 | 5 | MOLECULE: | HEMOGLOBIN |
| 97:00:00 | 1qxe-C | 4.4 | 3.4 | 85 | 141 | 4 | MOLECULE: | HEMOGLOBIN |
| 98:00:00 | 1ird-A | 4.4 | 3.6 | 88 | 141 | 3 | MOLECULE: | HEMOGLOBIN |
| 99:00:00 | 2qmb-A | 4.4 | 4.4 | 88 | 141 | 7 | MOLECULE: | HEMOGLOBIN |
| 0:00 | 2qu0-A | 4.4 | 3.6 | 88 | 141 | 3 | MOLECULE: | HEMOGLOBIN |
| 1:00 | 3ic0-C | 4.4 | 3.5 | 85 | 141 | 4 | MOLECULE: | HEMOGLOBIN |
| 2:00 | 6r2o-A | 4.4 | 3 | 84 | 141 | 4 | MOLECULE: | HEMOGLOBIN |
| 3:00 | 4yu4-C | 4.4 | 3.5 | 86 | 141 | 5 | MOLECULE: | HEMOGLOBIN |
| 4:00 | 1hv4-G | 4.4 | 3 | 83 | 141 | 6 | MOLECULE: | HEMOGLOBIN |
| 5:00 | 2zlt-A | 4.4 | 3.5 | 86 | 141 | 2 | MOLECULE: | HEMOGLOBIN |
| 6:00 | 3mjp-A | 4.4 | 3.4 | 86 | 141 | 5 | MOLECULE: | HEMOGLOBIN |
| 7:00 | 3ciu-C | 4.4 | 3.5 | 85 | 141 | 5 | MOLECULE: | HEMOGLOBIN |
| 8:00 | 1hbs-E | 4.4 | 2.9 | 84 | 141 | 4 | MOLECULE: | HEMOGLOBIN |
| 9:00 | 3d4x-A | 4.4 | 3.1 | 85 | 141 | 6 | MOLECULE: | HEMOGLOBIN |
| 10:00 | 1shr-A | 4.4 | 3.5 | 86 | 141 | 2 | MOLECULE: | HEMOGLOBIN |
| 11:00 | 3ic2-C | 4.4 | 3.5 | 85 | 141 | 4 | MOLECULE: | HEMOGLOBIN |
| 12:00 | 3a59-G | 4.4 | 3.6 | 87 | 141 | 5 | MOLECULE: | HEMOGLOBIN |
| 13:00 | 1qxd-C | 4.4 | 3 | 83 | 141 | 5 | MOLECULE: | HEMOGLOBIN |
| 14:00 | 2b7h-C | 4.4 | 3.5 | 87 | 141 | 7 | MOLECULE: | HEMOGLOBIN |
| 15:00 | 3d4x-C | 4.4 | 3.1 | 84 | 141 | 6 | MOLECULE: | HEMOGLOBIN |
| 16:00 | 7dy3-A | 4.4 | 3 | 84 | 141 | 6 | MOLECULE: | HEMOGLOBIN |
| 17:00 | 3fh9-A | 4.4 | 3.6 | 87 | 141 | 5 | MOLECULE: | HEMOGLOBIN |
| 18:00 | 1c40-A | 4.4 | 4.1 | 87 | 141 | 8 | MOLECULE: | PROTEIN (HE |
| 19:00 | 5u3i-C | 4.4 | 3.4 | 84 | 140 | 4 | MOLECULE: | HEMOGLOBIN |
| 20:00 | 3gys-A | 4.4 | 3.6 | 87 | 138 | 5 | MOLECULE: | HEMOGLOBIN |
| 21:00 | 4h2l-A | 4.4 | 3.5 | 87 | 141 | 8 | MOLECULE: | ALPHA-GLOBI |
| 22:00 | 3kmf-A | 4.4 | 3.1 | 85 | 141 | 6 | MOLECULE: | HEMOGLOBIN |
| 23:00 | 2ohb-A | 4.4 | 4.5 | 93 | 154 | 8 | MOLECULE: | MYOGLOBIN |
| 24:00:00 | 2mgj-A | 4.4 | 4.5 | 93 | 154 | 8 | MOLECULE: | MYOGLOBIN |
| 25:00:00 | 3pi1-B | 4.4 | 3.9 | 86 | 151 | 5 | MOLECULE: | HEMOGLOBIN |
| 26:00:00 | 2h8f-C | 4.4 | 3.3 | 85 | 142 | 7 | MOLECULE: | HEMOGLOBIN |
| 27:00:00 | 1ouu-B | 4.4 | 3.6 | 85 | 146 | 7 | MOLECULE: | HEMOGLOBIN |
| 28:00:00 | 3nfe-C | 4.4 | 3 | 84 | 142 | 7 | MOLECULE: | HEMOGLOBIN |
| 29:00:00 | 4g51-A | 4.4 | 3.3 | 85 | 142 | 7 | MOLECULE: | HEMOGLOBIN |
| 30:00:00 | 6zmy-D | 4.4 | 3.5 | 85 | 146 | 9 | MOLECULE: | HEMOGLOBIN |

| | | | | | | | | |
|---|---|---|---|---|---|---|---|---|
| 31:00:00 | 7qu4-H | 4.4 | 3.5 | 85 | 144 | 6 | MOLECULE: | HEMOGLOBIN |
| 32:00:00 | 6zmx-D | 4.4 | 3.5 | 85 | 146 | 9 | MOLECULE: | HEMOGLOBIN |
| 33:00:00 | 1j7s-D | 4.4 | 3.3 | 84 | 146 | 7 | MOLECULE: | HEMOGLOBIN |
| 34:00:00 | 6zmy-C | 4.4 | 3.3 | 84 | 137 | 6 | MOLECULE: | HEMOGLOBIN |
| 35:00:00 | 2g0s-A | 4.4 | 4.5 | 93 | 154 | 8 | MOLECULE: | MYOGLOBIN |
| 36:00:00 | 5x2t-E | 4.4 | 3.6 | 86 | 140 | 3 | MOLECULE: | HEMOGLOBIN |
| 37:00:00 | 1mz0-A | 4.4 | 4.5 | 93 | 154 | 8 | MOLECULE: | MYOGLOBIN |
| 38:00:00 | 2dc3-B | 4.4 | 4.2 | 98 | 168 | 9 | MOLECULE: | CYTOGLOBIN |
| 39:00:00 | 1faw-D | 4.4 | 3.5 | 85 | 146 | 11 | MOLECULE: | HEMOGLOBIN |
| 40:00:00 | 1s5x-A | 4.4 | 3.5 | 85 | 142 | 7 | MOLECULE: | HEMOGLOBIN |
| 41:00:00 | 1cio-A | 4.4 | 4.5 | 93 | 154 | 8 | MOLECULE: | PROTEIN (MY |
| 42:00:00 | 1ye2-D | 4.4 | 3.5 | 86 | 146 | 7 | MOLECULE: | HEMOGLOBIN |
| 43:00:00 | 6zmy-B | 4.4 | 3.5 | 85 | 146 | 9 | MOLECULE: | HEMOGLOBIN |
| 44:00:00 | 1y4b-D | 4.4 | 3.2 | 85 | 146 | 7 | MOLECULE: | HEMOGLOBIN |
| 45:00:00 | 1s5y-A | 4.4 | 3.5 | 85 | 142 | 7 | MOLECULE: | HEMOGLOBIN |
| 46:00:00 | 1c40-B | 4.4 | 3.5 | 85 | 146 | 11 | MOLECULE: | PROTEIN (HE |
| 47:00:00 | 5vmm-A | 4.4 | 3.6 | 87 | 139 | 3 | MOLECULE: | HEMOGLOBIN |
| 48:00:00 | 1yeu-B | 4.4 | 3.4 | 84 | 146 | 7 | MOLECULE: | HEMOGLOBIN |
| 49:00:00 | 1out-B | 4.4 | 3.8 | 87 | 146 | 7 | MOLECULE: | HEMOGLOBIN |
| 50:00:00 | 7ud7-D | 4.4 | 3.5 | 86 | 146 | 7 | MOLECULE: | HEMOGLOBIN |
| 51:00:00 | 6zmx-B | 4.4 | 3.5 | 85 | 146 | 9 | MOLECULE: | HEMOGLOBIN |
| 52:00:00 | 5lfg-C | 4.4 | 3.4 | 87 | 142 | 6 | MOLECULE: | HEMOGLOBIN |
| 53:00:00 | 1j7s-B | 4.4 | 3.3 | 84 | 146 | 7 | MOLECULE: | HEMOGLOBIN |
| 54:00:00 | 1ouu-D | 4.4 | 3.6 | 85 | 146 | 7 | MOLECULE: | HEMOGLOBIN |
| 55:00:00 | 5x2u-E | 4.4 | 3.2 | 85 | 140 | 4 | MOLECULE: | HEMOGLOBIN |
| 56:00:00 | 5x2u-I | 4.4 | 3.1 | 85 | 140 | 5 | MOLECULE: | HEMOGLOBIN |
| 57:00:00 | 1a01-B | 4.4 | 3.6 | 86 | 146 | 7 | MOLECULE: | HEMOGLOBIN |
| 58:00:00 | 1rqa-B | 4.4 | 3.5 | 86 | 146 | 7 | MOLECULE: | HEMOGLOBIN |
| 59:00:00 | 4hrt-E | 4.4 | 3.4 | 85 | 149 | 4 | MOLECULE: | GLOBIN-2 A |
| 60:00:00 | 1faw-B | 4.4 | 3.5 | 85 | 146 | 11 | MOLECULE: | HEMOGLOBIN |
| 61:00:00 | 1myz-A | 4.4 | 4.5 | 93 | 154 | 8 | MOLECULE: | MYOGLOBIN |
| 62:00:00 | 1y8w-A | 4.4 | 4.3 | 87 | 141 | 3 | MOLECULE: | HEMOGLOBIN |
| 63:00:00 | 1ch7-A | 4.4 | 4.4 | 93 | 154 | 8 | MOLECULE: | PROTEIN (MY |
| 64:00:00 | 1o1p-B | 4.4 | 3.3 | 84 | 146 | 7 | MOLECULE: | HEMOGLOBIN |
| 65:00:00 | 1dxu-B | 4.4 | 3.3 | 84 | 146 | 7 | MOLECULE: | HEMOGLOBIN |
| 66:00:00 | 1a00-D | 4.4 | 3.2 | 84 | 146 | 7 | MOLECULE: | HEMOGLOBIN |
| 67:00:00 | 1cik-A | 4.4 | 4.4 | 91 | 154 | 11 | MOLECULE: | PROTEIN (MY |
| 68:00:00 | 5x2r-A | 4.4 | 3.6 | 87 | 140 | 2 | MOLECULE: | HEMOGLOBIN |
| 69:00:00 | 1y4g-B | 4.4 | 3.5 | 86 | 146 | 7 | MOLECULE: | HEMOGLOBIN |
| 70:00:00 | 8fef-I | 4.3 | 6.6 | 101 | 253 | 9 | MOLECULE: | VIRULENCE F |
| 71:00:00 | 6myo-D | 4.3 | 8.1 | 83 | 102 | 10 | MOLECULE: | SUCCINATE D |
| 72:00:00 | 3sxk-A | 4.3 | 3.9 | 96 | 140 | 5 | MOLECULE: | TRANSCRIPTI |
| 73:00:00 | 7tn2-W | 4.3 | 19.7 | 95 | 895 | 6 | MOLECULE: | HISTONE H3 |
| 74:00:00 | 6vxm-A | 4.3 | 10.4 | 88 | 277 | 6 | MOLECULE: | MECHANOSENS |

| | | | | | | | | |
|---|---|---|---|---|---|---|---|---|
| 75:00:00 | 1o1n-D | 4.3 | 3.3 | 85 | 146 | 6 | MOLECULE: | HEMOGLOBIN |
| 76:00:00 | 3qm8-A | 4.3 | 4.2 | 84 | 145 | 7 | MOLECULE: | MYOGLOBIN |
| 77:00:00 | 4n7n-I | 4.3 | 3.6 | 88 | 141 | 3 | MOLECULE: | HEMOGLOBIN |
| 78:00:00 | 2hbs-B | 4.3 | 3.4 | 84 | 146 | 6 | MOLECULE: | HEMOGLOBIN |
| 79:00:00 | 1y8h-A | 4.3 | 3.5 | 85 | 141 | 4 | MOLECULE: | HEMOGLOBIN |
| 80:00:00 | 5me2-A | 4.3 | 3.5 | 85 | 140 | 4 | MOLECULE: | HEMOGLOBIN |
| 81:00:00 | 2hbs-F | 4.3 | 4.2 | 86 | 146 | 6 | MOLECULE: | HEMOGLOBIN |
| 82:00:00 | 1yeu-D | 4.3 | 3.7 | 87 | 146 | 6 | MOLECULE: | HEMOGLOBIN |
| 83:00:00 | 1a9w-F | 4.3 | 3.5 | 85 | 145 | 7 | MOLECULE: | HEMOGLOBIN |
| 84:00:00 | 2olp-A | 4.3 | 3.6 | 84 | 151 | 5 | MOLECULE: | HEMOGLOBIN |
| 85:00:00 | 2zlx-C | 4.3 | 3.1 | 84 | 139 | 4 | MOLECULE: | HEMOGLOBIN |
| 86:00:00 | 1hba-B | 4.3 | 3.4 | 84 | 146 | 6 | MOLECULE: | HEMOGLOBIN |
| 87:00:00 | 2zlw-C | 4.3 | 3.5 | 86 | 141 | 2 | MOLECULE: | HEMOGLOBIN |
| 88:00:00 | 2r80-A | 4.3 | 3.6 | 87 | 141 | 6 | MOLECULE: | HEMOGLOBIN |
| 89:00:00 | 1hv4-H | 4.3 | 3.6 | 85 | 146 | 11 | MOLECULE: | HEMOGLOBIN |
| 90:00:00 | 1yih-B | 4.3 | 4.1 | 86 | 146 | 6 | MOLECULE: | HEMOGLOBIN |
| 91:00:00 | 3eu1-A | 4.3 | 4 | 89 | 141 | 3 | MOLECULE: | HEMOGLOBIN |
| 92:00:00 | 1o1l-B | 4.3 | 3.3 | 85 | 146 | 6 | MOLECULE: | HEMOGLOBIN |
| 93:00:00 | 3fs4-A | 4.3 | 3.7 | 86 | 141 | 8 | MOLECULE: | HEMOGLOBIN |
| 94:00:00 | 7pcf-B | 4.3 | 3.3 | 84 | 146 | 10 | MOLECULE: | HEMOGLOBIN |
| 95:00:00 | 1hv4-F | 4.3 | 3.6 | 85 | 146 | 11 | MOLECULE: | HEMOGLOBIN |
| 96:00:00 | 2r80-B | 4.3 | 3.8 | 86 | 146 | 8 | MOLECULE: | HEMOGLOBIN |
| 97:00:00 | 4hrt-B | 4.3 | 3.8 | 89 | 146 | 9 | MOLECULE: | GLOBIN-2 A |
| 98:00:00 | 1hv4-B | 4.3 | 3.6 | 85 | 146 | 11 | MOLECULE: | HEMOGLOBIN |
| 99:00:00 | 4hrt-H | 4.3 | 4.1 | 90 | 148 | 8 | MOLECULE: | GLOBIN-2 A |
| 0:00 | 3a59-F | 4.3 | 3.6 | 85 | 146 | 8 | MOLECULE: | HEMOGLOBIN |
| 1:00 | 1hv4-D | 4.3 | 3.6 | 85 | 146 | 11 | MOLECULE: | HEMOGLOBIN |
| 2:00 | 1yie-B | 4.3 | 3.3 | 84 | 146 | 6 | MOLECULE: | HEMOGLOBIN |
| 3:00 | 4n7p-C | 4.3 | 3.1 | 84 | 141 | 5 | MOLECULE: | HEMOGLOBIN |
| 4:00 | 2dc3-A | 4.3 | 4.8 | 95 | 172 | 11 | MOLECULE: | CYTOGLOBIN |
| 5:00 | 1flp-A | 4.3 | 3.4 | 78 | 142 | 5 | MOLECULE: | HEMOGLOBIN |
| 6:00 | 4xs0-A | 4.3 | 3.3 | 83 | 141 | 5 | MOLECULE: | HEMOGLOBIN |
| 7:00 | 3hrw-A | 4.3 | 3.5 | 87 | 141 | 6 | MOLECULE: | HEMOGLOBIN |
| 8:00 | 2ri4-A | 4.3 | 3 | 84 | 139 | 5 | MOLECULE: | HEMOGLOBIN |
| 9:00 | 4n7p-A | 4.3 | 3.5 | 84 | 141 | 4 | MOLECULE: | HEMOGLOBIN |
| 10:00 | 5ker-A | 4.3 | 3.8 | 87 | 140 | 5 | MOLECULE: | ALPHA-GLOBI |
| 11:00 | 1hbs-C | 4.3 | 3.5 | 87 | 141 | 2 | MOLECULE: | HEMOGLOBIN |
| 12:00 | 3mju-A | 4.3 | 3.4 | 86 | 141 | 7 | MOLECULE: | HEMOGLOBIN |
| 13:00 | 6kai-E | 4.3 | 3 | 83 | 141 | 5 | MOLECULE: | HEMOGLOBIN |
| 14:00 | 3gdj-C | 4.3 | 3.5 | 86 | 141 | 6 | MOLECULE: | HEMOGLOBIN |
| 15:00 | 1hbs-A | 4.3 | 3.1 | 85 | 141 | 4 | MOLECULE: | HEMOGLOBIN |
| 16:00 | 2zfb-A | 4.3 | 3.6 | 87 | 141 | 6 | MOLECULE: | HEMOGLOBIN |
| 17:00 | 1o1m-A | 4.3 | 13.8 | 117 | 282 | 6 | MOLECULE: | HEMOGLOBIN |
| 18:00 | 3gqr-A | 4.3 | 4 | 89 | 141 | 6 | MOLECULE: | HEMOGLOBIN |

| | | | | | | | | |
|---|---|---|---|---|---|---|---|---|
| 19:00 | 2pgh-C | 4.3 | 3.1 | 84 | 141 | 6 | MOLECULE: | HEMOGLOBIN |
| 20:00 | 1hbs-H | 4.3 | 3.4 | 86 | 146 | 7 | MOLECULE: | HEMOGLOBIN |
| 21:00 | 3wtg-C | 4.3 | 3.4 | 85 | 138 | 8 | MOLECULE: | HEMOGLOBIN |
| 22:00 | 2zlx-A | 4.3 | 3.1 | 84 | 140 | 4 | MOLECULE: | HEMOGLOBIN |
| 23:00 | 5ker-G | 4.3 | 3.4 | 86 | 140 | 6 | MOLECULE: | ALPHA-GLOBI |
| 24:00:00 | 3gqp-A | 4.3 | 3.2 | 86 | 141 | 7 | MOLECULE: | HEMOGLOBIN |
| 25:00:00 | 5jgg-B | 4.3 | 4.1 | 87 | 144 | 6 | MOLECULE: | ALPHA CHAIN |
| 26:00:00 | 3k8b-D | 4.3 | 3.5 | 85 | 146 | 9 | MOLECULE: | HEMOGLOBIN |
| 27:00:00 | 3pi3-A | 4.3 | 3.7 | 83 | 151 | 4 | MOLECULE: | HEMOGLOBIN |
| 28:00:00 | 1yvq-C | 4.3 | 4 | 86 | 141 | 3 | MOLECULE: | HEMOGLOBIN |
| 29:00:00 | 3dhr-C | 4.3 | 3.1 | 84 | 141 | 7 | MOLECULE: | HEMOGLOBIN |
| 30:00:00 | 1o1n-B | 4.3 | 3.3 | 84 | 146 | 7 | MOLECULE: | HEMOGLOBIN |
| 31:00:00 | 1mtk-A | 4.3 | 4.5 | 93 | 154 | 8 | MOLECULE: | MYOGLOBIN |
| 32:00:00 | 1ch2-A | 4.3 | 4.5 | 93 | 154 | 8 | MOLECULE: | PROTEIN (MY |
| 33:00:00 | 1ye2-B | 4.3 | 3.5 | 86 | 146 | 7 | MOLECULE: | HEMOGLOBIN |
| 34:00:00 | 2oh8-A | 4.3 | 4.5 | 93 | 154 | 8 | MOLECULE: | MYOGLOBIN |
| 35:00:00 | 1y4g-D | 4.3 | 3.5 | 85 | 146 | 7 | MOLECULE: | HEMOGLOBIN |
| 36:00:00 | 1yie-D | 4.3 | 3.5 | 85 | 146 | 7 | MOLECULE: | HEMOGLOBIN |
| 37:00:00 | 1y4v-B | 4.3 | 3.4 | 84 | 146 | 7 | MOLECULE: | HEMOGLOBIN |
| 38:00:00 | 2oha-A | 4.3 | 4.4 | 93 | 154 | 8 | MOLECULE: | MYOGLOBIN |
| 39:00:00 | 2r80-D | 4.3 | 3.5 | 85 | 146 | 9 | MOLECULE: | HEMOGLOBIN |
| 40:00:00 | 1mlo-A | 4.3 | 4.4 | 92 | 154 | 8 | MOLECULE: | MYOGLOBIN |
| 41:00:00 | 1yev-D | 4.3 | 3.6 | 86 | 146 | 7 | MOLECULE: | HEMOGLOBIN |
| 42:00:00 | 5ut7-A | 4.3 | 4.5 | 92 | 154 | 9 | MOLECULE: | MYOGLOBIN |
| 43:00:00 | 1mls-A | 4.3 | 4.5 | 93 | 154 | 6 | MOLECULE: | MYOGLOBIN |
| 44:00:00 | 2qmb-B | 4.3 | 3.5 | 85 | 146 | 9 | MOLECULE: | HEMOGLOBIN |
| 45:00:00 | 1a9w-E | 4.3 | 3.5 | 85 | 145 | 7 | MOLECULE: | HEMOGLOBIN |
| 46:00:00 | 2mgl-A | 4.3 | 4.5 | 93 | 154 | 8 | MOLECULE: | MYOGLOBIN |
| 47:00:00 | 1ygd-B | 4.3 | 3.5 | 85 | 146 | 7 | MOLECULE: | HEMOGLOBIN |
| 48:00:00 | 1xye-C | 4.3 | 3.6 | 85 | 141 | 6 | MOLECULE: | HEMOGLOBIN |
| 49:00:00 | 1ofk-A | 4.3 | 4.5 | 93 | 154 | 8 | MOLECULE: | MYOGLOBIN |
| 50:00:00 | 1co8-A | 4.3 | 4.5 | 93 | 154 | 8 | MOLECULE: | PROTEIN (MY |
| 51:00:00 | 1y46-D | 4.3 | 3.2 | 84 | 146 | 7 | MOLECULE: | HEMOGLOBIN |
| 52:00:00 | 3pi2-B | 4.3 | 3.9 | 86 | 151 | 5 | MOLECULE: | HEMOGLOBIN |
| 53:00:00 | 1h1x-A | 4.3 | 4.5 | 92 | 154 | 9 | MOLECULE: | MYOGLOBIN |
| 54:00:00 | 1mcy-A | 4.3 | 4.1 | 89 | 154 | 10 | MOLECULE: | MYOGLOBIN ( |
| 55:00:00 | 4hrt-G | 4.3 | 3.5 | 86 | 149 | 3 | MOLECULE: | GLOBIN-2 A |
| 56:00:00 | 3gqp-D | 4.3 | 3.6 | 85 | 145 | 7 | MOLECULE: | HEMOGLOBIN |
| 57:00:00 | 3dhr-G | 4.3 | 3 | 82 | 141 | 6 | MOLECULE: | HEMOGLOBIN |
| 58:00:00 | 1buw-C | 4.3 | 4 | 86 | 141 | 6 | MOLECULE: | PROTEIN (HE |
| 59:00:00 | 6kav-C | 4.3 | 3.9 | 86 | 141 | 6 | MOLECULE: | HEMOGLOBIN |
| 60:00:00 | 4hrt-C | 4.3 | 3.4 | 85 | 149 | 4 | MOLECULE: | GLOBIN-2 A |
| 61:00:00 | 3k8b-C | 4.3 | 3.5 | 87 | 141 | 3 | MOLECULE: | HEMOGLOBIN |
| 62:00:00 | 1mlu-A | 4.3 | 4.5 | 93 | 154 | 8 | MOLECULE: | MYOGLOBIN |

| | | | | | | | |
|---|---|---|---|---|---|---|---|
| 63:00:00 | 3pi1-A | 4.3 | 3.8 | 85 | 151 | 4 | MOLECULE: HEMOGLOBIN |
| 64:00:00 | 2mgb-A | 4.3 | 4.5 | 93 | 154 | 8 | MOLECULE: MYOGLOBIN |
| 65:00:00 | 1ye0-C | 4.3 | 4.2 | 86 | 141 | 5 | MOLECULE: HEMOGLOBIN |
| 66:00:00 | 3dhr-B | 4.3 | 3.6 | 85 | 145 | 9 | MOLECULE: HEMOGLOBIN |
| 67:00:00 | 5hlq-A | 4.3 | 4.4 | 91 | 152 | 8 | MOLECULE: MYOGLOBIN |
| 68:00:00 | 1n9h-A | 4.3 | 4.5 | 92 | 154 | 9 | MOLECULE: MYOGLOBIN |
| 69:00:00 | 1yeo-D | 4.3 | 3.6 | 86 | 146 | 7 | MOLECULE: HEMOGLOBIN |
| 70:00:00 | 102m-A | 4.3 | 4.5 | 93 | 154 | 8 | MOLECULE: MYOGLOBIN |
| 71:00:00 | 1mym-A | 4.3 | 4.4 | 93 | 154 | 8 | MOLECULE: MYOGLOBIN |
| 72:00:00 | 1yg5-D | 4.3 | 3.5 | 86 | 146 | 7 | MOLECULE: HEMOGLOBIN |
| 73:00:00 | 1ufj-A | 4.3 | 4.1 | 91 | 154 | 8 | MOLECULE: MYOGLOBIN |
| 74:00:00 | 2mgc-A | 4.3 | 4.5 | 93 | 154 | 8 | MOLECULE: MYOGLOBIN |
| 75:00:00 | 3pi3-B | 4.3 | 3.8 | 84 | 151 | 5 | MOLECULE: HEMOGLOBIN |
| 76:00:00 | 1rqa-D | 4.3 | 3.5 | 86 | 146 | 7 | MOLECULE: HEMOGLOBIN |
| 77:00:00 | 1y46-B | 4.3 | 3.1 | 83 | 146 | 7 | MOLECULE: HEMOGLOBIN |
| 78:00:00 | 1co9-A | 4.3 | 4.5 | 93 | 154 | 8 | MOLECULE: PROTEIN (MY |
| 79:00:00 | 2mgh-A | 4.3 | 4.5 | 93 | 154 | 8 | MOLECULE: MYOGLOBIN |
| 80:00:00 | 1ch3-A | 4.3 | 4.5 | 93 | 154 | 6 | MOLECULE: PROTEIN (MY |
| 81:00:00 | 4n7n-A | 4.3 | 3.1 | 84 | 141 | 6 | MOLECULE: HEMOGLOBIN |
| 82:00:00 | 1mlk-A | 4.3 | 4.5 | 93 | 154 | 8 | MOLECULE: MYOGLOBIN |
| 83:00:00 | 1qi8-C | 4.3 | 3.4 | 85 | 141 | 5 | MOLECULE: HEMOGLOBIN |
| 84:00:00 | 1y0t-B | 4.3 | 3.4 | 84 | 146 | 7 | MOLECULE: HEMOGLOBIN |
| 85:00:00 | 2pgh-A | 4.3 | 3.1 | 84 | 141 | 5 | MOLECULE: HEMOGLOBIN |
| 86:00:00 | 2mge-A | 4.3 | 4.5 | 93 | 154 | 6 | MOLECULE: MYOGLOBIN |
| 87:00:00 | 2spl-A | 4.3 | 4.5 | 93 | 154 | 6 | MOLECULE: MYOGLOBIN |
| 88:00:00 | 1f63-A | 4.3 | 4.4 | 93 | 154 | 6 | MOLECULE: MYOGLOBIN |
| 89:00:00 | 4iro-A | 4.3 | 3.3 | 85 | 142 | 7 | MOLECULE: HEMOGLOBIN |
| 90:00:00 | 1y4f-B | 4.3 | 3.4 | 84 | 146 | 7 | MOLECULE: HEMOGLOBIN |
| 91:00:00 | 1j41-E | 4.3 | 3.3 | 86 | 141 | 3 | MOLECULE: HEMOGLOBIN |
| 92:00:00 | 6u8y-b | 4.2 | 6 | 69 | 79 | 7 | MOLECULE: MONOVALENT |
| 93:00:00 | 1cqx-A | 4.2 | 4.9 | 104 | 403 | 10 | MOLECULE: FLAVOHEMOPR |
| 94:00:00 | 7ol3-B | 4.2 | 11.6 | 92 | 425 | 7 | MOLECULE: ATLASTIN-1 |
| 95:00:00 | 7d3u-F | 4.2 | 6.9 | 75 | 85 | 8 | MOLECULE: MONOVALENT |
| 96:00:00 | 6wb9-6 | 4.2 | 4.7 | 79 | 98 | 4 | MOLECULE: ENDOPLASMIC |
| 97:00:00 | 2qmb-D | 4.2 | 3.6 | 85 | 146 | 9 | MOLECULE: HEMOGLOBIN |
| 98:00:00 | 3bcq-B | 4.2 | 3.6 | 86 | 146 | 9 | MOLECULE: ALPHA-CHAIN |
| 99:00:00 | 1myf-A | 4.2 | 3.8 | 87 | 153 | 10 | MOLECULE: MYOGLOBIN |
| 0:00 | 5x2r-K | 4.2 | 3.9 | 87 | 140 | 6 | MOLECULE: HEMOGLOBIN |
| 1:00 | 3gqp-B | 4.2 | 3.7 | 85 | 145 | 6 | MOLECULE: HEMOGLOBIN |
| 2:00 | 7uf7-A | 4.2 | 3.4 | 85 | 141 | 7 | MOLECULE: HEMOGLOBIN |
| 3:00 | 3pi4-B | 4.2 | 4.1 | 87 | 151 | 7 | MOLECULE: HEMOGLOBIN |
| 4:00 | 1i3e-B | 4.2 | 3.6 | 86 | 146 | 5 | MOLECULE: HEMOGLOBIN |
| 5:00 | 7pch-B | 4.2 | 3.6 | 85 | 146 | 6 | MOLECULE: HEMOGLOBIN |
| 6:00 | 1hds-A | 4.2 | 3.4 | 86 | 141 | 6 | MOLECULE: HEMOGLOBIN |

| | | | | | | | | |
|---|---|---|---|---|---|---|---|---|
| 7:00 | 2r1h-B | 4.2 | 3.6 | 86 | 147 | 9 | MOLECULE: | HEMOGLOBIN |
| 8:00 | 1a0u-D | 4.2 | 3.4 | 84 | 146 | 6 | MOLECULE: | HEMOGLOBIN |
| 9:00 | 3vrg-B | 4.2 | 3.6 | 85 | 146 | 6 | MOLECULE: | HEMOGLOBIN |
| 10:00 | 2qmb-C | 4.2 | 3.7 | 86 | 141 | 7 | MOLECULE: | HEMOGLOBIN |
| 11:00 | 3pt7-A | 4.2 | 3.8 | 84 | 151 | 5 | MOLECULE: | HEMOGLOBIN |
| 12:00 | 3bom-B | 4.2 | 3.6 | 86 | 147 | 9 | MOLECULE: | HEMOGLOBIN |
| 13:00 | 5x2s-E | 4.2 | 3.3 | 85 | 140 | 4 | MOLECULE: | HEMOGLOBIN |
| 14:00 | 3mju-B | 4.2 | 4 | 87 | 146 | 8 | MOLECULE: | HEMOGLOBIN |
| 15:00 | 1y0t-D | 4.2 | 3.3 | 85 | 146 | 6 | MOLECULE: | HEMOGLOBIN |
| 16:00 | 7ylk-A | 4.2 | 4.2 | 91 | 153 | 8 | MOLECULE: | MYOGLOBIN |
| 17:00 | 3k8b-A | 4.2 | 4 | 87 | 141 | 7 | MOLECULE: | HEMOGLOBIN |
| 18:00 | 3bcq-D | 4.2 | 3.5 | 85 | 146 | 9 | MOLECULE: | ALPHA-CHAIN |
| 19:00 | 3eok-B | 4.2 | 3.6 | 86 | 146 | 9 | MOLECULE: | HEMOGLOBIN |
| 20:00 | 1hv4-E | 4.2 | 4.3 | 87 | 141 | 8 | MOLECULE: | HEMOGLOBIN |
| 21:00 | 1coh-A | 4.2 | 3.4 | 84 | 141 | 5 | MOLECULE: | HEMOGLOBIN |
| 22:00 | 1a3o-B | 4.2 | 3.5 | 84 | 145 | 7 | MOLECULE: | HEMOGLOBIN |
| 23:00 | 1hv4-A | 4.2 | 4.3 | 87 | 141 | 8 | MOLECULE: | HEMOGLOBIN |
| 24:00:00 | 3lqd-C | 4.2 | 3.2 | 86 | 141 | 6 | MOLECULE: | HEMOGLOBIN |
| 25:00:00 | 3fs4-C | 4.2 | 4.4 | 88 | 141 | 7 | MOLECULE: | HEMOGLOBIN |
| 26:00:00 | 1faw-C | 4.2 | 3.7 | 86 | 141 | 8 | MOLECULE: | HEMOGLOBIN |
| 27:00:00 | 8egi-A | 4.2 | 3.4 | 84 | 141 | 6 | MOLECULE: | HEMOGLOBIN |
| 28:00:00 | 1a00-A | 4.2 | 3.6 | 84 | 141 | 6 | MOLECULE: | HEMOGLOBIN |
| 29:00:00 | 5urc-A | 4.2 | 3.7 | 86 | 141 | 7 | MOLECULE: | HEMOGLOBIN |
| 30:00:00 | 1b0b-A | 4.2 | 3.4 | 78 | 142 | 4 | MOLECULE: | HEMOGLOBIN |
| 31:00:00 | 1faw-A | 4.2 | 4.1 | 87 | 141 | 8 | MOLECULE: | HEMOGLOBIN |
| 32:00:00 | 4hrt-D | 4.2 | 4.1 | 90 | 151 | 8 | MOLECULE: | GLOBIN-2 A |
| 33:00:00 | 6xdt-A | 4.2 | 3.3 | 84 | 141 | 6 | MOLECULE: | HEMOGLOBIN |
| 34:00:00 | 1hv4-C | 4.2 | 4.2 | 87 | 141 | 8 | MOLECULE: | HEMOGLOBIN |
| 35:00:00 | 2zlw-A | 4.2 | 3.5 | 86 | 141 | 2 | MOLECULE: | HEMOGLOBIN |
| 36:00:00 | 6di4-A | 4.2 | 3.3 | 84 | 141 | 6 | MOLECULE: | HEMOGLOBIN |
| 37:00:00 | 3lqd-A | 4.2 | 3.5 | 86 | 141 | 5 | MOLECULE: | HEMOGLOBIN |
| 38:00:00 | 7xgy-C | 4.2 | 3.6 | 84 | 141 | 6 | MOLECULE: | HEMOGLOBIN |
| 39:00:00 | 1hbs-G | 4.2 | 3.6 | 87 | 141 | 2 | MOLECULE: | HEMOGLOBIN |
| 40:00:00 | 4hrt-F | 4.2 | 4.1 | 90 | 151 | 8 | MOLECULE: | GLOBIN-2 A |
| 41:00:00 | 7xgy-A | 4.2 | 3.6 | 85 | 141 | 5 | MOLECULE: | HEMOGLOBIN |
| 42:00:00 | 4n7p-K | 4.2 | 3.9 | 87 | 141 | 5 | MOLECULE: | HEMOGLOBIN |
| 43:00:00 | 3qm6-A | 4.2 | 4.3 | 84 | 145 | 7 | MOLECULE: | MYOGLOBIN |
| 44:00:00 | 5x2u-A | 4.2 | 4 | 86 | 140 | 6 | MOLECULE: | HEMOGLOBIN |
| 45:00:00 | 1nqp-C | 4.2 | 4 | 86 | 141 | 6 | MOLECULE: | HEMOGLOBIN |
| 46:00:00 | 1dxu-D | 4.2 | 3.8 | 86 | 146 | 5 | MOLECULE: | HEMOGLOBIN |
| 47:00:00 | 2ri4-K | 4.2 | 3 | 84 | 138 | 5 | MOLECULE: | HEMOGLOBIN |
| 48:00:00 | 4mqk-F | 4.2 | 4.1 | 86 | 145 | 6 | MOLECULE: | HEMOGLOBIN |
| 49:00:00 | 1j52-A | 4.2 | 4.2 | 91 | 154 | 10 | MOLECULE: | MYOGLOBIN |
| 50:00:00 | 1fhj-D | 4.2 | 3.5 | 84 | 146 | 6 | MOLECULE: | HEMOGLOBIN |

| | | | | | | | | |
|---|---|---|---|---|---|---|---|---|
| 51:00:00 | 6g5t-A | 4.2 | 4.5 | 93 | 155 | 6 | MOLECULE: | MYOGLOBIN |
| 52:00:00 | 109m-A | 4.2 | 4.5 | 93 | 154 | 6 | MOLECULE: | MYOGLOBIN |
| 53:00:00 | 8egi-D | 4.2 | 3.6 | 85 | 146 | 7 | MOLECULE: | HEMOGLOBIN |
| 54:00:00 | 1ch1-A | 4.2 | 4.5 | 93 | 154 | 6 | MOLECULE: | PROTEIN (MY |
| 55:00:00 | 3ovu-C | 4.2 | 3.7 | 87 | 139 | 2 | MOLECULE: | ALPHA-HEMOG |
| 56:00:00 | 2mgk-A | 4.2 | 4.5 | 93 | 154 | 6 | MOLECULE: | MYOGLOBIN |
| 57:00:00 | 3dhr-D | 4.2 | 3.6 | 85 | 145 | 9 | MOLECULE: | HEMOGLOBIN |
| 58:00:00 | 1gbu-B | 4.2 | 3.6 | 84 | 146 | 6 | MOLECULE: | HEMOGLOBIN |
| 59:00:00 | 1mlm-A | 4.2 | 4.4 | 92 | 154 | 8 | MOLECULE: | MYOGLOBIN |
| 60:00:00 | 2oh9-A | 4.2 | 4.5 | 93 | 154 | 6 | MOLECULE: | MYOGLOBIN |
| 61:00:00 | 3k8b-B | 4.2 | 3.5 | 85 | 146 | 9 | MOLECULE: | HEMOGLOBIN |
| 62:00:00 | 6otw-A | 4.2 | 3.7 | 84 | 151 | 5 | MOLECULE: | HEMOGLOBIN |
| 63:00:00 | 1lue-A | 4.2 | 4.5 | 93 | 154 | 6 | MOLECULE: | MYOGLOBIN |
| 64:00:00 | 2mgd-A | 4.2 | 4.5 | 93 | 154 | 6 | MOLECULE: | MYOGLOBIN |
| 65:00:00 | 1cp0-A | 4.2 | 4.4 | 93 | 154 | 6 | MOLECULE: | PROTEIN (MY |
| 66:00:00 | 2zfb-B | 4.2 | 3.6 | 85 | 146 | 9 | MOLECULE: | HEMOGLOBIN |
| 67:00:00 | 2dn1-B | 4.2 | 3.4 | 84 | 145 | 7 | MOLECULE: | HEMOGLOBIN |
| 68:00:00 | 1fhj-B | 4.2 | 3.5 | 84 | 146 | 6 | MOLECULE: | HEMOGLOBIN |
| 69:00:00 | 7jjq-B | 4.2 | 3.6 | 85 | 145 | 6 | MOLECULE: | HEMOGLOBIN |
| 70:00:00 | 1cpw-A | 4.2 | 4.5 | 93 | 154 | 6 | MOLECULE: | PROTEIN (MY |
| 71:00:00 | 2spn-A | 4.2 | 4.5 | 93 | 154 | 6 | MOLECULE: | MYOGLOBIN |
| 72:00:00 | 2dxm-A | 4.2 | 3.7 | 85 | 141 | 6 | MOLECULE: | HEMOGLOBIN |
| 73:00:00 | 5e83-D | 4.2 | 3.4 | 84 | 146 | 7 | MOLECULE: | HEMOGLOBIN |
| 74:00:00 | 112m-A | 4.2 | 4.5 | 93 | 154 | 6 | MOLECULE: | MYOGLOBIN |
| 75:00:00 | 4hrr-C | 4.2 | 3.5 | 86 | 149 | 3 | MOLECULE: | GLOBIN-2 A |
| 76:00:00 | 2spo-A | 4.2 | 4.4 | 93 | 154 | 6 | MOLECULE: | MYOGLOBIN |
| 77:00:00 | 1gli-D | 4.2 | 3.1 | 83 | 146 | 7 | MOLECULE: | DEOXYHEMOGL |
| 78:00:00 | 1yeq-B | 4.2 | 3.5 | 85 | 146 | 7 | MOLECULE: | HEMOGLOBIN |
| 79:00:00 | 1jy7-P | 4.2 | 4.6 | 88 | 141 | 6 | MOLECULE: | HEMOGLOBIN |
| 80:00:00 | 3pi4-A | 4.2 | 3.9 | 84 | 151 | 5 | MOLECULE: | HEMOGLOBIN |
| 81:00:00 | 1gli-B | 4.2 | 3.6 | 84 | 146 | 6 | MOLECULE: | DEOXYHEMOGL |
| 82:00:00 | 1ebt-A | 4.2 | 3.4 | 78 | 142 | 5 | MOLECULE: | HEMOGLOBIN |
| 83:00:00 | 7pcq-B | 4.2 | 3.6 | 85 | 146 | 6 | MOLECULE: | HEMOGLOBIN |
| 84:00:00 | 4hrr-E | 4.2 | 3.9 | 89 | 149 | 6 | MOLECULE: | GLOBIN-2 A |
| 85:00:00 | 2dxm-C | 4.2 | 3.7 | 86 | 141 | 6 | MOLECULE: | HEMOGLOBIN |
| 86:00:00 | 1cp5-A | 4.2 | 4.2 | 91 | 154 | 10 | MOLECULE: | PROTEIN (MY |
| 87:00:00 | 6oty-A | 4.2 | 4 | 85 | 151 | 5 | MOLECULE: | HEMOGLOBIN |
| 88:00:00 | 5e29-D | 4.2 | 3.1 | 83 | 145 | 7 | MOLECULE: | HEMOGLOBIN |
| 89:00:00 | 1mlf-A | 4.2 | 4.6 | 93 | 154 | 8 | MOLECULE: | MYOGLOBIN |
| 90:00:00 | 2spm-A | 4.2 | 4.5 | 93 | 154 | 6 | MOLECULE: | MYOGLOBIN |
| 91:00:00 | 3at6-B | 4.2 | 4 | 89 | 146 | 9 | MOLECULE: | ALPHAA-GLOB |
| 92:00:00 | 1hlb-A | 4.1 | 3.9 | 91 | 157 | 9 | MOLECULE: | HEMOGLOBIN |
| 93:00:00 | 6m5r-B | 4.1 | 17.2 | 101 | 689 | 6 | MOLECULE: | TRIPARTITE |
| 94:00:00 | 4zvb-A | 4.1 | 4.1 | 89 | 150 | 3 | MOLECULE: | DIGUANYLATE |

| | | | | | | | | |
|---|---|---|---|---|---|---|---|---|
| 95:00:00 | 2peg-B | 4.1 | 3.8 | 84 | 139 | 7 | MOLECULE: | HEMOGLOBIN |
| 96:00:00 | 4j19-A | 4.1 | 3.2 | 72 | 77 | 6 | MOLECULE: | HOMEOBOX-CO |
| 97:00:00 | 1dm1-A | 4.1 | 4.3 | 84 | 146 | 11 | MOLECULE: | MYOGLOBIN |
| 98:00:00 | 3gys-H | 4.1 | 3.6 | 85 | 145 | 7 | MOLECULE: | HEMOGLOBIN |
| 99:00:00 | 3uhk-D | 4.1 | 3.3 | 84 | 145 | 11 | MOLECULE: | GLOBIN-1 |
| 0:00 | 4sdh-A | 4.1 | 3.3 | 83 | 145 | 8 | MOLECULE: | HEMOGLOBIN |
| 1:00 | 3fs4-B | 4.1 | 3.8 | 86 | 146 | 6 | MOLECULE: | HEMOGLOBIN |
| 2:00 | 3dht-A | 4.1 | 3.4 | 84 | 139 | 8 | MOLECULE: | HEMOGLOBIN |
| 3:00 | 7xgy-B | 4.1 | 3.5 | 84 | 146 | 8 | MOLECULE: | HEMOGLOBIN |
| 4:00 | 4n7o-C | 4.1 | 3 | 82 | 141 | 4 | MOLECULE: | HEMOGLOBIN |
| 5:00 | 7pcq-D | 4.1 | 3.6 | 85 | 146 | 6 | MOLECULE: | HEMOGLOBIN |
| 6:00 | 2r1h-D | 4.1 | 3.6 | 86 | 147 | 9 | MOLECULE: | HEMOGLOBIN |
| 7:00 | 1y35-B | 4.1 | 3.6 | 82 | 146 | 7 | MOLECULE: | HEMOGLOBIN |
| 8:00 | 6hbi-B | 4.1 | 3.3 | 84 | 145 | 10 | MOLECULE: | HEMOGLOBIN |
| 9:00 | 3uhw-B | 4.1 | 3.8 | 86 | 145 | 10 | MOLECULE: | GLOBIN-1 |
| 10:00 | 5me2-C | 4.1 | 4 | 86 | 140 | 6 | MOLECULE: | HEMOGLOBIN |
| 11:00 | 3b75-C | 4.1 | 3.4 | 85 | 141 | 7 | MOLECULE: | HEMOGLOBIN |
| 12:00 | 1si4-C | 4.1 | 3.4 | 85 | 141 | 7 | MOLECULE: | HEMOGLOBIN |
| 13:00 | 5x2t-J | 4.1 | 3.5 | 84 | 145 | 6 | MOLECULE: | HEMOGLOBIN |
| 14:00 | 3bom-D | 4.1 | 3.7 | 86 | 147 | 7 | MOLECULE: | HEMOGLOBIN |
| 15:00 | 1y2z-B | 4.1 | 3.6 | 83 | 146 | 7 | MOLECULE: | HEMOGLOBIN |
| 16:00 | 7xgy-D | 4.1 | 3.4 | 84 | 146 | 8 | MOLECULE: | HEMOGLOBIN |
| 17:00 | 4mqj-B | 4.1 | 4.1 | 86 | 146 | 5 | MOLECULE: | HEMOGLOBIN |
| 18:00 | 3a59-E | 4.1 | 3.7 | 86 | 141 | 8 | MOLECULE: | HEMOGLOBIN |
| 19:00 | 1nxf-B | 4.1 | 3.5 | 85 | 145 | 9 | MOLECULE: | GLOBIN I |
| 20:00 | 3g4w-A | 4.1 | 3.4 | 85 | 145 | 9 | MOLECULE: | GLOBIN-1 |
| 21:00 | 3uhn-B | 4.1 | 3.4 | 85 | 145 | 9 | MOLECULE: | GLOBIN-1 |
| 22:00 | 3g4v-A | 4.1 | 3.5 | 86 | 145 | 9 | MOLECULE: | GLOBIN-1 |
| 23:00 | 8egi-C | 4.1 | 3.5 | 85 | 141 | 7 | MOLECULE: | HEMOGLOBIN |
| 24:00:00 | 5hy8-S | 4.1 | 3.3 | 84 | 141 | 6 | MOLECULE: | HEMOGLOBIN |
| 25:00:00 | 3mjp-C | 4.1 | 3.8 | 88 | 141 | 3 | MOLECULE: | HEMOGLOBIN |
| 26:00:00 | 1yff-E | 4.1 | 3.4 | 84 | 141 | 6 | MOLECULE: | HEMOGLOBIN |
| 27:00:00 | 5hy8-C | 4.1 | 3.4 | 85 | 141 | 7 | MOLECULE: | HEMOGLOBIN |
| 28:00:00 | 8vyl-C | 4.1 | 3.9 | 86 | 140 | 6 | MOLECULE: | HEMOGLOBIN |
| 29:00:00 | 1bij-A | 4.1 | 3.4 | 83 | 141 | 5 | MOLECULE: | HEMOGLOBIN |
| 30:00:00 | 6kat-C | 4.1 | 3.6 | 85 | 141 | 6 | MOLECULE: | HEMOGLOBIN |
| 31:00:00 | 6l5x-C | 4.1 | 3.9 | 86 | 141 | 6 | MOLECULE: | HEMOGLOBIN |
| 32:00:00 | 6kar-A | 4.1 | 4.1 | 87 | 141 | 7 | MOLECULE: | HEMOGLOBIN |
| 33:00:00 | 3g52-A | 4.1 | 3.5 | 86 | 145 | 9 | MOLECULE: | GLOBIN-1 |
| 34:00:00 | 6lcw-E | 4.1 | 4 | 88 | 141 | 6 | MOLECULE: | HEMOGLOBIN |
| 35:00:00 | 4n7p-I | 4.1 | 3.4 | 85 | 141 | 2 | MOLECULE: | HEMOGLOBIN |
| 36:00:00 | 7ud8-A | 4.1 | 3.3 | 84 | 141 | 6 | MOLECULE: | HEMOGLOBIN |
| 37:00:00 | 5hy8-A | 4.1 | 3.6 | 85 | 141 | 6 | MOLECULE: | HEMOGLOBIN |
| 38:00:00 | 4sdh-B | 4.1 | 3.3 | 84 | 145 | 8 | MOLECULE: | HEMOGLOBIN |

| | | | | | | | | |
|---|---|---|---|---|---|---|---|---|
| 39:00:00 | 5u3i-D | 4.1 | 3.5 | 84 | 145 | 7 | MOLECULE: | HEMOGLOBIN |
| 40:00:00 | 5hy8-G | 4.1 | 4.1 | 85 | 138 | 6 | MOLECULE: | HEMOGLOBIN |
| 41:00:00 | 1nwi-A | 4.1 | 3.3 | 85 | 145 | 9 | MOLECULE: | GLOBIN I |
| 42:00:00 | 6xd9-C | 4.1 | 3.4 | 86 | 141 | 6 | MOLECULE: | HEMOGLOBIN |
| 43:00:00 | 4n7n-E | 4.1 | 3.8 | 86 | 141 | 7 | MOLECULE: | HEMOGLOBIN |
| 44:00:00 | 4n7o-A | 4.1 | 4.1 | 88 | 141 | 6 | MOLECULE: | HEMOGLOBIN |
| 45:00:00 | 3hrw-C | 4.1 | 3.2 | 85 | 141 | 7 | MOLECULE: | HEMOGLOBIN |
| 46:00:00 | 3g4y-A | 4.1 | 3.5 | 86 | 145 | 9 | MOLECULE: | GLOBIN-1 |
| 47:00:00 | 4n7o-G | 4.1 | 4.1 | 87 | 141 | 6 | MOLECULE: | HEMOGLOBIN |
| 48:00:00 | 7vcf-W | 4.1 | 4.5 | 96 | 164 | 6 | MOLECULE: | TIC214 |
| 49:00:00 | 6xe7-C | 4.1 | 3.6 | 85 | 141 | 6 | MOLECULE: | HEMOGLOBIN |
| 50:00:00 | 3dhr-A | 4.1 | 4.1 | 87 | 141 | 8 | MOLECULE: | HEMOGLOBIN |
| 51:00:00 | 3uhk-A | 4.1 | 3.2 | 83 | 145 | 8 | MOLECULE: | GLOBIN-1 |
| 52:00:00 | 3uhs-B | 4.1 | 3.3 | 84 | 145 | 10 | MOLECULE: | GLOBIN-1 |
| 53:00:00 | 5x2t-C | 4.1 | 4 | 85 | 140 | 6 | MOLECULE: | HEMOGLOBIN |
| 54:00:00 | 1a01-D | 4.1 | 3.8 | 85 | 146 | 7 | MOLECULE: | HEMOGLOBIN |
| 55:00:00 | 3b75-A | 4.1 | 4 | 87 | 141 | 7 | MOLECULE: | HEMOGLOBIN |
| 56:00:00 | 1jzk-B | 4.1 | 3.5 | 84 | 145 | 13 | MOLECULE: | GLOBIN I - |
| 57:00:00 | 3g4u-A | 4.1 | 3.5 | 86 | 145 | 10 | MOLECULE: | GLOBIN-1 |
| 58:00:00 | 1yg5-B | 4.1 | 3.7 | 84 | 146 | 7 | MOLECULE: | HEMOGLOBIN |
| 59:00:00 | 1y4b-B | 4.1 | 3.6 | 83 | 146 | 7 | MOLECULE: | HEMOGLOBIN |
| 60:00:00 | 5x2r-I | 4.1 | 4.1 | 86 | 140 | 6 | MOLECULE: | HEMOGLOBIN |
| 61:00:00 | 5u3i-B | 4.1 | 3.5 | 84 | 146 | 7 | MOLECULE: | HEMOGLOBIN |
| 62:00:00 | 1y85-B | 4.1 | 3.6 | 83 | 145 | 7 | MOLECULE: | HEMOGLOBIN |
| 63:00:00 | 7pch-D | 4.1 | 3.6 | 84 | 146 | 6 | MOLECULE: | HEMOGLOBIN |
| 64:00:00 | 4hrr-B | 4.1 | 4.1 | 90 | 151 | 8 | MOLECULE: | GLOBIN-2 A |
| 65:00:00 | 1a3n-B | 4.1 | 3.6 | 83 | 145 | 7 | MOLECULE: | HEMOGLOBIN |
| 66:00:00 | 3uhx-B | 4.1 | 3.3 | 84 | 145 | 10 | MOLECULE: | GLOBIN-1 |
| 67:00:00 | 1nwi-D | 4.1 | 3.3 | 83 | 145 | 10 | MOLECULE: | GLOBIN I |
| 68:00:00 | 3gys-D | 4.1 | 3.6 | 85 | 145 | 7 | MOLECULE: | HEMOGLOBIN |
| 69:00:00 | 6jp1-B | 4.1 | 4.4 | 92 | 151 | 7 | MOLECULE: | MYOGLOBIN |
| 70:00:00 | 2qls-B | 4.1 | 3.5 | 84 | 146 | 6 | MOLECULE: | HEMOGLOBIN |
| 71:00:00 | 4hrt-A | 4.1 | 4 | 89 | 149 | 4 | MOLECULE: | GLOBIN-2 A |
| 72:00:00 | 1qpw-D | 4.1 | 3.5 | 84 | 146 | 8 | MOLECULE: | PORCINE HEM |
| 73:00:00 | 8vyl-D | 4.1 | 3.7 | 84 | 146 | 6 | MOLECULE: | HEMOGLOBIN |
| 74:00:00 | 8fdm-B | 4.1 | 3.6 | 84 | 146 | 6 | MOLECULE: | HEMOGLOBIN |
| 75:00:00 | 3wcp-B | 4.1 | 3 | 82 | 145 | 7 | MOLECULE: | HEMOGLOBIN |
| 76:00:00 | 2hbs-D | 4.1 | 4.2 | 85 | 146 | 7 | MOLECULE: | HEMOGLOBIN |
| 77:00:00 | 3pel-B | 4.1 | 3.8 | 86 | 146 | 5 | MOLECULE: | HEMOGLOBIN |
| 78:00:00 | 3a59-H | 4.1 | 3.6 | 85 | 146 | 8 | MOLECULE: | HEMOGLOBIN |
| 79:00:00 | 7dds-A | 4.1 | 4.3 | 90 | 154 | 11 | MOLECULE: | ANCESTRAL M |
| 80:00:00 | 8fdm-D | 4.1 | 3.5 | 84 | 146 | 7 | MOLECULE: | HEMOGLOBIN |
| 81:00:00 | 3pia-A | 4.1 | 4 | 85 | 141 | 8 | MOLECULE: | HEMOGLOBIN |
| 82:00:00 | 3g4r-A | 4.1 | 3.6 | 86 | 145 | 10 | MOLECULE: | GLOBIN-1 |

| | | | | | | | | |
|---|---|---|---|---|---|---|---|---|
| 83:00:00 | 4hrr-G | 4.1 | 4 | 90 | 149 | 6 | MOLECULE: | GLOBIN-2 A |
| 84:00:00 | 3fh9-B | 4.1 | 3.7 | 86 | 146 | 7 | MOLECULE: | HEMOGLOBIN |
| 85:00:00 | 2r80-C | 4.1 | 3.7 | 86 | 140 | 7 | MOLECULE: | HEMOGLOBIN |
| 86:00:00 | 7uf6-D | 4.1 | 3.6 | 84 | 146 | 6 | MOLECULE: | HEMOGLOBIN |
| 87:00:00 | 2b7h-D | 4.1 | 3.5 | 85 | 145 | 6 | MOLECULE: | HEMOGLOBIN |
| 88:00:00 | 8dov-B | 4.1 | 3.6 | 84 | 146 | 6 | MOLECULE: | HEMOGLOBIN |
| 89:00:00 | 5x2s-B | 4.1 | 3.5 | 84 | 145 | 7 | MOLECULE: | HEMOGLOBIN |
| 90:00:00 | 7ud8-D | 4.1 | 3.6 | 85 | 146 | 6 | MOLECULE: | HEMOGLOBIN |
| 91:00:00 | 5x2s-A | 4.1 | 4 | 87 | 140 | 6 | MOLECULE: | HEMOGLOBIN |
| 92:00:00 | 1hbs-B | 4.1 | 3.6 | 85 | 146 | 6 | MOLECULE: | HEMOGLOBIN |
| 93:00:00 | 1jzk-A | 4.1 | 3.5 | 85 | 145 | 11 | MOLECULE: | GLOBIN I - |
| 94:00:00 | 8dov-D | 4.1 | 3.6 | 85 | 145 | 6 | MOLECULE: | HEMOGLOBIN |
| 95:00:00 | 5xwp-A | 4 | 18 | 86 | 1125 | 3 | MOLECULE: | UNCHARACTER |
| 96:00:00 | 8cr1-C | 4 | 3.8 | 77 | 266 | 5 | MOLECULE: | ATPASE ASNA |
| 97:00:00 | 5m0n-A | 4 | 6.3 | 131 | 423 | 2 | MOLECULE: | TERMINAL OL |
| 98:00:00 | 6wig-A | 4 | 4.3 | 74 | 84 | 5 | MOLECULE: | STENOFOLIA |
| 99:00:00 | 4u8u-G | 4 | 3.6 | 86 | 151 | 5 | MOLECULE: | GLOBIN A CH |
| 0:00 | 5tsz-A | 4 | 6.8 | 86 | 130 | 9 | MOLECULE: | PV CELL-TRA |
| 1:00 | 8rjl-C | 4 | 19.8 | 94 | 273 | 5 | MOLECULE: | CITRATE SYN |
| 2:00 | 7wb4-b | 4 | 15.6 | 97 | 636 | 5 | MOLECULE: | OUTER NUP13 |
| 3:00 | 7xk3-E | 4 | 7.6 | 85 | 198 | 7 | MOLECULE: | NA(+)-TRANS |
| 4:00 | 1y0w-D | 4 | 3.6 | 83 | 146 | 7 | MOLECULE: | HEMOGLOBIN |
| 5:00 | 4hbi-B | 4 | 3.4 | 84 | 145 | 8 | MOLECULE: | HEMOGLOBIN |
| 6:00 | 2z85-A | 4 | 3.4 | 84 | 145 | 11 | MOLECULE: | GLOBIN-1 |
| 7:00 | 6hal-D | 4 | 3.8 | 85 | 145 | 7 | MOLECULE: | HEMOGLOBIN |
| 8:00 | 1hba-D | 4 | 3.6 | 83 | 146 | 7 | MOLECULE: | HEMOGLOBIN |
| 9:00 | 1y7c-B | 4 | 3.6 | 83 | 146 | 7 | MOLECULE: | HEMOGLOBIN |
| 10:00 | 3at5-B | 4 | 4.3 | 86 | 146 | 7 | MOLECULE: | ALPHAA-GLOB |
| 11:00 | 1jzm-B | 4 | 3.5 | 84 | 145 | 8 | MOLECULE: | GLOBIN I - |
| 12:00 | 1nwi-B | 4 | 3.3 | 84 | 145 | 11 | MOLECULE: | GLOBIN I |
| 13:00 | 1y4v-D | 4 | 3.6 | 83 | 146 | 7 | MOLECULE: | HEMOGLOBIN |
| 14:00 | 3vre-D | 4 | 3.7 | 83 | 146 | 7 | MOLECULE: | HEMOGLOBIN |
| 15:00 | 3uhd-A | 4 | 3.4 | 84 | 145 | 11 | MOLECULE: | GLOBIN-1 |
| 16:00 | 1jzk-C | 4 | 3.3 | 83 | 145 | 10 | MOLECULE: | GLOBIN I - |
| 17:00 | 3wr1-B | 4 | 4.3 | 87 | 146 | 10 | MOLECULE: | HEMOGLOBIN |
| 18:00 | 5x2s-L | 4 | 3.2 | 83 | 145 | 6 | MOLECULE: | HEMOGLOBIN |
| 19:00 | 1y7c-D | 4 | 3.6 | 82 | 146 | 7 | MOLECULE: | HEMOGLOBIN |
| 20:00 | 3a59-A | 4 | 3.7 | 86 | 141 | 7 | MOLECULE: | HEMOGLOBIN |
| 21:00 | 2hhe-D | 4 | 3.6 | 83 | 145 | 7 | MOLECULE: | HEMOGLOBIN |
| 22:00 | 1nxf-A | 4 | 3.3 | 82 | 145 | 12 | MOLECULE: | GLOBIN I |
| 23:00 | 1gbu-D | 4 | 3.6 | 83 | 146 | 7 | MOLECULE: | HEMOGLOBIN |
| 24:00:00 | 3a0g-B | 4 | 3.5 | 85 | 144 | 8 | MOLECULE: | HEMOGLOBIN |
| 25:00:00 | 3uhc-A | 4 | 3.3 | 84 | 145 | 11 | MOLECULE: | GLOBIN-1 |
| 26:00:00 | 3sdh-A | 4 | 3.3 | 84 | 145 | 11 | MOLECULE: | HEMOGLOBIN |

| | | | | | | | | |
|---|---|---|---|---|---|---|---|---|
| 27:00:00 | 7ud7-B | 4 | 3.8 | 86 | 146 | 6 | MOLECULE: | HEMOGLOBIN |
| 28:00:00 | 2r4x-B | 4 | 3.5 | 86 | 145 | 9 | MOLECULE: | GLOBIN-1 |
| 29:00:00 | 4f4o-B | 4 | 3.7 | 85 | 146 | 7 | MOLECULE: | HEMOGLOBIN |
| 30:00:00 | 3gys-F | 4 | 3.7 | 85 | 145 | 7 | MOLECULE: | HEMOGLOBIN |
| 31:00:00 | 1a3n-D | 4 | 3.6 | 82 | 145 | 7 | MOLECULE: | HEMOGLOBIN |
| 32:00:00 | 2av3-B | 4 | 3.5 | 84 | 145 | 8 | MOLECULE: | GLOBIN I |
| 33:00:00 | 3dhr-H | 4 | 4.2 | 85 | 145 | 9 | MOLECULE: | HEMOGLOBIN |
| 34:00:00 | 1y0w-B | 4 | 3.6 | 83 | 146 | 7 | MOLECULE: | HEMOGLOBIN |
| 35:00:00 | 2d5z-D | 4 | 3.6 | 82 | 145 | 7 | MOLECULE: | HEMOGLOBIN |
| 36:00:00 | 1ygd-D | 4 | 4.1 | 87 | 146 | 6 | MOLECULE: | HEMOGLOBIN |
| 37:00:00 | 3qob-A | 4 | 3.4 | 85 | 145 | 9 | MOLECULE: | GLOBIN-1 |
| 38:00:00 | 1o1l-D | 4 | 3.9 | 86 | 146 | 6 | MOLECULE: | HEMOGLOBIN |
| 39:00:00 | 4l7y-D | 4 | 3.6 | 82 | 145 | 7 | MOLECULE: | HEMOGLOBIN |
| 40:00:00 | 1yih-D | 4 | 3.8 | 86 | 146 | 6 | MOLECULE: | HEMOGLOBIN |
| 41:00:00 | 2auq-A | 4 | 3.4 | 84 | 145 | 11 | MOLECULE: | GLOBIN I |
| 42:00:00 | 3uhk-B | 4 | 3.3 | 82 | 145 | 12 | MOLECULE: | GLOBIN-1 |
| 43:00:00 | 1jwn-B | 4 | 3.5 | 83 | 145 | 12 | MOLECULE: | GLOBIN I - |
| 44:00:00 | 1fdh-H | 4 | 3.8 | 86 | 146 | 5 | MOLECULE: | HEMOGLOBIN |
| 45:00:00 | 1v5h-A | 4 | 4.1 | 86 | 151 | 10 | MOLECULE: | CYTOGLOBIN |
| 46:00:00 | 2aur-B | 4 | 3.5 | 85 | 145 | 8 | MOLECULE: | GLOBIN I |
| 47:00:00 | 2d60-B | 4 | 3.7 | 83 | 145 | 7 | MOLECULE: | HEMOGLOBIN |
| 48:00:00 | 1y4f-D | 4 | 3.6 | 83 | 146 | 7 | MOLECULE: | HEMOGLOBIN |
| 49:00:00 | 1a0u-B | 4 | 3.7 | 85 | 146 | 6 | MOLECULE: | HEMOGLOBIN |
| 50:00:00 | 1y85-D | 4 | 3.6 | 82 | 145 | 7 | MOLECULE: | HEMOGLOBIN |
| 51:00:00 | 3uht-B | 4 | 3.5 | 85 | 145 | 11 | MOLECULE: | GLOBIN-1 |
| 52:00:00 | 1fdh-G | 4 | 3.8 | 86 | 146 | 5 | MOLECULE: | HEMOGLOBIN |
| 53:00:00 | 3uhe-A | 4 | 3.3 | 84 | 145 | 10 | MOLECULE: | GLOBIN-1 |
| 54:00:00 | 1a0z-D | 4 | 3.6 | 82 | 146 | 7 | MOLECULE: | HEMOGLOBIN |
| 55:00:00 | 1y2z-D | 4 | 3.6 | 82 | 146 | 7 | MOLECULE: | HEMOGLOBIN |
| 56:00:00 | 1jzl-A | 4 | 3.3 | 84 | 145 | 11 | MOLECULE: | GLOBIN I - |
| 57:00:00 | 3ui0-B | 4 | 3.5 | 85 | 145 | 8 | MOLECULE: | GLOBIN-1 |
| 58:00:00 | 3uhz-A | 4 | 3.3 | 84 | 145 | 11 | MOLECULE: | GLOBIN-1 |
| 59:00:00 | 5x2r-F | 4 | 3.6 | 84 | 145 | 7 | MOLECULE: | HEMOGLOBIN |
| 60:00:00 | 3uhg-A | 4 | 3.4 | 84 | 145 | 11 | MOLECULE: | GLOBIN-1 |
| 61:00:00 | 3uht-A | 4 | 3.3 | 83 | 145 | 10 | MOLECULE: | GLOBIN-1 |
| 62:00:00 | 2z85-B | 4 | 3.3 | 83 | 145 | 11 | MOLECULE: | GLOBIN-1 |
| 63:00:00 | 1y4p-D | 4 | 3.8 | 86 | 146 | 6 | MOLECULE: | HEMOGLOBIN |
| 64:00:00 | 3uhz-B | 4 | 3.5 | 84 | 145 | 8 | MOLECULE: | GLOBIN-1 |
| 65:00:00 | 2d60-D | 4 | 3.6 | 82 | 145 | 7 | MOLECULE: | HEMOGLOBIN |
| 66:00:00 | 1a00-B | 4 | 3.7 | 85 | 146 | 6 | MOLECULE: | HEMOGLOBIN |
| 67:00:00 | 3uhi-A | 4 | 3.4 | 84 | 145 | 8 | MOLECULE: | GLOBIN-1 |
| 68:00:00 | 3vre-B | 4 | 3.3 | 83 | 146 | 6 | MOLECULE: | HEMOGLOBIN |
| 69:00:00 | 3uhy-B | 4 | 3.5 | 84 | 145 | 8 | MOLECULE: | GLOBIN-1 |
| 70:00:00 | 3uhk-C | 4 | 3.3 | 84 | 145 | 10 | MOLECULE: | GLOBIN-1 |

| | | | | | | | | |
|---|---|---|---|---|---|---|---|---|
| 71:00:00 | 1nwi-C | 4 | 3.4 | 85 | 145 | 9 | MOLECULE: | GLOBIN I |
| 72:00:00 | 3g4q-A | 4 | 3.5 | 86 | 145 | 9 | MOLECULE: | GLOBIN-1 |
| 73:00:00 | 4n7n-C | 4 | 4.1 | 86 | 141 | 6 | MOLECULE: | HEMOGLOBIN |
| 74:00:00 | 3g4w-B | 4 | 3.5 | 84 | 145 | 8 | MOLECULE: | GLOBIN-1 |
| 75:00:00 | 3g53-A | 4 | 3.3 | 84 | 145 | 10 | MOLECULE: | GLOBIN-1 |
| 76:00:00 | 7vuc-A | 4 | 4.9 | 77 | 153 | 12 | MOLECULE: | MYOGLOBIN |
| 77:00:00 | 7hbi-A | 4 | 3.3 | 83 | 145 | 8 | MOLECULE: | HEMOGLOBIN |
| 78:00:00 | 3gys-B | 4 | 3.7 | 85 | 145 | 7 | MOLECULE: | HEMOGLOBIN |
| 79:00:00 | 3uhx-A | 4 | 3.3 | 84 | 145 | 10 | MOLECULE: | GLOBIN-1 |
| 80:00:00 | 3ugy-A | 4 | 3.3 | 84 | 145 | 10 | MOLECULE: | GLOBIN-1 |
| 81:00:00 | 3uhw-A | 4 | 3.3 | 84 | 145 | 10 | MOLECULE: | GLOBIN-1 |
| 82:00:00 | 3uh7-A | 4 | 3.3 | 84 | 145 | 10 | MOLECULE: | GLOBIN-1 |
| 83:00:00 | 3uhr-B | 4 | 3.3 | 83 | 145 | 8 | MOLECULE: | GLOBIN-1 |
| 84:00:00 | 1jzk-D | 4 | 3.3 | 84 | 145 | 11 | MOLECULE: | GLOBIN I - |
| 85:00:00 | 3ag0-A | 4 | 4 | 89 | 158 | 10 | MOLECULE: | CYTOGLOBIN |
| 86:00:00 | 1jwn-D | 4 | 3.3 | 84 | 145 | 10 | MOLECULE: | GLOBIN I - |
| 87:00:00 | 3gqr-F | 4 | 3.5 | 84 | 145 | 8 | MOLECULE: | HEMOGLOBIN |
| 88:00:00 | 2hhe-B | 4 | 4.2 | 85 | 145 | 7 | MOLECULE: | HEMOGLOBIN |
| 89:00:00 | 5jdo-D | 4 | 3.5 | 84 | 145 | 6 | MOLECULE: | HAPTOGLOBIN |
| 90:00:00 | 1y35-D | 4 | 3.6 | 83 | 146 | 7 | MOLECULE: | HEMOGLOBIN |
| 91:00:00 | 1shr-B | 4 | 3.6 | 84 | 146 | 5 | MOLECULE: | HEMOGLOBIN |
| 92:00:00 | 1a3o-D | 4 | 3.7 | 83 | 145 | 7 | MOLECULE: | HEMOGLOBIN |
| 93:00:00 | 5e83-B | 4 | 3.7 | 84 | 146 | 6 | MOLECULE: | HEMOGLOBIN |
| 94:00:00 | 5hbi-A | 4 | 3.4 | 84 | 145 | 10 | MOLECULE: | HEMOGLOBIN |
| 95:00:00 | 3gou-D | 4 | 3.6 | 85 | 146 | 5 | MOLECULE: | HEMOGLOBIN |
| 96:00:00 | 3uhu-A | 4 | 3.3 | 83 | 145 | 8 | MOLECULE: | GLOBIN-1 |
| 97:00:00 | 3uhv-A | 4 | 3.3 | 84 | 145 | 10 | MOLECULE: | GLOBIN-1 |
| 98:00:00 | 3uhy-A | 4 | 3.3 | 82 | 146 | 12 | MOLECULE: | GLOBIN-1 |
| 99:00:00 | 3uhv-B | 4 | 3.5 | 86 | 145 | 10 | MOLECULE: | GLOBIN-1 |
| 0:00 | 3uh6-A | 4 | 3.4 | 85 | 145 | 11 | MOLECULE: | GLOBIN-1 |
| 1:00 | 8dov-F | 4 | 4.2 | 85 | 144 | 7 | MOLECULE: | HEMOGLOBIN |
| 2:00 | 5x2r-D | 4 | 3.7 | 85 | 145 | 6 | MOLECULE: | HEMOGLOBIN |
| 3:00 | 3uh3-A | 4 | 3.3 | 83 | 145 | 11 | MOLECULE: | GLOBIN-1 |
| 4:00 | 1ch4-D | 4 | 3.5 | 84 | 146 | 5 | MOLECULE: | MODULE-SUBS |
| 5:00 | 2aup-B | 4 | 3.5 | 84 | 145 | 12 | MOLECULE: | GLOBIN I |
| 6:00 | 3gqr-D | 4 | 3.7 | 84 | 145 | 7 | MOLECULE: | HEMOGLOBIN |
| 7:00 | 3g46-A | 4 | 3.5 | 85 | 146 | 8 | MOLECULE: | GLOBIN-1 |
| 8:00 | 1nej-D | 4 | 3.6 | 85 | 146 | 7 | MOLECULE: | HEMOGLOBIN |
| 9:00 | 2grf-A | 4 | 3.3 | 83 | 145 | 8 | MOLECULE: | GLOBIN-1 |
| 10:00 | 7uf6-B | 4 | 3.6 | 84 | 146 | 7 | MOLECULE: | HEMOGLOBIN |
| 11:00 | 5x2t-L | 4 | 3.4 | 84 | 145 | 7 | MOLECULE: | HEMOGLOBIN |
| 12:00 | 3uhu-B | 4 | 3.8 | 83 | 145 | 11 | MOLECULE: | GLOBIN-1 |
| 13:00 | 3uhr-A | 4 | 3.3 | 83 | 145 | 12 | MOLECULE: | GLOBIN-1 |
| 14:00 | 3uhn-A | 4 | 3.3 | 83 | 145 | 8 | MOLECULE: | GLOBIN-1 |

| | | | | | | | | |
|---|---|---|---|---|---|---|---|---|
| 15:00 | 6hbi-A | 4 | 3.6 | 85 | 145 | 9 | MOLECULE: | HEMOGLOBIN |
| 16:00 | 3uhs-A | 4 | 3.4 | 84 | 145 | 10 | MOLECULE: | GLOBIN-1 |
| 17:00 | 3ugz-A | 4 | 3.4 | 84 | 145 | 11 | MOLECULE: | GLOBIN-1 |
| 18:00 | 3ui0-A | 4 | 3.4 | 84 | 145 | 10 | MOLECULE: | GLOBIN-1 |
| 19:00 | 3gqr-H | 4 | 3.6 | 84 | 145 | 7 | MOLECULE: | HEMOGLOBIN |
| 20:00 | 1ch4-C | 4 | 3.5 | 84 | 146 | 5 | MOLECULE: | MODULE-SUBS |
| 21:00 | 3uhq-A | 4 | 3.3 | 83 | 145 | 8 | MOLECULE: | GLOBIN-1 |
| 22:00 | 6wg7-E | 4 | 8.3 | 102 | 228 | 8 | MOLECULE: | DNA (35-MER |
| 23:00 | 8qfs-A | 3.9 | 13 | 100 | 1183 | 12 | MOLECULE: | ELONGATION |
| 24:00:00 | 5y0r-A | 3.9 | 14.2 | 112 | 423 | 4 | MOLECULE: | TMTMCAL(SEM |
| 25:00:00 | 7m8i-A | 3.9 | 9.7 | 151 | 572 | 6 | MOLECULE: | ADRENODOXIN |
| 26:00:00 | 6m9a-C | 3.9 | 4.3 | 87 | 157 | 13 | MOLECULE: | SIGNALING P |
| 27:00:00 | 8q4h-C | 3.9 | 4.3 | 70 | 89 | 7 | MOLECULE: | TETRACHLORO |
| 28:00:00 | 3bhn-A | 3.9 | 5.5 | 81 | 214 | 9 | MOLECULE: | THIJ/PFPI D |
| 29:00:00 | 8q6o-2 | 3.9 | 6.5 | 92 | 267 | 9 | MOLECULE: | DNA REPLICA |
| 30:00:00 | 1or4-A | 3.9 | 5.5 | 106 | 169 | 8 | MOLECULE: | HEME-BASED |
| 31:00:00 | 6z16-d | 3.9 | 9.2 | 104 | 489 | 8 | MOLECULE: | MULTISUBUNI |
| 32:00:00 | 2p6v-A | 3.9 | 3.3 | 77 | 97 | 10 | MOLECULE: | TRANSCRIPTI |
| 33:00:00 | 2hvr-A | 3.9 | 11.5 | 90 | 319 | 6 | MOLECULE: | 5'- |
| 34:00:00 | 7ewp-A | 3.9 | 9.7 | 106 | 577 | 6 | MOLECULE: | PROBABLE G- |
| 35:00:00 | 3eqm-A | 3.9 | 4.5 | 147 | 452 | 4 | MOLECULE: | CYTOCHROME |
| 36:00:00 | 6zka-K | 3.9 | 8 | 81 | 98 | 7 | MOLECULE: | NADH-UBIQUI |
| 37:00:00 | 3mjp-B | 3.9 | 3.6 | 85 | 146 | 9 | MOLECULE: | HEMOGLOBIN |
| 38:00:00 | 4l7y-B | 3.9 | 3.6 | 83 | 145 | 7 | MOLECULE: | HEMOGLOBIN |
| 39:00:00 | 6hal-B | 3.9 | 3.9 | 85 | 145 | 7 | MOLECULE: | HEMOGLOBIN |
| 40:00:00 | 4f4o-E | 3.9 | 3.7 | 85 | 146 | 7 | MOLECULE: | HEMOGLOBIN |
| 41:00:00 | 3uhi-B | 3.9 | 3.4 | 83 | 145 | 10 | MOLECULE: | GLOBIN-1 |
| 42:00:00 | 5x2s-J | 3.9 | 3.8 | 83 | 145 | 7 | MOLECULE: | HEMOGLOBIN |
| 43:00:00 | 3uhq-B | 3.9 | 3.6 | 83 | 145 | 12 | MOLECULE: | GLOBIN-1 |
| 44:00:00 | 5e29-B | 3.9 | 3.6 | 82 | 145 | 7 | MOLECULE: | HEMOGLOBIN |
| 45:00:00 | 5x2s-C | 3.9 | 4.2 | 86 | 140 | 6 | MOLECULE: | HEMOGLOBIN |
| 46:00:00 | 5x2r-E | 3.9 | 4.1 | 86 | 140 | 6 | MOLECULE: | HEMOGLOBIN |
| 47:00:00 | 3ugy-B | 3.9 | 3.3 | 81 | 145 | 11 | MOLECULE: | GLOBIN-1 |
| 48:00:00 | 4f4o-H | 3.9 | 3.7 | 85 | 146 | 7 | MOLECULE: | HEMOGLOBIN |
| 49:00:00 | 2aup-A | 3.9 | 3.4 | 84 | 145 | 11 | MOLECULE: | GLOBIN I |
| 50:00:00 | 1a0z-B | 3.9 | 3.7 | 85 | 146 | 6 | MOLECULE: | HEMOGLOBIN |
| 51:00:00 | 7k4m-E | 3.9 | 3.9 | 86 | 142 | 6 | MOLECULE: | HEMOGLOBIN |
| 52:00:00 | 3sdh-B | 3.9 | 3.4 | 83 | 145 | 8 | MOLECULE: | HEMOGLOBIN |
| 53:00:00 | 3uhc-B | 3.9 | 3.4 | 81 | 145 | 12 | MOLECULE: | GLOBIN-1 |
| 54:00:00 | 3g4y-B | 3.9 | 3.4 | 82 | 145 | 12 | MOLECULE: | GLOBIN-1 |
| 55:00:00 | 3uh7-B | 3.9 | 3.4 | 83 | 145 | 8 | MOLECULE: | GLOBIN-1 |
| 56:00:00 | 3uhe-B | 3.9 | 3.4 | 83 | 145 | 8 | MOLECULE: | GLOBIN-1 |
| 57:00:00 | 3g52-B | 3.9 | 3.4 | 82 | 145 | 12 | MOLECULE: | GLOBIN-1 |
| 58:00:00 | 3uhi-C | 3.9 | 3.3 | 84 | 145 | 10 | MOLECULE: | GLOBIN-1 |

| | | | | | | | | |
|---|---|---|---|---|---|---|---|---|
| 59:00:00 | 3g4v-B | 3.9 | 3.4 | 82 | 145 | 12 | MOLECULE: | GLOBIN-1 |
| 60:00:00 | 1jwn-C | 3.9 | 3.4 | 84 | 145 | 11 | MOLECULE: | GLOBIN I - |
| 61:00:00 | 1bij-C | 3.9 | 3.8 | 86 | 141 | 7 | MOLECULE: | HEMOGLOBIN |
| 62:00:00 | 2aur-A | 3.9 | 3.3 | 82 | 145 | 12 | MOLECULE: | GLOBIN I |
| 63:00:00 | 3kmf-E | 3.9 | 3.8 | 86 | 141 | 7 | MOLECULE: | HEMOGLOBIN |
| 64:00:00 | 3uhb-B | 3.9 | 3.4 | 83 | 145 | 8 | MOLECULE: | GLOBIN-1 |
| 65:00:00 | 3uhh-B | 3.9 | 3.6 | 83 | 145 | 12 | MOLECULE: | GLOBIN-1 |
| 66:00:00 | 2auo-B | 3.9 | 3.4 | 82 | 145 | 12 | MOLECULE: | GLOBIN I |
| 67:00:00 | 2m6z-A | 3.9 | 3.2 | 85 | 141 | 5 | MOLECULE: | HEMOGLOBIN |
| 68:00:00 | 1hds-C | 3.9 | 3.6 | 86 | 141 | 6 | MOLECULE: | HEMOGLOBIN |
| 69:00:00 | 3g4q-B | 3.9 | 3.4 | 83 | 145 | 8 | MOLECULE: | GLOBIN-1 |
| 70:00:00 | 2grh-B | 3.9 | 3.5 | 84 | 145 | 8 | MOLECULE: | GLOBIN-1 |
| 71:00:00 | 7hbi-B | 3.9 | 3.4 | 82 | 145 | 12 | MOLECULE: | HEMOGLOBIN |
| 72:00:00 | 3uh6-B | 3.9 | 3.4 | 83 | 145 | 8 | MOLECULE: | GLOBIN-1 |
| 73:00:00 | 3uhg-B | 3.9 | 3.4 | 82 | 145 | 12 | MOLECULE: | GLOBIN-1 |
| 74:00:00 | 5x2u-J | 3.9 | 3.7 | 83 | 145 | 7 | MOLECULE: | HEMOGLOBIN |
| 75:00:00 | 3d4x-B | 3.9 | 4.1 | 85 | 145 | 7 | MOLECULE: | HEMOGLOBIN |
| 76:00:00 | 8vyl-B | 3.9 | 4.1 | 84 | 145 | 7 | MOLECULE: | HEMOGLOBIN |
| 77:00:00 | 3hyu-B | 3.9 | 3.7 | 86 | 146 | 8 | MOLECULE: | HEMOGLOBIN |
| 78:00:00 | 3d4x-D | 3.9 | 3.8 | 83 | 145 | 7 | MOLECULE: | HEMOGLOBIN |
| 79:00:00 | 3mjp-D | 3.9 | 3.6 | 85 | 146 | 9 | MOLECULE: | HEMOGLOBIN |
| 80:00:00 | 4h2l-B | 3.9 | 3.7 | 85 | 146 | 7 | MOLECULE: | ALPHA-GLOBI |
| 81:00:00 | 5x2u-L | 3.9 | 4.2 | 85 | 145 | 6 | MOLECULE: | HEMOGLOBIN |
| 82:00:00 | 2m6z-C | 3.9 | 3.2 | 85 | 141 | 5 | MOLECULE: | HEMOGLOBIN |
| 83:00:00 | 1jzm-A | 3.9 | 3.4 | 82 | 145 | 10 | MOLECULE: | GLOBIN I - |
| 84:00:00 | 7k4m-D | 3.9 | 3.5 | 84 | 147 | 7 | MOLECULE: | HEMOGLOBIN |
| 85:00:00 | 1jwn-A | 3.9 | 3.4 | 82 | 145 | 11 | MOLECULE: | GLOBIN I - |
| 86:00:00 | 5x2t-A | 3.9 | 4.1 | 87 | 140 | 7 | MOLECULE: | HEMOGLOBIN |
| 87:00:00 | 5x2s-H | 3.9 | 3.5 | 83 | 145 | 6 | MOLECULE: | HEMOGLOBIN |
| 88:00:00 | 3gqr-B | 3.9 | 3.7 | 85 | 145 | 7 | MOLECULE: | HEMOGLOBIN |
| 89:00:00 | 3gou-B | 3.9 | 4.2 | 85 | 146 | 6 | MOLECULE: | HEMOGLOBIN |
| 90:00:00 | 2r4y-B | 3.9 | 3.5 | 82 | 145 | 10 | MOLECULE: | GLOBIN-1 |
| 91:00:00 | 5ufj-B | 3.9 | 3.7 | 84 | 146 | 6 | MOLECULE: | HEMOGLOBIN |
| 92:00:00 | 7jjq-D | 3.9 | 4.1 | 85 | 144 | 6 | MOLECULE: | HEMOGLOBIN |
| 93:00:00 | 5ker-H | 3.9 | 3.7 | 86 | 146 | 8 | MOLECULE: | ALPHA-GLOBI |
| 94:00:00 | 1x3k-A | 3.8 | 3.8 | 81 | 152 | 9 | MOLECULE: | HEMOGLOBIN |
| 95:00:00 | 4bja-A | 3.8 | 5.3 | 99 | 182 | 7 | MOLECULE: | PROTEIN GLB |
| 96:00:00 | 8h17-A | 3.8 | 5 | 88 | 157 | 7 | MOLECULE: | TLR1989 PRO |
| 97:00:00 | 4m8m-B | 3.8 | 6.4 | 118 | 574 | 3 | MOLECULE: | GCN4 COILED |
| 98:00:00 | 2q0y-A | 3.8 | 3.8 | 57 | 153 | 4 | MOLECULE: | GCN5-RELATE |
| 99:00:00 | 5o5e-A | 3.8 | 10.7 | 129 | 381 | 1 | MOLECULE: | UDP-N-ACETY |
| 0:00 | 7xdi-D | 3.8 | 5.8 | 79 | 124 | 6 | MOLECULE: | VP1 |
| 1:00 | 2da6-A | 3.8 | 5 | 76 | 102 | 5 | MOLECULE: | HEPATOCYTE |
| 2:00 | 7w0y-k | 3.8 | 6.3 | 81 | 98 | 6 | MOLECULE: | NADH DEHYDR |

| | | | | | | | | |
|---|---|---|---|---|---|---|---|---|
| 3:00 | 5ex8-A | 3.8 | 5.9 | 143 | 414 | 8 | MOLECULE: | CYTOCHROME |
| 4:00 | 7uw5-A | 3.8 | 14 | 99 | 723 | 9 | MOLECULE: | MECHANOSENS |
| 5:00 | 3wft-A | 3.8 | 4 | 81 | 153 | 14 | MOLECULE: | MYOGLOBIN |
| 6:00 | 1shr-D | 3.8 | 4.8 | 85 | 146 | 4 | MOLECULE: | HEMOGLOBIN |
| 7:00 | 2d5z-B | 3.8 | 3.9 | 84 | 145 | 5 | MOLECULE: | HEMOGLOBIN |
| 8:00 | 5x2t-F | 3.8 | 3.7 | 84 | 145 | 6 | MOLECULE: | HEMOGLOBIN |
| 9:00 | 7spf-A | 3.8 | 3.8 | 84 | 152 | 11 | MOLECULE: | MYOGLOBIN |
| 10:00 | 6bmg-A | 3.8 | 4 | 85 | 154 | 9 | MOLECULE: | MYOGLOBIN |
| 11:00 | 3g4u-B | 3.8 | 4.1 | 84 | 145 | 15 | MOLECULE: | GLOBIN-1 |
| 12:00 | 2r4w-B | 3.8 | 3.6 | 83 | 145 | 11 | MOLECULE: | GLOBIN-1 |
| 13:00 | 3uh3-B | 3.8 | 3.6 | 82 | 145 | 13 | MOLECULE: | GLOBIN-1 |
| 14:00 | 6on4-A | 3.8 | 3.9 | 91 | 217 | 8 | MOLECULE: | HTH-TYPE TR |
| 15:00 | 3uh5-B | 3.8 | 3.5 | 84 | 145 | 8 | MOLECULE: | GLOBIN-1 |
| 16:00 | 2b7h-B | 3.8 | 4.6 | 86 | 144 | 6 | MOLECULE: | HEMOGLOBIN |
| 17:00 | 5ufj-D | 3.8 | 4.1 | 86 | 146 | 6 | MOLECULE: | HEMOGLOBIN |
| 18:00 | 7ud8-B | 3.8 | 4.1 | 85 | 146 | 6 | MOLECULE: | HEMOGLOBIN |
| 19:00 | 3g4r-B | 3.8 | 4 | 84 | 145 | 15 | MOLECULE: | GLOBIN-1 |
| 20:00 | 3uhd-B | 3.8 | 3.6 | 85 | 144 | 9 | MOLECULE: | GLOBIN-1 |
| 21:00 | 1nej-B | 3.8 | 3.6 | 84 | 146 | 7 | MOLECULE: | HEMOGLOBIN |
| 22:00 | 5x2u-B | 3.8 | 4.2 | 86 | 145 | 6 | MOLECULE: | HEMOGLOBIN |
| 23:00 | 5x2r-L | 3.8 | 4.2 | 86 | 145 | 7 | MOLECULE: | HEMOGLOBIN |
| 24:00:00 | 5x2r-J | 3.8 | 4.1 | 88 | 145 | 6 | MOLECULE: | HEMOGLOBIN |
| 25:00:00 | 5x2t-B | 3.8 | 4.2 | 86 | 145 | 6 | MOLECULE: | HEMOGLOBIN |
| 26:00:00 | 5x2u-F | 3.8 | 4.1 | 84 | 145 | 7 | MOLECULE: | HEMOGLOBIN |
| 27:00:00 | 5jdo-F | 3.8 | 4 | 84 | 144 | 7 | MOLECULE: | HAPTOGLOBIN |
| 28:00:00 | 1nwn-B | 3.8 | 3.9 | 84 | 145 | 12 | MOLECULE: | GLOBIN I |
| 29:00:00 | 6wfq-C | 3.8 | 8.1 | 107 | 228 | 7 | MOLECULE: | HTH-TYPE TR |
| 30:00:00 | 4g51-C | 3.8 | 3.7 | 86 | 142 | 6 | MOLECULE: | HEMOGLOBIN |
| 31:00:00 | 2pgh-D | 3.8 | 4 | 84 | 146 | 8 | MOLECULE: | HEMOGLOBIN |
| 32:00:00 | 5v7p-A | 3.7 | 6.2 | 101 | 281 | 7 | MOLECULE: | PROTEIN-S-I |
| 33:00:00 | 4noo-A | 3.7 | 3.7 | 77 | 200 | 6 | MOLECULE: | VGRG PROTEI |
| 34:00:00 | 7dpd-B | 3.7 | 4.3 | 80 | 273 | 4 | MOLECULE: | DNA HELICAS |
| 35:00:00 | 7lt2-A | 3.7 | 17.1 | 101 | 387 | 10 | MOLECULE: | MAB-21 DOMA |
| 36:00:00 | 1x46-A | 3.7 | 4 | 80 | 150 | 6 | MOLECULE: | HEMOGLOBIN |
| 37:00:00 | 8b6l-M | 3.7 | 6.4 | 74 | 110 | 3 | MOLECULE: | PROTEIN TRA |
| 38:00:00 | 5ca9-A | 3.7 | 25.6 | 116 | 656 | 7 | MOLECULE: | PROTEIN SEY |
| 39:00:00 | 7sjr-A | 3.7 | 7.7 | 99 | 911 | 10 | MOLECULE: | DNA HELICAS |
| 40:00:00 | 6ezn-B | 3.7 | 7.5 | 77 | 110 | 4 | MOLECULE: | DOLICHYL-DI |
| 41:00:00 | 8pi9-B | 3.7 | 10.3 | 92 | 177 | 8 | MOLECULE: | CHAINS: E |
| 42:00:00 | 2zfo-D | 3.7 | 3.8 | 83 | 145 | 8 | MOLECULE: | EXTRACELLUL |
| 43:00:00 | 8jem-B | 3.7 | 9 | 97 | 419 | 6 | MOLECULE: | TEICHOIC AC |
| 44:00:00 | 7rb4-A | 3.7 | 18.8 | 122 | 603 | 7 | MOLECULE: | DIPHTHERIA |
| 45:00:00 | 7k4m-B | 3.7 | 4.2 | 86 | 146 | 5 | MOLECULE: | HEMOGLOBIN |
| 46:00:00 | 3uhi-D | 3.7 | 3.8 | 83 | 145 | 12 | MOLECULE: | GLOBIN-1 |

| | | | | | | | | |
|---|---|---|---|---|---|---|---|---|
| 47:00:00 | 1hbi-A | 3.7 | 3.9 | 84 | 145 | 12 | MOLECULE: | HEMOGLOBIN |
| 48:00:00 | 5jom-A | 3.7 | 3.7 | 84 | 154 | 11 | MOLECULE: | MYOGLOBIN |
| 49:00:00 | 1ch4-B | 3.7 | 3.9 | 84 | 146 | 8 | MOLECULE: | MODULE-SUBS |
| 50:00:00 | 2r4z-A | 3.7 | 3.8 | 82 | 145 | 13 | MOLECULE: | GLOBIN-1 |
| 51:00:00 | 5ych-A | 3.7 | 3.8 | 84 | 153 | 12 | MOLECULE: | ANCESTRAL M |
| 52:00:00 | 2grh-A | 3.7 | 3.9 | 84 | 145 | 12 | MOLECULE: | GLOBIN-1 |
| 53:00:00 | 5x2t-H | 3.7 | 3.8 | 83 | 145 | 7 | MOLECULE: | HEMOGLOBIN |
| 54:00:00 | 1ch4-A | 3.7 | 3.9 | 84 | 146 | 8 | MOLECULE: | MODULE-SUBS |
| 55:00:00 | 4f4o-K | 3.7 | 4.3 | 85 | 146 | 8 | MOLECULE: | HEMOGLOBIN |
| 56:00:00 | 2grf-B | 3.7 | 3.9 | 85 | 145 | 12 | MOLECULE: | GLOBIN-1 |
| 57:00:00 | 1jzl-B | 3.7 | 4 | 84 | 145 | 15 | MOLECULE: | GLOBIN I - |
| 58:00:00 | 5m3r-A | 3.7 | 3.7 | 84 | 154 | 11 | MOLECULE: | MYOGLOBIN |
| 59:00:00 | 5hbi-B | 3.7 | 4 | 84 | 145 | 15 | MOLECULE: | HEMOGLOBIN |
| 60:00:00 | 7a45-A | 3.7 | 3.7 | 84 | 154 | 11 | MOLECULE: | MYOGLOBIN |
| 61:00:00 | 3uhh-A | 3.7 | 3.9 | 84 | 145 | 14 | MOLECULE: | GLOBIN-1 |
| 62:00:00 | 2auq-B | 3.7 | 4 | 84 | 145 | 15 | MOLECULE: | GLOBIN I |
| 63:00:00 | 4hbi-A | 3.7 | 3.8 | 83 | 145 | 12 | MOLECULE: | HEMOGLOBIN |
| 64:00:00 | 3g53-B | 3.7 | 4.1 | 84 | 145 | 15 | MOLECULE: | GLOBIN-1 |
| 65:00:00 | 2av0-A | 3.7 | 3.9 | 84 | 145 | 12 | MOLECULE: | GLOBIN I |
| 66:00:00 | 2pgh-B | 3.7 | 3.7 | 84 | 146 | 6 | MOLECULE: | HEMOGLOBIN |
| 67:00:00 | 3qob-B | 3.7 | 4 | 84 | 145 | 14 | MOLECULE: | GLOBIN-1 |
| 68:00:00 | 5x2u-H | 3.7 | 3.8 | 84 | 145 | 6 | MOLECULE: | HEMOGLOBIN |
| 69:00:00 | 6bb5-B | 3.7 | 3.8 | 83 | 145 | 6 | MOLECULE: | HEMOGLOBIN |
| 70:00:00 | 1mbs-A | 3.7 | 3.9 | 91 | 153 | 11 | MOLECULE: | MYOGLOBIN |
| 71:00:00 | 2av0-B | 3.7 | 4 | 84 | 145 | 15 | MOLECULE: | GLOBIN I |
| 72:00:00 | 1si4-D | 3.7 | 4.2 | 84 | 146 | 6 | MOLECULE: | HEMOGLOBIN |
| 73:00:00 | 6kye-H | 3.7 | 4.3 | 86 | 145 | 7 | MOLECULE: | HEMOGLOBIN |
| 74:00:00 | 7uvb-D | 3.7 | 4.1 | 84 | 145 | 7 | MOLECULE: | HEMOGLOBIN |
| 75:00:00 | 3ugz-B | 3.7 | 4.1 | 84 | 145 | 15 | MOLECULE: | GLOBIN-1 |
| 76:00:00 | 4ood-A | 3.7 | 3.5 | 80 | 154 | 11 | MOLECULE: | MYOGLOBIN |
| 77:00:00 | 2r4y-A | 3.7 | 3.9 | 84 | 145 | 14 | MOLECULE: | GLOBIN-1 |
| 78:00:00 | 3g46-B | 3.7 | 3.7 | 82 | 146 | 13 | MOLECULE: | GLOBIN-1 |
| 79:00:00 | 4x0l-B | 3.7 | 4.2 | 85 | 144 | 5 | MOLECULE: | HEMOGLOBIN |
| 80:00:00 | 2z8a-A | 3.7 | 3.9 | 85 | 145 | 11 | MOLECULE: | GLOBIN-1 |
| 81:00:00 | 5x2s-D | 3.7 | 4.2 | 85 | 145 | 6 | MOLECULE: | HEMOGLOBIN |
| 82:00:00 | 3hf4-B | 3.7 | 3.8 | 85 | 145 | 6 | MOLECULE: | HEMOGLOBIN |
| 83:00:00 | 5ker-D | 3.7 | 4.3 | 85 | 146 | 8 | MOLECULE: | ALPHA-GLOBI |
| 84:00:00 | 2auo-A | 3.7 | 3.9 | 85 | 145 | 12 | MOLECULE: | GLOBIN I |
| 85:00:00 | 3uh5-A | 3.7 | 3.9 | 85 | 145 | 12 | MOLECULE: | GLOBIN-1 |
| 86:00:00 | 6wg7-C | 3.7 | 8.1 | 107 | 228 | 7 | MOLECULE: | DNA (35-MER |
| 87:00:00 | 6wg7-F | 3.7 | 8.2 | 102 | 228 | 7 | MOLECULE: | DNA (35-MER |
| 88:00:00 | 3uhb-A | 3.7 | 3.9 | 85 | 145 | 12 | MOLECULE: | GLOBIN-1 |
| 89:00:00 | 3vw4-A | 3.6 | 18.9 | 85 | 122 | 7 | MOLECULE: | REP |
| 90:00:00 | 4ktp-A | 3.6 | 16.3 | 100 | 767 | 9 | MOLECULE: | GLYCOSIDE H |

| | | | | | | | | |
|---|---|---|---|---|---|---|---|---|
| 91:00:00 | 1us7-B | 3.6 | 20.6 | 100 | 193 | 2 | MOLECULE: | HEAT SHOCK |
| 92:00:00 | 6rd4-4 | 3.6 | 14 | 92 | 290 | 8 | MOLECULE: | ASA-10: POL |
| 93:00:00 | 4u6u-A | 3.6 | 4.5 | 59 | 67 | 7 | MOLECULE: | COG7 |
| 94:00:00 | 3bes-R | 3.6 | 11.8 | 88 | 250 | 9 | MOLECULE: | INTERFERON |
| 95:00:00 | 8slm-A | 3.6 | 8 | 96 | 248 | 10 | MOLECULE: | ZN DEPENDEN |
| 96:00:00 | 3v8d-A | 3.6 | 4.9 | 142 | 479 | 3 | MOLECULE: | CHOLESTEROL |
| 97:00:00 | 1lqs-L | 3.6 | 11.7 | 99 | 142 | 10 | MOLECULE: | INTERLEUKIN |
| 98:00:00 | 3zjh-B | 3.6 | 7.2 | 99 | 193 | 10 | MOLECULE: | PROTOGLOBIN |
| 99:00:00 | 1puf-A | 3.6 | 4.5 | 67 | 77 | 3 | MOLECULE: | 5'- |
| 0:00 | 8w20-A | 3.6 | 31.8 | 65 | 732 | 9 | MOLECULE: | SECRETED PR |
| 1:00 | 7etw-B | 3.6 | 22 | 108 | 483 | 6 | MOLECULE: | INSULIN-IND |
| 2:00 | 6s8h-F | 3.6 | 5 | 85 | 239 | 7 | MOLECULE: | LIPOPOLYSAC |
| 3:00 | 5eik-A | 3.6 | 9.3 | 87 | 234 | 6 | MOLECULE: | UNCHARACTER |
| 4:00 | 6iuy-A | 3.6 | 10.4 | 89 | 585 | 2 | MOLECULE: | GLYCEROL-3- |
| 5:00 | 6ygb-B | 3.6 | 19.2 | 106 | 725 | 5 | MOLECULE: | N-ALPHA-ACE |
| 6:00 | 7z8b-C | 3.6 | 26.3 | 131 | 1224 | 9 | MOLECULE: | CULLIN-7 |
| 7:00 | 1nek-D | 3.6 | 9.1 | 84 | 113 | 6 | MOLECULE: | SUCCINATE D |
| 8:00 | 1yvl-B | 3.6 | 13.9 | 107 | 653 | 7 | MOLECULE: | SIGNAL TRAN |
| 9:00 | 8hmc-B | 3.6 | 27.3 | 131 | 1195 | 4 | MOLECULE: | INTRAFLAGEL |
| 10:00 | 7uic-e | 3.6 | 23.8 | 92 | 729 | 12 | MOLECULE: | MEDIATOR OF |
| 11:00 | 7yug-A | 3.6 | 2.9 | 63 | 117 | 3 | MOLECULE: | PROTEIN BAN |
| 12:00 | 8fib-A | 3.6 | 5.2 | 134 | 418 | 10 | MOLECULE: | CYTOCHROME |
| 13:00 | 2z8a-B | 3.6 | 4.1 | 84 | 146 | 14 | MOLECULE: | GLOBIN-1 |
| 14:00 | 2av3-A | 3.6 | 3.9 | 84 | 145 | 12 | MOLECULE: | GLOBIN I |
| 15:00 | 1hbs-F | 3.6 | 3.7 | 84 | 146 | 6 | MOLECULE: | HEMOGLOBIN |
| 16:00 | 2r4z-B | 3.6 | 4 | 84 | 146 | 14 | MOLECULE: | GLOBIN-1 |
| 17:00 | 1hbi-B | 3.6 | 4.1 | 84 | 145 | 15 | MOLECULE: | HEMOGLOBIN |
| 18:00 | 5ycj-A | 3.6 | 4 | 81 | 154 | 12 | MOLECULE: | ANCESTRAL M |
| 19:00 | 8f9i-A | 3.6 | 3.7 | 80 | 154 | 11 | MOLECULE: | MYOGLOBIN |
| 20:00 | 1mnh-A | 3.6 | 3.7 | 80 | 153 | 15 | MOLECULE: | MYOGLOBIN |
| 21:00 | 7uvb-B | 3.6 | 4.3 | 86 | 146 | 7 | MOLECULE: | HEMOGLOBIN |
| 22:00 | 5x2r-H | 3.6 | 4.2 | 88 | 145 | 8 | MOLECULE: | HEMOGLOBIN |
| 23:00 | 1it3-B | 3.6 | 4 | 80 | 146 | 6 | MOLECULE: | HEMOGLOBIN |
| 24:00:00 | 7ddr-A | 3.6 | 4.6 | 90 | 154 | 10 | MOLECULE: | ANCESTRAL M |
| 25:00:00 | 1nwn-A | 3.6 | 3.9 | 84 | 145 | 12 | MOLECULE: | GLOBIN I |
| 26:00:00 | 2r4x-A | 3.6 | 3.9 | 84 | 145 | 12 | MOLECULE: | GLOBIN-1 |
| 27:00:00 | 1z8u-B | 3.6 | 3.6 | 81 | 135 | 7 | MOLECULE: | ALPHA-HEMOG |
| 28:00:00 | 1it3-D | 3.6 | 3.2 | 76 | 146 | 7 | MOLECULE: | HEMOGLOBIN |
| 29:00:00 | 1z8u-D | 3.6 | 3.7 | 81 | 135 | 7 | MOLECULE: | ALPHA-HEMOG |
| 30:00:00 | 3al0-B | 3.5 | 13.3 | 79 | 482 | 10 | MOLECULE: | GLUTAMYL-TR |
| 31:00:00 | 6rfl-I | 3.5 | 20.7 | 119 | 773 | 9 | MOLECULE: | DNA-DEPENDE |
| 32:00:00 | 4zi2-C | 3.5 | 4.1 | 80 | 131 | 6 | MOLECULE: | ADP-RIBOSYL |
| 33:00:00 | 7kzn-C | 3.5 | 34.3 | 121 | 965 | 7 | MOLECULE: | HEAVY CHAIN |
| 34:00:00 | 5o1l-A | 3.5 | 4.3 | 109 | 375 | 6 | MOLECULE: | RUBBER OXYG |

| | | | | | | | | |
|---|---|---|---|---|---|---|---|---|
| 35:00:00 | 8wu3-B | 3.5 | 24 | 121 | 592 | 7 | MOLECULE: | RNA POLYMER |
| 36:00:00 | 8ce1-c | 3.5 | 8.7 | 79 | 241 | 8 | MOLECULE: | CYTOCHROME |
| 37:00:00 | 1w9r-A | 3.5 | 12.3 | 88 | 119 | 2 | MOLECULE: | CHOLINE BIN |
| 38:00:00 | 5t8v-A | 3.5 | 11 | 103 | 1315 | 5 | MOLECULE: | PUTATIVE UN |
| 39:00:00 | 3l9w-B | 3.5 | 6.6 | 97 | 357 | 2 | MOLECULE: | GLUTATHIONE |
| 40:00:00 | 7obq-u | 3.5 | 15.6 | 76 | 441 | 3 | MOLECULE: | SRP RNA |
| 41:00:00 | 5tpm-A | 3.5 | 9 | 94 | 155 | 5 | MOLECULE: | PYRUVATE DE |
| 42:00:00 | 6csm-D | 3.5 | 11.4 | 129 | 277 | 8 | MOLECULE: | GTACR1 |
| 43:00:00 | 8xh6-A | 3.5 | 6.5 | 72 | 161 | 8 | MOLECULE: | LATENT MEMB |
| 44:00:00 | 7pkq-o | 3.5 | 7.1 | 76 | 163 | 3 | MOLECULE: | MS35 |
| 45:00:00 | 8p50-A | 3.5 | 24.6 | 95 | 1574 | 5 | MOLECULE: | TOXIN PROTE |
| 46:00:00 | 7qru-F | 3.5 | 7.9 | 80 | 91 | 6 | MOLECULE: | NA+/H+ ANTI |
| 47:00:00 | 7pik-C | 3.5 | 21.3 | 90 | 557 | 4 | MOLECULE: | TRANSPOSON |
| 48:00:00 | 6wg3-E | 3.5 | 10.2 | 97 | 1226 | 4 | MOLECULE: | STRUCTURAL |
| 49:00:00 | 1n9i-A | 3.5 | 3.8 | 83 | 154 | 10 | MOLECULE: | MYOGLOBIN |
| 50:00:00 | 3qma-A | 3.5 | 4.1 | 78 | 145 | 5 | MOLECULE: | MYOGLOBIN |
| 51:00:00 | 5ycg-A | 3.5 | 3.7 | 80 | 154 | 14 | MOLECULE: | ANCESTRAL M |
| 52:00:00 | 5hlu-A | 3.5 | 4 | 80 | 152 | 10 | MOLECULE: | MYOGLOBIN |
| 53:00:00 | 2w6x-A | 3.5 | 4.1 | 84 | 154 | 8 | MOLECULE: | MYOGLOBIN |
| 54:00:00 | 3h57-A | 3.5 | 3.8 | 83 | 154 | 10 | MOLECULE: | MYOGLOBIN |
| 55:00:00 | 5ojc-A | 3.5 | 3.8 | 84 | 154 | 10 | MOLECULE: | MYOGLOBIN |
| 56:00:00 | 3nml-A | 3.5 | 3.6 | 80 | 154 | 11 | MOLECULE: | MYOGLOBIN |
| 57:00:00 | 3hf4-F | 3.5 | 4.2 | 84 | 146 | 7 | MOLECULE: | HEMOGLOBIN |
| 58:00:00 | 1f65-A | 3.5 | 4.1 | 84 | 154 | 10 | MOLECULE: | MYOGLOBIN |
| 59:00:00 | 1n9x-A | 3.5 | 4.1 | 84 | 154 | 10 | MOLECULE: | MYOGLOBIN |
| 60:00:00 | 3dht-B | 3.5 | 4.2 | 84 | 145 | 7 | MOLECULE: | HEMOGLOBIN |
| 61:00:00 | 3a2g-A | 3.5 | 4.1 | 84 | 154 | 11 | MOLECULE: | MYOGLOBIN |
| 62:00:00 | 1mlh-A | 3.5 | 3.9 | 84 | 154 | 11 | MOLECULE: | MYOGLOBIN |
| 63:00:00 | 2e2y-A | 3.5 | 4.1 | 84 | 154 | 10 | MOLECULE: | MYOGLOBIN |
| 64:00:00 | 1n9f-A | 3.5 | 4.1 | 84 | 154 | 10 | MOLECULE: | MYOGLOBIN |
| 65:00:00 | 1naz-A | 3.5 | 3.7 | 79 | 154 | 10 | MOLECULE: | MYOGLOBIN |
| 66:00:00 | 2nx0-A | 3.5 | 3.9 | 78 | 146 | 5 | MOLECULE: | MYOGLOBIN |
| 67:00:00 | 3ia3-B | 3.5 | 3 | 79 | 135 | 8 | MOLECULE: | ALPHA-HEMOG |
| 68:00:00 | 6c94-A | 3.4 | 8.5 | 109 | 482 | 5 | MOLECULE: | CYTOCHROME |
| 69:00:00 | 8j5t-C | 3.4 | 4.1 | 100 | 288 | 12 | MOLECULE: | UNCHARACTER |
| 70:00:00 | 6z3w-F | 3.4 | 7.6 | 78 | 110 | 1 | MOLECULE: | ER MEMBRANE |
| 71:00:00 | 8iic-B | 3.4 | 6.5 | 128 | 425 | 5 | MOLECULE: | POLYMERASE |
| 72:00:00 | 7wji-A | 3.4 | 24.5 | 100 | 1763 | 11 | MOLECULE: | PROTEIN UNC |
| 73:00:00 | 4p96-A | 3.4 | 4.5 | 103 | 278 | 8 | MOLECULE: | FATTY ACID |
| 74:00:00 | 5zle-A | 3.4 | 5.8 | 98 | 225 | 7 | MOLECULE: | CYTOCHROME |
| 75:00:00 | 5yo8-A | 3.4 | 6.4 | 91 | 349 | 5 | MOLECULE: | TETRAPRENYL |
| 76:00:00 | 2mjm-A | 3.4 | 3.6 | 67 | 101 | 4 | MOLECULE: | PROTEIN NLR |
| 77:00:00 | 7v9g-D | 3.4 | 10.5 | 77 | 112 | 9 | MOLECULE: | BEN DOMAIN- |
| 78:00:00 | 7tt9-A | 3.4 | 4.7 | 71 | 116 | 8 | MOLECULE: | GROUP 1 TRU |

| | | | | | | | | |
|---|---|---|---|---|---|---|---|---|
| 79:00:00 | 6g2j-K | 3.4 | 6.1 | 82 | 98 | 6 | MOLECULE: | NADH-UBIQUI |
| 80:00:00 | 2pke-A | 3.4 | 5.2 | 82 | 233 | 5 | MOLECULE: | HALOACID DE |
| 81:00:00 | 7w5g-A | 3.4 | 16.2 | 69 | 355 | 1 | MOLECULE: | TERPENE CYC |
| 82:00:00 | 3jc6-5 | 3.4 | 4.3 | 81 | 254 | 5 | MOLECULE: | DNA REPLICA |
| 83:00:00 | 4q9v-B | 3.4 | 6.7 | 94 | 186 | 16 | MOLECULE: | TUMOR NECRO |
| 84:00:00 | 6uz3-A | 3.4 | 24.7 | 108 | 1126 | 6 | MOLECULE: | SODIUM CHAN |
| 85:00:00 | 4jle-B | 3.4 | 10.3 | 73 | 159 | 3 | MOLECULE: | PHIST |
| 86:00:00 | 6e8c-A | 3.4 | 5.6 | 65 | 135 | 6 | MOLECULE: | DOUBLE HOME |
| 87:00:00 | 2py8-C | 3.4 | 7.8 | 79 | 123 | 5 | MOLECULE: | HYPOTHETICA |
| 88:00:00 | 8e9g-L | 3.4 | 21.2 | 113 | 625 | 10 | MOLECULE: | TWO-COMPONE |
| 89:00:00 | 4h0b-A | 3.4 | 4.1 | 84 | 154 | 10 | MOLECULE: | MYOGLOBIN |
| 90:00:00 | 3hrw-B | 3.4 | 4 | 81 | 146 | 6 | MOLECULE: | HEMOGLOBIN |
| 91:00:00 | 3bj2-A | 3.4 | 3.5 | 80 | 142 | 10 | MOLECULE: | HEMOGLOBIN |
| 92:00:00 | 5eui-A | 3.4 | 3.2 | 76 | 134 | 7 | MOLECULE: | HBA PROTEIN |
| 93:00:00 | 1ury-B | 3.4 | 4.5 | 86 | 154 | 5 | MOLECULE: | CYTOGLOBIN |
| 94:00:00 | 4pnj-A | 3.4 | 4 | 84 | 154 | 10 | MOLECULE: | MYOGLOBIN |
| 95:00:00 | 3a59-D | 3.4 | 4.1 | 82 | 146 | 7 | MOLECULE: | HEMOGLOBIN |
| 96:00:00 | 6on4-B | 3.4 | 4 | 89 | 214 | 8 | MOLECULE: | HTH-TYPE TR |
| 97:00:00 | 5hy8-E | 3.4 | 3.7 | 80 | 137 | 6 | MOLECULE: | HEMOGLOBIN |
| 98:00:00 | 1f5o-E | 3.4 | 3.5 | 77 | 149 | 6 | MOLECULE: | HEMOGLOBIN |
| 99:00:00 | 1jeb-B | 3.4 | 4 | 82 | 143 | 7 | MOLECULE: | HEMOGLOBIN |
| 0:00 | 3lhb-A | 3.4 | 3.1 | 76 | 149 | 5 | MOLECULE: | PROTEIN (HE |
| 1:00 | 3lhb-F | 3.4 | 3.6 | 78 | 149 | 6 | MOLECULE: | PROTEIN (HE |
| 2:00 | 1f5p-F | 3.4 | 3.6 | 78 | 149 | 5 | MOLECULE: | HEMOGLOBIN |
| 3:00 | 3lhb-D | 3.4 | 3.6 | 78 | 149 | 6 | MOLECULE: | PROTEIN (HE |
| 4:00 | 1uc3-K | 3.4 | 3.6 | 78 | 149 | 5 | MOLECULE: | GLOBIN |
| 5:00 | 3lhb-H | 3.4 | 3.1 | 76 | 149 | 5 | MOLECULE: | PROTEIN (HE |
| 6:00 | 2h35-C | 3.4 | 3.4 | 82 | 141 | 6 | MOLECULE: | HEMOGLOBIN |
| 7:00 | 3lhb-G | 3.4 | 3.5 | 77 | 149 | 6 | MOLECULE: | PROTEIN (HE |
| 8:00 | 3lhb-K | 3.4 | 3.1 | 76 | 149 | 5 | MOLECULE: | PROTEIN (HE |
| 9:00 | 1f5p-A | 3.4 | 3.6 | 78 | 149 | 5 | MOLECULE: | HEMOGLOBIN |
| 10:00 | 3lhb-J | 3.4 | 3.6 | 78 | 149 | 6 | MOLECULE: | PROTEIN (HE |
| 11:00 | 3lhb-I | 3.4 | 3.1 | 76 | 149 | 5 | MOLECULE: | PROTEIN (HE |
| 12:00 | 3lhb-B | 3.4 | 3.6 | 78 | 149 | 6 | MOLECULE: | PROTEIN (HE |
| 13:00 | 1f5p-D | 3.4 | 3.6 | 78 | 149 | 5 | MOLECULE: | HEMOGLOBIN |
| 14:00 | 1f5p-C | 3.4 | 3.6 | 78 | 149 | 5 | MOLECULE: | HEMOGLOBIN |
| 15:00 | 1f5p-B | 3.4 | 3.6 | 78 | 149 | 5 | MOLECULE: | HEMOGLOBIN |
| 16:00 | 3lhb-L | 3.4 | 3.5 | 77 | 149 | 6 | MOLECULE: | PROTEIN (HE |
| 17:00 | 7yim-A | 3.3 | 15.7 | 84 | 592 | 4 | MOLECULE: | ALPHA-FETOP |
| 18:00 | 5nvu-A | 3.3 | 27.7 | 130 | 3169 | 0 | MOLECULE: | DYNEIN MOTO |
| 19:00 | 7uy5-K | 3.3 | 4 | 58 | 157 | 5 | MOLECULE: | TELOMERASE |
| 20:00 | 5w65-P | 3.3 | 7 | 78 | 481 | 10 | MOLECULE: | DNA-DIRECTE |
| 21:00 | 4pi0-A | 3.3 | 18.6 | 85 | 390 | 15 | MOLECULE: | UNKNOWN PEP |
| 22:00 | 6ewz-A | 3.3 | 6.6 | 71 | 202 | 3 | MOLECULE: | GTP PYROPHO |

| | | | | | | | | |
|---|---|---|---|---|---|---|---|---|
| 23:00 | 7exx-A | 3.3 | 12.1 | 74 | 440 | 4 | MOLECULE: | DNA PHOSPHO |
| 24:00:00 | 6vls-D | 3.3 | 32.5 | 131 | 963 | 8 | MOLECULE: | MALTOSE/MAL |
| 25:00:00 | 8arl-A | 3.3 | 14.2 | 84 | 232 | 8 | MOLECULE: | TRYPTOPHAN- |
| 26:00:00 | 7uic-b | 3.3 | 9.6 | 56 | 65 | 9 | MOLECULE: | MEDIATOR OF |
| 27:00:00 | 3g06-A | 3.3 | 20.7 | 87 | 601 | 7 | MOLECULE: | SSPH2 (LEUC |
| 28:00:00 | 8bpz-A | 3.3 | 16.3 | 68 | 322 | 9 | MOLECULE: | SPBETA PROP |
| 29:00:00 | 5gzb-A | 3.3 | 3.6 | 66 | 91 | 6 | MOLECULE: | TRANSCRIPTI |
| 30:00:00 | 6br8-A | 3.3 | 8.3 | 81 | 249 | 7 | MOLECULE: | PROTEIN A6 |
| 31:00:00 | 2f4q-A | 3.3 | 4.2 | 72 | 309 | 3 | MOLECULE: | TYPE I TOPO |
| 32:00:00 | 7dgu-A | 3.3 | 4.7 | 71 | 97 | 6 | MOLECULE: | A1 |
| 33:00:00 | 3p3o-A | 3.3 | 5.8 | 150 | 416 | 7 | MOLECULE: | CYTOCHROME |
| 34:00:00 | 6wq0-A | 3.3 | 7.4 | 73 | 131 | 3 | MOLECULE: | DNA (301-ME |
| 35:00:00 | 6xdc-A | 3.3 | 16.3 | 98 | 193 | 7 | MOLECULE: | PROTEIN 3A |
| 36:00:00 | 3ua3-A | 3.3 | 17.6 | 124 | 663 | 3 | MOLECULE: | PROTEIN ARG |
| 37:00:00 | 7bc6-A | 3.3 | 7 | 95 | 432 | 5 | MOLECULE: | PROTON-COUP |
| 38:00:00 | 7oqz-A | 3.3 | 12.6 | 67 | 237 | 4 | MOLECULE: | TRANSMEMBRA |
| 39:00:00 | 6zgd-A | 3.3 | 6.2 | 76 | 288 | 12 | MOLECULE: | PROTON-GATE |
| 40:00:00 | 8k49-T | 3.3 | 18.9 | 106 | 627 | 7 | MOLECULE: | VP2 |
| 41:00:00 | 7dgy-A | 3.3 | 3.8 | 70 | 99 | 3 | MOLECULE: | C2 |
| 42:00:00 | 3h36-A | 3.3 | 4.1 | 68 | 78 | 6 | MOLECULE: | POLYRIBONUC |
| 43:00:00 | 8b9z-K | 3.3 | 7.8 | 79 | 96 | 11 | MOLECULE: | NADH-UBIQUI |
| 44:00:00 | 7qe7-A | 3.3 | 17.9 | 135 | 1648 | 5 | MOLECULE: | ANAPHASE-PR |
| 45:00:00 | 7ar9-J | 3.3 | 8.6 | 87 | 145 | 3 | MOLECULE: | ND3 |
| 46:00:00 | 7mhz-A | 3.3 | 8.7 | 102 | 490 | 4 | MOLECULE: | SONIC HEDGE |
| 47:00:00 | 7mi6-A | 3.3 | 42.1 | 127 | 2419 | 7 | MOLECULE: | FUSION PROT |
| 48:00:00 | 2gb7-B | 3.3 | 19.1 | 79 | 295 | 4 | MOLECULE: | DNA STRAND |
| 49:00:00 | 1ux9-B | 3.3 | 4.5 | 86 | 154 | 5 | MOLECULE: | CYTOGLOBIN |
| 50:00:00 | 2h35-A | 3.3 | 3.3 | 80 | 141 | 5 | MOLECULE: | HEMOGLOBIN |
| 51:00:00 | 1f5o-C | 3.3 | 3.1 | 76 | 149 | 5 | MOLECULE: | HEMOGLOBIN |
| 52:00:00 | 1jeb-D | 3.3 | 4.6 | 84 | 146 | 7 | MOLECULE: | HEMOGLOBIN |
| 53:00:00 | 1f5o-A | 3.3 | 3.1 | 76 | 149 | 5 | MOLECULE: | HEMOGLOBIN |
| 54:00:00 | 1f5o-B | 3.3 | 3.1 | 76 | 149 | 5 | MOLECULE: | HEMOGLOBIN |
| 55:00:00 | 1uc3-I | 3.3 | 3.1 | 76 | 149 | 5 | MOLECULE: | GLOBIN |
| 56:00:00 | 1f5o-D | 3.3 | 3.5 | 77 | 149 | 6 | MOLECULE: | HEMOGLOBIN |
| 57:00:00 | 1uc3-H | 3.3 | 3.6 | 78 | 149 | 5 | MOLECULE: | GLOBIN |
| 58:00:00 | 3lhb-C | 3.3 | 3.6 | 78 | 149 | 6 | MOLECULE: | PROTEIN (HE |
| 59:00:00 | 5lcw-N | 3.3 | 17.4 | 90 | 703 | 11 | MOLECULE: | ANAPHASE-PR |
| 60:00:00 | 2nrm-A | 3.3 | 4 | 76 | 146 | 5 | MOLECULE: | MYOGLOBIN |
| 61:00:00 | 3k3u-B | 3.2 | 3.7 | 74 | 137 | 9 | MOLECULE: | DEHALOPEROX |
| 62:00:00 | 7a7c-A | 3.2 | 16.6 | 100 | 560 | 8 | MOLECULE: | CATALASE-PE |
| 63:00:00 | 1zs4-A | 3.2 | 3 | 45 | 82 | 7 | MOLECULE: | DNA - 27MER |
| 64:00:00 | 5oxf-C | 3.2 | 22.1 | 106 | 601 | 10 | MOLECULE: | GTP-BINDING |
| 65:00:00 | 6al9-B | 3.2 | 3.1 | 42 | 91 | 2 | MOLECULE: | CHORISMATE |
| 66:00:00 | 5k94-A | 3.2 | 13.8 | 99 | 506 | 6 | MOLECULE: | MALTOSE-BIN |

| | | | | | | | | |
|---|---|---|---|---|---|---|---|---|
| 67:00:00 | 6fwr-A | 3.2 | 21.8 | 89 | 699 | 3 | MOLECULE: | ATP-DEPENDE |
| 68:00:00 | 1kw2-A | 3.2 | 7.4 | 76 | 455 | 12 | MOLECULE: | VITAMIN D-B |
| 69:00:00 | 8v46-D | 3.2 | 3.6 | 87 | 305 | 6 | MOLECULE: | ARIA ANTITO |
| 70:00:00 | 8ieu-B | 3.2 | 14.9 | 75 | 336 | 9 | MOLECULE: | DUF2891 DOM |
| 71:00:00 | 2hs5-A | 3.2 | 4.7 | 97 | 207 | 7 | MOLECULE: | PUTATIVE TR |
| 72:00:00 | 8v3b-a | 3.2 | 12.3 | 79 | 416 | 3 | MOLECULE: | O43_129_+4 |
| 73:00:00 | 6w2r-B | 3.2 | 6.5 | 73 | 221 | 8 | MOLECULE: | JUNCTION 19 |
| 74:00:00 | 6ebu-A | 3.2 | 4.4 | 88 | 165 | 8 | MOLECULE: | LPXE |
| 75:00:00 | 1q2h-A | 3.2 | 4.3 | 57 | 63 | 7 | MOLECULE: | ADAPTOR PRO |
| 76:00:00 | 3a6m-A | 3.2 | 6.1 | 68 | 168 | 4 | MOLECULE: | PROTEIN GRP |
| 77:00:00 | 7ctp-A | 3.2 | 5 | 102 | 553 | 12 | MOLECULE: | PROTEIN NIB |
| 78:00:00 | 5z1n-A | 3.2 | 3.9 | 83 | 166 | 11 | MOLECULE: | G-PROTEIN I |
| 79:00:00 | 5jm8-A | 3.2 | 20.6 | 68 | 556 | 3 | MOLECULE: | AEROBACTIN |
| 80:00:00 | 5j6f-A | 3.2 | 16.6 | 59 | 352 | 7 | MOLECULE: | 3-DEOXY-D-A |
| 81:00:00 | 1r5i-D | 3.2 | 10 | 87 | 214 | 3 | MOLECULE: | HLA CLASS I |
| 82:00:00 | 6zyd-A | 3.2 | 21.9 | 121 | 323 | 5 | MOLECULE: | LOW CONDUCT |
| 83:00:00 | 4gr6-B | 3.2 | 3.9 | 71 | 105 | 7 | MOLECULE: | ATRBCX2 |
| 84:00:00 | 5cd4-I | 3.2 | 10 | 82 | 494 | 9 | MOLECULE: | CRISPR SYST |
| 85:00:00 | 3ed5-A | 3.2 | 6.1 | 96 | 232 | 7 | MOLECULE: | YFNB |
| 86:00:00 | 8j3r-B | 3.2 | 25.7 | 115 | 387 | 3 | MOLECULE: | TRANSPOSASE |
| 87:00:00 | 8uw3-A | 3.2 | 7.2 | 109 | 1265 | 6 | MOLECULE: | LINE-1 RETR |
| 88:00:00 | 4zsf-A | 3.2 | 17.6 | 99 | 272 | 5 | MOLECULE: | BSAWI ENDON |
| 89:00:00 | 3bzk-A | 3.2 | 13.4 | 92 | 728 | 7 | MOLECULE: | TEX |
| 90:00:00 | 3bni-A | 3.2 | 5.2 | 67 | 174 | 9 | MOLECULE: | PUTATIVE TE |
| 91:00:00 | 3bm3-A | 3.2 | 4.8 | 90 | 259 | 4 | MOLECULE: | DNA (5'-D(* |
| 92:00:00 | 8a3t-C | 3.2 | 26.7 | 134 | 1410 | 10 | MOLECULE: | ANAPHASE-PR |
| 93:00:00 | 5d2m-C | 3.2 | 4.6 | 83 | 158 | 17 | MOLECULE: | SUMO-CONJUG |
| 94:00:00 | 3dan-A | 3.2 | 6.7 | 106 | 473 | 2 | MOLECULE: | CYTOCHROME |
| 95:00:00 | 6dlu-P | 3.2 | 8.3 | 95 | 747 | 5 | MOLECULE: | DYNAMIN-1 |
| 96:00:00 | 6zyy-C | 3.2 | 8.4 | 90 | 2901 | 6 | MOLECULE: | DYNEIN HEAV |
| 97:00:00 | 4hkr-A | 3.2 | 8.8 | 89 | 165 | 2 | MOLECULE: | CALCIUM REL |
| 98:00:00 | 3vr8-D | 3.2 | 8.5 | 84 | 129 | 7 | MOLECULE: | FLAVOPROTEI |
| 99:00:00 | 8ity-V | 3.2 | 13.8 | 80 | 361 | 9 | MOLECULE: | SNRNA-ACTIV |
| 0:00 | 8ovw-Z | 3.2 | 21.5 | 93 | 151 | 8 | MOLECULE: | CENTROMERE- |
| 1:00 | 7ryq-B | 3.2 | 15.6 | 93 | 527 | 6 | MOLECULE: | KIF-BINDING |
| 2:00 | 7kfu-C | 3.2 | 19.3 | 137 | 909 | 7 | MOLECULE: | CAS2 |
| 3:00 | 6khj-E | 3.2 | 7.2 | 80 | 101 | 9 | MOLECULE: | NAD(P)H-QUI |
| 4:00 | 3w6j-A | 3.2 | 5.3 | 101 | 150 | 8 | MOLECULE: | SCPA |
| 5:00 | 5lbm-C | 3.2 | 5.7 | 70 | 90 | 7 | MOLECULE: | TRANSCRIPTI |
| 6:00 | 1uc3-J | 3.2 | 3.5 | 76 | 149 | 5 | MOLECULE: | GLOBIN |
| 7:00 | 3ia3-D | 3.2 | 3.6 | 82 | 135 | 5 | MOLECULE: | ALPHA-HEMOG |
| 8:00 | 6otx-A | 3.2 | 4.2 | 78 | 151 | 4 | MOLECULE: | HEMOGLOBIN |
| 9:00 | 4hrr-F | 3.2 | 4.1 | 87 | 151 | 7 | MOLECULE: | GLOBIN-2 A |
| 10:00 | 1s5y-C | 3.2 | 4.2 | 81 | 142 | 7 | MOLECULE: | HEMOGLOBIN |

| | | | | | | | | |
|---|---|---|---|---|---|---|---|---|
| 11:00 | 1uc3-D | 3.2 | 4.2 | 81 | 149 | 7 | MOLECULE: | GLOBIN |
| 12:00 | 3lhb-E | 3.2 | 4.1 | 80 | 149 | 8 | MOLECULE: | PROTEIN (HE |
| 13:00 | 6tlj-N | 3.2 | 19.8 | 99 | 703 | 7 | MOLECULE: | ANAPHASE-PR |
| 14:00 | 1f5o-F | 3.2 | 4.1 | 80 | 149 | 8 | MOLECULE: | HEMOGLOBIN |
| 15:00 | 1uc3-G | 3.2 | 3.6 | 78 | 149 | 6 | MOLECULE: | GLOBIN |
| 16:00 | 7z0o-G | 3.1 | 2.3 | 53 | 84 | 8 | MOLECULE: | HISTONE H3 |
| 17:00 | 7vvy-K | 3.1 | 31.3 | 90 | 237 | 8 | MOLECULE: | CHROMATIN M |
| 18:00 | 8pnj-A | 3.1 | 5.2 | 58 | 390 | 9 | MOLECULE: | CHORISMATE |
| 19:00 | 6zpa-A | 3.1 | 4.1 | 72 | 173 | 10 | MOLECULE: | DATZ |
| 20:00 | 7xdi-F | 3.1 | 6.6 | 69 | 456 | 6 | MOLECULE: | VP1 |
| 21:00 | 8ea3-W | 3.1 | 20.6 | 101 | 514 | 7 | MOLECULE: | TARGET_LE |
| 22:00 | 4fcy-B | 3.1 | 24.5 | 100 | 476 | 2 | MOLECULE: | TRANSPOSASE |
| 23:00 | 1ysm-A | 3.1 | 4.5 | 43 | 55 | 9 | MOLECULE: | CALCYCLIN-B |
| 24:00:00 | 3f1i-H | 3.1 | 19.9 | 77 | 98 | 13 | MOLECULE: | HEPATOCYTE |
| 25:00:00 | 8fuq-A | 3.1 | 4.1 | 40 | 51 | 10 | MOLECULE: | PHOSPHOPROT |
| 26:00:00 | 1m12-A | 3.1 | 2.9 | 57 | 84 | 7 | MOLECULE: | SAPOSIN C |
| 27:00:00 | 7ro2-A | 3.1 | 10.4 | 70 | 94 | 6 | MOLECULE: | MAJOR CAPSI |
| 28:00:00 | 1xdo-A | 3.1 | 20.3 | 103 | 687 | 8 | MOLECULE: | POLYPHOSPHA |
| 29:00:00 | 5szj-B | 3.1 | 9.5 | 85 | 145 | 5 | MOLECULE: | RAS-RELATED |
| 30:00:00 | 6o3s-A | 3.1 | 5.8 | 44 | 47 | 5 | MOLECULE: | RIBOSOME-IN |
| 31:00:00 | 7s00-c | 3.1 | 14.3 | 67 | 484 | 9 | MOLECULE: | DNA-DIRECTE |
| 32:00:00 | 7ymd-C | 3.1 | 11.4 | 78 | 285 | 9 | MOLECULE: | NON-STRUCTU |
| 33:00:00 | 7pq1-A | 3.1 | 5.8 | 130 | 395 | 8 | MOLECULE: | CYTOCHROME |
| 34:00:00 | 7pkq-C | 3.1 | 25.3 | 71 | 369 | 6 | MOLECULE: | MS35 |
| 35:00:00 | 6gys-B | 3.1 | 10.3 | 105 | 579 | 10 | MOLECULE: | CENTROMERE |
| 36:00:00 | 7emf-J | 3.1 | 16.2 | 67 | 122 | 6 | MOLECULE: | MEDIATOR OF |
| 37:00:00 | 1knz-A | 3.1 | 9.6 | 56 | 154 | 7 | MOLECULE: | 5'-R(*UP*GP |
| 38:00:00 | 6wc3-B | 3.1 | 8.3 | 67 | 94 | 15 | MOLECULE: | PROTEIN TRA |
| 39:00:00 | 7wst-A | 3.1 | 11.1 | 110 | 601 | 2 | MOLECULE: | IRON-PHYTOS |
| 40:00:00 | 7lma-A | 3.1 | 23.2 | 145 | 1012 | 6 | MOLECULE: | TELOMERASE |
| 41:00:00 | 8j2x-A | 3.1 | 4.1 | 77 | 337 | 10 | MOLECULE: | COBALAMIN-B |
| 42:00:00 | 6lum-D | 3.1 | 10.4 | 91 | 146 | 8 | MOLECULE: | SUCCINATE D |
| 43:00:00 | 5yhf-A | 3.1 | 5.6 | 107 | 734 | 9 | MOLECULE: | PROTEIN TRA |
| 44:00:00 | 7zmb-6 | 3.1 | 10.5 | 102 | 190 | 10 | MOLECULE: | NADH-UBIQUI |
| 45:00:00 | 7wuq-R | 3.1 | 9.7 | 84 | 278 | 4 | MOLECULE: | GUANINE NUC |
| 46:00:00 | 5e6g-B | 3.1 | 10.4 | 75 | 114 | 9 | MOLECULE: | DE NOVO DES |
| 47:00:00 | 7ot9-A | 3.1 | 11.9 | 84 | 221 | 7 | MOLECULE: | AI-2E MEMBE |
| 48:00:00 | 6k5s-A | 3.1 | 11 | 70 | 220 | 7 | MOLECULE: | TELOMERIC D |
| 49:00:00 | 7op8-A | 3.1 | 14.1 | 88 | 1118 | 3 | MOLECULE: | CATION-TRAN |
| 50:00:00 | 7mxb-A | 3.1 | 8.1 | 62 | 672 | 8 | MOLECULE: | SERINE/THRE |
| 51:00:00 | 6ysg-A | 3.1 | 17.8 | 126 | 1264 | 4 | MOLECULE: | MG-CHELATAS |
| 52:00:00 | 8uc3-C | 3.1 | 7.4 | 70 | 98 | 3 | MOLECULE: | ALBONOURSIN |
| 53:00:00 | 6zvh-R | 3.1 | 14.8 | 78 | 135 | 6 | MOLECULE: | 18S RRNA |
| 54:00:00 | 5iy7-M | 3.1 | 11.1 | 76 | 310 | 9 | MOLECULE: | DNA-DIRECTE |

| | | | | | | | | |
|---|---|---|---|---|---|---|---|---|
| 55:00:00 | 5w7g-B | 3.1 | 9.5 | 66 | 126 | 8 | MOLECULE: | ORF140 |
| 56:00:00 | 5ys9-A | 3.1 | 23.5 | 132 | 692 | 5 | MOLECULE: | ACYL-COENZY |
| 57:00:00 | 6d5f-A | 3.1 | 10.4 | 82 | 193 | 10 | MOLECULE: | FIMBRIAL PR |
| 58:00:00 | 7d3u-G | 3.1 | 8.8 | 87 | 111 | 7 | MOLECULE: | MONOVALENT |
| 59:00:00 | 6rao-I | 3.1 | 19.6 | 91 | 443 | 4 | MOLECULE: | AFP1 |
| 60:00:00 | 4y9j-B | 3.1 | 13.7 | 108 | 593 | 3 | MOLECULE: | PROTEIN ACD |
| 61:00:00 | 5w7g-A | 3.1 | 7.7 | 67 | 131 | 4 | MOLECULE: | ORF140 |
| 62:00:00 | 3waq-A | 3.1 | 6.9 | 78 | 134 | 8 | MOLECULE: | HEMERYTHRIN |
| 63:00:00 | 2wus-R | 3.1 | 2.9 | 60 | 87 | 5 | MOLECULE: | PUTATIVE UN |
| 64:00:00 | 8c3y-A | 3.1 | 5.7 | 105 | 929 | 2 | MOLECULE: | PFEMP1 |
| 65:00:00 | 2rld-C | 3.1 | 5 | 78 | 116 | 4 | MOLECULE: | UNCHARACTER |
| 66:00:00 | 3hc1-A | 3.1 | 4.9 | 80 | 298 | 10 | MOLECULE: | UNCHARACTER |
| 67:00:00 | 7vw0-B | 3.1 | 8.6 | 76 | 94 | 11 | MOLECULE: | DUF305 DOMA |
| 68:00:00 | 3pi2-A | 3.1 | 4.6 | 78 | 151 | 5 | MOLECULE: | HEMOGLOBIN |
| 69:00:00 | 2nrl-A | 3.1 | 4.3 | 76 | 145 | 5 | MOLECULE: | MYOGLOBIN |
| 70:00:00 | 1f6h-A | 3.1 | 4.1 | 82 | 153 | 11 | MOLECULE: | MYOGLOBIN ( |
| 71:00:00 | 7vde-D | 3.1 | 4.6 | 81 | 146 | 7 | MOLECULE: | HEMOGLOBIN |
| 72:00:00 | 7vde-B | 3.1 | 4.6 | 81 | 146 | 7 | MOLECULE: | HEMOGLOBIN |
| 73:00:00 | 1uc3-L | 3.1 | 3.6 | 78 | 149 | 6 | MOLECULE: | GLOBIN |
| 74:00:00 | 1yeq-D | 3.1 | 3.5 | 84 | 146 | 10 | MOLECULE: | HEMOGLOBIN |
| 75:00:00 | 8w6c-A | 3 | 17 | 92 | 530 | 11 | MOLECULE: | SOLUTE CARR |
| 76:00:00 | 6hqa-J | 3 | 3.3 | 45 | 51 | 4 | MOLECULE: | TAF2 |
| 77:00:00 | 5dw7-A | 3 | 4.1 | 79 | 309 | 8 | MOLECULE: | GERMACRADIE |
| 78:00:00 | 4wfc-B | 3 | 12.9 | 81 | 118 | 2 | MOLECULE: | EXOSOME COM |
| 79:00:00 | 2e87-A | 3 | 12.2 | 91 | 356 | 5 | MOLECULE: | HYPOTHETICA |
| 80:00:00 | 2btv-B | 3 | 25.2 | 86 | 885 | 8 | MOLECULE: | PROTEIN (VP |
| 81:00:00 | 6vz1-A | 3 | 11.1 | 90 | 411 | 13 | MOLECULE: | DIACYLGLYCE |
| 82:00:00 | 6kg7-A | 3 | 15.7 | 130 | 1817 | 5 | MOLECULE: | PIEZO-TYPE |
| 83:00:00 | 7u5d-A | 3 | 18.2 | 70 | 630 | 6 | MOLECULE: | CRRNA |
| 84:00:00 | 7x07-A | 3 | 11.9 | 122 | 633 | 5 | MOLECULE: | ATP-BINDING |
| 85:00:00 | 7x2q-A | 3 | 5.7 | 153 | 467 | 4 | MOLECULE: | SUGIOL SYNT |
| 86:00:00 | 6v4a-C | 3 | 14.4 | 63 | 593 | 5 | MOLECULE: | NEUR_CHAN_L |
| 87:00:00 | 4hts-A | 3 | 5.1 | 89 | 228 | 9 | MOLECULE: | SEC-INDEPEN |
| 88:00:00 | 8eja-A | 3 | 5.4 | 63 | 113 | 6 | MOLECULE: | ABAK |
| 89:00:00 | 8djk-A | 3 | 9.1 | 108 | 245 | 4 | MOLECULE: | 3-HYDROXY-3 |
| 90:00:00 | 8tr4-A | 3 | 13.8 | 60 | 276 | 5 | MOLECULE: | POLYKETIDE |
| 91:00:00 | 7ojn-L | 3 | 30.8 | 143 | 2010 | 7 | MOLECULE: | RNA-DIRECTE |
| 92:00:00 | 3tix-A | 3 | 6.3 | 84 | 153 | 8 | MOLECULE: | UBIQUITIN-L |
| 93:00:00 | 3mlg-A | 3 | 5.9 | 76 | 161 | 9 | MOLECULE: | 2X CHIMERA |
| 94:00:00 | 7n8x-A | 3 | 10.5 | 77 | 691 | 5 | MOLECULE: | CHONDROITIN |
| 95:00:00 | 7xr3-Z | 3 | 13.4 | 114 | 1320 | 5 | MOLECULE: | VP3 |
| 96:00:00 | 2ouw-B | 3 | 4.6 | 60 | 136 | 12 | MOLECULE: | ALKYLHYDROP |
| 97:00:00 | 8q85-a | 3 | 21.8 | 94 | 234 | 5 | MOLECULE: | KINETOCHORE |
| 98:00:00 | 8bou-A | 3 | 19.1 | 100 | 807 | 4 | MOLECULE: | N,N'-DIACET |

| | | | | | | | | |
|---|---|---|---|---|---|---|---|---|
| 99:00:00 | 2r6i-A | 3 | 16.4 | 66 | 268 | 8 | MOLECULE: | UNCHARACTER |
| 0:00 | 7lhf-A | 3 | 22.1 | 98 | 2123 | 4 | MOLECULE: | INOSITOL 1, |
| 1:00 | 6pvp-A | 3 | 17 | 78 | 661 | 6 | MOLECULE: | TRANSIENT R |
| 2:00 | 6cfw-A | 3 | 10.4 | 69 | 165 | 9 | MOLECULE: | MONOVALENT |
| 3:00 | 6hsy-A | 3 | 4.4 | 55 | 190 | 2 | MOLECULE: | TOLUENE TOL |
| 4:00 | 5e2q-A | 3 | 12.4 | 98 | 725 | 7 | MOLECULE: | DIPEPTIDYL |
| 5:00 | 1g8x-A | 3 | 18.5 | 103 | 1009 | 3 | MOLECULE: | MYOSIN II H |
| 6:00 | 5f3o-A | 3 | 6.3 | 82 | 194 | 9 | MOLECULE: | PUTATIVE UN |
| 7:00 | 5ean-A | 3 | 20.1 | 110 | 1051 | 7 | MOLECULE: | DNA REPLICA |
| 8:00 | 3dp7-A | 3 | 7.9 | 76 | 351 | 8 | MOLECULE: | SAM-DEPENDE |
| 9:00 | 8jpj-A | 3 | 14.7 | 125 | 733 | 7 | MOLECULE: | H(+)/CL(-) |
| 10:00 | 7lzh-A | 3 | 35.9 | 104 | 799 | 4 | MOLECULE: | GLUTAMATE R |
| 11:00 | 3i4m-D | 3 | 7 | 76 | 187 | 7 | MOLECULE: | DNA-DIRECTE |
| 12:00 | 7w6k-A | 3 | 28.3 | 119 | 410 | 4 | MOLECULE: | GMALMT12/QU |
| 13:00 | 6uak-A | 3 | 3.8 | 74 | 298 | 3 | MOLECULE: | SAM DEPENDE |
| 14:00 | 7vu7-A | 3 | 3.9 | 64 | 89 | 3 | MOLECULE: | FLAGELLIFOR |
| 15:00 | 7m5w-A | 3 | 9.5 | 69 | 138 | 12 | MOLECULE: | PROTEIN CAP |
| 16:00 | 8ifg-A | 3 | 16.3 | 90 | 879 | 11 | MOLECULE: | RBAP48-RELA |
| 17:00 | 3f7c-A | 3 | 4.3 | 64 | 199 | 8 | MOLECULE: | PROTEIN OF |
| 18:00 | 6cfw-G | 3 | 8.2 | 79 | 113 | 8 | MOLECULE: | MONOVALENT |
| 19:00 | 5a31-N | 3 | 12 | 72 | 703 | 13 | MOLECULE: | ANAPHASE-PR |
| 20:00 | 2q5z-B | 3 | 5.8 | 65 | 94 | 6 | MOLECULE: | HYPOTHETICA |
| 21:00 | 2qup-A | 3 | 8 | 79 | 119 | 5 | MOLECULE: | BH1478 PROT |
| 22:00 | 5lnk-J | 3 | 10.3 | 79 | 175 | 6 | MOLECULE: | MITOCHONDRI |
| 23:00 | 6xgx-A | 3 | 20.6 | 103 | 402 | 8 | MOLECULE: | MUTATOR FAM |
| 24:00:00 | 3mzy-A | 3 | 2.5 | 51 | 123 | 16 | MOLECULE: | RNA POLYMER |
| 25:00:00 | 7m2w-G | 3 | 10.4 | 89 | 712 | 7 | MOLECULE: | TUBULIN GAM |
| 26:00:00 | 1jog-D | 3 | 7.6 | 77 | 136 | 6 | MOLECULE: | HYPOTHETICA |
| 27:00:00 | 1gzs-B | 3 | 3.3 | 88 | 165 | 10 | MOLECULE: | SOPE |
| 28:00:00 | 8qq7-A | 3 | 10.8 | 100 | 399 | 9 | MOLECULE: | FAD-BINDING |
| 29:00:00 | 6r6h-O | 3 | 20.8 | 124 | 744 | 6 | MOLECULE: | COP9 SIGNAL |
| 30:00:00 | 8pm6-A | 3 | 15.5 | 88 | 872 | 8 | MOLECULE: | BILE SALT E |
| 31:00:00 | 2cqn-A | 3 | 3.6 | 62 | 77 | 0 | MOLECULE: | FORMIN-BIND |
| 32:00:00 | 7tmw-R | 3 | 11.4 | 113 | 499 | 9 | MOLECULE: | RELAXIN REC |
| 33:00:00 | 7woo-Z | 3 | 16.7 | 81 | 746 | 2 | MOLECULE: | NUCLEOPORIN |
| 34:00:00 | 1mti-A | 3 | 4.9 | 78 | 154 | 10 | MOLECULE: | MYOGLOBIN |
| 35:00:00 | 1mlg-A | 3 | 5 | 78 | 154 | 12 | MOLECULE: | MYOGLOBIN |
| 36:00:00 | 3wyo-D | 3 | 10.6 | 72 | 153 | 11 | MOLECULE: | MYOGLOBIN |
| 37:00:00 | 1i3d-B | 3 | 4.4 | 80 | 146 | 10 | MOLECULE: | HEMOGLOBIN |
| 38:00:00 | 3wtg-D | 3 | 4.7 | 83 | 146 | 13 | MOLECULE: | HEMOGLOBIN |
| 39:00:00 | 1ch5-A | 3 | 4.8 | 76 | 154 | 11 | MOLECULE: | PROTEIN (MY |
| 40:00:00 | 5g04-N | 3 | 28.1 | 113 | 631 | 5 | MOLECULE: | ANAPHASE-PR |
| 41:00:00 | 4hrr-D | 3 | 3.7 | 85 | 151 | 9 | MOLECULE: | GLOBIN-2 A |
| 42:00:00 | 2di3-B | 3 | 6.8 | 99 | 230 | 7 | MOLECULE: | BACTERIAL R |

| | | | | | | | | |
|---|---|---|---|---|---|---|---|---|
| 43:00:00 | 6q6g-N | 3 | 32 | 122 | 656 | 6 | MOLECULE: | CELL DIVISI |
| 44:00:00 | 7dgn-B | 3 | 10.1 | 72 | 153 | 7 | MOLECULE: | MYOGLOBIN |
| 45:00:00 | 7dgl-A | 3 | 13.3 | 60 | 153 | 7 | MOLECULE: | MYOGLOBIN |
| 46:00:00 | 3wyo-A | 3 | 7.1 | 54 | 153 | 2 | MOLECULE: | MYOGLOBIN |
| 47:00:00 | 1a4f-B | 3 | 3.6 | 85 | 146 | 11 | MOLECULE: | HEMOGLOBIN |
| 48:00:00 | 6wg7-G | 3 | 12.2 | 94 | 219 | 2 | MOLECULE: | DNA (35-MER |
| 49:00:00 | 7k4m-F | 3 | 3.7 | 85 | 146 | 8 | MOLECULE: | HEMOGLOBIN |
| 50:00:00 | 5x2r-B | 3 | 3.3 | 79 | 145 | 10 | MOLECULE: | HEMOGLOBIN |
| 51:00:00 | 7uf7-B | 3 | 3.5 | 84 | 146 | 8 | MOLECULE: | HEMOGLOBIN |
| 52:00:00 | 2vyw-A | 2.9 | 3.9 | 78 | 148 | 4 | MOLECULE: | HEMOGLOBIN |
| 53:00:00 | 5flv-M | 2.9 | 7.1 | 68 | 229 | 7 | MOLECULE: | HOMEOBOX PR |
| 54:00:00 | 8a5a-U | 2.9 | 19.8 | 88 | 690 | 6 | MOLECULE: | CHROMATIN-R |
| 55:00:00 | 6nd4-L | 2.9 | 13.5 | 91 | 473 | 4 | MOLECULE: | ETS RRNA |
| 56:00:00 | 7b1s-D | 2.9 | 3.7 | 93 | 593 | 3 | MOLECULE: | ETHYL-COENZ |
| 57:00:00 | 6h4c-D | 2.9 | 4.4 | 65 | 148 | 0 | MOLECULE: | DUTPASE |
| 58:00:00 | 4kp3-D | 2.9 | 12.8 | 52 | 82 | 10 | MOLECULE: | UNCONVENTIO |
| 59:00:00 | 3nmz-A | 2.9 | 23 | 89 | 406 | 7 | MOLECULE: | APC VARIANT |
| 60:00:00 | 5zr1-E | 2.9 | 26.5 | 70 | 460 | 6 | MOLECULE: | ORIGIN RECO |
| 61:00:00 | 4geh-D | 2.9 | 3.6 | 48 | 69 | 4 | MOLECULE: | PROGRAMMED |
| 62:00:00 | 7kpv-B | 2.9 | 9.3 | 80 | 290 | 10 | MOLECULE: | MEIOTIC MRN |
| 63:00:00 | 4wid-A | 2.9 | 7.3 | 66 | 353 | 5 | MOLECULE: | RHUL123 |
| 64:00:00 | 6nr8-6 | 2.9 | 3.6 | 41 | 102 | 15 | MOLECULE: | PREFOLDIN S |
| 65:00:00 | 8qrl-1 | 2.9 | 12 | 75 | 278 | 7 | MOLECULE: | 12S MITOCHO |
| 66:00:00 | 5wwl-N | 2.9 | 15.8 | 61 | 155 | 3 | MOLECULE: | CENTROMERE |
| 67:00:00 | 6ec8-A | 2.9 | 11.2 | 65 | 803 | 8 | MOLECULE: | LANTIBIOTIC |
| 68:00:00 | 1x8z-A | 2.9 | 5.3 | 81 | 151 | 9 | MOLECULE: | INVERTASE/P |
| 69:00:00 | 3lsg-A | 2.9 | 3.6 | 57 | 103 | 7 | MOLECULE: | TWO-COMPONE |
| 70:00:00 | 3hmj-A | 2.9 | 10 | 103 | 1750 | 10 | MOLECULE: | FATTY ACID |
| 71:00:00 | 6z6o-D | 2.9 | 19.9 | 73 | 542 | 4 | MOLECULE: | HISTONE DEA |
| 72:00:00 | 2d1l-A | 2.9 | 21.9 | 101 | 249 | 5 | MOLECULE: | METASTASIS |
| 73:00:00 | 6oap-A | 2.9 | 13.2 | 105 | 305 | 12 | MOLECULE: | DUAL SENSOR |
| 74:00:00 | 1whu-A | 2.9 | 6 | 72 | 104 | 10 | MOLECULE: | POLYNUCLEOT |
| 75:00:00 | 7o4i-1 | 2.9 | 12.9 | 68 | 522 | 6 | MOLECULE: | GENERAL TRA |
| 76:00:00 | 5j0l-A | 2.9 | 9.3 | 64 | 130 | 6 | MOLECULE: | DESIGNED PR |
| 77:00:00 | 7w7c-B | 2.9 | 4.9 | 73 | 159 | 10 | MOLECULE: | PUTATIVE AB |
| 78:00:00 | 7ml0-1 | 2.9 | 16.9 | 64 | 367 | 5 | MOLECULE: | DNA-DIRECTE |
| 79:00:00 | 8r33-A | 2.9 | 6.1 | 111 | 489 | 5 | MOLECULE: | PHO90 ISOFO |
| 80:00:00 | 6g18-y | 2.9 | 14.9 | 70 | 325 | 7 | MOLECULE: | PRE-18S RIB |
| 81:00:00 | 6e11-E | 2.9 | 11.3 | 103 | 210 | 5 | MOLECULE: | UNKNOWN (CL |
| 82:00:00 | 8hdu-B | 2.9 | 5.5 | 55 | 162 | 11 | MOLECULE: | DE NOVO DES |
| 83:00:00 | 7lt1-A | 2.9 | 12.6 | 100 | 403 | 4 | MOLECULE: | PROTEIN MB2 |
| 84:00:00 | 4v2o-B | 2.9 | 7.4 | 51 | 81 | 8 | MOLECULE: | SAPOSIN-B |
| 85:00:00 | 8th8-H | 2.9 | 12.1 | 61 | 820 | 5 | MOLECULE: | DYNEIN REGU |
| 86:00:00 | 1q1v-A | 2.9 | 3.8 | 63 | 70 | 3 | MOLECULE: | DEK PROTEIN |

| | | | | | | | | |
|---|---|---|---|---|---|---|---|---|
| 87:00:00 | 7elb-A | 2.9 | 29.3 | 121 | 1937 | 7 | MOLECULE: | RNA-DIRECTE |
| 88:00:00 | 8j22-C | 2.9 | 6.6 | 88 | 285 | 6 | MOLECULE: | GUANINE NUC |
| 89:00:00 | 8kee-C | 2.9 | 8.4 | 104 | 505 | 4 | MOLECULE: | SHEATH |
| 90:00:00 | 7kmt-B | 2.9 | 11.7 | 87 | 475 | 5 | MOLECULE: | TRAFFICKING |
| 91:00:00 | 7aqq-N | 2.9 | 12.6 | 92 | 488 | 3 | MOLECULE: | NADH-UBIQUI |
| 92:00:00 | 6nps-A | 2.9 | 18.9 | 102 | 968 | 5 | MOLECULE: | AXYAGU115A |
| 93:00:00 | 6csv-A | 2.9 | 11.9 | 74 | 90 | 5 | MOLECULE: | CENTROSOMAL |
| 94:00:00 | 6s6b-B | 2.9 | 3.6 | 76 | 154 | 7 | MOLECULE: | CRISPR-ASSO |
| 95:00:00 | 6ud5-D | 2.9 | 10.5 | 77 | 355 | 6 | MOLECULE: | TRYPTOPHAN |
| 96:00:00 | 5hzi-A | 2.9 | 9.8 | 96 | 476 | 5 | MOLECULE: | INTERSECTIN |
| 97:00:00 | 5fhp-C | 2.9 | 13.1 | 67 | 209 | 9 | MOLECULE: | NICR |
| 98:00:00 | 8d4y-B | 2.9 | 14.9 | 80 | 217 | 11 | MOLECULE: | CHROMODOMAI |
| 99:00:00 | 2qwt-A | 2.9 | 8.1 | 65 | 167 | 6 | MOLECULE: | TRANSCRIPTI |
| 0:00 | 8wt9-A | 2.9 | 11.2 | 86 | 319 | 9 | MOLECULE: | IS621 TRANS |
| 1:00 | 5oqj-W | 2.9 | 21.5 | 83 | 258 | 5 | MOLECULE: | DNA-DIRECTE |
| 2:00 | 7eeb-K | 2.9 | 15.9 | 91 | 143 | 5 | MOLECULE: | ENHANCED GR |
| 3:00 | 5c8j-L | 2.9 | 8.7 | 71 | 137 | 8 | MOLECULE: | ANTIBODY FR |
| 4:00 | 6ejq-B | 2.9 | 14.9 | 72 | 141 | 8 | MOLECULE: | TERMINASE S |
| 5:00 | 8ap7-j | 2.9 | 11 | 78 | 168 | 4 | MOLECULE: | ATP SYNTHAS |
| 6:00 | 8ttf-A | 2.9 | 12.6 | 89 | 388 | 8 | MOLECULE: | QUINOLONE R |
| 7:00 | 2dh4-A | 2.9 | 11.8 | 92 | 326 | 1 | MOLECULE: | YPL069C |
| 8:00 | 7oca-E | 2.9 | 10.8 | 93 | 158 | 6 | MOLECULE: | GLUTAMATE R |
| 9:00 | 7ody-C | 2.9 | 5.1 | 69 | 100 | 13 | MOLECULE: | MAZG-LIKE P |
| 10:00 | 6yxq-A | 2.9 | 17.6 | 64 | 188 | 6 | MOLECULE: | ACTIVATING |
| 11:00 | 8fed-J | 2.9 | 3.5 | 89 | 263 | 1 | MOLECULE: | VIRULENCE F |
| 12:00 | 7x3t-V | 2.9 | 14.6 | 93 | 792 | 8 | MOLECULE: | HISTONE H3 |
| 13:00 | 6w6x-A | 2.9 | 7.8 | 77 | 126 | 8 | MOLECULE: | DE NOVO DES |
| 14:00 | 6kex-A | 2.9 | 3.6 | 59 | 349 | 5 | MOLECULE: | PHOSPHORIBU |
| 15:00 | 5ajk-A | 2.9 | 5.7 | 72 | 148 | 4 | MOLECULE: | HOMOLOG OF |
| 16:00 | 6bhp-A | 2.9 | 4.7 | 78 | 170 | 8 | MOLECULE: | MEMBRANE PR |
| 17:00 | 7t7t-A | 2.9 | 10.4 | 84 | 466 | 4 | MOLECULE: | PROTEIN TON |
| 18:00 | 6wtw-A | 2.9 | 12.3 | 95 | 489 | 5 | MOLECULE: | DASS FAMILY |
| 19:00 | 8axa-A | 2.9 | 22 | 118 | 465 | 1 | MOLECULE: | CAS12K |
| 20:00 | 6swy-9 | 2.9 | 5.5 | 59 | 157 | 12 | MOLECULE: | VACUOLAR IM |
| 21:00 | 8p0w-A | 2.9 | 19.1 | 73 | 190 | 12 | MOLECULE: | COMM DOMAIN |
| 22:00 | 8ucq-A | 2.9 | 23.9 | 105 | 1232 | 9 | MOLECULE: | SEC7 DOMAIN |
| 23:00 | 7wb4-I | 2.9 | 33.6 | 118 | 796 | 3 | MOLECULE: | OUTER NUP13 |
| 24:00:00 | 4raf-A | 2.9 | 8.9 | 85 | 362 | 7 | MOLECULE: | PROTEIN PHO |
| 25:00:00 | 6elu-A | 2.9 | 5.5 | 74 | 193 | 12 | MOLECULE: | SERUM RESIS |
| 26:00:00 | 5z58-1 | 2.9 | 20.5 | 108 | 1038 | 6 | MOLECULE: | PRE-MRNA-PR |
| 27:00:00 | 5unk-A | 2.9 | 3.6 | 53 | 62 | 9 | MOLECULE: | SLEEPING BE |
| 28:00:00 | 5v2z-A | 2.9 | 10.3 | 64 | 350 | 2 | MOLECULE: | 2-OXOGLUTAR |
| 29:00:00 | 8hf3-A | 2.9 | 12.9 | 106 | 295 | 7 | MOLECULE: | PALMITOYLTR |
| 30:00:00 | 2z3r-E | 2.9 | 4.5 | 75 | 119 | 4 | MOLECULE: | INTERLEUKIN |

| | | | | | | | | |
|---|---|---|---|---|---|---|---|---|
| 31:00:00 | 4kha-A | 2.9 | 12.1 | 78 | 360 | 8 | MOLECULE: | SPT16M-HIST |
| 32:00:00 | 7jr7-A | 2.9 | 5.5 | 86 | 588 | 7 | MOLECULE: | ATP-BINDING |
| 33:00:00 | 4f9k-B | 2.9 | 3.2 | 46 | 58 | 2 | MOLECULE: | CAMP-DEPEND |
| 34:00:00 | 2hjm-A | 2.9 | 5.5 | 69 | 91 | 4 | MOLECULE: | HYPOTHETICA |
| 35:00:00 | 5jhf-D | 2.9 | 7.9 | 61 | 82 | 7 | MOLECULE: | KLTH0D11660 |
| 36:00:00 | 5do7-D | 2.9 | 4.5 | 74 | 571 | 9 | MOLECULE: | ATP-BINDING |
| 37:00:00 | 7bmh-A | 2.9 | 8.6 | 104 | 247 | 11 | MOLECULE: | OPSIN |
| 38:00:00 | 6whg-A | 2.9 | 19.5 | 121 | 523 | 7 | MOLECULE: | CALCIUM CHA |
| 39:00:00 | 7dgj-A | 2.9 | 8.3 | 55 | 153 | 4 | MOLECULE: | MYOGLOBIN |
| 40:00:00 | 7dgm-B | 2.9 | 8.1 | 57 | 153 | 4 | MOLECULE: | MYOGLOBIN |
| 41:00:00 | 3vm9-A | 2.9 | 11.5 | 57 | 153 | 5 | MOLECULE: | MYOGLOBIN |
| 42:00:00 | 4hrr-H | 2.9 | 4.1 | 75 | 150 | 8 | MOLECULE: | GLOBIN-2 A |
| 43:00:00 | 3dhr-F | 2.9 | 4.9 | 79 | 145 | 11 | MOLECULE: | HEMOGLOBIN |
| 44:00:00 | 7dgo-A | 2.9 | 8.3 | 54 | 153 | 2 | MOLECULE: | MYOGLOBIN |
| 45:00:00 | 4mqj-H | 2.9 | 4.8 | 83 | 146 | 10 | MOLECULE: | HEMOGLOBIN |
| 46:00:00 | 7v5q-A | 2.9 | 10.1 | 71 | 153 | 6 | MOLECULE: | MYOGLOBIN |
| 47:00:00 | 7dgm-A | 2.9 | 8.1 | 55 | 153 | 4 | MOLECULE: | MYOGLOBIN |
| 48:00:00 | 3fs4-D | 2.9 | 4.6 | 78 | 146 | 15 | MOLECULE: | HEMOGLOBIN |
| 49:00:00 | 7dgk-A | 2.9 | 10.5 | 65 | 153 | 3 | MOLECULE: | MYOGLOBIN |
| 50:00:00 | 7dgj-B | 2.9 | 9.9 | 72 | 153 | 6 | MOLECULE: | MYOGLOBIN |
| 51:00:00 | 3wyo-C | 2.9 | 11.4 | 56 | 153 | 2 | MOLECULE: | MYOGLOBIN |
| 52:00:00 | 6ltm-A | 2.9 | 8 | 56 | 153 | 4 | MOLECULE: | MYOGLOBIN |
| 53:00:00 | 7dgl-B | 2.9 | 10.1 | 72 | 153 | 7 | MOLECULE: | MYOGLOBIN |
| 54:00:00 | 7v5r-A | 2.9 | 9.6 | 63 | 153 | 5 | MOLECULE: | MYOGLOBIN |
| 55:00:00 | 2hbs-H | 2.9 | 4.6 | 82 | 146 | 10 | MOLECULE: | HEMOGLOBIN |
| 56:00:00 | 6ltl-A | 2.9 | 11.6 | 57 | 153 | 4 | MOLECULE: | MYOGLOBIN |
| 57:00:00 | 1y4p-B | 2.9 | 4.4 | 79 | 146 | 10 | MOLECULE: | HEMOGLOBIN |
| 58:00:00 | 8dov-H | 2.9 | 3.6 | 81 | 145 | 7 | MOLECULE: | HEMOGLOBIN |
| 59:00:00 | 7b1s-B | 2.8 | 10.1 | 105 | 466 | 10 | MOLECULE: | ETHYL-COENZ |
| 60:00:00 | 6zu0-C | 2.8 | 17.2 | 87 | 426 | 8 | MOLECULE: | CITRATE SYN |
| 61:00:00 | 6cgh-A | 2.8 | 4.2 | 47 | 89 | 6 | MOLECULE: | DNAJ HOMOLO |
| 62:00:00 | 2y7c-A | 2.8 | 3.8 | 63 | 464 | 10 | MOLECULE: | TYPE-1 REST |
| 63:00:00 | 7zgi-B | 2.8 | 6.1 | 58 | 329 | 7 | MOLECULE: | PEPTIDYLPRO |
| 64:00:00 | 8snl-B | 2.8 | 16.3 | 75 | 491 | 8 | MOLECULE: | DISINTEGRIN |
| 65:00:00 | 4f4h-A | 2.8 | 16.7 | 102 | 540 | 9 | MOLECULE: | GLUTAMINE D |
| 66:00:00 | 6j36-A | 2.8 | 4.7 | 55 | 459 | 4 | MOLECULE: | ENOLASE |
| 67:00:00 | 4jvy-A | 2.8 | 10.5 | 58 | 191 | 14 | MOLECULE: | FEMALE GERM |
| 68:00:00 | 6gyt-B | 2.8 | 6.2 | 52 | 168 | 8 | MOLECULE: | HISTONE ACE |
| 69:00:00 | 4cej-A | 2.8 | 22.4 | 129 | 1177 | 9 | MOLECULE: | ATP-DEPENDE |
| 70:00:00 | 6ffv-A | 2.8 | 13 | 59 | 194 | 3 | MOLECULE: | BTUM |
| 71:00:00 | 6gkw-A | 2.8 | 3.7 | 73 | 351 | 7 | MOLECULE: | PUTATIVE PH |
| 72:00:00 | 2yev-B | 2.8 | 17.3 | 92 | 319 | 4 | MOLECULE: | CYTOCHROME |
| 73:00:00 | 4nqj-A | 2.8 | 7.5 | 64 | 177 | 8 | MOLECULE: | E3 UBIQUITI |
| 74:00:00 | 5vp3-A | 2.8 | 11 | 72 | 195 | 6 | MOLECULE: | MNEMIOPSIN |

| | | | | | | | | |
|---|---|---|---|---|---|---|---|---|
| 75:00:00 | 6rfs-6 | 2.8 | 9.8 | 78 | 138 | 8 | MOLECULE: | SUBUNIT NUA |
| 76:00:00 | 5jhf-F | 2.8 | 24.9 | 95 | 405 | 12 | MOLECULE: | KLTH0D11660 |
| 77:00:00 | 7yoj-A | 2.8 | 28.3 | 103 | 867 | 5 | MOLECULE: | CASPI |
| 78:00:00 | 7asv-A | 2.8 | 10 | 71 | 155 | 7 | MOLECULE: | DNA-DIRECTE |
| 79:00:00 | 8vjn-A | 2.8 | 4.7 | 43 | 52 | 9 | MOLECULE: | ENCAPSULIN |
| 80:00:00 | 3stq-A | 2.8 | 6.4 | 51 | 86 | 6 | MOLECULE: | PUTATIVE UN |
| 81:00:00 | 6ql4-A | 2.8 | 10.6 | 114 | 660 | 6 | MOLECULE: | PUTATIVE MI |
| 82:00:00 | 7sbe-A | 2.8 | 6.3 | 96 | 228 | 8 | MOLECULE: | TELOMERASE |
| 83:00:00 | 4he8-D | 2.8 | 7.2 | 60 | 160 | 7 | MOLECULE: | NADH-QUINON |
| 84:00:00 | 2wtg-A | 2.8 | 4.3 | 76 | 158 | 5 | MOLECULE: | GLOBIN-LIKE |
| 85:00:00 | 6p18-Q | 2.8 | 7.2 | 73 | 156 | 8 | MOLECULE: | DNA (67-MER |
| 86:00:00 | 3wfw-A | 2.8 | 10.7 | 57 | 138 | 7 | MOLECULE: | HEMOGLOBIN- |
| 87:00:00 | 8izu-A | 2.8 | 4.5 | 79 | 137 | 5 | MOLECULE: | VIRION MORP |
| 88:00:00 | 6swg-B | 2.8 | 4.7 | 54 | 74 | 2 | MOLECULE: | PERIPHILIN- |
| 89:00:00 | 7ao3-A | 2.8 | 9.5 | 57 | 303 | 4 | MOLECULE: | CYCLOOCTAT- |
| 90:00:00 | 6jo0-A | 2.8 | 5.3 | 69 | 224 | 3 | MOLECULE: | VIRRDTS |
| 91:00:00 | 2wj9-B | 2.8 | 6.1 | 79 | 145 | 5 | MOLECULE: | INTERGENIC- |
| 92:00:00 | 1w0b-A | 2.8 | 6.2 | 72 | 102 | 10 | MOLECULE: | ALPHA-HEMOG |
| 93:00:00 | 7s5l-A | 2.8 | 4.5 | 87 | 378 | 9 | MOLECULE: | CEMBRENE A |
| 94:00:00 | 6jal-A | 2.8 | 11.6 | 58 | 416 | 10 | MOLECULE: | ABC TRANSPO |
| 95:00:00 | 6lum-G | 2.8 | 5.6 | 61 | 123 | 2 | MOLECULE: | SUCCINATE D |
| 96:00:00 | 3n1e-B | 2.8 | 7.8 | 83 | 141 | 2 | MOLECULE: | VACUOLAR PR |
| 97:00:00 | 7emf-P | 2.8 | 16 | 75 | 766 | 8 | MOLECULE: | MEDIATOR OF |
| 98:00:00 | 6rwy-g | 2.8 | 12.1 | 73 | 86 | 4 | MOLECULE: | INNER ROD P |
| 99:00:00 | 1it2-A | 2.8 | 3.9 | 76 | 146 | 12 | MOLECULE: | HEMOGLOBIN |
| 0:00 | 6idp-A | 2.8 | 13 | 103 | 438 | 7 | MOLECULE: | MATE FAMILY |
| 1:00 | 1akh-B | 2.8 | 3.6 | 66 | 78 | 14 | MOLECULE: | DNA (5'- |
| 2:00 | 6sum-A | 2.8 | 7.5 | 95 | 335 | 4 | MOLECULE: | AMICOUMACIN |
| 3:00 | 3cit-A | 2.8 | 9 | 64 | 155 | 6 | MOLECULE: | SENSOR HIST |
| 4:00 | 2r5z-A | 2.8 | 5.9 | 67 | 75 | 1 | MOLECULE: | HOMEOTIC PR |
| 5:00 | 6sz9-D | 2.8 | 19.3 | 69 | 282 | 4 | MOLECULE: | ICMO (DOTL) |
| 6:00 | 6wql-A | 2.8 | 5.4 | 44 | 49 | 11 | MOLECULE: | SEED PEPTID |
| 7:00 | 8ppr-P | 2.8 | 14.7 | 56 | 176 | 4 | MOLECULE: | KINETOCHORE |
| 8:00 | 3fx7-B | 2.8 | 13.1 | 73 | 87 | 3 | MOLECULE: | PUTATIVE UN |
| 9:00 | 4j41-B | 2.8 | 9.7 | 65 | 86 | 3 | MOLECULE: | SECRETED PR |
| 10:00 | 7eu3-G | 2.8 | 8.5 | 80 | 176 | 9 | MOLECULE: | NAD(P)H-QUI |
| 11:00 | 6ybw-y | 2.8 | 18.4 | 83 | 162 | 11 | MOLECULE: | 40S RIBOSOM |
| 12:00 | 2ilk-A | 2.8 | 17.1 | 85 | 155 | 9 | MOLECULE: | INTERLEUKIN |
| 13:00 | 4u7b-A | 2.8 | 19.4 | 75 | 342 | 3 | MOLECULE: | DNA (25-MER |
| 14:00 | 8if8-B | 2.8 | 14.2 | 97 | 620 | 5 | MOLECULE: | GALACTAN 5- |
| 15:00 | 7os0-A | 2.8 | 13.2 | 95 | 1130 | 5 | MOLECULE: | CAS13A |
| 16:00 | 6awl-A | 2.8 | 18 | 84 | 217 | 8 | MOLECULE: | UBIQUINONE |
| 17:00 | 2yvx-A | 2.8 | 7.1 | 99 | 442 | 8 | MOLECULE: | MG2+ TRANSP |
| 18:00 | 4gfq-A | 2.8 | 4.6 | 76 | 186 | 9 | MOLECULE: | RIBOSOME-RE |

| | | | | | | | | |
|---|---|---|---|---|---|---|---|---|
| 19:00 | 5abb-A | 2.8 | 9.2 | 90 | 443 | 8 | MOLECULE: | PROTEIN TRA |
| 20:00 | 7eqf-A | 2.8 | 5.8 | 77 | 184 | 12 | MOLECULE: | TETR/ACRR F |
| 21:00 | 6oqr-W | 2.8 | 8.7 | 89 | 173 | 7 | MOLECULE: | ATP SYNTHAS |
| 22:00 | 6k7l-A | 2.8 | 21.9 | 96 | 992 | 5 | MOLECULE: | PHOSPHOLIPI |
| 23:00 | 5o5j-O | 2.8 | 5.9 | 63 | 88 | 14 | MOLECULE: | 16S RRNA |
| 24:00:00 | 7ar9-K | 2.8 | 9.2 | 82 | 104 | 7 | MOLECULE: | ND3 |
| 25:00:00 | 6u8y-d | 2.8 | 6.3 | 67 | 89 | 1 | MOLECULE: | MONOVALENT |
| 26:00:00 | 3utk-A | 2.8 | 3.4 | 60 | 95 | 7 | MOLECULE: | LIPOPROTEIN |
| 27:00:00 | 1uzc-A | 2.8 | 2.8 | 55 | 69 | 7 | MOLECULE: | HYPOTHETICA |
| 28:00:00 | 7nhr-A | 2.8 | 20.8 | 85 | 663 | 4 | MOLECULE: | PUTATIVE TR |
| 29:00:00 | 2p6j-A | 2.8 | 2.3 | 44 | 52 | 0 | MOLECULE: | DESIGNED EN |
| 30:00:00 | 8j56-C | 2.8 | 3.9 | 65 | 160 | 6 | MOLECULE: | FLAGELLAR T |
| 31:00:00 | 8eau-e | 2.8 | 5.3 | 61 | 72 | 7 | MOLECULE: | V-TYPE PROT |
| 32:00:00 | 7lsv-B | 2.8 | 6.1 | 85 | 158 | 4 | MOLECULE: | CALMODULIN- |
| 33:00:00 | 5dn6-H | 2.8 | 4.4 | 69 | 110 | 3 | MOLECULE: | CHAIN A |
| 34:00:00 | 6z5s-W | 2.8 | 4 | 73 | 94 | 3 | MOLECULE: | LIGHT HARVE |
| 35:00:00 | 8ovt-A | 2.8 | 7.1 | 71 | 288 | 4 | MOLECULE: | YEGT GLYCOS |
| 36:00:00 | 5ch6-A | 2.8 | 8.3 | 81 | 286 | 6 | MOLECULE: | FRIGIDA |
| 37:00:00 | 6wxv-A | 2.8 | 17.3 | 121 | 1091 | 10 | MOLECULE: | DUAL OXIDAS |
| 38:00:00 | 3wvo-A | 2.8 | 4.6 | 76 | 543 | 12 | MOLECULE: | CRISPR-ASSO |
| 39:00:00 | 6wg3-C | 2.8 | 23.6 | 81 | 248 | 10 | MOLECULE: | STRUCTURAL |
| 40:00:00 | 4p9f-A | 2.8 | 7.8 | 102 | 212 | 8 | MOLECULE: | HTH-TYPE TR |
| 41:00:00 | 2k85-A | 2.8 | 3.2 | 58 | 65 | 9 | MOLECULE: | GLUCOCORTIC |
| 42:00:00 | 3pux-G | 2.8 | 5.2 | 88 | 293 | 5 | MOLECULE: | MALTOSE-BIN |
| 43:00:00 | 8eew-A | 2.8 | 4.7 | 112 | 766 | 3 | MOLECULE: | DYNAMIN-LIK |
| 44:00:00 | 6ekr-A | 2.8 | 6.6 | 89 | 303 | 10 | MOLECULE: | TYPE II SIT |
| 45:00:00 | 8q7e-J | 2.8 | 21.4 | 126 | 895 | 8 | MOLECULE: | CULLIN-9 |
| 46:00:00 | 7z1n-O | 2.8 | 22.7 | 96 | 570 | 5 | MOLECULE: | DNA-DIRECTE |
| 47:00:00 | 7yh5-B | 2.8 | 8.8 | 85 | 177 | 18 | MOLECULE: | NUCLEOSIDE |
| 48:00:00 | 5ero-A | 2.8 | 14.1 | 96 | 301 | 4 | MOLECULE: | FUSICOCCADI |
| 49:00:00 | 8hsb-A | 2.8 | 19.6 | 102 | 374 | 5 | MOLECULE: | CDNG |
| 50:00:00 | 1hbs-D | 2.8 | 3.4 | 77 | 146 | 8 | MOLECULE: | HEMOGLOBIN |
| 51:00:00 | 7v5p-B | 2.8 | 8.4 | 55 | 153 | 4 | MOLECULE: | MYOGLOBIN |
| 52:00:00 | 7dgk-B | 2.8 | 10.2 | 66 | 153 | 8 | MOLECULE: | MYOGLOBIN |
| 53:00:00 | 5x2s-F | 2.8 | 4.1 | 80 | 145 | 13 | MOLECULE: | HEMOGLOBIN |
| 54:00:00 | 6q6h-N | 2.8 | 30.9 | 120 | 653 | 4 | MOLECULE: | ANAPHASE-PR |
| 55:00:00 | 1qpw-B | 2.8 | 4.6 | 79 | 146 | 10 | MOLECULE: | PORCINE HEM |
| 56:00:00 | 7v5q-B | 2.8 | 10.1 | 65 | 153 | 5 | MOLECULE: | MYOGLOBIN |
| 57:00:00 | 7dgo-B | 2.8 | 10.1 | 66 | 153 | 3 | MOLECULE: | MYOGLOBIN |
| 58:00:00 | 3vrf-B | 2.8 | 4.8 | 79 | 146 | 8 | MOLECULE: | HEMOGLOBIN |
| 59:00:00 | 3wyo-B | 2.8 | 7.2 | 55 | 153 | 7 | MOLECULE: | MYOGLOBIN |
| 60:00:00 | 3wcp-D | 2.8 | 4.5 | 79 | 145 | 10 | MOLECULE: | HEMOGLOBIN |
| 61:00:00 | 7v5p-A | 2.8 | 11.3 | 56 | 153 | 5 | MOLECULE: | MYOGLOBIN |
| 62:00:00 | 1moh-A | 2.8 | 4.4 | 74 | 142 | 5 | MOLECULE: | MONOMERIC H |

| | | | | | | | | |
|---|---|---|---|---|---|---|---|---|
| 63:00:00 | 3w4u-B | 2.8 | 4.4 | 81 | 145 | 11 | MOLECULE: | HEMOGLOBIN |
| 64:00:00 | 5gko-A | 2.7 | 4.2 | 79 | 650 | 3 | MOLECULE: | MACROLIDE E |
| 65:00:00 | 6n7p-X | 2.7 | 12.2 | 83 | 826 | 2 | MOLECULE: | U1 SMALL NU |
| 66:00:00 | 6u4b-A | 2.7 | 11.5 | 122 | 570 | 4 | MOLECULE: | WBBM PROTEI |
| 67:00:00 | 6id0-L | 2.7 | 14 | 81 | 475 | 4 | MOLECULE: | PRE-MRNA-PR |
| 68:00:00 | 3ucq-A | 2.7 | 19.9 | 82 | 651 | 4 | MOLECULE: | AMYLOSUCRAS |
| 69:00:00 | 2rdc-B | 2.7 | 11.3 | 90 | 135 | 9 | MOLECULE: | UNCHARACTER |
| 70:00:00 | 6m36-I | 2.7 | 4.2 | 45 | 104 | 4 | MOLECULE: | SERINE-PROT |
| 71:00:00 | 6pij-A | 2.7 | 4.3 | 68 | 351 | 6 | MOLECULE: | CAS7 TYPE I |
| 72:00:00 | 3d1n-I | 2.7 | 13.7 | 67 | 149 | 7 | MOLECULE: | 5'- |
| 73:00:00 | 6xiv-A | 2.7 | 4.6 | 84 | 257 | 4 | MOLECULE: | REGULATORY |
| 74:00:00 | 7f52-A | 2.7 | 29.7 | 124 | 584 | 5 | MOLECULE: | NON-STRUCTU |
| 75:00:00 | 3etz-B | 2.7 | 14.9 | 76 | 115 | 3 | MOLECULE: | ADHESIN A |
| 76:00:00 | 3oov-A | 2.7 | 2.9 | 39 | 164 | 10 | MOLECULE: | METHYL-ACCE |
| 77:00:00 | 5gm2-B | 2.7 | 11 | 75 | 283 | 13 | MOLECULE: | O-METHYLRAN |
| 78:00:00 | 8bf9-G | 2.7 | 7.6 | 60 | 158 | 3 | MOLECULE: | RNA (1766) |
| 79:00:00 | 4j7o-A | 2.7 | 13.4 | 91 | 329 | 3 | MOLECULE: | PUTATIVE SU |
| 80:00:00 | 5w7d-A | 2.7 | 16.1 | 84 | 528 | 4 | MOLECULE: | ACYLOXYACYL |
| 81:00:00 | 6kp3-B | 2.7 | 9.5 | 42 | 112 | 2 | MOLECULE: | PROGRAMMED |
| 82:00:00 | 8bd7-A | 2.7 | 22.5 | 105 | 539 | 10 | MOLECULE: | IFT88 |
| 83:00:00 | 7z36-B | 2.7 | 9.5 | 62 | 446 | 5 | MOLECULE: | ENDOLYSIN,T |
| 84:00:00 | 6vmb-g | 2.7 | 4.5 | 42 | 323 | 10 | MOLECULE: | ATP SYNTHAS |
| 85:00:00 | 2ebi-A | 2.7 | 3.1 | 61 | 86 | 8 | MOLECULE: | DNA BINDING |
| 86:00:00 | 2pmq-B | 2.7 | 3.6 | 46 | 374 | 0 | MOLECULE: | MANDELATE R |
| 87:00:00 | 2g8l-B | 2.7 | 2.7 | 49 | 286 | 4 | MOLECULE: | 287AA LONG |
| 88:00:00 | 5yfp-B | 2.7 | 17.9 | 77 | 927 | 5 | MOLECULE: | EXOCYST COM |
| 89:00:00 | 8k2g-A | 2.7 | 20.7 | 75 | 505 | 8 | MOLECULE: | OLIGOSACCHA |
| 90:00:00 | 7e8k-B | 2.7 | 9.4 | 67 | 205 | 7 | MOLECULE: | RNA-FREE RI |
| 91:00:00 | 8sun-B | 2.7 | 16.3 | 116 | 743 | 4 | MOLECULE: | ANOCTAMIN-6 |
| 92:00:00 | 7bhq-B | 2.7 | 15 | 87 | 204 | 8 | MOLECULE: | BASAL-BODY |
| 93:00:00 | 2jvg-A | 2.7 | 4 | 64 | 72 | 11 | MOLECULE: | IGG-BINDING |
| 94:00:00 | 8ek4-B | 2.7 | 9.4 | 70 | 135 | 14 | MOLECULE: | ICE-BINDING |
| 95:00:00 | 7sbb-I | 2.7 | 7 | 111 | 680 | 6 | MOLECULE: | CAS7D |
| 96:00:00 | 8gwe-A | 2.7 | 24.7 | 121 | 931 | 5 | MOLECULE: | RNA-DIRECTE |
| 97:00:00 | 5k8o-B | 2.7 | 10.4 | 73 | 425 | 8 | MOLECULE: | 5-NITROANTH |
| 98:00:00 | 8cjf-B | 2.7 | 8.2 | 90 | 639 | 3 | MOLECULE: | AETF |
| 99:00:00 | 7ml7-A | 2.7 | 17.7 | 121 | 1216 | 5 | MOLECULE: | TOXIN B |
| 0:00 | 7tb3-A | 2.7 | 15.8 | 78 | 243 | 5 | MOLECULE: | ISOFORM 2 O |
| 1:00 | 5w10-A | 2.7 | 3.4 | 39 | 173 | 8 | MOLECULE: | CGMP-SPECIF |
| 2:00 | 3ps5-A | 2.7 | 5.2 | 90 | 529 | 8 | MOLECULE: | TYROSINE-PR |
| 3:00 | 1n93-X | 2.7 | 19 | 84 | 335 | 2 | MOLECULE: | P40 NUCLEOP |
| 4:00 | 7nza-A | 2.7 | 9.1 | 65 | 141 | 9 | MOLECULE: | ODORANT BIN |
| 5:00 | 5csk-A | 2.7 | 34.2 | 124 | 1996 | 7 | MOLECULE: | ACETYL-COA |
| 6:00 | 5d50-G | 2.7 | 9.7 | 55 | 77 | 9 | MOLECULE: | REPRESSOR |

| | | | | | | | | |
|---|---|---|---|---|---|---|---|---|
| 7:00 | 5im3-A | 2.7 | 15.8 | 77 | 874 | 5 | MOLECULE: | RIBONUCLEOS |
| 8:00 | 6wuc-H | 2.7 | 7.8 | 69 | 179 | 9 | MOLECULE: | INNER KINET |
| 9:00 | 8j5q-B | 2.7 | 2.8 | 42 | 325 | 10 | MOLECULE: | UNCHARACTER |
| 10:00 | 6idf-B | 2.7 | 8.2 | 98 | 312 | 9 | MOLECULE: | NICASTRIN |
| 11:00 | 2kjf-A | 2.7 | 2.6 | 50 | 60 | 14 | MOLECULE: | CARNOCYCLIN |
| 12:00 | 5t58-A | 2.7 | 13.2 | 71 | 233 | 4 | MOLECULE: | KLLA0F02343 |
| 13:00 | 4y2f-A | 2.7 | 9.1 | 61 | 143 | 10 | MOLECULE: | SENSOR PROT |
| 14:00 | 8wzb-A | 2.7 | 3 | 42 | 46 | 10 | MOLECULE: | DPY30 DOMAI |
| 15:00 | 8ear-A | 2.7 | 20.5 | 116 | 2389 | 6 | MOLECULE: | INOSITOL 1, |
| 16:00 | 7lvf-A | 2.7 | 6.4 | 48 | 57 | 4 | MOLECULE: | VICILIN JUG |
| 17:00 | 7zm7-2 | 2.7 | 15.7 | 100 | 558 | 4 | MOLECULE: | NADH-UBIQUI |
| 18:00 | 8h4p-A | 2.7 | 5.6 | 81 | 326 | 7 | MOLECULE: | LONGIBORNEO |
| 19:00 | 8fo9-F | 2.7 | 31.2 | 108 | 2289 | 6 | MOLECULE: | LEUCINE-RIC |
| 20:00 | 6r6b-F | 2.7 | 8.4 | 66 | 245 | 6 | MOLECULE: | SURFACE PRE |
| 21:00 | 3kdw-A | 2.7 | 5.3 | 76 | 206 | 8 | MOLECULE: | PUTATIVE SU |
| 22:00 | 6z70-A | 2.7 | 9.1 | 88 | 444 | 9 | MOLECULE: | AQ128 |
| 23:00 | 4n5a-A | 2.7 | 25.5 | 99 | 535 | 6 | MOLECULE: | PROTEIN EFR |
| 24:00:00 | 8tdl-A | 2.7 | 24.8 | 83 | 364 | 5 | MOLECULE: | MECHANOSENS |
| 25:00:00 | 4yo2-A | 2.7 | 4.7 | 67 | 186 | 9 | MOLECULE: | TRANSCRIPTI |
| 26:00:00 | 7kzn-B | 2.7 | 18.5 | 80 | 683 | 10 | MOLECULE: | HEAVY CHAIN |
| 27:00:00 | 6o3q-A | 2.7 | 3.7 | 46 | 49 | 7 | MOLECULE: | VICILIN |
| 28:00:00 | 2fbq-A | 2.7 | 11.7 | 70 | 213 | 7 | MOLECULE: | PROBABLE TR |
| 29:00:00 | 5n8k-C | 2.7 | 5.5 | 51 | 80 | 10 | MOLECULE: | GALACTOCERE |
| 30:00:00 | 3f0c-A | 2.7 | 6.7 | 74 | 193 | 4 | MOLECULE: | TRANSCRIPTI |
| 31:00:00 | 6bhc-A | 2.7 | 4 | 80 | 371 | 4 | MOLECULE: | PSEUDOPODIU |
| 32:00:00 | 2xco-A | 2.7 | 14.6 | 80 | 636 | 8 | MOLECULE: | DNA GYRASE |
| 33:00:00 | 2hcf-A | 2.7 | 6.3 | 63 | 225 | 8 | MOLECULE: | HYDROLASE, |
| 34:00:00 | 5mmi-Z | 2.7 | 10.8 | 77 | 101 | 3 | MOLECULE: | 50S RIBOSOM |
| 35:00:00 | 7zke-E | 2.7 | 22.4 | 132 | 1623 | 7 | MOLECULE: | DNA (36-MER |
| 36:00:00 | 2m2e-A | 2.7 | 3.7 | 56 | 70 | 11 | MOLECULE: | DNAJ HOMOLO |
| 37:00:00 | 4ejo-A | 2.7 | 4.3 | 53 | 112 | 13 | MOLECULE: | TRANSCRIPTI |
| 38:00:00 | 5xw4-B | 2.7 | 9.9 | 87 | 369 | 9 | MOLECULE: | TYROSINE-PR |
| 39:00:00 | 6ny2-Y | 2.7 | 29.7 | 137 | 915 | 7 | MOLECULE: | DNA TARGET |
| 40:00:00 | 7e40-A | 2.7 | 6.7 | 74 | 106 | 5 | MOLECULE: | PROTEIN PHO |
| 41:00:00 | 3lml-A | 2.7 | 3.9 | 77 | 440 | 9 | MOLECULE: | LIN1278 PRO |
| 42:00:00 | 6vrb-C | 2.7 | 7.3 | 64 | 229 | 5 | MOLECULE: | RNA (52-MER |
| 43:00:00 | 2ly9-A | 2.7 | 5.6 | 64 | 74 | 8 | MOLECULE: | ZINC FINGER |
| 44:00:00 | 8agg-A | 2.7 | 7.8 | 64 | 489 | 3 | MOLECULE: | PHOSPHOCHOL |
| 45:00:00 | 1a7d-A | 2.7 | 6.4 | 70 | 118 | 4 | MOLECULE: | MYOHEMERYTH |
| 46:00:00 | 6yxp-B | 2.7 | 11 | 76 | 272 | 4 | MOLECULE: | SWI/SNF COM |
| 47:00:00 | 2ijd-1 | 2.7 | 16.2 | 111 | 644 | 7 | MOLECULE: | PICORNAIN 3 |
| 48:00:00 | 5iz5-A | 2.7 | 9.2 | 113 | 757 | 4 | MOLECULE: | CYTOSOLIC P |
| 49:00:00 | 7way-A | 2.7 | 9 | 101 | 960 | 5 | MOLECULE: | DNA (33-MER |
| 50:00:00 | 4qvg-C | 2.7 | 7.6 | 104 | 349 | 8 | MOLECULE: | SIBL |

| | | | | | | | | |
|---|---|---|---|---|---|---|---|---|
| 51:00:00 | 5udb-4 | 2.7 | 8.6 | 111 | 751 | 9 | MOLECULE: | DNA REPLICA |
| 52:00:00 | 3nnq-A | 2.7 | 4.4 | 65 | 95 | 3 | MOLECULE: | N-TERMINAL |
| 53:00:00 | 2js9-A | 2.7 | 3.1 | 54 | 81 | 6 | MOLECULE: | SAPOSIN-LIK |
| 54:00:00 | 7bw0-R | 2.7 | 17.8 | 110 | 280 | 10 | MOLECULE: | SOLUBLE CYT |
| 55:00:00 | 8p62-D | 2.7 | 19 | 80 | 245 | 6 | MOLECULE: | DNA REPLICA |
| 56:00:00 | 5h72-A | 2.7 | 2.7 | 58 | 67 | 7 | MOLECULE: | FLAGELLAR B |
| 57:00:00 | 8r6y-A | 2.7 | 18.9 | 135 | 1991 | 7 | MOLECULE: | RNA-DIRECTE |
| 58:00:00 | 7c7e-A | 2.7 | 8.5 | 88 | 142 | 6 | MOLECULE: | PUTATIVE DN |
| 59:00:00 | 6l30-A | 2.7 | 13.2 | 81 | 518 | 9 | MOLECULE: | PROTEIN ECT |
| 60:00:00 | 8rf0-A | 2.7 | 16.4 | 111 | 2780 | 5 | MOLECULE: | CYCLIC BETA |
| 61:00:00 | 3trc-A | 2.7 | 4.7 | 74 | 168 | 9 | MOLECULE: | PHOSPHOENOL |
| 62:00:00 | 7yyg-A | 2.7 | 16.3 | 91 | 275 | 7 | MOLECULE: | PUTATIVE OU |
| 63:00:00 | 6gmh-M | 2.7 | 22.5 | 111 | 991 | 5 | MOLECULE: | RPB1 |
| 64:00:00 | 2be4-A | 2.7 | 7.1 | 93 | 270 | 5 | MOLECULE: | HYPOTHETICA |
| 65:00:00 | 3egw-C | 2.7 | 10.3 | 107 | 225 | 11 | MOLECULE: | RESPIRATORY |
| 66:00:00 | 5nzz-A | 2.7 | 15.2 | 101 | 378 | 11 | MOLECULE: | TGF-BETA-AC |
| 67:00:00 | 8iq4-R | 2.7 | 13.3 | 99 | 274 | 2 | MOLECULE: | PROSTAGLAND |
| 68:00:00 | 7jgr-D | 2.7 | 21.4 | 89 | 441 | 8 | MOLECULE: | ORIGIN RECO |
| 69:00:00 | 5zib-A | 2.7 | 26.6 | 107 | 579 | 5 | MOLECULE: | ALPHA-1,6-M |
| 70:00:00 | 3pde-B | 2.7 | 7.8 | 84 | 287 | 2 | MOLECULE: | FARNESYL-DI |
| 71:00:00 | 1hlv-A | 2.7 | 6.8 | 78 | 131 | 10 | MOLECULE: | CENP-B BOX |
| 72:00:00 | 7whm-A | 2.7 | 33.4 | 100 | 1214 | 6 | MOLECULE: | PPPDE DOMAI |
| 73:00:00 | 7aoa-A | 2.7 | 31.7 | 94 | 343 | 12 | MOLECULE: | METHYL-CPG- |
| 74:00:00 | 5zb3-A | 2.7 | 13.6 | 105 | 280 | 9 | MOLECULE: | ORF57 |
| 75:00:00 | 6thh-C | 2.7 | 13.5 | 79 | 816 | 6 | MOLECULE: | SIRV2 ACRID |
| 76:00:00 | 7p4a-A | 2.7 | 26.7 | 94 | 246 | 6 | MOLECULE: | STL |
| 77:00:00 | 5e6e-B | 2.7 | 4.5 | 75 | 146 | 11 | MOLECULE: | HEMOGLOBIN |
| 78:00:00 | 5x2u-D | 2.7 | 4.6 | 78 | 145 | 12 | MOLECULE: | HEMOGLOBIN |
| 79:00:00 | 1si4-B | 2.7 | 4.6 | 78 | 146 | 12 | MOLECULE: | HEMOGLOBIN |
| 80:00:00 | 3w4u-D | 2.7 | 4.5 | 79 | 145 | 10 | MOLECULE: | HEMOGLOBIN |
| 81:00:00 | 7dgn-A | 2.7 | 8.3 | 55 | 153 | 4 | MOLECULE: | MYOGLOBIN |
| 82:00:00 | 3vm9-B | 2.7 | 8.3 | 56 | 153 | 4 | MOLECULE: | MYOGLOBIN |
| 83:00:00 | 6ltl-B | 2.7 | 9.7 | 73 | 153 | 5 | MOLECULE: | MYOGLOBIN |
| 84:00:00 | 5ker-B | 2.7 | 4.5 | 79 | 146 | 10 | MOLECULE: | ALPHA-GLOBI |
| 85:00:00 | 6ltm-B | 2.7 | 8 | 57 | 153 | 4 | MOLECULE: | MYOGLOBIN |
| 86:00:00 | 7v5r-B | 2.7 | 8.1 | 57 | 153 | 4 | MOLECULE: | MYOGLOBIN |
| 87:00:00 | 2qls-D | 2.7 | 4.9 | 79 | 146 | 9 | MOLECULE: | HEMOGLOBIN |
| 88:00:00 | 7uf7-D | 2.7 | 4.6 | 76 | 146 | 11 | MOLECULE: | HEMOGLOBIN |
| 89:00:00 | 6tm5-N | 2.7 | 22.3 | 109 | 676 | 6 | MOLECULE: | ANAPHASE-PR |
| 90:00:00 | 8pkp-N | 2.7 | 8.5 | 121 | 652 | 5 | MOLECULE: | ANAPHASE-PR |
| 91:00:00 | 5hdi-A | 2.6 | 4.7 | 98 | 397 | 6 | MOLECULE: | CYTOCHROME |
| 92:00:00 | 7ubn-Q | 2.6 | 5.2 | 74 | 207 | 8 | MOLECULE: | DNA (53-MER |
| 93:00:00 | 2pyw-A | 2.6 | 7.9 | 77 | 417 | 1 | MOLECULE: | UNCHARACTER |
| 94:00:00 | 6em5-b | 2.6 | 16 | 88 | 421 | 6 | MOLECULE: | 5.8S RIBOSO |

| | | | | | | | | |
|---|---|---|---|---|---|---|---|---|
| 95:00:00 | 7cbb-A | 2.6 | 5.6 | 100 | 576 | 6 | MOLECULE: | IUCA/IUCC F |
| 96:00:00 | 7tjl-A | 2.6 | 5.4 | 58 | 85 | 5 | MOLECULE: | DE NOVO DES |
| 97:00:00 | 3zjc-E | 2.6 | 13.8 | 88 | 286 | 6 | MOLECULE: | GTPASE IMAP |
| 98:00:00 | 8c5s-A | 2.6 | 4.4 | 78 | 924 | 4 | MOLECULE: | MITOCHONDRI |
| 99:00:00 | 8t53-A | 2.6 | 15.2 | 106 | 398 | 6 | MOLECULE: | UNDECAPRENY |
| 0:00 | 5x3x-M | 2.6 | 6.9 | 66 | 207 | 6 | MOLECULE: | COBALT ABC |
| 1:00 | 3ut2-A | 2.6 | 18.9 | 108 | 735 | 6 | MOLECULE: | CATALASE-PE |
| 2:00 | 8sgi-A | 2.6 | 4 | 62 | 751 | 5 | MOLECULE: | SODIUM/CALC |
| 3:00 | 8es4-C | 2.6 | 4.9 | 74 | 181 | 5 | MOLECULE: | GP35 |
| 4:00 | 8hwa-A | 2.6 | 27.9 | 105 | 700 | 7 | MOLECULE: | PRIMASE D5 |
| 5:00 | 8wue-A | 2.6 | 12.7 | 70 | 226 | 7 | MOLECULE: | SIGMA NON-O |
| 6:00 | 7eld-A | 2.6 | 25.5 | 99 | 1137 | 8 | MOLECULE: | ENDORIBONUC |
| 7:00 | 8ahx-E | 2.6 | 9.2 | 76 | 213 | 9 | MOLECULE: | ION-TRANSLO |
| 8:00 | 8cws-A | 2.6 | 16.8 | 130 | 451 | 12 | MOLECULE: | F4132-2 CHA |
| 9:00 | 7e2d-H | 2.6 | 6.7 | 62 | 147 | 8 | MOLECULE: | TRAPP-ASSOC |
| 10:00 | 7pv1-D | 2.6 | 18.4 | 61 | 141 | 8 | MOLECULE: | MICOS COMPL |
| 11:00 | 6lk8-J | 2.6 | 19.3 | 83 | 1026 | 10 | MOLECULE: | MGC83295 PR |
| 12:00 | 5ve9-B | 2.6 | 3.7 | 67 | 91 | 7 | MOLECULE: | MICROTUBULE |
| 13:00 | 7ccl-C | 2.6 | 9.5 | 74 | 147 | 8 | MOLECULE: | BCL-2-LIKE |
| 14:00 | 8v46-A | 2.6 | 15.2 | 84 | 412 | 8 | MOLECULE: | ARIA ANTITO |
| 15:00 | 6dlm-B | 2.6 | 6 | 50 | 72 | 6 | MOLECULE: | DHD127_A |
| 16:00 | 7uy7-B | 2.6 | 16.3 | 82 | 291 | 9 | MOLECULE: | TELOMERASE- |
| 17:00 | 7xad-C | 2.6 | 6.9 | 55 | 96 | 5 | MOLECULE: | PROGRAMMED |
| 18:00 | 8tl7-A | 2.6 | 7.1 | 104 | 694 | 7 | MOLECULE: | COMPUTATION |
| 19:00 | 6ofb-A | 2.6 | 12.7 | 87 | 696 | 3 | MOLECULE: | GLUTAMINE-D |
| 20:00 | 6s3l-K | 2.6 | 7.2 | 80 | 193 | 5 | MOLECULE: | FLAGELLAR B |
| 21:00 | 8dmg-B | 2.6 | 10.5 | 144 | 1033 | 4 | MOLECULE: | BIFUNCTIONA |
| 22:00 | 7jts-s | 2.6 | 24.6 | 85 | 290 | 7 | MOLECULE: | RADIAL SPOK |
| 23:00 | 1dce-A | 2.6 | 11.1 | 110 | 567 | 6 | MOLECULE: | PROTEIN (RA |
| 24:00:00 | 2l7x-A | 2.6 | 2.8 | 47 | 77 | 2 | MOLECULE: | ENVELOPE GL |
| 25:00:00 | 5j8v-A | 2.6 | 19.4 | 76 | 3398 | 9 | MOLECULE: | RYANODINE R |
| 26:00:00 | 4qn1-A | 2.6 | 10.2 | 100 | 372 | 10 | MOLECULE: | E3 UBIQUITI |
| 27:00:00 | 7qoo-Q | 2.6 | 13.7 | 71 | 209 | 4 | MOLECULE: | CENTROMERE |
| 28:00:00 | 7wmp-A | 2.6 | 7.4 | 53 | 263 | 6 | MOLECULE: | PORTAL PROT |
| 29:00:00 | 8fm9-M | 2.6 | 4.8 | 94 | 333 | 7 | MOLECULE: | RNA-DIRECTE |
| 30:00:00 | 4hga-A | 2.6 | 20.5 | 104 | 207 | 5 | MOLECULE: | DEATH DOMAI |
| 31:00:00 | 3fey-A | 2.6 | 5.7 | 79 | 761 | 5 | MOLECULE: | NUCLEAR CAP |
| 32:00:00 | 7obq-y | 2.6 | 8.8 | 80 | 454 | 8 | MOLECULE: | SRP RNA |
| 33:00:00 | 8ovw-Q | 2.6 | 24.2 | 74 | 258 | 4 | MOLECULE: | CENTROMERE- |
| 34:00:00 | 8fed-K | 2.6 | 4.4 | 57 | 141 | 5 | MOLECULE: | VIRULENCE F |
| 35:00:00 | 2qf9-A | 2.6 | 6.7 | 75 | 162 | 8 | MOLECULE: | PUTATIVE SE |
| 36:00:00 | 3w5m-A | 2.6 | 17.6 | 103 | 1030 | 6 | MOLECULE: | PUTATIVE RH |
| 37:00:00 | 7ykr-A | 2.6 | 28.5 | 97 | 1035 | 12 | MOLECULE: | TRANSIENT R |
| 38:00:00 | 8d08-D | 2.6 | 6.6 | 58 | 67 | 7 | MOLECULE: | HALC4_135 |

| | | | | | | | | |
|---|---|---|---|---|---|---|---|---|
| 39:00:00 | 8h38-C | 2.6 | 22.7 | 117 | 407 | 5 | MOLECULE: | COP9 SIGNAL |
| 40:00:00 | 7t5p-B | 2.6 | 17.2 | 85 | 419 | 9 | MOLECULE: | SUMO-INTERA |
| 41:00:00 | 6yiz-A | 2.6 | 2.7 | 43 | 211 | 12 | MOLECULE: | TRANSCRIPTI |
| 42:00:00 | 6gyf-B | 2.6 | 11.8 | 89 | 366 | 7 | MOLECULE: | NICOTINAMID |
| 43:00:00 | 5izs-D | 2.6 | 10.2 | 56 | 82 | 4 | MOLECULE: | DESIGNED PR |
| 44:00:00 | 5m4s-A | 2.6 | 3.5 | 56 | 209 | 7 | MOLECULE: | TRANSCRIPTI |
| 45:00:00 | 6vjy-B | 2.6 | 10.6 | 63 | 280 | 5 | MOLECULE: | ERAD-ASSOCI |
| 46:00:00 | 8xj6-D | 2.6 | 19.1 | 95 | 589 | 9 | MOLECULE: | MONKEYPOX V |
| 47:00:00 | 8erc-A | 2.6 | 16.4 | 97 | 441 | 5 | MOLECULE: | LYSOPHOSPHO |
| 48:00:00 | 7zr6-C | 2.6 | 4.7 | 83 | 277 | 7 | MOLECULE: | HEAT SHOCK |
| 49:00:00 | 5t58-B | 2.6 | 11.1 | 55 | 190 | 5 | MOLECULE: | KLLA0F02343 |
| 50:00:00 | 2fiy-A | 2.6 | 8.2 | 106 | 285 | 16 | MOLECULE: | PROTEIN FDH |
| 51:00:00 | 6p8r-B | 2.6 | 6.5 | 64 | 168 | 5 | MOLECULE: | HORMA DOMAI |
| 52:00:00 | 6fkg-C | 2.6 | 4.8 | 50 | 111 | 10 | MOLECULE: | RV1989C (MB |
| 53:00:00 | 2maj-A | 2.6 | 9.7 | 61 | 82 | 11 | MOLECULE: | STROMAL INT |
| 54:00:00 | 8d8j-O | 2.6 | 12.3 | 60 | 210 | 5 | MOLECULE: | PROBABLE S- |
| 55:00:00 | 4mlb-B | 2.6 | 9.5 | 101 | 454 | 5 | MOLECULE: | PF0708 |
| 56:00:00 | 5vvh-D | 2.6 | 2.9 | 40 | 210 | 10 | MOLECULE: | OCTOPINE CA |
| 57:00:00 | 7f5v-A | 2.6 | 21.8 | 113 | 689 | 8 | MOLECULE: | DELTA-1-PYR |
| 58:00:00 | 5anz-A | 2.6 | 10.9 | 99 | 375 | 3 | MOLECULE: | SOLUBLE LYT |
| 59:00:00 | 7vcf-C | 2.6 | 12.2 | 74 | 217 | 8 | MOLECULE: | TIC214 |
| 60:00:00 | 7nur-A | 2.6 | 13.4 | 104 | 1044 | 12 | MOLECULE: | PROTEIN KIN |
| 61:00:00 | 7qd4-A | 2.6 | 14.4 | 108 | 908 | 6 | MOLECULE: | TRANSPOSASE |
| 62:00:00 | 7sp5-A | 2.6 | 9.9 | 122 | 433 | 11 | MOLECULE: | PHOSPHATE T |
| 63:00:00 | 4hzu-S | 2.6 | 5.5 | 91 | 164 | 5 | MOLECULE: | ENERGY-COUP |
| 64:00:00 | 2odm-B | 2.6 | 6.5 | 72 | 83 | 6 | MOLECULE: | UPF0358 PRO |
| 65:00:00 | 4mk6-A | 2.6 | 7.9 | 70 | 188 | 3 | MOLECULE: | PROBABLE DI |
| 66:00:00 | 7d3u-C | 2.6 | 7.6 | 88 | 129 | 9 | MOLECULE: | MONOVALENT |
| 67:00:00 | 6q6b-A | 2.6 | 6.7 | 76 | 132 | 8 | MOLECULE: | CYTOSOLIC C |
| 68:00:00 | 8d2z-A | 2.6 | 13.1 | 77 | 306 | 6 | MOLECULE: | METALLO-BET |
| 69:00:00 | 2khm-A | 2.6 | 11.8 | 89 | 140 | 11 | MOLECULE: | FIBROIN-3 |
| 70:00:00 | 4wbd-A | 2.6 | 21.2 | 63 | 540 | 8 | MOLECULE: | BSHC |
| 71:00:00 | 4avm-A | 2.6 | 14 | 112 | 230 | 7 | MOLECULE: | BRIDGING IN |
| 72:00:00 | 7yh3-D | 2.6 | 4.7 | 72 | 157 | 7 | MOLECULE: | TRAPPC3 FRO |
| 73:00:00 | 3dby-C | 2.6 | 12.6 | 91 | 267 | 7 | MOLECULE: | UNCHARACTER |
| 74:00:00 | 4x2c-A | 2.6 | 8.1 | 65 | 207 | 6 | MOLECULE: | FIC FAMILY |
| 75:00:00 | 3nrk-A | 2.6 | 9.1 | 89 | 312 | 10 | MOLECULE: | LIC12922 |
| 76:00:00 | 1e9s-A | 2.6 | 15.3 | 104 | 427 | 9 | MOLECULE: | CONJUGAL TR |
| 77:00:00 | 4uig-A | 2.6 | 7 | 67 | 91 | 7 | MOLECULE: | COPPER SENS |
| 78:00:00 | 3nrg-A | 2.6 | 8.4 | 76 | 217 | 5 | MOLECULE: | TETR FAMILY |
| 79:00:00 | 7emf-H | 2.6 | 7.9 | 75 | 181 | 9 | MOLECULE: | MEDIATOR OF |
| 80:00:00 | 2xl4-A | 2.6 | 9.4 | 81 | 147 | 6 | MOLECULE: | LMO0438 PRO |
| 81:00:00 | 8j0n-D | 2.6 | 35.8 | 95 | 156 | 4 | MOLECULE: | ER MEMBRANE |
| 82:00:00 | 3u8p-A | 2.6 | 16.2 | 97 | 336 | 10 | MOLECULE: | CYTOCHROME |

| | | | | | | | | |
|---|---|---|---|---|---|---|---|---|
| 83:00:00 | 3bey-D | 2.6 | 3 | 55 | 94 | 11 | MOLECULE: | CONSERVED P |
| 84:00:00 | 8twj-A | 2.6 | 5 | 90 | 303 | 7 | MOLECULE: | AVIRULENCE |
| 85:00:00 | 6nd4-M | 2.6 | 11.8 | 107 | 431 | 9 | MOLECULE: | ETS RRNA |
| 86:00:00 | 2w4m-A | 2.6 | 7.1 | 84 | 250 | 11 | MOLECULE: | N-ACYLNEURA |
| 87:00:00 | 5a5t-L | 2.6 | 23.9 | 87 | 372 | 6 | MOLECULE: | EUKARYOTIC |
| 88:00:00 | 7zol-B | 2.6 | 20.8 | 119 | 1702 | 7 | MOLECULE: | CAS7-11 |
| 89:00:00 | 8urw-Z | 2.6 | 8.2 | 122 | 1235 | 10 | MOLECULE: | DNA-DIRECTE |
| 90:00:00 | 5yfp-C | 2.6 | 15.7 | 101 | 790 | 7 | MOLECULE: | EXOCYST COM |
| 91:00:00 | 8vej-B | 2.6 | 3.1 | 53 | 242 | 4 | MOLECULE: | CHD_BUTTRES |
| 92:00:00 | 6hux-A | 2.6 | 14.2 | 85 | 369 | 7 | MOLECULE: | H(2)-FORMIN |
| 93:00:00 | 8ca1-A | 2.6 | 20.9 | 127 | 589 | 6 | MOLECULE: | VERY LONG-C |
| 94:00:00 | 3hug-A | 2.6 | 11.4 | 58 | 80 | 10 | MOLECULE: | RNA POLYMER |
| 95:00:00 | 1tjf-B | 2.6 | 8.9 | 86 | 186 | 3 | MOLECULE: | ADENYLYL CY |
| 96:00:00 | 6w2j-B | 2.6 | 21.3 | 96 | 1364 | 5 | MOLECULE: | CARBAMOYL-P |
| 97:00:00 | 3esw-A | 2.6 | 6.6 | 98 | 333 | 6 | MOLECULE: | PEPTIDE-N(4 |
| 98:00:00 | 5v5t-A | 2.6 | 7.5 | 107 | 306 | 7 | MOLECULE: | CONSERVED D |
| 99:00:00 | 7yca-O | 2.6 | 5 | 65 | 96 | 3 | MOLECULE: | LHCA1 |
| 0:00 | 6c66-I | 2.6 | 8.9 | 67 | 177 | 10 | MOLECULE: | CRISPR-ASSO |
| 1:00 | 6nsj-A | 2.6 | 9.4 | 86 | 182 | 6 | MOLECULE: | ACID-ACTIVA |
| 2:00 | 7v99-A | 2.6 | 23.8 | 152 | 991 | 9 | MOLECULE: | TELOMERASE |
| 3:00 | 6t58-A | 2.6 | 6.6 | 84 | 524 | 5 | MOLECULE: | CELLULAR TU |
| 4:00 | 8hmd-C | 2.6 | 10.2 | 81 | 1334 | 11 | MOLECULE: | INTRAFLAGEL |
| 5:00 | 6es9-A | 2.6 | 8.9 | 92 | 545 | 4 | MOLECULE: | ACYL-COA DE |
| 6:00 | 7jtk-A | 2.6 | 15.2 | 75 | 749 | 9 | MOLECULE: | FLAGELLAR R |
| 7:00 | 8vc8-A | 2.6 | 4 | 68 | 197 | 7 | MOLECULE: | HEM_3.C9 |
| 8:00 | 6qm5-A | 2.6 | 9.3 | 79 | 673 | 4 | MOLECULE: | PREDICTED P |
| 9:00 | 5eqw-B | 2.6 | 12.1 | 73 | 123 | 4 | MOLECULE: | PUTATIVE MA |
| 10:00 | 8foc-B | 2.6 | 11.2 | 81 | 450 | 7 | MOLECULE: | DNA PRIMASE |
| 11:00 | 8cqr-A | 2.6 | 6.4 | 71 | 103 | 3 | MOLECULE: | NINJURIN-1 |
| 12:00 | 8uq9-a | 2.6 | 14.8 | 83 | 433 | 7 | MOLECULE: | E3 UBIQUITI |
| 13:00 | 6cfw-C | 2.6 | 9.1 | 89 | 114 | 6 | MOLECULE: | MONOVALENT |
| 14:00 | 3c3w-A | 2.6 | 14.4 | 70 | 211 | 7 | MOLECULE: | TWO COMPONE |
| 15:00 | 8egi-B | 2.6 | 4.5 | 76 | 146 | 11 | MOLECULE: | HEMOGLOBIN |
| 16:00 | 5ker-F | 2.6 | 4.8 | 79 | 146 | 9 | MOLECULE: | ALPHA-GLOBI |
| 17:00 | 4ui9-N | 2.6 | 29.2 | 116 | 631 | 8 | MOLECULE: | ANAPHASE-PR |
| 18:00 | 3w4u-F | 2.6 | 4.4 | 78 | 145 | 10 | MOLECULE: | HEMOGLOBIN |
| 19:00 | 5x2t-D | 2.6 | 4.7 | 81 | 145 | 11 | MOLECULE: | HEMOGLOBIN |
| 20:00 | 8hki-a | 2.5 | 4.5 | 82 | 291 | 2 | MOLECULE: | T-COMPLEX P |
| 21:00 | 8t9f-K | 2.5 | 17.1 | 82 | 265 | 7 | MOLECULE: | HISTONE-LYS |
| 22:00 | 4uis-B | 2.5 | 20 | 90 | 241 | 1 | MOLECULE: | GAMMA-SECRE |
| 23:00 | 6psk-R | 2.5 | 3.3 | 46 | 72 | 2 | MOLECULE: | ANTIHOLIN |
| 24:00:00 | 8tkp-A | 2.5 | 26.7 | 125 | 614 | 8 | MOLECULE: | TRANSMEMBRA |
| 25:00:00 | 4hlq-C | 2.5 | 13.4 | 70 | 281 | 7 | MOLECULE: | TBC1 DOMAIN |
| 26:00:00 | 7qog-M | 2.5 | 4.1 | 50 | 126 | 6 | MOLECULE: | PORTAL PROT |

| | | | | | | | | |
|---|---|---|---|---|---|---|---|---|
| 27:00:00 | 2kob-A | 2.5 | 10 | 50 | 108 | 16 | MOLECULE: | UNCHARACTER |
| 28:00:00 | 2z0q-A | 2.5 | 12.8 | 104 | 321 | 10 | MOLECULE: | RHO GUANINE |
| 29:00:00 | 6qzh-A | 2.5 | 5 | 84 | 744 | 7 | MOLECULE: | HUMAN CHEMO |
| 30:00:00 | 5t58-N | 2.5 | 13.7 | 74 | 196 | 7 | MOLECULE: | KLLA0F02343 |
| 31:00:00 | 8ip4-A | 2.5 | 28.5 | 103 | 320 | 10 | MOLECULE: | MAGNESIUM T |
| 32:00:00 | 1wa8-A | 2.5 | 9.8 | 59 | 99 | 8 | MOLECULE: | ESAT-6 LIKE |
| 33:00:00 | 4q9u-A | 2.5 | 22.6 | 121 | 303 | 8 | MOLECULE: | RAB5 GDP/GT |
| 34:00:00 | 1vzs-A | 2.5 | 6.1 | 57 | 76 | 5 | MOLECULE: | ATP SYNTHAS |
| 35:00:00 | 7xpc-A | 2.5 | 13.3 | 66 | 337 | 5 | MOLECULE: | D-GLYCERATE |
| 36:00:00 | 4pfr-B | 2.5 | 4.1 | 63 | 229 | 5 | MOLECULE: | TRAP DICARB |
| 37:00:00 | 1y71-B | 2.5 | 8.1 | 53 | 112 | 8 | MOLECULE: | KINASE-ASSO |
| 38:00:00 | 2f9i-A | 2.5 | 11.8 | 64 | 309 | 3 | MOLECULE: | ACETYL-COEN |
| 39:00:00 | 7cun-G | 2.5 | 22 | 100 | 895 | 7 | MOLECULE: | INTEGRATOR |
| 40:00:00 | 8g3a-D | 2.5 | 17.4 | 66 | 334 | 3 | MOLECULE: | BACITRACIN |
| 41:00:00 | 2o57-A | 2.5 | 11.4 | 76 | 282 | 3 | MOLECULE: | PUTATIVE SA |
| 42:00:00 | 6ahp-A | 2.5 | 3.9 | 70 | 110 | 6 | MOLECULE: | FLAGELLAR P |
| 43:00:00 | 3bo0-A | 2.5 | 6.9 | 103 | 442 | 5 | MOLECULE: | 23S RIBOSOM |
| 44:00:00 | 4q2g-B | 2.5 | 6 | 96 | 269 | 11 | MOLECULE: | PHOSPHATIDA |
| 45:00:00 | 8iqi-C | 2.5 | 11.8 | 109 | 916 | 3 | MOLECULE: | PUTATIVE PR |
| 46:00:00 | 6ozu-A | 2.5 | 11.2 | 65 | 235 | 5 | MOLECULE: | EUKARYOTIC |
| 47:00:00 | 6hty-B | 2.5 | 4.7 | 71 | 302 | 11 | MOLECULE: | NUCLEAR REC |
| 48:00:00 | 2qqy-A | 2.5 | 6.6 | 73 | 139 | 3 | MOLECULE: | SIGMA B OPE |
| 49:00:00 | 6eqo-A | 2.5 | 16.4 | 127 | 1804 | 4 | MOLECULE: | ACETYL-COEN |
| 50:00:00 | 5che-A | 2.5 | 4.2 | 60 | 427 | 3 | MOLECULE: | GLUTAMYL-TR |
| 51:00:00 | 8pzp-A | 2.5 | 13.2 | 61 | 470 | 18 | MOLECULE: | NUCLEOPROTE |
| 52:00:00 | 7qh5-A | 2.5 | 8.3 | 81 | 313 | 11 | MOLECULE: | RNA POLYMER |
| 53:00:00 | 6ye4-A | 2.5 | 17.1 | 51 | 236 | 10 | MOLECULE: | BIOPOLYMER |
| 54:00:00 | 5way-A | 2.5 | 22.2 | 81 | 482 | 4 | MOLECULE: | TRANSCRIPTI |
| 55:00:00 | 7ywq-A | 2.5 | 2.8 | 51 | 79 | 14 | MOLECULE: | GENOME POLY |
| 56:00:00 | 7qcd-D | 2.5 | 12.9 | 94 | 326 | 9 | MOLECULE: | STRUCTURAL |
| 57:00:00 | 5x41-F | 2.5 | 10.2 | 82 | 244 | 6 | MOLECULE: | COBALT ABC |
| 58:00:00 | 4rp5-A | 2.5 | 4.7 | 57 | 94 | 11 | MOLECULE: | DISKS LARGE |
| 59:00:00 | 4knf-C | 2.5 | 4.9 | 81 | 232 | 5 | MOLECULE: | BLUE-LIGHT |
| 60:00:00 | 6xqk-A | 2.5 | 4.1 | 46 | 47 | 4 | MOLECULE: | CAMP-DEPEND |
| 61:00:00 | 8syn-A | 2.5 | 12.4 | 71 | 801 | 8 | MOLECULE: | VPS35 ENDOS |
| 62:00:00 | 7jv7-B | 2.5 | 6.4 | 80 | 310 | 10 | MOLECULE: | CTD KINASE |
| 63:00:00 | 8ity-W | 2.5 | 9.6 | 64 | 111 | 6 | MOLECULE: | SNRNA-ACTIV |
| 64:00:00 | 8cih-A | 2.5 | 6.4 | 63 | 188 | 6 | MOLECULE: | CYCLIN-DEPE |
| 65:00:00 | 6vzd-E | 2.5 | 8.1 | 61 | 85 | 5 | MOLECULE: | PULMONARY S |
| 66:00:00 | 8hfs-Y | 2.5 | 8.3 | 75 | 249 | 3 | MOLECULE: | MANNOSE-SPE |
| 67:00:00 | 8ts6-A | 2.5 | 17.4 | 66 | 500 | 5 | MOLECULE: | PORTAL PROT |
| 68:00:00 | 7xgr-A | 2.5 | 26.7 | 96 | 546 | 1 | MOLECULE: | GEM-ASSOCIA |
| 69:00:00 | 5xmw-A | 2.5 | 15.8 | 87 | 270 | 16 | MOLECULE: | ZEARALENONE |
| 70:00:00 | 5ip0-D | 2.5 | 10.3 | 65 | 108 | 11 | MOLECULE: | PHA GRANULE |

| | | | | | | | | |
|---|---|---|---|---|---|---|---|---|
| 71:00:00 | 6zu9-Q | 2.5 | 14.2 | 74 | 226 | 8 | MOLECULE: | 18S RIBOSOM |
| 72:00:00 | 7e5c-A | 2.5 | 16.1 | 78 | 439 | 8 | MOLECULE: | XAA-PRO DIP |
| 73:00:00 | 4zs9-B | 2.5 | 11 | 57 | 378 | 9 | MOLECULE: | SUGAR BINDI |
| 74:00:00 | 3r2x-C | 2.5 | 2.8 | 55 | 82 | 9 | MOLECULE: | HEMAGGLUTIN |
| 75:00:00 | 5yde-A | 2.5 | 5.4 | 56 | 111 | 9 | MOLECULE: | PARAFIBROMI |
| 76:00:00 | 8by2-A | 2.5 | 14.1 | 74 | 548 | 8 | MOLECULE: | GLUTATHIONE |
| 77:00:00 | 1q8c-A | 2.5 | 4 | 69 | 132 | 4 | MOLECULE: | HYPOTHETICA |
| 78:00:00 | 7y22-M | 2.5 | 8.3 | 81 | 215 | 4 | MOLECULE: | PHAGE CONNE |
| 79:00:00 | 6nd4-J | 2.5 | 23.8 | 83 | 493 | 11 | MOLECULE: | ETS RRNA |
| 80:00:00 | 1wpk-A | 2.5 | 11.7 | 74 | 146 | 7 | MOLECULE: | ADA REGULAT |
| 81:00:00 | 6giq-Q | 2.5 | 2.9 | 50 | 75 | 4 | MOLECULE: | CYTOCHROME |
| 82:00:00 | 7p2y-a | 2.5 | 10.7 | 82 | 277 | 6 | MOLECULE: | ATP SYNTHAS |
| 83:00:00 | 6mur-B | 2.5 | 9.6 | 56 | 162 | 9 | MOLECULE: | UNCHARACTER |
| 84:00:00 | 1ypy-A | 2.5 | 7.9 | 78 | 182 | 9 | MOLECULE: | VIRION MEMB |
| 85:00:00 | 4xig-M | 2.5 | 5.4 | 95 | 288 | 6 | MOLECULE: | ALGM1 |
| 86:00:00 | 7s0r-A | 2.5 | 2.8 | 45 | 85 | 7 | MOLECULE: | C PROTEIN B |
| 87:00:00 | 4kqt-A | 2.5 | 6 | 59 | 169 | 7 | MOLECULE: | PUTATIVE OU |
| 88:00:00 | 3cjh-I | 2.5 | 3.4 | 42 | 54 | 7 | MOLECULE: | MITOCHONDRI |
| 89:00:00 | 3ilk-A | 2.5 | 12 | 78 | 239 | 3 | MOLECULE: | UNCHARACTER |
| 90:00:00 | 3kfw-X | 2.5 | 8.7 | 74 | 234 | 3 | MOLECULE: | UNCHARACTER |
| 91:00:00 | 2owy-A | 2.5 | 20 | 82 | 306 | 6 | MOLECULE: | RECOMBINATI |
| 92:00:00 | 8hp7-A | 2.5 | 4.1 | 91 | 241 | 5 | MOLECULE: | (S)-2-HALOA |
| 93:00:00 | 8tsh-F | 2.5 | 4.1 | 46 | 265 | 4 | MOLECULE: | ABC TRANSPO |
| 94:00:00 | 8ppr-D | 2.5 | 15.5 | 80 | 248 | 9 | MOLECULE: | KINETOCHORE |
| 95:00:00 | 5kzb-A | 2.5 | 8.8 | 58 | 104 | 3 | MOLECULE: | VIRUS MATRI |
| 96:00:00 | 7ogp-D | 2.5 | 17.9 | 77 | 510 | 10 | MOLECULE: | PHIKZ055,PH |
| 97:00:00 | 2ehw-C | 2.5 | 17.5 | 61 | 117 | 16 | MOLECULE: | HYPOTHETICA |
| 98:00:00 | 8ijq-A | 2.5 | 7.6 | 105 | 264 | 4 | MOLECULE: | SPHINGOMYEL |
| 99:00:00 | 7jz2-A | 2.5 | 14.6 | 91 | 481 | 11 | MOLECULE: | SUCCINATE D |
| 0:00 | 5ojl-A | 2.5 | 6.2 | 78 | 284 | 9 | MOLECULE: | IMINE REDUC |
| 1:00 | 6wjv-A | 2.5 | 22.4 | 113 | 1951 | 4 | MOLECULE: | DNA POLYMER |
| 2:00 | 6swy-1 | 2.5 | 14.6 | 80 | 542 | 0 | MOLECULE: | VACUOLAR IM |
| 3:00 | 7udk-A | 2.5 | 12 | 62 | 172 | 8 | MOLECULE: | DESIGNED HE |
| 4:00 | 1zl8-B | 2.5 | 3.1 | 41 | 54 | 5 | MOLECULE: | LIN-7 |
| 5:00 | 7uv1-A | 2.5 | 3.1 | 49 | 56 | 8 | MOLECULE: | VICILIN-LIK |
| 6:00 | 7jrg-H | 2.5 | 2.9 | 50 | 65 | 2 | MOLECULE: | MITOCHONDRI |
| 7:00 | 4bzj-C | 2.5 | 12.6 | 91 | 693 | 12 | MOLECULE: | PROTEIN TRA |
| 8:00 | 6tqe-A | 2.5 | 17.9 | 103 | 634 | 9 | MOLECULE: | ABC TRANSPO |
| 9:00 | 5iwz-A | 2.5 | 6.9 | 78 | 372 | 8 | MOLECULE: | SYNAPTONEMA |
| 10:00 | 8pm4-A | 2.5 | 7.1 | 90 | 604 | 6 | MOLECULE: | TRANSPOSASE |
| 11:00 | 6zyx-K | 2.5 | 3.2 | 48 | 95 | 13 | MOLECULE: | DYNEIN HEAV |
| 12:00 | 5z62-B | 2.5 | 18.9 | 79 | 227 | 5 | MOLECULE: | CYTOCHROME |
| 13:00 | 7dl2-A | 2.5 | 24.3 | 98 | 1127 | 8 | MOLECULE: | HAMARTIN |
| 14:00 | 7nnh-X | 2.5 | 18.8 | 88 | 637 | 3 | MOLECULE: | ERYTHROCYTE |

| | | | | | | | | |
|---|---|---|---|---|---|---|---|---|
| 15:00 | 7bin-L | 2.5 | 10.6 | 57 | 75 | 11 | MOLECULE: | FLAGELLAR B |
| 16:00 | 6d5f-a | 2.5 | 9.8 | 81 | 131 | 11 | MOLECULE: | FIMBRIAL PR |
| 17:00 | 3k66-A | 2.5 | 19.2 | 85 | 217 | 5 | MOLECULE: | BETA-AMYLOI |
| 18:00 | 5b86-A | 2.5 | 30.9 | 102 | 579 | 6 | MOLECULE: | TUMOR NECRO |
| 19:00 | 1yo7-A | 2.5 | 5.2 | 46 | 120 | 0 | MOLECULE: | REGULATORY |
| 20:00 | 6jl7-A | 2.5 | 21.6 | 90 | 444 | 4 | MOLECULE: | TBC1 DOMAIN |
| 21:00 | 5jki-A | 2.5 | 3.8 | 82 | 201 | 7 | MOLECULE: | PUTATIVE LI |
| 22:00 | 1zxj-D | 2.5 | 8.2 | 56 | 193 | 9 | MOLECULE: | HYPOTHETICA |
| 23:00 | 5kbw-B | 2.5 | 8.7 | 59 | 171 | 7 | MOLECULE: | RIBOFLAVIN |
| 24:00:00 | 5h69-A | 2.5 | 6.3 | 49 | 252 | 8 | MOLECULE: | CHROMOSOME |
| 25:00:00 | 7xdi-A | 2.5 | 4.7 | 51 | 69 | 16 | MOLECULE: | VP1 |
| 26:00:00 | 6k9k-A | 2.5 | 11.9 | 83 | 2501 | 5 | MOLECULE: | SERINE-PROT |
| 27:00:00 | 2l3y-A | 2.5 | 4.6 | 79 | 184 | 5 | MOLECULE: | INTERLEUKIN |
| 28:00:00 | 1l9l-A | 2.5 | 6 | 41 | 74 | 2 | MOLECULE: | GRANULYSIN |
| 29:00:00 | 2mxx-A | 2.5 | 16.6 | 104 | 179 | 10 | MOLECULE: | AMYLASE-BIN |
| 30:00:00 | 6gdj-A | 2.5 | 5.5 | 47 | 71 | 11 | MOLECULE: | MTO2 |
| 31:00:00 | 4x5m-B | 2.5 | 10.2 | 58 | 93 | 10 | MOLECULE: | UNCHARACTER |
| 32:00:00 | 2ieq-B | 2.5 | 11.4 | 60 | 89 | 13 | MOLECULE: | SPIKE GLYCO |
| 33:00:00 | 5b0l-B | 2.5 | 8.2 | 102 | 334 | 8 | MOLECULE: | MOEN5,DNA-B |
| 34:00:00 | 7pkn-R | 2.5 | 5.7 | 60 | 80 | 7 | MOLECULE: | CENTROMERE |
| 35:00:00 | 5ir2-A | 2.5 | 4.6 | 58 | 222 | 5 | MOLECULE: | CELLULASE |
| 36:00:00 | 3cp8-A | 2.5 | 6.6 | 94 | 611 | 4 | MOLECULE: | TRNA URIDIN |
| 37:00:00 | 1sf9-A | 2.5 | 10.3 | 77 | 118 | 1 | MOLECULE: | YFHH HYPOTH |
| 38:00:00 | 7w0y-i | 2.5 | 8.6 | 95 | 347 | 6 | MOLECULE: | NADH DEHYDR |
| 39:00:00 | 8k9z-A | 2.5 | 17.8 | 122 | 405 | 5 | MOLECULE: | RDTND-RID C |
| 40:00:00 | 8psn-B | 2.5 | 10.9 | 117 | 515 | 9 | MOLECULE: | POLYMERASE |
| 41:00:00 | 3rip-A | 2.5 | 23.9 | 104 | 568 | 9 | MOLECULE: | GAMMA-TUBUL |
| 42:00:00 | 8y7g-B | 2.5 | 9.5 | 74 | 543 | 1 | MOLECULE: | CRISPR-ASSO |
| 43:00:00 | 7x1i-A | 2.5 | 10.8 | 94 | 678 | 5 | MOLECULE: | ISOFORM 1 O |
| 44:00:00 | 8hyj-D | 2.5 | 11.1 | 58 | 117 | 2 | MOLECULE: | DNA-DIRECTE |
| 45:00:00 | 3njc-A | 2.5 | 4.2 | 81 | 153 | 5 | MOLECULE: | YSLB PROTEI |
| 46:00:00 | 5eom-C | 2.5 | 6.2 | 83 | 353 | 7 | MOLECULE: | PROTEIN MAB |
| 47:00:00 | 4c0n-A | 2.5 | 4.9 | 78 | 150 | 6 | MOLECULE: | 2-ON-2 HEMO |
| 48:00:00 | 7f3t-A | 2.5 | 11 | 103 | 330 | 5 | MOLECULE: | TRANSMEMBRA |
| 49:00:00 | 7xsq-A | 2.5 | 11.2 | 108 | 1289 | 6 | MOLECULE: | RAMP SUPERF |
| 50:00:00 | 3nlc-A | 2.5 | 7.8 | 95 | 534 | 7 | MOLECULE: | UNCHARACTER |
| 51:00:00 | 5dy0-A | 2.5 | 3.2 | 57 | 220 | 4 | MOLECULE: | TETR FAMILY |
| 52:00:00 | 6zvh-y | 2.5 | 2.7 | 57 | 72 | 12 | MOLECULE: | 18S RRNA |
| 53:00:00 | 1fyz-F | 2.5 | 15.9 | 97 | 168 | 9 | MOLECULE: | METHANE MON |
| 54:00:00 | 7o3d-A | 2.5 | 16.1 | 120 | 709 | 10 | MOLECULE: | TRANSCRIPTI |
| 55:00:00 | 8xat-A | 2.5 | 4.7 | 70 | 187 | 4 | MOLECULE: | TWO-COMPONE |
| 56:00:00 | 8otx-A | 2.5 | 18.6 | 131 | 1054 | 10 | MOLECULE: | SPERM-SPECI |
| 57:00:00 | 8pmq-9 | 2.5 | 27.5 | 103 | 412 | 4 | MOLECULE: | E3 UBIQUITI |
| 58:00:00 | 7wwb-A | 2.5 | 6.4 | 86 | 333 | 9 | MOLECULE: | CHOLINE TRA |

| | | | | | | | | |
|---|---|---|---|---|---|---|---|---|
| 59:00:00 | 1qgi-A | 2.5 | 6.7 | 77 | 259 | 4 | MOLECULE: | PROTEIN (CH |
| 60:00:00 | 6h3w-A | 2.5 | 5.3 | 87 | 246 | 2 | MOLECULE: | ENVELOPMENT |
| 61:00:00 | 7e38-B | 2.5 | 9.8 | 92 | 305 | 1 | MOLECULE: | DEOXYPODOPH |
| 62:00:00 | 7k5c-A | 2.5 | 19.4 | 111 | 650 | 6 | MOLECULE: | INTERNAL VI |
| 63:00:00 | 3wa8-A | 2.5 | 6.4 | 63 | 179 | 6 | MOLECULE: | CRISPR-ASSO |
| 64:00:00 | 8e4g-0 | 2.5 | 7.5 | 71 | 196 | 7 | MOLECULE: | PORTAL PROT |
| 65:00:00 | 6zmq-A | 2.5 | 9.4 | 97 | 589 | 5 | MOLECULE: | CYTOCHROME |
| 66:00:00 | 8aw5-A | 2.5 | 10.4 | 96 | 266 | 9 | MOLECULE: | HEME O OXYG |
| 67:00:00 | 6ppl-B | 2.5 | 20.7 | 102 | 665 | 9 | MOLECULE: | N-ALPHA-ACE |
| 68:00:00 | 4ap2-B | 2.5 | 11.2 | 84 | 359 | 10 | MOLECULE: | KELCH-LIKE |
| 69:00:00 | 7awt-D | 2.5 | 7.5 | 105 | 589 | 9 | MOLECULE: | NADH-QUINON |
| 70:00:00 | 4jbw-F | 2.5 | 22.6 | 110 | 492 | 10 | MOLECULE: | MALTOSE TRA |
| 71:00:00 | 8xje-B | 2.5 | 4.2 | 59 | 149 | 5 | MOLECULE: | YQEY |
| 72:00:00 | 6wgv-A | 2.5 | 11.6 | 91 | 173 | 4 | MOLECULE: | CORRINOID A |
| 73:00:00 | 6qfd-B | 2.5 | 2.9 | 46 | 116 | 11 | MOLECULE: | DNA-BINDING |
| 74:00:00 | 6fkf-d | 2.5 | 8.1 | 86 | 179 | 5 | MOLECULE: | ATP SYNTHAS |
| 75:00:00 | 7z0s-G | 2.5 | 9.3 | 72 | 250 | 13 | MOLECULE: | FORMATE HYD |
| 76:00:00 | 7bss-A | 2.5 | 27.4 | 115 | 816 | 7 | MOLECULE: | ATP11C |
| 77:00:00 | 5xf9-D | 2.5 | 8 | 99 | 455 | 9 | MOLECULE: | NAD-REDUCIN |
| 78:00:00 | 3jac-A | 2.5 | 7 | 98 | 918 | 3 | MOLECULE: | PIEZO-TYPE |
| 79:00:00 | 8b17-A | 2.5 | 9.7 | 65 | 451 | 9 | MOLECULE: | DIPEPTIDE A |
| 80:00:00 | 7pds-A | 2.5 | 22.7 | 100 | 457 | 5 | MOLECULE: | SIMILAR TO |
| 81:00:00 | 8f2a-E | 2.5 | 9 | 76 | 117 | 4 | MOLECULE: | RECEPTOR AC |
| 82:00:00 | 7oci-C | 2.5 | 7.5 | 70 | 119 | 10 | MOLECULE: | DOLICHYL-DI |
| 83:00:00 | 3syk-A | 2.5 | 6 | 73 | 290 | 3 | MOLECULE: | PROTEIN CBB |
| 84:00:00 | 4n4w-A | 2.5 | 24.3 | 109 | 457 | 6 | MOLECULE: | CYTOCHROME |
| 85:00:00 | 3lss-B | 2.5 | 19.3 | 85 | 468 | 9 | MOLECULE: | SERYL-TRNA |
| 86:00:00 | 6w2q-A | 2.5 | 4.2 | 69 | 195 | 4 | MOLECULE: | JUNCTION 34 |
| 87:00:00 | 7arc-E | 2.5 | 4.9 | 87 | 235 | 9 | MOLECULE: | PSST |
| 88:00:00 | 4i1s-A | 2.5 | 7.5 | 71 | 238 | 8 | MOLECULE: | MELANOMA DI |
| 89:00:00 | 3s6j-A | 2.5 | 12.7 | 60 | 216 | 15 | MOLECULE: | HYDROLASE, |
| 90:00:00 | 6w38-A | 2.5 | 6.2 | 90 | 331 | 4 | MOLECULE: | TERMINAL NU |
| 91:00:00 | 2be3-A | 2.5 | 10.8 | 68 | 203 | 4 | MOLECULE: | GTP PYROPHO |
| 92:00:00 | 8gkh-P | 2.5 | 11.3 | 96 | 600 | 7 | MOLECULE: | DNA (5'-D(P |
| 93:00:00 | 3jyy-B | 2.5 | 7.5 | 90 | 266 | 4 | MOLECULE: | LINCOSAMIDE |
| 94:00:00 | 8odu-C | 2.5 | 21.3 | 114 | 232 | 7 | MOLECULE: | ATPASE GET3 |
| 95:00:00 | 4iwb-A | 2.5 | 7.8 | 81 | 165 | 4 | MOLECULE: | FLIC, FLIS |
| 96:00:00 | 8c81-B | 2.5 | 24.5 | 93 | 539 | 12 | MOLECULE: | PROTEIN ORM |
| 97:00:00 | 7qe5-A | 2.5 | 10.2 | 83 | 616 | 5 | MOLECULE: | SIALIC ACID |
| 98:00:00 | 2gtt-K | 2.5 | 15.2 | 92 | 420 | 7 | MOLECULE: | NUCLEOPROTE |
| 99:00:00 | 7jsr-A | 2.5 | 7.6 | 78 | 1496 | 5 | MOLECULE: | NAD-SPECIFI |
| 0:00 | 1jr5-A | 2.5 | 4.3 | 61 | 90 | 7 | MOLECULE: | 10 KDA ANTI |
| 1:00 | 2qeu-B | 2.5 | 4.9 | 58 | 133 | 7 | MOLECULE: | PUTATIVE CA |
| 2:00 | 7m5t-A | 2.5 | 7.4 | 66 | 100 | 3 | MOLECULE: | DE NOVO DES |

| | | | | | | | | |
|---|---|---|---|---|---|---|---|---|
| 3:00 | 1zp2-A | 2.5 | 3.9 | 74 | 227 | 5 | MOLECULE: | RNA POLYMER |
| 4:00 | 6ulo-A | 2.5 | 15.6 | 94 | 317 | 5 | MOLECULE: | UNCHARACTER |
| 5:00 | 4rng-C | 2.5 | 3.9 | 64 | 83 | 5 | MOLECULE: | MTN3/SALIVA |
| 6:00 | 2d7d-A | 2.5 | 17.5 | 87 | 621 | 3 | MOLECULE: | 5'-D(P*TP*T |
| 7:00 | 5kqj-A | 2.5 | 7.8 | 90 | 185 | 13 | MOLECULE: | 2''-AMINOGL |
| 8:00 | 7zkq-C | 2.5 | 22.9 | 97 | 416 | 9 | MOLECULE: | NADH DEHYDR |
| 9:00 | 3hrw-D | 2.5 | 4.6 | 79 | 146 | 9 | MOLECULE: | HEMOGLOBIN |
| 10:00 | 2nup-C | 2.4 | 4.2 | 52 | 137 | 10 | MOLECULE: | PROTEIN TRA |
| 11:00 | 7uv4-A | 2.4 | 3.5 | 44 | 60 | 7 | MOLECULE: | VICILIN PIS |
| 12:00 | 6n63-A | 2.4 | 9.9 | 77 | 141 | 9 | MOLECULE: | ENCAPSULIN |
| 13:00 | 8ebt-C | 2.4 | 19.4 | 107 | 191 | 6 | MOLECULE: | GENERAL TRA |
| 14:00 | 7kzn-F | 2.4 | 3.3 | 44 | 100 | 7 | MOLECULE: | HEAVY CHAIN |
| 15:00 | 7jqe-A | 2.4 | 18.9 | 88 | 357 | 9 | MOLECULE: | ESAT-6/WXG1 |
| 16:00 | 3c2g-A | 2.4 | 26.4 | 102 | 618 | 4 | MOLECULE: | SYS-1 PROTE |
| 17:00 | 5ztc-A | 2.4 | 16 | 63 | 197 | 2 | MOLECULE: | LMO2088 PRO |
| 18:00 | 8hel-A | 2.4 | 2.8 | 42 | 68 | 5 | MOLECULE: | ACRIE3 |
| 19:00 | 3sjb-D | 2.4 | 10 | 65 | 72 | 8 | MOLECULE: | ATPASE GET3 |
| 20:00 | 4i0x-F | 2.4 | 9.9 | 59 | 85 | 12 | MOLECULE: | ESAT-6-LIKE |
| 21:00 | 1uj8-A | 2.4 | 2.8 | 53 | 73 | 4 | MOLECULE: | HYPOTHETICA |
| 22:00 | 8env-A | 2.4 | 6.5 | 92 | 504 | 7 | MOLECULE: | SHEATH PROT |
| 23:00 | 1dek-A | 2.4 | 18.6 | 77 | 241 | 10 | MOLECULE: | DEOXYNUCLEO |
| 24:00:00 | 1bp3-A | 2.4 | 4.1 | 80 | 186 | 4 | MOLECULE: | PROTEIN (GR |
| 25:00:00 | 6b2z-M | 2.4 | 3.9 | 58 | 249 | 3 | MOLECULE: | ATP SYNTHAS |
| 26:00:00 | 8rc4-o | 2.4 | 17.7 | 73 | 372 | 5 | MOLECULE: | SERINE/THRE |
| 27:00:00 | 7r5s-T | 2.4 | 13.5 | 58 | 112 | 5 | MOLECULE: | CENTROMERE |
| 28:00:00 | 8eki-B | 2.4 | 6.7 | 48 | 85 | 8 | MOLECULE: | PROTEIN TRA |
| 29:00:00 | 8e4y-A | 2.4 | 9.4 | 89 | 651 | 8 | MOLECULE: | GLYCEROL-3- |
| 30:00:00 | 2fm8-C | 2.4 | 15.7 | 88 | 220 | 8 | MOLECULE: | SURFACE PRE |
| 31:00:00 | 6lpf-B | 2.4 | 22.7 | 82 | 1007 | 10 | MOLECULE: | LEUCINE--TR |
| 32:00:00 | 7dkh-I | 2.4 | 10.2 | 64 | 914 | 6 | MOLECULE: | RNA POLYMER |
| 33:00:00 | 6lo8-F | 2.4 | 3.7 | 55 | 84 | 9 | MOLECULE: | MITOCHONDRI |
| 34:00:00 | 2ew2-B | 2.4 | 6.2 | 82 | 314 | 7 | MOLECULE: | 2-DEHYDROPA |
| 35:00:00 | 5tgq-A | 2.4 | 9.1 | 102 | 226 | 6 | MOLECULE: | R.SWAI PROT |
| 36:00:00 | 5a9j-A | 2.4 | 15.6 | 96 | 805 | 7 | MOLECULE: | DNA POLYMER |
| 37:00:00 | 3t4r-A | 2.4 | 3.3 | 49 | 72 | 10 | MOLECULE: | PHOSPHOPROT |
| 38:00:00 | 3f8f-A | 2.4 | 10.5 | 57 | 114 | 11 | MOLECULE: | TRANSCRIPTI |
| 39:00:00 | 7evp-C | 2.4 | 4.6 | 36 | 43 | 17 | MOLECULE: | BETA SLIDIN |
| 40:00:00 | 5i1u-B | 2.4 | 10.6 | 80 | 294 | 8 | MOLECULE: | GERMACRADIE |
| 41:00:00 | 4fl2-A | 2.4 | 5.9 | 70 | 555 | 6 | MOLECULE: | TYROSINE-PR |
| 42:00:00 | 7l48-A | 2.4 | 11.5 | 93 | 522 | 9 | MOLECULE: | CAS12F |
| 43:00:00 | 6ulg-N | 2.4 | 13 | 66 | 483 | 3 | MOLECULE: | FOLLICULIN |
| 44:00:00 | 3pmd-A | 2.4 | 7.6 | 92 | 153 | 5 | MOLECULE: | CONSERVED D |
| 45:00:00 | 7wff-F | 2.4 | 7.4 | 105 | 677 | 9 | MOLECULE: | NAD(P)H-QUI |
| 46:00:00 | 7p2p-C | 2.4 | 20.6 | 84 | 174 | 6 | MOLECULE: | SIGNAL PEPT |

| | | | | | | | | |
|---|---|---|---|---|---|---|---|---|
| 47:00:00 | 2fna-A | 2.4 | 10.4 | 77 | 352 | 8 | MOLECULE: | CONSERVED H |
| 48:00:00 | 7l7v-A | 2.4 | 7.7 | 56 | 121 | 7 | MOLECULE: | PROBABLE DI |
| 49:00:00 | 6ero-A | 2.4 | 16.7 | 88 | 296 | 6 | MOLECULE: | DIMETHYLADE |
| 50:00:00 | 5eyb-A | 2.4 | 12.3 | 81 | 340 | 2 | MOLECULE: | DNA-BINDING |
| 51:00:00 | 7r3e-A | 2.4 | 18.3 | 96 | 545 | 3 | MOLECULE: | 2-AMINOBENZ |
| 52:00:00 | 2nb2-A | 2.4 | 1.9 | 34 | 38 | 12 | MOLECULE: | NIGELLIN-1. |
| 53:00:00 | 3aq0-C | 2.4 | 8 | 63 | 327 | 2 | MOLECULE: | GERANYL DIP |
| 54:00:00 | 1upt-B | 2.4 | 5.6 | 42 | 58 | 5 | MOLECULE: | ADP-RIBOSYL |
| 55:00:00 | 6vld-B | 2.4 | 5.4 | 57 | 470 | 7 | MOLECULE: | ALPHA-(1,6) |
| 56:00:00 | 7cgp-J | 2.4 | 13.3 | 69 | 83 | 7 | MOLECULE: | MITOCHONDRI |
| 57:00:00 | 6u05-A | 2.4 | 22.6 | 84 | 413 | 4 | MOLECULE: | TRNA LIGASE |
| 58:00:00 | 3jbr-F | 2.4 | 11.9 | 118 | 872 | 2 | MOLECULE: | VOLTAGE-DEP |
| 59:00:00 | 4ux3-B | 2.4 | 4.2 | 55 | 71 | 9 | MOLECULE: | STRUCTURAL |
| 60:00:00 | 5vvr-B | 2.4 | 28.4 | 95 | 1207 | 5 | MOLECULE: | DNA-DIRECTE |
| 61:00:00 | 2m7a-A | 2.4 | 4.2 | 63 | 86 | 3 | MOLECULE: | UNCHARACTER |
| 62:00:00 | 8etc-h | 2.4 | 18.7 | 80 | 121 | 5 | MOLECULE: | RNA (2151-M |
| 63:00:00 | 6f7s-C | 2.4 | 9.9 | 65 | 319 | 8 | MOLECULE: | SERRATE RNA |
| 64:00:00 | 7zcm-B | 2.4 | 6.6 | 55 | 66 | 9 | MOLECULE: | SENSORY RHO |
| 65:00:00 | 5gox-A | 2.4 | 7.9 | 58 | 181 | 12 | MOLECULE: | DNA REPAIR |
| 66:00:00 | 6zyx-L | 2.4 | 4.3 | 47 | 97 | 6 | MOLECULE: | DYNEIN HEAV |
| 67:00:00 | 8tdj-A | 2.4 | 32.2 | 109 | 467 | 9 | MOLECULE: | MECHANOSENS |
| 68:00:00 | 6xi6-A | 2.4 | 17.2 | 81 | 269 | 11 | MOLECULE: | HELICAL FUS |
| 69:00:00 | 7aqq-d | 2.4 | 5.3 | 54 | 75 | 15 | MOLECULE: | NADH-UBIQUI |
| 70:00:00 | 2khq-A | 2.4 | 15.7 | 50 | 110 | 6 | MOLECULE: | INTEGRASE |
| 71:00:00 | 7eu9-A | 2.4 | 21.6 | 117 | 1078 | 7 | MOLECULE: | CAS12I1 D64 |
| 72:00:00 | 4evv-A | 2.4 | 5.5 | 66 | 145 | 8 | MOLECULE: | METHYL-CPG- |
| 73:00:00 | 7ahd-A | 2.4 | 17.5 | 79 | 547 | 6 | MOLECULE: | ABC-TYPE PR |
| 74:00:00 | 7sn7-a | 2.4 | 15.5 | 83 | 546 | 4 | MOLECULE: | FLAGELLIN |
| 75:00:00 | 7o4i-W | 2.4 | 19.8 | 86 | 312 | 6 | MOLECULE: | GENERAL TRA |
| 76:00:00 | 7x29-B | 2.4 | 14.6 | 87 | 1190 | 7 | MOLECULE: | SPIKE GLYCO |
| 77:00:00 | 7mrw-C | 2.4 | 8.5 | 113 | 706 | 4 | MOLECULE: | CYTOADHEREN |
| 78:00:00 | 8k3h-A | 2.4 | 11.2 | 91 | 480 | 10 | MOLECULE: | PSEP |
| 79:00:00 | 4bk0-B | 2.4 | 11.4 | 75 | 97 | 9 | MOLECULE: | ATP-DEPENDE |
| 80:00:00 | 8ab6-Q | 2.4 | 2.9 | 49 | 71 | 0 | MOLECULE: | CYTOCHROME |
| 81:00:00 | 8bdk-A | 2.4 | 3.2 | 46 | 96 | 7 | MOLECULE: | DNA-BINDING |
| 82:00:00 | 4cc9-B | 2.4 | 6.4 | 48 | 98 | 8 | MOLECULE: | PROTEIN VPR |
| 83:00:00 | 4dvy-P | 2.4 | 16.6 | 110 | 656 | 4 | MOLECULE: | CYTOTOXICIT |
| 84:00:00 | 3eo8-A | 2.4 | 16.6 | 68 | 219 | 4 | MOLECULE: | BLUB-LIKE F |
| 85:00:00 | 3bqp-B | 2.4 | 6.7 | 42 | 80 | 7 | MOLECULE: | PROACTIVATO |
| 86:00:00 | 6nwo-D | 2.4 | 5.1 | 63 | 165 | 5 | MOLECULE: | TRANSCRIPTI |
| 87:00:00 | 4dlq-A | 2.4 | 9.3 | 74 | 353 | 8 | MOLECULE: | LATROPHILIN |
| 88:00:00 | 7np8-B | 2.4 | 13.6 | 77 | 620 | 8 | MOLECULE: | COENZYME F4 |
| 89:00:00 | 3zcj-A | 2.4 | 9.3 | 90 | 205 | 6 | MOLECULE: | CAGL |
| 90:00:00 | 6nbu-A | 2.4 | 4.5 | 73 | 126 | 8 | MOLECULE: | CRISPR-ASSO |

| | | | | | | | | |
|---|---|---|---|---|---|---|---|---|
| 91:00:00 | 5yfp-E | 2.4 | 30 | 119 | 776 | 7 | MOLECULE: | EXOCYST COM |
| 92:00:00 | 7mnn-A | 2.4 | 14.6 | 93 | 750 | 11 | MOLECULE: | E3 SUMO-PRO |
| 93:00:00 | 5l09-B | 2.4 | 2.4 | 38 | 164 | 3 | MOLECULE: | QUORUM-SENS |
| 94:00:00 | 3odn-A | 2.4 | 8.8 | 71 | 366 | 6 | MOLECULE: | DALLY-LIKE |
| 95:00:00 | 2ksr-A | 2.4 | 4.2 | 60 | 140 | 13 | MOLECULE: | NEURONAL AC |
| 96:00:00 | 4mt4-B | 2.4 | 4.4 | 99 | 473 | 7 | MOLECULE: | CMEC |
| 97:00:00 | 4djg-A | 2.4 | 5.5 | 40 | 49 | 5 | MOLECULE: | PLECTIN-REL |
| 98:00:00 | 7jzw-A | 2.4 | 15.1 | 61 | 427 | 3 | MOLECULE: | CRISPR TYPE |
| 99:00:00 | 1p8c-C | 2.4 | 6.7 | 63 | 119 | 8 | MOLECULE: | CONSERVED H |
| 0:00 | 6jly-G | 2.4 | 4.5 | 75 | 363 | 9 | MOLECULE: | TRANSLATION |
| 1:00 | 6ygu-C | 2.4 | 8.4 | 44 | 215 | 9 | MOLECULE: | ATP DEPENDE |
| 2:00 | 5muu-A | 2.4 | 6.1 | 118 | 766 | 6 | MOLECULE: | MAJOR INNER |
| 3:00 | 8idb-D | 2.4 | 6.6 | 84 | 295 | 13 | MOLECULE: | CELL DIVISI |
| 4:00 | 1f02-T | 2.4 | 2.8 | 42 | 66 | 7 | MOLECULE: | INTIMIN |
| 5:00 | 1yns-A | 2.4 | 4.5 | 83 | 254 | 11 | MOLECULE: | E-1 ENZYME |
| 6:00 | 2px0-A | 2.4 | 10.9 | 50 | 258 | 8 | MOLECULE: | FLAGELLAR B |
| 7:00 | 6ynw-H | 2.4 | 4.5 | 50 | 75 | 6 | MOLECULE: | SUBUNIT C |
| 8:00 | 8a9b-B | 2.4 | 22.8 | 95 | 809 | 9 | MOLECULE: | LIPID BINDI |
| 9:00 | 4lnb-A | 2.4 | 14.5 | 85 | 339 | 5 | MOLECULE: | CAAX FARNES |
| 10:00 | 7d3e-A | 2.4 | 17.5 | 80 | 1380 | 4 | MOLECULE: | DUAL OXIDAS |
| 11:00 | 4ql6-B | 2.4 | 3.4 | 66 | 509 | 11 | MOLECULE: | CARBOXY-TER |
| 12:00 | 1lj2-B | 2.4 | 15.6 | 64 | 110 | 6 | MOLECULE: | NONSTRUCTUR |
| 13:00 | 6g7c-C | 2.4 | 13 | 63 | 239 | 5 | MOLECULE: | IMPA-RELATE |
| 14:00 | 2kmd-A | 2.4 | 4.2 | 71 | 111 | 0 | MOLECULE: | PROTEIN C-E |
| 15:00 | 6gw6-B | 2.4 | 4.8 | 67 | 149 | 9 | MOLECULE: | RES TOXIN |
| 16:00 | 3kkb-B | 2.4 | 5.7 | 71 | 128 | 6 | MOLECULE: | SENSOR PROT |
| 17:00 | 8wm8-B | 2.4 | 6.1 | 91 | 265 | 9 | MOLECULE: | NITRATE TRA |
| 18:00 | 4djb-A | 2.4 | 3.9 | 76 | 118 | 11 | MOLECULE: | E4-ORF3 |
| 19:00 | 4kyw-A | 2.4 | 7.9 | 76 | 254 | 9 | MOLECULE: | TYPE-2 REST |
| 20:00 | 2a2f-X | 2.4 | 14.1 | 93 | 296 | 4 | MOLECULE: | EXOCYST COM |
| 21:00 | 7yzi-A | 2.4 | 12 | 103 | 378 | 8 | MOLECULE: | ADENYLATE C |
| 22:00 | 8i24-F | 2.4 | 12.3 | 68 | 233 | 10 | MOLECULE: | DNA-DIRECTE |
| 23:00 | 6wq2-A | 2.4 | 7.3 | 69 | 154 | 3 | MOLECULE: | A-DNA |
| 24:00:00 | 6vr4-A | 2.4 | 13.3 | 112 | 2166 | 5 | MOLECULE: | DNA-DEPENDE |
| 25:00:00 | 8gy3-A | 2.4 | 11.2 | 83 | 440 | 6 | MOLECULE: | CYTOCHROME |
| 26:00:00 | 6v6d-A | 2.4 | 20.6 | 82 | 209 | 11 | MOLECULE: | PANNEXIN-1 |
| 27:00:00 | 5dpl-A | 2.4 | 14.5 | 99 | 521 | 10 | MOLECULE: | PROTEIN LYS |
| 28:00:00 | 8f2r-E | 2.4 | 9.1 | 84 | 205 | 8 | MOLECULE: | COMM DOMAIN |
| 29:00:00 | 8t1l-I | 2.4 | 18.8 | 85 | 1111 | 13 | MOLECULE: | MEDIATOR OF |
| 30:00:00 | 2gta-A | 2.4 | 5.8 | 68 | 98 | 3 | MOLECULE: | HYPOTHETICA |
| 31:00:00 | 7eew-A | 2.4 | 20.9 | 112 | 611 | 8 | MOLECULE: | TYPE I REST |
| 32:00:00 | 7nj1-A | 2.4 | 20.2 | 99 | 1448 | 12 | MOLECULE: | SEPARIN |
| 33:00:00 | 2qby-B | 2.4 | 5.9 | 75 | 368 | 3 | MOLECULE: | CELL DIVISI |
| 34:00:00 | 6xss-A | 2.4 | 6.8 | 60 | 260 | 7 | MOLECULE: | C4_NAT_HFUS |

| | | | | | | | | |
|---|---|---|---|---|---|---|---|---|
| 35:00:00 | 2qv2-A | 2.4 | 15.2 | 90 | 305 | 10 | MOLECULE: | INOSITOL PO |
| 36:00:00 | 1i5p-A | 2.4 | 13.6 | 103 | 633 | 4 | MOLECULE: | PESTICIDIAL |
| 37:00:00 | 7oqe-K | 2.4 | 17.2 | 87 | 406 | 0 | MOLECULE: | PROTEIN NAM |
| 38:00:00 | 5f8c-B | 2.4 | 17 | 85 | 363 | 4 | MOLECULE: | METHYLTRANS |
| 39:00:00 | 5hiu-A | 2.4 | 15.5 | 108 | 407 | 7 | MOLECULE: | GTPASE ACTI |
| 40:00:00 | 2e0w-B | 2.4 | 12.1 | 103 | 500 | 9 | MOLECULE: | GAMMA-GLUTA |
| 41:00:00 | 4fhn-B | 2.4 | 9.1 | 66 | 1022 | 6 | MOLECULE: | NUCLEOPORIN |
| 42:00:00 | 6eud-A | 2.4 | 19 | 87 | 808 | 3 | MOLECULE: | ATP-DEPENDE |
| 43:00:00 | 6rjy-A | 2.4 | 7.2 | 101 | 415 | 11 | MOLECULE: | ARABINO-OLI |
| 44:00:00 | 8g01-A | 2.4 | 11.5 | 96 | 360 | 7 | MOLECULE: | PHOSPHO-N-A |
| 45:00:00 | 3qt2-C | 2.4 | 11.4 | 83 | 110 | 4 | MOLECULE: | INTERLEUKIN |
| 46:00:00 | 3sjr-A | 2.4 | 8.8 | 73 | 126 | 5 | MOLECULE: | UNCHARACTER |
| 47:00:00 | 4r29-B | 2.4 | 10.6 | 87 | 204 | 3 | MOLECULE: | UNCHARACTER |
| 48:00:00 | 8w15-A | 2.4 | 24.6 | 100 | 2431 | 5 | MOLECULE: | HUNTINGTIN |
| 49:00:00 | 6ibl-A | 2.4 | 13.5 | 102 | 397 | 6 | MOLECULE: | THIOREDOXIN |
| 50:00:00 | 7s37-P | 2.4 | 27 | 124 | 1033 | 10 | MOLECULE: | CRISPR-ASSO |
| 51:00:00 | 8adl-A | 2.4 | 20.4 | 111 | 633 | 6 | MOLECULE: | MTC5 ISOFOR |
| 52:00:00 | 7pp4-f | 2.4 | 8.9 | 57 | 142 | 7 | MOLECULE: | DNA-DIRECTE |
| 53:00:00 | 7vv6-R | 2.4 | 15.1 | 89 | 271 | 6 | MOLECULE: | MAS-RELATED |
| 54:00:00 | 6s7t-H | 2.4 | 8.3 | 73 | 112 | 4 | MOLECULE: | DOLICHYL-DI |
| 55:00:00 | 5yqz-R | 2.4 | 16.8 | 140 | 558 | 9 | MOLECULE: | GLUCAGON RE |
| 56:00:00 | 8qkd-A | 2.4 | 5.6 | 80 | 281 | 6 | MOLECULE: | SC-CC-5-24 |
| 57:00:00 | 4tq4-D | 2.4 | 6.7 | 91 | 290 | 3 | MOLECULE: | PRENYLTRANS |
| 58:00:00 | 4dwl-B | 2.4 | 4 | 66 | 110 | 11 | MOLECULE: | BBP7 |
| 59:00:00 | 7w72-U | 2.4 | 9 | 97 | 420 | 8 | MOLECULE: | PHOSPHATIDY |
| 60:00:00 | 3d5l-A | 2.4 | 5.3 | 68 | 203 | 3 | MOLECULE: | REGULATORY |
| 61:00:00 | 6yrf-A | 2.4 | 20.4 | 114 | 777 | 9 | MOLECULE: | VEGETATIVE |
| 62:00:00 | 5bs1-D | 2.4 | 5.2 | 76 | 121 | 1 | MOLECULE: | CRRBCX-IIA |
| 63:00:00 | 5xaz-A | 2.4 | 4.1 | 74 | 223 | 3 | MOLECULE: | GAMMA-BUTYR |
| 64:00:00 | 3h9p-A | 2.4 | 7.6 | 79 | 226 | 3 | MOLECULE: | PUTATIVE TR |
| 65:00:00 | 7nhr-C | 2.4 | 24.7 | 93 | 565 | 6 | MOLECULE: | PUTATIVE TR |
| 66:00:00 | 6ty9-A | 2.4 | 5.9 | 119 | 1208 | 8 | MOLECULE: | RNA-DEPENDE |
| 67:00:00 | 1sqf-A | 2.4 | 7.3 | 83 | 425 | 2 | MOLECULE: | SUN PROTEIN |
| 68:00:00 | 8age-H | 2.4 | 7.8 | 72 | 133 | 10 | MOLECULE: | DOLICHYL-DI |
| 69:00:00 | 7ljn-A | 2.4 | 6.8 | 74 | 342 | 8 | MOLECULE: | CD-NTASE |
| 70:00:00 | 8ojy-C | 2.4 | 5.4 | 106 | 952 | 4 | MOLECULE: | PHOSPHOENOL |
| 71:00:00 | 1knv-B | 2.4 | 8.9 | 92 | 291 | 9 | MOLECULE: | BSE634I RES |
| 72:00:00 | 6hqa-A | 2.4 | 19 | 108 | 887 | 3 | MOLECULE: | TAF2 |
| 73:00:00 | 7pp2-A | 2.4 | 21 | 100 | 442 | 4 | MOLECULE: | EXOCYST SUB |
| 74:00:00 | 8vc1-A | 2.4 | 7.5 | 101 | 391 | 8 | MOLECULE: | GUSTATORY R |
| 75:00:00 | 4qvz-B | 2.4 | 4 | 55 | 204 | 7 | MOLECULE: | FRAGILE X M |
| 76:00:00 | 2mab-A | 2.4 | 3.7 | 78 | 109 | 8 | MOLECULE: | ACINIFORM S |
| 77:00:00 | 9b8o-e | 2.4 | 6.2 | 59 | 80 | 5 | MOLECULE: | ATPASE H+-T |
| 78:00:00 | 5v8f-D | 2.4 | 5.2 | 75 | 456 | 9 | MOLECULE: | DNA REPLICA |

| | | | | | | | | |
|---|---|---|---|---|---|---|---|---|
| 79:00:00 | 7ye1-G | 2.4 | 13.2 | 60 | 103 | 5 | MOLECULE: | DNA-DIRECTE |
| 80:00:00 | 3k1z-A | 2.4 | 6.8 | 96 | 241 | 5 | MOLECULE: | HALOACID DE |
| 81:00:00 | 5b7j-A | 2.4 | 3.6 | 71 | 110 | 8 | MOLECULE: | SWITCH-ACTI |
| 82:00:00 | 7xr2-a | 2.4 | 7.5 | 62 | 274 | 6 | MOLECULE: | VP3 |
| 83:00:00 | 5hea-A | 2.4 | 10.7 | 57 | 276 | 5 | MOLECULE: | PUTATIVE GL |
| 84:00:00 | 4tkr-B | 2.4 | 4.2 | 79 | 186 | 5 | MOLECULE: | THIAMINE TR |
| 85:00:00 | 2gsc-B | 2.4 | 6.8 | 76 | 117 | 7 | MOLECULE: | PUTATIVE UN |
| 86:00:00 | 2v5d-A | 2.4 | 11.2 | 82 | 722 | 5 | MOLECULE: | O-GLCNACASE |
| 87:00:00 | 2huj-A | 2.4 | 10.5 | 78 | 125 | 10 | MOLECULE: | LIN2004 PRO |
| 88:00:00 | 4an8-A | 2.4 | 4.8 | 73 | 468 | 8 | MOLECULE: | CSE1 |
| 89:00:00 | 8tar-N | 2.4 | 13.6 | 123 | 682 | 4 | MOLECULE: | ANAPHASE-PR |
| 90:00:00 | 6tnt-N | 2.4 | 26.1 | 110 | 672 | 7 | MOLECULE: | ANAPHASE-PR |
| 91:00:00 | 8oqs-B | 2.3 | 5.9 | 83 | 734 | 4 | MOLECULE: | 3-HYDROXYAC |
| 92:00:00 | 1i2l-A | 2.3 | 14.4 | 63 | 270 | 6 | MOLECULE: | 4-AMINO-4-D |
| 93:00:00 | 5jw9-B | 2.3 | 9.2 | 67 | 115 | 6 | MOLECULE: | AF4/FMR2 FA |
| 94:00:00 | 3h0g-A | 2.3 | 25.7 | 117 | 1496 | 6 | MOLECULE: | DNA-DIRECTE |
| 95:00:00 | 7opl-A | 2.3 | 18.5 | 119 | 1070 | 8 | MOLECULE: | DNA POLYMER |
| 96:00:00 | 6lo8-D | 2.3 | 7.9 | 51 | 119 | 4 | MOLECULE: | MITOCHONDRI |
| 97:00:00 | 2iw3-B | 2.3 | 12.5 | 96 | 980 | 1 | MOLECULE: | ELONGATION |
| 98:00:00 | 8t05-A | 2.3 | 5 | 71 | 201 | 13 | MOLECULE: | MYOMAKER |
| 99:00:00 | 8dfu-0 | 2.3 | 6.7 | 54 | 84 | 7 | MOLECULE: | PILIN PROTE |
| 0:00 | 3dkq-A | 2.3 | 5.5 | 57 | 230 | 5 | MOLECULE: | PKHD-TYPE H |
| 1:00 | 3ni7-A | 2.3 | 4.7 | 72 | 174 | 11 | MOLECULE: | BACTERIAL R |
| 2:00 | 7bst-C | 2.3 | 15.2 | 102 | 992 | 8 | MOLECULE: | TYPE I REST |
| 3:00 | 7wji-B | 2.3 | 36.6 | 116 | 1663 | 4 | MOLECULE: | PROTEIN UNC |
| 4:00 | 6h08-B | 2.3 | 9.9 | 84 | 297 | 5 | MOLECULE: | CYTOCHROME |
| 5:00 | 7lve-A | 2.3 | 6.1 | 43 | 44 | 2 | MOLECULE: | VICILIN JUG |
| 6:00 | 5wcn-A | 2.3 | 14.3 | 89 | 382 | 4 | MOLECULE: | SIAD |
| 7:00 | 7auf-A | 2.3 | 3.1 | 47 | 89 | 6 | MOLECULE: | SIMILAR TO |
| 8:00 | 3f46-A | 2.3 | 7.9 | 86 | 345 | 3 | MOLECULE: | 5,10-METHEN |
| 9:00 | 1qdm-A | 2.3 | 19.4 | 88 | 430 | 10 | MOLECULE: | PROPHYTEPSI |
| 10:00 | 6ayi-A | 2.3 | 9.3 | 44 | 183 | 11 | MOLECULE: | HTH-TYPE TR |
| 11:00 | 7cun-D | 2.3 | 17.1 | 96 | 827 | 5 | MOLECULE: | INTEGRATOR |
| 12:00 | 4i0x-K | 2.3 | 8 | 59 | 77 | 10 | MOLECULE: | ESAT-6-LIKE |
| 13:00 | 7ag9-B | 2.3 | 16.7 | 80 | 384 | 5 | MOLECULE: | KAR9 |
| 14:00 | 3hjz-A | 2.3 | 8.8 | 91 | 333 | 7 | MOLECULE: | TRANSALDOLA |
| 15:00 | 7mq1-B | 2.3 | 9.8 | 59 | 83 | 8 | MOLECULE: | COPPER-SENS |
| 16:00 | 5n9j-D | 2.3 | 12.4 | 74 | 135 | 5 | MOLECULE: | MEDIATOR OF |
| 17:00 | 2e62-A | 2.3 | 3.4 | 46 | 61 | 9 | MOLECULE: | PROTEIN AT5 |
| 18:00 | 6uxu-A | 2.3 | 8.5 | 81 | 300 | 10 | MOLECULE: | CHLOROTHALO |
| 19:00 | 8jhk-A | 2.3 | 24.1 | 110 | 727 | 7 | MOLECULE: | ENGULFMENT |
| 20:00 | 1s56-B | 2.3 | 4.1 | 68 | 135 | 12 | MOLECULE: | HEMOGLOBIN- |
| 21:00 | 8tzk-E | 2.3 | 14.4 | 59 | 101 | 2 | MOLECULE: | CELL DIVISI |
| 22:00 | 7ppo-A | 2.3 | 7.7 | 111 | 500 | 6 | MOLECULE: | UBIQUITINAT |

| | | | | | | | | |
|---|---|---|---|---|---|---|---|---|
| 23:00 | 3ogi-B | 2.3 | 13.5 | 69 | 89 | 6 | MOLECULE: | PUTATIVE ES |
| 24:00:00 | 8c83-x | 2.3 | 17 | 55 | 259 | 9 | MOLECULE: | OTU DOMAIN- |
| 25:00:00 | 6fvb-A | 2.3 | 14.4 | 102 | 1001 | 8 | MOLECULE: | IMPORTIN BE |
| 26:00:00 | 7ab4-F | 2.3 | 8 | 90 | 340 | 6 | MOLECULE: | PREDICTED T |
| 27:00:00 | 5fgn-A | 2.3 | 10 | 98 | 536 | 9 | MOLECULE: | LIPOOLIGOSA |
| 28:00:00 | 7tjk-F | 2.3 | 24 | 60 | 262 | 5 | MOLECULE: | ORIGIN RECO |
| 29:00:00 | 3hr0-A | 2.3 | 11.6 | 74 | 250 | 4 | MOLECULE: | COG4 |
| 30:00:00 | 2m6u-A | 2.3 | 3.2 | 46 | 82 | 2 | MOLECULE: | CHOLINE BIN |
| 31:00:00 | 5xj5-A | 2.3 | 6.7 | 88 | 200 | 7 | MOLECULE: | GLYCEROL-3- |
| 32:00:00 | 8kg9-E | 2.3 | 19.1 | 72 | 568 | 6 | MOLECULE: | DNA REPLICA |
| 33:00:00 | 3wz4-G | 2.3 | 8.1 | 45 | 140 | 2 | MOLECULE: | DOTI |
| 34:00:00 | 2h7f-X | 2.3 | 11.2 | 79 | 314 | 5 | MOLECULE: | 5'-D(*TP*TP |
| 35:00:00 | 4ayb-Q | 2.3 | 3.5 | 40 | 50 | 10 | MOLECULE: | DNA-DIRECTE |
| 36:00:00 | 6mc9-A | 2.3 | 17.2 | 80 | 150 | 6 | MOLECULE: | SODIUM CHAN |
| 37:00:00 | 4cpg-A | 2.3 | 7.4 | 51 | 69 | 10 | MOLECULE: | SMALL GLUTA |
| 38:00:00 | 8ae6-W | 2.3 | 14.5 | 75 | 941 | 4 | MOLECULE: | NITROGEN PE |
| 39:00:00 | 6jhm-A | 2.3 | 13.2 | 82 | 494 | 11 | MOLECULE: | CHLOROPHENO |
| 40:00:00 | 2bsk-D | 2.3 | 3.8 | 48 | 90 | 4 | MOLECULE: | MITOCHONDRI |
| 41:00:00 | 2oc6-A | 2.3 | 9.3 | 45 | 124 | 2 | MOLECULE: | YDHG PROTEI |
| 42:00:00 | 2fzt-B | 2.3 | 7.5 | 63 | 79 | 14 | MOLECULE: | HYPOTHETICA |
| 43:00:00 | 8k0b-A | 2.3 | 16.7 | 100 | 689 | 3 | MOLECULE: | CALCIUM PER |
| 44:00:00 | 6juv-A | 2.3 | 15.9 | 66 | 192 | 15 | MOLECULE: | SEGREGATION |
| 45:00:00 | 8w8q-C | 2.3 | 8 | 98 | 287 | 9 | MOLECULE: | SOLUBLE CYT |
| 46:00:00 | 1urf-A | 2.3 | 12.4 | 69 | 81 | 10 | MOLECULE: | PROTEIN KIN |
| 47:00:00 | 5ham-A | 2.3 | 13.9 | 80 | 260 | 6 | MOLECULE: | RICKCE |
| 48:00:00 | 7b93-n | 2.3 | 8.6 | 83 | 178 | 0 | MOLECULE: | NADH-UBIQUI |
| 49:00:00 | 5k7l-A | 2.3 | 29.7 | 99 | 701 | 5 | MOLECULE: | POTASSIUM V |
| 50:00:00 | 8a3t-Q | 2.3 | 27.2 | 93 | 623 | 3 | MOLECULE: | ANAPHASE-PR |
| 51:00:00 | 5ydc-B | 2.3 | 6.6 | 65 | 122 | 5 | MOLECULE: | UNCHARACTER |
| 52:00:00 | 8ovw-U | 2.3 | 17.2 | 78 | 184 | 5 | MOLECULE: | CENTROMERE- |
| 53:00:00 | 3iuk-A | 2.3 | 20 | 88 | 552 | 3 | MOLECULE: | UNCHARACTER |
| 54:00:00 | 7k15-A | 2.3 | 4.6 | 96 | 455 | 2 | MOLECULE: | LEUKOTRIENE |
| 55:00:00 | 7el9-A | 2.3 | 33.4 | 132 | 1731 | 10 | MOLECULE: | RNA-DIRECTE |
| 56:00:00 | 5x3q-A | 2.3 | 20 | 75 | 313 | 3 | MOLECULE: | ENVELOPE GL |
| 57:00:00 | 6iy8-A | 2.3 | 17.7 | 86 | 469 | 3 | MOLECULE: | POSITIVE RE |
| 58:00:00 | 6c66-A | 2.3 | 16.2 | 82 | 511 | 4 | MOLECULE: | CRISPR-ASSO |
| 59:00:00 | 7d7n-A | 2.3 | 15.9 | 122 | 704 | 5 | MOLECULE: | ATP-BINDING |
| 60:00:00 | 2okx-A | 2.3 | 15.8 | 107 | 954 | 5 | MOLECULE: | RHAMNOSIDAS |
| 61:00:00 | 5wb2-A | 2.3 | 14.7 | 104 | 421 | 9 | MOLECULE: | ENVELOPE PR |
| 62:00:00 | 5nx9-D | 2.3 | 15.2 | 104 | 477 | 5 | MOLECULE: | ADENYLOSUCC |
| 63:00:00 | 8g8i-A | 2.3 | 21.8 | 78 | 465 | 5 | MOLECULE: | C3HR3_9R_SH |
| 64:00:00 | 1mje-A | 2.3 | 26.4 | 108 | 600 | 9 | MOLECULE: | 5'-D(P*TP*T |
| 65:00:00 | 3cfo-A | 2.3 | 10.1 | 73 | 906 | 8 | MOLECULE: | DNA POLYMER |
| 66:00:00 | 6xp5-D | 2.3 | 19.9 | 74 | 134 | 3 | MOLECULE: | MEDIATOR OF |

| | | | | | | | | |
|---|---|---|---|---|---|---|---|---|
| 67:00:00 | 8adb-A | 2.3 | 10.6 | 59 | 209 | 7 | MOLECULE: | WC-VDT1 |
| 68:00:00 | 1uuj-D | 2.3 | 5.2 | 64 | 79 | 5 | MOLECULE: | PLATELET-AC |
| 69:00:00 | 3b77-A | 2.3 | 10.6 | 55 | 188 | 9 | MOLECULE: | UNCHARACTER |
| 70:00:00 | 7ef9-A | 2.3 | 17.7 | 81 | 414 | 11 | MOLECULE: | ADENINE DNA |
| 71:00:00 | 8ghn-B | 2.3 | 11.7 | 95 | 800 | 8 | MOLECULE: | PROTEIN HIR |
| 72:00:00 | 1ej6-D | 2.3 | 13.2 | 106 | 417 | 8 | MOLECULE: | LAMBDA2 |
| 73:00:00 | 7amy-A | 2.3 | 23.4 | 77 | 353 | 3 | MOLECULE: | FLAGELLAR B |
| 74:00:00 | 7mbz-A | 2.3 | 5.7 | 87 | 222 | 13 | MOLECULE: | ABC TRANSPO |
| 75:00:00 | 7krw-A | 2.3 | 9 | 90 | 608 | 6 | MOLECULE: | CHAPERONE P |
| 76:00:00 | 6z2k-D | 2.3 | 18.8 | 87 | 157 | 9 | MOLECULE: | HISTONE DEA |
| 77:00:00 | 7rfq-A | 2.3 | 8.6 | 57 | 223 | 4 | MOLECULE: | BETA CELL E |
| 78:00:00 | 6ivh-A | 2.3 | 3 | 48 | 94 | 15 | MOLECULE: | SEGREGATION |
| 79:00:00 | 8e4v-A | 2.3 | 3.1 | 53 | 84 | 15 | MOLECULE: | ISOFORM 3 O |
| 80:00:00 | 7jmn-O | 2.3 | 13.5 | 66 | 164 | 6 | MOLECULE: | MEDIATOR OF |
| 81:00:00 | 1k32-A | 2.3 | 29.7 | 92 | 1023 | 7 | MOLECULE: | TRICORN PRO |
| 82:00:00 | 2ou3-A | 2.3 | 6.2 | 76 | 160 | 9 | MOLECULE: | TELLURITE R |
| 83:00:00 | 8tef-A | 2.3 | 27.7 | 99 | 516 | 4 | MOLECULE: | RESPONSE RE |
| 84:00:00 | 3v53-B | 2.3 | 3.4 | 55 | 112 | 16 | MOLECULE: | RNA-BINDING |
| 85:00:00 | 2uy1-A | 2.3 | 15.3 | 90 | 438 | 7 | MOLECULE: | CLEAVAGE ST |
| 86:00:00 | 7xuh-A | 2.3 | 25 | 88 | 707 | 5 | MOLECULE: | CHLORIDE AN |
| 87:00:00 | 8hvj-A | 2.3 | 5.5 | 66 | 281 | 9 | MOLECULE: | UPF0701 PRO |
| 88:00:00 | 6c6r-A | 2.3 | 10 | 76 | 451 | 4 | MOLECULE: | SQUALENE MO |
| 89:00:00 | 7khw-a | 2.3 | 9.6 | 52 | 174 | 2 | MOLECULE: | TRANSLOCON |
| 90:00:00 | 6mj1-A | 2.3 | 4.2 | 79 | 200 | 4 | MOLECULE: | PROBABLE HT |
| 91:00:00 | 3hhc-B | 2.3 | 11.3 | 60 | 157 | 5 | MOLECULE: | INTERLEUKIN |
| 92:00:00 | 8b9z-Y | 2.3 | 16.5 | 90 | 167 | 3 | MOLECULE: | NADH-UBIQUI |
| 93:00:00 | 7pp6-B | 2.3 | 23.3 | 101 | 1148 | 2 | MOLECULE: | MUCIN-2 |
| 94:00:00 | 5xfs-A | 2.3 | 4.8 | 47 | 78 | 4 | MOLECULE: | PE FAMILY P |
| 95:00:00 | 6c0f-w | 2.3 | 6.9 | 51 | 70 | 10 | MOLECULE: | SACCHAROMYC |
| 96:00:00 | 2khv-A | 2.3 | 3.9 | 48 | 106 | 0 | MOLECULE: | PHAGE INTEG |
| 97:00:00 | 8fcj-I | 2.3 | 13.2 | 97 | 494 | 7 | MOLECULE: | TYPE I-B CR |
| 98:00:00 | 8cqv-A | 2.3 | 16.1 | 76 | 202 | 3 | MOLECULE: | NITROREDUCT |
| 99:00:00 | 8ikj-R | 2.3 | 15.7 | 84 | 532 | 14 | MOLECULE: | ADHESION G |
| 0:00 | 7qoj-E | 2.3 | 3.6 | 49 | 238 | 12 | MOLECULE: | PORTAL PROT |
| 1:00 | 6ihr-A | 2.3 | 7.2 | 70 | 257 | 7 | MOLECULE: | PHOSPHORYLA |
| 2:00 | 6wlz-Y | 2.3 | 12.7 | 45 | 267 | 9 | MOLECULE: | V-TYPE PROT |
| 3:00 | 4xp7-A | 2.3 | 9.2 | 72 | 324 | 13 | MOLECULE: | TRNA-DIHYDR |
| 4:00 | 5l4k-S | 2.3 | 21.4 | 77 | 491 | 12 | MOLECULE: | 26S PROTEAS |
| 5:00 | 7ugc-A | 2.3 | 6.1 | 62 | 191 | 8 | MOLECULE: | PROTEIN VIA |
| 6:00 | 7cy6-A | 2.3 | 4.8 | 72 | 872 | 6 | MOLECULE: | MALTODEXTRI |
| 7:00 | 6urt-A | 2.3 | 17.5 | 91 | 331 | 4 | MOLECULE: | LOW CONDUCT |
| 8:00 | 7zmb-X | 2.3 | 13.6 | 65 | 187 | 5 | MOLECULE: | NADH-UBIQUI |
| 9:00 | 8ewi-A | 2.3 | 18.7 | 97 | 1775 | 10 | MOLECULE: | E3 UBIQUITI |
| 10:00 | 4bt9-B | 2.3 | 21.8 | 85 | 238 | 2 | MOLECULE: | PROLYL 4-HY |

| | | | | | | | | |
|---|---|---|---|---|---|---|---|---|
| 11:00 | 2fgy-A | 2.3 | 9.1 | 108 | 471 | 8 | MOLECULE: | CARBOXYSOME |
| 12:00 | 8cli-A | 2.3 | 12.4 | 109 | 542 | 6 | MOLECULE: | GENERAL TRA |
| 13:00 | 8btl-A | 2.3 | 9.7 | 78 | 353 | 6 | MOLECULE: | CDNA FLJ125 |
| 14:00 | 7upq-F | 2.3 | 10.6 | 59 | 77 | 5 | MOLECULE: | DHT03 PROTE |
| 15:00 | 8rhf-A | 2.3 | 16.7 | 97 | 327 | 6 | MOLECULE: | LYSM PEPTID |
| 16:00 | 6ird-B | 2.3 | 23.2 | 66 | 254 | 5 | MOLECULE: | 1-PHOSPHATI |
| 17:00 | 4oph-A | 2.3 | 14.1 | 77 | 202 | 6 | MOLECULE: | NONSTRUCTUR |
| 18:00 | 5oc0-A | 2.3 | 6.7 | 61 | 174 | 8 | MOLECULE: | CYTOCHROME |
| 19:00 | 8t8k-A | 2.3 | 4.2 | 61 | 183 | 2 | MOLECULE: | DUF507 FAMI |
| 20:00 | 7apd-B | 2.3 | 20.9 | 72 | 289 | 24 | MOLECULE: | REPLICATION |
| 21:00 | 5da8-K | 2.3 | 6.4 | 86 | 433 | 8 | MOLECULE: | 60 KDA CHAP |
| 22:00 | 3dee-A | 2.3 | 9 | 64 | 200 | 5 | MOLECULE: | PUTATIVE RE |
| 23:00 | 6l80-C | 2.3 | 10.1 | 74 | 107 | 3 | MOLECULE: | GAMMA-TUBUL |
| 24:00:00 | 7t62-A | 2.3 | 16.4 | 79 | 561 | 3 | MOLECULE: | GLYPICAN-2 |
| 25:00:00 | 4gmq-A | 2.3 | 6.3 | 47 | 92 | 4 | MOLECULE: | PUTATIVE RI |
| 26:00:00 | 3kfu-G | 2.3 | 7.9 | 48 | 86 | 4 | MOLECULE: | NON-DISCRIM |
| 27:00:00 | 8uz8-B | 2.3 | 4.8 | 52 | 60 | 8 | MOLECULE: | 2-OXOGLUTAR |
| 28:00:00 | 5cm8-A | 2.3 | 17 | 70 | 419 | 9 | MOLECULE: | RAL GUANINE |
| 29:00:00 | 6wqz-A | 2.3 | 10.2 | 112 | 536 | 4 | MOLECULE: | AUTOPHAGY-R |
| 30:00:00 | 2ixp-B | 2.3 | 9 | 90 | 316 | 7 | MOLECULE: | SERINE/THRE |
| 31:00:00 | 3wqy-A | 2.3 | 27.4 | 132 | 906 | 8 | MOLECULE: | ALANINE--TR |
| 32:00:00 | 3x38-B | 2.3 | 3.2 | 39 | 79 | 13 | MOLECULE: | MITOCHONDRI |
| 33:00:00 | 5xa5-A | 2.3 | 12 | 66 | 248 | 3 | MOLECULE: | ALPHA-CATEN |
| 34:00:00 | 5xsv-A | 2.3 | 25.8 | 81 | 486 | 6 | MOLECULE: | CHITINASE |
| 35:00:00 | 4wrp-A | 2.3 | 10.3 | 80 | 121 | 1 | MOLECULE: | UNCHARACTER |
| 36:00:00 | 6uek-A | 2.3 | 10.1 | 100 | 660 | 8 | MOLECULE: | UROCANATE H |
| 37:00:00 | 6gw5-A | 2.3 | 9 | 72 | 393 | 1 | MOLECULE: | PUTATIVE OU |
| 38:00:00 | 2oeb-A | 2.3 | 3.8 | 77 | 152 | 5 | MOLECULE: | HYPOTHETICA |
| 39:00:00 | 7zaw-A | 2.3 | 5.3 | 94 | 361 | 9 | MOLECULE: | GLYPICAN-3 |
| 40:00:00 | 2h1n-A | 2.3 | 17.6 | 109 | 566 | 4 | MOLECULE: | OLIGOENDOPE |
| 41:00:00 | 7vrc-C | 2.3 | 7.8 | 72 | 109 | 6 | MOLECULE: | TRANSCRIPTI |
| 42:00:00 | 2euf-A | 2.3 | 3 | 64 | 244 | 3 | MOLECULE: | VIRAL CYCLI |
| 43:00:00 | 3svi-A | 2.3 | 3.2 | 58 | 79 | 10 | MOLECULE: | TYPE III EF |
| 44:00:00 | 6qyi-B | 2.3 | 18.9 | 122 | 519 | 8 | MOLECULE: | 4-HYDROXYPH |
| 45:00:00 | 4z7f-B | 2.3 | 4.3 | 86 | 169 | 5 | MOLECULE: | FOLATE ECF |
| 46:00:00 | 6oiu-C | 2.3 | 20.1 | 119 | 900 | 7 | MOLECULE: | AMINOPEPTID |
| 47:00:00 | 7vmb-A | 2.3 | 18.3 | 99 | 359 | 6 | MOLECULE: | IQ MOTIF AN |
| 48:00:00 | 8yb7-C | 2.3 | 10.2 | 96 | 455 | 4 | MOLECULE: | PAPAIN-LIKE |
| 49:00:00 | 8egr-G | 2.3 | 5.8 | 59 | 313 | 7 | MOLECULE: | GP15, RECEP |
| 50:00:00 | 4hm9-A | 2.3 | 19.1 | 112 | 511 | 2 | MOLECULE: | BETA-CATENI |
| 51:00:00 | 2k9a-A | 2.3 | 8.9 | 95 | 136 | 7 | MOLECULE: | ADP-RIBOSYL |
| 52:00:00 | 5zi2-A | 2.3 | 6.6 | 77 | 339 | 6 | MOLECULE: | MALATE DEHY |
| 53:00:00 | 7d7c-F | 2.3 | 3.2 | 67 | 137 | 6 | MOLECULE: | DNA-DIRECTE |
| 54:00:00 | 8u53-A | 2.3 | 9.1 | 110 | 574 | 7 | MOLECULE: | CSC1-LIKE P |

| | | | | | | | | |
|---|---|---|---|---|---|---|---|---|
| 55:00:00 | 8emh-B | 2.3 | 20.3 | 105 | 676 | 11 | MOLECULE: | PROTEASE LO |
| 56:00:00 | 6x58-F | 2.3 | 12.3 | 70 | 135 | 3 | MOLECULE: | 10E8V4 FAB |
| 57:00:00 | 8evm-C | 2.3 | 10 | 71 | 237 | 8 | MOLECULE: | CHLOROPHYLL |
| 58:00:00 | 5yet-A | 2.3 | 10.1 | 87 | 397 | 3 | MOLECULE: | UNCHARACTER |
| 59:00:00 | 5tr1-A | 2.3 | 11.5 | 78 | 606 | 4 | MOLECULE: | CHLORIDE CH |
| 60:00:00 | 6xu2-A | 2.3 | 23.9 | 109 | 1091 | 9 | MOLECULE: | IMPORTIN-5 |
| 61:00:00 | 8qgy-B | 2.3 | 9.2 | 73 | 809 | 10 | MOLECULE: | MITOGEN-ACT |
| 62:00:00 | 4l3u-A | 2.3 | 6 | 71 | 123 | 8 | MOLECULE: | UNCHARACTER |
| 63:00:00 | 7e4n-A | 2.3 | 6.3 | 79 | 290 | 6 | MOLECULE: | SAT1646 |
| 64:00:00 | 4ysm-A | 2.3 | 7 | 78 | 475 | 8 | MOLECULE: | CALMODULIN- |
| 65:00:00 | 8bsb-B | 2.3 | 13.2 | 89 | 162 | 8 | MOLECULE: | METHYL-ACCE |
| 66:00:00 | 7njn-C | 2.3 | 18.8 | 89 | 536 | 10 | MOLECULE: | ATP SYNTHAS |
| 67:00:00 | 7y3f-1 | 2.3 | 11.8 | 77 | 466 | 8 | MOLECULE: | PHOTOSYSTEM |
| 68:00:00 | 6f42-W | 2.3 | 13.2 | 76 | 165 | 5 | MOLECULE: | DNA-DIRECTE |
| 69:00:00 | 1irx-B | 2.3 | 14.1 | 84 | 508 | 7 | MOLECULE: | LYSYL-TRNA |
| 70:00:00 | 7v9x-A | 2.3 | 15.1 | 74 | 314 | 11 | MOLECULE: | RNA-DIRECTE |
| 71:00:00 | 8xr6-q | 2.3 | 7.8 | 58 | 143 | 2 | MOLECULE: | PHOTOSYSTEM |
| 72:00:00 | 7kpo-A | 2.3 | 3.4 | 64 | 107 | 8 | MOLECULE: | RESPONSE RE |
| 73:00:00 | 8imi-0 | 2.3 | 27.3 | 121 | 1045 | 7 | MOLECULE: | APCE |
| 74:00:00 | 7x34-C | 2.3 | 3.1 | 56 | 100 | 5 | MOLECULE: | ZUOTIN |
| 75:00:00 | 6lb8-C | 2.3 | 11.9 | 94 | 483 | 1 | MOLECULE: | ENDOLYSIN,C |
| 76:00:00 | 2f07-A | 2.3 | 8.4 | 73 | 197 | 7 | MOLECULE: | YVDT |
| 77:00:00 | 8hwi-B | 2.3 | 4.5 | 67 | 177 | 12 | MOLECULE: | CD-NTASE-AS |
| 78:00:00 | 7wb4-g | 2.3 | 6.8 | 65 | 672 | 3 | MOLECULE: | OUTER NUP13 |
| 79:00:00 | 5oqm-n | 2.3 | 21.8 | 74 | 136 | 7 | MOLECULE: | DNA-DIRECTE |
| 80:00:00 | 6xwi-A | 2.3 | 3.5 | 57 | 66 | 7 | MOLECULE: | S0_2.126 |
| 81:00:00 | 3pxp-A | 2.3 | 10.4 | 74 | 292 | 11 | MOLECULE: | HELIX-TURN- |
| 82:00:00 | 5w99-A | 2.3 | 7.9 | 84 | 319 | 4 | MOLECULE: | PBTD |
| 83:00:00 | 6h02-A | 2.3 | 9.1 | 114 | 1334 | 8 | MOLECULE: | MEDIATOR OF |
| 84:00:00 | 7cgp-A | 2.3 | 2.6 | 61 | 147 | 10 | MOLECULE: | MITOCHONDRI |
| 85:00:00 | 8gxl-B | 2.3 | 9.9 | 64 | 122 | 6 | MOLECULE: | SURP AND G- |
| 86:00:00 | 7jmn-E | 2.3 | 8.1 | 79 | 584 | 5 | MOLECULE: | MEDIATOR OF |
| 87:00:00 | 2w96-A | 2.3 | 9.1 | 75 | 249 | 4 | MOLECULE: | CELL DIVISI |
| 88:00:00 | 6nf4-A | 2.3 | 13.3 | 90 | 421 | 4 | MOLECULE: | OTOPETRIN1 |
| 89:00:00 | 7k3z-A | 2.3 | 9.5 | 85 | 564 | 7 | MOLECULE: | 60 KDA CHAP |
| 90:00:00 | 2eul-A | 2.3 | 7.2 | 79 | 156 | 5 | MOLECULE: | ANTI-CLEAVA |
| 91:00:00 | 5u30-A | 2.3 | 18.1 | 81 | 1085 | 6 | MOLECULE: | CRISPR-ASSO |
| 92:00:00 | 4oe8-C | 2.3 | 8.8 | 74 | 87 | 15 | MOLECULE: | INTERLEUKIN |
| 93:00:00 | 7woo-F | 2.3 | 11.1 | 106 | 1622 | 8 | MOLECULE: | NUCLEOPORIN |
| 94:00:00 | 6qti-A | 2.3 | 19.3 | 105 | 1038 | 8 | MOLECULE: | NICOTINAMID |
| 95:00:00 | 5v8f-6 | 2.3 | 9.2 | 117 | 692 | 8 | MOLECULE: | DNA REPLICA |
| 96:00:00 | 6aqk-A | 2.3 | 14.3 | 69 | 216 | 7 | MOLECULE: | TOXIN PROTE |
| 97:00:00 | 1bmt-A | 2.3 | 11 | 97 | 246 | 6 | MOLECULE: | METHIONINE |
| 98:00:00 | 8wtc-B | 2.3 | 3.8 | 48 | 100 | 2 | MOLECULE: | PROTEIN-ARG |

| | | | | | | | | |
|---|---|---|---|---|---|---|---|---|
| 99:00:00 | 1b48-A | 2.3 | 5.1 | 79 | 221 | 11 | MOLECULE: | PROTEIN (GL |
| 0:00 | 6wb9-2 | 2.3 | 13.2 | 80 | 292 | 11 | MOLECULE: | ENDOPLASMIC |
| 1:00 | 4a01-A | 2.3 | 7 | 106 | 740 | 7 | MOLECULE: | PROTON PYRO |
| 2:00 | 6mit-G | 2.3 | 10.2 | 78 | 355 | 5 | MOLECULE: | LIPOPOLYSAC |
| 3:00 | 8dol-A | 2.3 | 19.3 | 106 | 540 | 2 | MOLECULE: | CAG PATHOGE |
| 4:00 | 3kwo-A | 2.3 | 7.5 | 77 | 149 | 5 | MOLECULE: | PUTATIVE BA |
| 5:00 | 8d00-A | 2.3 | 13.2 | 92 | 299 | 7 | MOLECULE: | MICROTUBULE |
| 6:00 | 6cp6-Y | 2.3 | 9.2 | 83 | 188 | 7 | MOLECULE: | ATP SYNTHAS |
| 7:00 | 8g1e-A | 2.3 | 6.5 | 102 | 1037 | 10 | MOLECULE: | ATP-CITRATE |
| 8:00 | 6hcy-A | 2.3 | 5.4 | 84 | 436 | 6 | MOLECULE: | METALLOREDU |
| 9:00 | 5y78-A | 2.3 | 11.3 | 89 | 305 | 8 | MOLECULE: | PUTATIVE HE |
| 10:00 | 8kem-A | 2.3 | 4.8 | 60 | 425 | 13 | MOLECULE: | SDR FAMILY |
| 11:00 | 6btm-F | 2.3 | 17.4 | 113 | 402 | 6 | MOLECULE: | ALTERNATIVE |
| 12:00 | 8p2m-A | 2.3 | 5.4 | 98 | 659 | 7 | MOLECULE: | NAD(+) HYDR |
| 13:00 | 6kn5-A | 2.3 | 8.2 | 78 | 225 | 9 | MOLECULE: | AF4/FMR2 FA |
| 14:00 | 2lp4-A | 2.3 | 20.1 | 110 | 225 | 5 | MOLECULE: | CHEMOTAXIS |
| 15:00 | 7k10-A | 2.3 | 15.2 | 121 | 1259 | 6 | MOLECULE: | DNA-DEPENDE |
| 16:00 | 2g7l-A | 2.3 | 7.4 | 68 | 201 | 9 | MOLECULE: | TETR-FAMILY |
| 17:00 | 6uhd-A | 2.3 | 4 | 86 | 322 | 3 | MOLECULE: | HISTONE ACE |
| 18:00 | 2gtv-X | 2.3 | 3.9 | 60 | 104 | 3 | MOLECULE: | CHORISMATE |
| 19:00 | 6cfz-J | 2.3 | 22.3 | 80 | 134 | 5 | MOLECULE: | ASK1 |
| 20:00 | 3mka-J | 2.3 | 13 | 65 | 252 | 6 | MOLECULE: | PROTEASOME |
| 21:00 | 8h77-F | 2.3 | 10.7 | 82 | 310 | 5 | MOLECULE: | HEAT SHOCK |
| 22:00 | 8cqn-B | 2.3 | 28.3 | 102 | 187 | 7 | MOLECULE: | LIPOPROTEIN |
| 23:00 | 6vek-A | 2.3 | 6.1 | 71 | 321 | 7 | MOLECULE: | CONTACT-DEP |
| 24:00:00 | 5dku-B | 2.3 | 16.4 | 88 | 581 | 8 | MOLECULE: | PREX DNA PO |
| 25:00:00 | 5wtk-A | 2.3 | 20.5 | 105 | 1215 | 7 | MOLECULE: | CRISPR-ASSO |
| 26:00:00 | 6wg7-H | 2.3 | 8.2 | 72 | 219 | 1 | MOLECULE: | DNA (35-MER |
| 27:00:00 | 6wg7-D | 2.3 | 8.2 | 72 | 219 | 1 | MOLECULE: | DNA (35-MER |
| 28:00:00 | 5wgg-A | 2.2 | 14.9 | 85 | 441 | 2 | MOLECULE: | RADICAL SAM |
| 29:00:00 | 3c2b-A | 2.2 | 13.6 | 63 | 200 | 6 | MOLECULE: | TRANSCRIPTI |
| 30:00:00 | 3euh-B | 2.2 | 8.9 | 104 | 321 | 6 | MOLECULE: | CHROMOSOME |
| 31:00:00 | 7thw-A | 2.2 | 8.6 | 72 | 158 | 11 | MOLECULE: | PUTATIVE OM |
| 32:00:00 | 6qeq-A | 2.2 | 8.9 | 67 | 111 | 7 | MOLECULE: | PCFF |
| 33:00:00 | 7y6t-A | 2.2 | 26.8 | 107 | 1124 | 8 | MOLECULE: | SPIKE GLYCO |
| 34:00:00 | 3cjh-B | 2.2 | 3.8 | 49 | 59 | 2 | MOLECULE: | MITOCHONDRI |
| 35:00:00 | 7z0o-D | 2.2 | 19.8 | 77 | 296 | 5 | MOLECULE: | HISTONE H3 |
| 36:00:00 | 6cc4-A | 2.2 | 12.8 | 75 | 608 | 7 | MOLECULE: | SOLUBLE CYT |
| 37:00:00 | 6hua-B | 2.2 | 4.3 | 57 | 229 | 9 | MOLECULE: | UNCHARACTER |
| 38:00:00 | 4x86-A | 2.2 | 3.9 | 38 | 51 | 8 | MOLECULE: | UBIQUITIN-L |
| 39:00:00 | 1jjo-A | 2.2 | 3.1 | 35 | 40 | 6 | MOLECULE: | NEUROSERPIN |
| 40:00:00 | 7bst-A | 2.2 | 3.7 | 48 | 384 | 8 | MOLECULE: | TYPE I REST |
| 41:00:00 | 2dsy-D | 2.2 | 3.5 | 49 | 81 | 4 | MOLECULE: | HYPOTHETICA |
| 42:00:00 | 2fdr-A | 2.2 | 6.5 | 54 | 222 | 11 | MOLECULE: | CONSERVED H |

| | | | | | | | | |
|---|---|---|---|---|---|---|---|---|
| 43:00:00 | 1snl-A | 2.2 | 12.1 | 61 | 99 | 3 | MOLECULE: | NUCLEOBINDI |
| 44:00:00 | 5chi-A | 2.2 | 9.3 | 69 | 230 | 7 | MOLECULE: | UNCHARACTER |
| 45:00:00 | 6wg3-A | 2.2 | 11.3 | 67 | 561 | 7 | MOLECULE: | STRUCTURAL |
| 46:00:00 | 7lqt-A | 2.2 | 15.8 | 64 | 173 | 8 | MOLECULE: | SERINE/THRE |
| 47:00:00 | 8eau-p | 2.2 | 4.3 | 40 | 56 | 3 | MOLECULE: | V-TYPE PROT |
| 48:00:00 | 5dfz-B | 2.2 | 15.4 | 90 | 1224 | 2 | MOLECULE: | VACUOLAR PR |
| 49:00:00 | 7oq4-Z | 2.2 | 4.4 | 52 | 118 | 0 | MOLECULE: | DNA-DIRECTE |
| 50:00:00 | 3rmi-A | 2.2 | 13.2 | 48 | 105 | 10 | MOLECULE: | CHORISMATE |
| 51:00:00 | 7qru-G | 2.2 | 10.2 | 85 | 113 | 7 | MOLECULE: | NA+/H+ ANTI |
| 52:00:00 | 5ncl-A | 2.2 | 10.3 | 69 | 418 | 7 | MOLECULE: | SERINE/THRE |
| 53:00:00 | 5b3f-A | 2.2 | 14.9 | 83 | 306 | 5 | MOLECULE: | PHOSPHORIBU |
| 54:00:00 | 2qq8-A | 2.2 | 8.8 | 72 | 288 | 7 | MOLECULE: | TBC1 DOMAIN |
| 55:00:00 | 2hh6-A | 2.2 | 9.1 | 50 | 112 | 8 | MOLECULE: | BH3980 PROT |
| 56:00:00 | 3cih-A | 2.2 | 19.2 | 102 | 714 | 5 | MOLECULE: | PUTATIVE AL |
| 57:00:00 | 3u2r-A | 2.2 | 3.6 | 44 | 135 | 2 | MOLECULE: | REGULATORY |
| 58:00:00 | 7xe4-F | 2.2 | 19.2 | 79 | 1503 | 5 | MOLECULE: | 1,3-BETA-GL |
| 59:00:00 | 7sox-A | 2.2 | 16.6 | 93 | 514 | 8 | MOLECULE: | EH DOMAIN-C |
| 60:00:00 | 6bs3-A | 2.2 | 13 | 61 | 330 | 8 | MOLECULE: | PUTATIVE AT |
| 61:00:00 | 8fvx-A | 2.2 | 2.7 | 42 | 63 | 10 | MOLECULE: | CBFD_NFYB_H |
| 62:00:00 | 3fay-A | 2.2 | 10 | 125 | 379 | 6 | MOLECULE: | RAS GTPASE- |
| 63:00:00 | 6xxv-C | 2.2 | 9 | 54 | 111 | 6 | MOLECULE: | ANTIBODY C5 |
| 64:00:00 | 3oao-A | 2.2 | 10.2 | 66 | 140 | 11 | MOLECULE: | UNCHARACTER |
| 65:00:00 | 4eba-A | 2.2 | 22.3 | 95 | 585 | 4 | MOLECULE: | MRNA 3'-END |
| 66:00:00 | 2azj-A | 2.2 | 3.7 | 55 | 276 | 11 | MOLECULE: | GERANYLGERA |
| 67:00:00 | 7x5e-B | 2.2 | 20 | 73 | 106 | 10 | MOLECULE: | TRANSCRIPTI |
| 68:00:00 | 3ufe-A | 2.2 | 13.7 | 51 | 109 | 4 | MOLECULE: | TRANSCRIPTI |
| 69:00:00 | 1ufi-A | 2.2 | 4.5 | 42 | 48 | 10 | MOLECULE: | MAJOR CENTR |
| 70:00:00 | 7t92-B | 2.2 | 9.4 | 83 | 359 | 7 | MOLECULE: | PEROXIN-12 |
| 71:00:00 | 1h0a-A | 2.2 | 7.1 | 70 | 158 | 6 | MOLECULE: | EPSIN |
| 72:00:00 | 7ocn-A | 2.2 | 11.2 | 98 | 690 | 4 | MOLECULE: | HAD HYDROLA |
| 73:00:00 | 5gmk-I | 2.2 | 5.8 | 61 | 102 | 3 | MOLECULE: | PRE-MRNA-SP |
| 74:00:00 | 2d3o-1 | 2.2 | 4.3 | 59 | 100 | 3 | MOLECULE: | 23S RIBOSOM |
| 75:00:00 | 6pc0-B | 2.2 | 15.7 | 73 | 484 | 1 | MOLECULE: | GUANOSINE P |
| 76:00:00 | 8pjn-b | 2.2 | 20.3 | 106 | 296 | 7 | MOLECULE: | E3 UBIQUITI |
| 77:00:00 | 5x62-B | 2.2 | 20.5 | 98 | 388 | 2 | MOLECULE: | CARNOSINE N |
| 78:00:00 | 8jzr-B | 2.2 | 11.5 | 108 | 456 | 10 | MOLECULE: | LYSOSOMAL T |
| 79:00:00 | 6rwt-A | 2.2 | 5.4 | 71 | 181 | 7 | MOLECULE: | UBIQUINOL-C |
| 80:00:00 | 8r8q-A | 2.2 | 10.1 | 80 | 391 | 6 | MOLECULE: | MAJOR FACIL |
| 81:00:00 | 9b8q-J | 2.2 | 11.5 | 66 | 149 | 0 | MOLECULE: | V-TYPE PROT |
| 82:00:00 | 8th8-P | 2.2 | 8.1 | 96 | 305 | 5 | MOLECULE: | DYNEIN REGU |
| 83:00:00 | 1kx5-A | 2.2 | 14 | 73 | 135 | 10 | MOLECULE: | DNA |
| 84:00:00 | 7uuy-A | 2.2 | 4.6 | 89 | 503 | 4 | MOLECULE: | SODIUM/IODI |
| 85:00:00 | 5mps-R | 2.2 | 6.9 | 56 | 108 | 7 | MOLECULE: | YEAST UBC4 |
| 86:00:00 | 6mdw-A | 2.2 | 14.8 | 80 | 185 | 3 | MOLECULE: | SPRT-LIKE D |

| | | | | | | | | |
|---|---|---|---|---|---|---|---|---|
| 87:00:00 | 5x55-A | 2.2 | 7.8 | 66 | 276 | 6 | MOLECULE: | PROBABLE UR |
| 88:00:00 | 6btm-D | 2.2 | 9.4 | 63 | 172 | 5 | MOLECULE: | ALTERNATIVE |
| 89:00:00 | 5e8b-A | 2.2 | 4.3 | 40 | 54 | 3 | MOLECULE: | RNA POLYMER |
| 90:00:00 | 3edv-A | 2.2 | 24.1 | 86 | 322 | 9 | MOLECULE: | SPECTRIN BE |
| 91:00:00 | 5w0p-C | 2.2 | 24 | 117 | 850 | 3 | MOLECULE: | ENDOLYSIN,R |
| 92:00:00 | 7te2-A | 2.2 | 9.4 | 62 | 205 | 10 | MOLECULE: | AERR |
| 93:00:00 | 2n72-A | 2.2 | 6.8 | 53 | 69 | 6 | MOLECULE: | GOLGI RESID |
| 94:00:00 | 7d6v-C | 2.2 | 4.8 | 57 | 264 | 5 | MOLECULE: | SUCCINATE D |
| 95:00:00 | 2lk2-A | 2.2 | 4.3 | 59 | 89 | 5 | MOLECULE: | HOMEOBOX PR |
| 96:00:00 | 7qy5-B | 2.2 | 10.7 | 67 | 320 | 3 | MOLECULE: | NURS COMPLE |
| 97:00:00 | 1ukf-A | 2.2 | 4.8 | 61 | 188 | 13 | MOLECULE: | AVIRULENCE |
| 98:00:00 | 3hj1-B | 2.2 | 16.7 | 71 | 387 | 11 | MOLECULE: | MINOR EDITO |
| 99:00:00 | 7zge-A | 2.2 | 12.6 | 67 | 199 | 9 | MOLECULE: | BRXA, A BRE |
| 0:00 | 6xf1-A | 2.2 | 23.2 | 88 | 230 | 9 | MOLECULE: | NESPRIN-2 |
| 1:00 | 7dtl-A | 2.2 | 6.8 | 53 | 61 | 6 | MOLECULE: | PSK |
| 2:00 | 8iqg-A | 2.2 | 14 | 76 | 213 | 11 | MOLECULE: | CHROMATIN A |
| 3:00 | 5mbv-A | 2.2 | 11.7 | 66 | 98 | 3 | MOLECULE: | RECBCD ENZY |
| 4:00 | 7wmv-A | 2.2 | 18.9 | 104 | 602 | 6 | MOLECULE: | SODIUM/GLUC |
| 5:00 | 4psn-C | 2.2 | 20.2 | 62 | 221 | 8 | MOLECULE: | SSDNA BINDI |
| 6:00 | 8teu-E | 2.2 | 34.4 | 73 | 495 | 10 | MOLECULE: | LARGE TEGUM |
| 7:00 | 2rq5-A | 2.2 | 5 | 75 | 121 | 13 | MOLECULE: | PROTEIN JUM |
| 8:00 | 8sl4-A | 2.2 | 28.4 | 89 | 970 | 7 | MOLECULE: | ADENYLATE C |
| 9:00 | 4abn-B | 2.2 | 18 | 89 | 426 | 7 | MOLECULE: | TETRATRICOP |
| 10:00 | 6y4b-A | 2.2 | 10.9 | 69 | 255 | 6 | MOLECULE: | RNA (77-MER |
| 11:00 | 7bin-A | 2.2 | 9 | 71 | 209 | 1 | MOLECULE: | FLAGELLAR B |
| 12:00 | 5htf-B | 2.2 | 5.3 | 45 | 260 | 13 | MOLECULE: | FOLDASE PRO |
| 13:00 | 8hnp-A | 2.2 | 6.9 | 62 | 141 | 10 | MOLECULE: | ARCHAEAL TR |
| 14:00 | 8ibw-C | 2.2 | 12.3 | 86 | 922 | 3 | MOLECULE: | DNA (60-MER |
| 15:00 | 6n04-B | 2.2 | 4.9 | 69 | 366 | 9 | MOLECULE: | ABSH3 |
| 16:00 | 8eki-A | 2.2 | 12.2 | 84 | 170 | 4 | MOLECULE: | PROTEIN TRA |
| 17:00 | 7kyp-F | 2.2 | 12.5 | 89 | 290 | 8 | MOLECULE: | MANGANESE A |
| 18:00 | 5yti-A | 2.2 | 5.4 | 42 | 330 | 2 | MOLECULE: | FLAGELLAR H |
| 19:00 | 2bf0-X | 2.2 | 4 | 67 | 131 | 12 | MOLECULE: | PCF11 |
| 20:00 | 4ily-B | 2.2 | 4.7 | 71 | 250 | 8 | MOLECULE: | CHITOSANASE |
| 21:00 | 3lph-A | 2.2 | 4.4 | 42 | 62 | 2 | MOLECULE: | PROTEIN REV |
| 22:00 | 7dm1-A | 2.2 | 3.3 | 66 | 334 | 6 | MOLECULE: | PHOSPHATE-B |
| 23:00 | 3wee-A | 2.2 | 23.1 | 81 | 428 | 5 | MOLECULE: | ACTIN-LIKE |
| 24:00:00 | 6iuq-B | 2.2 | 15 | 77 | 296 | 6 | MOLECULE: | PROLYL 4-HY |
| 25:00:00 | 6nd4-I | 2.2 | 29.8 | 85 | 487 | 5 | MOLECULE: | ETS RRNA |
| 26:00:00 | 1d2t-A | 2.2 | 5.2 | 89 | 222 | 8 | MOLECULE: | ACID PHOSPH |
| 27:00:00 | 8ppr-M | 2.2 | 15.2 | 63 | 205 | 5 | MOLECULE: | KINETOCHORE |
| 28:00:00 | 6tdw-C | 2.2 | 10.3 | 70 | 157 | 11 | MOLECULE: | ATPTB1 |
| 29:00:00 | 3veb-B | 2.2 | 13 | 58 | 150 | 16 | MOLECULE: | MACRODOMAIN |
| 30:00:00 | 2np5-A | 2.2 | 13.3 | 69 | 187 | 10 | MOLECULE: | TRANSCRIPTI |

| | | | | | | | | |
|---|---|---|---|---|---|---|---|---|
| 31:00:00 | 7vxs-m | 2.2 | 5.2 | 57 | 175 | 4 | MOLECULE: | NADH DEHYDR |
| 32:00:00 | 8p23-B | 2.2 | 12.1 | 110 | 715 | 6 | MOLECULE: | ANAEROBIC R |
| 33:00:00 | 1hs7-A | 2.2 | 5.3 | 58 | 97 | 5 | MOLECULE: | SYNTAXIN VA |
| 34:00:00 | 5mu4-A | 2.2 | 4.9 | 68 | 196 | 4 | MOLECULE: | TAIL TUBULA |
| 35:00:00 | 5ujc-A | 2.2 | 8 | 88 | 267 | 8 | MOLECULE: | MMACHC-LIKE |
| 36:00:00 | 6v4m-A | 2.2 | 4.6 | 58 | 163 | 7 | MOLECULE: | BCL-2 |
| 37:00:00 | 4okv-F | 2.2 | 3.1 | 43 | 66 | 12 | MOLECULE: | HEAVY CHAIN |
| 38:00:00 | 8w5z-A | 2.2 | 13.8 | 77 | 333 | 1 | MOLECULE: | PROTEIN-TYR |
| 39:00:00 | 1qhh-C | 2.2 | 10.9 | 57 | 102 | 7 | MOLECULE: | PROTEIN (PC |
| 40:00:00 | 1dj3-A | 2.2 | 13.2 | 78 | 432 | 6 | MOLECULE: | ADENYLOSUCC |
| 41:00:00 | 7kpx-C | 2.2 | 12.6 | 103 | 1352 | 3 | MOLECULE: | MEIOTIC MRN |
| 42:00:00 | 3c1d-B | 2.2 | 14.8 | 69 | 154 | 12 | MOLECULE: | REGULATORY |
| 43:00:00 | 5i0n-A | 2.2 | 10.8 | 86 | 489 | 13 | MOLECULE: | PHOSPHATIDY |
| 44:00:00 | 7qqe-A | 2.2 | 3.3 | 66 | 163 | 3 | MOLECULE: | NUCLEAR FAC |
| 45:00:00 | 6dkm-D | 2.2 | 9.5 | 59 | 79 | 3 | MOLECULE: | DHD131_A |
| 46:00:00 | 5gpa-A | 2.2 | 13.9 | 72 | 189 | 8 | MOLECULE: | TRANSCRIPTI |
| 47:00:00 | 5fqf-A | 2.2 | 5.5 | 48 | 583 | 2 | MOLECULE: | BETA-N-ACET |
| 48:00:00 | 5v8z-C | 2.2 | 8 | 65 | 98 | 11 | MOLECULE: | ENDOPLASMIC |
| 49:00:00 | 7alw-A | 2.2 | 15.6 | 71 | 344 | 4 | MOLECULE: | LOW CALCIUM |
| 50:00:00 | 2es4-E | 2.2 | 15.8 | 95 | 278 | 6 | MOLECULE: | LIPASE |
| 51:00:00 | 3bfm-A | 2.2 | 3.4 | 60 | 230 | 8 | MOLECULE: | BIOTIN PROT |
| 52:00:00 | 3d9d-A | 2.2 | 9.8 | 102 | 432 | 10 | MOLECULE: | NITROALKANE |
| 53:00:00 | 7amv-K | 2.2 | 19.5 | 80 | 708 | 8 | MOLECULE: | DNA-DIRECTE |
| 54:00:00 | 8jgw-A | 2.2 | 18.2 | 76 | 500 | 7 | MOLECULE: | EXOPOLYPHOS |
| 55:00:00 | 1nl3-A | 2.2 | 20.9 | 102 | 839 | 4 | MOLECULE: | PREPROTEIN |
| 56:00:00 | 5oqq-B | 2.2 | 24 | 111 | 824 | 3 | MOLECULE: | CONDENSIN C |
| 57:00:00 | 5j10-A | 2.2 | 9.3 | 53 | 70 | 11 | MOLECULE: | PEPTIDE DES |
| 58:00:00 | 2n8p-A | 2.2 | 7 | 41 | 53 | 15 | MOLECULE: | LACTICIN Q |
| 59:00:00 | 3jb9-r | 2.2 | 23.3 | 74 | 573 | 3 | MOLECULE: | PRE-MRNA-SP |
| 60:00:00 | 3g3z-A | 2.2 | 13.6 | 76 | 142 | 5 | MOLECULE: | TRANSCRIPTI |
| 61:00:00 | 7rsl-A | 2.2 | 3.3 | 46 | 235 | 9 | MOLECULE: | SEIPIN |
| 62:00:00 | 4hw5-A | 2.2 | 3.3 | 39 | 77 | 8 | MOLECULE: | COMPLEMENT |
| 63:00:00 | 5ndd-A | 2.2 | 9.1 | 103 | 567 | 9 | MOLECULE: | LYSOZYME,PR |
| 64:00:00 | 2g49-A | 2.2 | 17.3 | 108 | 970 | 7 | MOLECULE: | INSULIN-DEG |
| 65:00:00 | 5j2l-A | 2.2 | 9.3 | 59 | 76 | 7 | MOLECULE: | PROTEIN DES |
| 66:00:00 | 7upo-A | 2.2 | 8.2 | 57 | 74 | 2 | MOLECULE: | DHT03 PROTE |
| 67:00:00 | 6zyx-C | 2.2 | 23.4 | 103 | 278 | 5 | MOLECULE: | DYNEIN HEAV |
| 68:00:00 | 8ssk-D | 2.2 | 11.1 | 61 | 306 | 5 | MOLECULE: | CONTACT-DEP |
| 69:00:00 | 3txs-C | 2.2 | 7.9 | 53 | 91 | 4 | MOLECULE: | TERMINASE D |
| 70:00:00 | 6s8g-G | 2.2 | 9.3 | 92 | 244 | 14 | MOLECULE: | LIPOPOLYSAC |
| 71:00:00 | 3tek-A | 2.2 | 2.8 | 50 | 139 | 8 | MOLECULE: | THERMODBP-S |
| 72:00:00 | 2hi3-A | 2.2 | 4.1 | 59 | 73 | 2 | MOLECULE: | HOMEODOMAIN |
| 73:00:00 | 8bmw-E | 2.2 | 3.2 | 57 | 122 | 11 | MOLECULE: | CRISPR-ASSO |
| 74:00:00 | 7msk-A | 2.2 | 21.9 | 107 | 421 | 8 | MOLECULE: | GLYCO_TRANS |

| | | | | | | | | |
|---|---|---|---|---|---|---|---|---|
| 75:00:00 | 2gmf-A | 2.2 | 4.6 | 72 | 121 | 4 | MOLECULE: | GRANULOCYTE |
| 76:00:00 | 6zfb-d | 2.2 | 17 | 78 | 139 | 3 | MOLECULE: | DNA-DIRECTE |
| 77:00:00 | 7cth-E | 2.2 | 5.1 | 70 | 75 | 6 | MOLECULE: | CORE PROTEI |
| 78:00:00 | 6o0y-A | 2.2 | 6.6 | 105 | 1146 | 3 | MOLECULE: | CRISPR-ASSO |
| 79:00:00 | 6nvz-A | 2.2 | 2.7 | 47 | 60 | 6 | MOLECULE: | MYB FAMILY |
| 80:00:00 | 5nug-A | 2.2 | 9.6 | 92 | 2920 | 5 | MOLECULE: | CYTOPLASMIC |
| 81:00:00 | 3ajb-A | 2.2 | 6.4 | 82 | 293 | 6 | MOLECULE: | PEROXISOMAL |
| 82:00:00 | 8rd2-B | 2.2 | 3.7 | 59 | 400 | 8 | MOLECULE: | INVARIANT S |
| 83:00:00 | 8bb6-A | 2.2 | 6.3 | 69 | 476 | 4 | MOLECULE: | SUCROSE TRA |
| 84:00:00 | 7unk-A | 2.2 | 27.5 | 150 | 1047 | 7 | MOLECULE: | IMPORTIN-4 |
| 85:00:00 | 5a2v-B | 2.2 | 11.1 | 69 | 472 | 4 | MOLECULE: | MITOCHONDRI |
| 86:00:00 | 7qpg-B | 2.2 | 21.7 | 111 | 591 | 5 | MOLECULE: | PROTEIN ZWI |
| 87:00:00 | 8e1u-A | 2.2 | 7.9 | 85 | 1126 | 6 | MOLECULE: | PPI-DEPENDE |
| 88:00:00 | 7yxx-A | 2.2 | 27.1 | 125 | 1754 | 6 | MOLECULE: | PROBABLE UB |
| 89:00:00 | 7nj0-C | 2.2 | 13 | 69 | 270 | 3 | MOLECULE: | SECURIN,SEP |
| 90:00:00 | 3llw-D | 2.2 | 8.6 | 73 | 289 | 3 | MOLECULE: | GERANYLTRAN |
| 91:00:00 | 2ewg-A | 2.2 | 13.8 | 90 | 367 | 8 | MOLECULE: | FARNESYL PY |
| 92:00:00 | 3pvl-A | 2.2 | 26.4 | 89 | 597 | 10 | MOLECULE: | MYOSIN VIIA |
| 93:00:00 | 8h2h-D | 2.2 | 14.2 | 78 | 599 | 1 | MOLECULE: | LTRB |
| 94:00:00 | 7xn2-A | 2.2 | 3.3 | 69 | 148 | 7 | MOLECULE: | ALR3614 PRO |
| 95:00:00 | 2r44-A | 2.2 | 7.1 | 67 | 330 | 9 | MOLECULE: | UNCHARACTER |
| 96:00:00 | 7qba-D | 2.2 | 4.7 | 85 | 275 | 5 | MOLECULE: | PROBABLE AB |
| 97:00:00 | 8umd-B | 2.2 | 13.8 | 68 | 331 | 10 | MOLECULE: | FLAGELLAR M |
| 98:00:00 | 8d8l-8 | 2.2 | 5.8 | 86 | 452 | 2 | MOLECULE: | PROBABLE S- |
| 99:00:00 | 1l1l-A | 2.2 | 16.1 | 84 | 717 | 7 | MOLECULE: | RIBONUCLEOS |
| 0:00 | 8jo3-A | 2.2 | 7.2 | 85 | 350 | 5 | MOLECULE: | SUBSTRATE O |
| 1:00 | 3bve-A | 2.2 | 8.2 | 81 | 173 | 5 | MOLECULE: | FERRITIN |
| 2:00 | 3lpz-A | 2.2 | 8.8 | 95 | 314 | 3 | MOLECULE: | UNCHARACTER |
| 3:00 | 6p8v-G | 2.2 | 7.3 | 65 | 320 | 5 | MOLECULE: | ATPASE, AAA |
| 4:00 | 4w8j-A | 2.2 | 20.7 | 100 | 1017 | 8 | MOLECULE: | PESTICIDAL |
| 5:00 | 2rbd-A | 2.2 | 10.4 | 83 | 159 | 2 | MOLECULE: | BH2358 PROT |
| 6:00 | 2qkm-B | 2.2 | 6.1 | 84 | 266 | 4 | MOLECULE: | SPBC3B9.21 |
| 7:00 | 5z7c-A | 2.2 | 11.7 | 100 | 414 | 6 | MOLECULE: | 3'3'-CGAMP- |
| 8:00 | 2mn2-A | 2.2 | 9.3 | 77 | 122 | 4 | MOLECULE: | YMOB |
| 9:00 | 7ux8-A | 2.2 | 11.3 | 83 | 380 | 11 | MOLECULE: | MFNG |
| 10:00 | 4v19-2 | 2.2 | 13.8 | 89 | 178 | 6 | MOLECULE: | MITORIBOSOM |
| 11:00 | 2k3q-A | 2.2 | 4 | 68 | 118 | 12 | MOLECULE: | TUSP1 |
| 12:00 | 2nn4-A | 2.2 | 3.5 | 53 | 62 | 9 | MOLECULE: | HYPOTHETICA |
| 13:00 | 4i9e-A | 2.2 | 18.3 | 100 | 383 | 6 | MOLECULE: | RESPONSE RE |
| 14:00 | 3o77-A | 2.2 | 3.2 | 62 | 388 | 3 | MOLECULE: | MYOSIN LIGH |
| 15:00 | 3l9f-A | 2.2 | 6.6 | 67 | 170 | 4 | MOLECULE: | PUTATIVE UN |
| 16:00 | 6vrc-A | 2.2 | 12.4 | 103 | 1007 | 10 | MOLECULE: | CRISPR-ASSO |
| 17:00 | 4kis-A | 2.2 | 7.1 | 67 | 315 | 6 | MOLECULE: | PUTATIVE IN |
| 18:00 | 4uxv-A | 2.2 | 13.8 | 92 | 522 | 8 | MOLECULE: | SEPTATION R |

| | | | | | | | | |
|---|---|---|---|---|---|---|---|---|
| 19:00 | 5b4s-B | 2.2 | 5.1 | 75 | 310 | 3 | MOLECULE: | CHITOSANASE |
| 20:00 | 5lxf-B | 2.2 | 5.3 | 83 | 149 | 6 | MOLECULE: | MACROPHAGE |
| 21:00 | 2l6f-A | 2.2 | 8.6 | 92 | 176 | 5 | MOLECULE: | FOCAL ADHES |
| 22:00 | 7yzb-A | 2.2 | 3.6 | 56 | 125 | 7 | MOLECULE: | FORKHEAD BO |
| 23:00 | 7nyw-C | 2.2 | 11.2 | 68 | 335 | 10 | MOLECULE: | CHROMOSOME |
| 24:00:00 | 2hi4-A | 2.2 | 5 | 123 | 480 | 5 | MOLECULE: | CYTOCHROME |
| 25:00:00 | 2kmf-A | 2.2 | 9.1 | 68 | 115 | 4 | MOLECULE: | PHOTOSYSTEM |
| 26:00:00 | 2hxo-A | 2.2 | 5.2 | 57 | 207 | 11 | MOLECULE: | PUTATIVE TE |
| 27:00:00 | 6hgc-A | 2.2 | 12.6 | 81 | 304 | 6 | MOLECULE: | UBIQUITIN C |
| 28:00:00 | 6nmi-B | 2.2 | 9.3 | 101 | 760 | 8 | MOLECULE: | GENERAL TRA |
| 29:00:00 | 1mwk-A | 2.2 | 10.2 | 88 | 320 | 3 | MOLECULE: | PARM |
| 30:00:00 | 3ljl-A | 2.2 | 7.4 | 59 | 149 | 7 | MOLECULE: | TRANSCRIPTI |
| 31:00:00 | 5awf-A | 2.2 | 5.9 | 70 | 385 | 0 | MOLECULE: | FES CLUSTER |
| 32:00:00 | 7w7p-D | 2.2 | 17.7 | 96 | 284 | 6 | MOLECULE: | DNA HELICAS |
| 33:00:00 | 5vyl-A | 2.2 | 16.2 | 92 | 502 | 5 | MOLECULE: | INNER TEGUM |
| 34:00:00 | 6i3m-E | 2.2 | 7.3 | 88 | 354 | 8 | MOLECULE: | TRANSLATION |
| 35:00:00 | 2fdc-A | 2.2 | 17.4 | 89 | 505 | 4 | MOLECULE: | 5'-D(P*CP*G |
| 36:00:00 | 4cz8-A | 2.2 | 16.8 | 122 | 422 | 6 | MOLECULE: | NA+/H+ ANTI |
| 37:00:00 | 3fmt-A | 2.2 | 4.5 | 72 | 162 | 7 | MOLECULE: | PROTEIN SEQ |
| 38:00:00 | 7wvz-A | 2.2 | 12.9 | 110 | 1719 | 11 | MOLECULE: | BETA-KETOAC |
| 39:00:00 | 8hra-C | 2.2 | 6.9 | 76 | 852 | 12 | MOLECULE: | ARCHAEAL AT |
| 40:00:00 | 7etm-A | 2.2 | 22.4 | 92 | 498 | 5 | MOLECULE: | PORTAL PROT |
| 41:00:00 | 3h3m-A | 2.2 | 6.8 | 70 | 89 | 4 | MOLECULE: | FLAGELLAR P |
| 42:00:00 | 2js1-A | 2.2 | 4.3 | 63 | 80 | 8 | MOLECULE: | UNCHARACTER |
| 43:00:00 | 8q9t-E | 2.2 | 16.5 | 82 | 1060 | 7 | MOLECULE: | ANTIVIRAL H |
| 44:00:00 | 4bm5-A | 2.2 | 14.5 | 94 | 268 | 6 | MOLECULE: | SIMILAR TO |
| 45:00:00 | 6s6b-K | 2.2 | 13.9 | 102 | 1024 | 2 | MOLECULE: | CRISPR-ASSO |
| 46:00:00 | 4buj-A | 2.2 | 24 | 85 | 903 | 11 | MOLECULE: | SUPERKILLER |
| 47:00:00 | 1sc7-A | 2.2 | 7.2 | 88 | 567 | 5 | MOLECULE: | 5'-D(*AP*AP |
| 48:00:00 | 8thc-A | 2.2 | 4.4 | 55 | 479 | 9 | MOLECULE: | ELG1 ISOFOR |
| 49:00:00 | 8kde-D | 2.2 | 17.1 | 110 | 351 | 5 | MOLECULE: | PHOTOSYSTEM |
| 50:00:00 | 6ef3-A | 2.2 | 8.3 | 84 | 247 | 6 | MOLECULE: | PROTEASOME |
| 51:00:00 | 7vt4-A | 2.2 | 9.1 | 82 | 438 | 9 | MOLECULE: | ENDOGLUCANA |
| 52:00:00 | 5aqd-M | 2.2 | 7 | 83 | 184 | 6 | MOLECULE: | PHYCOERYTHR |
| 53:00:00 | 4otn-A | 2.2 | 11.9 | 54 | 123 | 11 | MOLECULE: | EUKARYOTIC |
| 54:00:00 | 8iwh-q | 2.2 | 8.1 | 58 | 156 | 9 | MOLECULE: | PHOTOSYSTEM |
| 55:00:00 | 7mi8-A | 2.2 | 12 | 97 | 868 | 5 | MOLECULE: | FUSION PROT |
| 56:00:00 | 8wr4-B | 2.2 | 31.2 | 103 | 1442 | 5 | MOLECULE: | CBCAS9 EFFE |
| 57:00:00 | 4ng2-E | 2.2 | 8.9 | 62 | 94 | 6 | MOLECULE: | TRANSCRIPTI |
| 58:00:00 | 5aor-A | 2.2 | 7.1 | 112 | 1009 | 6 | MOLECULE: | DOSAGE COMP |
| 59:00:00 | 5utg-A | 2.2 | 5.2 | 73 | 134 | 3 | MOLECULE: | EGG-LYSIN |
| 60:00:00 | 6jc4-A | 2.2 | 3.9 | 68 | 200 | 6 | MOLECULE: | UREASE ACCE |
| 61:00:00 | 7c6g-A | 2.2 | 3.8 | 55 | 420 | 11 | MOLECULE: | SUGAR ABC T |
| 62:00:00 | 6vqx-A | 2.2 | 3.6 | 65 | 97 | 11 | MOLECULE: | ACRF6 |

| | | | | | | | | |
|---|---|---|---|---|---|---|---|---|
| 63:00:00 | 6xpd-A | 2.2 | 18.6 | 106 | 303 | 5 | MOLECULE: | ZINC TRANSP |
| 64:00:00 | 2ia0-A | 2.2 | 20 | 59 | 156 | 8 | MOLECULE: | PUTATIVE HT |
| 65:00:00 | 6m9t-A | 2.2 | 12.2 | 77 | 444 | 6 | MOLECULE: | PROSTAGLAND |
| 66:00:00 | 8hrc-A | 2.2 | 21.8 | 102 | 776 | 6 | MOLECULE: | ADENOSINE D |
| 67:00:00 | 4hh0-A | 2.2 | 14.8 | 89 | 383 | 7 | MOLECULE: | APPA PROTEI |
| 68:00:00 | 7z12-A | 2.2 | 5.1 | 122 | 1853 | 10 | MOLECULE: | PAM1.4, HEA |
| 69:00:00 | 5jpw-A | 2.2 | 6.1 | 78 | 143 | 8 | MOLECULE: | DYNEIN LIGH |
| 70:00:00 | 8p0v-M | 2.2 | 14.7 | 92 | 357 | 5 | MOLECULE: | COILED-COIL |
| 71:00:00 | 5w5y-O | 2.2 | 10.4 | 102 | 640 | 7 | MOLECULE: | DNA-DIRECTE |
| 72:00:00 | 8hhe-B | 2.2 | 8.4 | 103 | 647 | 7 | MOLECULE: | CRYSTALINE |
| 73:00:00 | 6e6y-A | 2.2 | 9.9 | 98 | 268 | 8 | MOLECULE: | DIECKMANN C |
| 74:00:00 | 7wiu-B | 2.2 | 18.2 | 95 | 576 | 7 | MOLECULE: | MYCOBACTIN |
| 75:00:00 | 5b2o-A | 2.2 | 20 | 123 | 1455 | 7 | MOLECULE: | CRISPR-ASSO |
| 76:00:00 | 3cra-A | 2.2 | 7.6 | 94 | 239 | 12 | MOLECULE: | PROTEIN MAZ |
| 77:00:00 | 5kkl-B | 2.2 | 24.8 | 107 | 838 | 3 | MOLECULE: | PUTATIVE PO |
| 78:00:00 | 8j98-A | 2.2 | 4.1 | 75 | 184 | 7 | MOLECULE: | NEAR-INFRAR |
| 79:00:00 | 2wzk-A | 2.2 | 6 | 80 | 369 | 9 | MOLECULE: | CULLIN-5 |
| 80:00:00 | 5lq4-B | 2.2 | 22.2 | 89 | 465 | 10 | MOLECULE: | CYAGOX |
| 81:00:00 | 5oqj-1 | 2.2 | 28.2 | 99 | 491 | 6 | MOLECULE: | DNA-DIRECTE |
| 82:00:00 | 6upn-A | 2.2 | 6.7 | 78 | 245 | 4 | MOLECULE: | ENDOPHILIN- |
| 83:00:00 | 7zcv-B | 2.2 | 14.5 | 73 | 285 | 3 | MOLECULE: | TRANSCRIPTI |
| 84:00:00 | 8pda-A | 2.2 | 7 | 94 | 973 | 3 | MOLECULE: | ERAD-ASSOCI |
| 85:00:00 | 7sfj-A | 2.2 | 8.8 | 92 | 284 | 9 | MOLECULE: | CHRMINE |
| 86:00:00 | 7fbo-A | 2.2 | 10.3 | 82 | 283 | 9 | MOLECULE: | BEZA |
| 87:00:00 | 2q83-A | 2.2 | 6.8 | 65 | 332 | 6 | MOLECULE: | YTAA PROTEI |
| 88:00:00 | 1cf7-B | 2.2 | 4 | 69 | 82 | 4 | MOLECULE: | DNA (5'- |
| 89:00:00 | 7emf-G | 2.2 | 20.4 | 87 | 161 | 6 | MOLECULE: | MEDIATOR OF |
| 90:00:00 | 5lo9-A | 2.2 | 3.7 | 81 | 471 | 4 | MOLECULE: | CYTOCHROME |
| 91:00:00 | 6b85-A | 2.2 | 21.8 | 104 | 215 | 8 | MOLECULE: | TMHC4_R |
| 92:00:00 | 8p0w-J | 2.2 | 17.1 | 68 | 202 | 12 | MOLECULE: | COMM DOMAIN |
| 93:00:00 | 7jw3-C | 2.2 | 10.1 | 94 | 244 | 9 | MOLECULE: | EXONUCLEASE |
| 94:00:00 | 4lev-A | 2.2 | 4.2 | 72 | 369 | 4 | MOLECULE: | CYCLIC GMP- |
| 95:00:00 | 6lg3-A | 2.2 | 3.2 | 57 | 74 | 7 | MOLECULE: | PHIRV1 PHAG |
| 96:00:00 | 6wfq-D | 2.2 | 8.2 | 72 | 219 | 1 | MOLECULE: | HTH-TYPE TR |
| 97:00:00 | 8f2r-D | 2.1 | 17 | 60 | 199 | 7 | MOLECULE: | COMM DOMAIN |
| 98:00:00 | 6twr-A | 2.1 | 7.6 | 74 | 169 | 9 | MOLECULE: | BETA-GLUCOS |
| 99:00:00 | 8utk-A | 2.1 | 3.8 | 37 | 55 | 14 | MOLECULE: | 23R-B04DSLF |
| 0:00 | 1kr7-A | 2.1 | 4.2 | 61 | 110 | 13 | MOLECULE: | NEURAL GLOB |
| 1:00 | 6m3q-F | 2.1 | 20.6 | 108 | 301 | 10 | MOLECULE: | ANKYRIN-2 |
| 2:00 | 6lk8-a | 2.1 | 24.8 | 120 | 1272 | 8 | MOLECULE: | MGC83295 PR |
| 3:00 | 6s3f-A | 2.1 | 2.9 | 49 | 65 | 6 | MOLECULE: | 2S ALBUMIN |
| 4:00 | 4ryk-A | 2.1 | 15.7 | 71 | 295 | 3 | MOLECULE: | LMO0325 PRO |
| 5:00 | 4rvq-A | 2.1 | 5.7 | 69 | 122 | 14 | MOLECULE: | PRE-MRNA SP |
| 6:00 | 8a2n-B | 2.1 | 4.3 | 82 | 299 | 9 | MOLECULE: | CGND |

| | | | | | | | | |
|---|---|---|---|---|---|---|---|---|
| 7:00 | 3qnk-C | 2.1 | 25.3 | 84 | 511 | 6 | MOLECULE: | PUTATIVE LI |
| 8:00 | 6f2d-F | 2.1 | 14.9 | 65 | 258 | 2 | MOLECULE: | FLAGELLAR B |
| 9:00 | 7yul-A | 2.1 | 3.6 | 55 | 92 | 5 | MOLECULE: | BEN DOMAIN- |
| 10:00 | 6gmh-Q | 2.1 | 29.4 | 84 | 884 | 10 | MOLECULE: | RPB1 |
| 11:00 | 8d8j-5 | 2.1 | 7.7 | 76 | 270 | 7 | MOLECULE: | PROBABLE S- |
| 12:00 | 1gnc-A | 2.1 | 15.5 | 63 | 175 | 14 | MOLECULE: | GRANULOCYTE |
| 13:00 | 7nyw-B | 2.1 | 32.7 | 92 | 858 | 4 | MOLECULE: | CHROMOSOME |
| 14:00 | 6zbj-B | 2.1 | 26 | 124 | 501 | 2 | MOLECULE: | PRECURSOR O |
| 15:00 | 5lb7-B | 2.1 | 5.8 | 53 | 78 | 6 | MOLECULE: | KATANIN P80 |
| 16:00 | 8oix-b | 2.1 | 3.9 | 62 | 208 | 5 | MOLECULE: | FAMILY T1, |
| 17:00 | 7uqj-A | 2.1 | 10 | 62 | 585 | 6 | MOLECULE: | ATPASE HIST |
| 18:00 | 7v2c-p | 2.1 | 6.5 | 76 | 178 | 3 | MOLECULE: | NADH DEHYDR |
| 19:00 | 5fia-B | 2.1 | 4.9 | 79 | 397 | 8 | MOLECULE: | LPIR1 |
| 20:00 | 8th8-A | 2.1 | 16.4 | 71 | 290 | 10 | MOLECULE: | DYNEIN REGU |
| 21:00 | 6efn-A | 2.1 | 9.9 | 62 | 384 | 8 | MOLECULE: | SPORULATION |
| 22:00 | 8bw9-A | 2.1 | 3.6 | 54 | 88 | 4 | MOLECULE: | PROTEIN AVE |
| 23:00 | 1qoy-A | 2.1 | 7 | 77 | 303 | 5 | MOLECULE: | HEMOLYSIN E |
| 24:00:00 | 1rso-B | 2.1 | 3.1 | 40 | 56 | 3 | MOLECULE: | PRESYNAPTIC |
| 25:00:00 | 4k6j-A | 2.1 | 15.8 | 85 | 497 | 12 | MOLECULE: | WINGS APART |
| 26:00:00 | 7jqq-A | 2.1 | 12.4 | 59 | 327 | 5 | MOLECULE: | DNA PACKAGI |
| 27:00:00 | 5mpd-Z | 2.1 | 12.5 | 82 | 906 | 9 | MOLECULE: | 26S PROTEAS |
| 28:00:00 | 1nkl-A | 2.1 | 6.8 | 42 | 78 | 5 | MOLECULE: | NK-LYSIN |
| 29:00:00 | 8jar-B | 2.1 | 20 | 101 | 573 | 4 | MOLECULE: | AMYLOID PRO |
| 30:00:00 | 3mse-B | 2.1 | 7 | 56 | 168 | 2 | MOLECULE: | CALCIUM-DEP |
| 31:00:00 | 7qru-E | 2.1 | 7 | 58 | 158 | 14 | MOLECULE: | NA+/H+ ANTI |
| 32:00:00 | 7but-A | 2.1 | 7.7 | 59 | 142 | 7 | MOLECULE: | ACINIFORM S |
| 33:00:00 | 4z7k-A | 2.1 | 5 | 82 | 218 | 10 | MOLECULE: | UNCHARACTER |
| 34:00:00 | 3vej-B | 2.1 | 2.7 | 37 | 41 | 3 | MOLECULE: | UBIQUITIN-L |
| 35:00:00 | 3jcm-I | 2.1 | 21.6 | 92 | 416 | 7 | MOLECULE: | PRE-MRNA-SP |
| 36:00:00 | 1xnx-A | 2.1 | 4.5 | 78 | 232 | 14 | MOLECULE: | CONSTITUTIV |
| 37:00:00 | 7p37-A | 2.1 | 9.7 | 63 | 147 | 8 | MOLECULE: | TRANSCRIPTI |
| 38:00:00 | 7p2y-J | 2.1 | 9.5 | 54 | 80 | 13 | MOLECULE: | ATP SYNTHAS |
| 39:00:00 | 6f0x-B | 2.1 | 3.5 | 74 | 398 | 9 | MOLECULE: | PACHYTENE C |
| 40:00:00 | 8rz0-E | 2.1 | 5.6 | 63 | 113 | 3 | MOLECULE: | RH5-34EM |
| 41:00:00 | 7eaw-A | 2.1 | 15 | 83 | 560 | 10 | MOLECULE: | TREHALASE |
| 42:00:00 | 7ar9-d | 2.1 | 5.7 | 51 | 80 | 4 | MOLECULE: | ND3 |
| 43:00:00 | 7dkk-A | 2.1 | 2.4 | 42 | 89 | 5 | MOLECULE: | DE NOVO DES |
| 44:00:00 | 8osp-A | 2.1 | 6.9 | 62 | 174 | 6 | MOLECULE: | N-ACETYLTRA |
| 45:00:00 | 6k61-I | 2.1 | 5.1 | 32 | 34 | 3 | MOLECULE: | PHOTOSYSTEM |
| 46:00:00 | 4gd3-A | 2.1 | 12.9 | 67 | 179 | 4 | MOLECULE: | HYDROGENASE |
| 47:00:00 | 8ixk-E | 2.1 | 21.7 | 65 | 109 | 5 | MOLECULE: | ATTACHMENT |
| 48:00:00 | 7yov-E | 2.1 | 10.5 | 49 | 158 | 10 | MOLECULE: | NDV P PROTE |
| 49:00:00 | 3rh2-A | 2.1 | 11.7 | 78 | 210 | 4 | MOLECULE: | HYPOTHETICA |
| 50:00:00 | 4z8u-A | 2.1 | 13.6 | 112 | 332 | 4 | MOLECULE: | AVRRXO1-ORF |

| | | | | | | | | |
|---|---|---|---|---|---|---|---|---|
| 51:00:00 | 7et0-C | 2.1 | 16.9 | 113 | 684 | 3 | MOLECULE: | BACTERIA FA |
| 52:00:00 | 1xsx-A | 2.1 | 3.6 | 59 | 95 | 8 | MOLECULE: | SSO10A |
| 53:00:00 | 1g4a-E | 2.1 | 16 | 74 | 356 | 7 | MOLECULE: | ATP-DEPENDE |
| 54:00:00 | 7x5e-E | 2.1 | 21.9 | 70 | 101 | 6 | MOLECULE: | TRANSCRIPTI |
| 55:00:00 | 5jk9-A | 2.1 | 21.9 | 71 | 241 | 8 | MOLECULE: | IZUMO SPERM |
| 56:00:00 | 8dl7-A | 2.1 | 6.2 | 82 | 467 | 7 | MOLECULE: | SOLUTE CARR |
| 57:00:00 | 6jho-A | 2.1 | 7.8 | 74 | 200 | 8 | MOLECULE: | CAG PATHOGE |
| 58:00:00 | 2yml-A | 2.1 | 7 | 107 | 228 | 9 | MOLECULE: | L-HALOACID |
| 59:00:00 | 5iuy-C | 2.1 | 4.9 | 92 | 451 | 10 | MOLECULE: | MULTIDRUG E |
| 60:00:00 | 3eaf-A | 2.1 | 4.9 | 52 | 384 | 4 | MOLECULE: | ABC TRANSPO |
| 61:00:00 | 6jcc-A | 2.1 | 4.1 | 65 | 97 | 8 | MOLECULE: | COMPUTATION |
| 62:00:00 | 5hyc-A | 2.1 | 4.5 | 79 | 130 | 4 | MOLECULE: | UNCHARACTER |
| 63:00:00 | 6poo-A | 2.1 | 19.3 | 59 | 273 | 12 | MOLECULE: | BIBA |
| 64:00:00 | 1wa8-B | 2.1 | 7.5 | 54 | 95 | 7 | MOLECULE: | ESAT-6 LIKE |
| 65:00:00 | 5ea1-B | 2.1 | 3.8 | 49 | 128 | 2 | MOLECULE: | TRANSCRIPTI |
| 66:00:00 | 5dlq-B | 2.1 | 29.1 | 101 | 1052 | 9 | MOLECULE: | EXPORTIN-4 |
| 67:00:00 | 4gdf-A | 2.1 | 9.2 | 98 | 497 | 6 | MOLECULE: | LARGE T ANT |
| 68:00:00 | 5j4i-A | 2.1 | 10.5 | 103 | 437 | 6 | MOLECULE: | ARGININE/AG |
| 69:00:00 | 5y9o-A | 2.1 | 23.8 | 79 | 401 | 5 | MOLECULE: | WIPA |
| 70:00:00 | 4eyw-B | 2.1 | 11.8 | 104 | 634 | 9 | MOLECULE: | CARNITINE O |
| 71:00:00 | 6a5n-A | 2.1 | 4.6 | 77 | 502 | 3 | MOLECULE: | HISTONE-LYS |
| 72:00:00 | 3boy-A | 2.1 | 14.9 | 54 | 150 | 7 | MOLECULE: | 5'-R(*UP*UP |
| 73:00:00 | 5mz6-1 | 2.1 | 16.4 | 109 | 1018 | 6 | MOLECULE: | SEPARASE |
| 74:00:00 | 3s63-B | 2.1 | 7 | 44 | 90 | 5 | MOLECULE: | SAPOSIN-LIK |
| 75:00:00 | 6qld-Y | 2.1 | 21.1 | 68 | 223 | 4 | MOLECULE: | INNER KINET |
| 76:00:00 | 4ocu-A | 2.1 | 16.5 | 68 | 356 | 7 | MOLECULE: | N-ACETYLHEX |
| 77:00:00 | 7w1m-D | 2.1 | 19.1 | 107 | 967 | 10 | MOLECULE: | STRUCTURAL |
| 78:00:00 | 7nyw-E | 2.1 | 8.3 | 80 | 212 | 8 | MOLECULE: | CHROMOSOME |
| 79:00:00 | 5of3-C | 2.1 | 6.6 | 49 | 107 | 6 | MOLECULE: | DNA PRIMASE |
| 80:00:00 | 6s6v-A | 2.1 | 15.3 | 65 | 386 | 3 | MOLECULE: | NUCLEASE SB |
| 81:00:00 | 2mrl-A | 2.1 | 3.9 | 44 | 72 | 2 | MOLECULE: | UNCHARACTER |
| 82:00:00 | 5ek8-A | 2.1 | 7.4 | 103 | 668 | 7 | MOLECULE: | LIPOXYGENAS |
| 83:00:00 | 4hqo-B | 2.1 | 24.6 | 83 | 260 | 11 | MOLECULE: | SPOROZOITE |
| 84:00:00 | 5uju-A | 2.1 | 9.1 | 110 | 558 | 3 | MOLECULE: | NAD-DEPENDE |
| 85:00:00 | 5i6r-A | 2.1 | 25.2 | 78 | 419 | 6 | MOLECULE: | SLIT-ROBO R |
| 86:00:00 | 8hm3-B | 2.1 | 9.6 | 66 | 427 | 8 | MOLECULE: | PUTATIVE UB |
| 87:00:00 | 3mem-A | 2.1 | 5.3 | 68 | 453 | 10 | MOLECULE: | PUTATIVE SI |
| 88:00:00 | 2z3x-A | 2.1 | 4.3 | 48 | 56 | 10 | MOLECULE: | SMALL, ACID |
| 89:00:00 | 7b0p-A | 2.1 | 6.8 | 52 | 224 | 8 | MOLECULE: | HYPOTHETICA |
| 90:00:00 | 3ck9-B | 2.1 | 23.4 | 87 | 515 | 7 | MOLECULE: | SUSD |
| 91:00:00 | 7wo9-A | 2.1 | 30.5 | 99 | 1581 | 6 | MOLECULE: | NUCLEOPORIN |
| 92:00:00 | 4nwp-D | 2.1 | 10.6 | 54 | 149 | 9 | MOLECULE: | PUTATIVE UN |
| 93:00:00 | 7y88-A | 2.1 | 9.8 | 60 | 354 | 2 | MOLECULE: | PUTATIVE GL |
| 94:00:00 | 4nsm-A | 2.1 | 5.8 | 44 | 71 | 0 | MOLECULE: | COLLAGEN-LI |

| | | | | | | | | |
|---|---|---|---|---|---|---|---|---|
| 95:00:00 | 1uyv-A | 2.1 | 17.2 | 83 | 571 | 12 | MOLECULE: | ACETYL-COA |
| 96:00:00 | 7wkk-A | 2.1 | 28.9 | 103 | 1684 | 6 | MOLECULE: | MGC83295 PR |
| 97:00:00 | 6wq2-a | 2.1 | 7.5 | 73 | 202 | 15 | MOLECULE: | A-DNA |
| 98:00:00 | 6nr8-3 | 2.1 | 3.1 | 46 | 132 | 4 | MOLECULE: | PREFOLDIN S |
| 99:00:00 | 7rmy-A | 2.1 | 14.5 | 67 | 266 | 7 | MOLECULE: | DE NOVO DES |
| 0:00 | 5cqq-B | 2.1 | 3.2 | 53 | 75 | 6 | MOLECULE: | REGULATORY |
| 1:00 | 7p3x-A | 2.1 | 13.9 | 74 | 576 | 11 | MOLECULE: | AP-3 COMPLE |
| 2:00 | 5c4y-A | 2.1 | 3.2 | 55 | 136 | 11 | MOLECULE: | PUTATIVE TR |
| 3:00 | 8d0b-F | 2.1 | 9.7 | 69 | 915 | 3 | MOLECULE: | CST COMPLEX |
| 4:00 | 4wd9-A | 2.1 | 19.7 | 81 | 965 | 4 | MOLECULE: | NISIN BIOSY |
| 5:00 | 1nzb-A | 2.1 | 17.4 | 76 | 332 | 9 | MOLECULE: | LOXP DNA |
| 6:00 | 6yxu-H | 2.1 | 17.3 | 84 | 691 | 13 | MOLECULE: | DNA-DIRECTE |
| 7:00 | 8ga7-A | 2.1 | 15.9 | 107 | 503 | 7 | MOLECULE: | THR5 |
| 8:00 | 2iho-A | 2.1 | 4 | 83 | 292 | 8 | MOLECULE: | LECTIN |
| 9:00 | 2grm-A | 2.1 | 14.2 | 73 | 315 | 8 | MOLECULE: | PRGX |
| 10:00 | 1ku9-A | 2.1 | 7.9 | 72 | 151 | 8 | MOLECULE: | HYPOTHETICA |
| 11:00 | 8h8e-D | 2.1 | 4.5 | 41 | 293 | 7 | MOLECULE: | PROTON-ACTI |
| 12:00 | 1ung-D | 2.1 | 4 | 69 | 149 | 9 | MOLECULE: | CELL DIVISI |
| 13:00 | 7rpk-A | 2.1 | 18.9 | 123 | 953 | 11 | MOLECULE: | PROTEIN DIS |
| 14:00 | 2jps-A | 2.1 | 4.6 | 60 | 105 | 12 | MOLECULE: | NUCLEAR POL |
| 15:00 | 4bbp-A | 2.1 | 9.3 | 61 | 266 | 10 | MOLECULE: | ZINC ABC TR |
| 16:00 | 8cmk-A | 2.1 | 28.5 | 118 | 913 | 13 | MOLECULE: | TRANSPORTIN |
| 17:00 | 4akk-A | 2.1 | 16.7 | 76 | 368 | 7 | MOLECULE: | NITRATE REG |
| 18:00 | 4itr-A | 2.1 | 16.3 | 78 | 299 | 5 | MOLECULE: | ADENOSINE M |
| 19:00 | 8f2r-C | 2.1 | 24.6 | 78 | 195 | 8 | MOLECULE: | COMM DOMAIN |
| 20:00 | 8ibh-A | 2.1 | 8.9 | 55 | 75 | 9 | MOLECULE: | CENTROSOMAL |
| 21:00 | 7ek2-A | 2.1 | 16.5 | 95 | 341 | 7 | MOLECULE: | BESTROPHIN- |
| 22:00 | 5od9-A | 2.1 | 6.5 | 52 | 95 | 8 | MOLECULE: | MID1SC9 |
| 23:00 | 7yqh-E | 2.1 | 16.3 | 88 | 527 | 6 | MOLECULE: | STRUCTURAL |
| 24:00:00 | 7jqd-B | 2.1 | 3.5 | 35 | 38 | 3 | MOLECULE: | PITUITARY A |
| 25:00:00 | 8hqs-B | 2.1 | 21.5 | 80 | 408 | 8 | MOLECULE: | STRUCTURAL |
| 26:00:00 | 6ahf-C | 2.1 | 25.5 | 98 | 865 | 6 | MOLECULE: | HEAT SHOCK |
| 27:00:00 | 8f0x-A | 2.1 | 10.2 | 90 | 928 | 3 | MOLECULE: | IMPORTIN SU |
| 28:00:00 | 6dft-A | 2.1 | 6.5 | 113 | 384 | 3 | MOLECULE: | DEOXYHYPUSI |
| 29:00:00 | 7lhe-A | 2.1 | 22.5 | 101 | 2222 | 8 | MOLECULE: | INOSITOL 1, |
| 30:00:00 | 6n7p-H | 2.1 | 20.2 | 66 | 105 | 3 | MOLECULE: | U1 SMALL NU |
| 31:00:00 | 4kky-X | 2.1 | 5.4 | 114 | 411 | 11 | MOLECULE: | CAMPHOR 5-M |
| 32:00:00 | 5nf8-A | 2.1 | 13.8 | 73 | 158 | 4 | MOLECULE: | RESPIRATORY |
| 33:00:00 | 6f1t-f | 2.1 | 35.4 | 124 | 929 | 5 | MOLECULE: | ARP1 ACTIN |
| 34:00:00 | 6jo5-K | 2.1 | 2.9 | 39 | 45 | 10 | MOLECULE: | PHOTOSYSTEM |
| 35:00:00 | 6fht-B | 2.1 | 22.9 | 113 | 735 | 12 | MOLECULE: | BACTERIOPHY |
| 36:00:00 | 1xkp-A | 2.1 | 11.9 | 84 | 230 | 5 | MOLECULE: | PUTATIVE ME |
| 37:00:00 | 4rpu-A | 2.1 | 10 | 116 | 988 | 12 | MOLECULE: | PRESEQUENCE |
| 38:00:00 | 2p5g-B | 2.1 | 20.9 | 107 | 756 | 8 | MOLECULE: | TEMPLATE DN |

| | | | | | | | | |
|---|---|---|---|---|---|---|---|---|
| 39:00:00 | 3pf6-A | 2.1 | 3.9 | 44 | 60 | 11 | MOLECULE: | HYPOTHETICA |
| 40:00:00 | 7xcd-A | 2.1 | 5.5 | 52 | 178 | 2 | MOLECULE: | DISEASE SPE |
| 41:00:00 | 2c5z-A | 2.1 | 4.7 | 54 | 93 | 2 | MOLECULE: | SET DOMAIN |
| 42:00:00 | 6n7p-D | 2.1 | 25.8 | 111 | 544 | 5 | MOLECULE: | U1 SMALL NU |
| 43:00:00 | 2xe4-A | 2.1 | 13.9 | 60 | 721 | 5 | MOLECULE: | OLIGOPEPTID |
| 44:00:00 | 6k3b-B | 2.1 | 4.2 | 67 | 106 | 9 | MOLECULE: | LPG2147 |
| 45:00:00 | 4c2e-A | 2.1 | 18.3 | 69 | 435 | 12 | MOLECULE: | CARBOXY-TER |
| 46:00:00 | 7dbe-B | 2.1 | 4.6 | 64 | 332 | 6 | MOLECULE: | BRANCHED-CH |
| 47:00:00 | 6r6u-A | 2.1 | 7.8 | 114 | 458 | 11 | MOLECULE: | CIS-ACONITA |
| 48:00:00 | 3t6p-A | 2.1 | 15.1 | 62 | 330 | 8 | MOLECULE: | BACULOVIRAL |
| 49:00:00 | 7x8u-A | 2.1 | 9.2 | 73 | 227 | 7 | MOLECULE: | TOSPOVIRUS |
| 50:00:00 | 8t1l-V | 2.1 | 20.1 | 63 | 278 | 6 | MOLECULE: | MEDIATOR OF |
| 51:00:00 | 4qsz-A | 2.1 | 17.9 | 94 | 686 | 2 | MOLECULE: | MALTOSE-BIN |
| 52:00:00 | 7t5p-A | 2.1 | 15.6 | 82 | 403 | 6 | MOLECULE: | SUMO-INTERA |
| 53:00:00 | 5jfm-B | 2.1 | 9.4 | 114 | 452 | 10 | MOLECULE: | ALDEHYDE DE |
| 54:00:00 | 5w60-A | 2.1 | 10.7 | 55 | 179 | 5 | MOLECULE: | APOPTOSIS R |
| 55:00:00 | 8iaz-A | 2.1 | 18 | 77 | 384 | 3 | MOLECULE: | TRANSPOSASE |
| 56:00:00 | 6j3i-A | 2.1 | 4.4 | 84 | 583 | 12 | MOLECULE: | GAMMA-GLUTA |
| 57:00:00 | 1rw5-A | 2.1 | 5.1 | 89 | 199 | 4 | MOLECULE: | PROLACTIN |
| 58:00:00 | 6x1g-A | 2.1 | 6 | 65 | 216 | 12 | MOLECULE: | ULP_PROTEAS |
| 59:00:00 | 2b9z-A | 2.1 | 4.1 | 44 | 74 | 7 | MOLECULE: | B2 PROTEIN |
| 60:00:00 | 2r9i-A | 2.1 | 10.9 | 62 | 72 | 15 | MOLECULE: | PUTATIVE PH |
| 61:00:00 | 2zop-A | 2.1 | 3.9 | 64 | 113 | 8 | MOLECULE: | PUTATIVE UN |
| 62:00:00 | 8ke8-A | 2.1 | 11.7 | 60 | 201 | 3 | MOLECULE: | NALC |
| 63:00:00 | 2lck-A | 2.1 | 8.4 | 105 | 296 | 5 | MOLECULE: | MITOCHONDRI |
| 64:00:00 | 6xf9-D | 2.1 | 3.5 | 67 | 426 | 10 | MOLECULE: | PACKAGING P |
| 65:00:00 | 2o5h-A | 2.1 | 4.6 | 67 | 128 | 12 | MOLECULE: | HYPOTHETICA |
| 66:00:00 | 7ecd-A | 2.1 | 7 | 82 | 272 | 7 | MOLECULE: | PHOSPHATIDA |
| 67:00:00 | 5ool-v | 2.1 | 7.5 | 50 | 69 | 2 | MOLECULE: | 16S RIBOSOM |
| 68:00:00 | 8ui7-C | 2.1 | 8.3 | 76 | 339 | 11 | MOLECULE: | ATPASE FAMI |
| 69:00:00 | 5an6-A | 2.1 | 5.2 | 52 | 123 | 12 | MOLECULE: | CRISPR-ASSO |
| 70:00:00 | 6li2-A | 2.1 | 8.5 | 103 | 347 | 4 | MOLECULE: | CHIMERA OF |
| 71:00:00 | 8e9h-J | 2.1 | 24.2 | 74 | 231 | 9 | MOLECULE: | TWO-COMPONE |
| 72:00:00 | 4z7x-A | 2.1 | 6.8 | 82 | 211 | 16 | MOLECULE: | MDBA |
| 73:00:00 | 4mi2-B | 2.1 | 6.3 | 51 | 261 | 14 | MOLECULE: | PUTATIVE EN |
| 74:00:00 | 7z4w-a | 2.1 | 5.5 | 67 | 102 | 6 | MOLECULE: | PORTAL PROT |
| 75:00:00 | 6w8p-B | 2.1 | 9.3 | 108 | 426 | 5 | MOLECULE: | ENDOSOMAL/L |
| 76:00:00 | 7b4m-A | 2.1 | 13.4 | 109 | 459 | 6 | MOLECULE: | SODIUM/HYDR |
| 77:00:00 | 1fou-A | 2.1 | 8.5 | 75 | 257 | 5 | MOLECULE: | UPPER COLLA |
| 78:00:00 | 1d8b-A | 2.1 | 3.6 | 63 | 81 | 2 | MOLECULE: | SGS1 RECQ H |
| 79:00:00 | 3cuz-A | 2.1 | 7.4 | 66 | 529 | 6 | MOLECULE: | MALATE SYNT |
| 80:00:00 | 4ymt-B | 2.1 | 4.1 | 69 | 216 | 6 | MOLECULE: | ABC-TYPE PO |
| 81:00:00 | 6iqc-A | 2.1 | 6.4 | 70 | 103 | 11 | MOLECULE: | DNA-BINDING |
| 82:00:00 | 5ejq-A | 2.1 | 17.9 | 89 | 501 | 7 | MOLECULE: | MYOSIN-I HE |

| | | | | | | | | |
|---|---|---|---|---|---|---|---|---|
| 83:00:00 | 6n1c-A | 2.1 | 6.4 | 65 | 176 | 2 | MOLECULE: | INORGANIC P |
| 84:00:00 | 2juc-A | 2.1 | 3.2 | 49 | 55 | 2 | MOLECULE: | PRE-MRNA-SP |
| 85:00:00 | 7eeb-C | 2.1 | 13.8 | 91 | 278 | 4 | MOLECULE: | ENHANCED GR |
| 86:00:00 | 2kn6-A | 2.1 | 9.7 | 78 | 195 | 6 | MOLECULE: | APOPTOSIS-A |
| 87:00:00 | 7z6q-c | 2.1 | 20.7 | 110 | 121 | 6 | MOLECULE: | PHOTOSYSTEM |
| 88:00:00 | 7fap-A | 2.1 | 11.9 | 113 | 1236 | 9 | MOLECULE: | ERYTHROCYTE |
| 89:00:00 | 7ldk-B | 2.1 | 4.3 | 72 | 106 | 10 | MOLECULE: | NON-STRUCTU |
| 90:00:00 | 4zzl-A | 2.1 | 11.2 | 62 | 146 | 11 | MOLECULE: | MULTIDRUG R |
| 91:00:00 | 5m1m-A | 2.1 | 9.3 | 53 | 154 | 6 | MOLECULE: | MATRIX PROT |
| 92:00:00 | 6yj4-V | 2.1 | 12.4 | 97 | 126 | 3 | MOLECULE: | NADH-UBIQUI |
| 93:00:00 | 7qia-A | 2.1 | 5 | 89 | 401 | 12 | MOLECULE: | DIVALENT ME |
| 94:00:00 | 2nvu-A | 2.1 | 13.7 | 74 | 530 | 7 | MOLECULE: | NEDD8-ACTIV |
| 95:00:00 | 7dsq-B | 2.1 | 13.8 | 96 | 465 | 6 | MOLECULE: | 4F2 CELL-SU |
| 96:00:00 | 7y38-X | 2.1 | 16.8 | 129 | 611 | 5 | MOLECULE: | MRNA-CAPPIN |
| 97:00:00 | 5cx6-B | 2.1 | 18.4 | 139 | 665 | 7 | MOLECULE: | RNA-DEPENDE |
| 98:00:00 | 7f3x-A | 2.1 | 18.4 | 76 | 449 | 3 | MOLECULE: | LPCAT3 |
| 99:00:00 | 3taz-B | 2.1 | 9.9 | 104 | 433 | 5 | MOLECULE: | DNA DOUBLE- |
| 0:00 | 7kfk-C | 2.1 | 4.4 | 79 | 132 | 8 | MOLECULE: | ISOFORM 2 O |
| 1:00 | 8rc4-e | 2.1 | 14.1 | 103 | 873 | 4 | MOLECULE: | SERINE/THRE |
| 2:00 | 2p06-B | 2.1 | 5 | 72 | 88 | 4 | MOLECULE: | HYPOTHETICA |
| 3:00 | 8gi1-A | 2.1 | 6.8 | 72 | 144 | 3 | MOLECULE: | ACCESSORY P |
| 4:00 | 8j8j-A | 2.1 | 6.4 | 80 | 290 | 8 | MOLECULE: | DECAPRENYL- |
| 5:00 | 8bsh-C | 2.1 | 23.4 | 109 | 794 | 6 | MOLECULE: | PROTEIN TRA |
| 6:00 | 5dzt-A | 2.1 | 18.6 | 104 | 886 | 6 | MOLECULE: | CYLM |
| 7:00 | 3kd4-A | 2.1 | 17 | 103 | 505 | 8 | MOLECULE: | PUTATIVE PR |
| 8:00 | 4gp2-A | 2.1 | 8.4 | 89 | 342 | 7 | MOLECULE: | POLYPRENYL |
| 9:00 | 7k36-I | 2.1 | 17.3 | 102 | 599 | 5 | MOLECULE: | SERINE/THRE |
| 10:00 | 6mpb-A | 2.1 | 6.8 | 100 | 447 | 9 | MOLECULE: | NEUTRAL AMI |
| 11:00 | 4zht-A | 2.1 | 8.3 | 77 | 384 | 10 | MOLECULE: | BIFUNCTIONA |
| 12:00 | 7lma-H | 2.1 | 18.1 | 86 | 199 | 6 | MOLECULE: | TELOMERASE |
| 13:00 | 3zqg-A | 2.1 | 12.4 | 73 | 208 | 10 | MOLECULE: | TETRACYCLIN |
| 14:00 | 7pc1-A | 2.1 | 4.1 | 61 | 72 | 7 | MOLECULE: | STBA |
| 15:00 | 7roa-A | 2.1 | 3.3 | 69 | 117 | 3 | MOLECULE: | ENTV |
| 16:00 | 8d8j-0 | 2.1 | 9.8 | 78 | 493 | 4 | MOLECULE: | PROBABLE S- |
| 17:00 | 2c9e-A | 2.1 | 17.5 | 89 | 317 | 6 | MOLECULE: | PERIDININ-C |
| 18:00 | 3faj-A | 2.1 | 7 | 70 | 101 | 4 | MOLECULE: | PUTATIVE UN |
| 19:00 | 5x5y-F | 2.1 | 10.5 | 81 | 351 | 14 | MOLECULE: | PROBABLE AT |
| 20:00 | 6v3z-A | 2.1 | 10.2 | 76 | 178 | 7 | MOLECULE: | SEN1395 |
| 21:00 | 7qsd-N | 2.1 | 8.4 | 93 | 347 | 5 | MOLECULE: | NADH-UBIQUI |
| 22:00 | 6vzz-A | 2.1 | 5.4 | 85 | 374 | 9 | MOLECULE: | BAMAA.19900 |
| 23:00 | 6jkg-A | 2.1 | 7.2 | 79 | 216 | 4 | MOLECULE: | STEROL-4-AL |
| 24:00:00 | 7xpt-A | 2.1 | 7.6 | 79 | 375 | 11 | MOLECULE: | TRANSGLYCOS |
| 25:00:00 | 2aja-B | 2.1 | 18.2 | 83 | 345 | 4 | MOLECULE: | ANKYRIN REP |
| 26:00:00 | 8tfo-A | 2.1 | 13.5 | 89 | 390 | 9 | MOLECULE: | MEVALONATE |

| | | | | | | | | |
|---|---|---|---|---|---|---|---|---|
| 27:00:00 | 2dmn-A | 2.1 | 5.9 | 72 | 83 | 4 | MOLECULE: | HOMEOBOX PR |
| 28:00:00 | 5xn7-B | 2.1 | 9.3 | 103 | 551 | 7 | MOLECULE: | PUTATIVE RT |
| 29:00:00 | 5l7s-A | 2.1 | 17 | 83 | 267 | 12 | MOLECULE: | SECRETED RX |
| 30:00:00 | 5n9j-U | 2.1 | 11.1 | 94 | 198 | 5 | MOLECULE: | MEDIATOR OF |
| 31:00:00 | 5y7q-A | 2.1 | 11.7 | 105 | 546 | 10 | MOLECULE: | FANCONI-ASS |
| 32:00:00 | 2m64-A | 2.1 | 12.4 | 88 | 231 | 6 | MOLECULE: | PHLP5 |
| 33:00:00 | 6t1z-A | 2.1 | 15.6 | 82 | 393 | 5 | MOLECULE: | LMRP INTEGR |
| 34:00:00 | 2voh-A | 2.1 | 5 | 81 | 155 | 10 | MOLECULE: | BCL-2-RELAT |
| 35:00:00 | 8sfj-A | 2.1 | 19.1 | 116 | 1001 | 3 | MOLECULE: | CRISPR-ASSO |
| 36:00:00 | 4afi-A | 2.1 | 7.9 | 76 | 155 | 11 | MOLECULE: | AP-3 COMPLE |
| 37:00:00 | 2apl-A | 2.1 | 8 | 63 | 149 | 5 | MOLECULE: | HYPOTHETICA |
| 38:00:00 | 8fzb-A | 2.1 | 22.1 | 87 | 319 | 11 | MOLECULE: | PROTEIN-LYS |
| 39:00:00 | 7w78-B | 2.1 | 10.5 | 104 | 344 | 3 | MOLECULE: | PUTATIVE AB |
| 40:00:00 | 5y36-A | 2.1 | 25.8 | 156 | 1368 | 6 | MOLECULE: | CRISPR-ASSO |
| 41:00:00 | 6jql-A | 2.1 | 18.1 | 108 | 678 | 14 | MOLECULE: | BIFUNCTIONA |
| 42:00:00 | 6zfw-A | 2.1 | 4.8 | 54 | 70 | 4 | MOLECULE: | PEROXIN-14 |
| 43:00:00 | 3f75-P | 2.1 | 5.3 | 60 | 75 | 13 | MOLECULE: | CATHEPSIN L |
| 44:00:00 | 2ccy-A | 2.1 | 4.7 | 66 | 127 | 14 | MOLECULE: | CYTOCHROME |
| 45:00:00 | 7zkq-T | 2.1 | 20 | 89 | 351 | 6 | MOLECULE: | NADH DEHYDR |
| 46:00:00 | 2vqg-B | 2.1 | 4.2 | 65 | 77 | 9 | MOLECULE: | UNCHARACTER |
| 47:00:00 | 7t7r-A | 2.1 | 18.8 | 118 | 1132 | 9 | MOLECULE: | PROTEIN UNC |
| 48:00:00 | 7y7q-A | 2.1 | 13.1 | 108 | 951 | 6 | MOLECULE: | RNA-DEPENDE |
| 49:00:00 | 7mgq-A | 2.1 | 16 | 80 | 602 | 3 | MOLECULE: | 5-AMINOIMID |
| 50:00:00 | 3uit-D | 2.1 | 12.3 | 85 | 257 | 6 | MOLECULE: | INAD-LIKE P |
| 51:00:00 | 2ly1-A | 2.1 | 17.5 | 83 | 180 | 5 | MOLECULE: | TUDOR DOMAI |
| 52:00:00 | 5dcq-E | 2.1 | 7.4 | 59 | 200 | 5 | MOLECULE: | ARTIFICIAL |
| 53:00:00 | 7w01-A | 2.1 | 15.2 | 107 | 1598 | 8 | MOLECULE: | PHOSPHOLIPI |
| 54:00:00 | 7akx-A | 2.1 | 8.8 | 91 | 224 | 7 | MOLECULE: | VIRAL RHODO |
| 55:00:00 | 8tlt-E | 2.1 | 11.5 | 68 | 232 | 4 | MOLECULE: | DNA POLYMER |
| 56:00:00 | 6qum-N | 2.1 | 17.3 | 113 | 649 | 4 | MOLECULE: | V-TYPE ATP |
| 57:00:00 | 3iee-A | 2.1 | 10.4 | 79 | 256 | 4 | MOLECULE: | PUTATIVE EX |
| 58:00:00 | 3maj-A | 2.1 | 6.7 | 57 | 353 | 9 | MOLECULE: | DNA PROCESS |
| 59:00:00 | 6j8l-A | 2.1 | 3.6 | 72 | 132 | 13 | MOLECULE: | AVH240 |
| 60:00:00 | 8ap7-o | 2.1 | 11.2 | 78 | 96 | 6 | MOLECULE: | ATP SYNTHAS |
| 61:00:00 | 7rvb-A | 2.1 | 16.5 | 97 | 209 | 2 | MOLECULE: | FERRITIN |
| 62:00:00 | 7wj0-A | 2.1 | 9.1 | 42 | 49 | 12 | MOLECULE: | ANTITOXIN M |
| 63:00:00 | 8et6-A | 2.1 | 18.9 | 115 | 532 | 10 | MOLECULE: | 1-Oct |
| 64:00:00 | 7bu0-A | 2.1 | 5 | 77 | 407 | 10 | MOLECULE: | UNCHARACTER |
| 65:00:00 | 8tcw-A | 2.1 | 8.4 | 77 | 282 | 5 | MOLECULE: | PYRROLINE-5 |
| 66:00:00 | 7f4l-B | 2.1 | 5.5 | 75 | 130 | 11 | MOLECULE: | TRANSMEMBRA |
| 67:00:00 | 1jud-A | 2.1 | 6.1 | 102 | 220 | 9 | MOLECULE: | L-2-HALOACI |
| 68:00:00 | 3p8c-F | 2.1 | 28.3 | 99 | 156 | 4 | MOLECULE: | CYTOPLASMIC |
| 69:00:00 | 5wat-A | 2.1 | 6 | 76 | 310 | 8 | MOLECULE: | HOMOSERINE |
| 70:00:00 | 5usf-A | 2.1 | 23.2 | 96 | 683 | 9 | MOLECULE: | TYROSYL-TRN |

| | | | | | | | | |
|---|---|---|---|---|---|---|---|---|
| 71:00:00 | 4k2u-A | 2.1 | 8.4 | 65 | 230 | 6 | MOLECULE: | ERYTHROCYTE |
| 72:00:00 | 4elj-A | 2.1 | 8.5 | 86 | 590 | 6 | MOLECULE: | RETINOBLAST |
| 73:00:00 | 1ntc-A | 2.1 | 7.1 | 61 | 91 | 8 | MOLECULE: | PROTEIN (NI |
| 74:00:00 | 5tk6-A | 2.1 | 5.8 | 70 | 191 | 13 | MOLECULE: | OXSA PROTEI |
| 75:00:00 | 8t1l-S | 2.1 | 12.1 | 111 | 913 | 5 | MOLECULE: | MEDIATOR OF |
| 76:00:00 | 6wkr-A | 2.1 | 8.8 | 66 | 439 | 8 | MOLECULE: | UBIQUITIN |
| 77:00:00 | 1n81-A | 2.1 | 11 | 72 | 186 | 4 | MOLECULE: | PLASMODIUM |
| 78:00:00 | 6wge-C | 2.1 | 17.4 | 78 | 174 | 9 | MOLECULE: | STRUCTURAL |
| 79:00:00 | 6zie-A | 2.1 | 5 | 66 | 123 | 8 | MOLECULE: | CMPX-383B |
| 80:00:00 | 8izd-A | 2.1 | 12.5 | 95 | 315 | 6 | MOLECULE: | CERAMIDE SY |
| 81:00:00 | 7aft-F | 2.1 | 11 | 76 | 190 | 4 | MOLECULE: | PROTEIN TRA |
| 82:00:00 | 4acl-A | 2.1 | 8.2 | 73 | 155 | 7 | MOLECULE: | TSSL |
| 83:00:00 | 5uld-A | 2.1 | 5.5 | 83 | 445 | 5 | MOLECULE: | TRANSPORTER |
| 84:00:00 | 6tdy-M | 2.1 | 7.6 | 93 | 243 | 13 | MOLECULE: | ATP SYNTHAS |
| 85:00:00 | 8bhw-A | 2.1 | 2.9 | 58 | 332 | 3 | MOLECULE: | ECA POLYSAC |
| 86:00:00 | 8ck1-E | 2.1 | 4.6 | 71 | 206 | 4 | MOLECULE: | TAIL NOZZLE |
| 87:00:00 | 8xgr-R | 2.1 | 15.2 | 101 | 305 | 5 | MOLECULE: | GUANINE NUC |
| 88:00:00 | 7pw9-A | 2.1 | 19.6 | 113 | 1952 | 10 | MOLECULE: | SERINE/THRE |
| 89:00:00 | 6uzt-A | 2.1 | 25.2 | 113 | 587 | 6 | MOLECULE: | RECEPTOR-TY |
| 90:00:00 | 8hgu-A | 2.1 | 4.6 | 98 | 295 | 10 | MOLECULE: | ALPHA/BETA |
| 91:00:00 | 6icz-A | 2.1 | 20.3 | 134 | 2253 | 10 | MOLECULE: | PROTEIN MAG |
| 92:00:00 | 5lpc-A | 2.1 | 4.7 | 76 | 624 | 7 | MOLECULE: | VANADIUM-DE |
| 93:00:00 | 6xp5-H | 2.1 | 11.4 | 87 | 146 | 5 | MOLECULE: | MEDIATOR OF |
| 94:00:00 | 6m31-A | 2.1 | 14.2 | 95 | 280 | 5 | MOLECULE: | DIGERANYLGE |
| 95:00:00 | 3no4-A | 2.1 | 11 | 77 | 264 | 12 | MOLECULE: | CREATININE |
| 96:00:00 | 3fgg-A | 2.1 | 8.1 | 70 | 147 | 4 | MOLECULE: | UNCHARACTER |
| 97:00:00 | 3wfr-H | 2.1 | 20 | 109 | 463 | 3 | MOLECULE: | RNA (74-MER |
| 98:00:00 | 6rfl-C | 2.1 | 3.7 | 71 | 304 | 11 | MOLECULE: | DNA-DEPENDE |
| 99:00:00 | 6l1k-A | 2.1 | 8.3 | 68 | 384 | 12 | MOLECULE: | NADH-DEPEND |
| 0:00 | 6vk0-C | 2.1 | 6.7 | 93 | 180 | 4 | MOLECULE: | U1 SNP1-ASS |
| 1:00 | 6iu4-A | 2.1 | 5.7 | 77 | 225 | 5 | MOLECULE: | VIT1 |
| 2:00 | 8dgc-E | 2.1 | 20.5 | 98 | 538 | 9 | MOLECULE: | SEAVS3 |
| 3:00 | 5jb3-Q | 2.1 | 9.1 | 60 | 158 | 8 | MOLECULE: | 16S RIBOSOM |
| 4:00 | 6ofs-A | 2.1 | 8.5 | 73 | 905 | 8 | MOLECULE: | PROBABLE ZI |
| 5:00 | 8qrk-R | 2.1 | 12.8 | 82 | 295 | 4 | MOLECULE: | 28S RIBOSOM |
| 6:00 | 1x3w-B | 2.1 | 3.5 | 49 | 57 | 12 | MOLECULE: | PEPTIDE:N-G |
| 7:00 | 8osg-B | 2.1 | 7.7 | 75 | 323 | 8 | MOLECULE: | MAGNESIUM-C |
| 8:00 | 7k5c-B | 2.1 | 17.5 | 83 | 228 | 2 | MOLECULE: | INTERNAL VI |
| 9:00 | 5wue-A | 2.1 | 4.8 | 71 | 203 | 4 | MOLECULE: | UNCHARACTER |
| 10:00 | 2c5u-A | 2.1 | 14.3 | 77 | 375 | 4 | MOLECULE: | RNA LIGASE |
| 11:00 | 8jnt-A | 2.1 | 3.5 | 57 | 77 | 5 | MOLECULE: | RXLR19781 |
| 12:00 | 7z8v-D | 2.1 | 24.9 | 124 | 1158 | 7 | MOLECULE: | CULLIN-1 |
| 13:00 | 7jpp-E | 2 | 27.1 | 89 | 406 | 7 | MOLECULE: | ORIGIN RECO |
| 14:00 | 8ic6-B | 2 | 22.1 | 99 | 855 | 9 | MOLECULE: | EXO-BETA-D- |

| | | | | | | | | |
|---|---|---|---|---|---|---|---|---|
| 15:00 | 2lon-A | 2 | 15.2 | 74 | 99 | 5 | MOLECULE: | HIG1 DOMAIN |
| 16:00 | 7lxt-J | 2 | 3.8 | 63 | 215 | 14 | MOLECULE: | 20S PROTEAS |
| 17:00 | 1gk7-A | 2 | 3.3 | 39 | 39 | 8 | MOLECULE: | VIMENTIN |
| 18:00 | 6ugm-X | 2 | 10.1 | 58 | 73 | 12 | MOLECULE: | HISTONE H3 |
| 19:00 | 2a01-A | 2 | 13.3 | 74 | 243 | 7 | MOLECULE: | APOLIPOPROT |
| 20:00 | 3cs3-A | 2 | 8.1 | 68 | 266 | 4 | MOLECULE: | SUGAR-BINDI |
| 21:00 | 6mag-A | 2 | 10.4 | 82 | 282 | 13 | MOLECULE: | BBVCI ENDON |
| 22:00 | 5wvc-B | 2 | 3.9 | 57 | 103 | 7 | MOLECULE: | APOPTOTIC P |
| 23:00 | 2bdt-A | 2 | 3.6 | 48 | 172 | 13 | MOLECULE: | BH3686 |
| 24:00:00 | 3a1y-A | 2 | 2.6 | 44 | 58 | 9 | MOLECULE: | 50S RIBOSOM |
| 25:00:00 | 5y6r-A | 2 | 15.4 | 116 | 667 | 9 | MOLECULE: | GENOME POLY |
| 26:00:00 | 6jta-A | 2 | 17.5 | 81 | 1299 | 4 | MOLECULE: | PHOSPHORIBO |
| 27:00:00 | 8jhu-A | 2 | 20.2 | 99 | 811 | 4 | MOLECULE: | LEGIONELLA |
| 28:00:00 | 1zvr-A | 2 | 7.6 | 85 | 514 | 6 | MOLECULE: | MYOTUBULARI |
| 29:00:00 | 7twd-B | 2 | 3.2 | 41 | 45 | 15 | MOLECULE: | ALPHA- AND |
| 30:00:00 | 6cnb-R | 2 | 17.7 | 107 | 522 | 4 | MOLECULE: | DNA-DIRECTE |
| 31:00:00 | 8hr5-A | 2 | 28.2 | 94 | 466 | 9 | MOLECULE: | TRANSPOSASE |
| 32:00:00 | 1u4q-A | 2 | 38 | 122 | 318 | 8 | MOLECULE: | SPECTRIN AL |
| 33:00:00 | 6wut-C | 2 | 18.5 | 75 | 348 | 5 | MOLECULE: | SAM35 |
| 34:00:00 | 8sre-A | 2 | 25.9 | 103 | 1377 | 6 | MOLECULE: | TRPM2 CHANZ |
| 35:00:00 | 8q3v-e | 2 | 10.1 | 109 | 294 | 6 | MOLECULE: | TETRAHYDROM |
| 36:00:00 | 1u2m-C | 2 | 12.4 | 57 | 143 | 14 | MOLECULE: | HISTONE-LIK |
| 37:00:00 | 7px2-C | 2 | 3 | 50 | 85 | 6 | MOLECULE: | CONOTOXIN M |
| 38:00:00 | 4rs7-B | 2 | 5.1 | 58 | 88 | 5 | MOLECULE: | PARB-C |
| 39:00:00 | 4imj-A | 2 | 11.6 | 87 | 333 | 6 | MOLECULE: | SYMPLEKIN |
| 40:00:00 | 8h5s-A | 2 | 4.6 | 83 | 247 | 6 | MOLECULE: | BETA-PHOSPH |
| 41:00:00 | 1lsh-A | 2 | 22 | 98 | 955 | 3 | MOLECULE: | LIPOVITELLI |
| 42:00:00 | 7orl-A | 2 | 12.4 | 121 | 2183 | 7 | MOLECULE: | RNA (5'-D(* |
| 43:00:00 | 1hrk-A | 2 | 16.2 | 73 | 359 | 1 | MOLECULE: | FERROCHELAT |
| 44:00:00 | 7dsj-A | 2 | 6 | 56 | 363 | 11 | MOLECULE: | ANTHRANILAT |
| 45:00:00 | 6jxc-A | 2 | 22.2 | 100 | 2345 | 7 | MOLECULE: | SERINE/THRE |
| 46:00:00 | 8dwj-A | 2 | 7 | 64 | 339 | 8 | MOLECULE: | RNA(5'-PPP- |
| 47:00:00 | 6bzf-A | 2 | 19.1 | 62 | 197 | 5 | MOLECULE: | SPORULATION |
| 48:00:00 | 6p7r-A | 2 | 6 | 82 | 277 | 5 | MOLECULE: | TOXIN CO-RE |
| 49:00:00 | 5j08-A | 2 | 9 | 71 | 163 | 13 | MOLECULE: | EPSIN-5 |
| 50:00:00 | 7msw-A | 2 | 17.3 | 87 | 635 | 6 | MOLECULE: | NON-STRUCTU |
| 51:00:00 | 7qe7-O | 2 | 32.7 | 81 | 707 | 9 | MOLECULE: | ANAPHASE-PR |
| 52:00:00 | 6fes-A | 2 | 13.9 | 84 | 365 | 11 | MOLECULE: | D12_BRIC2, |
| 53:00:00 | 5erg-A | 2 | 8.5 | 69 | 340 | 6 | MOLECULE: | TRNA (ADENI |
| 54:00:00 | 5m48-A | 2 | 20.1 | 73 | 114 | 3 | MOLECULE: | REGULATOR O |
| 55:00:00 | 6lo8-E | 2 | 6 | 54 | 80 | 11 | MOLECULE: | MITOCHONDRI |
| 56:00:00 | 2o7g-A | 2 | 3.4 | 55 | 88 | 5 | MOLECULE: | PROBABLE RN |
| 57:00:00 | 8bgo-C | 2 | 9.2 | 64 | 271 | 8 | MOLECULE: | DIACETYLCHI |
| 58:00:00 | 4v1a-a | 2 | 20.1 | 92 | 393 | 7 | MOLECULE: | MITORIBOSOM |

| | | | | | | | | |
|---|---|---|---|---|---|---|---|---|
| 59:00:00 | 8kde-G | 2 | 8.8 | 62 | 145 | 10 | MOLECULE: | PHOTOSYSTEM |
| 60:00:00 | 7r07-F | 2 | 7.7 | 108 | 600 | 7 | MOLECULE: | ABIK |
| 61:00:00 | 6l1q-A | 2 | 14.1 | 70 | 267 | 7 | MOLECULE: | CBBQ PROTEI |
| 62:00:00 | 4pas-B | 2 | 4.7 | 39 | 39 | 3 | MOLECULE: | GAMMA-AMINO |
| 63:00:00 | 3wrn-A | 2 | 3.9 | 92 | 248 | 5 | MOLECULE: | NON-CAPSID |
| 64:00:00 | 1n1c-A | 2 | 14.4 | 92 | 199 | 8 | MOLECULE: | TORA SPECIF |
| 65:00:00 | 8hpo-C | 2 | 9.6 | 60 | 138 | 2 | MOLECULE: | TRANSCRIPTI |
| 66:00:00 | 3on2-B | 2 | 4.7 | 68 | 195 | 6 | MOLECULE: | PROBABLE TR |
| 67:00:00 | 6vej-A | 2 | 28.6 | 138 | 1022 | 3 | MOLECULE: | PROBABLE RE |
| 68:00:00 | 1zhc-A | 2 | 8.3 | 51 | 76 | 8 | MOLECULE: | HYPOTHETICA |
| 69:00:00 | 3dxj-D | 2 | 19.2 | 75 | 1504 | 3 | MOLECULE: | DNA-DIRECTE |
| 70:00:00 | 3n2o-A | 2 | 6.3 | 76 | 630 | 1 | MOLECULE: | BIOSYNTHETI |
| 71:00:00 | 1na6-B | 2 | 17.7 | 102 | 395 | 4 | MOLECULE: | RESTRICTION |
| 72:00:00 | 2jdi-H | 2 | 8.4 | 43 | 88 | 14 | MOLECULE: | ATP SYNTHAS |
| 73:00:00 | 3r6n-A | 2 | 37 | 112 | 450 | 6 | MOLECULE: | DESMOPLAKIN |
| 74:00:00 | 7pkq-N | 2 | 9.7 | 51 | 195 | 2 | MOLECULE: | MS35 |
| 75:00:00 | 1oks-A | 2 | 3.3 | 37 | 53 | 8 | MOLECULE: | RNA POLYMER |
| 76:00:00 | 8c1s-D | 2 | 10.9 | 61 | 155 | 7 | MOLECULE: | GLUTAMATE R |
| 77:00:00 | 3cwz-B | 2 | 16.1 | 59 | 310 | 5 | MOLECULE: | RAS-RELATED |
| 78:00:00 | 2xus-A | 2 | 3.3 | 41 | 41 | 10 | MOLECULE: | BREAST CANC |
| 79:00:00 | 6c8c-A | 2 | 5.1 | 71 | 119 | 8 | MOLECULE: | CHIMERIC PR |
| 80:00:00 | 3otv-C | 2 | 15.6 | 63 | 257 | 8 | MOLECULE: | PROBABLE CO |
| 81:00:00 | 8ozh-D | 2 | 21.5 | 84 | 405 | 7 | MOLECULE: | ENVELOPE GL |
| 82:00:00 | 5g49-A | 2 | 6.4 | 52 | 97 | 6 | MOLECULE: | NUCLEAR TRA |
| 83:00:00 | 1nxh-A | 2 | 9 | 66 | 124 | 8 | MOLECULE: | MTH396 PROT |
| 84:00:00 | 6its-A | 2 | 6 | 56 | 147 | 4 | MOLECULE: | METHYL-ACCE |
| 85:00:00 | 2n5n-A | 2 | 7.4 | 48 | 86 | 4 | MOLECULE: | CHROMODOMAI |
| 86:00:00 | 5jbr-A | 2 | 5.4 | 56 | 149 | 5 | MOLECULE: | UNCHARACTER |
| 87:00:00 | 6u1s-A | 2 | 13.8 | 73 | 310 | 7 | MOLECULE: | DE NOVO DES |
| 88:00:00 | 5a9e-A | 2 | 15.5 | 91 | 254 | 11 | MOLECULE: | DELTAMBD GA |
| 89:00:00 | 1f3m-A | 2 | 3.5 | 43 | 70 | 0 | MOLECULE: | SERINE/THRE |
| 90:00:00 | 4xig-N | 2 | 10.3 | 79 | 290 | 6 | MOLECULE: | ALGM1 |
| 91:00:00 | 5f0n-A | 2 | 13.3 | 125 | 1038 | 5 | MOLECULE: | COHESIN SUB |
| 92:00:00 | 6vwb-A | 2 | 4.5 | 52 | 90 | 15 | MOLECULE: | CROSSOVER J |
| 93:00:00 | 7lxk-A | 2 | 3.6 | 47 | 59 | 4 | MOLECULE: | ALLERGEN AR |
| 94:00:00 | 4dci-B | 2 | 25.1 | 80 | 148 | 11 | MOLECULE: | UNCHARACTER |
| 95:00:00 | 6sny-A | 2 | 4.7 | 61 | 107 | 2 | MOLECULE: | SYNTHETIC E |
| 96:00:00 | 5l1a-A | 2 | 6.6 | 48 | 108 | 8 | MOLECULE: | UNCHARACTER |
| 97:00:00 | 8a51-B | 2 | 3.6 | 39 | 39 | 8 | MOLECULE: | HEAT SHOCK |
| 98:00:00 | 4q2d-A | 2 | 5.2 | 128 | 927 | 4 | MOLECULE: | CRISPR-ASSO |
| 99:00:00 | 5ncn-B | 2 | 7.5 | 60 | 76 | 7 | MOLECULE: | DBF2 KINASE |
| 0:00 | 3ezh-A | 2 | 8.7 | 57 | 114 | 7 | MOLECULE: | NITRATE/NIT |
| 1:00 | 5b6b-G | 2 | 11 | 64 | 74 | 2 | MOLECULE: | MOB KINASE |
| 2:00 | 7yh8-A | 2 | 6.9 | 43 | 62 | 2 | MOLECULE: | L-19437 |

| | | | | | | | | |
|---|---|---|---|---|---|---|---|---|
| 3:00 | 7ktr-A | 2 | 22 | 90 | 3042 | 6 | MOLECULE: | TRANSFORMAT |
| 4:00 | 6wi5-A | 2 | 4.1 | 53 | 90 | 8 | MOLECULE: | DE NOVO DES |
| 5:00 | 6s18-A | 2 | 11.8 | 84 | 143 | 7 | MOLECULE: | AROMATIC AC |
| 6:00 | 1jwy-A | 2 | 18.3 | 117 | 1039 | 9 | MOLECULE: | MYOSIN-2 HE |
| 7:00 | 6o3c-A | 2 | 8.8 | 81 | 484 | 9 | MOLECULE: | SMOOTHENED |
| 8:00 | 6pqx-A | 2 | 8.7 | 82 | 480 | 4 | MOLECULE: | DNA-MEDIATE |
| 9:00 | 1tuz-A | 2 | 4.6 | 73 | 118 | 4 | MOLECULE: | DIACYLGLYCE |
| 10:00 | 4l6v-6 | 2 | 6.9 | 71 | 125 | 3 | MOLECULE: | PHOTOSYSTEM |
| 11:00 | 6lu0-A | 2 | 14.2 | 89 | 937 | 3 | MOLECULE: | CAS12I2 |
| 12:00 | 1puo-B | 2 | 7.3 | 54 | 142 | 2 | MOLECULE: | MAJOR ALLER |
| 13:00 | 6zyv-A | 2 | 14.8 | 77 | 238 | 4 | MOLECULE: | CIR PROTEIN |
| 14:00 | 8iue-P | 2 | 18.3 | 73 | 303 | 5 | MOLECULE: | DNA-DIRECTE |
| 15:00 | 6ei6-A | 2 | 4.2 | 59 | 242 | 8 | MOLECULE: | COILED-COIL |
| 16:00 | 5img-A | 2 | 14.9 | 75 | 467 | 4 | MOLECULE: | DIPEPTIDASE |
| 17:00 | 2guz-B | 2 | 5.4 | 53 | 65 | 8 | MOLECULE: | MITOCHONDRI |
| 18:00 | 8bpd-A | 2 | 11.7 | 75 | 370 | 8 | MOLECULE: | GLYCOSYL HY |
| 19:00 | 8irl-A | 2 | 19.2 | 106 | 540 | 6 | MOLECULE: | ADENINE/GUA |
| 20:00 | 4mi8-B | 2 | 6.7 | 71 | 136 | 7 | MOLECULE: | BCL-2 HOMOL |
| 21:00 | 2jes-A | 2 | 4.8 | 55 | 370 | 7 | MOLECULE: | PORTAL PROT |
| 22:00 | 4dra-H | 2 | 2.7 | 49 | 76 | 14 | MOLECULE: | CENTROMERE |
| 23:00 | 2oyy-A | 2 | 3.4 | 57 | 71 | 4 | MOLECULE: | HEXAMERIC C |
| 24:00:00 | 4gwp-C | 2 | 8.8 | 83 | 184 | 6 | MOLECULE: | MEDIATOR OF |
| 25:00:00 | 3f41-A | 2 | 19.4 | 95 | 590 | 11 | MOLECULE: | PHYTASE |
| 26:00:00 | 2mb9-A | 2 | 4.3 | 68 | 106 | 10 | MOLECULE: | B-CELL LYMP |
| 27:00:00 | 3sqn-A | 2 | 17.2 | 117 | 468 | 6 | MOLECULE: | CONSERVED D |
| 28:00:00 | 6kkk-A | 2 | 14.3 | 95 | 380 | 4 | MOLECULE: | SUGAR EFFLU |
| 29:00:00 | 3h20-A | 2 | 9.9 | 68 | 288 | 13 | MOLECULE: | REPLICATION |
| 30:00:00 | 5yfp-H | 2 | 10.2 | 66 | 518 | 8 | MOLECULE: | EXOCYST COM |
| 31:00:00 | 5okc-B | 2 | 5.9 | 99 | 377 | 5 | MOLECULE: | SISTER CHRO |
| 32:00:00 | 4nk2-A | 2 | 5.7 | 81 | 164 | 7 | MOLECULE: | HEMOGLOBIN- |
| 33:00:00 | 7ura-A | 2 | 16.6 | 98 | 432 | 6 | MOLECULE: | ISOFORM 2 O |
| 34:00:00 | 5k2a-A | 2 | 11.6 | 73 | 398 | 4 | MOLECULE: | ADENOSINE R |
| 35:00:00 | 7wb4-E | 2 | 10.3 | 120 | 1363 | 7 | MOLECULE: | OUTER NUP13 |
| 36:00:00 | 2dla-A | 2 | 5.2 | 83 | 222 | 7 | MOLECULE: | 397AA LONG |
| 37:00:00 | 8eg3-A | 2 | 27.6 | 94 | 554 | 6 | MOLECULE: | STERYL-SULF |
| 38:00:00 | 7n1j-B | 2 | 4.6 | 59 | 64 | 5 | MOLECULE: | FIBROBLAST |
| 39:00:00 | 5y77-A | 2 | 7.8 | 72 | 456 | 4 | MOLECULE: | KYNURENINE |
| 40:00:00 | 5mnt-B | 2 | 26.8 | 86 | 421 | 5 | MOLECULE: | A2 MATURATI |
| 41:00:00 | 7uqi-G | 2 | 13.3 | 83 | 197 | 4 | MOLECULE: | ATPASE HIST |
| 42:00:00 | 6l7r-A | 2 | 15 | 59 | 105 | 2 | MOLECULE: | PUTATIVE SP |
| 43:00:00 | 6lyy-A | 2 | 16.3 | 94 | 382 | 6 | MOLECULE: | MONOCARBOXY |
| 44:00:00 | 3icq-T | 2 | 13.8 | 98 | 949 | 7 | MOLECULE: | EXPORTIN-T |
| 45:00:00 | 6p5a-B | 2 | 4.7 | 74 | 118 | 4 | MOLECULE: | TRANSPOSABL |
| 46:00:00 | 7cww-A | 2 | 10.2 | 77 | 350 | 9 | MOLECULE: | TSRE |

| | | | | | | | | |
|---|---|---|---|---|---|---|---|---|
| 47:00:00 | 8exh-a | 2 | 11.4 | 63 | 69 | 5 | MOLECULE: | PROTEIN VIR |
| 48:00:00 | 6xtt-B | 2 | 2.9 | 59 | 116 | 2 | MOLECULE: | NTTA |
| 49:00:00 | 6exn-O | 2 | 18.8 | 88 | 320 | 6 | MOLECULE: | U2 SNRNA |
| 50:00:00 | 1yoz-B | 2 | 6.1 | 61 | 116 | 5 | MOLECULE: | HYPOTHETICA |
| 51:00:00 | 8jhq-A | 2 | 14.7 | 98 | 446 | 4 | MOLECULE: | SPHINGOSINE |
| 52:00:00 | 6f2d-G | 2 | 7.6 | 65 | 89 | 8 | MOLECULE: | FLAGELLAR B |
| 53:00:00 | 3hc7-A | 2 | 15.8 | 67 | 252 | 9 | MOLECULE: | GENE 12 PRO |
| 54:00:00 | 6drh-E | 2 | 5.1 | 79 | 366 | 6 | MOLECULE: | ADP-RIBOSYL |
| 55:00:00 | 8b4h-C | 2 | 19.9 | 86 | 362 | 8 | MOLECULE: | PUTATIVE TR |
| 56:00:00 | 7xno-B | 2 | 5.9 | 62 | 106 | 10 | MOLECULE: | BACTERIOCIN |
| 57:00:00 | 5w6l-A | 2 | 20.7 | 99 | 494 | 3 | MOLECULE: | RTX REPEAT- |
| 58:00:00 | 5t9f-B | 2 | 9.7 | 75 | 254 | 5 | MOLECULE: | PREPHENATE |
| 59:00:00 | 6rj9-C | 2 | 25 | 86 | 836 | 7 | MOLECULE: | ACRIIA6 |
| 60:00:00 | 7f47-A | 2 | 6.7 | 117 | 830 | 7 | MOLECULE: | HYPOTHETICA |
| 61:00:00 | 7dve-B | 2 | 3.7 | 71 | 299 | 6 | MOLECULE: | 6'''-HYDROX |
| 62:00:00 | 8e5t-6 | 2 | 18.9 | 80 | 514 | 8 | MOLECULE: | RIBOSOME BI |
| 63:00:00 | 7vf2-A | 2 | 20 | 108 | 1196 | 7 | MOLECULE: | PROTEIN VIR |
| 64:00:00 | 8e9b-G | 2 | 9.1 | 83 | 151 | 6 | MOLECULE: | ACTIN-RELAT |
| 65:00:00 | 4u7u-B | 2 | 5.9 | 64 | 157 | 8 | MOLECULE: | CRISPR SYST |
| 66:00:00 | 6p4o-F | 2 | 17.1 | 82 | 481 | 10 | MOLECULE: | DNA-DEPENDE |
| 67:00:00 | 7d60-B | 2 | 13.9 | 103 | 288 | 6 | MOLECULE: | CALCIUM HOM |
| 68:00:00 | 4y7k-A | 2 | 23.6 | 90 | 250 | 4 | MOLECULE: | LARGE CONDU |
| 69:00:00 | 7jhj-R | 2 | 10.6 | 79 | 277 | 5 | MOLECULE: | GUANINE NUC |
| 70:00:00 | 8h7g-C | 2 | 24.7 | 100 | 3261 | 6 | MOLECULE: | TRANSFORMAT |
| 71:00:00 | 4g33-A | 2 | 6.9 | 86 | 661 | 12 | MOLECULE: | 15S-LIPOXYG |
| 72:00:00 | 7y6b-A | 2 | 7.1 | 56 | 72 | 2 | MOLECULE: | ESCE/YSCE/S |
| 73:00:00 | 1g4u-S | 2 | 12.1 | 73 | 360 | 11 | MOLECULE: | PROTEIN TYR |
| 74:00:00 | 5tj5-A | 2 | 15.2 | 80 | 570 | 8 | MOLECULE: | V-TYPE PROT |
| 75:00:00 | 7ctq-A | 2 | 9.9 | 93 | 429 | 10 | MOLECULE: | PEPTIDYL TR |
| 76:00:00 | 2nnu-A | 2 | 8.1 | 88 | 201 | 13 | MOLECULE: | REGULATORY |
| 77:00:00 | 3ksy-A | 2 | 21.8 | 102 | 1007 | 3 | MOLECULE: | SON OF SEVE |
| 78:00:00 | 8eki-D | 2 | 12.6 | 96 | 684 | 8 | MOLECULE: | PROTEIN TRA |
| 79:00:00 | 8k4a-H | 2 | 4.4 | 74 | 301 | 8 | MOLECULE: | VP2 |
| 80:00:00 | 8s4g-N | 2 | 31 | 138 | 682 | 7 | MOLECULE: | ANAPHASE-PR |

**Query : s001 A**

**DALI results: MdA-1 Penton**

| No: | Chain | Z | rmsd | lali | nres | %id | PDB Description | |
|---|---|---|---|---|---|---|---|---|
| 1:00 | 6z7q-E | 12 | 3.8 | 224 | 416 | 13 | MOLECULE: | PENTON PROTEIN |
| 2:00 | 6z7q-B | 11.9 | 3.9 | 225 | 416 | 13 | MOLECULE: | PENTON PROTEIN |
| 3:00 | 2c6s-C | 11.8 | 3.9 | 224 | 443 | 14 | MOLECULE: | ADENOVIRUS 2,12 |
| 4:00 | 2c6s-I | 11.8 | 3.8 | 223 | 443 | 13 | MOLECULE: | ADENOVIRUS 2,12 |
| 5:00 | 2c6s-M | 11.8 | 3.9 | 224 | 443 | 14 | MOLECULE: | ADENOVIRUS 2,12 |
| 6:00 | 2c6s-L | 11.8 | 3.8 | 223 | 443 | 13 | MOLECULE: | ADENOVIRUS 2,12 |
| 7:00 | 6z7q-D | 11.8 | 3.9 | 225 | 416 | 13 | MOLECULE: | PENTON PROTEIN |
| 8:00 | 6z7q-A | 11.8 | 3.9 | 225 | 416 | 13 | MOLECULE: | PENTON PROTEIN |
| 9:00 | 2c6s-G | 11.8 | 3.8 | 223 | 443 | 14 | MOLECULE: | ADENOVIRUS 2,12 |
| 10:00 | 6z7q-C | 11.8 | 3.9 | 224 | 416 | 13 | MOLECULE: | PENTON PROTEIN |
| 11:00 | 2c6s-H | 11.8 | 3.9 | 224 | 443 | 14 | MOLECULE: | ADENOVIRUS 2,12 |
| 12:00 | 2c6s-O | 11.8 | 3.8 | 224 | 443 | 13 | MOLECULE: | ADENOVIRUS 2,12 |
| 13:00 | 2c6s-E | 11.8 | 3.9 | 224 | 443 | 14 | MOLECULE: | ADENOVIRUS 2,12 |
| 14:00 | 2c6s-D | 11.8 | 3.8 | 223 | 443 | 13 | MOLECULE: | ADENOVIRUS 2,12 |
| 15:00 | 2c6s-F | 11.8 | 3.8 | 223 | 443 | 13 | MOLECULE: | ADENOVIRUS 2,12 |
| 16:00 | 1x9p-A | 11.7 | 3.9 | 226 | 460 | 14 | MOLECULE: | PENTON PROTEIN |
| 17:00 | 2c6s-K | 11.7 | 3.8 | 223 | 443 | 13 | MOLECULE: | ADENOVIRUS 2,12 |
| 18:00 | 2c6s-A | 11.7 | 3.8 | 223 | 443 | 14 | MOLECULE: | ADENOVIRUS 2,12 |
| 19:00 | 2c6s-J | 11.7 | 3.8 | 223 | 443 | 13 | MOLECULE: | ADENOVIRUS 2,12 |
| 20:00 | 2c9g-B | 11.7 | 3.9 | 226 | 447 | 14 | MOLECULE: | PENTON PROTEIN |
| 21:00 | 2c6s-B | 11.7 | 3.8 | 224 | 443 | 14 | MOLECULE: | ADENOVIRUS 2,12 |
| 22:00 | 2c6s-N | 11.7 | 3.8 | 223 | 443 | 13 | MOLECULE: | ADENOVIRUS 2,12 |
| 23:00 | 2c9g-C | 11.7 | 3.9 | 222 | 447 | 14 | MOLECULE: | PENTON PROTEIN |
| 24:00:00 | 2c9g-E | 11.7 | 3.9 | 225 | 447 | 14 | MOLECULE: | PENTON PROTEIN |
| 25:00:00 | 2c9g-D | 11.7 | 3.9 | 222 | 447 | 14 | MOLECULE: | PENTON PROTEIN |
| 26:00:00 | 2c9g-A | 11.7 | 4 | 225 | 447 | 15 | MOLECULE: | PENTON PROTEIN |
| 27:00:00 | 1x9t-A | 11.6 | 3.9 | 222 | 440 | 14 | MOLECULE: | PENTON PROTEIN |
| 28:00:00 | 2c9f-A | 11.6 | 3.9 | 222 | 440 | 14 | MOLECULE: | PENTON PROTEIN |
| 29:00:00 | 2c9f-B | 11.6 | 3.9 | 222 | 440 | 13 | MOLECULE: | PENTON PROTEIN |
| 30:00:00 | 2c9f-D | 11.6 | 3.9 | 222 | 440 | 14 | MOLECULE: | PENTON PROTEIN |
| 31:00:00 | 2c9f-C | 11.6 | 3.9 | 222 | 440 | 14 | MOLECULE: | PENTON PROTEIN |
| 32:00:00 | 2c9f-E | 11.6 | 3.9 | 222 | 440 | 14 | MOLECULE: | PENTON PROTEIN |
| 33:00:00 | 4ar2-A | 11.4 | 3.9 | 226 | 455 | 13 | MOLECULE: | L2 PROTEIN III |
| 34:00:00 | 3izo-D | 11.4 | 3.9 | 227 | 455 | 13 | MOLECULE: | PENTON PROTEIN |
| 35:00:00 | 3izo-C | 11.4 | 3.9 | 226 | 455 | 13 | MOLECULE: | PENTON PROTEIN |
| 36:00:00 | 3izo-A | 11.4 | 3.9 | 227 | 455 | 13 | MOLECULE: | PENTON PROTEIN |
| 37:00:00 | 3izo-E | 11.4 | 3.9 | 226 | 455 | 13 | MOLECULE: | PENTON PROTEIN |
| 38:00:00 | 3izo-B | 11.4 | 3.9 | 227 | 455 | 13 | MOLECULE: | PENTON PROTEIN |
| 39:00:00 | 4aqq-A | 11.3 | 3.9 | 226 | 462 | 13 | MOLECULE: | L2 PROTEIN III |
| 40:00:00 | 6g41-F | 11.2 | 3.3 | 159 | 301 | 6 | MOLECULE: | MINOR CAPSID PR |
| 41:00:00 | 6g41-I | 11 | 3.3 | 154 | 301 | 6 | MOLECULE: | MINOR CAPSID PR |

| | | | | | | | | |
|---|---|---|---|---|---|---|---|---|
| 42:00:00 | 6g42-D | 11 | 3.3 | 155 | 302 | 6 | MOLECULE: | MINOR CAPSID PR |
| 43:00:00 | 6g42-E | 10.9 | 3.2 | 158 | 301 | 6 | MOLECULE: | MINOR CAPSID PR |
| 44:00:00 | 6g41-J | 10.8 | 3.1 | 153 | 301 | 6 | MOLECULE: | MINOR CAPSID PR |
| 45:00:00 | 6g41-C | 10.8 | 3.2 | 156 | 301 | 6 | MOLECULE: | MINOR CAPSID PR |
| 46:00:00 | 6g42-A | 10.8 | 3.4 | 161 | 301 | 6 | MOLECULE: | MINOR CAPSID PR |
| 47:00:00 | 6g42-C | 10.8 | 3.3 | 159 | 302 | 6 | MOLECULE: | MINOR CAPSID PR |
| 48:00:00 | 6g42-B | 10.8 | 3.6 | 162 | 301 | 6 | MOLECULE: | MINOR CAPSID PR |
| 49:00:00 | 6g41-A | 10.8 | 3.1 | 155 | 297 | 6 | MOLECULE: | MINOR CAPSID PR |
| 50:00:00 | 6g41-E | 10.7 | 3 | 150 | 301 | 6 | MOLECULE: | MINOR CAPSID PR |
| 51:00:00 | 6g41-B | 10.7 | 3.2 | 154 | 295 | 6 | MOLECULE: | MINOR CAPSID PR |
| 52:00:00 | 6g41-H | 10.7 | 3.5 | 162 | 301 | 6 | MOLECULE: | MINOR CAPSID PR |
| 53:00:00 | 6g41-D | 10.7 | 3.2 | 155 | 291 | 6 | MOLECULE: | MINOR CAPSID PR |
| 54:00:00 | 8rbt-S | 9.3 | 3.3 | 209 | 663 | 11 | MOLECULE: | MAJOR CAPSID PR |
| 55:00:00 | 8coi-I | 8.9 | 2.4 | 99 | 249 | 11 | MOLECULE: | ADDOBODY |
| 56:00:00 | 8coi-A | 8.4 | 2.3 | 95 | 245 | 11 | MOLECULE: | ADDOBODY |
| 57:00:00 | 8coi-J | 8.4 | 2.4 | 94 | 243 | 11 | MOLECULE: | ADDOBODY |
| 58:00:00 | 8coi-Q | 8.4 | 2.4 | 96 | 232 | 11 | MOLECULE: | ADDOBODY |
| 59:00:00 | 8coi-P | 8.4 | 2.3 | 95 | 243 | 12 | MOLECULE: | ADDOBODY |
| 60:00:00 | 3j26-N | 8.2 | 3.8 | 166 | 362 | 7 | MOLECULE: | CAPSID PROTEIN |
| 61:00:00 | 8coi-C | 8.2 | 2.5 | 99 | 249 | 11 | MOLECULE: | ADDOBODY |
| 62:00:00 | 8coi-E | 8.2 | 2.3 | 94 | 228 | 11 | MOLECULE: | ADDOBODY |
| 63:00:00 | 6g41-G | 8 | 3.3 | 156 | 287 | 6 | MOLECULE: | MINOR CAPSID PR |
| 64:00:00 | 8coi-N | 7.9 | 2.3 | 92 | 232 | 12 | MOLECULE: | ADDOBODY |
| 65:00:00 | 8coi-H | 7.7 | 2.3 | 96 | 243 | 11 | MOLECULE: | ADDOBODY |
| 66:00:00 | 8coi-F | 7.6 | 2.3 | 98 | 240 | 11 | MOLECULE: | ADDOBODY |
| 67:00:00 | 8coi-R | 7.1 | 2.3 | 93 | 227 | 11 | MOLECULE: | ADDOBODY |
| 68:00:00 | 8coi-D | 7.1 | 2.3 | 96 | 244 | 11 | MOLECULE: | ADDOBODY |
| 69:00:00 | 8coi-G | 7 | 2.3 | 96 | 243 | 11 | MOLECULE: | ADDOBODY |
| 70:00:00 | 8coi-M | 7 | 2.4 | 96 | 237 | 11 | MOLECULE: | ADDOBODY |
| 71:00:00 | 8coi-L | 7 | 2.4 | 95 | 244 | 12 | MOLECULE: | ADDOBODY |
| 72:00:00 | 8qb3-E | 6.9 | 2.7 | 89 | 233 | 10 | MOLECULE: | ADDOBODY |
| 73:00:00 | 8coi-S | 6.8 | 2.3 | 94 | 230 | 11 | MOLECULE: | ADDOBODY |
| 74:00:00 | 8coi-B | 6.7 | 2.3 | 93 | 246 | 12 | MOLECULE: | ADDOBODY |
| 75:00:00 | 8coi-O | 6.7 | 2.4 | 92 | 229 | 10 | MOLECULE: | ADDOBODY |
| 76:00:00 | 8coi-K | 6.6 | 2.3 | 92 | 250 | 12 | MOLECULE: | ADDOBODY |
| 77:00:00 | 8coi-T | 6.6 | 2.4 | 84 | 212 | 11 | MOLECULE: | ADDOBODY |
| 78:00:00 | 7cz6-C | 6.2 | 3.1 | 99 | 126 | 8 | MOLECULE: | CAPSID PROTEIN |
| 79:00:00 | 6ds5-E | 6.2 | 3.8 | 107 | 160 | 13 | MOLECULE: | SEIPIN |
| 80:00:00 | 6ds5-K | 6.2 | 3.8 | 107 | 160 | 13 | MOLECULE: | SEIPIN |
| 81:00:00 | 6ds5-B | 6.2 | 3.8 | 107 | 160 | 13 | MOLECULE: | SEIPIN |
| 82:00:00 | 6ds5-J | 6.2 | 3.8 | 107 | 160 | 13 | MOLECULE: | SEIPIN |
| 83:00:00 | 6ds5-D | 6.2 | 3.8 | 107 | 160 | 13 | MOLECULE: | SEIPIN |
| 84:00:00 | 6ds5-C | 6.2 | 3.8 | 107 | 160 | 13 | MOLECULE: | SEIPIN |
| 85:00:00 | 6ds5-G | 6.1 | 3.6 | 104 | 160 | 13 | MOLECULE: | SEIPIN |

| | | | | | | | | |
|---|---|---|---|---|---|---|---|---|
| 86:00:00 | 6ds5-H | 6.1 | 3.8 | 105 | 160 | 13 | MOLECULE: | SEIPIN |
| 87:00:00 | 6wpc-C | 6 | 3.4 | 113 | 588 | 9 | MOLECULE: | CRY1A.2 |
| 88:00:00 | 6ds5-A | 6 | 3.6 | 105 | 160 | 12 | MOLECULE: | SEIPIN |
| 89:00:00 | 6ds5-F | 6 | 3.7 | 104 | 160 | 13 | MOLECULE: | SEIPIN |
| 90:00:00 | 6ds5-I | 6 | 3.7 | 104 | 160 | 13 | MOLECULE: | SEIPIN |
| 91:00:00 | 6wpc-B | 5.9 | 3.4 | 109 | 588 | 11 | MOLECULE: | CRY1A.2 |
| 92:00:00 | 6wpc-A | 5.9 | 3.2 | 114 | 588 | 11 | MOLECULE: | CRY1A.2 |
| 93:00:00 | 8qb3-C | 5.9 | 2.6 | 89 | 216 | 10 | MOLECULE: | ADDOBODY |
| 94:00:00 | 6owk-A | 5.9 | 3.2 | 116 | 582 | 6 | MOLECULE: | PESTICIDAL CRYS |
| 95:00:00 | 7rsl-B | 5.7 | 3.3 | 108 | 235 | 11 | MOLECULE: | SEIPIN |
| 96:00:00 | 7rsl-D | 5.7 | 3.3 | 108 | 235 | 11 | MOLECULE: | SEIPIN |
| 97:00:00 | 7rsl-F | 5.7 | 3.4 | 109 | 235 | 11 | MOLECULE: | SEIPIN |
| 98:00:00 | 7rsl-H | 5.7 | 3.2 | 107 | 235 | 11 | MOLECULE: | SEIPIN |
| 99:00:00 | 1ciy-A | 5.7 | 3.9 | 115 | 577 | 8 | MOLECULE: | CRYIA(A) |
| 0:00 | 7rsl-J | 5.7 | 3.3 | 108 | 235 | 11 | MOLECULE: | SEIPIN |
| 1:00 | 9buv-A | 5.6 | 3.9 | 110 | 573 | 7 | MOLECULE: | PESTICIDAL CRYS |
| 2:00 | 8qb3-D | 5.6 | 2.5 | 84 | 206 | 11 | MOLECULE: | ADDOBODY |
| 3:00 | 7rsl-E | 5.5 | 5.2 | 111 | 235 | 13 | MOLECULE: | SEIPIN |
| 4:00 | 7rsl-G | 5.5 | 5.2 | 110 | 235 | 13 | MOLECULE: | SEIPIN |
| 5:00 | 7rsl-A | 5.5 | 5.2 | 110 | 235 | 13 | MOLECULE: | SEIPIN |
| 6:00 | 7rsl-I | 5.5 | 5.2 | 110 | 235 | 13 | MOLECULE: | SEIPIN |
| 7:00 | 7rsl-C | 5.5 | 5.2 | 112 | 235 | 13 | MOLECULE: | SEIPIN |
| 8:00 | 6mlu-A | 5.5 | 4.1 | 105 | 153 | 5 | MOLECULE: | SEIPIN |
| 9:00 | 6mlu-B | 5.5 | 4.1 | 105 | 153 | 5 | MOLECULE: | SEIPIN |
| 10:00 | 6za2-A | 5.5 | 3.7 | 104 | 1081 | 11 | MOLECULE: | POR SECRETION S |
| 11:00 | 8qb3-A | 5.5 | 2.4 | 85 | 233 | 11 | MOLECULE: | ADDOBODY |
| 12:00 | 3eb7-A | 5.4 | 3.8 | 109 | 589 | 8 | MOLECULE: | INSECTICIDAL DE |
| 13:00 | 3eb7-C | 5.4 | 3.3 | 105 | 588 | 9 | MOLECULE: | INSECTICIDAL DE |
| 14:00 | 1dlc-A | 5.4 | 4 | 114 | 584 | 4 | MOLECULE: | DELTA-ENDOTOXIN |
| 15:00 | 8qb3-B | 5.4 | 2.8 | 89 | 233 | 10 | MOLECULE: | ADDOBODY |
| 16:00 | 6dj4-A | 5.3 | 3.3 | 110 | 571 | 5 | MOLECULE: | CRY1A.105 |
| 17:00 | 6ovb-A | 5.3 | 4.1 | 110 | 566 | 11 | MOLECULE: | ACTIVE CORE CRY |
| 18:00 | 4qx3-A | 5.3 | 4 | 115 | 584 | 5 | MOLECULE: | PESTICIDAL CRYS |
| 19:00 | 4qx0-A | 5.3 | 4 | 115 | 584 | 5 | MOLECULE: | PESTICIDAL CRYS |
| 20:00 | 1ji6-A | 5.2 | 3.9 | 105 | 589 | 6 | MOLECULE: | PESTICIDIAL CRY |
| 21:00 | 4qx1-A | 5.2 | 4 | 116 | 584 | 4 | MOLECULE: | PESTICIDAL CRYS |
| 22:00 | 4qx2-A | 5.1 | 4 | 112 | 584 | 7 | MOLECULE: | PESTICIDAL CRYS |
| 23:00 | 7ear-A | 5.1 | 3.9 | 110 | 584 | 7 | MOLECULE: | CRYSTALINE ENTO |
| 24:00:00 | 3iii-A | 5 | 3.4 | 114 | 556 | 4 | MOLECULE: | COCE/NOND FAMIL |
| 25:00:00 | 4moa-A | 5 | 4 | 117 | 602 | 11 | MOLECULE: | PESTICIDAL CRYS |
| 26:00:00 | 6za2-B | 4.9 | 3.7 | 111 | 1083 | 10 | MOLECULE: | POR SECRETION S |
| 27:00:00 | 5hlz-A | 4.9 | 4.4 | 103 | 204 | 7 | MOLECULE: | INHIBIN BETA A |
| 28:00:00 | 4ary-B | 4.8 | 4.5 | 106 | 580 | 8 | MOLECULE: | PESTICIDAL CRYS |
| 29:00:00 | 4arx-C | 4.8 | 4.5 | 104 | 577 | 9 | MOLECULE: | PESTICIDAL CRYS |

| | | | | | | | | |
|---|---|---|---|---|---|---|---|---|
| 30:00:00 | 5hlz-C | 4.8 | 4.3 | 104 | 204 | 7 | MOLECULE: | INHIBIN BETA A |
| 31:00:00 | 5hly-A | 4.8 | 4.4 | 105 | 299 | 7 | MOLECULE: | INHIBIN BETA A |
| 32:00:00 | 4w8j-A | 4.7 | 4.3 | 103 | 1017 | 9 | MOLECULE: | PESTICIDAL CRYS |
| 33:00:00 | 4arx-B | 4.7 | 4.1 | 106 | 578 | 7 | MOLECULE: | PESTICIDAL CRYS |
| 34:00:00 | 5zi1-A | 4.7 | 4 | 115 | 590 | 7 | MOLECULE: | INSECTICIDAL CR |
| 35:00:00 | 4ary-C | 4.7 | 4.5 | 105 | 577 | 7 | MOLECULE: | PESTICIDAL CRYS |
| 36:00:00 | 5cvz-A | 4.6 | 3.6 | 104 | 141 | 6 | MOLECULE: | COAT PROTEIN |
| 37:00:00 | 3eb7-B | 4.6 | 4.5 | 97 | 589 | 7 | MOLECULE: | INSECTICIDAL DE |
| 38:00:00 | 5zi1-B | 4.6 | 3.5 | 115 | 590 | 7 | MOLECULE: | INSECTICIDAL CR |
| 39:00:00 | 3qd6-D | 4.6 | 3.7 | 100 | 135 | 11 | MOLECULE: | CD40 LIGAND |
| 40:00:00 | 1w99-A | 4.6 | 4.2 | 98 | 558 | 10 | MOLECULE: | PESTICIDIAL CRY |
| 41:00:00 | 1stm-A | 4.5 | 3.4 | 99 | 141 | 5 | MOLECULE: | SATELLITE PANIC |
| 42:00:00 | 1stm-C | 4.5 | 3.5 | 100 | 141 | 5 | MOLECULE: | SATELLITE PANIC |
| 43:00:00 | 1stm-B | 4.5 | 3.5 | 101 | 141 | 5 | MOLECULE: | SATELLITE PANIC |
| 44:00:00 | 4arx-D | 4.5 | 4.2 | 104 | 576 | 8 | MOLECULE: | PESTICIDAL CRYS |
| 45:00:00 | 1stm-D | 4.5 | 3.6 | 102 | 141 | 5 | MOLECULE: | SATELLITE PANIC |
| 46:00:00 | 1stm-E | 4.5 | 3.5 | 100 | 141 | 5 | MOLECULE: | SATELLITE PANIC |
| 47:00:00 | 4ary-D | 4.5 | 4.5 | 100 | 577 | 8 | MOLECULE: | PESTICIDAL CRYS |
| 48:00:00 | 3c7g-A | 4.4 | 3.7 | 101 | 488 | 7 | MOLECULE: | ENDO-1,4-BETA-X |
| 49:00:00 | 3ib3-B | 4.4 | 3.3 | 114 | 552 | 4 | MOLECULE: | COCE/NOND FAMIL |
| 50:00:00 | 1aly-A | 4.4 | 3.6 | 106 | 146 | 9 | MOLECULE: | CD40 LIGAND |
| 51:00:00 | 7r1m-A | 4.3 | 3.4 | 102 | 156 | 8 | MOLECULE: | EXOSPORIUM PROT |
| 52:00:00 | 8j3y-A | 4.3 | 4.2 | 98 | 392 | 5 | MOLECULE: | PUTATIVE POLYSA |
| 53:00:00 | 3ib3-A | 4.3 | 3.5 | 115 | 551 | 9 | MOLECULE: | COCE/NOND FAMIL |
| 54:00:00 | 6lfp-A | 4.3 | 4.5 | 95 | 582 | 6 | MOLECULE: | CRY3AA PROTEIN |
| 55:00:00 | 2c9k-A | 4.3 | 5 | 98 | 598 | 6 | MOLECULE: | PESTICIDAL CRYS |
| 56:00:00 | 4ind-A | 4.2 | 5.2 | 96 | 377 | 9 | MOLECULE: | C381 TURRET PRO |
| 57:00:00 | 3qd6-E | 4.2 | 3.8 | 95 | 135 | 11 | MOLECULE: | CD40 LIGAND |
| 58:00:00 | 8w7n-A | 4.2 | 5 | 96 | 1011 | 10 | MOLECULE: | PESTICIDAL CRYS |
| 59:00:00 | 4lr4-C | 4.1 | 3.7 | 96 | 346 | 6 | MOLECULE: | HYPOTHETICAL PR |
| 60:00:00 | 4qpw-A | 4.1 | 3 | 86 | 142 | 8 | MOLECULE: | GLYCOSYL HYDROL |
| 61:00:00 | 1i9r-A | 4.1 | 3.9 | 101 | 143 | 10 | MOLECULE: | CD40 LIGAND |
| 62:00:00 | 3qd6-B | 4.1 | 3.8 | 95 | 135 | 11 | MOLECULE: | CD40 LIGAND |
| 63:00:00 | 3qd6-F | 4.1 | 3.5 | 95 | 135 | 8 | MOLECULE: | CD40 LIGAND |
| 64:00:00 | 2rjl-A | 4.1 | 4 | 99 | 141 | 11 | MOLECULE: | TNF SUPERFAMILY |
| 65:00:00 | 1i9r-C | 4.1 | 3.8 | 99 | 143 | 9 | MOLECULE: | CD40 LIGAND |
| 66:00:00 | 6w9g-B | 4.1 | 3.5 | 95 | 137 | 9 | MOLECULE: | CD40 LIGAND |
| 67:00:00 | 3qd6-A | 4.1 | 3.3 | 99 | 135 | 8 | MOLECULE: | CD40 LIGAND |
| 68:00:00 | 4ary-A | 4.1 | 5.1 | 92 | 578 | 7 | MOLECULE: | PESTICIDAL CRYS |
| 69:00:00 | 4qhw-A | 4 | 3.2 | 91 | 400 | 3 | MOLECULE: | UNCHARACTERIZED |
| 70:00:00 | 6w9g-A | 4 | 3.6 | 93 | 137 | 11 | MOLECULE: | CD40 LIGAND |
| 71:00:00 | 4arx-A | 4 | 3.6 | 107 | 578 | 9 | MOLECULE: | PESTICIDAL CRYS |
| 72:00:00 | 7sgm-C | 4 | 3.7 | 96 | 137 | 9 | MOLECULE: | CD40 LIGAND |
| 73:00:00 | 3qd6-C | 4 | 3.5 | 100 | 135 | 11 | MOLECULE: | CD40 LIGAND |

| | | | | | | | | |
|---|---|---|---|---|---|---|---|---|
| 74:00:00 | 1i9r-B | 4 | 3.9 | 100 | 143 | 8 | MOLECULE: | CD40 LIGAND |
| 75:00:00 | 2rjk-A | 4 | 4.1 | 96 | 141 | 9 | MOLECULE: | TNF SUPERFAMILY |
| 76:00:00 | 5cw0-A | 4 | 3.8 | 100 | 141 | 6 | MOLECULE: | COAT PROTEIN |
| 77:00:00 | 7vt4-A | 3.9 | 4.3 | 86 | 438 | 7 | MOLECULE: | ENDOGLUCANASE H |
| 78:00:00 | 7kx0-A | 3.9 | 3.8 | 95 | 140 | 5 | MOLECULE: | CD70 ANTIGEN |
| 79:00:00 | 5jsd-A | 3.9 | 3.2 | 85 | 548 | 7 | MOLECULE: | PHIAB6 TAILSPIK |
| 80:00:00 | 8xcg-J | 3.9 | 6.8 | 106 | 598 | 5 | MOLECULE: | TAIL TIP PROTEI |
| 81:00:00 | 4mtm-A | 3.9 | 4.2 | 81 | 137 | 10 | MOLECULE: | PUTATIVE TAIL F |
| 82:00:00 | 6wpc-D | 3.9 | 3.3 | 114 | 588 | 11 | MOLECULE: | CRY1A.2 |
| 83:00:00 | 5hlz-G | 3.9 | 4.5 | 97 | 204 | 6 | MOLECULE: | INHIBIN BETA A |
| 84:00:00 | 3lkj-A | 3.9 | 3.8 | 97 | 133 | 11 | MOLECULE: | CD40 LIGAND |
| 85:00:00 | 6w9g-C | 3.9 | 3.5 | 98 | 139 | 8 | MOLECULE: | CD40 LIGAND |
| 86:00:00 | 4e4d-X | 3.9 | 3.9 | 96 | 154 | 3 | MOLECULE: | TUMOR NECROSIS |
| 87:00:00 | 2wys-B | 3.8 | 3.3 | 90 | 516 | 8 | MOLECULE: | ENDO-1,4-BETA-X |
| 88:00:00 | 6sih-A | 3.8 | 18.2 | 98 | 488 | 8 | MOLECULE: | FLAGELLAR HOOK- |
| 89:00:00 | 2hey-F | 3.8 | 3.8 | 87 | 135 | 10 | MOLECULE: | TUMOR NECROSIS |
| 90:00:00 | 2qe3-A | 3.8 | 4 | 96 | 140 | 10 | MOLECULE: | TNF SUPERFAMILY |
| 91:00:00 | 3lkj-C | 3.8 | 3.7 | 91 | 129 | 2 | MOLECULE: | CD40 LIGAND |
| 92:00:00 | 1jtz-X | 3.8 | 3.8 | 97 | 155 | 3 | MOLECULE: | TUMOR NECROSIS |
| 93:00:00 | 7dn2-1 | 3.7 | 3.2 | 89 | 126 | 8 | MOLECULE: | MAJOR STRUCTURA |
| 94:00:00 | 8des-A | 3.7 | 3.7 | 102 | 475 | 7 | MOLECULE: | MAJOR CAPSID PR |
| 95:00:00 | 4z11-A | 3.7 | 4.3 | 97 | 513 | 2 | MOLECULE: | AURONE SYNTHASE |
| 96:00:00 | 5hlz-E | 3.7 | 4.4 | 104 | 204 | 5 | MOLECULE: | INHIBIN BETA A |
| 97:00:00 | 4msv-A | 3.7 | 4.1 | 100 | 139 | 4 | MOLECULE: | TUMOR NECROSIS |
| 98:00:00 | 3wqb-A | 3.6 | 3.8 | 104 | 597 | 13 | MOLECULE: | EXTRACELLULAR S |
| 99:00:00 | 8ond-A | 3.6 | 3.5 | 103 | 344 | 5 | MOLECULE: | BD2133 |
| 0:00 | 4ht1-T | 3.6 | 3.8 | 87 | 131 | 5 | MOLECULE: | TUMOR NECROSIS |
| 1:00 | 6iwy-A | 3.6 | 2.6 | 75 | 366 | 12 | MOLECULE: | FLAGELLAR HOOK- |
| 2:00 | 5zru-A | 3.6 | 6.8 | 105 | 576 | 7 | MOLECULE: | ALPHA-1,3-GLUCA |
| 3:00 | 5cir-D | 3.6 | 4.3 | 102 | 150 | 8 | MOLECULE: | TUMOR NECROSIS |
| 4:00 | 5l19-A | 3.6 | 4.7 | 100 | 150 | 5 | MOLECULE: | TUMOR NECROSIS |
| 5:00 | 7ta6-B | 3.6 | 4.4 | 96 | 129 | 7 | MOLECULE: | TUMOR NECROSIS |
| 6:00 | 7sgm-A | 3.6 | 3.9 | 95 | 137 | 8 | MOLECULE: | CD40 LIGAND |
| 7:00 | 3lkj-B | 3.6 | 3.8 | 97 | 132 | 6 | MOLECULE: | CD40 LIGAND |
| 8:00 | 5cir-B | 3.6 | 4.1 | 101 | 151 | 9 | MOLECULE: | TUMOR NECROSIS |
| 9:00 | 4mxw-D | 3.6 | 4.4 | 104 | 135 | 6 | MOLECULE: | TUMOR NECROSIS |
| 10:00 | 5cir-A | 3.5 | 4.2 | 102 | 149 | 8 | MOLECULE: | TUMOR NECROSIS |
| 11:00 | 7ta6-A | 3.5 | 3.9 | 109 | 141 | 5 | MOLECULE: | TUMOR NECROSIS |
| 12:00 | 7ta6-H | 3.5 | 3.4 | 98 | 131 | 7 | MOLECULE: | TUMOR NECROSIS |
| 13:00 | 6w98-A | 3.4 | 9.3 | 141 | 1312 | 6 | MOLECULE: | F5/8 TYPE C DOM |
| 14:00 | 2vzs-A | 3.4 | 3.5 | 92 | 857 | 9 | MOLECULE: | EXO-BETA-D-GLUC |
| 15:00 | 1a6c-A | 3.4 | 7.1 | 96 | 513 | 4 | MOLECULE: | TOBACCO RINGSPO |
| 16:00 | 2gh8-A | 3.4 | 6.8 | 103 | 544 | 4 | MOLECULE: | CAPSID PROTEIN |
| 17:00 | 2wz8-A | 3.4 | 3.3 | 82 | 135 | 11 | MOLECULE: | CELLULOSOME PRO |

| | | | | | | | | |
|---|---|---|---|---|---|---|---|---|
| 18:00 | 7ta6-F | 3.4 | 3.8 | 105 | 138 | 8 | MOLECULE: | TUMOR NECROSIS |
| 19:00 | 7jra-B | 3.4 | 4.7 | 98 | 146 | 7 | MOLECULE: | TUMOR NECROSIS |
| 20:00 | 6x86-F | 3.4 | 3.8 | 103 | 144 | 7 | MOLECULE: | TUMOR NECROSIS |
| 21:00 | 6x85-F | 3.4 | 3.7 | 104 | 144 | 7 | MOLECULE: | TUMOR NECROSIS |
| 22:00 | 7sgm-B | 3.4 | 3.8 | 100 | 140 | 3 | MOLECULE: | CD40 LIGAND |
| 23:00 | 7ta6-G | 3.4 | 4.4 | 97 | 138 | 6 | MOLECULE: | TUMOR NECROSIS |
| 24:00:00 | 3k51-A | 3.4 | 3.7 | 97 | 143 | 3 | MOLECULE: | TUMOR NECROSIS |
| 25:00:00 | 7kp9-B | 3.4 | 3.9 | 97 | 139 | 10 | MOLECULE: | TUMOR NECROSIS |
| 26:00:00 | 7w63-B | 3.3 | 4.1 | 94 | 395 | 7 | MOLECULE: | TOXIN-COREGULAT |
| 27:00:00 | 4jdm-C | 3.3 | 4.2 | 102 | 269 | 7 | MOLECULE: | VIRULENCE PLASM |
| 28:00:00 | 7ta6-E | 3.3 | 3.9 | 102 | 136 | 7 | MOLECULE: | TUMOR NECROSIS |
| 29:00:00 | 2tun-D | 3.3 | 3.6 | 105 | 151 | 6 | MOLECULE: | TUMOR NECROSIS |
| 30:00:00 | 2tun-F | 3.3 | 3.9 | 103 | 151 | 10 | MOLECULE: | TUMOR NECROSIS |
| 31:00:00 | 2tun-A | 3.3 | 4 | 108 | 151 | 7 | MOLECULE: | TUMOR NECROSIS |
| 32:00:00 | 6ooy-B | 3.3 | 3.9 | 103 | 140 | 10 | MOLECULE: | TUMOR NECROSIS |
| 33:00:00 | 6ooy-A | 3.3 | 4 | 102 | 140 | 7 | MOLECULE: | TUMOR NECROSIS |
| 34:00:00 | 6ooz-C | 3.3 | 4 | 96 | 126 | 8 | MOLECULE: | TUMOR NECROSIS |
| 35:00:00 | 2zpx-B | 3.3 | 4.3 | 99 | 142 | 7 | MOLECULE: | TUMOR NECROSIS |
| 36:00:00 | 6rmj-C | 3.3 | 3.8 | 108 | 140 | 6 | MOLECULE: | TUMOR NECROSIS |
| 37:00:00 | 5mu8-f | 3.3 | 3.8 | 99 | 143 | 8 | MOLECULE: | TUMOR NECROSIS |
| 38:00:00 | 6ooz-B | 3.3 | 3.8 | 95 | 142 | 8 | MOLECULE: | TUMOR NECROSIS |
| 39:00:00 | 2o0o-B | 3.3 | 3.9 | 104 | 153 | 8 | MOLECULE: | TNF SUPERFAMILY |
| 40:00:00 | 6brb-A | 3.3 | 3.6 | 97 | 142 | 2 | MOLECULE: | CD40 LIGAND |
| 41:00:00 | 5mu8-L | 3.3 | 4 | 103 | 142 | 8 | MOLECULE: | TUMOR NECROSIS |
| 42:00:00 | 6x82-F | 3.3 | 3.6 | 95 | 140 | 6 | MOLECULE: | TUMOR NECROSIS |
| 43:00:00 | 2re9-A | 3.3 | 3.8 | 114 | 164 | 5 | MOLECULE: | TNF SUPERFAMILY |
| 44:00:00 | 8env-A | 3.2 | 15 | 87 | 504 | 9 | MOLECULE: | SHEATH PROTEIN |
| 45:00:00 | 8gld-A | 3.2 | 3.6 | 84 | 365 | 6 | MOLECULE: | SGNH HYDROLASE- |
| 46:00:00 | 1dg6-A | 3.2 | 4.2 | 95 | 149 | 9 | MOLECULE: | APO2L/TNF-RELAT |
| 47:00:00 | 5uui-A | 3.2 | 4.2 | 90 | 134 | 11 | MOLECULE: | TUMOR NECROSIS |
| 48:00:00 | 2zjc-A | 3.2 | 3.8 | 103 | 142 | 11 | MOLECULE: | TUMOR NECROSIS |
| 49:00:00 | 7ta6-D | 3.2 | 3.7 | 104 | 145 | 6 | MOLECULE: | TUMOR NECROSIS |
| 50:00:00 | 4mxw-Y | 3.2 | 3.5 | 95 | 131 | 8 | MOLECULE: | TUMOR NECROSIS |
| 51:00:00 | 1tnf-A | 3.2 | 3.8 | 102 | 152 | 8 | MOLECULE: | TUMOR NECROSIS |
| 52:00:00 | 2az5-B | 3.2 | 3.7 | 97 | 139 | 9 | MOLECULE: | TUMOR NECROSIS |
| 53:00:00 | 6x85-C | 3.2 | 3.9 | 100 | 144 | 7 | MOLECULE: | TUMOR NECROSIS |
| 54:00:00 | 7ta6-C | 3.2 | 4 | 102 | 138 | 8 | MOLECULE: | TUMOR NECROSIS |
| 55:00:00 | 3it8-G | 3.2 | 4 | 99 | 152 | 8 | MOLECULE: | TUMOR NECROSIS |
| 56:00:00 | 3it8-I | 3.2 | 4 | 107 | 152 | 5 | MOLECULE: | TUMOR NECROSIS |
| 57:00:00 | 6op0-B | 3.2 | 3.9 | 104 | 141 | 7 | MOLECULE: | TUMOR NECROSIS |
| 58:00:00 | 3it8-H | 3.2 | 4 | 107 | 152 | 5 | MOLECULE: | TUMOR NECROSIS |
| 59:00:00 | 5mu8-H | 3.2 | 4 | 101 | 150 | 7 | MOLECULE: | TUMOR NECROSIS |
| 60:00:00 | 5yoy-B | 3.2 | 3.8 | 98 | 154 | 9 | MOLECULE: | TUMOR NECROSIS |
| 61:00:00 | 1s55-A | 3.2 | 3.7 | 98 | 156 | 3 | MOLECULE: | TUMOR NECROSIS |

| | | | | | | | | |
|---|---|---|---|---|---|---|---|---|
| 62:00:00 | 7kp9-C | 3.2 | 3.6 | 100 | 131 | 7 | MOLECULE: | TUMOR NECROSIS |
| 63:00:00 | 6ooz-A | 3.2 | 4 | 101 | 147 | 7 | MOLECULE: | TUMOR NECROSIS |
| 64:00:00 | 3it8-C | 3.2 | 3.7 | 99 | 152 | 9 | MOLECULE: | TUMOR NECROSIS |
| 65:00:00 | 5yoy-C | 3.2 | 3.6 | 100 | 154 | 7 | MOLECULE: | TUMOR NECROSIS |
| 66:00:00 | 6x86-B | 3.2 | 3.7 | 95 | 130 | 6 | MOLECULE: | TUMOR NECROSIS |
| 67:00:00 | 5m2m-M | 3.2 | 3.8 | 102 | 148 | 8 | MOLECULE: | TUMOR NECROSIS |
| 68:00:00 | 1iqa-A | 3.2 | 3.8 | 99 | 156 | 4 | MOLECULE: | RECEPTOR ACTIVA |
| 69:00:00 | 1tnf-B | 3.2 | 3.8 | 102 | 152 | 9 | MOLECULE: | TUMOR NECROSIS |
| 70:00:00 | 6x85-A | 3.2 | 3.9 | 98 | 148 | 8 | MOLECULE: | TUMOR NECROSIS |
| 71:00:00 | 1tnf-C | 3.2 | 3.8 | 101 | 152 | 10 | MOLECULE: | TUMOR NECROSIS |
| 72:00:00 | 5yoy-L | 3.2 | 4.1 | 106 | 153 | 7 | MOLECULE: | TUMOR NECROSIS |
| 73:00:00 | 5mu8-b | 3.2 | 3.8 | 96 | 144 | 8 | MOLECULE: | TUMOR NECROSIS |
| 74:00:00 | 7kpa-B | 3.2 | 3.9 | 100 | 146 | 10 | MOLECULE: | TUMOR NECROSIS |
| 75:00:00 | 2zpx-C | 3.2 | 4.2 | 102 | 142 | 6 | MOLECULE: | TUMOR NECROSIS |
| 76:00:00 | 3wd5-A | 3.2 | 3.8 | 110 | 152 | 5 | MOLECULE: | TUMOR NECROSIS |
| 77:00:00 | 1a8m-B | 3.2 | 3.5 | 101 | 152 | 7 | MOLECULE: | TUMOR NECROSIS |
| 78:00:00 | 5l36-A | 3.2 | 3.7 | 90 | 138 | 6 | MOLECULE: | TUMOR NECROSIS |
| 79:00:00 | 6x83-E | 3.2 | 4.2 | 97 | 130 | 10 | MOLECULE: | TUMOR NECROSIS |
| 80:00:00 | 3mi8-A | 3.2 | 3.7 | 89 | 144 | 4 | MOLECULE: | TUMOR NECROSIS |
| 81:00:00 | 3it8-A | 3.2 | 4 | 107 | 152 | 5 | MOLECULE: | TUMOR NECROSIS |
| 82:00:00 | 6q5u-R | 3.1 | 3.4 | 81 | 125 | 12 | MOLECULE: | MAJOR CAPSID PR |
| 83:00:00 | 6x85-B | 3.1 | 3.9 | 98 | 129 | 8 | MOLECULE: | TUMOR NECROSIS |
| 84:00:00 | 3it8-B | 3.1 | 4.3 | 96 | 152 | 4 | MOLECULE: | TUMOR NECROSIS |
| 85:00:00 | 7kp8-C | 3.1 | 3.9 | 92 | 137 | 11 | MOLECULE: | TUMOR NECROSIS |
| 86:00:00 | 7kpb-C | 3.1 | 3.9 | 97 | 138 | 9 | MOLECULE: | TUMOR NECROSIS |
| 87:00:00 | 5yoy-A | 3.1 | 3.9 | 106 | 154 | 8 | MOLECULE: | TUMOR NECROSIS |
| 88:00:00 | 1iqa-C | 3.1 | 4.5 | 98 | 156 | 4 | MOLECULE: | RECEPTOR ACTIVA |
| 89:00:00 | 3alq-E | 3.1 | 4.2 | 96 | 141 | 9 | MOLECULE: | TUMOR NECROSIS |
| 90:00:00 | 1a8m-A | 3.1 | 3.9 | 98 | 152 | 8 | MOLECULE: | TUMOR NECROSIS |
| 91:00:00 | 5mu8-C | 3.1 | 4 | 100 | 146 | 9 | MOLECULE: | TUMOR NECROSIS |
| 92:00:00 | 7kp9-A | 3.1 | 4.1 | 105 | 148 | 9 | MOLECULE: | TUMOR NECROSIS |
| 93:00:00 | 3urf-A | 3.1 | 4.1 | 99 | 156 | 3 | MOLECULE: | TUMOR NECROSIS |
| 94:00:00 | 6x83-F | 3.1 | 4.1 | 101 | 138 | 9 | MOLECULE: | TUMOR NECROSIS |
| 95:00:00 | 5wux-G | 3.1 | 4.1 | 98 | 141 | 7 | MOLECULE: | HEAVY |
| 96:00:00 | 1s55-B | 3.1 | 4 | 95 | 156 | 4 | MOLECULE: | TUMOR NECROSIS |
| 97:00:00 | 5mu8-X | 3.1 | 3.8 | 101 | 150 | 6 | MOLECULE: | TUMOR NECROSIS |
| 98:00:00 | 5m2m-C | 3.1 | 4 | 98 | 150 | 6 | MOLECULE: | TUMOR NECROSIS |
| 99:00:00 | 6op0-C | 3.1 | 3.9 | 92 | 119 | 10 | MOLECULE: | TUMOR NECROSIS |
| 0:00 | 5mu8-I | 3.1 | 4 | 100 | 141 | 6 | MOLECULE: | TUMOR NECROSIS |
| 1:00 | 2tnf-B | 3.1 | 3.6 | 102 | 148 | 8 | MOLECULE: | PROTEIN (TUMOR |
| 2:00 | 7kpa-A | 3.1 | 4.2 | 102 | 147 | 8 | MOLECULE: | TUMOR NECROSIS |
| 3:00 | 1d0g-B | 3.1 | 4 | 105 | 151 | 4 | MOLECULE: | DEATH RECEPTOR- |
| 4:00 | 5mu8-d | 3.1 | 4.1 | 99 | 138 | 9 | MOLECULE: | TUMOR NECROSIS |
| 5:00 | 7kp8-A | 3.1 | 4 | 96 | 140 | 8 | MOLECULE: | TUMOR NECROSIS |

| | | | | | | | | |
|---|---|---|---|---|---|---|---|---|
| 6:00 | 1a8m-C | 3.1 | 4.1 | 101 | 152 | 7 | MOLECULE: | TUMOR NECROSIS |
| 7:00 | 2az5-A | 3.1 | 3.9 | 99 | 131 | 8 | MOLECULE: | TUMOR NECROSIS |
| 8:00 | 7kp7-B | 3.1 | 4 | 98 | 148 | 7 | MOLECULE: | TUMOR NECROSIS |
| 9:00 | 7kpb-B | 3.1 | 4 | 99 | 143 | 7 | MOLECULE: | TUMOR NECROSIS |
| 10:00 | 6x82-C | 3.1 | 3.3 | 98 | 143 | 7 | MOLECULE: | TUMOR NECROSIS |
| 11:00 | 6x83-C | 3.1 | 3.7 | 93 | 135 | 6 | MOLECULE: | TUMOR NECROSIS |
| 12:00 | 5m2m-G | 3.1 | 4 | 98 | 150 | 6 | MOLECULE: | TUMOR NECROSIS |
| 13:00 | 6x83-B | 3.1 | 4.1 | 99 | 138 | 10 | MOLECULE: | TUMOR NECROSIS |
| 14:00 | 2o0o-A | 3.1 | 4.2 | 102 | 153 | 4 | MOLECULE: | TNF SUPERFAMILY |
| 15:00 | 5mu8-i | 3.1 | 4.1 | 98 | 145 | 9 | MOLECULE: | TUMOR NECROSIS |
| 16:00 | 2e7a-B | 3.1 | 3.8 | 100 | 142 | 9 | MOLECULE: | TUMOR NECROSIS |
| 17:00 | 6x81-F | 3.1 | 3.8 | 102 | 144 | 7 | MOLECULE: | TUMOR NECROSIS |
| 18:00 | 4mxw-B | 3.1 | 3.5 | 87 | 126 | 6 | MOLECULE: | TUMOR NECROSIS |
| 19:00 | 6ooy-C | 3.1 | 4.1 | 91 | 134 | 5 | MOLECULE: | TUMOR NECROSIS |
| 20:00 | 7kpa-C | 3.1 | 3.9 | 97 | 133 | 7 | MOLECULE: | TUMOR NECROSIS |
| 21:00 | 5mu8-g | 3.1 | 4 | 102 | 145 | 9 | MOLECULE: | TUMOR NECROSIS |
| 22:00 | 4twt-D | 3.1 | 3.9 | 95 | 130 | 6 | MOLECULE: | TUMOR NECROSIS |
| 23:00 | 5m2m-I | 3.1 | 3.9 | 101 | 150 | 7 | MOLECULE: | TUMOR NECROSIS |
| 24:00:00 | 5yoy-K | 3.1 | 4 | 105 | 153 | 7 | MOLECULE: | TUMOR NECROSIS |
| 25:00:00 | 3qbq-A | 3.1 | 4.4 | 98 | 155 | 4 | MOLECULE: | TUMOR NECROSIS |
| 26:00:00 | 2zpx-A | 3.1 | 3.9 | 100 | 142 | 9 | MOLECULE: | TUMOR NECROSIS |
| 27:00:00 | 2az5-C | 3.1 | 3.6 | 96 | 128 | 8 | MOLECULE: | TUMOR NECROSIS |
| 28:00:00 | 7kp7-C | 3.1 | 3.8 | 102 | 148 | 10 | MOLECULE: | TUMOR NECROSIS |
| 29:00:00 | 2e7a-A | 3.1 | 3.7 | 103 | 150 | 6 | MOLECULE: | TUMOR NECROSIS |
| 30:00:00 | 6x81-D | 3.1 | 3.9 | 99 | 141 | 10 | MOLECULE: | TUMOR NECROSIS |
| 31:00:00 | 6x83-D | 3.1 | 3.9 | 98 | 127 | 11 | MOLECULE: | TUMOR NECROSIS |
| 32:00:00 | 5yoy-J | 3.1 | 3.7 | 108 | 153 | 6 | MOLECULE: | TUMOR NECROSIS |
| 33:00:00 | 2o14-A | 3 | 2.9 | 83 | 354 | 4 | MOLECULE: | HYPOTHETICAL PR |
| 34:00:00 | 1d2q-B | 3 | 3.5 | 84 | 134 | 6 | MOLECULE: | TNF-RELATED APO |
| 35:00:00 | 6x82-D | 3 | 4.1 | 95 | 142 | 9 | MOLECULE: | TUMOR NECROSIS |
| 36:00:00 | 5wux-F | 3 | 4.1 | 101 | 141 | 8 | MOLECULE: | HEAVY |
| 37:00:00 | 3alq-A | 3 | 4.3 | 97 | 141 | 11 | MOLECULE: | TUMOR NECROSIS |
| 38:00:00 | 6rmj-B | 3 | 4.1 | 100 | 147 | 6 | MOLECULE: | TUMOR NECROSIS |
| 39:00:00 | 5mu8-R | 3 | 4.1 | 101 | 138 | 7 | MOLECULE: | TUMOR NECROSIS |
| 40:00:00 | 5mu8-r | 3 | 3.9 | 100 | 144 | 9 | MOLECULE: | TUMOR NECROSIS |
| 41:00:00 | 8kb6-A | 3 | 3.9 | 98 | 148 | 6 | MOLECULE: | TUMOR NECROSIS |
| 42:00:00 | 6x86-D | 3 | 3.8 | 100 | 139 | 7 | MOLECULE: | TUMOR NECROSIS |
| 43:00:00 | 5mu8-j | 3 | 3.9 | 99 | 150 | 8 | MOLECULE: | TUMOR NECROSIS |
| 44:00:00 | 5mu8-k | 3 | 3.9 | 100 | 142 | 7 | MOLECULE: | TUMOR NECROSIS |
| 45:00:00 | 6x81-B | 3 | 4.4 | 95 | 146 | 6 | MOLECULE: | TUMOR NECROSIS |
| 46:00:00 | 6x86-A | 3 | 3.9 | 104 | 151 | 7 | MOLECULE: | TUMOR NECROSIS |
| 47:00:00 | 6x85-E | 3 | 4.1 | 98 | 135 | 7 | MOLECULE: | TUMOR NECROSIS |
| 48:00:00 | 1iqa-B | 3 | 4.6 | 103 | 156 | 5 | MOLECULE: | RECEPTOR ACTIVA |
| 49:00:00 | 5mu8-l | 3 | 3.7 | 97 | 143 | 8 | MOLECULE: | TUMOR NECROSIS |

| | | | | | | | | |
|---|---|---|---|---|---|---|---|---|
| 50:00:00 | 2o0o-C | 3 | 3.9 | 101 | 152 | 8 | MOLECULE: | TNF SUPERFAMILY |
| 51:00:00 | 6x82-B | 3 | 4.2 | 99 | 139 | 10 | MOLECULE: | TUMOR NECROSIS |
| 52:00:00 | 5mu8-q | 3 | 3.8 | 102 | 149 | 9 | MOLECULE: | TUMOR NECROSIS |
| 53:00:00 | 5mu8-F | 3 | 3.9 | 98 | 140 | 8 | MOLECULE: | TUMOR NECROSIS |
| 54:00:00 | 5m2i-C | 3 | 3.9 | 100 | 148 | 7 | MOLECULE: | TUMOR NECROSIS |
| 55:00:00 | 7kp7-A | 3 | 3.9 | 97 | 148 | 6 | MOLECULE: | TUMOR NECROSIS |
| 56:00:00 | 5m2m-B | 3 | 4.1 | 102 | 151 | 10 | MOLECULE: | TUMOR NECROSIS |
| 57:00:00 | 5mu8-P | 3 | 3.7 | 97 | 145 | 7 | MOLECULE: | TUMOR NECROSIS |
| 58:00:00 | 3alq-D | 3 | 3.7 | 101 | 141 | 6 | MOLECULE: | TUMOR NECROSIS |
| 59:00:00 | 7kpb-A | 3 | 3.8 | 104 | 153 | 9 | MOLECULE: | TUMOR NECROSIS |
| 60:00:00 | 5m2j-A | 3 | 3.8 | 104 | 139 | 7 | MOLECULE: | TUMOR NECROSIS |
| 61:00:00 | 2zjc-C | 3 | 4.1 | 100 | 142 | 11 | MOLECULE: | TUMOR NECROSIS |
| 62:00:00 | 5mu8-Z | 3 | 3.8 | 103 | 143 | 6 | MOLECULE: | TUMOR NECROSIS |
| 63:00:00 | 5mu8-M | 3 | 3.9 | 104 | 149 | 8 | MOLECULE: | TUMOR NECROSIS |
| 64:00:00 | 5mu8-e | 3 | 3.7 | 100 | 149 | 6 | MOLECULE: | TUMOR NECROSIS |
| 65:00:00 | 7jra-A | 3 | 3.9 | 102 | 153 | 9 | MOLECULE: | TUMOR NECROSIS |
| 66:00:00 | 5mu8-m | 3 | 4.1 | 96 | 147 | 8 | MOLECULE: | TUMOR NECROSIS |
| 67:00:00 | 5mu8-n | 3 | 3.7 | 97 | 147 | 8 | MOLECULE: | TUMOR NECROSIS |
| 68:00:00 | 3alq-B | 3 | 4.1 | 100 | 141 | 9 | MOLECULE: | TUMOR NECROSIS |
| 69:00:00 | 5m2m-A | 3 | 3.7 | 99 | 150 | 7 | MOLECULE: | TUMOR NECROSIS |
| 70:00:00 | 5m2i-F | 3 | 3.9 | 100 | 148 | 7 | MOLECULE: | TUMOR NECROSIS |
| 71:00:00 | 1d0g-D | 3 | 3.8 | 98 | 151 | 4 | MOLECULE: | DEATH RECEPTOR- |
| 72:00:00 | 5mu8-S | 3 | 4 | 101 | 149 | 7 | MOLECULE: | TUMOR NECROSIS |
| 73:00:00 | 6x81-E | 3 | 3.8 | 97 | 132 | 5 | MOLECULE: | TUMOR NECROSIS |
| 74:00:00 | 7kp8-B | 3 | 4 | 100 | 140 | 13 | MOLECULE: | TUMOR NECROSIS |
| 75:00:00 | 1s55-C | 3 | 4.4 | 100 | 156 | 4 | MOLECULE: | TUMOR NECROSIS |
| 76:00:00 | 3qbq-C | 3 | 4.6 | 98 | 155 | 3 | MOLECULE: | TUMOR NECROSIS |
| 77:00:00 | 5mu8-J | 3 | 4.1 | 97 | 138 | 9 | MOLECULE: | TUMOR NECROSIS |
| 78:00:00 | 1jtz-Z | 2.9 | 4.5 | 99 | 155 | 3 | MOLECULE: | TUMOR NECROSIS |
| 79:00:00 | 5bnq-A | 2.9 | 4.4 | 99 | 156 | 4 | MOLECULE: | TUMOR NECROSIS |
| 80:00:00 | 1jtz-Y | 2.9 | 4.1 | 98 | 155 | 3 | MOLECULE: | TUMOR NECROSIS |
| 81:00:00 | 4giq-A | 2.9 | 4.3 | 102 | 155 | 5 | MOLECULE: | TUMOR NECROSIS |
| 82:00:00 | 5wux-E | 2.9 | 4.5 | 102 | 136 | 9 | MOLECULE: | HEAVY |
| 83:00:00 | 4n90-B | 2.9 | 4.7 | 103 | 155 | 3 | MOLECULE: | TUMOR NECROSIS |
| 84:00:00 | 5mu8-a | 2.9 | 3.9 | 101 | 149 | 9 | MOLECULE: | TUMOR NECROSIS |
| 85:00:00 | 2tun-C | 2.9 | 4 | 103 | 151 | 10 | MOLECULE: | TUMOR NECROSIS |
| 86:00:00 | 1du3-L | 2.9 | 4.4 | 104 | 152 | 3 | MOLECULE: | DEATH RECEPTOR |
| 87:00:00 | 7ta3-D | 2.9 | 4.1 | 100 | 149 | 7 | MOLECULE: | ALPHA-PEPTIDE-3 |
| 88:00:00 | 1du3-E | 2.9 | 4.4 | 105 | 152 | 3 | MOLECULE: | DEATH RECEPTOR |
| 89:00:00 | 6x81-C | 2.9 | 3.7 | 97 | 144 | 6 | MOLECULE: | TUMOR NECROSIS |
| 90:00:00 | 2az5-D | 2.9 | 3.8 | 94 | 140 | 6 | MOLECULE: | TUMOR NECROSIS |
| 91:00:00 | 7jra-C | 2.9 | 4 | 99 | 142 | 6 | MOLECULE: | TUMOR NECROSIS |
| 92:00:00 | 5mu8-v | 2.9 | 3.8 | 97 | 144 | 9 | MOLECULE: | TUMOR NECROSIS |
| 93:00:00 | 5mu8-B | 2.9 | 3.9 | 94 | 145 | 10 | MOLECULE: | TUMOR NECROSIS |

| | | | | | | | | |
|---|---|---|---|---|---|---|---|---|
| 94:00:00 | 5mu8-s | 2.9 | 3.7 | 96 | 149 | 10 | MOLECULE: | TUMOR NECROSIS |
| 95:00:00 | 5m2i-A | 2.9 | 4.1 | 103 | 148 | 7 | MOLECULE: | TUMOR NECROSIS |
| 96:00:00 | 5mu8-G | 2.9 | 3.7 | 100 | 149 | 6 | MOLECULE: | TUMOR NECROSIS |
| 97:00:00 | 1du3-K | 2.9 | 4.2 | 100 | 152 | 4 | MOLECULE: | DEATH RECEPTOR |
| 98:00:00 | 4twt-A | 2.9 | 3.8 | 98 | 149 | 9 | MOLECULE: | TUMOR NECROSIS |
| 99:00:00 | 5mu8-V | 2.9 | 3.9 | 102 | 138 | 10 | MOLECULE: | TUMOR NECROSIS |
| 0:00 | 5mu8-O | 2.9 | 4.2 | 99 | 145 | 9 | MOLECULE: | TUMOR NECROSIS |
| 1:00 | 2tnf-C | 2.9 | 4.1 | 98 | 148 | 8 | MOLECULE: | PROTEIN (TUMOR |
| 2:00 | 5mu8-Q | 2.9 | 3.9 | 106 | 149 | 6 | MOLECULE: | TUMOR NECROSIS |
| 3:00 | 5mu8-u | 2.9 | 3.7 | 93 | 145 | 5 | MOLECULE: | TUMOR NECROSIS |
| 4:00 | 2zjc-B | 2.9 | 3.9 | 100 | 151 | 10 | MOLECULE: | TUMOR NECROSIS |
| 5:00 | 5mu8-h | 2.9 | 4.2 | 98 | 137 | 6 | MOLECULE: | TUMOR NECROSIS |
| 6:00 | 6x82-E | 2.9 | 4 | 100 | 136 | 9 | MOLECULE: | TUMOR NECROSIS |
| 7:00 | 6rmj-A | 2.9 | 4.2 | 99 | 149 | 7 | MOLECULE: | TUMOR NECROSIS |
| 8:00 | 1d2q-A | 2.9 | 3.4 | 85 | 134 | 5 | MOLECULE: | TNF-RELATED APO |
| 9:00 | 5mu8-U | 2.9 | 4.1 | 98 | 149 | 9 | MOLECULE: | TUMOR NECROSIS |
| 10:00 | 5mu8-o | 2.9 | 4 | 98 | 147 | 9 | MOLECULE: | TUMOR NECROSIS |
| 11:00 | 5m2i-B | 2.8 | 4.2 | 96 | 148 | 8 | MOLECULE: | TUMOR NECROSIS |
| 12:00 | 2e7a-C | 2.8 | 3.9 | 97 | 143 | 6 | MOLECULE: | TUMOR NECROSIS |
| 13:00 | 6x86-E | 2.8 | 4.1 | 98 | 135 | 8 | MOLECULE: | TUMOR NECROSIS |
| 14:00 | 5tsw-E | 2.8 | 4.3 | 95 | 148 | 9 | MOLECULE: | PROTEIN (TUMOR |
| 15:00 | 3l9j-T | 2.8 | 3.7 | 99 | 149 | 5 | MOLECULE: | TNFALPHA |
| 16:00 | 3me2-A | 2.8 | 4.5 | 99 | 156 | 4 | MOLECULE: | TUMOR NECROSIS |
| 17:00 | 1du3-J | 2.8 | 4 | 99 | 152 | 4 | MOLECULE: | DEATH RECEPTOR |
| 18:00 | 1du3-F | 2.8 | 4.5 | 105 | 152 | 3 | MOLECULE: | DEATH RECEPTOR |
| 19:00 | 2tun-E | 2.8 | 4.4 | 93 | 151 | 6 | MOLECULE: | TUMOR NECROSIS |
| 20:00 | 2tnf-A | 2.8 | 3.8 | 95 | 148 | 9 | MOLECULE: | PROTEIN (TUMOR |
| 21:00 | 5mu8-w | 2.8 | 3.9 | 98 | 145 | 6 | MOLECULE: | TUMOR NECROSIS |
| 22:00 | 1du3-D | 2.8 | 4.4 | 101 | 152 | 4 | MOLECULE: | DEATH RECEPTOR |
| 23:00 | 3alq-F | 2.8 | 4.2 | 100 | 141 | 10 | MOLECULE: | TUMOR NECROSIS |
| 24:00:00 | 5mu8-Y | 2.8 | 3.8 | 99 | 149 | 10 | MOLECULE: | TUMOR NECROSIS |
| 25:00:00 | 5mu8-A | 2.8 | 4.3 | 98 | 141 | 8 | MOLECULE: | TUMOR NECROSIS |
| 26:00:00 | 4mxw-Z | 2.8 | 3.8 | 92 | 134 | 9 | MOLECULE: | TUMOR NECROSIS |
| 27:00:00 | 5m2i-D | 2.8 | 4.1 | 94 | 148 | 6 | MOLECULE: | TUMOR NECROSIS |
| 28:00:00 | 6x83-A | 2.8 | 3.8 | 94 | 141 | 9 | MOLECULE: | TUMOR NECROSIS |
| 29:00:00 | 5mu8-W | 2.8 | 4 | 98 | 143 | 7 | MOLECULE: | TUMOR NECROSIS |
| 30:00:00 | 1d4v-B | 2.8 | 4.5 | 104 | 163 | 4 | MOLECULE: | TNF-RELATED APO |
| 31:00:00 | 4twt-C | 2.8 | 4.1 | 99 | 142 | 7 | MOLECULE: | TUMOR NECROSIS |
| 32:00:00 | 1d0g-A | 2.8 | 4 | 102 | 151 | 5 | MOLECULE: | DEATH RECEPTOR- |
| 33:00:00 | 5tsw-B | 2.8 | 4.2 | 101 | 148 | 11 | MOLECULE: | PROTEIN (TUMOR |
| 34:00:00 | 5mu8-t | 2.8 | 4.1 | 96 | 143 | 8 | MOLECULE: | TUMOR NECROSIS |
| 35:00:00 | 5tsw-C | 2.8 | 4 | 99 | 148 | 7 | MOLECULE: | PROTEIN (TUMOR |
| 36:00:00 | 5mu8-c | 2.8 | 3.9 | 93 | 145 | 9 | MOLECULE: | TUMOR NECROSIS |
| 37:00:00 | 5ubu-B | 2.7 | 4.8 | 116 | 331 | 7 | MOLECULE: | PUTATIVE ACETAM |

| | | | | | | | | |
|---|---|---|---|---|---|---|---|---|
| 38:00:00 | 6x81-A | 2.7 | 4.1 | 101 | 145 | 7 | MOLECULE: | TUMOR NECROSIS |
| 39:00:00 | 5mu8-N | 2.7 | 3.9 | 96 | 143 | 8 | MOLECULE: | TUMOR NECROSIS |
| 40:00:00 | 5mu8-T | 2.7 | 4.2 | 97 | 146 | 10 | MOLECULE: | TUMOR NECROSIS |
| 41:00:00 | 5mu8-D | 2.7 | 4 | 95 | 149 | 12 | MOLECULE: | TUMOR NECROSIS |
| 42:00:00 | 6x82-A | 2.6 | 4 | 101 | 143 | 4 | MOLECULE: | TUMOR NECROSIS |
| 43:00:00 | 4n90-A | 2.6 | 4.2 | 96 | 155 | 3 | MOLECULE: | TUMOR NECROSIS |
| 44:00:00 | 4n90-C | 2.6 | 3.9 | 98 | 154 | 7 | MOLECULE: | TUMOR NECROSIS |
| 45:00:00 | 5tsw-D | 2.6 | 3.7 | 102 | 148 | 9 | MOLECULE: | PROTEIN (TUMOR |
| 46:00:00 | 5tsw-A | 2.5 | 4 | 98 | 148 | 9 | MOLECULE: | PROTEIN (TUMOR |
| 47:00:00 | 5tsw-F | 2.5 | 4 | 98 | 148 | 7 | MOLECULE: | PROTEIN (TUMOR |
| 48:00:00 | 4g3y-C | 2.5 | 4 | 99 | 147 | 4 | MOLECULE: | INFLIXIMAB FAB |
| 49:00:00 | 3alq-C | 2.5 | 3.8 | 91 | 141 | 11 | MOLECULE: | TUMOR NECROSIS |
| 50:00:00 | 6kty-A | 2.4 | 2.7 | 63 | 176 | 6 | MOLECULE: | FLAGELLAR HOOK- |
| 51:00:00 | 6x80-C | 2.4 | 16.9 | 80 | 574 | 11 | MOLECULE: | FLAGELLIN A |
| 52:00:00 | 8slr-D | 2.2 | 4.1 | 89 | 155 | 4 | MOLECULE: | TUMOR NECROSIS |

**DALI results: MdA-1 Hexon**

| No: | Chain | Z | rmsd | lali | nres | %id | PDB | Description |
|---|---|---|---|---|---|---|---|---|
| 1:00 | 8rbt-A | 14.9 | 3.9 | 333 | 481 | 10 | MOLECULE: | MAJOR CAPSID PROTEIN |
| 2:00 | 6g43-A | 14.4 | 4.3 | 353 | 515 | 7 | MOLECULE: | PUTATIVE MAJOR CAPSID PROTEIN |
| 3:00 | 5j7o-F | 14.2 | 4.3 | 347 | 622 | 10 | MOLECULE: | MAJOR CAPSID PROTEIN |
| 4:00 | 3j26-A | 13.8 | 4.5 | 370 | 508 | 10 | MOLECULE: | CAPSID PROTEIN V20 |
| 5:00 | 5tip-A | 13.8 | 3.6 | 322 | 436 | 9 | MOLECULE: | MAJOR CAPSID PROTEIN |
| 6:00 | 3sam-B | 12.4 | 4 | 316 | 544 | 8 | MOLECULE: | RIFAMPICIN RESISTANCE PROTEIN |
| 7:00 | 7yjl-A | 11.4 | 4 | 335 | 459 | 6 | MOLECULE: | MAJOR CAPSID PROTEIN |
| 8:00 | 6l2t-C | 10.6 | 4.5 | 353 | 633 | 7 | MOLECULE: | B646L,MAJOR CAPSID PROTEIN |
| 9:00 | 2iny-A | 10.4 | 4.3 | 400 | 941 | 8 | MOLECULE: | HEXON PROTEIN |
| 10:00 | 6ojn-A | 10.4 | 4.2 | 313 | 423 | 7 | MOLECULE: | MAJOR CAPSID PROTEIN |
| 11:00 | 6q5u-C | 9.6 | 4.9 | 291 | 394 | 6 | MOLECULE: | MAJOR CAPSID PROTEIN (P3) |
| 12:00 | 9cls-K | 8.9 | 5.4 | 401 | 935 | 6 | MOLECULE: | HEXON PROTEIN |
| 13:00 | 2vvf-A | 7.5 | 4.2 | 207 | 269 | 7 | MOLECULE: | MAJOR CAPSID PROTEIN P2 |
| 14:00 | 3j31-A | 6.3 | 4.2 | 215 | 344 | 8 | MOLECULE: | A223 PENTON BASE |
| 15:00 | 5oac-A | 5.4 | 4.9 | 186 | 310 | 9 | MOLECULE: | MAJOR CAPSID PROTEIN |
| 16:00 | 7zzz-A | 4.4 | 4.2 | 172 | 239 | 7 | MOLECULE: | MAJOR CAPSID PROTEIN P5 |
| 17:00 | 7v1v-A | 3 | 4.7 | 205 | 447 | 4 | MOLECULE: | DIFRUCTOSE DIANHYDRIDE I SYNTHASE/HYDROLASE |

**DALI results: MdA-1 Adenain**

| No: | Chain | Z | rmsd | lali | nres | %id | PDB Description | |
|---|---|---|---|---|---|---|---|---|
| 1:00 | 4pis-A | 13.4 | 2.8 | 148 | 204 | 14 | MOLECULE: | PROTEASE |
| 2:00 | 4wx4-A | 13.3 | 2.7 | 146 | 201 | 14 | MOLECULE: | PROTEASE |
| 3:00 | 4piq-A | 13.3 | 2.7 | 146 | 204 | 12 | MOLECULE: | PROTEASE |
| 4:00 | 1nln-A | 13.2 | 2.9 | 148 | 203 | 16 | MOLECULE: | ADENAIN |
| 5:00 | 4wx7-C | 13.2 | 2.7 | 147 | 204 | 14 | MOLECULE: | PROTEASE |
| 6:00 | 4pie-A | 13.2 | 2.7 | 147 | 202 | 14 | MOLECULE: | PROTEASE |
| 7:00 | 4wx6-A | 13.2 | 2.7 | 147 | 204 | 14 | MOLECULE: | PROTEASE |
| 8:00 | 4wx6-C | 13.1 | 2.7 | 147 | 204 | 14 | MOLECULE: | PROTEASE |
| 9:00 | 4pid-A | 13.1 | 2.8 | 147 | 204 | 15 | MOLECULE: | PROTEASE |
| 10:00 | 4wx7-A | 13.1 | 2.8 | 147 | 204 | 12 | MOLECULE: | PROTEASE |
| 11:00 | 4ekf-A | 13.1 | 2.7 | 143 | 192 | 14 | MOLECULE: | ADENAIN |
| 12:00 | 5fgy-A | 12.9 | 2.8 | 146 | 203 | 14 | MOLECULE: | PROTEASE |
| 13:00 | 1avp-A | 12.8 | 2.8 | 146 | 204 | 14 | MOLECULE: | ADENOVIRAL PROTEINASE |
| 14:00 | 7fiu-A | 11.9 | 4.5 | 166 | 270 | 13 | MOLECULE: | ULP_PROTEASE DOMAIN-CONTAINING |
| 15:00 | 6ups-A | 11.6 | 3.6 | 147 | 245 | 15 | MOLECULE: | ULP_PROTEASE DOMAIN-CONTAINING |
| 16:00 | 6upu-E | 11.2 | 3.6 | 147 | 258 | 14 | MOLECULE: | ULP_PROTEASE DOMAIN-CONTAINING |
| 17:00 | 8efx-A | 11.1 | 3.9 | 150 | 259 | 13 | MOLECULE: | OTDUB |
| 18:00 | 6upu-M | 11 | 3.6 | 149 | 258 | 14 | MOLECULE: | ULP_PROTEASE DOMAIN-CONTAINING |
| 19:00 | 6upu-I | 10.8 | 3.5 | 148 | 258 | 14 | MOLECULE: | ULP_PROTEASE DOMAIN-CONTAINING |
| 20:00 | 5haf-A | 10.8 | 6.6 | 166 | 312 | 10 | MOLECULE: | DEUBIQUITINASE SSEL |
| 21:00 | 5ubw-B | 10.8 | 3.1 | 145 | 173 | 10 | MOLECULE: | DEUBIQUITINASE SSEL |
| 22:00 | 5ubw-A | 10.7 | 3.2 | 146 | 175 | 12 | MOLECULE: | DEUBIQUITINASE SSEL |
| 23:00 | 6upu-A | 10.7 | 3.4 | 146 | 259 | 14 | MOLECULE: | ULP_PROTEASE DOMAIN-CONTAINING |
| 24:00:00 | 6wtg-A | 10.6 | 3.6 | 147 | 169 | 10 | MOLECULE: | UBIQUITINATING/DEUBIQUITINATIN |
| 25:00:00 | 5cra-A | 10.4 | 3.5 | 143 | 172 | 10 | MOLECULE: | SDEA |
| 26:00:00 | 5ham-B | 10.3 | 2.9 | 136 | 260 | 13 | MOLECULE: | RICKCE |
| 27:00:00 | 1xt9-A | 10.3 | 3.4 | 143 | 208 | 13 | MOLECULE: | SENTRIN-SPECIFIC PROTEASE 8 |
| 28:00:00 | 5b5q-B | 10.3 | 3.5 | 140 | 244 | 16 | MOLECULE: | MEMBRANE THIOL PROTEASE |
| 29:00:00 | 5cra-B | 10.3 | 3.5 | 143 | 172 | 10 | MOLECULE: | SDEA |
| 30:00:00 | 5ham-A | 10.2 | 2.9 | 135 | 260 | 13 | MOLECULE: | RICKCE |
| 31:00:00 | 5haf-B | 10.1 | 9.3 | 169 | 327 | 10 | MOLECULE: | DEUBIQUITINASE SSEL |
| 32:00:00 | 2bkq-A | 10.1 | 3.5 | 146 | 210 | 12 | MOLECULE: | SENTRIN-SPECIFIC PROTEASE 8 |
| 33:00:00 | 2bkr-A | 10 | 3.4 | 142 | 211 | 12 | MOLECULE: | SENTRIN-SPECIFIC PROTEASE 8 |
| 34:00:00 | 6be0-A | 10 | 3.5 | 150 | 266 | 14 | MOLECULE: | AVRA |
| 35:00:00 | 2bkq-C | 9.9 | 3.6 | 142 | 207 | 13 | MOLECULE: | SENTRIN-SPECIFIC PROTEASE 8 |
| 36:00:00 | 5crb-B | 9.8 | 3.8 | 151 | 191 | 11 | MOLECULE: | SDEA |
| 37:00:00 | 6lj9-A | 9.8 | 6 | 139 | 272 | 14 | MOLECULE: | CYSTEINE PROTEASE S273R |
| 38:00:00 | 2bkq-B | 9.8 | 3.6 | 143 | 208 | 13 | MOLECULE: | SENTRIN-SPECIFIC PROTEASE 8 |
| 39:00:00 | 8efw-B | 9.7 | 4 | 153 | 183 | 12 | MOLECULE: | SDEA |
| 40:00:00 | 6fdu-A | 9.6 | 3.5 | 139 | 240 | 16 | MOLECULE: | DEUBIQUITINASE AND DENEDDYLASE |
| 41:00:00 | 6fdk-A | 9.6 | 3.5 | 139 | 242 | 16 | MOLECULE: | DEUBIQUITINASE AND DENEDDYLASE |
| 42:00:00 | 5crb-A | 9.5 | 3.7 | 148 | 192 | 10 | MOLECULE: | SDEA |

| | | | | | | | | |
|---|---|---|---|---|---|---|---|---|
| 43:00:00 | 5b5q-A | 9.5 | 3.4 | 139 | 243 | 14 | MOLECULE: | MEMBRANE THIOL PROTEASE |
| 44:00:00 | 6oam-B | 9.5 | 3.3 | 141 | 239 | 14 | MOLECULE: | DEUBIQUITINASE AND DENEDDYLASE |
| 45:00:00 | 6fdu-B | 9.5 | 3.4 | 137 | 240 | 13 | MOLECULE: | DEUBIQUITINASE AND DENEDDYLASE |
| 46:00:00 | 6fdq-A | 9.5 | 3.4 | 138 | 239 | 14 | MOLECULE: | DEUBIQUITINASE AND DENEDDYLASE |
| 47:00:00 | 6ljb-A | 9.5 | 6.1 | 138 | 267 | 14 | MOLECULE: | CYSTEINE PROTEASE S273R |
| 48:00:00 | 8oi3-A | 9.3 | 2.9 | 123 | 183 | 10 | MOLECULE: | TYPE III EFFECTOR |
| 49:00:00 | 6lj9-B | 9.3 | 6.1 | 130 | 259 | 15 | MOLECULE: | CYSTEINE PROTEASE S273R |
| 50:00:00 | 6fdq-B | 9.3 | 3.4 | 140 | 247 | 16 | MOLECULE: | DEUBIQUITINASE AND DENEDDYLASE |
| 51:00:00 | 5jp3-E | 9.3 | 3 | 133 | 193 | 11 | MOLECULE: | XANTHOMONAS OUTER PROTEIN D |
| 52:00:00 | 5klq-B | 9.2 | 3.8 | 153 | 320 | 12 | MOLECULE: | ORF34 |
| 53:00:00 | 5klq-C | 9.2 | 3.7 | 152 | 320 | 11 | MOLECULE: | ORF34 |
| 54:00:00 | 6oam-A | 9.2 | 3.3 | 141 | 239 | 14 | MOLECULE: | DEUBIQUITINASE AND DENEDDYLASE |
| 55:00:00 | 5klp-A | 9.2 | 4 | 153 | 309 | 10 | MOLECULE: | ORF34 |
| 56:00:00 | 2bkq-D | 9.1 | 3.6 | 138 | 209 | 12 | MOLECULE: | SENTRIN-SPECIFIC PROTEASE 8 |
| 57:00:00 | 6gzs-A | 9.1 | 3.5 | 142 | 250 | 13 | MOLECULE: | DEUBIQUITINASE AND DENEDDYLASE |
| 58:00:00 | 5klq-A | 9.1 | 3.8 | 153 | 324 | 12 | MOLECULE: | ORF34 |
| 59:00:00 | 5jp3-C | 9.1 | 3 | 130 | 196 | 11 | MOLECULE: | XANTHOMONAS OUTER PROTEIN D |
| 60:00:00 | 6mrn-A | 9.1 | 3.4 | 139 | 240 | 14 | MOLECULE: | DEUBIQUITINASE AND DENEDDYLASE |
| 61:00:00 | 5klp-B | 9.1 | 4 | 152 | 310 | 11 | MOLECULE: | ORF34 |
| 62:00:00 | 6gzt-A | 9 | 8.1 | 152 | 267 | 13 | MOLECULE: | DEUBIQUITINASE AND DENEDDYLASE |
| 63:00:00 | 6gzu-A | 9 | 2.9 | 130 | 241 | 14 | MOLECULE: | CONSERVED MEMBRANE PROTEIN |
| 64:00:00 | 5jp1-A | 9 | 3.1 | 129 | 199 | 12 | MOLECULE: | XANTHOMONAS OUTER PROTEIN D |
| 65:00:00 | 5hag-A | 9 | 7.5 | 150 | 266 | 13 | MOLECULE: | DEUBIQUITINASE AND DENEDDYLASE |
| 66:00:00 | 5klp-C | 9 | 4 | 155 | 315 | 10 | MOLECULE: | ORF34 |
| 67:00:00 | 8oi3-B | 9 | 3 | 125 | 183 | 10 | MOLECULE: | TYPE III EFFECTOR |
| 68:00:00 | 2oix-A | 9 | 2.8 | 122 | 178 | 11 | MOLECULE: | XANTHOMONAS OUTER PROTEIN D |
| 69:00:00 | 5aek-A | 8.9 | 3.5 | 129 | 223 | 10 | MOLECULE: | SENTRIN-SPECIFIC PROTEASE 2 |
| 70:00:00 | 5jp3-G | 8.9 | 3 | 129 | 196 | 12 | MOLECULE: | XANTHOMONAS OUTER PROTEIN D |
| 71:00:00 | 5jp3-A | 8.7 | 3 | 129 | 205 | 11 | MOLECULE: | XANTHOMONAS OUTER PROTEIN D |
| 72:00:00 | 7r2e-B | 8.7 | 4.5 | 144 | 243 | 12 | MOLECULE: | SENTRIN-SPECIFIC PROTEASE 7 |
| 73:00:00 | 2g4d-C | 8.7 | 3.3 | 127 | 205 | 8 | MOLECULE: | SENP1 PROTEIN |
| 74:00:00 | 5dgg-B | 8.7 | 3.3 | 125 | 176 | 15 | MOLECULE: | UNCHARACTERIZED PROTEIN |
| 75:00:00 | 7r2e-A | 8.7 | 4.5 | 148 | 244 | 11 | MOLECULE: | SENTRIN-SPECIFIC PROTEASE 7 |
| 76:00:00 | 5aek-M | 8.7 | 3.5 | 129 | 222 | 11 | MOLECULE: | SENTRIN-SPECIFIC PROTEASE 2 |
| 77:00:00 | 2g4d-A | 8.7 | 3.3 | 128 | 205 | 8 | MOLECULE: | SENP1 PROTEIN |
| 78:00:00 | 3eay-A | 8.6 | 4.4 | 146 | 247 | 12 | MOLECULE: | SENTRIN-SPECIFIC PROTEASE 7 |
| 79:00:00 | 1th0-B | 8.6 | 3.4 | 129 | 223 | 12 | MOLECULE: | SENTRIN-SPECIFIC PROTEASE 2 |
| 80:00:00 | 2hl9-A | 8.6 | 3.4 | 133 | 218 | 11 | MOLECULE: | UBIQUITIN-LIKE-SPECIFIC PROTEA |
| 81:00:00 | 2hkp-A | 8.6 | 3.3 | 133 | 219 | 11 | MOLECULE: | UBIQUITIN-LIKE-SPECIFIC PROTEA |
| 82:00:00 | 2hl8-A | 8.5 | 3.4 | 128 | 219 | 12 | MOLECULE: | UBIQUITIN-LIKE-SPECIFIC PROTEA |
| 83:00:00 | 1euv-A | 8.5 | 3.4 | 133 | 221 | 13 | MOLECULE: | ULP1 PROTEASE |
| 84:00:00 | 5aek-K | 8.4 | 3.2 | 127 | 223 | 9 | MOLECULE: | SENTRIN-SPECIFIC PROTEASE 2 |
| 85:00:00 | 5aek-U | 8.4 | 3.3 | 126 | 223 | 10 | MOLECULE: | SENTRIN-SPECIFIC PROTEASE 2 |
| 86:00:00 | 5lnb-B | 8.4 | 4.7 | 152 | 259 | 9 | MOLECULE: | UBIQUITIN-LIKE-SPECIFIC PROTEA |

| | | | | | | | | |
|---|---|---|---|---|---|---|---|---|
| 87:00:00 | 5dgg-A | 8.4 | 2.8 | 119 | 176 | 15 | MOLECULE: | UNCHARACTERIZED PROTEIN |
| 88:00:00 | 2iy1-A | 8.4 | 3.3 | 128 | 226 | 8 | MOLECULE: | SENTRIN-SPECIFIC PROTEASE 1 |
| 89:00:00 | 2xph-A | 8.4 | 3.5 | 130 | 227 | 9 | MOLECULE: | SENTRIN-SPECIFIC PROTEASE 1 |
| 90:00:00 | 6nnq-A | 8.4 | 3.3 | 128 | 224 | 9 | MOLECULE: | SENTRIN-SPECIFIC PROTEASE 1 |
| 91:00:00 | 5w40-D | 8.3 | 3.9 | 145 | 327 | 12 | MOLECULE: | POPP2 PROTEIN |
| 92:00:00 | 5aek-C | 8.3 | 3.5 | 129 | 224 | 10 | MOLECULE: | SENTRIN-SPECIFIC PROTEASE 2 |
| 93:00:00 | 2ckh-A | 8.3 | 3.5 | 128 | 225 | 9 | MOLECULE: | SENTRIN-SPECIFIC PROTEASE 1 |
| 94:00:00 | 2iyc-A | 8.3 | 3.4 | 128 | 226 | 9 | MOLECULE: | SENTRIN-SPECIFIC PROTEASE 1 |
| 95:00:00 | 5w3y-C | 8.3 | 3.8 | 143 | 335 | 12 | MOLECULE: | POPP2 PROTEIN |
| 96:00:00 | 2ckg-B | 8.3 | 3.5 | 130 | 225 | 9 | MOLECULE: | SENTRIN-SPECIFIC PROTEASE 1 |
| 97:00:00 | 2ckg-A | 8.3 | 3.4 | 128 | 225 | 9 | MOLECULE: | SENTRIN-SPECIFIC PROTEASE 1 |
| 98:00:00 | 2iy0-A | 8.3 | 3.4 | 128 | 226 | 8 | MOLECULE: | SENTRIN-SPECIFIC PROTEASE 1 |
| 99:00:00 | 2iyc-B | 8.3 | 3.4 | 128 | 226 | 9 | MOLECULE: | SENTRIN-SPECIFIC PROTEASE 1 |
| 0:00 | 5aek-G | 8.3 | 3.3 | 128 | 223 | 9 | MOLECULE: | SENTRIN-SPECIFIC PROTEASE 2 |
| 1:00 | 5aek-S | 8.3 | 3.3 | 128 | 224 | 9 | MOLECULE: | SENTRIN-SPECIFIC PROTEASE 2 |
| 2:00 | 1th0-A | 8.2 | 3.4 | 129 | 226 | 12 | MOLECULE: | SENTRIN-SPECIFIC PROTEASE 2 |
| 3:00 | 1tgz-A | 8.2 | 3.5 | 129 | 224 | 11 | MOLECULE: | SENTRIN-SPECIFIC PROTEASE 2 |
| 4:00 | 5aek-W | 8.2 | 3.5 | 131 | 222 | 11 | MOLECULE: | SENTRIN-SPECIFIC PROTEASE 2 |
| 5:00 | 5aek-I | 8.2 | 3.4 | 128 | 223 | 9 | MOLECULE: | SENTRIN-SPECIFIC PROTEASE 2 |
| 6:00 | 2iyd-A | 8.2 | 3.5 | 128 | 226 | 9 | MOLECULE: | SENTRIN-SPECIFIC PROTEASE 1 |
| 7:00 | 5w3t-B | 8.2 | 3.7 | 141 | 333 | 13 | MOLECULE: | POPP2 PROTEIN |
| 8:00 | 5aek-O | 8.2 | 3.4 | 128 | 223 | 11 | MOLECULE: | SENTRIN-SPECIFIC PROTEASE 2 |
| 9:00 | 2iy1-C | 8.2 | 3.4 | 125 | 226 | 7 | MOLECULE: | SENTRIN-SPECIFIC PROTEASE 1 |
| 10:00 | 5w40-B | 8.2 | 3.8 | 142 | 333 | 12 | MOLECULE: | POPP2 PROTEIN |
| 11:00 | 5aek-E | 8.2 | 3.5 | 131 | 224 | 11 | MOLECULE: | SENTRIN-SPECIFIC PROTEASE 2 |
| 12:00 | 2oiv-A | 8.2 | 2.8 | 116 | 170 | 11 | MOLECULE: | XANTHOMONAS OUTER PROTEIN D |
| 13:00 | 2xre-A | 8.2 | 3.4 | 128 | 227 | 9 | MOLECULE: | SENTRIN-SPECIFIC PROTEASE 1 |
| 14:00 | 2xph-B | 8.1 | 3.6 | 128 | 226 | 9 | MOLECULE: | SENTRIN-SPECIFIC PROTEASE 1 |
| 15:00 | 2io1-A | 8.1 | 3.3 | 128 | 223 | 9 | MOLECULE: | SENTRIN-SPECIFIC PROTEASE 2 |
| 16:00 | 2xre-B | 8 | 3.6 | 128 | 225 | 9 | MOLECULE: | SENTRIN-SPECIFIC PROTEASE 1 |
| 17:00 | 2io1-C | 8 | 3.5 | 127 | 224 | 11 | MOLECULE: | SENTRIN-SPECIFIC PROTEASE 2 |
| 18:00 | 6dg4-A | 8 | 3.5 | 131 | 253 | 11 | MOLECULE: | ULP1-LIKE SUMO PROTEASE |
| 19:00 | 2io1-E | 8 | 3.4 | 131 | 223 | 11 | MOLECULE: | SENTRIN-SPECIFIC PROTEASE 2 |
| 20:00 | 5ms8-A | 7.9 | 6.3 | 165 | 377 | 13 | MOLECULE: | LEGIONELLA PNEUMOPHILA EFFECTO |
| 21:00 | 2io2-A | 7.9 | 3.3 | 127 | 225 | 9 | MOLECULE: | SENTRIN-SPECIFIC PROTEASE 2 |
| 22:00 | 2io3-A | 7.9 | 3.3 | 129 | 224 | 9 | MOLECULE: | SENTRIN-SPECIFIC PROTEASE 2 |
| 23:00 | 3zo5-A | 7.9 | 3.4 | 131 | 230 | 10 | MOLECULE: | SENTRIN-SPECIFIC PROTEASE 2 |
| 24:00:00 | 5ms2-A | 7.8 | 7 | 160 | 405 | 14 | MOLECULE: | LEGIONELLA PNEUMOPHILA EFFECTO |
| 25:00:00 | 5io3-A | 7.8 | 7.1 | 158 | 385 | 11 | MOLECULE: | UNCHARACTERIZED PROTEIN RAVZ |
| 26:00:00 | 5w3y-B | 7.8 | 3.9 | 147 | 327 | 10 | MOLECULE: | POPP2 PROTEIN |
| 27:00:00 | 5aek-Q | 7.8 | 3.3 | 127 | 224 | 9 | MOLECULE: | SENTRIN-SPECIFIC PROTEASE 2 |
| 28:00:00 | 5izv-A | 7.8 | 6.8 | 165 | 376 | 12 | MOLECULE: | UNCHARACTERIZED PROTEIN RAVZ |
| 29:00:00 | 5w3y-D | 7.8 | 3.9 | 145 | 334 | 12 | MOLECULE: | POPP2 PROTEIN |
| 30:00:00 | 5w40-A | 7.8 | 3.9 | 146 | 336 | 12 | MOLECULE: | POPP2 PROTEIN |

| | | | | | | | | |
|---|---|---|---|---|---|---|---|---|
| 31:00:00 | 5w3x-A | 7.7 | 4 | 146 | 335 | 13 | MOLECULE: | POPP2 PROTEIN |
| 32:00:00 | 5w3t-D | 7.7 | 3.7 | 142 | 333 | 13 | MOLECULE: | POPP2 PROTEIN |
| 33:00:00 | 5w3t-A | 7.7 | 3.9 | 141 | 332 | 13 | MOLECULE: | POPP2 PROTEIN |
| 34:00:00 | 5w3t-C | 7.7 | 3.6 | 141 | 332 | 13 | MOLECULE: | POPP2 PROTEIN |
| 35:00:00 | 5w40-C | 7.7 | 3.8 | 140 | 333 | 12 | MOLECULE: | POPP2 PROTEIN |
| 36:00:00 | 5w3x-C | 7.7 | 3.9 | 144 | 342 | 13 | MOLECULE: | POPP2 PROTEIN |
| 37:00:00 | 5izv-B | 7.6 | 6.6 | 165 | 369 | 13 | MOLECULE: | UNCHARACTERIZED PROTEIN RAVZ |
| 38:00:00 | 5w3y-A | 7.6 | 3.9 | 141 | 336 | 12 | MOLECULE: | POPP2 PROTEIN |
| 39:00:00 | 2io0-A | 7.6 | 3.4 | 127 | 226 | 9 | MOLECULE: | SENTRIN-SPECIFIC PROTEASE 2 |
| 40:00:00 | 5hzy-A | 7.4 | 6.5 | 159 | 380 | 13 | MOLECULE: | UNCHARACTERIZED PROTEIN RAVZ |
| 41:00:00 | 5cqc-A | 7.4 | 6.3 | 150 | 352 | 12 | MOLECULE: | PUTATIVE RAVZ PROTEIN |
| 42:00:00 | 5ms7-A | 7.3 | 6.8 | 160 | 370 | 13 | MOLECULE: | LEGIONELLA PNEUMOPHILA EFFECTO |
| 43:00:00 | 7f3n-A | 6.8 | 3.8 | 125 | 247 | 14 | MOLECULE: | TYPE III EFFECTOR PROTEIN POPP |
| 44:00:00 | 5crc-A | 5.4 | 6.1 | 99 | 152 | 10 | MOLECULE: | SDEA |
| 45:00:00 | 5crc-B | 5.4 1 | 0.2 | 106 | 155 | 9 | MOLECULE: | SDEA |
| 46:00:00 | 6w9s-A | 5.3 | 8.1 | 106 | 178 | 8 | MOLECULE: | OTU DOMAIN-CONTAINING PROTEIN |
| 47:00:00 | 4g38-A | 5 | 5 | 103 | 449 | 8 | MOLECULE: | SULFITE REDUCTASE [NADPH] HEMO |
| 48:00:00 | 5gep-A | 4.8 1 | 2.3 | 107 | 456 | 7 | MOLECULE: | SULFITE REDUCTASE HEMOPROTEIN |
| 49:00:00 | 6gep-A | 4.8 1 | 2.3 | 107 | 456 | 7 | MOLECULE: | SULFITE REDUCTASE HEMOPROTEIN |
| 50:00:00 | 4gep-A | 4.7 1 | 2.2 | 108 | 456 | 6 | MOLECULE: | SULFITE REDUCTASE HEMOPROTEIN |
| 51:00:00 | 6c3z-A | 4.7 1 | 2.3 | 107 | 462 | 7 | MOLECULE: | SULFITE REDUCTASE [NADPH] HEMO |
| 52:00:00 | 3aop-A | 4.6 1 | 2.3 | 107 | 453 | 7 | MOLECULE: | SULFITE REDUCTASE HEMOPROTEIN |
| 53:00:00 | 7gep-A | 4.6 1 | 2.2 | 108 | 469 | 6 | MOLECULE: | SULFITE REDUCTASE HEMOPROTEIN |
| 54:00:00 | 4htr-A | 4.6 | 5 | 104 | 441 | 8 | MOLECULE: | SULFITE REDUCTASE [NADPH] HEMO |
| 55:00:00 | 5aop-A | 4.6 | 4.9 | 101 | 456 | 6 | MOLECULE: | SULFITE REDUCTASE HEMOPROTEIN |
| 56:00:00 | 6c3x-A | 4.5 1 | 2.2 | 108 | 464 | 6 | MOLECULE: | SULFITE REDUCTASE [NADPH] HEMO |
| 57:00:00 | 8gep-A | 4.5 | 5.1 | 104 | 456 | 8 | MOLECULE: | SULFITE REDUCTASE HEMOPROTEIN |
| 58:00:00 | 4aop-A | 4.4 1 | 2.2 | 108 | 453 | 6 | MOLECULE: | SULFITE REDUCTASE HEMOPROTEIN |
| 59:00:00 | 5h92-A | 4.4 1 | 3.7 | 105 | 567 | 11 | MOLECULE: | SULFITE REDUCTASE [FERREDOXIN] |
| 60:00:00 | 3geo-A | 4.4 1 | 2.2 | 107 | 455 | 7 | MOLECULE: | SULFITE REDUCTASE HEMOPROTEIN |
| 61:00:00 | 2aop-A | 4.4 1 | 2.3 | 107 | 456 | 7 | MOLECULE: | SULFITE REDUCTASE HEMOPROTEIN |
| 62:00:00 | 4g39-A | 4.4 1 | 2.2 | 108 | 465 | 5 | MOLECULE: | SULFITE REDUCTASE [NADPH] HEMO |
| 63:00:00 | 5h92-B | 4.3 1 | 3.8 | 106 | 572 | 11 | MOLECULE: | SULFITE REDUCTASE [FERREDOXIN] |
| 64:00:00 | 6c3m-A | 4.3 1 | 2.3 | 107 | 465 | 7 | MOLECULE: | SULFITE REDUCTASE [NADPH] HEMO |
| 65:00:00 | 5h8y-C | 4.3 1 | 4.6 | 107 | 563 | 11 | MOLECULE: | SULFITE REDUCTASE [FERREDOXIN] |
| 66:00:00 | 2gep-A | 4.2 1 | 2.2 | 107 | 472 | 5 | MOLECULE: | SULFITE REDUCTASE HEMOPROTEIN |
| 67:00:00 | 5h8v-B | 4.1 1 | 3.6 | 106 | 560 | 11 | MOLECULE: | SULFITE REDUCTASE [FERREDOXIN] |
| 68:00:00 | 1zj8-B | 4.1 | 5.2 | 103 | 546 | 4 | MOLECULE: | PROBABLE FERREDOXIN-DEPENDENT |
| 69:00:00 | 5h8v-A | 4.1 1 | 3.7 | 109 | 564 | 10 | MOLECULE: | SULFITE REDUCTASE [FERREDOXIN] |
| 70:00:00 | 1aop-A | 4.1 | 5 | 103 | 456 | 6 | MOLECULE: | SULFITE REDUCTASE HEMOPROTEIN |
| 71:00:00 | 7xql-A | 4 | 5.3 | 159 | 444 | 11 | MOLECULE: | ANKYRIN REPEAT-CONTAINING PROT |
| 72:00:00 | 5h8y-D | 4 | 5.2 | 105 | 554 | 12 | MOLECULE: | SULFITE REDUCTASE [FERREDOXIN] |
| 73:00:00 | 5h8y-A | 4.0 1 | 3.7 | 109 | 567 | 8 | MOLECULE: | SULFITE REDUCTASE [FERREDOXIN] |
| 74:00:00 | 1zj9-A | 3.8 1 | 0.4 | 110 | 546 | 5 | MOLECULE: | PROBABLE FERREDOXIN-DEPENDENT |

| | | | | | | | | |
|---|---|---|---|---|---|---|---|---|
| 75:00:00 | 1zj9-B | 3.8 1 | 0.3 | 110 | 546 | 5 | MOLECULE: | PROBABLE FERREDOXIN-DEPENDENT |
| 76:00:00 | 3b0n-A | 3.7 1 | 3.6 | 106 | 538 | 8 | MOLECULE: | NITRITE REDUCTASE |
| 77:00:00 | 3vm0-A | 3.7 1 | 3.7 | 107 | 538 | 7 | MOLECULE: | NITRITE REDUCTASE |
| 78:00:00 | 6mca-A | 3.6 1 | 1.8 | 148 | 460 | 12 | MOLECULE: | ANKYRIN REPEAT DOMAIN PROTEIN |
| 79:00:00 | 3vkr-A | 3.6 1 | 1.7 | 107 | 538 | 10 | MOLECULE: | NITRITE REDUCTASE |
| 80:00:00 | 3vks-A | 3.6 1 | 4.1 | 112 | 538 | 9 | MOLECULE: | NITRITE REDUCTASE |
| 81:00:00 | 3b0m-A | 3.6 1 | 2.9 | 107 | 538 | 8 | MOLECULE: | NITRITE REDUCTASE |
| 82:00:00 | 3vlz-A | 3.6 1 | 2.9 | 107 | 538 | 9 | MOLECULE: | NITRITE REDUCTASE |
| 83:00:00 | 5h8y-B | 3.5 1 | 3.1 | 110 | 565 | 12 | MOLECULE: | SULFITE REDUCTASE [FERREDOXIN] |
| 84:00:00 | 3b0l-A | 3.3 1 | 2.5 | 106 | 538 | 10 | MOLECULE: | NITRITE REDUCTASE |
| 85:00:00 | 4kqe-A | 3.2 1 | 7.2 | 93 | 597 | 5 | MOLECULE: | GLYCINE--TRNA LIGASE |
| 86:00:00 | 3vkt-A | 3.1 1 | 6.9 | 98 | 538 | 5 | MOLECULE: | NITRITE REDUCTASE |
| 87:00:00 | 2akj-A | 3.1 1 | 2.5 | 97 | 535 | 6 | MOLECULE: | FERREDOXIN--NITRITE REDUCTASE, |
| 88:00:00 | 5e6m-A | 3.0 1 | 7.1 | 95 | 520 | 6 | MOLECULE: | GLYCINE--TRNA LIGASE |
| 89:00:00 | 3b0j-A | 3.0 1 | 1.7 | 107 | 538 | 9 | MOLECULE: | NITRITE REDUCTASE |
| 90:00:00 | 3vkp-A | 3.0 1 | 3.7 | 107 | 538 | 7 | MOLECULE: | NITRITE REDUCTASE |
| 91:00:00 | 3b0h-A | 3.0 1 | 3.6 | 95 | 536 | 6 | MOLECULE: | NITRITE REDUCTASE |
| 92:00:00 | 3b0g-A | 3.0 1 | 4.8 | 94 | 538 | 5 | MOLECULE: | NITRITE REDUCTASE |
| 93:00:00 | 3vlx-A | 3.0 1 | 2.5 | 95 | 538 | 6 | MOLECULE: | NITRITE REDUCTASE |
| 94:00:00 | 3b0h-B | 3.0 1 | 3.1 | 83 | 535 | 7 | MOLECULE: | NITRITE REDUCTASE |
| 95:00:00 | 3vly-A | 3.0 1 | 2.5 | 95 | 538 | 6 | MOLECULE: | NITRITE REDUCTASE |
| 96:00:00 | 3vkq-A | 3.0 1 | 2.7 | 93 | 538 | 6 | MOLECULE: | NITRITE REDUCTASE |
| 97:00:00 | 1zj8-A | 2.9 | 5.2 | 103 | 546 | 4 | MOLECULE: | PROBABLE FERREDOXIN-DEPENDENT |
| 98:00:00 | 2zt7-A | 2.9 1 | 3.3 | 91 | 531 | 5 | MOLECULE: | GLYCYL-TRNA SYNTHETASE |
| 99:00:00 | 3vm1-A | 2.9 1 | 4.4 | 95 | 538 | 5 | MOLECULE: | NITRITE REDUCTASE |
| 0:00 | 6c3y-A | 2.9 1 | 1.9 | 107 | 464 | 7 | MOLECULE: | SULFITE REDUCTASE [NADPH] HEMO |
| 1:00 | 5uto-A | 2.8 | 3.6 | 95 | 288 | 4 | MOLECULE: | EDD DOMAIN PROTEIN, DEGV FAMIL |
| 2:00 | 2eo6-A | 2.6 | 3.7 | 85 | 141 | 7 | MOLECULE: | B-CELL LINKER PROTEIN |
| 3:00 | 7xrm-B | 2.6 | 3.7 | 100 | 253 | 8 | MOLECULE: | ETHANOLAMINE AMMONIA-LYASE LAR |
| 4:00 | 2j3l-B | 2.6 1 | 8.4 | 82 | 571 | 13 | MOLECULE: | PROLYL-TRNA SYNTHETASE |

**DALI results: MdA-1 Prim**

| No: | Chain | Z | rmsd l | al | i nres | %id | PDB Description |
|---|---|---|---|---|---|---|---|
| 1:00 | 5iok-A | 2.9 | 3.5 | 61 | 142 | 10 | MOLECULE: TRANSCRIPTION |
| 2:00 | 6miq-A | 2.5 | 3.4 | 61 | 141 | 10 | MOLECULE: TRANSCRIPTION |
| 3:00 | 5d7e-A | 2.5 | 3.5 | 61 | 140 | 10 | MOLECULE: TRANSCRIPTION |
| 4:00 | 7egm-H | 2.4 | 5.6 | 66 | 388 | 9 | MOLECULE: TRANSCRIPTION |
| 5:00 | 6mip-A | 2.3 | 4.2 | 62 | 139 | 10 | MOLECULE: TRANSCRIPTION |
| 6:00 | 6min-A | 2.3 | 3.6 | 61 | 139 | 10 | MOLECULE: TRANSCRIPTION |
| 7:00 | 2hcz-X | 2.2 | 4.3 | 66 | 242 | 3 | MOLECULE: BETA-EXPANSIN |
| 8:00 | 7f5m-A | 2.1 | 3.5 | 61 | 139 | 11 | MOLECULE: SOMETHING ABO |
| 9:00 | 6ans-B | 2.0 | 4.5 | 54 | 369 | 6 | MOLECULE: UNCHARACTERIZ |
| 10:00 | 6mio-A | 2.0 | 3.4 | 56 | 142 | 7 | MOLECULE: TRANSCRIPTION |

**DALI results: MdA-1 E1**

| No: | Chain | Z | rmsd | lali | nres | %id | PDB | Description |
|---|---|---|---|---|---|---|---|---|
| 1:00 | 7apd-B | 19.8 | 3.4 | 268 | 289 | 21 | MOLECULE: | REPLICATION PROTEIN E1 |
| 2:00 | 1tue-K | 17 | 2.4 | 185 | 192 | 22 | MOLECULE: | REPLICATION PROTEIN E1 |
| 3:00 | 4gdf-A | 16.5 | 5.7 | 334 | 497 | 16 | MOLECULE: | LARGE T ANTIGEN |
| 4:00 | 9bc5-F | 11.1 | 4.3 | 157 | 480 | 19 | MOLECULE: | PROTEIN REP68 |
| 5:00 | 7apd-G | 10.9 | 3.9 | 140 | 152 | 9 | MOLECULE: | REPLICATION PROTEIN E1 |
| 6:00 | 6k9c-A | 10.7 | 6.7 | 230 | 416 | 14 | MOLECULE: | PRIMASE |
| 7:00 | 4fb3-A | 10.3 | 2.6 | 112 | 115 | 9 | MOLECULE: | ORI DNA OLIGONUCLEOTIDE-CRICK ST |
| 8:00 | 1r9w-A | 9.5 | 3.4 | 124 | 138 | 11 | MOLECULE: | REPLICATION PROTEIN E1 |
| 9:00 | 7ola-A | 9.4 | 6.7 | 217 | 441 | 13 | MOLECULE: | DNA PRIMASE |
| 10:00 | 8wgy-E | 9.3 | 6 | 259 | 769 | 8 | MOLECULE: | UNCOATING FACTOR OPG117 |
| 11:00 | 8iqi-C | 9 | 4.3 | 262 | 916 | 11 | MOLECULE: | PUTATIVE PRIMASE C962R |
| 12:00 | 3dkx-A | 8 | 10.3 | 119 | 202 | 8 | MOLECULE: | REPLICATION PROTEIN REPB |
| 13:00 | 7mi8-A | 6.8 | 4.4 | 191 | 868 | 8 | MOLECULE: | FUSION PROTEIN OF DYNEIN AND END |
| 14:00 | 7tts-D | 6.8 | 13.6 | 177 | 463 | 11 | MOLECULE: | CASEINOLYTIC PEPTIDASE B PROTEIN |
| 15:00 | 4p6q-A | 6.6 | 10.9 | 121 | 286 | 6 | MOLECULE: | MSX2-INTERACTING PROTEIN |
| 16:00 | 5nvu-A | 6.4 | 16.3 | 197 | 3169 | 0 | MOLECULE: | DYNEIN MOTOR DOMAIN |
| 17:00 | 3bos-B | 6.4 | 3.6 | 136 | 231 | 13 | MOLECULE: | PUTATIVE DNA REPLICATION FACTOR |
| 18:00 | 7fd4-A | 6.3 | 15 | 235 | 779 | 10 | MOLECULE: | LON PROTEASE |
| 19:00 | 3j3r-A | 6.2 | 11.5 | 216 | 798 | 11 | MOLECULE: | ADAPTER PROTEIN MECA 1 |
| 20:00 | 5udb-4 | 6 | 8.3 | 227 | 751 | 12 | MOLECULE: | DNA REPLICATION LICENSING FACTOR |
| 21:00 | 5axk-B | 5.9 | 3.9 | 107 | 239 | 6 | MOLECULE: | TRNA(HIS)-5'-GUANYLYLTRANSFERASE |
| 22:00 | 3nbx-X | 5.8 | 4.1 | 173 | 481 | 9 | MOLECULE: | ATPASE RAVA |
| 23:00 | 8umy-A | 5.8 | 21.9 | 161 | 546 | 14 | MOLECULE: | CHROMOSOME TRANSMISSION FIDELITY |
| 24:00:00 | 7w42-B | 5.7 | 16.2 | 171 | 402 | 15 | MOLECULE: | UNCHARACTERIZED ATPASE YJOB |
| 25:00:00 | 8c0v-B | 5.7 | 12.2 | 213 | 1030 | 8 | MOLECULE: | PEROXISOMAL ATPASE PEX1 |
| 26:00:00 | 5nug-A | 5.7 | 22.2 | 214 | 2920 | 8 | MOLECULE: | CYTOPLASMIC DYNEIN 1 HEAVY CHAIN |
| 27:00:00 | 8xku-B | 5.7 | 12.3 | 186 | 845 | 13 | MOLECULE: | PROBABLE INACTIVE ATP-DEPENDENT |
| 28:00:00 | 8osg-B | 5.7 | 4.4 | 171 | 323 | 9 | MOLECULE: | MAGNESIUM-CHELATASE SUBUNIT CHLI |
| 29:00:00 | 4rh7-A | 5.6 | 22.9 | 222 | 3005 | 11 | MOLECULE: | GREEN FLUORESCENT PROTEIN/CYTOPL |
| 30:00:00 | 2hiy-D | 5.6 | 3 | 82 | 183 | 11 | MOLECULE: | HYPOTHETICAL PROTEIN |
| 31:00:00 | 3u5z-B | 5.6 | 11 | 138 | 320 | 17 | MOLECULE: | DNA POLYMERASE ACCESSORY PROTEIN |
| 32:00:00 | 8dr5-A | 5.5 | 12 | 154 | 646 | 11 | MOLECULE: | REPLICATION FACTOR C SUBUNIT 1 |
| 33:00:00 | 6f74-A | 5.5 | 14.6 | 132 | 572 | 4 | MOLECULE: | ALCOHOL OXIDASE |
| 34:00:00 | 5m7o-A | 5.5 | 14.5 | 162 | 448 | 12 | MOLECULE: | NITROGEN ASSIMILATION REGULATORY |
| 35:00:00 | 7b8s-C | 5.5 | 5.4 | 104 | 589 | 3 | MOLECULE: | MULTIDRUG EFFLUX PUMP SUBUNIT AC |
| 36:00:00 | 5dgk-A | 5.5 | 7.1 | 201 | 519 | 9 | MOLECULE: | ACTIVE HELICASE |
| 37:00:00 | 7uqj-A | 5.4 | 5.8 | 157 | 585 | 11 | MOLECULE: | ATPASE HISTONE CHAPERONE YTA7 |
| 38:00:00 | 7vcs-A | 5.4 | 9.1 | 210 | 766 | 11 | MOLECULE: | TRANSITIONAL ENDOPLASMIC RETICUL |
| 39:00:00 | 8k8v-A | 5.3 | 14.3 | 155 | 719 | 12 | MOLECULE: | ENDOPEPTIDASE LA |
| 40:00:00 | 7qo4-M | 5.3 | 10.4 | 183 | 421 | 13 | MOLECULE: | 26S PROTEASOME REGULATORY SUBUNI |
| 41:00:00 | 5udb-7 | 5.1 | 5.8 | 203 | 726 | 12 | MOLECULE: | DNA REPLICATION LICENSING FACTOR |
| 42:00:00 | 6fhs-C | 5.1 | 15.4 | 149 | 459 | 12 | MOLECULE: | RUVB-LIKE HELICASE |

| | | | | | | | | |
|---|---|---|---|---|---|---|---|---|
| 43:00:00 | 3tht-B | 5.1 | 15.3 | 118 | 312 | 5 | MOLECULE: | ALKYLATED DNA REPAIR PROTEIN ALK |
| 44:00:00 | 7ykk-A | 5.1 | 9.2 | 199 | 735 | 10 | MOLECULE: | ATPASE FAMILY GENE 2 PROTEIN |
| 45:00:00 | 6uen-A | 5 | 16.2 | 195 | 1392 | 6 | MOLECULE: | RNA-DIRECTED RNA POLYMERASE L |
| 46:00:00 | 3vf0-B | 5 | 5.3 | 105 | 283 | 4 | MOLECULE: | VINCULIN |
| 47:00:00 | 2qby-B | 5 | 7 | 137 | 368 | 9 | MOLECULE: | CELL DIVISION CONTROL PROTEIN 6 |
| 48:00:00 | 3f8t-A | 4.9 | 5.5 | 185 | 459 | 9 | MOLECULE: | PREDICTED ATPASE INVOLVED IN REP |
| 49:00:00 | 8xks-B | 4.9 | 34.3 | 144 | 603 | 13 | MOLECULE: | CTAP1 |
| 50:00:00 | 6dgd-A | 4.8 | 10.7 | 153 | 704 | 12 | MOLECULE: | PRIMOSOMAL PROTEIN N' |
| 51:00:00 | 3wbz-E | 4.8 | 4.5 | 115 | 268 | 4 | MOLECULE: | LIKELY HISTIDYL TRNA-SPECIFIC GU |
| 52:00:00 | 5yww-A | 4.8 | 13.7 | 175 | 485 | 11 | MOLECULE: | NUCLEOTIDE BINDING PROTEIN PINC |
| 53:00:00 | 5zr1-A | 4.8 | 17.9 | 168 | 494 | 8 | MOLECULE: | ORIGIN RECOGNITION COMPLEX SUBUN |
| 54:00:00 | 7crc-B | 4.8 | 13.1 | 186 | 1112 | 10 | MOLECULE: | NAD+ HYDROLASE (NADASE) |
| 55:00:00 | 7mjy-A | 4.7 | 15.9 | 121 | 449 | 10 | MOLECULE: | TRNA-2-METHYLTHIO-N(6)-DIMETHYLA |
| 56:00:00 | 8fwj-A | 4.7 | 10.1 | 140 | 552 | 11 | MOLECULE: | CIRCADIAN CLOCK PROTEIN KAIC |
| 57:00:00 | 3k1j-A | 4.7 | 12.2 | 177 | 566 | 8 | MOLECULE: | ATP-DEPENDENT PROTEASE LON |
| 58:00:00 | 8c0v-A | 4.7 | 11.9 | 186 | 823 | 6 | MOLECULE: | PEROXISOMAL ATPASE PEX1 |
| 59:00:00 | 8jx6-B | 4.6 | 11.1 | 122 | 451 | 9 | MOLECULE: | DEEP-SEA HELICASE 9 |
| 60:00:00 | 7qh2-C | 4.5 | 12.2 | 137 | 467 | 12 | MOLECULE: | LACTATE DEHYDROGENASE (NAD(+),FE |
| 61:00:00 | 8sk1-A | 4.5 | 3.3 | 91 | 162 | 11 | MOLECULE: | 2-AMINO-4-HYDROXY-6-HYDROXYMETHY |
| 62:00:00 | 6dhs-B | 4.5 | 6.8 | 93 | 176 | 4 | MOLECULE: | HETEROGENEOUS NUCLEAR RIBONUCLEO |
| 63:00:00 | 6fjw-A | 4.5 | 3.4 | 84 | 239 | 10 | MOLECULE: | CAS6 PROTEIN |
| 64:00:00 | 6f7s-C | 4.4 | 13.1 | 131 | 319 | 3 | MOLECULE: | SERRATE RNA EFFECTOR MOLECULE HO |
| 65:00:00 | 7dex-A | 4.4 | 9.2 | 80 | 460 | 4 | MOLECULE: | ANTHOCYANIN 5-AROMATIC ACYLTRANS |
| 66:00:00 | 7t8b-A | 4.4 | 8 | 170 | 518 | 9 | MOLECULE: | TWINKLE MTDNA HELICASE |
| 67:00:00 | 5e7p-A | 4.4 | 18.6 | 208 | 719 | 11 | MOLECULE: | CELL DIVISION CONTROL PROTEIN CD |
| 68:00:00 | 5aor-A | 4.4 | 18.1 | 167 | 1009 | 12 | MOLECULE: | DOSAGE COMPENSATION REGULATOR |
| 69:00:00 | 4kw3-A | 4.4 | 3.5 | 105 | 266 | 5 | MOLECULE: | NS1 |
| 70:00:00 | 8emh-B | 4.4 | 4.8 | 198 | 676 | 8 | MOLECULE: | PROTEASE LON-RELATED BREX SYSTEM |
| 71:00:00 | 4gwp-F | 4.3 | 3.5 | 81 | 209 | 4 | MOLECULE: | MEDIATOR OF RNA POLYMERASE II TR |
| 72:00:00 | 6v55-A | 4.3 | 21.7 | 156 | 746 | 6 | MOLECULE: | ADHESION G-PROTEIN COUPLED RECEP |
| 73:00:00 | 5che-A | 4.3 | 23.6 | 120 | 427 | 5 | MOLECULE: | GLUTAMYL-TRNA REDUCTASE 1, CHLOR |
| 74:00:00 | 5jpq-i | 4.3 | 7.2 | 120 | 659 | 13 | MOLECULE: | WD40 DOMAIN PROTEINS |
| 75:00:00 | 8tdl-A | 4.3 | 23.5 | 118 | 364 | 8 | MOLECULE: | MECHANOSENSITIVE ION CHANNEL PRO |
| 76:00:00 | 4ksr-A | 4.3 | 16.5 | 153 | 521 | 12 | MOLECULE: | TYPE II SECRETION SYSTEM PROTEIN |
| 77:00:00 | 8tzk-C | 4.3 | 22.4 | 118 | 304 | 6 | MOLECULE: | CELL DIVISION ATP-BINDING PROTEI |
| 78:00:00 | 8xku-D | 4.3 | 23.8 | 216 | 805 | 9 | MOLECULE: | PROBABLE INACTIVE ATP-DEPENDENT |
| 79:00:00 | 8fcv-R | 4.3 | 8.2 | 142 | 343 | 8 | MOLECULE: | DNA (60-MER) |
| 80:00:00 | 5wx8-A | 4.2 | 3.8 | 89 | 165 | 8 | MOLECULE: | IMMEDIATE-EARLY PROTEIN 2 |
| 81:00:00 | 1vr6-A | 4.2 | 14.7 | 109 | 343 | 7 | MOLECULE: | PHOSPHO-2-DEHYDRO-3-DEOXYHEPTONA |
| 82:00:00 | 1m1g-C | 4.2 | 7.6 | 101 | 244 | 10 | MOLECULE: | TRANSCRIPTION ANTITERMINATION PR |
| 83:00:00 | 6o16-A | 4.2 | 15.7 | 140 | 798 | 11 | MOLECULE: | DEAH (ASP-GLU-ALA-HIS) BOX POLYP |
| 84:00:00 | 8k4i-A | 4.1 | 2.8 | 68 | 163 | 1 | MOLECULE: | YAJQ |
| 85:00:00 | 8y6p-Q | 4.1 | 13.6 | 143 | 1228 | 11 | MOLECULE: | APAF-1 RELATED KILLER DARK |
| 86:00:00 | 4lgt-A | 4.1 | 16.6 | 98 | 251 | 5 | MOLECULE: | RIBOSOMAL LARGE SUBUNIT PSEUDOUR |

| | | | | | | | | |
|---|---|---|---|---|---|---|---|---|
| 87:00:00 | 4lya-A | 4.1 | 4.7 | 132 | 521 | 8 | MOLECULE: | UNCHARACTERIZED PROTEIN |
| 88:00:00 | 3cmu-A | 4.1 | 10.4 | 143 | 1937 | 13 | MOLECULE: | DNA (5'- |
| 89:00:00 | 6ted-Z | 4 | 5.4 | 131 | 510 | 5 | MOLECULE: | DNA-DIRECTED RNA POLYMERASE SUBU |
| 90:00:00 | 6hyp-A | 4 | 7.9 | 171 | 2272 | 8 | MOLECULE: | MIDASIN,MIDASIN |
| 91:00:00 | 3c19-A | 4 | 12 | 99 | 177 | 7 | MOLECULE: | UNCHARACTERIZED PROTEIN MK0293 |
| 92:00:00 | 2qv6-A | 4 | 11.3 | 89 | 253 | 7 | MOLECULE: | GTP CYCLOHYDROLASE III |
| 93:00:00 | 4m9s-B | 4 | 14.3 | 154 | 511 | 12 | MOLECULE: | CELL DEATH PROTEIN 4 |
| 94:00:00 | 8yha-B | 4 | 3.5 | 84 | 268 | 7 | MOLECULE: | CRISPR SYSTEM CASCADE SUBUNIT CA |
| 95:00:00 | 6mfv-A | 4 | 7.7 | 125 | 641 | 11 | MOLECULE: | TETRATRICOPEPTIDE REPEAT SENSOR |
| 96:00:00 | 3mtj-A | 4 | 9.7 | 113 | 431 | 12 | MOLECULE: | HOMOSERINE DEHYDROGENASE |
| 97:00:00 | 9esh-P | 3.9 | 16.6 | 110 | 271 | 6 | MOLECULE: | PRE-MRNA |
| 98:00:00 | 7v9x-A | 3.9 | 14.7 | 124 | 314 | 10 | MOLECULE: | RNA-DIRECTED DNA POLYMERASE FROM |
| 99:00:00 | 8th8-L | 3.9 | 11.7 | 174 | 862 | 8 | MOLECULE: | DYNEIN REGULATORY COMPLEX PROTEI |
| 0:00 | 1ahu-A | 3.9 | 13.8 | 127 | 555 | 12 | MOLECULE: | VANILLYL-ALCOHOL OXIDASE |
| 1:00 | 5o6b-A | 3.9 | 3.8 | 119 | 528 | 13 | MOLECULE: | ATP-DEPENDENT DNA HELICASE PIF1 |
| 2:00 | 3d5l-A | 3.8 | 4.1 | 58 | 203 | 12 | MOLECULE: | REGULATORY PROTEIN RECX |
| 3:00 | 7oky-Z | 3.8 | 7.8 | 145 | 374 | 7 | MOLECULE: | DNA-DIRECTED RNA POLYMERASE II S |
| 4:00 | 7epu-B | 3.8 | 16 | 134 | 748 | 10 | MOLECULE: | NON-IMMUNIZED HUMAN SCFV |
| 5:00 | 6d7y-B | 3.7 | 2.7 | 79 | 155 | 9 | MOLECULE: | HEMAGGLUTININ |
| 6:00 | 4nh0-A | 3.7 | 4.2 | 134 | 861 | 10 | MOLECULE: | CELL DIVISIONFTSK/SPOIIIE |
| 7:00 | 4znm-B | 3.7 | 12.6 | 91 | 444 | 10 | MOLECULE: | C-DOMAIN TYPE II PEPTIDE SYNTHET |
| 8:00 | 8peu-A | 3.7 | 9 | 164 | 419 | 12 | MOLECULE: | TRANSCRIPTION TERMINATION FACTOR |
| 9:00 | 5k8o-B | 3.7 | 12.2 | 104 | 425 | 11 | MOLECULE: | 5-NITROANTHRANILIC ACID AMINOHYD |
| 10:00 | 6pwn-A | 3.7 | 18.7 | 113 | 280 | 4 | MOLECULE: | SMALL-CONDUCTANCE MECHANOSENSITI |
| 11:00 | 8x5i-A | 3.7 | 13.4 | 146 | 477 | 12 | MOLECULE: | ENDONUCLEASE GAJA |
| 12:00 | 3zqj-F | 3.7 | 16.5 | 126 | 607 | 20 | MOLECULE: | UVRABC SYSTEM PROTEIN A |
| 13:00 | 3crv-A | 3.7 | 9.2 | 126 | 551 | 13 | MOLECULE: | XPD/RAD3 RELATED DNA HELICASE |
| 14:00 | 4d2i-A | 3.6 | 8.7 | 130 | 464 | 8 | MOLECULE: | HERA |
| 15:00 | 3gp8-A | 3.6 | 18.3 | 129 | 551 | 14 | MOLECULE: | EXODEOXYRIBONUCLEASE V, SUBUNIT |
| 16:00 | 2iai-A | 3.6 | 3.5 | 71 | 200 | 7 | MOLECULE: | PUTATIVE TRANSCRIPTIONAL REGULAT |
| 17:00 | 7sgr-A | 3.6 | 13.8 | 168 | 700 | 11 | MOLECULE: | ALPHA-HEMOLYSIN TRANSLOCATION AT |
| 18:00 | 7t5q-I | 3.6 | 22.3 | 90 | 283 | 9 | MOLECULE: | ACTIN-RELATED PROTEIN 3 |
| 19:00 | 9fzy-F | 3.6 | 13.4 | 121 | 442 | 5 | MOLECULE: | CORRINOID IRON-SULFUR PROTEIN LA |
| 20:00 | 2nyi-A | 3.6 | 10.8 | 92 | 171 | 7 | MOLECULE: | UNKNOWN PROTEIN |
| 21:00 | 3al0-B | 3.6 | 20.2 | 115 | 482 | 10 | MOLECULE: | GLUTAMYL-TRNA(GLN) AMIDOTRANSFER |
| 22:00 | 2pfd-A | 3.6 | 23.5 | 119 | 540 | 3 | MOLECULE: | FORMIMIDOYLTRANSFERASE-CYCLODEAM |
| 23:00 | 8oz7-A | 3.6 | 13 | 151 | 597 | 6 | MOLECULE: | ABIA |
| 24:00:00 | 8dgf-A | 3.6 | 12.8 | 188 | 1541 | 8 | MOLECULE: | ATP-BINDING PROTEIN AVS4 |
| 25:00:00 | 7t5q-J | 3.6 | 30.4 | 106 | 291 | 7 | MOLECULE: | ACTIN-RELATED PROTEIN 3 |
| 26:00:00 | 4a15-A | 3.6 | 10.7 | 118 | 596 | 12 | MOLECULE: | ATP-DEPENDENT DNA HELICASE TA005 |
| 27:00:00 | 8xks-E | 3.6 | 10.9 | 198 | 1458 | 10 | MOLECULE: | CTAP1 |
| 28:00:00 | 7cww-A | 3.6 | 6.8 | 116 | 350 | 10 | MOLECULE: | TSRE |
| 29:00:00 | 7pkq-g | 3.5 | 3.4 | 70 | 132 | 7 | MOLECULE: | MS35 |
| 30:00:00 | 5n9j-Y | 3.5 | 4.5 | 89 | 185 | 7 | MOLECULE: | MEDIATOR OF RNA POLYMERASE II TR |

| | | | | | | | | |
|---|---|---|---|---|---|---|---|---|
| 31:00:00 | 8uy1-A | 3.5 | 4.8 | 115 | 598 | 2 | MOLECULE: | METHYLENETETRAHYDROFOLATE REDUCT |
| 32:00:00 | 5ks8-F | 3.5 | 11.9 | 137 | 482 | 8 | MOLECULE: | PYRUVATE CARBOXYLASE SUBUNIT ALP |
| 33:00:00 | 5ijn-F | 3.5 | 17.8 | 118 | 335 | 8 | MOLECULE: | NUCLEAR PORE COMPLEX PROTEIN NUP |
| 34:00:00 | 2cg8-C | 3.5 | 8.5 | 96 | 248 | 11 | MOLECULE: | DIHYDRONEOPTERIN ALDOLASE 6-HYDR |
| 35:00:00 | 6fon-A | 3.5 | 7.2 | 78 | 252 | 5 | MOLECULE: | COPPER CHAPERONE FOR SUPEROXIDE |
| 36:00:00 | 6zyd-A | 3.5 | 12.6 | 99 | 323 | 11 | MOLECULE: | LOW CONDUCTANCE MECHANOSENSITIVE |
| 37:00:00 | 3tuz-C | 3.5 | 5.5 | 121 | 345 | 10 | MOLECULE: | D-METHIONINE TRANSPORT SYSTEM PE |
| 38:00:00 | 8fvu-A | 3.5 | 10.6 | 181 | 1361 | 10 | MOLECULE: | BACULOVIRAL IAP REPEAT-CONTAININ |
| 39:00:00 | 6bwo-A | 3.5 | 5.2 | 98 | 144 | 3 | MOLECULE: | PYRIDINIUM-3,5-BISTHIOCARBOXYLIC |
| 40:00:00 | 7wrx-C | 3.5 | 11.5 | 139 | 594 | 9 | MOLECULE: | HERA |
| 41:00:00 | 1wp9-A | 3.4 | 14.4 | 131 | 479 | 13 | MOLECULE: | ATP-DEPENDENT RNA HELICASE, PUTA |
| 42:00:00 | 6pif-G | 3.4 | 16.7 | 108 | 521 | 6 | MOLECULE: | CAS7, TYPE I-F CRISPR-ASSOCIATED |
| 43:00:00 | 7njn-C | 3.4 | 12.9 | 186 | 536 | 6 | MOLECULE: | ATP SYNTHASE SUBUNIT ALPHA |
| 44:00:00 | 9io5-J | 3.4 | 6.9 | 186 | 597 | 7 | MOLECULE: | G1-ATPASE SUBUNIT BETA |
| 45:00:00 | 8vx9-B | 3.4 | 20.4 | 170 | 1174 | 10 | MOLECULE: | HAMA |
| 46:00:00 | 6pe4-A | 3.4 | 5.8 | 88 | 758 | 3 | MOLECULE: | V-TYPE PROTON ATPASE SUBUNIT A, |
| 47:00:00 | 5gko-A | 3.4 | 10.9 | 122 | 650 | 11 | MOLECULE: | MACROLIDE EXPORT ATP-BINDING/PER |
| 48:00:00 | 6umm-A | 3.4 | 16.2 | 121 | 285 | 4 | MOLECULE: | ESX-3 SECRETION SYSTEM PROTEIN E |
| 49:00:00 | 7elb-A | 3.4 | 5.1 | 171 | 1937 | 3 | MOLECULE: | RNA-DIRECTED RNA POLYMERASE L |
| 50:00:00 | 6rfl-Y | 3.4 | 4.2 | 131 | 600 | 11 | MOLECULE: | DNA-DEPENDENT RNA POLYMERASE SUB |
| 51:00:00 | 8qvt-A | 3.4 | 12 | 127 | 514 | 11 | MOLECULE: | CYTOKININ DEHYDROGENASE |
| 52:00:00 | 2xgj-B | 3.3 | 8.5 | 126 | 773 | 6 | MOLECULE: | ATP-DEPENDENT RNA HELICASE DOB1 |
| 53:00:00 | 6ekk-B | 3.3 | 6.5 | 137 | 381 | 12 | MOLECULE: | DENN DOMAIN-CONTAINING PROTEIN 1 |
| 54:00:00 | 5okc-B | 3.3 | 14.4 | 105 | 377 | 6 | MOLECULE: | SISTER CHROMATID COHESION PROTEI |
| 55:00:00 | 3pfo-A | 3.3 | 14.2 | 115 | 426 | 10 | MOLECULE: | PUTATIVE ACETYLORNITHINE DEACETY |
| 56:00:00 | 6ner-E | 3.3 | 3.8 | 89 | 186 | 6 | MOLECULE: | BMC-H TANDEM FUSION PROTEIN |
| 57:00:00 | 5m86-A | 3.3 | 6.7 | 111 | 321 | 6 | MOLECULE: | TA1207 |
| 58:00:00 | 5a9j-A | 3.3 | 4.4 | 133 | 805 | 14 | MOLECULE: | DNA POLYMERASE THETA |
| 59:00:00 | 7r9x-A | 3.3 | 12.5 | 82 | 418 | 12 | MOLECULE: | AMBE |
| 60:00:00 | 3onq-B | 3.2 | 16.5 | 104 | 255 | 8 | MOLECULE: | REGULATOR OF POLYKETIDE SYNTHASE |
| 61:00:00 | 8hra-C | 3.2 | 14 | 141 | 852 | 9 | MOLECULE: | ARCHAEAL ATPASE |
| 62:00:00 | 7ou2-B | 3.2 | 12.4 | 170 | 650 | 8 | MOLECULE: | DNA MISMATCH REPAIR PROTEIN MUTS |
| 63:00:00 | 2oca-A | 3.2 | 9.5 | 134 | 494 | 6 | MOLECULE: | ATP-DEPENDENT DNA HELICASE UVSW |
| 64:00:00 | 6jyt-A | 3.2 | 21.2 | 143 | 597 | 10 | MOLECULE: | HELICASE |
| 65:00:00 | 3c6k-D | 3.2 | 14.3 | 92 | 348 | 13 | MOLECULE: | SPERMINE SYNTHASE |
| 66:00:00 | 9go3-A | 3.2 | 19.2 | 116 | 253 | 7 | MOLECULE: | MECHANOSENSITIVE CHANNEL PROTEIN |
| 67:00:00 | 7mi1-A | 3.2 | 16.1 | 107 | 2628 | 7 | MOLECULE: | CHIMERA PROTEIN OF DYNEIN AND EN |
| 68:00:00 | 5zxd-B | 3.2 | 13.3 | 116 | 491 | 16 | MOLECULE: | ATP-BINDING CASSETTE SUB-FAMILY |
| 69:00:00 | 6d9m-A | 3.2 | 13 | 103 | 326 | 6 | MOLECULE: | FUSION PROTEIN OF ENDOLYSIN,RESP |
| 70:00:00 | 1yoz-B | 3.2 | 2.8 | 55 | 116 | 4 | MOLECULE: | HYPOTHETICAL PROTEIN AF0941 |
| 71:00:00 | 9cvc-A | 3.2 | 16.8 | 98 | 570 | 8 | MOLECULE: | CODANIN-1 |
| 72:00:00 | 8k0b-A | 3.1 | 19.5 | 143 | 689 | 2 | MOLECULE: | CALCIUM PERMEABLE STRESS-GATED C |
| 73:00:00 | 6nmi-A | 3.1 | 19.2 | 145 | 653 | 10 | MOLECULE: | GENERAL TRANSCRIPTION AND DNA RE |
| 74:00:00 | 6njy-A | 3.1 | 7.4 | 85 | 237 | 4 | MOLECULE: | TYPE IV CRISPR ASSOCIATED CAS6 R |

| | | | | | | | | |
|---|---|---|---|---|---|---|---|---|
| 75:00:00 | 8jqb-E | 3.1 | 5.7 | 114 | 499 | 9 | MOLECULE: | ENDONUCLEASE GAJA |
| 76:00:00 | 5z79-C | 3.1 | 11 | 96 | 599 | 10 | MOLECULE: | HYDROXYMETHYLDIHYDROPTERIN PYROP |
| 77:00:00 | 7tmr-F | 3.1 | 6.7 | 151 | 478 | 9 | MOLECULE: | H(+)-TRANSPORTING TWO-SECTOR ATP |
| 78:00:00 | 6iy8-A | 3.1 | 10.8 | 96 | 469 | 7 | MOLECULE: | POSITIVE REGULATOR CAPR |
| 79:00:00 | 6xf9-D | 3.1 | 10 | 94 | 426 | 6 | MOLECULE: | PACKAGING PROTEIN UL32 |
| 80:00:00 | 3pyb-B | 3.1 | 4.9 | 97 | 689 | 7 | MOLECULE: | ENT-COPALYL DIPHOSPHATE SYNTHASE |
| 81:00:00 | 3dew-A | 3.1 | 3.5 | 77 | 192 | 3 | MOLECULE: | TRANSCRIPTIONAL REGULATOR, TETR |
| 82:00:00 | 4wd9-A | 3.1 | 16.5 | 85 | 965 | 7 | MOLECULE: | NISIN BIOSYNTHESIS PROTEIN NISB |
| 83:00:00 | 4qbu-A | 3.1 | 6 | 107 | 436 | 8 | MOLECULE: | ZMAA |
| 84:00:00 | 2pgc-A | 3 | 7.6 | 102 | 206 | 12 | MOLECULE: | UNCHARACTERIZED PROTEIN |
| 85:00:00 | 8wtk-A | 3 | 11.9 | 118 | 496 | 14 | MOLECULE: | CRISPR-ASSOCIATED HELICASE CAS3 |
| 86:00:00 | 7rgd-A | 3 | 10.2 | 142 | 519 | 9 | MOLECULE: | ISOFORM 5 OF INOSINE-5'-MONOPHOS |
| 87:00:00 | 6urt-A | 3 | 13.8 | 99 | 331 | 8 | MOLECULE: | LOW CONDUCTANCE MECHANOSENSITIVE |
| 88:00:00 | 1lfp-A | 3 | 4.9 | 94 | 243 | 6 | MOLECULE: | HYPOTHETICAL PROTEIN AQ_1575 |
| 89:00:00 | 1yb3-A | 3 | 5.5 | 85 | 166 | 8 | MOLECULE: | HYPOTHETICAL PROTEIN |
| 90:00:00 | 3a1y-G | 3 | 19.4 | 101 | 211 | 10 | MOLECULE: | 50S RIBOSOMAL PROTEIN P1 (L12P) |
| 91:00:00 | 4j8s-A | 3 | 11.3 | 66 | 180 | 9 | MOLECULE: | CCR4-NOT TRANSCRIPTION COMPLEX S |
| 92:00:00 | 8xj6-D | 3 | 15.9 | 132 | 589 | 11 | MOLECULE: | MONKEYPOX VIRUS E5 |
| 93:00:00 | 4cej-B | 2.9 | 15.7 | 146 | 1156 | 8 | MOLECULE: | ATP-DEPENDENT HELICASE/NUCLEASE |
| 94:00:00 | 8phk-P | 2.9 | 4.2 | 85 | 162 | 7 | MOLECULE: | TRANSCRIPTION ANTITERMINATION PR |
| 95:00:00 | 9gm7-C | 2.9 | 8.5 | 50 | 440 | 2 | MOLECULE: | CHROMOSOME PARTITION PROTEIN MUK |
| 96:00:00 | 5egf-C | 2.9 | 16.3 | 92 | 444 | 10 | MOLECULE: | TQAA |
| 97:00:00 | 8s7v-K | 2.9 | 14.6 | 119 | 521 | 15 | MOLECULE: | METHYL-COENZYME M REDUCTASE SUBU |
| 98:00:00 | 7z3h-A | 2.9 | 11.7 | 91 | 372 | 4 | MOLECULE: | TRANSFERASE-LIKE PROTEIN |
| 99:00:00 | 3on2-B | 2.9 | 3.5 | 76 | 195 | 7 | MOLECULE: | PROBABLE TRANSCRIPTIONAL REGULAT |
| 0:00 | 3gmg-A | 2.8 | 7 | 84 | 143 | 2 | MOLECULE: | UNCHARACTERIZED PROTEIN RV1825/M |
| 1:00 | 5dy0-A | 2.8 | 3.1 | 73 | 220 | 3 | MOLECULE: | TETR FAMILY TRANSCRIPTIONAL REGU |
| 2:00 | 6zca-H | 2.8 | 17.9 | 153 | 771 | 9 | MOLECULE: | PROBABLE DNA-DIRECTED RNA POLYME |
| 3:00 | 8j0s-d | 2.8 | 13.3 | 74 | 445 | 8 | MOLECULE: | ATP SYNTHASE SUBUNIT A |
| 4:00 | 2opl-B | 2.7 | 8.7 | 75 | 182 | 3 | MOLECULE: | HYPOTHETICAL PROTEIN |
| 5:00 | 5jb3-H | 2.7 | 3.2 | 69 | 214 | 9 | MOLECULE: | 16S RIBOSOMAL RNA |
| 6:00 | 7c0q-A | 2.7 | 3.2 | 57 | 279 | 7 | MOLECULE: | EFFECTOR LPG2505 |
| 7:00 | 7yqh-A | 2.6 | 4.1 | 125 | 1069 | 11 | MOLECULE: | STRUCTURAL MAINTENANCE OF CHROMO |
| 8:00 | 8xku-A | 2.6 | 9.8 | 112 | 730 | 12 | MOLECULE: | PROBABLE INACTIVE ATP-DEPENDENT |
| 9:00 | 9gm6-B | 2.6 | 4 | 111 | 1148 | 10 | MOLECULE: | CHROMOSOME PARTITION PROTEIN MUK |
| 10:00 | 7f52-A | 2.6 | 19.5 | 78 | 584 | 13 | MOLECULE: | NON-STRUCTURAL PROTEIN 2 |
| 11:00 | 2k5s-A | 2.6 | 4.3 | 45 | 73 | 7 | MOLECULE: | MODULATING PROTEIN YMOA |
| 12:00 | 7e7i-A | 2.6 | 13 | 147 | 2011 | 9 | MOLECULE: | RETINAL-SPECIFIC PHOSPHOLIPID-TR |
| 13:00 | 6w2r-B | 2.6 | 9 | 58 | 221 | 3 | MOLECULE: | JUNCTION 19 DHR54-DHR79 |
| 14:00 | 4x4r-A | 2.5 | 13.3 | 100 | 443 | 9 | MOLECULE: | CCA-ADDING ENZYME |
| 15:00 | 3j5s-D | 2.5 | 13.7 | 117 | 554 | 9 | MOLECULE: | 16S RIBOSOMAL RNA |
| 16:00 | 7roq-A | 2.5 | 12.9 | 154 | 1831 | 10 | MOLECULE: | PHOSPHOLIPID-TRANSPORTING ATPASE |
| 17:00 | 8wcn-A | 2.5 | 11.8 | 82 | 379 | 5 | MOLECULE: | DIGUANYLATE CYCLASE |
| 18:00 | 7pde-A | 2.5 | 21.9 | 122 | 846 | 6 | MOLECULE: | ADENYLATE CYCLASE 9 |

| | | | | | | | | |
|---|---|---|---|---|---|---|---|---|
| 19:00 | 7nyw-B | 2.5 | 3.9 | 110 | 858 | 10 | MOLECULE: | CHROMOSOME PARTITION PROTEIN MUK |
| 20:00 | 8s36-M | 2.5 | 12 | 124 | 519 | 10 | MOLECULE: | CRISPR TYPE AFERR-ASSOCIATED PRO |
| 21:00 | 1k8w-A | 2.5 | 3.5 | 81 | 303 | 10 | MOLECULE: | 5'-R(*GP*GP*CP*AP*AP*CP*GP*GP*UP |
| 22:00 | 7b0c-A | 2.4 | 3.4 | 57 | 144 | 7 | MOLECULE: | HTH-TYPE TRANSCRIPTIONAL REPRESS |
| 23:00 | 7sjr-A | 2.4 | 17.4 | 136 | 911 | 13 | MOLECULE: | DNA HELICASE |
| 24:00:00 | 4krr-A | 2.4 | 3 | 65 | 194 | 6 | MOLECULE: | WNT INHIBITOR OF DORSAL PROTEIN |
| 25:00:00 | 2hxi-B | 2.4 | 2.4 | 59 | 198 | 10 | MOLECULE: | PUTATIVE TRANSCRIPTIONAL REGULAT |
| 26:00:00 | 8f4x-0 | 2.4 | 3.2 | 43 | 67 | 19 | MOLECULE: | RC_I_1-H11 |
| 27:00:00 | 7q5b-B | 2.4 | 9.1 | 101 | 482 | 6 | MOLECULE: | DNA (56-MER) |
| 28:00:00 | 6r4p-A | 2.4 | 13.7 | 119 | 478 | 5 | MOLECULE: | ADENYLATE CYCLASE 9 |
| 29:00:00 | 5opt-h | 2.4 | 10.8 | 74 | 173 | 8 | MOLECULE: | ACTIVATED PROTEIN KINASE C RECEP |
| 30:00:00 | 2lc2-A | 2.3 | 3.2 | 47 | 108 | 9 | MOLECULE: | AVR3A4 |
| 31:00:00 | 7lcc-A | 2.2 | 25.2 | 110 | 1369 | 10 | MOLECULE: | HELRAISER K1068Q |
| 32:00:00 | 8xku-C | 2.1 | 16.6 | 105 | 921 | 13 | MOLECULE: | PROBABLE INACTIVE ATP-DEPENDENT |

**DALI results: MdA-1 RepE1**

| No: | Chain | Z | rmsd | lali | nres | %id | PDB | Description |
|---|---|---|---|---|---|---|---|---|
| 1:00 | 2gxa-I | 16.9 | 4.1 | 261 | 270 | 21 | MOLECULE: | 5'-D(*TP*TP*TP*TP*TP*TP*TP*TP*TP*TP*TP*TP*T)-3' |
| 2:00 | 2v9p-D | 16.9 | 3.8 | 254 | 274 | 21 | MOLECULE: | REPLICATION PROTEIN E1 |
| 3:00 | 5a9k-D | 16.8 | 3.8 | 254 | 274 | 21 | MOLECULE: | REPLICATION PROTEIN E1 |
| 4:00 | 2gxa-G | 16.8 | 3.8 | 251 | 270 | 21 | MOLECULE: | 5'-D(*TP*TP*TP*TP*TP*TP*TP*TP*TP*TP*TP*TP*T)-3' |
| 5:00 | 2gxa-D | 16.8 | 3.9 | 261 | 270 | 20 | MOLECULE: | 5'-D(*TP*TP*TP*TP*TP*TP*TP*TP*TP*TP*TP*TP*T)-3' |
| 6:00 | 2v9p-J | 16.7 | 3.9 | 256 | 272 | 21 | MOLECULE: | REPLICATION PROTEIN E1 |
| 7:00 | 2v9p-I | 16.7 | 3.5 | 238 | 270 | 21 | MOLECULE: | REPLICATION PROTEIN E1 |
| 8:00 | 2gxa-B | 16.7 | 4 | 258 | 270 | 21 | MOLECULE: | 5'-D(*TP*TP*TP*TP*TP*TP*TP*TP*TP*TP*TP*TP*T)-3' |
| 9:00 | 2v9p-H | 16.7 | 3.5 | 237 | 268 | 22 | MOLECULE: | REPLICATION PROTEIN E1 |
| 10:00 | 2gxa-C | 16.6 | 3.8 | 245 | 270 | 21 | MOLECULE: | 5'-D(*TP*TP*TP*TP*TP*TP*TP*TP*TP*TP*TP*TP*T)-3' |
| 11:00 | 2gxa-E | 16.6 | 4.2 | 262 | 274 | 21 | MOLECULE: | 5'-D(*TP*TP*TP*TP*TP*TP*TP*TP*TP*TP*TP*TP*T)-3' |
| 12:00 | 2gxa-J | 16.6 | 4.1 | 249 | 270 | 20 | MOLECULE: | 5'-D(*TP*TP*TP*TP*TP*TP*TP*TP*TP*TP*TP*TP*T)-3' |
| 13:00 | 2v9p-B | 16.6 | 3.5 | 237 | 268 | 22 | MOLECULE: | REPLICATION PROTEIN E1 |
| 14:00 | 2gxa-F | 16.5 | 4.2 | 257 | 270 | 21 | MOLECULE: | 5'-D(*TP*TP*TP*TP*TP*TP*TP*TP*TP*TP*TP*TP*T)-3' |
| 15:00 | 2gxa-H | 16.5 | 4.1 | 248 | 274 | 20 | MOLECULE: | 5'-D(*TP*TP*TP*TP*TP*TP*TP*TP*TP*TP*TP*TP*T)-3' |
| 16:00 | 5a9k-B | 16.5 | 3.5 | 236 | 268 | 21 | MOLECULE: | REPLICATION PROTEIN E1 |
| 17:00 | 5a9k-C | 16.5 | 3.5 | 236 | 267 | 22 | MOLECULE: | REPLICATION PROTEIN E1 |
| 18:00 | 2v9p-C | 16.5 | 3.5 | 236 | 267 | 22 | MOLECULE: | REPLICATION PROTEIN E1 |
| 19:00 | 2v9p-G | 16.4 | 3.9 | 254 | 270 | 21 | MOLECULE: | REPLICATION PROTEIN E1 |
| 20:00 | 2v9p-E | 16.3 | 3.7 | 255 | 270 | 22 | MOLECULE: | REPLICATION PROTEIN E1 |
| 21:00 | 7apd-E | 16.3 | 4.2 | 268 | 287 | 20 | MOLECULE: | REPLICATION PROTEIN E1 |
| 22:00 | 5a9k-E | 16.3 | 3.7 | 255 | 270 | 22 | MOLECULE: | REPLICATION PROTEIN E1 |
| 23:00 | 2gxa-K | 16.3 | 4.2 | 266 | 273 | 21 | MOLECULE: | 5'-D(*TP*TP*TP*TP*TP*TP*TP*TP*TP*TP*TP*TP*T)-3' |
| 24:00:00 | 5a9k-A | 16.3 | 4 | 256 | 272 | 21 | MOLECULE: | REPLICATION PROTEIN E1 |
| 25:00:00 | 2v9p-A | 16.3 | 3.9 | 240 | 272 | 22 | MOLECULE: | REPLICATION PROTEIN E1 |
| 26:00:00 | 2v9p-K | 16.2 | 3.6 | 253 | 269 | 22 | MOLECULE: | REPLICATION PROTEIN E1 |
| 27:00:00 | 2h1l-K | 16.1 | 5.4 | 272 | 362 | 17 | MOLECULE: | LARGE T ANTIGEN |
| 28:00:00 | 2gxa-A | 16 | 4.6 | 261 | 270 | 20 | MOLECULE: | 5'-D(*TP*TP*TP*TP*TP*TP*TP*TP*TP*TP*TP*TP*T)-3' |
| 29:00:00 | 1svm-F | 16 | 5.4 | 275 | 363 | 17 | MOLECULE: | LARGE T ANTIGEN |
| 30:00:00 | 2v9p-F | 15.9 | 4 | 253 | 269 | 21 | MOLECULE: | REPLICATION PROTEIN E1 |
| 31:00:00 | 2h1l-E | 15.9 | 5.2 | 269 | 362 | 17 | MOLECULE: | LARGE T ANTIGEN |
| 32:00:00 | 1svm-E | 15.9 | 5.1 | 268 | 362 | 17 | MOLECULE: | LARGE T ANTIGEN |
| 33:00:00 | 2h1l-L | 15.9 | 5.5 | 273 | 363 | 17 | MOLECULE: | LARGE T ANTIGEN |
| 34:00:00 | 1svm-C | 15.9 | 5.4 | 274 | 363 | 17 | MOLECULE: | LARGE T ANTIGEN |
| 35:00:00 | 2h1l-C | 15.9 | 5.5 | 274 | 363 | 17 | MOLECULE: | LARGE T ANTIGEN |
| 36:00:00 | 1svm-B | 15.8 | 5.2 | 268 | 362 | 17 | MOLECULE: | LARGE T ANTIGEN |
| 37:00:00 | 2h1l-F | 15.8 | 5.4 | 273 | 363 | 17 | MOLECULE: | LARGE T ANTIGEN |
| 38:00:00 | 1svm-A | 15.7 | 5.4 | 273 | 362 | 17 | MOLECULE: | LARGE T ANTIGEN |
| 39:00:00 | 2h1l-D | 15.7 | 5.4 | 270 | 362 | 17 | MOLECULE: | LARGE T ANTIGEN |
| 40:00:00 | 2h1l-B | 15.7 | 5.5 | 272 | 362 | 17 | MOLECULE: | LARGE T ANTIGEN |
| 41:00:00 | 5j4y-A | 15.7 | 4.8 | 258 | 363 | 16 | MOLECULE: | LARGE T ANTIGEN |
| 42:00:00 | 7apd-B | 15.6 | 3.4 | 270 | 289 | 21 | MOLECULE: | REPLICATION PROTEIN E1 |
| 43:00:00 | 2h1l-J | 15.6 | 5.1 | 266 | 362 | 17 | MOLECULE: | LARGE T ANTIGEN |
| 44:00:00 | 2v9p-L | 15.6 | 4.1 | 251 | 267 | 21 | MOLECULE: | REPLICATION PROTEIN E1 |
| 45:00:00 | 2h1l-H | 15.6 | 5.4 | 273 | 362 | 17 | MOLECULE: | LARGE T ANTIGEN |
| 46:00:00 | 7apd-D | 15.6 | 3.9 | 270 | 287 | 20 | MOLECULE: | REPLICATION PROTEIN E1 |
| 47:00:00 | 4e2i-J | 15.5 | 5.3 | 270 | 362 | 17 | MOLECULE: | LARGE T ANTIGEN |
| 48:00:00 | 4e2i-H | 15.5 | 5.3 | 270 | 362 | 17 | MOLECULE: | LARGE T ANTIGEN |
| 49:00:00 | 7apd-C | 15.5 | 3.8 | 271 | 287 | 20 | MOLECULE: | REPLICATION PROTEIN E1 |
| 50:00:00 | 4gdf-E | 15.5 | 7.2 | 308 | 497 | 16 | MOLECULE: | LARGE T ANTIGEN |
| 51:00:00 | 4e2i-A | 15.5 | 5.3 | 270 | 362 | 17 | MOLECULE: | LARGE T ANTIGEN |

| | | | | | | | | |
|---|---|---|---|---|---|---|---|---|
| 52:00:00 | 2h1l-G | 15.5 | 5.3 | 268 | 362 | 18 | MOLECULE: | LARGE T ANTIGEN |
| 53:00:00 | 2h1l-I | 15.5 | 5.4 | 270 | 363 | 17 | MOLECULE: | LARGE T ANTIGEN |
| 54:00:00 | 1svm-D | 15.5 | 5.2 | 269 | 362 | 17 | MOLECULE: | LARGE T ANTIGEN |
| 55:00:00 | 4e2i-E | 15.5 | 5.3 | 270 | 362 | 17 | MOLECULE: | LARGE T ANTIGEN |
| 56:00:00 | 4e2i-G | 15.5 | 5.3 | 270 | 362 | 17 | MOLECULE: | LARGE T ANTIGEN |
| 57:00:00 | 4e2i-D | 15.5 | 5.3 | 270 | 362 | 17 | MOLECULE: | LARGE T ANTIGEN |
| 58:00:00 | 4e2i-K | 15.5 | 5.3 | 270 | 362 | 17 | MOLECULE: | LARGE T ANTIGEN |
| 59:00:00 | 4e2i-I | 15.5 | 5.3 | 270 | 362 | 17 | MOLECULE: | LARGE T ANTIGEN |
| 60:00:00 | 4e2i-L | 15.5 | 5.3 | 270 | 362 | 17 | MOLECULE: | LARGE T ANTIGEN |
| 61:00:00 | 4e2i-F | 15.5 | 5.3 | 270 | 362 | 17 | MOLECULE: | LARGE T ANTIGEN |
| 62:00:00 | 2h1l-A | 15.5 | 5.2 | 270 | 362 | 17 | MOLECULE: | LARGE T ANTIGEN |
| 63:00:00 | 5tct-B | 15.5 | 5.7 | 277 | 362 | 17 | MOLECULE: | LARGE T ANTIGEN |
| 64:00:00 | 7apd-F | 15.4 | 4.5 | 271 | 286 | 20 | MOLECULE: | REPLICATION PROTEIN E1 |
| 65:00:00 | 4gdf-A | 15.4 | 7.2 | 307 | 497 | 16 | MOLECULE: | LARGE T ANTIGEN |
| 66:00:00 | 1svo-A | 15.4 | 4.9 | 253 | 362 | 19 | MOLECULE: | LARGE T ANTIGEN |
| 67:00:00 | 1n25-B | 15.4 | 4.8 | 257 | 362 | 18 | MOLECULE: | LARGE T ANTIGEN |
| 68:00:00 | 1svl-C | 15.3 | 5.7 | 280 | 362 | 16 | MOLECULE: | LARGE T ANTIGEN |
| 69:00:00 | 1svl-B | 15.3 | 4.4 | 252 | 362 | 18 | MOLECULE: | LARGE T ANTIGEN |
| 70:00:00 | 1n25-A | 15.3 | 4.7 | 250 | 362 | 18 | MOLECULE: | LARGE T ANTIGEN |
| 71:00:00 | 1svl-A | 15.3 | 5.2 | 253 | 363 | 17 | MOLECULE: | LARGE T ANTIGEN |
| 72:00:00 | 5a9k-F | 15.3 | 4 | 253 | 269 | 21 | MOLECULE: | REPLICATION PROTEIN E1 |
| 73:00:00 | 1svo-B | 15.3 | 4.5 | 252 | 362 | 19 | MOLECULE: | LARGE T ANTIGEN |
| 74:00:00 | 5j40-A | 15.3 | 4.7 | 251 | 358 | 16 | MOLECULE: | LARGE T ANTIGEN |
| 75:00:00 | 5tct-C | 15.2 | 5.2 | 260 | 362 | 18 | MOLECULE: | LARGE T ANTIGEN |
| 76:00:00 | 5tct-F | 15.2 | 4.7 | 255 | 362 | 19 | MOLECULE: | LARGE T ANTIGEN |
| 77:00:00 | 4e2i-C | 15.2 | 5.3 | 270 | 362 | 17 | MOLECULE: | LARGE T ANTIGEN |
| 78:00:00 | 4ij8-B | 15.2 | 2.3 | 124 | 151 | 18 | MOLECULE: | N-LYSINE METHYLTRANSFERASE SETD8 |
| 79:00:00 | 4e2i-B | 15.1 | 5.3 | 270 | 362 | 17 | MOLECULE: | LARGE T ANTIGEN |
| 80:00:00 | 5tct-D | 15.1 | 5 | 261 | 362 | 17 | MOLECULE: | LARGE T ANTIGEN |
| 81:00:00 | 1tue-A | 15.1 | 2.7 | 192 | 201 | 21 | MOLECULE: | REPLICATION PROTEIN E1 |
| 82:00:00 | 1tue-K | 15 | 2.4 | 184 | 192 | 22 | MOLECULE: | REPLICATION PROTEIN E1 |
| 83:00:00 | 3f9z-A | 15 | 2.6 | 125 | 157 | 18 | MOLECULE: | HISTONE-LYSINE N-METHYLTRANSFERASE SETD8 |
| 84:00:00 | 3f9y-A | 14.9 | 2.6 | 124 | 160 | 18 | MOLECULE: | HISTONE-LYSINE N-METHYLTRANSFERASE SETD8 |
| 85:00:00 | 5teg-B | 14.9 | 2.4 | 123 | 159 | 18 | MOLECULE: | N-LYSINE METHYLTRANSFERASE KMT5A |
| 86:00:00 | 5w1y-B | 14.9 | 2.2 | 120 | 151 | 18 | MOLECULE: | N-LYSINE METHYLTRANSFERASE KMT5A |
| 87:00:00 | 3f9w-B | 14.9 | 2.5 | 124 | 161 | 18 | MOLECULE: | HISTONE-LYSINE N-METHYLTRANSFERASE SETD8 |
| 88:00:00 | 5j47-A | 14.9 | 5 | 258 | 360 | 16 | MOLECULE: | LARGE T ANTIGEN |
| 89:00:00 | 5j4v-A | 14.9 | 5.6 | 264 | 358 | 16 | MOLECULE: | LARGE T ANTIGEN |
| 90:00:00 | 3f9y-B | 14.9 | 2.5 | 123 | 161 | 18 | MOLECULE: | HISTONE-LYSINE N-METHYLTRANSFERASE SETD8 |
| 91:00:00 | 5tct-A | 14.8 | 5.4 | 265 | 362 | 17 | MOLECULE: | LARGE T ANTIGEN |
| 92:00:00 | 1tue-D | 14.8 | 2.5 | 191 | 202 | 22 | MOLECULE: | REPLICATION PROTEIN E1 |
| 93:00:00 | 5tct-E | 14.8 | 5.2 | 260 | 362 | 18 | MOLECULE: | LARGE T ANTIGEN |
| 94:00:00 | 1zkk-D | 14.8 | 2.5 | 124 | 162 | 18 | MOLECULE: | HISTONE-LYSINE N-METHYLTRANSFERASE, H4 LYSINE-20 |
| 95:00:00 | 1zkk-B | 14.8 | 2.5 | 124 | 162 | 18 | MOLECULE: | HISTONE-LYSINE N-METHYLTRANSFERASE, H4 LYSINE-20 |
| 96:00:00 | 3f9w-D | 14.8 | 2.5 | 124 | 162 | 18 | MOLECULE: | HISTONE-LYSINE N-METHYLTRANSFERASE SETD8 |
| 97:00:00 | 3f9z-B | 14.8 | 2.5 | 123 | 162 | 19 | MOLECULE: | HISTONE-LYSINE N-METHYLTRANSFERASE SETD8 |
| 98:00:00 | 3f9z-D | 14.8 | 2.5 | 124 | 162 | 19 | MOLECULE: | HISTONE-LYSINE N-METHYLTRANSFERASE SETD8 |
| 99:00:00 | 3f9w-A | 14.8 | 2.5 | 123 | 161 | 18 | MOLECULE: | HISTONE-LYSINE N-METHYLTRANSFERASE SETD8 |
| 0:00 | 7apd-A | 14.7 | 4.7 | 271 | 284 | 20 | MOLECULE: | REPLICATION PROTEIN E1 |
| 1:00 | 1zkk-A | 14.7 | 2.5 | 123 | 161 | 18 | MOLECULE: | HISTONE-LYSINE N-METHYLTRANSFERASE, H4 LYSINE-20 |
| 2:00 | 5teg-A | 14.7 | 2.5 | 123 | 161 | 18 | MOLECULE: | N-LYSINE METHYLTRANSFERASE KMT5A |
| 3:00 | 3f9x-B | 14.7 | 2.5 | 123 | 161 | 18 | MOLECULE: | HISTONE-LYSINE N-METHYLTRANSFERASE SETD8 |
| 4:00 | 3f9x-D | 14.7 | 2.5 | 124 | 162 | 18 | MOLECULE: | HISTONE-LYSINE N-METHYLTRANSFERASE SETD8 |

| | | | | | | | | |
|---|---|---|---|---|---|---|---|---|
| 5:00 | 2bqz-A | 14.7 | 2.4 | 123 | 162 | 18 | MOLECULE: | SET8 PROTEIN |
| 6:00 | 3f9x-C | 14.7 | 2.5 | 123 | 161 | 18 | MOLECULE: | HISTONE-LYSINE N-METHYLTRANSFERASE SETD8 |
| 7:00 | 1tue-M | 14.7 | 2.5 | 187 | 196 | 21 | MOLECULE: | REPLICATION PROTEIN E1 |
| 8:00 | 1tue-H | 14.6 | 2.5 | 182 | 191 | 23 | MOLECULE: | REPLICATION PROTEIN E1 |
| 9:00 | 2bqz-E | 14.6 | 2.5 | 123 | 162 | 18 | MOLECULE: | SET8 PROTEIN |
| 10:00 | 3f9w-C | 14.6 | 2.6 | 124 | 161 | 18 | MOLECULE: | HISTONE-LYSINE N-METHYLTRANSFERASE SETD8 |
| 11:00 | 5w1y-A | 14.6 | 3 | 125 | 162 | 18 | MOLECULE: | N-LYSINE METHYLTRANSFERASE KMT5A |
| 12:00 | 3f9x-A | 14.6 | 2.5 | 123 | 161 | 18 | MOLECULE: | HISTONE-LYSINE N-METHYLTRANSFERASE SETD8 |
| 13:00 | 1zkk-C | 14.5 | 2.6 | 124 | 161 | 18 | MOLECULE: | HISTONE-LYSINE N-METHYLTRANSFERASE, H4 LYSINE-20 |
| 14:00 | 3f9z-C | 14.5 | 2.6 | 124 | 161 | 19 | MOLECULE: | HISTONE-LYSINE N-METHYLTRANSFERASE SETD8 |
| 15:00 | 5hq2-M | 14.4 | 2.4 | 123 | 160 | 18 | MOLECULE: | HISTONE H3.2 |
| 16:00 | 4ij8-A | 14.2 | 2.4 | 124 | 162 | 18 | MOLECULE: | N-LYSINE METHYLTRANSFERASE SETD8 |
| 17:00 | 6boz-B | 14.1 | 2.3 | 116 | 134 | 19 | MOLECULE: | N-LYSINE METHYLTRANSFERASE KMT5A |
| 18:00 | 2gxa-L | 14 | 3.7 | 225 | 270 | 20 | MOLECULE: | 5'-D(*TP*TP*TP*TP*TP*TP*TP*TP*TP*TP*TP*TP*T)-3' |
| 19:00 | 7d20-K | 14 | 2.5 | 124 | 166 | 18 | MOLECULE: | HISTONE H3-LIKE CENTROMERIC PROTEIN A |
| 20:00 | 5t5g-A | 14 | 2.4 | 118 | 143 | 19 | MOLECULE: | N-LYSINE METHYLTRANSFERASE KMT5A |
| 21:00 | 7u5v-A | 14 | 2.4 | 111 | 125 | 16 | MOLECULE: | HISTONE-LYSINE N-METHYLTRANSFERASE 2A |
| 22:00 | 1tue-F | 14 | 2.4 | 178 | 188 | 22 | MOLECULE: | REPLICATION PROTEIN E1 |
| 23:00 | 4ldg-A | 14 | 2 | 126 | 216 | 17 | MOLECULE: | PROTEIN WITH A SET DOMAIN WITHIN CARBOXY REGION |
| 24:00:00 | 4z4p-A | 13.9 | 2.2 | 119 | 162 | 24 | MOLECULE: | HISTONE-LYSINE N-METHYLTRANSFERASE 2D |
| 25:00:00 | 5v2n-A | 13.9 | 2.3 | 123 | 161 | 18 | MOLECULE: | N-LYSINE METHYLTRANSFERASE KMT5A |
| 26:00:00 | 6boz-A | 13.9 | 2.8 | 120 | 139 | 18 | MOLECULE: | N-LYSINE METHYLTRANSFERASE KMT5A |
| 27:00:00 | 6u9m-A | 13.8 | 2.3 | 118 | 148 | 17 | MOLECULE: | HISTONE-LYSINE N-METHYLTRANSFERASE |
| 28:00:00 | 6u9r-B | 13.8 | 2.3 | 118 | 148 | 17 | MOLECULE: | HISTONE-LYSINE N-METHYLTRANSFERASE |
| 29:00:00 | 6u9n-B | 13.8 | 2.3 | 118 | 147 | 17 | MOLECULE: | HISTONE-LYSINE N-METHYLTRANSFERASE |
| 30:00:00 | 6u9r-A | 13.8 | 2.4 | 119 | 149 | 17 | MOLECULE: | HISTONE-LYSINE N-METHYLTRANSFERASE |
| 31:00:00 | 6u9k-B | 13.7 | 2.3 | 118 | 149 | 17 | MOLECULE: | HISTONE-LYSINE N-METHYLTRANSFERASE |
| 32:00:00 | 7w6l-E | 13.7 | 2.3 | 119 | 152 | 20 | MOLECULE: | SET1/ASH2 HISTONE METHYLTRANSFERASE COMPLEX SUBUN |
| 33:00:00 | 6u9n-A | 13.7 | 2.3 | 117 | 147 | 17 | MOLECULE: | HISTONE-LYSINE N-METHYLTRANSFERASE |
| 34:00:00 | 5f6k-C | 13.7 | 2.2 | 116 | 151 | 20 | MOLECULE: | SET1/ASH2 HISTONE METHYLTRANSFERASE COMPLEX SUBUN |
| 35:00:00 | 5f59-A | 13.7 | 2.5 | 118 | 149 | 19 | MOLECULE: | HISTONE-LYSINE N-METHYLTRANSFERASE 2C |
| 36:00:00 | 7w67-C | 13.7 | 4.1 | 124 | 158 | 17 | MOLECULE: | SET1/ASH2 HISTONE METHYLTRANSFERASE COMPLEX SUBUN |
| 37:00:00 | 5f5e-A | 13.6 | 2.3 | 117 | 149 | 17 | MOLECULE: | HISTONE-LYSINE N-METHYLTRANSFERASE 2A |
| 38:00:00 | 6pww-C | 13.6 | 2.3 | 120 | 154 | 17 | MOLECULE: | RETINOBLASTOMA-BINDING PROTEIN 5 |
| 39:00:00 | 6u9k-A | 13.6 | 2.5 | 118 | 149 | 18 | MOLECULE: | HISTONE-LYSINE N-METHYLTRANSFERASE |
| 40:00:00 | 7w6i-C | 13.5 | 4.1 | 122 | 160 | 17 | MOLECULE: | SET1/ASH2 HISTONE METHYLTRANSFERASE COMPLEX SUBUN |
| 41:00:00 | 5f6k-E | 13.5 | 2.1 | 116 | 149 | 21 | MOLECULE: | SET1/ASH2 HISTONE METHYLTRANSFERASE COMPLEX SUBUN |
| 42:00:00 | 7w6j-C | 13.4 | 4.1 | 122 | 159 | 17 | MOLECULE: | SET1/ASH2 HISTONE METHYLTRANSFERASE COMPLEX SUBUN |
| 43:00:00 | 6u9m-B | 13.4 | 2.5 | 117 | 149 | 19 | MOLECULE: | HISTONE-LYSINE N-METHYLTRANSFERASE |
| 44:00:00 | 6w5n-C | 13.4 | 2.4 | 119 | 153 | 17 | MOLECULE: | RETINOBLASTOMA-BINDING PROTEIN 5 |
| 45:00:00 | 7xpx-K | 13.3 | 2.6 | 124 | 168 | 18 | MOLECULE: | HISTONE H3 |
| 46:00:00 | 5th7-B | 13.3 | 2.3 | 118 | 144 | 19 | MOLECULE: | N-LYSINE METHYLTRANSFERASE KMT5A |
| 47:00:00 | 6pwv-C | 13.3 | 2.4 | 119 | 155 | 17 | MOLECULE: | RETINOBLASTOMA-BINDING PROTEIN 5 |
| 48:00:00 | 7w6l-C | 13.3 | 3.2 | 124 | 159 | 19 | MOLECULE: | SET1/ASH2 HISTONE METHYLTRANSFERASE COMPLEX SUBUN |
| 49:00:00 | 6w5i-C | 13.3 | 2.3 | 116 | 152 | 17 | MOLECULE: | RETINOBLASTOMA-BINDING PROTEIN 5 |
| 50:00:00 | 7mbn-C | 13.3 | 2.4 | 119 | 153 | 17 | MOLECULE: | RETINOBLASTOMA-BINDING PROTEIN 5 |
| 51:00:00 | 6w5m-C | 13.3 | 2.2 | 117 | 148 | 17 | MOLECULE: | RETINOBLASTOMA-BINDING PROTEIN 5 |
| 52:00:00 | 7w6a-C | 13.3 | 2.4 | 121 | 160 | 17 | MOLECULE: | SET1/ASH2 HISTONE METHYLTRANSFERASE COMPLEX SUBUN |
| 53:00:00 | 5th7-A | 13.3 | 2.3 | 117 | 141 | 19 | MOLECULE: | N-LYSINE METHYLTRANSFERASE KMT5A |
| 54:00:00 | 5f6l-A | 13.2 | 2.5 | 120 | 154 | 17 | MOLECULE: | RETINOBLASTOMA-BINDING PROTEIN 5 |
| 55:00:00 | 7d1z-K | 13.1 | 2.5 | 124 | 166 | 18 | MOLECULE: | HISTONE H3.1 |
| 56:00:00 | 1s9h-A | 13.1 | 5.2 | 206 | 268 | 16 | MOLECULE: | REP 40 PROTEIN |
| 57:00:00 | 2w5y-A | 13 | 2.2 | 118 | 171 | 17 | MOLECULE: | HISTONE-LYSINE N-METHYLTRANSFERASE HRX |

| | | | | | | | |
|---|---|---|---|---|---|---|---|
| 58:00:00 | 7bre-B | 13 | 2.4 | 121 | 164 | 15 MOLECULE: | SET1/ASH2 HISTONE METHYLTRANSFERASE COMPLEX SUBUN |
| 59:00:00 | 5ht6-A | 12.9 | 2.6 | 110 | 123 | 19 MOLECULE: | HISTONE-LYSINE N-METHYLTRANSFERASE 2E |
| 60:00:00 | 1s9h-B | 12.9 | 4.3 | 203 | 267 | 16 MOLECULE: | REP 40 PROTEIN |
| 61:00:00 | 6chg-C | 12.8 | 2.1 | 114 | 144 | 18 MOLECULE: | KLLA0E24487P |
| 62:00:00 | 7bre-E | 12.8 | 2.4 | 120 | 164 | 15 MOLECULE: | SET1/ASH2 HISTONE METHYLTRANSFERASE COMPLEX SUBUN |
| 63:00:00 | 4gdf-B | 12.6 | 12.6 | 208 | 497 | 17 MOLECULE: | LARGE T ANTIGEN |
| 64:00:00 | 2w5z-A | 12.6 | 2.3 | 120 | 181 | 17 MOLECULE: | HISTONE-LYSINE N-METHYLTRANSFERASE HRX |
| 65:00:00 | 1u0j-A | 12.6 | 5.1 | 204 | 261 | 14 MOLECULE: | DNA REPLICATION PROTEIN |
| 66:00:00 | 4gdf-F | 12.2 | 11.7 | 210 | 497 | 18 MOLECULE: | LARGE T ANTIGEN |
| 67:00:00 | 5ht6-B | 12.2 | 2.5 | 104 | 109 | 20 MOLECULE: | HISTONE-LYSINE N-METHYLTRANSFERASE 2E |
| 68:00:00 | 6uh5-M | 12.1 | 2.4 | 114 | 222 | 18 MOLECULE: | HISTONE H3 |
| 69:00:00 | 5hyn-Q | 12.1 | 2.3 | 118 | 566 | 19 MOLECULE: | HISTONE-LYSINE N-METHYLTRANSFERASE EZH2 |
| 70:00:00 | 6kqq-B | 12 | 2.5 | 118 | 220 | 20 MOLECULE: | HISTONE-LYSINE N-METHYLTRANSFERASE, H3 LYSINE-36 |
| 71:00:00 | 5ij8-A | 12 | 2.1 | 116 | 464 | 20 MOLECULE: | ENHANCER OF ZESTE HOMOLOG 2 (EZH2),HISTONE-LYSINE |
| 72:00:00 | 6c24-K | 12 | 2.6 | 119 | 325 | 19 MOLECULE: | POLYCOMB PROTEIN SUZ12 |
| 73:00:00 | 6ven-N | 11.9 | 2.2 | 112 | 219 | 18 MOLECULE: | HISTONE H3.2 |
| 74:00:00 | 7jsg-F | 11.9 | 3.6 | 221 | 272 | 14 MOLECULE: | PROTEIN REP68 |
| 75:00:00 | 6kqp-A | 11.7 | 2.1 | 114 | 221 | 20 MOLECULE: | HISTONE-LYSINE N-METHYLTRANSFERASE, H3 LYSINE-36 |
| 76:00:00 | 4w2r-B | 11.7 | 2.6 | 120 | 465 | 19 MOLECULE: | ENHANCER OF ZESTE 2 POLYCOMB REPRESSIVE COMPLEX 2 |
| 77:00:00 | 7td5-A | 11.7 | 2.3 | 118 | 498 | 19 MOLECULE: | HISTONE-LYSINE N-METHYLTRANSFERASE EZH1 |
| 78:00:00 | 5ij8-B | 11.7 | 2.5 | 120 | 466 | 19 MOLECULE: | ENHANCER OF ZESTE HOMOLOG 2 (EZH2),HISTONE-LYSINE |
| 79:00:00 | 6b3w-B | 11.6 | 2.2 | 116 | 465 | 20 MOLECULE: | ENHANCER OF ZESTE 2 POLYCOMB REPRESSIVE COMPLEX 2 |
| 80:00:00 | 5ij7-A | 11.6 | 2.1 | 116 | 465 | 20 MOLECULE: | ENHANCER OF ZESTE HOMOLOG 2 (EZH2),HISTONE-LYSINE |
| 81:00:00 | 7jsi-E | 11.5 | 3.6 | 226 | 276 | 14 MOLECULE: | PROTEIN REP68 |
| 82:00:00 | 5kkl-B | 11.5 | 9 | 155 | 838 | 15 MOLECULE: | PUTATIVE POLYCOMB PROTEIN EED |
| 83:00:00 | 6cen-A | 11.4 | 2 | 114 | 227 | 19 MOLECULE: | HISTONE-LYSINE N-METHYLTRANSFERASE NSD3 |
| 84:00:00 | 4j7i-A | 11.4 | 2.6 | 114 | 244 | 18 MOLECULE: | HISTONE-LYSINE N-METHYLTRANSFERASE SETD7 |
| 85:00:00 | 1mt6-A | 11.4 | 2.7 | 116 | 281 | 18 MOLECULE: | SET9 |
| 86:00:00 | 5lt8-A | 11.4 | 1.9 | 113 | 228 | 19 MOLECULE: | HISTONE-LYSINE N-METHYLTRANSFERASE SETD2 |
| 87:00:00 | 7td5-F | 11.4 | 2.3 | 118 | 448 | 19 MOLECULE: | HISTONE-LYSINE N-METHYLTRANSFERASE EZH1 |
| 88:00:00 | 7jsf-E | 11.4 | 5.2 | 217 | 276 | 14 MOLECULE: | PROTEIN REP68 |
| 89:00:00 | 7e8d-K | 11.4 | 3.1 | 119 | 219 | 19 MOLECULE: | HISTONE H3.1 |
| 90:00:00 | 1h3i-A | 11.3 | 2.8 | 116 | 293 | 16 MOLECULE: | HISTONE H3 LYSINE 4 SPECIFIC METHYLTRANSFERASE |
| 91:00:00 | 5m5g-B | 11.3 | 6.7 | 156 | 796 | 18 MOLECULE: | PUTATIVE POLYCOMB PROTEIN EED |
| 92:00:00 | 7jsg-E | 11.3 | 4 | 223 | 272 | 13 MOLECULE: | PROTEIN REP68 |
| 93:00:00 | 6b3w-A | 11.3 | 2.6 | 120 | 463 | 19 MOLECULE: | ENHANCER OF ZESTE 2 POLYCOMB REPRESSIVE COMPLEX 2 |
| 94:00:00 | 1muf-A | 11.3 | 2.7 | 116 | 257 | 16 MOLECULE: | SET9 |
| 95:00:00 | 7jsh-B | 11.3 | 3.3 | 224 | 272 | 14 MOLECULE: | PROTEIN REP68 |
| 96:00:00 | 5tqr-B | 11.2 | 6.8 | 160 | 808 | 16 MOLECULE: | POLYCOMB PROTEIN EED |
| 97:00:00 | 5hyn-F | 11.2 | 2.4 | 118 | 569 | 19 MOLECULE: | HISTONE-LYSINE N-METHYLTRANSFERASE EZH2 |
| 98:00:00 | 4ynp-B | 11.2 | 2.2 | 116 | 194 | 17 MOLECULE: | HISTONE-LYSINE N-METHYLTRANSFERASE ASH1L |
| 99:00:00 | 7jsf-A | 11.2 | 5 | 213 | 276 | 14 MOLECULE: | PROTEIN REP68 |
| 0:00 | 7jsf-C | 11.2 | 5 | 213 | 276 | 14 MOLECULE: | PROTEIN REP68 |
| 1:00 | 7jsf-G | 11.2 | 5 | 211 | 276 | 14 MOLECULE: | PROTEIN REP68 |
| 2:00 | 7jsf-F | 11.2 | 5 | 213 | 276 | 14 MOLECULE: | PROTEIN REP68 |
| 3:00 | 7jsf-B | 11.2 | 5 | 213 | 276 | 14 MOLECULE: | PROTEIN REP68 |
| 4:00 | 7jsg-C | 11.2 | 3.4 | 214 | 272 | 14 MOLECULE: | PROTEIN REP68 |
| 5:00 | 1n6a-A | 11.1 | 2.8 | 116 | 235 | 18 MOLECULE: | SET DOMAIN-CONTAINING PROTEIN 7 |
| 6:00 | 4yz8-A | 11.1 | 2 | 113 | 210 | 19 MOLECULE: | HISTONE-LYSINE N-METHYLTRANSFERASE NSD3 |
| 7:00 | 4ynm-B | 11.1 | 2.2 | 116 | 197 | 17 MOLECULE: | HISTONE-LYSINE N-METHYLTRANSFERASE ASH1L |
| 8:00 | 4ypu-B | 11.1 | 2.3 | 116 | 191 | 17 MOLECULE: | HISTONE-LYSINE N-METHYLTRANSFERASE ASH1L |
| 9:00 | 7jsf-D | 11.1 | 5.1 | 215 | 276 | 14 MOLECULE: | PROTEIN REP68 |
| 10:00 | 7jsi-C | 11.1 | 3.5 | 228 | 276 | 14 MOLECULE: | PROTEIN REP68 |

| | | | | | | | | |
|---|---|---|---|---|---|---|---|---|
| 11:00 | 1h3i-B | 11 | 2.9 | 117 | 293 | 16 | MOLECULE: | HISTONE H3 LYSINE 4 SPECIFIC METHYLTRANSFERASE |
| 12:00 | 3vuz-A | 11 | 2.5 | 113 | 247 | 19 | MOLECULE: | HISTONE-LYSINE N-METHYLTRANSFERASE SETD7 |
| 13:00 | 3m56-A | 11 | 2.7 | 115 | 245 | 18 | MOLECULE: | HISTONE-LYSINE N-METHYLTRANSFERASE SETD7 |
| 14:00 | 3cbp-A | 11 | 3 | 120 | 240 | 17 | MOLECULE: | HISTONE-LYSINE N-METHYLTRANSFERASE SETD7 |
| 15:00 | 6kqq-A | 11 | 2.5 | 118 | 224 | 20 | MOLECULE: | HISTONE-LYSINE N-METHYLTRANSFERASE, H3 LYSINE-36 |
| 16:00 | 5lsu-A | 11 | 2.1 | 115 | 232 | 20 | MOLECULE: | HISTONE-LYSINE N-METHYLTRANSFERASE NSD2 |
| 17:00 | 5upd-A | 11 | 2 | 113 | 212 | 19 | MOLECULE: | HISTONE-LYSINE N-METHYLTRANSFERASE NSD3 |
| 18:00 | 3ope-B | 11 | 2.2 | 116 | 194 | 17 | MOLECULE: | PROBABLE HISTONE-LYSINE N-METHYLTRANSFERASE ASH1L |
| 19:00 | 6kiz-K | 11 | 3 | 121 | 180 | 17 | MOLECULE: | HISTONE H3 |
| 20:00 | 7lzb-A | 11 | 2.8 | 117 | 236 | 20 | MOLECULE: | HISTONE-LYSINE N-METHYLTRANSFERASE SETD2 |
| 21:00 | 5wfd-B | 11 | 7.6 | 156 | 772 | 14 | MOLECULE: | POLYCOMB PROTEIN EED |
| 22:00 | 7jsi-B | 11 | 3.5 | 226 | 276 | 14 | MOLECULE: | PROTEIN REP68 |
| 23:00 | 3m55-A | 10.9 | 3 | 120 | 245 | 17 | MOLECULE: | HISTONE-LYSINE N-METHYLTRANSFERASE SETD7 |
| 24:00:00 | 5ch2-B | 10.9 | 5.4 | 165 | 755 | 16 | MOLECULE: | PUTATIVE POLYCOMB PROTEIN EED |
| 25:00:00 | 4j83-A | 10.9 | 2.6 | 113 | 243 | 18 | MOLECULE: | HISTONE-LYSINE N-METHYLTRANSFERASE SETD7 |
| 26:00:00 | 3m54-A | 10.9 | 3 | 120 | 246 | 17 | MOLECULE: | HISTONE-LYSINE N-METHYLTRANSFERASE SETD7 |
| 27:00:00 | 3m57-A | 10.9 | 2.6 | 114 | 245 | 18 | MOLECULE: | HISTONE-LYSINE N-METHYLTRANSFERASE SETD7 |
| 28:00:00 | 4jds-B | 10.9 | 2.8 | 117 | 245 | 18 | MOLECULE: | HISTONE-LYSINE N-METHYLTRANSFERASE SETD7 |
| 29:00:00 | 4j7f-A | 10.9 | 3 | 119 | 242 | 17 | MOLECULE: | HISTONE-LYSINE N-METHYLTRANSFERASE SETD7 |
| 30:00:00 | 3m53-A | 10.9 | 2.6 | 113 | 247 | 18 | MOLECULE: | HISTONE-LYSINE N-METHYLTRANSFERASE SETD7 |
| 31:00:00 | 3cbo-A | 10.9 | 2.9 | 118 | 240 | 16 | MOLECULE: | HISTONE-LYSINE N-METHYLTRANSFERASE SETD7 |
| 32:00:00 | 5ls6-D | 10.9 | 2.2 | 115 | 570 | 20 | MOLECULE: | HISTONE-LYSINE N-METHYLTRANSFERASE EZH2,HISTONE-L |
| 33:00:00 | 5ylt-A | 10.9 | 3.7 | 123 | 248 | 18 | MOLECULE: | HISTONE-LYSINE N-METHYLTRANSFERASE SETD7 |
| 34:00:00 | 3cbm-A | 10.9 | 3 | 120 | 244 | 17 | MOLECULE: | HISTONE-LYSINE N-METHYLTRANSFERASE SETD7 |
| 35:00:00 | 5wf7-B | 10.9 | 7.5 | 163 | 764 | 15 | MOLECULE: | POLYCOMB PROTEIN EED |
| 36:00:00 | 4ype-B | 10.9 | 2.2 | 115 | 203 | 17 | MOLECULE: | HISTONE-LYSINE N-METHYLTRANSFERASE ASH1L |
| 37:00:00 | 5lsu-B | 10.9 | 2.1 | 115 | 223 | 20 | MOLECULE: | HISTONE-LYSINE N-METHYLTRANSFERASE NSD2 |
| 38:00:00 | 5eg2-A | 10.9 | 3 | 120 | 244 | 17 | MOLECULE: | HISTONE-LYSINE N-METHYLTRANSFERASE SETD7 |
| 39:00:00 | 4e47-A | 10.9 | 2.9 | 118 | 247 | 18 | MOLECULE: | HISTONE-LYSINE N-METHYLTRANSFERASE SETD7 |
| 40:00:00 | 7jsi-D | 10.9 | 3.6 | 226 | 276 | 15 | MOLECULE: | PROTEIN REP68 |
| 41:00:00 | 5wg6-A | 10.8 | 6 | 158 | 517 | 16 | MOLECULE: | HISTONE-LYSINE N-METHYLTRANSFERASE EZH2,POLYCOMB |
| 42:00:00 | 8sr6-A | 10.8 | 23.3 | 159 | 452 | 16 | MOLECULE: | EUKARYOTIC HUNTINGTIN INTERACTING PROTEIN B |
| 43:00:00 | 5jlb-A | 10.8 | 2.7 | 118 | 249 | 19 | MOLECULE: | HISTONE-LYSINE N-METHYLTRANSFERASE SETD2 |
| 44:00:00 | 4j8o-A | 10.8 | 2.9 | 118 | 246 | 17 | MOLECULE: | HISTONE-LYSINE N-METHYLTRANSFERASE SETD7 |
| 45:00:00 | 2f69-A | 10.8 | 3 | 119 | 245 | 17 | MOLECULE: | HISTONE-LYSINE N-METHYLTRANSFERASE, H3 LYSINE-4 |
| 46:00:00 | 3m5a-A | 10.8 | 3.1 | 120 | 247 | 17 | MOLECULE: | HISTONE-LYSINE N-METHYLTRANSFERASE SETD7 |
| 47:00:00 | 4mi5-A | 10.8 | 2.3 | 112 | 208 | 21 | MOLECULE: | HISTONE-LYSINE N-METHYLTRANSFERASE EZH2 |
| 48:00:00 | 4jds-A | 10.8 | 2.4 | 112 | 247 | 19 | MOLECULE: | HISTONE-LYSINE N-METHYLTRANSFERASE SETD7 |
| 49:00:00 | 3m59-A | 10.8 | 2.6 | 114 | 246 | 18 | MOLECULE: | HISTONE-LYSINE N-METHYLTRANSFERASE SETD7 |
| 50:00:00 | 8fbg-A | 10.8 | 2.1 | 114 | 218 | 20 | MOLECULE: | HISTONE-LYSINE N-METHYLTRANSFERASE, H3 LYSINE-36 |
| 51:00:00 | 6ine-A | 10.8 | 2.1 | 114 | 248 | 18 | MOLECULE: | HISTONE-LYSINE N-METHYLTRANSFERASE ASH1L |
| 52:00:00 | 5wfc-B | 10.8 | 7.2 | 150 | 768 | 16 | MOLECULE: | POLYCOMB PROTEIN EED |
| 53:00:00 | 8fyh-G | 10.8 | 2.1 | 116 | 514 | 20 | MOLECULE: | POLYCOMB PROTEIN SUZ12 |
| 54:00:00 | 4e47-B | 10.8 | 2.4 | 112 | 246 | 19 | MOLECULE: | HISTONE-LYSINE N-METHYLTRANSFERASE SETD7 |
| 55:00:00 | 5ayf-A | 10.8 | 2.8 | 117 | 245 | 16 | MOLECULE: | HISTONE-LYSINE N-METHYLTRANSFERASE SETD7 |
| 56:00:00 | 6ago-A | 10.8 | 2.1 | 116 | 243 | 17 | MOLECULE: | HISTONE-LYSINE N-METHYLTRANSFERASE ASH1L |
| 57:00:00 | 3os5-A | 10.7 | 4.1 | 124 | 250 | 18 | MOLECULE: | HISTONE-LYSINE N-METHYLTRANSFERASE SETD7 |
| 58:00:00 | 3m58-A | 10.7 | 2.6 | 114 | 254 | 18 | MOLECULE: | HISTONE-LYSINE N-METHYLTRANSFERASE SETD7 |
| 59:00:00 | 5ch1-B | 10.7 | 4.3 | 160 | 796 | 16 | MOLECULE: | PUTATIVE POLYCOMB PROTEIN EED |
| 60:00:00 | 5bjs-B | 10.7 | 7.2 | 156 | 799 | 16 | MOLECULE: | POLYCOMB PROTEIN EED |
| 61:00:00 | 4jds-C | 10.7 | 2.4 | 112 | 247 | 20 | MOLECULE: | HISTONE-LYSINE N-METHYLTRANSFERASE SETD7 |
| 62:00:00 | 4ypa-B | 10.7 | 2.2 | 116 | 213 | 17 | MOLECULE: | HISTONE-LYSINE N-METHYLTRANSFERASE ASH1L |
| 63:00:00 | 4ypa-D | 10.7 | 2.3 | 116 | 212 | 17 | MOLECULE: | HISTONE-LYSINE N-METHYLTRANSFERASE ASH1L |

| | | | | | | | | |
|---|---|---|---|---|---|---|---|---|
| 64:00:00 | 8fbg-B | 10.7 | 2.3 | 115 | 219 | 20 | MOLECULE: | HISTONE-LYSINE N-METHYLTRANSFERASE, H3 LYSINE-36 |
| 65:00:00 | 5lsz-A | 10.7 | 2.3 | 116 | 243 | 19 | MOLECULE: | HISTONE-LYSINE N-METHYLTRANSFERASE SETD2 |
| 66:00:00 | 6c23-K | 10.7 | 2.5 | 119 | 325 | 19 | MOLECULE: | POLYCOMB PROTEIN SUZ12 |
| 67:00:00 | 4e47-C | 10.7 | 2.7 | 115 | 246 | 18 | MOLECULE: | HISTONE-LYSINE N-METHYLTRANSFERASE SETD7 |
| 68:00:00 | 7jsh-G | 10.7 | 3.8 | 224 | 272 | 14 | MOLECULE: | PROTEIN REP68 |
| 69:00:00 | 1xqh-E | 10.6 | 4 | 118 | 251 | 18 | MOLECULE: | HISTONE-LYSINE N-METHYLTRANSFERASE, H3 LYSINE-4 |
| 70:00:00 | 1xqh-A | 10.6 | 4 | 119 | 251 | 19 | MOLECULE: | HISTONE-LYSINE N-METHYLTRANSFERASE, H3 LYSINE-4 |
| 71:00:00 | 1o9s-A | 10.6 | 4.2 | 124 | 251 | 18 | MOLECULE: | HISTONE-LYSINE N-METHYLTRANSFERASE, H3 LYSINE-4 |
| 72:00:00 | 5kji-B | 10.6 | 6.7 | 155 | 762 | 17 | MOLECULE: | PUTATIVE POLYCOMB PROTEIN EED |
| 73:00:00 | 3ooi-A | 10.6 | 2.3 | 114 | 233 | 20 | MOLECULE: | HISTONE-LYSINE N-METHYLTRANSFERASE, H3 LYSINE-36 |
| 74:00:00 | 4jds-D | 10.6 | 2.6 | 113 | 246 | 18 | MOLECULE: | HISTONE-LYSINE N-METHYLTRANSFERASE SETD7 |
| 75:00:00 | 4ynm-A | 10.6 | 2.2 | 116 | 216 | 17 | MOLECULE: | HISTONE-LYSINE N-METHYLTRANSFERASE ASH1L |
| 76:00:00 | 3ope-A | 10.6 | 2.2 | 116 | 214 | 17 | MOLECULE: | PROBABLE HISTONE-LYSINE N-METHYLTRANSFERASE ASH1L |
| 77:00:00 | 7jsi-F | 10.6 | 3.6 | 227 | 276 | 13 | MOLECULE: | PROTEIN REP68 |
| 78:00:00 | 7jsg-B | 10.6 | 3.7 | 217 | 272 | 15 | MOLECULE: | PROTEIN REP68 |
| 79:00:00 | 6kiv-K | 10.6 | 2.5 | 119 | 180 | 18 | MOLECULE: | HISTONE H3 |
| 80:00:00 | 1o9s-B | 10.6 | 3.3 | 121 | 251 | 17 | MOLECULE: | HISTONE-LYSINE N-METHYLTRANSFERASE, H3 LYSINE-4 |
| 81:00:00 | 6x0p-D | 10.6 | 2.2 | 117 | 221 | 17 | MOLECULE: | HISTONE-LYSINE N-METHYLTRANSFERASE ASH1L |
| 82:00:00 | 6x0p-B | 10.6 | 2.2 | 117 | 221 | 17 | MOLECULE: | HISTONE-LYSINE N-METHYLTRANSFERASE ASH1L |
| 83:00:00 | 1peg-B | 10.5 | 2.7 | 119 | 210 | 18 | MOLECULE: | HISTONE H3 METHYLTRANSFERASE DIM-5 |
| 84:00:00 | 4jlg-A | 10.5 | 3.6 | 121 | 249 | 16 | MOLECULE: | HISTONE-LYSINE N-METHYLTRANSFERASE SETD7 |
| 85:00:00 | 5h6z-A | 10.5 | 2.7 | 111 | 123 | 13 | MOLECULE: | SET DOMAIN-CONTAINING PROTEIN 7 |
| 86:00:00 | 4jlg-B | 10.5 | 3.6 | 117 | 248 | 19 | MOLECULE: | HISTONE-LYSINE N-METHYLTRANSFERASE SETD7 |
| 87:00:00 | 4ypa-C | 10.5 | 2.2 | 116 | 218 | 17 | MOLECULE: | HISTONE-LYSINE N-METHYLTRANSFERASE ASH1L |
| 88:00:00 | 4h12-A | 10.5 | 1.9 | 114 | 234 | 19 | MOLECULE: | HISTONE-LYSINE N-METHYLTRANSFERASE SETD2 |
| 89:00:00 | 4ynp-A | 10.5 | 2.1 | 115 | 214 | 17 | MOLECULE: | HISTONE-LYSINE N-METHYLTRANSFERASE ASH1L |
| 90:00:00 | 4ypu-A | 10.5 | 2.2 | 116 | 215 | 17 | MOLECULE: | HISTONE-LYSINE N-METHYLTRANSFERASE ASH1L |
| 91:00:00 | 5ww0-A | 10.5 | 2.6 | 111 | 123 | 13 | MOLECULE: | SET DOMAIN-CONTAINING PROTEIN 7 |
| 92:00:00 | 6x0p-A | 10.5 | 2.2 | 116 | 220 | 17 | MOLECULE: | HISTONE-LYSINE N-METHYLTRANSFERASE ASH1L |
| 93:00:00 | 7at8-A | 10.5 | 2.4 | 118 | 312 | 19 | MOLECULE: | ISOFORM 2 OF HISTONE-LYSINE N-METHYLTRANSFERASE E |
| 94:00:00 | 4ypa-A | 10.5 | 2.2 | 116 | 219 | 17 | MOLECULE: | HISTONE-LYSINE N-METHYLTRANSFERASE ASH1L |
| 95:00:00 | 4e47-D | 10.5 | 3.1 | 120 | 251 | 17 | MOLECULE: | HISTONE-LYSINE N-METHYLTRANSFERASE SETD7 |
| 96:00:00 | 6x0p-C | 10.5 | 2.2 | 116 | 221 | 17 | MOLECULE: | HISTONE-LYSINE N-METHYLTRANSFERASE ASH1L |
| 97:00:00 | 6wzw-A | 10.5 | 2.5 | 118 | 222 | 17 | MOLECULE: | HISTONE-LYSINE N-METHYLTRANSFERASE ASH1L |
| 98:00:00 | 8fbh-A | 10.4 | 2.3 | 114 | 233 | 20 | MOLECULE: | HISTONE-LYSINE N-METHYLTRANSFERASE, H3 LYSINE-36 |
| 99:00:00 | 6kiu-K | 10.4 | 2.5 | 116 | 182 | 17 | MOLECULE: | HISTONE H3 |
| 0:00 | 7lzd-A | 10.4 | 2.1 | 116 | 238 | 18 | MOLECULE: | HISTONE-LYSINE N-METHYLTRANSFERASE SETD2 |
| 1:00 | 5lsx-A | 10.4 | 2 | 114 | 231 | 19 | MOLECULE: | HISTONE-LYSINE N-METHYLTRANSFERASE SETD2 |
| 2:00 | 5jle-A | 10.4 | 2.9 | 118 | 233 | 19 | MOLECULE: | HISTONE-LYSINE N-METHYLTRANSFERASE SETD2 |
| 3:00 | 4ype-A | 10.4 | 2.2 | 115 | 218 | 17 | MOLECULE: | HISTONE-LYSINE N-METHYLTRANSFERASE ASH1L |
| 4:00 | 5vah-A | 10.4 | 3.1 | 123 | 214 | 15 | MOLECULE: | PROBABLE HISTONE-LYSINE N-METHYLTRANSFERASE ATXR5 |
| 5:00 | 5vsc-A | 10.4 | 5.4 | 120 | 270 | 17 | MOLECULE: | HISTONE-LYSINE N-METHYLTRANSFERASE EHMT2 |
| 6:00 | 7jsg-G | 10.4 | 4 | 215 | 272 | 13 | MOLECULE: | PROTEIN REP68 |
| 7:00 | 5vbc-B | 10.4 | 3.1 | 123 | 216 | 15 | MOLECULE: | PROBABLE HISTONE-LYSINE N-METHYLTRANSFERASE ATXR5 |
| 8:00 | 1n6c-A | 10.4 | 2.9 | 120 | 284 | 18 | MOLECULE: | SET DOMAIN-CONTAINING PROTEIN 7 |
| 9:00 | 7lzf-A | 10.3 | 2.5 | 118 | 240 | 19 | MOLECULE: | HISTONE-LYSINE N-METHYLTRANSFERASE SETD2 |
| 10:00 | 6vdb-A | 10.3 | 2.7 | 118 | 245 | 19 | MOLECULE: | HISTONE-LYSINE N-METHYLTRANSFERASE SETD2 |
| 11:00 | 4rz0-B | 10.3 | 2.3 | 99 | 110 | 12 | MOLECULE: | PFL0690C |
| 12:00 | 7ty3-A | 10.3 | 2.3 | 117 | 239 | 19 | MOLECULE: | HISTONE-LYSINE N-METHYLTRANSFERASE SETD2 |
| 13:00 | 5va6-B | 10.3 | 3.1 | 123 | 216 | 15 | MOLECULE: | PROBABLE HISTONE-LYSINE N-METHYLTRANSFERASE ATXR5 |
| 14:00 | 5vah-B | 10.3 | 3.1 | 123 | 215 | 15 | MOLECULE: | PROBABLE HISTONE-LYSINE N-METHYLTRANSFERASE ATXR5 |
| 15:00 | 7ty2-A | 10.3 | 2.3 | 117 | 235 | 18 | MOLECULE: | HISTONE-LYSINE N-METHYLTRANSFERASE SETD2 |
| 16:00 | 5vbc-A | 10.3 | 3.1 | 123 | 217 | 15 | MOLECULE: | PROBABLE HISTONE-LYSINE N-METHYLTRANSFERASE ATXR5 |

| | | | | | | | | |
|---|---|---|---|---|---|---|---|---|
| 17:00 | 4o30-B | 10.3 | 3.1 | 123 | 217 | 15 | MOLECULE: | HISTONE-LYSINE N-METHYLTRANSFERASE ATXR6, PUTATIV |
| 18:00 | 4mi0-A | 10.3 | 2.5 | 117 | 208 | 20 | MOLECULE: | HISTONE-LYSINE N-METHYLTRANSFERASE EZH2 |
| 19:00 | 5jjy-A | 10.2 | 2.7 | 118 | 248 | 19 | MOLECULE: | HISTONE-LYSINE N-METHYLTRANSFERASE SETD2 |
| 20:00 | 4o30-A | 10.2 | 3.1 | 124 | 218 | 15 | MOLECULE: | HISTONE-LYSINE N-METHYLTRANSFERASE ATXR6, PUTATIV |
| 21:00 | 5lsy-A | 10.2 | 2.3 | 116 | 246 | 19 | MOLECULE: | HISTONE-LYSINE N-METHYLTRANSFERASE SETD2 |
| 22:00 | 6j9j-A | 10.2 | 2.7 | 118 | 248 | 19 | MOLECULE: | HISTONE-LYSINE N-METHYLTRANSFERASE SETD2 |
| 23:00 | 5v21-A | 10.2 | 2.7 | 118 | 246 | 19 | MOLECULE: | HISTONE-LYSINE N-METHYLTRANSFERASE SETD2 |
| 24:00:00 | 5lss-A | 10.2 | 2.6 | 117 | 238 | 19 | MOLECULE: | HISTONE-LYSINE N-METHYLTRANSFERASE SETD2 |
| 25:00:00 | 8q5p-A | 10.2 | 2.8 | 118 | 246 | 19 | MOLECULE: | HISTONE-LYSINE N-METHYLTRANSFERASE SETD2 |
| 26:00:00 | 5lt7-A | 10.2 | 2.3 | 116 | 243 | 19 | MOLECULE: | HISTONE-LYSINE N-METHYLTRANSFERASE SETD2 |
| 27:00:00 | 5vac-A | 10.2 | 3.1 | 123 | 216 | 15 | MOLECULE: | PROBABLE HISTONE-LYSINE N-METHYLTRANSFERASE ATXR5 |
| 28:00:00 | 4fmu-A | 10.2 | 2.8 | 118 | 236 | 19 | MOLECULE: | HISTONE-LYSINE N-METHYLTRANSFERASE SETD2 |
| 29:00:00 | 5va6-A | 10.2 | 3.1 | 123 | 217 | 15 | MOLECULE: | PROBABLE HISTONE-LYSINE N-METHYLTRANSFERASE ATXR5 |
| 30:00:00 | 7xud-A | 10.2 | 2.5 | 121 | 269 | 17 | MOLECULE: | HISTONE-LYSINE N-METHYLTRANSFERASE EHMT2 |
| 31:00:00 | 3bo5-A | 10.2 | 3.1 | 128 | 270 | 20 | MOLECULE: | HISTONE-LYSINE N-METHYLTRANSFERASE SETMAR |
| 32:00:00 | 7jsi-A | 10.2 | 3.8 | 228 | 276 | 14 | MOLECULE: | PROTEIN REP68 |
| 33:00:00 | 7jsh-E | 10.2 | 3.5 | 224 | 272 | 14 | MOLECULE: | PROTEIN REP68 |
| 34:00:00 | 7jsg-D | 10.2 | 3.9 | 215 | 272 | 13 | MOLECULE: | PROTEIN REP68 |
| 35:00:00 | 5v22-A | 10.1 | 2.7 | 118 | 246 | 19 | MOLECULE: | HISTONE-LYSINE N-METHYLTRANSFERASE SETD2 |
| 36:00:00 | 5kjh-B | 10.1 | 6.6 | 155 | 806 | 17 | MOLECULE: | PUTATIVE POLYCOMB PROTEIN EED |
| 37:00:00 | 6ugm-M | 10.1 | 2.4 | 112 | 189 | 18 | MOLECULE: | HISTONE H3 |
| 38:00:00 | 6a5n-A | 10.1 | 2.9 | 122 | 502 | 16 | MOLECULE: | HISTONE-LYSINE N-METHYLTRANSFERASE, H3 LYSINE-9 S |
| 39:00:00 | 5ls6-G | 10 | 2.2 | 115 | 572 | 20 | MOLECULE: | HISTONE-LYSINE N-METHYLTRANSFERASE EZH2,HISTONE-L |
| 40:00:00 | 4w2r-A | 10 | 2.2 | 116 | 463 | 20 | MOLECULE: | ENHANCER OF ZESTE 2 POLYCOMB REPRESSIVE COMPLEX 2 |
| 41:00:00 | 5vsc-B | 10 | 5.6 | 121 | 273 | 18 | MOLECULE: | HISTONE-LYSINE N-METHYLTRANSFERASE EHMT2 |
| 42:00:00 | 7xub-A | 10 | 2.5 | 120 | 271 | 18 | MOLECULE: | HISTONE-LYSINE N-METHYLTRANSFERASE EHMT2 |
| 43:00:00 | 7om0-B | 10 | 7.2 | 216 | 439 | 14 | MOLECULE: | DNA PRIMASE |
| 44:00:00 | 7ola-B | 10 | 7.7 | 228 | 441 | 13 | MOLECULE: | DNA PRIMASE |
| 45:00:00 | 7mbm-C | 10 | 2.3 | 115 | 179 | 19 | MOLECULE: | RETINOBLASTOMA-BINDING PROTEIN 5 |
| 46:00:00 | 6kix-K | 10 | 2.4 | 116 | 182 | 19 | MOLECULE: | HISTONE H3 |
| 47:00:00 | 5czy-A | 9.9 | 23.4 | 151 | 450 | 19 | MOLECULE: | LEGIONELLA EFFECTOR LEGAS4 |
| 48:00:00 | 4rz0-A | 9.9 | 2.7 | 100 | 112 | 12 | MOLECULE: | PFL0690C |
| 49:00:00 | 3mo0-B | 9.9 | 2.2 | 117 | 243 | 19 | MOLECULE: | HISTONE-LYSINE N-METHYLTRANSFERASE, H3 LYSINE-9 S |
| 50:00:00 | 5ij7-B | 9.9 | 2.2 | 116 | 469 | 20 | MOLECULE: | ENHANCER OF ZESTE HOMOLOG 2 (EZH2),HISTONE-LYSINE |
| 51:00:00 | 5vk3-B | 9.9 | 7.6 | 152 | 792 | 16 | MOLECULE: | POLYCOMB PROTEIN EED |
| 52:00:00 | 5lt6-B | 9.9 | 2.3 | 113 | 227 | 19 | MOLECULE: | HISTONE-LYSINE N-METHYLTRANSFERASE SETD2 |
| 53:00:00 | 6ago-B | 9.9 | 2.1 | 112 | 224 | 18 | MOLECULE: | HISTONE-LYSINE N-METHYLTRANSFERASE ASH1L |
| 54:00:00 | 5tuz-B | 9.9 | 2.4 | 116 | 258 | 18 | MOLECULE: | HISTONE-LYSINE N-METHYLTRANSFERASE EHMT1 |
| 55:00:00 | 7jsg-A | 9.9 | 3.8 | 214 | 272 | 14 | MOLECULE: | PROTEIN REP68 |
| 56:00:00 | 8uh7-E | 9.9 | 4.4 | 127 | 314 | 20 | MOLECULE: | SLIDING-CLAMP-LOADER SMALL SUBUNIT |
| 57:00:00 | 7t7m-A | 9.9 | 2.8 | 121 | 260 | 19 | MOLECULE: | HISTONE-LYSINE N-METHYLTRANSFERASE EHMT1 |
| 58:00:00 | 7x73-A | 9.9 | 2.5 | 121 | 275 | 17 | MOLECULE: | HISTONE-LYSINE N-METHYLTRANSFERASE EHMT2 |
| 59:00:00 | 8xpt-C | 9.9 | 2.5 | 118 | 256 | 18 | MOLECULE: | HISTONE-LYSINE N-METHYLTRANSFERASE EHMT1 |
| 60:00:00 | 6wkr-C | 9.8 | 3.1 | 120 | 607 | 21 | MOLECULE: | UBIQUITIN |
| 61:00:00 | 5h6z-B | 9.8 | 2.6 | 102 | 120 | 14 | MOLECULE: | SET DOMAIN-CONTAINING PROTEIN 7 |
| 62:00:00 | 3swc-A | 9.8 | 2.5 | 121 | 262 | 18 | MOLECULE: | HISTONE-LYSINE N-METHYLTRANSFERASE EHMT1 |
| 63:00:00 | 5lt6-A | 9.8 | 2.3 | 113 | 234 | 19 | MOLECULE: | HISTONE-LYSINE N-METHYLTRANSFERASE SETD2 |
| 64:00:00 | 3kmt-C | 9.8 | 2.5 | 97 | 120 | 16 | MOLECULE: | A612L PROTEIN |
| 65:00:00 | 5vsf-A | 9.8 | 2.7 | 122 | 262 | 18 | MOLECULE: | HISTONE-LYSINE N-METHYLTRANSFERASE EHMT1 |
| 66:00:00 | 5v9j-B | 9.8 | 2.4 | 119 | 260 | 19 | MOLECULE: | HISTONE-LYSINE N-METHYLTRANSFERASE EHMT1 |
| 67:00:00 | 7t7m-C | 9.8 | 2.7 | 120 | 260 | 18 | MOLECULE: | HISTONE-LYSINE N-METHYLTRANSFERASE EHMT1 |
| 68:00:00 | 5ttf-B | 9.8 | 2.4 | 117 | 268 | 16 | MOLECULE: | HISTONE-LYSINE N-METHYLTRANSFERASE EHMT2 |
| 69:00:00 | 5ttf-D | 9.8 | 2.5 | 117 | 262 | 17 | MOLECULE: | HISTONE-LYSINE N-METHYLTRANSFERASE EHMT2 |

| | | | | | | | | |
|---|---|---|---|---|---|---|---|---|
| 70:00:00 | 8unh-E | 9.8 | 3.8 | 132 | 319 | 20 | MOLECULE: | SLIDING-CLAMP-LOADER LARGE SUBUNIT |
| 71:00:00 | 6mbp-A | 9.8 | 2.8 | 122 | 262 | 18 | MOLECULE: | HISTONE-LYSINE N-METHYLTRANSFERASE EHMT1 |
| 72:00:00 | 3kmt-A | 9.7 | 2.6 | 97 | 120 | 16 | MOLECULE: | A612L PROTEIN |
| 73:00:00 | 5hyn-A | 9.7 | 2.4 | 118 | 582 | 19 | MOLECULE: | HISTONE-LYSINE N-METHYLTRANSFERASE EZH2 |
| 74:00:00 | 8swi-A | 9.7 | 23.7 | 154 | 452 | 14 | MOLECULE: | EUKARYOTIC HUNTINGTIN INTERACTING PROTEIN B |
| 75:00:00 | 3kmt-B | 9.7 | 2.5 | 97 | 120 | 16 | MOLECULE: | A612L PROTEIN |
| 76:00:00 | 5ttf-A | 9.7 | 2.5 | 117 | 266 | 17 | MOLECULE: | HISTONE-LYSINE N-METHYLTRANSFERASE EHMT2 |
| 77:00:00 | 7x73-B | 9.7 | 2.4 | 121 | 274 | 17 | MOLECULE: | HISTONE-LYSINE N-METHYLTRANSFERASE EHMT2 |
| 78:00:00 | 7xud-B | 9.7 | 2.5 | 121 | 271 | 17 | MOLECULE: | HISTONE-LYSINE N-METHYLTRANSFERASE EHMT2 |
| 79:00:00 | 7jsh-A | 9.7 | 3.7 | 221 | 272 | 15 | MOLECULE: | PROTEIN REP68 |
| 80:00:00 | 7jsh-C | 9.7 | 3.9 | 216 | 272 | 13 | MOLECULE: | PROTEIN REP68 |
| 81:00:00 | 7jsh-F | 9.7 | 3.6 | 228 | 272 | 14 | MOLECULE: | PROTEIN REP68 |
| 82:00:00 | 3u61-C | 9.7 | 9.5 | 147 | 319 | 18 | MOLECULE: | DNA POLYMERASE ACCESSORY PROTEIN 44 |
| 83:00:00 | 8uh7-B | 9.7 | 9.6 | 139 | 319 | 18 | MOLECULE: | SLIDING-CLAMP-LOADER SMALL SUBUNIT |
| 84:00:00 | 3swc-B | 9.6 | 2.7 | 121 | 262 | 18 | MOLECULE: | HISTONE-LYSINE N-METHYLTRANSFERASE EHMT1 |
| 85:00:00 | 3sw9-A | 9.6 | 2.7 | 122 | 262 | 18 | MOLECULE: | HISTONE-LYSINE N-METHYLTRANSFERASE EHMT1 |
| 86:00:00 | 6a5m-A | 9.6 | 2.9 | 126 | 489 | 15 | MOLECULE: | HISTONE-LYSINE N-METHYLTRANSFERASE, H3 LYSINE-9 S |
| 87:00:00 | 5tuz-A | 9.6 | 2.5 | 119 | 260 | 18 | MOLECULE: | HISTONE-LYSINE N-METHYLTRANSFERASE EHMT1 |
| 88:00:00 | 7dcf-A | 9.6 | 2.3 | 118 | 261 | 17 | MOLECULE: | HISTONE-LYSINE N-METHYLTRANSFERASE EHMT2 |
| 89:00:00 | 5v9i-A | 9.6 | 2.3 | 115 | 266 | 17 | MOLECULE: | HISTONE-LYSINE N-METHYLTRANSFERASE EHMT2 |
| 90:00:00 | 7xuc-B | 9.6 | 2.5 | 121 | 270 | 17 | MOLECULE: | HISTONE-LYSINE N-METHYLTRANSFERASE EHMT2 |
| 91:00:00 | 5v9i-B | 9.6 | 2.3 | 115 | 268 | 17 | MOLECULE: | HISTONE-LYSINE N-METHYLTRANSFERASE EHMT2 |
| 92:00:00 | 6mbo-A | 9.6 | 2.7 | 121 | 262 | 18 | MOLECULE: | HISTONE-LYSINE N-METHYLTRANSFERASE EHMT1 |
| 93:00:00 | 6k9c-A | 9.6 | 5.6 | 229 | 416 | 16 | MOLECULE: | PRIMASE |
| 94:00:00 | 1peg-A | 9.6 | 2.7 | 120 | 249 | 18 | MOLECULE: | HISTONE H3 METHYLTRANSFERASE DIM-5 |
| 95:00:00 | 6mbp-B | 9.6 | 2.8 | 122 | 261 | 18 | MOLECULE: | HISTONE-LYSINE N-METHYLTRANSFERASE EHMT1 |
| 96:00:00 | 3mo0-A | 9.6 | 2.8 | 121 | 252 | 18 | MOLECULE: | HISTONE-LYSINE N-METHYLTRANSFERASE, H3 LYSINE-9 S |
| 97:00:00 | 5v9j-A | 9.6 | 2.4 | 117 | 258 | 19 | MOLECULE: | HISTONE-LYSINE N-METHYLTRANSFERASE EHMT1 |
| 98:00:00 | 5tuy-A | 9.6 | 2.5 | 121 | 268 | 17 | MOLECULE: | HISTONE-LYSINE N-METHYLTRANSFERASE EHMT2 |
| 99:00:00 | 8iqi-C | 9.5 | 5.1 | 246 | 916 | 10 | MOLECULE: | PUTATIVE PRIMASE C962R |
| 0:00 | 3kma-A | 9.5 | 2.3 | 93 | 105 | 17 | MOLECULE: | A612L PROTEIN |
| 1:00 | 7kso-A | 9.5 | 2.5 | 117 | 395 | 19 | MOLECULE: | HISTONE-LYSINE N-METHYLTRANSFERASE EZH1 |
| 2:00 | 4i51-A | 9.5 | 2.8 | 120 | 262 | 18 | MOLECULE: | HISTONE-LYSINE N-METHYLTRANSFERASE EHMT1 |
| 3:00 | 3sw9-B | 9.5 | 2.7 | 121 | 262 | 19 | MOLECULE: | HISTONE-LYSINE N-METHYLTRANSFERASE EHMT1 |
| 4:00 | 1mvx-A | 9.5 | 2.9 | 124 | 269 | 15 | MOLECULE: | CRYPTIC LOCI REGULATOR 4 |
| 5:00 | 2rfi-A | 9.5 | 2.8 | 120 | 263 | 18 | MOLECULE: | HISTONE-LYSINE N-METHYLTRANSFERASE, H3 LYSINE-9 |
| 6:00 | 3mo2-A | 9.5 | 2.5 | 118 | 255 | 19 | MOLECULE: | HISTONE-LYSINE N-METHYLTRANSFERASE, H3 LYSINE-9 S |
| 7:00 | 2igq-B | 9.5 | 2.5 | 117 | 255 | 18 | MOLECULE: | EUCHROMATIC HISTONE METHYLTRANSFERASE 1 |
| 8:00 | 3hna-B | 9.5 | 2.8 | 120 | 261 | 18 | MOLECULE: | HISTONE-LYSINE N-METHYLTRANSFERASE, H3 LYSINE-9 S |
| 9:00 | 2o8j-A | 9.5 | 2.6 | 118 | 255 | 17 | MOLECULE: | HISTONE-LYSINE N-METHYLTRANSFERASE, H3 LYSINE-9 |
| 10:00 | 5ww0-B | 9.5 | 2.6 | 102 | 120 | 14 | MOLECULE: | SET DOMAIN-CONTAINING PROTEIN 7 |
| 11:00 | 3kma-B | 9.5 | 2.3 | 93 | 105 | 17 | MOLECULE: | A612L PROTEIN |
| 12:00 | 6bx3-E | 9.5 | 2.6 | 115 | 192 | 17 | MOLECULE: | HISTONE-LYSINE N-METHYLTRANSFERASE, H3 LYSINE-4 S |
| 13:00 | 6xaz-B | 9.5 | 2.5 | 101 | 125 | 21 | MOLECULE: | PR DOMAIN ZINC FINGER PROTEIN 5 |
| 14:00 | 5ttg-A | 9.5 | 2.6 | 118 | 261 | 19 | MOLECULE: | HISTONE-LYSINE N-METHYLTRANSFERASE EHMT1 |
| 15:00 | 5ls6-J | 9.5 | 2.5 | 117 | 570 | 20 | MOLECULE: | HISTONE-LYSINE N-METHYLTRANSFERASE EZH2,HISTONE-L |
| 16:00 | 8xpt-B | 9.5 | 2.6 | 119 | 251 | 18 | MOLECULE: | HISTONE-LYSINE N-METHYLTRANSFERASE EHMT1 |
| 17:00 | 5ttg-B | 9.5 | 2.4 | 117 | 259 | 19 | MOLECULE: | HISTONE-LYSINE N-METHYLTRANSFERASE EHMT1 |
| 18:00 | 3kmj-A | 9.5 | 2.5 | 96 | 119 | 17 | MOLECULE: | A612L PROTEIN |
| 19:00 | 5vsd-A | 9.5 | 2.3 | 116 | 258 | 18 | MOLECULE: | HISTONE-LYSINE N-METHYLTRANSFERASE EHMT1 |
| 20:00 | 5vse-A | 9.5 | 5.5 | 121 | 270 | 17 | MOLECULE: | HISTONE-LYSINE N-METHYLTRANSFERASE EHMT2 |
| 21:00 | 7xuc-A | 9.5 | 2.5 | 121 | 273 | 17 | MOLECULE: | HISTONE-LYSINE N-METHYLTRANSFERASE EHMT2 |
| 22:00 | 7jsh-D | 9.5 | 3.7 | 221 | 272 | 13 | MOLECULE: | PROTEIN REP68 |

| | | | | | | | | |
|---|---|---|---|---|---|---|---|---|
| 23:00 | 8umy-A | 9.5 | 28.6 | 164 | 546 | 16 | MOLECULE: | CHROMOSOME TRANSMISSION FIDELITY PROTEIN 18 HOMOL |
| 24:00:00 | 5vsf-B | 9.5 | 2.6 | 118 | 259 | 19 | MOLECULE: | HISTONE-LYSINE N-METHYLTRANSFERASE EHMT1 |
| 25:00:00 | 5vsd-B | 9.5 | 2.6 | 118 | 259 | 19 | MOLECULE: | HISTONE-LYSINE N-METHYLTRANSFERASE EHMT1 |
| 26:00:00 | 3mo5-C | 9.5 | 2.6 | 118 | 259 | 19 | MOLECULE: | HISTONE-LYSINE N-METHYLTRANSFERASE, H3 LYSINE-9 S |
| 27:00:00 | 3mo5-B | 9.5 | 2.5 | 118 | 258 | 19 | MOLECULE: | HISTONE-LYSINE N-METHYLTRANSFERASE, H3 LYSINE-9 S |
| 28:00:00 | 8fyh-A | 9.5 | 2.1 | 116 | 514 | 20 | MOLECULE: | POLYCOMB PROTEIN SUZ12 |
| 29:00:00 | 6mbo-B | 9.5 | 2.6 | 119 | 259 | 18 | MOLECULE: | HISTONE-LYSINE N-METHYLTRANSFERASE EHMT1 |
| 30:00:00 | 3mo5-D | 9.4 | 2.5 | 117 | 257 | 18 | MOLECULE: | HISTONE-LYSINE N-METHYLTRANSFERASE, H3 LYSINE-9 S |
| 31:00:00 | 2o8j-D | 9.4 | 2.6 | 116 | 249 | 16 | MOLECULE: | HISTONE-LYSINE N-METHYLTRANSFERASE, H3 LYSINE-9 |
| 32:00:00 | 1mvh-A | 9.4 | 2.8 | 123 | 269 | 15 | MOLECULE: | CRYPTIC LOCI REGULATOR 4 |
| 33:00:00 | 3fpd-B | 9.4 | 2.8 | 121 | 261 | 18 | MOLECULE: | HISTONE-LYSINE N-METHYLTRANSFERASE, H3 LYSINE-9 |
| 34:00:00 | 5ls6-A | 9.4 | 2.2 | 115 | 570 | 20 | MOLECULE: | HISTONE-LYSINE N-METHYLTRANSFERASE EZH2,HISTONE-L |
| 35:00:00 | 2o8j-B | 9.4 | 2.6 | 121 | 272 | 17 | MOLECULE: | HISTONE-LYSINE N-METHYLTRANSFERASE, H3 LYSINE-9 |
| 36:00:00 | 3mo5-A | 9.4 | 2.5 | 118 | 255 | 19 | MOLECULE: | HISTONE-LYSINE N-METHYLTRANSFERASE, H3 LYSINE-9 S |
| 37:00:00 | 3rjw-A | 9.4 | 2.6 | 121 | 269 | 17 | MOLECULE: | HISTONE-LYSINE N-METHYLTRANSFERASE EHMT2 |
| 38:00:00 | 3fpd-A | 9.4 | 2.8 | 122 | 262 | 18 | MOLECULE: | HISTONE-LYSINE N-METHYLTRANSFERASE, H3 LYSINE-9 |
| 39:00:00 | 6bp4-A | 9.4 | 2.7 | 120 | 269 | 15 | MOLECULE: | HISTONE-LYSINE N-METHYLTRANSFERASE, H3 LYSINE-9 S |
| 40:00:00 | 5t0k-B | 9.4 | 5.1 | 125 | 266 | 17 | MOLECULE: | HISTONE-LYSINE N-METHYLTRANSFERASE EHMT2 |
| 41:00:00 | 7xua-A | 9.4 | 2.6 | 121 | 274 | 17 | MOLECULE: | HISTONE-LYSINE N-METHYLTRANSFERASE EHMT2 |
| 42:00:00 | 7xub-B | 9.4 | 2.4 | 117 | 267 | 17 | MOLECULE: | HISTONE-LYSINE N-METHYLTRANSFERASE EHMT2 |
| 43:00:00 | 1ml9-A | 9.4 | 2.7 | 121 | 250 | 17 | MOLECULE: | HISTONE H3 METHYLTRANSFERASE DIM-5 |
| 44:00:00 | 6p0r-B | 9.4 | 2.4 | 114 | 230 | 20 | MOLECULE: | HISTONE-LYSINE N-METHYLTRANSFERASE SUV39H2 |
| 45:00:00 | 6k9e-F | 9.4 | 5.3 | 215 | 404 | 16 | MOLECULE: | PRIMASE |
| 46:00:00 | 6k9e-B | 9.4 | 5.5 | 222 | 404 | 18 | MOLECULE: | PRIMASE |
| 47:00:00 | 6oax-D | 9.4 | 11.2 | 191 | 581 | 13 | MOLECULE: | HYPERACTIVE DISAGGREGASE CLPB |
| 48:00:00 | 3u61-D | 9.4 | 10.4 | 143 | 320 | 18 | MOLECULE: | DNA POLYMERASE ACCESSORY PROTEIN 44 |
| 49:00:00 | 8uk9-K | 9.4 | 10.4 | 135 | 319 | 19 | MOLECULE: | SLIDING-CLAMP-LOADER SMALL SUBUNIT |
| 50:00:00 | 8uh7-D | 9.4 | 9.2 | 135 | 319 | 20 | MOLECULE: | SLIDING-CLAMP-LOADER SMALL SUBUNIT |
| 51:00:00 | 6rn4-C | 9.4 | 10.6 | 189 | 689 | 13 | MOLECULE: | CHAPERONE PROTEIN CLPB |
| 52:00:00 | 3mo2-B | 9.4 | 2.5 | 118 | 258 | 18 | MOLECULE: | HISTONE-LYSINE N-METHYLTRANSFERASE, H3 LYSINE-9 S |
| 53:00:00 | 3mo2-C | 9.4 | 2.6 | 118 | 259 | 19 | MOLECULE: | HISTONE-LYSINE N-METHYLTRANSFERASE, H3 LYSINE-9 S |
| 54:00:00 | 5ecj-A | 9.3 | 7 | 125 | 263 | 14 | MOLECULE: | PR DOMAIN ZINC FINGER PROTEIN 14,PROTEIN CBFA2T2 |
| 55:00:00 | 6nzo-S | 9.3 | 2.4 | 117 | 233 | 20 | MOLECULE: | HISTONE H3 |
| 56:00:00 | 5wg6-C | 9.3 | 3.5 | 154 | 504 | 16 | MOLECULE: | HISTONE-LYSINE N-METHYLTRANSFERASE EZH2,POLYCOMB |
| 57:00:00 | 4i51-B | 9.3 | 2.3 | 115 | 259 | 17 | MOLECULE: | HISTONE-LYSINE N-METHYLTRANSFERASE EHMT1 |
| 58:00:00 | 2igq-A | 9.3 | 2.5 | 115 | 252 | 17 | MOLECULE: | EUCHROMATIC HISTONE METHYLTRANSFERASE 1 |
| 59:00:00 | 3vv0-A | 9.3 | 2.7 | 116 | 236 | 16 | MOLECULE: | HISTONE-LYSINE N-METHYLTRANSFERASE SETD7 |
| 60:00:00 | 3k5k-B | 9.3 | 5.6 | 122 | 270 | 16 | MOLECULE: | HISTONE-LYSINE N-METHYLTRANSFERASE, H3 LYSINE-9 S |
| 61:00:00 | 3mo2-D | 9.3 | 2.5 | 117 | 257 | 18 | MOLECULE: | HISTONE-LYSINE N-METHYLTRANSFERASE, H3 LYSINE-9 S |
| 62:00:00 | 7dcf-B | 9.3 | 2.1 | 114 | 260 | 18 | MOLECULE: | HISTONE-LYSINE N-METHYLTRANSFERASE EHMT2 |
| 63:00:00 | 6a5k-A | 9.3 | 2.6 | 119 | 467 | 17 | MOLECULE: | HISTONE-LYSINE N-METHYLTRANSFERASE, H3 LYSINE-9 S |
| 64:00:00 | 6px3-S | 9.3 | 2.4 | 117 | 233 | 20 | MOLECULE: | HISTONE H3 |
| 65:00:00 | 7t7l-C | 9.3 | 2.5 | 119 | 267 | 18 | MOLECULE: | HISTONE-LYSINE N-METHYLTRANSFERASE EHMT2 |
| 66:00:00 | 7t7l-B | 9.3 | 2.5 | 118 | 265 | 18 | MOLECULE: | HISTONE-LYSINE N-METHYLTRANSFERASE EHMT2 |
| 67:00:00 | 5t0m-A | 9.3 | 2.4 | 116 | 261 | 17 | MOLECULE: | HISTONE-LYSINE N-METHYLTRANSFERASE EHMT2 |
| 68:00:00 | 7btv-A | 9.3 | 2.5 | 120 | 272 | 18 | MOLECULE: | HISTONE-LYSINE N-METHYLTRANSFERASE EHMT2 |
| 69:00:00 | 8xpt-D | 9.3 | 2.4 | 115 | 248 | 19 | MOLECULE: | HISTONE-LYSINE N-METHYLTRANSFERASE EHMT1 |
| 70:00:00 | 5ttf-C | 9.3 | 2.4 | 117 | 265 | 16 | MOLECULE: | HISTONE-LYSINE N-METHYLTRANSFERASE EHMT2 |
| 71:00:00 | 7xua-B | 9.3 | 5.9 | 123 | 275 | 17 | MOLECULE: | HISTONE-LYSINE N-METHYLTRANSFERASE EHMT2 |
| 72:00:00 | 7buc-A | 9.3 | 5.7 | 125 | 274 | 17 | MOLECULE: | HISTONE-LYSINE N-METHYLTRANSFERASE EHMT2 |
| 73:00:00 | 3ray-A | 9.3 | 2.7 | 109 | 159 | 19 | MOLECULE: | PR DOMAIN-CONTAINING PROTEIN 11 |
| 74:00:00 | 7tts-B | 9.3 | 8.6 | 182 | 440 | 13 | MOLECULE: | CASEINOLYTIC PEPTIDASE B PROTEIN HOMOLOG |
| 75:00:00 | 3u61-B | 9.3 | 4.5 | 131 | 305 | 18 | MOLECULE: | DNA POLYMERASE ACCESSORY PROTEIN 44 |

| | | | | | | | | |
|---|---|---|---|---|---|---|---|---|
| 76:00:00 | 8unh-B | 9.3 | 4.1 | 128 | 319 | 20 | MOLECULE: | SLIDING-CLAMP-LOADER LARGE SUBUNIT |
| 77:00:00 | 8unf-B | 9.3 | 10.6 | 140 | 319 | 19 | MOLECULE: | PRIMER DNA |
| 78:00:00 | 8umu-A | 9.3 | 24 | 167 | 515 | 14 | MOLECULE: | CHROMOSOME TRANSMISSION FIDELITY PROTEIN 18 HOMOL |
| 79:00:00 | 2rfi-B | 9.2 | 2.3 | 114 | 252 | 18 | MOLECULE: | HISTONE-LYSINE N-METHYLTRANSFERASE, H3 LYSINE-9 |
| 80:00:00 | 4nvq-B | 9.2 | 5.6 | 118 | 264 | 16 | MOLECULE: | HISTONE-LYSINE N-METHYLTRANSFERASE EHMT2 |
| 81:00:00 | 2o8j-C | 9.2 | 2.6 | 117 | 263 | 17 | MOLECULE: | HISTONE-LYSINE N-METHYLTRANSFERASE, H3 LYSINE-9 |
| 82:00:00 | 3k5k-A | 9.2 | 2.5 | 119 | 273 | 17 | MOLECULE: | HISTONE-LYSINE N-METHYLTRANSFERASE, H3 LYSINE-9 S |
| 83:00:00 | 8xpt-A | 9.2 | 3.1 | 122 | 258 | 18 | MOLECULE: | HISTONE-LYSINE N-METHYLTRANSFERASE EHMT1 |
| 84:00:00 | 5jhn-A | 9.2 | 2.8 | 120 | 274 | 18 | MOLECULE: | HISTONE-LYSINE N-METHYLTRANSFERASE EHMT2 |
| 85:00:00 | 5jj0-A | 9.2 | 2.8 | 120 | 274 | 18 | MOLECULE: | HISTONE-LYSINE N-METHYLTRANSFERASE EHMT2 |
| 86:00:00 | 5jin-A | 9.2 | 2.8 | 120 | 274 | 18 | MOLECULE: | HISTONE-LYSINE N-METHYLTRANSFERASE EHMT2 |
| 87:00:00 | 5jhn-B | 9.2 | 5.5 | 120 | 268 | 18 | MOLECULE: | HISTONE-LYSINE N-METHYLTRANSFERASE EHMT2 |
| 88:00:00 | 5t0k-A | 9.2 | 2.4 | 116 | 265 | 17 | MOLECULE: | HISTONE-LYSINE N-METHYLTRANSFERASE EHMT2 |
| 89:00:00 | 5v9i-C | 9.2 | 2.3 | 115 | 265 | 17 | MOLECULE: | HISTONE-LYSINE N-METHYLTRANSFERASE EHMT2 |
| 90:00:00 | 7t7l-A | 9.2 | 2.7 | 120 | 267 | 16 | MOLECULE: | HISTONE-LYSINE N-METHYLTRANSFERASE EHMT2 |
| 91:00:00 | 5t0m-B | 9.2 | 2.8 | 120 | 272 | 18 | MOLECULE: | HISTONE-LYSINE N-METHYLTRANSFERASE EHMT2 |
| 92:00:00 | 5v9i-D | 9.2 | 2.3 | 115 | 265 | 17 | MOLECULE: | HISTONE-LYSINE N-METHYLTRANSFERASE EHMT2 |
| 93:00:00 | 6k9e-D | 9.2 | 5.7 | 222 | 404 | 16 | MOLECULE: | PRIMASE |
| 94:00:00 | 3u5z-O | 9.2 | 4.4 | 135 | 305 | 19 | MOLECULE: | DNA POLYMERASE ACCESSORY PROTEIN 44 |
| 95:00:00 | 3u5z-C | 9.2 | 10.9 | 139 | 320 | 19 | MOLECULE: | DNA POLYMERASE ACCESSORY PROTEIN 44 |
| 96:00:00 | 8uh7-C | 9.2 | 9.2 | 139 | 320 | 19 | MOLECULE: | SLIDING-CLAMP-LOADER SMALL SUBUNIT |
| 97:00:00 | 8unf-C | 9.2 | 10.5 | 140 | 319 | 19 | MOLECULE: | PRIMER DNA |
| 98:00:00 | 7ttr-D | 9.2 | 12.2 | 163 | 334 | 12 | MOLECULE: | CASEINOLYTIC PEPTIDASE B PROTEIN HOMOLOG |
| 99:00:00 | 7t7m-D | 9.2 | 2.5 | 117 | 262 | 19 | MOLECULE: | HISTONE-LYSINE N-METHYLTRANSFERASE EHMT1 |
| 0:00 | 5jiy-A | 9.1 | 2.8 | 120 | 274 | 18 | MOLECULE: | HISTONE-LYSINE N-METHYLTRANSFERASE EHMT2 |
| 1:00 | 3hna-A | 9.1 | 2.8 | 120 | 281 | 18 | MOLECULE: | HISTONE-LYSINE N-METHYLTRANSFERASE, H3 LYSINE-9 S |
| 2:00 | 4nvq-A | 9.1 | 2.6 | 117 | 268 | 18 | MOLECULE: | HISTONE-LYSINE N-METHYLTRANSFERASE EHMT2 |
| 3:00 | 3rjw-B | 9.1 | 11.4 | 122 | 271 | 16 | MOLECULE: | HISTONE-LYSINE N-METHYLTRANSFERASE EHMT2 |
| 4:00 | 6box-B | 9.1 | 2.6 | 118 | 268 | 15 | MOLECULE: | HISTONE-LYSINE N-METHYLTRANSFERASE, H3 LYSINE-9 S |
| 5:00 | 5tuy-B | 9.1 | 2.6 | 119 | 264 | 18 | MOLECULE: | HISTONE-LYSINE N-METHYLTRANSFERASE EHMT2 |
| 6:00 | 5jin-B | 9.1 | 5.5 | 120 | 268 | 18 | MOLECULE: | HISTONE-LYSINE N-METHYLTRANSFERASE EHMT2 |
| 7:00 | 7btv-B | 9.1 | 2.5 | 119 | 273 | 18 | MOLECULE: | HISTONE-LYSINE N-METHYLTRANSFERASE EHMT2 |
| 8:00 | 5jj0-B | 9.1 | 5.5 | 120 | 268 | 18 | MOLECULE: | HISTONE-LYSINE N-METHYLTRANSFERASE EHMT2 |
| 9:00 | 7ea5-K | 9.1 | 2.9 | 118 | 229 | 18 | MOLECULE: | HISTONE H3 |
| 10:00 | 7t7l-D | 9.1 | 2.5 | 118 | 270 | 17 | MOLECULE: | HISTONE-LYSINE N-METHYLTRANSFERASE EHMT2 |
| 11:00 | 5jiy-B | 9.1 | 5.4 | 120 | 268 | 18 | MOLECULE: | HISTONE-LYSINE N-METHYLTRANSFERASE EHMT2 |
| 12:00 | 5vse-B | 9.1 | 5.5 | 120 | 272 | 17 | MOLECULE: | HISTONE-LYSINE N-METHYLTRANSFERASE EHMT2 |
| 13:00 | 7buc-B | 9.1 | 2.4 | 116 | 270 | 17 | MOLECULE: | HISTONE-LYSINE N-METHYLTRANSFERASE EHMT2 |
| 14:00 | 7qnr-A | 9.1 | 3.8 | 121 | 427 | 14 | MOLECULE: | HISTONE-LYSINE N-METHYLTRANSFERASE SMYD3 |
| 15:00 | 6z2a-A | 9.1 | 2.8 | 122 | 284 | 15 | MOLECULE: | HISTONE-LYSINE N-METHYLTRANSFERASE, H3 LYSINE-9 S |
| 16:00 | 6z2a-B | 9.1 | 2.7 | 122 | 277 | 15 | MOLECULE: | HISTONE-LYSINE N-METHYLTRANSFERASE, H3 LYSINE-9 S |
| 17:00 | 6p0r-A | 9.1 | 2.8 | 122 | 254 | 20 | MOLECULE: | HISTONE-LYSINE N-METHYLTRANSFERASE SUV39H2 |
| 18:00 | 8dr5-A | 9.1 | 17.5 | 174 | 646 | 10 | MOLECULE: | REPLICATION FACTOR C SUBUNIT 1 |
| 19:00 | 7u1a-A | 9.1 | 13.8 | 166 | 617 | 12 | MOLECULE: | REPLICATION FACTOR C SUBUNIT 1 |
| 20:00 | 3u60-D | 9.1 | 8.6 | 135 | 319 | 19 | MOLECULE: | DNA POLYMERASE ACCESSORY PROTEIN 44 |
| 21:00 | 3u60-C | 9.1 | 9 | 138 | 320 | 19 | MOLECULE: | DNA POLYMERASE ACCESSORY PROTEIN 44 |
| 22:00 | 3u60-E | 9.1 | 4.2 | 130 | 305 | 19 | MOLECULE: | DNA POLYMERASE ACCESSORY PROTEIN 44 |
| 23:00 | 8uk9-B | 9.1 | 10 | 134 | 319 | 19 | MOLECULE: | SLIDING-CLAMP-LOADER SMALL SUBUNIT |
| 24:00:00 | 8unf-D | 9.1 | 10.8 | 140 | 319 | 19 | MOLECULE: | PRIMER DNA |
| 25:00:00 | 6rn3-E | 9.1 | 9.6 | 185 | 684 | 14 | MOLECULE: | CHAPERONE PROTEIN CLPB |
| 26:00:00 | 7t7m-B | 9.1 | 2.6 | 119 | 259 | 18 | MOLECULE: | HISTONE-LYSINE N-METHYLTRANSFERASE EHMT1 |
| 27:00:00 | 7ksr-A | 9 | 2 | 114 | 395 | 19 | MOLECULE: | HISTONE-LYSINE N-METHYLTRANSFERASE EZH1 |
| 28:00:00 | 5arg-A | 9 | 3.7 | 119 | 427 | 11 | MOLECULE: | N-LYSINE METHYLTRANSFERASE SMYD2 |

| | | | | | | | | |
|---|---|---|---|---|---|---|---|---|
| 29:00:00 | 5arf-A | 9 | 3.6 | 118 | 426 | 12 | MOLECULE: | N-LYSINE METHYLTRANSFERASE SMYD2 |
| 30:00:00 | 6xaz-A | 9 | 2.3 | 94 | 118 | 21 | MOLECULE: | PR DOMAIN ZINC FINGER PROTEIN 5 |
| 31:00:00 | 7pds-D | 9 | 4.4 | 237 | 454 | 11 | MOLECULE: | SIMILAR TO D. NODOSUS VAPE |
| 32:00:00 | 7pds-C | 9 | 4.4 | 239 | 451 | 11 | MOLECULE: | SIMILAR TO D. NODOSUS VAPE |
| 33:00:00 | 7tib-A | 9 | 11.1 | 151 | 495 | 14 | MOLECULE: | REPLICATION FACTOR C SUBUNIT 1 |
| 34:00:00 | 8dr1-A | 9 | 11.1 | 149 | 493 | 13 | MOLECULE: | REPLICATION FACTOR C SUBUNIT 1 |
| 35:00:00 | 8efy-N | 9 | 11.7 | 148 | 310 | 12 | MOLECULE: | HOLLIDAY JUNCTION ATP-DEPENDENT DNA HELICASE RUVB |
| 36:00:00 | 3u5z-D | 9 | 10.1 | 138 | 320 | 20 | MOLECULE: | DNA POLYMERASE ACCESSORY PROTEIN 44 |
| 37:00:00 | 8uk9-M | 9 | 9.4 | 135 | 317 | 18 | MOLECULE: | SLIDING-CLAMP-LOADER SMALL SUBUNIT |
| 38:00:00 | 9emc-F | 9 | 8.2 | 144 | 337 | 13 | MOLECULE: | RUVB-LIKE 1 |
| 39:00:00 | 7ktp-A | 9 | 1.9 | 114 | 395 | 19 | MOLECULE: | HISTONE-LYSINE N-METHYLTRANSFERASE EZH1 |
| 40:00:00 | 2r3a-A | 9 | 4.2 | 124 | 272 | 19 | MOLECULE: | HISTONE-LYSINE N-METHYLTRANSFERASE SUV39H2 |
| 41:00:00 | 6xau-A | 8.9 | 2.4 | 100 | 119 | 13 | MOLECULE: | HISTONE-LYSINE N-METHYLTRANSFERASE MECOM |
| 42:00:00 | 6kiw-K | 8.9 | 2.7 | 117 | 156 | 20 | MOLECULE: | HISTONE H3 |
| 43:00:00 | 7pds-B | 8.9 | 4 | 234 | 448 | 12 | MOLECULE: | SIMILAR TO D. NODOSUS VAPE |
| 44:00:00 | 7xc2-E | 8.9 | 26.7 | 172 | 858 | 13 | MOLECULE: | CNL9 |
| 45:00:00 | 8iqi-B | 8.9 | 6.4 | 254 | 915 | 10 | MOLECULE: | PUTATIVE PRIMASE C962R |
| 46:00:00 | 3u5z-B | 8.9 | 10.9 | 134 | 320 | 19 | MOLECULE: | DNA POLYMERASE ACCESSORY PROTEIN 44 |
| 47:00:00 | 3u5z-N | 8.9 | 10.2 | 138 | 320 | 20 | MOLECULE: | DNA POLYMERASE ACCESSORY PROTEIN 44 |
| 48:00:00 | 3u5z-L | 8.9 | 8.4 | 133 | 320 | 19 | MOLECULE: | DNA POLYMERASE ACCESSORY PROTEIN 44 |
| 49:00:00 | 3u5z-M | 8.9 | 9.4 | 137 | 320 | 19 | MOLECULE: | DNA POLYMERASE ACCESSORY PROTEIN 44 |
| 50:00:00 | 9emc-D | 8.9 | 4.7 | 141 | 337 | 13 | MOLECULE: | RUVB-LIKE 1 |
| 51:00:00 | 5hi7-A | 8.8 | 3.7 | 121 | 426 | 14 | MOLECULE: | HISTONE-LYSINE N-METHYLTRANSFERASE SMYD3 |
| 52:00:00 | 8iqi-D | 8.8 | 5 | 243 | 916 | 10 | MOLECULE: | PUTATIVE PRIMASE C962R |
| 53:00:00 | 3u61-E | 8.8 | 4 | 130 | 294 | 22 | MOLECULE: | DNA POLYMERASE ACCESSORY PROTEIN 44 |
| 54:00:00 | 3u60-B | 8.8 | 8.6 | 134 | 319 | 19 | MOLECULE: | DNA POLYMERASE ACCESSORY PROTEIN 44 |
| 55:00:00 | 8uk9-N | 8.8 | 4.5 | 127 | 305 | 19 | MOLECULE: | SLIDING-CLAMP-LOADER SMALL SUBUNIT |
| 56:00:00 | 8uk9-C | 8.8 | 11 | 141 | 320 | 18 | MOLECULE: | SLIDING-CLAMP-LOADER SMALL SUBUNIT |
| 57:00:00 | 8ui8-A | 8.8 | 17.8 | 168 | 643 | 11 | MOLECULE: | ATPASE FAMILY AAA DOMAIN-CONTAINING PROTEIN 5 |
| 58:00:00 | 8un0-A | 8.8 | 26.2 | 160 | 546 | 15 | MOLECULE: | CHROMOSOME TRANSMISSION FIDELITY PROTEIN 18 HOMOL |
| 59:00:00 | 7xc2-A | 8.7 | 25.5 | 163 | 858 | 13 | MOLECULE: | CNL9 |
| 60:00:00 | 7mi8-A | 8.7 | 4.9 | 190 | 868 | 8 | MOLECULE: | FUSION PROTEIN OF DYNEIN AND ENDOLYSIN |
| 61:00:00 | 6zrb-A | 8.7 | 3.8 | 121 | 426 | 14 | MOLECULE: | HISTONE-LYSINE N-METHYLTRANSFERASE SMYD3 |
| 62:00:00 | 7xc2-C | 8.7 | 24.5 | 171 | 858 | 13 | MOLECULE: | CNL9 |
| 63:00:00 | 7ttr-A | 8.7 | 8.8 | 147 | 304 | 12 | MOLECULE: | CASEINOLYTIC PEPTIDASE B PROTEIN HOMOLOG |
| 64:00:00 | 2qpw-A | 8.7 | 3.1 | 105 | 147 | 16 | MOLECULE: | PR DOMAIN ZINC FINGER PROTEIN 2 |
| 65:00:00 | 7om0-D | 8.7 | 6.5 | 217 | 438 | 12 | MOLECULE: | DNA PRIMASE |
| 66:00:00 | 7u1p-A | 8.7 | 13.3 | 161 | 617 | 12 | MOLECULE: | REPLICATION FACTOR C SUBUNIT 1 |
| 67:00:00 | 8iqi-A | 8.7 | 6.4 | 249 | 915 | 10 | MOLECULE: | PUTATIVE PRIMASE C962R |
| 68:00:00 | 7xc2-I | 8.7 | 26.4 | 172 | 858 | 13 | MOLECULE: | CNL9 |
| 69:00:00 | 8unf-E | 8.7 | 4.9 | 129 | 319 | 19 | MOLECULE: | PRIMER DNA |
| 70:00:00 | 8uk9-L | 8.7 | 9.2 | 137 | 320 | 18 | MOLECULE: | SLIDING-CLAMP-LOADER SMALL SUBUNIT |
| 71:00:00 | 6qs6-E | 8.7 | 15.2 | 222 | 577 | 11 | MOLECULE: | CHAPERONE PROTEIN CLPB |
| 72:00:00 | 6oax-B | 8.7 | 10.8 | 191 | 581 | 14 | MOLECULE: | HYPERACTIVE DISAGGREGASE CLPB |
| 73:00:00 | 4qep-A | 8.7 | 3.2 | 118 | 468 | 19 | MOLECULE: | HISTONE-LYSINE N-METHYLTRANSFERASE, H3 LYSINE-9 S |
| 74:00:00 | 3ep0-A | 8.6 | 2.2 | 89 | 113 | 15 | MOLECULE: | PR DOMAIN ZINC FINGER PROTEIN 12 |
| 75:00:00 | 7o2c-A | 8.6 | 3.8 | 121 | 428 | 14 | MOLECULE: | HISTONE-LYSINE N-METHYLTRANSFERASE SMYD3 |
| 76:00:00 | 4qeo-A | 8.6 | 3.4 | 120 | 473 | 18 | MOLECULE: | HISTONE-LYSINE N-METHYLTRANSFERASE, H3 LYSINE-9 S |
| 77:00:00 | 3ep0-B | 8.6 | 2.5 | 92 | 113 | 13 | MOLECULE: | PR DOMAIN ZINC FINGER PROTEIN 12 |
| 78:00:00 | 4qen-A | 8.6 | 3.3 | 120 | 473 | 18 | MOLECULE: | HISTONE-LYSINE N-METHYLTRANSFERASE, H3 LYSINE-9 S |
| 79:00:00 | 7ola-A | 8.6 | 6.9 | 216 | 441 | 13 | MOLECULE: | DNA PRIMASE |
| 80:00:00 | 8dr7-A | 8.6 | 13.5 | 149 | 497 | 13 | MOLECULE: | REPLICATION FACTOR C SUBUNIT 1 |
| 81:00:00 | 6oax-A | 8.6 | 14.6 | 189 | 581 | 14 | MOLECULE: | HYPERACTIVE DISAGGREGASE CLPB |

| | | | | | | | | |
|---|---|---|---|---|---|---|---|---|
| 82:00:00 | 6mfv-B | 8.6 | 8.8 | 131 | 641 | 11 | MOLECULE: | TETRATRICOPEPTIDE REPEAT SENSOR PH0952 |
| 83:00:00 | 8uk9-D | 8.6 | 10.4 | 138 | 318 | 19 | MOLECULE: | SLIDING-CLAMP-LOADER SMALL SUBUNIT |
| 84:00:00 | 3uk6-E | 8.6 | 3.1 | 139 | 290 | 12 | MOLECULE: | RUVB-LIKE 2 |
| 85:00:00 | 5fm7-B | 8.6 | 11.4 | 146 | 422 | 12 | MOLECULE: | RVB1 |
| 86:00:00 | 7crr-I | 8.5 | 3.2 | 116 | 223 | 21 | MOLECULE: | HISTONE H3 |
| 87:00:00 | 7o2b-A | 8.5 | 3.8 | 121 | 428 | 14 | MOLECULE: | HISTONE-LYSINE N-METHYLTRANSFERASE SMYD3 |
| 88:00:00 | 6paf-A | 8.5 | 3.7 | 121 | 424 | 15 | MOLECULE: | HISTONE-LYSINE N-METHYLTRANSFERASE SMYD3 |
| 89:00:00 | 3dal-B | 8.5 | 3.2 | 111 | 166 | 14 | MOLECULE: | PR DOMAIN ZINC FINGER PROTEIN 1 |
| 90:00:00 | 4au7-B | 8.5 | 3.1 | 98 | 209 | 10 | MOLECULE: | HISTONE-LYSINE N-METHYLTRANSFERASE SUV420H2 |
| 91:00:00 | 7qnu-A | 8.5 | 3.8 | 121 | 427 | 15 | MOLECULE: | HISTONE-LYSINE N-METHYLTRANSFERASE SMYD3 |
| 92:00:00 | 7om0-E | 8.5 | 6.2 | 215 | 439 | 13 | MOLECULE: | DNA PRIMASE |
| 93:00:00 | 7om0-A | 8.5 | 6.2 | 215 | 439 | 13 | MOLECULE: | DNA PRIMASE |
| 94:00:00 | 7om0-C | 8.5 | 6.5 | 225 | 439 | 13 | MOLECULE: | DNA PRIMASE |
| 95:00:00 | 7om0-F | 8.5 | 7.1 | 242 | 439 | 13 | MOLECULE: | DNA PRIMASE |
| 96:00:00 | 3u5z-E | 8.5 | 4.3 | 131 | 305 | 19 | MOLECULE: | DNA POLYMERASE ACCESSORY PROTEIN 44 |
| 97:00:00 | 3uk6-L | 8.5 | 8.6 | 128 | 232 | 12 | MOLECULE: | RUVB-LIKE 2 |
| 98:00:00 | 1s9h-C | 8.5 | 4.6 | 157 | 199 | 14 | MOLECULE: | REP 40 PROTEIN |
| 99:00:00 | 6o9o-A | 8.4 | 3.8 | 121 | 427 | 14 | MOLECULE: | HISTONE-LYSINE N-METHYLTRANSFERASE SMYD3 |
| 0:00 | 6z2r-A | 8.4 | 3.8 | 121 | 427 | 14 | MOLECULE: | HISTONE-LYSINE N-METHYLTRANSFERASE SMYD3 |
| 1:00 | 6p6g-A | 8.4 | 3.8 | 121 | 427 | 14 | MOLECULE: | HISTONE-LYSINE N-METHYLTRANSFERASE SMYD3 |
| 2:00 | 7las-A | 8.4 | 5.7 | 134 | 183 | 16 | MOLECULE: | ATP-DEPENDENT HELICASE REP |
| 3:00 | 5n7r-A | 8.4 | 14.9 | 143 | 380 | 10 | MOLECULE: | RUVB-LIKE 2 |
| 4:00 | 6rn2-D | 8.4 | 11.4 | 189 | 684 | 12 | MOLECULE: | CHAPERONE PROTEIN CLPB |
| 5:00 | 6vvo-A | 8.4 | 6.6 | 156 | 448 | 9 | MOLECULE: | REPLICATION FACTOR C SUBUNIT 1 |
| 6:00 | 3uk6-F | 8.4 | 11.8 | 144 | 301 | 13 | MOLECULE: | RUVB-LIKE 2 |
| 7:00 | 7qv9-a | 8.4 | 7.3 | 149 | 259 | 9 | MOLECULE: | DNA-DIRECTED RNA POLYMERASE SUBUNIT ALPHA |
| 8:00 | 9emc-E | 8.4 | 8.6 | 145 | 337 | 13 | MOLECULE: | RUVB-LIKE 1 |
| 9:00 | 7ud5-K | 8.3 | 2.7 | 114 | 176 | 18 | MOLECULE: | HISTONE H3 |
| 10:00 | 6bp4-B | 8.3 | 4.4 | 124 | 283 | 15 | MOLECULE: | HISTONE-LYSINE N-METHYLTRANSFERASE, H3 LYSINE-9 S |
| 11:00 | 7ln2-F | 8.3 | 14.6 | 185 | 700 | 9 | MOLECULE: | TRANSITIONAL ENDOPLASMIC RETICULUM ATPASE |
| 12:00 | 4bup-A | 8.3 | 3.2 | 105 | 233 | 10 | MOLECULE: | HISTONE-LYSINE N-METHYLTRANSFERASE SUV420H1 |
| 13:00 | 5v3h-A | 8.3 | 4.2 | 121 | 439 | 12 | MOLECULE: | N-LYSINE METHYLTRANSFERASE SMYD2 |
| 14:00 | 6oax-E | 8.3 | 10.5 | 184 | 579 | 14 | MOLECULE: | HYPERACTIVE DISAGGREGASE CLPB |
| 15:00 | 7ola-E | 8.3 | 4.3 | 198 | 344 | 14 | MOLECULE: | DNA PRIMASE |
| 16:00 | 8apl-D | 8.3 | 6.1 | 214 | 449 | 13 | MOLECULE: | PRIMASE D5 |
| 17:00 | 8hwe-D | 8.3 | 5 | 215 | 379 | 9 | MOLECULE: | PRIMASE D5 |
| 18:00 | 6u1y-F | 8.3 | 2.9 | 127 | 244 | 18 | MOLECULE: | MITOCHONDRIAL CHAPERONE BCS1 |
| 19:00 | 6mat-C | 8.3 | 12.4 | 159 | 578 | 11 | MOLECULE: | RIX7 MUTANT |
| 20:00 | 7jlv-A | 8.3 | 15.6 | 144 | 437 | 15 | MOLECULE: | DISEASE RESISTANCE PROTEIN ROQ1 |
| 21:00 | 7jlv-G | 8.3 | 13 | 136 | 437 | 15 | MOLECULE: | DISEASE RESISTANCE PROTEIN ROQ1 |
| 22:00 | 5oaf-D | 8.3 | 9.3 | 144 | 411 | 11 | MOLECULE: | RUVB-LIKE 1 |
| 23:00 | 5z6r-A | 8.3 | 11.5 | 143 | 273 | 11 | MOLECULE: | SPASTIN |
| 24:00:00 | 6mat-D | 8.3 | 18.9 | 188 | 578 | 9 | MOLECULE: | RIX7 MUTANT |
| 25:00:00 | 6og1-A | 8.3 | 14.2 | 220 | 581 | 15 | MOLECULE: | HYPERACTIVE DISAGGREGASE CLPB |
| 26:00:00 | 6box-A | 8.2 | 2.7 | 114 | 271 | 16 | MOLECULE: | HISTONE-LYSINE N-METHYLTRANSFERASE, H3 LYSINE-9 S |
| 27:00:00 | 4bup-B | 8.2 | 3.2 | 105 | 238 | 10 | MOLECULE: | HISTONE-LYSINE N-METHYLTRANSFERASE SUV420H1 |
| 28:00:00 | 3db5-B | 8.2 | 2.9 | 99 | 134 | 14 | MOLECULE: | PR DOMAIN ZINC FINGER PROTEIN 4 |
| 29:00:00 | 3ihx-C | 8.2 | 3 | 99 | 124 | 13 | MOLECULE: | PR DOMAIN ZINC FINGER PROTEIN 10 |
| 30:00:00 | 3db5-A | 8.2 | 2.5 | 93 | 124 | 14 | MOLECULE: | PR DOMAIN ZINC FINGER PROTEIN 4 |
| 31:00:00 | 8apl-C | 8.2 | 6.2 | 221 | 449 | 12 | MOLECULE: | PRIMASE D5 |
| 32:00:00 | 7lar-E | 8.2 | 5.8 | 136 | 183 | 16 | MOLECULE: | ATP-DEPENDENT HELICASE REP |
| 33:00:00 | 8apl-E | 8.2 | 6.1 | 217 | 449 | 12 | MOLECULE: | PRIMASE D5 |
| 34:00:00 | 8apl-B | 8.2 | 6.8 | 225 | 449 | 12 | MOLECULE: | PRIMASE D5 |

| | | | | | | | | |
|---|---|---|---|---|---|---|---|---|
| 35:00:00 | 8apm-D | 8.2 | 6.4 | 220 | 449 | 12 | MOLECULE: | PRIMASE D5 |
| 36:00:00 | 7las-E | 8.2 | 5.4 | 136 | 183 | 16 | MOLECULE: | ATP-DEPENDENT HELICASE REP |
| 37:00:00 | 7qo4-M | 8.2 | 11.4 | 178 | 421 | 11 | MOLECULE: | 26S PROTEASOME REGULATORY SUBUNIT RPN1 |
| 38:00:00 | 7las-C | 8.2 | 5.9 | 142 | 183 | 15 | MOLECULE: | ATP-DEPENDENT HELICASE REP |
| 39:00:00 | 8apl-F | 8.2 | 6.4 | 207 | 449 | 12 | MOLECULE: | PRIMASE D5 |
| 40:00:00 | 8apl-A | 8.2 | 6.2 | 214 | 449 | 12 | MOLECULE: | PRIMASE D5 |
| 41:00:00 | 5oaf-F | 8.2 | 8.7 | 147 | 407 | 12 | MOLECULE: | RUVB-LIKE 1 |
| 42:00:00 | 7jlv-B | 8.2 | 10.6 | 141 | 437 | 16 | MOLECULE: | DISEASE RESISTANCE PROTEIN ROQ1 |
| 43:00:00 | 3uk6-B | 8.2 | 3.4 | 139 | 300 | 13 | MOLECULE: | RUVB-LIKE 2 |
| 44:00:00 | 3uk6-J | 8.2 | 3.2 | 145 | 300 | 12 | MOLECULE: | RUVB-LIKE 2 |
| 45:00:00 | 3uk6-A | 8.2 | 3.1 | 139 | 298 | 12 | MOLECULE: | RUVB-LIKE 2 |
| 46:00:00 | 7ttr-C | 8.2 | 10.4 | 155 | 327 | 14 | MOLECULE: | CASEINOLYTIC PEPTIDASE B PROTEIN HOMOLOG |
| 47:00:00 | 5vy9-A | 8.2 | 10.2 | 211 | 849 | 9 | MOLECULE: | HEAT SHOCK PROTEIN 104 |
| 48:00:00 | 5z6q-A | 8.2 | 13.2 | 145 | 272 | 10 | MOLECULE: | SPASTIN |
| 49:00:00 | 3dal-A | 8.1 | 3.2 | 110 | 179 | 19 | MOLECULE: | PR DOMAIN ZINC FINGER PROTEIN 1 |
| 50:00:00 | 3s8p-B | 8.1 | 3.2 | 105 | 239 | 10 | MOLECULE: | HISTONE-LYSINE N-METHYLTRANSFERASE SUV420H1 |
| 51:00:00 | 7xc2-G | 8.1 | 26.8 | 172 | 858 | 13 | MOLECULE: | CNL9 |
| 52:00:00 | 7lar-C | 8.1 | 5.2 | 134 | 183 | 16 | MOLECULE: | ATP-DEPENDENT HELICASE REP |
| 53:00:00 | 8apm-E | 8.1 | 6 | 223 | 449 | 12 | MOLECULE: | PRIMASE D5 |
| 54:00:00 | 8hwf-C | 8.1 | 4.1 | 206 | 378 | 9 | MOLECULE: | PRIMASE D5 |
| 55:00:00 | 8hwd-B | 8.1 | 4.9 | 206 | 378 | 10 | MOLECULE: | PRIMASE D5 |
| 56:00:00 | 8xj7-B | 8.1 | 7 | 212 | 452 | 13 | MOLECULE: | MONKEYPOX VIRUS E5 |
| 57:00:00 | 8hwh-C | 8.1 | 4.2 | 204 | 361 | 9 | MOLECULE: | PRIMASE D5 |
| 58:00:00 | 7u19-A | 8.1 | 14.4 | 163 | 617 | 13 | MOLECULE: | REPLICATION FACTOR C SUBUNIT 1 |
| 59:00:00 | 6h7x-A | 8.1 | 13.9 | 144 | 371 | 11 | MOLECULE: | RUVB-LIKE 2 |
| 60:00:00 | 6e10-6 | 8.1 | 11.1 | 187 | 718 | 17 | MOLECULE: | HEAT SHOCK PROTEIN 101 |
| 61:00:00 | 5fm6-A | 8.1 | 3.4 | 145 | 426 | 16 | MOLECULE: | RVB1 |
| 62:00:00 | 5fm7-A | 8.1 | 3.5 | 146 | 418 | 13 | MOLECULE: | RVB1 |
| 63:00:00 | 6u1y-G | 8.1 | 3 | 128 | 244 | 18 | MOLECULE: | MITOCHONDRIAL CHAPERONE BCS1 |
| 64:00:00 | 7ttr-E | 8.1 | 11.5 | 164 | 332 | 12 | MOLECULE: | CASEINOLYTIC PEPTIDASE B PROTEIN HOMOLOG |
| 65:00:00 | 6rn2-E | 8.1 | 10.5 | 186 | 684 | 12 | MOLECULE: | CHAPERONE PROTEIN CLPB |
| 66:00:00 | 6qi9-E | 8.1 | 9.1 | 150 | 335 | 11 | MOLECULE: | RUVB-LIKE 1 |
| 67:00:00 | 7ea8-L | 8.1 | 3.1 | 119 | 246 | 18 | MOLECULE: | HISTONE-LYSINE N-METHYLTRANSFERASE SETD2 |
| 68:00:00 | 8y6p-Q | 8 | 8.5 | 152 | 1228 | 11 | MOLECULE: | APAF-1 RELATED KILLER DARK |
| 69:00:00 | 6hyp-A | 8 | 18.3 | 194 | 2272 | 9 | MOLECULE: | MIDASIN,MIDASIN |
| 70:00:00 | 7crp-I | 8 | 2.8 | 114 | 223 | 21 | MOLECULE: | HISTONE H3 |
| 71:00:00 | 4nj5-A | 8 | 2.6 | 114 | 482 | 16 | MOLECULE: | PROBABLE HISTONE-LYSINE N-METHYLTRANSFERASE, H3 L |
| 72:00:00 | 6p7z-A | 8 | 3.8 | 121 | 426 | 14 | MOLECULE: | HISTONE-LYSINE N-METHYLTRANSFERASE SMYD3 |
| 73:00:00 | 5ecj-B | 8 | 8.1 | 128 | 263 | 13 | MOLECULE: | PR DOMAIN ZINC FINGER PROTEIN 14,PROTEIN CBFA2T2 |
| 74:00:00 | 5wbv-A | 8 | 3.3 | 106 | 234 | 9 | MOLECULE: | HISTONE-LYSINE N-METHYLTRANSFERASE KMT5B |
| 75:00:00 | 6hz9-J | 8 | 3.7 | 137 | 284 | 11 | MOLECULE: | 5-METHYLCYTOSINE-SPECIFIC RESTRICTION ENZYME B |
| 76:00:00 | 8xj7-C | 8 | 5.6 | 207 | 454 | 10 | MOLECULE: | MONKEYPOX VIRUS E5 |
| 77:00:00 | 8hwf-A | 8 | 4.4 | 214 | 378 | 11 | MOLECULE: | PRIMASE D5 |
| 78:00:00 | 8hwf-B | 8 | 4.4 | 207 | 378 | 10 | MOLECULE: | PRIMASE D5 |
| 79:00:00 | 8hwa-B | 8 | 4.1 | 210 | 379 | 10 | MOLECULE: | PRIMASE D5 |
| 80:00:00 | 8xj7-A | 8 | 4.4 | 212 | 377 | 11 | MOLECULE: | MONKEYPOX VIRUS E5 |
| 81:00:00 | 6e10-1 | 8 | 5.6 | 184 | 719 | 15 | MOLECULE: | HEAT SHOCK PROTEIN 101 |
| 82:00:00 | 7tts-D | 8 | 11.4 | 177 | 463 | 12 | MOLECULE: | CASEINOLYTIC PEPTIDASE B PROTEIN HOMOLOG |
| 83:00:00 | 3uk6-I | 8 | 3.1 | 137 | 294 | 12 | MOLECULE: | RUVB-LIKE 2 |
| 84:00:00 | 3uk6-C | 8 | 3.2 | 146 | 321 | 12 | MOLECULE: | RUVB-LIKE 2 |
| 85:00:00 | 7xbk-H | 8 | 3.2 | 146 | 345 | 12 | MOLECULE: | ISOFORM 2 OF CASEINOLYTIC PEPTIDASE B PROTEIN HOM |
| 86:00:00 | 3s8p-A | 7.9 | 3.3 | 106 | 234 | 10 | MOLECULE: | HISTONE-LYSINE N-METHYLTRANSFERASE SUV420H1 |
| 87:00:00 | 5hq8-A | 7.9 | 3.8 | 121 | 427 | 13 | MOLECULE: | HISTONE-LYSINE N-METHYLTRANSFERASE SMYD3 |

| | | | | | | | | |
|---|---|---|---|---|---|---|---|---|
| 88:00:00 | 3rq4-A | 7.9 | 3.9 | 111 | 241 | 10 | MOLECULE: | HISTONE-LYSINE N-METHYLTRANSFERASE SUV420H2 |
| 89:00:00 | 4c1q-A | 7.9 | 2.6 | 107 | 174 | 16 | MOLECULE: | HISTONE-LYSINE N-METHYLTRANSFERASE PRDM9 |
| 90:00:00 | 7t0v-C | 7.9 | 11.4 | 155 | 570 | 11 | MOLECULE: | RIX7 |
| 91:00:00 | 6hz9-D | 7.9 | 3.7 | 137 | 284 | 11 | MOLECULE: | 5-METHYLCYTOSINE-SPECIFIC RESTRICTION ENZYME B |
| 92:00:00 | 7ola-F | 7.9 | 5.3 | 199 | 345 | 14 | MOLECULE: | DNA PRIMASE |
| 93:00:00 | 3syk-A | 7.9 | 11.3 | 146 | 290 | 11 | MOLECULE: | PROTEIN CBBX |
| 94:00:00 | 1g4a-E | 7.9 | 9.7 | 152 | 356 | 13 | MOLECULE: | ATP-DEPENDENT HSL PROTEASE ATP-BINDING SUBUNIT |
| 95:00:00 | 8hwf-D | 7.9 | 4.2 | 202 | 378 | 10 | MOLECULE: | PRIMASE D5 |
| 96:00:00 | 8hwe-C | 7.9 | 4.3 | 208 | 379 | 9 | MOLECULE: | PRIMASE D5 |
| 97:00:00 | 6po1-B | 7.9 | 12.5 | 164 | 352 | 14 | MOLECULE: | ATP-DEPENDENT CLP PROTEASE ATP-BINDING SUBUNIT CL |
| 98:00:00 | 8hwh-B | 7.9 | 4.6 | 209 | 357 | 11 | MOLECULE: | PRIMASE D5 |
| 99:00:00 | 8hwe-B | 7.9 | 4.4 | 209 | 379 | 10 | MOLECULE: | PRIMASE D5 |
| 0:00 | 6p07-D | 7.9 | 12.1 | 136 | 303 | 13 | MOLECULE: | SPASTIN |
| 1:00 | 6oay-D | 7.9 | 14.1 | 151 | 570 | 10 | MOLECULE: | HYPERACTIVE DISAGGREGASE CLPB |
| 2:00 | 3uk6-K | 7.9 | 3.1 | 136 | 283 | 12 | MOLECULE: | RUVB-LIKE 2 |
| 3:00 | 3uk6-H | 7.9 | 8.9 | 138 | 277 | 15 | MOLECULE: | RUVB-LIKE 2 |
| 4:00 | 3zuh-C | 7.9 | 12.2 | 143 | 286 | 11 | MOLECULE: | PROTEIN CBBX |
| 5:00 | 6e11-4 | 7.9 | 9.1 | 192 | 717 | 15 | MOLECULE: | UNKNOWN (CLAW) |
| 6:00 | 7crq-L | 7.8 | 2.8 | 115 | 223 | 21 | MOLECULE: | HISTONE H3 |
| 7:00 | 4au7-A | 7.8 | 3.8 | 109 | 244 | 12 | MOLECULE: | HISTONE-LYSINE N-METHYLTRANSFERASE SUV420H2 |
| 8:00 | 6fnd-C | 7.8 | 5.8 | 109 | 420 | 13 | MOLECULE: | APICAL COMPLEX LYSINE METHYLTRANSFERASE |
| 9:00 | 6mat-F | 7.8 | 9.8 | 160 | 568 | 12 | MOLECULE: | RIX7 MUTANT |
| 10:00 | 8umt-A | 7.8 | 24 | 162 | 537 | 15 | MOLECULE: | CHROMOSOME TRANSMISSION FIDELITY PROTEIN 18 HOMOL |
| 11:00 | 5wbv-B | 7.8 | 3.6 | 105 | 228 | 11 | MOLECULE: | HISTONE-LYSINE N-METHYLTRANSFERASE KMT5B |
| 12:00 | 5yww-A | 7.8 | 14.4 | 149 | 485 | 11 | MOLECULE: | NUCLEOTIDE BINDING PROTEIN PINC |
| 13:00 | 8hwb-B | 7.8 | 4.4 | 213 | 379 | 10 | MOLECULE: | PRIMASE D5 |
| 14:00 | 8hwb-C | 7.8 | 4.6 | 211 | 379 | 8 | MOLECULE: | PRIMASE D5 |
| 15:00 | 8hwb-A | 7.8 | 3.9 | 206 | 379 | 10 | MOLECULE: | PRIMASE D5 |
| 16:00 | 7xe0-C | 7.8 | 22.9 | 162 | 838 | 14 | MOLECULE: | CNL9 |
| 17:00 | 8xj8-C | 7.8 | 4.2 | 204 | 379 | 9 | MOLECULE: | DNA (70-MER) |
| 18:00 | 2qz4-A | 7.8 | 3.3 | 134 | 223 | 14 | MOLECULE: | PARAPLEGIN |
| 19:00 | 8umy-D | 7.8 | 15.1 | 147 | 329 | 12 | MOLECULE: | CHROMOSOME TRANSMISSION FIDELITY PROTEIN 18 HOMOL |
| 20:00 | 7crq-I | 7.8 | 2.8 | 116 | 223 | 20 | MOLECULE: | HISTONE H3 |
| 21:00 | 7fd4-A | 7.7 | 18.1 | 244 | 779 | 10 | MOLECULE: | LON PROTEASE |
| 22:00 | 8c0v-B | 7.7 | 11.9 | 204 | 1030 | 10 | MOLECULE: | PEROXISOMAL ATPASE PEX1 |
| 23:00 | 8hr8-H | 7.7 | 14.6 | 158 | 834 | 7 | MOLECULE: | ARCHAEAL ATPASE |
| 24:00:00 | 5hyn-K | 7.7 | 2.6 | 119 | 563 | 19 | MOLECULE: | HISTONE-LYSINE N-METHYLTRANSFERASE EZH2 |
| 25:00:00 | 5cpr-B | 7.7 | 4.4 | 107 | 233 | 11 | MOLECULE: | HISTONE-LYSINE N-METHYLTRANSFERASE SUV420H1 |
| 26:00:00 | 7yrd-K | 7.7 | 5.9 | 117 | 260 | 9 | MOLECULE: | HISTONE H3.2 |
| 27:00:00 | 7t3i-B | 7.7 | 13.1 | 165 | 578 | 10 | MOLECULE: | RIX7 |
| 28:00:00 | 3oxf-B | 7.7 | 3.8 | 121 | 426 | 15 | MOLECULE: | SET AND MYND DOMAIN-CONTAINING PROTEIN 3 |
| 29:00:00 | 7z11-D | 7.7 | 9.7 | 189 | 734 | 8 | MOLECULE: | ATPASE FAMILY GENE 2 PROTEIN |
| 30:00:00 | 7lar-F | 7.7 | 5.2 | 128 | 183 | 17 | MOLECULE: | ATP-DEPENDENT HELICASE REP |
| 31:00:00 | 7las-F | 7.7 | 5.2 | 124 | 183 | 16 | MOLECULE: | ATP-DEPENDENT HELICASE REP |
| 32:00:00 | 8apm-B | 7.7 | 6.2 | 221 | 449 | 12 | MOLECULE: | PRIMASE D5 |
| 33:00:00 | 8hwa-D | 7.7 | 4.2 | 202 | 379 | 9 | MOLECULE: | PRIMASE D5 |
| 34:00:00 | 8hwh-D | 7.7 | 4.6 | 204 | 363 | 11 | MOLECULE: | PRIMASE D5 |
| 35:00:00 | 8hwb-E | 7.7 | 4.7 | 223 | 700 | 11 | MOLECULE: | PRIMASE D5 |
| 36:00:00 | 8hwg-D | 7.7 | 4.8 | 213 | 378 | 9 | MOLECULE: | PRIMASE D5 |
| 37:00:00 | 8dr4-A | 7.7 | 10.2 | 144 | 497 | 14 | MOLECULE: | REPLICATION FACTOR C SUBUNIT 1 |
| 38:00:00 | 6fnd-B | 7.7 | 6.2 | 111 | 418 | 13 | MOLECULE: | APICAL COMPLEX LYSINE METHYLTRANSFERASE |
| 39:00:00 | 8xj8-A | 7.7 | 6.2 | 214 | 422 | 10 | MOLECULE: | DNA (70-MER) |
| 40:00:00 | 6opc-B | 7.7 | 9.4 | 166 | 741 | 10 | MOLECULE: | CELL DIVISION CONTROL PROTEIN 48 |

| | | | | | | | | |
|---|---|---|---|---|---|---|---|---|
| 41:00:00 | 6mat-B | 7.7 | 12.9 | 170 | 578 | 12 | MOLECULE: | RIX7 MUTANT |
| 42:00:00 | 3jc7-7 | 7.7 | 8.9 | 211 | 657 | 10 | MOLECULE: | DNA REPLICATION LICENSING FACTOR MCM2 |
| 43:00:00 | 2chq-A | 7.7 | 14.7 | 144 | 313 | 13 | MOLECULE: | REPLICATION FACTOR C SMALL SUBUNIT |
| 44:00:00 | 6z1f-2 | 7.7 | 11.3 | 145 | 281 | 11 | MOLECULE: | RIBULOSE BISPHOSPHATE CARBOXYLASE/OXYGENASE ACTIV |
| 45:00:00 | 7jlv-D | 7.7 | 12.4 | 131 | 437 | 17 | MOLECULE: | DISEASE RESISTANCE PROTEIN ROQ1 |
| 46:00:00 | 6igm-D | 7.7 | 4.6 | 135 | 396 | 13 | MOLECULE: | RUVB-LIKE 1 |
| 47:00:00 | 5fm6-B | 7.7 | 11.2 | 147 | 419 | 14 | MOLECULE: | RVB1 |
| 48:00:00 | 6epc-L | 7.7 | 19.5 | 159 | 389 | 10 | MOLECULE: | PROTEASOME SUBUNIT ALPHA TYPE-6 |
| 49:00:00 | 7xbk-A | 7.7 | 3.6 | 147 | 367 | 14 | MOLECULE: | ISOFORM 2 OF CASEINOLYTIC PEPTIDASE B PROTEIN HOM |
| 50:00:00 | 6oa9-B | 7.7 | 8.6 | 168 | 493 | 7 | MOLECULE: | CELL DIVISION CONTROL PROTEIN 48 |
| 51:00:00 | 5ji3-F | 7.7 | 9.6 | 154 | 353 | 14 | MOLECULE: | ATP-DEPENDENT PROTEASE SUBUNIT HSLV |
| 52:00:00 | 2fuf-A | 7.7 | 3.3 | 104 | 126 | 7 | MOLECULE: | LARGE T ANTIGEN |
| 53:00:00 | 2ipr-B | 7.7 | 3.3 | 105 | 124 | 7 | MOLECULE: | LARGE T ANTIGEN |
| 54:00:00 | 8hwa-A | 7.6 | 4.2 | 226 | 700 | 10 | MOLECULE: | PRIMASE D5 |
| 55:00:00 | 4ijd-A | 7.6 | 4.8 | 118 | 215 | 14 | MOLECULE: | HISTONE-LYSINE N-METHYLTRANSFERASE PRDM9 |
| 56:00:00 | 7cro-I | 7.6 | 2.6 | 113 | 220 | 22 | MOLECULE: | HISTONE H3 |
| 57:00:00 | 2h2e-C | 7.6 | 22.7 | 123 | 440 | 13 | MOLECULE: | RIBULOSE-1,5 BISPHOSPHATE CARBOXYLASE/OXYGENASE L |
| 58:00:00 | 8hr9-K | 7.6 | 15.3 | 150 | 836 | 10 | MOLECULE: | ARCHAEAL ATPASE |
| 59:00:00 | 3ihx-A | 7.6 | 2.7 | 96 | 126 | 15 | MOLECULE: | PR DOMAIN ZINC FINGER PROTEIN 10 |
| 60:00:00 | 2jv0-A | 7.6 | 3.4 | 108 | 163 | 16 | MOLECULE: | PR DOMAIN ZINC FINGER PROTEIN 2 |
| 61:00:00 | 6hz9-L | 7.6 | 3.7 | 137 | 285 | 11 | MOLECULE: | 5-METHYLCYTOSINE-SPECIFIC RESTRICTION ENZYME B |
| 62:00:00 | 8hwa-E | 7.6 | 6.2 | 208 | 379 | 10 | MOLECULE: | PRIMASE D5 |
| 63:00:00 | 7w46-A | 7.6 | 6.8 | 176 | 396 | 10 | MOLECULE: | UNCHARACTERIZED ATPASE YJOB |
| 64:00:00 | 7xe0-A | 7.6 | 24.6 | 165 | 838 | 15 | MOLECULE: | CNL9 |
| 65:00:00 | 6hz5-J | 7.6 | 3.7 | 138 | 284 | 12 | MOLECULE: | 5-METHYLCYTOSINE-SPECIFIC RESTRICTION ENZYME B |
| 66:00:00 | 7xe0-G | 7.6 | 22.8 | 163 | 838 | 15 | MOLECULE: | CNL9 |
| 67:00:00 | 4rh7-A | 7.6 | 18.3 | 217 | 3005 | 8 | MOLECULE: | GREEN FLUORESCENT PROTEIN/CYTOPLASMIC DYNEIN 2 HE |
| 68:00:00 | 5bk4-3 | 7.6 | 7.8 | 244 | 604 | 7 | MOLECULE: | DNA REPLICATION LICENSING FACTOR MCM2 |
| 69:00:00 | 7xe0-I | 7.6 | 24.5 | 165 | 838 | 15 | MOLECULE: | CNL9 |
| 70:00:00 | 6rn2-B | 7.6 | 5 | 187 | 689 | 16 | MOLECULE: | CHAPERONE PROTEIN CLPB |
| 71:00:00 | 8btg-B | 7.6 | 12.7 | 140 | 335 | 12 | MOLECULE: | CHROMOSOMAL REPLICATION INITIATOR PROTEIN DNAA |
| 72:00:00 | 3m6a-A | 7.6 | 10.6 | 166 | 496 | 14 | MOLECULE: | ATP-DEPENDENT PROTEASE LA 1 |
| 73:00:00 | 6qi9-A | 7.6 | 3.4 | 142 | 317 | 11 | MOLECULE: | RUVB-LIKE 1 |
| 74:00:00 | 6hz9-F | 7.6 | 3.7 | 137 | 285 | 11 | MOLECULE: | 5-METHYLCYTOSINE-SPECIFIC RESTRICTION ENZYME B |
| 75:00:00 | 2nl8-A | 7.6 | 3.2 | 99 | 117 | 8 | MOLECULE: | 18-NT PEN ELEMENT OF THE SV40 DNA ORIGIN |
| 76:00:00 | 3qn2-A | 7.6 | 3.3 | 105 | 126 | 9 | MOLECULE: | LARGE T ANTIGEN |
| 77:00:00 | 4fb3-B | 7.6 | 3.1 | 100 | 114 | 11 | MOLECULE: | ORI DNA OLIGONUCLEOTIDE-CRICK STRAND |
| 78:00:00 | 7crc-B | 7.5 | 8.9 | 177 | 1112 | 10 | MOLECULE: | NAD+ HYDROLASE (NADASE) |
| 79:00:00 | 2g46-A | 7.5 | 3 | 97 | 120 | 16 | MOLECULE: | PBCV-1 HISTONE H3-LYS 27 METHYLTRANSFERASE |
| 80:00:00 | 8v48-A | 7.5 | 3.5 | 109 | 412 | 15 | MOLECULE: | ARIA ANTITOXIN |
| 81:00:00 | 5v8f-6 | 7.5 | 5.9 | 231 | 692 | 10 | MOLECULE: | DNA REPLICATION LICENSING FACTOR MCM2 |
| 82:00:00 | 6uks-G | 7.5 | 4.3 | 153 | 362 | 15 | MOLECULE: | MITOCHONDRIAL CHAPERONE BCS1 |
| 83:00:00 | 6rn4-E | 7.5 | 10.5 | 190 | 684 | 13 | MOLECULE: | CHAPERONE PROTEIN CLPB |
| 84:00:00 | 6uks-B | 7.5 | 3.9 | 149 | 362 | 16 | MOLECULE: | MITOCHONDRIAL CHAPERONE BCS1 |
| 85:00:00 | 5udb-7 | 7.5 | 6.7 | 236 | 726 | 10 | MOLECULE: | DNA REPLICATION LICENSING FACTOR MCM2 |
| 86:00:00 | 7tfh-A | 7.5 | 19.1 | 181 | 628 | 9 | MOLECULE: | REPLICATION FACTOR C SUBUNIT 1 |
| 87:00:00 | 8hwa-F | 7.5 | 5 | 272 | 700 | 9 | MOLECULE: | PRIMASE D5 |
| 88:00:00 | 8hwb-D | 7.5 | 10.5 | 263 | 700 | 10 | MOLECULE: | PRIMASE D5 |
| 89:00:00 | 7w42-A | 7.5 | 7.2 | 182 | 397 | 13 | MOLECULE: | UNCHARACTERIZED ATPASE YJOB |
| 90:00:00 | 5grb-B | 7.5 | 4 | 128 | 212 | 8 | MOLECULE: | EV71 2C ATPASE |
| 91:00:00 | 7xe0-E | 7.5 | 24.5 | 167 | 838 | 14 | MOLECULE: | CNL9 |
| 92:00:00 | 8v45-F | 7.5 | 3.2 | 107 | 378 | 14 | MOLECULE: | ARIA ANTITOXIN |
| 93:00:00 | 6hz4-A | 7.5 | 3.8 | 137 | 290 | 12 | MOLECULE: | 5-METHYLCYTOSINE-SPECIFIC RESTRICTION ENZYME B |

| | | | | | | | | |
|---|---|---|---|---|---|---|---|---|
| 94:00:00 | 8ui7-A | 7.5 | 17.9 | 167 | 643 | 10 | MOLECULE: | ATPASE FAMILY AAA DOMAIN-CONTAINING PROTEIN 5 |
| 95:00:00 | 6rn2-A | 7.5 | 4.4 | 191 | 684 | 14 | MOLECULE: | CHAPERONE PROTEIN CLPB |
| 96:00:00 | 8dqx-A | 7.5 | 10.6 | 144 | 497 | 13 | MOLECULE: | REPLICATION FACTOR C SUBUNIT 1 |
| 97:00:00 | 1ixs-B | 7.5 | 10.1 | 146 | 315 | 13 | MOLECULE: | HOLLIDAY JUNCTION DNA HELICASE RUVA |
| 98:00:00 | 2xsz-D | 7.5 | 3.3 | 138 | 310 | 12 | MOLECULE: | RUVB-LIKE 1 |
| 99:00:00 | 2xsz-F | 7.5 | 3.2 | 139 | 311 | 14 | MOLECULE: | RUVB-LIKE 1 |
| 0:00 | 3uk6-G | 7.5 | 3.7 | 148 | 330 | 12 | MOLECULE: | RUVB-LIKE 2 |
| 1:00 | 3glf-D | 7.5 | 12.4 | 144 | 362 | 12 | MOLECULE: | DNA POLYMERASE III SUBUNIT DELTA |
| 2:00 | 3gli-I | 7.5 | 12.4 | 152 | 362 | 12 | MOLECULE: | DNA POLYMERASE III SUBUNIT DELTA |
| 3:00 | 6hz4-D | 7.5 | 3.7 | 138 | 284 | 12 | MOLECULE: | 5-METHYLCYTOSINE-SPECIFIC RESTRICTION ENZYME B |
| 4:00 | 6etx-B | 7.5 | 8.7 | 142 | 429 | 13 | MOLECULE: | RUVB-LIKE 1 |
| 5:00 | 6pek-A | 7.5 | 6.9 | 132 | 283 | 9 | MOLECULE: | SPASTIN |
| 6:00 | 6etx-F | 7.5 | 8.8 | 143 | 434 | 12 | MOLECULE: | RUVB-LIKE 1 |
| 7:00 | 8ui9-A | 7.5 | 23.7 | 169 | 643 | 9 | MOLECULE: | ATPASE FAMILY AAA DOMAIN-CONTAINING PROTEIN 5 |
| 8:00 | 6pen-A | 7.5 | 6.9 | 132 | 283 | 9 | MOLECULE: | SPASTIN |
| 9:00 | 6qs8-F | 7.5 | 15.3 | 213 | 573 | 12 | MOLECULE: | CHAPERONE PROTEIN CLPB |
| 10:00 | 2ntc-A | 7.5 | 3.4 | 106 | 127 | 8 | MOLECULE: | 21-NT PEN ELEMENT OF THE SV40 DNA ORIGIN |
| 11:00 | 4fb3-A | 7.5 | 3 | 97 | 115 | 11 | MOLECULE: | ORI DNA OLIGONUCLEOTIDE-CRICK STRAND |
| 12:00 | 2itl-A | 7.5 | 3.3 | 101 | 120 | 8 | MOLECULE: | 24-NT PEN ELEMENT OF THE SV40 DNA ORIGIN |
| 13:00 | 2ipr-A | 7.5 | 3.3 | 103 | 127 | 8 | MOLECULE: | LARGE T ANTIGEN |
| 14:00 | 4nbp-A | 7.5 | 3.6 | 108 | 129 | 9 | MOLECULE: | LARGE T ANTIGEN |
| 15:00 | 4fb3-E | 7.5 | 3.2 | 99 | 114 | 11 | MOLECULE: | ORI DNA OLIGONUCLEOTIDE-CRICK STRAND |
| 16:00 | 4lmd-B | 7.5 | 3.7 | 108 | 130 | 10 | MOLECULE: | LARGE T ANTIGEN |
| 17:00 | 4lmd-A | 7.5 | 3.7 | 110 | 128 | 9 | MOLECULE: | LARGE T ANTIGEN |
| 18:00 | 6fnd-A | 7.4 | 5.3 | 109 | 423 | 13 | MOLECULE: | APICAL COMPLEX LYSINE METHYLTRANSFERASE |
| 19:00 | 8hr9-E | 7.4 | 10.3 | 148 | 833 | 8 | MOLECULE: | ARCHAEAL ATPASE |
| 20:00 | 7wbb-A | 7.4 | 13.1 | 188 | 726 | 7 | MOLECULE: | AFG2 ISOFORM 1 |
| 21:00 | 3ihx-B | 7.4 | 3 | 100 | 129 | 14 | MOLECULE: | PR DOMAIN ZINC FINGER PROTEIN 10 |
| 22:00 | 8hr9-D | 7.4 | 15 | 146 | 836 | 8 | MOLECULE: | ARCHAEAL ATPASE |
| 23:00 | 7m9b-D | 7.4 | 9.2 | 135 | 257 | 10 | MOLECULE: | TNSC |
| 24:00:00 | 4ijd-B | 7.4 | 5.4 | 121 | 215 | 14 | MOLECULE: | HISTONE-LYSINE N-METHYLTRANSFERASE PRDM9 |
| 25:00:00 | 3tg4-A | 7.4 | 3.6 | 118 | 428 | 12 | MOLECULE: | N-LYSINE METHYLTRANSFERASE SMYD2 |
| 26:00:00 | 6hz7-F | 7.4 | 3.8 | 137 | 285 | 11 | MOLECULE: | 5-METHYLCYTOSINE-SPECIFIC RESTRICTION ENZYME B |
| 27:00:00 | 7swl-A | 7.4 | 8.9 | 159 | 553 | 11 | MOLECULE: | RIX7 |
| 28:00:00 | 6l1q-A | 7.4 | 6.7 | 138 | 267 | 11 | MOLECULE: | CBBQ PROTEIN |
| 29:00:00 | 5m7o-A | 7.4 | 13.3 | 162 | 448 | 9 | MOLECULE: | NITROGEN ASSIMILATION REGULATORY PROTEIN |
| 30:00:00 | 3pvs-D | 7.4 | 11.2 | 139 | 424 | 12 | MOLECULE: | REPLICATION-ASSOCIATED RECOMBINATION PROTEIN A |
| 31:00:00 | 8dr3-A | 7.4 | 15.5 | 168 | 643 | 11 | MOLECULE: | REPLICATION FACTOR C SUBUNIT 1 |
| 32:00:00 | 8hwh-A | 7.4 | 4.1 | 198 | 353 | 12 | MOLECULE: | PRIMASE D5 |
| 33:00:00 | 2uz3-B | 7.4 | 9.8 | 111 | 207 | 9 | MOLECULE: | THYMIDINE KINASE |
| 34:00:00 | 8hwe-E | 7.4 | 4 | 200 | 372 | 10 | MOLECULE: | PRIMASE D5 |
| 35:00:00 | 5vqa-A | 7.4 | 4.2 | 156 | 368 | 13 | MOLECULE: | PACHYTENE CHECKPOINT PROTEIN 2 HOMOLOG |
| 36:00:00 | 6hz7-L | 7.4 | 3.7 | 135 | 285 | 12 | MOLECULE: | 5-METHYLCYTOSINE-SPECIFIC RESTRICTION ENZYME B |
| 37:00:00 | 1sxj-B | 7.4 | 13.3 | 148 | 316 | 12 | MOLECULE: | ACTIVATOR 1 95 KDA SUBUNIT |
| 38:00:00 | 6rn2-C | 7.4 | 5.6 | 180 | 689 | 14 | MOLECULE: | CHAPERONE PROTEIN CLPB |
| 39:00:00 | 5bk4-7 | 7.4 | 5.5 | 219 | 689 | 10 | MOLECULE: | DNA REPLICATION LICENSING FACTOR MCM2 |
| 40:00:00 | 1iqp-A | 7.4 | 16 | 157 | 326 | 13 | MOLECULE: | RFCS |
| 41:00:00 | 4xgu-D | 7.4 | 4.5 | 168 | 377 | 13 | MOLECULE: | PUTATIVE PACHYTENE CHECKPOINT PROTEIN 2 |
| 42:00:00 | 2xsz-E | 7.4 | 3.2 | 139 | 315 | 14 | MOLECULE: | RUVB-LIKE 1 |
| 43:00:00 | 3gli-D | 7.4 | 9.8 | 150 | 362 | 11 | MOLECULE: | DNA POLYMERASE III SUBUNIT DELTA |
| 44:00:00 | 2x8a-A | 7.4 | 6.9 | 133 | 237 | 14 | MOLECULE: | NUCLEAR VALOSIN-CONTAINING PROTEIN-LIKE |
| 45:00:00 | 6hz5-L | 7.4 | 3.7 | 137 | 285 | 11 | MOLECULE: | 5-METHYLCYTOSINE-SPECIFIC RESTRICTION ENZYME B |
| 46:00:00 | 3bos-B | 7.4 | 3.5 | 133 | 231 | 13 | MOLECULE: | PUTATIVE DNA REPLICATION FACTOR |

| | | | | | | | | |
|---|---|---|---|---|---|---|---|---|
| 47:00:00 | 4fgn-B | 7.4 | 3.7 | 109 | 124 | 7 | MOLECULE: | LARGE T ANTIGEN |
| 48:00:00 | 5d9i-B | 7.4 | 3.4 | 105 | 132 | 7 | MOLECULE: | LARGE T ANTIGEN |
| 49:00:00 | 5cyn-A | 7.4 | 3.4 | 106 | 127 | 8 | MOLECULE: | LARGE T ANTIGEN |
| 50:00:00 | 2ntc-B | 7.4 | 3.4 | 106 | 126 | 8 | MOLECULE: | 21-NT PEN ELEMENT OF THE SV40 DNA ORIGIN |
| 51:00:00 | 2if9-A | 7.4 | 3.4 | 106 | 132 | 8 | MOLECULE: | LARGE T ANTIGEN |
| 52:00:00 | 5d9i-A | 7.4 | 3.5 | 106 | 133 | 8 | MOLECULE: | LARGE T ANTIGEN |
| 53:00:00 | 3qk2-A | 7.4 | 3.5 | 106 | 132 | 8 | MOLECULE: | LARGE T ANTIGEN |
| 54:00:00 | 6hz5-C | 7.3 | 3.7 | 139 | 290 | 11 | MOLECULE: | 5-METHYLCYTOSINE-SPECIFIC RESTRICTION ENZYME B |
| 55:00:00 | 7t0v-D | 7.3 | 9.5 | 143 | 570 | 13 | MOLECULE: | RIX7 |
| 56:00:00 | 6sh3-D | 7.3 | 14.6 | 170 | 376 | 11 | MOLECULE: | MITOCHONDRIAL CHAPERONE BCS1 |
| 57:00:00 | 6uks-C | 7.3 | 4.1 | 154 | 362 | 16 | MOLECULE: | MITOCHONDRIAL CHAPERONE BCS1 |
| 58:00:00 | 7upr-A | 7.3 | 9.4 | 134 | 235 | 13 | MOLECULE: | OUTER MITOCHONDRIAL TRANSMEMBRANE HELIX TRANSLOCA |
| 59:00:00 | 6hz4-F | 7.3 | 3.8 | 137 | 285 | 11 | MOLECULE: | 5-METHYLCYTOSINE-SPECIFIC RESTRICTION ENZYME B |
| 60:00:00 | 5j1s-A | 7.3 | 3.1 | 142 | 277 | 14 | MOLECULE: | TORSIN-1A |
| 61:00:00 | 6pod-A | 7.3 | 10.8 | 150 | 329 | 15 | MOLECULE: | ATP-DEPENDENT CLP PROTEASE ATP-BINDING SUBUNIT CL |
| 62:00:00 | 8hwb-F | 7.3 | 4.4 | 205 | 379 | 11 | MOLECULE: | PRIMASE D5 |
| 63:00:00 | 6hz6-J | 7.3 | 3.7 | 137 | 284 | 11 | MOLECULE: | 5-METHYLCYTOSINE-SPECIFIC RESTRICTION ENZYME B |
| 64:00:00 | 8hwc-A | 7.3 | 6.8 | 207 | 378 | 12 | MOLECULE: | PRIMASE D5 |
| 65:00:00 | 6rn3-C | 7.3 | 4.1 | 189 | 689 | 14 | MOLECULE: | CHAPERONE PROTEIN CLPB |
| 66:00:00 | 6l8d-A | 7.3 | 4.3 | 148 | 267 | 13 | MOLECULE: | MAGNESIUM-CHELATASE SUBUNIT CHLI |
| 67:00:00 | 6oay-C | 7.3 | 13.9 | 154 | 570 | 9 | MOLECULE: | HYPERACTIVE DISAGGREGASE CLPB |
| 68:00:00 | 4xgu-A | 7.3 | 5.7 | 162 | 376 | 14 | MOLECULE: | PUTATIVE PACHYTENE CHECKPOINT PROTEIN 2 |
| 69:00:00 | 5ifw-B | 7.3 | 17.5 | 175 | 702 | 9 | MOLECULE: | TETHER CONTAINING UBX DOMAIN FOR GLUT4 |
| 70:00:00 | 2rko-A | 7.3 | 14 | 144 | 280 | 13 | MOLECULE: | VACUOLAR PROTEIN SORTING-ASSOCIATED PROTEIN 4 |
| 71:00:00 | 3glg-C | 7.3 | 12 | 141 | 365 | 11 | MOLECULE: | DNA POLYMERASE III SUBUNIT DELTA |
| 72:00:00 | 3glf-I | 7.3 | 12 | 142 | 362 | 12 | MOLECULE: | DNA POLYMERASE III SUBUNIT DELTA |
| 73:00:00 | 3glg-I | 7.3 | 9.4 | 145 | 362 | 11 | MOLECULE: | DNA POLYMERASE III SUBUNIT DELTA |
| 74:00:00 | 6vvo-E | 7.3 | 3.2 | 127 | 346 | 9 | MOLECULE: | REPLICATION FACTOR C SUBUNIT 1 |
| 75:00:00 | 3glf-H | 7.3 | 9.8 | 140 | 365 | 11 | MOLECULE: | DNA POLYMERASE III SUBUNIT DELTA |
| 76:00:00 | 3gli-C | 7.3 | 11.8 | 143 | 365 | 11 | MOLECULE: | DNA POLYMERASE III SUBUNIT DELTA |
| 77:00:00 | 3gli-H | 7.3 | 11.6 | 143 | 365 | 12 | MOLECULE: | DNA POLYMERASE III SUBUNIT DELTA |
| 78:00:00 | 3glg-H | 7.3 | 11.8 | 140 | 365 | 12 | MOLECULE: | DNA POLYMERASE III SUBUNIT DELTA |
| 79:00:00 | 3glg-D | 7.3 | 12.7 | 141 | 362 | 11 | MOLECULE: | DNA POLYMERASE III SUBUNIT DELTA |
| 80:00:00 | 3glf-C | 7.3 | 11.6 | 141 | 365 | 12 | MOLECULE: | DNA POLYMERASE III SUBUNIT DELTA |
| 81:00:00 | 1sxj-A | 7.3 | 4.1 | 142 | 441 | 13 | MOLECULE: | ACTIVATOR 1 95 KDA SUBUNIT |
| 82:00:00 | 1w4r-A | 7.3 | 3.5 | 98 | 174 | 10 | MOLECULE: | THYMIDINE KINASE |
| 83:00:00 | 6pen-C | 7.3 | 10.1 | 137 | 288 | 11 | MOLECULE: | SPASTIN |
| 84:00:00 | 6pek-C | 7.3 | 10.1 | 137 | 288 | 11 | MOLECULE: | SPASTIN |
| 85:00:00 | 6pen-F | 7.3 | 11.8 | 140 | 272 | 11 | MOLECULE: | SPASTIN |
| 86:00:00 | 6p8v-F | 7.3 | 3.6 | 138 | 270 | 15 | MOLECULE: | ATPASE, AAA FAMILY |
| 87:00:00 | 6hz6-D | 7.3 | 3.7 | 137 | 284 | 11 | MOLECULE: | 5-METHYLCYTOSINE-SPECIFIC RESTRICTION ENZYME B |
| 88:00:00 | 4lif-A | 7.3 | 3.6 | 107 | 127 | 7 | MOLECULE: | LARGE T ANTIGEN |
| 89:00:00 | 2if9-B | 7.3 | 3.7 | 107 | 131 | 7 | MOLECULE: | LARGE T ANTIGEN |
| 90:00:00 | 2itj-A | 7.3 | 3.3 | 105 | 128 | 8 | MOLECULE: | LARGE T ANTIGEN |
| 91:00:00 | 2itl-B | 7.3 | 3.3 | 100 | 115 | 8 | MOLECULE: | 24-NT PEN ELEMENT OF THE SV40 DNA ORIGIN |
| 92:00:00 | 3qfq-A | 7.3 | 3.5 | 102 | 119 | 7 | MOLECULE: | LARGE T ANTIGEN |
| 93:00:00 | 7vcs-A | 7.2 | 7 | 180 | 766 | 8 | MOLECULE: | TRANSITIONAL ENDOPLASMIC RETICULUM ATPASE |
| 94:00:00 | 2h21-B | 7.2 | 16.4 | 123 | 441 | 12 | MOLECULE: | RIBULOSE-1,5 BISPHOSPHATE CARBOXYLASE/OXYGENASE |
| 95:00:00 | 8hr8-A | 7.2 | 15.6 | 149 | 836 | 9 | MOLECULE: | ARCHAEAL ATPASE |
| 96:00:00 | 6j6i-O | 7.2 | 21.9 | 150 | 790 | 11 | MOLECULE: | PROBABLE SERINE/THREONINE-PROTEIN KINASE PBL2 |
| 97:00:00 | 8hrb-B | 7.2 | 15.5 | 144 | 835 | 8 | MOLECULE: | ARCHAEAL ATPASE |
| 98:00:00 | 6hz6-F | 7.2 | 3.7 | 137 | 285 | 11 | MOLECULE: | 5-METHYLCYTOSINE-SPECIFIC RESTRICTION ENZYME B |
| 99:00:00 | 6sh4-F | 7.2 | 6.3 | 141 | 376 | 14 | MOLECULE: | MITOCHONDRIAL CHAPERONE BCS1 |

| | | | | | | | | |
|---|---|---|---|---|---|---|---|---|
| 0:00 | 5e7p-B | 7.2 | 16 | 165 | 704 | 11 | MOLECULE: | CELL DIVISION CONTROL PROTEIN CDC48 |
| 1:00 | 5bq5-B | 7.2 | 3.5 | 126 | 187 | 14 | MOLECULE: | INSERTION SEQUENCE IS5376 PUTATIVE ATP-BINDING PR |
| 2:00 | 7lar-A | 7.2 | 4.9 | 127 | 183 | 17 | MOLECULE: | ATP-DEPENDENT HELICASE REP |
| 3:00 | 5ex0-A | 7.2 | 4.4 | 125 | 430 | 14 | MOLECULE: | HISTONE-LYSINE N-METHYLTRANSFERASE SMYD3 |
| 4:00 | 6rn3-A | 7.2 | 12.7 | 153 | 684 | 8 | MOLECULE: | CHAPERONE PROTEIN CLPB |
| 5:00 | 8q6p-2 | 7.2 | 4.9 | 176 | 431 | 9 | MOLECULE: | DNA REPLICATION LICENSING FACTOR MCM2 |
| 6:00 | 1sxj-D | 7.2 | 14.5 | 140 | 328 | 11 | MOLECULE: | ACTIVATOR 1 95 KDA SUBUNIT |
| 7:00 | 3glg-B | 7.2 | 14.8 | 141 | 364 | 14 | MOLECULE: | DNA POLYMERASE III SUBUNIT DELTA |
| 8:00 | 8jx6-A | 7.2 | 6.8 | 128 | 447 | 10 | MOLECULE: | DEEP-SEA HELICASE 9 |
| 9:00 | 5ji3-E | 7.2 | 11.4 | 159 | 375 | 13 | MOLECULE: | ATP-DEPENDENT PROTEASE SUBUNIT HSLV |
| 10:00 | 6hz6-L | 7.2 | 3.7 | 137 | 285 | 11 | MOLECULE: | 5-METHYLCYTOSINE-SPECIFIC RESTRICTION ENZYME B |
| 11:00 | 6hz4-C | 7.2 | 3.8 | 139 | 290 | 11 | MOLECULE: | 5-METHYLCYTOSINE-SPECIFIC RESTRICTION ENZYME B |
| 12:00 | 6hz7-J | 7.2 | 3.7 | 137 | 284 | 11 | MOLECULE: | 5-METHYLCYTOSINE-SPECIFIC RESTRICTION ENZYME B |
| 13:00 | 7y53-C | 7.2 | 7.8 | 187 | 741 | 11 | MOLECULE: | DERLIN-1 |
| 14:00 | 7qv9-d | 7.2 | 7.3 | 148 | 256 | 9 | MOLECULE: | DNA-DIRECTED RNA POLYMERASE SUBUNIT ALPHA |
| 15:00 | 3zuh-D | 7.2 | 12.3 | 143 | 286 | 10 | MOLECULE: | PROTEIN CBBX |
| 16:00 | 4xgc-E | 7.2 | 8.6 | 136 | 402 | 10 | MOLECULE: | ORIGIN RECOGNITION COMPLEX SUBUNIT 2 |
| 17:00 | 6qs8-A | 7.2 | 7.9 | 210 | 568 | 14 | MOLECULE: | CHAPERONE PROTEIN CLPB |
| 18:00 | 3qfq-E | 7.2 | 3.3 | 100 | 117 | 6 | MOLECULE: | LARGE T ANTIGEN |
| 19:00 | 3qfq-B | 7.2 | 3.5 | 106 | 120 | 8 | MOLECULE: | LARGE T ANTIGEN |
| 20:00 | 7sh2-A | 7.1 | 11.3 | 163 | 405 | 12 | MOLECULE: | CHECKPOINT PROTEIN RAD24 |
| 21:00 | 8hr9-J | 7.1 | 15.7 | 147 | 835 | 7 | MOLECULE: | ARCHAEAL ATPASE |
| 22:00 | 7t3i-F | 7.1 | 9.5 | 167 | 552 | 12 | MOLECULE: | RIX7 |
| 23:00 | 8hr8-F | 7.1 | 15.6 | 145 | 835 | 10 | MOLECULE: | ARCHAEAL ATPASE |
| 24:00:00 | 8fcw-V | 7.1 | 7.9 | 140 | 343 | 9 | MOLECULE: | TNSC |
| 25:00:00 | 8fcx-U | 7.1 | 8.1 | 147 | 343 | 7 | MOLECULE: | TNSC |
| 26:00:00 | 7bj1-A | 7.1 | 3.8 | 121 | 427 | 14 | MOLECULE: | HISTONE-LYSINE N-METHYLTRANSFERASE SMYD3 |
| 27:00:00 | 6rn3-B | 7.1 | 9.1 | 195 | 689 | 12 | MOLECULE: | CHAPERONE PROTEIN CLPB |
| 28:00:00 | 3glf-G | 7.1 | 16.3 | 158 | 378 | 13 | MOLECULE: | DNA POLYMERASE III SUBUNIT DELTA |
| 29:00:00 | 7st9-E | 7.1 | 3.2 | 129 | 354 | 11 | MOLECULE: | CHECKPOINT PROTEIN RAD24 |
| 30:00:00 | 8xj7-D | 7.1 | 6.5 | 218 | 452 | 10 | MOLECULE: | MONKEYPOX VIRUS E5 |
| 31:00:00 | 6rn4-A | 7.1 | 12.9 | 151 | 675 | 11 | MOLECULE: | CHAPERONE PROTEIN CLPB |
| 32:00:00 | 6opc-F | 7.1 | 14.6 | 187 | 681 | 9 | MOLECULE: | CELL DIVISION CONTROL PROTEIN 48 |
| 33:00:00 | 5vca-P | 7.1 | 10.4 | 177 | 543 | 8 | MOLECULE: | VCP-LIKE ATPASE |
| 34:00:00 | 3glh-G | 7.1 | 11.1 | 145 | 364 | 10 | MOLECULE: | DNA POLYMERASE III SUBUNIT DELTA |
| 35:00:00 | 3glf-B | 7.1 | 13.6 | 146 | 364 | 12 | MOLECULE: | DNA POLYMERASE III SUBUNIT DELTA |
| 36:00:00 | 3glg-G | 7.1 | 16.4 | 159 | 378 | 13 | MOLECULE: | DNA POLYMERASE III SUBUNIT DELTA |
| 37:00:00 | 6pen-D | 7.1 | 9.2 | 140 | 293 | 11 | MOLECULE: | SPASTIN |
| 38:00:00 | 4w5w-A | 7.1 | 12.7 | 147 | 269 | 12 | MOLECULE: | RIBULOSE BISPHOSPHATE CARBOXYLASE/OXYGENASE ACTIV |
| 39:00:00 | 6pek-D | 7.1 | 9.2 | 140 | 293 | 11 | MOLECULE: | SPASTIN |
| 40:00:00 | 6igm-F | 7.1 | 4.7 | 141 | 366 | 13 | MOLECULE: | RUVB-LIKE 1 |
| 41:00:00 | 5bk4-5 | 7.1 | 5.6 | 199 | 661 | 9 | MOLECULE: | DNA REPLICATION LICENSING FACTOR MCM2 |
| 42:00:00 | 6hz5-D | 7.1 | 3.7 | 138 | 284 | 12 | MOLECULE: | 5-METHYLCYTOSINE-SPECIFIC RESTRICTION ENZYME B |
| 43:00:00 | 3zuh-B | 7.1 | 11.3 | 138 | 286 | 10 | MOLECULE: | PROTEIN CBBX |
| 44:00:00 | 2itj-B | 7.1 | 3.2 | 96 | 118 | 8 | MOLECULE: | LARGE T ANTIGEN |
| 45:00:00 | 3j3r-A | 7 | 19.1 | 227 | 798 | 13 | MOLECULE: | ADAPTER PROTEIN MECA 1 |
| 46:00:00 | 7stb-A | 7 | 7.6 | 153 | 515 | 15 | MOLECULE: | CHECKPOINT PROTEIN RAD24 |
| 47:00:00 | 8fcw-S | 7 | 6.5 | 141 | 343 | 9 | MOLECULE: | TNSC |
| 48:00:00 | 3ihx-D | 7 | 2.9 | 89 | 116 | 12 | MOLECULE: | PR DOMAIN ZINC FINGER PROTEIN 10 |
| 49:00:00 | 7y4w-C | 7 | 8 | 191 | 744 | 11 | MOLECULE: | DERLIN-1 |
| 50:00:00 | 6f0x-B | 7 | 3.9 | 155 | 398 | 13 | MOLECULE: | PACHYTENE CHECKPOINT PROTEIN 2 HOMOLOG |
| 51:00:00 | 6qi8-E | 7 | 8.7 | 150 | 335 | 11 | MOLECULE: | RUVB-LIKE 1 |
| 52:00:00 | 4ypl-C | 7 | 8.7 | 189 | 538 | 13 | MOLECULE: | LON PROTEASE |

| | | | | | | | | |
|---|---|---|---|---|---|---|---|---|
| 53:00:00 | 8osg-A | 7 | 4.8 | 162 | 283 | 10 | MOLECULE: | MAGNESIUM-CHELATASE SUBUNIT CHLI |
| 54:00:00 | 8fcv-R | 7 | 7.9 | 149 | 343 | 8 | MOLECULE: | DNA (60-MER) |
| 55:00:00 | 6j6i-F | 7 | 23.2 | 156 | 790 | 8 | MOLECULE: | PROBABLE SERINE/THREONINE-PROTEIN KINASE PBL2 |
| 56:00:00 | 6j5t-G | 7 | 24.8 | 140 | 808 | 9 | MOLECULE: | PROBABLE SERINE/THREONINE-PROTEIN KINASE PBL2 |
| 57:00:00 | 6oaa-D | 7 | 5.8 | 156 | 537 | 12 | MOLECULE: | CELL DIVISION CONTROL PROTEIN 48 |
| 58:00:00 | 6j6i-G | 7 | 23.9 | 155 | 790 | 8 | MOLECULE: | PROBABLE SERINE/THREONINE-PROTEIN KINASE PBL2 |
| 59:00:00 | 5vc7-F | 7 | 9.6 | 179 | 544 | 10 | MOLECULE: | VCP-LIKE ATPASE |
| 60:00:00 | 1sxj-C | 7 | 13 | 143 | 322 | 10 | MOLECULE: | ACTIVATOR 1 95 KDA SUBUNIT |
| 61:00:00 | 6oay-B | 7 | 11.7 | 134 | 570 | 10 | MOLECULE: | HYPERACTIVE DISAGGREGASE CLPB |
| 62:00:00 | 5vc7-C | 7 | 9.6 | 179 | 544 | 10 | MOLECULE: | VCP-LIKE ATPASE |
| 63:00:00 | 5vc7-D | 7 | 9.5 | 176 | 544 | 10 | MOLECULE: | VCP-LIKE ATPASE |
| 64:00:00 | 3cmw-A | 7 | 15.3 | 145 | 1609 | 12 | MOLECULE: | DNA (5'- |
| 65:00:00 | 3gli-B | 7 | 14.8 | 147 | 359 | 13 | MOLECULE: | DNA POLYMERASE III SUBUNIT DELTA |
| 66:00:00 | 3glh-M | 7 | 13.5 | 150 | 364 | 11 | MOLECULE: | DNA POLYMERASE III SUBUNIT DELTA |
| 67:00:00 | 1mo4-A | 7 | 10.4 | 147 | 324 | 12 | MOLECULE: | RECA |
| 68:00:00 | 6pen-E | 7 | 3.5 | 127 | 288 | 9 | MOLECULE: | SPASTIN |
| 69:00:00 | 6pxk-K | 7 | 11.4 | 159 | 429 | 12 | MOLECULE: | ATP-DEPENDENT PROTEASE ATPASE SUBUNIT HSLU |
| 70:00:00 | 6pek-E | 7 | 3.6 | 129 | 243 | 9 | MOLECULE: | SPASTIN |
| 71:00:00 | 3vfd-A | 7 | 11.1 | 135 | 266 | 12 | MOLECULE: | SPASTIN |
| 72:00:00 | 6qs8-B | 7 | 6.1 | 192 | 582 | 12 | MOLECULE: | CHAPERONE PROTEIN CLPB |
| 73:00:00 | 4fgn-A | 7 | 3.4 | 102 | 126 | 8 | MOLECULE: | LARGE T ANTIGEN |
| 74:00:00 | 6dgd-A | 6.9 | 21.2 | 166 | 704 | 10 | MOLECULE: | PRIMOSOMAL PROTEIN N' |
| 75:00:00 | 7m9c-G | 6.9 | 9.7 | 135 | 257 | 9 | MOLECULE: | TNSC |
| 76:00:00 | 8hr9-F | 6.9 | 14.9 | 144 | 835 | 8 | MOLECULE: | ARCHAEAL ATPASE |
| 77:00:00 | 7m9a-G | 6.9 | 9.7 | 136 | 257 | 9 | MOLECULE: | TNSC |
| 78:00:00 | 7m9a-H | 6.9 | 9.5 | 138 | 257 | 9 | MOLECULE: | TNSC |
| 79:00:00 | 8fcv-T | 6.9 | 8.6 | 150 | 343 | 9 | MOLECULE: | DNA (60-MER) |
| 80:00:00 | 7m9c-P | 6.9 | 9.7 | 135 | 257 | 9 | MOLECULE: | TNSC |
| 81:00:00 | 8hr8-E | 6.9 | 10 | 147 | 833 | 8 | MOLECULE: | ARCHAEAL ATPASE |
| 82:00:00 | 8hr8-G | 6.9 | 14.8 | 151 | 830 | 9 | MOLECULE: | ARCHAEAL ATPASE |
| 83:00:00 | 7m9b-G | 6.9 | 9.6 | 138 | 257 | 9 | MOLECULE: | TNSC |
| 84:00:00 | 7ykl-F | 6.9 | 9.4 | 207 | 735 | 9 | MOLECULE: | ATPASE FAMILY GENE 2 PROTEIN |
| 85:00:00 | 8owo-A | 6.9 | 3.8 | 121 | 427 | 14 | MOLECULE: | HISTONE-LYSINE N-METHYLTRANSFERASE SMYD3 |
| 86:00:00 | 6nyy-C | 6.9 | 3.4 | 128 | 491 | 12 | MOLECULE: | AFG3-LIKE PROTEIN 2 |
| 87:00:00 | 8apm-C | 6.9 | 6.2 | 221 | 449 | 11 | MOLECULE: | PRIMASE D5 |
| 88:00:00 | 6sh3-A | 6.9 | 14.8 | 164 | 376 | 10 | MOLECULE: | MITOCHONDRIAL CHAPERONE BCS1 |
| 89:00:00 | 6zyy-C | 6.9 | 20.5 | 216 | 2901 | 6 | MOLECULE: | DYNEIN HEAVY CHAIN, OUTER ARM PROTEIN |
| 90:00:00 | 3n70-A | 6.9 | 3.2 | 117 | 142 | 9 | MOLECULE: | TRANSPORT ACTIVATOR |
| 91:00:00 | 6sh3-F | 6.9 | 5.9 | 146 | 376 | 14 | MOLECULE: | MITOCHONDRIAL CHAPERONE BCS1 |
| 92:00:00 | 6hz8-F | 6.9 | 3.7 | 136 | 285 | 11 | MOLECULE: | 5-METHYLCYTOSINE-SPECIFIC RESTRICTION ENZYME B |
| 93:00:00 | 2is6-A | 6.9 | 16.6 | 136 | 654 | 6 | MOLECULE: | 5'-D(*CP*GP*AP*GP*CP*AP*CP*TP*GP*CP*AP*GP*TP*GP*C |
| 94:00:00 | 6rn3-D | 6.9 | 4.3 | 178 | 684 | 16 | MOLECULE: | CHAPERONE PROTEIN CLPB |
| 95:00:00 | 6og2-A | 6.9 | 7.3 | 192 | 548 | 13 | MOLECULE: | HYPERACTIVE DISAGGREGASE CLPB |
| 96:00:00 | 6j6i-L | 6.9 | 24 | 158 | 790 | 8 | MOLECULE: | PROBABLE SERINE/THREONINE-PROTEIN KINASE PBL2 |
| 97:00:00 | 6j5t-O | 6.9 | 24.5 | 134 | 808 | 9 | MOLECULE: | PROBABLE SERINE/THREONINE-PROTEIN KINASE PBL2 |
| 98:00:00 | 6j6i-C | 6.9 | 21.7 | 162 | 790 | 9 | MOLECULE: | PROBABLE SERINE/THREONINE-PROTEIN KINASE PBL2 |
| 99:00:00 | 6j5t-C | 6.9 | 24.5 | 137 | 808 | 9 | MOLECULE: | PROBABLE SERINE/THREONINE-PROTEIN KINASE PBL2 |
| 0:00 | 3j96-A | 6.9 | 17.6 | 183 | 678 | 11 | MOLECULE: | VESICLE-FUSING ATPASE |
| 1:00 | 5w0t-A | 6.9 | 5.5 | 144 | 298 | 15 | MOLECULE: | PROTEIN MSP1 |
| 2:00 | 3glh-I | 6.9 | 13.8 | 148 | 363 | 11 | MOLECULE: | DNA POLYMERASE III SUBUNIT DELTA |
| 3:00 | 3glh-H | 6.9 | 13.3 | 137 | 364 | 12 | MOLECULE: | DNA POLYMERASE III SUBUNIT DELTA |
| 4:00 | 4wvy-B | 6.9 | 11.1 | 144 | 408 | 12 | MOLECULE: | RUVB-LIKE 1 |
| 5:00 | 4hse-A | 6.9 | 11 | 148 | 369 | 11 | MOLECULE: | CHAPERONE PROTEIN CLPB |

| | | | | | | | | |
|---|---|---|---|---|---|---|---|---|
| 6:00 | 2c9o-A | 6.9 | 3.1 | 139 | 398 | 11 | MOLECULE: | RUVB-LIKE 1 |
| 7:00 | 1e94-E | 6.9 | 11.7 | 157 | 408 | 14 | MOLECULE: | HEAT SHOCK PROTEIN HSLV |
| 8:00 | 5oaf-B | 6.9 | 8.2 | 143 | 418 | 13 | MOLECULE: | RUVB-LIKE 1 |
| 9:00 | 6uks-E | 6.9 | 4.2 | 154 | 362 | 15 | MOLECULE: | MITOCHONDRIAL CHAPERONE BCS1 |
| 10:00 | 8t9f-K | 6.8 | 6.3 | 117 | 265 | 11 | MOLECULE: | HISTONE-LYSINE N-METHYLTRANSFERASE KMT5B |
| 11:00 | 3n71-A | 6.8 | 20.9 | 143 | 469 | 16 | MOLECULE: | HISTONE LYSINE METHYLTRANSFERASE SMYD1 |
| 12:00 | 6p6k-A | 6.8 | 3.8 | 121 | 425 | 15 | MOLECULE: | HISTONE-LYSINE N-METHYLTRANSFERASE SMYD3 |
| 13:00 | 7w42-B | 6.8 | 23.3 | 176 | 402 | 16 | MOLECULE: | UNCHARACTERIZED ATPASE YJOB |
| 14:00 | 7m9c-H | 6.8 | 8.9 | 133 | 257 | 11 | MOLECULE: | TNSC |
| 15:00 | 8osg-B | 6.8 | 4.3 | 167 | 323 | 10 | MOLECULE: | MAGNESIUM-CHELATASE SUBUNIT CHLI |
| 16:00 | 7ufi-4 | 6.8 | 10.5 | 149 | 311 | 7 | MOLECULE: | DNA (5'- |
| 17:00 | 7m9b-H | 6.8 | 9.5 | 138 | 257 | 9 | MOLECULE: | TNSC |
| 18:00 | 8fcx-R | 6.8 | 8 | 148 | 343 | 8 | MOLECULE: | TNSC |
| 19:00 | 7ufi-1 | 6.8 | 10.5 | 150 | 311 | 8 | MOLECULE: | DNA (5'- |
| 20:00 | 5hq8-B | 6.8 | 3.9 | 121 | 426 | 14 | MOLECULE: | HISTONE-LYSINE N-METHYLTRANSFERASE SMYD3 |
| 21:00 | 6fnd-D | 6.8 | 6.3 | 112 | 422 | 13 | MOLECULE: | APICAL COMPLEX LYSINE METHYLTRANSFERASE |
| 22:00 | 6p8v-C | 6.8 | 3.7 | 140 | 303 | 16 | MOLECULE: | ATPASE, AAA FAMILY |
| 23:00 | 7tts-C | 6.8 | 12.1 | 170 | 456 | 14 | MOLECULE: | CASEINOLYTIC PEPTIDASE B PROTEIN HOMOLOG |
| 24:00:00 | 6oab-D | 6.8 | 8.6 | 140 | 537 | 11 | MOLECULE: | CELL DIVISION CONTROL PROTEIN 48 |
| 25:00:00 | 7z6h-D | 6.8 | 14.7 | 136 | 324 | 10 | MOLECULE: | CELL CYCLE CHECKPOINT CONTROL PROTEIN RAD9A |
| 26:00:00 | 3j3t-F | 6.8 | 10.1 | 193 | 798 | 13 | MOLECULE: | ADAPTER PROTEIN MECA 1 |
| 27:00:00 | 5ftn-A | 6.8 | 11.6 | 192 | 737 | 12 | MOLECULE: | TRANSITIONAL ENDOPLASMIC RETICULUM ATPASE |
| 28:00:00 | 8hwg-C | 6.8 | 4.1 | 203 | 372 | 9 | MOLECULE: | PRIMASE D5 |
| 29:00:00 | 1sxj-E | 6.8 | 11.7 | 134 | 317 | 9 | MOLECULE: | ACTIVATOR 1 95 KDA SUBUNIT |
| 30:00:00 | 7tjj-I | 6.8 | 16.6 | 144 | 380 | 14 | MOLECULE: | ORIGIN RECOGNITION COMPLEX SUBUNIT 1 |
| 31:00:00 | 6j5t-L | 6.8 | 25.2 | 130 | 808 | 9 | MOLECULE: | PROBABLE SERINE/THREONINE-PROTEIN KINASE PBL2 |
| 32:00:00 | 8jx6-B | 6.8 | 6.7 | 124 | 451 | 10 | MOLECULE: | DEEP-SEA HELICASE 9 |
| 33:00:00 | 3b9p-A | 6.8 | 9 | 132 | 268 | 11 | MOLECULE: | CG5977-PA, ISOFORM A |
| 34:00:00 | 5ln3-J | 6.8 | 12.2 | 174 | 373 | 8 | MOLECULE: | 26S PROTEASOME NON-ATPASE REGULATORY SUBUNIT 2 |
| 35:00:00 | 3glh-C | 6.8 | 12.5 | 140 | 364 | 11 | MOLECULE: | DNA POLYMERASE III SUBUNIT DELTA |
| 36:00:00 | 2xsz-B | 6.8 | 3.3 | 142 | 319 | 11 | MOLECULE: | RUVB-LIKE 1 |
| 37:00:00 | 3glh-B | 6.8 | 14.8 | 144 | 364 | 10 | MOLECULE: | DNA POLYMERASE III SUBUNIT DELTA |
| 38:00:00 | 6p8v-A | 6.8 | 3.6 | 140 | 299 | 15 | MOLECULE: | ATPASE, AAA FAMILY |
| 39:00:00 | 3t15-A | 6.8 | 12.4 | 135 | 263 | 16 | MOLECULE: | RIBULOSE BISPHOSPHATE CARBOXYLASE/OXYGENASE ACTIV |
| 40:00:00 | 3s7d-A | 6.8 | 3.8 | 118 | 430 | 11 | MOLECULE: | N-LYSINE METHYLTRANSFERASE SMYD2 |
| 41:00:00 | 7ufi-5 | 6.7 | 10.2 | 150 | 311 | 8 | MOLECULE: | DNA (5'- |
| 42:00:00 | 3qwv-A | 6.7 | 3.6 | 119 | 431 | 9 | MOLECULE: | SET AND MYND DOMAIN-CONTAINING PROTEIN 2 |
| 43:00:00 | 8dgf-B | 6.7 | 15.5 | 165 | 1534 | 7 | MOLECULE: | ATP-BINDING PROTEIN AVS4 |
| 44:00:00 | 7mbw-C | 6.7 | 13.6 | 147 | 461 | 11 | MOLECULE: | TRANSPOSON TN7 TRANSPOSITION PROTEIN TNSC |
| 45:00:00 | 8fcx-S | 6.7 | 6.6 | 145 | 343 | 8 | MOLECULE: | TNSC |
| 46:00:00 | 8ff4-S | 6.7 | 6.6 | 145 | 343 | 8 | MOLECULE: | TYPE I-B CRISPR-ASSOCIATED PROTEIN CAS5 |
| 47:00:00 | 6oa9-D | 6.7 | 8 | 166 | 489 | 8 | MOLECULE: | CELL DIVISION CONTROL PROTEIN 48 |
| 48:00:00 | 1mlv-A | 6.7 | 16.3 | 116 | 425 | 13 | MOLECULE: | RIBULOSE-1,5 BIPHOSPHATE CARBOXYLASE/OXYGENASE LA |
| 49:00:00 | 2qby-B | 6.7 | 13.3 | 142 | 368 | 8 | MOLECULE: | CELL DIVISION CONTROL PROTEIN 6 HOMOLOG 1 |
| 50:00:00 | 5udb-4 | 6.7 | 6.9 | 243 | 751 | 12 | MOLECULE: | DNA REPLICATION LICENSING FACTOR MCM2 |
| 51:00:00 | 1g41-A | 6.7 | 9.9 | 154 | 334 | 15 | MOLECULE: | HEAT SHOCK PROTEIN HSLU |
| 52:00:00 | 6sh5-D | 6.7 | 4.5 | 148 | 334 | 13 | MOLECULE: | MITOCHONDRIAL CHAPERONE BCS1 |
| 53:00:00 | 6j5t-F | 6.7 | 25.9 | 132 | 808 | 11 | MOLECULE: | PROBABLE SERINE/THREONINE-PROTEIN KINASE PBL2 |
| 54:00:00 | 6sh5-B | 6.7 | 8.8 | 147 | 334 | 13 | MOLECULE: | MITOCHONDRIAL CHAPERONE BCS1 |
| 55:00:00 | 6sh5-C | 6.7 | 4.5 | 148 | 334 | 13 | MOLECULE: | MITOCHONDRIAL CHAPERONE BCS1 |
| 56:00:00 | 6sh5-A | 6.7 | 5.6 | 152 | 334 | 13 | MOLECULE: | MITOCHONDRIAL CHAPERONE BCS1 |
| 57:00:00 | 4d80-F | 6.7 | 10.1 | 139 | 274 | 13 | MOLECULE: | AAA ATPASE, CENTRAL DOMAIN PROTEIN |
| 58:00:00 | 4ww4-B | 6.7 | 11.9 | 148 | 435 | 11 | MOLECULE: | RUVB-LIKE 1 |

| | | | | | | | | |
|---|---|---|---|---|---|---|---|---|
| 59:00:00 | 2xsz-A | 6.7 | 3.3 | 143 | 318 | 11 | MOLECULE: | RUVB-LIKE 1 |
| 60:00:00 | 3ecc-A | 6.7 | 3.6 | 122 | 182 | 14 | MOLECULE: | DNA REPLICATION PROTEIN DNAC |
| 61:00:00 | 5xmi-A | 6.7 | 12.9 | 137 | 319 | 12 | MOLECULE: | VACUOLAR PROTEIN SORTING-ASSOCIATED PROTEIN 4 |
| 62:00:00 | 3glh-N | 6.7 | 8.4 | 147 | 363 | 12 | MOLECULE: | DNA POLYMERASE III SUBUNIT DELTA |
| 63:00:00 | 3gli-G | 6.7 | 16.4 | 158 | 373 | 12 | MOLECULE: | DNA POLYMERASE III SUBUNIT DELTA |
| 64:00:00 | 6p07-A | 6.7 | 12.1 | 131 | 270 | 12 | MOLECULE: | SPASTIN |
| 65:00:00 | 2awn-D | 6.7 | 8.6 | 119 | 301 | 15 | MOLECULE: | MALTOSE/MALTODEXTRIN IMPORT ATP-BINDING PROTEIN M |
| 66:00:00 | 3s7b-A | 6.7 | 3.9 | 119 | 430 | 10 | MOLECULE: | N-LYSINE METHYLTRANSFERASE SMYD2 |
| 67:00:00 | 7m9c-D | 6.6 | 8.5 | 132 | 257 | 10 | MOLECULE: | TNSC |
| 68:00:00 | 8hrb-P | 6.6 | 12 | 133 | 834 | 11 | MOLECULE: | ARCHAEAL ATPASE |
| 69:00:00 | 7ufi-2 | 6.6 | 10.1 | 150 | 311 | 8 | MOLECULE: | DNA (5'- |
| 70:00:00 | 8hr9-L | 6.6 | 14.5 | 143 | 833 | 8 | MOLECULE: | ARCHAEAL ATPASE |
| 71:00:00 | 8fcx-T | 6.6 | 7.9 | 148 | 343 | 9 | MOLECULE: | TNSC |
| 72:00:00 | 7m9a-F | 6.6 | 8.3 | 129 | 257 | 10 | MOLECULE: | TNSC |
| 73:00:00 | 7m9c-F | 6.6 | 9.4 | 138 | 257 | 9 | MOLECULE: | TNSC |
| 74:00:00 | 7m9a-I | 6.6 | 9.2 | 134 | 257 | 10 | MOLECULE: | TNSC |
| 75:00:00 | 7m99-C | 6.6 | 8.7 | 130 | 257 | 8 | MOLECULE: | TNSC |
| 76:00:00 | 7ufm-H | 6.6 | 9.2 | 143 | 311 | 8 | MOLECULE: | VCHTNSC |
| 77:00:00 | 7m9b-I | 6.6 | 9.2 | 134 | 257 | 10 | MOLECULE: | TNSC |
| 78:00:00 | 8fnt-G | 6.6 | 14.6 | 132 | 785 | 8 | MOLECULE: | ARCHAEAL ATPASE |
| 79:00:00 | 8ff4-T | 6.6 | 6.2 | 140 | 343 | 9 | MOLECULE: | TYPE I-B CRISPR-ASSOCIATED PROTEIN CAS5 |
| 80:00:00 | 2h21-A | 6.6 | 28.6 | 132 | 425 | 14 | MOLECULE: | RIBULOSE-1,5 BISPHOSPHATE CARBOXYLASE/OXYGENASE |
| 81:00:00 | 8fnt-F | 6.6 | 15 | 150 | 785 | 8 | MOLECULE: | ARCHAEAL ATPASE |
| 82:00:00 | 6oaa-C | 6.6 | 8.4 | 141 | 532 | 13 | MOLECULE: | CELL DIVISION CONTROL PROTEIN 48 |
| 83:00:00 | 3oxg-A | 6.6 | 3.8 | 121 | 425 | 13 | MOLECULE: | SET AND MYND DOMAIN-CONTAINING PROTEIN 3 |
| 84:00:00 | 5nvu-A | 6.6 | 17.1 | 221 | 3169 | 0 | MOLECULE: | DYNEIN MOTOR DOMAIN |
| 85:00:00 | 7ufi-6 | 6.6 | 10.5 | 149 | 311 | 8 | MOLECULE: | DNA (5'- |
| 86:00:00 | 8hwf-F | 6.6 | 4.4 | 202 | 371 | 11 | MOLECULE: | PRIMASE D5 |
| 87:00:00 | 5zr1-A | 6.6 | 19.8 | 169 | 494 | 8 | MOLECULE: | ORIGIN RECOGNITION COMPLEX SUBUNIT 1 |
| 88:00:00 | 5zr1-E | 6.6 | 12.9 | 143 | 460 | 8 | MOLECULE: | ORIGIN RECOGNITION COMPLEX SUBUNIT 1 |
| 89:00:00 | 6oa9-E | 6.6 | 8.5 | 166 | 496 | 8 | MOLECULE: | CELL DIVISION CONTROL PROTEIN 48 |
| 90:00:00 | 5mp9-H | 6.6 | 14.7 | 160 | 390 | 11 | MOLECULE: | PROTEASOME SUBUNIT ALPHA TYPE-1 |
| 91:00:00 | 4xgu-B | 6.6 | 6.9 | 165 | 382 | 13 | MOLECULE: | PUTATIVE PACHYTENE CHECKPOINT PROTEIN 2 |
| 92:00:00 | 6sh5-F | 6.6 | 5.6 | 152 | 334 | 13 | MOLECULE: | MITOCHONDRIAL CHAPERONE BCS1 |
| 93:00:00 | 6sh5-E | 6.6 | 4.5 | 149 | 334 | 13 | MOLECULE: | MITOCHONDRIAL CHAPERONE BCS1 |
| 94:00:00 | 5wc0-A | 6.6 | 10.9 | 137 | 275 | 12 | MOLECULE: | MEIOTIC SPINDLE FORMATION PROTEIN MEI-1 |
| 95:00:00 | 3glh-L | 6.6 | 12.9 | 150 | 364 | 11 | MOLECULE: | DNA POLYMERASE III SUBUNIT DELTA |
| 96:00:00 | 6etx-E | 6.6 | 4.5 | 139 | 443 | 12 | MOLECULE: | RUVB-LIKE 1 |
| 97:00:00 | 6rn4-B | 6.6 | 10.5 | 206 | 689 | 11 | MOLECULE: | CHAPERONE PROTEIN CLPB |
| 98:00:00 | 5jpq-i | 6.6 | 28.9 | 133 | 659 | 11 | MOLECULE: | WD40 DOMAIN PROTEINS |
| 99:00:00 | 2h23-C | 6.6 | 17.2 | 127 | 440 | 13 | MOLECULE: | RIBULOSE-1,5 BISPHOSPHATE CARBOXYLASE/OXYGENASE L |
| 0:00 | 3s7f-A | 6.6 | 4.4 | 121 | 430 | 12 | MOLECULE: | N-LYSINE METHYLTRANSFERASE SMYD2 |
| 1:00 | 8val-E | 6.6 | 13.7 | 139 | 337 | 11 | MOLECULE: | DNA POLYMERASE III SUBUNIT DELTA |
| 2:00 | 6etx-D | 6.6 | 5.5 | 140 | 434 | 14 | MOLECULE: | RUVB-LIKE 1 |
| 3:00 | 7t3i-C | 6.6 | 13.2 | 176 | 578 | 10 | MOLECULE: | RIX7 |
| 4:00 | 1z1d-B | 6.6 | 4 | 107 | 131 | 8 | MOLECULE: | REPLICATION PROTEIN A 32 KDA SUBUNIT |
| 5:00 | 8fcv-V | 6.5 | 8 | 147 | 343 | 9 | MOLECULE: | DNA (60-MER) |
| 6:00 | 7m9c-O | 6.5 | 9.1 | 133 | 257 | 10 | MOLECULE: | TNSC |
| 7:00 | 7m9a-A | 6.5 | 9.2 | 136 | 257 | 10 | MOLECULE: | TNSC |
| 8:00 | 8hra-B | 6.5 | 10.1 | 138 | 835 | 9 | MOLECULE: | ARCHAEAL ATPASE |
| 9:00 | 7st9-A | 6.5 | 11 | 168 | 522 | 14 | MOLECULE: | CHECKPOINT PROTEIN RAD24 |
| 10:00 | 8fcx-V | 6.5 | 8.5 | 153 | 343 | 9 | MOLECULE: | TNSC |
| 11:00 | 8fcw-T | 6.5 | 8.4 | 152 | 343 | 9 | MOLECULE: | TNSC |

| | | | | | | | | |
|---|---|---|---|---|---|---|---|---|
| 12:00 | 7m9b-A | 6.5 | 9.2 | 135 | 257 | 10 | MOLECULE: | TNSC |
| 13:00 | 8fnt-A | 6.5 | 14.5 | 137 | 785 | 9 | MOLECULE: | ARCHAEAL ATPASE |
| 14:00 | 7m9c-A | 6.5 | 9.1 | 135 | 257 | 10 | MOLECULE: | TNSC |
| 15:00 | 8c0v-C | 6.5 | 12 | 170 | 821 | 8 | MOLECULE: | PEROXISOMAL ATPASE PEX1 |
| 16:00 | 7ttr-B | 6.5 | 10.1 | 159 | 311 | 13 | MOLECULE: | CASEINOLYTIC PEPTIDASE B PROTEIN HOMOLOG |
| 17:00 | 6oa9-F | 6.5 | 7.2 | 168 | 533 | 10 | MOLECULE: | CELL DIVISION CONTROL PROTEIN 48 |
| 18:00 | 7ufi-3 | 6.5 | 10.1 | 149 | 311 | 8 | MOLECULE: | DNA (5'- |
| 19:00 | 7ykk-F | 6.5 | 7.9 | 208 | 735 | 9 | MOLECULE: | ATPASE FAMILY GENE 2 PROTEIN |
| 20:00 | 8hwa-C | 6.5 | 3.9 | 200 | 379 | 9 | MOLECULE: | PRIMASE D5 |
| 21:00 | 6epc-H | 6.5 | 12.7 | 158 | 396 | 8 | MOLECULE: | PROTEASOME SUBUNIT ALPHA TYPE-6 |
| 22:00 | 8osf-C | 6.5 | 4.4 | 168 | 321 | 11 | MOLECULE: | MAGNESIUM-CHELATASE SUBUNIT CHLI |
| 23:00 | 6sh3-C | 6.5 | 6 | 146 | 376 | 14 | MOLECULE: | MITOCHONDRIAL CHAPERONE BCS1 |
| 24:00:00 | 1lv7-A | 6.5 | 3.3 | 134 | 251 | 9 | MOLECULE: | FTSH |
| 25:00:00 | 1g64-B | 6.5 | 8.9 | 113 | 190 | 15 | MOLECULE: | COB(I)ALAMIN ADENOSYLTRANSFERASE |
| 26:00:00 | 4hut-A | 6.5 | 9.1 | 111 | 191 | 14 | MOLECULE: | COB(I)YRINIC ACID A,C-DIAMIDE ADENOSYLTRANSFERASE |
| 27:00:00 | 3glh-D | 6.5 | 11.3 | 138 | 363 | 12 | MOLECULE: | DNA POLYMERASE III SUBUNIT DELTA |
| 28:00:00 | 7yup-E | 6.5 | 11 | 156 | 773 | 12 | MOLECULE: | LON PROTEASE |
| 29:00:00 | 6pen-B | 6.5 | 10.8 | 138 | 288 | 10 | MOLECULE: | SPASTIN |
| 30:00:00 | 6qs6-F | 6.5 | 16.2 | 210 | 577 | 12 | MOLECULE: | CHAPERONE PROTEIN CLPB |
| 31:00:00 | 6pek-B | 6.5 | 10.8 | 138 | 288 | 10 | MOLECULE: | SPASTIN |
| 32:00:00 | 8emc-F | 6.5 | 6.8 | 188 | 666 | 9 | MOLECULE: | PROTEASE LON-RELATED BREX SYSTEM PROTEIN BRXL |
| 33:00:00 | 6xji-C | 6.5 | 7.2 | 130 | 290 | 12 | MOLECULE: | PHENOL-SOLUBLE MODULIN EXPORT ABC TRANSPORTER PER |
| 34:00:00 | 7mi1-A | 6.5 | 20.5 | 119 | 2628 | 8 | MOLECULE: | CHIMERA PROTEIN OF DYNEIN AND ENDOLYSIN |
| 35:00:00 | 6qs7-F | 6.5 | 12.6 | 199 | 573 | 11 | MOLECULE: | CHAPERONE PROTEIN CLPB |
| 36:00:00 | 6qs6-B | 6.5 | 8.7 | 193 | 582 | 11 | MOLECULE: | CHAPERONE PROTEIN CLPB |
| 37:00:00 | 1tbd-A | 6.5 | 4.2 | 111 | 134 | 9 | MOLECULE: | SV40 T-ANTIGEN |
| 38:00:00 | 6orb-A | 6.4 | 24.6 | 170 | 3352 | 9 | MOLECULE: | MIDASIN |
| 39:00:00 | 8fcv-Q | 6.4 | 7.5 | 142 | 310 | 9 | MOLECULE: | DNA (60-MER) |
| 40:00:00 | 8ff4-V | 6.4 | 8.2 | 148 | 343 | 9 | MOLECULE: | TYPE I-B CRISPR-ASSOCIATED PROTEIN CAS5 |
| 41:00:00 | 7n6i-C | 6.4 | 8.7 | 131 | 257 | 8 | MOLECULE: | TNIQ (HOMOLOGY MODEL) |
| 42:00:00 | 7n6i-E | 6.4 | 8.7 | 129 | 257 | 9 | MOLECULE: | TNIQ (HOMOLOGY MODEL) |
| 43:00:00 | 7m9b-F | 6.4 | 9.3 | 135 | 257 | 10 | MOLECULE: | TNSC |
| 44:00:00 | 8hra-E | 6.4 | 15.7 | 143 | 834 | 10 | MOLECULE: | ARCHAEAL ATPASE |
| 45:00:00 | 8fcu-Q | 6.4 | 7.5 | 142 | 310 | 9 | MOLECULE: | TYPE I-B CRISPR-ASSOCIATED PROTEIN CAS5 |
| 46:00:00 | 8ff4-R | 6.4 | 7.9 | 149 | 343 | 8 | MOLECULE: | TYPE I-B CRISPR-ASSOCIATED PROTEIN CAS5 |
| 47:00:00 | 7m9a-J | 6.4 | 9.2 | 133 | 257 | 10 | MOLECULE: | TNSC |
| 48:00:00 | 8fcv-S | 6.4 | 6.7 | 143 | 343 | 9 | MOLECULE: | DNA (60-MER) |
| 49:00:00 | 8fnt-B | 6.4 | 15 | 150 | 785 | 9 | MOLECULE: | ARCHAEAL ATPASE |
| 50:00:00 | 2h2j-A | 6.4 | 16.2 | 116 | 425 | 13 | MOLECULE: | RIBULOSE-1,5 BISPHOSPHATE CARBOXYLASE/OXYGENASE L |
| 51:00:00 | 8fnt-C | 6.4 | 13.1 | 139 | 785 | 9 | MOLECULE: | ARCHAEAL ATPASE |
| 52:00:00 | 6oaa-E | 6.4 | 5.9 | 174 | 442 | 9 | MOLECULE: | CELL DIVISION CONTROL PROTEIN 48 |
| 53:00:00 | 8bob-C | 6.4 | 8 | 125 | 412 | 17 | MOLECULE: | PROTEIN MALY |
| 54:00:00 | 6u1y-B | 6.4 | 3.2 | 131 | 245 | 18 | MOLECULE: | MITOCHONDRIAL CHAPERONE BCS1 |
| 55:00:00 | 4fw9-A | 6.4 | 14.4 | 156 | 602 | 9 | MOLECULE: | TTC1975 PEPTIDASE |
| 56:00:00 | 7tts-A | 6.4 | 9.9 | 156 | 433 | 13 | MOLECULE: | CASEINOLYTIC PEPTIDASE B PROTEIN HOMOLOG |
| 57:00:00 | 6oab-E | 6.4 | 8 | 136 | 442 | 10 | MOLECULE: | CELL DIVISION CONTROL PROTEIN 48 |
| 58:00:00 | 7jpp-E | 6.4 | 16.2 | 134 | 406 | 8 | MOLECULE: | ORIGIN RECOGNITION COMPLEX SUBUNIT 1 |
| 59:00:00 | 6p8v-D | 6.4 | 3.6 | 138 | 303 | 17 | MOLECULE: | ATPASE, AAA FAMILY |
| 60:00:00 | 6oay-E | 6.4 | 14.2 | 142 | 570 | 10 | MOLECULE: | HYPERACTIVE DISAGGREGASE CLPB |
| 61:00:00 | 6edo-A | 6.4 | 3.8 | 157 | 2031 | 15 | MOLECULE: | MIDASIN |
| 62:00:00 | 5c3c-B | 6.4 | 6.3 | 139 | 235 | 12 | MOLECULE: | CBBQ/NIRQ/NORQ DOMAIN PROTEIN |
| 63:00:00 | 5vca-O | 6.4 | 9.5 | 148 | 543 | 11 | MOLECULE: | VCP-LIKE ATPASE |
| 64:00:00 | 5vhm-F | 6.4 | 8.3 | 136 | 219 | 16 | MOLECULE: | 26S PROTEASOME NON-ATPASE REGULATORY SUBUNIT 10 |

| | | | | | | | | |
|---|---|---|---|---|---|---|---|---|
| 65:00:00 | 5vc7-A | 6.4 | 9.6 | 179 | 544 | 10 | MOLECULE: | VCP-LIKE ATPASE |
| 66:00:00 | 1g64-A | 6.4 | 3.4 | 103 | 169 | 15 | MOLECULE: | COB(I)ALAMIN ADENOSYLTRANSFERASE |
| 67:00:00 | 6p07-F | 6.4 | 11.6 | 139 | 300 | 13 | MOLECULE: | SPASTIN |
| 68:00:00 | 8osg-E | 6.4 | 4.5 | 167 | 323 | 10 | MOLECULE: | MAGNESIUM-CHELATASE SUBUNIT CHLI |
| 69:00:00 | 2tbd-A | 6.4 | 3.6 | 103 | 134 | 6 | MOLECULE: | SV40 T ANTIGEN |
| 70:00:00 | 6mdm-D | 6.3 | 10.5 | 162 | 713 | 13 | MOLECULE: | VESICLE-FUSING ATPASE |
| 71:00:00 | 7m9c-J | 6.3 | 9.2 | 133 | 257 | 10 | MOLECULE: | TNSC |
| 72:00:00 | 7n6i-J | 6.3 | 9.3 | 131 | 257 | 10 | MOLECULE: | TNIQ (HOMOLOGY MODEL) |
| 73:00:00 | 7n6i-I | 6.3 | 10.1 | 132 | 257 | 10 | MOLECULE: | TNIQ (HOMOLOGY MODEL) |
| 74:00:00 | 8fcw-R | 6.3 | 8.2 | 153 | 343 | 8 | MOLECULE: | TNSC |
| 75:00:00 | 7ufm-A | 6.3 | 9.2 | 143 | 311 | 8 | MOLECULE: | VCHTNSC |
| 76:00:00 | 7m9b-E | 6.3 | 8.2 | 132 | 257 | 8 | MOLECULE: | TNSC |
| 77:00:00 | 7m9b-L | 6.3 | 9.1 | 134 | 257 | 10 | MOLECULE: | TNSC |
| 78:00:00 | 8hrb-F | 6.3 | 15 | 150 | 851 | 9 | MOLECULE: | ARCHAEAL ATPASE |
| 79:00:00 | 7m9b-C | 6.3 | 9.4 | 133 | 257 | 10 | MOLECULE: | TNSC |
| 80:00:00 | 7m9a-L | 6.3 | 9.2 | 135 | 257 | 9 | MOLECULE: | TNSC |
| 81:00:00 | 8ff4-Q | 6.3 | 7.5 | 142 | 310 | 9 | MOLECULE: | TYPE I-B CRISPR-ASSOCIATED PROTEIN CAS5 |
| 82:00:00 | 7m9a-B | 6.3 | 9 | 132 | 257 | 11 | MOLECULE: | TNSC |
| 83:00:00 | 7m9c-I | 6.3 | 9.2 | 134 | 257 | 10 | MOLECULE: | TNSC |
| 84:00:00 | 7m9a-K | 6.3 | 9.1 | 132 | 257 | 11 | MOLECULE: | TNSC |
| 85:00:00 | 7m9c-B | 6.3 | 9.1 | 133 | 257 | 10 | MOLECULE: | TNSC |
| 86:00:00 | 8thb-A | 6.3 | 18.4 | 174 | 468 | 11 | MOLECULE: | ELG1 ISOFORM 1 |
| 87:00:00 | 7m9b-B | 6.3 | 9.1 | 133 | 257 | 10 | MOLECULE: | TNSC |
| 88:00:00 | 7m9c-L | 6.3 | 9.1 | 133 | 257 | 11 | MOLECULE: | TNSC |
| 89:00:00 | 7m9b-J | 6.3 | 9.2 | 133 | 257 | 10 | MOLECULE: | TNSC |
| 90:00:00 | 8hra-D | 6.3 | 15.5 | 141 | 848 | 10 | MOLECULE: | ARCHAEAL ATPASE |
| 91:00:00 | 7ufm-C | 6.3 | 10.1 | 146 | 311 | 8 | MOLECULE: | VCHTNSC |
| 92:00:00 | 7ufm-J | 6.3 | 10.3 | 150 | 311 | 8 | MOLECULE: | VCHTNSC |
| 93:00:00 | 7m9a-E | 6.3 | 8.2 | 131 | 257 | 9 | MOLECULE: | TNSC |
| 94:00:00 | 7m9a-C | 6.3 | 9.4 | 134 | 257 | 10 | MOLECULE: | TNSC |
| 95:00:00 | 7mcs-A | 6.3 | 3.7 | 129 | 249 | 12 | MOLECULE: | TRANSPOSON TN7 TRANSPOSITION PROTEIN TNSC |
| 96:00:00 | 7t3i-A | 6.3 | 9.4 | 142 | 577 | 15 | MOLECULE: | RIX7 |
| 97:00:00 | 1l8q-A | 6.3 | 15.9 | 137 | 321 | 9 | MOLECULE: | CHROMOSOMAL REPLICATION INITIATOR PROTEIN DNAA |
| 98:00:00 | 7rzy-7 | 6.3 | 10.5 | 147 | 311 | 8 | MOLECULE: | TN6677 VIBRIO CHOLERAE TRANSPOSON TNSC (VCHTNSC) |
| 99:00:00 | 6mat-A | 6.3 | 9 | 148 | 578 | 11 | MOLECULE: | RIX7 MUTANT |
| 0:00 | 6sh3-G | 6.3 | 14.1 | 161 | 376 | 10 | MOLECULE: | MITOCHONDRIAL CHAPERONE BCS1 |
| 1:00 | 6sh3-B | 6.3 | 12.2 | 153 | 376 | 10 | MOLECULE: | MITOCHONDRIAL CHAPERONE BCS1 |
| 2:00 | 6sh3-E | 6.3 | 12.2 | 154 | 376 | 10 | MOLECULE: | MITOCHONDRIAL CHAPERONE BCS1 |
| 3:00 | 5nug-A | 6.3 | 28.9 | 245 | 2920 | 8 | MOLECULE: | CYTOPLASMIC DYNEIN 1 HEAVY CHAIN 1 |
| 4:00 | 2xsz-C | 6.3 | 3.4 | 142 | 320 | 11 | MOLECULE: | RUVB-LIKE 1 |
| 5:00 | 2z4r-A | 6.3 | 7.9 | 127 | 240 | 14 | MOLECULE: | CHROMOSOMAL REPLICATION INITIATOR PROTEIN DNAA |
| 6:00 | 4ww4-A | 6.3 | 3.7 | 143 | 416 | 14 | MOLECULE: | RUVB-LIKE 1 |
| 7:00 | 8c0w-C | 6.3 | 18.2 | 211 | 1030 | 8 | MOLECULE: | PEROXISOMAL ATPASE PEX6 |
| 8:00 | 6qs6-A | 6.3 | 4.4 | 184 | 577 | 12 | MOLECULE: | CHAPERONE PROTEIN CLPB |
| 9:00 | 6u1y-A | 6.3 | 4 | 136 | 244 | 17 | MOLECULE: | MITOCHONDRIAL CHAPERONE BCS1 |
| 10:00 | 1w5s-B | 6.3 | 14.8 | 158 | 396 | 9 | MOLECULE: | ORIGIN RECOGNITION COMPLEX SUBUNIT 2 ORC2 |
| 11:00 | 1ng9-A | 6.2 | 16.1 | 183 | 794 | 7 | MOLECULE: | 5'-D(*AP*GP*CP*TP*GP*CP*CP*AP*GP*GP*CP*AP*CP*CP*A |
| 12:00 | 8hra-G | 6.2 | 15.9 | 140 | 835 | 10 | MOLECULE: | ARCHAEAL ATPASE |
| 13:00 | 7ufm-M | 6.2 | 9.2 | 143 | 311 | 8 | MOLECULE: | VCHTNSC |
| 14:00 | 7ufm-F | 6.2 | 9.2 | 143 | 311 | 8 | MOLECULE: | VCHTNSC |
| 15:00 | 7m9b-K | 6.2 | 9.3 | 131 | 257 | 10 | MOLECULE: | TNSC |
| 16:00 | 7m99-H | 6.2 | 8.7 | 129 | 257 | 9 | MOLECULE: | TNSC |
| 17:00 | 7ufi-7 | 6.2 | 9.9 | 148 | 311 | 7 | MOLECULE: | DNA (5'- |

| | | | | | | | | |
|---|---|---|---|---|---|---|---|---|
| 18:00 | 8hra-C | 6.2 | 9.8 | 143 | 852 | 9 | MOLECULE: | ARCHAEAL ATPASE |
| 19:00 | 8fcv-W | 6.2 | 8.2 | 148 | 343 | 9 | MOLECULE: | DNA (60-MER) |
| 20:00 | 7ufm-G | 6.2 | 9.1 | 143 | 311 | 8 | MOLECULE: | VCHTNSC |
| 21:00 | 7m9c-K | 6.2 | 9.3 | 132 | 257 | 10 | MOLECULE: | TNSC |
| 22:00 | 8hra-H | 6.2 | 14.6 | 136 | 834 | 10 | MOLECULE: | ARCHAEAL ATPASE |
| 23:00 | 8uii-A | 6.2 | 28.5 | 170 | 641 | 9 | MOLECULE: | ATPASE FAMILY AAA DOMAIN-CONTAINING PROTEIN 5 |
| 24:00:00 | 8hrb-Q | 6.2 | 15.1 | 144 | 835 | 9 | MOLECULE: | ARCHAEAL ATPASE |
| 25:00:00 | 7ufm-D | 6.2 | 9.1 | 143 | 311 | 8 | MOLECULE: | VCHTNSC |
| 26:00:00 | 7m9c-C | 6.2 | 9.4 | 133 | 257 | 10 | MOLECULE: | TNSC |
| 27:00:00 | 5ex3-A | 6.2 | 12.8 | 126 | 429 | 13 | MOLECULE: | HISTONE-LYSINE N-METHYLTRANSFERASE SMYD3 |
| 28:00:00 | 2r44-A | 6.2 | 4.8 | 165 | 330 | 8 | MOLECULE: | UNCHARACTERIZED PROTEIN |
| 29:00:00 | 1jr3-E | 6.2 | 14.7 | 129 | 334 | 11 | MOLECULE: | DNA POLYMERASE III SUBUNIT GAMMA |
| 30:00:00 | 7qo4-J | 6.2 | 13.2 | 181 | 405 | 9 | MOLECULE: | 26S PROTEASOME REGULATORY SUBUNIT RPN1 |
| 31:00:00 | 5vca-R | 6.2 | 8.8 | 143 | 543 | 12 | MOLECULE: | VCP-LIKE ATPASE |
| 32:00:00 | 8d8k-W | 6.2 | 27.6 | 137 | 395 | 9 | MOLECULE: | PROBABLE S-ADENOSYL-L-METHIONINE-DEPENDENT RNA |
| 33:00:00 | 6ef2-J | 6.2 | 3.5 | 128 | 262 | 12 | MOLECULE: | PROTEASOME SUBUNIT ALPHA TYPE-1 |
| 34:00:00 | 5g4f-B | 6.2 | 8.5 | 179 | 726 | 10 | MOLECULE: | VCP-LIKE ATPASE |
| 35:00:00 | 4kfu-B | 6.2 | 3.2 | 104 | 196 | 13 | MOLECULE: | GENOME PACKAGING NTPASE B204 |
| 36:00:00 | 5t0g-C | 6.2 | 11.3 | 164 | 384 | 12 | MOLECULE: | PROTEASOME SUBUNIT ALPHA TYPE-6 |
| 37:00:00 | 4wvy-A | 6.2 | 8.4 | 150 | 427 | 13 | MOLECULE: | RUVB-LIKE 1 |
| 38:00:00 | 8uk9-E | 6.2 | 4.4 | 130 | 318 | 18 | MOLECULE: | SLIDING-CLAMP-LOADER SMALL SUBUNIT |
| 39:00:00 | 1jbk-A | 6.2 | 3.8 | 129 | 189 | 12 | MOLECULE: | CLPB PROTEIN |
| 40:00:00 | 5irm-C | 6.2 | 11 | 159 | 770 | 8 | MOLECULE: | UNCHARACTERIZED PROTEIN |
| 41:00:00 | 5bk4-6 | 6.2 | 6.2 | 200 | 615 | 8 | MOLECULE: | DNA REPLICATION LICENSING FACTOR MCM2 |
| 42:00:00 | 8ag6-B | 6.1 | 14.4 | 161 | 975 | 9 | MOLECULE: | DNA MISMATCH REPAIR PROTEIN MSH2 |
| 43:00:00 | 3mek-A | 6.1 | 3.8 | 121 | 424 | 14 | MOLECULE: | SET AND MYND DOMAIN-CONTAINING PROTEIN 3 |
| 44:00:00 | 7m9c-E | 6.1 | 9.7 | 136 | 257 | 10 | MOLECULE: | TNSC |
| 45:00:00 | 7m9a-D | 6.1 | 9.4 | 139 | 257 | 9 | MOLECULE: | TNSC |
| 46:00:00 | 3qww-A | 6.1 | 3.6 | 120 | 431 | 9 | MOLECULE: | SET AND MYND DOMAIN-CONTAINING PROTEIN 2 |
| 47:00:00 | 8c0v-E | 6.1 | 9.2 | 172 | 821 | 9 | MOLECULE: | PEROXISOMAL ATPASE PEX1 |
| 48:00:00 | 8emc-K | 6.1 | 5.5 | 206 | 665 | 9 | MOLECULE: | PROTEASE LON-RELATED BREX SYSTEM PROTEIN BRXL |
| 49:00:00 | 8thd-A | 6.1 | 22.3 | 170 | 474 | 12 | MOLECULE: | ELG1 ISOFORM 1 |
| 50:00:00 | 8osh-A | 6.1 | 4.5 | 163 | 290 | 9 | MOLECULE: | MAGNESIUM-CHELATASE SUBUNIT CHLI |
| 51:00:00 | 7jgr-D | 6.1 | 17.4 | 152 | 441 | 9 | MOLECULE: | ORIGIN RECOGNITION COMPLEX SUBUNIT 2 |
| 52:00:00 | 5g4f-C | 6.1 | 8.9 | 180 | 726 | 11 | MOLECULE: | VCP-LIKE ATPASE |
| 53:00:00 | 6rn3-F | 6.1 | 8.1 | 192 | 675 | 14 | MOLECULE: | CHAPERONE PROTEIN CLPB |
| 54:00:00 | 6qem-L | 6.1 | 5.6 | 148 | 239 | 12 | MOLECULE: | REPLICATIVE DNA HELICASE |
| 55:00:00 | 4kfu-D | 6.1 | 3.7 | 109 | 195 | 14 | MOLECULE: | GENOME PACKAGING NTPASE B204 |
| 56:00:00 | 5wcb-A | 6.1 | 13.3 | 138 | 277 | 11 | MOLECULE: | MEIOTIC SPINDLE FORMATION PROTEIN MEI-1 |
| 57:00:00 | 4kft-C | 6.1 | 3.2 | 108 | 204 | 12 | MOLECULE: | GENOME PACKAGING NTPASE B204 |
| 58:00:00 | 5udb-A | 6.1 | 15.6 | 141 | 372 | 6 | MOLECULE: | DNA REPLICATION LICENSING FACTOR MCM2 |
| 59:00:00 | 7wrx-E | 6.1 | 9.7 | 137 | 590 | 7 | MOLECULE: | HERA |
| 60:00:00 | 2dhr-A | 6.1 | 9.1 | 153 | 458 | 12 | MOLECULE: | FTSH |
| 61:00:00 | 7m99-E | 6 | 8.7 | 131 | 257 | 8 | MOLECULE: | TNSC |
| 62:00:00 | 8ff4-W | 6 | 6.7 | 140 | 343 | 9 | MOLECULE: | TYPE I-B CRISPR-ASSOCIATED PROTEIN CAS5 |
| 63:00:00 | 7ufm-N | 6 | 9.1 | 144 | 311 | 8 | MOLECULE: | VCHTNSC |
| 64:00:00 | 7n6i-G | 6 | 9.4 | 133 | 257 | 10 | MOLECULE: | TNIQ (HOMOLOGY MODEL) |
| 65:00:00 | 8hr9-I | 6 | 15.5 | 143 | 836 | 10 | MOLECULE: | ARCHAEAL ATPASE |
| 66:00:00 | 8fnt-D | 6 | 15 | 150 | 784 | 9 | MOLECULE: | ARCHAEAL ATPASE |
| 67:00:00 | 7m99-D | 6 | 8.7 | 131 | 257 | 8 | MOLECULE: | TNSC |
| 68:00:00 | 7n6i-F | 6 | 8.7 | 129 | 257 | 9 | MOLECULE: | TNIQ (HOMOLOGY MODEL) |
| 69:00:00 | 7ufm-E | 6 | 9.1 | 141 | 311 | 9 | MOLECULE: | VCHTNSC |
| 70:00:00 | 8osh-B | 6 | 4.4 | 167 | 317 | 12 | MOLECULE: | MAGNESIUM-CHELATASE SUBUNIT CHLI |

| | | | | | | | | |
|---|---|---|---|---|---|---|---|---|
| 71:00:00 | 7plh-A | 6 | 8.5 | 128 | 260 | 10 | MOLECULE: | SHTNSC |
| 72:00:00 | 3zh9-B | 6 | 15 | 126 | 339 | 10 | MOLECULE: | DELTA |
| 73:00:00 | 5zr1-B | 6 | 12.5 | 133 | 374 | 11 | MOLECULE: | ORIGIN RECOGNITION COMPLEX SUBUNIT 1 |
| 74:00:00 | 4wia-A | 6 | 3.5 | 120 | 229 | 11 | MOLECULE: | PUTATIVE FLAGELLA-RELATED PROTEIN H |
| 75:00:00 | 4ai6-A | 6 | 15.8 | 102 | 2650 | 10 | MOLECULE: | GLUTATHIONE S-TRANSFERASE CLASS-MU 26 KDA ISOZYME |
| 76:00:00 | 2h23-A | 6 | 23.5 | 130 | 426 | 13 | MOLECULE: | RIBULOSE-1,5 BISPHOSPHATE CARBOXYLASE/OXYGENASE L |
| 77:00:00 | 6qi9-D | 6 | 10.6 | 140 | 268 | 9 | MOLECULE: | RUVB-LIKE 1 |
| 78:00:00 | 7rzy-1 | 6 | 7.1 | 142 | 311 | 8 | MOLECULE: | TN6677 VIBRIO CHOLERAE TRANSPOSON TNSC (VCHTNSC) |
| 79:00:00 | 6uks-D | 6 | 4.2 | 154 | 362 | 15 | MOLECULE: | MITOCHONDRIAL CHAPERONE BCS1 |
| 80:00:00 | 7ttr-F | 6 | 10.8 | 149 | 304 | 15 | MOLECULE: | CASEINOLYTIC PEPTIDASE B PROTEIN HOMOLOG |
| 81:00:00 | 5v8f-D | 6 | 14.9 | 148 | 456 | 11 | MOLECULE: | DNA REPLICATION LICENSING FACTOR MCM2 |
| 82:00:00 | 5vhr-B | 6 | 3.5 | 125 | 233 | 10 | MOLECULE: | 26S PROTEASOME NON-ATPASE REGULATORY SUBUNIT 10 |
| 83:00:00 | 4kfu-A | 6 | 3.5 | 105 | 205 | 13 | MOLECULE: | GENOME PACKAGING NTPASE B204 |
| 84:00:00 | 6oab-C | 6 | 8.5 | 139 | 532 | 12 | MOLECULE: | CELL DIVISION CONTROL PROTEIN 48 |
| 85:00:00 | 3co5-A | 6 | 4.2 | 109 | 134 | 7 | MOLECULE: | PUTATIVE TWO-COMPONENT SYSTEM TRANSCRIPTIONAL RES |
| 86:00:00 | 4kft-D | 6 | 3.6 | 109 | 194 | 14 | MOLECULE: | GENOME PACKAGING NTPASE B204 |
| 87:00:00 | 4kft-B | 6 | 3.6 | 110 | 197 | 14 | MOLECULE: | GENOME PACKAGING NTPASE B204 |
| 88:00:00 | 5x71-A | 5.9 | 6.1 | 129 | 796 | 6 | MOLECULE: | MRNA CAPPING ENZYME P5 |
| 89:00:00 | 8thc-A | 5.9 | 18.9 | 171 | 479 | 11 | MOLECULE: | ELG1 ISOFORM 1 |
| 90:00:00 | 8ff4-U | 5.9 | 8.5 | 152 | 343 | 9 | MOLECULE: | TYPE I-B CRISPR-ASSOCIATED PROTEIN CAS5 |
| 91:00:00 | 8hr9-G | 5.9 | 15.1 | 146 | 830 | 10 | MOLECULE: | ARCHAEAL ATPASE |
| 92:00:00 | 7f02-E | 5.9 | 3.3 | 100 | 202 | 8 | MOLECULE: | CYTOCHROME C BIOGENESIS ATP-BINDING EXPORT PROTEI |
| 93:00:00 | 7mcs-E | 5.9 | 4.2 | 149 | 402 | 12 | MOLECULE: | TRANSPOSON TN7 TRANSPOSITION PROTEIN TNSC |
| 94:00:00 | 7ufm-K | 5.9 | 9.1 | 142 | 311 | 8 | MOLECULE: | VCHTNSC |
| 95:00:00 | 8fcv-U | 5.9 | 8 | 149 | 343 | 8 | MOLECULE: | DNA (60-MER) |
| 96:00:00 | 8fcw-U | 5.9 | 8 | 145 | 343 | 9 | MOLECULE: | TNSC |
| 97:00:00 | 7mcs-F | 5.9 | 3.9 | 140 | 401 | 10 | MOLECULE: | TRANSPOSON TN7 TRANSPOSITION PROTEIN TNSC |
| 98:00:00 | 3k1j-A | 5.9 | 11.6 | 151 | 566 | 13 | MOLECULE: | ATP-DEPENDENT PROTEASE LON |
| 99:00:00 | 8p49-A | 5.9 | 4.9 | 148 | 377 | 7 | MOLECULE: | Q8U0N8 PROTEIN |
| 0:00 | 5ln3-M | 5.9 | 11.8 | 159 | 389 | 9 | MOLECULE: | 26S PROTEASOME NON-ATPASE REGULATORY SUBUNIT 2 |
| 1:00 | 7mbw-B | 5.9 | 10.6 | 155 | 469 | 12 | MOLECULE: | TRANSPOSON TN7 TRANSPOSITION PROTEIN TNSC |
| 2:00 | 6n8v-F | 5.9 | 10.9 | 235 | 710 | 9 | MOLECULE: | HEAT SHOCK PROTEIN 104 |
| 3:00 | 3cmw-C | 5.9 | 15.2 | 143 | 1608 | 12 | MOLECULE: | DNA (5'- |
| 4:00 | 4kft-A | 5.9 | 3.5 | 107 | 195 | 13 | MOLECULE: | GENOME PACKAGING NTPASE B204 |
| 5:00 | 2ce7-D | 5.9 | 8.6 | 145 | 413 | 12 | MOLECULE: | CELL DIVISION PROTEIN FTSH |
| 6:00 | 1xp8-A | 5.9 | 10.8 | 139 | 298 | 12 | MOLECULE: | RECA PROTEIN |
| 7:00 | 4kfs-A | 5.9 | 3.6 | 110 | 202 | 14 | MOLECULE: | GENOME PACKAGING NTPASE B204 |
| 8:00 | 7qh2-C | 5.8 | 12.5 | 138 | 467 | 9 | MOLECULE: | LACTATE DEHYDROGENASE (NAD(+),FERREDOXIN) SUBUNIT |
| 9:00 | 4cej-B | 5.8 | 15.6 | 136 | 1156 | 8 | MOLECULE: | ATP-DEPENDENT HELICASE/NUCLEASE SUBUNIT A |
| 10:00 | 7n6i-D | 5.8 | 8.7 | 129 | 257 | 9 | MOLECULE: | TNIQ (HOMOLOGY MODEL) |
| 11:00 | 8emh-G | 5.8 | 7.1 | 181 | 671 | 9 | MOLECULE: | PROTEASE LON-RELATED BREX SYSTEM PROTEIN BRXL |
| 12:00 | 8emh-I | 5.8 | 5.6 | 200 | 672 | 9 | MOLECULE: | PROTEASE LON-RELATED BREX SYSTEM PROTEIN BRXL |
| 13:00 | 8c0w-F | 5.8 | 11.4 | 187 | 823 | 9 | MOLECULE: | PEROXISOMAL ATPASE PEX6 |
| 14:00 | 7m99-B | 5.8 | 8.7 | 129 | 257 | 9 | MOLECULE: | TNSC |
| 15:00 | 1g8p-A | 5.8 | 4.4 | 176 | 321 | 11 | MOLECULE: | MAGNESIUM-CHELATASE 38 KDA SUBUNIT |
| 16:00 | 8th8-L | 5.8 | 11.9 | 206 | 862 | 11 | MOLECULE: | DYNEIN REGULATORY COMPLEX PROTEIN 1/2 N-TERMINAL |
| 17:00 | 7rzy-5 | 5.8 | 10.6 | 148 | 311 | 8 | MOLECULE: | TN6677 VIBRIO CHOLERAE TRANSPOSON TNSC (VCHTNSC) |
| 18:00 | 3b85-A | 5.8 | 3.4 | 114 | 187 | 10 | MOLECULE: | PHOSPHATE STARVATION-INDUCIBLE PROTEIN |
| 19:00 | 8iqh-J | 5.8 | 9 | 237 | 881 | 8 | MOLECULE: | PUTATIVE PRIMASE C962R |
| 20:00 | 4fwv-A | 5.8 | 3.2 | 115 | 340 | 10 | MOLECULE: | TTC1975 PEPTIDASE |
| 21:00 | 1xwi-A | 5.8 | 12.9 | 135 | 322 | 16 | MOLECULE: | SKD1 PROTEIN |
| 22:00 | 8gja-F | 5.8 | 5 | 143 | 342 | 5 | MOLECULE: | RAD51C |
| 23:00 | 4kfr-B | 5.8 | 3.5 | 112 | 208 | 13 | MOLECULE: | GENOME PACKAGING NTPASE B204 |

| | | | | | | | | |
|---|---|---|---|---|---|---|---|---|
| 24:00:00 | 5g4f-E | 5.8 | 8.9 | 197 | 726 | 11 | MOLECULE: | VCP-LIKE ATPASE |
| 25:00:00 | 5vhj-C | 5.8 | 9.1 | 133 | 213 | 13 | MOLECULE: | 26S PROTEASOME NON-ATPASE REGULATORY SUBUNIT 10 |
| 26:00:00 | 4kfr-A | 5.8 | 3.3 | 107 | 204 | 13 | MOLECULE: | GENOME PACKAGING NTPASE B204 |
| 27:00:00 | 6oab-B | 5.8 | 8.6 | 139 | 509 | 12 | MOLECULE: | CELL DIVISION CONTROL PROTEIN 48 |
| 28:00:00 | 3uk6-D | 5.8 | 8.5 | 143 | 299 | 10 | MOLECULE: | RUVB-LIKE 2 |
| 29:00:00 | 5og1-B | 5.8 | 9.4 | 174 | 572 | 11 | MOLECULE: | CHAPERONE PROTEIN CLPB,ATP-DEPENDENT CLP PROTEASE |
| 30:00:00 | 4kfs-B | 5.8 | 3.7 | 110 | 199 | 13 | MOLECULE: | GENOME PACKAGING NTPASE B204 |
| 31:00:00 | 6uks-F | 5.8 | 3.9 | 150 | 362 | 15 | MOLECULE: | MITOCHONDRIAL CHAPERONE BCS1 |
| 32:00:00 | 2oca-A | 5.8 | 6.1 | 122 | 494 | 7 | MOLECULE: | ATP-DEPENDENT DNA HELICASE UVSW |
| 33:00:00 | 8dgf-A | 5.7 | 10.2 | 168 | 1541 | 7 | MOLECULE: | ATP-BINDING PROTEIN AVS4 |
| 34:00:00 | 6o16-A | 5.7 | 10.7 | 138 | 798 | 11 | MOLECULE: | DEAH (ASP-GLU-ALA-HIS) BOX POLYPEPTIDE 37 |
| 35:00:00 | 7rzy-2 | 5.7 | 10 | 148 | 311 | 7 | MOLECULE: | TN6677 VIBRIO CHOLERAE TRANSPOSON TNSC (VCHTNSC) |
| 36:00:00 | 8c0w-B | 5.7 | 11.4 | 183 | 821 | 8 | MOLECULE: | PEROXISOMAL ATPASE PEX6 |
| 37:00:00 | 7m99-F | 5.7 | 8.7 | 129 | 257 | 9 | MOLECULE: | TNSC |
| 38:00:00 | 8fnt-E | 5.7 | 15.5 | 137 | 781 | 9 | MOLECULE: | ARCHAEAL ATPASE |
| 39:00:00 | 8c0w-D | 5.7 | 9.9 | 176 | 821 | 9 | MOLECULE: | PEROXISOMAL ATPASE PEX6 |
| 40:00:00 | 8hrb-K | 5.7 | 12.9 | 133 | 835 | 10 | MOLECULE: | ARCHAEAL ATPASE |
| 41:00:00 | 7n6i-H | 5.7 | 8.7 | 129 | 257 | 9 | MOLECULE: | TNIQ (HOMOLOGY MODEL) |
| 42:00:00 | 3rib-B | 5.7 | 3.8 | 120 | 419 | 10 | MOLECULE: | N-LYSINE METHYLTRANSFERASE SMYD2 |
| 43:00:00 | 6oaa-B | 5.7 | 12.1 | 147 | 509 | 13 | MOLECULE: | CELL DIVISION CONTROL PROTEIN 48 |
| 44:00:00 | 3f8t-A | 5.7 | 5.2 | 212 | 459 | 10 | MOLECULE: | PREDICTED ATPASE INVOLVED IN REPLICATION CONTROL, |
| 45:00:00 | 1jr3-D | 5.7 | 16.8 | 141 | 338 | 9 | MOLECULE: | DNA POLYMERASE III SUBUNIT GAMMA |
| 46:00:00 | 6eud-A | 5.7 | 14.1 | 138 | 808 | 14 | MOLECULE: | ATP-DEPENDENT RNA HELICASE HRPB |
| 47:00:00 | 7mcs-B | 5.7 | 4.1 | 144 | 402 | 11 | MOLECULE: | TRANSPOSON TN7 TRANSPOSITION PROTEIN TNSC |
| 48:00:00 | 5g4f-P | 5.7 | 8.7 | 181 | 726 | 10 | MOLECULE: | VCP-LIKE ATPASE |
| 49:00:00 | 2qgz-A | 5.7 | 3.9 | 124 | 183 | 14 | MOLECULE: | PUTATIVE PRIMOSOME COMPONENT |
| 50:00:00 | 5vhr-D | 5.7 | 6.9 | 128 | 236 | 15 | MOLECULE: | 26S PROTEASOME NON-ATPASE REGULATORY SUBUNIT 10 |
| 51:00:00 | 5vhn-D | 5.7 | 8.1 | 131 | 237 | 14 | MOLECULE: | 26S PROTEASOME NON-ATPASE REGULATORY SUBUNIT 10 |
| 52:00:00 | 5vhq-B | 5.7 | 8.8 | 131 | 233 | 11 | MOLECULE: | 26S PROTEASOME NON-ATPASE REGULATORY SUBUNIT 10 |
| 53:00:00 | 6jpu-A | 5.7 | 3.7 | 143 | 577 | 10 | MOLECULE: | UNCHARACTERIZED AAA DOMAIN-CONTAINING PROTEIN C31 |
| 54:00:00 | 5d4w-C | 5.7 | 4.7 | 179 | 688 | 12 | MOLECULE: | PUTATIVE HEAT SHOCK PROTEIN |
| 55:00:00 | 7ykk-A | 5.7 | 8.7 | 205 | 735 | 9 | MOLECULE: | ATPASE FAMILY GENE 2 PROTEIN |
| 56:00:00 | 4a15-A | 5.6 | 9.7 | 115 | 596 | 12 | MOLECULE: | ATP-DEPENDENT DNA HELICASE TA0057 |
| 57:00:00 | 8d8l-W | 5.6 | 5.5 | 127 | 395 | 10 | MOLECULE: | PROBABLE S-ADENOSYL-L-METHIONINE-DEPENDENT RNA |
| 58:00:00 | 8dgc-B | 5.6 | 15.4 | 165 | 2004 | 11 | MOLECULE: | SEAVS3 |
| 59:00:00 | 8fnu-B | 5.6 | 15.2 | 133 | 796 | 5 | MOLECULE: | KAP NTPASE DOMAIN-CONTAINING PROTEIN |
| 60:00:00 | 7yut-F | 5.6 | 21.8 | 191 | 773 | 13 | MOLECULE: | LON PROTEASE |
| 61:00:00 | 5a5b-M | 5.6 | 4 | 162 | 367 | 14 | MOLECULE: | PROTEASOME COMPONENT PRE3 |
| 62:00:00 | 7rzy-3 | 5.6 | 10.6 | 148 | 311 | 9 | MOLECULE: | TN6677 VIBRIO CHOLERAE TRANSPOSON TNSC (VCHTNSC) |
| 63:00:00 | 6og2-F | 5.6 | 10.5 | 133 | 558 | 11 | MOLECULE: | HYPERACTIVE DISAGGREGASE CLPB |
| 64:00:00 | 5g4f-D | 5.6 | 13.5 | 188 | 726 | 10 | MOLECULE: | VCP-LIKE ATPASE |
| 65:00:00 | 6mfv-D | 5.6 | 8.6 | 127 | 641 | 11 | MOLECULE: | TETRATRICOPEPTIDE REPEAT SENSOR PH0952 |
| 66:00:00 | 5vhm-D | 5.6 | 3.3 | 126 | 220 | 16 | MOLECULE: | 26S PROTEASOME NON-ATPASE REGULATORY SUBUNIT 10 |
| 67:00:00 | 4kfu-C | 5.6 | 3.7 | 109 | 195 | 14 | MOLECULE: | GENOME PACKAGING NTPASE B204 |
| 68:00:00 | 5og1-C | 5.6 | 9.8 | 187 | 624 | 10 | MOLECULE: | CHAPERONE PROTEIN CLPB,ATP-DEPENDENT CLP PROTEASE |
| 69:00:00 | 6qs7-B | 5.6 | 6.5 | 178 | 582 | 11 | MOLECULE: | CHAPERONE PROTEIN CLPB |
| 70:00:00 | 6ees-A | 5.6 | 7.5 | 157 | 2023 | 13 | MOLECULE: | MIDASIN |
| 71:00:00 | 6qs7-A | 5.6 | 8 | 213 | 568 | 14 | MOLECULE: | CHAPERONE PROTEIN CLPB |
| 72:00:00 | 8fnu-A | 5.5 | 13.6 | 137 | 796 | 7 | MOLECULE: | KAP NTPASE DOMAIN-CONTAINING PROTEIN |
| 73:00:00 | 7ufm-L | 5.5 | 9.3 | 142 | 311 | 8 | MOLECULE: | VCHTNSC |
| 74:00:00 | 8hrb-M | 5.5 | 15 | 136 | 849 | 11 | MOLECULE: | ARCHAEAL ATPASE |
| 75:00:00 | 7ufm-I | 5.5 | 9 | 143 | 311 | 8 | MOLECULE: | VCHTNSC |
| 76:00:00 | 3qxy-A | 5.5 | 18.5 | 129 | 421 | 12 | MOLECULE: | N-LYSINE METHYLTRANSFERASE SETD6 |

| | | | | | | | | |
|---|---|---|---|---|---|---|---|---|
| 77:00:00 | 8fnu-C | 5.5 | 13.7 | 139 | 796 | 5 | MOLECULE: | KAP NTPASE DOMAIN-CONTAINING PROTEIN |
| 78:00:00 | 6n9x-B | 5.5 | 21.6 | 174 | 500 | 9 | MOLECULE: | DNA PRIMASE/HELICASE |
| 79:00:00 | 6fhs-C | 5.5 | 9.6 | 141 | 459 | 15 | MOLECULE: | RUVB-LIKE HELICASE |
| 80:00:00 | 5vhf-D | 5.5 | 13.4 | 163 | 344 | 12 | MOLECULE: | 26S PROTEASOME NON-ATPASE REGULATORY SUBUNIT 10 |
| 81:00:00 | 3gp8-A | 5.5 | 14.3 | 141 | 551 | 14 | MOLECULE: | EXODEOXYRIBONUCLEASE V, SUBUNIT RECD, PUTATIVE |
| 82:00:00 | 7jgs-C | 5.5 | 18.9 | 171 | 614 | 9 | MOLECULE: | ORIGIN RECOGNITION COMPLEX SUBUNIT 2 |
| 83:00:00 | 5aor-A | 5.5 | 27.8 | 200 | 1009 | 10 | MOLECULE: | DOSAGE COMPENSATION REGULATOR |
| 84:00:00 | 5t0j-B | 5.5 | 4.2 | 144 | 348 | 11 | MOLECULE: | 26S PROTEASOME NON-ATPASE REGULATORY SUBUNIT 2 |
| 85:00:00 | 6igm-B | 5.5 | 4.8 | 141 | 398 | 11 | MOLECULE: | RUVB-LIKE 1 |
| 86:00:00 | 6oab-A | 5.5 | 14.3 | 180 | 493 | 12 | MOLECULE: | CELL DIVISION CONTROL PROTEIN 48 |
| 87:00:00 | 6fo1-D | 5.5 | 9.4 | 142 | 310 | 13 | MOLECULE: | RUVB-LIKE 1 |
| 88:00:00 | 6ppj-B | 5.5 | 15.6 | 119 | 848 | 13 | MOLECULE: | UVRD/REP HELICASE |
| 89:00:00 | 6qs4-A | 5.5 | 11.6 | 192 | 568 | 13 | MOLECULE: | CHAPERONE PROTEIN CLPB |
| 90:00:00 | 6ox1-B | 5.4 | 30 | 138 | 485 | 10 | MOLECULE: | ACTIN, CYTOPLASMIC 1 |
| 91:00:00 | 5e7p-A | 5.4 | 15.9 | 199 | 719 | 11 | MOLECULE: | CELL DIVISION CONTROL PROTEIN CDC48 |
| 92:00:00 | 2fna-A | 5.4 | 12.1 | 136 | 352 | 10 | MOLECULE: | CONSERVED HYPOTHETICAL PROTEIN |
| 93:00:00 | 6hz8-L | 5.4 | 3.7 | 136 | 285 | 11 | MOLECULE: | 5-METHYLCYTOSINE-SPECIFIC RESTRICTION ENZYME B |
| 94:00:00 | 7otj-B | 5.4 | 4.3 | 123 | 436 | 10 | MOLECULE: | ATP-DEPENDENT DNA HELICASE PIF1 |
| 95:00:00 | 3nbx-X | 5.4 | 5.5 | 175 | 481 | 10 | MOLECULE: | ATPASE RAVA |
| 96:00:00 | 5dgk-A | 5.4 | 4.8 | 202 | 519 | 9 | MOLECULE: | ACTIVE HELICASE |
| 97:00:00 | 7rzy-6 | 5.4 | 6.7 | 138 | 311 | 7 | MOLECULE: | TN6677 VIBRIO CHOLERAE TRANSPOSON TNSC (VCHTNSC) |
| 98:00:00 | 6ef1-M | 5.4 | 10.4 | 134 | 262 | 15 | MOLECULE: | PROTEASOME SUBUNIT ALPHA TYPE-1 |
| 99:00:00 | 5vhf-E | 5.4 | 10.2 | 151 | 345 | 10 | MOLECULE: | 26S PROTEASOME NON-ATPASE REGULATORY SUBUNIT 10 |
| 0:00 | 4tvs-A | 5.4 | 3.6 | 147 | 224 | 10 | MOLECULE: | TORSIN-1A-INTERACTING PROTEIN 1 |
| 1:00 | 5kne-F | 5.4 | 10.1 | 196 | 607 | 10 | MOLECULE: | HEAT SHOCK PROTEIN 104 |
| 2:00 | 5nss-N | 5.4 | 3.6 | 138 | 251 | 7 | MOLECULE: | DNA-DIRECTED RNA POLYMERASE SUBUNIT ALPHA |
| 3:00 | 3j16-B | 5.3 | 5.2 | 126 | 608 | 11 | MOLECULE: | DOM34P |
| 4:00 | 6nmi-A | 5.3 | 16.3 | 144 | 653 | 9 | MOLECULE: | GENERAL TRANSCRIPTION AND DNA REPAIR FACTOR IIH H |
| 5:00 | 7rzy-4 | 5.3 | 7.3 | 142 | 311 | 8 | MOLECULE: | TN6677 VIBRIO CHOLERAE TRANSPOSON TNSC (VCHTNSC) |
| 6:00 | 7pkq-x | 5.3 | 3.8 | 117 | 368 | 7 | MOLECULE: | MS35 |
| 7:00 | 8hr9-H | 5.3 | 15.4 | 139 | 834 | 8 | MOLECULE: | ARCHAEAL ATPASE |
| 8:00 | 7uqj-A | 5.3 | 9.2 | 202 | 585 | 13 | MOLECULE: | ATPASE HISTONE CHAPERONE YTA7 |
| 9:00 | 6oa9-C | 5.3 | 8.7 | 166 | 492 | 9 | MOLECULE: | CELL DIVISION CONTROL PROTEIN 48 |
| 10:00 | 7qy7-A | 5.3 | 15.8 | 160 | 342 | 8 | MOLECULE: | 26S PROTEASOME NON-ATPASE REGULATORY SUBUNIT 13 |
| 11:00 | 7mcs-C | 5.3 | 4 | 143 | 404 | 11 | MOLECULE: | TRANSPOSON TN7 TRANSPOSITION PROTEIN TNSC |
| 12:00 | 8giy-A | 5.3 | 14 | 130 | 343 | 11 | MOLECULE: | DNA POLYMERASE III SUBUNIT DELTA |
| 13:00 | 3io5-B | 5.3 | 4.3 | 142 | 272 | 8 | MOLECULE: | RECOMBINATION AND REPAIR PROTEIN |
| 14:00 | 5vhn-E | 5.3 | 6.4 | 121 | 241 | 12 | MOLECULE: | 26S PROTEASOME NON-ATPASE REGULATORY SUBUNIT 10 |
| 15:00 | 5vhh-E | 5.3 | 10.6 | 152 | 345 | 12 | MOLECULE: | 26S PROTEASOME REGULATORY SUBUNIT 7 |
| 16:00 | 6ahf-C | 5.3 | 15.5 | 215 | 865 | 10 | MOLECULE: | HEAT SHOCK PROTEIN 104 |
| 17:00 | 2xgj-B | 5.2 | 7.4 | 128 | 773 | 5 | MOLECULE: | ATP-DEPENDENT RNA HELICASE DOB1 |
| 18:00 | 8fnu-G | 5.2 | 15 | 142 | 796 | 6 | MOLECULE: | KAP NTPASE DOMAIN-CONTAINING PROTEIN |
| 19:00 | 8fnu-D | 5.2 | 15.7 | 129 | 796 | 6 | MOLECULE: | KAP NTPASE DOMAIN-CONTAINING PROTEIN |
| 20:00 | 5o6b-A | 5.2 | 4 | 121 | 528 | 12 | MOLECULE: | ATP-DEPENDENT DNA HELICASE PIF1 |
| 21:00 | 7z6h-K | 5.2 | 9.4 | 138 | 439 | 11 | MOLECULE: | CELL CYCLE CHECKPOINT CONTROL PROTEIN RAD9A |
| 22:00 | 8fnu-E | 5.2 | 13.5 | 137 | 796 | 5 | MOLECULE: | KAP NTPASE DOMAIN-CONTAINING PROTEIN |
| 23:00 | 7wi4-B | 5.2 | 11.6 | 132 | 410 | 11 | MOLECULE: | ATP-DEPENDENT ZINC METALLOPROTEASE FTSH |
| 24:00:00 | 6ef0-J | 5.2 | 8 | 139 | 272 | 12 | MOLECULE: | PROTEASOME SUBUNIT ALPHA TYPE-1 |
| 25:00:00 | 2f1h-A | 5.2 | 6.6 | 141 | 313 | 9 | MOLECULE: | DNA REPAIR AND RECOMBINATION PROTEIN RADA |
| 26:00:00 | 5vft-C | 5.2 | 7.8 | 151 | 384 | 12 | MOLECULE: | 26S PROTEASOME NON-ATPASE REGULATORY SUBUNIT 1 |
| 27:00:00 | 5udb-D | 5.2 | 11.6 | 143 | 447 | 10 | MOLECULE: | DNA REPLICATION LICENSING FACTOR MCM2 |
| 28:00:00 | 8emc-B | 5.2 | 6 | 177 | 671 | 10 | MOLECULE: | PROTEASE LON-RELATED BREX SYSTEM PROTEIN BRXL |
| 29:00:00 | 6acx-A | 5.1 | 16.7 | 173 | 1175 | 10 | MOLECULE: | MYCOBACTERIUM SMEGMATIS MFD |

| | | | | | | | | |
|---|---|---|---|---|---|---|---|---|
| 30:00:00 | 7ypj-E | 5.1 | 23.6 | 202 | 773 | 13 | MOLECULE: | LON PROTEASE |
| 31:00:00 | 7amv-W | 5.1 | 8.7 | 142 | 637 | 12 | MOLECULE: | DNA-DIRECTED RNA POLYMERASE 147 KDA POLYPEPTIDE |
| 32:00:00 | 8fnu-F | 5.1 | 13.6 | 139 | 796 | 6 | MOLECULE: | KAP NTPASE DOMAIN-CONTAINING PROTEIN |
| 33:00:00 | 2mpl-A | 5.1 | 7.2 | 91 | 127 | 5 | MOLECULE: | ZINC FINGER PROTEIN ZFPM1 |
| 34:00:00 | 4xgc-C | 5.1 | 12.4 | 157 | 567 | 9 | MOLECULE: | ORIGIN RECOGNITION COMPLEX SUBUNIT 2 |
| 35:00:00 | 6qem-I | 5.1 | 6.8 | 142 | 239 | 14 | MOLECULE: | REPLICATIVE DNA HELICASE |
| 36:00:00 | 3crv-A | 5.1 | 14.9 | 126 | 551 | 12 | MOLECULE: | XPD/RAD3 RELATED DNA HELICASE |
| 37:00:00 | 8faz-X | 5.1 | 3.4 | 111 | 236 | 9 | MOLECULE: | DNA REPAIR PROTEIN RAD51 HOMOLOG 2 |
| 38:00:00 | 8jqb-E | 5.1 | 5.7 | 112 | 499 | 9 | MOLECULE: | ENDONUCLEASE GAJA |
| 39:00:00 | 5vhi-B | 5.1 | 9.5 | 162 | 298 | 10 | MOLECULE: | 26S PROTEASOME REGULATORY SUBUNIT 7 |
| 40:00:00 | 6fhs-E | 5.1 | 14.3 | 146 | 441 | 11 | MOLECULE: | RUVB-LIKE HELICASE |
| 41:00:00 | 4fwh-A | 5.1 | 11.6 | 152 | 580 | 9 | MOLECULE: | TTC1975 PEPTIDASE |
| 42:00:00 | 7t1q-A | 5 | 5.8 | 105 | 377 | 4 | MOLECULE: | SUCCINYL-DIAMINOPIMELATE DESUCCINYLASE |
| 43:00:00 | 8fvu-A | 5 | 9.5 | 179 | 1361 | 10 | MOLECULE: | BACULOVIRAL IAP REPEAT-CONTAINING PROTEIN 1 |
| 44:00:00 | 7mqj-A | 5 | 16.1 | 140 | 773 | 10 | MOLECULE: | PROBABLE ATP-DEPENDENT RNA HELICASE DHR1 |
| 45:00:00 | 8hr8-D | 5 | 14.8 | 146 | 836 | 9 | MOLECULE: | ARCHAEAL ATPASE |
| 46:00:00 | 8osf-B | 5 | 4.3 | 166 | 321 | 10 | MOLECULE: | MAGNESIUM-CHELATASE SUBUNIT CHLI |
| 47:00:00 | 6sh5-G | 5 | 5.8 | 136 | 334 | 13 | MOLECULE: | MITOCHONDRIAL CHAPERONE BCS1 |
| 48:00:00 | 5vca-N | 5 | 9.6 | 147 | 543 | 9 | MOLECULE: | VCP-LIKE ATPASE |
| 49:00:00 | 8gbj-X | 5 | 3.5 | 116 | 232 | 9 | MOLECULE: | DNA REPAIR PROTEIN RAD51 HOMOLOG 2 |
| 50:00:00 | 5d4w-A | 5 | 13.5 | 191 | 688 | 10 | MOLECULE: | PUTATIVE HEAT SHOCK PROTEIN |
| 51:00:00 | 7ou2-B | 4.9 | 17 | 172 | 650 | 8 | MOLECULE: | DNA MISMATCH REPAIR PROTEIN MUTS |
| 52:00:00 | 4q2d-A | 4.9 | 11.8 | 163 | 927 | 7 | MOLECULE: | CRISPR-ASSOCIATED HELICASE CAS3 |
| 53:00:00 | 8c0v-A | 4.9 | 14.6 | 205 | 823 | 13 | MOLECULE: | PEROXISOMAL ATPASE PEX1 |
| 54:00:00 | 4cej-A | 4.9 | 11.8 | 119 | 1177 | 9 | MOLECULE: | ATP-DEPENDENT HELICASE/NUCLEASE SUBUNIT A |
| 55:00:00 | 3cmu-A | 4.9 | 10.8 | 140 | 1937 | 14 | MOLECULE: | DNA (5'- |
| 56:00:00 | 8dgc-A | 4.9 | 22 | 175 | 2028 | 9 | MOLECULE: | SEAVS3 |
| 57:00:00 | 7mbw-D | 4.9 | 10.3 | 154 | 467 | 11 | MOLECULE: | TRANSPOSON TN7 TRANSPOSITION PROTEIN TNSC |
| 58:00:00 | 7qy7-D | 4.9 | 5.6 | 169 | 378 | 9 | MOLECULE: | 26S PROTEASOME NON-ATPASE REGULATORY SUBUNIT 13 |
| 59:00:00 | 6npy-A | 4.9 | 9.8 | 147 | 798 | 12 | MOLECULE: | NACHT, LRR AND PYD DOMAINS-CONTAINING PROTEIN 3 |
| 60:00:00 | 8osf-D | 4.9 | 4.2 | 165 | 319 | 12 | MOLECULE: | MAGNESIUM-CHELATASE SUBUNIT CHLI |
| 61:00:00 | 7mcs-D | 4.9 | 4.1 | 146 | 403 | 11 | MOLECULE: | TRANSPOSON TN7 TRANSPOSITION PROTEIN TNSC |
| 62:00:00 | 6qem-K | 4.9 | 4.7 | 141 | 239 | 14 | MOLECULE: | REPLICATIVE DNA HELICASE |
| 63:00:00 | 6qem-H | 4.9 | 5.1 | 138 | 239 | 14 | MOLECULE: | REPLICATIVE DNA HELICASE |
| 64:00:00 | 5wcb-B | 4.9 | 10.3 | 121 | 277 | 12 | MOLECULE: | MEIOTIC SPINDLE FORMATION PROTEIN MEI-1 |
| 65:00:00 | 7jk5-A | 4.9 | 17.2 | 169 | 390 | 7 | MOLECULE: | ORIGIN RECOGNITION COMPLEX SUBUNIT 2 |
| 66:00:00 | 4xgc-A | 4.9 | 13.5 | 142 | 310 | 8 | MOLECULE: | ORIGIN RECOGNITION COMPLEX SUBUNIT 2 |
| 67:00:00 | 8ouz-D | 4.9 | 3.5 | 117 | 234 | 9 | MOLECULE: | DNA REPAIR PROTEIN RAD51 HOMOLOG 2 |
| 68:00:00 | 6qem-G | 4.9 | 15.3 | 153 | 239 | 10 | MOLECULE: | REPLICATIVE DNA HELICASE |
| 69:00:00 | 6rfl-Y | 4.8 | 4.3 | 131 | 600 | 11 | MOLECULE: | DNA-DEPENDENT RNA POLYMERASE SUBUNIT RPO132 |
| 70:00:00 | 6qem-J | 4.8 | 8.3 | 143 | 239 | 13 | MOLECULE: | REPLICATIVE DNA HELICASE |
| 71:00:00 | 6ef0-I | 4.8 | 10.4 | 137 | 271 | 14 | MOLECULE: | PROTEASOME SUBUNIT ALPHA TYPE-1 |
| 72:00:00 | 6mfv-C | 4.8 | 8.5 | 123 | 641 | 11 | MOLECULE: | TETRATRICOPEPTIDE REPEAT SENSOR PH0952 |
| 73:00:00 | 4ag5-B | 4.8 | 4 | 117 | 374 | 9 | MOLECULE: | TYPE IV SECRETORY PATHWAY VIRB4 COMPONENTS-LIKE P |
| 74:00:00 | 8odp-A | 4.7 | 4.5 | 110 | 505 | 5 | MOLECULE: | RNA-SPLICING LIGASE RTCB HOMOLOG |
| 75:00:00 | 7sjr-A | 4.7 | 16.9 | 140 | 911 | 11 | MOLECULE: | DNA HELICASE |
| 76:00:00 | 2wjy-A | 4.7 | 24.2 | 182 | 773 | 4 | MOLECULE: | REGULATOR OF NONSENSE TRANSCRIPTS 1 |
| 77:00:00 | 7mbw-A | 4.7 | 11.5 | 152 | 475 | 11 | MOLECULE: | TRANSPOSON TN7 TRANSPOSITION PROTEIN TNSC |
| 78:00:00 | 6ef1-J | 4.7 | 3.4 | 122 | 273 | 13 | MOLECULE: | PROTEASOME SUBUNIT ALPHA TYPE-1 |
| 79:00:00 | 2d7d-A | 4.7 | 16.7 | 138 | 621 | 9 | MOLECULE: | 5'-D(P*TP*TP*T)-3' |
| 80:00:00 | 4ag5-A | 4.7 | 3.8 | 116 | 374 | 11 | MOLECULE: | TYPE IV SECRETORY PATHWAY VIRB4 COMPONENTS-LIKE P |
| 81:00:00 | 4ag6-A | 4.7 | 4 | 118 | 373 | 8 | MOLECULE: | TYPE IV SECRETORY PATHWAY VIRB4 COMPONENTS-LIKE P |
| 82:00:00 | 4ag6-B | 4.7 | 3.8 | 117 | 376 | 9 | MOLECULE: | TYPE IV SECRETORY PATHWAY VIRB4 COMPONENTS-LIKE P |

| | | | | | | | | |
|---|---|---|---|---|---|---|---|---|
| 83:00:00 | 5vhp-D | 4.7 | 3.7 | 123 | 237 | 13 | MOLECULE: | 26S PROTEASOME NON-ATPASE REGULATORY SUBUNIT 10 |
| 84:00:00 | 7yux-B | 4.7 | 25.2 | 174 | 773 | 10 | MOLECULE: | LON PROTEASE |
| 85:00:00 | 7lcc-A | 4.7 | 26.6 | 121 | 1369 | 12 | MOLECULE: | HELRAISER K1068Q |
| 86:00:00 | 4zcf-C | 4.6 | 6.4 | 143 | 627 | 7 | MOLECULE: | RESTRICTION ENDONUCLEASE ECOP15I, MODIFICATION SU |
| 87:00:00 | 7lgp-B | 4.6 | 7.1 | 102 | 377 | 8 | MOLECULE: | SUCCINYL-DIAMINOPIMELATE DESUCCINYLASE |
| 88:00:00 | 2fdc-A | 4.6 | 4.3 | 123 | 505 | 10 | MOLECULE: | 5'-D(P*CP*GP*GP*CP*TP*CP*CP*AP*TP*CP*TP*CP*TP*AP* |
| 89:00:00 | 5vhn-A | 4.6 | 8.7 | 129 | 258 | 9 | MOLECULE: | 26S PROTEASOME NON-ATPASE REGULATORY SUBUNIT 10 |
| 90:00:00 | 4m9s-B | 4.6 | 12.3 | 153 | 511 | 10 | MOLECULE: | CELL DEATH PROTEIN 4 |
| 91:00:00 | 6qs4-B | 4.6 | 7 | 181 | 582 | 13 | MOLECULE: | CHAPERONE PROTEIN CLPB |
| 92:00:00 | 1gm5-A | 4.5 | 14.3 | 139 | 729 | 11 | MOLECULE: | RECG |
| 93:00:00 | 8b1r-B | 4.5 | 12.9 | 127 | 1170 | 9 | MOLECULE: | RECBCD ENZYME SUBUNIT RECB |
| 94:00:00 | 4ksr-A | 4.5 | 18.9 | 135 | 521 | 9 | MOLECULE: | TYPE II SECRETION SYSTEM PROTEIN E, HEMOLYSIN-COR |
| 95:00:00 | 7bpa-A | 4.5 | 17.7 | 200 | 742 | 10 | MOLECULE: | TRANSITIONAL ENDOPLASMIC RETICULUM ATPASE |
| 96:00:00 | 5vhq-A | 4.5 | 3.2 | 122 | 261 | 11 | MOLECULE: | 26S PROTEASOME NON-ATPASE REGULATORY SUBUNIT 10 |
| 97:00:00 | 6ef1-I | 4.5 | 6.2 | 128 | 271 | 13 | MOLECULE: | PROTEASOME SUBUNIT ALPHA TYPE-1 |
| 98:00:00 | 7yph-E | 4.5 | 13.6 | 74 | 773 | 22 | MOLECULE: | LON PROTEASE |
| 99:00:00 | 7mi6-A | 4.4 | 24 | 95 | 2419 | 14 | MOLECULE: | FUSION PROTEIN OF DYNEIN AND ENDOLYSIN |
| 0:00 | 7nyx-A | 4.4 | 3.7 | 112 | 1467 | 11 | MOLECULE: | CHROMOSOME PARTITION PROTEIN MUKB |
| 1:00 | 6fwr-A | 4.4 | 15.4 | 134 | 699 | 10 | MOLECULE: | ATP-DEPENDENT DNA HELICASE DING |
| 2:00 | 5j1s-B | 4.4 | 3.7 | 145 | 235 | 11 | MOLECULE: | TORSIN-1A |
| 3:00 | 5udb-9 | 4.4 | 16.7 | 147 | 376 | 7 | MOLECULE: | DNA REPLICATION LICENSING FACTOR MCM2 |
| 4:00 | 6sh4-B | 4.4 | 7.1 | 144 | 376 | 13 | MOLECULE: | MITOCHONDRIAL CHAPERONE BCS1 |
| 5:00 | 7mcs-G | 4.4 | 4.2 | 142 | 358 | 13 | MOLECULE: | TRANSPOSON TN7 TRANSPOSITION PROTEIN TNSC |
| 6:00 | 5t0i-B | 4.4 | 9.5 | 149 | 348 | 12 | MOLECULE: | 26S PROTEASE REGULATORY SUBUNIT 7 |
| 7:00 | 7b8s-C | 4.3 | 5 | 96 | 589 | 4 | MOLECULE: | MULTIDRUG EFFLUX PUMP SUBUNIT ACRB,MULTIDRUG EFFL |
| 8:00 | 3al0-B | 4.3 | 21.1 | 119 | 482 | 4 | MOLECULE: | GLUTAMYL-TRNA(GLN) AMIDOTRANSFERASE SUBUNIT A |
| 9:00 | 8t5s-A | 4.3 | 4.5 | 121 | 693 | 7 | MOLECULE: | DICER-RELATED HELICASE |
| 10:00 | 6slf-A | 4.3 | 12.8 | 111 | 398 | 4 | MOLECULE: | N-ALPHA-ACYL-GLUTAMINE AMINOACYLASE |
| 11:00 | 7w46-B | 4.3 | 7.8 | 156 | 402 | 13 | MOLECULE: | UNCHARACTERIZED ATPASE YJOB |
| 12:00 | 6sh4-A | 4.3 | 6.7 | 162 | 376 | 10 | MOLECULE: | MITOCHONDRIAL CHAPERONE BCS1 |
| 13:00 | 7wi4-C | 4.3 | 11.4 | 127 | 410 | 13 | MOLECULE: | ATP-DEPENDENT ZINC METALLOPROTEASE FTSH |
| 14:00 | 6sh4-G | 4.3 | 7.9 | 152 | 376 | 13 | MOLECULE: | MITOCHONDRIAL CHAPERONE BCS1 |
| 15:00 | 4m4w-O | 4.3 | 4.1 | 121 | 250 | 13 | MOLECULE: | REPLICATIVE HELICASE |
| 16:00 | 8oz7-A | 4.2 | 22.6 | 142 | 597 | 9 | MOLECULE: | ABIA |
| 17:00 | 8q1b-B | 4.2 | 7.4 | 124 | 406 | 10 | MOLECULE: | PROBABLE MITOCHONDRIAL-PROCESSING PEPTIDASE SUBUN |
| 18:00 | 6jpq-A | 4.2 | 8.2 | 196 | 577 | 11 | MOLECULE: | UNCHARACTERIZED AAA DOMAIN-CONTAINING PROTEIN C31 |
| 19:00 | 8osf-F | 4.2 | 4.2 | 115 | 226 | 10 | MOLECULE: | MAGNESIUM-CHELATASE SUBUNIT CHLI |
| 20:00 | 7wi4-A | 4.2 | 11.7 | 128 | 410 | 12 | MOLECULE: | ATP-DEPENDENT ZINC METALLOPROTEASE FTSH |
| 21:00 | 5vhn-B | 4.2 | 3.4 | 135 | 257 | 13 | MOLECULE: | 26S PROTEASOME NON-ATPASE REGULATORY SUBUNIT 10 |
| 22:00 | 6em8-B | 4.2 | 5.4 | 181 | 744 | 15 | MOLECULE: | ATP-DEPENDENT CLP PROTEASE ATP-BINDING SUBUNIT CL |
| 23:00 | 6bog-A | 4.1 | 5.1 | 174 | 967 | 10 | MOLECULE: | RNA POLYMERASE-ASSOCIATED PROTEIN RAPA |
| 24:00:00 | 4f4c-A | 4.1 | 18.7 | 97 | 1250 | 10 | MOLECULE: | MULTIDRUG RESISTANCE PROTEIN PGP-1 |
| 25:00:00 | 7ufm-B | 4.1 | 8.1 | 145 | 311 | 8 | MOLECULE: | VCHTNSC |
| 26:00:00 | 7wi4-F | 4.1 | 11.7 | 134 | 410 | 8 | MOLECULE: | ATP-DEPENDENT ZINC METALLOPROTEASE FTSH |
| 27:00:00 | 6ef0-M | 4.1 | 12.1 | 129 | 258 | 15 | MOLECULE: | PROTEASOME SUBUNIT ALPHA TYPE-1 |
| 28:00:00 | 4cr4-J | 4.1 | 10.5 | 164 | 373 | 10 | MOLECULE: | PROTEASOME COMPONENT PRE3 |
| 29:00:00 | 1w36-C | 4 | 17.8 | 127 | 1121 | 9 | MOLECULE: | EXODEOXYRIBONUCLEASE V BETA CHAIN |
| 30:00:00 | 8osh-D | 4 | 4.4 | 145 | 230 | 7 | MOLECULE: | MAGNESIUM-CHELATASE SUBUNIT CHLI |
| 31:00:00 | 4xgc-D | 4 | 15.6 | 143 | 440 | 8 | MOLECULE: | ORIGIN RECOGNITION COMPLEX SUBUNIT 2 |
| 32:00:00 | 6epc-K | 4 | 16.7 | 87 | 391 | 17 | MOLECULE: | PROTEASOME SUBUNIT ALPHA TYPE-6 |
| 33:00:00 | 3jb9-X | 3.9 | 27 | 199 | 1195 | 12 | MOLECULE: | PRE-MRNA-SPLICING FACTOR SPP42 |
| 34:00:00 | 3tuz-C | 3.9 | 7 | 110 | 345 | 7 | MOLECULE: | D-METHIONINE TRANSPORT SYSTEM PERMEASE PROTEIN ME |
| 35:00:00 | 6mfv-A | 3.9 | 8 | 123 | 641 | 11 | MOLECULE: | TETRATRICOPEPTIDE REPEAT SENSOR PH0952 |

| | | | | | | | | |
|---|---|---|---|---|---|---|---|---|
| 36:00:00 | 8iqh-C | 3.9 | 6.3 | 203 | 891 | 7 | MOLECULE: | PUTATIVE PRIMASE C962R |
| 37:00:00 | 5g4f-A | 3.9 | 7.5 | 173 | 726 | 9 | MOLECULE: | VCP-LIKE ATPASE |
| 38:00:00 | 8yax-A | 3.8 | 4.7 | 111 | 1314 | 9 | MOLECULE: | PAPAIN-LIKE PROTEASE NSP3 |
| 39:00:00 | 5cio-A | 3.8 | 5.6 | 176 | 770 | 7 | MOLECULE: | PYRROLOQUINOLINE QUINONE BIOSYNTHESIS PROTEIN PQQ |
| 40:00:00 | 8vx9-B | 3.8 | 9 | 165 | 1174 | 9 | MOLECULE: | HAMA |
| 41:00:00 | 4nh0-A | 3.8 | 4.2 | 130 | 861 | 12 | MOLECULE: | CELL DIVISIONFTSK/SPOIIIE |
| 42:00:00 | 8hr8-C | 3.8 | 15.2 | 143 | 835 | 10 | MOLECULE: | ARCHAEAL ATPASE |
| 43:00:00 | 8iqh-A | 3.8 | 5.4 | 202 | 882 | 8 | MOLECULE: | PUTATIVE PRIMASE C962R |
| 44:00:00 | 5kzf-K | 3.8 | 10.5 | 168 | 498 | 12 | MOLECULE: | PROTEASOME-ASSOCIATED ATPASE |
| 45:00:00 | 5wcb-E | 3.8 | 10.3 | 125 | 277 | 11 | MOLECULE: | MEIOTIC SPINDLE FORMATION PROTEIN MEI-1 |
| 46:00:00 | 5vho-A | 3.8 | 3.4 | 122 | 255 | 11 | MOLECULE: | 26S PROTEASOME NON-ATPASE REGULATORY SUBUNIT 10 |
| 47:00:00 | 7y8u-F | 3.7 | 8.4 | 96 | 347 | 10 | MOLECULE: | ALBE HOMOLOG |
| 48:00:00 | 7sr6-A | 3.7 | 12.7 | 166 | 566 | 5 | MOLECULE: | POLYMERASE |
| 49:00:00 | 6og1-F | 3.7 | 10.7 | 140 | 571 | 9 | MOLECULE: | HYPERACTIVE DISAGGREGASE CLPB |
| 50:00:00 | 2g49-A | 3.6 | 16 | 189 | 970 | 6 | MOLECULE: | INSULIN-DEGRADING ENZYME |
| 51:00:00 | 3c6k-D | 3.6 | 9.8 | 108 | 348 | 14 | MOLECULE: | SPERMINE SYNTHASE |
| 52:00:00 | 7yv9-A | 3.5 | 12.7 | 144 | 1067 | 9 | MOLECULE: | UNCONVENTIONAL MYOSIN-VA |
| 53:00:00 | 5b7i-A | 3.5 | 13.3 | 149 | 1044 | 11 | MOLECULE: | CRISPR-ASSOCIATED NUCLEASE/HELICASE CAS3 SUBTYPE |
| 54:00:00 | 4ke4-A | 3.5 | 3.5 | 86 | 437 | 5 | MOLECULE: | HYDROXYCINNAMOYL-COA:SHIKIMATE HYDROXYCINNAMOYL |
| 55:00:00 | 6jqh-B | 3.5 | 9.7 | 107 | 513 | 9 | MOLECULE: | MADA |
| 56:00:00 | 8iqh-E | 3.5 | 5.4 | 199 | 880 | 8 | MOLECULE: | PUTATIVE PRIMASE C962R |
| 57:00:00 | 7wi4-E | 3.5 | 10.3 | 134 | 410 | 10 | MOLECULE: | ATP-DEPENDENT ZINC METALLOPROTEASE FTSH |
| 58:00:00 | 8csp-X | 3.5 | 4.3 | 131 | 352 | 11 | MOLECULE: | 28S RIBOSOMAL PROTEIN S34, MITOCHONDRIAL |
| 59:00:00 | 4bj6-A | 3.5 | 6 | 132 | 293 | 5 | MOLECULE: | RAP1-INTERACTING FACTOR 2 |
| 60:00:00 | 7v2b-D | 3.5 | 7.5 | 107 | 376 | 9 | MOLECULE: | VPSR |
| 61:00:00 | 6ted-Z | 3.4 | 14.8 | 150 | 510 | 5 | MOLECULE: | DNA-DIRECTED RNA POLYMERASE SUBUNIT |
| 62:00:00 | 3ife-A | 3.4 | 10 | 92 | 416 | 7 | MOLECULE: | PEPTIDASE T |
| 63:00:00 | 7w01-A | 3.4 | 7.6 | 147 | 1598 | 12 | MOLECULE: | PHOSPHOLIPID-TRANSPORTING ATPASE ABCA3 |
| 64:00:00 | 3qmz-B | 3.4 | 21.8 | 87 | 2136 | 7 | MOLECULE: | CYTOPLASMIC DYNEIN HEAVY CHAIN |
| 65:00:00 | 7nfy-A | 3.4 | 18.8 | 101 | 776 | 17 | MOLECULE: | LON PROTEASE HOMOLOG, MITOCHONDRIAL |
| 66:00:00 | 8iqh-B | 3.4 | 6.7 | 202 | 892 | 6 | MOLECULE: | PUTATIVE PRIMASE C962R |
| 67:00:00 | 6sh4-D | 3.4 | 8.1 | 144 | 376 | 11 | MOLECULE: | MITOCHONDRIAL CHAPERONE BCS1 |
| 68:00:00 | 6sh4-E | 3.4 | 7.9 | 144 | 376 | 11 | MOLECULE: | MITOCHONDRIAL CHAPERONE BCS1 |
| 69:00:00 | 6sh4-C | 3.4 | 6.8 | 142 | 376 | 11 | MOLECULE: | MITOCHONDRIAL CHAPERONE BCS1 |
| 70:00:00 | 7oky-Z | 3.3 | 7.9 | 114 | 374 | 5 | MOLECULE: | DNA-DIRECTED RNA POLYMERASE II SUBUNIT RPB1 |
| 71:00:00 | 3fp0-A | 3.3 | 6.5 | 96 | 510 | 7 | MOLECULE: | 15-O-ACETYLTRANSFERASE |
| 72:00:00 | 6vh5-D | 3.3 | 4.6 | 88 | 283 | 7 | MOLECULE: | PREPHENATE DEHYDRATASE:AMINO ACID-BINDING ACT |
| 73:00:00 | 4pj3-A | 3.3 | 22.9 | 196 | 1335 | 8 | MOLECULE: | INTRON-BINDING PROTEIN AQUARIUS |
| 74:00:00 | 1g8x-A | 3.3 | 4.7 | 131 | 1009 | 4 | MOLECULE: | MYOSIN II HEAVY CHAIN FUSED TO ALPHA-ACTININ 3 |
| 75:00:00 | 7mdm-B | 3.3 | 8.1 | 101 | 701 | 12 | MOLECULE: | TRANSITIONAL ENDOPLASMIC RETICULUM ATPASE |
| 76:00:00 | 8iqh-K | 3.3 | 5.5 | 203 | 866 | 8 | MOLECULE: | PUTATIVE PRIMASE C962R |
| 77:00:00 | 8iqh-G | 3.3 | 5.7 | 209 | 863 | 7 | MOLECULE: | PUTATIVE PRIMASE C962R |
| 78:00:00 | 8iqh-D | 3.3 | 5.7 | 199 | 891 | 8 | MOLECULE: | PUTATIVE PRIMASE C962R |
| 79:00:00 | 4w8f-B | 3.3 | 26 | 138 | 2609 | 7 | MOLECULE: | DYNEIN HEAVY CHAIN LYSOZYME CHIMERA |
| 80:00:00 | 7wae-A | 3.2 | 16.9 | 136 | 877 | 6 | MOLECULE: | CYANOPHYCIN SYNTHASE |
| 81:00:00 | 8e5t-5 | 3.2 | 18.8 | 81 | 530 | 10 | MOLECULE: | RIBOSOME BIOGENESIS PROTEIN MAK21 |
| 82:00:00 | 5yvu-B | 3.2 | 21.7 | 129 | 601 | 6 | MOLECULE: | GENOME POLYPROTEIN |
| 83:00:00 | 7nhr-C | 3.2 | 18.2 | 115 | 565 | 7 | MOLECULE: | PUTATIVE TRANSMEMBRANE PROTEIN WZC |
| 84:00:00 | 7r9x-A | 3.2 | 14.3 | 91 | 418 | 9 | MOLECULE: | AMBE |
| 85:00:00 | 5ofo-A | 3.2 | 11.9 | 180 | 686 | 8 | MOLECULE: | CHAPERONE PROTEIN CLPB,ATP-DEPENDENT CLP PROTEASE |
| 86:00:00 | 7n7s-B | 3.1 | 18.3 | 123 | 441 | 8 | MOLECULE: | HYDROXYMETHYLGLUTARYL-COA REDUCTASE |
| 87:00:00 | 5k8o-B | 3.1 | 6.8 | 118 | 425 | 4 | MOLECULE: | 5-NITROANTHRANILIC ACID AMINOHYDROLASE |
| 88:00:00 | 7t5q-J | 3.1 | 21.9 | 87 | 291 | 7 | MOLECULE: | ACTIN-RELATED PROTEIN 3 |

| | | | | | | | | |
|---|---|---|---|---|---|---|---|---|
| 89:00:00 | 6c7n-D | 3.1 | 24.2 | 73 | 389 | 10 | MOLECULE: | MALIC ENZYME |
| 90:00:00 | 4ch7-A | 3.1 | 19.4 | 111 | 309 | 5 | MOLECULE: | NIRD-LIKE PROTEIN |
| 91:00:00 | 4qbu-A | 3.1 | 4.4 | 103 | 436 | 10 | MOLECULE: | ZMAA |
| 92:00:00 | 8iqh-F | 3.1 | 6.5 | 204 | 880 | 7 | MOLECULE: | PUTATIVE PRIMASE C962R |
| 93:00:00 | 8iqi-F | 3.1 | 5.5 | 201 | 910 | 6 | MOLECULE: | PUTATIVE PRIMASE C962R |
| 94:00:00 | 6s6b-J | 3 | 12.4 | 89 | 475 | 8 | MOLECULE: | CRISPR-ASSOCIATED PROTEIN, CMR5 FAMILY |
| 95:00:00 | 8dew-A | 3 | 23.1 | 150 | 1058 | 8 | MOLECULE: | EFFLUX PUMP MEMBRANE TRANSPORTER |
| 96:00:00 | 5yhf-A | 3 | 19.4 | 123 | 734 | 7 | MOLECULE: | PROTEIN TRANSLOCASE SUBUNIT SECDF |
| 97:00:00 | 6fsk-A | 3 | 3.6 | 85 | 484 | 6 | MOLECULE: | DYP-TYPE PEROXIDASE |
| 98:00:00 | 6mur-E | 3 | 3.5 | 88 | 286 | 6 | MOLECULE: | UNCHARACTERIZED PROTEIN |
| 99:00:00 | 8iqh-H | 3 | 5.4 | 201 | 872 | 9 | MOLECULE: | PUTATIVE PRIMASE C962R |
| 0:00 | 6qel-J | 3 | 4.4 | 86 | 237 | 16 | MOLECULE: | REPLICATIVE DNA HELICASE |
| 1:00 | 7uoi-A | 2.9 | 7 | 96 | 383 | 3 | MOLECULE: | SUCCINYL-DIAMINOPIMELATE DESUCCINYLASE |
| 2:00 | 8sg4-A | 2.9 | 21.1 | 90 | 1469 | 11 | MOLECULE: | METAL RESISTANCE PROTEIN YCF1 |
| 3:00 | 6cse-M | 2.9 | 13.5 | 98 | 432 | 7 | MOLECULE: | MONOCLONAL ANTIBODY FAB HEAVY CHAIN |
| 4:00 | 6em8-E | 2.9 | 7.3 | 216 | 664 | 8 | MOLECULE: | ATP-DEPENDENT CLP PROTEASE ATP-BINDING SUBUNIT CL |
| 5:00 | 6pwy-B | 2.8 | 8.8 | 125 | 513 | 7 | MOLECULE: | ZK177.8 |
| 6:00 | 6pns-A | 2.8 | 24.7 | 169 | 1291 | 6 | MOLECULE: | RNA-DIRECTED RNA POLYMERASE |
| 7:00 | 7nhr-A | 2.8 | 17.8 | 121 | 663 | 7 | MOLECULE: | PUTATIVE TRANSMEMBRANE PROTEIN WZC |
| 8:00 | 7r07-F | 2.8 | 16.9 | 142 | 600 | 8 | MOLECULE: | ABIK |
| 9:00 | 5mgu-A | 2.7 | 14.8 | 102 | 408 | 3 | MOLECULE: | PHENYLALANINE--TRNA LIGASE, MITOCHONDRIAL |
| 10:00 | 3nlc-A | 2.7 | 11.4 | 127 | 534 | 6 | MOLECULE: | UNCHARACTERIZED PROTEIN VP0956 |
| 11:00 | 4tx3-B | 2.7 | 15.8 | 108 | 457 | 13 | MOLECULE: | OXYB PROTEIN |
| 12:00 | 4yut-A | 2.7 | 14.9 | 91 | 351 | 9 | MOLECULE: | FAMILY 3 ADENYLATE CYCLASE |
| 13:00 | 6neq-g | 2.7 | 4.5 | 123 | 351 | 12 | MOLECULE: | 28S RIBOSOMAL RNA, MITOCHONDRIAL |
| 14:00 | 6qel-L | 2.7 | 4.4 | 78 | 237 | 17 | MOLECULE: | REPLICATIVE DNA HELICASE |
| 15:00 | 4xea-A | 2.6 | 4.6 | 119 | 423 | 8 | MOLECULE: | PEPTIDASE M16 DOMAIN PROTEIN |
| 16:00 | 6qel-H | 2.6 | 4.1 | 73 | 237 | 16 | MOLECULE: | REPLICATIVE DNA HELICASE |
| 17:00 | 7p04-A | 2.5 | 19.3 | 87 | 1353 | 14 | MOLECULE: | PLEIOTROPIC ABC EFFLUX TRANSPORTER OF MULTIPLE DR |
| 18:00 | 1q9j-B | 2.5 | 13.6 | 89 | 399 | 2 | MOLECULE: | POLYKETIDE SYNTHASE ASSOCIATED PROTEIN 5 |
| 19:00 | 8hwd-E | 2.5 | 4.7 | 139 | 247 | 10 | MOLECULE: | PRIMASE D5 |
| 20:00 | 6qel-I | 2.5 | 4.2 | 73 | 237 | 16 | MOLECULE: | REPLICATIVE DNA HELICASE |
| 21:00 | 5vlj-A | 2.5 | 25.6 | 86 | 2376 | 8 | MOLECULE: | DYNEIN HEAVY CHAIN, CYTOPLASMIC |
| 22:00 | 3d3y-A | 2.4 | 6.6 | 110 | 425 | 7 | MOLECULE: | UNCHARACTERIZED PROTEIN |
| 23:00 | 6qel-K | 2.4 | 4.4 | 68 | 237 | 15 | MOLECULE: | REPLICATIVE DNA HELICASE |
| 24:00:00 | 4ap2-B | 2.3 | 21.8 | 89 | 359 | 4 | MOLECULE: | KELCH-LIKE PROTEIN 11 |
| 25:00:00 | 5n8o-A | 2.3 | 19.2 | 80 | 1432 | 11 | MOLECULE: | DNA HELICASE I |
| 26:00:00 | 6ahf-F | 2.3 | 8.3 | 169 | 574 | 9 | MOLECULE: | HEAT SHOCK PROTEIN 104 |
| 27:00:00 | 3s4w-A | 2.2 | 11.6 | 84 | 1206 | 6 | MOLECULE: | FANCONI ANEMIA GROUP I PROTEIN HOMOLOG |
| 28:00:00 | 7c0q-A | 2.2 | 3.2 | 59 | 279 | 7 | MOLECULE: | EFFECTOR LPG2505 |
| 29:00:00 | 6qel-G | 2.2 | 4.3 | 87 | 229 | 18 | MOLECULE: | REPLICATIVE DNA HELICASE |
| 30:00:00 | 5t0j-C | 2.2 | 11.3 | 154 | 392 | 8 | MOLECULE: | 26S PROTEASOME NON-ATPASE REGULATORY SUBUNIT 2 |
| 31:00:00 | 5twv-B | 2.1 | 19.6 | 75 | 1375 | 11 | MOLECULE: | ATP-SENSITIVE INWARD RECTIFIER POTASSIUM CHANNEL |
| 32:00:00 | 8iqh-I | 2.1 | 7 | 199 | 866 | 7 | MOLECULE: | PUTATIVE PRIMASE C962R |
| 33:00:00 | 7jtk-A | 2.1 | 24.5 | 124 | 749 | 1 | MOLECULE: | FLAGELLAR RADIAL SPOKE PROTEIN 1 |

**DALI results: MdA-1 SET**

| No: | Chain | Z | rmsd | lali | nres | %id | PDB | Description |
|---|---|---|---|---|---|---|---|---|
| 1:00 | 4ldg-A | 13.8 | 2.1 | 125 | 216 | 17 | MOLECULE: | PROTEIN WITH A SET DOMAIN WITHIN CARBOXY REGION |
| 2:00 | 7xpx-K | 13.7 | 2.3 | 124 | 168 | 18 | MOLECULE: | HISTONE H3 |
| 3:00 | 6uh5-M | 12.1 | 3 | 116 | 222 | 18 | MOLECULE: | HISTONE H3 |
| 4:00 | 1h3i-A | 11.6 | 2.7 | 118 | 293 | 19 | MOLECULE: | HISTONE H3 LYSINE 4 SPECIFIC METHYLTRANSFERASE |
| 5:00 | 5kkl-B | 11.5 | 5.1 | 123 | 838 | 20 | MOLECULE: | PUTATIVE POLYCOMB PROTEIN EED |
| 6:00 | 8sr6-A | 10.8 | 2.4 | 110 | 452 | 18 | MOLECULE: | EUKARYOTIC HUNTINGTIN INTERACTING PROTEIN B |
| 7:00 | 8rzu-A | 10.7 | 2.3 | 121 | 251 | 17 | MOLECULE: | HISTONE-LYSINE N-METHYLTRANSFERASE SETD2 |
| 8:00 | 5wg6-A | 10.7 | 2.4 | 118 | 517 | 19 | MOLECULE: | HISTONE-LYSINE N-METHYLTRANSFERASE EZH2,POLYCOMB |
| 9:00 | 5h6z-A | 10.3 | 2.6 | 108 | 123 | 13 | MOLECULE: | SET DOMAIN-CONTAINING PROTEIN 7 |
| 10:00 | 6wkr-C | 10.1 | 2.7 | 122 | 607 | 19 | MOLECULE: | UBIQUITIN |
| 11:00 | 4o30-A | 10.1 | 3.2 | 129 | 218 | 16 | MOLECULE: | HISTONE-LYSINE N-METHYLTRANSFERASE ATXR6, PUTATIV |
| 12:00 | 6a5n-A | 10 | 2.7 | 125 | 502 | 17 | MOLECULE: | HISTONE-LYSINE N-METHYLTRANSFERASE, H3 LYSINE-9 S |
| 13:00 | 3kma-A | 9.8 | 2.3 | 94 | 105 | 17 | MOLECULE: | A612L PROTEIN |
| 14:00 | 5ecj-A | 9.1 | 3.1 | 117 | 263 | 15 | MOLECULE: | PR DOMAIN ZINC FINGER PROTEIN 14,PROTEIN CBFA2T2 |
| 15:00 | 4ijd-A | 7.8 | 2.7 | 108 | 215 | 17 | MOLECULE: | HISTONE-LYSINE N-METHYLTRANSFERASE PRDM9 |
| 16:00 | 3n71-A | 6.6 | 2.9 | 118 | 469 | 16 | MOLECULE: | HISTONE LYSINE METHYLTRANSFERASE SMYD1 |
| 17:00 | 8t9f-K | 6.4 | 3 | 106 | 265 | 11 | MOLECULE: | HISTONE-LYSINE N-METHYLTRANSFERASE KMT5B |
| 18:00 | 8xzv-I | 6.4 | 3.3 | 114 | 413 | 14 | MOLECULE: | DNA-DIRECTED RNA POLYMERASE SUBUNIT ALPHA |
| 19:00 | 6fnd-A | 6 | 2.9 | 98 | 423 | 13 | MOLECULE: | APICAL COMPLEX LYSINE METHYLTRANSFERASE |
| 20:00 | 6ox1-B | 5.5 | 3.7 | 112 | 485 | 12 | MOLECULE: | ACTIN, CYTOPLASMIC 1 |
| 21:00 | 2mpl-A | 4.2 | 3.9 | 77 | 127 | 8 | MOLECULE: | ZINC FINGER PROTEIN ZFPM1 |
| 22:00 | 2n1i-A | 3.8 | 3.2 | 87 | 176 | 9 | MOLECULE: | PR DOMAIN ZINC FINGER PROTEIN 16 |

**DALI results: MnA-1 Zifi**

| No: | Chain | Z | rmsd | lali | nres | %id | PDB Description | | | |
|---|---|---|---|---|---|---|---|---|---|---|
| 1:00 | 6k7e-A | 5.6 | 3.1 | 74 | 498 | 9 | MOLECULE: | MALTOSE | /MALTOD | EXTRIN-BINDING PERIPLASMIC PROTEIN, |
| 2:00 | 2ig7-B | 5.5 | 2.5 | 67 | 342 | 10 | MOLECULE: | CHOLINE | /ETHANO | LAMINE KINASE |
| 3:00 | 5eqp-A | 5.4 | 2.5 | 67 | 352 | 7 | MOLECULE: | CHOLINE | KINASE | ALPHA |
| 4:00 | 4r80-A | 5.4 | 2.8 | 68 | 76 | 9 | MOLECULE: | OR486 | | |
| 5:00 | 8bi6-A | 5.4 | 2.5 | 67 | 355 | 7 | MOLECULE: | CHOLINE | KINASE | ALPHA |
| 6:00 | 3g15-A | 5.4 | 2.5 | 67 | 354 | 7 | MOLECULE: | CHOLINE | KINASE | ALPHA |
| 7:00 | 8bi6-B | 5.4 | 2.5 | 67 | 354 | 7 | MOLECULE: | CHOLINE | KINASE | ALPHA |
| 8:00 | 5afv-B | 5.3 | 2.5 | 67 | 354 | 7 | MOLECULE: | CHOLINE | KINASE | ALPHA |
| 9:00 | 5afv-A | 5.3 | 2.5 | 67 | 354 | 7 | MOLECULE: | CHOLINE | KINASE | ALPHA |
| 10:00 | 3f2r-A | 5.3 | 2.5 | 66 | 348 | 8 | MOLECULE: | CHOLINE | KINASE | ALPHA |
| 11:00 | 5eqe-A | 5.3 | 2.5 | 68 | 353 | 7 | MOLECULE: | CHOLINE | KINASE | ALPHA |
| 12:00 | 3zm9-A | 5.3 | 2.9 | 69 | 354 | 9 | MOLECULE: | CHOLINE | KINASE | ALPHA |
| 13:00 | 4r80-B | 5.3 | 2.8 | 68 | 76 | 9 | MOLECULE: | OR486 | | |
| 14:00 | 7a04-B | 5.2 | 2.8 | 68 | 348 | 9 | MOLECULE: | CHOLINE | KINASE | ALPHA |
| 15:00 | 7sh3-B | 5.2 | 3.2 | 70 | 125 | 7 | MOLECULE: | SYNTHET | IC VIRB | 8 MINIPROTEIN BINDER |
| 16:00 | 7a04-A | 5.2 | 2.9 | 68 | 348 | 9 | MOLECULE: | CHOLINE | KINASE | ALPHA |
| 17:00 | 8bi5-B | 5.2 | 2.6 | 67 | 354 | 6 | MOLECULE: | CHOLINE | KINASE | ALPHA |
| 18:00 | 8bi5-A | 5.2 | 2.8 | 70 | 355 | 7 | MOLECULE: | CHOLINE | KINASE | ALPHA |
| 19:00 | 4cga-A | 5.2 | 2.8 | 69 | 354 | 9 | MOLECULE: | CHOLINE | KINASE | ALPHA |
| 20:00 | 4cg9-A | 5.2 | 2.9 | 68 | 353 | 7 | MOLECULE: | CHOLINE | KINASE | ALPHA |
| 21:00 | 3wz4-C | 5.2 | 3.5 | 73 | 139 | 10 | MOLECULE: | DOTI | | |
| 22:00 | 7a06-A | 5.2 | 3 | 70 | 354 | 9 | MOLECULE: | CHOLINE | KINASE | ALPHA |
| 23:00 | 5fut-A | 5.1 | 2.9 | 68 | 355 | 9 | MOLECULE: | CHOLINE | KINASE | ALPHA |
| 24:00:00 | 5w6o-A | 5.1 | 2.9 | 68 | 352 | 9 | MOLECULE: | CHOLINE | KINASE | ALPHA |
| 25:00:00 | 5kuy-G | 5.1 | 3.8 | 70 | 84 | 10 | MOLECULE: | HEMAGGL | UTININ | HA1 |
| 26:00:00 | 3wz4-D | 5.1 | 3 | 69 | 136 | 9 | MOLECULE: | DOTI | | |
| 27:00:00 | 4da5-A | 5.1 | 2.5 | 65 | 352 | 8 | MOLECULE: | CHOLINE | KINASE | ALPHA |
| 28:00:00 | 5eqy-B | 5.1 | 2.5 | 68 | 351 | 6 | MOLECULE: | CHOLINE | KINASE | ALPHA |
| 29:00:00 | 6k7f-A | 5 | 3.2 | 78 | 499 | 9 | MOLECULE: | MALTOSE | /MALTOD | EXTRIN-BINDING PERIPLASMIC PROTEIN, |
| 30:00:00 | 5ftg-A | 5 | 2.9 | 68 | 353 | 9 | MOLECULE: | CHOLINE | KINASE | ALPHA |
| 31:00:00 | 5kuy-H | 5 | 3.9 | 71 | 85 | 10 | MOLECULE: | HEMAGGL | UTININ | HA1 |
| 32:00:00 | 5w6o-B | 5 | 2.9 | 68 | 353 | 9 | MOLECULE: | CHOLINE | KINASE | ALPHA |
| 33:00:00 | 6k7d-A | 5 | 3.4 | 78 | 499 | 9 | MOLECULE: | MALTOSE | /MALTOD | EXTRIN-BINDING PERIPLASMIC PROTEIN, |
| 34:00:00 | 4lzi-A | 4.9 | 3.8 | 77 | 277 | 4 | MOLECULE: | MULTICY | STATIN | |
| 35:00:00 | 5eqp-B | 4.9 | 2.5 | 67 | 349 | 6 | MOLECULE: | CHOLINE | KINASE | ALPHA |
| 36:00:00 | 3fi8-A | 4.9 | 2.6 | 66 | 339 | 3 | MOLECULE: | CHOLINE | KINASE | |
| 37:00:00 | 5eqy-A | 4.9 | 2.8 | 69 | 352 | 7 | MOLECULE: | CHOLINE | KINASE | ALPHA |
| 38:00:00 | 3wz4-H | 4.9 | 3.6 | 73 | 139 | 10 | MOLECULE: | DOTI | | |
| 39:00:00 | 3h4z-C | 4.9 | 3 | 75 | 552 | 5 | MOLECULE: | MALTOSE | #NAME? | G PERIPLASMIC PROTEIN FUSED WITH AL |
| 40:00:00 | 6i1m-A | 4.8 | 3 | 69 | 93 | 9 | MOLECULE: | CYSTATI | N | |
| 41:00:00 | 3wz4-G | 4.8 | 3.5 | 73 | 140 | 10 | MOLECULE: | DOTI | | |
| 42:00:00 | 5kph-A | 4.8 | 2.7 | 68 | 85 | 12 | MOLECULE: | DE NOVO | BETA S | HEET DESIGN PROTEIN OR485 |
| 43:00:00 | 6l77-A | 4.8 | 4.3 | 75 | 214 | 8 | MOLECULE: | MS5A | | |
| 44:00:00 | 2w9p-G | 4.8 | 3.3 | 66 | 87 | 5 | MOLECULE: | MULTICY | STATIN | |
| 45:00:00 | 4n6t-A | 4.8 | 3.9 | 68 | 79 | 6 | MOLECULE: | ADHIRON | | |
| 46:00:00 | 3feg-A | 4.8 | 2.5 | 62 | 336 | 10 | MOLECULE: | CHOLINE | /ETHANO | LAMINE KINASE |
| 47:00:00 | 5kpe-A | 4.7 | 3.8 | 71 | 120 | 7 | MOLECULE: | DE NOVO | BETA S | HEET DESIGN PROTEIN OR664 |
| 48:00:00 | 2ig7-A | 4.7 | 2.7 | 68 | 335 | 10 | MOLECULE: | CHOLINE | /ETHANO | LAMINE KINASE |
| 49:00:00 | 4it7-C | 4.7 | 3.4 | 72 | 107 | 7 | MOLECULE: | CPI | | |
| 50:00:00 | 6qoz-C | 4.7 | 3.8 | 68 | 79 | 6 | MOLECULE: | RNA2 PO | LYPROTE | IN |
| 51:00:00 | 4it7-D | 4.7 | 3.3 | 72 | 107 | 8 | MOLECULE: | CPI | | |
| 52:00:00 | 3zm9-B | 4.7 | 2.6 | 64 | 346 | 8 | MOLECULE: | CHOLINE | KINASE | ALPHA |

| | | | | | | | | | | |
|---|---|---|---|---|---|---|---|---|---|---|
| 53:00:00 | 2w9p-F | 4.7 | 3.7 | 66 | 87 | 5 | MOLECULE: | MULTICY | STATIN | |
| 54:00:00 | 2w9p-M | 4.7 | 3.5 | 66 | 87 | 5 | MOLECULE: | MULTICY | STATIN | |
| 55:00:00 | 5eqe-B | 4.7 | 2.5 | 63 | 344 | 8 | MOLECULE: | CHOLINE | KINASE | ALPHA |
| 56:00:00 | 4it7-B | 4.7 | 3.3 | 72 | 107 | 8 | MOLECULE: | CPI | | |
| 57:00:00 | 4o3v-A | 4.7 | 3.3 | 69 | 138 | 9 | MOLECULE: | VIRB8-L | IKE PRO | TEIN OF TYPE IV SECRETION SYSTEM |
| 58:00:00 | 6rpv-A | 4.7 | 3.4 | 74 | 120 | 7 | MOLECULE: | CYSTATI | N-C | |
| 59:00:00 | 6w3g-B | 4.7 | 3 | 68 | 107 | 4 | MOLECULE: | RD1NTF2 | _04 | |
| 60:00:00 | 2w9p-I | 4.7 | 3.5 | 66 | 87 | 6 | MOLECULE: | MULTICY | STATIN | |
| 61:00:00 | 2w9p-K | 4.7 | 3.4 | 67 | 87 | 4 | MOLECULE: | MULTICY | STATIN | |
| 62:00:00 | 4r1k-B | 4.6 | 3.5 | 80 | 136 | 4 | MOLECULE: | UNCHARA | CTERIZE | D PROTEIN |
| 63:00:00 | 4it7-A | 4.6 | 3.3 | 71 | 107 | 7 | MOLECULE: | CPI | | |
| 64:00:00 | 2ckp-A | 4.6 | 2.8 | 64 | 316 | 8 | MOLECULE: | CHOLINE | KINASE | ALPHA |
| 65:00:00 | 2uug-D | 4.6 | 2.8 | 61 | 82 | 11 | MOLECULE: | URACIL- | DNA GLY | COSYLASE |
| 66:00:00 | 4n6v-6 | 4.6 | 3.3 | 68 | 91 | 9 | MOLECULE: | CYSTATI | N-B | |
| 67:00:00 | 4n6v-1 | 4.6 | 3.3 | 68 | 91 | 9 | MOLECULE: | CYSTATI | N-B | |
| 68:00:00 | 4n6u-A | 4.6 | 3.9 | 67 | 79 | 6 | MOLECULE: | ADHIRON | | |
| 69:00:00 | 3dc2-B | 4.6 | 3.2 | 75 | 524 | 11 | MOLECULE: | D-3-PHO | SPHOGLY | CERATE DEHYDROGENASE |
| 70:00:00 | 4n6v-7 | 4.6 | 3.3 | 68 | 91 | 9 | MOLECULE: | CYSTATI | N-B | |
| 71:00:00 | 4n6v-4 | 4.6 | 3.3 | 68 | 91 | 9 | MOLECULE: | CYSTATI | N-B | |
| 72:00:00 | 5ohm-J | 4.6 | 3.7 | 69 | 90 | 4 | MOLECULE: | POLYUBI | QUITIN- | C |
| 73:00:00 | 4br3-A | 4.6 | 2.8 | 69 | 354 | 9 | MOLECULE: | CHOLINE | KINASE | ALPHA |
| 74:00:00 | 2ugi-A | 4.6 | 2.6 | 60 | 83 | 10 | MOLECULE: | URACIL- | DNA GLY | COSYLASE INHIBITOR |
| 75:00:00 | 4n6v-3 | 4.6 | 3.3 | 68 | 91 | 9 | MOLECULE: | CYSTATI | N-B | |
| 76:00:00 | 4n6v-9 | 4.6 | 3.2 | 68 | 91 | 9 | MOLECULE: | CYSTATI | N-B | |
| 77:00:00 | 2w9p-A | 4.6 | 3.9 | 69 | 87 | 6 | MOLECULE: | MULTICY | STATIN | |
| 78:00:00 | 2w9p-E | 4.6 | 3.8 | 67 | 87 | 4 | MOLECULE: | MULTICY | STATIN | |
| 79:00:00 | 2w9p-D | 4.6 | 3.7 | 68 | 87 | 6 | MOLECULE: | MULTICY | STATIN | |
| 80:00:00 | 6hjl-D | 4.6 | 3.7 | 68 | 90 | 4 | MOLECULE: | BCL-2-L | IKE PRO | TEIN 1 |
| 81:00:00 | 2w9p-H | 4.6 | 3.6 | 67 | 87 | 4 | MOLECULE: | MULTICY | STATIN | |
| 82:00:00 | 5ohv-B | 4.6 | 3.5 | 68 | 89 | 4 | MOLECULE: | UBIQUIT | IN | |
| 83:00:00 | 3mes-A | 4.6 | 2.6 | 62 | 357 | 5 | MOLECULE: | CHOLINE | KINASE | |
| 84:00:00 | 2w9p-C | 4.6 | 3.7 | 68 | 87 | 6 | MOLECULE: | MULTICY | STATIN | |
| 85:00:00 | 4lyl-H | 4.5 | 2.9 | 61 | 82 | 11 | MOLECULE: | URACIL- | DNA GLY | COSYLASE |
| 86:00:00 | 2cko-B | 4.5 | 2.5 | 62 | 346 | 8 | MOLECULE: | CHOLINE | KINASE | ALPHA |
| 87:00:00 | 6k8q-A | 4.5 | 3 | 72 | 115 | 7 | MOLECULE: | MITOCHO | NDRIAL | IMPORT INNER MEMBRANE TRANSLOCASE S |
| 88:00:00 | 2ckq-A | 4.5 | 2.7 | 65 | 346 | 8 | MOLECULE: | CHOLINE | KINASE | ALPHA |
| 89:00:00 | 4n6v-8 | 4.5 | 3.3 | 68 | 91 | 9 | MOLECULE: | CYSTATI | N-B | |
| 90:00:00 | 1ugh-I | 4.5 | 2.3 | 56 | 82 | 11 | MOLECULE: | PROTEIN | (URACI | L-DNA GLYCOSYLASE) |
| 91:00:00 | 1roa-A | 4.5 | 2.8 | 65 | 111 | 2 | MOLECULE: | CYSTATI | N D | |
| 92:00:00 | 2owp-A | 4.5 | 3.4 | 76 | 129 | 7 | MOLECULE: | HYPOTHE | TICAL P | ROTEIN BXE_B1374 |
| 93:00:00 | 6hjl-H | 4.5 | 3.8 | 67 | 90 | 4 | MOLECULE: | BCL-2-L | IKE PRO | TEIN 1 |
| 94:00:00 | 4n6v-2 | 4.5 | 3.3 | 68 | 91 | 9 | MOLECULE: | CYSTATI | N-B | |
| 95:00:00 | 5ohm-F | 4.5 | 3.7 | 67 | 91 | 4 | MOLECULE: | POLYUBI | QUITIN- | C |
| 96:00:00 | 1ugi-A | 4.5 | 2.8 | 62 | 83 | 11 | MOLECULE: | URACIL- | DNA GLY | COSYLASE INHIBITOR |
| 97:00:00 | 3wz4-A | 4.5 | 3 | 69 | 137 | 9 | MOLECULE: | DOTI | | |
| 98:00:00 | 2w9p-B | 4.5 | 3.8 | 68 | 87 | 6 | MOLECULE: | MULTICY | STATIN | |
| 99:00:00 | 2w9p-J | 4.5 | 3.8 | 69 | 87 | 6 | MOLECULE: | MULTICY | STATIN | |
| 0:00 | 2mzv-A | 4.5 | 3.3 | 73 | 199 | 4 | MOLECULE: | CYSTATI | N | |
| 1:00 | 2w9p-N | 4.5 | 3.4 | 65 | 87 | 5 | MOLECULE: | MULTICY | STATIN | |
| 2:00 | 3dc2-A | 4.5 | 3.3 | 79 | 527 | 9 | MOLECULE: | D-3-PHO | SPHOGLY | CERATE DEHYDROGENASE |
| 3:00 | 3ima-D | 4.5 | 3.6 | 68 | 85 | 6 | MOLECULE: | PAPAIN | | |
| 4:00 | 1ugi-G | 4.5 | 2.7 | 61 | 83 | 11 | MOLECULE: | URACIL- | DNA GLY | COSYLASE INHIBITOR |
| 5:00 | 2q83-A | 4.4 | 2.7 | 57 | 332 | 7 | MOLECULE: | YTAA PR | OTEIN | |
| 6:00 | 1ylx-A | 4.4 | 3.3 | 68 | 100 | 6 | MOLECULE: | HYPOTHE | TICAL P | ROTEIN APC35702 |

| | | | | | | | | | | |
|---|---|---|---|---|---|---|---|---|---|---|
| 7:00 | 4ec6-A | 4.4 | 3 | 71 | 109 | 6 | MOLECULE: | PUTATIV | E UNCHA | RACTERIZED PROTEIN |
| 8:00 | 1rn7-A | 4.4 | 3 | 64 | 112 | 2 | MOLECULE: | CYSTATI | N D | |
| 9:00 | 6lg3-A | 4.4 | 2.8 | 60 | 74 | 10 | MOLECULE: | PHIRV1 | PHAGE P | ROTEIN |
| 10:00 | 2q83-B | 4.4 | 2.6 | 57 | 332 | 7 | MOLECULE: | YTAA PR | OTEIN | |
| 11:00 | 4cg8-A | 4.4 | 2.9 | 68 | 353 | 9 | MOLECULE: | CHOLINE | KINASE | ALPHA |
| 12:00 | 5elj-A | 4.4 | 3.8 | 69 | 90 | 4 | MOLECULE: | SUMO-AD | HIRON-S | 2D5 |
| 13:00 | 1ugi-H | 4.4 | 3 | 62 | 83 | 11 | MOLECULE: | URACIL- | DNA GLY | COSYLASE INHIBITOR |
| 14:00 | 4ec6-C | 4.4 | 2.9 | 69 | 109 | 6 | MOLECULE: | PUTATIV | E UNCHA | RACTERIZED PROTEIN |
| 15:00 | 2ciu-A | 4.4 | 3.7 | 71 | 123 | 8 | MOLECULE: | IMPORT | INNER M | EMBRANE TRANSLOCASE SUBUNIT TIM21 |
| 16:00 | 6yxs-A | 4.4 | 2.4 | 64 | 359 | 3 | MOLECULE: | CHOLINE | KINASE | |
| 17:00 | 3wz5-D | 4.4 | 3 | 69 | 135 | 9 | MOLECULE: | DOTI | | |
| 18:00 | 4ec6-F | 4.4 | 3 | 71 | 109 | 7 | MOLECULE: | PUTATIV | E UNCHA | RACTERIZED PROTEIN |
| 19:00 | 4ec6-D | 4.4 | 2.9 | 69 | 109 | 4 | MOLECULE: | PUTATIV | E UNCHA | RACTERIZED PROTEIN |
| 20:00 | 5ohv-D | 4.4 | 3.6 | 67 | 90 | 4 | MOLECULE: | UBIQUIT | IN | |
| 21:00 | 6stj-F | 4.4 | 3.6 | 65 | 90 | 5 | MOLECULE: | INDUCED | MYELOI | D LEUKEMIA CELL DIFFERENTIATION PRO |
| 22:00 | 7ds3-A | 4.4 | 3.3 | 71 | 266 | 4 | MOLECULE: | F-ACTIN | #NAME? | G PROTEIN SUBUNIT ALPHA-1 |
| 23:00 | 3h3h-A | 4.4 | 3.1 | 71 | 118 | 4 | MOLECULE: | UNCHARA | CTERIZE | D SNOAL-LIKE PROTEIN |
| 24:00:00 | 6pai-C | 4.4 | 3.3 | 74 | 401 | 7 | MOLECULE: | DNA DAM | AGE-BIN | DING PROTEIN 1 |
| 25:00:00 | 1nw1-A | 4.4 | 3.1 | 72 | 365 | 8 | MOLECULE: | CHOLINE | KINASE | (49.2 KD) |
| 26:00:00 | 2rcd-B | 4.4 | 3.4 | 74 | 128 | 11 | MOLECULE: | UNCHARA | CTERIZE | D PROTEIN |
| 27:00:00 | 2w9q-A | 4.4 | 3.7 | 68 | 87 | 6 | MOLECULE: | MULTICY | STATIN | |
| 28:00:00 | 8aim-G | 4.4 | 2.8 | 59 | 86 | 7 | MOLECULE: | URACIL- | DNA GLY | COSYLASE INHIBITOR |
| 29:00:00 | 3wz4-F | 4.4 | 3 | 69 | 136 | 9 | MOLECULE: | DOTI | | |
| 30:00:00 | 5ohm-B | 4.4 | 3.2 | 65 | 89 | 6 | MOLECULE: | POLYUBI | QUITIN- | C |
| 31:00:00 | 3ddn-A | 4.4 | 3.4 | 79 | 527 | 9 | MOLECULE: | D-3-PHO | SPHOGLY | CERATE DEHYDROGENASE |
| 32:00:00 | 5ohm-H | 4.4 | 3.6 | 66 | 89 | 5 | MOLECULE: | POLYUBI | QUITIN- | C |
| 33:00:00 | 8aim-B | 4.3 | 2.9 | 62 | 86 | 6 | MOLECULE: | URACIL- | DNA GLY | COSYLASE INHIBITOR |
| 34:00:00 | 4ccv-A | 4.3 | 3.4 | 70 | 115 | 7 | MOLECULE: | HISTIDI | NE-RICH | GLYCOPROTEIN |
| 35:00:00 | 3f14-A | 4.3 | 3.6 | 72 | 112 | 4 | MOLECULE: | UNCHARA | CTERIZE | D NTF2-LIKE PROTEIN |
| 36:00:00 | 1ygy-A | 4.3 | 3.3 | 79 | 527 | 9 | MOLECULE: | D-3-PHO | SPHOGLY | CERATE DEHYDROGENASE |
| 37:00:00 | 7eeb-H | 4.3 | 3.5 | 74 | 916 | 5 | MOLECULE: | ENHANCE | D GREEN | FLUORESCENT PROTEIN,CATION CHANNEL |
| 38:00:00 | 4br3-B | 4.3 | 2.9 | 67 | 349 | 7 | MOLECULE: | CHOLINE | KINASE | ALPHA |
| 39:00:00 | 5uxd-B | 4.3 | 2.5 | 57 | 299 | 7 | MOLECULE: | MACROLI | DE 2'-P | HOSPHOTRANSFERASE MPHH |
| 40:00:00 | 3fka-C | 4.3 | 3.5 | 70 | 119 | 7 | MOLECULE: | UNCHARA | CTERIZE | D NTF-2 LIKE PROTEIN |
| 41:00:00 | 2ckq-B | 4.3 | 2.8 | 68 | 348 | 9 | MOLECULE: | CHOLINE | KINASE | ALPHA |
| 42:00:00 | 5mn2-D | 4.3 | 3.7 | 65 | 90 | 5 | MOLECULE: | LOW AFF | INITY I | MMUNOGLOBULIN GAMMA FC REGION RECEP |
| 43:00:00 | 6q0w-C | 4.3 | 3 | 67 | 192 | 7 | MOLECULE: | DNA DAM | AGE-BIN | DING PROTEIN 1 |
| 44:00:00 | 7z8i-K | 4.3 | 3.5 | 74 | 278 | 3 | MOLECULE: | ARP1 AC | TIN REL | ATED PROTEIN 1 HOMOLOG A |
| 45:00:00 | 7ccc-C | 4.3 | 3.8 | 74 | 269 | 3 | MOLECULE: | ACTIN, | ALPHA S | KELETAL MUSCLE |
| 46:00:00 | 8ixi-B | 4.3 | 2.5 | 57 | 295 | 5 | MOLECULE: | ALDEHYD | E DEHYD | ROGENASE PROTEIN |
| 47:00:00 | 5uxb-B | 4.3 | 2.5 | 57 | 297 | 7 | MOLECULE: | MACROLI | DE 2'-P | HOSPHOTRANSFERASE MPHH |
| 48:00:00 | 4ec6-E | 4.3 | 2.8 | 69 | 109 | 4 | MOLECULE: | PUTATIV | E UNCHA | RACTERIZED PROTEIN |
| 49:00:00 | 6z0o-E | 4.3 | 3.6 | 67 | 90 | 4 | MOLECULE: | NUCLEOC | APSID | |
| 50:00:00 | 2cko-A | 4.3 | 2.6 | 63 | 344 | 10 | MOLECULE: | CHOLINE | KINASE | ALPHA |
| 51:00:00 | 5ohm-D | 4.3 | 3.5 | 64 | 89 | 5 | MOLECULE: | POLYUBI | QUITIN- | C |
| 52:00:00 | 6stj-G | 4.3 | 3.6 | 63 | 88 | 5 | MOLECULE: | INDUCED | MYELOI | D LEUKEMIA CELL DIFFERENTIATION PRO |
| 53:00:00 | 6stj-E | 4.3 | 3.6 | 64 | 89 | 5 | MOLECULE: | INDUCED | MYELOI | D LEUKEMIA CELL DIFFERENTIATION PRO |
| 54:00:00 | 3c5i-B | 4.3 | 3.4 | 67 | 345 | 4 | MOLECULE: | CHOLINE | KINASE | |
| 55:00:00 | 7ds4-A | 4.3 | 3.5 | 74 | 270 | 5 | MOLECULE: | F-ACTIN | #NAME? | G PROTEIN SUBUNIT ALPHA-1 |
| 56:00:00 | 3h4z-A | 4.3 | 2.9 | 75 | 568 | 5 | MOLECULE: | MALTOSE | #NAME? | G PERIPLASMIC PROTEIN FUSED WITH AL |
| 57:00:00 | 6gki-A | 4.3 | 3 | 68 | 185 | 1 | MOLECULE: | PROBABL | E PHOSP | HOLIPID-BINDING PROTEIN MLAC |
| 58:00:00 | 5igw-A | 4.3 | 2.5 | 56 | 296 | 9 | MOLECULE: | MACROLI | DE 2'-P | HOSPHOTRANSFERASE II |
| 59:00:00 | 1ylx-B | 4.3 | 3.2 | 70 | 100 | 6 | MOLECULE: | HYPOTHE | TICAL P | ROTEIN APC35702 |
| 60:00:00 | 3gax-A | 4.3 | 2.9 | 69 | 107 | 3 | MOLECULE: | CYSTATI | N-C | |

| | | | | | | | | | | | |
|---|---|---|---|---|---|---|---|---|---|---|---|
| 61:00:00 | 4r1k-A | 4.3 | 2.8 | 72 | 135 | 7 | MOLECULE: | UNCHARA | CTERIZE | D PROTEIN | |
| 62:00:00 | 3wz5-B | 4.3 | 2.9 | 69 | 135 | 9 | MOLECULE: | DOTI | | | |
| 63:00:00 | 9b8o-P | 4.3 | 2.9 | 76 | 204 | 16 | MOLECULE: | ATPASE | H+-TRAN | SPORTING V1 SUBUNIT D | |
| 64:00:00 | 3mwz-A | 4.3 | 4 | 72 | 115 | 6 | MOLECULE: | SIALOST | ATIN L2 | | |
| 65:00:00 | 3nx0-A | 4.3 | 3.6 | 72 | 109 | 7 | MOLECULE: | CYSTATI | N-C | | |
| 66:00:00 | 6roa-B | 4.3 | 3.6 | 72 | 109 | 7 | MOLECULE: | CYSTATI | N-C | | |
| 67:00:00 | 5ml9-B | 4.3 | 4 | 71 | 103 | 4 | MOLECULE: | LOW AFF | INITY I | MMUNOGLOBULIN GAMMA FC REGION RECEP | |
| 68:00:00 | 3ima-B | 4.3 | 3.6 | 67 | 84 | 6 | MOLECULE: | PAPAIN | | | |
| 69:00:00 | 6roa-A | 4.3 | 3.5 | 71 | 109 | 7 | MOLECULE: | CYSTATI | N-C | | |
| 70:00:00 | 2bo9-D | 4.3 | 4.1 | 75 | 217 | 7 | MOLECULE: | CARBOXY | PEPTIDA | SE A4 | |
| 71:00:00 | 6hjl-C | 4.3 | 3.8 | 69 | 90 | 6 | MOLECULE: | BCL-2-L | IKE PRO | TEIN 1 | |
| 72:00:00 | 2zhx-N | 4.3 | 2.6 | 58 | 82 | 12 | MOLECULE: | URACIL- | DNA GLY | COSYLASE | |
| 73:00:00 | 5igh-A | 4.2 | 2.7 | 59 | 301 | 14 | MOLECULE: | MACROLI | DE 2'-P | HOSPHOTRANSFERASE | |
| 74:00:00 | 3fka-A | 4.2 | 3.2 | 68 | 119 | 7 | MOLECULE: | UNCHARA | CTERIZE | D NTF-2 LIKE PROTEIN | |
| 75:00:00 | 5uxb-A | 4.2 | 2.3 | 55 | 290 | 7 | MOLECULE: | MACROLI | DE 2'-P | HOSPHOTRANSFERASE MPHH | |
| 76:00:00 | 2zhx-B | 4.2 | 3 | 59 | 82 | 10 | MOLECULE: | URACIL- | DNA GLY | COSYLASE | |
| 77:00:00 | 6yr8-B | 4.2 | 3.6 | 64 | 89 | 5 | MOLECULE: | GTPASE | KRAS | | |
| 78:00:00 | 5n9j-W | 4.2 | 4.1 | 77 | 463 | 9 | MOLECULE: | MEDIATO | R OF RN | A POLYMERASE II TRANSCRIPTION SUBUN | |
| 79:00:00 | 5ohm-L | 4.2 | 3.9 | 66 | 89 | 5 | MOLECULE: | POLYUBI | QUITIN- | C | |
| 80:00:00 | 5iwu-A | 4.2 | 2.6 | 58 | 299 | 9 | MOLECULE: | MACROLI | DE 2'-P | HOSPHOTRANSFERASE II | |
| 81:00:00 | 5u0s-Q | 4.2 | 4.2 | 76 | 500 | 9 | MOLECULE: | MEDIATO | R COMPL | EX SUBUNIT 6 | |
| 82:00:00 | 3c5i-D | 4.2 | 3.2 | 68 | 355 | 6 | MOLECULE: | CHOLINE | KINASE | | |
| 83:00:00 | 8ixi-A | 4.2 | 2.4 | 58 | 297 | 7 | MOLECULE: | ALDEHYD | E DEHYD | ROGENASE PROTEIN | |
| 84:00:00 | 6swt-B | 4.2 | 3.2 | 64 | 88 | 5 | MOLECULE: | ALPHA-A | CTININ- | | 2 |
| 85:00:00 | 6stj-H | 4.2 | 3.6 | 63 | 88 | 5 | MOLECULE: | INDUCED | MYELOI | D LEUKEMIA CELL DIFFERENTIATION PRO | |
| 86:00:00 | 5igv-A | 4.2 | 2.5 | 57 | 298 | 9 | MOLECULE: | MACROLI | DE 2'-P | HOSPHOTRANSFERASE II | |
| 87:00:00 | 2pun-B | 4.2 | 2.6 | 58 | 378 | 3 | MOLECULE: | METHYLT | HIORIBO | SE KINASE | |
| 88:00:00 | 5l33-A | 4.2 | 3.3 | 69 | 106 | 12 | MOLECULE: | DENOVO | NTF2 | | |
| 89:00:00 | 5elu-A | 4.2 | 3.6 | 65 | 91 | 5 | MOLECULE: | SUMO-AD | HIRON-S | 2B3 | |
| 90:00:00 | 3c5i-C | 4.2 | 3.5 | 69 | 353 | 6 | MOLECULE: | CHOLINE | KINASE | | |
| 91:00:00 | 6q0r-C | 4.2 | 3.5 | 72 | 183 | 7 | MOLECULE: | DNA DAM | AGE-BIN | DING PROTEIN 1 | |
| 92:00:00 | 5ih0-A | 4.2 | 2.6 | 58 | 299 | 9 | MOLECULE: | MACROLI | DE 2'-P | HOSPHOTRANSFERASE II | |
| 93:00:00 | 5igy-A | 4.2 | 2.3 | 55 | 295 | 9 | MOLECULE: | MACROLI | DE 2'-P | HOSPHOTRANSFERASE II | |
| 94:00:00 | 6uio-C | 4.2 | 2.9 | 69 | 112 | 7 | MOLECULE: | CYSTATI | N-8 | | |
| 95:00:00 | 6uio-B | 4.2 | 3 | 69 | 110 | 7 | MOLECULE: | CYSTATI | N-8 | | |
| 96:00:00 | 3wz4-B | 4.2 | 3.4 | 68 | 137 | 10 | MOLECULE: | DOTI | | | |
| 97:00:00 | 3c5i-A | 4.2 | 3.3 | 66 | 343 | 5 | MOLECULE: | CHOLINE | KINASE | | |
| 98:00:00 | 4ec6-B | 4.2 | 3.1 | 68 | 109 | 7 | MOLECULE: | PUTATIV | E UNCHA | RACTERIZED PROTEIN | |
| 99:00:00 | 8pk3-I | 4.2 | 3.2 | 66 | 87 | 6 | MOLECULE: | HEMAGGL | UTININ | HA1 CHAIN | |
| 0:00 | 1nb3-I | 4.2 | 2.9 | 66 | 98 | 5 | MOLECULE: | CATHEPS | IN H | | |
| 1:00 | 1kwi-A | 4.2 | 2.7 | 61 | 85 | 3 | MOLECULE: | PROTEGR | IN-3 PR | ECURSOR | |
| 2:00 | 1stf-I | 4.2 | 3.9 | 70 | 98 | 7 | MOLECULE: | PAPAIN | | | |
| 3:00 | 3k9m-D | 4.2 | 4.2 | 69 | 98 | 7 | MOLECULE: | CATHEPS | IN B | | |
| 4:00 | 1nb3-J | 4.2 | 4 | 71 | 98 | 8 | MOLECULE: | CATHEPS | IN H | | |
| 5:00 | 2mf7-A | 4.2 | 4.4 | 78 | 127 | 8 | MOLECULE: | MITOCHO | NDRIAL | IMPORT INNER MEMBRANE TRANSLOCASE S | |
| 6:00 | 1ygy-B | 4.2 | 3.1 | 73 | 527 | 11 | MOLECULE: | D-3-PHO | SPHOGLY | CERATE DEHYDROGENASE | |
| 7:00 | 6sj7-A | 4.2 | 3.2 | 74 | 355 | 8 | MOLECULE: | DDB1- A | ND CUL4 | -ASSOCIATED FACTOR 15 | |
| 8:00 | 3fka-B | 4.1 | 3.2 | 69 | 120 | 7 | MOLECULE: | UNCHARA | CTERIZE | D NTF-2 LIKE PROTEIN | |
| 9:00 | 6bjt-B | 4.1 | 3.1 | 71 | 130 | 6 | MOLECULE: | DUF4440 | DOMAIN | -CONTAINING PROTEIN | |
| 10:00 | 1nnv-A | 4.1 | 3.8 | 67 | 107 | 3 | MOLECULE: | HYPOTHE | TICAL P | ROTEIN HI1450 | |
| 11:00 | 8gqq-A | 4.1 | 3.1 | 66 | 99 | 3 | MOLECULE: | CYSTATI | N-A2 | | |
| 12:00 | 4i4k-A | 4.1 | 3.2 | 72 | 138 | 8 | MOLECULE: | UNCHARA | CTERIZE | D PROTEIN SGCJ | |
| 13:00 | 7pkw-A | 4.1 | 3.4 | 70 | 240 | 9 | MOLECULE: | PUTATIV | E TRANS | FER PROTEIN | |
| 14:00 | 4n6l-A | 4.1 | 2.8 | 67 | 114 | 3 | MOLECULE: | CYSTATI | N-M | | |

| | | | | | | | | | | |
|---|---|---|---|---|---|---|---|---|---|---|
| 15:00 | 3lh4-A | 4.1 | 4.3 | 73 | 115 | 7 | MOLECULE: | SECRETE | D CYSTA | TIN |
| 16:00 | 3b8l-A | 4.1 | 4.1 | 71 | 147 | 6 | MOLECULE: | UNCHARA | CTERIZE | D PROTEIN |
| 17:00 | 4pw1-B | 4.1 | 3.3 | 73 | 215 | 8 | MOLECULE: | UNCHARA | CTERIZE | D PROTEIN |
| 18:00 | 3fka-D | 4.1 | 3.2 | 68 | 119 | 7 | MOLECULE: | UNCHARA | CTERIZE | D NTF-2 LIKE PROTEIN |
| 19:00 | 8gt7-D | 4.1 | 3.7 | 70 | 98 | 9 | MOLECULE: | CYSTEIN | E PROTE | INASE FALCIPAIN 2A |
| 20:00 | 5a0o-B | 4.1 | 3.1 | 64 | 93 | 5 | MOLECULE: | ADHIRON | | |
| 21:00 | 6w3d-A | 4.1 | 2.8 | 69 | 114 | 7 | MOLECULE: | RD1NTF2 | _05 | |
| 22:00 | 5mn2-C | 4.1 | 3.6 | 63 | 88 | 5 | MOLECULE: | LOW AFF | INITY I | MMUNOGLOBULIN GAMMA FC REGION RECEP |
| 23:00 | 1spu-B | 4.1 | 3.3 | 67 | 720 | 6 | MOLECULE: | COPPER | AMINE O | XIDASE |
| 24:00:00 | 1nb3-K | 4.1 | 2.9 | 65 | 98 | 5 | MOLECULE: | CATHEPS | IN H | |
| 25:00:00 | 3kfq-D | 4.1 | 3 | 66 | 98 | 8 | MOLECULE: | CATHEPS | IN L2 | |
| 26:00:00 | 5zc1-C | 4.1 | 3.5 | 70 | 100 | 6 | MOLECULE: | CYSTATI | N-1 | |
| 27:00:00 | 4qvh-B | 4.1 | 3.1 | 74 | 595 | 1 | MOLECULE: | MALTOSE | #NAME? | G PERIPLASMIC PROTEIN, 4'-PHOSPHOPA |
| 28:00:00 | 3buu-B | 4.1 | 4 | 76 | 220 | 8 | MOLECULE: | UNCHARA | CTERIZE | D LOLA SUPERFAMILY PROTEIN NE2245 |
| 29:00:00 | 2olc-A | 4.1 | 1.8 | 51 | 376 | 4 | MOLECULE: | METHYLT | HIORIBO | SE KINASE |
| 30:00:00 | 5uxd-A | 4.1 | 2.5 | 57 | 300 | 7 | MOLECULE: | MACROLI | DE 2'-P | HOSPHOTRANSFERASE MPHH |
| 31:00:00 | 3ub1-A | 4.1 | 2.8 | 69 | 248 | 0 | MOLECULE: | ORF13-L | IKE PRO | TEIN |
| 32:00:00 | 7pk4-B | 4.1 | 3.2 | 58 | 81 | 5 | MOLECULE: | CATHEPS | IN L2 | |
| 33:00:00 | 5zc1-F | 4.1 | 3.4 | 67 | 98 | 4 | MOLECULE: | CYSTATI | N-1 | |
| 34:00:00 | 6uio-D | 4.1 | 2.8 | 67 | 109 | 7 | MOLECULE: | CYSTATI | N-8 | |
| 35:00:00 | 5uxa-A | 4.1 | 2.3 | 55 | 301 | 11 | MOLECULE: | MACROLI | DE 2'-P | HOSPHOTRANSFERASE II |
| 36:00:00 | 6yxw-D | 4.1 | 3.4 | 62 | 88 | 5 | MOLECULE: | GTPASE | KRAS | |
| 37:00:00 | 6qb2-A | 4.1 | 2.9 | 65 | 95 | 5 | MOLECULE: | MONOMEI | OF SQT | -1C |
| 38:00:00 | 3gax-B | 4.1 | 3.5 | 70 | 107 | 4 | MOLECULE: | CYSTATI | N-C | |
| 39:00:00 | 3wz5-A | 4.1 | 2.9 | 69 | 135 | 9 | MOLECULE: | DOTI | | |
| 40:00:00 | 3nx0-B | 4.1 | 3.6 | 68 | 109 | 6 | MOLECULE: | CYSTATI | N-C | |
| 41:00:00 | 1nb5-I | 4.1 | 4.1 | 69 | 98 | 7 | MOLECULE: | CATHEPS | IN H | |
| 42:00:00 | 3blz-A | 4.1 | 3.5 | 68 | 124 | 9 | MOLECULE: | NTF2-LI | KE PROT | EIN OF UNKNOWN FUNCTION |
| 43:00:00 | 7w1a-A | 4.1 | 2.6 | 56 | 287 | 5 | MOLECULE: | MACROLI | DE 2'-P | HOSPHOTRANSFERASE |
| 44:00:00 | 8pk3-G | 4.1 | 3.2 | 67 | 87 | 6 | MOLECULE: | HEMAGGL | UTININ | HA1 CHAIN |
| 45:00:00 | 7ny8-C | 4.1 | 4.2 | 69 | 91 | 4 | MOLECULE: | GTPASE | KRAS | |
| 46:00:00 | 6st2-C | 4.1 | 3.7 | 66 | 87 | 5 | MOLECULE: | BCL-2-L | IKE PRO | TEIN 1 |
| 47:00:00 | 1nb5-J | 4.1 | 3.1 | 67 | 98 | 4 | MOLECULE: | CATHEPS | IN H | |
| 48:00:00 | 6st2-D | 4.1 | 3.7 | 64 | 87 | 5 | MOLECULE: | BCL-2-L | IKE PRO | TEIN 1 |
| 49:00:00 | 6zgn-A | 4.1 | 3.2 | 63 | 101 | 5 | MOLECULE: | PUTATIV | E TRANS | FER PROTEIN |
| 50:00:00 | 3blz-H | 4.1 | 3.4 | 70 | 123 | 10 | MOLECULE: | NTF2-LI | KE PROT | EIN OF UNKNOWN FUNCTION |
| 51:00:00 | 5zc1-E | 4.1 | 3.2 | 70 | 98 | 7 | MOLECULE: | CYSTATI | N-1 | |
| 52:00:00 | 3blz-C | 4.1 | 3.3 | 68 | 124 | 9 | MOLECULE: | NTF2-LI | KE PROT | EIN OF UNKNOWN FUNCTION |
| 53:00:00 | 5o46-B | 4.1 | 4 | 70 | 114 | 3 | MOLECULE: | IRISTAT | IN | |
| 54:00:00 | 1nb5-K | 4.1 | 3 | 66 | 98 | 6 | MOLECULE: | CATHEPS | IN H | |
| 55:00:00 | 1n5h-A | 4 | 3 | 66 | 105 | 3 | MOLECULE: | PROTEGR | INS | |
| 56:00:00 | 7rag-A | 4 | 3.3 | 61 | 170 | 5 | MOLECULE: | LIPOPRO | TEIN | |
| 57:00:00 | 5ts4-B | 4 | 3.1 | 71 | 106 | 6 | MOLECULE: | DENOVO | NTF2 | |
| 58:00:00 | 3mes-B | 4 | 2.6 | 62 | 358 | 5 | MOLECULE: | CHOLINE | KINASE | |
| 59:00:00 | 1wnh-A | 4 | 3.7 | 68 | 220 | 4 | MOLECULE: | LATEXIN | | |
| 60:00:00 | 5ih1-A | 4 | 2.5 | 56 | 297 | 9 | MOLECULE: | MACROLI | DE 2'-P | HOSPHOTRANSFERASE II |
| 61:00:00 | 6yxw-B | 4 | 3.1 | 63 | 89 | 5 | MOLECULE: | GTPASE | KRAS | |
| 62:00:00 | 4n6m-B | 4 | 2.8 | 66 | 116 | 3 | MOLECULE: | CYSTATI | N-M | |
| 63:00:00 | 3kfq-C | 4 | 3.2 | 68 | 98 | 4 | MOLECULE: | CATHEPS | IN L2 | |
| 64:00:00 | 8f8q-G | 4 | 3.9 | 75 | 278 | 3 | MOLECULE: | ACTIN, | ALPHA S | KELETAL MUSCLE |
| 65:00:00 | 3kse-F | 4 | 3.6 | 72 | 98 | 4 | MOLECULE: | CATHEPS | IN L1 | |
| 66:00:00 | 5cnl-A | 4 | 3.3 | 72 | 139 | 13 | MOLECULE: | ICML-LI | KE | |
| 67:00:00 | 4akr-A | 4 | 3.2 | 75 | 269 | 4 | MOLECULE: | F-ACTIN | #NAME? | G PROTEIN SUBUNIT ALPHA |
| 68:00:00 | 3lq3-A | 4 | 2.5 | 63 | 339 | 10 | MOLECULE: | CHOLINE | /ETHANO | LAMINE KINASE |

| | | | | | | | | | | |
|---|---|---|---|---|---|---|---|---|---|---|
| 69:00:00 | 3wz4-E | 4 | 3.6 | 73 | 139 | 10 | MOLECULE: | DOTI | | |
| 70:00:00 | 2j8x-B | 4 | 2.5 | 58 | 83 | 12 | MOLECULE: | URACIL- | DNA GLY | COSYLASE |
| 71:00:00 | 8pk3-H | 4 | 3 | 64 | 87 | 5 | MOLECULE: | HEMAGGL | UTININ | HA1 CHAIN |
| 72:00:00 | 5a0o-S | 4 | 3.9 | 71 | 93 | 6 | MOLECULE: | ADHIRON | | |
| 73:00:00 | 7dsb-A | 4 | 3.5 | 73 | 264 | 5 | MOLECULE: | F-ACTIN | #NAME? | G PROTEIN SUBUNIT ALPHA-1 |
| 74:00:00 | 7ds8-A | 4 | 3.3 | 74 | 274 | 5 | MOLECULE: | F-ACTIN | #NAME? | G PROTEIN SUBUNIT ALPHA-1 |
| 75:00:00 | 5jou-A | 4 | 3.3 | 66 | 945 | 11 | MOLECULE: | ALPHA-X | YLOSIDA | SE BOGH31A |
| 76:00:00 | 1lqm-D | 4 | 2.5 | 59 | 83 | 14 | MOLECULE: | URACIL- | DNA GLY | COSYLASE |
| 77:00:00 | 3h4z-B | 4 | 3 | 74 | 563 | 5 | MOLECULE: | MALTOSE | #NAME? | G PERIPLASMIC PROTEIN FUSED WITH AL |
| 78:00:00 | 3blz-B | 4 | 3.1 | 66 | 124 | 9 | MOLECULE: | NTF2-LI | KE PROT | EIN OF UNKNOWN FUNCTION |
| 79:00:00 | 1cew-I | 4 | 3.6 | 69 | 108 | 4 | MOLECULE: | CYSTATI | N | |
| 80:00:00 | 1ugi-E | 4 | 2.8 | 60 | 83 | 13 | MOLECULE: | URACIL- | DNA GLY | COSYLASE INHIBITOR |
| 81:00:00 | 1nb3-L | 4 | 3 | 66 | 98 | 8 | MOLECULE: | CATHEPS | IN H | |
| 82:00:00 | 3ub1-E | 4 | 3 | 71 | 252 | 1 | MOLECULE: | ORF13-L | IKE PRO | TEIN |
| 83:00:00 | 7w1a-B | 4 | 2.6 | 56 | 287 | 5 | MOLECULE: | MACROLI | DE 2'-P | HOSPHOTRANSFERASE |
| 84:00:00 | 3kse-D | 4 | 3.2 | 69 | 98 | 4 | MOLECULE: | CATHEPS | IN L1 | |
| 85:00:00 | 5igp-A | 4 | 3.1 | 59 | 300 | 17 | MOLECULE: | MACROLI | DE 2'-P | HOSPHOTRANSFERASE |
| 86:00:00 | 5zc1-B | 4 | 3.4 | 68 | 101 | 9 | MOLECULE: | CYSTATI | N-1 | |
| 87:00:00 | 3ub1-F | 4 | 2.9 | 68 | 251 | 0 | MOLECULE: | ORF13-L | IKE PRO | TEIN |
| 88:00:00 | 7knc-A | 4 | 2.9 | 58 | 919 | 7 | MOLECULE: | ALPHA-X | YLOSIDA | SE |
| 89:00:00 | 8hbn-B | 3.9 | 3.2 | 71 | 165 | 11 | MOLECULE: | MRNA EX | PORT FA | CTOR MEX67 |
| 90:00:00 | 2lyx-A | 3.9 | 3.1 | 63 | 87 | 3 | MOLECULE: | UNCHARA | CTERIZE | D PROTEIN YQZG |
| 91:00:00 | 3ub1-D | 3.9 | 3.8 | 70 | 253 | 14 | MOLECULE: | ORF13-L | IKE PRO | TEIN |
| 92:00:00 | 6ud7-A | 3.9 | 3.4 | 73 | 426 | 5 | MOLECULE: | DDB1- A | ND CUL4 | -ASSOCIATED FACTOR 15 |
| 93:00:00 | 3buu-A | 3.9 | 3.7 | 75 | 224 | 7 | MOLECULE: | UNCHARA | CTERIZE | D LOLA SUPERFAMILY PROTEIN NE2245 |
| 94:00:00 | 6wlw-U | 3.9 | 3 | 76 | 205 | 12 | MOLECULE: | V-TYPE | PROTON | ATPASE 21 KDA PROTEOLIPID SUBUNIT |
| 95:00:00 | 3en8-A | 3.9 | 3.3 | 73 | 128 | 8 | MOLECULE: | UNCHARA | CTERIZE | D NTF-2 LIKE PROTEIN |
| 96:00:00 | 6w3g-A | 3.9 | 3.1 | 69 | 109 | 4 | MOLECULE: | RD1NTF2 | _04 | |
| 97:00:00 | 4k90-B | 3.9 | 3 | 71 | 207 | 6 | MOLECULE: | EXTRACE | LLULAR | METALLOPROTEINASE MEP |
| 98:00:00 | 7w15-A | 3.9 | 2.6 | 56 | 294 | 5 | MOLECULE: | MACROLI | DE 2'-P | HOSPHOTRANSFERASE |
| 99:00:00 | 4n6m-A | 3.9 | 2.8 | 66 | 117 | 3 | MOLECULE: | CYSTATI | N-M | |
| 0:00 | 5uxc-A | 3.9 | 2.5 | 57 | 296 | 7 | MOLECULE: | PREDICT | ED AMIN | OGLYCOSIDE PHOSPHOTRANSFERASE |
| 1:00 | 7w15-B | 3.9 | 2.5 | 56 | 293 | 5 | MOLECULE: | MACROLI | DE 2'-P | HOSPHOTRANSFERASE |
| 2:00 | 3ub1-C | 3.9 | 3.7 | 69 | 251 | 14 | MOLECULE: | ORF13-L | IKE PRO | TEIN |
| 3:00 | 6ihj-D | 3.9 | 3.7 | 74 | 122 | 3 | MOLECULE: | NUCLEAR | RNA EX | PORT FACTOR 1 |
| 4:00 | 6e5u-H | 3.9 | 2.6 | 69 | 138 | 12 | MOLECULE: | NUCLEAR | RNA EX | PORT FACTOR 1 |
| 5:00 | 3aa6-A | 3.9 | 3.3 | 73 | 270 | 5 | MOLECULE: | F-ACTIN | #NAME? | G PROTEIN SUBUNIT ALPHA-1 |
| 6:00 | 1lvn-B | 3.9 | 3.3 | 67 | 720 | 6 | MOLECULE: | COPPER | AMINE O | XIDASE |
| 7:00 | 1ugi-B | 3.9 | 2.7 | 60 | 83 | 13 | MOLECULE: | URACIL- | DNA GLY | COSYLASE INHIBITOR |
| 8:00 | 6ta7-D | 3.9 | 3.1 | 70 | 124 | 6 | MOLECULE: | RAS GTP | ASE-ACT | IVATING PROTEIN-BINDING PROTEIN 1 |
| 9:00 | 6q0v-C | 3.9 | 3.1 | 69 | 192 | 7 | MOLECULE: | DNA DAM | AGE-BIN | DING PROTEIN 1 |
| 10:00 | 4n4x-A | 3.9 | 3 | 71 | 535 | 10 | MOLECULE: | MALTOSE | #NAME? | G PERIPLASMIC/PALATE LUNG AND NASAL |
| 11:00 | 2r4i-B | 3.9 | 3.3 | 69 | 121 | 3 | MOLECULE: | UNCHARA | CTERIZE | D PROTEIN |
| 12:00 | 1ugi-C | 3.9 | 2.6 | 59 | 83 | 14 | MOLECULE: | URACIL- | DNA GLY | COSYLASE INHIBITOR |
| 13:00 | 2olc-B | 3.9 | 2.1 | 55 | 376 | 4 | MOLECULE: | METHYLT | HIORIBO | SE KINASE |
| 14:00 | 1udi-I | 3.9 | 2.7 | 59 | 83 | 14 | MOLECULE: | URACIL- | DNA GLY | COSYLASE |
| 15:00 | 3blz-G | 3.9 | 3.3 | 70 | 124 | 10 | MOLECULE: | NTF2-LI | KE PROT | EIN OF UNKNOWN FUNCTION |
| 16:00 | 1lqm-F | 3.9 | 2.8 | 59 | 82 | 14 | MOLECULE: | URACIL- | DNA GLY | COSYLASE |
| 17:00 | 5igr-A | 3.9 | 3.4 | 59 | 300 | 15 | MOLECULE: | MACROLI | DE 2'-P | HOSPHOTRANSFERASE |
| 18:00 | 3blz-K | 3.9 | 3 | 64 | 123 | 9 | MOLECULE: | NTF2-LI | KE PROT | EIN OF UNKNOWN FUNCTION |
| 19:00 | 5o46-A | 3.9 | 3.8 | 69 | 114 | 3 | MOLECULE: | IRISTAT | IN | |
| 20:00 | 4lyl-J | 3.9 | 2.5 | 58 | 82 | 14 | MOLECULE: | URACIL- | DNA GLY | COSYLASE |
| 21:00 | 5igj-A | 3.9 | 3.4 | 59 | 300 | 15 | MOLECULE: | MACROLI | DE 2'-P | HOSPHOTRANSFERASE |
| 22:00 | 7ny8-D | 3.9 | 3.3 | 62 | 97 | 5 | MOLECULE: | GTPASE | KRAS | |

| | | | | | | | | | | |
|---|---|---|---|---|---|---|---|---|---|---|
| 23:00 | 3blz-L | 3.9 | 3.4 | 70 | 123 | 10 | MOLECULE: | NTF2-LI | KE PROT | EIN OF UNKNOWN FUNCTION |
| 24:00:00 | 2w9p-L | 3.9 | 3.8 | 70 | 87 | 7 | MOLECULE: | MULTICY | STATIN | |
| 25:00:00 | 5igs-A | 3.9 | 3.3 | 59 | 300 | 15 | MOLECULE: | MACROLI | DE 2'-P | HOSPHOTRANSFERASE |
| 26:00:00 | 1ugi-F | 3.9 | 2.9 | 61 | 83 | 13 | MOLECULE: | URACIL- | DNA GLY | COSYLASE INHIBITOR |
| 27:00:00 | 1lvn-A | 3.9 | 3.2 | 63 | 718 | 6 | MOLECULE: | COPPER | AMINE O | XIDASE |
| 28:00:00 | 5zc1-A | 3.9 | 3.4 | 67 | 101 | 9 | MOLECULE: | CYSTATI | N-1 | |
| 29:00:00 | 3blz-F | 3.9 | 3.5 | 70 | 123 | 10 | MOLECULE: | NTF2-LI | KE PROT | EIN OF UNKNOWN FUNCTION |
| 30:00:00 | 5cnl-B | 3.9 | 3.3 | 71 | 139 | 13 | MOLECULE: | ICML-LI | KE | |
| 31:00:00 | 3kse-E | 3.9 | 3.6 | 71 | 98 | 4 | MOLECULE: | CATHEPS | IN L1 | |
| 32:00:00 | 2rgq-B | 3.8 | 3.3 | 72 | 134 | 10 | MOLECULE: | DOMAIN | OF UNKN | OWN FUNCTION WITH A CYSTATIN-LIKE F |
| 33:00:00 | 8cgm-A | 3.8 | 3.2 | 70 | 190 | 7 | MOLECULE: | OUTER M | EMBRANE | LIPOPROTEIN CARRIER PROTEIN LOLA |
| 34:00:00 | 1jkg-A | 3.8 | 2.9 | 71 | 139 | 11 | MOLECULE: | P15 | | |
| 35:00:00 | 2qiy-A | 3.8 | 2.9 | 67 | 134 | 10 | MOLECULE: | UBP3-AS | SOCIATE | D PROTEIN BRE5 |
| 36:00:00 | 3f7s-A | 3.8 | 3.5 | 68 | 142 | 7 | MOLECULE: | UNCHARA | CTERIZE | D NTF2-LIKE PROTEIN |
| 37:00:00 | 5uwa-A | 3.8 | 3.1 | 66 | 185 | 0 | MOLECULE: | PROBABL | E PHOSP | HOLIPID-BINDING PROTEIN MLAC |
| 38:00:00 | 7mhu-A | 3.8 | 3.4 | 61 | 374 | 11 | MOLECULE: | EXO-ALP | HA-SIAL | IDASE |
| 39:00:00 | 2gu3-A | 3.8 | 2.9 | 59 | 128 | 3 | MOLECULE: | YPMB PR | OTEIN | |
| 40:00:00 | 1lqm-H | 3.8 | 2.7 | 59 | 84 | 14 | MOLECULE: | URACIL- | DNA GLY | COSYLASE |
| 41:00:00 | 4k61-B | 3.8 | 2.8 | 61 | 138 | 8 | MOLECULE: | UNCHARA | CTERIZE | D PROTEIN |
| 42:00:00 | 2rgq-A | 3.8 | 3.3 | 72 | 133 | 10 | MOLECULE: | DOMAIN | OF UNKN | OWN FUNCTION WITH A CYSTATIN-LIKE F |
| 43:00:00 | 5ts4-A | 3.8 | 3.1 | 69 | 101 | 6 | MOLECULE: | DENOVO | NTF2 | |
| 44:00:00 | 4lyl-D | 3.8 | 3.1 | 62 | 82 | 13 | MOLECULE: | URACIL- | DNA GLY | COSYLASE |
| 45:00:00 | 5igz-A | 3.8 | 2.6 | 57 | 299 | 9 | MOLECULE: | MACROLI | DE 2'-P | HOSPHOTRANSFERASE II |
| 46:00:00 | 5tgn-C | 3.8 | 3.7 | 71 | 107 | 7 | MOLECULE: | UNCHARA | CTERIZE | D PROTEIN |
| 47:00:00 | 1nb5-L | 3.8 | 3 | 64 | 98 | 5 | MOLECULE: | CATHEPS | IN H | |
| 48:00:00 | 1yvb-I | 3.8 | 3.4 | 65 | 111 | 6 | MOLECULE: | FALCIPA | IN 2 | |
| 49:00:00 | 3ub1-B | 3.8 | 3.7 | 70 | 252 | 14 | MOLECULE: | ORF13-L | IKE PRO | TEIN |
| 50:00:00 | 1dvc-A | 3.8 | 3.2 | 62 | 98 | 5 | MOLECULE: | STEFIN | A | |
| 51:00:00 | 6hjl-G | 3.8 | 3.6 | 63 | 88 | 5 | MOLECULE: | BCL-2-L | IKE PRO | TEIN 1 |
| 52:00:00 | 4h63-Q | 3.8 | 4 | 67 | 410 | 7 | MOLECULE: | MEDIATO | R OF RN | A POLYMERASE II TRANSCRIPTION SUBUN |
| 53:00:00 | 3nv0-B | 3.8 | 3.2 | 72 | 136 | 3 | MOLECULE: | NUCLEAR | RNA EX | PORT FACTOR 2 |
| 54:00:00 | 6yxt-A | 3.8 | 2.6 | 63 | 359 | 5 | MOLECULE: | CHOLINE | KINASE | |
| 55:00:00 | 7ds2-A | 3.8 | 3.3 | 73 | 272 | 5 | MOLECULE: | F-ACTIN | #NAME? | G PROTEIN SUBUNIT ALPHA-1 |
| 56:00:00 | 7ds6-A | 3.8 | 3.4 | 74 | 275 | 5 | MOLECULE: | F-ACTIN | #NAME? | G PROTEIN SUBUNIT ALPHA-1 |
| 57:00:00 | 6ue5-A | 3.8 | 3.3 | 73 | 424 | 7 | MOLECULE: | DDB1- A | ND CUL4 | -ASSOCIATED FACTOR 15 |
| 58:00:00 | 6znl-K | 3.8 | 3.9 | 73 | 278 | 4 | MOLECULE: | ARP1 AC | TIN REL | ATED PROTEIN 1 HOMOLOG A |
| 59:00:00 | 2wo0-B | 3.8 | 3.3 | 67 | 720 | 7 | MOLECULE: | PRIMARY | AMINE | OXIDASE |
| 60:00:00 | 5uwa-B | 3.8 | 3.1 | 68 | 185 | 0 | MOLECULE: | PROBABL | E PHOSP | HOLIPID-BINDING PROTEIN MLAC |
| 61:00:00 | 1a67-A | 3.8 | 3.4 | 73 | 107 | 5 | MOLECULE: | CYSTATI | N | |
| 62:00:00 | 5ts4-C | 3.8 | 3.3 | 70 | 104 | 6 | MOLECULE: | DENOVO | NTF2 | |
| 63:00:00 | 7kmp-A | 3.8 | 2.6 | 57 | 920 | 7 | MOLECULE: | ALPHA-X | YLOSIDA | SE |
| 64:00:00 | 1lqm-B | 3.8 | 2.5 | 58 | 82 | 14 | MOLECULE: | URACIL- | DNA GLY | COSYLASE |
| 65:00:00 | 1eqk-A | 3.8 | 3.6 | 69 | 102 | 9 | MOLECULE: | ORYZACY | STATIN- | I |
| 66:00:00 | 6grr-A | 3.8 | 3.4 | 69 | 718 | 10 | MOLECULE: | AMINE O | XIDASE | |
| 67:00:00 | 8cgm-B | 3.8 | 3.6 | 71 | 190 | 8 | MOLECULE: | OUTER M | EMBRANE | LIPOPROTEIN CARRIER PROTEIN LOLA |
| 68:00:00 | 4lyl-F | 3.8 | 2.8 | 60 | 82 | 13 | MOLECULE: | URACIL- | DNA GLY | COSYLASE |
| 69:00:00 | 6e5u-F | 3.8 | 2.6 | 69 | 138 | 12 | MOLECULE: | NUCLEAR | RNA EX | PORT FACTOR 1 |
| 70:00:00 | 3wz3-A | 3.8 | 3.1 | 69 | 137 | 4 | MOLECULE: | TRAM PR | OTEIN | |
| 71:00:00 | 4z48-B | 3.8 | 4.4 | 73 | 240 | 5 | MOLECULE: | UNCHARA | CTERIZE | D PROTEIN |
| 72:00:00 | 5tgn-B | 3.8 | 3.5 | 71 | 108 | 7 | MOLECULE: | UNCHARA | CTERIZE | D PROTEIN |
| 73:00:00 | 4lyl-L | 3.8 | 2.7 | 60 | 82 | 13 | MOLECULE: | URACIL- | DNA GLY | COSYLASE |
| 74:00:00 | 4ebg-A | 3.8 | 3.4 | 71 | 97 | 4 | MOLECULE: | UNCHARA | CTERIZE | D PROTEIN |
| 75:00:00 | 5tgn-D | 3.8 | 3.7 | 69 | 109 | 9 | MOLECULE: | UNCHARA | CTERIZE | D PROTEIN |
| 76:00:00 | 4lyl-N | 3.8 | 2.6 | 58 | 82 | 14 | MOLECULE: | URACIL- | DNA GLY | COSYLASE |

| | | | | | | | | | | |
|---|---|---|---|---|---|---|---|---|---|---|
| 77:00:00 | 1qal-A | 3.8 | 3.1 | 64 | 718 | 11 | MOLECULE: | COPPER | AMINE O | XIDASE |
| 78:00:00 | 6uio-A | 3.8 | 3.4 | 62 | 112 | 8 | MOLECULE: | CYSTATI | N-8 | |
| 79:00:00 | 4wyk-D | 3.8 | 2.6 | 69 | 138 | 12 | MOLECULE: | NUCLEAR | RNA EX | PORT FACTOR 1 |
| 80:00:00 | 3blz-D | 3.8 | 3.3 | 73 | 124 | 11 | MOLECULE: | NTF2-LI | KE PROT | EIN OF UNKNOWN FUNCTION |
| 81:00:00 | 2bo9-B | 3.8 | 3.1 | 66 | 217 | 9 | MOLECULE: | CARBOXY | PEPTIDA | SE A4 |
| 82:00:00 | 1jrq-A | 3.8 | 3 | 64 | 718 | 11 | MOLECULE: | COPPER | AMINE O | XIDASE |
| 83:00:00 | 1ugi-D | 3.8 | 2.9 | 61 | 82 | 13 | MOLECULE: | URACIL- | DNA GLY | COSYLASE INHIBITOR |
| 84:00:00 | 6ezz-A | 3.8 | 3.1 | 66 | 718 | 9 | MOLECULE: | PRIMARY | AMINE | OXIDASE |
| 85:00:00 | 3blz-I | 3.8 | 3.3 | 67 | 123 | 9 | MOLECULE: | NTF2-LI | KE PROT | EIN OF UNKNOWN FUNCTION |
| 86:00:00 | 2m7o-A | 3.7 | 3.2 | 60 | 70 | 7 | MOLECULE: | UNCHARA | CTERIZE | D PROTEIN |
| 87:00:00 | 1oac-B | 3.7 | 3.2 | 66 | 723 | 8 | MOLECULE: | COPPER | AMINE O | XIDASE |
| 88:00:00 | 7zhu-B | 3.7 | 4.5 | 65 | 543 | 3 | MOLECULE: | GLUCOSE | #NAME? | PHATE 1-DEHYDROGENASE |
| 89:00:00 | 5tgn-A | 3.7 | 3.3 | 66 | 109 | 8 | MOLECULE: | UNCHARA | CTERIZE | D PROTEIN |
| 90:00:00 | 1nkg-A | 3.7 | 3 | 47 | 508 | 6 | MOLECULE: | RHAMNOG | ALACTUR | ONASE B |
| 91:00:00 | 3ke7-B | 3.7 | 3.9 | 73 | 132 | 7 | MOLECULE: | PUTATIV | E KETOS | TEROID ISOMERASE |
| 92:00:00 | 5jov-A | 3.7 | 2.9 | 65 | 949 | 12 | MOLECULE: | ALPHA-X | YLOSIDA | SE BOGH31A |
| 93:00:00 | 2pyw-A | 3.7 | 2.4 | 62 | 417 | 5 | MOLECULE: | UNCHARA | CTERIZE | D PROTEIN |
| 94:00:00 | 8wzb-G | 3.7 | 4.2 | 70 | 387 | 3 | MOLECULE: | DPY30 D | OMAIN C | ONTAINING 2 |
| 95:00:00 | 4exr-A | 3.7 | 3.4 | 59 | 152 | 12 | MOLECULE: | PUTATIV | E LIPOP | ROTEIN |
| 96:00:00 | 6lyd-I | 3.7 | 2.8 | 60 | 83 | 10 | MOLECULE: | PROBABL | E URACI | L-DNA GLYCOSYLASE |
| 97:00:00 | 3b8l-F | 3.7 | 4 | 70 | 143 | 9 | MOLECULE: | UNCHARA | CTERIZE | D PROTEIN |
| 98:00:00 | 4k61-A | 3.7 | 2.8 | 61 | 140 | 8 | MOLECULE: | UNCHARA | CTERIZE | D PROTEIN |
| 99:00:00 | 1dvd-A | 3.7 | 3.5 | 66 | 98 | 5 | MOLECULE: | STEFIN | A | |
| 0:00 | 5tph-A | 3.7 | 4.2 | 69 | 120 | 6 | MOLECULE: | DE NOVO | NTF2 H | OMODIMER |
| 1:00 | 8gt0-D | 3.7 | 3.6 | 71 | 98 | 3 | MOLECULE: | CYSTEIN | E PROTE | ASE FALCIPAIN-2 |
| 2:00 | 4wwu-I | 3.7 | 3.3 | 73 | 161 | 11 | MOLECULE: | MRNA EX | PORT FA | CTOR MEX67 |
| 3:00 | 4wwu-L | 3.7 | 3.3 | 73 | 162 | 11 | MOLECULE: | MRNA EX | PORT FA | CTOR MEX67 |
| 4:00 | 2uug-C | 3.7 | 2.5 | 58 | 82 | 14 | MOLECULE: | URACIL- | DNA GLY | COSYLASE |
| 5:00 | 2r4i-C | 3.7 | 3.5 | 69 | 120 | 3 | MOLECULE: | UNCHARA | CTERIZE | D PROTEIN |
| 6:00 | 1d6u-B | 3.7 | 2.8 | 63 | 720 | 8 | MOLECULE: | COPPER | AMINE O | XIDASE |
| 7:00 | 6z0o-F | 3.7 | 3.9 | 66 | 90 | 5 | MOLECULE: | NUCLEOC | APSID | |
| 8:00 | 2zhx-J | 3.7 | 2.6 | 59 | 82 | 12 | MOLECULE: | URACIL- | DNA GLY | COSYLASE |
| 9:00 | 3b8l-C | 3.7 | 3.8 | 71 | 144 | 8 | MOLECULE: | UNCHARA | CTERIZE | D PROTEIN |
| 10:00 | 3b8l-E | 3.7 | 3.9 | 71 | 144 | 8 | MOLECULE: | UNCHARA | CTERIZE | D PROTEIN |
| 11:00 | 3b8l-B | 3.7 | 3.6 | 71 | 144 | 8 | MOLECULE: | UNCHARA | CTERIZE | D PROTEIN |
| 12:00 | 3b8l-D | 3.7 | 3.8 | 71 | 144 | 8 | MOLECULE: | UNCHARA | CTERIZE | D PROTEIN |
| 13:00 | 2wyh-B | 3.7 | 3.1 | 76 | 891 | 5 | MOLECULE: | ALPHA-M | ANNOSID | ASE |
| 14:00 | 1ask-B | 3.7 | 3.3 | 70 | 120 | 10 | MOLECULE: | NUCLEAR | TRANSP | ORT FACTOR 2 |
| 15:00 | 4jf8-A | 3.7 | 3 | 63 | 144 | 6 | MOLECULE: | TRWG CO | MPONENT | OF TYPE IV SECRETION SYSTEM |
| 16:00 | 5zc1-D | 3.7 | 3.4 | 70 | 98 | 4 | MOLECULE: | CYSTATI | N-1 | |
| 17:00 | 9bry-c | 3.7 | 3.1 | 77 | 203 | 12 | MOLECULE: | V-TYPE | PROTON | ATPASE SUBUNIT S1 |
| 18:00 | 2vw2-A | 3.7 | 3.4 | 63 | 658 | 11 | MOLECULE: | SIALIDA | SE B | |
| 19:00 | 4ttg-B | 3.7 | 3 | 51 | 1015 | 4 | MOLECULE: | BETA-GA | LACTOSI | DASE |
| 20:00 | 2zhx-H | 3.7 | 2.8 | 60 | 82 | 12 | MOLECULE: | URACIL- | DNA GLY | COSYLASE |
| 21:00 | 2pul-B | 3.7 | 1.9 | 50 | 369 | 4 | MOLECULE: | METHYLT | HIORIBO | SE KINASE |
| 22:00 | 4fq4-A | 3.7 | 3.5 | 62 | 658 | 8 | MOLECULE: | SIALIDA | SE B | |
| 23:00 | 3k9m-C | 3.7 | 3 | 66 | 98 | 5 | MOLECULE: | CATHEPS | IN B | |
| 24:00:00 | 2qiy-B | 3.7 | 3.2 | 67 | 133 | 12 | MOLECULE: | UBP3-AS | SOCIATE | D PROTEIN BRE5 |
| 25:00:00 | 1uug-B | 3.7 | 2.6 | 58 | 82 | 12 | MOLECULE: | URACIL- | DNA GLY | COSYLASE |
| 26:00:00 | 1zx2-A | 3.7 | 3.5 | 73 | 130 | 12 | MOLECULE: | UBP3-AS | SOCIATE | D PROTEIN BRE5 |
| 27:00:00 | 6ihj-B | 3.7 | 3.8 | 73 | 128 | 3 | MOLECULE: | NUCLEAR | RNA EX | PORT FACTOR 1 |
| 28:00:00 | 6wm3-U | 3.7 | 3.1 | 78 | 204 | 10 | MOLECULE: | V-TYPE | PROTON | ATPASE 116 KDA SUBUNIT A ISOFORM 1 |
| 29:00:00 | 1qaf-A | 3.7 | 3.1 | 63 | 718 | 11 | MOLECULE: | PROTEIN | (COPPE | R AMINE OXIDASE) |
| 30:00:00 | 2pun-A | 3.7 | 2 | 51 | 380 | 4 | MOLECULE: | METHYLT | HIORIBO | SE KINASE |

| | | | | | | | | | | |
|---|---|---|---|---|---|---|---|---|---|---|
| 31:00:00 | 6grr-B | 3.7 | 3.2 | 65 | 720 | 8 | MOLECULE: | AMINE O | XIDASE | |
| 32:00:00 | 2ugi-B | 3.7 | 2.8 | 57 | 83 | 11 | MOLECULE: | URACIL- | DNA GLY | COSYLASE INHIBITOR |
| 33:00:00 | 3blz-J | 3.7 | 3.5 | 67 | 124 | 9 | MOLECULE: | NTF2-LI | KE PROT | EIN OF UNKNOWN FUNCTION |
| 34:00:00 | 5eql-A | 3.7 | 3.3 | 61 | 82 | 3 | MOLECULE: | SUMO-AD | HIRON-S | 2D5 |
| 35:00:00 | 6mrk-U | 3.7 | 3.4 | 70 | 132 | 10 | MOLECULE: | NUCLEAR | RNA EX | PORT FACTOR 2 |
| 36:00:00 | 4wyk-B | 3.7 | 2.7 | 69 | 138 | 12 | MOLECULE: | NUCLEAR | RNA EX | PORT FACTOR 1 |
| 37:00:00 | 6w90-A | 3.6 | 3.3 | 68 | 130 | 13 | MOLECULE: | NTF2 FO | LD PROT | EIN LOOP-HELIX-LOOP DESIGN NT-9 |
| 38:00:00 | 4hbr-A | 3.6 | 3.1 | 62 | 140 | 5 | MOLECULE: | PUTATIV | E PERIP | LASMIC PROTEIN |
| 39:00:00 | 4o3v-B | 3.6 | 3.2 | 69 | 140 | 9 | MOLECULE: | VIRB8-L | IKE PRO | TEIN OF TYPE IV SECRETION SYSTEM |
| 40:00:00 | 3u1w-C | 3.6 | 3.2 | 60 | 250 | 5 | MOLECULE: | HYPOTHE | TICAL P | ERIPLASMIC PROTEIN |
| 41:00:00 | 2l4v-A | 3.6 | 3.7 | 74 | 135 | 5 | MOLECULE: | CYSTATI | N | |
| 42:00:00 | 5u0p-Q | 3.6 | 4.1 | 75 | 508 | 9 | MOLECULE: | MEDIATO | R COMPL | EX SUBUNIT 14 |
| 43:00:00 | 4z48-A | 3.6 | 4.5 | 77 | 240 | 3 | MOLECULE: | UNCHARA | CTERIZE | D PROTEIN |
| 44:00:00 | 7a0h-A | 3.6 | 3.4 | 76 | 285 | 5 | MOLECULE: | F-ACTIN | #NAME? | G PROTEIN SUBUNIT ALPHA |
| 45:00:00 | 2r4i-A | 3.6 | 3.7 | 69 | 123 | 3 | MOLECULE: | UNCHARA | CTERIZE | D PROTEIN |
| 46:00:00 | 4ebg-B | 3.6 | 3.4 | 70 | 99 | 6 | MOLECULE: | UNCHARA | CTERIZE | D PROTEIN |
| 47:00:00 | 6myv-C | 3.6 | 3.8 | 64 | 526 | 6 | MOLECULE: | SIALIDA | SE26 | |
| 48:00:00 | 2wyh-A | 3.6 | 3.1 | 76 | 905 | 5 | MOLECULE: | ALPHA-M | ANNOSID | ASE |
| 49:00:00 | 3cu3-A | 3.6 | 3.4 | 71 | 162 | 3 | MOLECULE: | DOMAIN | OF UNKN | OWN FUNCTION WITH A CYSTATIN-LIKE F |
| 50:00:00 | 6s6z-A | 3.6 | 3.5 | 75 | 1083 | 4 | MOLECULE: | BETA-GA | LACTOSI | DASE |
| 51:00:00 | 8ppr-K | 3.6 | 4.2 | 63 | 246 | 8 | MOLECULE: | KINETOC | HORE-AS | SOCIATED PROTEIN DSN1 HOMOLOG |
| 52:00:00 | 3h3h-B | 3.6 | 3.1 | 71 | 120 | 4 | MOLECULE: | UNCHARA | CTERIZE | D SNOAL-LIKE PROTEIN |
| 53:00:00 | 7to3-A | 3.6 | 4.5 | 75 | 568 | 9 | MOLECULE: | CAP2 | | |
| 54:00:00 | 3bcy-A | 3.6 | 3.3 | 72 | 146 | 8 | MOLECULE: | PROTEIN | YER067 | W |
| 55:00:00 | 3dxq-A | 3.6 | 3.5 | 57 | 294 | 2 | MOLECULE: | CHOLINE | /ETHANO | LAMINE KINASE FAMILY PROTEIN |
| 56:00:00 | 3db7-A | 3.6 | 2.7 | 50 | 127 | 12 | MOLECULE: | PUTATIV | E CALCI | UM-REGULATED PERIPLASMIC PROTEIN |
| 57:00:00 | 3d00-A | 3.6 | 3.3 | 65 | 184 | 11 | MOLECULE: | TUNGSTE | N FORMY | LMETHANOFURAN DEHYDROGENASE SUBUNIT |
| 58:00:00 | 7e8h-I | 3.6 | 4.1 | 64 | 760 | 11 | MOLECULE: | DIPEPTI | DYL AMI | NOPEPTIDASE-LIKE PROTEIN 6 |
| 59:00:00 | 4hbr-D | 3.6 | 3.2 | 60 | 139 | 3 | MOLECULE: | PUTATIV | E PERIP | LASMIC PROTEIN |
| 60:00:00 | 3aae-A | 3.6 | 3.6 | 74 | 270 | 5 | MOLECULE: | F-ACTIN | #NAME? | G PROTEIN SUBUNIT ALPHA-1 |
| 61:00:00 | 4wwu-F | 3.6 | 2.9 | 70 | 164 | 9 | MOLECULE: | MRNA EX | PORT FA | CTOR MEX67 |
| 62:00:00 | 5u35-A | 3.6 | 4.2 | 73 | 124 | 5 | MOLECULE: | DE NOVO | NTF2 W | ITH LARGE CAVITY |
| 63:00:00 | 1zx2-B | 3.6 | 3.4 | 72 | 130 | 11 | MOLECULE: | UBP3-AS | SOCIATE | D PROTEIN BRE5 |
| 64:00:00 | 6bjt-A | 3.6 | 3.2 | 72 | 128 | 7 | MOLECULE: | DUF4440 | DOMAIN | -CONTAINING PROTEIN |
| 65:00:00 | 3lk4-M | 3.6 | 3.3 | 73 | 269 | 5 | MOLECULE: | F-ACTIN | #NAME? | G PROTEIN SUBUNIT ALPHA-1 |
| 66:00:00 | 3lk4-P | 3.6 | 3.3 | 73 | 269 | 5 | MOLECULE: | F-ACTIN | #NAME? | G PROTEIN SUBUNIT ALPHA-1 |
| 67:00:00 | 3lk4-G | 3.6 | 3.3 | 73 | 269 | 5 | MOLECULE: | F-ACTIN | #NAME? | G PROTEIN SUBUNIT ALPHA-1 |
| 68:00:00 | 2rcd-A | 3.6 | 3.3 | 73 | 128 | 4 | MOLECULE: | UNCHARA | CTERIZE | D PROTEIN |
| 69:00:00 | 6ta7-B | 3.6 | 3.4 | 67 | 129 | 6 | MOLECULE: | RAS GTP | ASE-ACT | IVATING PROTEIN-BINDING PROTEIN 1 |
| 70:00:00 | 2rcd-C | 3.6 | 3.3 | 73 | 128 | 4 | MOLECULE: | UNCHARA | CTERIZE | D PROTEIN |
| 71:00:00 | 6e5u-D | 3.6 | 3.1 | 73 | 138 | 5 | MOLECULE: | NUCLEAR | RNA EX | PORT FACTOR 1 |
| 72:00:00 | 3lk4-1 | 3.6 | 3.3 | 73 | 269 | 5 | MOLECULE: | F-ACTIN | #NAME? | G PROTEIN SUBUNIT ALPHA-1 |
| 73:00:00 | 1gy6-A | 3.6 | 3.5 | 72 | 125 | 11 | MOLECULE: | NUCLEAR | TRANSP | ORT FACTOR 2 |
| 74:00:00 | 1oun-A | 3.6 | 3.4 | 71 | 125 | 11 | MOLECULE: | NUCLEAR | TRANSP | ORT FACTOR 2 |
| 75:00:00 | 3lk4-A | 3.6 | 3.3 | 73 | 269 | 5 | MOLECULE: | F-ACTIN | #NAME? | G PROTEIN SUBUNIT ALPHA-1 |
| 76:00:00 | 3ke7-A | 3.6 | 4 | 72 | 131 | 7 | MOLECULE: | PUTATIV | E KETOS | TEROID ISOMERASE |
| 77:00:00 | 3lk4-Y | 3.6 | 3.3 | 73 | 269 | 5 | MOLECULE: | F-ACTIN | #NAME? | G PROTEIN SUBUNIT ALPHA-1 |
| 78:00:00 | 3lk4-J | 3.6 | 3.3 | 73 | 269 | 5 | MOLECULE: | F-ACTIN | #NAME? | G PROTEIN SUBUNIT ALPHA-1 |
| 79:00:00 | 6e5u-B | 3.6 | 3.2 | 73 | 138 | 5 | MOLECULE: | NUCLEAR | RNA EX | PORT FACTOR 1 |
| 80:00:00 | 1of5-B | 3.6 | 3.2 | 67 | 128 | 9 | MOLECULE: | MRNA EX | PORT FA | CTOR MEX67 |
| 81:00:00 | 2rcd-D | 3.6 | 3.3 | 73 | 128 | 4 | MOLECULE: | UNCHARA | CTERIZE | D PROTEIN |
| 82:00:00 | 3lk4-D | 3.6 | 3.3 | 73 | 269 | 5 | MOLECULE: | F-ACTIN | #NAME? | G PROTEIN SUBUNIT ALPHA-1 |
| 83:00:00 | 3lk4-V | 3.6 | 3.3 | 73 | 269 | 5 | MOLECULE: | F-ACTIN | #NAME? | G PROTEIN SUBUNIT ALPHA-1 |
| 84:00:00 | 2owp-B | 3.6 | 3.5 | 76 | 129 | 11 | MOLECULE: | HYPOTHE | TICAL P | ROTEIN BXE_B1374 |

| | | | | | | | | | | |
|---|---|---|---|---|---|---|---|---|---|---|
| 85:00:00 | 4ipb-A | 3.6 | 3.1 | 63 | 141 | 3 | MOLECULE: | UNCHARA | CTERIZE | D PROTEIN |
| 86:00:00 | 3lk4-7 | 3.6 | 3.3 | 73 | 269 | 5 | MOLECULE: | F-ACTIN | #NAME? | G PROTEIN SUBUNIT ALPHA-1 |
| 87:00:00 | 1ask-A | 3.6 | 3.4 | 72 | 125 | 11 | MOLECULE: | NUCLEAR | TRANSP | ORT FACTOR 2 |
| 88:00:00 | 1ar0-A | 3.6 | 3.4 | 71 | 125 | 11 | MOLECULE: | NUCLEAR | TRANSP | ORT FACTOR 2 |
| 89:00:00 | 1uug-D | 3.6 | 2.8 | 58 | 82 | 10 | MOLECULE: | URACIL- | DNA GLY | COSYLASE |
| 90:00:00 | 3lk4-S | 3.6 | 3.3 | 73 | 269 | 5 | MOLECULE: | F-ACTIN | #NAME? | G PROTEIN SUBUNIT ALPHA-1 |
| 91:00:00 | 2zhx-D | 3.6 | 2.6 | 59 | 82 | 12 | MOLECULE: | URACIL- | DNA GLY | COSYLASE |
| 92:00:00 | 3blz-E | 3.6 | 3 | 65 | 124 | 9 | MOLECULE: | NTF2-LI | KE PROT | EIN OF UNKNOWN FUNCTION |
| 93:00:00 | 2wyi-A | 3.6 | 3.1 | 76 | 896 | 5 | MOLECULE: | ALPHA-M | ANNOSID | ASE |
| 94:00:00 | 6wm4-U | 3.6 | 3 | 78 | 204 | 12 | MOLECULE: | V-TYPE | PROTON | ATPASE 116 KDA SUBUNIT A ISOFORM 1 |
| 95:00:00 | 4fpf-A | 3.6 | 3.4 | 62 | 658 | 8 | MOLECULE: | SIALIDA | SE B | |
| 96:00:00 | 7to3-B | 3.6 | 4.5 | 75 | 568 | 9 | MOLECULE: | CAP2 | | |
| 97:00:00 | 2j8x-D | 3.6 | 2.5 | 57 | 81 | 14 | MOLECULE: | URACIL- | DNA GLY | COSYLASE |
| 98:00:00 | 6s6z-B | 3.6 | 3.6 | 71 | 1083 | 4 | MOLECULE: | BETA-GA | LACTOSI | DASE |
| 99:00:00 | 5bxq-A | 3.6 | 3.3 | 70 | 124 | 10 | MOLECULE: | NUCLEAR | TRANSP | ORT FACTOR 2 |
| 0:00 | 1jz7-B | 3.6 | 3 | 51 | 1011 | 4 | MOLECULE: | BETA-GA | LACTOSI | DASE |
| 1:00 | 4lyl-P | 3.6 | 2.6 | 59 | 82 | 12 | MOLECULE: | URACIL- | DNA GLY | COSYLASE |
| 2:00 | 1oun-B | 3.6 | 3.3 | 70 | 121 | 10 | MOLECULE: | NUCLEAR | TRANSP | ORT FACTOR 2 |
| 3:00 | 1ar0-B | 3.6 | 3.3 | 70 | 121 | 10 | MOLECULE: | NUCLEAR | TRANSP | ORT FACTOR 2 |
| 4:00 | 5bxq-B | 3.6 | 3.3 | 70 | 124 | 10 | MOLECULE: | NUCLEAR | TRANSP | ORT FACTOR 2 |
| 5:00 | 1a2k-A | 3.6 | 3.3 | 70 | 124 | 10 | MOLECULE: | NUCLEAR | TRANSP | ORT FACTOR 2 |
| 6:00 | 6drv-B | 3.6 | 2.8 | 48 | 1024 | 4 | MOLECULE: | BETA-GA | LACTOSI | DASE |
| 7:00 | 2cc3-B | 3.6 | 2.5 | 65 | 144 | 9 | MOLECULE: | PROTEIN | VIRB8 | |
| 8:00 | 6wm2-U | 3.6 | 3 | 76 | 205 | 12 | MOLECULE: | V-TYPE | PROTON | ATPASE SUBUNIT E 1 |
| 9:00 | 4lyl-B | 3.6 | 2.7 | 59 | 82 | 12 | MOLECULE: | URACIL- | DNA GLY | COSYLASE |
| 10:00 | 1jb4-A | 3.6 | 3.4 | 70 | 123 | 9 | MOLECULE: | NUCLEAR | TRANSP | ORT FACTOR 2 |
| 11:00 | 5trv-A | 3.6 | 3.2 | 68 | 118 | 6 | MOLECULE: | DENOVO | NTF2 | |
| 12:00 | 6bju-C | 3.6 | 3 | 71 | 125 | 6 | MOLECULE: | ATZH | | |
| 13:00 | 7tqd-B | 3.6 | 4.5 | 74 | 513 | 11 | MOLECULE: | CAP2 | | |
| 14:00 | 6bju-A | 3.6 | 3.1 | 72 | 125 | 6 | MOLECULE: | ATZH | | |
| 15:00 | 4fpy-A | 3.6 | 3.5 | 62 | 658 | 6 | MOLECULE: | SIALIDA | SE B | |
| 16:00 | 8hbn-A | 3.5 | 3.3 | 72 | 371 | 3 | MOLECULE: | MRNA EX | PORT FA | CTOR MEX67 |
| 17:00 | 3w0s-A | 3.5 | 2.6 | 54 | 298 | 6 | MOLECULE: | HYGROMY | CIN-B 4 | -O-KINASE |
| 18:00 | 4gbf-B | 3.5 | 3.3 | 73 | 370 | 8 | MOLECULE: | PHIKZ13 | 1 | |
| 19:00 | 3a76-B | 3.5 | 3 | 69 | 153 | 9 | MOLECULE: | GAMMA-H | EXACHLO | ROCYCLOHEXANE DEHYDROCHLORINASE |
| 20:00 | 7zhm-C | 3.5 | 3.3 | 60 | 87 | 5 | MOLECULE: | RHS1 PR | OTEIN | |
| 21:00 | 2kxg-A | 3.5 | 3.7 | 64 | 95 | 6 | MOLECULE: | ASPARTI | C PROTE | ASE INHIBITOR |
| 22:00 | 7cd7-A | 3.5 | 3.3 | 64 | 99 | 6 | MOLECULE: | GREEN F | LUORESC | ENT PROTEIN |
| 23:00 | 8v1k-A | 3.5 | 3.2 | 65 | 185 | 5 | MOLECULE: | OUTER-M | EMBRANE | LIPOPROTEIN CARRIER PROTEIN |
| 24:00:00 | 7abw-B | 3.5 | 3.3 | 60 | 365 | 5 | MOLECULE: | PEPSY D | OMAIN-C | ONTAINING PROTEIN |
| 25:00:00 | 7kd9-A | 3.5 | 3.7 | 77 | 231 | 3 | MOLECULE: | GALLATE | DECARB | OXYLASE |
| 26:00:00 | 7bcz-A | 3.5 | 4 | 72 | 242 | 6 | MOLECULE: | 4'-PHOS | PHOPANT | ETHEINYL TRANSFERASE ENTD |
| 27:00:00 | 5iqc-A | 3.5 | 3.4 | 62 | 301 | 10 | MOLECULE: | BIFUNCT | IONAL A | AC/APH |
| 28:00:00 | 6ekt-A | 3.5 | 4.2 | 77 | 419 | 5 | MOLECULE: | P-47 PR | OTEIN | |
| 29:00:00 | 3cxj-A | 3.5 | 3.3 | 60 | 144 | 2 | MOLECULE: | UNCHARA | CTERIZE | D PROTEIN |
| 30:00:00 | 7sjy-A | 3.5 | 3.8 | 67 | 313 | 6 | MOLECULE: | ANTI-SI | GMA-I F | ACTOR RSGI9 |
| 31:00:00 | 3qv0-A | 3.5 | 3.3 | 59 | 179 | 10 | MOLECULE: | MITOCHO | NDRIAL | ACIDIC PROTEIN MAM33 |
| 32:00:00 | 1u5o-A | 3.5 | 3.5 | 73 | 125 | 11 | MOLECULE: | NUCLEAR | TRANSP | ORT FACTOR 2 |
| 33:00:00 | 7prq-B | 3.5 | 2.3 | 63 | 316 | 2 | MOLECULE: | PROBABL | E CHEMO | TAXIS TRANSDUCER |
| 34:00:00 | 3k0z-A | 3.5 | 3.6 | 75 | 149 | 9 | MOLECULE: | PUTATIV | E POLYK | ETIDE CYCLASE |
| 35:00:00 | 4std-A | 3.5 | 3 | 65 | 164 | 5 | MOLECULE: | SCYTALO | NE DEHY | DRATASE |
| 36:00:00 | 6ya7-A | 3.5 | 3.8 | 61 | 350 | 5 | MOLECULE: | CELL DI | VISION | CYCLE 7-RELATED PROTEIN KINASE,CELL |
| 37:00:00 | 3g16-A | 3.5 | 4.2 | 72 | 156 | 3 | MOLECULE: | UNCHARA | CTERIZE | D PROTEIN WITH CYSTATIN-LIKE FOLD |
| 38:00:00 | 8hsb-B | 3.5 | 3.7 | 69 | 159 | 17 | MOLECULE: | CDNG | | |

| | | | | | | | | | | | |
|---|---|---|---|---|---|---|---|---|---|---|---|
| 39:00:00 | 5cd6-A | 3.5 | 3 | 75 | 575 | 8 | MOLECULE: | TPR-DOM | AIN CON | TAINING PROTEIN | |
| 40:00:00 | 3ecf-A | 3.5 | 3.3 | 66 | 128 | 3 | MOLECULE: | NTF2-LI | KE PROT | EIN | |
| 41:00:00 | 2k54-A | 3.5 | 3.8 | 73 | 123 | 7 | MOLECULE: | PROTEIN | ATU074 | | 2 |
| 42:00:00 | 3f7w-A | 3.5 | 2.2 | 52 | 288 | 4 | MOLECULE: | PUTATIV | E FRUCT | OSAMINE-3-KINASE | |
| 43:00:00 | 5aiw-A | 3.5 | 4 | 71 | 127 | 7 | MOLECULE: | TRAH | | | |
| 44:00:00 | 2cc3-A | 3.5 | 2.3 | 61 | 144 | 10 | MOLECULE: | PROTEIN | VIRB8 | | |
| 45:00:00 | 5evh-A | 3.5 | 3.6 | 66 | 121 | 11 | MOLECULE: | UNCHARA | CTERIZE | D PROTEIN | |
| 46:00:00 | 3tdw-A | 3.5 | 2.1 | 53 | 302 | 6 | MOLECULE: | GENTAMI | CIN RES | ISTANCE PROTEIN | |
| 47:00:00 | 6w3f-A | 3.5 | 3.4 | 67 | 115 | 7 | MOLECULE: | RD1NTF2 | _05_I64 | F_A80G_T94P_D101K_L106W | |
| 48:00:00 | 4wwu-C | 3.5 | 2.9 | 70 | 165 | 9 | MOLECULE: | MRNA EX | PORT FA | CTOR MEX67 | |
| 49:00:00 | 1qma-A | 3.5 | 3.5 | 73 | 126 | 11 | MOLECULE: | NUCLEAR | TRANSP | ORT FACTOR 2 | |
| 50:00:00 | 3ddn-B | 3.5 | 3.2 | 77 | 525 | 9 | MOLECULE: | D-3-PHO | SPHOGLY | CERATE DEHYDROGENASE | |
| 51:00:00 | 1gy6-B | 3.5 | 3.5 | 72 | 123 | 11 | MOLECULE: | NUCLEAR | TRANSP | ORT FACTOR 2 | |
| 52:00:00 | 5nw4-Y | 3.5 | 3.6 | 74 | 275 | 4 | MOLECULE: | DYNEIN | HEAVY C | HAIN | |
| 53:00:00 | 5adx-K | 3.5 | 3.6 | 74 | 275 | 4 | MOLECULE: | ACTIN R | ELATED | PROTEIN 1 | |
| 54:00:00 | 4eyc-B | 3.5 | 3.5 | 63 | 99 | 5 | MOLECULE: | CATHELI | CIDIN A | NTIMICROBIAL PEPTIDE | |
| 55:00:00 | 1qak-A | 3.5 | 3.3 | 67 | 719 | 6 | MOLECULE: | COPPER | AMINE O | XIDASE | |
| 56:00:00 | 1d6y-B | 3.5 | 3.3 | 67 | 720 | 7 | MOLECULE: | COPPER | AMINE O | XIDASE | |
| 57:00:00 | 1n5p-A | 3.5 | 3.2 | 64 | 105 | 3 | MOLECULE: | PROTEGR | INS | | |
| 58:00:00 | 6w3f-B | 3.5 | 3.4 | 66 | 114 | 8 | MOLECULE: | RD1NTF2 | _05_I64 | F_A80G_T94P_D101K_L106W | |
| 59:00:00 | 1qma-C | 3.5 | 3.5 | 73 | 126 | 11 | MOLECULE: | NUCLEAR | TRANSP | ORT FACTOR 2 | |
| 60:00:00 | 4hbr-B | 3.5 | 3.4 | 60 | 140 | 3 | MOLECULE: | PUTATIV | E PERIP | LASMIC PROTEIN | |
| 61:00:00 | 4ovm-F | 3.5 | 3 | 70 | 129 | 9 | MOLECULE: | UNCHARA | CTERIZE | D PROTEIN SGCJ | |
| 62:00:00 | 8wzb-S | 3.5 | 4.3 | 69 | 392 | 7 | MOLECULE: | DPY30 D | OMAIN C | ONTAINING 2 | |
| 63:00:00 | 1u5o-B | 3.5 | 3.4 | 70 | 121 | 10 | MOLECULE: | NUCLEAR | TRANSP | ORT FACTOR 2 | |
| 64:00:00 | 4pw1-A | 3.5 | 3.3 | 71 | 214 | 8 | MOLECULE: | UNCHARA | CTERIZE | D PROTEIN | |
| 65:00:00 | 8u2w-A | 3.5 | 5.1 | 69 | 482 | 3 | MOLECULE: | ACETYL- | COENZYM | E A SYNTHETASE | |
| 66:00:00 | 6ztk-A | 3.5 | 3.2 | 62 | 108 | 5 | MOLECULE: | MIALOST | ATIN | | |
| 67:00:00 | 1jb2-A | 3.5 | 3.4 | 70 | 123 | 10 | MOLECULE: | NUCLEAR | TRANSP | ORT FACTOR 2 | |
| 68:00:00 | 5u35-B | 3.5 | 3.3 | 69 | 121 | 1 | MOLECULE: | DE NOVO | NTF2 W | ITH LARGE CAVITY | |
| 69:00:00 | 3l0r-B | 3.5 | 3.7 | 64 | 108 | 3 | MOLECULE: | CYSTATI | N-2 | | |
| 70:00:00 | 5kbp-B | 3.5 | 3.4 | 80 | 884 | 8 | MOLECULE: | GLYCOSY | L HYDRO | LASE, FAMILY 38 | |
| 71:00:00 | 2w0q-B | 3.5 | 3.3 | 66 | 719 | 8 | MOLECULE: | COPPER | AMINE O | XIDASE | |
| 72:00:00 | 1gy5-A | 3.5 | 3.1 | 66 | 123 | 11 | MOLECULE: | NUCLEAR | TRANSP | ORT FACTOR 2 | |
| 73:00:00 | 1jb5-A | 3.5 | 3.3 | 67 | 123 | 9 | MOLECULE: | NUCLEAR | TRANSP | ORT FACTOR 2 | |
| 74:00:00 | 1qma-D | 3.5 | 3.4 | 70 | 124 | 10 | MOLECULE: | NUCLEAR | TRANSP | ORT FACTOR 2 | |
| 75:00:00 | 6ztk-B | 3.5 | 3.5 | 65 | 110 | 5 | MOLECULE: | MIALOST | ATIN | | |
| 76:00:00 | 4ttg-C | 3.5 | 3.1 | 51 | 1015 | 4 | MOLECULE: | BETA-GA | LACTOSI | DASE | |
| 77:00:00 | 1a2k-B | 3.5 | 3.3 | 70 | 124 | 10 | MOLECULE: | NUCLEAR | TRANSP | ORT FACTOR 2 | |
| 78:00:00 | 3vd9-A | 3.5 | 3 | 50 | 1015 | 6 | MOLECULE: | BETA-GA | LACTOSI | DASE | |
| 79:00:00 | 3l0r-A | 3.5 | 3.6 | 64 | 108 | 3 | MOLECULE: | CYSTATI | N-2 | | |
| 80:00:00 | 2xvl-A | 3.5 | 2.7 | 63 | 944 | 11 | MOLECULE: | ALPHA-X | YLOSIDA | SE, PUTATIVE, XYL31A | |
| 81:00:00 | 3vd9-D | 3.5 | 3.1 | 51 | 1015 | 4 | MOLECULE: | BETA-GA | LACTOSI | DASE | |
| 82:00:00 | 1qma-B | 3.5 | 3.4 | 70 | 124 | 10 | MOLECULE: | NUCLEAR | TRANSP | ORT FACTOR 2 | |
| 83:00:00 | 9brz-c | 3.5 | 3.1 | 76 | 203 | 11 | MOLECULE: | V-TYPE | PROTON | ATPASE SUBUNIT S1 | |
| 84:00:00 | 2zhx-L | 3.5 | 2.9 | 62 | 82 | 11 | MOLECULE: | URACIL- | DNA GLY | COSYLASE | |
| 85:00:00 | 5igt-A | 3.5 | 3.4 | 59 | 300 | 15 | MOLECULE: | MACROLI | DE 2'-P | HOSPHOTRANSFERASE | |
| 86:00:00 | 4uvq-A | 3.4 | 3.2 | 75 | 263 | 12 | MOLECULE: | THIAZOL | INE OXI | DASE/SUBTILISIN-LIKE PROTEASE | |
| 87:00:00 | 2xf3-B | 3.4 | 3.1 | 68 | 428 | 6 | MOLECULE: | ORF12 | | | |
| 88:00:00 | 6p7l-A | 3.4 | 3.5 | 64 | 140 | 6 | MOLECULE: | ALN2 | | | |
| 89:00:00 | 3pgb-A | 3.4 | 3.7 | 83 | 740 | 4 | MOLECULE: | PUTATIV | E UNCHA | RACTERIZED PROTEIN | |
| 90:00:00 | 6e8a-A | 3.4 | 3.8 | 65 | 148 | 9 | MOLECULE: | DUF1795 | DOMAIN | -CONTAINING PROTEIN | |
| 91:00:00 | 4nlm-A | 3.4 | 4.5 | 65 | 326 | 3 | MOLECULE: | LMO1340 | PROTEI | N | |
| 92:00:00 | 8gm7-C | 3.4 | 3 | 61 | 170 | 7 | MOLECULE: | TK0353 | | | |

| | | | | | | | | | | |
|---|---|---|---|---|---|---|---|---|---|---|
| 93:00:00 | 3ef8-A | 3.4 | 3.2 | 67 | 149 | 9 | MOLECULE: | PUTATIV | E SCYAL | ONE DEHYDRATASE |
| 94:00:00 | 3nct-D | 3.4 | 3.7 | 73 | 140 | 7 | MOLECULE: | PROTEIN | PSIB | |
| 95:00:00 | 4cdb-A | 3.4 | 3.4 | 71 | 488 | 3 | MOLECULE: | LISTERI | OLYSIN | O |
| 96:00:00 | 5u6y-A | 3.4 | 3.7 | 69 | 459 | 1 | MOLECULE: | CALCIUM | /CALMOD | ULIN-DEPENDENT PROTEIN KINASE TYPE |
| 97:00:00 | 6e5u-A | 3.4 | 3.3 | 73 | 339 | 3 | MOLECULE: | NUCLEAR | RNA EX | PORT FACTOR 1 |
| 98:00:00 | 2ags-A | 3.4 | 3.9 | 67 | 634 | 6 | MOLECULE: | SIALIDA | SE | |
| 99:00:00 | 3d9r-B | 3.4 | 3.9 | 73 | 134 | 7 | MOLECULE: | KETOSTE | ROID IS | OMERASE-LIKE PROTEIN |
| 0:00 | 6w3w-A | 3.4 | 3.3 | 67 | 107 | 15 | MOLECULE: | DENOVO | NTF2 | |
| 1:00 | 4lmg-B | 3.4 | 2.9 | 59 | 121 | 5 | MOLECULE: | IRON-RE | GULATED | TRANSCRIPTIONAL ACTIVATOR AFT2 |
| 2:00 | 7vnx-A | 3.4 | 2.7 | 58 | 216 | 7 | MOLECULE: | TKARKI | | |
| 3:00 | 3os7-A | 3.4 | 3.5 | 76 | 338 | 8 | MOLECULE: | GALACTO | SE MUTA | ROTASE-LIKE PROTEIN |
| 4:00 | 4fr9-A | 3.4 | 3.7 | 66 | 141 | 11 | MOLECULE: | UNCHARA | CTERIZE | D PROTEIN |
| 5:00 | 1gyb-A | 3.4 | 3.5 | 74 | 122 | 5 | MOLECULE: | NUCLEAR | TRANSP | ORT FACTOR 2 |
| 6:00 | 4n6n-B | 3.4 | 3.4 | 62 | 117 | 3 | MOLECULE: | LEGUMAI | N | |
| 7:00 | 6mrk-V | 3.4 | 3.7 | 72 | 126 | 3 | MOLECULE: | NUCLEAR | RNA EX | PORT FACTOR 2 |
| 8:00 | 6d63-B | 3.4 | 3.2 | 72 | 128 | 13 | MOLECULE: | ATZH | | |
| 9:00 | 5tpj-A | 3.4 | 3.4 | 68 | 122 | 13 | MOLECULE: | DENOVO | NTF2 | |
| 10:00 | 1jn5-A | 3.4 | 2.8 | 67 | 136 | 12 | MOLECULE: | P15 | | |
| 11:00 | 8v1l-A | 3.4 | 3.8 | 71 | 126 | 8 | MOLECULE: | RAS GTP | ASE-ACT | IVATING PROTEIN-BINDING PROTEIN 1 |
| 12:00 | 6d63-I | 3.4 | 3.3 | 68 | 123 | 6 | MOLECULE: | ATZH | | |
| 13:00 | 5tph-B | 3.4 | 4.4 | 71 | 121 | 6 | MOLECULE: | DE NOVO | NTF2 H | OMODIMER |
| 14:00 | 6ta7-E | 3.4 | 3.6 | 72 | 125 | 7 | MOLECULE: | RAS GTP | ASE-ACT | IVATING PROTEIN-BINDING PROTEIN 1 |
| 15:00 | 6s5l-B | 3.4 | 3 | 61 | 124 | 10 | MOLECULE: | ALL4940 | PROTEI | N |
| 16:00 | 4n6o-B | 3.4 | 3.4 | 62 | 115 | 3 | MOLECULE: | LEGUMAI | N | |
| 17:00 | 6xby-s | 3.4 | 3.2 | 77 | 205 | 10 | MOLECULE: | V-TYPE | PROTON | ATPASE CATALYTIC SUBUNIT A |
| 18:00 | 6lye-I | 3.4 | 2.6 | 57 | 82 | 11 | MOLECULE: | PROBABL | E URACI | L-DNA GLYCOSYLASE |
| 19:00 | 6bju-B | 3.4 | 3.2 | 68 | 121 | 7 | MOLECULE: | ATZH | | |
| 20:00 | 4hbr-C | 3.4 | 2.9 | 62 | 140 | 2 | MOLECULE: | PUTATIV | E PERIP | LASMIC PROTEIN |
| 21:00 | 6d63-A | 3.4 | 2.9 | 70 | 127 | 6 | MOLECULE: | ATZH | | |
| 22:00 | 3njv-A | 3.4 | 3 | 47 | 508 | 6 | MOLECULE: | RHAMNOG | ALACTUR | ONASE B |
| 23:00 | 7toe-D | 3.4 | 4.1 | 65 | 467 | 3 | MOLECULE: | GLUCOSE | #NAME? | PHATE 1-DEHYDROGENASE |
| 24:00:00 | 4j9t-A | 3.4 | 3.2 | 63 | 355 | 16 | MOLECULE: | DESIGNE | D UNNAT | URAL AMINO ACID DEPENDENT METALLOPR |
| 25:00:00 | 7dmp-B | 3.4 | 3.5 | 71 | 336 | 3 | MOLECULE: | RADIAL | SPOKE H | EAD PROTEIN 4 HOMOLOG A |
| 26:00:00 | 1jb2-B | 3.4 | 3.5 | 70 | 123 | 10 | MOLECULE: | NUCLEAR | TRANSP | ORT FACTOR 2 |
| 27:00:00 | 1wcq-C | 3.4 | 3.4 | 66 | 599 | 15 | MOLECULE: | SIALIDA | SE | |
| 28:00:00 | 2zhx-F | 3.4 | 2.7 | 59 | 82 | 12 | MOLECULE: | URACIL- | DNA GLY | COSYLASE |
| 29:00:00 | 8feb-A | 3.4 | 3.3 | 65 | 501 | 5 | MOLECULE: | SIALIDA | SE | |
| 30:00:00 | 3njx-A | 3.4 | 3 | 47 | 508 | 6 | MOLECULE: | RHAMNOG | ALACTUR | ONASE B |
| 31:00:00 | 4ttg-D | 3.4 | 3 | 50 | 1015 | 6 | MOLECULE: | BETA-GA | LACTOSI | DASE |
| 32:00:00 | 6d63-G | 3.4 | 3.1 | 65 | 115 | 8 | MOLECULE: | ATZH | | |
| 33:00:00 | 1jb4-B | 3.4 | 3.4 | 70 | 123 | 10 | MOLECULE: | NUCLEAR | TRANSP | ORT FACTOR 2 |
| 34:00:00 | 1gy5-B | 3.4 | 3.4 | 70 | 124 | 10 | MOLECULE: | NUCLEAR | TRANSP | ORT FACTOR 2 |
| 35:00:00 | 4fpk-A | 3.4 | 3.4 | 60 | 658 | 10 | MOLECULE: | SIALIDA | SE B | |
| 36:00:00 | 4dux-C | 3.4 | 3.1 | 50 | 1015 | 6 | MOLECULE: | BETA-GA | LACTOSI | DASE |
| 37:00:00 | 2wof-A | 3.4 | 3 | 64 | 718 | 11 | MOLECULE: | PRIMARY | AMINE | OXIDASE |
| 38:00:00 | 7ual-D | 3.4 | 4.3 | 69 | 484 | 3 | MOLECULE: | GLUCOSE | #NAME? | PHATE 1-DEHYDROGENASE |
| 39:00:00 | 2pu8-A | 3.4 | 2.5 | 52 | 373 | 2 | MOLECULE: | METHYLT | HIORIBO | SE KINASE |
| 40:00:00 | 4orl-A | 3.3 | 3.2 | 66 | 110 | 11 | MOLECULE: | UNCHARA | CTERIZE | D PROTEIN |
| 41:00:00 | 4q51-B | 3.3 | 3.1 | 64 | 265 | 11 | MOLECULE: | UNCHARA | CTERIZE | D PROTEIN |
| 42:00:00 | 8t1l-V | 3.3 | 3.1 | 66 | 278 | 11 | MOLECULE: | MEDIATO | R OF RN | A POLYMERASE II TRANSCRIPTION SUBUN |
| 43:00:00 | 6mrk-B | 3.3 | 3.5 | 73 | 188 | 4 | MOLECULE: | NUCLEAR | RNA EX | PORT FACTOR 2 |
| 44:00:00 | 3flj-A | 3.3 | 3.5 | 71 | 141 | 8 | MOLECULE: | UNCHARA | CTERIZE | D PROTEIN CONSERVED IN BACTERIA WIT |
| 45:00:00 | 8frs-A | 3.3 | 3.1 | 66 | 317 | 8 | MOLECULE: | MAJOR S | TRUCTUR | AL PROTEIN |
| 46:00:00 | 4qvh-A | 3.3 | 3.2 | 69 | 596 | 4 | MOLECULE: | MALTOSE | #NAME? | G PERIPLASMIC PROTEIN, 4'-PHOSPHOPA |

| | | | | | | | | | | |
|---|---|---|---|---|---|---|---|---|---|---|
| 47:00:00 | 4kt3-B | 3.3 | 4 | 74 | 128 | 8 | MOLECULE: | UNCHARA | CTERIZE | D PROTEIN |
| 48:00:00 | 3h51-A | 3.3 | 3.6 | 67 | 142 | 9 | MOLECULE: | PUTATIV | E CALCI | UM/CALMODULIN DEPENDENT PROTEIN KIN |
| 49:00:00 | 2mc8-A | 3.3 | 2.7 | 59 | 114 | 10 | MOLECULE: | UNCHARA | CTERIZE | D PROTEIN |
| 50:00:00 | 3en1-B | 3.3 | 4.1 | 69 | 180 | 4 | MOLECULE: | BENZENE | 1,2-DI | OXYGENASE SUBUNIT ALPHA |
| 51:00:00 | 5k21-B | 3.3 | 3.3 | 58 | 123 | 3 | MOLECULE: | PYOCYAN | IN DEME | THYLASE |
| 52:00:00 | 3f8x-B | 3.3 | 3.1 | 61 | 132 | 10 | MOLECULE: | PUTATIV | E DELTA | -5-3-KETOSTEROID ISOMERASE |
| 53:00:00 | 6jzf-A | 3.3 | 4 | 68 | 153 | 4 | MOLECULE: | PLASTID | DIVISI | ON PROTEIN CDP1, CHLOROPLASTIC |
| 54:00:00 | 6w40-A | 3.3 | 3.7 | 69 | 120 | 13 | MOLECULE: | DENOVO | NTF2 | |
| 55:00:00 | 6e0a-B | 3.3 | 3 | 65 | 262 | 5 | MOLECULE: | METHYL- | ACCEPTI | NG CHEMOTAXIS PROTEIN TLPA |
| 56:00:00 | 5efv-B | 3.3 | 3.6 | 69 | 635 | 7 | MOLECULE: | PHI ETA | ORF 56 | -LIKE PROTEIN |
| 57:00:00 | 3aki-A | 3.3 | 3.2 | 68 | 448 | 7 | MOLECULE: | PUTATIV | E SECRE | TED ALPHA L-ARABINOFURANOSIDASE II |
| 58:00:00 | 5boi-A | 3.3 | 3.6 | 67 | 226 | 6 | MOLECULE: | GERMINA | TION PR | OTEIN YPEB |
| 59:00:00 | 2du7-A | 3.3 | 2.9 | 61 | 539 | 0 | MOLECULE: | O-PHOSP | HOSERYL | -TRNA SYNTHETASE |
| 60:00:00 | 2r41-C | 3.3 | 2.6 | 58 | 106 | 16 | MOLECULE: | UNCHARA | CTERIZE | D PROTEIN |
| 61:00:00 | 8t5t-A | 3.3 | 4.2 | 75 | 195 | 3 | MOLECULE: | OUTER M | EMBRANE | LIPOPROTEIN CARRIER PROTEIN (LOLA) |
| 62:00:00 | 3u1w-B | 3.3 | 3.2 | 60 | 248 | 5 | MOLECULE: | HYPOTHE | TICAL P | ERIPLASMIC PROTEIN |
| 63:00:00 | 6bju-D | 3.3 | 3.1 | 68 | 123 | 6 | MOLECULE: | ATZH | | |
| 64:00:00 | 6d63-L | 3.3 | 3.5 | 69 | 121 | 13 | MOLECULE: | ATZH | | |
| 65:00:00 | 7pdz-F | 3.3 | 3.6 | 73 | 272 | 4 | MOLECULE: | ISOFORM | 2 OF F | #NAME? |
| 66:00:00 | 1dyu-B | 3.3 | 3.3 | 67 | 720 | 7 | MOLECULE: | COPPER | AMINE O | XIDASE |
| 67:00:00 | 6d63-C | 3.3 | 3.2 | 68 | 125 | 7 | MOLECULE: | ATZH | | |
| 68:00:00 | 2wof-B | 3.3 | 3.3 | 67 | 720 | 7 | MOLECULE: | PRIMARY | AMINE | OXIDASE |
| 69:00:00 | 1jrq-B | 3.3 | 3.2 | 64 | 721 | 6 | MOLECULE: | COPPER | AMINE O | XIDASE |
| 70:00:00 | 8v1l-D | 3.3 | 3.2 | 69 | 128 | 7 | MOLECULE: | RAS GTP | ASE-ACT | IVATING PROTEIN-BINDING PROTEIN 1 |
| 71:00:00 | 7a0h-B | 3.3 | 3.3 | 73 | 255 | 3 | MOLECULE: | F-ACTIN | #NAME? | G PROTEIN SUBUNIT ALPHA |
| 72:00:00 | 2woh-B | 3.3 | 3.4 | 65 | 720 | 8 | MOLECULE: | PRIMARY | AMINE | OXIDASE |
| 73:00:00 | 4nfa-A | 3.3 | 4.2 | 63 | 204 | 11 | MOLECULE: | PROTEIN | CASC5 | |
| 74:00:00 | 7c03-A | 3.3 | 3.1 | 66 | 312 | 5 | MOLECULE: | POLARIS | ACT(T57 | S) |
| 75:00:00 | 4eyc-A | 3.3 | 3.3 | 61 | 103 | 5 | MOLECULE: | CATHELI | CIDIN A | NTIMICROBIAL PEPTIDE |
| 76:00:00 | 1d6z-B | 3.3 | 3.4 | 68 | 720 | 6 | MOLECULE: | COPPER | AMINE O | XIDASE |
| 77:00:00 | 2rgq-C | 3.3 | 3.5 | 65 | 134 | 5 | MOLECULE: | DOMAIN | OF UNKN | OWN FUNCTION WITH A CYSTATIN-LIKE F |
| 78:00:00 | 2pul-A | 3.3 | 2.4 | 54 | 381 | 4 | MOLECULE: | METHYLT | HIORIBO | SE KINASE |
| 79:00:00 | 7zhw-B | 3.3 | 4.3 | 68 | 535 | 3 | MOLECULE: | GLUCOSE | #NAME? | PHATE 1-DEHYDROGENASE |
| 80:00:00 | 6drv-D | 3.3 | 2.6 | 46 | 1024 | 4 | MOLECULE: | BETA-GA | LACTOSI | DASE |
| 81:00:00 | 6ezz-B | 3.3 | 3.3 | 66 | 720 | 8 | MOLECULE: | PRIMARY | AMINE | OXIDASE |
| 82:00:00 | 4dux-B | 3.3 | 3.1 | 50 | 1015 | 6 | MOLECULE: | BETA-GA | LACTOSI | DASE |
| 83:00:00 | 2wyi-B | 3.3 | 3.4 | 71 | 896 | 15 | MOLECULE: | ALPHA-M | ANNOSID | ASE |
| 84:00:00 | 7zvd-N | 3.3 | 4.1 | 64 | 484 | 3 | MOLECULE: | GLUCOSE | #NAME? | PHATE 1-DEHYDROGENASE |
| 85:00:00 | 5igi-A | 3.3 | 3.4 | 59 | 300 | 15 | MOLECULE: | MACROLI | DE 2'-P | HOSPHOTRANSFERASE |
| 86:00:00 | 1ms1-A | 3.3 | 3.9 | 67 | 622 | 7 | MOLECULE: | TRANS-S | IALIDAS | E |
| 87:00:00 | 1jb5-B | 3.3 | 3.5 | 69 | 123 | 10 | MOLECULE: | NUCLEAR | TRANSP | ORT FACTOR 2 |
| 88:00:00 | 7zhv-A | 3.3 | 5 | 67 | 533 | 3 | MOLECULE: | GLUCOSE | #NAME? | PHATE 1-DEHYDROGENASE |
| 89:00:00 | 3elg-A | 3.3 | 4.3 | 56 | 127 | 5 | MOLECULE: | UNCHARA | CTERIZE | D PERIPLASMIC PROTEIN |
| 90:00:00 | 1jz7-A | 3.3 | 3.2 | 51 | 1011 | 4 | MOLECULE: | BETA-GA | LACTOSI | DASE |
| 91:00:00 | 3due-A | 3.3 | 3.5 | 54 | 127 | 9 | MOLECULE: | PUTATIV | E PERIP | LASMIC PROTEIN |
| 92:00:00 | 1dyu-A | 3.3 | 3 | 64 | 718 | 11 | MOLECULE: | COPPER | AMINE O | XIDASE |
| 93:00:00 | 6d63-K | 3.3 | 3.3 | 68 | 120 | 7 | MOLECULE: | ATZH | | |
| 94:00:00 | 1spu-A | 3.3 | 3.2 | 64 | 718 | 11 | MOLECULE: | COPPER | AMINE O | XIDASE |
| 95:00:00 | 4lv8-A | 3.2 | 3.4 | 66 | 360 | 11 | MOLECULE: | RHOPTRY | PROTEI | N 5 C |
| 96:00:00 | 3dmc-A | 3.2 | 3.7 | 71 | 134 | 11 | MOLECULE: | NTF2-LI | KE PROT | EIN |
| 97:00:00 | 5hx0-A | 3.2 | 3.7 | 60 | 365 | 5 | MOLECULE: | UNCHARA | CTERIZE | D PROTEIN DFER_1899 |
| 98:00:00 | 2iut-A | 3.2 | 3.5 | 59 | 408 | 8 | MOLECULE: | DNA TRA | NSLOCAS | E FTSK |
| 99:00:00 | 6gvw-C | 3.2 | 3.7 | 69 | 383 | 4 | MOLECULE: | BRCA1-A | COMPLE | X SUBUNIT ABRAXAS 1 |
| 0:00 | 6qbe-A | 3.2 | 3.2 | 74 | 215 | 9 | MOLECULE: | NEP1-LI | KE PROT | EIN |

| | | | | | | | | | | |
|---|---|---|---|---|---|---|---|---|---|---|
| 1:00 | 6d8v-A | 3.2 | 2.8 | 60 | 270 | 7 | MOLECULE: | PROBABL | E CHEMO | RECEPTOR (METHYL-ACCEPTING CHEMOTAX |
| 2:00 | 8u50-A | 3.2 | 3.1 | 51 | 267 | 12 | MOLECULE: | KLEBSIE | LLA PNE | UMONIAE FAMILY 1 ENCAPSULIN SHELL |
| 3:00 | 7n34-A | 3.2 | 3.1 | 64 | 108 | 2 | MOLECULE: | CAP15 | | |
| 4:00 | 2d7d-A | 3.2 | 2.8 | 58 | 621 | 12 | MOLECULE: | 5'-D(P* | TP*TP*T | )-3' |
| 5:00 | 2h1t-A | 3.2 | 3.9 | 54 | 186 | 7 | MOLECULE: | HYPOTHE | TICAL P | ROTEIN |
| 6:00 | 6y2p-A | 3.2 | 2.7 | 54 | 354 | 9 | MOLECULE: | MRNA EN | DORIBON | UCLEASE TOXIN LS |
| 7:00 | 7f13-A | 3.2 | 3.6 | 71 | 154 | 6 | MOLECULE: | DCR3 | | |
| 8:00 | 4epa-A | 3.2 | 3.3 | 69 | 632 | 3 | MOLECULE: | PESTICI | N RECEP | TOR |
| 9:00 | 5hal-A | 3.2 | 3.4 | 65 | 103 | 9 | MOLECULE: | UNCHARA | CTERIZE | D PROTEIN |
| 10:00 | 1sa8-A | 3.2 | 3.5 | 60 | 106 | 8 | MOLECULE: | FATTY A | CID-BIN | DING PROTEIN, INTESTINAL |
| 11:00 | 4mjd-A | 3.2 | 3.8 | 66 | 113 | 12 | MOLECULE: | KETOSTE | ROID IS | OMERASE FOLD PROTEIN HMUK_0747 |
| 12:00 | 6s50-A | 3.2 | 3.7 | 53 | 248 | 4 | MOLECULE: | STREPTA | VIDIN | |
| 13:00 | 7fhr-A | 3.2 | 3.5 | 74 | 439 | 5 | MOLECULE: | PUTATIV | E PHTHA | LATE 4,5-DIOXYGENASE, SUBUNIT ALPHA |
| 14:00 | 7woo-D | 3.2 | 3.2 | 59 | 1398 | 3 | MOLECULE: | NUCLEOP | ORIN NI | C96 |
| 15:00 | 1jki-A | 3.2 | 4.1 | 59 | 525 | 10 | MOLECULE: | MYO-INO | SITOL-1 | -PHOSPHATE SYNTHASE |
| 16:00 | 8dml-B | 3.2 | 2.9 | 54 | 133 | 2 | MOLECULE: | VTRA | | |
| 17:00 | 7yls-A | 3.2 | 3.5 | 67 | 158 | 4 | MOLECULE: | AROMATI | C-RING- | HYDROXYLATING DIOXYGENASE BETA SUBU |
| 18:00 | 7z6e-C | 3.2 | 2.9 | 55 | 534 | 11 | MOLECULE: | SERINE/ | THREONI | NE-PROTEIN KINASE MRCK-1 |
| 19:00 | 6whp-A | 3.2 | 3.3 | 70 | 428 | 9 | MOLECULE: | CHOLINE | KINASE | |
| 20:00 | 6fhm-A | 3.2 | 3.7 | 59 | 193 | 5 | MOLECULE: | OUTER-M | EMBRANE | LIPOPROTEIN CARRIER PROTEIN |
| 21:00 | 7agv-F | 3.2 | 2.6 | 51 | 163 | 6 | MOLECULE: | K(+)/H( | +) ANTI | PORTER SUBUNIT KHTT |
| 22:00 | 8gt0-B | 3.2 | 3.5 | 71 | 98 | 4 | MOLECULE: | CYSTEIN | E PROTE | ASE FALCIPAIN-2 |
| 23:00 | 1gyb-D | 3.2 | 3.2 | 66 | 120 | 9 | MOLECULE: | NUCLEAR | TRANSP | ORT FACTOR 2 |
| 24:00:00 | 1gy7-D | 3.2 | 3.4 | 68 | 120 | 9 | MOLECULE: | NUCLEAR | TRANSP | ORT FACTOR 2 |
| 25:00:00 | 5drv-A | 3.2 | 3.7 | 72 | 131 | 7 | MOLECULE: | RAS GTP | ASE-ACT | IVATING PROTEIN-BINDING PROTEIN 2 |
| 26:00:00 | 8th5-D | 3.2 | 3.7 | 71 | 129 | 7 | MOLECULE: | RAS GTP | ASE-ACT | IVATING PROTEIN-BINDING PROTEIN 1 |
| 27:00:00 | 7xhf-A | 3.2 | 3.4 | 69 | 124 | 6 | MOLECULE: | RAS GTP | ASE-ACT | IVATING PROTEIN-BINDING PROTEIN 1 |
| 28:00:00 | 3fsd-A | 3.2 | 3.2 | 65 | 121 | 5 | MOLECULE: | NTF2-LI | KE PROT | EIN OF UNKNOWN FUNCTION IN NUTRIENT |
| 29:00:00 | 4nf9-A | 3.2 | 4.3 | 65 | 216 | 11 | MOLECULE: | PROTEIN | CASC5 | |
| 30:00:00 | 1izn-A | 3.2 | 3.6 | 71 | 275 | 7 | MOLECULE: | CAPZ AL | PHA-1 S | UBUNIT |
| 31:00:00 | 6d63-E | 3.2 | 3.7 | 68 | 119 | 7 | MOLECULE: | ATZH | | |
| 32:00:00 | 4ipb-B | 3.2 | 3 | 63 | 141 | 3 | MOLECULE: | UNCHARA | CTERIZE | D PROTEIN |
| 33:00:00 | 6d63-J | 3.2 | 3.3 | 68 | 129 | 7 | MOLECULE: | ATZH | | |
| 34:00:00 | 1gyb-C | 3.2 | 3.3 | 68 | 120 | 9 | MOLECULE: | NUCLEAR | TRANSP | ORT FACTOR 2 |
| 35:00:00 | 7qz3-B | 3.2 | 2.5 | 54 | 519 | 6 | MOLECULE: | BNR/ASP | #NAME? | PEAT PROTEIN |
| 36:00:00 | 4i4k-B | 3.2 | 3 | 70 | 131 | 10 | MOLECULE: | UNCHARA | CTERIZE | D PROTEIN SGCJ |
| 37:00:00 | 2w0q-A | 3.2 | 3.3 | 66 | 718 | 8 | MOLECULE: | COPPER | AMINE O | XIDASE |
| 38:00:00 | 1d6u-A | 3.2 | 3.1 | 65 | 718 | 8 | MOLECULE: | COPPER | AMINE O | XIDASE |
| 39:00:00 | 3u1w-A | 3.2 | 3.4 | 62 | 249 | 8 | MOLECULE: | HYPOTHE | TICAL P | ERIPLASMIC PROTEIN |
| 40:00:00 | 6d63-D | 3.2 | 3.1 | 66 | 115 | 8 | MOLECULE: | ATZH | | |
| 41:00:00 | 6s6z-G | 3.2 | 2.7 | 49 | 1083 | 10 | MOLECULE: | BETA-GA | LACTOSI | DASE |
| 42:00:00 | 1ms4-A | 3.2 | 3.2 | 62 | 623 | 6 | MOLECULE: | TRANS-S | IALIDAS | E |
| 43:00:00 | 1q42-A | 3.2 | 3.5 | 71 | 159 | 13 | MOLECULE: | MRNA TR | ANSPORT | REGULATOR MTR2 |
| 44:00:00 | 1a90-A | 3.2 | 3.3 | 62 | 107 | 5 | MOLECULE: | CYSTATI | N | |
| 45:00:00 | 7jr9-C | 3.2 | 4.3 | 69 | 280 | 4 | MOLECULE: | RADIAL | SPOKE P | ROTEIN 9 |
| 46:00:00 | 8agy-A | 3.1 | 3 | 55 | 346 | 7 | MOLECULE: | CORRAMY | CIN PHO | SPHOTRANSFERASE |
| 47:00:00 | 6szw-C | 3.1 | 2.8 | 53 | 186 | 8 | MOLECULE: | COMPLEM | ENT COM | PONENT 1 Q SUBCOMPONENT-BINDING PRO |
| 48:00:00 | 5u4z-A | 3.1 | 3.1 | 73 | 181 | 5 | MOLECULE: | REPRESS | OR OF R | NA POLYMERASE III TRANSCRIPTION |
| 49:00:00 | 4e72-A | 3.1 | 3.6 | 70 | 218 | 4 | MOLECULE: | UNCHARA | CTERIZE | D PROTEIN |
| 50:00:00 | 1vr7-A | 3.1 | 3.4 | 62 | 120 | 0 | MOLECULE: | S-ADENO | SYLMETH | IONINE DECARBOXYLASE PROENZYME |
| 51:00:00 | 2p4o-A | 3.1 | 3.1 | 61 | 302 | 13 | MOLECULE: | HYPOTHE | TICAL P | ROTEIN |
| 52:00:00 | 3ats-A | 3.1 | 3.4 | 62 | 352 | 5 | MOLECULE: | PUTATIV | E UNCHA | RACTERIZED PROTEIN |
| 53:00:00 | 3k1l-A | 3.1 | 3.8 | 61 | 376 | 5 | MOLECULE: | FANCL | | |
| 54:00:00 | 6iqt-A | 3.1 | 3.3 | 64 | 135 | 9 | MOLECULE: | CAG PAT | HOGENIC | ITY ISLAND PROTEIN (CAG10) |

| | | | | | | | | | | | |
|---|---|---|---|---|---|---|---|---|---|---|---|
| 55:00:00 | 6sy9-A | 3.1 | 4.1 | 61 | 307 | 10 | MOLECULE: | RESPONS | E REGUL | ATOR | |
| 56:00:00 | 3lid-A | 3.1 | 3.2 | 62 | 279 | 8 | MOLECULE: | PUTATIV | E SENSO | RY BOX/GGDEF FAMILY PROTEIN | |
| 57:00:00 | 5yvf-A | 3.1 | 3.9 | 71 | 322 | 8 | MOLECULE: | BFA1 | | | |
| 58:00:00 | 3elz-B | 3.1 | 3.9 | 54 | 135 | 7 | MOLECULE: | ILEAL B | ILE ACI | D-BINDING PROTEIN | |
| 59:00:00 | 6vxk-B | 3.1 | 3.8 | 73 | 747 | 5 | MOLECULE: | SEMAPHO | RIN-LIK | E PROTEIN 139 | |
| 60:00:00 | 1wzn-A | 3.1 | 5.9 | 49 | 245 | 6 | MOLECULE: | SAM-DEP | ENDENT | METHYLTRANSFERASE | |
| 61:00:00 | 1sgo-A | 3.1 | 4 | 69 | 139 | 7 | MOLECULE: | PROTEIN | C14ORF | | 129 |
| 62:00:00 | 3cwy-A | 3.1 | 3.6 | 66 | 130 | 9 | MOLECULE: | PROTEIN | CAGD | | |
| 63:00:00 | 6ck1-A | 3.1 | 3 | 59 | 375 | 7 | MOLECULE: | A1B2F4 | PROTEIN | | |
| 64:00:00 | 7vud-A | 3.1 | 3.5 | 59 | 535 | 7 | MOLECULE: | CAROTEN | OID CLE | AVAGE DIOXYGENASE 1 | |
| 65:00:00 | 7vwk-A | 3.1 | 2.9 | 54 | 260 | 6 | MOLECULE: | POLYKET | IDE SYN | THASE | |
| 66:00:00 | 6h3i-A | 3.1 | 4.5 | 70 | 2124 | 4 | MOLECULE: | PROTEIN | INVOLV | ED IN GLIDING MOTILITY SPRA | |
| 67:00:00 | 2h36-X | 3.1 | 3 | 55 | 108 | 4 | MOLECULE: | HYPOTHE | TICAL P | ROTEIN SIFV0014 | |
| 68:00:00 | 3eps-A | 3.1 | 4.5 | 79 | 566 | 3 | MOLECULE: | ISOCITR | ATE DEH | YDROGENASE KINASE/PHOSPHATASE | |
| 69:00:00 | 6zce-p | 3.1 | 3.8 | 67 | 646 | 4 | MOLECULE: | EUKARYO | TIC TRA | NSLATION INITIATION FACTOR 3 SUBUNI | |
| 70:00:00 | 3ff2-A | 3.1 | 4.1 | 65 | 117 | 8 | MOLECULE: | UNCHARA | CTERIZE | D CYSTATIN FOLD PROTEIN (YP_497570. | |
| 71:00:00 | 4pqg-A | 3.1 | 2.7 | 54 | 506 | 11 | MOLECULE: | GLYCOSY | LTRANSF | ERASE GTF1 | |
| 72:00:00 | 8ovw-N | 3.1 | 3.7 | 69 | 391 | 4 | MOLECULE: | CENTROM | ERE-BIN | DING PROTEIN 1 | |
| 73:00:00 | 7qh3-B | 3.1 | 4.5 | 63 | 148 | 5 | MOLECULE: | RSFG | | | |
| 74:00:00 | 4tqj-A | 3.1 | 3.5 | 66 | 403 | 5 | MOLECULE: | LECTIN | 2 | | |
| 75:00:00 | 8wxf-A | 3.1 | 3.9 | 54 | 730 | 13 | MOLECULE: | BIFUNCT | IONAL G | UANOSINE PENTAPHOSPHATE | |
| 76:00:00 | 3nqh-A | 3.1 | 4.3 | 57 | 439 | 4 | MOLECULE: | GLYCOSY | L HYDRO | LASE | |
| 77:00:00 | 3s25-A | 3.1 | 3.6 | 58 | 295 | 14 | MOLECULE: | HYPOTHE | TICAL 7 | #NAME? | |
| 78:00:00 | 2wgo-A | 3.1 | 4 | 74 | 98 | 4 | MOLECULE: | RANASPU | MIN-2 | | |
| 79:00:00 | 7jrj-C | 3.1 | 4.3 | 72 | 359 | 10 | MOLECULE: | RADIAL | SPOKE P | ROTEIN 10 | |
| 80:00:00 | 8th1-D | 3.1 | 3.5 | 69 | 137 | 6 | MOLECULE: | RAS GTP | ASE-ACT | IVATING PROTEIN-BINDING PROTEIN 1 | |
| 81:00:00 | 4fcj-A | 3.1 | 3.2 | 66 | 128 | 6 | MOLECULE: | RAS GTP | ASE-ACT | IVATING PROTEIN-BINDING PROTEIN 1 | |
| 82:00:00 | 8th5-A | 3.1 | 3.7 | 71 | 123 | 6 | MOLECULE: | RAS GTP | ASE-ACT | IVATING PROTEIN-BINDING PROTEIN 1 | |
| 83:00:00 | 7suo-B | 3.1 | 3.8 | 72 | 137 | 7 | MOLECULE: | RAS GTP | ASE-ACT | IVATING PROTEIN-BINDING PROTEIN 1 | |
| 84:00:00 | 1zo2-A | 3.1 | 3.2 | 67 | 124 | 10 | MOLECULE: | NUCLEAR | TRANSP | ORT FACTOR 2 | |
| 85:00:00 | 8x2u-G | 3.1 | 4.2 | 70 | 387 | 3 | MOLECULE: | DPY30 D | OMAIN C | ONTAINING 2 | |
| 86:00:00 | 8th1-A | 3.1 | 3.4 | 68 | 137 | 6 | MOLECULE: | RAS GTP | ASE-ACT | IVATING PROTEIN-BINDING PROTEIN 1 | |
| 87:00:00 | 2pup-B | 3.1 | 2.5 | 54 | 370 | 4 | MOLECULE: | METHYLT | HIORIBO | SE KINASE | |
| 88:00:00 | 6f38-K | 3.1 | 3.4 | 73 | 278 | 4 | MOLECULE: | ARP1 AC | TIN REL | ATED PROTEIN 1 HOMOLOG A | |
| 89:00:00 | 6fej-A | 3.1 | 3.3 | 68 | 124 | 10 | MOLECULE: | ALL4940 | PROTEI | N | |
| 90:00:00 | 4nf9-B | 3.1 | 4.2 | 65 | 207 | 11 | MOLECULE: | PROTEIN | CASC5 | | |
| 91:00:00 | 3ob8-D | 3.1 | 4.3 | 50 | 1024 | 8 | MOLECULE: | BETA-GA | LACTOSI | DASE | |
| 92:00:00 | 4xj9-A | 3.1 | 3.4 | 60 | 658 | 7 | MOLECULE: | SIALIDA | SE B | | |
| 93:00:00 | 4ovm-C | 3.1 | 3.1 | 71 | 129 | 10 | MOLECULE: | UNCHARA | CTERIZE | D PROTEIN SGCJ | |
| 94:00:00 | 2bzd-B | 3.1 | 3.1 | 64 | 601 | 16 | MOLECULE: | BACTERI | AL SIAL | IDASE | |
| 95:00:00 | 5fw5-B | 3.1 | 3.2 | 67 | 131 | 6 | MOLECULE: | RAS GTP | ASE-ACT | IVATING PROTEIN-BINDING PROTEIN 1 | |
| 96:00:00 | 3wz5-C | 3.1 | 3.9 | 70 | 136 | 7 | MOLECULE: | DOTI | | | |
| 97:00:00 | 1w0p-A | 3.1 | 3.3 | 73 | 753 | 7 | MOLECULE: | SIALIDA | SE | | |
| 98:00:00 | 7zhz-B | 3.1 | 4.6 | 70 | 527 | 3 | MOLECULE: | GLUCOSE | #NAME? | PHATE 1-DEHYDROGENASE | |
| 99:00:00 | 7w19-A | 3.1 | 2.5 | 55 | 293 | 5 | MOLECULE: | MACROLI | DE 2'-P | HOSPHOTRANSFERASE | |
| 0:00 | 1d6z-A | 3.1 | 3.1 | 65 | 718 | 8 | MOLECULE: | COPPER | AMINE O | XIDASE | |
| 1:00 | 3ujm-A | 3.1 | 3.1 | 64 | 117 | 5 | MOLECULE: | RASPUTI | N | | |
| 2:00 | 6s6z-F | 3.1 | 3.4 | 74 | 1083 | 4 | MOLECULE: | BETA-GA | LACTOSI | DASE | |
| 3:00 | 4fp3-A | 3.1 | 2.9 | 57 | 658 | 2 | MOLECULE: | SIALIDA | SE B | | |
| 4:00 | 4dux-D | 3.1 | 3 | 50 | 1015 | 6 | MOLECULE: | BETA-GA | LACTOSI | DASE | |
| 5:00 | 3lk3-A | 3.1 | 3.3 | 68 | 266 | 6 | MOLECULE: | F-ACTIN | #NAME? | G PROTEIN SUBUNIT ALPHA-1 | |
| 6:00 | 2ber-A | 3.1 | 3.3 | 65 | 601 | 15 | MOLECULE: | BACTERI | AL SIAL | IDASE | |
| 7:00 | 6s6z-H | 3.1 | 2.7 | 50 | 1083 | 10 | MOLECULE: | BETA-GA | LACTOSI | DASE | |
| 8:00 | 3aa1-A | 3.1 | 3.5 | 72 | 271 | 7 | MOLECULE: | F-ACTIN | #NAME? | G PROTEIN SUBUNIT ALPHA-1 | |

| | | | | | | | | | | |
|---|---|---|---|---|---|---|---|---|---|---|
| 9:00 | 3dxq-B | 3.1 | 2.5 | 49 | 292 | 6 | MOLECULE: | CHOLINE | /ETHANO | LAMINE KINASE FAMILY PROTEIN |
| 10:00 | 1jz7-C | 3.1 | 3.1 | 51 | 1011 | 4 | MOLECULE: | BETA-GA | LACTOSI | DASE |
| 11:00 | 6m1u-A | 3 | 3.5 | 63 | 122 | 3 | MOLECULE: | RNA N6- | ADENOSI | NE-METHYLTRANSFERASE METTL16,RNA N6 |
| 12:00 | 5wlz-C | 3 | 2.8 | 52 | 206 | 12 | MOLECULE: | DNA REP | AIR PRO | TEIN XRCC4,MYOSIN-7 |
| 13:00 | 8e1m-C | 3 | 3.5 | 69 | 264 | 6 | MOLECULE: | MITOCHO | NDRIAL | IMPORT INNER MEMBRANE TRANSLOCASE S |
| 14:00 | 4zsv-A | 3 | 3.6 | 60 | 294 | 5 | MOLECULE: | UNCHARA | CTERIZE | D PROTEIN |
| 15:00 | 8ju7-A | 3 | 3.8 | 60 | 104 | 8 | MOLECULE: | HISTIDI | NE KINA | SE |
| 16:00 | 2chc-A | 3 | 3.5 | 70 | 169 | 4 | MOLECULE: | PROTEIN | RV3472 | |
| 17:00 | 3duk-A | 3 | 3.4 | 70 | 125 | 9 | MOLECULE: | NTF2-LI | KE PROT | EIN OF UNKNOWN FUNCTION |
| 18:00 | 2glz-A | 3 | 3.8 | 70 | 149 | 3 | MOLECULE: | SIMILAR | TO FOR | MYLMETHANOFURAN DEHYDROGENASE SUBUN |
| 19:00 | 8pjn-2 | 3 | 3.8 | 64 | 156 | 6 | MOLECULE: | E3 UBIQ | UITIN-P | ROTEIN TRANSFERASE RMND5A |
| 20:00 | 4pj2-B | 3 | 2.4 | 61 | 123 | 10 | MOLECULE: | PUTATIV | E EXPOR | TED PROTEIN |
| 21:00 | 2rfr-A | 3 | 3.4 | 68 | 154 | 7 | MOLECULE: | UNCHARA | CTERIZE | D PROTEIN |
| 22:00 | 5fw5-A | 3 | 3.6 | 70 | 139 | 6 | MOLECULE: | RAS GTP | ASE-ACT | IVATING PROTEIN-BINDING PROTEIN 1 |
| 23:00 | 4zr7-D | 3 | 2.5 | 56 | 120 | 16 | MOLECULE: | SENSOR | HISTIDI | NE KINASE RESE |
| 24:00:00 | 6x1i-B | 3 | 3.6 | 72 | 146 | 7 | MOLECULE: | COB_ADE | NO_TRAN | S DOMAIN-CONTAINING PROTEIN PH0671 |
| 25:00:00 | 3lml-A | 3 | 4.1 | 68 | 440 | 4 | MOLECULE: | LIN1278 | PROTEI | N |
| 26:00:00 | 3soy-A | 3 | 3.2 | 67 | 142 | 13 | MOLECULE: | NTF2-LI | KE SUPE | RFAMILY PROTEIN |
| 27:00:00 | 6mni-A | 3 | 2.7 | 59 | 259 | 7 | MOLECULE: | METHYL- | ACCEPTI | NG CHEMOTAXIS PROTEIN |
| 28:00:00 | 5d16-A | 3 | 3.3 | 66 | 161 | 8 | MOLECULE: | TRANSPO | SON TN7 | TRANSPOSITION PROTEIN TNSE |
| 29:00:00 | 3k7c-A | 3 | 3.5 | 62 | 108 | 6 | MOLECULE: | PUTATIV | E NTF2- | LIKE TRANSPEPTIDASE |
| 30:00:00 | 2f98-C | 3 | 3.3 | 70 | 143 | 7 | MOLECULE: | AKLANON | IC ACID | METHYL ESTER CYCLASE, AKNH |
| 31:00:00 | 4lmi-B | 3 | 3.7 | 71 | 136 | 10 | MOLECULE: | UNCHARA | CTERIZE | D PROTEIN |
| 32:00:00 | 3h6j-A | 3 | 3.3 | 64 | 438 | 8 | MOLECULE: | NEURAMI | NIDASE | |
| 33:00:00 | 3i2n-A | 3 | 3.8 | 67 | 343 | 7 | MOLECULE: | WD REPE | AT-CONT | AINING PROTEIN 92 |
| 34:00:00 | 2gxf-A | 3 | 2.9 | 60 | 119 | 12 | MOLECULE: | HYPOTHE | TICAL P | ROTEIN YYBH |
| 35:00:00 | 6nd4-H | 3 | 3.9 | 59 | 834 | 5 | MOLECULE: | ETS RRN | A | |
| 36:00:00 | 5xq3-A | 3 | 2.8 | 52 | 901 | 8 | MOLECULE: | PCRGLX | PROTEIN | |
| 37:00:00 | 3g0k-A | 3 | 3.6 | 69 | 129 | 9 | MOLECULE: | PUTATIV | E MEMBR | ANE PROTEIN |
| 38:00:00 | 7fds-B | 3 | 3.7 | 54 | 118 | 4 | MOLECULE: | LIPOPRO | TEIN LP | QH |
| 39:00:00 | 2lqv-A | 3 | 3.5 | 64 | 99 | 9 | MOLECULE: | PROTEIN | YEBF | |
| 40:00:00 | 6p2k-B | 3 | 3.9 | 73 | 778 | 7 | MOLECULE: | FIBRONE | CTIN TY | PE III DOMAIN-CONTAINING PROTEIN |
| 41:00:00 | 8shj-A | 3 | 3.6 | 64 | 334 | 13 | MOLECULE: | WD REPE | AT-CONT | AINING PROTEIN 91 |
| 42:00:00 | 1pre-B | 3 | 3.7 | 75 | 451 | 7 | MOLECULE: | PROAERO | LYSIN | |
| 43:00:00 | 6qvi-A | 3 | 6.2 | 57 | 518 | 7 | MOLECULE: | COMZ | | |
| 44:00:00 | 1l0q-A | 3 | 3.4 | 60 | 391 | 12 | MOLECULE: | SURFACE | LAYER | PROTEIN |
| 45:00:00 | 6ht9-B | 3 | 3.9 | 67 | 316 | 4 | MOLECULE: | ASTACIN | | |
| 46:00:00 | 7pge-B | 3 | 4 | 60 | 198 | 13 | MOLECULE: | COPPER | RESISTA | NCE PROTEIN B |
| 47:00:00 | 3fh1-A | 3 | 4.1 | 70 | 122 | 7 | MOLECULE: | UNCHARA | CTERIZE | D NTF2-LIKE PROTEIN |
| 48:00:00 | 7aqc-R | 3 | 3.9 | 71 | 558 | 4 | MOLECULE: | 23S RIB | OSOMAL | RNA |
| 49:00:00 | 6igb-A | 3 | 3.5 | 67 | 363 | 9 | MOLECULE: | PERIPLA | SMIC GL | UCONOLACTONASE, PPGL |
| 50:00:00 | 2ch9-A | 3 | 4.4 | 68 | 126 | 10 | MOLECULE: | CYSTATI | N F | |
| 51:00:00 | 6b62-A | 3 | 3.5 | 64 | 256 | 6 | MOLECULE: | FRUCTOS | E-1,6-B | ISPHOSPHATASE/INOSITOL-1-MONOPHOSPH |
| 52:00:00 | 3f9s-A | 3 | 3.5 | 70 | 143 | 10 | MOLECULE: | PUTATIV | E POLYK | ETIDE CYCLASE |
| 53:00:00 | 6jp6-A | 3 | 3.3 | 66 | 998 | 5 | MOLECULE: | TRNA (G | UANOSIN | E(34)-2'-O)-METHYLTRANSFERASE NON-C |
| 54:00:00 | 4agi-A | 3 | 2.7 | 46 | 314 | 9 | MOLECULE: | FUCOSE- | SPECIFI | C LECTIN FLEA |
| 55:00:00 | 6z37-A | 3 | 7.1 | 69 | 418 | 1 | MOLECULE: | TODX | | |
| 56:00:00 | 7kbj-I | 3 | 3.1 | 61 | 153 | 5 | MOLECULE: | NEUTRAL | ALPHA- | GLUCOSIDASE AB TRYPSIN-CLEAVED FRAG |
| 57:00:00 | 6iy9-A | 3 | 2.7 | 58 | 325 | 7 | MOLECULE: | HYGROMY | CIN-B 7 | '-O-KINASE |
| 58:00:00 | 6mso-A | 3 | 3 | 63 | 540 | 3 | MOLECULE: | FUMARAT | E HYDRA | TASE |
| 59:00:00 | 7c5x-A | 3 | 3.6 | 71 | 506 | 1 | MOLECULE: | IOTA-CA | RBONIC | ANHYDRASE |
| 60:00:00 | 6mly-C | 3 | 3.7 | 71 | 774 | 10 | MOLECULE: | BIFUNCT | IONAL G | H43-CE PROTEIN |
| 61:00:00 | 8veh-C | 3 | 3.5 | 73 | 194 | 4 | MOLECULE: | OUTER M | EMBRANE | LIPOPROTEIN-SORTING PROTEIN LOLA |
| 62:00:00 | 1q40-A | 3 | 3.6 | 71 | 165 | 11 | MOLECULE: | MRNA TR | ANSPORT | REGULATOR MTR2 |

| | | | | | | | | | | |
|---|---|---|---|---|---|---|---|---|---|---|
| 63:00:00 | 3lyg-A | 3 | 3.2 | 64 | 120 | 13 | MOLECULE: | NTF2-LI | KE PROT | EIN OF UNKNOWN FUNCTION |
| 64:00:00 | 3ejv-A | 3 | 3.5 | 69 | 159 | 4 | MOLECULE: | UNCHARA | CTERIZE | D PROTEIN WITH CYSTATIN-LIKE FOLD |
| 65:00:00 | 4u13-A | 3 | 3.1 | 63 | 109 | 10 | MOLECULE: | PUTATIV | E POLYK | ETIDE CYCLASE SMA1630 |
| 66:00:00 | 4ghb-B | 3 | 3 | 53 | 257 | 2 | MOLECULE: | HYPOTHE | TICAL P | ROTEIN |
| 67:00:00 | 8th7-B | 3 | 3.9 | 70 | 139 | 6 | MOLECULE: | RAS GTP | ASE-ACT | IVATING PROTEIN-BINDING PROTEIN 1 |
| 68:00:00 | 4xju-A | 3 | 3.2 | 63 | 658 | 2 | MOLECULE: | SIALIDA | SE B | |
| 69:00:00 | 6f3a-K | 3 | 3.4 | 74 | 278 | 3 | MOLECULE: | ARP1 AC | TIN REL | ATED PROTEIN 1 HOMOLOG A |
| 70:00:00 | 4akr-C | 3 | 3.2 | 75 | 271 | 4 | MOLECULE: | F-ACTIN | #NAME? | G PROTEIN SUBUNIT ALPHA |
| 71:00:00 | 7suo-A | 3 | 3.7 | 71 | 136 | 6 | MOLECULE: | RAS GTP | ASE-ACT | IVATING PROTEIN-BINDING PROTEIN 1 |
| 72:00:00 | 3sil-A | 3 | 3.1 | 63 | 379 | 5 | MOLECULE: | SIALIDA | SE | |
| 73:00:00 | 7yth-B | 3 | 3.5 | 70 | 315 | 10 | MOLECULE: | KETOSTE | ROID IS | OMERASE-RELATED PROTEIN |
| 74:00:00 | 4fcm-B | 3 | 3.8 | 67 | 135 | 6 | MOLECULE: | RAS GTP | ASE-ACT | IVATING PROTEIN-BINDING PROTEIN 1 |
| 75:00:00 | 4xma-A | 3 | 3.3 | 63 | 658 | 2 | MOLECULE: | SIALIDA | SE B | |
| 76:00:00 | 8th6-B | 3 | 3.9 | 71 | 139 | 8 | MOLECULE: | RAS GTP | ASE-ACT | IVATING PROTEIN-BINDING PROTEIN 1 |
| 77:00:00 | 8th6-A | 3 | 3.5 | 72 | 138 | 7 | MOLECULE: | RAS GTP | ASE-ACT | IVATING PROTEIN-BINDING PROTEIN 1 |
| 78:00:00 | 6ta7-A | 3 | 3.9 | 71 | 131 | 7 | MOLECULE: | RAS GTP | ASE-ACT | IVATING PROTEIN-BINDING PROTEIN 1 |
| 79:00:00 | 3q90-B | 3 | 3.5 | 68 | 130 | 6 | MOLECULE: | RAS GTP | ASE-ACT | IVATING PROTEIN-BINDING PROTEIN 1 |
| 80:00:00 | 1qaf-B | 3 | 3.4 | 65 | 721 | 8 | MOLECULE: | PROTEIN | (COPPE | R AMINE OXIDASE) |
| 81:00:00 | 7xhg-C | 3 | 3.6 | 70 | 126 | 6 | MOLECULE: | RAS GTP | ASE-ACT | IVATING PROTEIN-BINDING PROTEIN 1 |
| 82:00:00 | 7vr6-A | 3 | 3.2 | 67 | 185 | 0 | MOLECULE: | INTERME | MBRANE | PHOSPHOLIPID TRANSPORT SYSTEM BINDI |
| 83:00:00 | 1zo2-B | 3 | 3.5 | 68 | 124 | 15 | MOLECULE: | NUCLEAR | TRANSP | ORT FACTOR 2 |
| 84:00:00 | 1mr5-A | 3 | 3.1 | 59 | 621 | 7 | MOLECULE: | TRANS-S | IALIDAS | E |
| 85:00:00 | 8v1l-C | 3 | 3.3 | 68 | 127 | 6 | MOLECULE: | RAS GTP | ASE-ACT | IVATING PROTEIN-BINDING PROTEIN 1 |
| 86:00:00 | 3ujm-B | 3 | 3 | 62 | 117 | 5 | MOLECULE: | RASPUTI | N | |
| 87:00:00 | 7uc2-C | 3 | 4.4 | 70 | 475 | 3 | MOLECULE: | GLUCOSE | #NAME? | PHATE 1-DEHYDROGENASE |
| 88:00:00 | 8th1-C | 3 | 3.9 | 70 | 133 | 7 | MOLECULE: | RAS GTP | ASE-ACT | IVATING PROTEIN-BINDING PROTEIN 1 |
| 89:00:00 | 6s6z-C | 3 | 2.7 | 50 | 1083 | 10 | MOLECULE: | BETA-GA | LACTOSI | DASE |
| 90:00:00 | 6d63-H | 3 | 3.1 | 66 | 121 | 8 | MOLECULE: | ATZH | | |
| 91:00:00 | 2ya7-B | 3 | 3.7 | 64 | 468 | 9 | MOLECULE: | NEURAMI | NIDASE | A |
| 92:00:00 | 1oac-A | 3 | 3.5 | 69 | 720 | 10 | MOLECULE: | COPPER | AMINE O | XIDASE |
| 93:00:00 | 1gy7-C | 3 | 3.2 | 62 | 119 | 6 | MOLECULE: | NUCLEAR | TRANSP | ORT FACTOR 2 |
| 94:00:00 | 1gy7-A | 3 | 3.3 | 63 | 121 | 8 | MOLECULE: | NUCLEAR | TRANSP | ORT FACTOR 2 |
| 95:00:00 | 1ms3-B | 3 | 3.1 | 62 | 623 | 6 | MOLECULE: | TRANS-S | IALIDAS | E |
| 96:00:00 | 4xjz-A | 3 | 3.3 | 63 | 658 | 2 | MOLECULE: | SIALIDA | SE B | |
| 97:00:00 | 6d63-F | 3 | 3.2 | 68 | 126 | 6 | MOLECULE: | ATZH | | |
| 98:00:00 | 7zhw-A | 3 | 4.4 | 67 | 534 | 3 | MOLECULE: | GLUCOSE | #NAME? | PHATE 1-DEHYDROGENASE |
| 99:00:00 | 7xhg-A | 3 | 3.2 | 67 | 133 | 4 | MOLECULE: | RAS GTP | ASE-ACT | IVATING PROTEIN-BINDING PROTEIN 1 |
| 0:00 | 8th5-C | 3 | 3.5 | 69 | 125 | 6 | MOLECULE: | RAS GTP | ASE-ACT | IVATING PROTEIN-BINDING PROTEIN 1 |
| 1:00 | 1q40-C | 3 | 3.6 | 71 | 164 | 11 | MOLECULE: | MRNA TR | ANSPORT | REGULATOR MTR2 |
| 2:00 | 1ms1-B | 3 | 3.7 | 61 | 622 | 7 | MOLECULE: | TRANS-S | IALIDAS | E |
| 3:00 | 4ovm-A | 3 | 3.1 | 69 | 128 | 9 | MOLECULE: | UNCHARA | CTERIZE | D PROTEIN SGCJ |
| 4:00 | 4bbw-A | 3 | 3 | 59 | 520 | 5 | MOLECULE: | SIALIDA | SE (NEU | RAMINIDASE) |
| 5:00 | 6ta7-C | 3 | 3.1 | 67 | 128 | 6 | MOLECULE: | RAS GTP | ASE-ACT | IVATING PROTEIN-BINDING PROTEIN 1 |
| 6:00 | 8t26-A | 3 | 3.3 | 65 | 504 | 5 | MOLECULE: | SIALIDA | SE, PUT | ATIVE |
| 7:00 | 2jkb-A | 3 | 3.4 | 60 | 661 | 8 | MOLECULE: | SIALIDA | SE B | |
| 8:00 | 6s6z-D | 3 | 2.7 | 50 | 1083 | 10 | MOLECULE: | BETA-GA | LACTOSI | DASE |
| 9:00 | 6baq-F | 2.9 | 2.8 | 72 | 211 | 10 | MOLECULE: | BPI FOL | D-CONTA | INING FAMILY A MEMBER 1 |
| 10:00 | 5d06-A | 2.9 | 3.7 | 71 | 1526 | 3 | MOLECULE: | UNCHARA | CTERIZE | D PROTEIN |
| 11:00 | 8rtd-S | 2.9 | 3.8 | 70 | 223 | 10 | MOLECULE: | TRWJ PR | OTEIN | |
| 12:00 | 3obu-A | 2.9 | 3.6 | 68 | 142 | 9 | MOLECULE: | TUMOR S | USCEPTI | BILITY GENE 101 PROTEIN |
| 13:00 | 3kkg-A | 2.9 | 4.2 | 72 | 144 | 10 | MOLECULE: | PUTATIV | E SNOAL | -LIKE POLYKETIDE CYCLASE |
| 14:00 | 3bb9-B | 2.9 | 3.3 | 64 | 125 | 8 | MOLECULE: | PUTATIV | E ORPHA | N PROTEIN |
| 15:00 | 3e29-B | 2.9 | 3.8 | 69 | 135 | 6 | MOLECULE: | UNCHARA | CTERIZE | D PROTEIN Q7WE92_BORBR |
| 16:00 | 1s5a-B | 2.9 | 3.5 | 65 | 143 | 5 | MOLECULE: | HYPOTHE | TICAL P | ROTEIN YESE |

| | | | | | | | | | | |
|---|---|---|---|---|---|---|---|---|---|---|
| 17:00 | 1tu1-A | 2.9 | 4.3 | 67 | 144 | 6 | MOLECULE: | HYPOTHE | TICAL P | ROTEIN PA0094 |
| 18:00 | 8a9b-B | 2.9 | 4 | 78 | 809 | 1 | MOLECULE: | LIPID B | INDING | PROTEIN P116 (MPN213) |
| 19:00 | 8jd7-A | 2.9 | 4.1 | 74 | 510 | 4 | MOLECULE: | ENDO-1, | 3-FUCAN | ASE |
| 20:00 | 3k5j-A | 2.9 | 3.8 | 70 | 182 | 13 | MOLECULE: | SUPPRES | SOR OF | FUSED FAMILY PROTEIN |
| 21:00 | 4ywz-A | 2.9 | 3 | 61 | 147 | 7 | MOLECULE: | SENSOR | PROTEIN | KINASE WALK |
| 22:00 | 4wp5-A | 2.9 | 3.2 | 73 | 199 | 4 | MOLECULE: | MRNA EX | PORT PR | OTEIN |
| 23:00 | 2k3d-A | 2.9 | 3.1 | 57 | 87 | 7 | MOLECULE: | LIN0334 | PROTEI | N |
| 24:00:00 | 7zei-A | 2.9 | 3.4 | 59 | 310 | 8 | MOLECULE: | CH_GAF1 | 59A | |
| 25:00:00 | 4gl6-B | 2.9 | 3.5 | 59 | 241 | 2 | MOLECULE: | HYPOTHE | TICAL P | ROTEIN |
| 26:00:00 | 1mdc-A | 2.9 | 3.6 | 60 | 131 | 5 | MOLECULE: | INSECT | FATTY A | CID BINDING PROTEIN |
| 27:00:00 | 2w5n-A | 2.9 | 3.9 | 67 | 365 | 12 | MOLECULE: | ALPHA-L | #NAME? | OFURANOSIDASE |
| 28:00:00 | 3ksp-A | 2.9 | 3.6 | 68 | 129 | 9 | MOLECULE: | CALCIUM | /CALMOD | ULIN-DEPENDENT KINASE II ASSOCIATIO |
| 29:00:00 | 5jk2-A | 2.9 | 3.3 | 71 | 128 | 6 | MOLECULE: | TP0751 | | |
| 30:00:00 | 4k17-A | 2.9 | 3.3 | 55 | 660 | 9 | MOLECULE: | LEUCINE | -RICH R | EPEAT-CONTAINING PROTEIN 16A |
| 31:00:00 | 8hpo-K | 2.9 | 3.2 | 55 | 413 | 15 | MOLECULE: | TRANSCR | IPTIONA | L REGULATORY PROTEIN UME1 |
| 32:00:00 | 1jp4-A | 2.9 | 5.3 | 72 | 302 | 3 | MOLECULE: | 3'(2'), | 5'-BISP | HOSPHATE NUCLEOTIDASE |
| 33:00:00 | 5uac-C | 2.9 | 6.2 | 51 | 1342 | 4 | MOLECULE: | DNA-DIR | ECTED R | NA POLYMERASE SUBUNIT ALPHA |
| 34:00:00 | 8ain-B | 2.9 | 3.2 | 59 | 107 | 7 | MOLECULE: | URACIL- | DNA GLY | COSYLASE |
| 35:00:00 | 6n8s-A | 2.9 | 3.5 | 72 | 834 | 4 | MOLECULE: | LETHAL( | 2) GIAN | T LARVAE PROTEIN HOMOLOG 2 |
| 36:00:00 | 5chs-B | 2.9 | 3.4 | 54 | 315 | 7 | MOLECULE: | RNA-DIR | ECTED R | NA POLYMERASE L |
| 37:00:00 | 7e85-A | 2.9 | 2.9 | 63 | 114 | 8 | MOLECULE: | SNOAL-L | IKE DOM | AIN-CONTAINING PROTEIN |
| 38:00:00 | 6pon-A | 2.9 | 4 | 71 | 255 | 1 | MOLECULE: | ADHEREN | CE AND | VIRULENCE PROTEIN A |
| 39:00:00 | 3eu7-A | 2.9 | 3.1 | 56 | 313 | 5 | MOLECULE: | PARTNER | AND LO | CALIZER OF BRCA2 |
| 40:00:00 | 8of7-A | 2.9 | 4.8 | 61 | 143 | 5 | MOLECULE: | RHS FAM | ILY PRO | TEIN |
| 41:00:00 | 2yzy-A | 2.9 | 3.5 | 59 | 163 | 8 | MOLECULE: | PUTATIV | E UNCHA | RACTERIZED PROTEIN TTHA1012 |
| 42:00:00 | 2ppq-A | 2.9 | 2.7 | 53 | 310 | 6 | MOLECULE: | HOMOSER | INE KIN | ASE |
| 43:00:00 | 4pr7-A | 2.9 | 3.5 | 52 | 171 | 12 | MOLECULE: | OLIGOGA | LACTURO | NATE-SPECIFIC PORIN KDGM |
| 44:00:00 | 4gfc-A | 2.9 | 3 | 55 | 165 | 4 | MOLECULE: | SPINDLE | ASSEMB | LY ABNORMAL PROTEIN 6 |
| 45:00:00 | 7r2x-A | 2.9 | 4.2 | 69 | 762 | 6 | MOLECULE: | MANNURC | NIC ACI | D SPECIFIC LYASE |
| 46:00:00 | 7apk-F | 2.9 | 3.3 | 58 | 337 | 16 | MOLECULE: | THO COM | PLEX SU | BUNIT 1 |
| 47:00:00 | 5hax-A | 2.9 | 3.3 | 60 | 698 | 10 | MOLECULE: | NUCLEOP | ORIN NU | P170 |
| 48:00:00 | 6vk3-B | 2.9 | 3.6 | 67 | 350 | 9 | MOLECULE: | HRD3 | | |
| 49:00:00 | 6sg8-B | 2.9 | 3.2 | 61 | 381 | 7 | MOLECULE: | HEMAGGL | UTININ- | NEURAMINIDASE |
| 50:00:00 | 7jj9-A | 2.9 | 3.3 | 72 | 461 | 6 | MOLECULE: | ZINC-BI | NDING L | IPOPROTEIN ADCA |
| 51:00:00 | 3i0y-A | 2.9 | 4.1 | 67 | 138 | 9 | MOLECULE: | PUTATIV | E POLYK | ETIDE CYCLASE |
| 52:00:00 | 3msw-A | 2.9 | 3.8 | 61 | 139 | 5 | MOLECULE: | UNCHARA | CTERIZE | D PROTEIN |
| 53:00:00 | 4gb5-A | 2.9 | 3.2 | 63 | 148 | 10 | MOLECULE: | UNCHARA | CTERIZE | D PROTEIN |
| 54:00:00 | 8cli-B | 2.9 | 3.1 | 65 | 689 | 0 | MOLECULE: | GENERAL | TRANSC | RIPTION FACTOR 3C POLYPEPTIDE 1 |
| 55:00:00 | 5yx4-A | 2.9 | 3.8 | 70 | 232 | 9 | MOLECULE: | CHALCON | E-FLAVO | NONE ISOMERASE FAMILY PROTEIN |
| 56:00:00 | 4keg-A | 2.9 | 3 | 71 | 535 | 10 | MOLECULE: | MALTOSE | #NAME? | G PERIPLASMIC/PALATE LUNG AND NASAL |
| 57:00:00 | 1qak-B | 2.9 | 3.4 | 64 | 722 | 8 | MOLECULE: | COPPER | AMINE O | XIDASE |
| 58:00:00 | 8qx7-A | 2.9 | 3 | 62 | 124 | 10 | MOLECULE: | ALL4940 | PROTEI | N |
| 59:00:00 | 6myv-B | 2.9 | 3.7 | 64 | 522 | 6 | MOLECULE: | SIALIDA | SE26 | |
| 60:00:00 | 8qx7-B | 2.9 | 3.1 | 62 | 126 | 10 | MOLECULE: | ALL4940 | PROTEI | N |
| 61:00:00 | 8v1l-E | 2.9 | 3.3 | 70 | 134 | 6 | MOLECULE: | RAS GTP | ASE-ACT | IVATING PROTEIN-BINDING PROTEIN 1 |
| 62:00:00 | 7jr9-D | 2.9 | 3.1 | 66 | 339 | 6 | MOLECULE: | RADIAL | SPOKE P | ROTEIN 9 |
| 63:00:00 | 4xja-A | 2.9 | 3.2 | 63 | 658 | 2 | MOLECULE: | SIALIDA | SE B | |
| 64:00:00 | 6eku-A | 2.9 | 3.1 | 71 | 755 | 7 | MOLECULE: | SIALIDA | SE | |
| 65:00:00 | 2pui-B | 2.9 | 1.9 | 49 | 367 | 4 | MOLECULE: | METHYLT | HIORIBO | SE KINASE |
| 66:00:00 | 1eui-D | 2.9 | 2.6 | 48 | 70 | 8 | MOLECULE: | URACIL- | DNA GLY | COSYLASE |
| 67:00:00 | 2r4i-D | 2.9 | 3.7 | 63 | 117 | 10 | MOLECULE: | UNCHARA | CTERIZE | D PROTEIN |
| 68:00:00 | 7w19-B | 2.9 | 2.4 | 55 | 291 | 5 | MOLECULE: | MACROLI | DE 2'-P | HOSPHOTRANSFERASE |
| 69:00:00 | 4ovm-B | 2.9 | 2.9 | 65 | 126 | 12 | MOLECULE: | UNCHARA | CTERIZE | D PROTEIN SGCJ |
| 70:00:00 | 3ob8-A | 2.9 | 3.4 | 48 | 1024 | 6 | MOLECULE: | BETA-GA | LACTOSI | DASE |

| | | | | | | | | | | |
|---|---|---|---|---|---|---|---|---|---|---|
| 71:00:00 | 7qy9-A | 2.9 | 3.8 | 63 | 519 | 5 | MOLECULE: | BNR/ASP | #NAME? | PEAT PROTEIN |
| 72:00:00 | 7xhg-B | 2.9 | 3.1 | 65 | 133 | 6 | MOLECULE: | RAS GTP | ASE-ACT | IVATING PROTEIN-BINDING PROTEIN 1 |
| 73:00:00 | 1eui-C | 2.9 | 2.6 | 48 | 70 | 8 | MOLECULE: | URACIL- | DNA GLY | COSYLASE |
| 74:00:00 | 4xyx-A | 2.9 | 2.9 | 55 | 660 | 0 | MOLECULE: | SIALIDA | SE B | |
| 75:00:00 | 7zht-D | 2.9 | 4.2 | 65 | 485 | 3 | MOLECULE: | GLUCOSE | #NAME? | PHATE 1-DEHYDROGENASE |
| 76:00:00 | 6sd0-C | 2.9 | 2.7 | 50 | 1083 | 10 | MOLECULE: | BETA-GA | LACTOSI | DASE |
| 77:00:00 | 1gd4-A | 2.9 | 3.7 | 66 | 98 | 5 | MOLECULE: | CYSTATI | N A | |
| 78:00:00 | 2chc-C | 2.9 | 3.4 | 63 | 159 | 10 | MOLECULE: | PROTEIN | RV3472 | |
| 79:00:00 | 5ohl-F | 2.9 | 3.7 | 58 | 88 | 5 | MOLECULE: | K6-SPEC | IFIC AF | FIMER |
| 80:00:00 | 2wgq-A | 2.9 | 3.1 | 65 | 720 | 11 | MOLECULE: | AMINE O | XIDASE | |
| 81:00:00 | 2sim-A | 2.9 | 2.9 | 62 | 381 | 5 | MOLECULE: | SIALIDA | SE | |
| 82:00:00 | 7zht-B | 2.9 | 4.3 | 65 | 525 | 3 | MOLECULE: | GLUCOSE | #NAME? | PHATE 1-DEHYDROGENASE |
| 83:00:00 | 3gzr-B | 2.9 | 3.7 | 68 | 142 | 9 | MOLECULE: | UNCHARA | CTERIZE | D PROTEIN WITH A NTF2-LIKE FOLD |
| 84:00:00 | 1lqg-D | 2.9 | 2.6 | 48 | 70 | 8 | MOLECULE: | URACIL- | DNA GLY | COSYLASE |
| 85:00:00 | 7a5x-B | 2.9 | 3.3 | 64 | 472 | 6 | MOLECULE: | SIALIDA | SE A | |
| 86:00:00 | 1gy7-B | 2.9 | 3.6 | 68 | 122 | 7 | MOLECULE: | NUCLEAR | TRANSP | ORT FACTOR 2 |
| 87:00:00 | 4fp2-A | 2.9 | 3.1 | 58 | 658 | 0 | MOLECULE: | SIALIDA | SE B | |
| 88:00:00 | 7zhy-B | 2.9 | 4.4 | 66 | 533 | 3 | MOLECULE: | GLUCOSE | #NAME? | PHATE 1-DEHYDROGENASE |
| 89:00:00 | 1izn-C | 2.9 | 3.6 | 73 | 273 | 5 | MOLECULE: | CAPZ AL | PHA-1 S | UBUNIT |
| 90:00:00 | 2imj-A | 2.8 | 3.1 | 71 | 158 | 4 | MOLECULE: | HYPOTHE | TICAL P | ROTEIN DUF1348 |
| 91:00:00 | 7k5n-A | 2.8 | 2.8 | 59 | 238 | 10 | MOLECULE: | SENSOR | DOMAIN- | CONTAINING DIGUANYLATE CYCLASE |
| 92:00:00 | 4i5q-A | 2.8 | 3.2 | 51 | 220 | 4 | MOLECULE: | THIOL:D | ISULFID | E INTERCHANGE PROTEIN DSBC |
| 93:00:00 | 4xzv-B | 2.8 | 3.9 | 64 | 142 | 5 | MOLECULE: | MALTOSE | #NAME? | G PERIPLASMIC PROTEIN,TP53-REGULATE |
| 94:00:00 | 3rob-A | 2.8 | 4.1 | 69 | 131 | 3 | MOLECULE: | UNCHARA | CTERIZE | D CONSERVED PROTEIN |
| 95:00:00 | 3bws-A | 2.8 | 3.3 | 61 | 407 | 15 | MOLECULE: | PROTEIN | LP49 | |
| 96:00:00 | 1vi8-B | 2.8 | 3.9 | 68 | 146 | 9 | MOLECULE: | HYPOTHE | TICAL P | ROTEIN YDII |
| 97:00:00 | 3hzp-A | 2.8 | 3.3 | 64 | 128 | 6 | MOLECULE: | NTF2-LI | KE PROT | EIN OF UNKNOWN FUNCTION |
| 98:00:00 | 8ghn-A | 2.8 | 3.6 | 59 | 667 | 5 | MOLECULE: | PROTEIN | HIR1 | |
| 99:00:00 | 2gu1-A | 2.8 | 3.7 | 62 | 324 | 11 | MOLECULE: | ZINC PE | PTIDASE | |
| 0:00 | 2ia7-A | 2.8 | 3.2 | 62 | 111 | 6 | MOLECULE: | TAIL LY | SOZYME, | PUTATIVE |
| 1:00 | 2gvi-A | 2.8 | 3.5 | 65 | 201 | 9 | MOLECULE: | CONSERV | ED HYPO | THETICAL PROTEIN |
| 2:00 | 2r5x-B | 2.8 | 3.2 | 55 | 118 | 5 | MOLECULE: | UNCHARA | CTERIZE | D CONSERVED PROTEIN |
| 3:00 | 7t5q-J | 2.8 | 3.6 | 72 | 291 | 3 | MOLECULE: | ACTIN-R | ELATED | PROTEIN 3 |
| 4:00 | 7xfk-C | 2.8 | 3.7 | 65 | 132 | 14 | MOLECULE: | SORDARI | N/HYPOX | YSORDARIN BIOSYNTHESIS CLUSTER PROT |
| 5:00 | 7aed-B | 2.8 | 3 | 58 | 138 | 5 | MOLECULE: | PRGL | | |
| 6:00 | 4hz9-B | 2.8 | 2.5 | 61 | 123 | 11 | MOLECULE: | PUTATIV | E CYTOP | LASMIC PROTEIN |
| 7:00 | 6xrb-A | 2.8 | 4.3 | 67 | 150 | 6 | MOLECULE: | SCIW | | |
| 8:00 | 8p32-D | 2.8 | 3.4 | 65 | 125 | 3 | MOLECULE: | BB0238 | | |
| 9:00 | 3put-B | 2.8 | 3.9 | 66 | 155 | 5 | MOLECULE: | HYPOTHE | TICAL C | ONSERVED PROTEIN |
| 10:00 | 4l8p-A | 2.8 | 3.3 | 73 | 168 | 3 | MOLECULE: | BILE AC | ID 7A-D | EHYDRATASE, BAIE |
| 11:00 | 6khi-V | 2.8 | 3 | 63 | 109 | 5 | MOLECULE: | NAD(P)H | #NAME? | E OXIDOREDUCTASE SUBUNIT 1 |
| 12:00 | 3by9-A | 2.8 | 2.7 | 54 | 259 | 7 | MOLECULE: | SENSOR | PROTEIN | |
| 13:00 | 8a9y-E | 2.8 | 3.5 | 64 | 497 | 3 | MOLECULE: | GLYCOSI | DE HYDR | OLASE FAMILY 32 |
| 14:00 | 2iv9-A | 2.8 | 3.1 | 57 | 236 | 0 | MOLECULE: | AP-2 CO | MPLEX S | UBUNIT BETA-2 |
| 15:00 | 5njb-B | 2.8 | 2.9 | 62 | 445 | 5 | MOLECULE: | METALLO | PROTEAS | E TLDD |
| 16:00 | 6moc-A | 2.8 | 2.8 | 57 | 363 | 5 | MOLECULE: | PEPTIDA | SE M23 | |
| 17:00 | 1nyc-A | 2.8 | 3.8 | 67 | 111 | 9 | MOLECULE: | CYSTEIN | E PROTE | ASE INHIBITOR |
| 18:00 | 5iw9-A | 2.8 | 3.2 | 60 | 126 | 10 | MOLECULE: | BASEPLA | TE WEDG | E PROTEIN GP25 |
| 19:00 | 8jxe-A | 2.8 | 3 | 62 | 1469 | 10 | MOLECULE: | LDL REC | EPTOR R | ELATED PROTEIN 2 |
| 20:00 | 3v7d-B | 2.8 | 2.5 | 46 | 450 | 11 | MOLECULE: | SUPPRES | SOR OF | KINETOCHORE PROTEIN 1 |
| 21:00 | 8t5j-A | 2.8 | 3.1 | 48 | 154 | 8 | MOLECULE: | OUTER M | EMBRANE | LIPOCARRIER LOLA FAMILY PROTEIN |
| 22:00 | 3ehc-B | 2.8 | 3.5 | 70 | 128 | 11 | MOLECULE: | SNOAL-L | IKE POL | YKETIDE CYCLASE |
| 23:00 | 3hk4-A | 2.8 | 3.5 | 67 | 118 | 12 | MOLECULE: | MLR7391 | PROTEI | N |
| 24:00:00 | 5edf-A | 2.8 | 3 | 54 | 224 | 4 | MOLECULE: | FRPC OP | ERON PR | OTEIN |

| | | | | | | | | | | |
|---|---|---|---|---|---|---|---|---|---|---|
| 25:00:00 | 6pny-A | 2.8 | 4 | 66 | 116 | 12 | MOLECULE: | FLPP3 | | |
| 26:00:00 | 7erl-A | 2.8 | 3.4 | 63 | 545 | 8 | MOLECULE: | BETA-XY | LANASE | |
| 27:00:00 | 2gey-A | 2.8 | 3.5 | 70 | 157 | 6 | MOLECULE: | ACLR PR | OTEIN | |
| 28:00:00 | 6h3i-F | 2.8 | 3.7 | 66 | 345 | 8 | MOLECULE: | PROTEIN | INVOLV | ED IN GLIDING MOTILITY SPRA |
| 29:00:00 | 3gzr-A | 2.8 | 3.6 | 64 | 143 | 9 | MOLECULE: | UNCHARA | CTERIZE | D PROTEIN WITH A NTF2-LIKE FOLD |
| 30:00:00 | 3f40-A | 2.8 | 3.8 | 62 | 112 | 10 | MOLECULE: | UNCHARA | CTERIZE | D NTF2-LIKE PROTEIN |
| 31:00:00 | 6i7s-G | 2.8 | 5.1 | 72 | 863 | 4 | MOLECULE: | PROTEIN | DISULF | IDE-ISOMERASE |
| 32:00:00 | 3vsf-C | 2.8 | 3.3 | 63 | 482 | 6 | MOLECULE: | RICIN B | LECTIN | |
| 33:00:00 | 6zlt-B | 2.8 | 7.1 | 65 | 933 | 3 | MOLECULE: | SUSD HO | MOLOG | |
| 34:00:00 | 8gex-A | 2.8 | 3.6 | 59 | 411 | 7 | MOLECULE: | DUF4374 | DOMAIN | -CONTAINING PROTEIN |
| 35:00:00 | 8ye0-C | 2.8 | 3.4 | 70 | 290 | 6 | MOLECULE: | LYNF/TR | UF/PATF | FAMILY PEPTIDE O-PRENYLTRANSFERASE |
| 36:00:00 | 3uv1-A | 2.8 | 3.2 | 71 | 190 | 6 | MOLECULE: | DER F 7 | ALLERG | EN |
| 37:00:00 | 7akv-A | 2.8 | 2.5 | 59 | 375 | 7 | MOLECULE: | PLASMA | PROTEAS | E C1 INHIBITOR |
| 38:00:00 | 8dwj-A | 2.8 | 4 | 58 | 339 | 10 | MOLECULE: | RNA(5'- | PPP-GCC | GA-3') |
| 39:00:00 | 6nhi-A | 2.8 | 4.2 | 64 | 448 | 6 | MOLECULE: | HISTIDI | NE--TRN | A LIGASE |
| 40:00:00 | 3df6-B | 2.8 | 3 | 56 | 99 | 13 | MOLECULE: | ORF99 | | |
| 41:00:00 | 3s5t-A | 2.8 | 3.7 | 69 | 253 | 7 | MOLECULE: | DUF3298 | FAMILY | PROTEIN |
| 42:00:00 | 4i0o-A | 2.8 | 2.6 | 47 | 463 | 11 | MOLECULE: | PROTEIN | ELYS | |
| 43:00:00 | 6yai-E | 2.8 | 3.8 | 66 | 726 | 5 | MOLECULE: | CLATHRI | N HEAVY | CHAIN |
| 44:00:00 | 6s9u-A | 2.8 | 3.9 | 69 | 520 | 9 | MOLECULE: | PUTATIV | E SUCRO | SE PHOSPHORYLASE |
| 45:00:00 | 5hi8-A | 2.8 | 3.4 | 73 | 136 | 5 | MOLECULE: | ANTENNA | PROTEI | N |
| 46:00:00 | 4v1a-i | 2.8 | 3.6 | 74 | 242 | 8 | MOLECULE: | MITORIB | OSOMAL | PROTEIN ML37, MRPL37 |
| 47:00:00 | 8bbg-B | 2.8 | 3.3 | 58 | 1426 | 9 | MOLECULE: | WD REPE | AT-CONT | AINING PROTEIN 19 |
| 48:00:00 | 7qpg-R | 2.8 | 3 | 55 | 2208 | 7 | MOLECULE: | PROTEIN | ZWILCH | HOMOLOG |
| 49:00:00 | 5hlb-A | 2.8 | 5 | 58 | 733 | 12 | MOLECULE: | PENICIL | LIN-BIN | DING PROTEIN 1B |
| 50:00:00 | 4ocu-A | 2.8 | 2.3 | 52 | 356 | 6 | MOLECULE: | N-ACETY | LHEXOSA | MINE 1-PHOSPHATE KINASE |
| 51:00:00 | 4irt-A | 2.8 | 3 | 57 | 398 | 7 | MOLECULE: | UNCHARA | CTERIZE | D PROTEIN |
| 52:00:00 | 6m36-I | 2.8 | 4.1 | 55 | 104 | 11 | MOLECULE: | SERINE- | PROTEIN | KINASE RSBW |
| 53:00:00 | 3b7c-A | 2.8 | 3.6 | 69 | 121 | 10 | MOLECULE: | UNCHARA | CTERIZE | D PROTEIN |
| 54:00:00 | 7eeb-F | 2.8 | 2.7 | 54 | 1042 | 7 | MOLECULE: | ENHANCE | D GREEN | FLUORESCENT PROTEIN,CATION CHANNEL |
| 55:00:00 | 5h2d-A | 2.8 | 7 | 62 | 418 | 8 | MOLECULE: | KLLA0C0 | 4147P | |
| 56:00:00 | 8rf0-A | 2.8 | 3 | 50 | 2780 | 12 | MOLECULE: | CYCLIC | BETA-(1 | ,2)-GLUCAN SYNTHASE NDVB |
| 57:00:00 | 5ogs-A | 2.8 | 3.3 | 59 | 403 | 7 | MOLECULE: | WD REPE | AT AND | HMG-BOX DNA-BINDING PROTEIN 1 |
| 58:00:00 | 3aae-G | 2.8 | 3.6 | 73 | 270 | 5 | MOLECULE: | F-ACTIN | #NAME? | G PROTEIN SUBUNIT ALPHA-1 |
| 59:00:00 | 4fpc-A | 2.8 | 3.3 | 63 | 658 | 2 | MOLECULE: | SIALIDA | SE B | |
| 60:00:00 | 6fej-B | 2.8 | 2.7 | 57 | 118 | 9 | MOLECULE: | ALL4940 | PROTEI | N |
| 61:00:00 | 7yth-C | 2.8 | 3.1 | 63 | 314 | 8 | MOLECULE: | KETOSTE | ROID IS | OMERASE-RELATED PROTEIN |
| 62:00:00 | 4fcm-A | 2.8 | 3 | 64 | 132 | 6 | MOLECULE: | RAS GTP | ASE-ACT | IVATING PROTEIN-BINDING PROTEIN 1 |
| 63:00:00 | 6f1u-K | 2.8 | 3.6 | 74 | 278 | 7 | MOLECULE: | ARP1 AC | TIN REL | ATED PROTEIN 1 HOMOLOG A |
| 64:00:00 | 2pup-A | 2.8 | 2.4 | 52 | 371 | 4 | MOLECULE: | METHYLT | HIORIBO | SE KINASE |
| 65:00:00 | 6w40-B | 2.8 | 3.2 | 62 | 115 | 16 | MOLECULE: | DENOVO | NTF2 | |
| 66:00:00 | 8th5-F | 2.8 | 3.7 | 69 | 130 | 7 | MOLECULE: | RAS GTP | ASE-ACT | IVATING PROTEIN-BINDING PROTEIN 1 |
| 67:00:00 | 2ya6-A | 2.8 | 3.1 | 60 | 470 | 10 | MOLECULE: | NEURAMI | NIDASE | A |
| 68:00:00 | 2woh-A | 2.8 | 3.3 | 68 | 718 | 10 | MOLECULE: | PRIMARY | AMINE | OXIDASE |
| 69:00:00 | 5ohl-E | 2.8 | 3.7 | 58 | 89 | 5 | MOLECULE: | K6-SPEC | IFIC AF | FIMER |
| 70:00:00 | 7zhu-A | 2.8 | 4.6 | 65 | 534 | 3 | MOLECULE: | GLUCOSE | #NAME? | PHATE 1-DEHYDROGENASE |
| 71:00:00 | 5kky-B | 2.8 | 3.6 | 71 | 470 | 4 | MOLECULE: | SIALIDA | SE A | |
| 72:00:00 | 7xhg-D | 2.8 | 3.5 | 65 | 133 | 6 | MOLECULE: | RAS GTP | ASE-ACT | IVATING PROTEIN-BINDING PROTEIN 1 |
| 73:00:00 | 6d24-A | 2.8 | 4.6 | 73 | 502 | 4 | MOLECULE: | GLUCOSE | #NAME? | PHATE 1-DEHYDROGENASE |
| 74:00:00 | 3pjq-A | 2.8 | 3.1 | 61 | 626 | 7 | MOLECULE: | TRANS-S | IALIDAS | E |
| 75:00:00 | 4xio-A | 2.8 | 3.4 | 60 | 658 | 7 | MOLECULE: | SIALIDA | SE B | |
| 76:00:00 | 1d6y-A | 2.8 | 3.1 | 65 | 718 | 8 | MOLECULE: | COPPER | AMINE O | XIDASE |
| 77:00:00 | 2xhn-A | 2.8 | 3 | 47 | 508 | 6 | MOLECULE: | RHAMNOG | ALACTUR | ONASE B |
| 78:00:00 | 6f1t-K | 2.8 | 3.6 | 74 | 278 | 7 | MOLECULE: | ARP1 AC | TIN REL | ATED PROTEIN 1 HOMOLOG A |

| | | | | | | | | | | |
|---|---|---|---|---|---|---|---|---|---|---|
| 79:00:00 | 1cyu-A | 2.8 | 3 | 59 | 98 | 5 | MOLECULE: | CYSTATI | N A | |
| 80:00:00 | 2xgl-A | 2.7 | 2.8 | 60 | 90 | 0 | MOLECULE: | COLICIN | #NAME? | NITY PROTEIN |
| 81:00:00 | 6sum-A | 2.7 | 3 | 57 | 335 | 2 | MOLECULE: | AMICOUM | ACIN KI | NASE |
| 82:00:00 | 7jsr-A | 2.7 | 3.8 | 78 | 1496 | 5 | MOLECULE: | NAD-SPE | CIFIC G | LUTAMATE DEHYDROGENASE |
| 83:00:00 | 2o3o-I | 2.7 | 3.2 | 59 | 244 | 5 | MOLECULE: | YYCI PR | OTEIN | |
| 84:00:00 | 4fvs-A | 2.7 | 3.7 | 81 | 214 | 6 | MOLECULE: | PUTATIV | E LIPOP | ROTEIN |
| 85:00:00 | 6zxf-z | 2.7 | 3.7 | 66 | 373 | 5 | MOLECULE: | PRE-18S | RIBOSO | MAL RNA |
| 86:00:00 | 3w9k-A | 2.7 | 3.8 | 61 | 135 | 7 | MOLECULE: | FATTY A | CID-BIN | DING PROTEIN |
| 87:00:00 | 1eq6-A | 2.7 | 3.4 | 72 | 189 | 8 | MOLECULE: | MOG1P | | |
| 88:00:00 | 6gn5-A | 2.7 | 3.8 | 67 | 177 | 6 | MOLECULE: | GRAM DO | MAIN-CO | NTAINING PROTEIN 1C |
| 89:00:00 | 4plp-A | 2.7 | 3 | 74 | 475 | 7 | MOLECULE: | HOMOSPE | RMIDINE | SYNTHASE |
| 90:00:00 | 4l9h-A | 2.7 | 4 | 69 | 155 | 4 | MOLECULE: | F-BOX O | NLY PRO | TEIN 7 |
| 91:00:00 | 6kac-P | 2.7 | 3.6 | 71 | 188 | 7 | MOLECULE: | PHOTOSY | STEM II | PROTEIN D1 |
| 92:00:00 | 3ub9-B | 2.7 | 2.5 | 57 | 165 | 7 | MOLECULE: | CHEMORE | CEPTOR | TLPB |
| 93:00:00 | 8qc6-A | 2.7 | 3.5 | 62 | 366 | 5 | MOLECULE: | OXIDORE | DUCTASE | |
| 94:00:00 | 1aom-B | 2.7 | 3.4 | 64 | 559 | 9 | MOLECULE: | NITRITE | REDUCT | ASE |
| 95:00:00 | 3g8z-A | 2.7 | 3.8 | 64 | 129 | 11 | MOLECULE: | PROTEIN | OF UNK | NOWN FUNCTION WITH CYSTATIN-LIKE FO |
| 96:00:00 | 3i0o-A | 2.7 | 3 | 58 | 329 | 9 | MOLECULE: | SPECTIN | OMYCIN | PHOSPHOTRANSFERASE |
| 97:00:00 | 2bmo-B | 2.7 | 3.5 | 70 | 194 | 10 | MOLECULE: | OXYGENA | SE-ALPH | A NBDO |
| 98:00:00 | 5g4y-A | 2.7 | 2.1 | 56 | 147 | 5 | MOLECULE: | CHEMOTA | XIS PRO | TEIN |
| 99:00:00 | 7qh5-A | 2.7 | 3.9 | 73 | 313 | 7 | MOLECULE: | RNA POL | YMERASE | SIGMA FACTOR |
| 0:00 | 2xlq-A | 2.7 | 3.7 | 67 | 316 | 3 | MOLECULE: | CLOQ | | |
| 1:00 | 6tlb-A | 2.7 | 4.1 | 69 | 185 | 13 | MOLECULE: | SERINE/ | THREONI | NE PROTEIN KINASE |
| 2:00 | 3byv-A | 2.7 | 3.9 | 63 | 345 | 8 | MOLECULE: | RHOPTRY | KINASE | |
| 3:00 | 3hw2-A | 2.7 | 4.8 | 56 | 308 | 9 | MOLECULE: | PROTEIN | SIFA | |
| 4:00 | 8bo9-A | 2.7 | 3.2 | 58 | 174 | 9 | MOLECULE: | NON STR | UCTURAL | POLYPROTEIN |
| 5:00 | 6eus-A | 2.7 | 3.9 | 56 | 346 | 5 | MOLECULE: | DCAP-LI | KE PROT | EIN |
| 6:00 | 6qp8-A | 2.7 | 3.6 | 66 | 622 | 3 | MOLECULE: | IP13724 | P | |
| 7:00 | 7q4p-C | 2.7 | 3 | 61 | 878 | 7 | MOLECULE: | SPLICIN | G FACTO | R 3A SUBUNIT 2 |
| 8:00 | 4r5o-A | 2.7 | 3.6 | 64 | 426 | 6 | MOLECULE: | QUINONP | ROTEIN | ALCOHOL DEHYDROGENASE-LIKE PROTEIN |
| 9:00 | 8hmc-B | 2.7 | 3.8 | 58 | 1195 | 5 | MOLECULE: | INTRAFL | AGELLAR | TRANSPORT PROTEIN 122 HOMOLOG |
| 10:00 | 6nd4-Q | 2.7 | 3.3 | 59 | 862 | 12 | MOLECULE: | ETS RRN | A | |
| 11:00 | 8wid-w | 2.7 | 3.4 | 60 | 126 | 10 | MOLECULE: | 16S RRN | A | |
| 12:00 | 3mlq-A | 2.7 | 4 | 60 | 186 | 5 | MOLECULE: | DNA-DIR | ECTED R | NA POLYMERASE SUBUNIT BETA |
| 13:00 | 6b7l-A | 2.7 | 8.4 | 51 | 290 | 4 | MOLECULE: | IMMUNE | MODULAT | OR A |
| 14:00 | 8alo-B | 2.7 | 3.9 | 60 | 148 | 12 | MOLECULE: | CHOLERA | TOXIN | TRANSCRIPTIONAL ACTIVATOR |
| 15:00 | 7jta-A | 2.7 | 3.2 | 48 | 56 | 2 | MOLECULE: | NTF2-LI | KE NUCL | EASE/ANTI-CRISPR |
| 16:00 | 3jr1-A | 2.7 | 2.1 | 53 | 296 | 8 | MOLECULE: | PUTATIV | E FRUCT | OSAMINE-3-KINASE |
| 17:00 | 8vwp-A | 2.7 | 4.1 | 65 | 502 | 9 | MOLECULE: | LANGYA | VIRUS A | TTACHMENT (G) PROTEIN |
| 18:00 | 6xp5-N | 2.7 | 3.4 | 61 | 544 | 2 | MOLECULE: | MEDIATO | R OF RN | A POLYMERASE II TRANSCRIPTION SUBUN |
| 19:00 | 3tkn-A | 2.7 | 4.2 | 58 | 451 | 9 | MOLECULE: | NUCLEOP | ORIN NU | P82 |
| 20:00 | 7nie-C | 2.7 | 5.3 | 61 | 294 | 10 | MOLECULE: | GLYCERO | L KINAS | E |
| 21:00 | 6m76-A | 2.7 | 3.3 | 64 | 923 | 5 | MOLECULE: | LPXTG-M | OTIF CE | LL WALL ANCHOR DOMAIN PROTEIN |
| 22:00 | 6f37-A | 2.7 | 2.7 | 55 | 113 | 5 | MOLECULE: | NANO3,F | UCOSE-B | INDING LECTIN PROTEIN |
| 23:00 | 5c9p-A | 2.7 | 3 | 59 | 354 | 10 | MOLECULE: | PLL LEC | TIN | |
| 24:00:00 | 4hxg-F | 2.7 | 4.1 | 62 | 614 | 5 | MOLECULE: | PUTATIV | E UNCHA | RACTERIZED PROTEIN PH0594 |
| 25:00:00 | 7n3v-A | 2.7 | 2.8 | 48 | 266 | 4 | MOLECULE: | LMCA | | |
| 26:00:00 | 2fkj-A | 2.7 | 4.4 | 63 | 361 | 5 | MOLECULE: | OUTER S | URFACE | PROTEIN A |
| 27:00:00 | 5x7o-A | 2.7 | 3.1 | 56 | 1247 | 7 | MOLECULE: | GLYCOSI | DE HYDR | OLASE FAMILY 31 ALPHA-GLUCOSIDASE |
| 28:00:00 | 6exp-F | 2.7 | 3.2 | 60 | 103 | 2 | MOLECULE: | SIRV3 A | CRID1 ( | GP02) ANTI-CRISPR PROTEIN |
| 29:00:00 | 5yje-B | 2.7 | 3.5 | 57 | 318 | 7 | MOLECULE: | PROTEIN | HIRA | |
| 30:00:00 | 6hxv-A | 2.7 | 1.5 | 46 | 163 | 4 | MOLECULE: | COILED- | COIL DO | MAIN-CONTAINING PROTEIN 61 |
| 31:00:00 | 6eu4-A | 2.7 | 4.5 | 57 | 587 | 7 | MOLECULE: | TAIL SP | IKE PRO | TEIN |
| 32:00:00 | 2p9w-A | 2.7 | 3.4 | 59 | 333 | 7 | MOLECULE: | MAL S 1 | ALLERG | ENIC PROTEIN |

| | | | | | | | | | | |
|---|---|---|---|---|---|---|---|---|---|---|
| 33:00:00 | 7ep9-A | 2.7 | 3.7 | 57 | 660 | 12 | MOLECULE: | S9 FAMI | LY PEPT | IDASE |
| 34:00:00 | 4ww9-A | 2.7 | 3.4 | 52 | 238 | 6 | MOLECULE: | EKC/KEO | PS COMP | LEX SUBUNIT BUD32 |
| 35:00:00 | 7r78-A | 2.7 | 5.7 | 65 | 1881 | 3 | MOLECULE: | DNA REP | AIR PRO | TEIN RAD8 |
| 36:00:00 | 6jdp-A | 2.7 | 4.5 | 70 | 241 | 6 | MOLECULE: | IMM52 F | AMILY P | ROTEIN |
| 37:00:00 | 2h0h-B | 2.7 | 3.1 | 54 | 230 | 9 | MOLECULE: | THIOL:D | ISULFID | E INTERCHANGE PROTEIN DSBG |
| 38:00:00 | 8uzl-A | 2.7 | 5.9 | 62 | 143 | 10 | MOLECULE: | DESIGNE | D TRANS | MEMBRANE BETA-BARREL TMB10_163 |
| 39:00:00 | 3ci0-K | 2.7 | 3.4 | 60 | 280 | 5 | MOLECULE: | PSEUDOP | ILIN GS | PI |
| 40:00:00 | 5gva-A | 2.7 | 3.1 | 56 | 305 | 5 | MOLECULE: | WD REPE | AT AND | HMG-BOX DNA-BINDING PROTEIN 1 |
| 41:00:00 | 4xmi-A | 2.7 | 3.3 | 63 | 657 | 2 | MOLECULE: | SIALIDA | SE B | |
| 42:00:00 | 2chc-B | 2.7 | 3.2 | 62 | 161 | 5 | MOLECULE: | PROTEIN | RV3472 | |
| 43:00:00 | 7mhu-B | 2.7 | 3.6 | 62 | 374 | 3 | MOLECULE: | EXO-ALP | HA-SIAL | IDASE |
| 44:00:00 | 6q0v-B | 2.7 | 3.9 | 66 | 208 | 11 | MOLECULE: | DNA DAM | AGE-BIN | DING PROTEIN 1 |
| 45:00:00 | 7dmp-b | 2.7 | 3.3 | 63 | 336 | 8 | MOLECULE: | RADIAL | SPOKE H | EAD PROTEIN 4 HOMOLOG A |
| 46:00:00 | 1gd3-A | 2.7 | 3.7 | 67 | 98 | 4 | MOLECULE: | CYSTATI | N A | |
| 47:00:00 | 4yw4-A | 2.7 | 3.3 | 56 | 659 | 2 | MOLECULE: | NEURAMI | NIDASE | C |
| 48:00:00 | 6eks-A | 2.7 | 2.6 | 66 | 749 | 8 | MOLECULE: | SIALIDA | SE | |
| 49:00:00 | 7yth-D | 2.7 | 3.5 | 68 | 311 | 10 | MOLECULE: | KETOSTE | ROID IS | OMERASE-RELATED PROTEIN |
| 50:00:00 | 2wo0-A | 2.7 | 3.3 | 67 | 718 | 9 | MOLECULE: | PRIMARY | AMINE | OXIDASE |
| 51:00:00 | 3vd9-B | 2.7 | 3.5 | 71 | 1016 | 8 | MOLECULE: | BETA-GA | LACTOSI | DASE |
| 52:00:00 | 8v1l-B | 2.7 | 3.2 | 66 | 133 | 6 | MOLECULE: | RAS GTP | ASE-ACT | IVATING PROTEIN-BINDING PROTEIN 1 |
| 53:00:00 | 7zht-A | 2.7 | 4.5 | 67 | 530 | 3 | MOLECULE: | GLUCOSE | #NAME? | PHATE 1-DEHYDROGENASE |
| 54:00:00 | 4ovm-D | 2.7 | 2.9 | 64 | 125 | 13 | MOLECULE: | UNCHARA | CTERIZE | D PROTEIN SGCJ |
| 55:00:00 | 5ohl-G | 2.7 | 3.8 | 56 | 89 | 5 | MOLECULE: | K6-SPEC | IFIC AF | FIMER |
| 56:00:00 | 5ohl-D | 2.7 | 3.8 | 58 | 89 | 5 | MOLECULE: | K6-SPEC | IFIC AF | FIMER |
| 57:00:00 | 4dux-A | 2.7 | 3.6 | 69 | 1015 | 9 | MOLECULE: | BETA-GA | LACTOSI | DASE |
| 58:00:00 | 2vw1-A | 2.7 | 3 | 57 | 658 | 0 | MOLECULE: | SIALIDA | SE B | |
| 59:00:00 | 2ya7-A | 2.7 | 3.5 | 67 | 469 | 6 | MOLECULE: | NEURAMI | NIDASE | A |
| 60:00:00 | 2wgq-B | 2.7 | 3.5 | 58 | 723 | 9 | MOLECULE: | AMINE O | XIDASE | |
| 61:00:00 | 8th1-B | 2.7 | 3.3 | 66 | 133 | 6 | MOLECULE: | RAS GTP | ASE-ACT | IVATING PROTEIN-BINDING PROTEIN 1 |
| 62:00:00 | 7xhf-B | 2.7 | 3.4 | 67 | 134 | 6 | MOLECULE: | RAS GTP | ASE-ACT | IVATING PROTEIN-BINDING PROTEIN 1 |
| 63:00:00 | 3opz-C | 2.7 | 3.1 | 59 | 625 | 7 | MOLECULE: | TRANS-S | IALIDAS | E |
| 64:00:00 | 4ovm-H | 2.7 | 3 | 64 | 126 | 13 | MOLECULE: | UNCHARA | CTERIZE | D PROTEIN SGCJ |
| 65:00:00 | 4xhb-A | 2.7 | 2.9 | 55 | 658 | 0 | MOLECULE: | SIALIDA | SE B | |
| 66:00:00 | 5f9t-B | 2.7 | 3.3 | 56 | 659 | 2 | MOLECULE: | NEURAMI | NIDASE | C |
| 67:00:00 | 4yw5-A | 2.7 | 2.7 | 46 | 659 | 2 | MOLECULE: | NEURAMI | NIDASE | C |
| 68:00:00 | 3ob8-C | 2.7 | 4.3 | 51 | 1024 | 8 | MOLECULE: | BETA-GA | LACTOSI | DASE |
| 69:00:00 | 1eur-A | 2.7 | 3.3 | 65 | 361 | 15 | MOLECULE: | SIALIDA | SE | |
| 70:00:00 | 3elg-B | 2.7 | 4.1 | 55 | 126 | 5 | MOLECULE: | UNCHARA | CTERIZE | D PERIPLASMIC PROTEIN |
| 71:00:00 | 4xil-A | 2.7 | 3 | 56 | 658 | 0 | MOLECULE: | SIALIDA | SE B | |
| 72:00:00 | 1gyb-B | 2.7 | 3.6 | 64 | 122 | 8 | MOLECULE: | NUCLEAR | TRANSP | ORT FACTOR 2 |
| 73:00:00 | 4yw4-B | 2.7 | 2.9 | 54 | 659 | 2 | MOLECULE: | NEURAMI | NIDASE | C |
| 74:00:00 | 4yz2-B | 2.7 | 3.3 | 56 | 656 | 2 | MOLECULE: | SIALIDA | SE NANC | |
| 75:00:00 | 3vd9-C | 2.7 | 3.1 | 64 | 1015 | 6 | MOLECULE: | BETA-GA | LACTOSI | DASE |
| 76:00:00 | 4fpo-B | 2.7 | 3.1 | 58 | 658 | 0 | MOLECULE: | SIALIDA | SE B | |
| 77:00:00 | 8p26-J | 2.6 | 2.4 | 51 | 147 | 12 | MOLECULE: | U2 SMAL | L NUCLE | AR RIBONUCLEOPROTEIN AUXILIARY FACT |
| 78:00:00 | 5ezu-A | 2.6 | 2.8 | 50 | 76 | 14 | MOLECULE: | PROTEIN | A46 | |
| 79:00:00 | 8abv-A | 2.6 | 3.3 | 71 | 244 | 7 | MOLECULE: | SNOAL-L | IKE DOM | AIN-CONTAINING PROTEIN |
| 80:00:00 | 3cnx-C | 2.6 | 3.1 | 61 | 148 | 3 | MOLECULE: | UNCHARA | CTERIZE | D PROTEIN |
| 81:00:00 | 2obd-A | 2.6 | 4 | 74 | 472 | 7 | MOLECULE: | CHOLEST | ERYL ES | TER TRANSFER PROTEIN |
| 82:00:00 | 4x2m-A | 2.6 | 3.3 | 67 | 177 | 3 | MOLECULE: | MTR2 | | |
| 83:00:00 | 2lnj-A | 2.6 | 4.9 | 74 | 170 | 1 | MOLECULE: | PUTATIV | E UNCHA | RACTERIZED PROTEIN SLL1418 |
| 84:00:00 | 6kaw-A | 2.6 | 3.6 | 70 | 381 | 6 | MOLECULE: | CGHA | | |
| 85:00:00 | 6hsy-A | 2.6 | 3.4 | 67 | 190 | 6 | MOLECULE: | TOLUENE | TOLERA | NCE PROTEIN TTG2D |
| 86:00:00 | 4y4v-A | 2.6 | 3 | 65 | 318 | 3 | MOLECULE: | CONSERV | ED HYPO | THETICAL SECRETED PROTEIN |

| | | | | | | | | | | |
|---|---|---|---|---|---|---|---|---|---|---|
| 87:00:00 | 5kku-B | 2.6 | 4 | 61 | 289 | 0 | MOLECULE: | POLYKET | IDE SYN | THASE TYPE I |
| 88:00:00 | 7w0w-A | 2.6 | 3.4 | 56 | 131 | 9 | MOLECULE: | METHYL- | ACCEPTI | NG CHEMOTAXIS PROTEIN |
| 89:00:00 | 8gqe-A | 2.6 | 3.5 | 59 | 348 | 8 | MOLECULE: | WD REPE | AT-CONT | AINING PROTEIN RUP2 |
| 90:00:00 | 2y3u-A | 2.6 | 3.9 | 53 | 679 | 2 | MOLECULE: | COLLAGE | NASE | |
| 91:00:00 | 7apk-m | 2.6 | 3.6 | 66 | 549 | 8 | MOLECULE: | THO COM | PLEX SU | BUNIT 1 |
| 92:00:00 | 4h3u-A | 2.6 | 3.3 | 65 | 131 | 6 | MOLECULE: | HYPOTHE | TICAL P | ROTEIN |
| 93:00:00 | 7yzi-A | 2.6 | 3.6 | 65 | 378 | 6 | MOLECULE: | ADENYLA | TE CYCL | ASE |
| 94:00:00 | 1s18-A | 2.6 | 3.2 | 64 | 317 | 6 | MOLECULE: | APYRASE | | |
| 95:00:00 | 2a8e-A | 2.6 | 3.9 | 77 | 210 | 8 | MOLECULE: | HYPOTHE | TICAL P | ROTEIN YKTB |
| 96:00:00 | 3vu4-A | 2.6 | 3.4 | 59 | 319 | 7 | MOLECULE: | KMHSV2 | | |
| 97:00:00 | 7pab-A | 2.6 | 3.8 | 62 | 415 | 6 | MOLECULE: | NUCLEAR | EGRESS | PROTEIN 2,NUCLEAR EGRESS PROTEIN 1 |
| 98:00:00 | 1k1x-A | 2.6 | 3.3 | 71 | 636 | 6 | MOLECULE: | 4-ALPHA | #NAME? | OTRANSFERASE |
| 99:00:00 | 2kcd-A | 2.6 | 3.4 | 69 | 120 | 6 | MOLECULE: | UNCHARA | CTERIZE | D PROTEIN SSP0047 |
| 0:00 | 5h3z-A | 2.6 | 6.7 | 55 | 1113 | 2 | MOLECULE: | UNCHARA | CTERIZE | D PROTEIN |
| 1:00 | 6kml-C | 2.6 | 3.7 | 52 | 105 | 4 | MOLECULE: | MRNA IN | TERFERA | SE TOXIN HIGB |
| 2:00 | 8t1l-L | 2.6 | 3.4 | 66 | 559 | 6 | MOLECULE: | MEDIATO | R OF RN | A POLYMERASE II TRANSCRIPTION SUBUN |
| 3:00 | 3dkq-A | 2.6 | 3.2 | 60 | 230 | 7 | MOLECULE: | PKHD-TY | PE HYDR | OXYLASE SBAL_3634 |
| 4:00 | 1pu1-A | 2.6 | 3.7 | 59 | 91 | 10 | MOLECULE: | HYPOTHE | TICAL P | ROTEIN MTH677 |
| 5:00 | 1r5m-A | 2.6 | 3.4 | 57 | 351 | 12 | MOLECULE: | SIR4-IN | TERACTI | NG PROTEIN SIF2 |
| 6:00 | 6nd4-O | 2.6 | 4.3 | 60 | 832 | 12 | MOLECULE: | ETS RRN | A | |
| 7:00 | 7uhy-C | 2.6 | 3.1 | 57 | 578 | 7 | MOLECULE: | GATOR C | OMPLEX | PROTEIN MIOS |
| 8:00 | 4o9d-B | 2.6 | 3.1 | 57 | 392 | 7 | MOLECULE: | RIK1-AS | SOCIATE | D FACTOR 1 |
| 9:00 | 2rk0-A | 2.6 | 3.5 | 51 | 135 | 2 | MOLECULE: | GLYOXAL | ASE/BLE | OMYCIN RESISTANCE PROTEIN/DIOXYGENA |
| 10:00 | 7zty-A | 2.6 | 3.6 | 54 | 345 | 4 | MOLECULE: | CNH DOM | AIN-CON | TAINING PROTEIN |
| 11:00 | 5mc1-B | 2.6 | 3.9 | 73 | 487 | 12 | MOLECULE: | XAA-PRO | DIPEPT | IDASE |
| 12:00 | 3cyg-A | 2.6 | 3.9 | 64 | 215 | 5 | MOLECULE: | UNCHARA | CTERIZE | D PROTEIN |
| 13:00 | 6rwb-A | 2.6 | 3.4 | 69 | 1873 | 7 | MOLECULE: | TOXIN,T | OXIN CO | MPLEX SUBUNIT TCAB,PUTATIVE TOXIN S |
| 14:00 | 4qtq-A | 2.6 | 5 | 54 | 209 | 4 | MOLECULE: | XAC2610 | PROTEI | N |
| 15:00 | 1fwx-A | 2.6 | 4.2 | 67 | 591 | 6 | MOLECULE: | NITROUS | OXIDE | REDUCTASE |
| 16:00 | 6mv2-A | 2.6 | 4.4 | 59 | 356 | 8 | MOLECULE: | CYTOCHR | OME B5 | REDUCTASE 4 |
| 17:00 | 6icl-A | 2.6 | 3.1 | 59 | 338 | 5 | MOLECULE: | METHYLX | ANTHINE | N3-DEMETHYLASE NDMB |
| 18:00 | 8q7n-F | 2.6 | 3.1 | 60 | 431 | 7 | MOLECULE: | U5 SNRN | A | |
| 19:00 | 8ih8-D | 2.6 | 4.4 | 60 | 130 | 3 | MOLECULE: | ANTI-SI | GMA-F F | ACTOR ANTAGONIST RSFB |
| 20:00 | 8r54-A | 2.6 | 4.6 | 63 | 1982 | 5 | MOLECULE: | TENEURI | N-3 | |
| 21:00 | 6e14-D | 2.6 | 7.3 | 64 | 805 | 6 | MOLECULE: | TYPE 1 | FIMBRIN | D-MANNOSE SPECIFIC ADHESIN |
| 22:00 | 5chx-A | 2.6 | 2.5 | 49 | 210 | 8 | MOLECULE: | XRCC4-M | YH7-159 | 0-1657 |
| 23:00 | 7btx-L | 2.6 | 6.1 | 59 | 314 | 7 | MOLECULE: | MITOCHO | NDRIAL | OUTER MEMBRANE BETA-BARREL PROTEIN |
| 24:00:00 | 6f3a-h | 2.6 | 3.8 | 70 | 401 | 10 | MOLECULE: | ARP1 AC | TIN REL | ATED PROTEIN 1 HOMOLOG A |
| 25:00:00 | 5ho2-A | 2.6 | 3.3 | 66 | 803 | 3 | MOLECULE: | EXTRACE | LLULAR | ARABINANASE |
| 26:00:00 | 7obm-A | 2.6 | 2.8 | 57 | 624 | 4 | MOLECULE: | PROLYL | ENDOPEP | TIDASE-LIKE |
| 27:00:00 | 3ldk-A | 2.6 | 3.5 | 56 | 634 | 5 | MOLECULE: | FRUCTOS | YLTRANS | FERASE |
| 28:00:00 | 7bkb-k | 2.6 | 3 | 47 | 386 | 13 | MOLECULE: | COB--CO | M HETER | ODISULFIDE REDUCTASE IRON-SULFUR SU |
| 29:00:00 | 7upn-C | 2.6 | 3.1 | 47 | 208 | 4 | MOLECULE: | PEPTIDY | L-PROLY | L CIS-TRANS ISOMERASE A |
| 30:00:00 | 6jq9-B | 2.6 | 2.6 | 58 | 483 | 10 | MOLECULE: | SHORT U | LVAN LY | ASE |
| 31:00:00 | 3lic-A | 2.6 | 2.8 | 59 | 265 | 12 | MOLECULE: | SENSOR | PROTEIN | |
| 32:00:00 | 3dcz-A | 2.6 | 3.6 | 67 | 170 | 12 | MOLECULE: | PUTATIV | E RNFG | SUBUNIT OF ELECTRON TRANSPORT COMPL |
| 33:00:00 | 7uea-U | 2.6 | 3.8 | 55 | 364 | 5 | MOLECULE: | PHOTOSY | STEM P8 | 40 REACTION CENTER, LARGE SUBUNIT |
| 34:00:00 | 2eyq-A | 2.6 | 3.1 | 57 | 1146 | 4 | MOLECULE: | TRANSCR | IPTION- | REPAIR COUPLING FACTOR |
| 35:00:00 | 5bw0-F | 2.6 | 7.4 | 49 | 91 | 4 | MOLECULE: | TYPE II | SECRET | ION SYSTEM PROTEIN J |
| 36:00:00 | 6k4e-B | 2.6 | 3.7 | 66 | 246 | 9 | MOLECULE: | HAMP DO | MAIN-CO | NTAINING PROTEIN |
| 37:00:00 | 7p3l-A | 2.6 | 2.9 | 66 | 332 | 8 | MOLECULE: | ISOPENI | CILLIN | N SYNTHASE |
| 38:00:00 | 3csl-A | 2.6 | 6.2 | 67 | 753 | 4 | MOLECULE: | HASR PR | OTEIN | |
| 39:00:00 | 1eut-A | 2.6 | 3.5 | 58 | 601 | 7 | MOLECULE: | SIALIDA | SE | |
| 40:00:00 | 7uic-n | 2.6 | 3.5 | 63 | 220 | 3 | MOLECULE: | MEDIATO | R OF RN | A POLYMERASE II TRANSCRIPTION SUBUN |

| | | | | | | | | | | |
|---|---|---|---|---|---|---|---|---|---|---|
| 41:00:00 | 5k1c-C | 2.6 | 2.8 | 53 | 407 | 8 | MOLECULE: | UBIQUIT | IN CARB | OXYL-TERMINAL HYDROLASE 12 |
| 42:00:00 | 6stx-C | 2.6 | 3.1 | 58 | 663 | 3 | MOLECULE: | KELCH D | OMAIN-C | ONTAINING PROTEIN |
| 43:00:00 | 8t8s-B | 2.6 | 3 | 57 | 664 | 4 | MOLECULE: | SORTILI | N | |
| 44:00:00 | 1cyv-A | 2.6 | 2.9 | 58 | 98 | 3 | MOLECULE: | CYSTATI | N A | |
| 45:00:00 | 2xhn-B | 2.6 | 2.9 | 67 | 508 | 4 | MOLECULE: | RHAMNOG | ALACTUR | ONASE B |
| 46:00:00 | 4bl9-C | 2.6 | 4.6 | 73 | 724 | 7 | MOLECULE: | MALTOSE | #NAME? | G PERIPLASMIC PROTEIN, SUPPRESSOR O |
| 47:00:00 | 6mnj-B | 2.6 | 2.9 | 58 | 511 | 7 | MOLECULE: | HZ136 | | |
| 48:00:00 | 1lqg-C | 2.6 | 3.1 | 52 | 73 | 8 | MOLECULE: | URACIL- | DNA GLY | COSYLASE |
| 49:00:00 | 2pu8-B | 2.6 | 2.4 | 48 | 364 | 4 | MOLECULE: | METHYLT | HIORIBO | SE KINASE |
| 50:00:00 | 7e8h-K | 2.6 | 4.3 | 72 | 760 | 14 | MOLECULE: | DIPEPTI | DYL AMI | NOPEPTIDASE-LIKE PROTEIN 6 |
| 51:00:00 | 4fpe-A | 2.6 | 3 | 56 | 658 | 0 | MOLECULE: | SIALIDA | SE B | |
| 52:00:00 | 2ya4-B | 2.6 | 3.2 | 61 | 471 | 8 | MOLECULE: | NEURAMI | NIDASE | A |
| 53:00:00 | 4ovm-J | 2.6 | 3.2 | 66 | 124 | 11 | MOLECULE: | UNCHARA | CTERIZE | D PROTEIN SGCJ |
| 54:00:00 | 4ovm-E | 2.6 | 3.1 | 67 | 126 | 9 | MOLECULE: | UNCHARA | CTERIZE | D PROTEIN SGCJ |
| 55:00:00 | 5ohl-B | 2.6 | 3.7 | 55 | 89 | 5 | MOLECULE: | K6-SPEC | IFIC AF | FIMER |
| 56:00:00 | 4fph-A | 2.6 | 2.9 | 56 | 658 | 0 | MOLECULE: | SIALIDA | SE B | |
| 57:00:00 | 2vvz-A | 2.6 | 4.1 | 73 | 470 | 4 | MOLECULE: | SIALIDA | SE A | |
| 58:00:00 | 4xa2-A | 2.6 | 3.7 | 62 | 490 | 5 | MOLECULE: | MALTOSE | #NAME? | G PERIPLASMIC PROTEIN,MBP-PILA: C |
| 59:00:00 | 7a5x-A | 2.6 | 3.1 | 60 | 472 | 7 | MOLECULE: | SIALIDA | SE A | |
| 60:00:00 | 3lvt-A | 2.6 | 3 | 70 | 833 | 4 | MOLECULE: | GLYCOSY | L HYDRO | LASE, FAMILY 38 |
| 61:00:00 | 3oba-B | 2.6 | 3.8 | 50 | 1024 | 8 | MOLECULE: | BETA-GA | LACTOSI | DASE |
| 62:00:00 | 4q0p-A | 2.5 | 3.6 | 68 | 249 | 4 | MOLECULE: | L-RIBOS | E ISOME | RASE |
| 63:00:00 | 2e3m-A | 2.5 | 4.3 | 73 | 237 | 10 | MOLECULE: | LIPID-T | RANSFER | PROTEIN CERT |
| 64:00:00 | 3kzt-A | 2.5 | 3.2 | 64 | 132 | 14 | MOLECULE: | UNCHARA | CTERIZE | D PROTEIN |
| 65:00:00 | 2bas-A | 2.5 | 2.7 | 60 | 402 | 10 | MOLECULE: | YKUI PR | OTEIN | |
| 66:00:00 | 2xu8-A | 2.5 | 5.1 | 52 | 116 | 10 | MOLECULE: | PA1645 | | |
| 67:00:00 | 5h7z-B | 2.5 | 5.4 | 57 | 302 | 9 | MOLECULE: | UNCHARA | CTERIZE | D PROTEIN |
| 68:00:00 | 2y32-C | 2.5 | 3.5 | 65 | 138 | 3 | MOLECULE: | BLR5658 | PROTEI | N |
| 69:00:00 | 5ku5-A | 2.5 | 2.9 | 60 | 148 | 7 | MOLECULE: | SENSOR | KINASE | CUSS |
| 70:00:00 | 8g1r-A | 2.5 | 3.5 | 55 | 331 | 5 | MOLECULE: | MAJOR H | EAD PRO | TEIN |
| 71:00:00 | 5lkh-A | 2.5 | 3.6 | 62 | 948 | 8 | MOLECULE: | TCDA1 | | |
| 72:00:00 | 2qhk-A | 2.5 | 2.6 | 56 | 148 | 11 | MOLECULE: | METHYL- | ACCEPTI | NG CHEMOTAXIS PROTEIN |
| 73:00:00 | 3te8-B | 2.5 | 3.3 | 57 | 125 | 14 | MOLECULE: | SENSOR | HISTIDI | NE KINASE RISS |
| 74:00:00 | 6p59-B | 2.5 | 4.5 | 63 | 196 | 13 | MOLECULE: | CORE-BI | NDING F | ACTOR SUBUNIT BETA |
| 75:00:00 | 7t5q-I | 2.5 | 3.6 | 76 | 283 | 1 | MOLECULE: | ACTIN-R | ELATED | PROTEIN 3 |
| 76:00:00 | 4eg9-A | 2.5 | 3.9 | 83 | 231 | 2 | MOLECULE: | UNCHARA | CTERIZE | D PROTEIN SAOUHSC_02783 |
| 77:00:00 | 7d9b-A | 2.5 | 3.6 | 65 | 588 | 5 | MOLECULE: | ALPHA-G | LYCOSID | ASE |
| 78:00:00 | 1oh1-A | 2.5 | 3 | 58 | 109 | 0 | MOLECULE: | STAPHOS | TATIN A | |
| 79:00:00 | 6z9u-D | 2.5 | 4.1 | 57 | 134 | 5 | MOLECULE: | TRNA-SP | LICING | ENDONUCLEASE SUBUNIT SEN34 |
| 80:00:00 | 1vl4-A | 2.5 | 3.7 | 65 | 428 | 8 | MOLECULE: | PMBA-RE | LATED P | ROTEIN |
| 81:00:00 | 8fg6-B | 2.5 | 3.7 | 54 | 151 | 4 | MOLECULE: | AMYLOID | OGENIC | PEPTIDE |
| 82:00:00 | 5nj5-A | 2.5 | 3.8 | 62 | 480 | 10 | MOLECULE: | METALLO | PROTEAS | E TLDD |
| 83:00:00 | 3nqz-A | 2.5 | 3.6 | 66 | 175 | 6 | MOLECULE: | SECRETE | D METAL | LOPROTEASE MCP02 |
| 84:00:00 | 5ul2-A | 2.5 | 3.1 | 59 | 731 | 3 | MOLECULE: | OXSB PR | OTEIN | |
| 85:00:00 | 6lk8-J | 2.5 | 3.7 | 84 | 1026 | 7 | MOLECULE: | MGC8329 | 5 PROTE | IN |
| 86:00:00 | 4lgq-A | 2.5 | 3.8 | 63 | 134 | 5 | MOLECULE: | PUTATIV | E POLYK | ETIDE CYCLASE |
| 87:00:00 | 4ouh-A | 2.5 | 3.4 | 62 | 149 | 6 | MOLECULE: | PROTEAS | OME INH | IBITOR PI31 SUBUNIT |
| 88:00:00 | 1pvj-A | 2.5 | 3.3 | 65 | 339 | 8 | MOLECULE: | PYROGEN | IC EXOT | OXIN B |
| 89:00:00 | 4ci8-A | 2.5 | 4.7 | 65 | 640 | 8 | MOLECULE: | ECHINOD | ERM MIC | ROTUBULE-ASSOCIATED PROTEIN-LIKE 1 |
| 90:00:00 | 1su3-B | 2.5 | 2.7 | 49 | 416 | 4 | MOLECULE: | INTERST | ITIAL C | OLLAGENASE |
| 91:00:00 | 6gxs-B | 2.5 | 3.3 | 56 | 363 | 11 | MOLECULE: | CV39L L | ECTIN | |
| 92:00:00 | 2x2h-A | 2.5 | 3.3 | 63 | 1025 | 6 | MOLECULE: | ALPHA-1 | ,4-GLUC | AN LYASE ISOZYME 1 |
| 93:00:00 | 5txc-A | 2.5 | 3.7 | 56 | 653 | 9 | MOLECULE: | ATXE2 | | |
| 94:00:00 | 3e99-A | 2.5 | 3.8 | 69 | 149 | 6 | MOLECULE: | BENZOAT | E 1,2-D | IOXYGENASE BETA SUBUNIT |

| | | | | | | | | | | | |
|---|---|---|---|---|---|---|---|---|---|---|---|
| 95:00:00 | 8i4w-A | 2.5 | 4.7 | 67 | 442 | 6 | MOLECULE: | STRUCTU | RAL MAI | NTENANCE OF CHROMOSOMES PROTEIN 5 | |
| 96:00:00 | 7esn-A | 2.5 | 3.7 | 64 | 435 | 8 | MOLECULE: | L-RHAMN | OSE-ALP | HA-1,4-D-GLUCURONATE LYASE | |
| 97:00:00 | 2dso-C | 2.5 | 3 | 57 | 323 | 4 | MOLECULE: | DRP35 | | | |
| 98:00:00 | 7epn-B | 2.5 | 3.5 | 61 | 137 | 15 | MOLECULE: | SNOAL-L | IKE DOM | AIN-CONTAINING PROTEIN | |
| 99:00:00 | 2n19-A | 2.5 | 4.3 | 53 | 87 | 6 | MOLECULE: | SERINE/ | THREONI | NE-PROTEIN KINASE PLK4 | |
| 0:00 | 3zpm-A | 2.5 | 3.3 | 71 | 211 | 8 | MOLECULE: | LATHERI | N | | |
| 1:00 | 6d7k-H | 2.5 | 2.4 | 48 | 68 | 8 | MOLECULE: | METHANE | MONOOX | YGENASE HYDROXYLASE, MMOX1 | |
| 2:00 | 6cl5-A | 2.5 | 3.3 | 54 | 374 | 11 | MOLECULE: | TAIL FI | BER PRO | TEIN | |
| 3:00 | 5iri-A | 2.5 | 3.3 | 58 | 105 | 10 | MOLECULE: | SERINE/ | THREONI | NE-PROTEIN KINASE BRSK1 | |
| 4:00 | 6xlf-C | 2.5 | 4.2 | 68 | 316 | 3 | MOLECULE: | ATP-DEP | ENDENT | MOLECULAR CHAPERONE HSC82 | |
| 5:00 | 7bkb-e | 2.5 | 2.6 | 43 | 411 | 9 | MOLECULE: | COB--CO | M HETER | ODISULFIDE REDUCTASE IRON-SULFUR SU | |
| 6:00 | 8qca-C | 2.5 | 3.4 | 64 | 384 | 11 | MOLECULE: | ANTIVIR | AL HELI | CASE SKI2 | |
| 7:00 | 7pi3-A | 2.5 | 4 | 67 | 328 | 4 | MOLECULE: | CYSTEIN | E-RICH | PROTECTIVE ANTIGEN | |
| 8:00 | 2pk8-A | 2.5 | 3.7 | 49 | 94 | 8 | MOLECULE: | UNCHARA | CTERIZE | D PROTEIN PF0899 | |
| 9:00 | 7azn-A | 2.5 | 4.6 | 68 | 177 | 3 | MOLECULE: | PROTEIN | ASTER- | C | |
| 10:00 | 5mqr-A | 2.5 | 3.8 | 64 | 1082 | 11 | MOLECULE: | BETA-L- | ARABINO | BIOSIDASE | |
| 11:00 | 2ich-B | 2.5 | 4 | 63 | 323 | 3 | MOLECULE: | PUTATIV | E ATTH | | |
| 12:00 | 7zb1-B | 2.5 | 2.8 | 51 | 725 | 4 | MOLECULE: | PROLYL | ENDOPEP | TIDASE | |
| 13:00 | 8hhv-A | 2.5 | 3.5 | 62 | 479 | 6 | MOLECULE: | ENDO-AL | PHA-D-A | RABINANASE | |
| 14:00 | 3dzm-A | 2.5 | 5.1 | 62 | 208 | 5 | MOLECULE: | HYPOTHE | TICAL C | ONSERVED PROTEIN | |
| 15:00 | 8ovo-A | 2.5 | 4.6 | 58 | 504 | 5 | MOLECULE: | PUTATIV | E PERIP | LASMIC BINDING TRANSPORT PROTEIN,GR | |
| 16:00 | 4gk9-A | 2.5 | 3.5 | 50 | 279 | 4 | MOLECULE: | AGGLUTI | NIN (BO | A) | |
| 17:00 | 2xe4-A | 2.5 | 3.6 | 63 | 721 | 13 | MOLECULE: | OLIGOPE | PTIDASE | B | |
| 18:00 | 6urg-A | 2.5 | 3.4 | 61 | 1202 | 7 | MOLECULE: | CLEAVAG | E AND P | OLYADENYLATION SPECIFICITY FACTOR S | |
| 19:00 | 1sli-A | 2.5 | 4.2 | 71 | 679 | 11 | MOLECULE: | INTRAMO | LECULAR | TRANS-SIALIDASE | |
| 20:00 | 1no3-A | 2.5 | 3.8 | 75 | 851 | 5 | MOLECULE: | LIPOXYG | ENASE-3 | | |
| 21:00 | 8tie-a | 2.5 | 3.5 | 65 | 1012 | 8 | MOLECULE: | NUCLEOP | ORIN NU | P120 | |
| 22:00 | 4wy9-A | 2.5 | 3.3 | 60 | 285 | 7 | MOLECULE: | PUTATIV | E MCP-T | YPE SIGNAL TRANSDUCTION PROTEIN | |
| 23:00 | 7kra-A | 2.5 | 4.3 | 65 | 694 | 8 | MOLECULE: | ER MEMB | RANE PR | OTEIN COMPLEX SUBUNIT 1 | |
| 24:00:00 | 7qe7-A | 2.5 | 3.9 | 63 | 1648 | 8 | MOLECULE: | ANAPHAS | E-PROMO | TING COMPLEX SUBUNIT 10 | |
| 25:00:00 | 6h6g-A | 2.5 | 2.7 | 56 | 2139 | 9 | MOLECULE: | TCDB2,T | CCC3 | | |
| 26:00:00 | 5l8s-A | 2.5 | 3.5 | 55 | 604 | 5 | MOLECULE: | AMINO A | CYL PEP | TIDASE | |
| 27:00:00 | 5mmj-f | 2.5 | 3.2 | 50 | 113 | 6 | MOLECULE: | 50S RIB | OSOMAL | PROTEIN L31 | |
| 28:00:00 | 8asw-A | 2.5 | 3.3 | 64 | 1255 | 11 | MOLECULE: | ELONGAT | OR COMP | LEX PROTEIN 1 | |
| 29:00:00 | 4h5j-B | 2.5 | 3.3 | 57 | 347 | 11 | MOLECULE: | GUANINE | NUCLEO | TIDE-EXCHANGE FACTOR SEC12 | |
| 30:00:00 | 5f75-C | 2.5 | 3.6 | 61 | 475 | 11 | MOLECULE: | THIOCYA | NATE DE | HYDROGENASE | |
| 31:00:00 | 5uam-A | 2.5 | 3.9 | 61 | 440 | 5 | MOLECULE: | ULVAN L | YASE-PL | | 25 |
| 32:00:00 | 3n7n-B | 2.5 | 3.3 | 51 | 164 | 8 | MOLECULE: | MONOPOL | IN COMP | LEX SUBUNIT CSM1 | |
| 33:00:00 | 7zkh-A | 2.5 | 6.4 | 51 | 267 | 6 | MOLECULE: | METHYLT | RANSFER | ASE | |
| 34:00:00 | 5ien-A | 2.5 | 3.7 | 63 | 129 | 6 | MOLECULE: | D30H_F1 | 4 | | |
| 35:00:00 | 6kyb-B | 2.5 | 3.1 | 60 | 330 | 3 | MOLECULE: | AUTOPHA | GY-RELA | TED PROTEIN 18 | |
| 36:00:00 | 4x49-A | 2.5 | 3 | 65 | 489 | 9 | MOLECULE: | ANHYDRO | SIALIDA | SE | |
| 37:00:00 | 7yth-A | 2.5 | 3.1 | 65 | 314 | 8 | MOLECULE: | KETOSTE | ROID IS | OMERASE-RELATED PROTEIN | |
| 38:00:00 | 8x2u-J | 2.5 | 3.9 | 69 | 392 | 9 | MOLECULE: | DPY30 D | OMAIN C | ONTAINING 2 | |
| 39:00:00 | 4fov-A | 2.5 | 3.2 | 63 | 658 | 5 | MOLECULE: | SIALIDA | SE B | | |
| 40:00:00 | 4bl9-A | 2.5 | 4.8 | 78 | 722 | 6 | MOLECULE: | MALTOSE | #NAME? | G PERIPLASMIC PROTEIN, SUPPRESSOR O | |
| 41:00:00 | 7brs-D | 2.5 | 3.2 | 73 | 1015 | 3 | MOLECULE: | BETA-GA | LACTOSI | DASE | |
| 42:00:00 | 4yz5-A | 2.5 | 3.3 | 56 | 656 | 2 | MOLECULE: | PUTATIV | E NEURA | MINIDASE | |
| 43:00:00 | 6s5l-F | 2.5 | 3.6 | 64 | 129 | 11 | MOLECULE: | ALL4940 | PROTEI | N | |
| 44:00:00 | 4iia-A | 2.5 | 3.5 | 68 | 131 | 4 | MOLECULE: | RAS GTP | ASE-ACT | IVATING PROTEIN-BINDING PROTEIN 1 | |
| 45:00:00 | 2sli-A | 2.5 | 3.5 | 62 | 679 | 6 | MOLECULE: | INTRAMO | LECULAR | TRANS-SIALIDASE | |
| 46:00:00 | 1qal-B | 2.5 | 3.5 | 58 | 721 | 9 | MOLECULE: | COPPER | AMINE O | XIDASE | |
| 47:00:00 | 4yw2-B | 2.5 | 3.3 | 56 | 659 | 2 | MOLECULE: | NEURAMI | NIDASE | C | |
| 48:00:00 | 4yz2-A | 2.5 | 3.3 | 57 | 655 | 2 | MOLECULE: | SIALIDA | SE NANC | | |

| | | | | | | | | | | |
|---|---|---|---|---|---|---|---|---|---|---|
| 49:00:00 | 1ms0-B | 2.5 | 3.2 | 62 | 623 | 6 | MOLECULE: | TRANS-S | IALIDAS | E |
| 50:00:00 | 1jz7-D | 2.5 | 3.6 | 69 | 1011 | 7 | MOLECULE: | BETA-GA | LACTOSI | DASE |
| 51:00:00 | 4xjw-A | 2.5 | 3.6 | 66 | 658 | 6 | MOLECULE: | SIALIDA | SE B | |
| 52:00:00 | 4yw5-B | 2.5 | 3.2 | 58 | 659 | 2 | MOLECULE: | NEURAMI | NIDASE | C |
| 53:00:00 | 3dec-A | 2.5 | 3.7 | 59 | 983 | 5 | MOLECULE: | BETA-GA | LACTOSI | DASE |
| 54:00:00 | 4yz1-B | 2.5 | 3.3 | 57 | 656 | 2 | MOLECULE: | PUTATIV | E NEURA | MINIDASE |
| 55:00:00 | 4xe9-A | 2.5 | 3.6 | 65 | 658 | 6 | MOLECULE: | SIALIDA | SE B | |
| 56:00:00 | 4ovm-G | 2.5 | 2.8 | 64 | 122 | 9 | MOLECULE: | UNCHARA | CTERIZE | D PROTEIN SGCJ |
| 57:00:00 | 1sll-A | 2.5 | 3.8 | 62 | 679 | 5 | MOLECULE: | SIALIDA | SE L | |
| 58:00:00 | 5tsp-A | 2.5 | 4 | 68 | 448 | 9 | MOLECULE: | SIALIDA | SE | |
| 59:00:00 | 1wcq-B | 2.5 | 3.5 | 68 | 601 | 4 | MOLECULE: | SIALIDA | SE | |
| 60:00:00 | 1eus-A | 2.5 | 4 | 70 | 358 | 4 | MOLECULE: | SIALIDA | SE | |
| 61:00:00 | 7lbu-A | 2.5 | 3.9 | 69 | 380 | 4 | MOLECULE: | EXO-ALP | HA-SIAL | IDASE |
| 62:00:00 | 2bzd-A | 2.5 | 3.8 | 68 | 601 | 13 | MOLECULE: | BACTERI | AL SIAL | IDASE |
| 63:00:00 | 7a54-B | 2.5 | 3.3 | 57 | 472 | 7 | MOLECULE: | SIALIDA | SE A | |
| 64:00:00 | 7bgh-A | 2.4 | 6 | 63 | 177 | 5 | MOLECULE: | OUTER E | NVELOPE | PORE PROTEIN 21, CHLOROPLASTIC |
| 65:00:00 | 6rw6-B | 2.4 | 3.4 | 65 | 2485 | 6 | MOLECULE: | TCDA1 | | |
| 66:00:00 | 4p07-A | 2.4 | 3.7 | 69 | 569 | 4 | MOLECULE: | ARYLSUL | FATE SU | LFOTRANSFERASE ASST |
| 67:00:00 | 7k5c-A | 2.4 | 3.2 | 52 | 650 | 8 | MOLECULE: | INTERNA | L VIRIO | N PROTEIN GP15 |
| 68:00:00 | 7fde-P | 2.4 | 4.2 | 72 | 181 | 10 | MOLECULE: | V-TYPE | PROTON | ATPASE SUBUNIT C |
| 69:00:00 | 6gcd-A | 2.4 | 4.6 | 63 | 159 | 5 | MOLECULE: | 5-METHY | LCYTOSI | NE-SPECIFIC RESTRICTION ENZYME B |
| 70:00:00 | 8ok9-C | 2.4 | 3.8 | 71 | 236 | 6 | MOLECULE: | PIWI PR | OTEIN A | F_1318 |
| 71:00:00 | 1twu-A | 2.4 | 2.8 | 50 | 137 | 10 | MOLECULE: | HYPOTHE | TICAL P | ROTEIN YYCE |
| 72:00:00 | 4x4r-A | 2.4 | 4.7 | 67 | 443 | 4 | MOLECULE: | CCA-ADD | ING ENZ | YME |
| 73:00:00 | 2ooj-B | 2.4 | 4 | 58 | 134 | 5 | MOLECULE: | HYPOTHE | TICAL P | ROTEIN |
| 74:00:00 | 8t0b-A | 2.4 | 4.7 | 62 | 121 | 10 | MOLECULE: | DUF1842 | DOMAIN | -CONTAINING PROTEIN |
| 75:00:00 | 8c47-B | 2.4 | 3.1 | 52 | 151 | 4 | MOLECULE: | PROFILI | N | |
| 76:00:00 | 8h8b-A | 2.4 | 3.4 | 57 | 1027 | 4 | MOLECULE: | PUTATIV | E RHS-F | AMILY PROTEIN |
| 77:00:00 | 5oa3-0 | 2.4 | 2.9 | 51 | 479 | 10 | MOLECULE: | EUKARYO | TIC TRA | NSLATION INITIATION FACTOR 2D |
| 78:00:00 | 8t2v-B | 2.4 | 3.6 | 63 | 718 | 3 | MOLECULE: | INTEGRI | N ALPHA | -IIB |
| 79:00:00 | 5mwv-A | 2.4 | 4.7 | 43 | 281 | 14 | MOLECULE: | OUTER M | EMBRANE | PROTEIN G |
| 80:00:00 | 6itw-A | 2.4 | 3.5 | 51 | 221 | 2 | MOLECULE: | TYPE VI | IMMUNI | TY PROTEIN ATU4351 |
| 81:00:00 | 7px8-A | 2.4 | 3.6 | 50 | 700 | 8 | MOLECULE: | ACYLAMI | NO-ACID | -RELEASING ENZYME |
| 82:00:00 | 1lsh-B | 2.4 | 3.1 | 50 | 174 | 6 | MOLECULE: | LIPOVIT | ELLIN ( | LV-1N, LV-1C) |
| 83:00:00 | 6rfl-E | 2.4 | 3.5 | 67 | 184 | 4 | MOLECULE: | DNA-DEP | ENDENT | RNA POLYMERASE SUBUNIT RPO132 |
| 84:00:00 | 7wb4-c | 2.4 | 3.1 | 59 | 346 | 5 | MOLECULE: | OUTER N | UP133 | |
| 85:00:00 | 6v04-A | 2.4 | 6.5 | 53 | 268 | 6 | MOLECULE: | UNCHARA | CTERIZE | D SRPBCC DOMAIN-CONTAINING PROTEIN |
| 86:00:00 | 7vcf-I | 2.4 | 5 | 61 | 363 | 7 | MOLECULE: | TIC214 | | |
| 87:00:00 | 4ffg-A | 2.4 | 3 | 63 | 480 | 5 | MOLECULE: | LEVAN F | RUCTOTR | ANSFERASE |
| 88:00:00 | 3j1w-A | 2.4 | 3.1 | 52 | 106 | 12 | MOLECULE: | PROTEIN | PRGH | |
| 89:00:00 | 6kme-A | 2.4 | 5.1 | 61 | 284 | 8 | MOLECULE: | PHYTOCH | ROMOBIL | IN SYNTHASE |
| 90:00:00 | 8feh-D | 2.4 | 5.9 | 59 | 268 | 7 | MOLECULE: | MINOR C | APSID P | ROTEIN A1 FUSION |
| 91:00:00 | 5jen-A | 2.4 | 3.4 | 62 | 210 | 8 | MOLECULE: | ANTI-SI | GMA-V F | ACTOR RSIV |
| 92:00:00 | 5o8o-A | 2.4 | 4.5 | 66 | 332 | 6 | MOLECULE: | MITOCHO | NDRIAL | IMPORT RECEPTOR SUBUNIT TOM40 |
| 93:00:00 | 2rjz-A | 2.4 | 5.1 | 62 | 135 | 15 | MOLECULE: | PILO PR | OTEIN | |
| 94:00:00 | 3c6k-D | 2.4 | 3.6 | 59 | 348 | 3 | MOLECULE: | SPERMIN | E SYNTH | ASE |
| 95:00:00 | 4hh4-C | 2.4 | 3.3 | 48 | 257 | 4 | MOLECULE: | CCBJ | | |
| 96:00:00 | 1mg2-A | 2.4 | 3.2 | 61 | 382 | 8 | MOLECULE: | METHYLA | MINE DE | HYDROGENASE, HEAVY CHAIN |
| 97:00:00 | 8orn-B | 2.4 | 3.4 | 54 | 180 | 11 | MOLECULE: | OUTER-M | EMBRANE | LIPOPROTEIN CARRIER PROTEIN |
| 98:00:00 | 6cd2-C | 2.4 | 7.1 | 62 | 727 | 2 | MOLECULE: | CHAPERO | NE PROT | EIN PAPD |
| 99:00:00 | 8wkl-G | 2.4 | 3.2 | 68 | 307 | 9 | MOLECULE: | STRICTO | SIDINE | SYNTHASE |
| 0:00 | 3lif-A | 2.4 | 3.2 | 56 | 243 | 13 | MOLECULE: | PUTATIV | E DIGUA | NYLATE CYCLASE (GGDEF) WITH PAS/PAC |
| 1:00 | 8fo9-F | 2.4 | 3.6 | 50 | 2289 | 12 | MOLECULE: | LEUCINE | -RICH R | EPEAT SERINE/THREONINE-PROTEIN KINA |
| 2:00 | 8eti-A | 2.4 | 4.9 | 62 | 254 | 5 | MOLECULE: | RNA (15 | 64-MER) | |

| | | | | | | | | | | |
|---|---|---|---|---|---|---|---|---|---|---|
| 3:00 | 5img-A | 2.4 | 3.2 | 57 | 467 | 7 | MOLECULE: | DIPEPTI | DASE | |
| 4:00 | 7ty0-C | 2.4 | 3.6 | 66 | 488 | 8 | MOLECULE: | GLYCOPR | OTEIN G | |
| 5:00 | 4dvy-P | 2.4 | 4.5 | 51 | 656 | 4 | MOLECULE: | CYTOTOX | ICITY-A | SSOCIATED IMMUNODOMINANT ANTIGEN |
| 6:00 | 8xpg-B | 2.4 | 3.4 | 62 | 224 | 8 | MOLECULE: | SERINE/ | THREONI | NE-PROTEIN KINASE PLK4 |
| 7:00 | 7eeb-G | 2.4 | 3 | 56 | 714 | 7 | MOLECULE: | ENHANCE | D GREEN | FLUORESCENT PROTEIN,CATION CHANNEL |
| 8:00 | 1jnw-A | 2.4 | 4.3 | 67 | 214 | 4 | MOLECULE: | PYRIDOX | INE 5'- | PHOSPHATE OXIDASE |
| 9:00 | 2i51-A | 2.4 | 4.3 | 65 | 194 | 8 | MOLECULE: | UNCHARA | CTERIZE | D CONSERVED PROTEIN OF COG5135 |
| 10:00 | 3qqz-A | 2.4 | 2.9 | 57 | 243 | 4 | MOLECULE: | PUTATIV | E UNCHA | RACTERIZED PROTEIN YJIK |
| 11:00 | 5m11-A | 2.4 | 3.2 | 60 | 748 | 10 | MOLECULE: | IMMUNOR | EACTIVE | 84KD ANTIGEN PG93 |
| 12:00 | 8u1o-o | 2.4 | 49.6 | 74 | 663 | 9 | MOLECULE: | TAIL SP | IKE PRO | TEIN |
| 13:00 | 7v2w-H | 2.4 | 3.3 | 63 | 379 | 8 | MOLECULE: | THO COM | PLEX SU | BUNIT HPR1 |
| 14:00 | 4uw8-A | 2.4 | 4.5 | 60 | 401 | 5 | MOLECULE: | L-SHAPE | D TAIL | FIBER PROTEIN |
| 15:00 | 8ptx-B | 2.4 | 3.3 | 58 | 800 | 7 | MOLECULE: | ELONGAT | OR COMP | LEX PROTEIN 1 |
| 16:00 | 8glo-A | 2.4 | 4.2 | 57 | 255 | 11 | MOLECULE: | HEMOPHI | LIN | |
| 17:00 | 7dl2-A | 2.4 | 4.7 | 63 | 1127 | 6 | MOLECULE: | HAMARTI | N | |
| 18:00 | 8chv-B | 2.4 | 2.8 | 47 | 147 | 6 | MOLECULE: | TRANSCR | IPTIONA | L ACTIVATOR PROTEIN PUR-ALPHA |
| 19:00 | 4u8u-M | 2.4 | 3 | 57 | 222 | 0 | MOLECULE: | GLOBIN | A CHAIN | |
| 20:00 | 3w7s-A | 2.4 | 3.8 | 60 | 760 | 8 | MOLECULE: | UNCHARA | CTERIZE | D PROTEIN YGJK |
| 21:00 | 8a3t-B | 2.4 | 2.8 | 52 | 440 | 10 | MOLECULE: | ANAPHAS | E-PROMO | TING COMPLEX SUBUNIT CDC27 |
| 22:00 | 8kg3-A | 2.4 | 3.5 | 63 | 311 | 5 | MOLECULE: | OS06G06 | 23700 P | ROTEIN |
| 23:00 | 4fhn-B | 2.4 | 3.3 | 62 | 1022 | 8 | MOLECULE: | NUCLEOP | ORIN NU | P37 |
| 24:00:00 | 5yyl-A | 2.4 | 4 | 63 | 405 | 10 | MOLECULE: | MAJOR R | OYAL JE | LLY PROTEIN 1 |
| 25:00:00 | 4d4o-A | 2.4 | 4.1 | 56 | 414 | 7 | MOLECULE: | PROTEIN | ATS1, | DIPHTHAMIDE BIOSYNTHESIS PROTEIN 3 |
| 26:00:00 | 4ui9-A | 2.4 | 3.1 | 58 | 1441 | 9 | MOLECULE: | ANAPHAS | E-PROMO | TING COMPLEX SUBUNIT 1 |
| 27:00:00 | 6s5l-A | 2.4 | 3.2 | 59 | 125 | 8 | MOLECULE: | ALL4940 | PROTEI | N |
| 28:00:00 | 3aaa-A | 2.4 | 3.5 | 74 | 267 | 5 | MOLECULE: | F-ACTIN | #NAME? | G PROTEIN SUBUNIT ALPHA-1 |
| 29:00:00 | 6mrx-A | 2.4 | 3.9 | 62 | 522 | 5 | MOLECULE: | SIALIDA | SE26 | |
| 30:00:00 | 3lk4-4 | 2.4 | 3.3 | 73 | 269 | 5 | MOLECULE: | F-ACTIN | #NAME? | G PROTEIN SUBUNIT ALPHA-1 |
| 31:00:00 | 3lk2-A | 2.4 | 3.5 | 74 | 270 | 5 | MOLECULE: | F-ACTIN | #NAME? | G PROTEIN SUBUNIT ALPHA-1 |
| 32:00:00 | 4ovm-I | 2.4 | 3.1 | 64 | 120 | 11 | MOLECULE: | UNCHARA | CTERIZE | D PROTEIN SGCJ |
| 33:00:00 | 4poi-B | 2.4 | 3.3 | 57 | 120 | 4 | MOLECULE: | PUTATIV | E PERIP | LASMIC PROTEIN |
| 34:00:00 | 3opz-A | 2.4 | 2.8 | 60 | 625 | 5 | MOLECULE: | TRANS-S | IALIDAS | E |
| 35:00:00 | 8x2u-R | 2.4 | 4.3 | 66 | 392 | 5 | MOLECULE: | DPY30 D | OMAIN C | ONTAINING 2 |
| 36:00:00 | 6d24-D | 2.4 | 4.5 | 70 | 494 | 4 | MOLECULE: | GLUCOSE | #NAME? | PHATE 1-DEHYDROGENASE |
| 37:00:00 | 4yz3-A | 2.4 | 3.2 | 56 | 654 | 2 | MOLECULE: | NEURAMI | NIDASE | |
| 38:00:00 | 6q0w-B | 2.4 | 3.7 | 66 | 208 | 6 | MOLECULE: | DNA DAM | AGE-BIN | DING PROTEIN 1 |
| 39:00:00 | 4yw1-B | 2.4 | 3.2 | 56 | 659 | 2 | MOLECULE: | NEURAMI | NIDASE | C |
| 40:00:00 | 6sd0-D | 2.4 | 2.7 | 50 | 1083 | 10 | MOLECULE: | BETA-GA | LACTOSI | DASE |
| 41:00:00 | 1ms5-A | 2.4 | 3.8 | 69 | 623 | 9 | MOLECULE: | TRANS-S | IALIDAS | E |
| 42:00:00 | 5ts4-D | 2.4 | 2.9 | 58 | 89 | 7 | MOLECULE: | DENOVO | NTF2 | |
| 43:00:00 | 4yw1-A | 2.4 | 3.3 | 59 | 659 | 2 | MOLECULE: | NEURAMI | NIDASE | C |
| 44:00:00 | 4yz3-B | 2.4 | 2.8 | 53 | 656 | 2 | MOLECULE: | NEURAMI | NIDASE | |
| 45:00:00 | 3b69-A | 2.4 | 3.6 | 65 | 626 | 8 | MOLECULE: | TRANS-S | IALIDAS | E |
| 46:00:00 | 4poi-A | 2.4 | 3.4 | 57 | 116 | 4 | MOLECULE: | PUTATIV | E PERIP | LASMIC PROTEIN |
| 47:00:00 | 4yw3-B | 2.4 | 3.3 | 56 | 659 | 2 | MOLECULE: | NEURAMI | NIDASE | C |
| 48:00:00 | 3sli-A | 2.4 | 3.3 | 61 | 679 | 8 | MOLECULE: | INTRAMO | LECULAR | TRANS-SIALIDASE |
| 49:00:00 | 3opz-B | 2.4 | 3.2 | 65 | 625 | 8 | MOLECULE: | TRANS-S | IALIDAS | E |
| 50:00:00 | 7brs-B | 2.4 | 3.6 | 69 | 1012 | 7 | MOLECULE: | BETA-GA | LACTOSI | DASE |
| 51:00:00 | 2lf2-A | 2.3 | 4 | 72 | 175 | 3 | MOLECULE: | UNCHARA | CTERIZE | D PROTEIN |
| 52:00:00 | 8d8l-C | 2.3 | 3.1 | 56 | 240 | 4 | MOLECULE: | PROBABL | E S-ADE | NOSYL-L-METHIONINE-DEPENDENT RNA |
| 53:00:00 | 7c8z-B | 2.3 | 3.6 | 70 | 161 | 11 | MOLECULE: | SALICYL | ATE 5-H | YDROXYLASE, LARGE OXYGENASE COMPONE |
| 54:00:00 | 6am0-F | 2.3 | 3.6 | 62 | 177 | 3 | MOLECULE: | KLLA0F2 | 3980P | |
| 55:00:00 | 7ofn-A | 2.3 | 4.2 | 68 | 163 | 3 | MOLECULE: | LIPOPRO | TEIN | |
| 56:00:00 | 2kd2-A | 2.3 | 3.3 | 56 | 94 | 5 | MOLECULE: | FAS APO | PTOTIC | INHIBITORY MOLECULE 1 |

| | | | | | | | | | | |
|---|---|---|---|---|---|---|---|---|---|---|
| 57:00:00 | 1ojg-A | 2.3 | 3.2 | 61 | 135 | 10 | MOLECULE: | SENSOR | PROTEIN | DCUS |
| 58:00:00 | 2y2l-B | 2.3 | 3.5 | 57 | 476 | 4 | MOLECULE: | PENICIL | LIN-BIN | DING PROTEIN 1B |
| 59:00:00 | 8x6f-C | 2.3 | 3.8 | 63 | 1153 | 11 | MOLECULE: | DNA-DIR | ECTED R | NA POLYMERASE SUBUNIT ALPHA |
| 60:00:00 | 7mp9-A | 2.3 | 4.9 | 65 | 414 | 5 | MOLECULE: | SERINE/ | THREONI | NE-PROTEIN KINASE PINK1, MITOCHONDR |
| 61:00:00 | 5dyv-A | 2.3 | 4.1 | 57 | 135 | 5 | MOLECULE: | YD REPE | AT-CONT | AINING PROTEIN |
| 62:00:00 | 3o2u-B | 2.3 | 4 | 71 | 185 | 1 | MOLECULE: | NEDD8-C | ONJUGAT | ING ENZYME UBC12 |
| 63:00:00 | 5x9j-B | 2.3 | 4.1 | 68 | 152 | 9 | MOLECULE: | PRHC | | |
| 64:00:00 | 3nqn-A | 2.3 | 2.9 | 60 | 151 | 2 | MOLECULE: | UNCHARA | CTERIZE | D PROTEIN |
| 65:00:00 | 3c7g-A | 2.3 | 3.4 | 59 | 488 | 5 | MOLECULE: | ENDO-1, | 4-BETA- | XYLANASE |
| 66:00:00 | 6o38-L | 2.3 | 4.3 | 63 | 130 | 2 | MOLECULE: | ACINETO | BACTER | SECRETED PROTEASE CPAA |
| 67:00:00 | 6r82-A | 2.3 | 4.4 | 73 | 181 | 3 | MOLECULE: | GTPASE- | ACTIVAT | ING PROTEIN SKYWALKER |
| 68:00:00 | 6ipv-A | 2.3 | 3.4 | 63 | 221 | 10 | MOLECULE: | CQSB2 | | |
| 69:00:00 | 4rny-A | 2.3 | 3.4 | 60 | 359 | 3 | MOLECULE: | CONSERV | ED HYPO | THETICAL SECRETED PROTEIN |
| 70:00:00 | 3kk7-A | 2.3 | 3.2 | 63 | 502 | 5 | MOLECULE: | PUTATIV | E CELL | INVASION PROTEIN WITH MAC/PERFORIN |
| 71:00:00 | 6jql-A | 2.3 | 3.2 | 58 | 678 | 3 | MOLECULE: | BIFUNCT | IONAL P | ROTEIN PAAZ |
| 72:00:00 | 4ped-A | 2.3 | 3.7 | 72 | 387 | 1 | MOLECULE: | CHAPERO | NE ACTI | VITY OF BC1 COMPLEX-LIKE, MITOCHOND |
| 73:00:00 | 7pd7-B | 2.3 | 9.3 | 57 | 251 | 9 | MOLECULE: | METHYLT | RANSFER | ASE |
| 74:00:00 | 6ns4-A | 2.3 | 4.9 | 68 | 685 | 4 | MOLECULE: | LIPOXYG | ENASE | |
| 75:00:00 | 7p2p-C | 2.3 | 3.6 | 61 | 174 | 5 | MOLECULE: | SIGNAL | PEPTIDA | SE COMPLEX CATALYTIC SUBUNIT SEC11A |
| 76:00:00 | 5yk1-A | 2.3 | 3.9 | 66 | 401 | 5 | MOLECULE: | PROBABL | E CONSE | RVED ATP-BINDING PROTEIN ABC TRANSP |
| 77:00:00 | 8ajm-B | 2.3 | 3.7 | 56 | 395 | 11 | MOLECULE: | DNA DAM | AGE-BIN | DING PROTEIN 1 |
| 78:00:00 | 7eu3-7 | 2.3 | 3.5 | 69 | 317 | 9 | MOLECULE: | NAD(P)H | #NAME? | E OXIDOREDUCTASE SUBUNIT 1, CHLOROP |
| 79:00:00 | 2o5p-A | 2.3 | 4.1 | 66 | 772 | 6 | MOLECULE: | FERRIPY | OVERDIN | E RECEPTOR |
| 80:00:00 | 7c3c-B | 2.3 | 3 | 58 | 346 | 7 | MOLECULE: | AOFLEA | | |
| 81:00:00 | 3zwl-D | 2.3 | 3.1 | 52 | 352 | 13 | MOLECULE: | EUKARYO | TIC TRA | NSLATION INITIATION FACTOR 3 SUBUNI |
| 82:00:00 | 7wb4-E | 2.3 | 3.8 | 71 | 1363 | 11 | MOLECULE: | OUTER N | UP133 | |
| 83:00:00 | 2gia-G | 2.3 | 3.7 | 52 | 155 | 13 | MOLECULE: | MITOCHO | NDRIAL | RNA-BINDING PROTEIN 2 |
| 84:00:00 | 2joz-A | 2.3 | 3.5 | 56 | 135 | 4 | MOLECULE: | HYPOTHE | TICAL P | ROTEIN YXEF |
| 85:00:00 | 1k32-A | 2.3 | 3.1 | 55 | 1023 | 7 | MOLECULE: | TRICORN | PROTEA | SE |
| 86:00:00 | 7oc9-A | 2.3 | 3.3 | 60 | 133 | 12 | MOLECULE: | BD0675 | | |
| 87:00:00 | 7r5z-B | 2.3 | 4.3 | 62 | 154 | 5 | MOLECULE: | DIRIGEN | T PROTE | IN |
| 88:00:00 | 4fnv-A | 2.3 | 3.8 | 69 | 659 | 9 | MOLECULE: | HEPARIN | ASE III | PROTEIN, HEPARITIN SULFATE LYASE |
| 89:00:00 | 6hzn-A | 2.3 | 3.3 | 59 | 742 | 7 | MOLECULE: | DERMATA | N-SULFA | TE EPIMERASE |
| 90:00:00 | 2qml-A | 2.3 | 4.9 | 47 | 193 | 11 | MOLECULE: | BH2621 | PROTEIN | |
| 91:00:00 | 3hi2-B | 2.3 | 3.2 | 62 | 97 | 2 | MOLECULE: | HTH-TYP | E TRANS | CRIPTIONAL REGULATOR MQSA(YGIT) |
| 92:00:00 | 4asc-A | 2.3 | 3 | 48 | 308 | 4 | MOLECULE: | KELCH R | EPEAT A | ND BTB DOMAIN-CONTAINING PROTEIN 5 |
| 93:00:00 | 8a3t-Q | 2.3 | 3.6 | 60 | 623 | 7 | MOLECULE: | ANAPHAS | E-PROMO | TING COMPLEX SUBUNIT CDC27 |
| 94:00:00 | 5h68-B | 2.3 | 3.6 | 61 | 332 | 11 | MOLECULE: | CHROMOS | OME PAR | TITION PROTEIN SMC |
| 95:00:00 | 7dtr-A | 2.3 | 2.5 | 46 | 163 | 11 | MOLECULE: | ACRIF24 | | |
| 96:00:00 | 6y86-A | 2.3 | 3.6 | 48 | 425 | 6 | MOLECULE: | MEMBRAN | E PROTE | IN INSERTASE YIDC |
| 97:00:00 | 1qjs-A | 2.3 | 2.5 | 43 | 408 | 2 | MOLECULE: | HEMOPEX | IN | |
| 98:00:00 | 8giu-A | 2.3 | 2.8 | 48 | 381 | 10 | MOLECULE: | GP_4 (C | APSID A | CCESSORY PROTEIN) |
| 99:00:00 | 3af5-A | 2.3 | 10.3 | 65 | 638 | 5 | MOLECULE: | PUTATIV | E UNCHA | RACTERIZED PROTEIN PH1404 |
| 0:00 | 5hl8-A | 2.3 | 3.6 | 52 | 81 | 12 | MOLECULE: | TYPE II | SECRET | ION SYSTEM PROTEIN L |
| 1:00 | 8beq-A | 2.3 | 3.3 | 61 | 534 | 7 | MOLECULE: | FRUCTOF | URANOSI | DASE FROM RHODOTORULA DAIRENENSIS |
| 2:00 | 6lnh-C | 2.3 | 3.4 | 60 | 228 | 7 | MOLECULE: | L-ISOLE | UCINE-4 | -HYDROXYLASE |
| 3:00 | 2crf-A | 2.3 | 3.4 | 59 | 150 | 5 | MOLECULE: | RAN BIN | DING PR | OTEIN 3 |
| 4:00 | 2pff-B | 2.3 | 7.1 | 50 | 2006 | 0 | MOLECULE: | FATTY A | CID SYN | THASE SUBUNIT ALPHA |
| 5:00 | 7aak-A | 2.3 | 3 | 59 | 345 | 5 | MOLECULE: | PORPHOB | ILINOGE | N DEAMINASE |
| 6:00 | 4doo-A | 2.3 | 3.1 | 64 | 203 | 2 | MOLECULE: | CHALCON | E-FLAVA | NONE ISOMERASE FAMILY PROTEIN |
| 7:00 | 6zrd-A | 2.3 | 2.9 | 51 | 411 | 10 | MOLECULE: | HISTONE | #NAME? | G PROTEIN RBBP4 |
| 8:00 | 8tar-R | 2.3 | 3.8 | 55 | 440 | 5 | MOLECULE: | ANAPHAS | E-PROMO | TING COMPLEX SUBUNIT 1 |
| 9:00 | 8wqr-B | 2.3 | 4.8 | 55 | 356 | 11 | MOLECULE: | ACTIVAT | ING MOL | ECULE IN BECN1-REGULATED AUTOPHAGY |
| 10:00 | 3dsb-A | 2.3 | 4.5 | 62 | 153 | 5 | MOLECULE: | PUTATIV | E ACETY | LTRANSFERASE |

| | | | | | | | | | | |
|---|---|---|---|---|---|---|---|---|---|---|
| 11:00 | 6n2g-A | 2.3 | 4.6 | 64 | 285 | 5 | MOLECULE: | NUCLEOS | OME ASS | EMBLY PROTEIN |
| 12:00 | 6dzk-Y | 2.3 | 3.2 | 57 | 103 | 4 | MOLECULE: | 16S RRN | A | |
| 13:00 | 8j0n-A | 2.3 | 3.7 | 62 | 949 | 3 | MOLECULE: | ER MEMB | RANE PR | OTEIN COMPLEX SUBUNIT 1 |
| 14:00 | 8ond-A | 2.3 | 4.6 | 55 | 344 | 5 | MOLECULE: | BD2133 | | |
| 15:00 | 6acx-A | 2.3 | 3 | 52 | 1175 | 8 | MOLECULE: | MYCOBAC | TERIUM | SMEGMATIS MFD |
| 16:00 | 5yvq-A | 2.3 | 7.3 | 54 | 358 | 2 | MOLECULE: | TAIL FI | BER PRO | TEIN S |
| 17:00 | 8bys-A | 2.3 | 3.7 | 59 | 323 | 3 | MOLECULE: | S-LAYER | HOMOLO | GY DOMAIN-CONTAINING PROTEIN |
| 18:00 | 6dn4-A | 2.3 | 10.1 | 60 | 275 | 7 | MOLECULE: | BETA-LA | CTAMASE | |
| 19:00 | 3e5d-A | 2.3 | 4 | 50 | 126 | 6 | MOLECULE: | PUTATIV | E GLYOX | ALASE I |
| 20:00 | 6zym-p | 2.3 | 3.9 | 62 | 325 | 5 | MOLECULE: | U2 SNRN | A | |
| 21:00 | 7mge-A | 2.3 | 2.9 | 52 | 349 | 8 | MOLECULE: | WD REPE | AT-CONT | AINING PROTEIN 41 |
| 22:00 | 8q66-A | 2.3 | 3.8 | 60 | 271 | 3 | MOLECULE: | EXONUCL | EASE MU | T-7 |
| 23:00 | 4aip-B | 2.3 | 8.1 | 65 | 665 | 12 | MOLECULE: | FE-REGU | LATED P | ROTEIN B |
| 24:00:00 | 2hhi-A | 2.3 | 3.1 | 67 | 204 | 6 | MOLECULE: | IMMUNOG | ENIC PR | OTEIN MPT64 |
| 25:00:00 | 8f0v-A | 2.3 | 3.9 | 57 | 155 | 5 | MOLECULE: | MILK PR | OTEIN | |
| 26:00:00 | 6vs7-A | 2.3 | 2.9 | 48 | 409 | 2 | MOLECULE: | ADHESIN | | |
| 27:00:00 | 7t3j-J | 2.3 | 2.5 | 46 | 228 | 11 | MOLECULE: | CRISPR- | ASSOCIA | TED PROTEIN CSY1 |
| 28:00:00 | 8chw-A | 2.3 | 2.9 | 44 | 67 | 7 | MOLECULE: | TRANSCR | IPTIONA | L ACTIVATOR PROTEIN PUR-ALPHA |
| 29:00:00 | 4ktp-A | 2.3 | 4.5 | 60 | 767 | 7 | MOLECULE: | GLYCOSI | DE HYDR | OLASE FAMILY 65 CENTRAL CATALYTIC |
| 30:00:00 | 8wo8-A | 2.3 | 3.7 | 68 | 463 | 3 | MOLECULE: | PROBABL | E RIBON | UCLEASE FAU-1 |
| 31:00:00 | 4e54-B | 2.3 | 2.9 | 51 | 402 | 14 | MOLECULE: | DNA DAM | AGE-BIN | DING PROTEIN 1 |
| 32:00:00 | 7svm-A | 2.3 | 3.4 | 51 | 852 | 10 | MOLECULE: | DIPEPTI | DYL PEP | TIDASE 8 |
| 33:00:00 | 1fj1-E | 2.3 | 5.5 | 58 | 251 | 3 | MOLECULE: | HYBRIDO | MA ANTI | BODY LA2 (LIGHT CHAIN) |
| 34:00:00 | 3aae-E | 2.3 | 3.5 | 75 | 270 | 5 | MOLECULE: | F-ACTIN | #NAME? | G PROTEIN SUBUNIT ALPHA-1 |
| 35:00:00 | 4x47-A | 2.3 | 3.4 | 69 | 489 | 7 | MOLECULE: | ANHYDRO | SIALIDA | SE |
| 36:00:00 | 4dsd-B | 2.3 | 4 | 58 | 126 | 10 | MOLECULE: | PUTATIV | E PERIP | LASMIC PROTEIN |
| 37:00:00 | 6q0r-B | 2.3 | 3.3 | 68 | 207 | 9 | MOLECULE: | DNA DAM | AGE-BIN | DING PROTEIN 1 |
| 38:00:00 | 4yw3-A | 2.3 | 3.4 | 60 | 659 | 2 | MOLECULE: | NEURAMI | NIDASE | C |
| 39:00:00 | 6drv-A | 2.3 | 2.9 | 62 | 1024 | 5 | MOLECULE: | BETA-GA | LACTOSI | DASE |
| 40:00:00 | 7e8h-J | 2.3 | 3 | 53 | 756 | 0 | MOLECULE: | DIPEPTI | DYL AMI | NOPEPTIDASE-LIKE PROTEIN 6 |
| 41:00:00 | 1dil-A | 2.3 | 4.1 | 70 | 381 | 6 | MOLECULE: | SIALIDA | SE | |
| 42:00:00 | 3oba-C | 2.3 | 3.6 | 73 | 1024 | 7 | MOLECULE: | BETA-GA | LACTOSI | DASE |
| 43:00:00 | 2vvz-B | 2.3 | 4.1 | 73 | 470 | 5 | MOLECULE: | SIALIDA | SE A | |
| 44:00:00 | 2ya8-B | 2.3 | 3.2 | 57 | 470 | 7 | MOLECULE: | NEURAMI | NIDASE | A |
| 45:00:00 | 2ya5-B | 2.3 | 3.2 | 57 | 470 | 7 | MOLECULE: | NEURAMI | NIDASE | A |
| 46:00:00 | 5kbp-A | 2.3 | 3.5 | 72 | 886 | 6 | MOLECULE: | GLYCOSY | L HYDRO | LASE, FAMILY 38 |
| 47:00:00 | 4foy-A | 2.3 | 2.9 | 55 | 658 | 0 | MOLECULE: | SIALIDA | SE B | |
| 48:00:00 | 8gn6-C | 2.3 | 3.5 | 62 | 494 | 10 | MOLECULE: | SIALIDA | SE | |
| 49:00:00 | 8gn6-B | 2.3 | 3.6 | 64 | 496 | 11 | MOLECULE: | SIALIDA | SE | |
| 50:00:00 | 4fj6-C | 2.3 | 3 | 69 | 519 | 10 | MOLECULE: | GLYCOSI | DE HYDR | OLASE FAMILY 33, CANDIDATE SIALIDAS |
| 51:00:00 | 4xog-A | 2.3 | 3.6 | 63 | 660 | 6 | MOLECULE: | SIALIDA | SE B | |
| 52:00:00 | 4q6k-D | 2.3 | 3.3 | 71 | 523 | 11 | MOLECULE: | BNR/ASP | #NAME? | PEAT PROTEIN |
| 53:00:00 | 5vg5-W | 2.3 | 4.4 | 64 | 486 | 5 | MOLECULE: | GLUCOSE | #NAME? | PHATE 1-DEHYDROGENASE |
| 54:00:00 | 5ohl-A | 2.3 | 2.9 | 50 | 89 | 6 | MOLECULE: | K6-SPEC | IFIC AF | FIMER |
| 55:00:00 | 7ual-A | 2.3 | 4.3 | 59 | 484 | 8 | MOLECULE: | GLUCOSE | #NAME? | PHATE 1-DEHYDROGENASE |
| 56:00:00 | 7a54-A | 2.3 | 3.5 | 70 | 472 | 4 | MOLECULE: | SIALIDA | SE A | |
| 57:00:00 | 4qa8-A | 2.2 | 3.8 | 73 | 210 | 10 | MOLECULE: | PUTATIV | E LIPOP | ROTEIN LPRF |
| 58:00:00 | 3cwf-A | 2.2 | 3.4 | 56 | 108 | 9 | MOLECULE: | ALKALIN | E PHOSP | HATASE SYNTHESIS SENSOR PROTEIN PHO |
| 59:00:00 | 1n3g-A | 2.2 | 3.7 | 60 | 113 | 5 | MOLECULE: | PROTEIN | YFIA | |
| 60:00:00 | 6pif-G | 2.2 | 3.2 | 61 | 521 | 5 | MOLECULE: | CAS7, T | YPE I-F | CRISPR-ASSOCIATED PROTEIN |
| 61:00:00 | 7e9d-A | 2.2 | 3.3 | 55 | 443 | 9 | MOLECULE: | IRG1 | | |
| 62:00:00 | 1tuh-A | 2.2 | 3.4 | 63 | 131 | 16 | MOLECULE: | HYPOTHE | TICAL P | ROTEIN EGC068 |
| 63:00:00 | 6hdv-A | 2.2 | 4.1 | 54 | 136 | 6 | MOLECULE: | AFIFAVI | DIN | |
| 64:00:00 | 7pkq-f | 2.2 | 4.2 | 57 | 103 | 7 | MOLECULE: | MS35 | | |

| | | | | | | | | | | |
|---|---|---|---|---|---|---|---|---|---|---|
| 65:00:00 | 7t69-A | 2.2 | 3.9 | 51 | 209 | 12 | MOLECULE: | AVR3 (S | IX1), S | ECRETED IN XYLEM 1 |
| 66:00:00 | 7s00-c | 2.2 | 3.3 | 60 | 484 | 5 | MOLECULE: | DNA-DIR | ECTED R | NA POLYMERASE BETA SUBUNIT |
| 67:00:00 | 6vqv-D | 2.2 | 3.9 | 65 | 306 | 2 | MOLECULE: | ACRF9 | | |
| 68:00:00 | 8f3h-A | 2.2 | 3.1 | 61 | 643 | 8 | MOLECULE: | PENICIL | LIN BIN | DING PROTEIN 5 |
| 69:00:00 | 3fo5-A | 2.2 | 3.4 | 71 | 239 | 10 | MOLECULE: | THIOEST | ERASE, | ADIPOSE ASSOCIATED, ISOFORM BFIT2 |
| 70:00:00 | 5zi2-A | 2.2 | 3.5 | 68 | 339 | 7 | MOLECULE: | MALATE | DEHYDRO | GENASE |
| 71:00:00 | 4i60-A | 2.2 | 4.7 | 56 | 128 | 5 | MOLECULE: | AVIDIN | | |
| 72:00:00 | 8ixw-A | 2.2 | 3 | 52 | 168 | 10 | MOLECULE: | CHAINS: | A,B | |
| 73:00:00 | 1zwy-B | 2.2 | 3.1 | 57 | 180 | 5 | MOLECULE: | HYPOTHE | TICAL U | PF0244 PROTEIN VC0702 |
| 74:00:00 | 8a0h-A | 2.2 | 3.2 | 63 | 324 | 5 | MOLECULE: | OCP N-T | ERMINAL | DOMAIN-CONTAINING PROTEIN |
| 75:00:00 | 8ttb-B | 2.2 | 3.8 | 66 | 426 | 9 | MOLECULE: | SERINE/ | THREONI | NE-PROTEIN PHOSPHATASE 2A 65 KDA RE |
| 76:00:00 | 6r1g-A | 2.2 | 3.4 | 65 | 148 | 8 | MOLECULE: | OUTER S | URFACE | 22 KDA LIPOPROTEIN |
| 77:00:00 | 8tl6-B | 2.2 | 3.7 | 52 | 353 | 8 | MOLECULE: | DNA DAM | AGE-BIN | DING PROTEIN 1 |
| 78:00:00 | 3c30-A | 2.2 | 3.5 | 61 | 219 | 8 | MOLECULE: | AUTOIND | UCER 2 | SENSOR KINASE/PHOSPHATASE LUXQ |
| 79:00:00 | 7vma-A | 2.2 | 3.7 | 62 | 620 | 5 | MOLECULE: | AMYLO-A | LPHA-1, | 6-GLUCOSIDASE, PUTATIVE ARCHAEAL TY |
| 80:00:00 | 7czf-B | 2.2 | 3 | 53 | 644 | 9 | MOLECULE: | EPHRIN | TYPE-A | RECEPTOR 2 |
| 81:00:00 | 6rte-A | 2.2 | 3.6 | 57 | 464 | 9 | MOLECULE: | CYTOCHR | OME C | |
| 82:00:00 | 5dky-A | 2.2 | 3.4 | 63 | 928 | 0 | MOLECULE: | ALPHA G | LUCOSID | ASE-LIKE PROTEIN |
| 83:00:00 | 5n9j-A | 2.2 | 4.7 | 64 | 566 | 5 | MOLECULE: | MEDIATO | R OF RN | A POLYMERASE II TRANSCRIPTION SUBUN |
| 84:00:00 | 1jc5-B | 2.2 | 4.6 | 54 | 147 | 9 | MOLECULE: | METHYLM | ALONYL- | COA EPIMERASE |
| 85:00:00 | 3b77-A | 2.2 | 3.4 | 52 | 188 | 13 | MOLECULE: | UNCHARA | CTERIZE | D PROTEIN |
| 86:00:00 | 7syf-A | 2.2 | 5 | 63 | 1391 | 10 | MOLECULE: | PHOSPHA | TIDYLIN | OSITOL 3,4,5-TRISPHOSPHATE-DEPENDEN |
| 87:00:00 | 7kpr-A | 2.2 | 4.8 | 65 | 403 | 8 | MOLECULE: | PROTEIN | PHOSPH | ATASE 1H |
| 88:00:00 | 2wjq-A | 2.2 | 3.6 | 52 | 205 | 4 | MOLECULE: | PROBABL | E N-ACE | TYLNEURAMINIC ACID OUTER MEMBRANE C |
| 89:00:00 | 3v89-A | 2.2 | 4.3 | 63 | 853 | 6 | MOLECULE: | TRANSFE | RRIN-BI | NDING PROTEIN A |
| 90:00:00 | 4hvt-A | 2.2 | 3.4 | 61 | 691 | 10 | MOLECULE: | POST-PR | OLINE C | LEAVING ENZYME |
| 91:00:00 | 6dm4-A | 2.2 | 3.4 | 60 | 119 | 2 | MOLECULE: | RAVO | | |
| 92:00:00 | 7st9-H | 2.2 | 4.1 | 53 | 316 | 4 | MOLECULE: | CHECKPO | INT PRO | TEIN RAD24 |
| 93:00:00 | 5fq6-M | 2.2 | 6.2 | 63 | 948 | 3 | MOLECULE: | PUTATIV | E LIPOP | ROTEIN |
| 94:00:00 | 7krw-A | 2.2 | 2.9 | 52 | 608 | 2 | MOLECULE: | CHAPERO | NE PROT | EIN DNAK FUSED WITH SUBSTRATE PEPTI |
| 95:00:00 | 4jpd-A | 2.2 | 4 | 60 | 109 | 8 | MOLECULE: | PROTEIN | CYAY | |
| 96:00:00 | 6z3b-A | 2.2 | 3.8 | 62 | 371 | 10 | MOLECULE: | GFO/IDH | /MOCA F | AMILY OXIDOREDUCTASE |
| 97:00:00 | 5if6-A | 2.2 | 4.2 | 60 | 134 | 10 | MOLECULE: | OHP9_1C | | |
| 98:00:00 | 3wpv-A | 2.2 | 3.7 | 70 | 538 | 6 | MOLECULE: | BETA-FR | UCTOFUR | ANOSIDASE |
| 99:00:00 | 5gzl-A | 2.2 | 4.8 | 63 | 358 | 6 | MOLECULE: | LYSINE | CYCLODE | AMINASE |
| 0:00 | 6nd4-T | 2.2 | 3.3 | 49 | 812 | 8 | MOLECULE: | ETS RRN | A | |
| 1:00 | 5a1u-E | 2.2 | 3.2 | 54 | 822 | 11 | MOLECULE: | ADP-RIB | OSYLATI | ON FACTOR 1 |
| 2:00 | 6abo-A | 2.2 | 2.7 | 43 | 213 | 5 | MOLECULE: | DNA REP | AIR PRO | TEIN XRCC4 |
| 3:00 | 5cvo-D | 2.2 | 3.8 | 61 | 612 | 7 | MOLECULE: | WD REPE | AT-CONT | AINING PROTEIN 48 |
| 4:00 | 8bmx-C | 2.2 | 4 | 68 | 633 | 1 | MOLECULE: | PUTATIV | E TONB- | LINKED OUTER MEMBRANE RECEPTOR |
| 5:00 | 7z8b-F | 2.2 | 3 | 57 | 457 | 2 | MOLECULE: | CULLIN- | 7 | |
| 6:00 | 8dgt-A | 2.2 | 4.6 | 54 | 431 | 7 | MOLECULE: | SERINE/ | THREONI | NE-PROTEIN KINASE B-RAF |
| 7:00 | 7zgr-A | 2.2 | 3.6 | 74 | 1253 | 1 | MOLECULE: | PROTEIN | CFT1 | |
| 8:00 | 7wkk-D | 2.2 | 4.2 | 53 | 1115 | 2 | MOLECULE: | MGC8329 | 5 PROTE | IN |
| 9:00 | 1jmx-B | 2.2 | 3.5 | 59 | 339 | 5 | MOLECULE: | AMINE D | EHYDROG | ENASE |
| 10:00 | 3u4y-A | 2.2 | 2.9 | 52 | 319 | 12 | MOLECULE: | UNCHARA | CTERIZE | D PROTEIN |
| 11:00 | 6r2q-B | 2.2 | 4.4 | 57 | 649 | 2 | MOLECULE: | CYSTATH | IONINE | BETA-SYNTHASE |
| 12:00 | 8s1p-V | 2.2 | 3.3 | 54 | 249 | 6 | MOLECULE: | 50S RIB | OSOMAL | PROTEIN L35 |
| 13:00 | 8an0-A | 2.2 | 3.4 | 54 | 282 | 7 | MOLECULE: | PUTATIV | E SECRE | TED PROTEIN |
| 14:00 | 3alx-B | 2.2 | 5.5 | 69 | 526 | 7 | MOLECULE: | HEMAGGL | UTININ, | LINKER,CDW150 |
| 15:00 | 7jrd-A | 2.2 | 7.3 | 63 | 566 | 2 | MOLECULE: | LACTOFE | RRIN-BI | NDING PROTEIN B |
| 16:00 | 8c7g-A | 2.2 | 3.8 | 65 | 607 | 14 | MOLECULE: | MIC1 DO | MAIN-CO | NTAINING PROTEIN |
| 17:00 | 7eeb-E | 2.2 | 3.1 | 50 | 1053 | 4 | MOLECULE: | ENHANCE | D GREEN | FLUORESCENT PROTEIN,CATION CHANNEL |
| 18:00 | 4pdy-A | 2.2 | 3.2 | 56 | 333 | 4 | MOLECULE: | AMINOGL | YCOSIDE | PHOSPHOTRANSFERASE |

| | | | | | | | | | | |
|---|---|---|---|---|---|---|---|---|---|---|
| 19:00 | 2prv-A | 2.2 | 4.9 | 52 | 153 | 4 | MOLECULE: | UNCHARA | CTERIZE | D PROTEIN YOBK |
| 20:00 | 7zet-A | 2.2 | 3.2 | 60 | 377 | 8 | MOLECULE: | CLUSTER | IN | |
| 21:00 | 3c5m-B | 2.2 | 3.8 | 66 | 378 | 11 | MOLECULE: | OLIGOGA | LACTURO | NATE LYASE |
| 22:00 | 7z4f-G | 2.2 | 3.9 | 60 | 922 | 3 | MOLECULE: | PUTATIV | E STRUC | TURAL PROTEIN |
| 23:00 | 8h93-B | 2.2 | 3 | 54 | 367 | 7 | MOLECULE: | NACHT, | LRR AND | PYD DOMAINS-CONTAINING PROTEIN 5 |
| 24:00:00 | 2qkd-A | 2.2 | 3.8 | 50 | 384 | 8 | MOLECULE: | ZINC FI | NGER PR | OTEIN ZPR1 |
| 25:00:00 | 7np8-B | 2.2 | 2.7 | 42 | 620 | 10 | MOLECULE: | COENZYM | E F420- | DEPENDENT SULFITE REDUCTASE |
| 26:00:00 | 8ptx-A | 2.2 | 2.8 | 58 | 1215 | 7 | MOLECULE: | ELONGAT | OR COMP | LEX PROTEIN 1 |
| 27:00:00 | 8ew9-A | 2.2 | 2.9 | 62 | 241 | 2 | MOLECULE: | ALTERED | INHERI | TANCE OF MITOCHONDRIA PROTEIN 46, |
| 28:00:00 | 3oan-A | 2.2 | 3 | 47 | 116 | 0 | MOLECULE: | ABR034W | P | |
| 29:00:00 | 6y9c-A | 2.2 | 3.8 | 66 | 365 | 9 | MOLECULE: | CARNITI | NE MONO | OXYGENASE OXYGENASE SUBUNIT |
| 30:00:00 | 6imj-A | 2.2 | 3.1 | 56 | 412 | 5 | MOLECULE: | DNA LIG | ASE | |
| 31:00:00 | 3wyb-A | 2.2 | 3.9 | 53 | 327 | 6 | MOLECULE: | MESO-DI | AMINOPI | MELATE D-DEHYDROGENASE |
| 32:00:00 | 3ong-A | 2.2 | 3.6 | 55 | 114 | 9 | MOLECULE: | UBIQUIT | IN-ACTI | VATING ENZYME E1-LIKE |
| 33:00:00 | 2l3a-A | 2.2 | 4.1 | 48 | 82 | 6 | MOLECULE: | UNCHARA | CTERIZE | D PROTEIN |
| 34:00:00 | 8gxj-A | 2.2 | 3.4 | 49 | 189 | 6 | MOLECULE: | N-ACETY | LTRANSF | ERASE DOMAIN-CONTAINING PROTEIN |
| 35:00:00 | 2mhg-A | 2.2 | 4.6 | 50 | 75 | 8 | MOLECULE: | UNCHARA | CTERIZE | D PROTEIN |
| 36:00:00 | 1q9u-A | 2.2 | 3.2 | 58 | 128 | 12 | MOLECULE: | UNCHARA | CTERIZE | D PROTEIN APC35924 |
| 37:00:00 | 6wl5-A | 2.2 | 3.5 | 52 | 158 | 6 | MOLECULE: | ECMRR T | RANSCRI | PTIONAL REGULATOR |
| 38:00:00 | 3a15-B | 2.2 | 5.3 | 67 | 362 | 9 | MOLECULE: | ALDOXIM | E DEHYD | RATASE |
| 39:00:00 | 7uic-p | 2.2 | 2.6 | 62 | 903 | 10 | MOLECULE: | MEDIATO | R OF RN | A POLYMERASE II TRANSCRIPTION SUBUN |
| 40:00:00 | 4zn4-B | 2.2 | 3.4 | 57 | 411 | 7 | MOLECULE: | SQT1 | | |
| 41:00:00 | 7t6a-A | 2.2 | 2.8 | 46 | 185 | 13 | MOLECULE: | AVR1 (S | IX4), A | VIRULENCE PROTEIN 1 |
| 42:00:00 | 3was-A | 2.2 | 3.6 | 59 | 389 | 5 | MOLECULE: | 4-O-BET | A-D-MAN | NOSYL-D-GLUCOSE PHOSPHORYLASE |
| 43:00:00 | 3ty1-A | 2.2 | 5 | 56 | 384 | 7 | MOLECULE: | HYPOTHE | TICAL A | LDOSE 1-EPIMERASE |
| 44:00:00 | 3hdp-A | 2.2 | 3.7 | 49 | 132 | 6 | MOLECULE: | GLYOXAL | ASE-I | |
| 45:00:00 | 4wvm-B | 2.2 | 4.8 | 51 | 616 | 2 | MOLECULE: | STONUST | OXIN SU | BUNIT ALPHA |
| 46:00:00 | 3d4e-A | 2.2 | 2.9 | 54 | 162 | 4 | MOLECULE: | PUTATIV | E BETA- | LACTAMASE INHIBITOR PROTEIN |
| 47:00:00 | 5nxh-A | 2.2 | 3.6 | 60 | 546 | 5 | MOLECULE: | LONG-TA | IL FIBE | R PROXIMAL SUBUNIT |
| 48:00:00 | 5tf2-A | 2.2 | 3.3 | 57 | 338 | 7 | MOLECULE: | PROLACT | IN REGU | LATORY ELEMENT-BINDING PROTEIN |
| 49:00:00 | 6q50-A | 2.2 | 3.7 | 63 | 318 | 6 | MOLECULE: | MPT-4 | | |
| 50:00:00 | 4ddt-A | 2.2 | 4.2 | 56 | 1102 | 4 | MOLECULE: | REVERSE | GYRASE | |
| 51:00:00 | 8jwu-C | 2.2 | 3.8 | 63 | 412 | 2 | MOLECULE: | PHD FIN | GER PRO | TEIN 7,UBIQUITIN-CONJUGATING ENZYME |
| 52:00:00 | 8tum-a | 2.2 | 3.1 | 52 | 143 | 0 | MOLECULE: | TYPE IV | MAJOR | PILIN PROTEIN PILA |
| 53:00:00 | 6zls-A | 2.2 | 3.7 | 60 | 292 | 12 | MOLECULE: | HISTIDI | NE KINA | SE |
| 54:00:00 | 4bla-A | 2.2 | 4.4 | 69 | 730 | 3 | MOLECULE: | MALTOSE | #NAME? | G PERIPLASMIC PROTEIN, SUPPRESSOR O |
| 55:00:00 | 4bla-C | 2.2 | 5.4 | 72 | 724 | 4 | MOLECULE: | MALTOSE | #NAME? | G PERIPLASMIC PROTEIN, SUPPRESSOR O |
| 56:00:00 | 4q6k-A | 2.2 | 3.9 | 62 | 523 | 5 | MOLECULE: | BNR/ASP | #NAME? | PEAT PROTEIN |
| 57:00:00 | 6mnj-A | 2.2 | 2.8 | 58 | 511 | 7 | MOLECULE: | HZ136 | | |
| 58:00:00 | 4dsd-A | 2.2 | 3.8 | 58 | 126 | 12 | MOLECULE: | PUTATIV | E PERIP | LASMIC PROTEIN |
| 59:00:00 | 2ya6-B | 2.2 | 3.8 | 70 | 470 | 3 | MOLECULE: | NEURAMI | NIDASE | A |
| 60:00:00 | 2ya7-D | 2.2 | 4.1 | 72 | 470 | 4 | MOLECULE: | NEURAMI | NIDASE | A |
| 61:00:00 | 1ms4-B | 2.2 | 3.1 | 65 | 623 | 8 | MOLECULE: | TRANS-S | IALIDAS | E |
| 62:00:00 | 1w8n-A | 2.2 | 4 | 67 | 601 | 4 | MOLECULE: | BACTERI | AL SIAL | IDASE |
| 63:00:00 | 2ah2-A | 2.2 | 3.2 | 65 | 624 | 8 | MOLECULE: | TRANS-S | IALIDAS | E |
| 64:00:00 | 4bl9-B | 2.2 | 4.8 | 71 | 729 | 6 | MOLECULE: | MALTOSE | #NAME? | G PERIPLASMIC PROTEIN, SUPPRESSOR O |
| 65:00:00 | 7lbv-A | 2.2 | 3.6 | 65 | 379 | 11 | MOLECULE: | EXO-ALP | HA-SIAL | IDASE |
| 66:00:00 | 8t1z-A | 2.2 | 3.7 | 62 | 502 | 10 | MOLECULE: | SIALIDA | SE | |
| 67:00:00 | 3ob8-B | 2.2 | 3.5 | 71 | 1024 | 7 | MOLECULE: | BETA-GA | LACTOSI | DASE |
| 68:00:00 | 5ihj-A | 2.2 | 3.9 | 59 | 484 | 2 | MOLECULE: | MALTOSE | #NAME? | G PERIPLASMIC PROTEIN,FIMBRIAL PROT |
| 69:00:00 | 6d23-B | 2.2 | 4.7 | 71 | 490 | 3 | MOLECULE: | GLUCOSE | #NAME? | PHATE 1-DEHYDROGENASE |
| 70:00:00 | 6mrx-D | 2.2 | 3.3 | 60 | 521 | 7 | MOLECULE: | SIALIDA | SE26 | |
| 71:00:00 | 5o6c-A | 2.1 | 3.4 | 71 | 252 | 7 | MOLECULE: | E3 UBIQ | UITIN-P | ROTEIN LIGASE MYCBP2 |
| 72:00:00 | 6s62-A | 2.1 | 3.7 | 54 | 485 | 11 | MOLECULE: | PROPION | ATE CAT | ABOLIC PROTEIN PRPD |

| | | | | | | | | | | |
|---|---|---|---|---|---|---|---|---|---|---|
| 73:00:00 | 6j6a-A | 2.1 | 3.8 | 68 | 442 | 7 | MOLECULE: | ZINC-DE | PENDENT | PROTEASE, TLDD/PMBA FAMILY |
| 74:00:00 | 5cl2-A | 2.1 | 3.8 | 74 | 246 | 7 | MOLECULE: | SPORULA | TION-CO | NTROL PROTEIN SPO0M |
| 75:00:00 | 4ebr-A | 2.1 | 3.5 | 58 | 157 | 9 | MOLECULE: | UBIQUIT | IN-LIKE | -CONJUGATING ENZYME ATG10 |
| 76:00:00 | 6t3x-A | 2.1 | 4.1 | 65 | 212 | 5 | MOLECULE: | NUCLEAR | EGRESS | PROTEIN 2,NUCLEAR EGRESS PROTEIN 1 |
| 77:00:00 | 4h5b-A | 2.1 | 3.3 | 61 | 152 | 10 | MOLECULE: | DR_1245 | PROTEI | N |
| 78:00:00 | 7mh2-B | 2.1 | 2.7 | 52 | 279 | 13 | MOLECULE: | T4GALA | ENGINEE | RED PROTEIN NANOCAGE |
| 79:00:00 | 5xta-A | 2.1 | 3.7 | 61 | 120 | 13 | MOLECULE: | VIRK PR | OTEIN | |
| 80:00:00 | 7e5c-A | 2.1 | 4.4 | 72 | 439 | 7 | MOLECULE: | XAA-PRO | DIPEPT | IDASE |
| 81:00:00 | 6bqi-A | 2.1 | 4.1 | 63 | 258 | 8 | MOLECULE: | PROTEIN | IMPACT | HOMOLOG |
| 82:00:00 | 3t8n-B | 2.1 | 4.5 | 70 | 130 | 7 | MOLECULE: | STEROID | DELTA- | ISOMERASE |
| 83:00:00 | 8w7p-A | 2.1 | 2.8 | 60 | 250 | 0 | MOLECULE: | EXTRACE | LLULAR | DOMAIN OF A SENSOR HISTIDINE KINASE |
| 84:00:00 | 8e16-A | 2.1 | 5.4 | 53 | 272 | 9 | MOLECULE: | MAJOR C | APSID P | ROTEIN, GP6 |
| 85:00:00 | 1x9y-A | 2.1 | 3.5 | 73 | 346 | 4 | MOLECULE: | CYSTEIN | E PROTE | INASE |
| 86:00:00 | 3wqy-A | 2.1 | 3.7 | 60 | 906 | 5 | MOLECULE: | ALANINE | #NAME? | LIGASE |
| 87:00:00 | 5npy-A | 2.1 | 5.7 | 73 | 472 | 3 | MOLECULE: | FLAGELL | AR BASA | L BODY PROTEIN |
| 88:00:00 | 4tq2-A | 2.1 | 3.5 | 64 | 177 | 2 | MOLECULE: | PUTATIV | E PHYCO | ERYTHRIN LYASE |
| 89:00:00 | 3nlc-A | 2.1 | 2.7 | 48 | 534 | 8 | MOLECULE: | UNCHARA | CTERIZE | D PROTEIN VP0956 |
| 90:00:00 | 1vjh-A | 2.1 | 4.1 | 60 | 120 | 10 | MOLECULE: | BET V I | ALLERG | EN FAMILY |
| 91:00:00 | 2gjv-A | 2.1 | 3.2 | 62 | 136 | 3 | MOLECULE: | PUTATIV | E CYTOP | LASMIC PROTEIN |
| 92:00:00 | 3uau-A | 2.1 | 4 | 77 | 344 | 6 | MOLECULE: | SURFACE | #NAME? | D LIPOPROTEIN |
| 93:00:00 | 4jgo-B | 2.1 | 3.5 | 61 | 210 | 2 | MOLECULE: | SPORULA | TION KI | NASE D |
| 94:00:00 | 4onw-A | 2.1 | 4.7 | 66 | 265 | 6 | MOLECULE: | SUCCINY | L-DIAMI | NOPIMELATE DESUCCINYLASE |
| 95:00:00 | 4o4s-A | 2.1 | 3.8 | 70 | 199 | 9 | MOLECULE: | PHYCOCY | ANOBILI | N LYASE CPCT |
| 96:00:00 | 4r9k-C | 2.1 | 4.8 | 68 | 160 | 4 | MOLECULE: | LIMONEN | E-1,2-E | POXIDE HYDROLASE |
| 97:00:00 | 2i9i-A | 2.1 | 3.8 | 65 | 221 | 8 | MOLECULE: | HYPOTHE | TICAL P | ROTEIN |
| 98:00:00 | 6xgq-A | 2.1 | 4.5 | 55 | 353 | 7 | MOLECULE: | YSD1_17 | | |
| 99:00:00 | 7x80-A | 2.1 | 3.5 | 56 | 148 | 7 | MOLECULE: | PLOI4 | | |
| 0:00 | 4xkz-A | 2.1 | 3.7 | 50 | 251 | 12 | MOLECULE: | VALINE- | TRNA LI | GASE |
| 1:00 | 5mmj-c | 2.1 | 3.4 | 59 | 216 | 7 | MOLECULE: | 50S RIB | OSOMAL | PROTEIN L31 |
| 2:00 | 5ux0-D | 2.1 | 4.9 | 69 | 639 | 1 | MOLECULE: | ARGONAU | TE PROT | EIN |
| 3:00 | 4rzm-B | 2.1 | 4.7 | 72 | 283 | 11 | MOLECULE: | EPOXIDE | HYDROL | ASE LASB |
| 4:00 | 8gmm-A | 2.1 | 3.8 | 51 | 232 | 8 | MOLECULE: | HEMOPHI | LIN | |
| 5:00 | 2k6p-A | 2.1 | 2.8 | 53 | 84 | 4 | MOLECULE: | UNCHARA | CTERIZE | D PROTEIN HP_1423 |
| 6:00 | 2h9f-A | 2.1 | 4.4 | 58 | 391 | 5 | MOLECULE: | HYPOTHE | TICAL P | ROTEIN |
| 7:00 | 7cqy-D | 2.1 | 3.2 | 50 | 464 | 6 | MOLECULE: | TETRATH | IONATE | HYDROLASE |
| 8:00 | 7cud-A | 2.1 | 3.6 | 56 | 116 | 4 | MOLECULE: | AMINO A | CID ABC | TRANSPORTER SUBSTRATE-BINDING PROT |
| 9:00 | 3u0s-A | 2.1 | 3.2 | 58 | 327 | 7 | MOLECULE: | DIISOPR | OPYL-FL | UOROPHOSPHATASE |
| 10:00 | 6rw8-B | 2.1 | 3.4 | 50 | 2337 | 2 | MOLECULE: | A COMPO | NENT OF | INSECTICIDAL TOXIN COMPLEX (TC) |
| 11:00 | 8xcj-F | 2.1 | 7.6 | 55 | 420 | 4 | MOLECULE: | MALTOPO | RIN | |
| 12:00 | 6mfi-A | 2.1 | 7.9 | 54 | 264 | 6 | MOLECULE: | METALLO | #NAME? | ACTAMASE |
| 13:00 | 1jyh-A | 2.1 | 4 | 54 | 155 | 2 | MOLECULE: | DNA GYR | ASE INH | IBITORY PROTEIN |
| 14:00 | 6e8e-A | 2.1 | 3.6 | 44 | 468 | 11 | MOLECULE: | BETA SL | IDING C | LAMP,DNA MISMATCH REPAIR PROTEIN MU |
| 15:00 | 8sy3-A | 2.1 | 3.6 | 58 | 209 | 3 | MOLECULE: | BURP DO | MAIN-CO | NTAINING PROTEIN |
| 16:00 | 6trq-C | 2.1 | 3.9 | 55 | 336 | 4 | MOLECULE: | M7GPPPX | DIPHOS | PHATASE |
| 17:00 | 8c8g-B | 2.1 | 3 | 55 | 1195 | 2 | MOLECULE: | PUTATIV | E BOTUL | INUM-LIKE TOXIN WO |
| 18:00 | 6goc-A | 2.1 | 3.2 | 49 | 444 | 8 | MOLECULE: | DUF3826 | DOMAIN | -CONTAINING PROTEIN |
| 19:00 | 5dl5-A | 2.1 | 5.3 | 59 | 416 | 7 | MOLECULE: | MEMBRAN | E PROTE | IN |
| 20:00 | 4ut1-A | 2.1 | 5 | 57 | 551 | 4 | MOLECULE: | FLAGELL | AR HOOK | -ASSOCIATED PROTEIN |
| 21:00 | 8iha-A | 2.1 | 4 | 58 | 129 | 10 | MOLECULE: | POLYKET | IDE CYC | LASE / DEHYDRASE AND LIPID TRANSPOR |
| 22:00 | 8y7e-3 | 2.1 | 6.6 | 60 | 1193 | 3 | MOLECULE: | PRE-MRN | A | |
| 23:00 | 7qe7-I | 2.1 | 5.8 | 57 | 742 | 7 | MOLECULE: | ANAPHAS | E-PROMO | TING COMPLEX SUBUNIT 10 |
| 24:00:00 | 8rt6-A | 2.1 | 3.5 | 58 | 221 | 3 | MOLECULE: | TRWE PR | OTEIN | |
| 25:00:00 | 6nd4-J | 2.1 | 3.3 | 55 | 493 | 7 | MOLECULE: | ETS RRN | A | |
| 26:00:00 | 7rtn-A | 2.1 | 4.4 | 63 | 270 | 8 | MOLECULE: | OUTER C | APSID P | ROTEIN VP5 |

| | | | | | | | | | | |
|---|---|---|---|---|---|---|---|---|---|---|
| 27:00:00 | 8jxc-A | 2.1 | 3.3 | 49 | 1337 | 14 | MOLECULE: | LDL REC | EPTOR R | ELATED PROTEIN 2 |
| 28:00:00 | 1ugj-A | 2.1 | 3.7 | 56 | 141 | 2 | MOLECULE: | RIKEN C | DNA 231 | 0057J16 PROTEIN |
| 29:00:00 | 8dit-C | 2.1 | 3.1 | 66 | 736 | 5 | MOLECULE: | VACUOLA | R PROTE | IN SORTING-ASSOCIATED PROTEIN 33 |
| 30:00:00 | 8a0j-A | 2.1 | 8 | 55 | 223 | 7 | MOLECULE: | UNCHARA | CTERIZE | D PROTEIN TCIL3000_11_11110 |
| 31:00:00 | 7fh6-A | 2.1 | 3.5 | 62 | 653 | 5 | MOLECULE: | CYLK | | |
| 32:00:00 | 8fs6-G | 2.1 | 3.2 | 44 | 298 | 9 | MOLECULE: | CHECKPO | INT PRO | TEIN RAD24 |
| 33:00:00 | 8ovw-O | 2.1 | 5.8 | 55 | 241 | 13 | MOLECULE: | CENTROM | ERE-BIN | DING PROTEIN 1 |
| 34:00:00 | 7pkq-N | 2.1 | 2.9 | 58 | 195 | 3 | MOLECULE: | MS35 | | |
| 35:00:00 | 7sg7-S | 2.1 | 3.3 | 56 | 471 | 4 | MOLECULE: | GENE 14 | PROTEI | N |
| 36:00:00 | 6mit-G | 2.1 | 4.6 | 58 | 355 | 3 | MOLECULE: | LIPOPOL | YSACCHA | RIDE EXPORT SYSTEM ATP-BINDING PROT |
| 37:00:00 | 7pkn-O | 2.1 | 4.1 | 62 | 210 | 6 | MOLECULE: | CENTROM | ERE PRO | TEIN H |
| 38:00:00 | 3no2-A | 2.1 | 3 | 57 | 274 | 2 | MOLECULE: | UNCHARA | CTERIZE | D PROTEIN |
| 39:00:00 | 1y8c-A | 2.1 | 8.7 | 58 | 246 | 0 | MOLECULE: | S-ADENO | SYLMETH | IONINE-DEPENDENT METHYLTRANSFERASE |
| 40:00:00 | 6tdy-M | 2.1 | 4.4 | 51 | 243 | 6 | MOLECULE: | ATP SYN | THASE S | UBUNIT ALPHA |
| 41:00:00 | 4jf7-A | 2.1 | 3.9 | 69 | 501 | 6 | MOLECULE: | HEMAGGL | UTININ- | NEURAMINIDASE |
| 42:00:00 | 6nd4-L | 2.1 | 3 | 43 | 473 | 5 | MOLECULE: | ETS RRN | A | |
| 43:00:00 | 5yjl-D | 2.1 | 4.1 | 64 | 253 | 6 | MOLECULE: | GLUTAMY | L-TRNA | REDUCTASE 1 |
| 44:00:00 | 7lc1-B | 2.1 | 4.1 | 56 | 218 | 7 | MOLECULE: | ISOFORM | 2B OF | GTPASE KRAS |
| 45:00:00 | 7bjt-A | 2.1 | 3.7 | 58 | 727 | 7 | MOLECULE: | ALGINAT | E LYASE | , FAMILY PL17 |
| 46:00:00 | 1vrq-D | 2.1 | 3.9 | 55 | 91 | 5 | MOLECULE: | SARCOSI | NE OXID | ASE ALPHA SUBUNIT |
| 47:00:00 | 6nd4-W | 2.1 | 2.7 | 51 | 385 | 14 | MOLECULE: | ETS RRN | A | |
| 48:00:00 | 8xcj-A | 2.1 | 4.4 | 64 | 422 | 9 | MOLECULE: | MALTOPO | RIN | |
| 49:00:00 | 6rh5-A | 2.1 | 2.9 | 51 | 138 | 6 | MOLECULE: | ADAPTIN | EAR-BI | NDING COAT-ASSOCIATED PROTEIN 1 |
| 50:00:00 | 3wy4-B | 2.1 | 2.9 | 59 | 421 | 8 | MOLECULE: | ALPHA-G | LUCOSID | ASE |
| 51:00:00 | 1bp1-A | 2.1 | 3.2 | 57 | 456 | 5 | MOLECULE: | BACTERI | CIDAL/P | ERMEABILITY-INCREASING PROTEIN |
| 52:00:00 | 5vf3-A | 2.1 | 2.4 | 52 | 456 | 8 | MOLECULE: | CAPSID | VERTEX | PROTEIN GP24 |
| 53:00:00 | 8k9z-A | 2.1 | 3.2 | 54 | 405 | 9 | MOLECULE: | RDTND-R | ID CBD | |
| 54:00:00 | 6yle-B | 2.1 | 2.8 | 53 | 415 | 6 | MOLECULE: | PRE-RRN | A-PROCE | SSING PROTEIN IPI3 |
| 55:00:00 | 7qf6-B | 2.1 | 4.1 | 61 | 440 | 5 | MOLECULE: | N(5)-HY | DROXYOR | NITHINE:CIS-ANHYDROMEVALONYL COENZY |
| 56:00:00 | 3csv-A | 2.1 | 3.2 | 53 | 322 | 6 | MOLECULE: | AMINOGL | YCOSIDE | PHOSPHOTRANSFERASE |
| 57:00:00 | 1g6o-A | 2.1 | 2.6 | 41 | 323 | 7 | MOLECULE: | CAG-ALP | HA | |
| 58:00:00 | 2ojh-A | 2.1 | 4.2 | 58 | 277 | 7 | MOLECULE: | UNCHARA | CTERIZE | D PROTEIN ATU1656/AGR_C_3050 |
| 59:00:00 | 4yhc-A | 2.1 | 3.7 | 56 | 439 | 9 | MOLECULE: | STEROL | REGULAT | ORY ELEMENT-BINDING PROTEIN CLEAVAG |
| 60:00:00 | 3lib-C | 2.1 | 3.1 | 56 | 268 | 5 | MOLECULE: | HYPOTHE | TICAL S | ENSORY TRANSDUCTION HISTIDINE KINAS |
| 61:00:00 | 7t4e-A | 2.1 | 3.7 | 51 | 307 | 10 | MOLECULE: | EPX1 | | |
| 62:00:00 | 4rqo-B | 2.1 | 2.9 | 57 | 448 | 7 | MOLECULE: | L-SERIN | E DEHYD | RATASE |
| 63:00:00 | 6em5-R | 2.1 | 3.3 | 52 | 120 | 8 | MOLECULE: | 5.8S RI | BOSOMAL | RNA |
| 64:00:00 | 8e2g-A | 2.1 | 3.8 | 67 | 523 | 7 | MOLECULE: | BACULOV | IRAL IA | P REPEAT-CONTAINING PROTEIN 6 |
| 65:00:00 | 7c06-A | 2.1 | 2.6 | 52 | 193 | 6 | MOLECULE: | SPLICIN | G FACTO | R U2AF 23 KDA SUBUNIT |
| 66:00:00 | 2nyk-A | 2.1 | 3.7 | 52 | 237 | 2 | MOLECULE: | M157 | | |
| 67:00:00 | 6nd4-U | 2.1 | 3.6 | 56 | 407 | 9 | MOLECULE: | ETS RRN | A | |
| 68:00:00 | 8bou-A | 2.1 | 3.2 | 69 | 807 | 10 | MOLECULE: | N,N'-DI | ACETYLC | HITOBIOSE PHOSPHORYLASE |
| 69:00:00 | 5wd6-A | 2.1 | 3.4 | 54 | 195 | 7 | MOLECULE: | SHORT P | ALATE, | LUNG AND NASAL EPITHELIUM CARCINOMA |
| 70:00:00 | 4upl-A | 2.1 | 6 | 55 | 555 | 7 | MOLECULE: | SULFATA | SE FAMI | LY PROTEIN |
| 71:00:00 | 1a87-A | 2.1 | 3.1 | 48 | 297 | 10 | MOLECULE: | COLICIN | N | |
| 72:00:00 | 5hzd-A | 2.1 | 2.6 | 46 | 476 | 4 | MOLECULE: | 3' TERM | INAL UR | IDYLYL TRANSFERASE |
| 73:00:00 | 8onx-A | 2.1 | 4.5 | 64 | 371 | 6 | MOLECULE: | METHION | INE AMI | NOPEPTIDASE 2 |
| 74:00:00 | 8adl-A | 2.1 | 3.1 | 51 | 633 | 8 | MOLECULE: | MTC5 IS | OFORM 1 | |
| 75:00:00 | 5hcc-C | 2.1 | 3.3 | 53 | 148 | 8 | MOLECULE: | COMPLEM | ENT C5 | |
| 76:00:00 | 8jxg-A | 2.1 | 3.3 | 61 | 1114 | 7 | MOLECULE: | RAT RAP | | |
| 77:00:00 | 4pk1-A | 2.1 | 2.9 | 56 | 494 | 9 | MOLECULE: | CHIMERA | PROTEI | N OF OUTER MEMBRANE PROTEIN ASSEMBL |
| 78:00:00 | 6k96-B | 2.1 | 4 | 55 | 343 | 9 | MOLECULE: | FIVE-ME | MBERED- | CYCLITOL-PHOSPHATE SYNTHASE |
| 79:00:00 | 8y6p-Q | 2.1 | 3 | 52 | 1228 | 8 | MOLECULE: | APAF-1 | RELATED | KILLER DARK |
| 80:00:00 | 5e4m-A | 2.1 | 4.4 | 63 | 177 | 5 | MOLECULE: | HYDROXY | NITRILE | LYASE |

| | | | | | | | | | | |
|---|---|---|---|---|---|---|---|---|---|---|
| 81:00:00 | 8kei-D | 2.1 | 3.8 | 50 | 164 | 16 | MOLECULE: | CYTOCHR | OME B-2 | 45 LIGHT CHAIN |
| 82:00:00 | 2y3w-A | 2.1 | 2.8 | 53 | 172 | 9 | MOLECULE: | SPINDLE | ASSEMB | LY ABNORMAL PROTEIN 6 HOMOLOG |
| 83:00:00 | 5c0p-A | 2.1 | 3.5 | 62 | 284 | 8 | MOLECULE: | ENDO-AR | ABINASE | |
| 84:00:00 | 8ovw-P | 2.1 | 3.6 | 56 | 257 | 4 | MOLECULE: | CENTROM | ERE-BIN | DING PROTEIN 1 |
| 85:00:00 | 4ntq-A | 2.1 | 2.9 | 49 | 76 | 2 | MOLECULE: | CONTACT | #NAME? | ENT INHIBITOR A |
| 86:00:00 | 2fgt-A | 2.1 | 4.4 | 52 | 373 | 12 | MOLECULE: | TWO-COM | PONENT | SYSTEM YYCF/YYCG REGULATORY PROTEIN |
| 87:00:00 | 8uci-A | 2.1 | 3.3 | 62 | 381 | 8 | MOLECULE: | ATP DEP | ENDENT | DNA LIGASE |
| 88:00:00 | 8du5-B | 2.1 | 3.6 | 63 | 366 | 3 | MOLECULE: | SIALIDA | SE-1 | |
| 89:00:00 | 5afu-K | 2.1 | 3.6 | 74 | 275 | 4 | MOLECULE: | DYNEIN | TAIL | |
| 90:00:00 | 3aae-C | 2.1 | 3.6 | 74 | 270 | 5 | MOLECULE: | F-ACTIN | #NAME? | G PROTEIN SUBUNIT ALPHA-1 |
| 91:00:00 | 1qki-A | 2.1 | 4.4 | 75 | 487 | 1 | MOLECULE: | GLUCOSE | #NAME? | PHATE 1-DEHYDROGENASE |
| 92:00:00 | 7qyp-A | 2.1 | 2.9 | 67 | 519 | 12 | MOLECULE: | BNR/ASP | #NAME? | PEAT PROTEIN |
| 93:00:00 | 2kxp-A | 2.1 | 3.8 | 74 | 275 | 5 | MOLECULE: | F-ACTIN | #NAME? | G PROTEIN SUBUNIT ALPHA-1 |
| 94:00:00 | 5tsp-B | 2.1 | 3.3 | 65 | 448 | 5 | MOLECULE: | SIALIDA | SE | |
| 95:00:00 | 8du5-A | 2.1 | 3.5 | 61 | 366 | 3 | MOLECULE: | SIALIDA | SE-1 | |
| 96:00:00 | 7qyj-A | 2.1 | 2.9 | 66 | 519 | 12 | MOLECULE: | BNR/ASP | #NAME? | PEAT PROTEIN |
| 97:00:00 | 1ms0-A | 2.1 | 2.9 | 64 | 623 | 11 | MOLECULE: | TRANS-S | IALIDAS | E |
| 98:00:00 | 4yz1-A | 2.1 | 3.2 | 59 | 656 | 3 | MOLECULE: | PUTATIV | E NEURA | MINIDASE |
| 99:00:00 | 6myv-D | 2.1 | 3.3 | 57 | 522 | 5 | MOLECULE: | SIALIDA | SE26 | |
| 0:00 | 7xhl-F | 2.1 | 4.1 | 75 | 462 | 7 | MOLECULE: | GLUCOSE | 6-PHOS | PHATE DEHYDROGENASE |
| 1:00 | 5cfv-A | 2.1 | 6.6 | 59 | 484 | 8 | MOLECULE: | MALTOSE | #NAME? | G PERIPLASMIC PROTEIN,MALTOSE-BINDI |
| 2:00 | 4xa2-B | 2.1 | 3.5 | 56 | 488 | 4 | MOLECULE: | MALTOSE | #NAME? | G PERIPLASMIC PROTEIN,MBP-PILA: C |
| 3:00 | 1w8o-A | 2.1 | 2.9 | 54 | 601 | 4 | MOLECULE: | BACTERI | AL SIAL | IDASE |
| 4:00 | 2vw0-A | 2.1 | 3.1 | 57 | 657 | 11 | MOLECULE: | SIALIDA | SE B | |
| 5:00 | 7qy8-B | 2.1 | 2.8 | 58 | 519 | 5 | MOLECULE: | BNR/ASP | #NAME? | PEAT PROTEIN |
| 6:00 | 2ya8-A | 2.1 | 3.2 | 64 | 470 | 6 | MOLECULE: | NEURAMI | NIDASE | A |
| 7:00 | 4q6k-C | 2.1 | 3.9 | 69 | 523 | 12 | MOLECULE: | BNR/ASP | #NAME? | PEAT PROTEIN |
| 8:00 | 4fow-A | 2.1 | 3.4 | 61 | 658 | 11 | MOLECULE: | SIALIDA | SE B | |
| 9:00 | 5u0p-F | 2 | 3.8 | 58 | 195 | 2 | MOLECULE: | MEDIATO | R COMPL | EX SUBUNIT 14 |
| 10:00 | 8e0f-B | 2 | 4 | 60 | 454 | 10 | MOLECULE: | DOUBLE- | STRANDE | D RNA-SPECIFIC EDITASE 1 |
| 11:00 | 6wub-f | 2 | 4.2 | 53 | 97 | 6 | MOLECULE: | 16S RRN | A | |
| 12:00 | 3mqz-A | 2 | 4 | 75 | 205 | 5 | MOLECULE: | UNCHARA | CTERIZE | D CONSERVED PROTEIN DUF1054 |
| 13:00 | 8bgm-D | 2 | 4.2 | 74 | 476 | 7 | MOLECULE: | TOXIN | | |
| 14:00 | 5dcx-A | 2 | 3.8 | 60 | 209 | 2 | MOLECULE: | PROTEIN | REP68 | |
| 15:00 | 7lzh-A | 2 | 4.8 | 78 | 799 | 8 | MOLECULE: | GLUTAMA | TE RECE | PTOR 3.4 |
| 16:00 | 3n72-B | 2 | 3.7 | 67 | 152 | 3 | MOLECULE: | PUTATIV | E ACTIV | ATOR OF HSP90 |
| 17:00 | 5z4g-A | 2 | 3.2 | 59 | 145 | 3 | MOLECULE: | SAHS4 | | |
| 18:00 | 7cn3-C | 2 | 4 | 77 | 349 | 4 | MOLECULE: | 2,5-DIH | YDROXYP | YRIDINE 5,6-DIOXYGENASE |
| 19:00 | 7b3a-A | 2 | 3.4 | 63 | 276 | 6 | MOLECULE: | PUTATIV | E ACYL- | TRANSFERASE DOMAIN PROTEIN |
| 20:00 | 5lal-B | 2 | 4.5 | 61 | 158 | 2 | MOLECULE: | DIRIGEN | T PROTE | IN 6 |
| 21:00 | 6tyd-V | 2 | 3.9 | 59 | 219 | 3 | MOLECULE: | LIM DOM | AIN-BIN | DING PROTEIN 1 |
| 22:00 | 7sn8-I | 2 | 2.6 | 47 | 634 | 11 | MOLECULE: | INTEGRA | TOR COM | PLEX SUBUNIT 4 |
| 23:00 | 1xko-B | 2 | 2.8 | 57 | 159 | 12 | MOLECULE: | CHEMOTA | XIS PRO | TEIN CHEX |
| 24:00:00 | 1dbf-A | 2 | 4.8 | 60 | 127 | 12 | MOLECULE: | PROTEIN | (CHORI | SMATE MUTASE) |
| 25:00:00 | 6lix-B | 2 | 3.6 | 74 | 196 | 7 | MOLECULE: | CHROMOF | HORE LY | ASE CRL, CHLOROPLASTIC |
| 26:00:00 | 2p2s-A | 2 | 3.6 | 67 | 333 | 9 | MOLECULE: | PUTATIV | E OXIDO | REDUCTASE |
| 27:00:00 | 7el9-A | 2 | 3.9 | 55 | 1731 | 5 | MOLECULE: | RNA-DIR | ECTED R | NA POLYMERASE L |
| 28:00:00 | 1ss4-A | 2 | 3.2 | 50 | 149 | 6 | MOLECULE: | GLYOXAL | ASE FAM | ILY PROTEIN |
| 29:00:00 | 3w15-A | 2 | 3.9 | 60 | 336 | 5 | MOLECULE: | PEROXIS | OMAL TA | RGETING SIGNAL 2 RECEPTOR |
| 30:00:00 | 7rw4-A | 2 | 4 | 45 | 321 | 0 | MOLECULE: | JUNCTOP | HILIN-1 | |
| 31:00:00 | 2hye-A | 2 | 3.6 | 55 | 1140 | 2 | MOLECULE: | DNA DAM | AGE-BIN | DING PROTEIN 1 |
| 32:00:00 | 5o51-A | 2 | 3 | 54 | 309 | 6 | MOLECULE: | RHO GUA | NYL NUC | LEOTIDE EXCHANGE FACTOR (ROM2), PUT |
| 33:00:00 | 4rki-A | 2 | 3.7 | 50 | 374 | 4 | MOLECULE: | DNA POL | YMERASE | III SUBUNIT BETA |
| 34:00:00 | 4yy8-B | 2 | 2.2 | 47 | 377 | 4 | MOLECULE: | KELCH P | ROTEIN | |

| | | | | | | | | | | | |
|---|---|---|---|---|---|---|---|---|---|---|---|
| 35:00:00 | 2opd-A | 2 | 3.3 | 59 | 121 | 5 | MOLECULE: | PILX | | | |
| 36:00:00 | 7st9-G | 2 | 3.6 | 53 | 340 | 6 | MOLECULE: | CHECKPO | INT PRO | TEIN RAD24 | |
| 37:00:00 | 3lp9-A | 2 | 3.1 | 49 | 227 | 6 | MOLECULE: | LS-24 | | | |
| 38:00:00 | 6l7x-A | 2 | 3.1 | 53 | 203 | 11 | MOLECULE: | MRNA_TR | IPASE D | OMAIN-CONTAINING PROTEIN | |
| 39:00:00 | 8suc-A | 2 | 2.6 | 52 | 282 | 8 | MOLECULE: | NHL (RI | NG FING | ER B-BOX COILED COIL) DOMAIN CONTAI | |
| 40:00:00 | 3r75-B | 2 | 4.8 | 60 | 622 | 7 | MOLECULE: | ANTHRAN | ILATE/P | ARA-AMINOBENZOATE SYNTHASES COMPONE | |
| 41:00:00 | 6fpt-A | 2 | 3.1 | 55 | 391 | 11 | MOLECULE: | E3 UBIQ | UITIN-P | ROTEIN LIGASE TRIM71 | |
| 42:00:00 | 2g7j-A | 2 | 3.1 | 55 | 112 | 5 | MOLECULE: | PUTATIV | E CYTOP | LASMIC PROTEIN | |
| 43:00:00 | 2ntk-A | 2 | 3.8 | 53 | 206 | 8 | MOLECULE: | IMP CYC | LOHYDRO | LASE | |
| 44:00:00 | 7l5m-A | 2 | 8.1 | 51 | 87 | 4 | MOLECULE: | LIPOCAL | IN FAMI | LY PROTEIN | |
| 45:00:00 | 6a5g-B | 2 | 3.9 | 66 | 157 | 9 | MOLECULE: | [4+2] A | ND [4+6 | ] CYCLASE STMD | |
| 46:00:00 | 5jrk-A | 2 | 3.2 | 57 | 695 | 12 | MOLECULE: | DIPEPTI | DYL AMI | NOPEPTIDASES/ACYLAMINOACYL-PEPTIDAS | |
| 47:00:00 | 6j6q-F | 2 | 3.8 | 56 | 161 | 2 | MOLECULE: | PRE-MRN | A-SPLIC | ING FACTOR 8 | |
| 48:00:00 | 3pqi-A | 2 | 3.7 | 47 | 202 | 2 | MOLECULE: | GENE PR | ODUCT 1 | | 38 |
| 49:00:00 | 5mq0-t | 2 | 3 | 48 | 438 | 8 | MOLECULE: | YEAST U | BC4 GEN | E FOR UBIQUITIN-CONJUGATING ENZYME | |
| 50:00:00 | 6gie-A | 2 | 4.8 | 60 | 218 | 5 | MOLECULE: | 33-36 K | DA OUTE | R MEMBRANE PROTEIN | |
| 51:00:00 | 7zao-A | 2 | 3.4 | 69 | 429 | 6 | MOLECULE: | SIALIDA | SE (NEU | RAMINIDASE) FAMILY PROTEIN-LIKE PRO | |
| 52:00:00 | 8ck1-A | 2 | 3.2 | 53 | 826 | 2 | MOLECULE: | TAIL NO | ZZLE | | |
| 53:00:00 | 8eec-A | 2 | 4.3 | 53 | 317 | 8 | MOLECULE: | ISOFORM | 2 OF M | ITOGEN-ACTIVATED PROTEIN KINASE KIN | |
| 54:00:00 | 3to3-B | 2 | 3.5 | 53 | 595 | 6 | MOLECULE: | PETROBA | CTIN BI | OSYNTHESIS PROTEIN ASBB | |
| 55:00:00 | 2fug-7 | 2 | 4.4 | 60 | 127 | 3 | MOLECULE: | NADH-QU | INONE O | XIDOREDUCTASE CHAIN 1 | |
| 56:00:00 | 1r0u-A | 2 | 4.6 | 50 | 142 | 8 | MOLECULE: | PROTEIN | YWIB | | |
| 57:00:00 | 8bux-A | 2 | 3.4 | 66 | 289 | 9 | MOLECULE: | GRANULE | ASSOCI | ATED RAC AND RHOG EFFECTOR PROTEIN | |
| 58:00:00 | 5vh9-B | 2 | 2.9 | 50 | 354 | 6 | MOLECULE: | DYNEIN | HEAVY C | HAIN, CYTOPLASMIC | |
| 59:00:00 | 5nqz-A | 2 | 3.7 | 52 | 267 | 2 | MOLECULE: | FACTOR | H BINDI | NG PROTEIN,MAJOR OUTER MEMBRANE PRO | |
| 60:00:00 | 5wru-E | 2 | 4.7 | 50 | 311 | 2 | MOLECULE: | PROBABL | E INORG | ANIC PYROPHOSPHATASE | |
| 61:00:00 | 6qam-A | 2 | 3.6 | 55 | 211 | 9 | MOLECULE: | OUTER M | EMBRANE | PROTEIN ALKL | |
| 62:00:00 | 3t4k-A | 2 | 3.8 | 54 | 268 | 11 | MOLECULE: | HISTIDI | NE KINA | SE 4 | |
| 63:00:00 | 1qqg-B | 2 | 4.3 | 51 | 210 | 4 | MOLECULE: | INSULIN | RECEPT | OR SUBSTRATE 1 | |
| 64:00:00 | 3kya-A | 2 | 3.6 | 61 | 471 | 10 | MOLECULE: | PUTATIV | E PHOSP | HATASE | |
| 65:00:00 | 6m48-A | 2 | 3.6 | 65 | 813 | 11 | MOLECULE: | SPAC | | | |
| 66:00:00 | 7wj9-A | 2 | 3.5 | 54 | 733 | 6 | MOLECULE: | ALPHA-X | YLOSIDA | SE | |
| 67:00:00 | 7snh-C | 2 | 3.6 | 62 | 476 | 3 | MOLECULE: | GLUCOSE | #NAME? | PHATE 1-DEHYDROGENASE | |
| 68:00:00 | 3aae-I | 2 | 3.5 | 75 | 270 | 5 | MOLECULE: | F-ACTIN | #NAME? | G PROTEIN SUBUNIT ALPHA-1 | |
| 69:00:00 | 2pyw-B | 2 | 2.4 | 55 | 411 | 5 | MOLECULE: | UNCHARA | CTERIZE | D PROTEIN | |
| 70:00:00 | 7qy9-B | 2 | 3.3 | 68 | 519 | 12 | MOLECULE: | BNR/ASP | #NAME? | PEAT PROTEIN | |
| 71:00:00 | 1ms3-A | 2 | 3.2 | 69 | 623 | 10 | MOLECULE: | TRANS-S | IALIDAS | E | |
| 72:00:00 | 5aq1-C | 2 | 4.3 | 69 | 484 | 4 | MOLECULE: | GLUCOSE | #NAME? | PHATE DEHYDROGENASE | |
| 73:00:00 | 4yz4-B | 2 | 3.6 | 59 | 656 | 7 | MOLECULE: | SIALIDA | SE, NAN | C | |
| 74:00:00 | 7sng-B | 2 | 3.4 | 71 | 467 | 7 | MOLECULE: | GLUCOSE | #NAME? | PHATE 1-DEHYDROGENASE | |
| 75:00:00 | 4yz5-B | 2 | 3.7 | 54 | 654 | 6 | MOLECULE: | PUTATIV | E NEURA | MINIDASE | |
| 76:00:00 | 7qyj-B | 2 | 3.2 | 67 | 519 | 12 | MOLECULE: | BNR/ASP | #NAME? | PEAT PROTEIN | |
| 77:00:00 | 3bk5-A | 2 | 3.2 | 53 | 235 | 2 | MOLECULE: | PUTATIV | E OUTER | MEMBRANE LIPOPROTEIN-SORTING PROTE | |

**DALI results: MnA-1 Col**

| No: | Chain | Z | rmsd | lali | nres | %id | PDB | Description |
|---|---|---|---|---|---|---|---|---|
| 1:00 | 8th8-J | 5.6 | 9 | 178 | 372 | 6 | MOLECULE: | DYNEIN REGULATORY COMPLEX PROTEIN 1/2 N-TERMINAL |
| 2:00 | 3o0z-D | 5.1 | 5.5 | 144 | 160 | 4 | MOLECULE: | RHO-ASSOCIATED PROTEIN KINASE 1 |
| 3:00 | 8q85-G | 5 | 6.9 | 152 | 178 | 8 | MOLECULE: | KINETOCHORE PROTEIN NDC80 |
| 4:00 | 4jpp-D | 5 | 9.4 | 128 | 129 | 3 | MOLECULE: | MINOR SPIKE PROTEIN H |
| 5:00 | 2fxo-A | 5 | 4.9 | 126 | 129 | 10 | MOLECULE: | MYOSIN HEAVY CHAIN, CARDIAC MUSCLE BETA ISOFORM |
| 6:00 | 7jg5-b | 4.9 | 8.1 | 136 | 145 | 2 | MOLECULE: | ATP SYNTHASE SUBUNIT ALPHA |
| 7:00 | 8txr-C | 4.8 | 3.9 | 167 | 431 | 5 | MOLECULE: | EXODEOXYRIBONUCLEASE 7 LARGE SUBUNIT |
| 8:00 | 4f61-I | 4.8 | 7.7 | 173 | 234 | 9 | MOLECULE: | TUBULIN ALPHA CHAIN |
| 9:00 | 9bw6-A | 4.8 | 16.6 | 159 | 783 | 9 | MOLECULE: | MAJOR VAULT PROTEIN |
| 10:00 | 8i3e-B | 4.8 | 7 | 128 | 135 | 8 | MOLECULE: | ELKS/RAB6-INTERACTING/CAST FAMILY MEMBER 1 |
| 11:00 | 2ocy-A | 4.5 | 9 | 145 | 149 | 6 | MOLECULE: | RAB GUANINE NUCLEOTIDE EXCHANGE FACTOR SEC2 |
| 12:00 | 6j5i-b | 4.4 | 12.3 | 154 | 209 | 4 | MOLECULE: | ATP SYNTHASE SUBUNIT ALPHA, MITOCHONDRIAL |
| 13:00 | 7znk-o | 4.4 | 8.7 | 134 | 164 | 9 | MOLECULE: | RNA |
| 14:00 | 5gai-B | 4.4 | 7.9 | 131 | 721 | 8 | MOLECULE: | PORTAL PROTEIN |
| 15:00 | 6fkf-p | 4.3 | 12.1 | 135 | 143 | 5 | MOLECULE: | ATP SYNTHASE SUBUNIT ALPHA, CHLOROPLASTIC |
| 16:00 | 8th8-A | 4.3 | 7.7 | 158 | 290 | 6 | MOLECULE: | DYNEIN REGULATORY COMPLEX PROTEIN 1/2 N-TERMINAL |
| 17:00 | 4xa6-A | 4.3 | 11.4 | 131 | 168 | 6 | MOLECULE: | GP7-MYH7(1777-1855)-EB1 CHIMERA PROTEIN |
| 18:00 | 3uf1-D | 4.3 | 3.4 | 112 | 112 | 10 | MOLECULE: | VIMENTIN |
| 19:00 | 6e2j-B | 4.3 | 3.1 | 103 | 104 | 2 | MOLECULE: | KERATIN, TYPE II CYTOSKELETAL 1 |
| 20:00 | 6fkh-b | 4.3 | 6.1 | 131 | 161 | 3 | MOLECULE: | ATP SYNTHASE SUBUNIT A, CHLOROPLASTIC |
| 21:00 | 7apk-m | 4.2 | 8 | 128 | 549 | 6 | MOLECULE: | THO COMPLEX SUBUNIT 1 |
| 22:00 | 8fef-B | 4.2 | 22.9 | 159 | 335 | 3 | MOLECULE: | VIRULENCE FACTOR MCE FAMILY PROTEIN |
| 23:00 | 8th8-D | 4.2 | 10.3 | 166 | 292 | 7 | MOLECULE: | DYNEIN REGULATORY COMPLEX PROTEIN 1/2 N-TERMINAL |
| 24:00:00 | 7fde-H | 4.2 | 10.8 | 112 | 112 | 5 | MOLECULE: | V-TYPE PROTON ATPASE SUBUNIT C |
| 25:00:00 | 6wm2-K | 4.1 | 10.9 | 114 | 114 | 9 | MOLECULE: | V-TYPE PROTON ATPASE SUBUNIT E 1 |
| 26:00:00 | 2no2-A | 4.1 | 3.4 | 102 | 102 | 8 | MOLECULE: | HUNTINGTIN-INTERACTING PROTEIN 1 |
| 27:00:00 | 6bl7-A | 4.1 | 2.8 | 99 | 99 | 6 | MOLECULE: | PROTEIN STU2 |
| 28:00:00 | 2oto-B | 4.1 | 6.5 | 134 | 138 | 8 | MOLECULE: | M PROTEIN |
| 29:00:00 | 8bd7-X | 4.1 | 6.7 | 103 | 106 | 7 | MOLECULE: | IFT88 |
| 30:00:00 | 5to5-A | 4.1 | 6.6 | 126 | 141 | 10 | MOLECULE: | NUCLEOPROTEIN TPR |
| 31:00:00 | 6ys4-A | 4.1 | 4.1 | 100 | 103 | 4 | MOLECULE: | SPINDLE ASSEMBLY ABNORMAL PROTEIN 6 HOMOLOG |
| 32:00:00 | 6iac-E | 4 | 27.4 | 131 | 146 | 3 | MOLECULE: | PORTAL PROTEIN |
| 33:00:00 | 4lin-A | 4 | 10.3 | 179 | 289 | 4 | MOLECULE: | TAIL NEEDLE PROTEIN GP26 |
| 34:00:00 | 1d7m-A | 4 | 4.4 | 99 | 101 | 10 | MOLECULE: | CORTEXILLIN I |
| 35:00:00 | 6uui-C | 4 | 6.1 | 107 | 107 | 6 | MOLECULE: | KERATIN, TYPE II CYTOSKELETAL 1 |
| 36:00:00 | 9ce3-C | 4 | 11.3 | 183 | 616 | 4 | MOLECULE: | ISOFORM 4 OF TUBERIN |
| 37:00:00 | 8tek-P | 3.9 | 8.4 | 128 | 138 | 5 | MOLECULE: | DYNEIN REGULATORY COMPLEX PROTEIN 1/2 N-TERMINAL |
| 38:00:00 | 8bd7-L | 3.9 | 41.1 | 151 | 303 | 7 | MOLECULE: | IFT88 |
| 39:00:00 | 5d80-G | 3.9 | 5 | 101 | 223 | 5 | MOLECULE: | V-TYPE PROTON ATPASE CATALYTIC SUBUNIT A |
| 40:00:00 | 8bd7-M | 3.9 | 22.9 | 137 | 164 | 2 | MOLECULE: | IFT88 |
| 41:00:00 | 8r1a-A | 3.9 | 5.2 | 167 | 587 | 5 | MOLECULE: | GUANYLATE BINDING PROTEIN 1 |
| 42:00:00 | 6eun-A | 3.9 | 3.2 | 112 | 147 | 3 | MOLECULE: | ADHESIN |
| 43:00:00 | 8tek-N | 3.9 | 5.6 | 158 | 187 | 5 | MOLECULE: | DYNEIN REGULATORY COMPLEX PROTEIN 1/2 N-TERMINAL |
| 44:00:00 | 6ff7-K | 3.9 | 33.4 | 123 | 213 | 3 | MOLECULE: | RNA-BINDING MOTIF PROTEIN, X-LINKED 2 |
| 45:00:00 | 8arf-B | 3.9 | 3.2 | 96 | 97 | 9 | MOLECULE: | PROTEIN SPINDLY |
| 46:00:00 | 8pqw-F | 3.9 | 13.9 | 139 | 150 | 9 | MOLECULE: | CYTOPLASMIC DYNEIN 1 HEAVY CHAIN 1 |
| 47:00:00 | 8i03-G | 3.9 | 4.1 | 112 | 166 | 13 | MOLECULE: | PAIRED AMPHIPATHIC HELIX PROTEIN PST1 |
| 48:00:00 | 4yto-B | 3.9 | 5.7 | 96 | 97 | 4 | MOLECULE: | SYNAPTONEMAL COMPLEX PROTEIN 1 |
| 49:00:00 | 6vq6-M | 3.9 | 9.8 | 111 | 114 | 7 | MOLECULE: | ATPASE H+TRANSPORTING V1 SUBUNIT A |
| 50:00:00 | 8p0v-L | 3.8 | 44.8 | 139 | 416 | 5 | MOLECULE: | COILED-COIL DOMAIN-CONTAINING PROTEIN 93 |

| | | | | | | | | |
|---|---|---|---|---|---|---|---|---|
| 51:00:00 | 8ppr-D | 3.8 | 13.9 | 152 | 248 | 5 | MOLECULE: | KINETOCHORE-ASSOCIATED PROTEIN DSN1 HOMOLOG |
| 52:00:00 | 8i03-E | 3.8 | 12.4 | 129 | 169 | 4 | MOLECULE: | PAIRED AMPHIPATHIC HELIX PROTEIN PST1 |
| 53:00:00 | 3oja-A | 3.8 | 32.1 | 146 | 482 | 7 | MOLECULE: | LEUCINE-RICH IMMUNE MOLECULE 1 |
| 54:00:00 | 3ghg-J | 3.8 | 8.7 | 129 | 186 | 3 | MOLECULE: | FIBRINOGEN ALPHA CHAIN |
| 55:00:00 | 3s4r-B | 3.8 | 3.9 | 91 | 91 | 3 | MOLECULE: | VIMENTIN |
| 56:00:00 | 6h9l-B | 3.8 | 7.5 | 114 | 129 | 11 | MOLECULE: | UNCHARACTERIZED PROTEIN |
| 57:00:00 | 8tek-B | 3.8 | 10 | 147 | 208 | 2 | MOLECULE: | DYNEIN REGULATORY COMPLEX PROTEIN 1/2 N-TERMINAL |
| 58:00:00 | 6e2j-A | 3.8 | 5.1 | 105 | 107 | 7 | MOLECULE: | KERATIN, TYPE II CYTOSKELETAL 1 |
| 59:00:00 | 7bjg-A | 3.8 | 5 | 97 | 98 | 6 | MOLECULE: | SD21996P |
| 60:00:00 | 8tek-A | 3.7 | 11.6 | 149 | 200 | 3 | MOLECULE: | DYNEIN REGULATORY COMPLEX PROTEIN 1/2 N-TERMINAL |
| 61:00:00 | 5hmo-A | 3.7 | 5.1 | 123 | 131 | 12 | MOLECULE: | UNCONVENTIONAL MYOSIN-X |
| 62:00:00 | 8fed-A | 3.7 | 25.9 | 152 | 392 | 5 | MOLECULE: | VIRULENCE FACTOR MCE FAMILY PROTEIN |
| 63:00:00 | 6r17-B | 3.7 | 3.9 | 91 | 94 | 2 | MOLECULE: | SYNAPTONEMAL COMPLEX CENTRAL ELEMENT PROTEIN 2 |
| 64:00:00 | 9b0z-A | 3.7 | 3.3 | 89 | 89 | 8 | MOLECULE: | OPTINEURIN |
| 65:00:00 | 6fia-D | 3.7 | 9.2 | 102 | 104 | 10 | MOLECULE: | LINE-1 RETROTRANSPOSABLE ELEMENT ORF1 PROTEIN |
| 66:00:00 | 8ovw-U | 3.6 | 15 | 128 | 184 | 2 | MOLECULE: | CENTROMERE-BINDING PROTEIN 1 |
| 67:00:00 | 9fn1-U | 3.6 | 7.9 | 120 | 158 | 8 | MOLECULE: | VPS20/32/60-LIKE PROTEIN (ESCRT-III) |
| 68:00:00 | 7woo-I | 3.6 | 25.4 | 109 | 187 | 4 | MOLECULE: | NUCLEOPORIN NIC96 |
| 69:00:00 | 3ni0-A | 3.6 | 5.2 | 92 | 95 | 4 | MOLECULE: | BONE MARROW STROMAL ANTIGEN 2 |
| 70:00:00 | 3oja-B | 3.6 | 6.6 | 109 | 534 | 10 | MOLECULE: | LEUCINE-RICH IMMUNE MOLECULE 1 |
| 71:00:00 | 5nvu-A | 3.6 | 33.4 | 163 | 3169 | 0 | MOLECULE: | DYNEIN MOTOR DOMAIN |
| 72:00:00 | 6u0t-A | 3.6 | 4.4 | 117 | 136 | 6 | MOLECULE: | TUBULIN ALPHA CHAIN |
| 73:00:00 | 6vzf-A | 3.6 | 3.2 | 86 | 86 | 7 | MOLECULE: | AUTOPHAGY-RELATED PROTEIN 11 |
| 74:00:00 | 8z22-C | 3.6 | 3.5 | 90 | 93 | 7 | MOLECULE: | REGULATING SYNAPTIC MEMBRANE EXOCYTOSIS 1 |
| 75:00:00 | 8ims-A | 3.6 | 3.7 | 92 | 93 | 3 | MOLECULE: | E3 UBIQUITIN-PROTEIN LIGASE TRAF7 |
| 76:00:00 | 8ppr-N | 3.5 | 15.2 | 148 | 217 | 7 | MOLECULE: | KINETOCHORE-ASSOCIATED PROTEIN DSN1 HOMOLOG |
| 77:00:00 | 8ovw-Z | 3.5 | 8.9 | 121 | 151 | 5 | MOLECULE: | CENTROMERE-BINDING PROTEIN 1 |
| 78:00:00 | 5mqf-L | 3.5 | 34.7 | 116 | 336 | 9 | MOLECULE: | PRE-MRNA-PROCESSING-SPLICING FACTOR 8 |
| 79:00:00 | 4zqa-A | 3.5 | 4.7 | 84 | 86 | 5 | MOLECULE: | SIN3 HISTONE DEACETYLASE COREPRESSOR COMPLEX COMP |
| 80:00:00 | 1x79-B | 3.5 | 3.7 | 88 | 90 | 6 | MOLECULE: | ADP-RIBOSYLATION FACTOR BINDING PROTEIN GGA1 |
| 81:00:00 | 6oqa-G | 3.5 | 2.7 | 87 | 87 | 6 | MOLECULE: | PEPTIDYL-PROLYL CIS-TRANS ISOMERASE FKBP1A |
| 82:00:00 | 6rd4-5 | 3.4 | 6.2 | 115 | 123 | 12 | MOLECULE: | ASA-10: POLYTOMELLA F-ATP SYNTHASE ASSOCIATED SUB |
| 83:00:00 | 2p22-A | 3.4 | 14 | 121 | 168 | 3 | MOLECULE: | SUPPRESSOR PROTEIN STP22 OF TEMPERATURE-SENSITIVE |
| 84:00:00 | 7qoo-H | 3.4 | 31.3 | 130 | 210 | 5 | MOLECULE: | CENTROMERE PROTEIN C |
| 85:00:00 | 2xnx-M | 3.4 | 18.6 | 106 | 107 | 6 | MOLECULE: | FIBRINOGEN ALPHA CHAIN |
| 86:00:00 | 6cfz-I | 3.4 | 13.2 | 104 | 106 | 6 | MOLECULE: | ASK1 |
| 87:00:00 | 9mhg-C | 3.4 | 10.4 | 124 | 304 | 2 | MOLECULE: | PHOSPHOINOSITIDE 3-KINASE REGULATORY SUBUNIT 4 |
| 88:00:00 | 8ppr-P | 3.4 | 9 | 124 | 176 | 8 | MOLECULE: | KINETOCHORE-ASSOCIATED PROTEIN DSN1 HOMOLOG |
| 89:00:00 | 4y66-C | 3.4 | 19.3 | 127 | 197 | 6 | MOLECULE: | MND1 |
| 90:00:00 | 6exn-D | 3.4 | 8.6 | 95 | 97 | 11 | MOLECULE: | U2 SNRNA |
| 91:00:00 | 5ijn-H | 3.4 | 20.9 | 98 | 169 | 6 | MOLECULE: | NUCLEAR PORE COMPLEX PROTEIN NUP155 |
| 92:00:00 | 6l5j-A | 3.4 | 7.3 | 150 | 208 | 7 | MOLECULE: | ROOTLETIN |
| 93:00:00 | 7yqh-A | 3.4 | 8.4 | 183 | 1069 | 4 | MOLECULE: | STRUCTURAL MAINTENANCE OF CHROMOSOMES PROTEIN 5 |
| 94:00:00 | 5bw9-g | 3.3 | 4.5 | 85 | 141 | 7 | MOLECULE: | V-TYPE PROTON ATPASE CATALYTIC SUBUNIT A |
| 95:00:00 | 5dfz-A | 3.3 | 21.9 | 157 | 343 | 4 | MOLECULE: | VACUOLAR PROTEIN SORTING-ASSOCIATED PROTEIN 38 |
| 96:00:00 | 6znl-n | 3.3 | 48.5 | 112 | 343 | 5 | MOLECULE: | ARP1 ACTIN RELATED PROTEIN 1 HOMOLOG A |
| 97:00:00 | 8th8-B | 3.3 | 30.2 | 166 | 276 | 8 | MOLECULE: | DYNEIN REGULATORY COMPLEX PROTEIN 1/2 N-TERMINAL |
| 98:00:00 | 3vp8-B | 3.3 | 4.5 | 79 | 80 | 5 | MOLECULE: | GENERAL TRANSCRIPTIONAL COREPRESSOR TUP1 |
| 99:00:00 | 5lxn-A | 3.3 | 3.6 | 81 | 81 | 9 | MOLECULE: | TRANSFORMING ACIDIC COILED-COIL-CONTAINING PROTEI |
| 0:00 | 8amr-A | 3.3 | 4.8 | 106 | 155 | 3 | MOLECULE: | TRIPARTITE MOTIF-CONTAINING PROTEIN 3 |
| 1:00 | 4h22-B | 3.3 | 4.1 | 82 | 83 | 6 | MOLECULE: | LEUCINE-RICH REPEAT FLIGHTLESS-INTERACTING PROTEI |
| 2:00 | 6fln-B | 3.3 | 47.9 | 127 | 367 | 7 | MOLECULE: | E3 UBIQUITIN/ISG15 LIGASE TRIM25 |

| | | | | | | | | |
|---|---|---|---|---|---|---|---|---|
| 3:00 | 7fde-G | 3.3 | 8.9 | 131 | 231 | 7 | MOLECULE: | V-TYPE PROTON ATPASE SUBUNIT C |
| 4:00 | 6b8h-b | 3.3 | 25.4 | 135 | 200 | 7 | MOLECULE: | ATP SYNTHASE SUBUNIT 9, MITOCHONDRIAL |
| 5:00 | 8fed-F | 3.2 | 21.3 | 153 | 399 | 5 | MOLECULE: | VIRULENCE FACTOR MCE FAMILY PROTEIN |
| 6:00 | 7puz-A | 3.2 | 4.5 | 101 | 148 | 9 | MOLECULE: | MICOS COMPLEX SUBUNIT MIC60 |
| 7:00 | 9na8-E | 3.2 | 4.9 | 84 | 244 | 7 | MOLECULE: | AUGMIN SUBUNIT 1 |
| 8:00 | 5nnv-D | 3.2 | 5.1 | 127 | 252 | 9 | MOLECULE: | CHROMOSOME PARTITION PROTEIN SMC,CHROMOSOME PARTI |
| 9:00 | 5hda-A | 3.2 | 8 | 97 | 121 | 8 | MOLECULE: | ZINC FINGER MYND DOMAIN-CONTAINING PROTEIN 11 |
| 10:00 | 4xa1-D | 3.2 | 7.3 | 103 | 122 | 9 | MOLECULE: | GP7-MYH7(1173-1238)-EB1 CHIMERA PROTEIN |
| 11:00 | 9bq2-A | 3.2 | 9.3 | 148 | 402 | 10 | MOLECULE: | FLOTILLIN-2 |
| 12:00 | 8xi2-K | 3.2 | 50.6 | 148 | 210 | 3 | MOLECULE: | MPN DOMAIN-CONTAINING PROTEIN |
| 13:00 | 6vq6-I | 3.2 | 4.6 | 101 | 225 | 6 | MOLECULE: | ATPASE H+TRANSPORTING V1 SUBUNIT A |
| 14:00 | 6y09-C | 3.2 | 5.1 | 83 | 86 | 7 | MOLECULE: | RAS-RELATED PROTEIN RAB-33B |
| 15:00 | 8wjo-B | 3.1 | 41.9 | 88 | 283 | 5 | MOLECULE: | STRUCTURAL MAINTENANCE OF CHROMOSOMES PROTEIN 5 |
| 16:00 | 5dfz-D | 3.1 | 13.4 | 135 | 341 | 7 | MOLECULE: | VACUOLAR PROTEIN SORTING-ASSOCIATED PROTEIN 38 |
| 17:00 | 7zr1-D | 3.1 | 7.9 | 146 | 780 | 8 | MOLECULE: | DOUBLE-STRAND BREAK REPAIR PROTEIN |
| 18:00 | 9dtr-D | 3.1 | 10.9 | 100 | 108 | 3 | MOLECULE: | U2 SNRNA |
| 19:00 | 8hpo-D | 3.1 | 11.6 | 109 | 165 | 4 | MOLECULE: | TRANSCRIPTIONAL REGULATORY PROTEIN UME1 |
| 20:00 | 1gk4-A | 3.1 | 2.9 | 78 | 79 | 6 | MOLECULE: | VIMENTIN |
| 21:00 | 2ch7-A | 3.1 | 6.2 | 150 | 309 | 5 | MOLECULE: | METHYL-ACCEPTING CHEMOTAXIS PROTEIN |
| 22:00 | 3iv1-A | 3.1 | 2.6 | 78 | 78 | 8 | MOLECULE: | TUMOR SUSCEPTIBILITY GENE 101 PROTEIN |
| 23:00 | 6lth-O | 3.1 | 16.9 | 111 | 325 | 8 | MOLECULE: | TRANSCRIPTION ACTIVATOR BRG1 |
| 24:00:00 | 3bas-B | 3.1 | 3.9 | 82 | 87 | 10 | MOLECULE: | MYOSIN HEAVY CHAIN, STRIATED MUSCLE/GENERAL CONTR |
| 25:00:00 | 1fav-A | 3.1 | 2.9 | 78 | 78 | 9 | MOLECULE: | HIV-1 ENVELOPE PROTEIN CHIMERA |
| 26:00:00 | 2xqh-A | 3 | 9.5 | 130 | 258 | 5 | MOLECULE: | IMMUNOGLOBULIN-BINDING PROTEIN EIBD |
| 27:00:00 | 8tek-M | 3 | 13 | 111 | 164 | 5 | MOLECULE: | DYNEIN REGULATORY COMPLEX PROTEIN 1/2 N-TERMINAL |
| 28:00:00 | 6oqr-X | 3 | 7.1 | 120 | 156 | 7 | MOLECULE: | ATP SYNTHASE SUBUNIT DELTA |
| 29:00:00 | 5h07-D | 3 | 3.9 | 78 | 78 | 8 | MOLECULE: | POLYUBIQUITIN-C |
| 30:00:00 | 8ppr-M | 3 | 9.3 | 129 | 205 | 7 | MOLECULE: | KINETOCHORE-ASSOCIATED PROTEIN DSN1 HOMOLOG |
| 31:00:00 | 8g0p-A | 3 | 6.8 | 101 | 143 | 10 | MOLECULE: | KINETOCHORE PROTEIN NDC80 HOMOLOG |
| 32:00:00 | 2jee-A | 3 | 4.2 | 77 | 78 | 6 | MOLECULE: | CELL DIVISION PROTEIN ZAPB |
| 33:00:00 | 7qoo-U | 3 | 4.4 | 110 | 186 | 4 | MOLECULE: | CENTROMERE PROTEIN C |
| 34:00:00 | 3v6i-A | 3 | 6.4 | 113 | 186 | 12 | MOLECULE: | V-TYPE ATP SYNTHASE SUBUNIT E |
| 35:00:00 | 6z6o-D | 3 | 65.4 | 162 | 542 | 7 | MOLECULE: | HISTONE DEACETYLASE HDA1 |
| 36:00:00 | 9bq2-B | 3 | 15.4 | 139 | 422 | 9 | MOLECULE: | FLOTILLIN-2 |
| 37:00:00 | 7vdv-A | 2.9 | 22.2 | 125 | 809 | 5 | MOLECULE: | HISTONE H4 |
| 38:00:00 | 9ftl-a | 2.9 | 29.5 | 116 | 170 | 6 | MOLECULE: | HEIMDALLARCHAEOTA ARCHAEON AB_125 ESCRT-IIIB |
| 39:00:00 | 8cqn-B | 2.9 | 7.7 | 122 | 187 | 6 | MOLECULE: | LIPOPROTEIN, PUTATIVE |
| 40:00:00 | 8u95-A | 2.9 | 9.8 | 180 | 1009 | 6 | MOLECULE: | MYOSIN HEAVY CHAIN, ISOFORM U |
| 41:00:00 | 9i8m-U | 2.9 | 5.6 | 74 | 75 | 3 | MOLECULE: | GAMMA-TUBULIN COMPLEX COMPONENT |
| 42:00:00 | 3trt-A | 2.9 | 2.8 | 75 | 75 | 5 | MOLECULE: | VIMENTIN |
| 43:00:00 | 3vem-C | 2.9 | 3.2 | 79 | 83 | 9 | MOLECULE: | HELICASE PROTEIN MOM1 |
| 44:00:00 | 2doq-D | 2.9 | 3.6 | 81 | 83 | 4 | MOLECULE: | CELL DIVISION CONTROL PROTEIN 31 |
| 45:00:00 | 4dt0-A | 2.9 | 7.9 | 98 | 181 | 8 | MOLECULE: | V-TYPE ATP SYNTHASE SUBUNIT E |
| 46:00:00 | 7sqk-E | 2.9 | 10.7 | 144 | 610 | 6 | MOLECULE: | HAUS AUGMIN-LIKE COMPLEX SUBUNIT 1 |
| 47:00:00 | 7jtk-F | 2.9 | 8.3 | 151 | 287 | 5 | MOLECULE: | FLAGELLAR RADIAL SPOKE PROTEIN 1 |
| 48:00:00 | 9kns-B | 2.9 | 5.1 | 74 | 75 | 9 | MOLECULE: | PROTEIN HOOK HOMOLOG 3 |
| 49:00:00 | 4wpe-A | 2.9 | 72.5 | 162 | 275 | 2 | MOLECULE: | CYTOKINESIS PROTEIN 2 |
| 50:00:00 | 8ovw-B | 2.9 | 5.8 | 90 | 107 | 7 | MOLECULE: | CENTROMERE-BINDING PROTEIN 1 |
| 51:00:00 | 8q85-U | 2.8 | 29.4 | 94 | 95 | 4 | MOLECULE: | KINETOCHORE PROTEIN NDC80 |
| 52:00:00 | 7sc0-A | 2.8 | 26.2 | 102 | 129 | 5 | MOLECULE: | CAVEOLIN-1 |
| 53:00:00 | 6wuc-K | 2.8 | 21.7 | 109 | 235 | 5 | MOLECULE: | INNER KINETOCHORE SUBUNIT MCM16 |
| 54:00:00 | 8d8k-V | 2.8 | 17.3 | 122 | 233 | 4 | MOLECULE: | PROBABLE S-ADENOSYL-L-METHIONINE-DEPENDENT RNA |

| | | | | | | | | |
|---|---|---|---|---|---|---|---|---|
| 55:00:00 | 4wpx-B | 2.8 | 8.8 | 77 | 81 | 9 | MOLECULE: | CELL DIVISION CONTROL PROTEIN 31-LIKE PROTEIN |
| 56:00:00 | 7sqk-H | 2.8 | 25.7 | 130 | 201 | 5 | MOLECULE: | HAUS AUGMIN-LIKE COMPLEX SUBUNIT 1 |
| 57:00:00 | 1m1j-B | 2.8 | 7.4 | 129 | 402 | 3 | MOLECULE: | FIBRINOGEN ALPHA SUBUNIT |
| 58:00:00 | 4mvd-B | 2.8 | 17.4 | 118 | 253 | 7 | MOLECULE: | CHOLINE-PHOSPHATE CYTIDYLYLTRANSFERASE A |
| 59:00:00 | 1m1j-D | 2.8 | 9.6 | 132 | 194 | 5 | MOLECULE: | FIBRINOGEN ALPHA SUBUNIT |
| 60:00:00 | 9esh-W | 2.8 | 33.6 | 144 | 526 | 6 | MOLECULE: | PRE-MRNA |
| 61:00:00 | 4aj5-B | 2.8 | 12 | 85 | 87 | 7 | MOLECULE: | SPINDLE AND KINETOCHORE-ASSOCIATED PROTEIN 1 |
| 62:00:00 | 8qbv-A | 2.8 | 12.9 | 128 | 217 | 8 | MOLECULE: | MEMBRANE-ASSOCIATED PROTEIN VIPP1 |
| 63:00:00 | 5wlq-A | 2.8 | 4.6 | 86 | 104 | 5 | MOLECULE: | CAPSID ASSEMBLY SCAFFOLDING PROTEIN,MYOSIN-7,MICR |
| 64:00:00 | 7x2e-A | 2.8 | 7.4 | 90 | 163 | 4 | MOLECULE: | HARMONIN |
| 65:00:00 | 3n7n-B | 2.7 | 2.6 | 84 | 164 | 2 | MOLECULE: | MONOPOLIN COMPLEX SUBUNIT CSM1 |
| 66:00:00 | 9cgi-C | 2.7 | 13.4 | 113 | 201 | 4 | MOLECULE: | RNA-DIRECTED RNA POLYMERASE L |
| 67:00:00 | 3wqa-A | 2.7 | 9.6 | 132 | 201 | 6 | MOLECULE: | TRIMERIC AUTOTRANSPORTER ADHESIN |
| 68:00:00 | 6oei-A | 2.7 | 7.4 | 94 | 243 | 10 | MOLECULE: | SPINDLE POLE BODY COMPONENT SPC42,SIGMA-54-DEPEND |
| 69:00:00 | 2ba2-A | 2.7 | 4.8 | 81 | 81 | 6 | MOLECULE: | HYPOTHETICAL UPF0134 PROTEIN MPN010 |
| 70:00:00 | 4dhx-A | 2.7 | 2.9 | 72 | 72 | 8 | MOLECULE: | 80 KDA MCM3-ASSOCIATED PROTEIN |
| 71:00:00 | 5wst-A | 2.7 | 2.1 | 71 | 71 | 10 | MOLECULE: | UNCONVENTIONAL MYOSIN-VIIA |
| 72:00:00 | 5eof-A | 2.7 | 6.6 | 74 | 74 | 11 | MOLECULE: | OPTINEURIN |
| 73:00:00 | 3k29-A | 2.7 | 5 | 98 | 162 | 6 | MOLECULE: | PUTATIVE UNCHARACTERIZED PROTEIN |
| 74:00:00 | 6djl-E | 2.7 | 56.2 | 106 | 226 | 8 | MOLECULE: | RAS-RELATED PROTEIN RAB-11A |
| 75:00:00 | 7v2w-J | 2.7 | 18.1 | 144 | 236 | 2 | MOLECULE: | THO COMPLEX SUBUNIT HPR1 |
| 76:00:00 | 7oac-A | 2.7 | 6.2 | 81 | 81 | 5 | MOLECULE: | GENERAL CONTROL TRANSCRIPTION FACTOR GCN4,CONSERV |
| 77:00:00 | 6n6s-D | 2.7 | 3.2 | 68 | 69 | 10 | MOLECULE: | TNFAIP3-INTERACTING PROTEIN 1 |
| 78:00:00 | 7yqh-B | 2.7 | 9.8 | 187 | 1066 | 6 | MOLECULE: | STRUCTURAL MAINTENANCE OF CHROMOSOMES PROTEIN 5 |
| 79:00:00 | 8q85-V | 2.7 | 9.6 | 92 | 117 | 2 | MOLECULE: | KINETOCHORE PROTEIN NDC80 |
| 80:00:00 | 5mq4-C | 2.7 | 8.1 | 92 | 117 | 7 | MOLECULE: | PROTEIN KINASE C-BINDING PROTEIN 1 |
| 81:00:00 | 5t58-B | 2.6 | 10.9 | 110 | 190 | 5 | MOLECULE: | KLLA0F02343P |
| 82:00:00 | 8q85-b | 2.6 | 6.2 | 68 | 70 | 4 | MOLECULE: | KINETOCHORE PROTEIN NDC80 |
| 83:00:00 | 9io5-G | 2.6 | 5.3 | 89 | 260 | 8 | MOLECULE: | G1-ATPASE SUBUNIT BETA |
| 84:00:00 | 6wcj-B | 2.6 | 12.4 | 87 | 109 | 7 | MOLECULE: | CLATHRIN HEAVY CHAIN 1 |
| 85:00:00 | 8qyd-A | 2.6 | 6.1 | 118 | 280 | 4 | MOLECULE: | ANTI-PHAGE DEFENSE ZORAB SYSTEM ZORA |
| 86:00:00 | 9cpo-B | 2.6 | 12.4 | 92 | 195 | 2 | MOLECULE: | RNA-DIRECTED RNA POLYMERASE NSP12 |
| 87:00:00 | 8qfc-D | 2.6 | 21.7 | 120 | 185 | 13 | MOLECULE: | 60S RIBOSOMAL PROTEIN L10A |
| 88:00:00 | 7emf-D | 2.6 | 11.5 | 109 | 158 | 6 | MOLECULE: | MEDIATOR OF RNA POLYMERASE II TRANSCRIPTION SUBUN |
| 89:00:00 | 6kn7-U | 2.6 | 9.4 | 108 | 170 | 6 | MOLECULE: | ACTIN, ALPHA SKELETAL MUSCLE |
| 90:00:00 | 8at3-B | 2.6 | 10.3 | 153 | 597 | 6 | MOLECULE: | HAUS AUGMIN-LIKE COMPLEX SUBUNIT 1 |
| 91:00:00 | 2nps-B | 2.6 | 3.8 | 66 | 68 | 2 | MOLECULE: | VESICLE-ASSOCIATED MEMBRANE PROTEIN 4 |
| 92:00:00 | 3zdo-F | 2.6 | 2.3 | 65 | 65 | 6 | MOLECULE: | PHOSPHOPROTEIN |
| 93:00:00 | 3b5n-C | 2.6 | 3.3 | 69 | 70 | 1 | MOLECULE: | SYNAPTOBREVIN HOMOLOG 1 |
| 94:00:00 | 5opt-P | 2.6 | 13.6 | 94 | 249 | 2 | MOLECULE: | ACTIVATED PROTEIN KINASE C RECEPTOR, PUTATIVE |
| 95:00:00 | 6b3o-A | 2.6 | 32 | 127 | 344 | 2 | MOLECULE: | SPIKE GLYCOPROTEIN |
| 96:00:00 | 5mq0-t | 2.6 | 17.8 | 96 | 438 | 6 | MOLECULE: | YEAST UBC4 GENE FOR UBIQUITIN-CONJUGATING ENZYME |
| 97:00:00 | 5toh-B | 2.6 | 5.3 | 69 | 70 | 1 | MOLECULE: | POLYMERASE COFACTOR VP35 |
| 98:00:00 | 4n21-C | 2.6 | 4 | 84 | 124 | 4 | MOLECULE: | GP2 ECTODOMAIN |
| 99:00:00 | 5ed9-B | 2.6 | 3.9 | 72 | 72 | 8 | MOLECULE: | SUN DOMAIN-CONTAINING PROTEIN 2 |
| 0:00 | 5kiu-A | 2.6 | 7.1 | 75 | 76 | 3 | MOLECULE: | SELENOPROTEIN S |
| 1:00 | 8a5p-G | 2.6 | 9.8 | 93 | 99 | 4 | MOLECULE: | INO80 ATPASE |
| 2:00 | 5oqm-h | 2.6 | 9.9 | 102 | 131 | 6 | MOLECULE: | DNA-DIRECTED RNA POLYMERASE II SUBUNIT RPB1 |
| 3:00 | 8j0s-d | 2.6 | 23.3 | 161 | 445 | 4 | MOLECULE: | ATP SYNTHASE SUBUNIT A |
| 4:00 | 2gv5-C | 2.6 | 2.4 | 69 | 73 | 4 | MOLECULE: | CELL DIVISION CONTROL PROTEIN 31 |
| 5:00 | 5zuv-A | 2.6 | 3.5 | 86 | 130 | 0 | MOLECULE: | SPIKE GLYCOPROTEIN,INHIBITOR EK1 |
| 6:00 | 9lc0-K | 2.5 | 12.2 | 134 | 298 | 4 | MOLECULE: | 60 KDA PROTEIN |

| | | | | | | | | |
|---|---|---|---|---|---|---|---|---|
| 7:00 | 8fed-E | 2.5 | 21.4 | 141 | 358 | 7 | MOLECULE: | VIRULENCE FACTOR MCE FAMILY PROTEIN |
| 8:00 | 6ird-B | 2.5 | 5.1 | 112 | 254 | 4 | MOLECULE: | 1-PHOSPHATIDYLINOSITOL 4,5-BISPHOSPHATE PHOSPHODI |
| 9:00 | 3jb9-i | 2.5 | 34.7 | 112 | 161 | 5 | MOLECULE: | PRE-MRNA-SPLICING FACTOR SPP42 |
| 10:00 | 4r8g-E | 2.5 | 20.2 | 118 | 324 | 3 | MOLECULE: | UNCONVENTIONAL MYOSIN-IC |
| 11:00 | 1joc-A | 2.5 | 7 | 75 | 123 | 0 | MOLECULE: | EARLY ENDOSOMAL AUTOANTIGEN 1 |
| 12:00 | 8ppr-F | 2.5 | 2.8 | 66 | 112 | 14 | MOLECULE: | KINETOCHORE-ASSOCIATED PROTEIN DSN1 HOMOLOG |
| 13:00 | 7woo-G | 2.5 | 6.3 | 91 | 200 | 3 | MOLECULE: | NUCLEOPORIN NIC96 |
| 14:00 | 6yvu-A | 2.5 | 61.7 | 146 | 1127 | 7 | MOLECULE: | STRUCTURAL MAINTENANCE OF CHROMOSOMES PROTEIN 2,S |
| 15:00 | 1gmj-A | 2.5 | 2.7 | 65 | 65 | 8 | MOLECULE: | ATPASE INHIBITOR |
| 16:00 | 4l2w-D | 2.5 | 3.7 | 67 | 68 | 10 | MOLECULE: | RHO-ASSOCIATED PROTEIN KINASE 1 |
| 17:00 | 5vpc-B | 2.5 | 3.9 | 68 | 68 | 4 | MOLECULE: | PROTEIN FOSB |
| 18:00 | 4e61-A | 2.5 | 2.5 | 68 | 91 | 9 | MOLECULE: | PROTEIN BIM1 |
| 19:00 | 4cg4-A | 2.5 | 30.3 | 95 | 376 | 12 | MOLECULE: | PYRIN |
| 20:00 | 2wz7-B | 2.5 | 3 | 67 | 67 | 3 | MOLECULE: | UNCHARACTERIZED PROTEIN YBGF |
| 21:00 | 3qh9-A | 2.5 | 3.8 | 66 | 66 | 9 | MOLECULE: | LIPRIN-BETA-2 |
| 22:00 | 5jhf-F | 2.5 | 48.7 | 160 | 405 | 1 | MOLECULE: | KLTH0D11660P |
| 23:00 | 5c9n-A | 2.5 | 2.3 | 64 | 64 | 6 | MOLECULE: | GEMININ COILED-COIL DOMAIN-CONTAINING PROTEIN 1 |
| 24:00:00 | 4aj5-T | 2.5 | 8.7 | 90 | 110 | 7 | MOLECULE: | SPINDLE AND KINETOCHORE-ASSOCIATED PROTEIN 1 |
| 25:00:00 | 4nad-A | 2.5 | 5.8 | 88 | 133 | 6 | MOLECULE: | REGULATION OF NUCLEAR PRE-MRNA DOMAIN-CONTAINING |
| 26:00:00 | 5apz-A | 2.5 | 6.5 | 104 | 112 | 4 | MOLECULE: | GENERAL CONTROL PROTEIN GCN4, NOR1 TCAR0761, GENE |
| 27:00:00 | 6otn-B | 2.5 | 3 | 66 | 67 | 5 | MOLECULE: | TROPOMYOSIN ALPHA-3 CHAIN |
| 28:00:00 | 8c5v-l | 2.4 | 5.9 | 154 | 516 | 3 | MOLECULE: | CHEMOTAXIS PROTEIN CHEA |
| 29:00:00 | 7jts-s | 2.4 | 20.8 | 103 | 290 | 4 | MOLECULE: | RADIAL SPOKE PROTEIN 3 |
| 30:00:00 | 2p22-C | 2.4 | 9.5 | 103 | 186 | 8 | MOLECULE: | SUPPRESSOR PROTEIN STP22 OF TEMPERATURE-SENSITIVE |
| 31:00:00 | 4r3z-A | 2.4 | 4 | 69 | 76 | 7 | MOLECULE: | AMINOACYL TRNA SYNTHASE COMPLEX-INTERACTING MULTI |
| 32:00:00 | 4orh-H | 2.4 | 2.8 | 74 | 142 | 8 | MOLECULE: | UBIQUITIN-CONJUGATING ENZYME E2 VARIANT 2 |
| 33:00:00 | 1aa0-A | 2.4 | 7.4 | 87 | 113 | 2 | MOLECULE: | FIBRITIN |
| 34:00:00 | 7bv6-B | 2.4 | 2.2 | 61 | 61 | 5 | MOLECULE: | VESICLE-ASSOCIATED MEMBRANE PROTEIN 8 |
| 35:00:00 | 3m0d-A | 2.4 | 3.6 | 63 | 63 | 8 | MOLECULE: | TNF RECEPTOR-ASSOCIATED FACTOR 2 |
| 36:00:00 | 4n78-E | 2.4 | 3.8 | 67 | 67 | 4 | MOLECULE: | CYTOPLASMIC FMR1-INTERACTING PROTEIN 1 |
| 37:00:00 | 3b5n-D | 2.4 | 3.5 | 64 | 64 | 6 | MOLECULE: | SYNAPTOBREVIN HOMOLOG 1 |
| 38:00:00 | 3lt6-F | 2.4 | 3 | 63 | 63 | 5 | MOLECULE: | ADHESIN YADA |
| 39:00:00 | 4c47-A | 2.4 | 6.2 | 84 | 191 | 5 | MOLECULE: | INNER MEMBRANE LIPOPROTEIN |
| 40:00:00 | 6w1s-Q | 2.4 | 28.6 | 101 | 131 | 8 | MOLECULE: | MEDIATOR OF RNA POLYMERASE II TRANSCRIPTION SUBUN |
| 41:00:00 | 7d2e-D | 2.4 | 2.5 | 63 | 63 | 6 | MOLECULE: | LIPRIN-ALPHA-2 |
| 42:00:00 | 8ia3-E | 2.4 | 7.4 | 87 | 111 | 1 | MOLECULE: | UPSTREAM STIMULATORY FACTOR 2 |
| 43:00:00 | 8p1u-C | 2.4 | 10.9 | 89 | 93 | 6 | MOLECULE: | CELL DIVISION PROTEIN FTSL |
| 44:00:00 | 5jvp-B | 2.4 | 6 | 68 | 90 | 12 | MOLECULE: | CHIMERA PROTEIN OF CENTROMERE-ASSOCIATED PROTEIN |
| 45:00:00 | 4pxj-C | 2.4 | 2.3 | 59 | 61 | 8 | MOLECULE: | C-JUN-AMINO-TERMINAL KINASE-INTERACTING PROTEIN 3 |
| 46:00:00 | 8q85-d | 2.4 | 17.8 | 87 | 108 | 7 | MOLECULE: | KINETOCHORE PROTEIN NDC80 |
| 47:00:00 | 2nps-A | 2.4 | 3.9 | 62 | 63 | 10 | MOLECULE: | VESICLE-ASSOCIATED MEMBRANE PROTEIN 4 |
| 48:00:00 | 5oqm-n | 2.4 | 16.1 | 97 | 136 | 6 | MOLECULE: | DNA-DIRECTED RNA POLYMERASE II SUBUNIT RPB1 |
| 49:00:00 | 7eeb-C | 2.3 | 4.1 | 80 | 278 | 6 | MOLECULE: | ENHANCED GREEN FLUORESCENT PROTEIN,CATION CHANNEL |
| 50:00:00 | 8q3v-g | 2.3 | 14.5 | 73 | 76 | 7 | MOLECULE: | TETRAHYDROMETHANOPTERIN S-METHYLTRANSFERASE SUBUN |
| 51:00:00 | 4y66-D | 2.3 | 47.7 | 115 | 208 | 11 | MOLECULE: | MND1 |
| 52:00:00 | 1go4-G | 2.3 | 16.1 | 88 | 100 | 9 | MOLECULE: | MITOTIC SPINDLE ASSEMBLY CHECKPOINT PROTEIN MAD2A |
| 53:00:00 | 5c1f-A | 2.3 | 43.2 | 129 | 298 | 5 | MOLECULE: | SEPTATION PROTEIN IMP2 |
| 54:00:00 | 8soi-B | 2.3 | 16.8 | 145 | 515 | 0 | MOLECULE: | RB1-INDUCIBLE COILED-COIL PROTEIN 1 |
| 55:00:00 | 3p8c-F | 2.3 | 11 | 105 | 156 | 3 | MOLECULE: | CYTOPLASMIC FMR1-INTERACTING PROTEIN 1 |
| 56:00:00 | 4cjd-A | 2.3 | 17.9 | 79 | 123 | 6 | MOLECULE: | NADA |
| 57:00:00 | 1z56-A | 2.3 | 4.8 | 75 | 77 | 4 | MOLECULE: | LIGASE INTERACTING FACTOR 1 |
| 58:00:00 | 5odw-D | 2.3 | 2.7 | 64 | 198 | 9 | MOLECULE: | FERRIPYOVERDINE RECEPTOR |

| | | | | | | | | |
|---|---|---|---|---|---|---|---|---|
| 59:00:00 | 1t2k-D | 2.3 | 2.8 | 60 | 61 | 3 | MOLECULE: | 31-MER |
| 60:00:00 | 6kn7-T | 2.3 | 15.7 | 82 | 138 | 10 | MOLECULE: | ACTIN, ALPHA SKELETAL MUSCLE |
| 61:00:00 | 3m0d-C | 2.3 | 2.6 | 63 | 63 | 5 | MOLECULE: | TNF RECEPTOR-ASSOCIATED FACTOR 2 |
| 62:00:00 | 2yo2-A | 2.3 | 5.8 | 106 | 160 | 3 | MOLECULE: | GENERAL CONTROL PROTEIN GCN4, PUTATIVE INNER MEMB |
| 63:00:00 | 1s1c-Y | 2.3 | 3.9 | 70 | 70 | 7 | MOLECULE: | TRANSFORMING PROTEIN RHOA |
| 64:00:00 | 2n64-A | 2.3 | 6 | 71 | 75 | 8 | MOLECULE: | SH3 DOMAIN-CONTAINING KINASE-BINDING PROTEIN 1 |
| 65:00:00 | 1h88-B | 2.3 | 5.3 | 69 | 71 | 4 | MOLECULE: | CCAAT/ENHANCER BINDING PROTEIN BETA |
| 66:00:00 | 7sqk-C | 2.3 | 32.2 | 164 | 600 | 5 | MOLECULE: | HAUS AUGMIN-LIKE COMPLEX SUBUNIT 1 |
| 67:00:00 | 4ke2-A | 2.3 | 38.6 | 107 | 196 | 5 | MOLECULE: | TYPE I HYPERACTIVE ANTIFREEZE PROTEIN |
| 68:00:00 | 3hrn-A | 2.3 | 3.6 | 63 | 63 | 8 | MOLECULE: | TRANSIENT RECEPTOR POTENTIAL (TRP) CHANNEL |
| 69:00:00 | 8at3-C | 2.3 | 19.8 | 144 | 353 | 10 | MOLECULE: | HAUS AUGMIN-LIKE COMPLEX SUBUNIT 1 |
| 70:00:00 | 7w5a-X | 2.3 | 11.3 | 86 | 87 | 6 | MOLECULE: | PRE-MRNA-PROCESSING-SPLICING FACTOR 8 |
| 71:00:00 | 7v2w-I | 2.3 | 17.5 | 137 | 239 | 9 | MOLECULE: | THO COMPLEX SUBUNIT HPR1 |
| 72:00:00 | 8t1l-I | 2.3 | 25.7 | 117 | 1111 | 3 | MOLECULE: | MEDIATOR OF RNA POLYMERASE II TRANSCRIPTION SUBUN |
| 73:00:00 | 5n9j-D | 2.3 | 16.9 | 95 | 135 | 7 | MOLECULE: | MEDIATOR OF RNA POLYMERASE II TRANSCRIPTION SUBUN |
| 74:00:00 | 8teu-E | 2.3 | 14.6 | 101 | 495 | 8 | MOLECULE: | LARGE TEGUMENT PROTEIN DENEDDYLASE |
| 75:00:00 | 7d2h-D | 2.3 | 3 | 59 | 60 | 15 | MOLECULE: | LIPRIN-ALPHA-2 |
| 76:00:00 | 7mge-B | 2.2 | 10.9 | 67 | 442 | 9 | MOLECULE: | WD REPEAT-CONTAINING PROTEIN 41 |
| 77:00:00 | 8hqo-x | 2.2 | 8.7 | 77 | 80 | 3 | MOLECULE: | PORTAL PROTEIN |
| 78:00:00 | 6em5-t | 2.2 | 15.7 | 77 | 290 | 9 | MOLECULE: | 5.8S RIBOSOMAL RNA |
| 79:00:00 | 6gmh-Q | 2.2 | 17.7 | 109 | 884 | 8 | MOLECULE: | RPB1 |
| 80:00:00 | 2m0q-A | 2.2 | 59.6 | 98 | 123 | 3 | MOLECULE: | POTASSIUM VOLTAGE-GATED CHANNEL SUBFAMILY E MEMBE |
| 81:00:00 | 4i1l-A | 2.2 | 3.7 | 61 | 62 | 3 | MOLECULE: | FORKHEAD BOX PROTEIN P3 |
| 82:00:00 | 7zcg-B | 2.2 | 12.5 | 104 | 147 | 5 | MOLECULE: | CHARGED MULTIVESICULAR BODY PROTEIN 2A |
| 83:00:00 | 4om3-D | 2.2 | 4.1 | 80 | 122 | 10 | MOLECULE: | TRANSDUCIN-LIKE ENHANCER PROTEIN 1 |
| 84:00:00 | 8idc-E | 2.2 | 42.3 | 106 | 300 | 4 | MOLECULE: | CELL DIVISION ATP-BINDING PROTEIN FTSE |
| 85:00:00 | 6gvw-F | 2.2 | 31 | 101 | 320 | 5 | MOLECULE: | BRCA1-A COMPLEX SUBUNIT ABRAXAS 1 |
| 86:00:00 | 7z47-D | 2.2 | 3.6 | 71 | 87 | 4 | MOLECULE: | ADAPTOR PROTEIN |
| 87:00:00 | 8q85-Y | 2.2 | 5.6 | 66 | 72 | 6 | MOLECULE: | KINETOCHORE PROTEIN NDC80 |
| 88:00:00 | 7zm8-W | 2.2 | 6.9 | 72 | 101 | 7 | MOLECULE: | NADH-UBIQUINONE OXIDOREDUCTASE CHAIN 1 |
| 89:00:00 | 9na9-A | 2.1 | 7.7 | 68 | 103 | 9 | MOLECULE: | AUGMIN SUBUNIT 1 |
| 90:00:00 | 5n9j-U | 2.1 | 25.2 | 100 | 198 | 9 | MOLECULE: | MEDIATOR OF RNA POLYMERASE II TRANSCRIPTION SUBUN |
| 91:00:00 | 2v0o-A | 2.1 | 54.5 | 122 | 273 | 4 | MOLECULE: | FCH DOMAIN ONLY PROTEIN 2 |
| 92:00:00 | 3hgf-A | 2.1 | 3.7 | 70 | 98 | 6 | MOLECULE: | RHOPTRY PROTEIN FRAGMENT |
| 93:00:00 | 7emf-G | 2.1 | 5.3 | 77 | 161 | 13 | MOLECULE: | MEDIATOR OF RNA POLYMERASE II TRANSCRIPTION SUBUN |
| 94:00:00 | 9f63-A | 2.1 | 42.5 | 88 | 422 | 1 | MOLECULE: | PROTEIN PNS1 |
| 95:00:00 | 4h8s-C | 2.1 | 14.1 | 122 | 382 | 8 | MOLECULE: | DCC-INTERACTING PROTEIN 13-BETA |
| 96:00:00 | 5yfp-C | 2.1 | 41.6 | 124 | 790 | 7 | MOLECULE: | EXOCYST COMPLEX COMPONENT SEC3 |
| 97:00:00 | 8ap7-c | 2.1 | 5.5 | 63 | 64 | 10 | MOLECULE: | ATP SYNTHASE SUBUNIT A |
| 98:00:00 | 8p0v-K | 2.1 | 29.9 | 145 | 304 | 5 | MOLECULE: | COILED-COIL DOMAIN-CONTAINING PROTEIN 93 |
| 99:00:00 | 5yfp-F | 2.1 | 15.3 | 165 | 725 | 4 | MOLECULE: | EXOCYST COMPLEX COMPONENT SEC3 |
| 0:00 | 2p2u-B | 2.1 | 23.4 | 80 | 153 | 6 | MOLECULE: | HOST-NUCLEASE INHIBITOR PROTEIN GAM, PUTATIVE |
| 1:00 | 1gl2-C | 2.1 | 3.3 | 58 | 59 | 3 | MOLECULE: | ENDOBREVIN |
| 2:00 | 2ahm-G | 2.1 | 9.1 | 80 | 191 | 3 | MOLECULE: | REPLICASE POLYPROTEIN 1AB, LIGHT CHAIN |
| 3:00 | 4bwd-A | 2.1 | 4.5 | 60 | 61 | 8 | MOLECULE: | SHORT COILED-COIL PROTEIN |
| 4:00 | 5gna-B | 2.1 | 3.9 | 55 | 56 | 5 | MOLECULE: | FLAGELLAR PROTEIN FLIT |
| 5:00 | 4njl-A | 2.1 | 2.7 | 81 | 129 | 5 | MOLECULE: | S PROTEIN |
| 6:00 | 8izl-C | 2.1 | 6.9 | 74 | 91 | 4 | MOLECULE: | PHOSPHOPROTEIN |
| 7:00 | 8cra-E | 2.1 | 13 | 77 | 96 | 5 | MOLECULE: | FLORAL HOMEOTIC PROTEIN AGAMOUS |
| 8:00 | 8q85-c | 2.1 | 4.7 | 62 | 64 | 3 | MOLECULE: | KINETOCHORE PROTEIN NDC80 |
| 9:00 | 8iss-D | 2 | 12 | 45 | 289 | 7 | MOLECULE: | TRNA-SPLICING ENDONUCLEASE SUBUNIT SEN15 |
| 10:00 | 6oee-A | 2 | 68.6 | 119 | 243 | 8 | MOLECULE: | TYPE IV SECRETION SYSTEM APPARATUS PROTEIN CAGT |

| | | | | | | | | |
|---|---|---|---|---|---|---|---|---|
| 11:00 | 5jeq-A | 2 | 5.1 | 88 | 227 | 1 | MOLECULE: | NITRATE/NITRITE SENSOR PROTEIN NARQ |
| 12:00 | 7zmg-n | 2 | 13.2 | 91 | 136 | 10 | MOLECULE: | NADH-UBIQUINONE OXIDOREDUCTASE CHAIN 1 |
| 13:00 | 8qby-L | 2 | 27.6 | 104 | 660 | 7 | MOLECULE: | NADH-QUINONE OXIDOREDUCTASE SUBUNIT K |
| 14:00 | 6od2-A | 2 | 2.2 | 53 | 53 | 6 | MOLECULE: | SPINDLE POLE BODY COMPONENT SPC42 |
| 15:00 | 8x5f-A | 2 | 30.8 | 89 | 616 | 4 | MOLECULE: | SOLUTE CARRIER FAMILY 53 MEMBER 1 |
| 16:00 | 7xyz-B | 2 | 17.5 | 115 | 456 | 7 | MOLECULE: | TRIPARTITE MOTIF-CONTAINING PROTEIN 72 |

**DALI results: MnA-1 Colalt**

| No: | Chain | Z | rmsd | lali | nres | %id | PDB Description | |
|---|---|---|---|---|---|---|---|---|
| 1:00 | 8cra-A | 2.7 | 14 | 77 | 96 | 8 | MOLECULE: | FLORAL HOMEOTIC PROTEIN AGAMOUS |
| 2:00 | 6y09-C | 2.7 | 5.7 | 66 | 86 | 9 | MOLECULE: | RAS-RELATED PROTEIN RAB-33B |
| 3:00 | 3f1i-H | 2.7 | 4.7 | 75 | 98 | 7 | MOLECULE: | HEPATOCYTE GROWTH FACTOR-REGULATED |
| 4:00 | 5vpc-B | 2.5 | 3.1 | 56 | 68 | 9 | MOLECULE: | PROTEIN FOSB |
| 5:00 | 5i50-B | 2.4 | 14.4 | 60 | 96 | 8 | MOLECULE: | MYC PROTO-ONCOGENE PROTEIN |
| 6:00 | 6c0f-8 | 2.4 | 4.5 | 66 | 98 | 6 | MOLECULE: | SACCHAROMYCES CEREVISIAE S288C 35S |
| 7:00 | 7x5e-E | 2.3 | 5.1 | 60 | 101 | 8 | MOLECULE: | TRANSCRIPTION FACTOR MAFG |
| 8:00 | 8pqw-F | 2.3 | 25.7 | 62 | 150 | 8 | MOLECULE: | CYTOPLASMIC DYNEIN 1 HEAVY CHAIN 1 |

**DALI results: MnA-1 Wasp**

| No: | Chain | Z | rmsd | lali | nres | %id | PDB | Description | |
|---|---|---|---|---|---|---|---|---|---|
| 1:00 | 2fzt-B | 5.9 | 4.6 | 62 | 79 | 10 | MOLECULE: | HYPOTHETICAL PROTEIN TM0693 | |
| 2:00 | 6cp3-7 | 5.3 | 34.7 | 111 | 171 | 7 | MOLECULE: | ATP SYNTHASE SUBUNIT 9, MITOCHONDRIAL | |
| 3:00 | 9b8o-U | 5.3 | 7.4 | 88 | 214 | 6 | MOLECULE: | ATPASE H+-TRANSPORTING V1 SUBUNIT D | |
| 4:00 | 6lk8-a | 5.1 | 22.3 | 115 | 1272 | 4 | MOLECULE: | MGC83295 PROTEIN | |
| 5:00 | 2maj-A | 5.1 | 4.8 | 64 | 82 | 6 | MOLECULE: | STROMAL INTERACTION MOLECULE 1 | |
| 6:00 | 9bpg-N | 5.1 | 2.8 | 57 | 219 | 2 | MOLECULE: | ATP SYNTHASE SUBUNIT C | |
| 7:00 | 8hmc-B | 5 | 7.4 | 71 | 1195 | 10 | MOLECULE: | INTRAFLAGELLAR TRANSPORT PROTEIN 122 HOMOLO | G |
| 8:00 | 7xzi-B | 5 | 8.9 | 57 | 166 | 7 | MOLECULE: | CTAP3 | |
| 9:00 | 7uic-b | 5 | 3.1 | 58 | 65 | 5 | MOLECULE: | MEDIATOR OF RNA POLYMERASE II TRANSCRIPTION | SUBUN |
| 10:00 | 7ryq-B | 4.9 | 21 | 100 | 527 | 4 | MOLECULE: | KIF-BINDING PROTEIN | |
| 11:00 | 8xij-A | 4.9 | 2.2 | 49 | 469 | 6 | MOLECULE: | GPI-ANCHORED WALL TRANSFER PROTEIN 1 | |
| 12:00 | 7bln-A | 4.9 | 13 | 85 | 769 | 5 | MOLECULE: | VACUOLAR PROTEIN SORTING-ASSOCIATED PROTEIN | 29 |
| 13:00 | 6xp5-G | 4.9 | 13.1 | 91 | 122 | 9 | MOLECULE: | MEDIATOR OF RNA POLYMERASE II TRANSCRIPTION | SUBUN |
| 14:00 | 5wlq-A | 4.9 | 17.4 | 66 | 104 | 6 | MOLECULE: | CAPSID ASSEMBLY SCAFFOLDING PROTEIN,MYOSIN- | 7,MICR |
| 15:00 | 6z0l-A | 4.9 | 2.7 | 47 | 48 | 0 | MOLECULE: | POSITIVE STRAND | |
| 16:00 | 8qkd-A | 4.8 | 6.7 | 67 | 281 | 12 | MOLECULE: | SC-CC-5-24 | |
| 17:00 | 5an6-A | 4.8 | 9.3 | 77 | 123 | 10 | MOLECULE: | CRISPR-ASSOCIATED PROTEIN, CSM2 FAMILY | |
| 18:00 | 4pi0-A | 4.8 | 26.9 | 94 | 390 | 5 | MOLECULE: | UNKNOWN PEPTIDE | |
| 19:00 | 5b49-B | 4.8 | 15.1 | 69 | 247 | 7 | MOLECULE: | UDP-2,3-DIACYLGLUCOSAMINE HYDROLASE | |
| 20:00 | 5j8v-A | 4.8 | 10.8 | 65 | 3398 | 9 | MOLECULE: | RYANODINE RECEPTOR 1 | |
| 21:00 | 4i0x-F | 4.8 | 4.8 | 61 | 85 | 8 | MOLECULE: | ESAT-6-LIKE PROTEIN MAB_3112 | |
| 22:00 | 5n9j-D | 4.7 | 14.7 | 95 | 135 | 7 | MOLECULE: | MEDIATOR OF RNA POLYMERASE II TRANSCRIPTION | SUBUN |
| 23:00 | 1lj2-B | 4.7 | 6.4 | 60 | 110 | 2 | MOLECULE: | NONSTRUCTURAL RNA-BINDING PROTEIN 34 | |
| 24:00:00 | 6w8p-B | 4.7 | 17.2 | 86 | 426 | 3 | MOLECULE: | ENDOSOMAL/LYSOSOMAL POTASSIUM CHANNEL TMEM1 | 75 |
| 25:00:00 | 6hgc-A | 4.7 | 4.7 | 97 | 304 | 1 | MOLECULE: | UBIQUITIN CARBOXYL-TERMINAL HYDROLASE CALYP | SO,UBI |
| 26:00:00 | 4ut1-A | 4.7 | 12.2 | 110 | 551 | 5 | MOLECULE: | FLAGELLAR HOOK-ASSOCIATED PROTEIN | |
| 27:00:00 | 3oa7-A | 4.7 | 29.8 | 104 | 193 | 5 | MOLECULE: | HEAD MORPHOGENESIS PROTEIN, CHAOTIC NUCLEAR | MIGRA |
| 28:00:00 | 6rie-A | 4.7 | 14.4 | 118 | 1285 | 7 | MOLECULE: | DNA-DEPENDENT RNA POLYMERASE SUBUNIT RPO147 | |
| 29:00:00 | 2spc-A | 4.7 | 5.4 | 70 | 107 | 4 | MOLECULE: | SPECTRIN | |
| 30:00:00 | 5izs-D | 4.7 | 2.4 | 62 | 82 | 6 | MOLECULE: | DESIGNED PROTEIN 5L6HC3_1 | |
| 31:00:00 | 4kqt-A | 4.7 | 7.4 | 82 | 169 | 4 | MOLECULE: | PUTATIVE OUTER MEMBRANE CHAPERONE (OMPH-LIK | E) |
| 32:00:00 | 8ap9-G | 4.7 | 3.8 | 61 | 279 | 0 | MOLECULE: | ATP SYNTHASE GAMMA SUBUNIT | |
| 33:00:00 | 2a01-A | 4.6 | 18.8 | 93 | 243 | 5 | MOLECULE: | APOLIPOPROTEIN A-I | |
| 34:00:00 | 8rsq-A | 4.6 | 17.7 | 104 | 389 | 8 | MOLECULE: | MICROBIAL RHODOPSIN - GTPASE CHIMERA,ADP-RI | BOSYLA |
| 35:00:00 | 8hc0-A | 4.6 | 7.3 | 60 | 360 | 13 | MOLECULE: | ADHESION G-PROTEIN COUPLED RECEPTOR F1 | |
| 36:00:00 | 3g80-A | 4.6 | 4 | 61 | 73 | 10 | MOLECULE: | PROTEIN B2 | |
| 37:00:00 | 8qfs-A | 4.6 | 22.7 | 75 | 1183 | 7 | MOLECULE: | ELONGATION FACTOR TU | |
| 38:00:00 | 6wg3-C | 4.6 | 33.2 | 99 | 248 | 7 | MOLECULE: | STRUCTURAL MAINTENANCE OF CHROMOSOMES PROTE | IN 1A |
| 39:00:00 | 7zw6-A | 4.6 | 17.3 | 109 | 793 | 3 | MOLECULE: | SLR0869 PROTEIN | |
| 40:00:00 | 7cbc-A | 4.5 | 26.9 | 123 | 319 | 8 | MOLECULE: | DE NOVO DESIGNED SWITCH PROTEIN CAGING A HE | MAGGLU |
| 41:00:00 | 8x5f-A | 4.5 | 22.8 | 103 | 616 | 10 | MOLECULE: | SOLUTE CARRIER FAMILY 53 MEMBER 1 | |
| 42:00:00 | 2r6i-A | 4.5 | 2.9 | 47 | 268 | 0 | MOLECULE: | UNCHARACTERIZED PROTEIN ATU1473 | |
| 43:00:00 | 7lzh-A | 4.5 | 19 | 99 | 799 | 5 | MOLECULE: | GLUTAMATE RECEPTOR 3.4 | |
| 44:00:00 | 6zyx-C | 4.5 | 8.8 | 55 | 278 | 4 | MOLECULE: | DYNEIN HEAVY CHAIN, OUTER ARM PROTEIN | |
| 45:00:00 | 6zyv-A | 4.5 | 3 | 48 | 238 | 4 | MOLECULE: | CIR PROTEIN | |
| 46:00:00 | 2v0p-A | 4.5 | 13.4 | 85 | 219 | 5 | MOLECULE: | TYPE 2A PHOSPHATASE-ASSOCIATED PROTEIN 42 | |
| 47:00:00 | 6nd4-T | 4.5 | 6.6 | 59 | 812 | 5 | MOLECULE: | ETS RRNA | |
| 48:00:00 | 7wkk-A | 4.5 | 7 | 54 | 1684 | 6 | MOLECULE: | MGC83295 PROTEIN | |
| 49:00:00 | 7pmk-X | 4.5 | 13.3 | 76 | 665 | 0 | MOLECULE: | DNA REPLICATION LICENSING FACTOR MCM2 | |
| 50:00:00 | 7tvz-A | 4.5 | 3.4 | 63 | 850 | 5 | MOLECULE: | BAND 3 ANION TRANSPORT PROTEIN | |
| 51:00:00 | 6upn-A | 4.5 | 25 | 113 | 245 | 8 | MOLECULE: | ENDOPHILIN-B1 | |
| 52:00:00 | 5xbt-A | 4.5 | 9 | 81 | 271 | 2 | MOLECULE: | PROBABLE TRANSCRIPTIONAL REGULATOR | |
| 53:00:00 | 5wwl-N | 4.5 | 7.7 | 73 | 155 | 11 | MOLECULE: | CENTROMERE PROTEIN MIS12 | |
| 54:00:00 | 6wc3-B | 4.5 | 3.3 | 57 | 94 | 2 | MOLECULE: | PROTEIN TRANSPORT PROTEIN TIP20 | |
| 55:00:00 | 8xku-C | 4.5 | 26.1 | 110 | 921 | 6 | MOLECULE: | PROBABLE INACTIVE ATP-DEPENDENT ZINC METALL | OPROTE |
| 56:00:00 | 2r9i-A | 4.5 | 4.3 | 61 | 72 | 13 | MOLECULE: | PUTATIVE PHAGE CAPSID PROTEIN | |
| 57:00:00 | 6irr-A | 4.4 | 33.9 | 103 | 133 | 6 | MOLECULE: | DISRUPTED IN SCHIZOPHRENIA 1 HOMOLOG,CYCLIC | AMP-D |
| 58:00:00 | 8y7m-I | 4.4 | 30.2 | 108 | 117 | 7 | MOLECULE: | POLYMERASE ACIDIC PROTEIN | |
| 59:00:00 | 6icz-A | 4.4 | 25.8 | 124 | 2253 | 4 | MOLECULE: | PROTEIN MAGO NASHI HOMOLOG 2 | |
| 60:00:00 | 4h54-A | 4.4 | 7.6 | 89 | 274 | 6 | MOLECULE: | DIGUANYLATE CYCLASE YDEH | |
| 61:00:00 | 5vj4-A | 4.4 | 9.5 | 80 | 275 | 10 | MOLECULE: | UNCHARACTERIZED PROTEIN | |
| 62:00:00 | 7wji-B | 4.4 | 12.7 | 83 | 1663 | 5 | MOLECULE: | PROTEIN UNC-80 HOMOLOG | |
| 63:00:00 | 8wjn-B | 4.4 | 29.1 | 140 | 481 | 6 | MOLECULE: | STRUCTURAL MAINTENANCE OF CHROMOSOMES PROTE | IN 6 |
| 64:00:00 | 7w01-A | 4.4 | 28.2 | 112 | 1598 | 6 | MOLECULE: | PHOSPHOLIPID-TRANSPORTING ATPASE ABCA3 | |
| 65:00:00 | 6dfp-A | 4.4 | 8.2 | 68 | 361 | 10 | MOLECULE: | VCA0883 | |
| 66:00:00 | 7vgr-A | 4.4 | 16.4 | 91 | 198 | 8 | MOLECULE: | YN7756_1 FAB LIGHT CHAIN | |
| 67:00:00 | 6uxe-B | 4.4 | 4 | 53 | 85 | 4 | MOLECULE: | CYSTEINE DESULFURASE, MITOCHONDRIAL | |
| 68:00:00 | 6h8q-A | 4.4 | 15.2 | 87 | 879 | 5 | MOLECULE: | COHESIN SUBUNIT SCC3 | |
| 69:00:00 | 4kpk-A | 4.4 | 4.1 | 50 | 265 | 0 | MOLECULE: | ENOYL-COA HYDRATASE/ISOMERASE | |
| 70:00:00 | 3rfw-A | 4.4 | 20.4 | 118 | 252 | 4 | MOLECULE: | CELL-BINDING FACTOR 2 | |
| 71:00:00 | 5j2l-A | 4.4 | 3.7 | 58 | 76 | 7 | MOLECULE: | PROTEIN DESIGN 2L4HC2_11 | |
| 72:00:00 | 6zyd-A | 4.4 | 28.4 | 119 | 323 | 2 | MOLECULE: | LOW CONDUCTANCE MECHANOSENSITIVE CHANNEL YN | AI,LOW |
| 73:00:00 | 4ux3-B | 4.4 | 2.7 | 54 | 71 | 9 | MOLECULE: | STRUCTURAL MAINTENANCE OF CHROMOSOMES PROTE | IN 3 |
| 74:00:00 | 6ql4-A | 4.4 | 3.4 | 49 | 660 | 12 | MOLECULE: | PUTATIVE MITOCHONDRIAL DYNAMIN PROTEIN | |
| 75:00:00 | 4lws-A | 4.4 | 3.5 | 68 | 100 | 7 | MOLECULE: | UNCHARACTERIZED PROTEIN | |
| 76:00:00 | 3fx7-B | 4.4 | 3.5 | 60 | 87 | 3 | MOLECULE: | PUTATIVE UNCHARACTERIZED PROTEIN | |
| 77:00:00 | 6gdj-A | 4.4 | 3 | 50 | 71 | 0 | MOLECULE: | MTO2 | |
| 78:00:00 | 5h11-A | 4.3 | 28.4 | 102 | 494 | 7 | MOLECULE: | UNCHARACTERIZED PROTEIN | |
| 79:00:00 | 7f3x-A | 4.3 | 7.7 | 90 | 449 | 6 | MOLECULE: | LPCAT3 | |
| 80:00:00 | 3u84-A | 4.3 | 12.1 | 72 | 506 | 7 | MOLECULE: | MENIN | |
| 81:00:00 | 5img-A | 4.3 | 3 | 63 | 467 | 13 | MOLECULE: | DIPEPTIDASE | |
| 82:00:00 | 6rax-N | 4.3 | 12.7 | 82 | 207 | 2 | MOLECULE: | DNA REPLICATION LICENSING FACTOR MCM2 | |
| 83:00:00 | 8ovw-Z | 4.3 | 29.4 | 99 | 151 | 8 | MOLECULE: | CENTROMERE-BINDING PROTEIN 1 | |
| 84:00:00 | 6cfz-J | 4.3 | 3.2 | 60 | 134 | 8 | MOLECULE: | ASK1 | |
| 85:00:00 | 4gwp-D | 4.3 | 5.9 | 91 | 121 | 4 | MOLECULE: | MEDIATOR OF RNA POLYMERASE II TRANSCRIPTION | SUBUN |
| 86:00:00 | 8g3a-D | 4.3 | 21.4 | 95 | 334 | 6 | MOLECULE: | BACITRACIN EXPORT PERMEASE PROTEIN BCEB | |
| 87:00:00 | 6arz-A | 4.3 | 5.6 | 68 | 96 | 4 | MOLECULE: | UNCHARACTERIZED PROTEIN | |
| 88:00:00 | 1kf6-C | 4.3 | 8.3 | 78 | 130 | 6 | MOLECULE: | FUMARATE REDUCTASE FLAVOPROTEIN | |
| 89:00:00 | 7wb4-b | 4.3 | 26.6 | 154 | 636 | 9 | MOLECULE: | OUTER NUP133 | |
| 90:00:00 | 1au1-A | 4.3 | 4.4 | 53 | 166 | 11 | MOLECULE: | INTERFERON-BETA | |
| 91:00:00 | 6c96-A | 4.2 | 25.1 | 140 | 723 | 4 | MOLECULE: | TWO PORE CALCIUM CHANNEL PROTEIN 1 | |
| 92:00:00 | 8pm6-A | 4.2 | 31.6 | 192 | 872 | 5 | MOLECULE: | BILE SALT EXPORT PUMP | |
| 93:00:00 | 7qoo-Q | 4.2 | 38.3 | 101 | 209 | 6 | MOLECULE: | CENTROMERE PROTEIN C | |
| 94:00:00 | 7mvv-A | 4.2 | 25.4 | 161 | 1542 | 7 | MOLECULE: | NUCLEOPORIN NUP192 | |
| 95:00:00 | 5yfp-B | 4.2 | 19.3 | 77 | 927 | 8 | MOLECULE: | EXOCYST COMPLEX COMPONENT SEC3 | |
| 96:00:00 | 1e7p-C | 4.2 | 7.9 | 78 | 254 | 8 | MOLECULE: | FUMARATE REDUCTASE FLAVOPROTEIN SUBUNIT | |
| 97:00:00 | 1xdo-A | 4.2 | 28 | 88 | 687 | 8 | MOLECULE: | POLYPHOSPHATE KINASE | |
| 98:00:00 | 5mbv-A | 4.2 | 12.8 | 87 | 98 | 10 | MOLECULE: | RECBCD ENZYME SUBUNIT RECB | |
| 99:00:00 | 5v5t-A | 4.2 | 2.5 | 47 | 306 | 15 | MOLECULE: | CONSERVED DOMAIN PROTEIN | |

| | | | | | | | | | | |
|---|---|---|---|---|---|---|---|---|---|---|
| 0:00 | 8tef-A | 4.2 | 8.1 | 78 | 516 | 5 | MOLECULE: | RESPONSE REGULATOR RECEIVER PROTEIN | | |
| 1:00 | 5sy1-A | 4.2 | 14.1 | 83 | 582 | 5 | MOLECULE: | CALMODULIN | | |
| 2:00 | 3zc0-B | 4.2 | 11.9 | 85 | 191 | 7 | MOLECULE: | AFTRAX | | |
| 3:00 | 8d8j-0 | 4.2 | 17.5 | 102 | 493 | 8 | MOLECULE: | PROBABLE S-ADENOSYL-L-METHIONINE-DEPENDENT | RNA | |
| 4:00 | 8eki-C | 4.2 | 6.8 | 97 | 701 | 7 | MOLECULE: | PROTEIN TRANSPORT PROTEIN SEC20 | | |
| 5:00 | 5c8j-L | 4.2 | 9.7 | 61 | 137 | 5 | MOLECULE: | ANTIBODY FRAGMENT, HEAVY CHAIN | | |
| 6:00 | 5oqj-1 | 4.2 | 22.6 | 100 | 491 | 2 | MOLECULE: | DNA-DIRECTED RNA POLYMERASE II SUBUNIT RPB1 | | |
| 7:00 | 7xjz-A | 4.2 | 24.3 | 81 | 983 | 4 | MOLECULE: | GLYCINE--TRNA LIGASE | | |
| 8:00 | 4gx2-B | 4.2 | 25.6 | 91 | 548 | 7 | MOLECULE: | TRKA DOMAIN PROTEIN | | |
| 9:00 | 3uum-B | 4.2 | 3.7 | 66 | 123 | 5 | MOLECULE: | UTROPHIN | | |
| 10:00 | 7qpg-W | 4.2 | 31.8 | 96 | 779 | 5 | MOLECULE: | PROTEIN ZWILCH HOMOLOG | | |
| 11:00 | 3he4-A | 4.2 | 1.6 | 42 | 45 | 10 | MOLECULE: | SYNZIP6 | | |
| 12:00 | 4j41-B | 4.2 | 3.3 | 62 | 86 | 10 | MOLECULE: | SECRETED PROTEIN ESXB | | |
| 13:00 | 2n72-A | 4.2 | 4.8 | 56 | 69 | 7 | MOLECULE: | GOLGI RESIDENT PROTEIN GCP60 | | |
| 14:00 | 3vop-C | 4.2 | 2.9 | 43 | 43 | 19 | MOLECULE: | PROTEIN A27 | | |
| 15:00 | 6zgd-A | 4.1 | 12.1 | 65 | 288 | 14 | MOLECULE: | PROTON-GATED ION CHANNEL | | |
| 16:00 | 8qqm-B | 4.1 | 28.8 | 107 | 180 | 7 | MOLECULE: | ACETYLCHOLINE RECEPTOR SUBUNIT ALPHA | | |
| 17:00 | 6vxm-A | 4.1 | 20.8 | 108 | 277 | 6 | MOLECULE: | MECHANOSENSITIVE ION CHANNEL PROTEIN 1, MIT | OCHOND | |
| 18:00 | 7egd-D | 4.1 | 17.3 | 91 | 197 | 8 | MOLECULE: | TRANSCRIPTION INITIATION FACTOR TFIID SUBUN | IT 1 | |
| 19:00 | 5n9j-V | 4.1 | 11.7 | 83 | 99 | 7 | MOLECULE: | MEDIATOR OF RNA POLYMERASE II TRANSCRIPTION | SUBUN | |
| 20:00 | 5dfz-B | 4.1 | 27 | 112 | 1224 | 6 | MOLECULE: | VACUOLAR PROTEIN SORTING-ASSOCIATED PROTEIN | | 38 |
| 21:00 | 6vjy-B | 4.1 | 9.1 | 94 | 280 | 9 | MOLECULE: | ERAD-ASSOCIATED E3 UBIQUITIN-PROTEIN LIGASE | HRD1 | |
| 22:00 | 8otx-A | 4.1 | 29.6 | 129 | 1054 | 5 | MOLECULE: | SPERM-SPECIFIC SODIUM PROTON EXCHANGER | | |
| 23:00 | 5j43-B | 4.1 | 3.2 | 57 | 101 | 5 | MOLECULE: | CYSTEINE SYNTHASE A | | |
| 24:00:00 | 4y7k-A | 4.1 | 18 | 111 | 250 | 7 | MOLECULE: | LARGE CONDUCTANCE MECHANOSENSITIVE CHANNEL | PROTEI | |
| 25:00:00 | 3rv0-B | 4.1 | 23.2 | 91 | 304 | 8 | MOLECULE: | K. POLYSPORUS DCR1 | | |
| 26:00:00 | 8h17-A | 4.1 | 2 | 49 | 157 | 2 | MOLECULE: | TLR1989 PROTEIN | | |
| 27:00:00 | 8tcw-A | 4.1 | 2.5 | 56 | 282 | 0 | MOLECULE: | PYRROLINE-5-CARBOXYLATE REDUCTASE 1, MITOCH | ONDRIA | |
| 28:00:00 | 8dfv-A | 4.1 | 27.6 | 118 | 1664 | 4 | MOLECULE: | ENDORIBONUCLEASE DCR-1 | | |
| 29:00:00 | 5ibw-C | 4.1 | 3.2 | 43 | 43 | 7 | MOLECULE: | CALCIUM-BINDING EF-HAND DOMAIN-CONTAINING P | ROTEIN | |
| 30:00:00 | 7ekn-F | 4.1 | 1.3 | 40 | 42 | 5 | MOLECULE: | IPEP | | |
| 31:00:00 | 2yev-B | 4.1 | 15.5 | 92 | 319 | 7 | MOLECULE: | CYTOCHROME C OXIDASE POLYPEPTIDE I+III | | |
| 32:00:00 | 7uw5-A | 4.1 | 22.8 | 107 | 723 | 6 | MOLECULE: | MECHANOSENSITIVE CHANNEL MSCK | | |
| 33:00:00 | 6al9-B | 4.1 | 3.2 | 49 | 91 | 10 | MOLECULE: | CHORISMATE MUTASE | | |
| 34:00:00 | 1q2h-A | 4.1 | 4.8 | 53 | 63 | 0 | MOLECULE: | ADAPTOR PROTEIN WITH PLECKSTRIN HOMOLOGY AN | D SRC | |
| 35:00:00 | 6b87-D | 4.1 | 3.8 | 63 | 102 | 10 | MOLECULE: | TMHC2_E | | |
| 36:00:00 | 7cgp-J | 4.1 | 4.1 | 58 | 83 | 5 | MOLECULE: | MITOCHONDRIAL IMPORT INNER MEMBRANE TRANSLO | CASE S | |
| 37:00:00 | 2kg7-A | 4.1 | 5.1 | 57 | 97 | 4 | MOLECULE: | UNCHARACTERIZED PROTEIN ESXG (PE FAMILY PRO | TEIN) | |
| 38:00:00 | 6cfw-G | 4.1 | 2.5 | 51 | 113 | 2 | MOLECULE: | MONOVALENT CATION/H+ ANTIPORTER SUBUNIT D | | |
| 39:00:00 | 6tj1-C | 4.1 | 3.3 | 49 | 74 | 8 | MOLECULE: | DE NOVO DESIGNED WSHC6 | | |
| 40:00:00 | 7drt-B | 4.1 | 7.5 | 121 | 496 | 5 | MOLECULE: | PROTEIN WNT-3A | | |
| 41:00:00 | 7cm3-A | 4 | 27.1 | 190 | 1278 | 7 | MOLECULE: | SODIUM LEAK CHANNEL NON-SELECTIVE PROTEIN | | |
| 42:00:00 | 5t58-N | 4 | 46.8 | 104 | 196 | 5 | MOLECULE: | KLLA0F02343P | | |
| 43:00:00 | 7wji-A | 4 | 31.7 | 129 | 1763 | 5 | MOLECULE: | PROTEIN UNC-80 HOMOLOG | | |
| 44:00:00 | 8ipr-C | 4 | 28.5 | 156 | 515 | 3 | MOLECULE: | COMPONENT LINKED WITH THE ASSEMBLY OF CYTOC | HROME' | |
| 45:00:00 | 7rmy-A | 4 | 29.1 | 97 | 266 | 6 | MOLECULE: | DE NOVO DESIGNED TUNABLE HOMODIMER, D_3-337 | | |
| 46:00:00 | 8t1l-I | 4 | 58.6 | 138 | 1111 | 5 | MOLECULE: | MEDIATOR OF RNA POLYMERASE II TRANSCRIPTION | SUBUN | |
| 47:00:00 | 5tgz-A | 4 | 8.7 | 127 | 439 | 6 | MOLECULE: | CANNABINOID RECEPTOR 1,FLAVODOXIN,CANNABINO | ID REC | |
| 48:00:00 | 6u8y-x | 4 | 28.4 | 108 | 603 | 4 | MOLECULE: | MONOVALENT CATION/H+ ANTIPORTER SUBUNIT E | | |
| 49:00:00 | 4hw5-A | 4 | 2.5 | 49 | 77 | 2 | MOLECULE: | COMPLEMENT C3 | | |
| 50:00:00 | 7xpt-A | 4 | 3.8 | 59 | 375 | 10 | MOLECULE: | TRANSGLYCOSYLSE | | |
| 51:00:00 | 8b6l-J | 4 | 6.9 | 90 | 121 | 4 | MOLECULE: | PROTEIN TRANSPORT PROTEIN SEC61 SUBUNIT ALP | HA ISO | |
| 52:00:00 | 7r5s-S | 4 | 11.4 | 76 | 120 | 5 | MOLECULE: | CENTROMERE PROTEIN H | | |
| 53:00:00 | 6gv8-A | 4 | 10.5 | 60 | 157 | 12 | MOLECULE: | HYPEROSMOLARITY RESISTANCE PROTEIN EMB | | |
| 54:00:00 | 6djs-A | 4 | 8.8 | 99 | 490 | 0 | MOLECULE: | SHORT TRANSIENT RECEPTOR POTENTIAL CHANNEL | | 3 |
| 55:00:00 | 7v2w-I | 4 | 28.5 | 128 | 239 | 10 | MOLECULE: | THO COMPLEX SUBUNIT HPR1 | | |
| 56:00:00 | 8j22-C | 4 | 5.4 | 77 | 285 | 6 | MOLECULE: | GUANINE NUCLEOTIDE-BINDING PROTEIN G(I)/G(S | )/G(T) | |
| 57:00:00 | 2be3-A | 4 | 5.6 | 59 | 203 | 8 | MOLECULE: | GTP PYROPHOSPHOKINASE | | |
| 58:00:00 | 8w5z-A | 4 | 14.4 | 75 | 333 | 8 | MOLECULE: | PROTEIN-TYROSINE SULFOTRANSFERASE (FRAGMENT | ) | |
| 59:00:00 | 3axj-B | 4 | 11.8 | 76 | 240 | 4 | MOLECULE: | GM27569P | | |
| 60:00:00 | 7w0b-A | 4 | 17.8 | 126 | 1586 | 5 | MOLECULE: | DICER-2, ISOFORM A | | |
| 61:00:00 | 5al6-A | 4 | 2.5 | 42 | 44 | 5 | MOLECULE: | ANASTRAL SPINDLE 2 | | |
| 62:00:00 | 3g9r-A | 4 | 1.8 | 40 | 42 | 13 | MOLECULE: | FUSION COMPLEX OF HIV-1 ENVELOPE GLYCOPROTE | IN AND | |
| 63:00:00 | 3mud-C | 4 | 2.2 | 40 | 41 | 0 | MOLECULE: | DNA REPAIR PROTEIN XRCC4,TROPOMYOSIN ALPHA- | 1 CHAI | |
| 64:00:00 | 2oqq-A | 4 | 1.9 | 41 | 42 | 5 | MOLECULE: | TRANSCRIPTION FACTOR HY5 | | |
| 65:00:00 | 6jxr-n | 4 | 3.7 | 42 | 287 | 12 | MOLECULE: | T-CELL SURFACE GLYCOPROTEIN CD3 ZETA CHAIN | | |
| 66:00:00 | 3he4-B | 4 | 1.3 | 40 | 44 | 10 | MOLECULE: | SYNZIP6 | | |
| 67:00:00 | 2dnx-A | 4 | 5.4 | 66 | 130 | 12 | MOLECULE: | SYNTAXIN-12 | | |
| 68:00:00 | 6c2j-A | 4 | 11.2 | 60 | 265 | 8 | MOLECULE: | RNAP1 | | |
| 69:00:00 | 3opc-B | 4 | 4 | 58 | 131 | 3 | MOLECULE: | UNCHARACTERIZED PROTEIN | | |
| 70:00:00 | 7dxb-A | 4 | 22.5 | 99 | 738 | 4 | MOLECULE: | SHORT TRANSIENT RECEPTOR POTENTIAL CHANNEL | | 3 |
| 71:00:00 | 6dkm-D | 4 | 4 | 57 | 79 | 4 | MOLECULE: | DHD131_A | | |
| 72:00:00 | 4kxr-A | 4 | 2.9 | 59 | 85 | 7 | MOLECULE: | PE25 | | |
| 73:00:00 | 3whj-A | 4 | 5.3 | 61 | 111 | 5 | MOLECULE: | PROBABLE 26S PROTEASOME REGULATORY SUBUNIT | P27 | |
| 74:00:00 | 5vyk-C | 4 | 4.1 | 53 | 203 | 4 | MOLECULE: | CHIMERA PROTEIN OF BRS DOMAIN OF BRAF AND C | C-SAM | |
| 75:00:00 | 4p96-A | 4 | 7.2 | 56 | 278 | 4 | MOLECULE: | FATTY ACID METABOLISM REGULATOR PROTEIN | | |
| 76:00:00 | 4y2f-A | 4 | 3.7 | 49 | 143 | 4 | MOLECULE: | SENSOR PROTEIN KDPD | | |
| 77:00:00 | 8gxl-B | 4 | 6.4 | 52 | 122 | 8 | MOLECULE: | SURP AND G-PATCH DOMAIN-CONTAINING PROTEIN | | 1 |
| 78:00:00 | 6oit-F | 4 | 3.9 | 62 | 342 | 8 | MOLECULE: | PROTEIN RDM1 | | |
| 79:00:00 | 6u1s-A | 4 | 2.9 | 62 | 310 | 3 | MOLECULE: | DE NOVO DESIGNED 16-HELIX TRANSMEMBRANE NAN | OPORE, | |
| 80:00:00 | 8rz0-E | 4 | 2.8 | 57 | 113 | 14 | MOLECULE: | RH5-34EM | | |
| 81:00:00 | 3h3m-A | 4 | 4 | 52 | 89 | 2 | MOLECULE: | FLAGELLAR PROTEIN FLIT | | |
| 82:00:00 | 5i9e-H | 4 | 2.9 | 44 | 44 | 11 | MOLECULE: | ACTIN-RELATED PROTEIN 4 | | |
| 83:00:00 | 7kzn-B | 3.9 | 28.1 | 148 | 683 | 5 | MOLECULE: | HEAVY CHAIN ALPHA | | |
| 84:00:00 | 8ppr-P | 3.9 | 25.7 | 114 | 176 | 4 | MOLECULE: | KINETOCHORE-ASSOCIATED PROTEIN DSN1 HOMOLOG | | |
| 85:00:00 | 5yi4-A | 3.9 | 19.5 | 73 | 145 | 4 | MOLECULE: | DISRUPTED IN SCHIZOPHRENIA 1 HOMOLOG,NUCLEA | R DIST | |
| 86:00:00 | 7jtk-A | 3.9 | 30.2 | 90 | 749 | 11 | MOLECULE: | FLAGELLAR RADIAL SPOKE PROTEIN 1 | | |
| 87:00:00 | 5wb2-A | 3.9 | 13.7 | 121 | 421 | 3 | MOLECULE: | ENVELOPE PROTEIN US28, NANOBODY 7 FUSION PR | OTEIN | |
| 88:00:00 | 7oqz-A | 3.9 | 6.4 | 94 | 237 | 6 | MOLECULE: | TRANSMEMBRANE PROTEIN 45A | | |
| 89:00:00 | 7tch-A | 3.9 | 14.2 | 108 | 645 | 4 | MOLECULE: | BACITRACIN EXPORT PERMEASE PROTEIN BCEB | | |
| 90:00:00 | 4xng-D | 3.9 | 3.8 | 61 | 144 | 5 | MOLECULE: | UNCHARACTERIZED PROTEIN MG218.1 | | |
| 91:00:00 | 5x62-B | 3.9 | 17.1 | 93 | 388 | 11 | MOLECULE: | CARNOSINE N-METHYLTRANSFERASE | | |
| 92:00:00 | 8odu-C | 3.9 | 15.3 | 98 | 232 | 6 | MOLECULE: | ATPASE GET3 | | |
| 93:00:00 | 7w72-A | 3.9 | 11 | 85 | 588 | 6 | MOLECULE: | PHOSPHATIDYLINOSITOL GLYCAN ANCHOR BIOSYNTH | ESIS C | |
| 94:00:00 | 3bhn-A | 3.9 | 2.7 | 49 | 214 | 0 | MOLECULE: | THIJ/PFPI DOMAIN PROTEIN | | |
| 95:00:00 | 2p7n-A | 3.9 | 5.3 | 62 | 335 | 11 | MOLECULE: | PATHOGENICITY ISLAND 1 EFFECTOR PROTEIN | | |
| 96:00:00 | 6its-A | 3.9 | 2.5 | 61 | 147 | 20 | MOLECULE: | METHYL-ACCEPTING CHEMOTAXIS SENSORY TRANSDU | CER | |
| 97:00:00 | 5x5y-F | 3.9 | 2.4 | 51 | 351 | 0 | MOLECULE: | PROBABLE ATP-BINDING COMPONENT OF ABC TRANS | PORTER | |
| 98:00:00 | 3p01-A | 3.9 | 6.8 | 70 | 178 | 3 | MOLECULE: | TWO-COMPONENT RESPONSE REGULATOR | | |
| 99:00:00 | 6xf1-A | 3.9 | 16.4 | 91 | 230 | 7 | MOLECULE: | NESPRIN-2 | | |
| 0:00 | 3he5-A | 3.9 | 3.2 | 44 | 47 | 9 | MOLECULE: | SYNZIP1 | | |

| | | | | | | | | | |
|---|---|---|---|---|---|---|---|---|---|
| 1:00 | 7edp-B | 3.9 | 2.8 | 40 | 40 | 15 | MOLECULE: | HISTONE-LYSINE N-METHYLTRANSFERASE, H3 LYSI | NE-79 |
| 2:00 | 5yr0-B | 3.9 | 2.6 | 42 | 44 | 10 | MOLECULE: | BECLIN-1 | |
| 3:00 | 1jek-A | 3.9 | 3 | 40 | 40 | 8 | MOLECULE: | ENV POLYPROTEIN | |
| 4:00 | 2xus-A | 3.9 | 1.7 | 40 | 41 | 5 | MOLECULE: | BREAST CANCER METASTASIS-SUPPRESSOR 1 | |
| 5:00 | 6iva-B | 3.9 | 2.9 | 41 | 42 | 0 | MOLECULE: | OXALOACETATE DECARBOXYLASE BETA CHAIN | |
| 6:00 | 6pl5-A | 3.9 | 3.5 | 52 | 336 | 0 | MOLECULE: | PEPTIDOGLYCAN GLYCOSYLTRANSFERASE RODA | |
| 7:00 | 1usd-A | 3.9 | 3.6 | 40 | 41 | 10 | MOLECULE: | VASODILATOR-STIMULATED PHOSPHOPROTEIN | |
| 8:00 | 6mit-G | 3.9 | 3.5 | 52 | 355 | 6 | MOLECULE: | LIPOPOLYSACCHARIDE EXPORT SYSTEM ATP-BINDIN | G PROT |
| 9:00 | 8eoi-K | 3.9 | 19.7 | 139 | 1116 | 3 | MOLECULE: | ER MEMBRANE PROTEIN COMPLEX SUBUNIT 1 | |
| 10:00 | 4gif-A | 3.9 | 2.7 | 42 | 45 | 7 | MOLECULE: | POLYCYSTIC KIDNEY DISEASE 2-LIKE 1 PROTEIN | |
| 11:00 | 6zbj-B | 3.8 | 24.3 | 136 | 501 | 7 | MOLECULE: | PRECURSOR OF THE MAJOR MEROZOITE SURFACE AN | TIGENS |
| 12:00 | 5oqm-n | 3.8 | 16.9 | 93 | 136 | 8 | MOLECULE: | DNA-DIRECTED RNA POLYMERASE II SUBUNIT RPB1 | |
| 13:00 | 7zkq-T | 3.8 | 5.2 | 77 | 351 | 6 | MOLECULE: | NADH DEHYDROGENASE SUBUNIT 2 | |
| 14:00 | 7xn9-A | 3.8 | 5.1 | 60 | 476 | 3 | MOLECULE: | SOMATOSTATIN RECEPTOR TYPE 2,ENDO-1,4-BETA- | XYLANA |
| 15:00 | 8vej-B | 3.8 | 12 | 62 | 242 | 5 | MOLECULE: | CHD_BUTTRESS | |
| 16:00 | 4kis-A | 3.8 | 10.1 | 69 | 315 | 4 | MOLECULE: | PUTATIVE INTEGRASE [BACTERIOPHAGE A118] | |
| 17:00 | 3tul-A | 3.8 | 3.9 | 86 | 136 | 6 | MOLECULE: | CELL INVASION PROTEIN SIPB | |
| 18:00 | 5yud-C | 3.8 | 5.6 | 67 | 75 | 7 | MOLECULE: | BACULOVIRAL IAP REPEAT-CONTAINING PROTEIN 1 | E |
| 19:00 | 8kfh-A | 3.8 | 2.5 | 49 | 160 | 16 | MOLECULE: | MYOGLOBIN | |
| 20:00 | 8ulg-A | 3.8 | 11.4 | 86 | 827 | 7 | MOLECULE: | ROD CGMP-SPECIFIC 3',5'-CYCLIC PHOSPHODIEST | ERASE |
| 21:00 | 3bo0-A | 3.8 | 12.7 | 95 | 442 | 4 | MOLECULE: | 23S RIBOSOMAL RNA | |
| 22:00 | 6gmh-M | 3.8 | 32.2 | 130 | 991 | 9 | MOLECULE: | RPB1 | |
| 23:00 | 5j6f-A | 3.8 | 12.8 | 73 | 352 | 10 | MOLECULE: | 3-DEOXY-D-ARABINO-HEPTULOSONATE 7-PHOSPHATE | SYNTH |
| 24:00:00 | 6s8h-F | 3.8 | 2.2 | 48 | 239 | 4 | MOLECULE: | LIPOPOLYSACCHARIDE ABC TRANSPORTER, ATP-BIN | DING P |
| 25:00:00 | 5xsv-A | 3.8 | 2.3 | 46 | 486 | 7 | MOLECULE: | CHITINASE | |
| 26:00:00 | 7k10-A | 3.8 | 24.3 | 77 | 1259 | 5 | MOLECULE: | DNA-DEPENDENT PROTEIN KINASE CATALYTIC SUBU | NIT |
| 27:00:00 | 8ug4-A | 3.8 | 29.6 | 102 | 486 | 6 | MOLECULE: | OTOPETRIN-2 | |
| 28:00:00 | 6sqw-A | 3.8 | 10.4 | 60 | 114 | 7 | MOLECULE: | DCTP PYROPHOSPHATASE 1 | |
| 29:00:00 | 2p2u-B | 3.8 | 17.9 | 96 | 153 | 3 | MOLECULE: | HOST-NUCLEASE INHIBITOR PROTEIN GAM, PUTATI | VE |
| 30:00:00 | 7t71-A | 3.8 | 8.7 | 57 | 348 | 4 | MOLECULE: | MEVALONATE 3,5-BISPHOSPHATE DECARBOXYLASE | |
| 31:00:00 | 4u6u-D | 3.8 | 19.8 | 110 | 283 | 8 | MOLECULE: | COG7 | |
| 32:00:00 | 4cgk-A | 3.8 | 15.3 | 153 | 351 | 4 | MOLECULE: | SECRETED 45 KDA PROTEIN | |
| 33:00:00 | 2cwo-A | 3.8 | 2.9 | 58 | 165 | 2 | MOLECULE: | RNA SILENCING SUPPRESSOR | |
| 34:00:00 | 5yfp-H | 3.8 | 7.6 | 70 | 518 | 7 | MOLECULE: | EXOCYST COMPLEX COMPONENT SEC3 | |
| 35:00:00 | 8op7-A | 3.8 | 20.6 | 128 | 1162 | 6 | MOLECULE: | CATION-TRANSPORTING ATPASE-LIKE PROTEIN | |
| 36:00:00 | 3oov-A | 3.8 | 2.8 | 47 | 164 | 11 | MOLECULE: | METHYL-ACCEPTING CHEMOTAXIS PROTEIN, PUTATI | VE |
| 37:00:00 | 5gw1-C | 3.8 | 2.8 | 43 | 176 | 12 | MOLECULE: | SORTING NEXIN-16 | |
| 38:00:00 | 4cgb-B | 3.8 | 2.5 | 42 | 45 | 2 | MOLECULE: | ECHINODERM MICROTUBULE-ASSOCIATED PROTEIN-L | IKE 2 |
| 39:00:00 | 3m9b-A | 3.8 | 4.6 | 64 | 186 | 9 | MOLECULE: | PROTEASOME-ASSOCIATED ATPASE | |
| 40:00:00 | 4r0r-A | 3.8 | 2.9 | 44 | 48 | 11 | MOLECULE: | EBOIZN21 | |
| 41:00:00 | 1n2d-C | 3.8 | 2.5 | 43 | 48 | 5 | MOLECULE: | MYOSIN LIGHT CHAIN | |
| 42:00:00 | 2z5h-A | 3.8 | 4.1 | 47 | 49 | 11 | MOLECULE: | GENERAL CONTROL PROTEIN GCN4 AND TROPOMYOSI | N ALPH |
| 43:00:00 | 7n15-A | 3.8 | 10.3 | 70 | 520 | 6 | MOLECULE: | CYCLIC NUCLEOTIDE-GATED CATION CHANNEL | |
| 44:00:00 | 8e4g-a | 3.8 | 3.2 | 65 | 127 | 3 | MOLECULE: | PORTAL PROTEIN | |
| 45:00:00 | 6ep3-B | 3.8 | 3.4 | 49 | 218 | 6 | MOLECULE: | LMO0651 PROTEIN | |
| 46:00:00 | 5fhp-C | 3.8 | 5.3 | 51 | 209 | 6 | MOLECULE: | NICR | |
| 47:00:00 | 7x29-A | 3.8 | 4.1 | 64 | 991 | 8 | MOLECULE: | SPIKE GLYCOPROTEIN | |
| 48:00:00 | 8zue-A | 3.7 | 25.1 | 95 | 531 | 3 | MOLECULE: | ANTI-BACTERIOPHAGE PROTEIN A | |
| 49:00:00 | 7vf2-A | 3.7 | 22.7 | 94 | 1196 | 9 | MOLECULE: | PROTEIN VIRILIZER HOMOLOG | |
| 50:00:00 | 7kmt-B | 3.7 | 25 | 111 | 475 | 8 | MOLECULE: | TRAFFICKING PROTEIN PARTICLE COMPLEX SUBUNI | T 23 |
| 51:00:00 | 8ek4-B | 3.7 | 13.8 | 69 | 135 | 7 | MOLECULE: | ICE-BINDING PROTEIN TIP-99A | |
| 52:00:00 | 5uz5-E | 3.7 | 18.9 | 95 | 583 | 6 | MOLECULE: | U1 SMALL NUCLEAR RIBONUCLEOPROTEIN 70 KDA H | OMOLOG |
| 53:00:00 | 9c59-d | 3.7 | 20.2 | 76 | 605 | 8 | MOLECULE: | AP-3 COMPLEX SUBUNIT DELTA-1 | |
| 54:00:00 | 8bf9-G | 3.7 | 9.3 | 74 | 158 | 7 | MOLECULE: | RNA (1766) | |
| 55:00:00 | 6s8g-G | 3.7 | 6.2 | 80 | 244 | 5 | MOLECULE: | LIPOPOLYSACCHARIDE ABC TRANSPORTER, ATP-BIN | DING P |
| 56:00:00 | 8ahx-E | 3.7 | 3.5 | 56 | 213 | 2 | MOLECULE: | ION-TRANSLOCATING OXIDOREDUCTASE COMPLEX SU | BUNIT |
| 57:00:00 | 6lo8-C | 3.7 | 2.9 | 56 | 108 | 5 | MOLECULE: | MITOCHONDRIAL IMPORT INNER MEMBRANE TRANSLO | CASE S |
| 58:00:00 | 3mhh-C | 3.7 | 9.1 | 64 | 93 | 5 | MOLECULE: | UBIQUITIN CARBOXYL-TERMINAL HYDROLASE 8 | |
| 59:00:00 | 8etc-F | 3.7 | 7.7 | 64 | 218 | 3 | MOLECULE: | RNA (2151-MER) | |
| 60:00:00 | 8jtv-A | 3.7 | 8.3 | 87 | 437 | 6 | MOLECULE: | SOLUTE CARRIER FAMILY 22 MEMBER 1 | |
| 61:00:00 | 3wfw-A | 3.7 | 2.6 | 49 | 138 | 10 | MOLECULE: | HEMOGLOBIN-LIKE FLAVOPROTEIN FUSED TO ROADB | LOCK/L |
| 62:00:00 | 5gox-A | 3.7 | 9 | 85 | 181 | 14 | MOLECULE: | DNA REPAIR PROTEIN RAD50 | |
| 63:00:00 | 7wkk-B | 3.7 | 7.6 | 83 | 1482 | 4 | MOLECULE: | MGC83295 PROTEIN | |
| 64:00:00 | 3dhi-A | 3.7 | 7.4 | 68 | 498 | 10 | MOLECULE: | TOLUENE 4-MONOOXYGENASE HYDROXYLASE ALPHA S | UBUNIT |
| 65:00:00 | 5e5w-B | 3.7 | 6.6 | 69 | 149 | 1 | MOLECULE: | HEMAGGLUTININ-ESTERASE | |
| 66:00:00 | 6q82-A | 3.7 | 26.9 | 108 | 1027 | 5 | MOLECULE: | IMPORTIN BETA-LIKE PROTEIN KAP122 | |
| 67:00:00 | 6w1s-Q | 3.7 | 10.3 | 118 | 131 | 6 | MOLECULE: | MEDIATOR OF RNA POLYMERASE II TRANSCRIPTION | SUBUN |
| 68:00:00 | 5vjx-H | 3.7 | 2.3 | 45 | 51 | 7 | MOLECULE: | CLOCK-INTERACTING PACEMAKER | |
| 69:00:00 | 1go4-G | 3.7 | 30.7 | 68 | 100 | 4 | MOLECULE: | MITOTIC SPINDLE ASSEMBLY CHECKPOINT PROTEIN | MAD2A |
| 70:00:00 | 3fga-D | 3.7 | 2.3 | 43 | 47 | 9 | MOLECULE: | SERINE/THREONINE-PROTEIN PHOSPHATASE 2A 65 | KDA RE |
| 71:00:00 | 7bhq-B | 3.7 | 7.9 | 86 | 204 | 7 | MOLECULE: | BASAL-BODY ROD MODIFICATION PROTEIN FLGD | |
| 72:00:00 | 5i50-B | 3.7 | 2.3 | 48 | 96 | 8 | MOLECULE: | MYC PROTO-ONCOGENE PROTEIN | |
| 73:00:00 | 5vr2-A | 3.7 | 3.3 | 44 | 50 | 2 | MOLECULE: | MYOCILIN | |
| 74:00:00 | 5y2h-A | 3.7 | 3 | 43 | 48 | 7 | MOLECULE: | NONSTRUCTURAL PROTEIN 4 | |
| 75:00:00 | 5hud-E | 3.7 | 4.5 | 54 | 84 | 6 | MOLECULE: | 3-DEOXY-D-ARABINO-HEPTULOSONATE 7-PHOSPHATE | (DAHP |
| 76:00:00 | 6dnq-E | 3.7 | 3.4 | 43 | 54 | 7 | MOLECULE: | BZIP FACTOR | |
| 77:00:00 | 3vn0-B | 3.7 | 3.8 | 42 | 49 | 7 | MOLECULE: | VOLTAGE-GATED HYDROGEN CHANNEL 1 | |
| 78:00:00 | 6ugm-X | 3.7 | 3.2 | 55 | 73 | 5 | MOLECULE: | HISTONE H3 | |
| 79:00:00 | 3ci9-B | 3.7 | 2.5 | 42 | 45 | 10 | MOLECULE: | HEAT SHOCK FACTOR-BINDING PROTEIN 1 | |
| 80:00:00 | 7exx-A | 3.7 | 4 | 57 | 440 | 4 | MOLECULE: | DNA PHOSPHOROTHIOATION-DEPENDENT RESTRICTIO | N PROT |
| 81:00:00 | 6akm-B | 3.7 | 3.7 | 44 | 46 | 5 | MOLECULE: | SUPPRESSOR OF IKBKE 1 | |
| 82:00:00 | 3txs-C | 3.7 | 3.3 | 54 | 91 | 9 | MOLECULE: | TERMINASE DNA PACKAGING ENZYME SMALL SUBUNI | T |
| 83:00:00 | 8qbv-A | 3.7 | 31 | 90 | 217 | 13 | MOLECULE: | MEMBRANE-ASSOCIATED PROTEIN VIPP1 | |
| 84:00:00 | 2v4h-A | 3.7 | 32.4 | 80 | 98 | 4 | MOLECULE: | NF-KAPPA-B ESSENTIAL MODULATOR | |
| 85:00:00 | 3ayf-A | 3.7 | 4.5 | 75 | 754 | 9 | MOLECULE: | NITRIC OXIDE REDUCTASE | |
| 86:00:00 | 6z6o-D | 3.7 | 28.4 | 102 | 542 | 7 | MOLECULE: | HISTONE DEACETYLASE HDA1 | |
| 87:00:00 | 6cv0-A | 3.7 | 3.9 | 63 | 993 | 8 | MOLECULE: | SPIKE GLYCOPROTEIN | |
| 88:00:00 | 5x41-F | 3.7 | 21.5 | 79 | 244 | 0 | MOLECULE: | COBALT ABC TRANSPORTER ATP-BINDING PROTEIN | |
| 89:00:00 | 3rmi-A | 3.7 | 2.2 | 50 | 105 | 10 | MOLECULE: | CHORISMATE MUTASE PROTEIN | |
| 90:00:00 | 3o1j-A | 3.6 | 27.9 | 93 | 273 | 8 | MOLECULE: | SENSOR PROTEIN TORS | |
| 91:00:00 | 6th1-R | 3.6 | 33 | 118 | 360 | 3 | MOLECULE: | IMMEDIATE EARLY PROTEIN 1 | |
| 92:00:00 | 6wuc-H | 3.6 | 17 | 95 | 179 | 6 | MOLECULE: | INNER KINETOCHORE SUBUNIT MCM16 | |
| 93:00:00 | 7sgr-A | 3.6 | 27.7 | 165 | 700 | 6 | MOLECULE: | ALPHA-HEMOLYSIN TRANSLOCATION ATP-BINDING P | ROTEIN |
| 94:00:00 | 6zie-A | 3.6 | 31.6 | 83 | 123 | 12 | MOLECULE: | CMPX-383B | |
| 95:00:00 | 8teu-A | 3.6 | 23.7 | 132 | 722 | 3 | MOLECULE: | LARGE TEGUMENT PROTEIN DENEDDYLASE | |
| 96:00:00 | 8yad-B | 3.6 | 85.7 | 171 | 1396 | 6 | MOLECULE: | SPATACSIN | |
| 97:00:00 | 6myo-D | 3.6 | 5.3 | 84 | 102 | 4 | MOLECULE: | SUCCINATE DEHYDROGENASE [UBIQUINONE] FLAVOP | ROTEIN |
| 98:00:00 | 8xr6-q | 3.6 | 11 | 71 | 143 | 6 | MOLECULE: | PHOTOSYSTEM II PROTEIN D1 | |
| 99:00:00 | 7a1g-z | 3.6 | 12.2 | 75 | 120 | 8 | MOLECULE: | 18S RIBOSOMAL RNA | |
| 0:00 | 7uxc-R | 3.6 | 7.3 | 65 | 107 | 3 | MOLECULE: | SERINE/THREONINE-PROTEIN KINASE MTOR | |
| 1:00 | 6wqz-A | 3.6 | 26.9 | 116 | 536 | 7 | MOLECULE: | AUTOPHAGY-RELATED PROTEIN 9A | |

| | | | | | | | | | | |
|---|---|---|---|---|---|---|---|---|---|---|
| 2:00 | 6swy-1 | 3.6 | 3.4 | 80 | 542 | 5 | MOLECULE: | VACUOLAR IMPORT AND DEGRADATION PROTEIN 28 | | |
| 3:00 | 7etm-A | 3.6 | 30 | 131 | 498 | 7 | MOLECULE: | PORTAL PROTEIN | | |
| 4:00 | 6kkk-A | 3.6 | 4.9 | 65 | 380 | 5 | MOLECULE: | SUGAR EFFLUX TRANSPORTER | | |
| 5:00 | 6a70-B | 3.6 | 6.2 | 128 | 704 | 4 | MOLECULE: | POLYCYSTIN-2 | | |
| 6:00 | 4wzx-A | 3.6 | 2.5 | 53 | 87 | 8 | MOLECULE: | SERINE/THREONINE-PROTEIN KINASE ULK3 | | |
| 7:00 | 8xks-S | 3.6 | 5 | 57 | 114 | 4 | MOLECULE: | CTAP1 | | |
| 8:00 | 1ej6-D | 3.6 | 5.3 | 60 | 417 | 13 | MOLECULE: | LAMBDA2 | | |
| 9:00 | 7y9n-A | 3.6 | 4.3 | 73 | 212 | 4 | MOLECULE: | SPIKE PROTEIN S2',5HB-H2 | | |
| 10:00 | 2ycd-A | 3.6 | 4.8 | 71 | 213 | 3 | MOLECULE: | GLUTATHIONE S-TRANSFERASE | | |
| 11:00 | 6wg3-B | 3.6 | 12.6 | 98 | 693 | 9 | MOLECULE: | STRUCTURAL MAINTENANCE OF CHROMOSOMES PROTE | IN 1A | |
| 12:00 | 7xdi-D | 3.6 | 11.6 | 91 | 124 | 3 | MOLECULE: | VP1 | | |
| 13:00 | 6uo8-B | 3.6 | 8.1 | 78 | 696 | 4 | MOLECULE: | GAMMA-AMINOBUTYRIC ACID TYPE B RECEPTOR SUB | UNIT 1 | |
| 14:00 | 1o7d-C | 3.6 | 9.8 | 62 | 150 | 6 | MOLECULE: | LYSOSOMAL ALPHA-MANNOSIDASE | | |
| 15:00 | 8ca1-A | 3.6 | 13.3 | 137 | 589 | 6 | MOLECULE: | VERY LONG-CHAIN SPECIFIC ACYL-COA DEHYDROGE | NASE, | |
| 16:00 | 6ye4-A | 3.6 | 22.5 | 108 | 236 | 7 | MOLECULE: | BIOPOLYMER TRANSPORT PROTEIN EXBB | | |
| 17:00 | 6dg6-A | 3.6 | 3.7 | 70 | 100 | 1 | MOLECULE: | NEOLEUKIN-2/15 | | |
| 18:00 | 2lon-A | 3.6 | 5.6 | 66 | 99 | 3 | MOLECULE: | HIG1 DOMAIN FAMILY MEMBER 1B | | |
| 19:00 | 9mny-A | 3.6 | 3.4 | 59 | 108 | 7 | MOLECULE: | MITOCHONDRIAL PYRUVATE CARRIER 1 | | |
| 20:00 | 9cpo-B | 3.6 | 7.7 | 93 | 195 | 6 | MOLECULE: | RNA-DIRECTED RNA POLYMERASE NSP12 | | |
| 21:00 | 7bst-A | 3.6 | 2.2 | 60 | 384 | 3 | MOLECULE: | TYPE I RESTRICTION ENZYME R PROTEIN | | |
| 22:00 | 6y1y-B | 3.6 | 4.4 | 64 | 128 | 3 | MOLECULE: | CHEA | | |
| 23:00 | 3nmd-A | 3.6 | 3.1 | 46 | 53 | 11 | MOLECULE: | CGMP DEPENDENT PROTEIN KINASE | | |
| 24:00:00 | 4r4l-C | 3.6 | 2.3 | 44 | 49 | 11 | MOLECULE: | CGMP-DEPENDENT PROTEIN KINASE 1 | | |
| 25:00:00 | 6akl-A | 3.6 | 2.3 | 41 | 52 | 5 | MOLECULE: | SUPPRESSOR OF IKBKE 1 | | |
| 26:00:00 | 4x01-C | 3.6 | 2.6 | 45 | 50 | 9 | MOLECULE: | DNA BINDING CTP1 | | |
| 27:00:00 | 6ku5-B | 3.6 | 12.7 | 66 | 207 | 5 | MOLECULE: | TRAF5 | | |
| 28:00:00 | 4gkg-A | 3.6 | 3.6 | 45 | 52 | 2 | MOLECULE: | C4-DICARBOXYLATE TRANSPORT SENSOR PROTEIN D | CTB | |
| 29:00:00 | 3he5-B | 3.6 | 2.2 | 41 | 48 | 12 | MOLECULE: | SYNZIP1 | | |
| 30:00:00 | 3efg-A | 3.6 | 4.1 | 44 | 51 | 5 | MOLECULE: | PROTEIN SLYX HOMOLOG | | |
| 31:00:00 | 6j6q-F | 3.6 | 3.2 | 42 | 161 | 2 | MOLECULE: | PRE-MRNA-SPLICING FACTOR 8 | | |
| 32:00:00 | 7tpr-A | 3.6 | 13.7 | 77 | 1145 | 4 | MOLECULE: | SPIKE GLYCOPROTEIN | | |
| 33:00:00 | 2m67-A | 3.6 | 3.6 | 59 | 81 | 3 | MOLECULE: | MERF | | |
| 34:00:00 | 5yij-A | 3.6 | 3 | 57 | 959 | 7 | MOLECULE: | SDEA | | |
| 35:00:00 | 3zid-A | 3.6 | 2.8 | 46 | 360 | 9 | MOLECULE: | TUBULIN/FTSZ, GTPASE | | |
| 36:00:00 | 7dkh-I | 3.6 | 2.8 | 65 | 914 | 8 | MOLECULE: | RNA POLYMERASE-ASSOCIATED PROTEIN CTR9 | | |
| 37:00:00 | 7yvb-A | 3.6 | 4.9 | 49 | 471 | 0 | MOLECULE: | FMRFAMIDE-GATED NA+ CHANNEL | | |
| 38:00:00 | 6hd5-t | 3.6 | 15 | 90 | 838 | 9 | MOLECULE: | N-TERMINAL ACETYLTRANSFERASE A COMPLEX SUBU | NIT NA | |
| 39:00:00 | 6xp5-H | 3.6 | 7.5 | 72 | 146 | 6 | MOLECULE: | MEDIATOR OF RNA POLYMERASE II TRANSCRIPTION | SUBUN | |
| 40:00:00 | 3u2r-A | 3.6 | 2.7 | 53 | 135 | 11 | MOLECULE: | REGULATORY PROTEIN MARR | | |
| 41:00:00 | 7bu0-A | 3.6 | 23.4 | 91 | 407 | 3 | MOLECULE: | UNCHARACTERIZED PROTEIN | | |
| 42:00:00 | 3vp8-B | 3.6 | 3.1 | 49 | 80 | 6 | MOLECULE: | GENERAL TRANSCRIPTIONAL COREPRESSOR TUP1 | | |
| 43:00:00 | 7znk-o | 3.5 | 41.4 | 108 | 164 | 8 | MOLECULE: | RNA | | |
| 44:00:00 | 6f1t-f | 3.5 | 24.4 | 125 | 929 | 5 | MOLECULE: | ARP1 ACTIN RELATED PROTEIN 1 HOMOLOG A | | |
| 45:00:00 | 7emf-K | 3.5 | 23 | 89 | 112 | 4 | MOLECULE: | MEDIATOR OF RNA POLYMERASE II TRANSCRIPTION | SUBUN | |
| 46:00:00 | 6ird-B | 3.5 | 26.7 | 97 | 254 | 6 | MOLECULE: | 1-PHOSPHATIDYLINOSITOL 4,5-BISPHOSPHATE PHO | SPHODI | |
| 47:00:00 | 7woo-D | 3.5 | 25.9 | 125 | 1398 | 6 | MOLECULE: | NUCLEOPORIN NIC96 | | |
| 48:00:00 | 5yfp-C | 3.5 | 41 | 177 | 790 | 7 | MOLECULE: | EXOCYST COMPLEX COMPONENT SEC3 | | |
| 49:00:00 | 8kg9-B | 3.5 | 6.5 | 58 | 193 | 10 | MOLECULE: | DNA REPLICATION LICENSING FACTOR MCM2 | | |
| 50:00:00 | 8ikj-R | 3.5 | 29.4 | 120 | 532 | 8 | MOLECULE: | ADHESION G PROTEIN-COUPLED RECEPTOR E5,SOLU | BLE CY | |
| 51:00:00 | 5eqz-A | 3.5 | 6.4 | 57 | 138 | 7 | MOLECULE: | REV PROTEIN | | |
| 52:00:00 | 8wrx-A | 3.5 | 4.7 | 50 | 86 | 4 | MOLECULE: | ANTI-CRISPR PROTEIN ACRIIA28 | | |
| 53:00:00 | 8vc1-A | 3.5 | 7 | 101 | 391 | 5 | MOLECULE: | GUSTATORY RECEPTOR | | |
| 54:00:00 | 6soy-A | 3.5 | 5.5 | 105 | 323 | 10 | MOLECULE: | ESAG6, SUBUNIT OF HETERODIMERIC TRANSFERRIN | RECEP | |
| 55:00:00 | 7xe4-F | 3.5 | 6.1 | 68 | 1503 | 3 | MOLECULE: | 1,3-BETA-GLUCAN SYNTHASE COMPONENT FKS1 | | |
| 56:00:00 | 8ave-A | 3.5 | 3.9 | 44 | 146 | 7 | MOLECULE: | LEPTIN | | |
| 57:00:00 | 1flc-B | 3.5 | 11.2 | 92 | 162 | 5 | MOLECULE: | HAEMAGGLUTININ-ESTERASE-FUSION GLYCOPROTEIN | | |
| 58:00:00 | 8hsi-A | 3.5 | 7.2 | 104 | 405 | 6 | MOLECULE: | TRANSMEMBRANE PROTEIN 87A | | |
| 59:00:00 | 6d03-E | 3.5 | 28.6 | 79 | 466 | 6 | MOLECULE: | TRANSFERRIN RECEPTOR PROTEIN 1 | | |
| 60:00:00 | 6r7x-A | 3.5 | 16.7 | 76 | 629 | 4 | MOLECULE: | ANOCTAMIN-10 | | |
| 61:00:00 | 8eg3-A | 3.5 | 23.3 | 86 | 554 | 2 | MOLECULE: | STERYL-SULFATASE | | |
| 62:00:00 | 6xpf-B | 3.5 | 7.3 | 68 | 285 | 1 | MOLECULE: | ZINC TRANSPORTER 8 | | |
| 63:00:00 | 8ip4-A | 3.5 | 17.3 | 139 | 320 | 4 | MOLECULE: | MAGNESIUM TRANSPORTER MRS2 HOMOLOG, MITOCHO | NDRIAL | |
| 64:00:00 | 5a42-A | 3.5 | 20.2 | 91 | 1597 | 7 | MOLECULE: | UNCHARACTERIZED LIPOPROTEIN YFHM | | |
| 65:00:00 | 5t58-B | 3.5 | 24.4 | 105 | 190 | 9 | MOLECULE: | KLLA0F02343P | | |
| 66:00:00 | 5a1v-K | 3.5 | 17.5 | 87 | 1125 | 5 | MOLECULE: | ADP-RIBOSYLATION FACTOR 1 | | |
| 67:00:00 | 2pms-C | 3.5 | 5.1 | 85 | 109 | 12 | MOLECULE: | LACTOTRANSFERRIN | | |
| 68:00:00 | 8pht-A | 3.5 | 3.6 | 64 | 111 | 2 | MOLECULE: | SCAFFOLD PROTEIN | | |
| 69:00:00 | 3anw-A | 3.5 | 9.4 | 70 | 171 | 6 | MOLECULE: | PUTATIVE UNCHARACTERIZED PROTEIN | | |
| 70:00:00 | 8q85-d | 3.5 | 37.9 | 104 | 108 | 3 | MOLECULE: | KINETOCHORE PROTEIN NDC80 | | |
| 71:00:00 | 7deg-A | 3.5 | 7.8 | 63 | 581 | 6 | MOLECULE: | CYTOCHROME C OXIDASE SUBUNIT I | | |
| 72:00:00 | 6wuh-A | 3.5 | 12.7 | 83 | 308 | 10 | MOLECULE: | SAM35 | | |
| 73:00:00 | 4mer-D | 3.5 | 4.7 | 50 | 97 | 4 | MOLECULE: | STREPTOCOCCAL HISTIDINE-RICH GLYCOPROTEIN I | NTERAC | |
| 74:00:00 | 3bey-D | 3.5 | 2.8 | 47 | 94 | 2 | MOLECULE: | CONSERVED PROTEIN O27018 | | |
| 75:00:00 | 7wff-F | 3.5 | 27.6 | 105 | 677 | 4 | MOLECULE: | NAD(P)H-QUINONE OXIDOREDUCTASE SUBUNIT 1, C | HLOROP | |
| 76:00:00 | 1sc7-A | 3.5 | 19.9 | 88 | 567 | 9 | MOLECULE: | 5'-D(*AP*AP*AP*AP*AP*GP*AP*CP*TP*T)-3' | | |
| 77:00:00 | 8eaw-a | 3.5 | 8.2 | 72 | 287 | 13 | MOLECULE: | CAPSID PROTEIN | | |
| 78:00:00 | 8y5f-B | 3.5 | 2.1 | 52 | 273 | 4 | MOLECULE: | SPERMIDINE/PUTRESCINE IMPORT ATP-BINDING PR | OTEIN | |
| 79:00:00 | 8t05-A | 3.5 | 5.1 | 55 | 201 | 2 | MOLECULE: | MYOMAKER | | |
| 80:00:00 | 8g52-A | 3.5 | 18.6 | 68 | 279 | 4 | MOLECULE: | TPR_REGION DOMAIN-CONTAINING PROTEIN | | |
| 81:00:00 | 4nnh-A | 3.5 | 3.1 | 48 | 164 | 10 | MOLECULE: | 30S RIBOSOMAL PROTEIN S1 | | |
| 82:00:00 | 7egm-H | 3.5 | 11.9 | 99 | 388 | 9 | MOLECULE: | TRANSCRIPTION REGULATORY PROTEIN SNF2 | | |
| 83:00:00 | 1dvo-A | 3.5 | 11 | 81 | 152 | 5 | MOLECULE: | FERTILITY INHIBITION PROTEIN O | | |
| 84:00:00 | 6gmh-Q | 3.5 | 16.7 | 80 | 884 | 10 | MOLECULE: | RPB1 | | |
| 85:00:00 | 3syk-A | 3.5 | 7 | 78 | 290 | 1 | MOLECULE: | PROTEIN CBBX | | |
| 86:00:00 | 8th8-K | 3.5 | 26.6 | 118 | 434 | 5 | MOLECULE: | DYNEIN REGULATORY COMPLEX PROTEIN 1/2 N-TER | MINAL | |
| 87:00:00 | 3n1e-B | 3.5 | 5.8 | 54 | 141 | 11 | MOLECULE: | VACUOLAR PROTEIN SORTING-ASSOCIATED PROTEIN | | 54 |
| 88:00:00 | 4um2-A | 3.5 | 24 | 85 | 510 | 4 | MOLECULE: | TELOMERASE-BINDING PROTEIN EST1A | | |
| 89:00:00 | 8vyb-A | 3.5 | 9.9 | 80 | 464 | 8 | MOLECULE: | ISOFORM 2 OF RAB3 GTPASE-ACTIVATING PROTEIN | CATAL | |
| 90:00:00 | 8fhc-B | 3.5 | 4 | 51 | 215 | 0 | MOLECULE: | TRNA-(MS[2]IO[6]A)-HYDROXYLASE | | |
| 91:00:00 | 6xwx-E | 3.5 | 7.6 | 84 | 566 | 8 | MOLECULE: | UNCHARACTERIZED PROTEIN,UNCHARACTERIZED PRO | TEIN | |
| 92:00:00 | 1u2m-C | 3.5 | 5.2 | 81 | 143 | 9 | MOLECULE: | HISTONE-LIKE PROTEIN HLP-1 | | |
| 93:00:00 | 8p0v-M | 3.5 | 3.2 | 48 | 357 | 8 | MOLECULE: | COILED-COIL DOMAIN-CONTAINING PROTEIN 93 | | |
| 94:00:00 | 7sqk-C | 3.5 | 36.9 | 138 | 600 | 6 | MOLECULE: | HAUS AUGMIN-LIKE COMPLEX SUBUNIT 1 | | |
| 95:00:00 | 5a31-N | 3.5 | 27.3 | 92 | 703 | 4 | MOLECULE: | ANAPHASE-PROMOTING COMPLEX SUBUNIT 1 | | |
| 96:00:00 | 5cx2-A | 3.5 | 2.9 | 47 | 55 | 11 | MOLECULE: | CORONIN | | |
| 97:00:00 | 1gl2-A | 3.5 | 3.6 | 43 | 53 | 9 | MOLECULE: | ENDOBREVIN | | |
| 98:00:00 | 2kes-A | 3.5 | 2.5 | 44 | 48 | 5 | MOLECULE: | SYNPHILIN-1 | | |
| 99:00:00 | 4b86-B | 3.5 | 3.8 | 46 | 51 | 7 | MOLECULE: | MALE-SPECIFIC LETHAL 1 HOMOLOG | | |
| 0:00 | 1deb-A | 3.5 | 2.1 | 44 | 54 | 11 | MOLECULE: | ADENOMATOUS POLYPOSIS COLI PROTEIN | | |
| 1:00 | 6cko-C | 3.5 | 4.7 | 44 | 50 | 16 | MOLECULE: | PROTEIN AF-10 | | |
| 2:00 | 1ifm-A | 3.5 | 3.4 | 43 | 46 | 5 | MOLECULE: | INOVIRUS | | |

| | | | | | | | | | |
|---|---|---|---|---|---|---|---|---|---|
| 3:00 | 6od2-A | 3.5 | 3 | 43 | 53 | 5 | MOLECULE: | SPINDLE POLE BODY COMPONENT SPC42 | |
| 4:00 | 2bsk-D | 3.5 | 6.3 | 64 | 90 | 13 | MOLECULE: | MITOCHONDRIAL IMPORT INNER MEMBRANE TRANSLO | CASE |
| 5:00 | 3dkq-A | 3.5 | 3.5 | 43 | 230 | 5 | MOLECULE: | PKHD-TYPE HYDROXYLASE SBAL_3634 | |
| 6:00 | 1ezj-A | 3.5 | 3.3 | 44 | 114 | 11 | MOLECULE: | NUCLEOCAPSID PHOSPHOPROTEIN | |
| 7:00 | 6s7t-A | 3.5 | 26.4 | 104 | 706 | 6 | MOLECULE: | DOLICHYL-DIPHOSPHOOLIGOSACCHARIDE--PROTEIN | |
| 8:00 | 3zta-A | 3.5 | 2.4 | 62 | 139 | 6 | MOLECULE: | ANTI-SIGMA-FACTOR ANTAGONIST (STAS) DOMAIN | PROTEI |
| 9:00 | 6lo8-E | 3.5 | 3.7 | 55 | 80 | 9 | MOLECULE: | MITOCHONDRIAL IMPORT INNER MEMBRANE TRANSLO | CASE S |
| 10:00 | 3bm3-A | 3.5 | 3.4 | 77 | 259 | 10 | MOLECULE: | DNA (5'-D(*CP*AP*TP*CP*CP*AP*GP*GP*TP*AP*C) | -3') |
| 11:00 | 5tj5-A | 3.5 | 27.6 | 109 | 570 | 2 | MOLECULE: | V-TYPE PROTON ATPASE SUBUNIT A | |
| 12:00 | 6hs5-A | 3.5 | 2.9 | 46 | 249 | 4 | MOLECULE: | TSSA | |
| 13:00 | 8th8-L | 3.5 | 14.2 | 108 | 862 | 8 | MOLECULE: | DYNEIN REGULATORY COMPLEX PROTEIN 1/2 N-TER | MINAL |
| 14:00 | 8b6l-P | 3.5 | 3.5 | 50 | 602 | 4 | MOLECULE: | PROTEIN TRANSPORT PROTEIN SEC61 SUBUNIT ALP | HA ISO |
| 15:00 | 6v4l-A | 3.4 | 27.9 | 99 | 479 | 7 | MOLECULE: | TRK SYSTEM POTASSIUM UPTAKE PROTEIN TRKH | |
| 16:00 | 3j9e-D | 3.4 | 19.2 | 112 | 520 | 13 | MOLECULE: | VP5 | |
| 17:00 | 8z9y-A | 3.4 | 25.3 | 125 | 781 | 3 | MOLECULE: | PROTEIN TIC 214 | |
| 18:00 | 8ppr-D | 3.4 | 69.6 | 147 | 248 | 13 | MOLECULE: | KINETOCHORE-ASSOCIATED PROTEIN DSN1 HOMOLOG | |
| 19:00 | 8sfo-A | 3.4 | 37.8 | 153 | 1240 | 7 | MOLECULE: | CRISPR-ASSOCIATED ENDONUCLEASE CAS12A | |
| 20:00 | 5j0l-A | 3.4 | 30.6 | 79 | 130 | 10 | MOLECULE: | DESIGNED PROTEIN 3L6HC2_2 | |
| 21:00 | 4fhn-B | 3.4 | 29.1 | 140 | 1022 | 9 | MOLECULE: | NUCLEOPORIN NUP37 | |
| 22:00 | 8gix-E | 3.4 | 22.7 | 102 | 991 | 6 | MOLECULE: | HISTONE TRANSCRIPTION REGULATOR 3 HOMOLOG | |
| 23:00 | 2o8p-A | 3.4 | 6.6 | 97 | 220 | 3 | MOLECULE: | 14-3-3 DOMAIN CONTAINING PROTEIN | |
| 24:00:00 | 5l1a-A | 3.4 | 29.5 | 67 | 108 | 10 | MOLECULE: | UNCHARACTERIZED PROTEIN | |
| 25:00:00 | 8ttf-A | 3.4 | 30.6 | 94 | 388 | 10 | MOLECULE: | QUINOLONE RESISTANCE PROTEIN NORA | |
| 26:00:00 | 8gb3-F | 3.4 | 2.8 | 39 | 90 | 3 | MOLECULE: | CHAPERONE PROTEIN DNAK | |
| 27:00:00 | 8bd7-M | 3.4 | 24 | 127 | 164 | 6 | MOLECULE: | IFT88 | |
| 28:00:00 | 8whn-A | 3.4 | 4.3 | 95 | 142 | 5 | MOLECULE: | RIBOPHORIN 1 SUPERFAMILY PROTEIN | |
| 29:00:00 | 3add-A | 3.4 | 2.5 | 45 | 251 | 11 | MOLECULE: | L-SERYL-TRNA(SEC) KINASE | |
| 30:00:00 | 2d6y-B | 3.4 | 2.3 | 43 | 188 | 5 | MOLECULE: | PUTATIVE TETR FAMILY REGULATORY PROTEIN | |
| 31:00:00 | 7z0l-B | 3.4 | 18.8 | 72 | 370 | 4 | MOLECULE: | INTERLEUKIN-6 RECEPTOR SUBUNIT BETA | |
| 32:00:00 | 7tmw-R | 3.4 | 17 | 113 | 499 | 4 | MOLECULE: | RELAXIN RECEPTOR 1, GUANINE NUCLEOTIDE-BIND | ING PR |
| 33:00:00 | 1llm-C | 3.4 | 2.5 | 45 | 87 | 7 | MOLECULE: | 5'-D(*TP*CP*CP*CP*AP*CP*GP*CP*GP*TP*GP*GP*G | )-3' |
| 34:00:00 | 7yzp-C | 3.4 | 27.7 | 115 | 448 | 5 | MOLECULE: | DNA HAIRPIN (59-MER) | |
| 35:00:00 | 4g6d-B | 3.4 | 3.4 | 48 | 198 | 10 | MOLECULE: | RNA POLYMERASE SIGMA FACTOR RPOD | |
| 36:00:00 | 8ij1-D | 3.4 | 20 | 75 | 627 | 11 | MOLECULE: | CULLIN-2 | |
| 37:00:00 | 7eu9-A | 3.4 | 4.9 | 72 | 1078 | 8 | MOLECULE: | CAS12I1 D647A MUTANT | |
| 38:00:00 | 7ewp-A | 3.4 | 14.5 | 110 | 577 | 2 | MOLECULE: | PROBABLE G-PROTEIN COUPLED RECEPTOR 158 | |
| 39:00:00 | 3j83-A | 3.4 | 4.3 | 60 | 276 | 2 | MOLECULE: | ESX-1 SECRETION-ASSOCIATED PROTEIN ESPB | |
| 40:00:00 | 9ivk-A | 3.4 | 4.1 | 67 | 278 | 7 | MOLECULE: | HBC599 MEMBRANE PROTEIN BINDER | |
| 41:00:00 | 4jio-B | 3.4 | 16.6 | 101 | 320 | 4 | MOLECULE: | BRO1 | |
| 42:00:00 | 7sjr-A | 3.4 | 10.2 | 88 | 911 | 2 | MOLECULE: | DNA HELICASE | |
| 43:00:00 | 4rp5-A | 3.4 | 6.5 | 67 | 94 | 10 | MOLECULE: | DISKS LARGE 1 TUMOR SUPPRESSOR PROTEIN | |
| 44:00:00 | 6zsi-D | 3.4 | 5.6 | 100 | 133 | 4 | MOLECULE: | RAS-RELATED PROTEIN RAB-8A | |
| 45:00:00 | 8d8k-V | 3.4 | 25.1 | 90 | 233 | 12 | MOLECULE: | PROBABLE S-ADENOSYL-L-METHIONINE-DEPENDENT | RNA |
| 46:00:00 | 8dqk-A | 3.4 | 25.8 | 111 | 733 | 5 | MOLECULE: | CELLULOSE SYNTHASE-LIKE CSLF6 | |
| 47:00:00 | 7e7i-A | 3.4 | 20.1 | 105 | 2011 | 7 | MOLECULE: | RETINAL-SPECIFIC PHOSPHOLIPID-TRANSPORTING | ATPASE |
| 48:00:00 | 6gy6-A | 3.4 | 12.4 | 90 | 365 | 6 | MOLECULE: | XAXA | |
| 49:00:00 | 7etw-A | 3.4 | 4.4 | 51 | 197 | 8 | MOLECULE: | INSULIN-INDUCED GENE 2 PROTEIN | |
| 50:00:00 | 8bh8-A | 3.4 | 10 | 93 | 523 | 3 | MOLECULE: | PCIF1_WW DOMAIN-CONTAINING PROTEIN | |
| 51:00:00 | 6wge-C | 3.4 | 32.8 | 76 | 174 | 13 | MOLECULE: | STRUCTURAL MAINTENANCE OF CHROMOSOMES PROTE | IN 1A |
| 52:00:00 | 4wpx-B | 3.4 | 10.2 | 56 | 81 | 14 | MOLECULE: | CELL DIVISION CONTROL PROTEIN 31-LIKE PROTE | IN |
| 53:00:00 | 6yvd-D | 3.4 | 21.8 | 89 | 414 | 6 | MOLECULE: | CONDENSIN COMPLEX SUBUNIT 2 | |
| 54:00:00 | 5bw9-g | 3.4 | 22.6 | 87 | 141 | 8 | MOLECULE: | V-TYPE PROTON ATPASE CATALYTIC SUBUNIT A | |
| 55:00:00 | 6xp5-D | 3.4 | 25.4 | 97 | 134 | 8 | MOLECULE: | MEDIATOR OF RNA POLYMERASE II TRANSCRIPTION | SUBUN |
| 56:00:00 | 5wah-A | 3.4 | 2.6 | 56 | 99 | 4 | MOLECULE: | IGA FC RECEPTOR | |
| 57:00:00 | 8j3r-B | 3.4 | 6.1 | 78 | 387 | 9 | MOLECULE: | TRANSPOSASE IS605 ORFB C-TERMINAL DOMAIN-CO | NTAINI |
| 58:00:00 | 6e1k-A | 3.4 | 4.5 | 66 | 536 | 5 | MOLECULE: | TWO PORE CALCIUM CHANNEL PROTEIN 1 | |
| 59:00:00 | 8wjg-A | 3.4 | 10.6 | 72 | 533 | 7 | MOLECULE: | SOLUTE CARRIER FAMILY 22 MEMBER 12 | |
| 60:00:00 | 7x07-A | 3.4 | 14.5 | 143 | 633 | 7 | MOLECULE: | ATP-BINDING CASSETTE SUB-FAMILY D MEMBER 1 | |
| 61:00:00 | 3h7x-E | 3.4 | 3.5 | 46 | 53 | 4 | MOLECULE: | ADHESIN YADA | |
| 62:00:00 | 6wbp-A | 3.4 | 2.7 | 47 | 56 | 9 | MOLECULE: | SEPTIN-6 | |
| 63:00:00 | 6hk5-B | 3.4 | 3.1 | 44 | 66 | 0 | MOLECULE: | COOJ | |
| 64:00:00 | 5mw9-H | 3.4 | 3.2 | 48 | 57 | 4 | MOLECULE: | CENTROSOMIN | |
| 65:00:00 | 6fpr-B | 3.4 | 3.4 | 49 | 58 | 4 | MOLECULE: | SIGNAL RECOGNITION PARTICLE RECEPTOR FTSY | |
| 66:00:00 | 1n73-C | 3.4 | 13 | 56 | 322 | 4 | MOLECULE: | FIBRIN ALPHA-1 CHAIN | |
| 67:00:00 | 6wkr-C | 3.4 | 21.7 | 85 | 607 | 9 | MOLECULE: | UBIQUITIN | |
| 68:00:00 | 1ifk-A | 3.4 | 3 | 44 | 51 | 7 | MOLECULE: | INOVIRUS | |
| 69:00:00 | 4yv4-C | 3.4 | 3.2 | 48 | 57 | 4 | MOLECULE: | SPINDLE ASSEMBLY ABNORMAL PROTEIN 5 | |
| 70:00:00 | 1gl2-D | 3.4 | 3.1 | 45 | 54 | 4 | MOLECULE: | ENDOBREVIN | |
| 71:00:00 | 3rk3-E | 3.4 | 2.2 | 41 | 50 | 10 | MOLECULE: | VAMP2 | |
| 72:00:00 | 3wmi-A | 3.4 | 3.2 | 50 | 57 | 4 | MOLECULE: | EIAV GP45 WILD TYPE | |
| 73:00:00 | 7a9w-A | 3.4 | 15.8 | 70 | 513 | 4 | MOLECULE: | PROTEIN RMD9, MITOCHONDRIAL | |
| 74:00:00 | 6lth-Q | 3.4 | 2.7 | 49 | 105 | 4 | MOLECULE: | TRANSCRIPTION ACTIVATOR BRG1 | |
| 75:00:00 | 3jb9-r | 3.4 | 2.4 | 40 | 573 | 0 | MOLECULE: | PRE-MRNA-SPLICING FACTOR SPP42 | |
| 76:00:00 | 3bhp-C | 3.4 | 5.7 | 50 | 54 | 8 | MOLECULE: | UPF0291 PROTEIN YNZC | |
| 77:00:00 | 4v1a-o | 3.4 | 3.6 | 49 | 94 | 8 | MOLECULE: | MITORIBOSOMAL PROTEIN ML37, MRPL37 | |
| 78:00:00 | 7asm-W | 3.4 | 4 | 51 | 66 | 12 | MOLECULE: | 50S RIBOSOMAL PROTEIN L19 | |
| 79:00:00 | 6nd4-Q | 3.4 | 6.6 | 57 | 862 | 11 | MOLECULE: | ETS RRNA | |
| 80:00:00 | 1eq7-A | 3.4 | 3.5 | 47 | 56 | 4 | MOLECULE: | OUTER MEMBRANE LIPOPROTEIN | |
| 81:00:00 | 7vv6-R | 3.3 | 24.7 | 71 | 271 | 6 | MOLECULE: | MAS-RELATED G-PROTEIN COUPLED RECEPTOR MEMB | ER X2 |
| 82:00:00 | 6ted-R | 3.3 | 23.7 | 88 | 244 | 6 | MOLECULE: | DNA-DIRECTED RNA POLYMERASE SUBUNIT | |
| 83:00:00 | 8ga7-A | 3.3 | 30.3 | 137 | 503 | 9 | MOLECULE: | THR5 | |
| 84:00:00 | 5j1f-A | 3.3 | 32.9 | 92 | 188 | 12 | MOLECULE: | PLECTIN,PLECTIN | |
| 85:00:00 | 8dl7-A | 3.3 | 29.4 | 129 | 467 | 7 | MOLECULE: | SOLUTE CARRIER FAMILY 40 MEMBER 1 | |
| 86:00:00 | 6m6z-A | 3.3 | 31.4 | 103 | 203 | 5 | MOLECULE: | TMH4C4 | |
| 87:00:00 | 7rz4-A | 3.3 | 18.7 | 98 | 993 | 3 | MOLECULE: | GLUTAMATE RECEPTOR 2 | |
| 88:00:00 | 6i3m-E | 3.3 | 29 | 102 | 354 | 4 | MOLECULE: | TRANSLATION INITIATION FACTOR EIF-2B SUBUNI | T ALPH |
| 89:00:00 | 4gd3-A | 3.3 | 6 | 106 | 179 | 5 | MOLECULE: | HYDROGENASE-1 SMALL CHAIN | |
| 90:00:00 | 1n1c-A | 3.3 | 7.7 | 79 | 199 | 8 | MOLECULE: | TORA SPECIFIC CHAPERONE | |
| 91:00:00 | 8th8-B | 3.3 | 37.2 | 126 | 276 | 10 | MOLECULE: | DYNEIN REGULATORY COMPLEX PROTEIN 1/2 N-TER | MINAL |
| 92:00:00 | 1oks-A | 3.3 | 1.7 | 39 | 53 | 8 | MOLECULE: | RNA POLYMERASE ALPHA SUBUNIT | |
| 93:00:00 | 8pnl-C | 3.3 | 9.7 | 80 | 486 | 5 | MOLECULE: | PUTATIVE TRIACYLGLYCERIDE TRANSPORTER | |
| 94:00:00 | 7nsb-7 | 3.3 | 17 | 117 | 531 | 9 | MOLECULE: | VACUOLAR IMPORT AND DEGRADATION PROTEIN 30 | |
| 95:00:00 | 5lbm-C | 3.3 | 6.1 | 77 | 90 | 10 | MOLECULE: | TRANSCRIPTIONAL REPRESSOR FRMR | |
| 96:00:00 | 9c7u-C | 3.3 | 7.5 | 70 | 205 | 4 | MOLECULE: | NICALIN | |
| 97:00:00 | 2oar-A | 3.3 | 18.2 | 86 | 125 | 6 | MOLECULE: | LARGE-CONDUCTANCE MECHANOSENSITIVE CHANNEL | |
| 98:00:00 | 8xh6-A | 3.3 | 4.7 | 75 | 161 | 1 | MOLECULE: | LATENT MEMBRANE PROTEIN 1 | |
| 99:00:00 | 6cfz-I | 3.3 | 30.3 | 86 | 106 | 2 | MOLECULE: | ASK1 | |
| 0:00 | 8rot-A | 3.3 | 29.4 | 86 | 897 | 1 | MOLECULE: | CHAT DOMAIN-CONTAINING PROTEIN | |
| 1:00 | 2n12-A | 3.3 | 4.3 | 57 | 82 | 5 | MOLECULE: | UNCONVENTIONAL MYOSIN-VI | |
| 2:00 | 1ux5-A | 3.3 | 21 | 105 | 411 | 3 | MOLECULE: | BNI1 PROTEIN | |
| 3:00 | 7byl-A | 3.3 | 29 | 114 | 354 | 8 | MOLECULE: | GREEN FLUORESCENT PROTEIN,POTASSIUM VOLTAGE | #NAME? |

| | | | | | | | | | |
|---|---|---|---|---|---|---|---|---|---|
| 4:00 | 8b6l-M | 3.3 | 6 | 86 | 110 | 6 | MOLECULE: | PROTEIN TRANSPORT PROTEIN SEC61 SUBUNIT ALP | HA ISO |
| 5:00 | 2p4w-B | 3.3 | 8.5 | 64 | 198 | 8 | MOLECULE: | TRANSCRIPTIONAL REGULATORY PROTEIN ARSR FAM | ILY |
| 6:00 | 6z2w-E | 3.3 | 24.7 | 87 | 2325 | 1 | MOLECULE: | DNA DAMAGE CHECKPOINT PROTEIN LCD1 | |
| 7:00 | 8ftk-A | 3.3 | 24.5 | 122 | 1584 | 4 | MOLECULE: | 5'-3' RNA HELICASE-LIKE PROTEIN | |
| 8:00 | 7u8g-A | 3.3 | 5.6 | 86 | 260 | 5 | MOLECULE: | EGFP, CYTOCHROME B-245 HEAVY CHAIN CHIMERA | |
| 9:00 | 3trc-A | 3.3 | 4.2 | 48 | 168 | 2 | MOLECULE: | PHOSPHOENOLPYRUVATE-PROTEIN PHOSPHOTRANSFEF | ASE |
| 10:00 | 3hef-B | 3.3 | 8 | 77 | 130 | 12 | MOLECULE: | GENE 1 PROTEIN | |
| 11:00 | 7mgq-A | 3.3 | 16.9 | 99 | 602 | 6 | MOLECULE: | 5-AMINOIMIDAZOLE-4-CARBOXAMIDE RIBONUCLEOTI | DE |
| 12:00 | 6b85-A | 3.3 | 7.4 | 101 | 215 | 5 | MOLECULE: | TMHC4_R | |
| 13:00 | 6xns-B | 3.3 | 9.1 | 85 | 333 | 8 | MOLECULE: | C3_CROWN-05 | |
| 14:00 | 3p8c-F | 3.3 | 9.5 | 104 | 156 | 3 | MOLECULE: | CYTOPLASMIC FMR1-INTERACTING PROTEIN 1 | |
| 15:00 | 5k47-A | 3.3 | 21.1 | 81 | 484 | 7 | MOLECULE: | POLYCYSTIN-2 | |
| 16:00 | 8smr-G | 3.3 | 20.3 | 133 | 312 | 5 | MOLECULE: | UBIQUINOL-CYTOCHROME C REDUCTASE IRON-SULFU | R SUBU |
| 17:00 | 5da9-B | 3.3 | 9.3 | 92 | 400 | 5 | MOLECULE: | PUTATIVE UNCHARACTERIZED PROTEIN,PUTATIVE U | NCHARA |
| 18:00 | 6mpb-A | 3.3 | 13.8 | 81 | 447 | 7 | MOLECULE: | NEUTRAL AMINO ACID TRANSPORTER B(0) | |
| 19:00 | 7zu0-F | 3.3 | 4.5 | 56 | 112 | 7 | MOLECULE: | E3 UBIQUITIN-PROTEIN LIGASE PEP5 | |
| 20:00 | 6nd4-P | 3.3 | 22.5 | 79 | 359 | 6 | MOLECULE: | ETS RRNA | |
| 21:00 | 1gnc-A | 3.3 | 5.5 | 68 | 175 | 6 | MOLECULE: | GRANULOCYTE COLONY-STIMULATING FACTOR | |
| 22:00 | 6whb-A | 3.3 | 10.8 | 71 | 284 | 8 | MOLECULE: | MITOGEN-ACTIVATED PROTEIN KINASE KINASE KIN | ASE 1 |
| 23:00 | 7d2h-D | 3.3 | 2.4 | 49 | 60 | 6 | MOLECULE: | LIPRIN-ALPHA-2 | |
| 24:00:00 | 6cfz-G | 3.3 | 2.9 | 45 | 56 | 4 | MOLECULE: | ASK1 | |
| 25:00:00 | 2x7r-C | 3.3 | 3.5 | 48 | 58 | 6 | MOLECULE: | TRANSMEMBRANE PROTEIN GP41 | |
| 26:00:00 | 3e7k-H | 3.3 | 3.8 | 46 | 56 | 2 | MOLECULE: | TRPM7 CHANNEL | |
| 27:00:00 | 3uf1-D | 3.3 | 3.7 | 48 | 112 | 10 | MOLECULE: | VIMENTIN | |
| 28:00:00 | 6fia-D | 3.3 | 4 | 50 | 104 | 12 | MOLECULE: | LINE-1 RETROTRANSPOSABLE ELEMENT ORF1 PROTE | IN |
| 29:00:00 | 4i1l-A | 3.3 | 3 | 51 | 62 | 2 | MOLECULE: | FORKHEAD BOX PROTEIN P3 | |
| 30:00:00 | 5oxf-A | 3.3 | 3.8 | 57 | 703 | 12 | MOLECULE: | GTP-BINDING PROTEIN | |
| 31:00:00 | 3swf-B | 3.3 | 3.5 | 48 | 58 | 4 | MOLECULE: | CGMP-GATED CATION CHANNEL ALPHA-1 | |
| 32:00:00 | 7bb3-A | 3.3 | 3.4 | 48 | 54 | 8 | MOLECULE: | SURVIVAL MOTOR NEURON-LIKE PROTEIN 1,SURVIV | AL MOT |
| 33:00:00 | 5xfs-A | 3.3 | 2.8 | 49 | 78 | 4 | MOLECULE: | PE FAMILY PROTEIN PE8 | |
| 34:00:00 | 1urq-A | 3.3 | 4.2 | 44 | 57 | 5 | MOLECULE: | M-TOMOSYN ISOFORM | |
| 35:00:00 | 4m3l-A | 3.3 | 2.5 | 44 | 60 | 9 | MOLECULE: | E3 UBIQUITIN-PROTEIN LIGASE TRIM63 | |
| 36:00:00 | 5mz6-1 | 3.3 | 9.7 | 71 | 1018 | 6 | MOLECULE: | SEPARASE | |
| 37:00:00 | 7xad-C | 3.3 | 2.6 | 53 | 96 | 15 | MOLECULE: | PROGRAMMED CELL DEATH 1 LIGAND 1 | |
| 38:00:00 | 7v2w-J | 3.3 | 41.3 | 113 | 236 | 4 | MOLECULE: | THO COMPLEX SUBUNIT HPR1 | |
| 39:00:00 | 3bvo-A | 3.3 | 4.1 | 48 | 197 | 19 | MOLECULE: | CO-CHAPERONE PROTEIN HSCB, MITOCHONDRIAL PR | ECURSO |
| 40:00:00 | 6jzt-A | 3.3 | 4 | 57 | 402 | 9 | MOLECULE: | FLAGELLAR HOOK PROTEIN FLGE | |
| 41:00:00 | 4tko-B | 3.3 | 26.9 | 98 | 326 | 13 | MOLECULE: | EMRA | |
| 42:00:00 | 2c5i-T | 3.3 | 3.3 | 51 | 94 | 16 | MOLECULE: | T-SNARE AFFECTING A LATE GOLGI COMPARTMENT | |
| 43:00:00 | 6kg7-A | 3.3 | 15.4 | 66 | 1817 | 2 | MOLECULE: | PIEZO-TYPE MECHANOSENSITIVE ION CHANNEL COM | PONENT |
| 44:00:00 | 6nr8-4 | 3.3 | 3.5 | 61 | 104 | 8 | MOLECULE: | PREFOLDIN SUBUNIT 1 | |
| 45:00:00 | 7z6q-c | 3.3 | 8.3 | 77 | 121 | 1 | MOLECULE: | PHOTOSYSTEM P840 REACTION CENTER, LARGE SUB | UNIT |
| 46:00:00 | 8hpo-H | 3.2 | 52.3 | 110 | 233 | 9 | MOLECULE: | TRANSCRIPTIONAL REGULATORY PROTEIN UME1 | |
| 47:00:00 | 9gm9-A | 3.2 | 48 | 180 | 999 | 4 | MOLECULE: | CHROMOSOME PARTITION PROTEIN MUKB | |
| 48:00:00 | 6h95-A | 3.2 | 30 | 91 | 225 | 4 | MOLECULE: | ALBICIDIN RESISTANCE PROTEIN | |
| 49:00:00 | 5v7p-A | 3.2 | 21.4 | 74 | 281 | 5 | MOLECULE: | PROTEIN-S-ISOPRENYLCYSTEINE O-METHYLTRANSFE | RASE |
| 50:00:00 | 5lnc-B | 3.2 | 20.6 | 102 | 367 | 9 | MOLECULE: | VACUOLAR TRANSPORTER CHAPERONE 4,CORE HISTO | NE MAC |
| 51:00:00 | 1e1d-A | 3.2 | 30 | 99 | 553 | 2 | MOLECULE: | HYDROXYLAMINE REDUCTASE | |
| 52:00:00 | 5n9j-A | 3.2 | 27.8 | 108 | 566 | 10 | MOLECULE: | MEDIATOR OF RNA POLYMERASE II TRANSCRIPTION | SUBUN |
| 53:00:00 | 8eki-D | 3.2 | 23.4 | 114 | 684 | 4 | MOLECULE: | PROTEIN TRANSPORT PROTEIN SEC20 | |
| 54:00:00 | 5n9j-Z | 3.2 | 22.6 | 111 | 119 | 5 | MOLECULE: | MEDIATOR OF RNA POLYMERASE II TRANSCRIPTION | SUBUN |
| 55:00:00 | 8qz7-A | 3.2 | 11.6 | 92 | 333 | 5 | MOLECULE: | ISOFORM 2 OF CERAMIDE SYNTHASE 6 | |
| 56:00:00 | 3hr0-A | 3.2 | 6.9 | 101 | 250 | 6 | MOLECULE: | COG4 | |
| 57:00:00 | 3nym-A | 3.2 | 4.2 | 69 | 124 | 3 | MOLECULE: | UNCHARACTERIZED PROTEIN | |
| 58:00:00 | 8h38-C | 3.2 | 19.9 | 72 | 407 | 8 | MOLECULE: | COP9 SIGNALOSOME COMPLEX SUBUNIT 1 | |
| 59:00:00 | 7tmr-F | 3.2 | 9 | 87 | 478 | 5 | MOLECULE: | H(+)-TRANSPORTING TWO-SECTOR ATPASE | |
| 60:00:00 | 3qz0-B | 3.2 | 1.3 | 41 | 79 | 2 | MOLECULE: | FACTOR H BINDING PROTEIN | |
| 61:00:00 | 8d0b-F | 3.2 | 32.8 | 111 | 915 | 3 | MOLECULE: | CST COMPLEX SUBUNIT CTC1 | |
| 62:00:00 | 7yjm-C | 3.2 | 3.4 | 41 | 48 | 5 | MOLECULE: | ATLCB1 | |
| 63:00:00 | 6r7n-B | 3.2 | 15.7 | 83 | 443 | 12 | MOLECULE: | COP9 SIGNALOSOME COMPLEX SUBUNIT 1 | |
| 64:00:00 | 1un8-A | 3.2 | 20.5 | 61 | 542 | 13 | MOLECULE: | DIHYDROXYACETONE KINASE | |
| 65:00:00 | 2xqh-A | 3.2 | 24.9 | 65 | 258 | 12 | MOLECULE: | IMMUNOGLOBULIN-BINDING PROTEIN EIBD | |
| 66:00:00 | 8iqg-A | 3.2 | 6.7 | 64 | 213 | 6 | MOLECULE: | CHROMATIN ASSEMBLY FACTOR 1 SUBUNIT A | |
| 67:00:00 | 4tsh-A | 3.2 | 4.4 | 57 | 121 | 12 | MOLECULE: | SURFACE PROTEIN ADHESIN | |
| 68:00:00 | 7pkn-R | 3.2 | 10.9 | 57 | 80 | 18 | MOLECULE: | CENTROMERE PROTEIN H | |
| 69:00:00 | 3dxq-A | 3.2 | 10.5 | 57 | 294 | 7 | MOLECULE: | CHOLINE/ETHANOLAMINE KINASE FAMILY PROTEIN | |
| 70:00:00 | 4dvy-P | 3.2 | 3.2 | 78 | 656 | 6 | MOLECULE: | CYTOTOXICITY-ASSOCIATED IMMUNODOMINANT ANTI | GEN |
| 71:00:00 | 8h1l-B | 3.2 | 4.5 | 43 | 423 | 7 | MOLECULE: | N-ACYLGLUCOSAMINE 2-EPIMERASE | |
| 72:00:00 | 6cj6-D | 3.2 | 4 | 51 | 166 | 10 | MOLECULE: | PROTEIN F9 | |
| 73:00:00 | 1kw2-A | 3.2 | 6.5 | 68 | 455 | 6 | MOLECULE: | VITAMIN D-BINDING PROTEIN | |
| 74:00:00 | 6ddq-A | 3.2 | 25.1 | 99 | 507 | 5 | MOLECULE: | CYTOSOLIC PURINE 5'-NUCLEOTIDASE | |
| 75:00:00 | 8qfc-C | 3.2 | 15.9 | 91 | 380 | 5 | MOLECULE: | 60S RIBOSOMAL PROTEIN L10A | |
| 76:00:00 | 8iht-N | 3.2 | 9 | 58 | 295 | 7 | MOLECULE: | HISTONE H3 | |
| 77:00:00 | 5vgz-D | 3.2 | 9.9 | 66 | 107 | 2 | MOLECULE: | 26S PROTEASOME REGULATORY SUBUNIT 7 | |
| 78:00:00 | 7cy6-A | 3.2 | 4.9 | 67 | 872 | 4 | MOLECULE: | MALTODEXTRIN-BINDING PROTEIN,5-METHYLCYTOSI | NE-MOD |
| 79:00:00 | 6rw3-B | 3.2 | 4.4 | 62 | 439 | 15 | MOLECULE: | HEXOSE TRANSPORTER 1 | |
| 80:00:00 | 6vq6-A | 3.2 | 2.6 | 51 | 600 | 6 | MOLECULE: | ATPASE H+-TRANSPORTING V1 SUBUNIT A | |
| 81:00:00 | 7o4i-1 | 3.2 | 5.1 | 60 | 522 | 3 | MOLECULE: | GENERAL TRANSCRIPTION AND DNA REPAIR FACTOR | IIH H |
| 82:00:00 | 7qhm-I | 3.2 | 14.7 | 98 | 135 | 7 | MOLECULE: | CYTOCHROME BC1 COMPLEX RIESKE IRON-SULFUR S | UBUNIT |
| 83:00:00 | 7v2y-B | 3.2 | 32 | 108 | 1226 | 6 | MOLECULE: | THO COMPLEX SUBUNIT HPR1 | |
| 84:00:00 | 8jh7-D | 3.2 | 4.1 | 89 | 431 | 4 | MOLECULE: | ANTI-BRIL FAB HEAVY CHAIN | |
| 85:00:00 | 2ets-A | 3.2 | 4.2 | 68 | 115 | 4 | MOLECULE: | HYPOTHETICAL PROTEIN | |
| 86:00:00 | 5doq-B | 3.2 | 10.7 | 88 | 330 | 6 | MOLECULE: | BD-TYPE QUINOL OXIDASE SUBUNIT I | |
| 87:00:00 | 8yjb-B | 3.2 | 13.3 | 86 | 1061 | 3 | MOLECULE: | 26S PROTEASOME COMPLEX SUBUNIT DSS1 | |
| 88:00:00 | 7wrg-A | 3.2 | 17.6 | 112 | 812 | 7 | MOLECULE: | KINESIN-LIKE PROTEIN | |
| 89:00:00 | 7pde-A | 3.2 | 18.1 | 125 | 846 | 6 | MOLECULE: | ADENYLATE CYCLASE 9 | |
| 90:00:00 | 8d8k-M | 3.2 | 3.8 | 50 | 83 | 10 | MOLECULE: | PROBABLE S-ADENOSYL-L-METHIONINE-DEPENDENT | RNA |
| 91:00:00 | 7jmn-O | 3.2 | 3.4 | 59 | 164 | 7 | MOLECULE: | MEDIATOR OF RNA POLYMERASE II TRANSCRIPTION | SUBUN |
| 92:00:00 | 8tww-A | 3.2 | 11.1 | 81 | 245 | 6 | MOLECULE: | AETD | |
| 93:00:00 | 6zxr-A | 3.2 | 3.3 | 52 | 212 | 8 | MOLECULE: | ER LUMEN PROTEIN-RETAINING RECEPTOR 2 | |
| 94:00:00 | 1qhh-C | 3.2 | 2.9 | 48 | 102 | 2 | MOLECULE: | PROTEIN (PCRA (SUBUNIT)) | |
| 95:00:00 | 7d7n-A | 3.2 | 16.6 | 156 | 704 | 6 | MOLECULE: | ATP-BINDING CASSETTE SUB-FAMILY B MEMBER 6, | MITOC |
| 96:00:00 | 7u5d-A | 3.2 | 24.1 | 73 | 630 | 5 | MOLECULE: | CRRNA | |
| 97:00:00 | 7ux0-A | 3.2 | 2.4 | 50 | 115 | 0 | MOLECULE: | SPERM-EGG FUSION PROTEIN TMEM95 | |
| 98:00:00 | 4orh-H | 3.2 | 5.6 | 76 | 142 | 9 | MOLECULE: | UBIQUITIN-CONJUGATING ENZYME E2 VARIANT 2 | |
| 99:00:00 | 4hkr-A | 3.2 | 8.7 | 94 | 165 | 9 | MOLECULE: | CALCIUM RELEASE-ACTIVATED CALCIUM CHANNEL P | ROTEIN |
| 0:00 | 6br8-A | 3.2 | 10.1 | 66 | 249 | 5 | MOLECULE: | PROTEIN A6 HOMOLOG | |
| 1:00 | 8tkp-A | 3.2 | 18.4 | 65 | 614 | 3 | MOLECULE: | TRANSMEMBRANE CHANNEL-LIKE PROTEIN 2 | |
| 2:00 | 7t7r-A | 3.2 | 23.2 | 115 | 1132 | 8 | MOLECULE: | PROTEIN UNC-13 HOMOLOG A | |
| 3:00 | 7dgy-A | 3.2 | 2.6 | 47 | 99 | 0 | MOLECULE: | C2 | |
| 4:00 | 6bpz-A | 3.2 | 4.8 | 66 | 900 | 2 | MOLECULE: | PIEZO-TYPE MECHANOSENSITIVE ION CHANNEL COM | PONENT |

| | | | | | | | | | | |
|---|---|---|---|---|---|---|---|---|---|---|
| 5:00 | 2w6a-A | 3.2 | 3.1 | 47 | 61 | 11 | MOLECULE: | ARF GTPASE-ACTIVATING PROTEIN GIT1 | | |
| 6:00 | 5vpe-A | 3.2 | 2.7 | 46 | 67 | 15 | MOLECULE: | PROTEIN FOSB | | |
| 7:00 | 1gmj-A | 3.2 | 3.2 | 50 | 65 | 10 | MOLECULE: | ATPASE INHIBITOR | | |
| 8:00 | 5ncn-B | 3.2 | 2.3 | 44 | 76 | 2 | MOLECULE: | DBF2 KINASE ACTIVATOR PROTEIN MOB1 | | |
| 9:00 | 6rfq-3 | 3.2 | 5.5 | 58 | 108 | 3 | MOLECULE: | SUBUNIT NUAM OF NADH:UBIQUINONE OXIDOREDUCT | ASE (C | |
| 10:00 | 4dl0-K | 3.2 | 14 | 56 | 97 | 18 | MOLECULE: | V-TYPE PROTON ATPASE SUBUNIT C | | |
| 11:00 | 8gkv-A | 3.2 | 2.4 | 41 | 44 | 5 | MOLECULE: | ANTI-ADAPTER PROTEIN IRAP | | |
| 12:00 | 5chx-A | 3.2 | 4.2 | 52 | 210 | 2 | MOLECULE: | XRCC4-MYH7-1590-1657 | | |
| 13:00 | 5dw7-A | 3.2 | 3.2 | 52 | 309 | 6 | MOLECULE: | GERMACRADIENOL/GEOSMIN SYNTHASE | | |
| 14:00 | 7jw1-e | 3.2 | 6.3 | 102 | 235 | 3 | MOLECULE: | CAPSID PROTEINS | | |
| 15:00 | 7x6g-D | 3.2 | 11.5 | 69 | 152 | 7 | MOLECULE: | QUORUM-SENSING REGULATOR PROTEIN G | | |
| 16:00 | 5j1i-A | 3.2 | 3.1 | 55 | 346 | 9 | MOLECULE: | PLECTIN | | |
| 17:00 | 3aei-A | 3.2 | 4.5 | 55 | 94 | 13 | MOLECULE: | PREFOLDIN BETA SUBUNIT 2 | | |
| 18:00 | 5i08-A | 3.2 | 7.8 | 72 | 958 | 6 | MOLECULE: | SPIKE GLYCOPROTEIN, ENVELOPE GLYCOPROTEIN C | HIMERA | |
| 19:00 | 8k9c-A | 3.2 | 8.2 | 74 | 207 | 7 | MOLECULE: | CAPRIN-2 | | |
| 20:00 | 7sfm-A | 3.1 | 26.2 | 93 | 472 | 5 | MOLECULE: | HIP1 | | |
| 21:00 | 7eew-A | 3.1 | 31.7 | 87 | 611 | 7 | MOLECULE: | TYPE I RESTRICTION-MODIFICATION SYSTEM METH | YLTRAN | |
| 22:00 | 6vej-A | 3.1 | 31.7 | 141 | 1022 | 9 | MOLECULE: | PROBABLE RESISTANCE-NODULATION-CELL DIVISIO | N (RND | |
| 23:00 | 6wb9-2 | 3.1 | 31.9 | 85 | 292 | 4 | MOLECULE: | ENDOPLASMIC RETICULUM MEMBRANE PROTEIN COMP | LEX SU | |
| 24:00:00 | 5t58-D | 3.1 | 36.6 | 113 | 201 | 5 | MOLECULE: | KLLA0F02343P | | |
| 25:00:00 | 4qfh-B | 3.1 | 26.8 | 121 | 611 | 7 | MOLECULE: | GLUCOSE-6-PHOSPHATE ISOMERASE | | |
| 26:00:00 | 7tpg-B | 3.1 | 32.5 | 128 | 403 | 7 | MOLECULE: | PUTATIVE CELL SURFACE POLYSACCHARIDE POLYME | RASE/L | |
| 27:00:00 | 5lc5-n | 3.1 | 26.4 | 78 | 166 | 9 | MOLECULE: | NADH-UBIQUINONE OXIDOREDUCTASE CHAIN 3 | | |
| 28:00:00 | 8at3-D | 3.1 | 31.8 | 144 | 666 | 5 | MOLECULE: | HAUS AUGMIN-LIKE COMPLEX SUBUNIT 1 | | |
| 29:00:00 | 6uz3-A | 3.1 | 29 | 137 | 1126 | 2 | MOLECULE: | SODIUM CHANNEL PROTEIN TYPE 5 SUBUNIT ALPHA | ,GREEN | |
| 30:00:00 | 6klb-A | 3.1 | 33.8 | 144 | 1118 | 10 | MOLECULE: | LBCAS12A | | |
| 31:00:00 | 8zxd-A | 3.1 | 10.7 | 64 | 388 | 5 | MOLECULE: | VANG-LIKE PROTEIN 1 | | |
| 32:00:00 | 1y2o-A | 3.1 | 6.4 | 98 | 248 | 3 | MOLECULE: | BAI1-ASSOCIATED PROTEIN 2 ISOFORM 1 | | |
| 33:00:00 | 7obq-y | 3.1 | 18.9 | 85 | 454 | 2 | MOLECULE: | SRP RNA | | |
| 34:00:00 | 7yh5-B | 3.1 | 8.4 | 60 | 177 | 8 | MOLECULE: | NUCLEOSIDE TRIPHOSPHATE PYROPHOSPHOHYDROLAS | E | |
| 35:00:00 | 2xra-A | 3.1 | 6.5 | 105 | 203 | 3 | MOLECULE: | TRANSMEMBRANE PROTEIN GP41 | | |
| 36:00:00 | 6lo8-D | 3.1 | 6.6 | 83 | 119 | 7 | MOLECULE: | MITOCHONDRIAL IMPORT INNER MEMBRANE TRANSLO | CASE S | |
| 37:00:00 | 6t9i-H | 3.1 | 20.3 | 63 | 227 | 5 | MOLECULE: | TRANSCRIPTION FACTOR SPT20 | | |
| 38:00:00 | 8syn-A | 3.1 | 18.3 | 69 | 801 | 0 | MOLECULE: | VPS35 ENDOSOMAL PROTEIN-SORTING FACTOR-LIKE | | |
| 39:00:00 | 6ypc-W | 3.1 | 5.5 | 53 | 69 | 4 | MOLECULE: | INNER KINETOCHORE SUBUNIT MCM22 | | |
| 40:00:00 | 2luh-A | 3.1 | 2.4 | 52 | 167 | 2 | MOLECULE: | VACUOLAR PROTEIN SORTING-ASSOCIATED PROTEIN | VTA1 | |
| 41:00:00 | 2nrk-A | 3.1 | 3 | 42 | 165 | 17 | MOLECULE: | HYPOTHETICAL PROTEIN GRPB | | |
| 42:00:00 | 7a6h-K | 3.1 | 2.1 | 49 | 107 | 10 | MOLECULE: | DNA-DIRECTED RNA POLYMERASE III SUBUNIT RPC | | 1 |
| 43:00:00 | 8z9y-B | 3.1 | 2.3 | 52 | 164 | 6 | MOLECULE: | PROTEIN TIC 214 | | |
| 44:00:00 | 9asc-A | 3.1 | 10.2 | 101 | 318 | 9 | MOLECULE: | UNDECAPRENYL-PHOSPHATE 4-DEOXY-4-FORMAMIDO- | L-ARAB | |
| 45:00:00 | 8d2s-A | 3.1 | 9.3 | 105 | 475 | 5 | MOLECULE: | SODIUM-DEPENDENT LYSOPHOSPHATIDYLCHOLINE SY | MPORTE | |
| 46:00:00 | 8kc4-A | 3.1 | 12.5 | 68 | 151 | 9 | MOLECULE: | DE NOVO DESIGN PROTEIN -NA05 | | |
| 47:00:00 | 3p3o-A | 3.1 | 6.1 | 74 | 416 | 4 | MOLECULE: | CYTOCHROME P450 | | |
| 48:00:00 | 6dfk-I | 3.1 | 10 | 62 | 236 | 8 | MOLECULE: | SUBUNIT OF PROTEASEOME ACTIVATOR COMPLEX,PU | TATIVE | |
| 49:00:00 | 6kko-A | 3.1 | 7 | 65 | 170 | 2 | MOLECULE: | PUTATIVE SERINE PHOSPHATASE | | |
| 50:00:00 | 8h2h-D | 3.1 | 16.5 | 69 | 599 | 4 | MOLECULE: | LTRB | | |
| 51:00:00 | 6n7p-X | 3.1 | 14.1 | 112 | 826 | 4 | MOLECULE: | U1 SMALL NUCLEAR RIBONUCLEOPROTEIN 70 KDA H | OMOLOG | |
| 52:00:00 | 3f46-A | 3.1 | 9.6 | 66 | 345 | 3 | MOLECULE: | 5,10-METHENYLTETRAHYDROMETHANOPTERIN HYDROG | ENASE | |
| 53:00:00 | 1k30-A | 3.1 | 21.2 | 82 | 363 | 2 | MOLECULE: | GLYCEROL-3-PHOSPHATE ACYLTRANSFERASE | | |
| 54:00:00 | 6qg0-G | 3.1 | 8.7 | 72 | 355 | 6 | MOLECULE: | TRANSLATION INITIATION FACTOR EIF-2B SUBUNI | T ALPH | |
| 55:00:00 | 6ln2-A | 3.1 | 8.4 | 76 | 437 | 11 | MOLECULE: | GLUCAGON-LIKE PEPTIDE 1 RECEPTOR,RUBREDOXIN | ,GLUCA | |
| 56:00:00 | 9na9-E | 3.1 | 6.9 | 80 | 153 | 5 | MOLECULE: | AUGMIN SUBUNIT 1 | | |
| 57:00:00 | 7bw0-R | 3.1 | 6.3 | 111 | 280 | 6 | MOLECULE: | SOLUBLE CYTOCHROME B562,G-PROTEIN COUPLED B | ILE AC | |
| 58:00:00 | 3s0x-A | 3.1 | 11.5 | 75 | 225 | 4 | MOLECULE: | PEPTIDASE A24B, FLAK DOMAIN PROTEIN | | |
| 59:00:00 | 5xyv-C | 3.1 | 4.3 | 45 | 52 | 2 | MOLECULE: | RHINO | | |
| 60:00:00 | 7y22-M | 3.1 | 12.4 | 95 | 215 | 5 | MOLECULE: | PHAGE CONNECTOR PROTEIN | | |
| 61:00:00 | 8hf3-A | 3.1 | 6.9 | 98 | 295 | 7 | MOLECULE: | PALMITOYLTRANSFERASE ZDHHC9 | | |
| 62:00:00 | 7nad-5 | 3.1 | 13.2 | 70 | 123 | 11 | MOLECULE: | 25S RRNA | | |
| 63:00:00 | 7nh9-A | 3.1 | 11.4 | 113 | 327 | 4 | MOLECULE: | CMAX PROTEIN | | |
| 64:00:00 | 3hh0-A | 3.1 | 5.9 | 65 | 134 | 8 | MOLECULE: | TRANSCRIPTIONAL REGULATOR, MERR FAMILY | | |
| 65:00:00 | 6h2x-A | 3.1 | 24.7 | 141 | 352 | 4 | MOLECULE: | CHROMOSOME PARTITION PROTEIN MUKB,CHROMOSOM | E PART | |
| 66:00:00 | 9bkv-A | 3.1 | 19.2 | 92 | 788 | 1 | MOLECULE: | OXYGEN SENSOR PROTEIN DOSP | | |
| 67:00:00 | 6qld-Y | 3.1 | 13.2 | 113 | 223 | 6 | MOLECULE: | INNER KINETOCHORE SUBUNIT MIF2 | | |
| 68:00:00 | 7mi8-A | 3.1 | 20.5 | 132 | 868 | 4 | MOLECULE: | FUSION PROTEIN OF DYNEIN AND ENDOLYSIN | | |
| 69:00:00 | 3vpq-A | 3.1 | 4.2 | 70 | 203 | 6 | MOLECULE: | GLUTATHIONE S-TRANSFERASE SIGMA | | |
| 70:00:00 | 2ymb-D | 3.1 | 7.1 | 69 | 231 | 3 | MOLECULE: | MIT DOMAIN-CONTAINING PROTEIN 1 | | |
| 71:00:00 | 6tlj-S | 3.1 | 18.7 | 66 | 293 | 3 | MOLECULE: | ANAPHASE-PROMOTING COMPLEX SUBUNIT 1 | | |
| 72:00:00 | 7wnq-A | 3.1 | 5.4 | 67 | 368 | 3 | MOLECULE: | GUARD CELL S-TYPE ANION CHANNEL SLAC1 | | |
| 73:00:00 | 6vm0-A | 3.1 | 6.6 | 97 | 363 | 3 | MOLECULE: | GLYCINE RECEPTOR SUBUNIT ALPHAZ1 | | |
| 74:00:00 | 8qbd-A | 3.1 | 12.5 | 110 | 272 | 2 | MOLECULE: | SPHINGOLIPID LONG CHAIN BASE-RESPONSIVE PRO | TEIN P | |
| 75:00:00 | 8uic-T | 3.1 | 22.5 | 114 | 803 | 7 | MOLECULE: | INTEGRATOR COMPLEX SUBUNIT 11 | | |
| 76:00:00 | 8bsh-C | 3.1 | 9.8 | 58 | 794 | 7 | MOLECULE: | PROTEIN TRANSPORT PROTEIN SEC23 | | |
| 77:00:00 | 2ilk-A | 3.1 | 2.3 | 40 | 155 | 13 | MOLECULE: | INTERLEUKIN-10 | | |
| 78:00:00 | 5d80-G | 3.1 | 24.9 | 87 | 223 | 5 | MOLECULE: | V-TYPE PROTON ATPASE CATALYTIC SUBUNIT A | | |
| 79:00:00 | 8dke-P | 3.1 | 7.4 | 62 | 333 | 3 | MOLECULE: | ISOFORM 2 OF CYSTINOSIN | | |
| 80:00:00 | 8szh-A | 3.1 | 6.7 | 103 | 836 | 5 | MOLECULE: | EXTRACELLULAR CALCIUM-SENSING RECEPTOR | | |
| 81:00:00 | 2lor-A | 3.1 | 5.4 | 60 | 108 | 3 | MOLECULE: | TRANSMEMBRANE PROTEIN 141 | | |
| 82:00:00 | 5xpd-A | 3.1 | 10.8 | 64 | 269 | 13 | MOLECULE: | SUGAR TRANSPORTER | | |
| 83:00:00 | 6pe4-A | 3.1 | 13.4 | 97 | 758 | 2 | MOLECULE: | V-TYPE PROTON ATPASE SUBUNIT A, VACUOLAR IS | OFORM | |
| 84:00:00 | 7r5s-T | 3.1 | 13 | 69 | 112 | 4 | MOLECULE: | CENTROMERE PROTEIN H | | |
| 85:00:00 | 7yil-B | 3.1 | 9.5 | 58 | 94 | 9 | MOLECULE: | GINS | | |
| 86:00:00 | 6xrx-A | 3.1 | 7.6 | 51 | 551 | 0 | MOLECULE: | MALTOSE/MALTODEXTRIN-BINDING PERIPLASMIC PR | OTEIN, | |
| 87:00:00 | 7qii-B | 3.1 | 8 | 63 | 76 | 2 | MOLECULE: | CHAPERONE PROTEIN YSCY | | |
| 88:00:00 | 9go3-A | 3.1 | 25.2 | 93 | 253 | 5 | MOLECULE: | MECHANOSENSITIVE CHANNEL PROTEIN | | |
| 89:00:00 | 1lqs-L | 3.1 | 7.9 | 69 | 142 | 3 | MOLECULE: | INTERLEUKIN-10 RECEPTOR ALPHA CHAIN | | |
| 90:00:00 | 7jmn-P | 3.1 | 22 | 79 | 810 | 5 | MOLECULE: | MEDIATOR OF RNA POLYMERASE II TRANSCRIPTION | SUBUN | |
| 91:00:00 | 6jql-A | 3.1 | 16.5 | 104 | 678 | 9 | MOLECULE: | BIFUNCTIONAL PROTEIN PAAZ | | |
| 92:00:00 | 4k2u-A | 3.1 | 10.5 | 80 | 230 | 6 | MOLECULE: | ERYTHROCYTE BINDING ANTIGEN 175 | | |
| 93:00:00 | 6poo-A | 3.1 | 9.6 | 102 | 273 | 8 | MOLECULE: | BIBA | | |
| 94:00:00 | 5odw-D | 3.1 | 4 | 65 | 198 | 6 | MOLECULE: | FERRIPYOVERDINE RECEPTOR | | |
| 95:00:00 | 7etw-B | 3.1 | 22.6 | 92 | 483 | 3 | MOLECULE: | INSULIN-INDUCED GENE 2 PROTEIN | | |
| 96:00:00 | 3cl3-D | 3.1 | 3.1 | 47 | 59 | 4 | MOLECULE: | ORF K13 | | |
| 97:00:00 | 4tt0-B | 3.1 | 17.4 | 82 | 131 | 4 | MOLECULE: | DENEDDYLASE | | |
| 98:00:00 | 2xco-A | 3.1 | 10.5 | 63 | 636 | 5 | MOLECULE: | DNA GYRASE SUBUNIT B, DNA GYRASE SUBUNIT A | | |
| 99:00:00 | 3wuv-Q | 3.1 | 4.8 | 47 | 62 | 4 | MOLECULE: | CENTROSOMAL PROTEIN OF 55 KDA | | |
| 0:00 | 5c9n-A | 3.1 | 3.9 | 53 | 64 | 6 | MOLECULE: | GEMININ COILED-COIL DOMAIN-CONTAINING PROTE | IN 1 | |
| 1:00 | 6wxu-B | 3.1 | 4.5 | 48 | 273 | 4 | MOLECULE: | DUAL OXIDASE 1 | | |
| 2:00 | 5fqf-A | 3.1 | 2.5 | 46 | 583 | 11 | MOLECULE: | BETA-N-ACETYLGALACTOSAMINIDASE | | |
| 3:00 | 7k36-D | 3.1 | 2.6 | 55 | 74 | 4 | MOLECULE: | SERINE/THREONINE-PROTEIN PHOSPHATASE 2A 65 | KDA RE | |
| 4:00 | 8cqn-B | 3.1 | 14.4 | 63 | 187 | 6 | MOLECULE: | LIPOPROTEIN, PUTATIVE | | |
| 5:00 | 7ug2-A | 3.1 | 1.6 | 45 | 49 | 2 | MOLECULE: | TRIPARTITE MOTIF-CONTAINING PROTEIN 75 | | |

| | | | | | | | | | | |
|---|---|---|---|---|---|---|---|---|---|---|
| 6:00 | 1zhc-A | 3.1 | 4.2 | 56 | 76 | 13 | MOLECULE: | HYPOTHETICAL PROTEIN HP1242 | | |
| 7:00 | 5l4k-V | 3.1 | 4.1 | 52 | 293 | 4 | MOLECULE: | 26S PROTEASOME NON-ATPASE REGULATORY SUBUNI | T 4 | |
| 8:00 | 3jcm-I | 3.1 | 21.6 | 91 | 416 | 5 | MOLECULE: | PRE-MRNA-SPLICING FACTOR 8 | | |
| 9:00 | 4mt8-A | 3.1 | 3.4 | 51 | 100 | 4 | MOLECULE: | ETHYLENE RESPONSE SENSOR 1 | | |
| 10:00 | 7y1a-p | 3.1 | 4.3 | 56 | 164 | 4 | MOLECULE: | B-PHYCOERYTHRIN BETA CHAIN | | |
| 11:00 | 8b9z-K | 3.1 | 5.6 | 51 | 96 | 8 | MOLECULE: | NADH-UBIQUINONE OXIDOREDUCTASE CHAIN 3 | | |
| 12:00 | 7qe5-A | 3.1 | 5.1 | 56 | 616 | 5 | MOLECULE: | SIALIC ACID TRAP TRANSPORTER PERMEASE PROTE | IN SIA | |
| 13:00 | 2nps-D | 3.1 | 3.1 | 50 | 63 | 2 | MOLECULE: | VESICLE-ASSOCIATED MEMBRANE PROTEIN 4 | | |
| 14:00 | 5dol-B | 3.1 | 3.6 | 47 | 62 | 4 | MOLECULE: | INITIATION-CONTROL PROTEIN YABA | | |
| 15:00 | 5ajs-B | 3.1 | 3.5 | 52 | 64 | 17 | MOLECULE: | THAP DOMAIN-CONTAINING PROTEIN 11 | | |
| 16:00 | 8iw0-A | 3 | 20.3 | 78 | 84 | 3 | MOLECULE: | LIPRIN-BETA-1,KN MOTIF AND ANKYRIN REPEAT D | OMAIN- | |
| 17:00 | 8c5v-I | 3 | 94.5 | 188 | 516 | 7 | MOLECULE: | CHEMOTAXIS PROTEIN CHEA | | |
| 18:00 | 5u1s-A | 3 | 16.6 | 122 | 1488 | 9 | MOLECULE: | SEPARIN | | |
| 19:00 | 1cm5-A | 3 | 24.6 | 104 | 759 | 6 | MOLECULE: | PROTEIN (PYRUVATE FORMATE-LYASE) | | |
| 20:00 | 8rt9-F | 3 | 16.8 | 72 | 272 | 4 | MOLECULE: | TRWJ PROTEIN | | |
| 21:00 | 7ni5-A | 3 | 28.7 | 112 | 2791 | 6 | MOLECULE: | SERINE-PROTEIN KINASE ATM | | |
| 22:00 | 8rhn-A | 3 | 27.1 | 102 | 198 | 5 | MOLECULE: | ATPASE FAMILY GENE 2 PROTEIN HOMOLOG A | | |
| 23:00 | 6g70-B | 3 | 28.4 | 111 | 510 | 7 | MOLECULE: | PRE-MRNA-PROCESSING FACTOR 39 | | |
| 24:00:00 | 3cwz-B | 3 | 6.1 | 87 | 310 | 8 | MOLECULE: | RAS-RELATED PROTEIN RAB-6A | | |
| 25:00:00 | 8h2m-A | 3 | 13.4 | 77 | 754 | 10 | MOLECULE: | TAPE TAIL MEASURE PROTEIN | | |
| 26:00:00 | 3ufe-A | 3 | 6.1 | 63 | 109 | 6 | MOLECULE: | TRANSCRIPTIONAL ANTITERMINATOR (BGLG FAMILY | ) | |
| 27:00:00 | 3wvo-A | 3 | 14 | 90 | 543 | 9 | MOLECULE: | CRISPR-ASSOCIATED PROTEIN, CSE1 FAMILY | | |
| 28:00:00 | 2m64-A | 3 | 3.4 | 50 | 231 | 10 | MOLECULE: | PHLP5 | | |
| 29:00:00 | 3r6n-A | 3 | 31.9 | 141 | 450 | 4 | MOLECULE: | DESMOPLAKIN | | |
| 30:00:00 | 7uqj-A | 3 | 2.3 | 43 | 585 | 7 | MOLECULE: | ATPASE HISTONE CHAPERONE YTA7 | | |
| 31:00:00 | 7v2c-d | 3 | 29.6 | 104 | 175 | 3 | MOLECULE: | NADH DEHYDROGENASE [UBIQUINONE] FLAVOPROTEI | N 1, | |
| 32:00:00 | 7w1m-D | 3 | 28.2 | 119 | 967 | 6 | MOLECULE: | STRUCTURAL MAINTENANCE OF CHROMOSOMES PROTE | IN 1A | |
| 33:00:00 | 6bzf-A | 3 | 11.2 | 77 | 197 | 4 | MOLECULE: | SPORULATION-SPECIFIC PROTEIN 16 | | |
| 34:00:00 | 5dlq-B | 3 | 27.8 | 118 | 1052 | 9 | MOLECULE: | EXPORTIN-4 | | |
| 35:00:00 | 3erm-A | 3 | 8.8 | 55 | 67 | 11 | MOLECULE: | UNCHARACTERIZED CONSERVED PROTEIN | | |
| 36:00:00 | 3kdw-A | 3 | 8 | 70 | 206 | 4 | MOLECULE: | PUTATIVE SUGAR BINDING PROTEIN | | |
| 37:00:00 | 4ql6-B | 3 | 2.7 | 43 | 509 | 2 | MOLECULE: | CARBOXY-TERMINAL PROCESSING PROTEASE | | |
| 38:00:00 | 7emf-J | 3 | 4.6 | 47 | 122 | 11 | MOLECULE: | MEDIATOR OF RNA POLYMERASE II TRANSCRIPTION | SUBUN | |
| 39:00:00 | 5vjx-J | 3 | 3.2 | 51 | 64 | 8 | MOLECULE: | CLOCK-INTERACTING PACEMAKER | | |
| 40:00:00 | 6zis-A | 3 | 8.3 | 68 | 569 | 4 | MOLECULE: | MALTOSE/MALTODEXTRIN-BINDING PERIPLASMIC PR | OTEIN, | |
| 41:00:00 | 3sng-A | 3 | 2.9 | 54 | 267 | 6 | MOLECULE: | NUCLEASE | | |
| 42:00:00 | 7pp4-f | 3 | 9.8 | 68 | 142 | 7 | MOLECULE: | DNA-DIRECTED RNA POLYMERASE SUBUNIT ALPHA | | |
| 43:00:00 | 7ae8-A | 3 | 6.4 | 59 | 152 | 3 | MOLECULE: | HEPN TOXIN | | |
| 44:00:00 | 4hh0-A | 3 | 8.7 | 65 | 383 | 6 | MOLECULE: | APPA PROTEIN | | |
| 45:00:00 | 4ejo-A | 3 | 6.1 | 66 | 112 | 8 | MOLECULE: | TRANSCRIPTIONAL REGULATOR, PADR-LIKE FAMILY | | |
| 46:00:00 | 3w0l-B | 3 | 7.3 | 67 | 598 | 6 | MOLECULE: | GLUCOKINASE | | |
| 47:00:00 | 8ou0-C | 3 | 5.6 | 63 | 153 | 11 | MOLECULE: | TUBULIN BETA-4B CHAIN | | |
| 48:00:00 | 2rdp-A | 3 | 10.7 | 82 | 140 | 7 | MOLECULE: | PUTATIVE TRANSCRIPTIONAL REGULATOR MARR | | |
| 49:00:00 | 7uh4-A | 3 | 3.4 | 84 | 110 | 4 | MOLECULE: | LXG-ASSOCIATED ALPHA-HELICAL PROTEIN D2 | | |
| 50:00:00 | 7ktr-E | 3 | 13.2 | 106 | 124 | 6 | MOLECULE: | TRANSFORMATION/TRANSCRIPTION DOMAIN-ASSOCIA | TED PR | |
| 51:00:00 | 6ekr-A | 3 | 4.4 | 51 | 303 | 8 | MOLECULE: | TYPE II SITE-SPECIFIC DEOXYRIBONUCLEASE | | |
| 52:00:00 | 5cm8-A | 3 | 6.7 | 97 | 419 | 2 | MOLECULE: | RAL GUANINE NUCLEOTIDE DISSOCIATION STIMULA | TOR-LI | |
| 53:00:00 | 4uj6-A | 3 | 10 | 124 | 711 | 3 | MOLECULE: | SURFACE LAYER PROTEIN | | |
| 54:00:00 | 8age-H | 3 | 8.6 | 95 | 133 | 5 | MOLECULE: | DOLICHYL-DIPHOSPHOOLIGOSACCHARIDE--PROTEIN | | |
| 55:00:00 | 6ffv-A | 3 | 11.5 | 75 | 194 | 7 | MOLECULE: | BTUM | | |
| 56:00:00 | 2iaz-B | 3 | 15 | 93 | 113 | 9 | MOLECULE: | HYPOTHETICAL PROTEIN SP1372 | | |
| 57:00:00 | 7mop-A | 3 | 30.5 | 102 | 2445 | 5 | MOLECULE: | E3 UBIQUITIN-PROTEIN LIGASE HUWE1 | | |
| 58:00:00 | 8cli-A | 3 | 3.2 | 40 | 542 | 8 | MOLECULE: | GENERAL TRANSCRIPTION FACTOR 3C POLYPEPTIDE | | 1 |
| 59:00:00 | 7xr2-a | 3 | 2.9 | 45 | 274 | 2 | MOLECULE: | VP3 | | |
| 60:00:00 | 6l5d-B | 3 | 7.6 | 84 | 244 | 8 | MOLECULE: | GAS VESICLE PROTEIN | | |
| 61:00:00 | 7zvw-E | 3 | 36.8 | 153 | 488 | 7 | MOLECULE: | TRANSCRIPTION-ASSOCIATED PROTEIN | | |
| 62:00:00 | 6vbv-7 | 3 | 14.4 | 89 | 706 | 6 | MOLECULE: | BARDET-BIEDL SYNDROME 18 PROTEIN | | |
| 63:00:00 | 8qyd-A | 3 | 21.9 | 140 | 280 | 4 | MOLECULE: | ANTI-PHAGE DEFENSE ZORAB SYSTEM ZORA | | |
| 64:00:00 | 7wjn-A | 3 | 24.7 | 115 | 808 | 4 | MOLECULE: | CHITIN SYNTHASE | | |
| 65:00:00 | 8q85-X | 3 | 14.5 | 54 | 68 | 15 | MOLECULE: | KINETOCHORE PROTEIN NDC80 | | |
| 66:00:00 | 7xr2-B | 3 | 17.4 | 84 | 846 | 7 | MOLECULE: | VP3 | | |
| 67:00:00 | 6z1b-B | 3 | 2.2 | 44 | 202 | 7 | MOLECULE: | HTH-TYPE TRANSCRIPTIONAL REGULATOR RUTR | | |
| 68:00:00 | 6m5r-B | 3 | 20.1 | 104 | 689 | 4 | MOLECULE: | TRIPARTITE TERMINASE SUBUNIT 3 | | |
| 69:00:00 | 6rx4-B | 3 | 7.2 | 85 | 379 | 4 | MOLECULE: | CYTOCHROME BD-I UBIQUINOL OXIDASE SUBUNIT 1 | | |
| 70:00:00 | 7rsl-A | 3 | 9.8 | 68 | 235 | 4 | MOLECULE: | SEIPIN | | |
| 71:00:00 | 8etq-A | 3 | 5.9 | 57 | 277 | 7 | MOLECULE: | K-CTRP5 | | |
| 72:00:00 | 7qh2-C | 3 | 6.4 | 61 | 467 | 5 | MOLECULE: | LACTATE DEHYDROGENASE (NAD(+),FERREDOXIN) S | UBUNIT | |
| 73:00:00 | 9l0d-B | 3 | 7 | 79 | 466 | 6 | MOLECULE: | VACUOLAR FUSION PROTEIN MON1 HOMOLOG A | | |
| 74:00:00 | 5f5p-A | 3 | 9.5 | 118 | 181 | 6 | MOLECULE: | PROTEIN SHROOM2 | | |
| 75:00:00 | 2dg6-A | 3 | 8.6 | 64 | 207 | 2 | MOLECULE: | PUTATIVE TRANSCRIPTIONAL REGULATOR | | |
| 76:00:00 | 8xi9-A | 3 | 5.3 | 87 | 201 | 7 | MOLECULE: | FRB-FKBP FUSION PROTEIN | | |
| 77:00:00 | 8p2m-A | 3 | 23.7 | 109 | 659 | 12 | MOLECULE: | NAD(+) HYDROLASE TIR-1 | | |
| 78:00:00 | 8vdq-A | 3 | 14.8 | 111 | 2192 | 4 | MOLECULE: | GREEN FLUORESCENT PROTEIN,TALIN-1 | | |
| 79:00:00 | 6sp2-A | 3 | 16.7 | 96 | 366 | 5 | MOLECULE: | MEMBRANE PROTEIN TMS1D | | |
| 80:00:00 | 3a8t-A | 3 | 15.6 | 92 | 289 | 4 | MOLECULE: | ADENYLATE ISOPENTENYLTRANSFERASE | | |
| 81:00:00 | 6ysf-F | 3 | 13.5 | 100 | 257 | 9 | MOLECULE: | CHEMOTAXIS MOTB PROTEIN | | |
| 82:00:00 | 5td8-C | 3 | 9.8 | 58 | 111 | 9 | MOLECULE: | KINETOCHORE PROTEIN NDC80 | | |
| 83:00:00 | 9cah-A | 3 | 7.7 | 51 | 718 | 8 | MOLECULE: | DNA TOPOISOMERASE 3-BETA-1 | | |
| 84:00:00 | 8jw0-b | 3 | 12.6 | 86 | 617 | 6 | MOLECULE: | PHOTOSYSTEM I PSAA | | |
| 85:00:00 | 3rx6-A | 3 | 7.5 | 89 | 187 | 6 | MOLECULE: | POLARITY SUPPRESSION PROTEIN | | |
| 86:00:00 | 9h9q-C | 3 | 28 | 130 | 735 | 5 | MOLECULE: | TUBULIN GAMMA CHAIN | | |
| 87:00:00 | 9ayc-A | 3 | 21.5 | 162 | 1481 | 6 | MOLECULE: | METAL RESISTANCE PROTEIN YCF1 | | |
| 88:00:00 | 5efv-B | 3 | 25.3 | 98 | 635 | 5 | MOLECULE: | PHI ETA ORF 56-LIKE PROTEIN | | |
| 89:00:00 | 8yw6-B | 3 | 11.5 | 67 | 126 | 6 | MOLECULE: | MITOCHONDRIAL PYRUVATE CARRIER 2 | | |
| 90:00:00 | 8rc0-D | 3 | 25.9 | 90 | 580 | 4 | MOLECULE: | CD2 ANTIGEN CYTOPLASMIC TAIL-BINDING PROTEI | N 2 | |
| 91:00:00 | 7vcs-A | 3 | 18.8 | 71 | 766 | 1 | MOLECULE: | TRANSITIONAL ENDOPLASMIC RETICULUM ATPASE | | |
| 92:00:00 | 7loi-A | 3 | 2.9 | 51 | 175 | 2 | MOLECULE: | TRANSMEMBRANE PROTEIN GP41 | | |
| 93:00:00 | 3jc5-A | 3 | 17.5 | 81 | 208 | 6 | MOLECULE: | DNA REPLICATION LICENSING FACTOR MCM2 | | |
| 94:00:00 | 4c47-A | 3 | 14.2 | 63 | 191 | 11 | MOLECULE: | INNER MEMBRANE LIPOPROTEIN | | |
| 95:00:00 | 5mps-R | 3 | 2.9 | 47 | 108 | 4 | MOLECULE: | YEAST UBC4 GENE FOR UBIQUITIN-CONJUGATING E | NZYME | |
| 96:00:00 | 5f3o-A | 3 | 3.7 | 51 | 194 | 6 | MOLECULE: | PUTATIVE UNCHARACTERIZED PROTEIN | | |
| 97:00:00 | 6r7f-E | 3 | 4.5 | 52 | 311 | 6 | MOLECULE: | COP9 SIGNALOSOME COMPLEX SUBUNIT 1 | | |
| 98:00:00 | 5cj1-A | 3 | 17.9 | 72 | 98 | 6 | MOLECULE: | GP7-MYH7-(1526-1571) CHIMERA PROTEIN | | |
| 99:00:00 | 4kxr-B | 3 | 3.3 | 90 | 174 | 9 | MOLECULE: | PE25 | | |
| 0:00 | 7f6j-C | 3 | 4 | 54 | 106 | 4 | MOLECULE: | RAS-RELATED PROTEIN RAB-7A | | |
| 1:00 | 8i8b-C | 3 | 7 | 61 | 229 | 8 | MOLECULE: | MAJOR VIRAL CAPSID PROTEIN | | |
| 2:00 | 5w10-A | 3 | 4.3 | 43 | 173 | 5 | MOLECULE: | CGMP-SPECIFIC PHOSPHODIESTERASE | | |
| 3:00 | 6otn-B | 3 | 3 | 51 | 67 | 6 | MOLECULE: | TROPOMYOSIN ALPHA-3 CHAIN | | |
| 4:00 | 1ifl-A | 3 | 3.9 | 46 | 53 | 4 | MOLECULE: | INOVIRUS | | |
| 5:00 | 1jad-A | 2.9 | 31.9 | 130 | 242 | 10 | MOLECULE: | PHOSPHOLIPASE C BETA | | |
| 6:00 | 6idp-A | 2.9 | 34.4 | 116 | 438 | 6 | MOLECULE: | MATE FAMILY EFFLUX TRANSPORTER | | |

| | | | | | | | | | | |
|---|---|---|---|---|---|---|---|---|---|---|
| 7:00 | 4nqi-D | 2.9 | 18.5 | 127 | 232 | 6 | MOLECULE: | SH3 DOMAIN-CONTAINING PROTEIN | | |
| 8:00 | 4n5c-D | 2.9 | 22.4 | 113 | 757 | 5 | MOLECULE: | CARGO-TRANSPORT PROTEIN YPP1 | | |
| 9:00 | 6fvb-A | 2.9 | 28.9 | 137 | 1001 | 7 | MOLECULE: | IMPORTIN BETA-LIKE PROTEIN KAP120 | | |
| 10:00 | 7qoo-U | 2.9 | 37.5 | 137 | 186 | 7 | MOLECULE: | CENTROMERE PROTEIN C | | |
| 11:00 | 4btf-A | 2.9 | 21.3 | 77 | 413 | 4 | MOLECULE: | MIXED LINEAGE KINASE DOMAIN-LIKE PROTEIN | | |
| 12:00 | 8ce1-c | 2.9 | 19.6 | 88 | 241 | 8 | MOLECULE: | CYTOCHROME C BIOGENESIS ATP-BINDING EXPORT | PROTEI | |
| 13:00 | 8ceg-B | 2.9 | 19.8 | 127 | 214 | 6 | MOLECULE: | CBY1-INTERACTING BAR DOMAIN-CONTAINING PROT | EIN 1 | |
| 14:00 | 6w17-F | 2.9 | 3.9 | 60 | 168 | 3 | MOLECULE: | ACTIN-RELATED PROTEIN 3 | | |
| 15:00 | 5ijo-J | 2.9 | 26.6 | 103 | 1256 | 4 | MOLECULE: | NUCLEAR PORE COMPLEX PROTEIN NUP155 | | |
| 16:00 | 8w20-A | 2.9 | 38 | 135 | 732 | 9 | MOLECULE: | SECRETED PROTEIN | | |
| 17:00 | 3zcj-A | 2.9 | 33.8 | 109 | 205 | 5 | MOLECULE: | CAGL | | |
| 18:00 | 6ptm-A | 2.9 | 33.8 | 106 | 726 | 6 | MOLECULE: | UNCHARACTERIZED PROTEIN | | |
| 19:00 | 8ka6-B | 2.9 | 12 | 74 | 148 | 8 | MOLECULE: | DE NOVO DESIGN PROTEIN -NA7 | | |
| 20:00 | 7wiu-B | 2.9 | 21.6 | 126 | 576 | 6 | MOLECULE: | MYCOBACTIN IMPORT ATP-BINDING/PERMEASE PROT | EIN IR | |
| 21:00 | 4hga-A | 2.9 | 24.4 | 96 | 207 | 5 | MOLECULE: | DEATH DOMAIN-ASSOCIATED PROTEIN 6 | | |
| 22:00 | 5b86-A | 2.9 | 35.2 | 124 | 579 | 6 | MOLECULE: | TUMOR NECROSIS FACTOR ALPHA-INDUCED PROTEIN | | 2 |
| 23:00 | 8tek-E | 2.9 | 51.9 | 151 | 219 | 7 | MOLECULE: | DYNEIN REGULATORY COMPLEX PROTEIN 1/2 N-TER | MINAL | |
| 24:00:00 | 8dc2-A | 2.9 | 19.2 | 93 | 673 | 3 | MOLECULE: | CASLAMBDA | | |
| 25:00:00 | 3iuk-A | 2.9 | 24.8 | 101 | 552 | 5 | MOLECULE: | UNCHARACTERIZED PROTEIN | | |
| 26:00:00 | 4jza-A | 2.9 | 33.1 | 101 | 817 | 6 | MOLECULE: | UNCHARACTERIZED PROTEIN | | |
| 27:00:00 | 5n9j-E | 2.9 | 14 | 106 | 201 | 10 | MOLECULE: | MEDIATOR OF RNA POLYMERASE II TRANSCRIPTION | SUBUN | |
| 28:00:00 | 7xzi-A | 2.9 | 49.2 | 166 | 1598 | 8 | MOLECULE: | CTAP3 | | |
| 29:00:00 | 5xjt-2 | 2.9 | 12.4 | 57 | 217 | 5 | MOLECULE: | GEM-ASSOCIATED PROTEIN 2 | | |
| 30:00:00 | 7vmc-B | 2.9 | 8.9 | 51 | 111 | 2 | MOLECULE: | ELONGATION FACTOR TU | | |
| 31:00:00 | 1nek-D | 2.9 | 4.6 | 90 | 113 | 10 | MOLECULE: | SUCCINATE DEHYDROGENASE FLAVOPROTEIN SUBUNI | T | |
| 32:00:00 | 8tlp-A | 2.9 | 5.2 | 80 | 226 | 5 | MOLECULE: | D_3_633_8X, NO PEPTIDE | | |
| 33:00:00 | 8erc-A | 2.9 | 5.5 | 56 | 441 | 5 | MOLECULE: | LYSOPHOSPHOLIPID ACYLTRANSFERASE 7 | | |
| 34:00:00 | 8yb7-C | 2.9 | 23.2 | 69 | 455 | 1 | MOLECULE: | PAPAIN-LIKE PROTEASE NSP3 | | |
| 35:00:00 | 5ed9-B | 2.9 | 20.3 | 69 | 72 | 7 | MOLECULE: | SUN DOMAIN-CONTAINING PROTEIN 2 | | |
| 36:00:00 | 8bbg-B | 2.9 | 7.1 | 81 | 1426 | 4 | MOLECULE: | WD REPEAT-CONTAINING PROTEIN 19 | | |
| 37:00:00 | 6qm5-A | 2.9 | 19.1 | 133 | 673 | 8 | MOLECULE: | PREDICTED PROTEIN | | |
| 38:00:00 | 6dcf-D | 2.9 | 23.5 | 133 | 1177 | 7 | MOLECULE: | RNA POLYMERASE-BINDING PROTEIN RBPA | | |
| 39:00:00 | 5n9y-A | 2.9 | 13 | 108 | 327 | 6 | MOLECULE: | ZINC TRANSPORT PROTEIN ZNTB | | |
| 40:00:00 | 3mzy-A | 2.9 | 9.5 | 64 | 123 | 9 | MOLECULE: | RNA POLYMERASE SIGMA-H FACTOR | | |
| 41:00:00 | 4uis-B | 2.9 | 9.9 | 84 | 241 | 2 | MOLECULE: | GAMMA-SECRETASE | | |
| 42:00:00 | 7m5f-A | 2.9 | 4.7 | 49 | 98 | 4 | MOLECULE: | CDII | | |
| 43:00:00 | 8ymk-A | 2.9 | 8.8 | 79 | 226 | 1 | MOLECULE: | ISOFORM S OF LARGE ENVELOPE PROTEIN | | |
| 44:00:00 | 1kx5-A | 2.9 | 7.6 | 67 | 135 | 3 | MOLECULE: | DNA | | |
| 45:00:00 | 8th8-J | 2.9 | 10.5 | 127 | 372 | 4 | MOLECULE: | DYNEIN REGULATORY COMPLEX PROTEIN 1/2 N-TER | MINAL | |
| 46:00:00 | 1joc-A | 2.9 | 31.3 | 88 | 123 | 6 | MOLECULE: | EARLY ENDOSOMAL AUTOANTIGEN 1 | | |
| 47:00:00 | 8of0-M | 2.9 | 4.7 | 62 | 140 | 6 | MOLECULE: | DNA-DIRECTED RNA POLYMERASE II SUBUNIT RPB1 | | |
| 48:00:00 | 7d6v-C | 2.9 | 6.2 | 89 | 264 | 1 | MOLECULE: | SUCCINATE DEHYDROGENASE SUBUNIT A | | |
| 49:00:00 | 8wg3-A | 2.9 | 9.9 | 83 | 548 | 4 | MOLECULE: | CSC1-LIKE PROTEIN 2,GREEN FLUORESCENT PROTE | IN | |
| 50:00:00 | 8tzk-C | 2.9 | 3.8 | 80 | 304 | 4 | MOLECULE: | CELL DIVISION ATP-BINDING PROTEIN FTSE | | |
| 51:00:00 | 7paf-A | 2.9 | 9.1 | 74 | 278 | 3 | MOLECULE: | NANOBODY | | |
| 52:00:00 | 7t6d-A | 2.9 | 14.3 | 60 | 389 | 8 | MOLECULE: | LIPOPOLYSACCHARIDE ASSEMBLY PROTEIN B | | |
| 53:00:00 | 4iyp-A | 2.9 | 10.7 | 84 | 205 | 4 | MOLECULE: | IMMUNOGLOBULIN-BINDING PROTEIN 1 | | |
| 54:00:00 | 2zy4-F | 2.9 | 2.6 | 48 | 535 | 4 | MOLECULE: | L-ASPARTATE BETA-DECARBOXYLASE | | |
| 55:00:00 | 1qv9-A | 2.9 | 6.7 | 96 | 282 | 8 | MOLECULE: | F420-DEPENDENT METHYLENETETRAHYDROMETHANOPT | ERIN | |
| 56:00:00 | 8yzk-A | 2.9 | 4.2 | 89 | 264 | 6 | MOLECULE: | SOLUBLE CYTOCHROME B562,G-PROTEIN COUPLED R | ECEPTO | |
| 57:00:00 | 5n3u-B | 2.9 | 11.4 | 59 | 170 | 3 | MOLECULE: | PHYCOCYANOBILIN LYASE SUBUNIT ALPHA | | |
| 58:00:00 | 8q7n-S | 2.9 | 22.8 | 86 | 171 | 10 | MOLECULE: | U5 SNRNA | | |
| 59:00:00 | 6urt-A | 2.9 | 11.3 | 130 | 331 | 12 | MOLECULE: | LOW CONDUCTANCE MECHANOSENSITIVE CHANNEL YN | AI | |
| 60:00:00 | 4bzj-C | 2.9 | 7.2 | 49 | 693 | 4 | MOLECULE: | PROTEIN TRANSPORT PROTEIN SEC31 | | |
| 61:00:00 | 8soi-B | 2.9 | 33.8 | 161 | 515 | 7 | MOLECULE: | RB1-INDUCIBLE COILED-COIL PROTEIN 1 | | |
| 62:00:00 | 8xuj-A | 2.9 | 9 | 73 | 424 | 4 | MOLECULE: | MEMBRANE PROTEIN | | |
| 63:00:00 | 8gs1-D | 2.9 | 3.4 | 55 | 337 | 5 | MOLECULE: | AZI28 | | |
| 64:00:00 | 8qae-A | 2.9 | 4.8 | 86 | 211 | 3 | MOLECULE: | SC-APCC-6-SLLA | | |
| 65:00:00 | 6tpk-A | 2.9 | 15.5 | 121 | 461 | 13 | MOLECULE: | OXYTOCIN RECEPTOR | | |
| 66:00:00 | 5hy3-A | 2.9 | 2.6 | 44 | 252 | 7 | MOLECULE: | MRNA ENDORIBONUCLEASE LSOA | | |
| 67:00:00 | 2ox6-C | 2.9 | 11.7 | 74 | 163 | 15 | MOLECULE: | HYPOTHETICAL PROTEIN SO3848 | | |
| 68:00:00 | 7jhj-R | 2.9 | 4.4 | 93 | 277 | 10 | MOLECULE: | GUANINE NUCLEOTIDE-BINDING PROTEIN G(I) SUB | UNIT A | |
| 69:00:00 | 8r5s-A | 2.9 | 5.5 | 111 | 237 | 4 | MOLECULE: | CHAIN A, B, C AND H OF THE SOLUBLE HOMOTETR | AMER | |
| 70:00:00 | 4g80-I | 2.9 | 3.9 | 82 | 139 | 5 | MOLECULE: | VOLTAGE-SENSOR CONTAINING PHOSPHATASE | | |
| 71:00:00 | 6yrf-A | 2.9 | 34.6 | 109 | 777 | 2 | MOLECULE: | VEGETATIVE INSECTICIDAL PROTEIN | | |
| 72:00:00 | 6ffh-A | 2.9 | 13 | 92 | 414 | 4 | MOLECULE: | METABOTROPIC GLUTAMATE RECEPTOR 5,ENDOLYSIN | | |
| 73:00:00 | 9j2f-M | 2.9 | 4.7 | 59 | 331 | 7 | MOLECULE: | PHOTOSYNTHETIC REACTION CENTER CYTOCHROME C | SUBUN | |
| 74:00:00 | 8j8y-A | 2.9 | 6.1 | 70 | 129 | 14 | MOLECULE: | IMMUNODOMINANT MEMBRANE PROTEIN | | |
| 75:00:00 | 1qgk-A | 2.9 | 6.6 | 55 | 876 | 11 | MOLECULE: | PROTEIN (IMPORTIN BETA SUBUNIT) | | |
| 76:00:00 | 8cmk-A | 2.9 | 8.6 | 80 | 913 | 4 | MOLECULE: | TRANSPORTIN-3 | | |
| 77:00:00 | 5fu7-A | 2.9 | 16 | 88 | 519 | 1 | MOLECULE: | CCR4-NOT TRANSCRIPTION COMPLEX SUBUNIT 1 | | |
| 78:00:00 | 8xks-O | 2.9 | 18.1 | 95 | 197 | 5 | MOLECULE: | CTAP1 | | |
| 79:00:00 | 8exh-a | 2.9 | 4 | 49 | 69 | 6 | MOLECULE: | PROTEIN VIRB2 | | |
| 80:00:00 | 4rh7-A | 2.9 | 41.5 | 96 | 3005 | 8 | MOLECULE: | GREEN FLUORESCENT PROTEIN/CYTOPLASMIC DYNEI | N 2 HE | |
| 81:00:00 | 6rgv-A | 2.9 | 3.7 | 62 | 406 | 5 | MOLECULE: | FLAGELLIN | | |
| 82:00:00 | 3zjc-E | 2.9 | 4.7 | 62 | 286 | 18 | MOLECULE: | GTPASE IMAP FAMILY MEMBER 7 | | |
| 83:00:00 | 5fiy-A | 2.9 | 3.2 | 57 | 78 | 9 | MOLECULE: | PRKC APOPTOSIS WT1 REGULATOR PROTEIN | | |
| 84:00:00 | 8uz8-B | 2.9 | 2.4 | 41 | 60 | 5 | MOLECULE: | 2-OXOGLUTARATE:ACCEPTOR OXIDOREDUCTASE | | |
| 85:00:00 | 3bas-B | 2.9 | 31 | 72 | 87 | 11 | MOLECULE: | MYOSIN HEAVY CHAIN, STRIATED MUSCLE/GENERAL | CONTR | |
| 86:00:00 | 7o4i-W | 2.9 | 33 | 109 | 312 | 8 | MOLECULE: | GENERAL TRANSCRIPTION AND DNA REPAIR FACTOR | IIH H | |
| 87:00:00 | 3ctw-B | 2.9 | 4 | 55 | 120 | 5 | MOLECULE: | RCDA | | |
| 88:00:00 | 5n8k-C | 2.9 | 5.6 | 57 | 80 | 12 | MOLECULE: | GALACTOCEREBROSIDASE | | |
| 89:00:00 | 8w6e-B | 2.9 | 3.2 | 53 | 161 | 11 | MOLECULE: | HBC599 IN COMPLEX WITH WFAP1.1 | | |
| 90:00:00 | 6tdv-f | 2.9 | 15 | 74 | 274 | 4 | MOLECULE: | ATPTB1 | | |
| 91:00:00 | 6sih-A | 2.9 | 24.5 | 81 | 488 | 7 | MOLECULE: | FLAGELLAR HOOK-ASSOCIATED PROTEIN 2 | | |
| 92:00:00 | 3k66-A | 2.8 | 29.6 | 109 | 217 | 5 | MOLECULE: | BETA-AMYLOID-LIKE PROTEIN | | |
| 93:00:00 | 8ywi-A | 2.8 | 28.9 | 121 | 980 | 10 | MOLECULE: | DNA POLYMERASE | | |
| 94:00:00 | 4wbd-A | 2.8 | 23.1 | 92 | 540 | 3 | MOLECULE: | BSHC | | |
| 95:00:00 | 4avm-A | 2.8 | 20.2 | 120 | 230 | 4 | MOLECULE: | BRIDGING INTEGRATOR 2 | | |
| 96:00:00 | 8xzv-D | 2.8 | 37 | 114 | 1216 | 4 | MOLECULE: | DNA-DIRECTED RNA POLYMERASE SUBUNIT ALPHA | | |
| 97:00:00 | 7xge-A | 2.8 | 8 | 89 | 168 | 4 | MOLECULE: | BCL-XL AND MCL-1 DUAL BINDER 2 | | |
| 98:00:00 | 8e2h-A | 2.8 | 16.4 | 97 | 656 | 3 | MOLECULE: | BACULOVIRAL IAP REPEAT-CONTAINING PROTEIN 6 | | |
| 99:00:00 | 1u4q-A | 2.8 | 16.6 | 85 | 318 | 6 | MOLECULE: | SPECTRIN ALPHA CHAIN, BRAIN | | |
| 0:00 | 6yjd-A | 2.8 | 17.3 | 91 | 104 | 3 | MOLECULE: | CAPSID ASSEMBLY SCAFFOLDING PROTEIN,PRELAMI | N-A/C | |
| 1:00 | 8r7n-D | 2.8 | 46.4 | 135 | 221 | 6 | MOLECULE: | U11 SNRNA | | |
| 2:00 | 6ikn-A | 2.8 | 13.8 | 96 | 305 | 7 | MOLECULE: | GROWTH ARREST-SPECIFIC PROTEIN 7 | | |
| 3:00 | 4gwp-C | 2.8 | 31.8 | 92 | 184 | 5 | MOLECULE: | MEDIATOR OF RNA POLYMERASE II TRANSCRIPTION | SUBUN | |
| 4:00 | 1s35-A | 2.8 | 27.2 | 128 | 211 | 5 | MOLECULE: | SPECTRIN BETA CHAIN, ERYTHROCYTE | | |
| 5:00 | 4v1a-w | 2.8 | 31.1 | 100 | 387 | 5 | MOLECULE: | MITORIBOSOMAL PROTEIN ML37, MRPL37 | | |
| 6:00 | 5mpd-Q | 2.8 | 36.3 | 127 | 434 | 8 | MOLECULE: | 26S PROTEASOME REGULATORY SUBUNIT RPN10 | | |
| 7:00 | 5jm8-A | 2.8 | 25 | 100 | 556 | 6 | MOLECULE: | AEROBACTIN SYNTHASE IUCA | | |

| | | | | | | | | | | |
|---|---|---|---|---|---|---|---|---|---|---|
| 8:00 | 2qe9-A | 2.8 | 33.7 | 86 | 165 | 6 | MOLECULE: | UNCHARACTERIZED PROTEIN YIZA | | |
| 9:00 | 9exz-A | 2.8 | 7.1 | 97 | 602 | 4 | MOLECULE: | DE NOVO PROTEIN P600 | | |
| 10:00 | 7kyp-F | 2.8 | 19.6 | 89 | 290 | 6 | MOLECULE: | MANGANESE ABC TRANSPORTER, ATP-BINDING PROT | EIN | |
| 11:00 | 6n88-A | 2.8 | 12 | 71 | 619 | 6 | MOLECULE: | MGC52556 PROTEIN | | |
| 12:00 | 7q4p-C | 2.8 | 5.5 | 43 | 878 | 9 | MOLECULE: | SPLICING FACTOR 3A SUBUNIT 2 | | |
| 13:00 | 6j5i-b | 2.8 | 19.7 | 87 | 209 | 8 | MOLECULE: | ATP SYNTHASE SUBUNIT ALPHA, MITOCHONDRIAL | | |
| 14:00 | 8cih-A | 2.8 | 5.6 | 103 | 188 | 7 | MOLECULE: | CYCLIN-DEPENDENT KINASE 2-INTERACTING PROTE | IN | |
| 15:00 | 7mwq-A | 2.8 | 8.1 | 57 | 204 | 12 | MOLECULE: | LHD29A53 | | |
| 16:00 | 3h36-A | 2.8 | 8.7 | 57 | 78 | 4 | MOLECULE: | POLYRIBONUCLEOTIDE NUCLEOTIDYLTRANSFERASE | | |
| 17:00 | 8qf8-A | 2.8 | 6.3 | 69 | 776 | 4 | MOLECULE: | GLYCOSYL HYDROLASE | | |
| 18:00 | 6yxq-B | 2.8 | 12.4 | 53 | 396 | 6 | MOLECULE: | ACTIVATING SIGNAL COINTEGRATOR 1 COMPLEX SU | BUNIT | |
| 19:00 | 7aqr-V | 2.8 | 13.7 | 60 | 140 | 3 | MOLECULE: | NADH DEHYDROGENASE [UBIQUINONE] IRON-SULFUR | PROTE | |
| 20:00 | 5kkl-B | 2.8 | 1.7 | 41 | 838 | 0 | MOLECULE: | PUTATIVE POLYCOMB PROTEIN EED | | |
| 21:00 | 4oph-A | 2.8 | 7.6 | 60 | 202 | 5 | MOLECULE: | NONSTRUCTURAL PROTEIN 1 | | |
| 22:00 | 4xb6-E | 2.8 | 6 | 64 | 149 | 3 | MOLECULE: | ALPHA-D-RIBOSE 1-METHYLPHOSPHONATE 5-TRIPHO | SPHATE | |
| 23:00 | 6zh3-A | 2.8 | 20.6 | 121 | 170 | 6 | MOLECULE: | VACUOLAR PROTEIN-SORTING-ASSOCIATED PROTEIN | | 24 |
| 24:00:00 | 6cp3-Z | 2.8 | 16.1 | 68 | 155 | 6 | MOLECULE: | ATP SYNTHASE SUBUNIT 9, MITOCHONDRIAL | | |
| 25:00:00 | 7v9g-D | 2.8 | 5.7 | 69 | 112 | 9 | MOLECULE: | BEN DOMAIN-CONTAINING PROTEIN 3 | | |
| 26:00:00 | 7vqo-A | 2.8 | 28.2 | 106 | 1180 | 8 | MOLECULE: | AMS1, NBR1 AND MALE FUSION PROTEIN | | |
| 27:00:00 | 7tt5-D | 2.8 | 6.1 | 57 | 153 | 7 | MOLECULE: | MALTOSE/MALTODEXTRIN-BINDING PERIPLASMIC PR | OTEIN, | |
| 28:00:00 | 7vf2-C | 2.8 | 38.4 | 119 | 184 | 10 | MOLECULE: | PROTEIN VIRILIZER HOMOLOG | | |
| 29:00:00 | 5t76-A | 2.8 | 6.1 | 74 | 208 | 7 | MOLECULE: | ALANINE--TRNA LIGASE, CYTOPLASMIC | | |
| 30:00:00 | 6xgx-A | 2.8 | 5.5 | 55 | 402 | 9 | MOLECULE: | MUTATOR FAMILY TRANSPOSASE | | |
| 31:00:00 | 9kqi-A | 2.8 | 6.2 | 74 | 378 | 3 | MOLECULE: | PHOSPHATIDYLSERINE SYNTHASE 1 | | |
| 32:00:00 | 8e1m-A | 2.8 | 12.5 | 94 | 133 | 5 | MOLECULE: | MITOCHONDRIAL IMPORT INNER MEMBRANE TRANSLO | CASE S | |
| 33:00:00 | 3err-A | 2.8 | 34.4 | 148 | 527 | 5 | MOLECULE: | FUSION PROTEIN OF MICROTUBULE BINDING DOMAI | N FROM | |
| 34:00:00 | 8at3-A | 2.8 | 41.6 | 175 | 286 | 4 | MOLECULE: | HAUS AUGMIN-LIKE COMPLEX SUBUNIT 1 | | |
| 35:00:00 | 8rwv-C | 2.8 | 29.7 | 94 | 678 | 5 | MOLECULE: | DNA REPLICATION LICENSING FACTOR MCM2 | | |
| 36:00:00 | 4qnd-A | 2.8 | 9.1 | 58 | 97 | 7 | MOLECULE: | CHEMICAL TRANSPORT PROTEIN | | |
| 37:00:00 | 8ket-B | 2.8 | 8.4 | 71 | 267 | 7 | MOLECULE: | E3 UBIQUITIN-PROTEIN LIGASE SYNOVIOLIN | | |
| 38:00:00 | 7mge-B | 2.8 | 29 | 124 | 442 | 6 | MOLECULE: | WD REPEAT-CONTAINING PROTEIN 41 | | |
| 39:00:00 | 8bou-A | 2.8 | 17.3 | 89 | 807 | 4 | MOLECULE: | N,N'-DIACETYLCHITOBIOSE PHOSPHORYLASE | | |
| 40:00:00 | 8qx8-A | 2.8 | 20.9 | 71 | 969 | 7 | MOLECULE: | VACUOLAR PROTEIN SORTING-ASSOCIATED PROTEIN | | 8 |
| 41:00:00 | 8a6m-A | 2.8 | 3.7 | 54 | 161 | 4 | MOLECULE: | ISOFORM 1B OF SYNAPTOGYRIN-1 | | |
| 42:00:00 | 8f7c-A | 2.8 | 6.5 | 103 | 287 | 7 | MOLECULE: | PANNEXIN-2, SOLUBLE CYTOCHROME B562 FUSION | | |
| 43:00:00 | 7rut-B | 2.8 | 9.7 | 67 | 187 | 7 | MOLECULE: | CORRINOID ADENOSYLTRANSFERASE | | |
| 44:00:00 | 2hsb-A | 2.8 | 3.9 | 50 | 126 | 4 | MOLECULE: | HYPOTHETICAL UPF0332 PROTEIN AF0298 | | |
| 45:00:00 | 7syf-A | 2.8 | 25.7 | 120 | 1391 | 8 | MOLECULE: | PHOSPHATIDYLINOSITOL 3,4,5-TRISPHOSPHATE-DE | PENDEN | |
| 46:00:00 | 6nsj-A | 2.8 | 4.3 | 80 | 182 | 5 | MOLECULE: | ACID-ACTIVATED UREA CHANNEL | | |
| 47:00:00 | 8vxq-A | 2.8 | 4.1 | 56 | 155 | 2 | MOLECULE: | GP72 | | |
| 48:00:00 | 6al2-A | 2.8 | 24.5 | 95 | 484 | 6 | MOLECULE: | MEMBRANE PROTEIN INSERTASE YIDC | | |
| 49:00:00 | 2y7c-A | 2.8 | 23.1 | 132 | 464 | 6 | MOLECULE: | TYPE-1 RESTRICTION ENZYME ECOKI SPECIFICITY | PROTE | |
| 50:00:00 | 7drj-A | 2.8 | 9.9 | 67 | 189 | 9 | MOLECULE: | CDP-DIACYLGLYCEROL--GLYCEROL-3-PHOSPHATE 3- | | |
| 51:00:00 | 1s3j-A | 2.8 | 9.7 | 79 | 143 | 6 | MOLECULE: | YUSO PROTEIN | | |
| 52:00:00 | 8afz-A | 2.8 | 20.6 | 111 | 381 | 8 | MOLECULE: | SORTING NEXIN-1 | | |
| 53:00:00 | 3kmi-A | 2.8 | 5.7 | 93 | 172 | 8 | MOLECULE: | PUTATIVE MEMBRANE PROTEIN COG4129 | | |
| 54:00:00 | 9crw-D | 2.8 | 7.9 | 83 | 228 | 8 | MOLECULE: | KINESIN-LIKE PROTEIN | | |
| 55:00:00 | 6lqo-L | 2.8 | 4.9 | 82 | 94 | 2 | MOLECULE: | CYTOPLASMIC ENVELOPMENT PROTEIN 1 | | |
| 56:00:00 | 6c6r-A | 2.8 | 4.8 | 83 | 451 | 5 | MOLECULE: | SQUALENE MONOOXYGENASE | | |
| 57:00:00 | 8t9l-A | 2.8 | 13.7 | 78 | 108 | 4 | MOLECULE: | NUCLEOPORIN POM152 | | |
| 58:00:00 | 6m3q-F | 2.8 | 12 | 116 | 301 | 6 | MOLECULE: | ANKYRIN-2 | | |
| 59:00:00 | 6ec8-A | 2.8 | 25.7 | 130 | 803 | 6 | MOLECULE: | LANTIBIOTIC DEHYDRATASE DOMAIN PROTEIN | | |
| 60:00:00 | 4rs7-B | 2.8 | 7.3 | 69 | 88 | 13 | MOLECULE: | PARB-C | | |
| 61:00:00 | 1cqx-A | 2.8 | 14.6 | 71 | 403 | 13 | MOLECULE: | FLAVOHEMOPROTEIN | | |
| 62:00:00 | 8ua7-A | 2.8 | 7.6 | 63 | 109 | 6 | MOLECULE: | HISTONE H3 | | |
| 63:00:00 | 3nr7-A | 2.8 | 4.2 | 62 | 81 | 5 | MOLECULE: | DNA-BINDING PROTEIN H-NS | | |
| 64:00:00 | 5b0l-B | 2.8 | 14.7 | 99 | 334 | 3 | MOLECULE: | MOEN5,DNA-BINDING PROTEIN 7D | | |
| 65:00:00 | 5oge-G | 2.8 | 4.3 | 68 | 306 | 6 | MOLECULE: | GDP-MANNOSE TRANSPORTER 1 | | |
| 66:00:00 | 5ixp-A | 2.8 | 3.3 | 40 | 393 | 3 | MOLECULE: | EXTRACELLULAR SOLUTE-BINDING PROTEIN FAMILY | | 1 |
| 67:00:00 | 5xsj-L | 2.8 | 4 | 75 | 122 | 11 | MOLECULE: | PERIPLASMIC BINDING PROTEIN/LACI TRANSCRIPT | IONAL | |
| 68:00:00 | 5mdx-A | 2.8 | 11.2 | 71 | 307 | 3 | MOLECULE: | PHOTOSYSTEM II PROTEIN D1 | | |
| 69:00:00 | 8h0m-A | 2.8 | 5.7 | 74 | 392 | 7 | MOLECULE: | VIOD | | |
| 70:00:00 | 4mnd-A | 2.8 | 8 | 63 | 408 | 8 | MOLECULE: | CTP L-MYO-INOSITOL-1-PHOSPHATE CYTIDYLYLTRA | NSFERA | |
| 71:00:00 | 3qt2-C | 2.8 | 4.4 | 50 | 110 | 16 | MOLECULE: | INTERLEUKIN-5 RECEPTOR SUBUNIT ALPHA | | |
| 72:00:00 | 5z25-A | 2.8 | 4.3 | 62 | 96 | 8 | MOLECULE: | CYTOCHROME C552 | | |
| 73:00:00 | 2rvq-D | 2.8 | 5.9 | 54 | 129 | 7 | MOLECULE: | HISTONE H2A TYPE 1-B/E | | |
| 74:00:00 | 7ty0-C | 2.8 | 33 | 89 | 488 | 11 | MOLECULE: | GLYCOPROTEIN G | | |
| 75:00:00 | 6m15-A | 2.8 | 26 | 88 | 965 | 3 | MOLECULE: | SPIKE GLYCOPROTEIN | | |
| 76:00:00 | 7rtn-A | 2.8 | 11.9 | 82 | 270 | 6 | MOLECULE: | OUTER CAPSID PROTEIN VP5 | | |
| 77:00:00 | 9c3f-A | 2.8 | 25.1 | 99 | 485 | 7 | MOLECULE: | ANHYDROMUROPEPTIDE PERMEASE,SOLUBLE CYTOCHR | OME B5 | |
| 78:00:00 | 8u54-B | 2.8 | 28.5 | 124 | 680 | 6 | MOLECULE: | INTEGRAL MEMBRANE PROTEIN GPR155 | | |
| 79:00:00 | 9gm6-B | 2.8 | 43.9 | 142 | 1148 | 5 | MOLECULE: | CHROMOSOME PARTITION PROTEIN MUKF | | |
| 80:00:00 | 8v1g-A | 2.8 | 14.4 | 111 | 990 | 11 | MOLECULE: | NIEMANN-PICK TYPE C1-RELATED PROTEIN | | |
| 81:00:00 | 7ojn-L | 2.8 | 17.8 | 87 | 2010 | 5 | MOLECULE: | RNA-DIRECTED RNA POLYMERASE L | | |
| 82:00:00 | 6pw5-B | 2.8 | 19.8 | 123 | 782 | 6 | MOLECULE: | TRP-LIKE ION CHANNEL | | |
| 83:00:00 | 8ezj-A | 2.8 | 3.6 | 48 | 1285 | 8 | MOLECULE: | ARF GUANINE-NUCLEOTIDE EXCHANGE FACTOR 2 | | |
| 84:00:00 | 4ap2-B | 2.8 | 10.7 | 55 | 359 | 4 | MOLECULE: | KELCH-LIKE PROTEIN 11 | | |
| 85:00:00 | 7y9x-A | 2.8 | 44.4 | 111 | 1458 | 10 | MOLECULE: | CHAT DOMAIN-CONTAINING PROTEIN | | |
| 86:00:00 | 1lgh-A | 2.8 | 4.8 | 45 | 56 | 7 | MOLECULE: | LIGHT HARVESTING COMPLEX II | | |
| 87:00:00 | 7t92-B | 2.8 | 22.7 | 117 | 359 | 5 | MOLECULE: | PEROXIN-12 | | |
| 88:00:00 | 6hd8-B | 2.8 | 5.8 | 73 | 232 | 7 | MOLECULE: | NANOBODY,MALTOSE/MALTODEXTRIN-BINDING PERIP | LASMIC | |
| 89:00:00 | 7lma-A | 2.8 | 6.5 | 96 | 1012 | 4 | MOLECULE: | TELOMERASE LA-RELATED PROTEIN P65 | | |
| 90:00:00 | 6bym-A | 2.8 | 5.2 | 56 | 200 | 13 | MOLECULE: | STEROL-BINDING PROTEIN | | |
| 91:00:00 | 7ktr-A | 2.8 | 23.2 | 104 | 3042 | 8 | MOLECULE: | TRANSFORMATION/TRANSCRIPTION DOMAIN-ASSOCIA | TED PR | |
| 92:00:00 | 7zjk-A | 2.8 | 5.3 | 100 | 221 | 6 | MOLECULE: | CSPZ | | |
| 93:00:00 | 8eki-A | 2.8 | 10.7 | 84 | 170 | 5 | MOLECULE: | PROTEIN TRANSPORT PROTEIN SEC20 | | |
| 94:00:00 | 6lcp-B | 2.8 | 22.9 | 109 | 376 | 9 | MOLECULE: | PHOSPHOLIPID-TRANSPORTING ATPASE | | |
| 95:00:00 | 7vrc-D | 2.8 | 5.2 | 48 | 51 | 8 | MOLECULE: | TRANSCRIPTION REGULATORY PROTEIN SNF11 | | |
| 96:00:00 | 2x3j-A | 2.8 | 19 | 105 | 586 | 4 | MOLECULE: | ACSD | | |
| 97:00:00 | 2odu-A | 2.8 | 6.8 | 92 | 217 | 9 | MOLECULE: | PLECTIN 1 | | |
| 98:00:00 | 7bii-B | 2.8 | 24.4 | 143 | 2144 | 6 | MOLECULE: | E3 UBIQUITIN-PROTEIN LIGASE HUWE1 | | |
| 99:00:00 | 6cgh-A | 2.8 | 2.4 | 44 | 89 | 16 | MOLECULE: | DNAJ HOMOLOG SUBFAMILY C MEMBER 2 | | |
| 0:00 | 5wst-A | 2.8 | 3 | 54 | 71 | 9 | MOLECULE: | UNCONVENTIONAL MYOSIN-VIIA | | |
| 1:00 | 6n6s-D | 2.8 | 3.5 | 55 | 69 | 11 | MOLECULE: | TNFAIP3-INTERACTING PROTEIN 1 | | |
| 2:00 | 8sjj-A | 2.8 | 3.5 | 58 | 74 | 7 | MOLECULE: | SEPTIN-14 | | |
| 3:00 | 5xau-E | 2.8 | 14.5 | 54 | 72 | 7 | MOLECULE: | LAMININ SUBUNIT ALPHA-5 | | |
| 4:00 | 6yip-A | 2.8 | 3.8 | 55 | 71 | 5 | MOLECULE: | KINESIN-LIKE PROTEIN KIF20A | | |
| 5:00 | 7eqb-A | 2.8 | 4.5 | 55 | 68 | 11 | MOLECULE: | KINESIN-LIKE PROTEIN | | |
| 6:00 | 6hk9-B | 2.8 | 4.7 | 59 | 78 | 7 | MOLECULE: | TESTIS-EXPRESSED PROTEIN 12 | | |
| 7:00 | 8fn2-a | 2.8 | 4.4 | 47 | 65 | 6 | MOLECULE: | 23S RIBOSOMAL RNA | | |
| 8:00 | 4akk-A | 2.8 | 11.3 | 68 | 368 | 6 | MOLECULE: | NITRATE REGULATORY PROTEIN | | |

| | | | | | | | | | |
|---|---|---|---|---|---|---|---|---|---|
| 9:00 | 5mnt-B | 2.8 | 10.2 | 87 | 421 | 3 | MOLECULE: | A2 MATURATION PROTEIN | |
| 10:00 | 9dva-G | 2.8 | 4.3 | 47 | 71 | 9 | MOLECULE: | ACTIN, ALPHA SKELETAL MUSCLE | |
| 11:00 | 8sl3-A | 2.8 | 5.8 | 66 | 578 | 11 | MOLECULE: | ADENYLATE CYCLASE TYPE 5 | |
| 12:00 | 2e5t-A | 2.8 | 3.5 | 41 | 46 | 5 | MOLECULE: | ATP SYNTHASE EPSILON CHAIN | |
| 13:00 | 6r5k-O | 2.7 | 22 | 88 | 447 | 9 | MOLECULE: | PAN2-PAN3 DEADENYLATION COMPLEX CATALYTIC S | UBUNIT |
| 14:00 | 6nd4-O | 2.7 | 7 | 64 | 832 | 13 | MOLECULE: | ETS RRNA | |
| 15:00 | 5yke-B | 2.7 | 20.4 | 171 | 845 | 7 | MOLECULE: | ATP-SENSITIVE INWARD RECTIFIER POTASSIUM CH | ANNEL |
| 16:00 | 8pr3-C | 2.7 | 34.4 | 86 | 106 | 1 | MOLECULE: | C-JUN-AMINO-TERMINAL KINASE-INTERACTING PRO | TEIN 3 |
| 17:00 | 6h9m-A | 2.7 | 22.3 | 73 | 94 | 4 | MOLECULE: | COILED-COIL DOMAIN-CONTAINING PROTEIN 90B, | MITOCH |
| 18:00 | 8b6l-O | 2.7 | 22.8 | 71 | 580 | 6 | MOLECULE: | PROTEIN TRANSPORT PROTEIN SEC61 SUBUNIT ALP | HA ISO |
| 19:00 | 7eld-A | 2.7 | 28.8 | 112 | 1137 | 6 | MOLECULE: | ENDORIBONUCLEASE DICER HOMOLOG 1 | |
| 20:00 | 7b93-L | 2.7 | 35.2 | 196 | 607 | 1 | MOLECULE: | NADH-UBIQUINONE OXIDOREDUCTASE CHAIN 3 | |
| 21:00 | 8j98-A | 2.7 | 16.5 | 59 | 184 | 8 | MOLECULE: | NEAR-INFRARED FLUORESCENT PROTEIN | |
| 22:00 | 6lul-A | 2.7 | 21.4 | 102 | 223 | 7 | MOLECULE: | RESPIRATORY SUPERCOMPLEX FACTOR 2, MITOCHON | DRIAL |
| 23:00 | 6tgz-F | 2.7 | 31.3 | 127 | 352 | 9 | MOLECULE: | 55 KDA IMMEDIATE-EARLY PROTEIN 1 | |
| 24:00:00 | 8tdl-A | 2.7 | 37 | 121 | 364 | 5 | MOLECULE: | MECHANOSENSITIVE ION CHANNEL PROTEIN 10 | |
| 25:00:00 | 2dh4-A | 2.7 | 13.9 | 59 | 326 | 10 | MOLECULE: | YPL069C | |
| 26:00:00 | 8p0s-A | 2.7 | 50.7 | 117 | 133 | 5 | MOLECULE: | RHO-ASSOCIATED PROTEIN KINASE 1 | |
| 27:00:00 | 4lin-A | 2.7 | 66.9 | 161 | 289 | 8 | MOLECULE: | TAIL NEEDLE PROTEIN GP26 | |
| 28:00:00 | 6hux-A | 2.7 | 16.1 | 85 | 369 | 7 | MOLECULE: | H(2)-FORMING METHYLENETETRAHYDROMETHANOPTER | IN |
| 29:00:00 | 8rg0-5 | 2.7 | 28.5 | 105 | 520 | 9 | MOLECULE: | EUKARYOTIC TRANSLATION INITIATION FACTOR 3 | SUBUNI |
| 30:00:00 | 5y5a-A | 2.7 | 37.2 | 147 | 514 | 3 | MOLECULE: | KLLA0F20702P | |
| 31:00:00 | 5cws-D | 2.7 | 27.4 | 130 | 180 | 5 | MOLECULE: | SAB-158 FAB LIGHT CHAIN | |
| 32:00:00 | 2huj-A | 2.7 | 30.2 | 80 | 125 | 1 | MOLECULE: | LIN2004 PROTEIN | |
| 33:00:00 | 6ysg-A | 2.7 | 30.4 | 131 | 1264 | 10 | MOLECULE: | MG-CHELATASE SUBUNIT CHLH | |
| 34:00:00 | 8t1l-S | 2.7 | 21.7 | 159 | 913 | 6 | MOLECULE: | MEDIATOR OF RNA POLYMERASE II TRANSCRIPTION | SUBUN |
| 35:00:00 | 6eqo-A | 2.7 | 26.4 | 122 | 1804 | 4 | MOLECULE: | ACETYL-COENZYME A SYNTHETASE | |
| 36:00:00 | 9ewy-A | 2.7 | 7.5 | 91 | 815 | 10 | MOLECULE: | [F-ACTIN]-MONOOXYGENASE MICAL1 | |
| 37:00:00 | 5x11-E | 2.7 | 3.9 | 64 | 181 | 5 | MOLECULE: | DNA (28-MER) | |
| 38:00:00 | 5lst-A | 2.7 | 6.3 | 66 | 617 | 3 | MOLECULE: | ATP-DEPENDENT DNA HELICASE Q4 | |
| 39:00:00 | 8fli-B | 2.7 | 3.8 | 51 | 458 | 14 | MOLECULE: | GROUP II INTRON | |
| 40:00:00 | 8egr-A | 2.7 | 31.6 | 131 | 609 | 9 | MOLECULE: | GP15, RECEPTOR-BINDING PROTEIN, TAIL FIBER | |
| 41:00:00 | 6em5-r | 2.7 | 12.6 | 78 | 176 | 6 | MOLECULE: | 5.8S RIBOSOMAL RNA | |
| 42:00:00 | 8e9h-J | 2.7 | 4.2 | 51 | 231 | 8 | MOLECULE: | TWO-COMPONENT SYSTEM RESPONSE REGULATOR | |
| 43:00:00 | 8xku-R | 2.7 | 8.9 | 91 | 267 | 7 | MOLECULE: | PROBABLE INACTIVE ATP-DEPENDENT ZINC METALL | OPROTE |
| 44:00:00 | 5b7j-A | 2.7 | 5.1 | 62 | 110 | 2 | MOLECULE: | SWITCH-ACTIVATING PROTEIN 1 | |
| 45:00:00 | 8gyz-B | 2.7 | 16.9 | 87 | 215 | 3 | MOLECULE: | TRANSCRIPTION FACTOR TGA7 | |
| 46:00:00 | 6a73-A | 2.7 | 15.6 | 90 | 290 | 10 | MOLECULE: | COP9 SIGNALOSOME COMPLEX SUBUNIT 2,ENDOLYSI | N |
| 47:00:00 | 6g3e-A | 2.7 | 11.9 | 133 | 497 | 5 | MOLECULE: | ARGININOSUCCINATE LYASE | |
| 48:00:00 | 8q3v-g | 2.7 | 4 | 54 | 76 | 4 | MOLECULE: | TETRAHYDROMETHANOPTERIN S-METHYLTRANSFERASE | SUBUN |
| 49:00:00 | 3dxp-A | 2.7 | 10.9 | 82 | 329 | 5 | MOLECULE: | PUTATIVE ACYL-COA DEHYDROGENASE | |
| 50:00:00 | 8yx6-A | 2.7 | 10.6 | 83 | 324 | 0 | MOLECULE: | LIGAND-GATED CATION CHANNEL ZACN | |
| 51:00:00 | 7dvr-A | 2.7 | 9.4 | 75 | 147 | 1 | MOLECULE: | HTH MARR-TYPE DOMAIN-CONTAINING PROTEIN | |
| 52:00:00 | 5lnf-A | 2.7 | 10.3 | 93 | 139 | 6 | MOLECULE: | PEROXISOMAL BIOGENESIS FACTOR 19 | |
| 53:00:00 | 8j0h-A | 2.7 | 4.6 | 92 | 134 | 4 | MOLECULE: | UNCHARACTERIZED PROTEIN C4H3.06 | |
| 54:00:00 | 3fgg-A | 2.7 | 7.2 | 67 | 147 | 12 | MOLECULE: | UNCHARACTERIZED PROTEIN BCE2196 | |
| 55:00:00 | 5ufl-A | 2.7 | 2.3 | 39 | 523 | 5 | MOLECULE: | PROTEIN CIP2A | |
| 56:00:00 | 6l4o-A | 2.7 | 7.9 | 57 | 429 | 5 | MOLECULE: | APOPTOSIS INHIBITOR 5 | |
| 57:00:00 | 6xm1-C | 2.7 | 29.8 | 128 | 570 | 0 | MOLECULE: | VPS45 | |
| 58:00:00 | 7nwu-B | 2.7 | 5.5 | 53 | 249 | 2 | MOLECULE: | REGULATOR OF NONSENSE TRANSCRIPTS 3B | |
| 59:00:00 | 5y58-A | 2.7 | 18.6 | 88 | 548 | 7 | MOLECULE: | ATP-DEPENDENT DNA HELICASE II SUBUNIT 1 | |
| 60:00:00 | 5y88-U | 2.7 | 20.4 | 104 | 488 | 11 | MOLECULE: | PRE-MRNA-SPLICING FACTOR 8 | |
| 61:00:00 | 9na8-C | 2.7 | 4 | 45 | 234 | 7 | MOLECULE: | AUGMIN SUBUNIT 1 | |
| 62:00:00 | 4l80-B | 2.7 | 17.1 | 76 | 347 | 7 | MOLECULE: | HPCH/HPAI ALDOLASE | |
| 63:00:00 | 9eto-A | 2.7 | 7.8 | 76 | 363 | 9 | MOLECULE: | 4-HYDROXYTRYPTAMINE KINASE | |
| 64:00:00 | 8y1x-A | 2.7 | 14.6 | 85 | 544 | 2 | MOLECULE: | ASPARTATE/ALANINE ANTIPORTER | |
| 65:00:00 | 4zi2-C | 2.7 | 8.1 | 59 | 131 | 7 | MOLECULE: | ADP-RIBOSYLATION FACTOR-LIKE PROTEIN 3 | |
| 66:00:00 | 7p2y-J | 2.7 | 3.3 | 61 | 80 | 7 | MOLECULE: | ATP SYNTHASE SUBUNIT ALPHA | |
| 67:00:00 | 8rd2-B | 2.7 | 13.1 | 124 | 400 | 6 | MOLECULE: | INVARIANT SURFACE GLYCOPROTEIN | |
| 68:00:00 | 7qj0-L | 2.7 | 5.1 | 71 | 566 | 7 | MOLECULE: | MITOTIC-SPINDLE ORGANIZING PROTEIN 1 | |
| 69:00:00 | 9h3q-C | 2.7 | 5.2 | 62 | 158 | 8 | MOLECULE: | LARGE RIBOSOMAL SUBUNIT PROTEIN BL34 | |
| 70:00:00 | 8hpo-D | 2.7 | 44.4 | 94 | 165 | 9 | MOLECULE: | TRANSCRIPTIONAL REGULATORY PROTEIN UME1 | |
| 71:00:00 | 6lum-G | 2.7 | 8.3 | 78 | 123 | 1 | MOLECULE: | SUCCINATE DEHYDROGENASE SUBUNIT C | |
| 72:00:00 | 8tdj-A | 2.7 | 14.2 | 156 | 467 | 4 | MOLECULE: | MECHANOSENSITIVE ION CHANNEL PROTEIN 10 | |
| 73:00:00 | 6h4b-B | 2.7 | 3.7 | 48 | 84 | 6 | MOLECULE: | ORF026 | |
| 74:00:00 | 6cnn-A | 2.7 | 7.7 | 86 | 360 | 6 | MOLECULE: | INTERMEDIATE CONDUCTANCE CALCIUM-ACTIVATED | POTASS |
| 75:00:00 | 7m7a-A | 2.7 | 4.2 | 72 | 538 | 7 | MOLECULE: | PHOSPHOINOSITIDE 3-KINASE MAVQ | |
| 76:00:00 | 5cd4-I | 2.7 | 9.5 | 90 | 494 | 6 | MOLECULE: | CRISPR SYSTEM CASCADE SUBUNIT CASE | |
| 77:00:00 | 5olk-A | 2.7 | 12.5 | 73 | 395 | 4 | MOLECULE: | RIBONUCLEOSIDE-DIPHOSPHATE REDUCTASE, BETA | SUBUNI |
| 78:00:00 | 8wt9-A | 2.7 | 2.8 | 40 | 319 | 8 | MOLECULE: | IS621 TRANSPOSASE | |
| 79:00:00 | 3gfi-A | 2.7 | 9.8 | 77 | 143 | 1 | MOLECULE: | 146AA LONG HYPOTHETICAL TRANSCRIPTIONAL REG | ULATOR |
| 80:00:00 | 4uig-A | 2.7 | 3.6 | 69 | 91 | 10 | MOLECULE: | COPPER SENSITIVE OPERON REPRESSOR | |
| 81:00:00 | 3cfo-A | 2.7 | 2.7 | 40 | 906 | 18 | MOLECULE: | DNA POLYMERASE | |
| 82:00:00 | 7oca-E | 2.7 | 8.6 | 91 | 158 | 2 | MOLECULE: | GLUTAMATE RECEPTOR 1 | |
| 83:00:00 | 8k58-G | 2.7 | 9 | 62 | 129 | 3 | MOLECULE: | DNA-DIRECTED RNA POLYMERASE SUBUNIT BETA | |
| 84:00:00 | 2xnx-M | 2.7 | 17.6 | 62 | 107 | 8 | MOLECULE: | FIBRINOGEN ALPHA CHAIN | |
| 85:00:00 | 5a63-D | 2.7 | 3.7 | 60 | 100 | 10 | MOLECULE: | NICASTRIN | |
| 86:00:00 | 2gd5-D | 2.7 | 4.9 | 88 | 162 | 11 | MOLECULE: | CHARGED MULTIVESICULAR BODY PROTEIN 3 | |
| 87:00:00 | 5iwz-A | 2.7 | 10.3 | 65 | 372 | 6 | MOLECULE: | SYNAPTONEMAL COMPLEX PROTEIN 2 | |
| 88:00:00 | 5oc0-A | 2.7 | 5.8 | 100 | 174 | 6 | MOLECULE: | CYTOCHROME B561 | |
| 89:00:00 | 6ian-A | 2.7 | 26 | 138 | 306 | 9 | MOLECULE: | INTRAFLAGELLAR TRANSPORT PROTEIN 74 | |
| 90:00:00 | 4w8j-A | 2.7 | 27.3 | 137 | 1017 | 4 | MOLECULE: | PESTICIDAL CRYSTAL PROTEIN CRY1AC | |
| 91:00:00 | 6v4m-A | 2.7 | 2.5 | 42 | 163 | 5 | MOLECULE: | BCL-2 | |
| 92:00:00 | 8z1y-A | 2.7 | 13 | 125 | 331 | 9 | MOLECULE: | DIPEPTIDE TRANSPORT SYSTEM PERMEASE PROTEIN | DPPB |
| 93:00:00 | 8evm-C | 2.7 | 4.9 | 85 | 237 | 7 | MOLECULE: | CHLOROPHYLL DIMER PROTEIN DESIGNS, SPECIAL | PAIR 3 |
| 94:00:00 | 7uic-c | 2.7 | 5.4 | 80 | 110 | 8 | MOLECULE: | MEDIATOR OF RNA POLYMERASE II TRANSCRIPTION | SUBUN |
| 95:00:00 | 6exn-D | 2.7 | 30.8 | 77 | 97 | 5 | MOLECULE: | U2 SNRNA | |
| 96:00:00 | 1dce-A | 2.7 | 7.7 | 73 | 567 | 7 | MOLECULE: | PROTEIN (RAB GERANYLGERANYLTRANSFERASE ALPH | A |
| 97:00:00 | 3vr8-D | 2.7 | 9.9 | 99 | 129 | 2 | MOLECULE: | FLAVOPROTEIN SUBUNIT OF COMPLEX II | |
| 98:00:00 | 6xp5-V | 2.7 | 25.2 | 107 | 112 | 9 | MOLECULE: | MEDIATOR OF RNA POLYMERASE II TRANSCRIPTION | SUBUN |
| 99:00:00 | 6jmg-A | 2.7 | 2.9 | 48 | 267 | 13 | MOLECULE: | DNAJ HOMOLOG SUBFAMILY C MEMBER 27-A | |
| 0:00 | 8hhf-B | 2.7 | 7.3 | 58 | 89 | 7 | MOLECULE: | CELL DIVISION PROTEIN FTSQ | |
| 1:00 | 8ifg-A | 2.7 | 26.1 | 88 | 879 | 8 | MOLECULE: | RBAP48-RELATED WD40 REPEAT-CONTAINING PROTE | IN PRW |
| 2:00 | 2pf4-E | 2.7 | 4.8 | 50 | 163 | 6 | MOLECULE: | SERINE/THREONINE-PROTEIN PHOSPHATASE 2A 65 | KDA RE |
| 3:00 | 6v35-A | 2.7 | 23 | 76 | 914 | 4 | MOLECULE: | CALCIUM-ACTIVATED POTASSIUM CHANNEL SUBUNIT | ALPHA |
| 4:00 | 6wvg-A | 2.7 | 21.9 | 84 | 443 | 7 | MOLECULE: | GREEN FLUORESCENT PROTEIN, LEUKOCYTE SURFAC | E ANTI |
| 5:00 | 1tjl-A | 2.7 | 4.5 | 68 | 145 | 9 | MOLECULE: | DNAK SUPPRESSOR PROTEIN | |
| 6:00 | 3jrt-A | 2.7 | 5.4 | 56 | 166 | 7 | MOLECULE: | INTEGRON CASSETTE PROTEIN VPC_CASS2 | |
| 7:00 | 4n9n-A | 2.7 | 28.2 | 79 | 428 | 5 | MOLECULE: | STEROL UPTAKE CONTROL PROTEIN 2, LYSOZYME | |
| 8:00 | 6v85-C | 2.7 | 3.3 | 58 | 74 | 5 | MOLECULE: | RNA-DIRECTED RNA POLYMERASE L | |
| 9:00 | 7bv6-H | 2.7 | 3.1 | 56 | 68 | 7 | MOLECULE: | VESICLE-ASSOCIATED MEMBRANE PROTEIN 8 | |

| | | | | | | | | | | |
|---|---|---|---|---|---|---|---|---|---|---|
| 10:00 | 6abo-B | 2.7 | 5.3 | 59 | 77 | 14 | MOLECULE: | DNA REPAIR PROTEIN XRCC4 | | |
| 11:00 | 6dm9-A | 2.7 | 3.3 | 54 | 78 | 6 | MOLECULE: | DHD15_EXTENDED_A | | |
| 12:00 | 8i3e-B | 2.7 | 30.8 | 61 | 135 | 8 | MOLECULE: | ELKS/RAB6-INTERACTING/CAST FAMILY MEMBER 1 | | |
| 13:00 | 5hzp-A | 2.7 | 2.9 | 54 | 71 | 7 | MOLECULE: | M PROTEIN, SEROTYPE 49 | | |
| 14:00 | 1nyh-A | 2.7 | 3.8 | 55 | 76 | 2 | MOLECULE: | REGULATORY PROTEIN SIR4 | | |
| 15:00 | 5vpc-B | 2.7 | 3.1 | 54 | 68 | 13 | MOLECULE: | PROTEIN FOSB | | |
| 16:00 | 9itj-X | 2.7 | 25.3 | 61 | 155 | 5 | MOLECULE: | ATP SYNTHASE SUBUNIT ALPHA | | |
| 17:00 | 5yyl-D | 2.7 | 5.5 | 41 | 43 | 10 | MOLECULE: | MAJOR ROYAL JELLY PROTEIN 1 | | |
| 18:00 | 7x5e-E | 2.7 | 3.7 | 58 | 101 | 9 | MOLECULE: | TRANSCRIPTION FACTOR MAFG | | |
| 19:00 | 2d3o-W | 2.7 | 4.4 | 50 | 66 | 6 | MOLECULE: | 23S RIBOSOMAL RNA | | |
| 20:00 | 8fxf-A | 2.7 | 4.9 | 57 | 78 | 4 | MOLECULE: | E3 UBIQUITIN-PROTEIN LIGASE TRIM56 | | |
| 21:00 | 4d7x-A | 2.7 | 2.8 | 46 | 86 | 13 | MOLECULE: | MEDIATOR OF RNA POLYMERASE II TRANSCRIPTION | SUBUN | |
| 22:00 | 8kbe-B | 2.7 | 5.4 | 54 | 131 | 4 | MOLECULE: | THOERIS ANTI-DEFENSE 1 | | |
| 23:00 | 5nen-A | 2.7 | 4.3 | 72 | 266 | 3 | MOLECULE: | LIPASE C | | |
| 24:00:00 | 7m2w-F | 2.7 | 6.1 | 71 | 674 | 4 | MOLECULE: | TUBULIN GAMMA CHAIN | | |
| 25:00:00 | 6ny2-Y | 2.7 | 12.7 | 84 | 915 | 7 | MOLECULE: | DNA TARGET STRAND | | |
| 26:00:00 | 7nvr-n | 2.7 | 13 | 76 | 132 | 8 | MOLECULE: | TFIIH BASAL TRANSCRIPTION FACTOR COMPLEX HE | LICASE | |
| 27:00:00 | 7twd-B | 2.7 | 4.3 | 41 | 45 | 7 | MOLECULE: | ALPHA- AND GAMMA-ADAPTIN-BINDING PROTEIN P3 | | 4 |
| 28:00:00 | 6whp-A | 2.6 | 33 | 100 | 428 | 4 | MOLECULE: | CHOLINE KINASE | | |
| 29:00:00 | 8k0b-A | 2.6 | 30.2 | 129 | 689 | 7 | MOLECULE: | CALCIUM PERMEABLE STRESS-GATED CATION CHANN | EL 1 | |
| 30:00:00 | 6bx3-F | 2.6 | 18.6 | 94 | 215 | 5 | MOLECULE: | HISTONE-LYSINE N-METHYLTRANSFERASE, H3 LYSI | NE-4 S | |
| 31:00:00 | 8urn-A | 2.6 | 26.9 | 69 | 152 | 9 | MOLECULE: | ESCI INNER ROD PROTEIN TYPE III SECRETION S | YSTEM, | |
| 32:00:00 | 2p0n-A | 2.6 | 33.5 | 97 | 161 | 7 | MOLECULE: | HYPOTHETICAL PROTEIN NMB1532 | | |
| 33:00:00 | 3nl9-A | 2.6 | 31.4 | 93 | 169 | 3 | MOLECULE: | PUTATIVE NTP PYROPHOSPHOHYDROLASE | | |
| 34:00:00 | 8gaa-A | 2.6 | 8.3 | 50 | 197 | 4 | MOLECULE: | C6HR1_4R | | |
| 35:00:00 | 9f63-A | 2.6 | 20.1 | 103 | 422 | 6 | MOLECULE: | PROTEIN PNS1 | | |
| 36:00:00 | 5yz0-A | 2.6 | 46.6 | 141 | 2362 | 6 | MOLECULE: | SERINE/THREONINE-PROTEIN KINASE ATR | | |
| 37:00:00 | 7sbe-A | 2.6 | 33.3 | 116 | 228 | 8 | MOLECULE: | TELOMERASE REVERSE TRANSCRIPTASE | | |
| 38:00:00 | 6zce-q | 2.6 | 26 | 83 | 636 | 7 | MOLECULE: | EUKARYOTIC TRANSLATION INITIATION FACTOR 3 | SUBUNI | |
| 39:00:00 | 9exk-A | 2.6 | 31.8 | 118 | 1000 | 6 | MOLECULE: | DE NOVO DESIGNED PROTEIN K12 | | |
| 40:00:00 | 8v29-A | 2.6 | 29.7 | 87 | 172 | 5 | MOLECULE: | ONCOSTATIN-M | | |
| 41:00:00 | 7sfj-A | 2.6 | 26.8 | 105 | 284 | 8 | MOLECULE: | CHRMINE | | |
| 42:00:00 | 8dt0-A | 2.6 | 15.3 | 54 | 140 | 7 | MOLECULE: | SCAFFOLDING PROTEIN FUNCTIONAL SITES | | |
| 43:00:00 | 8ssc-A | 2.6 | 28 | 83 | 1123 | 5 | MOLECULE: | METHIONINE SYNTHASE | | |
| 44:00:00 | 8a5a-U | 2.6 | 32.2 | 125 | 690 | 4 | MOLECULE: | CHROMATIN-REMODELING ATPASE INO80 | | |
| 45:00:00 | 8bd7-G | 2.6 | 28.1 | 162 | 205 | 8 | MOLECULE: | IFT88 | | |
| 46:00:00 | 2v71-A | 2.6 | 58.8 | 124 | 160 | 9 | MOLECULE: | NUCLEAR DISTRIBUTION PROTEIN NUDE-LIKE 1 | | |
| 47:00:00 | 8ovw-U | 2.6 | 33.8 | 99 | 184 | 6 | MOLECULE: | CENTROMERE-BINDING PROTEIN 1 | | |
| 48:00:00 | 6zpp-A | 2.6 | 33.3 | 90 | 157 | 3 | MOLECULE: | VIRULENCE FACTOR | | |
| 49:00:00 | 2d1l-A | 2.6 | 21.5 | 119 | 249 | 8 | MOLECULE: | METASTASIS SUPPRESSOR PROTEIN 1 | | |
| 50:00:00 | 8j2x-A | 2.6 | 22.2 | 87 | 337 | 0 | MOLECULE: | COBALAMIN-BINDING PROTEIN | | |
| 51:00:00 | 4wpc-B | 2.6 | 32.6 | 133 | 288 | 6 | MOLECULE: | RHO GTPASE-ACTIVATING PROTEIN RGD1 | | |
| 52:00:00 | 8ibw-C | 2.6 | 19.2 | 111 | 922 | 12 | MOLECULE: | DNA (60-MER) | | |
| 53:00:00 | 3ggz-D | 2.6 | 6.1 | 92 | 188 | 5 | MOLECULE: | INCREASED SODIUM TOLERANCE PROTEIN 1 | | |
| 54:00:00 | 4yif-B | 2.6 | 10 | 79 | 141 | 11 | MOLECULE: | MARR FAMILY PROTEIN RV0880 | | |
| 55:00:00 | 6x0a-a | 2.6 | 13.7 | 63 | 87 | 8 | MOLECULE: | PLASMID STABILIZATION SYSTEM | | |
| 56:00:00 | 8pjn-b | 2.6 | 31 | 104 | 296 | 9 | MOLECULE: | E3 UBIQUITIN-PROTEIN TRANSFERASE RMND5A | | |
| 57:00:00 | 6pwn-A | 2.6 | 18.5 | 147 | 280 | 7 | MOLECULE: | SMALL-CONDUCTANCE MECHANOSENSITIVE CHANNEL | | |
| 58:00:00 | 9knu-M | 2.6 | 8.3 | 76 | 140 | 7 | MOLECULE: | PORTAL PROTEIN | | |
| 59:00:00 | 8ppr-F | 2.6 | 17.4 | 77 | 112 | 5 | MOLECULE: | KINETOCHORE-ASSOCIATED PROTEIN DSN1 HOMOLOG | | |
| 60:00:00 | 8jj6-A | 2.6 | 13.1 | 101 | 554 | 5 | MOLECULE: | NEGATIVE ELONGATION FACTOR B | | |
| 61:00:00 | 8i6v-D | 2.6 | 10.5 | 78 | 678 | 4 | MOLECULE: | VACUOLAR TRANSPORTER CHAPERONE COMPLEX SUBU | NIT 1 | |
| 62:00:00 | 4mi2-B | 2.6 | 22.4 | 92 | 261 | 8 | MOLECULE: | PUTATIVE ENOYL-COA HYDRATASE/ISOMERASE | | |
| 63:00:00 | 5o5j-O | 2.6 | 5 | 69 | 88 | 4 | MOLECULE: | 16S RRNA | | |
| 64:00:00 | 5ajk-A | 2.6 | 3.4 | 41 | 148 | 7 | MOLECULE: | HOMOLOG OF VACCINIA VIRUS CDS F1L | | |
| 65:00:00 | 3dt5-A | 2.6 | 4.2 | 51 | 119 | 10 | MOLECULE: | UNCHARACTERIZED PROTEIN AF_0924 | | |
| 66:00:00 | 6wgv-A | 2.6 | 4 | 48 | 173 | 4 | MOLECULE: | CORRINOID ADENOSYLTRANSFERASE | | |
| 67:00:00 | 3g9g-A | 2.6 | 4.1 | 107 | 250 | 7 | MOLECULE: | SUPPRESSOR OF YEAST PROFILIN DELETION | | |
| 68:00:00 | 1j1n-A | 2.6 | 4.8 | 47 | 492 | 2 | MOLECULE: | ALGQ2 | | |
| 69:00:00 | 5abb-A | 2.6 | 34.2 | 114 | 443 | 5 | MOLECULE: | PROTEIN TRANSLOCASE SUBUNIT SECY | | |
| 70:00:00 | 6o0y-A | 2.6 | 17.9 | 105 | 1146 | 8 | MOLECULE: | CRISPR-ASSOCIATED ENDONUCLEASE CAS9/CSN1 | | |
| 71:00:00 | 8p0v-L | 2.6 | 10.4 | 61 | 416 | 10 | MOLECULE: | COILED-COIL DOMAIN-CONTAINING PROTEIN 93 | | |
| 72:00:00 | 6zz6-C | 2.6 | 27.6 | 52 | 83 | 12 | MOLECULE: | STRUCTURAL MAINTENANCE OF CHROMOSOMES PROTE | IN 1,S | |
| 73:00:00 | 6s6b-B | 2.6 | 3.6 | 61 | 154 | 7 | MOLECULE: | CRISPR-ASSOCIATED PROTEIN, CMR5 FAMILY | | |
| 74:00:00 | 8k0e-A | 2.6 | 17 | 122 | 646 | 4 | MOLECULE: | PROLYL 3-HYDROXYLASE 1 | | |
| 75:00:00 | 8zxc-A | 2.6 | 3 | 47 | 205 | 2 | MOLECULE: | ASH1L BROMODOMAIN AND PHD DOMAIN | | |
| 76:00:00 | 8ul9-A | 2.6 | 20.9 | 112 | 387 | 5 | MOLECULE: | CHOLINEPHOSPHOTRANSFERASE 1 | | |
| 77:00:00 | 7uuy-A | 2.6 | 8.4 | 106 | 503 | 7 | MOLECULE: | SODIUM/IODIDE COTRANSPORTER | | |
| 78:00:00 | 4akf-A | 2.6 | 19.9 | 105 | 560 | 8 | MOLECULE: | VIPD | | |
| 79:00:00 | 6rd4-1 | 2.6 | 17.5 | 106 | 595 | 7 | MOLECULE: | ASA-10: POLYTOMELLA F-ATP SYNTHASE ASSOCIAT | ED SUB | |
| 80:00:00 | 6lu0-A | 2.6 | 4.6 | 71 | 937 | 13 | MOLECULE: | CAS12I2 | | |
| 81:00:00 | 4xal-A | 2.6 | 7.7 | 68 | 90 | 1 | MOLECULE: | TEGUMENT PROTEIN VP22 | | |
| 82:00:00 | 5tuu-B | 2.6 | 5.8 | 50 | 106 | 6 | MOLECULE: | TRANSCRIPTION FACTOR DP1 | | |
| 83:00:00 | 7y7o-A | 2.6 | 3.2 | 39 | 147 | 5 | MOLECULE: | ENDORIBONUCLEASE YBEY | | |
| 84:00:00 | 8e4g-0 | 2.6 | 13.2 | 107 | 196 | 2 | MOLECULE: | PORTAL PROTEIN | | |
| 85:00:00 | 4aur-A | 2.6 | 5.5 | 94 | 564 | 2 | MOLECULE: | LEOA | | |
| 86:00:00 | 5t2x-C | 2.6 | 2.4 | 38 | 258 | 13 | MOLECULE: | UNCHARACTERIZED PROTEIN LPG1670 | | |
| 87:00:00 | 3kdq-C | 2.6 | 11.2 | 76 | 153 | 3 | MOLECULE: | UNCHARACTERIZED CONSERVED PROTEIN | | |
| 88:00:00 | 5ip0-D | 2.6 | 3.9 | 86 | 108 | 8 | MOLECULE: | PHA GRANULE-ASSOCIATED PROTEIN | | |
| 89:00:00 | 4wat-A | 2.6 | 5.4 | 67 | 332 | 9 | MOLECULE: | PFRH5 | | |
| 90:00:00 | 8xlr-A | 2.6 | 18.3 | 128 | 757 | 10 | MOLECULE: | ANOCTAMIN-1 | | |
| 91:00:00 | 6maf-A | 2.6 | 21.3 | 111 | 259 | 5 | MOLECULE: | BBVCI ENDONUCLEASE SUBUNIT 1 | | |
| 92:00:00 | 6uwm-A | 2.6 | 7.7 | 75 | 221 | 4 | MOLECULE: | VOLTAGE-GATED POTASSIUM CHANNEL | | |
| 93:00:00 | 2yo3-C | 2.6 | 32.1 | 106 | 257 | 5 | MOLECULE: | GENERAL CONTROL PROTEIN GCN4, PUTATIVE INNE | R MEMB | |
| 94:00:00 | 8xmd-A | 2.6 | 13.5 | 88 | 1159 | 6 | MOLECULE: | DNA-DIRECTED RNA POLYMERASE IV SUBUNIT 1 | | |
| 95:00:00 | 6cbr-A | 2.6 | 4.2 | 68 | 144 | 9 | MOLECULE: | DNA PRIMASE,SINGLE-STRANDED DNA-BINDING PRO | TEIN C | |
| 96:00:00 | 6pvp-A | 2.6 | 7.4 | 105 | 661 | 10 | MOLECULE: | TRANSIENT RECEPTOR POTENTIAL CATION CHANNEL | SUBFA | |
| 97:00:00 | 7ljn-A | 2.6 | 13.8 | 87 | 342 | 5 | MOLECULE: | CD-NTASE | | |
| 98:00:00 | 5mym-C | 2.6 | 4.9 | 53 | 205 | 6 | MOLECULE: | HTH-TYPE TRANSCRIPTIONAL REGULATOR ETHR | | |
| 99:00:00 | 8wpe-D | 2.6 | 6.7 | 48 | 69 | 10 | MOLECULE: | DNA POLYMERASE | | |
| 0:00 | 2fsf-B | 2.6 | 13.5 | 56 | 723 | 4 | MOLECULE: | PREPROTEIN TRANSLOCASE SECA SUBUNIT | | |
| 1:00 | 3p2c-A | 2.6 | 12.1 | 88 | 445 | 8 | MOLECULE: | PUTATIVE GLYCOSYL HYDROLASE | | |
| 2:00 | 7wze-A | 2.6 | 5.4 | 62 | 161 | 5 | MOLECULE: | UNCHARACTERIZED HTH-TYPE TRANSCRIPTIONAL RE | GULATO | |
| 3:00 | 7l7v-A | 2.6 | 4.7 | 49 | 121 | 10 | MOLECULE: | PROBABLE DISEASE RESISTANCE PROTEIN AT5G669 | | 0 |
| 4:00 | 8tvh-A | 2.6 | 5.3 | 53 | 86 | 6 | MOLECULE: | 4G5 LIGHT CHAIN | | |
| 5:00 | 6iu4-A | 2.6 | 7.6 | 72 | 225 | 4 | MOLECULE: | VIT1 | | |
| 6:00 | 7xe1-A | 2.6 | 9.3 | 65 | 751 | 5 | MOLECULE: | LYSINE-SPECIFIC HISTONE DEMETHYLASE 1B | | |
| 7:00 | 4gnk-E | 2.6 | 4.8 | 100 | 235 | 1 | MOLECULE: | GUANINE NUCLEOTIDE-BINDING PROTEIN G(Q) SUB | UNIT A | |
| 8:00 | 5a5t-E | 2.6 | 13.9 | 91 | 419 | 4 | MOLECULE: | EUKARYOTIC TRANSLATION INITIATION FACTOR 3 | SUBUNI | |
| 9:00 | 3a6m-A | 2.6 | 11.3 | 90 | 168 | 8 | MOLECULE: | PROTEIN GRPE | | |
| 10:00 | 8ih5-A | 2.6 | 5.1 | 75 | 689 | 5 | MOLECULE: | SYN-COPALYL DIPHOSPHATE SYNTHASE, CHLOROPLA | STIC | |

| | | | | | | | | | | |
|---|---|---|---|---|---|---|---|---|---|---|
| 11:00 | 9b9m-B | 2.6 | 9.6 | 59 | 329 | 3 | MOLECULE: | PYRROLOQUINOLINE QUINONE (COENZYME PQQ) BIO | SYNTHE | |
| 12:00 | 6w98-A | 2.6 | 16.2 | 90 | 1312 | 6 | MOLECULE: | F5/8 TYPE C DOMAIN-CONTAINING PROTEIN | | |
| 13:00 | 4di3-A | 2.6 | 17.6 | 66 | 273 | 9 | MOLECULE: | TP33 PROTEIN | | |
| 14:00 | 8tzj-C | 2.6 | 3.7 | 77 | 203 | 8 | MOLECULE: | CELL DIVISION ATP-BINDING PROTEIN FTSE | | |
| 15:00 | 5xfa-A | 2.6 | 9.9 | 94 | 581 | 7 | MOLECULE: | NAD-REDUCING HYDROGENASE | | |
| 16:00 | 6lcn-F | 2.6 | 10.1 | 72 | 309 | 4 | MOLECULE: | SERINE O-ACETYLTRANSFERASE | | |
| 17:00 | 7uus-Q | 2.6 | 34.2 | 109 | 170 | 6 | MOLECULE: | HYDROGENASE-2, LARGE SUBUNIT | | |
| 18:00 | 5vmn-A | 2.6 | 3.3 | 46 | 125 | 0 | MOLECULE: | BAK PROTEIN | | |
| 19:00 | 7xyb-G | 2.6 | 6.5 | 61 | 168 | 8 | MOLECULE: | DNA-DIRECTED RNA POLYMERASE SUBUNIT ALPHA | | |
| 20:00 | 8s37-B | 2.6 | 27.7 | 113 | 343 | 6 | MOLECULE: | CRISPR TYPE AFERR-ASSOCIATED PROTEIN CSF2 | | |
| 21:00 | 8p65-J | 2.6 | 3.2 | 51 | 64 | 6 | MOLECULE: | CYTOCHROME B-C1 COMPLEX SUBUNIT 1, MITOCHON | DRIAL | |
| 22:00 | 5k8o-B | 2.6 | 7 | 51 | 425 | 8 | MOLECULE: | 5-NITROANTHRANILIC ACID AMINOHYDROLASE | | |
| 23:00 | 8d7h-D | 2.6 | 8.8 | 66 | 178 | 3 | MOLECULE: | CILIARY NEUROTROPHIC FACTOR RECEPTOR SUBUNI | T ALPH | |
| 24:00:00 | 5tk6-A | 2.6 | 7.8 | 74 | 191 | 7 | MOLECULE: | OXSA PROTEIN | | |
| 25:00:00 | 5nvu-A | 2.6 | 29 | 127 | 3169 | 0 | MOLECULE: | DYNEIN MOTOR DOMAIN | | |
| 26:00:00 | 8yzt-C | 2.6 | 3.2 | 44 | 121 | 11 | MOLECULE: | PROTEIN BANP | | |
| 27:00:00 | 6xrw-B | 2.6 | 7.7 | 56 | 79 | 0 | MOLECULE: | PLASMID STABILISATION SYSTEM PROTEIN | | |
| 28:00:00 | 8hk0-C | 2.6 | 8.1 | 105 | 346 | 5 | MOLECULE: | DEHYDROGENASE | | |
| 29:00:00 | 8zl2-A | 2.6 | 6.7 | 54 | 542 | 6 | MOLECULE: | PROTEIN I'M NOT DEAD YET | | |
| 30:00:00 | 5opt-P | 2.6 | 18.3 | 62 | 249 | 8 | MOLECULE: | ACTIVATED PROTEIN KINASE C RECEPTOR, PUTATI | VE | |
| 31:00:00 | 6m9t-A | 2.6 | 4.3 | 101 | 444 | 8 | MOLECULE: | PROSTAGLANDIN E2 RECEPTOR EP3 SUBTYPE, ENDO | LYSIN | |
| 32:00:00 | 2x0l-A | 2.6 | 42.4 | 122 | 670 | 11 | MOLECULE: | LYSINE-SPECIFIC HISTONE DEMETHYLASE 1 | | |
| 33:00:00 | 2p5g-B | 2.6 | 2.7 | 40 | 756 | 18 | MOLECULE: | TEMPLATE DNA | | |
| 34:00:00 | 8i03-F | 2.6 | 25.1 | 94 | 228 | 7 | MOLECULE: | PAIRED AMPHIPATHIC HELIX PROTEIN PST1 | | |
| 35:00:00 | 8p0w-H | 2.6 | 10.6 | 76 | 183 | 7 | MOLECULE: | COMM DOMAIN-CONTAINING PROTEIN 1 | | |
| 36:00:00 | 6k9k-A | 2.6 | 14.5 | 125 | 2501 | 9 | MOLECULE: | SERINE-PROTEIN KINASE ATM | | |
| 37:00:00 | 3omb-A | 2.6 | 5.8 | 49 | 503 | 6 | MOLECULE: | EXTRACELLULAR SOLUTE-BINDING PROTEIN, FAMIL | Y 1 | |
| 38:00:00 | 2epo-A | 2.6 | 18.1 | 88 | 626 | 8 | MOLECULE: | N-ACETYL-BETA-D-GLUCOSAMINIDASE | | |
| 39:00:00 | 2qqy-A | 2.6 | 6.1 | 61 | 139 | 5 | MOLECULE: | SIGMA B OPERON | | |
| 40:00:00 | 8tcb-C | 2.6 | 2.6 | 54 | 80 | 2 | MOLECULE: | C4B-BINDING PROTEIN ALPHA CHAIN | | |
| 41:00:00 | 5djn-C | 2.6 | 3.5 | 59 | 81 | 7 | MOLECULE: | KINESIN-LIKE PROTEIN | | |
| 42:00:00 | 6wsm-A | 2.6 | 3.1 | 53 | 74 | 9 | MOLECULE: | SEPTIN-8 | | |
| 43:00:00 | 4dhx-A | 2.6 | 4.4 | 56 | 72 | 4 | MOLECULE: | 80 KDA MCM3-ASSOCIATED PROTEIN | | |
| 44:00:00 | 5hda-A | 2.6 | 17 | 63 | 121 | 10 | MOLECULE: | ZINC FINGER MYND DOMAIN-CONTAINING PROTEIN | | 11 |
| 45:00:00 | 2i7u-A | 2.6 | 3.5 | 50 | 62 | 8 | MOLECULE: | FOUR-ALPHA-HELIX BUNDLE | | |
| 46:00:00 | 6y09-C | 2.6 | 2.9 | 59 | 86 | 5 | MOLECULE: | RAS-RELATED PROTEIN RAB-33B | | |
| 47:00:00 | 3iv1-A | 2.6 | 3.9 | 58 | 78 | 7 | MOLECULE: | TUMOR SUSCEPTIBILITY GENE 101 PROTEIN | | |
| 48:00:00 | 4h22-B | 2.6 | 3.6 | 57 | 83 | 11 | MOLECULE: | LEUCINE-RICH REPEAT FLIGHTLESS-INTERACTING | PROTEI | |
| 49:00:00 | 5xmk-G | 2.6 | 5.6 | 49 | 54 | 6 | MOLECULE: | VACUOLAR PROTEIN SORTING-ASSOCIATED PROTEIN | | 4 |
| 50:00:00 | 2vrz-A | 2.6 | 4.6 | 59 | 98 | 8 | MOLECULE: | VIRULENCE FACTOR ESXA | | |
| 51:00:00 | 2nr5-E | 2.6 | 2.6 | 45 | 65 | 0 | MOLECULE: | HYPOTHETICAL PROTEIN SO2669 | | |
| 52:00:00 | 4djg-A | 2.6 | 4.7 | 44 | 49 | 0 | MOLECULE: | PLECTIN-RELATED PROTEIN | | |
| 53:00:00 | 3f0c-A | 2.6 | 2.4 | 44 | 193 | 5 | MOLECULE: | TRANSCRIPTIONAL REGULATOR | | |
| 54:00:00 | 7y3f-1 | 2.6 | 2.5 | 53 | 466 | 2 | MOLECULE: | PHOTOSYSTEM I P700 CHLOROPHYLL A APOPROTEIN | A1 | |
| 55:00:00 | 8djk-A | 2.6 | 4.1 | 54 | 245 | 2 | MOLECULE: | 3-HYDROXY-3-METHYLGLUTARYL-COENZYME A REDUC | TASE | |
| 56:00:00 | 8on7-A | 2.6 | 7.5 | 57 | 525 | 4 | MOLECULE: | FMRFAMIDE-GATED SODIUM CHANNEL 1 (FANAC1) | | |
| 57:00:00 | 3c98-B | 2.6 | 7.9 | 88 | 230 | 7 | MOLECULE: | SYNTAXIN-BINDING PROTEIN 1 | | |
| 58:00:00 | 2fup-A | 2.6 | 4.3 | 50 | 128 | 2 | MOLECULE: | HYPOTHETICAL PROTEIN PA3352 | | |
| 59:00:00 | 6yj4-L | 2.5 | 26.2 | 124 | 655 | 7 | MOLECULE: | NADH-UBIQUINONE OXIDOREDUCTASE CHAIN 3 | | |
| 60:00:00 | 4wpe-A | 2.5 | 32 | 127 | 275 | 8 | MOLECULE: | CYTOKINESIS PROTEIN 2 | | |
| 61:00:00 | 1wgz-A | 2.5 | 27.3 | 129 | 510 | 7 | MOLECULE: | CARBOXYPEPTIDASE 1 | | |
| 62:00:00 | 7khw-a | 2.5 | 20.2 | 68 | 174 | 3 | MOLECULE: | TRANSLOCON ESPA | | |
| 63:00:00 | 4ihq-A | 2.5 | 27 | 73 | 512 | 7 | MOLECULE: | FLAI ATPASE | | |
| 64:00:00 | 6tdv-H | 2.5 | 31.6 | 126 | 388 | 6 | MOLECULE: | ATPTB1 | | |
| 65:00:00 | 8bfi-B | 2.5 | 28.3 | 123 | 574 | 9 | MOLECULE: | CCR4-NOT TRANSCRIPTION COMPLEX SUBUNIT 1 | | |
| 66:00:00 | 4ymk-A | 2.5 | 13.3 | 73 | 322 | 5 | MOLECULE: | ACYL-COA DESATURASE 1 | | |
| 67:00:00 | 3ksy-A | 2.5 | 21.6 | 147 | 1007 | 11 | MOLECULE: | SON OF SEVENLESS HOMOLOG 1 | | |
| 68:00:00 | 4zuz-B | 2.5 | 28.5 | 110 | 858 | 6 | MOLECULE: | SIDC | | |
| 69:00:00 | 7mdy-B | 2.5 | 31.9 | 103 | 411 | 5 | MOLECULE: | LIPOPROTEIN TRANSPORTER SUBUNIT LOLE | | |
| 70:00:00 | 5h7b-A | 2.5 | 11 | 58 | 230 | 12 | MOLECULE: | IMMUNOGLOBULIN G-BINDING PROTEIN A | | |
| 71:00:00 | 5wgr-A | 2.5 | 29 | 116 | 664 | 6 | MOLECULE: | FLAVIN-DEPENDENT HALOGENASE | | |
| 72:00:00 | 7p5j-A | 2.5 | 35.1 | 87 | 399 | 6 | MOLECULE: | PROTEIN TWEETY HOMOLOG 1 | | |
| 73:00:00 | 8wcn-A | 2.5 | 29.8 | 123 | 379 | 7 | MOLECULE: | DIGUANYLATE CYCLASE | | |
| 74:00:00 | 8d8j-d | 2.5 | 26.1 | 90 | 660 | 6 | MOLECULE: | PROBABLE S-ADENOSYL-L-METHIONINE-DEPENDENT | RNA | |
| 75:00:00 | 7e40-D | 2.5 | 26.7 | 113 | 345 | 7 | MOLECULE: | PROTEIN PHOSPHATE STARVATION RESPONSE 2 | | |
| 76:00:00 | 4uxv-A | 2.5 | 35.7 | 116 | 522 | 7 | MOLECULE: | SEPTATION RING FORMATION REGULATOR EZRA | | |
| 77:00:00 | 7bv6-B | 2.5 | 23 | 45 | 61 | 4 | MOLECULE: | VESICLE-ASSOCIATED MEMBRANE PROTEIN 8 | | |
| 78:00:00 | 2pv4-A | 2.5 | 16.2 | 45 | 145 | 2 | MOLECULE: | UNCHARACTERIZED PROTEIN | | |
| 79:00:00 | 6yuf-D | 2.5 | 32.9 | 119 | 1272 | 5 | MOLECULE: | COHESIN SUBUNIT RAD21 | | |
| 80:00:00 | 7zqy-B | 2.5 | 12.6 | 71 | 177 | 10 | MOLECULE: | DH DOMAIN-CONTAINING PROTEIN | | |
| 81:00:00 | 6lkc-B | 2.5 | 17.8 | 65 | 533 | 9 | MOLECULE: | POLYUNSATURATED FATTY ACID SYNTHASE PFAD | | |
| 82:00:00 | 6xm1-D | 2.5 | 33.6 | 100 | 181 | 5 | MOLECULE: | VPS45 | | |
| 83:00:00 | 3etz-B | 2.5 | 17.7 | 72 | 115 | 7 | MOLECULE: | ADHESIN A | | |
| 84:00:00 | 7wji-C | 2.5 | 45.9 | 149 | 1394 | 3 | MOLECULE: | PROTEIN UNC-80 HOMOLOG | | |
| 85:00:00 | 4m9s-B | 2.5 | 20.2 | 102 | 511 | 5 | MOLECULE: | CELL DEATH PROTEIN 4 | | |
| 86:00:00 | 7tn2-W | 2.5 | 14.2 | 61 | 895 | 10 | MOLECULE: | HISTONE H3 | | |
| 87:00:00 | 1knz-A | 2.5 | 9.4 | 59 | 154 | 8 | MOLECULE: | 5'-R(*UP*GP*AP*CP*C)-3' | | |
| 88:00:00 | 8arl-A | 2.5 | 5.6 | 99 | 232 | 12 | MOLECULE: | TRYPTOPHAN-RICH ANTIGEN | | |
| 89:00:00 | 6v6d-A | 2.5 | 5.4 | 86 | 209 | 5 | MOLECULE: | PANNEXIN-1 | | |
| 90:00:00 | 5me8-A | 2.5 | 5.6 | 75 | 107 | 7 | MOLECULE: | INHIBITOR OF GROWTH PROTEIN 5 | | |
| 91:00:00 | 6o8b-B | 2.5 | 5.5 | 99 | 657 | 7 | MOLECULE: | STIMULATOR OF INTERFERON GENES PROTEIN | | |
| 92:00:00 | 6jfk-A | 2.5 | 22 | 109 | 428 | 4 | MOLECULE: | MITOFUSIN-2,CDNA FLJ57997, HIGHLY SIMILAR T | O TRAN | |
| 93:00:00 | 4w4u-B | 2.5 | 4 | 46 | 96 | 13 | MOLECULE: | UBIQUITIN CARBOXYL-TERMINAL HYDROLASE | | |
| 94:00:00 | 3f42-B | 2.5 | 2.1 | 40 | 84 | 5 | MOLECULE: | PROTEIN HP0035 | | |
| 95:00:00 | 8kg9-E | 2.5 | 24.5 | 100 | 568 | 7 | MOLECULE: | DNA REPLICATION LICENSING FACTOR MCM2 | | |
| 96:00:00 | 6hcy-A | 2.5 | 9.5 | 116 | 436 | 3 | MOLECULE: | METALLOREDUCTASE STEAP4 | | |
| 97:00:00 | 3j9t-P | 2.5 | 21.6 | 90 | 461 | 3 | MOLECULE: | V-TYPE PROTON ATPASE SUBUNIT D | | |
| 98:00:00 | 9d80-C | 2.5 | 21.7 | 93 | 275 | 6 | MOLECULE: | PORTAL PROTEIN | | |
| 99:00:00 | 6inw-B | 2.5 | 14 | 59 | 391 | 7 | MOLECULE: | O-METHYLTRANSFERASE LEPI | | |
| 0:00 | 6k3b-B | 2.5 | 3.9 | 50 | 106 | 8 | MOLECULE: | LPG2147 | | |
| 1:00 | 7zn7-D | 2.5 | 5.8 | 61 | 128 | 7 | MOLECULE: | DNA DAMAGE-BINDING PROTEIN 1 | | |
| 2:00 | 5c8d-E | 2.5 | 8.4 | 73 | 280 | 3 | MOLECULE: | LIGHT-DEPENDENT TRANSCRIPTIONAL REGULATOR C | ARH | |
| 3:00 | 7apk-m | 2.5 | 44.3 | 160 | 549 | 8 | MOLECULE: | THO COMPLEX SUBUNIT 1 | | |
| 4:00 | 3oop-A | 2.5 | 9.6 | 77 | 139 | 4 | MOLECULE: | LIN2960 PROTEIN | | |
| 5:00 | 5fia-B | 2.5 | 6.3 | 67 | 397 | 9 | MOLECULE: | LPIR1 | | |
| 6:00 | 1oxz-A | 2.5 | 4.1 | 88 | 132 | 7 | MOLECULE: | ADP-RIBOSYLATION FACTOR BINDING PROTEIN GGA | | 1 |
| 7:00 | 1t7s-A | 2.5 | 11.1 | 85 | 129 | 5 | MOLECULE: | BAG-1 COCHAPERONE | | |
| 8:00 | 7nhr-A | 2.5 | 11 | 74 | 663 | 3 | MOLECULE: | PUTATIVE TRANSMEMBRANE PROTEIN WZC | | |
| 9:00 | 2bb6-A | 2.5 | 18.4 | 80 | 414 | 9 | MOLECULE: | TRANSCOBALAMIN II | | |
| 10:00 | 6rd4-5 | 2.5 | 42.7 | 92 | 123 | 7 | MOLECULE: | ASA-10: POLYTOMELLA F-ATP SYNTHASE ASSOCIAT | ED SUB | |
| 11:00 | 7ugc-A | 2.5 | 17.3 | 71 | 191 | 1 | MOLECULE: | PROTEIN VIAA | | |

| | | | | | | | | | | |
|---|---|---|---|---|---|---|---|---|---|---|
| 12:00 | 2pjw-H | 2.5 | 4 | 48 | 88 | 6 | MOLECULE: | UNCHARACTERIZED PROTEIN YHL002W | | |
| 13:00 | 6txb-A | 2.5 | 6.4 | 65 | 263 | 3 | MOLECULE: | UBIQUITIN CARBOXYL-TERMINAL HYDROLASE MINDY | | -1 |
| 14:00 | 6thk-A | 2.5 | 9.7 | 115 | 466 | 6 | MOLECULE: | PYOCIN S5 | | |
| 15:00 | 6rjw-A | 2.5 | 5.9 | 72 | 161 | 10 | MOLECULE: | LYSM DOMAIN PROTEIN | | |
| 16:00 | 7c7e-A | 2.5 | 3.9 | 61 | 142 | 10 | MOLECULE: | PUTATIVE DNA-BINDING TRANSCRIPTIONAL REGULA | TOR | |
| 17:00 | 7emf-H | 2.5 | 10.1 | 106 | 181 | 7 | MOLECULE: | MEDIATOR OF RNA POLYMERASE II TRANSCRIPTION | SUBUN | |
| 18:00 | 3fcn-A | 2.5 | 6.1 | 62 | 158 | 6 | MOLECULE: | AN ALPHA-HELICAL PROTEIN OF UNKNOWN FUNCTIO | N (PFA | |
| 19:00 | 4nqw-B | 2.5 | 7 | 55 | 73 | 5 | MOLECULE: | ECF RNA POLYMERASE SIGMA FACTOR SIGK | | |
| 20:00 | 9f5y-I | 2.5 | 14 | 54 | 135 | 9 | MOLECULE: | NADH:UBIQUINONE OXIDOREDUCTASE 24 KD SUBUNI | T | |
| 21:00 | 8urw-Z | 2.5 | 8.3 | 69 | 1235 | 6 | MOLECULE: | DNA-DIRECTED RNA POLYMERASE SUBUNIT ALPHA | | |
| 22:00 | 6wgm-A | 2.5 | 11.8 | 74 | 305 | 5 | MOLECULE: | TRAP-TYPE C4-DICARBOXYLATE TRANSPORT SYSTEM | , PERI | |
| 23:00 | 7w0y-i | 2.5 | 7.5 | 92 | 347 | 8 | MOLECULE: | NADH DEHYDROGENASE [UBIQUINONE] FLAVOPROTEI | N 1, | |
| 24:00:00 | 6n39-A | 2.5 | 18.3 | 88 | 404 | 6 | MOLECULE: | DEPHOSPHO-COA KINASE | | |
| 25:00:00 | 8dft-A | 2.5 | 5 | 83 | 112 | 6 | MOLECULE: | PILIN PROTEIN | | |
| 26:00:00 | 8fn4-4 | 2.5 | 30.5 | 107 | 741 | 2 | MOLECULE: | RNA-EDITING SUBSTRATE-BINDING COMPLEX PROTE | IN 1 ( | |
| 27:00:00 | 7lhe-A | 2.5 | 39 | 155 | 2222 | 10 | MOLECULE: | INOSITOL 1,4,5-TRISPHOSPHATE RECEPTOR TYPE | | 1 |
| 28:00:00 | 4kk0-A | 2.5 | 11.1 | 83 | 403 | 4 | MOLECULE: | TUBEROUS SCLEROSIS 1 PROTEIN HOMOLOG | | |
| 29:00:00 | 9iz7-A | 2.5 | 48.4 | 162 | 2611 | 7 | MOLECULE: | SERINE/THREONINE-PROTEIN KINASE TEL1 | | |
| 30:00:00 | 2g0u-A | 2.5 | 4.2 | 67 | 92 | 4 | MOLECULE: | TYPE III SECRETION SYSTEM NEEDLE PROTEIN | | |
| 31:00:00 | 6n63-A | 2.5 | 5.8 | 86 | 141 | 3 | MOLECULE: | ENCAPSULIN CARGO PROTEIN | | |
| 32:00:00 | 3ajb-A | 2.5 | 2.8 | 52 | 293 | 2 | MOLECULE: | PEROXISOMAL BIOGENESIS FACTOR 3 | | |
| 33:00:00 | 9d80-B | 2.5 | 11.6 | 83 | 234 | 4 | MOLECULE: | PORTAL PROTEIN | | |
| 34:00:00 | 2pzi-A | 2.5 | 20.9 | 62 | 654 | 10 | MOLECULE: | PROBABLE SERINE/THREONINE-PROTEIN KINASE PK | NG | |
| 35:00:00 | 6qyi-B | 2.5 | 11.6 | 78 | 519 | 8 | MOLECULE: | 4-HYDROXYPHENYLACETATE 3-MONOOXYGENASE OXYG | ENASE | |
| 36:00:00 | 5cgz-A | 2.5 | 7.1 | 51 | 240 | 8 | MOLECULE: | 4-OXALMESACONATE HYDRATASE | | |
| 37:00:00 | 6v92-R | 2.5 | 20 | 156 | 325 | 5 | MOLECULE: | ACTIN-RELATED PROTEIN 7 | | |
| 38:00:00 | 5mx5-L | 2.5 | 5.4 | 86 | 214 | 7 | MOLECULE: | PROTEASOME ACTIVATOR COMPLEX SUBUNIT 1 | | |
| 39:00:00 | 2hsz-A | 2.5 | 6.6 | 59 | 225 | 10 | MOLECULE: | NOVEL PREDICTED PHOSPHATASE | | |
| 40:00:00 | 8c0v-B | 2.5 | 22 | 86 | 1030 | 10 | MOLECULE: | PEROXISOMAL ATPASE PEX1 | | |
| 41:00:00 | 6whg-A | 2.5 | 6.7 | 104 | 523 | 8 | MOLECULE: | CALCIUM CHANNEL YVC1 | | |
| 42:00:00 | 8pnu-A | 2.5 | 7.2 | 61 | 155 | 5 | MOLECULE: | STYRENE OXIDE ISOMERASE | | |
| 43:00:00 | 9im6-A | 2.5 | 22.6 | 84 | 1095 | 5 | MOLECULE: | IMPORTIN-5 | | |
| 44:00:00 | 6ezn-B | 2.5 | 5.8 | 80 | 110 | 1 | MOLECULE: | DOLICHYL-DIPHOSPHOOLIGOSACCHARIDE--PROTEIN | | |
| 45:00:00 | 2zqm-A | 2.5 | 2.8 | 76 | 114 | 7 | MOLECULE: | PREFOLDIN BETA SUBUNIT 1 | | |
| 46:00:00 | 2hh6-A | 2.5 | 2.3 | 42 | 112 | 10 | MOLECULE: | BH3980 PROTEIN | | |
| 47:00:00 | 8rc4-o | 2.5 | 5.7 | 78 | 372 | 5 | MOLECULE: | SERINE/THREONINE-PROTEIN PHOSPHATASE 2A 65 | KDA RE | |
| 48:00:00 | 7xsq-A | 2.5 | 24.7 | 94 | 1289 | 6 | MOLECULE: | RAMP SUPERFAMILY PROTEIN | | |
| 49:00:00 | 7mwa-A | 2.5 | 6.2 | 63 | 383 | 0 | MOLECULE: | 2-POLYPRENYL-6-METHOXYPHENOL 4-HYDROXYLASE | | |
| 50:00:00 | 7w9l-A | 2.5 | 20.8 | 120 | 1422 | 8 | MOLECULE: | SODIUM CHANNEL PROTEIN TYPE 9 SUBUNIT ALPHA | | |
| 51:00:00 | 9f5y-U | 2.5 | 7.6 | 82 | 105 | 1 | MOLECULE: | NADH:UBIQUINONE OXIDOREDUCTASE 24 KD SUBUNI | T | |
| 52:00:00 | 8rbz-j | 2.5 | 13.8 | 96 | 668 | 4 | MOLECULE: | UNK-UNK-UNK-UNK-UNK-UNK-UNK-UNK-UNK | | |
| 53:00:00 | 4ams-A | 2.5 | 8.3 | 86 | 367 | 7 | MOLECULE: | MG662 | | |
| 54:00:00 | 7zr6-C | 2.5 | 30 | 130 | 277 | 5 | MOLECULE: | HEAT SHOCK PROTEIN HSP 90-BETA | | |
| 55:00:00 | 3p8c-D | 2.5 | 21.9 | 121 | 204 | 5 | MOLECULE: | CYTOPLASMIC FMR1-INTERACTING PROTEIN 1 | | |
| 56:00:00 | 7qla-C | 2.5 | 18.7 | 76 | 441 | 9 | MOLECULE: | VACUOLAR FUSION PROTEIN MON1 | | |
| 57:00:00 | 3aai-A | 2.5 | 3.2 | 63 | 78 | 8 | MOLECULE: | COPPER HOMEOSTASIS OPERON REGULATORY PROTEI | N | |
| 58:00:00 | 5yfp-D | 2.5 | 18.8 | 95 | 954 | 6 | MOLECULE: | EXOCYST COMPLEX COMPONENT SEC3 | | |
| 59:00:00 | 8j1p-A | 2.5 | 32.5 | 95 | 1234 | 5 | MOLECULE: | UBIQUITIN FUSION DEGRADATION PROTEIN 4 | | |
| 60:00:00 | 1zgh-A | 2.5 | 10.3 | 47 | 227 | 13 | MOLECULE: | METHIONYL-TRNA FORMYLTRANSFERASE | | |
| 61:00:00 | 8g0l-B | 2.5 | 7 | 88 | 925 | 7 | MOLECULE: | N-ALPHA-ACETYLTRANSFERASE 20 | | |
| 62:00:00 | 6edw-B | 2.5 | 19.5 | 109 | 746 | 6 | MOLECULE: | ISOCITRATE LYASE 2 | | |
| 63:00:00 | 8e1m-C | 2.5 | 11.5 | 81 | 264 | 7 | MOLECULE: | MITOCHONDRIAL IMPORT INNER MEMBRANE TRANSLO | CASE S | |
| 64:00:00 | 4m75-A | 2.5 | 4.5 | 43 | 139 | 16 | MOLECULE: | U6 SNRNA-ASSOCIATED SM-LIKE PROTEIN LSM1 | | |
| 65:00:00 | 8gl3-A | 2.5 | 4.1 | 92 | 189 | 5 | MOLECULE: | PRLB-519 | | |
| 66:00:00 | 7bc6-A | 2.5 | 15.6 | 82 | 432 | 6 | MOLECULE: | PROTON-COUPLED FOLATE TRANSPORTER | | |
| 67:00:00 | 6gvw-F | 2.5 | 29.9 | 82 | 320 | 7 | MOLECULE: | BRCA1-A COMPLEX SUBUNIT ABRAXAS 1 | | |
| 68:00:00 | 8pmq-2 | 2.5 | 11.8 | 96 | 355 | 7 | MOLECULE: | E3 UBIQUITIN-PROTEIN LIGASE RMD5 | | |
| 69:00:00 | 7wwb-A | 2.5 | 14.2 | 53 | 333 | 0 | MOLECULE: | CHOLINE TRANSPORTER-LIKE PROTEIN 1 | | |
| 70:00:00 | 6rgn-A | 2.5 | 3.8 | 55 | 93 | 4 | MOLECULE: | DUF3120 DOMAIN-CONTAINING PROTEIN | | |
| 71:00:00 | 6qo1-A | 2.5 | 4.6 | 88 | 186 | 5 | MOLECULE: | PUTATIVE SURFACE PROTEIN | | |
| 72:00:00 | 6c0g-A | 2.5 | 5 | 48 | 114 | 4 | MOLECULE: | LYSINOALANINE SYNTHASE | | |
| 73:00:00 | 6wlw-0 | 2.5 | 3.8 | 69 | 204 | 3 | MOLECULE: | V-TYPE PROTON ATPASE 21 KDA PROTEOLIPID SUB | UNIT | |
| 74:00:00 | 8iff-A | 2.5 | 10.9 | 99 | 868 | 8 | MOLECULE: | PHYTOCHROME A | | |
| 75:00:00 | 7xzj-G | 2.5 | 4.4 | 55 | 73 | 2 | MOLECULE: | CTAP3 | | |
| 76:00:00 | 6mj1-A | 2.5 | 9.6 | 73 | 200 | 7 | MOLECULE: | PROBABLE HTH-TYPE TRANSCRIPTIONAL REGULATOR | YTTP | |
| 77:00:00 | 7d60-B | 2.5 | 20.9 | 101 | 288 | 7 | MOLECULE: | CALCIUM HOMEOSTASIS MODULATOR PROTEIN 5 | | |
| 78:00:00 | 6r6h-O | 2.5 | 30.5 | 90 | 744 | 3 | MOLECULE: | COP9 SIGNALOSOME COMPLEX SUBUNIT 1 | | |
| 79:00:00 | 5no8-A | 2.5 | 13.6 | 73 | 669 | 4 | MOLECULE: | BACCELL_00875 | | |
| 80:00:00 | 6s2e-A | 2.5 | 26.7 | 100 | 1006 | 7 | MOLECULE: | DNA POLYMERASE EPSILON CATALYTIC SUBUNIT A | | |
| 81:00:00 | 6tdw-C | 2.5 | 3.5 | 48 | 157 | 6 | MOLECULE: | ATPTB1 | | |
| 82:00:00 | 6fpg-F | 2.5 | 9.2 | 78 | 266 | 8 | MOLECULE: | CHROMOSOME 16, WHOLE GENOME SHOTGUN SEQUEN | E | |
| 83:00:00 | 2oap-1 | 2.5 | 3.1 | 38 | 498 | 5 | MOLECULE: | TYPE II SECRETION SYSTEM PROTEIN | | |
| 84:00:00 | 6n8a-B | 2.5 | 4.2 | 69 | 194 | 7 | MOLECULE: | TRANSCRIPTION REGULATOR ACAB | | |
| 85:00:00 | 2pgc-A | 2.5 | 15.6 | 58 | 206 | 2 | MOLECULE: | UNCHARACTERIZED PROTEIN | | |
| 86:00:00 | 5ex8-A | 2.5 | 5.4 | 71 | 414 | 13 | MOLECULE: | CYTOCHROME P450 | | |
| 87:00:00 | 8w8q-C | 2.5 | 7.6 | 79 | 287 | 4 | MOLECULE: | SOLUBLE CYTOCHROME B562,PROBABLE G-PROTEIN | COUPLE | |
| 88:00:00 | 3kkz-A | 2.5 | 7.8 | 58 | 257 | 3 | MOLECULE: | UNCHARACTERIZED PROTEIN Q5LES9 | | |
| 89:00:00 | 7edx-k | 2.5 | 8.7 | 69 | 98 | 7 | MOLECULE: | TRANSCRIPTION INITIATION FACTOR TFIID SUBUN | IT 1 | |
| 90:00:00 | 3jav-A | 2.5 | 32.7 | 152 | 868 | 8 | MOLECULE: | INOSITOL 1,4,5-TRISPHOSPHATE RECEPTOR TYPE | | 1 |
| 91:00:00 | 8ik3-A | 2.5 | 19.1 | 113 | 328 | 4 | MOLECULE: | STIMULATOR OF INTERFERON GENES PROTEIN,IMMU | NE PRO | |
| 92:00:00 | 7xl9-A | 2.5 | 11.7 | 88 | 164 | 6 | MOLECULE: | TRANSCRIPTIONAL REGULATOR, MARR FAMILY | | |
| 93:00:00 | 7ldg-A | 2.5 | 9.4 | 70 | 220 | 4 | MOLECULE: | HEAT SHOCK FACTOR 2-BINDING PROTEIN | | |
| 94:00:00 | 2oeq-C | 2.5 | 14.9 | 76 | 117 | 5 | MOLECULE: | PROTEIN OF UNKNOWN FUNCTION, DUF964 | | |
| 95:00:00 | 2o26-A | 2.5 | 6.5 | 73 | 144 | 5 | MOLECULE: | KIT LIGAND | | |
| 96:00:00 | 6nd4-M | 2.5 | 12.5 | 94 | 431 | 5 | MOLECULE: | ETS RRNA | | |
| 97:00:00 | 7nb6-A | 2.5 | 7.9 | 81 | 340 | 4 | MOLECULE: | AI-2 TRANSPORT PROTEIN TQSA | | |
| 98:00:00 | 6h3d-A | 2.5 | 5.3 | 48 | 429 | 4 | MOLECULE: | DUF2338 DOMAIN-CONTAINING PROTEIN | | |
| 99:00:00 | 3b8m-A | 2.5 | 7.4 | 68 | 255 | 6 | MOLECULE: | FERRIC ENTEROBACTIN (ENTEROCHELIN) TRANSPOR | T | |
| 0:00 | 2lf0-A | 2.5 | 8.6 | 71 | 120 | 7 | MOLECULE: | UNCHARACTERIZED PROTEIN YIBL | | |
| 1:00 | 4om3-D | 2.5 | 7.3 | 54 | 122 | 7 | MOLECULE: | TRANSDUCIN-LIKE ENHANCER PROTEIN 1 | | |
| 2:00 | 4zqa-A | 2.5 | 3.8 | 58 | 86 | 2 | MOLECULE: | SIN3 HISTONE DEACETYLASE COREPRESSOR COMPLE | X COMP | |
| 3:00 | 8sdi-A | 2.5 | 4.2 | 56 | 80 | 7 | MOLECULE: | TRIPARTITE MOTIF-CONTAINING PROTEIN 45 | | |
| 4:00 | 6oqa-G | 2.5 | 17 | 58 | 87 | 9 | MOLECULE: | PEPTIDYL-PROLYL CIS-TRANS ISOMERASE FKBP1A | | |
| 5:00 | 2jee-A | 2.5 | 3.4 | 55 | 78 | 9 | MOLECULE: | CELL DIVISION PROTEIN ZAPB | | |
| 6:00 | 4lnb-A | 2.5 | 2.7 | 39 | 339 | 3 | MOLECULE: | CAAX FARNESYLTRANSFERASE ALPHA SUBUNIT RAM2 | | |
| 7:00 | 5yfp-E | 2.5 | 15.1 | 110 | 776 | 8 | MOLECULE: | EXOCYST COMPLEX COMPONENT SEC3 | | |
| 8:00 | 5oi7-A | 2.5 | 3.5 | 56 | 88 | 4 | MOLECULE: | CENTROSOMAL PROTEIN OF 85 KDA | | |
| 9:00 | 8fck-F | 2.5 | 29.9 | 88 | 393 | 5 | MOLECULE: | HAUS AUGMIN-LIKE COMPLEX SUBUNIT 1 | | |
| 10:00 | 2wz7-B | 2.5 | 3.2 | 54 | 67 | 0 | MOLECULE: | UNCHARACTERIZED PROTEIN YBGF | | |
| 11:00 | 8gt9-A | 2.5 | 4.4 | 54 | 91 | 2 | MOLECULE: | BECLIN-2 | | |
| 12:00 | 6vhf-A | 2.5 | 3.7 | 61 | 140 | 7 | MOLECULE: | PHD-TYPE DOMAIN-CONTAINING PROTEIN | | |

| | | | | | | | | | | |
|---|---|---|---|---|---|---|---|---|---|---|
| 13:00 | 6nbx-C | 2.5 | 16.7 | 73 | 100 | 4 | MOLECULE: | NAD(P)H-QUINONE OXIDOREDUCTASE SUBUNIT 1 | | |
| 14:00 | 6t9i-T | 2.5 | 21.9 | 57 | 3513 | 11 | MOLECULE: | TRANSCRIPTION FACTOR SPT20 | | |
| 15:00 | 6m17-A | 2.5 | 11.1 | 72 | 606 | 6 | MOLECULE: | SODIUM-DEPENDENT NEUTRAL AMINO ACID TRANSPO | RTER B | |
| 16:00 | 7aqq-N | 2.5 | 10.4 | 94 | 488 | 3 | MOLECULE: | NADH-UBIQUINONE OXIDOREDUCTASE CHAIN 3 | | |
| 17:00 | 3vuq-C | 2.5 | 10.3 | 56 | 179 | 2 | MOLECULE: | TRANSCRIPTIONAL REGULATOR (TETR/ACRR FAMILY | ) | |
| 18:00 | 5xvj-A | 2.5 | 6.8 | 67 | 134 | 6 | MOLECULE: | PHD FINGER PROTEIN ALFIN-LIKE 7 | | |
| 19:00 | 7d3e-A | 2.5 | 18.6 | 86 | 1380 | 2 | MOLECULE: | DUAL OXIDASE 1 | | |
| 20:00 | 1fxk-C | 2.5 | 4.8 | 58 | 133 | 10 | MOLECULE: | PREFOLDIN | | |
| 21:00 | 5xbj-A | 2.4 | 32.2 | 126 | 500 | 9 | MOLECULE: | FLAGELLAR HOOK-ASSOCIATED PROTEIN FLGK | | |
| 22:00 | 9mty-B | 2.4 | 26.8 | 94 | 294 | 4 | MOLECULE: | TRANSPOSASE IS116/IS110/IS902 C-TERMINAL DO | MAIN-C | |
| 23:00 | 6t8d-X | 2.4 | 33.1 | 147 | 355 | 5 | MOLECULE: | MAKB | | |
| 24:00:00 | 4whj-A | 2.4 | 16.6 | 88 | 565 | 8 | MOLECULE: | INTERFERON-INDUCED GTP-BINDING PROTEIN MX2 | | |
| 25:00:00 | 8ti8-C | 2.4 | 11.6 | 72 | 259 | 7 | MOLECULE: | SHEDU PROTEIN SDUA | | |
| 26:00:00 | 7qpg-R | 2.4 | 33.6 | 145 | 2208 | 8 | MOLECULE: | PROTEIN ZWILCH HOMOLOG | | |
| 27:00:00 | 7s9w-A | 2.4 | 25.4 | 105 | 1238 | 5 | MOLECULE: | DRMA | | |
| 28:00:00 | 9fth-A | 2.4 | 27.7 | 82 | 601 | 10 | MOLECULE: | NEUROTRANSMITTER-GATED ION-CHANNEL LIGAND-B | INDING | |
| 29:00:00 | 7crc-E | 2.4 | 17.7 | 55 | 237 | 5 | MOLECULE: | NAD+ HYDROLASE (NADASE) | | |
| 30:00:00 | 6qum-N | 2.4 | 23.4 | 101 | 649 | 5 | MOLECULE: | V-TYPE ATP SYNTHASE ALPHA CHAIN | | |
| 31:00:00 | 5to5-A | 2.4 | 32.7 | 97 | 141 | 10 | MOLECULE: | NUCLEOPROTEIN TPR | | |
| 32:00:00 | 8u1a-A | 2.4 | 14.1 | 66 | 324 | 11 | MOLECULE: | DESIGNED NEAR INFRARED FLUORESCENT PROTEIN | MC7BP3 | |
| 33:00:00 | 8jzr-B | 2.4 | 34.3 | 131 | 456 | 5 | MOLECULE: | LYSOSOMAL TRANSPORTER,ALFA TAG | | |
| 34:00:00 | 4owt-A | 2.4 | 23.8 | 87 | 466 | 9 | MOLECULE: | INTEGRATOR COMPLEX SUBUNIT 3 | | |
| 35:00:00 | 7y8r-P | 2.4 | 31 | 101 | 385 | 6 | MOLECULE: | HISTONE H3 | | |
| 36:00:00 | 5c1f-A | 2.4 | 29.5 | 128 | 298 | 13 | MOLECULE: | SEPTATION PROTEIN IMP2 | | |
| 37:00:00 | 2ksr-A | 2.4 | 19.4 | 58 | 140 | 5 | MOLECULE: | NEURONAL ACETYLCHOLINE RECEPTOR SUBUNIT BET | A-2 | |
| 38:00:00 | 6ek4-A | 2.4 | 23.5 | 153 | 342 | 4 | MOLECULE: | PAXB | | |
| 39:00:00 | 8csp-8 | 2.4 | 27.1 | 97 | 326 | 6 | MOLECULE: | 28S RIBOSOMAL PROTEIN S34, MITOCHONDRIAL | | |
| 40:00:00 | 7c9m-C | 2.4 | 20.7 | 70 | 260 | 4 | MOLECULE: | D-HISTIDINE 2-AMINOBUTANOYLTRANSFERASE | | |
| 41:00:00 | 8pqw-F | 2.4 | 43.3 | 130 | 150 | 7 | MOLECULE: | CYTOPLASMIC DYNEIN 1 HEAVY CHAIN 1 | | |
| 42:00:00 | 7d2w-A | 2.4 | 15.2 | 58 | 125 | 7 | MOLECULE: | PRESAN DOMAIN-CONTAINING PROTEIN | | |
| 43:00:00 | 8gjg-A | 2.4 | 25.8 | 91 | 162 | 8 | MOLECULE: | GLUC_A04_0005 BINDER | | |
| 44:00:00 | 6jho-A | 2.4 | 32.8 | 113 | 200 | 6 | MOLECULE: | CAG PATHOGENICITY ISLAND PROTEIN (CAG6) | | |
| 45:00:00 | 5od9-A | 2.4 | 4 | 41 | 95 | 5 | MOLECULE: | MID1SC9 | | |
| 46:00:00 | 7ar9-p | 2.4 | 7.6 | 95 | 155 | 4 | MOLECULE: | ND3 | | |
| 47:00:00 | 5ve8-B | 2.4 | 26.1 | 119 | 1030 | 9 | MOLECULE: | KAP123 | | |
| 48:00:00 | 1s5j-A | 2.4 | 22.7 | 98 | 727 | 9 | MOLECULE: | DNA POLYMERASE I | | |
| 49:00:00 | 5b04-B | 2.4 | 15 | 56 | 319 | 7 | MOLECULE: | TRANSLATION INITIATION FACTOR EIF-2B SUBUNI | T ALPH | |
| 50:00:00 | 7lga-D | 2.4 | 3 | 48 | 91 | 8 | MOLECULE: | RETROTRANSPOSON-DERIVED PROTEIN PEG10 | | |
| 51:00:00 | 5d2s-A | 2.4 | 15.7 | 72 | 91 | 6 | MOLECULE: | FIBROIN-MODULATOR-BINDING PROTEIN-1 | | |
| 52:00:00 | 6nr8-2 | 2.4 | 4.6 | 72 | 103 | 6 | MOLECULE: | PREFOLDIN SUBUNIT 1 | | |
| 53:00:00 | 3dfu-A | 2.4 | 4.1 | 69 | 228 | 6 | MOLECULE: | UNCHARACTERIZED PROTEIN FROM 6-PHOSPHOGLUCO | NATE | |
| 54:00:00 | 2id3-A | 2.4 | 5.5 | 56 | 191 | 11 | MOLECULE: | PUTATIVE TRANSCRIPTIONAL REGULATOR | | |
| 55:00:00 | 3cqx-D | 2.4 | 2.9 | 65 | 84 | 6 | MOLECULE: | HEAT SHOCK COGNATE 71 KDA PROTEIN | | |
| 56:00:00 | 5iqc-A | 2.4 | 12.9 | 81 | 301 | 10 | MOLECULE: | BIFUNCTIONAL AAC/APH | | |
| 57:00:00 | 6yxj-B | 2.4 | 9 | 74 | 218 | 5 | MOLECULE: | NON-STRUCTURAL PROTEIN 3 | | |
| 58:00:00 | 9c2d-A | 2.4 | 10.9 | 83 | 366 | 5 | MOLECULE: | MAJOR CAPSID PROTEIN | | |
| 59:00:00 | 5tr1-A | 2.4 | 20.4 | 87 | 606 | 2 | MOLECULE: | CHLORIDE CHANNEL PROTEIN | | |
| 60:00:00 | 2h09-A | 2.4 | 5.1 | 59 | 127 | 10 | MOLECULE: | TRANSCRIPTIONAL REGULATOR MNTR | | |
| 61:00:00 | 3wxx-E | 2.4 | 8.6 | 70 | 151 | 7 | MOLECULE: | ACRH | | |
| 62:00:00 | 7vug-R | 2.4 | 5.2 | 107 | 282 | 3 | MOLECULE: | GUANINE NUCLEOTIDE-BINDING PROTEIN G(I) SUB | UNIT A | |
| 63:00:00 | 3k1m-B | 2.4 | 16 | 83 | 305 | 7 | MOLECULE: | HTH-TYPE TRANSCRIPTIONAL REGULATOR BENM | | |
| 64:00:00 | 3s84-B | 2.4 | 6.6 | 81 | 241 | 5 | MOLECULE: | APOLIPOPROTEIN A-IV | | |
| 65:00:00 | 1i49-A | 2.4 | 17.7 | 103 | 201 | 8 | MOLECULE: | ARFAPTIN 2 | | |
| 66:00:00 | 8ck1-E | 2.4 | 13.9 | 98 | 206 | 5 | MOLECULE: | TAIL NOZZLE | | |
| 67:00:00 | 6djl-E | 2.4 | 8.8 | 121 | 226 | 8 | MOLECULE: | RAS-RELATED PROTEIN RAB-11A | | |
| 68:00:00 | 9irz-A | 2.4 | 4.1 | 55 | 96 | 5 | MOLECULE: | PROBABLE HTH-TYPE TRANSCRIPTIONAL REGULATOR | YHAJ | |
| 69:00:00 | 7eu3-G | 2.4 | 8.2 | 74 | 176 | 1 | MOLECULE: | NAD(P)H-QUINONE OXIDOREDUCTASE SUBUNIT 1, C | HLOROP | |
| 70:00:00 | 2qup-A | 2.4 | 4.1 | 67 | 119 | 9 | MOLECULE: | BH1478 PROTEIN | | |
| 71:00:00 | 5kay-B | 2.4 | 5.2 | 80 | 200 | 6 | MOLECULE: | SPELTER | | |
| 72:00:00 | 6b4h-A | 2.4 | 12.5 | 80 | 318 | 3 | MOLECULE: | NUCLEOPORIN AMO1 | | |
| 73:00:00 | 4r3z-A | 2.4 | 3.8 | 53 | 76 | 4 | MOLECULE: | AMINOACYL TRNA SYNTHASE COMPLEX-INTERACTING | MULTI | |
| 74:00:00 | 7v1n-A | 2.4 | 29.1 | 108 | 2344 | 8 | MOLECULE: | TOXIN B | | |
| 75:00:00 | 3u9j-A | 2.4 | 7.1 | 65 | 157 | 6 | MOLECULE: | F-BOX/LRR-REPEAT PROTEIN 5 | | |
| 76:00:00 | 8h6h-A | 2.4 | 12.7 | 91 | 986 | 8 | MOLECULE: | CELLODEXTRIN PHOSPHORYLASE | | |
| 77:00:00 | 6bq1-E | 2.4 | 15.1 | 115 | 1510 | 7 | MOLECULE: | PHOSPHATIDYLINOSITOL 4-KINASE III ALPHA (PI | 4KA) | |
| 78:00:00 | 6db1-A | 2.4 | 3.7 | 69 | 137 | 10 | MOLECULE: | PUTATIVE METHYL-ACCEPTING CHEMOTAXIS PROTEI | N | |
| 79:00:00 | 2o34-A | 2.4 | 6.7 | 61 | 249 | 8 | MOLECULE: | HYPOTHETICAL PROTEIN | | |
| 80:00:00 | 7bi2-A | 2.4 | 25.9 | 94 | 1021 | 9 | MOLECULE: | PHOSPHATIDYLINOSITOL 4-PHOSPHATE 3-KINASE C | 2 DOMA | |
| 81:00:00 | 5hmo-A | 2.4 | 3.8 | 48 | 131 | 10 | MOLECULE: | UNCONVENTIONAL MYOSIN-X | | |
| 82:00:00 | 8jxo-A | 2.4 | 9.1 | 83 | 262 | 6 | MOLECULE: | BLCHR2 | | |
| 83:00:00 | 7pkq-w | 2.4 | 36.5 | 118 | 155 | 8 | MOLECULE: | MS35 | | |
| 84:00:00 | 8gju-F | 2.4 | 4.8 | 61 | 332 | 3 | MOLECULE: | METHYLMALONIC ACIDURIA TYPE A PROTEIN, MITO | CHONDR | |
| 85:00:00 | 8xej-X | 2.4 | 4.9 | 62 | 358 | 8 | MOLECULE: | ISOFORM 2 OF BASIGIN | | |
| 86:00:00 | 7nvr-3 | 2.4 | 16.9 | 73 | 214 | 12 | MOLECULE: | TFIIH BASAL TRANSCRIPTION FACTOR COMPLEX HE | LICASE | |
| 87:00:00 | 6j52-A | 2.4 | 8.2 | 50 | 94 | 4 | MOLECULE: | CASPASE RECRUITMENT DOMAIN-ONLY PROTEIN | | |
| 88:00:00 | 8eja-A | 2.4 | 5.8 | 75 | 113 | 5 | MOLECULE: | ABAK | | |
| 89:00:00 | 6xxv-C | 2.4 | 4.9 | 71 | 111 | 6 | MOLECULE: | ANTIBODY C57, HEAVY CHAIN | | |
| 90:00:00 | 2rkh-A | 2.4 | 5 | 60 | 164 | 0 | MOLECULE: | PUTATIVE APHA-LIKE TRANSCRIPTION FACTOR | | |
| 91:00:00 | 8jps-A | 2.4 | 9.9 | 71 | 257 | 4 | MOLECULE: | ATYPICAL CHEMOKINE RECEPTOR 1 | | |
| 92:00:00 | 1kf6-D | 2.4 | 5.6 | 96 | 119 | 6 | MOLECULE: | FUMARATE REDUCTASE FLAVOPROTEIN | | |
| 93:00:00 | 5mq4-C | 2.4 | 19 | 84 | 117 | 4 | MOLECULE: | PROTEIN KINASE C-BINDING PROTEIN 1 | | |
| 94:00:00 | 5xef-A | 2.4 | 4.3 | 75 | 123 | 3 | MOLECULE: | FLAGELLAR PROTEIN FLIS | | |
| 95:00:00 | 1na6-B | 2.4 | 5.7 | 75 | 395 | 4 | MOLECULE: | RESTRICTION ENDONUCLEASE ECORII | | |
| 96:00:00 | 8iq4-R | 2.4 | 4.2 | 95 | 274 | 9 | MOLECULE: | PROSTAGLANDIN F2-ALPHA RECEPTOR | | |
| 97:00:00 | 4dra-H | 2.4 | 7.5 | 63 | 76 | 2 | MOLECULE: | CENTROMERE PROTEIN S | | |
| 98:00:00 | 6xby-a | 2.4 | 7.3 | 77 | 494 | 4 | MOLECULE: | V-TYPE PROTON ATPASE CATALYTIC SUBUNIT A | | |
| 99:00:00 | 8hf2-A | 2.4 | 6.1 | 83 | 120 | 4 | MOLECULE: | PRA1 FAMILY PROTEIN | | |
| 0:00 | 4uw9-A | 2.4 | 5.9 | 71 | 228 | 6 | MOLECULE: | BETA-PHOSPHOGLUCOMUTASE | | |
| 1:00 | 8p62-D | 2.4 | 21 | 88 | 245 | 10 | MOLECULE: | DNA REPLICATION LICENSING FACTOR MCM2 | | |
| 2:00 | 8qx8-E | 2.4 | 5.3 | 46 | 216 | 13 | MOLECULE: | VACUOLAR PROTEIN SORTING-ASSOCIATED PROTEIN | | 8 |
| 3:00 | 7jrl-A | 2.4 | 20.4 | 96 | 457 | 4 | MOLECULE: | F5/8 TYPE C DOMAIN PROTEIN | | |
| 4:00 | 6ox1-B | 2.4 | 11.9 | 103 | 485 | 4 | MOLECULE: | ACTIN, CYTOPLASMIC 1 | | |
| 5:00 | 8vti-A | 2.4 | 6.5 | 84 | 358 | 4 | MOLECULE: | ISOFORM 4 OF ADHESION G PROTEIN-COUPLED REC | EPTOR | |
| 6:00 | 2xpl-A | 2.4 | 4.2 | 54 | 142 | 9 | MOLECULE: | IWS1 | | |
| 7:00 | 6cv9-A | 2.4 | 21.9 | 101 | 316 | 8 | MOLECULE: | SHORT TRANSIENT RECEPTOR POTENTIAL CHANNEL | | 6 |
| 8:00 | 8j0o-C | 2.4 | 29.6 | 80 | 258 | 5 | MOLECULE: | ER MEMBRANE PROTEIN COMPLEX SUBUNIT 1 | | |
| 9:00 | 8v14-A | 2.4 | 14.2 | 49 | 167 | 4 | MOLECULE: | NMDA RECEPTOR AUXILIARY PROTEIN | | |
| 10:00 | 7zet-A | 2.4 | 23.6 | 91 | 377 | 4 | MOLECULE: | CLUSTERIN | | |
| 11:00 | 5gqu-A | 2.4 | 5.8 | 60 | 755 | 7 | MOLECULE: | 1,4-ALPHA-GLUCAN BRANCHING ENZYME GLGB | | |
| 12:00 | 5j65-A | 2.4 | 15.4 | 101 | 402 | 5 | MOLECULE: | PESTICIDAL CRYSTAL PROTEIN CRY6AA | | |
| 13:00 | 6prk-A | 2.4 | 13.5 | 85 | 115 | 7 | MOLECULE: | RICF | | |

| | | | | | | | | | | |
|---|---|---|---|---|---|---|---|---|---|---|
| 14:00 | 7v7b-A | 2.4 | 25.3 | 118 | 1110 | 4 | MOLECULE: | DDB1- AND CUL4-ASSOCIATED FACTOR 1 | | |
| 15:00 | 8ovb-C | 2.4 | 4.6 | 100 | 266 | 10 | MOLECULE: | COMPLEMENT C3F FRAGMENT | | |
| 16:00 | 5myi-D | 2.4 | 6 | 62 | 165 | 8 | MOLECULE: | DUTPASE FROM DI S. AUREUS PHAGE | | |
| 17:00 | 8eki-E | 2.4 | 13.6 | 87 | 643 | 7 | MOLECULE: | PROTEIN TRANSPORT PROTEIN SEC20 | | |
| 18:00 | 2fo1-D | 2.4 | 3.5 | 44 | 63 | 5 | MOLECULE: | 5'- | | |
| 19:00 | 8p1l-A | 2.4 | 29.1 | 136 | 2036 | 3 | MOLECULE: | RNA-DIRECTED RNA POLYMERASE L | | |
| 20:00 | 7usd-A | 2.4 | 25.2 | 161 | 1185 | 4 | MOLECULE: | CYTOPLASMIC FMR1-INTERACTING PROTEIN 1 | | |
| 21:00 | 8ub7-B | 2.4 | 8.5 | 58 | 122 | 7 | MOLECULE: | REVERSE TRANSCRIPTASE | | |
| 22:00 | 5oqm-h | 2.4 | 27 | 93 | 131 | 6 | MOLECULE: | DNA-DIRECTED RNA POLYMERASE II SUBUNIT RPB1 | | |
| 23:00 | 5yx5-B | 2.4 | 10.8 | 71 | 270 | 7 | MOLECULE: | 49 PROTEIN | | |
| 24:00:00 | 5c0x-F | 2.4 | 3.7 | 40 | 215 | 5 | MOLECULE: | EXOSOME COMPLEX COMPONENT RRP45 | | |
| 25:00:00 | 6v0t-A | 2.4 | 22.2 | 81 | 402 | 6 | MOLECULE: | [PYRUVATE DEHYDROGENASE [ACETYL-TRANSFERRIN | G]]-PH | |
| 26:00:00 | 1wle-B | 2.4 | 24.2 | 137 | 470 | 7 | MOLECULE: | SERYL-TRNA SYNTHETASE | | |
| 27:00:00 | 6ch2-E | 2.4 | 3.9 | 64 | 147 | 6 | MOLECULE: | FLAGELLAR BIOSYNTHESIS PROTEIN FLHA | | |
| 28:00:00 | 6noy-A | 2.4 | 3.9 | 47 | 80 | 4 | MOLECULE: | MAINTENANCE OF CARBOXYSOME POSITIONING B PR | OTEIN, | |
| 29:00:00 | 9l2q-A | 2.4 | 10.4 | 59 | 105 | 14 | MOLECULE: | H2C2 | | |
| 30:00:00 | 4xwj-A | 2.4 | 4 | 71 | 153 | 4 | MOLECULE: | REGULATOR OF SIGMA D | | |
| 31:00:00 | 7z1n-O | 2.4 | 8.8 | 84 | 570 | 6 | MOLECULE: | DNA-DIRECTED RNA POLYMERASE III SUBUNIT RPC | | 1 |
| 32:00:00 | 7obq-u | 2.4 | 6 | 78 | 441 | 6 | MOLECULE: | SRP RNA | | |
| 33:00:00 | 5vkv-A | 2.4 | 15.3 | 62 | 224 | 3 | MOLECULE: | CYTOCHROME C-TYPE BIOGENESIS PROTEIN CCDA | | |
| 34:00:00 | 3edv-A | 2.4 | 17 | 101 | 322 | 7 | MOLECULE: | SPECTRIN BETA CHAIN, BRAIN 1 | | |
| 35:00:00 | 8xku-B | 2.4 | 26.3 | 95 | 845 | 7 | MOLECULE: | PROBABLE INACTIVE ATP-DEPENDENT ZINC METALL | OPROTE | |
| 36:00:00 | 8cbk-E | 2.4 | 4.2 | 38 | 470 | 3 | MOLECULE: | 3-HYDROXYACYL-COA DEHYDROGENASE TYPE-2 | | |
| 37:00:00 | 7f52-A | 2.4 | 16.3 | 123 | 584 | 9 | MOLECULE: | NON-STRUCTURAL PROTEIN 2 | | |
| 38:00:00 | 5lxj-A | 2.4 | 2.3 | 36 | 53 | 8 | MOLECULE: | PHOSPHOPROTEIN | | |
| 39:00:00 | 2q1f-A | 2.4 | 24.9 | 123 | 991 | 7 | MOLECULE: | CHONDROITINASE | | |
| 40:00:00 | 6t9i-I | 2.4 | 11.6 | 84 | 173 | 4 | MOLECULE: | TRANSCRIPTION FACTOR SPT20 | | |
| 41:00:00 | 6xz4-A | 2.4 | 9.9 | 87 | 304 | 5 | MOLECULE: | TALIN ROD DOMAIN-CONTAINING PROTEIN 1 | | |
| 42:00:00 | 4bt9-B | 2.4 | 22.7 | 107 | 238 | 7 | MOLECULE: | PROLYL 4-HYDROXYLASE SUBUNIT ALPHA-1 | | |
| 43:00:00 | 6vdq-A | 2.4 | 10.9 | 77 | 305 | 4 | MOLECULE: | 3-METHYL-L-TYROSINE PEROXYGENASE | | |
| 44:00:00 | 6wuc-K | 2.4 | 20.8 | 109 | 235 | 7 | MOLECULE: | INNER KINETOCHORE SUBUNIT MCM16 | | |
| 45:00:00 | 1ox3-A | 2.4 | 7.8 | 68 | 108 | 1 | MOLECULE: | FIBRITIN | | |
| 46:00:00 | 8pmq-9 | 2.4 | 28 | 114 | 412 | 7 | MOLECULE: | E3 UBIQUITIN-PROTEIN LIGASE RMD5 | | |
| 47:00:00 | 4k2p-A | 2.4 | 7.7 | 80 | 238 | 13 | MOLECULE: | T-LYMPHOMA INVASION AND METASTASIS-INDUCING | PROTE | |
| 48:00:00 | 3hai-A | 2.4 | 31 | 112 | 294 | 10 | MOLECULE: | HUMAN PACSIN1 F-BAR | | |
| 49:00:00 | 5z7g-C | 2.4 | 4.1 | 41 | 43 | 10 | MOLECULE: | TAX1-BINDING PROTEIN 1 | | |
| 50:00:00 | 6oap-A | 2.4 | 4.8 | 44 | 305 | 7 | MOLECULE: | DUAL SENSOR HISTIDINE KINASE | | |
| 51:00:00 | 5ool-v | 2.4 | 2.7 | 41 | 69 | 7 | MOLECULE: | 16S RIBOSOMAL RNA | | |
| 52:00:00 | 4abn-B | 2.4 | 14.3 | 86 | 426 | 2 | MOLECULE: | TETRATRICOPEPTIDE REPEAT PROTEIN 5 | | |
| 53:00:00 | 6vzf-A | 2.4 | 3.9 | 59 | 86 | 8 | MOLECULE: | AUTOPHAGY-RELATED PROTEIN 11 | | |
| 54:00:00 | 5h07-D | 2.4 | 13.4 | 57 | 78 | 9 | MOLECULE: | POLYUBIQUITIN-C | | |
| 55:00:00 | 1x79-B | 2.4 | 3.4 | 59 | 90 | 12 | MOLECULE: | ADP-RIBOSYLATION FACTOR BINDING PROTEIN GGA | | 1 |
| 56:00:00 | 1fav-A | 2.4 | 4.2 | 58 | 78 | 7 | MOLECULE: | HIV-1 ENVELOPE PROTEIN CHIMERA | | |
| 57:00:00 | 5cff-B | 2.4 | 3.3 | 55 | 88 | 9 | MOLECULE: | MIRANDA | | |
| 58:00:00 | 6pse-A | 2.4 | 4 | 58 | 78 | 7 | MOLECULE: | PROTEIN BICAUDAL D HOMOLOG 2 | | |
| 59:00:00 | 6k2j-B | 2.4 | 6.4 | 65 | 83 | 11 | MOLECULE: | UPF0335 PROTEIN CCNA_03428 | | |
| 60:00:00 | 8sjj-C | 2.4 | 18.7 | 56 | 86 | 13 | MOLECULE: | SEPTIN-14 | | |
| 61:00:00 | 7o97-A | 2.4 | 3.4 | 53 | 73 | 6 | MOLECULE: | HYPOTHETICAL PROTEIN UY81_C0065G0003 FROM C | ANDIDA | |
| 62:00:00 | 8sde-A | 2.4 | 3.8 | 56 | 89 | 11 | MOLECULE: | TRIPARTITE MOTIF-CONTAINING PROTEIN 29 | | |
| 63:00:00 | 5jxc-A | 2.4 | 19.2 | 56 | 86 | 5 | MOLECULE: | RAS/RAP GTPASE-ACTIVATING PROTEIN SYNGAP | | |
| 64:00:00 | 3a7p-A | 2.4 | 3.7 | 59 | 88 | 8 | MOLECULE: | AUTOPHAGY PROTEIN 16 | | |
| 65:00:00 | 2olt-C | 2.4 | 4.9 | 51 | 217 | 0 | MOLECULE: | HYPOTHETICAL PROTEIN | | |
| 66:00:00 | 6rwy-g | 2.4 | 5.2 | 59 | 86 | 7 | MOLECULE: | INNER ROD PROTEIN | | |
| 67:00:00 | 7zl9-A | 2.4 | 16.8 | 90 | 398 | 6 | MOLECULE: | HYDROXYCARBOXYLIC ACID RECEPTOR 2,SOLUBLE C | YTOCHR | |
| 68:00:00 | 6sl2-A | 2.4 | 19.5 | 96 | 619 | 2 | MOLECULE: | CALPONIN HOMOLOGY DOMAIN PROTEIN PUTATIVE | | |
| 69:00:00 | 7tds-A | 2.4 | 11.7 | 98 | 290 | 6 | MOLECULE: | 34K2 SALIVARY PROTEIN | | |
| 70:00:00 | 8srm-B | 2.4 | 34.6 | 109 | 235 | 6 | MOLECULE: | RB1-INDUCIBLE COILED-COIL PROTEIN 1 | | |
| 71:00:00 | 8egr-G | 2.4 | 17.5 | 61 | 313 | 8 | MOLECULE: | GP15, RECEPTOR-BINDING PROTEIN, TAIL FIBER | | |
| 72:00:00 | 8t1l-V | 2.4 | 12.8 | 104 | 278 | 7 | MOLECULE: | MEDIATOR OF RNA POLYMERASE II TRANSCRIPTION | SUBUN | |
| 73:00:00 | 7nyx-A | 2.3 | 98.3 | 194 | 1467 | 6 | MOLECULE: | CHROMOSOME PARTITION PROTEIN MUKB | | |
| 74:00:00 | 7z8b-C | 2.3 | 32.4 | 119 | 1224 | 4 | MOLECULE: | CULLIN-7 | | |
| 75:00:00 | 3k7d-A | 2.3 | 25.9 | 91 | 497 | 2 | MOLECULE: | GLUTAMATE-AMMONIA-LIGASE ADENYLYLTRANSFERAS | E | |
| 76:00:00 | 2ewg-A | 2.3 | 34.2 | 100 | 367 | 4 | MOLECULE: | FARNESYL PYROPHOSPHATE SYNTHASE | | |
| 77:00:00 | 3u24-A | 2.3 | 24.3 | 89 | 546 | 2 | MOLECULE: | PUTATIVE LIPOPROTEIN | | |
| 78:00:00 | 8tw7-1 | 2.3 | 23.9 | 94 | 258 | 5 | MOLECULE: | REPLICATION FACTOR C SUBUNIT 5 | | |
| 79:00:00 | 9bax-A | 2.3 | 22.6 | 101 | 1724 | 6 | MOLECULE: | PROTEIN EFR3 HOMOLOG A | | |
| 80:00:00 | 5jjh-A | 2.3 | 15.6 | 74 | 704 | 3 | MOLECULE: | 4-ALPHA-GLUCANOTRANSFERASE | | |
| 81:00:00 | 2vca-A | 2.3 | 31.1 | 95 | 886 | 6 | MOLECULE: | ALPHA-N-ACETYLGLUCOSAMINIDASE | | |
| 82:00:00 | 9j0n-W | 2.3 | 37 | 123 | 543 | 7 | MOLECULE: | DNA-DIRECTED RNA POLYMERASE SUBUNIT | | |
| 83:00:00 | 3jbz-A | 2.3 | 25.9 | 100 | 960 | 4 | MOLECULE: | SERINE/THREONINE-PROTEIN KINASE MTOR | | |
| 84:00:00 | 6aay-A | 2.3 | 23.2 | 131 | 1199 | 2 | MOLECULE: | BERGEYELLA ZOOHELCUM CAS13B (R1177A) MUTANT | | |
| 85:00:00 | 1qoy-A | 2.3 | 32.8 | 120 | 303 | 9 | MOLECULE: | HEMOLYSIN E | | |
| 86:00:00 | 8fti-A | 2.3 | 20.7 | 86 | 737 | 7 | MOLECULE: | INTEGRASE | | |
| 87:00:00 | 9dtr-S | 2.3 | 28.2 | 95 | 604 | 5 | MOLECULE: | U2 SNRNA | | |
| 88:00:00 | 6bfi-B | 2.3 | 31.9 | 179 | 806 | 7 | MOLECULE: | VIN1 | | |
| 89:00:00 | 2rd9-A | 2.3 | 35.6 | 96 | 189 | 5 | MOLECULE: | BH0186 PROTEIN | | |
| 90:00:00 | 5vhx-E | 2.3 | 32.8 | 89 | 179 | 6 | MOLECULE: | GLUTAMATE RECEPTOR 2,GERM CELL-SPECIFIC GEN | E 1-LI | |
| 91:00:00 | 4kpp-A | 2.3 | 29.8 | 141 | 395 | 6 | MOLECULE: | PUTATIVE UNCHARACTERIZED PROTEIN | | |
| 92:00:00 | 4yto-B | 2.3 | 33.3 | 87 | 97 | 9 | MOLECULE: | SYNAPTONEMAL COMPLEX PROTEIN 1 | | |
| 93:00:00 | 9fwf-D | 2.3 | 32.1 | 93 | 226 | 5 | MOLECULE: | N-VELCROVAX HBCAG WITH SUMO-AFFIMER INSERTE | D AT N | |
| 94:00:00 | 7b4m-A | 2.3 | 30.7 | 137 | 459 | 6 | MOLECULE: | SODIUM/HYDROGEN EXCHANGER 9B2 | | |
| 95:00:00 | 8ki9-A | 2.3 | 23.6 | 167 | 2238 | 8 | MOLECULE: | L PROTEIN | | |
| 96:00:00 | 5ys3-A | 2.3 | 28.4 | 85 | 188 | 8 | MOLECULE: | SUCCINATE-ACETATE PERMEASE | | |
| 97:00:00 | 8jhq-A | 2.3 | 31.6 | 137 | 446 | 4 | MOLECULE: | SPHINGOSINE-1-PHOSPHATE TRANSPORTER SPNS2,G | LGA GL | |
| 98:00:00 | 8cr1-C | 2.3 | 29 | 126 | 266 | 10 | MOLECULE: | ATPASE ASNA1 | | |
| 99:00:00 | 6csm-D | 2.3 | 8.9 | 95 | 277 | 5 | MOLECULE: | GTACR1 | | |
| 0:00 | 8ugc-A | 2.3 | 15.7 | 88 | 384 | 3 | MOLECULE: | FD15 | | |
| 1:00 | 7zke-E | 2.3 | 30.9 | 153 | 1623 | 10 | MOLECULE: | DNA (36-MER) | | |
| 2:00 | 6vls-D | 2.3 | 21.3 | 125 | 963 | 4 | MOLECULE: | MALTOSE/MALTODEXTRIN-BINDING PERIPLASMIC PR | OTEIN, | |
| 3:00 | 7zmg-d | 2.3 | 4.7 | 70 | 101 | 4 | MOLECULE: | NADH-UBIQUINONE OXIDOREDUCTASE CHAIN 1 | | |
| 4:00 | 7cgp-A | 2.3 | 11.5 | 75 | 147 | 7 | MOLECULE: | MITOCHONDRIAL IMPORT INNER MEMBRANE TRANSLO | CASE S | |
| 5:00 | 7zmb-3 | 2.3 | 19.7 | 103 | 130 | 6 | MOLECULE: | NADH-UBIQUINONE OXIDOREDUCTASE CHAIN 1 | | |
| 6:00 | 7dqk-B | 2.3 | 7.7 | 87 | 479 | 2 | MOLECULE: | PROTEIN DETOXIFICATION | | |
| 7:00 | 6iqc-A | 2.3 | 3.8 | 38 | 103 | 11 | MOLECULE: | DNA-BINDING PROTEIN SSO0352 | | |
| 8:00 | 8wcs-A | 2.3 | 14.6 | 98 | 1044 | 10 | MOLECULE: | CAS13H1 | | |
| 9:00 | 8v9u-A | 2.3 | 2 | 38 | 81 | 3 | MOLECULE: | DNA (CYTOSINE-5)-METHYLTRANSFERASE 1 | | |
| 10:00 | 8umy-A | 2.3 | 10.5 | 76 | 546 | 5 | MOLECULE: | CHROMOSOME TRANSMISSION FIDELITY PROTEIN 18 | HOMOL | |
| 11:00 | 9ce3-A | 2.3 | 29.6 | 104 | 1295 | 8 | MOLECULE: | ISOFORM 4 OF TUBERIN | | |
| 12:00 | 6wg3-A | 2.3 | 17.9 | 86 | 561 | 6 | MOLECULE: | STRUCTURAL MAINTENANCE OF CHROMOSOMES PROTE | IN 1A | |
| 13:00 | 9k9m-A | 2.3 | 4.4 | 66 | 216 | 2 | MOLECULE: | FLAGELLAR ASSOCIATED PROTEIN | | |
| 14:00 | 8upx-A | 2.3 | 21.9 | 97 | 906 | 6 | MOLECULE: | SPIKE GLYCOPROTEIN,SARS-COV-2 OMICRON SPIKE | WITH | |

| | | | | | | | | | | |
|---|---|---|---|---|---|---|---|---|---|---|
| 15:00 | 6k7x-C | 2.3 | 14 | 92 | 276 | 3 | MOLECULE: | CALCIUM UNIPORTER PROTEIN, MITOCHONDRIAL | | |
| 16:00 | 6hpn-A | 2.3 | 4 | 73 | 212 | 3 | MOLECULE: | ANTIGEN, P35 | | |
| 17:00 | 5xj5-A | 2.3 | 11.5 | 68 | 200 | 4 | MOLECULE: | GLYCEROL-3-PHOSPHATE ACYLTRANSFERASE | | |
| 18:00 | 8esv-B | 2.3 | 8.3 | 101 | 255 | 9 | MOLECULE: | DISINTEGRIN AND METALLOPROTEINASE DOMAIN-CO | NTAINI | |
| 19:00 | 7tjk-F | 2.3 | 2.3 | 39 | 262 | 3 | MOLECULE: | ORIGIN RECOGNITION COMPLEX SUBUNIT 1 | | |
| 20:00 | 8uwq-D | 2.3 | 12.9 | 64 | 280 | 13 | MOLECULE: | BT1282 | | |
| 21:00 | 5csk-A | 2.3 | 22.7 | 78 | 1996 | 5 | MOLECULE: | ACETYL-COA CARBOXYLASE | | |
| 22:00 | 7wgh-A | 2.3 | 11.2 | 83 | 350 | 6 | MOLECULE: | SQUALENE SYNTHASE | | |
| 23:00 | 9bgo-A | 2.3 | 15.7 | 93 | 395 | 8 | MOLECULE: | GP72 | | |
| 24:00:00 | 8i8b-H | 2.3 | 5.1 | 71 | 228 | 10 | MOLECULE: | MAJOR VIRAL CAPSID PROTEIN | | |
| 25:00:00 | 7ot9-A | 2.3 | 5.1 | 61 | 221 | 2 | MOLECULE: | AI-2E MEMBER YDIK | | |
| 26:00:00 | 6s3l-K | 2.3 | 11 | 83 | 193 | 6 | MOLECULE: | FLAGELLAR BIOSYNTHETIC PROTEIN FLIP | | |
| 27:00:00 | 8s1x-B | 2.3 | 3.7 | 44 | 61 | 2 | MOLECULE: | PEPTIDE DEFORMYLASE | | |
| 28:00:00 | 8ixk-A | 2.3 | 16.3 | 73 | 145 | 8 | MOLECULE: | ATTACHMENT PROTEIN G3P | | |
| 29:00:00 | 6vbu-2 | 2.3 | 4.5 | 78 | 659 | 6 | MOLECULE: | BARDET-BIEDL SYNDROME 18 PROTEIN | | |
| 30:00:00 | 6ukj-A | 2.3 | 8.8 | 83 | 350 | 8 | MOLECULE: | CHLOROQUINE RESISTANCE TRANSPORTER | | |
| 31:00:00 | 9b8v-E | 2.3 | 21.3 | 129 | 845 | 2 | MOLECULE: | CYCLIC DI-GMP-BINDING PROTEIN | | |
| 32:00:00 | 8yl8-A | 2.3 | 7.4 | 60 | 211 | 2 | MOLECULE: | DE NOVO PROTEIN | | |
| 33:00:00 | 8hpo-C | 2.3 | 7.6 | 103 | 138 | 5 | MOLECULE: | TRANSCRIPTIONAL REGULATORY PROTEIN UME1 | | |
| 34:00:00 | 8ap7-c | 2.3 | 7.4 | 57 | 64 | 9 | MOLECULE: | ATP SYNTHASE SUBUNIT A | | |
| 35:00:00 | 9j0x-B | 2.3 | 17.8 | 122 | 688 | 5 | MOLECULE: | POTASSIUM CHANNEL GORK | | |
| 36:00:00 | 3hgf-A | 2.3 | 13.9 | 74 | 98 | 9 | MOLECULE: | RHOPTRY PROTEIN FRAGMENT | | |
| 37:00:00 | 5ctq-A | 2.3 | 10.2 | 81 | 518 | 9 | MOLECULE: | SQUAMOUS CELL CARCINOMA ANTIGEN RECOGNIZED | BY T-C | |
| 38:00:00 | 7p2y-d | 2.3 | 8.4 | 63 | 174 | 6 | MOLECULE: | ATP SYNTHASE SUBUNIT ALPHA | | |
| 39:00:00 | 3let-B | 2.3 | 9.9 | 85 | 310 | 11 | MOLECULE: | ADENOSINE MONOPHOSPHATE-PROTEIN TRANSFERASE | VOPS | |
| 40:00:00 | 6e7e-A | 2.3 | 8.8 | 75 | 169 | 4 | MOLECULE: | INCLUSION MEMBRANE PROTEIN A | | |
| 41:00:00 | 7nuv-A | 2.3 | 6.1 | 60 | 76 | 10 | MOLECULE: | AUX2PLS20 | | |
| 42:00:00 | 7x1i-A | 2.3 | 10.3 | 114 | 678 | 6 | MOLECULE: | ISOFORM 1 OF SOLUTE CARRIER FAMILY 4 MEMBER | | 11 |
| 43:00:00 | 7zjr-A | 2.3 | 27.6 | 105 | 268 | 6 | MOLECULE: | PROTEIN PILJ | | |
| 44:00:00 | 3msv-A | 2.3 | 8.9 | 62 | 352 | 8 | MOLECULE: | NUCLEAR IMPORT ADAPTOR, NRO1 | | |
| 45:00:00 | 5y06-A | 2.3 | 4.3 | 88 | 235 | 6 | MOLECULE: | MSMEG_4306 | | |
| 46:00:00 | 9bw6-B | 2.3 | 3 | 49 | 136 | 8 | MOLECULE: | MAJOR VAULT PROTEIN | | |
| 47:00:00 | 3gp8-A | 2.3 | 7.5 | 71 | 551 | 7 | MOLECULE: | EXODEOXYRIBONUCLEASE V, SUBUNIT RECD, PUTAT | IVE | |
| 48:00:00 | 5hc9-A | 2.3 | 4.1 | 59 | 425 | 5 | MOLECULE: | TRNA NUCLEOTIDYL TRANSFERASE-RELATED PROTEI | N | |
| 49:00:00 | 7et3-1 | 2.3 | 7.2 | 76 | 285 | 4 | MOLECULE: | TRIPLEX CAPSID PROTEIN 2 | | |
| 50:00:00 | 6ait-A | 2.3 | 3.7 | 57 | 418 | 9 | MOLECULE: | BETA-BARREL ASSEMBLY-ENHANCING PROTEASE | | |
| 51:00:00 | 1szi-A | 2.3 | 7.6 | 85 | 194 | 6 | MOLECULE: | MANNOSE-6-PHOSPHATE RECEPTOR BINDING PROTEI | N 1 | |
| 52:00:00 | 7pgr-F | 2.3 | 26.6 | 107 | 2423 | 6 | MOLECULE: | NEUROFIBROMIN | | |
| 53:00:00 | 8wjo-B | 2.3 | 43 | 127 | 283 | 6 | MOLECULE: | STRUCTURAL MAINTENANCE OF CHROMOSOMES PROTE | IN 5 | |
| 54:00:00 | 2p01-A | 2.3 | 5.1 | 77 | 323 | 4 | MOLECULE: | ALPHA-2-MACROGLOBULIN RECEPTOR-ASSOCIATED | | |
| 55:00:00 | 6xy6-C | 2.3 | 2.6 | 41 | 138 | 15 | MOLECULE: | ANTI-APOPTOTIC MEMBRANE PROTEIN | | |
| 56:00:00 | 6ygh-D | 2.3 | 16.7 | 87 | 193 | 7 | MOLECULE: | CAPSID PROTEIN | | |
| 57:00:00 | 4it4-B | 2.3 | 12.6 | 89 | 210 | 2 | MOLECULE: | CG17282 | | |
| 58:00:00 | 7sq3-A | 2.3 | 7.8 | 51 | 146 | 4 | MOLECULE: | DESIGNED TREFOIL KNOT PROTEIN, VARIANT 1 | | |
| 59:00:00 | 6lum-D | 2.3 | 5.5 | 97 | 146 | 6 | MOLECULE: | SUCCINATE DEHYDROGENASE SUBUNIT C | | |
| 60:00:00 | 6mc9-A | 2.3 | 21.6 | 62 | 150 | 6 | MOLECULE: | SODIUM CHANNEL PROTEIN TYPE 4 SUBUNIT ALPHA | | |
| 61:00:00 | 6px3-C | 2.3 | 8.1 | 60 | 179 | 3 | MOLECULE: | HISTONE H3 | | |
| 62:00:00 | 4xqk-A | 2.3 | 14.1 | 105 | 1517 | 6 | MOLECULE: | LLABIII | | |
| 63:00:00 | 3gp4-A | 2.3 | 9.3 | 62 | 132 | 6 | MOLECULE: | TRANSCRIPTIONAL REGULATOR, MERR FAMILY | | |
| 64:00:00 | 3i3l-A | 2.3 | 8.7 | 84 | 550 | 2 | MOLECULE: | ALKYLHALIDASE CMLS | | |
| 65:00:00 | 3iv5-B | 2.3 | 3.2 | 52 | 98 | 2 | MOLECULE: | DNA-BINDING PROTEIN FIS | | |
| 66:00:00 | 7w7c-B | 2.3 | 4.7 | 72 | 159 | 1 | MOLECULE: | PUTATIVE ABC TRANSPORT SYSTEM, ATP-BINDING | PROTEI | |
| 67:00:00 | 8vwk-A | 2.3 | 6.7 | 70 | 430 | 4 | MOLECULE: | CYTOCHROME P450 | | |
| 68:00:00 | 3mnl-A | 2.3 | 6.5 | 60 | 188 | 10 | MOLECULE: | TRANSCRIPTIONAL REGULATORY PROTEIN (PROBABL | Y TETR | |
| 69:00:00 | 5vkq-A | 2.3 | 17.1 | 110 | 1499 | 5 | MOLECULE: | NO MECHANORECEPTOR POTENTIAL C ISOFORM L | | |
| 70:00:00 | 9b7y-A | 2.3 | 12.5 | 87 | 385 | 3 | MOLECULE: | TRANSCRIPTIONAL REPRESSOR MCE3R | | |
| 71:00:00 | 9ix4-A | 2.3 | 33.6 | 101 | 1178 | 5 | MOLECULE: | DDMD | | |
| 72:00:00 | 5dfz-D | 2.3 | 7.7 | 77 | 341 | 8 | MOLECULE: | VACUOLAR PROTEIN SORTING-ASSOCIATED PROTEIN | | 38 |
| 73:00:00 | 5jbr-A | 2.3 | 19.6 | 79 | 149 | 11 | MOLECULE: | UNCHARACTERIZED PROTEIN BCAV_2135 | | |
| 74:00:00 | 5xe3-E | 2.3 | 4.2 | 55 | 81 | 2 | MOLECULE: | ENDORIBONUCLEASE MAZF4 | | |
| 75:00:00 | 6tqe-A | 2.3 | 30.2 | 135 | 634 | 3 | MOLECULE: | ABC TRANSPORTER ATP-BINDING PROTEIN/PERMEAS | E | |
| 76:00:00 | 6btm-C | 2.3 | 6.7 | 74 | 457 | 7 | MOLECULE: | ALTERNATIVE COMPLEX III SUBUNIT A | | |
| 77:00:00 | 7yoj-A | 2.3 | 14 | 95 | 867 | 4 | MOLECULE: | CASPI | | |
| 78:00:00 | 8dr5-A | 2.3 | 26 | 62 | 646 | 11 | MOLECULE: | REPLICATION FACTOR C SUBUNIT 1 | | |
| 79:00:00 | 8qa2-A | 2.3 | 5.6 | 95 | 215 | 3 | MOLECULE: | GAP JUNCTION BETA-2 PROTEIN | | |
| 80:00:00 | 6wt8-A | 2.3 | 22.5 | 93 | 356 | 6 | MOLECULE: | STING-ASSOCIATED CDNE C-DI-GMP SYNTHASE | | |
| 81:00:00 | 5lm2-B | 2.3 | 26.3 | 124 | 340 | 11 | MOLECULE: | TYROSINE-PROTEIN PHOSPHATASE NON-RECEPTOR T | YPE 23 | |
| 82:00:00 | 5kqj-A | 2.3 | 3.3 | 39 | 185 | 5 | MOLECULE: | 2''-AMINOGLYCOSIDE NUCLEOTIDYLTRANSFERASE | | |
| 83:00:00 | 4d0n-B | 2.3 | 9.4 | 120 | 371 | 7 | MOLECULE: | TRANSFORMING PROTEIN RHOA | | |
| 84:00:00 | 7ttt-A | 2.3 | 21.1 | 92 | 351 | 3 | MOLECULE: | TUBULIN BETA CHAIN | | |
| 85:00:00 | 8oyv-A | 2.3 | 5.3 | 91 | 182 | 4 | MOLECULE: | DE NOVO DESIGNED SOLUBLE CLAUDIN | | |
| 86:00:00 | 5zki-B | 2.3 | 2.6 | 36 | 304 | 8 | MOLECULE: | NUCLEASE EXOG, MITOCHONDRIAL | | |
| 87:00:00 | 1bvp-1 | 2.3 | 3.5 | 42 | 349 | 7 | MOLECULE: | BLUETONGUE VIRUS COAT PROTEIN VP7 | | |
| 88:00:00 | 3ter-B | 2.3 | 5.6 | 70 | 121 | 4 | MOLECULE: | MAMMALIAN STROMAL INTERACTION MOLECULE-1 | | |
| 89:00:00 | 8kgn-A | 2.3 | 24.1 | 105 | 1190 | 7 | MOLECULE: | DNA TOPOISOMERASE 2 | | |
| 90:00:00 | 8ity-W | 2.3 | 4.4 | 42 | 111 | 5 | MOLECULE: | SNRNA-ACTIVATING PROTEIN COMPLEX SUBUNIT 1 | | |
| 91:00:00 | 2ipc-A | 2.3 | 29.1 | 140 | 939 | 8 | MOLECULE: | PREPROTEIN TRANSLOCASE SECA SUBUNIT | | |
| 92:00:00 | 8wag-B | 2.3 | 7.2 | 72 | 235 | 7 | MOLECULE: | PROTEIN CHUP1, CHLOROPLASTIC | | |
| 93:00:00 | 6pon-A | 2.3 | 5 | 50 | 255 | 8 | MOLECULE: | ADHERENCE AND VIRULENCE PROTEIN A | | |
| 94:00:00 | 5dyr-A | 2.3 | 7.6 | 57 | 139 | 4 | MOLECULE: | VIRULENCE-ASSOCIATED PROTEIN D | | |
| 95:00:00 | 5g49-A | 2.3 | 9 | 67 | 97 | 4 | MOLECULE: | NUCLEAR TRANSCRIPTION FACTOR Y SUBUNIT B-6 | | |
| 96:00:00 | 6mcp-A | 2.3 | 13.4 | 78 | 520 | 9 | MOLECULE: | LEGK7 | | |
| 97:00:00 | 6ka4-A | 2.3 | 9.5 | 86 | 534 | 7 | MOLECULE: | F22L4.1 PROTEIN | | |
| 98:00:00 | 4iuw-A | 2.3 | 25.4 | 115 | 630 | 10 | MOLECULE: | NEUTRAL ENDOPEPTIDASE | | |
| 99:00:00 | 8si6-A | 2.3 | 14.6 | 124 | 1107 | 7 | MOLECULE: | TRANSIENT RECEPTOR POTENTIAL CATION CHANNEL | SUBFA | |
| 0:00 | 3h0g-A | 2.3 | 10.2 | 96 | 1496 | 7 | MOLECULE: | DNA-DIRECTED RNA POLYMERASE II SUBUNIT RPB1 | | |
| 1:00 | 2e2e-A | 2.3 | 14.8 | 62 | 171 | 10 | MOLECULE: | FORMATE-DEPENDENT NITRITE REDUCTASE COMPLEX | NRFG | |
| 2:00 | 2win-M | 2.3 | 3.9 | 53 | 84 | 8 | MOLECULE: | COMPLEMENT C3 BETA CHAIN | | |
| 3:00 | 3n9m-A | 2.3 | 12.9 | 68 | 503 | 7 | MOLECULE: | PUTATIVE UNCHARACTERIZED PROTEIN | | |
| 4:00 | 7mrw-C | 2.3 | 24.7 | 95 | 706 | 7 | MOLECULE: | CYTOADHERENCE LINKED ASEXUAL PROTEIN 3.1 | | |
| 5:00 | 6c14-B | 2.3 | 5.9 | 88 | 164 | 2 | MOLECULE: | PROTOCADHERIN-15 | | |
| 6:00 | 4ztx-A | 2.3 | 11.4 | 82 | 768 | 9 | MOLECULE: | COBALAMIN-INDEPENDENT METHIONINE SYNTHASE | | |
| 7:00 | 6nf1-A | 2.3 | 7.4 | 104 | 550 | 4 | MOLECULE: | PROTO-ONCOGENE VAV | | |
| 8:00 | 4gr6-B | 2.3 | 7.4 | 57 | 105 | 4 | MOLECULE: | ATRBCX2 | | |
| 9:00 | 2qyw-A | 2.3 | 1.9 | 62 | 96 | 6 | MOLECULE: | VESICLE TRANSPORT THROUGH INTERACTION WITH | T-SNAR | |
| 10:00 | 1ypy-A | 2.3 | 9 | 62 | 182 | 3 | MOLECULE: | VIRION MEMBRANE PROTEIN | | |
| 11:00 | 6hyd-A | 2.3 | 20.1 | 108 | 1574 | 8 | MOLECULE: | MIDASIN,MIDASIN,MIDASIN | | |
| 12:00 | 1s7o-C | 2.3 | 8.7 | 68 | 108 | 4 | MOLECULE: | HYPOTHETICAL UPF0122 PROTEIN | | |
| 13:00 | 5jw9-B | 2.3 | 6.9 | 75 | 115 | 5 | MOLECULE: | AF4/FMR2 FAMILY MEMBER 4 | | |
| 14:00 | 5i1u-B | 2.3 | 14.8 | 75 | 294 | 7 | MOLECULE: | GERMACRADIEN-4-OL SYNTHASE | | |
| 15:00 | 6l82-A | 2.3 | 21 | 58 | 98 | 14 | MOLECULE: | SPINDLE POLE BODY COMPONENT | | |

| | | | | | | | | | | |
|---|---|---|---|---|---|---|---|---|---|---|
| 16:00 | 4f43-A | 2.3 | 8.9 | 64 | 320 | 6 | MOLECULE: | PROTELOMERASE | | |
| 17:00 | 5yx4-A | 2.3 | 24.1 | 64 | 232 | 9 | MOLECULE: | CHALCONE-FLAVONONE ISOMERASE FAMILY PROTEIN | | |
| 18:00 | 6pwy-B | 2.3 | 12.6 | 74 | 513 | 8 | MOLECULE: | ZK177.8 | | |
| 19:00 | 3vep-H | 2.3 | 6.9 | 53 | 71 | 4 | MOLECULE: | UNCHARACTERIZED PROTEIN RV3413C/MT3522 | | |
| 20:00 | 6ogd-A | 2.3 | 7.2 | 63 | 748 | 6 | MOLECULE: | TOXIN SUBUNIT YENA1 | | |
| 21:00 | 7arc-W | 2.3 | 3.3 | 45 | 127 | 7 | MOLECULE: | PSST | | |
| 22:00 | 8c06-A | 2.3 | 26.4 | 80 | 925 | 4 | MOLECULE: | E3 UBIQUITIN-PROTEIN LIGASE UBR5 | | |
| 23:00 | 7y17-C | 2.3 | 13.4 | 66 | 340 | 9 | MOLECULE: | POLYNUCLEOTIDE 5'-HYDROXYL-KINASE GRC3 | | |
| 24:00:00 | 7nnl-B | 2.3 | 23.5 | 97 | 682 | 5 | MOLECULE: | POTASSIUM-TRANSPORTING ATPASE POTASSIUM-BIN | DING S | |
| 25:00:00 | 1nek-C | 2.3 | 6.2 | 72 | 129 | 4 | MOLECULE: | SUCCINATE DEHYDROGENASE FLAVOPROTEIN SUBUNI | T | |
| 26:00:00 | 4zm1-B | 2.3 | 6 | 97 | 238 | 5 | MOLECULE: | CHAIN LENGTH DETERMINANT PROTEIN | | |
| 27:00:00 | 6bhc-A | 2.3 | 5.2 | 52 | 371 | 10 | MOLECULE: | PSEUDOPODIUM-ENRICHED ATYPICAL KINASE 1 | | |
| 28:00:00 | 6e6y-A | 2.3 | 8.4 | 61 | 268 | 3 | MOLECULE: | DIECKMANN CYCLASE, NCMC | | |
| 29:00:00 | 8sfn-A | 2.3 | 15.5 | 119 | 1302 | 8 | MOLECULE: | CRISPR-ASSOCIATED ENDONUCLEASE CAS12A | | |
| 30:00:00 | 8afz-B | 2.3 | 5.1 | 68 | 376 | 9 | MOLECULE: | SORTING NEXIN-1 | | |
| 31:00:00 | 6v8o-N | 2.3 | 15.2 | 105 | 412 | 9 | MOLECULE: | HIGH TEMPERATURE LETHAL PROTEIN 1 | | |
| 32:00:00 | 7s4m-C | 2.3 | 5.9 | 64 | 241 | 8 | MOLECULE: | PARTICULATE METHANE MONOOXYGENASE ALPHA SUB | UNIT | |
| 33:00:00 | 4h8s-C | 2.3 | 33.7 | 155 | 382 | 6 | MOLECULE: | DCC-INTERACTING PROTEIN 13-BETA | | |
| 34:00:00 | 4nad-A | 2.3 | 9.8 | 91 | 133 | 11 | MOLECULE: | REGULATION OF NUCLEAR PRE-MRNA DOMAIN-CONTA | INING | |
| 35:00:00 | 6g7c-C | 2.3 | 3.5 | 41 | 239 | 0 | MOLECULE: | IMPA-RELATED DOMAIN PROTEIN | | |
| 36:00:00 | 8d8k-Q | 2.3 | 4.1 | 50 | 205 | 4 | MOLECULE: | PROBABLE S-ADENOSYL-L-METHIONINE-DEPENDENT | RNA | |
| 37:00:00 | 3vou-A | 2.3 | 10.2 | 100 | 139 | 3 | MOLECULE: | ION TRANSPORT 2 DOMAIN PROTEIN, VOLTAGE-GAT | ED SOD | |
| 38:00:00 | 4me7-F | 2.3 | 8.1 | 55 | 79 | 4 | MOLECULE: | MRNA INTERFERASE ENDOA | | |
| 39:00:00 | 5o9e-B | 2.3 | 9.9 | 57 | 80 | 7 | MOLECULE: | PUTATIVE U3 SMALL NUCLEOLAR RIBONUCLEOPROTE | IN | |
| 40:00:00 | 3qwe-A | 2.3 | 22.5 | 117 | 260 | 5 | MOLECULE: | GEM-INTERACTING PROTEIN | | |
| 41:00:00 | 8j80-A | 2.3 | 16.6 | 96 | 290 | 10 | MOLECULE: | ZINC TRANSPORTER 7 | | |
| 42:00:00 | 3cit-A | 2.3 | 4.8 | 52 | 155 | 4 | MOLECULE: | SENSOR HISTIDINE KINASE | | |
| 43:00:00 | 7zkq-C | 2.3 | 15.3 | 77 | 416 | 5 | MOLECULE: | NADH DEHYDROGENASE SUBUNIT 2 | | |
| 44:00:00 | 6n7p-D | 2.3 | 22.9 | 99 | 544 | 10 | MOLECULE: | U1 SMALL NUCLEAR RIBONUCLEOPROTEIN 70 KDA H | OMOLOG | |
| 45:00:00 | 3j9t-O | 2.3 | 17.4 | 113 | 392 | 5 | MOLECULE: | V-TYPE PROTON ATPASE SUBUNIT D | | |
| 46:00:00 | 5vxv-A | 2.3 | 15.2 | 56 | 217 | 9 | MOLECULE: | PEROXISOMAL MEMBRANE PROTEIN PEX15 | | |
| 47:00:00 | 5l0w-B | 2.3 | 25.6 | 64 | 209 | 8 | MOLECULE: | SEC72 | | |
| 48:00:00 | 8au0-B | 2.3 | 4.9 | 38 | 41 | 5 | MOLECULE: | SUN DOMAIN-CONTAINING PROTEIN 1 | | |
| 49:00:00 | 7a48-B | 2.3 | 3.3 | 37 | 40 | 5 | MOLECULE: | NANOBODY 49 | | |
| 50:00:00 | 6q6e-A | 2.3 | 7.6 | 70 | 221 | 9 | MOLECULE: | CONDENSIN COMPLEX SUBUNIT 2,STRUCTURAL MAIN | TENANC | |
| 51:00:00 | 3c64-A | 2.3 | 4.8 | 54 | 152 | 7 | MOLECULE: | PFEMP1 VARIANT 2 OF STRAIN MC | | |
| 52:00:00 | 3pla-A | 2.3 | 9.7 | 71 | 375 | 4 | MOLECULE: | PRE MRNA SPLICING PROTEIN | | |
| 53:00:00 | 1i6z-A | 2.3 | 6.2 | 60 | 135 | 10 | MOLECULE: | BAG-FAMILY MOLECULAR CHAPERONE REGULATOR-1 | | |
| 54:00:00 | 8ims-A | 2.3 | 3.6 | 58 | 93 | 3 | MOLECULE: | E3 UBIQUITIN-PROTEIN LIGASE TRAF7 | | |
| 55:00:00 | 8hir-A | 2.3 | 24.3 | 97 | 918 | 9 | MOLECULE: | POTASSIUM CHANNEL SUBFAMILY T MEMBER 1 | | |
| 56:00:00 | 7c7y-C | 2.3 | 20 | 57 | 67 | 9 | MOLECULE: | UNCHARACTERIZED PROTEIN | | |
| 57:00:00 | 7bji-B | 2.3 | 3.7 | 56 | 92 | 11 | MOLECULE: | CENTROSOMAL PROTEIN OF 135 KDA | | |
| 58:00:00 | 4ytd-B | 2.3 | 3.1 | 55 | 99 | 4 | MOLECULE: | PROTEIN BICAUDAL D HOMOLOG 1 | | |
| 59:00:00 | 3wvz-A | 2.3 | 5.9 | 47 | 192 | 0 | MOLECULE: | PROTEIN HIKESHI | | |
| 60:00:00 | 6h9l-B | 2.3 | 5.2 | 55 | 129 | 11 | MOLECULE: | UNCHARACTERIZED PROTEIN | | |
| 61:00:00 | 6fkf-p | 2.3 | 34.1 | 89 | 143 | 8 | MOLECULE: | ATP SYNTHASE SUBUNIT ALPHA, CHLOROPLASTIC | | |
| 62:00:00 | 7xuz-A | 2.3 | 3.7 | 56 | 120 | 13 | MOLECULE: | HISTONE DEACETYLASE 4 | | |
| 63:00:00 | 7xfr-B | 2.3 | 4.1 | 52 | 69 | 0 | MOLECULE: | ISOFORM 2 OF WD REPEAT DOMAIN PHOSPHOINOSIT | IDE-IN | |
| 64:00:00 | 4nqf-B | 2.3 | 2.8 | 50 | 151 | 10 | MOLECULE: | NUCLEOTIDYLTRANSFERASE | | |
| 65:00:00 | 1izl-C | 2.3 | 18.7 | 70 | 349 | 0 | MOLECULE: | PHOTOSYSTEM II: SUBUNIT PSBA | | |
| 66:00:00 | 8vb0-H | 2.3 | 4.3 | 44 | 61 | 16 | MOLECULE: | MAJOR CAPSID PROTEIN (GP38) | | |
| 67:00:00 | 3m62-A | 2.3 | 19.1 | 108 | 955 | 6 | MOLECULE: | UBIQUITIN CONJUGATION FACTOR E4 | | |
| 68:00:00 | 7c1i-A | 2.3 | 4.3 | 49 | 115 | 2 | MOLECULE: | HISTIDINE KINASE | | |
| 69:00:00 | 6uak-A | 2.3 | 11 | 48 | 298 | 8 | MOLECULE: | SAM DEPENDENT METHYLTRANSFERASE LAHSB | | |
| 70:00:00 | 7vyx-A | 2.3 | 9.4 | 92 | 1211 | 4 | MOLECULE: | SELENOMETHIONINE (SEMET)-LABELED CAS12C1 D9 | 69A MU | |
| 71:00:00 | 6dql-A | 2.3 | 3.2 | 41 | 227 | 2 | MOLECULE: | REGULATOR OF PROTEINASE B ROPB | | |
| 72:00:00 | 6rd4-7 | 2.3 | 6 | 65 | 176 | 5 | MOLECULE: | ASA-10: POLYTOMELLA F-ATP SYNTHASE ASSOCIAT | ED SUB | |
| 73:00:00 | 6uuj-G | 2.3 | 3.5 | 49 | 74 | 4 | MOLECULE: | PE FAMILY IMMUNOMODULATOR PE5 | | |
| 74:00:00 | 7lhf-A | 2.2 | 38.3 | 166 | 2123 | 5 | MOLECULE: | INOSITOL 1,4,5-TRISPHOSPHATE RECEPTOR TYPE | | 1 |
| 75:00:00 | 8qby-L | 2.2 | 25.8 | 110 | 660 | 5 | MOLECULE: | NADH-QUINONE OXIDOREDUCTASE SUBUNIT K | | |
| 76:00:00 | 6hsy-A | 2.2 | 25.3 | 62 | 190 | 0 | MOLECULE: | TOLUENE TOLERANCE PROTEIN TTG2D | | |
| 77:00:00 | 8wge-A | 2.2 | 12.8 | 73 | 348 | 4 | MOLECULE: | ZAC | | |
| 78:00:00 | 6hu9-c | 2.2 | 13.1 | 89 | 269 | 4 | MOLECULE: | CYTOCHROME B-C1 COMPLEX SUBUNIT 1, MITOCHON | DRIAL | |
| 79:00:00 | 6rxd-A | 2.2 | 18.3 | 94 | 509 | 4 | MOLECULE: | HISTIDINE ACID PHOSPHATASE | | |
| 80:00:00 | 7ymo-A | 2.2 | 22.4 | 42 | 206 | 2 | MOLECULE: | RECOMBINATION PROTEIN RECO | | |
| 81:00:00 | 8yad-C | 2.2 | 29.5 | 127 | 1811 | 6 | MOLECULE: | SPATACSIN | | |
| 82:00:00 | 1gm5-A | 2.2 | 13.2 | 91 | 729 | 8 | MOLECULE: | RECG | | |
| 83:00:00 | 4ka7-A | 2.2 | 19.5 | 137 | 695 | 8 | MOLECULE: | OLIGOPEPTIDASE A | | |
| 84:00:00 | 2y44-A | 2.2 | 13.4 | 68 | 184 | 4 | MOLECULE: | GLUTAMIC ACID/ALANINE-RICH PROTEIN | | |
| 85:00:00 | 8ucj-c | 2.2 | 27.3 | 101 | 269 | 4 | MOLECULE: | SYNAPTIC VESICULAR AMINE TRANSPORTER | | |
| 86:00:00 | 8vlu-A | 2.2 | 27.4 | 117 | 550 | 8 | MOLECULE: | HEPARAN-ALPHA-GLUCOSAMINIDE N-ACETYLTRANSFE | RASE | |
| 87:00:00 | 7znk-b | 2.2 | 21.3 | 89 | 919 | 8 | MOLECULE: | RNA | | |
| 88:00:00 | 7zr1-D | 2.2 | 57.6 | 174 | 780 | 6 | MOLECULE: | DOUBLE-STRAND BREAK REPAIR PROTEIN | | |
| 89:00:00 | 6ibl-A | 2.2 | 5.9 | 121 | 397 | 2 | MOLECULE: | THIOREDOXIN 1,BETA-1 ADRENERGIC RECEPTOR | | |
| 90:00:00 | 7yn2-B | 2.2 | 29.8 | 94 | 349 | 5 | MOLECULE: | CCBD | | |
| 91:00:00 | 8rjj-A | 2.2 | 26.8 | 78 | 183 | 5 | MOLECULE: | GENOME POLYPROTEIN | | |
| 92:00:00 | 5wco-A | 2.2 | 15.6 | 62 | 196 | 8 | MOLECULE: | NS2 | | |
| 93:00:00 | 6fes-A | 2.2 | 21.6 | 91 | 365 | 4 | MOLECULE: | D12_BRIC2, A SYNTHETIC PROTEIN,D12_BRIC2, A | SYNTH | |
| 94:00:00 | 1iq0-A | 2.2 | 20.1 | 93 | 588 | 6 | MOLECULE: | ARGINYL-TRNA SYNTHETASE | | |
| 95:00:00 | 6hqa-A | 2.2 | 31.4 | 116 | 887 | 8 | MOLECULE: | TAF2 | | |
| 96:00:00 | 7lid-A | 2.2 | 35 | 119 | 381 | 8 | MOLECULE: | MHOR5 | | |
| 97:00:00 | 3eo8-A | 2.2 | 27.6 | 90 | 219 | 7 | MOLECULE: | BLUB-LIKE FLAVOPROTEIN | | |
| 98:00:00 | 2jqq-A | 2.2 | 14.9 | 65 | 154 | 6 | MOLECULE: | CONSERVED OLIGOMERIC GOLGI COMPLEX SUBUNIT | | 2 |
| 99:00:00 | 1yvl-B | 2.2 | 22.4 | 110 | 653 | 10 | MOLECULE: | SIGNAL TRANSDUCER AND ACTIVATOR OF TRANSCRI | PTION | |
| 0:00 | 3o0z-D | 2.2 | 53.6 | 134 | 160 | 8 | MOLECULE: | RHO-ASSOCIATED PROTEIN KINASE 1 | | |
| 1:00 | 6j36-A | 2.2 | 30.7 | 89 | 459 | 6 | MOLECULE: | ENOLASE | | |
| 2:00 | 2j68-A | 2.2 | 28.3 | 160 | 680 | 5 | MOLECULE: | BACTERIAL DYNAMIN-LIKE PROTEIN | | |
| 3:00 | 7wmp-m | 2.2 | 29.9 | 100 | 195 | 12 | MOLECULE: | PORTAL PROTEIN | | |
| 4:00 | 7ag9-B | 2.2 | 32.7 | 155 | 384 | 8 | MOLECULE: | KAR9 | | |
| 5:00 | 6s7t-H | 2.2 | 6.6 | 78 | 112 | 5 | MOLECULE: | DOLICHYL-DIPHOSPHOOLIGOSACCHARIDE--PROTEIN | | |
| 6:00 | 7qog-M | 2.2 | 3.7 | 58 | 126 | 10 | MOLECULE: | PORTAL PROTEIN GP20 | | |
| 7:00 | 6vtk-A | 2.2 | 3.7 | 61 | 446 | 5 | MOLECULE: | ACID-SENSING ION CHANNEL 1 | | |
| 8:00 | 3zhe-A | 2.2 | 21.9 | 110 | 399 | 4 | MOLECULE: | NONSENSE-MEDIATED MRNA DECAY PROTEIN | | |
| 9:00 | 7q21-f | 2.2 | 14.7 | 80 | 143 | 3 | MOLECULE: | CO-PURIFIED UNKNOWN TRANSMEMBRANE HELICES B | UILT A | |
| 10:00 | 3lvg-D | 2.2 | 15.2 | 64 | 180 | 9 | MOLECULE: | CLATHRIN HEAVY CHAIN 1 | | |
| 11:00 | 1x8z-A | 2.2 | 4.3 | 89 | 151 | 7 | MOLECULE: | INVERTASE/PECTIN METHYLESTERASE INHIBITOR F | AMILY | |
| 12:00 | 2oer-B | 2.2 | 11.8 | 70 | 189 | 6 | MOLECULE: | PROBABLE TRANSCRIPTIONAL REGULATOR | | |
| 13:00 | 6y92-A | 2.2 | 4.7 | 73 | 178 | 5 | MOLECULE: | B-LYMPHOCYTE ANTIGEN CD20 | | |
| 14:00 | 2uxx-B | 2.2 | 24.1 | 82 | 134 | 4 | MOLECULE: | LYSINE-SPECIFIC HISTONE DEMETHYLASE 1 | | |
| 15:00 | 1yx4-A | 2.2 | 16.4 | 69 | 112 | 6 | MOLECULE: | 26S PROTEASOME NON-ATPASE REGULATORY SUBUNI | T 4 | |
| 16:00 | 3k11-A | 2.2 | 13.9 | 82 | 439 | 6 | MOLECULE: | PUTATIVE GLYCOSYL HYDROLASE | | |

| | | | | | | | | | |
|---|---|---|---|---|---|---|---|---|---|
| 17:00 | 6jx7-A | 2.2 | 19.9 | 77 | 1245 | 8 | MOLECULE: | FELINE INFECTIOUS PERITONITIS VIRUS SPIKE P | ROTEIN |
| 18:00 | 8w97-A | 2.2 | 5.2 | 59 | 135 | 5 | MOLECULE: | DE NOVO DESIGN PROTEIN | |
| 19:00 | 9bq2-B | 2.2 | 22.4 | 82 | 422 | 6 | MOLECULE: | FLOTILLIN-2 | |
| 20:00 | 4boj-C | 2.2 | 7.9 | 66 | 344 | 2 | MOLECULE: | ALPHA-1,6-MANNANASE | |
| 21:00 | 4o8w-D | 2.2 | 7.8 | 43 | 120 | 7 | MOLECULE: | SPORE GERMINATION PROTEIN | |
| 22:00 | 3uau-A | 2.2 | 4.2 | 71 | 344 | 4 | MOLECULE: | SURFACE-EXPOSED LIPOPROTEIN | |
| 23:00 | 4cej-A | 2.2 | 30.4 | 106 | 1177 | 7 | MOLECULE: | ATP-DEPENDENT HELICASE/NUCLEASE SUBUNIT A | |
| 24:00:00 | 4gzu-A | 2.2 | 20 | 119 | 436 | 9 | MOLECULE: | FERM, RHOGEF AND PLECKSTRIN DOMAIN-CONTAINI | NG PRO |
| 25:00:00 | 8cq4-A | 2.2 | 13.5 | 90 | 393 | 7 | MOLECULE: | BIFUNCTIONAL CYCLOHEXADIENYL DEHYDRATASE/CH | ORISMA |
| 26:00:00 | 5fib-A | 2.2 | 18.7 | 125 | 534 | 10 | MOLECULE: | SPHINGOMYELIN PHOSPHODIESTERASE | |
| 27:00:00 | 2x3b-A | 2.2 | 7.2 | 69 | 295 | 3 | MOLECULE: | TOXIC EXTRACELLULAR ENDOPEPTIDASE | |
| 28:00:00 | 6em5-b | 2.2 | 13.5 | 132 | 421 | 8 | MOLECULE: | 5.8S RIBOSOMAL RNA | |
| 29:00:00 | 6xz6-A | 2.2 | 8.5 | 87 | 213 | 9 | MOLECULE: | GARP DOMAIN-CONTAINING PROTEIN | |
| 30:00:00 | 7mk3-B | 2.2 | 9.9 | 93 | 365 | 8 | MOLECULE: | REGULATORY PROTEIN NPR1 | |
| 31:00:00 | 8ofi-C | 2.2 | 10.3 | 91 | 415 | 10 | MOLECULE: | POTASSIUM/SODIUM HYPERPOLARIZATION-ACTIVATE | D CYCL |
| 32:00:00 | 8as8-E | 2.2 | 30.9 | 111 | 498 | 5 | MOLECULE: | JETC | |
| 33:00:00 | 8eld-L | 2.2 | 3.7 | 41 | 115 | 2 | MOLECULE: | GP37 | |
| 34:00:00 | 2fb5-A | 2.2 | 5.3 | 86 | 204 | 10 | MOLECULE: | HYPOTHETICAL MEMBRANE SPANNING PROTEIN | |
| 35:00:00 | 6s85-C | 2.2 | 22.9 | 89 | 374 | 7 | MOLECULE: | NUCLEASE SBCCD SUBUNIT C | |
| 36:00:00 | 4igg-B | 2.2 | 18.8 | 108 | 771 | 6 | MOLECULE: | CATENIN ALPHA-1 | |
| 37:00:00 | 8jpa-A | 2.2 | 9.1 | 70 | 139 | 11 | MOLECULE: | DE NOVO DESIGN CAVITATED PROTEIN | |
| 38:00:00 | 2yy5-B | 2.2 | 20.8 | 75 | 346 | 9 | MOLECULE: | TRYPTOPHANYL-TRNA SYNTHETASE | |
| 39:00:00 | 8wai-A | 2.2 | 8.8 | 83 | 277 | 6 | MOLECULE: | RVY_06210 | |
| 40:00:00 | 6wq0-A | 2.2 | 12.9 | 89 | 131 | 6 | MOLECULE: | DNA (301-MER) | |
| 41:00:00 | 6wq2-A | 2.2 | 9.8 | 97 | 154 | 5 | MOLECULE: | A-DNA | |
| 42:00:00 | 8hpp-C | 2.2 | 8.5 | 57 | 86 | 7 | MOLECULE: | INTEGRATOR COMPLEX SUBUNIT 3 | |
| 43:00:00 | 9na9-A | 2.2 | 5.5 | 74 | 103 | 5 | MOLECULE: | AUGMIN SUBUNIT 1 | |
| 44:00:00 | 2ocy-A | 2.2 | 31.5 | 81 | 149 | 12 | MOLECULE: | RAB GUANINE NUCLEOTIDE EXCHANGE FACTOR SEC2 | |
| 45:00:00 | 9jfl-A | 2.2 | 1.7 | 41 | 594 | 7 | MOLECULE: | DUF262 DOMAIN-CONTAINING PROTEIN | |
| 46:00:00 | 6ah0-w | 2.2 | 22.3 | 129 | 443 | 5 | MOLECULE: | U5SNRNA | |
| 47:00:00 | 5ue0-A | 2.2 | 8.3 | 76 | 280 | 4 | MOLECULE: | CT622 PROTEIN | |
| 48:00:00 | 7ykr-A | 2.2 | 14.1 | 119 | 1035 | 5 | MOLECULE: | TRANSIENT RECEPTOR POTENTIAL CATION CHANNEL | SUBFA |
| 49:00:00 | 3tvr-A | 2.2 | 2.7 | 51 | 172 | 2 | MOLECULE: | POLYKETIDE CYCLASE | |
| 50:00:00 | 4ckg-A | 2.2 | 28.4 | 132 | 368 | 7 | MOLECULE: | ARF-GAP WITH COILED-COIL, ANK REPEAT AND PH | DOMAI |
| 51:00:00 | 7bst-C | 2.2 | 6.8 | 60 | 992 | 2 | MOLECULE: | TYPE I RESTRICTION ENZYME R PROTEIN | |
| 52:00:00 | 6f0k-H | 2.2 | 6.5 | 84 | 156 | 2 | MOLECULE: | CYTOCHROME C FAMILY PROTEIN | |
| 53:00:00 | 7ypx-a | 2.2 | 3.9 | 58 | 162 | 14 | MOLECULE: | PAM3 TAIL FIBER PROREINS | |
| 54:00:00 | 9f4c-A | 2.2 | 12.5 | 52 | 141 | 13 | MOLECULE: | CYTOSKELETON-ASSOCIATED PROTEIN 5 | |
| 55:00:00 | 5c0x-K | 2.2 | 25.4 | 114 | 350 | 4 | MOLECULE: | EXOSOME COMPLEX COMPONENT RRP45 | |
| 56:00:00 | 6ulg-N | 2.2 | 18.8 | 88 | 483 | 9 | MOLECULE: | FOLLICULIN | |
| 57:00:00 | 2v0o-A | 2.2 | 29.3 | 123 | 273 | 6 | MOLECULE: | FCH DOMAIN ONLY PROTEIN 2 | |
| 58:00:00 | 6bhp-A | 2.2 | 4.8 | 89 | 170 | 6 | MOLECULE: | MEMBRANE PROTEIN | |
| 59:00:00 | 8fjg-A | 2.2 | 6.7 | 75 | 104 | 4 | MOLECULE: | H12 | |
| 60:00:00 | 8p23-B | 2.2 | 15 | 55 | 715 | 7 | MOLECULE: | ANAEROBIC RIBONUCLEOSIDE-TRIPHOSPHATE REDUC | TASE |
| 61:00:00 | 8bd7-L | 2.2 | 47.3 | 117 | 303 | 7 | MOLECULE: | IFT88 | |
| 62:00:00 | 6ar7-C | 2.2 | 4.8 | 102 | 207 | 7 | MOLECULE: | UNCHARACTERIZED PROTEIN | |
| 63:00:00 | 7but-A | 2.2 | 3.2 | 51 | 142 | 10 | MOLECULE: | ACINIFORM SPIDROIN | |
| 64:00:00 | 3cdh-B | 2.2 | 10.2 | 76 | 136 | 5 | MOLECULE: | TRANSCRIPTIONAL REGULATOR, MARR FAMILY | |
| 65:00:00 | 4xa6-A | 2.2 | 20.4 | 121 | 168 | 3 | MOLECULE: | GP7-MYH7(1777-1855)-EB1 CHIMERA PROTEIN | |
| 66:00:00 | 6tpn-A | 2.2 | 11 | 119 | 510 | 6 | MOLECULE: | OREXIN RECEPTOR TYPE 2,GLGA GLYCOGEN SYNTHA | SE,ORE |
| 67:00:00 | 8ap8-d | 2.2 | 17.1 | 87 | 270 | 2 | MOLECULE: | OSCP | |
| 68:00:00 | 8fed-E | 2.2 | 33 | 104 | 358 | 5 | MOLECULE: | VIRULENCE FACTOR MCE FAMILY PROTEIN | |
| 69:00:00 | 7k0i-C | 2.2 | 9.9 | 51 | 62 | 8 | MOLECULE: | SERINE PALMITOYLTRANSFERASE 1 | |
| 70:00:00 | 6h02-A | 2.2 | 25.5 | 129 | 1334 | 4 | MOLECULE: | MEDIATOR OF RNA POLYMERASE II TRANSCRIPTION | SUBUN |
| 71:00:00 | 8ua7-B | 2.2 | 9.1 | 62 | 81 | 15 | MOLECULE: | HISTONE H3 | |
| 72:00:00 | 6k6l-B | 2.2 | 10.1 | 68 | 263 | 7 | MOLECULE: | PSEUDO DEUBIQUITINASE | |
| 73:00:00 | 3woz-B | 2.2 | 29 | 73 | 229 | 8 | MOLECULE: | CLIP-ASSOCIATING PROTEIN 2 | |
| 74:00:00 | 8b1r-B | 2.2 | 18.3 | 101 | 1170 | 9 | MOLECULE: | RECBCD ENZYME SUBUNIT RECB | |
| 75:00:00 | 9g40-G | 2.2 | 13.3 | 118 | 797 | 7 | MOLECULE: | GAMMA-TUBULIN COMPLEX COMPONENT 3 | |
| 76:00:00 | 7q3e-B | 2.2 | 8.5 | 56 | 503 | 7 | MOLECULE: | WD REPEAT-CONTAINING AND PLANAR CELL POLARI | TY EFF |
| 77:00:00 | 8q4h-C | 2.2 | 6.6 | 75 | 89 | 3 | MOLECULE: | TETRACHLOROETHENE REDUCTIVE DEHALOGENASE | |
| 78:00:00 | 5ncl-A | 2.2 | 22.6 | 116 | 418 | 6 | MOLECULE: | SERINE/THREONINE-PROTEIN KINASE CBK1 | |
| 79:00:00 | 2guz-B | 2.2 | 3.5 | 45 | 65 | 7 | MOLECULE: | MITOCHONDRIAL IMPORT INNER MEMBRANE TRANSLO | CASE S |
| 80:00:00 | 7dhg-C | 2.2 | 5.4 | 60 | 470 | 7 | MOLECULE: | MITOCHONDRIAL IMPORT RECEPTOR SUBUNIT TOM70 | |
| 81:00:00 | 5tj4-H | 2.2 | 6.6 | 79 | 537 | 3 | MOLECULE: | SUGAR ABC TRANSPORTER SUBSTRATE-BINDING PRO | TEIN,G |
| 82:00:00 | 6t5a-A | 2.2 | 4.1 | 67 | 96 | 4 | MOLECULE: | TEGUMENT PROTEIN UL51 | |
| 83:00:00 | 2ia9-D | 2.2 | 3.9 | 42 | 99 | 2 | MOLECULE: | PUTATIVE SEPTATION PROTEIN SPOVG | |
| 84:00:00 | 6vz1-A | 2.2 | 12.9 | 115 | 411 | 7 | MOLECULE: | DIACYLGLYCEROL O-ACYLTRANSFERASE 1 | |
| 85:00:00 | 5guj-A | 2.2 | 2.3 | 37 | 324 | 11 | MOLECULE: | DNA PRIMASE | |
| 86:00:00 | 7pw5-B | 2.2 | 26.4 | 95 | 702 | 11 | MOLECULE: | SMG1,SERINE/THREONINE-PROTEIN KINASE SMG1,S | MG1, |
| 87:00:00 | 8hki-a | 2.2 | 8.8 | 70 | 291 | 4 | MOLECULE: | T-COMPLEX PROTEIN 1 SUBUNIT ALPHA | |
| 88:00:00 | 7ztb-A | 2.2 | 14.2 | 88 | 344 | 6 | MOLECULE: | RELA/SPOT FAMILY PROTEIN | |
| 89:00:00 | 8tn1-A | 2.2 | 3.5 | 76 | 147 | 5 | MOLECULE: | DE NOVO DESIGNED 4 HELIX BUNDLES | |
| 90:00:00 | 1q6u-A | 2.2 | 8 | 75 | 213 | 9 | MOLECULE: | FKBP-TYPE PEPTIDYL-PROLYL CIS-TRANS ISOMERA | SE FKP |
| 91:00:00 | 8ikg-R | 2.2 | 5.9 | 64 | 275 | 3 | MOLECULE: | GUANINE NUCLEOTIDE-BINDING PROTEIN G(I) SUB | UNIT A |
| 92:00:00 | 7c79-L | 2.2 | 3.3 | 73 | 131 | 1 | MOLECULE: | RIBONUCLEASE MRP RNA SUBUNIT NME1 | |
| 93:00:00 | 5lc5-p | 2.2 | 4.3 | 70 | 169 | 6 | MOLECULE: | NADH-UBIQUINONE OXIDOREDUCTASE CHAIN 3 | |
| 94:00:00 | 5oxf-C | 2.2 | 15.8 | 115 | 601 | 6 | MOLECULE: | GTP-BINDING PROTEIN | |
| 95:00:00 | 8r70-A | 2.2 | 23 | 96 | 618 | 5 | MOLECULE: | HEPARINASE | |
| 96:00:00 | 6nyi-A | 2.2 | 3.2 | 59 | 97 | 5 | MOLECULE: | DESIGN CONSTRUCT XXA | |
| 97:00:00 | 9dec-A | 2.2 | 2.1 | 38 | 82 | 8 | MOLECULE: | D9-THREADED | |
| 98:00:00 | 1miu-A | 2.2 | 16.1 | 84 | 671 | 8 | MOLECULE: | DELETED IN SPLIT HAND/SPLIT FOOT PROTEIN 1 | |
| 99:00:00 | 1yqg-A | 2.2 | 7.1 | 70 | 263 | 7 | MOLECULE: | PYRROLINE-5-CARBOXYLATE REDUCTASE | |
| 0:00 | 8q7n-X | 2.2 | 5.7 | 54 | 81 | 15 | MOLECULE: | U5 SNRNA | |
| 1:00 | 8ssn-B | 2.2 | 9.3 | 46 | 386 | 7 | MOLECULE: | TYROSINE-PROTEIN KINASE ABL1 | |
| 2:00 | 3dpn-A | 2.2 | 17 | 81 | 537 | 9 | MOLECULE: | PROTEIN CT_858 | |
| 3:00 | 7wvt-A | 2.2 | 2.8 | 71 | 364 | 8 | MOLECULE: | PHOSPHATIDYLINOSITOL TRANSFER PROTEIN CSR1 | |
| 4:00 | 7pkq-P | 2.2 | 9.7 | 75 | 337 | 8 | MOLECULE: | MS35 | |
| 5:00 | 7dwb-A | 2.2 | 13.6 | 124 | 421 | 5 | MOLECULE: | PANNEXIN-1 | |
| 6:00 | 8p62-C | 2.2 | 5.6 | 49 | 175 | 8 | MOLECULE: | DNA REPLICATION LICENSING FACTOR MCM2 | |
| 7:00 | 8a9b-B | 2.2 | 21.3 | 94 | 809 | 2 | MOLECULE: | LIPID BINDING PROTEIN P116 (MPN213) | |
| 8:00 | 5j1s-B | 2.2 | 6 | 60 | 235 | 5 | MOLECULE: | TORSIN-1A | |
| 9:00 | 7ag4-C | 2.2 | 10.5 | 90 | 425 | 8 | MOLECULE: | SULFOQUINOVOSE ISOMERASE | |
| 10:00 | 3ce2-A | 2.2 | 23.2 | 103 | 595 | 4 | MOLECULE: | PUTATIVE PEPTIDASE | |
| 11:00 | 8ccr-A | 2.2 | 3.7 | 63 | 105 | 5 | MOLECULE: | 4D2 (MUTANT T19D) | |
| 12:00 | 8vxq-J | 2.2 | 24.3 | 102 | 307 | 7 | MOLECULE: | GP72 | |
| 13:00 | 7bc4-B | 2.2 | 20.6 | 122 | 2054 | 6 | MOLECULE: | FATTY ACID SYNTHASE SUBUNIT ALPHA | |
| 14:00 | 8wzb-E | 2.2 | 11.3 | 66 | 201 | 5 | MOLECULE: | DPY30 DOMAIN CONTAINING 2 | |
| 15:00 | 6kn7-T | 2.2 | 47.1 | 119 | 138 | 6 | MOLECULE: | ACTIN, ALPHA SKELETAL MUSCLE | |
| 16:00 | 8bgo-C | 2.2 | 5.5 | 57 | 271 | 9 | MOLECULE: | DIACETYLCHITOBIOSE DEACETYLASE | |
| 17:00 | 3tdw-A | 2.2 | 15.5 | 85 | 302 | 5 | MOLECULE: | GENTAMICIN RESISTANCE PROTEIN | |

| | | | | | | | | | |
|---|---|---|---|---|---|---|---|---|---|
| 18:00 | 6gsi-J | 2.2 | 15.5 | 61 | 89 | 5 | MOLECULE: | CAPSID PROTEIN | |
| 19:00 | 7vw0-B | 2.2 | 5.8 | 82 | 94 | 5 | MOLECULE: | DUF305 DOMAIN-CONTAINING PROTEIN | |
| 20:00 | 4epz-A | 2.2 | 4.8 | 50 | 154 | 2 | MOLECULE: | TRANSCRIPTION ANTI-TERMINATOR ANTAGONIST UP | XZ |
| 21:00 | 1gqe-A | 2.2 | 12.3 | 102 | 362 | 1 | MOLECULE: | RELEASE FACTOR 2 | |
| 22:00 | 3hug-A | 2.2 | 8.5 | 51 | 80 | 12 | MOLECULE: | RNA POLYMERASE SIGMA FACTOR | |
| 23:00 | 8x2q-A | 2.2 | 3 | 79 | 109 | 11 | MOLECULE: | ADENOMATOUS POLYPOSIS COLI PROTEIN | |
| 24:00:00 | 8b9z-h | 2.2 | 10.2 | 99 | 145 | 7 | MOLECULE: | NADH-UBIQUINONE OXIDOREDUCTASE CHAIN 3 | |
| 25:00:00 | 6sc6-A | 2.2 | 12.3 | 75 | 365 | 7 | MOLECULE: | E3 UBIQUITIN-PROTEIN LIGASE RNF31 | |
| 26:00:00 | 7emf-D | 2.2 | 9.8 | 88 | 158 | 6 | MOLECULE: | MEDIATOR OF RNA POLYMERASE II TRANSCRIPTION | SUBUN |
| 27:00:00 | 1zbd-B | 2.2 | 5.1 | 80 | 124 | 5 | MOLECULE: | RABPHILIN-3A | |
| 28:00:00 | 8f2r-E | 2.2 | 2 | 41 | 205 | 10 | MOLECULE: | COMM DOMAIN-CONTAINING PROTEIN 1 | |
| 29:00:00 | 5xwp-A | 2.2 | 23.2 | 115 | 1125 | 6 | MOLECULE: | UNCHARACTERIZED PROTEIN | |
| 30:00:00 | 7qd4-A | 2.2 | 9.9 | 83 | 908 | 7 | MOLECULE: | TRANSPOSASE FOR TRANSPOSON TN4430 | |
| 31:00:00 | 6tt7-M | 2.2 | 26.5 | 70 | 156 | 11 | MOLECULE: | ATP SYNTHASE SUBUNIT ALPHA | |
| 32:00:00 | 8wzb-D | 2.2 | 4.4 | 41 | 46 | 5 | MOLECULE: | DPY30 DOMAIN CONTAINING 2 | |
| 33:00:00 | 7eqc-G | 2.2 | 48.2 | 75 | 107 | 11 | MOLECULE: | CYTOKINESIS DEFECT | |
| 34:00:00 | 6rx3-C | 2.2 | 3.4 | 48 | 99 | 8 | MOLECULE: | SYNCYTIN-2 | |
| 35:00:00 | 5w0p-C | 2.2 | 21.7 | 75 | 850 | 5 | MOLECULE: | ENDOLYSIN,RHODOPSIN,S-ARRESTIN | |
| 36:00:00 | 4bjm-A | 2.2 | 5.6 | 61 | 226 | 7 | MOLECULE: | AVRM | |
| 37:00:00 | 2fbq-A | 2.2 | 4.6 | 45 | 213 | 7 | MOLECULE: | PROBABLE TRANSCRIPTIONAL REGULATOR | |
| 38:00:00 | 8r5i-B | 2.2 | 3.2 | 40 | 43 | 10 | MOLECULE: | CORE PROTEIN A10 | |
| 39:00:00 | 8arf-B | 2.2 | 3.8 | 59 | 97 | 8 | MOLECULE: | PROTEIN SPINDLY | |
| 40:00:00 | 8i03-G | 2.2 | 21.2 | 88 | 166 | 7 | MOLECULE: | PAIRED AMPHIPATHIC HELIX PROTEIN PST1 | |
| 41:00:00 | 6z26-B | 2.2 | 21 | 61 | 103 | 7 | MOLECULE: | SPINDLE ASSEMBLY ABNORMAL PROTEIN 6 HOMOLOG | |
| 42:00:00 | 6r17-B | 2.2 | 30.6 | 65 | 94 | 9 | MOLECULE: | SYNAPTONEMAL COMPLEX CENTRAL ELEMENT PROTEI | N 2 |
| 43:00:00 | 4wij-B | 2.2 | 19.4 | 64 | 301 | 6 | MOLECULE: | SPLICING FACTOR, PROLINE- AND GLUTAMINE-RIC | H |
| 44:00:00 | 5oag-A | 2.2 | 3.6 | 55 | 98 | 13 | MOLECULE: | RAB-3A-INTERACTING PROTEIN | |
| 45:00:00 | 6zpm-A | 2.2 | 3.8 | 56 | 103 | 4 | MOLECULE: | TRYPANOSOMA CRUZI KKT4 117-218 | |
| 46:00:00 | 8fby-C | 2.2 | 3.6 | 56 | 93 | 9 | MOLECULE: | PLASMALEMMA VESICLE-ASSOCIATED PROTEIN | |
| 47:00:00 | 4bbp-A | 2.2 | 10.3 | 85 | 266 | 9 | MOLECULE: | ZINC ABC TRANSPORTER, PERIPLASMIC ZINC-BIND | ING PR |
| 48:00:00 | 7s0r-A | 2.2 | 2.2 | 44 | 85 | 11 | MOLECULE: | C PROTEIN BETA ANTIGEN | |
| 49:00:00 | 9cjl-A | 2.2 | 20.6 | 54 | 82 | 4 | MOLECULE: | TRANSMEMBRANE EMP24 DOMAIN-CONTAINING PROTE | IN 9 |
| 50:00:00 | 6jxc-A | 2.2 | 7.1 | 75 | 2345 | 9 | MOLECULE: | SERINE/THREONINE-PROTEIN KINASE TEL1 | |
| 51:00:00 | 5x3q-A | 2.2 | 9.9 | 97 | 313 | 8 | MOLECULE: | ENVELOPE GLYCOPROTEIN | |
| 52:00:00 | 1zw0-D | 2.2 | 2.8 | 45 | 66 | 11 | MOLECULE: | TYPE III SECRETION PROTEIN | |
| 53:00:00 | 7wb4-g | 2.2 | 8.3 | 57 | 672 | 11 | MOLECULE: | OUTER NUP133 | |
| 54:00:00 | 7yim-A | 2.2 | 3.1 | 39 | 592 | 8 | MOLECULE: | ALPHA-FETOPROTEIN | |
| 55:00:00 | 5fno-A | 2.2 | 3 | 50 | 568 | 2 | MOLECULE: | MANGANESE LIPOXYGENASE | |
| 56:00:00 | 1zs4-A | 2.2 | 8.7 | 61 | 82 | 5 | MOLECULE: | DNA - 27MER | |
| 57:00:00 | 7pgs-N | 2.2 | 10.9 | 67 | 1787 | 4 | MOLECULE: | NEUROFIBROMIN | |
| 58:00:00 | 6yhs-U | 2.2 | 4.5 | 46 | 61 | 4 | MOLECULE: | 23S RIBOSOMAL RNA | |
| 59:00:00 | 1g5z-A | 2.2 | 3.8 | 61 | 164 | 11 | MOLECULE: | OUTER SURFACE PROTEIN C | |
| 60:00:00 | 1z0x-A | 2.2 | 3.5 | 50 | 216 | 2 | MOLECULE: | TRANSCRIPTIONAL REGULATOR, TETR FAMILY | |
| 61:00:00 | 7nnt-C | 2.2 | 2.8 | 52 | 169 | 4 | MOLECULE: | ENERGY-COUPLING FACTOR TRANSPORTER ATP-BIND | ING PR |
| 62:00:00 | 2crb-A | 2.2 | 3.1 | 56 | 97 | 9 | MOLECULE: | NUCLEAR RECEPTOR BINDING FACTOR 2 | |
| 63:00:00 | 5tte-B | 2.2 | 11.2 | 103 | 442 | 5 | MOLECULE: | E3 UBIQUITIN-PROTEIN LIGASE ARIH1 | |
| 64:00:00 | 6h86-A | 2.1 | 22.5 | 60 | 76 | 10 | MOLECULE: | SYNAPTONEMAL COMPLEX CENTRAL ELEMENT PROTEI | N 3 |
| 65:00:00 | 6oht-A | 2.1 | 24 | 63 | 206 | 5 | MOLECULE: | 3-BETA-HYDROXYSTEROID-DELTA(8),DELTA(7)-ISO | MERASE |
| 66:00:00 | 9b3i-A | 2.1 | 34.4 | 122 | 944 | 6 | MOLECULE: | KAP114 ISOFORM 1 | |
| 67:00:00 | 8gji-A | 2.1 | 28.8 | 82 | 167 | 10 | MOLECULE: | GCG BINDER | |
| 68:00:00 | 1z7q-c | 2.1 | 18.4 | 105 | 230 | 4 | MOLECULE: | PROTEASOME COMPONENT C7-ALPHA | |
| 69:00:00 | 8dh9-D | 2.1 | 30 | 78 | 146 | 8 | MOLECULE: | LEPTIN RECEPTOR | |
| 70:00:00 | 8d8j-O | 2.1 | 37 | 78 | 210 | 6 | MOLECULE: | PROBABLE S-ADENOSYL-L-METHIONINE-DEPENDENT | RNA |
| 71:00:00 | 4acl-A | 2.1 | 25.8 | 58 | 155 | 0 | MOLECULE: | TSSL | |
| 72:00:00 | 6yj4-g | 2.1 | 12.6 | 74 | 198 | 7 | MOLECULE: | NADH-UBIQUINONE OXIDOREDUCTASE CHAIN 3 | |
| 73:00:00 | 8jme-A | 2.1 | 25.6 | 108 | 407 | 2 | MOLECULE: | GUSTATORY RECEPTOR FOR SUGAR TASTE 64A | |
| 74:00:00 | 7x3k-A | 2.1 | 11.5 | 91 | 262 | 12 | MOLECULE: | ZUOTIN | |
| 75:00:00 | 4ll7-A | 2.1 | 34.9 | 85 | 94 | 8 | MOLECULE: | SWI5-DEPENDENT HO EXPRESSION PROTEIN 3 | |
| 76:00:00 | 3hnw-A | 2.1 | 38.6 | 93 | 128 | 11 | MOLECULE: | UNCHARACTERIZED PROTEIN | |
| 77:00:00 | 7t3n-A | 2.1 | 28.1 | 113 | 452 | 9 | MOLECULE: | VESICULAR GLUTAMATE TRANSPORTER 2 | |
| 78:00:00 | 8upk-A | 2.1 | 40.7 | 100 | 739 | 7 | MOLECULE: | BACTERIOPHYTOCHROME (LIGHT-REGULATED SIGNAL | TRANS |
| 79:00:00 | 7met-B | 2.1 | 24.2 | 113 | 455 | 8 | MOLECULE: | ATP-DEPENDENT LIPID A-CORE FLIPPASE | |
| 80:00:00 | 1i5p-A | 2.1 | 24.6 | 74 | 633 | 5 | MOLECULE: | PESTICIDIAL CRYSTAL PROTEIN CRY2AA | |
| 81:00:00 | 8p1u-A | 2.1 | 24.3 | 87 | 370 | 2 | MOLECULE: | CELL DIVISION PROTEIN FTSL | |
| 82:00:00 | 5xis-D | 2.1 | 33.3 | 78 | 82 | 9 | MOLECULE: | E3 UBIQUITIN-PROTEIN LIGASE RNF168 | |
| 83:00:00 | 8ay2-A | 2.1 | 36.7 | 124 | 400 | 7 | MOLECULE: | EXOCYST COMPLEX COMPONENT 4 | |
| 84:00:00 | 8b17-A | 2.1 | 32.9 | 125 | 451 | 10 | MOLECULE: | DIPEPTIDE AND TRIPEPTIDE PERMEASE B | |
| 85:00:00 | 1ng9-A | 2.1 | 34.6 | 107 | 794 | 8 | MOLECULE: | 5'-D(*AP*GP*CP*TP*GP*CP*CP*AP*GP*GP*CP*AP*C | P*CP*A |
| 86:00:00 | 3caz-B | 2.1 | 22.4 | 138 | 210 | 9 | MOLECULE: | BAR PROTEIN | |
| 87:00:00 | 2f22-B | 2.1 | 30.7 | 74 | 143 | 9 | MOLECULE: | BH3987 | |
| 88:00:00 | 8i4t-R | 2.1 | 48 | 108 | 501 | 4 | MOLECULE: | ENVELOPMENT POLYPROTEIN | |
| 89:00:00 | 2gnx-A | 2.1 | 14.6 | 81 | 288 | 2 | MOLECULE: | HYPOTHETICAL PROTEIN | |
| 90:00:00 | 7wkk-I | 2.1 | 18.8 | 71 | 171 | 4 | MOLECULE: | MGC83295 PROTEIN | |
| 91:00:00 | 8q85-G | 2.1 | 61.5 | 113 | 178 | 6 | MOLECULE: | KINETOCHORE PROTEIN NDC80 | |
| 92:00:00 | 2qih-B | 2.1 | 43.1 | 119 | 137 | 5 | MOLECULE: | PROTEIN USPA1 | |
| 93:00:00 | 8xca-A | 2.1 | 22.3 | 83 | 668 | 7 | MOLECULE: | CASJ19 | |
| 94:00:00 | 7jm7-B | 2.1 | 27 | 82 | 218 | 2 | MOLECULE: | OSTEOPETROSIS-ASSOCIATED TRANSMEMBRANE PROT | EIN 1 |
| 95:00:00 | 7d3u-F | 2.1 | 4 | 67 | 85 | 10 | MOLECULE: | MONOVALENT NA+/H+ ANTIPORTER SUBUNIT D | |
| 96:00:00 | 9dtr-H | 2.1 | 18.6 | 104 | 460 | 8 | MOLECULE: | U2 SNRNA | |
| 97:00:00 | 8c0v-A | 2.1 | 5.9 | 58 | 823 | 5 | MOLECULE: | PEROXISOMAL ATPASE PEX1 | |
| 98:00:00 | 9h2d-A | 2.1 | 9.5 | 62 | 270 | 8 | MOLECULE: | INTRAFLAGELLAR TRANSPORT PROTEIN 172 HOMOLO | G |
| 99:00:00 | 3dby-C | 2.1 | 6.9 | 84 | 267 | 10 | MOLECULE: | UNCHARACTERIZED PROTEIN | |
| 0:00 | 8jc5-C | 2.1 | 5.2 | 53 | 196 | 4 | MOLECULE: | ADP-RIBOSYLATION FACTOR-LIKE PROTEIN 8B | |
| 1:00 | 8gel-A | 2.1 | 20.1 | 94 | 498 | 6 | MOLECULE: | SC4 | |
| 2:00 | 8t8k-A | 2.1 | 9 | 69 | 183 | 10 | MOLECULE: | DUF507 FAMILY PROTEIN | |
| 3:00 | 7s9y-A | 2.1 | 12.9 | 107 | 901 | 8 | MOLECULE: | CYTOCHROME C BIOGENESIS PROTEIN | |
| 4:00 | 4f52-E | 2.1 | 8.3 | 72 | 523 | 11 | MOLECULE: | CULLIN-1 | |
| 5:00 | 6u9f-A | 2.1 | 19.4 | 92 | 717 | 9 | MOLECULE: | PDPA | |
| 6:00 | 8gmh-A | 2.1 | 15.2 | 111 | 223 | 6 | MOLECULE: | LXG DOMAIN-CONTAINING PROTEIN | |
| 7:00 | 2nty-B | 2.1 | 13.2 | 95 | 337 | 6 | MOLECULE: | EMB\|CAB41934.1 | |
| 8:00 | 7oci-C | 2.1 | 4.8 | 74 | 119 | 5 | MOLECULE: | DOLICHYL-DIPHOSPHOOLIGOSACCHARIDE--PROTEIN | |
| 9:00 | 2efl-A | 2.1 | 28.2 | 131 | 281 | 6 | MOLECULE: | FORMIN-BINDING PROTEIN 1 | |
| 10:00 | 2eul-A | 2.1 | 6.8 | 66 | 156 | 8 | MOLECULE: | ANTI-CLEAVAGE ANTI-GREA TRANSCRIPTION FACTO | R GFH1 |
| 11:00 | 8p0u-A | 2.1 | 17.7 | 114 | 621 | 3 | MOLECULE: | POLYMERASE ACIDIC PROTEIN | |
| 12:00 | 2gyq-A | 2.1 | 7.4 | 81 | 162 | 6 | MOLECULE: | YCFI, PUTATIVE STRUCTURAL PROTEIN | |
| 13:00 | 5y3t-A | 2.1 | 13.7 | 42 | 135 | 12 | MOLECULE: | RANBP-TYPE AND C3HC4-TYPE ZINC FINGER-CONTA | INING |
| 14:00 | 7ktr-D | 2.1 | 9.9 | 83 | 219 | 8 | MOLECULE: | TRANSFORMATION/TRANSCRIPTION DOMAIN-ASSOCIA | TED PR |
| 15:00 | 4hwi-B | 2.1 | 2.8 | 70 | 177 | 11 | MOLECULE: | HEAT SHOCK COGNATE 71 KDA PROTEIN | |
| 16:00 | 6c94-A | 2.1 | 20.4 | 98 | 482 | 6 | MOLECULE: | CYTOCHROME P450 4B1 | |
| 17:00 | 6qrj-A | 2.1 | 21.8 | 87 | 463 | 5 | MOLECULE: | HYBRID KINASE | |
| 18:00 | 3b77-A | 2.1 | 7.1 | 86 | 188 | 6 | MOLECULE: | UNCHARACTERIZED PROTEIN | |

| | | | | | | | | | | |
|---|---|---|---|---|---|---|---|---|---|---|
| 19:00 | 8pnj-A | 2.1 | 5.5 | 68 | 390 | 6 | MOLECULE: | CHORISMATE MUTASE | | |
| 20:00 | 3bru-A | 2.1 | 10.3 | 77 | 191 | 5 | MOLECULE: | REGULATORY PROTEIN, TETR FAMILY | | |
| 21:00 | 6m3i-A | 2.1 | 15.7 | 68 | 316 | 9 | MOLECULE: | HISTONE PARYLATION FACTOR 1 | | |
| 22:00 | 6y37-C | 2.1 | 3.4 | 51 | 118 | 6 | MOLECULE: | HAPB | | |
| 23:00 | 2yfb-B | 2.1 | 12.4 | 110 | 238 | 10 | MOLECULE: | METHYL-ACCEPTING CHEMOTAXIS TRANSDUCER | | |
| 24:00:00 | 6xtv-B | 2.1 | 18.6 | 82 | 291 | 9 | MOLECULE: | LYSINE EXPORT TRANSCRIPTIONAL REGULATORY PR | OTEIN | |
| 25:00:00 | 4zrl-A | 2.1 | 2.7 | 41 | 313 | 5 | MOLECULE: | POLY(A) RNA POLYMERASE GLD-2 | | |
| 26:00:00 | 4drf-A | 2.1 | 7.2 | 107 | 408 | 7 | MOLECULE: | METALLOPHOSPHOESTERASE | | |
| 27:00:00 | 7pkn-O | 2.1 | 8.8 | 69 | 210 | 1 | MOLECULE: | CENTROMERE PROTEIN H | | |
| 28:00:00 | 4cti-A | 2.1 | 5.9 | 96 | 207 | 6 | MOLECULE: | OSMOLARITY SENSOR PROTEIN ENVZ, AF1503 | | |
| 29:00:00 | 6rmn-B | 2.1 | 8.6 | 68 | 222 | 4 | MOLECULE: | DNA MISMATCH REPAIR PROTEIN MLH1 | | |
| 30:00:00 | 7kak-E | 2.1 | 9.7 | 77 | 172 | 3 | MOLECULE: | PROTEIN TRANSPORT CHANNEL SEC61 COMPLEX, AL | PHA SU | |
| 31:00:00 | 8aw5-A | 2.1 | 7.5 | 97 | 266 | 5 | MOLECULE: | HEME O OXYGENASE | | |
| 32:00:00 | 8rt9-A | 2.1 | 33.1 | 93 | 207 | 10 | MOLECULE: | TRWJ PROTEIN | | |
| 33:00:00 | 8z4q-A | 2.1 | 8.5 | 86 | 463 | 5 | MOLECULE: | HYDROQUINONE DIOXYGENASE PAD | | |
| 34:00:00 | 7w72-U | 2.1 | 10 | 88 | 420 | 6 | MOLECULE: | PHOSPHATIDYLINOSITOL GLYCAN ANCHOR BIOSYNTH | ESIS C | |
| 35:00:00 | 7tbq-A | 2.1 | 6.1 | 88 | 375 | 7 | MOLECULE: | D7 LOV2-DARPIN FUSION | | |
| 36:00:00 | 2vwb-A | 2.1 | 2.6 | 43 | 519 | 5 | MOLECULE: | PUTATIVE O-SIALOGLYCOPROTEIN ENDOPEPTIDASE | | |
| 37:00:00 | 6wi1-A | 2.1 | 6.1 | 84 | 806 | 2 | MOLECULE: | IONOTROPIC GLUTAMATE RECEPTOR , NMDA RECEPT | OR GLU | |
| 38:00:00 | 5y69-A | 2.1 | 3.1 | 40 | 147 | 8 | MOLECULE: | CHAIN A | | |
| 39:00:00 | 9g8u-B | 2.1 | 15.4 | 65 | 285 | 8 | MOLECULE: | LACTONIZING LIPASE | | |
| 40:00:00 | 5da0-A | 2.1 | 15.6 | 80 | 467 | 1 | MOLECULE: | SULPHATE TRANSPORTER | | |
| 41:00:00 | 7zmb-9 | 2.1 | 7.7 | 74 | 103 | 3 | MOLECULE: | NADH-UBIQUINONE OXIDOREDUCTASE CHAIN 1 | | |
| 42:00:00 | 6ch3-B | 2.1 | 7.4 | 72 | 162 | 4 | MOLECULE: | FLAGELLAR BIOSYNTHESIS PROTEIN FLHA | | |
| 43:00:00 | 6grk-A | 2.1 | 7.5 | 106 | 354 | 6 | MOLECULE: | AHLB | | |
| 44:00:00 | 7xuh-A | 2.1 | 24.8 | 114 | 707 | 5 | MOLECULE: | CHLORIDE ANION EXCHANGER | | |
| 45:00:00 | 8t1l-Y | 2.1 | 16.7 | 93 | 154 | 8 | MOLECULE: | MEDIATOR OF RNA POLYMERASE II TRANSCRIPTION | SUBUN | |
| 46:00:00 | 8ear-A | 2.1 | 37 | 116 | 2389 | 3 | MOLECULE: | INOSITOL 1,4,5-TRISPHOSPHATE RECEPTOR TYPE | | 1 |
| 47:00:00 | 6krd-D | 2.1 | 10.5 | 71 | 429 | 3 | MOLECULE: | T-COMPLEX PROTEIN 1 SUBUNIT ALPHA | | |
| 48:00:00 | 2gtv-X | 2.1 | 3 | 46 | 104 | 11 | MOLECULE: | CHORISMATE MUTASE | | |
| 49:00:00 | 6nr8-1 | 2.1 | 2.9 | 70 | 107 | 10 | MOLECULE: | PREFOLDIN SUBUNIT 1 | | |
| 50:00:00 | 5t58-A | 2.1 | 33.4 | 122 | 233 | 7 | MOLECULE: | KLLA0F02343P | | |
| 51:00:00 | 5lj3-G | 2.1 | 3.2 | 58 | 97 | 10 | MOLECULE: | U5 SNRNA (SMALL NUCLEAR RNA) | | |
| 52:00:00 | 5bz1-A | 2.1 | 13.6 | 71 | 371 | 6 | MOLECULE: | SUPPRESSOR PROTEIN MPT5 | | |
| 53:00:00 | 8ppr-M | 2.1 | 36.1 | 114 | 205 | 4 | MOLECULE: | KINETOCHORE-ASSOCIATED PROTEIN DSN1 HOMOLOG | | |
| 54:00:00 | 1yf2-A | 2.1 | 4.3 | 88 | 425 | 6 | MOLECULE: | TYPE I RESTRICTION-MODIFICATION ENZYME, S S | UBUNIT | |
| 55:00:00 | 3eff-K | 2.1 | 11.3 | 96 | 139 | 2 | MOLECULE: | FAB | | |
| 56:00:00 | 5eqw-B | 2.1 | 4.3 | 69 | 123 | 6 | MOLECULE: | PUTATIVE MAJOR COAT PROTEIN | | |
| 57:00:00 | 3f3s-A | 2.1 | 6.2 | 52 | 312 | 4 | MOLECULE: | LAMBDA-CRYSTALLIN HOMOLOG | | |
| 58:00:00 | 5muu-A | 2.1 | 19.2 | 102 | 766 | 5 | MOLECULE: | MAJOR INNER PROTEIN P1 | | |
| 59:00:00 | 6o3v-B | 2.1 | 15.2 | 74 | 831 | 7 | MOLECULE: | PROTEIN VP3 | | |
| 60:00:00 | 6tv6-A | 2.1 | 2.7 | 44 | 362 | 9 | MOLECULE: | PROTEIN-ARGININE KINASE | | |
| 61:00:00 | 8khl-A | 2.1 | 10.8 | 62 | 275 | 16 | MOLECULE: | PROBABLE ACYL-COA LYASE BETA CHAIN | | |
| 62:00:00 | 8g8i-A | 2.1 | 15.1 | 80 | 465 | 6 | MOLECULE: | C3HR3_9R_SHIFT4 | | |
| 63:00:00 | 8ixk-E | 2.1 | 8.2 | 81 | 109 | 5 | MOLECULE: | ATTACHMENT PROTEIN G3P | | |
| 64:00:00 | 5uts-A | 2.1 | 13.6 | 82 | 447 | 6 | MOLECULE: | C-S LYASE EGT2 | | |
| 65:00:00 | 7xym-A | 2.1 | 22.3 | 104 | 488 | 4 | MOLECULE: | ENVELOPE GLYCOPROTEIN | | |
| 66:00:00 | 2mn2-A | 2.1 | 3.5 | 71 | 122 | 4 | MOLECULE: | YMOB | | |
| 67:00:00 | 7way-A | 2.1 | 30.9 | 136 | 960 | 7 | MOLECULE: | DNA (33-MER) | | |
| 68:00:00 | 7faw-A | 2.1 | 2.5 | 44 | 84 | 0 | MOLECULE: | TRANSCRIPTION ELONGATION FACTOR S-II | | |
| 69:00:00 | 2xeq-C | 2.1 | 15.5 | 79 | 241 | 6 | MOLECULE: | PAT1 HOMOLOG 1, | | |
| 70:00:00 | 8fef-D | 2.1 | 17.8 | 69 | 385 | 12 | MOLECULE: | VIRULENCE FACTOR MCE FAMILY PROTEIN | | |
| 71:00:00 | 7vpt-A | 2.1 | 23.9 | 90 | 469 | 10 | MOLECULE: | C. GLABRATA IMPORTIN ALPHA ARM DOMAIN - UPC | 2 NLS | |
| 72:00:00 | 5yo8-A | 2.1 | 17.6 | 77 | 349 | 9 | MOLECULE: | TETRAPRENYL-BETA-CURCUMENE SYNTHASE | | |
| 73:00:00 | 7vcf-C | 2.1 | 12.7 | 67 | 217 | 3 | MOLECULE: | TIC214 | | |
| 74:00:00 | 7wb4-E | 2.1 | 23.8 | 132 | 1363 | 4 | MOLECULE: | OUTER NUP133 | | |
| 75:00:00 | 1sms-A | 2.1 | 20.6 | 71 | 314 | 3 | MOLECULE: | RIBONUCLEOSIDE-DIPHOSPHATE REDUCTASE SMALL | CHAIN | |
| 76:00:00 | 4dnh-A | 2.1 | 3 | 35 | 386 | 0 | MOLECULE: | UNCHARACTERIZED PROTEIN | | |
| 77:00:00 | 4hts-A | 2.1 | 14.4 | 66 | 228 | 5 | MOLECULE: | SEC-INDEPENDENT PROTEIN TRANSLOCASE PROTEIN | TATC | |
| 78:00:00 | 2o6i-B | 2.1 | 11.4 | 76 | 430 | 7 | MOLECULE: | HD DOMAIN PROTEIN | | |
| 79:00:00 | 8ap7-a | 2.1 | 8.3 | 97 | 231 | 4 | MOLECULE: | ATP SYNTHASE SUBUNIT A | | |
| 80:00:00 | 7tuk-A | 2.1 | 17 | 80 | 142 | 6 | MOLECULE: | SAG PROTEIN | | |
| 81:00:00 | 5wrr-B | 2.1 | 7.2 | 85 | 438 | 13 | MOLECULE: | PSEUDOKINASE FAM20A | | |
| 82:00:00 | 7u1t-A | 2.1 | 3 | 40 | 176 | 18 | MOLECULE: | EPSTEIN-BARR NUCLEAR ANTIGEN 1 | | |
| 83:00:00 | 4u8u-G | 2.1 | 3 | 42 | 151 | 2 | MOLECULE: | GLOBIN A CHAIN | | |
| 84:00:00 | 6w4x-C | 2.1 | 17.7 | 112 | 375 | 4 | MOLECULE: | RIBONUCLEOSIDE-DIPHOSPHATE REDUCTASE 1 SUBU | NIT AL | |
| 85:00:00 | 7zw1-A | 2.1 | 8.6 | 102 | 317 | 4 | MOLECULE: | PERIPHERIN-2 | | |
| 86:00:00 | 5cow-A | 2.1 | 5.5 | 70 | 234 | 1 | MOLECULE: | PUTATIVE UNCHARACTERIZED PROTEIN | | |
| 87:00:00 | 8jbd-A | 2.1 | 9.8 | 56 | 316 | 7 | MOLECULE: | ADENYLOSUCCINATE LYASE | | |
| 88:00:00 | 2oqm-A | 2.1 | 4.4 | 50 | 180 | 8 | MOLECULE: | HYPOTHETICAL PROTEIN | | |
| 89:00:00 | 6zyx-Y | 2.1 | 32.4 | 111 | 1067 | 11 | MOLECULE: | DYNEIN HEAVY CHAIN, OUTER ARM PROTEIN | | |
| 90:00:00 | 9bi4-D | 2.1 | 8.7 | 91 | 459 | 9 | MOLECULE: | DNA REPAIR PROTEIN RAD50 | | |
| 91:00:00 | 7vmb-A | 2.1 | 1.5 | 38 | 359 | 3 | MOLECULE: | IQ MOTIF AND SEC7 DOMAIN-CONTAINING PROTEIN | | 1 |
| 92:00:00 | 5njx-A | 2.1 | 2.5 | 52 | 413 | 8 | MOLECULE: | PEPTIDYL-PROLYL CIS-TRANS ISOMERASE FKBP5 | | |
| 93:00:00 | 6mag-A | 2.1 | 14.2 | 78 | 282 | 9 | MOLECULE: | BBVCI ENDONUCLEASE SUBUNIT 2 | | |
| 94:00:00 | 7xqk-B | 2.1 | 8.8 | 46 | 276 | 4 | MOLECULE: | GLUTATHIONE S-TRANSFERASE CLASS-MU 26 KDA I | SOZYME | |
| 95:00:00 | 3gvd-A | 2.1 | 8.1 | 64 | 273 | 3 | MOLECULE: | SERINE ACETYLTRANSFERASE | | |
| 96:00:00 | 7rps-A | 2.1 | 9 | 88 | 154 | 5 | MOLECULE: | FIBRONECTIN-BINDING LIPOPROTEIN FBPB | | |
| 97:00:00 | 2qco-A | 2.1 | 4.2 | 44 | 202 | 7 | MOLECULE: | CMER | | |
| 98:00:00 | 4l6v-6 | 2.1 | 7.5 | 56 | 125 | 4 | MOLECULE: | PHOTOSYSTEM I P700 CHLOROPHYLL A APOPROTEIN | A1 | |
| 99:00:00 | 8qa9-B | 2.1 | 3.2 | 36 | 209 | 17 | MOLECULE: | TRANSCRIPTIONAL REPRESSOR PROTEIN KORB | | |
| 0:00 | 7qen-D | 2.1 | 16.6 | 124 | 1050 | 10 | MOLECULE: | DNA (35-MER) | | |
| 1:00 | 5dwd-B | 2.1 | 9.8 | 68 | 223 | 4 | MOLECULE: | ESTERASE | | |
| 2:00 | 4b6i-C | 2.1 | 3.5 | 43 | 102 | 5 | MOLECULE: | SMA2266 | | |
| 3:00 | 8ds9-A | 2.1 | 8 | 77 | 197 | 6 | MOLECULE: | VOLUME-REGULATED ANION CHANNEL SUBUNIT LRRC | 8A,SOL | |
| 4:00 | 8qat-C | 2.1 | 5 | 51 | 606 | 14 | MOLECULE: | PROTEIN HOOK HOMOLOG 3 | | |
| 5:00 | 6mzc-I | 2.1 | 23.8 | 105 | 338 | 9 | MOLECULE: | TRANSCRIPTION INITIATION FACTOR TFIID SUBUN | IT 2 | |
| 6:00 | 6wtw-A | 2.1 | 8.3 | 76 | 489 | 12 | MOLECULE: | DASS FAMILY SODIUM-COUPLED ANION SYMPORTER | | |
| 7:00 | 3lxu-X | 2.1 | 9.4 | 103 | 1217 | 13 | MOLECULE: | TRIPEPTIDYL-PEPTIDASE 2 | | |
| 8:00 | 6fze-A | 2.1 | 3.1 | 70 | 209 | 3 | MOLECULE: | PUTATIVE SURFACE PROTEIN | | |
| 9:00 | 4n2q-A | 2.1 | 22 | 70 | 198 | 1 | MOLECULE: | THA8 RNA BINDING PROTEIN | | |
| 10:00 | 7etr-A | 2.1 | 7.5 | 55 | 88 | 2 | MOLECULE: | TRANSCRIPTIONAL REGULATOR COPG FAMILY | | |
| 11:00 | 2yc2-C | 2.1 | 4.1 | 43 | 149 | 7 | MOLECULE: | INTRAFLAGELLAR TRANSPORT PROTEIN 25 | | |
| 12:00 | 7zcv-B | 2.1 | 11.9 | 81 | 285 | 6 | MOLECULE: | TRANSCRIPTIONAL REGULATOR | | |
| 13:00 | 7rcy-A | 2.1 | 26.2 | 75 | 829 | 13 | MOLECULE: | PENICILLIN-BINDING PROTEIN | | |
| 14:00 | 4dci-B | 2.1 | 8.8 | 87 | 148 | 7 | MOLECULE: | UNCHARACTERIZED PROTEIN | | |
| 15:00 | 6m31-A | 2.1 | 6.9 | 73 | 280 | 3 | MOLECULE: | DIGERANYLGERANYLGLYCERYL PHOSPHATE SYNTHASE | | |
| 16:00 | 1uus-A | 2.1 | 17 | 100 | 465 | 5 | MOLECULE: | STAT PROTEIN | | |
| 17:00 | 8tie-a | 2.1 | 17.2 | 112 | 1012 | 6 | MOLECULE: | NUCLEOPORIN NUP120 | | |
| 18:00 | 7ui4-A | 2.1 | 8.5 | 77 | 413 | 5 | MOLECULE: | DNA-GUANINE TRANSGLYCOSYLASE | | |
| 19:00 | 3iee-A | 2.1 | 9.3 | 93 | 256 | 8 | MOLECULE: | PUTATIVE EXPORTED PROTEIN | | |

| | | | | | | | | | | |
|---|---|---|---|---|---|---|---|---|---|---|
| 20:00 | 7pkq-O | 2.1 | 3.7 | 68 | 236 | 6 | MOLECULE: | MS35 | | |
| 21:00 | 9dof-D | 2.1 | 9 | 51 | 166 | 16 | MOLECULE: | GLYCOPROTEIN E | | |
| 22:00 | 6xdc-A | 2.1 | 9.9 | 83 | 193 | 5 | MOLECULE: | PROTEIN 3A | | |
| 23:00 | 5n9j-U | 2.1 | 17.4 | 120 | 198 | 10 | MOLECULE: | MEDIATOR OF RNA POLYMERASE II TRANSCRIPTION | SUBUN | |
| 24:00:00 | 6qd6-G | 2.1 | 36.5 | 120 | 469 | 8 | MOLECULE: | MB-CHOPQ-NB207,OUTER MEMBRANE PROTEIN,MB-CH | OPQ-NB | |
| 25:00:00 | 5v2c-b | 2.1 | 16.2 | 86 | 506 | 8 | MOLECULE: | PHOTOSYSTEM II PROTEIN D1 | | |
| 26:00:00 | 4m1p-A | 2.1 | 3.9 | 67 | 96 | 6 | MOLECULE: | COPPER-SENSITIVE OPERON REPRESSOR (CSOR) | | |
| 27:00:00 | 5ouz-A | 2.1 | 3.6 | 57 | 178 | 9 | MOLECULE: | FERRITIN | | |
| 28:00:00 | 6tmi-A | 2.1 | 7.8 | 60 | 109 | 7 | MOLECULE: | ATP SYNTHASE SUBUNIT ALPHA | | |
| 29:00:00 | 8tl7-A | 2.1 | 16.3 | 98 | 694 | 9 | MOLECULE: | COMPUTATIONALLY DESIGNED PROTEIN | | |
| 30:00:00 | 6yjj-A | 2.1 | 9.5 | 68 | 320 | 9 | MOLECULE: | CYRI-B (FAM49B) | | |
| 31:00:00 | 5i4q-B | 2.1 | 3.8 | 44 | 107 | 5 | MOLECULE: | CONTACT-DEPENDENT INHIBITOR A | | |
| 32:00:00 | 7ogt-A | 2.1 | 30.8 | 111 | 688 | 5 | MOLECULE: | STRUCTURAL MAINTENANCE OF CHROMOSOMES PROTE | IN 1 | |
| 33:00:00 | 3auy-A | 2.1 | 10.5 | 94 | 366 | 4 | MOLECULE: | DNA DOUBLE-STRAND BREAK REPAIR RAD50 ATPASE | | |
| 34:00:00 | 4r8g-E | 2.1 | 34.8 | 94 | 324 | 6 | MOLECULE: | UNCONVENTIONAL MYOSIN-IC | | |
| 35:00:00 | 4fgv-A | 2.1 | 22 | 82 | 1067 | 5 | MOLECULE: | CHROMOSOME REGION MAINTENANCE 1 (CRM1) OR E | XPORTI | |
| 36:00:00 | 5aww-G | 2.1 | 8 | 53 | 75 | 8 | MOLECULE: | PROTEIN TRANSLOCASE SUBUNIT SECY | | |
| 37:00:00 | 2no2-A | 2.1 | 13.1 | 60 | 102 | 7 | MOLECULE: | HUNTINGTIN-INTERACTING PROTEIN 1 | | |
| 38:00:00 | 8q85-F | 2.1 | 19.1 | 61 | 230 | 11 | MOLECULE: | KINETOCHORE PROTEIN NDC80 | | |
| 39:00:00 | 2qa7-C | 2.1 | 3.3 | 56 | 102 | 13 | MOLECULE: | HUNTINGTIN-INTERACTING PROTEIN 1 | | |
| 40:00:00 | 8th8-C | 2.1 | 5.4 | 91 | 533 | 7 | MOLECULE: | DYNEIN REGULATORY COMPLEX PROTEIN 1/2 N-TER | MINAL | |
| 41:00:00 | 5zak-A | 2.1 | 16.7 | 66 | 1314 | 8 | MOLECULE: | ENDORIBONUCLEASE DICER | | |
| 42:00:00 | 6c0f-w | 2.1 | 3 | 43 | 70 | 5 | MOLECULE: | SACCHAROMYCES CEREVISIAE S288C 35S PRE-RIBO | SOMAL | |
| 43:00:00 | 6m1h-B | 2.1 | 2.6 | 45 | 61 | 9 | MOLECULE: | PITUITARY ADENYLATE CYCLASE-ACTIVATING POLY | PEPTID | |
| 44:00:00 | 2db7-A | 2.1 | 2.5 | 37 | 57 | 3 | MOLECULE: | HAIRY/ENHANCER-OF-SPLIT RELATED WITH YRPW M | OTIF | |
| 45:00:00 | 1c02-A | 2.1 | 3.3 | 51 | 166 | 8 | MOLECULE: | PHOSPHOTRANSFERASE YPD1P | | |
| 46:00:00 | 7q83-D | 2.1 | 2.8 | 46 | 156 | 9 | MOLECULE: | EXOCYST COMPLEX COMPONENT SEC3 | | |
| 47:00:00 | 7zpo-M | 2.1 | 7.1 | 56 | 438 | 11 | MOLECULE: | KTR SYSTEM POTASSIUM UPTAKE PROTEIN A | | |
| 48:00:00 | 7bjg-A | 2.1 | 34 | 77 | 98 | 9 | MOLECULE: | SD21996P | | |
| 49:00:00 | 7k15-A | 2.1 | 7.1 | 65 | 455 | 6 | MOLECULE: | LEUKOTRIENE B4 RECEPTOR 1,FLAVODOXIN,LEUKOT | RIENE | |
| 50:00:00 | 6zbj-A | 2.1 | 6 | 75 | 666 | 4 | MOLECULE: | PRECURSOR OF THE MAJOR MEROZOITE SURFACE AN | TIGENS | |
| 51:00:00 | 3icq-T | 2 | 26.4 | 100 | 949 | 8 | MOLECULE: | EXPORTIN-T | | |
| 52:00:00 | 7p34-A | 2 | 24.8 | 146 | 391 | 5 | MOLECULE: | PEPTIDE ANTIBIOTIC TRANSPORTER SBMA | | |
| 53:00:00 | 9f3f-1 | 2 | 27.1 | 149 | 751 | 6 | MOLECULE: | NUCLEAR CAP BINDING COMPLEX SUBUNIT CBP110 | | |
| 54:00:00 | 6fln-B | 2 | 31.9 | 105 | 367 | 8 | MOLECULE: | E3 UBIQUITIN/ISG15 LIGASE TRIM25 | | |
| 55:00:00 | 2o2k-B | 2 | 16.2 | 59 | 333 | 5 | MOLECULE: | METHIONINE SYNTHASE | | |
| 56:00:00 | 7w7g-B | 2 | 31 | 134 | 1710 | 6 | MOLECULE: | PROTEIN UNC-79 HOMOLOG | | |
| 57:00:00 | 8uw3-A | 2 | 15.4 | 106 | 1265 | 6 | MOLECULE: | LINE-1 RETROTRANSPOSABLE ELEMENT ORF2 PROTE | IN | |
| 58:00:00 | 8hnc-A | 2 | 14.8 | 101 | 570 | 8 | MOLECULE: | SOLUTE CARRIER ORGANIC ANION TRANSPORTER FA | MILY M | |
| 59:00:00 | 8vb2-E | 2 | 46 | 160 | 844 | 4 | MOLECULE: | TETRAMERIC EJECTION PROTEIN (GP48) | | |
| 60:00:00 | 5eri-A | 2 | 23.8 | 73 | 153 | 8 | MOLECULE: | MARR FAMILY TRANSCRIPTIONAL REGULATOR | | |
| 61:00:00 | 8g0p-A | 2 | 28.9 | 106 | 143 | 5 | MOLECULE: | KINETOCHORE PROTEIN NDC80 HOMOLOG | | |
| 62:00:00 | 7bss-A | 2 | 33.7 | 145 | 816 | 7 | MOLECULE: | ATP11C | | |
| 63:00:00 | 8hhe-B | 2 | 11.4 | 99 | 647 | 3 | MOLECULE: | CRYSTALINE ENTOMOCIDAL PROTOXIN | | |
| 64:00:00 | 8agg-A | 2 | 2.7 | 35 | 489 | 9 | MOLECULE: | PHOSPHOCHOLINE HYDROLASE LEM3 | | |
| 65:00:00 | 7qpr-D | 2 | 25.5 | 92 | 689 | 5 | MOLECULE: | ACT DOMAIN PROTEIN | | |
| 66:00:00 | 7w0y-k | 2 | 8.7 | 77 | 98 | 4 | MOLECULE: | NADH DEHYDROGENASE [UBIQUINONE] FLAVOPROTEI | N 1, | |
| 67:00:00 | 9exv-C | 2 | 4.6 | 70 | 99 | 9 | MOLECULE: | NITROREDUCTASE | | |
| 68:00:00 | 3va9-A | 2 | 4.9 | 76 | 127 | 8 | MOLECULE: | SENSOR HISTIDINE KINASE | | |
| 69:00:00 | 5zle-A | 2 | 3.7 | 64 | 225 | 9 | MOLECULE: | CYTOCHROME B REDUCTASE 1 | | |
| 70:00:00 | 7xp9-A | 2 | 7.2 | 48 | 76 | 10 | MOLECULE: | RXLR EFFECTOR PROTEIN AVR-VNT11 | | |
| 71:00:00 | 3ezh-A | 2 | 4.1 | 69 | 114 | 3 | MOLECULE: | NITRATE/NITRITE SENSOR PROTEIN NARX | | |
| 72:00:00 | 4gcz-B | 2 | 11.1 | 84 | 378 | 4 | MOLECULE: | BLUE-LIGHT PHOTORECEPTOR, SENSOR PROTEIN FI | XL | |
| 73:00:00 | 6g49-A | 2 | 9.6 | 43 | 288 | 5 | MOLECULE: | PROTEIN-GLUTAMINE GAMMA-GLUTAMYLTRANSFERASE | | |
| 74:00:00 | 6h4j-A | 2 | 32.1 | 136 | 488 | 7 | MOLECULE: | UBIQUITIN CARBOXYL-TERMINAL HYDROLASE 25 | | |
| 75:00:00 | 4mk6-A | 2 | 7.1 | 80 | 188 | 5 | MOLECULE: | PROBABLE DIHYDROXYACETONE KINASE REGULATOR | DHSK_R | |
| 76:00:00 | 7qcd-C | 2 | 11.4 | 93 | 267 | 8 | MOLECULE: | STRUCTURAL MAINTENANCE OF CHROMOSOMES PROTE | IN 5 | |
| 77:00:00 | 6tg5-A | 2 | 6.7 | 63 | 111 | 3 | MOLECULE: | MACPD | | |
| 78:00:00 | 4wcx-A | 2 | 11.2 | 81 | 456 | 4 | MOLECULE: | BIOTIN AND THIAMIN SYNTHESIS ASSOCIATED | | |
| 79:00:00 | 6gts-B | 2 | 6 | 52 | 71 | 4 | MOLECULE: | ACETYLTRANSFERASE | | |
| 80:00:00 | 4iwb-A | 2 | 6.9 | 82 | 165 | 1 | MOLECULE: | FLIC, FLIS CHIMERA | | |
| 81:00:00 | 4mu6-A | 2 | 22.7 | 111 | 274 | 10 | MOLECULE: | KINECTIN 1 (KINESIN RECEPTOR) | | |
| 82:00:00 | 6s6b-I | 2 | 11 | 77 | 284 | 6 | MOLECULE: | CRISPR-ASSOCIATED PROTEIN, CMR5 FAMILY | | |
| 83:00:00 | 3mxz-A | 2 | 6.7 | 67 | 107 | 7 | MOLECULE: | TUBULIN-SPECIFIC CHAPERONE A | | |
| 84:00:00 | 3viq-A | 2 | 10.7 | 58 | 122 | 3 | MOLECULE: | SWI5-DEPENDENT RECOMBINATION DNA REPAIR PRO | TEIN 1 | |
| 85:00:00 | 8qem-M | 2 | 2.2 | 38 | 454 | 3 | MOLECULE: | PUTATIVE NECK PROTEIN | | |
| 86:00:00 | 8ec3-A | 2 | 9.2 | 90 | 155 | 11 | MOLECULE: | FIBRONECTIN-BINDING PROTEIN | | |
| 87:00:00 | 7ar9-M | 2 | 8.3 | 92 | 438 | 1 | MOLECULE: | ND3 | | |
| 88:00:00 | 6tdv-O | 2 | 19.7 | 84 | 100 | 2 | MOLECULE: | ATPTB1 | | |
| 89:00:00 | 9b8e-B | 2 | 8.4 | 57 | 225 | 7 | MOLECULE: | CEG10 | | |
| 90:00:00 | 7fh1-A | 2 | 7.3 | 96 | 874 | 6 | MOLECULE: | MECKELIN | | |
| 91:00:00 | 7y82-A | 2 | 23.7 | 90 | 1341 | 3 | MOLECULE: | RAMP SUPERFAMILY PROTEIN | | |
| 92:00:00 | 6giy-G | 2 | 6.4 | 103 | 444 | 3 | MOLECULE: | TSSF | | |
| 93:00:00 | 1wa8-A | 2 | 7.8 | 72 | 99 | 7 | MOLECULE: | ESAT-6 LIKE PROTEIN ESXB | | |
| 94:00:00 | 5y2v-A | 2 | 17.4 | 66 | 304 | 11 | MOLECULE: | RUBISCO OPERON TRANSCRIPTIONAL REGULATOR | | |
| 95:00:00 | 8fbi-A | 2 | 6.5 | 93 | 274 | 4 | MOLECULE: | KWOCA_39 | | |
| 96:00:00 | 6iiv-A | 2 | 4.8 | 97 | 461 | 1 | MOLECULE: | SOLUBLE CYTOCHROME B562,THROMBOXANE A2 RECE | PTOR,R | |
| 97:00:00 | 8z9a-A | 2 | 9.5 | 128 | 406 | 1 | MOLECULE: | ODORANT RECEPTOR, APISORCO | | |
| 98:00:00 | 7z0s-C | 2 | 13.3 | 118 | 604 | 6 | MOLECULE: | FORMATE HYDROGENLYASE SUBUNIT 3 | | |
| 99:00:00 | 6mev-A | 2 | 3.4 | 36 | 342 | 3 | MOLECULE: | BIFUNCTIONAL ARGININE DEMETHYLASE AND LYSYL | #NAME? | |
| 0:00 | 6z0c-A | 2 | 5.5 | 97 | 199 | 5 | MOLECULE: | MAQUETTE-3 | | |
| 1:00 | 3zqq-C | 2 | 12.2 | 82 | 134 | 9 | MOLECULE: | TERMINASE SMALL SUBUNIT | | |
| 2:00 | 6o1q-A | 2 | 4.5 | 74 | 119 | 4 | MOLECULE: | NEPHROCYSTIN-1 | | |
| 3:00 | 8d8j-5 | 2 | 9.2 | 54 | 270 | 4 | MOLECULE: | PROBABLE S-ADENOSYL-L-METHIONINE-DEPENDENT | RNA | |
| 4:00 | 8ver-B | 2 | 5.8 | 96 | 287 | 5 | MOLECULE: | ENDORIBONUCLEASE YICC | | |
| 5:00 | 5m1m-A | 2 | 4.4 | 65 | 154 | 9 | MOLECULE: | MATRIX PROTEIN 1 | | |
| 6:00 | 3l9f-A | 2 | 2.9 | 58 | 170 | 7 | MOLECULE: | PUTATIVE UNCHARACTERIZED PROTEIN SMU.1604C | | |
| 7:00 | 9c3i-S | 2 | 3.3 | 75 | 231 | 5 | MOLECULE: | CCR4-NOT TRANSCRIPTION COMPLEX SUBUNIT 3 | | |
| 8:00 | 2wtt-D | 2 | 2.6 | 43 | 47 | 5 | MOLECULE: | TUMOR PROTEIN P73 | | |
| 9:00 | 8qx8-D | 2 | 12.4 | 43 | 613 | 2 | MOLECULE: | VACUOLAR PROTEIN SORTING-ASSOCIATED PROTEIN | | 8 |
| 10:00 | 2k73-A | 2 | 11.5 | 94 | 183 | 5 | MOLECULE: | DISULFIDE BOND FORMATION PROTEIN B | | |
| 11:00 | 4bc2-A | 2 | 8 | 55 | 528 | 9 | MOLECULE: | XYLULOSE KINASE | | |
| 12:00 | 5yma-A | 2 | 7.9 | 77 | 179 | 5 | MOLECULE: | PUTATIVE RRNA PROCESSING PROTEIN | | |
| 13:00 | 6idy-A | 2 | 15.1 | 50 | 421 | 16 | MOLECULE: | LIPASE AFLB | | |
| 14:00 | 8rmm-E | 2 | 19.8 | 119 | 290 | 3 | MOLECULE: | CALCIUM HOMEOSTASIS MODULATOR PROTEIN 4 | | |
| 15:00 | 6umm-E | 2 | 5.6 | 87 | 343 | 11 | MOLECULE: | ESX-3 SECRETION SYSTEM PROTEIN ECCE3 | | |
| 16:00 | 6u8y-b | 2 | 4.9 | 65 | 79 | 6 | MOLECULE: | MONOVALENT CATION/H+ ANTIPORTER SUBUNIT E | | |
| 17:00 | 6iuq-B | 2 | 2.9 | 37 | 296 | 3 | MOLECULE: | PROLYL 4-HYDROXYLASE | | |
| 18:00 | 5cuf-C | 2 | 14.8 | 70 | 370 | 7 | MOLECULE: | SESTRIN-2 | | |
| 19:00 | 8z1e-A | 2 | 6.3 | 105 | 278 | 9 | MOLECULE: | UNCHARACTERIZED PROTEIN UL78 | | |
| 20:00 | 6tc0-C | 2 | 25.7 | 127 | 994 | 6 | MOLECULE: | PROBABLE CYTOSOLIC IRON-SULFUR PROTEIN ASSE | MBLY P | |

| | | | | | | | | | |
|---|---|---|---|---|---|---|---|---|---|
| 21:00 | 6m9a-C | 2 | 8.7 | 74 | 157 | 7 | MOLECULE: | SIGNALING PROTEIN | |
| 22:00 | 5nmo-A | 2 | 4.9 | 75 | 162 | 3 | MOLECULE: | CHROMOSOME PARTITION PROTEIN SMC,CHROMOSOME | PARTI |
| 23:00 | 6nr8-3 | 2 | 4.1 | 82 | 132 | 7 | MOLECULE: | PREFOLDIN SUBUNIT 1 | |
| 24:00:00 | 6cum-A | 2 | 5.4 | 48 | 51 | 4 | MOLECULE: | LAO/AO TRANSPORT SYSTEM ATPASE | |
| 25:00:00 | 8gi1-A | 2 | 4.7 | 72 | 144 | 6 | MOLECULE: | ACCESSORY PROTEIN CSX28 | |
| 26:00:00 | 5afr-A | 2 | 7.9 | 77 | 332 | 5 | MOLECULE: | DYNEIN HEAVY CHAIN, CYTOPLASMIC | |
| 27:00:00 | 4cpg-A | 2 | 2.5 | 40 | 69 | 5 | MOLECULE: | SMALL GLUTAMINE-RICH TETRATRICOPEPTIDE REPE | AT-CON |
| 28:00:00 | 6r24-A | 2 | 5.9 | 57 | 159 | 9 | MOLECULE: | TRANSPOSON TY3-I GAG-POL POLYPROTEIN | |
| 29:00:00 | 5w3s-A | 2 | 7.5 | 120 | 485 | 3 | MOLECULE: | MUCOLIPIN-3 ISOFORM 1 | |
| 30:00:00 | 6bk4-A | 2 | 5.1 | 60 | 106 | 3 | MOLECULE: | CAPRIN HOMOLOG | |
| 31:00:00 | 8qpk-S | 2 | 6 | 48 | 73 | 6 | MOLECULE: | PROBABLE ATP-DEPENDENT RNA HELICASE DDX23 | |
| 32:00:00 | 7pkq-C | 2 | 26.5 | 118 | 369 | 11 | MOLECULE: | MS35 | |
| 33:00:00 | 7k9y-A | 2 | 3.2 | 57 | 425 | 0 | MOLECULE: | TRT | |
| 34:00:00 | 8gup-A | 2 | 8.2 | 63 | 159 | 3 | MOLECULE: | TYPE VII SECRETION SYSTEM PROTEIN ESAG | |
| 35:00:00 | 2ptf-A | 2 | 7.8 | 51 | 200 | 8 | MOLECULE: | UNCHARACTERIZED PROTEIN MTH_863 | |
| 36:00:00 | 5ziy-A | 2 | 3.2 | 86 | 195 | 6 | MOLECULE: | FLAGELLAR HOOK-ASSOCIATED PROTEIN 3 | |
| 37:00:00 | 7woo-F | 2 | 19.3 | 120 | 1622 | 1 | MOLECULE: | NUCLEOPORIN NIC96 | |
| 38:00:00 | 8qt5-A | 2 | 15.5 | 83 | 248 | 5 | MOLECULE: | 14-3-3-LIKE PROTEIN G-BOX FACTOR 14 LAMBDA, | PROTEI |
| 39:00:00 | 8t5e-A | 2 | 4 | 42 | 130 | 2 | MOLECULE: | BIM_FULLDIFF | |
| 40:00:00 | 7jqe-A | 2 | 2.8 | 46 | 357 | 7 | MOLECULE: | ESAT-6/WXG100 SECRETION SYSTEM PROTEIN | |
| 41:00:00 | 4oe8-C | 2 | 3.7 | 41 | 87 | 15 | MOLECULE: | INTERLEUKIN-12 SUBUNIT BETA | |
| 42:00:00 | 8th8-s | 2 | 54 | 85 | 187 | 7 | MOLECULE: | DYNEIN REGULATORY COMPLEX PROTEIN 1/2 N-TER | MINAL |
| 43:00:00 | 6f62-A | 2 | 3.6 | 59 | 105 | 5 | MOLECULE: | SYNAPTONEMAL COMPLEX PROTEIN 1 | |
| 44:00:00 | 4ddg-A | 2 | 6.4 | 55 | 399 | 16 | MOLECULE: | UBIQUITIN-CONJUGATING ENZYME E2 D2, UBIQUIT | IN THI |
| 45:00:00 | 7upq-F | 2 | 4.3 | 51 | 77 | 10 | MOLECULE: | DHT03 PROTEIN A | |

**DALI results: MnA-1 Cah**

| No: | Chain | Z | rmsd | lali | nres | %id | PDB Description | |
|---|---|---|---|---|---|---|---|---|
| 1:00 | 2qih-A | 8.1 | 7.5 | 136 | 136 | 6 | MOLECULE: | PROTEIN USPA1 |
| 2:00 | 5jlh-J | 8 | 7 | 134 | 135 | 0 | MOLECULE: | ACTIN, CYTOPLASMIC 2 |
| 3:00 | 5jlh-H | 7.9 | 7 | 134 | 135 | 0 | MOLECULE: | ACTIN, CYTOPLASMIC 2 |
| 4:00 | 2b9c-B | 7.8 | 7.1 | 139 | 142 | 5 | MOLECULE: | STRIATED-MUSCLE ALPHA TROPOMYOSIN |
| 5:00 | 4cpc-H | 7.7 | 8.5 | 138 | 140 | 9 | MOLECULE: | SYNAPTONEMAL COMPLEX PROTEIN 3 |
| 6:00 | 4pxu-B | 7.7 | 6.8 | 136 | 136 | 6 | MOLECULE: | BIPOLAR KINESIN KRP-130 |
| 7:00 | 3j8a-G | 7.7 | 5.1 | 135 | 135 | 0 | MOLECULE: | TROPOMYOSIN ALPHA-1 |
| 8:00 | 4cpc-A | 7.5 | 7.7 | 141 | 143 | 9 | MOLECULE: | SYNAPTONEMAL COMPLEX PROTEIN 3 |
| 9:00 | 4cpc-C | 7.5 | 8 | 139 | 144 | 9 | MOLECULE: | SYNAPTONEMAL COMPLEX PROTEIN 3 |
| 10:00 | 6vol-J | 7.5 | 9.1 | 131 | 133 | 11 | MOLECULE: | ATP SYNTHASE SUBUNIT ALPHA, CHLOROPLASTIC |
| 11:00 | 2b9c-A | 7.5 | 7.9 | 135 | 136 | 7 | MOLECULE: | STRIATED-MUSCLE ALPHA TROPOMYOSIN |
| 12:00 | 2d3e-D | 7.5 | 5.2 | 130 | 130 | 12 | MOLECULE: | GENERAL CONTROL PROTEIN GCN4 AND TROPOMYOSIN |
| 13:00 | 2d3e-B | 7.5 | 5.2 | 130 | 130 | 12 | MOLECULE: | GENERAL CONTROL PROTEIN GCN4 AND TROPOMYOSIN |
| 14:00 | 2efr-C | 7.4 | 6.8 | 133 | 154 | 8 | MOLECULE: | GENERAL CONTROL PROTEIN GCN4 AND TROPOMYOSIN |
| 15:00 | 2efs-B | 7.4 | 5.4 | 133 | 154 | 8 | MOLECULE: | GENERAL CONTROL PROTEIN GCN4 AND TROPOMYOSIN |
| 16:00 | 6x5z-P | 7.4 | 8.6 | 151 | 166 | 6 | MOLECULE: | ACTIN, ALPHA SKELETAL MUSCLE |
| 17:00 | 4cpc-B | 7.4 | 7 | 130 | 144 | 6 | MOLECULE: | SYNAPTONEMAL COMPLEX PROTEIN 3 |
| 18:00 | 8h9g-K | 7.4 | 5.6 | 136 | 198 | 4 | MOLECULE: | ATP SYNTHASE F(0) COMPLEX SUBUNIT B1, MITOCH |
| 19:00 | 4cpc-F | 7.4 | 7 | 129 | 144 | 7 | MOLECULE: | SYNAPTONEMAL COMPLEX PROTEIN 3 |
| 20:00 | 6gap-B | 7.3 | 6.5 | 134 | 214 | 6 | MOLECULE: | OUTER CAPSID PROTEIN SIGMA-1 |
| 21:00 | 4cpc-E | 7.3 | 7.5 | 138 | 143 | 4 | MOLECULE: | SYNAPTONEMAL COMPLEX PROTEIN 3 |
| 22:00 | 5to7-C | 7.3 | 3.9 | 128 | 141 | 7 | MOLECULE: | NUCLEOPROTEIN TPR |
| 23:00 | 7z8k-W | 7.3 | 7.6 | 152 | 165 | 9 | MOLECULE: | CYTOPLASMIC DYNEIN 1 HEAVY CHAIN 1 |
| 24:00:00 | 7dl2-C | 7.2 | 4.3 | 134 | 224 | 8 | MOLECULE: | HAMARTIN |
| 25:00:00 | 6snz-D | 7.2 | 6.1 | 127 | 143 | 9 | MOLECULE: | PRELAMIN-A/C |
| 26:00:00 | 8enc-P | 7.2 | 9.6 | 145 | 166 | 6 | MOLECULE: | MYOSIN-7 |
| 27:00:00 | 8efi-P | 7.2 | 10 | 146 | 166 | 6 | MOLECULE: | MYOSIN-7 |
| 28:00:00 | 5to6-D | 7.2 | 5.4 | 128 | 140 | 6 | MOLECULE: | NUCLEOPROTEIN TPR |
| 29:00:00 | 6snz-C | 7.1 | 7 | 138 | 146 | 5 | MOLECULE: | PRELAMIN-A/C |
| 30:00:00 | 7qin-I | 7.1 | 5.2 | 118 | 118 | 0 | MOLECULE: | ACTIN, ALPHA SKELETAL MUSCLE |
| 31:00:00 | 7jg5-b | 7.1 | 8.9 | 142 | 145 | 8 | MOLECULE: | ATP SYNTHASE SUBUNIT ALPHA |
| 32:00:00 | 8aia-N | 7.1 | 7.8 | 150 | 156 | 5 | MOLECULE: | CRESCENTIN |
| 33:00:00 | 8aix-N | 7.1 | 7.8 | 150 | 156 | 5 | MOLECULE: | CRESCENTIN |
| 34:00:00 | 7ogt-A | 7.1 | 6.7 | 129 | 688 | 5 | MOLECULE: | STRUCTURAL MAINTENANCE OF CHROMOSOMES PROTEI |
| 35:00:00 | 8aix-X | 7.1 | 8 | 149 | 156 | 6 | MOLECULE: | CRESCENTIN |
| 36:00:00 | 5to6-B | 7.1 | 7 | 127 | 139 | 7 | MOLECULE: | NUCLEOPROTEIN TPR |
| 37:00:00 | 6vm4-I | 7 | 6.4 | 140 | 154 | 4 | MOLECULE: | ATP SYNTHASE SUBUNIT ALPHA, CHLOROPLASTIC |
| 38:00:00 | 6dd9-B | 7 | 6 | 123 | 127 | 5 | MOLECULE: | SYNAPTONEMAL COMPLEX PROTEIN 3 |
| 39:00:00 | 6dd8-A | 7 | 5.7 | 127 | 131 | 2 | MOLECULE: | SYNAPTONEMAL COMPLEX PROTEIN 3 |
| 40:00:00 | 8ajb-S | 7 | 9 | 148 | 155 | 5 | MOLECULE: | CRESCENTIN |
| 41:00:00 | 6gao-B | 7 | 7.7 | 145 | 225 | 8 | MOLECULE: | OUTER CAPSID PROTEIN SIGMA-1 |
| 42:00:00 | 5to7-B | 7 | 5.5 | 128 | 141 | 6 | MOLECULE: | NUCLEOPROTEIN TPR |
| 43:00:00 | 6snz-B | 7 | 6.9 | 136 | 155 | 4 | MOLECULE: | PRELAMIN-A/C |
| 44:00:00 | 7z8k-x | 7 | 4.7 | 131 | 135 | 10 | MOLECULE: | CYTOPLASMIC DYNEIN 1 HEAVY CHAIN 1 |
| 45:00:00 | 6f1u-X | 7 | 2.5 | 78 | 79 | 8 | MOLECULE: | ARP1 ACTIN RELATED PROTEIN 1 HOMOLOG A |
| 46:00:00 | 6wnq-Y | 7 | 9.8 | 131 | 156 | 8 | MOLECULE: | ATP SYNTHASE SUBUNIT DELTA |
| 47:00:00 | 6yvu-B | 7 | 11 | 150 | 1191 | 7 | MOLECULE: | STRUCTURAL MAINTENANCE OF CHROMOSOMES PROTEI |
| 48:00:00 | 5tvb-A | 7 | 5.6 | 127 | 140 | 7 | MOLECULE: | NUCLEOPROTEIN TPR |
| 49:00:00 | 5to6-A | 7 | 4.8 | 126 | 141 | 10 | MOLECULE: | NUCLEOPROTEIN TPR |

| | | | | | | | | |
|---|---|---|---|---|---|---|---|---|
| 50:00:00 | 6oqu-Y | 7 | 5 | 137 | 156 | 4 | MOLECULE: | ATP SYNTHASE SUBUNIT DELTA |
| 51:00:00 | 8tgt-A | 6.9 | 3.7 | 76 | 76 | 12 | MOLECULE: | M PROTEIN |
| 52:00:00 | 8ahl-G | 6.9 | 6.8 | 134 | 155 | 5 | MOLECULE: | CRESCENTIN |
| 53:00:00 | 8ga8-G | 6.9 | 4.6 | 79 | 97 | 6 | MOLECULE: | TRANSCRIPTIONAL REGULATORY PROTEIN SDS3 |
| 54:00:00 | 4wy4-C | 6.9 | 3.4 | 78 | 78 | 8 | MOLECULE: | VESICLE-ASSOCIATED MEMBRANE PROTEIN 8 |
| 55:00:00 | 5app-B | 6.9 | 8 | 126 | 128 | 10 | MOLECULE: | GENERAL CONTROL PROTEIN GCN4, OUTER MEMBRANE |
| 56:00:00 | 6vol-I | 6.9 | 8.4 | 143 | 150 | 8 | MOLECULE: | ATP SYNTHASE SUBUNIT ALPHA, CHLOROPLASTIC |
| 57:00:00 | 5fiy-E | 6.9 | 2.6 | 78 | 78 | 4 | MOLECULE: | PRKC APOPTOSIS WT1 REGULATOR PROTEIN |
| 58:00:00 | 5bu5-F | 6.9 | 3.2 | 81 | 174 | 9 | MOLECULE: | DNA STABILIZATION PROTEIN |
| 59:00:00 | 1gk4-D | 6.9 | 2.2 | 78 | 78 | 5 | MOLECULE: | VIMENTIN |
| 60:00:00 | 6hk9-B | 6.9 | 4.7 | 77 | 78 | 6 | MOLECULE: | TESTIS-EXPRESSED PROTEIN 12 |
| 61:00:00 | 5zvm-B | 6.9 | 4.8 | 76 | 76 | 3 | MOLECULE: | SPIKE GLYCOPROTEIN |
| 62:00:00 | 8csp-U | 6.9 | 10.5 | 127 | 174 | 11 | MOLECULE: | 28S RIBOSOMAL PROTEIN S34, MITOCHONDRIAL |
| 63:00:00 | 6z47-H | 6.9 | 5.1 | 135 | 257 | 13 | MOLECULE: | MYOSIN HEAVY CHAIN 11 |
| 64:00:00 | 2e7s-F | 6.9 | 6 | 124 | 124 | 10 | MOLECULE: | RAB GUANINE NUCLEOTIDE EXCHANGE FACTOR SEC2 |
| 65:00:00 | 2e7s-E | 6.9 | 3.6 | 124 | 124 | 8 | MOLECULE: | RAB GUANINE NUCLEOTIDE EXCHANGE FACTOR SEC2 |
| 66:00:00 | 8txr-C | 6.8 | 8.4 | 148 | 431 | 6 | MOLECULE: | EXODEOXYRIBONUCLEASE 7 LARGE SUBUNIT |
| 67:00:00 | 6vm4-J | 6.8 | 7.7 | 132 | 136 | 6 | MOLECULE: | ATP SYNTHASE SUBUNIT ALPHA, CHLOROPLASTIC |
| 68:00:00 | 6zpj-B | 6.8 | 4.2 | 77 | 77 | 9 | MOLECULE: | LEISHMANIA MEXICANA KKT4 |
| 69:00:00 | 8uzx-F | 6.8 | 2.7 | 82 | 82 | 5 | MOLECULE: | ACTIN, ALPHA CARDIAC MUSCLE 1 |
| 70:00:00 | 6voj-I | 6.8 | 7 | 140 | 154 | 5 | MOLECULE: | ATP SYNTHASE SUBUNIT ALPHA, CHLOROPLASTIC |
| 71:00:00 | 8afe-F | 6.8 | 4.9 | 79 | 79 | 10 | MOLECULE: | CRESCENTIN |
| 72:00:00 | 5ccg-C | 6.8 | 2.5 | 74 | 74 | 5 | MOLECULE: | VESICLE-ASSOCIATED MEMBRANE PROTEIN 2 |
| 73:00:00 | 8uzy-F | 6.8 | 2.8 | 82 | 82 | 5 | MOLECULE: | ACTIN, ALPHA CARDIAC MUSCLE 1 |
| 74:00:00 | 6fkf-p | 6.8 | 9.2 | 141 | 143 | 6 | MOLECULE: | ATP SYNTHASE SUBUNIT ALPHA, CHLOROPLASTIC |
| 75:00:00 | 6mi4-B | 6.8 | 3.1 | 121 | 123 | 10 | MOLECULE: | NF-KB ESSENTIAL MODULATOR |
| 76:00:00 | 8afh-F | 6.8 | 4.9 | 79 | 79 | 10 | MOLECULE: | CRESCENTIN |
| 77:00:00 | 6igv-A | 6.8 | 2.7 | 117 | 118 | 8 | MOLECULE: | KINESIN-1 HEAVY CHAIN |
| 78:00:00 | 5fiy-C | 6.8 | 4.9 | 78 | 78 | 3 | MOLECULE: | PRKC APOPTOSIS WT1 REGULATOR PROTEIN |
| 79:00:00 | 8afh-P | 6.8 | 4.9 | 79 | 79 | 10 | MOLECULE: | CRESCENTIN |
| 80:00:00 | 8v01-G | 6.8 | 4 | 81 | 82 | 6 | MOLECULE: | ACTIN, ALPHA CARDIAC MUSCLE 1 |
| 81:00:00 | 8v01-F | 6.8 | 4.8 | 81 | 82 | 6 | MOLECULE: | ACTIN, ALPHA CARDIAC MUSCLE 1 |
| 82:00:00 | 7bji-C | 6.8 | 5.1 | 79 | 80 | 3 | MOLECULE: | CENTROSOMAL PROTEIN OF 135 KDA |
| 83:00:00 | 8fcf-D | 6.8 | 3.6 | 77 | 79 | 6 | MOLECULE: | PLASMALEMMA VESICLE-ASSOCIATED PROTEIN |
| 84:00:00 | 8v0y-G | 6.8 | 4.3 | 81 | 82 | 6 | MOLECULE: | ACTIN, ALPHA CARDIAC MUSCLE 1 |
| 85:00:00 | 8uzy-G | 6.8 | 4 | 81 | 82 | 6 | MOLECULE: | ACTIN, ALPHA CARDIAC MUSCLE 1 |
| 86:00:00 | 7wxz-D | 6.8 | 4.8 | 76 | 187 | 3 | MOLECULE: | SPIKE PROTEIN S2' |
| 87:00:00 | 5xis-A | 6.8 | 4.8 | 79 | 79 | 4 | MOLECULE: | E3 UBIQUITIN-PROTEIN LIGASE RNF168 |
| 88:00:00 | 6h9l-C | 6.8 | 6.7 | 105 | 128 | 11 | MOLECULE: | UNCHARACTERIZED PROTEIN |
| 89:00:00 | 6l5h-A | 6.8 | 4.8 | 78 | 82 | 4 | MOLECULE: | ROOTLETIN |
| 90:00:00 | 5dn6-V | 6.8 | 3.7 | 78 | 78 | 0 | MOLECULE: | CHAIN A |
| 91:00:00 | 6l5h-B | 6.8 | 4.2 | 81 | 82 | 6 | MOLECULE: | ROOTLETIN |
| 92:00:00 | 8fcf-C | 6.8 | 3.4 | 81 | 83 | 7 | MOLECULE: | PLASMALEMMA VESICLE-ASSOCIATED PROTEIN |
| 93:00:00 | 6oqa-D | 6.8 | 4.8 | 83 | 83 | 13 | MOLECULE: | PEPTIDYL-PROLYL CIS-TRANS ISOMERASE FKBP1A |
| 94:00:00 | 5whf-F | 6.8 | 3.1 | 84 | 84 | 4 | MOLECULE: | VIMENTIN |
| 95:00:00 | 7jga-b | 6.8 | 9.9 | 133 | 141 | 9 | MOLECULE: | ATP SYNTHASE SUBUNIT ALPHA |
| 96:00:00 | 8ajb-H | 6.8 | 9 | 147 | 163 | 4 | MOLECULE: | CRESCENTIN |
| 97:00:00 | 8sjj-D | 6.8 | 3 | 81 | 86 | 6 | MOLECULE: | SEPTIN-14 |
| 98:00:00 | 8ahl-H | 6.8 | 9 | 147 | 163 | 4 | MOLECULE: | CRESCENTIN |
| 99:00:00 | 6qum-K | 6.8 | 3.5 | 85 | 98 | 5 | MOLECULE: | V-TYPE ATP SYNTHASE ALPHA CHAIN |
| 0:00 | 8jr0-b | 6.8 | 8.6 | 134 | 144 | 3 | MOLECULE: | ATP SYNTHASE SUBUNIT ALPHA |

| | | | | | | | | |
|---|---|---|---|---|---|---|---|---|
| 1:00 | 8j0t-b | 6.8 | 9.9 | 138 | 144 | 7 | MOLECULE: | ATP SYNTHASE SUBUNIT ALPHA |
| 2:00 | 5noj-F | 6.7 | 6.2 | 130 | 136 | 0 | MOLECULE: | ACTIN, ALPHA SKELETAL MUSCLE |
| 3:00 | 6vmg-I | 6.7 | 6.8 | 135 | 148 | 8 | MOLECULE: | ATP SYNTHASE SUBUNIT A, CHLOROPLASTIC |
| 4:00 | 5to7-A | 6.7 | 5.5 | 131 | 141 | 10 | MOLECULE: | NUCLEOPROTEIN TPR |
| 5:00 | 5adx-z | 6.7 | 22.1 | 119 | 419 | 0 | MOLECULE: | ACTIN RELATED PROTEIN 1 |
| 6:00 | 5to7-D | 6.7 | 4.4 | 130 | 141 | 11 | MOLECULE: | NUCLEOPROTEIN TPR |
| 7:00 | 6aoz-A | 6.7 | 3.6 | 74 | 74 | 5 | MOLECULE: | CASP8-ASSOCIATED PROTEIN 2 |
| 8:00 | 4yv3-A | 6.7 | 4.2 | 77 | 77 | 6 | MOLECULE: | VIMENTIN |
| 9:00 | 7rzt-A | 6.7 | 4.7 | 71 | 71 | 10 | MOLECULE: | SARS-COV-2 HR1 S940F LINKED TO A SCAFFOLD,SP |
| 10:00 | 7rzv-B | 6.7 | 4.7 | 71 | 71 | 11 | MOLECULE: | SARS-COV-2 HR1 LINKED TO A SCAFFOLD,SPIKE PR |
| 11:00 | 7tik-A | 6.7 | 4.8 | 71 | 71 | 11 | MOLECULE: | FERRITIN, DPS FAMILY PROTEIN AND SPIKE PROTE |
| 12:00 | 7tik-C | 6.7 | 4.8 | 71 | 71 | 11 | MOLECULE: | FERRITIN, DPS FAMILY PROTEIN AND SPIKE PROTE |
| 13:00 | 8v0y-F | 6.7 | 2.7 | 82 | 82 | 4 | MOLECULE: | ACTIN, ALPHA CARDIAC MUSCLE 1 |
| 14:00 | 6vmb-I | 6.7 | 8.9 | 143 | 153 | 7 | MOLECULE: | ATP SYNTHASE SUBUNIT ALPHA, CHLOROPLASTIC |
| 15:00 | 1gk4-B | 6.7 | 4.3 | 79 | 79 | 8 | MOLECULE: | VIMENTIN |
| 16:00 | 3bat-D | 6.7 | 4.6 | 78 | 80 | 12 | MOLECULE: | MYOSIN HEAVY CHAIN, STRIATED MUSCLE/GENERAL |
| 17:00 | 7tj7-V | 6.7 | 5 | 128 | 135 | 0 | MOLECULE: | CARDIAC ACTIN |
| 18:00 | 8fxf-A | 6.7 | 4.1 | 78 | 78 | 4 | MOLECULE: | E3 UBIQUITIN-PROTEIN LIGASE TRIM56 |
| 19:00 | 2fxo-A | 6.7 | 3.6 | 127 | 129 | 6 | MOLECULE: | MYOSIN HEAVY CHAIN, CARDIAC MUSCLE BETA ISOF |
| 20:00 | 5cci-C | 6.7 | 2.5 | 73 | 73 | 5 | MOLECULE: | VESICLE-ASSOCIATED MEMBRANE PROTEIN 2 |
| 21:00 | 5ccg-I | 6.7 | 2.6 | 74 | 74 | 5 | MOLECULE: | VESICLE-ASSOCIATED MEMBRANE PROTEIN 2 |
| 22:00 | 7jh7-J | 6.7 | 5 | 128 | 135 | 0 | MOLECULE: | ACTIN, ALPHA CARDIAC MUSCLE 1 |
| 23:00 | 5cch-C | 6.7 | 2.6 | 74 | 74 | 5 | MOLECULE: | VESICLE-ASSOCIATED MEMBRANE PROTEIN 2 |
| 24:00:00 | 2fxo-C | 6.7 | 4.5 | 126 | 127 | 6 | MOLECULE: | MYOSIN HEAVY CHAIN, CARDIAC MUSCLE BETA ISOF |
| 25:00:00 | 8i3e-B | 6.7 | 4.2 | 107 | 135 | 10 | MOLECULE: | ELKS/RAB6-INTERACTING/CAST FAMILY MEMBER 1 |
| 26:00:00 | 7uw5-B | 6.7 | 24.7 | 132 | 723 | 8 | MOLECULE: | MECHANOSENSITIVE CHANNEL MSCK |
| 27:00:00 | 8uzx-G | 6.7 | 4.2 | 81 | 82 | 6 | MOLECULE: | ACTIN, ALPHA CARDIAC MUSCLE 1 |
| 28:00:00 | 8uwx-G | 6.7 | 4.1 | 81 | 82 | 6 | MOLECULE: | ACTIN, ALPHA CARDIAC MUSCLE 1 |
| 29:00:00 | 8v0k-G | 6.7 | 4 | 81 | 82 | 6 | MOLECULE: | ACTIN, ALPHA CARDIAC MUSCLE 1 |
| 30:00:00 | 8uz6-G | 6.7 | 4.1 | 81 | 82 | 6 | MOLECULE: | ACTIN, ALPHA CARDIAC MUSCLE 1 |
| 31:00:00 | 8uyd-F | 6.7 | 4.9 | 81 | 82 | 6 | MOLECULE: | ACTIN, ALPHA CARDIAC MUSCLE 1 |
| 32:00:00 | 8v0i-F | 6.7 | 4.8 | 81 | 82 | 6 | MOLECULE: | ACTIN, ALPHA CARDIAC MUSCLE 1 |
| 33:00:00 | 8uyd-G | 6.7 | 4.1 | 81 | 82 | 6 | MOLECULE: | ACTIN, ALPHA CARDIAC MUSCLE 1 |
| 34:00:00 | 8v0k-F | 6.7 | 4.9 | 81 | 82 | 6 | MOLECULE: | ACTIN, ALPHA CARDIAC MUSCLE 1 |
| 35:00:00 | 8uww-G | 6.7 | 4.1 | 81 | 82 | 6 | MOLECULE: | ACTIN, ALPHA CARDIAC MUSCLE 1 |
| 36:00:00 | 3bas-A | 6.7 | 4.4 | 79 | 80 | 11 | MOLECULE: | MYOSIN HEAVY CHAIN, STRIATED MUSCLE/GENERAL |
| 37:00:00 | 5jxc-C | 6.7 | 3.8 | 78 | 78 | 4 | MOLECULE: | RAS/RAP GTPASE-ACTIVATING PROTEIN SYNGAP |
| 38:00:00 | 6dm9-C | 6.7 | 4.5 | 78 | 78 | 5 | MOLECULE: | DHD15_EXTENDED_A |
| 39:00:00 | 2bez-C | 6.7 | 2.5 | 77 | 77 | 6 | MOLECULE: | E2 GLYCOPROTEIN |
| 40:00:00 | 7tk9-U | 6.7 | 5.6 | 128 | 155 | 9 | MOLECULE: | ATP SYNTHASE SUBUNIT 9, MITOCHONDRIAL |
| 41:00:00 | 8uwy-F | 6.7 | 3.5 | 80 | 82 | 8 | MOLECULE: | ACTIN, ALPHA CARDIAC MUSCLE 1 |
| 42:00:00 | 2nps-C | 6.7 | 4.1 | 79 | 79 | 9 | MOLECULE: | VESICLE-ASSOCIATED MEMBRANE PROTEIN 4 |
| 43:00:00 | 8uwx-F | 6.7 | 3.5 | 80 | 82 | 8 | MOLECULE: | ACTIN, ALPHA CARDIAC MUSCLE 1 |
| 44:00:00 | 8uz5-F | 6.7 | 3.5 | 80 | 82 | 8 | MOLECULE: | ACTIN, ALPHA CARDIAC MUSCLE 1 |
| 45:00:00 | 5xis-D | 6.7 | 4.6 | 81 | 82 | 6 | MOLECULE: | E3 UBIQUITIN-PROTEIN LIGASE RNF168 |
| 46:00:00 | 8uww-F | 6.7 | 3.5 | 80 | 82 | 8 | MOLECULE: | ACTIN, ALPHA CARDIAC MUSCLE 1 |
| 47:00:00 | 2jee-A | 6.7 | 2.4 | 77 | 78 | 6 | MOLECULE: | CELL DIVISION PROTEIN ZAPB |
| 48:00:00 | 2jee-B | 6.7 | 2.4 | 77 | 78 | 6 | MOLECULE: | CELL DIVISION PROTEIN ZAPB |
| 49:00:00 | 2jee-C | 6.7 | 2.4 | 77 | 78 | 6 | MOLECULE: | CELL DIVISION PROTEIN ZAPB |
| 50:00:00 | 2jee-D | 6.7 | 2.4 | 77 | 78 | 6 | MOLECULE: | CELL DIVISION PROTEIN ZAPB |
| 51:00:00 | 4zqa-A | 6.7 | 4.4 | 83 | 86 | 12 | MOLECULE: | SIN3 HISTONE DEACETYLASE COREPRESSOR COMPLEX |

| | | | | | | | | |
|---|---|---|---|---|---|---|---|---|
| 52:00:00 | 8sdi-D | 6.7 | 3.8 | 79 | 80 | 10 | MOLECULE: | TRIPARTITE MOTIF-CONTAINING PROTEIN 45 |
| 53:00:00 | 8sjj-C | 6.7 | 3.4 | 83 | 86 | 6 | MOLECULE: | SEPTIN-14 |
| 54:00:00 | 2e7s-B | 6.7 | 3.6 | 121 | 121 | 7 | MOLECULE: | RAB GUANINE NUCLEOTIDE EXCHANGE FACTOR SEC2 |
| 55:00:00 | 2e7s-D | 6.7 | 3.3 | 124 | 124 | 10 | MOLECULE: | RAB GUANINE NUCLEOTIDE EXCHANGE FACTOR SEC2 |
| 56:00:00 | 8th8-D | 6.6 | 9.9 | 148 | 292 | 9 | MOLECULE: | DYNEIN REGULATORY COMPLEX PROTEIN 1/2 N-TERM |
| 57:00:00 | 6mi3-A | 6.6 | 3.7 | 119 | 122 | 8 | MOLECULE: | NF-KB ESSENTIAL MODULATOR,NF-KAPPA-B ESSENTI |
| 58:00:00 | 6jlb-A | 6.6 | 8 | 148 | 202 | 5 | MOLECULE: | LAMIN A/C |
| 59:00:00 | 2ocy-A | 6.6 | 9.6 | 144 | 149 | 9 | MOLECULE: | RAB GUANINE NUCLEOTIDE EXCHANGE FACTOR SEC2 |
| 60:00:00 | 6ign-A | 6.6 | 5.4 | 122 | 123 | 6 | MOLECULE: | KINESIN-1 HEAVY CHAIN |
| 61:00:00 | 7p2y-p | 6.6 | 8.9 | 146 | 155 | 4 | MOLECULE: | ATP SYNTHASE SUBUNIT ALPHA |
| 62:00:00 | 5xau-B | 6.6 | 4.1 | 71 | 71 | 4 | MOLECULE: | LAMININ SUBUNIT ALPHA-5 |
| 63:00:00 | 8cst-U | 6.6 | 10.9 | 126 | 174 | 13 | MOLECULE: | 28S RIBOSOMAL PROTEIN S34, MITOCHONDRIAL |
| 64:00:00 | 6vof-I | 6.6 | 6.7 | 138 | 151 | 4 | MOLECULE: | ATP SYNTHASE SUBUNIT ALPHA, CHLOROPLASTIC |
| 65:00:00 | 4mvd-B | 6.6 | 6.8 | 97 | 253 | 7 | MOLECULE: | CHOLINE-PHOSPHATE CYTIDYLYLTRANSFERASE A |
| 66:00:00 | 1kil-C | 6.6 | 1.8 | 72 | 72 | 7 | MOLECULE: | SYNAPTOBREVIN SNARE MOTIF |
| 67:00:00 | 6von-J | 6.6 | 8.8 | 129 | 130 | 7 | MOLECULE: | ATP SYNTHASE SUBUNIT ALPHA, CHLOROPLASTIC |
| 68:00:00 | 5djn-C | 6.6 | 4.8 | 80 | 81 | 6 | MOLECULE: | KINESIN-LIKE PROTEIN |
| 69:00:00 | 7tj7-T | 6.6 | 4.5 | 129 | 135 | 0 | MOLECULE: | CARDIAC ACTIN |
| 70:00:00 | 7tit-N | 6.6 | 4.9 | 128 | 135 | 0 | MOLECULE: | CARDIAC ACTIN |
| 71:00:00 | 4mvd-F | 6.6 | 4.5 | 96 | 253 | 7 | MOLECULE: | CHOLINE-PHOSPHATE CYTIDYLYLTRANSFERASE A |
| 72:00:00 | 5djn-A | 6.6 | 4.5 | 77 | 77 | 6 | MOLECULE: | KINESIN-LIKE PROTEIN |
| 73:00:00 | 4a7f-B | 6.6 | 6.4 | 133 | 136 | 5 | MOLECULE: | ACTIN, ALPHA SKELETAL MUSCLE |
| 74:00:00 | 8uz5-G | 6.6 | 4.1 | 81 | 82 | 6 | MOLECULE: | ACTIN, ALPHA CARDIAC MUSCLE 1 |
| 75:00:00 | 6ano-A | 6.6 | 3.8 | 75 | 75 | 4 | MOLECULE: | CASP8-ASSOCIATED PROTEIN 2 |
| 76:00:00 | 8qrl-U | 6.6 | 19.3 | 134 | 176 | 10 | MOLECULE: | 12S MITOCHONDRIAL RRNA |
| 77:00:00 | 1fav-A | 6.6 | 4.3 | 78 | 78 | 3 | MOLECULE: | HIV-1 ENVELOPE PROTEIN CHIMERA |
| 78:00:00 | 7bji-A | 6.6 | 4.5 | 80 | 82 | 3 | MOLECULE: | CENTROSOMAL PROTEIN OF 135 KDA |
| 79:00:00 | 5lxn-A | 6.6 | 4.2 | 80 | 81 | 5 | MOLECULE: | TRANSFORMING ACIDIC COILED-COIL-CONTAINING P |
| 80:00:00 | 6oqa-H | 6.6 | 3.3 | 84 | 85 | 6 | MOLECULE: | PEPTIDYL-PROLYL CIS-TRANS ISOMERASE FKBP1A |
| 81:00:00 | 8uz6-F | 6.6 | 3.5 | 80 | 82 | 8 | MOLECULE: | ACTIN, ALPHA CARDIAC MUSCLE 1 |
| 82:00:00 | 5k7b-C | 6.6 | 3.5 | 83 | 86 | 6 | MOLECULE: | BECLIN-2 |
| 83:00:00 | 8afh-E | 6.6 | 3.7 | 78 | 79 | 8 | MOLECULE: | CRESCENTIN |
| 84:00:00 | 8v0i-G | 6.6 | 4.9 | 81 | 82 | 6 | MOLECULE: | ACTIN, ALPHA CARDIAC MUSCLE 1 |
| 85:00:00 | 7tk4-U | 6.6 | 7.2 | 140 | 155 | 4 | MOLECULE: | ATP SYNTHASE SUBUNIT 9, MITOCHONDRIAL |
| 86:00:00 | 4jpn-J | 6.6 | 2.1 | 72 | 72 | 6 | MOLECULE: | MINOR SPIKE PROTEIN H |
| 87:00:00 | 5k7b-A | 6.6 | 2 | 78 | 87 | 13 | MOLECULE: | BECLIN-2 |
| 88:00:00 | 8gt9-B | 6.6 | 2.5 | 87 | 88 | 9 | MOLECULE: | BECLIN-2 |
| 89:00:00 | 3bas-B | 6.6 | 3.4 | 82 | 87 | 4 | MOLECULE: | MYOSIN HEAVY CHAIN, STRIATED MUSCLE/GENERAL |
| 90:00:00 | 7qin-J | 6.6 | 3.4 | 87 | 88 | 0 | MOLECULE: | ACTIN, ALPHA SKELETAL MUSCLE |
| 91:00:00 | 6vzf-A | 6.6 | 3.4 | 82 | 86 | 5 | MOLECULE: | AUTOPHAGY-RELATED PROTEIN 11 |
| 92:00:00 | 5d3a-A | 6.6 | 2.8 | 79 | 82 | 6 | MOLECULE: | KINESIN-LIKE PROTEIN KIF21A |
| 93:00:00 | 5jxc-E | 6.6 | 1.7 | 82 | 84 | 7 | MOLECULE: | RAS/RAP GTPASE-ACTIVATING PROTEIN SYNGAP |
| 94:00:00 | 6ys4-D | 6.6 | 4.9 | 85 | 87 | 7 | MOLECULE: | SPINDLE ASSEMBLY ABNORMAL PROTEIN 6 HOMOLOG |
| 95:00:00 | 5yl9-A | 6.6 | 3 | 86 | 88 | 8 | MOLECULE: | SPIKE GLYCOPROTEIN |
| 96:00:00 | 7eqc-C | 6.6 | 5.9 | 102 | 106 | 6 | MOLECULE: | CYTOKINESIS DEFECT |
| 97:00:00 | 2e7s-C | 6.6 | 2.3 | 118 | 118 | 3 | MOLECULE: | RAB GUANINE NUCLEOTIDE EXCHANGE FACTOR SEC2 |
| 98:00:00 | 5hmo-A | 6.5 | 3.9 | 124 | 131 | 10 | MOLECULE: | UNCONVENTIONAL MYOSIN-X |
| 99:00:00 | 4zry-A | 6.5 | 2.4 | 110 | 110 | 5 | MOLECULE: | KERATIN, TYPE I CYTOSKELETAL 10 |
| 0:00 | 6wvw-G | 6.5 | 4.2 | 74 | 74 | 5 | MOLECULE: | VESICLE-ASSOCIATED MEMBRANE PROTEIN 2 |
| 1:00 | 5zvk-A | 6.5 | 4 | 70 | 70 | 6 | MOLECULE: | HUMAN CORONAVIRUS MERS HR1 MOTIF |
| 2:00 | 6klq-A | 6.5 | 5.7 | 132 | 135 | 0 | MOLECULE: | TROPOMYOSIN |

| | | | | | | | | |
|---|---|---|---|---|---|---|---|---|
| 3:00 | 5mq4-D | 6.5 | 4.8 | 74 | 111 | 7 | MOLECULE: | PROTEIN KINASE C-BINDING PROTEIN 1 |
| 4:00 | 6wsm-A | 6.5 | 4.4 | 74 | 74 | 7 | MOLECULE: | SEPTIN-8 |
| 5:00 | 5nog-G | 6.5 | 6.3 | 131 | 135 | 0 | MOLECULE: | CARDIAC MUSCLE ALPHA ACTIN 1 |
| 6:00 | 8fxf-C | 6.5 | 4.3 | 78 | 78 | 4 | MOLECULE: | E3 UBIQUITIN-PROTEIN LIGASE TRIM56 |
| 7:00 | 8fxf-D | 6.5 | 4.2 | 78 | 78 | 3 | MOLECULE: | E3 UBIQUITIN-PROTEIN LIGASE TRIM56 |
| 8:00 | 7eqc-B | 6.5 | 4.5 | 101 | 101 | 6 | MOLECULE: | CYTOKINESIS DEFECT |
| 9:00 | 8fxf-B | 6.5 | 4.1 | 78 | 78 | 3 | MOLECULE: | E3 UBIQUITIN-PROTEIN LIGASE TRIM56 |
| 10:00 | 5ydk-F | 6.5 | 5.7 | 80 | 82 | 10 | MOLECULE: | E3 UBIQUITIN-PROTEIN LIGASE RNF168 |
| 11:00 | 5xit-E | 6.5 | 3.3 | 75 | 75 | 7 | MOLECULE: | UBIQUITIN-40S RIBOSOMAL PROTEIN S27A |
| 12:00 | 5whf-D | 6.5 | 3.6 | 85 | 85 | 5 | MOLECULE: | VIMENTIN |
| 13:00 | 6cp3-Z | 6.5 | 6.6 | 138 | 155 | 4 | MOLECULE: | ATP SYNTHASE SUBUNIT 9, MITOCHONDRIAL |
| 14:00 | 7tkp-U | 6.5 | 11.5 | 146 | 155 | 8 | MOLECULE: | ATP SYNTHASE SUBUNIT 9 |
| 15:00 | 6abo-B | 6.5 | 4.2 | 75 | 77 | 3 | MOLECULE: | DNA REPAIR PROTEIN XRCC4 |
| 16:00 | 6ap0-B | 6.5 | 4.3 | 88 | 88 | 8 | MOLECULE: | CASP8-ASSOCIATED PROTEIN 2 |
| 17:00 | 2e7s-K | 6.5 | 4.2 | 116 | 117 | 5 | MOLECULE: | RAB GUANINE NUCLEOTIDE EXCHANGE FACTOR SEC2 |
| 18:00 | 7uw5-A | 6.4 | 36.1 | 111 | 723 | 8 | MOLECULE: | MECHANOSENSITIVE CHANNEL MSCK |
| 19:00 | 4f61-I | 6.4 | 5.2 | 146 | 234 | 3 | MOLECULE: | TUBULIN ALPHA CHAIN |
| 20:00 | 5oag-B | 6.4 | 2.7 | 97 | 97 | 11 | MOLECULE: | RAB-3A-INTERACTING PROTEIN |
| 21:00 | 5jvd-E | 6.4 | 4.3 | 104 | 122 | 3 | MOLECULE: | TUBULIN ALPHA-1B CHAIN |
| 22:00 | 4a7h-B | 6.4 | 6.3 | 130 | 136 | 7 | MOLECULE: | ACTIN, ALPHA SKELETAL MUSCLE |
| 23:00 | 5zvk-C | 6.4 | 4.1 | 70 | 70 | 6 | MOLECULE: | HUMAN CORONAVIRUS MERS HR1 MOTIF |
| 24:00:00 | 8a0l-E | 6.4 | 4.4 | 104 | 123 | 3 | MOLECULE: | TUBULIN ALPHA-1B CHAIN |
| 25:00:00 | 5wst-A | 6.4 | 4.2 | 71 | 71 | 4 | MOLECULE: | UNCONVENTIONAL MYOSIN-VIIA |
| 26:00:00 | 5zvk-B | 6.4 | 4.2 | 70 | 70 | 6 | MOLECULE: | HUMAN CORONAVIRUS MERS HR1 MOTIF |
| 27:00:00 | 7db9-E | 6.4 | 4.5 | 105 | 124 | 3 | MOLECULE: | TUBULIN ALPHA-1B CHAIN |
| 28:00:00 | 7dbb-E | 6.4 | 1.6 | 100 | 123 | 3 | MOLECULE: | TUBULIN ALPHA-1B CHAIN |
| 29:00:00 | 6znn-S | 6.4 | 3.3 | 68 | 68 | 0 | MOLECULE: | ARP1 ACTIN RELATED PROTEIN 1 HOMOLOG A |
| 30:00:00 | 4bl6-A | 6.4 | 4.6 | 82 | 82 | 7 | MOLECULE: | PROTEIN BICAUDAL D |
| 31:00:00 | 1nkn-A | 6.4 | 4.7 | 73 | 74 | 4 | MOLECULE: | S2N51-GCN4 |
| 32:00:00 | 1sfc-K | 6.4 | 1.7 | 73 | 73 | 5 | MOLECULE: | PROTEIN (SYNAPTOBREVIN 2) |
| 33:00:00 | 1jth-C | 6.4 | 2.4 | 69 | 69 | 6 | MOLECULE: | SNAP25 |
| 34:00:00 | 6gbo-F | 6.4 | 4.8 | 73 | 73 | 0 | MOLECULE: | POLYMERASE COFACTOR VP35 |
| 35:00:00 | 2n1t-C | 6.4 | 3.8 | 75 | 77 | 7 | MOLECULE: | VESICLE-ASSOCIATED MEMBRANE PROTEIN 2 |
| 36:00:00 | 7bv6-G | 6.4 | 3.8 | 74 | 74 | 5 | MOLECULE: | VESICLE-ASSOCIATED MEMBRANE PROTEIN 8 |
| 37:00:00 | 4mvd-H | 6.4 | 6.5 | 99 | 253 | 10 | MOLECULE: | CHOLINE-PHOSPHATE CYTIDYLYLTRANSFERASE A |
| 38:00:00 | 6vq6-O | 6.4 | 8.1 | 113 | 114 | 12 | MOLECULE: | ATPASE H+-TRANSPORTING V1 SUBUNIT A |
| 39:00:00 | 5ydk-A | 6.4 | 4.2 | 80 | 81 | 9 | MOLECULE: | E3 UBIQUITIN-PROTEIN LIGASE RNF168 |
| 40:00:00 | 7y5c-b | 6.4 | 9 | 134 | 145 | 6 | MOLECULE: | ATP SYNTHASE SUBUNIT ALPHA |
| 41:00:00 | 7znk-o | 6.4 | 8.4 | 134 | 164 | 8 | MOLECULE: | RNA |
| 42:00:00 | 6pqv-Y | 6.4 | 5.6 | 129 | 156 | 9 | MOLECULE: | ATP SYNTHASE SUBUNIT DELTA |
| 43:00:00 | 7y5b-b | 6.4 | 10.3 | 140 | 145 | 6 | MOLECULE: | ATP SYNTHASE SUBUNIT ALPHA |
| 44:00:00 | 5fiy-D | 6.4 | 3.7 | 77 | 77 | 10 | MOLECULE: | PRKC APOPTOSIS WT1 REGULATOR PROTEIN |
| 45:00:00 | 6qum-L | 6.4 | 4.1 | 81 | 186 | 7 | MOLECULE: | V-TYPE ATP SYNTHASE ALPHA CHAIN |
| 46:00:00 | 7tkb-U | 6.4 | 7.2 | 139 | 155 | 4 | MOLECULE: | ATP SYNTHASE SUBUNIT 9, MITOCHONDRIAL |
| 47:00:00 | 5whf-H | 6.4 | 3.2 | 84 | 84 | 5 | MOLECULE: | VIMENTIN |
| 48:00:00 | 4jpn-B | 6.4 | 3.6 | 74 | 74 | 9 | MOLECULE: | MINOR SPIKE PROTEIN H |
| 49:00:00 | 3swk-B | 6.4 | 2.6 | 85 | 85 | 6 | MOLECULE: | VIMENTIN |
| 50:00:00 | 7tk2-U | 6.4 | 7.7 | 138 | 155 | 4 | MOLECULE: | ATP SYNTHASE SUBUNIT 9, MITOCHONDRIAL |
| 51:00:00 | 4jpn-A | 6.4 | 3.6 | 74 | 75 | 8 | MOLECULE: | MINOR SPIKE PROTEIN H |
| 52:00:00 | 8sde-A | 6.4 | 4 | 87 | 89 | 8 | MOLECULE: | TRIPARTITE MOTIF-CONTAINING PROTEIN 29 |
| 53:00:00 | 7tkg-U | 6.4 | 11.1 | 145 | 155 | 7 | MOLECULE: | ATP SYNTHASE SUBUNIT 9 |

| | | | | | | | | |
|---|---|---|---|---|---|---|---|---|
| 54:00:00 | 7jg8-b | 6.4 | 9.3 | 129 | 138 | 9 | MOLECULE: | ATP SYNTHASE SUBUNIT ALPHA |
| 55:00:00 | 6y5k-B | 6.4 | 3.3 | 87 | 133 | 9 | MOLECULE: | X-31 INFLUENZA HAEMAGGLUTININ HA1 |
| 56:00:00 | 6y5k-D | 6.4 | 3.3 | 87 | 133 | 9 | MOLECULE: | X-31 INFLUENZA HAEMAGGLUTININ HA1 |
| 57:00:00 | 6y5k-F | 6.4 | 3.3 | 87 | 133 | 7 | MOLECULE: | X-31 INFLUENZA HAEMAGGLUTININ HA1 |
| 58:00:00 | 4ll8-B | 6.4 | 4.2 | 112 | 113 | 10 | MOLECULE: | MYOSIN-4 |
| 59:00:00 | 5d3a-B | 6.4 | 4.3 | 82 | 82 | 4 | MOLECULE: | KINESIN-LIKE PROTEIN KIF21A |
| 60:00:00 | 8afh-O | 6.4 | 2.4 | 79 | 79 | 9 | MOLECULE: | CRESCENTIN |
| 61:00:00 | 8j0s-b | 6.4 | 6.5 | 127 | 144 | 6 | MOLECULE: | ATP SYNTHASE SUBUNIT A |
| 62:00:00 | 2e7s-Q | 6.4 | 2.3 | 111 | 111 | 3 | MOLECULE: | RAB GUANINE NUCLEOTIDE EXCHANGE FACTOR SEC2 |
| 63:00:00 | 6oqv-X | 6.4 | 9.1 | 132 | 156 | 10 | MOLECULE: | ATP SYNTHASE SUBUNIT DELTA |
| 64:00:00 | 2e7s-A | 6.4 | 2.2 | 113 | 114 | 7 | MOLECULE: | RAB GUANINE NUCLEOTIDE EXCHANGE FACTOR SEC2 |
| 65:00:00 | 2e7s-M | 6.4 | 3.4 | 117 | 117 | 6 | MOLECULE: | RAB GUANINE NUCLEOTIDE EXCHANGE FACTOR SEC2 |
| 66:00:00 | 6vmg-J | 6.4 | 5.9 | 112 | 131 | 6 | MOLECULE: | ATP SYNTHASE SUBUNIT A, CHLOROPLASTIC |
| 67:00:00 | 6fkj-E | 6.3 | 1.9 | 100 | 122 | 2 | MOLECULE: | TUBULIN ALPHA-1B CHAIN |
| 68:00:00 | 5s4v-E | 6.3 | 1.6 | 100 | 123 | 3 | MOLECULE: | TUBULIN ALPHA-1B CHAIN |
| 69:00:00 | 7cbz-E | 6.3 | 2.1 | 100 | 123 | 3 | MOLECULE: | TUBULIN ALPHA-1B CHAIN |
| 70:00:00 | 7yhn-E | 6.3 | 1.7 | 100 | 123 | 2 | MOLECULE: | TUBULIN ALPHA-1B CHAIN |
| 71:00:00 | 5d80-G | 6.3 | 5.2 | 103 | 223 | 7 | MOLECULE: | V-TYPE PROTON ATPASE CATALYTIC SUBUNIT A |
| 72:00:00 | 7dae-E | 6.3 | 1.9 | 101 | 124 | 2 | MOLECULE: | TUBULIN ALPHA-1B CHAIN |
| 73:00:00 | 7xqx-E | 6.3 | 2 | 100 | 123 | 5 | MOLECULE: | TUBULIN ALPHA-1B CHAIN |
| 74:00:00 | 6a9p-C | 6.3 | 5.3 | 104 | 104 | 9 | MOLECULE: | GLIAL FIBRILLARY ACIDIC PROTEIN |
| 75:00:00 | 6y6d-E | 6.3 | 4.3 | 104 | 122 | 3 | MOLECULE: | TUBULIN ALPHA-1B CHAIN |
| 76:00:00 | 5s4t-E | 6.3 | 4.4 | 104 | 123 | 3 | MOLECULE: | TUBULIN ALPHA-1B CHAIN |
| 77:00:00 | 5s5g-E | 6.3 | 4.4 | 104 | 123 | 3 | MOLECULE: | TUBULIN ALPHA-1B CHAIN |
| 78:00:00 | 6fkl-E | 6.3 | 1.6 | 100 | 122 | 3 | MOLECULE: | TUBULIN ALPHA-1B CHAIN |
| 79:00:00 | 5ov7-E | 6.3 | 2 | 99 | 123 | 5 | MOLECULE: | TUBULIN ALPHA-1B CHAIN |
| 80:00:00 | 7z2p-E | 6.3 | 2 | 100 | 123 | 5 | MOLECULE: | TUBULIN ALPHA-1B CHAIN |
| 81:00:00 | 5sb5-E | 6.3 | 2 | 100 | 123 | 5 | MOLECULE: | TUBULIN ALPHA-1B CHAIN |
| 82:00:00 | 5vpc-B | 6.3 | 4.2 | 68 | 68 | 10 | MOLECULE: | PROTEIN FOSB |
| 83:00:00 | 7ucc-J | 6.3 | 4.3 | 68 | 68 | 10 | MOLECULE: | PROTEIN FOSB |
| 84:00:00 | 7emj-E | 6.3 | 4.4 | 104 | 123 | 3 | MOLECULE: | TUBULIN ALPHA-1B CHAIN |
| 85:00:00 | 8c0f-E | 6.3 | 2 | 100 | 122 | 5 | MOLECULE: | TUBULIN ALPHA-1B CHAIN |
| 86:00:00 | 7vmk-E | 6.3 | 4.4 | 104 | 123 | 3 | MOLECULE: | TUBULIN ALPHA-1B CHAIN |
| 87:00:00 | 6n6s-D | 6.3 | 3.6 | 69 | 69 | 12 | MOLECULE: | TNFAIP3-INTERACTING PROTEIN 1 |
| 88:00:00 | 3cve-A | 6.3 | 4.3 | 68 | 68 | 7 | MOLECULE: | HOMER PROTEIN HOMOLOG 1 |
| 89:00:00 | 5sb8-E | 6.3 | 4.4 | 104 | 123 | 3 | MOLECULE: | TUBULIN ALPHA-1B CHAIN |
| 90:00:00 | 7xr1-E | 6.3 | 4.4 | 104 | 123 | 3 | MOLECULE: | TUBULIN ALPHA-1B CHAIN |
| 91:00:00 | 5wq4-C | 6.3 | 4.5 | 72 | 72 | 8 | MOLECULE: | UBIQUITIN |
| 92:00:00 | 5nnv-A | 6.3 | 5.9 | 119 | 233 | 8 | MOLECULE: | CHROMOSOME PARTITION PROTEIN SMC,CHROMOSOME |
| 93:00:00 | 1gk4-C | 6.3 | 3 | 70 | 70 | 0 | MOLECULE: | VIMENTIN |
| 94:00:00 | 6ltj-I | 6.3 | 21.4 | 118 | 551 | 7 | MOLECULE: | HISTONE H3.3 |
| 95:00:00 | 6gbo-L | 6.3 | 5 | 72 | 72 | 0 | MOLECULE: | POLYMERASE COFACTOR VP35 |
| 96:00:00 | 7bv6-W | 6.3 | 4.3 | 72 | 72 | 7 | MOLECULE: | VESICLE-ASSOCIATED MEMBRANE PROTEIN 8 |
| 97:00:00 | 6gbo-B | 6.3 | 5.2 | 73 | 73 | 4 | MOLECULE: | POLYMERASE COFACTOR VP35 |
| 98:00:00 | 5vpe-A | 6.3 | 2.9 | 67 | 67 | 7 | MOLECULE: | PROTEIN FOSB |
| 99:00:00 | 8b8d-D | 6.3 | 4.7 | 103 | 103 | 7 | MOLECULE: | PHOSPHOPROTEIN |
| 0:00 | 6y4n-E | 6.3 | 3.9 | 104 | 123 | 6 | MOLECULE: | TUBULIN ALPHA-1B CHAIN |
| 1:00 | 5s5x-E | 6.3 | 3.5 | 107 | 123 | 11 | MOLECULE: | TUBULIN ALPHA-1B CHAIN |
| 2:00 | 5s5q-E | 6.3 | 3.5 | 106 | 123 | 11 | MOLECULE: | TUBULIN ALPHA-1B CHAIN |
| 3:00 | 5s51-E | 6.3 | 3.6 | 109 | 123 | 12 | MOLECULE: | TUBULIN ALPHA-1B CHAIN |
| 4:00 | 5s4y-E | 6.3 | 3.5 | 106 | 123 | 11 | MOLECULE: | TUBULIN ALPHA-1B CHAIN |

| | | | | | | | | |
|---|---|---|---|---|---|---|---|---|
| 5:00 | 5s4p-E | 6.3 | 3.5 | 106 | 123 | 11 | MOLECULE: | TUBULIN ALPHA-1B CHAIN |
| 6:00 | 5s4u-E | 6.3 | 3.3 | 107 | 123 | 12 | MOLECULE: | TUBULIN ALPHA-1B CHAIN |
| 7:00 | 5s67-E | 6.3 | 3.4 | 107 | 123 | 12 | MOLECULE: | TUBULIN ALPHA-1B CHAIN |
| 8:00 | 5s5t-E | 6.3 | 3.4 | 108 | 123 | 12 | MOLECULE: | TUBULIN ALPHA-1B CHAIN |
| 9:00 | 5s4l-E | 6.3 | 3.4 | 106 | 123 | 11 | MOLECULE: | TUBULIN ALPHA-1B CHAIN |
| 10:00 | 5s55-E | 6.3 | 3.5 | 106 | 123 | 11 | MOLECULE: | TUBULIN ALPHA-1B CHAIN |
| 11:00 | 6tiu-E | 6.3 | 3.9 | 100 | 127 | 8 | MOLECULE: | TUBULIN ALPHA-1 CHAIN |
| 12:00 | 5hzp-A | 6.3 | 4.4 | 71 | 71 | 8 | MOLECULE: | M PROTEIN, SEROTYPE 49 |
| 13:00 | 3cvf-C | 6.3 | 1.6 | 74 | 74 | 12 | MOLECULE: | HOMER PROTEIN HOMOLOG 3 |
| 14:00 | 6gbp-F | 6.3 | 5 | 73 | 73 | 0 | MOLECULE: | POLYMERASE COFACTOR VP35 |
| 15:00 | 6hk9-A | 6.3 | 4.9 | 74 | 74 | 7 | MOLECULE: | TESTIS-EXPRESSED PROTEIN 12 |
| 16:00 | 6r0y-L | 6.3 | 4.6 | 81 | 186 | 7 | MOLECULE: | V-TYPE ATP SYNTHASE ALPHA CHAIN |
| 17:00 | 4jpn-G | 6.3 | 3.5 | 74 | 75 | 8 | MOLECULE: | MINOR SPIKE PROTEIN H |
| 18:00 | 5mq4-B | 6.3 | 2.8 | 74 | 111 | 8 | MOLECULE: | PROTEIN KINASE C-BINDING PROTEIN 1 |
| 19:00 | 4jpn-F | 6.3 | 3.4 | 72 | 72 | 8 | MOLECULE: | MINOR SPIKE PROTEIN H |
| 20:00 | 7tkc-U | 6.3 | 7.7 | 140 | 155 | 4 | MOLECULE: | ATP SYNTHASE SUBUNIT 9, MITOCHONDRIAL |
| 21:00 | 3swk-A | 6.3 | 4 | 85 | 86 | 5 | MOLECULE: | VIMENTIN |
| 22:00 | 4lhy-E | 6.3 | 3.8 | 76 | 76 | 9 | MOLECULE: | RAS-RELATED PROTEIN RAB-8A |
| 23:00 | 4lhz-E | 6.3 | 3.9 | 76 | 76 | 9 | MOLECULE: | RAS-RELATED PROTEIN RAB-8A |
| 24:00:00 | 7tkf-U | 6.3 | 12.4 | 147 | 155 | 9 | MOLECULE: | ATP SYNTHASE SUBUNIT 9 |
| 25:00:00 | 5nfz-E | 6.3 | 4 | 102 | 125 | 9 | MOLECULE: | TUBULIN ALPHA-1B CHAIN |
| 26:00:00 | 7jg6-b | 6.3 | 9 | 129 | 138 | 9 | MOLECULE: | ATP SYNTHASE SUBUNIT ALPHA |
| 27:00:00 | 6ses-E | 6.3 | 4.1 | 100 | 123 | 8 | MOLECULE: | TUBULIN ALPHA-1B CHAIN |
| 28:00:00 | 6qlf-U | 6.3 | 14.6 | 121 | 171 | 7 | MOLECULE: | INNER KINETOCHORE SUBUNIT IML3 |
| 29:00:00 | 6vq7-N | 6.3 | 8.7 | 113 | 114 | 7 | MOLECULE: | ATPASE H+-TRANSPORTING V1 SUBUNIT A |
| 30:00:00 | 2msd-C | 6.3 | 44.7 | 116 | 198 | 4 | MOLECULE: | APOLIPOPROTEIN A-I |
| 31:00:00 | 6zay-D | 6.3 | 3.6 | 85 | 86 | 7 | MOLECULE: | RAS-RELATED PROTEIN RAB-33B |
| 32:00:00 | 6wnr-X | 6.3 | 9.3 | 135 | 153 | 5 | MOLECULE: | ATP SYNTHASE SUBUNIT DELTA |
| 33:00:00 | 6lk8-M | 6.3 | 4.6 | 73 | 73 | 0 | MOLECULE: | MGC83295 PROTEIN |
| 34:00:00 | 4c47-A | 6.2 | 8.3 | 81 | 191 | 10 | MOLECULE: | INNER MEMBRANE LIPOPROTEIN |
| 35:00:00 | 7au5-E | 6.2 | 1.8 | 100 | 124 | 2 | MOLECULE: | TUBULIN ALPHA-1B CHAIN |
| 36:00:00 | 6th4-E | 6.2 | 6.2 | 103 | 128 | 2 | MOLECULE: | TUBULIN ALPHA CHAIN |
| 37:00:00 | 5s5o-E | 6.2 | 1.9 | 100 | 123 | 2 | MOLECULE: | TUBULIN ALPHA-1B CHAIN |
| 38:00:00 | 5s4x-E | 6.2 | 1.9 | 100 | 123 | 2 | MOLECULE: | TUBULIN ALPHA-1B CHAIN |
| 39:00:00 | 7e4z-E | 6.2 | 1.8 | 99 | 123 | 2 | MOLECULE: | TUBULIN ALPHA-1B CHAIN |
| 40:00:00 | 5s58-E | 6.2 | 1.7 | 99 | 123 | 2 | MOLECULE: | TUBULIN ALPHA-1B CHAIN |
| 41:00:00 | 5sb4-E | 6.2 | 1.8 | 99 | 123 | 2 | MOLECULE: | TUBULIN ALPHA-1B CHAIN |
| 42:00:00 | 5s62-E | 6.2 | 1.9 | 100 | 123 | 2 | MOLECULE: | TUBULIN ALPHA-1B CHAIN |
| 43:00:00 | 5s5e-E | 6.2 | 1.9 | 100 | 123 | 2 | MOLECULE: | TUBULIN ALPHA-1B CHAIN |
| 44:00:00 | 5s5j-E | 6.2 | 1.9 | 100 | 123 | 2 | MOLECULE: | TUBULIN ALPHA-1B CHAIN |
| 45:00:00 | 5s5h-E | 6.2 | 1.8 | 99 | 123 | 2 | MOLECULE: | TUBULIN ALPHA-1B CHAIN |
| 46:00:00 | 5sbd-E | 6.2 | 2.6 | 98 | 121 | 6 | MOLECULE: | TUBULIN ALPHA-1B CHAIN |
| 47:00:00 | 5s4r-E | 6.2 | 1.8 | 99 | 123 | 2 | MOLECULE: | TUBULIN ALPHA-1B CHAIN |
| 48:00:00 | 7daf-E | 6.2 | 1.9 | 100 | 123 | 2 | MOLECULE: | TUBULIN ALPHA-1B CHAIN |
| 49:00:00 | 5s4q-E | 6.2 | 1.8 | 100 | 123 | 2 | MOLECULE: | TUBULIN ALPHA-1B CHAIN |
| 50:00:00 | 5s4w-E | 6.2 | 1.8 | 100 | 123 | 2 | MOLECULE: | TUBULIN ALPHA-1B CHAIN |
| 51:00:00 | 7lz7-E | 6.2 | 1.7 | 98 | 121 | 3 | MOLECULE: | TUBULIN ALPHA-1B CHAIN |
| 52:00:00 | 8dbs-Y | 6.2 | 11.8 | 130 | 156 | 4 | MOLECULE: | ATP SYNTHASE SUBUNIT ALPHA |
| 53:00:00 | 7cnn-E | 6.2 | 4.5 | 103 | 122 | 3 | MOLECULE: | TUBULIN ALPHA-1B CHAIN |
| 54:00:00 | 7ogn-E | 6.2 | 4.4 | 102 | 120 | 4 | MOLECULE: | TUBULIN ALPHA-1B CHAIN |
| 55:00:00 | 6fjm-E | 6.2 | 1.8 | 99 | 123 | 2 | MOLECULE: | TUBULIN ALPHA-1B CHAIN |

| | | | | | | | | |
|---|---|---|---|---|---|---|---|---|
| 56:00:00 | 5xkg-E | 6.2 | 2 | 98 | 121 | 5 | MOLECULE: | TUBULIN ALPHA-1B CHAIN |
| 57:00:00 | 7e4y-E | 6.2 | 2.7 | 100 | 123 | 7 | MOLECULE: | TUBULIN ALPHA-1B CHAIN |
| 58:00:00 | 6v8o-J | 6.2 | 8 | 86 | 115 | 3 | MOLECULE: | HIGH TEMPERATURE LETHAL PROTEIN 1 |
| 59:00:00 | 6kw3-W | 6.2 | 3.9 | 69 | 69 | 6 | MOLECULE: | HISTONE H4 |
| 60:00:00 | 3cve-B | 6.2 | 4.1 | 66 | 66 | 8 | MOLECULE: | HOMER PROTEIN HOMOLOG 1 |
| 61:00:00 | 6n6r-D | 6.2 | 3.4 | 67 | 67 | 12 | MOLECULE: | UBIQUITIN |
| 62:00:00 | 5sba-E | 6.2 | 1.6 | 98 | 121 | 3 | MOLECULE: | TUBULIN ALPHA-1B CHAIN |
| 63:00:00 | 5sbb-E | 6.2 | 1.6 | 97 | 120 | 2 | MOLECULE: | TUBULIN ALPHA-1B CHAIN |
| 64:00:00 | 5sbc-E | 6.2 | 4.4 | 102 | 121 | 3 | MOLECULE: | TUBULIN ALPHA-1B CHAIN |
| 65:00:00 | 8amr-A | 6.2 | 3.7 | 79 | 155 | 5 | MOLECULE: | TRIPARTITE MOTIF-CONTAINING PROTEIN 3 |
| 66:00:00 | 6wvw-F | 6.2 | 2.1 | 66 | 66 | 6 | MOLECULE: | VESICLE-ASSOCIATED MEMBRANE PROTEIN 2 |
| 67:00:00 | 7bv6-S | 6.2 | 4.2 | 74 | 74 | 9 | MOLECULE: | VESICLE-ASSOCIATED MEMBRANE PROTEIN 8 |
| 68:00:00 | 6v86-C | 6.2 | 3.9 | 74 | 74 | 8 | MOLECULE: | RNA-DIRECTED RNA POLYMERASE L |
| 69:00:00 | 7rsc-D | 6.2 | 30.1 | 138 | 187 | 10 | MOLECULE: | GTPASE KRAS |
| 70:00:00 | 1jth-B | 6.2 | 2.7 | 67 | 67 | 4 | MOLECULE: | SNAP25 |
| 71:00:00 | 1urq-C | 6.2 | 1.6 | 68 | 68 | 6 | MOLECULE: | M-TOMOSYN ISOFORM |
| 72:00:00 | 5vpd-B | 6.2 | 4.2 | 67 | 67 | 10 | MOLECULE: | PROTEIN FOSB |
| 73:00:00 | 1gk4-E | 6.2 | 2.8 | 69 | 70 | 0 | MOLECULE: | VIMENTIN |
| 74:00:00 | 1sfc-B | 6.2 | 3.6 | 72 | 72 | 3 | MOLECULE: | PROTEIN (SYNAPTOBREVIN 2) |
| 75:00:00 | 1sfc-G | 6.2 | 2 | 72 | 72 | 6 | MOLECULE: | PROTEIN (SYNAPTOBREVIN 2) |
| 76:00:00 | 1gk4-F | 6.2 | 3.4 | 74 | 74 | 0 | MOLECULE: | VIMENTIN |
| 77:00:00 | 6cxi-T | 6.2 | 5.1 | 125 | 127 | 0 | MOLECULE: | ACTIN, CYTOPLASMIC 2 |
| 78:00:00 | 6cxj-T | 6.2 | 5 | 125 | 127 | 0 | MOLECULE: | ACTIN, CYTOPLASMIC 2 |
| 79:00:00 | 8pqw-F | 6.2 | 14.3 | 138 | 150 | 9 | MOLECULE: | CYTOPLASMIC DYNEIN 1 HEAVY CHAIN 1 |
| 80:00:00 | 3b5n-G | 6.2 | 1.7 | 69 | 69 | 6 | MOLECULE: | SYNAPTOBREVIN HOMOLOG 1 |
| 81:00:00 | 7c7y-C | 6.2 | 2.5 | 67 | 67 | 4 | MOLECULE: | UNCHARACTERIZED PROTEIN |
| 82:00:00 | 5s5k-E | 6.2 | 3.4 | 106 | 123 | 11 | MOLECULE: | TUBULIN ALPHA-1B CHAIN |
| 83:00:00 | 5s64-E | 6.2 | 3.5 | 106 | 123 | 11 | MOLECULE: | TUBULIN ALPHA-1B CHAIN |
| 84:00:00 | 5vpb-D | 6.2 | 4 | 66 | 66 | 11 | MOLECULE: | PROTEIN FOSB |
| 85:00:00 | 5s59-E | 6.2 | 3.5 | 106 | 123 | 11 | MOLECULE: | TUBULIN ALPHA-1B CHAIN |
| 86:00:00 | 7yg4-E | 6.2 | 4 | 67 | 67 | 7 | MOLECULE: | PROTEIN VIRILIZER HOMOLOG |
| 87:00:00 | 5s4n-E | 6.2 | 3.3 | 106 | 123 | 12 | MOLECULE: | TUBULIN ALPHA-1B CHAIN |
| 88:00:00 | 5vpe-D | 6.2 | 2.9 | 67 | 67 | 12 | MOLECULE: | PROTEIN FOSB |
| 89:00:00 | 6v85-E | 6.2 | 4.8 | 74 | 74 | 4 | MOLECULE: | RNA-DIRECTED RNA POLYMERASE L |
| 90:00:00 | 6vag-A | 6.2 | 4.8 | 74 | 74 | 4 | MOLECULE: | PHOSPHOPROTEIN |
| 91:00:00 | 6vqj-O | 6.2 | 4 | 67 | 67 | 1 | MOLECULE: | V-TYPE PROTON ATPASE SUBUNIT C 1 |
| 92:00:00 | 3cvf-D | 6.2 | 2.3 | 71 | 72 | 14 | MOLECULE: | HOMER PROTEIN HOMOLOG 3 |
| 93:00:00 | 4l2w-A | 6.2 | 2.1 | 67 | 67 | 4 | MOLECULE: | RHO-ASSOCIATED PROTEIN KINASE 1 |
| 94:00:00 | 6kpp-E | 6.2 | 3.9 | 107 | 123 | 11 | MOLECULE: | TUBULIN ALPHA-1B CHAIN |
| 95:00:00 | 4l2w-D | 6.2 | 1.6 | 68 | 68 | 7 | MOLECULE: | RHO-ASSOCIATED PROTEIN KINASE 1 |
| 96:00:00 | 6xx0-B | 6.2 | 4.5 | 75 | 75 | 9 | MOLECULE: | INHIBITOR OF KAPPA LIGHT POLYPEPTIDE GENE EN |
| 97:00:00 | 7njo-b | 6.2 | 9.2 | 132 | 141 | 5 | MOLECULE: | ATP SYNTHASE SUBUNIT ALPHA |
| 98:00:00 | 6ff7-J | 6.2 | 3.8 | 77 | 135 | 9 | MOLECULE: | RNA-BINDING MOTIF PROTEIN, X-LINKED 2 |
| 99:00:00 | 6r10-K | 6.2 | 3.8 | 98 | 98 | 7 | MOLECULE: | V-TYPE ATP SYNTHASE ALPHA CHAIN |
| 0:00 | 6vq6-N | 6.2 | 8.1 | 113 | 114 | 9 | MOLECULE: | ATPASE H+-TRANSPORTING V1 SUBUNIT A |
| 1:00 | 6vq8-M | 6.2 | 9.5 | 114 | 114 | 5 | MOLECULE: | ATPASE H+-TRANSPORTING V1 SUBUNIT A |
| 2:00 | 6vq7-M | 6.2 | 8.7 | 113 | 114 | 9 | MOLECULE: | ATPASE H+-TRANSPORTING V1 SUBUNIT A |
| 3:00 | 3uf1-C | 6.2 | 5.4 | 107 | 109 | 5 | MOLECULE: | VIMENTIN |
| 4:00 | 5hmo-C | 6.2 | 5.6 | 111 | 122 | 11 | MOLECULE: | UNCONVENTIONAL MYOSIN-X |
| 5:00 | 7ttf-E | 6.2 | 5 | 113 | 123 | 12 | MOLECULE: | TUBULIN ALPHA-1B CHAIN |
| 6:00 | 7y5d-b | 6.2 | 7.1 | 130 | 145 | 8 | MOLECULE: | ATP SYNTHASE SUBUNIT A |

| | | | | | | | | |
|---|---|---|---|---|---|---|---|---|
| 7:00 | 2e7s-L | 6.2 | 3.1 | 114 | 114 | 3 | MOLECULE: | RAB GUANINE NUCLEOTIDE EXCHANGE FACTOR SEC2 |
| 8:00 | 5t4q-J | 6.2 | 9.5 | 148 | 155 | 10 | MOLECULE: | ATP SYNTHASE SUBUNIT ALPHA |
| 9:00 | 7jts-s | 6.1 | 18.4 | 104 | 290 | 7 | MOLECULE: | RADIAL SPOKE PROTEIN 3 |
| 10:00 | 8q85-b | 6.1 | 2.7 | 68 | 70 | 6 | MOLECULE: | KINETOCHORE PROTEIN NDC80 |
| 11:00 | 6eun-A | 6.1 | 3.8 | 112 | 147 | 4 | MOLECULE: | ADHESIN |
| 12:00 | 8q85-G | 6.1 | 7 | 126 | 178 | 6 | MOLECULE: | KINETOCHORE PROTEIN NDC80 |
| 13:00 | 5bmv-E | 6.1 | 2.1 | 99 | 123 | 2 | MOLECULE: | TUBULIN ALPHA-1B CHAIN |
| 14:00 | 4zhq-E | 6.1 | 1.9 | 99 | 123 | 2 | MOLECULE: | TUBULIN ALPHA-1B CHAIN |
| 15:00 | 5s5n-E | 6.1 | 1.9 | 100 | 122 | 2 | MOLECULE: | TUBULIN ALPHA-1B CHAIN |
| 16:00 | 6xes-E | 6.1 | 5.9 | 101 | 125 | 3 | MOLECULE: | TUBULIN ALPHA-1B CHAIN |
| 17:00 | 5s4s-E | 6.1 | 1.7 | 97 | 120 | 2 | MOLECULE: | TUBULIN ALPHA-1B CHAIN |
| 18:00 | 6br1-E | 6.1 | 2 | 98 | 121 | 2 | MOLECULE: | TUBULIN ALPHA-1B CHAIN |
| 19:00 | 8ahl-D | 6.1 | 6.6 | 145 | 234 | 6 | MOLECULE: | CRESCENTIN |
| 20:00 | 7cda-E | 6.1 | 4.5 | 102 | 121 | 3 | MOLECULE: | TUBULIN ALPHA-1B CHAIN |
| 21:00 | 6nng-E | 6.1 | 2 | 98 | 121 | 2 | MOLECULE: | TUBULIN ALPHA-1B CHAIN |
| 22:00 | 8ajb-D | 6.1 | 9.4 | 154 | 234 | 9 | MOLECULE: | CRESCENTIN |
| 23:00 | 8ajb-P | 6.1 | 6.5 | 146 | 234 | 12 | MOLECULE: | CRESCENTIN |
| 24:00:00 | 6h9b-E | 6.1 | 6.4 | 110 | 136 | 4 | MOLECULE: | TUBULIN ALPHA CHAIN |
| 25:00:00 | 6s9e-E | 6.1 | 1.9 | 98 | 120 | 2 | MOLECULE: | TUBULIN ALPHA-1B CHAIN |
| 26:00:00 | 4zi7-E | 6.1 | 2 | 100 | 123 | 2 | MOLECULE: | TUBULIN ALPHA-1B CHAIN |
| 27:00:00 | 5h7o-E | 6.1 | 2.1 | 98 | 121 | 2 | MOLECULE: | TUBULIN ALPHA-1B CHAIN |
| 28:00:00 | 6k9v-E | 6.1 | 2 | 98 | 121 | 2 | MOLECULE: | TUBULIN ALPHA-1B CHAIN |
| 29:00:00 | 6y4m-E | 6.1 | 2.2 | 100 | 123 | 2 | MOLECULE: | TUBULIN ALPHA-1B CHAIN |
| 30:00:00 | 8ahl-B | 6.1 | 8.4 | 154 | 238 | 6 | MOLECULE: | CRESCENTIN |
| 31:00:00 | 5kx5-E | 6.1 | 4.5 | 102 | 121 | 3 | MOLECULE: | TUBULIN ALPHA CHAIN |
| 32:00:00 | 6d88-E | 6.1 | 4.5 | 102 | 121 | 3 | MOLECULE: | TUBULIN ALPHA-1B CHAIN |
| 33:00:00 | 8ajb-N | 6.1 | 8.4 | 154 | 238 | 6 | MOLECULE: | CRESCENTIN |
| 34:00:00 | 5yl4-E | 6.1 | 1.7 | 98 | 120 | 3 | MOLECULE: | TUBULIN ALPHA CHAIN |
| 35:00:00 | 8asn-E | 6.1 | 2.3 | 101 | 123 | 2 | MOLECULE: | TUBULIN ALPHA-1B CHAIN |
| 36:00:00 | 7en3-E | 6.1 | 2 | 98 | 121 | 5 | MOLECULE: | TUBULIN ALPHA-1B CHAIN |
| 37:00:00 | 6wm2-H | 6.1 | 13.2 | 100 | 225 | 6 | MOLECULE: | V-TYPE PROTON ATPASE SUBUNIT E 1 |
| 38:00:00 | 8clc-E | 6.1 | 1.7 | 100 | 123 | 3 | MOLECULE: | TUBULIN ALPHA-1B CHAIN |
| 39:00:00 | 5wjb-C | 6.1 | 3.7 | 97 | 112 | 4 | MOLECULE: | CAPSID ASSEMBLY SCAFFOLDING PROTEIN,MYOSIN-7 |
| 40:00:00 | 7bjs-A | 6.1 | 4.6 | 69 | 69 | 6 | MOLECULE: | KINESIN HEAVY CHAIN |
| 41:00:00 | 1wt6-B | 6.1 | 3.9 | 67 | 67 | 7 | MOLECULE: | MYOTONIN-PROTEIN KINASE |
| 42:00:00 | 7c7y-A | 6.1 | 4.7 | 66 | 66 | 6 | MOLECULE: | UNCHARACTERIZED PROTEIN |
| 43:00:00 | 8b8b-B | 6.1 | 3 | 95 | 95 | 12 | MOLECULE: | MUNIA BORNAVIRUS 1 PHOSPHOPROTEIN |
| 44:00:00 | 7say-D | 6.1 | 3.2 | 67 | 67 | 6 | MOLECULE: | GENERAL CONTROL TRANSCRIPTION FACTOR GCN4/M |
| 45:00:00 | 5sb9-E | 6.1 | 4.4 | 102 | 123 | 3 | MOLECULE: | TUBULIN ALPHA-1B CHAIN |
| 46:00:00 | 7bv6-O | 6.1 | 4.3 | 73 | 73 | 8 | MOLECULE: | VESICLE-ASSOCIATED MEMBRANE PROTEIN 8 |
| 47:00:00 | 1sfc-F | 6.1 | 4.7 | 73 | 73 | 4 | MOLECULE: | PROTEIN (SYNAPTOBREVIN 2) |
| 48:00:00 | 6g2t-P | 6.1 | 4.9 | 129 | 135 | 0 | MOLECULE: | ACTIN, CYTOPLASMIC 2 |
| 49:00:00 | 3gn4-A | 6.1 | 4.5 | 89 | 129 | 3 | MOLECULE: | MYOSIN-VI |
| 50:00:00 | 3b5n-F | 6.1 | 3.6 | 68 | 68 | 4 | MOLECULE: | SYNAPTOBREVIN HOMOLOG 1 |
| 51:00:00 | 7bv6-K | 6.1 | 4.4 | 73 | 73 | 7 | MOLECULE: | VESICLE-ASSOCIATED MEMBRANE PROTEIN 8 |
| 52:00:00 | 7eqb-A | 6.1 | 4.5 | 68 | 68 | 9 | MOLECULE: | KINESIN-LIKE PROTEIN |
| 53:00:00 | 6oqt-X | 6.1 | 8.7 | 135 | 156 | 7 | MOLECULE: | ATP SYNTHASE SUBUNIT DELTA |
| 54:00:00 | 5bw9-i | 6.1 | 9 | 126 | 217 | 7 | MOLECULE: | V-TYPE PROTON ATPASE CATALYTIC SUBUNIT A |
| 55:00:00 | 7e4r-E | 6.1 | 6.8 | 119 | 121 | 5 | MOLECULE: | TUBULIN ALPHA-1B CHAIN |
| 56:00:00 | 5s66-E | 6.1 | 3.3 | 105 | 123 | 11 | MOLECULE: | TUBULIN ALPHA-1B CHAIN |
| 57:00:00 | 5s52-E | 6.1 | 3.3 | 106 | 123 | 12 | MOLECULE: | TUBULIN ALPHA-1B CHAIN |

| | | | | | | | | |
|---|---|---|---|---|---|---|---|---|
| 58:00:00 | 4bwn-B | 6.1 | 4.9 | 81 | 86 | 6 | MOLECULE: | NF-KAPPA-B ESSENTIAL MODULATOR |
| 59:00:00 | 5ylj-E | 6.1 | 4.2 | 98 | 121 | 7 | MOLECULE: | TUBULIN ALPHA-1B CHAIN |
| 60:00:00 | 1sfc-C | 6.1 | 4 | 77 | 77 | 5 | MOLECULE: | PROTEIN (SYNAPTOBREVIN 2) |
| 61:00:00 | 6gbq-A | 6.1 | 4.3 | 70 | 70 | 3 | MOLECULE: | POLYMERASE COFACTOR VP35 |
| 62:00:00 | 3lj5-C | 6.1 | 8 | 131 | 692 | 8 | MOLECULE: | PORTAL PROTEIN |
| 63:00:00 | 3lj5-F | 6.1 | 8 | 131 | 692 | 8 | MOLECULE: | PORTAL PROTEIN |
| 64:00:00 | 5gai-G | 6.1 | 11.5 | 130 | 721 | 7 | MOLECULE: | PORTAL PROTEIN |
| 65:00:00 | 7syg-H | 6.1 | 7.4 | 72 | 237 | 8 | MOLECULE: | 18S RRNA |
| 66:00:00 | 6gbp-L | 6.1 | 5.5 | 73 | 73 | 4 | MOLECULE: | POLYMERASE COFACTOR VP35 |
| 67:00:00 | 8fby-A | 6.1 | 3.5 | 83 | 89 | 6 | MOLECULE: | PLASMALEMMA VESICLE-ASSOCIATED PROTEIN |
| 68:00:00 | 7njl-b | 6.1 | 9.3 | 131 | 145 | 5 | MOLECULE: | ATP SYNTHASE SUBUNIT ALPHA |
| 69:00:00 | 7tkj-U | 6.1 | 12.5 | 146 | 155 | 9 | MOLECULE: | ATP SYNTHASE SUBUNIT 9 |
| 70:00:00 | 7tmr-H | 6.1 | 8.7 | 112 | 112 | 5 | MOLECULE: | H(+)-TRANSPORTING TWO-SECTOR ATPASE |
| 71:00:00 | 7njn-b | 6.1 | 9.3 | 132 | 145 | 5 | MOLECULE: | ATP SYNTHASE SUBUNIT ALPHA |
| 72:00:00 | 5gon-E | 6.1 | 2 | 98 | 121 | 5 | MOLECULE: | TUBULIN ALPHA-1B CHAIN |
| 73:00:00 | 7l05-E | 6.1 | 5.8 | 111 | 121 | 8 | MOLECULE: | TUBULIN ALPHA-1B CHAIN |
| 74:00:00 | 3v6i-X | 6.1 | 4.4 | 99 | 99 | 6 | MOLECULE: | V-TYPE ATP SYNTHASE SUBUNIT E |
| 75:00:00 | 6pc4-E | 6.1 | 4.1 | 98 | 121 | 7 | MOLECULE: | TUBULIN ALPHA-1B CHAIN |
| 76:00:00 | 6vq8-O | 6.1 | 8.6 | 112 | 113 | 9 | MOLECULE: | ATPASE H+-TRANSPORTING V1 SUBUNIT A |
| 77:00:00 | 5lov-E | 6.1 | 1.9 | 97 | 117 | 5 | MOLECULE: | TUBULIN ALPHA-1B CHAIN |
| 78:00:00 | 4lhy-C | 6.1 | 4.2 | 72 | 72 | 6 | MOLECULE: | RAS-RELATED PROTEIN RAB-8A |
| 79:00:00 | 4cpc-D | 6.1 | 5.1 | 112 | 141 | 8 | MOLECULE: | SYNAPTONEMAL COMPLEX PROTEIN 3 |
| 80:00:00 | 5vpe-C | 6.1 | 2.6 | 66 | 66 | 8 | MOLECULE: | PROTEIN FOSB |
| 81:00:00 | 6eb8-E | 6.1 | 3.9 | 75 | 103 | 8 | MOLECULE: | PHOSPHOPROTEIN |
| 82:00:00 | 2e7s-S | 6.1 | 4.1 | 116 | 117 | 7 | MOLECULE: | RAB GUANINE NUCLEOTIDE EXCHANGE FACTOR SEC2 |
| 83:00:00 | 2e7s-R | 6.1 | 2.9 | 111 | 111 | 8 | MOLECULE: | RAB GUANINE NUCLEOTIDE EXCHANGE FACTOR SEC2 |
| 84:00:00 | 4xa6-A | 6 | 10.5 | 129 | 168 | 9 | MOLECULE: | GP7-MYH7(1777-1855)-EB1 CHIMERA PROTEIN |
| 85:00:00 | 8bd7-M | 6 | 17.7 | 131 | 164 | 8 | MOLECULE: | IFT88 |
| 86:00:00 | 6oei-A | 6 | 10.8 | 100 | 243 | 8 | MOLECULE: | SPINDLE POLE BODY COMPONENT SPC42,SIGMA-54-D |
| 87:00:00 | 8clf-E | 6 | 2 | 98 | 121 | 2 | MOLECULE: | TUBULIN ALPHA-1B CHAIN |
| 88:00:00 | 5xke-E | 6 | 2 | 98 | 121 | 2 | MOLECULE: | TUBULIN ALPHA-1B CHAIN |
| 89:00:00 | 7s5u-A | 6 | 6.3 | 139 | 179 | 7 | MOLECULE: | KINESIN-LIKE PROTEIN KLP61F |
| 90:00:00 | 4zol-E | 6 | 2.3 | 99 | 122 | 2 | MOLECULE: | TUBULIN ALPHA-1B CHAIN |
| 91:00:00 | 8cld-E | 6 | 2 | 98 | 121 | 2 | MOLECULE: | DETYROSINATED TUBULIN ALPHA-1B CHAIN |
| 92:00:00 | 6o61-E | 6 | 2.1 | 97 | 121 | 2 | MOLECULE: | TUBULIN ALPHA-1B CHAIN |
| 93:00:00 | 8i03-G | 6 | 7.7 | 112 | 166 | 8 | MOLECULE: | PAIRED AMPHIPATHIC HELIX PROTEIN PST1 |
| 94:00:00 | 8tek-N | 6 | 7.2 | 144 | 187 | 10 | MOLECULE: | DYNEIN REGULATORY COMPLEX PROTEIN 1/2 N-TERM |
| 95:00:00 | 6gze-E | 6 | 4.3 | 100 | 116 | 2 | MOLECULE: | TUBULIN ALPHA-1B CHAIN |
| 96:00:00 | 6jcj-E | 6 | 4.5 | 102 | 120 | 3 | MOLECULE: | TUBULIN ALPHA-1B CHAIN |
| 97:00:00 | 5xaf-E | 6 | 2 | 98 | 121 | 5 | MOLECULE: | TUBULIN ALPHA-1B CHAIN |
| 98:00:00 | 6bs2-E | 6 | 2 | 98 | 121 | 2 | MOLECULE: | TUBULIN ALPHA-1B CHAIN |
| 99:00:00 | 5wq4-E | 6 | 4.7 | 69 | 69 | 4 | MOLECULE: | UBIQUITIN |
| 0:00 | 6uci-D | 6 | 4 | 67 | 67 | 6 | MOLECULE: | PROTEIN FOSB |
| 1:00 | 7c7y-B | 6 | 4.3 | 66 | 66 | 6 | MOLECULE: | UNCHARACTERIZED PROTEIN |
| 2:00 | 3cve-C | 6 | 4.1 | 66 | 66 | 8 | MOLECULE: | HOMER PROTEIN HOMOLOG 1 |
| 3:00 | 7yer-E | 6 | 4.6 | 67 | 67 | 3 | MOLECULE: | RNA-DIRECTED RNA POLYMERASE L |
| 4:00 | 6lk8-K | 6 | 3.8 | 68 | 69 | 0 | MOLECULE: | MGC83295 PROTEIN |
| 5:00 | 5wwl-N | 6 | 5.9 | 73 | 155 | 7 | MOLECULE: | CENTROMERE PROTEIN MIS12 |
| 6:00 | 7yes-E | 6 | 4.7 | 67 | 67 | 3 | MOLECULE: | RNA-DIRECTED RNA POLYMERASE L |
| 7:00 | 6iak-F | 6 | 3.5 | 71 | 71 | 8 | MOLECULE: | UNCHARACTERIZED PROTEIN |
| 8:00 | 7bv6-C | 6 | 3.6 | 72 | 72 | 8 | MOLECULE: | VESICLE-ASSOCIATED MEMBRANE PROTEIN 8 |

| | | | | | | | | |
|---|---|---|---|---|---|---|---|---|
| 9:00 | 1n7s-B | 6 | 3.6 | 68 | 68 | 7 | MOLECULE: | VESICLE-ASSOCIATED MEMBRANE PROTEIN 2 |
| 10:00 | 5eof-B | 6 | 4.2 | 73 | 74 | 1 | MOLECULE: | OPTINEURIN |
| 11:00 | 5d80-J | 6 | 6 | 104 | 104 | 8 | MOLECULE: | V-TYPE PROTON ATPASE CATALYTIC SUBUNIT A |
| 12:00 | 6ptw-C | 6 | 35.8 | 150 | 198 | 9 | MOLECULE: | APOLIPOPROTEIN A-I |
| 13:00 | 3wmi-A | 6 | 2.4 | 56 | 57 | 4 | MOLECULE: | EIAV GP45 WILD TYPE |
| 14:00 | 6v85-D | 6 | 4.7 | 71 | 72 | 4 | MOLECULE: | RNA-DIRECTED RNA POLYMERASE L |
| 15:00 | 5bw9-m | 6 | 10.6 | 94 | 181 | 4 | MOLECULE: | V-TYPE PROTON ATPASE CATALYTIC SUBUNIT A |
| 16:00 | 5toi-C | 6 | 3.1 | 70 | 70 | 9 | MOLECULE: | POLYMERASE COFACTOR VP35 |
| 17:00 | 5toh-B | 6 | 4.8 | 70 | 70 | 9 | MOLECULE: | POLYMERASE COFACTOR VP35 |
| 18:00 | 6v86-E | 6 | 4.9 | 74 | 74 | 5 | MOLECULE: | RNA-DIRECTED RNA POLYMERASE L |
| 19:00 | 6vag-B | 6 | 2.9 | 71 | 72 | 10 | MOLECULE: | PHOSPHOPROTEIN |
| 20:00 | 1gu5-B | 6 | 3.1 | 67 | 67 | 16 | MOLECULE: | CAAT/ENHANCER BINDING PROTEIN BETA |
| 21:00 | 1gu4-B | 6 | 3.2 | 67 | 67 | 16 | MOLECULE: | CAAT/ENHANCER BINDING PROTEIN BETA |
| 22:00 | 6qld-Q | 6 | 18.6 | 102 | 204 | 5 | MOLECULE: | INNER KINETOCHORE SUBUNIT MIF2 |
| 23:00 | 7c4j-H | 6 | 22.4 | 132 | 242 | 8 | MOLECULE: | TRANSCRIPTION REGULATORY PROTEIN SNF12 |
| 24:00:00 | 6gbp-B | 6 | 5.4 | 73 | 73 | 4 | MOLECULE: | POLYMERASE COFACTOR VP35 |
| 25:00:00 | 6n6s-B | 6 | 2.5 | 65 | 65 | 17 | MOLECULE: | TNFAIP3-INTERACTING PROTEIN 1 |
| 26:00:00 | 6v85-B | 6 | 4.7 | 71 | 72 | 4 | MOLECULE: | RNA-DIRECTED RNA POLYMERASE L |
| 27:00:00 | 5whf-B | 6 | 4.7 | 82 | 82 | 9 | MOLECULE: | VIMENTIN |
| 28:00:00 | 4l2w-C | 6 | 3.3 | 67 | 67 | 6 | MOLECULE: | RHO-ASSOCIATED PROTEIN KINASE 1 |
| 29:00:00 | 5wq4-D | 6 | 4 | 72 | 72 | 7 | MOLECULE: | UBIQUITIN |
| 30:00:00 | 6gbq-C | 6 | 3.9 | 69 | 69 | 4 | MOLECULE: | POLYMERASE COFACTOR VP35 |
| 31:00:00 | 4bry-A | 6 | 3.5 | 69 | 69 | 7 | MOLECULE: | GEMININ |
| 32:00:00 | 7a5p-J | 6 | 8.6 | 80 | 110 | 6 | MOLECULE: | U2 SNRNA |
| 33:00:00 | 7njv-b | 6 | 3.9 | 62 | 63 | 10 | MOLECULE: | ATP SYNTHASE SUBUNIT C |
| 34:00:00 | 6fki-p | 6 | 7.7 | 132 | 143 | 7 | MOLECULE: | ATP SYNTHASE SUBUNIT A, CHLOROPLASTIC |
| 35:00:00 | 8b8b-A | 6 | 2.9 | 99 | 99 | 6 | MOLECULE: | MUNIA BORNAVIRUS 1 PHOSPHOPROTEIN |
| 36:00:00 | 6bry-E | 6 | 4.2 | 98 | 121 | 7 | MOLECULE: | TUBULIN ALPHA-1B CHAIN |
| 37:00:00 | 5bw9-j | 6 | 4.8 | 101 | 101 | 2 | MOLECULE: | V-TYPE PROTON ATPASE CATALYTIC SUBUNIT A |
| 38:00:00 | 5mqf-J | 6 | 3.5 | 74 | 135 | 8 | MOLECULE: | PRE-MRNA-PROCESSING-SPLICING FACTOR 8 |
| 39:00:00 | 7njr-b | 6 | 7.2 | 113 | 145 | 6 | MOLECULE: | ATP SYNTHASE SUBUNIT ALPHA |
| 40:00:00 | 6vm1-J | 6 | 7.5 | 123 | 131 | 11 | MOLECULE: | ATP SYNTHASE SUBUNIT ALPHA, CHLOROPLASTIC |
| 41:00:00 | 7njq-b | 6 | 7.7 | 113 | 144 | 5 | MOLECULE: | ATP SYNTHASE SUBUNIT ALPHA |
| 42:00:00 | 6j5i-b | 5.9 | 13.4 | 145 | 209 | 7 | MOLECULE: | ATP SYNTHASE SUBUNIT ALPHA, MITOCHONDRIAL |
| 43:00:00 | 8fef-B | 5.9 | 23.9 | 149 | 335 | 13 | MOLECULE: | VIRULENCE FACTOR MCE FAMILY PROTEIN |
| 44:00:00 | 7apk-m | 5.9 | 6.6 | 121 | 549 | 4 | MOLECULE: | THO COMPLEX SUBUNIT 1 |
| 45:00:00 | 5bw9-g | 5.9 | 3.3 | 87 | 141 | 6 | MOLECULE: | V-TYPE PROTON ATPASE CATALYTIC SUBUNIT A |
| 46:00:00 | 7zyw-E | 5.9 | 1.9 | 96 | 119 | 2 | MOLECULE: | TUBULIN ALPHA-1B CHAIN |
| 47:00:00 | 8clb-E | 5.9 | 2.1 | 98 | 121 | 2 | MOLECULE: | TUBULIN ALPHA-1B CHAIN |
| 48:00:00 | 5nvu-A | 5.9 | 10.3 | 146 | 3169 | 0 | MOLECULE: | DYNEIN MOTOR DOMAIN |
| 49:00:00 | 8wd0-E | 5.9 | 1.9 | 97 | 121 | 2 | MOLECULE: | TUBULIN ALPHA-1B CHAIN |
| 50:00:00 | 8cle-E | 5.9 | 2.1 | 97 | 120 | 5 | MOLECULE: | TUBULIN ALPHA-1B CHAIN |
| 51:00:00 | 6cch-C | 5.9 | 33.5 | 144 | 198 | 4 | MOLECULE: | APOLIPOPROTEIN A-I |
| 52:00:00 | 7c7y-D | 5.9 | 4 | 64 | 64 | 6 | MOLECULE: | UNCHARACTERIZED PROTEIN |
| 53:00:00 | 7aw9-E | 5.9 | 2.1 | 65 | 65 | 9 | MOLECULE: | HAPB |
| 54:00:00 | 1jth-A | 5.9 | 1.5 | 62 | 62 | 8 | MOLECULE: | SNAP25 |
| 55:00:00 | 5bw9-L | 5.9 | 4.8 | 68 | 68 | 9 | MOLECULE: | V-TYPE PROTON ATPASE CATALYTIC SUBUNIT A |
| 56:00:00 | 3b5n-K | 5.9 | 2.9 | 65 | 65 | 5 | MOLECULE: | SYNAPTOBREVIN HOMOLOG 1 |
| 57:00:00 | 6kw4-V | 5.9 | 4.1 | 69 | 69 | 6 | MOLECULE: | HISTONE H3.2 |
| 58:00:00 | 5vpf-D | 5.9 | 3.7 | 65 | 65 | 12 | MOLECULE: | PROTEIN FOSB |
| 59:00:00 | 7upz-B | 5.9 | 4.6 | 68 | 68 | 6 | MOLECULE: | CCAAT/ENHANCER-BINDING PROTEIN BETA |

| | | | | | | | | |
|---|---|---|---|---|---|---|---|---|
| 60:00:00 | 1s1c-X | 5.9 | 3.5 | 69 | 69 | 4 | MOLECULE: | TRANSFORMING PROTEIN RHOA |
| 61:00:00 | 6wvw-D | 5.9 | 4.9 | 63 | 63 | 8 | MOLECULE: | VESICLE-ASSOCIATED MEMBRANE PROTEIN 2 |
| 62:00:00 | 8rtd-B | 5.9 | 7.6 | 73 | 197 | 10 | MOLECULE: | TRWJ PROTEIN |
| 63:00:00 | 5toi-B | 5.9 | 5.1 | 70 | 70 | 9 | MOLECULE: | POLYMERASE COFACTOR VP35 |
| 64:00:00 | 3cvf-B | 5.9 | 1.3 | 70 | 70 | 6 | MOLECULE: | HOMER PROTEIN HOMOLOG 3 |
| 65:00:00 | 6zr2-Z | 5.9 | 13 | 95 | 141 | 8 | MOLECULE: | NADH-UBIQUINONE OXIDOREDUCTASE CHAIN 3 |
| 66:00:00 | 7n84-Z | 5.9 | 3.4 | 63 | 63 | 0 | MOLECULE: | NUCLEOPORIN NUP188 |
| 67:00:00 | 7tmp-J | 5.9 | 9 | 108 | 108 | 6 | MOLECULE: | H(+)-TRANSPORTING TWO-SECTOR ATPASE |
| 68:00:00 | 8xwx-D | 5.9 | 4.8 | 65 | 65 | 8 | MOLECULE: | B-CELL RECEPTOR-ASSOCIATED PROTEIN 31 |
| 69:00:00 | 7l4v-B | 5.9 | 2.6 | 69 | 69 | 9 | MOLECULE: | CCAAT/ENHANCER-BINDING PROTEIN BETA |
| 70:00:00 | 1gtw-B | 5.9 | 3 | 67 | 67 | 16 | MOLECULE: | CAAT/ENHANCER BINDING PROTEIN BETA |
| 71:00:00 | 5toi-A | 5.9 | 5.1 | 68 | 68 | 6 | MOLECULE: | POLYMERASE COFACTOR VP35 |
| 72:00:00 | 7bv6-P | 5.9 | 2.1 | 65 | 65 | 12 | MOLECULE: | VESICLE-ASSOCIATED MEMBRANE PROTEIN 8 |
| 73:00:00 | 5cch-D | 5.9 | 4.1 | 64 | 64 | 6 | MOLECULE: | VESICLE-ASSOCIATED MEMBRANE PROTEIN 2 |
| 74:00:00 | 6mti-D | 5.9 | 4.1 | 64 | 64 | 5 | MOLECULE: | SYNAPTOTAGMIN-1 |
| 75:00:00 | 7tmq-J | 5.9 | 10.6 | 108 | 108 | 6 | MOLECULE: | H(+)-TRANSPORTING TWO-SECTOR ATPASE |
| 76:00:00 | 6hx8-E | 5.9 | 4.2 | 100 | 123 | 8 | MOLECULE: | TUBULIN ALPHA-1B CHAIN |
| 77:00:00 | 6wm2-M | 5.9 | 8.8 | 112 | 114 | 9 | MOLECULE: | V-TYPE PROTON ATPASE SUBUNIT E 1 |
| 78:00:00 | 4lhx-C | 5.9 | 4.7 | 72 | 72 | 4 | MOLECULE: | RAS-RELATED PROTEIN RAB-8A |
| 79:00:00 | 5bw9-J | 5.9 | 5.1 | 103 | 104 | 4 | MOLECULE: | V-TYPE PROTON ATPASE CATALYTIC SUBUNIT A |
| 80:00:00 | 7qin-K | 5.9 | 4.6 | 88 | 88 | 0 | MOLECULE: | ACTIN, ALPHA SKELETAL MUSCLE |
| 81:00:00 | 4lhz-C | 5.9 | 4.8 | 72 | 72 | 4 | MOLECULE: | RAS-RELATED PROTEIN RAB-8A |
| 82:00:00 | 4ytd-A | 5.9 | 3.4 | 95 | 96 | 13 | MOLECULE: | PROTEIN BICAUDAL D HOMOLOG 1 |
| 83:00:00 | 6vmb-J | 5.9 | 6.5 | 112 | 136 | 5 | MOLECULE: | ATP SYNTHASE SUBUNIT ALPHA, CHLOROPLASTIC |
| 84:00:00 | 8fed-A | 5.8 | 20.1 | 134 | 392 | 7 | MOLECULE: | VIRULENCE FACTOR MCE FAMILY PROTEIN |
| 85:00:00 | 1joc-A | 5.8 | 27.3 | 81 | 123 | 7 | MOLECULE: | EARLY ENDOSOMAL AUTOANTIGEN 1 |
| 86:00:00 | 5jcb-E | 5.8 | 2.8 | 96 | 121 | 3 | MOLECULE: | TUBULIN ALPHA-1B CHAIN |
| 87:00:00 | 1x79-B | 5.8 | 4.1 | 90 | 90 | 6 | MOLECULE: | ADP-RIBOSYLATION FACTOR BINDING PROTEIN GGA1 |
| 88:00:00 | 3tnu-B | 5.8 | 2.1 | 95 | 95 | 6 | MOLECULE: | KERATIN, TYPE I CYTOSKELETAL 14 |
| 89:00:00 | 7tmm-I | 5.8 | 10.3 | 91 | 220 | 7 | MOLECULE: | H(+)-TRANSPORTING TWO-SECTOR ATPASE |
| 90:00:00 | 5vpc-D | 5.8 | 4.6 | 63 | 63 | 3 | MOLECULE: | PROTEIN FOSB |
| 91:00:00 | 8k8d-A | 5.8 | 2.1 | 68 | 68 | 3 | MOLECULE: | CCAAT/ENHANCER-BINDING PROTEIN BETA |
| 92:00:00 | 7bv6-T | 5.8 | 4.6 | 67 | 67 | 6 | MOLECULE: | VESICLE-ASSOCIATED MEMBRANE PROTEIN 8 |
| 93:00:00 | 3cve-D | 5.8 | 2.7 | 61 | 61 | 7 | MOLECULE: | HOMER PROTEIN HOMOLOG 1 |
| 94:00:00 | 1l2p-A | 5.8 | 2.9 | 61 | 61 | 10 | MOLECULE: | ATP SYNTHASE B CHAIN |
| 95:00:00 | 7eao-B | 5.8 | 4.2 | 63 | 63 | 5 | MOLECULE: | POLYUBIQUITIN-C |
| 96:00:00 | 7bv6-D | 5.8 | 4.6 | 67 | 67 | 6 | MOLECULE: | VESICLE-ASSOCIATED MEMBRANE PROTEIN 8 |
| 97:00:00 | 6vqi-I | 5.8 | 4.1 | 64 | 64 | 5 | MOLECULE: | V-TYPE PROTON ATPASE SUBUNIT C 1 |
| 98:00:00 | 7eal-E | 5.8 | 4.3 | 64 | 65 | 5 | MOLECULE: | UBIQUITIN |
| 99:00:00 | 6iak-E | 5.8 | 3.8 | 67 | 67 | 9 | MOLECULE: | UNCHARACTERIZED PROTEIN |
| 0:00 | 6j6g-c | 5.8 | 19.5 | 125 | 436 | 3 | MOLECULE: | PRE-MRNA-SPLICING FACTOR 8 |
| 1:00 | 6vqk-I | 5.8 | 4.3 | 64 | 64 | 5 | MOLECULE: | V-TYPE PROTON ATPASE SUBUNIT C 1 |
| 2:00 | 6j6q-c | 5.8 | 31.3 | 133 | 436 | 3 | MOLECULE: | PRE-MRNA-SPLICING FACTOR 8 |
| 3:00 | 6j6h-c | 5.8 | 31.3 | 133 | 436 | 3 | MOLECULE: | PRE-MRNA-SPLICING FACTOR 8 |
| 4:00 | 1kil-D | 5.8 | 3.4 | 66 | 66 | 2 | MOLECULE: | SYNAPTOBREVIN SNARE MOTIF |
| 5:00 | 1urq-B | 5.8 | 4.4 | 63 | 63 | 3 | MOLECULE: | M-TOMOSYN ISOFORM |
| 6:00 | 5low-D | 5.8 | 4.3 | 67 | 67 | 6 | MOLECULE: | RABPHILIN-3A |
| 7:00 | 7dp8-E | 5.8 | 5 | 112 | 123 | 4 | MOLECULE: | TUBULIN ALPHA-1B CHAIN |
| 8:00 | 1n7s-D | 5.8 | 3.6 | 66 | 66 | 2 | MOLECULE: | VESICLE-ASSOCIATED MEMBRANE PROTEIN 2 |
| 9:00 | 3b5n-H | 5.8 | 2.7 | 63 | 63 | 8 | MOLECULE: | SYNAPTOBREVIN HOMOLOG 1 |
| 10:00 | 6j6n-c | 5.8 | 31.3 | 133 | 436 | 3 | MOLECULE: | PRE-MRNA-SPLICING FACTOR 8 |

| | | | | | | | | |
|---|---|---|---|---|---|---|---|---|
| 11:00 | 6ucm-C | 5.8 | 4.9 | 65 | 65 | 14 | MOLECULE: | PROTEIN FOSB |
| 12:00 | 4yv3-C | 5.8 | 4 | 63 | 63 | 8 | MOLECULE: | VIMENTIN |
| 13:00 | 1wt6-A | 5.8 | 4.7 | 65 | 65 | 5 | MOLECULE: | MYOTONIN-PROTEIN KINASE |
| 14:00 | 7xfr-D | 5.8 | 3.7 | 62 | 62 | 5 | MOLECULE: | ISOFORM 2 OF WD REPEAT DOMAIN PHOSPHOINOSITI |
| 15:00 | 8rtd-A | 5.8 | 4.6 | 68 | 197 | 9 | MOLECULE: | TRWJ PROTEIN |
| 16:00 | 6v92-J | 5.8 | 3.9 | 80 | 115 | 6 | MOLECULE: | ACTIN-RELATED PROTEIN 7 |
| 17:00 | 6gbr-A | 5.8 | 3.5 | 67 | 67 | 3 | MOLECULE: | POLYMERASE COFACTOR VP35 |
| 18:00 | 2gv5-C | 5.8 | 3.2 | 72 | 73 | 10 | MOLECULE: | CELL DIVISION CONTROL PROTEIN 31 |
| 19:00 | 6sct-D | 5.8 | 11.8 | 85 | 104 | 12 | MOLECULE: | CLATHRIN HEAVY CHAIN |
| 20:00 | 6mg2-B | 5.8 | 2.4 | 68 | 69 | 9 | MOLECULE: | CCAAT/ENHANCER-BINDING PROTEIN BETA |
| 21:00 | 1hjb-D | 5.8 | 2.6 | 68 | 68 | 3 | MOLECULE: | CCAAT/ENHANCER BINDING PROTEIN BETA |
| 22:00 | 4jzl-C | 5.8 | 3 | 62 | 62 | 8 | MOLECULE: | B-CELL RECEPTOR-ASSOCIATED PROTEIN 31 |
| 23:00 | 2e42-B | 5.8 | 2.1 | 65 | 67 | 5 | MOLECULE: | CCAAT/ENHANCER-BINDING PROTEIN BETA |
| 24:00:00 | 4wy4-D | 5.8 | 4.6 | 64 | 64 | 6 | MOLECULE: | VESICLE-ASSOCIATED MEMBRANE PROTEIN 8 |
| 25:00:00 | 5cci-D | 5.8 | 4.1 | 64 | 64 | 5 | MOLECULE: | VESICLE-ASSOCIATED MEMBRANE PROTEIN 2 |
| 26:00:00 | 6gbq-D | 5.8 | 4.7 | 68 | 68 | 4 | MOLECULE: | POLYMERASE COFACTOR VP35 |
| 27:00:00 | 6pph-l | 5.8 | 6.9 | 83 | 83 | 6 | MOLECULE: | CAPSID VERTEX COMPONENT 1 |
| 28:00:00 | 4bry-B | 5.8 | 5 | 69 | 69 | 13 | MOLECULE: | GEMININ |
| 29:00:00 | 2e43-B | 5.8 | 2.1 | 64 | 64 | 5 | MOLECULE: | CCAAT/ENHANCER-BINDING PROTEIN BETA |
| 30:00:00 | 6mti-H | 5.8 | 4.1 | 64 | 64 | 5 | MOLECULE: | SYNAPTOTAGMIN-1 |
| 31:00:00 | 6mti-L | 5.8 | 4.1 | 64 | 64 | 5 | MOLECULE: | SYNAPTOTAGMIN-1 |
| 32:00:00 | 6mti-P | 5.8 | 4.1 | 64 | 64 | 5 | MOLECULE: | SYNAPTOTAGMIN-1 |
| 33:00:00 | 6r2f-B | 5.8 | 4.6 | 64 | 64 | 6 | MOLECULE: | TESTIS-EXPRESSED PROTEIN 12 |
| 34:00:00 | 6h9l-B | 5.8 | 4.3 | 110 | 129 | 8 | MOLECULE: | UNCHARACTERIZED PROTEIN |
| 35:00:00 | 6mti-T | 5.8 | 4.1 | 64 | 64 | 5 | MOLECULE: | SYNAPTOTAGMIN-1 |
| 36:00:00 | 7fde-H | 5.8 | 8.5 | 112 | 112 | 5 | MOLECULE: | V-TYPE PROTON ATPASE SUBUNIT C |
| 37:00:00 | 6r2f-A | 5.8 | 4.6 | 64 | 64 | 6 | MOLECULE: | TESTIS-EXPRESSED PROTEIN 12 |
| 38:00:00 | 5vpd-D | 5.8 | 2.3 | 63 | 63 | 13 | MOLECULE: | PROTEIN FOSB |
| 39:00:00 | 7n84-Y | 5.8 | 4.4 | 63 | 63 | 0 | MOLECULE: | NUCLEOPORIN NUP188 |
| 40:00:00 | 6mti-X | 5.8 | 4.1 | 64 | 64 | 5 | MOLECULE: | SYNAPTOTAGMIN-1 |
| 41:00:00 | 4o9b-C | 5.8 | 3.6 | 92 | 92 | 12 | MOLECULE: | STROMAL INTERACTION MOLECULE 1 |
| 42:00:00 | 5cff-B | 5.8 | 4.3 | 88 | 88 | 3 | MOLECULE: | MIRANDA |
| 43:00:00 | 5cff-D | 5.8 | 5.3 | 88 | 88 | 3 | MOLECULE: | MIRANDA |
| 44:00:00 | 8b8b-D | 5.8 | 4.7 | 97 | 97 | 5 | MOLECULE: | MUNIA BORNAVIRUS 1 PHOSPHOPROTEIN |
| 45:00:00 | 8b8d-C | 5.8 | 4.3 | 101 | 101 | 9 | MOLECULE: | PHOSPHOPROTEIN |
| 46:00:00 | 5wsu-D | 5.8 | 4.9 | 91 | 91 | 8 | MOLECULE: | CALMODULIN |
| 47:00:00 | 7a5p-I | 5.8 | 3.1 | 70 | 133 | 10 | MOLECULE: | U2 SNRNA |
| 48:00:00 | 6ian-A | 5.7 | 36.4 | 121 | 306 | 9 | MOLECULE: | INTRAFLAGELLAR TRANSPORT PROTEIN 74 |
| 49:00:00 | 8r1a-A | 5.7 | 6.6 | 143 | 587 | 7 | MOLECULE: | GUANYLATE BINDING PROTEIN 1 |
| 50:00:00 | 8fed-E | 5.7 | 20.1 | 119 | 358 | 8 | MOLECULE: | VIRULENCE FACTOR MCE FAMILY PROTEIN |
| 51:00:00 | 7wkk-L | 5.7 | 4.8 | 91 | 156 | 4 | MOLECULE: | MGC83295 PROTEIN |
| 52:00:00 | 5cws-C | 5.7 | 4 | 91 | 169 | 5 | MOLECULE: | SAB-158 FAB LIGHT CHAIN |
| 53:00:00 | 5low-F | 5.7 | 4.5 | 67 | 68 | 4 | MOLECULE: | RABPHILIN-3A |
| 54:00:00 | 5cws-I | 5.7 | 3.6 | 91 | 169 | 11 | MOLECULE: | SAB-158 FAB LIGHT CHAIN |
| 55:00:00 | 7l4v-A | 5.7 | 3.9 | 64 | 66 | 8 | MOLECULE: | CCAAT/ENHANCER-BINDING PROTEIN BETA |
| 56:00:00 | 7d2e-A | 5.7 | 1.1 | 62 | 62 | 8 | MOLECULE: | LIPRIN-ALPHA-2 |
| 57:00:00 | 6mti-U | 5.7 | 4.2 | 62 | 62 | 6 | MOLECULE: | SYNAPTOTAGMIN-1 |
| 58:00:00 | 7tmm-H | 5.7 | 11.1 | 107 | 109 | 6 | MOLECULE: | H(+)-TRANSPORTING TWO-SECTOR ATPASE |
| 59:00:00 | 6vqi-J | 5.7 | 3.9 | 64 | 64 | 11 | MOLECULE: | V-TYPE PROTON ATPASE SUBUNIT C 1 |
| 60:00:00 | 5wsu-C | 5.7 | 3 | 90 | 90 | 8 | MOLECULE: | CALMODULIN |
| 61:00:00 | 5gmk-c | 5.7 | 31.5 | 133 | 436 | 2 | MOLECULE: | PRE-MRNA-SPLICING FACTOR 8 |

| | | | | | | | | |
|---|---|---|---|---|---|---|---|---|
| 62:00:00 | 8ovw-K | 5.7 | 16.4 | 82 | 219 | 10 | MOLECULE: | CENTROMERE-BINDING PROTEIN 1 |
| 63:00:00 | 1gd2-F | 5.7 | 4.5 | 64 | 64 | 5 | MOLECULE: | DNA (5'- |
| 64:00:00 | 3b5n-J | 5.7 | 3.1 | 64 | 64 | 5 | MOLECULE: | SYNAPTOBREVIN HOMOLOG 1 |
| 65:00:00 | 2n9b-B | 5.7 | 4 | 68 | 69 | 6 | MOLECULE: | UNCONVENTIONAL MYOSIN-X, GENERAL CONTROL PRO |
| 66:00:00 | 1hvv-B | 5.7 | 2.6 | 62 | 62 | 3 | MOLECULE: | SYNTAXIN 1A |
| 67:00:00 | 9b8q-M | 5.7 | 6.2 | 102 | 102 | 4 | MOLECULE: | V-TYPE PROTON ATPASE SUBUNIT C 1 |
| 68:00:00 | 5vox-L | 5.7 | 7 | 103 | 104 | 7 | MOLECULE: | V-TYPE PROTON ATPASE CATALYTIC SUBUNIT A,V-T |
| 69:00:00 | 1n7s-A | 5.7 | 4.6 | 63 | 63 | 6 | MOLECULE: | VESICLE-ASSOCIATED MEMBRANE PROTEIN 2 |
| 70:00:00 | 4om2-A | 5.7 | 3.9 | 80 | 112 | 8 | MOLECULE: | TRANSDUCIN-LIKE ENHANCER PROTEIN 1 |
| 71:00:00 | 5vox-G | 5.7 | 7.1 | 117 | 217 | 6 | MOLECULE: | V-TYPE PROTON ATPASE CATALYTIC SUBUNIT A,V-T |
| 72:00:00 | 5wsg-c | 5.7 | 31.5 | 133 | 436 | 2 | MOLECULE: | PRE-MRNA-SPLICING FACTOR 8 |
| 73:00:00 | 6wvw-E | 5.7 | 4.1 | 63 | 63 | 10 | MOLECULE: | VESICLE-ASSOCIATED MEMBRANE PROTEIN 2 |
| 74:00:00 | 2n1t-A | 5.7 | 2.4 | 69 | 69 | 6 | MOLECULE: | VESICLE-ASSOCIATED MEMBRANE PROTEIN 2 |
| 75:00:00 | 3lj5-A | 5.7 | 9.7 | 129 | 692 | 5 | MOLECULE: | PORTAL PROTEIN |
| 76:00:00 | 6yai-D | 5.7 | 6.8 | 76 | 104 | 14 | MOLECULE: | CLATHRIN HEAVY CHAIN |
| 77:00:00 | 3lj5-K | 5.7 | 8 | 131 | 692 | 8 | MOLECULE: | PORTAL PROTEIN |
| 78:00:00 | 5gai-E | 5.7 | 11.7 | 126 | 721 | 9 | MOLECULE: | PORTAL PROTEIN |
| 79:00:00 | 5tsj-H | 5.7 | 11 | 101 | 182 | 8 | MOLECULE: | V-TYPE ATP SYNTHASE ALPHA CHAIN |
| 80:00:00 | 6sbz-D | 5.7 | 10.8 | 82 | 104 | 12 | MOLECULE: | CLATHRIN HEAVY CHAIN |
| 81:00:00 | 5gai-F | 5.7 | 11.6 | 130 | 721 | 7 | MOLECULE: | PORTAL PROTEIN |
| 82:00:00 | 5d80-l | 5.7 | 5.4 | 68 | 68 | 6 | MOLECULE: | V-TYPE PROTON ATPASE CATALYTIC SUBUNIT A |
| 83:00:00 | 6tt7-K | 5.7 | 5.6 | 140 | 208 | 5 | MOLECULE: | ATP SYNTHASE SUBUNIT ALPHA |
| 84:00:00 | 3lj5-J | 5.7 | 7.8 | 130 | 692 | 8 | MOLECULE: | PORTAL PROTEIN |
| 85:00:00 | 5gmk-o | 5.7 | 8.6 | 79 | 126 | 9 | MOLECULE: | PRE-MRNA-SPLICING FACTOR 8 |
| 86:00:00 | 1gtw-A | 5.7 | 2.5 | 65 | 65 | 3 | MOLECULE: | CAAT/ENHANCER BINDING PROTEIN BETA |
| 87:00:00 | 6sbz-N | 5.7 | 10.7 | 83 | 104 | 12 | MOLECULE: | CLATHRIN HEAVY CHAIN |
| 88:00:00 | 2gv5-F | 5.7 | 4.1 | 73 | 73 | 7 | MOLECULE: | CELL DIVISION CONTROL PROTEIN 31 |
| 89:00:00 | 6sbz-I | 5.7 | 10.8 | 82 | 104 | 12 | MOLECULE: | CLATHRIN HEAVY CHAIN |
| 90:00:00 | 1h8a-B | 5.7 | 3.8 | 65 | 67 | 18 | MOLECULE: | CAAT/ENHANCER BINDING PROTEIN BETA |
| 91:00:00 | 2e42-A | 5.7 | 2.6 | 65 | 65 | 3 | MOLECULE: | CCAAT/ENHANCER-BINDING PROTEIN BETA |
| 92:00:00 | 1hjb-E | 5.7 | 3.6 | 67 | 68 | 7 | MOLECULE: | CCAAT/ENHANCER BINDING PROTEIN BETA |
| 93:00:00 | 5kj7-G | 5.7 | 4.6 | 63 | 63 | 10 | MOLECULE: | VESICLE-ASSOCIATED MEMBRANE PROTEIN 3 |
| 94:00:00 | 6m3w-A | 5.7 | 35.5 | 135 | 344 | 2 | MOLECULE: | SPIKE GLYCOPROTEIN |
| 95:00:00 | 1h88-A | 5.7 | 3.3 | 68 | 70 | 7 | MOLECULE: | CCAAT/ENHANCER BINDING PROTEIN BETA |
| 96:00:00 | 1hjb-B | 5.7 | 2.2 | 67 | 67 | 3 | MOLECULE: | CCAAT/ENHANCER BINDING PROTEIN BETA |
| 97:00:00 | 7znl-M | 5.7 | 6.4 | 122 | 549 | 6 | MOLECULE: | THO COMPLEX SUBUNIT 1 |
| 98:00:00 | 6wvw-H | 5.7 | 4.8 | 62 | 62 | 6 | MOLECULE: | VESICLE-ASSOCIATED MEMBRANE PROTEIN 2 |
| 99:00:00 | 5ccg-D | 5.7 | 4.6 | 64 | 64 | 5 | MOLECULE: | VESICLE-ASSOCIATED MEMBRANE PROTEIN 2 |
| 0:00 | 5low-K | 5.7 | 4.8 | 72 | 72 | 1 | MOLECULE: | RABPHILIN-3A |
| 1:00 | 6kn7-T | 5.7 | 15.7 | 80 | 138 | 9 | MOLECULE: | ACTIN, ALPHA SKELETAL MUSCLE |
| 2:00 | 5mw9-F | 5.7 | 4.3 | 64 | 64 | 3 | MOLECULE: | CENTROSOMIN |
| 3:00 | 5cff-A | 5.7 | 4.9 | 87 | 87 | 3 | MOLECULE: | MIRANDA |
| 4:00 | 5cff-C | 5.7 | 4.9 | 87 | 87 | 3 | MOLECULE: | MIRANDA |
| 5:00 | 5dol-A | 5.7 | 2.8 | 61 | 61 | 7 | MOLECULE: | INITIATION-CONTROL PROTEIN YABA |
| 6:00 | 6v86-D | 5.7 | 2.1 | 71 | 72 | 14 | MOLECULE: | RNA-DIRECTED RNA POLYMERASE L |
| 7:00 | 5d60-D | 5.7 | 3.2 | 61 | 61 | 7 | MOLECULE: | PUTATIVE TRANSCRIPTION FACTOR |
| 8:00 | 4lin-L | 5.7 | 7.1 | 139 | 289 | 12 | MOLECULE: | TAIL NEEDLE PROTEIN GP26 |
| 9:00 | 8ovw-U | 5.6 | 13.8 | 132 | 184 | 8 | MOLECULE: | CENTROMERE-BINDING PROTEIN 1 |
| 10:00 | 7woo-I | 5.6 | 35 | 131 | 187 | 10 | MOLECULE: | NUCLEOPORIN NIC96 |
| 11:00 | 3oja-A | 5.6 | 8.7 | 118 | 482 | 6 | MOLECULE: | LEUCINE-RICH IMMUNE MOLECULE 1 |
| 12:00 | 6fia-D | 5.6 | 8.9 | 104 | 104 | 7 | MOLECULE: | LINE-1 RETROTRANSPOSABLE ELEMENT ORF1 PROTEI |

| | | | | | | | | |
|---|---|---|---|---|---|---|---|---|
| 13:00 | 6wuc-K | 5.6 | 20.8 | 121 | 235 | 5 | MOLECULE: | INNER KINETOCHORE SUBUNIT MCM16 |
| 14:00 | 8tek-P | 5.6 | 4.9 | 120 | 138 | 6 | MOLECULE: | DYNEIN REGULATORY COMPLEX PROTEIN 1/2 N-TERM |
| 15:00 | 3opc-B | 5.6 | 2.1 | 72 | 131 | 6 | MOLECULE: | UNCHARACTERIZED PROTEIN |
| 16:00 | 7znk-M | 5.6 | 9.9 | 128 | 549 | 9 | MOLECULE: | RNA |
| 17:00 | 5ajs-B | 5.6 | 4.6 | 64 | 64 | 11 | MOLECULE: | THAP DOMAIN-CONTAINING PROTEIN 11 |
| 18:00 | 7bv6-U | 5.6 | 4.8 | 66 | 66 | 9 | MOLECULE: | VESICLE-ASSOCIATED MEMBRANE PROTEIN 8 |
| 19:00 | 1qbz-C | 5.6 | 4.2 | 63 | 120 | 2 | MOLECULE: | PROTEIN (SIV GP41 ECTODOMAIN) |
| 20:00 | 1wt6-D | 5.6 | 3.4 | 63 | 63 | 6 | MOLECULE: | MYOTONIN-PROTEIN KINASE |
| 21:00 | 1x79-C | 5.6 | 4.2 | 88 | 88 | 6 | MOLECULE: | ADP-RIBOSYLATION FACTOR BINDING PROTEIN GGA1 |
| 22:00 | 6vqj-J | 5.6 | 4.1 | 63 | 64 | 6 | MOLECULE: | V-TYPE PROTON ATPASE SUBUNIT C 1 |
| 23:00 | 7bv6-M | 5.6 | 4.6 | 66 | 66 | 9 | MOLECULE: | VESICLE-ASSOCIATED MEMBRANE PROTEIN 8 |
| 24:00:00 | 1h89-A | 5.6 | 2.3 | 64 | 64 | 5 | MOLECULE: | CAAT/ENHANCER BINDING PROTEIN BETA |
| 25:00:00 | 1gmj-C | 5.6 | 2.7 | 59 | 59 | 8 | MOLECULE: | ATPASE INHIBITOR |
| 26:00:00 | 3jb9-T | 5.6 | 10.8 | 73 | 134 | 11 | MOLECULE: | PRE-MRNA-SPLICING FACTOR SPP42 |
| 27:00:00 | 3b5n-L | 5.6 | 3 | 62 | 62 | 8 | MOLECULE: | SYNAPTOBREVIN HOMOLOG 1 |
| 28:00:00 | 3gn4-E | 5.6 | 4 | 83 | 117 | 2 | MOLECULE: | MYOSIN-VI |
| 29:00:00 | 6pse-B | 5.6 | 2.2 | 67 | 68 | 6 | MOLECULE: | PROTEIN BICAUDAL D HOMOLOG 2 |
| 30:00:00 | 4m3l-A | 5.6 | 2.9 | 60 | 60 | 5 | MOLECULE: | E3 UBIQUITIN-PROTEIN LIGASE TRIM63 |
| 31:00:00 | 9b8q-I | 5.6 | 7.2 | 117 | 148 | 7 | MOLECULE: | V-TYPE PROTON ATPASE SUBUNIT C 1 |
| 32:00:00 | 6o7x-L | 5.6 | 7.4 | 105 | 105 | 7 | MOLECULE: | V-TYPE PROTON ATPASE SUBUNIT C |
| 33:00:00 | 8t9d-B | 5.6 | 9.1 | 110 | 158 | 6 | MOLECULE: | MEDIATOR OF RNA POLYMERASE II TRANSCRIPTION |
| 34:00:00 | 5gai-D | 5.6 | 6.7 | 114 | 721 | 4 | MOLECULE: | PORTAL PROTEIN |
| 35:00:00 | 6qle-U | 5.6 | 14.2 | 124 | 172 | 9 | MOLECULE: | CENTRAL KINETOCHORE SUBUNIT MCM16,CENTRAL KI |
| 36:00:00 | 5gai-H | 5.6 | 10.8 | 126 | 721 | 4 | MOLECULE: | PORTAL PROTEIN |
| 37:00:00 | 3lj5-E | 5.6 | 8 | 131 | 692 | 8 | MOLECULE: | PORTAL PROTEIN |
| 38:00:00 | 7fdc-H | 5.6 | 7.9 | 109 | 111 | 7 | MOLECULE: | YEAST VACUOLAR ATPASE A SUBUNIT |
| 39:00:00 | 2e43-A | 5.6 | 2 | 62 | 62 | 3 | MOLECULE: | CCAAT/ENHANCER-BINDING PROTEIN BETA |
| 40:00:00 | 7saf-A | 5.6 | 3.3 | 62 | 62 | 6 | MOLECULE: | GENERAL CONTROL TRANSCRIPTION FACTOR GCN4/M |
| 41:00:00 | 6wvw-A | 5.6 | 4.6 | 62 | 62 | 10 | MOLECULE: | VESICLE-ASSOCIATED MEMBRANE PROTEIN 2 |
| 42:00:00 | 7bv6-X | 5.6 | 4.6 | 65 | 65 | 6 | MOLECULE: | VESICLE-ASSOCIATED MEMBRANE PROTEIN 8 |
| 43:00:00 | 6iak-A | 5.6 | 3.5 | 68 | 68 | 9 | MOLECULE: | UNCHARACTERIZED PROTEIN |
| 44:00:00 | 1h89-B | 5.6 | 2.9 | 64 | 64 | 3 | MOLECULE: | CAAT/ENHANCER BINDING PROTEIN BETA |
| 45:00:00 | 7eao-C | 5.6 | 4.3 | 62 | 62 | 5 | MOLECULE: | POLYUBIQUITIN-C |
| 46:00:00 | 5mqf-L | 5.6 | 36.3 | 116 | 336 | 8 | MOLECULE: | PRE-MRNA-PROCESSING-SPLICING FACTOR 8 |
| 47:00:00 | 5dn6-W | 5.6 | 6.1 | 109 | 124 | 0 | MOLECULE: | CHAIN A |
| 48:00:00 | 6fki-b | 5.6 | 10.3 | 125 | 161 | 6 | MOLECULE: | ATP SYNTHASE SUBUNIT A, CHLOROPLASTIC |
| 49:00:00 | 3oja-B | 5.6 | 4.9 | 110 | 534 | 12 | MOLECULE: | LEUCINE-RICH IMMUNE MOLECULE 1 |
| 50:00:00 | 5d80-n | 5.6 | 7.8 | 105 | 105 | 2 | MOLECULE: | V-TYPE PROTON ATPASE CATALYTIC SUBUNIT A |
| 51:00:00 | 5w5c-C | 5.6 | 4.7 | 65 | 65 | 2 | MOLECULE: | VESICLE-ASSOCIATED MEMBRANE PROTEIN 2 |
| 52:00:00 | 6gbr-D | 5.6 | 2.2 | 67 | 67 | 4 | MOLECULE: | POLYMERASE COFACTOR VP35 |
| 53:00:00 | 8dbs-X | 5.6 | 8.5 | 137 | 156 | 6 | MOLECULE: | ATP SYNTHASE SUBUNIT ALPHA |
| 54:00:00 | 7qoo-H | 5.5 | 18.9 | 121 | 210 | 7 | MOLECULE: | CENTROMERE PROTEIN C |
| 55:00:00 | 2xnx-M | 5.5 | 4.3 | 106 | 107 | 8 | MOLECULE: | FIBRINOGEN ALPHA CHAIN |
| 56:00:00 | 8ap7-L | 5.5 | 3.6 | 63 | 65 | 5 | MOLECULE: | ATP SYNTHASE SUBUNIT A |
| 57:00:00 | 7bjs-B | 5.5 | 4.4 | 62 | 62 | 5 | MOLECULE: | KINESIN HEAVY CHAIN |
| 58:00:00 | 6zkl-q | 5.5 | 11 | 89 | 139 | 1 | MOLECULE: | NADH DEHYDROGENASE [UBIQUINONE] FLAVOPROTEIN |
| 59:00:00 | 1gd2-H | 5.5 | 2.7 | 63 | 64 | 5 | MOLECULE: | DNA (5'- |
| 60:00:00 | 6fia-F | 5.5 | 4.1 | 88 | 88 | 9 | MOLECULE: | LINE-1 RETROTRANSPOSABLE ELEMENT ORF1 PROTEI |
| 61:00:00 | 5gox-A | 5.5 | 6.9 | 95 | 181 | 6 | MOLECULE: | DNA REPAIR PROTEIN RAD50 |
| 62:00:00 | 1hvv-D | 5.5 | 3.4 | 61 | 61 | 3 | MOLECULE: | SYNTAXIN 1A |
| 63:00:00 | 7fda-H | 5.5 | 7.1 | 111 | 111 | 8 | MOLECULE: | YEAST VACUOLAR ATPASE A SUBUNIT |

| | | | | | | | | |
|---|---|---|---|---|---|---|---|---|
| 64:00:00 | 1urq-D | 5.5 | 4 | 62 | 62 | 6 | MOLECULE: | M-TOMOSYN ISOFORM |
| 65:00:00 | 6id1-t | 5.5 | 3.2 | 67 | 67 | 12 | MOLECULE: | PRE-MRNA-PROCESSING-SPLICING FACTOR 8 |
| 66:00:00 | 5jvp-B | 5.5 | 5.7 | 68 | 90 | 6 | MOLECULE: | CHIMERA PROTEIN OF CENTROMERE-ASSOCIATED PRO |
| 67:00:00 | 6ip1-D | 5.5 | 4.6 | 61 | 62 | 3 | MOLECULE: | VESICLE-ASSOCIATED MEMBRANE PROTEIN 2 |
| 68:00:00 | 1sfc-L | 5.5 | 2.7 | 68 | 73 | 4 | MOLECULE: | PROTEIN (SYNAPTOBREVIN 2) |
| 69:00:00 | 8t1l-B | 5.5 | 8.8 | 105 | 158 | 8 | MOLECULE: | MEDIATOR OF RNA POLYMERASE II TRANSCRIPTION |
| 70:00:00 | 8uwy-G | 5.5 | 2.2 | 81 | 82 | 10 | MOLECULE: | ACTIN, ALPHA CARDIAC MUSCLE 1 |
| 71:00:00 | 6pfp-B | 5.5 | 5.5 | 124 | 156 | 7 | MOLECULE: | MYOSIN-7 FUSED TO GP7 AND EB1 |
| 72:00:00 | 1h8a-A | 5.5 | 2.4 | 66 | 68 | 9 | MOLECULE: | CAAT/ENHANCER BINDING PROTEIN BETA |
| 73:00:00 | 1io4-B | 5.5 | 3.9 | 68 | 70 | 6 | MOLECULE: | CSF-1R PROMOTER |
| 74:00:00 | 8aia-H | 5.5 | 7.6 | 138 | 179 | 6 | MOLECULE: | CRESCENTIN |
| 75:00:00 | 5ccg-J | 5.5 | 4.1 | 63 | 63 | 5 | MOLECULE: | VESICLE-ASSOCIATED MEMBRANE PROTEIN 2 |
| 76:00:00 | 2n1t-D | 5.5 | 2 | 67 | 74 | 12 | MOLECULE: | VESICLE-ASSOCIATED MEMBRANE PROTEIN 2 |
| 77:00:00 | 5ylz-t | 5.5 | 5.4 | 69 | 128 | 3 | MOLECULE: | PRE-MRNA-SPLICING FACTOR 8 |
| 78:00:00 | 2zxx-A | 5.5 | 6.7 | 69 | 70 | 7 | MOLECULE: | GEMININ |
| 79:00:00 | 6id1-q | 5.5 | 11.6 | 90 | 132 | 10 | MOLECULE: | PRE-MRNA-PROCESSING-SPLICING FACTOR 8 |
| 80:00:00 | 6vqj-I | 5.5 | 2.4 | 63 | 64 | 8 | MOLECULE: | V-TYPE PROTON ATPASE SUBUNIT C 1 |
| 81:00:00 | 4bl6-C | 5.5 | 5 | 86 | 86 | 7 | MOLECULE: | PROTEIN BICAUDAL D |
| 82:00:00 | 6icz-L | 5.5 | 39.3 | 124 | 454 | 8 | MOLECULE: | PROTEIN MAGO NASHI HOMOLOG 2 |
| 83:00:00 | 7w59-L | 5.5 | 37.9 | 124 | 437 | 7 | MOLECULE: | PRE-MRNA-PROCESSING-SPLICING FACTOR 8 |
| 84:00:00 | 5whf-E | 5.5 | 5.3 | 85 | 85 | 5 | MOLECULE: | VIMENTIN |
| 85:00:00 | 4ll7-G | 5.5 | 4.6 | 87 | 87 | 7 | MOLECULE: | SWI5-DEPENDENT HO EXPRESSION PROTEIN 3 |
| 86:00:00 | 7njx-d | 5.5 | 3 | 59 | 61 | 5 | MOLECULE: | ATP SYNTHASE SUBUNIT C |
| 87:00:00 | 6mg3-A | 5.5 | 3.9 | 64 | 64 | 6 | MOLECULE: | CCAAT/ENHANCER-BINDING PROTEIN BETA |
| 88:00:00 | 6mti-M | 5.5 | 4.5 | 62 | 62 | 10 | MOLECULE: | SYNAPTOTAGMIN-1 |
| 89:00:00 | 8qrm-U | 5.5 | 16.4 | 137 | 176 | 5 | MOLECULE: | 12S MITOCHONDRIAL RRNA |
| 90:00:00 | 5w5d-C | 5.5 | 4.6 | 61 | 61 | 2 | MOLECULE: | VESICLE-ASSOCIATED MEMBRANE PROTEIN 2 |
| 91:00:00 | 1uii-A | 5.5 | 4.7 | 61 | 61 | 3 | MOLECULE: | GEMININ |
| 92:00:00 | 5y88-r | 5.5 | 5.4 | 69 | 129 | 3 | MOLECULE: | PRE-MRNA-SPLICING FACTOR 8 |
| 93:00:00 | 7usd-C | 5.5 | 6 | 69 | 191 | 6 | MOLECULE: | CYTOPLASMIC FMR1-INTERACTING PROTEIN 1 |
| 94:00:00 | 6o7v-J | 5.5 | 5.2 | 105 | 105 | 5 | MOLECULE: | V-TYPE PROTON ATPASE SUBUNIT D |
| 95:00:00 | 6cfz-A | 5.5 | 4.8 | 66 | 66 | 6 | MOLECULE: | ASK1 |
| 96:00:00 | 6fkh-p | 5.5 | 7.5 | 116 | 143 | 10 | MOLECULE: | ATP SYNTHASE SUBUNIT A, CHLOROPLASTIC |
| 97:00:00 | 7p3w-b | 5.5 | 7.3 | 117 | 150 | 7 | MOLECULE: | ATP SYNTHASE SUBUNIT ALPHA |
| 98:00:00 | 4wpx-B | 5.4 | 8.1 | 78 | 81 | 10 | MOLECULE: | CELL DIVISION CONTROL PROTEIN 31-LIKE PROTEI |
| 99:00:00 | 6exn-D | 5.4 | 7.7 | 96 | 97 | 4 | MOLECULE: | U2 SNRNA |
| 0:00 | 6iac-E | 5.4 | 23.5 | 128 | 146 | 5 | MOLECULE: | PORTAL PROTEIN |
| 1:00 | 8th8-A | 5.4 | 3.3 | 134 | 290 | 6 | MOLECULE: | DYNEIN REGULATORY COMPLEX PROTEIN 1/2 N-TERM |
| 2:00 | 6qaj-B | 5.4 | 10 | 123 | 441 | 6 | MOLECULE: | ENDOLYSIN,TRANSCRIPTION INTERMEDIARY FACTOR |
| 3:00 | 7d2e-B | 5.4 | 3.1 | 58 | 58 | 7 | MOLECULE: | LIPRIN-ALPHA-2 |
| 4:00 | 6iak-D | 5.4 | 3.5 | 62 | 62 | 10 | MOLECULE: | UNCHARACTERIZED PROTEIN |
| 5:00 | 4n3y-C | 5.4 | 3.8 | 83 | 83 | 12 | MOLECULE: | RAB5 GDP/GTP EXCHANGE FACTOR |
| 6:00 | 6fia-E | 5.4 | 4.5 | 88 | 88 | 9 | MOLECULE: | LINE-1 RETROTRANSPOSABLE ELEMENT ORF1 PROTEI |
| 7:00 | 7pkq-w | 5.4 | 12.1 | 137 | 155 | 9 | MOLECULE: | MS35 |
| 8:00 | 1sfc-H | 5.4 | 2.9 | 67 | 72 | 4 | MOLECULE: | PROTEIN (SYNAPTOBREVIN 2) |
| 9:00 | 1io4-A | 5.4 | 3.7 | 63 | 63 | 8 | MOLECULE: | CSF-1R PROMOTER |
| 10:00 | 5afu-5 | 5.4 | 4.9 | 139 | 275 | 0 | MOLECULE: | DYNEIN TAIL |
| 11:00 | 5voy-L | 5.4 | 6.6 | 104 | 104 | 4 | MOLECULE: | V-TYPE PROTON ATPASE CATALYTIC SUBUNIT A,V-T |
| 12:00 | 5gmk-p | 5.4 | 5.2 | 69 | 128 | 7 | MOLECULE: | PRE-MRNA-SPLICING FACTOR 8 |
| 13:00 | 8x7z-A | 5.4 | 1.5 | 83 | 83 | 8 | MOLECULE: | HR1 |
| 14:00 | 8pr4-x | 5.4 | 2.6 | 58 | 92 | 2 | MOLECULE: | ARP11 |

| | | | | | | | | |
|---|---|---|---|---|---|---|---|---|
| 15:00 | 4m3l-B | 5.4 | 1.8 | 58 | 58 | 5 | MOLECULE: | E3 UBIQUITIN-PROTEIN LIGASE TRIM63 |
| 16:00 | 6id0-t | 5.4 | 4.4 | 67 | 67 | 3 | MOLECULE: | PRE-MRNA-PROCESSING-SPLICING FACTOR 8 |
| 17:00 | 7fdc-J | 5.4 | 7.2 | 109 | 111 | 5 | MOLECULE: | YEAST VACUOLAR ATPASE A SUBUNIT |
| 18:00 | 5ajs-C | 5.4 | 4.5 | 61 | 61 | 11 | MOLECULE: | THAP DOMAIN-CONTAINING PROTEIN 11 |
| 19:00 | 6za9-T | 5.4 | 6.8 | 69 | 71 | 7 | MOLECULE: | ATP SYNTHASE F(0) COMPLEX SUBUNIT C1, MITOCH |
| 20:00 | 5c9n-B | 5.4 | 3.4 | 61 | 61 | 0 | MOLECULE: | GEMININ COILED-COIL DOMAIN-CONTAINING PROTEI |
| 21:00 | 6j6g-p | 5.4 | 5.4 | 69 | 128 | 3 | MOLECULE: | PRE-MRNA-SPLICING FACTOR 8 |
| 22:00 | 5i7c-A | 5.4 | 2.7 | 58 | 59 | 2 | MOLECULE: | CENTROSOMIN |
| 23:00 | 8aix-H | 5.4 | 7.6 | 138 | 179 | 6 | MOLECULE: | CRESCENTIN |
| 24:00:00 | 7nju-d | 5.4 | 4.5 | 61 | 61 | 3 | MOLECULE: | ATP SYNTHASE SUBUNIT C |
| 25:00:00 | 6h9l-A | 5.4 | 5.9 | 100 | 127 | 5 | MOLECULE: | UNCHARACTERIZED PROTEIN |
| 26:00:00 | 6oqa-G | 5.4 | 2.7 | 87 | 87 | 8 | MOLECULE: | PEPTIDYL-PROLYL CIS-TRANS ISOMERASE FKBP1A |
| 27:00:00 | 5bvz-C | 5.4 | 5.8 | 101 | 180 | 4 | MOLECULE: | DNA STABILIZATION PROTEIN |
| 28:00:00 | 5i7c-C | 5.4 | 2.8 | 58 | 59 | 2 | MOLECULE: | CENTROSOMIN |
| 29:00:00 | 6r0z-L | 5.4 | 5.3 | 112 | 185 | 10 | MOLECULE: | V-TYPE ATP SYNTHASE ALPHA CHAIN |
| 30:00:00 | 1m1j-B | 5.4 | 8.2 | 128 | 402 | 10 | MOLECULE: | FIBRINOGEN ALPHA SUBUNIT |
| 31:00:00 | 5mq0-u | 5.4 | 4.7 | 64 | 116 | 3 | MOLECULE: | YEAST UBC4 GENE FOR UBIQUITIN-CONJUGATING EN |
| 32:00:00 | 6gap-C | 5.4 | 9.5 | 144 | 213 | 8 | MOLECULE: | OUTER CAPSID PROTEIN SIGMA-1 |
| 33:00:00 | 2w4u-U | 5.4 | 9.4 | 153 | 277 | 5 | MOLECULE: | TROPONIN C, SKELETAL MUSCLE |
| 34:00:00 | 6id1-L | 5.4 | 39.6 | 124 | 475 | 8 | MOLECULE: | PRE-MRNA-PROCESSING-SPLICING FACTOR 8 |
| 35:00:00 | 5fil-T | 5.4 | 7.4 | 128 | 174 | 3 | MOLECULE: | ATP SYNTHASE SUBUNIT ALPHA, MITOCHONDRIAL |
| 36:00:00 | 5low-M | 5.4 | 3.5 | 70 | 72 | 3 | MOLECULE: | RABPHILIN-3A |
| 37:00:00 | 7njs-b | 5.4 | 6.6 | 105 | 144 | 3 | MOLECULE: | ATP SYNTHASE SUBUNIT ALPHA |
| 38:00:00 | 8fl8-Z | 5.4 | 6.7 | 125 | 155 | 4 | MOLECULE: | ATP SYNTHASE PROTEIN 8 |
| 39:00:00 | 8cqn-B | 5.3 | 5.7 | 123 | 187 | 7 | MOLECULE: | LIPOPROTEIN, PUTATIVE |
| 40:00:00 | 8fed-F | 5.3 | 21.8 | 149 | 399 | 5 | MOLECULE: | VIRULENCE FACTOR MCE FAMILY PROTEIN |
| 41:00:00 | 7nl9-d | 5.3 | 3.9 | 60 | 61 | 8 | MOLECULE: | ATP SYNTHASE SUBUNIT ALPHA |
| 42:00:00 | 1gl2-C | 5.3 | 4.1 | 59 | 59 | 7 | MOLECULE: | ENDOBREVIN |
| 43:00:00 | 6iak-B | 5.3 | 4.3 | 62 | 62 | 6 | MOLECULE: | UNCHARACTERIZED PROTEIN |
| 44:00:00 | 6wcj-O | 5.3 | 3.4 | 59 | 59 | 2 | MOLECULE: | CLATHRIN HEAVY CHAIN 1 |
| 45:00:00 | 7fde-G | 5.3 | 6.4 | 111 | 231 | 8 | MOLECULE: | V-TYPE PROTON ATPASE SUBUNIT C |
| 46:00:00 | 7njy-d | 5.3 | 4 | 60 | 61 | 8 | MOLECULE: | ATP SYNTHASE SUBUNIT C |
| 47:00:00 | 5gas-I | 5.3 | 6.4 | 98 | 99 | 5 | MOLECULE: | V-TYPE ATP SYNTHASE ALPHA CHAIN |
| 48:00:00 | 6wcj-N | 5.3 | 3.1 | 59 | 59 | 5 | MOLECULE: | CLATHRIN HEAVY CHAIN 1 |
| 49:00:00 | 3fwc-B | 5.3 | 3.4 | 83 | 83 | 10 | MOLECULE: | CELL DIVISION CONTROL PROTEIN 31 |
| 50:00:00 | 4n3y-B | 5.3 | 3.2 | 83 | 83 | 0 | MOLECULE: | RAB5 GDP/GTP EXCHANGE FACTOR |
| 51:00:00 | 6id0-L | 5.3 | 43.5 | 143 | 475 | 6 | MOLECULE: | PRE-MRNA-PROCESSING-SPLICING FACTOR 8 |
| 52:00:00 | 3b5n-E | 5.3 | 3.4 | 58 | 58 | 9 | MOLECULE: | SYNAPTOBREVIN HOMOLOG 1 |
| 53:00:00 | 7bv6-Q | 5.3 | 4.3 | 61 | 61 | 3 | MOLECULE: | VESICLE-ASSOCIATED MEMBRANE PROTEIN 8 |
| 54:00:00 | 1hvv-C | 5.3 | 4.1 | 66 | 66 | 14 | MOLECULE: | SYNTAXIN 1A |
| 55:00:00 | 5adx-Z | 5.3 | 29 | 129 | 419 | 0 | MOLECULE: | ACTIN RELATED PROTEIN 1 |
| 56:00:00 | 6v85-C | 5.3 | 3.1 | 74 | 74 | 7 | MOLECULE: | RNA-DIRECTED RNA POLYMERASE L |
| 57:00:00 | 5gai-C | 5.3 | 9.1 | 125 | 721 | 3 | MOLECULE: | PORTAL PROTEIN |
| 58:00:00 | 8ppr-P | 5.3 | 4.6 | 86 | 176 | 10 | MOLECULE: | KINETOCHORE-ASSOCIATED PROTEIN DSN1 HOMOLOG |
| 59:00:00 | 6vqi-a | 5.3 | 13.4 | 83 | 309 | 11 | MOLECULE: | V-TYPE PROTON ATPASE SUBUNIT C 1 |
| 60:00:00 | 5apv-B | 5.3 | 4 | 67 | 67 | 6 | MOLECULE: | GENERAL CONTROL PROTEIN GCN4 |
| 61:00:00 | 4m3l-C | 5.3 | 1.6 | 57 | 57 | 5 | MOLECULE: | E3 UBIQUITIN-PROTEIN LIGASE TRIM63 |
| 62:00:00 | 5f5p-E | 5.3 | 3.8 | 57 | 57 | 11 | MOLECULE: | PROTEIN SHROOM2 |
| 63:00:00 | 6gbq-H | 5.3 | 4.5 | 63 | 63 | 3 | MOLECULE: | POLYMERASE COFACTOR VP35 |
| 64:00:00 | 4wij-B | 5.3 | 3.6 | 118 | 301 | 7 | MOLECULE: | SPLICING FACTOR, PROLINE- AND GLUTAMINE-RICH |
| 65:00:00 | 6r0y-J | 5.3 | 6.4 | 115 | 185 | 8 | MOLECULE: | V-TYPE ATP SYNTHASE ALPHA CHAIN |

| | | | | | | | | |
|---|---|---|---|---|---|---|---|---|
| 66:00:00 | 5oi7-A | 5.3 | 3.5 | 86 | 88 | 10 | MOLECULE: | CENTROSOMAL PROTEIN OF 85 KDA |
| 67:00:00 | 3wuv-A | 5.3 | 2.7 | 54 | 56 | 17 | MOLECULE: | CENTROSOMAL PROTEIN OF 55 KDA |
| 68:00:00 | 4jpp-C | 5.3 | 6.3 | 101 | 122 | 5 | MOLECULE: | MINOR SPIKE PROTEIN H |
| 69:00:00 | 5jlh-I | 5.3 | 9.2 | 120 | 135 | 0 | MOLECULE: | ACTIN, CYTOPLASMIC 2 |
| 70:00:00 | 5jvp-C | 5.3 | 4 | 68 | 90 | 4 | MOLECULE: | CHIMERA PROTEIN OF CENTROMERE-ASSOCIATED PRO |
| 71:00:00 | 5t4q-I | 5.3 | 13.5 | 117 | 155 | 7 | MOLECULE: | ATP SYNTHASE SUBUNIT ALPHA |
| 72:00:00 | 8txr-A | 5.3 | 8.9 | 142 | 427 | 8 | MOLECULE: | EXODEOXYRIBONUCLEASE 7 LARGE SUBUNIT |
| 73:00:00 | 4n5b-A | 5.2 | 5.2 | 81 | 104 | 5 | MOLECULE: | PHOSPHOPROTEIN |
| 74:00:00 | 2fo1-D | 5.2 | 5.6 | 63 | 63 | 8 | MOLECULE: | 5'- |
| 75:00:00 | 8th8-J | 5.2 | 48.9 | 144 | 372 | 9 | MOLECULE: | DYNEIN REGULATORY COMPLEX PROTEIN 1/2 N-TERM |
| 76:00:00 | 6j9r-B | 5.2 | 5.4 | 102 | 126 | 7 | MOLECULE: | BRAIN TUMOR PROTEIN |
| 77:00:00 | 5hda-A | 5.2 | 6.7 | 95 | 121 | 6 | MOLECULE: | ZINC FINGER MYND DOMAIN-CONTAINING PROTEIN 1 |
| 78:00:00 | 8p0v-K | 5.2 | 27.9 | 128 | 304 | 9 | MOLECULE: | COILED-COIL DOMAIN-CONTAINING PROTEIN 93 |
| 79:00:00 | 7puz-A | 5.2 | 3.4 | 102 | 148 | 9 | MOLECULE: | MICOS COMPLEX SUBUNIT MIC60 |
| 80:00:00 | 5cws-D | 5.2 | 4.8 | 90 | 180 | 7 | MOLECULE: | SAB-158 FAB LIGHT CHAIN |
| 81:00:00 | 8iah-W | 5.2 | 8.3 | 142 | 248 | 10 | MOLECULE: | ADDUCIN 1 |
| 82:00:00 | 5aj3-d | 5.2 | 15.5 | 139 | 177 | 9 | MOLECULE: | MITORIBOSOMAL 12S RRNA |
| 83:00:00 | 3bat-C | 5.2 | 4.1 | 83 | 85 | 7 | MOLECULE: | MYOSIN HEAVY CHAIN, STRIATED MUSCLE/GENERAL |
| 84:00:00 | 3b5n-I | 5.2 | 4.4 | 58 | 58 | 9 | MOLECULE: | SYNAPTOBREVIN HOMOLOG 1 |
| 85:00:00 | 5mw9-H | 5.2 | 3.4 | 57 | 57 | 7 | MOLECULE: | CENTROSOMIN |
| 86:00:00 | 5nw4-5 | 5.2 | 8 | 154 | 275 | 0 | MOLECULE: | DYNEIN HEAVY CHAIN |
| 87:00:00 | 6wcj-J | 5.2 | 3.3 | 59 | 59 | 2 | MOLECULE: | CLATHRIN HEAVY CHAIN 1 |
| 88:00:00 | 7qoo-K | 5.2 | 15.7 | 132 | 253 | 7 | MOLECULE: | CENTROMERE PROTEIN C |
| 89:00:00 | 7qix-P | 5.2 | 7.8 | 69 | 237 | 10 | MOLECULE: | 18S RRNA BODY |
| 90:00:00 | 7oz3-A | 5.2 | 10.5 | 78 | 202 | 9 | MOLECULE: | GNTR FAMILY TRANSCRIPTIONAL REGULATOR |
| 91:00:00 | 7woo-H | 5.2 | 19.4 | 91 | 246 | 7 | MOLECULE: | NUCLEOPORIN NIC96 |
| 92:00:00 | 6wm2-K | 5.2 | 10 | 113 | 114 | 4 | MOLECULE: | V-TYPE PROTON ATPASE SUBUNIT E 1 |
| 93:00:00 | 5nt2-N | 5.2 | 3.5 | 96 | 173 | 9 | MOLECULE: | NON-STRUCTURAL PROTEIN 1 |
| 94:00:00 | 7p3n-p | 5.2 | 10.4 | 126 | 155 | 6 | MOLECULE: | ATP SYNTHASE SUBUNIT ALPHA |
| 95:00:00 | 7vf5-C | 5.2 | 7.8 | 109 | 184 | 6 | MOLECULE: | PROTEIN VIRILIZER HOMOLOG |
| 96:00:00 | 6r0w-J | 5.2 | 18.4 | 107 | 185 | 9 | MOLECULE: | V-TYPE ATP SYNTHASE ALPHA CHAIN |
| 97:00:00 | 6ip1-B | 5.2 | 3.7 | 62 | 62 | 5 | MOLECULE: | VESICLE-ASSOCIATED MEMBRANE PROTEIN 2 |
| 98:00:00 | 8x5f-B | 5.2 | 28.7 | 91 | 616 | 4 | MOLECULE: | SOLUTE CARRIER FAMILY 53 MEMBER 1 |
| 99:00:00 | 2e7s-O | 5.2 | 7.5 | 112 | 124 | 7 | MOLECULE: | RAB GUANINE NUCLEOTIDE EXCHANGE FACTOR SEC2 |
| 0:00 | 8bd7-L | 5.1 | 34.6 | 155 | 303 | 7 | MOLECULE: | IFT88 |
| 1:00 | 7woo-G | 5.1 | 3.5 | 91 | 200 | 12 | MOLECULE: | NUCLEOPORIN NIC96 |
| 2:00 | 8u95-A | 5.1 | 7 | 143 | 1009 | 8 | MOLECULE: | MYOSIN HEAVY CHAIN, ISOFORM U |
| 3:00 | 7d7n-A | 5.1 | 10.4 | 93 | 704 | 11 | MOLECULE: | ATP-BINDING CASSETTE SUB-FAMILY B MEMBER 6, |
| 4:00 | 7o3x-B | 5.1 | 26.3 | 133 | 215 | 8 | MOLECULE: | PROTEIN SLL0617 |
| 5:00 | 6cfz-I | 5.1 | 12.4 | 106 | 106 | 13 | MOLECULE: | ASK1 |
| 6:00 | 8a5a-G | 5.1 | 7.9 | 104 | 110 | 7 | MOLECULE: | CHROMATIN-REMODELING ATPASE INO80 |
| 7:00 | 4yv3-B | 5.1 | 2.8 | 75 | 75 | 12 | MOLECULE: | VIMENTIN |
| 8:00 | 5jxc-D | 5.1 | 4 | 78 | 78 | 8 | MOLECULE: | RAS/RAP GTPASE-ACTIVATING PROTEIN SYNGAP |
| 9:00 | 5jxc-B | 5.1 | 3 | 78 | 78 | 1 | MOLECULE: | RAS/RAP GTPASE-ACTIVATING PROTEIN SYNGAP |
| 10:00 | 7bv6-B | 5.1 | 1.8 | 61 | 61 | 15 | MOLECULE: | VESICLE-ASSOCIATED MEMBRANE PROTEIN 8 |
| 11:00 | 2zvo-D | 5.1 | 2.1 | 87 | 88 | 9 | MOLECULE: | UBC PROTEIN |
| 12:00 | 2n5e-B | 5.1 | 25.9 | 110 | 167 | 8 | MOLECULE: | APOLIPOPROTEIN A-I |
| 13:00 | 3lj5-H | 5.1 | 8.5 | 136 | 692 | 8 | MOLECULE: | PORTAL PROTEIN |
| 14:00 | 6von-I | 5.1 | 14.4 | 128 | 153 | 9 | MOLECULE: | ATP SYNTHASE SUBUNIT ALPHA, CHLOROPLASTIC |
| 15:00 | 5f5p-F | 5.1 | 3.2 | 58 | 58 | 10 | MOLECULE: | PROTEIN SHROOM2 |
| 16:00 | 6ian-C | 5.1 | 8.6 | 105 | 432 | 12 | MOLECULE: | INTRAFLAGELLAR TRANSPORT PROTEIN 74 |

| | | | | | | | | |
|---|---|---|---|---|---|---|---|---|
| 17:00 | 6wm3-K | 5.1 | 9.5 | 111 | 114 | 5 | MOLECULE: | V-TYPE PROTON ATPASE 116 KDA SUBUNIT A ISOFO |
| 18:00 | 8fby-D | 5.1 | 5.2 | 86 | 86 | 8 | MOLECULE: | PLASMALEMMA VESICLE-ASSOCIATED PROTEIN |
| 19:00 | 8k8a-A | 5.1 | 3.2 | 61 | 62 | 2 | MOLECULE: | NUCLEAR FACTOR INTERLEUKIN-3-REGULATED PROTE |
| 20:00 | 3wut-D | 5.1 | 2.9 | 53 | 54 | 17 | MOLECULE: | CENTROSOMAL PROTEIN OF 55 KDA |
| 21:00 | 2e7s-H | 5.1 | 4.7 | 88 | 107 | 11 | MOLECULE: | RAB GUANINE NUCLEOTIDE EXCHANGE FACTOR SEC2 |
| 22:00 | 4m8m-B | 5 | 17.4 | 92 | 574 | 7 | MOLECULE: | GCN4 COILED-COIL FUSED ZEBRAFISH PLEXINC1 |
| 23:00 | 6vq8-H | 5 | 3.6 | 94 | 214 | 10 | MOLECULE: | ATPASE H+-TRANSPORTING V1 SUBUNIT A |
| 24:00:00 | 6aoz-C | 5 | 3.2 | 74 | 74 | 9 | MOLECULE: | CASP8-ASSOCIATED PROTEIN 2 |
| 25:00:00 | 7rzr-C | 5 | 0.9 | 71 | 71 | 11 | MOLECULE: | SARS-COV-2 HR1 D936Y LINKED TO A SCAFFOLD,SP |
| 26:00:00 | 7rzv-C | 5 | 0.9 | 71 | 71 | 11 | MOLECULE: | SARS-COV-2 HR1 LINKED TO A SCAFFOLD,SPIKE PR |
| 27:00:00 | 7rzr-A | 5 | 0.9 | 71 | 71 | 11 | MOLECULE: | SARS-COV-2 HR1 D936Y LINKED TO A SCAFFOLD,SP |
| 28:00:00 | 7rzq-A | 5 | 0.9 | 71 | 71 | 11 | MOLECULE: | SARS-COV-2 HR1 LINKED TO A SCAFFOLD,SPIKE PR |
| 29:00:00 | 7rzt-C | 5 | 0.9 | 71 | 71 | 11 | MOLECULE: | SARS-COV-2 HR1 S940F LINKED TO A SCAFFOLD,SP |
| 30:00:00 | 7rzq-B | 5 | 0.9 | 71 | 71 | 11 | MOLECULE: | SARS-COV-2 HR1 LINKED TO A SCAFFOLD,SPIKE PR |
| 31:00:00 | 1kil-B | 5 | 2 | 59 | 59 | 12 | MOLECULE: | SYNAPTOBREVIN SNARE MOTIF |
| 32:00:00 | 6f3a-5 | 5 | 8 | 154 | 275 | 0 | MOLECULE: | ARP1 ACTIN RELATED PROTEIN 1 HOMOLOG A |
| 33:00:00 | 5fiy-A | 5 | 2.3 | 77 | 78 | 1 | MOLECULE: | PRKC APOPTOSIS WT1 REGULATOR PROTEIN |
| 34:00:00 | 5xit-A | 5 | 2.8 | 76 | 76 | 11 | MOLECULE: | UBIQUITIN-40S RIBOSOMAL PROTEIN S27A |
| 35:00:00 | 6fpr-B | 5 | 2.4 | 58 | 58 | 9 | MOLECULE: | SIGNAL RECOGNITION PARTICLE RECEPTOR FTSY |
| 36:00:00 | 5ip0-M | 5 | 3.8 | 57 | 107 | 5 | MOLECULE: | PHA GRANULE-ASSOCIATED PROTEIN |
| 37:00:00 | 8sdi-A | 5 | 3.4 | 80 | 80 | 5 | MOLECULE: | TRIPARTITE MOTIF-CONTAINING PROTEIN 45 |
| 38:00:00 | 2xqh-A | 4.9 | 8.5 | 118 | 258 | 10 | MOLECULE: | IMMUNOGLOBULIN-BINDING PROTEIN EIBD |
| 39:00:00 | 4dl0-K | 4.9 | 5.8 | 97 | 97 | 8 | MOLECULE: | V-TYPE PROTON ATPASE SUBUNIT C |
| 40:00:00 | 8tek-A | 4.9 | 8.8 | 141 | 200 | 11 | MOLECULE: | DYNEIN REGULATORY COMPLEX PROTEIN 1/2 N-TERM |
| 41:00:00 | 5dfz-D | 4.9 | 11.5 | 128 | 341 | 8 | MOLECULE: | VACUOLAR PROTEIN SORTING-ASSOCIATED PROTEIN |
| 42:00:00 | 4r8g-E | 4.9 | 17.7 | 110 | 324 | 7 | MOLECULE: | UNCONVENTIONAL MYOSIN-IC |
| 43:00:00 | 8wjo-B | 4.9 | 5.8 | 113 | 283 | 7 | MOLECULE: | STRUCTURAL MAINTENANCE OF CHROMOSOMES PROTEI |
| 44:00:00 | 3v6i-A | 4.9 | 5.7 | 114 | 186 | 10 | MOLECULE: | V-TYPE ATP SYNTHASE SUBUNIT E |
| 45:00:00 | 8p0v-L | 4.9 | 32.8 | 93 | 416 | 9 | MOLECULE: | COILED-COIL DOMAIN-CONTAINING PROTEIN 93 |
| 46:00:00 | 8fwj-A | 4.9 | 7.9 | 95 | 552 | 11 | MOLECULE: | CIRCADIAN CLOCK PROTEIN KAIC |
| 47:00:00 | 8g0q-B | 4.9 | 14.7 | 88 | 230 | 9 | MOLECULE: | KINETOCHORE PROTEIN NDC80 |
| 48:00:00 | 2eqb-C | 4.9 | 4 | 91 | 93 | 5 | MOLECULE: | RAS-RELATED PROTEIN SEC4 |
| 49:00:00 | 7tik-B | 4.9 | 0.9 | 71 | 71 | 11 | MOLECULE: | FERRITIN, DPS FAMILY PROTEIN AND SPIKE PROTE |
| 50:00:00 | 6f5x-A | 4.9 | 3.6 | 76 | 76 | 13 | MOLECULE: | SYNAPTONEMAL COMPLEX PROTEIN 1 |
| 51:00:00 | 6mti-C | 4.9 | 3.2 | 73 | 73 | 7 | MOLECULE: | SYNAPTOTAGMIN-1 |
| 52:00:00 | 1gmj-D | 4.9 | 1.9 | 56 | 56 | 7 | MOLECULE: | ATPASE INHIBITOR |
| 53:00:00 | 6mti-O | 4.9 | 3.2 | 73 | 73 | 7 | MOLECULE: | SYNAPTOTAGMIN-1 |
| 54:00:00 | 6mti-W | 4.9 | 3.2 | 73 | 73 | 7 | MOLECULE: | SYNAPTOTAGMIN-1 |
| 55:00:00 | 6mti-K | 4.9 | 3.2 | 73 | 73 | 7 | MOLECULE: | SYNAPTOTAGMIN-1 |
| 56:00:00 | 7y4s-A | 4.9 | 10 | 93 | 343 | 3 | MOLECULE: | TRIPARTITE MOTIF-CONTAINING PROTEIN 72 |
| 57:00:00 | 6mti-G | 4.9 | 3.2 | 73 | 73 | 7 | MOLECULE: | SYNAPTOTAGMIN-1 |
| 58:00:00 | 6mti-S | 4.9 | 3.2 | 73 | 73 | 7 | MOLECULE: | SYNAPTOTAGMIN-1 |
| 59:00:00 | 5nt1-A | 4.9 | 5.2 | 105 | 173 | 8 | MOLECULE: | E3 UBIQUITIN/ISG15 LIGASE TRIM25 |
| 60:00:00 | 3lj5-L | 4.9 | 8 | 131 | 692 | 8 | MOLECULE: | PORTAL PROTEIN |
| 61:00:00 | 3lj5-B | 4.9 | 8 | 131 | 692 | 8 | MOLECULE: | PORTAL PROTEIN |
| 62:00:00 | 3lj5-D | 4.9 | 8 | 131 | 692 | 8 | MOLECULE: | PORTAL PROTEIN |
| 63:00:00 | 5gai-J | 4.9 | 8.9 | 125 | 721 | 3 | MOLECULE: | PORTAL PROTEIN |
| 64:00:00 | 7z47-F | 4.9 | 5.9 | 66 | 68 | 5 | MOLECULE: | ADAPTOR PROTEIN |
| 65:00:00 | 3wuv-J | 4.9 | 4.3 | 56 | 56 | 4 | MOLECULE: | CENTROSOMAL PROTEIN OF 55 KDA |
| 66:00:00 | 4m3l-D | 4.9 | 2.4 | 53 | 53 | 6 | MOLECULE: | E3 UBIQUITIN-PROTEIN LIGASE TRIM63 |
| 67:00:00 | 5low-N | 4.9 | 4.5 | 60 | 60 | 8 | MOLECULE: | RABPHILIN-3A |

| | | | | | | | | |
|---|---|---|---|---|---|---|---|---|
| 68:00:00 | 6yjd-A | 4.9 | 3.8 | 87 | 104 | 8 | MOLECULE: | CAPSID ASSEMBLY SCAFFOLDING PROTEIN,PRELAMIN |
| 69:00:00 | 1nyh-A | 4.9 | 3.3 | 76 | 76 | 11 | MOLECULE: | REGULATORY PROTEIN SIR4 |
| 70:00:00 | 5opt-P | 4.8 | 8.7 | 87 | 249 | 6 | MOLECULE: | ACTIVATED PROTEIN KINASE C RECEPTOR, PUTATIV |
| 71:00:00 | 5ijn-F | 4.8 | 29.7 | 130 | 335 | 10 | MOLECULE: | NUCLEAR PORE COMPLEX PROTEIN NUP155 |
| 72:00:00 | 6znl-n | 4.8 | 13.7 | 93 | 343 | 10 | MOLECULE: | ARP1 ACTIN RELATED PROTEIN 1 HOMOLOG A |
| 73:00:00 | 5chx-A | 4.8 | 10.5 | 118 | 210 | 14 | MOLECULE: | XRCC4-MYH7-1590-1657 |
| 74:00:00 | 1aa0-A | 4.8 | 3.8 | 85 | 113 | 9 | MOLECULE: | FIBRITIN |
| 75:00:00 | 8srm-B | 4.8 | 9.7 | 85 | 235 | 5 | MOLECULE: | RB1-INDUCIBLE COILED-COIL PROTEIN 1 |
| 76:00:00 | 7qoo-U | 4.8 | 15.5 | 132 | 186 | 6 | MOLECULE: | CENTROMERE PROTEIN C |
| 77:00:00 | 4lin-A | 4.8 | 10.2 | 144 | 289 | 5 | MOLECULE: | TAIL NEEDLE PROTEIN GP26 |
| 78:00:00 | 4q9u-C | 4.8 | 3.9 | 83 | 83 | 6 | MOLECULE: | RAB5 GDP/GTP EXCHANGE FACTOR |
| 79:00:00 | 7r5v-U | 4.8 | 7.2 | 111 | 165 | 5 | MOLECULE: | CENTROMERE PROTEIN H |
| 80:00:00 | 4q9u-G | 4.8 | 5 | 84 | 84 | 0 | MOLECULE: | RAB5 GDP/GTP EXCHANGE FACTOR |
| 81:00:00 | 2yo3-B | 4.8 | 9 | 134 | 258 | 9 | MOLECULE: | GENERAL CONTROL PROTEIN GCN4, PUTATIVE INNER |
| 82:00:00 | 5tby-A | 4.8 | 36.5 | 76 | 954 | 7 | MOLECULE: | MYOSIN-7 |
| 83:00:00 | 3uia-A | 4.8 | 3.4 | 53 | 53 | 0 | MOLECULE: | GLYCOPROTEIN 41 |
| 84:00:00 | 7tkd-U | 4.8 | 15.5 | 130 | 155 | 11 | MOLECULE: | ATP SYNTHASE SUBUNIT 9, MITOCHONDRIAL |
| 85:00:00 | 8th8-E | 4.8 | 10.1 | 140 | 291 | 5 | MOLECULE: | DYNEIN REGULATORY COMPLEX PROTEIN 1/2 N-TERM |
| 86:00:00 | 4jpn-C | 4.8 | 5 | 77 | 77 | 1 | MOLECULE: | MINOR SPIKE PROTEIN H |
| 87:00:00 | 4mvd-E | 4.8 | 15 | 110 | 252 | 12 | MOLECULE: | CHOLINE-PHOSPHATE CYTIDYLYLTRANSFERASE A |
| 88:00:00 | 5oqm-h | 4.7 | 8.8 | 101 | 131 | 8 | MOLECULE: | DNA-DIRECTED RNA POLYMERASE II SUBUNIT RPB1 |
| 89:00:00 | 2p2u-B | 4.7 | 23.3 | 83 | 153 | 8 | MOLECULE: | HOST-NUCLEASE INHIBITOR PROTEIN GAM, PUTATIV |
| 90:00:00 | 8q85-V | 4.7 | 6.8 | 96 | 117 | 8 | MOLECULE: | KINETOCHORE PROTEIN NDC80 |
| 91:00:00 | 5dfz-A | 4.7 | 23.1 | 130 | 343 | 7 | MOLECULE: | VACUOLAR PROTEIN SORTING-ASSOCIATED PROTEIN |
| 92:00:00 | 2p22-A | 4.7 | 13.5 | 121 | 168 | 9 | MOLECULE: | SUPPRESSOR PROTEIN STP22 OF TEMPERATURE-SENS |
| 93:00:00 | 4q9u-H | 4.7 | 4.1 | 81 | 82 | 11 | MOLECULE: | RAB5 GDP/GTP EXCHANGE FACTOR |
| 94:00:00 | 4mvd-A | 4.7 | 17.8 | 100 | 252 | 8 | MOLECULE: | CHOLINE-PHOSPHATE CYTIDYLYLTRANSFERASE A |
| 95:00:00 | 4mvd-C | 4.7 | 18.6 | 100 | 253 | 5 | MOLECULE: | CHOLINE-PHOSPHATE CYTIDYLYLTRANSFERASE A |
| 96:00:00 | 8amr-C | 4.7 | 3.8 | 95 | 147 | 11 | MOLECULE: | TRIPARTITE MOTIF-CONTAINING PROTEIN 3 |
| 97:00:00 | 4jpn-H | 4.7 | 4.1 | 74 | 74 | 4 | MOLECULE: | MINOR SPIKE PROTEIN H |
| 98:00:00 | 5yfp-C | 4.7 | 39 | 127 | 790 | 6 | MOLECULE: | EXOCYST COMPLEX COMPONENT SEC3 |
| 99:00:00 | 4lin-C | 4.7 | 9.7 | 137 | 289 | 6 | MOLECULE: | TAIL NEEDLE PROTEIN GP26 |
| 0:00 | 5n9j-D | 4.6 | 17 | 97 | 135 | 3 | MOLECULE: | MEDIATOR OF RNA POLYMERASE II TRANSCRIPTION |
| 1:00 | 8tek-B | 4.6 | 9 | 145 | 208 | 8 | MOLECULE: | DYNEIN REGULATORY COMPLEX PROTEIN 1/2 N-TERM |
| 2:00 | 8ppr-N | 4.6 | 6.1 | 103 | 217 | 9 | MOLECULE: | KINETOCHORE-ASSOCIATED PROTEIN DSN1 HOMOLOG |
| 3:00 | 6b85-A | 4.6 | 5.1 | 66 | 215 | 3 | MOLECULE: | TMHC4_R |
| 4:00 | 6h9m-A | 4.6 | 3.9 | 84 | 94 | 8 | MOLECULE: | COILED-COIL DOMAIN-CONTAINING PROTEIN 90B, M |
| 5:00 | 5mq4-C | 4.6 | 7.2 | 95 | 117 | 16 | MOLECULE: | PROTEIN KINASE C-BINDING PROTEIN 1 |
| 6:00 | 2eqb-B | 4.6 | 8.3 | 76 | 93 | 11 | MOLECULE: | RAS-RELATED PROTEIN SEC4 |
| 7:00 | 4mvd-G | 4.6 | 17.5 | 100 | 252 | 8 | MOLECULE: | CHOLINE-PHOSPHATE CYTIDYLYLTRANSFERASE A |
| 8:00 | 6n6r-B | 4.6 | 2.5 | 68 | 68 | 3 | MOLECULE: | UBIQUITIN |
| 9:00 | 7cld-E | 4.6 | 10.2 | 78 | 123 | 6 | MOLECULE: | TUBULIN ALPHA-1B CHAIN |
| 10:00 | 3f1i-H | 4.6 | 5.4 | 76 | 98 | 9 | MOLECULE: | HEPATOCYTE GROWTH FACTOR-REGULATED TYROSINE |
| 11:00 | 8dd0-O | 4.6 | 1.4 | 70 | 70 | 6 | MOLECULE: | ACTIN, ALPHA CARDIAC MUSCLE 1 |
| 12:00 | 1kfm-A | 4.6 | 3.3 | 50 | 50 | 6 | MOLECULE: | MAJOR OUTER MEMBRANE LIPOPROTEIN |
| 13:00 | 1ik7-A | 4.6 | 2.6 | 52 | 52 | 8 | MOLECULE: | PROBABLE SERINE/THREONINE-PROTEIN KINASE PEL |
| 14:00 | 3htk-B | 4.6 | 3.7 | 73 | 73 | 10 | MOLECULE: | STRUCTURAL MAINTENANCE OF CHROMOSOMES PROTEI |
| 15:00 | 6aoz-D | 4.6 | 1.8 | 68 | 68 | 9 | MOLECULE: | CASP8-ASSOCIATED PROTEIN 2 |
| 16:00 | 8q5h-B | 4.6 | 8.3 | 75 | 174 | 12 | MOLECULE: | KINETOCHORE PROTEIN SPC24 |
| 17:00 | 6wko-A | 4.6 | 2.6 | 60 | 90 | 5 | MOLECULE: | HEMAGGLUTININ-ESTERASE-FUSION GLYCOPROTEIN |
| 18:00 | 3wqa-B | 4.6 | 6.3 | 118 | 201 | 10 | MOLECULE: | TRIMERIC AUTOTRANSPORTER ADHESIN |

| | | | | | | | | |
|---|---|---|---|---|---|---|---|---|
| 19:00 | 2yo3-A | 4.6 | 7.4 | 96 | 257 | 8 | MOLECULE: | GENERAL CONTROL PROTEIN GCN4, PUTATIVE INNER |
| 20:00 | 8at3-D | 4.5 | 36.2 | 107 | 666 | 7 | MOLECULE: | HAUS AUGMIN-LIKE COMPLEX SUBUNIT 1 |
| 21:00 | 8etc-F | 4.5 | 17.3 | 55 | 218 | 13 | MOLECULE: | RNA (2151-MER) |
| 22:00 | 1lj2-B | 4.5 | 7.3 | 66 | 110 | 2 | MOLECULE: | NONSTRUCTURAL RNA-BINDING PROTEIN 34 |
| 23:00 | 8hpo-D | 4.5 | 11.5 | 112 | 165 | 9 | MOLECULE: | TRANSCRIPTIONAL REGULATORY PROTEIN UME1 |
| 24:00:00 | 4y66-C | 4.5 | 19.2 | 115 | 197 | 6 | MOLECULE: | MND1 |
| 25:00:00 | 8hd0-E | 4.5 | 26.8 | 99 | 380 | 8 | MOLECULE: | CELL DIVISION ATP-BINDING PROTEIN FTSE |
| 26:00:00 | 6wm2-I | 4.5 | 7.4 | 111 | 225 | 5 | MOLECULE: | V-TYPE PROTON ATPASE SUBUNIT E 1 |
| 27:00:00 | 4xa6-B | 4.5 | 12.8 | 124 | 165 | 7 | MOLECULE: | GP7-MYH7(1777-1855)-EB1 CHIMERA PROTEIN |
| 28:00:00 | 5wst-B | 4.5 | 2.4 | 70 | 70 | 9 | MOLECULE: | UNCONVENTIONAL MYOSIN-VIIA |
| 29:00:00 | 5hzp-C | 4.5 | 1.5 | 71 | 71 | 3 | MOLECULE: | M PROTEIN, SEROTYPE 49 |
| 30:00:00 | 6ano-B | 4.5 | 3.6 | 70 | 70 | 6 | MOLECULE: | CASP8-ASSOCIATED PROTEIN 2 |
| 31:00:00 | 6cfz-D | 4.5 | 3.3 | 72 | 73 | 1 | MOLECULE: | ASK1 |
| 32:00:00 | 6aoz-B | 4.5 | 3.7 | 69 | 69 | 6 | MOLECULE: | CASP8-ASSOCIATED PROTEIN 2 |
| 33:00:00 | 6eb9-A | 4.5 | 8.1 | 74 | 101 | 9 | MOLECULE: | PHOSPHOPROTEIN |
| 34:00:00 | 5kj8-B | 4.5 | 3.6 | 66 | 66 | 2 | MOLECULE: | VESICLE-ASSOCIATED MEMBRANE PROTEIN 3 |
| 35:00:00 | 3lj5-I | 4.5 | 8 | 131 | 692 | 8 | MOLECULE: | PORTAL PROTEIN |
| 36:00:00 | 5zuv-C | 4.5 | 1.3 | 84 | 119 | 8 | MOLECULE: | SPIKE GLYCOPROTEIN,INHIBITOR EK1 |
| 37:00:00 | 2ba2-A | 4.5 | 3.8 | 81 | 81 | 5 | MOLECULE: | HYPOTHETICAL UPF0134 PROTEIN MPN010 |
| 38:00:00 | 7x5e-A | 4.5 | 2.9 | 80 | 99 | 9 | MOLECULE: | TRANSCRIPTION FACTOR MAFG |
| 39:00:00 | 4aj5-B | 4.5 | 11.5 | 86 | 87 | 5 | MOLECULE: | SPINDLE AND KINETOCHORE-ASSOCIATED PROTEIN 1 |
| 40:00:00 | 8iw0-A | 4.5 | 3.3 | 63 | 84 | 8 | MOLECULE: | LIPRIN-BETA-1,KN MOTIF AND ANKYRIN REPEAT DO |
| 41:00:00 | 6tda-L | 4.5 | 57.3 | 120 | 596 | 6 | MOLECULE: | HISTONE H3.2 |
| 42:00:00 | 6vy1-g | 4.5 | 2.8 | 46 | 121 | 4 | MOLECULE: | PREFOLDIN SUBUNIT ALPHA 2 |
| 43:00:00 | 6vy1-e | 4.5 | 3.5 | 47 | 121 | 4 | MOLECULE: | PREFOLDIN SUBUNIT ALPHA 2 |
| 44:00:00 | 8j0s-d | 4.5 | 15.7 | 149 | 445 | 7 | MOLECULE: | ATP SYNTHASE SUBUNIT A |
| 45:00:00 | 4egx-C | 4.5 | 4.2 | 66 | 174 | 11 | MOLECULE: | KINESIN-LIKE PROTEIN KIF1A |
| 46:00:00 | 3wpr-C | 4.5 | 7.4 | 106 | 218 | 6 | MOLECULE: | TRIMERIC AUTOTRANSPORTER ADHESIN |
| 47:00:00 | 6h9m-C | 4.5 | 4.1 | 84 | 94 | 7 | MOLECULE: | COILED-COIL DOMAIN-CONTAINING PROTEIN 90B, M |
| 48:00:00 | 3hgf-A | 4.4 | 4.1 | 69 | 98 | 3 | MOLECULE: | RHOPTRY PROTEIN FRAGMENT |
| 49:00:00 | 6gvw-F | 4.4 | 23.1 | 86 | 320 | 14 | MOLECULE: | BRCA1-A COMPLEX SUBUNIT ABRAXAS 1 |
| 50:00:00 | 8at3-A | 4.4 | 43.1 | 73 | 286 | 7 | MOLECULE: | HAUS AUGMIN-LIKE COMPLEX SUBUNIT 1 |
| 51:00:00 | 3n7n-B | 4.4 | 7.8 | 93 | 164 | 5 | MOLECULE: | MONOPOLIN COMPLEX SUBUNIT CSM1 |
| 52:00:00 | 8akr-R | 4.4 | 12.8 | 132 | 196 | 14 | MOLECULE: | CHLOROPLAST MEMBRANE-ASSOCIATED 30 KD PROTEI |
| 53:00:00 | 4q9u-D | 4.4 | 3.8 | 75 | 75 | 11 | MOLECULE: | RAB5 GDP/GTP EXCHANGE FACTOR |
| 54:00:00 | 6vqk-M | 4.4 | 4.1 | 67 | 67 | 4 | MOLECULE: | V-TYPE PROTON ATPASE SUBUNIT C 1 |
| 55:00:00 | 5d80-k | 4.4 | 9.3 | 91 | 178 | 4 | MOLECULE: | V-TYPE PROTON ATPASE CATALYTIC SUBUNIT A |
| 56:00:00 | 7ucd-J | 4.4 | 2.6 | 68 | 68 | 7 | MOLECULE: | PROTEIN FOSB |
| 57:00:00 | 6vqk-O | 4.4 | 1.9 | 66 | 66 | 9 | MOLECULE: | V-TYPE PROTON ATPASE SUBUNIT C 1 |
| 58:00:00 | 6k15-H | 4.4 | 21.3 | 123 | 393 | 7 | MOLECULE: | CHROMATIN STRUCTURE-REMODELING COMPLEX SUBUN |
| 59:00:00 | 5eoa-A | 4.4 | 5.3 | 72 | 72 | 7 | MOLECULE: | OPTINEURIN |
| 60:00:00 | 6vqj-M | 4.4 | 4.1 | 67 | 67 | 4 | MOLECULE: | V-TYPE PROTON ATPASE SUBUNIT C 1 |
| 61:00:00 | 6w2d-x | 4.4 | 3.5 | 68 | 68 | 6 | MOLECULE: | MAJOR CAPSID PROTEIN |
| 62:00:00 | 7uo7-B | 4.4 | 10.8 | 90 | 186 | 4 | MOLECULE: | RNA-DIRECTED RNA POLYMERASE |
| 63:00:00 | 8pr3-B | 4.4 | 3.3 | 84 | 105 | 11 | MOLECULE: | C-JUN-AMINO-TERMINAL KINASE-INTERACTING PROT |
| 64:00:00 | 2b9b-B | 4.4 | 9.1 | 97 | 482 | 7 | MOLECULE: | FUSION GLYCOPROTEIN F0 |
| 65:00:00 | 6wcj-F | 4.4 | 7 | 77 | 109 | 16 | MOLECULE: | CLATHRIN HEAVY CHAIN 1 |
| 66:00:00 | 4lin-F | 4.4 | 9.7 | 95 | 289 | 7 | MOLECULE: | TAIL NEEDLE PROTEIN GP26 |
| 67:00:00 | 5djo-A | 4.4 | 3.9 | 57 | 158 | 5 | MOLECULE: | KINESIN-LIKE PROTEIN |
| 68:00:00 | 6h9m-B | 4.4 | 4.9 | 84 | 94 | 7 | MOLECULE: | COILED-COIL DOMAIN-CONTAINING PROTEIN 90B, M |
| 69:00:00 | 2yny-A | 4.4 | 9.7 | 71 | 106 | 11 | MOLECULE: | GENERAL CONTROL PROTEIN GCN4, PUTATIVE INNER |

| | | | | | | | | |
|---|---|---|---|---|---|---|---|---|
| 70:00:00 | 7ty0-C | 4.3 | 38.3 | 70 | 488 | 6 | MOLECULE: | GLYCOPROTEIN G |
| 71:00:00 | 8tek-T | 4.3 | 18.6 | 110 | 137 | 4 | MOLECULE: | DYNEIN REGULATORY COMPLEX PROTEIN 1/2 N-TERM |
| 72:00:00 | 7mge-B | 4.3 | 10.2 | 68 | 442 | 7 | MOLECULE: | WD REPEAT-CONTAINING PROTEIN 41 |
| 73:00:00 | 3ub0-D | 4.3 | 8 | 70 | 194 | 7 | MOLECULE: | NON-STRUCTURAL PROTEIN 6, NSP6, |
| 74:00:00 | 7eeb-C | 4.3 | 24 | 88 | 278 | 8 | MOLECULE: | ENHANCED GREEN FLUORESCENT PROTEIN,CATION CH |
| 75:00:00 | 6wcj-B | 4.3 | 7.5 | 78 | 109 | 6 | MOLECULE: | CLATHRIN HEAVY CHAIN 1 |
| 76:00:00 | 8ppr-M | 4.3 | 8.9 | 116 | 205 | 5 | MOLECULE: | KINETOCHORE-ASSOCIATED PROTEIN DSN1 HOMOLOG |
| 77:00:00 | 7emf-D | 4.3 | 9.4 | 107 | 158 | 16 | MOLECULE: | MEDIATOR OF RNA POLYMERASE II TRANSCRIPTION |
| 78:00:00 | 7c4j-F | 4.3 | 20 | 114 | 383 | 9 | MOLECULE: | TRANSCRIPTION REGULATORY PROTEIN SNF12 |
| 79:00:00 | 7oqe-K | 4.3 | 23.2 | 120 | 406 | 8 | MOLECULE: | PROTEIN NAM8 |
| 80:00:00 | 8q85-X | 4.3 | 32.1 | 68 | 68 | 10 | MOLECULE: | KINETOCHORE PROTEIN NDC80 |
| 81:00:00 | 7a5p-K | 4.3 | 12.7 | 96 | 138 | 5 | MOLECULE: | U2 SNRNA |
| 82:00:00 | 4efa-E | 4.3 | 9.1 | 127 | 225 | 6 | MOLECULE: | V-TYPE PROTON ATPASE SUBUNIT C |
| 83:00:00 | 7yg4-B | 4.3 | 4 | 67 | 67 | 3 | MOLECULE: | PROTEIN VIRILIZER HOMOLOG |
| 84:00:00 | 7vdv-X | 4.3 | 32.6 | 120 | 252 | 5 | MOLECULE: | HISTONE H4 |
| 85:00:00 | 6wcj-E | 4.3 | 9 | 80 | 109 | 8 | MOLECULE: | CLATHRIN HEAVY CHAIN 1 |
| 86:00:00 | 5zuv-B | 4.3 | 2.2 | 85 | 129 | 9 | MOLECULE: | SPIKE GLYCOPROTEIN,INHIBITOR EK1 |
| 87:00:00 | 5zuv-A | 4.3 | 2.4 | 86 | 130 | 10 | MOLECULE: | SPIKE GLYCOPROTEIN,INHIBITOR EK1 |
| 88:00:00 | 6w2e-x | 4.3 | 3.4 | 68 | 68 | 4 | MOLECULE: | MAJOR CAPSID PROTEIN |
| 89:00:00 | 3wpr-A | 4.3 | 7.7 | 97 | 225 | 6 | MOLECULE: | TRIMERIC AUTOTRANSPORTER ADHESIN |
| 90:00:00 | 7aej-B | 4.3 | 4.2 | 80 | 145 | 9 | MOLECULE: | ENVELOPE GLYCOPROTEIN GP160,ENVELOPE GLYCOPR |
| 91:00:00 | 6abo-A | 4.2 | 9 | 108 | 213 | 7 | MOLECULE: | DNA REPAIR PROTEIN XRCC4 |
| 92:00:00 | 8ovw-Z | 4.2 | 6.6 | 99 | 151 | 13 | MOLECULE: | CENTROMERE-BINDING PROTEIN 1 |
| 93:00:00 | 8q3v-b | 4.2 | 3.9 | 57 | 70 | 2 | MOLECULE: | TETRAHYDROMETHANOPTERIN S-METHYLTRANSFERASE |
| 94:00:00 | 7zmg-n | 4.2 | 9.2 | 69 | 136 | 13 | MOLECULE: | NADH-UBIQUINONE OXIDOREDUCTASE CHAIN 1 |
| 95:00:00 | 5tsj-K | 4.2 | 4.3 | 94 | 209 | 2 | MOLECULE: | V-TYPE ATP SYNTHASE ALPHA CHAIN |
| 96:00:00 | 5y05-A | 4.2 | 9.3 | 99 | 233 | 14 | MOLECULE: | MSMEG_4306 |
| 97:00:00 | 6j5j-b | 4.2 | 14.7 | 140 | 209 | 9 | MOLECULE: | ATP SYNTHASE SUBUNIT ALPHA, MITOCHONDRIAL |
| 98:00:00 | 8u95-B | 4.2 | 6.5 | 145 | 1009 | 7 | MOLECULE: | MYOSIN HEAVY CHAIN, ISOFORM U |
| 99:00:00 | 2d3e-C | 4.2 | 8.6 | 102 | 130 | 5 | MOLECULE: | GENERAL CONTROL PROTEIN GCN4 AND TROPOMYOSIN |
| 0:00 | 5vox-M | 4.2 | 44 | 76 | 209 | 13 | MOLECULE: | V-TYPE PROTON ATPASE CATALYTIC SUBUNIT A,V-T |
| 1:00 | 5gai-I | 4.2 | 10.5 | 126 | 721 | 3 | MOLECULE: | PORTAL PROTEIN |
| 2:00 | 6ztq-Z | 4.2 | 13 | 95 | 141 | 8 | MOLECULE: | NADH-UBIQUINONE OXIDOREDUCTASE CHAIN 3 |
| 3:00 | 7tkk-U | 4.2 | 12.7 | 119 | 155 | 6 | MOLECULE: | ATP SYNTHASE SUBUNIT 9 |
| 4:00 | 8j9s-B | 4.2 | 4.8 | 67 | 67 | 12 | MOLECULE: | AMINOACYL TRNA SYNTHASE COMPLEX-INTERACTING |
| 5:00 | 6qld-U | 4.2 | 17.2 | 111 | 171 | 7 | MOLECULE: | INNER KINETOCHORE SUBUNIT MIF2 |
| 6:00 | 6zkp-q | 4.2 | 10.8 | 88 | 139 | 1 | MOLECULE: | NADH DEHYDROGENASE [UBIQUINONE] FLAVOPROTEIN |
| 7:00 | 6zkg-q | 4.2 | 11.3 | 88 | 139 | 1 | MOLECULE: | NADH DEHYDROGENASE [UBIQUINONE] FLAVOPROTEIN |
| 8:00 | 6zks-q | 4.2 | 10.7 | 88 | 139 | 1 | MOLECULE: | NADH DEHYDROGENASE [UBIQUINONE] FLAVOPROTEIN |
| 9:00 | 4jpp-B | 4.2 | 9.3 | 94 | 127 | 11 | MOLECULE: | MINOR SPIKE PROTEIN H |
| 10:00 | 7mf3-H | 4.2 | 4.5 | 101 | 211 | 6 | MOLECULE: | MYOSIN-11 |
| 11:00 | 7vhq-G | 4.2 | 14.6 | 106 | 299 | 3 | MOLECULE: | ATP-DEPENDENT ZINC METALLOPROTEASE FTSH |
| 12:00 | 6zke-q | 4.2 | 10.8 | 88 | 139 | 1 | MOLECULE: | NADH DEHYDROGENASE [UBIQUINONE] FLAVOPROTEIN |
| 13:00 | 6z47-G | 4.2 | 49.8 | 135 | 257 | 8 | MOLECULE: | MYOSIN HEAVY CHAIN 11 |
| 14:00 | 2yo1-C | 4.2 | 6 | 109 | 273 | 7 | MOLECULE: | GENERAL CONTROL PROTEIN GCN4, PUTATIVE INNER |
| 15:00 | 3wpr-B | 4.2 | 6.7 | 104 | 207 | 4 | MOLECULE: | TRIMERIC AUTOTRANSPORTER ADHESIN |
| 16:00 | 2yny-B | 4.2 | 4.6 | 82 | 104 | 7 | MOLECULE: | GENERAL CONTROL PROTEIN GCN4, PUTATIVE INNER |
| 17:00 | 5t58-B | 4.1 | 11.4 | 112 | 190 | 8 | MOLECULE: | KLLA0F02343P |
| 18:00 | 8rt9-A | 4.1 | 6.9 | 91 | 207 | 0 | MOLECULE: | TRWJ PROTEIN |
| 19:00 | 3p8c-D | 4.1 | 42.8 | 107 | 204 | 5 | MOLECULE: | CYTOPLASMIC FMR1-INTERACTING PROTEIN 1 |
| 20:00 | 7tch-A | 4.1 | 16 | 72 | 645 | 6 | MOLECULE: | BACITRACIN EXPORT PERMEASE PROTEIN BCEB |

| | | | | | | | | |
|---|---|---|---|---|---|---|---|---|
| 21:00 | 5ed9-B | 4.1 | 4 | 72 | 72 | 6 | MOLECULE: | SUN DOMAIN-CONTAINING PROTEIN 2 |
| 22:00 | 7egm-H | 4.1 | 11.8 | 82 | 388 | 7 | MOLECULE: | TRANSCRIPTION REGULATORY PROTEIN SNF2 |
| 23:00 | 8ppr-F | 4.1 | 2.7 | 70 | 112 | 4 | MOLECULE: | KINETOCHORE-ASSOCIATED PROTEIN DSN1 HOMOLOG |
| 24:00:00 | 8i21-A | 4.1 | 61.5 | 133 | 256 | 10 | MOLECULE: | STRUCTURAL MAINTENANCE OF CHROMOSOMES PROTEI |
| 25:00:00 | 7vc4-C | 4.1 | 15.2 | 84 | 107 | 5 | MOLECULE: | MITOCHONDRIAL IMPORT RECEPTOR SUBUNIT TOM6 H |
| 26:00:00 | 4r3z-A | 4.1 | 2.1 | 71 | 76 | 11 | MOLECULE: | AMINOACYL TRNA SYNTHASE COMPLEX-INTERACTING |
| 27:00:00 | 5j6f-A | 4.1 | 13.8 | 59 | 352 | 10 | MOLECULE: | 3-DEOXY-D-ARABINO-HEPTULOSONATE 7-PHOSPHATE |
| 28:00:00 | 8qfc-C | 4.1 | 8.2 | 97 | 380 | 9 | MOLECULE: | 60S RIBOSOMAL PROTEIN L10A |
| 29:00:00 | 8ap7-c | 4.1 | 4.9 | 60 | 64 | 3 | MOLECULE: | ATP SYNTHASE SUBUNIT A |
| 30:00:00 | 7tmq-M | 4.1 | 4.9 | 93 | 214 | 8 | MOLECULE: | H(+)-TRANSPORTING TWO-SECTOR ATPASE |
| 31:00:00 | 5y88-J | 4.1 | 30.8 | 133 | 431 | 3 | MOLECULE: | PRE-MRNA-SPLICING FACTOR 8 |
| 32:00:00 | 5bw9-l | 4.1 | 4.4 | 66 | 66 | 9 | MOLECULE: | V-TYPE PROTON ATPASE CATALYTIC SUBUNIT A |
| 33:00:00 | 3zx6-B | 4.1 | 8.5 | 112 | 303 | 12 | MOLECULE: | HAMP, METHYL-ACCEPTING CHEMOTAXIS PROTEIN I |
| 34:00:00 | 5lhg-A | 4.1 | 50.7 | 84 | 666 | 10 | MOLECULE: | LYSINE-SPECIFIC HISTONE DEMETHYLASE 1A |
| 35:00:00 | 8akr-u | 4.1 | 8.7 | 121 | 196 | 9 | MOLECULE: | CHLOROPLAST MEMBRANE-ASSOCIATED 30 KD PROTEI |
| 36:00:00 | 8at4-E | 4.1 | 22.4 | 108 | 217 | 7 | MOLECULE: | HAUS AUGMIN-LIKE COMPLEX SUBUNIT 1 |
| 37:00:00 | 6hgc-A | 4.1 | 9.2 | 66 | 304 | 6 | MOLECULE: | UBIQUITIN CARBOXYL-TERMINAL HYDROLASE CALYPS |
| 38:00:00 | 7b93-Z | 4.1 | 11.2 | 97 | 143 | 5 | MOLECULE: | NADH-UBIQUINONE OXIDOREDUCTASE CHAIN 3 |
| 39:00:00 | 3jbh-B | 4.1 | 25.7 | 104 | 964 | 12 | MOLECULE: | MYOSIN 2 HEAVY CHAIN STRIATED MUSCLE |
| 40:00:00 | 7uus-Q | 4 | 8.3 | 64 | 170 | 8 | MOLECULE: | HYDROGENASE-2, LARGE SUBUNIT |
| 41:00:00 | 2pjw-H | 4 | 7 | 65 | 88 | 6 | MOLECULE: | UNCHARACTERIZED PROTEIN YHL002W |
| 42:00:00 | 4orh-H | 4 | 2.6 | 77 | 142 | 6 | MOLECULE: | UBIQUITIN-CONJUGATING ENZYME E2 VARIANT 2 |
| 43:00:00 | 7v2w-J | 4 | 13.9 | 126 | 236 | 6 | MOLECULE: | THO COMPLEX SUBUNIT HPR1 |
| 44:00:00 | 4lws-A | 4 | 5 | 61 | 100 | 3 | MOLECULE: | UNCHARACTERIZED PROTEIN |
| 45:00:00 | 8dft-A | 4 | 2.3 | 46 | 112 | 15 | MOLECULE: | PILIN PROTEIN |
| 46:00:00 | 7qog-A | 4 | 9.9 | 86 | 657 | 2 | MOLECULE: | PORTAL PROTEIN GP20 |
| 47:00:00 | 3jb9-i | 4 | 30.1 | 107 | 161 | 7 | MOLECULE: | PRE-MRNA-SPLICING FACTOR SPP42 |
| 48:00:00 | 8aud-A | 4 | 3.8 | 94 | 350 | 13 | MOLECULE: | CELL WALL-ASSOCIATED HYDROLASES (INVASION-AS |
| 49:00:00 | 8i03-E | 4 | 16.6 | 115 | 169 | 9 | MOLECULE: | PAIRED AMPHIPATHIC HELIX PROTEIN PST1 |
| 50:00:00 | 1y9b-A | 4 | 6.4 | 66 | 81 | 8 | MOLECULE: | CONSERVED HYPOTHETICAL PROTEIN |
| 51:00:00 | 5o31-Z | 4 | 12.4 | 88 | 137 | 6 | MOLECULE: | NADH-UBIQUINONE OXIDOREDUCTASE CHAIN 3 |
| 52:00:00 | 6zls-A | 4 | 15.2 | 105 | 292 | 7 | MOLECULE: | HISTIDINE KINASE |
| 53:00:00 | 7sqk-C | 4 | 8 | 131 | 600 | 3 | MOLECULE: | HAUS AUGMIN-LIKE COMPLEX SUBUNIT 1 |
| 54:00:00 | 7tmo-M | 4 | 4.3 | 94 | 214 | 7 | MOLECULE: | H(+)-TRANSPORTING TWO-SECTOR ATPASE |
| 55:00:00 | 8gwk-B | 4 | 13.2 | 90 | 187 | 4 | MOLECULE: | RNA-DIRECTED RNA POLYMERASE |
| 56:00:00 | 8c5v-S | 4 | 5.3 | 117 | 516 | 6 | MOLECULE: | CHEMOTAXIS PROTEIN CHEA |
| 57:00:00 | 6m3w-B | 4 | 40.4 | 132 | 344 | 2 | MOLECULE: | SPIKE GLYCOPROTEIN |
| 58:00:00 | 7z47-D | 4 | 3.8 | 72 | 87 | 6 | MOLECULE: | ADAPTOR PROTEIN |
| 59:00:00 | 7apk-E | 4 | 14.6 | 111 | 523 | 5 | MOLECULE: | THO COMPLEX SUBUNIT 1 |
| 60:00:00 | 6pse-A | 4 | 9.1 | 78 | 78 | 9 | MOLECULE: | PROTEIN BICAUDAL D HOMOLOG 2 |
| 61:00:00 | 2e50-P | 4 | 10.7 | 84 | 186 | 13 | MOLECULE: | PROTEIN SET |
| 62:00:00 | 4g2k-B | 4 | 2.2 | 76 | 108 | 8 | MOLECULE: | GENERAL CONTROL PROTEIN GCN4, ENVELOPE GLYCO |
| 63:00:00 | 3lvg-D | 3.9 | 10.8 | 88 | 180 | 8 | MOLECULE: | CLATHRIN HEAVY CHAIN 1 |
| 64:00:00 | 6r7l-E | 3.9 | 14.4 | 72 | 98 | 3 | MOLECULE: | SECG |
| 65:00:00 | 5nnv-D | 3.9 | 5.1 | 120 | 252 | 7 | MOLECULE: | CHROMOSOME PARTITION PROTEIN SMC,CHROMOSOME |
| 66:00:00 | 4cjd-A | 3.9 | 17.4 | 80 | 123 | 8 | MOLECULE: | NADA |
| 67:00:00 | 7x2e-A | 3.9 | 10.3 | 96 | 163 | 9 | MOLECULE: | HARMONIN |
| 68:00:00 | 6cv9-A | 3.9 | 11.9 | 57 | 316 | 11 | MOLECULE: | SHORT TRANSIENT RECEPTOR POTENTIAL CHANNEL 6 |
| 69:00:00 | 5xbt-A | 3.9 | 10.4 | 75 | 271 | 11 | MOLECULE: | PROBABLE TRANSCRIPTIONAL REGULATOR |
| 70:00:00 | 8x5f-A | 3.9 | 32.1 | 90 | 616 | 10 | MOLECULE: | SOLUTE CARRIER FAMILY 53 MEMBER 1 |
| 71:00:00 | 8hmy-B | 3.9 | 13.8 | 77 | 264 | 4 | MOLECULE: | TRNA-SPLICING ENDONUCLEASE SUBUNIT SEN2 |

| | | | | | | | | |
|---|---|---|---|---|---|---|---|---|
| 72:00:00 | 2p4w-B | 3.9 | 13.8 | 64 | 198 | 16 | MOLECULE: | TRANSCRIPTIONAL REGULATORY PROTEIN ARSR FAMI |
| 73:00:00 | 8fck-E | 3.9 | 35.3 | 142 | 222 | 4 | MOLECULE: | HAUS AUGMIN-LIKE COMPLEX SUBUNIT 1 |
| 74:00:00 | 6ly8-G | 3.9 | 4.4 | 94 | 210 | 7 | MOLECULE: | V-TYPE ATP SYNTHASE ALPHA CHAIN |
| 75:00:00 | 5nt1-E | 3.9 | 5 | 85 | 173 | 8 | MOLECULE: | E3 UBIQUITIN/ISG15 LIGASE TRIM25 |
| 76:00:00 | 3lj5-G | 3.9 | 8.4 | 134 | 692 | 7 | MOLECULE: | PORTAL PROTEIN |
| 77:00:00 | 7yo1-B | 3.9 | 18.1 | 72 | 263 | 10 | MOLECULE: | CALCIUM-ACTIVATED POTASSIUM CHANNEL SUBUNIT |
| 78:00:00 | 8jsm-D | 3.9 | 3.3 | 68 | 68 | 9 | MOLECULE: | RNA-DIRECTED RNA POLYMERASE L |
| 79:00:00 | 4lin-K | 3.9 | 34.1 | 123 | 289 | 10 | MOLECULE: | TAIL NEEDLE PROTEIN GP26 |
| 80:00:00 | 3wpo-A | 3.9 | 6.7 | 102 | 193 | 5 | MOLECULE: | TRIMERIC AUTOTRANSPORTER ADHESIN |
| 81:00:00 | 7emf-I | 3.8 | 6.2 | 67 | 73 | 3 | MOLECULE: | MEDIATOR OF RNA POLYMERASE II TRANSCRIPTION |
| 82:00:00 | 7qoo-Q | 3.8 | 17.2 | 97 | 209 | 6 | MOLECULE: | CENTROMERE PROTEIN C |
| 83:00:00 | 6gmh-M | 3.8 | 14.5 | 53 | 991 | 8 | MOLECULE: | RPB1 |
| 84:00:00 | 7emf-G | 3.8 | 4.1 | 79 | 161 | 9 | MOLECULE: | MEDIATOR OF RNA POLYMERASE II TRANSCRIPTION |
| 85:00:00 | 7uxc-R | 3.8 | 18.8 | 57 | 107 | 7 | MOLECULE: | SERINE/THREONINE-PROTEIN KINASE MTOR |
| 86:00:00 | 1z56-A | 3.8 | 2.7 | 76 | 77 | 5 | MOLECULE: | LIGASE INTERACTING FACTOR 1 |
| 87:00:00 | 6yvu-A | 3.8 | 57.1 | 149 | 1127 | 9 | MOLECULE: | STRUCTURAL MAINTENANCE OF CHROMOSOMES PROTEI |
| 88:00:00 | 4wpe-A | 3.8 | 4.7 | 113 | 275 | 6 | MOLECULE: | CYTOKINESIS PROTEIN 2 |
| 89:00:00 | 6gmh-Q | 3.8 | 9.5 | 101 | 884 | 13 | MOLECULE: | RPB1 |
| 90:00:00 | 3eff-K | 3.8 | 3.8 | 81 | 139 | 10 | MOLECULE: | FAB |
| 91:00:00 | 6qld-Y | 3.8 | 14.2 | 129 | 223 | 9 | MOLECULE: | INNER KINETOCHORE SUBUNIT MIF2 |
| 92:00:00 | 5n9j-E | 3.8 | 6.9 | 85 | 201 | 9 | MOLECULE: | MEDIATOR OF RNA POLYMERASE II TRANSCRIPTION |
| 93:00:00 | 7p3w-p | 3.8 | 11.8 | 107 | 155 | 9 | MOLECULE: | ATP SYNTHASE SUBUNIT ALPHA |
| 94:00:00 | 5w9a-A | 3.8 | 10.3 | 96 | 193 | 5 | MOLECULE: | TRIPARTITE MOTIF-CONTAINING PROTEIN 5 |
| 95:00:00 | 6gbq-G | 3.8 | 4.7 | 66 | 66 | 6 | MOLECULE: | POLYMERASE COFACTOR VP35 |
| 96:00:00 | 5lki-D | 3.8 | 11.5 | 146 | 1427 | 5 | MOLECULE: | TCDA1 |
| 97:00:00 | 5vwl-A | 3.8 | 9.4 | 82 | 105 | 7 | MOLECULE: | CYTOPLASMIC TAIL OF HIV-1 GP41 PROTEIN |
| 98:00:00 | 4lin-E | 3.8 | 10.3 | 139 | 289 | 6 | MOLECULE: | TAIL NEEDLE PROTEIN GP26 |
| 99:00:00 | 5t96-B | 3.7 | 5.4 | 68 | 330 | 4 | MOLECULE: | HE PROTEIN |
| 0:00 | 5y06-A | 3.7 | 12 | 101 | 235 | 16 | MOLECULE: | MSMEG_4306 |
| 1:00 | 4aur-A | 3.7 | 10.3 | 71 | 564 | 8 | MOLECULE: | LEOA |
| 2:00 | 2ehw-C | 3.7 | 2.1 | 47 | 117 | 9 | MOLECULE: | HYPOTHETICAL PROTEIN TTHB059 |
| 3:00 | 8q85-Z | 3.7 | 5.8 | 80 | 94 | 5 | MOLECULE: | KINETOCHORE PROTEIN NDC80 |
| 4:00 | 6kn7-U | 3.7 | 7.8 | 108 | 170 | 8 | MOLECULE: | ACTIN, ALPHA SKELETAL MUSCLE |
| 5:00 | 8x2q-A | 3.7 | 5 | 63 | 109 | 6 | MOLECULE: | ADENOMATOUS POLYPOSIS COLI PROTEIN |
| 6:00 | 4om3-D | 3.7 | 13.5 | 84 | 122 | 1 | MOLECULE: | TRANSDUCIN-LIKE ENHANCER PROTEIN 1 |
| 7:00 | 2ve7-D | 3.7 | 8 | 98 | 242 | 4 | MOLECULE: | KINETOCHORE PROTEIN HEC1, KINETOCHORE PROTEI |
| 8:00 | 6wuc-H | 3.7 | 16.2 | 83 | 179 | 8 | MOLECULE: | INNER KINETOCHORE SUBUNIT MCM16 |
| 9:00 | 8e4c-C | 3.7 | 9.6 | 73 | 142 | 10 | MOLECULE: | ISOFORM 2 OF IMMUNOGLOBULIN HEAVY CONSTANT M |
| 10:00 | 2p22-C | 3.7 | 8.5 | 107 | 186 | 5 | MOLECULE: | SUPPRESSOR PROTEIN STP22 OF TEMPERATURE-SENS |
| 11:00 | 8t1l-W | 3.7 | 15.3 | 71 | 119 | 3 | MOLECULE: | MEDIATOR OF RNA POLYMERASE II TRANSCRIPTION |
| 12:00 | 4tt0-B | 3.7 | 7.3 | 87 | 131 | 3 | MOLECULE: | DENEDDYLASE |
| 13:00 | 8th8-B | 3.7 | 37 | 145 | 276 | 8 | MOLECULE: | DYNEIN REGULATORY COMPLEX PROTEIN 1/2 N-TERM |
| 14:00 | 7sc0-A | 3.7 | 16.5 | 78 | 129 | 10 | MOLECULE: | CAVEOLIN-1 |
| 15:00 | 8ixk-E | 3.7 | 3 | 61 | 109 | 8 | MOLECULE: | ATTACHMENT PROTEIN G3P |
| 16:00 | 5xe3-E | 3.7 | 10.3 | 65 | 81 | 9 | MOLECULE: | ENDORIBONUCLEASE MAZF4 |
| 17:00 | 5ip0-D | 3.7 | 1.5 | 51 | 108 | 2 | MOLECULE: | PHA GRANULE-ASSOCIATED PROTEIN |
| 18:00 | 3vou-A | 3.7 | 5 | 68 | 139 | 7 | MOLECULE: | ION TRANSPORT 2 DOMAIN PROTEIN, VOLTAGE-GATE |
| 19:00 | 8tkp-A | 3.7 | 29.1 | 68 | 614 | 9 | MOLECULE: | TRANSMEMBRANE CHANNEL-LIKE PROTEIN 2 |
| 20:00 | 4mu6-A | 3.7 | 4 | 98 | 274 | 6 | MOLECULE: | KINECTIN 1 (KINESIN RECEPTOR) |
| 21:00 | 6z6o-C | 3.7 | 11.7 | 109 | 548 | 6 | MOLECULE: | HISTONE DEACETYLASE HDA1 |
| 22:00 | 6f5d-G | 3.7 | 7.2 | 68 | 277 | 4 | MOLECULE: | ATP SYNTHASE ALPHA CHAIN, MITOCHONDRIAL |

| | | | | | | | | |
|---|---|---|---|---|---|---|---|---|
| 23:00 | 8tdj-A | 3.7 | 31.8 | 70 | 467 | 11 | MOLECULE: | MECHANOSENSITIVE ION CHANNEL PROTEIN 10 |
| 24:00:00 | 7kon-T | 3.7 | 16.2 | 99 | 126 | 5 | MOLECULE: | ACTIN, ALPHA SKELETAL MUSCLE |
| 25:00:00 | 6vy1-F | 3.7 | 3.3 | 47 | 121 | 4 | MOLECULE: | PREFOLDIN SUBUNIT ALPHA 2 |
| 26:00:00 | 7e9t-A | 3.7 | 10.5 | 111 | 371 | 3 | MOLECULE: | SPIKE PROTEIN S2 |
| 27:00:00 | 5uhq-D | 3.7 | 5.6 | 56 | 81 | 5 | MOLECULE: | SUGAR TRANSPORTER SEMISWEET |
| 28:00:00 | 3zmf-C | 3.7 | 6.6 | 85 | 111 | 6 | MOLECULE: | GENERAL CONTROL PROTEIN GCN4, PUTATIVE INNER |
| 29:00:00 | 2yny-C | 3.7 | 6 | 80 | 105 | 9 | MOLECULE: | GENERAL CONTROL PROTEIN GCN4, PUTATIVE INNER |
| 30:00:00 | 2yo1-A | 3.7 | 6.8 | 103 | 273 | 11 | MOLECULE: | GENERAL CONTROL PROTEIN GCN4, PUTATIVE INNER |
| 31:00:00 | 3zmf-B | 3.7 | 4.8 | 81 | 111 | 7 | MOLECULE: | GENERAL CONTROL PROTEIN GCN4, PUTATIVE INNER |
| 32:00:00 | 2efl-A | 3.6 | 3.7 | 121 | 281 | 2 | MOLECULE: | FORMIN-BINDING PROTEIN 1 |
| 33:00:00 | 7p64-L | 3.6 | 4.8 | 84 | 606 | 4 | MOLECULE: | NADH-QUINONE OXIDOREDUCTASE SUBUNIT F |
| 34:00:00 | 8q3v-g | 3.6 | 7.4 | 71 | 76 | 7 | MOLECULE: | TETRAHYDROMETHANOPTERIN S-METHYLTRANSFERASE |
| 35:00:00 | 5oxf-A | 3.6 | 9.6 | 120 | 703 | 5 | MOLECULE: | GTP-BINDING PROTEIN |
| 36:00:00 | 9f63-A | 3.6 | 24 | 82 | 422 | 5 | MOLECULE: | PROTEIN PNS1 |
| 37:00:00 | 5c1f-A | 3.6 | 4.6 | 123 | 298 | 7 | MOLECULE: | SEPTATION PROTEIN IMP2 |
| 38:00:00 | 7jtk-C | 3.6 | 12.2 | 122 | 427 | 8 | MOLECULE: | FLAGELLAR RADIAL SPOKE PROTEIN 1 |
| 39:00:00 | 5odw-D | 3.6 | 2.9 | 65 | 198 | 9 | MOLECULE: | FERRIPYOVERDINE RECEPTOR |
| 40:00:00 | 4gcz-B | 3.6 | 7.2 | 82 | 378 | 7 | MOLECULE: | BLUE-LIGHT PHOTORECEPTOR, SENSOR PROTEIN FIX |
| 41:00:00 | 8qfc-D | 3.6 | 13.5 | 103 | 185 | 10 | MOLECULE: | 60S RIBOSOMAL PROTEIN L10A |
| 42:00:00 | 7tmp-M | 3.6 | 4.4 | 94 | 214 | 7 | MOLECULE: | H(+)-TRANSPORTING TWO-SECTOR ATPASE |
| 43:00:00 | 7tmq-H | 3.6 | 7.1 | 51 | 51 | 6 | MOLECULE: | H(+)-TRANSPORTING TWO-SECTOR ATPASE |
| 44:00:00 | 6sct-O | 3.6 | 2.6 | 59 | 59 | 5 | MOLECULE: | CLATHRIN HEAVY CHAIN |
| 45:00:00 | 6sct-J | 3.6 | 2.6 | 59 | 59 | 5 | MOLECULE: | CLATHRIN HEAVY CHAIN |
| 46:00:00 | 2yo1-B | 3.6 | 8.7 | 114 | 273 | 10 | MOLECULE: | GENERAL CONTROL PROTEIN GCN4, PUTATIVE INNER |
| 47:00:00 | 2yo2-A | 3.6 | 22.5 | 95 | 160 | 7 | MOLECULE: | GENERAL CONTROL PROTEIN GCN4, PUTATIVE INNER |
| 48:00:00 | 3wqa-C | 3.6 | 28.6 | 86 | 201 | 7 | MOLECULE: | TRIMERIC AUTOTRANSPORTER ADHESIN |
| 49:00:00 | 7zr1-D | 3.5 | 10 | 130 | 780 | 6 | MOLECULE: | DOUBLE-STRAND BREAK REPAIR PROTEIN |
| 50:00:00 | 2v0o-A | 3.5 | 5.2 | 115 | 273 | 6 | MOLECULE: | FCH DOMAIN ONLY PROTEIN 2 |
| 51:00:00 | 7nvr-n | 3.5 | 12.1 | 93 | 132 | 11 | MOLECULE: | TFIIH BASAL TRANSCRIPTION FACTOR COMPLEX HEL |
| 52:00:00 | 8wjn-A | 3.5 | 38.2 | 103 | 491 | 10 | MOLECULE: | STRUCTURAL MAINTENANCE OF CHROMOSOMES PROTEI |
| 53:00:00 | 6ted-R | 3.5 | 12.8 | 82 | 244 | 5 | MOLECULE: | DNA-DIRECTED RNA POLYMERASE SUBUNIT |
| 54:00:00 | 7vf2-C | 3.5 | 35.1 | 102 | 184 | 7 | MOLECULE: | PROTEIN VIRILIZER HOMOLOG |
| 55:00:00 | 8d8k-V | 3.5 | 15.5 | 113 | 233 | 11 | MOLECULE: | PROBABLE S-ADENOSYL-L-METHIONINE-DEPENDENT R |
| 56:00:00 | 3jac-A | 3.5 | 39.9 | 103 | 918 | 3 | MOLECULE: | PIEZO-TYPE MECHANOSENSITIVE ION CHANNEL COMP |
| 57:00:00 | 8rhn-A | 3.5 | 3.9 | 64 | 198 | 13 | MOLECULE: | ATPASE FAMILY GENE 2 PROTEIN HOMOLOG A |
| 58:00:00 | 6sp2-A | 3.5 | 9.7 | 59 | 366 | 7 | MOLECULE: | MEMBRANE PROTEIN TMS1D |
| 59:00:00 | 8cbk-F | 3.5 | 34.3 | 87 | 344 | 6 | MOLECULE: | 3-HYDROXYACYL-COA DEHYDROGENASE TYPE-2 |
| 60:00:00 | 6vq7-H | 3.5 | 3.7 | 94 | 214 | 10 | MOLECULE: | ATPASE H+-TRANSPORTING V1 SUBUNIT A |
| 61:00:00 | 4iff-C | 3.5 | 2.3 | 66 | 82 | 8 | MOLECULE: | FUSION OF PHAGE PHI29 GP7 PROTEIN AND CELL D |
| 62:00:00 | 4mh6-A | 3.5 | 3.6 | 85 | 159 | 6 | MOLECULE: | PUTATIVE TYPE III SECRETION PROTEIN YSCO |
| 63:00:00 | 6g72-Z | 3.5 | 11.9 | 91 | 139 | 7 | MOLECULE: | NADH-UBIQUINONE OXIDOREDUCTASE CHAIN 3 |
| 64:00:00 | 1jch-C | 3.5 | 15.3 | 82 | 468 | 12 | MOLECULE: | COLICIN E3 |
| 65:00:00 | 6w1s-W | 3.5 | 20.2 | 89 | 118 | 10 | MOLECULE: | MEDIATOR OF RNA POLYMERASE II TRANSCRIPTION |
| 66:00:00 | 5lki-E | 3.5 | 11.5 | 145 | 1427 | 6 | MOLECULE: | TCDA1 |
| 67:00:00 | 7pgg-A | 3.5 | 3.3 | 77 | 147 | 12 | MOLECULE: | ION TRANSPORT PROTEIN |
| 68:00:00 | 8ia3-E | 3.5 | 7.6 | 87 | 111 | 5 | MOLECULE: | UPSTREAM STIMULATORY FACTOR 2 |
| 69:00:00 | 6sct-E | 3.5 | 1.7 | 59 | 59 | 8 | MOLECULE: | CLATHRIN HEAVY CHAIN |
| 70:00:00 | 5ylz-r | 3.5 | 4.7 | 63 | 125 | 3 | MOLECULE: | PRE-MRNA-SPLICING FACTOR 8 |
| 71:00:00 | 6xl0-P | 3.5 | 9.4 | 82 | 271 | 7 | MOLECULE: | FLAGELLIN |
| 72:00:00 | 8ebt-K | 3.4 | 10.6 | 84 | 172 | 10 | MOLECULE: | GENERAL TRANSCRIPTION AND DNA REPAIR FACTOR |
| 73:00:00 | 7zu0-F | 3.4 | 8.5 | 59 | 112 | 8 | MOLECULE: | E3 UBIQUITIN-PROTEIN LIGASE PEP5 |

| | | | | | | | | |
|---|---|---|---|---|---|---|---|---|
| 74:00:00 | 5oqm-n | 3.4 | 15.1 | 93 | 136 | 5 | MOLECULE: | DNA-DIRECTED RNA POLYMERASE II SUBUNIT RPB1 |
| 75:00:00 | 8gb3-F | 3.4 | 4.3 | 70 | 90 | 13 | MOLECULE: | CHAPERONE PROTEIN DNAK |
| 76:00:00 | 8cr1-C | 3.4 | 7.8 | 73 | 266 | 3 | MOLECULE: | ATPASE ASNA1 |
| 77:00:00 | 3p8c-F | 3.4 | 10.6 | 105 | 156 | 8 | MOLECULE: | CYTOPLASMIC FMR1-INTERACTING PROTEIN 1 |
| 78:00:00 | 8b9z-h | 3.4 | 8.7 | 84 | 145 | 12 | MOLECULE: | NADH-UBIQUINONE OXIDOREDUCTASE CHAIN 3 |
| 79:00:00 | 7jrg-G | 3.4 | 9.4 | 64 | 70 | 5 | MOLECULE: | MITOCHONDRIAL-PROCESSING PEPTIDASE SUBUNIT B |
| 80:00:00 | 5wkq-A | 3.4 | 3.7 | 85 | 166 | 7 | MOLECULE: | INVASIN IPAB |
| 81:00:00 | 8b9z-Z | 3.4 | 33.6 | 100 | 146 | 7 | MOLECULE: | NADH-UBIQUINONE OXIDOREDUCTASE CHAIN 3 |
| 82:00:00 | 5n9j-U | 3.4 | 23 | 103 | 198 | 3 | MOLECULE: | MEDIATOR OF RNA POLYMERASE II TRANSCRIPTION |
| 83:00:00 | 8h36-G | 3.4 | 7.8 | 79 | 379 | 4 | MOLECULE: | E3 UBIQUITIN-PROTEIN LIGASE RBX1 |
| 84:00:00 | 4cgk-A | 3.4 | 2.4 | 84 | 351 | 5 | MOLECULE: | SECRETED 45 KDA PROTEIN |
| 85:00:00 | 4cq4-A | 3.4 | 3.7 | 78 | 106 | 10 | MOLECULE: | ENGINEERED VERSION OF TRANSMEMBRANE RECEPTOR |
| 86:00:00 | 4zwt-N | 3.4 | 3.6 | 61 | 125 | 10 | MOLECULE: | RECOMBINATION PROTEIN UVSY |
| 87:00:00 | 6qum-G | 3.4 | 3.8 | 93 | 207 | 9 | MOLECULE: | V-TYPE ATP SYNTHASE ALPHA CHAIN |
| 88:00:00 | 7fde-M | 3.4 | 5.6 | 104 | 218 | 7 | MOLECULE: | V-TYPE PROTON ATPASE SUBUNIT C |
| 89:00:00 | 5sxi-B | 3.4 | 3.3 | 75 | 209 | 3 | MOLECULE: | PHOSPHATIDYLINOSITOL 4,5-BISPHOSPHATE 3-KINA |
| 90:00:00 | 5sxf-B | 3.4 | 3.6 | 72 | 183 | 13 | MOLECULE: | PHOSPHATIDYLINOSITOL 4,5-BISPHOSPHATE 3-KINA |
| 91:00:00 | 6gol-A | 3.4 | 2.7 | 45 | 65 | 2 | MOLECULE: | HEMAGGLUTININ TRI-STALK |
| 92:00:00 | 8h37-J | 3.4 | 5.1 | 79 | 168 | 8 | MOLECULE: | CULLIN-3 |
| 93:00:00 | 8pjn-i | 3.4 | 6 | 60 | 395 | 3 | MOLECULE: | E3 UBIQUITIN-PROTEIN TRANSFERASE RMND5A |
| 94:00:00 | 8t1i-B | 3.4 | 11.4 | 92 | 158 | 8 | MOLECULE: | MEDIATOR OF RNA POLYMERASE II TRANSCRIPTION |
| 95:00:00 | 5b83-B | 3.4 | 2.4 | 58 | 59 | 3 | MOLECULE: | TETRA UBIQUITIN |
| 96:00:00 | 5ip0-A | 3.4 | 3.7 | 57 | 107 | 5 | MOLECULE: | PHA GRANULE-ASSOCIATED PROTEIN |
| 97:00:00 | 8h37-G | 3.4 | 3.4 | 79 | 168 | 11 | MOLECULE: | CULLIN-3 |
| 98:00:00 | 6h2f-B | 3.4 | 5.5 | 112 | 344 | 7 | MOLECULE: | AHLB |
| 99:00:00 | 6e95-A | 3.4 | 16.4 | 74 | 240 | 14 | MOLECULE: | STAPHYLOCOCCUS AUREUS AGRC HISTIDINE KINASE |
| 0:00 | 7aej-A | 3.4 | 7.3 | 72 | 129 | 8 | MOLECULE: | ENVELOPE GLYCOPROTEIN GP160,ENVELOPE GLYCOPR |
| 1:00 | 5jeq-A | 3.3 | 4.9 | 88 | 227 | 10 | MOLECULE: | NITRATE/NITRITE SENSOR PROTEIN NARQ |
| 2:00 | 8afz-B | 3.3 | 2.4 | 74 | 376 | 9 | MOLECULE: | SORTING NEXIN-1 |
| 3:00 | 7wkk-l | 3.3 | 4.9 | 88 | 171 | 10 | MOLECULE: | MGC83295 PROTEIN |
| 4:00 | 5n9j-A | 3.3 | 24.9 | 124 | 566 | 4 | MOLECULE: | MEDIATOR OF RNA POLYMERASE II TRANSCRIPTION |
| 5:00 | 6ikn-A | 3.3 | 4.1 | 115 | 305 | 6 | MOLECULE: | GROWTH ARREST-SPECIFIC PROTEIN 7 |
| 6:00 | 8gju-F | 3.3 | 12.9 | 81 | 332 | 4 | MOLECULE: | METHYLMALONIC ACIDURIA TYPE A PROTEIN, MITOC |
| 7:00 | 3mhh-C | 3.3 | 10.3 | 65 | 93 | 6 | MOLECULE: | UBIQUITIN CARBOXYL-TERMINAL HYDROLASE 8 |
| 8:00 | 6bpz-A | 3.3 | 39.8 | 107 | 900 | 5 | MOLECULE: | PIEZO-TYPE MECHANOSENSITIVE ION CHANNEL COMP |
| 9:00 | 8tek-E | 3.3 | 15.8 | 116 | 219 | 7 | MOLECULE: | DYNEIN REGULATORY COMPLEX PROTEIN 1/2 N-TERM |
| 10:00 | 7w5a-X | 3.3 | 10.3 | 87 | 87 | 0 | MOLECULE: | PRE-MRNA-PROCESSING-SPLICING FACTOR 8 |
| 11:00 | 8fef-D | 3.3 | 25.2 | 139 | 385 | 6 | MOLECULE: | VIRULENCE FACTOR MCE FAMILY PROTEIN |
| 12:00 | 5j9u-G | 3.3 | 9.9 | 59 | 297 | 8 | MOLECULE: | HISTONE ACETYLTRANSFERASE ESA1 |
| 13:00 | 7xzj-G | 3.3 | 2.6 | 53 | 73 | 9 | MOLECULE: | CTAP3 |
| 14:00 | 8i4t-R | 3.3 | 12.3 | 103 | 501 | 2 | MOLECULE: | ENVELOPMENT POLYPROTEIN |
| 15:00 | 8pmq-9 | 3.3 | 10 | 68 | 412 | 4 | MOLECULE: | E3 UBIQUITIN-PROTEIN LIGASE RMD5 |
| 16:00 | 6cnn-A | 3.3 | 38.8 | 68 | 360 | 3 | MOLECULE: | INTERMEDIATE CONDUCTANCE CALCIUM-ACTIVATED P |
| 17:00 | 8t1i-F | 3.3 | 5 | 60 | 73 | 13 | MOLECULE: | MEDIATOR OF RNA POLYMERASE II TRANSCRIPTION |
| 18:00 | 7emf-K | 3.3 | 14.2 | 82 | 112 | 6 | MOLECULE: | MEDIATOR OF RNA POLYMERASE II TRANSCRIPTION |
| 19:00 | 5yfp-F | 3.3 | 16.3 | 137 | 725 | 8 | MOLECULE: | EXOCYST COMPLEX COMPONENT SEC3 |
| 20:00 | 7aqw-m | 3.3 | 8.8 | 60 | 70 | 7 | MOLECULE: | NADH-UBIQUINONE OXIDOREDUCTASE CHAIN 5 |
| 21:00 | 6oee-A | 3.3 | 36.4 | 110 | 243 | 7 | MOLECULE: | TYPE IV SECRETION SYSTEM APPARATUS PROTEIN C |
| 22:00 | 6rd4-5 | 3.3 | 5.8 | 99 | 123 | 5 | MOLECULE: | ASA-10: POLYTOMELLA F-ATP SYNTHASE ASSOCIATE |
| 23:00 | 6lom-A | 3.3 | 19.5 | 98 | 295 | 5 | MOLECULE: | CALCIUM HOMEOSTASIS MODULATOR PROTEIN |
| 24:00:00 | 7zqy-B | 3.3 | 4 | 74 | 177 | 9 | MOLECULE: | DH DOMAIN-CONTAINING PROTEIN |

| | | | | | | | | |
|---|---|---|---|---|---|---|---|---|
| 25:00:00 | 6r0w-G | 3.3 | 3.7 | 93 | 207 | 9 | MOLECULE: | V-TYPE ATP SYNTHASE ALPHA CHAIN |
| 26:00:00 | 5sxc-B | 3.3 | 4.5 | 77 | 208 | 10 | MOLECULE: | PHOSPHATIDYLINOSITOL 4,5-BISPHOSPHATE 3-KINA |
| 27:00:00 | 6wm2-G | 3.3 | 4.4 | 97 | 213 | 11 | MOLECULE: | V-TYPE PROTON ATPASE SUBUNIT E 1 |
| 28:00:00 | 5knb-G | 3.3 | 4 | 79 | 164 | 8 | MOLECULE: | V-TYPE SODIUM ATPASE CATALYTIC SUBUNIT A |
| 29:00:00 | 6r0y-G | 3.3 | 3.2 | 91 | 206 | 9 | MOLECULE: | V-TYPE ATP SYNTHASE ALPHA CHAIN |
| 30:00:00 | 6wm4-G | 3.3 | 4.2 | 97 | 213 | 11 | MOLECULE: | V-TYPE PROTON ATPASE 116 KDA SUBUNIT A ISOFO |
| 31:00:00 | 5sxe-B | 3.3 | 3.3 | 73 | 225 | 3 | MOLECULE: | PHOSPHATIDYLINOSITOL 4,5-BISPHOSPHATE 3-KINA |
| 32:00:00 | 6w1s-Q | 3.3 | 8.4 | 78 | 131 | 12 | MOLECULE: | MEDIATOR OF RNA POLYMERASE II TRANSCRIPTION |
| 33:00:00 | 6o7v-a | 3.3 | 36.2 | 83 | 625 | 4 | MOLECULE: | V-TYPE PROTON ATPASE SUBUNIT D |
| 34:00:00 | 1ujw-B | 3.3 | 4.2 | 62 | 115 | 8 | MOLECULE: | VITAMIN B12 RECEPTOR |
| 35:00:00 | 6m3w-C | 3.3 | 40.4 | 132 | 344 | 2 | MOLECULE: | SPIKE GLYCOPROTEIN |
| 36:00:00 | 5j9t-L | 3.3 | 4.2 | 60 | 119 | 10 | MOLECULE: | HISTONE ACETYLTRANSFERASE ESA1 |
| 37:00:00 | 5lgt-A | 3.3 | 51.8 | 86 | 666 | 10 | MOLECULE: | LYSINE-SPECIFIC HISTONE DEMETHYLASE 1A |
| 38:00:00 | 8pmq-2 | 3.3 | 7.2 | 84 | 355 | 8 | MOLECULE: | E3 UBIQUITIN-PROTEIN LIGASE RMD5 |
| 39:00:00 | 6ip1-A | 3.3 | 5 | 63 | 63 | 5 | MOLECULE: | VESICLE-ASSOCIATED MEMBRANE PROTEIN 2 |
| 40:00:00 | 8bjw-F | 3.2 | 5.2 | 57 | 97 | 12 | MOLECULE: | PROTEIN C |
| 41:00:00 | 5wlz-C | 3.2 | 12 | 100 | 206 | 7 | MOLECULE: | DNA REPAIR PROTEIN XRCC4,MYOSIN-7 |
| 42:00:00 | 4p1m-B | 3.2 | 2.9 | 61 | 107 | 10 | MOLECULE: | CELL DIVISION PROTEIN ZAPA |
| 43:00:00 | 1kmi-Z | 3.2 | 9.5 | 97 | 177 | 8 | MOLECULE: | CHEMOTAXIS PROTEIN CHEY |
| 44:00:00 | 4h8s-C | 3.2 | 12 | 122 | 382 | 3 | MOLECULE: | DCC-INTERACTING PROTEIN 13-BETA |
| 45:00:00 | 5vjx-J | 3.2 | 4.3 | 52 | 64 | 12 | MOLECULE: | CLOCK-INTERACTING PACEMAKER |
| 46:00:00 | 3na7-A | 3.2 | 12.8 | 100 | 237 | 13 | MOLECULE: | HP0958 |
| 47:00:00 | 1llm-C | 3.2 | 5.2 | 54 | 87 | 4 | MOLECULE: | 5'-D(*TP*CP*CP*CP*AP*CP*GP*CP*GP*TP*GP*GP*G) |
| 48:00:00 | 4f52-E | 3.2 | 23.6 | 99 | 523 | 7 | MOLECULE: | CULLIN-1 |
| 49:00:00 | 5vk5-B | 3.2 | 4.1 | 61 | 282 | 5 | MOLECULE: | POTASSIUM CHANNEL SUBFAMILY K MEMBER 2 |
| 50:00:00 | 7x6g-D | 3.2 | 7.2 | 79 | 152 | 0 | MOLECULE: | QUORUM-SENSING REGULATOR PROTEIN G |
| 51:00:00 | 7w5a-2 | 3.2 | 1.8 | 61 | 123 | 18 | MOLECULE: | PRE-MRNA-PROCESSING-SPLICING FACTOR 8 |
| 52:00:00 | 4jio-B | 3.2 | 5.5 | 59 | 320 | 7 | MOLECULE: | BRO1 |
| 53:00:00 | 7nb6-A | 3.2 | 26.6 | 70 | 340 | 9 | MOLECULE: | AI-2 TRANSPORT PROTEIN TQSA |
| 54:00:00 | 5l4k-S | 3.2 | 10.4 | 83 | 491 | 12 | MOLECULE: | 26S PROTEASOME NON-ATPASE REGULATORY SUBUNIT |
| 55:00:00 | 8hhf-B | 3.2 | 3.9 | 68 | 89 | 6 | MOLECULE: | CELL DIVISION PROTEIN FTSQ |
| 56:00:00 | 6djl-C | 3.2 | 28.7 | 109 | 224 | 5 | MOLECULE: | RAS-RELATED PROTEIN RAB-11A |
| 57:00:00 | 6ek8-A | 3.2 | 4.7 | 82 | 249 | 9 | MOLECULE: | YAXB |
| 58:00:00 | 4ke2-B | 3.2 | 5.3 | 91 | 196 | 8 | MOLECULE: | TYPE I HYPERACTIVE ANTIFREEZE PROTEIN |
| 59:00:00 | 6qum-J | 3.2 | 11.3 | 93 | 185 | 12 | MOLECULE: | V-TYPE ATP SYNTHASE ALPHA CHAIN |
| 60:00:00 | 1ebo-A | 3.2 | 5.1 | 77 | 112 | 6 | MOLECULE: | EBOLA VIRUS ENVELOPE PROTEIN CHIMERA CONSIST |
| 61:00:00 | 8v8h-D | 3.2 | 3.5 | 81 | 239 | 6 | MOLECULE: | PHOSPHATIDYLINOSITOL 4,5-BISPHOSPHATE 3-KINA |
| 62:00:00 | 4egx-A | 3.2 | 5.3 | 67 | 180 | 10 | MOLECULE: | KINESIN-LIKE PROTEIN KIF1A |
| 63:00:00 | 3zmf-A | 3.2 | 3.7 | 79 | 111 | 5 | MOLECULE: | GENERAL CONTROL PROTEIN GCN4, PUTATIVE INNER |
| 64:00:00 | 6mdm-H | 3.1 | 20.4 | 79 | 147 | 8 | MOLECULE: | VESICLE-FUSING ATPASE |
| 65:00:00 | 8c5v-I | 3.1 | 8.7 | 141 | 516 | 13 | MOLECULE: | CHEMOTAXIS PROTEIN CHEA |
| 66:00:00 | 5yfp-A | 3.1 | 36.1 | 93 | 672 | 8 | MOLECULE: | EXOCYST COMPLEX COMPONENT SEC3 |
| 67:00:00 | 5twv-B | 3.1 | 11 | 86 | 1375 | 5 | MOLECULE: | ATP-SENSITIVE INWARD RECTIFIER POTASSIUM CHA |
| 68:00:00 | 8ftk-A | 3.1 | 26.9 | 106 | 1584 | 8 | MOLECULE: | 5'-3' RNA HELICASE-LIKE PROTEIN |
| 69:00:00 | 1esx-A | 3.1 | 16.6 | 72 | 96 | 7 | MOLECULE: | VPR PROTEIN |
| 70:00:00 | 4dyl-A | 3.1 | 4.4 | 115 | 376 | 8 | MOLECULE: | TYROSINE-PROTEIN KINASE FES/FPS |
| 71:00:00 | 8ikg-R | 3.1 | 8.4 | 61 | 275 | 5 | MOLECULE: | GUANINE NUCLEOTIDE-BINDING PROTEIN G(I) SUBU |
| 72:00:00 | 6v8o-C | 3.1 | 8.6 | 53 | 60 | 9 | MOLECULE: | HIGH TEMPERATURE LETHAL PROTEIN 1 |
| 73:00:00 | 8b6j-k | 3.1 | 6.3 | 57 | 58 | 9 | MOLECULE: | PEPTIDASE M16 INACTIVE DOMAIN PROTEIN |
| 74:00:00 | 8e1m-C | 3.1 | 17.4 | 70 | 264 | 6 | MOLECULE: | MITOCHONDRIAL IMPORT INNER MEMBRANE TRANSLOC |
| 75:00:00 | 7oi8-q | 3.1 | 16.5 | 109 | 164 | 4 | MOLECULE: | 39S RIBOSOMAL PROTEIN L2, MITOCHONDRIAL |

| | | | | | | | | |
|---|---|---|---|---|---|---|---|---|
| 76:00:00 | 8ppr-D | 3.1 | 16.4 | 129 | 248 | 8 | MOLECULE: | KINETOCHORE-ASSOCIATED PROTEIN DSN1 HOMOLOG |
| 77:00:00 | 7m2w-K | 3.1 | 15.4 | 88 | 95 | 9 | MOLECULE: | TUBULIN GAMMA CHAIN |
| 78:00:00 | 1ezj-A | 3.1 | 7.3 | 72 | 114 | 7 | MOLECULE: | NUCLEOCAPSID PHOSPHOPROTEIN |
| 79:00:00 | 8j8p-R | 3.1 | 10.7 | 60 | 69 | 2 | MOLECULE: | CTR9-LIKE PROTEIN |
| 80:00:00 | 8u7i-A | 3.1 | 54.6 | 94 | 626 | 12 | MOLECULE: | ENDONUCLEASE GAJA |
| 81:00:00 | 3f6n-B | 3.1 | 21.9 | 70 | 74 | 7 | MOLECULE: | VIRION-ASSOCIATED PROTEIN |
| 82:00:00 | 6nct-B | 3.1 | 3.2 | 72 | 225 | 3 | MOLECULE: | PHOSPHATIDYLINOSITOL 4,5-BISPHOSPHATE 3-KINA |
| 83:00:00 | 5nen-B | 3.1 | 3.7 | 91 | 213 | 15 | MOLECULE: | LIPASE C |
| 84:00:00 | 6el1-Q | 3.1 | 5 | 112 | 332 | 6 | MOLECULE: | YAXA |
| 85:00:00 | 2yo3-C | 3.1 | 11.5 | 125 | 257 | 6 | MOLECULE: | GENERAL CONTROL PROTEIN GCN4, PUTATIVE INNER |
| 86:00:00 | 6grj-A | 3.1 | 5.7 | 108 | 338 | 7 | MOLECULE: | AHLB |
| 87:00:00 | 5lm1-A | 3.1 | 11.6 | 118 | 339 | 6 | MOLECULE: | TYROSINE-PROTEIN PHOSPHATASE NON-RECEPTOR TY |
| 88:00:00 | 5n9j-G | 3.1 | 18.2 | 110 | 163 | 5 | MOLECULE: | MEDIATOR OF RNA POLYMERASE II TRANSCRIPTION |
| 89:00:00 | 1fzb-A | 3.1 | 2.3 | 55 | 81 | 9 | MOLECULE: | FIBRINOGEN |
| 90:00:00 | 7yo1-H | 3.1 | 21.2 | 68 | 263 | 7 | MOLECULE: | CALCIUM-ACTIVATED POTASSIUM CHANNEL SUBUNIT |
| 91:00:00 | 5lki-C | 3.1 | 11.5 | 145 | 1427 | 6 | MOLECULE: | TCDA1 |
| 92:00:00 | 7tke-U | 3.1 | 8.3 | 100 | 155 | 11 | MOLECULE: | ATP SYNTHASE SUBUNIT 9 |
| 93:00:00 | 2hod-D | 3.1 | 3.7 | 46 | 71 | 7 | MOLECULE: | FIBRINOGEN ALPHA CHAIN |
| 94:00:00 | 5vox-J | 3.1 | 10.8 | 82 | 104 | 9 | MOLECULE: | V-TYPE PROTON ATPASE CATALYTIC SUBUNIT A,V-T |
| 95:00:00 | 5ip0-P | 3.1 | 3.3 | 55 | 108 | 5 | MOLECULE: | PHA GRANULE-ASSOCIATED PROTEIN |
| 96:00:00 | 6el1-O | 3.1 | 61 | 116 | 332 | 9 | MOLECULE: | YAXA |
| 97:00:00 | 6vy2-D | 3.1 | 3.9 | 44 | 121 | 9 | MOLECULE: | GLYCOPROTEIN 120 |
| 98:00:00 | 6vy2-F | 3.1 | 3.9 | 44 | 121 | 9 | MOLECULE: | GLYCOPROTEIN 120 |
| 99:00:00 | 6vy2-B | 3.1 | 3.9 | 44 | 121 | 9 | MOLECULE: | GLYCOPROTEIN 120 |
| 0:00 | 1y2o-A | 3 | 5.6 | 99 | 248 | 5 | MOLECULE: | BAI1-ASSOCIATED PROTEIN 2 ISOFORM 1 |
| 1:00 | 4nqi-D | 3 | 5 | 97 | 232 | 7 | MOLECULE: | SH3 DOMAIN-CONTAINING PROTEIN |
| 2:00 | 4tko-B | 3 | 37.6 | 112 | 326 | 6 | MOLECULE: | EMRA |
| 3:00 | 6em5-t | 3 | 16.9 | 77 | 290 | 8 | MOLECULE: | 5.8S RIBOSOMAL RNA |
| 4:00 | 8ap7-m | 3 | 9.8 | 59 | 129 | 5 | MOLECULE: | ATP SYNTHASE SUBUNIT A |
| 5:00 | 8hqo-x | 3 | 7.3 | 77 | 80 | 8 | MOLECULE: | PORTAL PROTEIN |
| 6:00 | 6djl-E | 3 | 28.9 | 89 | 226 | 3 | MOLECULE: | RAS-RELATED PROTEIN RAB-11A |
| 7:00 | 2kn8-A | 3 | 10 | 68 | 68 | 7 | MOLECULE: | DNA CLEAVAGE AND PACKAGING PROTEIN LARGE SUB |
| 8:00 | 7b93-L | 3 | 22.2 | 94 | 607 | 9 | MOLECULE: | NADH-UBIQUINONE OXIDOREDUCTASE CHAIN 3 |
| 9:00 | 6c0f-8 | 3 | 9 | 71 | 98 | 10 | MOLECULE: | SACCHAROMYCES CEREVISIAE S288C 35S PRE-RIBOS |
| 10:00 | 8q85-d | 3 | 21.1 | 72 | 108 | 11 | MOLECULE: | KINETOCHORE PROTEIN NDC80 |
| 11:00 | 2py8-C | 3 | 7.2 | 61 | 123 | 8 | MOLECULE: | HYPOTHETICAL PROTEIN RBCX |
| 12:00 | 8t1l-X | 3 | 11.2 | 72 | 127 | 13 | MOLECULE: | MEDIATOR OF RNA POLYMERASE II TRANSCRIPTION |
| 13:00 | 8p1u-C | 3 | 9.9 | 82 | 93 | 5 | MOLECULE: | CELL DIVISION PROTEIN FTSL |
| 14:00 | 1ox3-A | 3 | 3.1 | 60 | 108 | 12 | MOLECULE: | FIBRITIN |
| 15:00 | 8kde-3 | 3 | 11 | 90 | 97 | 7 | MOLECULE: | PHOTOSYSTEM II CP47 REACTION CENTER PROTEIN |
| 16:00 | 8i4u-A | 3 | 7.7 | 124 | 453 | 6 | MOLECULE: | STRUCTURAL MAINTENANCE OF CHROMOSOMES PROTEI |
| 17:00 | 6el1-S | 3 | 5 | 112 | 332 | 6 | MOLECULE: | YAXA |
| 18:00 | 6r0z-J | 3 | 11.5 | 88 | 185 | 14 | MOLECULE: | V-TYPE ATP SYNTHASE ALPHA CHAIN |
| 19:00 | 1jad-A | 3 | 3.7 | 95 | 242 | 7 | MOLECULE: | PHOSPHOLIPASE C BETA |
| 20:00 | 4mvd-D | 3 | 5 | 100 | 253 | 6 | MOLECULE: | CHOLINE-PHOSPHATE CYTIDYLYLTRANSFERASE A |
| 21:00 | 3uum-B | 3 | 2.3 | 51 | 123 | 10 | MOLECULE: | UTROPHIN |
| 22:00 | 1qce-B | 3 | 3.6 | 62 | 123 | 15 | MOLECULE: | PROTEIN (GP41) |
| 23:00 | 1qce-C | 3 | 3.6 | 62 | 123 | 15 | MOLECULE: | PROTEIN (GP41) |
| 24:00:00 | 6el1-N | 3 | 5 | 112 | 332 | 6 | MOLECULE: | YAXA |
| 25:00:00 | 1qce-A | 3 | 3.6 | 62 | 123 | 15 | MOLECULE: | PROTEIN (GP41) |
| 26:00:00 | 6el1-K | 3 | 5.1 | 112 | 332 | 6 | MOLECULE: | YAXA |

| | | | | | | | | |
|---|---|---|---|---|---|---|---|---|
| 27:00:00 | 5uky-A | 3 | 2.7 | 43 | 74 | 5 | MOLECULE: | ATP-BINDING PROTEIN |
| 28:00:00 | 6vo3-B | 3 | 4.2 | 44 | 124 | 7 | MOLECULE: | PGV04 HEAVY CHAIN |
| 29:00:00 | 6ot1-D | 3 | 4.8 | 47 | 132 | 4 | MOLECULE: | ENVELOPE GLYCOPROTEIN GP41 |
| 30:00:00 | 3wpo-C | 3 | 6.9 | 98 | 193 | 8 | MOLECULE: | TRIMERIC AUTOTRANSPORTER ADHESIN |
| 31:00:00 | 4lin-B | 3 | 51.5 | 111 | 289 | 4 | MOLECULE: | TAIL NEEDLE PROTEIN GP26 |
| 32:00:00 | 6vo3-E | 3 | 4.2 | 44 | 124 | 7 | MOLECULE: | PGV04 HEAVY CHAIN |
| 33:00:00 | 1llm-D | 3 | 4.9 | 56 | 85 | 7 | MOLECULE: | 5'-D(*TP*CP*CP*CP*AP*CP*GP*CP*GP*TP*GP*GP*G) |
| 34:00:00 | 6vo3-F | 3 | 4.2 | 44 | 124 | 7 | MOLECULE: | PGV04 HEAVY CHAIN |
| 35:00:00 | 5vo5-A | 2.9 | 3.3 | 76 | 179 | 7 | MOLECULE: | COILED-COIL AND C2 DOMAIN-CONTAINING PROTEIN |
| 36:00:00 | 6zz8-A | 2.9 | 4.6 | 91 | 215 | 10 | MOLECULE: | CENTRIOLE PROTEIN |
| 37:00:00 | 8cpz-B | 2.9 | 8.1 | 146 | 2122 | 11 | MOLECULE: | TCDA1 |
| 38:00:00 | 8f7n-A | 2.9 | 7.5 | 111 | 364 | 10 | MOLECULE: | METHYL-ACCEPTING CHEMOTAXIS PROTEIN |
| 39:00:00 | 6ah0-w | 2.9 | 17.4 | 86 | 443 | 10 | MOLECULE: | U5SNRNA |
| 40:00:00 | 5c21-A | 2.9 | 5.1 | 92 | 267 | 8 | MOLECULE: | CHROMOSOMAL HEMOLYSIN D |
| 41:00:00 | 8auc-B | 2.9 | 33.5 | 123 | 493 | 7 | MOLECULE: | CELL WALL-ASSOCIATED HYDROLASES (INVASION-AS |
| 42:00:00 | 5td8-D | 2.9 | 3.6 | 65 | 128 | 11 | MOLECULE: | KINETOCHORE PROTEIN NDC80 |
| 43:00:00 | 4ilo-A | 2.9 | 18.4 | 102 | 236 | 8 | MOLECULE: | CT398 |
| 44:00:00 | 3wpa-A | 2.9 | 6.6 | 112 | 346 | 4 | MOLECULE: | TRIMERIC AUTOTRANSPORTER ADHESIN |
| 45:00:00 | 4hkr-A | 2.9 | 10.9 | 67 | 165 | 6 | MOLECULE: | CALCIUM RELEASE-ACTIVATED CALCIUM CHANNEL PR |
| 46:00:00 | 8eki-C | 2.9 | 29.5 | 109 | 701 | 7 | MOLECULE: | PROTEIN TRANSPORT PROTEIN SEC20 |
| 47:00:00 | 7sn7-a | 2.9 | 7 | 104 | 546 | 10 | MOLECULE: | FLAGELLIN |
| 48:00:00 | 6fj3-A | 2.9 | 9.7 | 56 | 562 | 5 | MOLECULE: | PARATHYROID HORMONE/PARATHYROID HORMONE-RELA |
| 49:00:00 | 4he8-D | 2.9 | 26.5 | 72 | 160 | 8 | MOLECULE: | NADH-QUINONE OXIDOREDUCTASE SUBUNIT 7 |
| 50:00:00 | 5nen-A | 2.9 | 3.7 | 91 | 266 | 3 | MOLECULE: | LIPASE C |
| 51:00:00 | 8f7c-A | 2.9 | 9.6 | 65 | 287 | 6 | MOLECULE: | PANNEXIN-2, SOLUBLE CYTOCHROME B562 FUSION |
| 52:00:00 | 6tpi-A | 2.9 | 7.4 | 105 | 380 | 10 | MOLECULE: | MUREIN HYDROLASE ACTIVATOR ENVC |
| 53:00:00 | 7xzj-A | 2.9 | 3.7 | 49 | 423 | 10 | MOLECULE: | CTAP3 |
| 54:00:00 | 8av6-H | 2.9 | 8.1 | 78 | 98 | 12 | MOLECULE: | RUVB-LIKE HELICASE |
| 55:00:00 | 4gwp-D | 2.9 | 8.7 | 77 | 121 | 6 | MOLECULE: | MEDIATOR OF RNA POLYMERASE II TRANSCRIPTION |
| 56:00:00 | 8hpo-C | 2.9 | 13 | 84 | 138 | 4 | MOLECULE: | TRANSCRIPTIONAL REGULATORY PROTEIN UME1 |
| 57:00:00 | 7xym-A | 2.9 | 15.4 | 114 | 488 | 10 | MOLECULE: | ENVELOPE GLYCOPROTEIN |
| 58:00:00 | 8xud-D | 2.9 | 10.7 | 64 | 651 | 8 | MOLECULE: | LIPOPROTEIN NLPI |
| 59:00:00 | 6zh3-A | 2.9 | 16.6 | 88 | 170 | 8 | MOLECULE: | VACUOLAR PROTEIN-SORTING-ASSOCIATED PROTEIN |
| 60:00:00 | 3j9v-M | 2.9 | 4.1 | 96 | 210 | 6 | MOLECULE: | V-TYPE PROTON ATPASE SUBUNIT A, VACUOLAR ISO |
| 61:00:00 | 2ezp-B | 2.9 | 3.8 | 64 | 123 | 11 | MOLECULE: | GP41 |
| 62:00:00 | 8ap9-G | 2.9 | 2.7 | 54 | 279 | 6 | MOLECULE: | ATP SYNTHASE GAMMA SUBUNIT |
| 63:00:00 | 2ezs-B | 2.9 | 3.9 | 63 | 123 | 11 | MOLECULE: | GP41 |
| 64:00:00 | 7yo1-D | 2.9 | 19 | 66 | 263 | 9 | MOLECULE: | CALCIUM-ACTIVATED POTASSIUM CHANNEL SUBUNIT |
| 65:00:00 | 2ezr-A | 2.9 | 3.9 | 63 | 123 | 11 | MOLECULE: | GP41 |
| 66:00:00 | 2ezq-B | 2.9 | 3.9 | 64 | 123 | 11 | MOLECULE: | GP41 |
| 67:00:00 | 2ezr-C | 2.9 | 3.9 | 63 | 123 | 11 | MOLECULE: | GP41 |
| 68:00:00 | 2ezp-C | 2.9 | 3.8 | 64 | 123 | 11 | MOLECULE: | GP41 |
| 69:00:00 | 2ezr-B | 2.9 | 3.9 | 63 | 123 | 11 | MOLECULE: | GP41 |
| 70:00:00 | 2ezs-A | 2.9 | 3.9 | 63 | 123 | 11 | MOLECULE: | GP41 |
| 71:00:00 | 2ezs-C | 2.9 | 3.9 | 63 | 123 | 11 | MOLECULE: | GP41 |
| 72:00:00 | 7jh6-D | 2.9 | 5.9 | 48 | 174 | 13 | MOLECULE: | TWO-DOMAIN DI-ZN(II) AND PORPHYRIN-BINDING P |
| 73:00:00 | 6o58-E | 2.9 | 16.4 | 106 | 270 | 6 | MOLECULE: | CALCIUM UNIPORTER PROTEIN, MITOCHONDRIAL |
| 74:00:00 | 5lhh-A | 2.9 | 50.5 | 84 | 666 | 11 | MOLECULE: | LYSINE-SPECIFIC HISTONE DEMETHYLASE 1A |
| 75:00:00 | 5t8f-B | 2.9 | 6.6 | 82 | 169 | 9 | MOLECULE: | PHOSPHATIDYLINOSITOL 4,5-BISPHOSPHATE 3-KINA |
| 76:00:00 | 7jh6-C | 2.9 | 5.9 | 48 | 174 | 13 | MOLECULE: | TWO-DOMAIN DI-ZN(II) AND PORPHYRIN-BINDING P |
| 77:00:00 | 7khw-D | 2.9 | 3.2 | 62 | 174 | 3 | MOLECULE: | TRANSLOCON ESPA |

| | | | | | | | | |
|---|---|---|---|---|---|---|---|---|
| 78:00:00 | 5k7b-D | 2.9 | 11 | 59 | 88 | 12 | MOLECULE: | BECLIN-2 |
| 79:00:00 | 6ot1-B | 2.9 | 3.8 | 44 | 132 | 5 | MOLECULE: | ENVELOPE GLYCOPROTEIN GP41 |
| 80:00:00 | 6ot1-O | 2.9 | 3.8 | 44 | 132 | 5 | MOLECULE: | ENVELOPE GLYCOPROTEIN GP41 |
| 81:00:00 | 7qen-B | 2.9 | 11.8 | 75 | 419 | 4 | MOLECULE: | DNA (35-MER) |
| 82:00:00 | 7nuv-A | 2.8 | 5.5 | 52 | 76 | 4 | MOLECULE: | AUX2PLS20 |
| 83:00:00 | 3qwe-A | 2.8 | 5.4 | 93 | 260 | 9 | MOLECULE: | GEM-INTERACTING PROTEIN |
| 84:00:00 | 6tdv-M | 2.8 | 19.2 | 89 | 166 | 12 | MOLECULE: | ATPTB1 |
| 85:00:00 | 4o8w-D | 2.8 | 9.4 | 67 | 120 | 10 | MOLECULE: | SPORE GERMINATION PROTEIN |
| 86:00:00 | 6gsi-J | 2.8 | 17.4 | 70 | 89 | 6 | MOLECULE: | CAPSID PROTEIN |
| 87:00:00 | 9b8o-U | 2.8 | 3.8 | 56 | 214 | 5 | MOLECULE: | ATPASE H+-TRANSPORTING V1 SUBUNIT D |
| 88:00:00 | 6m3q-F | 2.8 | 6.4 | 98 | 301 | 5 | MOLECULE: | ANKYRIN-2 |
| 89:00:00 | 7y22-e | 2.8 | 4.2 | 64 | 172 | 5 | MOLECULE: | PHAGE CONNECTOR PROTEIN |
| 90:00:00 | 7zmg-j | 2.8 | 12.2 | 65 | 75 | 5 | MOLECULE: | NADH-UBIQUINONE OXIDOREDUCTASE CHAIN 1 |
| 91:00:00 | 4nnh-A | 2.8 | 3.1 | 42 | 164 | 7 | MOLECULE: | 30S RIBOSOMAL PROTEIN S1 |
| 92:00:00 | 7bqx-G | 2.8 | 8 | 73 | 84 | 3 | MOLECULE: | TRIPLEX CAPSID PROTEIN 1 |
| 93:00:00 | 7c4j-C | 2.8 | 37.4 | 98 | 154 | 8 | MOLECULE: | TRANSCRIPTION REGULATORY PROTEIN SNF12 |
| 94:00:00 | 8q85-a | 2.8 | 7.3 | 78 | 234 | 4 | MOLECULE: | KINETOCHORE PROTEIN NDC80 |
| 95:00:00 | 3viq-A | 2.8 | 16.3 | 75 | 122 | 9 | MOLECULE: | SWI5-DEPENDENT RECOMBINATION DNA REPAIR PROT |
| 96:00:00 | 7wtt-z | 2.8 | 25.1 | 59 | 82 | 3 | MOLECULE: | 18S RRNA |
| 97:00:00 | 2wwb-B | 2.8 | 10.1 | 57 | 68 | 9 | MOLECULE: | PROTEIN TRANSPORT PROTEIN SEC61 SUBUNIT ALPH |
| 98:00:00 | 7qo4-J | 2.8 | 18.7 | 101 | 405 | 4 | MOLECULE: | 26S PROTEASOME REGULATORY SUBUNIT RPN1 |
| 99:00:00 | 6yvd-D | 2.8 | 8.3 | 73 | 414 | 8 | MOLECULE: | CONDENSIN COMPLEX SUBUNIT 2 |
| 0:00 | 6h2x-A | 2.8 | 8 | 126 | 352 | 10 | MOLECULE: | CHROMOSOME PARTITION PROTEIN MUKB,CHROMOSOME |
| 1:00 | 6rw8-B | 2.8 | 58 | 136 | 2337 | 5 | MOLECULE: | A COMPONENT OF INSECTICIDAL TOXIN COMPLEX (T |
| 2:00 | 5knd-G | 2.8 | 4.8 | 84 | 178 | 7 | MOLECULE: | V-TYPE SODIUM ATPASE CATALYTIC SUBUNIT A |
| 3:00 | 5y5y-G | 2.8 | 4.6 | 95 | 210 | 13 | MOLECULE: | V-TYPE ATP SYNTHASE ALPHA CHAIN |
| 4:00 | 2ezo-B | 2.8 | 3.8 | 63 | 123 | 11 | MOLECULE: | GP41 |
| 5:00 | 6o58-C | 2.8 | 16.4 | 112 | 268 | 4 | MOLECULE: | CALCIUM UNIPORTER PROTEIN, MITOCHONDRIAL |
| 6:00 | 2ezq-C | 2.8 | 3.7 | 63 | 123 | 11 | MOLECULE: | GP41 |
| 7:00 | 2ezq-A | 2.8 | 3.7 | 63 | 123 | 11 | MOLECULE: | GP41 |
| 8:00 | 2ezp-A | 2.8 | 4 | 63 | 123 | 11 | MOLECULE: | GP41 |
| 9:00 | 6wdo-C | 2.8 | 15.8 | 119 | 268 | 3 | MOLECULE: | CALCIUM UNIPORTER PROTEIN, MITOCHONDRIAL |
| 10:00 | 6ud4-H | 2.8 | 4.6 | 41 | 134 | 5 | MOLECULE: | GLUTAMATE RECEPTOR 2 |
| 11:00 | 6ud4-F | 2.8 | 4.6 | 41 | 134 | 5 | MOLECULE: | GLUTAMATE RECEPTOR 2 |
| 12:00 | 5j9t-D | 2.8 | 4.5 | 58 | 119 | 7 | MOLECULE: | HISTONE ACETYLTRANSFERASE ESA1 |
| 13:00 | 6vqa-N | 2.8 | 6.7 | 46 | 47 | 4 | MOLECULE: | ATPASE H+-TRANSPORTING V1 SUBUNIT A |
| 14:00 | 4r61-A | 2.8 | 2.1 | 52 | 140 | 19 | MOLECULE: | GP41-BASED CONSTRUCT COVNHR3-ABC |
| 15:00 | 7llk-B | 2.8 | 4.2 | 47 | 126 | 19 | MOLECULE: | ENVELOPE GLYCOPROTEIN GP120 |
| 16:00 | 7rai-B | 2.8 | 4.1 | 47 | 128 | 4 | MOLECULE: | ENVELOPE GLYCOPROTEIN GP160 |
| 17:00 | 8tni-F | 2.8 | 4.1 | 49 | 126 | 2 | MOLECULE: | HIV-1 BG505 DS-SOSIP GP120 |
| 18:00 | 4wbd-A | 2.7 | 7 | 65 | 540 | 8 | MOLECULE: | BSHC |
| 19:00 | 8odu-C | 2.7 | 4.8 | 70 | 232 | 7 | MOLECULE: | ATPASE GET3 |
| 20:00 | 8w8q-C | 2.7 | 8.5 | 53 | 287 | 6 | MOLECULE: | SOLUBLE CYTOCHROME B562,PROBABLE G-PROTEIN C |
| 21:00 | 5lc5-Z | 2.7 | 14.7 | 91 | 138 | 5 | MOLECULE: | NADH-UBIQUINONE OXIDOREDUCTASE CHAIN 3 |
| 22:00 | 1lgh-A | 2.7 | 5.1 | 47 | 56 | 9 | MOLECULE: | LIGHT HARVESTING COMPLEX II |
| 23:00 | 8b6j-i | 2.7 | 16.7 | 73 | 117 | 3 | MOLECULE: | PEPTIDASE M16 INACTIVE DOMAIN PROTEIN |
| 24:00:00 | 2mi2-A | 2.7 | 15.2 | 84 | 104 | 8 | MOLECULE: | SEC-INDEPENDENT PROTEIN TRANSLOCASE PROTEIN |
| 25:00:00 | 5lm2-B | 2.7 | 13.7 | 113 | 340 | 4 | MOLECULE: | TYROSINE-PROTEIN PHOSPHATASE NON-RECEPTOR TY |
| 26:00:00 | 5w82-B | 2.7 | 6.3 | 55 | 101 | 11 | MOLECULE: | PROTEIN DELTA |
| 27:00:00 | 8i3j-A | 2.7 | 5.1 | 96 | 277 | 7 | MOLECULE: | DYNEIN AXONEMAL HEAVY CHAIN 1 |
| 28:00:00 | 5n9j-Z | 2.7 | 6.9 | 75 | 119 | 4 | MOLECULE: | MEDIATOR OF RNA POLYMERASE II TRANSCRIPTION |

| | | | | | | | | |
|---|---|---|---|---|---|---|---|---|
| 29:00:00 | 6yj4-i | 2.7 | 8.6 | 61 | 86 | 10 | MOLECULE: | NADH-UBIQUINONE OXIDOREDUCTASE CHAIN 3 |
| 30:00:00 | 5efv-B | 2.7 | 27.1 | 93 | 635 | 8 | MOLECULE: | PHI ETA ORF 56-LIKE PROTEIN |
| 31:00:00 | 7xe4-F | 2.7 | 27.8 | 95 | 1503 | 6 | MOLECULE: | 1,3-BETA-GLUCAN SYNTHASE COMPONENT FKS1 |
| 32:00:00 | 8bf9-G | 2.7 | 9.2 | 64 | 158 | 5 | MOLECULE: | RNA (1766) |
| 33:00:00 | 6wg3-C | 2.7 | 31.6 | 58 | 248 | 10 | MOLECULE: | STRUCTURAL MAINTENANCE OF CHROMOSOMES PROTEI |
| 34:00:00 | 4wxa-E | 2.7 | 5.6 | 50 | 69 | 14 | MOLECULE: | EKC/KEOPS COMPLEX SUBUNIT PCC1 |
| 35:00:00 | 7w76-D | 2.7 | 5.4 | 64 | 168 | 2 | MOLECULE: | UBIQUITIN-CONJUGATING ENZYME E2 2 |
| 36:00:00 | 6szw-C | 2.7 | 13.7 | 57 | 186 | 9 | MOLECULE: | COMPLEMENT COMPONENT 1 Q SUBCOMPONENT-BINDIN |
| 37:00:00 | 6foc-G | 2.7 | 3.8 | 71 | 193 | 8 | MOLECULE: | ATP SYNTHASE SUBUNIT ALPHA,ATP SYNTHASE SUBU |
| 38:00:00 | 5sxa-B | 2.7 | 3.2 | 70 | 218 | 7 | MOLECULE: | PHOSPHATIDYLINOSITOL 4,5-BISPHOSPHATE 3-KINA |
| 39:00:00 | 7zqy-A | 2.7 | 3.5 | 73 | 175 | 4 | MOLECULE: | DH DOMAIN-CONTAINING PROTEIN |
| 40:00:00 | 3j9t-M | 2.7 | 4.8 | 97 | 210 | 5 | MOLECULE: | V-TYPE PROTON ATPASE SUBUNIT D |
| 41:00:00 | 6xj1-B | 2.7 | 6.8 | 124 | 277 | 3 | MOLECULE: | CELL DIVISION CONTROL PROTEIN 15 |
| 42:00:00 | 7tmo-I | 2.7 | 10 | 91 | 217 | 8 | MOLECULE: | H(+)-TRANSPORTING TWO-SECTOR ATPASE |
| 43:00:00 | 7fda-L | 2.7 | 14.2 | 91 | 111 | 7 | MOLECULE: | YEAST VACUOLAR ATPASE A SUBUNIT |
| 44:00:00 | 4zwr-C | 2.7 | 6 | 58 | 137 | 9 | MOLECULE: | RECOMBINATION PROTEIN UVSY |
| 45:00:00 | 8dk2-C | 2.7 | 15.9 | 144 | 723 | 3 | MOLECULE: | JETA |
| 46:00:00 | 2ezo-C | 2.7 | 3.5 | 61 | 123 | 15 | MOLECULE: | GP41 |
| 47:00:00 | 7khw-k | 2.7 | 3.2 | 62 | 174 | 3 | MOLECULE: | TRANSLOCON ESPA |
| 48:00:00 | 7khw-A | 2.7 | 3.2 | 62 | 174 | 3 | MOLECULE: | TRANSLOCON ESPA |
| 49:00:00 | 7khw-C | 2.7 | 3.2 | 62 | 174 | 3 | MOLECULE: | TRANSLOCON ESPA |
| 50:00:00 | 6wdn-G | 2.7 | 11.8 | 88 | 174 | 13 | MOLECULE: | CALCIUM UPTAKE PROTEIN 2, MITOCHONDRIAL |
| 51:00:00 | 7khw-v | 2.7 | 3.2 | 62 | 174 | 3 | MOLECULE: | TRANSLOCON ESPA |
| 52:00:00 | 7khw-V | 2.7 | 3.2 | 62 | 174 | 3 | MOLECULE: | TRANSLOCON ESPA |
| 53:00:00 | 7khw-x | 2.7 | 3.2 | 62 | 174 | 3 | MOLECULE: | TRANSLOCON ESPA |
| 54:00:00 | 7khw-Q | 2.7 | 3.2 | 62 | 174 | 3 | MOLECULE: | TRANSLOCON ESPA |
| 55:00:00 | 5lgn-A | 2.7 | 50.4 | 84 | 670 | 10 | MOLECULE: | LYSINE-SPECIFIC HISTONE DEMETHYLASE 1A |
| 56:00:00 | 7khw-J | 2.7 | 3.2 | 62 | 174 | 3 | MOLECULE: | TRANSLOCON ESPA |
| 57:00:00 | 7khw-e | 2.7 | 3.2 | 62 | 174 | 3 | MOLECULE: | TRANSLOCON ESPA |
| 58:00:00 | 6vqb-M | 2.7 | 6.2 | 47 | 47 | 0 | MOLECULE: | ATPASE H+-TRANSPORTING V1 SUBUNIT A |
| 59:00:00 | 7khw-w | 2.7 | 3.2 | 62 | 174 | 3 | MOLECULE: | TRANSLOCON ESPA |
| 60:00:00 | 7khw-p | 2.7 | 3.2 | 62 | 174 | 3 | MOLECULE: | TRANSLOCON ESPA |
| 61:00:00 | 7rai-D | 2.7 | 3.4 | 47 | 128 | 9 | MOLECULE: | ENVELOPE GLYCOPROTEIN GP160 |
| 62:00:00 | 6u59-B | 2.7 | 3.6 | 45 | 120 | 4 | MOLECULE: | SOSIP.664 GP120,SOSIP.664 GP120 |
| 63:00:00 | 8euu-D | 2.7 | 3.9 | 44 | 130 | 5 | MOLECULE: | ENVELOPE GLYCOPROTEIN GP120 |
| 64:00:00 | 7l7u-D | 2.7 | 4.3 | 47 | 145 | 11 | MOLECULE: | BG505 SOSIP.V5.2(7S) - GP120 |
| 65:00:00 | 7ur6-I | 2.7 | 4.3 | 45 | 132 | 4 | MOLECULE: | GP120 |
| 66:00:00 | 8ovw-Q | 2.6 | 19.1 | 110 | 258 | 4 | MOLECULE: | CENTROMERE-BINDING PROTEIN 1 |
| 67:00:00 | 7jw1-e | 2.6 | 9.6 | 100 | 235 | 4 | MOLECULE: | CAPSID PROTEINS |
| 68:00:00 | 8t8k-A | 2.6 | 3.3 | 45 | 183 | 13 | MOLECULE: | DUF507 FAMILY PROTEIN |
| 69:00:00 | 4u6u-D | 2.6 | 11.1 | 90 | 283 | 7 | MOLECULE: | COG7 |
| 70:00:00 | 6jfk-A | 2.6 | 4.5 | 81 | 428 | 9 | MOLECULE: | MITOFUSIN-2,CDNA FLJ57997, HIGHLY SIMILAR TO |
| 71:00:00 | 5a31-N | 2.6 | 28.7 | 78 | 703 | 3 | MOLECULE: | ANAPHASE-PROMOTING COMPLEX SUBUNIT 1 |
| 72:00:00 | 5t58-A | 2.6 | 25.6 | 95 | 233 | 9 | MOLECULE: | KLLA0F02343P |
| 73:00:00 | 4ad8-A | 2.6 | 9.1 | 104 | 452 | 11 | MOLECULE: | DNA REPAIR PROTEIN RECN |
| 74:00:00 | 1yx4-A | 2.6 | 37.7 | 83 | 112 | 6 | MOLECULE: | 26S PROTEASOME NON-ATPASE REGULATORY SUBUNIT |
| 75:00:00 | 6r1j-J | 2.6 | 4.3 | 90 | 264 | 4 | MOLECULE: | UNCHARACTERIZED PROTEIN |
| 76:00:00 | 8soi-B | 2.6 | 6.4 | 99 | 515 | 3 | MOLECULE: | RB1-INDUCIBLE COILED-COIL PROTEIN 1 |
| 77:00:00 | 8eup-3 | 2.6 | 11.1 | 61 | 194 | 7 | MOLECULE: | RNA (1422-MER) |
| 78:00:00 | 2gf4-A | 2.6 | 2.5 | 47 | 89 | 4 | MOLECULE: | PROTEIN VNG1086C |
| 79:00:00 | 5ncl-A | 2.6 | 25.2 | 69 | 418 | 7 | MOLECULE: | SERINE/THREONINE-PROTEIN KINASE CBK1 |

| | | | | | | | | |
|---|---|---|---|---|---|---|---|---|
| 80:00:00 | 8ixk-A | 2.6 | 4.7 | 43 | 145 | 5 | MOLECULE: | ATTACHMENT PROTEIN G3P |
| 81:00:00 | 7loi-A | 2.6 | 18.3 | 81 | 175 | 9 | MOLECULE: | TRANSMEMBRANE PROTEIN GP41 |
| 82:00:00 | 8i8b-H | 2.6 | 10.2 | 65 | 228 | 6 | MOLECULE: | MAJOR VIRAL CAPSID PROTEIN |
| 83:00:00 | 5oxf-C | 2.6 | 21 | 107 | 601 | 6 | MOLECULE: | GTP-BINDING PROTEIN |
| 84:00:00 | 6tmi-B | 2.6 | 8.4 | 97 | 246 | 5 | MOLECULE: | ATP SYNTHASE SUBUNIT ALPHA |
| 85:00:00 | 1flc-B | 2.6 | 3 | 57 | 162 | 5 | MOLECULE: | HAEMAGGLUTININ-ESTERASE-FUSION GLYCOPROTEIN |
| 86:00:00 | 6gy6-A | 2.6 | 6 | 99 | 365 | 10 | MOLECULE: | XAXA |
| 87:00:00 | 5c0w-K | 2.6 | 38.5 | 99 | 576 | 6 | MOLECULE: | EXOSOME COMPLEX COMPONENT RRP45 |
| 88:00:00 | 7mjp-A | 2.6 | 6.8 | 61 | 366 | 3 | MOLECULE: | ATP-SENSITIVE INWARD RECTIFIER POTASSIUM CHA |
| 89:00:00 | 8b9z-m | 2.6 | 12.2 | 79 | 108 | 5 | MOLECULE: | NADH-UBIQUINONE OXIDOREDUCTASE CHAIN 3 |
| 90:00:00 | 6xky-A | 2.6 | 5.3 | 101 | 271 | 8 | MOLECULE: | FLAGELLIN |
| 91:00:00 | 6xf1-A | 2.6 | 6.1 | 78 | 230 | 5 | MOLECULE: | NESPRIN-2 |
| 92:00:00 | 6v22-E | 2.6 | 3.4 | 65 | 199 | 3 | MOLECULE: | CALCIUM-ACTIVATED POTASSIUM CHANNEL SUBUNIT |
| 93:00:00 | 8h2j-A | 2.6 | 6.9 | 49 | 92 | 8 | MOLECULE: | P26 |
| 94:00:00 | 7w6x-A | 2.6 | 7.2 | 47 | 447 | 6 | MOLECULE: | REGULATOR OF SIGMA-E PROTEASE RSEP |
| 95:00:00 | 7yzp-C | 2.6 | 7.5 | 92 | 448 | 8 | MOLECULE: | DNA HAIRPIN (59-MER) |
| 96:00:00 | 5f5p-A | 2.6 | 3.1 | 56 | 181 | 9 | MOLECULE: | PROTEIN SHROOM2 |
| 97:00:00 | 6zka-p | 2.6 | 21.6 | 89 | 128 | 2 | MOLECULE: | NADH-UBIQUINONE OXIDOREDUCTASE CHAIN 3 |
| 98:00:00 | 5knc-G | 2.6 | 6.6 | 88 | 195 | 10 | MOLECULE: | V-TYPE SODIUM ATPASE CATALYTIC SUBUNIT A |
| 99:00:00 | 6ixv-B | 2.6 | 33.8 | 102 | 226 | 5 | MOLECULE: | SH3 DOMAIN-BINDING PROTEIN 5 |
| 0:00 | 7jtk-D | 2.6 | 15.3 | 127 | 427 | 5 | MOLECULE: | FLAGELLAR RADIAL SPOKE PROTEIN 1 |
| 1:00 | 6sih-E | 2.6 | 6.8 | 108 | 488 | 5 | MOLECULE: | FLAGELLAR HOOK-ASSOCIATED PROTEIN 2 |
| 2:00 | 3etz-B | 2.6 | 2.8 | 52 | 115 | 6 | MOLECULE: | ADHESIN A |
| 3:00 | 8at4-B | 2.6 | 10.4 | 149 | 597 | 7 | MOLECULE: | HAUS AUGMIN-LIKE COMPLEX SUBUNIT 1 |
| 4:00 | 6lth-I | 2.6 | 13.1 | 84 | 120 | 8 | MOLECULE: | TRANSCRIPTION ACTIVATOR BRG1 |
| 5:00 | 7khw-d | 2.6 | 3.2 | 62 | 174 | 3 | MOLECULE: | TRANSLOCON ESPA |
| 6:00 | 7khw-t | 2.6 | 3.2 | 62 | 174 | 3 | MOLECULE: | TRANSLOCON ESPA |
| 7:00 | 7khw-R | 2.6 | 3.2 | 62 | 174 | 3 | MOLECULE: | TRANSLOCON ESPA |
| 8:00 | 7khw-I | 2.6 | 3.2 | 62 | 174 | 3 | MOLECULE: | TRANSLOCON ESPA |
| 9:00 | 6elu-G | 2.6 | 8.6 | 49 | 168 | 0 | MOLECULE: | SERUM RESISTANCE ASSOCIATED VSG PROTEIN |
| 10:00 | 7khw-T | 2.6 | 3.2 | 62 | 174 | 3 | MOLECULE: | TRANSLOCON ESPA |
| 11:00 | 7khw-K | 2.6 | 3.2 | 62 | 174 | 3 | MOLECULE: | TRANSLOCON ESPA |
| 12:00 | 7khw-m | 2.6 | 3.2 | 62 | 174 | 3 | MOLECULE: | TRANSLOCON ESPA |
| 13:00 | 7khw-f | 2.6 | 3.2 | 62 | 174 | 3 | MOLECULE: | TRANSLOCON ESPA |
| 14:00 | 7khw-X | 2.6 | 3.2 | 62 | 174 | 3 | MOLECULE: | TRANSLOCON ESPA |
| 15:00 | 7khw-o | 2.6 | 3.2 | 62 | 174 | 3 | MOLECULE: | TRANSLOCON ESPA |
| 16:00 | 7khw-h | 2.6 | 3.2 | 62 | 174 | 3 | MOLECULE: | TRANSLOCON ESPA |
| 17:00 | 7khw-q | 2.6 | 3.2 | 62 | 174 | 3 | MOLECULE: | TRANSLOCON ESPA |
| 18:00 | 7khw-M | 2.6 | 3.2 | 62 | 174 | 3 | MOLECULE: | TRANSLOCON ESPA |
| 19:00 | 7khw-s | 2.6 | 3.2 | 62 | 174 | 3 | MOLECULE: | TRANSLOCON ESPA |
| 20:00 | 7khw-O | 2.6 | 3.2 | 62 | 174 | 3 | MOLECULE: | TRANSLOCON ESPA |
| 21:00 | 7khw-Z | 2.6 | 3.2 | 62 | 174 | 3 | MOLECULE: | TRANSLOCON ESPA |
| 22:00 | 6vq9-N | 2.6 | 6.8 | 46 | 47 | 9 | MOLECULE: | ATPASE H+-TRANSPORTING V1 SUBUNIT A |
| 23:00 | 7khw-G | 2.6 | 3.2 | 62 | 174 | 3 | MOLECULE: | TRANSLOCON ESPA |
| 24:00:00 | 7khw-E | 2.6 | 3.2 | 62 | 174 | 3 | MOLECULE: | TRANSLOCON ESPA |
| 25:00:00 | 7khw-H | 2.6 | 3.2 | 62 | 174 | 3 | MOLECULE: | TRANSLOCON ESPA |
| 26:00:00 | 7khw-j | 2.6 | 3.2 | 62 | 174 | 3 | MOLECULE: | TRANSLOCON ESPA |
| 27:00:00 | 7khw-c | 2.6 | 3.2 | 62 | 174 | 3 | MOLECULE: | TRANSLOCON ESPA |
| 28:00:00 | 7khw-B | 2.6 | 3.2 | 62 | 174 | 3 | MOLECULE: | TRANSLOCON ESPA |
| 29:00:00 | 7khw-S | 2.6 | 3.2 | 62 | 174 | 3 | MOLECULE: | TRANSLOCON ESPA |
| 30:00:00 | 7khw-l | 2.6 | 3.2 | 62 | 174 | 3 | MOLECULE: | TRANSLOCON ESPA |

| | | | | | | | | |
|---|---|---|---|---|---|---|---|---|
| 31:00:00 | 7khw-u | 2.6 | 3.2 | 62 | 174 | 3 | MOLECULE: | TRANSLOCON ESPA |
| 32:00:00 | 6wdo-G | 2.6 | 22.7 | 109 | 255 | 9 | MOLECULE: | CALCIUM UNIPORTER PROTEIN, MITOCHONDRIAL |
| 33:00:00 | 7khw-g | 2.6 | 3.2 | 62 | 174 | 3 | MOLECULE: | TRANSLOCON ESPA |
| 34:00:00 | 7khw-L | 2.6 | 3.2 | 62 | 174 | 3 | MOLECULE: | TRANSLOCON ESPA |
| 35:00:00 | 7khw-n | 2.6 | 3.2 | 62 | 174 | 3 | MOLECULE: | TRANSLOCON ESPA |
| 36:00:00 | 7khw-U | 2.6 | 3.2 | 62 | 174 | 3 | MOLECULE: | TRANSLOCON ESPA |
| 37:00:00 | 7khw-N | 2.6 | 3.2 | 62 | 174 | 3 | MOLECULE: | TRANSLOCON ESPA |
| 38:00:00 | 7khw-P | 2.6 | 3.2 | 62 | 174 | 3 | MOLECULE: | TRANSLOCON ESPA |
| 39:00:00 | 7khw-b | 2.6 | 3.2 | 62 | 174 | 3 | MOLECULE: | TRANSLOCON ESPA |
| 40:00:00 | 7khw-i | 2.6 | 3.2 | 62 | 174 | 3 | MOLECULE: | TRANSLOCON ESPA |
| 41:00:00 | 7khw-F | 2.6 | 3.2 | 62 | 174 | 3 | MOLECULE: | TRANSLOCON ESPA |
| 42:00:00 | 7khw-Y | 2.6 | 3.2 | 62 | 174 | 3 | MOLECULE: | TRANSLOCON ESPA |
| 43:00:00 | 8q0e-B | 2.6 | 3.2 | 49 | 241 | 8 | MOLECULE: | VARIANT SURFACE GLYCOPROTEIN 545 |
| 44:00:00 | 7khw-W | 2.6 | 3.2 | 62 | 174 | 3 | MOLECULE: | TRANSLOCON ESPA |
| 45:00:00 | 7khw-r | 2.6 | 3.2 | 62 | 174 | 3 | MOLECULE: | TRANSLOCON ESPA |
| 46:00:00 | 6x5b-E | 2.6 | 3.8 | 50 | 149 | 12 | MOLECULE: | ENVELOPE GLYCOPROTEIN GP120 |
| 47:00:00 | 6x9r-B | 2.6 | 3.6 | 42 | 121 | 5 | MOLECULE: | HIV-1 ENVELOPE GLYCOPROTEIN BG505 SOSIP.664 |
| 48:00:00 | 6x9t-B | 2.6 | 3.5 | 42 | 121 | 17 | MOLECULE: | HIV-1 ENVELOPE GLYCOPROTEIN BG505 SOSIP.664 |
| 49:00:00 | 8gpj-U | 2.6 | 3.8 | 43 | 128 | 9 | MOLECULE: | 8ANC195 FAB HEAVY CHAIN |
| 50:00:00 | 6vo0-E | 2.6 | 4.2 | 47 | 122 | 4 | MOLECULE: | 43A2 LIGHT CHAIN |
| 51:00:00 | 6nc3-M | 2.6 | 5.1 | 47 | 128 | 6 | MOLECULE: | HIV-1 ENV AMC011 V4.2 SOSIP GP120 |
| 52:00:00 | 6nc2-N | 2.6 | 12 | 45 | 129 | 7 | MOLECULE: | AMC011 V4.2 SOSIP GP120 |
| 53:00:00 | 6nc3-K | 2.6 | 3.5 | 44 | 128 | 9 | MOLECULE: | HIV-1 ENV AMC011 V4.2 SOSIP GP120 |
| 54:00:00 | 6vo1-B | 2.6 | 3.8 | 44 | 129 | 7 | MOLECULE: | ENVELOPE GLYCOPROTEIN GP120 |
| 55:00:00 | 8tni-D | 2.6 | 3.7 | 46 | 126 | 11 | MOLECULE: | HIV-1 BG505 DS-SOSIP GP120 |
| 56:00:00 | 7lpn-E | 2.6 | 4 | 43 | 121 | 9 | MOLECULE: | HIV-1 ENVELOPE GLYCOPROTEIN BG505 SOSIP.664 |
| 57:00:00 | 7l6o-f | 2.6 | 3.8 | 43 | 126 | 5 | MOLECULE: | CH848.3.D0949.10.17CHIM.6R.DS.SOSIP.664 - GP |
| 58:00:00 | 6nc3-B | 2.6 | 3.5 | 44 | 128 | 9 | MOLECULE: | HIV-1 ENV AMC011 V4.2 SOSIP GP120 |
| 59:00:00 | 6vo0-F | 2.6 | 4.2 | 47 | 122 | 4 | MOLECULE: | 43A2 LIGHT CHAIN |
| 60:00:00 | 8g4m-F | 2.6 | 4.1 | 45 | 131 | 11 | MOLECULE: | ENVELOPE GLYCOPROTEIN GP120 |
| 61:00:00 | 6vo0-B | 2.6 | 4.2 | 47 | 122 | 4 | MOLECULE: | 43A2 LIGHT CHAIN |
| 62:00:00 | 6vo1-F | 2.6 | 3.8 | 44 | 129 | 7 | MOLECULE: | ENVELOPE GLYCOPROTEIN GP120 |
| 63:00:00 | 2vky-B | 2.6 | 8.6 | 59 | 152 | 12 | MOLECULE: | TAIL PROTEIN, PIIGCN4 |
| 64:00:00 | 4gnk-E | 2.5 | 42.2 | 72 | 235 | 11 | MOLECULE: | GUANINE NUCLEOTIDE-BINDING PROTEIN G(Q) SUBU |
| 65:00:00 | 8ceg-B | 2.5 | 2.7 | 79 | 214 | 4 | MOLECULE: | CBY1-INTERACTING BAR DOMAIN-CONTAINING PROTE |
| 66:00:00 | 6a6x-C | 2.5 | 9.3 | 61 | 75 | 5 | MOLECULE: | PROBABLE ENDORIBONUCLEASE MAZF7 |
| 67:00:00 | 6f7j-A | 2.5 | 8.9 | 59 | 97 | 8 | MOLECULE: | SERRATE RNA EFFECTOR MOLECULE HOMOLOG |
| 68:00:00 | 5yfp-B | 2.5 | 24.9 | 144 | 927 | 6 | MOLECULE: | EXOCYST COMPLEX COMPONENT SEC3 |
| 69:00:00 | 7p5j-A | 2.5 | 38.4 | 59 | 399 | 7 | MOLECULE: | PROTEIN TWEETY HOMOLOG 1 |
| 70:00:00 | 6y1y-B | 2.5 | 5.5 | 68 | 128 | 10 | MOLECULE: | CHEA |
| 71:00:00 | 7o6t-A | 2.5 | 3.1 | 46 | 73 | 4 | MOLECULE: | PROTEIN VERNALIZATION INSENSITIVE 3 |
| 72:00:00 | 7jve-A | 2.5 | 14.2 | 85 | 253 | 4 | MOLECULE: | DSBA FAMILY PROTEIN |
| 73:00:00 | 8k32-C | 2.5 | 30.6 | 64 | 333 | 5 | MOLECULE: | KETOL-ACID REDUCTOISOMERASE (NADP(+)) |
| 74:00:00 | 8hpo-H | 2.5 | 27.7 | 99 | 233 | 7 | MOLECULE: | TRANSCRIPTIONAL REGULATORY PROTEIN UME1 |
| 75:00:00 | 5sva-V | 2.5 | 6.9 | 58 | 85 | 12 | MOLECULE: | DNA-DIRECTED RNA POLYMERASE II SUBUNIT RPB1 |
| 76:00:00 | 5wjt-A | 2.5 | 6.1 | 105 | 302 | 3 | MOLECULE: | FLAGELLIN |
| 77:00:00 | 7nyx-A | 2.5 | 12.7 | 127 | 1467 | 11 | MOLECULE: | CHROMOSOME PARTITION PROTEIN MUKB |
| 78:00:00 | 6zyy-C | 2.5 | 41.4 | 142 | 2901 | 6 | MOLECULE: | DYNEIN HEAVY CHAIN, OUTER ARM PROTEIN |
| 79:00:00 | 2yjf-M | 2.5 | 15.6 | 78 | 121 | 8 | MOLECULE: | ACTIN, ALPHA SKELETAL MUSCLE |
| 80:00:00 | 5n0c-A | 2.5 | 34.6 | 99 | 1293 | 5 | MOLECULE: | TETANUS TOXIN |
| 81:00:00 | 2yfb-B | 2.5 | 4.8 | 71 | 238 | 7 | MOLECULE: | METHYL-ACCEPTING CHEMOTAXIS TRANSDUCER |

| | | | | | | | | |
|---|---|---|---|---|---|---|---|---|
| 82:00:00 | 4abx-A | 2.5 | 6.6 | 82 | 167 | 7 | MOLECULE: | DNA REPAIR PROTEIN RECN |
| 83:00:00 | 7nad-8 | 2.5 | 8.3 | 58 | 58 | 5 | MOLECULE: | 25S RRNA |
| 84:00:00 | 2lzs-A | 2.5 | 11 | 47 | 48 | 13 | MOLECULE: | SEC-INDEPENDENT PROTEIN TRANSLOCASE PROTEIN |
| 85:00:00 | 7vcf-C | 2.5 | 7.1 | 75 | 217 | 8 | MOLECULE: | TIC214 |
| 86:00:00 | 1few-A | 2.5 | 4 | 75 | 173 | 8 | MOLECULE: | SECOND MITOCHONDRIA-DERIVED ACTIVATOR OF CAS |
| 87:00:00 | 6x0a-a | 2.5 | 41 | 66 | 87 | 6 | MOLECULE: | PLASMID STABILIZATION SYSTEM |
| 88:00:00 | 7qtt-Y | 2.5 | 47.1 | 110 | 306 | 8 | MOLECULE: | SPLICING FACTOR 3B SUBUNIT 3 |
| 89:00:00 | 1ej6-D | 2.5 | 12.4 | 60 | 417 | 7 | MOLECULE: | LAMBDA2 |
| 90:00:00 | 6uz8-A | 2.5 | 51.8 | 119 | 734 | 9 | MOLECULE: | SHORT TRANSIENT RECEPTOR POTENTIAL CHANNEL 6 |
| 91:00:00 | 7jrj-G | 2.5 | 11.2 | 78 | 148 | 9 | MOLECULE: | RADIAL SPOKE PROTEIN 10 |
| 92:00:00 | 6h7w-O | 2.5 | 4.4 | 81 | 220 | 7 | MOLECULE: | VACUOLAR PROTEIN SORTING-ASSOCIATED PROTEIN |
| 93:00:00 | 5wwl-M | 2.5 | 24.3 | 99 | 214 | 2 | MOLECULE: | CENTROMERE PROTEIN MIS12 |
| 94:00:00 | 8fbi-A | 2.5 | 8.6 | 48 | 274 | 2 | MOLECULE: | KWOCA_39 |
| 95:00:00 | 7egm-A | 2.5 | 5.8 | 68 | 109 | 4 | MOLECULE: | TRANSCRIPTION REGULATORY PROTEIN SNF2 |
| 96:00:00 | 8eja-A | 2.5 | 3.6 | 39 | 113 | 10 | MOLECULE: | ABAK |
| 97:00:00 | 6xl0-B | 2.5 | 9.4 | 82 | 271 | 7 | MOLECULE: | FLAGELLIN |
| 98:00:00 | 6xl0-D | 2.5 | 9.4 | 82 | 271 | 7 | MOLECULE: | FLAGELLIN |
| 99:00:00 | 6xl0-G | 2.5 | 9.4 | 82 | 271 | 7 | MOLECULE: | FLAGELLIN |
| 0:00 | 6xl0-I | 2.5 | 9.4 | 82 | 271 | 7 | MOLECULE: | FLAGELLIN |
| 1:00 | 5xi5-E | 2.5 | 10.9 | 61 | 120 | 5 | MOLECULE: | TUBULIN ALPHA CHAIN |
| 2:00 | 6xl0-R | 2.5 | 9.4 | 82 | 271 | 7 | MOLECULE: | FLAGELLIN |
| 3:00 | 8v8j-B | 2.5 | 4.4 | 84 | 239 | 5 | MOLECULE: | PHOSPHATIDYLINOSITOL 4,5-BISPHOSPHATE 3-KINA |
| 4:00 | 6xl0-F | 2.5 | 9.4 | 82 | 271 | 7 | MOLECULE: | FLAGELLIN |
| 5:00 | 6xl0-H | 2.5 | 9.4 | 82 | 271 | 7 | MOLECULE: | FLAGELLIN |
| 6:00 | 6vqb-O | 2.5 | 6.8 | 46 | 47 | 9 | MOLECULE: | ATPASE H+-TRANSPORTING V1 SUBUNIT A |
| 7:00 | 6xl0-J | 2.5 | 9.4 | 82 | 271 | 7 | MOLECULE: | FLAGELLIN |
| 8:00 | 8g6u-B | 2.5 | 2.9 | 46 | 120 | 11 | MOLECULE: | CRF01_AE T/F100 HIV-1 GP120 |
| 9:00 | 3wpo-B | 2.5 | 9.1 | 54 | 196 | 4 | MOLECULE: | TRIMERIC AUTOTRANSPORTER ADHESIN |
| 10:00 | 6ce0-B | 2.5 | 3.7 | 43 | 137 | 9 | MOLECULE: | ENVELOPE GLYCOPROTEIN GP160 |
| 11:00 | 6x5c-G | 2.5 | 3 | 50 | 148 | 6 | MOLECULE: | ENVELOPE GLYCOPROTEIN GP160 |
| 12:00 | 7rai-F | 2.5 | 3.9 | 46 | 128 | 4 | MOLECULE: | ENVELOPE GLYCOPROTEIN GP160 |
| 13:00 | 6nc3-J | 2.5 | 3.4 | 43 | 128 | 9 | MOLECULE: | HIV-1 ENV AMC011 V4.2 SOSIP GP120 |
| 14:00 | 8d5c-Y | 2.5 | 4.4 | 50 | 129 | 4 | MOLECULE: | ENVELOPE GLYCOPROTEIN GP41 |
| 15:00 | 7mdt-D | 2.5 | 4.3 | 44 | 122 | 7 | MOLECULE: | SURFACE PROTEIN GP120 |
| 16:00 | 8g6u-F | 2.5 | 2.9 | 46 | 120 | 11 | MOLECULE: | CRF01_AE T/F100 HIV-1 GP120 |
| 17:00 | 8saq-F | 2.5 | 4.6 | 50 | 127 | 2 | MOLECULE: | CH848.0526.25 GP120 |
| 18:00 | 6u0n-Z | 2.5 | 3.8 | 50 | 124 | 2 | MOLECULE: | ENVELOPE GLYCOPROTEIN GP120 |
| 19:00 | 6opq-B | 2.5 | 2.9 | 47 | 128 | 9 | MOLECULE: | ENVELOPE GLYCOPROTEIN GP160 |
| 20:00 | 6nc3-I | 2.5 | 3.4 | 43 | 128 | 9 | MOLECULE: | HIV-1 ENV AMC011 V4.2 SOSIP GP120 |
| 21:00 | 6vo1-E | 2.5 | 4 | 44 | 129 | 9 | MOLECULE: | ENVELOPE GLYCOPROTEIN GP120 |
| 22:00 | 6um6-J | 2.5 | 4.4 | 45 | 132 | 7 | MOLECULE: | CH848 10.17DT GP120 |
| 23:00 | 6x9s-B | 2.5 | 3.8 | 42 | 121 | 5 | MOLECULE: | HIV-1 ENVELOPE GLYCOPROTEIN BG505 SOSIP.664 |
| 24:00:00 | 8g6u-J | 2.5 | 2.9 | 46 | 120 | 11 | MOLECULE: | CRF01_AE T/F100 HIV-1 GP120 |
| 25:00:00 | 6nc3-N | 2.5 | 3.4 | 43 | 128 | 9 | MOLECULE: | HIV-1 ENV AMC011 V4.2 SOSIP GP120 |
| 26:00:00 | 6um6-F | 2.5 | 4.2 | 44 | 132 | 7 | MOLECULE: | CH848 10.17DT GP120 |
| 27:00:00 | 7nnl-B | 2.4 | 17 | 70 | 682 | 1 | MOLECULE: | POTASSIUM-TRANSPORTING ATPASE POTASSIUM-BIND |
| 28:00:00 | 4d6k-A | 2.4 | 7 | 46 | 76 | 4 | MOLECULE: | DEOXYNUCLEOTIDYLTRANSFERASE TERMINAL-INTERAC |
| 29:00:00 | 7wb4-E | 2.4 | 24 | 60 | 1363 | 5 | MOLECULE: | OUTER NUP133 |
| 30:00:00 | 6sl1-A | 2.4 | 18.2 | 77 | 2651 | 5 | MOLECULE: | SERINE/THREONINE-PROTEIN KINASE TEL1 |
| 31:00:00 | 6zbj-A | 2.4 | 36.7 | 130 | 666 | 5 | MOLECULE: | PRECURSOR OF THE MAJOR MEROZOITE SURFACE ANT |
| 32:00:00 | 6nr8-4 | 2.4 | 2.8 | 50 | 104 | 2 | MOLECULE: | PREFOLDIN SUBUNIT 1 |

| | | | | | | | | |
|---|---|---|---|---|---|---|---|---|
| 33:00:00 | 6s3l-K | 2.4 | 5.1 | 58 | 193 | 5 | MOLECULE: | FLAGELLAR BIOSYNTHETIC PROTEIN FLIP |
| 34:00:00 | 1fxk-C | 2.4 | 5.7 | 60 | 133 | 7 | MOLECULE: | PREFOLDIN |
| 35:00:00 | 5jr9-A | 2.4 | 7.4 | 58 | 188 | 14 | MOLECULE: | NEQ131 |
| 36:00:00 | 6qum-N | 2.4 | 8.5 | 65 | 649 | 9 | MOLECULE: | V-TYPE ATP SYNTHASE ALPHA CHAIN |
| 37:00:00 | 7xdi-D | 2.4 | 4.9 | 51 | 124 | 4 | MOLECULE: | VP1 |
| 38:00:00 | 5d2s-A | 2.4 | 16 | 58 | 91 | 16 | MOLECULE: | FIBROIN-MODULATOR-BINDING PROTEIN-1 |
| 39:00:00 | 4uy3-A | 2.4 | 4.5 | 71 | 194 | 7 | MOLECULE: | SEPTATION RING FORMATION REGULATOR EZRA |
| 40:00:00 | 6xby-a | 2.4 | 46.8 | 82 | 494 | 7 | MOLECULE: | V-TYPE PROTON ATPASE CATALYTIC SUBUNIT A |
| 41:00:00 | 5z25-A | 2.4 | 6.1 | 51 | 96 | 8 | MOLECULE: | CYTOCHROME C552 |
| 42:00:00 | 3edv-A | 2.4 | 9.7 | 88 | 322 | 6 | MOLECULE: | SPECTRIN BETA CHAIN, BRAIN 1 |
| 43:00:00 | 6ye4-A | 2.4 | 3.9 | 83 | 236 | 8 | MOLECULE: | BIOPOLYMER TRANSPORT PROTEIN EXBB |
| 44:00:00 | 5j1i-A | 2.4 | 6.6 | 105 | 346 | 10 | MOLECULE: | PLECTIN |
| 45:00:00 | 5i4q-B | 2.4 | 2.7 | 45 | 107 | 9 | MOLECULE: | CONTACT-DEPENDENT INHIBITOR A |
| 46:00:00 | 3caz-B | 2.4 | 6.1 | 82 | 210 | 5 | MOLECULE: | BAR PROTEIN |
| 47:00:00 | 2kog-A | 2.4 | 21.9 | 76 | 116 | 7 | MOLECULE: | VESICLE-ASSOCIATED MEMBRANE PROTEIN 2 |
| 48:00:00 | 2iaz-B | 2.4 | 13.6 | 55 | 113 | 5 | MOLECULE: | HYPOTHETICAL PROTEIN SP1372 |
| 49:00:00 | 8z9a-A | 2.4 | 8.8 | 64 | 406 | 14 | MOLECULE: | ODORANT RECEPTOR, APISORCO |
| 50:00:00 | 4yxw-G | 2.4 | 4.1 | 77 | 213 | 5 | MOLECULE: | ATP SYNTHASE SUBUNIT ALPHA, MITOCHONDRIAL |
| 51:00:00 | 3g3z-A | 2.4 | 12.3 | 58 | 142 | 2 | MOLECULE: | TRANSCRIPTIONAL REGULATOR, MARR FAMILY |
| 52:00:00 | 7uh4-A | 2.4 | 2.3 | 63 | 110 | 5 | MOLECULE: | LXG-ASSOCIATED ALPHA-HELICAL PROTEIN D2 |
| 53:00:00 | 4uig-A | 2.4 | 5.3 | 44 | 91 | 7 | MOLECULE: | COPPER SENSITIVE OPERON REPRESSOR |
| 54:00:00 | 6uz3-A | 2.4 | 30.6 | 67 | 1126 | 4 | MOLECULE: | SODIUM CHANNEL PROTEIN TYPE 5 SUBUNIT ALPHA, |
| 55:00:00 | 7pde-A | 2.4 | 12.6 | 86 | 846 | 3 | MOLECULE: | ADENYLATE CYCLASE 9 |
| 56:00:00 | 5z7b-B | 2.4 | 10 | 49 | 197 | 4 | MOLECULE: | PADR FAMILY TRANSCRIPTIONAL REGULATOR |
| 57:00:00 | 5td8-C | 2.4 | 9.2 | 73 | 111 | 8 | MOLECULE: | KINETOCHORE PROTEIN NDC80 |
| 58:00:00 | 2nb1-A | 2.4 | 7.5 | 46 | 60 | 4 | MOLECULE: | TUMOR PROTEIN 63 |
| 59:00:00 | 7qj0-L | 2.4 | 10.4 | 64 | 566 | 5 | MOLECULE: | MITOTIC-SPINDLE ORGANIZING PROTEIN 1 |
| 60:00:00 | 8ulg-A | 2.4 | 7.2 | 60 | 827 | 7 | MOLECULE: | ROD CGMP-SPECIFIC 3',5'-CYCLIC PHOSPHODIESTE |
| 61:00:00 | 1x04-A | 2.4 | 2.7 | 72 | 200 | 8 | MOLECULE: | SH3-CONTAINING GRB2-LIKE PROTEIN 2 |
| 62:00:00 | 6gje-C | 2.4 | 7.8 | 64 | 83 | 5 | MOLECULE: | PROTEIN AMNIONLESS |
| 63:00:00 | 3r6n-A | 2.4 | 5.1 | 123 | 450 | 11 | MOLECULE: | DESMOPLAKIN |
| 64:00:00 | 3rx6-A | 2.4 | 8.7 | 73 | 187 | 4 | MOLECULE: | POLARITY SUPPRESSION PROTEIN |
| 65:00:00 | 4tql-B | 2.4 | 3.5 | 83 | 235 | 11 | MOLECULE: | THREE HELIX BUNDLE |
| 66:00:00 | 6sl2-A | 2.4 | 4.5 | 121 | 619 | 5 | MOLECULE: | CALPONIN HOMOLOGY DOMAIN PROTEIN PUTATIVE |
| 67:00:00 | 5j9q-D | 2.4 | 5.6 | 63 | 120 | 8 | MOLECULE: | HISTONE ACETYLTRANSFERASE ESA1 |
| 68:00:00 | 2zdi-C | 2.4 | 11.9 | 71 | 148 | 8 | MOLECULE: | PREFOLDIN SUBUNIT BETA |
| 69:00:00 | 1sc7-A | 2.4 | 5.9 | 55 | 567 | 7 | MOLECULE: | 5'-D(*AP*AP*AP*AP*AP*GP*AP*CP*TP*T)-3' |
| 70:00:00 | 8i4u-B | 2.4 | 7.1 | 129 | 458 | 9 | MOLECULE: | STRUCTURAL MAINTENANCE OF CHROMOSOMES PROTEI |
| 71:00:00 | 6sih-G | 2.4 | 7.6 | 102 | 488 | 8 | MOLECULE: | FLAGELLAR HOOK-ASSOCIATED PROTEIN 2 |
| 72:00:00 | 6h7w-M | 2.4 | 3.3 | 85 | 220 | 6 | MOLECULE: | VACUOLAR PROTEIN SORTING-ASSOCIATED PROTEIN |
| 73:00:00 | 6nyk-B | 2.4 | 3.1 | 51 | 90 | 8 | MOLECULE: | DESIGN CONSTRUCT XAX_GGDQ |
| 74:00:00 | 6kn7-b | 2.4 | 40 | 93 | 170 | 11 | MOLECULE: | ACTIN, ALPHA SKELETAL MUSCLE |
| 75:00:00 | 6el1-I | 2.4 | 8.1 | 109 | 351 | 9 | MOLECULE: | YAXA |
| 76:00:00 | 7lic-A | 2.4 | 5.2 | 51 | 377 | 4 | MOLECULE: | MHOR5 |
| 77:00:00 | 7lic-C | 2.4 | 5 | 51 | 377 | 2 | MOLECULE: | MHOR5 |
| 78:00:00 | 7lic-D | 2.4 | 5 | 51 | 377 | 2 | MOLECULE: | MHOR5 |
| 79:00:00 | 5xiw-E | 2.4 | 6.1 | 55 | 121 | 5 | MOLECULE: | TUBULIN ALPHA-1B CHAIN |
| 80:00:00 | 8eav-F | 2.4 | 2.4 | 44 | 159 | 0 | MOLECULE: | YAR027W OR YAR028W |
| 81:00:00 | 6v8x-J | 2.4 | 4.6 | 54 | 128 | 9 | MOLECULE: | ENVELOPE GLYCOPROTEIN GP120 |
| 82:00:00 | 2mk3-A | 2.4 | 12.3 | 56 | 68 | 5 | MOLECULE: | TRANSMEMBRANE GLYCOPROTEIN, CHIMERIC CONSTRU |
| 83:00:00 | 8euu-F | 2.4 | 3.9 | 44 | 130 | 5 | MOLECULE: | ENVELOPE GLYCOPROTEIN GP120 |

| | | | | | | | | |
|---|---|---|---|---|---|---|---|---|
| 84:00:00 | 7mep-E | 2.4 | 3.4 | 44 | 123 | 16 | MOLECULE: | RM19R MAB LIGHT CHAIN |
| 85:00:00 | 6osy-A | 2.4 | 4.5 | 44 | 132 | 7 | MOLECULE: | BG505 GP120 |
| 86:00:00 | 7tcn-F | 2.4 | 10.1 | 49 | 122 | 2 | MOLECULE: | ENVELOPE GLYCOPROTEIN GP160 |
| 87:00:00 | 6cue-1 | 2.4 | 4.2 | 45 | 132 | 2 | MOLECULE: | ENVELOPE GLYCOPROTEIN GP120 |
| 88:00:00 | 6v8x-B | 2.4 | 4.6 | 54 | 128 | 9 | MOLECULE: | ENVELOPE GLYCOPROTEIN GP120 |
| 89:00:00 | 5i8h-B | 2.4 | 4.9 | 45 | 132 | 7 | MOLECULE: | BG505 SOSIP.664 GP120 |
| 90:00:00 | 7uoj-B | 2.4 | 4.7 | 45 | 131 | 7 | MOLECULE: | ENVELOPE GLYCOPROTEIN GP120 |
| 91:00:00 | 7mxe-C | 2.4 | 5.7 | 49 | 124 | 8 | MOLECULE: | 8ANC195 G52K5 FAB HEAVY CHAIN |
| 92:00:00 | 8dok-B | 2.4 | 3.4 | 44 | 121 | 7 | MOLECULE: | CRF-1_AE T/F100 ENV GP120 |
| 93:00:00 | 6u0n-X | 2.4 | 2.1 | 41 | 129 | 5 | MOLECULE: | ENVELOPE GLYCOPROTEIN GP120 |
| 94:00:00 | 6v8x-F | 2.4 | 4.6 | 54 | 128 | 9 | MOLECULE: | ENVELOPE GLYCOPROTEIN GP120 |
| 95:00:00 | 6x9v-B | 2.4 | 3.9 | 42 | 121 | 5 | MOLECULE: | HIV-1 ENVELOPE GLYCOPROTEIN BG505 SOSIP.664 |
| 96:00:00 | 1qo0-E | 2.3 | 5.7 | 72 | 194 | 7 | MOLECULE: | AMIC |
| 97:00:00 | 7kzn-E | 2.3 | 18.9 | 75 | 474 | 8 | MOLECULE: | HEAVY CHAIN ALPHA |
| 98:00:00 | 6rwb-A | 2.3 | 29.3 | 153 | 1873 | 10 | MOLECULE: | TOXIN,TOXIN COMPLEX SUBUNIT TCAB,PUTATIVE TO |
| 99:00:00 | 2aze-A | 2.3 | 1.5 | 52 | 149 | 13 | MOLECULE: | TRANSCRIPTION FACTOR DP-1 |
| 0:00 | 6vy1-A | 2.3 | 5.1 | 58 | 121 | 9 | MOLECULE: | PREFOLDIN SUBUNIT ALPHA 2 |
| 1:00 | 6ogd-E | 2.3 | 10.4 | 119 | 1078 | 6 | MOLECULE: | TOXIN SUBUNIT YENA1 |
| 2:00 | 5dfz-C | 2.3 | 38.6 | 77 | 783 | 6 | MOLECULE: | VACUOLAR PROTEIN SORTING-ASSOCIATED PROTEIN |
| 3:00 | 7z7s-A | 2.3 | 21 | 63 | 131 | 5 | MOLECULE: | NADH-QUINONE OXIDOREDUCTASE SUBUNIT F |
| 4:00 | 3lss-B | 2.3 | 35.4 | 58 | 468 | 2 | MOLECULE: | SERYL-TRNA SYNTHETASE |
| 5:00 | 3euh-B | 2.3 | 21.5 | 78 | 321 | 9 | MOLECULE: | CHROMOSOME PARTITION PROTEIN MUKF |
| 6:00 | 6zz6-C | 2.3 | 29.4 | 40 | 83 | 8 | MOLECULE: | STRUCTURAL MAINTENANCE OF CHROMOSOMES PROTEI |
| 7:00 | 6x80-C | 2.3 | 5.1 | 115 | 574 | 6 | MOLECULE: | FLAGELLIN A |
| 8:00 | 6thk-A | 2.3 | 9.6 | 108 | 466 | 9 | MOLECULE: | PYOCIN S5 |
| 9:00 | 8anq-C | 2.3 | 8.2 | 45 | 227 | 7 | MOLECULE: | BACTERIORHODOPSIN |
| 10:00 | 7etr-A | 2.3 | 10.1 | 50 | 88 | 8 | MOLECULE: | TRANSCRIPTIONAL REGULATOR COPG FAMILY |
| 11:00 | 2d1l-A | 2.3 | 4.6 | 87 | 249 | 7 | MOLECULE: | METASTASIS SUPPRESSOR PROTEIN 1 |
| 12:00 | 8k49-T | 2.3 | 16.9 | 59 | 627 | 12 | MOLECULE: | VP2 |
| 13:00 | 1u4q-A | 2.3 | 6.5 | 102 | 318 | 11 | MOLECULE: | SPECTRIN ALPHA CHAIN, BRAIN |
| 14:00 | 8t9l-B | 2.3 | 16.4 | 70 | 110 | 4 | MOLECULE: | NUCLEOPORIN POM152 |
| 15:00 | 2xzb-B | 2.3 | 13.3 | 60 | 175 | 8 | MOLECULE: | POTASSIUM-TRANSPORTING ATPASE ALPHA CHAIN 1 |
| 16:00 | 5lc5-h | 2.3 | 20.5 | 95 | 134 | 1 | MOLECULE: | NADH-UBIQUINONE OXIDOREDUCTASE CHAIN 3 |
| 17:00 | 5me8-A | 2.3 | 2.6 | 51 | 107 | 6 | MOLECULE: | INHIBITOR OF GROWTH PROTEIN 5 |
| 18:00 | 6cfw-I | 2.3 | 44.5 | 95 | 114 | 6 | MOLECULE: | MONOVALENT CATION/H+ ANTIPORTER SUBUNIT D |
| 19:00 | 5n9j-V | 2.3 | 10.6 | 67 | 99 | 7 | MOLECULE: | MEDIATOR OF RNA POLYMERASE II TRANSCRIPTION |
| 20:00 | 3b0z-A | 2.3 | 4.1 | 39 | 41 | 5 | MOLECULE: | FLAGELLAR BIOSYNTHETIC PROTEIN FLHB |
| 21:00 | 8fck-F | 2.3 | 12.5 | 100 | 393 | 6 | MOLECULE: | HAUS AUGMIN-LIKE COMPLEX SUBUNIT 1 |
| 22:00 | 2zqm-A | 2.3 | 3.6 | 58 | 114 | 3 | MOLECULE: | PREFOLDIN BETA SUBUNIT 1 |
| 23:00 | 1jfi-B | 2.3 | 18.7 | 78 | 135 | 13 | MOLECULE: | 5'- |
| 24:00:00 | 4gwp-A | 2.3 | 9.9 | 71 | 112 | 6 | MOLECULE: | MEDIATOR OF RNA POLYMERASE II TRANSCRIPTION |
| 25:00:00 | 8bd7-G | 2.3 | 37.8 | 89 | 205 | 10 | MOLECULE: | IFT88 |
| 26:00:00 | 3pwx-A | 2.3 | 2.7 | 66 | 224 | 20 | MOLECULE: | PUTATIVE FLAGELLAR HOOK-ASSOCIATED PROTEIN |
| 27:00:00 | 4wid-A | 2.3 | 3.2 | 67 | 353 | 10 | MOLECULE: | RHUL123 |
| 28:00:00 | 1ylf-C | 2.3 | 2.8 | 38 | 119 | 8 | MOLECULE: | RRF2 FAMILY PROTEIN |
| 29:00:00 | 6tgz-F | 2.3 | 5.6 | 94 | 352 | 5 | MOLECULE: | 55 KDA IMMEDIATE-EARLY PROTEIN 1 |
| 30:00:00 | 6hb4-A | 2.3 | 26.9 | 60 | 197 | 8 | MOLECULE: | TRANSCRIPTION FACTOR A, MITOCHONDRIAL |
| 31:00:00 | 5g06-P | 2.3 | 6 | 56 | 95 | 7 | MOLECULE: | EXOSOME COMPLEX COMPONENT RRP45 |
| 32:00:00 | 6wg3-B | 2.3 | 14.9 | 85 | 693 | 7 | MOLECULE: | STRUCTURAL MAINTENANCE OF CHROMOSOMES PROTEI |
| 33:00:00 | 7v7b-A | 2.3 | 39.6 | 96 | 1110 | 6 | MOLECULE: | DDB1- AND CUL4-ASSOCIATED FACTOR 1 |
| 34:00:00 | 6xp5-D | 2.3 | 5.5 | 93 | 134 | 5 | MOLECULE: | MEDIATOR OF RNA POLYMERASE II TRANSCRIPTION |

| | | | | | | | | |
|---|---|---|---|---|---|---|---|---|
| 35:00:00 | 6n7p-H | 2.3 | 10.8 | 57 | 105 | 2 | MOLECULE: | U1 SMALL NUCLEAR RIBONUCLEOPROTEIN 70 KDA HO |
| 36:00:00 | 5s5z-E | 2.3 | 6.9 | 56 | 123 | 5 | MOLECULE: | TUBULIN ALPHA-1B CHAIN |
| 37:00:00 | 5t1y-E | 2.3 | 8.1 | 49 | 110 | 6 | MOLECULE: | MLA10 |
| 38:00:00 | 6c5w-B | 2.3 | 7.7 | 89 | 217 | 2 | MOLECULE: | CALCIUM UNIPORTER |
| 39:00:00 | 6nrb-5 | 2.3 | 2.8 | 50 | 127 | 8 | MOLECULE: | T-COMPLEX PROTEIN 1 SUBUNIT ALPHA |
| 40:00:00 | 6xjx-E | 2.3 | 6.3 | 71 | 153 | 3 | MOLECULE: | CALCIUM UNIPORTER PROTEIN, MITOCHONDRIAL |
| 41:00:00 | 7dad-E | 2.3 | 10.6 | 60 | 121 | 8 | MOLECULE: | TUBULIN ALPHA-1B CHAIN |
| 42:00:00 | 5tj5-J | 2.3 | 4 | 46 | 150 | 7 | MOLECULE: | V-TYPE PROTON ATPASE SUBUNIT A |
| 43:00:00 | 8jtd-B | 2.3 | 4.4 | 46 | 126 | 7 | MOLECULE: | GP120 PROTEIN OF HIV ENVELOPE TRIMER |
| 44:00:00 | 6nnj-B | 2.3 | 3.5 | 45 | 127 | 11 | MOLECULE: | ENVELOPE GLYCOPROTEIN GP41 |
| 45:00:00 | 7tfn-Z | 2.3 | 3.3 | 50 | 125 | 12 | MOLECULE: | ENVELOPE GLYCOPROTEIN BG505 SOSIP.664 - GP12 |
| 46:00:00 | 8g9y-D | 2.3 | 4.5 | 45 | 132 | 7 | MOLECULE: | ENVELOPE GLYCOPROTEIN GP41 |
| 47:00:00 | 8tni-B | 2.3 | 3.6 | 46 | 127 | 7 | MOLECULE: | HIV-1 BG505 DS-SOSIP GP120 |
| 48:00:00 | 8dok-J | 2.3 | 3.3 | 43 | 120 | 7 | MOLECULE: | CRF-1_AE T/F100 ENV GP120 |
| 49:00:00 | 6ohy-E | 2.3 | 3.5 | 44 | 120 | 11 | MOLECULE: | ENVELOPE GLYCOPROTEIN GP160 |
| 50:00:00 | 8sb5-G | 2.3 | 3.6 | 43 | 132 | 9 | MOLECULE: | CH848.10.17.SOSIP GP120 |
| 51:00:00 | 6x5b-B | 2.3 | 4 | 49 | 149 | 10 | MOLECULE: | ENVELOPE GLYCOPROTEIN GP120 |
| 52:00:00 | 6uda-F | 2.3 | 4.7 | 47 | 126 | 2 | MOLECULE: | CH505.N279K.G458Y.SOSIP.664 GP120 |
| 53:00:00 | 6h4j-A | 2.2 | 23.4 | 101 | 488 | 7 | MOLECULE: | UBIQUITIN CARBOXYL-TERMINAL HYDROLASE 25 |
| 54:00:00 | 8bbt-A | 2.2 | 4.2 | 58 | 230 | 7 | MOLECULE: | MOBP |
| 55:00:00 | 5c0w-B | 2.2 | 6.6 | 52 | 244 | 12 | MOLECULE: | EXOSOME COMPLEX COMPONENT RRP45 |
| 56:00:00 | 6j6q-F | 2.2 | 12.7 | 54 | 161 | 7 | MOLECULE: | PRE-MRNA-SPLICING FACTOR 8 |
| 57:00:00 | 1t7s-A | 2.2 | 3.2 | 41 | 129 | 0 | MOLECULE: | BAG-1 COCHAPERONE |
| 58:00:00 | 7r6k-w | 2.2 | 26.1 | 82 | 214 | 5 | MOLECULE: | 25S RRNA |
| 59:00:00 | 5w0p-C | 2.2 | 7.7 | 54 | 850 | 7 | MOLECULE: | ENDOLYSIN,RHODOPSIN,S-ARRESTIN |
| 60:00:00 | 5i6r-A | 2.2 | 2.6 | 73 | 419 | 10 | MOLECULE: | SLIT-ROBO RHO GTPASE-ACTIVATING PROTEIN 2 |
| 61:00:00 | 2kjy-A | 2.2 | 4.1 | 46 | 60 | 4 | MOLECULE: | PROTEIN PHOSPHATASE 1 REGULATORY SUBUNIT 12A |
| 62:00:00 | 7e84-A | 2.2 | 22.6 | 65 | 460 | 8 | MOLECULE: | POTASSIUM VOLTAGE-GATED CHANNEL SUBFAMILY D |
| 63:00:00 | 6xz6-A | 2.2 | 4.3 | 67 | 213 | 7 | MOLECULE: | GARP DOMAIN-CONTAINING PROTEIN |
| 64:00:00 | 2pih-A | 2.2 | 27.2 | 66 | 123 | 6 | MOLECULE: | PROTEIN YMCA |
| 65:00:00 | 8t1l-Y | 2.2 | 33.6 | 93 | 154 | 6 | MOLECULE: | MEDIATOR OF RNA POLYMERASE II TRANSCRIPTION |
| 66:00:00 | 8gkh-P | 2.2 | 20.6 | 79 | 600 | 4 | MOLECULE: | DNA (5'-D(P*CP*GP*GP*TP*AP*CP*CP*CP*GP*GP*GP |
| 67:00:00 | 7cun-B | 2.2 | 22.5 | 77 | 1058 | 3 | MOLECULE: | INTEGRATOR COMPLEX SUBUNIT 1 |
| 68:00:00 | 7wb4-b | 2.2 | 28.2 | 91 | 636 | 4 | MOLECULE: | OUTER NUP133 |
| 69:00:00 | 8cwo-T | 2.2 | 3.5 | 45 | 87 | 9 | MOLECULE: | 16S RIBOSOMAL RNA |
| 70:00:00 | 5vtl-A | 2.2 | 3.7 | 61 | 210 | 11 | MOLECULE: | TB427.07.360- PUTATIVE UNCHARACTERIZED METAC |
| 71:00:00 | 3whj-A | 2.2 | 5.8 | 47 | 111 | 11 | MOLECULE: | PROBABLE 26S PROTEASOME REGULATORY SUBUNIT P |
| 72:00:00 | 7byl-A | 2.2 | 16.9 | 69 | 354 | 7 | MOLECULE: | GREEN FLUORESCENT PROTEIN,POTASSIUM VOLTAGE- |
| 73:00:00 | 8v8b-A | 2.2 | 33.3 | 75 | 476 | 7 | MOLECULE: | PORTAL PROTEIN |
| 74:00:00 | 6y6k-A | 2.2 | 27.7 | 80 | 1299 | 8 | MOLECULE: | RNA-DEPENDENT RNA POLYMERASE |
| 75:00:00 | 6nr8-3 | 2.2 | 11.3 | 68 | 132 | 12 | MOLECULE: | PREFOLDIN SUBUNIT 1 |
| 76:00:00 | 7lid-A | 2.2 | 6.8 | 67 | 381 | 9 | MOLECULE: | MHOR5 |
| 77:00:00 | 3cqx-D | 2.2 | 4.8 | 47 | 84 | 2 | MOLECULE: | HEAT SHOCK COGNATE 71 KDA PROTEIN |
| 78:00:00 | 1s35-A | 2.2 | 3.1 | 71 | 211 | 14 | MOLECULE: | SPECTRIN BETA CHAIN, ERYTHROCYTE |
| 79:00:00 | 7vxs-o | 2.2 | 23.3 | 93 | 128 | 6 | MOLECULE: | NADH DEHYDROGENASE [UBIQUINONE] IRON-SULFUR |
| 80:00:00 | 3on0-A | 2.2 | 12.4 | 67 | 121 | 4 | MOLECULE: | PROTEIN TRAM |
| 81:00:00 | 8ip4-A | 2.2 | 10.6 | 106 | 320 | 4 | MOLECULE: | MAGNESIUM TRANSPORTER MRS2 HOMOLOG, MITOCHON |
| 82:00:00 | 8cli-B | 2.2 | 7.9 | 53 | 689 | 11 | MOLECULE: | GENERAL TRANSCRIPTION FACTOR 3C POLYPEPTIDE |
| 83:00:00 | 7x29-A | 2.2 | 24 | 87 | 991 | 9 | MOLECULE: | SPIKE GLYCOPROTEIN |
| 84:00:00 | 2w0c-S | 2.2 | 22.4 | 72 | 84 | 3 | MOLECULE: | MAJOR CAPSID PROTEIN P2 |
| 85:00:00 | 6epg-C | 2.2 | 9.8 | 51 | 56 | 6 | MOLECULE: | EPSILON_1 ANTITOXIN |

| | | | | | | | | |
|---|---|---|---|---|---|---|---|---|
| 86:00:00 | 2qri-A | 2.2 | 18.5 | 60 | 398 | 8 | MOLECULE: | H-2 CLASS I HISTOCOMPATIBILITY ANTIGEN K-B A |
| 87:00:00 | 8wjn-B | 2.2 | 45.5 | 67 | 481 | 9 | MOLECULE: | STRUCTURAL MAINTENANCE OF CHROMOSOMES PROTEI |
| 88:00:00 | 7pv1-D | 2.2 | 6 | 56 | 141 | 14 | MOLECULE: | MICOS COMPLEX SUBUNIT MIC60 FUSED TO MIC19 |
| 89:00:00 | 7zu0-E | 2.2 | 6.3 | 47 | 343 | 2 | MOLECULE: | E3 UBIQUITIN-PROTEIN LIGASE PEP5 |
| 90:00:00 | 5hdi-A | 2.2 | 15.6 | 66 | 397 | 6 | MOLECULE: | CYTOCHROME P450 144 |
| 91:00:00 | 6a68-A | 2.2 | 12.3 | 80 | 162 | 3 | MOLECULE: | CALCIUM-DEPENDENT SECRETION ACTIVATOR 1 |
| 92:00:00 | 6tdx-O | 2.2 | 3.7 | 40 | 81 | 10 | MOLECULE: | ATP SYNTHASE F1 SUBUNIT GAMMA |
| 93:00:00 | 6xl0-N | 2.2 | 4.7 | 98 | 271 | 8 | MOLECULE: | FLAGELLIN |
| 94:00:00 | 6xl0-O | 2.2 | 4.7 | 98 | 271 | 8 | MOLECULE: | FLAGELLIN |
| 95:00:00 | 6xky-E | 2.2 | 5.4 | 97 | 271 | 8 | MOLECULE: | FLAGELLIN |
| 96:00:00 | 6xl0-K | 2.2 | 4.9 | 98 | 271 | 9 | MOLECULE: | FLAGELLIN |
| 97:00:00 | 6xky-I | 2.2 | 4.1 | 97 | 271 | 7 | MOLECULE: | FLAGELLIN |
| 98:00:00 | 6xky-M | 2.2 | 4.2 | 95 | 271 | 6 | MOLECULE: | FLAGELLIN |
| 99:00:00 | 6xky-D | 2.2 | 5.5 | 97 | 271 | 7 | MOLECULE: | FLAGELLIN |
| 0:00 | 6xky-L | 2.2 | 5.5 | 97 | 271 | 7 | MOLECULE: | FLAGELLIN |
| 1:00 | 6xky-K | 2.2 | 5.6 | 96 | 271 | 6 | MOLECULE: | FLAGELLIN |
| 2:00 | 3k8v-A | 2.2 | 8.1 | 97 | 323 | 3 | MOLECULE: | FLAGELLIN HOMOLOG |
| 3:00 | 6xky-G | 2.2 | 5.5 | 98 | 271 | 7 | MOLECULE: | FLAGELLIN |
| 4:00 | 6xky-C | 2.2 | 5.6 | 96 | 271 | 6 | MOLECULE: | FLAGELLIN |
| 5:00 | 6x80-I | 2.2 | 7.1 | 108 | 574 | 10 | MOLECULE: | FLAGELLIN A |
| 6:00 | 6xky-F | 2.2 | 4.1 | 97 | 271 | 7 | MOLECULE: | FLAGELLIN |
| 7:00 | 6ek7-A | 2.2 | 3.3 | 83 | 364 | 4 | MOLECULE: | YAXA |
| 8:00 | 4zwq-E | 2.2 | 3.2 | 50 | 135 | 10 | MOLECULE: | RECOMBINATION PROTEIN UVSY |
| 9:00 | 1fxk-A | 2.2 | 2.8 | 48 | 107 | 2 | MOLECULE: | PREFOLDIN |
| 10:00 | 8jsn-C | 2.2 | 12.3 | 67 | 99 | 6 | MOLECULE: | RNA-DIRECTED RNA POLYMERASE L |
| 11:00 | 5jsb-L | 2.2 | 2.6 | 44 | 116 | 9 | MOLECULE: | INDUCED MYELOID LEUKEMIA CELL DIFFERENTIATIO |
| 12:00 | 6u59-I | 2.2 | 3.5 | 44 | 120 | 5 | MOLECULE: | SOSIP.664 GP120,SOSIP.664 GP120 |
| 13:00 | 6w03-B | 2.2 | 4.9 | 45 | 124 | 7 | MOLECULE: | ENVELOPE GLYCOPROTEIN GP41 |
| 14:00 | 5wdu-J | 2.2 | 3.9 | 44 | 126 | 0 | MOLECULE: | ENVELOPE GLYCOPROTEIN GP160 |
| 15:00 | 6opn-B | 2.2 | 3.4 | 42 | 124 | 19 | MOLECULE: | ENVELOPE GLYCOPROTEIN GP160 |
| 16:00 | 6ieq-B | 2.2 | 4.1 | 47 | 131 | 4 | MOLECULE: | ENVELOPE GLYCOPROTEIN GP160 |
| 17:00 | 6mug-B | 2.2 | 3.4 | 44 | 123 | 9 | MOLECULE: | ENVELOPE GLYCOPROTEIN GP160 |
| 18:00 | 6opq-F | 2.2 | 3.3 | 47 | 128 | 2 | MOLECULE: | ENVELOPE GLYCOPROTEIN GP160 |
| 19:00 | 8sb1-B | 2.2 | 3.5 | 48 | 124 | 8 | MOLECULE: | CH848.10.17 GP120 |
| 20:00 | 6opq-G | 2.2 | 3.3 | 47 | 128 | 2 | MOLECULE: | ENVELOPE GLYCOPROTEIN GP160 |
| 21:00 | 8sb1-G | 2.2 | 3.5 | 48 | 124 | 8 | MOLECULE: | CH848.10.17 GP120 |
| 22:00 | 6nf2-I | 2.2 | 15 | 51 | 132 | 4 | MOLECULE: | ENVELOPE GLYCOPROTEIN GP120 |
| 23:00 | 8sax-B | 2.2 | 10.5 | 56 | 132 | 0 | MOLECULE: | CH848.10.17.SOSIP GP120 |
| 24:00:00 | 7lpn-B | 2.2 | 5.1 | 44 | 121 | 5 | MOLECULE: | HIV-1 ENVELOPE GLYCOPROTEIN BG505 SOSIP.664 |
| 25:00:00 | 6oz4-B | 2.2 | 4 | 44 | 122 | 2 | MOLECULE: | ENVELOPE GLYCOPROTEIN GP160 |
| 26:00:00 | 6muf-B | 2.2 | 3.4 | 44 | 122 | 9 | MOLECULE: | ENVELOPE GLYCOPROTEIN GP160 |
| 27:00:00 | 6nf5-B | 2.2 | 4 | 44 | 122 | 5 | MOLECULE: | HIV-1 ENV BG505 MD64 N332-GT5 SOSIP GP120 |
| 28:00:00 | 8g9w-O | 2.2 | 17.2 | 48 | 132 | 0 | MOLECULE: | ENVELOPE GLYCOPROTEIN GP41 |
| 29:00:00 | 8tnh-D | 2.2 | 9.3 | 47 | 126 | 9 | MOLECULE: | HIV-1 BG505 DS-SOSIP GP120 |
| 30:00:00 | 8d5c-Z | 2.2 | 7.5 | 43 | 129 | 7 | MOLECULE: | ENVELOPE GLYCOPROTEIN GP41 |
| 31:00:00 | 7sq1-B | 2.2 | 4.2 | 42 | 121 | 5 | MOLECULE: | TRANSMEMBRANE PROTEIN GP41 |
| 32:00:00 | 6uda-J | 2.2 | 4.7 | 45 | 126 | 4 | MOLECULE: | CH505.N279K.G458Y.SOSIP.664 GP120 |
| 33:00:00 | 8sav-F | 2.2 | 3.8 | 46 | 136 | 9 | MOLECULE: | CH848.0526.25 GP120 |
| 34:00:00 | 5t3s-B | 2.2 | 3.7 | 40 | 122 | 5 | MOLECULE: | ENVELOPE GLYCOPROTEIN GP160 |
| 35:00:00 | 6opn-E | 2.2 | 3.4 | 42 | 124 | 19 | MOLECULE: | ENVELOPE GLYCOPROTEIN GP160 |
| 36:00:00 | 8dok-F | 2.2 | 3.4 | 44 | 117 | 7 | MOLECULE: | CRF-1_AE T/F100 ENV GP120 |

| | | | | | | | | |
|---|---|---|---|---|---|---|---|---|
| 37:00:00 | 4gfq-A | 2.1 | 2.9 | 70 | 186 | 6 | MOLECULE: | RIBOSOME-RECYCLING FACTOR |
| 38:00:00 | 8f2r-E | 2.1 | 3.1 | 43 | 205 | 12 | MOLECULE: | COMM DOMAIN-CONTAINING PROTEIN 1 |
| 39:00:00 | 7x5e-B | 2.1 | 12.6 | 64 | 106 | 6 | MOLECULE: | TRANSCRIPTION FACTOR MAFG |
| 40:00:00 | 3g5o-A | 2.1 | 8.4 | 52 | 92 | 2 | MOLECULE: | UNCHARACTERIZED PROTEIN RV2865 |
| 41:00:00 | 8p65-J | 2.1 | 7.1 | 54 | 64 | 7 | MOLECULE: | CYTOCHROME B-C1 COMPLEX SUBUNIT 1, MITOCHOND |
| 42:00:00 | 6j72-A | 2.1 | 3.3 | 80 | 561 | 8 | MOLECULE: | ISONIAZID INDUCIBLE GENE PROTEIN INIA |
| 43:00:00 | 5szj-B | 2.1 | 2.2 | 57 | 145 | 5 | MOLECULE: | RAS-RELATED PROTEIN RAB-10 |
| 44:00:00 | 7e40-D | 2.1 | 18.9 | 87 | 345 | 5 | MOLECULE: | PROTEIN PHOSPHATE STARVATION RESPONSE 2 |
| 45:00:00 | 5yud-C | 2.1 | 2.9 | 45 | 75 | 2 | MOLECULE: | BACULOVIRAL IAP REPEAT-CONTAINING PROTEIN 1E |
| 46:00:00 | 8eki-A | 2.1 | 3.7 | 71 | 170 | 3 | MOLECULE: | PROTEIN TRANSPORT PROTEIN SEC20 |
| 47:00:00 | 6ww6-B | 2.1 | 3.6 | 64 | 192 | 9 | MOLECULE: | RESPONSE REGULATOR |
| 48:00:00 | 6qeq-A | 2.1 | 2.5 | 37 | 111 | 5 | MOLECULE: | PCFF |
| 49:00:00 | 7qlr-A | 2.1 | 17.2 | 92 | 603 | 2 | MOLECULE: | CDHS1_22 PUTATIVE TAIL FIBER PROTEIN |
| 50:00:00 | 2oar-A | 2.1 | 20.9 | 62 | 125 | 6 | MOLECULE: | LARGE-CONDUCTANCE MECHANOSENSITIVE CHANNEL |
| 51:00:00 | 8ox6-A | 2.1 | 13.7 | 55 | 575 | 5 | MOLECULE: | PHOSPHOLIPID-TRANSPORTING ATPASE IC |
| 52:00:00 | 7nyw-A | 2.1 | 15.1 | 102 | 685 | 7 | MOLECULE: | CHROMOSOME PARTITION PROTEIN MUKB |
| 53:00:00 | 6th1-R | 2.1 | 5.3 | 91 | 360 | 7 | MOLECULE: | IMMEDIATE EARLY PROTEIN 1 |
| 54:00:00 | 7f6j-C | 2.1 | 1.8 | 40 | 106 | 8 | MOLECULE: | RAS-RELATED PROTEIN RAB-7A |
| 55:00:00 | 6cum-A | 2.1 | 4.9 | 39 | 51 | 3 | MOLECULE: | LAO/AO TRANSPORT SYSTEM ATPASE |
| 56:00:00 | 4rp5-A | 2.1 | 5.9 | 48 | 94 | 8 | MOLECULE: | DISKS LARGE 1 TUMOR SUPPRESSOR PROTEIN |
| 57:00:00 | 3cf4-A | 2.1 | 34.9 | 73 | 766 | 7 | MOLECULE: | ACETYL-COA DECARBOXYLASE/SYNTHASE ALPHA SUBU |
| 58:00:00 | 4y66-D | 2.1 | 17.5 | 93 | 208 | 4 | MOLECULE: | MND1 |
| 59:00:00 | 5che-A | 2.1 | 7.6 | 66 | 427 | 2 | MOLECULE: | GLUTAMYL-TRNA REDUCTASE 1, CHLOROPLASTIC |
| 60:00:00 | 2odu-A | 2.1 | 3.1 | 77 | 217 | 13 | MOLECULE: | PLECTIN 1 |
| 61:00:00 | 6yj4-g | 2.1 | 7.5 | 79 | 198 | 5 | MOLECULE: | NADH-UBIQUINONE OXIDOREDUCTASE CHAIN 3 |
| 62:00:00 | 6t9i-K | 2.1 | 11.5 | 68 | 279 | 3 | MOLECULE: | TRANSCRIPTION FACTOR SPT20 |
| 63:00:00 | 6yj4-o | 2.1 | 6.1 | 46 | 83 | 4 | MOLECULE: | NADH-UBIQUINONE OXIDOREDUCTASE CHAIN 3 |
| 64:00:00 | 5lbm-C | 2.1 | 3.3 | 41 | 90 | 12 | MOLECULE: | TRANSCRIPTIONAL REPRESSOR FRMR |
| 65:00:00 | 2ox6-C | 2.1 | 11.1 | 46 | 163 | 11 | MOLECULE: | HYPOTHETICAL PROTEIN SO3848 |
| 66:00:00 | 6rlr-A | 2.1 | 4.8 | 46 | 80 | 2 | MOLECULE: | CD9 ANTIGEN |
| 67:00:00 | 3av0-B | 2.1 | 12 | 74 | 365 | 11 | MOLECULE: | DNA DOUBLE-STRAND BREAK REPAIR PROTEIN MRE11 |
| 68:00:00 | 6swy-1 | 2.1 | 21.8 | 68 | 542 | 1 | MOLECULE: | VACUOLAR IMPORT AND DEGRADATION PROTEIN 28 |
| 69:00:00 | 2q83-A | 2.1 | 27.1 | 63 | 332 | 16 | MOLECULE: | YTAA PROTEIN |
| 70:00:00 | 6xl0-A | 2.1 | 4.6 | 98 | 271 | 9 | MOLECULE: | FLAGELLIN |
| 71:00:00 | 6xky-B | 2.1 | 5.6 | 102 | 271 | 8 | MOLECULE: | FLAGELLIN |
| 72:00:00 | 6xky-R | 2.1 | 5.3 | 98 | 271 | 8 | MOLECULE: | FLAGELLIN |
| 73:00:00 | 6xky-S | 2.1 | 5.5 | 97 | 271 | 7 | MOLECULE: | FLAGELLIN |
| 74:00:00 | 6xl0-L | 2.1 | 4.7 | 98 | 271 | 8 | MOLECULE: | FLAGELLIN |
| 75:00:00 | 6xl0-Q | 2.1 | 4.2 | 92 | 271 | 9 | MOLECULE: | FLAGELLIN |
| 76:00:00 | 5ik2-G | 2.1 | 4 | 82 | 285 | 10 | MOLECULE: | ATP SYNTHASE SUBUNIT ALPHA |
| 77:00:00 | 6xl0-T | 2.1 | 4.2 | 92 | 271 | 10 | MOLECULE: | FLAGELLIN |
| 78:00:00 | 6xky-Q | 2.1 | 5.3 | 98 | 271 | 8 | MOLECULE: | FLAGELLIN |
| 79:00:00 | 6xl0-M | 2.1 | 4.2 | 92 | 271 | 10 | MOLECULE: | FLAGELLIN |
| 80:00:00 | 6xl0-C | 2.1 | 4.2 | 92 | 271 | 10 | MOLECULE: | FLAGELLIN |
| 81:00:00 | 2vrz-A | 2.1 | 6.8 | 55 | 98 | 9 | MOLECULE: | VIRULENCE FACTOR ESXA |
| 82:00:00 | 5j9w-H | 2.1 | 5.7 | 63 | 118 | 13 | MOLECULE: | HISTONE ACETYLTRANSFERASE ESA1 |
| 83:00:00 | 6cnm-A | 2.1 | 19.1 | 98 | 359 | 3 | MOLECULE: | INTERMEDIATE CONDUCTANCE CALCIUM-ACTIVATED P |
| 84:00:00 | 5t1y-A | 2.1 | 8 | 49 | 108 | 6 | MOLECULE: | MLA10 |
| 85:00:00 | 7u1s-E | 2.1 | 12.1 | 68 | 1440 | 3 | MOLECULE: | ATP-SENSITIVE INWARD RECTIFIER POTASSIUM CHA |
| 86:00:00 | 1i4l-A | 2.1 | 4 | 60 | 185 | 10 | MOLECULE: | ARFAPTIN 2 |
| 87:00:00 | 7yo0-H | 2.1 | 22.7 | 59 | 262 | 8 | MOLECULE: | CALCIUM-ACTIVATED POTASSIUM CHANNEL SUBUNIT |

| | | | | | | | | |
|---|---|---|---|---|---|---|---|---|
| 88:00:00 | 1i4d-A | 2.1 | 3.9 | 60 | 188 | 5 | MOLECULE: | ARFAPTIN 2 |
| 89:00:00 | 5da9-A | 2.1 | 13.6 | 75 | 433 | 4 | MOLECULE: | PUTATIVE UNCHARACTERIZED PROTEIN,PUTATIVE UN |
| 90:00:00 | 1i4t-B | 2.1 | 4.6 | 57 | 179 | 4 | MOLECULE: | ARFAPTIN 2 |
| 91:00:00 | 5vox-b | 2.1 | 24.2 | 100 | 627 | 9 | MOLECULE: | V-TYPE PROTON ATPASE CATALYTIC SUBUNIT A,V-T |
| 92:00:00 | 7sb2-A | 2.1 | 5.4 | 45 | 299 | 2 | MOLECULE: | GLDM |
| 93:00:00 | 6cnm-C | 2.1 | 25.9 | 84 | 359 | 6 | MOLECULE: | INTERMEDIATE CONDUCTANCE CALCIUM-ACTIVATED P |
| 94:00:00 | 6ysl-C | 2.1 | 6.6 | 70 | 255 | 9 | MOLECULE: | MOTILITY PROTEIN A |
| 95:00:00 | 6nfc-I | 2.1 | 3.7 | 42 | 120 | 5 | MOLECULE: | BASE-BINDING RM20A3 FRAGMENT ANTIGEN BINDING |
| 96:00:00 | 7l7u-F | 2.1 | 4.3 | 47 | 145 | 15 | MOLECULE: | BG505 SOSIP.V5.2(7S) - GP120 |
| 97:00:00 | 6x9u-B | 2.1 | 3.2 | 42 | 121 | 10 | MOLECULE: | HIV-1 ENVELOPE GLYCOPROTEIN BG505 SOSIP.664 |
| 98:00:00 | 7lu9-b | 2.1 | 9.9 | 49 | 135 | 10 | MOLECULE: | DH851.3 LIGHT CHAIN |
| 99:00:00 | 8g9w-D | 2.1 | 4.3 | 47 | 132 | 4 | MOLECULE: | ENVELOPE GLYCOPROTEIN GP41 |
| 0:00 | 8g9x-O | 2.1 | 15.4 | 50 | 132 | 4 | MOLECULE: | ENVELOPE GLYCOPROTEIN GP41 |
| 1:00 | 7mxd-B | 2.1 | 4.9 | 47 | 136 | 9 | MOLECULE: | ENVELOPE GLYCOPROTEIN GP120 |
| 2:00 | 8czz-B | 2.1 | 7.9 | 43 | 125 | 2 | MOLECULE: | CRF01_AE T/F100 HIV-1 GP120 |
| 3:00 | 7sq1-D | 2.1 | 4.2 | 42 | 121 | 5 | MOLECULE: | TRANSMEMBRANE PROTEIN GP41 |
| 4:00 | 8tnh-B | 2.1 | 3.5 | 47 | 127 | 9 | MOLECULE: | HIV-1 BG505 DS-SOSIP GP120 |
| 5:00 | 7lpn-A | 2.1 | 4.6 | 43 | 121 | 5 | MOLECULE: | HIV-1 ENVELOPE GLYCOPROTEIN BG505 SOSIP.664 |
| 6:00 | 6nfc-G | 2.1 | 4.1 | 41 | 120 | 7 | MOLECULE: | BASE-BINDING RM20A3 FRAGMENT ANTIGEN BINDING |
| 7:00 | 5v8l-B | 2.1 | 4.5 | 46 | 126 | 2 | MOLECULE: | GP120 |
| 8:00 | 6x5b-K | 2.1 | 3.5 | 49 | 149 | 4 | MOLECULE: | ENVELOPE GLYCOPROTEIN GP120 |
| 9:00 | 6ohy-F | 2.1 | 7 | 49 | 120 | 10 | MOLECULE: | ENVELOPE GLYCOPROTEIN GP160 |
| 10:00 | 8ttw-J | 2.1 | 4.9 | 44 | 123 | 5 | MOLECULE: | ENVELOPE GLYCOPROTEIN GP160 |
| 11:00 | 8flw-B | 2.1 | 3 | 45 | 123 | 7 | MOLECULE: | PGT145 DU303 HEAVY |
| 12:00 | 8dow-J | 2.1 | 10.9 | 48 | 132 | 6 | MOLECULE: | ENVELOPE GLYCOPROTEIN GP120 |
| 13:00 | 5g06-J | 2 | 27.1 | 80 | 993 | 8 | MOLECULE: | EXOSOME COMPLEX COMPONENT RRP45 |
| 14:00 | 8ffz-A | 2 | 10.2 | 46 | 314 | 4 | MOLECULE: | TRANSCRIPTION FACTOR IIIA |
| 15:00 | 1sjj-A | 2 | 14.5 | 139 | 863 | 11 | MOLECULE: | ACTININ |
| 16:00 | 4mer-D | 2 | 7.1 | 60 | 97 | 10 | MOLECULE: | STREPTOCOCCAL HISTIDINE-RICH GLYCOPROTEIN IN |
| 17:00 | 6ezn-E | 2 | 7.4 | 52 | 85 | 4 | MOLECULE: | DOLICHYL-DIPHOSPHOOLIGOSACCHARIDE--PROTEIN |
| 18:00 | 6lcn-F | 2 | 23.7 | 81 | 309 | 2 | MOLECULE: | SERINE O-ACETYLTRANSFERASE |
| 19:00 | 7x29-B | 2 | 12.9 | 83 | 1190 | 6 | MOLECULE: | SPIKE GLYCOPROTEIN |
| 20:00 | 6q0x-A | 2 | 5.6 | 92 | 373 | 4 | MOLECULE: | SORTING NEXIN MVP1 |
| 21:00 | 6wg3-A | 2 | 21.8 | 79 | 561 | 5 | MOLECULE: | STRUCTURAL MAINTENANCE OF CHROMOSOMES PROTEI |
| 22:00 | 4drf-A | 2 | 4.5 | 54 | 408 | 11 | MOLECULE: | METALLOPHOSPHOESTERASE |
| 23:00 | 6zsi-D | 2 | 1.6 | 34 | 133 | 21 | MOLECULE: | RAS-RELATED PROTEIN RAB-8A |
| 24:00:00 | 8ap7-j | 2 | 40 | 81 | 168 | 1 | MOLECULE: | ATP SYNTHASE SUBUNIT A |
| 25:00:00 | 8pjn-b | 2 | 6.1 | 80 | 296 | 6 | MOLECULE: | E3 UBIQUITIN-PROTEIN TRANSFERASE RMND5A |
| 26:00:00 | 7o4i-1 | 2 | 8.5 | 44 | 522 | 11 | MOLECULE: | GENERAL TRANSCRIPTION AND DNA REPAIR FACTOR |
| 27:00:00 | 7rz4-A | 2 | 9.5 | 54 | 993 | 9 | MOLECULE: | GLUTAMATE RECEPTOR 2 |
| 28:00:00 | 8ebw-H | 2 | 5 | 75 | 532 | 12 | MOLECULE: | TFIIH BASAL TRANSCRIPTION FACTOR COMPLEX HEL |
| 29:00:00 | 6al9-B | 2 | 3.5 | 41 | 91 | 12 | MOLECULE: | CHORISMATE MUTASE |
| 30:00:00 | 7nyw-B | 2 | 16.8 | 97 | 858 | 5 | MOLECULE: | CHROMOSOME PARTITION PROTEIN MUKB |
| 31:00:00 | 7cgp-C | 2 | 32.2 | 98 | 223 | 11 | MOLECULE: | MITOCHONDRIAL IMPORT INNER MEMBRANE TRANSLOC |
| 32:00:00 | 6zyd-A | 2 | 29.3 | 50 | 323 | 2 | MOLECULE: | LOW CONDUCTANCE MECHANOSENSITIVE CHANNEL YNA |
| 33:00:00 | 3o1j-A | 2 | 4.4 | 86 | 273 | 9 | MOLECULE: | SENSOR PROTEIN TORS |
| 34:00:00 | 1pug-A | 2 | 3.5 | 43 | 94 | 12 | MOLECULE: | HYPOTHETICAL UPF0133 PROTEIN YBAB |
| 35:00:00 | 6xky-N | 2 | 4.1 | 97 | 271 | 7 | MOLECULE: | FLAGELLIN |
| 36:00:00 | 3zgx-B | 2 | 34.6 | 77 | 351 | 8 | MOLECULE: | CHROMOSOME PARTITION PROTEIN SMC |
| 37:00:00 | 5j9u-K | 2 | 5.8 | 61 | 120 | 16 | MOLECULE: | HISTONE ACETYLTRANSFERASE ESA1 |
| 38:00:00 | 8fef-F | 2 | 34.8 | 72 | 399 | 1 | MOLECULE: | VIRULENCE FACTOR MCE FAMILY PROTEIN |

| | | | | | | | | |
|---|---|---|---|---|---|---|---|---|
| 39:00:00 | 3jac-B | 2 | 37.6 | 94 | 918 | 1 | MOLECULE: | PIEZO-TYPE MECHANOSENSITIVE ION CHANNEL COMP |
| 40:00:00 | 5j9u-D | 2 | 5.5 | 63 | 120 | 8 | MOLECULE: | HISTONE ACETYLTRANSFERASE ESA1 |
| 41:00:00 | 7nh9-A | 2 | 6 | 91 | 327 | 11 | MOLECULE: | CMAX PROTEIN |
| 42:00:00 | 2x2v-B | 2 | 1.9 | 37 | 69 | 5 | MOLECULE: | ATP SYNTHASE SUBUNIT C |
| 43:00:00 | 7ur6-B | 2 | 11.3 | 49 | 132 | 6 | MOLECULE: | GP120 |
| 44:00:00 | 6um5-B | 2 | 4.1 | 48 | 131 | 2 | MOLECULE: | CH848 10.17 DT GP120 |
| 45:00:00 | 7ucg-M | 2 | 4.4 | 44 | 134 | 7 | MOLECULE: | ENVELOPE GLYCOPROTEIN GP41 |
| 46:00:00 | 6mu6-B | 2 | 3.5 | 45 | 128 | 11 | MOLECULE: | ENVELOPE GLYCOPROTEIN GP160 |
| 47:00:00 | 6u59-D | 2 | 4.2 | 44 | 120 | 7 | MOLECULE: | SOSIP.664 GP120,SOSIP.664 GP120 |
| 48:00:00 | 6vtt-B | 2 | 5.1 | 46 | 129 | 7 | MOLECULE: | ENVELOPE GLYCOPROTEIN GP120 |
| 49:00:00 | 6udj-F | 2 | 3.3 | 44 | 123 | 9 | MOLECULE: | 10-1074 FAB HEAVY CHAIN |

**DALI results: MnA-1 Penton**

| No: | Chain | Z | rmsd | lali | nres | %id | PDB Description |
|---|---|---|---|---|---|---|---|
| 1:00 | 6z7q-B | 11.8 | 3.9 | 224 | 416 | 14 | MOLECULE: PENTON PROTEIN |
| 2:00 | 6z7q-C | 11.7 | 3.9 | 224 | 416 | 14 | MOLECULE: PENTON PROTEIN |
| 3:00 | 6z7q-D | 11.7 | 3.9 | 224 | 416 | 14 | MOLECULE: PENTON PROTEIN |
| 4:00 | 6z7q-E | 11.7 | 3.9 | 224 | 416 | 14 | MOLECULE: PENTON PROTEIN |
| 5:00 | 2c6s-K | 11.6 | 3.9 | 222 | 443 | 15 | MOLECULE: ADENOVIRUS 2,12 PENTON BASE CHIMERA |
| 6:00 | 2c6s-H | 11.6 | 3.9 | 223 | 443 | 15 | MOLECULE: ADENOVIRUS 2,12 PENTON BASE CHIMERA |
| 7:00 | 2c6s-D | 11.6 | 3.9 | 222 | 443 | 15 | MOLECULE: ADENOVIRUS 2,12 PENTON BASE CHIMERA |
| 8:00 | 2c6s-G | 11.6 | 3.9 | 223 | 443 | 15 | MOLECULE: ADENOVIRUS 2,12 PENTON BASE CHIMERA |
| 9:00 | 2c6s-C | 11.6 | 3.9 | 223 | 443 | 15 | MOLECULE: ADENOVIRUS 2,12 PENTON BASE CHIMERA |
| 10:00 | 2c6s-L | 11.6 | 3.9 | 222 | 443 | 15 | MOLECULE: ADENOVIRUS 2,12 PENTON BASE CHIMERA |
| 11:00 | 2c6s-F | 11.6 | 3.9 | 222 | 443 | 15 | MOLECULE: ADENOVIRUS 2,12 PENTON BASE CHIMERA |
| 12:00 | 2c6s-A | 11.6 | 3.9 | 222 | 443 | 15 | MOLECULE: ADENOVIRUS 2,12 PENTON BASE CHIMERA |
| 13:00 | 6z7q-A | 11.6 | 3.9 | 222 | 416 | 14 | MOLECULE: PENTON PROTEIN |
| 14:00 | 2c6s-J | 11.6 | 3.9 | 223 | 443 | 15 | MOLECULE: ADENOVIRUS 2,12 PENTON BASE CHIMERA |
| 15:00 | 2c6s-N | 11.6 | 3.9 | 224 | 443 | 15 | MOLECULE: ADENOVIRUS 2,12 PENTON BASE CHIMERA |
| 16:00 | 2c6s-O | 11.6 | 3.9 | 222 | 443 | 15 | MOLECULE: ADENOVIRUS 2,12 PENTON BASE CHIMERA |
| 17:00 | 2c6s-E | 11.6 | 3.9 | 223 | 443 | 15 | MOLECULE: ADENOVIRUS 2,12 PENTON BASE CHIMERA |
| 18:00 | 2c6s-I | 11.6 | 3.9 | 222 | 443 | 15 | MOLECULE: ADENOVIRUS 2,12 PENTON BASE CHIMERA |
| 19:00 | 2c6s-B | 11.6 | 3.9 | 223 | 443 | 15 | MOLECULE: ADENOVIRUS 2,12 PENTON BASE CHIMERA |
| 20:00 | 2c6s-M | 11.5 | 3.9 | 222 | 443 | 14 | MOLECULE: ADENOVIRUS 2,12 PENTON BASE CHIMERA |
| 21:00 | 2c9g-B | 11.5 | 4 | 223 | 447 | 15 | MOLECULE: PENTON PROTEIN |
| 22:00 | 2c9g-A | 11.5 | 4 | 223 | 447 | 15 | MOLECULE: PENTON PROTEIN |
| 23:00 | 2c9g-D | 11.5 | 4 | 223 | 447 | 15 | MOLECULE: PENTON PROTEIN |
| 24:00:00 | 2c9g-C | 11.5 | 4 | 223 | 447 | 15 | MOLECULE: PENTON PROTEIN |
| 25:00:00 | 2c9g-E | 11.5 | 4 | 223 | 447 | 15 | MOLECULE: PENTON PROTEIN |
| 26:00:00 | 1x9p-A | 11.4 | 4 | 222 | 460 | 15 | MOLECULE: PENTON PROTEIN |
| 27:00:00 | 2c9f-B | 11.4 | 4 | 222 | 440 | 15 | MOLECULE: PENTON PROTEIN |
| 28:00:00 | 2c9f-D | 11.4 | 4.1 | 223 | 440 | 15 | MOLECULE: PENTON PROTEIN |
| 29:00:00 | 1x9t-A | 11.4 | 4.1 | 223 | 440 | 15 | MOLECULE: PENTON PROTEIN |
| 30:00:00 | 2c9f-A | 11.4 | 4 | 222 | 440 | 15 | MOLECULE: PENTON PROTEIN |
| 31:00:00 | 2c9f-E | 11.4 | 4.1 | 223 | 440 | 15 | MOLECULE: PENTON PROTEIN |
| 32:00:00 | 2c9f-C | 11.4 | 4.1 | 223 | 440 | 15 | MOLECULE: PENTON PROTEIN |
| 33:00:00 | 3izo-A | 11.2 | 4 | 224 | 455 | 14 | MOLECULE: PENTON PROTEIN |
| 34:00:00 | 3izo-E | 11.2 | 4 | 224 | 455 | 14 | MOLECULE: PENTON PROTEIN |
| 35:00:00 | 3izo-D | 11.2 | 4 | 224 | 455 | 14 | MOLECULE: PENTON PROTEIN |
| 36:00:00 | 3izo-C | 11.2 | 4 | 224 | 455 | 14 | MOLECULE: PENTON PROTEIN |
| 37:00:00 | 6g41-F | 11.2 | 3.2 | 153 | 301 | 7 | MOLECULE: MINOR CAPSID PR OTEIN |
| 38:00:00 | 4aqq-A | 11.1 | 4.1 | 226 | 462 | 14 | MOLECULE: L2 PROTEIN III (PENTON BASE ) |
| 39:00:00 | 3izo-B | 11.1 | 3.9 | 222 | 455 | 14 | MOLECULE: PENTON PROTEIN |
| 40:00:00 | 4ar2-A | 11 | 4.2 | 226 | 455 | 13 | MOLECULE: L2 PROTEIN III (PENTON BASE ) |
| 41:00:00 | 6g42-D | 10.9 | 3.2 | 155 | 302 | 6 | MOLECULE: MINOR CAPSID PR OTEIN |
| 42:00:00 | 6g41-I | 10.9 | 3.2 | 154 | 301 | 6 | MOLECULE: MINOR CAPSID PR OTEIN |

| | | | | | | | | | |
|---|---|---|---|---|---|---|---|---|---|
| 43:00:00 | 6g42-E | 10.8 | 3.1 | 154 | 301 | 6 | MOLECULE: | MINOR CAPSID PR | OTEIN |
| 44:00:00 | 6g41-J | 10.8 | 3.2 | 154 | 301 | 6 | MOLECULE: | MINOR CAPSID PR | OTEIN |
| 45:00:00 | 6g42-C | 10.7 | 3.2 | 155 | 302 | 6 | MOLECULE: | MINOR CAPSID PR | OTEIN |
| 46:00:00 | 6g41-A | 10.7 | 3.5 | 155 | 297 | 6 | MOLECULE: | MINOR CAPSID PR | OTEIN |
| 47:00:00 | 6g42-B | 10.7 | 3.2 | 154 | 301 | 6 | MOLECULE: | MINOR CAPSID PR | OTEIN |
| 48:00:00 | 6g42-A | 10.7 | 3.1 | 153 | 301 | 7 | MOLECULE: | MINOR CAPSID PR | OTEIN |
| 49:00:00 | 6g41-C | 10.7 | 3.2 | 154 | 301 | 6 | MOLECULE: | MINOR CAPSID PR | OTEIN |
| 50:00:00 | 6g41-E | 10.7 | 3.1 | 152 | 301 | 7 | MOLECULE: | MINOR CAPSID PR | OTEIN |
| 51:00:00 | 6g41-H | 10.7 | 3.1 | 152 | 301 | 7 | MOLECULE: | MINOR CAPSID PR | OTEIN |
| 52:00:00 | 6g41-D | 10.6 | 3.2 | 153 | 291 | 7 | MOLECULE: | MINOR CAPSID PR | OTEIN |
| 53:00:00 | 6g41-B | 10.6 | 3.1 | 152 | 295 | 7 | MOLECULE: | MINOR CAPSID PR | OTEIN |
| 54:00:00 | 8rbt-S | 8.9 | 3.3 | 209 | 663 | 12 | MOLECULE: | MAJOR CAPSID PR | OTEIN |
| 55:00:00 | 8coi-I | 8.6 | 2.6 | 99 | 249 | 15 | MOLECULE: | ADDOBODY | |
| 56:00:00 | 6g41-G | 8.1 | 3.1 | 152 | 287 | 7 | MOLECULE: | MINOR CAPSID PR | OTEIN |
| 57:00:00 | 8coi-A | 8 | 2.7 | 96 | 245 | 15 | MOLECULE: | ADDOBODY | |
| 58:00:00 | 8coi-N | 8 | 2.6 | 95 | 232 | 16 | MOLECULE: | ADDOBODY | |
| 59:00:00 | 3j26-N | 8 | 4 | 165 | 362 | 7 | MOLECULE: | CAPSID PROTEIN | V20 |
| 60:00:00 | 8coi-C | 7.9 | 2.7 | 99 | 249 | 15 | MOLECULE: | ADDOBODY | |
| 61:00:00 | 8coi-Q | 7.9 | 2.6 | 96 | 232 | 16 | MOLECULE: | ADDOBODY | |
| 62:00:00 | 8coi-J | 7.9 | 2.6 | 94 | 243 | 15 | MOLECULE: | ADDOBODY | |
| 63:00:00 | 8coi-E | 7.8 | 2.5 | 94 | 228 | 15 | MOLECULE: | ADDOBODY | |
| 64:00:00 | 8coi-P | 7.6 | 2.6 | 92 | 243 | 16 | MOLECULE: | ADDOBODY | |
| 65:00:00 | 8coi-H | 7.5 | 2.5 | 96 | 243 | 16 | MOLECULE: | ADDOBODY | |
| 66:00:00 | 8coi-F | 7.2 | 2.6 | 98 | 240 | 15 | MOLECULE: | ADDOBODY | |
| 67:00:00 | 8coi-B | 6.7 | 2.5 | 96 | 246 | 16 | MOLECULE: | ADDOBODY | |
| 68:00:00 | 8coi-G | 6.7 | 2.6 | 96 | 243 | 16 | MOLECULE: | ADDOBODY | |
| 69:00:00 | 8coi-M | 6.7 | 2.7 | 96 | 237 | 16 | MOLECULE: | ADDOBODY | |
| 70:00:00 | 8coi-R | 6.7 | 2.6 | 93 | 227 | 15 | MOLECULE: | ADDOBODY | |
| 71:00:00 | 8coi-D | 6.7 | 2.6 | 96 | 244 | 16 | MOLECULE: | ADDOBODY | |
| 72:00:00 | 8coi-K | 6.6 | 2.6 | 95 | 250 | 16 | MOLECULE: | ADDOBODY | |
| 73:00:00 | 8qb3-E | 6.5 | 2.9 | 89 | 233 | 15 | MOLECULE: | ADDOBODY | |
| 74:00:00 | 8coi-S | 6.5 | 2.6 | 94 | 230 | 15 | MOLECULE: | ADDOBODY | |
| 75:00:00 | 8coi-L | 6.5 | 2.6 | 95 | 244 | 16 | MOLECULE: | ADDOBODY | |
| 76:00:00 | 8coi-T | 6.3 | 2.7 | 85 | 212 | 15 | MOLECULE: | ADDOBODY | |
| 77:00:00 | 8coi-O | 6.3 | 2.5 | 91 | 229 | 14 | MOLECULE: | ADDOBODY | |
| 78:00:00 | 6wpc-C | 6 | 3.3 | 112 | 588 | 8 | MOLECULE: | CRY1A.2 | |
| 79:00:00 | 6ds5-E | 5.9 | 3.8 | 105 | 160 | 13 | MOLECULE: | SEIPIN | |
| 80:00:00 | 6ds5-G | 5.9 | 3.8 | 105 | 160 | 13 | MOLECULE: | SEIPIN | |
| 81:00:00 | 7cz6-C | 5.9 | 3.2 | 99 | 126 | 7 | MOLECULE: | CAPSID PROTEIN | |
| 82:00:00 | 7rsl-A | 5.8 | 5.3 | 108 | 235 | 12 | MOLECULE: | SEIPIN | |
| 83:00:00 | 7rsl-F | 5.8 | 3.2 | 106 | 235 | 12 | MOLECULE: | SEIPIN | |
| 84:00:00 | 7rsl-D | 5.8 | 3.2 | 106 | 235 | 12 | MOLECULE: | SEIPIN | |
| 85:00:00 | 1ciy-A | 5.8 | 3.8 | 107 | 577 | 7 | MOLECULE: | CRYIA(A) | |
| 86:00:00 | 7rsl-G | 5.8 | 5.2 | 109 | 235 | 13 | MOLECULE: | SEIPIN | |

| | | | | | | | | | | |
|---|---|---|---|---|---|---|---|---|---|---|
| 87:00:00 | 6ds5-F | 5.8 | 3.8 | 105 | 160 | 13 | MOLECULE: | SEIPIN | | | |
| 88:00:00 | 6ds5-C | 5.8 | 3.8 | 105 | 160 | 13 | MOLECULE: | SEIPIN | | | |
| 89:00:00 | 6ds5-H | 5.8 | 3.8 | 105 | 160 | 13 | MOLECULE: | SEIPIN | | | |
| 90:00:00 | 6ds5-K | 5.8 | 3.8 | 105 | 160 | 13 | MOLECULE: | SEIPIN | | | |
| 91:00:00 | 6ds5-D | 5.8 | 3.8 | 105 | 160 | 13 | MOLECULE: | SEIPIN | | | |
| 92:00:00 | 6ds5-A | 5.8 | 3.8 | 105 | 160 | 13 | MOLECULE: | SEIPIN | | | |
| 93:00:00 | 6ds5-B | 5.8 | 3.8 | 105 | 160 | 13 | MOLECULE: | SEIPIN | | | |
| 94:00:00 | 6ds5-I | 5.8 | 3.8 | 105 | 160 | 13 | MOLECULE: | SEIPIN | | | |
| 95:00:00 | 6ds5-J | 5.8 | 3.6 | 103 | 160 | 13 | MOLECULE: | SEIPIN | | | |
| 96:00:00 | 7rsl-I | 5.8 | 5.2 | 110 | 235 | 13 | MOLECULE: | SEIPIN | | | |
| 97:00:00 | 7rsl-B | 5.8 | 3.1 | 102 | 235 | 13 | MOLECULE: | SEIPIN | | | |
| 98:00:00 | 7rsl-C | 5.8 | 5.2 | 111 | 235 | 13 | MOLECULE: | SEIPIN | | | |
| 99:00:00 | 6owk-A | 5.7 | 3.2 | 106 | 582 | 7 | MOLECULE: | PESTICIDAL CRYS | TAL PRO | TEIN | CRY1BE, CI |
| 0:00 | 6za2-A | 5.7 | 3.9 | 102 | 1081 | 7 | MOLECULE: | POR SECRETION S | YSTEM P | ROTEI | N PORU |
| 1:00 | 7rsl-E | 5.7 | 5.3 | 110 | 235 | 12 | MOLECULE: | SEIPIN | | | |
| 2:00 | 6wpc-B | 5.7 | 3.8 | 110 | 588 | 10 | MOLECULE: | CRY1A.2 | | | |
| 3:00 | 9buv-A | 5.6 | 4 | 106 | 573 | 8 | MOLECULE: | PESTICIDAL CRYS | TAL PRO | TEIN | CRY1FA |
| 4:00 | 6wpc-A | 5.6 | 3.9 | 111 | 588 | 9 | MOLECULE: | CRY1A.2 | | | |
| 5:00 | 8qb3-C | 5.5 | 2.7 | 88 | 216 | 15 | MOLECULE: | ADDOBODY | | | |
| 6:00 | 2c9k-A | 5.5 | 4 | 108 | 598 | 7 | MOLECULE: | PESTICIDAL CRYS | TAL PRO | TEIN | CRY4AA |
| 7:00 | 7rsl-J | 5.4 | 3.2 | 106 | 235 | 10 | MOLECULE: | SEIPIN | | | |
| 8:00 | 7rsl-H | 5.4 | 3.2 | 106 | 235 | 10 | MOLECULE: | SEIPIN | | | |
| 9:00 | 3eb7-A | 5.3 | 3.5 | 106 | 589 | 8 | MOLECULE: | INSECTICIDAL DE | LTA-END | OTOXI | N CRY8EA1 |
| 10:00 | 3eb7-C | 5.3 | 3.2 | 104 | 588 | 10 | MOLECULE: | INSECTICIDAL DE | LTA-END | OTOXI | N CRY8EA1 |
| 11:00 | 8qb3-D | 5.3 | 2.8 | 85 | 206 | 15 | MOLECULE: | ADDOBODY | | | |
| 12:00 | 8qb3-A | 5.2 | 2.7 | 87 | 233 | 15 | MOLECULE: | ADDOBODY | | | |
| 13:00 | 3eb7-B | 5.2 | 3.5 | 112 | 589 | 13 | MOLECULE: | INSECTICIDAL DE | LTA-END | OTOXI | N CRY8EA1 |
| 14:00 | 4qx0-A | 5.2 | 3.7 | 113 | 584 | 5 | MOLECULE: | PESTICIDAL CRYS | TAL PRO | TEIN | CRY3AA |
| 15:00 | 6za2-B | 5.2 | 3.9 | 100 | 1083 | 7 | MOLECULE: | POR SECRETION S | YSTEM P | ROTEI | N PORU |
| 16:00 | 4qx2-A | 5.2 | 3.6 | 113 | 584 | 4 | MOLECULE: | PESTICIDAL CRYS | TAL PRO | TEIN | CRY3AA |
| 17:00 | 4qx3-A | 5.2 | 3.5 | 112 | 584 | 5 | MOLECULE: | PESTICIDAL CRYS | TAL PRO | TEIN | CRY3AA |
| 18:00 | 6ovb-A | 5.1 | 4.4 | 110 | 566 | 10 | MOLECULE: | ACTIVE CORE CRY | STAL TO | XIN P | ROTEIN 1D |
| 19:00 | 8j3x-B | 5.1 | 3.3 | 91 | 390 | 7 | MOLECULE: | PUTATIVE POLYSA | CCHARID | E-BIN | DING PROT |
| 20:00 | 6lfp-A | 5.1 | 3.6 | 111 | 582 | 7 | MOLECULE: | CRY3AA PROTEIN | | | |
| 21:00 | 8qb3-B | 5.1 | 3 | 89 | 233 | 15 | MOLECULE: | ADDOBODY | | | |
| 22:00 | 8j3y-A | 5 | 4.2 | 93 | 392 | 5 | MOLECULE: | PUTATIVE POLYSA | CCHARID | E-BIN | DING PROT |
| 23:00 | 7ear-A | 5 | 3.3 | 108 | 584 | 5 | MOLECULE: | CRYSTALINE ENTO | MOCIDAL | PROT | OXIN |
| 24:00:00 | 4qx1-A | 5 | 3.9 | 116 | 584 | 5 | MOLECULE: | PESTICIDAL CRYS | TAL PRO | TEIN | CRY3AA |
| 25:00:00 | 4arx-D | 5 | 4 | 105 | 576 | 8 | MOLECULE: | PESTICIDAL CRYS | TAL PRO | TEIN | CRY1AC |
| 26:00:00 | 1dlc-A | 5 | 3.3 | 109 | 584 | 6 | MOLECULE: | DELTA-ENDOTOXIN | CRYIII | A | |
| 27:00:00 | 6mlu-B | 5 | 4.2 | 97 | 153 | 6 | MOLECULE: | SEIPIN | | | |
| 28:00:00 | 4arx-A | 5 | 4.1 | 106 | 578 | 8 | MOLECULE: | PESTICIDAL CRYS | TAL PRO | TEIN | CRY1AC |
| 29:00:00 | 4arx-C | 4.9 | 4 | 106 | 577 | 8 | MOLECULE: | PESTICIDAL CRYS | TAL PRO | TEIN | CRY1AC |
| 30:00:00 | 4arx-B | 4.9 | 4 | 106 | 578 | 8 | MOLECULE: | PESTICIDAL CRYS | TAL PRO | TEIN | CRY1AC |

| | | | | | | | | | | | |
|---|---|---|---|---|---|---|---|---|---|---|---|
| 31:00:00 | 8j3x-A | 4.9 | 4.2 | 92 | 389 | 5 | MOLECULE: | PUTATIVE POLYSA | CCHARID | E-BIN | DING PROT |
| 32:00:00 | 4ary-B | 4.9 | 4 | 105 | 580 | 9 | MOLECULE: | PESTICIDAL CRYS | TAL PRO | TEIN | CRY1AC |
| 33:00:00 | 4ary-A | 4.9 | 4 | 106 | 578 | 8 | MOLECULE: | PESTICIDAL CRYS | TAL PRO | TEIN | CRY1AC |
| 34:00:00 | 4ary-C | 4.9 | 4 | 106 | 577 | 8 | MOLECULE: | PESTICIDAL CRYS | TAL PRO | TEIN | CRY1AC |
| 35:00:00 | 4w8j-A | 4.8 | 3.9 | 103 | 1017 | 8 | MOLECULE: | PESTICIDAL CRYS | TAL PRO | TEIN | CRY1AC |
| 36:00:00 | 5hly-A | 4.8 | 4.3 | 101 | 299 | 10 | MOLECULE: | INHIBIN BETA A | CHAIN | | |
| 37:00:00 | 5hlz-A | 4.8 | 4.3 | 104 | 204 | 11 | MOLECULE: | INHIBIN BETA A | CHAIN | | |
| 38:00:00 | 5hlz-C | 4.7 | 4.4 | 100 | 204 | 8 | MOLECULE: | INHIBIN BETA A | CHAIN | | |
| 39:00:00 | 6dj4-A | 4.7 | 3.7 | 112 | 571 | 5 | MOLECULE: | CRY1A.105 | | | |
| 40:00:00 | 1ji6-A | 4.6 | 3.5 | 110 | 589 | 7 | MOLECULE: | PESTICIDIAL CRY | STAL PR | OTEIN | CRY3BB |
| 41:00:00 | 5g56-A | 4.6 | 4.4 | 100 | 706 | 12 | MOLECULE: | CARBOHYDRATE BI | NDING F | AMILY | 6 |
| 42:00:00 | 4ary-D | 4.6 | 4 | 99 | 577 | 8 | MOLECULE: | PESTICIDAL CRYS | TAL PRO | TEIN | CRY1AC |
| 43:00:00 | 1w99-A | 4.6 | 4.4 | 100 | 558 | 10 | MOLECULE: | PESTICIDIAL CRY | STAL PR | OTEIN | CRY4BA |
| 44:00:00 | 5zi1-A | 4.5 | 3.5 | 114 | 590 | 7 | MOLECULE: | INSECTICIDAL CR | YSTAL P | ROTEI | N CRY7CAI |
| 45:00:00 | 5la2-B | 4.5 | 4.5 | 97 | 465 | 12 | MOLECULE: | CARBOHYDRATE BI | NDING F | AMILY | 6 |
| 46:00:00 | 6mlu-A | 4.5 | 4.3 | 93 | 153 | 8 | MOLECULE: | SEIPIN | | | |
| 47:00:00 | 3iii-A | 4.4 | 3.3 | 110 | 556 | 10 | MOLECULE: | COCE/NOND FAMIL | Y HYDRO | LASE | |
| 48:00:00 | 1stm-A | 4.4 | 3.6 | 100 | 141 | 5 | MOLECULE: | SATELLITE PANIC | UM MOSA | IC VI | RUS |
| 49:00:00 | 5zi1-B | 4.4 | 3.5 | 114 | 590 | 7 | MOLECULE: | INSECTICIDAL CR | YSTAL P | ROTEI | N CRY7CAI |
| 50:00:00 | 5hlz-G | 4.4 | 4.4 | 98 | 204 | 9 | MOLECULE: | INHIBIN BETA A | CHAIN | | |
| 51:00:00 | 3wnm-A | 4.4 | 3.4 | 92 | 701 | 9 | MOLECULE: | CYCLOISOMALTOOL | IGOSACC | HARID | E GLUCAN( |
| 52:00:00 | 8w7n-A | 4.4 | 4.8 | 98 | 1011 | 10 | MOLECULE: | PESTICIDAL CRYS | TAL PRO | TEIN | CRY1AA |
| 53:00:00 | 1pnf-A | 4.3 | 3.7 | 97 | 314 | 8 | MOLECULE: | PEPTIDE-N(4)-(N | #NAME? | #### | -D-GLUCOS |
| 54:00:00 | 2y8k-A | 4.3 | 4.4 | 101 | 480 | 9 | MOLECULE: | CARBOHYDRATE BI | NDING F | AMILY | 6 |
| 55:00:00 | 3wnl-A | 4.3 | 3.4 | 91 | 701 | 8 | MOLECULE: | CYCLOISOMALTOOL | IGOSACC | HARID | E GLUCAN( |
| 56:00:00 | 4jdm-C | 4.2 | 3.5 | 96 | 269 | 7 | MOLECULE: | VIRULENCE PLASM | ID PROT | EIN P | GP3-D |
| 57:00:00 | 6i56-A | 4.2 | 4.8 | 89 | 279 | 4 | MOLECULE: | PHAGE-LIKE ELEM | ENT PBS | X PRO | TEIN XEPA |
| 58:00:00 | 3wnk-A | 4.2 | 3.1 | 88 | 712 | 8 | MOLECULE: | CYCLOISOMALTOOL | IGOSACC | HARID | E GLUCAN( |
| 59:00:00 | 3wno-A | 4.2 | 3.2 | 87 | 700 | 8 | MOLECULE: | CYCLOISOMALTOOL | IGOSACC | HARID | E GLUCAN( |
| 60:00:00 | 5jsd-A | 4.1 | 3.8 | 87 | 548 | 7 | MOLECULE: | PHIAB6 TAILSPIK | E | | |
| 61:00:00 | 8erm-B | 4.1 | 2.6 | 72 | 203 | 8 | MOLECULE: | B-TYPE FLAGELLI | N | | |
| 62:00:00 | 8j3y-B | 4.1 | 4 | 92 | 392 | 4 | MOLECULE: | PUTATIVE POLYSA | CCHARID | E-BIN | DING PROT |
| 63:00:00 | 7r1m-A | 4 | 3.6 | 98 | 156 | 8 | MOLECULE: | EXOSPORIUM PROT | EIN | | |
| 64:00:00 | 3qd6-D | 4 | 3.8 | 95 | 135 | 9 | MOLECULE: | CD40 LIGAND | | | |
| 65:00:00 | 7vt4-A | 4 | 4.5 | 83 | 438 | 5 | MOLECULE: | ENDOGLUCANASE H | | | |
| 66:00:00 | 5la0-A | 4 | 4.6 | 96 | 476 | 9 | MOLECULE: | CARBOHYDRATE BI | NDING F | AMILY | 6 |
| 67:00:00 | 4mtm-A | 3.9 | 4.4 | 81 | 137 | 10 | MOLECULE: | PUTATIVE TAIL F | IBER PR | OTEIN | |
| 68:00:00 | 8xcg-J | 3.9 | 6.5 | 106 | 598 | 7 | MOLECULE: | TAIL TIP PROTEI | N M | | |
| 69:00:00 | 1yq8-A | 3.9 | 3.6 | 95 | 191 | 6 | MOLECULE: | MINOR CAPSID PR | OTEIN | | |
| 70:00:00 | 5ak1-A | 3.9 | 4.1 | 100 | 842 | 12 | MOLECULE: | CARBOHYDRATE BI | NDING F | AMILY | 6 |
| 71:00:00 | 3wnn-A | 3.9 | 3.1 | 79 | 699 | 11 | MOLECULE: | CYCLOISOMALTOOL | IGOSACC | HARID | E GLUCAN( |
| 72:00:00 | 4moa-A | 3.9 | 4 | 99 | 602 | 10 | MOLECULE: | PESTICIDAL CRYS | TAL PRO | TEIN | CRY4BA |
| 73:00:00 | 7crc-B | 3.8 | 3.3 | 91 | 1112 | 5 | MOLECULE: | NAD+ HYDROLASE | (NADASE | ) | |
| 74:00:00 | 6s44-A | 3.8 | 3.4 | 92 | 146 | 3 | MOLECULE: | CAPSID PROTEIN | | | |

| | | | | | | | | | | | | |
|---|---|---|---|---|---|---|---|---|---|---|---|---|
| 75:00:00 | 3a21-A | 3.8 | 3.3 | 89 | 614 | 6 | MOLECULE: | PUTATIVE SECRET | ED ALPH | A-GAL | ACTOSIDAS | |
| 76:00:00 | 3wno-B | 3.8 | 3.1 | 78 | 699 | 10 | MOLECULE: | CYCLOISOMALTOOL | IGOSACC | HARID | E GLUCAN | |
| 77:00:00 | 8yzo-A | 3.7 | 7.3 | 101 | 506 | 12 | MOLECULE: | LIPASE | | | | |
| 78:00:00 | 7pug-A | 3.7 | 4.7 | 106 | 829 | 4 | MOLECULE: | XYLAN ALPHA-1,2 | #NAME? | ONIDA | SE | |
| 79:00:00 | 5zru-A | 3.7 | 5.2 | 103 | 576 | 6 | MOLECULE: | ALPHA-1,3-GLUCA | NASE | | | |
| 80:00:00 | 6wpc-D | 3.7 | 3.6 | 115 | 588 | 10 | MOLECULE: | CRY1A.2 | | | | |
| 81:00:00 | 5la2-A | 3.7 | 4.4 | 100 | 464 | 10 | MOLECULE: | CARBOHYDRATE BI | NDING F | AMILY | | 6 |
| 82:00:00 | 5hlz-E | 3.7 | 4.5 | 105 | 204 | 7 | MOLECULE: | INHIBIN BETA A | CHAIN | | | |
| 83:00:00 | 6nff-A | 3.6 | 7.3 | 102 | 794 | 10 | MOLECULE: | XAA-PRO DIPEPTI | DYL-PEP | TIDAS | E | |
| 84:00:00 | 2hey-F | 3.6 | 4.1 | 96 | 135 | 11 | MOLECULE: | TUMOR NECROSIS | FACTOR | LIGAN | D SUPERFA | |
| 85:00:00 | 5e7t-H | 3.6 | 4.5 | 97 | 173 | 7 | MOLECULE: | MINOR STRUCTURA | L PROTE | IN 4 | | |
| 86:00:00 | 7ufg-A | 3.6 | 4.3 | 103 | 1151 | 6 | MOLECULE: | PAPPALYSIN-1 | | | | |
| 87:00:00 | 2wz8-A | 3.6 | 3.5 | 87 | 135 | 10 | MOLECULE: | CELLULOSOME PRO | TEIN DO | CKERI | N TYPE I | |
| 88:00:00 | 3wnn-B | 3.6 | 3.4 | 86 | 698 | 8 | MOLECULE: | CYCLOISOMALTOOL | IGOSACC | HARID | E GLUCAN | |
| 89:00:00 | 3wqb-A | 3.5 | 3.9 | 104 | 597 | 13 | MOLECULE: | EXTRACELLULAR S | ERINE P | ROTEA | SE | |
| 90:00:00 | 7a02-A | 3.5 | 6 | 83 | 112 | 10 | MOLECULE: | DUF3992 DOMAIN- | CONTAIN | ING P | ROTEIN | |
| 91:00:00 | 3c7g-A | 3.5 | 4.1 | 101 | 488 | 3 | MOLECULE: | ENDO-1,4-BETA-X | YLANASE | | | |
| 92:00:00 | 4ind-A | 3.4 | 4.6 | 89 | 377 | 7 | MOLECULE: | C381 TURRET PRO | TEIN | | | |
| 93:00:00 | 4r4x-A | 3.3 | 4.2 | 109 | 544 | 13 | MOLECULE: | PNGF-II | | | | |
| 94:00:00 | 8gld-A | 3.3 | 3.3 | 85 | 365 | 8 | MOLECULE: | SGNH HYDROLASE- | TYPE ES | TERAS | E DOMAIN- | |
| 95:00:00 | 2wys-B | 3.2 | 3.8 | 91 | 516 | 7 | MOLECULE: | ENDO-1,4-BETA-X | YLANASE | Y | | |
| 96:00:00 | 7kx0-A | 3.2 | 4 | 93 | 140 | 5 | MOLECULE: | CD70 ANTIGEN | | | | |
| 97:00:00 | 5la1-A | 3.2 | 5.8 | 93 | 474 | 13 | MOLECULE: | CARBOHYDRATE BI | NDING F | AMILY | | 6 |
| 98:00:00 | 4qpw-A | 3.1 | 3.6 | 90 | 142 | 9 | MOLECULE: | GLYCOSYL HYDROL | ASE FAM | ILY 1 | | 0 |
| 99:00:00 | 7w63-B | 3.1 | 4.2 | 94 | 395 | 10 | MOLECULE: | TOXIN-COREGULAT | ED PILU | S BIO | SYNTHESIS | |
| 0:00 | 1a6c-A | 3.1 | 6.8 | 91 | 513 | 4 | MOLECULE: | TOBACCO RINGSPO | T VIRUS | CAPS | ID PROTEIN | |
| 1:00 | 6iwy-A | 3 | 2.7 | 73 | 366 | 11 | MOLECULE: | FLAGELLAR HOOK- | ASSOCIA | TED P | ROTEIN 2 | |
| 2:00 | 3b9t-A | 3 | 5.5 | 118 | 419 | 5 | MOLECULE: | TWIN-ARGININE T | RANSLOC | ATION | PATHWAY S | |
| 3:00 | 6q5u-R | 2.9 | 3.2 | 87 | 125 | 7 | MOLECULE: | MAJOR CAPSID PR | OTEIN ( | P3) | | |
| 4:00 | 5ubu-B | 2.7 | 5.2 | 116 | 331 | 7 | MOLECULE: | PUTATIVE ACETAM | IDASE/F | ORMA | IDASE | |
| 5:00 | 8hqo-P | 2.5 | 4 | 68 | 170 | 6 | MOLECULE: | PORTAL PROTEIN | | | | |
| 6:00 | 6x80-C | 2.5 | 16.8 | 80 | 574 | 14 | MOLECULE: | FLAGELLIN A | | | | |

**DALI results: MnA-1 Macc**

| No: | Chain | Z | rmsd | lali | nres | %id | PDB Description | | | |
|---|---|---|---|---|---|---|---|---|---|---|
| 1:00 | 5jdo-C | 5.9 | 3 | 84 | 141 | 7 | MOLECULE: | HAPTOGLOBI | N-HAEMOG | LOBIN RECEPTOR |
| 2:00 | 7kyr-A | 5.7 | 3.9 | 88 | 151 | 11 | MOLECULE: | MYOGLOBIN | | |
| 3:00 | 3e5o-A | 5.6 | 4.1 | 88 | 153 | 11 | MOLECULE: | MYOGLOBIN | | |
| 4:00 | 6g5b-A | 5.6 | 4 | 88 | 154 | 11 | MOLECULE: | MYOGLOBIN | | |
| 5:00 | 7sph-A | 5.6 | 4 | 91 | 154 | 12 | MOLECULE: | MYOGLOBIN | | |
| 6:00 | 7a44-A | 5.5 | 4 | 91 | 154 | 12 | MOLECULE: | MYOGLOBIN | | |
| 7:00 | 1jw8-A | 5.4 | 4.1 | 90 | 154 | 11 | MOLECULE: | MYOGLOBIN | | |
| 8:00 | 6e03-A | 5.4 | 4.1 | 88 | 153 | 11 | MOLECULE: | MYOGLOBIN | | |
| 9:00 | 4tyx-A | 5.4 | 4 | 88 | 153 | 11 | MOLECULE: | MYOGLOBIN | | |
| 10:00 | 5m3s-A | 5.4 | 4.2 | 90 | 154 | 11 | MOLECULE: | MYOGLOBIN | | |
| 11:00 | 1ch9-A | 5.4 | 4.1 | 90 | 154 | 12 | MOLECULE: | PROTEIN (MYOGLOBIN) | | |
| 12:00 | 5b84-A | 5.4 | 4.1 | 88 | 153 | 11 | MOLECULE: | MYOGLOBIN | | |
| 13:00 | 7spe-A | 5.3 | 4.1 | 90 | 154 | 11 | MOLECULE: | MYOGLOBIN | | |
| 14:00 | 2r4w-A | 5.3 | 3.3 | 84 | 145 | 8 | MOLECULE: | GLOBIN-1 | | |
| 15:00 | 2blh-A | 5.3 | 4 | 90 | 153 | 12 | MOLECULE: | MYOGLOBIN | | |
| 16:00 | 7l3u-A | 5.3 | 4.1 | 91 | 153 | 11 | MOLECULE: | MYOGLOBIN | | |
| 17:00 | 8eko-A | 5.3 | 4.1 | 88 | 153 | 11 | MOLECULE: | MYOGLOBIN | | |
| 18:00 | 2ri4-C | 5.3 | 3 | 83 | 138 | 7 | MOLECULE: | HEMOGLOBIN | SUBUNIT | ALPHA-1/2 |
| 19:00 | 6az6-A | 5.2 | 8.3 | 105 | 209 | 8 | MOLECULE: | GNTR FAMIL | Y TRANSC | RIPTIONAL REGULATOR |
| 20:00 | 1myt-A | 5.2 | 4.1 | 85 | 145 | 9 | MOLECULE: | MYOGLOBIN | | |
| 21:00 | 5zzf-A | 5.2 | 4.2 | 88 | 153 | 11 | MOLECULE: | MYOGLOBIN | | |
| 22:00 | 7xcq-A | 5.2 | 4.1 | 88 | 153 | 11 | MOLECULE: | MYOGLOBIN | | |
| 23:00 | 7xc9-A | 5.2 | 4.1 | 88 | 153 | 11 | MOLECULE: | MYOGLOBIN | | |
| 24:00:00 | 1dtm-A | 5.2 | 4.1 | 91 | 152 | 11 | MOLECULE: | RECOMBINAN | T SPERM | WHALE MYOGLOBIN VARIANT H93G |
| 25:00:00 | 8j4k-A | 5.2 | 4.1 | 88 | 153 | 11 | MOLECULE: | MYOGLOBIN | | |
| 26:00:00 | 5yzf-A | 5.2 | 4 | 89 | 154 | 12 | MOLECULE: | SPERM WHAL | E MYOGLO | BIN |
| 27:00:00 | 7xzi-B | 5.1 | 4.4 | 88 | 166 | 2 | MOLECULE: | CTAP3 | | |
| 28:00:00 | 5hav-A | 5.1 | 4.1 | 90 | 153 | 9 | MOLECULE: | MYOGLOBIN | | |
| 29:00:00 | 6krc-A | 5.1 | 4.1 | 90 | 153 | 9 | MOLECULE: | MYOGLOBIN | | |
| 30:00:00 | 7l3y-A | 5.1 | 4 | 88 | 153 | 11 | MOLECULE: | MYOGLOBIN | | |
| 31:00:00 | 3qm9-A | 5.1 | 4 | 84 | 144 | 7 | MOLECULE: | MYOGLOBIN | | |
| 32:00:00 | 3a59-B | 5.1 | 3.5 | 85 | 146 | 9 | MOLECULE: | HEMOGLOBIN | SUBUNIT | ALPHA-A |
| 33:00:00 | 5yce-A | 5.1 | 4.1 | 88 | 151 | 11 | MOLECULE: | MYOGLOBIN | | |
| 34:00:00 | 5yci-B | 5 | 4.1 | 93 | 157 | 16 | MOLECULE: | ANCESTRAL | MYOGLOBI | N AMBWB' OF BASILOSAURUS RELATI |
| 35:00:00 | 7spf-A | 5 | 4.1 | 92 | 152 | 11 | MOLECULE: | MYOGLOBIN | | |
| 36:00:00 | 6ls8-I | 5 | 4 | 91 | 153 | 13 | MOLECULE: | MYOGLOBIN | | |
| 37:00:00 | 5zzg-A | 5 | 4.1 | 88 | 153 | 11 | MOLECULE: | MYOGLOBIN | | |
| 38:00:00 | 2nrm-A | 5 | 4 | 85 | 146 | 7 | MOLECULE: | MYOGLOBIN | | |
| 39:00:00 | 8oup-D | 5 | 4.1 | 91 | 151 | 5 | MOLECULE: | NERVE HEMO | GLOBIN | |
| 40:00:00 | 1v75-A | 5 | 3.9 | 89 | 141 | 8 | MOLECULE: | HEMOGLOBIN | D ALPHA | CHAIN |
| 41:00:00 | 3w4u-A | 5 | 3.6 | 86 | 141 | 3 | MOLECULE: | HEMOGLOBIN | SUBUNIT | ZETA |
| 42:00:00 | 3sxy-B | 5 | 4.4 | 105 | 201 | 6 | MOLECULE: | TRANSCRIPT | IONAL RE | GULATOR, GNTR FAMILY |
| 43:00:00 | 8oup-B | 5 | 3.9 | 91 | 151 | 5 | MOLECULE: | NERVE HEMO | GLOBIN | |
| 44:00:00 | 1wmu-A | 5 | 3.9 | 88 | 141 | 8 | MOLECULE: | HEMOGLOBIN | D ALPHA | CHAIN |
| 45:00:00 | 6az6-B | 5 | 7.6 | 105 | 205 | 6 | MOLECULE: | GNTR FAMIL | Y TRANSC | RIPTIONAL REGULATOR |
| 46:00:00 | 3w4u-E | 5 | 3.1 | 85 | 138 | 6 | MOLECULE: | HEMOGLOBIN | SUBUNIT | ZETA |
| 47:00:00 | 3ecx-A | 5 | 4.1 | 88 | 153 | 11 | MOLECULE: | MYOGLOBIN | | |
| 48:00:00 | 7spg-A | 5 | 4.2 | 92 | 152 | 11 | MOLECULE: | MYOGLOBIN | | |
| 49:00:00 | 6zca-H | 4.9 | 10.3 | 123 | 771 | 7 | MOLECULE: | PROBABLE D | NA-DIREC | TED RNA POLYMERASE SUBUNIT DELT |
| 50:00:00 | 3pt7-B | 4.9 | 4.1 | 87 | 152 | 6 | MOLECULE: | HEMOGLOBIN | II | |
| 51:00:00 | 2bnl-C | 4.9 | 3.7 | 83 | 134 | 10 | MOLECULE: | MODULATOR | PROTEIN | RSBR |
| 52:00:00 | 3sxk-A | 4.9 | 4.4 | 100 | 140 | 10 | MOLECULE: | TRANSCRIPT | IONAL RE | GULATOR, GNTR FAMILY |

| | | | | | | | | | | |
|---|---|---|---|---|---|---|---|---|---|---|
| 53:00:00 | 6bmg-A | 4.9 | 4.1 | 90 | 154 | 14 | MOLECULE: | MYOGLOBIN | | |
| 54:00:00 | 6ls8-G | 4.9 | 4.1 | 89 | 153 | 17 | MOLECULE: | MYOGLOBIN | | |
| 55:00:00 | 6f1a-A | 4.9 | 4.2 | 90 | 154 | 9 | MOLECULE: | MYOGLOBIN | | |
| 56:00:00 | 1wvp-A | 4.9 | 4.1 | 89 | 151 | 12 | MOLECULE: | MYOGLOBIN | | |
| 57:00:00 | 3qm5-A | 4.9 | 4 | 84 | 145 | 7 | MOLECULE: | MYOGLOBIN | | |
| 58:00:00 | 3w4u-C | 4.9 | 3.6 | 86 | 141 | 3 | MOLECULE: | HEMOGLOBIN | SUBUNIT | ZETA |
| 59:00:00 | 2w6x-A | 4.9 | 4.2 | 90 | 154 | 13 | MOLECULE: | MYOGLOBIN | | |
| 60:00:00 | 3s1j-A | 4.9 | 3.6 | 85 | 131 | 9 | MOLECULE: | HEMOGLOBIN | #NAME? | AVOPROTEIN |
| 61:00:00 | 3s1i-A | 4.9 | 3.6 | 85 | 131 | 9 | MOLECULE: | HEMOGLOBIN | #NAME? | AVOPROTEIN |
| 62:00:00 | 6oty-B | 4.9 | 4.2 | 88 | 152 | 6 | MOLECULE: | HEMOGLOBIN | II | |
| 63:00:00 | 3s1i-C | 4.9 | 3.7 | 84 | 131 | 8 | MOLECULE: | HEMOGLOBIN | #NAME? | AVOPROTEIN |
| 64:00:00 | 3ubc-D | 4.9 | 3.7 | 85 | 131 | 9 | MOLECULE: | HEMOGLOBIN | #NAME? | AVOPROTEIN |
| 65:00:00 | 8oup-C | 4.9 | 4.1 | 91 | 151 | 5 | MOLECULE: | NERVE HEMO | GLOBIN | |
| 66:00:00 | 8oup-A | 4.9 | 4 | 90 | 150 | 6 | MOLECULE: | NERVE HEMO | GLOBIN | |
| 67:00:00 | 5ycj-B | 4.9 | 4.2 | 94 | 157 | 15 | MOLECULE: | ANCESTRAL | MYOGLOBI | N AMBWB' OF BASILOSAURUS RELATI |
| 68:00:00 | 2z6n-A | 4.9 | 3.9 | 88 | 141 | 8 | MOLECULE: | HEMOGLOBIN | D SUBUN | IT ALPHA |
| 69:00:00 | 6ls8-C | 4.9 | 4.2 | 90 | 153 | 17 | MOLECULE: | MYOGLOBIN | | |
| 70:00:00 | 3ogb-A | 4.9 | 4 | 91 | 154 | 12 | MOLECULE: | MYOGLOBIN | | |
| 71:00:00 | 3sxm-A | 4.9 | 4.1 | 96 | 137 | 8 | MOLECULE: | TRANSCRIPT | IONAL RE | GULATOR, GNTR FAMILY |
| 72:00:00 | 1nz3-A | 4.9 | 4.2 | 89 | 153 | 17 | MOLECULE: | MYOGLOBIN | | |
| 73:00:00 | 1jeb-A | 4.9 | 3.4 | 86 | 141 | 6 | MOLECULE: | HEMOGLOBIN | ZETA CH | AIN |
| 74:00:00 | 4nxa-A | 4.9 | 4 | 89 | 153 | 12 | MOLECULE: | MYOGLOBIN | | |
| 75:00:00 | 1ebc-A | 4.9 | 4.1 | 90 | 153 | 12 | MOLECULE: | PROTEIN (M | YOGLOBIN | ) |
| 76:00:00 | 5m3r-A | 4.9 | 4 | 91 | 154 | 12 | MOLECULE: | MYOGLOBIN | | |
| 77:00:00 | 1l2k-A | 4.9 | 4.1 | 88 | 151 | 11 | MOLECULE: | MYOGLOBIN | | |
| 78:00:00 | 3rgk-A | 4.9 | 3.4 | 85 | 149 | 15 | MOLECULE: | MYOGLOBIN | | |
| 79:00:00 | 1lhs-A | 4.9 | 3.9 | 89 | 153 | 9 | MOLECULE: | MYOGLOBIN | | |
| 80:00:00 | 4h07-A | 4.8 | 4.2 | 90 | 154 | 12 | MOLECULE: | MYOGLOBIN | | |
| 81:00:00 | 1emy-A | 4.8 | 4 | 89 | 153 | 11 | MOLECULE: | MYOGLOBIN | | |
| 82:00:00 | 2frk-X | 4.8 | 4.2 | 91 | 152 | 15 | MOLECULE: | MYOGLOBIN | | |
| 83:00:00 | 6n03-A | 4.8 | 4 | 91 | 154 | 12 | MOLECULE: | MYOGLOBIN, | CO-BOUN | D |
| 84:00:00 | 6ls8-A | 4.8 | 4.1 | 91 | 153 | 13 | MOLECULE: | MYOGLOBIN | | |
| 85:00:00 | 2w6y-A | 4.8 | 4.1 | 88 | 153 | 11 | MOLECULE: | MYOGLOBIN | | |
| 86:00:00 | 8f9i-B | 4.8 | 4.1 | 91 | 154 | 12 | MOLECULE: | MYOGLOBIN | | |
| 87:00:00 | 6ls8-E | 4.8 | 4.2 | 91 | 153 | 15 | MOLECULE: | MYOGLOBIN | | |
| 88:00:00 | 1n9f-A | 4.8 | 4.3 | 90 | 154 | 12 | MOLECULE: | MYOGLOBIN | | |
| 89:00:00 | 5xl0-A | 4.8 | 4.1 | 88 | 151 | 11 | MOLECULE: | MYOGLOBIN | | |
| 90:00:00 | 2zsz-A | 4.8 | 4.1 | 88 | 153 | 11 | MOLECULE: | MYOGLOBIN | | |
| 91:00:00 | 3rj6-B | 4.8 | 4.1 | 90 | 152 | 13 | MOLECULE: | MYOGLOBIN | | |
| 92:00:00 | 1hbr-C | 4.8 | 3.1 | 86 | 140 | 5 | MOLECULE: | PROTEIN (H | EMOGLOBI | N D) |
| 93:00:00 | 4rol-C | 4.8 | 2.9 | 84 | 141 | 7 | MOLECULE: | HEMOGLOBIN | SUBUNIT | ALPHA |
| 94:00:00 | 3qm6-A | 4.8 | 4 | 83 | 145 | 6 | MOLECULE: | MYOGLOBIN | | |
| 95:00:00 | 1mlq-A | 4.8 | 4.2 | 90 | 154 | 11 | MOLECULE: | MYOGLOBIN | | |
| 96:00:00 | 2r1h-C | 4.8 | 3.3 | 85 | 142 | 8 | MOLECULE: | HEMOGLOBIN | SUBUNIT | ALPHA-4 |
| 97:00:00 | 8wf5-A | 4.8 | 4.1 | 90 | 152 | 13 | MOLECULE: | MYOGLOBIN | | |
| 98:00:00 | 4pqb-A | 4.8 | 4.2 | 89 | 153 | 13 | MOLECULE: | MYOGLOBIN | | |
| 99:00:00 | 3rj6-A | 4.8 | 4.2 | 90 | 152 | 16 | MOLECULE: | MYOGLOBIN | | |
| 0:00 | 3hxn-C | 4.8 | 3 | 84 | 141 | 7 | MOLECULE: | HEMOGLOBIN | SUBUNIT | ALPHA |
| 1:00 | 8ess-A | 4.8 | 4 | 89 | 154 | 11 | MOLECULE: | MYOGLOBIN | | |
| 2:00 | 5ilp-A | 4.8 | 4.1 | 88 | 151 | 11 | MOLECULE: | MYOGLOBIN | | |
| 3:00 | 2ekt-A | 4.8 | 4.2 | 90 | 153 | 13 | MOLECULE: | MYOGLOBIN | | |
| 4:00 | 1m6m-A | 4.8 | 4 | 90 | 153 | 13 | MOLECULE: | PROTEIN (M | YOGLOBIN | ) |
| 5:00 | 5iks-A | 4.8 | 4.1 | 88 | 151 | 11 | MOLECULE: | MYOGLOBIN | | |
| 6:00 | 1n9x-A | 4.8 | 4.1 | 90 | 154 | 11 | MOLECULE: | MYOGLOBIN | | |

| | | | | | | | | | | |
|---|---|---|---|---|---|---|---|---|---|---|
| 7:00 | 3pt8-B | 4.8 | 4.1 | 87 | 152 | 6 | MOLECULE: | HEMOGLOBIN | II | |
| 8:00 | 7sli-A | 4.8 | 4.2 | 89 | 152 | 11 | MOLECULE: | MYOGLOBIN | | |
| 9:00 | 1n9h-A | 4.8 | 4.3 | 90 | 154 | 12 | MOLECULE: | MYOGLOBIN | | |
| 10:00 | 1fcs-A | 4.8 | 4.1 | 90 | 154 | 11 | MOLECULE: | MYOGLOBIN | | |
| 11:00 | 1iop-A | 4.8 | 4.1 | 91 | 153 | 11 | MOLECULE: | MYOGLOBIN | | |
| 12:00 | 112m-A | 4.8 | 4 | 90 | 154 | 11 | MOLECULE: | MYOGLOBIN | | |
| 13:00 | 3ase-A | 4.8 | 4.2 | 90 | 154 | 11 | MOLECULE: | MYOGLOBIN | | |
| 14:00 | 3bom-A | 4.8 | 2.8 | 83 | 142 | 8 | MOLECULE: | HEMOGLOBIN | SUBUNIT | ALPHA-4 |
| 15:00 | 1mlk-A | 4.8 | 4 | 90 | 154 | 11 | MOLECULE: | MYOGLOBIN | | |
| 16:00 | 3m3b-A | 4.8 | 4 | 91 | 153 | 11 | MOLECULE: | MYOGLOBIN | | |
| 17:00 | 3a2g-A | 4.8 | 4.1 | 90 | 154 | 11 | MOLECULE: | MYOGLOBIN | | |
| 18:00 | 4twv-A | 4.8 | 4.1 | 90 | 153 | 13 | MOLECULE: | MYOGLOBIN | | |
| 19:00 | 1mob-A | 4.8 | 4 | 90 | 154 | 11 | MOLECULE: | MYOGLOBIN | | |
| 20:00 | 1mtk-A | 4.8 | 4 | 90 | 154 | 11 | MOLECULE: | MYOGLOBIN | | |
| 21:00 | 4rol-A | 4.8 | 3 | 84 | 141 | 7 | MOLECULE: | HEMOGLOBIN | SUBUNIT | ALPHA |
| 22:00 | 1mlr-A | 4.8 | 4.2 | 90 | 154 | 11 | MOLECULE: | MYOGLOBIN | | |
| 23:00 | 1yog-A | 4.8 | 4.1 | 89 | 153 | 12 | MOLECULE: | MYOGLOBIN | | |
| 24:00:00 | 2nx0-A | 4.8 | 3.9 | 84 | 146 | 7 | MOLECULE: | MYOGLOBIN | | |
| 25:00:00 | 2spl-A | 4.8 | 4.1 | 90 | 154 | 11 | MOLECULE: | MYOGLOBIN | | |
| 26:00:00 | 3s1j-C | 4.8 | 3.7 | 84 | 131 | 8 | MOLECULE: | HEMOGLOBIN | #NAME? | AVOPROTEIN |
| 27:00:00 | 7vdn-A | 4.8 | 4.1 | 88 | 151 | 11 | MOLECULE: | MYOGLOBIN | | |
| 28:00:00 | 3h57-A | 4.8 | 4.1 | 90 | 154 | 11 | MOLECULE: | MYOGLOBIN | | |
| 29:00:00 | 5ile-A | 4.8 | 4.2 | 90 | 154 | 11 | MOLECULE: | MYOGLOBIN | | |
| 30:00:00 | 1j3f-A | 4.8 | 3.6 | 88 | 152 | 13 | MOLECULE: | MYOGLOBIN | | |
| 31:00:00 | 5kkk-A | 4.8 | 4.2 | 90 | 154 | 12 | MOLECULE: | MYOGLOBIN | | |
| 32:00:00 | 2mb5-A | 4.8 | 4.1 | 89 | 153 | 12 | MOLECULE: | MYOGLOBIN | | |
| 33:00:00 | 1spe-A | 4.8 | 4.2 | 90 | 153 | 12 | MOLECULE: | MYOGLOBIN | | |
| 34:00:00 | 6otw-B | 4.8 | 4.1 | 87 | 152 | 6 | MOLECULE: | HEMOGLOBIN | II | |
| 35:00:00 | 1h1x-A | 4.8 | 4.2 | 90 | 154 | 12 | MOLECULE: | MYOGLOBIN | | |
| 36:00:00 | 1hbr-A | 4.8 | 3.6 | 87 | 141 | 3 | MOLECULE: | PROTEIN (H | EMOGLOBI | N D) |
| 37:00:00 | 1xzu-C | 4.8 | 3 | 84 | 141 | 7 | MOLECULE: | HEMOGLOBIN | ALPHA C | HAIN |
| 38:00:00 | 2mga-A | 4.8 | 4.1 | 90 | 154 | 11 | MOLECULE: | MYOGLOBIN | | |
| 39:00:00 | 3bom-C | 4.8 | 3.3 | 85 | 142 | 8 | MOLECULE: | HEMOGLOBIN | SUBUNIT | ALPHA-4 |
| 40:00:00 | 2r1h-A | 4.8 | 3.3 | 85 | 142 | 8 | MOLECULE: | HEMOGLOBIN | SUBUNIT | ALPHA-4 |
| 41:00:00 | 3ubc-G | 4.8 | 3.8 | 84 | 131 | 8 | MOLECULE: | HEMOGLOBIN | #NAME? | AVOPROTEIN |
| 42:00:00 | 3h58-A | 4.8 | 4.1 | 90 | 154 | 11 | MOLECULE: | MYOGLOBIN | | |
| 43:00:00 | 8esu-A | 4.8 | 4.2 | 90 | 154 | 11 | MOLECULE: | MYOGLOBIN | | |
| 44:00:00 | 1abs-A | 4.8 | 4.1 | 90 | 154 | 11 | MOLECULE: | MYOGLOBIN | | |
| 45:00:00 | 3sxz-A | 4.8 | 3.9 | 97 | 139 | 7 | MOLECULE: | TRANSCRIPT | IONAL RE | GULATOR, GNTR FAMILY |
| 46:00:00 | 2oha-A | 4.8 | 4.1 | 90 | 154 | 11 | MOLECULE: | MYOGLOBIN | | |
| 47:00:00 | 6m8f-A | 4.8 | 4.2 | 90 | 154 | 11 | MOLECULE: | MYOGLOBIN | | |
| 48:00:00 | 1n9i-A | 4.8 | 4.2 | 90 | 154 | 11 | MOLECULE: | MYOGLOBIN | | |
| 49:00:00 | 2o5m-X | 4.8 | 4.2 | 91 | 152 | 13 | MOLECULE: | MYOGLOBIN | | |
| 50:00:00 | 1jeb-C | 4.8 | 3.4 | 86 | 141 | 6 | MOLECULE: | HEMOGLOBIN | ZETA CH | AIN |
| 51:00:00 | 2v1j-A | 4.8 | 4.1 | 90 | 153 | 13 | MOLECULE: | MYOGLOBIN | | |
| 52:00:00 | 2o5t-X | 4.8 | 4.1 | 90 | 152 | 13 | MOLECULE: | MYOGLOBIN | | |
| 53:00:00 | 2o5o-X | 4.8 | 4.2 | 91 | 152 | 15 | MOLECULE: | MYOGLOBIN | | |
| 54:00:00 | 4twu-A | 4.8 | 4.1 | 88 | 153 | 16 | MOLECULE: | MYOGLOBIN | | |
| 55:00:00 | 6mv0-A | 4.8 | 4 | 88 | 154 | 11 | MOLECULE: | MYOGLOBIN | | |
| 56:00:00 | 2o58-X | 4.8 | 4.2 | 91 | 152 | 13 | MOLECULE: | MYOGLOBIN | | |
| 57:00:00 | 5vzq-A | 4.8 | 4.1 | 91 | 153 | 11 | MOLECULE: | MYOGLOBIN | | |
| 58:00:00 | 2o5q-X | 4.8 | 4.2 | 91 | 152 | 15 | MOLECULE: | MYOGLOBIN | | |
| 59:00:00 | 1ufp-A | 4.8 | 4.1 | 88 | 154 | 11 | MOLECULE: | MYOGLOBIN | | |
| 60:00:00 | 1bzp-A | 4.8 | 4.1 | 89 | 153 | 12 | MOLECULE: | PROTEIN (M | YOGLOBIN | ) |

| | | | | | | | | | | |
|---|---|---|---|---|---|---|---|---|---|---|
| 61:00:00 | 2oh9-A | 4.8 | 4.1 | 90 | 154 | 11 | MOLECULE: | MYOGLOBIN | | |
| 62:00:00 | 1c7c-A | 4.8 | 9.4 | 114 | 283 | 7 | MOLECULE: | PROTEIN (D | EOXYHEMO | GLOBIN (ALPHA CHAIN)) |
| 63:00:00 | 2mgj-A | 4.8 | 4.2 | 90 | 154 | 11 | MOLECULE: | MYOGLOBIN | | |
| 64:00:00 | 1cpw-A | 4.8 | 4.2 | 90 | 154 | 11 | MOLECULE: | PROTEIN (M | YOGLOBIN | ) |
| 65:00:00 | 1vxc-A | 4.8 | 4.1 | 89 | 153 | 12 | MOLECULE: | MYOGLOBIN | | |
| 66:00:00 | 1mlj-A | 4.8 | 4.1 | 90 | 154 | 11 | MOLECULE: | MYOGLOBIN | | |
| 67:00:00 | 1cp0-A | 4.8 | 4.2 | 90 | 154 | 11 | MOLECULE: | PROTEIN (M | YOGLOBIN | ) |
| 68:00:00 | 1moc-A | 4.8 | 4.2 | 90 | 154 | 11 | MOLECULE: | MYOGLOBIN | | |
| 69:00:00 | 1ofj-A | 4.8 | 4.1 | 90 | 154 | 11 | MOLECULE: | MYOGLOBIN | | |
| 70:00:00 | 104m-A | 4.8 | 4.1 | 89 | 153 | 12 | MOLECULE: | MYOGLOBIN | | |
| 71:00:00 | 103m-A | 4.8 | 4.2 | 91 | 154 | 12 | MOLECULE: | MYOGLOBIN | | |
| 72:00:00 | 2mya-A | 4.8 | 4.1 | 89 | 153 | 12 | MOLECULE: | MYOGLOBIN | (ETHYL I | SOCYANIDE) |
| 73:00:00 | 2spo-A | 4.8 | 4.1 | 90 | 154 | 11 | MOLECULE: | MYOGLOBIN | | |
| 74:00:00 | 1dti-A | 4.8 | 4.1 | 91 | 154 | 12 | MOLECULE: | PROTEIN (M | YOGLOBIN | ) |
| 75:00:00 | 2oh8-A | 4.8 | 4.1 | 90 | 154 | 11 | MOLECULE: | MYOGLOBIN | | |
| 76:00:00 | 1vxe-A | 4.8 | 4.1 | 89 | 153 | 12 | MOLECULE: | MYOGLOBIN | | |
| 77:00:00 | 1co9-A | 4.8 | 4.2 | 90 | 154 | 11 | MOLECULE: | PROTEIN (M | YOGLOBIN | ) |
| 78:00:00 | 1cio-A | 4.8 | 4.1 | 90 | 154 | 11 | MOLECULE: | PROTEIN (M | YOGLOBIN | ) |
| 79:00:00 | 5zze-A | 4.8 | 4.1 | 90 | 152 | 13 | MOLECULE: | MYOGLOBIN | | |
| 80:00:00 | 1mlm-A | 4.8 | 4.3 | 90 | 154 | 11 | MOLECULE: | MYOGLOBIN | | |
| 81:00:00 | 1bz6-A | 4.8 | 4.1 | 89 | 153 | 12 | MOLECULE: | PROTEIN (M | YOGLOBIN | ) |
| 82:00:00 | 4it8-A | 4.8 | 4.1 | 88 | 153 | 11 | MOLECULE: | MYOGLOBIN | | |
| 83:00:00 | 1j52-A | 4.8 | 4.2 | 90 | 154 | 11 | MOLECULE: | MYOGLOBIN | | |
| 84:00:00 | 1ch1-A | 4.8 | 4.2 | 90 | 154 | 11 | MOLECULE: | PROTEIN (M | YOGLOBIN | ) |
| 85:00:00 | 1mln-A | 4.8 | 4.3 | 90 | 154 | 11 | MOLECULE: | MYOGLOBIN | | |
| 86:00:00 | 1o16-A | 4.8 | 4.2 | 90 | 154 | 11 | MOLECULE: | MYOGLOBIN | | |
| 87:00:00 | 1myz-A | 4.8 | 4.1 | 90 | 154 | 11 | MOLECULE: | MYOGLOBIN | | |
| 88:00:00 | 1vxd-A | 4.8 | 4.1 | 88 | 153 | 11 | MOLECULE: | MYOGLOBIN | | |
| 89:00:00 | 1o1l-A | 4.8 | 10.2 | 115 | 283 | 7 | MOLECULE: | HEMOGLOBIN | ALPHA C | HAIN |
| 90:00:00 | 1mll-A | 4.8 | 4.1 | 90 | 154 | 11 | MOLECULE: | MYOGLOBIN | | |
| 91:00:00 | 4mbn-A | 4.8 | 4.1 | 90 | 153 | 12 | MOLECULE: | MYOGLOBIN | | |
| 92:00:00 | 1mz0-A | 4.8 | 4.1 | 90 | 154 | 11 | MOLECULE: | MYOGLOBIN | | |
| 93:00:00 | 102m-A | 4.8 | 4.2 | 90 | 154 | 11 | MOLECULE: | MYOGLOBIN | | |
| 94:00:00 | 1ltw-A | 4.8 | 4.2 | 90 | 154 | 11 | MOLECULE: | MYOGLOBIN | | |
| 95:00:00 | 1ch7-A | 4.8 | 4.1 | 90 | 154 | 11 | MOLECULE: | PROTEIN (M | YOGLOBIN | ) |
| 96:00:00 | 1tes-A | 4.8 | 4.2 | 90 | 154 | 11 | MOLECULE: | MYOGLOBIN | | |
| 97:00:00 | 1cik-A | 4.8 | 4.1 | 90 | 154 | 11 | MOLECULE: | PROTEIN (M | YOGLOBIN | ) |
| 98:00:00 | 106m-A | 4.8 | 4.2 | 90 | 154 | 11 | MOLECULE: | MYOGLOBIN | | |
| 99:00:00 | 2mge-A | 4.8 | 4.1 | 90 | 154 | 11 | MOLECULE: | MYOGLOBIN | | |
| 0:00 | 1mlu-A | 4.8 | 4.2 | 90 | 154 | 11 | MOLECULE: | MYOGLOBIN | | |
| 1:00 | 2mgm-A | 4.8 | 4.1 | 90 | 154 | 11 | MOLECULE: | MYOGLOBIN | | |
| 2:00 | 108m-A | 4.8 | 4.2 | 90 | 154 | 11 | MOLECULE: | MYOGLOBIN | | |
| 3:00 | 2spm-A | 4.8 | 4.2 | 90 | 154 | 11 | MOLECULE: | MYOGLOBIN | | |
| 4:00 | 1mlf-A | 4.8 | 4.1 | 90 | 154 | 11 | MOLECULE: | MYOGLOBIN | | |
| 5:00 | 1a6g-A | 4.8 | 4.1 | 88 | 151 | 11 | MOLECULE: | MYOGLOBIN | | |
| 6:00 | 1nz4-A | 4.8 | 4.2 | 89 | 153 | 17 | MOLECULE: | MYOGLOBIN | | |
| 7:00 | 6q6p-A | 4.8 | 4.1 | 90 | 156 | 2 | MOLECULE: | CYTOGLOBIN | | |
| 8:00 | 111m-A | 4.8 | 4.2 | 90 | 154 | 11 | MOLECULE: | MYOGLOBIN | | |
| 9:00 | 2mgb-A | 4.8 | 4.2 | 90 | 154 | 11 | MOLECULE: | MYOGLOBIN | | |
| 10:00 | 1mlo-A | 4.8 | 4.3 | 90 | 154 | 11 | MOLECULE: | MYOGLOBIN | | |
| 11:00 | 5cn4-A | 4.8 | 4.1 | 89 | 152 | 15 | MOLECULE: | MYOGLOBIN | | |
| 12:00 | 2mgk-A | 4.8 | 4.1 | 90 | 154 | 11 | MOLECULE: | MYOGLOBIN | | |
| 13:00 | 101m-A | 4.8 | 4.2 | 90 | 154 | 11 | MOLECULE: | MYOGLOBIN | | |
| 14:00 | 1cq2-A | 4.8 | 4.1 | 89 | 153 | 12 | MOLECULE: | MYOGLOBIN | | |

| | | | | | | | | | | |
|---|---|---|---|---|---|---|---|---|---|---|
| 15:00 | 2v1g-A | 4.8 | 4.1 | 90 | 153 | 13 | MOLECULE: | MYOGLOBIN | | |
| 16:00 | 2ohb-A | 4.8 | 4.2 | 90 | 154 | 11 | MOLECULE: | MYOGLOBIN | | |
| 17:00 | 1o1p-A | 4.8 | 10.6 | 115 | 283 | 8 | MOLECULE: | HEMOGLOBIN | ALPHA C | HAIN |
| 18:00 | 109m-A | 4.8 | 4.2 | 90 | 154 | 11 | MOLECULE: | MYOGLOBIN | | |
| 19:00 | 5mbn-A | 4.8 | 4.1 | 90 | 153 | 12 | MOLECULE: | MYOGLOBIN | | |
| 20:00 | 1mls-A | 4.8 | 4.1 | 90 | 154 | 11 | MOLECULE: | MYOGLOBIN | | |
| 21:00 | 1cp5-A | 4.8 | 4.1 | 90 | 154 | 11 | MOLECULE: | PROTEIN (M | YOGLOBIN | ) |
| 22:00 | 1o1j-A | 4.8 | 9.5 | 112 | 283 | 8 | MOLECULE: | HEMOGLOBIN | ALPHA C | HAIN |
| 23:00 | 5cn5-A | 4.8 | 4.1 | 90 | 152 | 13 | MOLECULE: | MYOGLOBIN | | |
| 24:00:00 | 1yoi-A | 4.8 | 4.1 | 89 | 153 | 12 | MOLECULE: | MYOGLOBIN | | |
| 25:00:00 | 2mgc-A | 4.8 | 4.1 | 90 | 154 | 11 | MOLECULE: | MYOGLOBIN | | |
| 26:00:00 | 2mgl-A | 4.8 | 4.2 | 90 | 154 | 11 | MOLECULE: | MYOGLOBIN | | |
| 27:00:00 | 1ofk-A | 4.8 | 4.1 | 90 | 154 | 11 | MOLECULE: | MYOGLOBIN | | |
| 28:00:00 | 2mye-A | 4.8 | 4.1 | 89 | 153 | 12 | MOLECULE: | MYOGLOBIN | (ETHYL I | SOCYANIDE) |
| 29:00:00 | 1lht-A | 4.8 | 4 | 89 | 153 | 9 | MOLECULE: | MYOGLOBIN | | |
| 30:00:00 | 2g0s-A | 4.8 | 4.1 | 90 | 154 | 11 | MOLECULE: | MYOGLOBIN | | |
| 31:00:00 | 1lue-A | 4.8 | 4.2 | 90 | 154 | 11 | MOLECULE: | MYOGLOBIN | | |
| 32:00:00 | 1vxh-A | 4.8 | 4.1 | 89 | 153 | 12 | MOLECULE: | MYOGLOBIN | | |
| 33:00:00 | 1ufj-A | 4.8 | 4.1 | 88 | 154 | 11 | MOLECULE: | MYOGLOBIN | | |
| 34:00:00 | 1f63-A | 4.8 | 4.1 | 90 | 154 | 11 | MOLECULE: | MYOGLOBIN | | |
| 35:00:00 | 1mym-A | 4.8 | 4.2 | 90 | 154 | 11 | MOLECULE: | MYOGLOBIN | | |
| 36:00:00 | 1swm-A | 4.8 | 4.1 | 89 | 153 | 12 | MOLECULE: | MYOGLOBIN | | |
| 37:00:00 | 1mbc-A | 4.8 | 4.1 | 88 | 153 | 11 | MOLECULE: | MYOGLOBIN | | |
| 38:00:00 | 2mgi-A | 4.8 | 4.2 | 90 | 154 | 11 | MOLECULE: | MYOGLOBIN | | |
| 39:00:00 | 2mgf-A | 4.8 | 4.2 | 90 | 154 | 11 | MOLECULE: | MYOGLOBIN | | |
| 40:00:00 | 6q6p-B | 4.8 | 4.2 | 91 | 157 | 2 | MOLECULE: | CYTOGLOBIN | | |
| 41:00:00 | 1obm-A | 4.8 | 4.2 | 90 | 154 | 11 | MOLECULE: | MYOGLOBIN | | |
| 42:00:00 | 6ird-B | 4.7 | 6.5 | 90 | 254 | 4 | MOLECULE: | 1-PHOSPHAT | IDYLINOS | ITOL 4,5-BISPHOSPHATE PHOSPHODI |
| 43:00:00 | 1abw-A | 4.7 | 9.3 | 111 | 287 | 7 | MOLECULE: | HEMOGLOBIN | #NAME? | LOOD SUBSTITUTE |
| 44:00:00 | 1mwd-A | 4.7 | 4 | 90 | 153 | 13 | MOLECULE: | PROTEIN (M | YOGLOBIN | ) |
| 45:00:00 | 2v1i-A | 4.7 | 4.1 | 90 | 153 | 13 | MOLECULE: | MYOGLOBIN | | |
| 46:00:00 | 1myj-B | 4.7 | 4.1 | 90 | 153 | 13 | MOLECULE: | MYOGLOBIN | | |
| 47:00:00 | 1mnk-B | 4.7 | 4.2 | 90 | 151 | 13 | MOLECULE: | MYOGLOBIN | | |
| 48:00:00 | 5ilm-A | 4.7 | 4.1 | 88 | 153 | 11 | MOLECULE: | MYOGLOBIN | | |
| 49:00:00 | 2w6w-A | 4.7 | 4.1 | 88 | 153 | 11 | MOLECULE: | MYOGLOBIN | | |
| 50:00:00 | 2d6c-B | 4.7 | 4 | 89 | 151 | 12 | MOLECULE: | MYOGLOBIN | | |
| 51:00:00 | 8f9i-A | 4.7 | 4.2 | 92 | 154 | 11 | MOLECULE: | MYOGLOBIN | | |
| 52:00:00 | 1lfl-P | 4.7 | 3 | 84 | 141 | 7 | MOLECULE: | HEMOGLOBIN | ALPHA C | HAIN |
| 53:00:00 | 1j3z-C | 4.7 | 3 | 84 | 141 | 7 | MOLECULE: | HEMOGLOBIN | ALPHA C | HAIN |
| 54:00:00 | 2dn2-A | 4.7 | 3 | 84 | 141 | 7 | MOLECULE: | HEMOGLOBIN | ALPHA S | UBUNIT |
| 55:00:00 | 1hgb-A | 4.7 | 3 | 84 | 141 | 7 | MOLECULE: | HEMOGLOBIN | (AQUO M | ET) (ALPHA CHAIN) |
| 56:00:00 | 6e04-A | 4.7 | 4.1 | 88 | 153 | 11 | MOLECULE: | MYOGLOBIN | | |
| 57:00:00 | 6f18-A | 4.7 | 4 | 88 | 154 | 11 | MOLECULE: | MYOGLOBIN | | |
| 58:00:00 | 3szk-A | 4.7 | 3.4 | 85 | 140 | 5 | MOLECULE: | HEMOGLOBIN | SUBUNIT | ALPHA |
| 59:00:00 | 1j41-G | 4.7 | 3 | 84 | 141 | 7 | MOLECULE: | HEMOGLOBIN | ALPHA C | HAIN |
| 60:00:00 | 1dxv-C | 4.7 | 3 | 84 | 141 | 7 | MOLECULE: | HEMOGLOBIN | (DEOXY) | (ALPHA CHAIN) |
| 61:00:00 | 3m39-A | 4.7 | 4 | 88 | 153 | 11 | MOLECULE: | MYOGLOBIN | | |
| 62:00:00 | 2dn2-C | 4.7 | 3 | 84 | 141 | 7 | MOLECULE: | HEMOGLOBIN | ALPHA S | UBUNIT |
| 63:00:00 | 1a3o-C | 4.7 | 3 | 84 | 141 | 7 | MOLECULE: | HEMOGLOBIN | (ALPHA | CHAIN) |
| 64:00:00 | 1do1-A | 4.7 | 4.1 | 88 | 153 | 11 | MOLECULE: | MYOGLOBIN | | |
| 65:00:00 | 1hjt-A | 4.7 | 4.1 | 91 | 153 | 11 | MOLECULE: | MYOGLOBIN | | |
| 66:00:00 | 6f17-A | 4.7 | 4.1 | 89 | 153 | 11 | MOLECULE: | MYOGLOBIN | | |
| 67:00:00 | 1y7g-C | 4.7 | 3 | 84 | 141 | 7 | MOLECULE: | HEMOGLOBIN | ALPHA C | HAIN |
| 68:00:00 | 3qje-A | 4.7 | 3 | 84 | 141 | 7 | MOLECULE: | HEMOGLOBIN | SUBUNIT | ALPHA |

| | | | | | | | | | | |
|---|---|---|---|---|---|---|---|---|---|---|
| 69:00:00 | 1myg-B | 4.7 | 4.1 | 90 | 153 | 13 | MOLECULE: | MYOGLOBIN | | |
| 70:00:00 | 5ut7-A | 4.7 | 4.1 | 90 | 154 | 11 | MOLECULE: | MYOGLOBIN | | |
| 71:00:00 | 1do7-A | 4.7 | 4.1 | 87 | 153 | 11 | MOLECULE: | MYOGLOBIN | | |
| 72:00:00 | 6nbd-A | 4.7 | 3 | 84 | 140 | 7 | MOLECULE: | HEMOGLOBIN | SUBUNIT | ALPHA |
| 73:00:00 | 2d5z-A | 4.7 | 3 | 84 | 141 | 7 | MOLECULE: | HEMOGLOBIN | ALPHA S | UBUNIT |
| 74:00:00 | 1yh9-C | 4.7 | 3 | 84 | 141 | 7 | MOLECULE: | HEMOGLOBIN | ALPHA C | HAIN |
| 75:00:00 | 1y0t-C | 4.7 | 3 | 84 | 141 | 7 | MOLECULE: | HEMOGLOBIN | ALPHA C | HAIN |
| 76:00:00 | 3ecz-A | 4.7 | 4.1 | 88 | 153 | 11 | MOLECULE: | MYOGLOBIN | | |
| 77:00:00 | 5ut9-A | 4.7 | 4.2 | 88 | 153 | 13 | MOLECULE: | MYOGLOBIN | | |
| 78:00:00 | 8f9j-A | 4.7 | 4.2 | 88 | 153 | 11 | MOLECULE: | MYOGLOBIN | | |
| 79:00:00 | 3at5-A | 4.7 | 2.9 | 84 | 141 | 8 | MOLECULE: | ALPHAA-GLO | BIN | |
| 80:00:00 | 1a01-A | 4.7 | 3 | 84 | 141 | 7 | MOLECULE: | HEMOGLOBIN | (ALPHA | CHAIN) |
| 81:00:00 | 3d17-C | 4.7 | 3.5 | 87 | 141 | 6 | MOLECULE: | HEMOGLOBIN | SUBUNIT | ALPHA |
| 82:00:00 | 3s1j-B | 4.7 | 3.7 | 85 | 131 | 9 | MOLECULE: | HEMOGLOBIN | #NAME? | AVOPROTEIN |
| 83:00:00 | 1c7b-C | 4.7 | 3 | 84 | 141 | 7 | MOLECULE: | PROTEIN (D | EOXYHEMO | GLOBIN (ALPHA CHAIN)) |
| 84:00:00 | 1ch5-A | 4.7 | 4.1 | 90 | 154 | 12 | MOLECULE: | PROTEIN (M | YOGLOBIN | ) |
| 85:00:00 | 1bz0-C | 4.7 | 3 | 84 | 141 | 7 | MOLECULE: | PROTEIN (H | EMOGLOBI | N ALPHA CHAIN) |
| 86:00:00 | 1j3y-C | 4.7 | 3 | 84 | 141 | 7 | MOLECULE: | HEMOGLOBIN | ALPHA C | HAIN |
| 87:00:00 | 1xz4-A | 4.7 | 3 | 84 | 141 | 8 | MOLECULE: | HEMOGLOBIN | ALPHA C | HAIN |
| 88:00:00 | 1a00-A | 4.7 | 3 | 84 | 141 | 7 | MOLECULE: | HEMOGLOBIN | (ALPHA | CHAIN) |
| 89:00:00 | 6kye-K | 4.7 | 3.1 | 84 | 140 | 7 | MOLECULE: | HEMOGLOBIN | SUBUNIT | ALPHA |
| 90:00:00 | 5kdq-A | 4.7 | 3 | 84 | 141 | 7 | MOLECULE: | HEMOGLOBIN | SUBUNIT | ALPHA |
| 91:00:00 | 1gli-A | 4.7 | 3 | 84 | 141 | 7 | MOLECULE: | DEOXYHEMOG | LOBIN | |
| 92:00:00 | 1vwt-C | 4.7 | 3 | 84 | 141 | 7 | MOLECULE: | HEMOGLOBIN | | |
| 93:00:00 | 1a6n-A | 4.7 | 4.1 | 88 | 151 | 11 | MOLECULE: | MYOGLOBIN | | |
| 94:00:00 | 1hgc-C | 4.7 | 3.1 | 85 | 141 | 6 | MOLECULE: | HEMOGLOBIN | (OXY) ( | ALPHA CHAIN) |
| 95:00:00 | 2zso-A | 4.7 | 4.1 | 88 | 153 | 11 | MOLECULE: | MYOGLOBIN | | |
| 96:00:00 | 1j7y-A | 4.7 | 3 | 84 | 141 | 7 | MOLECULE: | HEMOGLOBIN | | |
| 97:00:00 | 1xz2-A | 4.7 | 3 | 84 | 141 | 7 | MOLECULE: | HEMOGLOBIN | ALPHA C | HAIN |
| 98:00:00 | 3d7o-A | 4.7 | 3.4 | 85 | 141 | 7 | MOLECULE: | HEMOGLOBIN | SUBUNIT | ALPHA |
| 99:00:00 | 1y0d-C | 4.7 | 3 | 84 | 140 | 7 | MOLECULE: | HEMOGLOBIN | ALPHA C | HAIN |
| 0:00 | 1gbu-C | 4.7 | 3 | 84 | 141 | 7 | MOLECULE: | HEMOGLOBIN | | |
| 1:00 | 1y5k-C | 4.7 | 3 | 84 | 141 | 7 | MOLECULE: | HEMOGLOBIN | ALPHA C | HAIN |
| 2:00 | 2zt2-A | 4.7 | 4.1 | 88 | 153 | 11 | MOLECULE: | MYOGLOBIN | | |
| 3:00 | 3dut-A | 4.7 | 3 | 84 | 141 | 7 | MOLECULE: | HEMOGLOBIN | SUBUNIT | ALPHA |
| 4:00 | 1y5j-C | 4.7 | 3 | 84 | 141 | 7 | MOLECULE: | HEMOGLOBIN | ALPHA C | HAIN |
| 5:00 | 1gzx-C | 4.7 | 3 | 84 | 141 | 7 | MOLECULE: | HEMOGLOBIN | SUBUNIT | ALPHA |
| 6:00 | 2d60-C | 4.7 | 3 | 84 | 141 | 7 | MOLECULE: | HEMOGLOBIN | ALPHA S | UBUNIT |
| 7:00 | 2hbd-A | 4.7 | 2.9 | 83 | 141 | 7 | MOLECULE: | HEMOGLOBIN | A (METH | YL ISOCYANIDE) (ALPHA CHAIN) |
| 8:00 | 1yhr-A | 4.7 | 3 | 84 | 141 | 7 | MOLECULE: | HEMOGLOBIN | ALPHA C | HAIN |
| 9:00 | 1coh-A | 4.7 | 3 | 84 | 141 | 7 | MOLECULE: | HEMOGLOBIN | (FERROU | S CARBONMONOXY) (ALPHA CHAIN) |
| 10:00 | 3s1i-B | 4.7 | 3.8 | 85 | 131 | 9 | MOLECULE: | HEMOGLOBIN | #NAME? | AVOPROTEIN |
| 11:00 | 1a0u-C | 4.7 | 3 | 84 | 141 | 7 | MOLECULE: | HEMOGLOBIN | (ALPHA | CHAIN) |
| 12:00 | 1o1n-A | 4.7 | 9.8 | 115 | 282 | 6 | MOLECULE: | HEMOGLOBIN | ALPHA C | HAIN |
| 13:00 | 2mgh-A | 4.7 | 4.2 | 90 | 154 | 11 | MOLECULE: | MYOGLOBIN | | |
| 14:00 | 2spn-A | 4.7 | 4.1 | 89 | 154 | 11 | MOLECULE: | MYOGLOBIN | | |
| 15:00 | 1y09-C | 4.7 | 3 | 84 | 141 | 7 | MOLECULE: | HEMOGLOBIN | ALPHA C | HAIN |
| 16:00 | 1xz7-C | 4.7 | 3 | 84 | 141 | 7 | MOLECULE: | HEMOGLOBIN | ALPHA C | HAIN |
| 17:00 | 6cf0-A | 4.7 | 4.2 | 88 | 153 | 13 | MOLECULE: | MYOGLOBIN | | |
| 18:00 | 1mwc-A | 4.7 | 4.1 | 90 | 153 | 13 | MOLECULE: | PROTEIN (M | YOGLOBIN | ) |
| 19:00 | 1dxv-A | 4.7 | 3 | 84 | 141 | 7 | MOLECULE: | HEMOGLOBIN | (DEOXY) | (ALPHA CHAIN) |
| 20:00 | 3wft-A | 4.7 | 4.1 | 90 | 153 | 13 | MOLECULE: | MYOGLOBIN | | |
| 21:00 | 3ba2-A | 4.7 | 4.2 | 88 | 152 | 14 | MOLECULE: | MYOGLOBIN | | |
| 22:00 | 2hbc-A | 4.7 | 3 | 84 | 141 | 7 | MOLECULE: | HEMOGLOBIN | A (ETHY | L ISOCYANIDE) (ALPHA CHAIN) |

| | | | | | | | | | | |
|---|---|---|---|---|---|---|---|---|---|---|
| 23:00 | 8fdl-A | 4.7 | 3 | 84 | 141 | 7 | MOLECULE: | HEMOGLOBIN | SUBUNIT | ALPHA |
| 24:00:00 | 1gbv-A | 4.7 | 3 | 84 | 141 | 7 | MOLECULE: | HEMOGLOBIN | | |
| 25:00:00 | 1xz2-C | 4.7 | 3 | 84 | 141 | 7 | MOLECULE: | HEMOGLOBIN | ALPHA C | HAIN |
| 26:00:00 | 3mn0-A | 4.7 | 3.9 | 88 | 153 | 11 | MOLECULE: | MYOGLOBIN | | |
| 27:00:00 | 2hhe-C | 4.7 | 3 | 84 | 141 | 7 | MOLECULE: | HEMOGLOBIN | (DEOXY) | (ALPHA CHAIN) |
| 28:00:00 | 1dxu-C | 4.7 | 3 | 84 | 141 | 7 | MOLECULE: | HEMOGLOBIN | (DEOXY) | (ALPHA CHAIN) |
| 29:00:00 | 2hbs-C | 4.7 | 3 | 84 | 141 | 7 | MOLECULE: | HEMOGLOBIN | S (DEOX | Y), ALPHA CHAIN |
| 30:00:00 | 1y4v-C | 4.7 | 3 | 84 | 141 | 7 | MOLECULE: | HEMOGLOBIN | ALPHA C | HAIN |
| 31:00:00 | 4n7n-K | 4.7 | 3 | 84 | 141 | 7 | MOLECULE: | HEMOGLOBIN | SUBUNIT | ALPHA |
| 32:00:00 | 2mgg-A | 4.7 | 4.2 | 90 | 154 | 11 | MOLECULE: | MYOGLOBIN | | |
| 33:00:00 | 8dov-G | 4.7 | 3.4 | 85 | 140 | 7 | MOLECULE: | HEMOGLOBIN | SUBUNIT | ALPHA |
| 34:00:00 | 3ed9-A | 4.7 | 4.1 | 88 | 153 | 11 | MOLECULE: | MYOGLOBIN | | |
| 35:00:00 | 1dxt-A | 4.7 | 3 | 84 | 141 | 7 | MOLECULE: | HEMOGLOBIN | (DEOXY) | (ALPHA CHAIN) |
| 36:00:00 | 1irc-A | 4.7 | 4 | 89 | 152 | 12 | MOLECULE: | MYOGLOBIN | (METAQUO | ) |
| 37:00:00 | 1sdk-A | 4.7 | 3 | 84 | 141 | 7 | MOLECULE: | HEMOGLOBIN | A | |
| 38:00:00 | 1ajh-A | 4.7 | 4.1 | 88 | 153 | 11 | MOLECULE: | MYOGLOBIN | | |
| 39:00:00 | 3hhb-C | 4.7 | 3 | 84 | 141 | 7 | MOLECULE: | HEMOGLOBIN | (DEOXY) | (ALPHA CHAIN) |
| 40:00:00 | 1y4r-C | 4.7 | 3 | 84 | 141 | 7 | MOLECULE: | HEMOGLOBIN | ALPHA C | HAIN |
| 41:00:00 | 1hbb-C | 4.7 | 3 | 84 | 141 | 7 | MOLECULE: | HEMOGLOBIN | A (DEOX | Y) (ALPHA CHAIN) |
| 42:00:00 | 3ecl-A | 4.7 | 4.1 | 88 | 153 | 11 | MOLECULE: | MYOGLOBIN | | |
| 43:00:00 | 1yhe-C | 4.7 | 3 | 84 | 141 | 7 | MOLECULE: | HEMOGLOBIN | ALPHA C | HAIN |
| 44:00:00 | 2eku-A | 4.7 | 4.1 | 88 | 153 | 11 | MOLECULE: | MYOGLOBIN | | |
| 45:00:00 | 3fms-A | 4.7 | 4.2 | 101 | 209 | 7 | MOLECULE: | TRANSCRIPT | IONAL RE | GULATOR, GNTR FAMILY |
| 46:00:00 | 1dxu-A | 4.7 | 3 | 84 | 141 | 7 | MOLECULE: | HEMOGLOBIN | (DEOXY) | (ALPHA CHAIN) |
| 47:00:00 | 1qsh-A | 4.7 | 3 | 84 | 141 | 7 | MOLECULE: | PROTEIN (H | EMOGLOBI | N ALPHA CHAIN) |
| 48:00:00 | 1mno-A | 4.7 | 4 | 90 | 153 | 13 | MOLECULE: | PROTEIN (M | YOGLOBIN | ) |
| 49:00:00 | 2zt3-A | 4.7 | 4.1 | 88 | 153 | 11 | MOLECULE: | MYOGLOBIN | | |
| 50:00:00 | 3eda-A | 4.7 | 4.1 | 88 | 153 | 11 | MOLECULE: | MYOGLOBIN | | |
| 51:00:00 | 8f9n-A | 4.7 | 4.2 | 89 | 153 | 11 | MOLECULE: | MYOGLOBIN | | |
| 52:00:00 | 1mnj-B | 4.7 | 4.2 | 90 | 151 | 13 | MOLECULE: | MYOGLOBIN | | |
| 53:00:00 | 3k9z-A | 4.7 | 4 | 88 | 153 | 11 | MOLECULE: | MYOGLOBIN | | |
| 54:00:00 | 4h0b-A | 4.7 | 4 | 89 | 154 | 11 | MOLECULE: | MYOGLOBIN | | |
| 55:00:00 | 1do4-A | 4.7 | 4.1 | 88 | 153 | 11 | MOLECULE: | MYOGLOBIN | | |
| 56:00:00 | 3m38-A | 4.7 | 4.1 | 89 | 153 | 12 | MOLECULE: | MYOGLOBIN | | |
| 57:00:00 | 6kae-G | 4.7 | 3 | 84 | 141 | 7 | MOLECULE: | HEMOGLOBIN | SUBUNIT | ALPHA |
| 58:00:00 | 1dxd-A | 4.7 | 4.1 | 88 | 153 | 11 | MOLECULE: | MYOGLOBIN | | |
| 59:00:00 | 1j3y-G | 4.7 | 3 | 84 | 141 | 7 | MOLECULE: | HEMOGLOBIN | ALPHA C | HAIN |
| 60:00:00 | 4m4a-A | 4.7 | 3.4 | 85 | 140 | 7 | MOLECULE: | HEMOGLOBIN | SUBUNIT | ALPHA |
| 61:00:00 | 1r1y-A | 4.7 | 3.3 | 85 | 141 | 7 | MOLECULE: | HEMOGLOBIN | ALPHA C | HAIN |
| 62:00:00 | 1vwt-A | 4.7 | 3 | 84 | 141 | 7 | MOLECULE: | HEMOGLOBIN | | |
| 63:00:00 | 1k1k-A | 4.7 | 3 | 84 | 141 | 7 | MOLECULE: | HEMOGLOBIN | ALPHA C | HAIN |
| 64:00:00 | 6hit-A | 4.7 | 3.3 | 86 | 142 | 5 | MOLECULE: | HEMOGLOBIN | ALPHA 2 | CHAIN |
| 65:00:00 | 3e4n-A | 4.7 | 4.1 | 88 | 153 | 11 | MOLECULE: | MYOGLOBIN | | |
| 66:00:00 | 1yen-C | 4.7 | 2.9 | 84 | 138 | 6 | MOLECULE: | HEMOGLOBIN | ALPHA C | HAIN |
| 67:00:00 | 1do3-A | 4.7 | 4.2 | 88 | 153 | 13 | MOLECULE: | MYOGLOBIN | | |
| 68:00:00 | 1ye2-C | 4.7 | 3 | 84 | 141 | 7 | MOLECULE: | HEMOGLOBIN | ALPHA C | HAIN |
| 69:00:00 | 1y7z-C | 4.7 | 3 | 84 | 141 | 7 | MOLECULE: | HEMOGLOBIN | ALPHA C | HAIN |
| 70:00:00 | 3e55-A | 4.7 | 4.1 | 88 | 153 | 11 | MOLECULE: | MYOGLOBIN | | |
| 71:00:00 | 3ubv-D | 4.7 | 3.7 | 85 | 131 | 9 | MOLECULE: | HEMOGLOBIN | #NAME? | AVOPROTEIN |
| 72:00:00 | 4rom-A | 4.7 | 3 | 84 | 141 | 7 | MOLECULE: | HEMOGLOBIN | SUBUNIT | ALPHA |
| 73:00:00 | 1qsh-C | 4.7 | 3 | 84 | 141 | 7 | MOLECULE: | PROTEIN (H | EMOGLOBI | N ALPHA CHAIN) |
| 74:00:00 | 1duo-A | 4.7 | 4.1 | 89 | 152 | 12 | MOLECULE: | SPERM WHAL | E METAQU | OMYOGLOBIN VARIANT H93G |
| 75:00:00 | 7ud7-A | 4.7 | 3 | 84 | 141 | 7 | MOLECULE: | HEMOGLOBIN | SUBUNIT | ALPHA |
| 76:00:00 | 1xxt-C | 4.7 | 3 | 84 | 141 | 7 | MOLECULE: | HEMOGLOBIN | ALPHA C | HAIN |

| | | | | | | | | | | |
|---|---|---|---|---|---|---|---|---|---|---|
| 77:00:00 | 1y22-C | 4.7 | 3 | 84 | 141 | 7 | MOLECULE: | HEMOGLOBIN | ALPHA C | HAIN |
| 78:00:00 | 6nq5-A | 4.7 | 3.1 | 85 | 140 | 7 | MOLECULE: | HEMOGLOBIN | SUBUNIT | ALPHA |
| 79:00:00 | 1bzz-C | 4.7 | 3 | 84 | 141 | 7 | MOLECULE: | PROTEIN (H | EMOGLOBI | N ALPHA CHAIN) |
| 80:00:00 | 5yci-A | 4.7 | 4.2 | 92 | 154 | 15 | MOLECULE: | ANCESTRAL | MYOGLOBI | N AMBWB' OF BASILOSAURUS RELATI |
| 81:00:00 | 2zt0-A | 4.7 | 4.1 | 88 | 153 | 11 | MOLECULE: | MYOGLOBIN | | |
| 82:00:00 | 1ye0-A | 4.7 | 3 | 84 | 141 | 7 | MOLECULE: | HEMOGLOBIN | ALPHA C | HAIN |
| 83:00:00 | 3sdn-A | 4.7 | 4.1 | 89 | 152 | 12 | MOLECULE: | MYOGLOBIN | | |
| 84:00:00 | 3oo5-A | 4.7 | 3.3 | 85 | 140 | 7 | MOLECULE: | HEMOGLOBIN | SUBUNIT | ALPHA |
| 85:00:00 | 6g5a-A | 4.7 | 4 | 89 | 153 | 12 | MOLECULE: | MYOGLOBIN | | |
| 86:00:00 | 1y45-A | 4.7 | 3 | 84 | 141 | 7 | MOLECULE: | HEMOGLOBIN | ALPHA C | HAIN |
| 87:00:00 | 1xy0-C | 4.7 | 3 | 84 | 141 | 7 | MOLECULE: | HEMOGLOBIN | ALPHA C | HAIN |
| 88:00:00 | 1g9v-A | 4.7 | 3 | 84 | 141 | 7 | MOLECULE: | HEMOGLOBIN | ALPHA C | HAIN |
| 89:00:00 | 1y0d-A | 4.7 | 3 | 84 | 140 | 7 | MOLECULE: | HEMOGLOBIN | ALPHA C | HAIN |
| 90:00:00 | 4pqc-A | 4.7 | 4 | 88 | 153 | 11 | MOLECULE: | MYOGLOBIN | | |
| 91:00:00 | 1coh-C | 4.7 | 3 | 84 | 141 | 7 | MOLECULE: | HEMOGLOBIN | (FERROU | S CARBONMONOXY) (ALPHA CHAIN) |
| 92:00:00 | 1pmb-B | 4.7 | 4.1 | 89 | 153 | 15 | MOLECULE: | MYOGLOBIN | | |
| 93:00:00 | 4lpi-A | 4.7 | 4.1 | 88 | 153 | 11 | MOLECULE: | MYOGLOBIN | | |
| 94:00:00 | 1vxa-A | 4.7 | 4.1 | 89 | 153 | 12 | MOLECULE: | MYOGLOBIN | | |
| 95:00:00 | 2ef2-A | 4.7 | 3.6 | 87 | 153 | 11 | MOLECULE: | MYOGLOBIN | | |
| 96:00:00 | 4m4b-A | 4.7 | 3.4 | 85 | 140 | 7 | MOLECULE: | HEMOGLOBIN | SUBUNIT | ALPHA |
| 97:00:00 | 3p5q-A | 4.7 | 3.4 | 85 | 139 | 7 | MOLECULE: | HEMOGLOBIN | SUBUNIT | ALPHA |
| 98:00:00 | 5c6y-A | 4.7 | 4.1 | 88 | 153 | 11 | MOLECULE: | MYOGLOBIN | | |
| 99:00:00 | 1y0c-C | 4.7 | 3 | 84 | 141 | 7 | MOLECULE: | HEMOGLOBIN | ALPHA C | HAIN |
| 0:00 | 1yih-C | 4.7 | 3 | 84 | 141 | 7 | MOLECULE: | HEMOGLOBIN | ALPHA C | HAIN |
| 1:00 | 1dxt-C | 4.7 | 3 | 84 | 141 | 7 | MOLECULE: | HEMOGLOBIN | (DEOXY) | (ALPHA CHAIN) |
| 2:00 | 4n7o-I | 4.7 | 3.1 | 85 | 141 | 7 | MOLECULE: | HEMOGLOBIN | SUBUNIT | ALPHA |
| 3:00 | 1bz0-A | 4.7 | 3 | 84 | 141 | 7 | MOLECULE: | PROTEIN (H | EMOGLOBI | N ALPHA CHAIN) |
| 4:00 | 1bab-A | 4.7 | 3 | 84 | 142 | 7 | MOLECULE: | HEMOGLOBIN | THIONVI | LLE (DEOXY) (ALPHA CHAIN) |
| 5:00 | 1ydz-C | 4.7 | 3 | 84 | 141 | 7 | MOLECULE: | HEMOGLOBIN | ALPHA C | HAIN |
| 6:00 | 5vzp-A | 4.7 | 4.2 | 89 | 153 | 11 | MOLECULE: | MYOGLOBIN | | |
| 7:00 | 1a3n-C | 4.7 | 3 | 84 | 141 | 7 | MOLECULE: | HEMOGLOBIN | (ALPHA | CHAIN) |
| 8:00 | 2d5z-C | 4.7 | 3 | 84 | 141 | 7 | MOLECULE: | HEMOGLOBIN | ALPHA S | UBUNIT |
| 9:00 | 1o1o-C | 4.7 | 3 | 84 | 141 | 7 | MOLECULE: | HEMOGLOBIN | ALPHA C | HAIN |
| 10:00 | 6otx-B | 4.7 | 4.1 | 88 | 152 | 6 | MOLECULE: | HEMOGLOBIN | II | |
| 11:00 | 1a6m-A | 4.7 | 4.1 | 88 | 151 | 11 | MOLECULE: | MYOGLOBIN | | |
| 12:00 | 1rvw-A | 4.7 | 3.3 | 85 | 141 | 7 | MOLECULE: | HEMOGLOBIN | | |
| 13:00 | 1j3z-G | 4.7 | 3 | 84 | 141 | 7 | MOLECULE: | HEMOGLOBIN | ALPHA C | HAIN |
| 14:00 | 1y83-C | 4.7 | 3 | 84 | 141 | 7 | MOLECULE: | HEMOGLOBIN | ALPHA C | HAIN |
| 15:00 | 1bzz-A | 4.7 | 3 | 84 | 141 | 7 | MOLECULE: | PROTEIN (H | EMOGLOBI | N ALPHA CHAIN) |
| 16:00 | 2hhd-A | 4.7 | 3.3 | 85 | 141 | 7 | MOLECULE: | HEMOGLOBIN | (DEOXY) | (ALPHA CHAIN) |
| 17:00 | 1hsy-A | 4.7 | 4.2 | 90 | 153 | 16 | MOLECULE: | MYOGLOBIN | | |
| 18:00 | 1qsi-C | 4.7 | 3 | 84 | 141 | 7 | MOLECULE: | PROTEIN (H | EMOGLOBI | N ALPHA CHAIN) |
| 19:00 | 1y4g-C | 4.7 | 3 | 84 | 141 | 7 | MOLECULE: | HEMOGLOBIN | ALPHA C | HAIN |
| 20:00 | 1c7b-A | 4.7 | 3 | 84 | 141 | 7 | MOLECULE: | PROTEIN (D | EOXYHEMO | GLOBIN (ALPHA CHAIN)) |
| 21:00 | 1myj-A | 4.7 | 4.1 | 89 | 153 | 15 | MOLECULE: | MYOGLOBIN | | |
| 22:00 | 1xz7-A | 4.7 | 3 | 84 | 141 | 7 | MOLECULE: | HEMOGLOBIN | ALPHA C | HAIN |
| 23:00 | 1y8w-C | 4.7 | 3.1 | 85 | 141 | 6 | MOLECULE: | HEMOGLOBIN | ALPHA C | HAIN |
| 24:00:00 | 8fdm-C | 4.7 | 3.4 | 85 | 139 | 7 | MOLECULE: | HEMOGLOBIN | SUBUNIT | ALPHA |
| 25:00:00 | 3szk-D | 4.7 | 3.4 | 85 | 140 | 5 | MOLECULE: | HEMOGLOBIN | SUBUNIT | ALPHA |
| 26:00:00 | 2zsn-A | 4.7 | 4.1 | 88 | 153 | 11 | MOLECULE: | MYOGLOBIN | | |
| 27:00:00 | 1hdb-C | 4.7 | 3 | 84 | 141 | 7 | MOLECULE: | HEMOGLOBIN | (DEOXY) | BETA-V67T |
| 28:00:00 | 1a0z-C | 4.7 | 3 | 84 | 141 | 7 | MOLECULE: | HEMOGLOBIN | (ALPHA | CHAIN) |
| 29:00:00 | 3wfu-A | 4.7 | 4.1 | 90 | 152 | 13 | MOLECULE: | MYOGLOBIN | | |
| 30:00:00 | 1lfl-A | 4.7 | 3.1 | 85 | 141 | 6 | MOLECULE: | HEMOGLOBIN | ALPHA C | HAIN |

| | | | | | | | | | | |
|---|---|---|---|---|---|---|---|---|---|---|
| 31:00:00 | 2zt1-A | 4.7 | 4.1 | 88 | 153 | 11 | MOLECULE: | MYOGLOBIN | | |
| 32:00:00 | 6hbw-A | 4.7 | 3 | 84 | 141 | 7 | MOLECULE: | PROTEIN (H | EMOGLOBI | N ALPHA 1) |
| 33:00:00 | 1yhr-C | 4.7 | 3 | 85 | 141 | 6 | MOLECULE: | HEMOGLOBIN | ALPHA C | HAIN |
| 34:00:00 | 1a0z-A | 4.7 | 3 | 84 | 141 | 7 | MOLECULE: | HEMOGLOBIN | (ALPHA | CHAIN) |
| 35:00:00 | 1y35-C | 4.7 | 3 | 84 | 141 | 7 | MOLECULE: | HEMOGLOBIN | ALPHA C | HAIN |
| 36:00:00 | 5e6e-A | 4.7 | 3.4 | 85 | 141 | 7 | MOLECULE: | HEMOGLOBIN | SUBUNIT | ALPHA |
| 37:00:00 | 1k0y-A | 4.7 | 3 | 84 | 141 | 7 | MOLECULE: | HEMOGLOBIN | ALPHA C | HAIN |
| 38:00:00 | 3qjd-A | 4.7 | 3 | 84 | 141 | 7 | MOLECULE: | HEMOGLOBIN | SUBUNIT | ALPHA |
| 39:00:00 | 1y0w-C | 4.7 | 3 | 84 | 141 | 7 | MOLECULE: | HEMOGLOBIN | ALPHA C | HAIN |
| 40:00:00 | 2zsq-A | 4.7 | 4.1 | 88 | 153 | 11 | MOLECULE: | MYOGLOBIN | | |
| 41:00:00 | 3v2v-A | 4.7 | 4.1 | 89 | 152 | 17 | MOLECULE: | MYOGLOBIN | | |
| 42:00:00 | 1mbn-A | 4.7 | 4.1 | 90 | 153 | 12 | MOLECULE: | MYOGLOBIN | | |
| 43:00:00 | 3v2z-A | 4.7 | 4.2 | 88 | 152 | 16 | MOLECULE: | MYOGLOBIN | | |
| 44:00:00 | 7ddt-A | 4.7 | 4.2 | 90 | 153 | 12 | MOLECULE: | ANCESTRAL | MYOGLOBI | N AMBSE |
| 45:00:00 | 1y4q-C | 4.7 | 3 | 84 | 141 | 7 | MOLECULE: | HEMOGLOBIN | ALPHA C | HAIN |
| 46:00:00 | 3hhb-A | 4.7 | 3 | 84 | 141 | 7 | MOLECULE: | HEMOGLOBIN | (DEOXY) | (ALPHA CHAIN) |
| 47:00:00 | 1g08-C | 4.7 | 3.1 | 85 | 141 | 7 | MOLECULE: | HEMOGLOBIN | ALPHA C | HAIN |
| 48:00:00 | 5vzo-A | 4.7 | 4.2 | 88 | 153 | 13 | MOLECULE: | MYOGLOBIN | | |
| 49:00:00 | 4of9-A | 4.7 | 4.1 | 88 | 153 | 11 | MOLECULE: | MYOGLOBIN | | |
| 50:00:00 | 1y0a-C | 4.7 | 3 | 84 | 141 | 7 | MOLECULE: | HEMOGLOBIN | ALPHA C | HAIN |
| 51:00:00 | 8fb0-A | 4.7 | 4.1 | 88 | 153 | 11 | MOLECULE: | MYOGLOBIN | | |
| 52:00:00 | 2zsr-A | 4.7 | 4.1 | 88 | 153 | 11 | MOLECULE: | MYOGLOBIN | | |
| 53:00:00 | 1k0y-C | 4.7 | 3 | 84 | 141 | 7 | MOLECULE: | HEMOGLOBIN | ALPHA C | HAIN |
| 54:00:00 | 1qsi-A | 4.7 | 3 | 84 | 141 | 7 | MOLECULE: | PROTEIN (H | EMOGLOBI | N ALPHA CHAIN) |
| 55:00:00 | 5ut8-A | 4.7 | 4.2 | 89 | 153 | 11 | MOLECULE: | MYOGLOBIN | | |
| 56:00:00 | 1bz1-C | 4.7 | 3 | 84 | 142 | 7 | MOLECULE: | PROTEIN (H | EMOGLOBI | N ALPHA CHAIN) |
| 57:00:00 | 3onz-A | 4.7 | 3.4 | 85 | 140 | 7 | MOLECULE: | HEMOGLOBIN | SUBUNIT | ALPHA |
| 58:00:00 | 7k4m-G | 4.7 | 3 | 85 | 140 | 6 | MOLECULE: | HEMOGLOBIN | SUBUNIT | ALPHA |
| 59:00:00 | 1bz1-A | 4.7 | 3 | 84 | 142 | 7 | MOLECULE: | PROTEIN (H | EMOGLOBI | N ALPHA CHAIN) |
| 60:00:00 | 5kd1-A | 4.7 | 4.1 | 88 | 153 | 11 | MOLECULE: | MYOGLOBIN | | |
| 61:00:00 | 1y7d-C | 4.7 | 3 | 84 | 141 | 7 | MOLECULE: | HEMOGLOBIN | ALPHA C | HAIN |
| 62:00:00 | 1lfl-C | 4.7 | 3.1 | 85 | 141 | 6 | MOLECULE: | HEMOGLOBIN | ALPHA C | HAIN |
| 63:00:00 | 1sdl-C | 4.7 | 3 | 84 | 141 | 7 | MOLECULE: | HEMOGLOBIN | A | |
| 64:00:00 | 8fdm-A | 4.7 | 3.4 | 85 | 139 | 7 | MOLECULE: | HEMOGLOBIN | SUBUNIT | ALPHA |
| 65:00:00 | 2z6t-A | 4.7 | 4.1 | 88 | 151 | 11 | MOLECULE: | MYOGLOBIN | | |
| 66:00:00 | 6ka9-C | 4.7 | 3 | 84 | 141 | 7 | MOLECULE: | HEMOGLOBIN | SUBUNIT | ALPHA |
| 67:00:00 | 3edb-A | 4.7 | 4.1 | 88 | 153 | 11 | MOLECULE: | MYOGLOBIN | | |
| 68:00:00 | 1mod-A | 4.7 | 4.2 | 90 | 154 | 11 | MOLECULE: | MYOGLOBIN | | |
| 69:00:00 | 1yev-C | 4.7 | 3 | 84 | 138 | 7 | MOLECULE: | HEMOGLOBIN | ALPHA C | HAIN |
| 70:00:00 | 1gli-C | 4.7 | 3 | 84 | 141 | 7 | MOLECULE: | DEOXYHEMOG | LOBIN | |
| 71:00:00 | 2zss-A | 4.7 | 4.1 | 88 | 153 | 11 | MOLECULE: | MYOGLOBIN | | |
| 72:00:00 | 1y4p-C | 4.7 | 3 | 84 | 141 | 7 | MOLECULE: | HEMOGLOBIN | ALPHA C | HAIN |
| 73:00:00 | 1rq3-C | 4.7 | 3 | 84 | 141 | 7 | MOLECULE: | HEMOGLOBIN | ALPHA C | HAIN |
| 74:00:00 | 2zt4-A | 4.7 | 4.1 | 88 | 153 | 11 | MOLECULE: | MYOGLOBIN | | |
| 75:00:00 | 1o1o-A | 4.7 | 3 | 84 | 141 | 7 | MOLECULE: | HEMOGLOBIN | ALPHA C | HAIN |
| 76:00:00 | 1ajg-A | 4.7 | 4.1 | 88 | 153 | 11 | MOLECULE: | MYOGLOBIN | | |
| 77:00:00 | 3m3a-A | 4.7 | 4 | 89 | 153 | 12 | MOLECULE: | MYOGLOBIN | | |
| 78:00:00 | 1xzv-C | 4.7 | 3 | 84 | 141 | 7 | MOLECULE: | HEMOGLOBIN | ALPHA C | HAIN |
| 79:00:00 | 7slh-A | 4.7 | 4.2 | 89 | 152 | 11 | MOLECULE: | MYOGLOBIN | | |
| 80:00:00 | 1mnk-A | 4.7 | 4.1 | 90 | 151 | 13 | MOLECULE: | MYOGLOBIN | | |
| 81:00:00 | 1xz4-C | 4.7 | 3.6 | 86 | 141 | 5 | MOLECULE: | HEMOGLOBIN | ALPHA C | HAIN |
| 82:00:00 | 5uta-A | 4.7 | 4.2 | 88 | 153 | 13 | MOLECULE: | MYOGLOBIN | | |
| 83:00:00 | 2eb9-A | 4.7 | 3.6 | 86 | 150 | 12 | MOLECULE: | MYOGLOBIN | | |
| 84:00:00 | 1y5f-C | 4.7 | 3 | 84 | 141 | 7 | MOLECULE: | HEMOGLOBIN | ALPHA C | HAIN |

| | | | | | | | | | | |
|---|---|---|---|---|---|---|---|---|---|---|
| 85:00:00 | 4l7y-A | 4.7 | 3 | 84 | 141 | 7 | MOLECULE: | HEMOGLOBIN | SUBUNIT | ALPHA |
| 86:00:00 | 2jho-A | 4.7 | 4.1 | 89 | 153 | 12 | MOLECULE: | MYOGLOBIN | | |
| 87:00:00 | 6e02-A | 4.7 | 4.2 | 89 | 153 | 12 | MOLECULE: | MYOGLOBIN | | |
| 88:00:00 | 5ilr-A | 4.7 | 4.1 | 88 | 151 | 11 | MOLECULE: | MYOGLOBIN | | |
| 89:00:00 | 6zmx-A | 4.7 | 3 | 85 | 138 | 6 | MOLECULE: | HEMOGLOBIN | SUBUNIT | ALPHA-A |
| 90:00:00 | 3e5i-A | 4.7 | 4.1 | 88 | 153 | 11 | MOLECULE: | MYOGLOBIN | | |
| 91:00:00 | 4fwx-A | 4.7 | 4 | 88 | 153 | 11 | MOLECULE: | MYOGLOBIN | | |
| 92:00:00 | 1gbv-C | 4.7 | 3 | 84 | 141 | 7 | MOLECULE: | HEMOGLOBIN | | |
| 93:00:00 | 2zsp-A | 4.7 | 4.1 | 88 | 153 | 11 | MOLECULE: | MYOGLOBIN | | |
| 94:00:00 | 5vrt-A | 4.7 | 4.1 | 88 | 153 | 11 | MOLECULE: | MYOGLOBIN | | |
| 95:00:00 | 1dke-C | 4.7 | 2.9 | 84 | 141 | 6 | MOLECULE: | HEMOGLOBIN | : ALPHA | CHAIN |
| 96:00:00 | 2o5b-X | 4.7 | 4.1 | 89 | 152 | 16 | MOLECULE: | MYOGLOBIN | | |
| 97:00:00 | 5ych-A | 4.7 | 4.1 | 90 | 153 | 13 | MOLECULE: | ANCESTRAL | MYOGLOBI | N AMBWB OF BASILOSAURUS RELATIV |
| 98:00:00 | 5yl3-A | 4.7 | 4.1 | 90 | 151 | 13 | MOLECULE: | MYOGLOBIN | | |
| 99:00:00 | 1nz2-A | 4.7 | 4.2 | 90 | 153 | 16 | MOLECULE: | MYOGLOBIN | | |
| 0:00 | 1mbd-A | 4.7 | 4.2 | 90 | 153 | 12 | MOLECULE: | MYOGLOBIN | | |
| 1:00 | 2yrs-I | 4.7 | 3.3 | 85 | 141 | 7 | MOLECULE: | HEMOGLOBIN | SUBUNIT | ALPHA |
| 2:00 | 1mcy-A | 4.7 | 4.1 | 89 | 154 | 11 | MOLECULE: | MYOGLOBIN | (CARBONM | ONOXY) |
| 3:00 | 5jnz-A | 4.7 | 3.1 | 85 | 142 | 1 | MOLECULE: | ALPHA CHAI | N | |
| 4:00 | 1cls-A | 4.7 | 3 | 84 | 141 | 7 | MOLECULE: | HEMOGLOBIN | | |
| 5:00 | 4pq6-A | 4.7 | 4 | 88 | 153 | 11 | MOLECULE: | MYOGLOBIN | | |
| 6:00 | 6krf-A | 4.7 | 4.1 | 88 | 153 | 11 | MOLECULE: | MYOGLOBIN | | |
| 7:00 | 2hbs-G | 4.7 | 3 | 84 | 141 | 7 | MOLECULE: | HEMOGLOBIN | S (DEOX | Y), ALPHA CHAIN |
| 8:00 | 1mtj-A | 4.7 | 4.2 | 90 | 154 | 11 | MOLECULE: | MYOGLOBIN | | |
| 9:00 | 2zsy-A | 4.7 | 4.1 | 88 | 153 | 11 | MOLECULE: | MYOGLOBIN | | |
| 10:00 | 8qbc-A | 4.7 | 4.1 | 89 | 156 | 11 | MOLECULE: | MYOGLOBIN | | |
| 11:00 | 4mxk-A | 4.7 | 4 | 88 | 153 | 11 | MOLECULE: | MYOGLOBIN | | |
| 12:00 | 3ubv-A | 4.7 | 3.7 | 84 | 131 | 8 | MOLECULE: | HEMOGLOBIN | #NAME? | AVOPROTEIN |
| 13:00 | 2mgd-A | 4.7 | 4.2 | 90 | 154 | 11 | MOLECULE: | MYOGLOBIN | | |
| 14:00 | 1ch2-A | 4.7 | 4.1 | 89 | 154 | 11 | MOLECULE: | PROTEIN (M | YOGLOBIN | ) |
| 15:00 | 1nz5-A | 4.7 | 4.2 | 90 | 153 | 16 | MOLECULE: | MYOGLOBIN | | |
| 16:00 | 1myg-A | 4.7 | 4.1 | 89 | 153 | 15 | MOLECULE: | MYOGLOBIN | | |
| 17:00 | 107m-A | 4.7 | 4.2 | 90 | 154 | 11 | MOLECULE: | MYOGLOBIN | | |
| 18:00 | 4ns2-A | 4.7 | 4 | 88 | 153 | 14 | MOLECULE: | MYOGLOBIN | | |
| 19:00 | 1jpb-A | 4.7 | 4.1 | 88 | 151 | 11 | MOLECULE: | MYOGLOBIN | | |
| 20:00 | 2bwh-A | 4.7 | 4 | 88 | 153 | 13 | MOLECULE: | MYOGLOBIN | | |
| 21:00 | 6kai-G | 4.7 | 3 | 84 | 141 | 7 | MOLECULE: | HEMOGLOBIN | SUBUNIT | ALPHA |
| 22:00 | 5wjk-A | 4.7 | 4.2 | 89 | 153 | 11 | MOLECULE: | MYOGLOBIN | | |
| 23:00 | 4qau-A | 4.7 | 4.1 | 88 | 153 | 11 | MOLECULE: | MYOGLOBIN | | |
| 24:00:00 | 8fdj-A | 4.7 | 4.1 | 88 | 151 | 11 | MOLECULE: | MYOGLOBIN | | |
| 25:00:00 | 3at6-A | 4.7 | 2.9 | 84 | 141 | 8 | MOLECULE: | ALPHAA-GLO | BIN | |
| 26:00:00 | 7xcf-A | 4.7 | 4.1 | 88 | 153 | 11 | MOLECULE: | MYOGLOBIN | | |
| 27:00:00 | 3nmm-C | 4.7 | 3 | 84 | 141 | 7 | MOLECULE: | HEMOGLOBIN | SUBUNIT | ALPHA |
| 28:00:00 | 3ubc-A | 4.7 | 3.8 | 84 | 131 | 8 | MOLECULE: | HEMOGLOBIN | #NAME? | AVOPROTEIN |
| 29:00:00 | 6g5t-A | 4.7 | 4.1 | 89 | 155 | 11 | MOLECULE: | MYOGLOBIN | | |
| 30:00:00 | 5vnu-A | 4.7 | 4 | 88 | 153 | 11 | MOLECULE: | MYOGLOBIN | | |
| 31:00:00 | 1a01-C | 4.7 | 3 | 84 | 141 | 7 | MOLECULE: | HEMOGLOBIN | (ALPHA | CHAIN) |
| 32:00:00 | 1y46-C | 4.7 | 3 | 84 | 141 | 7 | MOLECULE: | HEMOGLOBIN | ALPHA C | HAIN |
| 33:00:00 | 6kah-G | 4.7 | 3 | 84 | 141 | 7 | MOLECULE: | HEMOGLOBIN | SUBUNIT | ALPHA |
| 34:00:00 | 1ch3-A | 4.7 | 4.1 | 89 | 154 | 11 | MOLECULE: | PROTEIN (M | YOGLOBIN | ) |
| 35:00:00 | 2bli-A | 4.7 | 4 | 89 | 153 | 11 | MOLECULE: | MYOGLOBIN | | |
| 36:00:00 | 2zsx-A | 4.7 | 4.1 | 88 | 153 | 11 | MOLECULE: | MYOGLOBIN | | |
| 37:00:00 | 1moa-A | 4.7 | 4.1 | 89 | 154 | 11 | MOLECULE: | MYOGLOBIN | | |
| 38:00:00 | 1j7w-A | 4.7 | 3 | 84 | 141 | 7 | MOLECULE: | HEMOGLOBIN | | |

| | | | | | | | | | | |
|---|---|---|---|---|---|---|---|---|---|---|
| 39:00:00 | 5xkw-A | 4.7 | 4.1 | 89 | 153 | 11 | MOLECULE: | MYOGLOBIN | | |
| 40:00:00 | 6iyi-A | 4.7 | 3 | 84 | 142 | 2 | MOLECULE: | ALPHA CHAI | N | |
| 41:00:00 | 1xy0-A | 4.7 | 3 | 84 | 141 | 7 | MOLECULE: | HEMOGLOBIN | ALPHA C | HAIN |
| 42:00:00 | 4mxl-A | 4.7 | 4.1 | 89 | 153 | 12 | MOLECULE: | MYOGLOBIN | | |
| 43:00:00 | 1r1y-C | 4.7 | 3.3 | 85 | 141 | 7 | MOLECULE: | HEMOGLOBIN | ALPHA C | HAIN |
| 44:00:00 | 1c7d-A | 4.7 | 11.4 | 111 | 284 | 9 | MOLECULE: | PROTEIN (D | EOXYHEMO | GLOBIN (ALPHA CHAIN)) |
| 45:00:00 | 1hba-C | 4.7 | 3 | 84 | 141 | 7 | MOLECULE: | HEMOGLOBIN | ROTHSCH | ILD (DEOXY) (ALPHA CHAIN) |
| 46:00:00 | 2d6c-A | 4.7 | 4.2 | 88 | 153 | 11 | MOLECULE: | MYOGLOBIN | | |
| 47:00:00 | 6n02-A | 4.7 | 4 | 89 | 154 | 12 | MOLECULE: | MYOGLOBIN, | CO-BOUN | D |
| 48:00:00 | 1j40-G | 4.7 | 3 | 84 | 141 | 7 | MOLECULE: | HEMOGLOBIN | ALPHA C | HAIN |
| 49:00:00 | 2hbs-E | 4.7 | 3 | 84 | 141 | 7 | MOLECULE: | HEMOGLOBIN | S (DEOX | Y), ALPHA CHAIN |
| 50:00:00 | 1rse-A | 4.7 | 4.2 | 90 | 153 | 16 | MOLECULE: | HORSE HEAR | T MYOGLO | BIN |
| 51:00:00 | 5oja-A | 4.7 | 4.1 | 90 | 152 | 12 | MOLECULE: | MYOGLOBIN | | |
| 52:00:00 | 4fwy-A | 4.7 | 4.1 | 89 | 153 | 12 | MOLECULE: | MYOGLOBIN | | |
| 53:00:00 | 2zst-A | 4.7 | 4.1 | 88 | 153 | 11 | MOLECULE: | MYOGLOBIN | | |
| 54:00:00 | 2nss-A | 4.7 | 4.1 | 90 | 152 | 16 | MOLECULE: | MYOGLOBIN | | |
| 55:00:00 | 2hbf-A | 4.7 | 2.9 | 83 | 141 | 7 | MOLECULE: | HEMOGLOBIN | A (N-PR | OPYL ISOCYANIDE) (ALPHA CHAIN) |
| 56:00:00 | 7cen-A | 4.7 | 4.1 | 88 | 153 | 11 | MOLECULE: | MYOGLOBIN | | |
| 57:00:00 | 1bje-A | 4.7 | 4.2 | 90 | 153 | 16 | MOLECULE: | MYOGLOBIN | | |
| 58:00:00 | 1mbo-A | 4.7 | 4.1 | 90 | 153 | 12 | MOLECULE: | MYOGLOBIN | | |
| 59:00:00 | 1g9v-C | 4.7 | 3 | 84 | 141 | 7 | MOLECULE: | HEMOGLOBIN | ALPHA C | HAIN |
| 60:00:00 | 1xye-C | 4.7 | 3 | 84 | 141 | 7 | MOLECULE: | HEMOGLOBIN | ALPHA C | HAIN |
| 61:00:00 | 6bwp-A | 4.7 | 3 | 84 | 141 | 7 | MOLECULE: | HEMOGLOBIN | SUBUNIT | ALPHA |
| 62:00:00 | 2myc-A | 4.7 | 4.1 | 89 | 153 | 12 | MOLECULE: | MYOGLOBIN | (N-BUTYL | ISOCYANIDE) |
| 63:00:00 | 1d8u-A | 4.6 | 4.8 | 95 | 165 | 9 | MOLECULE: | NON-SYMBIO | TIC HEMO | GLOBIN |
| 64:00:00 | 5ca9-A | 4.6 | 20.1 | 110 | 656 | 6 | MOLECULE: | PROTEIN SE | Y1 | |
| 65:00:00 | 1mwd-B | 4.6 | 4.3 | 90 | 153 | 16 | MOLECULE: | PROTEIN (M | YOGLOBIN | ) |
| 66:00:00 | 7pcf-A | 4.6 | 3.6 | 87 | 141 | 6 | MOLECULE: | HEMOGLOBIN | SUBUNIT | ALPHA |
| 67:00:00 | 5ojb-A | 4.6 | 3.6 | 89 | 154 | 12 | MOLECULE: | MYOGLOBIN | | |
| 68:00:00 | 1mno-B | 4.6 | 4.2 | 90 | 153 | 16 | MOLECULE: | PROTEIN (M | YOGLOBIN | ) |
| 69:00:00 | 1ycb-B | 4.6 | 4.2 | 89 | 153 | 15 | MOLECULE: | MYOGLOBIN | | |
| 70:00:00 | 6lcw-C | 4.6 | 3 | 84 | 141 | 7 | MOLECULE: | HEMOGLOBIN | SUBUNIT | ALPHA |
| 71:00:00 | 1spg-A | 4.6 | 3.4 | 86 | 142 | 6 | MOLECULE: | HEMOGLOBIN | | |
| 72:00:00 | 1kd2-C | 4.6 | 3.1 | 85 | 141 | 7 | MOLECULE: | HEMOGLOBIN | ALPHA C | HAIN |
| 73:00:00 | 1g09-A | 4.6 | 3.1 | 85 | 141 | 7 | MOLECULE: | HEMOGLOBIN | ALPHA C | HAIN |
| 74:00:00 | 4rom-C | 4.6 | 3 | 84 | 141 | 7 | MOLECULE: | HEMOGLOBIN | SUBUNIT | ALPHA |
| 75:00:00 | 1hab-C | 4.6 | 3.3 | 85 | 141 | 7 | MOLECULE: | HEMOGLOBIN | A | |
| 76:00:00 | 6bnr-A | 4.6 | 2.9 | 83 | 141 | 7 | MOLECULE: | HEMOGLOBIN | SUBUNIT | ALPHA |
| 77:00:00 | 3sxz-B | 4.6 | 3.8 | 94 | 140 | 7 | MOLECULE: | TRANSCRIPT | IONAL RE | GULATOR, GNTR FAMILY |
| 78:00:00 | 6kat-A | 4.6 | 3.1 | 85 | 141 | 6 | MOLECULE: | HEMOGLOBIN | SUBUNIT | ALPHA |
| 79:00:00 | 1yeq-C | 4.6 | 2.9 | 84 | 141 | 6 | MOLECULE: | HEMOGLOBIN | ALPHA C | HAIN |
| 80:00:00 | 1b86-C | 4.6 | 3.1 | 85 | 141 | 6 | MOLECULE: | PROTEIN (H | EMOGLOBI | N ALPHA CHAIN) |
| 81:00:00 | 1yie-A | 4.6 | 3.3 | 85 | 141 | 7 | MOLECULE: | HEMOGLOBIN | ALPHA C | HAIN |
| 82:00:00 | 1xxt-A | 4.6 | 3.1 | 85 | 141 | 7 | MOLECULE: | HEMOGLOBIN | ALPHA C | HAIN |
| 83:00:00 | 3odq-C | 4.6 | 3.4 | 86 | 141 | 6 | MOLECULE: | HEMOGLOBIN | SUBUNIT | ALPHA |
| 84:00:00 | 1ye0-C | 4.6 | 2.9 | 83 | 141 | 7 | MOLECULE: | HEMOGLOBIN | ALPHA C | HAIN |
| 85:00:00 | 2qsp-C | 4.6 | 3.1 | 84 | 141 | 7 | MOLECULE: | HEMOGLOBIN | SUBUNIT | ALPHA |
| 86:00:00 | 6kau-C | 4.6 | 3 | 84 | 141 | 6 | MOLECULE: | HEMOGLOBIN | SUBUNIT | ALPHA |
| 87:00:00 | 5jdo-E | 4.6 | 3.4 | 85 | 140 | 7 | MOLECULE: | HAPTOGLOBI | N-HAEMOG | LOBIN RECEPTOR |
| 88:00:00 | 6ka9-G | 4.6 | 3 | 84 | 141 | 7 | MOLECULE: | HEMOGLOBIN | SUBUNIT | ALPHA |
| 89:00:00 | 1hda-C | 4.6 | 3 | 84 | 141 | 7 | MOLECULE: | HEMOGLOBIN | (DEOXY) | (ALPHA CHAIN) |
| 90:00:00 | 1thb-A | 4.6 | 3 | 84 | 141 | 7 | MOLECULE: | HEMOGLOBIN | A (OXY) | (ALPHA CHAIN) |
| 91:00:00 | 5ee4-E | 4.6 | 3.5 | 85 | 141 | 5 | MOLECULE: | HPUA | | |
| 92:00:00 | 2w72-C | 4.6 | 2.8 | 83 | 141 | 7 | MOLECULE: | HUMAN HEMO | GLOBIN A | |

| | | | | | | | | | | |
|---|---|---|---|---|---|---|---|---|---|---|
| 93:00:00 | 2hco-A | 4.6 | 3 | 84 | 141 | 7 | MOLECULE: | HEMOGLOBIN | (CARBON | MONOXY) (ALPHA CHAIN) |
| 94:00:00 | 3oo4-A | 4.6 | 3.4 | 85 | 141 | 7 | MOLECULE: | HEMOGLOBIN | SUBUNIT | ALPHA |
| 95:00:00 | 1j3y-E | 4.6 | 3 | 84 | 141 | 7 | MOLECULE: | HEMOGLOBIN | ALPHA C | HAIN |
| 96:00:00 | 3d17-A | 4.6 | 3.5 | 86 | 141 | 6 | MOLECULE: | HEMOGLOBIN | SUBUNIT | ALPHA |
| 97:00:00 | 4ni0-A | 4.6 | 3.4 | 86 | 141 | 6 | MOLECULE: | HEMOGLOBIN | SUBUNIT | ALPHA |
| 98:00:00 | 7dy4-E | 4.6 | 3.1 | 85 | 141 | 7 | MOLECULE: | HEMOGLOBIN | SUBUNIT | ALPHA |
| 99:00:00 | 3dut-C | 4.6 | 3 | 84 | 141 | 7 | MOLECULE: | HEMOGLOBIN | SUBUNIT | ALPHA |
| 0:00 | 5zeo-A | 4.6 | 4.1 | 89 | 153 | 11 | MOLECULE: | MYOGLOBIN | | |
| 1:00 | 6l5y-C | 4.6 | 3.1 | 85 | 141 | 6 | MOLECULE: | HEMOGLOBIN | SUBUNIT | ALPHA |
| 2:00 | 3wcp-C | 4.6 | 3 | 84 | 141 | 7 | MOLECULE: | HEMOGLOBIN | SUBUNIT | ALPHA |
| 3:00 | 5kdq-C | 4.6 | 3 | 85 | 141 | 7 | MOLECULE: | HEMOGLOBIN | SUBUNIT | ALPHA |
| 4:00 | 3wtg-A | 4.6 | 3 | 85 | 138 | 9 | MOLECULE: | HEMOGLOBIN | SUBUNIT | ALPHA-A |
| 5:00 | 1yeo-A | 4.6 | 3 | 84 | 141 | 7 | MOLECULE: | HEMOGLOBIN | ALPHA C | HAIN |
| 6:00 | 3qje-C | 4.6 | 3 | 84 | 141 | 7 | MOLECULE: | HEMOGLOBIN | SUBUNIT | ALPHA |
| 7:00 | 1yeo-C | 4.6 | 3 | 84 | 141 | 7 | MOLECULE: | HEMOGLOBIN | ALPHA C | HAIN |
| 8:00 | 6kao-A | 4.6 | 3.4 | 85 | 141 | 7 | MOLECULE: | HEMOGLOBIN | SUBUNIT | ALPHA |
| 9:00 | 6l5y-A | 4.6 | 3 | 84 | 141 | 6 | MOLECULE: | HEMOGLOBIN | SUBUNIT | ALPHA |
| 10:00 | 7dy4-A | 4.6 | 3 | 84 | 141 | 7 | MOLECULE: | HEMOGLOBIN | SUBUNIT | ALPHA |
| 11:00 | 4n8t-A | 4.6 | 3.4 | 85 | 140 | 7 | MOLECULE: | HEMOGLOBIN | SUBUNIT | ALPHA |
| 12:00 | 6di4-A | 4.6 | 3.5 | 85 | 141 | 5 | MOLECULE: | HEMOGLOBIN | SUBUNIT | ALPHA |
| 13:00 | 1xzu-A | 4.6 | 3.1 | 85 | 141 | 7 | MOLECULE: | HEMOGLOBIN | ALPHA C | HAIN |
| 14:00 | 3bj3-A | 4.6 | 3.3 | 86 | 142 | 3 | MOLECULE: | HEMOGLOBIN | ALPHA | |
| 15:00 | 6ihx-A | 4.6 | 3.1 | 84 | 140 | 7 | MOLECULE: | HEMOGLOBIN | SUBUNIT | ALPHA |
| 16:00 | 5hlx-A | 4.6 | 4.2 | 88 | 153 | 11 | MOLECULE: | MYOGLOBIN | | |
| 17:00 | 4b3w-B | 4.6 | 4.2 | 89 | 154 | 9 | MOLECULE: | CYTOGLOBIN | | |
| 18:00 | 1hgc-A | 4.6 | 2.9 | 84 | 141 | 7 | MOLECULE: | HEMOGLOBIN | (OXY) ( | ALPHA CHAIN) |
| 19:00 | 1hab-A | 4.6 | 3 | 84 | 141 | 7 | MOLECULE: | HEMOGLOBIN | A | |
| 20:00 | 1qi8-C | 4.6 | 3 | 84 | 141 | 7 | MOLECULE: | HEMOGLOBIN | | |
| 21:00 | 6ihx-C | 4.6 | 3.1 | 84 | 140 | 7 | MOLECULE: | HEMOGLOBIN | SUBUNIT | ALPHA |
| 22:00 | 3vrf-A | 4.6 | 3.2 | 86 | 141 | 7 | MOLECULE: | HEMOGLOBIN | SUBUNIT | ALPHA |
| 23:00 | 4hhb-A | 4.6 | 3 | 84 | 141 | 7 | MOLECULE: | HEMOGLOBIN | SUBUNIT | ALPHA |
| 24:00:00 | 1bbb-A | 4.6 | 3.4 | 85 | 141 | 5 | MOLECULE: | HEMOGLOBIN | A (CARB | ONMONOXY) (ALPHA CHAIN) |
| 25:00:00 | 2hbe-A | 4.6 | 2.9 | 83 | 141 | 7 | MOLECULE: | HEMOGLOBIN | A (N-BU | TYL ISOCYANIDE) (ALPHA CHAIN) |
| 26:00:00 | 1yen-A | 4.6 | 3.1 | 85 | 138 | 6 | MOLECULE: | HEMOGLOBIN | ALPHA C | HAIN |
| 27:00:00 | 1y45-C | 4.6 | 3 | 84 | 141 | 7 | MOLECULE: | HEMOGLOBIN | ALPHA C | HAIN |
| 28:00:00 | 3qjd-C | 4.6 | 3 | 84 | 141 | 7 | MOLECULE: | HEMOGLOBIN | SUBUNIT | ALPHA |
| 29:00:00 | 1y4p-A | 4.6 | 3 | 84 | 141 | 7 | MOLECULE: | HEMOGLOBIN | ALPHA C | HAIN |
| 30:00:00 | 4hhb-C | 4.6 | 3 | 84 | 141 | 7 | MOLECULE: | HEMOGLOBIN | SUBUNIT | ALPHA |
| 31:00:00 | 1mni-B | 4.6 | 4.2 | 90 | 153 | 13 | MOLECULE: | MYOGLOBIN | | |
| 32:00:00 | 6kai-C | 4.6 | 3 | 84 | 141 | 7 | MOLECULE: | HEMOGLOBIN | SUBUNIT | ALPHA |
| 33:00:00 | 1y4q-A | 4.6 | 3.6 | 86 | 141 | 5 | MOLECULE: | HEMOGLOBIN | ALPHA C | HAIN |
| 34:00:00 | 8dov-A | 4.6 | 3.4 | 86 | 140 | 7 | MOLECULE: | HEMOGLOBIN | SUBUNIT | ALPHA |
| 35:00:00 | 6xdt-A | 4.6 | 2.9 | 83 | 141 | 7 | MOLECULE: | HEMOGLOBIN | SUBUNIT | ALPHA |
| 36:00:00 | 6kat-C | 4.6 | 3.1 | 85 | 141 | 6 | MOLECULE: | HEMOGLOBIN | SUBUNIT | ALPHA |
| 37:00:00 | 1y22-A | 4.6 | 3 | 84 | 141 | 7 | MOLECULE: | HEMOGLOBIN | ALPHA C | HAIN |
| 38:00:00 | 1qxd-A | 4.6 | 2.9 | 83 | 141 | 7 | MOLECULE: | HEMOGLOBIN | ALPHA C | HAIN |
| 39:00:00 | 6kas-C | 4.6 | 3.1 | 85 | 141 | 6 | MOLECULE: | HEMOGLOBIN | SUBUNIT | ALPHA |
| 40:00:00 | 1y09-A | 4.6 | 3 | 84 | 141 | 7 | MOLECULE: | HEMOGLOBIN | ALPHA C | HAIN |
| 41:00:00 | 6kav-C | 4.6 | 3 | 84 | 141 | 6 | MOLECULE: | HEMOGLOBIN | SUBUNIT | ALPHA |
| 42:00:00 | 1fdh-B | 4.6 | 3 | 84 | 141 | 7 | MOLECULE: | HEMOGLOBIN | F (DEOX | Y) (ALPHA CHAIN) |
| 43:00:00 | 1hac-C | 4.6 | 3.4 | 86 | 141 | 6 | MOLECULE: | HEMOGLOBIN | A | |
| 44:00:00 | 7uvb-C | 4.6 | 2.9 | 83 | 141 | 7 | MOLECULE: | HEMOGLOBIN | SUBUNIT | ALPHA |
| 45:00:00 | 7jy3-A | 4.6 | 2.9 | 83 | 141 | 7 | MOLECULE: | HEMOGLOBIN | SUBUNIT | ALPHA |
| 46:00:00 | 6l5w-A | 4.6 | 3.4 | 85 | 141 | 7 | MOLECULE: | HEMOGLOBIN | SUBUNIT | ALPHA |

| | | | | | | | | | | |
|---|---|---|---|---|---|---|---|---|---|---|
| 47:00:00 | 1y0w-A | 4.6 | 3 | 84 | 141 | 7 | MOLECULE: | HEMOGLOBIN | ALPHA C | HAIN |
| 48:00:00 | 2evp-A | 4.6 | 4.1 | 89 | 153 | 11 | MOLECULE: | MYOGLOBIN | | |
| 49:00:00 | 1y83-A | 4.6 | 3 | 84 | 141 | 7 | MOLECULE: | HEMOGLOBIN | ALPHA C | HAIN |
| 50:00:00 | 2w6v-C | 4.6 | 3 | 84 | 141 | 7 | MOLECULE: | HEMOGLOBIN | SUBUNIT | ALPHA |
| 51:00:00 | 5ucu-A | 4.6 | 3.4 | 85 | 140 | 7 | MOLECULE: | HEMOGLOBIN | SUBUNIT | ALPHA |
| 52:00:00 | 1rq4-C | 4.6 | 3 | 84 | 141 | 7 | MOLECULE: | HEMOGLOBIN | ALPHA C | HAIN |
| 53:00:00 | 1sdk-C | 4.6 | 3.1 | 84 | 141 | 7 | MOLECULE: | HEMOGLOBIN | A | |
| 54:00:00 | 6bwu-A | 4.6 | 3.4 | 85 | 140 | 7 | MOLECULE: | HEMOGLOBIN | SUBUNIT | ALPHA |
| 55:00:00 | 1ye1-C | 4.6 | 3 | 84 | 141 | 7 | MOLECULE: | HEMOGLOBIN | ALPHA C | HAIN |
| 56:00:00 | 1pmb-A | 4.6 | 4.1 | 89 | 153 | 15 | MOLECULE: | MYOGLOBIN | | |
| 57:00:00 | 6kap-A | 4.6 | 3.4 | 85 | 141 | 7 | MOLECULE: | HEMOGLOBIN | SUBUNIT | ALPHA |
| 58:00:00 | 6xdt-C | 4.6 | 3.5 | 86 | 141 | 3 | MOLECULE: | HEMOGLOBIN | SUBUNIT | ALPHA |
| 59:00:00 | 1y7c-C | 4.6 | 3 | 84 | 141 | 7 | MOLECULE: | HEMOGLOBIN | ALPHA C | HAIN |
| 60:00:00 | 1lft-A | 4.6 | 3 | 84 | 141 | 7 | MOLECULE: | HEMOGLOBIN | ALPHA C | HAIN |
| 61:00:00 | 7pch-A | 4.6 | 3.4 | 85 | 141 | 7 | MOLECULE: | HEMOGLOBIN | SUBUNIT | ALPHA |
| 62:00:00 | 1v4w-C | 4.6 | 3.3 | 85 | 143 | 5 | MOLECULE: | HEMOGLOBIN | ALPHA C | HAIN |
| 63:00:00 | 1j7s-C | 4.6 | 3 | 84 | 141 | 7 | MOLECULE: | HEMOGLOBIN | | |
| 64:00:00 | 3qjb-A | 4.6 | 3.4 | 85 | 141 | 7 | MOLECULE: | HEMOGLOBIN | SUBUNIT | ALPHA |
| 65:00:00 | 4n7n-G | 4.6 | 3.1 | 86 | 141 | 6 | MOLECULE: | HEMOGLOBIN | SUBUNIT | ALPHA |
| 66:00:00 | 1gbu-A | 4.6 | 3 | 84 | 141 | 7 | MOLECULE: | HEMOGLOBIN | | |
| 67:00:00 | 1v4x-A | 4.6 | 3.3 | 87 | 143 | 5 | MOLECULE: | HEMOGLOBIN | ALPHA C | HAIN |
| 68:00:00 | 2hhe-A | 4.6 | 3 | 84 | 141 | 7 | MOLECULE: | HEMOGLOBIN | (DEOXY) | (ALPHA CHAIN) |
| 69:00:00 | 3fs4-A | 4.6 | 3.6 | 87 | 141 | 5 | MOLECULE: | HEMOGLOBIN | SUBUNIT | ALPHA-A |
| 70:00:00 | 6kar-A | 4.6 | 3.4 | 85 | 141 | 7 | MOLECULE: | HEMOGLOBIN | SUBUNIT | ALPHA |
| 71:00:00 | 6kas-A | 4.6 | 3 | 84 | 141 | 7 | MOLECULE: | HEMOGLOBIN | SUBUNIT | ALPHA |
| 72:00:00 | 1j7y-C | 4.6 | 3 | 84 | 141 | 7 | MOLECULE: | HEMOGLOBIN | | |
| 73:00:00 | 1xz5-A | 4.6 | 3 | 84 | 141 | 7 | MOLECULE: | HEMOGLOBIN | ALPHA C | HAIN |
| 74:00:00 | 6hbw-C | 4.6 | 2.9 | 84 | 141 | 7 | MOLECULE: | PROTEIN (H | EMOGLOBI | N ALPHA 1) |
| 75:00:00 | 1ye2-A | 4.6 | 3.6 | 86 | 141 | 5 | MOLECULE: | HEMOGLOBIN | ALPHA C | HAIN |
| 76:00:00 | 1kd2-A | 4.6 | 3 | 84 | 141 | 7 | MOLECULE: | HEMOGLOBIN | ALPHA C | HAIN |
| 77:00:00 | 1yeu-A | 4.6 | 3 | 84 | 141 | 7 | MOLECULE: | HEMOGLOBIN | ALPHA C | HAIN |
| 78:00:00 | 3sxy-A | 4.6 | 4.4 | 103 | 201 | 6 | MOLECULE: | TRANSCRIPT | IONAL RE | GULATOR, GNTR FAMILY |
| 79:00:00 | 7uf6-C | 4.6 | 3 | 84 | 141 | 6 | MOLECULE: | HEMOGLOBIN | SUBUNIT | ALPHA |
| 80:00:00 | 5sw7-A | 4.6 | 3.4 | 85 | 142 | 7 | MOLECULE: | HEMOGLOBIN | SUBUNIT | ALPHA |
| 81:00:00 | 1hgb-C | 4.6 | 3 | 84 | 141 | 7 | MOLECULE: | HEMOGLOBIN | (AQUO M | ET) (ALPHA CHAIN) |
| 82:00:00 | 1y85-A | 4.6 | 3.1 | 85 | 141 | 7 | MOLECULE: | HEMOGLOBIN | ALPHA C | HAIN |
| 83:00:00 | 3bcq-A | 4.6 | 3.3 | 85 | 142 | 8 | MOLECULE: | ALPHA-CHAI | N HEMOGL | OBIN |
| 84:00:00 | 7dy4-G | 4.6 | 3 | 84 | 141 | 7 | MOLECULE: | HEMOGLOBIN | SUBUNIT | ALPHA |
| 85:00:00 | 1y5j-A | 4.6 | 3.6 | 86 | 141 | 5 | MOLECULE: | HEMOGLOBIN | ALPHA C | HAIN |
| 86:00:00 | 1qi8-A | 4.6 | 3 | 84 | 141 | 7 | MOLECULE: | HEMOGLOBIN | | |
| 87:00:00 | 1yvq-A | 4.6 | 2.9 | 83 | 141 | 7 | MOLECULE: | HEMOGLOBIN | ALPHA C | HAIN |
| 88:00:00 | 6ii1-C | 4.6 | 3.1 | 84 | 138 | 7 | MOLECULE: | HEMOGLOBIN | SUBUNIT | ALPHA |
| 89:00:00 | 6lcw-E | 4.6 | 3 | 84 | 141 | 7 | MOLECULE: | HEMOGLOBIN | SUBUNIT | ALPHA |
| 90:00:00 | 3wtg-C | 4.6 | 3.3 | 86 | 138 | 9 | MOLECULE: | HEMOGLOBIN | SUBUNIT | ALPHA-A |
| 91:00:00 | 1cls-C | 4.6 | 3 | 84 | 141 | 7 | MOLECULE: | HEMOGLOBIN | | |
| 92:00:00 | 3whm-E | 4.6 | 3 | 84 | 140 | 7 | MOLECULE: | HEMOGLOBIN | SUBUNIT | ALPHA |
| 93:00:00 | 5e29-A | 4.6 | 3 | 84 | 141 | 7 | MOLECULE: | HEMOGLOBIN | SUBUNIT | ALPHA |
| 94:00:00 | 1j7w-C | 4.6 | 3 | 84 | 141 | 7 | MOLECULE: | HEMOGLOBIN | | |
| 95:00:00 | 6ka9-E | 4.6 | 3 | 84 | 141 | 7 | MOLECULE: | HEMOGLOBIN | SUBUNIT | ALPHA |
| 96:00:00 | 1a0u-A | 4.6 | 3 | 84 | 141 | 7 | MOLECULE: | HEMOGLOBIN | (ALPHA | CHAIN) |
| 97:00:00 | 6z4t-A | 4.6 | 4.1 | 88 | 154 | 11 | MOLECULE: | MYOGLOBIN | | |
| 98:00:00 | 1yvt-A | 4.6 | 3.4 | 85 | 141 | 7 | MOLECULE: | HEMOGLOBIN | ALPHA C | HAIN |
| 99:00:00 | 6lcx-C | 4.6 | 3 | 84 | 141 | 7 | MOLECULE: | HEMOGLOBIN | SUBUNIT | ALPHA |
| 0:00 | 1fsx-C | 4.6 | 3 | 84 | 141 | 7 | MOLECULE: | HEMOGLOBIN | ALPHA C | HAIN |

| | | | | | | | | | | |
|---|---|---|---|---|---|---|---|---|---|---|
| 1:00 | 1y4g-A | 4.6 | 3 | 84 | 141 | 7 | MOLECULE: | HEMOGLOBIN | ALPHA C | HAIN |
| 2:00 | 1xye-A | 4.6 | 3 | 84 | 141 | 7 | MOLECULE: | HEMOGLOBIN | ALPHA C | HAIN |
| 3:00 | 2hbs-A | 4.6 | 3 | 84 | 141 | 7 | MOLECULE: | HEMOGLOBIN | S (DEOX | Y), ALPHA CHAIN |
| 4:00 | 6ka9-A | 4.6 | 3 | 84 | 141 | 7 | MOLECULE: | HEMOGLOBIN | SUBUNIT | ALPHA |
| 5:00 | 1y7d-A | 4.6 | 3 | 84 | 141 | 7 | MOLECULE: | HEMOGLOBIN | ALPHA C | HAIN |
| 6:00 | 6lcx-E | 4.6 | 3 | 84 | 141 | 7 | MOLECULE: | HEMOGLOBIN | SUBUNIT | ALPHA |
| 7:00 | 8fdk-C | 4.6 | 3.1 | 84 | 141 | 7 | MOLECULE: | HEMOGLOBIN | SUBUNIT | ALPHA |
| 8:00 | 4mqc-A | 4.6 | 3.1 | 85 | 141 | 7 | MOLECULE: | HEMOGLOBIN | SUBUNIT | ALPHA |
| 9:00 | 1ygd-A | 4.6 | 3 | 84 | 141 | 7 | MOLECULE: | HEMOGLOBIN | ALPHA C | HAIN |
| 10:00 | 4mqh-A | 4.6 | 3.2 | 86 | 139 | 6 | MOLECULE: | HEMOGLOBIN | SUBUNIT | ALPHA |
| 11:00 | 1y85-C | 4.6 | 3.6 | 86 | 141 | 5 | MOLECULE: | HEMOGLOBIN | ALPHA C | HAIN |
| 12:00 | 7cue-C | 4.6 | 3.5 | 86 | 141 | 6 | MOLECULE: | HEMOGLOBIN | SUBUNIT | ALPHA |
| 13:00 | 1yhe-A | 4.6 | 3 | 84 | 141 | 7 | MOLECULE: | HEMOGLOBIN | ALPHA C | HAIN |
| 14:00 | 1yg5-C | 4.6 | 3 | 84 | 141 | 7 | MOLECULE: | HEMOGLOBIN | ALPHA C | HAIN |
| 15:00 | 1j3z-E | 4.6 | 3 | 84 | 141 | 7 | MOLECULE: | HEMOGLOBIN | ALPHA C | HAIN |
| 16:00 | 2yrs-C | 4.6 | 3.3 | 86 | 141 | 7 | MOLECULE: | HEMOGLOBIN | SUBUNIT | ALPHA |
| 17:00 | 1rqa-C | 4.6 | 3 | 84 | 141 | 7 | MOLECULE: | HEMOGLOBIN | ALPHA C | HAIN |
| 18:00 | 2hhb-A | 4.6 | 3 | 84 | 141 | 7 | MOLECULE: | HEMOGLOBIN | (DEOXY) | (ALPHA CHAIN) |
| 19:00 | 7ehx-A | 4.6 | 4.1 | 88 | 153 | 11 | MOLECULE: | MYOGLOBIN | | |
| 20:00 | 1y2z-C | 4.6 | 3 | 84 | 141 | 7 | MOLECULE: | HEMOGLOBIN | ALPHA C | HAIN |
| 21:00 | 6hit-E | 4.6 | 3.4 | 86 | 142 | 5 | MOLECULE: | HEMOGLOBIN | ALPHA 2 | CHAIN |
| 22:00 | 1buw-A | 4.6 | 3.4 | 85 | 141 | 5 | MOLECULE: | PROTEIN (H | EMOGLOBI | N) |
| 23:00 | 1y7z-A | 4.6 | 3 | 84 | 141 | 7 | MOLECULE: | HEMOGLOBIN | ALPHA C | HAIN |
| 24:00:00 | 1y31-A | 4.6 | 3.6 | 86 | 141 | 5 | MOLECULE: | HEMOGLOBIN | ALPHA C | HAIN |
| 25:00:00 | 8fdk-A | 4.6 | 3.4 | 85 | 141 | 7 | MOLECULE: | HEMOGLOBIN | SUBUNIT | ALPHA |
| 26:00:00 | 6kye-I | 4.6 | 3.6 | 86 | 141 | 5 | MOLECULE: | HEMOGLOBIN | SUBUNIT | ALPHA |
| 27:00:00 | 2hhd-C | 4.6 | 3 | 85 | 141 | 7 | MOLECULE: | HEMOGLOBIN | (DEOXY) | (ALPHA CHAIN) |
| 28:00:00 | 1vxb-A | 4.6 | 4.1 | 88 | 153 | 11 | MOLECULE: | MYOGLOBIN | | |
| 29:00:00 | 1ye1-A | 4.6 | 3 | 84 | 141 | 7 | MOLECULE: | HEMOGLOBIN | ALPHA C | HAIN |
| 30:00:00 | 5ee4-C | 4.6 | 2.9 | 83 | 141 | 7 | MOLECULE: | HPUA | | |
| 31:00:00 | 1yie-C | 4.6 | 3 | 84 | 141 | 7 | MOLECULE: | HEMOGLOBIN | ALPHA C | HAIN |
| 32:00:00 | 8dov-C | 4.6 | 3.4 | 86 | 140 | 7 | MOLECULE: | HEMOGLOBIN | SUBUNIT | ALPHA |
| 33:00:00 | 1y35-A | 4.6 | 3 | 84 | 141 | 7 | MOLECULE: | HEMOGLOBIN | ALPHA C | HAIN |
| 34:00:00 | 5hu6-A | 4.6 | 2.9 | 83 | 141 | 7 | MOLECULE: | HEMOGLOBIN | SUBUNIT | ALPHA |
| 35:00:00 | 1y4b-A | 4.6 | 3 | 84 | 141 | 7 | MOLECULE: | HEMOGLOBIN | ALPHA C | HAIN |
| 36:00:00 | 5vmm-A | 4.6 | 3.5 | 88 | 139 | 6 | MOLECULE: | HEMOGLOBIN | SUBUNIT | ALPHA |
| 37:00:00 | 3bj1-C | 4.6 | 3.4 | 86 | 142 | 3 | MOLECULE: | HEMOGLOBIN | ALPHA | |
| 38:00:00 | 1yeq-A | 4.6 | 2.9 | 83 | 141 | 7 | MOLECULE: | HEMOGLOBIN | ALPHA C | HAIN |
| 39:00:00 | 1xzv-A | 4.6 | 3 | 84 | 141 | 7 | MOLECULE: | HEMOGLOBIN | ALPHA C | HAIN |
| 40:00:00 | 1nqp-A | 4.6 | 2.9 | 84 | 141 | 6 | MOLECULE: | HEMOGLOBIN | ALPHA C | HAIN |
| 41:00:00 | 1rq4-A | 4.6 | 3 | 84 | 141 | 7 | MOLECULE: | HEMOGLOBIN | ALPHA C | HAIN |
| 42:00:00 | 6kye-C | 4.6 | 3.1 | 85 | 140 | 6 | MOLECULE: | HEMOGLOBIN | SUBUNIT | ALPHA |
| 43:00:00 | 7jy0-C | 4.6 | 2.9 | 83 | 141 | 7 | MOLECULE: | HEMOGLOBIN | SUBUNIT | ALPHA |
| 44:00:00 | 1lfl-R | 4.6 | 3 | 85 | 141 | 6 | MOLECULE: | HEMOGLOBIN | ALPHA C | HAIN |
| 45:00:00 | 2evk-A | 4.6 | 4.1 | 88 | 153 | 11 | MOLECULE: | MYOGLOBIN | | |
| 46:00:00 | 7jy1-A | 4.6 | 3 | 84 | 141 | 7 | MOLECULE: | HEMOGLOBIN | SUBUNIT | ALPHA |
| 47:00:00 | 6kau-A | 4.6 | 3.1 | 85 | 141 | 6 | MOLECULE: | HEMOGLOBIN | SUBUNIT | ALPHA |
| 48:00:00 | 6kye-A | 4.6 | 3.5 | 87 | 140 | 6 | MOLECULE: | HEMOGLOBIN | SUBUNIT | ALPHA |
| 49:00:00 | 1hdb-A | 4.6 | 3 | 84 | 141 | 7 | MOLECULE: | HEMOGLOBIN | (DEOXY) | BETA-V67T |
| 50:00:00 | 6l5x-A | 4.6 | 3 | 84 | 141 | 7 | MOLECULE: | HEMOGLOBIN | SUBUNIT | ALPHA |
| 51:00:00 | 1hga-A | 4.6 | 3.3 | 85 | 141 | 7 | MOLECULE: | HEMOGLOBIN | (DEOXY) | (ALPHA CHAIN) |
| 52:00:00 | 1y5k-A | 4.6 | 3 | 84 | 141 | 7 | MOLECULE: | HEMOGLOBIN | ALPHA C | HAIN |
| 53:00:00 | 7uvb-A | 4.6 | 3.1 | 85 | 141 | 6 | MOLECULE: | HEMOGLOBIN | SUBUNIT | ALPHA |
| 54:00:00 | 2yrs-M | 4.6 | 3.3 | 85 | 141 | 7 | MOLECULE: | HEMOGLOBIN | SUBUNIT | ALPHA |

| | | | | | | | | | | |
|---|---|---|---|---|---|---|---|---|---|---|
| 55:00:00 | 5u3i-A | 4.6 | 3 | 84 | 141 | 6 | MOLECULE: | HEMOGLOBIN | SUBUNIT | ALPHA |
| 56:00:00 | 3fs4-C | 4.6 | 3.1 | 85 | 141 | 7 | MOLECULE: | HEMOGLOBIN | SUBUNIT | ALPHA-A |
| 57:00:00 | 1yeu-C | 4.6 | 3 | 84 | 141 | 7 | MOLECULE: | HEMOGLOBIN | ALPHA C | HAIN |
| 58:00:00 | 1y7c-A | 4.6 | 3 | 84 | 141 | 7 | MOLECULE: | HEMOGLOBIN | ALPHA C | HAIN |
| 59:00:00 | 1y31-C | 4.6 | 3 | 84 | 141 | 7 | MOLECULE: | HEMOGLOBIN | ALPHA C | HAIN |
| 60:00:00 | 6lcw-G | 4.6 | 2.9 | 84 | 141 | 7 | MOLECULE: | HEMOGLOBIN | SUBUNIT | ALPHA |
| 61:00:00 | 8fdn-A | 4.6 | 3.1 | 84 | 141 | 7 | MOLECULE: | HEMOGLOBIN | SUBUNIT | ALPHA |
| 62:00:00 | 6bwp-C | 4.6 | 2.9 | 84 | 141 | 7 | MOLECULE: | HEMOGLOBIN | SUBUNIT | ALPHA |
| 63:00:00 | 5hlq-A | 4.6 | 4.2 | 88 | 152 | 11 | MOLECULE: | MYOGLOBIN | | |
| 64:00:00 | 1nih-A | 4.6 | 3 | 85 | 141 | 7 | MOLECULE: | HEMOGLOBIN | (NICKEL | OUS DEOXY) (ALPHA CHAIN) |
| 65:00:00 | 6l5v-A | 4.6 | 3.4 | 85 | 141 | 7 | MOLECULE: | HEMOGLOBIN | SUBUNIT | ALPHA |
| 66:00:00 | 1y5f-A | 4.6 | 3 | 84 | 141 | 7 | MOLECULE: | HEMOGLOBIN | ALPHA C | HAIN |
| 67:00:00 | 1yih-A | 4.6 | 3 | 84 | 141 | 7 | MOLECULE: | HEMOGLOBIN | ALPHA C | HAIN |
| 68:00:00 | 7jjq-A | 4.6 | 3.4 | 85 | 140 | 7 | MOLECULE: | HEMOGLOBIN | SUBUNIT | ALPHA |
| 69:00:00 | 1g08-A | 4.6 | 3.6 | 87 | 141 | 5 | MOLECULE: | HEMOGLOBIN | ALPHA C | HAIN |
| 70:00:00 | 3bj1-A | 4.6 | 3.4 | 86 | 142 | 3 | MOLECULE: | HEMOGLOBIN | ALPHA | |
| 71:00:00 | 4esa-A | 4.6 | 3 | 84 | 142 | 5 | MOLECULE: | HEMOGLOBIN | ALPHA C | HAIN |
| 72:00:00 | 1y0a-A | 4.6 | 3 | 85 | 141 | 7 | MOLECULE: | HEMOGLOBIN | ALPHA C | HAIN |
| 73:00:00 | 4x0i-A | 4.6 | 2.9 | 83 | 141 | 7 | MOLECULE: | HEMOGLOBIN | SUBUNIT | ALPHA |
| 74:00:00 | 1r1x-A | 4.6 | 3.1 | 84 | 141 | 7 | MOLECULE: | HEMOGLOBIN | ALPHA C | HAIN |
| 75:00:00 | 3vrg-A | 4.6 | 3.2 | 85 | 141 | 7 | MOLECULE: | HEMOGLOBIN | SUBUNIT | ALPHA |
| 76:00:00 | 6kye-E | 4.6 | 3.4 | 86 | 140 | 6 | MOLECULE: | HEMOGLOBIN | SUBUNIT | ALPHA |
| 77:00:00 | 6kah-C | 4.6 | 3 | 84 | 141 | 7 | MOLECULE: | HEMOGLOBIN | SUBUNIT | ALPHA |
| 78:00:00 | 1ycb-A | 4.6 | 4.1 | 90 | 153 | 13 | MOLECULE: | MYOGLOBIN | | |
| 79:00:00 | 7k4m-A | 4.6 | 3 | 84 | 141 | 7 | MOLECULE: | HEMOGLOBIN | SUBUNIT | ALPHA |
| 80:00:00 | 1gzx-A | 4.6 | 3 | 84 | 141 | 7 | MOLECULE: | HEMOGLOBIN | SUBUNIT | ALPHA |
| 81:00:00 | 7ud8-A | 4.6 | 3 | 84 | 141 | 6 | MOLECULE: | HEMOGLOBIN | SUBUNIT | ALPHA |
| 82:00:00 | 4fwz-A | 4.6 | 4 | 88 | 153 | 11 | MOLECULE: | MYOGLOBIN | | |
| 83:00:00 | 7vw4-A | 4.6 | 4.2 | 88 | 152 | 11 | MOLECULE: | MYOGLOBIN | | |
| 84:00:00 | 6kae-C | 4.6 | 3 | 84 | 141 | 7 | MOLECULE: | HEMOGLOBIN | SUBUNIT | ALPHA |
| 85:00:00 | 1ljw-A | 4.6 | 3.1 | 84 | 141 | 7 | MOLECULE: | HEMOGLOBIN | ALPHA C | HAIN |
| 86:00:00 | 6kaq-A | 4.6 | 3.4 | 85 | 141 | 7 | MOLECULE: | HEMOGLOBIN | SUBUNIT | ALPHA |
| 87:00:00 | 2hhb-C | 4.6 | 3.6 | 86 | 141 | 5 | MOLECULE: | HEMOGLOBIN | (DEOXY) | (ALPHA CHAIN) |
| 88:00:00 | 7uf7-A | 4.6 | 3.5 | 85 | 141 | 5 | MOLECULE: | HEMOGLOBIN | SUBUNIT | ALPHA |
| 89:00:00 | 1bab-C | 4.6 | 3 | 84 | 142 | 7 | MOLECULE: | HEMOGLOBIN | THIONVI | LLE (DEOXY) (ALPHA CHAIN) |
| 90:00:00 | 1yev-A | 4.6 | 3 | 84 | 138 | 7 | MOLECULE: | HEMOGLOBIN | ALPHA C | HAIN |
| 91:00:00 | 2w6v-A | 4.6 | 3 | 85 | 141 | 7 | MOLECULE: | HEMOGLOBIN | SUBUNIT | ALPHA |
| 92:00:00 | 1j41-C | 4.6 | 3.1 | 85 | 141 | 7 | MOLECULE: | HEMOGLOBIN | ALPHA C | HAIN |
| 93:00:00 | 1y0t-A | 4.6 | 3 | 84 | 141 | 7 | MOLECULE: | HEMOGLOBIN | ALPHA C | HAIN |
| 94:00:00 | 1j3z-A | 4.6 | 3 | 84 | 141 | 7 | MOLECULE: | HEMOGLOBIN | ALPHA C | HAIN |
| 95:00:00 | 5ksi-C | 4.6 | 3 | 84 | 141 | 7 | MOLECULE: | HEMOGLOBIN | SUBUNIT | ALPHA |
| 96:00:00 | 1fsx-A | 4.6 | 3.5 | 85 | 141 | 5 | MOLECULE: | HEMOGLOBIN | ALPHA C | HAIN |
| 97:00:00 | 4ni1-A | 4.6 | 3.4 | 86 | 141 | 7 | MOLECULE: | HEMOGLOBIN | SUBUNIT | ALPHA |
| 98:00:00 | 1y4v-A | 4.6 | 3 | 84 | 141 | 7 | MOLECULE: | HEMOGLOBIN | ALPHA C | HAIN |
| 99:00:00 | 3ic2-A | 4.6 | 3 | 84 | 141 | 6 | MOLECULE: | HEMOGLOBIN | SUBUNIT | ALPHA |
| 0:00 | 8fdn-C | 4.6 | 3.4 | 85 | 141 | 7 | MOLECULE: | HEMOGLOBIN | SUBUNIT | ALPHA |
| 1:00 | 4xs0-A | 4.6 | 2.9 | 83 | 141 | 7 | MOLECULE: | HEMOGLOBIN | SUBUNIT | ALPHA |
| 2:00 | 7dy4-C | 4.6 | 3 | 84 | 141 | 7 | MOLECULE: | HEMOGLOBIN | SUBUNIT | ALPHA |
| 3:00 | 1xz5-C | 4.6 | 3 | 84 | 141 | 7 | MOLECULE: | HEMOGLOBIN | ALPHA C | HAIN |
| 4:00 | 1hbb-A | 4.6 | 3.1 | 85 | 141 | 7 | MOLECULE: | HEMOGLOBIN | A (DEOX | Y) (ALPHA CHAIN) |
| 5:00 | 1buw-C | 4.6 | 3.5 | 86 | 141 | 3 | MOLECULE: | PROTEIN (H | EMOGLOBI | N) |
| 6:00 | 1bvd-A | 4.6 | 4.1 | 88 | 153 | 11 | MOLECULE: | APOMYOGLOB | IN | |
| 7:00 | 1y4b-C | 4.6 | 3.5 | 85 | 141 | 5 | MOLECULE: | HEMOGLOBIN | ALPHA C | HAIN |
| 8:00 | 5ksi-A | 4.6 | 3 | 84 | 141 | 7 | MOLECULE: | HEMOGLOBIN | SUBUNIT | ALPHA |

| | | | | | | | | | | |
|---|---|---|---|---|---|---|---|---|---|---|
| 9:00 | 1thb-C | 4.6 | 3 | 84 | 141 | 7 | MOLECULE: | HEMOGLOBIN | A (OXY) | (ALPHA CHAIN) |
| 10:00 | 1y2z-A | 4.6 | 3.6 | 86 | 141 | 5 | MOLECULE: | HEMOGLOBIN | ALPHA C | HAIN |
| 11:00 | 1ygd-C | 4.6 | 3.1 | 85 | 141 | 6 | MOLECULE: | HEMOGLOBIN | ALPHA C | HAIN |
| 12:00 | 1hda-A | 4.6 | 3 | 84 | 141 | 7 | MOLECULE: | HEMOGLOBIN | (DEOXY) | (ALPHA CHAIN) |
| 13:00 | 1ygf-C | 4.6 | 3 | 84 | 141 | 7 | MOLECULE: | HEMOGLOBIN | ALPHA C | HAIN |
| 14:00 | 4mqg-A | 4.6 | 3.4 | 85 | 141 | 7 | MOLECULE: | HEMOGLOBIN | SUBUNIT | ALPHA |
| 15:00 | 7dy3-E | 4.6 | 3.1 | 85 | 141 | 7 | MOLECULE: | HEMOGLOBIN | SUBUNIT | ALPHA |
| 16:00 | 3vre-C | 4.6 | 3.1 | 86 | 141 | 7 | MOLECULE: | HEMOGLOBIN | SUBUNIT | ALPHA |
| 17:00 | 5ufj-A | 4.6 | 3 | 84 | 141 | 7 | MOLECULE: | HEMOGLOBIN | SUBUNIT | ALPHA |
| 18:00 | 1y4f-A | 4.6 | 3 | 84 | 141 | 7 | MOLECULE: | HEMOGLOBIN | ALPHA C | HAIN |
| 19:00 | 1y8w-A | 4.6 | 3 | 84 | 141 | 7 | MOLECULE: | HEMOGLOBIN | ALPHA C | HAIN |
| 20:00 | 1rps-A | 4.6 | 3 | 84 | 141 | 7 | MOLECULE: | HEMOGLOBIN | ALPHA C | HAIN |
| 21:00 | 1yh9-A | 4.6 | 3.6 | 86 | 141 | 5 | MOLECULE: | HEMOGLOBIN | ALPHA C | HAIN |
| 22:00 | 1o1i-A | 4.6 | 3 | 83 | 141 | 7 | MOLECULE: | HEMOGLOBIN | ALPHA C | HAIN |
| 23:00 | 3cy5-A | 4.6 | 3.6 | 86 | 141 | 5 | MOLECULE: | HEMOGLOBIN | SUBUNIT | ALPHA-2 |
| 24:00:00 | 1yca-B | 4.6 | 4.1 | 90 | 153 | 13 | MOLECULE: | MYOGLOBIN | | |
| 25:00:00 | 3bj3-C | 4.6 | 3.3 | 86 | 142 | 3 | MOLECULE: | HEMOGLOBIN | ALPHA | |
| 26:00:00 | 1a00-C | 4.6 | 3 | 85 | 141 | 7 | MOLECULE: | HEMOGLOBIN | (ALPHA | CHAIN) |
| 27:00:00 | 2w72-A | 4.6 | 3 | 84 | 141 | 7 | MOLECULE: | HUMAN HEMO | GLOBIN A | |
| 28:00:00 | 1yca-A | 4.6 | 4.1 | 90 | 153 | 13 | MOLECULE: | MYOGLOBIN | | |
| 29:00:00 | 1a3n-A | 4.6 | 3 | 84 | 141 | 7 | MOLECULE: | HEMOGLOBIN | (ALPHA | CHAIN) |
| 30:00:00 | 8fdl-C | 4.6 | 3 | 85 | 141 | 7 | MOLECULE: | HEMOGLOBIN | SUBUNIT | ALPHA |
| 31:00:00 | 1bbb-C | 4.6 | 3.5 | 85 | 141 | 5 | MOLECULE: | HEMOGLOBIN | A (CARB | ONMONOXY) (ALPHA CHAIN) |
| 32:00:00 | 3nl7-A | 4.6 | 3.4 | 85 | 141 | 7 | MOLECULE: | HEMOGLOBIN | SUBUNIT | ALPHA |
| 33:00:00 | 1uiw-C | 4.6 | 3 | 84 | 141 | 7 | MOLECULE: | HEMOGLOBIN | ALPHA C | HAIN |
| 34:00:00 | 1v4w-A | 4.6 | 3.3 | 85 | 143 | 5 | MOLECULE: | HEMOGLOBIN | ALPHA C | HAIN |
| 35:00:00 | 1nqp-C | 4.6 | 2.9 | 83 | 141 | 7 | MOLECULE: | HEMOGLOBIN | ALPHA C | HAIN |
| 36:00:00 | 3nmm-A | 4.6 | 3 | 84 | 141 | 7 | MOLECULE: | HEMOGLOBIN | SUBUNIT | ALPHA |
| 37:00:00 | 1rps-C | 4.6 | 3 | 84 | 141 | 7 | MOLECULE: | HEMOGLOBIN | ALPHA C | HAIN |
| 38:00:00 | 3qjc-A | 4.6 | 3.1 | 84 | 141 | 7 | MOLECULE: | HEMOGLOBIN | SUBUNIT | ALPHA |
| 39:00:00 | 1hba-A | 4.6 | 3.5 | 85 | 141 | 5 | MOLECULE: | HEMOGLOBIN | ROTHSCH | ILD (DEOXY) (ALPHA CHAIN) |
| 40:00:00 | 2qss-C | 4.6 | 3.4 | 85 | 141 | 7 | MOLECULE: | HEMOGLOBIN | SUBUNIT | ALPHA |
| 41:00:00 | 6lcx-G | 4.6 | 2.9 | 84 | 141 | 7 | MOLECULE: | HEMOGLOBIN | SUBUNIT | ALPHA |
| 42:00:00 | 1y0c-A | 4.6 | 3.1 | 85 | 141 | 7 | MOLECULE: | HEMOGLOBIN | ALPHA C | HAIN |
| 43:00:00 | 1y4f-C | 4.6 | 3 | 84 | 141 | 7 | MOLECULE: | HEMOGLOBIN | ALPHA C | HAIN |
| 44:00:00 | 1b86-A | 4.6 | 3 | 84 | 141 | 7 | MOLECULE: | PROTEIN (H | EMOGLOBI | N ALPHA CHAIN) |
| 45:00:00 | 1rq3-A | 4.6 | 3 | 84 | 141 | 7 | MOLECULE: | HEMOGLOBIN | ALPHA C | HAIN |
| 46:00:00 | 6kae-E | 4.6 | 3 | 84 | 141 | 7 | MOLECULE: | HEMOGLOBIN | SUBUNIT | ALPHA |
| 47:00:00 | 4esa-C | 4.6 | 3.3 | 85 | 142 | 5 | MOLECULE: | HEMOGLOBIN | ALPHA C | HAIN |
| 48:00:00 | 7jjq-C | 4.6 | 3.4 | 86 | 139 | 7 | MOLECULE: | HEMOGLOBIN | SUBUNIT | ALPHA |
| 49:00:00 | 5wog-A | 4.6 | 3.1 | 85 | 137 | 6 | MOLECULE: | HEMOGLOBIN | SUBUNIT | ALPHA |
| 50:00:00 | 2nsr-A | 4.6 | 4.4 | 91 | 152 | 15 | MOLECULE: | MYOGLOBIN | | |
| 51:00:00 | 1yvq-C | 4.6 | 3 | 84 | 141 | 6 | MOLECULE: | HEMOGLOBIN | ALPHA C | HAIN |
| 52:00:00 | 5woh-A | 4.6 | 3.1 | 85 | 137 | 6 | MOLECULE: | HEMOGLOBIN | SUBUNIT | ALPHA |
| 53:00:00 | 7dy3-C | 4.6 | 3 | 84 | 141 | 7 | MOLECULE: | HEMOGLOBIN | SUBUNIT | ALPHA |
| 54:00:00 | 3ubv-G | 4.6 | 3.5 | 83 | 131 | 8 | MOLECULE: | HEMOGLOBIN | #NAME? | AVOPROTEIN |
| 55:00:00 | 1j40-E | 4.6 | 3 | 84 | 141 | 7 | MOLECULE: | HEMOGLOBIN | ALPHA C | HAIN |
| 56:00:00 | 4l7y-C | 4.6 | 3 | 84 | 141 | 7 | MOLECULE: | HEMOGLOBIN | SUBUNIT | ALPHA |
| 57:00:00 | 1y4r-A | 4.6 | 3 | 84 | 141 | 7 | MOLECULE: | HEMOGLOBIN | ALPHA C | HAIN |
| 58:00:00 | 4b3w-A | 4.6 | 4.2 | 89 | 154 | 9 | MOLECULE: | CYTOGLOBIN | | |
| 59:00:00 | 7ddu-A | 4.6 | 4.1 | 88 | 154 | 10 | MOLECULE: | MYOGLOBIN | | |
| 60:00:00 | 6f19-A | 4.6 | 4.1 | 88 | 154 | 11 | MOLECULE: | MYOGLOBIN | | |
| 61:00:00 | 1a3o-A | 4.6 | 3 | 84 | 141 | 7 | MOLECULE: | HEMOGLOBIN | (ALPHA | CHAIN) |
| 62:00:00 | 5b85-A | 4.6 | 4.2 | 88 | 153 | 13 | MOLECULE: | MYOGLOBIN | | |

| | | | | | | | | | | |
|---|---|---|---|---|---|---|---|---|---|---|
| 63:00:00 | 1o1k-A | 4.6 | 3.1 | 84 | 140 | 7 | MOLECULE: | HEMOGLOBIN | ALPHA C | HAIN |
| 64:00:00 | 1mnh-A | 4.6 | 4.1 | 89 | 153 | 17 | MOLECULE: | MYOGLOBIN | | |
| 65:00:00 | 1o1k-C | 4.6 | 3.1 | 84 | 140 | 7 | MOLECULE: | HEMOGLOBIN | ALPHA C | HAIN |
| 66:00:00 | 6ii1-A | 4.6 | 3.1 | 84 | 137 | 7 | MOLECULE: | HEMOGLOBIN | SUBUNIT | ALPHA |
| 67:00:00 | 7uf6-A | 4.6 | 2.9 | 83 | 141 | 7 | MOLECULE: | HEMOGLOBIN | SUBUNIT | ALPHA |
| 68:00:00 | 5jgg-A | 4.6 | 3.5 | 87 | 142 | 1 | MOLECULE: | ALPHA CHAI | N | |
| 69:00:00 | 1urv-B | 4.6 | 4.2 | 90 | 154 | 10 | MOLECULE: | CYTOGLOBIN | | |
| 70:00:00 | 7dy3-G | 4.6 | 2.9 | 83 | 141 | 7 | MOLECULE: | HEMOGLOBIN | SUBUNIT | ALPHA |
| 71:00:00 | 1aby-A | 4.6 | 9.3 | 105 | 283 | 8 | MOLECULE: | HEMOGLOBIN | | |
| 72:00:00 | 6kae-A | 4.6 | 3 | 84 | 141 | 7 | MOLECULE: | HEMOGLOBIN | SUBUNIT | ALPHA |
| 73:00:00 | 4hrt-E | 4.6 | 3.4 | 86 | 149 | 3 | MOLECULE: | GLOBIN-2 A | CHAIN | |
| 74:00:00 | 1myh-A | 4.6 | 4 | 89 | 153 | 15 | MOLECULE: | MYOGLOBIN | | |
| 75:00:00 | 6zmx-C | 4.6 | 3.1 | 86 | 138 | 6 | MOLECULE: | HEMOGLOBIN | SUBUNIT | ALPHA-A |
| 76:00:00 | 5ycj-A | 4.6 | 4.2 | 89 | 154 | 17 | MOLECULE: | ANCESTRAL | MYOGLOBI | N AMBWB' OF BASILOSAURUS RELATI |
| 77:00:00 | 1y46-A | 4.6 | 3 | 84 | 141 | 7 | MOLECULE: | HEMOGLOBIN | ALPHA C | HAIN |
| 78:00:00 | 3ciu-A | 4.6 | 3 | 84 | 140 | 7 | MOLECULE: | HEMOGLOBIN | SUBUNIT | ALPHA |
| 79:00:00 | 3hxn-A | 4.6 | 3.3 | 85 | 141 | 7 | MOLECULE: | HEMOGLOBIN | SUBUNIT | ALPHA |
| 80:00:00 | 1ux9-B | 4.6 | 4.2 | 90 | 154 | 9 | MOLECULE: | CYTOGLOBIN | | |
| 81:00:00 | 1m6c-A | 4.6 | 4.1 | 90 | 153 | 13 | MOLECULE: | PROTEIN (M | YOGLOBIN | ) |
| 82:00:00 | 6iyh-A | 4.6 | 3.4 | 85 | 142 | 2 | MOLECULE: | ALPHA CHAI | N | |
| 83:00:00 | 2bnl-D | 4.6 | 4 | 79 | 129 | 8 | MOLECULE: | MODULATOR | PROTEIN | RSBR |
| 84:00:00 | 1ygf-A | 4.6 | 3 | 84 | 141 | 7 | MOLECULE: | HEMOGLOBIN | ALPHA C | HAIN |
| 85:00:00 | 2bnl-E | 4.6 | 3.6 | 77 | 129 | 8 | MOLECULE: | MODULATOR | PROTEIN | RSBR |
| 86:00:00 | 1j7s-A | 4.6 | 3 | 84 | 141 | 7 | MOLECULE: | HEMOGLOBIN | | |
| 87:00:00 | 6kai-A | 4.6 | 3 | 84 | 141 | 7 | MOLECULE: | HEMOGLOBIN | SUBUNIT | ALPHA |
| 88:00:00 | 2bnl-A | 4.6 | 4 | 79 | 129 | 8 | MOLECULE: | MODULATOR | PROTEIN | RSBR |
| 89:00:00 | 1ut0-B | 4.6 | 4.2 | 90 | 154 | 10 | MOLECULE: | CYTOGLOBIN | | |
| 90:00:00 | 6zmy-C | 4.6 | 3.1 | 85 | 137 | 6 | MOLECULE: | HEMOGLOBIN | SUBUNIT | ALPHA-A |
| 91:00:00 | 6lcw-A | 4.6 | 3 | 84 | 141 | 7 | MOLECULE: | HEMOGLOBIN | SUBUNIT | ALPHA |
| 92:00:00 | 8j4l-A | 4.6 | 4.1 | 88 | 153 | 11 | MOLECULE: | MYOGLOBIN | | |
| 93:00:00 | 7pcq-C | 4.6 | 3.4 | 85 | 141 | 7 | MOLECULE: | HEMOGLOBIN | SUBUNIT | ALPHA |
| 94:00:00 | 1uiw-A | 4.6 | 3 | 84 | 141 | 7 | MOLECULE: | HEMOGLOBIN | ALPHA C | HAIN |
| 95:00:00 | 1yma-A | 4.6 | 4.3 | 90 | 153 | 14 | MOLECULE: | MYOGLOBIN | | |
| 96:00:00 | 6lcx-A | 4.6 | 3 | 84 | 141 | 7 | MOLECULE: | HEMOGLOBIN | SUBUNIT | ALPHA |
| 97:00:00 | 1fdh-A | 4.6 | 3 | 84 | 141 | 7 | MOLECULE: | HEMOGLOBIN | F (DEOX | Y) (ALPHA CHAIN) |
| 98:00:00 | 1hbg-A | 4.5 | 3.5 | 87 | 147 | 8 | MOLECULE: | HEMOGLOBIN | (CARBON | MONOXY) |
| 99:00:00 | 4hrr-A | 4.5 | 3.3 | 87 | 149 | 3 | MOLECULE: | GLOBIN-2 A | CHAIN | |
| 0:00 | 1yhu-U | 4.5 | 3.3 | 84 | 145 | 12 | MOLECULE: | HEMOGLOBIN | A1 CHAI | N |
| 1:00 | 3bj2-C | 4.5 | 3.9 | 88 | 142 | 2 | MOLECULE: | HEMOGLOBIN | ALPHA | |
| 2:00 | 8u5a-B | 4.5 | 3.7 | 84 | 149 | 7 | MOLECULE: | DESIGNED M | YOGLOBIN | |
| 3:00 | 8u5a-A | 4.5 | 3.6 | 85 | 149 | 8 | MOLECULE: | DESIGNED M | YOGLOBIN | |
| 4:00 | 1hco-A | 4.5 | 3.7 | 88 | 141 | 3 | MOLECULE: | HEMOGLOBIN | (CARBON | MONOXY) (ALPHA CHAIN) |
| 5:00 | 1ymb-A | 4.5 | 4.1 | 90 | 153 | 13 | MOLECULE: | METMYOGLOB | IN | |
| 6:00 | 1sdl-A | 4.5 | 3.4 | 86 | 141 | 7 | MOLECULE: | HEMOGLOBIN | A | |
| 7:00 | 3eu1-C | 4.5 | 3.5 | 87 | 141 | 6 | MOLECULE: | HEMOGLOBIN | SUBUNIT | ALPHA-1/2 |
| 8:00 | 1g0a-A | 4.5 | 3.5 | 85 | 141 | 5 | MOLECULE: | HEMOGLOBIN | ALPHA C | HAIN |
| 9:00 | 5hy8-C | 4.5 | 3.5 | 85 | 141 | 5 | MOLECULE: | HEMOGLOBIN | SUBUNIT | ALPHA |
| 10:00 | 6hk2-A | 4.5 | 3.4 | 85 | 141 | 7 | MOLECULE: | HEMOGLOBIN | SUBUNIT | ALPHA |
| 11:00 | 1g09-C | 4.5 | 3.6 | 85 | 141 | 5 | MOLECULE: | HEMOGLOBIN | ALPHA C | HAIN |
| 12:00 | 1aj9-A | 4.5 | 3.4 | 86 | 141 | 7 | MOLECULE: | HEMOGLOBIN | (ALPHA | CHAIN) |
| 13:00 | 1y7g-A | 4.5 | 3.1 | 85 | 141 | 7 | MOLECULE: | HEMOGLOBIN | ALPHA C | HAIN |
| 14:00 | 6kah-E | 4.5 | 3.1 | 85 | 141 | 7 | MOLECULE: | HEMOGLOBIN | SUBUNIT | ALPHA |
| 15:00 | 7jy1-C | 4.5 | 2.9 | 83 | 141 | 7 | MOLECULE: | HEMOGLOBIN | SUBUNIT | ALPHA |
| 16:00 | 8egi-C | 4.5 | 3.5 | 86 | 141 | 3 | MOLECULE: | HEMOGLOBIN | SUBUNIT | ALPHA |

| | | | | | | | | | |
|---|---|---|---|---|---|---|---|---|---|
| 17:00 | 1yzi-A | 4.5 | 3.4 | 86 | 141 | 6 | MOLECULE: | HEMOGLOBIN | ALPHA C | HAIN |
| 18:00 | 1fhj-C | 4.5 | 3.6 | 86 | 141 | 5 | MOLECULE: | HEMOGLOBIN | (ALPHA | CHAIN) |
| 19:00 | 1xq5-C | 4.5 | 3.4 | 86 | 142 | 6 | MOLECULE: | HEMOGLOBIN | ALPHA-1 | CHAIN |
| 20:00 | 5e29-C | 4.5 | 3.1 | 85 | 141 | 7 | MOLECULE: | HEMOGLOBIN | SUBUNIT | ALPHA |
| 21:00 | 5e83-A | 4.5 | 2.9 | 83 | 141 | 7 | MOLECULE: | HEMOGLOBIN | SUBUNIT | ALPHA |
| 22:00 | 7jy0-A | 4.5 | 3 | 84 | 141 | 7 | MOLECULE: | HEMOGLOBIN | SUBUNIT | ALPHA |
| 23:00 | 6r2o-C | 4.5 | 2.9 | 83 | 141 | 7 | MOLECULE: | HEMOGLOBIN | SUBUNIT | ALPHA |
| 24:00:00 | 1a9w-A | 4.5 | 3.5 | 85 | 141 | 5 | MOLECULE: | HEMOGLOBIN | (ALPHA | CHAIN) |
| 25:00:00 | 4f4o-G | 4.5 | 3.1 | 84 | 141 | 7 | MOLECULE: | HEMOGLOBIN | SUBUNIT | ALPHA |
| 26:00:00 | 1yff-C | 4.5 | 3.5 | 85 | 141 | 5 | MOLECULE: | HEMOGLOBIN | ALPHA C | HAIN |
| 27:00:00 | 1yff-A | 4.5 | 3.1 | 84 | 141 | 7 | MOLECULE: | HEMOGLOBIN | ALPHA C | HAIN |
| 28:00:00 | 3gqg-C | 4.5 | 3.3 | 85 | 142 | 6 | MOLECULE: | HEMOGLOBIN | SUBUNIT | ALPHA |
| 29:00:00 | 1dke-A | 4.5 | 3.1 | 85 | 141 | 7 | MOLECULE: | HEMOGLOBIN | : ALPHA | CHAIN |
| 30:00:00 | 7jxz-C | 4.5 | 3.5 | 85 | 141 | 5 | MOLECULE: | HEMOGLOBIN | SUBUNIT | ALPHA |
| 31:00:00 | 3gdj-A | 4.5 | 3.2 | 86 | 141 | 7 | MOLECULE: | HEMOGLOBIN | SUBUNIT | ALPHA |
| 32:00:00 | 6kye-G | 4.5 | 3 | 84 | 140 | 7 | MOLECULE: | HEMOGLOBIN | SUBUNIT | ALPHA |
| 33:00:00 | 1jy7-A | 4.5 | 2.9 | 83 | 141 | 7 | MOLECULE: | HEMOGLOBIN | ALPHA C | HAIN |
| 34:00:00 | 3gys-G | 4.5 | 3.1 | 85 | 138 | 8 | MOLECULE: | HEMOGLOBIN | SUBUNIT | ALPHA |
| 35:00:00 | 7cue-A | 4.5 | 3.4 | 85 | 141 | 7 | MOLECULE: | HEMOGLOBIN | SUBUNIT | ALPHA |
| 36:00:00 | 1ird-A | 4.5 | 3.4 | 85 | 141 | 7 | MOLECULE: | HEMOGLOBIN | ALPHA C | HAIN |
| 37:00:00 | 4fc3-A | 4.5 | 2.9 | 83 | 140 | 7 | MOLECULE: | HEMOGLOBIN | SUBUNIT | ALPHA |
| 38:00:00 | 1yff-E | 4.5 | 3.5 | 85 | 141 | 5 | MOLECULE: | HEMOGLOBIN | ALPHA C | HAIN |
| 39:00:00 | 2dn3-A | 4.5 | 3.4 | 85 | 141 | 7 | MOLECULE: | HEMOGLOBIN | ALPHA S | UBUNIT |
| 40:00:00 | 1jy7-C | 4.5 | 3.5 | 85 | 141 | 5 | MOLECULE: | HEMOGLOBIN | ALPHA C | HAIN |
| 41:00:00 | 4wjg-Z | 4.5 | 3.6 | 85 | 141 | 5 | MOLECULE: | HEMOGLOBIN | SUBUNIT | ALPHA |
| 42:00:00 | 3pia-A | 4.5 | 3.5 | 85 | 141 | 5 | MOLECULE: | HEMOGLOBIN | SUBUNIT | ALPHA |
| 43:00:00 | 1hho-A | 4.5 | 3.1 | 84 | 141 | 7 | MOLECULE: | HEMOGLOBIN | A (OXY) | (ALPHA CHAIN) |
| 44:00:00 | 6hit-C | 4.5 | 3.1 | 85 | 142 | 5 | MOLECULE: | HEMOGLOBIN | ALPHA 2 | CHAIN |
| 45:00:00 | 6hit-G | 4.5 | 3 | 84 | 142 | 6 | MOLECULE: | HEMOGLOBIN | ALPHA 2 | CHAIN |
| 46:00:00 | 4mqi-A | 4.5 | 3.1 | 85 | 140 | 7 | MOLECULE: | HEMOGLOBIN | SUBUNIT | ALPHA |
| 47:00:00 | 1jy7-P | 4.5 | 3.5 | 85 | 141 | 5 | MOLECULE: | HEMOGLOBIN | ALPHA C | HAIN |
| 48:00:00 | 6xd9-A | 4.5 | 3.5 | 85 | 141 | 5 | MOLECULE: | HEMOGLOBIN | SUBUNIT | ALPHA |
| 49:00:00 | 1bij-C | 4.5 | 2.9 | 83 | 141 | 7 | MOLECULE: | HEMOGLOBIN | A | |
| 50:00:00 | 3s65-C | 4.5 | 3.5 | 86 | 141 | 3 | MOLECULE: | HEMOGLOBIN | SUBUNIT | ALPHA |
| 51:00:00 | 3ic0-A | 4.5 | 3 | 84 | 141 | 6 | MOLECULE: | HEMOGLOBIN | SUBUNIT | ALPHA |
| 52:00:00 | 5hlu-A | 4.5 | 4.2 | 89 | 152 | 11 | MOLECULE: | MYOGLOBIN | | |
| 53:00:00 | 1j40-C | 4.5 | 3.1 | 85 | 141 | 7 | MOLECULE: | HEMOGLOBIN | ALPHA C | HAIN |
| 54:00:00 | 1jy7-W | 4.5 | 2.9 | 83 | 141 | 7 | MOLECULE: | HEMOGLOBIN | ALPHA C | HAIN |
| 55:00:00 | 5ksj-C | 4.5 | 3.5 | 85 | 141 | 5 | MOLECULE: | HEMOGLOBIN | SUBUNIT | ALPHA |
| 56:00:00 | 6bnr-C | 4.5 | 3 | 84 | 141 | 6 | MOLECULE: | HEMOGLOBIN | SUBUNIT | ALPHA |
| 57:00:00 | 4mqk-A | 4.5 | 3.5 | 85 | 141 | 7 | MOLECULE: | HEMOGLOBIN | SUBUNIT | ALPHA |
| 58:00:00 | 1j3y-A | 4.5 | 3.1 | 85 | 141 | 7 | MOLECULE: | HEMOGLOBIN | ALPHA C | HAIN |
| 59:00:00 | 3gqg-A | 4.5 | 3.3 | 85 | 142 | 6 | MOLECULE: | HEMOGLOBIN | SUBUNIT | ALPHA |
| 60:00:00 | 7pch-C | 4.5 | 3.9 | 87 | 141 | 5 | MOLECULE: | HEMOGLOBIN | SUBUNIT | ALPHA |
| 61:00:00 | 3pi8-C | 4.5 | 2.9 | 83 | 141 | 7 | MOLECULE: | HEMOGLOBIN | SUBUNIT | ALPHA |
| 62:00:00 | 6xe7-A | 4.5 | 3.5 | 85 | 141 | 5 | MOLECULE: | HEMOGLOBIN | SUBUNIT | ALPHA |
| 63:00:00 | 1bij-A | 4.5 | 3.5 | 85 | 141 | 5 | MOLECULE: | HEMOGLOBIN | A | |
| 64:00:00 | 7uf7-C | 4.5 | 3.5 | 86 | 141 | 3 | MOLECULE: | HEMOGLOBIN | SUBUNIT | ALPHA |
| 65:00:00 | 3whm-A | 4.5 | 2.9 | 83 | 140 | 7 | MOLECULE: | HEMOGLOBIN | SUBUNIT | ALPHA |
| 66:00:00 | 6hk2-C | 4.5 | 3.5 | 86 | 141 | 7 | MOLECULE: | HEMOGLOBIN | SUBUNIT | ALPHA |
| 67:00:00 | 3ng6-A | 4.5 | 3.3 | 85 | 142 | 6 | MOLECULE: | HEMOGLOBIN | SUBUNIT | ALPHA-1 |
| 68:00:00 | 2qsp-A | 4.5 | 3.5 | 85 | 141 | 5 | MOLECULE: | HEMOGLOBIN | SUBUNIT | ALPHA |
| 69:00:00 | 1hbh-A | 4.5 | 3.3 | 85 | 142 | 6 | MOLECULE: | HEMOGLOBIN | (DEOXY) | (ALPHA CHAIN) |
| 70:00:00 | 6kai-E | 4.5 | 3.1 | 85 | 141 | 7 | MOLECULE: | HEMOGLOBIN | SUBUNIT | ALPHA |

| | | | | | | | | | | |
|---|---|---|---|---|---|---|---|---|---|---|
| 71:00:00 | 2h8d-A | 4.5 | 3.3 | 85 | 142 | 6 | MOLECULE: | HEMOGLOBIN | ALPHA S | UBUNIT |
| 72:00:00 | 6nbc-C | 4.5 | 3 | 84 | 140 | 7 | MOLECULE: | HEMOGLOBIN | SUBUNIT | ALPHA |
| 73:00:00 | 1cmy-A | 4.5 | 3 | 84 | 141 | 7 | MOLECULE: | HEMOGLOBIN | YPSILAN | TI (CARBONMONOXY) (ALPHA |
| 74:00:00 | 2aa1-C | 4.5 | 3 | 84 | 142 | 6 | MOLECULE: | HEMOGLOBIN | ALPHA-1 | CHAIN |
| 75:00:00 | 1mko-A | 4.5 | 3.1 | 84 | 141 | 7 | MOLECULE: | HEMOGLOBIN | ALPHA C | HAIN |
| 76:00:00 | 1faw-C | 4.5 | 3.1 | 85 | 141 | 8 | MOLECULE: | HEMOGLOBIN | (ALPHA | SUBUNIT) |
| 77:00:00 | 3cy5-C | 4.5 | 3.4 | 86 | 141 | 7 | MOLECULE: | HEMOGLOBIN | SUBUNIT | ALPHA-2 |
| 78:00:00 | 2yrs-A | 4.5 | 3 | 85 | 141 | 7 | MOLECULE: | HEMOGLOBIN | SUBUNIT | ALPHA |
| 79:00:00 | 7qu4-A | 4.5 | 3.5 | 86 | 142 | 3 | MOLECULE: | HEMOGLOBIN | SUBUNIT | GAMMA-2 |
| 80:00:00 | 4n7n-E | 4.5 | 3 | 84 | 141 | 7 | MOLECULE: | HEMOGLOBIN | SUBUNIT | ALPHA |
| 81:00:00 | 3s48-D | 4.5 | 2.9 | 83 | 140 | 7 | MOLECULE: | IRON-REGUL | ATED SUR | FACE DETERMINANT PROTEIN H |
| 82:00:00 | 5ksj-A | 4.5 | 3 | 84 | 141 | 7 | MOLECULE: | HEMOGLOBIN | SUBUNIT | ALPHA |
| 83:00:00 | 3pia-C | 4.5 | 3.5 | 85 | 141 | 5 | MOLECULE: | HEMOGLOBIN | SUBUNIT | ALPHA |
| 84:00:00 | 6l5x-C | 4.5 | 3.1 | 85 | 141 | 6 | MOLECULE: | HEMOGLOBIN | SUBUNIT | ALPHA |
| 85:00:00 | 7qu4-B | 4.5 | 3.1 | 84 | 141 | 7 | MOLECULE: | HEMOGLOBIN | SUBUNIT | GAMMA-2 |
| 86:00:00 | 3vre-A | 4.5 | 3.1 | 85 | 141 | 7 | MOLECULE: | HEMOGLOBIN | SUBUNIT | ALPHA |
| 87:00:00 | 1jy7-U | 4.5 | 2.9 | 83 | 141 | 7 | MOLECULE: | HEMOGLOBIN | ALPHA C | HAIN |
| 88:00:00 | 4hrr-E | 4.5 | 3.3 | 87 | 149 | 3 | MOLECULE: | GLOBIN-2 A | CHAIN | |
| 89:00:00 | 1v4u-C | 4.5 | 3 | 84 | 143 | 5 | MOLECULE: | HEMOGLOBIN | ALPHA C | HAIN |
| 90:00:00 | 2qss-A | 4.5 | 3.1 | 84 | 141 | 7 | MOLECULE: | HEMOGLOBIN | SUBUNIT | ALPHA |
| 91:00:00 | 4mqj-C | 4.5 | 3.5 | 85 | 141 | 7 | MOLECULE: | HEMOGLOBIN | SUBUNIT | ALPHA |
| 92:00:00 | 1lfq-A | 4.5 | 3.5 | 85 | 141 | 5 | MOLECULE: | HEMOGLOBIN | ALPHA C | HAIN |
| 93:00:00 | 1cmy-C | 4.5 | 3 | 84 | 141 | 7 | MOLECULE: | HEMOGLOBIN | YPSILAN | TI (CARBONMONOXY) (ALPHA |
| 94:00:00 | 1nej-C | 4.5 | 2.9 | 83 | 141 | 7 | MOLECULE: | HEMOGLOBIN | ALPHA C | HAIN |
| 95:00:00 | 3bcq-C | 4.5 | 3.3 | 85 | 142 | 8 | MOLECULE: | ALPHA-CHAI | N HEMOGL | OBIN |
| 96:00:00 | 1j40-A | 4.5 | 3.1 | 85 | 141 | 7 | MOLECULE: | HEMOGLOBIN | ALPHA C | HAIN |
| 97:00:00 | 7ud7-C | 4.5 | 2.9 | 84 | 141 | 7 | MOLECULE: | HEMOGLOBIN | SUBUNIT | ALPHA |
| 98:00:00 | 3s66-A | 4.5 | 3.1 | 85 | 139 | 7 | MOLECULE: | HEMOGLOBIN | SUBUNIT | ALPHA |
| 99:00:00 | 6kav-A | 4.5 | 3 | 84 | 141 | 6 | MOLECULE: | HEMOGLOBIN | SUBUNIT | ALPHA |
| 0:00 | 1uiw-G | 4.5 | 3 | 84 | 141 | 7 | MOLECULE: | HEMOGLOBIN | ALPHA C | HAIN |
| 1:00 | 4n7o-G | 4.5 | 3.4 | 86 | 141 | 7 | MOLECULE: | HEMOGLOBIN | SUBUNIT | ALPHA |
| 2:00 | 1ouu-A | 4.5 | 3.3 | 85 | 142 | 6 | MOLECULE: | HEMOGLOBIN | I | |
| 3:00 | 1a9w-C | 4.5 | 2.9 | 83 | 141 | 7 | MOLECULE: | HEMOGLOBIN | (ALPHA | CHAIN) |
| 4:00 | 6zmy-A | 4.5 | 3.1 | 85 | 139 | 6 | MOLECULE: | HEMOGLOBIN | SUBUNIT | ALPHA-A |
| 5:00 | 6xe7-C | 4.5 | 3 | 84 | 141 | 6 | MOLECULE: | HEMOGLOBIN | SUBUNIT | ALPHA |
| 6:00 | 3wcp-A | 4.5 | 3.1 | 85 | 141 | 7 | MOLECULE: | HEMOGLOBIN | SUBUNIT | ALPHA |
| 7:00 | 7vde-A | 4.5 | 3 | 84 | 141 | 7 | MOLECULE: | HEMOGLOBIN | SUBUNIT | ALPHA |
| 8:00 | 4mqj-A | 4.5 | 3.5 | 86 | 141 | 6 | MOLECULE: | HEMOGLOBIN | SUBUNIT | ALPHA |
| 9:00 | 7vde-C | 4.5 | 3 | 84 | 141 | 7 | MOLECULE: | HEMOGLOBIN | SUBUNIT | ALPHA |
| 10:00 | 1m9p-C | 4.5 | 3.5 | 85 | 141 | 5 | MOLECULE: | HEMOGLOBIN | ALPHA C | HAIN |
| 11:00 | 4f4o-J | 4.5 | 3.1 | 85 | 141 | 6 | MOLECULE: | HEMOGLOBIN | SUBUNIT | ALPHA |
| 12:00 | 2qmb-A | 4.5 | 3.6 | 87 | 141 | 3 | MOLECULE: | HEMOGLOBIN | SUBUNIT | ALPHA-A |
| 13:00 | 2dxm-C | 4.5 | 3.7 | 88 | 141 | 3 | MOLECULE: | HEMOGLOBIN | SUBUNIT | ALPHA |
| 14:00 | 2aa1-A | 4.5 | 3.3 | 85 | 142 | 6 | MOLECULE: | HEMOGLOBIN | ALPHA-1 | CHAIN |
| 15:00 | 3r5i-A | 4.5 | 3 | 84 | 141 | 7 | MOLECULE: | HEMOGLOBIN | SUBUNIT | ALPHA |
| 16:00 | 3bj2-A | 4.5 | 3.4 | 86 | 142 | 3 | MOLECULE: | HEMOGLOBIN | ALPHA | |
| 17:00 | 1pbx-A | 4.5 | 3.3 | 85 | 142 | 6 | MOLECULE: | HEMOGLOBIN | (CARBON | MONOXY) (ALPHA CHAIN) |
| 18:00 | 1lfv-A | 4.5 | 3.1 | 84 | 141 | 7 | MOLECULE: | HEMOGLOBIN | ALPHA C | HAIN |
| 19:00 | 1m9p-A | 4.5 | 3.5 | 85 | 141 | 5 | MOLECULE: | HEMOGLOBIN | ALPHA C | HAIN |
| 20:00 | 4mqk-E | 4.5 | 3.3 | 86 | 138 | 6 | MOLECULE: | HEMOGLOBIN | SUBUNIT | ALPHA |
| 21:00 | 1g0a-C | 4.5 | 3.5 | 85 | 141 | 5 | MOLECULE: | HEMOGLOBIN | ALPHA C | HAIN |
| 22:00 | 2qls-C | 4.5 | 3.1 | 85 | 141 | 7 | MOLECULE: | HEMOGLOBIN | SUBUNIT | ALPHA |
| 23:00 | 8egi-A | 4.5 | 3.5 | 85 | 141 | 5 | MOLECULE: | HEMOGLOBIN | SUBUNIT | ALPHA |
| 24:00:00 | 1j41-A | 4.5 | 3.1 | 85 | 141 | 7 | MOLECULE: | HEMOGLOBIN | ALPHA C | HAIN |

| | | | | | | | | | | | |
|---|---|---|---|---|---|---|---|---|---|---|---|
| 25:00:00 | 1nih-C | 4.5 | 3 | 84 | 141 | 7 | MOLECULE: | HEMOGLOBIN | (NICKEL | OUS DEOXY) (ALPHA CHAIN) | |
| 26:00:00 | 2dxm-A | 4.5 | 3 | 84 | 141 | 7 | MOLECULE: | HEMOGLOBIN | SUBUNIT | ALPHA | |
| 27:00:00 | 6di4-C | 4.5 | 2.9 | 83 | 141 | 7 | MOLECULE: | HEMOGLOBIN | SUBUNIT | ALPHA | |
| 28:00:00 | 1v4x-C | 4.5 | 3.3 | 85 | 143 | 5 | MOLECULE: | HEMOGLOBIN | ALPHA C | HAIN | |
| 29:00:00 | 1si4-A | 4.5 | 3 | 84 | 141 | 6 | MOLECULE: | HEMOGLOBIN | ALPHA C | HAIN | |
| 30:00:00 | 5ker-E | 4.5 | 3.2 | 86 | 140 | 7 | MOLECULE: | ALPHA-GLOB | IN | | |
| 31:00:00 | 5urc-C | 4.5 | 3.5 | 86 | 141 | 3 | MOLECULE: | HEMOGLOBIN | SUBUNIT | ALPHA | |
| 32:00:00 | 1hac-A | 4.5 | 3.1 | 86 | 141 | 6 | MOLECULE: | HEMOGLOBIN | A | | |
| 33:00:00 | 4n7o-K | 4.5 | 3.1 | 86 | 141 | 6 | MOLECULE: | HEMOGLOBIN | SUBUNIT | ALPHA | |
| 34:00:00 | 3pi8-A | 4.5 | 3.5 | 85 | 141 | 5 | MOLECULE: | HEMOGLOBIN | SUBUNIT | ALPHA | |
| 35:00:00 | 7jy3-C | 4.5 | 3 | 84 | 141 | 7 | MOLECULE: | HEMOGLOBIN | SUBUNIT | ALPHA | |
| 36:00:00 | 6nbd-C | 4.5 | 3.4 | 86 | 140 | 7 | MOLECULE: | HEMOGLOBIN | SUBUNIT | ALPHA | |
| 37:00:00 | 8pur-A | 4.5 | 3.4 | 85 | 140 | 8 | MOLECULE: | HEMOGLOBIN | , ALPHA | | 1 |
| 38:00:00 | 1yg5-A | 4.5 | 3 | 84 | 141 | 7 | MOLECULE: | HEMOGLOBIN | ALPHA C | HAIN | |
| 39:00:00 | 1v4u-A | 4.5 | 3.3 | 85 | 143 | 5 | MOLECULE: | HEMOGLOBIN | ALPHA C | HAIN | |
| 40:00:00 | 2d60-A | 4.5 | 3.1 | 85 | 141 | 7 | MOLECULE: | HEMOGLOBIN | ALPHA S | UBUNIT | |
| 41:00:00 | 3pi9-C | 4.5 | 3.5 | 85 | 141 | 5 | MOLECULE: | HEMOGLOBIN | SUBUNIT | ALPHA | |
| 42:00:00 | 5ufj-C | 4.5 | 2.9 | 83 | 141 | 7 | MOLECULE: | HEMOGLOBIN | SUBUNIT | ALPHA | |
| 43:00:00 | 6kah-A | 4.5 | 3.1 | 85 | 141 | 7 | MOLECULE: | HEMOGLOBIN | SUBUNIT | ALPHA | |
| 44:00:00 | 1rqa-A | 4.5 | 3.5 | 85 | 141 | 5 | MOLECULE: | HEMOGLOBIN | ALPHA C | HAIN | |
| 45:00:00 | 6nbc-A | 4.5 | 3 | 84 | 140 | 7 | MOLECULE: | HEMOGLOBIN | SUBUNIT | ALPHA | |
| 46:00:00 | 5hy8-S | 4.5 | 2.9 | 83 | 141 | 7 | MOLECULE: | HEMOGLOBIN | SUBUNIT | ALPHA | |
| 47:00:00 | 7pcq-A | 4.5 | 3.6 | 86 | 141 | 5 | MOLECULE: | HEMOGLOBIN | SUBUNIT | ALPHA | |
| 48:00:00 | 1j41-E | 4.5 | 3.1 | 85 | 141 | 7 | MOLECULE: | HEMOGLOBIN | ALPHA C | HAIN | |
| 49:00:00 | 1jy7-R | 4.5 | 2.9 | 83 | 141 | 7 | MOLECULE: | HEMOGLOBIN | ALPHA C | HAIN | |
| 50:00:00 | 7k4m-C | 4.5 | 3 | 84 | 140 | 7 | MOLECULE: | HEMOGLOBIN | SUBUNIT | ALPHA | |
| 51:00:00 | 5e83-C | 4.5 | 3 | 84 | 141 | 6 | MOLECULE: | HEMOGLOBIN | SUBUNIT | ALPHA | |
| 52:00:00 | 4n7o-E | 4.5 | 3.1 | 84 | 141 | 7 | MOLECULE: | HEMOGLOBIN | SUBUNIT | ALPHA | |
| 53:00:00 | 1ydz-A | 4.5 | 3.3 | 85 | 141 | 7 | MOLECULE: | HEMOGLOBIN | ALPHA C | HAIN | |
| 54:00:00 | 1qxe-A | 4.5 | 3 | 84 | 141 | 6 | MOLECULE: | HEMOGLOBIN | ALPHA C | HAIN | |
| 55:00:00 | 1uiw-E | 4.5 | 2.9 | 83 | 141 | 7 | MOLECULE: | HEMOGLOBIN | ALPHA C | HAIN | |
| 56:00:00 | 1lfy-A | 4.5 | 3.1 | 84 | 141 | 7 | MOLECULE: | HEMOGLOBIN | ALPHA C | HAIN | |
| 57:00:00 | 1hga-C | 4.5 | 3 | 85 | 141 | 6 | MOLECULE: | HEMOGLOBIN | (DEOXY) | (ALPHA CHAIN) | |
| 58:00:00 | 1yff-G | 4.5 | 3.5 | 85 | 141 | 5 | MOLECULE: | HEMOGLOBIN | ALPHA C | HAIN | |
| 59:00:00 | 4mqk-C | 4.5 | 3.5 | 86 | 141 | 6 | MOLECULE: | HEMOGLOBIN | SUBUNIT | ALPHA | |
| 60:00:00 | 3wr1-A | 4.5 | 3.6 | 87 | 141 | 6 | MOLECULE: | HEMOGLOBIN | SUBUNIT | ALPHA-A | |
| 61:00:00 | 5hy8-A | 4.5 | 3.5 | 85 | 141 | 5 | MOLECULE: | HEMOGLOBIN | SUBUNIT | ALPHA | |
| 62:00:00 | 1mnj-A | 4.5 | 4.3 | 90 | 153 | 13 | MOLECULE: | MYOGLOBIN | | | |
| 63:00:00 | 3gqr-G | 4.5 | 3.1 | 85 | 141 | 8 | MOLECULE: | HEMOGLOBIN | SUBUNIT | ALPHA | |
| 64:00:00 | 8f9h-A | 4.5 | 4 | 88 | 151 | 11 | MOLECULE: | MYOGLOBIN | | | |
| 65:00:00 | 5vzn-A | 4.5 | 4 | 88 | 151 | 11 | MOLECULE: | MYOGLOBIN | | | |
| 66:00:00 | 1a4f-A | 4.5 | 2.9 | 84 | 141 | 8 | MOLECULE: | HEMOGLOBIN | (ALPHA | CHAIN) | |
| 67:00:00 | 5x2r-G | 4.5 | 3.4 | 85 | 140 | 7 | MOLECULE: | HEMOGLOBIN | SUBUNIT | ALPHA | |
| 68:00:00 | 1ux9-A | 4.5 | 4.1 | 90 | 154 | 9 | MOLECULE: | CYTOGLOBIN | | | |
| 69:00:00 | 4mqk-H | 4.5 | 3.3 | 85 | 145 | 6 | MOLECULE: | HEMOGLOBIN | SUBUNIT | ALPHA | |
| 70:00:00 | 5x2s-G | 4.5 | 3.5 | 86 | 140 | 6 | MOLECULE: | HEMOGLOBIN | SUBUNIT | ALPHA | |
| 71:00:00 | 1s5y-C | 4.5 | 3.1 | 84 | 142 | 6 | MOLECULE: | HEMOGLOBIN | ALPHA C | HAIN | |
| 72:00:00 | 8dov-E | 4.5 | 3.4 | 85 | 138 | 7 | MOLECULE: | HEMOGLOBIN | SUBUNIT | ALPHA | |
| 73:00:00 | 105m-A | 4.5 | 4.2 | 90 | 153 | 9 | MOLECULE: | MYOGLOBIN | | | |
| 74:00:00 | 4mqk-D | 4.5 | 3.4 | 85 | 145 | 6 | MOLECULE: | HEMOGLOBIN | SUBUNIT | ALPHA | |
| 75:00:00 | 5x2u-G | 4.5 | 3.5 | 86 | 140 | 6 | MOLECULE: | HEMOGLOBIN | SUBUNIT | ALPHA | |
| 76:00:00 | 2ri4-I | 4.5 | 3 | 84 | 137 | 6 | MOLECULE: | HEMOGLOBIN | SUBUNIT | ALPHA-1/2 | |
| 77:00:00 | 2dn1-A | 4.5 | 3.4 | 85 | 140 | 7 | MOLECULE: | HEMOGLOBIN | ALPHA S | UBUNIT | |
| 78:00:00 | 3d1k-A | 4.5 | 3.4 | 85 | 142 | 6 | MOLECULE: | HEMOGLOBIN | SUBUNIT | ALPHA-1 | |

| | | | | | | | | | | |
|---|---|---|---|---|---|---|---|---|---|---|
| 79:00:00 | 6hal-A | 4.5 | 3.1 | 84 | 139 | 7 | MOLECULE: | HEMOGLOBIN | SUBUNIT | ALPHA |
| 80:00:00 | 8vyl-C | 4.5 | 3 | 84 | 140 | 7 | MOLECULE: | HEMOGLOBIN | SUBUNIT | ALPHA |
| 81:00:00 | 6bb5-A | 4.5 | 3.4 | 85 | 139 | 7 | MOLECULE: | HEMOGLOBIN | SUBUNIT | ALPHA |
| 82:00:00 | 5ycg-A | 4.5 | 4.1 | 88 | 154 | 14 | MOLECULE: | ANCESTRAL | MYOGLOBI | N AMBWP OF PAKICETUS RELATIVE |
| 83:00:00 | 5wog-B | 4.5 | 3.4 | 85 | 137 | 7 | MOLECULE: | HEMOGLOBIN | SUBUNIT | ALPHA |
| 84:00:00 | 5x2s-I | 4.5 | 3.1 | 85 | 140 | 6 | MOLECULE: | HEMOGLOBIN | SUBUNIT | ALPHA |
| 85:00:00 | 1la6-A | 4.5 | 3.4 | 85 | 142 | 6 | MOLECULE: | HEMOGLOBIN | ALPHA-1 | CHAIN |
| 86:00:00 | 1fhj-A | 4.5 | 3.1 | 85 | 141 | 7 | MOLECULE: | HEMOGLOBIN | (ALPHA | CHAIN) |
| 87:00:00 | 1i3d-A | 4.5 | 3.4 | 86 | 146 | 6 | MOLECULE: | HEMOGLOBIN | GAMMA C | HAINS |
| 88:00:00 | 4hrt-G | 4.5 | 3.4 | 85 | 149 | 4 | MOLECULE: | GLOBIN-2 A | CHAIN | |
| 89:00:00 | 1i3d-B | 4.5 | 3.4 | 86 | 146 | 6 | MOLECULE: | HEMOGLOBIN | GAMMA C | HAINS |
| 90:00:00 | 6rp5-A | 4.5 | 3.5 | 85 | 142 | 5 | MOLECULE: | HEMOGLOBIN | SUBUNIT | ALPHA 1 |
| 91:00:00 | 5x2t-I | 4.5 | 3.1 | 84 | 140 | 7 | MOLECULE: | HEMOGLOBIN | SUBUNIT | ALPHA |
| 92:00:00 | 3gqp-C | 4.5 | 3.2 | 85 | 141 | 8 | MOLECULE: | HEMOGLOBIN | SUBUNIT | ALPHA |
| 93:00:00 | 2bnl-B | 4.5 | 3.6 | 76 | 128 | 8 | MOLECULE: | MODULATOR | PROTEIN | RSBR |
| 94:00:00 | 4mqj-F | 4.5 | 3.4 | 86 | 146 | 6 | MOLECULE: | HEMOGLOBIN | SUBUNIT | ALPHA |
| 95:00:00 | 3a59-C | 4.5 | 3.4 | 86 | 141 | 7 | MOLECULE: | HEMOGLOBIN | SUBUNIT | ALPHA-A |
| 96:00:00 | 5x2u-C | 4.5 | 3.5 | 86 | 140 | 6 | MOLECULE: | HEMOGLOBIN | SUBUNIT | ALPHA |
| 97:00:00 | 2r80-C | 4.5 | 3.1 | 85 | 140 | 8 | MOLECULE: | HEMOGLOBIN | SUBUNIT | ALPHA-A |
| 98:00:00 | 4mqj-D | 4.5 | 3.3 | 85 | 145 | 6 | MOLECULE: | HEMOGLOBIN | SUBUNIT | ALPHA |
| 99:00:00 | 5woh-C | 4.5 | 3.4 | 85 | 137 | 7 | MOLECULE: | HEMOGLOBIN | SUBUNIT | ALPHA |
| 0:00 | 5x2u-K | 4.5 | 3 | 83 | 140 | 7 | MOLECULE: | HEMOGLOBIN | SUBUNIT | ALPHA |
| 1:00 | 4hrt-C | 4.5 | 3.5 | 86 | 149 | 3 | MOLECULE: | GLOBIN-2 A | CHAIN | |
| 2:00 | 5ziq-A | 4.4 | 4.4 | 91 | 155 | 8 | MOLECULE: | GLOBIN PRO | TEIN | |
| 3:00 | 7m8i-A | 4.4 | 7.8 | 145 | 572 | 9 | MOLECULE: | ADRENODOXI | N, CYTOC | HROME P450 11B2, MITOCHONDRIAL |
| 4:00 | 6p5b-A | 4.4 | 10.9 | 94 | 388 | 4 | MOLECULE: | MAVC | | |
| 5:00 | 7d3u-F | 4.4 | 7.5 | 76 | 85 | 9 | MOLECULE: | MONOVALENT | NA+/H+ | ANTIPORTER SUBUNIT D |
| 6:00 | 6vxm-A | 4.4 | 14.8 | 100 | 277 | 6 | MOLECULE: | MECHANOSEN | SITIVE I | ON CHANNEL PROTEIN 1, MITOCHOND |
| 7:00 | 5utd-A | 4.4 | 4.5 | 92 | 151 | 8 | MOLECULE: | MYOGLOBIN | | |
| 8:00 | 6ls8-K | 4.4 | 4.2 | 91 | 153 | 14 | MOLECULE: | MYOGLOBIN | | |
| 9:00 | 3pi2-B | 4.4 | 4 | 86 | 151 | 7 | MOLECULE: | HEMOGLOBIN | II | |
| 10:00 | 1bvc-A | 4.4 | 4.3 | 90 | 153 | 10 | MOLECULE: | APOMYOGLOB | IN | |
| 11:00 | 2eb8-A | 4.4 | 3.8 | 90 | 154 | 8 | MOLECULE: | MYOGLOBIN | | |
| 12:00 | 2olp-B | 4.4 | 4 | 84 | 151 | 5 | MOLECULE: | HEMOGLOBIN | II | |
| 13:00 | 1u7s-A | 4.4 | 4.3 | 91 | 153 | 10 | MOLECULE: | MYOGLOBIN | | |
| 14:00 | 3pi1-B | 4.4 | 4 | 85 | 151 | 6 | MOLECULE: | HEMOGLOBIN | II | |
| 15:00 | 3pt8-A | 4.4 | 3.8 | 84 | 151 | 7 | MOLECULE: | HEMOGLOBIN | II | |
| 16:00 | 3a59-G | 4.4 | 3.1 | 85 | 141 | 7 | MOLECULE: | HEMOGLOBIN | SUBUNIT | ALPHA-A |
| 17:00 | 4n7p-I | 4.4 | 3.4 | 85 | 141 | 7 | MOLECULE: | HEMOGLOBIN | SUBUNIT | ALPHA |
| 18:00 | 1ouu-C | 4.4 | 3.4 | 86 | 142 | 6 | MOLECULE: | HEMOGLOBIN | I | |
| 19:00 | 1si4-C | 4.4 | 3.5 | 85 | 141 | 5 | MOLECULE: | HEMOGLOBIN | ALPHA C | HAIN |
| 20:00 | 4wjg-F | 4.4 | 3.1 | 84 | 141 | 7 | MOLECULE: | HEMOGLOBIN | SUBUNIT | ALPHA |
| 21:00 | 3eok-A | 4.4 | 3 | 83 | 141 | 7 | MOLECULE: | HEMOGLOBIN | SUBUNIT | ALPHA-A |
| 22:00 | 3b75-C | 4.4 | 2.9 | 83 | 141 | 7 | MOLECULE: | HEMOGLOBIN | SUBUNIT | ALPHA |
| 23:00 | 4wjg-P | 4.4 | 3.6 | 85 | 141 | 5 | MOLECULE: | HEMOGLOBIN | SUBUNIT | ALPHA |
| 24:00:00 | 4wjg-U | 4.4 | 3.1 | 84 | 141 | 7 | MOLECULE: | HEMOGLOBIN | SUBUNIT | ALPHA |
| 25:00:00 | 3dhr-C | 4.4 | 3.6 | 87 | 141 | 6 | MOLECULE: | HEMOGLOBIN | SUBUNIT | ALPHA-A |
| 26:00:00 | 2qu0-A | 4.4 | 3.7 | 88 | 141 | 3 | MOLECULE: | HEMOGLOBIN | SUBUNIT | ALPHA-1/2 |
| 27:00:00 | 3gys-C | 4.4 | 3.6 | 87 | 139 | 6 | MOLECULE: | HEMOGLOBIN | SUBUNIT | ALPHA |
| 28:00:00 | 1ns9-A | 4.4 | 3.2 | 85 | 141 | 7 | MOLECULE: | HEMOGLOBIN | ALPHA S | UBUNIT |
| 29:00:00 | 6xd9-C | 4.4 | 3.5 | 86 | 141 | 3 | MOLECULE: | HEMOGLOBIN | SUBUNIT | ALPHA |
| 30:00:00 | 4wjg-A | 4.4 | 3.6 | 85 | 141 | 5 | MOLECULE: | HEMOGLOBIN | SUBUNIT | ALPHA |
| 31:00:00 | 3d1a-C | 4.4 | 3.4 | 85 | 141 | 7 | MOLECULE: | HEMOGLOBIN | SUBUNIT | ALPHA-1/2 |
| 32:00:00 | 1lfz-A | 4.4 | 3.4 | 86 | 141 | 6 | MOLECULE: | HEMOGLOBIN | ALPHA C | HAIN |

| | | | | | | | | | | |
|---|---|---|---|---|---|---|---|---|---|---|
| 33:00:00 | 2zlu-A | 4.4 | 3.1 | 84 | 140 | 7 | MOLECULE: | HEMOGLOBIN | SUBUNIT | ALPHA |
| 34:00:00 | 3d4x-C | 4.4 | 3 | 83 | 141 | 10 | MOLECULE: | HEMOGLOBIN | SUBUNIT | ALPHA |
| 35:00:00 | 1qpw-C | 4.4 | 3.2 | 85 | 141 | 7 | MOLECULE: | PORCINE HE | MOGLOBIN | (ALPHA SUBUNIT) |
| 36:00:00 | 1y8i-C | 4.4 | 3 | 83 | 141 | 7 | MOLECULE: | HEMOGLOBIN | ALPHA C | HAINS |
| 37:00:00 | 6tb2-A | 4.4 | 3.6 | 85 | 141 | 5 | MOLECULE: | HEMOGLOBIN | SUBUNIT | ALPHA |
| 38:00:00 | 1g0b-A | 4.4 | 3.1 | 84 | 141 | 7 | MOLECULE: | HEMOGLOBIN | ALPHA C | HAIN |
| 39:00:00 | 1y8k-C | 4.4 | 2.9 | 83 | 141 | 7 | MOLECULE: | HEMOGLOBIN | ALPHA C | HAINS |
| 40:00:00 | 3ic2-C | 4.4 | 3.5 | 85 | 141 | 5 | MOLECULE: | HEMOGLOBIN | SUBUNIT | ALPHA |
| 41:00:00 | 1out-A | 4.4 | 3 | 84 | 142 | 6 | MOLECULE: | HEMOGLOBIN | I | |
| 42:00:00 | 1qxe-C | 4.4 | 3 | 83 | 141 | 7 | MOLECULE: | HEMOGLOBIN | ALPHA C | HAIN |
| 43:00:00 | 3dht-A | 4.4 | 3.2 | 86 | 139 | 7 | MOLECULE: | HEMOGLOBIN | SUBUNIT | ALPHA-1/2 |
| 44:00:00 | 7jxz-A | 4.4 | 3 | 84 | 141 | 7 | MOLECULE: | HEMOGLOBIN | SUBUNIT | ALPHA |
| 45:00:00 | 3kmf-E | 4.4 | 3 | 84 | 141 | 7 | MOLECULE: | HEMOGLOBIN | SUBUNIT | ALPHA |
| 46:00:00 | 4wjg-K | 4.4 | 3.6 | 85 | 141 | 5 | MOLECULE: | HEMOGLOBIN | SUBUNIT | ALPHA |
| 47:00:00 | 3d1a-A | 4.4 | 3.8 | 88 | 141 | 3 | MOLECULE: | HEMOGLOBIN | SUBUNIT | ALPHA-1/2 |
| 48:00:00 | 4mqj-G | 4.4 | 3.2 | 84 | 141 | 7 | MOLECULE: | HEMOGLOBIN | SUBUNIT | ALPHA |
| 49:00:00 | 3dhr-A | 4.4 | 3.1 | 85 | 141 | 8 | MOLECULE: | HEMOGLOBIN | SUBUNIT | ALPHA-A |
| 50:00:00 | 2zlt-A | 4.4 | 3.1 | 84 | 141 | 7 | MOLECULE: | HEMOGLOBIN | SUBUNIT | ALPHA |
| 51:00:00 | 3gou-A | 4.4 | 3.2 | 86 | 141 | 7 | MOLECULE: | HEMOGLOBIN | SUBUNIT | ALPHA |
| 52:00:00 | 1qxd-C | 4.4 | 3.1 | 84 | 141 | 7 | MOLECULE: | HEMOGLOBIN | ALPHA C | HAIN |
| 53:00:00 | 4mqk-G | 4.4 | 3.3 | 86 | 140 | 6 | MOLECULE: | HEMOGLOBIN | SUBUNIT | ALPHA |
| 54:00:00 | 4n7o-A | 4.4 | 3.8 | 87 | 141 | 5 | MOLECULE: | HEMOGLOBIN | SUBUNIT | ALPHA |
| 55:00:00 | 3b75-E | 4.4 | 3.3 | 86 | 141 | 6 | MOLECULE: | HEMOGLOBIN | SUBUNIT | ALPHA |
| 56:00:00 | 3r5i-C | 4.4 | 3.1 | 84 | 141 | 7 | MOLECULE: | HEMOGLOBIN | SUBUNIT | ALPHA |
| 57:00:00 | 3gys-E | 4.4 | 2.9 | 83 | 138 | 10 | MOLECULE: | HEMOGLOBIN | SUBUNIT | ALPHA |
| 58:00:00 | 2qu0-C | 4.4 | 3.1 | 84 | 141 | 7 | MOLECULE: | HEMOGLOBIN | SUBUNIT | ALPHA-1/2 |
| 59:00:00 | 3pel-A | 4.4 | 3.1 | 85 | 141 | 7 | MOLECULE: | HEMOGLOBIN | SUBUNIT | ALPHA |
| 60:00:00 | 3dhr-G | 4.4 | 3.5 | 86 | 141 | 6 | MOLECULE: | HEMOGLOBIN | SUBUNIT | ALPHA-A |
| 61:00:00 | 3dhr-E | 4.4 | 3 | 84 | 141 | 8 | MOLECULE: | HEMOGLOBIN | SUBUNIT | ALPHA-A |
| 62:00:00 | 1nej-A | 4.4 | 2.9 | 83 | 141 | 7 | MOLECULE: | HEMOGLOBIN | ALPHA C | HAIN |
| 63:00:00 | 5c6e-A | 4.4 | 3.1 | 84 | 141 | 7 | MOLECULE: | HEMOGLOBIN | SUBUNIT | ALPHA |
| 64:00:00 | 1mko-C | 4.4 | 3.1 | 85 | 141 | 6 | MOLECULE: | HEMOGLOBIN | ALPHA C | HAIN |
| 65:00:00 | 2bw9-M | 4.4 | 4.2 | 90 | 153 | 9 | MOLECULE: | MYOGLOBIN | | |
| 66:00:00 | 3d4x-A | 4.4 | 3.1 | 85 | 141 | 9 | MOLECULE: | HEMOGLOBIN | SUBUNIT | ALPHA |
| 67:00:00 | 3gqr-E | 4.4 | 3.1 | 85 | 141 | 8 | MOLECULE: | HEMOGLOBIN | SUBUNIT | ALPHA |
| 68:00:00 | 3gou-C | 4.4 | 3.4 | 85 | 141 | 8 | MOLECULE: | HEMOGLOBIN | SUBUNIT | ALPHA |
| 69:00:00 | 4h2l-A | 4.4 | 3.2 | 85 | 141 | 8 | MOLECULE: | ALPHA-GLOB | IN | |
| 70:00:00 | 1iwh-A | 4.4 | 3.1 | 84 | 141 | 7 | MOLECULE: | HEMOGLOBIN | ALPHA C | HAIN |
| 71:00:00 | 3b75-S | 4.4 | 3.5 | 86 | 141 | 3 | MOLECULE: | HEMOGLOBIN | SUBUNIT | ALPHA |
| 72:00:00 | 3s65-A | 4.4 | 2.9 | 83 | 141 | 7 | MOLECULE: | HEMOGLOBIN | SUBUNIT | ALPHA |
| 73:00:00 | 2rao-A | 4.4 | 3.5 | 85 | 141 | 6 | MOLECULE: | HEMOGLOBIN | SUBUNIT | ALPHA-1/2 |
| 74:00:00 | 1shr-C | 4.4 | 3 | 83 | 141 | 7 | MOLECULE: | HEMOGLOBIN | ALPHA C | HAIN |
| 75:00:00 | 4n7p-C | 4.4 | 3.1 | 84 | 141 | 7 | MOLECULE: | HEMOGLOBIN | SUBUNIT | ALPHA |
| 76:00:00 | 3pi9-A | 4.4 | 3.5 | 85 | 141 | 5 | MOLECULE: | HEMOGLOBIN | SUBUNIT | ALPHA |
| 77:00:00 | 3b75-G | 4.4 | 3.5 | 86 | 141 | 7 | MOLECULE: | HEMOGLOBIN | SUBUNIT | ALPHA |
| 78:00:00 | 3a59-A | 4.4 | 3.6 | 87 | 141 | 5 | MOLECULE: | HEMOGLOBIN | SUBUNIT | ALPHA-A |
| 79:00:00 | 4yu4-A | 4.4 | 3.5 | 85 | 141 | 7 | MOLECULE: | HEMOGLOBIN | | |
| 80:00:00 | 1a01-B | 4.4 | 3.6 | 86 | 146 | 5 | MOLECULE: | HEMOGLOBIN | (ALPHA | CHAIN) |
| 81:00:00 | 3nfe-A | 4.4 | 3 | 84 | 142 | 6 | MOLECULE: | HEMOGLOBIN | SUBUNIT | ALPHA-1 |
| 82:00:00 | 7ud8-C | 4.4 | 3.1 | 84 | 141 | 7 | MOLECULE: | HEMOGLOBIN | SUBUNIT | ALPHA |
| 83:00:00 | 1fn3-A | 4.4 | 3.1 | 85 | 141 | 7 | MOLECULE: | HEMOGLOBIN | ALPHA C | HAIN |
| 84:00:00 | 4g51-A | 4.4 | 3.3 | 85 | 142 | 6 | MOLECULE: | HEMOGLOBIN | SUBUNIT | ALPHA |
| 85:00:00 | 4mqj-E | 4.4 | 3.2 | 84 | 141 | 7 | MOLECULE: | HEMOGLOBIN | SUBUNIT | ALPHA |
| 86:00:00 | 2h8d-C | 4.4 | 3.3 | 85 | 142 | 6 | MOLECULE: | HEMOGLOBIN | ALPHA S | UBUNIT |

| | | | | | | | | | | |
|---|---|---|---|---|---|---|---|---|---|---|
| 87:00:00 | 2mhb-A | 4.4 | 3.4 | 85 | 141 | 7 | MOLECULE: | HEMOGLOBIN | (AQUO M | ET) (ALPHA CHAIN) |
| 88:00:00 | 2d5x-A | 4.4 | 3.4 | 85 | 141 | 7 | MOLECULE: | HEMOGLOBIN | ALPHA S | UBUNIT |
| 89:00:00 | 1fn3-C | 4.4 | 3.1 | 85 | 141 | 7 | MOLECULE: | HEMOGLOBIN | ALPHA C | HAIN |
| 90:00:00 | 1hbs-E | 4.4 | 2.8 | 83 | 141 | 7 | MOLECULE: | HEMOGLOBIN | S (DEOX | Y) (ALPHA CHAIN) |
| 91:00:00 | 1ibe-A | 4.4 | 3.3 | 85 | 141 | 8 | MOLECULE: | HEMOGLOBIN | (DEOXY) | |
| 92:00:00 | 2h8f-C | 4.4 | 3.3 | 85 | 142 | 6 | MOLECULE: | HEMOGLOBIN | ALPHA S | UBUNIT |
| 93:00:00 | 2h8f-A | 4.4 | 3.3 | 85 | 142 | 6 | MOLECULE: | HEMOGLOBIN | ALPHA S | UBUNIT |
| 94:00:00 | 2qls-A | 4.4 | 3.6 | 87 | 141 | 5 | MOLECULE: | HEMOGLOBIN | SUBUNIT | ALPHA |
| 95:00:00 | 1hbh-C | 4.4 | 2.8 | 83 | 142 | 6 | MOLECULE: | HEMOGLOBIN | (DEOXY) | (ALPHA CHAIN) |
| 96:00:00 | 3b75-A | 4.4 | 3 | 84 | 141 | 6 | MOLECULE: | HEMOGLOBIN | SUBUNIT | ALPHA |
| 97:00:00 | 4f4o-A | 4.4 | 3.6 | 86 | 141 | 3 | MOLECULE: | HEMOGLOBIN | SUBUNIT | ALPHA |
| 98:00:00 | 3gqr-C | 4.4 | 3.1 | 84 | 141 | 10 | MOLECULE: | HEMOGLOBIN | SUBUNIT | ALPHA |
| 99:00:00 | 3a0g-A | 4.4 | 3.6 | 86 | 139 | 5 | MOLECULE: | HEMOGLOBIN | SUBUNIT | ALPHA |
| 0:00 | 3hyu-A | 4.4 | 3.6 | 86 | 141 | 5 | MOLECULE: | HEMOGLOBIN | SUBUNIT | ALPHA |
| 1:00 | 7dy3-A | 4.4 | 3.1 | 85 | 141 | 7 | MOLECULE: | HEMOGLOBIN | SUBUNIT | ALPHA |
| 2:00 | 3gys-A | 4.4 | 3.1 | 85 | 138 | 8 | MOLECULE: | HEMOGLOBIN | SUBUNIT | ALPHA |
| 3:00 | 4f4o-D | 4.4 | 3.1 | 84 | 141 | 7 | MOLECULE: | HEMOGLOBIN | SUBUNIT | ALPHA |
| 4:00 | 4yu4-C | 4.4 | 3.5 | 85 | 141 | 7 | MOLECULE: | HEMOGLOBIN | | |
| 5:00 | 3kmf-A | 4.4 | 3.1 | 85 | 141 | 7 | MOLECULE: | HEMOGLOBIN | SUBUNIT | ALPHA |
| 6:00 | 2zlv-A | 4.4 | 3 | 83 | 141 | 7 | MOLECULE: | HEMOGLOBIN | SUBUNIT | ALPHA |
| 7:00 | 3nfe-C | 4.4 | 3 | 84 | 142 | 6 | MOLECULE: | HEMOGLOBIN | SUBUNIT | ALPHA-1 |
| 8:00 | 1y8i-A | 4.4 | 2.9 | 83 | 141 | 7 | MOLECULE: | HEMOGLOBIN | ALPHA C | HAINS |
| 9:00 | 3ng6-C | 4.4 | 3 | 84 | 142 | 6 | MOLECULE: | HEMOGLOBIN | SUBUNIT | ALPHA-1 |
| 10:00 | 2b7h-C | 4.4 | 3.4 | 86 | 141 | 8 | MOLECULE: | HEMOGLOBIN | ALPHA C | HAIN |
| 11:00 | 4iro-C | 4.4 | 3.3 | 85 | 142 | 6 | MOLECULE: | HEMOGLOBIN | SUBUNIT | ALPHA |
| 12:00 | 1qpw-A | 4.4 | 3 | 84 | 141 | 7 | MOLECULE: | PORCINE HE | MOGLOBIN | (ALPHA SUBUNIT) |
| 13:00 | 4n7p-K | 4.4 | 3.4 | 85 | 141 | 7 | MOLECULE: | HEMOGLOBIN | SUBUNIT | ALPHA |
| 14:00 | 1ouu-B | 4.4 | 3.5 | 85 | 146 | 7 | MOLECULE: | HEMOGLOBIN | I | |
| 15:00 | 1mdn-A | 4.4 | 4.4 | 91 | 153 | 13 | MOLECULE: | PROTEIN (M | YOGLOBIN | ) |
| 16:00 | 1o1p-D | 4.4 | 3.3 | 85 | 146 | 6 | MOLECULE: | HEMOGLOBIN | ALPHA C | HAIN |
| 17:00 | 5vmm-C | 4.4 | 3.1 | 84 | 141 | 7 | MOLECULE: | HEMOGLOBIN | SUBUNIT | ALPHA |
| 18:00 | 5x2u-I | 4.4 | 3.1 | 84 | 140 | 7 | MOLECULE: | HEMOGLOBIN | SUBUNIT | ALPHA |
| 19:00 | 5x2r-I | 4.4 | 3.1 | 84 | 140 | 7 | MOLECULE: | HEMOGLOBIN | SUBUNIT | ALPHA |
| 20:00 | 1a01-D | 4.4 | 3.2 | 85 | 146 | 6 | MOLECULE: | HEMOGLOBIN | (ALPHA | CHAIN) |
| 21:00 | 1c40-B | 4.4 | 3.5 | 85 | 146 | 9 | MOLECULE: | PROTEIN (H | EMOGLOBI | N (ALPHA CHAIN)) |
| 22:00 | 1faw-D | 4.4 | 3.5 | 85 | 146 | 9 | MOLECULE: | HEMOGLOBIN | (ALPHA | SUBUNIT) |
| 23:00 | 1i3e-A | 4.4 | 3.4 | 86 | 146 | 6 | MOLECULE: | HEMOGLOBIN | GAMMA C | HAINS |
| 24:00:00 | 1yeo-B | 4.4 | 3.5 | 86 | 146 | 6 | MOLECULE: | HEMOGLOBIN | ALPHA C | HAIN |
| 25:00:00 | 1out-B | 4.4 | 3.6 | 85 | 146 | 7 | MOLECULE: | HEMOGLOBIN | I | |
| 26:00:00 | 1j7s-B | 4.4 | 3.1 | 84 | 146 | 6 | MOLECULE: | HEMOGLOBIN | | |
| 27:00:00 | 3pi3-B | 4.4 | 3.9 | 84 | 151 | 7 | MOLECULE: | HEMOGLOBIN | II | |
| 28:00:00 | 5lfg-A | 4.4 | 3.3 | 85 | 142 | 6 | MOLECULE: | HEMOGLOBIN | SUBUNIT | ALPHA-1 |
| 29:00:00 | 1faw-B | 4.4 | 3.5 | 85 | 146 | 9 | MOLECULE: | HEMOGLOBIN | (ALPHA | SUBUNIT) |
| 30:00:00 | 5x2t-G | 4.4 | 3.4 | 85 | 140 | 7 | MOLECULE: | HEMOGLOBIN | SUBUNIT | ALPHA |
| 31:00:00 | 4nxc-A | 4.4 | 4.4 | 92 | 153 | 9 | MOLECULE: | MYOGLOBIN | | |
| 32:00:00 | 3s48-C | 4.4 | 3.4 | 86 | 138 | 6 | MOLECULE: | IRON-REGUL | ATED SUR | FACE DETERMINANT PROTEIN H |
| 33:00:00 | 1y4b-D | 4.4 | 3.2 | 85 | 146 | 6 | MOLECULE: | HEMOGLOBIN | ALPHA C | HAIN |
| 34:00:00 | 4n7p-E | 4.4 | 3.2 | 86 | 141 | 6 | MOLECULE: | HEMOGLOBIN | SUBUNIT | ALPHA |
| 35:00:00 | 1y4g-D | 4.4 | 3.5 | 86 | 146 | 6 | MOLECULE: | HEMOGLOBIN | ALPHA C | HAIN |
| 36:00:00 | 1ns6-A | 4.4 | 3.2 | 85 | 141 | 7 | MOLECULE: | HEMOGLOBIN | ALPHA S | UBUNIT |
| 37:00:00 | 2blj-M | 4.4 | 4.7 | 91 | 153 | 10 | MOLECULE: | MYOGLOBIN | | |
| 38:00:00 | 5hy8-G | 4.4 | 3.1 | 83 | 138 | 7 | MOLECULE: | HEMOGLOBIN | SUBUNIT | ALPHA |
| 39:00:00 | 5ni1-C | 4.4 | 3.1 | 85 | 141 | 6 | MOLECULE: | HEMOGLOBIN | SUBUNIT | ALPHA |
| 40:00:00 | 1ouu-D | 4.4 | 3.6 | 85 | 146 | 7 | MOLECULE: | HEMOGLOBIN | I | |

| | | | | | | | | | | |
|---|---|---|---|---|---|---|---|---|---|---|
| 41:00:00 | 1yev-B | 4.4 | 3.5 | 86 | 146 | 6 | MOLECULE: | HEMOGLOBIN | ALPHA C | HAIN |
| 42:00:00 | 1yeu-D | 4.4 | 3.5 | 86 | 146 | 6 | MOLECULE: | HEMOGLOBIN | ALPHA C | HAIN |
| 43:00:00 | 7qu4-H | 4.4 | 3.4 | 84 | 144 | 6 | MOLECULE: | HEMOGLOBIN | SUBUNIT | GAMMA-2 |
| 44:00:00 | 5u3i-C | 4.4 | 2.9 | 82 | 140 | 7 | MOLECULE: | HEMOGLOBIN | SUBUNIT | ALPHA |
| 45:00:00 | 1t1n-A | 4.4 | 3.3 | 85 | 142 | 6 | MOLECULE: | PROTEIN (H | EMOGLOBI | N) |
| 46:00:00 | 1m6c-B | 4.4 | 4.1 | 89 | 153 | 13 | MOLECULE: | PROTEIN (M | YOGLOBIN | ) |
| 47:00:00 | 1yih-D | 4.4 | 3.5 | 85 | 146 | 6 | MOLECULE: | HEMOGLOBIN | ALPHA C | HAIN |
| 48:00:00 | 5ni1-A | 4.4 | 3.1 | 85 | 141 | 6 | MOLECULE: | HEMOGLOBIN | SUBUNIT | ALPHA |
| 49:00:00 | 1j7s-D | 4.4 | 3.3 | 85 | 146 | 6 | MOLECULE: | HEMOGLOBIN | | |
| 50:00:00 | 1rqa-B | 4.4 | 3.5 | 86 | 146 | 6 | MOLECULE: | HEMOGLOBIN | ALPHA C | HAIN |
| 51:00:00 | 6hal-C | 4.4 | 3.1 | 84 | 139 | 7 | MOLECULE: | HEMOGLOBIN | SUBUNIT | ALPHA |
| 52:00:00 | 1a0z-D | 4.4 | 3.5 | 86 | 146 | 6 | MOLECULE: | HEMOGLOBIN | (ALPHA | CHAIN) |
| 53:00:00 | 3gkv-A | 4.4 | 3.4 | 85 | 142 | 6 | MOLECULE: | HEMOGLOBIN | SUBUNIT | ALPHA |
| 54:00:00 | 1yeu-B | 4.4 | 3.5 | 86 | 146 | 6 | MOLECULE: | HEMOGLOBIN | ALPHA C | HAIN |
| 55:00:00 | 1o1p-B | 4.4 | 3.3 | 85 | 146 | 6 | MOLECULE: | HEMOGLOBIN | ALPHA C | HAIN |
| 56:00:00 | 1a00-D | 4.4 | 3.1 | 84 | 146 | 6 | MOLECULE: | HEMOGLOBIN | (ALPHA | CHAIN) |
| 57:00:00 | 1v9q-A | 4.4 | 4.6 | 92 | 154 | 9 | MOLECULE: | MYOGLOBIN | | |
| 58:00:00 | 1ye2-D | 4.4 | 3.5 | 86 | 146 | 6 | MOLECULE: | HEMOGLOBIN | ALPHA C | HAIN |
| 59:00:00 | 6sva-A | 4.4 | 3.1 | 84 | 139 | 7 | MOLECULE: | HEMOGLOBIN | SUBUNIT | ALPHA |
| 60:00:00 | 4n7n-I | 4.4 | 3.2 | 86 | 141 | 7 | MOLECULE: | HEMOGLOBIN | SUBUNIT | ALPHA |
| 61:00:00 | 2peg-A | 4.4 | 3.4 | 85 | 142 | 6 | MOLECULE: | HEMOGLOBIN | SUBUNIT | ALPHA |
| 62:00:00 | 5lfg-C | 4.4 | 3.1 | 84 | 142 | 6 | MOLECULE: | HEMOGLOBIN | SUBUNIT | ALPHA-1 |
| 63:00:00 | 1y4g-B | 4.4 | 3.5 | 86 | 146 | 6 | MOLECULE: | HEMOGLOBIN | ALPHA C | HAIN |
| 64:00:00 | 4n7p-A | 4.4 | 3.4 | 85 | 141 | 7 | MOLECULE: | HEMOGLOBIN | SUBUNIT | ALPHA |
| 65:00:00 | 5hy8-E | 4.4 | 3 | 82 | 137 | 7 | MOLECULE: | HEMOGLOBIN | SUBUNIT | ALPHA |
| 66:00:00 | 4odc-A | 4.4 | 3.4 | 85 | 142 | 6 | MOLECULE: | HEMOGLOBIN | SUBUNIT | ALPHA |
| 67:00:00 | 1dm1-A | 4.3 | 3.5 | 81 | 146 | 7 | MOLECULE: | MYOGLOBIN | | |
| 68:00:00 | 6myo-D | 4.3 | 8.1 | 84 | 102 | 7 | MOLECULE: | SUCCINATE | DEHYDROG | ENASE [UBIQUINONE] FLAVOPROTEIN |
| 69:00:00 | 5hv1-A | 4.3 | 11.2 | 108 | 847 | 8 | MOLECULE: | PHOSPHOENO | LPYRUVAT | E SYNTHASE |
| 70:00:00 | 1f65-A | 4.3 | 4.4 | 91 | 154 | 9 | MOLECULE: | MYOGLOBIN | | |
| 71:00:00 | 6oty-A | 4.3 | 4 | 86 | 151 | 6 | MOLECULE: | HEMOGLOBIN | II | |
| 72:00:00 | 5oj9-A | 4.3 | 4.1 | 90 | 154 | 9 | MOLECULE: | MYOGLOBIN | | |
| 73:00:00 | 7cez-A | 4.3 | 4.3 | 90 | 153 | 10 | MOLECULE: | MYOGLOBIN | | |
| 74:00:00 | 1hbs-A | 4.3 | 3.5 | 86 | 141 | 5 | MOLECULE: | HEMOGLOBIN | S (DEOX | Y) (ALPHA CHAIN) |
| 75:00:00 | 3pt7-A | 4.3 | 3.9 | 85 | 151 | 6 | MOLECULE: | HEMOGLOBIN | II | |
| 76:00:00 | 3pi2-A | 4.3 | 3.9 | 85 | 151 | 8 | MOLECULE: | HEMOGLOBIN | II | |
| 77:00:00 | 8qba-A | 4.3 | 4.3 | 91 | 156 | 8 | MOLECULE: | MYOGLOBIN | | |
| 78:00:00 | 6bmg-B | 4.3 | 4.4 | 91 | 154 | 8 | MOLECULE: | MYOGLOBIN | | |
| 79:00:00 | 6zfb-h | 4.3 | 19.6 | 126 | 771 | 7 | MOLECULE: | DNA-DIRECT | ED RNA P | OLYMERASE SUBUNIT DELTA |
| 80:00:00 | 1hrm-A | 4.3 | 4.7 | 92 | 153 | 14 | MOLECULE: | MYOGLOBIN | | |
| 81:00:00 | 3wtg-B | 4.3 | 3.5 | 85 | 146 | 11 | MOLECULE: | HEMOGLOBIN | SUBUNIT | ALPHA-A |
| 82:00:00 | 1flp-A | 4.3 | 3.4 | 78 | 142 | 5 | MOLECULE: | HEMOGLOBIN | I (AQUO | MET) |
| 83:00:00 | 6d45-A | 4.3 | 4.3 | 90 | 153 | 10 | MOLECULE: | MYOGLOBIN | | |
| 84:00:00 | 1moh-A | 4.3 | 3.4 | 78 | 142 | 5 | MOLECULE: | MONOMERIC | HEMOGLOB | IN I |
| 85:00:00 | 2e2y-A | 4.3 | 4.2 | 91 | 154 | 8 | MOLECULE: | MYOGLOBIN | | |
| 86:00:00 | 5ker-C | 4.3 | 3 | 83 | 140 | 8 | MOLECULE: | ALPHA-GLOB | IN | |
| 87:00:00 | 1shr-A | 4.3 | 3.5 | 85 | 141 | 5 | MOLECULE: | HEMOGLOBIN | ALPHA C | HAIN |
| 88:00:00 | 4x0l-A | 4.3 | 3.6 | 85 | 141 | 5 | MOLECULE: | HEMOGLOBIN | SUBUNIT | ALPHA |
| 89:00:00 | 1y4f-B | 4.3 | 3.5 | 86 | 146 | 5 | MOLECULE: | HEMOGLOBIN | ALPHA C | HAIN |
| 90:00:00 | 1dxu-B | 4.3 | 3.6 | 86 | 146 | 5 | MOLECULE: | HEMOGLOBIN | (DEOXY) | (ALPHA CHAIN) |
| 91:00:00 | 4hrt-F | 4.3 | 3.9 | 88 | 151 | 9 | MOLECULE: | GLOBIN-2 A | CHAIN | |
| 92:00:00 | 1o1l-B | 4.3 | 3.6 | 86 | 146 | 5 | MOLECULE: | HEMOGLOBIN | ALPHA C | HAIN |
| 93:00:00 | 4n7n-C | 4.3 | 3.1 | 85 | 141 | 7 | MOLECULE: | HEMOGLOBIN | SUBUNIT | ALPHA |
| 94:00:00 | 1c40-A | 4.3 | 3.5 | 85 | 141 | 7 | MOLECULE: | PROTEIN (H | EMOGLOBI | N (ALPHA CHAIN)) |

| | | | | | | | | | | |
|---|---|---|---|---|---|---|---|---|---|---|
| 95:00:00 | 3a59-E | 4.3 | 3.4 | 86 | 141 | 7 | MOLECULE: | HEMOGLOBIN | SUBUNIT | ALPHA-A |
| 96:00:00 | 3k8b-C | 4.3 | 3.5 | 87 | 141 | 3 | MOLECULE: | HEMOGLOBIN | SUBUNIT | ALPHA-A |
| 97:00:00 | 3sxm-B | 4.3 | 3.8 | 93 | 137 | 8 | MOLECULE: | TRANSCRIPT | IONAL RE | GULATOR, GNTR FAMILY |
| 98:00:00 | 1yih-B | 4.3 | 3.4 | 84 | 146 | 5 | MOLECULE: | HEMOGLOBIN | ALPHA C | HAIN |
| 99:00:00 | 1y8k-A | 4.3 | 2.9 | 83 | 141 | 7 | MOLECULE: | HEMOGLOBIN | ALPHA C | HAINS |
| 0:00 | 6r2o-A | 4.3 | 2.9 | 83 | 141 | 7 | MOLECULE: | HEMOGLOBIN | SUBUNIT | ALPHA |
| 1:00 | 1rqa-D | 4.3 | 3.6 | 86 | 146 | 5 | MOLECULE: | HEMOGLOBIN | ALPHA C | HAIN |
| 2:00 | 1y8h-A | 4.3 | 3 | 83 | 141 | 7 | MOLECULE: | HEMOGLOBIN | ALPHA C | HAINS |
| 3:00 | 5ker-G | 4.3 | 3.1 | 84 | 140 | 8 | MOLECULE: | ALPHA-GLOB | IN | |
| 4:00 | 2zlx-C | 4.3 | 3 | 84 | 139 | 6 | MOLECULE: | HEMOGLOBIN | SUBUNIT | ALPHA |
| 5:00 | 2hbs-H | 4.3 | 3.4 | 84 | 146 | 5 | MOLECULE: | HEMOGLOBIN | S (DEOX | Y), ALPHA CHAIN |
| 6:00 | 2r80-A | 4.3 | 3.2 | 86 | 141 | 8 | MOLECULE: | HEMOGLOBIN | SUBUNIT | ALPHA-A |
| 7:00 | 3fh9-A | 4.3 | 3.7 | 86 | 141 | 6 | MOLECULE: | HEMOGLOBIN | ALPHA C | HAIN |
| 8:00 | 7pcf-B | 4.3 | 3.3 | 84 | 146 | 8 | MOLECULE: | HEMOGLOBIN | SUBUNIT | ALPHA |
| 9:00 | 3mju-A | 4.3 | 3.4 | 86 | 141 | 8 | MOLECULE: | HEMOGLOBIN | SUBUNIT | ALPHA-A |
| 10:00 | 2pgh-C | 4.3 | 3.2 | 85 | 141 | 7 | MOLECULE: | HEMOGLOBIN | (AQUO M | ET) (ALPHA CHAIN) |
| 11:00 | 3ic0-C | 4.3 | 3 | 83 | 141 | 7 | MOLECULE: | HEMOGLOBIN | SUBUNIT | ALPHA |
| 12:00 | 1y4p-D | 4.3 | 3.6 | 86 | 146 | 5 | MOLECULE: | HEMOGLOBIN | ALPHA C | HAIN |
| 13:00 | 7k4m-E | 4.3 | 3.5 | 85 | 142 | 5 | MOLECULE: | HEMOGLOBIN | SUBUNIT | ALPHA |
| 14:00 | 1o1n-D | 4.3 | 3.6 | 86 | 146 | 5 | MOLECULE: | HEMOGLOBIN | ALPHA C | HAIN |
| 15:00 | 2b7h-A | 4.3 | 3.5 | 86 | 141 | 9 | MOLECULE: | HEMOGLOBIN | ALPHA C | HAIN |
| 16:00 | 4g51-C | 4.3 | 3.4 | 85 | 142 | 6 | MOLECULE: | HEMOGLOBIN | SUBUNIT | ALPHA |
| 17:00 | 1y46-B | 4.3 | 3.5 | 86 | 146 | 5 | MOLECULE: | HEMOGLOBIN | ALPHA C | HAIN |
| 18:00 | 1o1n-B | 4.3 | 3.6 | 86 | 146 | 5 | MOLECULE: | HEMOGLOBIN | ALPHA C | HAIN |
| 19:00 | 3gqr-A | 4.3 | 3.5 | 87 | 141 | 8 | MOLECULE: | HEMOGLOBIN | SUBUNIT | ALPHA |
| 20:00 | 5ker-A | 4.3 | 3.6 | 85 | 140 | 6 | MOLECULE: | ALPHA-GLOB | IN | |
| 21:00 | 3eu1-A | 4.3 | 3.6 | 87 | 141 | 6 | MOLECULE: | HEMOGLOBIN | SUBUNIT | ALPHA-1/2 |
| 22:00 | 3gqp-A | 4.3 | 3.2 | 85 | 141 | 9 | MOLECULE: | HEMOGLOBIN | SUBUNIT | ALPHA |
| 23:00 | 1ye2-B | 4.3 | 3.5 | 86 | 146 | 5 | MOLECULE: | HEMOGLOBIN | ALPHA C | HAIN |
| 24:00:00 | 4yu3-A | 4.3 | 3.6 | 85 | 141 | 7 | MOLECULE: | HEMOGLOBIN | | |
| 25:00:00 | 1yg5-B | 4.3 | 3.5 | 86 | 146 | 5 | MOLECULE: | HEMOGLOBIN | ALPHA C | HAIN |
| 26:00:00 | 2dhb-A | 4.3 | 2.9 | 83 | 141 | 6 | MOLECULE: | HEMOGLOBIN | (DEOXY) | (ALPHA CHAIN) |
| 27:00:00 | 3mjp-A | 4.3 | 3.4 | 87 | 141 | 6 | MOLECULE: | HEMOGLOBIN | SUBUNIT | ALPHA-A |
| 28:00:00 | 2pgh-A | 4.3 | 3.1 | 84 | 141 | 7 | MOLECULE: | HEMOGLOBIN | (AQUO M | ET) (ALPHA CHAIN) |
| 29:00:00 | 1dxu-D | 4.3 | 3.6 | 86 | 146 | 5 | MOLECULE: | HEMOGLOBIN | (DEOXY) | (ALPHA CHAIN) |
| 30:00:00 | 1y8h-C | 4.3 | 3 | 83 | 141 | 7 | MOLECULE: | HEMOGLOBIN | ALPHA C | HAINS |
| 31:00:00 | 1y4f-D | 4.3 | 3.6 | 86 | 146 | 5 | MOLECULE: | HEMOGLOBIN | ALPHA C | HAIN |
| 32:00:00 | 2ri4-A | 4.3 | 3 | 84 | 139 | 7 | MOLECULE: | HEMOGLOBIN | SUBUNIT | ALPHA-1/2 |
| 33:00:00 | 1o1l-D | 4.3 | 3.6 | 86 | 146 | 5 | MOLECULE: | HEMOGLOBIN | ALPHA C | HAIN |
| 34:00:00 | 1s0h-A | 4.3 | 3.1 | 84 | 141 | 8 | MOLECULE: | HEMOGLOBIN | ALPHA C | HAIN |
| 35:00:00 | 1fdh-G | 4.3 | 3.6 | 86 | 146 | 5 | MOLECULE: | HEMOGLOBIN | F (DEOX | Y) (ALPHA CHAIN) |
| 36:00:00 | 1hbs-C | 4.3 | 3.1 | 85 | 141 | 6 | MOLECULE: | HEMOGLOBIN | S (DEOX | Y) (ALPHA CHAIN) |
| 37:00:00 | 1y4p-B | 4.3 | 3.5 | 86 | 146 | 5 | MOLECULE: | HEMOGLOBIN | ALPHA C | HAIN |
| 38:00:00 | 1dxc-A | 4.3 | 4.7 | 92 | 154 | 9 | MOLECULE: | MYOGLOBIN | | |
| 39:00:00 | 1yie-D | 4.3 | 3.5 | 85 | 146 | 6 | MOLECULE: | HEMOGLOBIN | ALPHA C | HAIN |
| 40:00:00 | 1a9w-F | 4.3 | 3.5 | 85 | 145 | 5 | MOLECULE: | HEMOGLOBIN | (ALPHA | CHAIN) |
| 41:00:00 | 6zmx-B | 4.3 | 3.5 | 85 | 146 | 8 | MOLECULE: | HEMOGLOBIN | SUBUNIT | ALPHA-A |
| 42:00:00 | 1a9w-E | 4.3 | 3.5 | 85 | 145 | 5 | MOLECULE: | HEMOGLOBIN | (ALPHA | CHAIN) |
| 43:00:00 | 3wr1-B | 4.3 | 3.5 | 86 | 146 | 9 | MOLECULE: | HEMOGLOBIN | SUBUNIT | ALPHA-A |
| 44:00:00 | 1naz-A | 4.3 | 4.7 | 92 | 154 | 9 | MOLECULE: | MYOGLOBIN | | |
| 45:00:00 | 3hf4-E | 4.3 | 3 | 84 | 141 | 7 | MOLECULE: | HEMOGLOBIN | SUBUNIT | ALPHA-1/2 |
| 46:00:00 | 1yev-D | 4.3 | 3.5 | 86 | 146 | 6 | MOLECULE: | HEMOGLOBIN | ALPHA C | HAIN |
| 47:00:00 | 3nml-A | 4.3 | 4.5 | 91 | 154 | 9 | MOLECULE: | MYOGLOBIN | | |
| 48:00:00 | 1mdn-B | 4.3 | 4.3 | 89 | 153 | 16 | MOLECULE: | PROTEIN (M | YOGLOBIN | ) |

| | | | | | | | | | | |
|---|---|---|---|---|---|---|---|---|---|---|
| 49:00:00 | 1yeo-D | 4.3 | 3.5 | 86 | 146 | 6 | MOLECULE: | HEMOGLOBIN | ALPHA C | HAIN |
| 50:00:00 | 1gli-D | 4.3 | 3.3 | 85 | 146 | 6 | MOLECULE: | DEOXYHEMOG | LOBIN | |
| 51:00:00 | 5me2-C | 4.3 | 2.9 | 83 | 140 | 7 | MOLECULE: | HEMOGLOBIN | SUBUNIT | ALPHA |
| 52:00:00 | 4hrr-G | 4.3 | 3.3 | 87 | 149 | 3 | MOLECULE: | GLOBIN-2 A | CHAIN | |
| 53:00:00 | 1yie-B | 4.3 | 3.4 | 84 | 146 | 6 | MOLECULE: | HEMOGLOBIN | ALPHA C | HAIN |
| 54:00:00 | 1hbr-D | 4.3 | 3.6 | 86 | 146 | 8 | MOLECULE: | PROTEIN (H | EMOGLOBI | N D) |
| 55:00:00 | 1yg5-D | 4.3 | 3.5 | 86 | 146 | 6 | MOLECULE: | HEMOGLOBIN | ALPHA C | HAIN |
| 56:00:00 | 2hbs-B | 4.3 | 3.1 | 83 | 146 | 6 | MOLECULE: | HEMOGLOBIN | S (DEOX | Y), ALPHA CHAIN |
| 57:00:00 | 1y4v-D | 4.3 | 3.5 | 86 | 146 | 6 | MOLECULE: | HEMOGLOBIN | ALPHA C | HAIN |
| 58:00:00 | 1o1m-A | 4.3 | 9.7 | 102 | 282 | 9 | MOLECULE: | HEMOGLOBIN | ALPHA C | HAIN |
| 59:00:00 | 2cmm-A | 4.3 | 4.4 | 91 | 153 | 9 | MOLECULE: | MYOGLOBIN | | |
| 60:00:00 | 3k8b-D | 4.3 | 3.5 | 86 | 146 | 8 | MOLECULE: | HEMOGLOBIN | SUBUNIT | ALPHA-A |
| 61:00:00 | 5me2-A | 4.3 | 2.9 | 83 | 140 | 7 | MOLECULE: | HEMOGLOBIN | SUBUNIT | ALPHA |
| 62:00:00 | 1s5y-A | 4.3 | 3.5 | 85 | 142 | 6 | MOLECULE: | HEMOGLOBIN | ALPHA C | HAIN |
| 63:00:00 | 1yeq-D | 4.3 | 3.5 | 86 | 146 | 6 | MOLECULE: | HEMOGLOBIN | ALPHA C | HAIN |
| 64:00:00 | 1dxt-B | 4.3 | 3.1 | 83 | 147 | 6 | MOLECULE: | HEMOGLOBIN | (DEOXY) | (ALPHA CHAIN) |
| 65:00:00 | 8puq-A | 4.3 | 3.2 | 85 | 139 | 6 | MOLECULE: | HEMOGLOBIN | SUBUNIT | ALPHA |
| 66:00:00 | 1y0t-D | 4.3 | 3.5 | 86 | 146 | 6 | MOLECULE: | HEMOGLOBIN | ALPHA C | HAIN |
| 67:00:00 | 1hbs-H | 4.3 | 3.4 | 86 | 146 | 6 | MOLECULE: | HEMOGLOBIN | S (DEOX | Y) (ALPHA CHAIN) |
| 68:00:00 | 1co8-A | 4.3 | 4.5 | 92 | 154 | 8 | MOLECULE: | PROTEIN (M | YOGLOBIN | ) |
| 69:00:00 | 1y0t-B | 4.3 | 3 | 83 | 146 | 6 | MOLECULE: | HEMOGLOBIN | ALPHA C | HAIN |
| 70:00:00 | 1y01-B | 4.3 | 3.2 | 85 | 126 | 7 | MOLECULE: | ALPHA-HEMO | GLOBIN S | TABILIZING PROTEIN |
| 71:00:00 | 2hbs-F | 4.3 | 2.9 | 82 | 146 | 6 | MOLECULE: | HEMOGLOBIN | S (DEOX | Y), ALPHA CHAIN |
| 72:00:00 | 2r80-B | 4.3 | 3.6 | 86 | 146 | 8 | MOLECULE: | HEMOGLOBIN | SUBUNIT | ALPHA-A |
| 73:00:00 | 1y4b-B | 4.3 | 3.2 | 85 | 146 | 6 | MOLECULE: | HEMOGLOBIN | ALPHA C | HAIN |
| 74:00:00 | 3pi1-A | 4.3 | 3.9 | 86 | 151 | 7 | MOLECULE: | HEMOGLOBIN | II | |
| 75:00:00 | 1s5x-A | 4.3 | 3.1 | 84 | 142 | 6 | MOLECULE: | HEMOGLOBIN | ALPHA C | HAIN |
| 76:00:00 | 5x2r-E | 4.3 | 3.1 | 84 | 140 | 7 | MOLECULE: | HEMOGLOBIN | SUBUNIT | ALPHA |
| 77:00:00 | 3ciu-C | 4.3 | 3 | 83 | 141 | 7 | MOLECULE: | HEMOGLOBIN | SUBUNIT | ALPHA |
| 78:00:00 | 1ygd-B | 4.3 | 3.5 | 86 | 146 | 6 | MOLECULE: | HEMOGLOBIN | ALPHA C | HAIN |
| 79:00:00 | 1y2z-B | 4.3 | 3.1 | 83 | 146 | 6 | MOLECULE: | HEMOGLOBIN | ALPHA C | HAIN |
| 80:00:00 | 2qmb-B | 4.3 | 3.5 | 85 | 146 | 8 | MOLECULE: | HEMOGLOBIN | SUBUNIT | ALPHA-A |
| 81:00:00 | 1gli-B | 4.3 | 3.3 | 85 | 146 | 6 | MOLECULE: | DEOXYHEMOG | LOBIN | |
| 82:00:00 | 2rao-C | 4.3 | 3 | 83 | 141 | 8 | MOLECULE: | HEMOGLOBIN | SUBUNIT | ALPHA-1/2 |
| 83:00:00 | 1y35-B | 4.3 | 3 | 83 | 146 | 6 | MOLECULE: | HEMOGLOBIN | ALPHA C | HAIN |
| 84:00:00 | 1y46-D | 4.3 | 3.2 | 85 | 146 | 6 | MOLECULE: | HEMOGLOBIN | ALPHA C | HAIN |
| 85:00:00 | 1y7c-D | 4.3 | 3.5 | 86 | 146 | 6 | MOLECULE: | HEMOGLOBIN | ALPHA C | HAIN |
| 86:00:00 | 1a0u-D | 4.3 | 3.5 | 86 | 146 | 6 | MOLECULE: | HEMOGLOBIN | (ALPHA | CHAIN) |
| 87:00:00 | 5x2t-A | 4.3 | 3.5 | 85 | 140 | 7 | MOLECULE: | HEMOGLOBIN | SUBUNIT | ALPHA |
| 88:00:00 | 2olp-A | 4.3 | 3.9 | 84 | 151 | 6 | MOLECULE: | HEMOGLOBIN | II | |
| 89:00:00 | 1y4v-B | 4.3 | 3 | 83 | 146 | 6 | MOLECULE: | HEMOGLOBIN | ALPHA C | HAIN |
| 90:00:00 | 3pi3-A | 4.3 | 3.9 | 85 | 151 | 7 | MOLECULE: | HEMOGLOBIN | II | |
| 91:00:00 | 3at5-B | 4.3 | 3.5 | 86 | 146 | 9 | MOLECULE: | ALPHAA-GLO | BIN | |
| 92:00:00 | 1a4f-B | 4.3 | 3.5 | 85 | 146 | 9 | MOLECULE: | HEMOGLOBIN | (ALPHA | CHAIN) |
| 93:00:00 | 7wb4-b | 4.2 | 24 | 137 | 636 | 4 | MOLECULE: | OUTER NUP1 | 33 | |
| 94:00:00 | 8fef-I | 4.2 | 10.3 | 97 | 253 | 6 | MOLECULE: | VIRULENCE | FACTOR M | CE FAMILY PROTEIN |
| 95:00:00 | 6u8y-b | 4.2 | 6.2 | 70 | 79 | 7 | MOLECULE: | MONOVALENT | CATION/ | H+ ANTIPORTER SUBUNIT E |
| 96:00:00 | 3hen-A | 4.2 | 4.2 | 90 | 153 | 12 | MOLECULE: | MYOGLOBIN | | |
| 97:00:00 | 1ebt-A | 4.2 | 3.4 | 78 | 142 | 5 | MOLECULE: | HEMOGLOBIN | | |
| 98:00:00 | 1hv4-A | 4.2 | 3.6 | 85 | 141 | 7 | MOLECULE: | HEMOGLOBIN | ALPHA-A | CHAIN |
| 99:00:00 | 7vuc-A | 4.2 | 4.4 | 91 | 153 | 9 | MOLECULE: | MYOGLOBIN | | |
| 0:00 | 2in4-A | 4.2 | 4.1 | 90 | 153 | 12 | MOLECULE: | MYOGLOBIN | | |
| 1:00 | 3hep-A | 4.2 | 4.3 | 90 | 153 | 14 | MOLECULE: | MYOGLOBIN | | |
| 2:00 | 5ojc-A | 4.2 | 4.2 | 91 | 154 | 9 | MOLECULE: | MYOGLOBIN | | |

| | | | | | | | | | | |
|---|---|---|---|---|---|---|---|---|---|---|
| 3:00 | 1hv4-C | 4.2 | 3.6 | 85 | 141 | 7 | MOLECULE: | HEMOGLOBIN | ALPHA-A | CHAIN |
| 4:00 | 4pnj-A | 4.2 | 4.4 | 91 | 154 | 9 | MOLECULE: | MYOGLOBIN | | |
| 5:00 | 3rjn-B | 4.2 | 4.2 | 90 | 152 | 12 | MOLECULE: | MYOGLOBIN | | |
| 6:00 | 3u3e-A | 4.2 | 4.4 | 90 | 153 | 10 | MOLECULE: | MYOGLOBIN | | |
| 7:00 | 3heo-A | 4.2 | 4.2 | 90 | 153 | 14 | MOLECULE: | MYOGLOBIN | | |
| 8:00 | 6jp1-A | 4.2 | 4.5 | 91 | 151 | 9 | MOLECULE: | MYOGLOBIN | | |
| 9:00 | 3g52-A | 4.2 | 3.3 | 85 | 145 | 8 | MOLECULE: | GLOBIN-1 | | |
| 10:00 | 7ylk-A | 4.2 | 4.6 | 91 | 153 | 8 | MOLECULE: | MYOGLOBIN | | |
| 11:00 | 1jp8-A | 4.2 | 3.9 | 88 | 152 | 9 | MOLECULE: | MYOGLOBIN | | |
| 12:00 | 1y35-D | 4.2 | 3.6 | 86 | 146 | 5 | MOLECULE: | HEMOGLOBIN | ALPHA C | HAIN |
| 13:00 | 5urc-A | 4.2 | 4.2 | 87 | 141 | 7 | MOLECULE: | HEMOGLOBIN | SUBUNIT | ALPHA |
| 14:00 | 3hc9-A | 4.2 | 4.3 | 90 | 153 | 14 | MOLECULE: | MYOGLOBIN | | |
| 15:00 | 1gbu-B | 4.2 | 3.6 | 86 | 146 | 5 | MOLECULE: | HEMOGLOBIN | | |
| 16:00 | 1y0w-D | 4.2 | 3.6 | 86 | 146 | 5 | MOLECULE: | HEMOGLOBIN | ALPHA C | HAIN |
| 17:00 | 1a00-B | 4.2 | 3.5 | 84 | 146 | 5 | MOLECULE: | HEMOGLOBIN | (ALPHA | CHAIN) |
| 18:00 | 6hbi-B | 4.2 | 3.3 | 85 | 145 | 8 | MOLECULE: | HEMOGLOBIN | | |
| 19:00 | 3hrw-A | 4.2 | 3.1 | 85 | 141 | 7 | MOLECULE: | HEMOGLOBIN | SUBUNIT | ALPHA |
| 20:00 | 3k8b-A | 4.2 | 3.5 | 85 | 141 | 6 | MOLECULE: | HEMOGLOBIN | SUBUNIT | ALPHA-A |
| 21:00 | 3gdj-C | 4.2 | 3.6 | 86 | 141 | 7 | MOLECULE: | HEMOGLOBIN | SUBUNIT | ALPHA |
| 22:00 | 4sdh-B | 4.2 | 3.3 | 84 | 145 | 11 | MOLECULE: | HEMOGLOBIN | I (DEOX | Y) |
| 23:00 | 3uhs-B | 4.2 | 3.2 | 84 | 145 | 11 | MOLECULE: | GLOBIN-1 | | |
| 24:00:00 | 2zlx-A | 4.2 | 3.2 | 85 | 140 | 6 | MOLECULE: | HEMOGLOBIN | SUBUNIT | ALPHA |
| 25:00:00 | 2zlw-C | 4.2 | 3.1 | 83 | 141 | 7 | MOLECULE: | HEMOGLOBIN | SUBUNIT | ALPHA |
| 26:00:00 | 1nwn-B | 4.2 | 3.3 | 85 | 145 | 9 | MOLECULE: | GLOBIN I | | |
| 27:00:00 | 3ui0-B | 4.2 | 3.4 | 84 | 145 | 11 | MOLECULE: | GLOBIN-1 | | |
| 28:00:00 | 1jp9-A | 4.2 | 4 | 90 | 151 | 9 | MOLECULE: | MYOGLOBIN | | |
| 29:00:00 | 1a6k-A | 4.2 | 4.2 | 89 | 151 | 9 | MOLECULE: | MYOGLOBIN | | |
| 30:00:00 | 3uhw-B | 4.2 | 3.3 | 85 | 145 | 8 | MOLECULE: | GLOBIN-1 | | |
| 31:00:00 | 3uhx-B | 4.2 | 3.3 | 85 | 145 | 8 | MOLECULE: | GLOBIN-1 | | |
| 32:00:00 | 3lqd-A | 4.2 | 3 | 83 | 141 | 8 | MOLECULE: | HEMOGLOBIN | SUBUNIT | ALPHA |
| 33:00:00 | 1hba-B | 4.2 | 3.4 | 84 | 146 | 5 | MOLECULE: | HEMOGLOBIN | ROTHSCH | ILD (DEOXY) (ALPHA CHAIN) |
| 34:00:00 | 4hrt-A | 4.2 | 3.7 | 88 | 149 | 5 | MOLECULE: | GLOBIN-2 A | CHAIN | |
| 35:00:00 | 2zfb-A | 4.2 | 3.1 | 85 | 141 | 8 | MOLECULE: | HEMOGLOBIN | SUBUNIT | ALPHA |
| 36:00:00 | 1gbu-D | 4.2 | 3.5 | 84 | 146 | 5 | MOLECULE: | HEMOGLOBIN | | |
| 37:00:00 | 3g4y-A | 4.2 | 3.4 | 86 | 145 | 9 | MOLECULE: | GLOBIN-1 | | |
| 38:00:00 | 1hba-D | 4.2 | 3.5 | 84 | 146 | 5 | MOLECULE: | HEMOGLOBIN | ROTHSCH | ILD (DEOXY) (ALPHA CHAIN) |
| 39:00:00 | 4hbi-B | 4.2 | 3.3 | 84 | 145 | 11 | MOLECULE: | HEMOGLOBIN | | |
| 40:00:00 | 3uhv-B | 4.2 | 3.3 | 84 | 145 | 11 | MOLECULE: | GLOBIN-1 | | |
| 41:00:00 | 6jp1-B | 4.2 | 4.2 | 89 | 151 | 10 | MOLECULE: | MYOGLOBIN | | |
| 42:00:00 | 6z4r-A | 4.2 | 4.6 | 91 | 154 | 9 | MOLECULE: | MYOGLOBIN | | |
| 43:00:00 | 3uhr-B | 4.2 | 3.4 | 84 | 145 | 11 | MOLECULE: | GLOBIN-1 | | |
| 44:00:00 | 3o89-A | 4.2 | 4.7 | 91 | 153 | 10 | MOLECULE: | MYOGLOBIN | | |
| 45:00:00 | 3vre-B | 4.2 | 3.3 | 84 | 146 | 7 | MOLECULE: | HEMOGLOBIN | SUBUNIT | ALPHA |
| 46:00:00 | 1jzk-B | 4.2 | 3.3 | 84 | 145 | 12 | MOLECULE: | GLOBIN I - | ARK SHE | LL |
| 47:00:00 | 5jgg-B | 4.2 | 3.7 | 85 | 144 | 7 | MOLECULE: | ALPHA CHAI | N | |
| 48:00:00 | 3uhi-A | 4.2 | 3.3 | 84 | 145 | 11 | MOLECULE: | GLOBIN-1 | | |
| 49:00:00 | 1yeq-B | 4.2 | 3.3 | 85 | 146 | 6 | MOLECULE: | HEMOGLOBIN | ALPHA C | HAIN |
| 50:00:00 | 1jzm-B | 4.2 | 3.3 | 84 | 145 | 11 | MOLECULE: | GLOBIN I - | ARK SHE | LL |
| 51:00:00 | 2grf-B | 4.2 | 3.1 | 83 | 145 | 11 | MOLECULE: | GLOBIN-1 | | |
| 52:00:00 | 3uhz-B | 4.2 | 3.3 | 84 | 145 | 11 | MOLECULE: | GLOBIN-1 | | |
| 53:00:00 | 2zfb-B | 4.2 | 3.6 | 85 | 146 | 8 | MOLECULE: | HEMOGLOBIN | SUBUNIT | ALPHA |
| 54:00:00 | 3uhn-B | 4.2 | 3.4 | 84 | 145 | 11 | MOLECULE: | GLOBIN-1 | | |
| 55:00:00 | 3uht-B | 4.2 | 3.4 | 84 | 145 | 12 | MOLECULE: | GLOBIN-1 | | |
| 56:00:00 | 7xgy-C | 4.2 | 3.5 | 84 | 141 | 7 | MOLECULE: | HEMOGLOBIN | SUBUNIT | ALPHA |

| | | | | | | | | | | |
|---|---|---|---|---|---|---|---|---|---|---|
| 57:00:00 | 7dy4-B | 4.2 | 3.2 | 84 | 146 | 6 | MOLECULE: | HEMOGLOBIN | SUBUNIT | ALPHA |
| 58:00:00 | 7xgy-A | 4.2 | 3.6 | 84 | 141 | 7 | MOLECULE: | HEMOGLOBIN | SUBUNIT | ALPHA |
| 59:00:00 | 3eok-B | 4.2 | 3.5 | 85 | 146 | 8 | MOLECULE: | HEMOGLOBIN | SUBUNIT | ALPHA-A |
| 60:00:00 | 1jzk-A | 4.2 | 3.3 | 85 | 145 | 9 | MOLECULE: | GLOBIN I - | ARK SHE | LL |
| 61:00:00 | 7pcq-B | 4.2 | 3.5 | 84 | 146 | 6 | MOLECULE: | HEMOGLOBIN | SUBUNIT | ALPHA |
| 62:00:00 | 1hv4-H | 4.2 | 3.5 | 85 | 146 | 9 | MOLECULE: | HEMOGLOBIN | ALPHA-A | CHAIN |
| 63:00:00 | 1nwi-A | 4.2 | 3.3 | 85 | 145 | 8 | MOLECULE: | GLOBIN I | | |
| 64:00:00 | 1b0b-A | 4.2 | 3.4 | 78 | 142 | 5 | MOLECULE: | HEMOGLOBIN | | |
| 65:00:00 | 3uhy-B | 4.2 | 3.3 | 84 | 145 | 11 | MOLECULE: | GLOBIN-1 | | |
| 66:00:00 | 4iro-A | 4.2 | 3.3 | 85 | 142 | 6 | MOLECULE: | HEMOGLOBIN | SUBUNIT | ALPHA |
| 67:00:00 | 3g4q-A | 4.2 | 3.4 | 85 | 145 | 8 | MOLECULE: | GLOBIN-1 | | |
| 68:00:00 | 1y0w-B | 4.2 | 3.1 | 83 | 146 | 6 | MOLECULE: | HEMOGLOBIN | ALPHA C | HAIN |
| 69:00:00 | 4hrr-C | 4.2 | 3.7 | 87 | 149 | 3 | MOLECULE: | GLOBIN-2 A | CHAIN | |
| 70:00:00 | 7pch-B | 4.2 | 3.4 | 84 | 146 | 6 | MOLECULE: | HEMOGLOBIN | SUBUNIT | ALPHA |
| 71:00:00 | 3a59-F | 4.2 | 3.5 | 85 | 146 | 9 | MOLECULE: | HEMOGLOBIN | SUBUNIT | ALPHA-A |
| 72:00:00 | 1y7c-B | 4.2 | 3.4 | 84 | 146 | 6 | MOLECULE: | HEMOGLOBIN | ALPHA C | HAIN |
| 73:00:00 | 1hv4-B | 4.2 | 3.5 | 85 | 146 | 9 | MOLECULE: | HEMOGLOBIN | ALPHA-A | CHAIN |
| 74:00:00 | 1hv4-F | 4.2 | 3.6 | 86 | 146 | 9 | MOLECULE: | HEMOGLOBIN | ALPHA-A | CHAIN |
| 75:00:00 | 7dy4-F | 4.2 | 2.9 | 82 | 146 | 6 | MOLECULE: | HEMOGLOBIN | SUBUNIT | ALPHA |
| 76:00:00 | 3fs4-D | 4.2 | 3.5 | 85 | 146 | 9 | MOLECULE: | HEMOGLOBIN | SUBUNIT | ALPHA-A |
| 77:00:00 | 1i3e-B | 4.2 | 3.5 | 86 | 146 | 6 | MOLECULE: | HEMOGLOBIN | GAMMA C | HAINS |
| 78:00:00 | 1hds-A | 4.2 | 3.4 | 85 | 141 | 6 | MOLECULE: | HEMOGLOBIN | S (DEOX | Y) (ALPHA CHAIN) |
| 79:00:00 | 4ij2-A | 4.2 | 3.5 | 84 | 138 | 5 | MOLECULE: | HEMOGLOBIN | SUBUNIT | ALPHA |
| 80:00:00 | 7dy3-F | 4.2 | 3.1 | 83 | 146 | 6 | MOLECULE: | HEMOGLOBIN | SUBUNIT | ALPHA |
| 81:00:00 | 3mju-B | 4.2 | 3.5 | 85 | 146 | 8 | MOLECULE: | HEMOGLOBIN | SUBUNIT | ALPHA-A |
| 82:00:00 | 3bcq-B | 4.2 | 3.6 | 86 | 146 | 7 | MOLECULE: | ALPHA-CHAI | N HEMOGL | OBIN |
| 83:00:00 | 8egi-D | 4.2 | 3.6 | 86 | 146 | 6 | MOLECULE: | HEMOGLOBIN | SUBUNIT | ALPHA |
| 84:00:00 | 3uhu-B | 4.2 | 3.4 | 84 | 145 | 11 | MOLECULE: | GLOBIN-1 | | |
| 85:00:00 | 3uhk-A | 4.2 | 3.2 | 84 | 145 | 8 | MOLECULE: | GLOBIN-1 | | |
| 86:00:00 | 4ij2-C | 4.2 | 3.5 | 84 | 138 | 5 | MOLECULE: | HEMOGLOBIN | SUBUNIT | ALPHA |
| 87:00:00 | 3g4v-A | 4.2 | 3.3 | 84 | 145 | 11 | MOLECULE: | GLOBIN-1 | | |
| 88:00:00 | 6xdt-D | 4.2 | 3.5 | 86 | 146 | 6 | MOLECULE: | HEMOGLOBIN | SUBUNIT | ALPHA |
| 89:00:00 | 3gqp-B | 4.2 | 3.5 | 85 | 145 | 7 | MOLECULE: | HEMOGLOBIN | SUBUNIT | ALPHA |
| 90:00:00 | 7dy4-D | 4.2 | 3 | 82 | 146 | 6 | MOLECULE: | HEMOGLOBIN | SUBUNIT | ALPHA |
| 91:00:00 | 3ovu-C | 4.2 | 3.6 | 86 | 139 | 7 | MOLECULE: | ALPHA-HEMO | GLOBIN-S | TABILIZING PROTEIN |
| 92:00:00 | 7dds-A | 4.2 | 4.5 | 92 | 154 | 12 | MOLECULE: | ANCESTRAL | MYOGLOBI | N AMBSP |
| 93:00:00 | 2zlw-A | 4.2 | 3 | 83 | 141 | 7 | MOLECULE: | HEMOGLOBIN | SUBUNIT | ALPHA |
| 94:00:00 | 3vre-D | 4.2 | 3.5 | 84 | 146 | 7 | MOLECULE: | HEMOGLOBIN | SUBUNIT | ALPHA |
| 95:00:00 | 3dhr-D | 4.2 | 3.6 | 85 | 145 | 8 | MOLECULE: | HEMOGLOBIN | SUBUNIT | ALPHA-A |
| 96:00:00 | 5x2s-C | 4.2 | 3.1 | 84 | 140 | 6 | MOLECULE: | HEMOGLOBIN | SUBUNIT | ALPHA |
| 97:00:00 | 1a0z-B | 4.2 | 3.1 | 83 | 146 | 6 | MOLECULE: | HEMOGLOBIN | (ALPHA | CHAIN) |
| 98:00:00 | 3uhk-D | 4.2 | 3.3 | 84 | 145 | 11 | MOLECULE: | GLOBIN-1 | | |
| 99:00:00 | 1hlb-A | 4.1 | 4.4 | 91 | 157 | 11 | MOLECULE: | HEMOGLOBIN | (DEOXY) | |
| 0:00 | 6wb9-6 | 4.1 | 3.8 | 74 | 98 | 4 | MOLECULE: | ENDOPLASMI | C RETICU | LUM MEMBRANE PROTEIN COMPLEX SU |
| 1:00 | 7ol3-B | 4.1 | 12.6 | 88 | 425 | 5 | MOLECULE: | ATLASTIN-1 | | |
| 2:00 | 1mwc-B | 4.1 | 4.3 | 90 | 153 | 13 | MOLECULE: | PROTEIN (M | YOGLOBIN | ) |
| 3:00 | 7ddr-A | 4.1 | 4.3 | 91 | 154 | 13 | MOLECULE: | ANCESTRAL | MYOGLOBI | N AMBSP |
| 4:00 | 1m6m-B | 4.1 | 4.3 | 90 | 153 | 13 | MOLECULE: | PROTEIN (M | YOGLOBIN | ) |
| 5:00 | 4sdh-A | 4.1 | 3.3 | 84 | 145 | 8 | MOLECULE: | HEMOGLOBIN | I (DEOX | Y) |
| 6:00 | 1hv4-G | 4.1 | 3.6 | 85 | 141 | 7 | MOLECULE: | HEMOGLOBIN | ALPHA-A | CHAIN |
| 7:00 | 4hrr-B | 4.1 | 3.9 | 88 | 151 | 9 | MOLECULE: | GLOBIN-2 A | CHAIN | |
| 8:00 | 1hbi-A | 4.1 | 3.3 | 84 | 145 | 8 | MOLECULE: | HEMOGLOBIN | (OXY) | |
| 9:00 | 2z85-A | 4.1 | 3.3 | 84 | 145 | 8 | MOLECULE: | GLOBIN-1 | | |
| 10:00 | 3g53-A | 4.1 | 3.3 | 85 | 145 | 9 | MOLECULE: | GLOBIN-1 | | |

| | | | | | | | | | | |
|---|---|---|---|---|---|---|---|---|---|---|
| 11:00 | 3uhz-A | 4.1 | 3.3 | 84 | 145 | 8 | MOLECULE: | GLOBIN-1 | | |
| 12:00 | 3g4r-A | 4.1 | 3.3 | 84 | 145 | 8 | MOLECULE: | GLOBIN-1 | | |
| 13:00 | 1nwi-D | 4.1 | 3.3 | 84 | 145 | 8 | MOLECULE: | GLOBIN I | | |
| 14:00 | 3lqd-C | 4.1 | 3 | 83 | 141 | 8 | MOLECULE: | HEMOGLOBIN | SUBUNIT | ALPHA |
| 15:00 | 2qmb-C | 4.1 | 3.5 | 85 | 141 | 6 | MOLECULE: | HEMOGLOBIN | SUBUNIT | ALPHA-A |
| 16:00 | 3uhw-A | 4.1 | 3.3 | 84 | 145 | 11 | MOLECULE: | GLOBIN-1 | | |
| 17:00 | 4hrt-H | 4.1 | 3.4 | 87 | 148 | 9 | MOLECULE: | GLOBIN-2 A | CHAIN | |
| 18:00 | 1nwi-C | 4.1 | 3.2 | 84 | 145 | 8 | MOLECULE: | GLOBIN I | | |
| 19:00 | 3g4u-A | 4.1 | 3.3 | 85 | 145 | 9 | MOLECULE: | GLOBIN-1 | | |
| 20:00 | 3ugy-A | 4.1 | 3.3 | 85 | 145 | 9 | MOLECULE: | GLOBIN-1 | | |
| 21:00 | 3uhe-A | 4.1 | 3.3 | 84 | 145 | 8 | MOLECULE: | GLOBIN-1 | | |
| 22:00 | 2auq-A | 4.1 | 3.3 | 84 | 145 | 8 | MOLECULE: | GLOBIN I | | |
| 23:00 | 3uh7-A | 4.1 | 3.3 | 85 | 145 | 9 | MOLECULE: | GLOBIN-1 | | |
| 24:00:00 | 2r4x-B | 4.1 | 3.3 | 84 | 145 | 8 | MOLECULE: | GLOBIN-1 | | |
| 25:00:00 | 2r4y-B | 4.1 | 3.3 | 84 | 145 | 8 | MOLECULE: | GLOBIN-1 | | |
| 26:00:00 | 1hv4-E | 4.1 | 3.6 | 85 | 141 | 7 | MOLECULE: | HEMOGLOBIN | ALPHA-A | CHAIN |
| 27:00:00 | 6hbi-A | 4.1 | 3.3 | 84 | 145 | 8 | MOLECULE: | HEMOGLOBIN | | |
| 28:00:00 | 3uhv-A | 4.1 | 3.3 | 84 | 145 | 8 | MOLECULE: | GLOBIN-1 | | |
| 29:00:00 | 4n7o-C | 4.1 | 3 | 83 | 141 | 6 | MOLECULE: | HEMOGLOBIN | SUBUNIT | ALPHA |
| 30:00:00 | 3ui0-A | 4.1 | 3.3 | 84 | 145 | 8 | MOLECULE: | GLOBIN-1 | | |
| 31:00:00 | 1nxf-B | 4.1 | 3.5 | 85 | 145 | 9 | MOLECULE: | GLOBIN I | | |
| 32:00:00 | 3uhg-A | 4.1 | 3.3 | 84 | 145 | 8 | MOLECULE: | GLOBIN-1 | | |
| 33:00:00 | 3uhn-A | 4.1 | 3.3 | 84 | 145 | 8 | MOLECULE: | GLOBIN-1 | | |
| 34:00:00 | 3uhu-A | 4.1 | 3.3 | 84 | 145 | 8 | MOLECULE: | GLOBIN-1 | | |
| 35:00:00 | 3uhq-B | 4.1 | 3.3 | 84 | 145 | 8 | MOLECULE: | GLOBIN-1 | | |
| 36:00:00 | 2z85-B | 4.1 | 3.5 | 85 | 145 | 9 | MOLECULE: | GLOBIN-1 | | |
| 37:00:00 | 4hrr-F | 4.1 | 3.7 | 87 | 151 | 10 | MOLECULE: | GLOBIN-2 A | CHAIN | |
| 38:00:00 | 3g4w-A | 4.1 | 3.6 | 87 | 145 | 9 | MOLECULE: | GLOBIN-1 | | |
| 39:00:00 | 3sdh-A | 4.1 | 3.3 | 84 | 145 | 8 | MOLECULE: | HEMOGLOBIN | I (CARB | ONMONOXY) |
| 40:00:00 | 1faw-A | 4.1 | 3.6 | 85 | 141 | 7 | MOLECULE: | HEMOGLOBIN | (ALPHA | SUBUNIT) |
| 41:00:00 | 2av3-B | 4.1 | 3.5 | 85 | 145 | 8 | MOLECULE: | GLOBIN I | | |
| 42:00:00 | 2grf-A | 4.1 | 3.3 | 84 | 145 | 8 | MOLECULE: | GLOBIN-1 | | |
| 43:00:00 | 2z8a-A | 4.1 | 3.6 | 86 | 145 | 8 | MOLECULE: | GLOBIN-1 | | |
| 44:00:00 | 5xkv-A | 4.1 | 3.6 | 80 | 153 | 10 | MOLECULE: | MYOGLOBIN | | |
| 45:00:00 | 1jzk-D | 4.1 | 3.3 | 84 | 145 | 8 | MOLECULE: | GLOBIN I - | ARK SHE | LL |
| 46:00:00 | 2auo-A | 4.1 | 3.3 | 84 | 145 | 8 | MOLECULE: | GLOBIN I | | |
| 47:00:00 | 1jwn-A | 4.1 | 3.3 | 84 | 145 | 8 | MOLECULE: | GLOBIN I - | ARK SHE | LL |
| 48:00:00 | 3mjp-C | 4.1 | 3.9 | 88 | 141 | 2 | MOLECULE: | HEMOGLOBIN | SUBUNIT | ALPHA-A |
| 49:00:00 | 1jzm-A | 4.1 | 3.3 | 84 | 145 | 8 | MOLECULE: | GLOBIN I - | ARK SHE | LL |
| 50:00:00 | 3uhr-A | 4.1 | 3.2 | 82 | 145 | 11 | MOLECULE: | GLOBIN-1 | | |
| 51:00:00 | 2av0-A | 4.1 | 3.5 | 85 | 145 | 8 | MOLECULE: | GLOBIN I | | |
| 52:00:00 | 2aup-B | 4.1 | 3.4 | 84 | 145 | 11 | MOLECULE: | GLOBIN I | | |
| 53:00:00 | 3uhs-A | 4.1 | 3.3 | 84 | 145 | 8 | MOLECULE: | GLOBIN-1 | | |
| 54:00:00 | 4hbi-A | 4.1 | 3.3 | 84 | 145 | 8 | MOLECULE: | HEMOGLOBIN | | |
| 55:00:00 | 3qob-A | 4.1 | 3.4 | 85 | 145 | 8 | MOLECULE: | GLOBIN-1 | | |
| 56:00:00 | 3uh6-A | 4.1 | 3.3 | 85 | 145 | 9 | MOLECULE: | GLOBIN-1 | | |
| 57:00:00 | 3a59-D | 4.1 | 3.5 | 85 | 146 | 9 | MOLECULE: | HEMOGLOBIN | SUBUNIT | ALPHA-A |
| 58:00:00 | 2aur-B | 4.1 | 3.4 | 84 | 145 | 11 | MOLECULE: | GLOBIN I | | |
| 59:00:00 | 1jwn-B | 4.1 | 3.2 | 83 | 145 | 12 | MOLECULE: | GLOBIN I - | ARK SHE | LL |
| 60:00:00 | 3g46-B | 4.1 | 3.2 | 83 | 146 | 11 | MOLECULE: | GLOBIN-1 | | |
| 61:00:00 | 7xgy-B | 4.1 | 3.4 | 84 | 146 | 7 | MOLECULE: | HEMOGLOBIN | SUBUNIT | ALPHA |
| 62:00:00 | 2ri4-K | 4.1 | 3 | 84 | 138 | 6 | MOLECULE: | HEMOGLOBIN | SUBUNIT | ALPHA-1/2 |
| 63:00:00 | 3g4u-B | 4.1 | 3.4 | 84 | 145 | 8 | MOLECULE: | GLOBIN-1 | | |
| 64:00:00 | 1qpw-B | 4.1 | 3.6 | 86 | 146 | 5 | MOLECULE: | PORCINE HE | MOGLOBIN | (ALPHA SUBUNIT) |

| | | | | | | | | | | |
|---|---|---|---|---|---|---|---|---|---|---|
| 65:00:00 | 7xgy-D | 4.1 | 3.4 | 84 | 146 | 7 | MOLECULE: | HEMOGLOBIN | SUBUNIT | ALPHA |
| 66:00:00 | 3g4w-B | 4.1 | 3.4 | 85 | 145 | 8 | MOLECULE: | GLOBIN-1 | | |
| 67:00:00 | 7ud8-D | 4.1 | 3.5 | 84 | 146 | 6 | MOLECULE: | HEMOGLOBIN | SUBUNIT | ALPHA |
| 68:00:00 | 3uhk-C | 4.1 | 3.3 | 85 | 145 | 8 | MOLECULE: | GLOBIN-1 | | |
| 69:00:00 | 1fhj-B | 4.1 | 3.5 | 84 | 146 | 7 | MOLECULE: | HEMOGLOBIN | (ALPHA | CHAIN) |
| 70:00:00 | 2qmb-D | 4.1 | 3.5 | 85 | 146 | 8 | MOLECULE: | HEMOGLOBIN | SUBUNIT | ALPHA-A |
| 71:00:00 | 3uhy-A | 4.1 | 3.4 | 85 | 146 | 8 | MOLECULE: | GLOBIN-1 | | |
| 72:00:00 | 3fh9-B | 4.1 | 3.6 | 86 | 146 | 6 | MOLECULE: | HEMOGLOBIN | ALPHA C | HAIN |
| 73:00:00 | 5e83-D | 4.1 | 3.4 | 84 | 146 | 6 | MOLECULE: | HEMOGLOBIN | SUBUNIT | ALPHA |
| 74:00:00 | 2pgh-D | 4.1 | 3.5 | 85 | 146 | 5 | MOLECULE: | HEMOGLOBIN | (AQUO M | ET) (ALPHA CHAIN) |
| 75:00:00 | 8dov-B | 4.1 | 3.5 | 84 | 146 | 6 | MOLECULE: | HEMOGLOBIN | SUBUNIT | ALPHA |
| 76:00:00 | 8fdm-B | 4.1 | 3.4 | 84 | 146 | 6 | MOLECULE: | HEMOGLOBIN | SUBUNIT | ALPHA |
| 77:00:00 | 7dy3-H | 4.1 | 3.5 | 84 | 146 | 6 | MOLECULE: | HEMOGLOBIN | SUBUNIT | ALPHA |
| 78:00:00 | 5hbi-A | 4.1 | 3.4 | 84 | 145 | 8 | MOLECULE: | HEMOGLOBIN | | |
| 79:00:00 | 3gou-D | 4.1 | 3.5 | 84 | 146 | 7 | MOLECULE: | HEMOGLOBIN | SUBUNIT | ALPHA |
| 80:00:00 | 7hbi-A | 4.1 | 3.3 | 84 | 145 | 8 | MOLECULE: | HEMOGLOBIN | | |
| 81:00:00 | 3g46-A | 4.1 | 3.3 | 84 | 146 | 8 | MOLECULE: | GLOBIN-1 | | |
| 82:00:00 | 3uh5-A | 4.1 | 3.3 | 84 | 145 | 8 | MOLECULE: | GLOBIN-1 | | |
| 83:00:00 | 7ddu-C | 4.1 | 4.6 | 92 | 154 | 10 | MOLECULE: | MYOGLOBIN | | |
| 84:00:00 | 2r1h-B | 4.1 | 3.6 | 86 | 147 | 6 | MOLECULE: | HEMOGLOBIN | SUBUNIT | ALPHA-4 |
| 85:00:00 | 3uhb-A | 4.1 | 3.4 | 85 | 145 | 8 | MOLECULE: | GLOBIN-1 | | |
| 86:00:00 | 3uh3-A | 4.1 | 3.4 | 85 | 145 | 9 | MOLECULE: | GLOBIN-1 | | |
| 87:00:00 | 7jy1-D | 4.1 | 3.5 | 84 | 146 | 6 | MOLECULE: | HEMOGLOBIN | SUBUNIT | ALPHA |
| 88:00:00 | 2grh-A | 4.1 | 3.3 | 84 | 145 | 8 | MOLECULE: | GLOBIN-1 | | |
| 89:00:00 | 3uhq-A | 4.1 | 3.3 | 84 | 145 | 8 | MOLECULE: | GLOBIN-1 | | |
| 90:00:00 | 3hrw-C | 4.1 | 3.3 | 86 | 141 | 7 | MOLECULE: | HEMOGLOBIN | SUBUNIT | ALPHA |
| 91:00:00 | 6xe7-D | 4.1 | 3.6 | 86 | 146 | 6 | MOLECULE: | HEMOGLOBIN | SUBUNIT | ALPHA |
| 92:00:00 | 8fdm-D | 4.1 | 3.4 | 84 | 146 | 6 | MOLECULE: | HEMOGLOBIN | SUBUNIT | ALPHA |
| 93:00:00 | 5e6e-B | 4.1 | 3.4 | 84 | 146 | 6 | MOLECULE: | HEMOGLOBIN | SUBUNIT | ALPHA |
| 94:00:00 | 1jwn-D | 4.1 | 3.5 | 86 | 145 | 10 | MOLECULE: | GLOBIN I - | ARK SHE | LL |
| 95:00:00 | 3bcq-D | 4.1 | 3.6 | 87 | 146 | 7 | MOLECULE: | ALPHA-CHAI | N HEMOGL | OBIN |
| 96:00:00 | 3g4v-B | 4.1 | 3.4 | 84 | 145 | 8 | MOLECULE: | GLOBIN-1 | | |
| 97:00:00 | 7dy3-D | 4.1 | 3.5 | 84 | 146 | 6 | MOLECULE: | HEMOGLOBIN | SUBUNIT | ALPHA |
| 98:00:00 | 7jjq-D | 4.1 | 3.4 | 83 | 144 | 6 | MOLECULE: | HEMOGLOBIN | SUBUNIT | ALPHA |
| 99:00:00 | 5x2s-E | 4.1 | 3.2 | 83 | 140 | 7 | MOLECULE: | HEMOGLOBIN | SUBUNIT | ALPHA |
| 0:00 | 3uhx-A | 4.1 | 3.4 | 85 | 145 | 8 | MOLECULE: | GLOBIN-1 | | |
| 1:00 | 7rb4-A | 4 | 10.8 | 116 | 603 | 9 | MOLECULE: | DIPHTHERIA | TOXIN, | C DOMAIN |
| 2:00 | 5tsz-A | 4 | 4.8 | 86 | 130 | 9 | MOLECULE: | PV CELL-TR | AVERSAL | PROTEIN |
| 3:00 | 2peg-B | 4 | 3.9 | 84 | 139 | 6 | MOLECULE: | HEMOGLOBIN | SUBUNIT | ALPHA |
| 4:00 | 7ewp-A | 4 | 15.3 | 109 | 577 | 4 | MOLECULE: | PROBABLE G | #NAME? | COUPLED RECEPTOR 158 |
| 5:00 | 7uw5-A | 4 | 20.6 | 123 | 723 | 7 | MOLECULE: | MECHANOSEN | SITIVE C | HANNEL MSCK |
| 6:00 | 5o5e-A | 4 | 6.2 | 118 | 381 | 2 | MOLECULE: | UDP-N-ACET | YLGLUCOS | AMINE--DOLICHYL-PHOSPHATE N- |
| 7:00 | 6z16-d | 4 | 13.7 | 105 | 489 | 6 | MOLECULE: | MULTISUBUN | IT NA+/H | + ANTIPORTER, A SUBUNIT |
| 8:00 | 1nwi-B | 4 | 3.3 | 84 | 145 | 8 | MOLECULE: | GLOBIN I | | |
| 9:00 | 2auo-B | 4 | 3.5 | 85 | 145 | 8 | MOLECULE: | GLOBIN I | | |
| 10:00 | 3hf4-A | 4 | 4.3 | 86 | 141 | 7 | MOLECULE: | HEMOGLOBIN | SUBUNIT | ALPHA-1/2 |
| 11:00 | 1nxf-A | 4 | 3.3 | 84 | 145 | 8 | MOLECULE: | GLOBIN I | | |
| 12:00 | 3uhd-A | 4 | 3.3 | 85 | 145 | 9 | MOLECULE: | GLOBIN-1 | | |
| 13:00 | 3uhi-D | 4 | 3.3 | 84 | 145 | 8 | MOLECULE: | GLOBIN-1 | | |
| 14:00 | 3g4y-B | 4 | 3.5 | 85 | 145 | 8 | MOLECULE: | GLOBIN-1 | | |
| 15:00 | 1a0u-B | 4 | 3.7 | 83 | 146 | 5 | MOLECULE: | HEMOGLOBIN | (ALPHA | CHAIN) |
| 16:00 | 3g4q-B | 4 | 3.5 | 85 | 145 | 8 | MOLECULE: | GLOBIN-1 | | |
| 17:00 | 3uh7-B | 4 | 3.5 | 85 | 145 | 8 | MOLECULE: | GLOBIN-1 | | |
| 18:00 | 2auq-B | 4 | 3.5 | 85 | 145 | 8 | MOLECULE: | GLOBIN I | | |

| | | | | | | | | | | |
|---|---|---|---|---|---|---|---|---|---|---|
| 19:00 | 1jzk-C | 4 | 3.3 | 84 | 145 | 8 | MOLECULE: | GLOBIN I - | ARK SHE | LL |
| 20:00 | 3uhi-C | 4 | 3.3 | 85 | 145 | 9 | MOLECULE: | GLOBIN-1 | | |
| 21:00 | 4mqj-B | 4 | 4.4 | 87 | 146 | 2 | MOLECULE: | HEMOGLOBIN | SUBUNIT | ALPHA |
| 22:00 | 1hbr-B | 4 | 3.9 | 84 | 143 | 6 | MOLECULE: | PROTEIN (H | EMOGLOBI | N D) |
| 23:00 | 4n7n-A | 4 | 4.1 | 87 | 141 | 7 | MOLECULE: | HEMOGLOBIN | SUBUNIT | ALPHA |
| 24:00:00 | 3g52-B | 4 | 3.5 | 85 | 145 | 8 | MOLECULE: | GLOBIN-1 | | |
| 25:00:00 | 3uhb-B | 4 | 3.5 | 85 | 145 | 8 | MOLECULE: | GLOBIN-1 | | |
| 26:00:00 | 1nwn-A | 4 | 3.3 | 85 | 145 | 9 | MOLECULE: | GLOBIN I | | |
| 27:00:00 | 2r4x-A | 4 | 3.3 | 84 | 145 | 8 | MOLECULE: | GLOBIN-1 | | |
| 28:00:00 | 3uhc-B | 4 | 3.5 | 85 | 145 | 8 | MOLECULE: | GLOBIN-1 | | |
| 29:00:00 | 3uhi-B | 4 | 3.4 | 84 | 145 | 8 | MOLECULE: | GLOBIN-1 | | |
| 30:00:00 | 3uht-A | 4 | 3.3 | 84 | 145 | 10 | MOLECULE: | GLOBIN-1 | | |
| 31:00:00 | 1jwn-C | 4 | 3.3 | 84 | 145 | 8 | MOLECULE: | GLOBIN I - | ARK SHE | LL |
| 32:00:00 | 2av3-A | 4 | 3.3 | 84 | 145 | 8 | MOLECULE: | GLOBIN I | | |
| 33:00:00 | 1jzl-A | 4 | 3.3 | 84 | 145 | 8 | MOLECULE: | GLOBIN I - | ARK SHE | LL |
| 34:00:00 | 3uhh-A | 4 | 3.3 | 84 | 145 | 8 | MOLECULE: | GLOBIN-1 | | |
| 35:00:00 | 4hrt-D | 4 | 3.4 | 87 | 151 | 9 | MOLECULE: | GLOBIN-2 A | CHAIN | |
| 36:00:00 | 2aup-A | 4 | 3.4 | 85 | 145 | 8 | MOLECULE: | GLOBIN I | | |
| 37:00:00 | 8vyl-A | 4 | 3.3 | 84 | 137 | 7 | MOLECULE: | HEMOGLOBIN | SUBUNIT | ALPHA |
| 38:00:00 | 3uhk-B | 4 | 3.3 | 84 | 145 | 8 | MOLECULE: | GLOBIN-1 | | |
| 39:00:00 | 2aur-A | 4 | 3.3 | 84 | 145 | 8 | MOLECULE: | GLOBIN I | | |
| 40:00:00 | 3ugz-A | 4 | 3.3 | 84 | 145 | 8 | MOLECULE: | GLOBIN-1 | | |
| 41:00:00 | 3g4r-B | 4 | 3.5 | 85 | 145 | 8 | MOLECULE: | GLOBIN-1 | | |
| 42:00:00 | 3uhc-A | 4 | 3.3 | 85 | 145 | 9 | MOLECULE: | GLOBIN-1 | | |
| 43:00:00 | 7k4m-I | 4 | 4.1 | 86 | 140 | 7 | MOLECULE: | HEMOGLOBIN | SUBUNIT | ALPHA |
| 44:00:00 | 2r4y-A | 4 | 3.3 | 84 | 145 | 8 | MOLECULE: | GLOBIN-1 | | |
| 45:00:00 | 5hbi-B | 4 | 3.5 | 85 | 145 | 8 | MOLECULE: | HEMOGLOBIN | | |
| 46:00:00 | 2qls-B | 4 | 3.5 | 84 | 146 | 7 | MOLECULE: | HEMOGLOBIN | SUBUNIT | ALPHA |
| 47:00:00 | 1hbi-B | 4 | 3.4 | 84 | 145 | 8 | MOLECULE: | HEMOGLOBIN | (OXY) | |
| 48:00:00 | 3a0g-B | 4 | 3.6 | 86 | 144 | 8 | MOLECULE: | HEMOGLOBIN | SUBUNIT | ALPHA |
| 49:00:00 | 4x0l-B | 4 | 3.5 | 84 | 144 | 6 | MOLECULE: | HEMOGLOBIN | SUBUNIT | ALPHA |
| 50:00:00 | 6wg7-F | 4 | 8.3 | 106 | 228 | 10 | MOLECULE: | DNA (35-ME | R) | |
| 51:00:00 | 5jdo-F | 4 | 3.5 | 84 | 144 | 6 | MOLECULE: | HAPTOGLOBI | N-HAEMOG | LOBIN RECEPTOR |
| 52:00:00 | 3gys-D | 4 | 3.6 | 85 | 145 | 7 | MOLECULE: | HEMOGLOBIN | SUBUNIT | ALPHA |
| 53:00:00 | 3gqr-F | 4 | 3.5 | 84 | 145 | 7 | MOLECULE: | HEMOGLOBIN | SUBUNIT | ALPHA |
| 54:00:00 | 1qpw-D | 4 | 3.5 | 84 | 146 | 5 | MOLECULE: | PORCINE HE | MOGLOBIN | (ALPHA SUBUNIT) |
| 55:00:00 | 1hbs-B | 4 | 3.1 | 83 | 146 | 6 | MOLECULE: | HEMOGLOBIN | S (DEOX | Y) (ALPHA CHAIN) |
| 56:00:00 | 5x2s-A | 4 | 3.9 | 86 | 140 | 6 | MOLECULE: | HEMOGLOBIN | SUBUNIT | ALPHA |
| 57:00:00 | 2bnl-F | 4 | 4.4 | 80 | 129 | 8 | MOLECULE: | MODULATOR | PROTEIN | RSBR |
| 58:00:00 | 7ud8-B | 4 | 3.5 | 84 | 146 | 6 | MOLECULE: | HEMOGLOBIN | SUBUNIT | ALPHA |
| 59:00:00 | 1hbs-D | 4 | 3.4 | 85 | 146 | 6 | MOLECULE: | HEMOGLOBIN | S (DEOX | Y) (ALPHA CHAIN) |
| 60:00:00 | 2m6z-C | 4 | 3.6 | 88 | 141 | 8 | MOLECULE: | HEMOGLOBIN | SUBUNIT | ALPHA |
| 61:00:00 | 7jy1-B | 4 | 3.6 | 85 | 146 | 6 | MOLECULE: | HEMOGLOBIN | SUBUNIT | ALPHA |
| 62:00:00 | 7vcf-W | 4 | 4.1 | 87 | 164 | 6 | MOLECULE: | TIC214 | | |
| 63:00:00 | 6wg7-C | 4 | 8 | 103 | 228 | 11 | MOLECULE: | DNA (35-ME | R) | |
| 64:00:00 | 7xk3-E | 3.9 | 6.6 | 93 | 198 | 6 | MOLECULE: | NA(+)-TRAN | SLOCATIN | G NADH-QUINONE REDUCTASE SUBUNI |
| 65:00:00 | 8a5y-T | 3.9 | 22.2 | 108 | 650 | 5 | MOLECULE: | ANAPHASE-P | ROMOTING | COMPLEX SUBUNIT CDC27 |
| 66:00:00 | 1or4-A | 3.9 | 6.3 | 96 | 169 | 13 | MOLECULE: | HEME-BASED | AEROTAC | TIC TRANSDUCER HEMAT |
| 67:00:00 | 6v3f-A | 3.9 | 16.3 | 131 | 1234 | 9 | MOLECULE: | NPC1-LIKE | INTRACEL | LULAR CHOLESTEROL TRANSPORTER 1 |
| 68:00:00 | 8q6o-2 | 3.9 | 14 | 98 | 267 | 4 | MOLECULE: | DNA REPLIC | ATION LI | CENSING FACTOR MCM2 |
| 69:00:00 | 8q4h-C | 3.9 | 4.1 | 70 | 89 | 4 | MOLECULE: | TETRACHLOR | OETHENE | REDUCTIVE DEHALOGENASE |
| 70:00:00 | 5y0r-A | 3.9 | 12.3 | 101 | 423 | 5 | MOLECULE: | TMTMCAL(SE | MET) | |
| 71:00:00 | 8xh6-A | 3.9 | 10.6 | 80 | 161 | 9 | MOLECULE: | LATENT MEM | BRANE PR | OTEIN 1 |
| 72:00:00 | 5ex8-A | 3.9 | 11.4 | 140 | 414 | 9 | MOLECULE: | CYTOCHROME | P450 | |

| | | | | | | | | | | |
|---|---|---|---|---|---|---|---|---|---|---|
| 73:00:00 | 6ygb-B | 3.9 | 23.2 | 143 | 725 | 6 | MOLECULE: | N-ALPHA-AC | ETYLTRAN | SFERASE 30 |
| 74:00:00 | 5azr-A | 3.9 | 3.7 | 81 | 153 | 12 | MOLECULE: | MYOGLOBIN | | |
| 75:00:00 | 1dwr-A | 3.9 | 4.1 | 82 | 152 | 12 | MOLECULE: | MYOGLOBIN | | |
| 76:00:00 | 2vlz-A | 3.9 | 4.1 | 82 | 153 | 12 | MOLECULE: | MYOGLOBIN | | |
| 77:00:00 | 1xch-A | 3.9 | 3.7 | 81 | 153 | 11 | MOLECULE: | MYOGLOBIN | | |
| 78:00:00 | 6zfb-H | 3.9 | 19.6 | 126 | 771 | 7 | MOLECULE: | DNA-DIRECT | ED RNA P | OLYMERASE SUBUNIT DELTA |
| 79:00:00 | 3lr7-A | 3.9 | 4 | 82 | 151 | 12 | MOLECULE: | MYOGLOBIN | | |
| 80:00:00 | 5z7f-A | 3.9 | 4 | 82 | 152 | 12 | MOLECULE: | MYOGLOBIN | | |
| 81:00:00 | 5x2u-E | 3.9 | 4.5 | 86 | 140 | 7 | MOLECULE: | HEMOGLOBIN | SUBUNIT | ALPHA |
| 82:00:00 | 3ugy-B | 3.9 | 3.5 | 85 | 145 | 8 | MOLECULE: | GLOBIN-1 | | |
| 83:00:00 | 1y85-D | 3.9 | 3.5 | 82 | 145 | 5 | MOLECULE: | HEMOGLOBIN | ALPHA C | HAIN |
| 84:00:00 | 5x2r-C | 3.9 | 4.4 | 86 | 140 | 7 | MOLECULE: | HEMOGLOBIN | SUBUNIT | ALPHA |
| 85:00:00 | 3uhh-B | 3.9 | 3.5 | 85 | 145 | 8 | MOLECULE: | GLOBIN-1 | | |
| 86:00:00 | 7dy4-H | 3.9 | 3.4 | 82 | 146 | 4 | MOLECULE: | HEMOGLOBIN | SUBUNIT | ALPHA |
| 87:00:00 | 5x2r-K | 3.9 | 4.3 | 86 | 140 | 7 | MOLECULE: | HEMOGLOBIN | SUBUNIT | ALPHA |
| 88:00:00 | 3uhe-B | 3.9 | 3.5 | 85 | 145 | 8 | MOLECULE: | GLOBIN-1 | | |
| 89:00:00 | 2r4w-B | 3.9 | 3.4 | 84 | 145 | 8 | MOLECULE: | GLOBIN-1 | | |
| 90:00:00 | 3qob-B | 3.9 | 3.4 | 85 | 145 | 9 | MOLECULE: | GLOBIN-1 | | |
| 91:00:00 | 5x2t-C | 3.9 | 4.5 | 86 | 140 | 7 | MOLECULE: | HEMOGLOBIN | SUBUNIT | ALPHA |
| 92:00:00 | 1ygd-D | 3.9 | 4.1 | 86 | 146 | 5 | MOLECULE: | HEMOGLOBIN | ALPHA C | HAIN |
| 93:00:00 | 3odq-A | 3.9 | 4 | 86 | 141 | 7 | MOLECULE: | HEMOGLOBIN | SUBUNIT | ALPHA |
| 94:00:00 | 2av0-B | 3.9 | 3.5 | 85 | 145 | 8 | MOLECULE: | GLOBIN I | | |
| 95:00:00 | 3uh5-B | 3.9 | 3.5 | 85 | 145 | 8 | MOLECULE: | GLOBIN-1 | | |
| 96:00:00 | 3uh6-B | 3.9 | 3.5 | 85 | 145 | 8 | MOLECULE: | GLOBIN-1 | | |
| 97:00:00 | 7ud7-D | 3.9 | 3.9 | 83 | 146 | 5 | MOLECULE: | HEMOGLOBIN | SUBUNIT | ALPHA |
| 98:00:00 | 2grh-B | 3.9 | 3.5 | 85 | 145 | 8 | MOLECULE: | GLOBIN-1 | | |
| 99:00:00 | 1jzl-B | 3.9 | 3.5 | 85 | 145 | 8 | MOLECULE: | GLOBIN I - | ARK SHE | LL |
| 0:00 | 2r80-D | 3.9 | 4.1 | 86 | 146 | 7 | MOLECULE: | HEMOGLOBIN | SUBUNIT | ALPHA-A |
| 1:00 | 1fdh-H | 3.9 | 3.9 | 86 | 146 | 2 | MOLECULE: | HEMOGLOBIN | F (DEOX | Y) (ALPHA CHAIN) |
| 2:00 | 3g53-B | 3.9 | 3.5 | 85 | 145 | 8 | MOLECULE: | GLOBIN-1 | | |
| 3:00 | 7hbi-B | 3.9 | 3.5 | 85 | 145 | 8 | MOLECULE: | HEMOGLOBIN | | |
| 4:00 | 3sdh-B | 3.9 | 3.5 | 85 | 145 | 8 | MOLECULE: | HEMOGLOBIN | I (CARB | ONMONOXY) |
| 5:00 | 1hds-C | 3.9 | 3.5 | 86 | 141 | 5 | MOLECULE: | HEMOGLOBIN | S (DEOX | Y) (ALPHA CHAIN) |
| 6:00 | 7dy3-B | 3.9 | 3.8 | 83 | 146 | 5 | MOLECULE: | HEMOGLOBIN | SUBUNIT | ALPHA |
| 7:00 | 3uhg-B | 3.9 | 3.5 | 85 | 145 | 8 | MOLECULE: | GLOBIN-1 | | |
| 8:00 | 1y2z-D | 3.9 | 4.4 | 86 | 146 | 5 | MOLECULE: | HEMOGLOBIN | ALPHA C | HAIN |
| 9:00 | 3uh3-B | 3.9 | 3.5 | 85 | 145 | 9 | MOLECULE: | GLOBIN-1 | | |
| 10:00 | 4n7p-G | 3.9 | 4.4 | 87 | 141 | 7 | MOLECULE: | HEMOGLOBIN | SUBUNIT | ALPHA |
| 11:00 | 3lr9-A | 3.9 | 4 | 82 | 152 | 12 | MOLECULE: | MYOGLOBIN | | |
| 12:00 | 3ugz-B | 3.9 | 3.5 | 85 | 145 | 8 | MOLECULE: | GLOBIN-1 | | |
| 13:00 | 2o5s-X | 3.9 | 4.1 | 82 | 152 | 12 | MOLECULE: | MYOGLOBIN | | |
| 14:00 | 7vde-B | 3.9 | 3.6 | 85 | 146 | 6 | MOLECULE: | HEMOGLOBIN | SUBUNIT | ALPHA |
| 15:00 | 7vde-D | 3.9 | 3.6 | 85 | 146 | 6 | MOLECULE: | HEMOGLOBIN | SUBUNIT | ALPHA |
| 16:00 | 4h2l-B | 3.9 | 3.5 | 84 | 146 | 6 | MOLECULE: | ALPHA-GLOB | IN | |
| 17:00 | 5ufj-B | 3.9 | 3.6 | 84 | 146 | 6 | MOLECULE: | HEMOGLOBIN | SUBUNIT | ALPHA |
| 18:00 | 5ker-F | 3.9 | 3.5 | 84 | 146 | 5 | MOLECULE: | ALPHA-GLOB | IN | |
| 19:00 | 2v1k-A | 3.9 | 4.1 | 82 | 153 | 12 | MOLECULE: | MYOGLOBIN | | |
| 20:00 | 5ker-D | 3.9 | 3.6 | 84 | 146 | 5 | MOLECULE: | ALPHA-GLOB | IN | |
| 21:00 | 5ker-H | 3.9 | 3.7 | 86 | 146 | 5 | MOLECULE: | ALPHA-GLOB | IN | |
| 22:00 | 7jy0-D | 3.9 | 3.6 | 84 | 146 | 6 | MOLECULE: | HEMOGLOBIN | SUBUNIT | ALPHA |
| 23:00 | 2m6z-A | 3.9 | 3.5 | 88 | 141 | 7 | MOLECULE: | HEMOGLOBIN | SUBUNIT | ALPHA |
| 24:00:00 | 3gys-B | 3.9 | 3.5 | 84 | 145 | 10 | MOLECULE: | HEMOGLOBIN | SUBUNIT | ALPHA |
| 25:00:00 | 4f4o-E | 3.9 | 3.7 | 86 | 146 | 5 | MOLECULE: | HEMOGLOBIN | SUBUNIT | ALPHA |
| 26:00:00 | 4f4o-K | 3.9 | 3.6 | 85 | 146 | 5 | MOLECULE: | HEMOGLOBIN | SUBUNIT | ALPHA |

| | | | | | | | | | | |
|---|---|---|---|---|---|---|---|---|---|---|
| 27:00:00 | 3mjp-D | 3.9 | 3.6 | 85 | 146 | 8 | MOLECULE: | HEMOGLOBIN | SUBUNIT | ALPHA-A |
| 28:00:00 | 2hvr-A | 3.8 | 13.9 | 77 | 319 | 5 | MOLECULE: | 5'- | | |
| 29:00:00 | 1it2-A | 3.8 | 3.2 | 77 | 146 | 5 | MOLECULE: | HEMOGLOBIN | | |
| 30:00:00 | 2vwa-A | 3.8 | 2.9 | 70 | 99 | 13 | MOLECULE: | PUTATIVE U | NCHARACT | ERIZED PROTEIN PF13_0012 |
| 31:00:00 | 8cr1-C | 3.8 | 9.8 | 114 | 266 | 4 | MOLECULE: | ATPASE ASN | A1 | |
| 32:00:00 | 6rmg-A | 3.8 | 7 | 118 | 1026 | 8 | MOLECULE: | PROTEIN PA | TCHED HO | MOLOG 1,GFP-LIKE FLUORESCENT |
| 33:00:00 | 7ctp-A | 3.8 | 4.3 | 113 | 553 | 6 | MOLECULE: | PROTEIN NI | BAN 2 | |
| 34:00:00 | 7dgy-A | 3.8 | 3.5 | 61 | 99 | 3 | MOLECULE: | C2 | | |
| 35:00:00 | 4bja-A | 3.8 | 5.2 | 99 | 182 | 6 | MOLECULE: | PROTEIN GL | B-12 | |
| 36:00:00 | 5lbm-C | 3.8 | 4 | 72 | 90 | 6 | MOLECULE: | TRANSCRIPT | IONAL RE | PRESSOR FRMR |
| 37:00:00 | 7etm-A | 3.8 | 9.1 | 105 | 498 | 12 | MOLECULE: | PORTAL PRO | TEIN | |
| 38:00:00 | 1cqx-A | 3.8 | 9 | 122 | 403 | 7 | MOLECULE: | FLAVOHEMOP | ROTEIN | |
| 39:00:00 | 7w0y-k | 3.8 | 6.6 | 82 | 98 | 5 | MOLECULE: | NADH DEHYD | ROGENASE | [UBIQUINONE] FLAVOPROTEIN 1, |
| 40:00:00 | 1x46-A | 3.8 | 4.1 | 82 | 150 | 10 | MOLECULE: | HEMOGLOBIN | COMPONE | NT VII |
| 41:00:00 | 8fib-A | 3.8 | 4.7 | 123 | 418 | 7 | MOLECULE: | CYTOCHROME | P450 | |
| 42:00:00 | 8b6l-M | 3.8 | 6.7 | 76 | 110 | 1 | MOLECULE: | PROTEIN TR | ANSPORT | PROTEIN SEC61 SUBUNIT ALPHA ISO |
| 43:00:00 | 2p6v-A | 3.8 | 3.2 | 76 | 97 | 11 | MOLECULE: | TRANSCRIPT | ION INIT | IATION FACTOR TFIID SUBUNIT 4 |
| 44:00:00 | 1azi-A | 3.8 | 3.7 | 81 | 153 | 12 | MOLECULE: | MYOGLOBIN | | |
| 45:00:00 | 2vlx-A | 3.8 | 3.9 | 85 | 153 | 12 | MOLECULE: | MYOGLOBIN | | |
| 46:00:00 | 3wtg-D | 3.8 | 3.9 | 84 | 146 | 8 | MOLECULE: | HEMOGLOBIN | SUBUNIT | ALPHA-A |
| 47:00:00 | 5x2u-A | 3.8 | 4.5 | 86 | 140 | 7 | MOLECULE: | HEMOGLOBIN | SUBUNIT | ALPHA |
| 48:00:00 | 3uhd-B | 3.8 | 3.5 | 84 | 144 | 7 | MOLECULE: | GLOBIN-1 | | |
| 49:00:00 | 3at6-B | 3.8 | 4.2 | 88 | 146 | 7 | MOLECULE: | ALPHAA-GLO | BIN | |
| 50:00:00 | 5x2s-K | 3.8 | 4.3 | 86 | 140 | 7 | MOLECULE: | HEMOGLOBIN | SUBUNIT | ALPHA |
| 51:00:00 | 3pel-B | 3.8 | 3.6 | 82 | 146 | 6 | MOLECULE: | HEMOGLOBIN | SUBUNIT | ALPHA |
| 52:00:00 | 7pch-D | 3.8 | 3.8 | 84 | 146 | 4 | MOLECULE: | HEMOGLOBIN | SUBUNIT | ALPHA |
| 53:00:00 | 7cue-D | 3.8 | 3.6 | 82 | 146 | 5 | MOLECULE: | HEMOGLOBIN | SUBUNIT | ALPHA |
| 54:00:00 | 3dht-B | 3.8 | 3.7 | 85 | 145 | 6 | MOLECULE: | HEMOGLOBIN | SUBUNIT | ALPHA-1/2 |
| 55:00:00 | 1nej-B | 3.8 | 3.6 | 84 | 146 | 6 | MOLECULE: | HEMOGLOBIN | ALPHA C | HAIN |
| 56:00:00 | 1xq5-A | 3.8 | 3.3 | 85 | 142 | 7 | MOLECULE: | HEMOGLOBIN | ALPHA-1 | CHAIN |
| 57:00:00 | 6wfq-C | 3.8 | 8 | 103 | 228 | 11 | MOLECULE: | HTH-TYPE T | RANSCRIP | TIONAL REPRESSOR NANR |
| 58:00:00 | 5x2t-E | 3.8 | 3.4 | 84 | 140 | 8 | MOLECULE: | HEMOGLOBIN | SUBUNIT | ALPHA |
| 59:00:00 | 6wg7-E | 3.8 | 8 | 103 | 228 | 11 | MOLECULE: | DNA (35-ME | R) | |
| 60:00:00 | 1hbs-G | 3.8 | 3.4 | 85 | 141 | 4 | MOLECULE: | HEMOGLOBIN | S (DEOX | Y) (ALPHA CHAIN) |
| 61:00:00 | 5cne-A | 3.8 | 4 | 82 | 152 | 12 | MOLECULE: | MYOGLOBIN | | |
| 62:00:00 | 1nek-D | 3.7 | 8.7 | 84 | 113 | 7 | MOLECULE: | SUCCINATE | DEHYDROG | ENASE FLAVOPROTEIN SUBUNIT |
| 63:00:00 | 8egr-G | 3.7 | 12.5 | 94 | 313 | 4 | MOLECULE: | GP15, RECE | PTOR-BIN | DING PROTEIN, TAIL FIBER |
| 64:00:00 | 7qru-F | 3.7 | 7.5 | 76 | 91 | 5 | MOLECULE: | NA+/H+ ANT | IPORTER | SUBUNIT D |
| 65:00:00 | 7vgh-B | 3.7 | 24.2 | 146 | 1182 | 3 | MOLECULE: | CELL CYCLE | CONTROL | PROTEIN 50B |
| 66:00:00 | 7qe7-A | 3.7 | 21.6 | 122 | 1648 | 6 | MOLECULE: | ANAPHASE-P | ROMOTING | COMPLEX SUBUNIT 10 |
| 67:00:00 | 4zi2-C | 3.7 | 7.8 | 102 | 131 | 4 | MOLECULE: | ADP-RIBOSY | LATION F | ACTOR-LIKE PROTEIN 3 |
| 68:00:00 | 8jem-B | 3.7 | 13.4 | 112 | 419 | 4 | MOLECULE: | TEICHOIC A | CID D-AL | ANYLTRANSFERASE |
| 69:00:00 | 6zka-K | 3.7 | 6.5 | 84 | 98 | 6 | MOLECULE: | NADH-UBIQU | INONE OX | IDOREDUCTASE CHAIN 3 |
| 70:00:00 | 3a59-H | 3.7 | 4 | 85 | 146 | 7 | MOLECULE: | HEMOGLOBIN | SUBUNIT | ALPHA-A |
| 71:00:00 | 2myd-A | 3.7 | 3.6 | 80 | 153 | 10 | MOLECULE: | MYOGLOBIN | (N-PROPY | L ISOCYANIDE) |
| 72:00:00 | 1y85-B | 3.7 | 3.8 | 82 | 145 | 2 | MOLECULE: | HEMOGLOBIN | ALPHA C | HAIN |
| 73:00:00 | 1myh-B | 3.7 | 4.5 | 89 | 153 | 7 | MOLECULE: | MYOGLOBIN | | |
| 74:00:00 | 1mni-A | 3.7 | 4.8 | 90 | 153 | 7 | MOLECULE: | MYOGLOBIN | | |
| 75:00:00 | 7pcq-D | 3.7 | 3.9 | 83 | 146 | 2 | MOLECULE: | HEMOGLOBIN | SUBUNIT | ALPHA |
| 76:00:00 | 7ud7-B | 3.7 | 3.9 | 84 | 146 | 4 | MOLECULE: | HEMOGLOBIN | SUBUNIT | ALPHA |
| 77:00:00 | 1mbi-A | 3.7 | 3.7 | 80 | 153 | 10 | MOLECULE: | MYOGLOBIN | | |
| 78:00:00 | 3hf4-B | 3.7 | 3.7 | 84 | 145 | 6 | MOLECULE: | HEMOGLOBIN | SUBUNIT | ALPHA-1/2 |
| 79:00:00 | 2r4z-B | 3.7 | 3.8 | 82 | 146 | 12 | MOLECULE: | GLOBIN-1 | | |
| 80:00:00 | 2r4z-A | 3.7 | 3.7 | 82 | 145 | 11 | MOLECULE: | GLOBIN-1 | | |

| | | | | | | | | | | |
|---|---|---|---|---|---|---|---|---|---|---|
| 81:00:00 | 2myb-A | 3.7 | 3.6 | 80 | 153 | 10 | MOLECULE: | MYOGLOBIN | (METHYL | ISOCYANIDE) |
| 82:00:00 | 1vxg-A | 3.7 | 3.6 | 80 | 153 | 10 | MOLECULE: | MYOGLOBIN | | |
| 83:00:00 | 3gou-B | 3.7 | 3.8 | 84 | 146 | 5 | MOLECULE: | HEMOGLOBIN | SUBUNIT | ALPHA |
| 84:00:00 | 1yoh-A | 3.7 | 3.6 | 80 | 153 | 10 | MOLECULE: | MYOGLOBIN | | |
| 85:00:00 | 1ch4-B | 3.7 | 4 | 85 | 146 | 6 | MOLECULE: | MODULE-SUB | STITUTED | CHIMERA HEMOGLOBIN BETA-ALPHA |
| 86:00:00 | 2z6s-A | 3.7 | 3.6 | 80 | 151 | 10 | MOLECULE: | MYOGLOBIN | | |
| 87:00:00 | 3eqm-A | 3.6 | 4.8 | 155 | 452 | 6 | MOLECULE: | CYTOCHROME | P450 19 | A1 |
| 88:00:00 | 8djk-A | 3.6 | 12.3 | 119 | 245 | 8 | MOLECULE: | 3-HYDROXY- | 3-METHYL | GLUTARYL-COENZYME A REDUCTASE |
| 89:00:00 | 7yug-A | 3.6 | 3 | 64 | 117 | 3 | MOLECULE: | PROTEIN BA | NP | |
| 90:00:00 | 6wu0-B | 3.6 | 6.1 | 114 | 854 | 4 | MOLECULE: | HOPANOID B | IOSYNTHE | SIS ASSOCIATED RND TRANSPORTER |
| 91:00:00 | 6uak-A | 3.6 | 17.9 | 96 | 298 | 11 | MOLECULE: | SAM DEPEND | ENT METH | YLTRANSFERASE LAHSB |
| 92:00:00 | 3zjh-B | 3.6 | 5.5 | 98 | 193 | 8 | MOLECULE: | PROTOGLOBI | N | |
| 93:00:00 | 5yo8-A | 3.6 | 11.3 | 95 | 349 | 4 | MOLECULE: | TETRAPRENY | L-BETA-C | URCUMENE SYNTHASE |
| 94:00:00 | 2hjm-A | 3.6 | 5.4 | 73 | 91 | 5 | MOLECULE: | HYPOTHETIC | AL PROTE | IN PF1176 |
| 95:00:00 | 8h17-A | 3.6 | 4.9 | 91 | 157 | 8 | MOLECULE: | TLR1989 PR | OTEIN | |
| 96:00:00 | 6s8h-F | 3.6 | 14 | 93 | 239 | 8 | MOLECULE: | LIPOPOLYSA | CCHARIDE | ABC TRANSPORTER, ATP-BINDING P |
| 97:00:00 | 5jki-A | 3.6 | 5.2 | 99 | 201 | 14 | MOLECULE: | PUTATIVE L | IPID PHO | SPHATE PHOSPHATASE YODM |
| 98:00:00 | 3p3o-A | 3.6 | 5 | 142 | 416 | 5 | MOLECULE: | CYTOCHROME | P450 | |
| 99:00:00 | 4zvb-A | 3.6 | 4.1 | 92 | 150 | 5 | MOLECULE: | DIGUANYLAT | E CYCLAS | E DOSC |
| 0:00 | 2gb7-B | 3.6 | 16 | 85 | 295 | 5 | MOLECULE: | DNA STRAND | 1 | |
| 1:00 | 1it3-B | 3.6 | 3.1 | 76 | 146 | 7 | MOLECULE: | HEMOGLOBIN | | |
| 2:00 | 1it3-D | 3.6 | 3.2 | 76 | 146 | 7 | MOLECULE: | HEMOGLOBIN | | |
| 3:00 | 4f4o-B | 3.6 | 3.8 | 85 | 146 | 1 | MOLECULE: | HEMOGLOBIN | SUBUNIT | ALPHA |
| 4:00 | 2mbw-A | 3.6 | 3.6 | 80 | 154 | 10 | MOLECULE: | MYOGLOBIN | | |
| 5:00 | 7k4m-F | 3.6 | 4.3 | 85 | 146 | 4 | MOLECULE: | HEMOGLOBIN | SUBUNIT | ALPHA |
| 6:00 | 110m-A | 3.6 | 3.8 | 84 | 154 | 10 | MOLECULE: | MYOGLOBIN | | |
| 7:00 | 1myi-B | 3.6 | 4.4 | 89 | 153 | 8 | MOLECULE: | MYOGLOBIN | | |
| 8:00 | 1bzr-A | 3.6 | 4.4 | 89 | 153 | 7 | MOLECULE: | PROTEIN (M | YOGLOBIN | ) |
| 9:00 | 1duk-A | 3.6 | 4 | 81 | 152 | 10 | MOLECULE: | WILD-TYPE | RECOMBIN | ANT SPERM WHALE |
| 10:00 | 2g0r-A | 3.6 | 3.8 | 84 | 154 | 10 | MOLECULE: | MYOGLOBIN | | |
| 11:00 | 1u7r-A | 3.6 | 3.7 | 80 | 153 | 10 | MOLECULE: | MYOGLOBIN | | |
| 12:00 | 1shr-D | 3.6 | 4.4 | 83 | 146 | 2 | MOLECULE: | HEMOGLOBIN | ALPHA C | HAIN |
| 13:00 | 1mti-A | 3.6 | 3.8 | 84 | 154 | 10 | MOLECULE: | MYOGLOBIN | | |
| 14:00 | 1vxf-A | 3.6 | 3.6 | 80 | 153 | 10 | MOLECULE: | MYOGLOBIN | | |
| 15:00 | 5utc-A | 3.6 | 4.5 | 89 | 152 | 8 | MOLECULE: | MYOGLOBIN | | |
| 16:00 | 1ch4-A | 3.6 | 4 | 85 | 146 | 6 | MOLECULE: | MODULE-SUB | STITUTED | CHIMERA HEMOGLOBIN BETA-ALPHA |
| 17:00 | 1mgn-A | 3.6 | 3.8 | 84 | 154 | 10 | MOLECULE: | METMYOGLOB | IN | |
| 18:00 | 2z8a-B | 3.6 | 3.8 | 82 | 146 | 12 | MOLECULE: | GLOBIN-1 | | |
| 19:00 | 5v7p-A | 3.5 | 6.2 | 108 | 281 | 6 | MOLECULE: | PROTEIN-S- | ISOPRENY | LCYSTEINE O-METHYLTRANSFERASE |
| 20:00 | 3vw4-A | 3.5 | 13 | 91 | 122 | 7 | MOLECULE: | REP | | |
| 21:00 | 5eik-A | 3.5 | 5 | 76 | 234 | 5 | MOLECULE: | UNCHARACTE | RIZED PR | OTEIN Y57A10A.28 |
| 22:00 | 7bc6-A | 3.5 | 17.5 | 114 | 432 | 6 | MOLECULE: | PROTON-COU | PLED FOL | ATE TRANSPORTER |
| 23:00 | 1us7-B | 3.5 | 14.8 | 94 | 193 | 7 | MOLECULE: | HEAT SHOCK | PROTEIN | HSP82 |
| 24:00:00 | 5zle-A | 3.5 | 5.8 | 98 | 225 | 8 | MOLECULE: | CYTOCHROME | B REDUC | TASE 1 |
| 25:00:00 | 7nnt-C | 3.5 | 5 | 87 | 169 | 3 | MOLECULE: | ENERGY-COU | PLING FA | CTOR TRANSPORTER ATP-BINDING PR |
| 26:00:00 | 3jc6-5 | 3.5 | 12.3 | 86 | 254 | 5 | MOLECULE: | DNA REPLIC | ATION LI | CENSING FACTOR MCM2 |
| 27:00:00 | 8slm-A | 3.5 | 4.4 | 89 | 248 | 11 | MOLECULE: | ZN DEPENDE | NT HYDRO | LASE FUSED TO HTH DOMAIN, IRRE |
| 28:00:00 | 4p9f-A | 3.5 | 4.5 | 94 | 212 | 9 | MOLECULE: | HTH-TYPE T | RANSCRIP | TIONAL REGULATOR MCBR |
| 29:00:00 | 8wu3-B | 3.5 | 12.6 | 131 | 592 | 7 | MOLECULE: | RNA POLYME | RASE | |
| 30:00:00 | 4m8m-B | 3.5 | 16.3 | 104 | 574 | 5 | MOLECULE: | GCN4 COILE | D-COIL F | USED ZEBRAFISH PLEXINC1 |
| 31:00:00 | 5t8v-A | 3.5 | 19.2 | 109 | 1315 | 6 | MOLECULE: | PUTATIVE U | NCHARACT | ERIZED PROTEIN |
| 32:00:00 | 7tch-A | 3.5 | 13.2 | 104 | 645 | 3 | MOLECULE: | BACITRACIN | EXPORT | PERMEASE PROTEIN BCEB |
| 33:00:00 | 6g2j-K | 3.5 | 6.6 | 81 | 98 | 5 | MOLECULE: | NADH-UBIQU | INONE OX | IDOREDUCTASE CHAIN 3 |
| 34:00:00 | 6csm-D | 3.5 | 9.6 | 112 | 277 | 10 | MOLECULE: | GTACR1 | | |

| | | | | | | | | | | |
|---|---|---|---|---|---|---|---|---|---|---|
| 35:00:00 | 7obq-u | 3.5 | 16.4 | 88 | 441 | 6 | MOLECULE: | SRP RNA | | |
| 36:00:00 | 6wg3-E | 3.5 | 16.3 | 103 | 1226 | 6 | MOLECULE: | STRUCTURAL | MAINTEN | ANCE OF CHROMOSOMES PROTEIN 1A |
| 37:00:00 | 6iuy-A | 3.5 | 18.1 | 93 | 585 | 8 | MOLECULE: | GLYCEROL-3 | #NAME? | TE DEHYDROGENASE [NAD(+)] |
| 38:00:00 | 4dwl-B | 3.5 | 4.8 | 77 | 110 | 10 | MOLECULE: | BBP7 | | |
| 39:00:00 | 7dpd-B | 3.5 | 15.3 | 91 | 273 | 4 | MOLECULE: | DNA HELICA | SE MCM9 | |
| 40:00:00 | 1jdo-A | 3.5 | 4 | 85 | 154 | 9 | MOLECULE: | MYOGLOBIN | | |
| 41:00:00 | 2pgh-B | 3.5 | 4.1 | 85 | 146 | 2 | MOLECULE: | HEMOGLOBIN | (AQUO M | ET) (ALPHA CHAIN) |
| 42:00:00 | 1mlg-A | 3.5 | 3.8 | 84 | 154 | 10 | MOLECULE: | MYOGLOBIN | | |
| 43:00:00 | 1myi-A | 3.5 | 3.7 | 80 | 153 | 14 | MOLECULE: | MYOGLOBIN | | |
| 44:00:00 | 4f4o-H | 3.5 | 4.2 | 87 | 146 | 1 | MOLECULE: | HEMOGLOBIN | SUBUNIT | ALPHA |
| 45:00:00 | 1hbs-F | 3.5 | 3.8 | 84 | 146 | 4 | MOLECULE: | HEMOGLOBIN | S (DEOX | Y) (ALPHA CHAIN) |
| 46:00:00 | 6on4-B | 3.5 | 13 | 102 | 214 | 10 | MOLECULE: | HTH-TYPE T | RANSCRIP | TIONAL REPRESSOR NANR |
| 47:00:00 | 3qma-A | 3.5 | 4 | 78 | 145 | 5 | MOLECULE: | MYOGLOBIN | | |
| 48:00:00 | 2nrl-A | 3.5 | 4 | 78 | 145 | 5 | MOLECULE: | MYOGLOBIN | | |
| 49:00:00 | 1mlh-A | 3.5 | 3.8 | 84 | 154 | 10 | MOLECULE: | MYOGLOBIN | | |
| 50:00:00 | 1z8u-B | 3.5 | 3.7 | 84 | 135 | 8 | MOLECULE: | ALPHA-HEMO | GLOBIN S | TABILIZING PROTEIN |
| 51:00:00 | 4k0f-A | 3.4 | 10.6 | 146 | 531 | 8 | MOLECULE: | LANOSTEROL | 14-ALPH | A DEMETHYLASE |
| 52:00:00 | 4p96-A | 3.4 | 4.7 | 115 | 278 | 10 | MOLECULE: | FATTY ACID | METABOL | ISM REGULATOR PROTEIN |
| 53:00:00 | 7x2q-A | 3.4 | 16.3 | 97 | 467 | 7 | MOLECULE: | SUGIOL SYN | THASE | |
| 54:00:00 | 7b1s-D | 3.4 | 16.5 | 122 | 593 | 11 | MOLECULE: | ETHYL-COEN | ZYME M R | EDUCTASE ALPHA SUBUNIT |
| 55:00:00 | 1lqs-L | 3.4 | 12.1 | 118 | 142 | 12 | MOLECULE: | INTERLEUKI | N-10 REC | EPTOR ALPHA CHAIN |
| 56:00:00 | 8j5t-C | 3.4 | 10 | 107 | 288 | 12 | MOLECULE: | UNCHARACTE | RIZED PR | OTEIN RV1280C |
| 57:00:00 | 7tt9-A | 3.4 | 3.8 | 66 | 116 | 6 | MOLECULE: | GROUP 1 TR | UNCATED | HEMOGLOBIN |
| 58:00:00 | 6m9a-C | 3.4 | 4.4 | 87 | 157 | 8 | MOLECULE: | SIGNALING | PROTEIN | |
| 59:00:00 | 7z8b-C | 3.4 | 14.3 | 109 | 1224 | 13 | MOLECULE: | CULLIN-7 | | |
| 60:00:00 | 7eu3-G | 3.4 | 7 | 98 | 176 | 5 | MOLECULE: | NAD(P)H-QU | INONE OX | IDOREDUCTASE SUBUNIT 1, CHLOROP |
| 61:00:00 | 7pkq-o | 3.4 | 13.5 | 88 | 163 | 2 | MOLECULE: | MS35 | | |
| 62:00:00 | 2pke-A | 3.4 | 5.4 | 95 | 233 | 2 | MOLECULE: | HALOACID D | ELAHOGEN | ASE-LIKE FAMILY HYDROLASE |
| 63:00:00 | 7lt2-A | 3.4 | 20.7 | 96 | 387 | 7 | MOLECULE: | MAB-21 DOM | AIN-CONT | AINING PROTEIN |
| 64:00:00 | 6ezn-B | 3.4 | 7.4 | 77 | 110 | 3 | MOLECULE: | DOLICHYL-D | IPHOSPHO | OLIGOSACCHARIDE--PROTEIN |
| 65:00:00 | 3l9w-B | 3.4 | 9 | 103 | 357 | 4 | MOLECULE: | GLUTATHION | E-REGULA | TED POTASSIUM-EFFLUX SYSTEM PRO |
| 66:00:00 | 8dl7-A | 3.4 | 16 | 115 | 467 | 5 | MOLECULE: | SOLUTE CAR | RIER FAM | ILY 40 MEMBER 1 |
| 67:00:00 | 7ar9-J | 3.4 | 7.1 | 83 | 145 | 2 | MOLECULE: | ND3 | | |
| 68:00:00 | 2dc3-B | 3.4 | 5 | 91 | 168 | 4 | MOLECULE: | CYTOGLOBIN | | |
| 69:00:00 | 1ury-B | 3.4 | 4.2 | 84 | 154 | 5 | MOLECULE: | CYTOGLOBIN | | |
| 70:00:00 | 5x2t-K | 3.4 | 3.6 | 80 | 140 | 9 | MOLECULE: | HEMOGLOBIN | SUBUNIT | ALPHA |
| 71:00:00 | 5cnc-A | 3.4 | 4.8 | 81 | 152 | 10 | MOLECULE: | MYOGLOBIN | | |
| 72:00:00 | 3wi8-A | 3.4 | 4.5 | 79 | 151 | 10 | MOLECULE: | MYOGLOBIN | | |
| 73:00:00 | 3qm8-A | 3.4 | 4.1 | 81 | 145 | 5 | MOLECULE: | MYOGLOBIN | | |
| 74:00:00 | 6otw-A | 3.4 | 4.9 | 86 | 151 | 7 | MOLECULE: | HEMOGLOBIN | II | |
| 75:00:00 | 4hrt-B | 3.4 | 4.1 | 86 | 146 | 12 | MOLECULE: | GLOBIN-2 A | CHAIN | |
| 76:00:00 | 3qm7-A | 3.4 | 4.3 | 77 | 144 | 5 | MOLECULE: | MYOGLOBIN | | |
| 77:00:00 | 5cnf-A | 3.4 | 4.8 | 81 | 152 | 10 | MOLECULE: | MYOGLOBIN | | |
| 78:00:00 | 5x2r-A | 3.4 | 3.6 | 79 | 140 | 9 | MOLECULE: | HEMOGLOBIN | SUBUNIT | ALPHA |
| 79:00:00 | 3lhb-H | 3.4 | 3 | 76 | 149 | 3 | MOLECULE: | PROTEIN (H | EMOGLOBI | N) |
| 80:00:00 | 1uc3-K | 3.4 | 3.1 | 76 | 149 | 3 | MOLECULE: | GLOBIN | | |
| 81:00:00 | 1f5p-F | 3.4 | 3.2 | 77 | 149 | 3 | MOLECULE: | HEMOGLOBIN | V | |
| 82:00:00 | 3lhb-B | 3.4 | 3.2 | 77 | 149 | 3 | MOLECULE: | PROTEIN (H | EMOGLOBI | N) |
| 83:00:00 | 3lhb-C | 3.4 | 3.2 | 77 | 149 | 3 | MOLECULE: | PROTEIN (H | EMOGLOBI | N) |
| 84:00:00 | 1f5p-D | 3.4 | 3.2 | 77 | 149 | 3 | MOLECULE: | HEMOGLOBIN | V | |
| 85:00:00 | 3lhb-A | 3.4 | 3.1 | 76 | 149 | 3 | MOLECULE: | PROTEIN (H | EMOGLOBI | N) |
| 86:00:00 | 3lhb-D | 3.4 | 3.2 | 77 | 149 | 3 | MOLECULE: | PROTEIN (H | EMOGLOBI | N) |
| 87:00:00 | 1dwt-A | 3.4 | 4.6 | 81 | 152 | 10 | MOLECULE: | MYOGLOBIN | | |
| 88:00:00 | 3lhb-F | 3.4 | 3.2 | 77 | 149 | 3 | MOLECULE: | PROTEIN (H | EMOGLOBI | N) |

| | | | | | | | | | | | |
|---|---|---|---|---|---|---|---|---|---|---|---|
| 89:00:00 | 3lhb-I | 3.4 | 3.2 | 77 | 149 | 3 | MOLECULE: | PROTEIN (H | EMOGLOBI | N) | |
| 90:00:00 | 1f5p-B | 3.4 | 3.2 | 77 | 149 | 3 | MOLECULE: | HEMOGLOBIN | V | | |
| 91:00:00 | 1f5p-A | 3.4 | 3.2 | 77 | 149 | 3 | MOLECULE: | HEMOGLOBIN | V | | |
| 92:00:00 | 3lhb-L | 3.4 | 3.2 | 77 | 149 | 3 | MOLECULE: | PROTEIN (H | EMOGLOBI | N) | |
| 93:00:00 | 3lhb-G | 3.4 | 3.2 | 77 | 149 | 3 | MOLECULE: | PROTEIN (H | EMOGLOBI | N) | |
| 94:00:00 | 3lhb-K | 3.4 | 3.1 | 76 | 149 | 3 | MOLECULE: | PROTEIN (H | EMOGLOBI | N) | |
| 95:00:00 | 3hrw-D | 3.4 | 3.7 | 83 | 146 | 7 | MOLECULE: | HEMOGLOBIN | SUBUNIT | ALPHA | |
| 96:00:00 | 1f5p-C | 3.4 | 3.6 | 78 | 149 | 3 | MOLECULE: | HEMOGLOBIN | V | | |
| 97:00:00 | 3lhb-J | 3.4 | 3.2 | 77 | 149 | 3 | MOLECULE: | PROTEIN (H | EMOGLOBI | N) | |
| 98:00:00 | 1f5p-E | 3.4 | 3.2 | 77 | 149 | 3 | MOLECULE: | HEMOGLOBIN | V | | |
| 99:00:00 | 3lhb-E | 3.4 | 3.1 | 76 | 149 | 3 | MOLECULE: | PROTEIN (H | EMOGLOBI | N) | |
| 0:00 | 6wq0-A | 3.3 | 14.2 | 80 | 131 | 5 | MOLECULE: | DNA (301-M | ER) | | |
| 1:00 | 6z3w-F | 3.3 | 14.1 | 76 | 110 | 8 | MOLECULE: | ER MEMBRAN | E PROTEI | N COMPLEX SUBUNIT 1,ER MEMBRANE | |
| 2:00 | 2cqn-A | 3.3 | 4.2 | 66 | 77 | 2 | MOLECULE: | FORMIN-BIN | DING PRO | TEIN 3 | |
| 3:00 | 5k94-A | 3.3 | 19.3 | 98 | 506 | 8 | MOLECULE: | MALTOSE-BI | NDING PE | RIPLASMIC PROTEIN,PROTEIN TRANS | |
| 4:00 | 7a7c-A | 3.3 | 17.4 | 127 | 560 | 6 | MOLECULE: | CATALASE-P | EROXIDAS | E | |
| 5:00 | 2mjm-A | 3.3 | 3.6 | 67 | 101 | 3 | MOLECULE: | PROTEIN NL | RC5 | | |
| 6:00 | 3al0-B | 3.3 | 16.7 | 90 | 482 | 7 | MOLECULE: | GLUTAMYL-T | RNA(GLN) | AMIDOTRANSFERASE SUBUNIT A | |
| 7:00 | 7c79-L | 3.3 | 9.3 | 85 | 131 | 4 | MOLECULE: | RIBONUCLEA | SE MRP R | NA SUBUNIT NME1 | |
| 8:00 | 4gr6-B | 3.3 | 14.8 | 79 | 105 | 3 | MOLECULE: | ATRBCX2 | | | |
| 9:00 | 8ce1-c | 3.3 | 15.7 | 99 | 241 | 3 | MOLECULE: | CYTOCHROME | C BIOGE | NESIS ATP-BINDING EXPORT PROTEI | |
| 10:00 | 6zgd-A | 3.3 | 16.6 | 90 | 288 | 10 | MOLECULE: | PROTON-GAT | ED ION C | HANNEL | |
| 11:00 | 3ua3-A | 3.3 | 12.1 | 105 | 663 | 2 | MOLECULE: | PROTEIN AR | GININE N | -METHYLTRANSFERASE 5 | |
| 12:00 | 7etw-B | 3.3 | 13.3 | 99 | 483 | 6 | MOLECULE: | INSULIN-IN | DUCED GE | NE 2 PROTEIN | |
| 13:00 | 1q1v-A | 3.3 | 6.4 | 60 | 70 | 3 | MOLECULE: | DEK PROTEI | N | | |
| 14:00 | 3dan-A | 3.3 | 6.1 | 100 | 473 | 2 | MOLECULE: | CYTOCHROME | P450 74 | A2 | |
| 15:00 | 2zop-A | 3.3 | 3.3 | 72 | 113 | 6 | MOLECULE: | PUTATIVE U | NCHARACT | ERIZED PROTEIN TTHB164 | |
| 16:00 | 6c94-A | 3.3 | 18.1 | 106 | 482 | 8 | MOLECULE: | CYTOCHROME | P450 4B | | 1 |
| 17:00 | 7dgu-A | 3.3 | 4.6 | 71 | 97 | 7 | MOLECULE: | A1 | | | |
| 18:00 | 7mhz-A | 3.3 | 10.9 | 92 | 490 | 3 | MOLECULE: | SONIC HEDG | EHOG PRO | TEIN N-PRODUCT PEPTIDE | |
| 19:00 | 7pq1-A | 3.3 | 7 | 82 | 395 | 4 | MOLECULE: | CYTOCHROME | P450 PR | OTEIN | |
| 20:00 | 1gzs-B | 3.3 | 4.1 | 85 | 165 | 8 | MOLECULE: | SOPE | | | |
| 21:00 | 8p50-A | 3.3 | 25.6 | 111 | 1574 | 5 | MOLECULE: | TOXIN PROT | EIN | | |
| 22:00 | 1dws-A | 3.3 | 4.8 | 81 | 152 | 10 | MOLECULE: | MYOGLOBIN | | | |
| 23:00 | 5jom-A | 3.3 | 4.4 | 79 | 154 | 9 | MOLECULE: | MYOGLOBIN | | | |
| 24:00:00 | 1npf-A | 3.3 | 4.8 | 81 | 152 | 10 | MOLECULE: | MYOGLOBIN | | | |
| 25:00:00 | 2frj-X | 3.3 | 4.8 | 81 | 152 | 10 | MOLECULE: | MYOGLOBIN | | | |
| 26:00:00 | 5o41-A | 3.3 | 4.4 | 78 | 154 | 9 | MOLECULE: | MYOGLOBIN | | | |
| 27:00:00 | 5cn8-A | 3.3 | 4.8 | 81 | 152 | 10 | MOLECULE: | MYOGLOBIN | | | |
| 28:00:00 | 5cnd-A | 3.3 | 4.8 | 81 | 152 | 10 | MOLECULE: | MYOGLOBIN | | | |
| 29:00:00 | 1gjn-A | 3.3 | 4.8 | 81 | 153 | 10 | MOLECULE: | MYOGLOBIN | | | |
| 30:00:00 | 1wla-A | 3.3 | 5.4 | 82 | 153 | 10 | MOLECULE: | MYOGLOBIN | | | |
| 31:00:00 | 2vly-A | 3.3 | 4.9 | 81 | 153 | 10 | MOLECULE: | MYOGLOBIN | | | |
| 32:00:00 | 1z8u-D | 3.3 | 3.3 | 80 | 135 | 6 | MOLECULE: | ALPHA-HEMO | GLOBIN S | TABILIZING PROTEIN | |
| 33:00:00 | 1npg-A | 3.3 | 4.8 | 81 | 152 | 10 | MOLECULE: | MYOGLOBIN | | | |
| 34:00:00 | 2frf-A | 3.3 | 4.8 | 81 | 152 | 10 | MOLECULE: | MYOGLOBIN | | | |
| 35:00:00 | 5cn9-A | 3.3 | 4.8 | 81 | 152 | 10 | MOLECULE: | MYOGLOBIN | | | |
| 36:00:00 | 2v1f-A | 3.3 | 4.8 | 81 | 153 | 10 | MOLECULE: | MYOGLOBIN | | | |
| 37:00:00 | 4dc7-A | 3.3 | 4.8 | 81 | 152 | 10 | MOLECULE: | MYOGLOBIN | | | |
| 38:00:00 | 4dc8-A | 3.3 | 4.8 | 81 | 152 | 10 | MOLECULE: | MYOGLOBIN | | | |
| 39:00:00 | 5z7e-A | 3.3 | 4.8 | 81 | 153 | 10 | MOLECULE: | MYOGLOBIN | | | |
| 40:00:00 | 5cnb-A | 3.3 | 4.8 | 81 | 152 | 10 | MOLECULE: | MYOGLOBIN | | | |
| 41:00:00 | 5cng-A | 3.3 | 4.5 | 79 | 152 | 10 | MOLECULE: | MYOGLOBIN | | | |
| 42:00:00 | 2v1e-A | 3.3 | 4.8 | 81 | 153 | 10 | MOLECULE: | MYOGLOBIN | | | |

| | | | | | | | | | | |
|---|---|---|---|---|---|---|---|---|---|---|
| 43:00:00 | 5cmv-A | 3.3 | 4.8 | 81 | 152 | 10 | MOLECULE: | MYOGLOBIN | | |
| 44:00:00 | 2o5l-X | 3.3 | 4.9 | 81 | 152 | 10 | MOLECULE: | MYOGLOBIN | | |
| 45:00:00 | 2fri-X | 3.3 | 4.8 | 81 | 152 | 10 | MOLECULE: | MYOGLOBIN | | |
| 46:00:00 | 2vm0-A | 3.3 | 4.8 | 81 | 153 | 10 | MOLECULE: | MYOGLOBIN | | |
| 47:00:00 | 5azq-A | 3.3 | 5.6 | 84 | 153 | 10 | MOLECULE: | MYOGLOBIN | | |
| 48:00:00 | 3vau-A | 3.3 | 4.8 | 81 | 151 | 10 | MOLECULE: | MYOGLOBIN | | |
| 49:00:00 | 2v1h-A | 3.3 | 4.8 | 81 | 153 | 10 | MOLECULE: | MYOGLOBIN | | |
| 50:00:00 | 1uc3-D | 3.3 | 3.2 | 77 | 149 | 3 | MOLECULE: | GLOBIN | | |
| 51:00:00 | 2h35-C | 3.3 | 4 | 86 | 141 | 8 | MOLECULE: | HEMOGLOBIN | ALPHA S | UBUNIT |
| 52:00:00 | 1uc3-G | 3.3 | 3.4 | 78 | 149 | 3 | MOLECULE: | GLOBIN | | |
| 53:00:00 | 2h35-A | 3.3 | 4 | 86 | 141 | 8 | MOLECULE: | HEMOGLOBIN | ALPHA S | UBUNIT |
| 54:00:00 | 1uc3-H | 3.3 | 3.6 | 78 | 149 | 3 | MOLECULE: | GLOBIN | | |
| 55:00:00 | 1uc3-A | 3.3 | 3.2 | 77 | 149 | 3 | MOLECULE: | GLOBIN | | |
| 56:00:00 | 1qoy-A | 3.2 | 4.5 | 93 | 303 | 5 | MOLECULE: | HEMOLYSIN | E | |
| 57:00:00 | 4cru-A | 3.2 | 10.3 | 103 | 242 | 7 | MOLECULE: | CCR4-NOT T | RANSCRIP | TION COMPLEX SUBUNIT 1 |
| 58:00:00 | 7xdi-F | 3.2 | 9.3 | 83 | 456 | 5 | MOLECULE: | VP1 | | |
| 59:00:00 | 2hs5-A | 3.2 | 4.5 | 98 | 207 | 15 | MOLECULE: | PUTATIVE T | RANSCRIP | TIONAL REGULATOR GNTR |
| 60:00:00 | 6dk9-F | 3.2 | 4.9 | 122 | 232 | 8 | MOLECULE: | DNA DAMAGE | #NAME? | LE PROTEIN |
| 61:00:00 | 6btx-A | 3.2 | 19.8 | 106 | 377 | 15 | MOLECULE: | SOLUTE CAR | RIER FAM | ILY 39 (IRON-REGULATED TRANSPOR |
| 62:00:00 | 4cej-B | 3.2 | 18.6 | 125 | 1156 | 8 | MOLECULE: | ATP-DEPEND | ENT HELI | CASE/NUCLEASE SUBUNIT A |
| 63:00:00 | 5w65-P | 3.2 | 13.4 | 86 | 481 | 6 | MOLECULE: | DNA-DIRECT | ED RNA P | OLYMERASE I SUBUNIT RPA190 |
| 64:00:00 | 5ajk-A | 3.2 | 3.8 | 66 | 148 | 2 | MOLECULE: | HOMOLOG OF | VACCINI | A VIRUS CDS F1L |
| 65:00:00 | 5tpm-A | 3.2 | 5.3 | 102 | 155 | 9 | MOLECULE: | PYRUVATE D | EHYDROGE | NASE COMPLEX REPRESSOR |
| 66:00:00 | 3bni-A | 3.2 | 4.9 | 62 | 174 | 11 | MOLECULE: | PUTATIVE T | ETR-FAMI | LY TRANSCRIPTIONAL REGULATOR |
| 67:00:00 | 7v9g-D | 3.2 | 15.6 | 80 | 112 | 5 | MOLECULE: | BEN DOMAIN | #NAME? | ING PROTEIN 3 |
| 68:00:00 | 6khj-E | 3.2 | 6.3 | 80 | 101 | 3 | MOLECULE: | NAD(P)H-QU | INONE OX | IDOREDUCTASE SUBUNIT 1 |
| 69:00:00 | 7dqk-B | 3.2 | 8.3 | 84 | 479 | 4 | MOLECULE: | PROTEIN DE | TOXIFICA | TION |
| 70:00:00 | 8uw3-A | 3.2 | 11.2 | 122 | 1265 | 5 | MOLECULE: | LINE-1 RET | ROTRANSP | OSABLE ELEMENT ORF2 PROTEIN |
| 71:00:00 | 8q7h-A | 3.2 | 17.9 | 155 | 1403 | 9 | MOLECULE: | CULLIN-9 | | |
| 72:00:00 | 5z1n-A | 3.2 | 3.8 | 83 | 166 | 12 | MOLECULE: | G-PROTEIN | INTERACT | ING PROTEIN 1 |
| 73:00:00 | 8j2x-A | 3.2 | 11.1 | 74 | 337 | 7 | MOLECULE: | COBALAMIN- | BINDING | PROTEIN |
| 74:00:00 | 6zyy-C | 3.2 | 26.7 | 126 | 2901 | 7 | MOLECULE: | DYNEIN HEA | VY CHAIN | , OUTER ARM PROTEIN |
| 75:00:00 | 8q3v-e | 3.2 | 11.8 | 111 | 294 | 8 | MOLECULE: | TETRAHYDRO | METHANOP | TERIN S-METHYLTRANSFERASE SUBUN |
| 76:00:00 | 5d2m-C | 3.2 | 4.7 | 76 | 158 | 16 | MOLECULE: | SUMO-CONJU | GATING E | NZYME UBC9 |
| 77:00:00 | 3h36-A | 3.2 | 3.9 | 67 | 78 | 0 | MOLECULE: | POLYRIBONU | CLEOTIDE | NUCLEOTIDYLTRANSFERASE |
| 78:00:00 | 4zsf-A | 3.2 | 5.1 | 68 | 272 | 6 | MOLECULE: | BSAWI ENDO | NUCLEASE | |
| 79:00:00 | 7kfu-C | 3.2 | 14.5 | 147 | 909 | 5 | MOLECULE: | CAS2 | | |
| 80:00:00 | 4y9j-B | 3.2 | 25.6 | 125 | 593 | 7 | MOLECULE: | PROTEIN AC | DH-11, I | SOFORM B |
| 81:00:00 | 8p1u-A | 3.2 | 7.8 | 88 | 370 | 10 | MOLECULE: | CELL DIVIS | ION PROT | EIN FTSL |
| 82:00:00 | 6xdc-A | 3.2 | 12.7 | 98 | 193 | 9 | MOLECULE: | PROTEIN 3A | | |
| 83:00:00 | 2yvx-A | 3.2 | 10.8 | 96 | 442 | 10 | MOLECULE: | MG2+ TRANS | PORTER M | GTE |
| 84:00:00 | 6rao-I | 3.2 | 20.8 | 110 | 443 | 3 | MOLECULE: | AFP1 | | |
| 85:00:00 | 2khm-A | 3.2 | 10.6 | 82 | 140 | 7 | MOLECULE: | FIBROIN-3 | | |
| 86:00:00 | 7pik-C | 3.2 | 24.7 | 87 | 557 | 1 | MOLECULE: | TRANSPOSON | TN7 TRA | NSPOSITION PROTEIN TNSB |
| 87:00:00 | 4hkr-A | 3.2 | 9.2 | 97 | 165 | 2 | MOLECULE: | CALCIUM RE | LEASE-AC | TIVATED CALCIUM CHANNEL PROTEIN |
| 88:00:00 | 3ia3-B | 3.2 | 3.6 | 82 | 135 | 6 | MOLECULE: | ALPHA-HEMO | GLOBIN-S | TABILIZING PROTEIN |
| 89:00:00 | 1ymc-A | 3.2 | 5.6 | 82 | 153 | 10 | MOLECULE: | CYANOMET-S | ULFMYOGL | OBIN |
| 90:00:00 | 3ia3-D | 3.2 | 3.6 | 82 | 135 | 6 | MOLECULE: | ALPHA-HEMO | GLOBIN-S | TABILIZING PROTEIN |
| 91:00:00 | 1umo-B | 3.2 | 4.7 | 88 | 154 | 5 | MOLECULE: | CYTOGLOBIN | | |
| 92:00:00 | 7a45-A | 3.2 | 4.4 | 80 | 154 | 9 | MOLECULE: | MYOGLOBIN | | |
| 93:00:00 | 6otx-A | 3.2 | 4.2 | 76 | 151 | 5 | MOLECULE: | HEMOGLOBIN | II | |
| 94:00:00 | 7dgo-A | 3.2 | 11.3 | 87 | 153 | 3 | MOLECULE: | MYOGLOBIN | | |
| 95:00:00 | 7uvb-B | 3.2 | 3.3 | 81 | 146 | 11 | MOLECULE: | HEMOGLOBIN | SUBUNIT | ALPHA |
| 96:00:00 | 3hyu-B | 3.2 | 3.6 | 84 | 146 | 11 | MOLECULE: | HEMOGLOBIN | SUBUNIT | ALPHA |

| | | | | | | | | | | | |
|---|---|---|---|---|---|---|---|---|---|---|---|
| 97:00:00 | 3hrw-B | 3.2 | 4.2 | 83 | 146 | 6 | MOLECULE: | HEMOGLOBIN | SUBUNIT | ALPHA | |
| 98:00:00 | 8q7e-J | 3.1 | 18.3 | 126 | 895 | 10 | MOLECULE: | CULLIN-9 | | | |
| 99:00:00 | 6wtw-A | 3.1 | 7.7 | 107 | 489 | 7 | MOLECULE: | DASS FAMIL | Y SODIUM | -COUPLED ANION SYMPORTER | |
| 0:00 | 7c7e-A | 3.1 | 4.7 | 97 | 142 | 5 | MOLECULE: | PUTATIVE D | NA-BINDI | NG TRANSCRIPTIONAL REGULATOR | |
| 1:00 | 3h9p-A | 3.1 | 4.3 | 82 | 226 | 9 | MOLECULE: | PUTATIVE T | RIPHOSPH | ORIBOSYL-DEPHOSPHO-COA SYNTHASE | |
| 2:00 | 5do7-D | 3.1 | 12.5 | 101 | 571 | 7 | MOLECULE: | ATP-BINDIN | G CASSET | TE SUB-FAMILY G MEMBER 5 | |
| 3:00 | 9b8q-J | 3.1 | 14.5 | 74 | 149 | 11 | MOLECULE: | V-TYPE PRO | TON ATPA | SE SUBUNIT C 1 | |
| 4:00 | 2ilk-A | 3.1 | 14.7 | 118 | 155 | 10 | MOLECULE: | INTERLEUKI | N-10 | | |
| 5:00 | 4q9v-B | 3.1 | 16.4 | 93 | 186 | 14 | MOLECULE: | TUMOR NECR | OSIS FAC | TOR ALPHA-INDUCED PROTEIN 8-LIK | |
| 6:00 | 7ccl-C | 3.1 | 9.2 | 72 | 147 | 6 | MOLECULE: | BCL-2-LIKE | PROTEIN | | 15 |
| 7:00 | 8v46-D | 3.1 | 4.4 | 89 | 305 | 8 | MOLECULE: | ARIA ANTIT | OXIN | | |
| 8:00 | 4mu6-A | 3.1 | 4.7 | 85 | 274 | 7 | MOLECULE: | KINECTIN 1 | (KINESI | N RECEPTOR) | |
| 9:00 | 6w6x-A | 3.1 | 8.2 | 78 | 126 | 9 | MOLECULE: | DE NOVO DE | SIGNED A | BLE PROTEIN | |
| 10:00 | 6vjy-B | 3.1 | 12.7 | 104 | 280 | 8 | MOLECULE: | ERAD-ASSOC | IATED E3 | UBIQUITIN-PROTEIN LIGASE HRD1 | |
| 11:00 | 3v8d-A | 3.1 | 11.2 | 92 | 479 | 5 | MOLECULE: | CHOLESTERO | L 7-ALPH | A-MONOOXYGENASE | |
| 12:00 | 8j98-A | 3.1 | 8.3 | 79 | 184 | 6 | MOLECULE: | NEAR-INFRA | RED FLUO | RESCENT PROTEIN | |
| 13:00 | 7z0o-G | 3.1 | 4 | 58 | 84 | 7 | MOLECULE: | HISTONE H3 | | | |
| 14:00 | 1r5i-D | 3.1 | 17.7 | 79 | 214 | 4 | MOLECULE: | HLA CLASS | II HISTO | COMPATIBILITY ANTIGEN, DR | |
| 15:00 | 7vw0-B | 3.1 | 9.1 | 76 | 94 | 9 | MOLECULE: | DUF305 DOM | AIN-CONT | AINING PROTEIN | |
| 16:00 | 8uc3-C | 3.1 | 7.4 | 70 | 98 | 3 | MOLECULE: | ALBONOURSI | N SYNTHA | SE | |
| 17:00 | 5e2q-A | 3.1 | 20.6 | 105 | 725 | 7 | MOLECULE: | DIPEPTIDYL | PEPTIDA | SE 3 | |
| 18:00 | 8w6c-A | 3.1 | 10.8 | 112 | 530 | 8 | MOLECULE: | SOLUTE CAR | RIER FAM | ILY 13 MEMBER 2 | |
| 19:00 | 1w9r-A | 3.1 | 10.9 | 93 | 119 | 6 | MOLECULE: | CHOLINE BI | NDING PR | OTEIN A | |
| 20:00 | 8tzk-C | 3.1 | 6.2 | 81 | 304 | 7 | MOLECULE: | CELL DIVIS | ION ATP- | BINDING PROTEIN FTSE | |
| 21:00 | 4uis-B | 3.1 | 4.2 | 88 | 241 | 7 | MOLECULE: | GAMMA-SECR | ETASE | | |
| 22:00 | 7yet-A | 3.1 | 29 | 131 | 999 | 6 | MOLECULE: | RNA-DIRECT | ED RNA P | OLYMERASE L | |
| 23:00 | 2qup-A | 3.1 | 7.9 | 78 | 119 | 4 | MOLECULE: | BH1478 PRO | TEIN | | |
| 24:00:00 | 8pm6-A | 3.1 | 23.1 | 108 | 872 | 6 | MOLECULE: | BILE SALT | EXPORT P | UMP | |
| 25:00:00 | 3rh2-A | 3.1 | 12 | 77 | 210 | 6 | MOLECULE: | HYPOTHETIC | AL TETR- | LIKE TRANSCRIPTIONAL REGULATOR | |
| 26:00:00 | 3bm3-A | 3.1 | 4.8 | 95 | 259 | 7 | MOLECULE: | DNA (5'-D( | *CP*AP*T | P*CP*CP*AP*GP*GP*TP*AP*C)-3') | |
| 27:00:00 | 6lum-G | 3.1 | 7.3 | 77 | 123 | 8 | MOLECULE: | SUCCINATE | DEHYDROG | ENASE SUBUNIT C | |
| 28:00:00 | 7wuq-R | 3.1 | 8.9 | 94 | 278 | 4 | MOLECULE: | GUANINE NU | CLEOTIDE | #NAME? | |
| 29:00:00 | 2f4q-A | 3.1 | 14.5 | 74 | 309 | 7 | MOLECULE: | TYPE I TOP | OISOMERA | SE, PUTATIVE | |
| 30:00:00 | 6cfw-G | 3.1 | 8.4 | 80 | 113 | 6 | MOLECULE: | MONOVALENT | CATION/ | H+ ANTIPORTER SUBUNIT D | |
| 31:00:00 | 3ps5-A | 3.1 | 7.6 | 103 | 529 | 7 | MOLECULE: | TYROSINE-P | ROTEIN P | HOSPHATASE NON-RECEPTOR TYPE 6 | |
| 32:00:00 | 5yhf-A | 3.1 | 7.8 | 121 | 734 | 7 | MOLECULE: | PROTEIN TR | ANSLOCAS | E SUBUNIT SECDF | |
| 33:00:00 | 8if8-B | 3.1 | 9.7 | 99 | 620 | 6 | MOLECULE: | GALACTAN 5 | #NAME? | NOFURANOSYLTRANSFERASE | |
| 34:00:00 | 7d3u-G | 3.1 | 6.1 | 79 | 111 | 6 | MOLECULE: | MONOVALENT | NA+/H+ | ANTIPORTER SUBUNIT D | |
| 35:00:00 | 7op8-A | 3.1 | 21.7 | 111 | 1118 | 5 | MOLECULE: | CATION-TRA | NSPORTIN | G ATPASE | |
| 36:00:00 | 6lum-D | 3.1 | 8.9 | 82 | 146 | 5 | MOLECULE: | SUCCINATE | DEHYDROG | ENASE SUBUNIT C | |
| 37:00:00 | 7ar9-K | 3.1 | 6.5 | 72 | 104 | 10 | MOLECULE: | ND3 | | | |
| 38:00:00 | 7tmw-R | 3.1 | 16.7 | 100 | 499 | 7 | MOLECULE: | RELAXIN RE | CEPTOR 1 | , GUANINE NUCLEOTIDE-BINDING PR | |
| 39:00:00 | 4ap2-B | 3.1 | 20.1 | 125 | 359 | 7 | MOLECULE: | KELCH-LIKE | PROTEIN | | 11 |
| 40:00:00 | 8fm9-M | 3.1 | 12.5 | 108 | 333 | 8 | MOLECULE: | RNA-DIRECT | ED RNA P | OLYMERASE | |
| 41:00:00 | 2wus-R | 3.1 | 2.9 | 60 | 87 | 7 | MOLECULE: | PUTATIVE U | NCHARACT | ERIZED PROTEIN | |
| 42:00:00 | 6zmx-D | 3.1 | 4 | 77 | 146 | 17 | MOLECULE: | HEMOGLOBIN | SUBUNIT | ALPHA-A | |
| 43:00:00 | 4hrr-D | 3.1 | 4.2 | 87 | 151 | 7 | MOLECULE: | GLOBIN-2 A | CHAIN | | |
| 44:00:00 | 6zmy-B | 3.1 | 3.9 | 77 | 146 | 17 | MOLECULE: | HEMOGLOBIN | SUBUNIT | ALPHA-A | |
| 45:00:00 | 5eui-A | 3.1 | 4.1 | 75 | 134 | 11 | MOLECULE: | HBA PROTEI | N | | |
| 46:00:00 | 1jp6-A | 3.1 | 4.9 | 80 | 152 | 9 | MOLECULE: | MYOGLOBIN | | | |
| 47:00:00 | 4ood-A | 3.1 | 4.9 | 88 | 154 | 6 | MOLECULE: | MYOGLOBIN | | | |
| 48:00:00 | 4mqk-B | 3.1 | 4.4 | 80 | 145 | 14 | MOLECULE: | HEMOGLOBIN | SUBUNIT | ALPHA | |
| 49:00:00 | 1hv4-D | 3.1 | 4 | 76 | 146 | 17 | MOLECULE: | HEMOGLOBIN | ALPHA-A | CHAIN | |
| 50:00:00 | 3pi4-A | 3.1 | 4.2 | 77 | 151 | 5 | MOLECULE: | HEMOGLOBIN | II | | |

| | | | | | | | | | | |
|---|---|---|---|---|---|---|---|---|---|---|
| 51:00:00 | 1ch4-D | 3.1 | 3.5 | 77 | 146 | 8 | MOLECULE: | MODULE-SUB | STITUTED | CHIMERA HEMOGLOBIN BETA-ALPHA |
| 52:00:00 | 7k4m-D | 3.1 | 3.4 | 81 | 147 | 11 | MOLECULE: | HEMOGLOBIN | SUBUNIT | ALPHA |
| 53:00:00 | 5ufj-D | 3.1 | 3.3 | 82 | 146 | 11 | MOLECULE: | HEMOGLOBIN | SUBUNIT | ALPHA |
| 54:00:00 | 1ch4-C | 3.1 | 3.6 | 78 | 146 | 9 | MOLECULE: | MODULE-SUB | STITUTED | CHIMERA HEMOGLOBIN BETA-ALPHA |
| 55:00:00 | 3fs4-B | 3.1 | 3.6 | 85 | 146 | 12 | MOLECULE: | HEMOGLOBIN | SUBUNIT | ALPHA-A |
| 56:00:00 | 1myf-A | 3.1 | 4.2 | 79 | 153 | 9 | MOLECULE: | MYOGLOBIN | | |
| 57:00:00 | 7ot9-A | 3 | 8.2 | 79 | 221 | 3 | MOLECULE: | AI-2E MEMB | ER YDIK | |
| 58:00:00 | 6zy2-E | 3 | 17 | 93 | 259 | 8 | MOLECULE: | YRBD PROTE | IN | |
| 59:00:00 | 8gkh-P | 3 | 23.2 | 114 | 600 | 7 | MOLECULE: | DNA (5'-D( | P*CP*GP* | GP*TP*AP*CP*CP*CP*GP*GP*GP*CP*A |
| 60:00:00 | 6gkw-A | 3 | 4.8 | 76 | 351 | 8 | MOLECULE: | PUTATIVE P | HAGE XKD | K-LIKE PROTEIN |
| 61:00:00 | 7nur-A | 3 | 15.7 | 145 | 1044 | 10 | MOLECULE: | PROTEIN KI | NASE DOM | AIN-CONTAINING PROTEIN |
| 62:00:00 | 7rpk-A | 3 | 19.8 | 147 | 953 | 7 | MOLECULE: | PROTEIN DI | SPATCHED | HOMOLOG 1 |
| 63:00:00 | 2z3r-E | 3 | 3.5 | 70 | 119 | 4 | MOLECULE: | INTERLEUKI | N-15 | |
| 64:00:00 | 7lhe-A | 3 | 13.3 | 107 | 2222 | 5 | MOLECULE: | INOSITOL 1 | ,4,5-TRI | SPHOSPHATE RECEPTOR TYPE 1 |
| 65:00:00 | 7vu7-A | 3 | 4.2 | 64 | 89 | 11 | MOLECULE: | FLAGELLIFO | RM FIBRO | IN |
| 66:00:00 | 7d3u-C | 3 | 7.9 | 90 | 129 | 10 | MOLECULE: | MONOVALENT | NA+/H+ | ANTIPORTER SUBUNIT D |
| 67:00:00 | 2oeb-A | 3 | 3.5 | 79 | 152 | 5 | MOLECULE: | HYPOTHETIC | AL PROTE | IN |
| 68:00:00 | 1iie-A | 3 | 5.2 | 51 | 75 | 6 | MOLECULE: | PROTEIN (H | LA-DR AN | TIGENS ASSOCIATED INVARIANT |
| 69:00:00 | 6zvh-R | 3 | 14.4 | 80 | 135 | 3 | MOLECULE: | 18S RRNA | | |
| 70:00:00 | 5yi4-A | 3 | 12.5 | 83 | 145 | 7 | MOLECULE: | DISRUPTED | IN SCHIZ | OPHRENIA 1 HOMOLOG,NUCLEAR DIST |
| 71:00:00 | 7awt-D | 3 | 20.4 | 114 | 589 | 11 | MOLECULE: | NADH-QUINO | NE OXIDO | REDUCTASE SUBUNIT B |
| 72:00:00 | 8j3r-B | 3 | 9.8 | 91 | 387 | 7 | MOLECULE: | TRANSPOSAS | E IS605 | ORFB C-TERMINAL DOMAIN-CONTAINI |
| 73:00:00 | 7sbe-A | 3 | 4 | 90 | 228 | 7 | MOLECULE: | TELOMERASE | REVERSE | TRANSCRIPTASE |
| 74:00:00 | 1xsz-A | 3 | 14.9 | 69 | 356 | 4 | MOLECULE: | GUANINE NU | CLEOTIDE | EXCHANGE PROTEIN |
| 75:00:00 | 7wst-A | 3 | 17.4 | 103 | 601 | 6 | MOLECULE: | IRON-PHYTO | SIDEROPH | ORE TRANSPORTER |
| 76:00:00 | 6ebu-A | 3 | 4.9 | 91 | 165 | 8 | MOLECULE: | LPXE | | |
| 77:00:00 | 5mps-R | 3 | 11.2 | 56 | 108 | 0 | MOLECULE: | YEAST UBC4 | GENE FO | R UBIQUITIN-CONJUGATING ENZYME |
| 78:00:00 | 8tl7-A | 3 | 8 | 106 | 694 | 6 | MOLECULE: | COMPUTATIO | NALLY DE | SIGNED PROTEIN |
| 79:00:00 | 1knz-A | 3 | 6.1 | 79 | 154 | 3 | MOLECULE: | 5'-R(*UP*G | P*AP*CP* | C)-3' |
| 80:00:00 | 5m0n-A | 3 | 13.5 | 132 | 423 | 7 | MOLECULE: | TERMINAL O | LEFIN-FO | RMING FATTY ACID DECARBOXYLASE |
| 81:00:00 | 1tu9-A | 3 | 4.5 | 76 | 131 | 9 | MOLECULE: | HYPOTHETIC | AL PROTE | IN PA3967 |
| 82:00:00 | 3k3u-B | 3 | 4.6 | 72 | 137 | 4 | MOLECULE: | DEHALOPERO | XIDASE A | |
| 83:00:00 | 8kdb-A | 3 | 11.7 | 91 | 2117 | 3 | MOLECULE: | RNA-DIRECT | ED RNA P | OLYMERASE L |
| 84:00:00 | 6gys-B | 3 | 27.1 | 112 | 579 | 4 | MOLECULE: | CENTROMERE | DNA-BIN | DING PROTEIN COMPLEX CBF3 SUBUN |
| 85:00:00 | 3f7c-A | 3 | 5.7 | 67 | 199 | 9 | MOLECULE: | PROTEIN OF | UNKNOWN | FUNCTION (DUF416) |
| 86:00:00 | 4raf-A | 3 | 17.9 | 88 | 362 | 10 | MOLECULE: | PROTEIN PH | OSPHATAS | E 1A |
| 87:00:00 | 8a3t-C | 3 | 31.5 | 117 | 1410 | 9 | MOLECULE: | ANAPHASE-P | ROMOTING | COMPLEX SUBUNIT CDC27 |
| 88:00:00 | 8qq7-A | 3 | 19.1 | 120 | 399 | 8 | MOLECULE: | FAD-BINDIN | G FR-TYP | E DOMAIN-CONTAINING PROTEIN |
| 89:00:00 | 7zmb-6 | 3 | 7 | 87 | 190 | 11 | MOLECULE: | NADH-UBIQU | INONE OX | IDOREDUCTASE CHAIN 1 |
| 90:00:00 | 7ryq-B | 3 | 13.9 | 101 | 527 | 8 | MOLECULE: | KIF-BINDIN | G PROTEI | N |
| 91:00:00 | 4jbw-F | 3 | 15.8 | 99 | 492 | 3 | MOLECULE: | MALTOSE TR | ANSPORT | SYSTEM PERMEASE PROTEIN MALF |
| 92:00:00 | 8ca1-A | 3 | 12.9 | 129 | 589 | 8 | MOLECULE: | VERY LONG- | CHAIN SP | ECIFIC ACYL-COA DEHYDROGENASE, |
| 93:00:00 | 3tix-A | 3 | 10.3 | 75 | 153 | 12 | MOLECULE: | UBIQUITIN- | LIKE PRO | TEIN SMT3,RNA-INDUCED TRANSCRIP |
| 94:00:00 | 8c3y-A | 3 | 17.3 | 114 | 929 | 4 | MOLECULE: | PFEMP1 | | |
| 95:00:00 | 6ysg-A | 3 | 19.1 | 133 | 1264 | 7 | MOLECULE: | MG-CHELATA | SE SUBUN | IT CHLH |
| 96:00:00 | 5w7g-B | 3 | 9.5 | 66 | 126 | 9 | MOLECULE: | ORF140 | | |
| 97:00:00 | 7yh5-B | 3 | 9.1 | 91 | 177 | 15 | MOLECULE: | NUCLEOSIDE | TRIPHOS | PHATE PYROPHOSPHOHYDROLASE |
| 98:00:00 | 7mxb-A | 3 | 10.7 | 75 | 672 | 12 | MOLECULE: | SERINE/THR | EONINE P | ROTEIN KINASES |
| 99:00:00 | 5w7g-A | 3 | 10.7 | 72 | 131 | 6 | MOLECULE: | ORF140 | | |
| 0:00 | 1uc3-J | 3 | 3.9 | 79 | 149 | 4 | MOLECULE: | GLOBIN | | |
| 1:00 | 2lhb-A | 3 | 3.9 | 78 | 149 | 4 | MOLECULE: | HEMOGLOBIN | V (CYAN | O MET) |
| 2:00 | 2di3-B | 3 | 9.5 | 101 | 230 | 6 | MOLECULE: | BACTERIAL | REGULATO | RY PROTEINS, GNTR FAMILY |
| 3:00 | 3pi4-B | 3 | 4.3 | 76 | 151 | 4 | MOLECULE: | HEMOGLOBIN | II | |
| 4:00 | 4mqj-H | 3 | 4.5 | 79 | 146 | 11 | MOLECULE: | HEMOGLOBIN | SUBUNIT | ALPHA |

| | | | | | | | | | |
|---|---|---|---|---|---|---|---|---|---|
| 5:00 | 4mqk-F | 3 | 4.5 | 79 | 145 | 14 | MOLECULE: | HEMOGLOBIN | SUBUNIT | ALPHA |
| 6:00 | 6zmy-D | 3 | 4.7 | 79 | 146 | 15 | MOLECULE: | HEMOGLOBIN | SUBUNIT | ALPHA-A |
| 7:00 | 3d4x-D | 3 | 3.9 | 76 | 145 | 14 | MOLECULE: | HEMOGLOBIN | SUBUNIT | ALPHA |
| 8:00 | 3gys-F | 3 | 4.2 | 77 | 145 | 13 | MOLECULE: | HEMOGLOBIN | SUBUNIT | ALPHA |
| 9:00 | 3dhr-B | 3 | 4.6 | 79 | 145 | 16 | MOLECULE: | HEMOGLOBIN | SUBUNIT | ALPHA-A |
| 10:00 | 3dhr-F | 3 | 4.8 | 79 | 145 | 15 | MOLECULE: | HEMOGLOBIN | SUBUNIT | ALPHA-A |
| 11:00 | 3d4x-B | 3 | 4.5 | 79 | 145 | 14 | MOLECULE: | HEMOGLOBIN | SUBUNIT | ALPHA |
| 12:00 | 2hbs-D | 3 | 4.1 | 76 | 146 | 13 | MOLECULE: | HEMOGLOBIN | S (DEOX | Y), ALPHA CHAIN |
| 13:00 | 1dxt-D | 3 | 4.2 | 80 | 147 | 13 | MOLECULE: | HEMOGLOBIN | (DEOXY) | (ALPHA CHAIN) |
| 14:00 | 3dhr-H | 3 | 4.8 | 79 | 145 | 15 | MOLECULE: | HEMOGLOBIN | SUBUNIT | ALPHA-A |
| 15:00 | 3gys-H | 3 | 3.7 | 78 | 145 | 10 | MOLECULE: | HEMOGLOBIN | SUBUNIT | ALPHA |
| 16:00 | 3gqr-D | 3 | 4 | 77 | 145 | 14 | MOLECULE: | HEMOGLOBIN | SUBUNIT | ALPHA |
| 17:00 | 7jy3-B | 3 | 3.4 | 82 | 146 | 10 | MOLECULE: | HEMOGLOBIN | SUBUNIT | ALPHA |
| 18:00 | 1si4-B | 3 | 3.5 | 84 | 146 | 10 | MOLECULE: | HEMOGLOBIN | ALPHA C | HAIN |
| 19:00 | 7qu4-G | 3 | 4 | 77 | 143 | 12 | MOLECULE: | HEMOGLOBIN | SUBUNIT | GAMMA-2 |
| 20:00 | 7uf7-D | 3 | 3.6 | 79 | 146 | 11 | MOLECULE: | HEMOGLOBIN | SUBUNIT | ALPHA |
| 21:00 | 2di3-A | 2.9 | 7 | 104 | 231 | 8 | MOLECULE: | BACTERIAL | REGULATO | RY PROTEINS, GNTR FAMILY |
| 22:00 | 7kpv-B | 2.9 | 10 | 92 | 290 | 8 | MOLECULE: | MEIOTIC MR | NA STABI | LITY PROTEIN KINASE SSN3 |
| 23:00 | 6mpb-A | 2.9 | 7.4 | 110 | 447 | 6 | MOLECULE: | NEUTRAL AM | INO ACID | TRANSPORTER B(0) |
| 24:00:00 | 7kzn-B | 2.9 | 22.5 | 110 | 683 | 5 | MOLECULE: | HEAVY CHAI | N ALPHA | |
| 25:00:00 | 2wtg-A | 2.9 | 4.2 | 76 | 158 | 5 | MOLECULE: | GLOBIN-LIK | E PROTEI | N |
| 26:00:00 | 7ro2-A | 2.9 | 8.4 | 61 | 94 | 10 | MOLECULE: | MAJOR CAPS | ID PROTE | IN |
| 27:00:00 | 2be4-A | 2.9 | 8.2 | 103 | 270 | 9 | MOLECULE: | HYPOTHETIC | AL PROTE | IN LOC449832 |
| 28:00:00 | 5cd4-I | 2.9 | 18.5 | 83 | 494 | 7 | MOLECULE: | CRISPR SYS | TEM CASC | ADE SUBUNIT CASE |
| 29:00:00 | 7qru-G | 2.9 | 9.2 | 93 | 113 | 4 | MOLECULE: | NA+/H+ ANT | IPORTER | SUBUNIT D |
| 30:00:00 | 8eki-E | 2.9 | 18.1 | 115 | 643 | 6 | MOLECULE: | PROTEIN TR | ANSPORT | PROTEIN SEC20 |
| 31:00:00 | 6i3m-E | 2.9 | 6.2 | 94 | 354 | 10 | MOLECULE: | TRANSLATIO | N INITIA | TION FACTOR EIF-2B SUBUNIT ALPH |
| 32:00:00 | 3qbg-A | 2.9 | 10.3 | 103 | 261 | 8 | MOLECULE: | HALORHODOP | SIN | |
| 33:00:00 | 4ymt-B | 2.9 | 4 | 83 | 216 | 10 | MOLECULE: | ABC-TYPE P | OLAR AMI | NO ACID TRANSPORT SYSTEM, ATPAS |
| 34:00:00 | 5lnk-J | 2.9 | 6.7 | 80 | 175 | 5 | MOLECULE: | MITOCHONDR | IAL COMP | LEX I, 51 KDA SUBUNIT |
| 35:00:00 | 1whu-A | 2.9 | 5.5 | 74 | 104 | 9 | MOLECULE: | POLYNUCLEO | TIDE PHO | SPHORYLASE 3'-5' RNA |
| 36:00:00 | 4c0n-A | 2.9 | 4.7 | 86 | 150 | 8 | MOLECULE: | 2-ON-2 HEM | OGLOBIN | |
| 37:00:00 | 8qa2-A | 2.9 | 10.6 | 72 | 215 | 7 | MOLECULE: | GAP JUNCTI | ON BETA- | 2 PROTEIN |
| 38:00:00 | 6vm0-A | 2.9 | 18.1 | 85 | 363 | 4 | MOLECULE: | GLYCINE RE | CEPTOR S | UBUNIT ALPHAZ1 |
| 39:00:00 | 8wm8-B | 2.9 | 5.3 | 89 | 265 | 10 | MOLECULE: | NITRATE TR | ANSPORT | ATP-BINDING PROTEIN |
| 40:00:00 | 3h3m-A | 2.9 | 5.6 | 73 | 89 | 12 | MOLECULE: | FLAGELLAR | PROTEIN | FLIT |
| 41:00:00 | 7xdi-D | 2.9 | 6.1 | 79 | 124 | 5 | MOLECULE: | VP1 | | |
| 42:00:00 | 8odu-C | 2.9 | 23.2 | 90 | 232 | 10 | MOLECULE: | ATPASE GET | 3 | |
| 43:00:00 | 3nnq-A | 2.9 | 4.4 | 66 | 95 | 8 | MOLECULE: | N-TERMINAL | DOMAIN | OF MOLONEY MURINE LEUKEMIA VIRU |
| 44:00:00 | 3c1d-B | 2.9 | 16.7 | 76 | 154 | 8 | MOLECULE: | REGULATORY | PROTEIN | RECX |
| 45:00:00 | 6zyd-A | 2.9 | 21.6 | 106 | 323 | 4 | MOLECULE: | LOW CONDUC | TANCE ME | CHANOSENSITIVE CHANNEL YNAI,LOW |
| 46:00:00 | 2q5z-B | 2.9 | 15.7 | 69 | 94 | 7 | MOLECULE: | HYPOTHETIC | AL PROTE | IN |
| 47:00:00 | 5o5j-O | 2.9 | 5.9 | 63 | 88 | 16 | MOLECULE: | 16S RRNA | | |
| 48:00:00 | 7woo-Z | 2.9 | 30 | 109 | 746 | 4 | MOLECULE: | NUCLEOPORI | N NIC96 | |
| 49:00:00 | 3i4m-D | 2.9 | 10.4 | 84 | 187 | 7 | MOLECULE: | DNA-DIRECT | ED RNA P | OLYMERASE II SUBUNIT RPB1 |
| 50:00:00 | 4cej-A | 2.9 | 30 | 128 | 1177 | 7 | MOLECULE: | ATP-DEPEND | ENT HELI | CASE/NUCLEASE SUBUNIT A |
| 51:00:00 | 8t53-A | 2.9 | 5.6 | 122 | 398 | 6 | MOLECULE: | UNDECAPREN | YL-PHOSP | HATE GALACTOSE PHOSPHOTRANSFERA |
| 52:00:00 | 6rfl-I | 2.9 | 8.6 | 79 | 773 | 4 | MOLECULE: | DNA-DEPEND | ENT RNA | POLYMERASE SUBUNIT RPO132 |
| 53:00:00 | 7way-A | 2.9 | 33.4 | 131 | 960 | 5 | MOLECULE: | DNA (33-ME | R) | |
| 54:00:00 | 8kee-C | 2.9 | 13.9 | 89 | 505 | 7 | MOLECULE: | SHEATH | | |
| 55:00:00 | 1jog-D | 2.9 | 7.5 | 77 | 136 | 6 | MOLECULE: | HYPOTHETIC | AL PROTE | IN HI0074 |
| 56:00:00 | 2k85-A | 2.9 | 3.1 | 58 | 65 | 5 | MOLECULE: | GLUCOCORTI | COID REC | EPTOR DNA-BINDING FACTOR 1 |
| 57:00:00 | 6u8y-d | 2.9 | 6.5 | 72 | 89 | 3 | MOLECULE: | MONOVALENT | CATION/ | H+ ANTIPORTER SUBUNIT E |
| 58:00:00 | 8arl-A | 2.9 | 4.6 | 80 | 232 | 5 | MOLECULE: | TRYPTOPHAN | #NAME? | TIGEN |

| | | | | | | | | | | | |
|---|---|---|---|---|---|---|---|---|---|---|---|
| 59:00:00 | 8eau-e | 2.9 | 13.9 | 67 | 72 | 4 | MOLECULE: | V-TYPE PRO | TON ATPA | SE SUBUNIT A, VACUOLAR ISOFORM | |
| 60:00:00 | 5d67-D | 2.9 | 7.6 | 74 | 95 | 5 | MOLECULE: | EF-HAND CA | LCIUM-BI | NDING DOMAIN-CONTAINING PROTEIN | |
| 61:00:00 | 8iqi-C | 2.9 | 19.7 | 115 | 916 | 3 | MOLECULE: | PUTATIVE P | RIMASE C | 962R | |
| 62:00:00 | 1x8z-A | 2.9 | 8.9 | 92 | 151 | 8 | MOLECULE: | INVERTASE/ | PECTIN M | ETHYLESTERASE INHIBITOR FAMILY | |
| 63:00:00 | 3vr8-D | 2.9 | 14.7 | 83 | 129 | 10 | MOLECULE: | FLAVOPROTE | IN SUBUN | IT OF COMPLEX II | |
| 64:00:00 | 3mjp-B | 2.9 | 4.3 | 84 | 146 | 5 | MOLECULE: | HEMOGLOBIN | SUBUNIT | ALPHA-A | |
| 65:00:00 | 5u3i-D | 2.9 | 4 | 77 | 145 | 14 | MOLECULE: | HEMOGLOBIN | SUBUNIT | ALPHA | |
| 66:00:00 | 2hhe-B | 2.9 | 4.7 | 79 | 145 | 13 | MOLECULE: | HEMOGLOBIN | (DEOXY) | (ALPHA CHAIN) | |
| 67:00:00 | 3k8b-B | 2.9 | 4 | 77 | 146 | 17 | MOLECULE: | HEMOGLOBIN | SUBUNIT | ALPHA-A | |
| 68:00:00 | 8vyl-D | 2.9 | 4.5 | 78 | 146 | 14 | MOLECULE: | HEMOGLOBIN | SUBUNIT | ALPHA | |
| 69:00:00 | 3gqp-D | 2.9 | 4.4 | 78 | 145 | 14 | MOLECULE: | HEMOGLOBIN | SUBUNIT | ALPHA | |
| 70:00:00 | 3vrf-B | 2.9 | 4.5 | 79 | 146 | 13 | MOLECULE: | HEMOGLOBIN | SUBUNIT | ALPHA | |
| 71:00:00 | 2hhe-D | 2.9 | 4.7 | 80 | 145 | 13 | MOLECULE: | HEMOGLOBIN | (DEOXY) | (ALPHA CHAIN) | |
| 72:00:00 | 1fhj-D | 2.9 | 4 | 76 | 146 | 16 | MOLECULE: | HEMOGLOBIN | (ALPHA | CHAIN) | |
| 73:00:00 | 3vrg-B | 2.9 | 4 | 77 | 146 | 14 | MOLECULE: | HEMOGLOBIN | SUBUNIT | ALPHA | |
| 74:00:00 | 6wg7-D | 2.9 | 11.8 | 104 | 219 | 7 | MOLECULE: | DNA (35-ME | R) | | |
| 75:00:00 | 7uvb-D | 2.9 | 3.3 | 79 | 145 | 11 | MOLECULE: | HEMOGLOBIN | SUBUNIT | ALPHA | |
| 76:00:00 | 6on4-A | 2.9 | 15.5 | 93 | 217 | 4 | MOLECULE: | HTH-TYPE T | RANSCRIP | TIONAL REPRESSOR NANR | |
| 77:00:00 | 6wg7-H | 2.9 | 11.8 | 104 | 219 | 7 | MOLECULE: | DNA (35-ME | R) | | |
| 78:00:00 | 3kdw-A | 2.8 | 15.1 | 100 | 206 | 3 | MOLECULE: | PUTATIVE S | UGAR BIN | DING PROTEIN | |
| 79:00:00 | 4pi0-A | 2.8 | 27.6 | 99 | 390 | 7 | MOLECULE: | UNKNOWN PE | PTIDE | | |
| 80:00:00 | 7lsy-Y | 2.8 | 10.4 | 108 | 855 | 7 | MOLECULE: | X-RAY REPA | IR CROSS | -COMPLEMENTING PROTEIN 6 | |
| 81:00:00 | 7zw6-A | 2.8 | 26 | 115 | 793 | 5 | MOLECULE: | SLR0869 PR | OTEIN | | |
| 82:00:00 | 1uzc-A | 2.8 | 2.7 | 54 | 69 | 7 | MOLECULE: | HYPOTHETIC | AL PROTE | IN FLJ21157 | |
| 83:00:00 | 2lor-A | 2.8 | 7.2 | 64 | 108 | 5 | MOLECULE: | TRANSMEMBR | ANE PROT | EIN 141 | |
| 84:00:00 | 5gm2-B | 2.8 | 22.8 | 90 | 283 | 8 | MOLECULE: | O-METHYLRA | NSFERASE | | |
| 85:00:00 | 5hdi-A | 2.8 | 4.9 | 123 | 397 | 2 | MOLECULE: | CYTOCHROME | P450 14 | | 4 |
| 86:00:00 | 7s5l-A | 2.8 | 5.6 | 90 | 378 | 12 | MOLECULE: | CEMBRENE A | SYNTHAS | E | |
| 87:00:00 | 5udb-4 | 2.8 | 22.7 | 95 | 751 | 9 | MOLECULE: | DNA REPLIC | ATION LI | CENSING FACTOR MCM2 | |
| 88:00:00 | 4uig-A | 2.8 | 5.4 | 56 | 91 | 7 | MOLECULE: | COPPER SEN | SITIVE O | PERON REPRESSOR | |
| 89:00:00 | 7zpo-M | 2.8 | 5.8 | 124 | 438 | 5 | MOLECULE: | KTR SYSTEM | POTASSI | UM UPTAKE PROTEIN A | |
| 90:00:00 | 6zz6-B | 2.8 | 26.1 | 93 | 423 | 8 | MOLECULE: | STRUCTURAL | MAINTEN | ANCE OF CHROMOSOMES PROTEIN 1,S | |
| 91:00:00 | 8iic-B | 2.8 | 12.7 | 122 | 425 | 7 | MOLECULE: | POLYMERASE | POLYPRO | TEIN | |
| 92:00:00 | 2ijd-1 | 2.8 | 14.8 | 119 | 644 | 6 | MOLECULE: | PICORNAIN | 3C, RNA- | DIRECTED RNA POLYMERASE | |
| 93:00:00 | 7r5s-I | 2.8 | 25.3 | 121 | 622 | 6 | MOLECULE: | CENTROMERE | PROTEIN | H | |
| 94:00:00 | 6s6b-B | 2.8 | 3.7 | 79 | 154 | 5 | MOLECULE: | CRISPR-ASS | OCIATED | PROTEIN, CMR5 FAMILY | |
| 95:00:00 | 7e38-B | 2.8 | 8.3 | 96 | 305 | 4 | MOLECULE: | DEOXYPODOP | HYLLOTOX | IN SYNTHASE | |
| 96:00:00 | 6tmi-B | 2.8 | 19.5 | 104 | 246 | 8 | MOLECULE: | ATP SYNTHA | SE SUBUN | IT ALPHA | |
| 97:00:00 | 6e11-E | 2.8 | 15.9 | 99 | 210 | 6 | MOLECULE: | UNKNOWN (C | LAW) | | |
| 98:00:00 | 2wj9-B | 2.8 | 6 | 79 | 145 | 6 | MOLECULE: | INTERGENIC | #NAME? | PROTEIN | |
| 99:00:00 | 3bjd-C | 2.8 | 14.8 | 90 | 324 | 3 | MOLECULE: | PUTATIVE 3 | #NAME? | -(ACYL-CARRIER-PROTEIN) SYNTHAS | |
| 0:00 | 5abb-A | 2.8 | 18.7 | 72 | 443 | 7 | MOLECULE: | PROTEIN TR | ANSLOCAS | E SUBUNIT SECY | |
| 1:00 | 8tsh-C | 2.8 | 5.5 | 97 | 259 | 5 | MOLECULE: | ABC TRANSP | ORTER AT | P-BINDING PROTEIN | |
| 2:00 | 8hra-C | 2.8 | 15.7 | 109 | 852 | 6 | MOLECULE: | ARCHAEAL A | TPASE | | |
| 3:00 | 8ciw-A | 2.8 | 14.5 | 105 | 555 | 9 | MOLECULE: | (2S)-METHY | LSUCCINY | L-COA DEHYDROGENASE | |
| 4:00 | 6lo8-D | 2.8 | 8 | 83 | 119 | 11 | MOLECULE: | MITOCHONDR | IAL IMPO | RT INNER MEMBRANE TRANSLOCASE S | |
| 5:00 | 7zr6-C | 2.8 | 16.8 | 90 | 277 | 6 | MOLECULE: | HEAT SHOCK | PROTEIN | HSP 90-BETA | |
| 6:00 | 8ipr-A | 2.8 | 24.2 | 120 | 504 | 8 | MOLECULE: | COMPONENT | LINKED W | ITH THE ASSEMBLY OF CYTOCHROME' | |
| 7:00 | 7bmh-A | 2.8 | 8.4 | 92 | 247 | 4 | MOLECULE: | OPSIN | | | |
| 8:00 | 6zvh-y | 2.8 | 2.7 | 57 | 72 | 11 | MOLECULE: | 18S RRNA | | | |
| 9:00 | 6n39-A | 2.8 | 11.3 | 76 | 404 | 3 | MOLECULE: | DEPHOSPHO- | COA KINA | SE | |
| 10:00 | 7v99-A | 2.8 | 20.7 | 156 | 991 | 6 | MOLECULE: | TELOMERASE | REVERSE | TRANSCRIPTASE | |
| 11:00 | 6k7l-A | 2.8 | 26 | 92 | 992 | 10 | MOLECULE: | PHOSPHOLIP | ID-TRANS | PORTING ATPASE | |
| 12:00 | 5b7j-A | 2.8 | 3.7 | 82 | 110 | 5 | MOLECULE: | SWITCH-ACT | IVATING | PROTEIN 1 | |

| | | | | | | | | | | |
|---|---|---|---|---|---|---|---|---|---|---|
| 13:00 | 6whg-A | 2.8 | 21.1 | 140 | 523 | 8 | MOLECULE: | CALCIUM CH | ANNEL YV | C1 |
| 14:00 | 8esv-B | 2.8 | 4.5 | 71 | 255 | 6 | MOLECULE: | DISINTEGRI | N AND ME | TALLOPROTEINASE DOMAIN-CONTAINI |
| 15:00 | 6dlu-P | 2.8 | 24.6 | 121 | 747 | 6 | MOLECULE: | DYNAMIN-1 | | |
| 16:00 | 6kex-A | 2.8 | 7.3 | 66 | 349 | 5 | MOLECULE: | PHOSPHORIB | ULOKINAS | E |
| 17:00 | 5dn6-H | 2.8 | 4.3 | 68 | 110 | 3 | MOLECULE: | CHAIN A | | |
| 18:00 | 6urt-A | 2.8 | 18.7 | 125 | 331 | 8 | MOLECULE: | LOW CONDUC | TANCE ME | CHANOSENSITIVE CHANNEL YNAI |
| 19:00 | 4kha-A | 2.8 | 16.1 | 75 | 360 | 7 | MOLECULE: | SPT16M-HIS | TONE H2B | 1.1 CHIMERA |
| 20:00 | 7bw0-R | 2.8 | 14.6 | 108 | 280 | 8 | MOLECULE: | SOLUBLE CY | TOCHROME | B562,G-PROTEIN COUPLED BILE AC |
| 21:00 | 6ql4-A | 2.8 | 25.5 | 106 | 660 | 7 | MOLECULE: | PUTATIVE M | ITOCHOND | RIAL DYNAMIN PROTEIN |
| 22:00 | 5iy7-M | 2.8 | 16.1 | 82 | 310 | 5 | MOLECULE: | DNA-DIRECT | ED RNA P | OLYMERASE II SUBUNIT RPB1 |
| 23:00 | 7jr7-A | 2.8 | 16.9 | 119 | 588 | 7 | MOLECULE: | ATP-BINDIN | G CASSET | TE SUB-FAMILY G MEMBER 5 |
| 24:00:00 | 8anq-C | 2.8 | 9.3 | 96 | 227 | 5 | MOLECULE: | BACTERIORH | ODOPSIN | |
| 25:00:00 | 5ys9-A | 2.8 | 33.7 | 130 | 692 | 5 | MOLECULE: | ACYL-COENZ | YME A OX | IDASE 3 |
| 26:00:00 | 8otx-A | 2.8 | 18.6 | 132 | 1054 | 6 | MOLECULE: | SPERM-SPEC | IFIC SOD | IUM PROTON EXCHANGER |
| 27:00:00 | 3pux-G | 2.8 | 10.3 | 85 | 293 | 2 | MOLECULE: | MALTOSE-BI | NDING PE | RIPLASMIC PROTEIN |
| 28:00:00 | 7eeb-K | 2.8 | 7.9 | 87 | 143 | 2 | MOLECULE: | ENHANCED G | REEN FLU | ORESCENT PROTEIN,CATION CHANNEL |
| 29:00:00 | 8qfs-A | 2.8 | 30.1 | 140 | 1183 | 7 | MOLECULE: | ELONGATION | FACTOR | TU |
| 30:00:00 | 6wgv-A | 2.8 | 13.1 | 92 | 173 | 3 | MOLECULE: | CORRINOID | ADENOSYL | TRANSFERASE |
| 31:00:00 | 7o3d-A | 2.8 | 11.7 | 112 | 709 | 11 | MOLECULE: | TRANSCRIPT | ION ELON | GATION FACTOR SPT6 |
| 32:00:00 | 3k1z-A | 2.8 | 5.6 | 105 | 241 | 5 | MOLECULE: | HALOACID D | EHALOGEN | ASE-LIKE HYDROLASE DOMAIN-CONTA |
| 33:00:00 | 8axa-A | 2.8 | 8.4 | 88 | 465 | 3 | MOLECULE: | CAS12K | | |
| 34:00:00 | 1zp2-A | 2.8 | 15.5 | 84 | 227 | 6 | MOLECULE: | RNA POLYME | RASE II | HOLOENZYME CYCLIN-LIKE SUBUNIT |
| 35:00:00 | 3ed5-A | 2.8 | 5.3 | 89 | 232 | 8 | MOLECULE: | YFNB | | |
| 36:00:00 | 8yb7-C | 2.8 | 15 | 90 | 455 | 7 | MOLECULE: | PAPAIN-LIK | E PROTEA | SE NSP3 |
| 37:00:00 | 5xj5-A | 2.8 | 9.9 | 86 | 200 | 7 | MOLECULE: | GLYCEROL-3 | #NAME? | TE ACYLTRANSFERASE |
| 38:00:00 | 5edl-A | 2.8 | 12 | 95 | 197 | 9 | MOLECULE: | PUTATIVE H | MP/THIAM | INE PERMEASE PROTEIN YKOE |
| 39:00:00 | 7drt-B | 2.8 | 18.4 | 121 | 496 | 4 | MOLECULE: | PROTEIN WN | T-3A | |
| 40:00:00 | 8e4y-A | 2.8 | 17 | 100 | 651 | 5 | MOLECULE: | GLYCEROL-3 | #NAME? | TE ACYLTRANSFERASE 1, MITOCHOND |
| 41:00:00 | 1gak-A | 2.8 | 2.9 | 65 | 137 | 8 | MOLECULE: | FERTILIZAT | ION PROT | EIN |
| 42:00:00 | 8j56-C | 2.8 | 4 | 74 | 160 | 7 | MOLECULE: | FLAGELLAR | TRANSCRI | PTIONAL REGULATOR FLHD |
| 43:00:00 | 8fed-J | 2.8 | 7.4 | 98 | 263 | 1 | MOLECULE: | VIRULENCE | FACTOR M | CE FAMILY PROTEIN |
| 44:00:00 | 2voh-A | 2.8 | 4.5 | 76 | 155 | 9 | MOLECULE: | BCL-2-RELA | TED PROT | EIN A1 |
| 45:00:00 | 8ovw-Q | 2.8 | 8.2 | 83 | 258 | 8 | MOLECULE: | CENTROMERE | #NAME? | PROTEIN 1 |
| 46:00:00 | 2rld-C | 2.8 | 4.9 | 77 | 116 | 5 | MOLECULE: | UNCHARACTE | RIZED PR | OTEIN |
| 47:00:00 | 7w6k-A | 2.8 | 25.7 | 113 | 410 | 3 | MOLECULE: | GMALMT12/Q | UAC1 | |
| 48:00:00 | 7m2w-G | 2.8 | 19.5 | 108 | 712 | 4 | MOLECULE: | TUBULIN GA | MMA CHAI | N |
| 49:00:00 | 5ean-A | 2.8 | 25.1 | 135 | 1051 | 8 | MOLECULE: | DNA REPLIC | ATION AT | P-DEPENDENT HELICASE/NUCLEASE D |
| 50:00:00 | 7k4m-B | 2.8 | 4.2 | 75 | 146 | 13 | MOLECULE: | HEMOGLOBIN | SUBUNIT | ALPHA |
| 51:00:00 | 5e83-B | 2.8 | 4 | 77 | 146 | 14 | MOLECULE: | HEMOGLOBIN | SUBUNIT | ALPHA |
| 52:00:00 | 5u3i-B | 2.8 | 4.1 | 77 | 146 | 14 | MOLECULE: | HEMOGLOBIN | SUBUNIT | ALPHA |
| 53:00:00 | 1ut0-A | 2.8 | 5.1 | 89 | 154 | 8 | MOLECULE: | CYTOGLOBIN | | |
| 54:00:00 | 3bom-D | 2.8 | 4.5 | 80 | 147 | 13 | MOLECULE: | HEMOGLOBIN | SUBUNIT | ALPHA-4 |
| 55:00:00 | 7jy0-B | 2.8 | 4.6 | 79 | 146 | 14 | MOLECULE: | HEMOGLOBIN | SUBUNIT | ALPHA |
| 56:00:00 | 1mbs-A | 2.8 | 4.6 | 79 | 153 | 8 | MOLECULE: | MYOGLOBIN | | |
| 57:00:00 | 1shr-B | 2.8 | 4.1 | 77 | 146 | 13 | MOLECULE: | HEMOGLOBIN | ALPHA C | HAIN |
| 58:00:00 | 7uf6-D | 2.8 | 4.5 | 76 | 146 | 13 | MOLECULE: | HEMOGLOBIN | SUBUNIT | ALPHA |
| 59:00:00 | 3w4u-B | 2.8 | 4 | 77 | 145 | 14 | MOLECULE: | HEMOGLOBIN | SUBUNIT | ZETA |
| 60:00:00 | 3w4u-F | 2.8 | 3.9 | 76 | 145 | 14 | MOLECULE: | HEMOGLOBIN | SUBUNIT | ZETA |
| 61:00:00 | 7jxz-D | 2.8 | 4.1 | 77 | 146 | 14 | MOLECULE: | HEMOGLOBIN | SUBUNIT | ALPHA |
| 62:00:00 | 2qls-D | 2.8 | 4.5 | 75 | 146 | 15 | MOLECULE: | HEMOGLOBIN | SUBUNIT | ALPHA |
| 63:00:00 | 7jxz-B | 2.8 | 4.1 | 77 | 146 | 14 | MOLECULE: | HEMOGLOBIN | SUBUNIT | ALPHA |
| 64:00:00 | 8egi-B | 2.8 | 3.9 | 76 | 146 | 14 | MOLECULE: | HEMOGLOBIN | SUBUNIT | ALPHA |
| 65:00:00 | 7uf6-B | 2.8 | 4 | 77 | 146 | 14 | MOLECULE: | HEMOGLOBIN | SUBUNIT | ALPHA |
| 66:00:00 | 3w4u-D | 2.8 | 4.4 | 79 | 145 | 14 | MOLECULE: | HEMOGLOBIN | SUBUNIT | ZETA |

| | | | | | | | | | | |
|---|---|---|---|---|---|---|---|---|---|---|
| 67:00:00 | 2b7h-B | 2.8 | 4 | 76 | 144 | 14 | MOLECULE: | HEMOGLOBIN | ALPHA C | HAIN |
| 68:00:00 | 3gqr-B | 2.8 | 4 | 76 | 145 | 14 | MOLECULE: | HEMOGLOBIN | SUBUNIT | ALPHA |
| 69:00:00 | 7cue-B | 2.8 | 4.6 | 79 | 146 | 14 | MOLECULE: | HEMOGLOBIN | SUBUNIT | ALPHA |
| 70:00:00 | 1urv-A | 2.8 | 5.1 | 89 | 154 | 8 | MOLECULE: | CYTOGLOBIN | | |
| 71:00:00 | 6xe7-B | 2.8 | 4.1 | 77 | 146 | 14 | MOLECULE: | HEMOGLOBIN | SUBUNIT | ALPHA |
| 72:00:00 | 1nej-D | 2.8 | 4.6 | 78 | 146 | 14 | MOLECULE: | HEMOGLOBIN | ALPHA C | HAIN |
| 73:00:00 | 3bom-B | 2.8 | 4.1 | 78 | 147 | 13 | MOLECULE: | HEMOGLOBIN | SUBUNIT | ALPHA-4 |
| 74:00:00 | 8dov-F | 2.8 | 4 | 76 | 144 | 14 | MOLECULE: | HEMOGLOBIN | SUBUNIT | ALPHA |
| 75:00:00 | 6wg7-G | 2.8 | 12.6 | 106 | 219 | 8 | MOLECULE: | DNA (35-ME | R) | |
| 76:00:00 | 6wfq-D | 2.8 | 12.6 | 106 | 219 | 8 | MOLECULE: | HTH-TYPE T | RANSCRIP | TIONAL REPRESSOR NANR |
| 77:00:00 | 6u4b-A | 2.7 | 19.8 | 120 | 570 | 8 | MOLECULE: | WBBM PROTE | IN | |
| 78:00:00 | 3bhn-A | 2.7 | 10.1 | 64 | 214 | 5 | MOLECULE: | THIJ/PFPI | DOMAIN P | ROTEIN |
| 79:00:00 | 3f0c-A | 2.7 | 3.7 | 66 | 193 | 9 | MOLECULE: | TRANSCRIPT | IONAL RE | GULATOR |
| 80:00:00 | 7yim-A | 2.7 | 6.3 | 116 | 592 | 8 | MOLECULE: | ALPHA-FETO | PROTEIN | |
| 81:00:00 | 5nzz-A | 2.7 | 12.1 | 103 | 378 | 10 | MOLECULE: | TGF-BETA-A | CTIVATED | KINASE 1 AND MAP3K7-BINDING PR |
| 82:00:00 | 6tgb-A | 2.7 | 22.6 | 99 | 1450 | 3 | MOLECULE: | DEDICATOR | OF CYTOK | INESIS PROTEIN 2 |
| 83:00:00 | 6ppi-A | 2.7 | 18.2 | 104 | 413 | 8 | MOLECULE: | PORTAL PRO | TEIN | |
| 84:00:00 | 6vy1-A | 2.7 | 10.4 | 71 | 121 | 7 | MOLECULE: | PREFOLDIN | SUBUNIT | ALPHA 2 |
| 85:00:00 | 5hzi-A | 2.7 | 7 | 85 | 476 | 8 | MOLECULE: | INTERSECTI | N-1,NPH1 | -1,INTERSECTIN-1 |
| 86:00:00 | 2gnx-A | 2.7 | 13 | 80 | 288 | 5 | MOLECULE: | HYPOTHETIC | AL PROTE | IN |
| 87:00:00 | 8tdl-A | 2.7 | 18 | 116 | 364 | 4 | MOLECULE: | MECHANOSEN | SITIVE I | ON CHANNEL PROTEIN 10 |
| 88:00:00 | 2p01-A | 2.7 | 11.3 | 93 | 323 | 5 | MOLECULE: | ALPHA-2-MA | CROGLOBU | LIN RECEPTOR-ASSOCIATED |
| 89:00:00 | 2vyw-A | 2.7 | 4.4 | 66 | 148 | 6 | MOLECULE: | HEMOGLOBIN | | |
| 90:00:00 | 2xwu-B | 2.7 | 15.3 | 123 | 910 | 10 | MOLECULE: | SUMO-CONJU | GATING E | NZYME UBC9 |
| 91:00:00 | 7roa-A | 2.7 | 4.8 | 69 | 117 | 6 | MOLECULE: | ENTV | | |
| 92:00:00 | 8osg-B | 2.7 | 13.4 | 89 | 323 | 7 | MOLECULE: | MAGNESIUM- | CHELATAS | E SUBUNIT CHLI |
| 93:00:00 | 2js9-A | 2.7 | 3.2 | 57 | 81 | 4 | MOLECULE: | SAPOSIN-LI | KE PROTE | IN FAMILY PROTEIN 5 |
| 94:00:00 | 3lwj-A | 2.7 | 4.9 | 71 | 193 | 10 | MOLECULE: | PUTATIVE T | ETR-FAMI | LY TRANSCRIPTIONAL REGULATOR |
| 95:00:00 | 3ajb-A | 2.7 | 6.6 | 104 | 293 | 5 | MOLECULE: | PEROXISOMA | L BIOGEN | ESIS FACTOR 3 |
| 96:00:00 | 8eki-A | 2.7 | 20.7 | 82 | 170 | 6 | MOLECULE: | PROTEIN TR | ANSPORT | PROTEIN SEC20 |
| 97:00:00 | 8ppr-D | 2.7 | 24.3 | 88 | 248 | 11 | MOLECULE: | KINETOCHOR | E-ASSOCI | ATED PROTEIN DSN1 HOMOLOG |
| 98:00:00 | 5v8f-6 | 2.7 | 18.1 | 100 | 692 | 8 | MOLECULE: | DNA REPLIC | ATION LI | CENSING FACTOR MCM2 |
| 99:00:00 | 7eew-A | 2.7 | 17.2 | 89 | 611 | 9 | MOLECULE: | TYPE I RES | TRICTION | #NAME? |
| 0:00 | 2qkm-B | 2.7 | 9.7 | 87 | 266 | 1 | MOLECULE: | SPBC3B9.21 | PROTEIN | |
| 1:00 | 7p4a-A | 2.7 | 12.2 | 98 | 246 | 3 | MOLECULE: | STL | | |
| 2:00 | 7vf2-A | 2.7 | 23.3 | 121 | 1196 | 12 | MOLECULE: | PROTEIN VI | RILIZER | HOMOLOG |
| 3:00 | 7sjr-A | 2.7 | 29.2 | 96 | 911 | 13 | MOLECULE: | DNA HELICA | SE | |
| 4:00 | 6c66-I | 2.7 | 3.8 | 71 | 177 | 8 | MOLECULE: | CRISPR-ASS | OCIATED | HELICASE, CAS3 FAMILY |
| 5:00 | 2e87-A | 2.7 | 15.2 | 102 | 356 | 5 | MOLECULE: | HYPOTHETIC | AL PROTE | IN PH1320 |
| 6:00 | 7yyg-A | 2.7 | 7.7 | 87 | 275 | 7 | MOLECULE: | PUTATIVE O | UTER PRO | TEIN N |
| 7:00 | 8ucq-A | 2.7 | 23.2 | 114 | 1232 | 6 | MOLECULE: | SEC7 DOMAI | N-CONTAI | NING PROTEIN |
| 8:00 | 2fbq-A | 2.7 | 14.9 | 64 | 213 | 6 | MOLECULE: | PROBABLE T | RANSCRIP | TIONAL REGULATOR |
| 9:00 | 7pgr-F | 2.7 | 22 | 151 | 2423 | 6 | MOLECULE: | NEUROFIBRO | MIN | |
| 10:00 | 2k3q-A | 2.7 | 4.1 | 81 | 118 | 12 | MOLECULE: | TUSP1 | | |
| 11:00 | 5wrr-B | 2.7 | 6.5 | 64 | 438 | 8 | MOLECULE: | PSEUDOKINA | SE FAM20 | A |
| 12:00 | 6vrb-C | 2.7 | 7.4 | 73 | 229 | 4 | MOLECULE: | RNA (52-ME | R) | |
| 13:00 | 7pds-A | 2.7 | 15.8 | 88 | 457 | 6 | MOLECULE: | SIMILAR TO | D. NODO | SUS VAPE |
| 14:00 | 5dca-A | 2.7 | 15.1 | 134 | 1948 | 7 | MOLECULE: | PRE-MRNA-S | PLICING | HELICASE BRR2 |
| 15:00 | 7bss-A | 2.7 | 23.4 | 84 | 816 | 8 | MOLECULE: | ATP11C | | |
| 16:00 | 7obq-y | 2.7 | 14 | 82 | 454 | 7 | MOLECULE: | SRP RNA | | |
| 17:00 | 1sf9-A | 2.7 | 9.1 | 80 | 118 | 3 | MOLECULE: | YFHH HYPOT | HETICAL | PROTEIN |
| 18:00 | 7ody-C | 2.7 | 11.6 | 73 | 100 | 5 | MOLECULE: | MAZG-LIKE | PYROPHOS | PHOHYDROLASE (MAZZ) |
| 19:00 | 6nf4-A | 2.7 | 19.3 | 98 | 421 | 6 | MOLECULE: | OTOPETRIN1 | | |
| 20:00 | 1tjf-B | 2.7 | 10 | 89 | 186 | 2 | MOLECULE: | ADENYLYL C | YCLASE-A | SSOCIATED PROTEIN |

| | | | | | | | | | | |
|---|---|---|---|---|---|---|---|---|---|---|
| 21:00 | 8ovt-A | 2.7 | 5 | 70 | 288 | 4 | MOLECULE: | YEGT GLYCO | SYLTRANS | FERASE |
| 22:00 | 6cfw-C | 2.7 | 10 | 86 | 114 | 7 | MOLECULE: | MONOVALENT | CATION/ | H+ ANTIPORTER SUBUNIT D |
| 23:00 | 2xl4-A | 2.7 | 9.4 | 81 | 147 | 5 | MOLECULE: | LMO0438 PR | OTEIN | |
| 24:00:00 | 7exx-A | 2.7 | 28.3 | 117 | 440 | 7 | MOLECULE: | DNA PHOSPH | OROTHIOA | TION-DEPENDENT RESTRICTION PROT |
| 25:00:00 | 7t5p-B | 2.7 | 9.6 | 82 | 419 | 7 | MOLECULE: | SUMO-INTER | ACTING M | OTIF-CONTAINING PROTEIN 1 |
| 26:00:00 | 6d5f-A | 2.7 | 13.4 | 93 | 193 | 9 | MOLECULE: | FIMBRIAL P | ROTEIN | |
| 27:00:00 | 5kkl-B | 2.7 | 9.4 | 87 | 838 | 2 | MOLECULE: | PUTATIVE P | OLYCOMB | PROTEIN EED |
| 28:00:00 | 5h72-A | 2.7 | 2.7 | 58 | 67 | 9 | MOLECULE: | FLAGELLAR | BIOSYNTH | ETIC PROTEIN FLIP |
| 29:00:00 | 8rf0-A | 2.7 | 30 | 136 | 2780 | 4 | MOLECULE: | CYCLIC BET | A-(1,2)- | GLUCAN SYNTHASE NDVB |
| 30:00:00 | 5t58-A | 2.7 | 13.5 | 93 | 233 | 10 | MOLECULE: | KLLA0F0234 | 3P | |
| 31:00:00 | 5htf-B | 2.7 | 7.9 | 91 | 260 | 7 | MOLECULE: | FOLDASE PR | OTEIN PR | SA 1 |
| 32:00:00 | 5yet-A | 2.7 | 16.3 | 93 | 397 | 4 | MOLECULE: | UNCHARACTE | RIZED PR | OTEIN R354 |
| 33:00:00 | 5eqw-B | 2.7 | 12.2 | 75 | 123 | 3 | MOLECULE: | PUTATIVE M | AJOR COA | T PROTEIN |
| 34:00:00 | 6w2q-A | 2.7 | 4.2 | 66 | 195 | 5 | MOLECULE: | JUNCTION 3 | 4 | |
| 35:00:00 | 2dh4-A | 2.7 | 10.8 | 113 | 326 | 3 | MOLECULE: | YPL069C | | |
| 36:00:00 | 5yfp-C | 2.7 | 13.4 | 89 | 790 | 8 | MOLECULE: | EXOCYST CO | MPLEX CO | MPONENT SEC3 |
| 37:00:00 | 7nhr-A | 2.7 | 13.1 | 88 | 663 | 6 | MOLECULE: | PUTATIVE T | RANSMEMB | RANE PROTEIN WZC |
| 38:00:00 | 8j0h-A | 2.7 | 8.4 | 82 | 134 | 6 | MOLECULE: | UNCHARACTE | RIZED PR | OTEIN C4H3.06 |
| 39:00:00 | 2ly9-A | 2.7 | 5.2 | 61 | 74 | 8 | MOLECULE: | ZINC FINGE | RS AND H | OMEOBOXES PROTEIN 1 |
| 40:00:00 | 8c81-B | 2.7 | 17.6 | 95 | 539 | 9 | MOLECULE: | PROTEIN OR | M1 | |
| 41:00:00 | 3wvo-A | 2.7 | 17.4 | 89 | 543 | 10 | MOLECULE: | CRISPR-ASS | OCIATED | PROTEIN, CSE1 FAMILY |
| 42:00:00 | 7os0-A | 2.7 | 9.2 | 111 | 1130 | 5 | MOLECULE: | CAS13A | | |
| 43:00:00 | 3egw-C | 2.7 | 17.1 | 104 | 225 | 9 | MOLECULE: | RESPIRATOR | Y NITRAT | E REDUCTASE 1 ALPHA CHAIN |
| 44:00:00 | 3mw6-A | 2.7 | 6.4 | 80 | 122 | 13 | MOLECULE: | UNCHARACTE | RIZED PR | OTEIN NMB1681 |
| 45:00:00 | 6ekr-A | 2.7 | 10.8 | 91 | 303 | 11 | MOLECULE: | TYPE II SI | TE-SPECI | FIC DEOXYRIBONUCLEASE |
| 46:00:00 | 3rgc-B | 2.7 | 4.8 | 85 | 249 | 5 | MOLECULE: | POSSIBLE P | ERIPLASM | IC PROTEIN |
| 47:00:00 | 7x07-A | 2.7 | 24 | 89 | 633 | 10 | MOLECULE: | ATP-BINDIN | G CASSET | TE SUB-FAMILY D MEMBER 1 |
| 48:00:00 | 8hp7-A | 2.7 | 4.6 | 97 | 241 | 9 | MOLECULE: | (S)-2-HALO | ACID DEH | ALOGENASE |
| 49:00:00 | 6xpd-A | 2.7 | 27.1 | 115 | 303 | 8 | MOLECULE: | ZINC TRANS | PORTER 8 | |
| 50:00:00 | 7zke-E | 2.7 | 19.7 | 112 | 1623 | 6 | MOLECULE: | DNA (36-ME | R) | |
| 51:00:00 | 7xzj-A | 2.7 | 30.3 | 97 | 423 | 9 | MOLECULE: | CTAP3 | | |
| 52:00:00 | 3utk-A | 2.7 | 3.4 | 60 | 95 | 5 | MOLECULE: | LIPOPROTEI | N OUTS | |
| 53:00:00 | 6xdt-B | 2.7 | 4 | 74 | 146 | 14 | MOLECULE: | HEMOGLOBIN | SUBUNIT | ALPHA |
| 54:00:00 | 2b7h-D | 2.7 | 4 | 74 | 145 | 15 | MOLECULE: | HEMOGLOBIN | ALPHA C | HAIN |
| 55:00:00 | 1si4-D | 2.7 | 4.7 | 79 | 146 | 13 | MOLECULE: | HEMOGLOBIN | ALPHA C | HAIN |
| 56:00:00 | 7uf7-B | 2.7 | 4.5 | 75 | 146 | 15 | MOLECULE: | HEMOGLOBIN | SUBUNIT | ALPHA |
| 57:00:00 | 5ker-B | 2.7 | 4.3 | 75 | 146 | 13 | MOLECULE: | ALPHA-GLOB | IN | |
| 58:00:00 | 2r1h-D | 2.7 | 4.5 | 76 | 147 | 13 | MOLECULE: | HEMOGLOBIN | SUBUNIT | ALPHA-4 |
| 59:00:00 | 3hf4-F | 2.7 | 3.9 | 73 | 146 | 14 | MOLECULE: | HEMOGLOBIN | SUBUNIT | ALPHA-1/2 |
| 60:00:00 | 3gqr-H | 2.7 | 4.4 | 75 | 145 | 15 | MOLECULE: | HEMOGLOBIN | SUBUNIT | ALPHA |
| 61:00:00 | 1v5h-A | 2.7 | 4.6 | 78 | 151 | 8 | MOLECULE: | CYTOGLOBIN | | |
| 62:00:00 | 5aqd-M | 2.6 | 4.4 | 104 | 184 | 4 | MOLECULE: | PHYCOERYTH | RIN ALPH | A SUBUNIT |
| 63:00:00 | 8agg-A | 2.6 | 9.2 | 60 | 489 | 5 | MOLECULE: | PHOSPHOCHO | LINE HYD | ROLASE LEM3 |
| 64:00:00 | 5lpa-A | 2.6 | 3.9 | 70 | 267 | 9 | MOLECULE: | STREPTOMYC | IN 3''-A | DENYLYLTRANSFERASE |
| 65:00:00 | 5e6g-B | 2.6 | 8 | 72 | 114 | 10 | MOLECULE: | DE NOVO DE | SIGNED P | ROTEIN CA01 |
| 66:00:00 | 4me9-A | 2.6 | 4.3 | 67 | 188 | 4 | MOLECULE: | TRANSCRIPT | IONAL RE | GULATOR, TETR FAMILY |
| 67:00:00 | 7f5v-A | 2.6 | 16.3 | 134 | 689 | 7 | MOLECULE: | DELTA-1-PY | RROLINE- | 5-CARBOXYLATE SYNTHASE |
| 68:00:00 | 3ut2-A | 2.6 | 16.7 | 117 | 735 | 4 | MOLECULE: | CATALASE-P | EROXIDAS | E 2 |
| 69:00:00 | 7uic-b | 2.6 | 5.3 | 49 | 65 | 6 | MOLECULE: | MEDIATOR O | F RNA PO | LYMERASE II TRANSCRIPTION SUBUN |
| 70:00:00 | 8t1l-X | 2.6 | 20.3 | 80 | 127 | 9 | MOLECULE: | MEDIATOR O | F RNA PO | LYMERASE II TRANSCRIPTION SUBUN |
| 71:00:00 | 7m5t-A | 2.6 | 8.6 | 73 | 100 | 8 | MOLECULE: | DE NOVO DE | SIGNED P | ROTEIN 0515 |
| 72:00:00 | 8dmg-B | 2.6 | 10.3 | 187 | 1033 | 7 | MOLECULE: | BIFUNCTION | AL CYTOC | HROME P450/NADPH--P450 REDUCTAS |
| 73:00:00 | 7w7c-B | 2.6 | 4.6 | 82 | 159 | 9 | MOLECULE: | PUTATIVE A | BC TRANS | PORT SYSTEM, ATP-BINDING PROTEI |
| 74:00:00 | 8d0b-F | 2.6 | 18.2 | 110 | 915 | 7 | MOLECULE: | CST COMPLE | X SUBUNI | T CTC1 |

| | | | | | | | | | | |
|---|---|---|---|---|---|---|---|---|---|---|
| 75:00:00 | 4a01-A | 2.6 | 7.4 | 132 | 740 | 3 | MOLECULE: | PROTON PYR | OPHOSPHA | TASE |
| 76:00:00 | 2a6o-A | 2.6 | 5.4 | 78 | 152 | 6 | MOLECULE: | 5'-D(*CP*C | P*CP*CP* | TP*AP*GP*CP*TP*TP*TP*AP*GP*CP*T |
| 77:00:00 | 8cws-A | 2.6 | 12.3 | 94 | 451 | 5 | MOLECULE: | F4132-2 CH | AIN A | |
| 78:00:00 | 6xjh-A | 2.6 | 5.2 | 93 | 251 | 6 | MOLECULE: | PHENOL-SOL | UBLE MOD | ULIN EXPORT ABC TRANSPORTER PER |
| 79:00:00 | 8d4a-A | 2.6 | 18.7 | 113 | 1207 | 8 | MOLECULE: | ORFB_ZN_RI | BBON DOM | AIN-CONTAINING PROTEIN |
| 80:00:00 | 4hts-A | 2.6 | 4.7 | 98 | 228 | 7 | MOLECULE: | SEC-INDEPE | NDENT PR | OTEIN TRANSLOCASE PROTEIN TATC |
| 81:00:00 | 8r33-A | 2.6 | 12.5 | 102 | 489 | 3 | MOLECULE: | PHO90 ISOF | ORM 1 | |
| 82:00:00 | 6d0j-B | 2.6 | 4.9 | 99 | 396 | 5 | MOLECULE: | CLC-TYPE F | LUORIDE/ | PROTON ANTIPORTER |
| 83:00:00 | 8ap9-G | 2.6 | 7.2 | 62 | 279 | 5 | MOLECULE: | ATP SYNTHA | SE GAMMA | SUBUNIT |
| 84:00:00 | 6hgc-A | 2.6 | 19.1 | 81 | 304 | 2 | MOLECULE: | UBIQUITIN | CARBOXYL | #NAME? |
| 85:00:00 | 4i1s-A | 2.6 | 8.7 | 87 | 238 | 11 | MOLECULE: | MELANOMA D | IFFERENT | IATION ASSOCIATED PROTEIN-5 |
| 86:00:00 | 7s8u-A | 2.6 | 8.5 | 97 | 667 | 4 | MOLECULE: | SOLUTE CAR | RIER FAM | ILY 15 MEMBER 1 |
| 87:00:00 | 8ea3-W | 2.6 | 24.5 | 96 | 514 | 3 | MOLECULE: | TARGET_LE | | |
| 88:00:00 | 8csp-F | 2.6 | 11.8 | 86 | 208 | 1 | MOLECULE: | 28S RIBOSO | MAL PROT | EIN S34, MITOCHONDRIAL |
| 89:00:00 | 8cqv-A | 2.6 | 5 | 96 | 202 | 6 | MOLECULE: | NITROREDUC | TASE | |
| 90:00:00 | 3e00-D | 2.6 | 13.8 | 123 | 363 | 2 | MOLECULE: | RETINOIC A | CID RECE | PTOR RXR-ALPHA |
| 91:00:00 | 8jpj-A | 2.6 | 30.2 | 137 | 733 | 4 | MOLECULE: | H(+)/CL(-) | EXCHANG | E TRANSPORTER 6 |
| 92:00:00 | 2qeu-B | 2.6 | 3.1 | 54 | 133 | 6 | MOLECULE: | PUTATIVE C | ARBOXYMU | CONOLACTONE DECARBOXYLASE |
| 93:00:00 | 8es4-C | 2.6 | 10.6 | 81 | 181 | 6 | MOLECULE: | GP35 | | |
| 94:00:00 | 6zxr-A | 2.6 | 6.4 | 89 | 212 | 6 | MOLECULE: | ER LUMEN P | ROTEIN-R | ETAINING RECEPTOR 2 |
| 95:00:00 | 3o7q-A | 2.6 | 7 | 107 | 414 | 7 | MOLECULE: | L-FUCOSE-P | ROTON SY | MPORTER |
| 96:00:00 | 4xom-A | 2.6 | 5.7 | 98 | 206 | 4 | MOLECULE: | COENZYME F | 420:L-GL | UTAMATE LIGASE |
| 97:00:00 | 7y7q-A | 2.6 | 18.8 | 138 | 951 | 10 | MOLECULE: | RNA-DEPEND | ENT RNA | POLYMERASE |
| 98:00:00 | 7wkk-A | 2.6 | 23.2 | 116 | 1684 | 4 | MOLECULE: | MGC83295 P | ROTEIN | |
| 99:00:00 | 6csv-A | 2.6 | 6.9 | 58 | 90 | 9 | MOLECULE: | CENTROSOMA | L PROTEI | N OF 63 KDA,CENTROSOMAL PROTEIN |
| 0:00 | 7mdy-B | 2.6 | 5.9 | 78 | 411 | 4 | MOLECULE: | LIPOPROTEI | N TRANSP | ORTER SUBUNIT LOLE |
| 1:00 | 1nek-C | 2.6 | 5.7 | 79 | 129 | 4 | MOLECULE: | SUCCINATE | DEHYDROG | ENASE FLAVOPROTEIN SUBUNIT |
| 2:00 | 8b9z-K | 2.6 | 8.1 | 78 | 96 | 8 | MOLECULE: | NADH-UBIQU | INONE OX | IDOREDUCTASE CHAIN 3 |
| 3:00 | 6hux-A | 2.6 | 25.9 | 113 | 369 | 9 | MOLECULE: | H(2)-FORMI | NG METHY | LENETETRAHYDROMETHANOPTERIN |
| 4:00 | 8h2m-A | 2.6 | 28 | 109 | 754 | 8 | MOLECULE: | TAPE TAIL | MEASURE | PROTEIN |
| 5:00 | 2da3-A | 2.6 | 6.7 | 70 | 80 | 9 | MOLECULE: | ALPHA-FETO | PROTEIN | ENHANCER BINDING PROTEIN |
| 6:00 | 6rfl-C | 2.6 | 11.2 | 66 | 304 | 8 | MOLECULE: | DNA-DEPEND | ENT RNA | POLYMERASE SUBUNIT RPO132 |
| 7:00 | 6thh-C | 2.6 | 18.7 | 98 | 816 | 7 | MOLECULE: | SIRV2 ACRI | D1 (GP02 | ) ANTI-CRISPR PROTEIN |
| 8:00 | 3cit-A | 2.6 | 4 | 72 | 155 | 11 | MOLECULE: | SENSOR HIS | TIDINE K | INASE |
| 9:00 | 6ahp-A | 2.6 | 3.7 | 68 | 110 | 6 | MOLECULE: | FLAGELLAR | PROTEIN | FLIL |
| 10:00 | 2xyk-B | 2.6 | 3.8 | 72 | 130 | 10 | MOLECULE: | 2-ON-2 HEM | OGLOBIN | |
| 11:00 | 8syn-A | 2.6 | 26 | 117 | 801 | 9 | MOLECULE: | VPS35 ENDO | SOMAL PR | OTEIN-SORTING FACTOR-LIKE |
| 12:00 | 1dek-A | 2.6 | 8.7 | 96 | 241 | 5 | MOLECULE: | DEOXYNUCLE | OSIDE MO | NOPHOSPHATE KINASE |
| 13:00 | 2py8-C | 2.6 | 16 | 84 | 123 | 8 | MOLECULE: | HYPOTHETIC | AL PROTE | IN RBCX |
| 14:00 | 1ypy-A | 2.6 | 6.4 | 86 | 182 | 8 | MOLECULE: | VIRION MEM | BRANE PR | OTEIN |
| 15:00 | 7w6x-A | 2.6 | 11.5 | 82 | 447 | 12 | MOLECULE: | REGULATOR | OF SIGMA | -E PROTEASE RSEP |
| 16:00 | 6idp-A | 2.6 | 12.1 | 94 | 438 | 9 | MOLECULE: | MATE FAMIL | Y EFFLUX | TRANSPORTER |
| 17:00 | 2d7d-A | 2.6 | 27.7 | 100 | 621 | 6 | MOLECULE: | 5'-D(P*TP* | TP*T)-3' | |
| 18:00 | 6yxp-B | 2.6 | 8.1 | 75 | 272 | 0 | MOLECULE: | SWI/SNF CO | MPLEX SU | BUNIT SMARCC1 |
| 19:00 | 5dw7-A | 2.6 | 14.1 | 94 | 309 | 3 | MOLECULE: | GERMACRADI | ENOL/GEO | SMIN SYNTHASE |
| 20:00 | 6xlp-A | 2.6 | 13.8 | 98 | 586 | 8 | MOLECULE: | LPS BINDIN | G PROTEI | N |
| 21:00 | 7f3t-A | 2.6 | 16 | 107 | 330 | 5 | MOLECULE: | TRANSMEMBR | ANE PROT | EIN 120A |
| 22:00 | 6v4a-C | 2.6 | 16.3 | 88 | 593 | 3 | MOLECULE: | NEUR_CHAN_ | LBD DOMA | IN-CONTAINING PROTEIN |
| 23:00 | 8psn-B | 2.6 | 14.3 | 111 | 515 | 11 | MOLECULE: | POLYMERASE | ACIDIC | PROTEIN (PA-LIKE) |
| 24:00:00 | 6vrc-A | 2.6 | 13.8 | 105 | 1007 | 6 | MOLECULE: | CRISPR-ASS | OCIATED | ENDORIBONUCLEASE CAS13A |
| 25:00:00 | 7znk-b | 2.6 | 16.1 | 109 | 919 | 6 | MOLECULE: | RNA | | |
| 26:00:00 | 1jud-A | 2.6 | 4.8 | 95 | 220 | 7 | MOLECULE: | L-2-HALOAC | ID DEHAL | OGENASE |
| 27:00:00 | 6lo8-C | 2.6 | 4 | 61 | 108 | 10 | MOLECULE: | MITOCHONDR | IAL IMPO | RT INNER MEMBRANE TRANSLOCASE S |
| 28:00:00 | 5gko-A | 2.6 | 11.8 | 98 | 650 | 5 | MOLECULE: | MACROLIDE | EXPORT A | TP-BINDING/PERMEASE PROTEIN MAC |

| | | | | | | | | | | | |
|---|---|---|---|---|---|---|---|---|---|---|---|
| 29:00:00 | 3fay-A | 2.6 | 24.7 | 139 | 379 | 9 | MOLECULE: | RAS GTPASE | #NAME? | ING-LIKE PROTEIN IQGAP1 | |
| 30:00:00 | 8sgi-A | 2.6 | 10.7 | 100 | 751 | 9 | MOLECULE: | SODIUM/CAL | CIUM EXC | HANGER 1 | |
| 31:00:00 | 6w2r-B | 2.6 | 8 | 81 | 221 | 15 | MOLECULE: | JUNCTION 1 | 9 DHR54- | DHR79 | |
| 32:00:00 | 5csk-A | 2.6 | 23.1 | 100 | 1996 | 6 | MOLECULE: | ACETYL-COA | CARBOXY | LASE | |
| 33:00:00 | 7nnh-X | 2.6 | 7.7 | 122 | 637 | 11 | MOLECULE: | ERYTHROCYT | E MEMBRA | NE PROTEIN 1 | |
| 34:00:00 | 8iq4-R | 2.6 | 12 | 99 | 274 | 5 | MOLECULE: | PROSTAGLAN | DIN F2-A | LPHA RECEPTOR | |
| 35:00:00 | 7yca-O | 2.6 | 4.2 | 63 | 96 | 3 | MOLECULE: | LHCA1 | | | |
| 36:00:00 | 6zpa-A | 2.6 | 4.5 | 65 | 173 | 2 | MOLECULE: | DATZ | | | |
| 37:00:00 | 6ny2-Y | 2.6 | 22.1 | 135 | 915 | 8 | MOLECULE: | DNA TARGET | STRAND | | |
| 38:00:00 | 8hdu-B | 2.6 | 4 | 85 | 162 | 12 | MOLECULE: | DE NOVO DE | SIGN CAV | ITATED PROTEIN | |
| 39:00:00 | 6rfq-2 | 2.6 | 12.8 | 99 | 469 | 5 | MOLECULE: | SUBUNIT NU | AM OF NA | DH:UBIQUINONE OXIDOREDUCTASE (C | |
| 40:00:00 | 5ir2-A | 2.6 | 10.4 | 67 | 222 | 9 | MOLECULE: | CELLULASE | | | |
| 41:00:00 | 2mab-A | 2.6 | 3.6 | 76 | 109 | 7 | MOLECULE: | ACINIFORM | SPIDROIN | | |
| 42:00:00 | 5jeq-A | 2.6 | 32.8 | 110 | 227 | 5 | MOLECULE: | NITRATE/NI | TRITE SE | NSOR PROTEIN NARQ | |
| 43:00:00 | 8cqr-A | 2.6 | 6.2 | 70 | 103 | 4 | MOLECULE: | NINJURIN-1 | | | |
| 44:00:00 | 7lsv-B | 2.6 | 6.2 | 83 | 158 | 4 | MOLECULE: | CALMODULIN | #NAME? | NT PROTEIN KINASE | |
| 45:00:00 | 5k8o-B | 2.6 | 7.8 | 70 | 425 | 10 | MOLECULE: | 5-NITROANT | HRANILIC | ACID AMINOHYDROLASE | |
| 46:00:00 | 4btp-A | 2.6 | 12.3 | 114 | 748 | 5 | MOLECULE: | P1 | | | |
| 47:00:00 | 8xje-B | 2.6 | 9.3 | 64 | 149 | 2 | MOLECULE: | YQEY | | | |
| 48:00:00 | 5z58-1 | 2.6 | 21.5 | 93 | 1038 | 5 | MOLECULE: | PRE-MRNA-P | ROCESSIN | G-SPLICING FACTOR 8 | |
| 49:00:00 | 7z1n-O | 2.6 | 21 | 91 | 570 | 4 | MOLECULE: | DNA-DIRECT | ED RNA P | OLYMERASE III SUBUNIT RPC1 | |
| 50:00:00 | 2odm-B | 2.6 | 6.4 | 72 | 83 | 4 | MOLECULE: | UPF0358 PR | OTEIN MW | | 995 |
| 51:00:00 | 2nnu-A | 2.6 | 15 | 99 | 201 | 13 | MOLECULE: | REGULATORY | PROTEIN | E2 | |
| 52:00:00 | 3u8p-A | 2.6 | 9.6 | 78 | 336 | 5 | MOLECULE: | CYTOCHROME | B562 IN | TEGRAL FUSION WITH ENHANCED GRE | |
| 53:00:00 | 1jeb-B | 2.6 | 3.8 | 72 | 143 | 14 | MOLECULE: | HEMOGLOBIN | ZETA CH | AIN | |
| 54:00:00 | 7jy3-D | 2.6 | 4.6 | 76 | 146 | 13 | MOLECULE: | HEMOGLOBIN | SUBUNIT | ALPHA | |
| 55:00:00 | 1jeb-D | 2.6 | 4 | 73 | 146 | 14 | MOLECULE: | HEMOGLOBIN | ZETA CH | AIN | |
| 56:00:00 | 6nq5-B | 2.6 | 4.2 | 78 | 145 | 14 | MOLECULE: | HEMOGLOBIN | SUBUNIT | ALPHA | |
| 57:00:00 | 8p23-B | 2.5 | 14.8 | 103 | 715 | 6 | MOLECULE: | ANAEROBIC | RIBONUCL | EOSIDE-TRIPHOSPHATE REDUCTASE | |
| 58:00:00 | 4x2c-A | 2.5 | 8 | 75 | 207 | 5 | MOLECULE: | FIC FAMILY | PROTEIN | PUTATIVE FILAMENTATION INDUCED | |
| 59:00:00 | 8g01-A | 2.5 | 7.7 | 121 | 360 | 7 | MOLECULE: | PHOSPHO-N- | ACETYLMU | RAMOYL-PENTAPEPTIDE-TRANSFERASE | |
| 60:00:00 | 7e8k-B | 2.5 | 12.4 | 79 | 205 | 8 | MOLECULE: | RNA-FREE R | IBONUCLE | ASE P | |
| 61:00:00 | 2lk2-A | 2.5 | 7.5 | 67 | 89 | 7 | MOLECULE: | HOMEOBOX P | ROTEIN T | GIF1 | |
| 62:00:00 | 3ezh-A | 2.5 | 4.6 | 71 | 114 | 6 | MOLECULE: | NITRATE/NI | TRITE SE | NSOR PROTEIN NARX | |
| 63:00:00 | 8iaz-A | 2.5 | 10.4 | 104 | 384 | 5 | MOLECULE: | TRANSPOSAS | E | | |
| 64:00:00 | 7ef9-A | 2.5 | 4.9 | 80 | 414 | 6 | MOLECULE: | ADENINE DN | A GLYCOS | YLASE | |
| 65:00:00 | 5muu-A | 2.5 | 14.1 | 108 | 766 | 8 | MOLECULE: | MAJOR INNE | R PROTEI | N P1 | |
| 66:00:00 | 8jhq-A | 2.5 | 23.2 | 123 | 446 | 2 | MOLECULE: | SPHINGOSIN | E-1-PHOS | PHATE TRANSPORTER SPNS2,GLGA GL | |
| 67:00:00 | 7upq-F | 2.5 | 7.3 | 57 | 77 | 9 | MOLECULE: | DHT03 PROT | EIN A | | |
| 68:00:00 | 2oer-B | 2.5 | 6.3 | 80 | 189 | 13 | MOLECULE: | PROBABLE T | RANSCRIP | TIONAL REGULATOR | |
| 69:00:00 | 1fyz-F | 2.5 | 8.9 | 91 | 168 | 8 | MOLECULE: | METHANE MO | NOOXYGEN | ASE COMPONENT A, ALPHA CHAIN | |
| 70:00:00 | 8ppr-M | 2.5 | 17.2 | 74 | 205 | 3 | MOLECULE: | KINETOCHOR | E-ASSOCI | ATED PROTEIN DSN1 HOMOLOG | |
| 71:00:00 | 8d2q-A | 2.5 | 18 | 100 | 906 | 7 | MOLECULE: | CRISPR-ASS | OCIATED | ENDONUCLEASE, CSN1 FAMILY | |
| 72:00:00 | 8cqn-B | 2.5 | 16.8 | 71 | 187 | 13 | MOLECULE: | LIPOPROTEI | N, PUTAT | IVE | |
| 73:00:00 | 6rfs-6 | 2.5 | 6.5 | 77 | 138 | 10 | MOLECULE: | SUBUNIT NU | AM OF NA | DH:UBIQUINONE OXIDOREDUCTASE (C | |
| 74:00:00 | 7n7s-B | 2.5 | 29.5 | 128 | 441 | 5 | MOLECULE: | HYDROXYMET | HYLGLUTA | RYL-COA REDUCTASE | |
| 75:00:00 | 3lml-A | 2.5 | 4.4 | 76 | 440 | 8 | MOLECULE: | LIN1278 PR | OTEIN | | |
| 76:00:00 | 5lnk-M | 2.5 | 5.6 | 93 | 459 | 9 | MOLECULE: | MITOCHONDR | IAL COMP | LEX I, 51 KDA SUBUNIT | |
| 77:00:00 | 3taz-B | 2.5 | 16 | 122 | 433 | 7 | MOLECULE: | DNA DOUBLE | #NAME? | BREAK REPAIR PROTEIN NURA | |
| 78:00:00 | 6bhc-A | 2.5 | 4 | 80 | 371 | 6 | MOLECULE: | PSEUDOPODI | UM-ENRIC | HED ATYPICAL KINASE 1 | |
| 79:00:00 | 2qqy-A | 2.5 | 6.5 | 73 | 139 | 3 | MOLECULE: | SIGMA B OP | ERON | | |
| 80:00:00 | 7mrw-B | 2.5 | 18.3 | 98 | 972 | 5 | MOLECULE: | CYTOADHERE | NCE LINK | ED ASEXUAL PROTEIN 3.1 | |
| 81:00:00 | 6g2a-A | 2.5 | 4.3 | 89 | 361 | 7 | MOLECULE: | [PROTEIN A | DP-RIBOS | YLARGININE] HYDROLASE | |
| 82:00:00 | 8j0n-A | 2.5 | 18.1 | 108 | 949 | 5 | MOLECULE: | ER MEMBRAN | E PROTEI | N COMPLEX SUBUNIT 1 | |

| | | | | | | | | | | | |
|---|---|---|---|---|---|---|---|---|---|---|---|
| 83:00:00 | 6ajf-A | 2.5 | 24.4 | 127 | 901 | 6 | MOLECULE: | DRUG EXPOR | TERS OF | THE RND SUPERFAMILY-LIKE PROTEI | |
| 84:00:00 | 2mfz-A | 2.5 | 3.7 | 78 | 125 | 8 | MOLECULE: | MINOR AMPU | LLATE SP | IDROIN | |
| 85:00:00 | 5xef-A | 2.5 | 10.5 | 73 | 123 | 4 | MOLECULE: | FLAGELLAR | PROTEIN | FLIS | |
| 86:00:00 | 7ose-A | 2.5 | 6.2 | 74 | 503 | 5 | MOLECULE: | CYTOCHROME | BD-II U | BIQUINOL OXIDASE SUBUNIT 1 | |
| 87:00:00 | 5kbw-B | 2.5 | 8.6 | 81 | 171 | 9 | MOLECULE: | RIBOFLAVIN | TRANSPO | RTER RIBU | |
| 88:00:00 | 6qld-Y | 2.5 | 20.5 | 90 | 223 | 3 | MOLECULE: | INNER KINE | TOCHORE | SUBUNIT MIF2 | |
| 89:00:00 | 6oqr-W | 2.5 | 7.1 | 101 | 173 | 8 | MOLECULE: | ATP SYNTHA | SE SUBUN | IT DELTA | |
| 90:00:00 | 6vr4-A | 2.5 | 7.6 | 122 | 2166 | 6 | MOLECULE: | DNA-DEPEND | ENT RNA | POLYMERASE | |
| 91:00:00 | 6h08-B | 2.5 | 7.3 | 108 | 297 | 6 | MOLECULE: | CYTOCHROME | C PEROX | IDASE, MITOCHONDRIAL | |
| 92:00:00 | 5o1l-A | 2.5 | 9.8 | 116 | 375 | 7 | MOLECULE: | RUBBER OXY | GENASE | | |
| 93:00:00 | 2gta-A | 2.5 | 11.7 | 68 | 98 | 3 | MOLECULE: | HYPOTHETIC | AL PROTE | IN YPJD | |
| 94:00:00 | 3au4-A | 2.5 | 8.4 | 87 | 524 | 3 | MOLECULE: | MYOSIN-X | | | |
| 95:00:00 | 4n4w-A | 2.5 | 12.6 | 86 | 457 | 7 | MOLECULE: | CYTOCHROME | B(562), | SMOOTHENED HOMOLOG | |
| 96:00:00 | 8t9f-K | 2.5 | 5.2 | 94 | 265 | 7 | MOLECULE: | HISTONE-LY | SINE N-M | ETHYLTRANSFERASE KMT5B | |
| 97:00:00 | 4gfq-A | 2.5 | 4.4 | 73 | 186 | 10 | MOLECULE: | RIBOSOME-R | ECYCLING | FACTOR | |
| 98:00:00 | 7xqk-B | 2.5 | 3.9 | 78 | 276 | 5 | MOLECULE: | GLUTATHION | E S-TRAN | SFERASE CLASS-MU 26 KDA ISOZYME | |
| 99:00:00 | 8k9z-A | 2.5 | 16.9 | 114 | 405 | 9 | MOLECULE: | RDTND-RID | CBD | | |
| 0:00 | 4tq4-D | 2.5 | 11.1 | 93 | 290 | 4 | MOLECULE: | PRENYLTRAN | SFERASE | | |
| 1:00 | 8snl-B | 2.5 | 7.8 | 86 | 491 | 9 | MOLECULE: | DISINTEGRI | N AND ME | TALLOPROTEINASE DOMAIN-CONTAINI | |
| 2:00 | 5tgz-A | 2.5 | 11.6 | 95 | 439 | 4 | MOLECULE: | CANNABINOI | D RECEPT | OR 1,FLAVODOXIN,CANNABINOID REC | |
| 3:00 | 8pi9-B | 2.5 | 6.3 | 76 | 177 | 9 | MOLECULE: | CHAINS: E | | | |
| 4:00 | 4djb-A | 2.5 | 3.8 | 71 | 118 | 11 | MOLECULE: | E4-ORF3 | | | |
| 5:00 | 7xsq-A | 2.5 | 19.2 | 99 | 1289 | 5 | MOLECULE: | RAMP SUPER | FAMILY P | ROTEIN | |
| 6:00 | 2gtv-X | 2.5 | 4.1 | 67 | 104 | 3 | MOLECULE: | CHORISMATE | MUTASE | | |
| 7:00 | 8p2m-A | 2.5 | 10 | 104 | 659 | 9 | MOLECULE: | NAD(+) HYD | ROLASE T | IR-1 | |
| 8:00 | 7qd4-A | 2.5 | 19.6 | 106 | 908 | 14 | MOLECULE: | TRANSPOSAS | E FOR TR | ANSPOSON TN4430 | |
| 9:00 | 7aft-F | 2.5 | 17.4 | 84 | 190 | 2 | MOLECULE: | PROTEIN TR | ANSPORT | PROTEIN SEC61 | |
| 10:00 | 6u0o-A | 2.5 | 5.1 | 88 | 248 | 6 | MOLECULE: | LYSOSTAPHI | N RESIST | ANCE PROTEIN A | |
| 11:00 | 7apd-B | 2.5 | 14.2 | 78 | 289 | 3 | MOLECULE: | REPLICATIO | N PROTEI | N E1 | |
| 12:00 | 7sbb-I | 2.5 | 17.4 | 90 | 680 | 3 | MOLECULE: | CAS7D | | | |
| 13:00 | 1e9s-A | 2.5 | 5.9 | 94 | 427 | 13 | MOLECULE: | CONJUGAL T | RANSFER | PROTEIN TRWB | |
| 14:00 | 7e5c-A | 2.5 | 12.4 | 84 | 439 | 6 | MOLECULE: | XAA-PRO DI | PEPTIDAS | E | |
| 15:00 | 4qvg-C | 2.5 | 10.1 | 100 | 349 | 8 | MOLECULE: | SIBL | | | |
| 16:00 | 4qvz-B | 2.5 | 13.5 | 66 | 204 | 5 | MOLECULE: | FRAGILE X | MENTAL R | ETARDATION PROTEIN 1 | |
| 17:00 | 6rw3-B | 2.5 | 15.9 | 100 | 439 | 6 | MOLECULE: | HEXOSE TRA | NSPORTER | | 1 |
| 18:00 | 8u53-A | 2.5 | 16.2 | 104 | 574 | 9 | MOLECULE: | CSC1-LIKE | PROTEIN | ERD4 | |
| 19:00 | 6ukj-A | 2.5 | 21.7 | 109 | 350 | 7 | MOLECULE: | CHLOROQUIN | E RESIST | ANCE TRANSPORTER | |
| 20:00 | 4cc9-B | 2.5 | 6 | 74 | 98 | 9 | MOLECULE: | PROTEIN VP | RBP | | |
| 21:00 | 2fm8-C | 2.5 | 15.9 | 76 | 220 | 4 | MOLECULE: | SURFACE PR | ESENTATI | ON OF ANTIGENS PROTEIN SPAK | |
| 22:00 | 5xw4-B | 2.5 | 8.1 | 84 | 369 | 5 | MOLECULE: | TYROSINE-P | ROTEIN P | HOSPHATASE CDC14 | |
| 23:00 | 7etr-A | 2.5 | 12.7 | 52 | 88 | 0 | MOLECULE: | TRANSCRIPT | IONAL RE | GULATOR COPG FAMILY | |
| 24:00:00 | 8vej-B | 2.5 | 3 | 53 | 242 | 6 | MOLECULE: | CHD_BUTTRE | SS | | |
| 25:00:00 | 6mdw-A | 2.5 | 10.1 | 79 | 185 | 5 | MOLECULE: | SPRT-LIKE | DOMAIN-C | ONTAINING PROTEIN SPARTAN | |
| 26:00:00 | 2iu5-A | 2.5 | 12.3 | 77 | 179 | 8 | MOLECULE: | HTH-TYPE D | HAKLM OP | ERON TRANSCRIPTIONAL ACTIVATOR | |
| 27:00:00 | 8vc8-A | 2.5 | 5.7 | 93 | 197 | 12 | MOLECULE: | HEM_3.C9 | | | |
| 28:00:00 | 8e9g-L | 2.5 | 26.2 | 116 | 625 | 8 | MOLECULE: | TWO-COMPON | ENT SYST | EM RESPONSE REGULATOR | |
| 29:00:00 | 8d8j-O | 2.5 | 10.1 | 91 | 210 | 12 | MOLECULE: | PROBABLE S | #NAME? | L-L-METHIONINE-DEPENDENT RNA | |
| 30:00:00 | 1td6-A | 2.5 | 15.9 | 87 | 286 | 6 | MOLECULE: | HYPOTHETIC | AL PROTE | IN MG237 HOMOLOG | |
| 31:00:00 | 8e9h-J | 2.5 | 20.1 | 91 | 231 | 10 | MOLECULE: | TWO-COMPON | ENT SYST | EM RESPONSE REGULATOR | |
| 32:00:00 | 6g7c-C | 2.5 | 14.2 | 75 | 239 | 9 | MOLECULE: | IMPA-RELAT | ED DOMAI | N PROTEIN | |
| 33:00:00 | 6yrf-A | 2.5 | 13.2 | 97 | 777 | 6 | MOLECULE: | VEGETATIVE | INSECTI | CIDAL PROTEIN | |
| 34:00:00 | 5utg-A | 2.5 | 4.2 | 71 | 134 | 3 | MOLECULE: | EGG-LYSIN | | | |
| 35:00:00 | 5anz-A | 2.5 | 12 | 125 | 375 | 7 | MOLECULE: | SOLUBLE LY | TIC TRAN | SGLYCOSYLASE B3 | |
| 36:00:00 | 3syk-A | 2.5 | 12.3 | 106 | 290 | 4 | MOLECULE: | PROTEIN CB | BX | | |

| | | | | | | | | | | | |
|---|---|---|---|---|---|---|---|---|---|---|---|
| 37:00:00 | 2gtt-K | 2.5 | 25.8 | 87 | 420 | 3 | MOLECULE: | NUCLEOPROT | EIN | | |
| 38:00:00 | 7jgr-D | 2.5 | 10.6 | 80 | 441 | 6 | MOLECULE: | ORIGIN REC | OGNITION | COMPLEX SUBUNIT 2 | |
| 39:00:00 | 7w0y-i | 2.5 | 13.2 | 89 | 347 | 9 | MOLECULE: | NADH DEHYD | ROGENASE | [UBIQUINONE] FLAVOPROTEIN 1, | |
| 40:00:00 | 8in1-A | 2.5 | 21.9 | 112 | 952 | 10 | MOLECULE: | BETA-GLUCO | SIDASE | | |
| 41:00:00 | 6tpn-A | 2.5 | 19 | 101 | 510 | 4 | MOLECULE: | OREXIN REC | EPTOR TY | PE 2,GLGA GLYCOGEN SYNTHASE,ORE | |
| 42:00:00 | 7cun-G | 2.5 | 20.5 | 131 | 895 | 7 | MOLECULE: | INTEGRATOR | COMPLEX | SUBUNIT 1 | |
| 43:00:00 | 6qti-A | 2.5 | 24.9 | 119 | 1038 | 3 | MOLECULE: | NICOTINAMI | DE NUCLE | OTIDE TRANSHYDROGENASE | |
| 44:00:00 | 6g2d-C | 2.5 | 26.5 | 102 | 2121 | 9 | MOLECULE: | ACETYL-COA | CARBOXY | LASE 1 | |
| 45:00:00 | 5vkq-A | 2.5 | 21.2 | 136 | 1499 | 10 | MOLECULE: | NO MECHANO | RECEPTOR | POTENTIAL C ISOFORM L | |
| 46:00:00 | 6mur-B | 2.5 | 3.9 | 67 | 162 | 4 | MOLECULE: | UNCHARACTE | RIZED PR | OTEIN | |
| 47:00:00 | 8urw-Z | 2.5 | 13.9 | 120 | 1235 | 7 | MOLECULE: | DNA-DIRECT | ED RNA P | OLYMERASE SUBUNIT ALPHA | |
| 48:00:00 | 8hki-a | 2.4 | 4.1 | 84 | 291 | 6 | MOLECULE: | T-COMPLEX | PROTEIN | 1 SUBUNIT ALPHA | |
| 49:00:00 | 2o2e-B | 2.4 | 5 | 117 | 316 | 11 | MOLECULE: | TRYPTOPHAN | SYNTHAS | E BETA CHAIN | |
| 50:00:00 | 5yx5-B | 2.4 | 4.6 | 65 | 270 | 3 | MOLECULE: | 49 PROTEIN | | | |
| 51:00:00 | 5h9c-A | 2.4 | 6.8 | 55 | 78 | 7 | MOLECULE: | ENVELOPE G | LYCOPROT | EIN GP95 | |
| 52:00:00 | 4tkr-B | 2.4 | 3.7 | 77 | 186 | 6 | MOLECULE: | THIAMINE T | RANSPORT | ER THIA | |
| 53:00:00 | 7t92-C | 2.4 | 5.9 | 114 | 317 | 5 | MOLECULE: | PEROXIN-12 | | | |
| 54:00:00 | 8h36-G | 2.4 | 20.2 | 94 | 379 | 6 | MOLECULE: | E3 UBIQUIT | IN-PROTE | IN LIGASE RBX1 | |
| 55:00:00 | 6zg3-C | 2.4 | 6.1 | 101 | 207 | 4 | MOLECULE: | ENERGY-COU | PLING FA | CTOR TRANSPORTER ATP-BINDING PR | |
| 56:00:00 | 7qh5-A | 2.4 | 5.7 | 84 | 313 | 12 | MOLECULE: | RNA POLYME | RASE SIG | MA FACTOR | |
| 57:00:00 | 7yqh-A | 2.4 | 16.8 | 106 | 1069 | 5 | MOLECULE: | STRUCTURAL | MAINTEN | ANCE OF CHROMOSOMES PROTEIN 5 | |
| 58:00:00 | 3fx7-B | 2.4 | 9 | 65 | 87 | 8 | MOLECULE: | PUTATIVE U | NCHARACT | ERIZED PROTEIN | |
| 59:00:00 | 4kw3-A | 2.4 | 4.6 | 102 | 266 | 7 | MOLECULE: | NS1 | | | |
| 60:00:00 | 5ip0-D | 2.4 | 8.3 | 64 | 108 | 5 | MOLECULE: | PHA GRANUL | E-ASSOCI | ATED PROTEIN | |
| 61:00:00 | 7ura-A | 2.4 | 10 | 91 | 432 | 8 | MOLECULE: | ISOFORM 2 | OF PROTE | IN-SERINE O-PALMITOLEOYLTRANSFE | |
| 62:00:00 | 6l85-B | 2.4 | 8.5 | 98 | 401 | 7 | MOLECULE: | PHOSPHATE | TRANSPOR | TER | |
| 63:00:00 | 4u8u-G | 2.4 | 4.7 | 74 | 151 | 11 | MOLECULE: | GLOBIN A C | HAIN | | |
| 64:00:00 | 2hi4-A | 2.4 | 12.8 | 96 | 480 | 4 | MOLECULE: | CYTOCHROME | P450 1A | | 2 |
| 65:00:00 | 8i6v-D | 2.4 | 23.9 | 95 | 678 | 8 | MOLECULE: | VACUOLAR T | RANSPORT | ER CHAPERONE COMPLEX SUBUNIT 1 | |
| 66:00:00 | 2z0q-A | 2.4 | 13.8 | 88 | 321 | 7 | MOLECULE: | RHO GUANIN | E NUCLEO | TIDE EXCHANGE FACTOR 3 | |
| 67:00:00 | 6tqe-A | 2.4 | 21.1 | 122 | 634 | 9 | MOLECULE: | ABC TRANSP | ORTER AT | P-BINDING PROTEIN/PERMEASE | |
| 68:00:00 | 8ahx-E | 2.4 | 5.6 | 103 | 213 | 8 | MOLECULE: | ION-TRANSL | OCATING | OXIDOREDUCTASE COMPLEX SUBUNIT | |
| 69:00:00 | 6btm-C | 2.4 | 8.4 | 104 | 457 | 10 | MOLECULE: | ALTERNATIV | E COMPLE | X III SUBUNIT A | |
| 70:00:00 | 5bs1-D | 2.4 | 9.5 | 83 | 121 | 5 | MOLECULE: | CRRBCX-IIA | | | |
| 71:00:00 | 2a6b-A | 2.4 | 4.9 | 110 | 219 | 5 | MOLECULE: | HYPOTHETIC | AL PROTE | IN SPR0628 | |
| 72:00:00 | 7bin-L | 2.4 | 6.8 | 56 | 75 | 5 | MOLECULE: | FLAGELLAR | BIOSYNTH | ETIC PROTEIN FLIP | |
| 73:00:00 | 8gxl-B | 2.4 | 8.1 | 58 | 122 | 3 | MOLECULE: | SURP AND G | #NAME? | OMAIN-CONTAINING PROTEIN 1 | |
| 74:00:00 | 5j6f-A | 2.4 | 10.6 | 81 | 352 | 7 | MOLECULE: | 3-DEOXY-D- | ARABINO- | HEPTULOSONATE 7-PHOSPHATE SYNTH | |
| 75:00:00 | 8hr5-A | 2.4 | 15.1 | 93 | 466 | 8 | MOLECULE: | TRANSPOSAS | E | | |
| 76:00:00 | 5wd6-A | 2.4 | 4.1 | 94 | 195 | 4 | MOLECULE: | SHORT PALA | TE, LUNG | AND NASAL EPITHELIUM CARCINOMA | |
| 77:00:00 | 7y22-M | 2.4 | 12.5 | 87 | 215 | 3 | MOLECULE: | PHAGE CONN | ECTOR PR | OTEIN | |
| 78:00:00 | 6j7k-A | 2.4 | 8.7 | 79 | 385 | 3 | MOLECULE: | PSEUDOMONA | S AERUGI | NOSA EARP | |
| 79:00:00 | 3h0g-A | 2.4 | 28.4 | 130 | 1496 | 12 | MOLECULE: | DNA-DIRECT | ED RNA P | OLYMERASE II SUBUNIT RPB1 | |
| 80:00:00 | 6b87-D | 2.4 | 8.5 | 67 | 102 | 6 | MOLECULE: | TMHC2_E | | | |
| 81:00:00 | 6upn-A | 2.4 | 20.4 | 88 | 245 | 9 | MOLECULE: | ENDOPHILIN | -B1 | | |
| 82:00:00 | 2x0l-A | 2.4 | 35.9 | 115 | 670 | 5 | MOLECULE: | LYSINE-SPE | CIFIC HI | STONE DEMETHYLASE 1 | |
| 83:00:00 | 7lhf-A | 2.4 | 13 | 90 | 2123 | 7 | MOLECULE: | INOSITOL 1 | ,4,5-TRI | SPHOSPHATE RECEPTOR TYPE 1 | |
| 84:00:00 | 8ek4-B | 2.4 | 8.4 | 70 | 135 | 6 | MOLECULE: | ICE-BINDIN | G PROTEI | N TIP-99A | |
| 85:00:00 | 5xpd-A | 2.4 | 19.4 | 85 | 269 | 5 | MOLECULE: | SUGAR TRAN | SPORTER | | |
| 86:00:00 | 7oqz-A | 2.4 | 6.9 | 83 | 237 | 8 | MOLECULE: | TRANSMEMBR | ANE PROT | EIN 45A | |
| 87:00:00 | 6m5r-B | 2.4 | 18.4 | 107 | 689 | 6 | MOLECULE: | TRIPARTITE | TERMINA | SE SUBUNIT 3 | |
| 88:00:00 | 8b9z-Y | 2.4 | 12.9 | 105 | 167 | 6 | MOLECULE: | NADH-UBIQU | INONE OX | IDOREDUCTASE CHAIN 3 | |
| 89:00:00 | 8q7n-T | 2.4 | 15.1 | 81 | 412 | 6 | MOLECULE: | U5 SNRNA | | | |
| 90:00:00 | 5yqz-R | 2.4 | 17.1 | 131 | 558 | 8 | MOLECULE: | GLUCAGON R | ECEPTOR, | ENDOLYSIN,GLUCAGON RECEPTOR | |

| | | | | | | | | | | |
|---|---|---|---|---|---|---|---|---|---|---|
| 91:00:00 | 6cgh-A | 2.4 | 5.4 | 52 | 89 | 8 | MOLECULE: | DNAJ HOMOL | OG SUBFA | MILY C MEMBER 2 |
| 92:00:00 | 7jrg-F | 2.4 | 10.5 | 82 | 116 | 10 | MOLECULE: | MITOCHONDR | IAL-PROC | ESSING PEPTIDASE SUBUNIT BETA, |
| 93:00:00 | 8soj-B | 2.4 | 14 | 91 | 428 | 10 | MOLECULE: | CST COMPLE | X SUBUNI | T CTC1 |
| 94:00:00 | 5xf9-D | 2.4 | 17.3 | 95 | 455 | 7 | MOLECULE: | NAD-REDUCI | NG HYDRO | GENASE |
| 95:00:00 | 8d2z-A | 2.4 | 13.8 | 71 | 306 | 7 | MOLECULE: | METALLO-BE | TA-LACTA | MASE SUPERFAMILY PROTEIN |
| 96:00:00 | 8uq9-a | 2.4 | 16.4 | 91 | 433 | 7 | MOLECULE: | E3 UBIQUIT | IN-PROTE | IN LIGASE RNF168,UBIQUITIN-CONJ |
| 97:00:00 | 3qwe-A | 2.4 | 7.2 | 85 | 260 | 8 | MOLECULE: | GEM-INTERA | CTING PR | OTEIN |
| 98:00:00 | 6elu-A | 2.4 | 11.9 | 70 | 193 | 7 | MOLECULE: | SERUM RESI | STANCE A | SSOCIATED VSG PROTEIN |
| 99:00:00 | 7jw1-e | 2.4 | 33.2 | 98 | 235 | 8 | MOLECULE: | CAPSID PRO | TEINS | |
| 0:00 | 4y2f-A | 2.4 | 4 | 68 | 143 | 6 | MOLECULE: | SENSOR PRO | TEIN KDP | D |
| 1:00 | 2g7l-A | 2.4 | 8 | 80 | 201 | 10 | MOLECULE: | TETR-FAMIL | Y TRANSC | RIPTIONAL REGULATOR |
| 2:00 | 5xe3-E | 2.4 | 4.8 | 44 | 81 | 5 | MOLECULE: | ENDORIBONU | CLEASE M | AZF4 |
| 3:00 | 4fhn-B | 2.4 | 12.8 | 73 | 1022 | 5 | MOLECULE: | NUCLEOPORI | N NUP37 | |
| 4:00 | 6uzt-A | 2.4 | 7.5 | 87 | 587 | 7 | MOLECULE: | RECEPTOR-T | YPE TYRO | SINE-PROTEIN PHOSPHATASE ALPHA |
| 5:00 | 7tb3-A | 2.4 | 9.6 | 87 | 243 | 7 | MOLECULE: | ISOFORM 2 | OF CREB- | BINDING PROTEIN,FERRITIN HEAVY |
| 6:00 | 2huj-A | 2.4 | 10.3 | 75 | 125 | 8 | MOLECULE: | LIN2004 PR | OTEIN | |
| 7:00 | 1sqf-A | 2.4 | 7.9 | 90 | 425 | 6 | MOLECULE: | SUN PROTEI | N | |
| 8:00 | 5aor-A | 2.4 | 21.9 | 109 | 1009 | 6 | MOLECULE: | DOSAGE COM | PENSATIO | N REGULATOR |
| 9:00 | 7t5p-A | 2.4 | 10.5 | 89 | 403 | 11 | MOLECULE: | SUMO-INTER | ACTING M | OTIF-CONTAINING PROTEIN 1 |
| 10:00 | 6o0y-A | 2.4 | 22.3 | 127 | 1146 | 6 | MOLECULE: | CRISPR-ASS | OCIATED | ENDONUCLEASE CAS9/CSN1 |
| 11:00 | 9b8o-e | 2.4 | 8.7 | 64 | 80 | 5 | MOLECULE: | ATPASE H+- | TRANSPOR | TING V1 SUBUNIT D |
| 12:00 | 3n1e-B | 2.4 | 4.6 | 72 | 141 | 3 | MOLECULE: | VACUOLAR P | ROTEIN S | ORTING-ASSOCIATED PROTEIN 54 |
| 13:00 | 6q6b-A | 2.4 | 5.9 | 71 | 132 | 7 | MOLECULE: | CYTOSOLIC | COPPER S | TORAGE PROTEIN |
| 14:00 | 7pp4-f | 2.4 | 8.2 | 59 | 142 | 3 | MOLECULE: | DNA-DIRECT | ED RNA P | OLYMERASE SUBUNIT ALPHA |
| 15:00 | 4r29-B | 2.4 | 14 | 77 | 204 | 4 | MOLECULE: | UNCHARACTE | RIZED PR | OTEIN |
| 16:00 | 6v6d-A | 2.4 | 13.7 | 78 | 209 | 13 | MOLECULE: | PANNEXIN-1 | | |
| 17:00 | 7udk-A | 2.4 | 5.8 | 89 | 172 | 7 | MOLECULE: | DESIGNED H | ELICAL R | EPEAT PROTEIN (DHR) RPB_LRP2_R4 |
| 18:00 | 7z12-A | 2.4 | 20.6 | 136 | 1853 | 5 | MOLECULE: | PAM1.4, HE | AVY CHAI | N |
| 19:00 | 7akx-A | 2.4 | 10 | 102 | 224 | 7 | MOLECULE: | VIRAL RHOD | OPSIN OL | PVR1 |
| 20:00 | 8eg3-A | 2.4 | 18.1 | 81 | 554 | 9 | MOLECULE: | STERYL-SUL | FATASE | |
| 21:00 | 7b4m-A | 2.4 | 23.7 | 132 | 459 | 5 | MOLECULE: | SODIUM/HYD | ROGEN EX | CHANGER 9B2 |
| 22:00 | 6h3w-A | 2.4 | 3.6 | 74 | 246 | 4 | MOLECULE: | ENVELOPMEN | T POLYPR | OTEIN |
| 23:00 | 4hga-A | 2.4 | 22.9 | 82 | 207 | 9 | MOLECULE: | DEATH DOMA | IN-ASSOC | IATED PROTEIN 6 |
| 24:00:00 | 7oci-C | 2.4 | 7.5 | 69 | 119 | 12 | MOLECULE: | DOLICHYL-D | IPHOSPHO | OLIGOSACCHARIDE--PROTEIN |
| 25:00:00 | 1x3k-A | 2.4 | 3 | 67 | 152 | 7 | MOLECULE: | HEMOGLOBIN | COMPONE | NT V |
| 26:00:00 | 3qt2-C | 2.4 | 11.2 | 80 | 110 | 6 | MOLECULE: | INTERLEUKI | N-5 RECE | PTOR SUBUNIT ALPHA |
| 27:00:00 | 6s7t-H | 2.4 | 7.7 | 77 | 112 | 4 | MOLECULE: | DOLICHYL-D | IPHOSPHO | OLIGOSACCHARIDE--PROTEIN |
| 28:00:00 | 8y7g-B | 2.4 | 14.5 | 104 | 543 | 2 | MOLECULE: | CRISPR-ASS | OCIATED | PROTEIN |
| 29:00:00 | 2hxo-A | 2.4 | 15.7 | 65 | 207 | 9 | MOLECULE: | PUTATIVE T | ETR-FAMI | LY TRANSCRIPTIONAL REGULATOR |
| 30:00:00 | 7oqe-K | 2.4 | 12.1 | 74 | 406 | 3 | MOLECULE: | PROTEIN NA | M8 | |
| 31:00:00 | 7qba-D | 2.4 | 5.3 | 88 | 275 | 5 | MOLECULE: | PROBABLE A | BC TRANS | PORTER BINDING PROTEIN NOSD |
| 32:00:00 | 6wuc-H | 2.4 | 15.7 | 83 | 179 | 4 | MOLECULE: | INNER KINE | TOCHORE | SUBUNIT MCM16 |
| 33:00:00 | 3nlc-A | 2.4 | 7.5 | 106 | 534 | 9 | MOLECULE: | UNCHARACTE | RIZED PR | OTEIN VP0956 |
| 34:00:00 | 2mn2-A | 2.4 | 8.9 | 75 | 122 | 4 | MOLECULE: | YMOB | | |
| 35:00:00 | 8age-H | 2.4 | 7.5 | 75 | 133 | 8 | MOLECULE: | DOLICHYL-D | IPHOSPHO | OLIGOSACCHARIDE--PROTEIN |
| 36:00:00 | 3rkl-A | 2.4 | 7 | 66 | 80 | 5 | MOLECULE: | STIV-A81 | | |
| 37:00:00 | 5h11-A | 2.4 | 17.2 | 111 | 494 | 8 | MOLECULE: | UNCHARACTE | RIZED PR | OTEIN |
| 38:00:00 | 6gw6-B | 2.4 | 5.5 | 63 | 149 | 8 | MOLECULE: | RES TOXIN | | |
| 39:00:00 | 4txa-A | 2.4 | 11.1 | 83 | 367 | 12 | MOLECULE: | ROQUIN-1 | | |
| 40:00:00 | 5jwy-A | 2.4 | 12.9 | 72 | 259 | 3 | MOLECULE: | PHOSPHATID | YLGLYCER | OPHOSPHATASE B |
| 41:00:00 | 8ck1-E | 2.4 | 17.1 | 83 | 206 | 4 | MOLECULE: | TAIL NOZZL | E | |
| 42:00:00 | 1w0b-A | 2.4 | 4.6 | 72 | 102 | 11 | MOLECULE: | ALPHA-HEMO | GLOBIN S | TABILIZING PROTEIN |
| 43:00:00 | 6rf0-C | 2.4 | 11.9 | 95 | 275 | 8 | MOLECULE: | SODIUM PUM | PING RHO | DOPSIN |
| 44:00:00 | 6qeq-A | 2.4 | 16.4 | 71 | 111 | 3 | MOLECULE: | PCFF | | |

| | | | | | | | | | | |
|---|---|---|---|---|---|---|---|---|---|---|
| 45:00:00 | 4ql6-B | 2.4 | 15 | 82 | 509 | 9 | MOLECULE: | CARBOXY-TE | RMINAL P | ROCESSING PROTEASE |
| 46:00:00 | 8g3a-D | 2.4 | 15.5 | 80 | 334 | 6 | MOLECULE: | BACITRACIN | EXPORT | PERMEASE PROTEIN BCEB |
| 47:00:00 | 7emf-H | 2.4 | 21.4 | 78 | 181 | 9 | MOLECULE: | MEDIATOR O | F RNA PO | LYMERASE II TRANSCRIPTION SUBUN |
| 48:00:00 | 7ljn-A | 2.4 | 17 | 75 | 342 | 8 | MOLECULE: | CD-NTASE | | |
| 49:00:00 | 6p18-Q | 2.4 | 22.1 | 71 | 156 | 1 | MOLECULE: | DNA (67-ME | R) FRAGM | ENT CARRYING PHAGE-21 PR' PROMO |
| 50:00:00 | 6zu0-C | 2.4 | 15.7 | 96 | 426 | 5 | MOLECULE: | CITRATE SY | NTHASE | |
| 51:00:00 | 3dby-C | 2.4 | 12.5 | 104 | 267 | 7 | MOLECULE: | UNCHARACTE | RIZED PR | OTEIN |
| 52:00:00 | 8bsh-C | 2.4 | 7.3 | 126 | 794 | 7 | MOLECULE: | PROTEIN TR | ANSPORT | PROTEIN SEC23 |
| 53:00:00 | 6r6h-O | 2.4 | 20.8 | 126 | 744 | 7 | MOLECULE: | COP9 SIGNA | LOSOME C | OMPLEX SUBUNIT 1 |
| 54:00:00 | 1yvl-B | 2.4 | 10.4 | 84 | 653 | 4 | MOLECULE: | SIGNAL TRA | NSDUCER | AND ACTIVATOR OF TRANSCRIPTION |
| 55:00:00 | 8bsb-B | 2.4 | 19.5 | 96 | 162 | 8 | MOLECULE: | METHYL-ACC | EPTING C | HEMOTAXIS PROTEIN |
| 56:00:00 | 8hsb-A | 2.4 | 12.4 | 113 | 374 | 4 | MOLECULE: | CDNG | | |
| 57:00:00 | 2bf0-X | 2.4 | 3.9 | 68 | 131 | 12 | MOLECULE: | PCF11 | | |
| 58:00:00 | 2kd1-A | 2.4 | 3.5 | 77 | 118 | 3 | MOLECULE: | DNA INTEGR | ATION/RE | COMBINATION/INVERTION PROTEIN |
| 59:00:00 | 5mlq-A | 2.4 | 16.3 | 85 | 226 | 12 | MOLECULE: | CDPS | | |
| 60:00:00 | 8u12-A | 2.4 | 5.2 | 41 | 50 | 2 | MOLECULE: | ANTITOXIN | RV0298 | |
| 61:00:00 | 5m4s-A | 2.4 | 4.8 | 91 | 209 | 2 | MOLECULE: | TRANSCRIPT | ION INIT | IATION FACTOR IIA SUBUNIT 2,TRA |
| 62:00:00 | 7wb4-I | 2.4 | 14.6 | 127 | 796 | 7 | MOLECULE: | OUTER NUP1 | 33 | |
| 63:00:00 | 5c8j-L | 2.4 | 12.8 | 90 | 137 | 9 | MOLECULE: | ANTIBODY F | RAGMENT, | HEAVY CHAIN |
| 64:00:00 | 6ypq-A | 2.4 | 4.2 | 89 | 164 | 7 | MOLECULE: | C-PHYCOCYA | NIN ALPH | A CHAIN |
| 65:00:00 | 4e6k-G | 2.4 | 2.5 | 48 | 57 | 10 | MOLECULE: | BACTERIOFE | RRITIN | |
| 66:00:00 | 2ig3-A | 2.4 | 5.4 | 73 | 127 | 7 | MOLECULE: | GROUP III | TRUNCATE | D HAEMOGLOBIN |
| 67:00:00 | 3njc-A | 2.4 | 4.3 | 79 | 153 | 4 | MOLECULE: | YSLB PROTE | IN | |
| 68:00:00 | 3llw-D | 2.4 | 7.4 | 80 | 289 | 4 | MOLECULE: | GERANYLTRA | NSTRANSF | ERASE (ISPA) |
| 69:00:00 | 6kwy-c | 2.4 | 25.3 | 120 | 1811 | 4 | MOLECULE: | PROTEASOME | SUBUNIT | ALPHA TYPE-2 |
| 70:00:00 | 8e4g-0 | 2.4 | 9.9 | 79 | 196 | 5 | MOLECULE: | PORTAL PRO | TEIN | |
| 71:00:00 | 5a5t-L | 2.4 | 25.3 | 95 | 372 | 5 | MOLECULE: | EUKARYOTIC | TRANSLA | TION INITIATION FACTOR 3 SUBUNI |
| 72:00:00 | 8ts6-A | 2.4 | 5.6 | 77 | 500 | 10 | MOLECULE: | PORTAL PRO | TEIN | |
| 73:00:00 | 8pda-A | 2.4 | 26.7 | 117 | 973 | 3 | MOLECULE: | ERAD-ASSOC | IATED E3 | UBIQUITIN-PROTEIN LIGASE DOA10 |
| 74:00:00 | 5mmi-Z | 2.4 | 9.2 | 67 | 101 | 6 | MOLECULE: | 50S RIBOSO | MAL PROT | EIN L31 |
| 75:00:00 | 6t58-A | 2.4 | 18.1 | 90 | 524 | 7 | MOLECULE: | CELLULAR T | UMOR ANT | IGEN P53,PROTEIN S100-A4,PROTEI |
| 76:00:00 | 8j5q-B | 2.4 | 8.5 | 97 | 325 | 5 | MOLECULE: | UNCHARACTE | RIZED PR | OTEIN RV1280C |
| 77:00:00 | 7nj1-A | 2.4 | 13.2 | 88 | 1448 | 10 | MOLECULE: | SEPARIN | | |
| 78:00:00 | 6ibl-A | 2.4 | 16.1 | 109 | 397 | 6 | MOLECULE: | THIOREDOXI | N 1,BETA | -1 ADRENERGIC RECEPTOR |
| 79:00:00 | 8r6y-A | 2.4 | 17.9 | 138 | 1991 | 7 | MOLECULE: | RNA-DIRECT | ED RNA P | OLYMERASE L |
| 80:00:00 | 8p0u-A | 2.4 | 15 | 95 | 621 | 9 | MOLECULE: | POLYMERASE | ACIDIC | PROTEIN |
| 81:00:00 | 6vqx-A | 2.4 | 3.9 | 65 | 97 | 11 | MOLECULE: | ACRF6 | | |
| 82:00:00 | 4dvg-B | 2.4 | 8.1 | 95 | 305 | 7 | MOLECULE: | RHO-LIKE S | MALL GTP | ASE |
| 83:00:00 | 7zol-B | 2.4 | 24.9 | 122 | 1702 | 6 | MOLECULE: | CAS7-11 | | |
| 84:00:00 | 6ly9-M | 2.4 | 12.5 | 83 | 321 | 11 | MOLECULE: | V-TYPE ATP | SYNTHAS | E SUBUNIT I |
| 85:00:00 | 6fkf-d | 2.4 | 6.7 | 96 | 179 | 5 | MOLECULE: | ATP SYNTHA | SE SUBUN | IT ALPHA, CHLOROPLASTIC |
| 86:00:00 | 1uc3-B | 2.4 | 3.5 | 59 | 149 | 7 | MOLECULE: | GLOBIN | | |
| 87:00:00 | 2is4-B | 2.4 | 24.8 | 95 | 632 | 6 | MOLECULE: | 25-MER | | |
| 88:00:00 | 1uc3-I | 2.4 | 3.5 | 59 | 149 | 7 | MOLECULE: | GLOBIN | | |
| 89:00:00 | 3du5-A | 2.3 | 10.4 | 102 | 596 | 5 | MOLECULE: | TELOMERASE | REVERSE | TRANSCRIPTASE |
| 90:00:00 | 8hwi-B | 2.3 | 4.4 | 67 | 177 | 9 | MOLECULE: | CD-NTASE-A | SSOCIATE | D PROTEIN 12 |
| 91:00:00 | 1szi-A | 2.3 | 8.5 | 84 | 194 | 7 | MOLECULE: | MANNOSE-6- | PHOSPHAT | E RECEPTOR BINDING PROTEIN 1 |
| 92:00:00 | 4tko-B | 2.3 | 40.7 | 106 | 326 | 8 | MOLECULE: | EMRA | | |
| 93:00:00 | 2ieq-B | 2.3 | 13 | 60 | 89 | 5 | MOLECULE: | SPIKE GLYC | OPROTEIN | |
| 94:00:00 | 4dve-A | 2.3 | 3.6 | 72 | 189 | 10 | MOLECULE: | BIOTIN TRA | NSPORTER | BIOY |
| 95:00:00 | 7d7c-F | 2.3 | 3.1 | 66 | 137 | 9 | MOLECULE: | DNA-DIRECT | ED RNA P | OLYMERASE SUBUNIT ALPHA |
| 96:00:00 | 8dc2-A | 2.3 | 8.5 | 106 | 673 | 1 | MOLECULE: | CASLAMBDA | | |
| 97:00:00 | 2ox6-C | 2.3 | 5.7 | 52 | 163 | 8 | MOLECULE: | HYPOTHETIC | AL PROTE | IN SO3848 |
| 98:00:00 | 2a2f-X | 2.3 | 13.5 | 101 | 296 | 6 | MOLECULE: | EXOCYST CO | MPLEX CO | MPONENT SEC15 |

| | | | | | | | | | | |
|---|---|---|---|---|---|---|---|---|---|---|
| 99:00:00 | 3m1m-A | 2.3 | 15.2 | 65 | 320 | 2 | MOLECULE: | ORF904 | | |
| 0:00 | 3k1t-A | 2.3 | 8.5 | 99 | 424 | 12 | MOLECULE: | GLUTAMATE- | #NAME? | E LIGASE GSHA |
| 1:00 | 6c48-D | 2.3 | 3.9 | 55 | 87 | 4 | MOLECULE: | PROTEIN LI | N-9 HOMO | LOG |
| 2:00 | 7ppo-A | 2.3 | 10.2 | 106 | 500 | 5 | MOLECULE: | UBIQUITINA | TING/DEU | BIQUITINATING ENZYME SDEA |
| 3:00 | 5tgq-A | 2.3 | 11.5 | 96 | 226 | 4 | MOLECULE: | R.SWAI PRO | TEIN | |
| 4:00 | 2mxx-A | 2.3 | 16.5 | 101 | 179 | 8 | MOLECULE: | AMYLASE-BI | NDING PR | OTEIN ABPA |
| 5:00 | 7kpx-C | 2.3 | 35.1 | 120 | 1352 | 6 | MOLECULE: | MEIOTIC MR | NA STABI | LITY PROTEIN KINASE SSN3 |
| 6:00 | 6uhd-A | 2.3 | 12.1 | 82 | 322 | 5 | MOLECULE: | HISTONE AC | ETYLTRAN | SFERASE TYPE B CATALYTIC SUBUNI |
| 7:00 | 8pmq-2 | 2.3 | 18.1 | 99 | 355 | 8 | MOLECULE: | E3 UBIQUIT | IN-PROTE | IN LIGASE RMD5 |
| 8:00 | 6ar7-C | 2.3 | 5.4 | 86 | 207 | 5 | MOLECULE: | UNCHARACTE | RIZED PR | OTEIN |
| 9:00 | 6hrz-B | 2.3 | 14.3 | 70 | 202 | 3 | MOLECULE: | PROBABLE T | RANSCRIP | TIONAL REGULATORY PROTEIN |
| 10:00 | 8etq-A | 2.3 | 14 | 75 | 277 | 4 | MOLECULE: | K-CTRP5 | | |
| 11:00 | 5j0l-A | 2.3 | 7.3 | 69 | 130 | 6 | MOLECULE: | DESIGNED P | ROTEIN 3 | L6HC2_2 |
| 12:00 | 8gjg-A | 2.3 | 11.9 | 73 | 162 | 4 | MOLECULE: | GLUC_A04_0 | 005 BIND | ER |
| 13:00 | 8eew-A | 2.3 | 22.1 | 123 | 766 | 3 | MOLECULE: | DYNAMIN-LI | KE 120 K | DA PROTEIN, FORM S1 |
| 14:00 | 7nj0-C | 2.3 | 5 | 80 | 270 | 8 | MOLECULE: | SECURIN,SE | PARIN | |
| 15:00 | 4l3u-A | 2.3 | 9.5 | 78 | 123 | 5 | MOLECULE: | UNCHARACTE | RIZED PR | OTEIN |
| 16:00 | 6vmb-g | 2.3 | 6.9 | 64 | 323 | 3 | MOLECULE: | ATP SYNTHA | SE SUBUN | IT ALPHA, CHLOROPLASTIC |
| 17:00 | 4uoj-A | 2.3 | 23.4 | 87 | 917 | 6 | MOLECULE: | BETA-MANNO | SIDASE G | H2 |
| 18:00 | 8hir-A | 2.3 | 22.1 | 103 | 918 | 8 | MOLECULE: | POTASSIUM | CHANNEL | SUBFAMILY T MEMBER 1 |
| 19:00 | 3jac-A | 2.3 | 15.3 | 105 | 918 | 5 | MOLECULE: | PIEZO-TYPE | MECHANO | SENSITIVE ION CHANNEL COMPONENT |
| 20:00 | 1s56-B | 2.3 | 3.9 | 66 | 135 | 11 | MOLECULE: | HEMOGLOBIN | #NAME? | OTEIN HBN |
| 21:00 | 6tmi-A | 2.3 | 5.5 | 63 | 109 | 13 | MOLECULE: | ATP SYNTHA | SE SUBUN | IT ALPHA |
| 22:00 | 3k17-A | 2.3 | 20.5 | 111 | 355 | 10 | MOLECULE: | LIN0012 PR | OTEIN | |
| 23:00 | 6eyu-A | 2.3 | 9 | 92 | 232 | 9 | MOLECULE: | BACTERIORH | ODOPSIN | |
| 24:00:00 | 6its-A | 2.3 | 6 | 89 | 147 | 9 | MOLECULE: | METHYL-ACC | EPTING C | HEMOTAXIS SENSORY TRANSDUCER |
| 25:00:00 | 6tt7-M | 2.3 | 11.9 | 76 | 156 | 9 | MOLECULE: | ATP SYNTHA | SE SUBUN | IT ALPHA |
| 26:00:00 | 8dql-A | 2.3 | 16.5 | 88 | 257 | 6 | MOLECULE: | SECRETION | SYSTEM P | ROTEIN |
| 27:00:00 | 5x3x-M | 2.3 | 3.9 | 88 | 207 | 11 | MOLECULE: | COBALT ABC | TRANSPO | RTER ATP-BINDING PROTEIN |
| 28:00:00 | 4h8s-C | 2.3 | 15.8 | 117 | 382 | 3 | MOLECULE: | DCC-INTERA | CTING PR | OTEIN 13-BETA |
| 29:00:00 | 7mbz-A | 2.3 | 4 | 80 | 222 | 15 | MOLECULE: | ABC TRANSP | ORTER, P | ERMEASE PROTEIN |
| 30:00:00 | 8bhw-A | 2.3 | 33.7 | 99 | 332 | 5 | MOLECULE: | ECA POLYSA | CCHARIDE | CHAIN LENGTH MODULATION PROTEI |
| 31:00:00 | 2l6f-A | 2.3 | 7.8 | 80 | 176 | 11 | MOLECULE: | FOCAL ADHE | SION KIN | ASE 1, LINKER1, PAXILLIN, LINKE |
| 32:00:00 | 3tul-A | 2.3 | 8.1 | 62 | 136 | 3 | MOLECULE: | CELL INVAS | ION PROT | EIN SIPB |
| 33:00:00 | 3zcj-A | 2.3 | 10.8 | 93 | 205 | 3 | MOLECULE: | CAGL | | |
| 34:00:00 | 8x2q-A | 2.3 | 7.9 | 67 | 109 | 9 | MOLECULE: | ADENOMATOU | S POLYPO | SIS COLI PROTEIN |
| 35:00:00 | 6s3l-K | 2.3 | 11.8 | 86 | 193 | 7 | MOLECULE: | FLAGELLAR | BIOSYNTH | ETIC PROTEIN FLIP |
| 36:00:00 | 1g8x-A | 2.3 | 17.9 | 108 | 1009 | 7 | MOLECULE: | MYOSIN II | HEAVY CH | AIN FUSED TO ALPHA-ACTININ 3 |
| 37:00:00 | 8k9t-A | 2.3 | 12.2 | 66 | 961 | 5 | MOLECULE: | GPI INOSIT | OL-DEACY | LASE,MCHERRY PROTEIN |
| 38:00:00 | 7kah-D | 2.3 | 18.6 | 123 | 493 | 8 | MOLECULE: | PROTEIN TR | ANSPORT | PROTEIN SEC61 |
| 39:00:00 | 7f02-B | 2.3 | 6 | 81 | 220 | 7 | MOLECULE: | CYTOCHROME | C BIOGE | NESIS ATP-BINDING EXPORT PROTEI |
| 40:00:00 | 8ity-V | 2.3 | 27.6 | 122 | 361 | 6 | MOLECULE: | SNRNA-ACTI | VATING P | ROTEIN COMPLEX SUBUNIT 1 |
| 41:00:00 | 2r6i-A | 2.3 | 6 | 58 | 268 | 7 | MOLECULE: | UNCHARACTE | RIZED PR | OTEIN ATU1473 |
| 42:00:00 | 7w01-A | 2.3 | 23.6 | 121 | 1598 | 3 | MOLECULE: | PHOSPHOLIP | ID-TRANS | PORTING ATPASE ABCA3 |
| 43:00:00 | 7lt1-A | 2.3 | 13 | 72 | 403 | 6 | MOLECULE: | PROTEIN MB | 21D2 | |
| 44:00:00 | 2eul-A | 2.3 | 7.4 | 70 | 156 | 6 | MOLECULE: | ANTI-CLEAV | AGE ANTI | #NAME? |
| 45:00:00 | 8imi-0 | 2.3 | 18.3 | 86 | 1045 | 6 | MOLECULE: | APCE | | |
| 46:00:00 | 4hh0-A | 2.3 | 16.7 | 88 | 383 | 6 | MOLECULE: | APPA PROTE | IN | |
| 47:00:00 | 3zjc-E | 2.3 | 24.2 | 81 | 286 | 9 | MOLECULE: | GTPASE IMA | P FAMILY | MEMBER 7 |
| 48:00:00 | 4elj-A | 2.3 | 21.9 | 107 | 590 | 6 | MOLECULE: | RETINOBLAS | TOMA-ASS | OCIATED PROTEIN |
| 49:00:00 | 7tvz-A | 2.3 | 23.8 | 134 | 850 | 4 | MOLECULE: | BAND 3 ANI | ON TRANS | PORT PROTEIN |
| 50:00:00 | 3bo0-A | 2.3 | 4.7 | 96 | 442 | 6 | MOLECULE: | 23S RIBOSO | MAL RNA | |
| 51:00:00 | 3uit-D | 2.3 | 6.3 | 79 | 257 | 8 | MOLECULE: | INAD-LIKE | PROTEIN, | MAGUK P55 SUBFAMILY MEMBER 5, |
| 52:00:00 | 2hi3-A | 2.3 | 3.8 | 57 | 73 | 0 | MOLECULE: | HOMEODOMAI | N-ONLY P | ROTEIN |

| | | | | | | | | | | |
|---|---|---|---|---|---|---|---|---|---|---|
| 53:00:00 | 3bve-A | 2.3 | 8.1 | 80 | 173 | 4 | MOLECULE: | FERRITIN | | |
| 54:00:00 | 7mi8-A | 2.3 | 14.5 | 113 | 868 | 4 | MOLECULE: | FUSION PRO | TEIN OF | DYNEIN AND ENDOLYSIN |
| 55:00:00 | 6hqa-A | 2.3 | 27.8 | 120 | 887 | 2 | MOLECULE: | TAF2 | | |
| 56:00:00 | 5lox-1 | 2.3 | 11.9 | 94 | 241 | 9 | MOLECULE: | PEPTIDASE | | |
| 57:00:00 | 8j8j-A | 2.3 | 13.7 | 88 | 290 | 9 | MOLECULE: | DECAPRENYL | #NAME? | TE PHOSPHORIBOSYLTRANSFERASE |
| 58:00:00 | 6e6y-A | 2.3 | 8.2 | 94 | 268 | 10 | MOLECULE: | DIECKMANN | CYCLASE, | NCMC |
| 59:00:00 | 3va9-A | 2.3 | 7.9 | 70 | 127 | 4 | MOLECULE: | SENSOR HIS | TIDINE K | INASE |
| 60:00:00 | 5chs-B | 2.3 | 17.7 | 87 | 315 | 13 | MOLECULE: | RNA-DIRECT | ED RNA P | OLYMERASE L |
| 61:00:00 | 8adl-A | 2.3 | 23.3 | 120 | 633 | 6 | MOLECULE: | MTC5 ISOFO | RM 1 | |
| 62:00:00 | 2m6u-A | 2.3 | 4.8 | 56 | 82 | 4 | MOLECULE: | CHOLINE BI | NDING PR | OTEIN A |
| 63:00:00 | 4ocu-A | 2.3 | 11.4 | 88 | 356 | 5 | MOLECULE: | N-ACETYLHE | XOSAMINE | 1-PHOSPHATE KINASE |
| 64:00:00 | 2np9-A | 2.3 | 3.8 | 113 | 423 | 12 | MOLECULE: | DPGC | | |
| 65:00:00 | 6gw5-A | 2.3 | 9 | 72 | 393 | 3 | MOLECULE: | PUTATIVE O | UTER MEM | BRANE PROTEIN |
| 66:00:00 | 6kbh-A | 2.3 | 13.4 | 159 | 765 | 9 | MOLECULE: | CYTOCHROME | P450 MO | NOOXYGENASE |
| 67:00:00 | 7wf6-A | 2.3 | 10.4 | 85 | 145 | 6 | MOLECULE: | SORTING NE | XIN-13 | |
| 68:00:00 | 7cjn-B | 2.3 | 3.5 | 65 | 115 | 3 | MOLECULE: | INTERLEUKI | N | |
| 69:00:00 | 7ocn-A | 2.3 | 17.6 | 91 | 690 | 1 | MOLECULE: | HAD HYDROL | ASE, FAM | ILY IA, VARIANT 3 |
| 70:00:00 | 8hwa-A | 2.3 | 17.4 | 86 | 700 | 10 | MOLECULE: | PRIMASE D5 | | |
| 71:00:00 | 6nsj-A | 2.3 | 8.5 | 86 | 182 | 13 | MOLECULE: | ACID-ACTIV | ATED URE | A CHANNEL |
| 72:00:00 | 3o77-A | 2.3 | 3.2 | 61 | 388 | 3 | MOLECULE: | MYOSIN LIG | HT CHAIN | KINASE, SMOOTH MUSCLE,GREEN FL |
| 73:00:00 | 7qe5-A | 2.3 | 8.8 | 97 | 616 | 4 | MOLECULE: | SIALIC ACI | D TRAP T | RANSPORTER PERMEASE PROTEIN SIA |
| 74:00:00 | 2nn4-A | 2.3 | 3.7 | 52 | 62 | 13 | MOLECULE: | HYPOTHETIC | AL PROTE | IN YQGQ |
| 75:00:00 | 1cf7-B | 2.3 | 3.9 | 69 | 82 | 7 | MOLECULE: | DNA (5'- | | |
| 76:00:00 | 3v53-B | 2.3 | 4.2 | 65 | 112 | 5 | MOLECULE: | RNA-BINDIN | G PROTEI | N 25 |
| 77:00:00 | 6r6b-F | 2.3 | 9.6 | 72 | 245 | 8 | MOLECULE: | SURFACE PR | ESENTATI | ON OF ANTIGENS PROTEIN SPAP |
| 78:00:00 | 1au1-A | 2.3 | 6.6 | 63 | 166 | 6 | MOLECULE: | INTERFERON | -BETA | |
| 79:00:00 | 8evm-C | 2.3 | 15.4 | 75 | 237 | 7 | MOLECULE: | CHLOROPHYL | L DIMER | PROTEIN DESIGNS, SPECIAL PAIR 3 |
| 80:00:00 | 6xss-A | 2.3 | 9.4 | 74 | 260 | 9 | MOLECULE: | C4_NAT_HFU | SE-7900 | |
| 81:00:00 | 3pvl-A | 2.3 | 21.4 | 115 | 597 | 6 | MOLECULE: | MYOSIN VII | A ISOFOR | M 1 |
| 82:00:00 | 5wtk-A | 2.3 | 25.7 | 124 | 1215 | 4 | MOLECULE: | CRISPR-ASS | OCIATED | ENDORIBONUCLEASE C2C2 |
| 83:00:00 | 7smo-C | 2.3 | 18.4 | 85 | 639 | 6 | MOLECULE: | DEAD/DEAH | BOX RNA | HELICASE |
| 84:00:00 | 8eki-C | 2.3 | 18.4 | 115 | 701 | 7 | MOLECULE: | PROTEIN TR | ANSPORT | PROTEIN SEC20 |
| 85:00:00 | 8hrc-A | 2.3 | 7.3 | 115 | 776 | 3 | MOLECULE: | ADENOSINE | DEAMINAS | E |
| 86:00:00 | 3oao-A | 2.3 | 6.7 | 68 | 140 | 6 | MOLECULE: | UNCHARACTE | RIZED PR | OTEIN FROM DUF2059 FAMILY |
| 87:00:00 | 2dmn-A | 2.3 | 6.2 | 72 | 83 | 6 | MOLECULE: | HOMEOBOX P | ROTEIN T | GIF2LX |
| 88:00:00 | 8dgc-A | 2.3 | 18.8 | 114 | 2028 | 5 | MOLECULE: | SEAVS3 | | |
| 89:00:00 | 2h1n-A | 2.3 | 13.8 | 130 | 566 | 5 | MOLECULE: | OLIGOENDOP | EPTIDASE | F |
| 90:00:00 | 6jc4-A | 2.3 | 3.6 | 68 | 200 | 7 | MOLECULE: | UREASE ACC | ESSORY P | ROTEIN UREF |
| 91:00:00 | 8ojy-C | 2.3 | 5.5 | 95 | 952 | 2 | MOLECULE: | PHOSPHOENO | LPYRUVAT | E CARBOXYLASE 1 |
| 92:00:00 | 6swy-1 | 2.3 | 10.9 | 87 | 542 | 7 | MOLECULE: | VACUOLAR I | MPORT AN | D DEGRADATION PROTEIN 28 |
| 93:00:00 | 3wfw-A | 2.3 | 13.7 | 73 | 138 | 5 | MOLECULE: | HEMOGLOBIN | #NAME? | AVOPROTEIN FUSED TO ROADBLOCK/L |
| 94:00:00 | 7b1s-B | 2.3 | 13.2 | 100 | 466 | 4 | MOLECULE: | ETHYL-COEN | ZYME M R | EDUCTASE ALPHA SUBUNIT |
| 95:00:00 | 6z5s-W | 2.3 | 4 | 70 | 94 | 1 | MOLECULE: | LIGHT HARV | ESTING C | OMPLEX 1 PROTEIN W |
| 96:00:00 | 7t7r-A | 2.3 | 13.2 | 105 | 1132 | 8 | MOLECULE: | PROTEIN UN | C-13 HOM | OLOG A |
| 97:00:00 | 7jsr-A | 2.3 | 12.8 | 94 | 1496 | 5 | MOLECULE: | NAD-SPECIF | IC GLUTA | MATE DEHYDROGENASE |
| 98:00:00 | 3trc-A | 2.3 | 4.2 | 67 | 168 | 7 | MOLECULE: | PHOSPHOENO | LPYRUVAT | E-PROTEIN PHOSPHOTRANSFERASE |
| 99:00:00 | 7arc-E | 2.3 | 15.3 | 81 | 235 | 6 | MOLECULE: | PSST | | |
| 0:00 | 3pde-B | 2.3 | 9.1 | 79 | 287 | 5 | MOLECULE: | FARNESYL-D | IPHOSPHA | TE SYNTHASE |
| 1:00 | 5jfm-B | 2.3 | 6.5 | 120 | 452 | 10 | MOLECULE: | ALDEHYDE D | EHYDROGE | NASE |
| 2:00 | 3kwo-A | 2.3 | 7.6 | 79 | 149 | 6 | MOLECULE: | PUTATIVE B | ACTERIOF | ERRITIN |
| 3:00 | 6eud-A | 2.3 | 20.5 | 94 | 808 | 7 | MOLECULE: | ATP-DEPEND | ENT RNA | HELICASE HRPB |
| 4:00 | 8gy3-A | 2.3 | 13.7 | 75 | 440 | 7 | MOLECULE: | CYTOCHROME | C SUBUN | IT OF ALDEHYDE DEHYDROGENASE |
| 5:00 | 2lp4-A | 2.3 | 20.3 | 105 | 225 | 4 | MOLECULE: | CHEMOTAXIS | PROTEIN | CHEA |
| 6:00 | 2ksr-A | 2.3 | 8.9 | 79 | 140 | 5 | MOLECULE: | NEURONAL A | CETYLCHO | LINE RECEPTOR SUBUNIT BETA-2 |

| | | | | | | | | | | | |
|---|---|---|---|---|---|---|---|---|---|---|---|
| 7:00 | 2gsc-B | 2.3 | 6.9 | 75 | 117 | 4 | MOLECULE: | PUTATIVE U | NCHARACT | ERIZED PROTEIN XCC0516 | |
| 8:00 | 7w72-A | 2.3 | 15.7 | 82 | 588 | 9 | MOLECULE: | PHOSPHATID | YLINOSIT | OL GLYCAN ANCHOR BIOSYNTHESIS C | |
| 9:00 | 4itr-A | 2.3 | 16.8 | 98 | 299 | 1 | MOLECULE: | ADENOSINE | MONOPHOS | PHATE-PROTEIN TRANSFERASE AND C | |
| 10:00 | 7m5w-A | 2.3 | 9.2 | 59 | 138 | 5 | MOLECULE: | PROTEIN CA | PICUA HO | MOLOG | |
| 11:00 | 6k7x-C | 2.3 | 12.9 | 73 | 276 | 5 | MOLECULE: | CALCIUM UN | IPORTER | PROTEIN, MITOCHONDRIAL | |
| 12:00 | 7bu0-A | 2.3 | 16.2 | 78 | 407 | 6 | MOLECULE: | UNCHARACTE | RIZED PR | OTEIN | |
| 13:00 | 5b4s-B | 2.3 | 6 | 78 | 310 | 5 | MOLECULE: | CHITOSANAS | E | | |
| 14:00 | 2v5d-A | 2.3 | 17.6 | 119 | 722 | 8 | MOLECULE: | O-GLCNACAS | E NAGJ | | |
| 15:00 | 4m9s-B | 2.3 | 18.6 | 88 | 511 | 8 | MOLECULE: | CELL DEATH | PROTEIN | | 4 |
| 16:00 | 1bmt-A | 2.3 | 17.1 | 94 | 246 | 9 | MOLECULE: | METHIONINE | SYNTHAS | E | |
| 17:00 | 3pxp-A | 2.3 | 19.3 | 89 | 292 | 6 | MOLECULE: | HELIX-TURN | #NAME? | OMAIN PROTEIN | |
| 18:00 | 6agb-K | 2.3 | 9 | 59 | 128 | 8 | MOLECULE: | CHROMOSOME | V, COMP | LETE SEQUENCE | |
| 19:00 | 1knv-B | 2.3 | 4.8 | 84 | 291 | 10 | MOLECULE: | BSE634I RE | STRICTIO | N ENDONUCLEASE | |
| 20:00 | 6vek-A | 2.3 | 6.7 | 66 | 321 | 9 | MOLECULE: | CONTACT-DE | PENDENT | TOXIN CDIA | |
| 21:00 | 7qia-A | 2.3 | 9.5 | 90 | 401 | 18 | MOLECULE: | DIVALENT M | ETAL CAT | ION TRANSPORTER | |
| 22:00 | 8ttf-A | 2.3 | 7 | 98 | 388 | 5 | MOLECULE: | QUINOLONE | RESISTAN | CE PROTEIN NORA | |
| 23:00 | 7xpt-A | 2.3 | 15.3 | 91 | 375 | 10 | MOLECULE: | TRANSGLYCO | SYLSE | | |
| 24:00:00 | 6qyi-B | 2.3 | 16.7 | 126 | 519 | 7 | MOLECULE: | 4-HYDROXYP | HENYLACE | TATE 3-MONOOXYGENASE OXYGENASE | |
| 25:00:00 | 6cp6-Y | 2.3 | 6.5 | 97 | 188 | 5 | MOLECULE: | ATP SYNTHA | SE SUBUN | IT 9, MITOCHONDRIAL | |
| 26:00:00 | 8oyy-A | 2.3 | 11.7 | 103 | 233 | 5 | MOLECULE: | DE NOVO DE | SIGNED S | OLUBLE GPCR-LIKE PROTEIN | |
| 27:00:00 | 7cun-D | 2.3 | 20.3 | 111 | 827 | 5 | MOLECULE: | INTEGRATOR | COMPLEX | SUBUNIT 1 | |
| 28:00:00 | 4fcy-B | 2.3 | 21.3 | 104 | 476 | 10 | MOLECULE: | TRANSPOSAS | E | | |
| 29:00:00 | 5che-A | 2.3 | 22.9 | 69 | 427 | 1 | MOLECULE: | GLUTAMYL-T | RNA REDU | CTASE 1, CHLOROPLASTIC | |
| 30:00:00 | 8pm4-A | 2.3 | 16.5 | 98 | 604 | 11 | MOLECULE: | TRANSPOSAS | E | | |
| 31:00:00 | 5zi2-A | 2.3 | 7 | 73 | 339 | 7 | MOLECULE: | MALATE DEH | YDROGENA | SE | |
| 32:00:00 | 7nhr-C | 2.3 | 6 | 78 | 565 | 6 | MOLECULE: | PUTATIVE T | RANSMEMB | RANE PROTEIN WZC | |
| 33:00:00 | 7nyw-C | 2.3 | 16.7 | 70 | 335 | 13 | MOLECULE: | CHROMOSOME | PARTITI | ON PROTEIN MUKB | |
| 34:00:00 | 8eki-D | 2.3 | 13 | 92 | 684 | 10 | MOLECULE: | PROTEIN TR | ANSPORT | PROTEIN SEC20 | |
| 35:00:00 | 5hyc-A | 2.3 | 4.3 | 77 | 130 | 3 | MOLECULE: | UNCHARACTE | RIZED PR | OTEIN | |
| 36:00:00 | 6cfz-J | 2.3 | 12.9 | 95 | 134 | 8 | MOLECULE: | ASK1 | | | |
| 37:00:00 | 8i24-F | 2.3 | 17 | 79 | 233 | 8 | MOLECULE: | DNA-DIRECT | ED RNA P | OLYMERASE SUBUNIT ALPHA | |
| 38:00:00 | 3cra-A | 2.3 | 15.5 | 71 | 239 | 4 | MOLECULE: | PROTEIN MA | ZG | | |
| 39:00:00 | 4hwi-B | 2.3 | 15.7 | 86 | 177 | 9 | MOLECULE: | HEAT SHOCK | COGNATE | 71 KDA PROTEIN | |
| 40:00:00 | 7x34-C | 2.3 | 3.4 | 57 | 100 | 4 | MOLECULE: | ZUOTIN | | | |
| 41:00:00 | 2euf-A | 2.3 | 3 | 64 | 244 | 3 | MOLECULE: | VIRAL CYCL | IN | | |
| 42:00:00 | 5fd4-A | 2.3 | 14.3 | 78 | 309 | 4 | MOLECULE: | COMR | | | |
| 43:00:00 | 6usf-A | 2.3 | 16.1 | 75 | 464 | 3 | MOLECULE: | CHIMERA OF | SOLUBLE | CYTOCHROME B562 (BRIL) AND NEU | |
| 44:00:00 | 5utb-A | 2.3 | 5.9 | 71 | 152 | 6 | MOLECULE: | MYOGLOBIN | | | |
| 45:00:00 | 1f5o-E | 2.3 | 4.3 | 64 | 149 | 5 | MOLECULE: | HEMOGLOBIN | V | | |
| 46:00:00 | 1f5o-A | 2.3 | 3.5 | 61 | 149 | 5 | MOLECULE: | HEMOGLOBIN | V | | |
| 47:00:00 | 2is1-A | 2.3 | 24.1 | 99 | 649 | 9 | MOLECULE: | 5'-D(*GP*C | P*AP*GP* | TP*GP*CP*TP*CP*GP*TP*TP*TP*TP*T | |
| 48:00:00 | 1f5o-D | 2.3 | 3.2 | 60 | 149 | 5 | MOLECULE: | HEMOGLOBIN | V | | |
| 49:00:00 | 3wyo-B | 2.3 | 5.5 | 63 | 153 | 8 | MOLECULE: | MYOGLOBIN | | | |
| 50:00:00 | 1f5o-F | 2.3 | 3.3 | 59 | 149 | 5 | MOLECULE: | HEMOGLOBIN | V | | |
| 51:00:00 | 1f5o-B | 2.3 | 4.3 | 64 | 149 | 5 | MOLECULE: | HEMOGLOBIN | V | | |
| 52:00:00 | 8ftk-A | 2.2 | 19.1 | 115 | 1584 | 10 | MOLECULE: | 5'-3' RNA | HELICASE | -LIKE PROTEIN | |
| 53:00:00 | 8rt9-F | 2.2 | 4 | 66 | 272 | 9 | MOLECULE: | TRWJ PROTE | IN | | |
| 54:00:00 | 7sba-I | 2.2 | 15.2 | 89 | 899 | 7 | MOLECULE: | CAS7D | | | |
| 55:00:00 | 2i1k-A | 2.2 | 30.1 | 107 | 545 | 7 | MOLECULE: | MOESIN | | | |
| 56:00:00 | 3bhq-A | 2.2 | 3.2 | 78 | 202 | 5 | MOLECULE: | TRANSCRIPT | IONAL RE | GULATOR | |
| 57:00:00 | 6dlm-B | 2.2 | 3.6 | 50 | 72 | 14 | MOLECULE: | DHD127_A | | | |
| 58:00:00 | 5lnf-A | 2.2 | 12.1 | 91 | 139 | 2 | MOLECULE: | PEROXISOMA | L BIOGEN | ESIS FACTOR 19 | |
| 59:00:00 | 2ehw-C | 2.2 | 11.1 | 70 | 117 | 9 | MOLECULE: | HYPOTHETIC | AL PROTE | IN TTHB059 | |
| 60:00:00 | 8eki-B | 2.2 | 7.9 | 60 | 85 | 5 | MOLECULE: | PROTEIN TR | ANSPORT | PROTEIN SEC20 | |

| | | | | | | | | | | |
|---|---|---|---|---|---|---|---|---|---|---|
| 61:00:00 | 8bw9-B | 2.2 | 13 | 102 | 288 | 7 | MOLECULE: | PROTEIN AV | EUGLE | |
| 62:00:00 | 2fiy-A | 2.2 | 8.2 | 116 | 285 | 12 | MOLECULE: | PROTEIN FD | HE HOMOL | OG |
| 63:00:00 | 8gmh-A | 2.2 | 31.2 | 81 | 223 | 17 | MOLECULE: | LXG DOMAIN | #NAME? | ING PROTEIN |
| 64:00:00 | 5ebz-A | 2.2 | 27.6 | 98 | 655 | 0 | MOLECULE: | INHIBITOR | OF NUCLE | AR FACTOR KAPPA-B KINASE SUBUNI |
| 65:00:00 | 4gx2-B | 2.2 | 19.9 | 89 | 548 | 4 | MOLECULE: | TRKA DOMAI | N PROTEI | N |
| 66:00:00 | 4knf-C | 2.2 | 4.1 | 83 | 232 | 5 | MOLECULE: | BLUE-LIGHT | ABSORBI | NG PROTEORHODOPSIN |
| 67:00:00 | 2w96-A | 2.2 | 4.6 | 75 | 249 | 3 | MOLECULE: | CELL DIVIS | ION PROT | EIN KINASE 4 |
| 68:00:00 | 5mu4-A | 2.2 | 4.8 | 70 | 196 | 3 | MOLECULE: | TAIL TUBUL | AR PROTE | IN A |
| 69:00:00 | 5da8-K | 2.2 | 6.1 | 82 | 433 | 4 | MOLECULE: | 60 KDA CHA | PERONIN | |
| 70:00:00 | 7v2w-J | 2.2 | 15 | 80 | 236 | 11 | MOLECULE: | THO COMPLE | X SUBUNI | T HPR1 |
| 71:00:00 | 1bp3-A | 2.2 | 4.8 | 86 | 186 | 6 | MOLECULE: | PROTEIN (G | ROWTH HO | RMONE) |
| 72:00:00 | 5lo9-A | 2.2 | 19.5 | 103 | 471 | 6 | MOLECULE: | CYTOCHROME | C | |
| 73:00:00 | 5uju-A | 2.2 | 5.8 | 126 | 558 | 3 | MOLECULE: | NAD-DEPEND | ENT ALDE | HYDE DEHYDROGENASE |
| 74:00:00 | 5usf-A | 2.2 | 18.3 | 95 | 683 | 9 | MOLECULE: | TYROSYL-TR | NA SYNTH | ETASE, PUTATIVE |
| 75:00:00 | 8z9a-A | 2.2 | 11.7 | 90 | 406 | 6 | MOLECULE: | ODORANT RE | CEPTOR, | APISORCO |
| 76:00:00 | 8ojc-A | 2.2 | 3.3 | 58 | 395 | 9 | MOLECULE: | DNA (47-ME | R) | |
| 77:00:00 | 8tzk-E | 2.2 | 8.5 | 63 | 101 | 5 | MOLECULE: | CELL DIVIS | ION ATP- | BINDING PROTEIN FTSE |
| 78:00:00 | 5h1q-A | 2.2 | 7.4 | 97 | 360 | 2 | MOLECULE: | INNEXIN-6 | | |
| 79:00:00 | 8wjn-A | 2.2 | 14.3 | 101 | 491 | 9 | MOLECULE: | STRUCTURAL | MAINTEN | ANCE OF CHROMOSOMES PROTEIN 6 |
| 80:00:00 | 5j2l-A | 2.2 | 4.4 | 52 | 76 | 8 | MOLECULE: | PROTEIN DE | SIGN 2L4 | HC2_11 |
| 81:00:00 | 8k5p-M | 2.2 | 16.7 | 129 | 744 | 11 | MOLECULE: | DNA-DIRECT | ED RNA P | OLYMERASE II SUBUNIT RPB1 |
| 82:00:00 | 8fhc-B | 2.2 | 6.2 | 78 | 215 | 8 | MOLECULE: | TRNA-(MS[2 | ]IO[6]A) | -HYDROXYLASE |
| 83:00:00 | 2wcq-A | 2.2 | 3.4 | 75 | 153 | 3 | MOLECULE: | TNF-ALPHA | INDUCER | PROTEIN |
| 84:00:00 | 6s8g-G | 2.2 | 11.5 | 94 | 244 | 3 | MOLECULE: | LIPOPOLYSA | CCHARIDE | ABC TRANSPORTER, ATP-BINDING P |
| 85:00:00 | 1g5z-A | 2.2 | 8.5 | 78 | 164 | 9 | MOLECULE: | OUTER SURF | ACE PROT | EIN C |
| 86:00:00 | 3l8i-C | 2.2 | 13.3 | 101 | 212 | 5 | MOLECULE: | PROGRAMMED | CELL DE | ATH PROTEIN 10 |
| 87:00:00 | 4jle-B | 2.2 | 10 | 71 | 159 | 7 | MOLECULE: | PHIST | | |
| 88:00:00 | 3ub0-D | 2.2 | 13.8 | 74 | 194 | 9 | MOLECULE: | NON-STRUCT | URAL PRO | TEIN 6, NSP6, |
| 89:00:00 | 6zsi-D | 2.2 | 13.3 | 68 | 133 | 6 | MOLECULE: | RAS-RELATE | D PROTEI | N RAB-8A |
| 90:00:00 | 4acl-A | 2.2 | 8.2 | 76 | 155 | 5 | MOLECULE: | TSSL | | |
| 91:00:00 | 6irr-A | 2.2 | 8 | 70 | 133 | 3 | MOLECULE: | DISRUPTED | IN SCHIZ | OPHRENIA 1 HOMOLOG,CYCLIC AMP-D |
| 92:00:00 | 6m9t-A | 2.2 | 12.5 | 106 | 444 | 10 | MOLECULE: | PROSTAGLAN | DIN E2 R | ECEPTOR EP3 SUBTYPE, ENDOLYSIN |
| 93:00:00 | 6z6o-C | 2.2 | 24.1 | 103 | 548 | 7 | MOLECULE: | HISTONE DE | ACETYLAS | E HDA1 |
| 94:00:00 | 3hiu-D | 2.2 | 4.9 | 101 | 152 | 5 | MOLECULE: | UNCHARACTE | RIZED PR | OTEIN |
| 95:00:00 | 6c66-G | 2.2 | 19 | 127 | 832 | 5 | MOLECULE: | CRISPR-ASS | OCIATED | HELICASE, CAS3 FAMILY |
| 96:00:00 | 4i5s-B | 2.2 | 17.4 | 82 | 395 | 4 | MOLECULE: | PUTATIVE H | ISTIDINE | KINASE COVS VICK-LIKE PROTEIN |
| 97:00:00 | 5eke-C | 2.2 | 15.4 | 83 | 307 | 4 | MOLECULE: | UNCHARACTE | RIZED GL | YCOSYLTRANSFERASE SLL0501 |
| 98:00:00 | 3flo-B | 2.2 | 4.1 | 67 | 180 | 6 | MOLECULE: | DNA POLYME | RASE ALP | HA SUBUNIT B |
| 99:00:00 | 3mzo-B | 2.2 | 5.5 | 106 | 211 | 3 | MOLECULE: | LIN2634 PR | OTEIN | |
| 0:00 | 6pie-A | 2.2 | 8.5 | 79 | 162 | 5 | MOLECULE: | HEMERYTHRI | N HHE CA | TION BINDING DOMAIN PROTEIN |
| 1:00 | 6c66-A | 2.2 | 10.9 | 88 | 511 | 6 | MOLECULE: | CRISPR-ASS | OCIATED | HELICASE, CAS3 FAMILY |
| 2:00 | 8h9g-M | 2.2 | 11.4 | 77 | 156 | 5 | MOLECULE: | ATP SYNTHA | SE F(0) | COMPLEX SUBUNIT B1, MITOCHONDRI |
| 3:00 | 8h6u-A | 2.2 | 11.6 | 99 | 338 | 6 | MOLECULE: | PRESILPHIP | ERFOLAN- | 8-BETA-OL SYNTHASE |
| 4:00 | 6orb-A | 2.2 | 37.8 | 93 | 3352 | 5 | MOLECULE: | MIDASIN | | |
| 5:00 | 8jw3-A | 2.2 | 10.7 | 96 | 275 | 4 | MOLECULE: | TERPENOID | SYNTHASE | |
| 6:00 | 8xr6-q | 2.2 | 9.2 | 69 | 143 | 4 | MOLECULE: | PHOTOSYSTE | M II PRO | TEIN D1 |
| 7:00 | 8th8-A | 2.2 | 16.9 | 78 | 290 | 12 | MOLECULE: | DYNEIN REG | ULATORY | COMPLEX PROTEIN 1/2 N-TERMINAL |
| 8:00 | 6c14-B | 2.2 | 6.4 | 58 | 164 | 7 | MOLECULE: | PROTOCADHE | RIN-15 | |
| 9:00 | 7yul-A | 2.2 | 3.2 | 54 | 92 | 6 | MOLECULE: | BEN DOMAIN | #NAME? | ING PROTEIN 6 |
| 10:00 | 6zgs-A | 2.2 | 22.3 | 112 | 404 | 6 | MOLECULE: | L-LACTATE | TRANSPOR | TER |
| 11:00 | 3jbz-A | 2.2 | 19.8 | 89 | 960 | 7 | MOLECULE: | SERINE/THR | EONINE-P | ROTEIN KINASE MTOR |
| 12:00 | 4ilo-A | 2.2 | 35.8 | 108 | 236 | 5 | MOLECULE: | CT398 | | |
| 13:00 | 6h7d-A | 2.2 | 19.9 | 119 | 487 | 6 | MOLECULE: | SUGAR TRAN | SPORT PR | OTEIN 10 |
| 14:00 | 4eyz-A | 2.2 | 3.5 | 92 | 246 | 5 | MOLECULE: | CELLULOSOM | E-RELATE | D PROTEIN MODULE FROM RUMINOCOC |

| | | | | | | | | | | |
|---|---|---|---|---|---|---|---|---|---|---|
| 15:00 | 8p03-L | 2.2 | 12.2 | 73 | 188 | 8 | MOLECULE: | INITIATOR | METHIONY | LATED TRNA |
| 16:00 | 3col-A | 2.2 | 7.8 | 65 | 180 | 11 | MOLECULE: | PUTATIVE T | RANSCRIP | TION REGULATOR |
| 17:00 | 6n63-A | 2.2 | 4.1 | 68 | 141 | 6 | MOLECULE: | ENCAPSULIN | CARGO P | ROTEIN |
| 18:00 | 5yfp-G | 2.2 | 11.3 | 98 | 615 | 9 | MOLECULE: | EXOCYST CO | MPLEX CO | MPONENT SEC3 |
| 19:00 | 2h7f-X | 2.2 | 6.3 | 82 | 314 | 2 | MOLECULE: | 5'-D(*TP*T | P*GP*TP* | CP*GP*CP*CP*CP*TP*T)-3' |
| 20:00 | 4u3e-A | 2.2 | 20.7 | 99 | 637 | 7 | MOLECULE: | RIBONUCLEO | SIDE TRI | PHOSPHATE REDUCTASE |
| 21:00 | 3p8c-F | 2.2 | 15.5 | 69 | 156 | 9 | MOLECULE: | CYTOPLASMI | C FMR1-I | NTERACTING PROTEIN 1 |
| 22:00 | 2kmd-A | 2.2 | 4 | 65 | 111 | 3 | MOLECULE: | PROTEIN C- | ETS-1 | |
| 23:00 | 5an6-A | 2.2 | 7.8 | 69 | 123 | 4 | MOLECULE: | CRISPR-ASS | OCIATED | PROTEIN, CSM2 FAMILY |
| 24:00:00 | 6plm-B | 2.2 | 24.7 | 122 | 752 | 8 | MOLECULE: | SIDJ PROTE | IN | |
| 25:00:00 | 8ave-A | 2.2 | 4.6 | 73 | 146 | 11 | MOLECULE: | LEPTIN | | |
| 26:00:00 | 6nd4-M | 2.2 | 12.9 | 101 | 431 | 10 | MOLECULE: | ETS RRNA | | |
| 27:00:00 | 7y8r-L | 2.2 | 11.6 | 81 | 521 | 4 | MOLECULE: | HISTONE H3 | | |
| 28:00:00 | 8hmc-B | 2.2 | 14.6 | 83 | 1195 | 6 | MOLECULE: | INTRAFLAGE | LLAR TRA | NSPORT PROTEIN 122 HOMOLOG |
| 29:00:00 | 7z8v-D | 2.2 | 16.4 | 124 | 1158 | 7 | MOLECULE: | CULLIN-1 | | |
| 30:00:00 | 8rc4-e | 2.2 | 13.5 | 112 | 873 | 4 | MOLECULE: | SERINE/THR | EONINE-P | ROTEIN PHOSPHATASE 2A 65 KDA RE |
| 31:00:00 | 5xn7-B | 2.2 | 13.2 | 130 | 551 | 8 | MOLECULE: | PUTATIVE R | TX-TOXIN | |
| 32:00:00 | 6j8l-A | 2.2 | 3.9 | 70 | 132 | 9 | MOLECULE: | AVH240 | | |
| 33:00:00 | 8jjr-l | 2.2 | 11.3 | 82 | 253 | 4 | MOLECULE: | PCPI-7 | | |
| 34:00:00 | 8j0n-D | 2.2 | 20.6 | 88 | 156 | 7 | MOLECULE: | ER MEMBRAN | E PROTEI | N COMPLEX SUBUNIT 1 |
| 35:00:00 | 1ku9-A | 2.2 | 7.1 | 63 | 151 | 6 | MOLECULE: | HYPOTHETIC | AL PROTE | IN MJ223 |
| 36:00:00 | 8ox6-A | 2.2 | 12.6 | 79 | 575 | 10 | MOLECULE: | PHOSPHOLIP | ID-TRANS | PORTING ATPASE IC |
| 37:00:00 | 5yfp-H | 2.2 | 12.3 | 81 | 518 | 5 | MOLECULE: | EXOCYST CO | MPLEX CO | MPONENT SEC3 |
| 38:00:00 | 3h20-A | 2.2 | 10.3 | 72 | 288 | 10 | MOLECULE: | REPLICATIO | N PROTEI | N B |
| 39:00:00 | 6btc-A | 2.2 | 3.1 | 52 | 85 | 2 | MOLECULE: | LP1413 - S | CCMEC TY | PE IV-ENCODED DNA BINDING PROTE |
| 40:00:00 | 8pnj-A | 2.2 | 5.8 | 119 | 390 | 7 | MOLECULE: | CHORISMATE | MUTASE | |
| 41:00:00 | 8iwh-q | 2.2 | 13.3 | 71 | 156 | 4 | MOLECULE: | PHOTOSYSTE | M II PRO | TEIN D1 |
| 42:00:00 | 8kde-D | 2.2 | 14 | 104 | 351 | 6 | MOLECULE: | PHOTOSYSTE | M II CP4 | 7 REACTION CENTER PROTEIN |
| 43:00:00 | 7emf-G | 2.2 | 25.5 | 99 | 161 | 7 | MOLECULE: | MEDIATOR O | F RNA PO | LYMERASE II TRANSCRIPTION SUBUN |
| 44:00:00 | 7y38-X | 2.2 | 27.5 | 144 | 611 | 7 | MOLECULE: | MRNA-CAPPI | NG ENZYM | E NSP1,AFFINITY-TAG (STREPII-3X |
| 45:00:00 | 6ynw-H | 2.2 | 5 | 58 | 75 | 5 | MOLECULE: | SUBUNIT C | | |
| 46:00:00 | 7zcv-B | 2.2 | 15.6 | 73 | 285 | 0 | MOLECULE: | TRANSCRIPT | IONAL RE | GULATOR |
| 47:00:00 | 6ig2-C | 2.2 | 3.9 | 86 | 278 | 5 | MOLECULE: | PHOSPHATID | ATE CYTI | DYLYLTRANSFERASE, MITOCHONDRIAL |
| 48:00:00 | 7eld-A | 2.2 | 29.9 | 110 | 1137 | 9 | MOLECULE: | ENDORIBONU | CLEASE D | ICER HOMOLOG 1 |
| 49:00:00 | 7jzw-A | 2.2 | 10.2 | 87 | 427 | 2 | MOLECULE: | CRISPR TYP | E I-F/YP | EST-ASSOCIATED PROTEIN CSY1 |
| 50:00:00 | 7xn2-A | 2.2 | 3.1 | 70 | 148 | 9 | MOLECULE: | ALR3614 PR | OTEIN | |
| 51:00:00 | 6qum-N | 2.2 | 13.7 | 125 | 649 | 4 | MOLECULE: | V-TYPE ATP | SYNTHAS | E ALPHA CHAIN |
| 52:00:00 | 7uic-e | 2.2 | 17.3 | 107 | 729 | 13 | MOLECULE: | MEDIATOR O | F RNA PO | LYMERASE II TRANSCRIPTION SUBUN |
| 53:00:00 | 3faj-A | 2.2 | 6.8 | 69 | 101 | 4 | MOLECULE: | PUTATIVE U | NCHARACT | ERIZED PROTEIN |
| 54:00:00 | 7c6g-A | 2.2 | 3.5 | 53 | 420 | 8 | MOLECULE: | SUGAR ABC | TRANSPOR | TER, PERIPLASMIC SUGAR-BINDING |
| 55:00:00 | 4hlq-C | 2.2 | 20.1 | 94 | 281 | 3 | MOLECULE: | TBC1 DOMAI | N FAMILY | MEMBER 20 |
| 56:00:00 | 4xig-M | 2.2 | 13.3 | 116 | 288 | 5 | MOLECULE: | ALGM1 | | |
| 57:00:00 | 7tve-G | 2.2 | 15.6 | 96 | 307 | 6 | MOLECULE: | DNA (68-ME | R) | |
| 58:00:00 | 3ljl-A | 2.2 | 4.8 | 63 | 149 | 8 | MOLECULE: | TRANSCRIPT | IONAL RE | GULATOR LUXT |
| 59:00:00 | 8twj-A | 2.2 | 5.7 | 99 | 303 | 8 | MOLECULE: | AVIRULENCE | PROTEIN | B |
| 60:00:00 | 8hel-A | 2.2 | 3.5 | 50 | 68 | 4 | MOLECULE: | ACRIE3 | | |
| 61:00:00 | 7pc1-A | 2.2 | 3.2 | 59 | 72 | 5 | MOLECULE: | STBA | | |
| 62:00:00 | 1hlv-A | 2.2 | 13.5 | 58 | 131 | 3 | MOLECULE: | CENP-B BOX | DNA | |
| 63:00:00 | 7xc2-B | 2.2 | 10.2 | 97 | 431 | 2 | MOLECULE: | CNL9 | | |
| 64:00:00 | 5dy0-A | 2.2 | 10.1 | 70 | 220 | 4 | MOLECULE: | TETR FAMIL | Y TRANSC | RIPTIONAL REGULATOR |
| 65:00:00 | 5jpw-A | 2.2 | 5.8 | 73 | 143 | 10 | MOLECULE: | DYNEIN LIG | HT CHAIN | TCTEX-TYPE 1,CYTOPLASMIC DYNEI |
| 66:00:00 | 7cbb-A | 2.2 | 10.8 | 97 | 576 | 8 | MOLECULE: | IUCA/IUCC | FAMILY S | IDEROPHORE BIOSYNTHESIS PROTEIN |
| 67:00:00 | 3p01-A | 2.2 | 8.6 | 53 | 178 | 2 | MOLECULE: | TWO-COMPON | ENT RESP | ONSE REGULATOR |
| 68:00:00 | 1l1l-A | 2.2 | 12.2 | 103 | 717 | 6 | MOLECULE: | RIBONUCLEO | SIDE TRI | PHOSPHATE REDUCTASE |

| | | | | | | | | | | |
|---|---|---|---|---|---|---|---|---|---|---|
| 69:00:00 | 4b8b-B | 2.2 | 15.4 | 92 | 529 | 4 | MOLECULE: | GENERAL NE | GATIVE R | EGULATOR OF TRANSCRIPTION SUBUN |
| 70:00:00 | 8qkd-A | 2.2 | 5.3 | 77 | 281 | 6 | MOLECULE: | SC-CC-5-24 | | |
| 71:00:00 | 6kbn-C | 2.2 | 16 | 116 | 504 | 10 | MOLECULE: | VACUOLAR P | ROTEIN 8 | |
| 72:00:00 | 6wti-D | 2.2 | 6.8 | 83 | 99 | 7 | MOLECULE: | CYTOCHROME | O UBIQU | INOL OXIDASE, SUBUNIT I |
| 73:00:00 | 5okc-B | 2.2 | 12.4 | 101 | 377 | 6 | MOLECULE: | SISTER CHR | OMATID C | OHESION PROTEIN DCC1 |
| 74:00:00 | 8ap7-j | 2.2 | 8.4 | 76 | 168 | 7 | MOLECULE: | ATP SYNTHA | SE SUBUN | IT A |
| 75:00:00 | 6e8c-A | 2.2 | 6.7 | 64 | 135 | 8 | MOLECULE: | DOUBLE HOM | EOBOX PR | OTEIN 4 |
| 76:00:00 | 4evv-A | 2.2 | 3.6 | 59 | 145 | 2 | MOLECULE: | METHYL-CPG | #NAME? | DOMAIN PROTEIN 4 |
| 77:00:00 | 5loi-A | 2.2 | 11 | 86 | 391 | 5 | MOLECULE: | RAD26 | | |
| 78:00:00 | 3wfr-H | 2.2 | 22.9 | 87 | 463 | 7 | MOLECULE: | RNA (74-ME | R) | |
| 79:00:00 | 5of3-C | 2.2 | 2.7 | 52 | 107 | 4 | MOLECULE: | DNA PRIMAS | E SMALL | SUBUNIT PRIS |
| 80:00:00 | 5n5e-A | 2.2 | 9.1 | 73 | 98 | 5 | MOLECULE: | PFC_05175 | | |
| 81:00:00 | 7ldg-A | 2.2 | 5.8 | 90 | 220 | 10 | MOLECULE: | HEAT SHOCK | FACTOR | 2-BINDING PROTEIN |
| 82:00:00 | 5t2x-C | 2.2 | 12.2 | 74 | 258 | 3 | MOLECULE: | UNCHARACTE | RIZED PR | OTEIN LPG1670 |
| 83:00:00 | 8xgr-R | 2.2 | 14.9 | 90 | 305 | 7 | MOLECULE: | GUANINE NU | CLEOTIDE | #NAME? |
| 84:00:00 | 6db1-A | 2.2 | 10.3 | 91 | 137 | 7 | MOLECULE: | PUTATIVE M | ETHYL-AC | CEPTING CHEMOTAXIS PROTEIN |
| 85:00:00 | 3jyy-B | 2.2 | 10.8 | 87 | 266 | 3 | MOLECULE: | LINCOSAMID | E NUCLEO | TIDYLTRANSFERASE |
| 86:00:00 | 5oqj-1 | 2.2 | 29.3 | 94 | 491 | 9 | MOLECULE: | DNA-DIRECT | ED RNA P | OLYMERASE II SUBUNIT RPB1 |
| 87:00:00 | 6xwi-A | 2.2 | 3.5 | 57 | 66 | 2 | MOLECULE: | S0_2.126 | | |
| 88:00:00 | 8e1u-A | 2.2 | 14.7 | 77 | 1126 | 8 | MOLECULE: | PPI-DEPEND | ENT PEPC | K |
| 89:00:00 | 7s37-P | 2.2 | 16.3 | 119 | 1033 | 4 | MOLECULE: | CRISPR-ASS | OCIATED | ENDONUCLEASE CAS9/CSN1 |
| 90:00:00 | 4oe8-C | 2.2 | 9.2 | 73 | 87 | 12 | MOLECULE: | INTERLEUKI | N-12 SUB | UNIT BETA |
| 91:00:00 | 3l0m-A | 2.2 | 12.4 | 82 | 328 | 6 | MOLECULE: | DRRA | | |
| 92:00:00 | 8q9t-E | 2.2 | 17.3 | 122 | 1060 | 5 | MOLECULE: | ANTIVIRAL | HELICASE | SKI2 |
| 93:00:00 | 6fht-B | 2.2 | 15.2 | 114 | 735 | 6 | MOLECULE: | BACTERIOPH | YTOCHROM | E,ADENYLATE CYCLASE |
| 94:00:00 | 2ou3-A | 2.2 | 4.3 | 60 | 160 | 8 | MOLECULE: | TELLURITE | RESISTAN | CE PROTEIN OF COG3793 |
| 95:00:00 | 8k4a-H | 2.2 | 4.7 | 65 | 301 | 9 | MOLECULE: | VP2 | | |
| 96:00:00 | 5nug-A | 2.2 | 23.8 | 128 | 2920 | 6 | MOLECULE: | CYTOPLASMI | C DYNEIN | 1 HEAVY CHAIN 1 |
| 97:00:00 | 4kis-A | 2.2 | 15.2 | 88 | 315 | 6 | MOLECULE: | PUTATIVE I | NTEGRASE | [BACTERIOPHAGE A118] |
| 98:00:00 | 7wb4-E | 2.2 | 21.9 | 114 | 1363 | 6 | MOLECULE: | OUTER NUP1 | 33 | |
| 99:00:00 | 8gi1-A | 2.2 | 17.3 | 93 | 144 | 5 | MOLECULE: | ACCESSORY | PROTEIN | CSX28 |
| 0:00 | 7f66-J | 2.2 | 12.4 | 95 | 591 | 4 | MOLECULE: | TRANSLATIO | N INITIA | TION FACTOR EIF-2B SUBUNIT ALPH |
| 1:00 | 8har-A | 2.2 | 11.2 | 96 | 358 | 3 | MOLECULE: | FUR6 | | |
| 2:00 | 4g3v-A | 2.2 | 4.3 | 72 | 162 | 8 | MOLECULE: | TRANSCRIPT | IONAL RE | GULATOR NLH2 |
| 3:00 | 8jnt-A | 2.2 | 3.4 | 54 | 77 | 6 | MOLECULE: | RXLR19781 | | |
| 4:00 | 4g33-A | 2.2 | 18.7 | 131 | 661 | 10 | MOLECULE: | 15S-LIPOXY | GENASE | |
| 5:00 | 6sny-A | 2.2 | 6.8 | 79 | 107 | 5 | MOLECULE: | SYNTHETIC | EPCR BIN | DING PROTEIN |
| 6:00 | 8hm3-B | 2.2 | 12.3 | 103 | 427 | 6 | MOLECULE: | PUTATIVE U | BIQUITIN | |
| 7:00 | 4jw1-A | 2.2 | 19.3 | 98 | 515 | 7 | MOLECULE: | EFFECTOR P | ROTEIN B | |
| 8:00 | 5a2v-B | 2.2 | 7.4 | 84 | 472 | 4 | MOLECULE: | MITOCHONDR | IAL PROT | EIN |
| 9:00 | 4m1p-A | 2.2 | 9 | 73 | 96 | 3 | MOLECULE: | COPPER-SEN | SITIVE O | PERON REPRESSOR (CSOR) |
| 10:00 | 7jw3-C | 2.2 | 4 | 75 | 244 | 11 | MOLECULE: | EXONUCLEAS | E MUT-7 | HOMOLOG |
| 11:00 | 4gmq-A | 2.2 | 3.1 | 57 | 92 | 11 | MOLECULE: | PUTATIVE R | IBOSOME | ASSOCIATED PROTEIN |
| 12:00 | 5w99-A | 2.2 | 5 | 84 | 319 | 5 | MOLECULE: | PBTD | | |
| 13:00 | 7y3f-1 | 2.2 | 17.3 | 95 | 466 | 6 | MOLECULE: | PHOTOSYSTE | M I P700 | CHLOROPHYLL A APOPROTEIN A1 |
| 14:00 | 7sol-A | 2.2 | 19.3 | 90 | 1011 | 3 | MOLECULE: | UBIQUITIN- | LIKE MOD | IFIER-ACTIVATING ENZYME 6 |
| 15:00 | 5awf-A | 2.2 | 6.3 | 65 | 385 | 9 | MOLECULE: | FES CLUSTE | R ASSEMB | LY PROTEIN SUFB |
| 16:00 | 8d8j-5 | 2.2 | 16.7 | 74 | 270 | 5 | MOLECULE: | PROBABLE S | #NAME? | L-L-METHIONINE-DEPENDENT RNA |
| 17:00 | 8hdw-M | 2.2 | 15.7 | 84 | 382 | 7 | MOLECULE: | PAM3 TUBE | | |
| 18:00 | 8h2h-D | 2.2 | 15.9 | 100 | 599 | 6 | MOLECULE: | LTRB | | |
| 19:00 | 8h77-F | 2.2 | 15.8 | 97 | 310 | 5 | MOLECULE: | HEAT SHOCK | PROTEIN | HSP 90-BETA |
| 20:00 | 3l9f-A | 2.2 | 6.6 | 65 | 170 | 11 | MOLECULE: | PUTATIVE U | NCHARACT | ERIZED PROTEIN SMU.1604C |
| 21:00 | 5i1u-B | 2.2 | 6.5 | 76 | 294 | 3 | MOLECULE: | GERMACRADI | EN-4-OL | SYNTHASE |
| 22:00 | 7lma-H | 2.2 | 16.3 | 75 | 199 | 7 | MOLECULE: | TELOMERASE | LA-RELA | TED PROTEIN P65 |

| | | | | | | | | | | | |
|---|---|---|---|---|---|---|---|---|---|---|---|
| 23:00 | 8bb6-A | 2.2 | 6.1 | 77 | 476 | 5 | MOLECULE: | SUCROSE TR | ANSPORT | PROTEIN SUC1 | |
| 24:00:00 | 8vx9-B | 2.2 | 18.7 | 106 | 1174 | 2 | MOLECULE: | HAMA | | | |
| 25:00:00 | 4i9e-A | 2.2 | 19.2 | 117 | 383 | 8 | MOLECULE: | RESPONSE R | EGULATOR | ASPARTATE PHOSPHATASE F | |
| 26:00:00 | 8ap7-e | 2.2 | 11.7 | 113 | 383 | 8 | MOLECULE: | ATP SYNTHA | SE SUBUN | IT A | |
| 27:00:00 | 6igx-D | 2.2 | 9.4 | 84 | 731 | 8 | MOLECULE: | CONDENSIN | COMPLEX | SUBUNIT 3 | |
| 28:00:00 | 7w78-B | 2.2 | 12.2 | 97 | 344 | 7 | MOLECULE: | PUTATIVE A | BC TRANS | PORT SYSTEM, ATP-BINDING PROTEI | |
| 29:00:00 | 2qv2-A | 2.2 | 15.1 | 83 | 305 | 8 | MOLECULE: | INOSITOL P | OLYPHOSP | HATE 5-PHOSPHATASE OCRL-1 | |
| 30:00:00 | 3waq-A | 2.2 | 7 | 75 | 134 | 7 | MOLECULE: | HEMERYTHRI | N-LIKE D | OMAIN PROTEIN DCRH | |
| 31:00:00 | 6p8v-G | 2.2 | 4.2 | 65 | 320 | 6 | MOLECULE: | ATPASE, AA | A FAMILY | | |
| 32:00:00 | 2cwe-A | 2.2 | 10.8 | 78 | 191 | 5 | MOLECULE: | HYPOTHETIC | AL TRANS | CRIPTION REGULATOR PROTEIN, PH1 | |
| 33:00:00 | 7rvb-A | 2.2 | 8.8 | 97 | 209 | 1 | MOLECULE: | FERRITIN | | | |
| 34:00:00 | 6h4j-A | 2.2 | 14.2 | 88 | 488 | 8 | MOLECULE: | UBIQUITIN | CARBOXYL | -TERMINAL HYDROLASE 25 | |
| 35:00:00 | 6t1z-A | 2.2 | 5.7 | 103 | 393 | 4 | MOLECULE: | LMRP INTEG | RAL MEMB | RANE PROTEIN | |
| 36:00:00 | 2klq-A | 2.2 | 3.8 | 63 | 114 | 5 | MOLECULE: | DNA REPLIC | ATION LI | CENSING FACTOR MCM6 | |
| 37:00:00 | 2js1-A | 2.2 | 4.1 | 64 | 80 | 8 | MOLECULE: | UNCHARACTE | RIZED PR | OTEIN YVFG | |
| 38:00:00 | 4cth-A | 2.2 | 19.6 | 95 | 698 | 2 | MOLECULE: | NEPRILYSIN | | | |
| 39:00:00 | 2kmf-A | 2.2 | 9 | 68 | 115 | 4 | MOLECULE: | PHOTOSYSTE | M II 11 | KDA PROTEIN | |
| 40:00:00 | 4k2p-A | 2.2 | 9 | 74 | 238 | 4 | MOLECULE: | T-LYMPHOMA | INVASIO | N AND METASTASIS-INDUCING PROTE | |
| 41:00:00 | 2ols-A | 2.2 | 12.3 | 97 | 725 | 8 | MOLECULE: | PHOSPHOENO | LPYRUVAT | E SYNTHASE | |
| 42:00:00 | 8dol-A | 2.2 | 31.3 | 124 | 540 | 3 | MOLECULE: | CAG PATHOG | ENICITY | ISLAND PROTEIN (CAG5) | |
| 43:00:00 | 5way-A | 2.2 | 22.3 | 107 | 482 | 8 | MOLECULE: | TRANSCRIPT | IONAL AC | TIVATOR | |
| 44:00:00 | 5vyl-A | 2.2 | 17 | 80 | 502 | 5 | MOLECULE: | INNER TEGU | MENT PRO | TEIN | |
| 45:00:00 | 1wte-A | 2.2 | 4 | 80 | 272 | 5 | MOLECULE: | 5'-D(*AP*C | P*CP*GP* | GP*GP*CP*CP*CP*TP*GP*CP*C)-3' | |
| 46:00:00 | 5w5y-O | 2.2 | 17.9 | 121 | 640 | 4 | MOLECULE: | DNA-DIRECT | ED RNA P | OLYMERASE I SUBUNIT RPA190 | |
| 47:00:00 | 8hmd-C | 2.2 | 18.9 | 88 | 1334 | 7 | MOLECULE: | INTRAFLAGE | LLAR TRA | NSPORT PROTEIN 122 HOMOLOG | |
| 48:00:00 | 7fbo-A | 2.2 | 8.1 | 78 | 283 | 5 | MOLECULE: | BEZA | | | |
| 49:00:00 | 7cgp-A | 2.2 | 11.1 | 70 | 147 | 7 | MOLECULE: | MITOCHONDR | IAL IMPO | RT INNER MEMBRANE TRANSLOCASE S | |
| 50:00:00 | 8env-A | 2.2 | 14.5 | 89 | 504 | 8 | MOLECULE: | SHEATH PRO | TEIN GP3 | | 1 |
| 51:00:00 | 5w6l-A | 2.2 | 21.2 | 104 | 494 | 7 | MOLECULE: | RTX REPEAT | #NAME? | ING CYTOTOXIN | |
| 52:00:00 | 3ufe-A | 2.2 | 12.8 | 70 | 109 | 7 | MOLECULE: | TRANSCRIPT | IONAL AN | TITERMINATOR (BGLG FAMILY) | |
| 53:00:00 | 6irb-A | 2.2 | 22.4 | 81 | 211 | 7 | MOLECULE: | 1-PHOSPHAT | IDYLINOS | ITOL 4,5-BISPHOSPHATE PHOSPHODI | |
| 54:00:00 | 6irb-B | 2.2 | 20.8 | 75 | 207 | 4 | MOLECULE: | 1-PHOSPHAT | IDYLINOS | ITOL 4,5-BISPHOSPHATE PHOSPHODI | |
| 55:00:00 | 3wyo-A | 2.2 | 5.7 | 62 | 153 | 8 | MOLECULE: | MYOGLOBIN | | | |
| 56:00:00 | 3ag0-A | 2.2 | 4 | 72 | 158 | 6 | MOLECULE: | CYTOGLOBIN | | | |
| 57:00:00 | 5b7i-C | 2.1 | 4.9 | 76 | 138 | 8 | MOLECULE: | CRISPR-ASS | OCIATED | NUCLEASE/HELICASE CAS3 SUBTYPE | |
| 58:00:00 | 4kqt-A | 2.1 | 11.1 | 72 | 169 | 6 | MOLECULE: | PUTATIVE O | UTER MEM | BRANE CHAPERONE (OMPH-LIKE) | |
| 59:00:00 | 2w7y-A | 2.1 | 15.2 | 83 | 389 | 11 | MOLECULE: | PROBABLE S | UGAR ABC | TRANSPORTER, SUGAR-BINDING PRO | |
| 60:00:00 | 7kdx-B | 2.1 | 13.6 | 106 | 665 | 5 | MOLECULE: | METHYLTRAN | SFERASE | TOKK | |
| 61:00:00 | 6cmv-B | 2.1 | 10.3 | 64 | 113 | 9 | MOLECULE: | TRANSCRIPT | IONAL RE | GULATOR LRS14-LIKE PROTEIN | |
| 62:00:00 | 3ggz-D | 2.1 | 4.8 | 80 | 188 | 4 | MOLECULE: | INCREASED | SODIUM T | OLERANCE PROTEIN 1 | |
| 63:00:00 | 3s0x-A | 2.1 | 15.7 | 69 | 225 | 6 | MOLECULE: | PEPTIDASE | A24B, FL | AK DOMAIN PROTEIN | |
| 64:00:00 | 5z62-B | 2.1 | 15.3 | 73 | 227 | 5 | MOLECULE: | CYTOCHROME | C OXIDA | SE SUBUNIT 1 | |
| 65:00:00 | 7nyw-E | 2.1 | 12.4 | 69 | 212 | 6 | MOLECULE: | CHROMOSOME | PARTITI | ON PROTEIN MUKB | |
| 66:00:00 | 3zc0-B | 2.1 | 12.8 | 91 | 191 | 2 | MOLECULE: | AFTRAX | | | |
| 67:00:00 | 6jo0-A | 2.1 | 7.2 | 68 | 224 | 12 | MOLECULE: | VIRRDTS | | | |
| 68:00:00 | 7yh3-D | 2.1 | 4.3 | 75 | 157 | 5 | MOLECULE: | TRAPPC3 FR | OM THORA | RCHAEOTA SMTZ1-45 | |
| 69:00:00 | 7qog-B | 2.1 | 19.3 | 88 | 226 | 6 | MOLECULE: | PORTAL PRO | TEIN GP2 | | 0 |
| 70:00:00 | 2r9i-A | 2.1 | 7.2 | 56 | 72 | 11 | MOLECULE: | PUTATIVE P | HAGE CAP | SID PROTEIN | |
| 71:00:00 | 7o4i-1 | 2.1 | 16.2 | 99 | 522 | 7 | MOLECULE: | GENERAL TR | ANSCRIPT | ION AND DNA REPAIR FACTOR IIH H | |
| 72:00:00 | 8f7n-A | 2.1 | 31.3 | 93 | 364 | 12 | MOLECULE: | METHYL-ACC | EPTING C | HEMOTAXIS PROTEIN | |
| 73:00:00 | 7dm1-A | 2.1 | 3.3 | 68 | 334 | 6 | MOLECULE: | PHOSPHATE- | BINDING | PROTEIN PSTS 1 | |
| 74:00:00 | 3lwu-A | 2.1 | 6.9 | 119 | 372 | 8 | MOLECULE: | SUCCINYLGL | UTAMATE | DESUCCINYLASE/ASPARTOACYLASE | |
| 75:00:00 | 6dft-A | 2.1 | 19.9 | 114 | 384 | 6 | MOLECULE: | DEOXYHYPUS | INE SYNT | HASE | |
| 76:00:00 | 5y6r-A | 2.1 | 12.1 | 129 | 667 | 5 | MOLECULE: | GENOME POL | YPROTEIN | | |

| | | | | | | | | | | | |
|---|---|---|---|---|---|---|---|---|---|---|---|
| 77:00:00 | 3sae-A | 2.1 | 17.2 | 86 | 780 | 5 | MOLECULE: | ALPHA-BISA | BOLENE S | YNTHASE | |
| 78:00:00 | 2rbd-A | 2.1 | 11.7 | 76 | 159 | 9 | MOLECULE: | BH2358 PRO | TEIN | | |
| 79:00:00 | 3aqt-B | 2.1 | 7.4 | 84 | 196 | 5 | MOLECULE: | BACTERIAL | REGULATO | RY PROTEINS, TETR FAMILY | |
| 80:00:00 | 8vc1-A | 2.1 | 10 | 133 | 391 | 9 | MOLECULE: | GUSTATORY | RECEPTOR | | |
| 81:00:00 | 8ok8-C | 2.1 | 8.9 | 75 | 335 | 12 | MOLECULE: | VARIANT SU | RFACE GL | YCOPROTEIN 615 | |
| 82:00:00 | 7nyw-B | 2.1 | 32.3 | 129 | 858 | 9 | MOLECULE: | CHROMOSOME | PARTITI | ON PROTEIN MUKB | |
| 83:00:00 | 7qtt-W | 2.1 | 11.6 | 91 | 459 | 9 | MOLECULE: | SPLICING F | ACTOR 3B | SUBUNIT 3 | |
| 84:00:00 | 5j10-A | 2.1 | 4.2 | 49 | 70 | 6 | MOLECULE: | PEPTIDE DE | SIGN 2L4 | HC2_24 | |
| 85:00:00 | 6cnn-A | 2.1 | 21 | 102 | 360 | 3 | MOLECULE: | INTERMEDIA | TE CONDU | CTANCE CALCIUM-ACTIVATED POTASS | |
| 86:00:00 | 5olk-A | 2.1 | 10.9 | 80 | 395 | 4 | MOLECULE: | RIBONUCLEO | SIDE-DIP | HOSPHATE REDUCTASE, BETA SUBUNI | |
| 87:00:00 | 8pmq-9 | 2.1 | 21.1 | 98 | 412 | 2 | MOLECULE: | E3 UBIQUIT | IN-PROTE | IN LIGASE RMD5 | |
| 88:00:00 | 8ibh-A | 2.1 | 6.9 | 58 | 75 | 3 | MOLECULE: | CENTROSOMA | L PROTEI | N OF 57 KDA | |
| 89:00:00 | 6kg7-A | 2.1 | 22.7 | 131 | 1817 | 9 | MOLECULE: | PIEZO-TYPE | MECHANO | SENSITIVE ION CHANNEL COMPONENT | |
| 90:00:00 | 4mt4-B | 2.1 | 6.3 | 109 | 473 | 8 | MOLECULE: | CMEC | | | |
| 91:00:00 | 2yev-B | 2.1 | 13.7 | 79 | 319 | 6 | MOLECULE: | CYTOCHROME | C OXIDA | SE POLYPEPTIDE I+III | |
| 92:00:00 | 7csp-A | 2.1 | 13.5 | 91 | 455 | 11 | MOLECULE: | RHO GUANIN | E NUCLEO | TIDE EXCHANGE FACTOR 16 | |
| 93:00:00 | 7wff-F | 2.1 | 12.5 | 126 | 677 | 3 | MOLECULE: | NAD(P)H-QU | INONE OX | IDOREDUCTASE SUBUNIT 1, CHLOROP | |
| 94:00:00 | 6bq1-E | 2.1 | 22.6 | 89 | 1510 | 7 | MOLECULE: | PHOSPHATID | YLINOSIT | OL 4-KINASE III ALPHA (PI4KA) | |
| 95:00:00 | 6dkm-D | 2.1 | 6 | 55 | 79 | 7 | MOLECULE: | DHD131_A | | | |
| 96:00:00 | 1urj-B | 2.1 | 22.4 | 116 | 1030 | 5 | MOLECULE: | MAJOR DNA- | BINDING | PROTEIN | |
| 97:00:00 | 6ch3-B | 2.1 | 12.7 | 77 | 162 | 6 | MOLECULE: | FLAGELLAR | BIOSYNTH | ESIS PROTEIN FLHA | |
| 98:00:00 | 4nxt-A | 2.1 | 18.2 | 96 | 337 | 9 | MOLECULE: | MITOCHONDR | IAL DYNA | MIC PROTEIN MID51 | |
| 99:00:00 | 6s8t-A | 2.1 | 19.8 | 106 | 442 | 5 | MOLECULE: | ERYTHROCYT | E MEMBRA | NE PROTEIN 1 | |
| 0:00 | 8i21-A | 2.1 | 37.4 | 103 | 256 | 4 | MOLECULE: | STRUCTURAL | MAINTEN | ANCE OF CHROMOSOMES PROTEIN 6 | |
| 1:00 | 5l4k-V | 2.1 | 12.6 | 88 | 293 | 7 | MOLECULE: | 26S PROTEA | SOME NON | -ATPASE REGULATORY SUBUNIT 4 | |
| 2:00 | 8gju-F | 2.1 | 11.1 | 83 | 332 | 6 | MOLECULE: | METHYLMALO | NIC ACID | URIA TYPE A PROTEIN, MITOCHONDR | |
| 3:00 | 3a8e-C | 2.1 | 5.1 | 74 | 159 | 3 | MOLECULE: | CELLULOSE | SYNTHASE | OPERON PROTEIN D | |
| 4:00 | 7pde-A | 2.1 | 21.3 | 110 | 846 | 5 | MOLECULE: | ADENYLATE | CYCLASE | | 9 |
| 5:00 | 7z6q-A | 2.1 | 12.7 | 125 | 667 | 6 | MOLECULE: | PHOTOSYSTE | M P840 R | EACTION CENTER, LARGE SUBUNIT | |
| 6:00 | 3td7-A | 2.1 | 12.7 | 83 | 253 | 6 | MOLECULE: | FAD-LINKED | SULFHYD | RYL OXIDASE R596 | |
| 7:00 | 7k3z-A | 2.1 | 5.4 | 83 | 564 | 4 | MOLECULE: | 60 KDA CHA | PERONIN | | |
| 8:00 | 2ygu-F | 2.1 | 4.1 | 69 | 118 | 9 | MOLECULE: | VENOM ALLE | RGEN 2 | | |
| 9:00 | 7et0-C | 2.1 | 8.3 | 108 | 684 | 4 | MOLECULE: | BACTERIA F | ACTOR B | | |
| 10:00 | 7p64-L | 2.1 | 9.5 | 111 | 606 | 6 | MOLECULE: | NADH-QUINO | NE OXIDO | REDUCTASE SUBUNIT F | |
| 11:00 | 8d2s-A | 2.1 | 23.5 | 115 | 475 | 6 | MOLECULE: | SODIUM-DEP | ENDENT L | YSOPHOSPHATIDYLCHOLINE SYMPORTE | |
| 12:00 | 4kvq-A | 2.1 | 6.3 | 78 | 223 | 6 | MOLECULE: | ALDEHYDE D | ECARBONY | LASE | |
| 13:00 | 6dlm-A | 2.1 | 5 | 49 | 74 | 4 | MOLECULE: | DHD127_A | | | |
| 14:00 | 3na7-A | 2.1 | 29.3 | 93 | 237 | 3 | MOLECULE: | HP0958 | | | |
| 15:00 | 5tj5-A | 2.1 | 14.2 | 106 | 570 | 5 | MOLECULE: | V-TYPE PRO | TON ATPA | SE SUBUNIT A | |
| 16:00 | 3okq-A | 2.1 | 17.9 | 74 | 125 | 3 | MOLECULE: | BUD SITE S | ELECTION | PROTEIN 6 | |
| 17:00 | 8etc-h | 2.1 | 18.1 | 69 | 121 | 14 | MOLECULE: | RNA (2151- | MER) | | |
| 18:00 | 6wt4-A | 2.1 | 3.4 | 63 | 147 | 8 | MOLECULE: | BACTERIAL | STING | | |
| 19:00 | 8dh3-E | 2.1 | 11.4 | 132 | 825 | 7 | MOLECULE: | T7 RNA POL | YMERASE | | |
| 20:00 | 7y3w-A | 2.1 | 15.2 | 127 | 440 | 7 | MOLECULE: | QUESTIN OX | IDASE | | |
| 21:00 | 8eup-3 | 2.1 | 18.3 | 62 | 194 | 6 | MOLECULE: | RNA (1422- | MER) | | |
| 22:00 | 7x5e-E | 2.1 | 15.1 | 52 | 101 | 6 | MOLECULE: | TRANSCRIPT | ION FACT | OR MAFG | |
| 23:00 | 6cp3-7 | 2.1 | 9.4 | 80 | 171 | 8 | MOLECULE: | ATP SYNTHA | SE SUBUN | IT 9, MITOCHONDRIAL | |
| 24:00:00 | 8srm-B | 2.1 | 12.5 | 75 | 235 | 4 | MOLECULE: | RB1-INDUCI | BLE COIL | ED-COIL PROTEIN 1 | |
| 25:00:00 | 3iee-A | 2.1 | 5.5 | 77 | 256 | 4 | MOLECULE: | PUTATIVE E | XPORTED | PROTEIN | |
| 26:00:00 | 4gzu-A | 2.1 | 15.1 | 103 | 436 | 10 | MOLECULE: | FERM, RHOG | EF AND P | LECKSTRIN DOMAIN-CONTAINING PRO | |
| 27:00:00 | 4lws-A | 2.1 | 8.6 | 67 | 100 | 4 | MOLECULE: | UNCHARACTE | RIZED PR | OTEIN | |
| 28:00:00 | 8gyz-B | 2.1 | 6.1 | 78 | 215 | 12 | MOLECULE: | TRANSCRIPT | ION FACT | OR TGA7 | |
| 29:00:00 | 8e4g-a | 2.1 | 7.8 | 63 | 127 | 13 | MOLECULE: | PORTAL PRO | TEIN | | |
| 30:00:00 | 7x1i-A | 2.1 | 18 | 107 | 678 | 4 | MOLECULE: | ISOFORM 1 | OF SOLUT | E CARRIER FAMILY 4 MEMBER 11 | |

| | | | | | | | | | |
|---|---|---|---|---|---|---|---|---|---|
| 31:00:00 | 4wbd-A | 2.1 | 23.4 | 86 | 540 | 2 MOLECULE: | BSHC | | |
| 32:00:00 | 5vvh-D | 2.1 | 4.4 | 74 | 210 | 7 MOLECULE: | OCTOPINE C | ATABOLIS | M/UPTAKE OPERON REGULATORY PROT |
| 33:00:00 | 6d5f-a | 2.1 | 14.1 | 72 | 131 | 6 MOLECULE: | FIMBRIAL P | ROTEIN | |
| 34:00:00 | 6h02-A | 2.1 | 25.2 | 110 | 1334 | 7 MOLECULE: | MEDIATOR O | F RNA PO | LYMERASE II TRANSCRIPTION SUBUN |
| 35:00:00 | 6nf1-A | 2.1 | 29.9 | 128 | 550 | 2 MOLECULE: | PROTO-ONCO | GENE VAV | |
| 36:00:00 | 7y6b-A | 2.1 | 7.7 | 61 | 72 | 5 MOLECULE: | ESCE/YSCE/ | SSAE FAM | ILY TYPE III SECRETION SYSTEM N |
| 37:00:00 | 8wlj-A | 2.1 | 6.4 | 90 | 409 | 8 MOLECULE: | SYNAPTIC V | ESICULAR | AMINE TRANSPORTER |
| 38:00:00 | 2xeq-C | 2.1 | 12.7 | 93 | 241 | 3 MOLECULE: | PAT1 HOMOL | OG 1, | |
| 39:00:00 | 4w4u-B | 2.1 | 4.4 | 66 | 96 | 5 MOLECULE: | UBIQUITIN | CARBOXYL | -TERMINAL HYDROLASE |
| 40:00:00 | 2ebi-A | 2.1 | 2.9 | 59 | 86 | 3 MOLECULE: | DNA BINDIN | G PROTEI | N GT-1 |
| 41:00:00 | 7wss-A | 2.1 | 15.8 | 101 | 533 | 8 MOLECULE: | MICROBIAL | COLLAGEN | ASE |
| 42:00:00 | 6whb-A | 2.1 | 15.9 | 91 | 284 | 4 MOLECULE: | MITOGEN-AC | TIVATED | PROTEIN KINASE KINASE KINASE 1 |
| 43:00:00 | 9auc-E | 2.1 | 16.2 | 78 | 117 | 9 MOLECULE: | GUANINE NU | CLEOTIDE | #NAME? |
| 44:00:00 | 8p0v-M | 2.1 | 16.6 | 74 | 357 | 7 MOLECULE: | COILED-COI | L DOMAIN | -CONTAINING PROTEIN 93 |
| 45:00:00 | 5oqm-n | 2.1 | 10.8 | 77 | 136 | 6 MOLECULE: | DNA-DIRECT | ED RNA P | OLYMERASE II SUBUNIT RPB1 |
| 46:00:00 | 2is6-A | 2.1 | 19.7 | 80 | 654 | 6 MOLECULE: | 5'-D(*CP*G | P*AP*GP* | CP*AP*CP*TP*GP*CP*AP*GP*TP*GP*C |
| 47:00:00 | 5dwa-A | 2.1 | 11.6 | 84 | 278 | 5 MOLECULE: | TYPE-2 RES | TRICTION | ENZYME AGEI |
| 48:00:00 | 2r44-A | 2.1 | 6.7 | 62 | 330 | 6 MOLECULE: | UNCHARACTE | RIZED PR | OTEIN |
| 49:00:00 | 7rsl-A | 2.1 | 3.8 | 55 | 235 | 11 MOLECULE: | SEIPIN | | |
| 50:00:00 | 4w8j-A | 2.1 | 16.4 | 106 | 1017 | 8 MOLECULE: | PESTICIDAL | CRYSTAL | PROTEIN CRY1AC |
| 51:00:00 | 6prk-A | 2.1 | 4.3 | 58 | 115 | 7 MOLECULE: | RICF | | |
| 52:00:00 | 8b17-A | 2.1 | 11.7 | 84 | 451 | 7 MOLECULE: | DIPEPTIDE | AND TRIP | EPTIDE PERMEASE B |
| 53:00:00 | 8ebt-C | 2.1 | 13.3 | 81 | 191 | 7 MOLECULE: | GENERAL TR | ANSCRIPT | ION AND DNA REPAIR FACTOR IIH H |
| 54:00:00 | 7njn-C | 2.1 | 19.4 | 97 | 536 | 7 MOLECULE: | ATP SYNTHA | SE SUBUN | IT ALPHA |
| 55:00:00 | 3svi-A | 2.1 | 3 | 58 | 79 | 3 MOLECULE: | TYPE III E | FFECTOR | HOPAB2 |
| 56:00:00 | 8e9b-G | 2.1 | 13.1 | 89 | 151 | 3 MOLECULE: | ACTIN-RELA | TED PROT | EIN 3 |
| 57:00:00 | 5xgc-A | 2.1 | 14.8 | 125 | 473 | 8 MOLECULE: | RAP1 GTPAS | E-GDP DI | SSOCIATION STIMULATOR 1 |
| 58:00:00 | 6pqx-A | 2.1 | 11.8 | 86 | 480 | 7 MOLECULE: | DNA-MEDIAT | ED TRANS | POSASE |
| 59:00:00 | 7wwb-A | 2.1 | 5.3 | 99 | 333 | 5 MOLECULE: | CHOLINE TR | ANSPORTE | R-LIKE PROTEIN 1 |
| 60:00:00 | 1eku-B | 2.1 | 5.9 | 87 | 254 | 8 MOLECULE: | INTERFERON | GAMMA | |
| 61:00:00 | 7bqn-A | 2.1 | 5.2 | 69 | 121 | 4 MOLECULE: | REI | | |
| 62:00:00 | 5j0n-F | 2.1 | 14.7 | 109 | 356 | 6 MOLECULE: | ATTP(-117 | TO +79) | |
| 63:00:00 | 5dzt-A | 2.1 | 18.7 | 70 | 886 | 6 MOLECULE: | CYLM | | |
| 64:00:00 | 3icq-T | 2.1 | 17.2 | 82 | 949 | 7 MOLECULE: | EXPORTIN-T | | |
| 65:00:00 | 4iwb-A | 2.1 | 4.3 | 74 | 165 | 9 MOLECULE: | FLIC, FLIS | CHIMERA | |
| 66:00:00 | 7fap-A | 2.1 | 25.3 | 114 | 1236 | 6 MOLECULE: | ERYTHROCYT | E MEMBRA | NE PROTEIN 1, PFEMP1 |
| 67:00:00 | 5y36-A | 2.1 | 16.8 | 116 | 1368 | 9 MOLECULE: | CRISPR-ASS | OCIATED | ENDONUCLEASE CAS9/CSN1 |
| 68:00:00 | 2ofy-A | 2.1 | 8.3 | 60 | 82 | 5 MOLECULE: | PUTATIVE X | RE-FAMIL | Y TRANSCRIPTIONAL REGULATOR |
| 69:00:00 | 2wzk-A | 2.1 | 14.5 | 81 | 369 | 7 MOLECULE: | CULLIN-5 | | |
| 70:00:00 | 6k3b-B | 2.1 | 4.1 | 61 | 106 | 7 MOLECULE: | LPG2147 | | |
| 71:00:00 | 6jql-A | 2.1 | 9.4 | 81 | 678 | 14 MOLECULE: | BIFUNCTION | AL PROTE | IN PAAZ |
| 72:00:00 | 5zdz-C | 2.1 | 5.1 | 74 | 623 | 8 MOLECULE: | MOUSE RAG1 | | |
| 73:00:00 | 4l6v-6 | 2.1 | 7.7 | 71 | 125 | 4 MOLECULE: | PHOTOSYSTE | M I P700 | CHLOROPHYLL A APOPROTEIN A1 |
| 74:00:00 | 6xu2-A | 2.1 | 20.8 | 94 | 1091 | 7 MOLECULE: | IMPORTIN-5 | | |
| 75:00:00 | 6eld-A | 2.1 | 4.5 | 67 | 107 | 6 MOLECULE: | NUCLEOLYSI | N TIA-1 | ISOFORM P40,U1 SMALL NUCLEAR |
| 76:00:00 | 5dcq-E | 2.1 | 12.2 | 59 | 200 | 3 MOLECULE: | ARTIFICIAL | REPEAT | PROTEINS (ALPHAREP3) |
| 77:00:00 | 8t1l-S | 2.1 | 20.7 | 115 | 913 | 5 MOLECULE: | MEDIATOR O | F RNA PO | LYMERASE II TRANSCRIPTION SUBUN |
| 78:00:00 | 8v9u-A | 2.1 | 3.9 | 55 | 81 | 5 MOLECULE: | DNA (CYTOS | INE-5)-M | ETHYLTRANSFERASE 1 |
| 79:00:00 | 7mex-A | 2.1 | 21.9 | 109 | 1737 | 6 MOLECULE: | UBIQUITIN | | |
| 80:00:00 | 4uxv-A | 2.1 | 12.5 | 89 | 522 | 9 MOLECULE: | SEPTATION | RING FOR | MATION REGULATOR EZRA |
| 81:00:00 | 5ed8-A | 2.1 | 13.2 | 83 | 245 | 5 MOLECULE: | MKIAA0668 | PROTEIN | |
| 82:00:00 | 7eqf-A | 2.1 | 6.5 | 70 | 184 | 10 MOLECULE: | TETR/ACRR | FAMILY T | RANSCRIPTIONAL REGULATOR |
| 83:00:00 | 2c9e-A | 2.1 | 12.5 | 84 | 317 | 6 MOLECULE: | PERIDININ- | CHLOROPH | YLL A PROTEIN |
| 84:00:00 | 7unk-A | 2.1 | 20.6 | 139 | 1047 | 11 MOLECULE: | IMPORTIN-4 | | |

| | | | | | | | | | | |
|---|---|---|---|---|---|---|---|---|---|---|
| 85:00:00 | 8thc-A | 2.1 | 24.6 | 71 | 479 | 4 | MOLECULE: | ELG1 ISOFO | RM 1 | |
| 86:00:00 | 1e7p-C | 2.1 | 10.5 | 96 | 254 | 9 | MOLECULE: | FUMARATE R | EDUCTASE | FLAVOPROTEIN SUBUNIT |
| 87:00:00 | 5u30-A | 2.1 | 21.1 | 94 | 1085 | 6 | MOLECULE: | CRISPR-ASS | OCIATED | ENDONUCLEASE C2C1 |
| 88:00:00 | 8srz-A | 2.1 | 3.5 | 61 | 88 | 10 | MOLECULE: | PROBABLE S | ERINE PR | OTEASE FE772_23065 |
| 89:00:00 | 6l1k-A | 2.1 | 9.5 | 69 | 384 | 10 | MOLECULE: | NADH-DEPEN | DENT BUT | ANOL DEHYDROGENASE A |
| 90:00:00 | 7wiu-B | 2.1 | 26.1 | 103 | 576 | 8 | MOLECULE: | MYCOBACTIN | IMPORT | ATP-BINDING/PERMEASE PROTEIN IR |
| 91:00:00 | 2rae-A | 2.1 | 5.1 | 75 | 193 | 7 | MOLECULE: | TRANSCRIPT | IONAL RE | GULATOR, ACRR FAMILY PROTEIN |
| 92:00:00 | 6nq8-A | 2.1 | 11.1 | 87 | 208 | 8 | MOLECULE: | UNCHARACTE | RIZED PR | OTEIN YETJ |
| 93:00:00 | 6t8d-X | 2.1 | 12 | 96 | 355 | 6 | MOLECULE: | MAKB | | |
| 94:00:00 | 7mi6-A | 2.1 | 22 | 117 | 2419 | 3 | MOLECULE: | FUSION PRO | TEIN OF | DYNEIN AND ENDOLYSIN |
| 95:00:00 | 2qyu-A | 2.1 | 20.2 | 133 | 620 | 8 | MOLECULE: | SECRETED E | FFECTOR | PROTEIN |
| 96:00:00 | 8d00-A | 2.1 | 9.1 | 82 | 299 | 9 | MOLECULE: | MICROTUBUL | E-ASSOCI | ATED PROTEIN TORTIFOLIA1 |
| 97:00:00 | 3sjr-A | 2.1 | 4.6 | 62 | 126 | 6 | MOLECULE: | UNCHARACTE | RIZED PR | OTEIN |
| 98:00:00 | 3cvj-A | 2.1 | 8.8 | 99 | 243 | 5 | MOLECULE: | PUTATIVE P | HOSPHOHE | PTOSE ISOMERASE |
| 99:00:00 | 4he8-D | 2.1 | 5.5 | 86 | 160 | 3 | MOLECULE: | NADH-QUINO | NE OXIDO | REDUCTASE SUBUNIT 7 |
| 0:00 | 5zx1-A | 2.1 | 3.8 | 58 | 84 | 10 | MOLECULE: | ENT | | |
| 1:00 | 3bkh-A | 2.1 | 9.9 | 75 | 268 | 8 | MOLECULE: | LYTIC TRAN | SGLYCOSY | LASE |
| 2:00 | 6tdz-C | 2.1 | 19.5 | 91 | 530 | 8 | MOLECULE: | OLIGOMYCIN | SENSITI | VITY CONFERRING PROTEIN (OSCP) |
| 3:00 | 6w8p-B | 2.1 | 20.8 | 113 | 426 | 4 | MOLECULE: | ENDOSOMAL/ | LYSOSOMA | L POTASSIUM CHANNEL TMEM175 |
| 4:00 | 3crl-B | 2.1 | 16.4 | 105 | 383 | 5 | MOLECULE: | PYRUVATE D | EHYDROGE | NASE [LIPOAMIDE] KINASE ISOZYME |
| 5:00 | 3cny-A | 2.1 | 17.9 | 81 | 300 | 10 | MOLECULE: | INOSITOL C | ATABOLIS | M PROTEIN IOLE |
| 6:00 | 3s1s-A | 2.1 | 18.4 | 98 | 871 | 10 | MOLECULE: | RESTRICTIO | N ENDONU | CLEASE BPUSI |
| 7:00 | 5azb-A | 2.1 | 15.8 | 75 | 284 | 3 | MOLECULE: | PROLIPOPRO | TEIN DIA | CYLGLYCERYL TRANSFERASE |
| 8:00 | 3euh-B | 2.1 | 12.5 | 73 | 321 | 10 | MOLECULE: | CHROMOSOME | PARTITI | ON PROTEIN MUKF |
| 9:00 | 5b86-A | 2.1 | 15.4 | 108 | 579 | 5 | MOLECULE: | TUMOR NECR | OSIS FAC | TOR ALPHA-INDUCED PROTEIN 2 |
| 10:00 | 7cth-E | 2.1 | 4.5 | 61 | 75 | 5 | MOLECULE: | CORE PROTE | IN | |
| 11:00 | 5iwz-A | 2.1 | 6.6 | 86 | 372 | 9 | MOLECULE: | SYNAPTONEM | AL COMPL | EX PROTEIN 2 |
| 12:00 | 6yj4-V | 2.1 | 11 | 82 | 126 | 2 | MOLECULE: | NADH-UBIQU | INONE OX | IDOREDUCTASE CHAIN 3 |
| 13:00 | 2rkh-A | 2.1 | 8.2 | 69 | 164 | 4 | MOLECULE: | PUTATIVE A | PHA-LIKE | TRANSCRIPTION FACTOR |
| 14:00 | 2gmf-A | 2.1 | 4.7 | 72 | 121 | 10 | MOLECULE: | GRANULOCYT | E-MACROP | HAGE COLONY-STIMULATING FACTOR |
| 15:00 | 3fgg-A | 2.1 | 5.2 | 65 | 147 | 3 | MOLECULE: | UNCHARACTE | RIZED PR | OTEIN BCE2196 |
| 16:00 | 6xgt-C | 2.1 | 5.2 | 81 | 160 | 9 | MOLECULE: | CYANATE HY | DRATASE | |
| 17:00 | 8bmw-E | 2.1 | 4.6 | 60 | 122 | 10 | MOLECULE: | CRISPR-ASS | OCIATED | SMALL SUBUNIT PROTEIN (TYPE III |
| 18:00 | 7sfj-A | 2.1 | 10.2 | 98 | 284 | 10 | MOLECULE: | CHRMINE | | |
| 19:00 | 4bwc-A | 2.1 | 10.3 | 74 | 170 | 5 | MOLECULE: | PHOSPHOLIP | ASE B-LI | KE 1 |
| 20:00 | 3aai-A | 2.1 | 9.1 | 70 | 78 | 3 | MOLECULE: | COPPER HOM | EOSTASIS | OPERON REGULATORY PROTEIN |
| 21:00 | 4cz8-A | 2.1 | 18 | 114 | 422 | 4 | MOLECULE: | NA+/H+ ANT | IPORTER, | PUTATIVE |
| 22:00 | 7dl2-A | 2.1 | 19 | 111 | 1127 | 4 | MOLECULE: | HAMARTIN | | |
| 23:00 | 5ch6-A | 2.1 | 17.2 | 77 | 286 | 4 | MOLECULE: | FRIGIDA | | |
| 24:00:00 | 4zht-A | 2.1 | 14.3 | 68 | 384 | 10 | MOLECULE: | BIFUNCTION | AL UDP-N | -ACETYLGLUCOSAMINE 2-EPIMERASE/ |
| 25:00:00 | 7khw-a | 2.1 | 7.8 | 76 | 174 | 9 | MOLECULE: | TRANSLOCON | ESPA | |
| 26:00:00 | 7v7b-A | 2.1 | 15.5 | 119 | 1110 | 8 | MOLECULE: | DDB1- AND | CUL4-ASS | OCIATED FACTOR 1 |
| 27:00:00 | 6zfw-A | 2.1 | 3.7 | 52 | 70 | 2 | MOLECULE: | PEROXIN-14 | | |
| 28:00:00 | 5tr1-A | 2.1 | 27.7 | 92 | 606 | 8 | MOLECULE: | CHLORIDE C | HANNEL P | ROTEIN |
| 29:00:00 | 2juc-A | 2.1 | 3.1 | 49 | 55 | 4 | MOLECULE: | PRE-MRNA-S | PLICING | FACTOR URN1 |
| 30:00:00 | 8b4h-C | 2.1 | 18.7 | 82 | 362 | 5 | MOLECULE: | PUTATIVE T | RANSPOSA | SE FOR INSERTION SEQUENCE ELEME |
| 31:00:00 | 6ep3-B | 2.1 | 8.3 | 71 | 218 | 4 | MOLECULE: | LMO0651 PR | OTEIN | |
| 32:00:00 | 4nqa-I | 2.1 | 12.8 | 114 | 385 | 6 | MOLECULE: | RETINOIC A | CID RECE | PTOR RXR-ALPHA |
| 33:00:00 | 8hhe-B | 2.1 | 17.5 | 101 | 647 | 14 | MOLECULE: | CRYSTALINE | ENTOMOC | IDAL PROTOXIN |
| 34:00:00 | 6bpz-A | 2.1 | 15.4 | 121 | 900 | 6 | MOLECULE: | PIEZO-TYPE | MECHANO | SENSITIVE ION CHANNEL COMPONENT |
| 35:00:00 | 3f41-A | 2.1 | 13.5 | 90 | 590 | 7 | MOLECULE: | PHYTASE | | |
| 36:00:00 | 1kw2-A | 2.1 | 18.5 | 64 | 455 | 2 | MOLECULE: | VITAMIN D- | BINDING | PROTEIN |
| 37:00:00 | 3mka-J | 2.1 | 5.1 | 70 | 252 | 9 | MOLECULE: | PROTEASOME | SUBUNIT | ALPHA |
| 38:00:00 | 1ntc-A | 2.1 | 8.1 | 70 | 91 | 9 | MOLECULE: | PROTEIN (N | ITROGEN | REGULATION PROTEIN (NTRC)) |

| | | | | | | | | | | |
|---|---|---|---|---|---|---|---|---|---|---|
| 39:00:00 | 5y78-A | 2.1 | 6.7 | 99 | 305 | 9 | MOLECULE: | PUTATIVE H | EXOSE PH | OSPHATE TRANSLOCATOR |
| 40:00:00 | 8et6-A | 2.1 | 18.6 | 110 | 532 | 5 | MOLECULE: | 1-Oct | | |
| 41:00:00 | 6c6r-A | 2.1 | 14.9 | 87 | 451 | 7 | MOLECULE: | SQUALENE M | ONOOXYGE | NASE |
| 42:00:00 | 7fax-A | 2.1 | 3.9 | 73 | 103 | 3 | MOLECULE: | TBLW | | |
| 43:00:00 | 7pw9-A | 2.1 | 19.9 | 103 | 1952 | 7 | MOLECULE: | SERINE/THR | EONINE-P | ROTEIN KINASE SMG1,SERINE/THREO |
| 44:00:00 | 4v2x-A | 2.1 | 19.1 | 101 | 534 | 6 | MOLECULE: | ENDO-BETA- | 1,4-GLUC | ANASE (CELLULASE B) |
| 45:00:00 | 7ojn-L | 2.1 | 13.9 | 94 | 2010 | 5 | MOLECULE: | RNA-DIRECT | ED RNA P | OLYMERASE L |
| 46:00:00 | 4fp9-B | 2.1 | 12.6 | 80 | 245 | 4 | MOLECULE: | METHYLTRAN | SFERASE | NSUN4 |
| 47:00:00 | 1n81-A | 2.1 | 4.1 | 68 | 186 | 3 | MOLECULE: | PLASMODIUM | FALCIPA | RUM GAMETE ANTIGEN 27/25 |
| 48:00:00 | 5fhp-C | 2.1 | 5.4 | 88 | 209 | 3 | MOLECULE: | NICR | | |
| 49:00:00 | 8e2h-A | 2.1 | 10.7 | 115 | 656 | 9 | MOLECULE: | BACULOVIRA | L IAP RE | PEAT-CONTAINING PROTEIN 6 |
| 50:00:00 | 7wvz-A | 2.1 | 28.3 | 108 | 1719 | 12 | MOLECULE: | BETA-KETOA | CYL-ACYL | -CARRIER-PROTEIN SYNTHASE I |
| 51:00:00 | 8wcs-A | 2.1 | 10.1 | 92 | 1044 | 5 | MOLECULE: | CAS13H1 | | |
| 52:00:00 | 7z4w-a | 2.1 | 5.6 | 68 | 102 | 9 | MOLECULE: | PORTAL PRO | TEIN | |
| 53:00:00 | 6fpg-F | 2.1 | 5.3 | 89 | 266 | 2 | MOLECULE: | CHROMOSOME | 16, WHO | LE GENOME SHOTGUN SEQUENCE |
| 54:00:00 | 7xe4-F | 2.1 | 7.8 | 98 | 1503 | 8 | MOLECULE: | 1,3-BETA-G | LUCAN SY | NTHASE COMPONENT FKS1 |
| 55:00:00 | 6xi6-A | 2.1 | 15.4 | 83 | 269 | 10 | MOLECULE: | HELICAL FU | SION DES | IGN |
| 56:00:00 | 7loi-A | 2.1 | 12.5 | 70 | 175 | 6 | MOLECULE: | TRANSMEMBR | ANE PROT | EIN GP41 |
| 57:00:00 | 6mcp-A | 2.1 | 11.5 | 101 | 520 | 6 | MOLECULE: | LEGK7 | | |
| 58:00:00 | 8t8k-A | 2.1 | 10.1 | 71 | 183 | 1 | MOLECULE: | DUF507 FAM | ILY PROT | EIN |
| 59:00:00 | 6lu0-A | 2.1 | 16.5 | 126 | 937 | 3 | MOLECULE: | CAS12I2 | | |
| 60:00:00 | 6vls-D | 2.1 | 26.2 | 122 | 963 | 5 | MOLECULE: | MALTOSE/MA | LTODEXTR | IN-BINDING PERIPLASMIC PROTEIN, |
| 61:00:00 | 7ctq-A | 2.1 | 13.2 | 88 | 429 | 9 | MOLECULE: | PEPTIDYL T | RYPTOPHA | N DIHYDROXYLASE |
| 62:00:00 | 7qru-E | 2.1 | 11.8 | 74 | 158 | 7 | MOLECULE: | NA+/H+ ANT | IPORTER | SUBUNIT D |
| 63:00:00 | 5ul2-A | 2.1 | 21.6 | 112 | 731 | 3 | MOLECULE: | OXSB PROTE | IN | |
| 64:00:00 | 4k2u-A | 2.1 | 8.8 | 65 | 230 | 5 | MOLECULE: | ERYTHROCYT | E BINDIN | G ANTIGEN 175 |
| 65:00:00 | 2aml-A | 2.1 | 8.9 | 104 | 366 | 6 | MOLECULE: | SIS DOMAIN | PROTEIN | |
| 66:00:00 | 7o4i-W | 2.1 | 18.4 | 93 | 312 | 2 | MOLECULE: | GENERAL TR | ANSCRIPT | ION AND DNA REPAIR FACTOR IIH H |
| 67:00:00 | 2bma-A | 2.1 | 18.6 | 96 | 467 | 6 | MOLECULE: | GLUTAMATE | DEHYDROG | ENASE (NADP+) |
| 68:00:00 | 7w2q-B | 2.1 | 6.1 | 74 | 201 | 7 | MOLECULE: | PROBABLE T | RANSCRIP | TIONAL REGULATOR |
| 69:00:00 | 8hf3-B | 2.1 | 7.9 | 66 | 124 | 2 | MOLECULE: | PALMITOYLT | RANSFERA | SE ZDHHC9 |
| 70:00:00 | 2kiw-A | 2.1 | 6.9 | 76 | 111 | 8 | MOLECULE: | INT PROTEI | N | |
| 71:00:00 | 2fdc-A | 2.1 | 15.9 | 87 | 505 | 5 | MOLECULE: | 5'-D(P*CP* | GP*GP*CP | *TP*CP*CP*AP*TP*CP*TP*CP*TP*AP* |
| 72:00:00 | 6wb9-2 | 2.1 | 13.9 | 91 | 292 | 7 | MOLECULE: | ENDOPLASMI | C RETICU | LUM MEMBRANE PROTEIN COMPLEX SU |
| 73:00:00 | 2iho-A | 2.1 | 3.9 | 73 | 292 | 8 | MOLECULE: | LECTIN | | |
| 74:00:00 | 1m5y-D | 2.1 | 3.9 | 73 | 389 | 5 | MOLECULE: | SURVIVAL P | ROTEIN S | URA |
| 75:00:00 | 4rxx-A | 2.1 | 12.7 | 92 | 426 | 9 | MOLECULE: | UBIQUITIN | CARBOXYL | -TERMINAL HYDROLASE 38 |
| 76:00:00 | 4ng2-E | 2.1 | 3.3 | 54 | 94 | 7 | MOLECULE: | TRANSCRIPT | IONAL AC | TIVATOR PROTEIN LASR |
| 77:00:00 | 6vk0-C | 2.1 | 4.8 | 83 | 180 | 2 | MOLECULE: | U1 SNP1-AS | SOCIATIN | G PROTEIN 1 |
| 78:00:00 | 8by2-A | 2.1 | 11.2 | 104 | 548 | 10 | MOLECULE: | GLUTATHION | E-REGULA | TED POTASSIUM-EFFLUX SYSTEM PRO |
| 79:00:00 | 2m64-A | 2.1 | 21.6 | 91 | 231 | 5 | MOLECULE: | PHLP5 | | |
| 80:00:00 | 4xp7-A | 2.1 | 8.3 | 61 | 324 | 10 | MOLECULE: | TRNA-DIHYD | ROURIDIN | E(20) SYNTHASE [NAD(P)+]-LIKE |
| 81:00:00 | 7yzp-C | 2.1 | 34.8 | 81 | 448 | 6 | MOLECULE: | DNA HAIRPI | N (59-ME | R) |
| 82:00:00 | 6wkr-A | 2.1 | 9.2 | 68 | 439 | 9 | MOLECULE: | UBIQUITIN | | |
| 83:00:00 | 6rwb-A | 2.1 | 19.3 | 105 | 1873 | 6 | MOLECULE: | TOXIN,TOXI | N COMPLE | X SUBUNIT TCAB,PUTATIVE TOXIN S |
| 84:00:00 | 4hm9-A | 2.1 | 14.9 | 105 | 511 | 5 | MOLECULE: | BETA-CATEN | IN-LIKE | PROTEIN 1 |
| 85:00:00 | 7kbf-K | 2.1 | 3 | 57 | 78 | 7 | MOLECULE: | HISTONE H3 | 0.2 | |
| 86:00:00 | 2is2-B | 2.1 | 26 | 103 | 642 | 11 | MOLECULE: | 33-MER | | |
| 87:00:00 | 3wyo-C | 2.1 | 6.6 | 65 | 153 | 8 | MOLECULE: | MYOGLOBIN | | |
| 88:00:00 | 2dc3-A | 2.1 | 5 | 88 | 172 | 7 | MOLECULE: | CYTOGLOBIN | | |
| 89:00:00 | 8rt9-A | 2 | 23.5 | 104 | 207 | 7 | MOLECULE: | TRWJ PROTE | IN | |
| 90:00:00 | 8afz-B | 2 | 15.9 | 87 | 376 | 5 | MOLECULE: | SORTING NE | XIN-1 | |
| 91:00:00 | 5f5p-A | 2 | 7.3 | 90 | 181 | 13 | MOLECULE: | PROTEIN SH | ROOM2 | |
| 92:00:00 | 2q6h-A | 2 | 7.4 | 110 | 513 | 6 | MOLECULE: | TRANSPORTE | R | |

| | | | | | | | | | | | |
|---|---|---|---|---|---|---|---|---|---|---|---|
| 93:00:00 | 2bte-A | 2 | 19.6 | 122 | 878 | 5 | MOLECULE: | AMINOACYL- | TRNA SYN | THETASE | |
| 94:00:00 | 6teq-A | 2 | 17.5 | 94 | 418 | 10 | MOLECULE: | GALACTOKIN | ASE | | |
| 95:00:00 | 2w2h-A | 2 | 6.1 | 80 | 263 | 4 | MOLECULE: | CYCLIN-T1 | | | |
| 96:00:00 | 7eu3-D | 2 | 8.6 | 105 | 499 | 9 | MOLECULE: | NAD(P)H-QU | INONE OX | IDOREDUCTASE SUBUNIT 1, CHLOROP | |
| 97:00:00 | 6lp3-E | 2 | 11.5 | 63 | 122 | 5 | MOLECULE: | UNCHARACTE | RIZED PR | OTEIN YMR124W | |
| 98:00:00 | 5xa5-A | 2 | 12.2 | 96 | 248 | 9 | MOLECULE: | ALPHA-CATE | NIN-LIKE | PROTEIN HMP-1 | |
| 99:00:00 | 2e0w-B | 2 | 17.1 | 109 | 500 | 6 | MOLECULE: | GAMMA-GLUT | AMYLTRAN | SPEPTIDASE | |
| 0:00 | 3g06-A | 2 | 6.6 | 96 | 601 | 5 | MOLECULE: | SSPH2 (LEU | CINE-RIC | H REPEAT PROTEIN) | |
| 1:00 | 3tmb-B | 2 | 7.8 | 117 | 318 | 5 | MOLECULE: | UNCHARACTE | RIZED PR | OTEIN BD1817 | |
| 2:00 | 5tk6-A | 2 | 5.2 | 99 | 191 | 6 | MOLECULE: | OXSA PROTE | IN | | |
| 3:00 | 8hqs-B | 2 | 14.2 | 82 | 408 | 5 | MOLECULE: | STRUCTURAL | MAINTEN | ANCE OF CHROMOSOMES PROTEIN 5 | |
| 4:00 | 6xp5-H | 2 | 17.4 | 76 | 146 | 1 | MOLECULE: | MEDIATOR O | F RNA PO | LYMERASE II TRANSCRIPTION SUBUN | |
| 5:00 | 4oh3-B | 2 | 8.3 | 104 | 537 | 7 | MOLECULE: | NITRATE TR | ANSPORTE | R 1.1 | |
| 6:00 | 6fln-B | 2 | 34.7 | 97 | 367 | 5 | MOLECULE: | E3 UBIQUIT | IN/ISG15 | LIGASE TRIM25 | |
| 7:00 | 6su3-A | 2 | 10 | 90 | 256 | 10 | MOLECULE: | 48C12 HELI | ORHODOPS | IN | |
| 8:00 | 7jts-s | 2 | 18.4 | 93 | 290 | 8 | MOLECULE: | RADIAL SPO | KE PROTE | IN 3 | |
| 9:00 | 2pih-A | 2 | 20.5 | 70 | 123 | 4 | MOLECULE: | PROTEIN YM | CA | | |
| 10:00 | 8erc-A | 2 | 6.4 | 143 | 441 | 6 | MOLECULE: | LYSOPHOSPH | OLIPID A | CYLTRANSFERASE 7 | |
| 11:00 | 2zqm-A | 2 | 5.9 | 63 | 114 | 5 | MOLECULE: | PREFOLDIN | BETA SUB | UNIT 1 | |
| 12:00 | 1t33-A | 2 | 4.8 | 74 | 220 | 16 | MOLECULE: | PUTATIVE T | RANSCRIP | TIONAL REPRESSOR (TETR/ACRR FAM | |
| 13:00 | 6x58-F | 2 | 8.8 | 67 | 135 | 7 | MOLECULE: | 10E8V4 FAB | HEAVY C | HAIN | |
| 14:00 | 4k6m-A | 2 | 22.4 | 151 | 889 | 3 | MOLECULE: | POLYPROTEI | N | | |
| 15:00 | 8pjn-b | 2 | 15.5 | 85 | 296 | 6 | MOLECULE: | E3 UBIQUIT | IN-PROTE | IN TRANSFERASE RMND5A | |
| 16:00 | 2dla-A | 2 | 9.6 | 79 | 222 | 5 | MOLECULE: | 397AA LONG | HYPOTHE | TICAL PROTEIN | |
| 17:00 | 4q2d-A | 2 | 18 | 128 | 927 | 3 | MOLECULE: | CRISPR-ASS | OCIATED | HELICASE CAS3 | |
| 18:00 | 3go9-A | 2 | 8.9 | 96 | 458 | 7 | MOLECULE: | INSULINASE | FAMILY | PROTEASE | |
| 19:00 | 5b2o-A | 2 | 25.4 | 103 | 1455 | 9 | MOLECULE: | CRISPR-ASS | OCIATED | ENDONUCLEASE CAS9 | |
| 20:00 | 4g9s-A | 2 | 3.8 | 81 | 187 | 7 | MOLECULE: | GOOSE-TYPE | LYSOZYM | E | |
| 21:00 | 7oca-E | 2 | 9.9 | 93 | 158 | 4 | MOLECULE: | GLUTAMATE | RECEPTOR | | 1 |
| 22:00 | 3sjb-D | 2 | 7.4 | 56 | 72 | 4 | MOLECULE: | ATPASE GET | 3 | | |
| 23:00 | 6tc0-C | 2 | 22.5 | 114 | 994 | 11 | MOLECULE: | PROBABLE C | YTOSOLIC | IRON-SULFUR PROTEIN ASSEMBLY P | |
| 24:00:00 | 7o3x-B | 2 | 22.1 | 83 | 215 | 8 | MOLECULE: | PROTEIN SL | L0617 | | |
| 25:00:00 | 8ap8-d | 2 | 16.6 | 113 | 270 | 6 | MOLECULE: | OSCP | | | |
| 26:00:00 | 8hf2-A | 2 | 6.6 | 72 | 120 | 10 | MOLECULE: | PRA1 FAMIL | Y PROTEI | N | |
| 27:00:00 | 3mxz-A | 2 | 4.3 | 58 | 107 | 5 | MOLECULE: | TUBULIN-SP | ECIFIC C | HAPERONE A | |
| 28:00:00 | 2zy4-F | 2 | 25.6 | 108 | 535 | 7 | MOLECULE: | L-ASPARTAT | E BETA-D | ECARBOXYLASE | |
| 29:00:00 | 1cii-A | 2 | 25.1 | 111 | 602 | 8 | MOLECULE: | COLICIN IA | | | |
| 30:00:00 | 8k3b-A | 2 | 11.3 | 111 | 281 | 14 | MOLECULE: | MURR/RPIR | FAMILY T | RANSCRIPTIONAL REGULATOR | |
| 31:00:00 | 8kde-G | 2 | 4.6 | 64 | 145 | 6 | MOLECULE: | PHOTOSYSTE | M II CP4 | 7 REACTION CENTER PROTEIN | |
| 32:00:00 | 7asv-A | 2 | 6.3 | 71 | 155 | 15 | MOLECULE: | DNA-DIRECT | ED RNA P | OLYMERASE III SUBUNIT RPC5 | |
| 33:00:00 | 6h3s-A | 2 | 10.9 | 86 | 408 | 6 | MOLECULE: | ENVELOPMEN | T POLYPR | OTEIN | |
| 34:00:00 | 6u9w-A | 2 | 19.2 | 102 | 562 | 3 | MOLECULE: | P2X PURINO | CEPTOR 7 | | |
| 35:00:00 | 6wq2-A | 2 | 14.5 | 67 | 154 | 6 | MOLECULE: | A-DNA | | | |
| 36:00:00 | 7qoo-Q | 2 | 35 | 105 | 209 | 4 | MOLECULE: | CENTROMERE | PROTEIN | C | |
| 37:00:00 | 2hsb-A | 2 | 4.4 | 68 | 126 | 4 | MOLECULE: | HYPOTHETIC | AL UPF03 | 32 PROTEIN AF0298 | |
| 38:00:00 | 8tfo-A | 2 | 15.9 | 83 | 390 | 8 | MOLECULE: | MEVALONATE | KINASE | | |
| 39:00:00 | 2uxx-B | 2 | 12.7 | 80 | 134 | 11 | MOLECULE: | LYSINE-SPE | CIFIC HI | STONE DEMETHYLASE 1 | |
| 40:00:00 | 6uek-A | 2 | 16.8 | 102 | 660 | 7 | MOLECULE: | UROCANATE | HYDRATAS | E | |
| 41:00:00 | 1tuz-A | 2 | 7.5 | 83 | 118 | 6 | MOLECULE: | DIACYLGLYC | EROL KIN | ASE ALPHA | |
| 42:00:00 | 6xtt-B | 2 | 14.6 | 67 | 116 | 4 | MOLECULE: | NTTA | | | |
| 43:00:00 | 1zvr-A | 2 | 10.7 | 88 | 514 | 7 | MOLECULE: | MYOTUBULAR | IN-RELAT | ED PROTEIN 2 | |
| 44:00:00 | 3l6a-A | 2 | 19 | 85 | 350 | 8 | MOLECULE: | EUKARYOTIC | TRANSLA | TION INITIATION FACTOR 4 GAMMA | |
| 45:00:00 | 6zie-A | 2 | 9.6 | 83 | 123 | 5 | MOLECULE: | CMPX-383B | | | |
| 46:00:00 | 5n0c-A | 2 | 26.7 | 115 | 1293 | 6 | MOLECULE: | TETANUS TO | XIN | | |

| | | | | | | | | | | | |
|---|---|---|---|---|---|---|---|---|---|---|---|
| 47:00:00 | 4ysm-A | 2 | 15.6 | 87 | 475 | 6 | MOLECULE: | CALMODULIN | #NAME? | MAIN PROTEIN KINASE | |
| 48:00:00 | 2a3v-B | 2 | 15.3 | 71 | 320 | 11 | MOLECULE: | DNA (31-ME | R) | | |
| 49:00:00 | 8a3t-Q | 2 | 33.6 | 117 | 623 | 3 | MOLECULE: | ANAPHASE-P | ROMOTING | COMPLEX SUBUNIT CDC27 | |
| 50:00:00 | 2c5z-A | 2 | 3.5 | 55 | 93 | 4 | MOLECULE: | SET DOMAIN | PROTEIN | | 2 |
| 51:00:00 | 7t6d-A | 2 | 21.2 | 112 | 389 | 6 | MOLECULE: | LIPOPOLYSA | CCHARIDE | ASSEMBLY PROTEIN B | |
| 52:00:00 | 1fx2-A | 2 | 4.3 | 64 | 235 | 6 | MOLECULE: | RECEPTOR-T | YPE ADEN | YLATE CYCLASE GRESAG 4.1 | |
| 53:00:00 | 5cx6-B | 2 | 28.4 | 150 | 665 | 3 | MOLECULE: | RNA-DEPEND | ENT RNA | POLYMERASE | |
| 54:00:00 | 4u7u-B | 2 | 5.1 | 69 | 157 | 7 | MOLECULE: | CRISPR SYS | TEM CASC | ADE SUBUNIT CASA | |
| 55:00:00 | 7ubn-Q | 2 | 4.7 | 88 | 207 | 9 | MOLECULE: | DNA (53-ME | R) | | |
| 56:00:00 | 8p0w-A | 2 | 16.4 | 84 | 190 | 7 | MOLECULE: | COMM DOMAI | N-CONTAI | NING PROTEIN 1 | |
| 57:00:00 | 6maf-A | 2 | 14.2 | 94 | 259 | 4 | MOLECULE: | BBVCI ENDO | NUCLEASE | SUBUNIT 1 | |
| 58:00:00 | 6x80-C | 2 | 6.8 | 59 | 574 | 3 | MOLECULE: | FLAGELLIN | A | | |
| 59:00:00 | 8k0b-A | 2 | 18.2 | 89 | 689 | 6 | MOLECULE: | CALCIUM PE | RMEABLE | STRESS-GATED CATION CHANNEL 1 | |
| 60:00:00 | 7s01-A | 2 | 10.4 | 82 | 463 | 1 | MOLECULE: | DNA-DIRECT | ED RNA P | OLYMERASE SUBUNIT | |
| 61:00:00 | 6icz-A | 2 | 22.2 | 127 | 2253 | 6 | MOLECULE: | PROTEIN MA | GO NASHI | HOMOLOG 2 | |
| 62:00:00 | 5iz5-A | 2 | 18.3 | 99 | 757 | 10 | MOLECULE: | CYTOSOLIC | PHOSPHOL | IPASE A2 DELTA | |
| 63:00:00 | 7qj2-G | 2 | 13.9 | 110 | 640 | 7 | MOLECULE: | GAMMA-TUBU | LIN COMP | LEX COMPONENT 5 | |
| 64:00:00 | 2jes-A | 2 | 13.1 | 64 | 370 | 5 | MOLECULE: | PORTAL PRO | TEIN | | |
| 65:00:00 | 1eyy-A | 2 | 6.7 | 119 | 504 | 6 | MOLECULE: | ALDEHYDE D | EHYDROGE | NASE | |
| 66:00:00 | 7t5q-I | 2 | 8.1 | 70 | 283 | 7 | MOLECULE: | ACTIN-RELA | TED PROT | EIN 3 | |
| 67:00:00 | 6q82-A | 2 | 24.7 | 124 | 1027 | 7 | MOLECULE: | IMPORTIN B | ETA-LIKE | PROTEIN KAP122 | |
| 68:00:00 | 5eom-C | 2 | 14.8 | 75 | 353 | 8 | MOLECULE: | PROTEIN MA | B-21-LIK | E 1 | |
| 69:00:00 | 6uxu-A | 2 | 3.6 | 63 | 300 | 10 | MOLECULE: | CHLOROTHAL | ONIL HYD | ROLYTIC DEHALOGENASE | |
| 70:00:00 | 2kj5-A | 2 | 3.5 | 74 | 116 | 8 | MOLECULE: | PHAGE INTE | GRASE | | |
| 71:00:00 | 6kp3-B | 2 | 4.1 | 66 | 112 | 9 | MOLECULE: | PROGRAMMED | CELL DE | ATH 6-INTERACTING PROTEIN | |
| 72:00:00 | 7ml0-1 | 2 | 23.5 | 81 | 367 | 9 | MOLECULE: | DNA-DIRECT | ED RNA P | OLYMERASE SUBUNIT | |
| 73:00:00 | 6s18-A | 2 | 10.2 | 80 | 143 | 8 | MOLECULE: | AROMATIC A | CID CHEM | ORECEPTOR | |
| 74:00:00 | 6hty-B | 2 | 8.1 | 98 | 302 | 5 | MOLECULE: | NUCLEAR RE | CEPTOR S | UBFAMILY 1 GROUP I MEMBER 2,NUC | |
| 75:00:00 | 7vrc-C | 2 | 7.8 | 73 | 109 | 8 | MOLECULE: | TRANSCRIPT | ION REGU | LATORY PROTEIN SNF11 | |
| 76:00:00 | 6swy-8 | 2 | 6.3 | 88 | 325 | 7 | MOLECULE: | VACUOLAR I | MPORT AN | D DEGRADATION PROTEIN 28 | |
| 77:00:00 | 6tdy-M | 2 | 8.2 | 91 | 243 | 5 | MOLECULE: | ATP SYNTHA | SE SUBUN | IT ALPHA | |
| 78:00:00 | 3add-A | 2 | 12.2 | 74 | 251 | 7 | MOLECULE: | L-SERYL-TR | NA(SEC) | KINASE | |
| 79:00:00 | 3veb-B | 2 | 10.2 | 91 | 150 | 2 | MOLECULE: | MACRODOMAI | N TER PR | OTEIN | |
| 80:00:00 | 6pw5-B | 2 | 18.1 | 128 | 782 | 7 | MOLECULE: | TRP-LIKE I | ON CHANN | EL | |
| 81:00:00 | 4faa-A | 2 | 18.2 | 119 | 535 | 4 | MOLECULE: | CYTOCHROME | C OXIDA | SE SUBUNIT 1 | |
| 82:00:00 | 6iqc-A | 2 | 7.6 | 77 | 103 | 10 | MOLECULE: | DNA-BINDIN | G PROTEI | N SSO0352 | |
| 83:00:00 | 6ulo-A | 2 | 4.1 | 66 | 317 | 6 | MOLECULE: | UNCHARACTE | RIZED PR | OTEIN | |
| 84:00:00 | 2rre-A | 2 | 3 | 55 | 74 | 4 | MOLECULE: | PUTATIVE U | NCHARACT | ERIZED PROTEIN | |
| 85:00:00 | 4kyw-A | 2 | 12.8 | 70 | 254 | 7 | MOLECULE: | TYPE-2 RES | TRICTION | ENZYME DPNI | |
| 86:00:00 | 5b6e-A | 2 | 14.1 | 77 | 319 | 6 | MOLECULE: | CMP 5-HYDR | OXYMETHY | LASE | |
| 87:00:00 | 6ks0-A | 2 | 14.7 | 80 | 286 | 9 | MOLECULE: | ADIPONECTI | N RECEPT | OR PROTEIN 1 | |
| 88:00:00 | 7ktr-D | 2 | 22.2 | 94 | 219 | 5 | MOLECULE: | TRANSFORMA | TION/TRA | NSCRIPTION DOMAIN-ASSOCIATED PR | |
| 89:00:00 | 4u7b-A | 2 | 6.3 | 68 | 342 | 6 | MOLECULE: | DNA (25-ME | R) | | |
| 90:00:00 | 3lf9-A | 2 | 7.3 | 74 | 120 | 7 | MOLECULE: | 4E10_D0_1I | S1A_001_ | C (T161) | |
| 91:00:00 | 8t1l-V | 2 | 14.1 | 96 | 278 | 6 | MOLECULE: | MEDIATOR O | F RNA PO | LYMERASE II TRANSCRIPTION SUBUN | |
| 92:00:00 | 6n04-B | 2 | 9.9 | 70 | 366 | 10 | MOLECULE: | ABSH3 | | | |
| 93:00:00 | 4nwp-D | 2 | 12.4 | 87 | 149 | 7 | MOLECULE: | PUTATIVE U | NCHARACT | ERIZED PROTEIN | |
| 94:00:00 | 7qog-C | 2 | 4 | 60 | 225 | 10 | MOLECULE: | PORTAL PRO | TEIN GP2 | | 0 |
| 95:00:00 | 3zqg-A | 2 | 14.1 | 88 | 208 | 7 | MOLECULE: | TETRACYCLI | NE REPRE | SSOR PROTEIN CLASS B FROM TRANS | |
| 96:00:00 | 5xaz-A | 2 | 5.5 | 63 | 223 | 6 | MOLECULE: | GAMMA-BUTY | ROLACTON | E RECEPTOR PROTEIN | |
| 97:00:00 | 7m2w-F | 2 | 16.3 | 89 | 674 | 6 | MOLECULE: | TUBULIN GA | MMA CHAI | N | |
| 98:00:00 | 3qhq-A | 2 | 26.1 | 84 | 222 | 4 | MOLECULE: | SAG0897 FA | MILY CRI | SPR-ASSOCIATED PROTEIN | |
| 99:00:00 | 8h5s-A | 2 | 3.7 | 75 | 247 | 4 | MOLECULE: | BETA-PHOSP | HOGLUCOM | UTASE | |
| 0:00 | 3zni-A | 2 | 23.5 | 95 | 390 | 7 | MOLECULE: | E3 UBIQUIT | IN-PROTE | IN LIGASE CBL-B | |

| | | | | | | | | | | |
|---|---|---|---|---|---|---|---|---|---|---|
| 1:00 | 7a6h-O | 2 | 7.7 | 70 | 512 | 6 | MOLECULE: | DNA-DIRECT | ED RNA P | OLYMERASE III SUBUNIT RPC1 |
| 2:00 | 6vzz-A | 2 | 14.3 | 96 | 374 | 10 | MOLECULE: | BAMAA.1990 | 0.A | |
| 3:00 | 1i5p-A | 2 | 27.1 | 126 | 633 | 10 | MOLECULE: | PESTICIDIA | L CRYSTA | L PROTEIN CRY2AA |
| 4:00 | 8agy-A | 2 | 11.4 | 93 | 346 | 6 | MOLECULE: | CORRAMYCIN | PHOSPHO | TRANSFERASE |
| 5:00 | 4mk6-A | 2 | 11.8 | 73 | 188 | 5 | MOLECULE: | PROBABLE D | IHYDROXY | ACETONE KINASE REGULATOR DHSK_R |
| 6:00 | 6idf-B | 2 | 4.2 | 94 | 312 | 6 | MOLECULE: | NICASTRIN | | |
| 7:00 | 4mi2-B | 2 | 9.8 | 64 | 261 | 16 | MOLECULE: | PUTATIVE E | NOYL-COA | HYDRATASE/ISOMERASE |
| 8:00 | 7y82-A | 2 | 24.1 | 109 | 1341 | 2 | MOLECULE: | RAMP SUPER | FAMILY P | ROTEIN |
| 9:00 | 7mrw-A | 2 | 19.1 | 125 | 1209 | 6 | MOLECULE: | CYTOADHERE | NCE LINK | ED ASEXUAL PROTEIN 3.1 |
| 10:00 | 5img-A | 2 | 9.6 | 75 | 467 | 5 | MOLECULE: | DIPEPTIDAS | E | |
| 11:00 | 5lpc-A | 2 | 17.5 | 86 | 624 | 8 | MOLECULE: | VANADIUM-D | EPENDENT | BROMOPEROXIDASE |
| 12:00 | 1jr5-A | 2 | 3.6 | 61 | 90 | 7 | MOLECULE: | 10 KDA ANT | I-SIGMA | FACTOR |
| 13:00 | 4qfh-B | 2 | 6 | 92 | 611 | 9 | MOLECULE: | GLUCOSE-6- | PHOSPHAT | E ISOMERASE |
| 14:00 | 1qha-A | 2 | 16.8 | 129 | 903 | 3 | MOLECULE: | PROTEIN (H | EXOKINAS | E) |
| 15:00 | 3wyo-D | 2 | 8 | 76 | 153 | 7 | MOLECULE: | MYOGLOBIN | | |
| 16:00 | 4zs8-B | 2 | 4.2 | 64 | 231 | 9 | MOLECULE: | HTH-TYPE T | RANSCRIP | TIONAL REPRESSOR DASR |

**DALI results: MnA-1 Hexon**

| No: | Chain | Z | rmsd | lali | nres | %id | PDB | Description |
|---|---|---|---|---|---|---|---|---|
| 1:00 | 8rbt-A | 17.3 | 4.4 | 333 | 481 | 10 | MOLECULE: | MAJOR CAPSID PROTEIN |
| 2:00 | 5tip-A | 16.2 | 3.6 | 318 | 436 | 10 | MOLECULE: | MAJOR CAPSID PROTEIN |
| 3:00 | 6g43-A | 15.6 | 4.3 | 352 | 515 | 6 | MOLECULE: | PUTATIVE MAJOR CAPSID PROTEIN |
| 4:00 | 3j26-A | 14.9 | 3.9 | 360 | 508 | 9 | MOLECULE: | CAPSID PROTEIN V20 |
| 5:00 | 3sam-B | 14.8 | 4.4 | 311 | 544 | 9 | MOLECULE: | RIFAMPICIN RESISTANCE PROTEIN |
| 6:00 | 7yjl-A | 12.4 | 3.9 | 329 | 459 | 6 | MOLECULE: | MAJOR CAPSID PROTEIN |
| 7:00 | 6l2t-C | 11.7 | 4.7 | 338 | 633 | 7 | MOLECULE: | B646L,MAJOR CAPSID PROTEIN |
| 8:00 | 2iny-A | 11.3 | 4.7 | 400 | 941 | 7 | MOLECULE: | HEXON PROTEIN |
| 9:00 | 6q5u-C | 10.1 | 4.8 | 293 | 394 | 6 | MOLECULE: | MAJOR CAPSID PROTEIN (P3) |
| 10:00 | 2vvf-A | 7.6 | 4.3 | 205 | 269 | 6 | MOLECULE: | MAJOR CAPSID PROTEIN P2 |
| 11:00 | 5oac-A | 7.1 | 6.8 | 206 | 310 | 11 | MOLECULE: | MAJOR CAPSID PROTEIN |
| 12:00 | 3j31-A | 7 | 4.7 | 199 | 344 | 9 | MOLECULE: | A223 PENTON BASE |
| 13:00 | 6mw4-A | 6.6 | 5.5 | 158 | 551 | 6 | MOLECULE: | PUTATIVE GERMINATION-SPECIFIC PROTEASE |
| 14:00 | 7zzz-A | 6.1 | 4.3 | 161 | 239 | 11 | MOLECULE: | MAJOR CAPSID PROTEIN P5 |
| 15:00 | 4kh9-A | 5.4 | 17.6 | 123 | 361 | 8 | MOLECULE: | HYPOTHETICAL PROTEIN |
| 16:00 | 6wpx-B | 5.2 | 8.8 | 118 | 451 | 8 | MOLECULE: | BLEST2 |
| 17:00 | 4aqs-A | 4.9 | 11.8 | 140 | 453 | 10 | MOLECULE: | LAMININ SUBUNIT BETA-1 |
| 18:00 | 7drt-B | 4.5 | 10.2 | 136 | 496 | 5 | MOLECULE: | PROTEIN WNT-3A |
| 19:00 | 8e2f-A | 4.3 | 10.4 | 141 | 771 | 9 | MOLECULE: | BACULOVIRAL IAP REPEAT-CONTAINING PROTEIN 6 |
| 20:00 | 7ula-A | 3.9 | 11 | 123 | 430 | 7 | MOLECULE: | ALGINATE BIOSYNTHESIS PROTEIN ALGK |
| 21:00 | 2w0c-L | 3.8 | 9 | 87 | 335 | 7 | MOLECULE: | MAJOR CAPSID PROTEIN P2 |
| 22:00 | 8vhx-H | 3.4 | 15.3 | 101 | 375 | 5 | MOLECULE: | NECK 1 |
| 23:00 | 8beg-A | 3.4 | 3.3 | 86 | 650 | 8 | MOLECULE: | PRGB |
| 24:00:00 | 8a3t-A | 3.1 | 8 | 110 | 219 | 5 | MOLECULE: | ANAPHASE-PROMOTING COMPLEX SUBUNIT CDC27 |
| 25:00:00 | 1wky-A | 3 | 20.5 | 99 | 446 | 4 | MOLECULE: | ENDO-BETA-1,4-MANNANASE |
| 26:00:00 | 8x7n-A | 3 | 3.6 | 84 | 118 | 12 | MOLECULE: | CONJUGATIVE TRANSFER: AGGREGATE STABILITY |
| 27:00:00 | 7zqp-b | 3 | 14.3 | 113 | 719 | 5 | MOLECULE: | PROBABLE BASEPLATE HUB PROTEIN |
| 28:00:00 | 9b8i-C | 2.9 | 18 | 121 | 719 | 6 | MOLECULE: | CELLULOSE SYNTHASE BCSB-BCSC FUSION PROTEIN |
| 29:00:00 | 7wrk-A | 2.9 | 4.3 | 100 | 160 | 8 | MOLECULE: | HYPOTHETICAL PROTEIN TTHA1873 |
| 30:00:00 | 4pxq-A | 2.8 | 15 | 119 | 516 | 9 | MOLECULE: | D-GLUCURONYL C5 EPIMERASE B |

**DALI results: MnA-1 Adenain**

| No: | Chain | Z | rmsd | lali | nres | %id | PDB Description | |
|---|---|---|---|---|---|---|---|---|
| 1:00 | 4pis-A | 13.6 | 2.9 | 148 | 204 | 15 | MOLECULE: | PROTEASE |
| 2:00 | 4wx4-A | 13.5 | 2.7 | 146 | 201 | 14 | MOLECULE: | PROTEASE |
| 3:00 | 4ekf-A | 13.5 | 2.9 | 147 | 192 | 14 | MOLECULE: | ADENAIN |
| 4:00 | 4wx7-C | 13.4 | 2.8 | 147 | 204 | 16 | MOLECULE: | PROTEASE |
| 5:00 | 4pie-A | 13.4 | 2.7 | 147 | 202 | 14 | MOLECULE: | PROTEASE |
| 6:00 | 4wx6-A | 13.4 | 2.6 | 146 | 204 | 14 | MOLECULE: | PROTEASE |
| 7:00 | 1nln-A | 13.4 | 2.8 | 147 | 203 | 14 | MOLECULE: | ADENAIN |
| 8:00 | 4piq-A | 13.4 | 2.8 | 147 | 204 | 16 | MOLECULE: | PROTEASE |
| 9:00 | 4wx7-A | 13.3 | 2.7 | 146 | 204 | 14 | MOLECULE: | PROTEASE |
| 10:00 | 4pid-A | 13.3 | 2.8 | 147 | 204 | 14 | MOLECULE: | PROTEASE |
| 11:00 | 4wx6-C | 13.2 | 2.7 | 146 | 204 | 13 | MOLECULE: | PROTEASE |
| 12:00 | 5fgy-A | 13.1 | 2.8 | 145 | 203 | 14 | MOLECULE: | PROTEASE |
| 13:00 | 1avp-A | 12.9 | 2.9 | 147 | 204 | 14 | MOLECULE: | ADENOVIRAL PROTEINASE |
| 14:00 | 7fiu-A | 12.3 | 3.9 | 166 | 270 | 13 | MOLECULE: | ULP_PROTEASE DOMAIN-CONTAINING |
| 15:00 | 6ups-A | 11.9 | 4.1 | 150 | 245 | 11 | MOLECULE: | ULP_PROTEASE DOMAIN-CONTAINING |
| 16:00 | 8efx-A | 11.3 | 4.3 | 149 | 259 | 11 | MOLECULE: | OTDUB |
| 17:00 | 6upu-E | 11.1 | 3.8 | 151 | 258 | 11 | MOLECULE: | ULP_PROTEASE DOMAIN-CONTAINING |
| 18:00 | 6upu-M | 11.1 | 3.9 | 153 | 258 | 12 | MOLECULE: | ULP_PROTEASE DOMAIN-CONTAINING |
| 19:00 | 6upu-I | 11 | 3.8 | 153 | 258 | 12 | MOLECULE: | ULP_PROTEASE DOMAIN-CONTAINING |
| 20:00 | 6upu-A | 11 | 3.8 | 152 | 259 | 12 | MOLECULE: | ULP_PROTEASE DOMAIN-CONTAINING |
| 21:00 | 5b5q-B | 10.9 | 3.4 | 137 | 244 | 15 | MOLECULE: | MEMBRANE THIOL PROTEASE |
| 22:00 | 5ubw-B | 10.9 | 3.1 | 145 | 173 | 10 | MOLECULE: | DEUBIQUITINASE SSEL |
| 23:00 | 5ubw-A | 10.8 | 3.1 | 144 | 175 | 10 | MOLECULE: | DEUBIQUITINASE SSEL |
| 24:00:00 | 5cra-A | 10.7 | 3.7 | 148 | 172 | 11 | MOLECULE: | SDEA |
| 25:00:00 | 5cra-B | 10.7 | 3.7 | 148 | 172 | 11 | MOLECULE: | SDEA |
| 26:00:00 | 5haf-A | 10.6 | 4.3 | 162 | 312 | 10 | MOLECULE: | DEUBIQUITINASE SSEL |
| 27:00:00 | 6wtg-A | 10.4 | 3.6 | 146 | 169 | 10 | MOLECULE: | UBIQUITINATING/DEUBIQUITINATIN |
| 28:00:00 | 6be0-A | 10.3 | 3.9 | 151 | 266 | 12 | MOLECULE: | AVRA |
| 29:00:00 | 1xt9-A | 10.2 | 3.8 | 144 | 208 | 13 | MOLECULE: | SENTRIN-SPECIFIC PROTEASE 8 |
| 30:00:00 | 5haf-B | 10.1 | 4.7 | 164 | 327 | 10 | MOLECULE: | DEUBIQUITINASE SSEL |
| 31:00:00 | 6fdu-A | 10.1 | 3.2 | 138 | 240 | 14 | MOLECULE: | DEUBIQUITINASE AND DENEDDYLASE |
| 32:00:00 | 5ham-A | 10.1 | 3 | 136 | 260 | 11 | MOLECULE: | RICKCE |
| 33:00:00 | 6fdq-A | 10 | 3.3 | 138 | 239 | 14 | MOLECULE: | DEUBIQUITINASE AND DENEDDYLASE |
| 34:00:00 | 6fdu-B | 10 | 3.3 | 138 | 240 | 15 | MOLECULE: | DEUBIQUITINASE AND DENEDDYLASE |
| 35:00:00 | 5b5q-A | 10 | 3.3 | 139 | 243 | 14 | MOLECULE: | MEMBRANE THIOL PROTEASE |
| 36:00:00 | 2bkr-A | 10 | 3.4 | 139 | 211 | 13 | MOLECULE: | SENTRIN-SPECIFIC PROTEASE 8 |
| 37:00:00 | 2bkq-A | 10 | 4 | 147 | 210 | 12 | MOLECULE: | SENTRIN-SPECIFIC PROTEASE 8 |
| 38:00:00 | 6fdk-A | 9.9 | 3.2 | 136 | 242 | 15 | MOLECULE: | DEUBIQUITINASE AND DENEDDYLASE |
| 39:00:00 | 6fdq-B | 9.9 | 3.3 | 140 | 247 | 14 | MOLECULE: | DEUBIQUITINASE AND DENEDDYLASE |
| 40:00:00 | 6gzs-A | 9.8 | 3.4 | 140 | 250 | 14 | MOLECULE: | DEUBIQUITINASE AND DENEDDYLASE |
| 41:00:00 | 5crb-B | 9.8 | 3.7 | 151 | 191 | 11 | MOLECULE: | SDEA |
| 42:00:00 | 6lj9-A | 9.8 | 8.4 | 143 | 272 | 13 | MOLECULE: | CYSTEINE PROTEASE S273R |

| | | | | | | | | |
|---|---|---|---|---|---|---|---|---|
| 43:00:00 | 6ljb-A | 9.7 | 5.7 | 139 | 267 | 14 | MOLECULE: | CYSTEINE PROTEASE S273R |
| 44:00:00 | 2bkq-C | 9.7 | 3.6 | 141 | 207 | 12 | MOLECULE: | SENTRIN-SPECIFIC PROTEASE 8 |
| 45:00:00 | 8oi3-A | 9.7 | 3 | 126 | 183 | 12 | MOLECULE: | TYPE III EFFECTOR |
| 46:00:00 | 5crb-A | 9.7 | 3.7 | 149 | 192 | 11 | MOLECULE: | SDEA |
| 47:00:00 | 6oam-B | 9.7 | 3.3 | 142 | 239 | 17 | MOLECULE: | DEUBIQUITINASE AND DENEDDYLASE |
| 48:00:00 | 6lj9-B | 9.7 | 6.5 | 137 | 259 | 15 | MOLECULE: | CYSTEINE PROTEASE S273R |
| 49:00:00 | 8oi3-B | 9.7 | 3 | 124 | 183 | 10 | MOLECULE: | TYPE III EFFECTOR |
| 50:00:00 | 5hag-A | 9.6 | 4.4 | 149 | 266 | 15 | MOLECULE: | DEUBIQUITINASE AND DENEDDYLASE |
| 51:00:00 | 5ham-B | 9.6 | 2.8 | 128 | 260 | 13 | MOLECULE: | RICKCE |
| 52:00:00 | 8efw-B | 9.6 | 4.1 | 152 | 183 | 11 | MOLECULE: | SDEA |
| 53:00:00 | 6oam-A | 9.6 | 3.3 | 143 | 239 | 17 | MOLECULE: | DEUBIQUITINASE AND DENEDDYLASE |
| 54:00:00 | 6gzu-A | 9.6 | 2.9 | 130 | 241 | 14 | MOLECULE: | CONSERVED MEMBRANE PROTEIN |
| 55:00:00 | 2bkq-B | 9.6 | 4 | 143 | 208 | 12 | MOLECULE: | SENTRIN-SPECIFIC PROTEASE 8 |
| 56:00:00 | 6gzt-A | 9.5 | 4.2 | 147 | 267 | 14 | MOLECULE: | DEUBIQUITINASE AND DENEDDYLASE |
| 57:00:00 | 5aek-A | 9.4 | 3.5 | 128 | 223 | 8 | MOLECULE: | SENTRIN-SPECIFIC PROTEASE 2 |
| 58:00:00 | 6mrn-A | 9.4 | 3.3 | 139 | 240 | 17 | MOLECULE: | DEUBIQUITINASE AND DENEDDYLASE |
| 59:00:00 | 2oix-A | 9.4 | 2.9 | 122 | 178 | 11 | MOLECULE: | XANTHOMONAS OUTER PROTEIN D |
| 60:00:00 | 5jp3-E | 9.3 | 3.2 | 134 | 193 | 12 | MOLECULE: | XANTHOMONAS OUTER PROTEIN D |
| 61:00:00 | 5jp1-A | 9.3 | 3 | 127 | 199 | 11 | MOLECULE: | XANTHOMONAS OUTER PROTEIN D |
| 62:00:00 | 5jp3-C | 9.3 | 3.3 | 132 | 196 | 11 | MOLECULE: | XANTHOMONAS OUTER PROTEIN D |
| 63:00:00 | 5klp-A | 9.2 | 4.6 | 159 | 309 | 11 | MOLECULE: | ORF34 |
| 64:00:00 | 5klq-C | 9.1 | 4.4 | 158 | 320 | 11 | MOLECULE: | ORF34 |
| 65:00:00 | 5jp3-G | 9.1 | 3.5 | 136 | 196 | 12 | MOLECULE: | XANTHOMONAS OUTER PROTEIN D |
| 66:00:00 | 5klp-C | 9.1 | 4.6 | 154 | 315 | 10 | MOLECULE: | ORF34 |
| 67:00:00 | 5klq-B | 9.1 | 4.4 | 160 | 320 | 13 | MOLECULE: | ORF34 |
| 68:00:00 | 5klp-B | 9 | 4.3 | 158 | 310 | 10 | MOLECULE: | ORF34 |
| 69:00:00 | 5klq-A | 9 | 4.4 | 158 | 324 | 11 | MOLECULE: | ORF34 |
| 70:00:00 | 5aek-M | 9 | 3.9 | 131 | 222 | 8 | MOLECULE: | SENTRIN-SPECIFIC PROTEASE 2 |
| 71:00:00 | 2bkq-D | 8.9 | 4 | 145 | 209 | 10 | MOLECULE: | SENTRIN-SPECIFIC PROTEASE 8 |
| 72:00:00 | 5jp3-A | 8.8 | 4.9 | 142 | 205 | 11 | MOLECULE: | XANTHOMONAS OUTER PROTEIN D |
| 73:00:00 | 2g4d-C | 8.8 | 3.7 | 126 | 205 | 10 | MOLECULE: | SENP1 PROTEIN |
| 74:00:00 | 2g4d-A | 8.8 | 3.7 | 127 | 205 | 9 | MOLECULE: | SENP1 PROTEIN |
| 75:00:00 | 1th0-B | 8.8 | 3.8 | 129 | 223 | 9 | MOLECULE: | SENTRIN-SPECIFIC PROTEASE 2 |
| 76:00:00 | 5dgg-B | 8.8 | 3.6 | 130 | 176 | 13 | MOLECULE: | UNCHARACTERIZED PROTEIN |
| 77:00:00 | 5dgg-A | 8.7 | 3.1 | 125 | 176 | 17 | MOLECULE: | UNCHARACTERIZED PROTEIN |
| 78:00:00 | 5aek-K | 8.7 | 3.7 | 128 | 223 | 9 | MOLECULE: | SENTRIN-SPECIFIC PROTEASE 2 |
| 79:00:00 | 2oiv-A | 8.7 | 2.9 | 117 | 170 | 11 | MOLECULE: | XANTHOMONAS OUTER PROTEIN D |
| 80:00:00 | 5aek-U | 8.6 | 3.5 | 128 | 223 | 8 | MOLECULE: | SENTRIN-SPECIFIC PROTEASE 2 |
| 81:00:00 | 2hl9-A | 8.6 | 3.9 | 129 | 218 | 12 | MOLECULE: | UBIQUITIN-LIKE-SPECIFIC PROTEA |
| 82:00:00 | 2io1-C | 8.6 | 3.5 | 129 | 224 | 9 | MOLECULE: | SENTRIN-SPECIFIC PROTEASE 2 |
| 83:00:00 | 5aek-S | 8.6 | 4.1 | 131 | 224 | 8 | MOLECULE: | SENTRIN-SPECIFIC PROTEASE 2 |
| 84:00:00 | 2iy1-C | 8.6 | 4 | 133 | 226 | 10 | MOLECULE: | SENTRIN-SPECIFIC PROTEASE 1 |
| 85:00:00 | 2xph-A | 8.6 | 3.9 | 137 | 227 | 11 | MOLECULE: | SENTRIN-SPECIFIC PROTEASE 1 |
| 86:00:00 | 5aek-G | 8.6 | 4 | 131 | 223 | 8 | MOLECULE: | SENTRIN-SPECIFIC PROTEASE 2 |

| | | | | | | | |
|---|---|---|---|---|---|---|---|
| 87:00:00 | 2hl8-A | 8.5 | 3.7 | 129 | 219 | 11 | MOLECULE: UBIQUITIN-LIKE-SPECIFIC PROTEA |
| 88:00:00 | 5aek-C | 8.5 | 3.7 | 127 | 224 | 8 | MOLECULE: SENTRIN-SPECIFIC PROTEASE 2 |
| 89:00:00 | 2hkp-A | 8.5 | 3.7 | 129 | 219 | 10 | MOLECULE: UBIQUITIN-LIKE-SPECIFIC PROTEA |
| 90:00:00 | 6nnq-A | 8.5 | 3.8 | 132 | 224 | 11 | MOLECULE: SENTRIN-SPECIFIC PROTEASE 1 |
| 91:00:00 | 1euv-A | 8.5 | 4 | 132 | 221 | 11 | MOLECULE: ULP1 PROTEASE |
| 92:00:00 | 5aek-I | 8.5 | 3.9 | 132 | 223 | 8 | MOLECULE: SENTRIN-SPECIFIC PROTEASE 2 |
| 93:00:00 | 2ckg-B | 8.5 | 4 | 135 | 225 | 11 | MOLECULE: SENTRIN-SPECIFIC PROTEASE 1 |
| 94:00:00 | 1th0-A | 8.4 | 3.7 | 132 | 226 | 8 | MOLECULE: SENTRIN-SPECIFIC PROTEASE 2 |
| 95:00:00 | 1tgz-A | 8.4 | 3.5 | 128 | 224 | 9 | MOLECULE: SENTRIN-SPECIFIC PROTEASE 2 |
| 96:00:00 | 2ckh-A | 8.4 | 4.3 | 135 | 225 | 10 | MOLECULE: SENTRIN-SPECIFIC PROTEASE 1 |
| 97:00:00 | 5aek-E | 8.4 | 3.5 | 130 | 224 | 8 | MOLECULE: SENTRIN-SPECIFIC PROTEASE 2 |
| 98:00:00 | 2iy1-A | 8.4 | 3.8 | 131 | 226 | 9 | MOLECULE: SENTRIN-SPECIFIC PROTEASE 1 |
| 99:00:00 | 7r2e-A | 8.4 | 3.9 | 142 | 244 | 9 | MOLECULE: SENTRIN-SPECIFIC PROTEASE 7 |
| 0:00 | 2iy0-A | 8.4 | 4 | 132 | 226 | 9 | MOLECULE: SENTRIN-SPECIFIC PROTEASE 1 |
| 1:00 | 5w40-D | 8.4 | 5.5 | 151 | 327 | 12 | MOLECULE: POPP2 PROTEIN |
| 2:00 | 2iyd-A | 8.4 | 3.8 | 135 | 226 | 10 | MOLECULE: SENTRIN-SPECIFIC PROTEASE 1 |
| 3:00 | 2iyc-A | 8.3 | 3.9 | 133 | 226 | 10 | MOLECULE: SENTRIN-SPECIFIC PROTEASE 1 |
| 4:00 | 5aek-O | 8.3 | 3.6 | 128 | 223 | 8 | MOLECULE: SENTRIN-SPECIFIC PROTEASE 2 |
| 5:00 | 5w3y-C | 8.3 | 5 | 152 | 335 | 9 | MOLECULE: POPP2 PROTEIN |
| 6:00 | 5aek-W | 8.3 | 3.5 | 128 | 222 | 8 | MOLECULE: SENTRIN-SPECIFIC PROTEASE 2 |
| 7:00 | 2iyc-B | 8.3 | 3.9 | 133 | 226 | 11 | MOLECULE: SENTRIN-SPECIFIC PROTEASE 1 |
| 8:00 | 7r2e-B | 8.3 | 3.9 | 140 | 243 | 10 | MOLECULE: SENTRIN-SPECIFIC PROTEASE 7 |
| 9:00 | 2ckg-A | 8.3 | 4.1 | 133 | 225 | 9 | MOLECULE: SENTRIN-SPECIFIC PROTEASE 1 |
| 10:00 | 2xre-A | 8.3 | 4.1 | 134 | 227 | 11 | MOLECULE: SENTRIN-SPECIFIC PROTEASE 1 |
| 11:00 | 3eay-A | 8.3 | 4.1 | 142 | 247 | 10 | MOLECULE: SENTRIN-SPECIFIC PROTEASE 7 |
| 12:00 | 5w3t-B | 8.2 | 5.4 | 155 | 333 | 11 | MOLECULE: POPP2 PROTEIN |
| 13:00 | 5w40-B | 8.2 | 5.4 | 155 | 333 | 10 | MOLECULE: POPP2 PROTEIN |
| 14:00 | 5lnb-B | 8.2 | 6.7 | 156 | 259 | 12 | MOLECULE: UBIQUITIN-LIKE-SPECIFIC PROTEA |
| 15:00 | 2xph-B | 8.1 | 3.7 | 129 | 226 | 9 | MOLECULE: SENTRIN-SPECIFIC PROTEASE 1 |
| 16:00 | 2io1-A | 8.1 | 3.5 | 128 | 223 | 8 | MOLECULE: SENTRIN-SPECIFIC PROTEASE 2 |
| 17:00 | 2io1-E | 8.1 | 3.5 | 128 | 223 | 8 | MOLECULE: SENTRIN-SPECIFIC PROTEASE 2 |
| 18:00 | 6dg4-A | 8 | 3.9 | 137 | 253 | 9 | MOLECULE: ULP1-LIKE SUMO PROTEASE |
| 19:00 | 3zo5-A | 8 | 3.6 | 131 | 230 | 8 | MOLECULE: SENTRIN-SPECIFIC PROTEASE 2 |
| 20:00 | 2io2-A | 8 | 3.8 | 131 | 225 | 8 | MOLECULE: SENTRIN-SPECIFIC PROTEASE 2 |
| 21:00 | 2io3-A | 8 | 4.2 | 131 | 224 | 8 | MOLECULE: SENTRIN-SPECIFIC PROTEASE 2 |
| 22:00 | 5izv-B | 8 | 4.4 | 167 | 369 | 10 | MOLECULE: UNCHARACTERIZED PROTEIN RAVZ |
| 23:00 | 2xre-B | 7.9 | 3.7 | 130 | 225 | 9 | MOLECULE: SENTRIN-SPECIFIC PROTEASE 1 |
| 24:00:00 | 5aek-Q | 7.9 | 3.5 | 128 | 224 | 8 | MOLECULE: SENTRIN-SPECIFIC PROTEASE 2 |
| 25:00:00 | 2io0-A | 7.9 | 3.6 | 130 | 226 | 7 | MOLECULE: SENTRIN-SPECIFIC PROTEASE 2 |
| 26:00:00 | 5w3t-D | 7.9 | 5.4 | 155 | 333 | 10 | MOLECULE: POPP2 PROTEIN |
| 27:00:00 | 5w3y-B | 7.8 | 5.5 | 157 | 327 | 10 | MOLECULE: POPP2 PROTEIN |
| 28:00:00 | 5izv-A | 7.8 | 4.4 | 163 | 376 | 10 | MOLECULE: UNCHARACTERIZED PROTEIN RAVZ |
| 29:00:00 | 5w40-A | 7.8 | 5.4 | 154 | 336 | 9 | MOLECULE: POPP2 PROTEIN |
| 30:00:00 | 5w3y-D | 7.8 | 5.4 | 156 | 334 | 10 | MOLECULE: POPP2 PROTEIN |

| | | | | | | | | |
|---|---|---|---|---|---|---|---|---|
| 31:00:00 | 5ms8-A | 7.8 | 4.6 | 166 | 377 | 10 | MOLECULE: | LEGIONELLA PNEUMOPHILA EFFECTO |
| 32:00:00 | 5io3-A | 7.8 | 4.8 | 169 | 385 | 9 | MOLECULE: | UNCHARACTERIZED PROTEIN RAVZ |
| 33:00:00 | 5w3t-C | 7.8 | 5.4 | 157 | 332 | 11 | MOLECULE: | POPP2 PROTEIN |
| 34:00:00 | 5w3x-C | 7.7 | 5.4 | 156 | 342 | 10 | MOLECULE: | POPP2 PROTEIN |
| 35:00:00 | 5w3y-A | 7.7 | 5 | 152 | 336 | 11 | MOLECULE: | POPP2 PROTEIN |
| 36:00:00 | 5w3x-A | 7.7 | 5.3 | 155 | 335 | 10 | MOLECULE: | POPP2 PROTEIN |
| 37:00:00 | 5w3t-A | 7.7 | 5.4 | 153 | 332 | 12 | MOLECULE: | POPP2 PROTEIN |
| 38:00:00 | 5w40-C | 7.7 | 5.4 | 154 | 333 | 10 | MOLECULE: | POPP2 PROTEIN |
| 39:00:00 | 5hzy-A | 7.6 | 5.1 | 173 | 380 | 10 | MOLECULE: | UNCHARACTERIZED PROTEIN RAVZ |
| 40:00:00 | 5ms2-A | 7.5 | 4.7 | 165 | 405 | 9 | MOLECULE: | LEGIONELLA PNEUMOPHILA EFFECTO |
| 41:00:00 | 5cqc-A | 7.4 | 4.4 | 154 | 352 | 10 | MOLECULE: | PUTATIVE RAVZ PROTEIN |
| 42:00:00 | 5ms7-A | 7.3 | 5.1 | 166 | 370 | 9 | MOLECULE: | LEGIONELLA PNEUMOPHILA EFFECTO |
| 43:00:00 | 7f3n-A | 6.9 | 5.1 | 139 | 247 | 13 | MOLECULE: | TYPE III EFFECTOR PROTEIN POPP |
| 44:00:00 | 5crc-B | 6 | 9.7 | 104 | 155 | 11 | MOLECULE: | SDEA |
| 45:00:00 | 5crc-A | 5.5 | 6.3 | 99 | 152 | 12 | MOLECULE: | SDEA |
| 46:00:00 | 6w9s-A | 5.1 | 3.8 | 103 | 178 | 9 | MOLECULE: | OTU DOMAIN-CONTAINING PROTEIN |
| 47:00:00 | 7np8-D | 4.6 | 9.1 | 118 | 620 | 8 | MOLECULE: | COENZYME F420-DEPENDENT SULFIT |
| 48:00:00 | 7np8-B | 4.6 | 9.7 | 116 | 620 | 9 | MOLECULE: | COENZYME F420-DEPENDENT SULFIT |
| 49:00:00 | 7np8-C | 4.6 | 8.9 | 117 | 619 | 9 | MOLECULE: | COENZYME F420-DEPENDENT SULFIT |
| 50:00:00 | 7npa-L | 4.4 | 8.9 | 117 | 617 | 7 | MOLECULE: | COENZYME F420-DEPENDENT SULFIT |
| 51:00:00 | 7npa-O | 4.4 | 9.5 | 118 | 617 | 7 | MOLECULE: | COENZYME F420-DEPENDENT SULFIT |
| 52:00:00 | 7npa-K | 4.4 | 9 | 116 | 617 | 7 | MOLECULE: | COENZYME F420-DEPENDENT SULFIT |
| 53:00:00 | 7npa-I | 4.4 | 8.3 | 117 | 618 | 7 | MOLECULE: | COENZYME F420-DEPENDENT SULFIT |
| 54:00:00 | 7npa-N | 4.1 | 8.6 | 117 | 616 | 7 | MOLECULE: | COENZYME F420-DEPENDENT SULFIT |
| 55:00:00 | 7npa-M | 4.1 | 8.3 | 117 | 617 | 7 | MOLECULE: | COENZYME F420-DEPENDENT SULFIT |
| 56:00:00 | 7npa-P | 4.1 | 8 | 117 | 617 | 7 | MOLECULE: | COENZYME F420-DEPENDENT SULFIT |
| 57:00:00 | 7npa-C | 4.1 1 | 1.7 | 108 | 618 | 9 | MOLECULE: | COENZYME F420-DEPENDENT SULFIT |
| 58:00:00 | 2xsj-D | 3.9 1 | 5.6 | 121 | 436 | 3 | MOLECULE: | SULFITE REDUCTASE ALPHA SUBUNI |
| 59:00:00 | 6mca-A | 3.8 | 6.1 | 144 | 460 | 8 | MOLECULE: | ANKYRIN REPEAT DOMAIN PROTEIN |
| 60:00:00 | 7xql-A | 3.7 | 5.3 | 154 | 444 | 9 | MOLECULE: | ANKYRIN REPEAT-CONTAINING PROT |
| 61:00:00 | 7npa-A | 3.7 | 8.4 | 115 | 618 | 9 | MOLECULE: | COENZYME F420-DEPENDENT SULFIT |
| 62:00:00 | 3mmb-D | 3.6 | 5.4 | 104 | 417 | 5 | MOLECULE: | SULFITE REDUCTASE, DISSIMILATO |
| 63:00:00 | 7npa-G | 3.6 | 8.6 | 117 | 618 | 7 | MOLECULE: | COENZYME F420-DEPENDENT SULFIT |
| 64:00:00 | 2v4j-A | 3.6 1 | 4.5 | 116 | 436 | 3 | MOLECULE: | SULFITE REDUCTASE, DISSIMILATO |
| 65:00:00 | 3or1-A | 3.6 1 | 4.4 | 116 | 435 | 3 | MOLECULE: | SULFITE REDUCTASE ALPHA |
| 66:00:00 | 5h92-B | 3.5 1 | 7.6 | 110 | 572 | 8 | MOLECULE: | SULFITE REDUCTASE [FERREDOXIN] |
| 67:00:00 | 7np8-A | 3.5 | 9.5 | 117 | 619 | 9 | MOLECULE: | COENZYME F420-DEPENDENT SULFIT |
| 68:00:00 | 3or1-D | 3.5 | 5.9 | 114 | 435 | 3 | MOLECULE: | SULFITE REDUCTASE ALPHA |
| 69:00:00 | 2akj-A | 3.5 | 4 | 103 | 535 | 6 | MOLECULE: | FERREDOXIN--NITRITE REDUCTASE, |
| 70:00:00 | 2v4j-D | 3.5 1 | 4.5 | 115 | 436 | 3 | MOLECULE: | SULFITE REDUCTASE, DISSIMILATO |
| 71:00:00 | 3or2-A | 3.5 | 4.6 | 107 | 435 | 4 | MOLECULE: | SULFITE REDCUTASE SUBUNIT ALPH |
| 72:00:00 | 7npa-J | 3.3 | 8.4 | 115 | 618 | 9 | MOLECULE: | COENZYME F420-DEPENDENT SULFIT |
| 73:00:00 | 3or2-D | 3.3 | 5.6 | 113 | 435 | 4 | MOLECULE: | SULFITE REDCUTASE SUBUNIT ALPH |
| 74:00:00 | 2xsj-A | 3.3 1 | 5.3 | 119 | 436 | 3 | MOLECULE: | SULFITE REDUCTASE ALPHA SUBUNI |

| | | | | | | | | |
|---|---|---|---|---|---|---|---|---|
| 75:00:00 | 3mm5-D | 3.2 | 5.4 | 103 | 417 | 4 | MOLECULE: | SULFITE REDUCTASE, DISSIMILATO |
| 76:00:00 | 3mm5-A | 3.1 | 5.4 | 104 | 417 | 5 | MOLECULE: | SULFITE REDUCTASE, DISSIMILATO |
| 77:00:00 | 3mma-D | 3.1 | 5.4 | 105 | 417 | 5 | MOLECULE: | SULFITE REDUCTASE, DISSIMILATO |
| 78:00:00 | 3mmc-D | 3.1 | 4.4 | 103 | 417 | 5 | MOLECULE: | SULFITE REDUCTASE, DISSIMILATO |
| 79:00:00 | 3mm8-A | 3.1 | 5.5 | 104 | 417 | 5 | MOLECULE: | SULFITE REDUCTASE, DISSIMILATO |
| 80:00:00 | 1zj8-A | 3 | 4.6 | 108 | 546 | 6 | MOLECULE: | PROBABLE FERREDOXIN-DEPENDENT |
| 81:00:00 | 4kqe-A | 3.0 1 | 1.7 | 90 | 597 | 7 | MOLECULE: | GLYCINE--TRNA LIGASE |
| 82:00:00 | 3mm6-D | 3 | 5.4 | 105 | 417 | 5 | MOLECULE: | SULFITE REDUCTASE, DISSIMILATO |
| 83:00:00 | 3mm7-D | 3 | 5.5 | 104 | 417 | 5 | MOLECULE: | SULFITE REDUCTASE, DISSIMILATO |
| 84:00:00 | 3mm7-A | 3 | 5.4 | 103 | 417 | 5 | MOLECULE: | SULFITE REDUCTASE, DISSIMILATO |
| 85:00:00 | 3mm8-D | 3 | 5.4 | 103 | 417 | 5 | MOLECULE: | SULFITE REDUCTASE, DISSIMILATO |
| 86:00:00 | 3mma-A | 3 | 5.4 | 104 | 417 | 5 | MOLECULE: | SULFITE REDUCTASE, DISSIMILATO |
| 87:00:00 | 3mmc-A | 3 | 5.4 | 104 | 417 | 5 | MOLECULE: | SULFITE REDUCTASE, DISSIMILATO |
| 88:00:00 | 3mm6-A | 3 | 5.4 | 104 | 417 | 5 | MOLECULE: | SULFITE REDUCTASE, DISSIMILATO |
| 89:00:00 | 3mmb-A | 3 | 5.5 | 103 | 417 | 4 | MOLECULE: | SULFITE REDUCTASE, DISSIMILATO |
| 90:00:00 | 3mm9-D | 3 | 5.4 | 103 | 417 | 5 | MOLECULE: | SULFITE REDUCTASE, DISSIMILATO |
| 91:00:00 | 1v95-A | 2.9 | 3.5 | 77 | 130 | 6 | MOLECULE: | NUCLEAR RECEPTOR COACTIVATOR 5 |
| 92:00:00 | 7npa-E | 2.9 | 8.1 | 117 | 617 | 7 | MOLECULE: | COENZYME F420-DEPENDENT SULFIT |
| 93:00:00 | 5e6m-A | 2.8 1 | 4 | 93 | 520 | 6 | MOLECULE: | GLYCINE--TRNA LIGASE |
| 94:00:00 | 6vr7-A | 2.7 1 | 1.6 | 114 | 485 | 10 | MOLECULE: | MUR LIGASE MIDDLE DOMAIN PROTE |
| 95:00:00 | 2zt7-A | 2.7 1 | 1.9 | 85 | 531 | 7 | MOLECULE: | GLYCYL-TRNA SYNTHETASE |
| 96:00:00 | 3mm9-A | 2.5 | 5.2 | 102 | 417 | 6 | MOLECULE: | SULFITE REDUCTASE, DISSIMILATO |
| 97:00:00 | 2l82-A | 2.2 | 2.9 | 77 | 162 | 19 | MOLECULE: | DESIGNED PROTEIN OR32 |
| 98:00:00 | 7npa-F | 2.2 | 8.6 | 117 | 618 | 7 | MOLECULE: | COENZYME F420-DEPENDENT SULFIT |
| 99:00:00 | 7npa-H | 2.2 | 8.3 | 117 | 617 | 7 | MOLECULE: | COENZYME F420-DEPENDENT SULFIT |
| 0:00 | 7npa-B | 2.2 | 8.4 | 117 | 618 | 8 | MOLECULE: | COENZYME F420-DEPENDENT SULFIT |
| 1:00 | 7npa-D | 2.2 | 8.3 | 117 | 617 | 7 | MOLECULE: | COENZYME F420-DEPENDENT SULFIT |

**DALI results: MnA-1 Prim**

| No: | Chain | Z | rmsd l | ali | nres | %id | PDB Description |
|---|---|---|---|---|---|---|---|
| 1:00 | 3f1y-A | 2.5 | 11.4 | 65 | 318 | 5 | MOLECULE: MANNOSYL-3-PHOSPHOGLYCERATE SYNTHASE |
| 2:00 | 8pk5-A | 2.4 | 4.1 | 48 | 535 | 13 | MOLECULE: INTEGRATOR COMPLEX SUBUNIT 13 |
| 3:00 | 7m0q-B | 2.3 | 3.5 | 50 | 97 | 4 | MOLECULE: NETWORK HALLUCINATED PROTEIN 0738_MOD |
| 4:00 | 7zsb-4 | 2.3 | 2.8 | 45 | 306 | 7 | MOLECULE: DNA-DIRECTED RNA POLYMERASE II SUBUNIT RPB1 |
| 5:00 | 6msp-A | 2.3 | 4.7 | 48 | 97 | 15 | MOLECULE: DE NOVO DESIGNED PROTEIN FOLDIT3 |
| 6:00 | 4gsl-A | 2.2 | 4 | 67 | 598 | 1 | MOLECULE: UBIQUITIN-LIKE MODIFIER-ACTIVATING ENZYME |
| 7:00 | 4m0m-A | 2.2 | 4.1 | 63 | 721 | 5 | MOLECULE: PUTATIVE UNCHARACTERIZED PROTEIN |
| 8:00 | 3oyz-A | 2.1 | 6.6 | 60 | 376 | 5 | MOLECULE: MALATE SYNTHASE |
| 9:00 | 6t8d-X | 2.1 | 13.7 | 66 | 355 | 12 | MOLECULE: MAKB |
| 10:00 | 7y5y-A | 2.0 | 3.5 | 73 | 152 | 10 | MOLECULE: STAYGREEN DOMAIN-CONTAINING PROTEIN |
| 11:00 | 4xo1-A | 2.0 | 4.3 | 45 | 60 | 2 | MOLECULE: PROTEIN GNSA |

**DALI results: MnA-1 RepE1**

| *No:* | *Chain* | *Z* | *rmsd* | *lali* | *nres* | *%id* | *PDB Description* | |
|---|---|---|---|---|---|---|---|---|
| 1:00 | 2gxa-D | 19.2 | 3.4 | 260 | 270 | 19 | MOLECULE: | 5'-D(*TP*TP*TP*TP*TP*TP*TP*TP*TP*TP*TP*TP*T)-3' |
| 2:00 | 2gxa-G | 19.1 | 3.5 | 257 | 270 | 19 | MOLECULE: | 5'-D(*TP*TP*TP*TP*TP*TP*TP*TP*TP*TP*TP*TP*T)-3' |
| 3:00 | 5a9k-D | 19.1 | 3.4 | 230 | 274 | 19 | MOLECULE: | REPLICATION PROTEIN E1 |
| 4:00 | 2gxa-C | 19 | 3.4 | 257 | 270 | 19 | MOLECULE: | 5'-D(*TP*TP*TP*TP*TP*TP*TP*TP*TP*TP*TP*TP*T)-3' |
| 5:00 | 2gxa-I | 19 | 3.4 | 252 | 270 | 20 | MOLECULE: | 5'-D(*TP*TP*TP*TP*TP*TP*TP*TP*TP*TP*TP*TP*T)-3' |
| 6:00 | 2gxa-B | 19 | 3.5 | 256 | 270 | 18 | MOLECULE: | 5'-D(*TP*TP*TP*TP*TP*TP*TP*TP*TP*TP*TP*TP*T)-3' |
| 7:00 | 2gxa-J | 19 | 3.7 | 257 | 270 | 19 | MOLECULE: | 5'-D(*TP*TP*TP*TP*TP*TP*TP*TP*TP*TP*TP*TP*T)-3' |
| 8:00 | 2gxa-E | 18.9 | 3.8 | 259 | 274 | 19 | MOLECULE: | 5'-D(*TP*TP*TP*TP*TP*TP*TP*TP*TP*TP*TP*TP*T)-3' |
| 9:00 | 5a9k-E | 18.9 | 3.8 | 254 | 270 | 20 | MOLECULE: | REPLICATION PROTEIN E1 |
| 10:00 | 2v9p-J | 18.9 | 3.5 | 254 | 272 | 18 | MOLECULE: | REPLICATION PROTEIN E1 |
| 11:00 | 2v9p-D | 18.9 | 3.5 | 257 | 274 | 19 | MOLECULE: | REPLICATION PROTEIN E1 |
| 12:00 | 2v9p-E | 18.9 | 3.8 | 254 | 270 | 20 | MOLECULE: | REPLICATION PROTEIN E1 |
| 13:00 | 2gxa-A | 18.8 | 3.5 | 254 | 270 | 19 | MOLECULE: | 5'-D(*TP*TP*TP*TP*TP*TP*TP*TP*TP*TP*TP*TP*T)-3' |
| 14:00 | 5a9k-A | 18.8 | 3.6 | 255 | 272 | 19 | MOLECULE: | REPLICATION PROTEIN E1 |
| 15:00 | 2v9p-G | 18.8 | 3.7 | 254 | 270 | 19 | MOLECULE: | REPLICATION PROTEIN E1 |
| 16:00 | 2v9p-A | 18.8 | 3.6 | 255 | 272 | 19 | MOLECULE: | REPLICATION PROTEIN E1 |
| 17:00 | 2v9p-H | 18.8 | 3.3 | 252 | 268 | 19 | MOLECULE: | REPLICATION PROTEIN E1 |
| 18:00 | 5a9k-B | 18.8 | 3.3 | 252 | 268 | 19 | MOLECULE: | REPLICATION PROTEIN E1 |
| 19:00 | 2v9p-B | 18.8 | 3.3 | 252 | 268 | 19 | MOLECULE: | REPLICATION PROTEIN E1 |
| 20:00 | 2v9p-I | 18.8 | 3.3 | 253 | 270 | 18 | MOLECULE: | REPLICATION PROTEIN E1 |
| 21:00 | 2v9p-K | 18.7 | 3.7 | 254 | 269 | 20 | MOLECULE: | REPLICATION PROTEIN E1 |
| 22:00 | 5a9k-C | 18.6 | 3.4 | 250 | 267 | 18 | MOLECULE: | REPLICATION PROTEIN E1 |
| 23:00 | 2v9p-C | 18.6 | 3.4 | 250 | 267 | 18 | MOLECULE: | REPLICATION PROTEIN E1 |
| 24:00:00 | 2v9p-F | 18.6 | 4.4 | 254 | 269 | 19 | MOLECULE: | REPLICATION PROTEIN E1 |
| 25:00:00 | 5a9k-F | 18.5 | 3.7 | 217 | 269 | 21 | MOLECULE: | REPLICATION PROTEIN E1 |
| 26:00:00 | 2gxa-F | 18.4 | 4.7 | 227 | 270 | 21 | MOLECULE: | 5'-D(*TP*TP*TP*TP*TP*TP*TP*TP*TP*TP*TP*TP*T)-3' |
| 27:00:00 | 2gxa-H | 18.4 | 3.4 | 256 | 274 | 18 | MOLECULE: | 5'-D(*TP*TP*TP*TP*TP*TP*TP*TP*TP*TP*TP*TP*T)-3' |
| 28:00:00 | 2gxa-K | 18.2 | 9.9 | 237 | 273 | 19 | MOLECULE: | 5'-D(*TP*TP*TP*TP*TP*TP*TP*TP*TP*TP*TP*TP*T)-3' |
| 29:00:00 | 7apd-E | 18.2 | 5.1 | 241 | 287 | 19 | MOLECULE: | REPLICATION PROTEIN E1 |
| 30:00:00 | 2v9p-L | 18.1 | 4.6 | 252 | 267 | 19 | MOLECULE: | REPLICATION PROTEIN E1 |
| 31:00:00 | 7apd-A | 18 | 5.1 | 234 | 284 | 19 | MOLECULE: | REPLICATION PROTEIN E1 |
| 32:00:00 | 7apd-C | 17.7 | 3.8 | 268 | 287 | 19 | MOLECULE: | REPLICATION PROTEIN E1 |
| 33:00:00 | 7apd-D | 17.6 | 4.7 | 240 | 287 | 19 | MOLECULE: | REPLICATION PROTEIN E1 |
| 34:00:00 | 7apd-F | 17.5 | 4.9 | 232 | 286 | 19 | MOLECULE: | REPLICATION PROTEIN E1 |
| 35:00:00 | 7apd-B | 17.3 | 4 | 267 | 289 | 19 | MOLECULE: | REPLICATION PROTEIN E1 |
| 36:00:00 | 1tue-D | 17.1 | 2.5 | 192 | 202 | 20 | MOLECULE: | REPLICATION PROTEIN E1 |
| 37:00:00 | 1tue-M | 16.9 | 2.4 | 185 | 196 | 19 | MOLECULE: | REPLICATION PROTEIN E1 |
| 38:00:00 | 1tue-A | 16.8 | 2.6 | 191 | 201 | 19 | MOLECULE: | REPLICATION PROTEIN E1 |
| 39:00:00 | 1tue-K | 16.7 | 2.4 | 181 | 192 | 20 | MOLECULE: | REPLICATION PROTEIN E1 |
| 40:00:00 | 1tue-H | 16.4 | 2.4 | 179 | 191 | 21 | MOLECULE: | REPLICATION PROTEIN E1 |
| 41:00:00 | 1tue-F | 16.3 | 2.4 | 177 | 188 | 21 | MOLECULE: | REPLICATION PROTEIN E1 |
| 42:00:00 | 2h1l-C | 16 | 5 | 284 | 363 | 19 | MOLECULE: | LARGE T ANTIGEN |
| 43:00:00 | 2h1l-D | 15.9 | 5 | 281 | 362 | 19 | MOLECULE: | LARGE T ANTIGEN |
| 44:00:00 | 5j4y-A | 15.9 | 5.6 | 261 | 363 | 17 | MOLECULE: | LARGE T ANTIGEN |
| 45:00:00 | 1svm-E | 15.8 | 5.1 | 281 | 362 | 19 | MOLECULE: | LARGE T ANTIGEN |
| 46:00:00 | 1svm-B | 15.8 | 5.1 | 281 | 362 | 19 | MOLECULE: | LARGE T ANTIGEN |
| 47:00:00 | 2h1l-E | 15.8 | 5.1 | 282 | 362 | 19 | MOLECULE: | LARGE T ANTIGEN |
| 48:00:00 | 2h1l-K | 15.8 | 4.9 | 241 | 362 | 19 | MOLECULE: | LARGE T ANTIGEN |
| 49:00:00 | 4e2i-B | 15.7 | 5 | 280 | 362 | 19 | MOLECULE: | LARGE T ANTIGEN |
| 50:00:00 | 2h1l-F | 15.7 | 4.2 | 242 | 363 | 19 | MOLECULE: | LARGE T ANTIGEN |
| 51:00:00 | 1svm-C | 15.7 | 4.5 | 241 | 363 | 19 | MOLECULE: | LARGE T ANTIGEN |

| | | | | | | | | |
|---|---|---|---|---|---|---|---|---|
| 52:00:00 | 1svm-A | 15.7 | 4.5 | 242 | 362 | 19 | MOLECULE: | LARGE T ANTIGEN |
| 53:00:00 | 2h1l-L | 15.7 | 5.5 | 289 | 363 | 19 | MOLECULE: | LARGE T ANTIGEN |
| 54:00:00 | 4gdf-B | 15.7 | 13.6 | 311 | 497 | 16 | MOLECULE: | LARGE T ANTIGEN |
| 55:00:00 | 4gdf-F | 15.6 | 16.2 | 336 | 497 | 18 | MOLECULE: | LARGE T ANTIGEN |
| 56:00:00 | 2h1l-B | 15.6 | 5.1 | 281 | 362 | 19 | MOLECULE: | LARGE T ANTIGEN |
| 57:00:00 | 1svm-F | 15.6 | 5.2 | 262 | 363 | 18 | MOLECULE: | LARGE T ANTIGEN |
| 58:00:00 | 2gxa-L | 15.5 | 4.7 | 232 | 270 | 18 | MOLECULE: | 5'-D(*TP*TP*TP*TP*TP*TP*TP*TP*TP*TP*TP*TP*T)-3' |
| 59:00:00 | 4gdf-A | 15.5 | 8.8 | 278 | 497 | 16 | MOLECULE: | LARGE T ANTIGEN |
| 60:00:00 | 2h1l-H | 15.5 | 5.1 | 282 | 362 | 19 | MOLECULE: | LARGE T ANTIGEN |
| 61:00:00 | 2h1l-I | 15.5 | 5.1 | 286 | 363 | 19 | MOLECULE: | LARGE T ANTIGEN |
| 62:00:00 | 4gdf-E | 15.5 | 8.8 | 266 | 497 | 18 | MOLECULE: | LARGE T ANTIGEN |
| 63:00:00 | 1svl-A | 15.4 | 5.5 | 280 | 363 | 18 | MOLECULE: | LARGE T ANTIGEN |
| 64:00:00 | 1svl-C | 15.4 | 5.5 | 280 | 362 | 19 | MOLECULE: | LARGE T ANTIGEN |
| 65:00:00 | 1svl-B | 15.4 | 5.2 | 279 | 362 | 19 | MOLECULE: | LARGE T ANTIGEN |
| 66:00:00 | 2h1l-A | 15.4 | 5.2 | 285 | 362 | 19 | MOLECULE: | LARGE T ANTIGEN |
| 67:00:00 | 2h1l-G | 15.4 | 5.1 | 279 | 362 | 19 | MOLECULE: | LARGE T ANTIGEN |
| 68:00:00 | 5j40-A | 15.4 | 5.3 | 276 | 358 | 17 | MOLECULE: | LARGE T ANTIGEN |
| 69:00:00 | 2h1l-J | 15.4 | 4.4 | 242 | 362 | 19 | MOLECULE: | LARGE T ANTIGEN |
| 70:00:00 | 1svm-D | 15.4 | 4.5 | 243 | 362 | 19 | MOLECULE: | LARGE T ANTIGEN |
| 71:00:00 | 1n25-B | 15.4 | 5.8 | 255 | 362 | 20 | MOLECULE: | LARGE T ANTIGEN |
| 72:00:00 | 5tct-B | 15.4 | 5.1 | 251 | 362 | 20 | MOLECULE: | LARGE T ANTIGEN |
| 73:00:00 | 5tct-F | 15.3 | 5.5 | 282 | 362 | 19 | MOLECULE: | LARGE T ANTIGEN |
| 74:00:00 | 5j4v-A | 15.3 | 5 | 267 | 358 | 17 | MOLECULE: | LARGE T ANTIGEN |
| 75:00:00 | 1svo-A | 15.3 | 5.2 | 245 | 362 | 20 | MOLECULE: | LARGE T ANTIGEN |
| 76:00:00 | 1svo-B | 15.3 | 5.8 | 256 | 362 | 20 | MOLECULE: | LARGE T ANTIGEN |
| 77:00:00 | 1n25-A | 15.3 | 5.2 | 244 | 362 | 20 | MOLECULE: | LARGE T ANTIGEN |
| 78:00:00 | 5tct-D | 15.2 | 5.5 | 281 | 362 | 19 | MOLECULE: | LARGE T ANTIGEN |
| 79:00:00 | 5j47-A | 15.2 | 5.1 | 257 | 360 | 17 | MOLECULE: | LARGE T ANTIGEN |
| 80:00:00 | 4e2i-F | 15.2 | 4.3 | 247 | 362 | 19 | MOLECULE: | LARGE T ANTIGEN |
| 81:00:00 | 4e2i-G | 15.2 | 4.7 | 251 | 362 | 19 | MOLECULE: | LARGE T ANTIGEN |
| 82:00:00 | 4e2i-A | 15.2 | 4.9 | 255 | 362 | 18 | MOLECULE: | LARGE T ANTIGEN |
| 83:00:00 | 4e2i-L | 15.2 | 5.1 | 259 | 362 | 18 | MOLECULE: | LARGE T ANTIGEN |
| 84:00:00 | 4e2i-K | 15.2 | 4.5 | 250 | 362 | 19 | MOLECULE: | LARGE T ANTIGEN |
| 85:00:00 | 4e2i-C | 15.1 | 5 | 280 | 362 | 19 | MOLECULE: | LARGE T ANTIGEN |
| 86:00:00 | 4e2i-H | 15.1 | 4.7 | 255 | 362 | 18 | MOLECULE: | LARGE T ANTIGEN |
| 87:00:00 | 4e2i-I | 15.1 | 4.6 | 253 | 362 | 19 | MOLECULE: | LARGE T ANTIGEN |
| 88:00:00 | 4e2i-D | 15.1 | 4.5 | 250 | 362 | 19 | MOLECULE: | LARGE T ANTIGEN |
| 89:00:00 | 4e2i-E | 15.1 | 4.3 | 236 | 362 | 19 | MOLECULE: | LARGE T ANTIGEN |
| 90:00:00 | 5tct-C | 15.1 | 4.6 | 239 | 362 | 20 | MOLECULE: | LARGE T ANTIGEN |
| 91:00:00 | 4e2i-J | 15 | 5 | 279 | 362 | 19 | MOLECULE: | LARGE T ANTIGEN |
| 92:00:00 | 1s9h-A | 15 | 5.6 | 220 | 268 | 13 | MOLECULE: | REP 40 PROTEIN |
| 93:00:00 | 5tct-A | 14.8 | 7 | 250 | 362 | 20 | MOLECULE: | LARGE T ANTIGEN |
| 94:00:00 | 5tct-E | 14.8 | 6.8 | 244 | 362 | 20 | MOLECULE: | LARGE T ANTIGEN |
| 95:00:00 | 1s9h-B | 14.8 | 3.5 | 206 | 267 | 14 | MOLECULE: | REP 40 PROTEIN |
| 96:00:00 | 1u0j-A | 14.6 | 3.2 | 197 | 261 | 14 | MOLECULE: | DNA REPLICATION PROTEIN |
| 97:00:00 | 7jsh-B | 13.7 | 3.8 | 236 | 272 | 16 | MOLECULE: | PROTEIN REP68 |
| 98:00:00 | 7jsi-E | 13.3 | 3.9 | 222 | 276 | 13 | MOLECULE: | PROTEIN REP68 |
| 99:00:00 | 7jsg-C | 13.2 | 4.4 | 229 | 272 | 14 | MOLECULE: | PROTEIN REP68 |
| 0:00 | 7jsg-E | 13.2 | 4.1 | 230 | 272 | 15 | MOLECULE: | PROTEIN REP68 |
| 1:00 | 7jsf-E | 13.1 | 4.7 | 215 | 276 | 13 | MOLECULE: | PROTEIN REP68 |
| 2:00 | 7jsg-F | 13 | 4 | 215 | 272 | 15 | MOLECULE: | PROTEIN REP68 |
| 3:00 | 7jsf-B | 12.8 | 4.9 | 216 | 276 | 13 | MOLECULE: | PROTEIN REP68 |
| 4:00 | 7jsh-E | 12.8 | 4.2 | 237 | 272 | 14 | MOLECULE: | PROTEIN REP68 |

| | | | | | | | | |
|---|---|---|---|---|---|---|---|---|
| 5:00 | 7jsf-A | 12.7 | 4.7 | 215 | 276 | 13 | MOLECULE: | PROTEIN REP68 |
| 6:00 | 7jsf-C | 12.7 | 4.8 | 216 | 276 | 13 | MOLECULE: | PROTEIN REP68 |
| 7:00 | 7jsf-G | 12.7 | 4.9 | 216 | 276 | 13 | MOLECULE: | PROTEIN REP68 |
| 8:00 | 7jsf-F | 12.7 | 4.9 | 216 | 276 | 13 | MOLECULE: | PROTEIN REP68 |
| 9:00 | 7jsg-B | 12.7 | 4.4 | 225 | 272 | 14 | MOLECULE: | PROTEIN REP68 |
| 10:00 | 7jsi-C | 12.7 | 4 | 225 | 276 | 14 | MOLECULE: | PROTEIN REP68 |
| 11:00 | 7jsi-D | 12.6 | 3.8 | 222 | 276 | 14 | MOLECULE: | PROTEIN REP68 |
| 12:00 | 7jsf-D | 12.5 | 4.9 | 216 | 276 | 13 | MOLECULE: | PROTEIN REP68 |
| 13:00 | 7jsi-B | 12.4 | 4.2 | 225 | 276 | 15 | MOLECULE: | PROTEIN REP68 |
| 14:00 | 7jsg-D | 12.3 | 4.6 | 229 | 272 | 12 | MOLECULE: | PROTEIN REP68 |
| 15:00 | 7jsi-F | 12.2 | 4 | 227 | 276 | 14 | MOLECULE: | PROTEIN REP68 |
| 16:00 | 7jsh-G | 12.1 | 3.9 | 227 | 272 | 14 | MOLECULE: | PROTEIN REP68 |
| 17:00 | 7jsh-C | 11.9 | 4.5 | 235 | 272 | 14 | MOLECULE: | PROTEIN REP68 |
| 18:00 | 7jsg-G | 11.8 | 4.5 | 222 | 272 | 14 | MOLECULE: | PROTEIN REP68 |
| 19:00 | 7jsh-A | 11.7 | 4.3 | 226 | 272 | 12 | MOLECULE: | PROTEIN REP68 |
| 20:00 | 7jsi-A | 11.6 | 4.1 | 229 | 276 | 13 | MOLECULE: | PROTEIN REP68 |
| 21:00 | 6k9c-B | 11.6 | 4.8 | 215 | 414 | 16 | MOLECULE: | PRIMASE |
| 22:00 | 6k9c-A | 11.4 | 5.9 | 208 | 416 | 16 | MOLECULE: | PRIMASE |
| 23:00 | 7jsh-F | 11.3 | 4.2 | 233 | 272 | 13 | MOLECULE: | PROTEIN REP68 |
| 24:00:00 | 6lrb-F | 11.3 | 5.9 | 209 | 408 | 17 | MOLECULE: | PRIMASE |
| 25:00:00 | 6k9e-A | 11.3 | 4.1 | 201 | 402 | 15 | MOLECULE: | PRIMASE |
| 26:00:00 | 6k9e-C | 11.2 | 4.1 | 204 | 401 | 16 | MOLECULE: | PRIMASE |
| 27:00:00 | 6lrb-B | 11.2 | 7 | 206 | 408 | 16 | MOLECULE: | PRIMASE |
| 28:00:00 | 6lrb-A | 11.2 | 5.8 | 207 | 410 | 17 | MOLECULE: | PRIMASE |
| 29:00:00 | 7jsh-D | 11.2 | 4.2 | 215 | 272 | 14 | MOLECULE: | PROTEIN REP68 |
| 30:00:00 | 6lrb-C | 11.2 | 5.9 | 208 | 410 | 17 | MOLECULE: | PRIMASE |
| 31:00:00 | 6k9e-F | 11 | 4.3 | 202 | 404 | 16 | MOLECULE: | PRIMASE |
| 32:00:00 | 6k9e-D | 11 | 4 | 196 | 404 | 16 | MOLECULE: | PRIMASE |
| 33:00:00 | 6k9e-B | 11 | 4.1 | 201 | 404 | 16 | MOLECULE: | PRIMASE |
| 34:00:00 | 7ola-C | 10.8 | 11.5 | 218 | 426 | 12 | MOLECULE: | DNA PRIMASE |
| 35:00:00 | 7jsg-A | 10.8 | 4.2 | 211 | 272 | 14 | MOLECULE: | PROTEIN REP68 |
| 36:00:00 | 6lrb-D | 10.8 | 3.4 | 193 | 409 | 17 | MOLECULE: | PRIMASE |
| 37:00:00 | 7om0-E | 10.6 | 11.1 | 208 | 439 | 13 | MOLECULE: | DNA PRIMASE |
| 38:00:00 | 7om0-C | 10.5 | 9 | 221 | 439 | 12 | MOLECULE: | DNA PRIMASE |
| 39:00:00 | 1s9h-C | 10.5 | 4.4 | 157 | 199 | 12 | MOLECULE: | REP 40 PROTEIN |
| 40:00:00 | 8uh7-D | 10.4 | 10.5 | 144 | 319 | 17 | MOLECULE: | SLIDING-CLAMP-LOADER SMALL SUBUNIT |
| 41:00:00 | 8unf-C | 10.4 | 10.5 | 144 | 319 | 17 | MOLECULE: | PRIMER DNA |
| 42:00:00 | 4fb3-A | 10.4 | 2.8 | 110 | 115 | 10 | MOLECULE: | ORI DNA OLIGONUCLEOTIDE-CRICK STRAND |
| 43:00:00 | 8hwh-C | 10.2 | 3.6 | 200 | 361 | 10 | MOLECULE: | PRIMASE D5 |
| 44:00:00 | 7ola-F | 10.2 | 7.1 | 197 | 345 | 12 | MOLECULE: | DNA PRIMASE |
| 45:00:00 | 3u5z-C | 10.2 | 11.7 | 144 | 320 | 16 | MOLECULE: | DNA POLYMERASE ACCESSORY PROTEIN 44 |
| 46:00:00 | 5c3c-A | 10.2 | 6.9 | 140 | 231 | 14 | MOLECULE: | CBBQ/NIRQ/NORQ DOMAIN PROTEIN |
| 47:00:00 | 2nl8-A | 10.2 | 2.9 | 111 | 117 | 8 | MOLECULE: | 18-NT PEN ELEMENT OF THE SV40 DNA ORIGIN |
| 48:00:00 | 3qfq-E | 10.2 | 3 | 112 | 117 | 8 | MOLECULE: | LARGE T ANTIGEN |
| 49:00:00 | 4fb3-E | 10.2 | 2.8 | 110 | 114 | 10 | MOLECULE: | ORI DNA OLIGONUCLEOTIDE-CRICK STRAND |
| 50:00:00 | 8apm-F | 10.1 | 4 | 202 | 359 | 11 | MOLECULE: | PRIMASE D5 |
| 51:00:00 | 3u60-E | 10.1 | 4.8 | 133 | 305 | 16 | MOLECULE: | DNA POLYMERASE ACCESSORY PROTEIN 44 |
| 52:00:00 | 8uk9-K | 10.1 | 8.1 | 139 | 319 | 17 | MOLECULE: | SLIDING-CLAMP-LOADER SMALL SUBUNIT |
| 53:00:00 | 2itl-A | 10.1 | 2.9 | 112 | 120 | 8 | MOLECULE: | 24-NT PEN ELEMENT OF THE SV40 DNA ORIGIN |
| 54:00:00 | 4fb3-B | 10.1 | 2.7 | 109 | 114 | 10 | MOLECULE: | ORI DNA OLIGONUCLEOTIDE-CRICK STRAND |
| 55:00:00 | 2fuf-A | 10.1 | 3.1 | 111 | 126 | 8 | MOLECULE: | LARGE T ANTIGEN |
| 56:00:00 | 7om0-D | 10 | 8.6 | 215 | 438 | 13 | MOLECULE: | DNA PRIMASE |
| 57:00:00 | 8unf-B | 10 | 10 | 143 | 319 | 15 | MOLECULE: | PRIMER DNA |

| | | | | | | | | |
|---|---|---|---|---|---|---|---|---|
| 58:00:00 | 7pds-D | 10 | 5.6 | 219 | 454 | 12 | MOLECULE: | SIMILAR TO D. NODOSUS VAPE |
| 59:00:00 | 2ipr-A | 10 | 3.5 | 117 | 127 | 8 | MOLECULE: | LARGE T ANTIGEN |
| 60:00:00 | 3qfq-B | 10 | 3 | 113 | 120 | 9 | MOLECULE: | LARGE T ANTIGEN |
| 61:00:00 | 6lrb-E | 10 | 3.7 | 199 | 409 | 17 | MOLECULE: | PRIMASE |
| 62:00:00 | 2ipr-B | 10 | 2.9 | 112 | 124 | 8 | MOLECULE: | LARGE T ANTIGEN |
| 63:00:00 | 8apm-B | 9.9 | 5.5 | 209 | 449 | 11 | MOLECULE: | PRIMASE D5 |
| 64:00:00 | 6k9e-E | 9.9 | 4.1 | 203 | 402 | 16 | MOLECULE: | PRIMASE |
| 65:00:00 | 8apm-C | 9.9 | 5.7 | 209 | 449 | 11 | MOLECULE: | PRIMASE D5 |
| 66:00:00 | 8unf-E | 9.9 | 10.9 | 142 | 319 | 15 | MOLECULE: | PRIMER DNA |
| 67:00:00 | 8uk9-M | 9.9 | 9.6 | 139 | 317 | 17 | MOLECULE: | SLIDING-CLAMP-LOADER SMALL SUBUNIT |
| 68:00:00 | 4lmd-B | 9.9 | 3.4 | 116 | 130 | 9 | MOLECULE: | LARGE T ANTIGEN |
| 69:00:00 | 4nbp-A | 9.9 | 3.4 | 115 | 129 | 9 | MOLECULE: | LARGE T ANTIGEN |
| 70:00:00 | 5cyn-A | 9.9 | 3.6 | 116 | 127 | 9 | MOLECULE: | LARGE T ANTIGEN |
| 71:00:00 | 3qn2-A | 9.9 | 2.9 | 111 | 126 | 8 | MOLECULE: | LARGE T ANTIGEN |
| 72:00:00 | 4fgn-B | 9.9 | 3.5 | 116 | 124 | 8 | MOLECULE: | LARGE T ANTIGEN |
| 73:00:00 | 4lmd-A | 9.9 | 3.6 | 117 | 128 | 9 | MOLECULE: | LARGE T ANTIGEN |
| 74:00:00 | 2ntc-B | 9.9 | 3.5 | 116 | 126 | 8 | MOLECULE: | 21-NT PEN ELEMENT OF THE SV40 DNA ORIGIN |
| 75:00:00 | 2itj-B | 9.9 | 3.3 | 110 | 118 | 8 | MOLECULE: | LARGE T ANTIGEN |
| 76:00:00 | 7om0-A | 9.8 | 7.5 | 209 | 439 | 12 | MOLECULE: | DNA PRIMASE |
| 77:00:00 | 8unh-E | 9.8 | 7.3 | 139 | 319 | 17 | MOLECULE: | SLIDING-CLAMP-LOADER LARGE SUBUNIT |
| 78:00:00 | 8efy-C | 9.8 | 9 | 140 | 302 | 14 | MOLECULE: | HOLLIDAY JUNCTION ATP-DEPENDENT DNA HELICASE RUVB |
| 79:00:00 | 4lif-A | 9.8 | 3.6 | 116 | 127 | 9 | MOLECULE: | LARGE T ANTIGEN |
| 80:00:00 | 7pbt-C | 9.8 | 9.8 | 141 | 309 | 10 | MOLECULE: | HOLLIDAY JUNCTION ATP-DEPENDENT DNA HELICASE RUVB |
| 81:00:00 | 2if9-A | 9.8 | 3.7 | 117 | 132 | 8 | MOLECULE: | LARGE T ANTIGEN |
| 82:00:00 | 7pbs-D | 9.8 | 10.2 | 145 | 313 | 10 | MOLECULE: | HOLLIDAY JUNCTION ATP-DEPENDENT DNA HELICASE RUVB |
| 83:00:00 | 5d9i-B | 9.8 | 3.7 | 118 | 132 | 8 | MOLECULE: | LARGE T ANTIGEN |
| 84:00:00 | 3qfq-A | 9.8 | 3.1 | 112 | 119 | 8 | MOLECULE: | LARGE T ANTIGEN |
| 85:00:00 | 7om0-B | 9.7 | 7.9 | 213 | 439 | 13 | MOLECULE: | DNA PRIMASE |
| 86:00:00 | 7tts-C | 9.7 | 16.8 | 190 | 456 | 12 | MOLECULE: | CASEINOLYTIC PEPTIDASE B PROTEIN HOMOLOG |
| 87:00:00 | 6j7e-A | 9.7 | 3.6 | 138 | 251 | 7 | MOLECULE: | NITROGEN ASSIMILATION REGULATORY PROTEIN |
| 88:00:00 | 8efv-A | 9.7 | 9.6 | 139 | 313 | 14 | MOLECULE: | HOLLIDAY JUNCTION ATP-DEPENDENT DNA HELICASE RUVB |
| 89:00:00 | 7pbs-C | 9.7 | 10.1 | 140 | 309 | 11 | MOLECULE: | HOLLIDAY JUNCTION ATP-DEPENDENT DNA HELICASE RUVB |
| 90:00:00 | 2itj-A | 9.7 | 3.5 | 115 | 128 | 8 | MOLECULE: | LARGE T ANTIGEN |
| 91:00:00 | 2itl-B | 9.7 | 2.7 | 108 | 115 | 8 | MOLECULE: | 24-NT PEN ELEMENT OF THE SV40 DNA ORIGIN |
| 92:00:00 | 2if9-B | 9.7 | 3.4 | 117 | 131 | 8 | MOLECULE: | LARGE T ANTIGEN |
| 93:00:00 | 2ntc-A | 9.7 | 3 | 112 | 127 | 8 | MOLECULE: | 21-NT PEN ELEMENT OF THE SV40 DNA ORIGIN |
| 94:00:00 | 3qk2-A | 9.7 | 3.3 | 115 | 132 | 8 | MOLECULE: | LARGE T ANTIGEN |
| 95:00:00 | 5d9i-A | 9.7 | 3.5 | 116 | 133 | 9 | MOLECULE: | LARGE T ANTIGEN |
| 96:00:00 | 7om0-F | 9.6 | 8.6 | 218 | 439 | 13 | MOLECULE: | DNA PRIMASE |
| 97:00:00 | 3m0e-C | 9.6 | 3.7 | 141 | 247 | 9 | MOLECULE: | TRANSCRIPTIONAL REGULATOR (NTRC FAMILY) |
| 98:00:00 | 7ola-E | 9.6 | 3.7 | 197 | 344 | 15 | MOLECULE: | DNA PRIMASE |
| 99:00:00 | 6on2-E | 9.6 | 13.2 | 185 | 523 | 14 | MOLECULE: | ATP-DEPENDENT PROTEASE LA |
| 0:00 | 8efv-B | 9.6 | 8.9 | 136 | 314 | 14 | MOLECULE: | HOLLIDAY JUNCTION ATP-DEPENDENT DNA HELICASE RUVB |
| 1:00 | 8efy-A | 9.6 | 9.8 | 142 | 313 | 13 | MOLECULE: | HOLLIDAY JUNCTION ATP-DEPENDENT DNA HELICASE RUVB |
| 2:00 | 1in7-A | 9.6 | 9.5 | 140 | 298 | 14 | MOLECULE: | HOLLIDAY JUNCTION DNA HELICASE RUVB |
| 3:00 | 7pbs-A | 9.6 | 9.6 | 140 | 312 | 11 | MOLECULE: | HOLLIDAY JUNCTION ATP-DEPENDENT DNA HELICASE RUVB |
| 4:00 | 8umy-A | 9.5 | 21.9 | 165 | 546 | 16 | MOLECULE: | CHROMOSOME TRANSMISSION FIDELITY PROTEIN 18 HOMOL |
| 5:00 | 8apm-A | 9.5 | 4.5 | 210 | 359 | 11 | MOLECULE: | PRIMASE D5 |
| 6:00 | 3m0e-E | 9.5 | 3.7 | 141 | 247 | 9 | MOLECULE: | TRANSCRIPTIONAL REGULATOR (NTRC FAMILY) |
| 7:00 | 2c96-A | 9.5 | 3.5 | 138 | 237 | 9 | MOLECULE: | PSP OPERON TRANSCRIPTIONAL ACTIVATOR |
| 8:00 | 3m0e-F | 9.5 | 3.7 | 141 | 247 | 9 | MOLECULE: | TRANSCRIPTIONAL REGULATOR (NTRC FAMILY) |
| 9:00 | 3m0e-G | 9.5 | 3.7 | 141 | 247 | 9 | MOLECULE: | TRANSCRIPTIONAL REGULATOR (NTRC FAMILY) |
| 10:00 | 3m0e-B | 9.5 | 3.7 | 141 | 247 | 9 | MOLECULE: | TRANSCRIPTIONAL REGULATOR (NTRC FAMILY) |

| | | | | | | | | |
|---|---|---|---|---|---|---|---|---|
| 11:00 | 7ttr-E | 9.5 | 11.1 | 157 | 332 | 12 | MOLECULE: | CASEINOLYTIC PEPTIDASE B PROTEIN HOMOLOG |
| 12:00 | 3pfi-A | 9.5 | 4.7 | 144 | 301 | 12 | MOLECULE: | HOLLIDAY JUNCTION ATP-DEPENDENT DNA HELICASE RUVB |
| 13:00 | 3m0e-D | 9.5 | 3.7 | 141 | 247 | 9 | MOLECULE: | TRANSCRIPTIONAL REGULATOR (NTRC FAMILY) |
| 14:00 | 3m0e-A | 9.5 | 3.7 | 141 | 247 | 9 | MOLECULE: | TRANSCRIPTIONAL REGULATOR (NTRC FAMILY) |
| 15:00 | 8efv-E | 9.5 | 10.7 | 139 | 298 | 13 | MOLECULE: | HOLLIDAY JUNCTION ATP-DEPENDENT DNA HELICASE RUVB |
| 16:00 | 3pfi-B | 9.5 | 6.9 | 135 | 283 | 11 | MOLECULE: | HOLLIDAY JUNCTION ATP-DEPENDENT DNA HELICASE RUVB |
| 17:00 | 8efv-C | 9.5 | 11.9 | 146 | 301 | 14 | MOLECULE: | HOLLIDAY JUNCTION ATP-DEPENDENT DNA HELICASE RUVB |
| 18:00 | 8efy-B | 9.5 | 8.4 | 139 | 310 | 14 | MOLECULE: | HOLLIDAY JUNCTION ATP-DEPENDENT DNA HELICASE RUVB |
| 19:00 | 8efy-F | 9.5 | 10.1 | 140 | 308 | 14 | MOLECULE: | HOLLIDAY JUNCTION ATP-DEPENDENT DNA HELICASE RUVB |
| 20:00 | 8umu-A | 9.5 | 14.7 | 155 | 515 | 15 | MOLECULE: | CHROMOSOME TRANSMISSION FIDELITY PROTEIN 18 HOMOL |
| 21:00 | 1z1d-B | 9.5 | 3.1 | 111 | 131 | 9 | MOLECULE: | REPLICATION PROTEIN A 32 KDA SUBUNIT |
| 22:00 | 7pbs-B | 9.5 | 9.9 | 141 | 312 | 11 | MOLECULE: | HOLLIDAY JUNCTION ATP-DEPENDENT DNA HELICASE RUVB |
| 23:00 | 7pbl-C | 9.5 | 9.8 | 140 | 309 | 10 | MOLECULE: | HOLLIDAY JUNCTION ATP-DEPENDENT DNA HELICASE RUVB |
| 24:00:00 | 4fgn-A | 9.5 | 3 | 112 | 126 | 8 | MOLECULE: | LARGE T ANTIGEN |
| 25:00:00 | 7pbq-C | 9.5 | 10.1 | 142 | 309 | 11 | MOLECULE: | HOLLIDAY JUNCTION ATP-DEPENDENT DNA HELICASE RUVB |
| 26:00:00 | 7pbp-C | 9.5 | 10.2 | 139 | 309 | 11 | MOLECULE: | HOLLIDAY JUNCTION ATP-DEPENDENT DNA HELICASE RUVB |
| 27:00:00 | 7pbt-A | 9.5 | 9.6 | 139 | 312 | 11 | MOLECULE: | HOLLIDAY JUNCTION ATP-DEPENDENT DNA HELICASE RUVB |
| 28:00:00 | 7pbm-C | 9.5 | 10.3 | 141 | 309 | 12 | MOLECULE: | HOLLIDAY JUNCTION ATP-DEPENDENT DNA HELICASE RUVB |
| 29:00:00 | 8dzz-A | 9.5 | 18.7 | 179 | 2363 | 15 | MOLECULE: | DYNEIN HEAVY CHAIN, CYTOPLASMIC |
| 30:00:00 | 7mi8-A | 9.4 | 10.5 | 215 | 868 | 7 | MOLECULE: | FUSION PROTEIN OF DYNEIN AND ENDOLYSIN |
| 31:00:00 | 8hwf-E | 9.4 | 4.2 | 208 | 368 | 13 | MOLECULE: | PRIMASE D5 |
| 32:00:00 | 7ola-B | 9.4 | 8.7 | 217 | 441 | 14 | MOLECULE: | DNA PRIMASE |
| 33:00:00 | 8hwf-C | 9.4 | 3.7 | 204 | 378 | 10 | MOLECULE: | PRIMASE D5 |
| 34:00:00 | 8hwf-A | 9.4 | 4.5 | 215 | 378 | 11 | MOLECULE: | PRIMASE D5 |
| 35:00:00 | 4lzz-N | 9.4 | 3.7 | 138 | 247 | 9 | MOLECULE: | TRANSCRIPTIONAL REGULATOR (NTRC FAMILY) |
| 36:00:00 | 1ny6-N | 9.4 | 6.8 | 144 | 247 | 9 | MOLECULE: | TRANSCRIPTIONAL REGULATOR (NTRC FAMILY) |
| 37:00:00 | 8uh7-E | 9.4 | 4.9 | 133 | 314 | 17 | MOLECULE: | SLIDING-CLAMP-LOADER SMALL SUBUNIT |
| 38:00:00 | 7ttr-C | 9.4 | 12.7 | 161 | 327 | 12 | MOLECULE: | CASEINOLYTIC PEPTIDASE B PROTEIN HOMOLOG |
| 39:00:00 | 8efv-F | 9.4 | 10.2 | 145 | 299 | 14 | MOLECULE: | HOLLIDAY JUNCTION ATP-DEPENDENT DNA HELICASE RUVB |
| 40:00:00 | 8efy-K | 9.4 | 8.7 | 140 | 304 | 14 | MOLECULE: | HOLLIDAY JUNCTION ATP-DEPENDENT DNA HELICASE RUVB |
| 41:00:00 | 8efv-D | 9.4 | 9.9 | 140 | 296 | 13 | MOLECULE: | HOLLIDAY JUNCTION ATP-DEPENDENT DNA HELICASE RUVB |
| 42:00:00 | 7pbp-E | 9.4 | 10.3 | 142 | 312 | 10 | MOLECULE: | HOLLIDAY JUNCTION ATP-DEPENDENT DNA HELICASE RUVB |
| 43:00:00 | 7pbl-E | 9.4 | 9.4 | 139 | 312 | 11 | MOLECULE: | HOLLIDAY JUNCTION ATP-DEPENDENT DNA HELICASE RUVB |
| 44:00:00 | 7pbn-C | 9.4 | 10.6 | 141 | 309 | 11 | MOLECULE: | HOLLIDAY JUNCTION ATP-DEPENDENT DNA HELICASE RUVB |
| 45:00:00 | 7pbp-F | 9.4 | 9.3 | 139 | 312 | 11 | MOLECULE: | HOLLIDAY JUNCTION ATP-DEPENDENT DNA HELICASE RUVB |
| 46:00:00 | 8apm-E | 9.3 | 6.3 | 213 | 449 | 10 | MOLECULE: | PRIMASE D5 |
| 47:00:00 | 8apm-D | 9.3 | 5.8 | 210 | 449 | 10 | MOLECULE: | PRIMASE D5 |
| 48:00:00 | 4ly6-K | 9.3 | 3.6 | 142 | 247 | 11 | MOLECULE: | TRANSCRIPTIONAL REGULATOR (NTRC FAMILY) |
| 49:00:00 | 4ly6-N | 9.3 | 6.8 | 143 | 247 | 11 | MOLECULE: | TRANSCRIPTIONAL REGULATOR (NTRC FAMILY) |
| 50:00:00 | 1ny6-M | 9.3 | 6.9 | 146 | 247 | 10 | MOLECULE: | TRANSCRIPTIONAL REGULATOR (NTRC FAMILY) |
| 51:00:00 | 6n2i-A | 9.3 | 10.4 | 169 | 338 | 14 | MOLECULE: | DNA-BINDING ATP-DEPENDENT PROTEASE LA |
| 52:00:00 | 3u61-D | 9.3 | 10.9 | 146 | 320 | 16 | MOLECULE: | DNA POLYMERASE ACCESSORY PROTEIN 44 |
| 53:00:00 | 6jdl-A | 9.3 | 3.4 | 141 | 252 | 9 | MOLECULE: | NITROGEN ASSIMILATION REGULATORY PROTEIN |
| 54:00:00 | 7pds-B | 9.3 | 4.6 | 211 | 448 | 12 | MOLECULE: | SIMILAR TO D. NODOSUS VAPE |
| 55:00:00 | 7pbo-A | 9.3 | 10.2 | 141 | 312 | 10 | MOLECULE: | HOLLIDAY JUNCTION ATP-DEPENDENT DNA HELICASE RUVB |
| 56:00:00 | 6blb-A | 9.3 | 9.6 | 138 | 311 | 10 | MOLECULE: | HOLLIDAY JUNCTION ATP-DEPENDENT DNA HELICASE RUVB |
| 57:00:00 | 7x5b-A | 9.3 | 11.2 | 141 | 312 | 12 | MOLECULE: | HOLLIDAY JUNCTION ATP-DEPENDENT DNA HELICASE RUVB |
| 58:00:00 | 7pbn-E | 9.3 | 10.3 | 141 | 312 | 9 | MOLECULE: | HOLLIDAY JUNCTION ATP-DEPENDENT DNA HELICASE RUVB |
| 59:00:00 | 7pbo-C | 9.3 | 11.2 | 151 | 309 | 11 | MOLECULE: | HOLLIDAY JUNCTION ATP-DEPENDENT DNA HELICASE RUVB |
| 60:00:00 | 7pbq-D | 9.3 | 10.8 | 143 | 313 | 10 | MOLECULE: | HOLLIDAY JUNCTION ATP-DEPENDENT DNA HELICASE RUVB |
| 61:00:00 | 7pbq-A | 9.3 | 9.8 | 141 | 312 | 11 | MOLECULE: | HOLLIDAY JUNCTION ATP-DEPENDENT DNA HELICASE RUVB |
| 62:00:00 | 7pbo-B | 9.3 | 10 | 140 | 312 | 10 | MOLECULE: | HOLLIDAY JUNCTION ATP-DEPENDENT DNA HELICASE RUVB |
| 63:00:00 | 7ola-A | 9.2 | 7.1 | 213 | 441 | 12 | MOLECULE: | DNA PRIMASE |

| | | | | | | | | |
|---|---|---|---|---|---|---|---|---|
| 64:00:00 | 8efy-N | 9.2 | 10 | 145 | 310 | 13 | MOLECULE: | HOLLIDAY JUNCTION ATP-DEPENDENT DNA HELICASE RUVB |
| 65:00:00 | 8xj7-B | 9.2 | 6.7 | 207 | 452 | 10 | MOLECULE: | MONKEYPOX VIRUS E5 |
| 66:00:00 | 8hwa-B | 9.2 | 4.2 | 209 | 379 | 10 | MOLECULE: | PRIMASE D5 |
| 67:00:00 | 4lzz-R | 9.2 | 3.7 | 142 | 247 | 10 | MOLECULE: | TRANSCRIPTIONAL REGULATOR (NTRC FAMILY) |
| 68:00:00 | 4ly6-M | 9.2 | 3.4 | 140 | 247 | 11 | MOLECULE: | TRANSCRIPTIONAL REGULATOR (NTRC FAMILY) |
| 69:00:00 | 4lzz-K | 9.2 | 3.7 | 141 | 247 | 10 | MOLECULE: | TRANSCRIPTIONAL REGULATOR (NTRC FAMILY) |
| 70:00:00 | 4ly6-A | 9.2 | 3.5 | 141 | 247 | 11 | MOLECULE: | TRANSCRIPTIONAL REGULATOR (NTRC FAMILY) |
| 71:00:00 | 4lzz-W | 9.2 | 3.9 | 140 | 250 | 9 | MOLECULE: | TRANSCRIPTIONAL REGULATOR (NTRC FAMILY) |
| 72:00:00 | 4ly6-S | 9.2 | 6.9 | 144 | 247 | 11 | MOLECULE: | TRANSCRIPTIONAL REGULATOR (NTRC FAMILY) |
| 73:00:00 | 5ep3-A | 9.2 | 3.4 | 136 | 253 | 10 | MOLECULE: | PUTATIVE REPRESSOR PROTEIN LUXO |
| 74:00:00 | 4lzz-T | 9.2 | 3.6 | 138 | 247 | 10 | MOLECULE: | TRANSCRIPTIONAL REGULATOR (NTRC FAMILY) |
| 75:00:00 | 1ny6-K | 9.2 | 6.7 | 145 | 248 | 9 | MOLECULE: | TRANSCRIPTIONAL REGULATOR (NTRC FAMILY) |
| 76:00:00 | 1ny6-J | 9.2 | 6.9 | 142 | 247 | 9 | MOLECULE: | TRANSCRIPTIONAL REGULATOR (NTRC FAMILY) |
| 77:00:00 | 3u60-D | 9.2 | 13.8 | 144 | 319 | 18 | MOLECULE: | DNA POLYMERASE ACCESSORY PROTEIN 44 |
| 78:00:00 | 3u60-C | 9.2 | 12 | 144 | 320 | 16 | MOLECULE: | DNA POLYMERASE ACCESSORY PROTEIN 44 |
| 79:00:00 | 8xj8-B | 9.2 | 3.7 | 201 | 372 | 8 | MOLECULE: | DNA (70-MER) |
| 80:00:00 | 3u5z-N | 9.2 | 13.5 | 148 | 320 | 16 | MOLECULE: | DNA POLYMERASE ACCESSORY PROTEIN 44 |
| 81:00:00 | 3u61-B | 9.2 | 13.3 | 140 | 305 | 17 | MOLECULE: | DNA POLYMERASE ACCESSORY PROTEIN 44 |
| 82:00:00 | 3u61-C | 9.2 | 13.5 | 152 | 319 | 16 | MOLECULE: | DNA POLYMERASE ACCESSORY PROTEIN 44 |
| 83:00:00 | 3u5z-D | 9.2 | 12.7 | 144 | 320 | 17 | MOLECULE: | DNA POLYMERASE ACCESSORY PROTEIN 44 |
| 84:00:00 | 8uh7-C | 9.2 | 9 | 138 | 320 | 18 | MOLECULE: | SLIDING-CLAMP-LOADER SMALL SUBUNIT |
| 85:00:00 | 7thv-A | 9.2 | 6.8 | 164 | 477 | 14 | MOLECULE: | REPLICATION FACTOR C SUBUNIT 1 |
| 86:00:00 | 8unf-D | 9.2 | 11 | 146 | 319 | 16 | MOLECULE: | PRIMER DNA |
| 87:00:00 | 7ksm-A | 9.2 | 10.2 | 181 | 526 | 13 | MOLECULE: | LON PROTEASE HOMOLOG, MITOCHONDRIAL |
| 88:00:00 | 7tts-B | 9.2 | 15.5 | 191 | 440 | 15 | MOLECULE: | CASEINOLYTIC PEPTIDASE B PROTEIN HOMOLOG |
| 89:00:00 | 6jdi-A | 9.2 | 3.5 | 142 | 252 | 9 | MOLECULE: | NITROGEN ASSIMILATION REGULATORY PROTEIN |
| 90:00:00 | 8efy-J | 9.2 | 9.4 | 136 | 314 | 13 | MOLECULE: | HOLLIDAY JUNCTION ATP-DEPENDENT DNA HELICASE RUVB |
| 91:00:00 | 4ly6-E | 9.2 | 7 | 145 | 247 | 10 | MOLECULE: | TRANSCRIPTIONAL REGULATOR (NTRC FAMILY) |
| 92:00:00 | 8efy-D | 9.2 | 9.1 | 139 | 309 | 14 | MOLECULE: | HOLLIDAY JUNCTION ATP-DEPENDENT DNA HELICASE RUVB |
| 93:00:00 | 7pbm-A | 9.2 | 7.2 | 139 | 312 | 13 | MOLECULE: | HOLLIDAY JUNCTION ATP-DEPENDENT DNA HELICASE RUVB |
| 94:00:00 | 7pbl-A | 9.2 | 8.8 | 139 | 312 | 12 | MOLECULE: | HOLLIDAY JUNCTION ATP-DEPENDENT DNA HELICASE RUVB |
| 95:00:00 | 7tts-D | 9.2 | 12.8 | 183 | 463 | 12 | MOLECULE: | CASEINOLYTIC PEPTIDASE B PROTEIN HOMOLOG |
| 96:00:00 | 7pbo-F | 9.2 | 11.1 | 144 | 312 | 11 | MOLECULE: | HOLLIDAY JUNCTION ATP-DEPENDENT DNA HELICASE RUVB |
| 97:00:00 | 7pbr-B | 9.2 | 11.5 | 144 | 312 | 10 | MOLECULE: | HOLLIDAY JUNCTION ATP-DEPENDENT DNA HELICASE RUVB |
| 98:00:00 | 7pbs-E | 9.2 | 9.1 | 139 | 312 | 11 | MOLECULE: | HOLLIDAY JUNCTION ATP-DEPENDENT DNA HELICASE RUVB |
| 99:00:00 | 1in8-A | 9.2 | 8.8 | 146 | 298 | 14 | MOLECULE: | HOLLIDAY JUNCTION DNA HELICASE RUVB |
| 0:00 | 7pbl-F | 9.2 | 9.9 | 141 | 312 | 11 | MOLECULE: | HOLLIDAY JUNCTION ATP-DEPENDENT DNA HELICASE RUVB |
| 1:00 | 1in4-A | 9.2 | 9.5 | 138 | 298 | 14 | MOLECULE: | HOLLIDAY JUNCTION DNA HELICASE RUVB |
| 2:00 | 7pbq-F | 9.2 | 9.7 | 141 | 312 | 11 | MOLECULE: | HOLLIDAY JUNCTION ATP-DEPENDENT DNA HELICASE RUVB |
| 3:00 | 7pbp-B | 9.2 | 10 | 139 | 312 | 10 | MOLECULE: | HOLLIDAY JUNCTION ATP-DEPENDENT DNA HELICASE RUVB |
| 4:00 | 8fdt-A | 9.2 | 12.8 | 200 | 2736 | 8 | MOLECULE: | CYTOPLASMIC DYNEIN 1 HEAVY CHAIN 1 |
| 5:00 | 7pbm-D | 9.2 | 10.4 | 143 | 313 | 11 | MOLECULE: | HOLLIDAY JUNCTION ATP-DEPENDENT DNA HELICASE RUVB |
| 6:00 | 8dr7-A | 9.2 | 9.6 | 143 | 497 | 13 | MOLECULE: | REPLICATION FACTOR C SUBUNIT 1 |
| 7:00 | 7pbr-D | 9.2 | 11.7 | 151 | 313 | 10 | MOLECULE: | HOLLIDAY JUNCTION ATP-DEPENDENT DNA HELICASE RUVB |
| 8:00 | 7ola-D | 9.1 | 7.4 | 208 | 372 | 11 | MOLECULE: | DNA PRIMASE |
| 9:00 | 3sc3-A | 9.1 | 3.7 | 122 | 212 | 13 | MOLECULE: | PUTATIVE DNA REPLICATION REGULATOR HDA |
| 10:00 | 8hwf-D | 9.1 | 4.2 | 207 | 378 | 11 | MOLECULE: | PRIMASE D5 |
| 11:00 | 5exp-A | 9.1 | 3.4 | 136 | 250 | 8 | MOLECULE: | TRANSCRIPTIONAL REGULATOR FLEQ |
| 12:00 | 5ep2-A | 9.1 | 3.5 | 136 | 252 | 10 | MOLECULE: | PUTATIVE REPRESSOR PROTEIN LUXO |
| 13:00 | 4lzz-Q | 9.1 | 3.8 | 141 | 247 | 9 | MOLECULE: | TRANSCRIPTIONAL REGULATOR (NTRC FAMILY) |
| 14:00 | 4ly6-G | 9.1 | 6.7 | 145 | 247 | 11 | MOLECULE: | TRANSCRIPTIONAL REGULATOR (NTRC FAMILY) |
| 15:00 | 4ly6-R | 9.1 | 3.6 | 140 | 247 | 10 | MOLECULE: | TRANSCRIPTIONAL REGULATOR (NTRC FAMILY) |
| 16:00 | 4ly6-W | 9.1 | 6.7 | 146 | 247 | 10 | MOLECULE: | TRANSCRIPTIONAL REGULATOR (NTRC FAMILY) |

| | | | | | | | | |
|---|---|---|---|---|---|---|---|---|
| 17:00 | 4lzz-F | 9.1 | 3.6 | 140 | 247 | 11 | MOLECULE: | TRANSCRIPTIONAL REGULATOR (NTRC FAMILY) |
| 18:00 | 4lzz-H | 9.1 | 3.7 | 140 | 247 | 9 | MOLECULE: | TRANSCRIPTIONAL REGULATOR (NTRC FAMILY) |
| 19:00 | 4ly6-H | 9.1 | 6.9 | 143 | 247 | 10 | MOLECULE: | TRANSCRIPTIONAL REGULATOR (NTRC FAMILY) |
| 20:00 | 3u61-E | 9.1 | 4.7 | 136 | 294 | 16 | MOLECULE: | DNA POLYMERASE ACCESSORY PROTEIN 44 |
| 21:00 | 3u5z-M | 9.1 | 12.1 | 143 | 320 | 16 | MOLECULE: | DNA POLYMERASE ACCESSORY PROTEIN 44 |
| 22:00 | 8uk9-C | 9.1 | 10 | 141 | 320 | 17 | MOLECULE: | SLIDING-CLAMP-LOADER SMALL SUBUNIT |
| 23:00 | 8uh7-B | 9.1 | 9.5 | 139 | 319 | 17 | MOLECULE: | SLIDING-CLAMP-LOADER SMALL SUBUNIT |
| 24:00:00 | 8unh-B | 9.1 | 4.4 | 131 | 319 | 18 | MOLECULE: | SLIDING-CLAMP-LOADER LARGE SUBUNIT |
| 25:00:00 | 8giy-D | 9.1 | 12.3 | 148 | 360 | 14 | MOLECULE: | DNA POLYMERASE III SUBUNIT DELTA |
| 26:00:00 | 6rn2-E | 9.1 | 10.3 | 207 | 684 | 10 | MOLECULE: | CHAPERONE PROTEIN CLPB |
| 27:00:00 | 1j7k-A | 9.1 | 9 | 136 | 299 | 15 | MOLECULE: | HOLLIDAY JUNCTION DNA HELICASE RUVB |
| 28:00:00 | 7vbw-A | 9.1 | 3.5 | 138 | 238 | 12 | MOLECULE: | SIGMA-54 DEPENDENT TRANCSRIPTIONAL REGULATOR |
| 29:00:00 | 7vbs-A | 9.1 | 3.7 | 137 | 238 | 12 | MOLECULE: | SIGMA-54 DEPENDENT TRANCSRIPTIONAL REGULATOR |
| 30:00:00 | 1in5-A | 9.1 | 10.3 | 146 | 301 | 14 | MOLECULE: | HOLLIDAY JUNCTION DNA HELICASE RUVB |
| 31:00:00 | 4ly6-B | 9.1 | 7.1 | 145 | 247 | 10 | MOLECULE: | TRANSCRIPTIONAL REGULATOR (NTRC FAMILY) |
| 32:00:00 | 7pbn-A | 9.1 | 11.8 | 143 | 312 | 12 | MOLECULE: | HOLLIDAY JUNCTION ATP-DEPENDENT DNA HELICASE RUVB |
| 33:00:00 | 4lzz-O | 9.1 | 3.7 | 140 | 247 | 10 | MOLECULE: | TRANSCRIPTIONAL REGULATOR (NTRC FAMILY) |
| 34:00:00 | 4qnm-A | 9.1 | 3.7 | 131 | 238 | 9 | MOLECULE: | PSP OPERON TRANSCRIPTIONAL ACTIVATOR |
| 35:00:00 | 7pbm-E | 9.1 | 9.9 | 142 | 312 | 11 | MOLECULE: | HOLLIDAY JUNCTION ATP-DEPENDENT DNA HELICASE RUVB |
| 36:00:00 | 7pbm-F | 9.1 | 9.7 | 142 | 312 | 11 | MOLECULE: | HOLLIDAY JUNCTION ATP-DEPENDENT DNA HELICASE RUVB |
| 37:00:00 | 8dr3-A | 9.1 | 12.2 | 177 | 643 | 10 | MOLECULE: | REPLICATION FACTOR C SUBUNIT 1 |
| 38:00:00 | 7pbn-F | 9.1 | 9.6 | 142 | 312 | 11 | MOLECULE: | HOLLIDAY JUNCTION ATP-DEPENDENT DNA HELICASE RUVB |
| 39:00:00 | 7pbl-B | 9.1 | 9.9 | 139 | 312 | 10 | MOLECULE: | HOLLIDAY JUNCTION ATP-DEPENDENT DNA HELICASE RUVB |
| 40:00:00 | 7u1p-A | 9.1 | 11.3 | 170 | 617 | 12 | MOLECULE: | REPLICATION FACTOR C SUBUNIT 1 |
| 41:00:00 | 8hwg-B | 9 | 3.6 | 200 | 369 | 9 | MOLECULE: | PRIMASE D5 |
| 42:00:00 | 8hwh-A | 9 | 4.4 | 197 | 353 | 11 | MOLECULE: | PRIMASE D5 |
| 43:00:00 | 8iqi-A | 9 | 13.1 | 251 | 915 | 10 | MOLECULE: | PUTATIVE PRIMASE C962R |
| 44:00:00 | 8hwe-B | 9 | 3.7 | 202 | 379 | 9 | MOLECULE: | PRIMASE D5 |
| 45:00:00 | 8iqi-C | 9 | 17.7 | 209 | 916 | 11 | MOLECULE: | PUTATIVE PRIMASE C962R |
| 46:00:00 | 4lzz-U | 9 | 3.7 | 139 | 247 | 10 | MOLECULE: | TRANSCRIPTIONAL REGULATOR (NTRC FAMILY) |
| 47:00:00 | 4ly6-V | 9 | 6.7 | 145 | 247 | 10 | MOLECULE: | TRANSCRIPTIONAL REGULATOR (NTRC FAMILY) |
| 48:00:00 | 4qos-A | 9 | 3.7 | 134 | 240 | 10 | MOLECULE: | PSP OPERON TRANSCRIPTIONAL ACTIVATOR |
| 49:00:00 | 4ly6-Q | 9 | 3.7 | 142 | 247 | 10 | MOLECULE: | TRANSCRIPTIONAL REGULATOR (NTRC FAMILY) |
| 50:00:00 | 4lzz-C | 9 | 3.7 | 140 | 247 | 10 | MOLECULE: | TRANSCRIPTIONAL REGULATOR (NTRC FAMILY) |
| 51:00:00 | 2c99-A | 9 | 3.5 | 136 | 243 | 10 | MOLECULE: | PSP OPERON TRANSCRIPTIONAL ACTIVATOR |
| 52:00:00 | 3dzd-B | 9 | 6.3 | 176 | 368 | 10 | MOLECULE: | TRANSCRIPTIONAL REGULATOR (NTRC FAMILY) |
| 53:00:00 | 4ly6-D | 9 | 6.8 | 145 | 247 | 10 | MOLECULE: | TRANSCRIPTIONAL REGULATOR (NTRC FAMILY) |
| 54:00:00 | 4lj8-A | 9 | 12.3 | 169 | 322 | 15 | MOLECULE: | CHAPERONE PROTEIN CLPB |
| 55:00:00 | 4lj4-A | 9 | 11.7 | 172 | 319 | 14 | MOLECULE: | CHAPERONE PROTEIN CLPB |
| 56:00:00 | 4ypn-A | 9 | 5.7 | 161 | 250 | 13 | MOLECULE: | LON PROTEASE |
| 57:00:00 | 3u60-B | 9 | 14 | 148 | 319 | 16 | MOLECULE: | DNA POLYMERASE ACCESSORY PROTEIN 44 |
| 58:00:00 | 8uk9-D | 9 | 10.6 | 146 | 318 | 16 | MOLECULE: | SLIDING-CLAMP-LOADER SMALL SUBUNIT |
| 59:00:00 | 7tts-A | 9 | 12.8 | 181 | 433 | 10 | MOLECULE: | CASEINOLYTIC PEPTIDASE B PROTEIN HOMOLOG |
| 60:00:00 | 8var-D | 9 | 12.4 | 148 | 360 | 11 | MOLECULE: | DNA POLYMERASE III SUBUNIT DELTA |
| 61:00:00 | 6e10-1 | 9 | 14.7 | 220 | 719 | 14 | MOLECULE: | HEAT SHOCK PROTEIN 101 |
| 62:00:00 | 7ttr-D | 9 | 11.9 | 154 | 334 | 12 | MOLECULE: | CASEINOLYTIC PEPTIDASE B PROTEIN HOMOLOG |
| 63:00:00 | 8pb9-D | 9 | 7.8 | 157 | 390 | 8 | MOLECULE: | TRANSCRIPTIONAL REGULATOR FLEQ |
| 64:00:00 | 8dr6-A | 9 | 13.2 | 147 | 493 | 14 | MOLECULE: | REPLICATION FACTOR C SUBUNIT 1 |
| 65:00:00 | 1in6-A | 9 | 9.7 | 144 | 300 | 16 | MOLECULE: | HOLLIDAY JUNCTION DNA HELICASE RUVB |
| 66:00:00 | 7pbn-B | 9 | 11.9 | 143 | 312 | 11 | MOLECULE: | HOLLIDAY JUNCTION ATP-DEPENDENT DNA HELICASE RUVB |
| 67:00:00 | 3u5z-O | 9 | 4.7 | 132 | 305 | 17 | MOLECULE: | DNA POLYMERASE ACCESSORY PROTEIN 44 |
| 68:00:00 | 6rla-B | 9 | 12.1 | 204 | 2750 | 8 | MOLECULE: | O6-ALKYLGUANINE-DNA ALKYLTRANSFERASE MUTANT,DYNC2 |
| 69:00:00 | 2tbd-A | 9 | 3.5 | 113 | 134 | 8 | MOLECULE: | SV40 T ANTIGEN |

| | | | | | | | | |
|---|---|---|---|---|---|---|---|---|
| 70:00:00 | 7u1a-A | 9 | 12.1 | 169 | 617 | 14 | MOLECULE: | REPLICATION FACTOR C SUBUNIT 1 |
| 71:00:00 | 8umu-B | 9 | 10.5 | 136 | 316 | 13 | MOLECULE: | CHROMOSOME TRANSMISSION FIDELITY PROTEIN 18 HOMOL |
| 72:00:00 | 8dr1-A | 9 | 11.7 | 149 | 493 | 13 | MOLECULE: | REPLICATION FACTOR C SUBUNIT 1 |
| 73:00:00 | 7ti8-A | 9 | 11.1 | 149 | 491 | 13 | MOLECULE: | REPLICATION FACTOR C SUBUNIT 1 |
| 74:00:00 | 8dr0-A | 9 | 10.9 | 153 | 492 | 13 | MOLECULE: | REPLICATION FACTOR C SUBUNIT 1 |
| 75:00:00 | 7u19-A | 9 | 12.8 | 172 | 617 | 13 | MOLECULE: | REPLICATION FACTOR C SUBUNIT 1 |
| 76:00:00 | 8apl-B | 8.9 | 6.3 | 216 | 449 | 11 | MOLECULE: | PRIMASE D5 |
| 77:00:00 | 8apl-F | 8.9 | 6.6 | 217 | 449 | 11 | MOLECULE: | PRIMASE D5 |
| 78:00:00 | 8apl-C | 8.9 | 5.9 | 213 | 449 | 10 | MOLECULE: | PRIMASE D5 |
| 79:00:00 | 4ly6-J | 8.9 | 7 | 145 | 247 | 10 | MOLECULE: | TRANSCRIPTIONAL REGULATOR (NTRC FAMILY) |
| 80:00:00 | 4lzz-G | 8.9 | 6.9 | 145 | 247 | 10 | MOLECULE: | TRANSCRIPTIONAL REGULATOR (NTRC FAMILY) |
| 81:00:00 | 4ly6-C | 8.9 | 3.8 | 140 | 247 | 9 | MOLECULE: | TRANSCRIPTIONAL REGULATOR (NTRC FAMILY) |
| 82:00:00 | 4lzz-D | 8.9 | 3.6 | 137 | 247 | 11 | MOLECULE: | TRANSCRIPTIONAL REGULATOR (NTRC FAMILY) |
| 83:00:00 | 5ext-A | 8.9 | 3.5 | 141 | 252 | 7 | MOLECULE: | TRANSCRIPTIONAL REGULATOR FLEQ |
| 84:00:00 | 4qnr-A | 8.9 | 3.6 | 134 | 243 | 9 | MOLECULE: | PSP OPERON TRANSCRIPTIONAL ACTIVATOR |
| 85:00:00 | 4ly6-I | 8.9 | 3.6 | 139 | 247 | 10 | MOLECULE: | TRANSCRIPTIONAL REGULATOR (NTRC FAMILY) |
| 86:00:00 | 5exs-A | 8.9 | 3.4 | 141 | 252 | 9 | MOLECULE: | TRANSCRIPTIONAL REGULATOR FLEQ |
| 87:00:00 | 6pod-A | 8.9 | 12.7 | 155 | 329 | 15 | MOLECULE: | ATP-DEPENDENT CLP PROTEASE ATP-BINDING SUBUNIT CL |
| 88:00:00 | 3bos-B | 8.9 | 7.7 | 136 | 231 | 15 | MOLECULE: | PUTATIVE DNA REPLICATION FACTOR |
| 89:00:00 | 4lj6-A | 8.9 | 11.9 | 173 | 322 | 14 | MOLECULE: | CHAPERONE PROTEIN CLPB |
| 90:00:00 | 6on2-B | 8.9 | 11.7 | 183 | 524 | 13 | MOLECULE: | ATP-DEPENDENT PROTEASE LA |
| 91:00:00 | 7tfi-A | 8.9 | 14 | 149 | 391 | 13 | MOLECULE: | REPLICATION FACTOR C SUBUNIT 1 |
| 92:00:00 | 7p0m-F | 8.9 | 11.7 | 191 | 520 | 10 | MOLECULE: | LON PROTEASE HOMOLOG, MITOCHONDRIAL |
| 93:00:00 | 7ttr-A | 8.9 | 11 | 157 | 304 | 11 | MOLECULE: | CASEINOLYTIC PEPTIDASE B PROTEIN HOMOLOG |
| 94:00:00 | 6l1q-C | 8.9 | 6.7 | 143 | 267 | 11 | MOLECULE: | CBBQ PROTEIN |
| 95:00:00 | 7p6x-A | 8.9 | 3.2 | 142 | 320 | 12 | MOLECULE: | RUVB-LIKE 1 |
| 96:00:00 | 6z1e-B | 8.9 | 15.5 | 157 | 259 | 13 | MOLECULE: | RIBULOSE BISPHOSPHATE CARBOXYLASE/OXYGENASE ACTIV |
| 97:00:00 | 7pbs-F | 8.9 | 9.1 | 136 | 312 | 12 | MOLECULE: | HOLLIDAY JUNCTION ATP-DEPENDENT DNA HELICASE RUVB |
| 98:00:00 | 3u5z-B | 8.9 | 11.5 | 144 | 320 | 16 | MOLECULE: | DNA POLYMERASE ACCESSORY PROTEIN 44 |
| 99:00:00 | 7xbk-H | 8.9 | 8.1 | 153 | 345 | 12 | MOLECULE: | ISOFORM 2 OF CASEINOLYTIC PEPTIDASE B PROTEIN HOM |
| 0:00 | 7xbk-A | 8.9 | 11 | 158 | 367 | 12 | MOLECULE: | ISOFORM 2 OF CASEINOLYTIC PEPTIDASE B PROTEIN HOM |
| 1:00 | 8fd6-A | 8.9 | 13 | 256 | 2747 | 9 | MOLECULE: | CYTOPLASMIC DYNEIN 1 HEAVY CHAIN 1 |
| 2:00 | 7tid-A | 8.9 | 10.3 | 147 | 486 | 12 | MOLECULE: | REPLICATION FACTOR C SUBUNIT 1 |
| 3:00 | 7u19-C | 8.9 | 14.5 | 153 | 328 | 13 | MOLECULE: | REPLICATION FACTOR C SUBUNIT 1 |
| 4:00 | 7thv-C | 8.9 | 14.2 | 141 | 322 | 13 | MOLECULE: | REPLICATION FACTOR C SUBUNIT 1 |
| 5:00 | 8umy-C | 8.9 | 11.6 | 150 | 325 | 14 | MOLECULE: | CHROMOSOME TRANSMISSION FIDELITY PROTEIN 18 HOMOL |
| 6:00 | 7tib-A | 8.9 | 11.3 | 156 | 495 | 13 | MOLECULE: | REPLICATION FACTOR C SUBUNIT 1 |
| 7:00 | 8fs5-C | 8.9 | 12.3 | 150 | 327 | 11 | MOLECULE: | CHECKPOINT PROTEIN RAD24 |
| 8:00 | 8e00-A | 8.9 | 20.6 | 184 | 2363 | 14 | MOLECULE: | DYNEIN HEAVY CHAIN, CYTOPLASMIC |
| 9:00 | 7tic-A | 8.9 | 11.8 | 172 | 477 | 15 | MOLECULE: | REPLICATION FACTOR C SUBUNIT 1 |
| 10:00 | 7tfi-B | 8.9 | 14.4 | 158 | 320 | 11 | MOLECULE: | REPLICATION FACTOR C SUBUNIT 1 |
| 11:00 | 8apl-E | 8.8 | 6.3 | 212 | 449 | 11 | MOLECULE: | PRIMASE D5 |
| 12:00 | 8apl-D | 8.8 | 6.6 | 215 | 449 | 10 | MOLECULE: | PRIMASE D5 |
| 13:00 | 8apl-A | 8.8 | 6 | 211 | 449 | 10 | MOLECULE: | PRIMASE D5 |
| 14:00 | 8hwg-E | 8.8 | 4.3 | 209 | 369 | 12 | MOLECULE: | PRIMASE D5 |
| 15:00 | 6hz8-J | 8.8 | 3.4 | 136 | 284 | 11 | MOLECULE: | 5-METHYLCYTOSINE-SPECIFIC RESTRICTION ENZYME B |
| 16:00 | 8xj7-A | 8.8 | 3.6 | 200 | 377 | 10 | MOLECULE: | MONKEYPOX VIRUS E5 |
| 17:00 | 2vii-A | 8.8 | 3.6 | 138 | 247 | 9 | MOLECULE: | PSP OPERON TRANSCRIPTIONAL ACTIVATOR |
| 18:00 | 4lzz-P | 8.8 | 3.7 | 140 | 247 | 11 | MOLECULE: | TRANSCRIPTIONAL REGULATOR (NTRC FAMILY) |
| 19:00 | 4ly6-U | 8.8 | 3.8 | 141 | 247 | 10 | MOLECULE: | TRANSCRIPTIONAL REGULATOR (NTRC FAMILY) |
| 20:00 | 4ly6-T | 8.8 | 6.8 | 143 | 247 | 11 | MOLECULE: | TRANSCRIPTIONAL REGULATOR (NTRC FAMILY) |
| 21:00 | 4ly6-P | 8.8 | 3.4 | 136 | 247 | 10 | MOLECULE: | TRANSCRIPTIONAL REGULATOR (NTRC FAMILY) |
| 22:00 | 4ly6-X | 8.8 | 3.5 | 139 | 247 | 10 | MOLECULE: | TRANSCRIPTIONAL REGULATOR (NTRC FAMILY) |

| | | | | | | | | |
|---|---|---|---|---|---|---|---|---|
| 23:00 | 6hz7-G | 8.8 | 3.8 | 137 | 290 | 10 | MOLECULE: | 5-METHYLCYTOSINE-SPECIFIC RESTRICTION ENZYME B |
| 24:00:00 | 3m6a-A | 8.8 | 13.3 | 176 | 496 | 13 | MOLECULE: | ATP-DEPENDENT PROTEASE LA 1 |
| 25:00:00 | 6hz9-G | 8.8 | 3.7 | 138 | 290 | 12 | MOLECULE: | 5-METHYLCYTOSINE-SPECIFIC RESTRICTION ENZYME B |
| 26:00:00 | 4fct-A | 8.8 | 9.7 | 155 | 292 | 14 | MOLECULE: | CHAPERONE PROTEIN CLPB |
| 27:00:00 | 3m6a-B | 8.8 | 12.2 | 177 | 496 | 13 | MOLECULE: | ATP-DEPENDENT PROTEASE LA 1 |
| 28:00:00 | 6hz9-A | 8.8 | 3.7 | 137 | 290 | 12 | MOLECULE: | 5-METHYLCYTOSINE-SPECIFIC RESTRICTION ENZYME B |
| 29:00:00 | 3u5z-L | 8.8 | 12.9 | 144 | 320 | 16 | MOLECULE: | DNA POLYMERASE ACCESSORY PROTEIN 44 |
| 30:00:00 | 3pvs-C | 8.8 | 7.2 | 138 | 420 | 12 | MOLECULE: | REPLICATION-ASSOCIATED RECOMBINATION PROTEIN A |
| 31:00:00 | 6oax-C | 8.8 | 15.7 | 179 | 581 | 16 | MOLECULE: | HYPERACTIVE DISAGGREGASE CLPB |
| 32:00:00 | 8dqz-A | 8.8 | 11.6 | 147 | 492 | 14 | MOLECULE: | REPLICATION FACTOR C SUBUNIT 1 |
| 33:00:00 | 4qht-C | 8.8 | 3.6 | 137 | 248 | 13 | MOLECULE: | FLAGELLAR REGULATORY PROTEIN C |
| 34:00:00 | 6luf-G | 8.8 | 3.8 | 139 | 246 | 12 | MOLECULE: | FLAGELLAR REGULATORY PROTEIN C |
| 35:00:00 | 6luf-A | 8.8 | 3.6 | 137 | 247 | 13 | MOLECULE: | FLAGELLAR REGULATORY PROTEIN C |
| 36:00:00 | 3pvs-D | 8.8 | 6.9 | 136 | 424 | 13 | MOLECULE: | REPLICATION-ASSOCIATED RECOMBINATION PROTEIN A |
| 37:00:00 | 8efy-L | 8.8 | 10 | 140 | 309 | 14 | MOLECULE: | HOLLIDAY JUNCTION ATP-DEPENDENT DNA HELICASE RUVB |
| 38:00:00 | 8efy-M | 8.8 | 9.4 | 141 | 306 | 13 | MOLECULE: | HOLLIDAY JUNCTION ATP-DEPENDENT DNA HELICASE RUVB |
| 39:00:00 | 7pbt-E | 8.8 | 8.9 | 136 | 312 | 10 | MOLECULE: | HOLLIDAY JUNCTION ATP-DEPENDENT DNA HELICASE RUVB |
| 40:00:00 | 7x7p-O | 8.8 | 7.7 | 146 | 309 | 11 | MOLECULE: | DNA |
| 41:00:00 | 8un0-A | 8.8 | 15.8 | 158 | 546 | 15 | MOLECULE: | CHROMOSOME TRANSMISSION FIDELITY PROTEIN 18 HOMOL |
| 42:00:00 | 8fs3-C | 8.8 | 14.9 | 154 | 328 | 12 | MOLECULE: | CHECKPOINT PROTEIN RAD24 |
| 43:00:00 | 3pvs-B | 8.8 | 15.6 | 143 | 417 | 12 | MOLECULE: | REPLICATION-ASSOCIATED RECOMBINATION PROTEIN A |
| 44:00:00 | 1tbd-A | 8.8 | 3.2 | 112 | 134 | 9 | MOLECULE: | SV40 T-ANTIGEN |
| 45:00:00 | 8dr7-B | 8.8 | 14.9 | 150 | 318 | 11 | MOLECULE: | REPLICATION FACTOR C SUBUNIT 1 |
| 46:00:00 | 8fs7-C | 8.8 | 13.6 | 143 | 327 | 13 | MOLECULE: | CHECKPOINT PROTEIN RAD24 |
| 47:00:00 | 2chq-A | 8.8 | 14.2 | 141 | 313 | 14 | MOLECULE: | REPLICATION FACTOR C SMALL SUBUNIT |
| 48:00:00 | 7tfh-A | 8.8 | 8.5 | 164 | 628 | 12 | MOLECULE: | REPLICATION FACTOR C SUBUNIT 1 |
| 49:00:00 | 8dr4-B | 8.8 | 12.5 | 148 | 318 | 11 | MOLECULE: | REPLICATION FACTOR C SUBUNIT 1 |
| 50:00:00 | 8fs8-C | 8.8 | 12.4 | 151 | 327 | 11 | MOLECULE: | CHECKPOINT PROTEIN RAD24 |
| 51:00:00 | 8dqx-C | 8.8 | 12 | 158 | 330 | 11 | MOLECULE: | REPLICATION FACTOR C SUBUNIT 1 |
| 52:00:00 | 7tku-B | 8.8 | 14.2 | 147 | 318 | 11 | MOLECULE: | REPLICATION FACTOR C SUBUNIT 1 |
| 53:00:00 | 8dqz-C | 8.8 | 12.7 | 160 | 329 | 11 | MOLECULE: | REPLICATION FACTOR C SUBUNIT 1 |
| 54:00:00 | 7pbo-E | 8.8 | 10.5 | 140 | 312 | 10 | MOLECULE: | HOLLIDAY JUNCTION ATP-DEPENDENT DNA HELICASE RUVB |
| 55:00:00 | 7tfh-C | 8.8 | 10.8 | 145 | 330 | 12 | MOLECULE: | REPLICATION FACTOR C SUBUNIT 1 |
| 56:00:00 | 6hz5-B | 8.7 | 3.5 | 135 | 290 | 11 | MOLECULE: | 5-METHYLCYTOSINE-SPECIFIC RESTRICTION ENZYME B |
| 57:00:00 | 4lja-A | 8.7 | 11.6 | 168 | 320 | 14 | MOLECULE: | CHAPERONE PROTEIN CLPB |
| 58:00:00 | 4lzz-A | 8.7 | 7.2 | 145 | 247 | 10 | MOLECULE: | TRANSCRIPTIONAL REGULATOR (NTRC FAMILY) |
| 59:00:00 | 4lzz-M | 8.7 | 6.8 | 145 | 247 | 10 | MOLECULE: | TRANSCRIPTIONAL REGULATOR (NTRC FAMILY) |
| 60:00:00 | 2bjw-A | 8.7 | 7.3 | 136 | 242 | 9 | MOLECULE: | PSP OPERON TRANSCRIPTIONAL ACTIVATOR |
| 61:00:00 | 2c9c-A | 8.7 | 3.7 | 137 | 244 | 9 | MOLECULE: | PSP OPERON TRANSCRIPTIONAL ACTIVATOR |
| 62:00:00 | 4lzz-E | 8.7 | 4.4 | 144 | 258 | 10 | MOLECULE: | TRANSCRIPTIONAL REGULATOR (NTRC FAMILY) |
| 63:00:00 | 4qht-F | 8.7 | 3.4 | 136 | 245 | 13 | MOLECULE: | FLAGELLAR REGULATORY PROTEIN C |
| 64:00:00 | 4l4u-A | 8.7 | 9.7 | 165 | 388 | 10 | MOLECULE: | TRANSCRIPTIONAL REGULATOR (NTRC FAMILY) |
| 65:00:00 | 3uk6-I | 8.7 | 3.1 | 138 | 294 | 12 | MOLECULE: | RUVB-LIKE 2 |
| 66:00:00 | 6hz9-D | 8.7 | 3.4 | 136 | 284 | 11 | MOLECULE: | 5-METHYLCYTOSINE-SPECIFIC RESTRICTION ENZYME B |
| 67:00:00 | 3m6a-E | 8.7 | 14.3 | 178 | 496 | 14 | MOLECULE: | ATP-DEPENDENT PROTEASE LA 1 |
| 68:00:00 | 3m6a-C | 8.7 | 12.2 | 177 | 496 | 14 | MOLECULE: | ATP-DEPENDENT PROTEASE LA 1 |
| 69:00:00 | 4ypl-F | 8.7 | 13.4 | 185 | 538 | 14 | MOLECULE: | LON PROTEASE |
| 70:00:00 | 4fcw-F | 8.7 | 9.6 | 150 | 306 | 17 | MOLECULE: | CHAPERONE PROTEIN CLPB |
| 71:00:00 | 7ksm-E | 8.7 | 10.7 | 168 | 524 | 10 | MOLECULE: | LON PROTEASE HOMOLOG, MITOCHONDRIAL |
| 72:00:00 | 7tku-A | 8.7 | 10.3 | 147 | 486 | 12 | MOLECULE: | REPLICATION FACTOR C SUBUNIT 1 |
| 73:00:00 | 8uk9-B | 8.7 | 10.1 | 137 | 319 | 16 | MOLECULE: | SLIDING-CLAMP-LOADER SMALL SUBUNIT |
| 74:00:00 | 8giz-D | 8.7 | 12.3 | 142 | 360 | 12 | MOLECULE: | DNA POLYMERASE III SUBUNIT DELTA |
| 75:00:00 | 6l1q-B | 8.7 | 7 | 140 | 267 | 11 | MOLECULE: | CBBQ PROTEIN |

| | | | | | | | | |
|---|---|---|---|---|---|---|---|---|
| 76:00:00 | 8vap-D | 8.7 | 11.5 | 135 | 358 | 14 | MOLECULE: | DNA POLYMERASE III SUBUNIT DELTA |
| 77:00:00 | 4qhs-G | 8.7 | 3.7 | 136 | 243 | 13 | MOLECULE: | FLAGELLAR REGULATORY PROTEIN C |
| 78:00:00 | 6fo1-A | 8.7 | 11.5 | 147 | 323 | 12 | MOLECULE: | RUVB-LIKE 1 |
| 79:00:00 | 1ojl-D | 8.7 | 12.2 | 156 | 297 | 12 | MOLECULE: | TRANSCRIPTIONAL REGULATORY PROTEIN ZRAR |
| 80:00:00 | 6luf-C | 8.7 | 3.8 | 141 | 246 | 11 | MOLECULE: | FLAGELLAR REGULATORY PROTEIN C |
| 81:00:00 | 1ojl-A | 8.7 | 13.3 | 150 | 292 | 9 | MOLECULE: | TRANSCRIPTIONAL REGULATORY PROTEIN ZRAR |
| 82:00:00 | 8umt-A | 8.7 | 4.3 | 144 | 537 | 16 | MOLECULE: | CHROMOSOME TRANSMISSION FIDELITY PROTEIN 18 HOMOL |
| 83:00:00 | 2chg-A | 8.7 | 3.1 | 131 | 223 | 11 | MOLECULE: | REPLICATION FACTOR C SMALL SUBUNIT |
| 84:00:00 | 2chg-C | 8.7 | 3 | 127 | 221 | 12 | MOLECULE: | REPLICATION FACTOR C SMALL SUBUNIT |
| 85:00:00 | 4lzz-B | 8.7 | 6.8 | 142 | 247 | 11 | MOLECULE: | TRANSCRIPTIONAL REGULATOR (NTRC FAMILY) |
| 86:00:00 | 3uk6-E | 8.7 | 3.1 | 138 | 290 | 12 | MOLECULE: | RUVB-LIKE 2 |
| 87:00:00 | 6z1e-A | 8.7 | 7 | 151 | 274 | 13 | MOLECULE: | RIBULOSE BISPHOSPHATE CARBOXYLASE/OXYGENASE ACTIV |
| 88:00:00 | 1hqc-B | 8.7 | 8.8 | 138 | 314 | 12 | MOLECULE: | RUVB |
| 89:00:00 | 1ixs-B | 8.7 | 10.1 | 144 | 315 | 13 | MOLECULE: | HOLLIDAY JUNCTION DNA HELICASE RUVA |
| 90:00:00 | 2c9o-C | 8.7 | 3.1 | 139 | 311 | 13 | MOLECULE: | RUVB-LIKE 1 |
| 91:00:00 | 8pqw-A | 8.7 | 20.2 | 208 | 2892 | 8 | MOLECULE: | CYTOPLASMIC DYNEIN 1 HEAVY CHAIN 1 |
| 92:00:00 | 7tic-C | 8.7 | 14.3 | 142 | 322 | 13 | MOLECULE: | REPLICATION FACTOR C SUBUNIT 1 |
| 93:00:00 | 7u1p-B | 8.7 | 14.5 | 146 | 319 | 13 | MOLECULE: | REPLICATION FACTOR C SUBUNIT 1 |
| 94:00:00 | 8dzz-D | 8.7 | 8.1 | 222 | 2363 | 7 | MOLECULE: | DYNEIN HEAVY CHAIN, CYTOPLASMIC |
| 95:00:00 | 1iqp-C | 8.7 | 14.2 | 151 | 319 | 13 | MOLECULE: | RFCS |
| 96:00:00 | 7pbt-D | 8.7 | 11.3 | 146 | 313 | 10 | MOLECULE: | HOLLIDAY JUNCTION ATP-DEPENDENT DNA HELICASE RUVB |
| 97:00:00 | 8thd-B | 8.7 | 11.7 | 140 | 318 | 12 | MOLECULE: | ELG1 ISOFORM 1 |
| 98:00:00 | 8dr5-B | 8.7 | 15.5 | 145 | 318 | 12 | MOLECULE: | REPLICATION FACTOR C SUBUNIT 1 |
| 99:00:00 | 8dr6-B | 8.7 | 14.4 | 139 | 318 | 13 | MOLECULE: | REPLICATION FACTOR C SUBUNIT 1 |
| 0:00 | 7tfj-B | 8.7 | 14.7 | 144 | 320 | 12 | MOLECULE: | REPLICATION FACTOR C SUBUNIT 1 |
| 1:00 | 8dqw-C | 8.7 | 14.3 | 146 | 327 | 12 | MOLECULE: | REPLICATION FACTOR C SUBUNIT 4 |
| 2:00 | 8dr1-C | 8.7 | 11.1 | 148 | 330 | 12 | MOLECULE: | REPLICATION FACTOR C SUBUNIT 1 |
| 3:00 | 8thd-C | 8.7 | 12.5 | 145 | 325 | 13 | MOLECULE: | ELG1 ISOFORM 1 |
| 4:00 | 7thj-A | 8.7 | 10.5 | 167 | 482 | 12 | MOLECULE: | REPLICATION FACTOR C SUBUNIT 1 |
| 5:00 | 8umy-B | 8.7 | 12.1 | 142 | 319 | 13 | MOLECULE: | CHROMOSOME TRANSMISSION FIDELITY PROTEIN 18 HOMOL |
| 6:00 | 7pbn-D | 8.7 | 9.5 | 140 | 313 | 10 | MOLECULE: | HOLLIDAY JUNCTION ATP-DEPENDENT DNA HELICASE RUVB |
| 7:00 | 7tfj-C | 8.7 | 12.5 | 144 | 330 | 10 | MOLECULE: | REPLICATION FACTOR C SUBUNIT 1 |
| 8:00 | 8ui7-B | 8.7 | 11.7 | 161 | 319 | 12 | MOLECULE: | ATPASE FAMILY AAA DOMAIN-CONTAINING PROTEIN 5 |
| 9:00 | 7tfh-B | 8.7 | 13.1 | 154 | 320 | 12 | MOLECULE: | REPLICATION FACTOR C SUBUNIT 1 |
| 10:00 | 2chv-B | 8.7 | 13.5 | 141 | 313 | 13 | MOLECULE: | REPLICATION FACTOR C SMALL SUBUNIT |
| 11:00 | 8unj-C | 8.7 | 11.1 | 142 | 325 | 15 | MOLECULE: | CHROMOSOME TRANSMISSION FIDELITY PROTEIN 18 HOMOL |
| 12:00 | 8dqx-B | 8.7 | 13.1 | 147 | 315 | 12 | MOLECULE: | REPLICATION FACTOR C SUBUNIT 1 |
| 13:00 | 7tfi-C | 8.7 | 10.8 | 144 | 330 | 13 | MOLECULE: | REPLICATION FACTOR C SUBUNIT 1 |
| 14:00 | 8dr5-C | 8.7 | 11.9 | 151 | 330 | 11 | MOLECULE: | REPLICATION FACTOR C SUBUNIT 1 |
| 15:00 | 8dr1-B | 8.7 | 12.9 | 141 | 318 | 11 | MOLECULE: | REPLICATION FACTOR C SUBUNIT 1 |
| 16:00 | 8hwa-A | 8.6 | 17.4 | 231 | 700 | 9 | MOLECULE: | PRIMASE D5 |
| 17:00 | 6z1f-3 | 8.6 | 7 | 148 | 284 | 14 | MOLECULE: | RIBULOSE BISPHOSPHATE CARBOXYLASE/OXYGENASE ACTIV |
| 18:00 | 8hwg-A | 8.6 | 3.6 | 200 | 369 | 11 | MOLECULE: | PRIMASE D5 |
| 19:00 | 8hwd-B | 8.6 | 4.7 | 206 | 378 | 9 | MOLECULE: | PRIMASE D5 |
| 20:00 | 8hwe-D | 8.6 | 3.7 | 202 | 379 | 9 | MOLECULE: | PRIMASE D5 |
| 21:00 | 2qz4-A | 8.6 | 3.3 | 136 | 223 | 13 | MOLECULE: | PARAPLEGIN |
| 22:00 | 3sc3-B | 8.6 | 10.4 | 133 | 211 | 14 | MOLECULE: | PUTATIVE DNA REPLICATION REGULATOR HDA |
| 23:00 | 6mat-C | 8.6 | 14.1 | 148 | 578 | 14 | MOLECULE: | RIX7 MUTANT |
| 24:00:00 | 1ojl-B | 8.6 | 3.6 | 137 | 251 | 11 | MOLECULE: | TRANSCRIPTIONAL REGULATORY PROTEIN ZRAR |
| 25:00:00 | 4qht-B | 8.6 | 3.5 | 139 | 245 | 11 | MOLECULE: | FLAGELLAR REGULATORY PROTEIN C |
| 26:00:00 | 4qht-E | 8.6 | 3.5 | 139 | 246 | 11 | MOLECULE: | FLAGELLAR REGULATORY PROTEIN C |
| 27:00:00 | 4qht-G | 8.6 | 3.6 | 139 | 246 | 11 | MOLECULE: | FLAGELLAR REGULATORY PROTEIN C |
| 28:00:00 | 4qht-D | 8.6 | 3.6 | 139 | 246 | 12 | MOLECULE: | FLAGELLAR REGULATORY PROTEIN C |

| | | | | | | | | |
|---|---|---|---|---|---|---|---|---|
| 29:00:00 | 4ly6-O | 8.6 | 6.8 | 143 | 247 | 10 | MOLECULE: | TRANSCRIPTIONAL REGULATOR (NTRC FAMILY) |
| 30:00:00 | 4lzz-S | 8.6 | 6.7 | 143 | 247 | 11 | MOLECULE: | TRANSCRIPTIONAL REGULATOR (NTRC FAMILY) |
| 31:00:00 | 4qht-A | 8.6 | 3.6 | 140 | 246 | 11 | MOLECULE: | FLAGELLAR REGULATORY PROTEIN C |
| 32:00:00 | 5ep1-A | 8.6 | 3.5 | 136 | 252 | 10 | MOLECULE: | PUTATIVE REPRESSOR PROTEIN LUXO |
| 33:00:00 | 4lj5-A | 8.6 | 10.3 | 164 | 320 | 16 | MOLECULE: | CHAPERONE PROTEIN CLPB |
| 34:00:00 | 5bq5-A | 8.6 | 3.5 | 126 | 180 | 16 | MOLECULE: | INSERTION SEQUENCE IS5376 PUTATIVE ATP-BINDING PR |
| 35:00:00 | 6wqh-B | 8.6 | 11.9 | 185 | 537 | 10 | MOLECULE: | LON PROTEASE |
| 36:00:00 | 3u5z-E | 8.6 | 3.7 | 128 | 305 | 17 | MOLECULE: | DNA POLYMERASE ACCESSORY PROTEIN 44 |
| 37:00:00 | 6v11-E | 8.6 | 14.4 | 150 | 464 | 15 | MOLECULE: | LON PROTEASE |
| 38:00:00 | 7krz-E | 8.6 | 11.8 | 182 | 517 | 9 | MOLECULE: | LON PROTEASE HOMOLOG, MITOCHONDRIAL |
| 39:00:00 | 8unj-B | 8.6 | 12 | 139 | 319 | 12 | MOLECULE: | CHROMOSOME TRANSMISSION FIDELITY PROTEIN 18 HOMOL |
| 40:00:00 | 6luf-E | 8.6 | 3.6 | 138 | 246 | 11 | MOLECULE: | FLAGELLAR REGULATORY PROTEIN C |
| 41:00:00 | 6luf-F | 8.6 | 3.7 | 137 | 247 | 14 | MOLECULE: | FLAGELLAR REGULATORY PROTEIN C |
| 42:00:00 | 1ojl-F | 8.6 | 3.5 | 133 | 247 | 9 | MOLECULE: | TRANSCRIPTIONAL REGULATORY PROTEIN ZRAR |
| 43:00:00 | 6fo1-B | 8.6 | 3.2 | 144 | 313 | 12 | MOLECULE: | RUVB-LIKE 1 |
| 44:00:00 | 9emc-A | 8.6 | 3.7 | 141 | 322 | 13 | MOLECULE: | RUVB-LIKE 1 |
| 45:00:00 | 6l1q-A | 8.6 | 6.9 | 142 | 267 | 11 | MOLECULE: | CBBQ PROTEIN |
| 46:00:00 | 7qv9-f | 8.6 | 8.8 | 151 | 259 | 8 | MOLECULE: | DNA-DIRECTED RNA POLYMERASE SUBUNIT ALPHA |
| 47:00:00 | 4qhs-F | 8.6 | 3.6 | 132 | 242 | 14 | MOLECULE: | FLAGELLAR REGULATORY PROTEIN C |
| 48:00:00 | 3uk6-J | 8.6 | 3.3 | 143 | 300 | 12 | MOLECULE: | RUVB-LIKE 2 |
| 49:00:00 | 4qhs-A | 8.6 | 3.5 | 131 | 244 | 14 | MOLECULE: | FLAGELLAR REGULATORY PROTEIN C |
| 50:00:00 | 1hqc-A | 8.6 | 10.8 | 153 | 314 | 12 | MOLECULE: | RUVB |
| 51:00:00 | 1ixr-C | 8.6 | 3.3 | 140 | 308 | 11 | MOLECULE: | HOLLIDAY JUNCTION DNA HELICASE RUVA |
| 52:00:00 | 2c9o-B | 8.6 | 3 | 138 | 393 | 12 | MOLECULE: | RUVB-LIKE 1 |
| 53:00:00 | 7x7p-N | 8.6 | 11.3 | 156 | 309 | 8 | MOLECULE: | DNA |
| 54:00:00 | 7mgm-A | 8.6 | 8.6 | 220 | 2274 | 8 | MOLECULE: | DYNEIN AAA3-WALKERB MUTANT (E2488Q) |
| 55:00:00 | 2chg-B | 8.6 | 3.2 | 130 | 222 | 12 | MOLECULE: | REPLICATION FACTOR C SMALL SUBUNIT |
| 56:00:00 | 7u1a-C | 8.6 | 13.8 | 148 | 327 | 14 | MOLECULE: | REPLICATION FACTOR C SUBUNIT 1 |
| 57:00:00 | 7thv-B | 8.6 | 12.7 | 145 | 314 | 12 | MOLECULE: | REPLICATION FACTOR C SUBUNIT 1 |
| 58:00:00 | 8dr0-C | 8.6 | 13.8 | 145 | 330 | 12 | MOLECULE: | REPLICATION FACTOR C SUBUNIT 1 |
| 59:00:00 | 6vvo-A | 8.6 | 6.1 | 162 | 448 | 10 | MOLECULE: | REPLICATION FACTOR C SUBUNIT 1 |
| 60:00:00 | 2chq-C | 8.6 | 15 | 143 | 313 | 15 | MOLECULE: | REPLICATION FACTOR C SMALL SUBUNIT |
| 61:00:00 | 8dr3-B | 8.6 | 16.5 | 146 | 318 | 12 | MOLECULE: | REPLICATION FACTOR C SUBUNIT 1 |
| 62:00:00 | 8uii-C | 8.6 | 14.5 | 147 | 324 | 14 | MOLECULE: | ATPASE FAMILY AAA DOMAIN-CONTAINING PROTEIN 5 |
| 63:00:00 | 7stb-C | 8.6 | 15.4 | 153 | 327 | 13 | MOLECULE: | CHECKPOINT PROTEIN RAD24 |
| 64:00:00 | 8dr3-C | 8.6 | 12.8 | 158 | 330 | 12 | MOLECULE: | REPLICATION FACTOR C SUBUNIT 1 |
| 65:00:00 | 6vvo-D | 8.6 | 15.6 | 136 | 304 | 11 | MOLECULE: | REPLICATION FACTOR C SUBUNIT 1 |
| 66:00:00 | 8fs6-C | 8.6 | 11.7 | 149 | 327 | 11 | MOLECULE: | CHECKPOINT PROTEIN RAD24 |
| 67:00:00 | 8fs5-B | 8.6 | 13.7 | 155 | 317 | 13 | MOLECULE: | CHECKPOINT PROTEIN RAD24 |
| 68:00:00 | 8dyu-A | 8.6 | 5.9 | 192 | 2562 | 8 | MOLECULE: | CYTOPLASMIC DYNEIN 1 HEAVY CHAIN 1 |
| 69:00:00 | 7tib-C | 8.6 | 12.4 | 151 | 327 | 12 | MOLECULE: | REPLICATION FACTOR C SUBUNIT 1 |
| 70:00:00 | 2chq-B | 8.6 | 12 | 139 | 313 | 14 | MOLECULE: | REPLICATION FACTOR C SMALL SUBUNIT |
| 71:00:00 | 8thb-B | 8.6 | 12.2 | 146 | 318 | 12 | MOLECULE: | ELG1 ISOFORM 1 |
| 72:00:00 | 7tic-B | 8.6 | 11.9 | 144 | 314 | 13 | MOLECULE: | REPLICATION FACTOR C SUBUNIT 1 |
| 73:00:00 | 7sgz-C | 8.6 | 10.6 | 144 | 325 | 13 | MOLECULE: | CHECKPOINT PROTEIN RAD24 |
| 74:00:00 | 7ti8-B | 8.6 | 14.3 | 151 | 318 | 11 | MOLECULE: | REPLICATION FACTOR C SUBUNIT 1 |
| 75:00:00 | 7pbr-F | 8.6 | 9 | 139 | 312 | 10 | MOLECULE: | HOLLIDAY JUNCTION ATP-DEPENDENT DNA HELICASE RUVB |
| 76:00:00 | 7thj-B | 8.6 | 9.5 | 150 | 315 | 13 | MOLECULE: | REPLICATION FACTOR C SUBUNIT 1 |
| 77:00:00 | 8thb-C | 8.6 | 12.5 | 141 | 325 | 11 | MOLECULE: | ELG1 ISOFORM 1 |
| 78:00:00 | 8dqx-A | 8.6 | 9.8 | 145 | 497 | 12 | MOLECULE: | REPLICATION FACTOR C SUBUNIT 1 |
| 79:00:00 | 6vvo-B | 8.6 | 11.4 | 147 | 316 | 12 | MOLECULE: | REPLICATION FACTOR C SUBUNIT 1 |
| 80:00:00 | 8dr6-C | 8.6 | 14.5 | 154 | 330 | 11 | MOLECULE: | REPLICATION FACTOR C SUBUNIT 1 |
| 81:00:00 | 7t3i-B | 8.5 | 12.6 | 163 | 578 | 13 | MOLECULE: | RIX7 |

| | | | | | | | | |
|---|---|---|---|---|---|---|---|---|
| 82:00:00 | 7stb-A | 8.5 | 9.9 | 155 | 515 | 11 | MOLECULE: | CHECKPOINT PROTEIN RAD24 |
| 83:00:00 | 8hwa-D | 8.5 | 4.3 | 210 | 379 | 11 | MOLECULE: | PRIMASE D5 |
| 84:00:00 | 8hwb-B | 8.5 | 4.3 | 210 | 379 | 10 | MOLECULE: | PRIMASE D5 |
| 85:00:00 | 8xj7-C | 8.5 | 7.1 | 208 | 454 | 9 | MOLECULE: | MONKEYPOX VIRUS E5 |
| 86:00:00 | 3bos-A | 8.5 | 7.9 | 135 | 228 | 16 | MOLECULE: | PUTATIVE DNA REPLICATION FACTOR |
| 87:00:00 | 6hz8-A | 8.5 | 3.5 | 135 | 290 | 12 | MOLECULE: | 5-METHYLCYTOSINE-SPECIFIC RESTRICTION ENZYME B |
| 88:00:00 | 6hz8-G | 8.5 | 3.5 | 135 | 290 | 12 | MOLECULE: | 5-METHYLCYTOSINE-SPECIFIC RESTRICTION ENZYME B |
| 89:00:00 | 2c98-A | 8.5 | 7.2 | 138 | 244 | 10 | MOLECULE: | PSP OPERON TRANSCRIPTIONAL ACTIVATOR |
| 90:00:00 | 4ly6-L | 8.5 | 3.5 | 139 | 247 | 10 | MOLECULE: | TRANSCRIPTIONAL REGULATOR (NTRC FAMILY) |
| 91:00:00 | 4qhs-E | 8.5 | 3.7 | 139 | 244 | 11 | MOLECULE: | FLAGELLAR REGULATORY PROTEIN C |
| 92:00:00 | 1ny6-L | 8.5 | 3.4 | 137 | 246 | 10 | MOLECULE: | TRANSCRIPTIONAL REGULATOR (NTRC FAMILY) |
| 93:00:00 | 5x06-E | 8.5 | 8.3 | 139 | 220 | 17 | MOLECULE: | DNA POLYMERASE III SUBUNIT BETA |
| 94:00:00 | 5x06-G | 8.5 | 7.7 | 138 | 205 | 18 | MOLECULE: | DNA POLYMERASE III SUBUNIT BETA |
| 95:00:00 | 5bq5-B | 8.5 | 3.5 | 126 | 187 | 16 | MOLECULE: | INSERTION SEQUENCE IS5376 PUTATIVE ATP-BINDING PR |
| 96:00:00 | 4fcw-C | 8.5 | 9.5 | 149 | 306 | 17 | MOLECULE: | CHAPERONE PROTEIN CLPB |
| 97:00:00 | 5bq5-C | 8.5 | 3.5 | 126 | 180 | 16 | MOLECULE: | INSERTION SEQUENCE IS5376 PUTATIVE ATP-BINDING PR |
| 98:00:00 | 1iqp-F | 8.5 | 13.5 | 148 | 326 | 9 | MOLECULE: | RFCS |
| 99:00:00 | 4w5w-A | 8.5 | 17.9 | 152 | 269 | 13 | MOLECULE: | RIBULOSE BISPHOSPHATE CARBOXYLASE/OXYGENASE ACTIV |
| 0:00 | 8uk9-E | 8.5 | 4.8 | 135 | 318 | 18 | MOLECULE: | SLIDING-CLAMP-LOADER SMALL SUBUNIT |
| 1:00 | 7krz-D | 8.5 | 12.3 | 173 | 520 | 10 | MOLECULE: | LON PROTEASE HOMOLOG, MITOCHONDRIAL |
| 2:00 | 7p09-F | 8.5 | 11.3 | 180 | 520 | 11 | MOLECULE: | LON PROTEASE HOMOLOG, MITOCHONDRIAL |
| 3:00 | 8un0-B | 8.5 | 13.8 | 144 | 319 | 12 | MOLECULE: | CHROMOSOME TRANSMISSION FIDELITY PROTEIN 18 HOMOL |
| 4:00 | 7w42-A | 8.5 | 10.3 | 197 | 397 | 13 | MOLECULE: | UNCHARACTERIZED ATPASE YJOB |
| 5:00 | 8dqz-B | 8.5 | 15 | 146 | 318 | 12 | MOLECULE: | REPLICATION FACTOR C SUBUNIT 1 |
| 6:00 | 5oaf-F | 8.5 | 9.7 | 148 | 407 | 12 | MOLECULE: | RUVB-LIKE 1 |
| 7:00 | 7ksl-C | 8.5 | 10.2 | 179 | 489 | 9 | MOLECULE: | LON PROTEASE HOMOLOG, MITOCHONDRIAL |
| 8:00 | 6hz4-A | 8.5 | 3.7 | 135 | 290 | 13 | MOLECULE: | 5-METHYLCYTOSINE-SPECIFIC RESTRICTION ENZYME B |
| 9:00 | 6oay-E | 8.5 | 5.3 | 212 | 570 | 12 | MOLECULE: | HYPERACTIVE DISAGGREGASE CLPB |
| 10:00 | 6oax-A | 8.5 | 10.2 | 194 | 581 | 13 | MOLECULE: | HYPERACTIVE DISAGGREGASE CLPB |
| 11:00 | 6hz4-D | 8.5 | 3.4 | 135 | 284 | 11 | MOLECULE: | 5-METHYLCYTOSINE-SPECIFIC RESTRICTION ENZYME B |
| 12:00 | 6luf-D | 8.5 | 3.7 | 135 | 246 | 13 | MOLECULE: | FLAGELLAR REGULATORY PROTEIN C |
| 13:00 | 6luf-B | 8.5 | 3.6 | 139 | 247 | 12 | MOLECULE: | FLAGELLAR REGULATORY PROTEIN C |
| 14:00 | 7tfj-A | 8.5 | 12 | 151 | 396 | 14 | MOLECULE: | REPLICATION FACTOR C SUBUNIT 1 |
| 15:00 | 4qhs-C | 8.5 | 3.7 | 135 | 243 | 13 | MOLECULE: | FLAGELLAR REGULATORY PROTEIN C |
| 16:00 | 3uk6-F | 8.5 | 12.5 | 145 | 301 | 12 | MOLECULE: | RUVB-LIKE 2 |
| 17:00 | 5j1s-A | 8.5 | 3.1 | 141 | 277 | 15 | MOLECULE: | TORSIN-1A |
| 18:00 | 6z1f-6 | 8.5 | 7.3 | 151 | 276 | 12 | MOLECULE: | RIBULOSE BISPHOSPHATE CARBOXYLASE/OXYGENASE ACTIV |
| 19:00 | 1ny6-I | 8.5 | 3.6 | 137 | 247 | 9 | MOLECULE: | TRANSCRIPTIONAL REGULATOR (NTRC FAMILY) |
| 20:00 | 1ht2-H | 8.5 | 11.2 | 160 | 408 | 12 | MOLECULE: | HEAT SHOCK LOCUS HSLV |
| 21:00 | 7qv9-a | 8.5 | 7 | 147 | 259 | 10 | MOLECULE: | DNA-DIRECTED RNA POLYMERASE SUBUNIT ALPHA |
| 22:00 | 7x7p-M | 8.5 | 8.4 | 141 | 309 | 10 | MOLECULE: | DNA |
| 23:00 | 8fs3-B | 8.5 | 14.9 | 147 | 319 | 12 | MOLECULE: | CHECKPOINT PROTEIN RAD24 |
| 24:00:00 | 1iqp-A | 8.5 | 14.8 | 155 | 326 | 13 | MOLECULE: | RFCS |
| 25:00:00 | 7tfk-C | 8.5 | 14.4 | 149 | 329 | 13 | MOLECULE: | REPLICATION FACTOR C SUBUNIT 1 |
| 26:00:00 | 1iqp-D | 8.5 | 15.9 | 146 | 326 | 9 | MOLECULE: | RFCS |
| 27:00:00 | 4w8f-A | 8.5 | 20.8 | 209 | 2608 | 11 | MOLECULE: | DYNEIN HEAVY CHAIN LYSOZYME CHIMERA |
| 28:00:00 | 8dr4-C | 8.5 | 13 | 143 | 330 | 12 | MOLECULE: | REPLICATION FACTOR C SUBUNIT 1 |
| 29:00:00 | 8ui9-B | 8.5 | 11.3 | 140 | 319 | 12 | MOLECULE: | ATPASE FAMILY AAA DOMAIN-CONTAINING PROTEIN 5 |
| 30:00:00 | 7tid-C | 8.5 | 14.2 | 144 | 327 | 13 | MOLECULE: | REPLICATION FACTOR C SUBUNIT 1 |
| 31:00:00 | 7u19-B | 8.5 | 13.7 | 154 | 319 | 12 | MOLECULE: | REPLICATION FACTOR C SUBUNIT 1 |
| 32:00:00 | 8ui8-B | 8.5 | 12.8 | 157 | 319 | 13 | MOLECULE: | ATPASE FAMILY AAA DOMAIN-CONTAINING PROTEIN 5 |
| 33:00:00 | 8thc-B | 8.5 | 10.1 | 145 | 318 | 12 | MOLECULE: | ELG1 ISOFORM 1 |
| 34:00:00 | 8dr7-C | 8.5 | 12.4 | 141 | 330 | 11 | MOLECULE: | REPLICATION FACTOR C SUBUNIT 1 |

| | | | | | | | | |
|---|---|---|---|---|---|---|---|---|
| 35:00:00 | 6vvo-C | 8.5 | 14.8 | 147 | 312 | 15 | MOLECULE: | REPLICATION FACTOR C SUBUNIT 1 |
| 36:00:00 | 2chg-D | 8.5 | 3.3 | 127 | 221 | 10 | MOLECULE: | REPLICATION FACTOR C SMALL SUBUNIT |
| 37:00:00 | 8ui7-C | 8.5 | 14.2 | 143 | 339 | 13 | MOLECULE: | ATPASE FAMILY AAA DOMAIN-CONTAINING PROTEIN 5 |
| 38:00:00 | 7tku-C | 8.5 | 14 | 148 | 329 | 14 | MOLECULE: | REPLICATION FACTOR C SUBUNIT 1 |
| 39:00:00 | 8fs6-B | 8.5 | 14.3 | 155 | 319 | 12 | MOLECULE: | CHECKPOINT PROTEIN RAD24 |
| 40:00:00 | 7t3i-F | 8.4 | 9.4 | 165 | 552 | 14 | MOLECULE: | RIX7 |
| 41:00:00 | 8hwe-C | 8.4 | 3.6 | 201 | 379 | 9 | MOLECULE: | PRIMASE D5 |
| 42:00:00 | 8hwb-A | 8.4 | 4 | 208 | 379 | 10 | MOLECULE: | PRIMASE D5 |
| 43:00:00 | 8hwf-B | 8.4 | 3.6 | 201 | 378 | 8 | MOLECULE: | PRIMASE D5 |
| 44:00:00 | 8hwb-C | 8.4 | 3.9 | 205 | 379 | 10 | MOLECULE: | PRIMASE D5 |
| 45:00:00 | 2xsz-C | 8.4 | 3.2 | 143 | 320 | 13 | MOLECULE: | RUVB-LIKE 1 |
| 46:00:00 | 4bs1-B | 8.4 | 3.6 | 128 | 173 | 11 | MOLECULE: | TRANSCRIPTIONAL REGULATOR (NTRC FAMILY) |
| 47:00:00 | 4bt1-B | 8.4 | 3.6 | 128 | 173 | 11 | MOLECULE: | TRANSCRIPTIONAL REGULATOR |
| 48:00:00 | 1ny5-A | 8.4 | 8.7 | 170 | 384 | 9 | MOLECULE: | TRANSCRIPTIONAL REGULATOR (NTRC FAMILY) |
| 49:00:00 | 8q3w-B | 8.4 | 8.8 | 159 | 246 | 13 | MOLECULE: | INSERTION SEQUENCE IS5376 PUTATIVE ATP-BINDING PR |
| 50:00:00 | 3uk6-A | 8.4 | 3.2 | 140 | 298 | 12 | MOLECULE: | RUVB-LIKE 2 |
| 51:00:00 | 3m6a-F | 8.4 | 13.9 | 181 | 496 | 14 | MOLECULE: | ATP-DEPENDENT PROTEASE LA 1 |
| 52:00:00 | 4lj7-C | 8.4 | 9.5 | 155 | 294 | 15 | MOLECULE: | CHAPERONE PROTEIN CLPB |
| 53:00:00 | 6hz6-G | 8.4 | 3.4 | 135 | 290 | 12 | MOLECULE: | 5-METHYLCYTOSINE-SPECIFIC RESTRICTION ENZYME B |
| 54:00:00 | 2chv-D | 8.4 | 13.7 | 144 | 313 | 13 | MOLECULE: | REPLICATION FACTOR C SMALL SUBUNIT |
| 55:00:00 | 2xsz-B | 8.4 | 3.2 | 142 | 319 | 13 | MOLECULE: | RUVB-LIKE 1 |
| 56:00:00 | 2chv-C | 8.4 | 13.6 | 142 | 313 | 13 | MOLECULE: | REPLICATION FACTOR C SMALL SUBUNIT |
| 57:00:00 | 6on2-D | 8.4 | 12.8 | 174 | 518 | 13 | MOLECULE: | ATP-DEPENDENT PROTEASE LA |
| 58:00:00 | 6hz7-A | 8.4 | 3.9 | 137 | 290 | 11 | MOLECULE: | 5-METHYLCYTOSINE-SPECIFIC RESTRICTION ENZYME B |
| 59:00:00 | 1iqp-B | 8.4 | 15 | 151 | 326 | 11 | MOLECULE: | RFCS |
| 60:00:00 | 7ng5-F | 8.4 | 30.1 | 219 | 776 | 11 | MOLECULE: | LON PROTEASE HOMOLOG, MITOCHONDRIAL |
| 61:00:00 | 8umt-B | 8.4 | 13 | 142 | 316 | 11 | MOLECULE: | CHROMOSOME TRANSMISSION FIDELITY PROTEIN 18 HOMOL |
| 62:00:00 | 6hz5-A | 8.4 | 3.7 | 136 | 290 | 13 | MOLECULE: | 5-METHYLCYTOSINE-SPECIFIC RESTRICTION ENZYME B |
| 63:00:00 | 8dqw-B | 8.4 | 14 | 144 | 315 | 12 | MOLECULE: | REPLICATION FACTOR C SUBUNIT 4 |
| 64:00:00 | 6hz5-G | 8.4 | 3.7 | 136 | 290 | 13 | MOLECULE: | 5-METHYLCYTOSINE-SPECIFIC RESTRICTION ENZYME B |
| 65:00:00 | 8gj0-D | 8.4 | 11 | 149 | 360 | 11 | MOLECULE: | DNA POLYMERASE III SUBUNIT DELTA |
| 66:00:00 | 7w46-A | 8.4 | 10.7 | 192 | 396 | 14 | MOLECULE: | UNCHARACTERIZED ATPASE YJOB |
| 67:00:00 | 6qi9-A | 8.4 | 13.4 | 150 | 317 | 12 | MOLECULE: | RUVB-LIKE 1 |
| 68:00:00 | 6lua-B | 8.4 | 3.8 | 134 | 247 | 13 | MOLECULE: | FLAGELLAR REGULATORY PROTEIN C |
| 69:00:00 | 4qhs-D | 8.4 | 3.8 | 134 | 244 | 13 | MOLECULE: | FLAGELLAR REGULATORY PROTEIN C |
| 70:00:00 | 6iy8-A | 8.4 | 35.6 | 184 | 469 | 8 | MOLECULE: | POSITIVE REGULATOR CAPR |
| 71:00:00 | 3cf3-A | 8.4 | 7.1 | 182 | 723 | 10 | MOLECULE: | TRANSITIONAL ENDOPLASMIC RETICULUM ATPASE |
| 72:00:00 | 3cf1-A | 8.4 | 4.7 | 181 | 718 | 11 | MOLECULE: | TRANSITIONAL ENDOPLASMIC RETICULUM ATPASE |
| 73:00:00 | 8dr0-B | 8.4 | 14 | 142 | 318 | 12 | MOLECULE: | REPLICATION FACTOR C SUBUNIT 1 |
| 74:00:00 | 1iqp-E | 8.4 | 14.8 | 150 | 326 | 11 | MOLECULE: | RFCS |
| 75:00:00 | 8uii-B | 8.4 | 15.1 | 147 | 319 | 13 | MOLECULE: | ATPASE FAMILY AAA DOMAIN-CONTAINING PROTEIN 5 |
| 76:00:00 | 7sh2-B | 8.4 | 13.2 | 144 | 317 | 10 | MOLECULE: | CHECKPOINT PROTEIN RAD24 |
| 77:00:00 | 8ui9-D | 8.4 | 16.6 | 145 | 328 | 12 | MOLECULE: | ATPASE FAMILY AAA DOMAIN-CONTAINING PROTEIN 5 |
| 78:00:00 | 8umy-D | 8.4 | 14.3 | 143 | 329 | 13 | MOLECULE: | CHROMOSOME TRANSMISSION FIDELITY PROTEIN 18 HOMOL |
| 79:00:00 | 1sxj-B | 8.4 | 11.7 | 147 | 316 | 12 | MOLECULE: | ACTIVATOR 1 95 KDA SUBUNIT |
| 80:00:00 | 8umt-C | 8.4 | 14.7 | 147 | 322 | 13 | MOLECULE: | CHROMOSOME TRANSMISSION FIDELITY PROTEIN 18 HOMOL |
| 81:00:00 | 7tid-B | 8.4 | 14.3 | 147 | 319 | 12 | MOLECULE: | REPLICATION FACTOR C SUBUNIT 1 |
| 82:00:00 | 8ui8-C | 8.4 | 13.1 | 143 | 339 | 13 | MOLECULE: | ATPASE FAMILY AAA DOMAIN-CONTAINING PROTEIN 5 |
| 83:00:00 | 8uii-D | 8.4 | 16.2 | 146 | 328 | 12 | MOLECULE: | ATPASE FAMILY AAA DOMAIN-CONTAINING PROTEIN 5 |
| 84:00:00 | 7sh2-C | 8.4 | 11.2 | 144 | 325 | 11 | MOLECULE: | CHECKPOINT PROTEIN RAD24 |
| 85:00:00 | 8hwh-D | 8.3 | 4 | 205 | 363 | 11 | MOLECULE: | PRIMASE D5 |
| 86:00:00 | 8hwg-D | 8.3 | 4.1 | 206 | 378 | 10 | MOLECULE: | PRIMASE D5 |
| 87:00:00 | 6hz7-H | 8.3 | 3.5 | 138 | 290 | 12 | MOLECULE: | 5-METHYLCYTOSINE-SPECIFIC RESTRICTION ENZYME B |

| | | | | | | | | |
|---|---|---|---|---|---|---|---|---|
| 88:00:00 | 6hz8-C | 8.3 | 3.4 | 136 | 290 | 13 | MOLECULE: | 5-METHYLCYTOSINE-SPECIFIC RESTRICTION ENZYME B |
| 89:00:00 | 6hz9-B | 8.3 | 3.5 | 138 | 290 | 11 | MOLECULE: | 5-METHYLCYTOSINE-SPECIFIC RESTRICTION ENZYME B |
| 90:00:00 | 6hz8-I | 8.3 | 3.3 | 136 | 290 | 12 | MOLECULE: | 5-METHYLCYTOSINE-SPECIFIC RESTRICTION ENZYME B |
| 91:00:00 | 6hz7-B | 8.3 | 3.5 | 138 | 290 | 12 | MOLECULE: | 5-METHYLCYTOSINE-SPECIFIC RESTRICTION ENZYME B |
| 92:00:00 | 4w8f-B | 8.3 | 25.8 | 212 | 2609 | 9 | MOLECULE: | DYNEIN HEAVY CHAIN LYSOZYME CHIMERA |
| 93:00:00 | 8xj8-C | 8.3 | 3.6 | 200 | 379 | 9 | MOLECULE: | DNA (70-MER) |
| 94:00:00 | 3zw6-A | 8.3 | 13.7 | 144 | 263 | 13 | MOLECULE: | RIBULOSE BISPHOSPHATE CARBOXYLASE/OXYGENASE ACTIV |
| 95:00:00 | 4bt0-B | 8.3 | 3.6 | 128 | 173 | 11 | MOLECULE: | TRANSCRIPTIONAL REGULATOR |
| 96:00:00 | 4lzz-X | 8.3 | 7.9 | 139 | 246 | 10 | MOLECULE: | TRANSCRIPTIONAL REGULATOR (NTRC FAMILY) |
| 97:00:00 | 7ejw-C | 8.3 | 3.6 | 141 | 252 | 9 | MOLECULE: | TRANSCRIPTIONAL ANTIACTIVATOR FLEN |
| 98:00:00 | 4lj7-A | 8.3 | 11.5 | 156 | 304 | 15 | MOLECULE: | CHAPERONE PROTEIN CLPB |
| 99:00:00 | 3m6a-D | 8.3 | 12.2 | 175 | 496 | 13 | MOLECULE: | ATP-DEPENDENT PROTEASE LA 1 |
| 0:00 | 2chv-E | 8.3 | 15.3 | 141 | 313 | 13 | MOLECULE: | REPLICATION FACTOR C SMALL SUBUNIT |
| 1:00 | 1sxj-A | 8.3 | 5.4 | 150 | 441 | 12 | MOLECULE: | ACTIVATOR 1 95 KDA SUBUNIT |
| 2:00 | 3glf-D | 8.3 | 11.9 | 143 | 362 | 11 | MOLECULE: | DNA POLYMERASE III SUBUNIT DELTA |
| 3:00 | 8uk9-N | 8.3 | 4.1 | 126 | 305 | 21 | MOLECULE: | SLIDING-CLAMP-LOADER SMALL SUBUNIT |
| 4:00 | 2chv-F | 8.3 | 13.7 | 142 | 313 | 13 | MOLECULE: | REPLICATION FACTOR C SMALL SUBUNIT |
| 5:00 | 7ksm-C | 8.3 | 11.1 | 184 | 526 | 10 | MOLECULE: | LON PROTEASE HOMOLOG, MITOCHONDRIAL |
| 6:00 | 6on2-C | 8.3 | 12.1 | 186 | 523 | 11 | MOLECULE: | ATP-DEPENDENT PROTEASE LA |
| 7:00 | 7ksm-B | 8.3 | 13 | 189 | 520 | 10 | MOLECULE: | LON PROTEASE HOMOLOG, MITOCHONDRIAL |
| 8:00 | 6u5z-A | 8.3 | 12.8 | 190 | 529 | 13 | MOLECULE: | LON PROTEASE |
| 9:00 | 7krz-C | 8.3 | 11.6 | 176 | 527 | 11 | MOLECULE: | LON PROTEASE HOMOLOG, MITOCHONDRIAL |
| 10:00 | 7krz-F | 8.3 | 11.5 | 182 | 512 | 9 | MOLECULE: | LON PROTEASE HOMOLOG, MITOCHONDRIAL |
| 11:00 | 7ksm-D | 8.3 | 11.9 | 171 | 525 | 11 | MOLECULE: | LON PROTEASE HOMOLOG, MITOCHONDRIAL |
| 12:00 | 7z6h-F | 8.3 | 13.5 | 146 | 315 | 10 | MOLECULE: | CELL CYCLE CHECKPOINT CONTROL PROTEIN RAD9A |
| 13:00 | 7ste-B | 8.3 | 12.6 | 144 | 315 | 10 | MOLECULE: | REPLICATION FACTOR C SUBUNIT 4 |
| 14:00 | 6oax-E | 8.3 | 15.4 | 208 | 579 | 14 | MOLECULE: | HYPERACTIVE DISAGGREGASE CLPB |
| 15:00 | 8p53-E | 8.3 | 4.9 | 169 | 390 | 9 | MOLECULE: | ANTIACTIVATOR FLEN |
| 16:00 | 6h7x-A | 8.3 | 14.8 | 145 | 371 | 11 | MOLECULE: | RUVB-LIKE 2 |
| 17:00 | 6hz5-D | 8.3 | 3.4 | 135 | 284 | 11 | MOLECULE: | 5-METHYLCYTOSINE-SPECIFIC RESTRICTION ENZYME B |
| 18:00 | 7ejw-D | 8.3 | 3.5 | 139 | 254 | 9 | MOLECULE: | TRANSCRIPTIONAL ANTIACTIVATOR FLEN |
| 19:00 | 6lua-E | 8.3 | 3.7 | 133 | 246 | 14 | MOLECULE: | FLAGELLAR REGULATORY PROTEIN C |
| 20:00 | 6lua-G | 8.3 | 3.6 | 133 | 246 | 14 | MOLECULE: | FLAGELLAR REGULATORY PROTEIN C |
| 21:00 | 6lua-F | 8.3 | 3.7 | 133 | 247 | 14 | MOLECULE: | FLAGELLAR REGULATORY PROTEIN C |
| 22:00 | 7p6x-B | 8.3 | 3.4 | 144 | 313 | 11 | MOLECULE: | RUVB-LIKE 1 |
| 23:00 | 3hu1-C | 8.3 | 9.3 | 161 | 451 | 14 | MOLECULE: | TRANSITIONAL ENDOPLASMIC RETICULUM ATPASE |
| 24:00:00 | 6z1d-B | 8.3 | 16 | 160 | 259 | 13 | MOLECULE: | RIBULOSE BISPHOSPHATE CARBOXYLASE/OXYGENASE ACTIV |
| 25:00:00 | 3uk6-H | 8.3 | 3.2 | 134 | 277 | 14 | MOLECULE: | RUVB-LIKE 2 |
| 26:00:00 | 3hu1-E | 8.3 | 11 | 165 | 451 | 13 | MOLECULE: | TRANSITIONAL ENDOPLASMIC RETICULUM ATPASE |
| 27:00:00 | 3cf3-B | 8.3 | 5.7 | 182 | 723 | 10 | MOLECULE: | TRANSITIONAL ENDOPLASMIC RETICULUM ATPASE |
| 28:00:00 | 3cf1-B | 8.3 | 7.3 | 192 | 723 | 9 | MOLECULE: | TRANSITIONAL ENDOPLASMIC RETICULUM ATPASE |
| 29:00:00 | 3hu2-C | 8.3 | 11.8 | 165 | 452 | 13 | MOLECULE: | TRANSITIONAL ENDOPLASMIC RETICULUM ATPASE |
| 30:00:00 | 7x7p-P | 8.3 | 10.2 | 142 | 309 | 11 | MOLECULE: | DNA |
| 31:00:00 | 3cf3-C | 8.3 | 6.3 | 187 | 723 | 9 | MOLECULE: | TRANSITIONAL ENDOPLASMIC RETICULUM ATPASE |
| 32:00:00 | 8ui7-D | 8.3 | 16.6 | 142 | 328 | 11 | MOLECULE: | ATPASE FAMILY AAA DOMAIN-CONTAINING PROTEIN 5 |
| 33:00:00 | 2chv-A | 8.3 | 11.7 | 134 | 313 | 14 | MOLECULE: | REPLICATION FACTOR C SMALL SUBUNIT |
| 34:00:00 | 8fs7-B | 8.3 | 12.3 | 137 | 319 | 12 | MOLECULE: | CHECKPOINT PROTEIN RAD24 |
| 35:00:00 | 7tfl-C | 8.3 | 15 | 149 | 326 | 12 | MOLECULE: | REPLICATION FACTOR C SUBUNIT 1 |
| 36:00:00 | 7ste-C | 8.3 | 15.5 | 145 | 324 | 12 | MOLECULE: | REPLICATION FACTOR C SUBUNIT 4 |
| 37:00:00 | 8fs4-B | 8.3 | 13 | 149 | 316 | 11 | MOLECULE: | CHECKPOINT PROTEIN RAD24 |
| 38:00:00 | 8un0-D | 8.3 | 13.8 | 150 | 329 | 11 | MOLECULE: | CHROMOSOME TRANSMISSION FIDELITY PROTEIN 18 HOMOL |
| 39:00:00 | 8dr5-A | 8.2 | 15.2 | 165 | 646 | 12 | MOLECULE: | REPLICATION FACTOR C SUBUNIT 1 |
| 40:00:00 | 7fd4-A | 8.2 | 27.3 | 233 | 779 | 11 | MOLECULE: | LON PROTEASE |

| | | | | | | | | |
|---|---|---|---|---|---|---|---|---|
| 41:00:00 | 5nvu-A | 8.2 | 19.6 | 210 | 3169 | 0 | MOLECULE: | DYNEIN MOTOR DOMAIN |
| 42:00:00 | 8xj7-D | 8.2 | 6.1 | 203 | 452 | 10 | MOLECULE: | MONKEYPOX VIRUS E5 |
| 43:00:00 | 5vy9-A | 8.2 | 25.4 | 236 | 849 | 11 | MOLECULE: | HEAT SHOCK PROTEIN 104 |
| 44:00:00 | 6mat-D | 8.2 | 13 | 160 | 578 | 13 | MOLECULE: | RIX7 MUTANT |
| 45:00:00 | 8xj8-A | 8.2 | 5.8 | 204 | 422 | 11 | MOLECULE: | DNA (70-MER) |
| 46:00:00 | 5n7r-A | 8.2 | 15.1 | 144 | 380 | 11 | MOLECULE: | RUVB-LIKE 2 |
| 47:00:00 | 5c3c-B | 8.2 | 10.1 | 144 | 235 | 12 | MOLECULE: | CBBQ/NIRQ/NORQ DOMAIN PROTEIN |
| 48:00:00 | 4ly6-F | 8.2 | 3.6 | 141 | 247 | 11 | MOLECULE: | TRANSCRIPTIONAL REGULATOR (NTRC FAMILY) |
| 49:00:00 | 2bjv-A | 8.2 | 3.2 | 128 | 236 | 12 | MOLECULE: | PSP OPERON TRANSCRIPTIONAL ACTIVATOR |
| 50:00:00 | 5x06-H | 8.2 | 3.8 | 135 | 191 | 16 | MOLECULE: | DNA POLYMERASE III SUBUNIT BETA |
| 51:00:00 | 4qhs-B | 8.2 | 3.6 | 131 | 243 | 14 | MOLECULE: | FLAGELLAR REGULATORY PROTEIN C |
| 52:00:00 | 4lj7-B | 8.2 | 9.1 | 153 | 300 | 16 | MOLECULE: | CHAPERONE PROTEIN CLPB |
| 53:00:00 | 3uk6-K | 8.2 | 3.2 | 136 | 283 | 13 | MOLECULE: | RUVB-LIKE 2 |
| 54:00:00 | 4ypl-D | 8.2 | 13.7 | 183 | 537 | 14 | MOLECULE: | LON PROTEASE |
| 55:00:00 | 7ksm-F | 8.2 | 11.3 | 186 | 516 | 10 | MOLECULE: | LON PROTEASE HOMOLOG, MITOCHONDRIAL |
| 56:00:00 | 1do0-A | 8.2 | 11.4 | 157 | 406 | 11 | MOLECULE: | PROTEIN (HEAT SHOCK LOCUS U) |
| 57:00:00 | 7krz-A | 8.2 | 11.3 | 173 | 525 | 9 | MOLECULE: | LON PROTEASE HOMOLOG, MITOCHONDRIAL |
| 58:00:00 | 7ksl-A | 8.2 | 10.7 | 178 | 520 | 11 | MOLECULE: | LON PROTEASE HOMOLOG, MITOCHONDRIAL |
| 59:00:00 | 3gli-D | 8.2 | 11.1 | 146 | 362 | 12 | MOLECULE: | DNA POLYMERASE III SUBUNIT DELTA |
| 60:00:00 | 7ngc-F | 8.2 | 30.7 | 213 | 776 | 9 | MOLECULE: | LON PROTEASE HOMOLOG, MITOCHONDRIAL |
| 61:00:00 | 7ngl-C | 8.2 | 27.8 | 210 | 776 | 10 | MOLECULE: | LON PROTEASE HOMOLOG, MITOCHONDRIAL |
| 62:00:00 | 7ngl-F | 8.2 | 31.6 | 224 | 776 | 8 | MOLECULE: | LON PROTEASE HOMOLOG, MITOCHONDRIAL |
| 63:00:00 | 3gli-I | 8.2 | 11.1 | 138 | 362 | 12 | MOLECULE: | DNA POLYMERASE III SUBUNIT DELTA |
| 64:00:00 | 8p53-F | 8.2 | 4.5 | 166 | 390 | 9 | MOLECULE: | ANTIACTIVATOR FLEN |
| 65:00:00 | 7stb-B | 8.2 | 14.1 | 144 | 315 | 12 | MOLECULE: | CHECKPOINT PROTEIN RAD24 |
| 66:00:00 | 6sh3-G | 8.2 | 16.8 | 167 | 376 | 11 | MOLECULE: | MITOCHONDRIAL CHAPERONE BCS1 |
| 67:00:00 | 6hz9-L | 8.2 | 3.5 | 135 | 285 | 11 | MOLECULE: | 5-METHYLCYTOSINE-SPECIFIC RESTRICTION ENZYME B |
| 68:00:00 | 6lua-C | 8.2 | 3.8 | 137 | 247 | 12 | MOLECULE: | FLAGELLAR REGULATORY PROTEIN C |
| 69:00:00 | 6k0r-B | 8.2 | 3.4 | 145 | 318 | 12 | MOLECULE: | RUVB-LIKE 1,RUVB-LIKE 1 |
| 70:00:00 | 9emc-B | 8.2 | 3.7 | 141 | 322 | 13 | MOLECULE: | RUVB-LIKE 1 |
| 71:00:00 | 6ut5-C | 8.2 | 11.1 | 136 | 393 | 15 | MOLECULE: | GTPASE SUBUNIT OF RESTRICTION ENDONUCLEASE |
| 72:00:00 | 8fs8-B | 8.2 | 14.4 | 147 | 319 | 9 | MOLECULE: | CHECKPOINT PROTEIN RAD24 |
| 73:00:00 | 3hu2-A | 8.2 | 11.6 | 165 | 452 | 14 | MOLECULE: | TRANSITIONAL ENDOPLASMIC RETICULUM ATPASE |
| 74:00:00 | 6hd3-A | 8.2 | 6.5 | 162 | 450 | 10 | MOLECULE: | CELL DIVISION CONTROL PROTEIN 48 HOMOLOG A |
| 75:00:00 | 6z1f-1 | 8.2 | 16.1 | 158 | 280 | 13 | MOLECULE: | RIBULOSE BISPHOSPHATE CARBOXYLASE/OXYGENASE ACTIV |
| 76:00:00 | 3cf2-D | 8.2 | 9 | 194 | 659 | 10 | MOLECULE: | TRANSITIONAL ENDOPLASMIC RETICULUM ATPASE |
| 77:00:00 | 3cf2-B | 8.2 | 9 | 195 | 659 | 11 | MOLECULE: | TRANSITIONAL ENDOPLASMIC RETICULUM ATPASE |
| 78:00:00 | 3cf2-A | 8.2 | 7.4 | 175 | 659 | 11 | MOLECULE: | TRANSITIONAL ENDOPLASMIC RETICULUM ATPASE |
| 79:00:00 | 6z1d-A | 8.2 | 8 | 153 | 274 | 12 | MOLECULE: | RIBULOSE BISPHOSPHATE CARBOXYLASE/OXYGENASE ACTIV |
| 80:00:00 | 8hrz-I | 8.2 | 4.3 | 152 | 436 | 11 | MOLECULE: | TRANSITIONAL ENDOPLASMIC RETICULUM ATPASE |
| 81:00:00 | 5j1t-A | 8.2 | 3.3 | 147 | 277 | 16 | MOLECULE: | TORSIN-1A |
| 82:00:00 | 3cf2-C | 8.2 | 8.5 | 200 | 659 | 11 | MOLECULE: | TRANSITIONAL ENDOPLASMIC RETICULUM ATPASE |
| 83:00:00 | 4kln-B | 8.2 | 11.2 | 162 | 451 | 14 | MOLECULE: | TRANSITIONAL ENDOPLASMIC RETICULUM ATPASE |
| 84:00:00 | 7z11-D | 8.2 | 10.8 | 181 | 734 | 9 | MOLECULE: | ATPASE FAMILY GENE 2 PROTEIN |
| 85:00:00 | 6pxl-F | 8.2 | 11.5 | 155 | 410 | 12 | MOLECULE: | ATP-DEPENDENT PROTEASE ATPASE SUBUNIT HSLU |
| 86:00:00 | 8fcy-A | 8.2 | 27.4 | 214 | 2864 | 8 | MOLECULE: | CYTOPLASMIC DYNEIN 1 HEAVY CHAIN 1 |
| 87:00:00 | 8dr4-A | 8.2 | 13.1 | 154 | 497 | 11 | MOLECULE: | REPLICATION FACTOR C SUBUNIT 1 |
| 88:00:00 | 7w42-B | 8.2 | 23.3 | 189 | 402 | 13 | MOLECULE: | UNCHARACTERIZED ATPASE YJOB |
| 89:00:00 | 7z6h-G | 8.2 | 9.7 | 135 | 317 | 16 | MOLECULE: | CELL CYCLE CHECKPOINT CONTROL PROTEIN RAD9A |
| 90:00:00 | 8umu-D | 8.2 | 14.5 | 139 | 327 | 9 | MOLECULE: | CHROMOSOME TRANSMISSION FIDELITY PROTEIN 18 HOMOL |
| 91:00:00 | 7t3i-D | 8.1 | 13 | 150 | 578 | 15 | MOLECULE: | RIX7 |
| 92:00:00 | 8iqi-D | 8.1 | 16.8 | 191 | 916 | 12 | MOLECULE: | PUTATIVE PRIMASE C962R |
| 93:00:00 | 8hwa-E | 8.1 | 4 | 206 | 379 | 10 | MOLECULE: | PRIMASE D5 |

| | | | | | | | | |
|---|---|---|---|---|---|---|---|---|
| 94:00:00 | 6hz5-H | 8.1 | 3.5 | 136 | 290 | 12 | MOLECULE: | 5-METHYLCYTOSINE-SPECIFIC RESTRICTION ENZYME B |
| 95:00:00 | 6hz9-F | 8.1 | 3.5 | 135 | 285 | 11 | MOLECULE: | 5-METHYLCYTOSINE-SPECIFIC RESTRICTION ENZYME B |
| 96:00:00 | 3zw6-F | 8.1 | 11 | 134 | 263 | 15 | MOLECULE: | RIBULOSE BISPHOSPHATE CARBOXYLASE/OXYGENASE ACTIV |
| 97:00:00 | 5ep0-A | 8.1 | 6.6 | 167 | 380 | 8 | MOLECULE: | PUTATIVE REPRESSOR PROTEIN LUXO |
| 98:00:00 | 5fm6-A | 8.1 | 3.7 | 144 | 426 | 14 | MOLECULE: | RVB1 |
| 99:00:00 | 5x06-F | 8.1 | 8.3 | 141 | 228 | 16 | MOLECULE: | DNA POLYMERASE III SUBUNIT BETA |
| 0:00 | 6po1-B | 8.1 | 9.3 | 153 | 352 | 14 | MOLECULE: | ATP-DEPENDENT CLP PROTEASE ATP-BINDING SUBUNIT CL |
| 1:00 | 3glg-H | 8.1 | 11.5 | 138 | 365 | 10 | MOLECULE: | DNA POLYMERASE III SUBUNIT DELTA |
| 2:00 | 3gli-C | 8.1 | 11.8 | 144 | 365 | 11 | MOLECULE: | DNA POLYMERASE III SUBUNIT DELTA |
| 3:00 | 3glg-C | 8.1 | 11.8 | 140 | 365 | 10 | MOLECULE: | DNA POLYMERASE III SUBUNIT DELTA |
| 4:00 | 5ji3-E | 8.1 | 11.4 | 155 | 375 | 11 | MOLECULE: | ATP-DEPENDENT PROTEASE SUBUNIT HSLV |
| 5:00 | 3gli-H | 8.1 | 13.8 | 140 | 365 | 11 | MOLECULE: | DNA POLYMERASE III SUBUNIT DELTA |
| 6:00 | 5vqa-A | 8.1 | 4.5 | 154 | 368 | 14 | MOLECULE: | PACHYTENE CHECKPOINT PROTEIN 2 HOMOLOG |
| 7:00 | 3glf-H | 8.1 | 12.6 | 147 | 365 | 11 | MOLECULE: | DNA POLYMERASE III SUBUNIT DELTA |
| 8:00 | 3glg-I | 8.1 | 10.4 | 150 | 362 | 11 | MOLECULE: | DNA POLYMERASE III SUBUNIT DELTA |
| 9:00 | 3glg-D | 8.1 | 12 | 145 | 362 | 12 | MOLECULE: | DNA POLYMERASE III SUBUNIT DELTA |
| 10:00 | 6u5z-D | 8.1 | 13.2 | 189 | 529 | 14 | MOLECULE: | LON PROTEASE |
| 11:00 | 3glf-I | 8.1 | 12.1 | 146 | 362 | 12 | MOLECULE: | DNA POLYMERASE III SUBUNIT DELTA |
| 12:00 | 6u5z-C | 8.1 | 13.8 | 185 | 529 | 15 | MOLECULE: | LON PROTEASE |
| 13:00 | 6u5z-F | 8.1 | 14.1 | 185 | 529 | 14 | MOLECULE: | LON PROTEASE |
| 14:00 | 6u5z-E | 8.1 | 13.4 | 188 | 529 | 13 | MOLECULE: | LON PROTEASE |
| 15:00 | 6hz8-K | 8.1 | 3.8 | 137 | 285 | 12 | MOLECULE: | 5-METHYLCYTOSINE-SPECIFIC RESTRICTION ENZYME B |
| 16:00 | 7st9-B | 8.1 | 13.1 | 143 | 315 | 14 | MOLECULE: | CHECKPOINT PROTEIN RAD24 |
| 17:00 | 7tfl-B | 8.1 | 13 | 139 | 319 | 11 | MOLECULE: | REPLICATION FACTOR C SUBUNIT 1 |
| 18:00 | 1do0-D | 8.1 | 12.1 | 160 | 406 | 12 | MOLECULE: | PROTEIN (HEAT SHOCK LOCUS U) |
| 19:00 | 1d2n-A | 8.1 | 7.8 | 135 | 246 | 13 | MOLECULE: | N-ETHYLMALEIMIDE-SENSITIVE FUSION PROTEIN |
| 20:00 | 3glf-C | 8.1 | 11.8 | 143 | 365 | 11 | MOLECULE: | DNA POLYMERASE III SUBUNIT DELTA |
| 21:00 | 6e11-4 | 8.1 | 4.8 | 206 | 717 | 15 | MOLECULE: | UNKNOWN (CLAW) |
| 22:00 | 7ttr-B | 8.1 | 11.8 | 162 | 311 | 13 | MOLECULE: | CASEINOLYTIC PEPTIDASE B PROTEIN HOMOLOG |
| 23:00 | 7v2b-D | 8.1 | 7.9 | 164 | 376 | 11 | MOLECULE: | VPSR |
| 24:00:00 | 6rn4-F | 8.1 | 13.1 | 201 | 680 | 11 | MOLECULE: | CHAPERONE PROTEIN CLPB |
| 25:00:00 | 6ut3-F | 8.1 | 3.4 | 115 | 235 | 12 | MOLECULE: | GTPASE SUBUNIT OF RESTRICTION ENDONUCLEASE |
| 26:00:00 | 6hz8-E | 8.1 | 3.8 | 137 | 285 | 12 | MOLECULE: | 5-METHYLCYTOSINE-SPECIFIC RESTRICTION ENZYME B |
| 27:00:00 | 6igm-D | 8.1 | 3.7 | 140 | 396 | 12 | MOLECULE: | RUVB-LIKE 1 |
| 28:00:00 | 3hu2-E | 8.1 | 11.5 | 167 | 452 | 14 | MOLECULE: | TRANSITIONAL ENDOPLASMIC RETICULUM ATPASE |
| 29:00:00 | 6qi8-A | 8.1 | 3.7 | 145 | 317 | 11 | MOLECULE: | RUVB-LIKE 1 |
| 30:00:00 | 2c9o-A | 8.1 | 3.2 | 139 | 398 | 12 | MOLECULE: | RUVB-LIKE 1 |
| 31:00:00 | 4kln-E | 8.1 | 11 | 166 | 451 | 11 | MOLECULE: | TRANSITIONAL ENDOPLASMIC RETICULUM ATPASE |
| 32:00:00 | 6lua-D | 8.1 | 3.7 | 134 | 246 | 13 | MOLECULE: | FLAGELLAR REGULATORY PROTEIN C |
| 33:00:00 | 9emc-C | 8.1 | 3.7 | 141 | 322 | 13 | MOLECULE: | RUVB-LIKE 1 |
| 34:00:00 | 6ut5-D | 8.1 | 14.8 | 146 | 395 | 14 | MOLECULE: | GTPASE SUBUNIT OF RESTRICTION ENDONUCLEASE |
| 35:00:00 | 6qi8-B | 8.1 | 3.6 | 145 | 310 | 11 | MOLECULE: | RUVB-LIKE 1 |
| 36:00:00 | 3hu2-F | 8.1 | 11.5 | 167 | 452 | 14 | MOLECULE: | TRANSITIONAL ENDOPLASMIC RETICULUM ATPASE |
| 37:00:00 | 4kln-D | 8.1 | 10.9 | 162 | 451 | 14 | MOLECULE: | TRANSITIONAL ENDOPLASMIC RETICULUM ATPASE |
| 38:00:00 | 4kod-E | 8.1 | 12.1 | 154 | 439 | 10 | MOLECULE: | TRANSITIONAL ENDOPLASMIC RETICULUM ATPASE |
| 39:00:00 | 3dzd-A | 8.1 | 11.5 | 171 | 368 | 11 | MOLECULE: | TRANSCRIPTIONAL REGULATOR (NTRC FAMILY) |
| 40:00:00 | 3hu3-A | 8.1 | 10.9 | 169 | 453 | 13 | MOLECULE: | TRANSITIONAL ENDOPLASMIC RETICULUM ATPASE |
| 41:00:00 | 7pbr-C | 8.1 | 9.9 | 140 | 309 | 11 | MOLECULE: | HOLLIDAY JUNCTION ATP-DEPENDENT DNA HELICASE RUVB |
| 42:00:00 | 4kod-H | 8.1 | 5.9 | 156 | 438 | 12 | MOLECULE: | TRANSITIONAL ENDOPLASMIC RETICULUM ATPASE |
| 43:00:00 | 5ji3-F | 8.1 | 10 | 152 | 353 | 11 | MOLECULE: | ATP-DEPENDENT PROTEASE SUBUNIT HSLV |
| 44:00:00 | 4kod-I | 8.1 | 3.7 | 154 | 440 | 11 | MOLECULE: | TRANSITIONAL ENDOPLASMIC RETICULUM ATPASE |
| 45:00:00 | 1sxj-C | 8.1 | 13.1 | 147 | 322 | 12 | MOLECULE: | ACTIVATOR 1 95 KDA SUBUNIT |
| 46:00:00 | 8ui9-C | 8.1 | 11.8 | 151 | 339 | 14 | MOLECULE: | ATPASE FAMILY AAA DOMAIN-CONTAINING PROTEIN 5 |

| | | | | | | | | |
|---|---|---|---|---|---|---|---|---|
| 47:00:00 | 7st9-C | 8.1 | 11 | 144 | 327 | 13 | MOLECULE: | CHECKPOINT PROTEIN RAD24 |
| 48:00:00 | 5m7o-A | 8 | 12.7 | 181 | 448 | 11 | MOLECULE: | NITROGEN ASSIMILATION REGULATORY PROTEIN |
| 49:00:00 | 7t0v-D | 8 | 8.9 | 142 | 570 | 15 | MOLECULE: | RIX7 |
| 50:00:00 | 7sh2-A | 8 | 10.2 | 151 | 405 | 13 | MOLECULE: | CHECKPOINT PROTEIN RAD24 |
| 51:00:00 | 7qv9-c | 8 | 3.6 | 139 | 256 | 12 | MOLECULE: | DNA-DIRECTED RNA POLYMERASE SUBUNIT ALPHA |
| 52:00:00 | 7t3i-C | 8 | 15.5 | 152 | 578 | 14 | MOLECULE: | RIX7 |
| 53:00:00 | 8hwa-C | 8 | 3.4 | 199 | 379 | 9 | MOLECULE: | PRIMASE D5 |
| 54:00:00 | 8hwc-A | 8 | 3.7 | 202 | 378 | 10 | MOLECULE: | PRIMASE D5 |
| 55:00:00 | 5ofo-A | 8 | 15.3 | 222 | 686 | 10 | MOLECULE: | CHAPERONE PROTEIN CLPB,ATP-DEPENDENT CLP PROTEASE |
| 56:00:00 | 6hz6-H | 8 | 3.5 | 138 | 290 | 11 | MOLECULE: | 5-METHYLCYTOSINE-SPECIFIC RESTRICTION ENZYME B |
| 57:00:00 | 4xgu-E | 8 | 7.7 | 159 | 379 | 13 | MOLECULE: | PUTATIVE PACHYTENE CHECKPOINT PROTEIN 2 |
| 58:00:00 | 3t15-A | 8 | 12.7 | 140 | 263 | 14 | MOLECULE: | RIBULOSE BISPHOSPHATE CARBOXYLASE/OXYGENASE ACTIV |
| 59:00:00 | 5nss-K | 8 | 3.8 | 130 | 248 | 12 | MOLECULE: | DNA-DIRECTED RNA POLYMERASE SUBUNIT ALPHA |
| 60:00:00 | 3k1j-B | 8 | 11.3 | 156 | 561 | 12 | MOLECULE: | ATP-DEPENDENT PROTEASE LON |
| 61:00:00 | 1ojl-C | 8 | 3.4 | 136 | 252 | 12 | MOLECULE: | TRANSCRIPTIONAL REGULATORY PROTEIN ZRAR |
| 62:00:00 | 4ypl-A | 8 | 11 | 190 | 537 | 12 | MOLECULE: | LON PROTEASE |
| 63:00:00 | 3ja8-7 | 8 | 6.8 | 211 | 689 | 12 | MOLECULE: | MINICHROMOSOME MAINTENANCE 2 |
| 64:00:00 | 8q3w-I | 8 | 8.8 | 161 | 246 | 13 | MOLECULE: | INSERTION SEQUENCE IS5376 PUTATIVE ATP-BINDING PR |
| 65:00:00 | 3glf-B | 8 | 13.3 | 140 | 364 | 11 | MOLECULE: | DNA POLYMERASE III SUBUNIT DELTA |
| 66:00:00 | 6wqh-C | 8 | 12.6 | 186 | 538 | 12 | MOLECULE: | LON PROTEASE |
| 67:00:00 | 6wqh-A | 8 | 10.7 | 183 | 537 | 12 | MOLECULE: | LON PROTEASE |
| 68:00:00 | 7nfy-D | 8 | 24.9 | 209 | 776 | 10 | MOLECULE: | LON PROTEASE HOMOLOG, MITOCHONDRIAL |
| 69:00:00 | 7ng5-A | 8 | 34 | 214 | 776 | 12 | MOLECULE: | LON PROTEASE HOMOLOG, MITOCHONDRIAL |
| 70:00:00 | 7p0m-D | 8 | 11.7 | 179 | 531 | 11 | MOLECULE: | LON PROTEASE HOMOLOG, MITOCHONDRIAL |
| 71:00:00 | 7p0m-B | 8 | 11 | 175 | 536 | 10 | MOLECULE: | LON PROTEASE HOMOLOG, MITOCHONDRIAL |
| 72:00:00 | 6sh3-C | 8 | 16 | 175 | 376 | 9 | MOLECULE: | MITOCHONDRIAL CHAPERONE BCS1 |
| 73:00:00 | 6sh4-B | 8 | 22.2 | 172 | 376 | 10 | MOLECULE: | MITOCHONDRIAL CHAPERONE BCS1 |
| 74:00:00 | 6e10-6 | 8 | 12.1 | 225 | 718 | 15 | MOLECULE: | HEAT SHOCK PROTEIN 101 |
| 75:00:00 | 6hz5-J | 8 | 3.4 | 135 | 284 | 11 | MOLECULE: | 5-METHYLCYTOSINE-SPECIFIC RESTRICTION ENZYME B |
| 76:00:00 | 4r7z-N | 8 | 4.9 | 162 | 316 | 9 | MOLECULE: | CELL DIVISION CONTROL PROTEIN 21 |
| 77:00:00 | 6hz6-B | 8 | 3.9 | 140 | 290 | 12 | MOLECULE: | 5-METHYLCYTOSINE-SPECIFIC RESTRICTION ENZYME B |
| 78:00:00 | 1ojl-E | 8 | 3.6 | 132 | 285 | 9 | MOLECULE: | TRANSCRIPTIONAL REGULATORY PROTEIN ZRAR |
| 79:00:00 | 6fo1-C | 8 | 3.2 | 142 | 319 | 12 | MOLECULE: | RUVB-LIKE 1 |
| 80:00:00 | 6ut7-I | 8 | 11.3 | 141 | 392 | 13 | MOLECULE: | GTPASE SUBUNIT OF RESTRICTION ENDONUCLEASE |
| 81:00:00 | 6ut7-C | 8 | 13.3 | 139 | 390 | 14 | MOLECULE: | GTPASE SUBUNIT OF RESTRICTION ENDONUCLEASE |
| 82:00:00 | 6vfx-E | 8 | 11.9 | 160 | 341 | 12 | MOLECULE: | ATP-DEPENDENT CLP PROTEASE ATP-BINDING SUBUNIT CL |
| 83:00:00 | 3j99-C | 8 | 16.8 | 188 | 676 | 14 | MOLECULE: | VESICLE-FUSING ATPASE |
| 84:00:00 | 6k0r-A | 8 | 3.3 | 142 | 316 | 13 | MOLECULE: | RUVB-LIKE 1,RUVB-LIKE 1 |
| 85:00:00 | 6k0r-H | 8 | 3.3 | 144 | 318 | 13 | MOLECULE: | RUVB-LIKE 1,RUVB-LIKE 1 |
| 86:00:00 | 6ut5-F | 8 | 10.1 | 139 | 395 | 12 | MOLECULE: | GTPASE SUBUNIT OF RESTRICTION ENDONUCLEASE |
| 87:00:00 | 6ut7-J | 8 | 13.3 | 139 | 390 | 14 | MOLECULE: | GTPASE SUBUNIT OF RESTRICTION ENDONUCLEASE |
| 88:00:00 | 3j94-C | 8 | 9.6 | 144 | 488 | 14 | MOLECULE: | VESICLE-FUSING ATPASE |
| 89:00:00 | 8osg-A | 8 | 4.3 | 156 | 283 | 8 | MOLECULE: | MAGNESIUM-CHELATASE SUBUNIT CHLI |
| 90:00:00 | 3hu1-F | 8 | 11.1 | 163 | 451 | 12 | MOLECULE: | TRANSITIONAL ENDOPLASMIC RETICULUM ATPASE |
| 91:00:00 | 4lcb-A | 8 | 11.7 | 132 | 252 | 14 | MOLECULE: | CELL DIVISION PROTEIN CDVC, VPS4 |
| 92:00:00 | 3hu2-D | 8 | 11.1 | 166 | 452 | 14 | MOLECULE: | TRANSITIONAL ENDOPLASMIC RETICULUM ATPASE |
| 93:00:00 | 5w0t-A | 8 | 4.7 | 143 | 298 | 17 | MOLECULE: | PROTEIN MSP1 |
| 94:00:00 | 4kod-C | 8 | 3.7 | 153 | 438 | 12 | MOLECULE: | TRANSITIONAL ENDOPLASMIC RETICULUM ATPASE |
| 95:00:00 | 3hu3-B | 8 | 12.1 | 167 | 446 | 12 | MOLECULE: | TRANSITIONAL ENDOPLASMIC RETICULUM ATPASE |
| 96:00:00 | 4kod-D | 8 | 3.7 | 154 | 439 | 11 | MOLECULE: | TRANSITIONAL ENDOPLASMIC RETICULUM ATPASE |
| 97:00:00 | 4kod-L | 8 | 3.9 | 159 | 442 | 9 | MOLECULE: | TRANSITIONAL ENDOPLASMIC RETICULUM ATPASE |
| 98:00:00 | 3hu1-B | 8 | 10.9 | 165 | 451 | 12 | MOLECULE: | TRANSITIONAL ENDOPLASMIC RETICULUM ATPASE |
| 99:00:00 | 8hrz-E | 8 | 4 | 153 | 436 | 12 | MOLECULE: | TRANSITIONAL ENDOPLASMIC RETICULUM ATPASE |

| | | | | | | | | |
|---|---|---|---|---|---|---|---|---|
| 0:00 | 8hrz-J | 8 | 4.4 | 155 | 436 | 12 | MOLECULE: | TRANSITIONAL ENDOPLASMIC RETICULUM ATPASE |
| 1:00 | 8hrz-D | 8 | 4.3 | 157 | 436 | 11 | MOLECULE: | TRANSITIONAL ENDOPLASMIC RETICULUM ATPASE |
| 2:00 | 8hrz-B | 8 | 10.9 | 157 | 436 | 12 | MOLECULE: | TRANSITIONAL ENDOPLASMIC RETICULUM ATPASE |
| 3:00 | 8hrz-H | 8 | 4.9 | 153 | 436 | 12 | MOLECULE: | TRANSITIONAL ENDOPLASMIC RETICULUM ATPASE |
| 4:00 | 8efy-E | 8 | 10.7 | 146 | 309 | 14 | MOLECULE: | HOLLIDAY JUNCTION ATP-DEPENDENT DNA HELICASE RUVB |
| 5:00 | 3j99-D | 8 | 21.5 | 159 | 673 | 13 | MOLECULE: | VESICLE-FUSING ATPASE |
| 6:00 | 7pbm-B | 8 | 11.7 | 146 | 312 | 10 | MOLECULE: | HOLLIDAY JUNCTION ATP-DEPENDENT DNA HELICASE RUVB |
| 7:00 | 4ko8-A | 8 | 4.1 | 154 | 456 | 10 | MOLECULE: | TRANSITIONAL ENDOPLASMIC RETICULUM ATPASE |
| 8:00 | 4kod-B | 8 | 3.7 | 154 | 439 | 11 | MOLECULE: | TRANSITIONAL ENDOPLASMIC RETICULUM ATPASE |
| 9:00 | 1ht1-F | 8 | 11.4 | 161 | 408 | 11 | MOLECULE: | HEAT SHOCK LOCUS HSLV |
| 10:00 | 6pxk-I | 8 | 11.6 | 156 | 411 | 11 | MOLECULE: | ATP-DEPENDENT PROTEASE ATPASE SUBUNIT HSLU |
| 11:00 | 5nug-B | 8 | 15.6 | 217 | 2920 | 11 | MOLECULE: | CYTOPLASMIC DYNEIN 1 HEAVY CHAIN 1 |
| 12:00 | 8thc-C | 8 | 14.2 | 143 | 325 | 13 | MOLECULE: | ELG1 ISOFORM 1 |
| 13:00 | 7pbr-A | 8 | 12.9 | 144 | 312 | 11 | MOLECULE: | HOLLIDAY JUNCTION ATP-DEPENDENT DNA HELICASE RUVB |
| 14:00 | 7sgz-B | 8 | 11.8 | 137 | 317 | 12 | MOLECULE: | CHECKPOINT PROTEIN RAD24 |
| 15:00 | 4aki-B | 8 | 18.4 | 198 | 2650 | 10 | MOLECULE: | GLUTATHIONE S-TRANSFERASE CLASS-MU 26 KDA ISOZYME |
| 16:00 | 8ui8-D | 8 | 16.6 | 142 | 328 | 13 | MOLECULE: | ATPASE FAMILY AAA DOMAIN-CONTAINING PROTEIN 5 |
| 17:00 | 7tib-B | 8 | 15.3 | 141 | 319 | 13 | MOLECULE: | REPLICATION FACTOR C SUBUNIT 1 |
| 18:00 | 4d80-A | 8 | 12.2 | 153 | 269 | 14 | MOLECULE: | AAA ATPASE, CENTRAL DOMAIN PROTEIN |
| 19:00 | 4d80-C | 8 | 12 | 147 | 272 | 14 | MOLECULE: | AAA ATPASE, CENTRAL DOMAIN PROTEIN |
| 20:00 | 8umu-C | 8 | 10.9 | 142 | 322 | 14 | MOLECULE: | CHROMOSOME TRANSMISSION FIDELITY PROTEIN 18 HOMOL |
| 21:00 | 7tfk-B | 8 | 14.2 | 153 | 319 | 10 | MOLECULE: | REPLICATION FACTOR C SUBUNIT 1 |
| 22:00 | 4d80-E | 8 | 11.4 | 150 | 262 | 15 | MOLECULE: | AAA ATPASE, CENTRAL DOMAIN PROTEIN |
| 23:00 | 5nug-A | 7.9 | 15.3 | 224 | 2920 | 9 | MOLECULE: | CYTOPLASMIC DYNEIN 1 HEAVY CHAIN 1 |
| 24:00:00 | 1g4a-E | 7.9 | 11.9 | 154 | 356 | 13 | MOLECULE: | ATP-DEPENDENT HSL PROTEASE ATP-BINDING SUBUNIT |
| 25:00:00 | 8hwe-A | 7.9 | 3.5 | 180 | 329 | 10 | MOLECULE: | PRIMASE D5 |
| 26:00:00 | 7t0v-C | 7.9 | 12.9 | 160 | 570 | 13 | MOLECULE: | RIX7 |
| 27:00:00 | 8hwg-C | 7.9 | 3.7 | 201 | 372 | 10 | MOLECULE: | PRIMASE D5 |
| 28:00:00 | 6hz8-B | 7.9 | 3.5 | 137 | 290 | 12 | MOLECULE: | 5-METHYLCYTOSINE-SPECIFIC RESTRICTION ENZYME B |
| 29:00:00 | 8hwb-F | 7.9 | 4 | 204 | 379 | 11 | MOLECULE: | PRIMASE D5 |
| 30:00:00 | 8hwf-F | 7.9 | 4 | 207 | 371 | 11 | MOLECULE: | PRIMASE D5 |
| 31:00:00 | 8hwh-B | 7.9 | 3.7 | 197 | 357 | 10 | MOLECULE: | PRIMASE D5 |
| 32:00:00 | 3uk6-G | 7.9 | 4.1 | 148 | 330 | 12 | MOLECULE: | RUVB-LIKE 2 |
| 33:00:00 | 6hz8-H | 7.9 | 3.9 | 140 | 290 | 13 | MOLECULE: | 5-METHYLCYTOSINE-SPECIFIC RESTRICTION ENZYME B |
| 34:00:00 | 3zw6-D | 7.9 | 13.2 | 140 | 263 | 14 | MOLECULE: | RIBULOSE BISPHOSPHATE CARBOXYLASE/OXYGENASE ACTIV |
| 35:00:00 | 4lzz-J | 7.9 | 3.7 | 140 | 247 | 11 | MOLECULE: | TRANSCRIPTIONAL REGULATOR (NTRC FAMILY) |
| 36:00:00 | 4lzz-I | 7.9 | 3.9 | 141 | 248 | 10 | MOLECULE: | TRANSCRIPTIONAL REGULATOR (NTRC FAMILY) |
| 37:00:00 | 4lzz-V | 7.9 | 3.7 | 140 | 247 | 11 | MOLECULE: | TRANSCRIPTIONAL REGULATOR (NTRC FAMILY) |
| 38:00:00 | 7lmz-C | 7.9 | 7.3 | 169 | 752 | 10 | MOLECULE: | TRANSITIONAL ENDOPLASMIC RETICULUM ATPASE |
| 39:00:00 | 8q3w-E | 7.9 | 18 | 167 | 246 | 13 | MOLECULE: | INSERTION SEQUENCE IS5376 PUTATIVE ATP-BINDING PR |
| 40:00:00 | 3glg-G | 7.9 | 18.3 | 145 | 378 | 12 | MOLECULE: | DNA POLYMERASE III SUBUNIT DELTA |
| 41:00:00 | 4ypl-B | 7.9 | 16.5 | 192 | 537 | 13 | MOLECULE: | LON PROTEASE |
| 42:00:00 | 8q3w-H | 7.9 | 8.7 | 152 | 246 | 13 | MOLECULE: | INSERTION SEQUENCE IS5376 PUTATIVE ATP-BINDING PR |
| 43:00:00 | 8q3w-F | 7.9 | 9 | 160 | 246 | 14 | MOLECULE: | INSERTION SEQUENCE IS5376 PUTATIVE ATP-BINDING PR |
| 44:00:00 | 3glg-B | 7.9 | 13.3 | 135 | 364 | 13 | MOLECULE: | DNA POLYMERASE III SUBUNIT DELTA |
| 45:00:00 | 7p09-C | 7.9 | 12.5 | 178 | 532 | 10 | MOLECULE: | LON PROTEASE HOMOLOG, MITOCHONDRIAL |
| 46:00:00 | 7ngf-A | 7.9 | 26.9 | 225 | 776 | 8 | MOLECULE: | LON PROTEASE HOMOLOG, MITOCHONDRIAL |
| 47:00:00 | 7ngq-F | 7.9 | 26.5 | 210 | 776 | 9 | MOLECULE: | LON PROTEASE HOMOLOG, MITOCHONDRIAL |
| 48:00:00 | 6og1-A | 7.9 | 11.1 | 214 | 581 | 12 | MOLECULE: | HYPERACTIVE DISAGGREGASE CLPB |
| 49:00:00 | 8thc-A | 7.9 | 21.5 | 158 | 479 | 14 | MOLECULE: | ELG1 ISOFORM 1 |
| 50:00:00 | 8thd-A | 7.9 | 11.3 | 154 | 474 | 12 | MOLECULE: | ELG1 ISOFORM 1 |
| 51:00:00 | 1ht1-E | 7.9 | 11.5 | 157 | 408 | 12 | MOLECULE: | HEAT SHOCK LOCUS HSLV |
| 52:00:00 | 4r7z-B | 7.9 | 4.7 | 159 | 316 | 9 | MOLECULE: | CELL DIVISION CONTROL PROTEIN 21 |

| | | | | | | | |
|---|---|---|---|---|---|---|---|
| 53:00:00 | 6ut7-B | 7.9 | 11.3 | 141 | 392 | 13 MOLECULE: | GTPASE SUBUNIT OF RESTRICTION ENDONUCLEASE |
| 54:00:00 | 7aho-A | 7.9 | 3.1 | 138 | 305 | 12 MOLECULE: | RUVB-LIKE 1 |
| 55:00:00 | 3glf-G | 7.9 | 20.1 | 150 | 378 | 11 MOLECULE: | DNA POLYMERASE III SUBUNIT DELTA |
| 56:00:00 | 4kod-G | 7.9 | 10.8 | 162 | 437 | 10 MOLECULE: | TRANSITIONAL ENDOPLASMIC RETICULUM ATPASE |
| 57:00:00 | 3j96-C | 7.9 | 13.2 | 191 | 676 | 13 MOLECULE: | VESICLE-FUSING ATPASE |
| 58:00:00 | 4kln-A | 7.9 | 10.8 | 163 | 451 | 12 MOLECULE: | TRANSITIONAL ENDOPLASMIC RETICULUM ATPASE |
| 59:00:00 | 5txv-L | 7.9 | 11.5 | 158 | 371 | 11 MOLECULE: | ATP-DEPENDENT PROTEASE ATPASE SUBUNIT HSLU |
| 60:00:00 | 7v2v-A | 7.9 | 11.3 | 156 | 374 | 11 MOLECULE: | VPSR |
| 61:00:00 | 6chs-E | 7.9 | 12.1 | 178 | 550 | 10 MOLECULE: | NPL4 |
| 62:00:00 | 8hrz-G | 7.9 | 4.2 | 156 | 436 | 12 MOLECULE: | TRANSITIONAL ENDOPLASMIC RETICULUM ATPASE |
| 63:00:00 | 8hrz-C | 7.9 | 10.4 | 157 | 436 | 11 MOLECULE: | TRANSITIONAL ENDOPLASMIC RETICULUM ATPASE |
| 64:00:00 | 3j94-D | 7.9 | 10.4 | 148 | 485 | 13 MOLECULE: | VESICLE-FUSING ATPASE |
| 65:00:00 | 3j97-C | 7.9 | 13.5 | 204 | 676 | 12 MOLECULE: | VESICLE-FUSING ATPASE |
| 66:00:00 | 3j96-D | 7.9 | 18.4 | 150 | 673 | 13 MOLECULE: | VESICLE-FUSING ATPASE |
| 67:00:00 | 4ww0-A | 7.9 | 11.7 | 168 | 420 | 11 MOLECULE: | ATP-DEPENDENT ZINC METALLOPROTEASE FTSH |
| 68:00:00 | 6pxk-L | 7.9 | 11.1 | 159 | 394 | 11 MOLECULE: | ATP-DEPENDENT PROTEASE ATPASE SUBUNIT HSLU |
| 69:00:00 | 6pxl-L | 7.9 | 11.1 | 160 | 396 | 11 MOLECULE: | ATP-DEPENDENT PROTEASE ATPASE SUBUNIT HSLU |
| 70:00:00 | 7st9-A | 7.9 | 10.4 | 158 | 522 | 13 MOLECULE: | CHECKPOINT PROTEIN RAD24 |
| 71:00:00 | 4akh-B | 7.9 | 9.6 | 236 | 2650 | 7 MOLECULE: | GLUTATHIONE S-TRANSFERASE CLASS-MU 26 KDA ISOZYME |
| 72:00:00 | 8dyv-A | 7.9 | 4.6 | 190 | 2549 | 8 MOLECULE: | CYTOPLASMIC DYNEIN 1 HEAVY CHAIN 1 |
| 73:00:00 | 6pdy-B | 7.9 | 8.7 | 153 | 282 | 13 MOLECULE: | MEMBRANE-SPANNING ATPASE-LIKE PROTEIN |
| 74:00:00 | 4d80-F | 7.9 | 10.4 | 144 | 274 | 15 MOLECULE: | AAA ATPASE, CENTRAL DOMAIN PROTEIN |
| 75:00:00 | 7ti8-C | 7.9 | 12.6 | 152 | 329 | 12 MOLECULE: | REPLICATION FACTOR C SUBUNIT 1 |
| 76:00:00 | 4ai6-B | 7.9 | 10.4 | 224 | 2650 | 8 MOLECULE: | GLUTATHIONE S-TRANSFERASE CLASS-MU 26 KDA ISOZYME |
| 77:00:00 | 8ui8-A | 7.9 | 8.1 | 170 | 643 | 11 MOLECULE: | ATPASE FAMILY AAA DOMAIN-CONTAINING PROTEIN 5 |
| 78:00:00 | 4aki-A | 7.9 | 11.3 | 249 | 2650 | 8 MOLECULE: | GLUTATHIONE S-TRANSFERASE CLASS-MU 26 KDA ISOZYME |
| 79:00:00 | 8hwd-C | 7.8 | 4 | 186 | 344 | 11 MOLECULE: | PRIMASE D5 |
| 80:00:00 | 8hwh-E | 7.8 | 3.9 | 180 | 329 | 10 MOLECULE: | PRIMASE D5 |
| 81:00:00 | 7m99-G | 7.8 | 7.4 | 130 | 257 | 9 MOLECULE: | TNSC |
| 82:00:00 | 8hwe-E | 7.8 | 3.6 | 194 | 372 | 9 MOLECULE: | PRIMASE D5 |
| 83:00:00 | 5ji2-F | 7.8 | 11.2 | 157 | 353 | 11 MOLECULE: | ATP-DEPENDENT PROTEASE SUBUNIT HSLV |
| 84:00:00 | 4tvs-A | 7.8 | 3.5 | 143 | 224 | 11 MOLECULE: | TORSIN-1A-INTERACTING PROTEIN 1 |
| 85:00:00 | 1fnn-A | 7.8 | 13.8 | 148 | 379 | 10 MOLECULE: | CELL DIVISION CONTROL PROTEIN 6 |
| 86:00:00 | 5ftn-A | 7.8 | 7.6 | 174 | 737 | 12 MOLECULE: | TRANSITIONAL ENDOPLASMIC RETICULUM ATPASE |
| 87:00:00 | 5ep4-A | 7.8 | 3.6 | 135 | 253 | 10 MOLECULE: | PUTATIVE REPRESSOR PROTEIN LUXO |
| 88:00:00 | 5nss-G | 7.8 | 7.8 | 137 | 251 | 15 MOLECULE: | DNA-DIRECTED RNA POLYMERASE SUBUNIT ALPHA |
| 89:00:00 | 7cad-C | 7.8 | 6 | 134 | 384 | 10 MOLECULE: | ABC TRANSPORTER, ATP-BINDING PROTEIN SUGC |
| 90:00:00 | 5fm6-B | 7.8 | 11.3 | 148 | 419 | 15 MOLECULE: | RVB1 |
| 91:00:00 | 6on2-F | 7.8 | 10.8 | 177 | 512 | 16 MOLECULE: | ATP-DEPENDENT PROTEASE LA |
| 92:00:00 | 3hte-E | 7.8 | 10.7 | 140 | 302 | 14 MOLECULE: | ATP-DEPENDENT CLP PROTEASE ATP-BINDING SUBUNIT CL |
| 93:00:00 | 7ksl-B | 7.8 | 11 | 173 | 520 | 9 MOLECULE: | LON PROTEASE HOMOLOG, MITOCHONDRIAL |
| 94:00:00 | 8uk9-L | 7.8 | 10 | 140 | 320 | 17 MOLECULE: | SLIDING-CLAMP-LOADER SMALL SUBUNIT |
| 95:00:00 | 7p0m-E | 7.8 | 8.7 | 171 | 532 | 10 MOLECULE: | LON PROTEASE HOMOLOG, MITOCHONDRIAL |
| 96:00:00 | 7ngc-E | 7.8 | 29.2 | 210 | 776 | 10 MOLECULE: | LON PROTEASE HOMOLOG, MITOCHONDRIAL |
| 97:00:00 | 7p0m-C | 7.8 | 11.9 | 169 | 532 | 11 MOLECULE: | LON PROTEASE HOMOLOG, MITOCHONDRIAL |
| 98:00:00 | 5ifw-B | 7.8 | 13 | 170 | 702 | 11 MOLECULE: | TETHER CONTAINING UBX DOMAIN FOR GLUT4 |
| 99:00:00 | 7p09-A | 7.8 | 11.1 | 187 | 534 | 10 MOLECULE: | LON PROTEASE HOMOLOG, MITOCHONDRIAL |
| 0:00 | 1do0-C | 7.8 | 11.2 | 150 | 406 | 13 MOLECULE: | PROTEIN (HEAT SHOCK LOCUS U) |
| 1:00 | 6sh5-B | 7.8 | 19.3 | 155 | 334 | 11 MOLECULE: | MITOCHONDRIAL CHAPERONE BCS1 |
| 2:00 | 6z1f-2 | 7.8 | 7.4 | 147 | 281 | 14 MOLECULE: | RIBULOSE BISPHOSPHATE CARBOXYLASE/OXYGENASE ACTIV |
| 3:00 | 6u1y-C | 7.8 | 2.9 | 126 | 242 | 19 MOLECULE: | MITOCHONDRIAL CHAPERONE BCS1 |
| 4:00 | 1ofi-A | 7.8 | 10.5 | 152 | 299 | 15 MOLECULE: | ATP-DEPENDENT HSL PROTEASE ATP-BINDING SUBUNIT |
| 5:00 | 6u1y-E | 7.8 | 3.1 | 128 | 246 | 20 MOLECULE: | MITOCHONDRIAL CHAPERONE BCS1 |

| | | | | | | | | |
|---|---|---|---|---|---|---|---|---|
| 6:00 | 6sh5-A | 7.8 | 19.6 | 157 | 334 | 11 | MOLECULE: | MITOCHONDRIAL CHAPERONE BCS1 |
| 7:00 | 7mi6-A | 7.8 | 27.6 | 215 | 2419 | 9 | MOLECULE: | FUSION PROTEIN OF DYNEIN AND ENDOLYSIN |
| 8:00 | 6hz8-D | 7.8 | 3.4 | 137 | 284 | 9 | MOLECULE: | 5-METHYLCYTOSINE-SPECIFIC RESTRICTION ENZYME B |
| 9:00 | 6sh5-D | 7.8 | 19.3 | 158 | 334 | 11 | MOLECULE: | MITOCHONDRIAL CHAPERONE BCS1 |
| 10:00 | 6sh5-F | 7.8 | 7.1 | 157 | 334 | 11 | MOLECULE: | MITOCHONDRIAL CHAPERONE BCS1 |
| 11:00 | 3whk-A | 7.8 | 3.1 | 127 | 248 | 13 | MOLECULE: | PROTEASOME-ACTIVATING NUCLEOTIDASE, 26S PROTEASE |
| 12:00 | 6oa9-B | 7.8 | 6.2 | 146 | 493 | 9 | MOLECULE: | CELL DIVISION CONTROL PROTEIN 48 |
| 13:00 | 4r7z-M | 7.8 | 4.7 | 158 | 316 | 9 | MOLECULE: | CELL DIVISION CONTROL PROTEIN 21 |
| 14:00 | 4kod-A | 7.8 | 10.7 | 163 | 438 | 10 | MOLECULE: | TRANSITIONAL ENDOPLASMIC RETICULUM ATPASE |
| 15:00 | 6ut3-E | 7.8 | 15.6 | 141 | 384 | 12 | MOLECULE: | GTPASE SUBUNIT OF RESTRICTION ENDONUCLEASE |
| 16:00 | 7aho-C | 7.8 | 3.2 | 139 | 306 | 13 | MOLECULE: | RUVB-LIKE 1 |
| 17:00 | 6k0r-C | 7.8 | 8.5 | 146 | 313 | 13 | MOLECULE: | RUVB-LIKE 1,RUVB-LIKE 1 |
| 18:00 | 4kod-F | 7.8 | 3.8 | 155 | 438 | 12 | MOLECULE: | TRANSITIONAL ENDOPLASMIC RETICULUM ATPASE |
| 19:00 | 3cf1-C | 7.8 | 7.9 | 192 | 723 | 9 | MOLECULE: | TRANSITIONAL ENDOPLASMIC RETICULUM ATPASE |
| 20:00 | 6hz9-K | 7.8 | 3.8 | 136 | 285 | 10 | MOLECULE: | 5-METHYLCYTOSINE-SPECIFIC RESTRICTION ENZYME B |
| 21:00 | 3hu1-A | 7.8 | 11.6 | 161 | 451 | 12 | MOLECULE: | TRANSITIONAL ENDOPLASMIC RETICULUM ATPASE |
| 22:00 | 2x8a-A | 7.8 | 9.4 | 134 | 237 | 13 | MOLECULE: | NUCLEAR VALOSIN-CONTAINING PROTEIN-LIKE |
| 23:00 | 6chs-F | 7.8 | 11.4 | 177 | 550 | 10 | MOLECULE: | NPL4 |
| 24:00:00 | 6chs-D | 7.8 | 7 | 171 | 550 | 10 | MOLECULE: | NPL4 |
| 25:00:00 | 2xsz-F | 7.8 | 3.2 | 138 | 311 | 14 | MOLECULE: | RUVB-LIKE 1 |
| 26:00:00 | 6chs-I | 7.8 | 11.7 | 176 | 550 | 10 | MOLECULE: | NPL4 |
| 27:00:00 | 1nsf-A | 7.8 | 7.3 | 138 | 247 | 14 | MOLECULE: | N-ETHYLMALEIMIDE SENSITIVE FACTOR |
| 28:00:00 | 8hrz-F | 7.8 | 4.6 | 152 | 436 | 11 | MOLECULE: | TRANSITIONAL ENDOPLASMIC RETICULUM ATPASE |
| 29:00:00 | 8hrz-K | 7.8 | 3.7 | 149 | 436 | 13 | MOLECULE: | TRANSITIONAL ENDOPLASMIC RETICULUM ATPASE |
| 30:00:00 | 3j95-D | 7.8 | 12.1 | 150 | 479 | 17 | MOLECULE: | VESICLE-FUSING ATPASE |
| 31:00:00 | 7pbt-F | 7.8 | 10.1 | 143 | 312 | 10 | MOLECULE: | HOLLIDAY JUNCTION ATP-DEPENDENT DNA HELICASE RUVB |
| 32:00:00 | 4kod-J | 7.8 | 14.1 | 161 | 445 | 11 | MOLECULE: | TRANSITIONAL ENDOPLASMIC RETICULUM ATPASE |
| 33:00:00 | 4ko8-B | 7.8 | 11.1 | 165 | 444 | 12 | MOLECULE: | TRANSITIONAL ENDOPLASMIC RETICULUM ATPASE |
| 34:00:00 | 6pxk-B | 7.8 | 11.1 | 158 | 397 | 11 | MOLECULE: | ATP-DEPENDENT PROTEASE ATPASE SUBUNIT HSLU |
| 35:00:00 | 8unj-D | 7.8 | 15.8 | 141 | 329 | 10 | MOLECULE: | CHROMOSOME TRANSMISSION FIDELITY PROTEIN 18 HOMOL |
| 36:00:00 | 8un0-C | 7.8 | 11.2 | 144 | 325 | 15 | MOLECULE: | CHROMOSOME TRANSMISSION FIDELITY PROTEIN 18 HOMOL |
| 37:00:00 | 3pvs-A | 7.8 | 12.1 | 132 | 414 | 11 | MOLECULE: | REPLICATION-ASSOCIATED RECOMBINATION PROTEIN A |
| 38:00:00 | 8umt-D | 7.8 | 15 | 133 | 325 | 9 | MOLECULE: | CHROMOSOME TRANSMISSION FIDELITY PROTEIN 18 HOMOL |
| 39:00:00 | 6pdw-D | 7.8 | 9.6 | 155 | 299 | 13 | MOLECULE: | MEMBRANE-SPANNING ATPASE-LIKE PROTEIN |
| 40:00:00 | 8ui9-A | 7.8 | 12.7 | 171 | 643 | 13 | MOLECULE: | ATPASE FAMILY AAA DOMAIN-CONTAINING PROTEIN 5 |
| 41:00:00 | 8hwg-F | 7.7 | 4.2 | 193 | 352 | 11 | MOLECULE: | PRIMASE D5 |
| 42:00:00 | 1ofh-A | 7.7 | 14 | 162 | 309 | 10 | MOLECULE: | ATP-DEPENDENT HSL PROTEASE ATP-BINDING SUBUNIT |
| 43:00:00 | 3hws-A | 7.7 | 11.9 | 151 | 320 | 16 | MOLECULE: | ATP-DEPENDENT CLP PROTEASE ATP-BINDING SUBUNIT CL |
| 44:00:00 | 6oay-C | 7.7 | 12.3 | 206 | 570 | 12 | MOLECULE: | HYPERACTIVE DISAGGREGASE CLPB |
| 45:00:00 | 6opc-B | 7.7 | 9.1 | 156 | 741 | 13 | MOLECULE: | CELL DIVISION CONTROL PROTEIN 48 |
| 46:00:00 | 2xsz-A | 7.7 | 3.2 | 142 | 318 | 13 | MOLECULE: | RUVB-LIKE 1 |
| 47:00:00 | 3gli-B | 7.7 | 13.4 | 139 | 359 | 12 | MOLECULE: | DNA POLYMERASE III SUBUNIT DELTA |
| 48:00:00 | 8q3w-C | 7.7 | 8.7 | 152 | 246 | 13 | MOLECULE: | INSERTION SEQUENCE IS5376 PUTATIVE ATP-BINDING PR |
| 49:00:00 | 1e32-A | 7.7 | 6 | 156 | 438 | 11 | MOLECULE: | P97 |
| 50:00:00 | 4xgu-B | 7.7 | 8.1 | 157 | 382 | 11 | MOLECULE: | PUTATIVE PACHYTENE CHECKPOINT PROTEIN 2 |
| 51:00:00 | 1e94-E | 7.7 | 11.6 | 151 | 408 | 15 | MOLECULE: | HEAT SHOCK PROTEIN HSLV |
| 52:00:00 | 7nfy-C | 7.7 | 32.2 | 215 | 776 | 11 | MOLECULE: | LON PROTEASE HOMOLOG, MITOCHONDRIAL |
| 53:00:00 | 7ksl-F | 7.7 | 10.5 | 179 | 520 | 8 | MOLECULE: | LON PROTEASE HOMOLOG, MITOCHONDRIAL |
| 54:00:00 | 7p09-D | 7.7 | 8.8 | 182 | 531 | 9 | MOLECULE: | LON PROTEASE HOMOLOG, MITOCHONDRIAL |
| 55:00:00 | 7ngp-E | 7.7 | 39.8 | 222 | 776 | 8 | MOLECULE: | LON PROTEASE HOMOLOG, MITOCHONDRIAL |
| 56:00:00 | 7ngf-B | 7.7 | 32.7 | 221 | 776 | 10 | MOLECULE: | LON PROTEASE HOMOLOG, MITOCHONDRIAL |
| 57:00:00 | 1g4b-E | 7.7 | 10.3 | 149 | 393 | 15 | MOLECULE: | ATP-DEPENDENT HSL PROTEASE ATP-BINDING SUBUNIT |
| 58:00:00 | 7ngf-C | 7.7 | 31.1 | 201 | 776 | 12 | MOLECULE: | LON PROTEASE HOMOLOG, MITOCHONDRIAL |

| | | | | | | | | |
|---|---|---|---|---|---|---|---|---|
| 59:00:00 | 6sh5-C | 7.7 | 19.3 | 155 | 334 | 11 | MOLECULE: | MITOCHONDRIAL CHAPERONE BCS1 |
| 60:00:00 | 7tfk-A | 7.7 | 13.6 | 137 | 360 | 12 | MOLECULE: | REPLICATION FACTOR C SUBUNIT 1 |
| 61:00:00 | 1g3i-C | 7.7 | 11.8 | 156 | 319 | 13 | MOLECULE: | ATP-DEPENDENT HSLU PROTEASE ATP-BINDING SUBUNIT |
| 62:00:00 | 1do2-C | 7.7 | 9.1 | 143 | 407 | 15 | MOLECULE: | PROTEIN (HEAT SHOCK LOCUS U) |
| 63:00:00 | 6rn4-A | 7.7 | 13.2 | 196 | 675 | 13 | MOLECULE: | CHAPERONE PROTEIN CLPB |
| 64:00:00 | 6sh5-G | 7.7 | 19.3 | 157 | 334 | 11 | MOLECULE: | MITOCHONDRIAL CHAPERONE BCS1 |
| 65:00:00 | 6sh5-E | 7.7 | 7.1 | 158 | 334 | 11 | MOLECULE: | MITOCHONDRIAL CHAPERONE BCS1 |
| 66:00:00 | 6ut4-C | 7.7 | 14.3 | 143 | 384 | 15 | MOLECULE: | GTPASE SUBUNIT OF RESTRICTION ENDONUCLEASE |
| 67:00:00 | 1do0-F | 7.7 | 11.8 | 155 | 406 | 13 | MOLECULE: | PROTEIN (HEAT SHOCK LOCUS U) |
| 68:00:00 | 4r7z-G | 7.7 | 4.6 | 153 | 316 | 8 | MOLECULE: | CELL DIVISION CONTROL PROTEIN 21 |
| 69:00:00 | 4d80-D | 7.7 | 12.4 | 142 | 273 | 15 | MOLECULE: | AAA ATPASE, CENTRAL DOMAIN PROTEIN |
| 70:00:00 | 7qo4-J | 7.7 | 23 | 198 | 405 | 8 | MOLECULE: | 26S PROTEASOME REGULATORY SUBUNIT RPN1 |
| 71:00:00 | 3j95-C | 7.7 | 12.2 | 155 | 485 | 17 | MOLECULE: | VESICLE-FUSING ATPASE |
| 72:00:00 | 3j98-D | 7.7 | 23.6 | 171 | 673 | 13 | MOLECULE: | VESICLE-FUSING ATPASE |
| 73:00:00 | 6pxk-K | 7.7 | 11.4 | 157 | 429 | 11 | MOLECULE: | ATP-DEPENDENT PROTEASE ATPASE SUBUNIT HSLU |
| 74:00:00 | 3j97-D | 7.7 | 21.6 | 155 | 673 | 14 | MOLECULE: | VESICLE-FUSING ATPASE |
| 75:00:00 | 7st9-E | 7.7 | 8.1 | 132 | 354 | 11 | MOLECULE: | CHECKPOINT PROTEIN RAD24 |
| 76:00:00 | 3hu1-D | 7.7 | 10.9 | 163 | 451 | 12 | MOLECULE: | TRANSITIONAL ENDOPLASMIC RETICULUM ATPASE |
| 77:00:00 | 6chs-H | 7.7 | 7 | 169 | 550 | 10 | MOLECULE: | NPL4 |
| 78:00:00 | 5vc7-D | 7.7 | 5.2 | 175 | 544 | 14 | MOLECULE: | VCP-LIKE ATPASE |
| 79:00:00 | 6mdp-F | 7.7 | 3.6 | 132 | 243 | 13 | MOLECULE: | VESICLE-FUSING ATPASE |
| 80:00:00 | 6pxi-E | 7.7 | 11.6 | 154 | 374 | 12 | MOLECULE: | ATP-DEPENDENT PROTEASE SUBUNIT HSLV |
| 81:00:00 | 8hrz-L | 7.7 | 4.3 | 155 | 436 | 11 | MOLECULE: | TRANSITIONAL ENDOPLASMIC RETICULUM ATPASE |
| 82:00:00 | 4cr2-J | 7.7 | 20 | 168 | 373 | 13 | MOLECULE: | PROTEASOME COMPONENT PRE3 |
| 83:00:00 | 8hrz-A | 7.7 | 4.4 | 155 | 436 | 12 | MOLECULE: | TRANSITIONAL ENDOPLASMIC RETICULUM ATPASE |
| 84:00:00 | 4kln-C | 7.7 | 10.9 | 164 | 451 | 11 | MOLECULE: | TRANSITIONAL ENDOPLASMIC RETICULUM ATPASE |
| 85:00:00 | 7pbp-A | 7.7 | 11.6 | 141 | 312 | 10 | MOLECULE: | HOLLIDAY JUNCTION ATP-DEPENDENT DNA HELICASE RUVB |
| 86:00:00 | 7pbr-E | 7.7 | 9.2 | 140 | 312 | 10 | MOLECULE: | HOLLIDAY JUNCTION ATP-DEPENDENT DNA HELICASE RUVB |
| 87:00:00 | 5bk4-6 | 7.7 | 9.2 | 204 | 615 | 9 | MOLECULE: | DNA REPLICATION LICENSING FACTOR MCM2 |
| 88:00:00 | 7v3w-D | 7.7 | 5.7 | 166 | 376 | 13 | MOLECULE: | VPSR |
| 89:00:00 | 4xgu-A | 7.7 | 10.5 | 160 | 376 | 13 | MOLECULE: | PUTATIVE PACHYTENE CHECKPOINT PROTEIN 2 |
| 90:00:00 | 1ht2-E | 7.7 | 11.4 | 155 | 408 | 12 | MOLECULE: | HEAT SHOCK LOCUS HSLV |
| 91:00:00 | 1e94-F | 7.7 | 10 | 152 | 408 | 12 | MOLECULE: | HEAT SHOCK PROTEIN HSLV |
| 92:00:00 | 7w46-B | 7.7 | 25 | 181 | 402 | 15 | MOLECULE: | UNCHARACTERIZED ATPASE YJOB |
| 93:00:00 | 7u1a-B | 7.7 | 13.9 | 138 | 319 | 12 | MOLECULE: | REPLICATION FACTOR C SUBUNIT 1 |
| 94:00:00 | 4d82-A | 7.7 | 11.8 | 143 | 267 | 15 | MOLECULE: | AAA ATPASE, CENTRAL DOMAIN PROTEIN |
| 95:00:00 | 7pbp-D | 7.7 | 9.9 | 140 | 313 | 11 | MOLECULE: | HOLLIDAY JUNCTION ATP-DEPENDENT DNA HELICASE RUVB |
| 96:00:00 | 7thj-C | 7.7 | 14.4 | 143 | 322 | 13 | MOLECULE: | REPLICATION FACTOR C SUBUNIT 1 |
| 97:00:00 | 6pe0-F | 7.7 | 9 | 143 | 248 | 15 | MOLECULE: | MEMBRANE-SPANNING ATPASE-LIKE PROTEIN |
| 98:00:00 | 7xc2-A | 7.6 | 28.1 | 167 | 858 | 14 | MOLECULE: | CNL9 |
| 99:00:00 | 6orb-A | 7.6 | 7 | 198 | 3352 | 10 | MOLECULE: | MIDASIN |
| 0:00 | 8iqh-D | 7.6 | 27.7 | 230 | 891 | 10 | MOLECULE: | PUTATIVE PRIMASE C962R |
| 1:00 | 7ufi-7 | 7.6 | 9.9 | 147 | 311 | 7 | MOLECULE: | DNA (5'- |
| 2:00 | 5e7p-B | 7.6 | 8.8 | 168 | 704 | 10 | MOLECULE: | CELL DIVISION CONTROL PROTEIN CDC48 |
| 3:00 | 1xwi-A | 7.6 | 13.6 | 144 | 322 | 13 | MOLECULE: | SKD1 PROTEIN |
| 4:00 | 5ji2-E | 7.6 | 11.2 | 153 | 375 | 12 | MOLECULE: | ATP-DEPENDENT PROTEASE SUBUNIT HSLV |
| 5:00 | 6p07-D | 7.6 | 3.6 | 132 | 303 | 14 | MOLECULE: | SPASTIN |
| 6:00 | 6rn3-A | 7.6 | 12.2 | 159 | 684 | 9 | MOLECULE: | CHAPERONE PROTEIN CLPB |
| 7:00 | 2ce7-C | 7.6 | 10.1 | 162 | 421 | 13 | MOLECULE: | CELL DIVISION PROTEIN FTSH |
| 8:00 | 4ypl-C | 7.6 | 10.9 | 195 | 538 | 14 | MOLECULE: | LON PROTEASE |
| 9:00 | 3jc7-7 | 7.6 | 5.7 | 217 | 657 | 12 | MOLECULE: | DNA REPLICATION LICENSING FACTOR MCM2 |
| 10:00 | 8q3w-J | 7.6 | 8.7 | 152 | 246 | 13 | MOLECULE: | INSERTION SEQUENCE IS5376 PUTATIVE ATP-BINDING PR |
| 11:00 | 5fm7-A | 7.6 | 3.8 | 146 | 418 | 14 | MOLECULE: | RVB1 |

| | | | | | | | | |
|---|---|---|---|---|---|---|---|---|
| 12:00 | 1iy2-A | 7.6 | 3.2 | 132 | 245 | 13 | MOLECULE: | ATP-DEPENDENT METALLOPROTEASE FTSH |
| 13:00 | 6on2-A | 7.6 | 11 | 190 | 523 | 13 | MOLECULE: | ATP-DEPENDENT PROTEASE LA |
| 14:00 | 7ng4-B | 7.6 | 34 | 218 | 776 | 9 | MOLECULE: | LON PROTEASE HOMOLOG, MITOCHONDRIAL |
| 15:00 | 7nfy-B | 7.6 | 29.9 | 219 | 776 | 10 | MOLECULE: | LON PROTEASE HOMOLOG, MITOCHONDRIAL |
| 16:00 | 5vc7-A | 7.6 | 3.9 | 166 | 544 | 14 | MOLECULE: | VCP-LIKE ATPASE |
| 17:00 | 7ngf-D | 7.6 | 30.2 | 203 | 776 | 11 | MOLECULE: | LON PROTEASE HOMOLOG, MITOCHONDRIAL |
| 18:00 | 7p0m-A | 7.6 | 11.4 | 185 | 534 | 10 | MOLECULE: | LON PROTEASE HOMOLOG, MITOCHONDRIAL |
| 19:00 | 7lar-B | 7.6 | 3.7 | 126 | 183 | 17 | MOLECULE: | ATP-DEPENDENT HELICASE REP |
| 20:00 | 1g3i-E | 7.6 | 10.3 | 155 | 317 | 12 | MOLECULE: | ATP-DEPENDENT HSLU PROTEASE ATP-BINDING SUBUNIT |
| 21:00 | 1do0-B | 7.6 | 10.2 | 151 | 406 | 13 | MOLECULE: | PROTEIN (HEAT SHOCK LOCUS U) |
| 22:00 | 6uks-C | 7.6 | 5.2 | 157 | 362 | 18 | MOLECULE: | MITOCHONDRIAL CHAPERONE BCS1 |
| 23:00 | 6u1y-B | 7.6 | 2.9 | 126 | 245 | 19 | MOLECULE: | MITOCHONDRIAL CHAPERONE BCS1 |
| 24:00:00 | 6u1y-D | 7.6 | 2.9 | 126 | 244 | 19 | MOLECULE: | MITOCHONDRIAL CHAPERONE BCS1 |
| 25:00:00 | 3j3t-F | 7.6 | 13.5 | 227 | 798 | 11 | MOLECULE: | ADAPTER PROTEIN MECA 1 |
| 26:00:00 | 5vc7-C | 7.6 | 8.4 | 182 | 544 | 9 | MOLECULE: | VCP-LIKE ATPASE |
| 27:00:00 | 6etx-E | 7.6 | 3.9 | 141 | 443 | 13 | MOLECULE: | RUVB-LIKE 1 |
| 28:00:00 | 6u1y-A | 7.6 | 2.9 | 126 | 244 | 19 | MOLECULE: | MITOCHONDRIAL CHAPERONE BCS1 |
| 29:00:00 | 6ef2-J | 7.6 | 11.1 | 142 | 262 | 11 | MOLECULE: | PROTEASOME SUBUNIT ALPHA TYPE-1 |
| 30:00:00 | 1do0-E | 7.6 | 10.2 | 153 | 406 | 12 | MOLECULE: | PROTEIN (HEAT SHOCK LOCUS U) |
| 31:00:00 | 8osh-C | 7.6 | 4.5 | 168 | 315 | 10 | MOLECULE: | MAGNESIUM-CHELATASE SUBUNIT CHLI |
| 32:00:00 | 7las-B | 7.6 | 3.7 | 126 | 183 | 17 | MOLECULE: | ATP-DEPENDENT HELICASE REP |
| 33:00:00 | 7lar-D | 7.6 | 3.8 | 125 | 183 | 17 | MOLECULE: | ATP-DEPENDENT HELICASE REP |
| 34:00:00 | 4r7z-D | 7.6 | 4.8 | 156 | 316 | 11 | MOLECULE: | CELL DIVISION CONTROL PROTEIN 21 |
| 35:00:00 | 4r7z-P | 7.6 | 4.8 | 156 | 316 | 11 | MOLECULE: | CELL DIVISION CONTROL PROTEIN 21 |
| 36:00:00 | 6qi8-C | 7.6 | 3.6 | 144 | 316 | 10 | MOLECULE: | RUVB-LIKE 1 |
| 37:00:00 | 6k0r-G | 7.6 | 3.3 | 140 | 311 | 13 | MOLECULE: | RUVB-LIKE 1,RUVB-LIKE 1 |
| 38:00:00 | 6ut7-M | 7.6 | 14 | 138 | 392 | 15 | MOLECULE: | GTPASE SUBUNIT OF RESTRICTION ENDONUCLEASE |
| 39:00:00 | 6ut8-B | 7.6 | 11.3 | 141 | 392 | 13 | MOLECULE: | GTPASE SUBUNIT OF RESTRICTION ENDONUCLEASE |
| 40:00:00 | 2qby-B | 7.6 | 3.7 | 132 | 368 | 8 | MOLECULE: | CELL DIVISION CONTROL PROTEIN 6 HOMOLOG 1 |
| 41:00:00 | 3j95-F | 7.6 | 8.8 | 181 | 473 | 12 | MOLECULE: | VESICLE-FUSING ATPASE |
| 42:00:00 | 3hu2-B | 7.6 | 11.1 | 162 | 452 | 12 | MOLECULE: | TRANSITIONAL ENDOPLASMIC RETICULUM ATPASE |
| 43:00:00 | 7las-D | 7.6 | 3.8 | 128 | 183 | 17 | MOLECULE: | ATP-DEPENDENT HELICASE REP |
| 44:00:00 | 3j98-C | 7.6 | 21.3 | 165 | 676 | 13 | MOLECULE: | VESICLE-FUSING ATPASE |
| 45:00:00 | 4lgm-A | 7.6 | 12.8 | 138 | 251 | 16 | MOLECULE: | VPS4 AAA ATPASE |
| 46:00:00 | 5g4f-C | 7.6 | 8.6 | 190 | 726 | 9 | MOLECULE: | VCP-LIKE ATPASE |
| 47:00:00 | 3cf0-F | 7.6 | 10.1 | 138 | 281 | 12 | MOLECULE: | TRANSITIONAL ENDOPLASMIC RETICULUM ATPASE |
| 48:00:00 | 1ht1-G | 7.6 | 11.4 | 156 | 408 | 13 | MOLECULE: | HEAT SHOCK LOCUS HSLV |
| 49:00:00 | 7pbl-D | 7.6 | 11.2 | 139 | 313 | 12 | MOLECULE: | HOLLIDAY JUNCTION ATP-DEPENDENT DNA HELICASE RUVB |
| 50:00:00 | 8osg-E | 7.6 | 4.2 | 158 | 323 | 10 | MOLECULE: | MAGNESIUM-CHELATASE SUBUNIT CHLI |
| 51:00:00 | 6pxk-A | 7.6 | 11.3 | 156 | 420 | 13 | MOLECULE: | ATP-DEPENDENT PROTEASE ATPASE SUBUNIT HSLU |
| 52:00:00 | 6pxk-H | 7.6 | 11.3 | 157 | 392 | 11 | MOLECULE: | ATP-DEPENDENT PROTEASE ATPASE SUBUNIT HSLU |
| 53:00:00 | 4d80-B | 7.6 | 12.3 | 137 | 257 | 13 | MOLECULE: | AAA ATPASE, CENTRAL DOMAIN PROTEIN |
| 54:00:00 | 6pdw-B | 7.6 | 11.4 | 158 | 294 | 13 | MOLECULE: | MEMBRANE-SPANNING ATPASE-LIKE PROTEIN |
| 55:00:00 | 7t3i-A | 7.5 | 9.1 | 143 | 577 | 16 | MOLECULE: | RIX7 |
| 56:00:00 | 5og1-B | 7.5 | 10.6 | 209 | 572 | 10 | MOLECULE: | CHAPERONE PROTEIN CLPB,ATP-DEPENDENT CLP PROTEASE |
| 57:00:00 | 7las-E | 7.5 | 3.6 | 123 | 183 | 17 | MOLECULE: | ATP-DEPENDENT HELICASE REP |
| 58:00:00 | 6rn3-B | 7.5 | 12.5 | 180 | 689 | 8 | MOLECULE: | CHAPERONE PROTEIN CLPB |
| 59:00:00 | 4ww4-B | 7.5 | 12.4 | 152 | 435 | 11 | MOLECULE: | RUVB-LIKE 1 |
| 60:00:00 | 6oax-B | 7.5 | 13.5 | 155 | 581 | 10 | MOLECULE: | HYPERACTIVE DISAGGREGASE CLPB |
| 61:00:00 | 4eiw-E | 7.5 | 10 | 157 | 457 | 13 | MOLECULE: | ATP-DEPENDENT ZINC METALLOPROTEASE FTSH |
| 62:00:00 | 4ai6-A | 7.5 | 28.2 | 224 | 2650 | 9 | MOLECULE: | GLUTATHIONE S-TRANSFERASE CLASS-MU 26 KDA ISOZYME |
| 63:00:00 | 4lzz-L | 7.5 | 3.6 | 133 | 246 | 11 | MOLECULE: | TRANSCRIPTIONAL REGULATOR (NTRC FAMILY) |
| 64:00:00 | 1ht2-G | 7.5 | 10 | 155 | 408 | 10 | MOLECULE: | HEAT SHOCK LOCUS HSLV |

| | | | | | | | | |
|---|---|---|---|---|---|---|---|---|
| 65:00:00 | 3n70-A | 7.5 | 4.2 | 117 | 142 | 9 | MOLECULE: | TRANSPORT ACTIVATOR |
| 66:00:00 | 8q3w-A | 7.5 | 15.9 | 149 | 246 | 13 | MOLECULE: | INSERTION SEQUENCE IS5376 PUTATIVE ATP-BINDING PR |
| 67:00:00 | 6wqh-E | 7.5 | 12.2 | 188 | 537 | 13 | MOLECULE: | LON PROTEASE |
| 68:00:00 | 1njf-C | 7.5 | 3.5 | 127 | 239 | 12 | MOLECULE: | DNA POLYMERASE III SUBUNIT GAMMA |
| 69:00:00 | 6u5z-B | 7.5 | 13.2 | 186 | 529 | 15 | MOLECULE: | LON PROTEASE |
| 70:00:00 | 6v11-D | 7.5 | 12.8 | 169 | 465 | 13 | MOLECULE: | LON PROTEASE |
| 71:00:00 | 7ng5-E | 7.5 | 25.6 | 195 | 776 | 8 | MOLECULE: | LON PROTEASE HOMOLOG, MITOCHONDRIAL |
| 72:00:00 | 7ngl-E | 7.5 | 32.3 | 209 | 776 | 9 | MOLECULE: | LON PROTEASE HOMOLOG, MITOCHONDRIAL |
| 73:00:00 | 1kyi-A | 7.5 | 10.3 | 153 | 321 | 14 | MOLECULE: | ATP-DEPENDENT HSL PROTEASE ATP-BINDING SUBUNIT HS |
| 74:00:00 | 6uks-B | 7.5 | 5.7 | 159 | 362 | 18 | MOLECULE: | MITOCHONDRIAL CHAPERONE BCS1 |
| 75:00:00 | 1hqy-E | 7.5 | 9.9 | 152 | 408 | 11 | MOLECULE: | HEAT SHOCK LOCUS HSLV |
| 76:00:00 | 1g3i-B | 7.5 | 10.4 | 148 | 326 | 14 | MOLECULE: | ATP-DEPENDENT HSLU PROTEASE ATP-BINDING SUBUNIT |
| 77:00:00 | 1g3i-W | 7.5 | 11.6 | 157 | 312 | 11 | MOLECULE: | ATP-DEPENDENT HSLU PROTEASE ATP-BINDING SUBUNIT |
| 78:00:00 | 6j6i-C | 7.5 | 26.5 | 161 | 790 | 8 | MOLECULE: | PROBABLE SERINE/THREONINE-PROTEIN KINASE PBL2 |
| 79:00:00 | 1do2-A | 7.5 | 12.4 | 154 | 407 | 13 | MOLECULE: | PROTEIN (HEAT SHOCK LOCUS U) |
| 80:00:00 | 1g4b-K | 7.5 | 10.1 | 152 | 393 | 11 | MOLECULE: | ATP-DEPENDENT HSL PROTEASE ATP-BINDING SUBUNIT |
| 81:00:00 | 6u1y-G | 7.5 | 2.9 | 126 | 244 | 19 | MOLECULE: | MITOCHONDRIAL CHAPERONE BCS1 |
| 82:00:00 | 6edo-A | 7.5 | 23.4 | 187 | 2031 | 10 | MOLECULE: | MIDASIN |
| 83:00:00 | 8pb9-E | 7.5 | 11.2 | 149 | 380 | 11 | MOLECULE: | TRANSCRIPTIONAL REGULATOR FLEQ |
| 84:00:00 | 3j97-F | 7.5 | 14.9 | 183 | 654 | 15 | MOLECULE: | VESICLE-FUSING ATPASE |
| 85:00:00 | 6igm-E | 7.5 | 9.8 | 140 | 380 | 14 | MOLECULE: | RUVB-LIKE 1 |
| 86:00:00 | 2x31-J | 7.5 | 5.5 | 160 | 321 | 13 | MOLECULE: | MAGNESIUM-CHELATASE 60 KDA SUBUNIT |
| 87:00:00 | 6mdo-F | 7.5 | 8 | 133 | 241 | 14 | MOLECULE: | VESICLE-FUSING ATPASE |
| 88:00:00 | 4kln-F | 7.5 | 10.7 | 161 | 451 | 14 | MOLECULE: | TRANSITIONAL ENDOPLASMIC RETICULUM ATPASE |
| 89:00:00 | 4wvy-B | 7.5 | 11.1 | 141 | 408 | 13 | MOLECULE: | RUVB-LIKE 1 |
| 90:00:00 | 1ny5-B | 7.5 | 8.8 | 165 | 385 | 8 | MOLECULE: | TRANSCRIPTIONAL REGULATOR (NTRC FAMILY) |
| 91:00:00 | 1g3i-A | 7.5 | 11.5 | 158 | 326 | 11 | MOLECULE: | ATP-DEPENDENT HSLU PROTEASE ATP-BINDING SUBUNIT |
| 92:00:00 | 3cf0-I | 7.5 | 10.7 | 138 | 281 | 12 | MOLECULE: | TRANSITIONAL ENDOPLASMIC RETICULUM ATPASE |
| 93:00:00 | 8hr8-H | 7.5 | 15 | 151 | 834 | 8 | MOLECULE: | ARCHAEAL ATPASE |
| 94:00:00 | 3h4m-B | 7.5 | 8.3 | 138 | 261 | 12 | MOLECULE: | PROTEASOME-ACTIVATING NUCLEOTIDASE |
| 95:00:00 | 2rko-A | 7.5 | 14.1 | 141 | 280 | 13 | MOLECULE: | VACUOLAR PROTEIN SORTING-ASSOCIATED PROTEIN 4 |
| 96:00:00 | 8ff4-T | 7.5 | 8.4 | 151 | 343 | 8 | MOLECULE: | TYPE I-B CRISPR-ASSOCIATED PROTEIN CAS5 |
| 97:00:00 | 8thb-A | 7.5 | 24.4 | 168 | 468 | 12 | MOLECULE: | ELG1 ISOFORM 1 |
| 98:00:00 | 6pxk-E | 7.5 | 11.6 | 158 | 426 | 12 | MOLECULE: | ATP-DEPENDENT PROTEASE ATPASE SUBUNIT HSLU |
| 99:00:00 | 1ht1-I | 7.5 | 10.1 | 154 | 408 | 12 | MOLECULE: | HEAT SHOCK LOCUS HSLV |
| 0:00 | 7u1p-C | 7.5 | 13.6 | 143 | 328 | 13 | MOLECULE: | REPLICATION FACTOR C SUBUNIT 1 |
| 1:00 | 8fs4-C | 7.5 | 14.4 | 147 | 328 | 14 | MOLECULE: | CHECKPOINT PROTEIN RAD24 |
| 2:00 | 4d82-C | 7.5 | 8.5 | 138 | 272 | 15 | MOLECULE: | AAA ATPASE, CENTRAL DOMAIN PROTEIN |
| 3:00 | 7uqj-A | 7.4 | 5.5 | 157 | 585 | 12 | MOLECULE: | ATPASE HISTONE CHAPERONE YTA7 |
| 4:00 | 8fwj-A | 7.4 | 3.7 | 132 | 552 | 14 | MOLECULE: | CIRCADIAN CLOCK PROTEIN KAIC |
| 5:00 | 7vcs-A | 7.4 | 4.7 | 153 | 766 | 10 | MOLECULE: | TRANSITIONAL ENDOPLASMIC RETICULUM ATPASE |
| 6:00 | 6hyp-A | 7.4 | 14 | 221 | 2272 | 6 | MOLECULE: | MIDASIN,MIDASIN |
| 7:00 | 5j1s-B | 7.4 | 5.5 | 156 | 235 | 11 | MOLECULE: | TORSIN-1A |
| 8:00 | 3syk-A | 7.4 | 11.7 | 147 | 290 | 14 | MOLECULE: | PROTEIN CBBX |
| 9:00 | 7ufm-A | 7.4 | 10 | 146 | 311 | 8 | MOLECULE: | VCHTNSC |
| 10:00 | 7ufm-K | 7.4 | 10 | 146 | 311 | 8 | MOLECULE: | VCHTNSC |
| 11:00 | 1um8-A | 7.4 | 10.8 | 154 | 327 | 15 | MOLECULE: | ATP-DEPENDENT CLP PROTEASE ATP-BINDING SUBUNIT CL |
| 12:00 | 7lar-F | 7.4 | 4.8 | 136 | 183 | 15 | MOLECULE: | ATP-DEPENDENT HELICASE REP |
| 13:00 | 6qel-I | 7.4 | 26.2 | 166 | 237 | 14 | MOLECULE: | REPLICATIVE DNA HELICASE |
| 14:00 | 6j5t-L | 7.4 | 27 | 156 | 808 | 8 | MOLECULE: | PROBABLE SERINE/THREONINE-PROTEIN KINASE PBL2 |
| 15:00 | 5ln3-J | 7.4 | 20.5 | 169 | 373 | 11 | MOLECULE: | 26S PROTEASOME NON-ATPASE REGULATORY SUBUNIT 2 |
| 16:00 | 5bk4-7 | 7.4 | 15.2 | 229 | 689 | 10 | MOLECULE: | DNA REPLICATION LICENSING FACTOR MCM2 |
| 17:00 | 6mfv-B | 7.4 | 10.2 | 131 | 641 | 11 | MOLECULE: | TETRATRICOPEPTIDE REPEAT SENSOR PH0952 |

| | | | | | | | | |
|---|---|---|---|---|---|---|---|---|
| 18:00 | 6p07-E | 7.4 | 3.8 | 133 | 303 | 14 | MOLECULE: | SPASTIN |
| 19:00 | 4eiw-A | 7.4 | 9.2 | 158 | 457 | 12 | MOLECULE: | ATP-DEPENDENT ZINC METALLOPROTEASE FTSH |
| 20:00 | 5z6r-A | 7.4 | 13.5 | 145 | 273 | 10 | MOLECULE: | SPASTIN |
| 21:00 | 3zw6-B | 7.4 | 13.8 | 146 | 263 | 13 | MOLECULE: | RIBULOSE BISPHOSPHATE CARBOXYLASE/OXYGENASE ACTIV |
| 22:00 | 3zw6-E | 7.4 | 13.8 | 145 | 263 | 13 | MOLECULE: | RIBULOSE BISPHOSPHATE CARBOXYLASE/OXYGENASE ACTIV |
| 23:00 | 1sxj-D | 7.4 | 14.2 | 146 | 328 | 12 | MOLECULE: | ACTIVATOR 1 95 KDA SUBUNIT |
| 24:00:00 | 3zw6-C | 7.4 | 14.1 | 135 | 263 | 15 | MOLECULE: | RIBULOSE BISPHOSPHATE CARBOXYLASE/OXYGENASE ACTIV |
| 25:00:00 | 3d8b-A | 7.4 | 8.8 | 136 | 281 | 10 | MOLECULE: | FIDGETIN-LIKE PROTEIN 1 |
| 26:00:00 | 5exx-A | 7.4 | 7.4 | 142 | 256 | 11 | MOLECULE: | TRANSCRIPTIONAL REGULATOR FLEQ |
| 27:00:00 | 5fm7-B | 7.4 | 11.3 | 144 | 422 | 13 | MOLECULE: | RVB1 |
| 28:00:00 | 6j6i-F | 7.4 | 26.1 | 156 | 790 | 9 | MOLECULE: | PROBABLE SERINE/THREONINE-PROTEIN KINASE PBL2 |
| 29:00:00 | 1xxh-D | 7.4 | 13.9 | 147 | 364 | 13 | MOLECULE: | DNA POLYMERASE III, DELTA SUBUNIT |
| 30:00:00 | 2zr0-A | 7.4 | 8.7 | 145 | 316 | 12 | MOLECULE: | PROTEIN RECA |
| 31:00:00 | 1xxh-G | 7.4 | 11.2 | 145 | 364 | 10 | MOLECULE: | DNA POLYMERASE III, DELTA SUBUNIT |
| 32:00:00 | 3gli-G | 7.4 | 19.9 | 148 | 373 | 11 | MOLECULE: | DNA POLYMERASE III SUBUNIT DELTA |
| 33:00:00 | 7ng4-D | 7.4 | 30.3 | 225 | 776 | 11 | MOLECULE: | LON PROTEASE HOMOLOG, MITOCHONDRIAL |
| 34:00:00 | 7nfy-E | 7.4 | 24.5 | 187 | 776 | 10 | MOLECULE: | LON PROTEASE HOMOLOG, MITOCHONDRIAL |
| 35:00:00 | 3glh-C | 7.4 | 15.1 | 151 | 364 | 10 | MOLECULE: | DNA POLYMERASE III SUBUNIT DELTA |
| 36:00:00 | 6epc-L | 7.4 | 18.3 | 162 | 389 | 10 | MOLECULE: | PROTEASOME SUBUNIT ALPHA TYPE-6 |
| 37:00:00 | 1g3i-D | 7.4 | 11.5 | 154 | 318 | 16 | MOLECULE: | ATP-DEPENDENT HSLU PROTEASE ATP-BINDING SUBUNIT |
| 38:00:00 | 1g3i-F | 7.4 | 10.1 | 155 | 320 | 12 | MOLECULE: | ATP-DEPENDENT HSLU PROTEASE ATP-BINDING SUBUNIT |
| 39:00:00 | 1g4b-F | 7.4 | 10.2 | 151 | 393 | 14 | MOLECULE: | ATP-DEPENDENT HSL PROTEASE ATP-BINDING SUBUNIT |
| 40:00:00 | 8gj2-D | 7.4 | 12.5 | 148 | 360 | 13 | MOLECULE: | DNA POLYMERASE III SUBUNIT DELTA |
| 41:00:00 | 1g4b-L | 7.4 | 11.6 | 158 | 393 | 11 | MOLECULE: | ATP-DEPENDENT HSL PROTEASE ATP-BINDING SUBUNIT |
| 42:00:00 | 1g3i-V | 7.4 | 10.2 | 151 | 313 | 13 | MOLECULE: | ATP-DEPENDENT HSLU PROTEASE ATP-BINDING SUBUNIT |
| 43:00:00 | 1g3i-U | 7.4 | 10.1 | 155 | 322 | 12 | MOLECULE: | ATP-DEPENDENT HSLU PROTEASE ATP-BINDING SUBUNIT |
| 44:00:00 | 8pb9-C | 7.4 | 3.9 | 135 | 252 | 8 | MOLECULE: | TRANSCRIPTIONAL REGULATOR FLEQ |
| 45:00:00 | 1g3i-S | 7.4 | 10 | 147 | 317 | 12 | MOLECULE: | ATP-DEPENDENT HSLU PROTEASE ATP-BINDING SUBUNIT |
| 46:00:00 | 6rn4-B | 7.4 | 9.4 | 198 | 689 | 12 | MOLECULE: | CHAPERONE PROTEIN CLPB |
| 47:00:00 | 6uks-E | 7.4 | 5.2 | 158 | 362 | 18 | MOLECULE: | MITOCHONDRIAL CHAPERONE BCS1 |
| 48:00:00 | 6ut3-D | 7.4 | 15.2 | 142 | 381 | 14 | MOLECULE: | GTPASE SUBUNIT OF RESTRICTION ENDONUCLEASE |
| 49:00:00 | 1hqy-F | 7.4 | 10 | 158 | 408 | 11 | MOLECULE: | HEAT SHOCK LOCUS HSLV |
| 50:00:00 | 2qp9-X | 7.4 | 14 | 144 | 288 | 14 | MOLECULE: | VACUOLAR PROTEIN SORTING-ASSOCIATED PROTEIN 4 |
| 51:00:00 | 3j94-B | 7.4 | 9.5 | 151 | 484 | 13 | MOLECULE: | VESICLE-FUSING ATPASE |
| 52:00:00 | 7qv9-b | 7.4 | 7.7 | 150 | 258 | 11 | MOLECULE: | DNA-DIRECTED RNA POLYMERASE SUBUNIT ALPHA |
| 53:00:00 | 6qi9-B | 7.4 | 3.6 | 145 | 310 | 12 | MOLECULE: | RUVB-LIKE 1 |
| 54:00:00 | 2r65-A | 7.4 | 9 | 134 | 249 | 11 | MOLECULE: | CELL DIVISION PROTEASE FTSH HOMOLOG |
| 55:00:00 | 3j98-F | 7.4 | 13.3 | 180 | 654 | 11 | MOLECULE: | VESICLE-FUSING ATPASE |
| 56:00:00 | 6sh3-F | 7.4 | 16.9 | 169 | 376 | 10 | MOLECULE: | MITOCHONDRIAL CHAPERONE BCS1 |
| 57:00:00 | 3cf0-K | 7.4 | 10.9 | 139 | 281 | 13 | MOLECULE: | TRANSITIONAL ENDOPLASMIC RETICULUM ATPASE |
| 58:00:00 | 2qpa-A | 7.4 | 13.9 | 138 | 305 | 18 | MOLECULE: | VACUOLAR PROTEIN SORTING-ASSOCIATED PROTEIN 4 |
| 59:00:00 | 1cr0-A | 7.4 | 5.4 | 121 | 239 | 9 | MOLECULE: | DNA PRIMASE/HELICASE |
| 60:00:00 | 2cea-C | 7.4 | 9.5 | 152 | 421 | 15 | MOLECULE: | CELL DIVISION PROTEIN FTSH |
| 61:00:00 | 4r7y-B | 7.4 | 9.1 | 212 | 591 | 12 | MOLECULE: | MINICHROMOSOME MAINTENANCE PROTEIN MCM, CELL DIVI |
| 62:00:00 | 8osf-C | 7.4 | 4.4 | 162 | 321 | 9 | MOLECULE: | MAGNESIUM-CHELATASE SUBUNIT CHLI |
| 63:00:00 | 8osg-C | 7.4 | 4.3 | 156 | 323 | 12 | MOLECULE: | MAGNESIUM-CHELATASE SUBUNIT CHLI |
| 64:00:00 | 7pbo-D | 7.4 | 10.6 | 147 | 313 | 10 | MOLECULE: | HOLLIDAY JUNCTION ATP-DEPENDENT DNA HELICASE RUVB |
| 65:00:00 | 3b9p-A | 7.4 | 9.2 | 135 | 268 | 10 | MOLECULE: | CG5977-PA, ISOFORM A |
| 66:00:00 | 3j99-B | 7.4 | 22.2 | 168 | 672 | 13 | MOLECULE: | VESICLE-FUSING ATPASE |
| 67:00:00 | 7z8h-A | 7.4 | 15.7 | 160 | 1083 | 10 | MOLECULE: | CYTOPLASMIC DYNEIN 1 HEAVY CHAIN 1 |
| 68:00:00 | 4akg-B | 7.4 | 16.1 | 184 | 2650 | 10 | MOLECULE: | GLUTATHIONE S-TRANSFERASE CLASS-MU 26 KDA ISOZYME |
| 69:00:00 | 6mdm-D | 7.3 | 19.6 | 196 | 713 | 10 | MOLECULE: | VESICLE-FUSING ATPASE |
| 70:00:00 | 7ufm-H | 7.3 | 10 | 147 | 311 | 7 | MOLECULE: | VCHTNSC |

| | | | | | | | | |
|---|---|---|---|---|---|---|---|---|
| 71:00:00 | 7m9b-F | 7.3 | 7.1 | 130 | 257 | 8 | MOLECULE: | TNSC |
| 72:00:00 | 7n6i-F | 7.3 | 7.4 | 132 | 257 | 9 | MOLECULE: | TNIQ (HOMOLOGY MODEL) |
| 73:00:00 | 7m9c-D | 7.3 | 7.1 | 136 | 257 | 9 | MOLECULE: | TNSC |
| 74:00:00 | 8hwd-A | 7.3 | 3.7 | 172 | 308 | 11 | MOLECULE: | PRIMASE D5 |
| 75:00:00 | 6oaa-D | 7.3 | 8.4 | 142 | 537 | 13 | MOLECULE: | CELL DIVISION CONTROL PROTEIN 48 |
| 76:00:00 | 7las-A | 7.3 | 4 | 129 | 183 | 16 | MOLECULE: | ATP-DEPENDENT HELICASE REP |
| 77:00:00 | 7las-F | 7.3 | 4.8 | 131 | 183 | 16 | MOLECULE: | ATP-DEPENDENT HELICASE REP |
| 78:00:00 | 1jbk-A | 7.3 | 4.2 | 129 | 189 | 11 | MOLECULE: | CLPB PROTEIN |
| 79:00:00 | 1lv7-A | 7.3 | 8.7 | 136 | 251 | 10 | MOLECULE: | FTSH |
| 80:00:00 | 2x31-L | 7.3 | 5.6 | 161 | 321 | 11 | MOLECULE: | MAGNESIUM-CHELATASE 60 KDA SUBUNIT |
| 81:00:00 | 5og1-C | 7.3 | 13.9 | 230 | 624 | 10 | MOLECULE: | CHAPERONE PROTEIN CLPB,ATP-DEPENDENT CLP PROTEASE |
| 82:00:00 | 4xgc-E | 7.3 | 13.8 | 150 | 402 | 12 | MOLECULE: | ORIGIN RECOGNITION COMPLEX SUBUNIT 2 |
| 83:00:00 | 4wvy-A | 7.3 | 3.4 | 143 | 427 | 14 | MOLECULE: | RUVB-LIKE 1 |
| 84:00:00 | 4eiw-C | 7.3 | 9.1 | 155 | 457 | 14 | MOLECULE: | ATP-DEPENDENT ZINC METALLOPROTEASE FTSH |
| 85:00:00 | 1njf-D | 7.3 | 3.6 | 127 | 239 | 13 | MOLECULE: | DNA POLYMERASE III SUBUNIT GAMMA |
| 86:00:00 | 4ww4-A | 7.3 | 3.7 | 145 | 416 | 14 | MOLECULE: | RUVB-LIKE 1 |
| 87:00:00 | 1njf-A | 7.3 | 3.5 | 125 | 239 | 14 | MOLECULE: | DNA POLYMERASE III SUBUNIT GAMMA |
| 88:00:00 | 7jlv-D | 7.3 | 14.7 | 144 | 437 | 17 | MOLECULE: | DISEASE RESISTANCE PROTEIN ROQ1 |
| 89:00:00 | 7sxo-A | 7.3 | 12.5 | 186 | 544 | 11 | MOLECULE: | LON PROTEASE HOMOLOG, MITOCHONDRIAL |
| 90:00:00 | 7jlv-G | 7.3 | 15 | 145 | 437 | 17 | MOLECULE: | DISEASE RESISTANCE PROTEIN ROQ1 |
| 91:00:00 | 1njg-A | 7.3 | 4.4 | 130 | 240 | 13 | MOLECULE: | DNA POLYMERASE III SUBUNIT GAMMA |
| 92:00:00 | 2x31-H | 7.3 | 5.6 | 161 | 321 | 11 | MOLECULE: | MAGNESIUM-CHELATASE 60 KDA SUBUNIT |
| 93:00:00 | 6uks-G | 7.3 | 5.1 | 158 | 362 | 16 | MOLECULE: | MITOCHONDRIAL CHAPERONE BCS1 |
| 94:00:00 | 6qel-K | 7.3 | 27.1 | 163 | 237 | 13 | MOLECULE: | REPLICATIVE DNA HELICASE |
| 95:00:00 | 1szp-A | 7.3 | 7.5 | 137 | 294 | 10 | MOLECULE: | DNA REPAIR PROTEIN RAD51 |
| 96:00:00 | 6n8v-E | 7.3 | 18.3 | 234 | 709 | 12 | MOLECULE: | HEAT SHOCK PROTEIN 104 |
| 97:00:00 | 1g3i-X | 7.3 | 11.6 | 157 | 322 | 11 | MOLECULE: | ATP-DEPENDENT HSLU PROTEASE ATP-BINDING SUBUNIT |
| 98:00:00 | 5kne-F | 7.3 | 9.5 | 223 | 607 | 10 | MOLECULE: | HEAT SHOCK PROTEIN 104 |
| 99:00:00 | 6p8v-C | 7.3 | 6.8 | 143 | 303 | 16 | MOLECULE: | ATPASE, AAA FAMILY |
| 0:00 | 2gno-A | 7.3 | 17.4 | 130 | 296 | 8 | MOLECULE: | DNA POLYMERASE III, GAMMA SUBUNIT-RELATED PROTEIN |
| 1:00 | 5grb-B | 7.3 | 4 | 131 | 212 | 8 | MOLECULE: | EV71 2C ATPASE |
| 2:00 | 4r7z-K | 7.3 | 4.8 | 156 | 316 | 11 | MOLECULE: | CELL DIVISION CONTROL PROTEIN 21 |
| 3:00 | 6ut4-D | 7.3 | 14.1 | 146 | 383 | 14 | MOLECULE: | GTPASE SUBUNIT OF RESTRICTION ENDONUCLEASE |
| 4:00 | 3zuh-F | 7.3 | 11.4 | 134 | 286 | 13 | MOLECULE: | PROTEIN CBBX |
| 5:00 | 3eih-A | 7.3 | 10.2 | 136 | 319 | 17 | MOLECULE: | VACUOLAR PROTEIN SORTING-ASSOCIATED PROTEIN 4 |
| 6:00 | 6ut3-C | 7.3 | 13.2 | 141 | 375 | 14 | MOLECULE: | GTPASE SUBUNIT OF RESTRICTION ENDONUCLEASE |
| 7:00 | 4ww0-C | 7.3 | 7.7 | 151 | 412 | 13 | MOLECULE: | ATP-DEPENDENT ZINC METALLOPROTEASE FTSH |
| 8:00 | 3j99-F | 7.3 | 16.5 | 196 | 654 | 13 | MOLECULE: | VESICLE-FUSING ATPASE |
| 9:00 | 6ut5-A | 7.3 | 13 | 141 | 395 | 13 | MOLECULE: | GTPASE SUBUNIT OF RESTRICTION ENDONUCLEASE |
| 10:00 | 3syl-B | 7.3 | 12.9 | 140 | 286 | 13 | MOLECULE: | PROTEIN CBBX |
| 11:00 | 1w5t-B | 7.3 | 12.6 | 152 | 396 | 10 | MOLECULE: | ORC2 |
| 12:00 | 8fcx-T | 7.3 | 7.7 | 154 | 343 | 8 | MOLECULE: | TNSC |
| 13:00 | 8hr9-K | 7.3 | 9.3 | 139 | 836 | 7 | MOLECULE: | ARCHAEAL ATPASE |
| 14:00 | 7y53-C | 7.3 | 5.5 | 208 | 741 | 11 | MOLECULE: | DERLIN-1 |
| 15:00 | 2qpa-B | 7.3 | 13 | 130 | 303 | 18 | MOLECULE: | VACUOLAR PROTEIN SORTING-ASSOCIATED PROTEIN 4 |
| 16:00 | 2qpa-C | 7.3 | 13.5 | 130 | 296 | 17 | MOLECULE: | VACUOLAR PROTEIN SORTING-ASSOCIATED PROTEIN 4 |
| 17:00 | 3h4m-C | 7.3 | 9.2 | 138 | 261 | 13 | MOLECULE: | PROTEASOME-ACTIVATING NUCLEOTIDASE |
| 18:00 | 3j98-E | 7.3 | 17.4 | 161 | 670 | 14 | MOLECULE: | VESICLE-FUSING ATPASE |
| 19:00 | 3j99-E | 7.3 | 13.2 | 145 | 670 | 14 | MOLECULE: | VESICLE-FUSING ATPASE |
| 20:00 | 1cr1-A | 7.3 | 5.5 | 123 | 245 | 9 | MOLECULE: | DNA PRIMASE/HELICASE |
| 21:00 | 1im2-A | 7.3 | 10.2 | 154 | 346 | 12 | MOLECULE: | ATP-DEPENDENT HSL PROTEASE ATP-BINDING SUBUNIT |
| 22:00 | 6pdy-A | 7.3 | 11.9 | 165 | 262 | 12 | MOLECULE: | MEMBRANE-SPANNING ATPASE-LIKE PROTEIN |
| 23:00 | 5zr1-A | 7.2 | 23.7 | 185 | 494 | 7 | MOLECULE: | ORIGIN RECOGNITION COMPLEX SUBUNIT 1 |

| | | | | | | | |
|---|---|---|---|---|---|---|---|
| 24:00:00 | 3nbx-X | 7.2 | 4.8 | 177 | 481 | 8 MOLECULE: | ATPASE RAVA |
| 25:00:00 | 7m9b-E | 7.2 | 7.1 | 130 | 257 | 8 MOLECULE: | TNSC |
| 26:00:00 | 7z6h-D | 7.2 | 14.8 | 142 | 324 | 11 MOLECULE: | CELL CYCLE CHECKPOINT CONTROL PROTEIN RAD9A |
| 27:00:00 | 7m9c-O | 7.2 | 7.4 | 139 | 257 | 9 MOLECULE: | TNSC |
| 28:00:00 | 8btg-B | 7.2 | 12.9 | 146 | 335 | 11 MOLECULE: | CHROMOSOMAL REPLICATION INITIATOR PROTEIN DNAA |
| 29:00:00 | 7m9a-D | 7.2 | 7.1 | 135 | 257 | 9 MOLECULE: | TNSC |
| 30:00:00 | 7m9c-K | 7.2 | 7 | 132 | 257 | 9 MOLECULE: | TNSC |
| 31:00:00 | 7m9b-A | 7.2 | 7.4 | 137 | 257 | 9 MOLECULE: | TNSC |
| 32:00:00 | 8hwh-F | 7.2 | 3.8 | 174 | 316 | 10 MOLECULE: | PRIMASE D5 |
| 33:00:00 | 7m9a-A | 7.2 | 8 | 137 | 257 | 9 MOLECULE: | TNSC |
| 34:00:00 | 7m9c-A | 7.2 | 8 | 137 | 257 | 9 MOLECULE: | TNSC |
| 35:00:00 | 8emc-K | 7.2 | 5.9 | 208 | 665 | 9 MOLECULE: | PROTEASE LON-RELATED BREX SYSTEM PROTEIN BRXL |
| 36:00:00 | 6qel-H | 7.2 | 23.9 | 160 | 237 | 13 MOLECULE: | REPLICATIVE DNA HELICASE |
| 37:00:00 | 7lar-E | 7.2 | 3.4 | 122 | 183 | 18 MOLECULE: | ATP-DEPENDENT HELICASE REP |
| 38:00:00 | 3cf0-J | 7.2 | 10.7 | 136 | 281 | 15 MOLECULE: | TRANSITIONAL ENDOPLASMIC RETICULUM ATPASE |
| 39:00:00 | 6oaa-B | 7.2 | 10.8 | 152 | 509 | 14 MOLECULE: | CELL DIVISION CONTROL PROTEIN 48 |
| 40:00:00 | 6uks-D | 7.2 | 4.6 | 154 | 362 | 14 MOLECULE: | MITOCHONDRIAL CHAPERONE BCS1 |
| 41:00:00 | 6uks-F | 7.2 | 4.7 | 155 | 362 | 14 MOLECULE: | MITOCHONDRIAL CHAPERONE BCS1 |
| 42:00:00 | 3cf0-G | 7.2 | 11.2 | 143 | 281 | 13 MOLECULE: | TRANSITIONAL ENDOPLASMIC RETICULUM ATPASE |
| 43:00:00 | 2dhr-A | 7.2 | 8.7 | 160 | 458 | 13 MOLECULE: | FTSH |
| 44:00:00 | 3kds-G | 7.2 | 11 | 143 | 427 | 15 MOLECULE: | CELL DIVISION PROTEIN FTSH |
| 45:00:00 | 3h4m-A | 7.2 | 8.3 | 139 | 261 | 12 MOLECULE: | PROTEASOME-ACTIVATING NUCLEOTIDASE |
| 46:00:00 | 6j5t-C | 7.2 | 27 | 160 | 808 | 9 MOLECULE: | PROBABLE SERINE/THREONINE-PROTEIN KINASE PBL2 |
| 47:00:00 | 7jlv-B | 7.2 | 15.9 | 143 | 437 | 14 MOLECULE: | DISEASE RESISTANCE PROTEIN ROQ1 |
| 48:00:00 | 5vho-C | 7.2 | 3.2 | 133 | 237 | 12 MOLECULE: | 26S PROTEASOME NON-ATPASE REGULATORY SUBUNIT 10 |
| 49:00:00 | 6qi9-D | 7.2 | 13.4 | 132 | 268 | 11 MOLECULE: | RUVB-LIKE 1 |
| 50:00:00 | 7jlv-A | 7.2 | 16 | 145 | 437 | 16 MOLECULE: | DISEASE RESISTANCE PROTEIN ROQ1 |
| 51:00:00 | 1sxj-E | 7.2 | 8.7 | 132 | 317 | 10 MOLECULE: | ACTIVATOR 1 95 KDA SUBUNIT |
| 52:00:00 | 3glh-D | 7.2 | 11.7 | 137 | 363 | 10 MOLECULE: | DNA POLYMERASE III SUBUNIT DELTA |
| 53:00:00 | 1xxh-B | 7.2 | 15.7 | 138 | 364 | 10 MOLECULE: | DNA POLYMERASE III, DELTA SUBUNIT |
| 54:00:00 | 3glh-I | 7.2 | 11.8 | 135 | 363 | 13 MOLECULE: | DNA POLYMERASE III SUBUNIT DELTA |
| 55:00:00 | 1jr3-A | 7.2 | 12.7 | 143 | 366 | 10 MOLECULE: | DNA POLYMERASE III SUBUNIT GAMMA |
| 56:00:00 | 3glh-B | 7.2 | 11.8 | 138 | 364 | 11 MOLECULE: | DNA POLYMERASE III SUBUNIT DELTA |
| 57:00:00 | 3glh-M | 7.2 | 11.3 | 145 | 364 | 11 MOLECULE: | DNA POLYMERASE III SUBUNIT DELTA |
| 58:00:00 | 5uj7-A | 7.2 | 17.5 | 158 | 337 | 8 MOLECULE: | ORIGIN RECOGNITION COMPLEX SUBUNIT 1 |
| 59:00:00 | 5ujm-A | 7.2 | 17.6 | 155 | 337 | 7 MOLECULE: | ORIGIN RECOGNITION COMPLEX SUBUNIT 1 |
| 60:00:00 | 3glh-N | 7.2 | 12.1 | 140 | 363 | 11 MOLECULE: | DNA POLYMERASE III SUBUNIT DELTA |
| 61:00:00 | 1g3i-T | 7.2 | 11.6 | 157 | 331 | 11 MOLECULE: | ATP-DEPENDENT HSLU PROTEASE ATP-BINDING SUBUNIT |
| 62:00:00 | 5t0g-C | 7.2 | 22.7 | 180 | 384 | 12 MOLECULE: | PROTEASOME SUBUNIT ALPHA TYPE-6 |
| 63:00:00 | 1g4a-F | 7.2 | 9.8 | 153 | 356 | 12 MOLECULE: | ATP-DEPENDENT HSL PROTEASE ATP-BINDING SUBUNIT |
| 64:00:00 | 6ut3-B | 7.2 | 14.1 | 146 | 365 | 12 MOLECULE: | GTPASE SUBUNIT OF RESTRICTION ENDONUCLEASE |
| 65:00:00 | 6oay-A | 7.2 | 11.2 | 200 | 548 | 14 MOLECULE: | HYPERACTIVE DISAGGREGASE CLPB |
| 66:00:00 | 6sh4-C | 7.2 | 18 | 175 | 376 | 9 MOLECULE: | MITOCHONDRIAL CHAPERONE BCS1 |
| 67:00:00 | 4r7z-F | 7.2 | 4.8 | 168 | 316 | 10 MOLECULE: | CELL DIVISION CONTROL PROTEIN 21 |
| 68:00:00 | 4r7z-A | 7.2 | 4.7 | 166 | 316 | 10 MOLECULE: | CELL DIVISION CONTROL PROTEIN 21 |
| 69:00:00 | 4r7z-O | 7.2 | 4.7 | 162 | 316 | 9 MOLECULE: | CELL DIVISION CONTROL PROTEIN 21 |
| 70:00:00 | 7wbb-A | 7.2 | 6.5 | 190 | 726 | 8 MOLECULE: | AFG2 ISOFORM 1 |
| 71:00:00 | 6ut7-F | 7.2 | 14.2 | 138 | 392 | 14 MOLECULE: | GTPASE SUBUNIT OF RESTRICTION ENDONUCLEASE |
| 72:00:00 | 3vfd-A | 7.2 | 12.6 | 135 | 266 | 10 MOLECULE: | SPASTIN |
| 73:00:00 | 7jgr-D | 7.2 | 16.7 | 155 | 441 | 7 MOLECULE: | ORIGIN RECOGNITION COMPLEX SUBUNIT 2 |
| 74:00:00 | 6jpq-A | 7.2 | 4.6 | 151 | 577 | 11 MOLECULE: | UNCHARACTERIZED AAA DOMAIN-CONTAINING PROTEIN C31 |
| 75:00:00 | 3uk6-D | 7.2 | 3.3 | 139 | 299 | 13 MOLECULE: | RUVB-LIKE 2 |
| 76:00:00 | 4xgu-D | 7.2 | 6.2 | 165 | 377 | 12 MOLECULE: | PUTATIVE PACHYTENE CHECKPOINT PROTEIN 2 |

| | | | | | | | |
|---|---|---|---|---|---|---|---|
| 77:00:00 | 5zr1-E | 7.2 | 14.1 | 134 | 460 | 9 MOLECULE: | ORIGIN RECOGNITION COMPLEX SUBUNIT 1 |
| 78:00:00 | 3cf0-H | 7.2 | 10.3 | 135 | 281 | 13 MOLECULE: | TRANSITIONAL ENDOPLASMIC RETICULUM ATPASE |
| 79:00:00 | 3cf0-A | 7.2 | 10.5 | 136 | 281 | 13 MOLECULE: | TRANSITIONAL ENDOPLASMIC RETICULUM ATPASE |
| 80:00:00 | 4ww0-B | 7.2 | 8.7 | 166 | 425 | 14 MOLECULE: | ATP-DEPENDENT ZINC METALLOPROTEASE FTSH |
| 81:00:00 | 8osg-D | 7.2 | 5.1 | 161 | 323 | 10 MOLECULE: | MAGNESIUM-CHELATASE SUBUNIT CHLI |
| 82:00:00 | 4kod-K | 7.2 | 3.5 | 150 | 438 | 11 MOLECULE: | TRANSITIONAL ENDOPLASMIC RETICULUM ATPASE |
| 83:00:00 | 3j97-B | 7.2 | 21.8 | 151 | 672 | 15 MOLECULE: | VESICLE-FUSING ATPASE |
| 84:00:00 | 3j94-E | 7.2 | 8.4 | 142 | 482 | 14 MOLECULE: | VESICLE-FUSING ATPASE |
| 85:00:00 | 7v2b-A | 7.2 | 11.6 | 166 | 375 | 11 MOLECULE: | VPSR |
| 86:00:00 | 1l8q-A | 7.2 | 14.6 | 141 | 321 | 11 MOLECULE: | CHROMOSOMAL REPLICATION INITIATOR PROTEIN DNAA |
| 87:00:00 | 4akh-A | 7.2 | 12.1 | 237 | 2650 | 7 MOLECULE: | GLUTATHIONE S-TRANSFERASE CLASS-MU 26 KDA ISOZYME |
| 88:00:00 | 6rla-A | 7.2 | 11.9 | 200 | 2750 | 8 MOLECULE: | O6-ALKYLGUANINE-DNA ALKYLTRANSFERASE MUTANT,DYNC2 |
| 89:00:00 | 4d82-B | 7.2 | 8.2 | 137 | 274 | 16 MOLECULE: | AAA ATPASE, CENTRAL DOMAIN PROTEIN |
| 90:00:00 | 4d81-A | 7.2 | 11.5 | 146 | 263 | 16 MOLECULE: | AAA ATPASE, CENTRAL DOMAIN PROTEIN |
| 91:00:00 | 4akg-A | 7.2 | 18.9 | 184 | 2650 | 11 MOLECULE: | GLUTATHIONE S-TRANSFERASE CLASS-MU 26 KDA ISOZYME |
| 92:00:00 | 8uii-A | 7.2 | 6.4 | 165 | 641 | 13 MOLECULE: | ATPASE FAMILY AAA DOMAIN-CONTAINING PROTEIN 5 |
| 93:00:00 | 6pdy-E | 7.2 | 10.4 | 164 | 279 | 14 MOLECULE: | MEMBRANE-SPANNING ATPASE-LIKE PROTEIN |
| 94:00:00 | 6dgd-A | 7.1 | 11.1 | 166 | 704 | 8 MOLECULE: | PRIMOSOMAL PROTEIN N' |
| 95:00:00 | 5yww-A | 7.1 | 6.4 | 167 | 485 | 12 MOLECULE: | NUCLEOTIDE BINDING PROTEIN PINC |
| 96:00:00 | 8jx6-B | 7.1 | 11.2 | 121 | 451 | 8 MOLECULE: | DEEP-SEA HELICASE 9 |
| 97:00:00 | 7m9a-H | 7.1 | 6.9 | 133 | 257 | 9 MOLECULE: | TNSC |
| 98:00:00 | 7m9b-K | 7.1 | 7.2 | 131 | 257 | 9 MOLECULE: | TNSC |
| 99:00:00 | 7m9c-H | 7.1 | 6.9 | 133 | 257 | 9 MOLECULE: | TNSC |
| 0:00 | 7m9c-F | 7.1 | 7.1 | 131 | 257 | 8 MOLECULE: | TNSC |
| 1:00 | 7m9b-H | 7.1 | 7 | 135 | 257 | 9 MOLECULE: | TNSC |
| 2:00 | 7m9a-K | 7.1 | 7 | 130 | 257 | 9 MOLECULE: | TNSC |
| 3:00 | 3cf0-C | 7.1 | 10.4 | 132 | 281 | 14 MOLECULE: | TRANSITIONAL ENDOPLASMIC RETICULUM ATPASE |
| 4:00 | 8iqi-B | 7.1 | 22.2 | 141 | 915 | 11 MOLECULE: | PUTATIVE PRIMASE C962R |
| 5:00 | 7m9b-D | 7.1 | 7.1 | 136 | 257 | 9 MOLECULE: | TNSC |
| 6:00 | 6oab-C | 7.1 | 9.4 | 178 | 532 | 11 MOLECULE: | CELL DIVISION CONTROL PROTEIN 48 |
| 7:00 | 6qem-I | 7.1 | 22.7 | 164 | 239 | 12 MOLECULE: | REPLICATIVE DNA HELICASE |
| 8:00 | 6rn2-A | 7.1 | 14.2 | 179 | 684 | 9 MOLECULE: | CHAPERONE PROTEIN CLPB |
| 9:00 | 3cf0-M | 7.1 | 10.7 | 136 | 281 | 14 MOLECULE: | TRANSITIONAL ENDOPLASMIC RETICULUM ATPASE |
| 10:00 | 3cf0-B | 7.1 | 10.2 | 130 | 281 | 15 MOLECULE: | TRANSITIONAL ENDOPLASMIC RETICULUM ATPASE |
| 11:00 | 6qel-J | 7.1 | 24.7 | 166 | 237 | 14 MOLECULE: | REPLICATIVE DNA HELICASE |
| 12:00 | 6hz9-H | 7.1 | 3.5 | 138 | 290 | 12 MOLECULE: | 5-METHYLCYTOSINE-SPECIFIC RESTRICTION ENZYME B |
| 13:00 | 6mat-B | 7.1 | 9.7 | 148 | 578 | 14 MOLECULE: | RIX7 MUTANT |
| 14:00 | 6uks-A | 7.1 | 4.5 | 156 | 362 | 14 MOLECULE: | MITOCHONDRIAL CHAPERONE BCS1 |
| 15:00 | 3qmz-A | 7.1 | 23.1 | 204 | 2136 | 7 MOLECULE: | CYTOPLASMIC DYNEIN HEAVY CHAIN |
| 16:00 | 2cea-B | 7.1 | 6.1 | 146 | 411 | 14 MOLECULE: | CELL DIVISION PROTEIN FTSH |
| 17:00 | 2cea-F | 7.1 | 7.4 | 146 | 412 | 14 MOLECULE: | CELL DIVISION PROTEIN FTSH |
| 18:00 | 1xxh-H | 7.1 | 13.8 | 139 | 366 | 13 MOLECULE: | DNA POLYMERASE III, DELTA SUBUNIT |
| 19:00 | 7ngp-A | 7.1 | 29.6 | 203 | 776 | 9 MOLECULE: | LON PROTEASE HOMOLOG, MITOCHONDRIAL |
| 20:00 | 1xxh-I | 7.1 | 11 | 133 | 364 | 13 MOLECULE: | DNA POLYMERASE III, DELTA SUBUNIT |
| 21:00 | 5xf8-4 | 7.1 | 9.3 | 211 | 716 | 11 MOLECULE: | DNA REPLICATION LICENSING FACTOR MCM2 |
| 22:00 | 1xxh-C | 7.1 | 14.2 | 143 | 366 | 10 MOLECULE: | DNA POLYMERASE III, DELTA SUBUNIT |
| 23:00 | 7ttr-F | 7.1 | 12 | 155 | 304 | 13 MOLECULE: | CASEINOLYTIC PEPTIDASE B PROTEIN HOMOLOG |
| 24:00:00 | 6igm-F | 7.1 | 3.4 | 135 | 366 | 15 MOLECULE: | RUVB-LIKE 1 |
| 25:00:00 | 4r7z-L | 7.1 | 4.8 | 163 | 316 | 9 MOLECULE: | CELL DIVISION CONTROL PROTEIN 21 |
| 26:00:00 | 6lua-A | 7.1 | 3.7 | 137 | 247 | 12 MOLECULE: | FLAGELLAR REGULATORY PROTEIN C |
| 27:00:00 | 3zuh-A | 7.1 | 13.2 | 138 | 286 | 14 MOLECULE: | PROTEIN CBBX |
| 28:00:00 | 5wc2-A | 7.1 | 4.3 | 155 | 364 | 12 MOLECULE: | PACHYTENE CHECKPOINT PROTEIN 2 HOMOLOG |
| 29:00:00 | 8fcv-T | 7.1 | 8.4 | 151 | 343 | 8 MOLECULE: | DNA (60-MER) |

| | | | | | | | | |
|---|---|---|---|---|---|---|---|---|
| 30:00:00 | 7y4w-C | 7.1 | 5.2 | 187 | 744 | 12 | MOLECULE: | DERLIN-1 |
| 31:00:00 | 7jpp-E | 7.1 | 16.6 | 139 | 406 | 10 | MOLECULE: | ORIGIN RECOGNITION COMPLEX SUBUNIT 1 |
| 32:00:00 | 8fcw-S | 7.1 | 7.7 | 150 | 343 | 8 | MOLECULE: | TNSC |
| 33:00:00 | 6chs-J | 7.1 | 11.5 | 177 | 550 | 11 | MOLECULE: | NPL4 |
| 34:00:00 | 5vc7-E | 7.1 | 5.2 | 178 | 544 | 13 | MOLECULE: | VCP-LIKE ATPASE |
| 35:00:00 | 7tjj-I | 7.1 | 15.4 | 140 | 380 | 14 | MOLECULE: | ORIGIN RECOGNITION COMPLEX SUBUNIT 1 |
| 36:00:00 | 8c0v-C | 7.1 | 11.9 | 187 | 821 | 6 | MOLECULE: | PEROXISOMAL ATPASE PEX1 |
| 37:00:00 | 3kds-E | 7.1 | 12.3 | 143 | 428 | 15 | MOLECULE: | CELL DIVISION PROTEIN FTSH |
| 38:00:00 | 7ln2-F | 7.1 | 12.1 | 188 | 700 | 12 | MOLECULE: | TRANSITIONAL ENDOPLASMIC RETICULUM ATPASE |
| 39:00:00 | 2zrj-A | 7.1 | 8.6 | 142 | 314 | 13 | MOLECULE: | PROTEIN RECA |
| 40:00:00 | 6p07-C | 7.1 | 11.6 | 140 | 301 | 12 | MOLECULE: | SPASTIN |
| 41:00:00 | 6fhs-C | 7.1 | 18.6 | 151 | 459 | 13 | MOLECULE: | RUVB-LIKE HELICASE |
| 42:00:00 | 6pdw-E | 7.1 | 11.5 | 164 | 298 | 13 | MOLECULE: | MEMBRANE-SPANNING ATPASE-LIKE PROTEIN |
| 43:00:00 | 8y6p-Q | 7 | 18.1 | 154 | 1228 | 12 | MOLECULE: | APAF-1 RELATED KILLER DARK |
| 44:00:00 | 7m99-E | 7 | 8.1 | 136 | 257 | 9 | MOLECULE: | TNSC |
| 45:00:00 | 8iqh-G | 7 | 29.6 | 228 | 863 | 12 | MOLECULE: | PUTATIVE PRIMASE C962R |
| 46:00:00 | 7n6i-J | 7 | 8.1 | 136 | 257 | 9 | MOLECULE: | TNIQ (HOMOLOGY MODEL) |
| 47:00:00 | 7m9c-E | 7 | 7.1 | 132 | 257 | 8 | MOLECULE: | TNSC |
| 48:00:00 | 7n6i-G | 7 | 8.5 | 135 | 257 | 9 | MOLECULE: | TNIQ (HOMOLOGY MODEL) |
| 49:00:00 | 7n6i-E | 7 | 7.4 | 130 | 257 | 9 | MOLECULE: | TNIQ (HOMOLOGY MODEL) |
| 50:00:00 | 7ufi-1 | 7 | 10.3 | 145 | 311 | 10 | MOLECULE: | DNA (5'- |
| 51:00:00 | 7m9c-I | 7 | 7.2 | 132 | 257 | 9 | MOLECULE: | TNSC |
| 52:00:00 | 7n6i-C | 7 | 7.4 | 131 | 257 | 9 | MOLECULE: | TNIQ (HOMOLOGY MODEL) |
| 53:00:00 | 7m99-C | 7 | 8.1 | 134 | 257 | 9 | MOLECULE: | TNSC |
| 54:00:00 | 7m99-H | 7 | 7.4 | 131 | 257 | 9 | MOLECULE: | TNSC |
| 55:00:00 | 7m99-A | 7 | 8.6 | 135 | 257 | 9 | MOLECULE: | TNSC |
| 56:00:00 | 7m9a-F | 7 | 7 | 129 | 257 | 9 | MOLECULE: | TNSC |
| 57:00:00 | 7m9a-J | 7 | 7 | 132 | 257 | 9 | MOLECULE: | TNSC |
| 58:00:00 | 7ufi-4 | 7 | 10.1 | 147 | 311 | 8 | MOLECULE: | DNA (5'- |
| 59:00:00 | 7m9a-E | 7 | 7.1 | 132 | 257 | 8 | MOLECULE: | TNSC |
| 60:00:00 | 3r8f-B | 7 | 11.9 | 135 | 319 | 10 | MOLECULE: | CHROMOSOMAL REPLICATION INITIATOR PROTEIN DNAA |
| 61:00:00 | 6rn4-D | 7 | 13.1 | 162 | 684 | 10 | MOLECULE: | CHAPERONE PROTEIN CLPB |
| 62:00:00 | 7ufi-2 | 7 | 10.3 | 145 | 311 | 10 | MOLECULE: | DNA (5'- |
| 63:00:00 | 6j5t-O | 7 | 29.1 | 159 | 808 | 8 | MOLECULE: | PROBABLE SERINE/THREONINE-PROTEIN KINASE PBL2 |
| 64:00:00 | 6rn2-B | 7 | 12.5 | 162 | 689 | 9 | MOLECULE: | CHAPERONE PROTEIN CLPB |
| 65:00:00 | 6qem-H | 7 | 14.4 | 158 | 239 | 13 | MOLECULE: | REPLICATIVE DNA HELICASE |
| 66:00:00 | 2hcb-A | 7 | 13.5 | 131 | 317 | 11 | MOLECULE: | CHROMOSOMAL REPLICATION INITIATOR PROTEIN DNAA |
| 67:00:00 | 7m9c-P | 7 | 9.6 | 136 | 257 | 10 | MOLECULE: | TNSC |
| 68:00:00 | 5lkm-C | 7 | 4.2 | 132 | 373 | 11 | MOLECULE: | DNA REPAIR PROTEIN RADA |
| 69:00:00 | 3r8f-A | 7 | 12.2 | 136 | 319 | 10 | MOLECULE: | CHROMOSOMAL REPLICATION INITIATOR PROTEIN DNAA |
| 70:00:00 | 3cf0-E | 7 | 10.8 | 136 | 281 | 14 | MOLECULE: | TRANSITIONAL ENDOPLASMIC RETICULUM ATPASE |
| 71:00:00 | 6qel-L | 7 | 21.2 | 162 | 237 | 15 | MOLECULE: | REPLICATIVE DNA HELICASE |
| 72:00:00 | 6oab-E | 7 | 9.4 | 144 | 442 | 14 | MOLECULE: | CELL DIVISION CONTROL PROTEIN 48 |
| 73:00:00 | 3r8f-C | 7 | 11.8 | 135 | 319 | 10 | MOLECULE: | CHROMOSOMAL REPLICATION INITIATOR PROTEIN DNAA |
| 74:00:00 | 2hcb-D | 7 | 13.3 | 131 | 320 | 10 | MOLECULE: | CHROMOSOMAL REPLICATION INITIATOR PROTEIN DNAA |
| 75:00:00 | 3r8f-D | 7 | 12 | 136 | 319 | 10 | MOLECULE: | CHROMOSOMAL REPLICATION INITIATOR PROTEIN DNAA |
| 76:00:00 | 3kds-F | 7 | 10.9 | 144 | 426 | 14 | MOLECULE: | CELL DIVISION PROTEIN FTSH |
| 77:00:00 | 2cea-E | 7 | 4.2 | 144 | 405 | 14 | MOLECULE: | CELL DIVISION PROTEIN FTSH |
| 78:00:00 | 5zr1-B | 7 | 8.6 | 126 | 374 | 9 | MOLECULE: | ORIGIN RECOGNITION COMPLEX SUBUNIT 1 |
| 79:00:00 | 2ce7-F | 7 | 8.1 | 150 | 412 | 11 | MOLECULE: | CELL DIVISION PROTEIN FTSH |
| 80:00:00 | 4eiw-D | 7 | 9.9 | 154 | 446 | 14 | MOLECULE: | ATP-DEPENDENT ZINC METALLOPROTEASE FTSH |
| 81:00:00 | 2ce7-B | 7 | 10 | 157 | 411 | 12 | MOLECULE: | CELL DIVISION PROTEIN FTSH |
| 82:00:00 | 3ja8-6 | 7 | 11 | 215 | 616 | 9 | MOLECULE: | MINICHROMOSOME MAINTENANCE 2 |

| | | | | | | | | |
|---|---|---|---|---|---|---|---|---|
| 83:00:00 | 6zyy-C | 7 | 18.8 | 235 | 2901 | 7 | MOLECULE: | DYNEIN HEAVY CHAIN, OUTER ARM PROTEIN |
| 84:00:00 | 5f4h-A | 7 | 16.9 | 158 | 437 | 9 | MOLECULE: | NUCLEOTIDE BINDING PROTEIN PINC |
| 85:00:00 | 1njf-B | 7 | 3.7 | 126 | 239 | 13 | MOLECULE: | DNA POLYMERASE III SUBUNIT GAMMA |
| 86:00:00 | 7p09-B | 7 | 11.1 | 176 | 536 | 11 | MOLECULE: | LON PROTEASE HOMOLOG, MITOCHONDRIAL |
| 87:00:00 | 7ngq-E | 7 | 32.2 | 208 | 776 | 8 | MOLECULE: | LON PROTEASE HOMOLOG, MITOCHONDRIAL |
| 88:00:00 | 6etx-D | 7 | 3.6 | 136 | 434 | 13 | MOLECULE: | RUVB-LIKE 1 |
| 89:00:00 | 6opc-F | 7 | 11.4 | 188 | 681 | 12 | MOLECULE: | CELL DIVISION CONTROL PROTEIN 48 |
| 90:00:00 | 2zr9-A | 7 | 8.7 | 143 | 312 | 13 | MOLECULE: | PROTEIN RECA |
| 91:00:00 | 1do2-D | 7 | 10.8 | 153 | 407 | 14 | MOLECULE: | PROTEIN (HEAT SHOCK LOCUS U) |
| 92:00:00 | 1do2-B | 7 | 12.1 | 151 | 407 | 16 | MOLECULE: | PROTEIN (HEAT SHOCK LOCUS U) |
| 93:00:00 | 5oaf-D | 7 | 3.7 | 139 | 411 | 12 | MOLECULE: | RUVB-LIKE 1 |
| 94:00:00 | 8van-D | 7 | 10.6 | 144 | 359 | 9 | MOLECULE: | DNA POLYMERASE III SUBUNIT DELTA |
| 95:00:00 | 6rn2-C | 7 | 9.7 | 192 | 689 | 12 | MOLECULE: | CHAPERONE PROTEIN CLPB |
| 96:00:00 | 4r7z-H | 7 | 4.8 | 168 | 316 | 10 | MOLECULE: | CELL DIVISION CONTROL PROTEIN 21 |
| 97:00:00 | 6ut7-E | 7 | 14.2 | 144 | 392 | 14 | MOLECULE: | GTPASE SUBUNIT OF RESTRICTION ENDONUCLEASE |
| 98:00:00 | 8fnt-G | 7 | 15.5 | 137 | 785 | 8 | MOLECULE: | ARCHAEAL ATPASE |
| 99:00:00 | 8osf-A | 7 | 4.8 | 166 | 321 | 10 | MOLECULE: | MAGNESIUM-CHELATASE SUBUNIT CHLI |
| 0:00 | 3glf-E | 7 | 14.4 | 129 | 334 | 14 | MOLECULE: | DNA POLYMERASE III SUBUNIT DELTA |
| 1:00 | 3glf-J | 7 | 14.4 | 129 | 334 | 14 | MOLECULE: | DNA POLYMERASE III SUBUNIT DELTA |
| 2:00 | 5txv-C | 7 | 11.3 | 154 | 366 | 11 | MOLECULE: | ATP-DEPENDENT PROTEASE ATPASE SUBUNIT HSLU |
| 3:00 | 3uk6-C | 7 | 3.2 | 143 | 321 | 13 | MOLECULE: | RUVB-LIKE 2 |
| 4:00 | 4z8x-C | 7 | 4.5 | 142 | 423 | 13 | MOLECULE: | ATP-DEPENDENT ZINC METALLOPROTEASE FTSH |
| 5:00 | 3cf0-D | 7 | 10 | 137 | 281 | 13 | MOLECULE: | TRANSITIONAL ENDOPLASMIC RETICULUM ATPASE |
| 6:00 | 8efy-I | 7 | 9.6 | 141 | 313 | 13 | MOLECULE: | HOLLIDAY JUNCTION ATP-DEPENDENT DNA HELICASE RUVB |
| 7:00 | 3j94-F | 7 | 10 | 155 | 466 | 17 | MOLECULE: | VESICLE-FUSING ATPASE |
| 8:00 | 3j98-B | 7 | 16.4 | 171 | 671 | 16 | MOLECULE: | VESICLE-FUSING ATPASE |
| 9:00 | 3j94-A | 7 | 11.8 | 149 | 490 | 15 | MOLECULE: | VESICLE-FUSING ATPASE |
| 10:00 | 3j99-A | 7 | 13.3 | 146 | 678 | 14 | MOLECULE: | VESICLE-FUSING ATPASE |
| 11:00 | 1w5s-B | 7 | 11.6 | 155 | 396 | 10 | MOLECULE: | ORIGIN RECOGNITION COMPLEX SUBUNIT 2 ORC2 |
| 12:00 | 6nyy-C | 7 | 3.5 | 142 | 491 | 10 | MOLECULE: | AFG3-LIKE PROTEIN 2 |
| 13:00 | 6pdy-C | 7 | 12.8 | 164 | 298 | 14 | MOLECULE: | MEMBRANE-SPANNING ATPASE-LIKE PROTEIN |
| 14:00 | 8ui7-A | 7 | 10.9 | 168 | 643 | 12 | MOLECULE: | ATPASE FAMILY AAA DOMAIN-CONTAINING PROTEIN 5 |
| 15:00 | 7pbt-B | 7 | 11.2 | 146 | 312 | 10 | MOLECULE: | HOLLIDAY JUNCTION ATP-DEPENDENT DNA HELICASE RUVB |
| 16:00 | 6pdy-F | 7 | 9.7 | 144 | 236 | 14 | MOLECULE: | MEMBRANE-SPANNING ATPASE-LIKE PROTEIN |
| 17:00 | 7ufi-5 | 6.9 | 10.4 | 145 | 311 | 10 | MOLECULE: | DNA (5'- |
| 18:00 | 7m9c-J | 6.9 | 7.1 | 132 | 257 | 8 | MOLECULE: | TNSC |
| 19:00 | 7ufm-M | 6.9 | 10 | 147 | 311 | 7 | MOLECULE: | VCHTNSC |
| 20:00 | 7n6i-D | 6.9 | 7.4 | 130 | 257 | 8 | MOLECULE: | TNIQ (HOMOLOGY MODEL) |
| 21:00 | 7m9a-B | 6.9 | 7.2 | 133 | 257 | 9 | MOLECULE: | TNSC |
| 22:00 | 7ufm-F | 6.9 | 10 | 146 | 311 | 8 | MOLECULE: | VCHTNSC |
| 23:00 | 7ufm-E | 6.9 | 10 | 146 | 311 | 8 | MOLECULE: | VCHTNSC |
| 24:00:00 | 7m9a-I | 6.9 | 7.2 | 132 | 257 | 9 | MOLECULE: | TNSC |
| 25:00:00 | 7n6i-I | 6.9 | 7.4 | 131 | 257 | 9 | MOLECULE: | TNIQ (HOMOLOGY MODEL) |
| 26:00:00 | 7ufm-D | 6.9 | 10 | 146 | 311 | 8 | MOLECULE: | VCHTNSC |
| 27:00:00 | 7m9b-I | 6.9 | 7.2 | 134 | 257 | 9 | MOLECULE: | TNSC |
| 28:00:00 | 7m9c-B | 6.9 | 7.2 | 133 | 257 | 9 | MOLECULE: | TNSC |
| 29:00:00 | 7m99-B | 6.9 | 7.4 | 130 | 257 | 8 | MOLECULE: | TNSC |
| 30:00:00 | 7m9b-J | 6.9 | 7.1 | 132 | 257 | 9 | MOLECULE: | TNSC |
| 31:00:00 | 8iqh-E | 6.9 | 17.5 | 207 | 880 | 8 | MOLECULE: | PUTATIVE PRIMASE C962R |
| 32:00:00 | 6oab-B | 6.9 | 8.3 | 138 | 509 | 13 | MOLECULE: | CELL DIVISION CONTROL PROTEIN 48 |
| 33:00:00 | 6qem-J | 6.9 | 9.9 | 148 | 239 | 14 | MOLECULE: | REPLICATIVE DNA HELICASE |
| 34:00:00 | 7lar-C | 6.9 | 3.9 | 120 | 183 | 18 | MOLECULE: | ATP-DEPENDENT HELICASE REP |
| 35:00:00 | 7las-C | 6.9 | 3.8 | 118 | 183 | 19 | MOLECULE: | ATP-DEPENDENT HELICASE REP |

| | | | | | | | | |
|---|---|---|---|---|---|---|---|---|
| 36:00:00 | 2qgz-A | 6.9 | 4 | 124 | 183 | 13 | MOLECULE: | PUTATIVE PRIMOSOME COMPONENT |
| 37:00:00 | 6p8v-E | 6.9 | 3.5 | 140 | 298 | 16 | MOLECULE: | ATPASE, AAA FAMILY |
| 38:00:00 | 6j6i-L | 6.9 | 28.4 | 152 | 790 | 9 | MOLECULE: | PROBABLE SERINE/THREONINE-PROTEIN KINASE PBL2 |
| 39:00:00 | 6rn3-C | 6.9 | 13 | 176 | 689 | 10 | MOLECULE: | CHAPERONE PROTEIN CLPB |
| 40:00:00 | 2ce7-D | 6.9 | 7.4 | 147 | 413 | 14 | MOLECULE: | CELL DIVISION PROTEIN FTSH |
| 41:00:00 | 6j6i-O | 6.9 | 27 | 154 | 790 | 8 | MOLECULE: | PROBABLE SERINE/THREONINE-PROTEIN KINASE PBL2 |
| 42:00:00 | 4ciu-A | 6.9 | 10.3 | 157 | 657 | 10 | MOLECULE: | CHAPERONE PROTEIN CLPB |
| 43:00:00 | 3uk6-L | 6.9 | 3.1 | 124 | 232 | 12 | MOLECULE: | RUVB-LIKE 2 |
| 44:00:00 | 5nss-L | 6.9 | 3.7 | 134 | 250 | 14 | MOLECULE: | DNA-DIRECTED RNA POLYMERASE SUBUNIT ALPHA |
| 45:00:00 | 5nss-F | 6.9 | 6.3 | 137 | 248 | 14 | MOLECULE: | DNA-DIRECTED RNA POLYMERASE SUBUNIT ALPHA |
| 46:00:00 | 5vc7-F | 6.9 | 9.1 | 169 | 544 | 12 | MOLECULE: | VCP-LIKE ATPASE |
| 47:00:00 | 3glh-L | 6.9 | 13 | 145 | 364 | 12 | MOLECULE: | DNA POLYMERASE III SUBUNIT DELTA |
| 48:00:00 | 5bk4-2 | 6.9 | 13.4 | 213 | 603 | 10 | MOLECULE: | DNA REPLICATION LICENSING FACTOR MCM2 |
| 49:00:00 | 7ng5-C | 6.9 | 25 | 195 | 776 | 9 | MOLECULE: | LON PROTEASE HOMOLOG, MITOCHONDRIAL |
| 50:00:00 | 3glh-H | 6.9 | 15.1 | 148 | 364 | 11 | MOLECULE: | DNA POLYMERASE III SUBUNIT DELTA |
| 51:00:00 | 7p09-E | 6.9 | 8.7 | 171 | 532 | 10 | MOLECULE: | LON PROTEASE HOMOLOG, MITOCHONDRIAL |
| 52:00:00 | 7ngp-B | 6.9 | 34.1 | 229 | 776 | 9 | MOLECULE: | LON PROTEASE HOMOLOG, MITOCHONDRIAL |
| 53:00:00 | 7ngq-C | 6.9 | 27.7 | 211 | 776 | 9 | MOLECULE: | LON PROTEASE HOMOLOG, MITOCHONDRIAL |
| 54:00:00 | 6rn3-D | 6.9 | 13.9 | 172 | 684 | 9 | MOLECULE: | CHAPERONE PROTEIN CLPB |
| 55:00:00 | 6jpu-A | 6.9 | 6.6 | 152 | 577 | 13 | MOLECULE: | UNCHARACTERIZED AAA DOMAIN-CONTAINING PROTEIN C31 |
| 56:00:00 | 5vhj-C | 6.9 | 8.6 | 130 | 213 | 14 | MOLECULE: | 26S PROTEASOME NON-ATPASE REGULATORY SUBUNIT 10 |
| 57:00:00 | 4z8x-A | 6.9 | 6.7 | 146 | 425 | 12 | MOLECULE: | ATP-DEPENDENT ZINC METALLOPROTEASE FTSH |
| 58:00:00 | 6k0r-I | 6.9 | 3.8 | 142 | 313 | 12 | MOLECULE: | RUVB-LIKE 1,RUVB-LIKE 1 |
| 59:00:00 | 6ut4-F | 6.9 | 15.2 | 137 | 390 | 15 | MOLECULE: | GTPASE SUBUNIT OF RESTRICTION ENDONUCLEASE |
| 60:00:00 | 2p65-A | 6.9 | 3.5 | 117 | 185 | 12 | MOLECULE: | HYPOTHETICAL PROTEIN PF08_0063 |
| 61:00:00 | 4hse-A | 6.9 | 10.1 | 153 | 369 | 13 | MOLECULE: | CHAPERONE PROTEIN CLPB |
| 62:00:00 | 3gli-E | 6.9 | 12.2 | 127 | 334 | 13 | MOLECULE: | DNA POLYMERASE III SUBUNIT DELTA |
| 63:00:00 | 3glg-E | 6.9 | 13.9 | 133 | 334 | 13 | MOLECULE: | DNA POLYMERASE III SUBUNIT DELTA |
| 64:00:00 | 3gli-J | 6.9 | 14 | 129 | 334 | 12 | MOLECULE: | DNA POLYMERASE III SUBUNIT DELTA |
| 65:00:00 | 3glg-J | 6.9 | 14 | 131 | 334 | 13 | MOLECULE: | DNA POLYMERASE III SUBUNIT DELTA |
| 66:00:00 | 2qby-A | 6.9 | 13.5 | 139 | 366 | 9 | MOLECULE: | CELL DIVISION CONTROL PROTEIN 6 HOMOLOG 1 |
| 67:00:00 | 6f0x-F | 6.9 | 5.7 | 145 | 282 | 12 | MOLECULE: | PACHYTENE CHECKPOINT PROTEIN 2 HOMOLOG |
| 68:00:00 | 8osf-E | 6.9 | 4.7 | 169 | 321 | 8 | MOLECULE: | MAGNESIUM-CHELATASE SUBUNIT CHLI |
| 69:00:00 | 5v8f-D | 6.9 | 11.1 | 151 | 456 | 9 | MOLECULE: | DNA REPLICATION LICENSING FACTOR MCM2 |
| 70:00:00 | 7qv9-d | 6.9 | 7.1 | 145 | 256 | 8 | MOLECULE: | DNA-DIRECTED RNA POLYMERASE SUBUNIT ALPHA |
| 71:00:00 | 3jc5-6 | 6.9 | 9.6 | 218 | 671 | 10 | MOLECULE: | DNA REPLICATION LICENSING FACTOR MCM2 |
| 72:00:00 | 1xx6-A | 6.9 | 3.9 | 109 | 178 | 9 | MOLECULE: | THYMIDINE KINASE |
| 73:00:00 | 5v8f-6 | 6.9 | 7.3 | 241 | 692 | 10 | MOLECULE: | DNA REPLICATION LICENSING FACTOR MCM2 |
| 74:00:00 | 4cr2-I | 6.9 | 5.2 | 169 | 362 | 14 | MOLECULE: | PROTEASOME COMPONENT PRE3 |
| 75:00:00 | 7qv9-e | 6.9 | 7.4 | 147 | 259 | 11 | MOLECULE: | DNA-DIRECTED RNA POLYMERASE SUBUNIT ALPHA |
| 76:00:00 | 6amn-A | 6.9 | 6.2 | 159 | 349 | 13 | MOLECULE: | HEAT SHOCK PROTEIN 104 |
| 77:00:00 | 8val-E | 6.9 | 14 | 134 | 337 | 10 | MOLECULE: | DNA POLYMERASE III SUBUNIT DELTA |
| 78:00:00 | 6igm-C | 6.9 | 4 | 139 | 370 | 11 | MOLECULE: | RUVB-LIKE 1 |
| 79:00:00 | 7pbq-B | 6.9 | 9.7 | 137 | 312 | 10 | MOLECULE: | HOLLIDAY JUNCTION ATP-DEPENDENT DNA HELICASE RUVB |
| 80:00:00 | 4l15-A | 6.9 | 12.1 | 143 | 298 | 8 | MOLECULE: | FIDGETIN-LIKE PROTEIN 1 |
| 81:00:00 | 3j95-E | 6.9 | 10.9 | 156 | 482 | 15 | MOLECULE: | VESICLE-FUSING ATPASE |
| 82:00:00 | 5txv-A | 6.9 | 11.5 | 153 | 330 | 12 | MOLECULE: | ATP-DEPENDENT PROTEASE ATPASE SUBUNIT HSLU |
| 83:00:00 | 5vh9-A | 6.9 | 13.2 | 248 | 2376 | 7 | MOLECULE: | DYNEIN HEAVY CHAIN, CYTOPLASMIC |
| 84:00:00 | 3qmz-B | 6.9 | 20.4 | 205 | 2136 | 6 | MOLECULE: | CYTOPLASMIC DYNEIN HEAVY CHAIN |
| 85:00:00 | 7mi1-A | 6.9 | 25.2 | 211 | 2628 | 10 | MOLECULE: | CHIMERA PROTEIN OF DYNEIN AND ENDOLYSIN |
| 86:00:00 | 8emh-B | 6.8 | 5 | 210 | 676 | 7 | MOLECULE: | PROTEASE LON-RELATED BREX SYSTEM PROTEIN BRXL |
| 87:00:00 | 5e7p-A | 6.8 | 14 | 205 | 719 | 12 | MOLECULE: | CELL DIVISION CONTROL PROTEIN CDC48 |
| 88:00:00 | 8iqh-B | 6.8 | 28.6 | 243 | 892 | 12 | MOLECULE: | PUTATIVE PRIMASE C962R |

| | | | | | | | |
|---|---|---|---|---|---|---|---|
| 89:00:00 | 7m9a-L | 6.8 | 7.2 | 131 | 257 | 8 | MOLECULE: TNSC |
| 90:00:00 | 7swl-A | 6.8 | 10.7 | 190 | 553 | 13 | MOLECULE: RIX7 |
| 91:00:00 | 7m9a-G | 6.8 | 8.7 | 134 | 257 | 10 | MOLECULE: TNSC |
| 92:00:00 | 7ufm-C | 6.8 | 10 | 146 | 311 | 8 | MOLECULE: VCHTNSC |
| 93:00:00 | 7m99-F | 6.8 | 8.5 | 133 | 257 | 9 | MOLECULE: TNSC |
| 94:00:00 | 7m9c-L | 6.8 | 7.3 | 131 | 257 | 8 | MOLECULE: TNSC |
| 95:00:00 | 7ufm-I | 6.8 | 10 | 146 | 311 | 8 | MOLECULE: VCHTNSC |
| 96:00:00 | 7m9c-G | 6.8 | 7.7 | 134 | 257 | 10 | MOLECULE: TNSC |
| 97:00:00 | 7ufm-L | 6.8 | 10 | 146 | 311 | 8 | MOLECULE: VCHTNSC |
| 98:00:00 | 7m9b-L | 6.8 | 7.2 | 130 | 257 | 8 | MOLECULE: TNSC |
| 99:00:00 | 7m9b-B | 6.8 | 7.2 | 133 | 257 | 9 | MOLECULE: TNSC |
| 0:00 | 7n6i-H | 6.8 | 7.4 | 131 | 257 | 9 | MOLECULE: TNIQ (HOMOLOGY MODEL) |
| 1:00 | 7ufm-J | 6.8 | 10 | 146 | 311 | 8 | MOLECULE: VCHTNSC |
| 2:00 | 7m9b-G | 6.8 | 7.7 | 134 | 257 | 10 | MOLECULE: TNSC |
| 3:00 | 2hcb-B | 6.8 | 12.2 | 141 | 318 | 10 | MOLECULE: CHROMOSOMAL REPLICATION INITIATOR PROTEIN DNAA |
| 4:00 | 6qem-G | 6.8 | 20.7 | 167 | 239 | 12 | MOLECULE: REPLICATIVE DNA HELICASE |
| 5:00 | 8emh-H | 6.8 | 5.5 | 213 | 671 | 8 | MOLECULE: PROTEASE LON-RELATED BREX SYSTEM PROTEIN BRXL |
| 6:00 | 5he8-I | 6.8 | 3.7 | 111 | 165 | 16 | MOLECULE: HELICASE LOADER |
| 7:00 | 2z4r-B | 6.8 | 9 | 124 | 240 | 12 | MOLECULE: CHROMOSOMAL REPLICATION INITIATOR PROTEIN DNAA |
| 8:00 | 3cf0-L | 6.8 | 11.3 | 139 | 281 | 14 | MOLECULE: TRANSITIONAL ENDOPLASMIC RETICULUM ATPASE |
| 9:00 | 6rn3-F | 6.8 | 15.3 | 175 | 675 | 10 | MOLECULE: CHAPERONE PROTEIN CLPB |
| 10:00 | 3zuh-E | 6.8 | 11.5 | 130 | 286 | 15 | MOLECULE: PROTEIN CBBX |
| 11:00 | 6ef0-J | 6.8 | 10.6 | 144 | 272 | 9 | MOLECULE: PROTEASOME SUBUNIT ALPHA TYPE-1 |
| 12:00 | 2z4s-A | 6.8 | 8.6 | 126 | 240 | 13 | MOLECULE: CHROMOSOMAL REPLICATION INITIATOR PROTEIN DNAA |
| 13:00 | 2cea-D | 6.8 | 8.6 | 146 | 413 | 13 | MOLECULE: CELL DIVISION PROTEIN FTSH |
| 14:00 | 3j96-A | 6.8 | 13.7 | 207 | 678 | 14 | MOLECULE: VESICLE-FUSING ATPASE |
| 15:00 | 3te6-A | 6.8 | 14.6 | 148 | 304 | 5 | MOLECULE: REGULATORY PROTEIN SIR3 |
| 16:00 | 5he9-A | 6.8 | 3.7 | 110 | 168 | 17 | MOLECULE: HELICASE LOADER |
| 17:00 | 4r7y-A | 6.8 | 5.3 | 204 | 591 | 13 | MOLECULE: MINICHROMOSOME MAINTENANCE PROTEIN MCM, CELL DIVI |
| 18:00 | 6j6i-G | 6.8 | 26.6 | 151 | 790 | 9 | MOLECULE: PROBABLE SERINE/THREONINE-PROTEIN KINASE PBL2 |
| 19:00 | 6j5t-F | 6.8 | 27.2 | 155 | 808 | 10 | MOLECULE: PROBABLE SERINE/THREONINE-PROTEIN KINASE PBL2 |
| 20:00 | 3ja8-5 | 6.8 | 11 | 207 | 633 | 11 | MOLECULE: MINICHROMOSOME MAINTENANCE 2 |
| 21:00 | 3glh-G | 6.8 | 10.4 | 138 | 364 | 12 | MOLECULE: DNA POLYMERASE III SUBUNIT DELTA |
| 22:00 | 7ngc-C | 6.8 | 13.8 | 181 | 776 | 10 | MOLECULE: LON PROTEASE HOMOLOG, MITOCHONDRIAL |
| 23:00 | 5t0h-C | 6.8 | 20 | 171 | 384 | 12 | MOLECULE: 26S PROTEASE REGULATORY SUBUNIT 7 |
| 24:00:00 | 5bk4-3 | 6.8 | 12.3 | 234 | 604 | 9 | MOLECULE: DNA REPLICATION LICENSING FACTOR MCM2 |
| 25:00:00 | 5bk4-5 | 6.8 | 10.7 | 215 | 661 | 9 | MOLECULE: DNA REPLICATION LICENSING FACTOR MCM2 |
| 26:00:00 | 6og2-A | 6.8 | 12.7 | 205 | 548 | 14 | MOLECULE: HYPERACTIVE DISAGGREGASE CLPB |
| 27:00:00 | 5udb-E | 6.8 | 16.1 | 134 | 417 | 10 | MOLECULE: DNA REPLICATION LICENSING FACTOR MCM2 |
| 28:00:00 | 6p07-A | 6.8 | 13.1 | 134 | 270 | 14 | MOLECULE: SPASTIN |
| 29:00:00 | 5vft-C | 6.8 | 19.8 | 171 | 384 | 12 | MOLECULE: 26S PROTEASOME NON-ATPASE REGULATORY SUBUNIT 1 |
| 30:00:00 | 4hyy-A | 6.8 | 3 | 121 | 241 | 8 | MOLECULE: MEIOTIC RECOMBINATION PROTEIN DMC1/LIM15 HOMOLOG |
| 31:00:00 | 6ut7-K | 6.8 | 11.4 | 140 | 391 | 16 | MOLECULE: GTPASE SUBUNIT OF RESTRICTION ENDONUCLEASE |
| 32:00:00 | 1jr3-E | 6.8 | 14 | 146 | 334 | 12 | MOLECULE: DNA POLYMERASE III SUBUNIT GAMMA |
| 33:00:00 | 8emh-I | 6.8 | 4.9 | 204 | 672 | 7 | MOLECULE: PROTEASE LON-RELATED BREX SYSTEM PROTEIN BRXL |
| 34:00:00 | 6ut7-A | 6.8 | 13.8 | 149 | 390 | 12 | MOLECULE: GTPASE SUBUNIT OF RESTRICTION ENDONUCLEASE |
| 35:00:00 | 6ut7-D | 6.8 | 14.9 | 142 | 391 | 17 | MOLECULE: GTPASE SUBUNIT OF RESTRICTION ENDONUCLEASE |
| 36:00:00 | 6igm-A | 6.8 | 9.7 | 140 | 405 | 11 | MOLECULE: RUVB-LIKE 1 |
| 37:00:00 | 8osg-B | 6.8 | 4.9 | 171 | 323 | 11 | MOLECULE: MAGNESIUM-CHELATASE SUBUNIT CHLI |
| 38:00:00 | 1xp8-A | 6.8 | 5.6 | 129 | 298 | 16 | MOLECULE: RECA PROTEIN |
| 39:00:00 | 3j97-A | 6.8 | 21.8 | 156 | 678 | 15 | MOLECULE: VESICLE-FUSING ATPASE |
| 40:00:00 | 8osh-A | 6.8 | 4.3 | 155 | 290 | 9 | MOLECULE: MAGNESIUM-CHELATASE SUBUNIT CHLI |
| 41:00:00 | 6f0x-B | 6.8 | 4.1 | 150 | 398 | 14 | MOLECULE: PACHYTENE CHECKPOINT PROTEIN 2 HOMOLOG |

| | | | | | | | | |
|---|---|---|---|---|---|---|---|---|
| 42:00:00 | 8fcw-V | 6.8 | 6.4 | 142 | 343 | 10 | MOLECULE: | TNSC |
| 43:00:00 | 3j98-A | 6.8 | 19 | 158 | 676 | 15 | MOLECULE: | VESICLE-FUSING ATPASE |
| 44:00:00 | 3j96-F | 6.8 | 13 | 174 | 654 | 14 | MOLECULE: | VESICLE-FUSING ATPASE |
| 45:00:00 | 6pxl-D | 6.8 | 10.3 | 152 | 401 | 11 | MOLECULE: | ATP-DEPENDENT PROTEASE ATPASE SUBUNIT HSLU |
| 46:00:00 | 6pe0-A | 6.8 | 11.5 | 162 | 262 | 12 | MOLECULE: | MEMBRANE-SPANNING ATPASE-LIKE PROTEIN |
| 47:00:00 | 7t8b-A | 6.7 | 6.9 | 153 | 518 | 10 | MOLECULE: | TWINKLE MTDNA HELICASE |
| 48:00:00 | 7ykk-A | 6.7 | 9.1 | 207 | 735 | 8 | MOLECULE: | ATPASE FAMILY GENE 2 PROTEIN |
| 49:00:00 | 6qi8-E | 6.7 | 3.7 | 147 | 335 | 12 | MOLECULE: | RUVB-LIKE 1 |
| 50:00:00 | 7m9b-C | 6.7 | 7.3 | 133 | 257 | 9 | MOLECULE: | TNSC |
| 51:00:00 | 7rzy-2 | 6.7 | 9.9 | 144 | 311 | 8 | MOLECULE: | TN6677 VIBRIO CHOLERAE TRANSPOSON TNSC (VCHTNSC) |
| 52:00:00 | 6mat-A | 6.7 | 10.3 | 149 | 578 | 16 | MOLECULE: | RIX7 MUTANT |
| 53:00:00 | 5xmi-A | 6.7 | 13.7 | 142 | 319 | 13 | MOLECULE: | VACUOLAR PROTEIN SORTING-ASSOCIATED PROTEIN 4 |
| 54:00:00 | 5he8-D | 6.7 | 4 | 113 | 170 | 17 | MOLECULE: | HELICASE LOADER |
| 55:00:00 | 2hcb-C | 6.7 | 13.3 | 132 | 317 | 11 | MOLECULE: | CHROMOSOMAL REPLICATION INITIATOR PROTEIN DNAA |
| 56:00:00 | 6s2p-N | 6.7 | 4.8 | 107 | 252 | 13 | MOLECULE: | NRC1 |
| 57:00:00 | 5vc7-G | 6.7 | 12.1 | 173 | 544 | 10 | MOLECULE: | VCP-LIKE ATPASE |
| 58:00:00 | 6qel-G | 6.7 | 21.3 | 153 | 229 | 13 | MOLECULE: | REPLICATIVE DNA HELICASE |
| 59:00:00 | 7jk6-A | 6.7 | 16.6 | 155 | 373 | 8 | MOLECULE: | ORIGIN RECOGNITION COMPLEX SUBUNIT 4 |
| 60:00:00 | 6vvo-E | 6.7 | 10.6 | 131 | 346 | 10 | MOLECULE: | REPLICATION FACTOR C SUBUNIT 1 |
| 61:00:00 | 3ecc-A | 6.7 | 3.5 | 121 | 182 | 17 | MOLECULE: | DNA REPLICATION PROTEIN DNAC |
| 62:00:00 | 2z4r-C | 6.7 | 8.4 | 124 | 240 | 12 | MOLECULE: | CHROMOSOMAL REPLICATION INITIATOR PROTEIN DNAA |
| 63:00:00 | 3co5-A | 6.7 | 3.9 | 114 | 134 | 8 | MOLECULE: | PUTATIVE TWO-COMPONENT SYSTEM TRANSCRIPTIONAL RES |
| 64:00:00 | 3co5-B | 6.7 | 3.9 | 111 | 132 | 10 | MOLECULE: | PUTATIVE TWO-COMPONENT SYSTEM TRANSCRIPTIONAL RES |
| 65:00:00 | 5he8-B | 6.7 | 3.5 | 112 | 164 | 18 | MOLECULE: | HELICASE LOADER |
| 66:00:00 | 7ngf-E | 6.7 | 30 | 211 | 776 | 9 | MOLECULE: | LON PROTEASE HOMOLOG, MITOCHONDRIAL |
| 67:00:00 | 1nlz-C | 6.7 | 15.2 | 132 | 278 | 7 | MOLECULE: | VIRB11 HOMOLOG |
| 68:00:00 | 5he8-A | 6.7 | 3.5 | 109 | 163 | 14 | MOLECULE: | HELICASE LOADER |
| 69:00:00 | 6epe-K | 6.7 | 17.9 | 192 | 391 | 9 | MOLECULE: | PROTEASOME SUBUNIT ALPHA TYPE-6 |
| 70:00:00 | 4r7z-C | 6.7 | 4.6 | 151 | 316 | 11 | MOLECULE: | CELL DIVISION CONTROL PROTEIN 21 |
| 71:00:00 | 6ut8-D | 6.7 | 11.4 | 140 | 391 | 15 | MOLECULE: | GTPASE SUBUNIT OF RESTRICTION ENDONUCLEASE |
| 72:00:00 | 6ut8-E | 6.7 | 14.6 | 143 | 392 | 17 | MOLECULE: | GTPASE SUBUNIT OF RESTRICTION ENDONUCLEASE |
| 73:00:00 | 6ut4-A | 6.7 | 14.3 | 144 | 387 | 14 | MOLECULE: | GTPASE SUBUNIT OF RESTRICTION ENDONUCLEASE |
| 74:00:00 | 2w58-A | 6.7 | 4.1 | 126 | 188 | 15 | MOLECULE: | PRIMOSOME COMPONENT (HELICASE LOADER) |
| 75:00:00 | 6ut7-L | 6.7 | 14.6 | 143 | 392 | 17 | MOLECULE: | GTPASE SUBUNIT OF RESTRICTION ENDONUCLEASE |
| 76:00:00 | 6ut8-F | 6.7 | 14.2 | 135 | 392 | 15 | MOLECULE: | GTPASE SUBUNIT OF RESTRICTION ENDONUCLEASE |
| 77:00:00 | 6ut5-E | 6.7 | 14.3 | 144 | 395 | 15 | MOLECULE: | GTPASE SUBUNIT OF RESTRICTION ENDONUCLEASE |
| 78:00:00 | 6ut8-C | 6.7 | 14.9 | 141 | 390 | 13 | MOLECULE: | GTPASE SUBUNIT OF RESTRICTION ENDONUCLEASE |
| 79:00:00 | 1mo4-A | 6.7 | 11.9 | 151 | 324 | 13 | MOLECULE: | RECA |
| 80:00:00 | 1xxh-J | 6.7 | 15.6 | 127 | 334 | 12 | MOLECULE: | DNA POLYMERASE III, DELTA SUBUNIT |
| 81:00:00 | 1q57-A | 6.7 | 18.6 | 163 | 483 | 6 | MOLECULE: | DNA PRIMASE/HELICASE |
| 82:00:00 | 8fcv-W | 6.7 | 8.6 | 150 | 343 | 8 | MOLECULE: | DNA (60-MER) |
| 83:00:00 | 7yph-E | 6.7 | 23.6 | 223 | 773 | 11 | MOLECULE: | LON PROTEASE |
| 84:00:00 | 1w4r-A | 6.7 | 4.1 | 100 | 174 | 11 | MOLECULE: | THYMIDINE KINASE |
| 85:00:00 | 5ltj-A | 6.7 | 4.1 | 133 | 703 | 11 | MOLECULE: | PRE-MRNA-SPLICING FACTOR ATP-DEPENDENT RNA HELICA |
| 86:00:00 | 8ff4-S | 6.7 | 6.2 | 143 | 343 | 8 | MOLECULE: | TYPE I-B CRISPR-ASSOCIATED PROTEIN CAS5 |
| 87:00:00 | 7pbq-E | 6.7 | 9.3 | 140 | 312 | 10 | MOLECULE: | HOLLIDAY JUNCTION ATP-DEPENDENT DNA HELICASE RUVB |
| 88:00:00 | 3j95-B | 6.7 | 13.2 | 158 | 481 | 16 | MOLECULE: | VESICLE-FUSING ATPASE |
| 89:00:00 | 4nl4-H | 6.7 | 19.9 | 165 | 698 | 10 | MOLECULE: | PRIMOSOME ASSEMBLY PROTEIN PRIA |
| 90:00:00 | 4cr2-K | 6.7 | 26.2 | 164 | 381 | 12 | MOLECULE: | PROTEASOME COMPONENT PRE3 |
| 91:00:00 | 7crc-B | 6.6 | 8.6 | 176 | 1112 | 9 | MOLECULE: | NAD+ HYDROLASE (NADASE) |
| 92:00:00 | 7m9a-C | 6.6 | 7.2 | 135 | 257 | 9 | MOLECULE: | TNSC |
| 93:00:00 | 7ufm-N | 6.6 | 10 | 146 | 311 | 8 | MOLECULE: | VCHTNSC |
| 94:00:00 | 6l8d-A | 6.6 | 4.3 | 142 | 267 | 15 | MOLECULE: | MAGNESIUM-CHELATASE SUBUNIT CHLI |

| | | | | | | | | |
|---|---|---|---|---|---|---|---|---|
| 95:00:00 | 8emh-K | 6.6 | 5.9 | 203 | 664 | 10 | MOLECULE: | PROTEASE LON-RELATED BREX SYSTEM PROTEIN BRXL |
| 96:00:00 | 8q6p-2 | 6.6 | 4.7 | 179 | 431 | 9 | MOLECULE: | DNA REPLICATION LICENSING FACTOR MCM2 |
| 97:00:00 | 5lta-A | 6.6 | 4.2 | 128 | 698 | 11 | MOLECULE: | PRE-MRNA-SPLICING FACTOR ATP-DEPENDENT RNA HELICA |
| 98:00:00 | 2z4r-A | 6.6 | 9 | 125 | 240 | 12 | MOLECULE: | CHROMOSOMAL REPLICATION INITIATOR PROTEIN DNAA |
| 99:00:00 | 2ce7-E | 6.6 | 4.5 | 143 | 405 | 15 | MOLECULE: | CELL DIVISION PROTEIN FTSH |
| 0:00 | 5nss-J | 6.6 | 8 | 136 | 251 | 11 | MOLECULE: | DNA-DIRECTED RNA POLYMERASE SUBUNIT ALPHA |
| 1:00 | 1g6o-A | 6.6 | 17.8 | 143 | 323 | 6 | MOLECULE: | CAG-ALPHA |
| 2:00 | 5vhm-B | 6.6 | 8.8 | 136 | 232 | 10 | MOLECULE: | 26S PROTEASOME NON-ATPASE REGULATORY SUBUNIT 10 |
| 3:00 | 1xxh-E | 6.6 | 14.8 | 127 | 334 | 13 | MOLECULE: | DNA POLYMERASE III, DELTA SUBUNIT |
| 4:00 | 7nfy-A | 6.6 | 34.4 | 207 | 776 | 12 | MOLECULE: | LON PROTEASE HOMOLOG, MITOCHONDRIAL |
| 5:00 | 7ngp-C | 6.6 | 31.5 | 217 | 776 | 9 | MOLECULE: | LON PROTEASE HOMOLOG, MITOCHONDRIAL |
| 6:00 | 8fcv-Q | 6.6 | 6.3 | 145 | 310 | 10 | MOLECULE: | DNA (60-MER) |
| 7:00 | 7p6u-F | 6.6 | 33 | 221 | 772 | 11 | MOLECULE: | LON PROTEASE |
| 8:00 | 6rn4-C | 6.6 | 11.9 | 163 | 689 | 10 | MOLECULE: | CHAPERONE PROTEIN CLPB |
| 9:00 | 2v1u-A | 6.6 | 14.9 | 146 | 382 | 11 | MOLECULE: | CELL DIVISION CONTROL PROTEIN 6 HOMOLOG |
| 10:00 | 8hr9-E | 6.6 | 14 | 145 | 833 | 9 | MOLECULE: | ARCHAEAL ATPASE |
| 11:00 | 4fdg-E | 6.6 | 5.2 | 231 | 592 | 9 | MOLECULE: | MINICHROMOSOME MAINTENANCE PROTEIN MCM |
| 12:00 | 4fdg-D | 6.6 | 4.9 | 225 | 592 | 9 | MOLECULE: | MINICHROMOSOME MAINTENANCE PROTEIN MCM |
| 13:00 | 4fdg-B | 6.6 | 5.1 | 230 | 592 | 10 | MOLECULE: | MINICHROMOSOME MAINTENANCE PROTEIN MCM |
| 14:00 | 6ut4-B | 6.6 | 14.9 | 142 | 385 | 13 | MOLECULE: | GTPASE SUBUNIT OF RESTRICTION ENDONUCLEASE |
| 15:00 | 6ut4-E | 6.6 | 14.7 | 140 | 386 | 15 | MOLECULE: | GTPASE SUBUNIT OF RESTRICTION ENDONUCLEASE |
| 16:00 | 7plh-A | 6.6 | 7.6 | 133 | 260 | 9 | MOLECULE: | SHTNSC |
| 17:00 | 5vhr-D | 6.6 | 6.7 | 127 | 236 | 15 | MOLECULE: | 26S PROTEASOME NON-ATPASE REGULATORY SUBUNIT 10 |
| 18:00 | 5txv-O | 6.6 | 11.5 | 158 | 348 | 12 | MOLECULE: | ATP-DEPENDENT PROTEASE ATPASE SUBUNIT HSLU |
| 19:00 | 8fcx-U | 6.6 | 7.7 | 149 | 343 | 8 | MOLECULE: | TNSC |
| 20:00 | 3f8t-A | 6.5 | 5.9 | 186 | 459 | 11 | MOLECULE: | PREDICTED ATPASE INVOLVED IN REPLICATION CONTROL, |
| 21:00 | 7ufm-G | 6.5 | 10 | 146 | 311 | 8 | MOLECULE: | VCHTNSC |
| 22:00 | 7m99-D | 6.5 | 8.1 | 136 | 257 | 9 | MOLECULE: | TNSC |
| 23:00 | 7m9c-C | 6.5 | 7.3 | 133 | 257 | 9 | MOLECULE: | TNSC |
| 24:00:00 | 5wcb-A | 6.5 | 3.2 | 124 | 277 | 13 | MOLECULE: | MEIOTIC SPINDLE FORMATION PROTEIN MEI-1 |
| 25:00:00 | 6p07-B | 6.5 | 3.7 | 132 | 303 | 14 | MOLECULE: | SPASTIN |
| 26:00:00 | 7rzy-4 | 6.5 | 10.3 | 148 | 311 | 10 | MOLECULE: | TN6677 VIBRIO CHOLERAE TRANSPOSON TNSC (VCHTNSC) |
| 27:00:00 | 8jx6-A | 6.5 | 11.3 | 125 | 447 | 8 | MOLECULE: | DEEP-SEA HELICASE 9 |
| 28:00:00 | 6j5t-G | 6.5 | 28.8 | 159 | 808 | 8 | MOLECULE: | PROBABLE SERINE/THREONINE-PROTEIN KINASE PBL2 |
| 29:00:00 | 5he8-H | 6.5 | 3.8 | 109 | 165 | 17 | MOLECULE: | HELICASE LOADER |
| 30:00:00 | 8emh-F | 6.5 | 5.4 | 207 | 666 | 10 | MOLECULE: | PROTEASE LON-RELATED BREX SYSTEM PROTEIN BRXL |
| 31:00:00 | 8fnt-B | 6.5 | 16 | 137 | 785 | 8 | MOLECULE: | ARCHAEAL ATPASE |
| 32:00:00 | 8fnt-C | 6.5 | 14.9 | 140 | 785 | 8 | MOLECULE: | ARCHAEAL ATPASE |
| 33:00:00 | 6epc-H | 6.5 | 20.2 | 161 | 396 | 6 | MOLECULE: | PROTEASOME SUBUNIT ALPHA TYPE-6 |
| 34:00:00 | 1xu4-A | 6.5 | 5.6 | 145 | 318 | 10 | MOLECULE: | DNA REPAIR AND RECOMBINATION PROTEIN RADA |
| 35:00:00 | 5he8-J | 6.5 | 3.5 | 106 | 163 | 14 | MOLECULE: | HELICASE LOADER |
| 36:00:00 | 5he8-L | 6.5 | 3.6 | 108 | 163 | 14 | MOLECULE: | HELICASE LOADER |
| 37:00:00 | 2f1h-A | 6.5 | 5.9 | 144 | 313 | 10 | MOLECULE: | DNA REPAIR AND RECOMBINATION PROTEIN RADA |
| 38:00:00 | 5vhp-B | 6.5 | 3.6 | 127 | 227 | 9 | MOLECULE: | 26S PROTEASOME NON-ATPASE REGULATORY SUBUNIT 10 |
| 39:00:00 | 7ng4-A | 6.5 | 33.3 | 212 | 776 | 9 | MOLECULE: | LON PROTEASE HOMOLOG, MITOCHONDRIAL |
| 40:00:00 | 7ngq-A | 6.5 | 28.3 | 207 | 776 | 12 | MOLECULE: | LON PROTEASE HOMOLOG, MITOCHONDRIAL |
| 41:00:00 | 5t0i-C | 6.5 | 16.8 | 175 | 384 | 11 | MOLECULE: | 26S PROTEASE REGULATORY SUBUNIT 7 |
| 42:00:00 | 5d4w-B | 6.5 | 4.8 | 196 | 688 | 10 | MOLECULE: | PUTATIVE HEAT SHOCK PROTEIN |
| 43:00:00 | 7jk5-A | 6.5 | 15.3 | 150 | 390 | 6 | MOLECULE: | ORIGIN RECOGNITION COMPLEX SUBUNIT 2 |
| 44:00:00 | 6ees-A | 6.5 | 16.9 | 169 | 2023 | 11 | MOLECULE: | MIDASIN |
| 45:00:00 | 4fdg-C | 6.5 | 5 | 223 | 592 | 10 | MOLECULE: | MINICHROMOSOME MAINTENANCE PROTEIN MCM |
| 46:00:00 | 8van-C | 6.5 | 12 | 142 | 365 | 12 | MOLECULE: | DNA POLYMERASE III SUBUNIT DELTA |
| 47:00:00 | 8fcw-T | 6.5 | 7.7 | 150 | 343 | 8 | MOLECULE: | TNSC |

| | | | | | | | | |
|---|---|---|---|---|---|---|---|---|
| 48:00:00 | 1v5w-A | 6.5 | 2.9 | 114 | 239 | 9 | MOLECULE: | MEIOTIC RECOMBINATION PROTEIN DMC1/LIM15 HOMOLOG |
| 49:00:00 | 5tsh-A | 6.5 | 13.6 | 146 | 374 | 9 | MOLECULE: | TYPE IV PILUS BIOGENESIS ATPASE PILB |
| 50:00:00 | 5vq9-D | 6.5 | 4.3 | 152 | 367 | 14 | MOLECULE: | PACHYTENE CHECKPOINT PROTEIN 2 HOMOLOG |
| 51:00:00 | 8fcv-R | 6.5 | 7.2 | 148 | 343 | 8 | MOLECULE: | DNA (60-MER) |
| 52:00:00 | 7yup-E | 6.5 | 24.4 | 226 | 773 | 11 | MOLECULE: | LON PROTEASE |
| 53:00:00 | 8ff4-R | 6.5 | 7.2 | 148 | 343 | 8 | MOLECULE: | TYPE I-B CRISPR-ASSOCIATED PROTEIN CAS5 |
| 54:00:00 | 2orw-B | 6.5 | 5.6 | 108 | 173 | 12 | MOLECULE: | THYMIDINE KINASE |
| 55:00:00 | 8fcx-R | 6.5 | 7.7 | 150 | 343 | 8 | MOLECULE: | TNSC |
| 56:00:00 | 8c0w-F | 6.5 | 8.4 | 204 | 823 | 6 | MOLECULE: | PEROXISOMAL ATPASE PEX6 |
| 57:00:00 | 2qen-A | 6.5 | 10.1 | 131 | 350 | 18 | MOLECULE: | WALKER-TYPE ATPASE |
| 58:00:00 | 7ypj-E | 6.5 | 23.9 | 214 | 773 | 12 | MOLECULE: | LON PROTEASE |
| 59:00:00 | 5jpq-i | 6.4 | 8.5 | 135 | 659 | 10 | MOLECULE: | WD40 DOMAIN PROTEINS |
| 60:00:00 | 4p6q-A | 6.4 | 23.6 | 155 | 286 | 4 | MOLECULE: | MSX2-INTERACTING PROTEIN |
| 61:00:00 | 5udb-7 | 6.4 | 7.5 | 214 | 726 | 13 | MOLECULE: | DNA REPLICATION LICENSING FACTOR MCM2 |
| 62:00:00 | 6o16-A | 6.4 | 16.3 | 134 | 798 | 11 | MOLECULE: | DEAH (ASP-GLU-ALA-HIS) BOX POLYPEPTIDE 37 |
| 63:00:00 | 2o0h-A | 6.4 | 8.1 | 151 | 357 | 13 | MOLECULE: | DNA PACKAGING PROTEIN GP17 |
| 64:00:00 | 8gja-F | 6.4 | 13.9 | 145 | 342 | 6 | MOLECULE: | RAD51C |
| 65:00:00 | 8v48-A | 6.4 | 3.8 | 116 | 412 | 14 | MOLECULE: | ARIA ANTITOXIN |
| 66:00:00 | 5he8-K | 6.4 | 3.6 | 108 | 164 | 16 | MOLECULE: | HELICASE LOADER |
| 67:00:00 | 5kwa-A | 6.4 | 12.7 | 162 | 454 | 12 | MOLECULE: | PROTEASOME-ASSOCIATED ATPASE |
| 68:00:00 | 7upr-A | 6.4 | 9.1 | 131 | 235 | 12 | MOLECULE: | OUTER MITOCHONDRIAL TRANSMEMBRANE HELIX TRANSLOCA |
| 69:00:00 | 5he8-G | 6.4 | 3.8 | 107 | 165 | 16 | MOLECULE: | HELICASE LOADER |
| 70:00:00 | 5he8-C | 6.4 | 3.8 | 111 | 171 | 17 | MOLECULE: | HELICASE LOADER |
| 71:00:00 | 5he8-F | 6.4 | 3.5 | 106 | 164 | 18 | MOLECULE: | HELICASE LOADER |
| 72:00:00 | 7ng4-E | 6.4 | 27 | 204 | 776 | 10 | MOLECULE: | LON PROTEASE HOMOLOG, MITOCHONDRIAL |
| 73:00:00 | 7krz-B | 6.4 | 10.9 | 182 | 526 | 12 | MOLECULE: | LON PROTEASE HOMOLOG, MITOCHONDRIAL |
| 74:00:00 | 3cmv-F | 6.4 | 15.1 | 132 | 1175 | 12 | MOLECULE: | PROTEIN RECA |
| 75:00:00 | 7ngc-A | 6.4 | 32.9 | 220 | 776 | 9 | MOLECULE: | LON PROTEASE HOMOLOG, MITOCHONDRIAL |
| 76:00:00 | 5vhm-D | 6.4 | 3.3 | 124 | 220 | 15 | MOLECULE: | 26S PROTEASOME NON-ATPASE REGULATORY SUBUNIT 10 |
| 77:00:00 | 5vhr-B | 6.4 | 3.5 | 124 | 233 | 9 | MOLECULE: | 26S PROTEASOME NON-ATPASE REGULATORY SUBUNIT 10 |
| 78:00:00 | 8c0w-B | 6.4 | 9.5 | 204 | 821 | 7 | MOLECULE: | PEROXISOMAL ATPASE PEX6 |
| 79:00:00 | 3j96-B | 6.4 | 24.8 | 155 | 672 | 14 | MOLECULE: | VESICLE-FUSING ATPASE |
| 80:00:00 | 6ut8-A | 6.4 | 15 | 145 | 390 | 12 | MOLECULE: | GTPASE SUBUNIT OF RESTRICTION ENDONUCLEASE |
| 81:00:00 | 4cr4-I | 6.4 | 17.3 | 170 | 362 | 11 | MOLECULE: | PROTEASOME COMPONENT PRE3 |
| 82:00:00 | 6ut7-H | 6.4 | 14.8 | 140 | 390 | 12 | MOLECULE: | GTPASE SUBUNIT OF RESTRICTION ENDONUCLEASE |
| 83:00:00 | 4fdg-A | 6.4 | 6.7 | 230 | 592 | 10 | MOLECULE: | MINICHROMOSOME MAINTENANCE PROTEIN MCM |
| 84:00:00 | 4r7z-J | 6.4 | 4.8 | 162 | 316 | 9 | MOLECULE: | CELL DIVISION CONTROL PROTEIN 21 |
| 85:00:00 | 7lcc-A | 6.4 | 24.2 | 196 | 1369 | 7 | MOLECULE: | HELRAISER K1068Q |
| 86:00:00 | 5txv-Q | 6.4 | 11.1 | 154 | 370 | 12 | MOLECULE: | ATP-DEPENDENT PROTEASE ATPASE SUBUNIT HSLU |
| 87:00:00 | 2cvf-A | 6.4 | 3.2 | 124 | 220 | 19 | MOLECULE: | DNA REPAIR AND RECOMBINATION PROTEIN RADB |
| 88:00:00 | 8hr8-E | 6.4 | 12.9 | 140 | 833 | 7 | MOLECULE: | ARCHAEAL ATPASE |
| 89:00:00 | 8fcx-S | 6.4 | 7.1 | 149 | 343 | 8 | MOLECULE: | TNSC |
| 90:00:00 | 8osf-B | 6.4 | 4.4 | 166 | 321 | 9 | MOLECULE: | MAGNESIUM-CHELATASE SUBUNIT CHLI |
| 91:00:00 | 3j96-E | 6.4 | 21.7 | 161 | 670 | 14 | MOLECULE: | VESICLE-FUSING ATPASE |
| 92:00:00 | 5txv-B | 6.4 | 11.2 | 152 | 337 | 12 | MOLECULE: | ATP-DEPENDENT PROTEASE ATPASE SUBUNIT HSLU |
| 93:00:00 | 8c0v-A | 6.3 | 13.2 | 185 | 823 | 6 | MOLECULE: | PEROXISOMAL ATPASE PEX1 |
| 94:00:00 | 5vca-P | 6.3 | 9.1 | 149 | 543 | 13 | MOLECULE: | VCP-LIKE ATPASE |
| 95:00:00 | 5vhq-B | 6.3 | 9.7 | 133 | 233 | 10 | MOLECULE: | 26S PROTEASOME NON-ATPASE REGULATORY SUBUNIT 10 |
| 96:00:00 | 6oa9-C | 6.3 | 8.3 | 141 | 492 | 9 | MOLECULE: | CELL DIVISION CONTROL PROTEIN 48 |
| 97:00:00 | 1n0w-A | 6.3 | 3.1 | 113 | 210 | 8 | MOLECULE: | DNA REPAIR PROTEIN RAD51 HOMOLOG 1 |
| 98:00:00 | 6qem-L | 6.3 | 23.5 | 166 | 239 | 13 | MOLECULE: | REPLICATIVE DNA HELICASE |
| 99:00:00 | 4eiw-B | 6.3 | 8.4 | 154 | 446 | 13 | MOLECULE: | ATP-DEPENDENT ZINC METALLOPROTEASE FTSH |
| 0:00 | 5vca-N | 6.3 | 10.4 | 140 | 543 | 12 | MOLECULE: | VCP-LIKE ATPASE |

| | | | | | | | | |
|---|---|---|---|---|---|---|---|---|
| 1:00 | 3ec2-A | 6.3 | 3.4 | 115 | 177 | 16 | MOLECULE: | DNA REPLICATION PROTEIN DNAC |
| 2:00 | 4eiw-F | 6.3 | 9.8 | 155 | 446 | 13 | MOLECULE: | ATP-DEPENDENT ZINC METALLOPROTEASE FTSH |
| 3:00 | 2r44-A | 6.3 | 4.3 | 161 | 330 | 11 | MOLECULE: | UNCHARACTERIZED PROTEIN |
| 4:00 | 5irm-C | 6.3 | 11.5 | 161 | 770 | 9 | MOLECULE: | UNCHARACTERIZED PROTEIN |
| 5:00 | 5mp9-H | 6.3 | 18.9 | 168 | 390 | 10 | MOLECULE: | PROTEASOME SUBUNIT ALPHA TYPE-1 |
| 6:00 | 3cmv-E | 6.3 | 14 | 132 | 1165 | 12 | MOLECULE: | PROTEIN RECA |
| 7:00 | 3cmv-D | 6.3 | 14.4 | 138 | 1175 | 12 | MOLECULE: | PROTEIN RECA |
| 8:00 | 3cmv-G | 6.3 | 14.6 | 136 | 1167 | 13 | MOLECULE: | PROTEIN RECA |
| 9:00 | 1z6t-C | 6.3 | 11.3 | 158 | 586 | 11 | MOLECULE: | APOPTOTIC PROTEASE ACTIVATING FACTOR 1 |
| 10:00 | 5oe8-A | 6.3 | 17.2 | 124 | 407 | 8 | MOLECULE: | LARGE SUBUNIT TERMINASE |
| 11:00 | 6fhs-E | 6.3 | 13.8 | 149 | 441 | 13 | MOLECULE: | RUVB-LIKE HELICASE |
| 12:00 | 6em8-B | 6.3 | 16.3 | 213 | 744 | 11 | MOLECULE: | ATP-DEPENDENT CLP PROTEASE ATP-BINDING SUBUNIT CL |
| 13:00 | 8ff4-Q | 6.3 | 5.8 | 137 | 310 | 9 | MOLECULE: | TYPE I-B CRISPR-ASSOCIATED PROTEIN CAS5 |
| 14:00 | 8fnu-E | 6.3 | 16 | 146 | 796 | 8 | MOLECULE: | KAP NTPASE DOMAIN-CONTAINING PROTEIN |
| 15:00 | 6rn2-D | 6.3 | 13.7 | 164 | 684 | 12 | MOLECULE: | CHAPERONE PROTEIN CLPB |
| 16:00 | 8fcu-Q | 6.3 | 7.1 | 142 | 310 | 10 | MOLECULE: | TYPE I-B CRISPR-ASSOCIATED PROTEIN CAS5 |
| 17:00 | 5vhn-D | 6.3 | 9.2 | 130 | 237 | 12 | MOLECULE: | 26S PROTEASOME NON-ATPASE REGULATORY SUBUNIT 10 |
| 18:00 | 8fcw-R | 6.3 | 7.3 | 149 | 343 | 8 | MOLECULE: | TNSC |
| 19:00 | 6sh4-G | 6.3 | 22.2 | 182 | 376 | 10 | MOLECULE: | MITOCHONDRIAL CHAPERONE BCS1 |
| 20:00 | 4r7z-I | 6.3 | 4.7 | 153 | 316 | 11 | MOLECULE: | CELL DIVISION CONTROL PROTEIN 21 |
| 21:00 | 8bob-C | 6.3 | 8.7 | 126 | 412 | 16 | MOLECULE: | PROTEIN MALY |
| 22:00 | 8ff4-V | 6.3 | 8.3 | 145 | 343 | 8 | MOLECULE: | TYPE I-B CRISPR-ASSOCIATED PROTEIN CAS5 |
| 23:00 | 6ut5-B | 6.3 | 12.1 | 141 | 395 | 14 | MOLECULE: | GTPASE SUBUNIT OF RESTRICTION ENDONUCLEASE |
| 24:00:00 | 8c0v-E | 6.3 | 10.1 | 201 | 821 | 7 | MOLECULE: | PEROXISOMAL ATPASE PEX1 |
| 25:00:00 | 6em8-I | 6.3 | 4.4 | 158 | 436 | 13 | MOLECULE: | ATP-DEPENDENT CLP PROTEASE ATP-BINDING SUBUNIT CL |
| 26:00:00 | 5nss-N | 6.3 | 3.7 | 138 | 251 | 12 | MOLECULE: | DNA-DIRECTED RNA POLYMERASE SUBUNIT ALPHA |
| 27:00:00 | 5txv-H | 6.3 | 11.6 | 156 | 371 | 12 | MOLECULE: | ATP-DEPENDENT PROTEASE ATPASE SUBUNIT HSLU |
| 28:00:00 | 8hr9-G | 6.3 | 14.1 | 139 | 830 | 9 | MOLECULE: | ARCHAEAL ATPASE |
| 29:00:00 | 8fcv-S | 6.3 | 7.6 | 150 | 343 | 8 | MOLECULE: | DNA (60-MER) |
| 30:00:00 | 8faz-D | 6.3 | 7.6 | 147 | 303 | 12 | MOLECULE: | DNA REPAIR PROTEIN RAD51 HOMOLOG 2 |
| 31:00:00 | 6pdw-F | 6.3 | 12.4 | 148 | 261 | 11 | MOLECULE: | MEMBRANE-SPANNING ATPASE-LIKE PROTEIN |
| 32:00:00 | 8hwc-F | 6.2 | 4.1 | 170 | 308 | 11 | MOLECULE: | PRIMASE D5 |
| 33:00:00 | 2uz3-B | 6.2 | 9.6 | 113 | 207 | 10 | MOLECULE: | THYMIDINE KINASE |
| 34:00:00 | 6qem-K | 6.2 | 10.2 | 146 | 239 | 12 | MOLECULE: | REPLICATIVE DNA HELICASE |
| 35:00:00 | 5vhm-A | 6.2 | 3.5 | 126 | 258 | 10 | MOLECULE: | 26S PROTEASOME NON-ATPASE REGULATORY SUBUNIT 10 |
| 36:00:00 | 1g64-A | 6.2 | 3.9 | 102 | 169 | 15 | MOLECULE: | COB(I)ALAMIN ADENOSYLTRANSFERASE |
| 37:00:00 | 6uge-A | 6.2 | 13.6 | 145 | 304 | 12 | MOLECULE: | MEIOTIC SPINDLE FORMATION PROTEIN MEI-1 |
| 38:00:00 | 3b85-A | 6.2 | 3.4 | 112 | 187 | 10 | MOLECULE: | PHOSPHATE STARVATION-INDUCIBLE PROTEIN |
| 39:00:00 | 6rn3-E | 6.2 | 13 | 168 | 684 | 13 | MOLECULE: | CHAPERONE PROTEIN CLPB |
| 40:00:00 | 5vhp-C | 6.2 | 6.1 | 131 | 236 | 13 | MOLECULE: | 26S PROTEASOME NON-ATPASE REGULATORY SUBUNIT 10 |
| 41:00:00 | 5ln3-M | 6.2 | 13.7 | 171 | 389 | 9 | MOLECULE: | 26S PROTEASOME NON-ATPASE REGULATORY SUBUNIT 2 |
| 42:00:00 | 2zpa-B | 6.2 | 9.7 | 141 | 662 | 10 | MOLECULE: | UNCHARACTERIZED PROTEIN YPFI |
| 43:00:00 | 1g5r-A | 6.2 | 3.5 | 99 | 157 | 15 | MOLECULE: | COB(I)ALAMIN ADENOSYLTRANSFERASE |
| 44:00:00 | 6wqh-F | 6.2 | 11.7 | 188 | 537 | 13 | MOLECULE: | LON PROTEASE |
| 45:00:00 | 5vho-D | 6.2 | 8.3 | 119 | 207 | 10 | MOLECULE: | 26S PROTEASOME NON-ATPASE REGULATORY SUBUNIT 10 |
| 46:00:00 | 3cmv-B | 6.2 | 14.5 | 136 | 1163 | 13 | MOLECULE: | PROTEIN RECA |
| 47:00:00 | 3cmv-H | 6.2 | 13.8 | 136 | 1173 | 13 | MOLECULE: | PROTEIN RECA |
| 48:00:00 | 6sh4-F | 6.2 | 22.3 | 185 | 376 | 11 | MOLECULE: | MITOCHONDRIAL CHAPERONE BCS1 |
| 49:00:00 | 8p49-A | 6.2 | 4 | 149 | 377 | 6 | MOLECULE: | Q8U0N8 PROTEIN |
| 50:00:00 | 6sh4-E | 6.2 | 18 | 179 | 376 | 9 | MOLECULE: | MITOCHONDRIAL CHAPERONE BCS1 |
| 51:00:00 | 5vhm-C | 6.2 | 5.4 | 129 | 229 | 10 | MOLECULE: | 26S PROTEASOME NON-ATPASE REGULATORY SUBUNIT 10 |
| 52:00:00 | 6sh4-D | 6.2 | 18.2 | 180 | 376 | 10 | MOLECULE: | MITOCHONDRIAL CHAPERONE BCS1 |
| 53:00:00 | 6sh4-A | 6.2 | 22.4 | 184 | 376 | 10 | MOLECULE: | MITOCHONDRIAL CHAPERONE BCS1 |

| | | | | | | | | |
|---|---|---|---|---|---|---|---|---|
| 54:00:00 | 8hrb-B | 6.2 | 11 | 128 | 835 | 10 | MOLECULE: | ARCHAEAL ATPASE |
| 55:00:00 | 4iei-A | 6.2 | 15.9 | 137 | 449 | 13 | MOLECULE: | GENE 2 PROTEIN |
| 56:00:00 | 5a5b-I | 6.2 | 9.9 | 163 | 362 | 11 | MOLECULE: | PROTEASOME COMPONENT PRE3 |
| 57:00:00 | 8hr9-I | 6.2 | 14.6 | 136 | 836 | 10 | MOLECULE: | ARCHAEAL ATPASE |
| 58:00:00 | 4r7z-E | 6.2 | 4.7 | 163 | 316 | 10 | MOLECULE: | CELL DIVISION CONTROL PROTEIN 21 |
| 59:00:00 | 5vhp-D | 6.2 | 7.2 | 126 | 237 | 11 | MOLECULE: | 26S PROTEASOME NON-ATPASE REGULATORY SUBUNIT 10 |
| 60:00:00 | 8ff4-U | 6.2 | 7.7 | 149 | 343 | 8 | MOLECULE: | TYPE I-B CRISPR-ASSOCIATED PROTEIN CAS5 |
| 61:00:00 | 4cr3-I | 6.2 | 9.7 | 166 | 362 | 11 | MOLECULE: | PROTEASOME COMPONENT PRE3 |
| 62:00:00 | 8fcv-U | 6.2 | 7.7 | 149 | 343 | 8 | MOLECULE: | DNA (60-MER) |
| 63:00:00 | 5uj7-B | 6.2 | 17.2 | 151 | 337 | 6 | MOLECULE: | ORIGIN RECOGNITION COMPLEX SUBUNIT 1 |
| 64:00:00 | 5vhr-C | 6.2 | 7.8 | 137 | 242 | 12 | MOLECULE: | 26S PROTEASOME NON-ATPASE REGULATORY SUBUNIT 10 |
| 65:00:00 | 8pqz-J | 6.2 | 19.9 | 208 | 2892 | 9 | MOLECULE: | CYTOPLASMIC DYNEIN 1 HEAVY CHAIN 1 |
| 66:00:00 | 7tfl-A | 6.2 | 10.4 | 127 | 338 | 12 | MOLECULE: | REPLICATION FACTOR C SUBUNIT 1 |
| 67:00:00 | 4ksr-A | 6.1 | 25.3 | 161 | 521 | 9 | MOLECULE: | TYPE II SECRETION SYSTEM PROTEIN E, HEMOLYSIN-COR |
| 68:00:00 | 5dgk-A | 6.1 | 5.1 | 186 | 519 | 9 | MOLECULE: | ACTIVE HELICASE |
| 69:00:00 | 7otj-B | 6.1 | 4.3 | 123 | 436 | 10 | MOLECULE: | ATP-DEPENDENT DNA HELICASE PIF1 |
| 70:00:00 | 8iqh-J | 6.1 | 28.8 | 236 | 881 | 12 | MOLECULE: | PUTATIVE PRIMASE C962R |
| 71:00:00 | 4xgc-A | 6.1 | 13 | 136 | 310 | 7 | MOLECULE: | ORIGIN RECOGNITION COMPLEX SUBUNIT 2 |
| 72:00:00 | 5aor-A | 6.1 | 24.2 | 186 | 1009 | 7 | MOLECULE: | DOSAGE COMPENSATION REGULATOR |
| 73:00:00 | 5vca-R | 6.1 | 10 | 140 | 543 | 12 | MOLECULE: | VCP-LIKE ATPASE |
| 74:00:00 | 5vhf-B | 6.1 | 11.6 | 160 | 330 | 10 | MOLECULE: | 26S PROTEASOME NON-ATPASE REGULATORY SUBUNIT 10 |
| 75:00:00 | 5g4g-F | 6.1 | 5.8 | 167 | 721 | 10 | MOLECULE: | VCP-LIKE ATPASE |
| 76:00:00 | 5udb-A | 6.1 | 17 | 150 | 372 | 9 | MOLECULE: | DNA REPLICATION LICENSING FACTOR MCM2 |
| 77:00:00 | 6p07-F | 6.1 | 10.1 | 138 | 300 | 12 | MOLECULE: | SPASTIN |
| 78:00:00 | 3sfz-A | 6.1 | 15.1 | 135 | 1133 | 12 | MOLECULE: | APOPTOTIC PEPTIDASE ACTIVATING FACTOR 1 |
| 79:00:00 | 4hut-A | 6.1 | 7.8 | 118 | 191 | 15 | MOLECULE: | COB(I)YRINIC ACID A,C-DIAMIDE ADENOSYLTRANSFERASE |
| 80:00:00 | 4a8j-B | 6.1 | 3.2 | 113 | 220 | 12 | MOLECULE: | ELONGATOR COMPLEX PROTEIN 4 |
| 81:00:00 | 6mfv-D | 6.1 | 8.6 | 128 | 641 | 11 | MOLECULE: | TETRATRICOPEPTIDE REPEAT SENSOR PH0952 |
| 82:00:00 | 5kne-D | 6.1 | 12.6 | 168 | 770 | 10 | MOLECULE: | HEAT SHOCK PROTEIN 104 |
| 83:00:00 | 1g64-B | 6.1 | 7.9 | 115 | 190 | 14 | MOLECULE: | COB(I)ALAMIN ADENOSYLTRANSFERASE |
| 84:00:00 | 7ngc-B | 6.1 | 12.2 | 196 | 776 | 10 | MOLECULE: | LON PROTEASE HOMOLOG, MITOCHONDRIAL |
| 85:00:00 | 4a8j-E | 6.1 | 3 | 113 | 221 | 12 | MOLECULE: | ELONGATOR COMPLEX PROTEIN 4 |
| 86:00:00 | 7ngf-F | 6.1 | 32.6 | 219 | 776 | 9 | MOLECULE: | LON PROTEASE HOMOLOG, MITOCHONDRIAL |
| 87:00:00 | 5vhf-D | 6.1 | 16 | 162 | 344 | 10 | MOLECULE: | 26S PROTEASOME NON-ATPASE REGULATORY SUBUNIT 10 |
| 88:00:00 | 6rn2-F | 6.1 | 12.3 | 168 | 684 | 11 | MOLECULE: | CHAPERONE PROTEIN CLPB |
| 89:00:00 | 4iee-A | 6.1 | 16.1 | 138 | 452 | 12 | MOLECULE: | GENE 2 PROTEIN |
| 90:00:00 | 7wi4-B | 6.1 | 11 | 128 | 410 | 15 | MOLECULE: | ATP-DEPENDENT ZINC METALLOPROTEASE FTSH |
| 91:00:00 | 3f9v-A | 6.1 | 6.8 | 216 | 595 | 6 | MOLECULE: | MINICHROMOSOME MAINTENANCE PROTEIN MCM |
| 92:00:00 | 5vhs-F | 6.1 | 8.7 | 152 | 339 | 14 | MOLECULE: | 26S PROTEASOME REGULATORY SUBUNIT 7 |
| 93:00:00 | 1jr3-D | 6.1 | 15.8 | 131 | 338 | 11 | MOLECULE: | DNA POLYMERASE III SUBUNIT GAMMA |
| 94:00:00 | 8fcv-V | 6.1 | 8.2 | 149 | 343 | 8 | MOLECULE: | DNA (60-MER) |
| 95:00:00 | 7qy7-A | 6.1 | 17.3 | 158 | 342 | 9 | MOLECULE: | 26S PROTEASOME NON-ATPASE REGULATORY SUBUNIT 13 |
| 96:00:00 | 8fcx-V | 6.1 | 8.3 | 147 | 343 | 7 | MOLECULE: | TNSC |
| 97:00:00 | 7qy7-D | 6.1 | 14.6 | 174 | 378 | 9 | MOLECULE: | 26S PROTEASOME NON-ATPASE REGULATORY SUBUNIT 13 |
| 98:00:00 | 2vl7-A | 6.1 | 13.3 | 121 | 459 | 12 | MOLECULE: | XPD |
| 99:00:00 | 5he8-E | 6.1 | 3.4 | 105 | 165 | 15 | MOLECULE: | HELICASE LOADER |
| 0:00 | 3ja8-2 | 6.1 | 12 | 210 | 603 | 10 | MOLECULE: | MINICHROMOSOME MAINTENANCE 2 |
| 1:00 | 4z8x-B | 6.1 | 4.4 | 147 | 425 | 14 | MOLECULE: | ATP-DEPENDENT ZINC METALLOPROTEASE FTSH |
| 2:00 | 8pqz-A | 6.1 | 15.5 | 211 | 2892 | 8 | MOLECULE: | CYTOPLASMIC DYNEIN 1 HEAVY CHAIN 1 |
| 3:00 | 5vhi-B | 6.1 | 9.4 | 147 | 298 | 10 | MOLECULE: | 26S PROTEASOME REGULATORY SUBUNIT 7 |
| 4:00 | 6n9x-B | 6 | 9.3 | 155 | 500 | 6 | MOLECULE: | DNA PRIMASE/HELICASE |
| 5:00 | 8x5i-A | 6 | 10.5 | 135 | 477 | 13 | MOLECULE: | ENDONUCLEASE GAJA |
| 6:00 | 7jgs-C | 6 | 21.6 | 176 | 614 | 11 | MOLECULE: | ORIGIN RECOGNITION COMPLEX SUBUNIT 2 |

| | | | | | | | | |
|---|---|---|---|---|---|---|---|---|
| 7:00 | 3k1j-A | 6 | 12 | 163 | 566 | 12 | MOLECULE: | ATP-DEPENDENT PROTEASE LON |
| 8:00 | 3io5-B | 6 | 5.7 | 139 | 272 | 6 | MOLECULE: | RECOMBINATION AND REPAIR PROTEIN |
| 9:00 | 8ouz-C | 6 | 13.9 | 127 | 308 | 11 | MOLECULE: | DNA REPAIR PROTEIN RAD51 HOMOLOG 2 |
| 10:00 | 5ftc-A | 6 | 3.4 | 112 | 431 | 12 | MOLECULE: | TPR DOMAIN PROTEIN |
| 11:00 | 8hr8-G | 6 | 11.6 | 145 | 830 | 8 | MOLECULE: | ARCHAEAL ATPASE |
| 12:00 | 5g4g-C | 6 | 12.4 | 184 | 721 | 11 | MOLECULE: | VCP-LIKE ATPASE |
| 13:00 | 8hr9-J | 6 | 14.4 | 153 | 835 | 8 | MOLECULE: | ARCHAEAL ATPASE |
| 14:00 | 4fwv-A | 6 | 3.4 | 119 | 340 | 12 | MOLECULE: | TTC1975 PEPTIDASE |
| 15:00 | 5udb-D | 6 | 14.1 | 144 | 447 | 10 | MOLECULE: | DNA REPLICATION LICENSING FACTOR MCM2 |
| 16:00 | 3cmv-C | 6 | 14.5 | 136 | 1175 | 13 | MOLECULE: | PROTEIN RECA |
| 17:00 | 1g8p-A | 6 | 4.3 | 166 | 321 | 12 | MOLECULE: | MAGNESIUM-CHELATASE 38 KDA SUBUNIT |
| 18:00 | 8hrb-Q | 6 | 15 | 140 | 835 | 9 | MOLECULE: | ARCHAEAL ATPASE |
| 19:00 | 4kft-D | 6 | 3.5 | 105 | 194 | 12 | MOLECULE: | GENOME PACKAGING NTPASE B204 |
| 20:00 | 7yux-B | 6 | 14.3 | 203 | 773 | 10 | MOLECULE: | LON PROTEASE |
| 21:00 | 4m9s-B | 6 | 15.6 | 175 | 511 | 14 | MOLECULE: | CELL DEATH PROTEIN 4 |
| 22:00 | 5vhh-D | 6 | 22.7 | 170 | 344 | 11 | MOLECULE: | 26S PROTEASOME REGULATORY SUBUNIT 7 |
| 23:00 | 5axk-B | 5.9 | 5.7 | 120 | 239 | 8 | MOLECULE: | TRNA(HIS)-5'-GUANYLYLTRANSFERASE (THG1) LIKE PROT |
| 24:00:00 | 2is6-A | 5.9 | 16.2 | 130 | 654 | 5 | MOLECULE: | 5'-D(*CP*GP*AP*GP*CP*AP*CP*TP*GP*CP*AP*GP*TP*GP*C |
| 25:00:00 | 8hwc-B | 5.9 | 4.2 | 175 | 320 | 10 | MOLECULE: | PRIMASE D5 |
| 26:00:00 | 8dgf-B | 5.9 | 16.5 | 169 | 1534 | 7 | MOLECULE: | ATP-BINDING PROTEIN AVS4 |
| 27:00:00 | 6oab-A | 5.9 | 10.4 | 143 | 493 | 11 | MOLECULE: | CELL DIVISION CONTROL PROTEIN 48 |
| 28:00:00 | 4fwh-A | 5.9 | 11.2 | 152 | 580 | 9 | MOLECULE: | TTC1975 PEPTIDASE |
| 29:00:00 | 5t0j-C | 5.9 | 21.2 | 161 | 392 | 9 | MOLECULE: | 26S PROTEASOME NON-ATPASE REGULATORY SUBUNIT 2 |
| 30:00:00 | 8hr9-F | 5.9 | 15.2 | 143 | 835 | 9 | MOLECULE: | ARCHAEAL ATPASE |
| 31:00:00 | 7ksl-E | 5.9 | 7.9 | 170 | 502 | 14 | MOLECULE: | LON PROTEASE HOMOLOG, MITOCHONDRIAL |
| 32:00:00 | 5g4g-B | 5.9 | 6 | 166 | 721 | 8 | MOLECULE: | VCP-LIKE ATPASE |
| 33:00:00 | 3ux8-A | 5.9 | 13.4 | 133 | 564 | 13 | MOLECULE: | EXCINUCLEASE ABC, A SUBUNIT |
| 34:00:00 | 5kne-B | 5.9 | 7.8 | 221 | 711 | 10 | MOLECULE: | HEAT SHOCK PROTEIN 104 |
| 35:00:00 | 6igm-B | 5.9 | 8.7 | 137 | 398 | 12 | MOLECULE: | RUVB-LIKE 1 |
| 36:00:00 | 6og2-F | 5.9 | 10.2 | 136 | 558 | 13 | MOLECULE: | HYPERACTIVE DISAGGREGASE CLPB |
| 37:00:00 | 8gj3-D | 5.9 | 10.8 | 148 | 360 | 9 | MOLECULE: | DNA POLYMERASE III SUBUNIT DELTA |
| 38:00:00 | 8emc-F | 5.9 | 21.5 | 212 | 666 | 9 | MOLECULE: | PROTEASE LON-RELATED BREX SYSTEM PROTEIN BRXL |
| 39:00:00 | 4pht-B | 5.9 | 7.2 | 171 | 442 | 8 | MOLECULE: | TYPE II SECRETION SYSTEM PROTEIN L |
| 40:00:00 | 7ykl-F | 5.9 | 9.5 | 200 | 735 | 10 | MOLECULE: | ATPASE FAMILY GENE 2 PROTEIN |
| 41:00:00 | 4cr4-K | 5.9 | 12.7 | 162 | 381 | 12 | MOLECULE: | PROTEASOME COMPONENT PRE3 |
| 42:00:00 | 8fcw-U | 5.9 | 6.6 | 145 | 343 | 9 | MOLECULE: | TNSC |
| 43:00:00 | 3tuz-C | 5.8 | 10.5 | 135 | 345 | 12 | MOLECULE: | D-METHIONINE TRANSPORT SYSTEM PERMEASE PROTEIN ME |
| 44:00:00 | 3cmu-A | 5.8 | 15.7 | 138 | 1937 | 12 | MOLECULE: | DNA (5'- |
| 45:00:00 | 5udb-4 | 5.8 | 10.1 | 227 | 751 | 14 | MOLECULE: | DNA REPLICATION LICENSING FACTOR MCM2 |
| 46:00:00 | 6acx-A | 5.8 | 7.6 | 171 | 1175 | 10 | MOLECULE: | MYCOBACTERIUM SMEGMATIS MFD |
| 47:00:00 | 8dgf-D | 5.8 | 19 | 165 | 1534 | 6 | MOLECULE: | ATP-BINDING PROTEIN AVS4 |
| 48:00:00 | 1jqj-C | 5.8 | 15.9 | 129 | 328 | 12 | MOLECULE: | DNA POLYMERASE III, BETA CHAIN |
| 49:00:00 | 4xgc-C | 5.8 | 13.5 | 159 | 567 | 6 | MOLECULE: | ORIGIN RECOGNITION COMPLEX SUBUNIT 2 |
| 50:00:00 | 6epc-K | 5.8 | 25 | 159 | 391 | 11 | MOLECULE: | PROTEASOME SUBUNIT ALPHA TYPE-6 |
| 51:00:00 | 4wia-A | 5.8 | 3.5 | 116 | 229 | 10 | MOLECULE: | PUTATIVE FLAGELLA-RELATED PROTEIN H |
| 52:00:00 | 6wqh-D | 5.8 | 13.2 | 179 | 537 | 13 | MOLECULE: | LON PROTEASE |
| 53:00:00 | 1jql-B | 5.8 | 3.2 | 105 | 140 | 12 | MOLECULE: | DNA POLYMERASE III, BETA CHAIN |
| 54:00:00 | 6ef1-J | 5.8 | 9.2 | 136 | 273 | 9 | MOLECULE: | PROTEASOME SUBUNIT ALPHA TYPE-1 |
| 55:00:00 | 4kfu-D | 5.8 | 3.3 | 106 | 195 | 12 | MOLECULE: | GENOME PACKAGING NTPASE B204 |
| 56:00:00 | 8fnu-D | 5.8 | 14.1 | 138 | 796 | 9 | MOLECULE: | KAP NTPASE DOMAIN-CONTAINING PROTEIN |
| 57:00:00 | 8fnu-G | 5.8 | 15.8 | 134 | 796 | 7 | MOLECULE: | KAP NTPASE DOMAIN-CONTAINING PROTEIN |
| 58:00:00 | 7ykk-F | 5.8 | 7.3 | 215 | 735 | 12 | MOLECULE: | ATPASE FAMILY GENE 2 PROTEIN |
| 59:00:00 | 4kfu-A | 5.8 | 3.2 | 105 | 205 | 12 | MOLECULE: | GENOME PACKAGING NTPASE B204 |

| | | | | | | | |
|---|---|---|---|---|---|---|---|
| 60:00:00 | 8osf-D | 5.8 | 4.4 | 163 | 319 | 9 MOLECULE: | MAGNESIUM-CHELATASE SUBUNIT CHLI |
| 61:00:00 | 6ef1-M | 5.8 | 10.8 | 139 | 262 | 15 MOLECULE: | PROTEASOME SUBUNIT ALPHA TYPE-1 |
| 62:00:00 | 5vhq-C | 5.8 | 8.3 | 131 | 234 | 8 MOLECULE: | 26S PROTEASOME NON-ATPASE REGULATORY SUBUNIT 10 |
| 63:00:00 | 8faz-X | 5.8 | 3.3 | 112 | 236 | 8 MOLECULE: | DNA REPAIR PROTEIN RAD51 HOMOLOG 2 |
| 64:00:00 | 7ufi-6 | 5.8 | 10.3 | 147 | 311 | 10 MOLECULE: | DNA (5'- |
| 65:00:00 | 7ufi-3 | 5.8 | 10.3 | 146 | 311 | 10 MOLECULE: | DNA (5'- |
| 66:00:00 | 4a15-A | 5.7 | 10.8 | 118 | 596 | 11 MOLECULE: | ATP-DEPENDENT DNA HELICASE TA0057 |
| 67:00:00 | 8hra-C | 5.7 | 11.2 | 145 | 852 | 8 MOLECULE: | ARCHAEAL ATPASE |
| 68:00:00 | 7b8s-C | 5.7 | 4.3 | 103 | 589 | 4 MOLECULE: | MULTIDRUG EFFLUX PUMP SUBUNIT ACRB,MULTIDRUG EFFL |
| 69:00:00 | 8oz7-A | 5.7 | 22.5 | 135 | 597 | 10 MOLECULE: | ABIA |
| 70:00:00 | 3zh9-B | 5.7 | 4 | 118 | 339 | 11 MOLECULE: | DELTA |
| 71:00:00 | 4xgc-B | 5.7 | 3.3 | 114 | 281 | 12 MOLECULE: | ORIGIN RECOGNITION COMPLEX SUBUNIT 2 |
| 72:00:00 | 5vhr-A | 5.7 | 6.1 | 133 | 258 | 9 MOLECULE: | 26S PROTEASOME NON-ATPASE REGULATORY SUBUNIT 10 |
| 73:00:00 | 5wc0-B | 5.7 | 11.1 | 136 | 277 | 13 MOLECULE: | MEIOTIC SPINDLE FORMATION PROTEIN MEI-1 |
| 74:00:00 | 3cmw-C | 5.7 | 15.3 | 144 | 1608 | 11 MOLECULE: | DNA (5'- |
| 75:00:00 | 4yds-A | 5.7 | 3.4 | 120 | 226 | 12 MOLECULE: | FLAGELLA-RELATED PROTEIN H |
| 76:00:00 | 4kfr-B | 5.7 | 3.5 | 111 | 208 | 13 MOLECULE: | GENOME PACKAGING NTPASE B204 |
| 77:00:00 | 4ypl-E | 5.7 | 10.6 | 188 | 537 | 13 MOLECULE: | LON PROTEASE |
| 78:00:00 | 6rn4-E | 5.7 | 13.3 | 175 | 684 | 11 MOLECULE: | CHAPERONE PROTEIN CLPB |
| 79:00:00 | 6mfv-C | 5.7 | 8.2 | 131 | 641 | 11 MOLECULE: | TETRATRICOPEPTIDE REPEAT SENSOR PH0952 |
| 80:00:00 | 5wc0-A | 5.7 | 11.6 | 134 | 275 | 11 MOLECULE: | MEIOTIC SPINDLE FORMATION PROTEIN MEI-1 |
| 81:00:00 | 8fnu-F | 5.7 | 13.7 | 133 | 796 | 6 MOLECULE: | KAP NTPASE DOMAIN-CONTAINING PROTEIN |
| 82:00:00 | 8fnu-B | 5.7 | 13.8 | 136 | 796 | 8 MOLECULE: | KAP NTPASE DOMAIN-CONTAINING PROTEIN |
| 83:00:00 | 8fnu-C | 5.7 | 15.2 | 138 | 796 | 7 MOLECULE: | KAP NTPASE DOMAIN-CONTAINING PROTEIN |
| 84:00:00 | 6ef0-M | 5.7 | 11.9 | 133 | 258 | 14 MOLECULE: | PROTEASOME SUBUNIT ALPHA TYPE-1 |
| 85:00:00 | 7mbw-B | 5.7 | 13.5 | 159 | 469 | 12 MOLECULE: | TRANSPOSON TN7 TRANSPOSITION PROTEIN TNSC |
| 86:00:00 | 4r2i-A | 5.7 | 3 | 103 | 207 | 13 MOLECULE: | STIV B204 ATPASE |
| 87:00:00 | 1nlf-A | 5.7 | 6.4 | 127 | 254 | 12 MOLECULE: | REGULATORY PROTEIN REPA |
| 88:00:00 | 1f3o-A | 5.7 | 3.1 | 112 | 232 | 10 MOLECULE: | HYPOTHETICAL ABC TRANSPORTER ATP-BINDING PROTEIN |
| 89:00:00 | 4kfs-A | 5.7 | 3.4 | 107 | 202 | 12 MOLECULE: | GENOME PACKAGING NTPASE B204 |
| 90:00:00 | 7rzy-1 | 5.7 | 6.9 | 132 | 311 | 9 MOLECULE: | TN6677 VIBRIO CHOLERAE TRANSPOSON TNSC (VCHTNSC) |
| 91:00:00 | 1ewq-A | 5.7 | 31.9 | 191 | 759 | 9 MOLECULE: | DNA (5'- |
| 92:00:00 | 7z6h-K | 5.7 | 14.2 | 140 | 439 | 9 MOLECULE: | CELL CYCLE CHECKPOINT CONTROL PROTEIN RAD9A |
| 93:00:00 | 7mcs-A | 5.7 | 3.7 | 128 | 249 | 11 MOLECULE: | TRANSPOSON TN7 TRANSPOSITION PROTEIN TNSC |
| 94:00:00 | 7mqj-A | 5.6 | 9.4 | 140 | 773 | 9 MOLECULE: | PROBABLE ATP-DEPENDENT RNA HELICASE DHR1 |
| 95:00:00 | 8fvu-A | 5.6 | 11.7 | 195 | 1361 | 8 MOLECULE: | BACULOVIRAL IAP REPEAT-CONTAINING PROTEIN 1 |
| 96:00:00 | 5gko-A | 5.6 | 7 | 135 | 650 | 13 MOLECULE: | MACROLIDE EXPORT ATP-BINDING/PERMEASE PROTEIN MAC |
| 97:00:00 | 6yxu-H | 5.6 | 14.4 | 149 | 691 | 8 MOLECULE: | DNA-DIRECTED RNA POLYMERASE SUBUNIT ALPHA |
| 98:00:00 | 6ef0-I | 5.6 | 7 | 128 | 271 | 13 MOLECULE: | PROTEASOME SUBUNIT ALPHA TYPE-1 |
| 99:00:00 | 3cmw-A | 5.6 | 15.6 | 140 | 1609 | 13 MOLECULE: | DNA (5'- |
| 0:00 | 5vhh-B | 5.6 | 9.7 | 158 | 330 | 9 MOLECULE: | 26S PROTEASOME REGULATORY SUBUNIT 7 |
| 1:00 | 3kx2-A | 5.6 | 21.8 | 164 | 755 | 10 MOLECULE: | PRE-MRNA-SPLICING FACTOR ATP-DEPENDENT RNA |
| 2:00 | 7mcs-F | 5.6 | 3.6 | 136 | 401 | 12 MOLECULE: | TRANSPOSON TN7 TRANSPOSITION PROTEIN TNSC |
| 3:00 | 5vhi-F | 5.6 | 8.8 | 156 | 351 | 13 MOLECULE: | 26S PROTEASOME REGULATORY SUBUNIT 7 |
| 4:00 | 5og1-F | 5.6 | 5.4 | 187 | 686 | 14 MOLECULE: | CHAPERONE PROTEIN CLPB,ATP-DEPENDENT CLP PROTEASE |
| 5:00 | 8hr8-A | 5.6 | 14.8 | 140 | 836 | 8 MOLECULE: | ARCHAEAL ATPASE |
| 6:00 | 2fna-A | 5.6 | 12.1 | 132 | 352 | 11 MOLECULE: | CONSERVED HYPOTHETICAL PROTEIN |
| 7:00 | 5a5b-K | 5.6 | 13.3 | 165 | 381 | 12 MOLECULE: | PROTEASOME COMPONENT PRE3 |
| 8:00 | 8hrb-P | 5.6 | 17 | 138 | 834 | 9 MOLECULE: | ARCHAEAL ATPASE |
| 9:00 | 4kfu-B | 5.6 | 3.4 | 105 | 196 | 12 MOLECULE: | GENOME PACKAGING NTPASE B204 |
| 10:00 | 5ylz-W | 5.6 | 27.4 | 149 | 703 | 9 MOLECULE: | PRE-MRNA-SPLICING FACTOR 8 |
| 11:00 | 7wi4-D | 5.6 | 11.1 | 128 | 410 | 13 MOLECULE: | ATP-DEPENDENT ZINC METALLOPROTEASE FTSH |
| 12:00 | 7qh2-C | 5.5 | 20.2 | 130 | 467 | 7 MOLECULE: | LACTATE DEHYDROGENASE (NAD(+),FERREDOXIN) SUBUNIT |

| | | | | | | | | |
|---|---|---|---|---|---|---|---|---|
| 13:00 | 8hwc-E | 5.5 | 4.3 | 160 | 284 | 11 | MOLECULE: | PRIMASE D5 |
| 14:00 | 8giy-A | 5.5 | 14 | 141 | 343 | 11 | MOLECULE: | DNA POLYMERASE III SUBUNIT DELTA |
| 15:00 | 8gbj-X | 5.5 | 3.4 | 116 | 232 | 9 | MOLECULE: | DNA REPAIR PROTEIN RAD51 HOMOLOG 2 |
| 16:00 | 5udb-B | 5.5 | 18.9 | 128 | 319 | 10 | MOLECULE: | DNA REPLICATION LICENSING FACTOR MCM2 |
| 17:00 | 8emh-G | 5.5 | 5.3 | 204 | 671 | 8 | MOLECULE: | PROTEASE LON-RELATED BREX SYSTEM PROTEIN BRXL |
| 18:00 | 7f02-E | 5.5 | 3.2 | 97 | 202 | 9 | MOLECULE: | CYTOCHROME C BIOGENESIS ATP-BINDING EXPORT PROTEI |
| 19:00 | 2d7d-A | 5.5 | 17.4 | 139 | 621 | 10 | MOLECULE: | 5'-D(P*TP*TP*T)-3' |
| 20:00 | 8osh-B | 5.5 | 4.4 | 163 | 317 | 13 | MOLECULE: | MAGNESIUM-CHELATASE SUBUNIT CHLI |
| 21:00 | 6eud-A | 5.5 | 14 | 133 | 808 | 13 | MOLECULE: | ATP-DEPENDENT RNA HELICASE HRPB |
| 22:00 | 5vca-O | 5.5 | 10.1 | 151 | 543 | 9 | MOLECULE: | VCP-LIKE ATPASE |
| 23:00 | 8hra-B | 5.5 | 16.5 | 134 | 835 | 9 | MOLECULE: | ARCHAEAL ATPASE |
| 24:00:00 | 4cr3-K | 5.5 | 13.4 | 162 | 381 | 14 | MOLECULE: | PROTEASOME COMPONENT PRE3 |
| 25:00:00 | 5kzf-K | 5.5 | 12.6 | 167 | 498 | 11 | MOLECULE: | PROTEASOME-ASSOCIATED ATPASE |
| 26:00:00 | 6ef1-I | 5.5 | 6 | 126 | 271 | 13 | MOLECULE: | PROTEASOME SUBUNIT ALPHA TYPE-1 |
| 27:00:00 | 8ouy-D | 5.5 | 3.4 | 117 | 234 | 9 | MOLECULE: | DNA REPAIR PROTEIN RAD51 HOMOLOG 2 |
| 28:00:00 | 8ouz-D | 5.5 | 3.3 | 119 | 234 | 8 | MOLECULE: | DNA REPAIR PROTEIN RAD51 HOMOLOG 2 |
| 29:00:00 | 6oay-F | 5.5 | 12.3 | 139 | 558 | 14 | MOLECULE: | HYPERACTIVE DISAGGREGASE CLPB |
| 30:00:00 | 8c0w-D | 5.5 | 18.3 | 193 | 821 | 15 | MOLECULE: | PEROXISOMAL ATPASE PEX6 |
| 31:00:00 | 8osh-D | 5.5 | 4.7 | 147 | 230 | 7 | MOLECULE: | MAGNESIUM-CHELATASE SUBUNIT CHLI |
| 32:00:00 | 4cr4-J | 5.5 | 24.3 | 162 | 373 | 10 | MOLECULE: | PROTEASOME COMPONENT PRE3 |
| 33:00:00 | 2is4-B | 5.4 | 14.6 | 124 | 632 | 6 | MOLECULE: | 25-MER |
| 34:00:00 | 5udb-9 | 5.4 | 15.7 | 152 | 376 | 14 | MOLECULE: | DNA REPLICATION LICENSING FACTOR MCM2 |
| 35:00:00 | 2x31-K | 5.4 | 4.9 | 165 | 321 | 11 | MOLECULE: | MAGNESIUM-CHELATASE 60 KDA SUBUNIT |
| 36:00:00 | 7mcs-E | 5.4 | 3.6 | 140 | 402 | 11 | MOLECULE: | TRANSPOSON TN7 TRANSPOSITION PROTEIN TNSC |
| 37:00:00 | 8v45-F | 5.4 | 3.6 | 112 | 378 | 15 | MOLECULE: | ARIA ANTITOXIN |
| 38:00:00 | 5oea-E | 5.4 | 18.9 | 129 | 408 | 9 | MOLECULE: | LARGE SUBUNIT TERMINASE |
| 39:00:00 | 6n8v-F | 5.4 | 22.7 | 244 | 710 | 9 | MOLECULE: | HEAT SHOCK PROTEIN 104 |
| 40:00:00 | 8hr8-F | 5.4 | 14 | 139 | 835 | 9 | MOLECULE: | ARCHAEAL ATPASE |
| 41:00:00 | 1d2m-A | 5.4 | 4.4 | 120 | 552 | 10 | MOLECULE: | EXCINUCLEASE ABC SUBUNIT B |
| 42:00:00 | 2x31-I | 5.4 | 4.5 | 164 | 321 | 11 | MOLECULE: | MAGNESIUM-CHELATASE 60 KDA SUBUNIT |
| 43:00:00 | 8ff4-W | 5.4 | 7.2 | 145 | 343 | 7 | MOLECULE: | TYPE I-B CRISPR-ASSOCIATED PROTEIN CAS5 |
| 44:00:00 | 4m4w-O | 5.4 | 12.7 | 135 | 250 | 13 | MOLECULE: | REPLICATIVE HELICASE |
| 45:00:00 | 5vhm-E | 5.4 | 9.2 | 116 | 209 | 12 | MOLECULE: | 26S PROTEASOME NON-ATPASE REGULATORY SUBUNIT 10 |
| 46:00:00 | 5vlj-A | 5.4 | 18.5 | 180 | 2376 | 9 | MOLECULE: | DYNEIN HEAVY CHAIN, CYTOPLASMIC |
| 47:00:00 | 7dex-A | 5.3 | 7.2 | 93 | 460 | 5 | MOLECULE: | ANTHOCYANIN 5-AROMATIC ACYLTRANSFERASE |
| 48:00:00 | 4buj-A | 5.3 | 18.6 | 151 | 903 | 9 | MOLECULE: | SUPERKILLER PROTEIN 3 |
| 49:00:00 | 3gp8-A | 5.3 | 9.3 | 128 | 551 | 12 | MOLECULE: | EXODEOXYRIBONUCLEASE V, SUBUNIT RECD, PUTATIVE |
| 50:00:00 | 3crv-A | 5.3 | 13.7 | 122 | 551 | 11 | MOLECULE: | XPD/RAD3 RELATED DNA HELICASE |
| 51:00:00 | 7epu-B | 5.3 | 15.7 | 153 | 748 | 8 | MOLECULE: | NON-IMMUNIZED HUMAN SCFV |
| 52:00:00 | 8th8-L | 5.3 | 11.3 | 175 | 862 | 7 | MOLECULE: | DYNEIN REGULATORY COMPLEX PROTEIN 1/2 N-TERMINAL |
| 53:00:00 | 1ng9-A | 5.3 | 15.7 | 184 | 794 | 8 | MOLECULE: | 5'-D(*AP*GP*CP*TP*GP*CP*CP*AP*GP*GP*CP*AP*CP*CP*A |
| 54:00:00 | 4fw9-A | 5.3 | 11.7 | 156 | 602 | 5 | MOLECULE: | TTC1975 PEPTIDASE |
| 55:00:00 | 8ag6-B | 5.3 | 31.1 | 169 | 975 | 8 | MOLECULE: | DNA MISMATCH REPAIR PROTEIN MSH2 |
| 56:00:00 | 4bj1-A | 5.3 | 6.2 | 139 | 261 | 8 | MOLECULE: | PROTEIN RIF2 |
| 57:00:00 | 7rzy-3 | 5.3 | 9.7 | 145 | 311 | 8 | MOLECULE: | TN6677 VIBRIO CHOLERAE TRANSPOSON TNSC (VCHTNSC) |
| 58:00:00 | 7wi4-A | 5.3 | 11.5 | 129 | 410 | 13 | MOLECULE: | ATP-DEPENDENT ZINC METALLOPROTEASE FTSH |
| 59:00:00 | 3n70-B | 5.3 | 2.9 | 89 | 108 | 7 | MOLECULE: | TRANSPORT ACTIVATOR |
| 60:00:00 | 5jji-C | 5.3 | 5.5 | 174 | 417 | 14 | MOLECULE: | TRANSCRIPTION TERMINATION FACTOR RHO |
| 61:00:00 | 6j17-A | 5.3 | 3.9 | 116 | 254 | 16 | MOLECULE: | ESX-3 SECRETION SYSTEM PROTEIN ECCC3 |
| 62:00:00 | 5vhj-A | 5.3 | 3.9 | 122 | 212 | 11 | MOLECULE: | 26S PROTEASOME NON-ATPASE REGULATORY SUBUNIT 10 |
| 63:00:00 | 5vca-Q | 5.3 | 9 | 144 | 543 | 12 | MOLECULE: | VCP-LIKE ATPASE |
| 64:00:00 | 7mcs-D | 5.3 | 3.7 | 143 | 403 | 11 | MOLECULE: | TRANSPOSON TN7 TRANSPOSITION PROTEIN TNSC |
| 65:00:00 | 6ahf-F | 5.3 | 11.7 | 209 | 574 | 9 | MOLECULE: | HEAT SHOCK PROTEIN 104 |

| | | | | | | | | |
|---|---|---|---|---|---|---|---|---|
| 66:00:00 | 7mcs-C | 5.3 | 3.6 | 137 | 404 | 12 | MOLECULE: | TRANSPOSON TN7 TRANSPOSITION PROTEIN TNSC |
| 67:00:00 | 8hr8-D | 5.3 | 12.7 | 147 | 836 | 7 | MOLECULE: | ARCHAEAL ATPASE |
| 68:00:00 | 8emc-C | 5.3 | 7.6 | 180 | 668 | 9 | MOLECULE: | PROTEASE LON-RELATED BREX SYSTEM PROTEIN BRXL |
| 69:00:00 | 1ye8-A | 5.3 | 3.5 | 100 | 172 | 10 | MOLECULE: | HYPOTHETICAL UPF0334 KINASE-LIKE PROTEIN AQ_1292 |
| 70:00:00 | 7yut-F | 5.3 | 26.2 | 190 | 773 | 12 | MOLECULE: | LON PROTEASE |
| 71:00:00 | 8hra-H | 5.3 | 15.4 | 133 | 834 | 8 | MOLECULE: | ARCHAEAL ATPASE |
| 72:00:00 | 4nph-A | 5.3 | 4.6 | 142 | 324 | 13 | MOLECULE: | PROBABLE SECRETION SYSTEM APPARATUS ATP SYNTHASE |
| 73:00:00 | 8dgf-C | 5.3 | 16.7 | 180 | 1541 | 5 | MOLECULE: | ATP-BINDING PROTEIN AVS4 |
| 74:00:00 | 5vho-F | 5.3 | 8.8 | 135 | 238 | 14 | MOLECULE: | 26S PROTEASOME NON-ATPASE REGULATORY SUBUNIT 10 |
| 75:00:00 | 8c0v-B | 5.3 | 26.3 | 173 | 1030 | 8 | MOLECULE: | PEROXISOMAL ATPASE PEX1 |
| 76:00:00 | 8dgf-A | 5.2 | 14.7 | 158 | 1541 | 7 | MOLECULE: | ATP-BINDING PROTEIN AVS4 |
| 77:00:00 | 7njn-C | 5.2 | 4.8 | 175 | 536 | 6 | MOLECULE: | ATP SYNTHASE SUBUNIT ALPHA |
| 78:00:00 | 2xgj-B | 5.2 | 7.8 | 131 | 773 | 7 | MOLECULE: | ATP-DEPENDENT RNA HELICASE DOB1 |
| 79:00:00 | 6uen-A | 5.2 | 11 | 164 | 1392 | 6 | MOLECULE: | RNA-DIRECTED RNA POLYMERASE L |
| 80:00:00 | 3pxi-C | 5.2 | 21.7 | 153 | 711 | 9 | MOLECULE: | ADAPTER PROTEIN MECA 1 |
| 81:00:00 | 5vhq-A | 5.2 | 9.5 | 137 | 261 | 9 | MOLECULE: | 26S PROTEASOME NON-ATPASE REGULATORY SUBUNIT 10 |
| 82:00:00 | 5vhp-E | 5.2 | 12.3 | 123 | 209 | 11 | MOLECULE: | 26S PROTEASOME NON-ATPASE REGULATORY SUBUNIT 10 |
| 83:00:00 | 5vhf-E | 5.2 | 13.9 | 164 | 345 | 9 | MOLECULE: | 26S PROTEASOME NON-ATPASE REGULATORY SUBUNIT 10 |
| 84:00:00 | 4kfs-B | 5.2 | 3.3 | 103 | 199 | 9 | MOLECULE: | GENOME PACKAGING NTPASE B204 |
| 85:00:00 | 6em8-D | 5.2 | 17.5 | 216 | 436 | 14 | MOLECULE: | ATP-DEPENDENT CLP PROTEASE ATP-BINDING SUBUNIT CL |
| 86:00:00 | 5oea-F | 5.2 | 18.7 | 130 | 406 | 7 | MOLECULE: | LARGE SUBUNIT TERMINASE |
| 87:00:00 | 7wrx-E | 5.2 | 16.6 | 147 | 590 | 7 | MOLECULE: | HERA |
| 88:00:00 | 8jqb-E | 5.1 | 5.6 | 115 | 499 | 8 | MOLECULE: | ENDONUCLEASE GAJA |
| 89:00:00 | 3wbz-E | 5.1 | 7.4 | 115 | 268 | 8 | MOLECULE: | LIKELY HISTIDYL TRNA-SPECIFIC GUANYLYLTRANSFERASE |
| 90:00:00 | 2oca-A | 5.1 | 8.6 | 115 | 494 | 6 | MOLECULE: | ATP-DEPENDENT DNA HELICASE UVSW |
| 91:00:00 | 8hwd-D | 5.1 | 3.7 | 157 | 299 | 10 | MOLECULE: | PRIMASE D5 |
| 92:00:00 | 5aj3-g | 5.1 | 4.3 | 123 | 346 | 12 | MOLECULE: | MITORIBOSOMAL 12S RRNA |
| 93:00:00 | 7mbw-C | 5.1 | 15 | 147 | 461 | 9 | MOLECULE: | TRANSPOSON TN7 TRANSPOSITION PROTEIN TNSC |
| 94:00:00 | 7cg3-E | 5.1 | 14.1 | 161 | 686 | 9 | MOLECULE: | HEAT SHOCK PROTEIN 104 |
| 95:00:00 | 5b0o-B | 5.1 | 11.7 | 172 | 455 | 9 | MOLECULE: | FLAGELLUM-SPECIFIC ATP SYNTHASE |
| 96:00:00 | 5t0j-B | 5.1 | 13.1 | 156 | 348 | 12 | MOLECULE: | 26S PROTEASOME NON-ATPASE REGULATORY SUBUNIT 2 |
| 97:00:00 | 7mcs-B | 5.1 | 3.6 | 138 | 402 | 11 | MOLECULE: | TRANSPOSON TN7 TRANSPOSITION PROTEIN TNSC |
| 98:00:00 | 5wcb-E | 5.1 | 9.8 | 130 | 277 | 10 | MOLECULE: | MEIOTIC SPINDLE FORMATION PROTEIN MEI-1 |
| 99:00:00 | 4q47-A | 5.1 | 5.5 | 133 | 513 | 8 | MOLECULE: | DNA HELICASE RECQ |
| 0:00 | 5ujm-C | 5.1 | 7.3 | 157 | 553 | 7 | MOLECULE: | ORIGIN RECOGNITION COMPLEX SUBUNIT 1 |
| 1:00 | 1gm5-A | 5 | 17.5 | 163 | 729 | 10 | MOLECULE: | RECG |
| 2:00 | 4cej-B | 5 | 17.6 | 141 | 1156 | 11 | MOLECULE: | ATP-DEPENDENT HELICASE/NUCLEASE SUBUNIT A |
| 3:00 | 6fws-B | 5 | 15.9 | 130 | 683 | 12 | MOLECULE: | ATP-DEPENDENT DNA HELICASE DING |
| 4:00 | 5g4g-E | 5 | 5.7 | 165 | 721 | 12 | MOLECULE: | VCP-LIKE ATPASE |
| 5:00 | 2x31-G | 5 | 4.3 | 156 | 321 | 11 | MOLECULE: | MAGNESIUM-CHELATASE 60 KDA SUBUNIT |
| 6:00 | 7ng5-B | 5 | 35.2 | 223 | 776 | 9 | MOLECULE: | LON PROTEASE HOMOLOG, MITOCHONDRIAL |
| 7:00 | 5t0i-B | 5 | 7.8 | 143 | 348 | 14 | MOLECULE: | 26S PROTEASE REGULATORY SUBUNIT 7 |
| 8:00 | 7wi4-C | 5 | 11.4 | 134 | 410 | 13 | MOLECULE: | ATP-DEPENDENT ZINC METALLOPROTEASE FTSH |
| 9:00 | 6ppj-B | 5 | 19.8 | 120 | 848 | 11 | MOLECULE: | UVRD/REP HELICASE |
| 10:00 | 5f9f-I | 5 | 4.5 | 113 | 674 | 9 | MOLECULE: | PROBABLE ATP-DEPENDENT RNA HELICASE DDX58 |
| 11:00 | 6npy-A | 5 | 12.1 | 145 | 798 | 11 | MOLECULE: | NACHT, LRR AND PYD DOMAINS-CONTAINING PROTEIN 3 |
| 12:00 | 1jnv-A | 5 | 10.1 | 165 | 487 | 0 | MOLECULE: | ATP SYNTHASE ALPHA CHAIN |
| 13:00 | 4bj5-B | 5 | 5.6 | 137 | 291 | 7 | MOLECULE: | PROTEIN RIF2 |
| 14:00 | 4bj5-A | 5 | 5.7 | 136 | 292 | 9 | MOLECULE: | PROTEIN RIF2 |
| 15:00 | 6p74-A | 4.9 | 10.1 | 134 | 512 | 14 | MOLECULE: | PUTATIVE ATP-DEPENDENT ENDONUCLEASE OF THE OLD FA |
| 16:00 | 5x71-A | 4.9 | 8.3 | 143 | 796 | 7 | MOLECULE: | MRNA CAPPING ENZYME P5 |
| 17:00 | 4ihq-A | 4.9 | 21 | 187 | 512 | 7 | MOLECULE: | FLAI ATPASE |
| 18:00 | 4gwp-F | 4.9 | 3.8 | 85 | 209 | 8 | MOLECULE: | MEDIATOR OF RNA POLYMERASE II TRANSCRIPTION SUBUN |

| | | | | | | | | |
|---|---|---|---|---|---|---|---|---|
| 19:00 | 6tdz-C | 4.9 | 12.4 | 183 | 530 | 8 | MOLECULE: | OLIGOMYCIN SENSITIVITY CONFERRING PROTEIN (OSCP) |
| 20:00 | 5zxd-B | 4.9 | 4.6 | 117 | 491 | 13 | MOLECULE: | ATP-BINDING CASSETTE SUB-FAMILY F MEMBER 1 |
| 21:00 | 5o6b-A | 4.9 | 3.4 | 114 | 528 | 13 | MOLECULE: | ATP-DEPENDENT DNA HELICASE PIF1 |
| 22:00 | 4ag6-A | 4.9 | 3.8 | 114 | 373 | 9 | MOLECULE: | TYPE IV SECRETORY PATHWAY VIRB4 COMPONENTS-LIKE P |
| 23:00 | 8dgc-B | 4.9 | 12.8 | 161 | 2004 | 10 | MOLECULE: | SEAVS3 |
| 24:00:00 | 7amv-W | 4.9 | 9.5 | 140 | 637 | 11 | MOLECULE: | DNA-DIRECTED RNA POLYMERASE 147 KDA POLYPEPTIDE |
| 25:00:00 | 7ng4-C | 4.9 | 23.2 | 204 | 776 | 10 | MOLECULE: | LON PROTEASE HOMOLOG, MITOCHONDRIAL |
| 26:00:00 | 7ngl-A | 4.9 | 30.7 | 206 | 776 | 10 | MOLECULE: | LON PROTEASE HOMOLOG, MITOCHONDRIAL |
| 27:00:00 | 5wcb-B | 4.9 | 11 | 130 | 277 | 12 | MOLECULE: | MEIOTIC SPINDLE FORMATION PROTEIN MEI-1 |
| 28:00:00 | 8hra-G | 4.9 | 14.8 | 142 | 835 | 10 | MOLECULE: | ARCHAEAL ATPASE |
| 29:00:00 | 5uj8-C | 4.9 | 12.3 | 167 | 553 | 6 | MOLECULE: | ORIGIN RECOGNITION COMPLEX SUBUNIT 3 |
| 30:00:00 | 5vhq-D | 4.9 | 9 | 121 | 236 | 13 | MOLECULE: | 26S PROTEASOME NON-ATPASE REGULATORY SUBUNIT 10 |
| 31:00:00 | 7wi4-E | 4.9 | 10.2 | 141 | 410 | 11 | MOLECULE: | ATP-DEPENDENT ZINC METALLOPROTEASE FTSH |
| 32:00:00 | 2fdc-A | 4.9 | 4.4 | 123 | 505 | 7 | MOLECULE: | 5'-D(P*CP*GP*GP*CP*TP*CP*CP*AP*TP*CP*TP*CP*TP*AP* |
| 33:00:00 | 1ji0-A | 4.9 | 4 | 115 | 240 | 11 | MOLECULE: | ABC TRANSPORTER |
| 34:00:00 | 5wcb-D | 4.9 | 10.9 | 126 | 277 | 10 | MOLECULE: | MEIOTIC SPINDLE FORMATION PROTEIN MEI-1 |
| 35:00:00 | 5vhn-B | 4.9 | 3.3 | 125 | 257 | 14 | MOLECULE: | 26S PROTEASOME NON-ATPASE REGULATORY SUBUNIT 10 |
| 36:00:00 | 1u98-A | 4.9 | 8.4 | 131 | 311 | 11 | MOLECULE: | RECA PROTEIN |
| 37:00:00 | 5w99-A | 4.8 | 5.9 | 108 | 319 | 6 | MOLECULE: | PBTD |
| 38:00:00 | 1wp9-A | 4.8 | 14.1 | 139 | 479 | 10 | MOLECULE: | ATP-DEPENDENT RNA HELICASE, PUTATIVE |
| 39:00:00 | 7jr7-A | 4.8 | 8.7 | 148 | 588 | 11 | MOLECULE: | ATP-BINDING CASSETTE SUB-FAMILY G MEMBER 5 |
| 40:00:00 | 8hwb-D | 4.8 | 16.9 | 117 | 700 | 16 | MOLECULE: | PRIMASE D5 |
| 41:00:00 | 4ag5-A | 4.8 | 3.7 | 116 | 374 | 11 | MOLECULE: | TYPE IV SECRETORY PATHWAY VIRB4 COMPONENTS-LIKE P |
| 42:00:00 | 4ag6-B | 4.8 | 3.8 | 116 | 376 | 9 | MOLECULE: | TYPE IV SECRETORY PATHWAY VIRB4 COMPONENTS-LIKE P |
| 43:00:00 | 4xgc-D | 4.8 | 16.7 | 157 | 440 | 10 | MOLECULE: | ORIGIN RECOGNITION COMPLEX SUBUNIT 2 |
| 44:00:00 | 5vhs-E | 4.8 | 12.6 | 156 | 313 | 11 | MOLECULE: | 26S PROTEASOME REGULATORY SUBUNIT 7 |
| 45:00:00 | 7mbw-D | 4.8 | 11.3 | 145 | 467 | 12 | MOLECULE: | TRANSPOSON TN7 TRANSPOSITION PROTEIN TNSC |
| 46:00:00 | 4d2i-A | 4.8 | 8.5 | 142 | 464 | 8 | MOLECULE: | HERA |
| 47:00:00 | 5g4g-A | 4.8 | 9.1 | 168 | 721 | 11 | MOLECULE: | VCP-LIKE ATPASE |
| 48:00:00 | 5vhh-E | 4.8 | 13.3 | 163 | 345 | 9 | MOLECULE: | 26S PROTEASOME REGULATORY SUBUNIT 7 |
| 49:00:00 | 5lb3-B | 4.8 | 11 | 138 | 439 | 8 | MOLECULE: | ATP-DEPENDENT DNA HELICASE Q5 |
| 50:00:00 | 8d8k-W | 4.8 | 5.4 | 121 | 395 | 9 | MOLECULE: | PROBABLE S-ADENOSYL-L-METHIONINE-DEPENDENT RNA |
| 51:00:00 | 5xc6-A | 4.8 | 3.8 | 95 | 451 | 8 | MOLECULE: | NS3 HELICASE |
| 52:00:00 | 1sky-B | 4.8 | 11.2 | 159 | 479 | 11 | MOLECULE: | F1-ATPASE |
| 53:00:00 | 3pxg-D | 4.8 | 27.3 | 146 | 434 | 11 | MOLECULE: | ADAPTER PROTEIN MECA 1 |
| 54:00:00 | 1xpo-A | 4.8 | 5.9 | 170 | 408 | 11 | MOLECULE: | 5'-R(*CP*UP*CP*UP*CP*UP*CP*U)-3' |
| 55:00:00 | 3pxg-E | 4.8 | 23.7 | 141 | 434 | 13 | MOLECULE: | ADAPTER PROTEIN MECA 1 |
| 56:00:00 | 4xqk-A | 4.7 | 9.4 | 157 | 1517 | 8 | MOLECULE: | LLABIII |
| 57:00:00 | 6mfv-A | 4.7 | 8.9 | 132 | 641 | 11 | MOLECULE: | TETRATRICOPEPTIDE REPEAT SENSOR PH0952 |
| 58:00:00 | 4lya-A | 4.7 | 4.6 | 129 | 521 | 8 | MOLECULE: | UNCHARACTERIZED PROTEIN |
| 59:00:00 | 7e7i-A | 4.7 | 34.3 | 176 | 2011 | 9 | MOLECULE: | RETINAL-SPECIFIC PHOSPHOLIPID-TRANSPORTING ATPASE |
| 60:00:00 | 8puw-B | 4.7 | 15.8 | 137 | 531 | 9 | MOLECULE: | LAS1 |
| 61:00:00 | 5n9j-Y | 4.7 | 3.9 | 90 | 185 | 4 | MOLECULE: | MEDIATOR OF RNA POLYMERASE II TRANSCRIPTION SUBUN |
| 62:00:00 | 8vx9-B | 4.7 | 6.5 | 150 | 1174 | 8 | MOLECULE: | HAMA |
| 63:00:00 | 8hwa-F | 4.7 | 18.8 | 148 | 700 | 10 | MOLECULE: | PRIMASE D5 |
| 64:00:00 | 6rfl-Y | 4.7 | 3.9 | 126 | 600 | 11 | MOLECULE: | DNA-DEPENDENT RNA POLYMERASE SUBUNIT RPO132 |
| 65:00:00 | 8spg-A | 4.7 | 6.6 | 161 | 852 | 11 | MOLECULE: | RNA POLYMERASE-ASSOCIATED PROTEIN RAPA |
| 66:00:00 | 5uj8-A | 4.7 | 12.8 | 169 | 553 | 7 | MOLECULE: | ORIGIN RECOGNITION COMPLEX SUBUNIT 3 |
| 67:00:00 | 5vhn-C | 4.7 | 9.1 | 129 | 224 | 14 | MOLECULE: | 26S PROTEASOME NON-ATPASE REGULATORY SUBUNIT 10 |
| 68:00:00 | 7wi4-F | 4.7 | 11.6 | 134 | 410 | 12 | MOLECULE: | ATP-DEPENDENT ZINC METALLOPROTEASE FTSH |
| 69:00:00 | 5vhj-E | 4.7 | 9.3 | 121 | 211 | 9 | MOLECULE: | 26S PROTEASOME NON-ATPASE REGULATORY SUBUNIT 10 |
| 70:00:00 | 5vhi-E | 4.7 | 14.3 | 149 | 311 | 11 | MOLECULE: | 26S PROTEASOME REGULATORY SUBUNIT 7 |
| 71:00:00 | 7pkq-x | 4.7 | 3.9 | 125 | 368 | 11 | MOLECULE: | MS35 |

| | | | | | | | | |
|---|---|---|---|---|---|---|---|---|
| 72:00:00 | 1c9k-B | 4.7 | 3.1 | 94 | 180 | 19 | MOLECULE: | ADENOSYLCOBINAMIDE KINASE |
| 73:00:00 | 4q2d-A | 4.6 | 15.6 | 176 | 927 | 11 | MOLECULE: | CRISPR-ASSOCIATED HELICASE CAS3 |
| 74:00:00 | 8j5q-D | 4.6 | 3.9 | 128 | 611 | 14 | MOLECULE: | UNCHARACTERIZED PROTEIN RV1280C |
| 75:00:00 | 2oap-1 | 4.6 | 23.4 | 180 | 498 | 7 | MOLECULE: | TYPE II SECRETION SYSTEM PROTEIN |
| 76:00:00 | 8hwb-E | 4.6 | 23.5 | 129 | 700 | 13 | MOLECULE: | PRIMASE D5 |
| 77:00:00 | 2kjq-A | 4.6 | 6.9 | 99 | 149 | 11 | MOLECULE: | DNAA-RELATED PROTEIN |
| 78:00:00 | 8d8l-W | 4.6 | 5.6 | 124 | 395 | 8 | MOLECULE: | PROBABLE S-ADENOSYL-L-METHIONINE-DEPENDENT RNA |
| 79:00:00 | 5vhn-E | 4.6 | 11.3 | 115 | 241 | 12 | MOLECULE: | 26S PROTEASOME NON-ATPASE REGULATORY SUBUNIT 10 |
| 80:00:00 | 8sk1-A | 4.5 | 3.5 | 94 | 162 | 12 | MOLECULE: | 2-AMINO-4-HYDROXY-6-HYDROXYMETHYLDIHYDROPTERIDINE |
| 81:00:00 | 8fnu-A | 4.5 | 16.4 | 144 | 796 | 10 | MOLECULE: | KAP NTPASE DOMAIN-CONTAINING PROTEIN |
| 82:00:00 | 4yut-A | 4.5 | 16.2 | 95 | 351 | 11 | MOLECULE: | FAMILY 3 ADENYLATE CYCLASE |
| 83:00:00 | 7nyw-B | 4.5 | 3.9 | 111 | 858 | 11 | MOLECULE: | CHROMOSOME PARTITION PROTEIN MUKB |
| 84:00:00 | 8hwe-F | 4.5 | 4.2 | 146 | 274 | 14 | MOLECULE: | PRIMASE D5 |
| 85:00:00 | 6fws-A | 4.5 | 7.4 | 129 | 686 | 10 | MOLECULE: | ATP-DEPENDENT DNA HELICASE DING |
| 86:00:00 | 2r2a-A | 4.5 | 5.1 | 107 | 191 | 8 | MOLECULE: | UNCHARACTERIZED PROTEIN |
| 87:00:00 | 7ngq-B | 4.5 | 29.5 | 109 | 776 | 15 | MOLECULE: | LON PROTEASE HOMOLOG, MITOCHONDRIAL |
| 88:00:00 | 7ngl-B | 4.5 | 26.7 | 110 | 776 | 14 | MOLECULE: | LON PROTEASE HOMOLOG, MITOCHONDRIAL |
| 89:00:00 | 6og1-F | 4.5 | 13 | 139 | 571 | 11 | MOLECULE: | HYPERACTIVE DISAGGREGASE CLPB |
| 90:00:00 | 2fwr-A | 4.5 | 7.4 | 155 | 434 | 5 | MOLECULE: | DNA REPAIR PROTEIN RAD25 |
| 91:00:00 | 1t5l-A | 4.5 | 4.5 | 126 | 595 | 8 | MOLECULE: | UVRABC SYSTEM PROTEIN B |
| 92:00:00 | 3jbl-A | 4.5 | 13.9 | 145 | 909 | 11 | MOLECULE: | NLR FAMILY CARD DOMAIN-CONTAINING PROTEIN 4 |
| 93:00:00 | 5vho-E | 4.5 | 8.9 | 119 | 209 | 12 | MOLECULE: | 26S PROTEASOME NON-ATPASE REGULATORY SUBUNIT 10 |
| 94:00:00 | 4f4c-A | 4.4 | 33.9 | 164 | 1250 | 10 | MOLECULE: | MULTIDRUG RESISTANCE PROTEIN PGP-1 |
| 95:00:00 | 1ahu-A | 4.4 | 20.7 | 138 | 555 | 8 | MOLECULE: | VANILLYL-ALCOHOL OXIDASE |
| 96:00:00 | 8dgc-A | 4.4 | 29.8 | 201 | 2028 | 10 | MOLECULE: | SEAVS3 |
| 97:00:00 | 3h1t-A | 4.4 | 10.2 | 131 | 515 | 11 | MOLECULE: | TYPE I SITE-SPECIFIC RESTRICTION-MODIFICATION |
| 98:00:00 | 7roq-A | 4.4 | 41.7 | 164 | 1831 | 11 | MOLECULE: | PHOSPHOLIPID-TRANSPORTING ATPASE ABCA1 |
| 99:00:00 | 5ld2-D | 4.4 | 15.7 | 127 | 605 | 8 | MOLECULE: | RECBCD ENZYME SUBUNIT RECB,RECBCD ENZYME SUBUNIT |
| 0:00 | 3j16-B | 4.4 | 6.9 | 122 | 608 | 8 | MOLECULE: | DOM34P |
| 1:00 | 8fmv-D | 4.4 | 27.6 | 169 | 665 | 13 | MOLECULE: | PROBABLE MULTIDRUG RESISTANCE ABC TRANSPORTER ATP |
| 2:00 | 6pwn-A | 4.4 | 29.6 | 139 | 280 | 8 | MOLECULE: | SMALL-CONDUCTANCE MECHANOSENSITIVE CHANNEL |
| 3:00 | 8hwc-C | 4.4 | 4.1 | 157 | 280 | 11 | MOLECULE: | PRIMASE D5 |
| 4:00 | 6ahf-C | 4.4 | 12.7 | 185 | 865 | 6 | MOLECULE: | HEAT SHOCK PROTEIN 104 |
| 5:00 | 6ppu-B | 4.4 | 17.9 | 131 | 752 | 11 | MOLECULE: | UVRD/REP HELICASE |
| 6:00 | 7ngq-D | 4.4 | 27.8 | 146 | 776 | 10 | MOLECULE: | LON PROTEASE HOMOLOG, MITOCHONDRIAL |
| 7:00 | 7wrx-C | 4.3 | 15.6 | 143 | 594 | 6 | MOLECULE: | HERA |
| 8:00 | 3c19-A | 4.3 | 4.8 | 100 | 177 | 6 | MOLECULE: | UNCHARACTERIZED PROTEIN MK0293 |
| 9:00 | 4nh0-A | 4.3 | 5.3 | 136 | 861 | 8 | MOLECULE: | CELL DIVISIONFTSK/SPOIIIE |
| 10:00 | 7bst-C | 4.3 | 12.6 | 157 | 992 | 5 | MOLECULE: | TYPE I RESTRICTION ENZYME R PROTEIN |
| 11:00 | 6wb2-A | 4.3 | 11 | 125 | 540 | 6 | MOLECULE: | HIV-1 VIRAL RNA GENOME FRAGMENT |
| 12:00 | 3mtj-A | 4.3 | 12 | 124 | 431 | 8 | MOLECULE: | HOMOSERINE DEHYDROGENASE |
| 13:00 | 3af5-A | 4.3 | 8.4 | 115 | 638 | 7 | MOLECULE: | PUTATIVE UNCHARACTERIZED PROTEIN PH1404 |
| 14:00 | 8csp-X | 4.3 | 4.5 | 129 | 352 | 11 | MOLECULE: | 28S RIBOSOMAL PROTEIN S34, MITOCHONDRIAL |
| 15:00 | 5wcb-F | 4.3 | 12.8 | 121 | 277 | 12 | MOLECULE: | MEIOTIC SPINDLE FORMATION PROTEIN MEI-1 |
| 16:00 | 5vhi-A | 4.3 | 11.3 | 148 | 321 | 9 | MOLECULE: | 26S PROTEASOME REGULATORY SUBUNIT 7 |
| 17:00 | 4lgt-A | 4.2 | 13.7 | 91 | 251 | 5 | MOLECULE: | RIBOSOMAL LARGE SUBUNIT PSEUDOURIDINE SYNTHASE B |
| 18:00 | 7nyw-A | 4.2 | 3.5 | 104 | 685 | 11 | MOLECULE: | CHROMOSOME PARTITION PROTEIN MUKB |
| 19:00 | 7nhr-A | 4.2 | 5.4 | 125 | 663 | 10 | MOLECULE: | PUTATIVE TRANSMEMBRANE PROTEIN WZC |
| 20:00 | 8b1r-B | 4.2 | 20.3 | 140 | 1170 | 11 | MOLECULE: | RECBCD ENZYME SUBUNIT RECB |
| 21:00 | 7nhr-C | 4.2 | 5.4 | 121 | 565 | 11 | MOLECULE: | PUTATIVE TRANSMEMBRANE PROTEIN WZC |
| 22:00 | 4cej-A | 4.2 | 12.3 | 122 | 1177 | 9 | MOLECULE: | ATP-DEPENDENT HELICASE/NUCLEASE SUBUNIT A |
| 23:00 | 5wwp-A | 4.2 | 25.8 | 134 | 515 | 7 | MOLECULE: | ORF1AB |
| 24:00:00 | 7r7j-B | 4.2 | 5.2 | 124 | 574 | 13 | MOLECULE: | PUTATIVE DNA REPAIR HELICASE RADD |

| | | | | | | | | |
|---|---|---|---|---|---|---|---|---|
| 25:00:00 | 7ngp-F | 4.2 | 28.3 | 133 | 776 | 13 | MOLECULE: | LON PROTEASE HOMOLOG, MITOCHONDRIAL |
| 26:00:00 | 4znm-B | 4.1 | 11.2 | 98 | 444 | 8 | MOLECULE: | C-DOMAIN TYPE II PEPTIDE SYNTHETASE |
| 27:00:00 | 7mbw-A | 4.1 | 11.4 | 148 | 475 | 9 | MOLECULE: | TRANSPOSON TN7 TRANSPOSITION PROTEIN TNSC |
| 28:00:00 | 7r9g-A | 4.1 | 3.8 | 91 | 386 | 10 | MOLECULE: | TRNA PSEUDOURIDINE SYNTHASE 1 |
| 29:00:00 | 6k0w-B | 4.1 | 19.3 | 143 | 500 | 6 | MOLECULE: | ADENINE SPECIFIC DNA METHYLTRANSFERASE (MOD) |
| 30:00:00 | 7ou2-B | 4.1 | 8.9 | 141 | 650 | 12 | MOLECULE: | DNA MISMATCH REPAIR PROTEIN MUTS |
| 31:00:00 | 7d7n-A | 4.1 | 24.5 | 184 | 704 | 10 | MOLECULE: | ATP-BINDING CASSETTE SUB-FAMILY B MEMBER 6, MITOC |
| 32:00:00 | 4n5c-D | 4.1 | 21.7 | 161 | 757 | 4 | MOLECULE: | CARGO-TRANSPORT PROTEIN YPP1 |
| 33:00:00 | 7sjr-A | 4.1 | 17.7 | 136 | 911 | 11 | MOLECULE: | DNA HELICASE |
| 34:00:00 | 8dgc-E | 4.1 | 6.4 | 112 | 538 | 9 | MOLECULE: | SEAVS3 |
| 35:00:00 | 7xg3-L | 4.1 | 12.7 | 124 | 594 | 7 | MOLECULE: | CSF1 |
| 36:00:00 | 4bj6-A | 4.1 | 5.5 | 134 | 293 | 7 | MOLECULE: | RAP1-INTERACTING FACTOR 2 |
| 37:00:00 | 8c0w-C | 4.1 | 22.8 | 110 | 1030 | 9 | MOLECULE: | PEROXISOMAL ATPASE PEX6 |
| 38:00:00 | 8p53-D | 4.1 | 5.3 | 88 | 377 | 10 | MOLECULE: | ANTIACTIVATOR FLEN |
| 39:00:00 | 7lar-A | 4.1 | 5.6 | 118 | 183 | 16 | MOLECULE: | ATP-DEPENDENT HELICASE REP |
| 40:00:00 | 6ir9-W | 4 | 10.7 | 104 | 275 | 4 | MOLECULE: | DNA-DIRECTED RNA POLYMERASE SUBUNIT |
| 41:00:00 | 2fsf-B | 4 | 11.9 | 121 | 723 | 9 | MOLECULE: | PREPROTEIN TRANSLOCASE SECA SUBUNIT |
| 42:00:00 | 5cio-A | 4 | 18.5 | 175 | 770 | 7 | MOLECULE: | PYRROLOQUINOLINE QUINONE BIOSYNTHESIS PROTEIN PQQ |
| 43:00:00 | 5b7i-A | 4 | 6.7 | 161 | 1044 | 6 | MOLECULE: | CRISPR-ASSOCIATED NUCLEASE/HELICASE CAS3 SUBTYPE |
| 44:00:00 | 8rtd-T | 4 | 23.9 | 184 | 808 | 7 | MOLECULE: | TRWJ PROTEIN |
| 45:00:00 | 6nmi-A | 4 | 11.1 | 154 | 653 | 7 | MOLECULE: | GENERAL TRANSCRIPTION AND DNA REPAIR FACTOR IIH H |
| 46:00:00 | 3d3y-A | 4 | 10.4 | 128 | 425 | 9 | MOLECULE: | UNCHARACTERIZED PROTEIN |
| 47:00:00 | 1k8w-A | 4 | 4.6 | 86 | 303 | 10 | MOLECULE: | 5'-R(*GP*GP*CP*AP*AP*CP*GP*GP*UP*(FHU) |
| 48:00:00 | 7nfy-F | 4 | 27.2 | 97 | 776 | 16 | MOLECULE: | LON PROTEASE HOMOLOG, MITOCHONDRIAL |
| 49:00:00 | 5vho-A | 4 | 3.5 | 123 | 255 | 11 | MOLECULE: | 26S PROTEASOME NON-ATPASE REGULATORY SUBUNIT 10 |
| 50:00:00 | 8umd-B | 3.9 | 24.7 | 99 | 331 | 11 | MOLECULE: | FLAGELLAR M-RING PROTEIN |
| 51:00:00 | 6zca-H | 3.9 | 7.7 | 117 | 771 | 8 | MOLECULE: | PROBABLE DNA-DIRECTED RNA POLYMERASE SUBUNIT DELT |
| 52:00:00 | 5lkl-A | 3.9 | 13.9 | 133 | 704 | 7 | MOLECULE: | GENOME POLYPROTEIN |
| 53:00:00 | 7oky-Z | 3.9 | 15.7 | 107 | 374 | 6 | MOLECULE: | DNA-DIRECTED RNA POLYMERASE II SUBUNIT RPB1 |
| 54:00:00 | 3al0-B | 3.9 | 21.7 | 119 | 482 | 5 | MOLECULE: | GLUTAMYL-TRNA(GLN) AMIDOTRANSFERASE SUBUNIT A |
| 55:00:00 | 6urt-A | 3.9 | 22.3 | 119 | 331 | 7 | MOLECULE: | LOW CONDUCTANCE MECHANOSENSITIVE CHANNEL YNAI |
| 56:00:00 | 5do7-D | 3.9 | 6.7 | 134 | 571 | 14 | MOLECULE: | ATP-BINDING CASSETTE SUB-FAMILY G MEMBER 5 |
| 57:00:00 | 6wg3-A | 3.9 | 3.5 | 110 | 561 | 11 | MOLECULE: | STRUCTURAL MAINTENANCE OF CHROMOSOMES PROTEIN 1A |
| 58:00:00 | 6z6d-A | 3.9 | 9 | 129 | 479 | 10 | MOLECULE: | TERMINASE LARGE SUBUNIT |
| 59:00:00 | 3wrn-A | 3.9 | 3.2 | 104 | 248 | 5 | MOLECULE: | NON-CAPSID PROTEIN NS-1 |
| 60:00:00 | 6dhs-B | 3.9 | 10.5 | 93 | 176 | 4 | MOLECULE: | HETEROGENEOUS NUCLEAR RIBONUCLEOPROTEIN H |
| 61:00:00 | 2wjy-A | 3.9 | 28.3 | 161 | 773 | 5 | MOLECULE: | REGULATOR OF NONSENSE TRANSCRIPTS 1 |
| 62:00:00 | 7mjy-A | 3.9 | 6.7 | 127 | 449 | 7 | MOLECULE: | TRNA-2-METHYLTHIO-N(6)-DIMETHYLALLYLADENOSINE SYN |
| 63:00:00 | 6pns-A | 3.9 | 16.9 | 202 | 1291 | 5 | MOLECULE: | RNA-DIRECTED RNA POLYMERASE |
| 64:00:00 | 8osf-F | 3.9 | 4.3 | 117 | 226 | 8 | MOLECULE: | MAGNESIUM-CHELATASE SUBUNIT CHLI |
| 65:00:00 | 7mcs-G | 3.9 | 3.7 | 136 | 358 | 11 | MOLECULE: | TRANSPOSON TN7 TRANSPOSITION PROTEIN TNSC |
| 66:00:00 | 5k8o-B | 3.8 | 11.3 | 118 | 425 | 9 | MOLECULE: | 5-NITROANTHRANILIC ACID AMINOHYDROLASE |
| 67:00:00 | 5juy-B | 3.8 | 15 | 161 | 1234 | 9 | MOLECULE: | APOPTOTIC PROTEASE-ACTIVATING FACTOR 1 |
| 68:00:00 | 7x17-C | 3.8 | 14.7 | 109 | 455 | 6 | MOLECULE: | AMB ANTIMETABOLITE SYNTHASE AMBB |
| 69:00:00 | 5d4w-A | 3.8 | 5.4 | 172 | 688 | 8 | MOLECULE: | PUTATIVE HEAT SHOCK PROTEIN |
| 70:00:00 | 7ole-C | 3.8 | 13.1 | 138 | 427 | 9 | MOLECULE: | RUVB-LIKE 1 |
| 71:00:00 | 4etx-A | 3.7 | 13.3 | 94 | 300 | 6 | MOLECULE: | PELD |
| 72:00:00 | 7nyx-A | 3.7 | 4 | 112 | 1467 | 13 | MOLECULE: | CHROMOSOME PARTITION PROTEIN MUKB |
| 73:00:00 | 4eze-A | 3.7 | 18.7 | 93 | 291 | 8 | MOLECULE: | HALOACID DEHALOGENASE-LIKE HYDROLASE |
| 74:00:00 | 2nyi-A | 3.7 | 3.4 | 80 | 171 | 4 | MOLECULE: | UNKNOWN PROTEIN |
| 75:00:00 | 3pfo-A | 3.7 | 5.3 | 114 | 426 | 8 | MOLECULE: | PUTATIVE ACETYLORNITHINE DEACETYLASE |
| 76:00:00 | 6fon-A | 3.7 | 3.3 | 81 | 252 | 5 | MOLECULE: | COPPER CHAPERONE FOR SUPEROXIDE DISMUTASE |
| 77:00:00 | 6fwr-A | 3.7 | 13.4 | 139 | 699 | 10 | MOLECULE: | ATP-DEPENDENT DNA HELICASE DING |

| | | | | | | | | |
|---|---|---|---|---|---|---|---|---|
| 78:00:00 | 8iqh-C | 3.7 | 16.3 | 200 | 891 | 8 | MOLECULE: | PUTATIVE PRIMASE C962R |
| 79:00:00 | 8iqh-F | 3.7 | 16.4 | 195 | 880 | 9 | MOLECULE: | PUTATIVE PRIMASE C962R |
| 80:00:00 | 8q3w-D | 3.7 | 11.7 | 117 | 246 | 15 | MOLECULE: | INSERTION SEQUENCE IS5376 PUTATIVE ATP-BINDING PR |
| 81:00:00 | 7ng4-F | 3.7 | 27.6 | 97 | 776 | 16 | MOLECULE: | LON PROTEASE HOMOLOG, MITOCHONDRIAL |
| 82:00:00 | 7ngc-D | 3.7 | 27.5 | 102 | 776 | 14 | MOLECULE: | LON PROTEASE HOMOLOG, MITOCHONDRIAL |
| 83:00:00 | 6b03-A | 3.6 | 21.4 | 149 | 896 | 5 | MOLECULE: | PUTATIVE ZINC PROTEASE |
| 84:00:00 | 7r9x-A | 3.6 | 10.3 | 97 | 418 | 7 | MOLECULE: | AMBE |
| 85:00:00 | 7vdv-A | 3.6 | 28.5 | 179 | 809 | 6 | MOLECULE: | HISTONE H4 |
| 86:00:00 | 6slf-A | 3.6 | 4.3 | 109 | 398 | 6 | MOLECULE: | N-ALPHA-ACYL-GLUTAMINE AMINOACYLASE |
| 87:00:00 | 1fx2-A | 3.6 | 5.4 | 79 | 235 | 1 | MOLECULE: | RECEPTOR-TYPE ADENYLATE CYCLASE GRESAG 4.1 |
| 88:00:00 | 7n7s-B | 3.6 | 7.6 | 145 | 441 | 8 | MOLECULE: | HYDROXYMETHYLGLUTARYL-COA REDUCTASE |
| 89:00:00 | 3i5x-A | 3.6 | 13.2 | 114 | 509 | 8 | MOLECULE: | ATP-DEPENDENT RNA HELICASE MSS116 |
| 90:00:00 | 7nad-x | 3.6 | 17.1 | 114 | 539 | 11 | MOLECULE: | 25S RRNA |
| 91:00:00 | 4rh7-A | 3.6 | 22.4 | 132 | 3005 | 13 | MOLECULE: | GREEN FLUORESCENT PROTEIN/CYTOPLASMIC DYNEIN 2 HE |
| 92:00:00 | 1e2f-A | 3.6 | 3.7 | 89 | 210 | 15 | MOLECULE: | THYMIDYLATE KINASE |
| 93:00:00 | 1e9s-A | 3.6 | 4.5 | 115 | 427 | 10 | MOLECULE: | CONJUGAL TRANSFER PROTEIN TRWB |
| 94:00:00 | 4cgz-A | 3.5 | 10.5 | 128 | 629 | 8 | MOLECULE: | BLOOM'S SYNDROME HELICASE |
| 95:00:00 | 5u9m-D | 3.5 | 15.7 | 81 | 234 | 4 | MOLECULE: | SUPEROXIDE DISMUTASE [CU-ZN] |
| 96:00:00 | 7uoi-A | 3.5 | 9.5 | 115 | 383 | 7 | MOLECULE: | SUCCINYL-DIAMINOPIMELATE DESUCCINYLASE |
| 97:00:00 | 8tzk-C | 3.5 | 27.3 | 133 | 304 | 7 | MOLECULE: | CELL DIVISION ATP-BINDING PROTEIN FTSE |
| 98:00:00 | 7ns9-A | 3.5 | 3.6 | 74 | 178 | 7 | MOLECULE: | TRIPHOSPHATE TUNNEL METALLOENZYME SACI_0718 |
| 99:00:00 | 7w01-A | 3.5 | 17.2 | 162 | 1598 | 9 | MOLECULE: | PHOSPHOLIPID-TRANSPORTING ATPASE ABCA3 |
| 0:00 | 4ddt-A | 3.5 | 5.6 | 149 | 1102 | 9 | MOLECULE: | REVERSE GYRASE |
| 1:00 | 2iai-A | 3.5 | 7.4 | 84 | 200 | 4 | MOLECULE: | PUTATIVE TRANSCRIPTIONAL REGULATOR SCO3833 |
| 2:00 | 3luy-A | 3.5 | 7.5 | 104 | 326 | 10 | MOLECULE: | PROBABLE CHORISMATE MUTASE |
| 3:00 | 4ney-B | 3.5 | 5.6 | 95 | 173 | 8 | MOLECULE: | ENGINEERED PROTEIN OR277 |
| 4:00 | 8iqh-K | 3.5 | 28.4 | 196 | 866 | 8 | MOLECULE: | PUTATIVE PRIMASE C962R |
| 5:00 | 8p53-C | 3.5 | 5 | 83 | 378 | 10 | MOLECULE: | ANTIACTIVATOR FLEN |
| 6:00 | 7eih-A | 3.4 | 7.1 | 114 | 588 | 7 | MOLECULE: | FAD DEPENDENT ENZYME |
| 7:00 | 2qv6-A | 3.4 | 17.2 | 88 | 253 | 7 | MOLECULE: | GTP CYCLOHYDROLASE III |
| 8:00 | 5d08-A | 3.4 | 12.5 | 119 | 384 | 6 | MOLECULE: | EPOXYQUEUOSINE REDUCTASE |
| 9:00 | 6bua-A | 3.4 | 7 | 116 | 942 | 9 | MOLECULE: | DICER-2, ISOFORM A |
| 10:00 | 4zcf-C | 3.4 | 7.2 | 135 | 627 | 7 | MOLECULE: | RESTRICTION ENDONUCLEASE ECOP15I, MODIFICATION SU |
| 11:00 | 6s6y-L | 3.4 | 4.5 | 121 | 311 | 5 | MOLECULE: | FORMYLMETHANOFURAN DEHYDROGENASE SUBUNIT A |
| 12:00 | 5yvu-B | 3.4 | 15.6 | 106 | 601 | 8 | MOLECULE: | GENOME POLYPROTEIN |
| 13:00 | 5egf-C | 3.4 | 14.1 | 110 | 444 | 9 | MOLECULE: | TQAA |
| 14:00 | 8y65-A | 3.4 | 13.4 | 82 | 467 | 11 | MOLECULE: | SOLUTE CARRIER FAMILY 2, FACILITATED GLUCOSE TRAN |
| 15:00 | 2y08-B | 3.4 | 16.5 | 137 | 500 | 8 | MOLECULE: | TAML |
| 16:00 | 3j3r-A | 3.4 | 15.8 | 110 | 798 | 15 | MOLECULE: | ADAPTER PROTEIN MECA 1 |
| 17:00 | 7ngp-D | 3.4 | 26.2 | 100 | 776 | 14 | MOLECULE: | LON PROTEASE HOMOLOG, MITOCHONDRIAL |
| 18:00 | 5vhi-C | 3.4 | 22.3 | 132 | 350 | 10 | MOLECULE: | 26S PROTEASOME REGULATORY SUBUNIT 7 |
| 19:00 | 6neq-g | 3.4 | 4.3 | 117 | 351 | 15 | MOLECULE: | 28S RIBOSOMAL RNA, MITOCHONDRIAL |
| 20:00 | 8unj-A | 3.4 | 16.4 | 80 | 172 | 8 | MOLECULE: | CHROMOSOME TRANSMISSION FIDELITY PROTEIN 18 HOMOL |
| 21:00 | 7fse-A | 3.3 | 5.4 | 142 | 1040 | 11 | MOLECULE: | REVERSE GYRASE |
| 22:00 | 6ner-E | 3.3 | 3.3 | 73 | 186 | 5 | MOLECULE: | BMC-H TANDEM FUSION PROTEIN |
| 23:00 | 3rc8-A | 3.3 | 20.4 | 102 | 609 | 10 | MOLECULE: | ATP-DEPENDENT RNA HELICASE SUPV3L1, MITOCHONDRIAL |
| 24:00:00 | 7t5q-J | 3.3 | 33.8 | 86 | 291 | 3 | MOLECULE: | ACTIN-RELATED PROTEIN 3 |
| 25:00:00 | 1b0p-A | 3.3 | 6.2 | 123 | 1231 | 8 | MOLECULE: | PROTEIN (PYRUVATE-FERREDOXIN OXIDOREDUCTASE) |
| 26:00:00 | 8hwc-D | 3.3 | 4 | 138 | 266 | 9 | MOLECULE: | PRIMASE D5 |
| 27:00:00 | 2qnd-B | 3.2 | 3.7 | 72 | 143 | 11 | MOLECULE: | FMR1 PROTEIN |
| 28:00:00 | 6eg2-A | 3.2 | 18.8 | 156 | 619 | 6 | MOLECULE: | MALTOSE/MALTODEXTRIN-BINDING PERIPLASMIC PROTEIN, |
| 29:00:00 | 7z8v-D | 3.2 | 8.4 | 74 | 1158 | 9 | MOLECULE: | CULLIN-1 |
| 30:00:00 | 7yv9-A | 3.2 | 6 | 141 | 1067 | 6 | MOLECULE: | UNCONVENTIONAL MYOSIN-VA |

| | | | | | | | | |
|---|---|---|---|---|---|---|---|---|
| 31:00:00 | 4gcl-B | 3.2 | 10.7 | 73 | 196 | 7 | MOLECULE: | NUCLEOID OCCLUSION FACTOR SLMA |
| 32:00:00 | 7r08-E | 3.2 | 19.2 | 114 | 488 | 5 | MOLECULE: | REVERSE TRANSCRIPTASE |
| 33:00:00 | 6d97-A | 3.1 | 11.8 | 96 | 522 | 8 | MOLECULE: | ALDEHYDE DEHYDROGENASE 12 |
| 34:00:00 | 7x4c-A | 3.1 | 8.2 | 104 | 306 | 7 | MOLECULE: | EFCDNE |
| 35:00:00 | 1yb3-A | 3.1 | 3.5 | 79 | 166 | 10 | MOLECULE: | HYPOTHETICAL PROTEIN |
| 36:00:00 | 3on2-B | 3.1 | 11.3 | 82 | 195 | 2 | MOLECULE: | PROBABLE TRANSCRIPTIONAL REGULATOR |
| 37:00:00 | 7ea3-J | 3.1 | 29.5 | 133 | 883 | 7 | MOLECULE: | TRAPP-ASSOCIATED PROTEIN TCA17 |
| 38:00:00 | 7zr1-D | 3.1 | 3.6 | 114 | 780 | 10 | MOLECULE: | DOUBLE-STRAND BREAK REPAIR PROTEIN |
| 39:00:00 | 6zyd-A | 3.1 | 19.7 | 102 | 323 | 3 | MOLECULE: | LOW CONDUCTANCE MECHANOSENSITIVE CHANNEL YNAI,LOW |
| 40:00:00 | 6iy8-C | 3.1 | 5.6 | 81 | 468 | 10 | MOLECULE: | POSITIVE REGULATOR CAPR |
| 41:00:00 | 3him-B | 3 | 3.8 | 76 | 192 | 1 | MOLECULE: | PROBABLE TRANSCRIPTIONAL REGULATOR |
| 42:00:00 | 3wdo-A | 3 | 4 | 71 | 453 | 6 | MOLECULE: | MFS TRANSPORTER |
| 43:00:00 | 7yqh-A | 3 | 3.7 | 119 | 1069 | 11 | MOLECULE: | STRUCTURAL MAINTENANCE OF CHROMOSOMES PROTEIN 5 |
| 44:00:00 | 6yvu-B | 3 | 3.9 | 109 | 1191 | 7 | MOLECULE: | STRUCTURAL MAINTENANCE OF CHROMOSOMES PROTEIN 2,S |
| 45:00:00 | 1g8x-A | 3 | 5.3 | 136 | 1009 | 4 | MOLECULE: | MYOSIN II HEAVY CHAIN FUSED TO ALPHA-ACTININ 3 |
| 46:00:00 | 8iqh-I | 3 | 19.2 | 144 | 866 | 11 | MOLECULE: | PUTATIVE PRIMASE C962R |
| 47:00:00 | 5w7g-A | 2.9 | 9.5 | 66 | 131 | 12 | MOLECULE: | ORF140 |
| 48:00:00 | 7k9y-A | 2.9 | 21.4 | 136 | 425 | 8 | MOLECULE: | TRT |
| 49:00:00 | 5zh8-A | 2.9 | 7.4 | 75 | 358 | 9 | MOLECULE: | PROTEIN FMTA |
| 50:00:00 | 1jwy-A | 2.9 | 6.1 | 137 | 1039 | 3 | MOLECULE: | MYOSIN-2 HEAVY CHAIN,DYNAMIN-A |
| 51:00:00 | 3bjd-C | 2.8 | 8.4 | 54 | 324 | 9 | MOLECULE: | PUTATIVE 3-OXOACYL-(ACYL-CARRIER-PROTEIN) SYNTHAS |
| 52:00:00 | 6xf9-D | 2.8 | 6.4 | 102 | 426 | 7 | MOLECULE: | PACKAGING PROTEIN UL32 |
| 53:00:00 | 8g7w-B | 2.8 | 11.3 | 129 | 1059 | 4 | MOLECULE: | TYPE I PKS MODULE 4, MODULE 5 |
| 54:00:00 | 7qo4-M | 2.8 | 24.8 | 180 | 421 | 7 | MOLECULE: | 26S PROTEASOME REGULATORY SUBUNIT RPN1 |
| 55:00:00 | 4j8s-A | 2.7 | 15.6 | 59 | 180 | 10 | MOLECULE: | CCR4-NOT TRANSCRIPTION COMPLEX SUBUNIT 1 |
| 56:00:00 | 7b1s-D | 2.7 | 10.2 | 127 | 593 | 9 | MOLECULE: | ETHYL-COENZYME M REDUCTASE ALPHA SUBUNIT |
| 57:00:00 | 8h6s-A | 2.7 | 7 | 106 | 306 | 6 | MOLECULE: | MALONYL-COA-[ACYL-CARRIER-PROTEIN] TRANSACYLASE |
| 58:00:00 | 7opl-A | 2.7 | 7.9 | 59 | 1070 | 12 | MOLECULE: | DNA POLYMERASE ALPHA CATALYTIC SUBUNIT |
| 59:00:00 | 4mzv-A | 2.7 | 7.2 | 93 | 243 | 10 | MOLECULE: | EPITHELIAL CELL ADHESION MOLECULE |
| 60:00:00 | 8hwd-F | 2.7 | 3.8 | 128 | 258 | 9 | MOLECULE: | PRIMASE D5 |
| 61:00:00 | 8q3w-G | 2.7 | 20.5 | 102 | 246 | 17 | MOLECULE: | INSERTION SEQUENCE IS5376 PUTATIVE ATP-BINDING PR |
| 62:00:00 | 6r5k-H | 2.6 | 7.6 | 85 | 394 | 1 | MOLECULE: | PAN2-PAN3 DEADENYLATION COMPLEX CATALYTIC SUBUNIT |
| 63:00:00 | 3nlc-A | 2.6 | 18.8 | 103 | 534 | 8 | MOLECULE: | UNCHARACTERIZED PROTEIN VP0956 |
| 64:00:00 | 8ibw-C | 2.6 | 14.3 | 165 | 922 | 8 | MOLECULE: | DNA (60-MER) |
| 65:00:00 | 8sg4-A | 2.6 | 20.3 | 101 | 1469 | 12 | MOLECULE: | METAL RESISTANCE PROTEIN YCF1 |
| 66:00:00 | 8hwd-E | 2.6 | 3.7 | 117 | 247 | 9 | MOLECULE: | PRIMASE D5 |
| 67:00:00 | 8twj-A | 2.5 | 4.1 | 57 | 303 | 9 | MOLECULE: | AVIRULENCE PROTEIN B |
| 68:00:00 | 5i5n-A | 2.5 | 6.2 | 52 | 136 | 8 | MOLECULE: | MATRIX PROTEIN |
| 69:00:00 | 7zge-A | 2.5 | 8.4 | 72 | 199 | 4 | MOLECULE: | BRXA, A BREX PHAGE DEFENCE PROTEIN |
| 70:00:00 | 4ch7-A | 2.5 | 13 | 92 | 309 | 9 | MOLECULE: | NIRD-LIKE PROTEIN |
| 71:00:00 | 3jck-F | 2.5 | 12.8 | 86 | 388 | 6 | MOLECULE: | 26S PROTEASOME REGULATORY SUBUNIT RPN3 |
| 72:00:00 | 1zr6-A | 2.4 | 15 | 136 | 479 | 7 | MOLECULE: | GLUCOOLIGOSACCHARIDE OXIDASE |
| 73:00:00 | 2xhc-A | 2.4 | 4.2 | 86 | 341 | 9 | MOLECULE: | TRANSCRIPTION ANTITERMINATION PROTEIN NUSG |
| 74:00:00 | 5h78-A | 2.4 | 9.8 | 64 | 192 | 11 | MOLECULE: | CAMP-DEPENDENT PROTEIN KINASE TYPE II-ALPHA REGUL |
| 75:00:00 | 4g1i-B | 2.4 | 3.8 | 82 | 196 | 6 | MOLECULE: | PROTEIN PRGH |
| 76:00:00 | 7qla-A | 2.4 | 15.8 | 109 | 433 | 7 | MOLECULE: | VACUOLAR FUSION PROTEIN MON1 |
| 77:00:00 | 3kiz-B | 2.4 | 10.8 | 82 | 371 | 9 | MOLECULE: | PHOSPHORIBOSYLFORMYLGLYCINAMIDINE CYCLO-LIGASE |
| 78:00:00 | 2i9d-A | 2.4 | 4.3 | 71 | 214 | 11 | MOLECULE: | CHLORAMPHENICOL ACETYLTRANSFERASE |
| 79:00:00 | 3ezu-A | 2.3 | 10.9 | 90 | 336 | 3 | MOLECULE: | GGDEF DOMAIN PROTEIN |
| 80:00:00 | 7w26-A | 2.3 | 7.6 | 94 | 440 | 6 | MOLECULE: | FERULATE MONOLIGNOL TRANSFERASE |
| 81:00:00 | 7oxo-F | 2.3 | 31.5 | 94 | 776 | 19 | MOLECULE: | LON PROTEASE HOMOLOG, MITOCHONDRIAL |
| 82:00:00 | 5abb-A | 2.2 | 17.2 | 114 | 443 | 8 | MOLECULE: | PROTEIN TRANSLOCASE SUBUNIT SECY |
| 83:00:00 | 5m1m-A | 2.2 | 12 | 47 | 154 | 9 | MOLECULE: | MATRIX PROTEIN 1 |

| | | | | | | | | |
|---|---|---|---|---|---|---|---|---|
| 84:00:00 | 4zi2-C | 2.2 | 9.5 | 62 | 131 | 6 | MOLECULE: | ADP-RIBOSYLATION FACTOR-LIKE PROTEIN 3 |
| 85:00:00 | 5vhh-C | 2.2 | 13.2 | 123 | 342 | 15 | MOLECULE: | 26S PROTEASOME REGULATORY SUBUNIT 7 |
| 86:00:00 | 7ng5-D | 2.2 | 24.4 | 97 | 776 | 14 | MOLECULE: | LON PROTEASE HOMOLOG, MITOCHONDRIAL |
| 87:00:00 | 6u9d-L | 2.1 | 18.8 | 88 | 255 | 10 | MOLECULE: | ACETOLACTATE SYNTHASE CATALYTIC SUBUNIT, MITOCHON |
| 88:00:00 | 5vhf-C | 2.1 | 19.4 | 109 | 342 | 15 | MOLECULE: | 26S PROTEASOME NON-ATPASE REGULATORY SUBUNIT 10 |
| 89:00:00 | 7p04-A | 2 | 24.1 | 107 | 1353 | 13 | MOLECULE: | PLEIOTROPIC ABC EFFLUX TRANSPORTER OF MULTIPLE DR |

**DALI results: MnA-1 SET**

| No: | Chain | Z | rmsd | lali | nres | %id | PDB | Description |
|---|---|---|---|---|---|---|---|---|
| 1:00 | 3f9z-A | 15.2 | 2.4 | 127 | 157 | 18 | MOLECULE: | HISTONE-LYSINE N-METHYLTRANSFERASE SETD8 |
| 2:00 | 5teg-B | 15.1 | 2.2 | 125 | 159 | 18 | MOLECULE: | N-LYSINE METHYLTRANSFERASE KMT5A |
| 3:00 | 5w1y-B | 15.1 | 2.3 | 123 | 151 | 18 | MOLECULE: | N-LYSINE METHYLTRANSFERASE KMT5A |
| 4:00 | 2bqz-A | 15 | 2.1 | 124 | 162 | 18 | MOLECULE: | SET8 PROTEIN |
| 5:00 | 3f9y-A | 15 | 2.3 | 126 | 160 | 17 | MOLECULE: | HISTONE-LYSINE N-METHYLTRANSFERASE SETD8 |
| 6:00 | 2bqz-E | 14.9 | 2.3 | 125 | 162 | 18 | MOLECULE: | SET8 PROTEIN |
| 7:00 | 3f9w-B | 14.9 | 2.4 | 126 | 161 | 17 | MOLECULE: | HISTONE-LYSINE N-METHYLTRANSFERASE SETD8 |
| 8:00 | 1zkk-D | 14.9 | 2.3 | 126 | 162 | 17 | MOLECULE: | HISTONE-LYSINE N-METHYLTRANSFERASE, H4 LYSINE-20 |
| 9:00 | 1zkk-C | 14.9 | 2.2 | 124 | 161 | 18 | MOLECULE: | HISTONE-LYSINE N-METHYLTRANSFERASE, H4 LYSINE-20 |
| 10:00 | 4ij8-B | 14.9 | 2.8 | 128 | 151 | 18 | MOLECULE: | N-LYSINE METHYLTRANSFERASE SETD8 |
| 11:00 | 3f9w-C | 14.9 | 2.2 | 124 | 161 | 18 | MOLECULE: | HISTONE-LYSINE N-METHYLTRANSFERASE SETD8 |
| 12:00 | 3f9z-C | 14.9 | 2.2 | 124 | 161 | 19 | MOLECULE: | HISTONE-LYSINE N-METHYLTRANSFERASE SETD8 |
| 13:00 | 5teg-A | 14.9 | 2.2 | 124 | 161 | 18 | MOLECULE: | N-LYSINE METHYLTRANSFERASE KMT5A |
| 14:00 | 3f9y-B | 14.9 | 2.2 | 124 | 161 | 18 | MOLECULE: | HISTONE-LYSINE N-METHYLTRANSFERASE SETD8 |
| 15:00 | 3f9z-D | 14.9 | 2.3 | 126 | 162 | 18 | MOLECULE: | HISTONE-LYSINE N-METHYLTRANSFERASE SETD8 |
| 16:00 | 3f9w-D | 14.9 | 2.3 | 126 | 162 | 17 | MOLECULE: | HISTONE-LYSINE N-METHYLTRANSFERASE SETD8 |
| 17:00 | 3f9w-A | 14.9 | 2.3 | 125 | 161 | 18 | MOLECULE: | HISTONE-LYSINE N-METHYLTRANSFERASE SETD8 |
| 18:00 | 1zkk-B | 14.8 | 2.4 | 126 | 162 | 17 | MOLECULE: | HISTONE-LYSINE N-METHYLTRANSFERASE, H4 LYSINE-20 |
| 19:00 | 3f9x-C | 14.8 | 2.2 | 124 | 161 | 18 | MOLECULE: | HISTONE-LYSINE N-METHYLTRANSFERASE SETD8 |
| 20:00 | 1zkk-A | 14.8 | 2.3 | 125 | 161 | 18 | MOLECULE: | HISTONE-LYSINE N-METHYLTRANSFERASE, H4 LYSINE-20 |
| 21:00 | 3f9x-B | 14.8 | 2.3 | 125 | 161 | 18 | MOLECULE: | HISTONE-LYSINE N-METHYLTRANSFERASE SETD8 |
| 22:00 | 3f9z-B | 14.8 | 2.3 | 125 | 162 | 18 | MOLECULE: | HISTONE-LYSINE N-METHYLTRANSFERASE SETD8 |
| 23:00 | 3f9x-D | 14.8 | 2.3 | 126 | 162 | 17 | MOLECULE: | HISTONE-LYSINE N-METHYLTRANSFERASE SETD8 |
| 24:00:00 | 3f9x-A | 14.7 | 2.4 | 126 | 161 | 17 | MOLECULE: | HISTONE-LYSINE N-METHYLTRANSFERASE SETD8 |
| 25:00:00 | 5hq2-M | 14.5 | 2.4 | 126 | 160 | 17 | MOLECULE: | HISTONE H3.2 |
| 26:00:00 | 5w1y-A | 14.5 | 3 | 127 | 162 | 17 | MOLECULE: | N-LYSINE METHYLTRANSFERASE KMT5A |
| 27:00:00 | 6boz-A | 14.4 | 2.4 | 122 | 139 | 18 | MOLECULE: | N-LYSINE METHYLTRANSFERASE KMT5A |
| 28:00:00 | 6boz-B | 14.4 | 2.1 | 117 | 134 | 19 | MOLECULE: | N-LYSINE METHYLTRANSFERASE KMT5A |
| 29:00:00 | 7d20-K | 14.3 | 2.4 | 126 | 166 | 17 | MOLECULE: | HISTONE H3-LIKE CENTROMERIC PROTEIN A |
| 30:00:00 | 4z4p-A | 14.2 | 2.3 | 124 | 162 | 23 | MOLECULE: | HISTONE-LYSINE N-METHYLTRANSFERASE 2D |
| 31:00:00 | 7u5v-A | 14.1 | 2.2 | 111 | 125 | 16 | MOLECULE: | HISTONE-LYSINE N-METHYLTRANSFERASE 2A |
| 32:00:00 | 4ij8-A | 14 | 2.7 | 126 | 162 | 17 | MOLECULE: | N-LYSINE METHYLTRANSFERASE SETD8 |
| 33:00:00 | 6u9n-B | 14 | 2.4 | 121 | 147 | 17 | MOLECULE: | HISTONE-LYSINE N-METHYLTRANSFERASE |
| 34:00:00 | 6u9k-A | 14 | 2.6 | 123 | 149 | 16 | MOLECULE: | HISTONE-LYSINE N-METHYLTRANSFERASE |
| 35:00:00 | 6u9r-A | 14 | 2.5 | 122 | 149 | 16 | MOLECULE: | HISTONE-LYSINE N-METHYLTRANSFERASE |
| 36:00:00 | 7w6l-E | 13.9 | 2.6 | 123 | 152 | 20 | MOLECULE: | SET1/ASH2 HISTONE METHYLTRANSFERASE COMPLEX SUBUN |
| 37:00:00 | 6u9r-B | 13.9 | 2.4 | 121 | 148 | 17 | MOLECULE: | HISTONE-LYSINE N-METHYLTRANSFERASE |
| 38:00:00 | 6u9m-A | 13.9 | 2.4 | 121 | 148 | 17 | MOLECULE: | HISTONE-LYSINE N-METHYLTRANSFERASE |
| 39:00:00 | 5t5g-A | 13.9 | 2.3 | 118 | 143 | 19 | MOLECULE: | N-LYSINE METHYLTRANSFERASE KMT5A |
| 40:00:00 | 4ldg-A | 13.8 | 2.1 | 125 | 216 | 17 | MOLECULE: | PROTEIN WITH A SET DOMAIN WITHIN CARBOXY REGION |
| 41:00:00 | 5f5e-A | 13.8 | 2.5 | 121 | 149 | 17 | MOLECULE: | HISTONE-LYSINE N-METHYLTRANSFERASE 2A |
| 42:00:00 | 7w67-C | 13.8 | 2.4 | 122 | 158 | 17 | MOLECULE: | SET1/ASH2 HISTONE METHYLTRANSFERASE COMPLEX SUBUN |
| 43:00:00 | 6u9n-A | 13.8 | 2.5 | 121 | 147 | 17 | MOLECULE: | HISTONE-LYSINE N-METHYLTRANSFERASE |
| 44:00:00 | 6u9k-B | 13.8 | 2.4 | 121 | 149 | 17 | MOLECULE: | HISTONE-LYSINE N-METHYLTRANSFERASE |
| 45:00:00 | 7xpx-K | 13.7 | 2.3 | 124 | 168 | 18 | MOLECULE: | HISTONE H3 |
| 46:00:00 | 7w6j-C | 13.7 | 2.5 | 124 | 159 | 17 | MOLECULE: | SET1/ASH2 HISTONE METHYLTRANSFERASE COMPLEX SUBUN |
| 47:00:00 | 7w6i-C | 13.7 | 2.8 | 126 | 160 | 17 | MOLECULE: | SET1/ASH2 HISTONE METHYLTRANSFERASE COMPLEX SUBUN |
| 48:00:00 | 5f6k-E | 13.6 | 2.4 | 120 | 149 | 20 | MOLECULE: | SET1/ASH2 HISTONE METHYLTRANSFERASE COMPLEX SUBUN |
| 49:00:00 | 6u9m-B | 13.6 | 2.5 | 121 | 149 | 17 | MOLECULE: | HISTONE-LYSINE N-METHYLTRANSFERASE |

| | | | | | | | | |
|---|---|---|---|---|---|---|---|---|
| 50:00:00 | 7w6a-C | 13.5 | 2.7 | 125 | 160 | 17 | MOLECULE: | SET1/ASH2 HISTONE METHYLTRANSFERASE COMPLEX SUBUN |
| 51:00:00 | 6pww-C | 13.5 | 2.6 | 124 | 154 | 16 | MOLECULE: | RETINOBLASTOMA-BINDING PROTEIN 5 |
| 52:00:00 | 5f6k-C | 13.5 | 2.5 | 121 | 151 | 21 | MOLECULE: | SET1/ASH2 HISTONE METHYLTRANSFERASE COMPLEX SUBUN |
| 53:00:00 | 5f59-A | 13.5 | 2.7 | 121 | 149 | 19 | MOLECULE: | HISTONE-LYSINE N-METHYLTRANSFERASE 2C |
| 54:00:00 | 5th7-A | 13.4 | 2 | 114 | 141 | 19 | MOLECULE: | N-LYSINE METHYLTRANSFERASE KMT5A |
| 55:00:00 | 5th7-B | 13.4 | 2.4 | 119 | 144 | 18 | MOLECULE: | N-LYSINE METHYLTRANSFERASE KMT5A |
| 56:00:00 | 6w5n-C | 13.4 | 2.5 | 123 | 153 | 16 | MOLECULE: | RETINOBLASTOMA-BINDING PROTEIN 5 |
| 57:00:00 | 7w6l-C | 13.4 | 2.6 | 122 | 159 | 20 | MOLECULE: | SET1/ASH2 HISTONE METHYLTRANSFERASE COMPLEX SUBUN |
| 58:00:00 | 2w5y-A | 13.3 | 2.7 | 123 | 171 | 16 | MOLECULE: | HISTONE-LYSINE N-METHYLTRANSFERASE HRX |
| 59:00:00 | 5v2n-A | 13.3 | 2.8 | 120 | 161 | 19 | MOLECULE: | N-LYSINE METHYLTRANSFERASE KMT5A |
| 60:00:00 | 5f6l-A | 13.3 | 2.4 | 119 | 154 | 17 | MOLECULE: | RETINOBLASTOMA-BINDING PROTEIN 5 |
| 61:00:00 | 6w5i-C | 13.3 | 2.4 | 120 | 152 | 17 | MOLECULE: | RETINOBLASTOMA-BINDING PROTEIN 5 |
| 62:00:00 | 7mbn-C | 13.3 | 2.5 | 122 | 153 | 16 | MOLECULE: | RETINOBLASTOMA-BINDING PROTEIN 5 |
| 63:00:00 | 6pwv-C | 13.3 | 2.5 | 123 | 155 | 16 | MOLECULE: | RETINOBLASTOMA-BINDING PROTEIN 5 |
| 64:00:00 | 7d1z-K | 13.2 | 2.3 | 125 | 166 | 18 | MOLECULE: | HISTONE H3.1 |
| 65:00:00 | 6w5m-C | 13.2 | 2.1 | 117 | 148 | 17 | MOLECULE: | RETINOBLASTOMA-BINDING PROTEIN 5 |
| 66:00:00 | 6kqp-A | 13.1 | 2.3 | 124 | 221 | 19 | MOLECULE: | HISTONE-LYSINE N-METHYLTRANSFERASE, H3 LYSINE-36 |
| 67:00:00 | 7bre-B | 13.1 | 2.4 | 121 | 164 | 15 | MOLECULE: | SET1/ASH2 HISTONE METHYLTRANSFERASE COMPLEX SUBUN |
| 68:00:00 | 6kqq-B | 13 | 2.3 | 124 | 220 | 19 | MOLECULE: | HISTONE-LYSINE N-METHYLTRANSFERASE, H3 LYSINE-36 |
| 69:00:00 | 6chg-C | 12.9 | 2.1 | 115 | 144 | 18 | MOLECULE: | KLLA0E24487P |
| 70:00:00 | 5ht6-A | 12.9 | 3.4 | 114 | 123 | 18 | MOLECULE: | HISTONE-LYSINE N-METHYLTRANSFERASE 2E |
| 71:00:00 | 2w5z-A | 12.9 | 2.6 | 124 | 181 | 16 | MOLECULE: | HISTONE-LYSINE N-METHYLTRANSFERASE HRX |
| 72:00:00 | 7bre-E | 12.9 | 2.4 | 121 | 164 | 15 | MOLECULE: | SET1/ASH2 HISTONE METHYLTRANSFERASE COMPLEX SUBUN |
| 73:00:00 | 6c24-K | 12.4 | 2.4 | 121 | 325 | 19 | MOLECULE: | POLYCOMB PROTEIN SUZ12 |
| 74:00:00 | 5hyn-Q | 12.3 | 2.7 | 123 | 566 | 19 | MOLECULE: | HISTONE-LYSINE N-METHYLTRANSFERASE EZH2 |
| 75:00:00 | 4ynp-B | 12.2 | 2.3 | 125 | 194 | 17 | MOLECULE: | HISTONE-LYSINE N-METHYLTRANSFERASE ASH1L |
| 76:00:00 | 6uh5-M | 12.1 | 3 | 116 | 222 | 18 | MOLECULE: | HISTONE H3 |
| 77:00:00 | 5lsu-A | 12.1 | 2.3 | 124 | 232 | 19 | MOLECULE: | HISTONE-LYSINE N-METHYLTRANSFERASE NSD2 |
| 78:00:00 | 5ij8-A | 12.1 | 2.5 | 121 | 464 | 21 | MOLECULE: | ENHANCER OF ZESTE HOMOLOG 2 (EZH2),HISTONE-LYSINE |
| 79:00:00 | 5lsu-B | 12 | 2.3 | 124 | 223 | 19 | MOLECULE: | HISTONE-LYSINE N-METHYLTRANSFERASE NSD2 |
| 80:00:00 | 6ven-N | 12 | 2.3 | 113 | 219 | 18 | MOLECULE: | HISTONE H3.2 |
| 81:00:00 | 3bo5-A | 12 | 2.6 | 137 | 270 | 19 | MOLECULE: | HISTONE-LYSINE N-METHYLTRANSFERASE SETMAR |
| 82:00:00 | 3ope-B | 12 | 2.4 | 125 | 194 | 17 | MOLECULE: | PROBABLE HISTONE-LYSINE N-METHYLTRANSFERASE ASH1L |
| 83:00:00 | 6kqq-A | 12 | 2.3 | 124 | 224 | 19 | MOLECULE: | HISTONE-LYSINE N-METHYLTRANSFERASE, H3 LYSINE-36 |
| 84:00:00 | 4ynm-B | 12 | 2.5 | 126 | 197 | 17 | MOLECULE: | HISTONE-LYSINE N-METHYLTRANSFERASE ASH1L |
| 85:00:00 | 8fbg-A | 12 | 2.3 | 124 | 218 | 19 | MOLECULE: | HISTONE-LYSINE N-METHYLTRANSFERASE, H3 LYSINE-36 |
| 86:00:00 | 5ht6-B | 11.9 | 2.2 | 103 | 109 | 20 | MOLECULE: | HISTONE-LYSINE N-METHYLTRANSFERASE 2E |
| 87:00:00 | 8fbg-B | 11.9 | 2.4 | 124 | 219 | 19 | MOLECULE: | HISTONE-LYSINE N-METHYLTRANSFERASE, H3 LYSINE-36 |
| 88:00:00 | 4h12-A | 11.8 | 2.2 | 124 | 234 | 18 | MOLECULE: | HISTONE-LYSINE N-METHYLTRANSFERASE SETD2 |
| 89:00:00 | 5tqr-B | 11.8 | 2.4 | 122 | 808 | 19 | MOLECULE: | POLYCOMB PROTEIN EED |
| 90:00:00 | 7td5-A | 11.8 | 2.7 | 123 | 498 | 18 | MOLECULE: | HISTONE-LYSINE N-METHYLTRANSFERASE EZH1 |
| 91:00:00 | 5ij8-B | 11.7 | 2.6 | 121 | 466 | 21 | MOLECULE: | ENHANCER OF ZESTE HOMOLOG 2 (EZH2),HISTONE-LYSINE |
| 92:00:00 | 5lt8-A | 11.7 | 2.3 | 118 | 228 | 19 | MOLECULE: | HISTONE-LYSINE N-METHYLTRANSFERASE SETD2 |
| 93:00:00 | 4ype-B | 11.7 | 2.5 | 125 | 203 | 16 | MOLECULE: | HISTONE-LYSINE N-METHYLTRANSFERASE ASH1L |
| 94:00:00 | 4w2r-B | 11.7 | 2.5 | 121 | 465 | 21 | MOLECULE: | ENHANCER OF ZESTE 2 POLYCOMB REPRESSIVE COMPLEX 2 |
| 95:00:00 | 6cen-A | 11.7 | 2.4 | 119 | 227 | 19 | MOLECULE: | HISTONE-LYSINE N-METHYLTRANSFERASE NSD3 |
| 96:00:00 | 1h3i-A | 11.6 | 2.7 | 118 | 293 | 19 | MOLECULE: | HISTONE H3 LYSINE 4 SPECIFIC METHYLTRANSFERASE |
| 97:00:00 | 3ooi-A | 11.6 | 2.5 | 123 | 233 | 19 | MOLECULE: | HISTONE-LYSINE N-METHYLTRANSFERASE, H3 LYSINE-36 |
| 98:00:00 | 4ypa-B | 11.6 | 2.4 | 125 | 213 | 16 | MOLECULE: | HISTONE-LYSINE N-METHYLTRANSFERASE ASH1L |
| 99:00:00 | 5wfd-B | 11.6 | 2.1 | 116 | 772 | 19 | MOLECULE: | POLYCOMB PROTEIN EED |
| 0:00 | 4ypa-D | 11.6 | 2.4 | 125 | 212 | 16 | MOLECULE: | HISTONE-LYSINE N-METHYLTRANSFERASE ASH1L |

| | | | | | | | |
|---|---|---|---|---|---|---|---|
| 1:00 | 5ij7-A | 11.6 | 2.5 | 121 | 465 | 21 MOLECULE: | ENHANCER OF ZESTE HOMOLOG 2 (EZH2),HISTONE-LYSINE |
| 2:00 | 6b3w-B | 11.6 | 2.5 | 121 | 465 | 21 MOLECULE: | ENHANCER OF ZESTE 2 POLYCOMB REPRESSIVE COMPLEX 2 |
| 3:00 | 8fbh-A | 11.6 | 2.5 | 123 | 233 | 19 MOLECULE: | HISTONE-LYSINE N-METHYLTRANSFERASE, H3 LYSINE-36 |
| 4:00 | 3ope-A | 11.5 | 2.2 | 123 | 214 | 18 MOLECULE: | PROBABLE HISTONE-LYSINE N-METHYLTRANSFERASE ASH1L |
| 5:00 | 4j7i-A | 11.5 | 3.3 | 121 | 244 | 18 MOLECULE: | HISTONE-LYSINE N-METHYLTRANSFERASE SETD7 |
| 6:00 | 4ypu-B | 11.5 | 2.3 | 120 | 191 | 17 MOLECULE: | HISTONE-LYSINE N-METHYLTRANSFERASE ASH1L |
| 7:00 | 4yz8-A | 11.5 | 2.2 | 118 | 210 | 19 MOLECULE: | HISTONE-LYSINE N-METHYLTRANSFERASE NSD3 |
| 8:00 | 4ynm-A | 11.5 | 2.5 | 125 | 216 | 18 MOLECULE: | HISTONE-LYSINE N-METHYLTRANSFERASE ASH1L |
| 9:00 | 5hyn-F | 11.5 | 2.7 | 123 | 569 | 19 MOLECULE: | HISTONE-LYSINE N-METHYLTRANSFERASE EZH2 |
| 10:00 | 7lzb-A | 11.5 | 2.4 | 119 | 236 | 18 MOLECULE: | HISTONE-LYSINE N-METHYLTRANSFERASE SETD2 |
| 11:00 | 4ypa-A | 11.5 | 2.4 | 125 | 219 | 16 MOLECULE: | HISTONE-LYSINE N-METHYLTRANSFERASE ASH1L |
| 12:00 | 7td5-F | 11.5 | 2.7 | 123 | 448 | 18 MOLECULE: | HISTONE-LYSINE N-METHYLTRANSFERASE EZH1 |
| 13:00 | 6b3w-A | 11.5 | 2.5 | 121 | 463 | 21 MOLECULE: | ENHANCER OF ZESTE 2 POLYCOMB REPRESSIVE COMPLEX 2 |
| 14:00 | 4ypa-C | 11.5 | 2.4 | 125 | 218 | 16 MOLECULE: | HISTONE-LYSINE N-METHYLTRANSFERASE ASH1L |
| 15:00 | 5kkl-B | 11.5 | 5.1 | 123 | 838 | 20 MOLECULE: | PUTATIVE POLYCOMB PROTEIN EED |
| 16:00 | 1h3i-B | 11.4 | 2.8 | 118 | 293 | 19 MOLECULE: | HISTONE H3 LYSINE 4 SPECIFIC METHYLTRANSFERASE |
| 17:00 | 5m5g-B | 11.4 | 2.7 | 124 | 796 | 19 MOLECULE: | PUTATIVE POLYCOMB PROTEIN EED |
| 18:00 | 6x0p-C | 11.4 | 2.4 | 125 | 221 | 16 MOLECULE: | HISTONE-LYSINE N-METHYLTRANSFERASE ASH1L |
| 19:00 | 5wfc-B | 11.4 | 2.3 | 120 | 768 | 19 MOLECULE: | POLYCOMB PROTEIN EED |
| 20:00 | 6x0p-D | 11.4 | 2.4 | 125 | 221 | 17 MOLECULE: | HISTONE-LYSINE N-METHYLTRANSFERASE ASH1L |
| 21:00 | 5upd-A | 11.4 | 2.2 | 118 | 212 | 19 MOLECULE: | HISTONE-LYSINE N-METHYLTRANSFERASE NSD3 |
| 22:00 | 6x0p-B | 11.4 | 2.5 | 126 | 221 | 17 MOLECULE: | HISTONE-LYSINE N-METHYLTRANSFERASE ASH1L |
| 23:00 | 5wf7-B | 11.4 | 2.5 | 122 | 764 | 19 MOLECULE: | POLYCOMB PROTEIN EED |
| 24:00:00 | 6x0p-A | 11.4 | 2.4 | 125 | 220 | 16 MOLECULE: | HISTONE-LYSINE N-METHYLTRANSFERASE ASH1L |
| 25:00:00 | 1n6a-A | 11.4 | 3 | 118 | 235 | 19 MOLECULE: | SET DOMAIN-CONTAINING PROTEIN 7 |
| 26:00:00 | 1mt6-A | 11.3 | 2.6 | 111 | 281 | 19 MOLECULE: | SET9 |
| 27:00:00 | 6ago-A | 11.3 | 2.6 | 124 | 243 | 17 MOLECULE: | HISTONE-LYSINE N-METHYLTRANSFERASE ASH1L |
| 28:00:00 | 4ypu-A | 11.3 | 2.2 | 123 | 215 | 17 MOLECULE: | HISTONE-LYSINE N-METHYLTRANSFERASE ASH1L |
| 29:00:00 | 4ynp-A | 11.3 | 2.3 | 123 | 214 | 19 MOLECULE: | HISTONE-LYSINE N-METHYLTRANSFERASE ASH1L |
| 30:00:00 | 6kiz-K | 11.3 | 2 | 117 | 180 | 17 MOLECULE: | HISTONE H3 |
| 31:00:00 | 6wzw-A | 11.3 | 2.4 | 125 | 222 | 17 MOLECULE: | HISTONE-LYSINE N-METHYLTRANSFERASE ASH1L |
| 32:00:00 | 3vuz-A | 11.2 | 3.3 | 121 | 247 | 17 MOLECULE: | HISTONE-LYSINE N-METHYLTRANSFERASE SETD7 |
| 33:00:00 | 1muf-A | 11.1 | 2.6 | 111 | 257 | 17 MOLECULE: | SET9 |
| 34:00:00 | 3m5a-A | 11.1 | 3.2 | 119 | 247 | 17 MOLECULE: | HISTONE-LYSINE N-METHYLTRANSFERASE SETD7 |
| 35:00:00 | 5eg2-A | 11.1 | 3.2 | 120 | 244 | 17 MOLECULE: | HISTONE-LYSINE N-METHYLTRANSFERASE SETD7 |
| 36:00:00 | 3m55-A | 11.1 | 3.3 | 121 | 245 | 17 MOLECULE: | HISTONE-LYSINE N-METHYLTRANSFERASE SETD7 |
| 37:00:00 | 3m56-A | 11.1 | 3.2 | 120 | 245 | 17 MOLECULE: | HISTONE-LYSINE N-METHYLTRANSFERASE SETD7 |
| 38:00:00 | 4j7f-A | 11.1 | 3.3 | 121 | 242 | 17 MOLECULE: | HISTONE-LYSINE N-METHYLTRANSFERASE SETD7 |
| 39:00:00 | 3m54-A | 11.1 | 3.2 | 120 | 246 | 17 MOLECULE: | HISTONE-LYSINE N-METHYLTRANSFERASE SETD7 |
| 40:00:00 | 4ype-A | 11.1 | 2.6 | 125 | 218 | 16 MOLECULE: | HISTONE-LYSINE N-METHYLTRANSFERASE ASH1L |
| 41:00:00 | 6ine-A | 11.1 | 2.3 | 119 | 248 | 18 MOLECULE: | HISTONE-LYSINE N-METHYLTRANSFERASE ASH1L |
| 42:00:00 | 8fyh-G | 11.1 | 2.5 | 125 | 514 | 20 MOLECULE: | POLYCOMB PROTEIN SUZ12 |
| 43:00:00 | 3cbo-A | 11.1 | 3.3 | 120 | 240 | 16 MOLECULE: | HISTONE-LYSINE N-METHYLTRANSFERASE SETD7 |
| 44:00:00 | 2f69-A | 11 | 3.2 | 119 | 245 | 17 MOLECULE: | HISTONE-LYSINE N-METHYLTRANSFERASE, H3 LYSINE-4 |
| 45:00:00 | 4j8o-A | 11 | 3.2 | 120 | 246 | 17 MOLECULE: | HISTONE-LYSINE N-METHYLTRANSFERASE SETD7 |
| 46:00:00 | 3m57-A | 11 | 3.3 | 121 | 245 | 17 MOLECULE: | HISTONE-LYSINE N-METHYLTRANSFERASE SETD7 |
| 47:00:00 | 3m59-A | 11 | 3.2 | 119 | 246 | 17 MOLECULE: | HISTONE-LYSINE N-METHYLTRANSFERASE SETD7 |
| 48:00:00 | 4j83-A | 11 | 3.3 | 120 | 243 | 19 MOLECULE: | HISTONE-LYSINE N-METHYLTRANSFERASE SETD7 |
| 49:00:00 | 3m53-A | 11 | 3.2 | 120 | 247 | 17 MOLECULE: | HISTONE-LYSINE N-METHYLTRANSFERASE SETD7 |
| 50:00:00 | 4jds-A | 11 | 3 | 119 | 247 | 18 MOLECULE: | HISTONE-LYSINE N-METHYLTRANSFERASE SETD7 |
| 51:00:00 | 3cbp-A | 11 | 3 | 118 | 240 | 17 MOLECULE: | HISTONE-LYSINE N-METHYLTRANSFERASE SETD7 |

| | | | | | | | | |
|---|---|---|---|---|---|---|---|---|
| 52:00:00 | 4e47-B | 11 | 3 | 119 | 246 | 18 | MOLECULE: | HISTONE-LYSINE N-METHYLTRANSFERASE SETD7 |
| 53:00:00 | 4jds-B | 11 | 2.8 | 117 | 245 | 16 | MOLECULE: | HISTONE-LYSINE N-METHYLTRANSFERASE SETD7 |
| 54:00:00 | 4e47-A | 11 | 3 | 119 | 247 | 18 | MOLECULE: | HISTONE-LYSINE N-METHYLTRANSFERASE SETD7 |
| 55:00:00 | 5vsc-A | 11 | 2.8 | 126 | 270 | 17 | MOLECULE: | HISTONE-LYSINE N-METHYLTRANSFERASE EHMT2 |
| 56:00:00 | 3cbm-A | 11 | 3.3 | 120 | 244 | 17 | MOLECULE: | HISTONE-LYSINE N-METHYLTRANSFERASE SETD7 |
| 57:00:00 | 1peg-B | 10.9 | 2.8 | 123 | 210 | 19 | MOLECULE: | HISTONE H3 METHYLTRANSFERASE DIM-5 |
| 58:00:00 | 3os5-A | 10.9 | 3.2 | 119 | 250 | 17 | MOLECULE: | HISTONE-LYSINE N-METHYLTRANSFERASE SETD7 |
| 59:00:00 | 5ch2-B | 10.9 | 2.4 | 120 | 755 | 19 | MOLECULE: | PUTATIVE POLYCOMB PROTEIN EED |
| 60:00:00 | 5ylt-A | 10.9 | 3.1 | 117 | 248 | 18 | MOLECULE: | HISTONE-LYSINE N-METHYLTRANSFERASE SETD7 |
| 61:00:00 | 6kiu-K | 10.9 | 2.2 | 117 | 182 | 17 | MOLECULE: | HISTONE H3 |
| 62:00:00 | 5jle-A | 10.9 | 2.4 | 122 | 233 | 18 | MOLECULE: | HISTONE-LYSINE N-METHYLTRANSFERASE SETD2 |
| 63:00:00 | 6c23-K | 10.9 | 2.6 | 123 | 325 | 19 | MOLECULE: | POLYCOMB PROTEIN SUZ12 |
| 64:00:00 | 3m58-A | 10.9 | 3.2 | 119 | 254 | 18 | MOLECULE: | HISTONE-LYSINE N-METHYLTRANSFERASE SETD7 |
| 65:00:00 | 4jds-C | 10.9 | 3.1 | 119 | 247 | 18 | MOLECULE: | HISTONE-LYSINE N-METHYLTRANSFERASE SETD7 |
| 66:00:00 | 8sr6-A | 10.8 | 2.4 | 110 | 452 | 18 | MOLECULE: | EUKARYOTIC HUNTINGTIN INTERACTING PROTEIN B |
| 67:00:00 | 5ch1-B | 10.8 | 2.6 | 121 | 796 | 19 | MOLECULE: | PUTATIVE POLYCOMB PROTEIN EED |
| 68:00:00 | 6vdb-A | 10.8 | 2.3 | 120 | 245 | 18 | MOLECULE: | HISTONE-LYSINE N-METHYLTRANSFERASE SETD2 |
| 69:00:00 | 4jds-D | 10.8 | 3.1 | 119 | 246 | 18 | MOLECULE: | HISTONE-LYSINE N-METHYLTRANSFERASE SETD7 |
| 70:00:00 | 1o9s-A | 10.8 | 3.3 | 121 | 251 | 17 | MOLECULE: | HISTONE-LYSINE N-METHYLTRANSFERASE, H3 LYSINE-4 |
| 71:00:00 | 5lt6-A | 10.8 | 2.4 | 121 | 234 | 18 | MOLECULE: | HISTONE-LYSINE N-METHYLTRANSFERASE SETD2 |
| 72:00:00 | 5lsx-A | 10.8 | 2.1 | 118 | 231 | 19 | MOLECULE: | HISTONE-LYSINE N-METHYLTRANSFERASE SETD2 |
| 73:00:00 | 4e47-C | 10.8 | 3.1 | 119 | 246 | 18 | MOLECULE: | HISTONE-LYSINE N-METHYLTRANSFERASE SETD7 |
| 74:00:00 | 1xqh-A | 10.7 | 3.2 | 119 | 251 | 17 | MOLECULE: | HISTONE-LYSINE N-METHYLTRANSFERASE, H3 LYSINE-4 |
| 75:00:00 | 5lss-A | 10.7 | 2.4 | 122 | 238 | 17 | MOLECULE: | HISTONE-LYSINE N-METHYLTRANSFERASE SETD2 |
| 76:00:00 | 1xqh-E | 10.7 | 3.2 | 119 | 251 | 17 | MOLECULE: | HISTONE-LYSINE N-METHYLTRANSFERASE, H3 LYSINE-4 |
| 77:00:00 | 5lsy-A | 10.7 | 2.4 | 122 | 246 | 17 | MOLECULE: | HISTONE-LYSINE N-METHYLTRANSFERASE SETD2 |
| 78:00:00 | 5lt7-A | 10.7 | 2.4 | 122 | 243 | 17 | MOLECULE: | HISTONE-LYSINE N-METHYLTRANSFERASE SETD2 |
| 79:00:00 | 5lsz-A | 10.7 | 2.4 | 122 | 243 | 17 | MOLECULE: | HISTONE-LYSINE N-METHYLTRANSFERASE SETD2 |
| 80:00:00 | 8q5p-A | 10.7 | 2.3 | 120 | 246 | 18 | MOLECULE: | HISTONE-LYSINE N-METHYLTRANSFERASE SETD2 |
| 81:00:00 | 7e8d-K | 10.7 | 2.3 | 119 | 219 | 19 | MOLECULE: | HISTONE H3.1 |
| 82:00:00 | 5kji-B | 10.7 | 2.6 | 123 | 762 | 19 | MOLECULE: | PUTATIVE POLYCOMB PROTEIN EED |
| 83:00:00 | 6kiv-K | 10.7 | 2.7 | 121 | 180 | 17 | MOLECULE: | HISTONE H3 |
| 84:00:00 | 4jlg-A | 10.7 | 3.3 | 119 | 249 | 18 | MOLECULE: | HISTONE-LYSINE N-METHYLTRANSFERASE SETD7 |
| 85:00:00 | 4e47-D | 10.7 | 3.3 | 120 | 251 | 17 | MOLECULE: | HISTONE-LYSINE N-METHYLTRANSFERASE SETD7 |
| 86:00:00 | 1o9s-B | 10.7 | 3.1 | 119 | 251 | 18 | MOLECULE: | HISTONE-LYSINE N-METHYLTRANSFERASE, H3 LYSINE-4 |
| 87:00:00 | 5wg6-A | 10.7 | 2.4 | 118 | 517 | 19 | MOLECULE: | HISTONE-LYSINE N-METHYLTRANSFERASE EZH2,POLYCOMB |
| 88:00:00 | 5jjy-A | 10.6 | 2.3 | 120 | 248 | 18 | MOLECULE: | HISTONE-LYSINE N-METHYLTRANSFERASE SETD2 |
| 89:00:00 | 6j9j-A | 10.6 | 2.3 | 120 | 248 | 18 | MOLECULE: | HISTONE-LYSINE N-METHYLTRANSFERASE SETD2 |
| 90:00:00 | 1n6c-A | 10.6 | 2.7 | 116 | 284 | 18 | MOLECULE: | SET DOMAIN-CONTAINING PROTEIN 7 |
| 91:00:00 | 4mi0-A | 10.6 | 2.4 | 118 | 208 | 19 | MOLECULE: | HISTONE-LYSINE N-METHYLTRANSFERASE EZH2 |
| 92:00:00 | 5jlb-A | 10.6 | 2.3 | 120 | 249 | 18 | MOLECULE: | HISTONE-LYSINE N-METHYLTRANSFERASE SETD2 |
| 93:00:00 | 5v21-A | 10.6 | 2.3 | 120 | 246 | 18 | MOLECULE: | HISTONE-LYSINE N-METHYLTRANSFERASE SETD2 |
| 94:00:00 | 5v22-A | 10.6 | 2.3 | 120 | 246 | 18 | MOLECULE: | HISTONE-LYSINE N-METHYLTRANSFERASE SETD2 |
| 95:00:00 | 4fmu-A | 10.6 | 2.4 | 121 | 236 | 17 | MOLECULE: | HISTONE-LYSINE N-METHYLTRANSFERASE SETD2 |
| 96:00:00 | 7ty3-A | 10.6 | 2.3 | 119 | 239 | 18 | MOLECULE: | HISTONE-LYSINE N-METHYLTRANSFERASE SETD2 |
| 97:00:00 | 4jlg-B | 10.6 | 2.9 | 117 | 248 | 18 | MOLECULE: | HISTONE-LYSINE N-METHYLTRANSFERASE SETD7 |
| 98:00:00 | 3swc-A | 10.6 | 2.8 | 129 | 262 | 19 | MOLECULE: | HISTONE-LYSINE N-METHYLTRANSFERASE EHMT1 |
| 99:00:00 | 7lzd-A | 10.5 | 2.1 | 118 | 238 | 19 | MOLECULE: | HISTONE-LYSINE N-METHYLTRANSFERASE SETD2 |
| 0:00 | 7lzf-A | 10.5 | 2.1 | 118 | 240 | 19 | MOLECULE: | HISTONE-LYSINE N-METHYLTRANSFERASE SETD2 |
| 1:00 | 7ty2-A | 10.5 | 2.1 | 118 | 235 | 19 | MOLECULE: | HISTONE-LYSINE N-METHYLTRANSFERASE SETD2 |
| 2:00 | 3kmj-A | 10.5 | 2.8 | 103 | 119 | 16 | MOLECULE: | A612L PROTEIN |

| | | | | | | | | |
|---|---|---|---|---|---|---|---|---|
| 3:00 | 5czy-A | 10.5 | 2.8 | 114 | 450 | 18 | MOLECULE: | LEGIONELLA EFFECTOR LEGAS4 |
| 4:00 | 7t7m-C | 10.5 | 2.8 | 127 | 260 | 17 | MOLECULE: | HISTONE-LYSINE N-METHYLTRANSFERASE EHMT1 |
| 5:00 | 6z2a-A | 10.5 | 2.8 | 133 | 284 | 15 | MOLECULE: | HISTONE-LYSINE N-METHYLTRANSFERASE, H3 LYSINE-9 S |
| 6:00 | 5vsf-A | 10.5 | 2.9 | 130 | 262 | 18 | MOLECULE: | HISTONE-LYSINE N-METHYLTRANSFERASE EHMT1 |
| 7:00 | 5v9j-B | 10.5 | 2.9 | 129 | 260 | 19 | MOLECULE: | HISTONE-LYSINE N-METHYLTRANSFERASE EHMT1 |
| 8:00 | 3kmt-A | 10.4 | 2.7 | 102 | 120 | 17 | MOLECULE: | A612L PROTEIN |
| 9:00 | 4mi5-A | 10.4 | 2.5 | 118 | 208 | 19 | MOLECULE: | HISTONE-LYSINE N-METHYLTRANSFERASE EZH2 |
| 10:00 | 5vk3-B | 10.4 | 2.4 | 121 | 792 | 19 | MOLECULE: | POLYCOMB PROTEIN EED |
| 11:00 | 3mo0-B | 10.4 | 2.9 | 126 | 243 | 19 | MOLECULE: | HISTONE-LYSINE N-METHYLTRANSFERASE, H3 LYSINE-9 S |
| 12:00 | 3kmt-C | 10.4 | 2.7 | 103 | 120 | 16 | MOLECULE: | A612L PROTEIN |
| 13:00 | 3kmt-B | 10.4 | 2.8 | 103 | 120 | 16 | MOLECULE: | A612L PROTEIN |
| 14:00 | 7dcf-A | 10.4 | 2.6 | 126 | 261 | 17 | MOLECULE: | HISTONE-LYSINE N-METHYLTRANSFERASE EHMT2 |
| 15:00 | 7t7m-A | 10.4 | 2.9 | 129 | 260 | 18 | MOLECULE: | HISTONE-LYSINE N-METHYLTRANSFERASE EHMT1 |
| 16:00 | 7xud-B | 10.4 | 3 | 130 | 271 | 17 | MOLECULE: | HISTONE-LYSINE N-METHYLTRANSFERASE EHMT2 |
| 17:00 | 8fyh-A | 10.4 | 2.5 | 125 | 514 | 20 | MOLECULE: | POLYCOMB PROTEIN SUZ12 |
| 18:00 | 6mbo-A | 10.4 | 2.9 | 129 | 262 | 19 | MOLECULE: | HISTONE-LYSINE N-METHYLTRANSFERASE EHMT1 |
| 19:00 | 5v9j-A | 10.4 | 2.9 | 127 | 258 | 19 | MOLECULE: | HISTONE-LYSINE N-METHYLTRANSFERASE EHMT1 |
| 20:00 | 3swc-B | 10.4 | 3 | 129 | 262 | 19 | MOLECULE: | HISTONE-LYSINE N-METHYLTRANSFERASE EHMT1 |
| 21:00 | 7xud-A | 10.4 | 2.9 | 129 | 269 | 17 | MOLECULE: | HISTONE-LYSINE N-METHYLTRANSFERASE EHMT2 |
| 22:00 | 5tuz-A | 10.4 | 2.9 | 127 | 260 | 19 | MOLECULE: | HISTONE-LYSINE N-METHYLTRANSFERASE EHMT1 |
| 23:00 | 5ttf-D | 10.4 | 2.8 | 124 | 262 | 17 | MOLECULE: | HISTONE-LYSINE N-METHYLTRANSFERASE EHMT2 |
| 24:00:00 | 5h6z-A | 10.3 | 2.6 | 108 | 123 | 13 | MOLECULE: | SET DOMAIN-CONTAINING PROTEIN 7 |
| 25:00:00 | 2rfi-A | 10.3 | 3.1 | 128 | 263 | 19 | MOLECULE: | HISTONE-LYSINE N-METHYLTRANSFERASE, H3 LYSINE-9 |
| 26:00:00 | 3mo0-A | 10.3 | 3.1 | 131 | 252 | 18 | MOLECULE: | HISTONE-LYSINE N-METHYLTRANSFERASE, H3 LYSINE-9 S |
| 27:00:00 | 4rz0-B | 10.3 | 2.3 | 98 | 110 | 12 | MOLECULE: | PFL0690C |
| 28:00:00 | 5vah-A | 10.3 | 3.2 | 129 | 214 | 15 | MOLECULE: | PROBABLE HISTONE-LYSINE N-METHYLTRANSFERASE ATXR5 |
| 29:00:00 | 3hna-B | 10.3 | 3.2 | 130 | 261 | 18 | MOLECULE: | HISTONE-LYSINE N-METHYLTRANSFERASE, H3 LYSINE-9 S |
| 30:00:00 | 5vah-B | 10.3 | 3.2 | 129 | 215 | 16 | MOLECULE: | PROBABLE HISTONE-LYSINE N-METHYLTRANSFERASE ATXR5 |
| 31:00:00 | 4i51-A | 10.3 | 3.2 | 130 | 262 | 18 | MOLECULE: | HISTONE-LYSINE N-METHYLTRANSFERASE EHMT1 |
| 32:00:00 | 7mbm-C | 10.3 | 2.3 | 117 | 179 | 17 | MOLECULE: | RETINOBLASTOMA-BINDING PROTEIN 5 |
| 33:00:00 | 7xub-A | 10.3 | 2.7 | 127 | 271 | 17 | MOLECULE: | HISTONE-LYSINE N-METHYLTRANSFERASE EHMT2 |
| 34:00:00 | 7x73-B | 10.3 | 2.7 | 128 | 274 | 17 | MOLECULE: | HISTONE-LYSINE N-METHYLTRANSFERASE EHMT2 |
| 35:00:00 | 7xuc-B | 10.3 | 2.9 | 129 | 270 | 17 | MOLECULE: | HISTONE-LYSINE N-METHYLTRANSFERASE EHMT2 |
| 36:00:00 | 5kjh-B | 10.3 | 2.5 | 120 | 806 | 19 | MOLECULE: | PUTATIVE POLYCOMB PROTEIN EED |
| 37:00:00 | 7at8-A | 10.3 | 2.7 | 123 | 312 | 19 | MOLECULE: | ISOFORM 2 OF HISTONE-LYSINE N-METHYLTRANSFERASE E |
| 38:00:00 | 3mo5-B | 10.3 | 2.8 | 127 | 258 | 17 | MOLECULE: | HISTONE-LYSINE N-METHYLTRANSFERASE, H3 LYSINE-9 S |
| 39:00:00 | 5ttg-B | 10.3 | 2.6 | 125 | 259 | 18 | MOLECULE: | HISTONE-LYSINE N-METHYLTRANSFERASE EHMT1 |
| 40:00:00 | 3fpd-A | 10.3 | 3 | 130 | 262 | 18 | MOLECULE: | HISTONE-LYSINE N-METHYLTRANSFERASE, H3 LYSINE-9 |
| 41:00:00 | 3sw9-B | 10.3 | 3 | 129 | 262 | 19 | MOLECULE: | HISTONE-LYSINE N-METHYLTRANSFERASE EHMT1 |
| 42:00:00 | 3mo5-C | 10.3 | 2.9 | 127 | 259 | 19 | MOLECULE: | HISTONE-LYSINE N-METHYLTRANSFERASE, H3 LYSINE-9 S |
| 43:00:00 | 5vsd-B | 10.3 | 2.9 | 126 | 259 | 19 | MOLECULE: | HISTONE-LYSINE N-METHYLTRANSFERASE EHMT1 |
| 44:00:00 | 6mbp-A | 10.3 | 2.9 | 128 | 262 | 19 | MOLECULE: | HISTONE-LYSINE N-METHYLTRANSFERASE EHMT1 |
| 45:00:00 | 5tuz-B | 10.3 | 2.8 | 126 | 258 | 18 | MOLECULE: | HISTONE-LYSINE N-METHYLTRANSFERASE EHMT1 |
| 46:00:00 | 5tuy-A | 10.3 | 2.9 | 130 | 268 | 17 | MOLECULE: | HISTONE-LYSINE N-METHYLTRANSFERASE EHMT2 |
| 47:00:00 | 5vsf-B | 10.3 | 2.8 | 126 | 259 | 19 | MOLECULE: | HISTONE-LYSINE N-METHYLTRANSFERASE EHMT1 |
| 48:00:00 | 4i51-B | 10.3 | 2.8 | 125 | 259 | 18 | MOLECULE: | HISTONE-LYSINE N-METHYLTRANSFERASE EHMT1 |
| 49:00:00 | 6mbp-B | 10.3 | 2.9 | 128 | 261 | 19 | MOLECULE: | HISTONE-LYSINE N-METHYLTRANSFERASE EHMT1 |
| 50:00:00 | 6mbo-B | 10.3 | 2.8 | 126 | 259 | 19 | MOLECULE: | HISTONE-LYSINE N-METHYLTRANSFERASE EHMT1 |
| 51:00:00 | 5vsd-A | 10.3 | 2.8 | 126 | 258 | 18 | MOLECULE: | HISTONE-LYSINE N-METHYLTRANSFERASE EHMT1 |
| 52:00:00 | 4o30-B | 10.2 | 3.2 | 128 | 217 | 15 | MOLECULE: | HISTONE-LYSINE N-METHYLTRANSFERASE ATXR6, PUTATIV |
| 53:00:00 | 5bjs-B | 10.2 | 2.5 | 122 | 799 | 19 | MOLECULE: | POLYCOMB PROTEIN EED |

| | | | | | | | | |
|---|---|---|---|---|---|---|---|---|
| 54:00:00 | 3sw9-A | 10.2 | 3 | 130 | 262 | 18 | MOLECULE: | HISTONE-LYSINE N-METHYLTRANSFERASE EHMT1 |
| 55:00:00 | 5vbc-A | 10.2 | 3.2 | 129 | 217 | 16 | MOLECULE: | PROBABLE HISTONE-LYSINE N-METHYLTRANSFERASE ATXR5 |
| 56:00:00 | 5ttg-A | 10.2 | 3.1 | 129 | 261 | 19 | MOLECULE: | HISTONE-LYSINE N-METHYLTRANSFERASE EHMT1 |
| 57:00:00 | 5vbc-B | 10.2 | 3.2 | 128 | 216 | 15 | MOLECULE: | PROBABLE HISTONE-LYSINE N-METHYLTRANSFERASE ATXR5 |
| 58:00:00 | 5va6-B | 10.2 | 3.2 | 129 | 216 | 15 | MOLECULE: | PROBABLE HISTONE-LYSINE N-METHYLTRANSFERASE ATXR5 |
| 59:00:00 | 6kix-K | 10.2 | 2.3 | 117 | 182 | 17 | MOLECULE: | HISTONE H3 |
| 60:00:00 | 5va6-A | 10.2 | 3.2 | 129 | 217 | 16 | MOLECULE: | PROBABLE HISTONE-LYSINE N-METHYLTRANSFERASE ATXR5 |
| 61:00:00 | 3mo2-D | 10.2 | 2.9 | 126 | 257 | 18 | MOLECULE: | HISTONE-LYSINE N-METHYLTRANSFERASE, H3 LYSINE-9 S |
| 62:00:00 | 3mo2-C | 10.2 | 2.7 | 125 | 259 | 18 | MOLECULE: | HISTONE-LYSINE N-METHYLTRANSFERASE, H3 LYSINE-9 S |
| 63:00:00 | 5lt6-B | 10.2 | 2.5 | 117 | 227 | 19 | MOLECULE: | HISTONE-LYSINE N-METHYLTRANSFERASE SETD2 |
| 64:00:00 | 5ww0-A | 10.2 | 2.6 | 108 | 123 | 13 | MOLECULE: | SET DOMAIN-CONTAINING PROTEIN 7 |
| 65:00:00 | 7xuc-A | 10.2 | 2.9 | 129 | 273 | 17 | MOLECULE: | HISTONE-LYSINE N-METHYLTRANSFERASE EHMT2 |
| 66:00:00 | 5t0k-B | 10.2 | 3.1 | 128 | 266 | 17 | MOLECULE: | HISTONE-LYSINE N-METHYLTRANSFERASE EHMT2 |
| 67:00:00 | 5vse-A | 10.2 | 2.8 | 127 | 270 | 17 | MOLECULE: | HISTONE-LYSINE N-METHYLTRANSFERASE EHMT2 |
| 68:00:00 | 3fpd-B | 10.2 | 3 | 129 | 261 | 19 | MOLECULE: | HISTONE-LYSINE N-METHYLTRANSFERASE, H3 LYSINE-9 |
| 69:00:00 | 3mo2-B | 10.2 | 2.9 | 125 | 258 | 18 | MOLECULE: | HISTONE-LYSINE N-METHYLTRANSFERASE, H3 LYSINE-9 S |
| 70:00:00 | 3mo5-A | 10.2 | 2.8 | 125 | 255 | 19 | MOLECULE: | HISTONE-LYSINE N-METHYLTRANSFERASE, H3 LYSINE-9 S |
| 71:00:00 | 5ttf-A | 10.2 | 2.8 | 125 | 266 | 17 | MOLECULE: | HISTONE-LYSINE N-METHYLTRANSFERASE EHMT2 |
| 72:00:00 | 6wkr-C | 10.1 | 2.7 | 122 | 607 | 19 | MOLECULE: | UBIQUITIN |
| 73:00:00 | 4o30-A | 10.1 | 3.2 | 129 | 218 | 16 | MOLECULE: | HISTONE-LYSINE N-METHYLTRANSFERASE ATXR6, PUTATIV |
| 74:00:00 | 2r3a-A | 10.1 | 2.7 | 130 | 272 | 18 | MOLECULE: | HISTONE-LYSINE N-METHYLTRANSFERASE SUV39H2 |
| 75:00:00 | 6ugm-M | 10.1 | 2.3 | 112 | 189 | 18 | MOLECULE: | HISTONE H3 |
| 76:00:00 | 3mo5-D | 10.1 | 2.9 | 127 | 257 | 18 | MOLECULE: | HISTONE-LYSINE N-METHYLTRANSFERASE, H3 LYSINE-9 S |
| 77:00:00 | 7btv-A | 10.1 | 2.6 | 127 | 272 | 17 | MOLECULE: | HISTONE-LYSINE N-METHYLTRANSFERASE EHMT2 |
| 78:00:00 | 5ij7-B | 10.1 | 2.5 | 121 | 469 | 21 | MOLECULE: | ENHANCER OF ZESTE HOMOLOG 2 (EZH2),HISTONE-LYSINE |
| 79:00:00 | 5t0m-A | 10.1 | 2.9 | 126 | 261 | 17 | MOLECULE: | HISTONE-LYSINE N-METHYLTRANSFERASE EHMT2 |
| 80:00:00 | 5v9i-B | 10.1 | 2.8 | 125 | 268 | 16 | MOLECULE: | HISTONE-LYSINE N-METHYLTRANSFERASE EHMT2 |
| 81:00:00 | 8xpt-C | 10.1 | 2.9 | 126 | 256 | 17 | MOLECULE: | HISTONE-LYSINE N-METHYLTRANSFERASE EHMT1 |
| 82:00:00 | 7xub-B | 10.1 | 2.6 | 124 | 267 | 17 | MOLECULE: | HISTONE-LYSINE N-METHYLTRANSFERASE EHMT2 |
| 83:00:00 | 7x73-A | 10.1 | 2.8 | 128 | 275 | 17 | MOLECULE: | HISTONE-LYSINE N-METHYLTRANSFERASE EHMT2 |
| 84:00:00 | 2igq-B | 10.1 | 2.8 | 125 | 255 | 18 | MOLECULE: | EUCHROMATIC HISTONE METHYLTRANSFERASE 1 |
| 85:00:00 | 5vac-A | 10.1 | 3.3 | 125 | 216 | 15 | MOLECULE: | PROBABLE HISTONE-LYSINE N-METHYLTRANSFERASE ATXR5 |
| 86:00:00 | 7t7l-C | 10.1 | 2.8 | 127 | 267 | 17 | MOLECULE: | HISTONE-LYSINE N-METHYLTRANSFERASE EHMT2 |
| 87:00:00 | 7t7l-A | 10.1 | 2.7 | 126 | 267 | 17 | MOLECULE: | HISTONE-LYSINE N-METHYLTRANSFERASE EHMT2 |
| 88:00:00 | 7buc-A | 10.1 | 2.9 | 130 | 274 | 17 | MOLECULE: | HISTONE-LYSINE N-METHYLTRANSFERASE EHMT2 |
| 89:00:00 | 3mo2-A | 10.1 | 2.9 | 125 | 255 | 19 | MOLECULE: | HISTONE-LYSINE N-METHYLTRANSFERASE, H3 LYSINE-9 S |
| 90:00:00 | 5vsc-B | 10.1 | 2.9 | 127 | 273 | 17 | MOLECULE: | HISTONE-LYSINE N-METHYLTRANSFERASE EHMT2 |
| 91:00:00 | 6a5n-A | 10 | 2.7 | 125 | 502 | 17 | MOLECULE: | HISTONE-LYSINE N-METHYLTRANSFERASE, H3 LYSINE-9 S |
| 92:00:00 | 2rfi-B | 10 | 2.6 | 122 | 252 | 16 | MOLECULE: | HISTONE-LYSINE N-METHYLTRANSFERASE, H3 LYSINE-9 |
| 93:00:00 | 2o8j-B | 10 | 2.8 | 128 | 272 | 17 | MOLECULE: | HISTONE-LYSINE N-METHYLTRANSFERASE, H3 LYSINE-9 |
| 94:00:00 | 4nvq-B | 10 | 2.8 | 125 | 264 | 16 | MOLECULE: | HISTONE-LYSINE N-METHYLTRANSFERASE EHMT2 |
| 95:00:00 | 7xua-A | 10 | 2.8 | 128 | 274 | 17 | MOLECULE: | HISTONE-LYSINE N-METHYLTRANSFERASE EHMT2 |
| 96:00:00 | 5jin-B | 10 | 2.9 | 126 | 268 | 17 | MOLECULE: | HISTONE-LYSINE N-METHYLTRANSFERASE EHMT2 |
| 97:00:00 | 7dcf-B | 10 | 2.8 | 126 | 260 | 17 | MOLECULE: | HISTONE-LYSINE N-METHYLTRANSFERASE EHMT2 |
| 98:00:00 | 6ago-B | 10 | 1.9 | 112 | 224 | 17 | MOLECULE: | HISTONE-LYSINE N-METHYLTRANSFERASE ASH1L |
| 99:00:00 | 5jhn-B | 10 | 2.9 | 126 | 268 | 17 | MOLECULE: | HISTONE-LYSINE N-METHYLTRANSFERASE EHMT2 |
| 0:00 | 5ttf-C | 10 | 2.9 | 127 | 265 | 16 | MOLECULE: | HISTONE-LYSINE N-METHYLTRANSFERASE EHMT2 |
| 1:00 | 7t7l-B | 10 | 2.6 | 125 | 265 | 18 | MOLECULE: | HISTONE-LYSINE N-METHYLTRANSFERASE EHMT2 |
| 2:00 | 5t0k-A | 10 | 2.9 | 126 | 265 | 17 | MOLECULE: | HISTONE-LYSINE N-METHYLTRANSFERASE EHMT2 |
| 3:00 | 4w2r-A | 10 | 2.5 | 121 | 463 | 21 | MOLECULE: | ENHANCER OF ZESTE 2 POLYCOMB REPRESSIVE COMPLEX 2 |
| 4:00 | 5v9i-D | 10 | 2.8 | 125 | 265 | 16 | MOLECULE: | HISTONE-LYSINE N-METHYLTRANSFERASE EHMT2 |

| | | | | | | | | |
|---|---|---|---|---|---|---|---|---|
| 5:00 | 7t7m-B | 10 | 2.7 | 126 | 259 | 17 | MOLECULE: | HISTONE-LYSINE N-METHYLTRANSFERASE EHMT1 |
| 6:00 | 7xua-B | 10 | 2.6 | 125 | 275 | 18 | MOLECULE: | HISTONE-LYSINE N-METHYLTRANSFERASE EHMT2 |
| 7:00 | 3k5k-B | 10 | 2.6 | 125 | 270 | 17 | MOLECULE: | HISTONE-LYSINE N-METHYLTRANSFERASE, H3 LYSINE-9 S |
| 8:00 | 5tuy-B | 10 | 2.9 | 127 | 264 | 17 | MOLECULE: | HISTONE-LYSINE N-METHYLTRANSFERASE EHMT2 |
| 9:00 | 5hyn-A | 9.9 | 2.7 | 123 | 582 | 19 | MOLECULE: | HISTONE-LYSINE N-METHYLTRANSFERASE EZH2 |
| 10:00 | 4rz0-A | 9.9 | 2.7 | 98 | 112 | 12 | MOLECULE: | PFL0690C |
| 11:00 | 5h6z-B | 9.9 | 2.4 | 102 | 120 | 14 | MOLECULE: | SET DOMAIN-CONTAINING PROTEIN 7 |
| 12:00 | 5jiy-A | 9.9 | 3.3 | 130 | 274 | 17 | MOLECULE: | HISTONE-LYSINE N-METHYLTRANSFERASE EHMT2 |
| 13:00 | 3hna-A | 9.9 | 3.2 | 130 | 281 | 18 | MOLECULE: | HISTONE-LYSINE N-METHYLTRANSFERASE, H3 LYSINE-9 S |
| 14:00 | 1peg-A | 9.9 | 2.4 | 120 | 249 | 18 | MOLECULE: | HISTONE H3 METHYLTRANSFERASE DIM-5 |
| 15:00 | 8xpt-A | 9.9 | 3 | 127 | 258 | 19 | MOLECULE: | HISTONE-LYSINE N-METHYLTRANSFERASE EHMT1 |
| 16:00 | 5jin-A | 9.9 | 3.1 | 128 | 274 | 17 | MOLECULE: | HISTONE-LYSINE N-METHYLTRANSFERASE EHMT2 |
| 17:00 | 5jj0-B | 9.9 | 2.9 | 126 | 268 | 17 | MOLECULE: | HISTONE-LYSINE N-METHYLTRANSFERASE EHMT2 |
| 18:00 | 5jhn-A | 9.9 | 3.2 | 130 | 274 | 17 | MOLECULE: | HISTONE-LYSINE N-METHYLTRANSFERASE EHMT2 |
| 19:00 | 5jiy-B | 9.9 | 2.9 | 125 | 268 | 17 | MOLECULE: | HISTONE-LYSINE N-METHYLTRANSFERASE EHMT2 |
| 20:00 | 5t0m-B | 9.9 | 3.1 | 128 | 272 | 17 | MOLECULE: | HISTONE-LYSINE N-METHYLTRANSFERASE EHMT2 |
| 21:00 | 5jj0-A | 9.9 | 3.1 | 128 | 274 | 17 | MOLECULE: | HISTONE-LYSINE N-METHYLTRANSFERASE EHMT2 |
| 22:00 | 5v9i-C | 9.9 | 2.6 | 123 | 265 | 16 | MOLECULE: | HISTONE-LYSINE N-METHYLTRANSFERASE EHMT2 |
| 23:00 | 6bp4-A | 9.9 | 2.6 | 125 | 269 | 15 | MOLECULE: | HISTONE-LYSINE N-METHYLTRANSFERASE, H3 LYSINE-9 S |
| 24:00:00 | 5ttf-B | 9.9 | 2.9 | 127 | 268 | 16 | MOLECULE: | HISTONE-LYSINE N-METHYLTRANSFERASE EHMT2 |
| 25:00:00 | 5v9i-A | 9.9 | 2.7 | 124 | 266 | 16 | MOLECULE: | HISTONE-LYSINE N-METHYLTRANSFERASE EHMT2 |
| 26:00:00 | 3k5k-A | 9.9 | 2.6 | 126 | 273 | 17 | MOLECULE: | HISTONE-LYSINE N-METHYLTRANSFERASE, H3 LYSINE-9 S |
| 27:00:00 | 5vse-B | 9.9 | 2.7 | 125 | 272 | 17 | MOLECULE: | HISTONE-LYSINE N-METHYLTRANSFERASE EHMT2 |
| 28:00:00 | 7btv-B | 9.9 | 2.9 | 127 | 273 | 17 | MOLECULE: | HISTONE-LYSINE N-METHYLTRANSFERASE EHMT2 |
| 29:00:00 | 2igq-A | 9.9 | 2.7 | 121 | 252 | 17 | MOLECULE: | EUCHROMATIC HISTONE METHYLTRANSFERASE 1 |
| 30:00:00 | 7t7m-D | 9.9 | 3.1 | 129 | 262 | 19 | MOLECULE: | HISTONE-LYSINE N-METHYLTRANSFERASE EHMT1 |
| 31:00:00 | 7t7l-D | 9.9 | 2.8 | 126 | 270 | 17 | MOLECULE: | HISTONE-LYSINE N-METHYLTRANSFERASE EHMT2 |
| 32:00:00 | 3kma-A | 9.8 | 2.3 | 94 | 105 | 17 | MOLECULE: | A612L PROTEIN |
| 33:00:00 | 6a5m-A | 9.8 | 2.8 | 127 | 489 | 17 | MOLECULE: | HISTONE-LYSINE N-METHYLTRANSFERASE, H3 LYSINE-9 S |
| 34:00:00 | 3rjw-A | 9.8 | 2.9 | 129 | 269 | 17 | MOLECULE: | HISTONE-LYSINE N-METHYLTRANSFERASE EHMT2 |
| 35:00:00 | 3rjw-B | 9.8 | 2.7 | 126 | 271 | 17 | MOLECULE: | HISTONE-LYSINE N-METHYLTRANSFERASE EHMT2 |
| 36:00:00 | 2o8j-C | 9.8 | 2.7 | 123 | 263 | 17 | MOLECULE: | HISTONE-LYSINE N-METHYLTRANSFERASE, H3 LYSINE-9 |
| 37:00:00 | 2o8j-A | 9.8 | 2.7 | 123 | 255 | 16 | MOLECULE: | HISTONE-LYSINE N-METHYLTRANSFERASE, H3 LYSINE-9 |
| 38:00:00 | 3kma-B | 9.8 | 2.3 | 94 | 105 | 17 | MOLECULE: | A612L PROTEIN |
| 39:00:00 | 4nvq-A | 9.8 | 2.8 | 124 | 268 | 17 | MOLECULE: | HISTONE-LYSINE N-METHYLTRANSFERASE EHMT2 |
| 40:00:00 | 1mvx-A | 9.8 | 2.7 | 126 | 269 | 15 | MOLECULE: | CRYPTIC LOCI REGULATOR 4 |
| 41:00:00 | 6a5k-A | 9.8 | 2.5 | 122 | 467 | 19 | MOLECULE: | HISTONE-LYSINE N-METHYLTRANSFERASE, H3 LYSINE-9 S |
| 42:00:00 | 1mvh-A | 9.8 | 2.8 | 127 | 269 | 15 | MOLECULE: | CRYPTIC LOCI REGULATOR 4 |
| 43:00:00 | 7buc-B | 9.8 | 2.6 | 125 | 270 | 17 | MOLECULE: | HISTONE-LYSINE N-METHYLTRANSFERASE EHMT2 |
| 44:00:00 | 1ml9-A | 9.7 | 2.5 | 122 | 250 | 18 | MOLECULE: | HISTONE H3 METHYLTRANSFERASE DIM-5 |
| 45:00:00 | 8swi-A | 9.7 | 2.9 | 115 | 452 | 19 | MOLECULE: | EUKARYOTIC HUNTINGTIN INTERACTING PROTEIN B |
| 46:00:00 | 6p0r-B | 9.7 | 2.5 | 118 | 230 | 19 | MOLECULE: | HISTONE-LYSINE N-METHYLTRANSFERASE SUV39H2 |
| 47:00:00 | 5ww0-B | 9.7 | 2.4 | 102 | 120 | 14 | MOLECULE: | SET DOMAIN-CONTAINING PROTEIN 7 |
| 48:00:00 | 6z2a-B | 9.7 | 2.6 | 127 | 277 | 14 | MOLECULE: | HISTONE-LYSINE N-METHYLTRANSFERASE, H3 LYSINE-9 S |
| 49:00:00 | 2o8j-D | 9.7 | 2.6 | 120 | 249 | 16 | MOLECULE: | HISTONE-LYSINE N-METHYLTRANSFERASE, H3 LYSINE-9 |
| 50:00:00 | 6nzo-S | 9.6 | 2.2 | 118 | 233 | 19 | MOLECULE: | HISTONE H3 |
| 51:00:00 | 6px3-S | 9.6 | 2.2 | 118 | 233 | 19 | MOLECULE: | HISTONE H3 |
| 52:00:00 | 6box-B | 9.6 | 2.6 | 124 | 268 | 16 | MOLECULE: | HISTONE-LYSINE N-METHYLTRANSFERASE, H3 LYSINE-9 S |
| 53:00:00 | 6bp4-B | 9.6 | 2.6 | 126 | 283 | 14 | MOLECULE: | HISTONE-LYSINE N-METHYLTRANSFERASE, H3 LYSINE-9 S |
| 54:00:00 | 6box-A | 9.6 | 2.8 | 124 | 271 | 15 | MOLECULE: | HISTONE-LYSINE N-METHYLTRANSFERASE, H3 LYSINE-9 S |
| 55:00:00 | 8xpt-B | 9.6 | 2.4 | 119 | 251 | 18 | MOLECULE: | HISTONE-LYSINE N-METHYLTRANSFERASE EHMT1 |

| | | | | | | | | |
|---|---|---|---|---|---|---|---|---|
| 56:00:00 | 5ls6-A | 9.5 | 2.4 | 117 | 570 | 20 | MOLECULE: | HISTONE-LYSINE N-METHYLTRANSFERASE EZH2,HISTONE-L |
| 57:00:00 | 5ls6-D | 9.5 | 2.4 | 117 | 570 | 20 | MOLECULE: | HISTONE-LYSINE N-METHYLTRANSFERASE EZH2,HISTONE-L |
| 58:00:00 | 7ea5-K | 9.5 | 2.2 | 118 | 229 | 16 | MOLECULE: | HISTONE H3 |
| 59:00:00 | 8xpt-D | 9.5 | 2.3 | 116 | 248 | 19 | MOLECULE: | HISTONE-LYSINE N-METHYLTRANSFERASE EHMT1 |
| 60:00:00 | 3vv0-A | 9.4 | 2.9 | 117 | 236 | 16 | MOLECULE: | HISTONE-LYSINE N-METHYLTRANSFERASE SETD7 |
| 61:00:00 | 3ray-A | 9.4 | 2.4 | 107 | 159 | 20 | MOLECULE: | PR DOMAIN-CONTAINING PROTEIN 11 |
| 62:00:00 | 6p0r-A | 9.3 | 3.4 | 124 | 254 | 19 | MOLECULE: | HISTONE-LYSINE N-METHYLTRANSFERASE SUV39H2 |
| 63:00:00 | 7kso-A | 9.2 | 2.1 | 116 | 395 | 19 | MOLECULE: | HISTONE-LYSINE N-METHYLTRANSFERASE EZH1 |
| 64:00:00 | 6bx3-E | 9.2 | 2.6 | 115 | 192 | 17 | MOLECULE: | HISTONE-LYSINE N-METHYLTRANSFERASE, H3 LYSINE-4 S |
| 65:00:00 | 7ksr-A | 9.2 | 2.3 | 116 | 395 | 19 | MOLECULE: | HISTONE-LYSINE N-METHYLTRANSFERASE EZH1 |
| 66:00:00 | 6xaz-B | 9.2 | 2.6 | 102 | 125 | 22 | MOLECULE: | PR DOMAIN ZINC FINGER PROTEIN 5 |
| 67:00:00 | 5ecj-A | 9.1 | 3.1 | 117 | 263 | 15 | MOLECULE: | PR DOMAIN ZINC FINGER PROTEIN 14,PROTEIN CBFA2T2 |
| 68:00:00 | 5ayf-A | 9.1 | 3 | 116 | 245 | 19 | MOLECULE: | HISTONE-LYSINE N-METHYLTRANSFERASE SETD7 |
| 69:00:00 | 7ktp-A | 9.1 | 2.1 | 116 | 395 | 19 | MOLECULE: | HISTONE-LYSINE N-METHYLTRANSFERASE EZH1 |
| 70:00:00 | 5ls6-G | 9.1 | 2.5 | 120 | 572 | 19 | MOLECULE: | HISTONE-LYSINE N-METHYLTRANSFERASE EZH2,HISTONE-L |
| 71:00:00 | 5arf-A | 9.1 | 2.7 | 111 | 426 | 11 | MOLECULE: | N-LYSINE METHYLTRANSFERASE SMYD2 |
| 72:00:00 | 4qen-A | 9 | 2.8 | 119 | 473 | 18 | MOLECULE: | HISTONE-LYSINE N-METHYLTRANSFERASE, H3 LYSINE-9 S |
| 73:00:00 | 5ls6-J | 9 | 2.4 | 117 | 570 | 20 | MOLECULE: | HISTONE-LYSINE N-METHYLTRANSFERASE EZH2,HISTONE-L |
| 74:00:00 | 4qep-A | 9 | 2.7 | 118 | 468 | 19 | MOLECULE: | HISTONE-LYSINE N-METHYLTRANSFERASE, H3 LYSINE-9 S |
| 75:00:00 | 4qeo-A | 8.9 | 2.8 | 118 | 473 | 19 | MOLECULE: | HISTONE-LYSINE N-METHYLTRANSFERASE, H3 LYSINE-9 S |
| 76:00:00 | 6xaz-A | 8.9 | 2.7 | 99 | 118 | 21 | MOLECULE: | PR DOMAIN ZINC FINGER PROTEIN 5 |
| 77:00:00 | 6mon-A | 8.9 | 2.6 | 108 | 430 | 12 | MOLECULE: | N-LYSINE METHYLTRANSFERASE SMYD2 |
| 78:00:00 | 7qa6-A | 8.9 | 2.6 | 111 | 427 | 14 | MOLECULE: | HISTONE-LYSINE N-METHYLTRANSFERASE SMYD3 |
| 79:00:00 | 6zrb-A | 8.8 | 2.4 | 109 | 426 | 16 | MOLECULE: | HISTONE-LYSINE N-METHYLTRANSFERASE SMYD3 |
| 80:00:00 | 7crr-I | 8.8 | 2.5 | 117 | 223 | 18 | MOLECULE: | HISTONE H3 |
| 81:00:00 | 6kiw-K | 8.7 | 2.6 | 116 | 156 | 20 | MOLECULE: | HISTONE H3 |
| 82:00:00 | 6xau-A | 8.7 | 2.7 | 100 | 119 | 14 | MOLECULE: | HISTONE-LYSINE N-METHYLTRANSFERASE MECOM |
| 83:00:00 | 2qpw-A | 8.7 | 3.1 | 105 | 147 | 16 | MOLECULE: | PR DOMAIN ZINC FINGER PROTEIN 2 |
| 84:00:00 | 4bup-A | 8.7 | 3.1 | 107 | 233 | 12 | MOLECULE: | HISTONE-LYSINE N-METHYLTRANSFERASE SUV420H1 |
| 85:00:00 | 3dal-B | 8.7 | 2.9 | 111 | 166 | 16 | MOLECULE: | PR DOMAIN ZINC FINGER PROTEIN 1 |
| 86:00:00 | 4au7-B | 8.6 | 2.8 | 99 | 209 | 10 | MOLECULE: | HISTONE-LYSINE N-METHYLTRANSFERASE SUV420H2 |
| 87:00:00 | 4bup-B | 8.6 | 3.3 | 107 | 238 | 12 | MOLECULE: | HISTONE-LYSINE N-METHYLTRANSFERASE SUV420H1 |
| 88:00:00 | 3ep0-B | 8.5 | 2.5 | 92 | 113 | 14 | MOLECULE: | PR DOMAIN ZINC FINGER PROTEIN 12 |
| 89:00:00 | 7ea8-L | 8.5 | 2.7 | 120 | 246 | 18 | MOLECULE: | HISTONE-LYSINE N-METHYLTRANSFERASE SETD2 |
| 90:00:00 | 7ud5-K | 8.5 | 2.9 | 119 | 176 | 18 | MOLECULE: | HISTONE H3 |
| 91:00:00 | 3ep0-A | 8.5 | 2.5 | 92 | 113 | 14 | MOLECULE: | PR DOMAIN ZINC FINGER PROTEIN 12 |
| 92:00:00 | 3ihx-C | 8.5 | 2.8 | 102 | 124 | 14 | MOLECULE: | PR DOMAIN ZINC FINGER PROTEIN 10 |
| 93:00:00 | 7crp-I | 8.5 | 2.5 | 116 | 223 | 18 | MOLECULE: | HISTONE H3 |
| 94:00:00 | 3db5-B | 8.4 | 2.4 | 98 | 134 | 12 | MOLECULE: | PR DOMAIN ZINC FINGER PROTEIN 4 |
| 95:00:00 | 3dal-A | 8.4 | 3 | 112 | 179 | 15 | MOLECULE: | PR DOMAIN ZINC FINGER PROTEIN 1 |
| 96:00:00 | 6nm4-B | 8.3 | 2.8 | 111 | 184 | 15 | MOLECULE: | HISTONE-LYSINE N-METHYLTRANSFERASE PRDM9 |
| 97:00:00 | 4nj5-A | 8.3 | 2.3 | 116 | 482 | 16 | MOLECULE: | PROBABLE HISTONE-LYSINE N-METHYLTRANSFERASE, H3 L |
| 98:00:00 | 4c1q-A | 8.2 | 3.1 | 115 | 174 | 13 | MOLECULE: | HISTONE-LYSINE N-METHYLTRANSFERASE PRDM9 |
| 99:00:00 | 5wg6-C | 8.2 | 4.8 | 122 | 504 | 19 | MOLECULE: | HISTONE-LYSINE N-METHYLTRANSFERASE EZH2,POLYCOMB |
| 0:00 | 3db5-A | 8.2 | 2.5 | 94 | 124 | 14 | MOLECULE: | PR DOMAIN ZINC FINGER PROTEIN 4 |
| 1:00 | 5kjn-A | 8.2 | 2.7 | 112 | 430 | 11 | MOLECULE: | N-LYSINE METHYLTRANSFERASE SMYD2 |
| 2:00 | 7crq-L | 8.2 | 2.7 | 118 | 223 | 18 | MOLECULE: | HISTONE H3 |
| 3:00 | 7cro-I | 8.1 | 2.6 | 118 | 220 | 20 | MOLECULE: | HISTONE H3 |
| 4:00 | 7crq-I | 8.1 | 2.6 | 117 | 223 | 18 | MOLECULE: | HISTONE H3 |
| 5:00 | 2jv0-A | 8 | 3.3 | 110 | 163 | 15 | MOLECULE: | PR DOMAIN ZINC FINGER PROTEIN 2 |
| 6:00 | 6nm4-A | 8 | 2.6 | 108 | 186 | 16 | MOLECULE: | HISTONE-LYSINE N-METHYLTRANSFERASE PRDM9 |

| | | | | | | | | |
|---|---|---|---|---|---|---|---|---|
| 7:00 | 3ihx-A | 8 | 2.7 | 99 | 126 | 14 | MOLECULE: | PR DOMAIN ZINC FINGER PROTEIN 10 |
| 8:00 | 7yrg-L | 8 | 3.3 | 110 | 241 | 11 | MOLECULE: | HISTONE H3.2 |
| 9:00 | 5ecj-B | 7.9 | 3.3 | 116 | 263 | 15 | MOLECULE: | PR DOMAIN ZINC FINGER PROTEIN 14,PROTEIN CBFA2T2 |
| 10:00 | 5hq8-A | 7.9 | 2.4 | 110 | 427 | 15 | MOLECULE: | HISTONE-LYSINE N-METHYLTRANSFERASE SMYD3 |
| 11:00 | 4ijd-A | 7.8 | 2.7 | 108 | 215 | 17 | MOLECULE: | HISTONE-LYSINE N-METHYLTRANSFERASE PRDM9 |
| 12:00 | 4ijd-B | 7.8 | 2.8 | 109 | 215 | 16 | MOLECULE: | HISTONE-LYSINE N-METHYLTRANSFERASE PRDM9 |
| 13:00 | 3s8p-B | 7.8 | 3.1 | 107 | 239 | 12 | MOLECULE: | HISTONE-LYSINE N-METHYLTRANSFERASE SUV420H1 |
| 14:00 | 5hyn-K | 7.8 | 2.7 | 121 | 563 | 20 | MOLECULE: | HISTONE-LYSINE N-METHYLTRANSFERASE EZH2 |
| 15:00 | 5cpr-B | 7.8 | 3.5 | 107 | 233 | 10 | MOLECULE: | HISTONE-LYSINE N-METHYLTRANSFERASE SUV420H1 |
| 16:00 | 3s8p-A | 7.7 | 3.1 | 107 | 234 | 12 | MOLECULE: | HISTONE-LYSINE N-METHYLTRANSFERASE SUV420H1 |
| 17:00 | 3ihx-B | 7.7 | 2.8 | 100 | 129 | 14 | MOLECULE: | PR DOMAIN ZINC FINGER PROTEIN 10 |
| 18:00 | 2g46-A | 7.6 | 3.4 | 99 | 120 | 16 | MOLECULE: | PBCV-1 HISTONE H3-LYS 27 METHYLTRANSFERASE |
| 19:00 | 2g46-B | 7.6 | 3.1 | 99 | 120 | 16 | MOLECULE: | PBCV-1 HISTONE H3-LYS 27 METHYLTRANSFERASE |
| 20:00 | 5hi7-A | 7.6 | 2.6 | 111 | 426 | 14 | MOLECULE: | HISTONE-LYSINE N-METHYLTRANSFERASE SMYD3 |
| 21:00 | 7yrg-K | 7.6 | 3.1 | 108 | 260 | 11 | MOLECULE: | HISTONE H3.2 |
| 22:00 | 5kjk-A | 7.6 | 2.6 | 108 | 430 | 12 | MOLECULE: | N-LYSINE METHYLTRANSFERASE SMYD2 |
| 23:00 | 5wbv-B | 7.6 | 3.4 | 109 | 228 | 12 | MOLECULE: | HISTONE-LYSINE N-METHYLTRANSFERASE KMT5B |
| 24:00:00 | 2h2e-C | 7.5 | 3.3 | 109 | 440 | 13 | MOLECULE: | RIBULOSE-1,5 BISPHOSPHATE CARBOXYLASE/OXYGENASE L |
| 25:00:00 | 7qnr-A | 7.5 | 2.5 | 110 | 427 | 15 | MOLECULE: | HISTONE-LYSINE N-METHYLTRANSFERASE SMYD3 |
| 26:00:00 | 5arg-A | 7.4 | 2.5 | 109 | 427 | 12 | MOLECULE: | N-LYSINE METHYLTRANSFERASE SMYD2 |
| 27:00:00 | 5kjm-A | 7.4 | 2.7 | 109 | 430 | 13 | MOLECULE: | N-LYSINE METHYLTRANSFERASE SMYD2 |
| 28:00:00 | 6cby-A | 7.4 | 2.9 | 109 | 428 | 11 | MOLECULE: | N-LYSINE METHYLTRANSFERASE SMYD2 |
| 29:00:00 | 3ihx-D | 7.3 | 2.5 | 90 | 116 | 12 | MOLECULE: | PR DOMAIN ZINC FINGER PROTEIN 10 |
| 30:00:00 | 3ru0-B | 7.3 | 2.4 | 110 | 420 | 15 | MOLECULE: | SET AND MYND DOMAIN-CONTAINING PROTEIN 3 |
| 31:00:00 | 5v3h-A | 7.3 | 2.8 | 114 | 439 | 11 | MOLECULE: | N-LYSINE METHYLTRANSFERASE SMYD2 |
| 32:00:00 | 6mon-B | 7.3 | 2.8 | 109 | 430 | 13 | MOLECULE: | N-LYSINE METHYLTRANSFERASE SMYD2 |
| 33:00:00 | 5xxj-A | 7.3 | 2.6 | 110 | 424 | 15 | MOLECULE: | SMYD3 |
| 34:00:00 | 6cby-B | 7.3 | 2.7 | 110 | 422 | 11 | MOLECULE: | N-LYSINE METHYLTRANSFERASE SMYD2 |
| 35:00:00 | 6n3g-A | 7.3 | 2.9 | 112 | 431 | 13 | MOLECULE: | N-LYSINE METHYLTRANSFERASE SMYD2 |
| 36:00:00 | 5ccl-A | 7.1 | 2.5 | 109 | 426 | 16 | MOLECULE: | HISTONE-LYSINE N-METHYLTRANSFERASE SMYD3 |
| 37:00:00 | 7o2c-A | 7.1 | 2.9 | 110 | 428 | 14 | MOLECULE: | HISTONE-LYSINE N-METHYLTRANSFERASE SMYD3 |
| 38:00:00 | 3qwp-A | 7.1 | 2.4 | 109 | 426 | 16 | MOLECULE: | SET AND MYND DOMAIN-CONTAINING PROTEIN 3 |
| 39:00:00 | 5xxd-A | 7.1 | 2.6 | 110 | 424 | 15 | MOLECULE: | SMYD3 METHYLTRANSFERASE |
| 40:00:00 | 2h21-B | 7 | 3.6 | 112 | 441 | 13 | MOLECULE: | RIBULOSE-1,5 BISPHOSPHATE CARBOXYLASE/OXYGENASE |
| 41:00:00 | 7o2b-A | 7 | 2.7 | 110 | 428 | 15 | MOLECULE: | HISTONE-LYSINE N-METHYLTRANSFERASE SMYD3 |
| 42:00:00 | 6paf-A | 7 | 2.7 | 111 | 424 | 14 | MOLECULE: | HISTONE-LYSINE N-METHYLTRANSFERASE SMYD3 |
| 43:00:00 | 3oxf-B | 7 | 2.4 | 110 | 426 | 15 | MOLECULE: | SET AND MYND DOMAIN-CONTAINING PROTEIN 3 |
| 44:00:00 | 5wcg-A | 7 | 2.7 | 110 | 430 | 11 | MOLECULE: | N-LYSINE METHYLTRANSFERASE SMYD2 |
| 45:00:00 | 5yjo-A | 7 | 2.6 | 111 | 424 | 14 | MOLECULE: | HISTONE-LYSINE N-METHYLTRANSFERASE SMYD3 |
| 46:00:00 | 6z2r-A | 6.9 | 2.4 | 110 | 427 | 15 | MOLECULE: | HISTONE-LYSINE N-METHYLTRANSFERASE SMYD3 |
| 47:00:00 | 6p6k-A | 6.9 | 2.4 | 109 | 425 | 16 | MOLECULE: | HISTONE-LYSINE N-METHYLTRANSFERASE SMYD3 |
| 48:00:00 | 6o9o-A | 6.9 | 2.4 | 109 | 427 | 16 | MOLECULE: | HISTONE-LYSINE N-METHYLTRANSFERASE SMYD3 |
| 49:00:00 | 6p6g-A | 6.9 | 2.4 | 109 | 427 | 16 | MOLECULE: | HISTONE-LYSINE N-METHYLTRANSFERASE SMYD3 |
| 50:00:00 | 3oxf-A | 6.9 | 2.9 | 111 | 427 | 14 | MOLECULE: | SET AND MYND DOMAIN-CONTAINING PROTEIN 3 |
| 51:00:00 | 7qnu-A | 6.9 | 2.6 | 111 | 427 | 14 | MOLECULE: | HISTONE-LYSINE N-METHYLTRANSFERASE SMYD3 |
| 52:00:00 | 4au7-A | 6.8 | 3 | 109 | 244 | 11 | MOLECULE: | HISTONE-LYSINE N-METHYLTRANSFERASE SUV420H2 |
| 53:00:00 | 1mlv-B | 6.8 | 3.3 | 108 | 441 | 13 | MOLECULE: | RIBULOSE-1,5 BIPHOSPHATE CARBOXYLASE/OXYGENASE LA |
| 54:00:00 | 5v37-A | 6.8 | 2.6 | 109 | 425 | 15 | MOLECULE: | HISTONE-LYSINE N-METHYLTRANSFERASE SMYD3 |
| 55:00:00 | 6cbx-B | 6.8 | 3.2 | 112 | 433 | 11 | MOLECULE: | N-LYSINE METHYLTRANSFERASE SMYD2 |
| 56:00:00 | 3rq4-A | 6.7 | 3 | 108 | 241 | 11 | MOLECULE: | HISTONE-LYSINE N-METHYLTRANSFERASE SUV420H2 |
| 57:00:00 | 6p7z-A | 6.7 | 2.4 | 109 | 426 | 16 | MOLECULE: | HISTONE-LYSINE N-METHYLTRANSFERASE SMYD3 |

| | | | | | | | |
|---|---|---|---|---|---|---|---|
| 58:00:00 | 5wbv-A | 6.7 | 3.2 | 108 | 234 | 12 | MOLECULE: HISTONE-LYSINE N-METHYLTRANSFERASE KMT5B |
| 59:00:00 | 3n71-A | 6.6 | 2.9 | 118 | 469 | 16 | MOLECULE: HISTONE LYSINE METHYLTRANSFERASE SMYD1 |
| 60:00:00 | 2h21-A | 6.6 | 3.6 | 102 | 425 | 14 | MOLECULE: RIBULOSE-1,5 BISPHOSPHATE CARBOXYLASE/OXYGENASE |
| 61:00:00 | 1ozv-B | 6.6 | 3.8 | 112 | 442 | 13 | MOLECULE: RIBULOSE-1,5 BISPHOSPHATE CARBOXYLASE/OXYGENASE L |
| 62:00:00 | 1mlv-A | 6.6 | 3.5 | 103 | 425 | 15 | MOLECULE: RIBULOSE-1,5 BIPHOSPHATE CARBOXYLASE/OXYGENASE LA |
| 63:00:00 | 3s7d-A | 6.6 | 2.9 | 110 | 430 | 13 | MOLECULE: N-LYSINE METHYLTRANSFERASE SMYD2 |
| 64:00:00 | 4ynd-A | 6.6 | 2.9 | 112 | 429 | 11 | MOLECULE: N-LYSINE METHYLTRANSFERASE SMYD2 |
| 65:00:00 | 2h23-C | 6.5 | 3.3 | 108 | 440 | 13 | MOLECULE: RIBULOSE-1,5 BISPHOSPHATE CARBOXYLASE/OXYGENASE L |
| 66:00:00 | 4o6f-A | 6.5 | 2.9 | 111 | 431 | 13 | MOLECULE: N-LYSINE METHYLTRANSFERASE SMYD2 |
| 67:00:00 | 3qwv-A | 6.5 | 2.5 | 109 | 431 | 9 | MOLECULE: SET AND MYND DOMAIN-CONTAINING PROTEIN 2 |
| 68:00:00 | 8t9f-K | 6.4 | 3 | 106 | 265 | 11 | MOLECULE: HISTONE-LYSINE N-METHYLTRANSFERASE KMT5B |
| 69:00:00 | 2h2j-A | 6.4 | 3.4 | 103 | 425 | 14 | MOLECULE: RIBULOSE-1,5 BISPHOSPHATE CARBOXYLASE/OXYGENASE L |
| 70:00:00 | 5ccm-A | 6.4 | 2.7 | 110 | 424 | 15 | MOLECULE: HISTONE-LYSINE N-METHYLTRANSFERASE SMYD3 |
| 71:00:00 | 5hq8-B | 6.4 | 2.4 | 110 | 426 | 15 | MOLECULE: HISTONE-LYSINE N-METHYLTRANSFERASE SMYD3 |
| 72:00:00 | 3s7f-A | 6.4 | 2.6 | 109 | 430 | 11 | MOLECULE: N-LYSINE METHYLTRANSFERASE SMYD2 |
| 73:00:00 | 3s7j-A | 6.4 | 2.9 | 110 | 429 | 11 | MOLECULE: N-LYSINE METHYLTRANSFERASE SMYD2 |
| 74:00:00 | 5kjl-A | 6.4 | 2.8 | 109 | 429 | 13 | MOLECULE: N-LYSINE METHYLTRANSFERASE SMYD2 |
| 75:00:00 | 6cbx-A | 6.4 | 2.9 | 109 | 428 | 11 | MOLECULE: N-LYSINE METHYLTRANSFERASE SMYD2 |
| 76:00:00 | 6ijl-A | 6.4 | 2.6 | 111 | 424 | 14 | MOLECULE: HISTONE-LYSINE N-METHYLTRANSFERASE SMYD3 |
| 77:00:00 | 7yrd-K | 6.3 | 3.1 | 108 | 260 | 11 | MOLECULE: HISTONE H3.2 |
| 78:00:00 | 6fnd-C | 6.3 | 2.9 | 98 | 420 | 13 | MOLECULE: APICAL COMPLEX LYSINE METHYLTRANSFERASE |
| 79:00:00 | 8t9h-K | 6.3 | 3.1 | 109 | 249 | 10 | MOLECULE: HISTONE H3.2 |
| 80:00:00 | 3s7b-A | 6.3 | 2.6 | 109 | 430 | 11 | MOLECULE: N-LYSINE METHYLTRANSFERASE SMYD2 |
| 81:00:00 | 6yuh-A | 6.3 | 2.5 | 109 | 426 | 15 | MOLECULE: HISTONE-LYSINE N-METHYLTRANSFERASE SMYD3 |
| 82:00:00 | 6fnd-B | 6.2 | 3.1 | 99 | 418 | 13 | MOLECULE: APICAL COMPLEX LYSINE METHYLTRANSFERASE |
| 83:00:00 | 2h23-B | 6.2 | 3.4 | 111 | 442 | 14 | MOLECULE: RIBULOSE-1,5 BISPHOSPHATE CARBOXYLASE/OXYGENASE L |
| 84:00:00 | 2h2e-B | 6.2 | 3.4 | 110 | 442 | 13 | MOLECULE: RIBULOSE-1,5 BISPHOSPHATE CARBOXYLASE/OXYGENASE L |
| 85:00:00 | 5ex0-A | 6.2 | 2.6 | 114 | 430 | 15 | MOLECULE: HISTONE-LYSINE N-METHYLTRANSFERASE SMYD3 |
| 86:00:00 | 1ozv-C | 6.2 | 3.3 | 110 | 441 | 15 | MOLECULE: RIBULOSE-1,5 BISPHOSPHATE CARBOXYLASE/OXYGENASE L |
| 87:00:00 | 7qlb-A | 6.2 | 2.5 | 110 | 427 | 15 | MOLECULE: HISTONE-LYSINE N-METHYLTRANSFERASE SMYD3 |
| 88:00:00 | 6fnd-A | 6 | 2.9 | 98 | 423 | 13 | MOLECULE: APICAL COMPLEX LYSINE METHYLTRANSFERASE |
| 89:00:00 | 6fnd-D | 6 | 2.9 | 101 | 422 | 12 | MOLECULE: APICAL COMPLEX LYSINE METHYLTRANSFERASE |
| 90:00:00 | 1mlv-C | 6 | 3.1 | 108 | 439 | 14 | MOLECULE: RIBULOSE-1,5 BIPHOSPHATE CARBOXYLASE/OXYGENASE LA |
| 91:00:00 | 2h2j-C | 6 | 3.4 | 109 | 440 | 13 | MOLECULE: RIBULOSE-1,5 BISPHOSPHATE CARBOXYLASE/OXYGENASE L |
| 92:00:00 | 2h2j-B | 6 | 3.9 | 114 | 441 | 13 | MOLECULE: RIBULOSE-1,5 BISPHOSPHATE CARBOXYLASE/OXYGENASE L |
| 93:00:00 | 5ex3-A | 6 | 2.6 | 113 | 429 | 15 | MOLECULE: HISTONE-LYSINE N-METHYLTRANSFERASE SMYD3 |
| 94:00:00 | 1ozv-A | 5.9 | 3.1 | 103 | 433 | 15 | MOLECULE: RIBULOSE-1,5 BISPHOSPHATE CARBOXYLASE/OXYGENASE L |
| 95:00:00 | 3ru0-A | 5.9 | 2.4 | 109 | 426 | 16 | MOLECULE: SET AND MYND DOMAIN-CONTAINING PROTEIN 3 |
| 96:00:00 | 3qxy-A | 5.9 | 3.4 | 112 | 421 | 11 | MOLECULE: N-LYSINE METHYLTRANSFERASE SETD6 |
| 97:00:00 | 3rc0-A | 5.9 | 3.4 | 112 | 421 | 11 | MOLECULE: N-LYSINE METHYLTRANSFERASE SETD6 |
| 98:00:00 | 7o2a-A | 5.9 | 2.5 | 110 | 428 | 15 | MOLECULE: HISTONE-LYSINE N-METHYLTRANSFERASE SMYD3 |
| 99:00:00 | 3qww-A | 5.9 | 2.6 | 110 | 431 | 9 | MOLECULE: SET AND MYND DOMAIN-CONTAINING PROTEIN 2 |
| 0:00 | 8owo-A | 5.9 | 2.5 | 110 | 427 | 15 | MOLECULE: HISTONE-LYSINE N-METHYLTRANSFERASE SMYD3 |
| 1:00 | 3tg5-A | 5.8 | 2.5 | 110 | 426 | 12 | MOLECULE: N-LYSINE METHYLTRANSFERASE SMYD2 |
| 2:00 | 3oxl-A | 5.8 | 2.4 | 110 | 427 | 15 | MOLECULE: SET AND MYND DOMAIN-CONTAINING PROTEIN 3 |
| 3:00 | 3tg4-A | 5.8 | 2.6 | 111 | 428 | 12 | MOLECULE: N-LYSINE METHYLTRANSFERASE SMYD2 |
| 4:00 | 3qxy-B | 5.8 | 3.2 | 107 | 415 | 10 | MOLECULE: N-LYSINE METHYLTRANSFERASE SETD6 |
| 5:00 | 5xxg-A | 5.8 | 2.6 | 110 | 424 | 15 | MOLECULE: SMYD3 |
| 6:00 | 7bj1-A | 5.8 | 2.7 | 110 | 427 | 15 | MOLECULE: HISTONE-LYSINE N-METHYLTRANSFERASE SMYD3 |
| 7:00 | 3pdn-A | 5.8 | 2.6 | 109 | 428 | 15 | MOLECULE: SET AND MYND DOMAIN-CONTAINING PROTEIN 3 |
| 8:00 | 3mek-A | 5.7 | 2.7 | 109 | 424 | 14 | MOLECULE: SET AND MYND DOMAIN-CONTAINING PROTEIN 3 |

| | | | | | | | | |
|---|---|---|---|---|---|---|---|---|
| 9:00 | 4wuy-A | 5.7 | 2.8 | 108 | 415 | 11 | MOLECULE: | N-LYSINE METHYLTRANSFERASE SMYD2 |
| 10:00 | 2h23-A | 5.7 | 3.1 | 100 | 426 | 14 | MOLECULE: | RIBULOSE-1,5 BISPHOSPHATE CARBOXYLASE/OXYGENASE L |
| 11:00 | 2h21-C | 5.7 | 3.4 | 109 | 439 | 13 | MOLECULE: | RIBULOSE-1,5 BISPHOSPHATE CARBOXYLASE/OXYGENASE |
| 12:00 | 3rc0-B | 5.7 | 3.1 | 108 | 415 | 10 | MOLECULE: | N-LYSINE METHYLTRANSFERASE SETD6 |
| 13:00 | 3oxg-A | 5.6 | 2.7 | 110 | 425 | 15 | MOLECULE: | SET AND MYND DOMAIN-CONTAINING PROTEIN 3 |
| 14:00 | 6ox1-B | 5.5 | 3.7 | 112 | 485 | 12 | MOLECULE: | ACTIN, CYTOPLASMIC 1 |
| 15:00 | 3rib-B | 5.5 | 2.6 | 108 | 419 | 11 | MOLECULE: | N-LYSINE METHYLTRANSFERASE SMYD2 |
| 16:00 | 3rib-A | 5.4 | 3 | 109 | 431 | 11 | MOLECULE: | N-LYSINE METHYLTRANSFERASE SMYD2 |
| 17:00 | 2h2e-A | 5.4 | 3.2 | 100 | 426 | 13 | MOLECULE: | RIBULOSE-1,5 BISPHOSPHATE CARBOXYLASE/OXYGENASE L |
| 18:00 | 3smt-A | 5.4 | 3.9 | 115 | 466 | 10 | MOLECULE: | HISTONE-LYSINE N-METHYLTRANSFERASE SETD3 |
| 19:00 | 1p0y-B | 5.3 | 3.5 | 111 | 442 | 14 | MOLECULE: | RIBULOSE-1,5 BISPHOSPHATE CARBOXYLASE/OXYGENASE |
| 20:00 | 1p0y-A | 5 | 3.7 | 107 | 431 | 14 | MOLECULE: | RIBULOSE-1,5 BISPHOSPHATE CARBOXYLASE/OXYGENASE |
| 21:00 | 8jhf-K | 5 | 3.3 | 104 | 257 | 11 | MOLECULE: | HISTONE-LYSINE N-METHYLTRANSFERASE KMT5B |
| 22:00 | 2mpl-A | 4.2 | 3.9 | 77 | 127 | 8 | MOLECULE: | ZINC FINGER PROTEIN ZFPM1 |
| 23:00 | 2n1i-A | 3.8 | 3.2 | 87 | 176 | 9 | MOLECULE: | PR DOMAIN ZINC FINGER PROTEIN 16 |
| 24:00:00 | 1n3j-B | 3.7 | 3.8 | 88 | 119 | 16 | MOLECULE: | HISTONE H3 LYSINE METHYLTRANSFERASE |
| 25:00:00 | 1n3j-A | 3.5 | 3.8 | 86 | 119 | 16 | MOLECULE: | HISTONE H3 LYSINE METHYLTRANSFERASE |
