## Supplementary material for "Novel Adomaviruses Associated with Blotchy Bass Syndrome in Black Basses (*Micropterus spp.)*": S1 Text

***Fish collection***

Smallmouth bass (*Micropterus dolomieu*)

Smallmouth bass observed in Lake St. Clair were collected by trap net between 2002 and 2015, and a combination of trap net or electrofishing in subsequent years. These fish were all collected as part of annual efforts by the Michigan Department of Natural Resources to assess size-structure. During 2021 one or 2 scales were removed with sterilized forceps from a subsample of fish with hyperpigmented melanistic lesions (HPMLs; S2 Table). Samples were stabilized in RNA*Later* (Thermo Fisher Scientific, Waltham, MA). Only HMPLs were sampled from these fish to confirm the presence of virus. When possible, melanistic lesions on fin margins were sampled as an alternative to scales. During 2022 non-lethal sampling included the use of swabs and collection tubes containing DNA/ RNA Shield (Zymo Research, Irvine, CA). The HPMLs or normal skin were sampled by rubbing the swab back and forth with gentle, yet firm pressure for 10-15 seconds. The swab head was then rotated such that the entire surface contacted the sample area. Transfer of melanin to the swab confirmed tissue transfer in the case of HPMLs. Swabs were then inserted in the collection tube for subsequent nucleic acid extraction. A subset bass with clinical blotchy bass syndrome and an equal number of clinically normal fish were samples for this effort to evaluate if this virus was simply part of the normal mucosal microbiome.

Smallmouth bass sampled from all other locations were collected via electroshocking. Fish sampled in 2017 and 2019 were collected as part of other ongoing projects and included skin scrapings or scales within HPMLs and stabilized in RNALater [1, 2]. Bass samples in 2022 were collected using swabs and collection tubes containing DNA/ RNA Shield as described above. HPMLs and clinically normal skin from clinically affected fish as well as paired samples from clinically normal fish were collected from the Susquehanna River, PA site (S1 Table).

Largemouth bass (*Micropterus nigricans*)

Largemouth bass were sampled from environmental settings or aquatic exhibits at Bass Pro Shops and Cabela’s. All wild-caught bass were collected via boat DC electrofishing. Largemouth bass from VA during 2021 were sampled by the Virginia Department of Wildlife Resources as part of a mark-recapture survey which also included recording the observation of HPMLs. Scales or fin clips were collected from a subset of these and stabilized in RNAlater. Samples from bass samples in 2022 were collected using swabs and collection tubes containing DNA/ RNA Shield as described above. HPMLs and clinically normal skin from clinically affected bass as well as paired samples from clinically normal fish were collected from the bass in VA using the same experimental design as that for smallmouth bass described above (S1 Table). Largemouth bass expressing clinical signs of blotchy bass syndrome that were maintained in Bass Pro Shops and Cabela’s exhibits across the United States were non-lethally sampled using proprietary in-house veterinarian-approved protocols. HPMLs were sampled using swabs and collection tubes containing DNA/ RNA Shield as described above and fish were returned to the live exhibits. The origin of these fish was often unknown, but in many cases were collected from local waterbodies.

***RNA extraction and sequencing***

Total RNA was extracted from RNA stabilized skin tissues from HPMLs on smallmouth bass collected in 2017 using the E.Z.N.A. Total RNA Kit I (Omega Bio-tek) according to the manufacturer protocol. Due to the limited tissue available from the HPMLs, tandem DNA extractions were not possible. RNA integrity was determined using an Agilent RNA 6000 Nano Kit on the Agillent 2100 Bioanalyzer (Agilent Technologies, Santa Clara, CA). Total RNA (RIN>6) from five melanistic lesions was the Penn State Genomics Core Facility, University Park, PA for massively parallel sequencing. Sequencing libraries were prepared using the TrueSeq Stranded mRNA Library Preparation Kit (Illumina, San Diego, CA), and was carried out as 150 nt single end reads in duplicate per sample on an Illumina HiSeq2500 platform (Illumina, San Diego, CA).

RNA was extracted from DNA/ RNA Shield preserved swab samples of HPMLs and normal skin sampled from largemouth bass and smallmouth bass sampled in 2022 (S1 Table). Extraction was conducted using the *Quick*-DNA/RNA Microprep Plus Kits (Zymo Research) following manufacturer protocols. Samples were diluted to 15 ng/ μl and shipped to the Oklahoma Medical Research Foundation NGS Core for library construction and sequencing. Libraries were constructed with and IDT xGen RNA Library Kit and preparation included polyA-selection. Libraries were sequenced on an Illumina NovaSeq – S4 – PE 150cycle. Runs yielded ~20M in each direction.

Bioinformatic assembly of MdA-1 from RNA utilized the Iterative Virus Assembler [3].

***DNA extraction and sequencing***

DNA extractions differed given the sample collection type. Sample from RNAlater stabilized tissues were extracted using a DNeasy Blood and Tissue Kit (Qiagen, CA, USA), following manufacturer instructions for Purification of Total DNA from Animal Tissues (Spin-Column Protocol; DNeasy Blood & Tissue Handbook version 07/2020). As scales were dentinous, collected tissues were consumptively digested overnight (~10 hours) in 180µl of lysis buffer and 20µl proteinase K (600 mAU/ml) prior to proceeding with the specified protocol. Extracted DNA was quantified using a Qubit dsDNA HS Assay Kit and a Qubit 4.0 Fluorometer (Invitrogen, CA, USA). Swab samples preserved in DNA/ RNA Shield were extracted using *Quick*-DNA/RNA Microprep Plus Kits (Zymo Research) following manufacturer protocols.

***Random Circle Amplification***

All DNA samples used for genome sequencing were pre-amplified using random circle amplification. Samples collected prior to 2022 utilized a rolling circle amplification (RCA) method utilizing the EquiPhi29 polymerase. In short, template DNA was added to EquiPhi29 buffer (ThermoFisher Scientific) containing 20 µM of Exo-Resistant Random Primers (ThermoFisher Scientific). Template was denatured by heating to 95°C for 3min. The reaction was then cooled in a stepwise manner (50°C for 1 min, 30°C for 1 min and 4°C for 30s). To this an equal volume of amplification cycling mix was added such that the final reaction included 10 µM of exo-resistant random primers, 2 ng/ml of BSA, 15 mM of dNTPs and 2 U/ul of EquiPhi29. The reaction was incubated at 30°C for 18h, 65°C for 10 min and held at 4°C forever. The resulting product was purified using a 3x bead clean up. The purified DNA was used as template for sequencing on the Illumina MiSeq platform.

For samples collected during 2022 or later RCA was conducted using an Illustra TempliPhi 100 Amplification Kit (Cytiva), in accordance with manufacturer instructions specified for M13 Phage DNA, as it was the nearest DNA virus analog. The process was initiated by combining 0.5 µl template DNA (10ng/µl) with the included sample buffer. Using a thermocycler, the mix was denatured for 3 minutes at 95°C and immediately cooled to 4°C. A TempliPhi™ reaction master mix was made by combining 5μl of reaction buffer and 0.2μl enzyme mix, with a total of 5μl of the cocktail added to the denatured samples upon cooling. The combined mixture was then incubated on a thermocycler at 30°C for 18 hours, followed by heat-inactivating the enzyme at 65°C for 10 minutes. The resultant RCA product was diluted with 40µl of nuclease free water before preparation for on the Illumina MiSeq platform sequencing.

***Nextera XT and MiSeq prep***

The diluted amplification products from the RCA were quantified using a Qubit dsDNA HS Assay Kit and a Qubit 4.0 Fluorometer. The product was normalized to 0.2 ng/µL using nuclease free water. A total of 1ng (5µL) of normalized product was then used as starting material for next generation library preparation. An Illumina Nextera XT Library Preparation Kit (Illumina, CA, USA) was used in accordance with Nextera XT Library Preparation Reference Guide (Doc # 15031942 ver. 5) for MiSeq preparation. The final library was normalized with Illumina’s Bead-Based Normalization (BBN) method and pooled as described in the BBN Loading Concentrations Exceptions Table 2 (MiSeq System Denature and Dilute Libraries Guide (Doc # 15039740 ver. 10). Pooled libraries were then sequenced for 2 x 301 cycles and loaded with a 15% PhiX (12.5pM) spike.

Assembly of short reads was conducted in CLC Genomic Workbench (v.23.0.3).

***Oxford Nanopore (ONT) sequencing of MnA-1***

The Col ORF of MnA-1 includes long repetitive repeats. We were concerned that that even the 2 x 301 sequence reads yielded by the MiSeq may not be sufficient to accurately sequence this region of the genome. We used the MnA-1 draft genome as template to design long-range PCR primers to amplify the complete Col ORF. 1. In short, 50µL lrPCR reactions were conducted using the TaKaRa LA PCR™ Kit Ver.2.1 (Takara Bio Incorporation, Kusatsu, Shiga, Japan). Reactions included 5µL of template, 1µL of 10µM forward primer (MnA-1F_2600; TTGTCACTCAACCAGATCCGG), 1µL of 10µM reverse primer (MnA-1R_2600; GTCGTTCTTTTGTGCGCACT), and 29.5µL of nuclease free water, and 8µL of dNTP mixture, 5µL of 10X LA PCR Buffer II (Mg2+ plus), and 0.5µL of TaKaRa LA Taq DNA polymerase. Estimated amplicon size was 2600 bp. Viral DNA was amplified via long-rang PCR (TaKaRa LA PCR™ Kit Ver.2.1; Takara Bio Incorporation). PCR product was purified using a QIAquick PCR Purification Kit and the purified amplicon concentration was normalized to 15uM with nuclease free water and shipped to Plasmidsaurus (Eugene, OR, USA) for ONT sequencing.
